# Comparative pathogenesis of two lineages of Powassan virus reveals distinct clinical outcome, neuropathology, and inflammation

**DOI:** 10.1101/2023.08.01.551588

**Authors:** Erin S. Reynolds, Charles E. Hart, Jacob T. Nelson, Brandon J. Marzullo, Allen T. Esterly, Dakota N. Paine, Jessica Crooker, Paul T. Massa, Saravanan Thangamani

**Affiliations:** Department of Microbiology and Immunology, SUNY Upstate Medical University, Syracuse, New York, USA; SUNY Center for Vector-Borne Diseases, SUNY Upstate Medical University, Syracuse, New York, USA; Institute for Global Health and Translational Science, SUNY Upstate Medical University, Syracuse, New York, USA; Molecular Biology Section, Laboratory Sciences Directorate, Defense Centers for Public Health Aberdeen, Aberdeen Proving Grounds, MD, USA; Lovelace Biomedical Research Institute, Albuquerque, New Mexico, USA; Department of Biochemistry, Jacobs School of Medicine and Biomedical Sciences, SUNY Buffalo, Buffalo, New York, USA; Genomics and Bioinformatics Core, New York State Center of Excellence Bioinformatics & Life Sciences, SUNY Buffalo, Buffalo, New York, USA; Section of Infectious Diseases, Department of Internal Medicine, Yale School of Medicine, New Haven, CT, USA; Department of Neurology, SUNY Upstate Medical University, Syracuse, New York, USA

## Abstract

Tick-borne flaviviruses (TBFV) can cause severe neuroinvasive disease which may result in death or long-term neurological deficit in over 50% of survivors. Multiple mechanisms for invasion of the central nervous system (CNS) by flaviviruses have been proposed including axonal transport, transcytosis, endothelial infection, and Trojan horse routes. Flaviviruses may utilize different or multiple mechanisms of neuroinvasion depending on the specific virus, infection site, and host variability. In this work we have shown that infection of BALB/cJ mice with either Powassan virus lineage I (Powassan virus) or lineage II (deer tick virus) results in distinct spatial tropism of infection in the CNS which correlated with unique clinical presentation for each lineage. Comparative transcriptomics of infected brains demonstrates activation of different immune pathways and downstream host responses. Ultimately the comparative pathology and transcriptomics are congruent with different clinical signs in a murine model. These results suggest that different disease presentations would be occur in clinical cases due to the innate differences in the two lineages of Powassan virus.

**Author Summary:** Powassan virus causes a nationally notifiable disease which can cause severe neurological disease in humans and has no approved vaccines or therapeutics. Although two distinct lineages circulate in North America, clinical differentiation is not typically performed, and pathology has been assumed to be similar between lineages. In this work, a direct comparison of lineage I (Powassan virus) and lineage II (deer tick virus) demonstrated distinct differences in the clinical presentation, pathology of the central nervous system, and immune response in immunocompetent mice. These differences suggest that deer tick virus and Powassan virus do not utilize the same mechanisms for neuroinvasion and dissemination within the CNS. This is clinically relevant as the development of treatment plans and therapeutics need to be evaluated for these virus lineages.

## Introduction

Powassan virus is the only known TBFV circulating in North America and has two distinct genetic lineages. Prototypical Powassan virus, lineage I (POWV), was isolated in 1958 and known to be vectored by *Ixodes cookei* ticks ^1,2^. Lineage II Powassan virus is also known as deer tick virus (DTV) and was initially isolated in 1995 and is vectored primarily by *Ixodes scapularis* ticks ^3,4^. Powassan virus and DTV share approximately 84% nucleotide sequence identity and 94% amino acid identity but are serologically indistinguishable ^5–8^. The number of cases and geographic range for both POWV and DTV have increased, coinciding with the expansion of *Ixodes* tick and increased surveillance. In addition, recent studies have demonstrated that Powassan lineages may not only be vectored by *Ixodes* ticks but may also be transmitted by some species of *Dermacentor*, *Amblyomma*, and *Haemaphysalis* ticks ^9–11^.

Infection with POWV is usually asymptomatic; however, some individuals will experience a febrile illness that can progress to severe neuroinvasive disease that may present with meningoencephalitis or paralysis. Case fatality rates are estimated to be between 10% to 15%, and over 50% of POWV survivors experience long-term neurological sequelae ^5,12^. While there have been advances in the study of POWV neuroinvasive disease, the underlying pathology is not completely defined, especially in relation to the similarly pathogenic DTV. Powassan virus disease is a nationally notifiable condition but differentiation of lineages is not required ^13^. When lineage differentiation is performed for human cases it is most often done through sequencing of viral RNA as methods such as neutralization and immunohistochemistry use antibodies which are cross reactive ^14–17^.

The neuropathology seen in POWV is currently believed to be mediated by a combination of inflammatory cytokines breaching the integrity of the blood-brain barrier (BBB) and intravasating leukocytes. Of possible relevance to differential POWV and DTV pathogenesis, infections by clinically important TBFV may provide important clues on what determines particular disease outcomes. For instance, the levels of TNFα, IL-1β, IL-6, and IL-8 have been associated with the proinflammatory breakdown of the BBB through the loss of tight junctions, with MMP-9 being crucial for neuroinvasive disease with tick-borne encephalitis virus (TBEV) ^18,19^. Once the integrity of the BBB has been compromised, the cellular destruction and pathology associated with TBEV infection are driven by the migration of CD8+ T-lymphocytes and NK cells ^20^. Besides immune-mediated neuronal destruction, neuronal demise can occur throughout the CNS with deficits related to the area in the brain damaged, as evidenced by mapping a neuronal network with brains infected by the related Langat virus (LGTV) ^21^. Noting the differences in CNS pathogenesis of TBFV, we describe here essential differences in the pathogenicity of POWV and DTV which are of relevance as historically they were assumed to be interchangeable, which makes these findings novel and clinically relevant. As such, we propose that the present studies support a reconsideration of distinct pathogenetic outcomes of infection by POWV lineages in humans.

## Results

### Immunocompetent mice inoculated with DTV and POWV had different clinical disease presentations

We investigated whether equivalent doses of either DTV or POWV resulted in a different clinical presentation in six-week-old BALB/cJ mice. The first outward signs of illness occurred at 5 days post infection (dpi) when weight loss was observed in both infected groups (Fig. 1A, B). Although weight loss occurred across both infected groups, it was more severe in the DTV-infected mice. Seventy-five percent of DTV-infected mice (n=9) and 8% of POWV-infected mice (n=1) reached loss of 20% or greater from their baseline weight and required humane euthanasia (Fig. 1D). Starting at 6dpi generalized signs of febrile illness occurred in both groups including ruffled coat, hunched posture, lethargy, and rapid or labored respiration. Ocular discharge and closure of one or both eyes occurred in both infected groups between 6dpi and 8dpi but was more common in DTV-infected mice (50%, n=6) than POWV-infected mice (33%, n=4.) Onset of neurological illness occurred at 6dpi in the POWV infected mice and at 7dpi for the DTV infected mice and was markedly different between groups. Forty-two percent of DTV-infected mice presented with neurological illness, including seizure (n=4) (Fig. 1D) and weak grip which progressed to paralysis (n=1) (Fig. 1D). In contrast, neurological illness was present in 83% POWV-infected mice, with 25% mice exhibiting multiple signs. Observed signs in POWV-infected mice included weak grip (n=3), paresis (n=3), loss of righting reflex/prostration (n=2), seizure (n=1), and paralysis (n=5) (Fig. 1D).

**Fig. 1.**
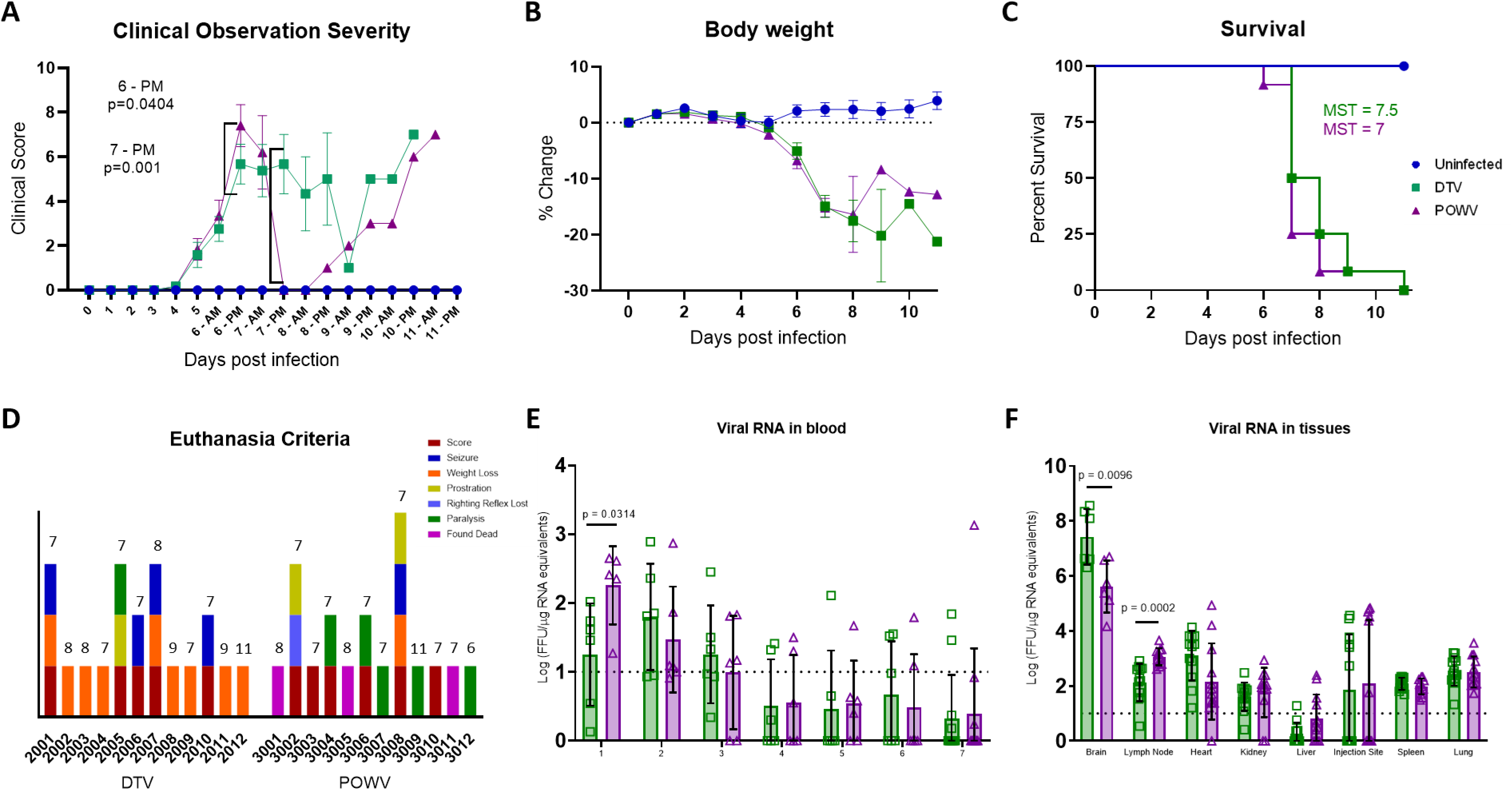
Clinical presentation of DTV and POWV infection. Mice (n = 12 per group) were inoculated via the footpad with 10^3^ FFU of media (blue), DTV (green), or POWV (purple) and monitored for signs of disease for up to 11 dpi. (A) Clinical signs of disease were categorized and scored by severity (Table S1). Significant differences in DTV and POWV mice clinical scores occurred at the PM observations on 6dpi and 7dpi. (B) Percent weight change from pre-infection. Comparisons on (A) and (B) were made using one-way ANOVA with Šίdάk multiple comparisons test. Error bars indicate SEM. (C) Survival of mice across all treatments were evaluated using the Mantel-Cox log-rank test and median survival times (MST) were calculated. (D) Humane euthanasia was performed when mice met specific criteria and are categorized to demonstrate differences between DTV and POWV with date of death shown above each bar. (E, F) Blood was collected from alternating cohorts of male (n=3) and female (n=3) mice from 0 to 7dpi and at termination blood and tissues (brain, popliteal lymph node, heart, kidney, liver, injection site, spleen, and lung) were collected from all animals to determine viral loads via rt-qPCR. For brain analysis, six animals per group (3M and 3F) were tested and for all other tissues twelve animals (6M and 6F) were tested. Viral loads are expressed as Log FFU equivalents per microgram of RNA after normalization to a standard curve. Comparisons on (D) and (E) were performed using two-tailed Student’s t-test with Welch’s correction.

Mean clinical disease scores and percent weight change from baseline were compared between all groups until 8dpi, after which sample sizes were too small for statistical comparison. Between 5dpi and 8dpi, there were statistically significant differences in the clinical disease scores between uninfected mice and each infected group and between DTV and POWV infected mice at the second daily observations on 6dpi [p=0.0404] and 7dpi [p=0.001] (Fig. 1A). Body weight changes from baseline were significantly different between uninfected and each infected group starting at 6dpi but not between DTV and POWV infected groups (Fig. 1B).

Differences in mean clinical disease scores and percent weight change from baseline were also compared between male and female mice within a group. The DTV infected male and female mice had significantly different clinical scores during the first daily observation at 7dpi [p=0.0298] (Fig. S1A) and no significant differences were found in the POWV infected male and female mice (Fig. S1F). In addition, trends in weight loss were consistent between male and female DTV and POWV infected mice (Fig. S1B, G). Overall, DTV infected mice experienced a higher degree of weight loss than POWV infected mice while POWV infected mice experienced more loss of ambulation than DTV infected mice.

### Immunocompetent mice inoculated with DTV and POWV had the same time to death

Despite differences in clinical disease, all mice inoculated via footpad with 10^3^ FFU of DTV or POWV succumbed to disease between 6 and 11 dpi. Log-rank test (Mantel-Cox) did not show significant differences in survival between DTV and POWV infected mice (Fig. 1C), which had median survival times (MST) of 7.5 and 7 days, respectively. In addition, no significant differences were present between male and female mice within a group (Fig. S1C, H) using the same analyses. MST for DTV-infected mice were 7 days for males and 8.5 days for females, while MST for POWV-infected mice were 7 days for both males and females.

### Viral titers differ in blood and tissues of DTV and POWV infected mice

Viral RNA was detected in blood samples from all DTV-infected mice and all except one POWV-infected mouse between 1dpi and 3dpi (Fig. 1E). Viral RNA declined following 3dpi and was not detected in blood samples after 9dpi. Mean viremia for the DTV group peaked on 2dpi at 1.798 Log FFU/µg RNA equivalents, while the POWV group peaked on 1dpi at 2.258 Log FFU/µg RNA equivalents (Fig. 1E). At 1dpi, there was higher viremia in POWV-infected mice than in DTV-infected mice [p=0.0314].

Detection of viral RNA in organs demonstrated that DTV and POWV disseminate throughout the body. Viral RNA was present in the brain, popliteal lymph node from the injected leg, spleen, and lung of all infected mice (Fig. 1F). There was viral RNA detected in the heart and kidney of all DTV-infected mice and 92% POWV-infected mice (n=11.) Viral RNA was present at the injection site for 50% of the mice in each infected group (n=6.) Limited viral RNA was found in the liver with either no viral RNA or RNA below the detection limit in 92% DTV-infected mice (n=11) and 67% POWV-infected mice (n=8.)

The highest amounts of viral RNA were detected in the brains of both infected groups, with an average of 7.43 Log FFU/µg RNA equivalents present in the DTV group and 5.62 Log FFU/µg RNA equivalents present in the POWV group (Fig. 1F). The brains of DTV-infected mice had higher viral loads than POWV-infected mice [p=0.0096], while the POWV-infected mice had higher viral loads in the lymph node [p=0.0002.] A comparison of viral RNA present in organs was also performed between infected male and female mice and the only significant difference was found in the livers of POWV-infected mice (Fig. S1I).

### Comparative Histopathology of Deer Tick Virus and Powassan Virus Infected Brains and Spinal Cords

Histopathological changes were observed in POWV and DTV infected mouse brains across multiple regions including the olfactory bulb (OB), cerebral cortex (CTX), hippocampal formation (HPF), hypothalamus (HY), thalamus (TH), midbrain (MB), pons (P), medulla (MY), and cerebellum (Purkinje layer: CB – P, granular layer: CB-G, molecular layer: CB – M) and scored by severity (Table S2). Infected mice exhibited microgliosis and neuronal necrosis in the cerebellum (Fig. 2A-C) and olfactory bulb (Fig. 2J-L), degenerating neurons and vacuolation of neuropil were observed Ammon’s horn of the hippocampal formation (Fig. 2D-F), and the isocortex showed perivascular cuffing and meningoencephalitis (Fig. 2G-I). Deer tick virus infected mice had higher lesion severity scores in the hippocampal formation, thalamus, and midbrain regions compared to POWV infected mice (Fig. 3E,F).

**Fig. 2.**
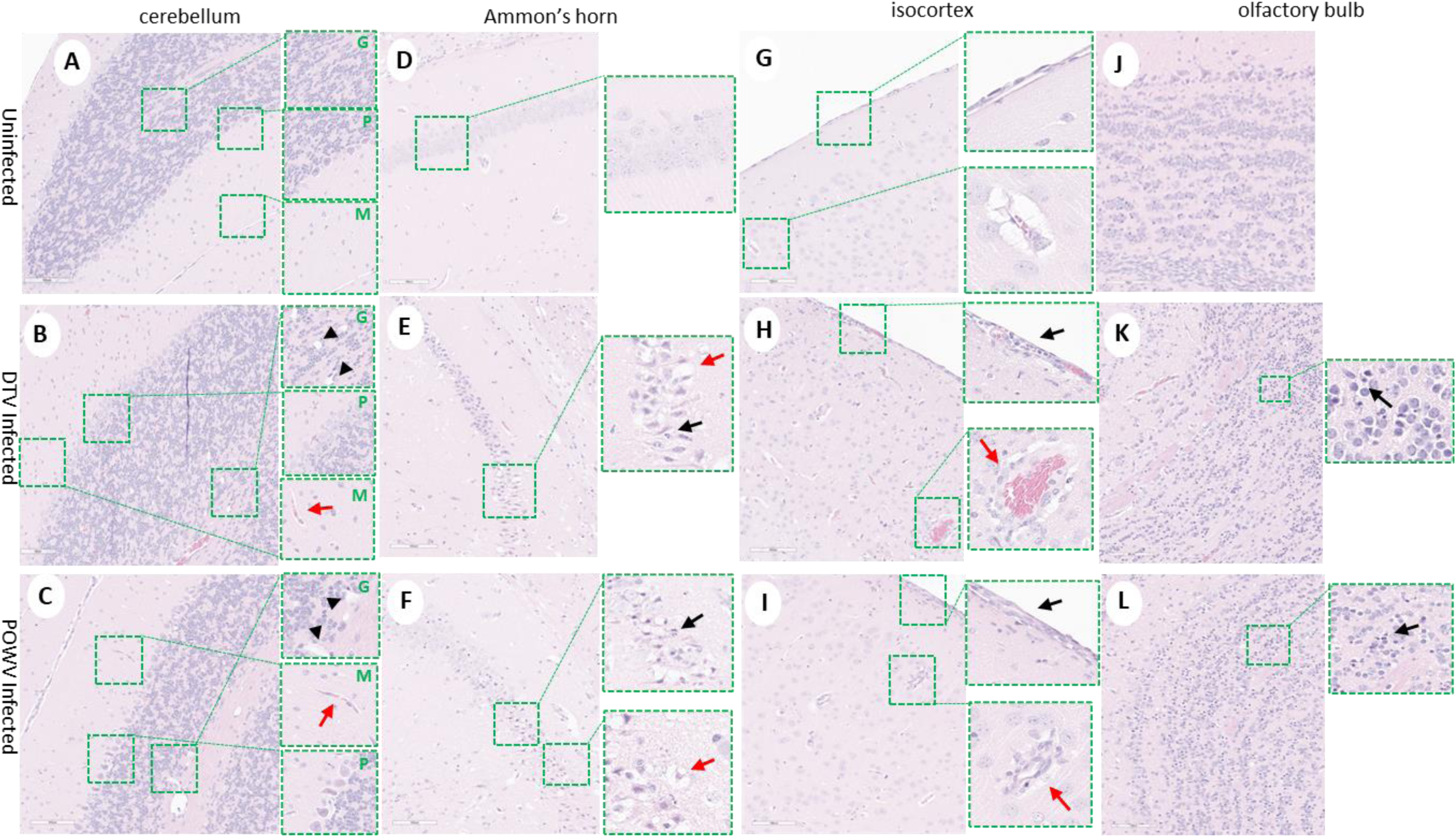
Neuropathological examination of DTV and POWV infected mouse brains. Uninfected mouse with normal structures shown in (A) cerebellum with insets showing molecular cell layer – M, Purkinje cell layer – P, and granular cell layers – G, (D) Ammon’s horn of the hippocampus, (G) isocortex, and (J) olfactory bulb. Cerebellum of DTV (B) and POWV (C) infected mice with microgliosis (inset, red arrow) and neuronal cell necrosis (inset, black arrowheads). Ammon’s horn of the hippocampus from DTV (E) and POWV (F) infected mice with neuropil vacuolation (inset, red arrow) and hyper eosinophilic degenerating neurons (inset, black arrow). Isocortex of DTV (H) and POWV (I) with infiltration of leptomeninges (inset, black arrow) and perivascular cuffing (inset, red arrow). Olfactory bulb of DTV (K) and POWV (L) infected mice with neuronal cell necrosis in granular cell layers (black arrow).

**Fig. 3.**
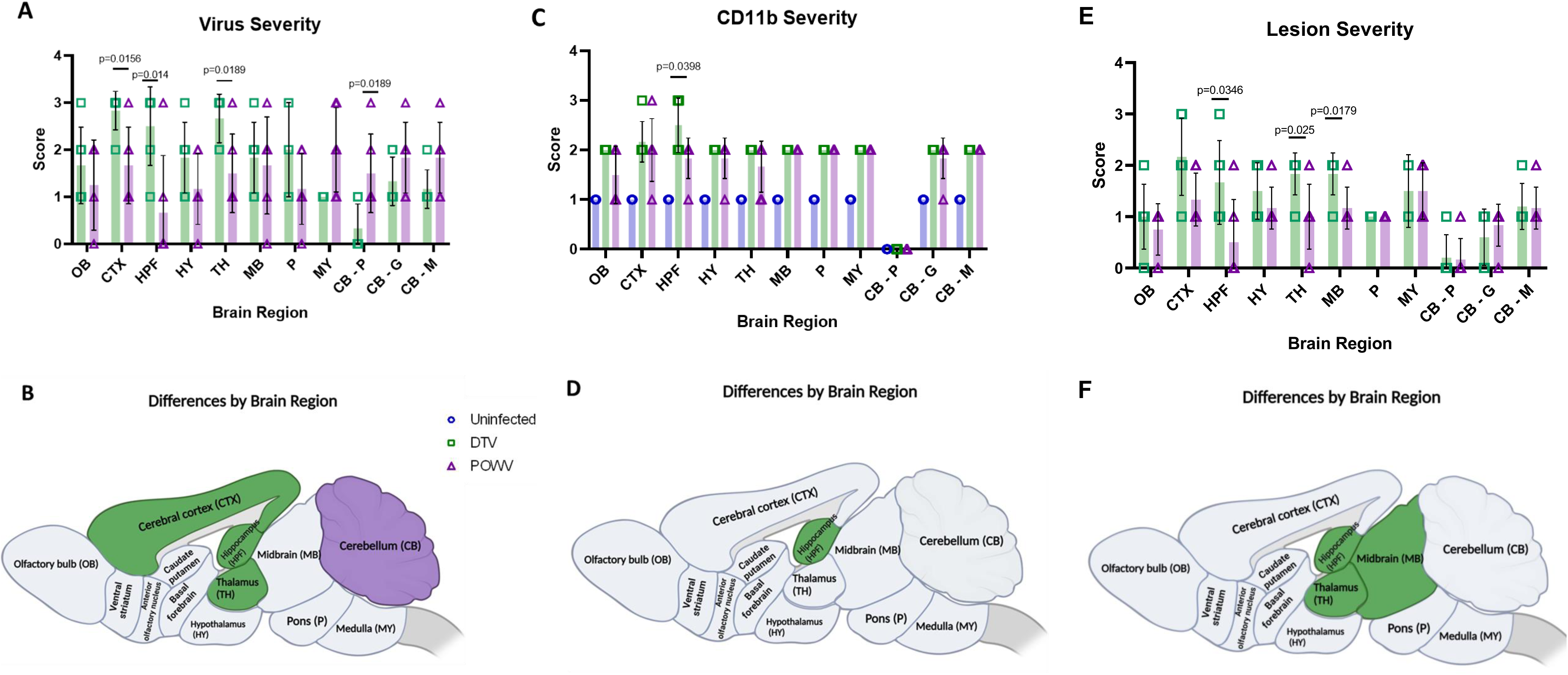
Severity of virus,CD11b, and lesions in DTV and POWV-infected mouse brains by region. (A) Virus severity individual mice by region and (B) brain regions of DTV infected mice had statistically more virus severity in CTX, HPF, and TH while POWV infected mice had more virus severity in CB. (C) CD11b severity of individual mice by region. (D) Brain regions of DTV infected mice had statistically significant increases in CD11b positive cells in HPF. (E) Severity of individual mice by region. (F) Brain regions of DTV infected mice had statistically significant differences in severity in the HPF, MB, and TH regions. All comparisons performed using unpaired two-tailed Student’s t-test with Welch’s correction.

Powassan virus infected brains showed meningoencephalitis (Fig. 2I) that had progressed into the cervical spinal cord (Fig. 4C), while the meninges of the spinal cord appeared normal in DTV infected samples (Fig. 4B). Cervical spinal cords of POWV infected mice exhibited considerable histopathology in which nuclei of ventral gray matter motoneurons were either shrunken with irregular cell bodies or had entirely degenerated (Fig. 4F). There was also conspicuous extravasation of inflammatory cells in spinal cord gray matter of POWV infected mice, especially in the cervical spinal cord (Fig. 4C, F, I, L). In contrast, neurons in the cerebellum and spinal cord of DTV infected mice appeared normal in numbers and morphology (Fig. 2B, Fig. 4B, E).

**Fig. 4.**
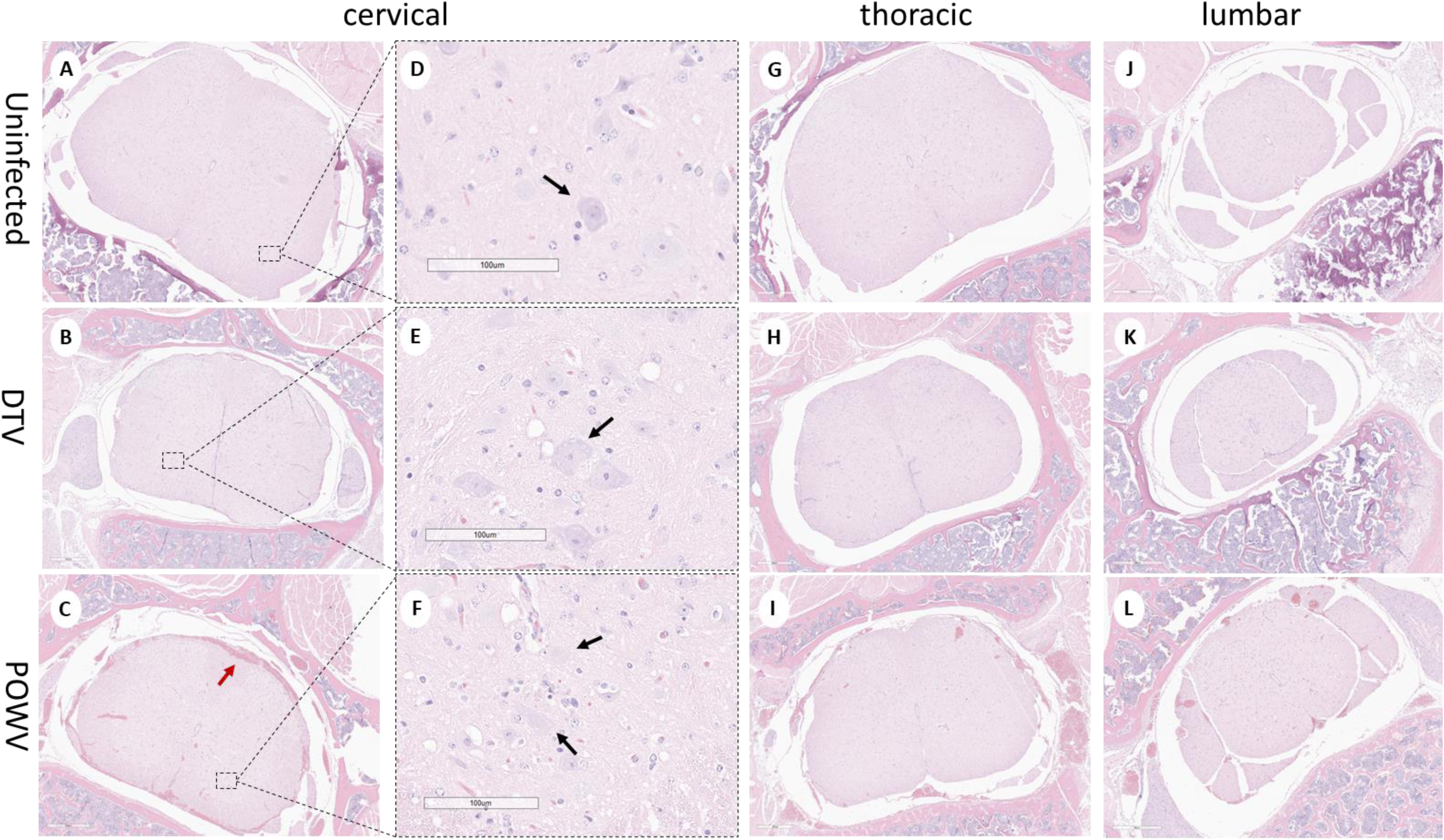
Neuropathological examination of DTV and POWV infected mouse spinal cords. Uninfected mouse with normal structures shown in (A, D) cervical region (black arrow shows motor neuron in ventral horn), (G) thoracic region, and (J) lumbar region. DTV infected mouse (B) cervical region, (E) motor neuron (black arrow) in dorsal horn, (H) thoracic region, and (K) lumbar region. POWV infected mouse with (C) meningoencephalitis (red arrow) in cervical region, (F) absent and degenerating motor neuron (black arrow) in dorsal horn, and infiltrations in (I) thoracic and (L) lumbar regions.

Brain sections stained for viral RNA and CD11b+ cells were examined and scored by severity in a manner consistent with histopathological analysis (Table S3). Powassan virus and DTV infection of neurons were noted throughout all infected animal brains; however, the regions infected differed between viruses. The brains infected with POWV exhibited staining of viral RNA in all regions but were concentrated primarily in the cerebellum and brainstem (Fig. 5C). Powassan virus only rarely infected the hippocampus, isocortex, and olfactory bulb in most animals (Fig. 5G, H, K, L) compared to relatively high levels of infection of Purkinje cells and granule cells in in the cerebellum (Fig. 5C).

**Fig. 5.**
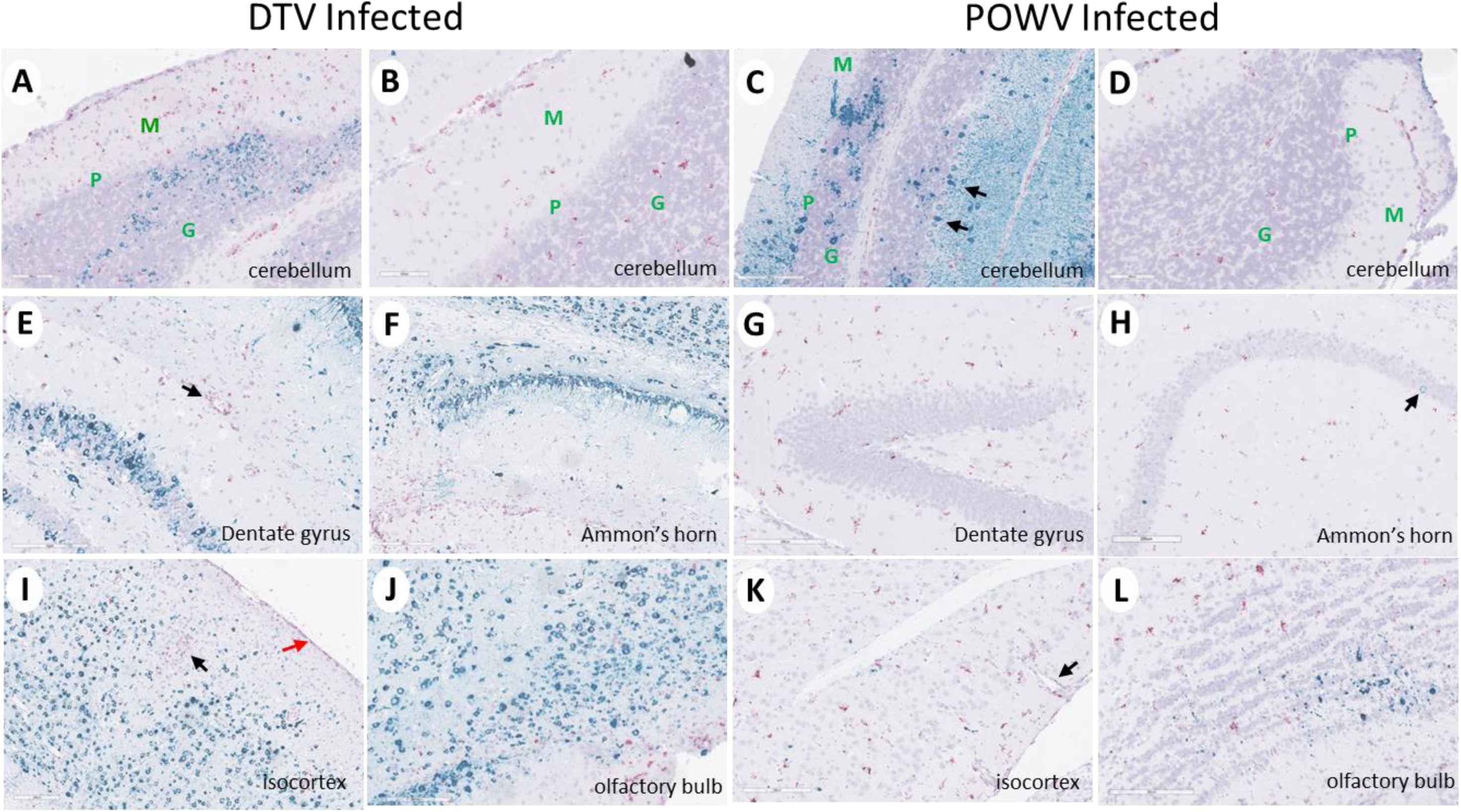
Divergent patterns of infection in DTV and POWV infected mouse brains. Representative images of DTV infected (A, B, E, F, I, J) and POWV infected (C, D, G, H, K, L) brains stained for viral RNA (teal) and CD11b^+^ cells (pink). (A – D) Molecular (green M), Purkinje (green P), and granular (green G) cell layers of the cerebellum and (C) POWV infected Purkinje cells (black arrow). (E) CD11b positive cells (black arrow) around a vessel in the hippocampal region of a DTV infected mouse brain. (H) Minimal infected cells (black arrow) in the hippocampal region of a POWV infected mouse brain. (I) CD11b positive cells around a vessel (black arrow) and in the leptomeninges (red arrow) of the isocortex of a DTV infected mouse brain. (K) CD11b positive cells in the perivascular (Virchow-Robin) space (black arrow) of a POWV infected mouse brain.

Deer tick virus-infected brains showed a markedly different pattern of viral infection. There was reduced infection in the cerebellum and brainstem (Fig. 5A, B) in comparison to POWV infected brains but a pronounced increase of viral RNA found in neurons in the isocortex, olfactory bulb, and hippocampal formation including the dentate gyrus granule cells and pyramidal neurons in Ammon’s horn (Fig. 5E, F, I, J). Virus severity scores show increased virus in the cortex, hippocampal formation, and thalamus of DTV infected mice and increased virus in the cerebellum of POWV infected mice (Fig. 3A, B). CD11b severity scores show an increased positive cells in the hippocampal formation of DTV-compared to POWV-infected mice (Figure 3C).

Neuron specific staining differed in the spinal cords of POWV and DTV infected mice. In DTV infected mice minimal viral RNA was present across the cervical, thoracic, and lumbar regions (Fig. 6B, E, H, K). Prominent infection of cervical spinal cord neurons of POWV infected mice was consistent with contiguous brainstem, cerebellum and midbrain regions that were also positive for POWV. The presence of viral RNA-containing neurons decreased considerably from cervical to thoracic and lumbar regions in several animals indicating that the cervical spinal cord was particularly affected at this stage of infection (Fig. 6C, F, I, L). One POWV-infected mouse had prominent infection of the hippocampus and isocortex (Table S3) however, this animal also had the characteristic lower brain stem and spinal cord infection seen in the other POWV-infected mice.

**Fig. 6.**
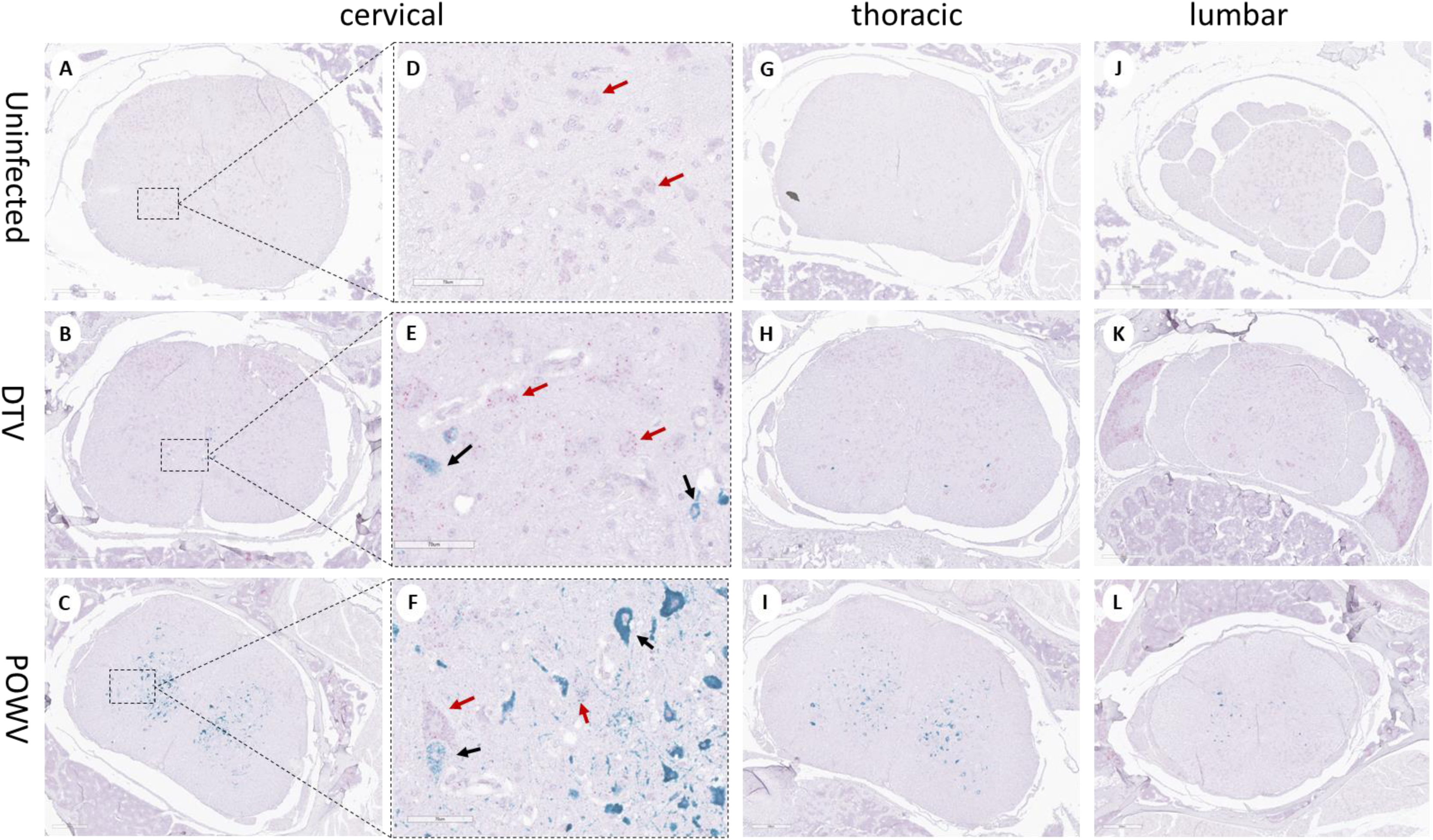
Divergent patterns of infection in DTV and POWV-infected mouse spinal cords. Representative images of uninfected, DTV infected, and POWV infected mouse spinal cord with teal staining of viral RNA and pink staining of neuronal nuclei (NeuN.) Uninfected cervical (A, D), thoracic (G), and lumbar (J) regions with normal motor neurons shown (D, red arrows). DTV infected mouse spinal cord cervical (B, E), thoracic (H), and lumbar (K) regions with minimal infected neurons. POWV infected mouse spinal cord cervical (C, F), thoracic (I), and lumbar (L) regions with decreasing infection across the cord and degenerating motor neurons (F, black arrows.)

In summary, infection of the brainstem, cerebellum, and contiguous cervical spinal cord by POWV correlates with the observed clinical signs seen during POWV infection, in which animals exhibited substantial paralysis and motor deficits while infection with DTV was concentrated mainly to the cerebral cortex, olfactory bulbs, and midbrain with less lower brainstem and spinal cord involvement. The distinct regional patterns of infection by DTV and POWV are depicted schematically in Figure 3B.

### Comparative Transcriptomics of POWV and DTV

Transcripts from either DTV or POWV-infected brains were generated and compared against the uninfected controls and represented increased levels of inflammatory cytokine and chemokine genes (Fig. S3A, B; Fig. S4). The smallest p-adjusted value for either group was STAT1, with genes involved in immunological signaling, inflammation, and interferon response following infection (Fig. S3C, D). The 25 genes with the smallest p-values in POWV-infected brains are involved with interferon signaling, antigen presentation and viral response (Fig. S6 C, D). There was a significant upregulation in genes responsible for controlling viral infection in both samples, including interferon-stimulated genes and innate immune signaling genes associated with MyD88 and RIG-I like receptors. Individually, both POWV and DTV appear to activate the JAK-STAT pathway through TLR stimulation and interferon production producing an anti-viral state.

The transcript libraries generated from POWV- and DTV-infected brains were then compared (Fig. 7A) and smaller libraries were generated based on GO enrichment analysis terms: inflammatory response (Fig. 7B), immune system process (Fig. 7C), and defense response (Fig. 7D). The brain tissues infected with POWV had significantly higher levels of inflammatory pathway genes including NLRP6 and chemokines associated with hematopoietic inflammatory cell recruitment including CX3CR1. The tissues infected with DTV displayed higher levels of Cav1, which induces NF-κB through T-cell activation. Furthermore, there is a strong difference in the levels of CXCL3, with DTV-infected tissues having significantly higher levels of the neutrophil chemotactic cytokine. The results from this comparison suggest that while there is a similar pattern to the interferon response and establishment of the anti-viral state, DTV and POWV infections of the CNS involve the recruitment of different cell types and activation of different inflammatory pathways.

**Fig. 7.**
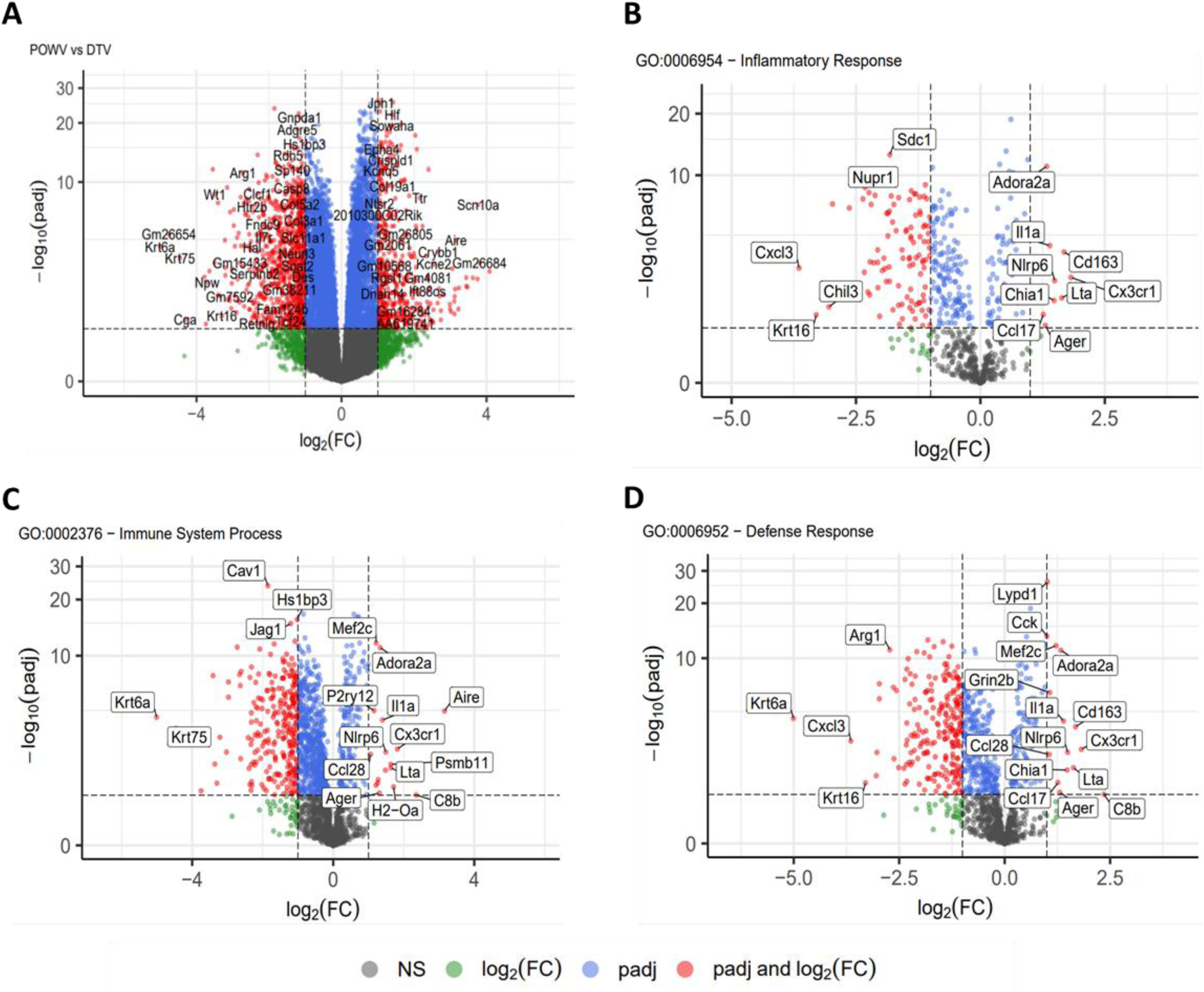
RNA sequencing analysis of brain tissue. Brains were harvested at time of euthanasia and stored in trizol before homogenization and RNA extraction followed by RNA-Seq. Transcript libraries were analyzed using R with (A) POWV compared to DTV. Genes were separated into smaller gene ontology sets and compared for (B) inflammatory response (C) immune system process and (D) defense response.

### Cytokine and chemokine analysis of terminal serum

Levels of 23 analytes were measured in terminal blood serum and evaluated to determine differences between uninfected (n=12), DTV infected (n=11), and POWV infected (n=8) mice (Fig. 8, Fig. S2). There were significantly higher levels of six analytes in DTV-infected groups compared to uninfected groups: IL-10 [p=0.0321](Fig. 8D), IL-12(p40) [p=0.006](Fig. 8E), MCP-1 [p=0.031](Fig. 8G), MIP-1α [p=0.0059](Fig. 8H), MIP-1β [p=0.0237](Fig. 8I), and RANTES [p=<0.0001](Fig. 8J). In contrast, there were only significantly lower levels in four analytes in POWV-infected animals compared to controls: GM-CSF [p=0.0112](Fig. 8A), IL-2 [p=0.0249](Fig. 8B), IL-5 [p=0.0121](Fig. 8C), and IL-13 [p=0.0004](Fig. 8F). The only difference between DTV and POWV was in IL-13 [p=0.0451](Fig. 8F). These data indicate an entirely different proinflammatory response in the periphery following infection by these two viruses. Analytes with no significant differences are shown in Fig. S2.

**Fig. 8.**
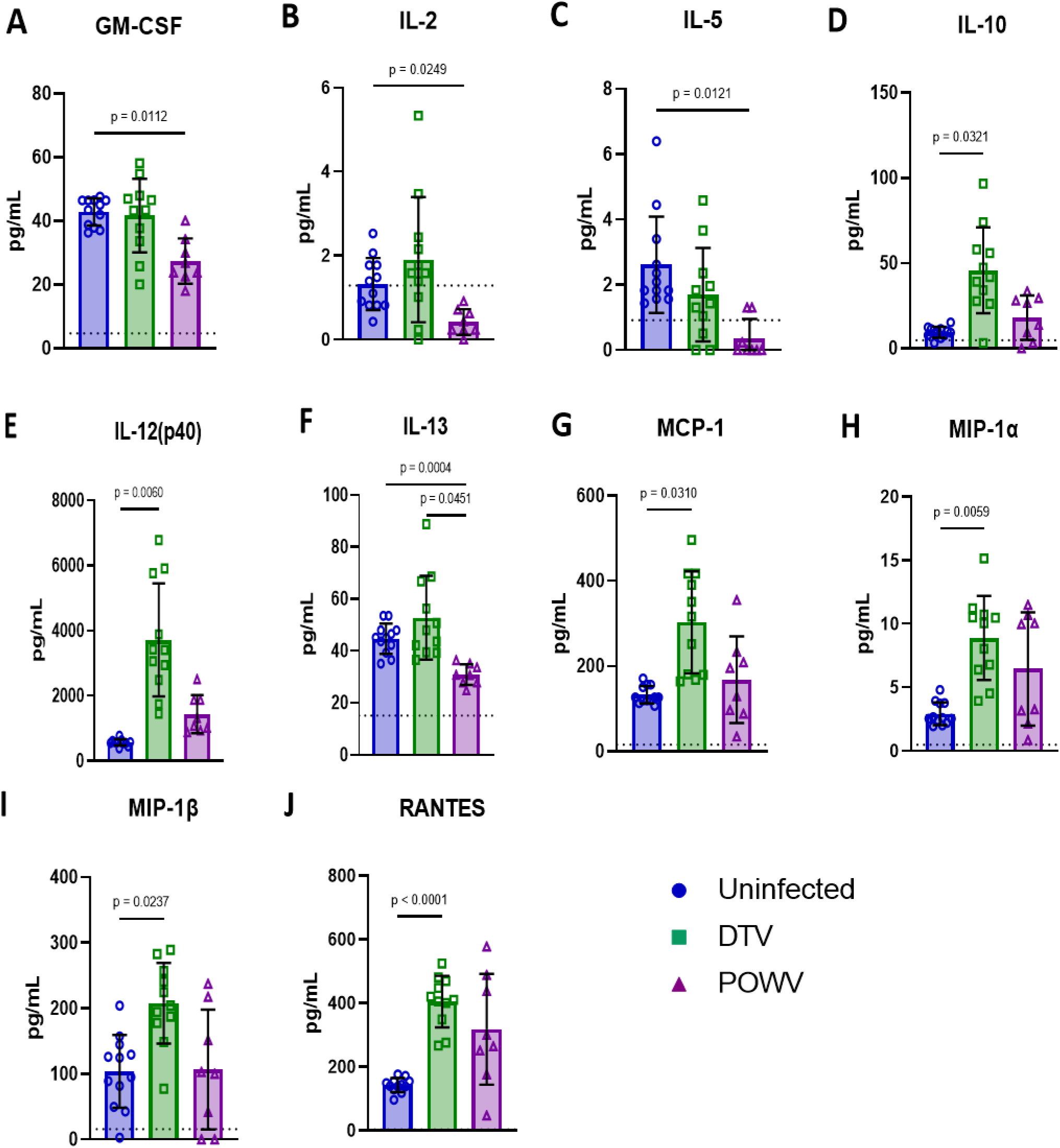
Cytokine and chemokine levels were measured in terminally collected serum from uninfected (n = 6M/6F), DTV infected (n = 5M/6F), and POWV infected (n = 3M/5F) mice by Bio-Plex Mouse Cytokine 23-plex panel. Significant differences were present between uninfected and POWV-infected mice for four parameters (A, B, C, F), between uninfected and DTV-infected mice for six parameters (D, E, G, H, I, J), and between DTV-and POWV infected mice for one parameter (F). Comparisons were made using Brown-Forsythe and Welch’s ANOVA test with Dunnett’s T3 multiple comparisons test.

## Discussion

Flaviviruses are endemic across much of the globe and cause millions of cases of human illness each year ^22^. Severity of neurotropic flaviviruses such as West Nile virus (WNV), Japanese encephalitis virus (JEV), TBEV, and Powassan virus can range from mild to fatal and can cause severe neuroinvasive disease with long-term sequelae in a high percentage of survivors. For example, disease severity, fatality rate, and long-term sequelae associated with TBEV infection has been shown to vary depending on the subtype despite the high sequence similarity between these viruses ^23,24^. Similarly, DTV and POWV share approximately 94% amino acid identity and 84% nucleotide sequence similarity and are serologically indistinguishable ^6–8^. Coupled with the lack of a vaccine or approved treatment and overlapping circulation in the United States, testing to differentiate lineages is typically not performed for human cases.

### Immunocompetent mice inoculated with Powassan virus and Deer tick virus have different clinical disease presentations

In this report, we have shown that DTV and POWV do not have the same clinical presentation when inoculated via the same route and dosage. Common clinical signs in both virus groups included ruffled fur, hunched posture, ocular discharge, and weight loss. The degree of weight loss experienced by DTV-infected mice was more severe than POWV-infected mice. Seizure was the most common neurological symptom present in DTV-infected mice and was consistent with prominent infection and inflammation in the hippocampus. Mice infected with POWV experienced neurological signs more consistent with infection and inflammation of the lower brainstem, cerebellum and spinal cord including paresis and paralysis.

### Powassan virus and Deer tick virus induce neuronal death in differing regions of the brain

The reduced levels of CD11b^+^ cells seen in the histopathology of POWV-infected brains as compared to the DTV-infected brains are congruent with the histopathology and spatial distribution of the virus. The spatial distribution of POWV in the brain is concentrated in the lower brain and spinal cord, while DTV preferentially infected and induced inflammation in the hippocampus and isocortices, which suggests a different neuroinvasive route and/or spread of infection in the brain. The neurologic clinical signs exhibited during late-stage infections are dissimilar as well with DTV eliciting seizures while POWV neuropathology is manifested as motor deficits. The comparative neuropathology of POWV and DTV reveals a stark contrast which is recapitulated in chemokines, histopathology, and observed clinical signs; DTV infects the hippocampus and isocortex, while POWV infection is concentrated within the cerebellum, brain stem, and spinal cord. Interestingly, acute seizures have been linked to loss of the pyramidal cell layer and neuronal loss in the hippocampus in Theiler’s murine encephalomyelitis virus-infected mice, which is consistent with pathology findings for mice that experienced seizures in DTV-infected mice in the present work ^25,26^. Published human cases of fatal DTV infection have shown severe inflammation and infection of the cerebellum and brainstem which is more consistent with neuropathology present in POWV-infected mice ^14,15^. In these human cases limb weakness and difficulty ambulating were noted and are consistent with the deficits we observed with mice with heavy infection in the cerebellum, brainstem, and cervical spinal cord.

Changes in the clinical presentation of disease and pattern of infection in the CNS could be attributed to a variety of factors, including person-to-person variability, site of infection, and genomic diversity of viruses. Other flaviviruses have demonstrated that genetic diversity in the E and NS5 regions change the phenotype of the virus including immunogenicity and pathogenicity ^27^. The strain of POWV or DTV used for inoculation, route and method of inoculation and time spent replicating in the periphery could affect the clinical signs and response to infection ^21,23,28–30^. This is underscored by our findings in that the vectored route of inoculation can cause a diverse set of clinical signs but ultimately be relegated to either a cortical region meningoencephalitis or involve the cerebellum and spinal cord, resulting in descending myelitis.

### Host Chemokine and Inflammatory Response to Powassan virus is different compared to Deer tick virus

Neuroinvasion of flaviviruses has been shown to be a result of inflammatory breakdown of the blood-brain barrier; however, other mechanisms may result in CNS infection as well. Our findings are consistent with previous studies showing an upregulation of STAT1 and genes associated with inflammation and interferon in the brain of moribund animals. The histopathological differences seen between the regions of the brain infected by DTV and POWV suggest a spatial tropism which may be affected by a region-specific neuroinvasion and dissemination. The olfactory bulb of DTV-infected animals had high levels of virus present which suggests that this area was significant for viral replication and consequent spreading to cerebral cortex and to the underlying midbrain structures. The clinical signs in POWV-infected animals combined with the histopathology clearly show myelitis and destruction of neurons in the ventral horns of the cervical spinal cord, contiguous lower brain stem, and cerebellum, illustrating a differing spatial tropism compared to DTV. The differences in these spatial tropisms could be explained by differential routes of neuroinvasion, with DTV for instance preferentially crossing the cribiform plate from olfactory neurons to infect the olfactory bulb, while POWV may utilize retrograde axonal transport of virus from the periphery into the spinal cord by the neural route or perhaps by a unique viremic route initiated by preferential inflammatory degradation of the BBB in the spinal cord.

The comparative RNA-seq data suggests that there are different innate immune pathways responding to DTV and POWV during CNS infection. While both viruses elicit a strong interferon response, there is a significant increase in NLRP6 and IL1α transcripts in tissues infected with POWV relative to tissues infected with DTV. There is also a significant increase in CX3CR1 and AIRE genes which indicate the recruitment of CD8+ T-cells in POWV infected tissues. The DTV-infected tissues conversely exhibit significant increases in CXCL3 CD177 and CAV1, suggesting neutrophilic recruitment combined with T-cell dependent NF-κB transcription. The spatial tropism exhibited in the histopathology is supported by the levels of Krt6a seen in DTV-infected tissues, which exhibit anti-microbial properties in the olfactory bulb, an area with extensive DTV infection. These findings suggest that the pathology seen in POWV is driven by CD8+ cytotoxic T-lymphocytes, while DTV elicits repetitive NF-κB-mediated transcription and inflammation. These findings are congruent with recent studies that suggest POWV is less inflammatory in neuronal cell cultures and that the pathogenesis of POWV, like TBEV, is driven by the extravasation of cytotoxic T-lymphocytes ^20,31^. The response to POWV has recently been expanded with experiments examining the role of TRIM5 in POWV infection ^32^. We have also noted in this report that TRIM5 is upregulated in both DTV and POWV infection compared to uninfected controls; however, TRIM5 is also significantly upregulated in response to POWV as contrasted to the response to DTV infection. This comparative view of RNA transcriptomics in DTV and POWV infection illustrates a difference in cellular and inflammatory response to infection.

In conclusion, a direct comparison of POWV (lineage I) and DTV (lineage II) demonstrated distinct differences in clinical presentation and neuro-immunopathology. These differences suggest these two viral lineages do not utilize the same routes of neuroinvasion and stimulate distinct neuroimmunological responses.

## Methods

### Ethics Statement

Mouse experiments were conducted in an American Association for the Accreditation of Laboratory Animal Care (AAALAC) approved facility within Animal Biosafety Level 3 (ABSL3) containment. All procedures were performed in accordance with SUNY Upstate Medical University Institutional Animal Care and Use Committee approved protocol (#455) which adhered to the Guide for the Care and Use of Laboratory Animals ^33^.

### Viruses, animals, and infection

Deer tick virus (Spooner strain) and POWV (LB strain) stocks were provided by the World Reference Center for Emerging Viruses and Arboviruses at University of Texas Medical Branch and had been previously passaged once in suckling mouse brains. Stocks were then passaged six times on Vero E6 cells. Vero E6 cells (CRL-1586; American Type Culture Collection) were cultured using Modified Eagle’s Medium (MEM; Corning) and supplemented with fetal bovine serum (FBS), non-essential amino acids, and penicillin/streptomycin antibiotic mixture and maintained at 37°C with 5% CO_2_. Virus titers were determined by focus-forming assay as previously described ^34^.

Next generation sequencing demonstrated that the consensus nucleotide sequence of our DTV stock was 99.84% identical to the DTV Spooner, Wisconsin isolate (GenBank: <u>HM440560.1</u>) and our POWV stock was 99.96% identical to the POWV LB, Ontario isolate (GenBank: <u>NC_003687.1</u>). Of 17 nucleotide differences between our DTV stock and DTV Spooner isolate there were three resulting amino acid sequence differences in M (N213K), NS4B (S2368T), and NS5 (K2903R). There were three nucleotide differences between our POWV stock and POWV LB isolate resulting in one amino acid sequence difference in E (Q1436R) as well as one nucleotide gap which may have resulted in one amino acid sequence difference in NS5 (W8255 to L or S). Thirty-six 5-week-old male (n=18) and female (n=18) BALB/cJ mice were purchased from The Jackson Laboratory (Bar Harbor, ME). Upon receipt, mice were housed three per cage in individually ventilated cages with HEPA-filtered supply and exhaust air. Food and water were provided *ad libitum*. Animals were provided a 12h light/12h dark cycle within a temperature and humidity-controlled room. Mice were acclimated to the facilities for one week. On the day of study initiation, prior to infection, animals were observed and weighed to ensure there were no apparent physical abnormalities and that each group had a similar weight distribution. Uninfected control mice had an average weight of 21.43g (± 2.09), DTV-infected mice had an average weight of 20.74g (± 1.98), and POWV-infected mice had an average weight of 21.06g (± 2.16).

All mice received a single footpad injection of 30µL under isoflurane anesthesia. Control mice received serum-free Dulbecco’s Modified Eagle Medium (DMEM, Corning), and infected mice received 10^3^ focus forming units (FFU) of either POWV Lineage I (POWV LB) or Lineage II (DTV Spooner). Each group contained six male and six female mice. These numbers were selected to provide the minimum number of animals required to determine statistical significance between sexes within a treatment group. Bodyweight measurements and detailed clinical observations were performed daily to assess the animal appearance, behavior, respiration, and neurological signs of disease (Table S1). Submandibular blood collection was performed from Days 0 to 7 using alternating cohorts of mice and alternating sites. Terminal blood was collected via cardiac puncture following euthanasia.

When animals reached the criteria for humane euthanasia or at 11 days post-infection (dpi) euthanasia was performed by CO_2_ inhalation followed by cervical dislocation. Whole body perfusion was performed using 10mL of sterile 0.9% sodium chloride, USP. Tissues collected at necropsy were stored in either Buffer RLT (Qiagen) or 10% Neutral Buffered Formalin (NBF).

Blood was stored in Buffer RLT. Samples stored in Buffer RLT were maintained in ABSL3 facilities for a minimum of 24 hours to allow inactivation. Samples stored in 10% NBF were fixed for a minimum of 72 hours, with one change of NBF occurring 24 hours after collection.

No specific methods were used to minimize confounders or blind staff to group allocation during the experiment. Each day procedures were performed with uninfected animals handled first, followed by DTV-infected animals, and POWV-infected animals handled last. Equipment and PPE were cleaned or changed between infected groups to prevent cross-contamination.

### Viral RNA detection via RT-qPCR

Total RNA was extracted from blood and tissue samples using RNeasy Mini Kits (Qiagen) using previously established modifications to the kit protocol ^35^. RNA was quantified using a DS-11+ Spectrophotometer (Denovix) and tested for the presence of viral RNA by RT-qPCR on a Bio-Rad CFX96 Touch Real-Time PCR System. Viral RNA was detected with forward (CCGAGCCARRGTGAGGATGT), reverse (TCTTTTGCYGARCTCCACTT), and probe (TTCATAGCGAAGGTKAGRTCCAACG) primers which have been previously validated to detect a region of the NS5 gene conserved between POWV and DTV ^36^. Standard curves for POWV and DTV were generated from 10-fold dilutions of a known quantity of viral RNA and used to estimate viral burden in experimental samples.

### Histology and RNA in situ hybridization

Inactivated tissues from three male and three female mice per group were transferred to 70% ethanol and shipped to Histowiz, Inc. for embedding and sectioning. Brains were sectioned at 5µM thickness and mounted on glass slides. Hematoxylin and eosin staining was performed at Histowiz, Inc in accordance with their established methods. RNA in situ hybridization was performed using RNAscope 2.5 HD Duplex kits (catalog 322430, Advanced Cell Diagnostics) per manufacturer recommendations. Brain sections underwent dual staining protocol for POWV positive-sense RNA probe (catalog 415641, Advanced Cell Diagnostics) and *Mus musculus* integrin alpha M/CD11b (catalog 311491, Advanced Cell Diagnostics) to identify infected cells and innate leukocytes. Spine sections underwent dual staining protocol using the same viral RNA probe used on brain sections and *Mus musculus* RNA binding protein, fox-1 homolog (*C. elegans*) 3/NeuN (catalog 481707, Advanced Cell Diagnostics), to identify infected cells including NeuN^+^ neurons. All slides were counterstained with a 50% solution of Gill’s hematoxylin I. Positive control samples were processed using *Mus musculus* Duplex Ppib and Polr2a (catalog 321651, Advanced Cell Diagnostics) for process verification. Both Hematoxylin and eosin staining and RNA in situ hybridization was performed on all brains sectioned.

Sagittal brain sections were examined in a rostro-caudal direction, including the olfactory bulb, cerebral cortex, hippocampus, thalamus, hypothalamus, midbrain, pons, medulla oblongata, and Purkinje, granular, and molecular layers of the cerebellum. Microscopic lesions, including microgliosis, perivascular cuffing, and neuronal degeneration were graded as: 0 (absence of lesions); 1 (minimal); 2 (mild); 3 (marked.) The presence of meningitis, encephalitis, and meningoencephalitis were noted as M, E, and ME, respectively. RNAscope stained cells were graded a: 0 (absence of staining); 1 (very few to low); 2 (moderate); 3 (numerous). Areas that were not present were noted as NP.

### RNA Sequencing and Bioinformatics

RNA sequencing and bioinformatics was performed for the same three male and three female mice per group which had H&E and RNA in situ hybridization performed. Total RNA was quality checked using an Agilent Fragment Analyzer to assess RNA quality and Qubit Fluorescence (Invitrogen) to measure concentration. RNA libraries (200ng total starting material) were prepared following the Illumina RiboZero Total Stranded RNA library prep kit using the ribosomal removal protocol. Following library preparation, concentration and quality control, final libraries were pooled to 10nM and the concentration of the pool was determined using the QuantaBio Universal qPCR reaction kit. After dilution and denaturing, the pooled library was loaded onto an Illumina NovaSeq6000 (PE100) for sequencing at 250pM as final concentration.

Per-cycle base call (BCL) files generated by the Illumina NovaSeq were converted to per-read FASTQ files using bcl2fastq version 2.20.0.422 using default parameters. The quality of the sequencing was reviewed using FastQC version 0.11.6. Detection of potential contamination was done using FastQ Screen version 0.14.1. Finally, FastQC and FastQ Screen quality reports were summarized using MultiQC version 1.9.0.

Adapter trimming was not performed. Genomic alignments were performed using HISAT2 version 2.2.1 using default parameters. Ensembl reference GRCm38 was used for the reference genome and gene annotation set. Sequence alignments were compressed and sorted into binary alignment map (BAM) files using sam tools version 1.3. Counting of mapped reads for genomic features was performed using Subread feature Counts version 2.0.0 using the parameters -p -s 2 –g gene_name –t exon –Q 60 -B -C, the annotation file specified with –a was the Ensembl GRCm38 reference provided by Illumina’s iGenomes. Alignment and feature assignment statistics were summarized using MultiQC.

Differentially expressed genes were detected using the Bioconductor package DESeq2 version 1.38.1. Significance was set to an adjusted p-value less than or equal to 0.05. DESeq2 tests were run for differential expression using a negative binomial generalized linear models, dispersion estimates, and logarithmic fold changes. DESeq2 calculated log2 fold changes and Wald test p-values as well as performing independent filtering and adjusted for multiple testing using the Benjamini-Hochberg procedure to control the false discovery rate (FDR).

Gene ontology analysis was performed using the GOSeq Bioconductor package version 1.50.0. GOSeq performs Gene Ontology analysis while addressing biases present in RNA-seq data not found using other techniques, namely that expected read counts for a transcript are based on both, the gene level of expression, and the length of the transcript. While other tools are commonly used for GO term analysis, they do not account for this effect leading to results that are biased toward GO terms with longer transcripts. The Wallenius approximation used by GOSeq approximate GO term enrichment and calculated p-values for each GO category being overrepresented among genes that were differentially expressed. These p-values were corrected for multiple testing using the Benjamini-Hochberg Procedure.

### Cytokine and chemokine analysis of terminal serum

Terminal serum samples were tested in duplicate for cytokine and chemokine levels on a Bio-plex 200 (Bio-Rad) using a Bio-plex Pro Mouse Cytokine 23-plex assay according to kit instructions. Cytokines detected were IL-1α, IL-1β, IL-2, IL-3, IL-4, IL-5, IL-6, IL-9, IL-10, IL-12 (p40), IL-12 (p70), IL-13, IL-17A, eotaxin, G-CSF, GM-CSF, IFN-γ, KC, MCP-1 (MCAF), MIP-1α, MIP-1β, RANTES, and TNF-α. Serum was diluted 1:4 in the kit supplied diluent, incubated with cytokine-specific labeled beads and detected with biotinylated antibody and streptavidin-PE.

### Statistical analysis

Statistical analysis was conducted with GraphPad Prism software version 9.4.1 (GraphPad.) No data were excluded from the analysis.

