## Supplementary figures and images for "Comparative pathogenesis of two lineages of Powassan virus reveals distinct clinical outcome, neuropathology, and inflammation"

### Fig S1

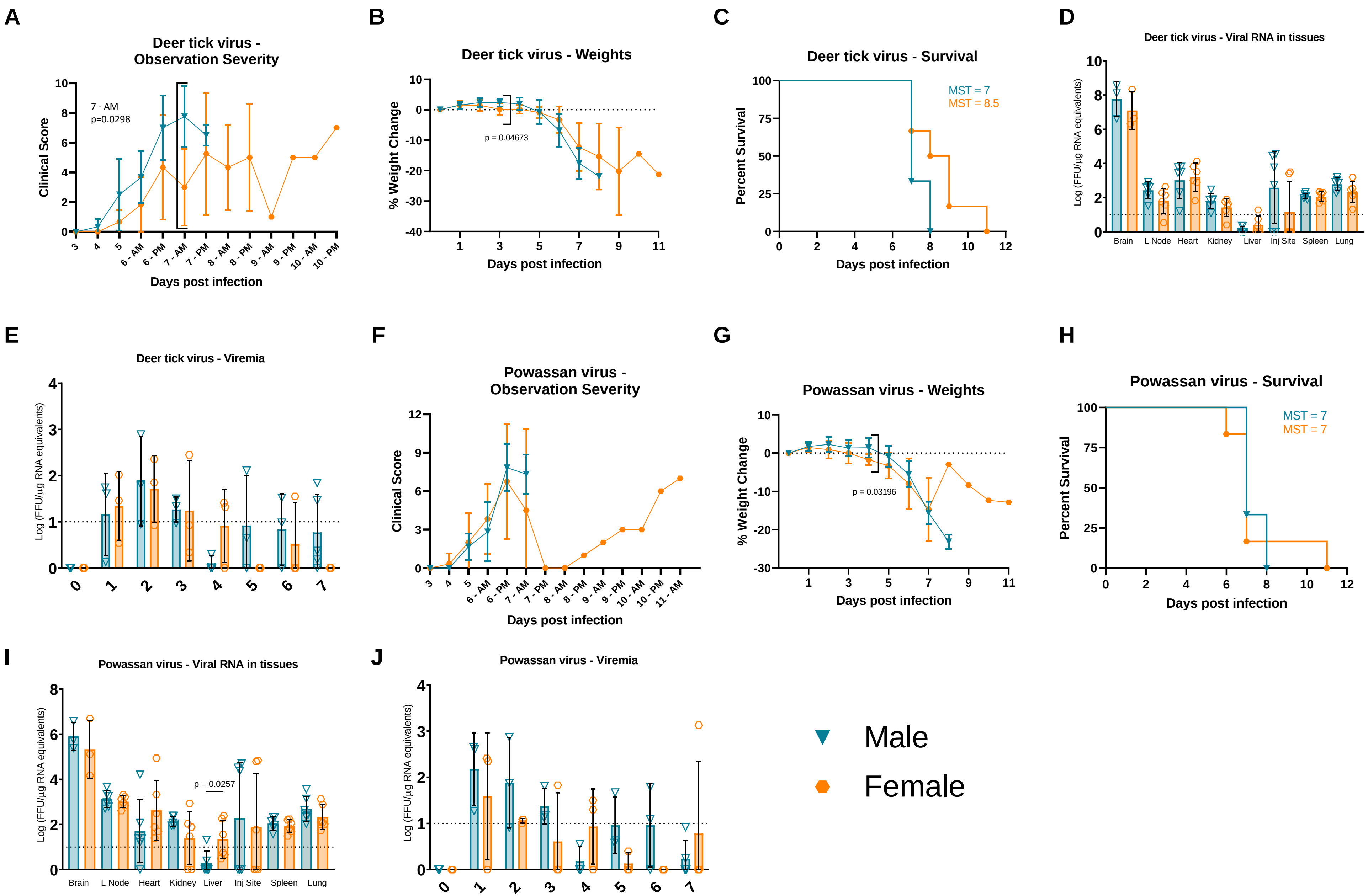

### Fig S2

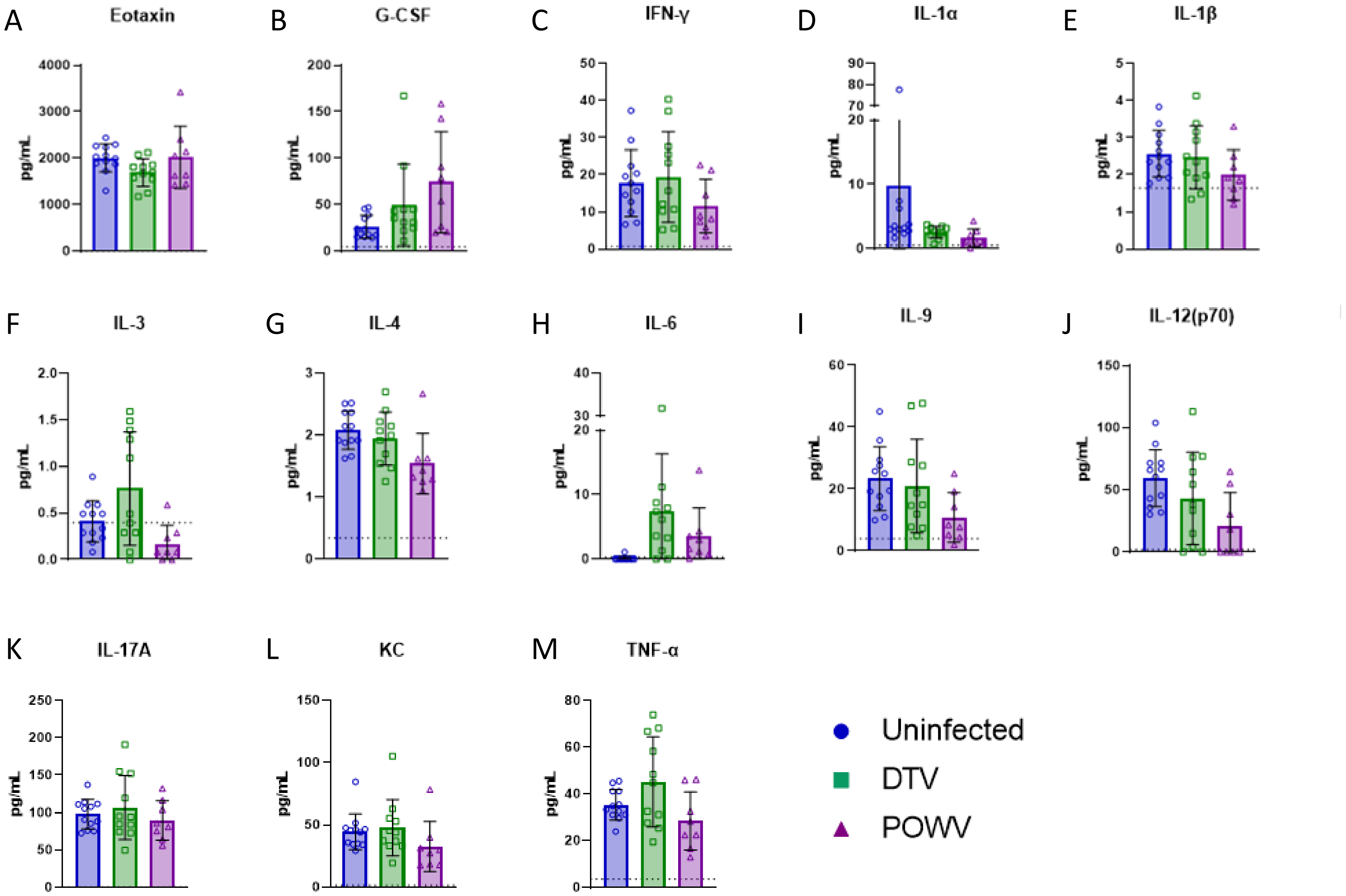

### Fig S3

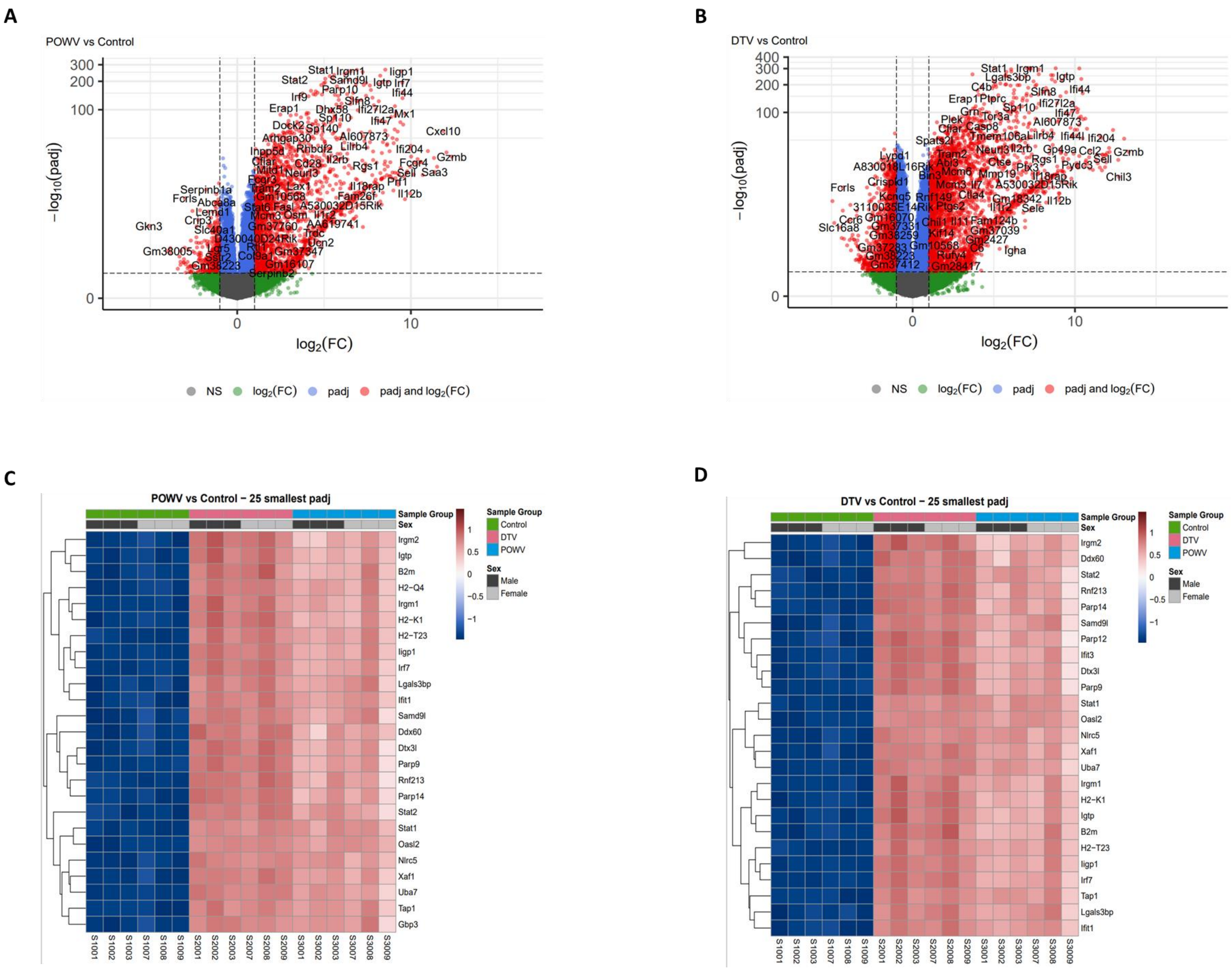

### Fig S4

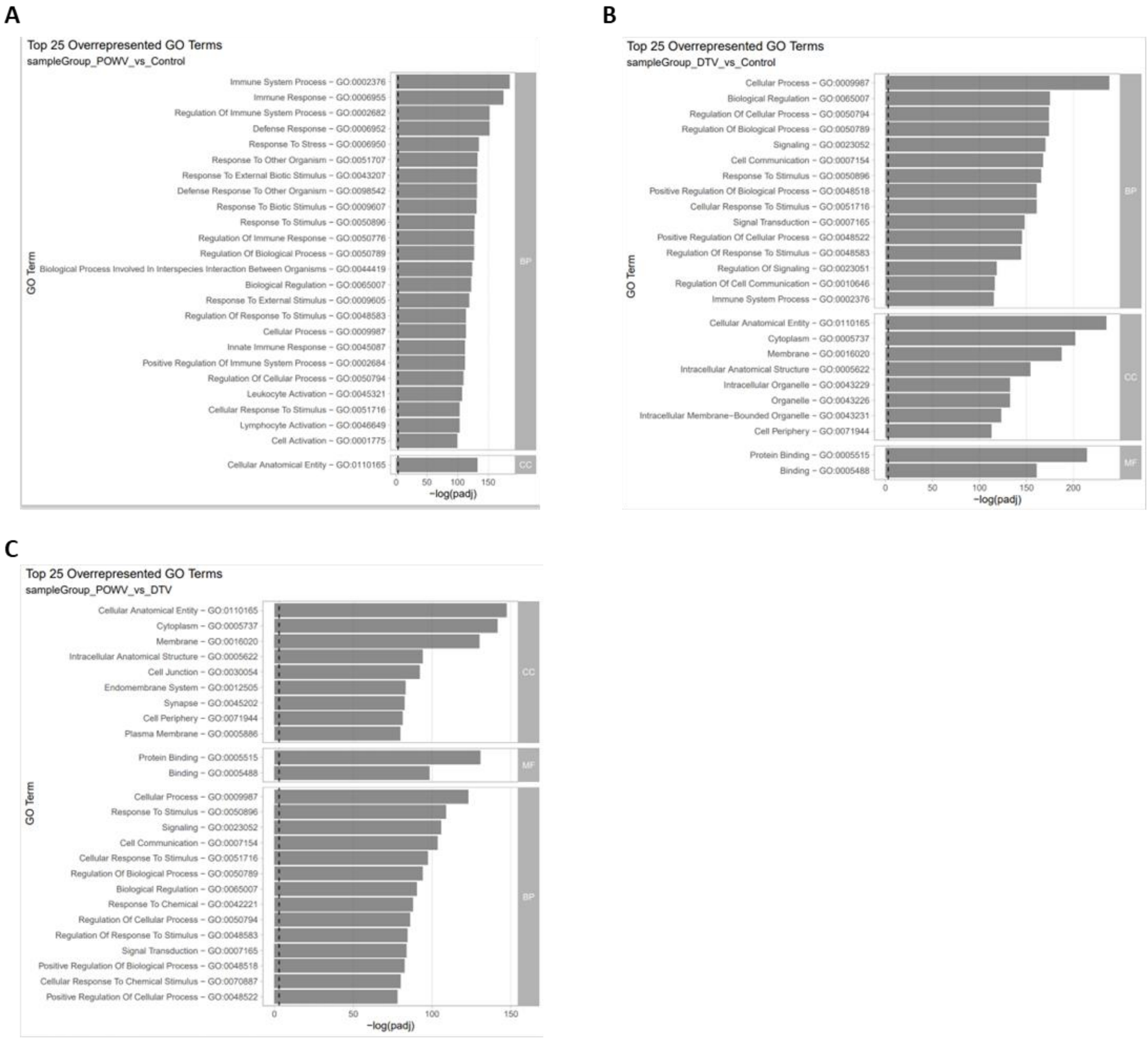
