## Supplementary material for "Comparative pathogenesis of two lineages of Powassan virus reveals distinct clinical outcome, neuropathology, and inflammation": Table S1

Clinical Scoring Chart

| Parameter | Degree of Parameter | Possible Score |
|---|---|---|
| **Weight** | Loss of up to 4.99% | 0 |
| | Loss of 5 to 9.99% | 1 |
| | Loss of 10 to 19.99% | 2 |
| | Loss ≥20%* | 3 |
| **Appearance** | Normal (smooth coat, eyes/nose clear) | 0 |
| | Reduced grooming, slightly ruffled | 1 |
| | Ruffled coat, ocular/nasal discharge, eye(s) partially closed, warm to touch | 2 |
| | No grooming, eye(s) closed, hunched posture, pale, piloerection, cold to touch | 3 |
| **Neurological Signs of Disease** | Normal | 0 |
| | Weak grip, reduced limb usage | 1 |
| | Paresis, ataxia, tremors, head tilt | 2 |
| | Paralysis*, seizures*, loss of righting reflex | 3 |
| **Provoked Behavior** | Normal | 0 |
| | Subdued but normal when stimulated | 1 |
| | Subdued even when stimulated, lethargic | 2 |
| | Unresponsive when stimulated*, moribund*, prostrate* | 3 |
| **Respiration** | Normal | 0 |
| | Rapid, shallow | 1 |
| | Diaphragmatic, labored | 2 |
| | Gasping* | 3 |

| | |
|---|---|
| **Cumulative Score** | Score of 0 - 3 = No intervention |
| | Score of ≥ 4 = More frequent monitoring |
| | Score of 6 to 8 = Contact PI and/or DLAR Veterinarian |
| | Score of ≥ 9 = Euthanasia |

* Observation requires humane euthanasia
