## Supplementary material for "Comparative pathogenesis of two lineages of Powassan virus reveals distinct clinical outcome, neuropathology, and inflammation": Table S1

| Group | Animal | Sex | Olfactory Bulb | Cerebral Cortex | Hippocampal formation | Thalamus | Hypothalamus | Midbrain | Pons | Medulla | Cerebellum | | | Meningitis/ encephalitis |
|---|---|---|---|---|---|---|---|---|---|---|---|---|---|---|
| | | | | | | | | | | | Purkinje cells | Granule cells | Molecular cells | |
| Uninfected | 1001 | M | - | - | - | - | - | - | - | - | - | - | - | - |
| | 1002 | M | - | - | - | - | - | - | - | - | - | - | - | - |
| | 1003 | M | - | - | - | - | - | - | - | - | - | - | - | - |
| | 1007 | F | NP | - | - | - | - | - | - | - | - | - | - | - |
| | 1008 | F | NP | - | - | - | - | - | - | - | - | - | - | - |
| | 1009 | F | - | - | - | - | - | - | - | - | - | - | - | - |
| DTV | 2001 | M | +<br>MG, NN | ++<br>MG, PC, NN | ++<br>MG, PC, NN | ++<br>MG, PC | +<br>MG | ++<br>MG, PC | NP | NP | - | +<br>NN | +<br>MG | ME |
| | 2002 | M | +<br>MG, NN | ++<br>MG, PC, NN | +<br>MG, PC, NN | ++<br>MG, PC | +<br>MG, NN | ++<br>MG | NP | NP | NP | NP | NP | ME |
| | 2003 | M | - | ++<br>MG, PC, NN | +<br>MG, PC, NN | ++<br>MG, PC, NN | ++<br>MG, PC, NN | ++<br>MG, PC, NN | +<br>MG, PC | ++<br>MG, PC | +<br>NN | +<br>NN | ++<br>MG, PC | ME |
| | 2007 | F | ++<br>MG, NN | +++<br>MG, PC, NN | +++<br>MG, PC, NN | ++<br>MG, PC, NN | ++<br>MG, PC, NN | ++<br>MG, PC, NN | +<br>MG, PC, NN | NP | - | +<br>NN | +<br>MG | ME |
| | 2008 | F | +<br>MG, NN | +++<br>MG, PC, NN | ++<br>MG, PC, NN | ++<br>MG, PC, NN | ++<br>MG, PC, NN | ++<br>MG, PC | +<br>MG, NN | +<br>MG, PC | - | - | +<br>MG, PC | ME |
| | 2009 | F | +<br>MG | +<br>MG, PC, NN | +<br>MG, PC, NN | +<br>MG, PC | +<br>MG, PC | +<br>MG, PC | NP | NP | - | - | +<br>MG, PC | ME |
| POWV | 3001 | M | NP | +<br>MG, PC | - | +<br>MG | +<br>MG, PC | +<br>MG, PC | +<br>MG, PC | ++<br>MG, PC | +<br>NN | +<br>NN | ++<br>MG | ME |
| | 3002 | M | +<br>MG | +<br>MG | - | +<br>MG, PC | +<br>MG, PC | +<br>MG, PC, NN | +<br>MG, PC | ++<br>MG, PC, NN | - | +<br>NN | +<br>MG | ME |
| | 3003 | M | NP | +<br>MG, PC, NN | - | +<br>MG, PC, NN | +<br>MG, PC | +<br>MG, PC, NN | +<br>MG | +<br>MG, NN | - | +<br>NN | +<br>MG | ME |
| | 3007 | F | +<br>MG, NN | +<br>MG, PC, NN | - | - | +<br>MG | +<br>MG | +<br>PC | +<br>MG, PC | - | +<br>NN | +<br>MG | ME |
| | 3008 | F | +<br>MG, NN | ++<br>MG, PC, NN | ++<br>MG, PC, NN | ++<br>MG, PC, NN | ++<br>MG, PC, NN | ++<br>MG, PC, NN | +<br>MG, PC | ++<br>MG, NN | - | +<br>NN | +<br>MG | ME |
| | 3009 | F | - | ++<br>MG, PC, NN | +<br>MG, PC | +<br>MG | +<br>MG, PC | +<br>MG | +<br>MG | +<br>MG, PC | - | - | +<br>MG, PC | ME |
