## Supplementary material for "Comparative pathogenesis of two lineages of Powassan virus reveals distinct clinical outcome, neuropathology, and inflammation": Table S1

| Group | Animal | Sex | Olfactory Bulb | Cerebral Cortex | Hippocampal formation | Thalamus | Hypothalamus | Midbrain | Pons | Medulla | Cerebellum: Purkinje cells | Cerebellum: Granule cells | Cerebellum: Molecular cells | Meningitis/ encephalitis |
|---|---|---|---|---|---|---|---|---|---|---|---|---|---|---|
| Uninfected | 1001 | M | - | - | - | - | - | - | - | - | - | - | - | - |
| | | | + | + | + | + | + | + | + | + | - | + | + | |
| | 1002 | M | - | - | - | - | - | - | - | - | - | - | - | - |
| | | | + | + | + | + | + | + | + | + | - | + | + | |
| | 1003 | M | - | - | - | - | - | - | - | - | - | - | - | - |
| | | | + | + | + | + | + | + | + | + | - | + | + | |
| | 1007 | F | - | - | - | - | - | - | - | - | - | - | - | - |
| | | | + | + | + | + | + | + | + | + | - | + | + | |
| | 1008 | F | NP | - | - | - | - | - | - | - | - | - | - | - |
| | | | | + | + | + | + | + | + | + | - | + | + | |
| | 1009 | F | - | - | - | - | - | - | - | - | - | - | - | - |
| | | | + | + | + | + | + | + | + | + | - | + | + | |
| DTV | 2001 | M | +++ | +++ | +++ | +++ | ++ | ++ | NP | NP | + | ++ | + | ME |
| | | | ++ | ++ | +++ | ++ | ++ | ++ | | | - | ++ | ++ | |
| | 2002 | M | ++ | +++ | +++ | +++ | ++ | ++ | NP | NP | - | ++ | + | ME |
| | | | ++ | ++ | ++ | ++ | ++ | ++ | | | - | ++ | ++ | |
| | 2003 | M | + | +++ | ++ | ++ | ++ | + | ++ | + | + | + | ++ | ME |
| | | | ++ | +++ | ++ | ++ | ++ | ++ | ++ | ++ | - | ++ | ++ | |
| | 2007 | F | ++ | +++ | +++ | +++ | +++ | +++ | +++ | NP | - | + | + | ME |
| | | | ++ | ++ | +++ | ++ | ++ | ++ | ++ | | - | ++ | ++ | |
| | 2008 | F | + | +++ | + | +++ | + | ++ | + | + | - | + | + | ME |
| | | | ++ | ++ | ++ | ++ | ++ | ++ | ++ | ++ | - | ++ | ++ | |
| | 2009 | F | + | ++ | +++ | ++ | + | + | NP | NP | - | + | + | ME |
| | | | ++ | ++ | +++ | ++ | ++ | ++ | | | - | ++ | ++ | |
| POWV | 3001 | M | NP | + | - | + | + | + | + | + | + | + | ++ | ME |
| | | | | + | + | + | ++ | ++ | ++ | ++ | - | + | ++ | |
| | 3002 | M | ++ | ++ | - | ++ | ++ | +++ | ++ | +++ | +++ | +++ | +++ | ME |
| | | | ++ | ++ | ++ | ++ | ++ | ++ | ++ | ++ | - | ++ | ++ | |
| | 3003 | M | NP | ++ | - | + | + | ++ | + | ++ | + | ++ | + | ME |
| | | | | ++ | ++ | ++ | ++ | ++ | ++ | ++ | - | ++ | ++ | |
| | 3007 | F | + | + | + | + | + | ++ | + | ++ | ++ | ++ | ++ | ME |
| | | | ++ | ++ | ++ | + | ++ | ++ | ++ | ++ | - | ++ | ++ | |
| | 3008 | F | ++ | +++ | +++ | +++ | ++ | ++ | ++ | +++ | + | ++ | ++ | ME |
| | | | + | ++ | ++ | ++ | + | ++ | ++ | ++ | - | ++ | ++ | |
| | 3009 | F | - | + | - | + | - | - | - | + | + | + | + | ME |
| | | | + | +++ | ++ | ++ | ++ | ++ | ++ | ++ | - | ++ | ++ | |
