## Supplementary material for "Comparative pathogenesis of two lineages of Powassan virus reveals distinct clinical outcome, neuropathology, and inflammation": Table S1

| Gene | baseMean | log2FoldChange | lfcSE | stat |
|---|---|---|---|---|
| 4933401J01Rik | 0 | NA | NA | NA |
| Gm26206 | 0 | NA | NA | NA |
| Xkr4 | 1236.723264 | 0.839366087 | 0.099232743 | 8.458559728 |
| Gm18956 | 0.257781492 | 1.475868219 | 2.50666619 | 0.588777327 |
| Gm37180 | 183.2012087 | 0.572810098 | 0.243356518 | 2.353789835 |
| Gm37363 | 156.7123429 | 0.803465212 | 0.242261484 | 3.316520642 |
| Gm37686 | 55.26816088 | 0.837496087 | 0.287117778 | 2.916907803 |
| Gm1992 | 0 | NA | NA | NA |
| Gm37329 | 74.22829483 | 0.632754959 | 0.244101202 | 2.592182894 |
| Gm7341 | 0.472128458 | -1.085619329 | 1.531169846 | -0.709013002 |
| Gm38148 | 143.524156 | 0.708383683 | 0.271276604 | 2.611296636 |
| Gm19938 | 181.8288108 | 0.713542293 | 0.257309833 | 2.773085987 |
| Gm10568 | 17.47471811 | 1.182893378 | 0.286107292 | 4.134439816 |
| Gm38385 | 0 | NA | NA | NA |
| Gm27396 | 0 | NA | NA | NA |
| Gm37381 | 0.042716945 | 0.037062677 | 3.724945049 | 0.009949859 |
| Gm6101 | 0.038981395 | -0.443816764 | 3.724945049 | -0.119147198 |
| Rp1 | 1.19669966 | 1.637609498 | 1.06567115 | 1.536693095 |
| Gm37483 | 0 | NA | NA | NA |
| Sox17 | 14.01852502 | 0.918698863 | 0.314319317 | 2.922820242 |
| Gm37587 | 0.590622753 | 0.01607133 | 1.363104756 | 0.011790239 |
| Gm7357 | 0.431020911 | 0.079633132 | 1.491702371 | 0.053384062 |
| Gm22307 | 0.205539566 | -1.28460531 | 2.601261962 | -0.493839271 |
| Gm38076 | 0 | NA | NA | NA |
| Gm37323 | 0.424160584 | 0.037077311 | 1.52868388 | 0.024254401 |
| Gm7369 | 0.062426873 | 0.517959912 | 3.724945049 | 0.139051692 |
| Gm6085 | 7.026365196 | -0.361122986 | 0.438896948 | -0.82279676 |
| Gm6119 | 1.007284973 | -0.037681575 | 1.093564618 | -0.034457566 |
| Gm25493 | 0.04393175 | 0.037062677 | 3.724945049 | 0.009949859 |
| Gm2053 | 0.509840029 | -0.443818745 | 1.49354531 | -0.297157871 |
| Gm6123 | 6.523572364 | -0.130398727 | 0.379097846 | -0.343971164 |
| Mrpl15 | 339.6237885 | -0.243341084 | 0.09965222 | -2.441903283 |
| Gm37144 | 10.84448109 | -0.115826652 | 0.434033141 | -0.266861309 |
| Lypla1 | 301.4400782 | -0.19793214 | 0.105767192 | -1.871394492 |
| Gm37988 | 0.099106094 | -0.93550548 | 3.724188147 | -0.251197158 |
| Tcea1 | 660.8047814 | 0.063848162 | 0.09766048 | 0.653776859 |
| Gm6104 | 7.397713871 | -0.020051845 | 0.383950468 | -0.052225082 |
| Gm37277 | 41.2031627 | 0.20306938 | 0.173930692 | 1.167530454 |
| Rgs20 | 298.6651195 | -0.453484928 | 0.103294312 | -4.39022167 |
| Gm17100 | 2.600587843 | -0.074727993 | 0.679424236 | -0.109987236 |
| Gm37079 | 17.87741427 | 0.06594537 | 0.35902948 | 0.183676755 |
| Gm16041 | 0.448921586 | -1.875220217 | 1.415597091 | -1.324684989 |
| Gm17101 | 1.407567115 | -0.428649352 | 0.913663053 | -0.469154741 |
| Atp6v1h | 1710.406335 | -0.13474914 | 0.09227176 | -1.460350821 |
| Gm7182 | 0 | NA | NA | NA |
| Gm37567 | 5.040767136 | 1.163623822 | 0.580149784 | 2.005729992 |

| | | | | |
|---|---|---|---|---|
| Gm38264 | 0.54045421 | 1.698276827 | 1.317488612 | 1.289025811 |
| Oprk1 | 98.2787742 | 0.739873124 | 0.157257215 | 4.704859653 |
| Gm36965 | 0 | NA | NA | NA |
| Gm22463 | 0 | NA | NA | NA |
| Gm37429 | 0 | NA | NA | NA |
| Npbwr1 | 11.64878932 | 0.29048174 | 0.321551037 | 0.903376781 |
| Gm19214 | 0.066050875 | -0.443818988 | 3.724945049 | -0.119147795 |
| Rb1cc1 | 2300.690646 | 0.128502568 | 0.05402501 | 2.378575539 |
| 4732440D04Rik | 117.8176336 | -0.388320676 | 0.171362594 | -2.26607608 |
| Fam150a | 0.409266314 | 0.847619893 | 1.425607067 | 0.594567685 |
| Gm2147 | 0.280898025 | 0.477135039 | 1.621170667 | 0.29431512 |
| Gm7417 | 0 | NA | NA | NA |
| Gm19026 | 17.7062556 | 0.030799775 | 0.263375966 | 0.11694224 |
| St18 | 573.9728649 | -0.030181126 | 0.131542062 | -0.229440877 |
| Gm37405 | 0 | NA | NA | NA |
| Gm7449 | 0.090074004 | 0.517955464 | 3.724945049 | 0.139050498 |
| Gm37108 | 10.32889666 | -0.476705456 | 0.313753354 | -1.519363697 |
| Gm37275 | 0 | NA | NA | NA |
| Gm37225 | 0 | NA | NA | NA |
| Gm37489 | 0 | NA | NA | NA |
| Gm5694 | 0 | NA | NA | NA |
| Pcmtd1 | 1844.737977 | 0.065395714 | 0.081494647 | 0.802454111 |
| Gm26983 | 0 | NA | NA | NA |
| Gm38372 | 224.9946318 | -0.213249049 | 0.237704369 | -0.89711876 |
| Gm9826 | 6.912980464 | 0.242114438 | 0.39759771 | 0.608943239 |
| Nras-ps2 | 0 | NA | NA | NA |
| Gm23274 | 0 | NA | NA | NA |
| Gm19002 | 0.085327104 | -0.905414734 | 3.723500899 | -0.243162217 |
| Gm26901 | 31.62112565 | 0.541966618 | 0.313647047 | 1.727950647 |
| Gm18984 | 0 | NA | NA | NA |
| Gm30414 | 0 | NA | NA | NA |
| Gm37791 | 0 | NA | NA | NA |
| Gm7470 | 0.066050875 | -0.443818988 | 3.724945049 | -0.119147795 |
| Rps2-ps2 | 0 | NA | NA | NA |
| Gm38216 | 0 | NA | NA | NA |
| Sntg1 | 622.888287 | 0.648560069 | 0.113789424 | 5.699651578 |
| Gm38024 | 20.53816256 | 0.92384983 | 0.399690276 | 2.311414328 |
| Gm16284 | 4.895741945 | 1.712018131 | 0.591368227 | 2.895012029 |
| Gm38259 | 46.04207795 | 1.124145625 | 0.261264824 | 4.302705611 |
| Gm23358 | 1.395067866 | 0.591861816 | 0.921270079 | 0.642441158 |
| Gm37005 | 8.035261982 | 0.700329929 | 0.393858168 | 1.778127217 |
| Gm15452 | 1.564680506 | 0.423758667 | 0.849829132 | 0.498639845 |
| Gm36964 | 0.112396584 | -0.917692504 | 3.724434845 | -0.246397787 |
| Gm24276 | 0.547110457 | -0.751378452 | 1.607894986 | -0.467305675 |
| Gm6152 | 0.180148697 | 1.020111592 | 2.703307731 | 0.377356814 |
| Gm7445 | 0.326379708 | 0.069252218 | 3.697733777 | 0.018728287 |
| Gm7493 | 10.60824579 | 0.221564487 | 0.50769971 | 0.436408535 |

| | | | | |
|---|---|---|---|---|
| Gm37629 | 21.84260411 | 0.523235587 | 0.308929041 | 1.693707999 |
| Gm38008 | 8.876683074 | 0.05310988 | 0.436727585 | 0.121608715 |
| Gm37143 | 0 | NA | NA | NA |
| Gm6187 | 0.1709406 | 0.51795796 | 3.385131608 | 0.153009697 |
| Gm18299 | 0 | NA | NA | NA |
| Gm7512 | 0 | NA | NA | NA |
| Gm24765 | 0 | NA | NA | NA |
| Gm18300 | 1.926091363 | -0.073674472 | 0.75418272 | -0.097687828 |
| Rrs1 | 218.6880439 | -0.406013159 | 0.122557728 | -3.312831977 |
| Adhfe1 | 178.1879691 | 0.608081743 | 0.144905344 | 4.1964066 |
| 2610203C22Rik | 5.506898143 | 0.294937423 | 0.490441196 | 0.601371634 |
| Gm6161 | 3.14279713 | 0.683078468 | 0.613070081 | 1.114193124 |
| 3110035E14Rik | 1247.772309 | 0.971760564 | 0.182741247 | 5.317685964 |
| Gm29520 | 0 | NA | NA | NA |
| Mybl1 | 100.6319816 | -0.148832133 | 0.165421491 | -0.899714612 |
| Vcpip1 | 2191.495749 | -0.06469258 | 0.077414883 | -0.835660757 |
| 1700034P13Rik | 2.716773814 | 0.427400965 | 0.654560432 | 0.652958756 |
| Sgk3 | 387.7590365 | -0.000350125 | 0.10926653 | -0.003204319 |
| Gm6195 | 2.037763675 | 0.336083743 | 0.889266089 | 0.377933835 |
| Gm22607 | 0 | NA | NA | NA |
| Mcmdc2 | 2.919172438 | -0.277632118 | 0.590917089 | -0.46983261 |
| Snhg6 | 57.04786074 | 0.111697356 | 0.196983798 | 0.567038291 |
| Snord87 | 1.794878382 | 0.225109532 | 0.794925799 | 0.283183075 |
| Tcf24 | 4.573727287 | -1.408858873 | 0.541206977 | -2.603179439 |
| E330040D14Rik | 0.234971822 | 0.517974441 | 2.12154591 | 0.244149532 |
| Ppp1r42 | 4.644333044 | 0.410135707 | 0.508052943 | 0.807269622 |
| Gm15818 | 0 | NA | NA | NA |
| Gm24674 | 0 | NA | NA | NA |
| Cops5 | 641.9122966 | -0.146331943 | 0.098909909 | -1.479446744 |
| Cspp1 | 856.404923 | 0.146182108 | 0.169870257 | 0.86055152 |
| 1700047N06Rik | 0 | NA | NA | NA |
| Arfgef1 | 2686.960645 | -0.050831768 | 0.050402977 | -1.008507266 |
| Gm26348 | 0 | NA | NA | NA |
| Gm37569 | 39.97338864 | 0.743057028 | 0.30556449 | 2.431751894 |
| Cpa6 | 18.52223898 | -0.585981257 | 0.532792462 | -1.099830233 |
| Gm38005 | 2.757436252 | -2.202176663 | 1.155508327 | -1.905807697 |
| Gm15604 | 1.883799777 | -1.552219789 | 1.100653383 | -1.410271219 |
| Gm25253 | 0.347911031 | -0.854719034 | 3.247678021 | -0.263178501 |
| Gm15603 | 0.042716945 | 0.037062677 | 3.724945049 | 0.009949859 |
| Gm37133 | 0 | NA | NA | NA |
| Gm5522 | 0.046345709 | -0.443818988 | 3.724945049 | -0.119147795 |
| Gm28659 | 0 | NA | NA | NA |
| Gm22963 | 0 | NA | NA | NA |
| Prex2 | 1191.400917 | 0.463428626 | 0.15724724 | 2.947133603 |
| Gm24173 | 0 | NA | NA | NA |
| Gm28686 | 0 | NA | NA | NA |
| Gm37410 | 0.060174537 | 0.517957688 | 3.724945049 | 0.139051095 |

| | | | | |
|---|---|---|---|---|
| A830018L16Rik | 657.3086267 | 0.91501361 | 0.107533783 | 8.509080443 |
| Gm38178 | 28.55896715 | 0.752088367 | 0.319584325 | 2.353333091 |
| Gm29273 | 0 | NA | NA | NA |
| Gm38069 | 9.696124784 | 0.965278986 | 0.478672604 | 2.016574542 |
| Gm7560 | 0.344328494 | 1.456618836 | 1.672981588 | 0.870672365 |
| Gm22633 | 0 | NA | NA | NA |
| Mir6341 | 0 | NA | NA | NA |
| Gm2383 | 0.481037375 | 1.001999486 | 1.448396742 | 0.691799047 |
| Gm6216 | 0 | NA | NA | NA |
| Gm17644 | 0.118371171 | 0.037073798 | 3.724945049 | 0.009952844 |
| Gm29663 | 0 | NA | NA | NA |
| Sulf1 | 425.5919252 | -0.63780741 | 0.128650739 | -4.957666138 |
| Slco5a1 | 118.4898984 | -0.550325282 | 0.150086984 | -3.666708921 |
| Gm37162 | 0 | NA | NA | NA |
| Gm5250 | 0 | NA | NA | NA |
| Gm29283 | 0 | NA | NA | NA |
| Prdm14 | 0.178837887 | -0.891757593 | 2.696671676 | -0.330688233 |
| Ncoa2 | 2049.993035 | -0.012796522 | 0.046549008 | -0.274904288 |
| Gm38376 | 24.50757996 | 0.671204296 | 0.387949316 | 1.730133986 |
| Gm37409 | 22.73165489 | 0.262248234 | 0.336809477 | 0.778624865 |
| Gm27881 | 0 | NA | NA | NA |
| Gm38223 | 5.5083264 | 0.167902337 | 0.494925826 | 0.339247475 |
| Gm37702 | 18.1938087 | 0.367059698 | 0.315639263 | 1.162908867 |
| Gm23169 | 4.580170235 | 0.198850125 | 0.615107268 | 0.323277151 |
| Gm38120 | 65.50301172 | 0.171688424 | 0.257753075 | 0.666096512 |
| Gm38380 | 51.36537569 | 0.016916191 | 0.307188317 | 0.055067819 |
| Gm29570 | 1.379700947 | 0.140163467 | 0.918745562 | 0.152559612 |
| Gm7593 | 0.038981395 | -0.443816764 | 3.724945049 | -0.119147198 |
| Tram1 | 601.6593407 | -0.451622353 | 0.101186089 | -4.463284981 |
| Lactb2 | 271.8818834 | -0.281230646 | 0.097681096 | -2.879069303 |
| Gm38319 | 9.470931737 | 0.541563311 | 0.527936976 | 1.025810533 |
| Xkr9 | 0.562487033 | 0.260482293 | 1.292375828 | 0.201553052 |
| Gm26273 | 0 | NA | NA | NA |
| Gm5523 | 0.58821479 | -0.869014867 | 1.196971373 | -0.726011404 |
| Gm36947 | 0.09689823 | 0.037062677 | 3.724945049 | 0.009949859 |
| Gm25491 | 0 | NA | NA | NA |
| Gm37400 | 0 | NA | NA | NA |
| Eya1 | 126.1615464 | 0.551004623 | 0.132893918 | 4.146198954 |
| Gm22616 | 0 | NA | NA | NA |
| Gm37444 | 2.632415511 | 2.263514579 | 0.968656782 | 2.336756032 |
| Gm9947 | 16.04594573 | 0.706877729 | 0.299888798 | 2.357132826 |
| Msc | 6.16192073 | -0.355832647 | 0.444071316 | -0.801296176 |
| Gm37412 | 7.68759231 | 1.179372772 | 0.40858492 | 2.886481398 |
| Smt3h2-ps4 | 2.907781834 | 0.883839915 | 0.577157009 | 1.531368244 |
| Rpl5-ps1 | 8.684837921 | -0.361251474 | 0.377561621 | -0.956801364 |
| Trpa1 | 3.284363746 | 0.848509662 | 0.585245151 | 1.449836295 |
| Gm26345 | 0 | NA | NA | NA |

| | | | | |
|---|---|---|---|---|
| Gm10566 | 6.613104402 | 0.006590946 | 0.399862001 | 0.016483053 |
| Kcnb2 | 762.7240551 | 0.429911087 | 0.10583834 | 4.061959844 |
| Gm38116 | 50.40214469 | 0.166346069 | 0.35806602 | 0.464568151 |
| Gm37138 | 26.46315223 | 0.45425943 | 0.290445992 | 1.564006537 |
| Gm25227 | 0 | NA | NA | NA |
| Gm25168 | 0.155005853 | 0.03706712 | 2.830519028 | 0.01309552 |
| Gm6060 | 0.066050875 | -0.443818988 | 3.724945049 | -0.119147795 |
| Gm28669 | 0.095435502 | 0.037073798 | 3.724945049 | 0.009952844 |
| Terf1 | 191.9488994 | 0.286549774 | 0.130635868 | 2.193499986 |
| Gm37509 | 44.48996355 | 0.331795657 | 0.326979769 | 1.014728396 |
| Sbspon | 3.624262949 | -0.070727585 | 0.620241482 | -0.114032336 |
| Gm23259 | 0.061920613 | -0.443816764 | 3.724945049 | -0.119147198 |
| Gm7634 | 0 | NA | NA | NA |
| 4930444P10Rik | 4.660125666 | -0.179298963 | 0.476527829 | -0.376261264 |
| Rpl7 | 1699.234633 | -0.189322773 | 0.113641324 | -1.665967688 |
| Rdh10 | 199.9939912 | -0.794994498 | 0.158117544 | -5.02787025 |
| Gm38249 | 4.612676316 | -0.671659517 | 0.505258126 | -1.329339368 |
| Gm28095 | 0 | NA | NA | NA |
| Stau2 | 1742.3238 | 0.4112672 | 0.076169242 | 5.399386786 |
| Gm7568 | 0.218065207 | 0.504347713 | 2.536190695 | 0.198860328 |
| Ube2w | 820.6053904 | 0.102783931 | 0.058932309 | 1.744101556 |
| Gm38342 | 3.77249287 | 0.094940041 | 0.47909208 | 0.198166585 |
| Tceb1 | 826.0456009 | 0.1850123 | 0.084003272 | 2.202441603 |
| D030040B21Rik | 0.250367046 | 0.517972245 | 2.484698329 | 0.208464842 |
| Gm7654 | 0.171036444 | -0.443823548 | 2.777278927 | -0.159805176 |
| Tmem70 | 222.9577256 | -0.330027288 | 0.107762061 | -3.062555455 |
| Ly96 | 21.7470159 | -0.484892797 | 0.249337981 | -1.944720955 |
| Gm29649 | 0 | NA | NA | NA |
| Gm5828 | 11.26054785 | -0.470341703 | 0.295018037 | -1.594281175 |
| Gm28376 | 0.229484813 | 0.998854299 | 3.114678805 | 0.320692553 |
| 4921511E07Rik | 0 | NA | NA | NA |
| Jph1 | 403.3093643 | 1.089325376 | 0.096021365 | 11.34461458 |
| Gm28783 | 0.484663642 | 0.486139371 | 1.315328391 | 0.369595436 |
| Gdap1 | 1402.76127 | 0.24637053 | 0.061793181 | 3.987018064 |
| Gm28784 | 4.433742602 | -0.332390296 | 0.516116927 | -0.644021304 |
| Gm28785 | 0.049553047 | -0.443818988 | 3.724945049 | -0.119147795 |
| Gm5251 | 0.15956642 | 0.036118649 | 3.418731032 | 0.010564929 |
| Rbm6-ps1 | 6.916369844 | -0.134833302 | 0.421868167 | -0.31961004 |
| Gm6075 | 0.038981395 | -0.443816764 | 3.724945049 | -0.119147198 |
| Gm25166 | 0 | NA | NA | NA |
| Pi15 | 14.69231755 | -0.41879861 | 0.287561965 | -1.456376923 |
| Gm28154 | 8.974735632 | 1.124579047 | 0.406191311 | 2.768594545 |
| Gm16070 | 24.47860644 | 1.016849254 | 0.261340994 | 3.890890751 |
| Crispld1 | 57.83831356 | 1.360605004 | 0.168580749 | 8.070939384 |
| Gm28153 | 1.598622982 | 0.980374497 | 0.915953365 | 1.070332327 |
| Gm7690 | 0 | NA | NA | NA |
| Gm18775 | 13.00822407 | 0.20149837 | 0.310280129 | 0.649407909 |

| | | | | |
|---|---|---|---|---|
| Gm17969 | 0 | NA | NA | NA |
| Gm28755 | 1.336776882 | 0.076913585 | 0.914745472 | 0.084081952 |
| Gm28756 | 10.18614479 | 0.880419506 | 0.397773617 | 2.213368276 |
| Crisp4 | 0 | NA | NA | NA |
| Defb44-ps | 0.047948698 | 0.037064902 | 3.724945049 | 0.009950456 |
| Defb18 | 0 | NA | NA | NA |
| Gm15386 | 0 | NA | NA | NA |
| Gm17837 | 0.172125443 | 0.517973273 | 2.321733368 | 0.223097656 |
| Gm29006 | 0 | NA | NA | NA |
| Gm5252 | 0 | NA | NA | NA |
| Gm24075 | 0 | NA | NA | NA |
| Gm28343 | 0 | NA | NA | NA |
| Gm28344 | 0 | NA | NA | NA |
| Gm15825 | 1.225430203 | 1.166367557 | 0.896499873 | 1.301023673 |
| Tfap2d | 23.18306498 | 0.796129894 | 0.278367329 | 2.859997602 |
| Gm10075 | 1.913880918 | 0.26584631 | 0.709121635 | 0.374895219 |
| Tfap2b | 140.1785569 | 0.125836013 | 0.156126507 | 0.805987501 |
| Gm28340 | 0.106520246 | 0.037071574 | 3.724945049 | 0.009952247 |
| Gm28341 | 0 | NA | NA | NA |
| Gm4849 | 4.195570756 | -0.813154239 | 0.484402021 | -1.678676396 |
| Gm24633 | 0 | NA | NA | NA |
| Gm38382 | 0.088383099 | 0.037058229 | 3.724945049 | 0.009948665 |
| Pkhd1 | 2.807822083 | -0.38387076 | 0.724980824 | -0.529490914 |
| Gm37315 | 0 | NA | NA | NA |
| 4930486I03Rik | 0.427990203 | 1.406301512 | 1.751356532 | 0.802978426 |
| Gm15795 | 0.244548831 | 0.559688928 | 2.102040361 | 0.266259839 |
| Gm24162 | 0 | NA | NA | NA |
| Mir206 | 0 | NA | NA | NA |
| Gm28653 | 1.730304082 | 0.256793103 | 0.848702544 | 0.302571383 |
| Mir133b | 0 | NA | NA | NA |
| Il17a | 0 | NA | NA | NA |
| Il17f | 0.328867834 | -1.355993497 | 2.0739129 | -0.653833387 |
| Mcm3 | 85.70957654 | -0.723034362 | 0.258445496 | -2.797628026 |
| Gm20587 | 0 | NA | NA | NA |
| Gm28065 | 0.068097359 | 0.037064902 | 3.724945049 | 0.009950456 |
| Gm24723 | 0 | NA | NA | NA |
| 6720483E21Rik | 3.174408629 | 0.721729482 | 0.721889369 | 0.999778516 |
| Paqr8 | 1990.654714 | 0.186162451 | 0.112734986 | 1.651328107 |
| Gm24171 | 0 | NA | NA | NA |
| Gm28064 | 5.60213491 | 0.871763563 | 0.460525875 | 1.89297412 |
| Efhc1 | 14.45078694 | -0.09782058 | 0.282389116 | -0.346403505 |
| Tram2 | 100.5415102 | -0.799154516 | 0.170778576 | -4.679477563 |
| Gm28287 | 0.218494371 | 0.037076968 | 2.166806342 | 0.017111344 |
| Gm2693 | 2.478342139 | 1.311551398 | 0.708517837 | 1.851119803 |
| Tmem14a | 140.3427173 | 0.894348645 | 0.157935315 | 5.662752778 |
| Gm38224 | 3.730747895 | -0.643410946 | 0.578798863 | -1.111631323 |
| Gsta3 | 15.33756154 | 0.929618843 | 0.368072714 | 2.525639116 |

| | | | | |
|---|---|---|---|---|
| Gm28836 | 1.863919813 | 0.699450339 | 0.810755385 | 0.862714392 |
| Gm7094 | 1.910334788 | 0.612995952 | 0.794477148 | 0.771571534 |
| Gm4956 | 0.357620736 | -1.173832251 | 1.524107575 | -0.770176771 |
| Gm21909 | 0.066050875 | -0.443818988 | 3.724945049 | -0.119147795 |
| Khdc1a | 0.075043205 | 0.517957688 | 3.724945049 | 0.139051095 |
| Gm16329 | 0 | NA | NA | NA |
| Gm16327 | 0 | NA | NA | NA |
| Khdc1c | 0.097139547 | -0.443816764 | 3.724945049 | -0.119147198 |
| Gm2709 | 0 | NA | NA | NA |
| Gm16533 | 0 | NA | NA | NA |
| Khdc1b | 0 | NA | NA | NA |
| Gm16328 | 0 | NA | NA | NA |
| Kcnq5 | 996.2770014 | 1.094875613 | 0.144750476 | 7.563882658 |
| Gm36949 | 26.09595903 | 1.364961762 | 0.333863046 | 4.088388268 |
| Gm7658 | 100.9699313 | 0.086301869 | 0.16178377 | 0.533439598 |
| Gm38243 | 62.35611887 | 0.74032138 | 0.351112113 | 2.108504242 |
| Gm26580 | 1.456296832 | 1.175751633 | 0.84150354 | 1.39720343 |
| Gm5696 | 0 | NA | NA | NA |
| Gm28239 | 0 | NA | NA | NA |
| Rims1 | 3306.597451 | 0.539517338 | 0.095311337 | 5.660578836 |
| Gm26866 | 0.048949532 | 0.037064902 | 3.724945049 | 0.009950456 |
| Gm7761 | 0.676990728 | 0.357541483 | 1.276891951 | 0.280009192 |
| Gm29506 | 0.481916195 | -0.75323141 | 1.568451902 | -0.480238768 |
| 4933415F23Rik | 0.213028513 | 1.797852896 | 2.12463566 | 0.846193505 |
| Gm24176 | 0 | NA | NA | NA |
| Gm6420 | 0.044191549 | 0.037062677 | 3.724945049 | 0.009949859 |
| Mir30a | 2.056819932 | -0.90899265 | 0.765574171 | -1.187334533 |
| Gm27028 | 24.51280018 | -0.211338419 | 0.235620217 | -0.896945186 |
| Mir30c-2 | 0 | NA | NA | NA |
| Gm29107 | 3.847956237 | -0.792456391 | 0.593726149 | -1.334717011 |
| Gm20954 | 0.386554245 | 0.349142726 | 3.239696621 | 0.107770192 |
| Ogfrl1 | 2085.851771 | 0.464571998 | 0.074774725 | 6.212954978 |
| Gm7784 | 0.046345709 | -0.443818988 | 3.724945049 | -0.119147795 |
| Gm28822 | 1.28195289 | -0.153503415 | 0.839921665 | -0.182759204 |
| B3gat2 | 201.421169 | 0.529376883 | 0.108975064 | 4.857779968 |
| Smap1 | 968.2258995 | -0.112279461 | 0.062208303 | -1.804895089 |
| Sdhaf4 | 66.50993585 | 0.071921259 | 0.177349275 | 0.405534553 |
| Gm26524 | 0.087261593 | 0.51795324 | 3.724945049 | 0.139049901 |
| Fam135a | 882.3529123 | 0.024493409 | 0.075555447 | 0.324177937 |
| Gm37580 | 2.220335296 | -0.070459313 | 0.72017215 | -0.097836765 |
| Gm26607 | 0 | NA | NA | NA |
| Col9a1 | 27.58382149 | 0.125588939 | 0.30253549 | 0.415121343 |
| Col19a1 | 256.8225277 | 1.40917105 | 0.204159565 | 6.902302362 |
| Gm26418 | 0.607053417 | 1.860687561 | 1.39255623 | 1.336166914 |
| Tubb4b-ps2 | 1.899816927 | -0.852454179 | 0.766801505 | -1.111701233 |
| Gm28439 | 270.7744438 | 0.356682044 | 0.163592168 | 2.180312472 |
| Gm10222 | 81.6198627 | 0.305533898 | 0.186857783 | 1.635114653 |

| | | | | |
|---|---|---|---|---|
| Gm28438 | 233.7213687 | 0.109975328 | 0.186488005 | 0.589717968 |
| Gm28437 | 654.6201061 | 0.271702502 | 0.18610974 | 1.459904795 |
| Gm10925 | 568.5256472 | 0.410696502 | 0.192143581 | 2.137445865 |
| Gm28661 | 361.8513258 | 0.393333656 | 0.141192793 | 2.785791309 |
| Gm29216 | 452.3609317 | -0.040453166 | 0.179166511 | -0.225785306 |
| Lmbrd1 | 1834.054128 | 0.289197172 | 0.058758779 | 4.921769561 |
| Gm28594 | 0 | NA | NA | NA |
| Gm28237 | 0.181654796 | -1.405197014 | 3.311020957 | -0.424399915 |
| Gm9443 | 0.049553047 | -0.443818988 | 3.724945049 | -0.119147795 |
| Gm29414 | 0.167977605 | 0.503907071 | 2.756630395 | 0.182798199 |
| Adgrb3 | 2707.39029 | 0.606735987 | 0.121774347 | 4.982461413 |
| Gm15702 | 0.267089184 | -0.712414163 | 1.786686398 | -0.398734867 |
| Gm20172 | 56.45506021 | 0.402614478 | 0.294620209 | 1.36655418 |
| Gm22260 | 10.97169895 | 0.532499198 | 0.387895236 | 1.372791283 |
| Gm37543 | 46.77977661 | 0.646954498 | 0.342731134 | 1.887644375 |
| Gm37331 | 188.7440919 | 0.428605429 | 0.206293112 | 2.077652642 |
| Gm37211 | 123.4678815 | 0.421152505 | 0.247098393 | 1.704391922 |
| Gm37862 | 145.3838556 | 0.365368265 | 0.240731614 | 1.5177411 |
| Gm38286 | 155.9317435 | 0.626454927 | 0.2354131 | 2.661087799 |
| Gm37880 | 213.8342296 | 0.417295167 | 0.300588931 | 1.388258598 |
| Gm25294 | 0 | NA | NA | NA |
| Gm37011 | 137.8405266 | 0.407925882 | 0.188452079 | 2.16461333 |
| Gm29214 | 43.40613233 | 0.521882018 | 0.237195207 | 2.200221603 |
| Gm37051 | 172.0940709 | 0.474941279 | 0.221176713 | 2.147338539 |
| Gm38260 | 342.1641184 | 0.598058651 | 0.177477911 | 3.369763866 |
| Gm37628 | 460.1944707 | 0.536187243 | 0.248017127 | 2.161896027 |
| 8030445P17Rik | 222.3518729 | 0.40929985 | 0.247937671 | 1.650817514 |
| Gm38198 | 365.9544784 | 0.422956681 | 0.29620599 | 1.427914004 |
| Gm37077 | 292.4722539 | 0.332899135 | 0.189031521 | 1.761077374 |
| Gm37895 | 566.8750103 | 0.457815678 | 0.183378477 | 2.496561669 |
| Gm36952 | 499.6234323 | 0.428068388 | 0.210193008 | 2.036549135 |
| Gm37602 | 362.6410622 | 0.379524126 | 0.152651105 | 2.486219315 |
| 2900022M07Rik | 83.257532 | 0.439690927 | 0.279090429 | 1.57544251 |
| 2900002M20Rik | 64.13250466 | 0.601329126 | 0.258963844 | 2.322058233 |
| Gm9884 | 9.752194563 | 0.10960257 | 0.394688064 | 0.27769416 |
| Gm18443 | 0 | NA | NA | NA |
| Gm19123 | 0 | NA | NA | NA |
| Gm24064 | 0 | NA | NA | NA |
| 4931408C20Rik | 0 | NA | NA | NA |
| n-R5s209 | 0 | NA | NA | NA |
| Gm19641 | 0 | NA | NA | NA |
| Gm5524 | 0.04393175 | 0.037062677 | 3.724945049 | 0.009949859 |
| Gm18776 | 0 | NA | NA | NA |
| Gm19028 | 0.124853745 | 1.05357957 | 3.721447096 | 0.283110183 |
| Gm38089 | 0 | NA | NA | NA |
| Gm7846 | 6.28322175 | -0.244872394 | 0.401145702 | -0.61043255 |
| Gm597 | 0.068097359 | 0.037064902 | 3.724945049 | 0.009950456 |

| | | | | |
|---|---|---|---|---|
| Gm5697 | 0.151010504 | -0.443820511 | 2.854938278 | -0.15545713 |
| Gm6462 | 0 | NA | NA | NA |
| Gm18033 | 0 | NA | NA | NA |
| Gm25586 | 0 | NA | NA | NA |
| Gm22679 | 0 | NA | NA | NA |
| Gm23785 | 0 | NA | NA | NA |
| Mir6342 | 0.042716945 | 0.037062677 | 3.724945049 | 0.009949859 |
| Gm5525 | 5.440920671 | -0.467666638 | 0.407342886 | -1.148090845 |
| Gm7858 | 0 | NA | NA | NA |
| Gm23771 | 0 | NA | NA | NA |
| Gm27430 | 0.117046891 | 0.037062677 | 3.724945049 | 0.009949859 |
| Gm9898 | 0.52260535 | 1.235334911 | 1.387474396 | 0.890347897 |
| Gm6473 | 7.058970522 | 0.56990739 | 0.404065964 | 1.410431564 |
| Gm2914 | 0.311886478 | 0.849699245 | 1.733757653 | 0.490091128 |
| Phf3 | 1888.771435 | 0.003125995 | 0.059183777 | 0.052818441 |
| Gm37522 | 62.34479475 | 0.376024248 | 0.241755653 | 1.555389682 |
| Gm37906 | 105.3489438 | 0.510980369 | 0.228293296 | 2.2382627 |
| Ptp4a1 | 625.4291984 | -0.456123064 | 0.115294325 | -3.956162327 |
| Gm5698 | 0 | NA | NA | NA |
| Gm9458 | 0 | NA | NA | NA |
| Gm5699 | 0.061920613 | -0.443816764 | 3.724945049 | -0.119147198 |
| Gm29669 | 0 | NA | NA | NA |
| Gm28644 | 0 | NA | NA | NA |
| Gm27362 | 0 | NA | NA | NA |
| 4931428L18Rik | 0.048949532 | 0.037064902 | 3.724945049 | 0.009950456 |
| Gm29128 | 0 | NA | NA | NA |
| Gm7893 | 0 | NA | NA | NA |
| Lgsn | 0.247864911 | -0.3352147 | 2.504674553 | -0.133835631 |
| Pih1d3 | 0 | NA | NA | NA |
| Gm26145 | 0 | NA | NA | NA |
| Gm4850 | 0 | NA | NA | NA |
| Gm28645 | 0.066050875 | -0.443818988 | 3.724945049 | -0.119147795 |
| Gm6489 | 0 | NA | NA | NA |
| Gm10307 | 0 | NA | NA | NA |
| Gm19680 | 8.167021386 | -0.274205115 | 0.361075965 | -0.759411155 |
| Gm23032 | 0 | NA | NA | NA |
| Gm7910 | 0 | NA | NA | NA |
| Khdrbs2 | 245.3494503 | 0.365114248 | 0.139896034 | 2.609897065 |
| Gm25814 | 0.068097359 | 0.037064902 | 3.724945049 | 0.009950456 |
| Gm37999 | 19.71845274 | 0.741182876 | 0.366965723 | 2.019760513 |
| Gm37110 | 1.003269596 | 1.055746694 | 1.139926502 | 0.926153303 |
| Gm37911 | 37.09896806 | 0.381645312 | 0.305902544 | 1.247604243 |
| Gm9839 | 0.136963818 | 0.037074342 | 3.599544405 | 0.010299732 |
| Gm37792 | 46.79344322 | 0.792551692 | 0.249415327 | 3.177638281 |
| Gm5415 | 0.486008019 | 0.914532954 | 1.294841417 | 0.706289544 |
| Gm24732 | 0 | NA | NA | NA |
| Gm37426 | 6.59084447 | 0.505899426 | 0.474522508 | 1.066123139 |

| | | | | |
|---|---|---|---|---|
| Gm23024 | 0.184258256 | 1.308934158 | 2.68866045 | 0.486835055 |
| Gm37151 | 1.344706689 | 1.780256408 | 1.066437181 | 1.669349531 |
| Gm28624 | 0.510604427 | 1.473961297 | 1.592515853 | 0.92555518 |
| Gm37724 | 1.002429578 | -0.095758161 | 1.092457263 | -0.08765392 |
| Gm7114 | 3.168220283 | 0.497618514 | 0.555171562 | 0.896332859 |
| Gm4785 | 0.341712212 | 1.016239099 | 1.572012736 | 0.646457294 |
| Gm24901 | 0 | NA | NA | NA |
| Gm37591 | 0.710426184 | 2.173613078 | 1.327744275 | 1.637072077 |
| Gm2987 | 0 | NA | NA | NA |
| Gm25792 | 0 | NA | NA | NA |
| Gm18799 | 2.333919785 | 0.579184675 | 0.75727525 | 0.764827156 |
| Gm7933 | 0.088383099 | 0.037058229 | 3.724945049 | 0.009948665 |
| Gm23453 | 0 | NA | NA | NA |
| Prim2 | 97.92590214 | -0.062795678 | 0.129295662 | -0.485675057 |
| Gm29228 | 0 | NA | NA | NA |
| Gm37618 | 13.20070015 | 0.011559808 | 0.341228739 | 0.033877007 |
| Gm17813 | 0 | NA | NA | NA |
| 1700001G17Rik | 1.688998503 | -0.851914469 | 0.874651824 | -0.974004108 |
| Gm38173 | 0.528487055 | 1.750354048 | 1.387695837 | 1.261338401 |
| Rab23 | 419.6998972 | -0.056259255 | 0.061160194 | -0.91986717 |
| Bag2 | 54.75618287 | -0.054102058 | 0.170716256 | -0.31691216 |
| Gm37905 | 0.392625049 | 0.033916234 | 1.540436226 | 0.022017292 |
| Zfp451 | 1940.361675 | 0.015126847 | 0.069287615 | 0.218319635 |
| Gm15455 | 3.94145278 | 1.438120205 | 0.607949508 | 2.365525734 |
| Bend6 | 1529.651517 | 0.40316452 | 0.129797817 | 3.106096308 |
| Gm37067 | 37.0527924 | 0.31983603 | 0.381059935 | 0.839332611 |
| Dst | 13187.49407 | 0.216815127 | 0.062167421 | 3.487600501 |
| Gm37354 | 31.03348255 | 0.370328314 | 0.339022884 | 1.092340166 |
| Gm28631 | 0 | NA | NA | NA |
| Gm23792 | 0 | NA | NA | NA |
| Gm38157 | 47.50224476 | 0.275635644 | 0.332476432 | 0.829038144 |
| mmu-mir-6896 | 1.465090628 | 0.995946575 | 1.064203511 | 0.935861012 |
| Gm26788 | 4.676338759 | 0.554962855 | 0.458920385 | 1.209279154 |
| D630036G22Rik | 69.39949552 | 0.403408851 | 0.198824748 | 2.028966996 |
| A930005N03Rik | 0.933986728 | 0.630266202 | 1.130144336 | 0.557686467 |
| Gm37958 | 0 | NA | NA | NA |
| Gm37385 | 0.204111068 | 0.538732764 | 2.597989528 | 0.207365256 |
| Gm5266 | 0 | NA | NA | NA |
| Gm3052 | 10.39265568 | 0.137166596 | 0.385529451 | 0.355787595 |
| Gm37233 | 84.28593999 | 0.161045342 | 0.276076754 | 0.583335394 |
| Mir5103 | 0 | NA | NA | NA |
| Ccdc115 | 140.6358465 | 0.055905012 | 0.119623875 | 0.467339916 |
| Gm28306 | 0 | NA | NA | NA |
| Imp4 | 379.5447932 | -0.435739396 | 0.085205453 | -5.113984855 |
| Ptpn18 | 38.27817853 | -0.547948484 | 0.238892628 | -2.29370194 |
| Gm28417 | 1.241452806 | -0.767100277 | 0.811426369 | -0.945372626 |
| 4930568A12Rik | 0.934086492 | 0.234510218 | 0.961412847 | 0.243922492 |

| | | | | |
|---|---|---|---|---|
| Prss39 | 0.301806123 | -1.763305685 | 1.704209074 | -1.034676855 |
| Gm28416 | 0 | NA | NA | NA |
| Cfc1 | 0.191490881 | 1.024132856 | 2.702348728 | 0.378978792 |
| Gm28419 | 0 | NA | NA | NA |
| Prss40 | 0 | NA | NA | NA |
| 1700101I19Rik | 0.559566189 | 0.690320814 | 3.64189201 | 0.18955005 |
| Amer3 | 415.894219 | 0.65035823 | 0.094430466 | 6.887165332 |
| Gm28418 | 0.045882455 | 0.517959912 | 3.724945049 | 0.139051692 |
| Gm36984 | 0.129408498 | 0.50354616 | 2.990398536 | 0.168387643 |
| Gm29730 | 0.628347421 | 2.044811954 | 1.196052662 | 1.709633713 |
| AA619741 | 4.444019085 | 1.761113323 | 0.693340406 | 2.540041384 |
| Arhgef4 | 1837.032545 | 0.511705958 | 0.051159367 | 10.00219481 |
| AC107766.1 | 0 | NA | NA | NA |
| Gm37224 | 4.445654744 | 0.444507979 | 0.504694858 | 0.880746003 |
| Gm37845 | 4.660398994 | 0.178188066 | 0.665508411 | 0.267747278 |
| Gm38336 | 10.69144326 | -0.513445783 | 0.438567653 | -1.170733361 |
| Fam168b | 3287.598993 | 0.053322646 | 0.048338526 | 1.103108642 |
| Gm37127 | 1.263832448 | 0.830618244 | 1.115726064 | 0.744464319 |
| Plekhb2 | 2551.489122 | -0.006008712 | 0.072235776 | -0.083181934 |
| Gm19430 | 0.332159689 | -1.613390556 | 1.924908899 | -0.83816463 |
| Gm37670 | 0.517156391 | 0.947297377 | 1.352325262 | 0.700495217 |
| Gm8009 | 0.062426873 | 0.517959912 | 3.724945049 | 0.139051692 |
| Gm38027 | 0 | NA | NA | NA |
| Gm25096 | 0 | NA | NA | NA |
| Gm37068 | 0.62070299 | 0.416861183 | 1.168449858 | 0.356764289 |
| Gm25634 | 0 | NA | NA | NA |
| Gm28415 | 0 | NA | NA | NA |
| 1110002O04Rik | 0.877137428 | -1.843932044 | 1.177214102 | -1.566352323 |
| Gm8022 | 0 | NA | NA | NA |
| Gm37146 | 0 | NA | NA | NA |
| Gm33222 | 0.331946588 | 0.037078315 | 1.728827035 | 0.021447093 |
| 4930535G08Rik | 0 | NA | NA | NA |
| Gm38049 | 0.068097359 | 0.037064902 | 3.724945049 | 0.009950456 |
| Hs6st1 | 777.2491163 | 0.075388795 | 0.075853426 | 0.993874613 |
| Gm37335 | 0.364365233 | 0.517975206 | 1.592798299 | 0.325198241 |
| Gm33280 | 0.253548292 | 0.492388018 | 1.82916605 | 0.269187162 |
| Uggt1 | 1213.9728 | -0.209167874 | 0.060017074 | -3.4851395 |
| Mir6897 | 0.539626406 | -0.436514127 | 1.477751519 | -0.295390748 |
| Neurl3 | 71.81858134 | -1.235046942 | 0.273563839 | -4.514657154 |
| Arid5a | 533.3445611 | -0.162302787 | 0.245516789 | -0.661065942 |
| 4930403P22Rik | 0.909075398 | 2.168731894 | 2.130234901 | 1.018071713 |
| Kansl3 | 1110.948051 | 0.023594258 | 0.070668473 | 0.333872472 |
| Mir6898 | 0.044191549 | 0.037062677 | 3.724945049 | 0.009949859 |
| Fer1l5 | 1.367163391 | -0.589249851 | 0.826536492 | -0.712914501 |
| Lman2l | 453.4304442 | -0.206327442 | 0.092931543 | -2.220208941 |
| Cnnm4 | 233.328149 | 0.018841621 | 0.109078288 | 0.172734839 |
| Gm37909 | 10.35866414 | 0.401670979 | 0.410471853 | 0.978559128 |

| | | | | |
|---|---|---|---|---|
| AC084391.1 | 0 | NA | NA | NA |
| Cnnm3 | 394.0709263 | 0.347520851 | 0.081478989 | 4.265159085 |
| Gm38033 | 0.17077006 | 0.486095726 | 2.737003846 | 0.177601404 |
| Ankrd23 | 35.9631227 | -0.23995546 | 0.194729709 | -1.23224885 |
| Ankrd39 | 86.73678357 | 0.180752809 | 0.141442835 | 1.277921286 |
| Sema4c | 276.9910831 | -0.226262437 | 0.116918818 | -1.935209748 |
| D430040D24Rik | 19.3189591 | 0.931600895 | 0.244217946 | 3.814629142 |
| Fam178b | 2.063825205 | -0.865785258 | 0.732551797 | -1.181875823 |
| Gm6428 | 1.366437448 | -0.719588722 | 0.93382606 | -0.77058111 |
| Gm24133 | 0 | NA | NA | NA |
| Cox5b | 1330.17609 | 0.14909439 | 0.118654978 | 1.256537167 |
| Actr1b | 2273.379145 | 0.016691488 | 0.040527655 | 0.411854277 |
| 4933424G06Rik | 1.049948633 | -1.121241275 | 1.136148522 | -0.986879139 |
| Gm33533 | 12.17216712 | 1.331149503 | 0.476536103 | 2.793386472 |
| RP23-247A1.6 | 1.790960893 | 2.033267259 | 0.964648959 | 2.107779457 |
| Zap70 | 42.79799715 | -0.480487568 | 0.26193633 | -1.834367793 |
| Gm18828 | 0 | NA | NA | NA |
| Tmem131 | 1521.534467 | -0.066211482 | 0.07156652 | -0.925173979 |
| Gm37506 | 4.051657186 | 1.211179623 | 0.607653602 | 1.993207345 |
| Gm38115 | 35.47402398 | 0.043898781 | 0.319897877 | 0.137227484 |
| F830112A20Rik | 59.60486576 | -0.011812999 | 0.261737158 | -0.045133061 |
| Rpl12-ps1 | 0.348012853 | 1.082097679 | 1.954522599 | 0.553637845 |
| n-R5s210 | 0 | NA | NA | NA |
| Vwa3b | 17.13175727 | 0.065063902 | 0.314226139 | 0.207060756 |
| Gm37020 | 0.834730223 | 2.17794748 | 1.302087143 | 1.672658771 |
| Cnga3 | 3.017996072 | -0.407333053 | 0.681786482 | -0.597449588 |
| 1700074A21Rik | 0 | NA | NA | NA |
| Inpp4a | 3160.431147 | 0.448270404 | 0.061321657 | 7.310148288 |
| 4930439A04Rik | 0.093831153 | 0.517957688 | 3.724945049 | 0.139051095 |
| Gm37135 | 0 | NA | NA | NA |
| Coa5 | 899.7403321 | -0.204669739 | 0.106961746 | -1.913485403 |
| Unc50 | 302.5469516 | -0.076342746 | 0.10251636 | -0.744688419 |
| Mgat4a | 840.1121684 | 0.309014917 | 0.05143316 | 6.008087311 |
| 4930594C11Rik | 47.43194901 | 0.758151841 | 0.326515641 | 2.321946475 |
| 2010300C02Rik | 1765.094823 | 1.003652939 | 0.173927925 | 5.770510615 |
| 4930556I23Rik | 0.066050875 | -0.443818988 | 3.724945049 | -0.119147795 |
| 4930470B04Rik | 0.188205214 | -1.274234033 | 2.673954082 | -0.476535495 |
| Gm19863 | 38.19950049 | 1.229974628 | 0.429028581 | 2.866882727 |
| Tsga10 | 318.6599432 | -0.153383903 | 0.125258178 | -1.22454203 |
| Gm26805 | 21.25672272 | 1.763510056 | 0.343921915 | 5.127646645 |
| Lipt1 | 33.80092537 | -0.132331285 | 0.195903242 | -0.67549308 |
| Mitd1 | 202.445721 | -0.567483455 | 0.170765441 | -3.323175056 |
| Mrpl30 | 491.4665078 | -0.145792555 | 0.103793739 | -1.404637269 |
| Lyg2 | 0.741337718 | 1.833676122 | 1.380833237 | 1.327949004 |
| Gm15457 | 0.21650593 | -1.791584846 | 2.13065922 | -0.840859406 |
| Lyg1 | 0.136120942 | -0.876404342 | 3.595683781 | -0.24373788 |
| Gm23409 | 0 | NA | NA | NA |

| | | | | |
|---|---|---|---|---|
| Txndc9 | 544.4411505 | -0.256330712 | 0.088551278 | -2.89471498 |
| Eif5b | 954.1717507 | -0.089210331 | 0.116695794 | -0.764469124 |
| Rev1 | 696.4174186 | 0.335735752 | 0.112645104 | 2.980473541 |
| Gm37062 | 35.92510916 | 0.718841246 | 0.232248388 | 3.095139876 |
| Gm5099 | 0.141094079 | 0.037072457 | 3.563850528 | 0.01040236 |
| Aff3 | 992.3491355 | 0.429359915 | 0.075562 | 5.682220085 |
| Gm16150 | 1.482000077 | 0.052757784 | 0.867905123 | 0.060787501 |
| Gm16151 | 0 | NA | NA | NA |
| Gm16152 | 1.435447043 | -3.002644392 | 1.040635487 | -2.885394963 |
| Gm34727 | 0 | NA | NA | NA |
| Gm38135 | 0 | NA | NA | NA |
| 2610300A13Rik | 0 | NA | NA | NA |
| Lonrf2 | 2520.190482 | 0.105140137 | 0.079808824 | 1.317399897 |
| Chst10 | 707.3698007 | 0.188597289 | 0.057722698 | 3.267298568 |
| Gm37707 | 0 | NA | NA | NA |
| Gm6621 | 0.169398393 | 1.480600686 | 3.393596657 | 0.436292475 |
| Nms | 1.222583469 | -1.607199921 | 0.918550389 | -1.749713397 |
| Pdcl3 | 219.8699854 | -0.338733532 | 0.129490148 | -2.615901955 |
| Gm37309 | 20.51143474 | 0.154727911 | 0.360836929 | 0.428802871 |
| Gm37821 | 0.54509763 | 1.168377088 | 1.230361895 | 0.949620671 |
| Gm37091 | 0.205567437 | 1.019593774 | 2.592100261 | 0.393346581 |
| Mir6349 | 0 | NA | NA | NA |
| Npas2 | 366.6556838 | 0.741108408 | 0.16034147 | 4.622063197 |
| Gm20428 | 0.263795272 | -0.898096705 | 1.814983137 | -0.494823719 |
| Gm3617 | 6.543142137 | 0.085644793 | 0.612653665 | 0.139793161 |
| Rpl31 | 1558.50621 | -0.110696855 | 0.125505845 | -0.882005574 |
| Tbc1d8 | 368.5922422 | 0.410339828 | 0.107902972 | 3.802859358 |
| Gm37265 | 6.813130323 | 0.426769305 | 0.519850457 | 0.820946293 |
| Gm24826 | 0 | NA | NA | NA |
| ACEA_U3 | 3.059522033 | -1.614139763 | 0.644319257 | -2.505186281 |
| Cnot11 | 371.6623297 | 0.014766408 | 0.098902902 | 0.149302066 |
| Gm15832 | 37.25351844 | -0.259915656 | 0.304831492 | -0.852653556 |
| Snord89 | 3.528574751 | -0.670668502 | 0.556889238 | -1.204312197 |
| Rnf149 | 381.317406 | -0.564374447 | 0.174766796 | -3.229300191 |
| Gm23722 | 10.82063883 | -0.314066135 | 0.371864659 | -0.844571074 |
| Gm5100 | 0.671689889 | -1.506795894 | 1.303446651 | -1.156008874 |
| Creg2 | 1118.864952 | 0.309165868 | 0.150385131 | 2.05582736 |
| D930019O06Rik | 42.32280546 | 0.224155441 | 0.226567726 | 0.989352918 |
| Rfx8 | 1.59726488 | 1.110832093 | 0.86618718 | 1.282438852 |
| Gm3646 | 0.042716945 | 0.037062677 | 3.724945049 | 0.009949859 |
| 1700066B17Rik | 0.337370548 | 0.951763055 | 2.007107691 | 0.474196307 |
| Map4k4 | 1991.375243 | -0.237964815 | 0.047559682 | -5.003498875 |
| 4930456G14Rik | 5.719135214 | 0.866733592 | 0.53440196 | 1.621875772 |
| Il1r2 | 32.73985817 | -0.392125485 | 0.683189043 | -0.573963369 |
| Gm16894 | 21.14797858 | 0.625655472 | 0.273357635 | 2.288779937 |
| Gm37347 | 2.446674312 | 0.178503689 | 0.595214429 | 0.299898121 |
| Il1r1 | 110.9761096 | -0.211826729 | 0.152078284 | -1.392879528 |

| | | | | |
|---|---|---|---|---|
| Gm38070 | 5.586992304 | -0.167902182 | 0.419655814 | -0.400094974 |
| Il1rl2 | 14.87794066 | -0.740468792 | 0.32279191 | -2.293950896 |
| Gm17970 | 0.509421185 | -0.406407176 | 1.431831616 | -0.283837269 |
| Il1rl1 | 3.917850035 | -1.566706751 | 0.590964043 | -2.651103345 |
| Il18r1 | 13.77975115 | -0.55812725 | 0.321801205 | -1.734385207 |
| Il18rap | 55.22165596 | -0.951856192 | 0.263313827 | -3.614911538 |
| Gm35801 | 0.072025462 | 0.517959912 | 3.724945049 | 0.139051692 |
| Slc9a4 | 29.46342029 | 0.032824689 | 0.233357667 | 0.14066257 |
| Gm37435 | 0.132101749 | -0.93550548 | 3.724188147 | -0.251197158 |
| Slc9a2 | 85.94852822 | 0.713506724 | 0.174611625 | 4.086249825 |
| Gm37623 | 0 | NA | NA | NA |
| Gm37915 | 0.583248015 | 0.372948301 | 1.287808429 | 0.289599208 |
| Mfsd9 | 54.36389778 | -0.405269765 | 0.196558456 | -2.061828186 |
| Tmem182 | 1.853692403 | -0.84744857 | 1.042744097 | -0.812710014 |
| Gm5973 | 0 | NA | NA | NA |
| 4930448I06Rik | 0.044544648 | 0.517957688 | 3.724945049 | 0.139051095 |
| Gm28634 | 0.143022002 | 0.03610674 | 3.547735707 | 0.010177404 |
| Gm29260 | 4.366569511 | -0.601920426 | 0.4779481 | -1.259384495 |
| Gm5267 | 0 | NA | NA | NA |
| Gm8176 | 0 | NA | NA | NA |
| Gm9915 | 0.24066848 | -0.438188839 | 2.104550651 | -0.208210165 |
| Gm29664 | 0 | NA | NA | NA |
| Gm28175 | 3.438849202 | 0.488226219 | 0.585799067 | 0.833436322 |
| Gm28140 | 0.100902008 | -0.876201588 | 3.72093227 | -0.235479048 |
| Gm37047 | 0.333556256 | 0.987447516 | 1.589607944 | 0.621189344 |
| Pantr1 | 157.9407554 | 0.620175301 | 0.162888998 | 3.807349237 |
| 4930420N18Rik | 0 | NA | NA | NA |
| Gm20646 | 38.88834011 | 0.803994853 | 0.208178684 | 3.862042148 |
| Pou3f3 | 1367.814843 | 0.069496341 | 0.090230301 | 0.770210675 |
| Pantr2 | 14.1555736 | 0.093353148 | 0.3842796 | 0.242930273 |
| Dalir | 16.94524304 | 1.543475436 | 0.38160892 | 4.044652409 |
| Mrps9 | 398.9447759 | 0.276416024 | 0.086845632 | 3.182843143 |
| Gm23126 | 0 | NA | NA | NA |
| Gpr45 | 153.2010998 | -0.10985205 | 0.139473679 | -0.787618502 |
| Gm24001 | 0.318202806 | 0.037076871 | 2.018121239 | 0.018371974 |
| Tgfbrap1 | 851.4724595 | 0.054305122 | 0.057199065 | 0.949405769 |
| Gm28782 | 1.198752213 | -2.022261524 | 1.283346663 | -1.575771833 |
| 8430432A02Rik | 7.526368882 | 1.094436704 | 0.483113393 | 2.265382659 |
| AI597479 | 352.8821411 | -0.035549511 | 0.082804199 | -0.429320155 |
| Fhl2 | 255.1157235 | 0.052496451 | 0.119511623 | 0.439258119 |
| Gm8210 | 95.32441064 | -0.16463052 | 0.187835637 | -0.876460521 |
| Gm29040 | 0.048949532 | 0.037064902 | 3.724945049 | 0.009950456 |
| Gm29610 | 0 | NA | NA | NA |
| Gm29041 | 0 | NA | NA | NA |
| Nck2 | 373.3238872 | 0.107602821 | 0.083090731 | 1.295003911 |
| Gm37536 | 17.59062222 | 0.421666922 | 0.366714287 | 1.149851361 |
| Gm38228 | 18.03991603 | 0.314524352 | 0.389312481 | 0.807896914 |

| | | | | |
|---|---|---|---|---|
| 1500015O10Rik | 35.22777033 | 0.002115856 | 0.287684019 | 0.007354793 |
| Gm29155 | 11.99474671 | 0.580468772 | 0.363807815 | 1.595536839 |
| Gm29157 | 0.378942021 | 1.002441346 | 1.603163396 | 0.625289567 |
| Gm29156 | 12.6916125 | 0.4945292 | 0.349516904 | 1.414893513 |
| Uxs1 | 508.6202783 | 0.062777537 | 0.08343946 | 0.752372278 |
| Gm16103 | 0 | NA | NA | NA |
| Gm19587 | 1.101887088 | 1.521374752 | 0.921782744 | 1.650469985 |
| Gm8241 | 0 | NA | NA | NA |
| Gm26352 | 0 | NA | NA | NA |
| Gm24180 | 0.220091977 | 0.036348664 | 2.162314232 | 0.016810075 |
| Tpp2 | 1340.114203 | 0.113175362 | 0.083869737 | 1.349418346 |
| Mettl21c | 0.903763266 | 1.216842532 | 1.05177353 | 1.156943484 |
| Gm15838 | 0 | NA | NA | NA |
| Gm8251 | 2.971797373 | -0.286940953 | 0.587715307 | -0.488231206 |
| Tex30 | 39.96396815 | 0.082765434 | 0.23262042 | 0.355796079 |
| Gm29247 | 0 | NA | NA | NA |
| Kdelc1 | 74.7610229 | -0.102929443 | 0.135642754 | -0.758827433 |
| Bivm | 338.2242815 | 0.052649602 | 0.074846401 | 0.703435323 |
| Ercc5 | 350.2066079 | 0.266566725 | 0.061823238 | 4.311756146 |
| Mettl21e | 0.63532651 | 2.829823704 | 1.234738659 | 2.291840208 |
| Gm15833 | 0.607153663 | 0.815459804 | 1.348533869 | 0.604701018 |
| Gm5700 | 0 | NA | NA | NA |
| Gm28893 | 0 | NA | NA | NA |
| Txn-ps1 | 3.667605082 | -0.399898579 | 0.548265597 | -0.729388422 |
| Gulp1 | 83.35613432 | -0.041870607 | 0.154442436 | -0.271108173 |
| Gm37626 | 20.77771962 | 0.236483092 | 0.31423876 | 0.752558636 |
| Gm16102 | 1.145880322 | 0.893514184 | 1.188239958 | 0.751964431 |
| 4930521E06Rik | 0.181654796 | -1.405197014 | 3.311020957 | -0.424399915 |
| Gm38110 | 3.191642098 | 0.755787289 | 0.617470585 | 1.224005332 |
| Gm3489 | 0.12092566 | 0.998850474 | 3.724945049 | 0.268151734 |
| Gm19480 | 0 | NA | NA | NA |
| Gm3496 | 0 | NA | NA | NA |
| Gm5268 | 0 | NA | NA | NA |
| Col3a1 | 232.7682847 | -1.039270252 | 0.186152995 | -5.582882267 |
| AC125167.1 | 0 | NA | NA | NA |
| Col5a2 | 180.1321098 | -1.148108629 | 0.1855071 | -6.18902796 |
| Gm38211 | 9.579960985 | -1.434021035 | 0.4119725 | -3.480865921 |
| Gm8296 | 0.048949532 | 0.037064902 | 3.724945049 | 0.009950456 |
| Gm7063 | 0.241951302 | -0.512996056 | 2.113262818 | -0.242750713 |
| Gm8304 | 0 | NA | NA | NA |
| Gm23216 | 0 | NA | NA | NA |
| Gm23974 | 0 | NA | NA | NA |
| Gm8307 | 0.148708979 | -0.443201667 | 2.868467393 | -0.154508177 |
| Gm23984 | 0 | NA | NA | NA |
| Wdr75 | 417.7461979 | -0.19488926 | 0.067117941 | -2.903683539 |
| Gm28906 | 0 | NA | NA | NA |
| Gm5526 | 49.19564311 | -0.279987027 | 0.218703887 | -1.280210565 |

| | | | | |
|---|---|---|---|---|
| Gm5269 | 0.154938607 | 0.037075974 | 2.832095776 | 0.013091356 |
| Gm18303 | 0.136201742 | -0.443823494 | 2.946222509 | -0.150641539 |
| Slc40a1 | 53.73490417 | -0.103336257 | 0.272587025 | -0.379094555 |
| Gm9496 | 0 | NA | NA | NA |
| Rpl23a-ps1 | 0 | NA | NA | NA |
| Gm3852 | 0.218641893 | -0.515041282 | 2.176122945 | -0.236678393 |
| Stk-ps2 | 0 | NA | NA | NA |
| Gm18849 | 0 | NA | NA | NA |
| Dnah7b | 166.4217659 | 0.17829765 | 0.152654026 | 1.167985241 |
| n-R5s211 | 0.044191549 | 0.037062677 | 3.724945049 | 0.009949859 |
| Gm8326 | 1.175435645 | 1.896240357 | 1.023346412 | 1.852979925 |
| Dnah7c | 35.29196087 | 0.561677659 | 0.244949333 | 2.29303608 |
| Slc39a10 | 1977.950324 | 0.437288661 | 0.116933463 | 3.739636615 |
| Gm28151 | 62.07798039 | 0.176824784 | 0.234612016 | 0.753690231 |
| Gm37393 | 2.121468972 | 1.884149655 | 1.277242078 | 1.475170359 |
| Gm28527 | 0.844069991 | 2.190571035 | 1.202085134 | 1.822309396 |
| Gm28825 | 0.476041195 | -1.853763939 | 1.438379201 | -1.288786668 |
| Gm28826 | 0.77428951 | 0.297040945 | 1.010552562 | 0.293939134 |
| Gm4852 | 3.57865669 | 0.457957822 | 0.515175409 | 0.888935718 |
| Gm38090 | 0 | NA | NA | NA |
| Mir6350 | 0 | NA | NA | NA |
| Gm18117 | 0.885425998 | 0.193266066 | 1.328459891 | 0.145481295 |
| Gm37196 | 1.287564413 | -1.295400726 | 1.089716459 | -1.188750261 |
| Gm8337 | 21.49979967 | -0.945636731 | 0.297795434 | -3.175457458 |
| Gm22419 | 0 | NA | NA | NA |
| Gm23829 | 0 | NA | NA | NA |
| Gm19206 | 0 | NA | NA | NA |
| Gm5974 | 0 | NA | NA | NA |
| Gm23240 | 0 | NA | NA | NA |
| Gm28830 | 0 | NA | NA | NA |
| Gm8354 | 0 | NA | NA | NA |
| Gm19124 | 0 | NA | NA | NA |
| Gm8357 | 0 | NA | NA | NA |
| Gm22693 | 0 | NA | NA | NA |
| Gm28740 | 0 | NA | NA | NA |
| Gm8367 | 0 | NA | NA | NA |
| C230029F24Rik | 0 | NA | NA | NA |
| Gm6757 | 0 | NA | NA | NA |
| Gm3605 | 0 | NA | NA | NA |
| Gm28388 | 0 | NA | NA | NA |
| Gm8384 | 0 | NA | NA | NA |
| 4933411E06Rik | 0.068097359 | 0.037064902 | 3.724945049 | 0.009950456 |
| Gm8391 | 0 | NA | NA | NA |
| Gm27330 | 0 | NA | NA | NA |
| Gm29665 | 0 | NA | NA | NA |
| Gm20118 | 0 | NA | NA | NA |
| Gm25843 | 0 | NA | NA | NA |

| | | | | |
|---|---|---|---|---|
| Gm28321 | 0.092691418 | -0.908687078 | 3.724560878 | -0.2439716 |
| Tmeff2 | 912.3017426 | -0.091208381 | 0.074502606 | -1.224230754 |
| Gm28322 | 2.545969209 | -1.013390648 | 0.873197894 | -1.160550953 |
| Gm25736 | 12.09809119 | -0.228919591 | 0.380992436 | -0.600850749 |
| Gm28319 | 0.231621402 | 0.446609028 | 2.519475245 | 0.177262717 |
| Gm28320 | 83.01456838 | 0.132469604 | 0.230988311 | 0.57349051 |
| Gm29325 | 37.76672809 | 0.455396601 | 0.27307503 | 1.667661085 |
| 9330175M20Rik | 4.588839173 | -0.150919209 | 0.539530171 | -0.279723391 |
| Sdpr | 92.2652614 | -0.112039117 | 0.145326403 | -0.770948119 |
| Gm5253 | 0.982135049 | -1.496388651 | 1.189199831 | -1.25831556 |
| 1700072G22Rik | 0.174149299 | -0.429800891 | 3.359371215 | -0.127940874 |
| Nabp1 | 240.5774001 | -1.037357207 | 0.194355384 | -5.337424599 |
| Gm17767 | 3.178614829 | -0.148527508 | 0.549315838 | -0.270386357 |
| Gm37535 | 0.140088945 | 0.037059971 | 3.605664301 | 0.010278264 |
| Gm28055 | 0.781209455 | -1.350381146 | 1.298219267 | -1.04017956 |
| Myo1b | 450.2003702 | 0.332752118 | 0.131614009 | 2.528242398 |
| Gm29453 | 3.059357988 | 0.499895559 | 0.651914871 | 0.7668111 |
| Gm28323 | 3.764028883 | -0.681494699 | 0.630232474 | -1.081338596 |
| Gm8419 | 0 | NA | NA | NA |
| Gm27607 | 0 | NA | NA | NA |
| Gm8420 | 0.200957741 | -1.699401354 | 3.22964838 | -0.526187731 |
| Stat4 | 19.2643103 | -0.133744089 | 0.245071132 | -0.545735797 |
| Gm28177 | 2.000698716 | -0.555811371 | 0.627014694 | -0.886440744 |
| Gm3940 | 0.598154365 | -0.866182043 | 1.356754267 | -0.638422199 |
| Stat1 | 2376.29098 | -0.174947891 | 0.10255783 | -1.705846263 |
| Gm18301 | 0.092228164 | 0.037073798 | 3.724945049 | 0.009952844 |
| Gls | 3738.856511 | 0.089533403 | 0.06892184 | 1.299057063 |
| Gm5975 | 0.042716945 | 0.037062677 | 3.724945049 | 0.009949859 |
| Gm23460 | 0 | NA | NA | NA |
| Nab1 | 590.3308437 | -0.43608816 | 0.114955025 | -3.793554553 |
| Gm5976 | 0.038981395 | -0.443816764 | 3.724945049 | -0.119147198 |
| Rps27a-ps1 | 3.694411362 | 0.190591274 | 0.660436282 | 0.288583893 |
| Gm553 | 3.129775993 | 0.86696428 | 0.624696696 | 1.387816337 |
| Gm28178 | 0.452230046 | -2.065951209 | 1.296981003 | -1.592892421 |
| Gm5527 | 9.620821135 | 0.024789178 | 0.369722892 | 0.067047992 |
| Tmem194b | 84.55902287 | 0.025242266 | 0.187157687 | 0.134871647 |
| Gm37242 | 11.17186488 | 0.307153051 | 0.337460479 | 0.910189696 |
| Mfsd6 | 1706.135027 | 0.147335336 | 0.078913318 | 1.867052861 |
| Gm29670 | 25.59481537 | 0.645176092 | 0.316332571 | 2.039549988 |
| Gm28650 | 0.805100971 | 0.075635668 | 1.171389803 | 0.064569171 |
| Inpp1 | 600.5457346 | 0.081968779 | 0.105354491 | 0.778028328 |
| Hibch | 147.5834649 | -0.534644474 | 0.168486961 | -3.173209782 |
| Gm16107 | 1.228146034 | 1.960070329 | 0.915464674 | 2.141066044 |
| 1700019D03Rik | 237.2523422 | 0.071232143 | 0.09031842 | 0.788677907 |
| Mstn | 4.314567637 | 0.382715351 | 0.471150911 | 0.812298866 |
| Gm24349 | 0 | NA | NA | NA |
| 1700019A02Rik | 5.126622909 | 0.072558099 | 0.5781026 | 0.125510764 |

| | | | | |
|---|---|---|---|---|
| Pms1 | 234.6633514 | 0.13107784 | 0.095341045 | 1.374831162 |
| Gm28777 | 0 | NA | NA | NA |
| Ormdl1 | 92.39055438 | -0.392414916 | 0.154935238 | -2.532767379 |
| Gm28778 | 0.044191549 | 0.037062677 | 3.724945049 | 0.009949859 |
| Osgepl1 | 175.8191109 | 0.081461347 | 0.111134254 | 0.732999448 |
| Gm28551 | 1.277925253 | 0.804054002 | 0.938517113 | 0.85672812 |
| Gm18800 | 2.110212028 | 0.312188817 | 0.76241515 | 0.409473522 |
| Asnsd1 | 39.89699009 | -0.525406504 | 0.183493953 | -2.863345061 |
| Gm28779 | 0.097139547 | -0.443816764 | 3.724945049 | -0.119147198 |
| Gm6799 | 0 | NA | NA | NA |
| Gm6801 | 0 | NA | NA | NA |
| Dnah7a | 93.9630857 | 0.455508459 | 0.173707774 | 2.622268698 |
| Gm25240 | 0.108123235 | 0.517955464 | 3.724945049 | 0.139050498 |
| Stk17b | 400.9618622 | -0.655607633 | 0.142124406 | -4.612913787 |
| Gm37863 | 0 | NA | NA | NA |
| Hecw2 | 1282.188723 | 0.271043898 | 0.077317263 | 3.505606462 |
| Mir7681 | 0.068097359 | 0.037064902 | 3.724945049 | 0.009950456 |
| Gm24251 | 2.00647346 | 1.501440396 | 0.844626094 | 1.777639131 |
| Gm37633 | 69.5255469 | 0.693350109 | 0.355763015 | 1.948910035 |
| Gm25829 | 0.15586508 | 0.037074716 | 2.826809592 | 0.013115392 |
| Ccdc150 | 3.346209518 | 1.514027136 | 0.724768843 | 2.088979336 |
| Gtf3c3 | 606.9969703 | 0.062430212 | 0.065719772 | 0.949945654 |
| Pgap1 | 1020.253931 | 0.106207152 | 0.081156459 | 1.308671589 |
| Gm36955 | 30.15051617 | 0.084315495 | 0.264200431 | 0.319134587 |
| Gm19637 | 0.11197992 | 0.037073798 | 3.724945049 | 0.009952844 |
| Ankrd44 | 326.5543126 | 0.079287125 | 0.096067304 | 0.825328925 |
| 4930444A19Rik | 0.477440416 | 0.545819953 | 1.481194802 | 0.368499776 |
| Gm37370 | 15.65747994 | 0.824155678 | 0.298116308 | 2.764544089 |
| Gm24053 | 0 | NA | NA | NA |
| A130048G24Rik | 23.27167967 | 0.273273832 | 0.322012648 | 0.848643163 |
| Sf3b1 | 4398.794146 | 0.132895473 | 0.072703791 | 1.827902922 |
| E330011M16Rik | 2.812432955 | -0.310210975 | 0.619693801 | -0.500587507 |
| Coq10b | 351.6495304 | -0.288556184 | 0.132039884 | -2.185371386 |
| Gm17971 | 11.17290908 | 0.750545545 | 0.537706805 | 1.395826755 |
| Hspd1 | 1608.718732 | -0.575985716 | 0.102374875 | -5.626240949 |
| Hspe1 | 313.2638214 | -0.372416242 | 0.149423284 | -2.492357497 |
| Gm5147 | 0.483827566 | 1.517596466 | 1.418529972 | 1.069837434 |
| Gm16581 | 13.32551551 | -0.045443362 | 0.361012197 | -0.125877637 |
| Mob4 | 696.0879921 | -0.142508773 | 0.090973635 | -1.566484325 |
| Gm6822 | 0.477893029 | -0.878128209 | 1.359916118 | -0.645722333 |
| Rftn2 | 338.1297183 | 0.279083809 | 0.128594821 | 2.17025698 |
| Gm10561 | 20.71569524 | 0.052479726 | 0.268787751 | 0.195245973 |
| Mars2 | 418.0161997 | 0.317352815 | 0.113101816 | 2.805903797 |
| Boll | 0.988924806 | -0.289690779 | 1.056287214 | -0.274253797 |
| Gm8292 | 51.67441755 | -0.284134307 | 0.190126962 | -1.4944451 |
| Gm29138 | 0 | NA | NA | NA |
| Gm8531 | 0.047948698 | 0.037064902 | 3.724945049 | 0.009950456 |

| | | | | |
|---|---|---|---|---|
| Plcl1 | 931.6952418 | 0.187520345 | 0.094107812 | 1.992611895 |
| Gm38056 | 155.7691939 | 0.604249177 | 0.368596262 | 1.639325299 |
| Gm6644 | 4.857481711 | -0.131328489 | 0.443203561 | -0.296316413 |
| Gm37382 | 65.4083701 | 0.474543447 | 0.267463598 | 1.774235641 |
| 9130227L01Rik | 1.675708655 | -0.092518596 | 0.897921763 | -0.103036366 |
| Gm36986 | 0 | NA | NA | NA |
| 1700003I22Rik | 0 | NA | NA | NA |
| Gm28900 | 0.298162442 | -1.505552873 | 1.690851424 | -0.890411098 |
| Gm28240 | 0 | NA | NA | NA |
| Hsfy2 | 0 | NA | NA | NA |
| Satb2 | 608.4974551 | 0.610646719 | 0.198311601 | 3.07922842 |
| 9130024F11Rik | 127.5850177 | 1.737917811 | 0.242223473 | 7.174852995 |
| Gm4081 | 9.66177043 | 2.383180163 | 0.626857202 | 3.801791152 |
| Gm23943 | 0 | NA | NA | NA |
| 1700126A01Rik | 0.044544648 | 0.517957688 | 3.724945049 | 0.139051095 |
| BC055402 | 2.819616475 | -0.858359501 | 0.653875776 | -1.312725646 |
| Gm5254 | 0 | NA | NA | NA |
| 4930558J18Rik | 2.344527321 | -0.545186401 | 0.735505294 | -0.74124062 |
| 1700066M21Rik | 269.4751219 | 0.227736505 | 0.102579805 | 2.22009104 |
| Tyw5 | 138.2526831 | -0.053428385 | 0.091208915 | -0.585780295 |
| 9430016H08Rik | 289.7250437 | 0.08949221 | 0.114494139 | 0.781631362 |
| Gm22371 | 0 | NA | NA | NA |
| Gm24014 | 0 | NA | NA | NA |
| Gm8581 | 0 | NA | NA | NA |
| Gm37373 | 0.354472284 | 0.517953661 | 3.236003702 | 0.160059663 |
| Gm17929 | 0.044544648 | 0.517957688 | 3.724945049 | 0.139051095 |
| Spats2l | 531.7349419 | -0.429231503 | 0.085858397 | -4.999295558 |
| Gm17234 | 0.08486385 | 0.037730769 | 3.724898049 | 0.010129343 |
| Kctd18 | 231.8874669 | 0.184335344 | 0.125536317 | 1.468382605 |
| Sgol2a | 34.52874643 | 0.214308415 | 0.269679145 | 0.794679227 |
| Aox1 | 275.5358162 | -0.470668041 | 0.117793791 | -3.995694833 |
| Aox3 | 36.77314627 | -0.529924542 | 0.229063741 | -2.313437033 |
| Gm24548 | 0 | NA | NA | NA |
| Aox4 | 64.4352454 | -0.107177896 | 0.227373968 | -0.471372764 |
| Gm15759 | 0.15100991 | 0.51796601 | 3.527597456 | 0.146832516 |
| Aox2 | 2.732211411 | -0.404786306 | 0.709369236 | -0.570628503 |
| Gm37607 | 38.38692177 | -0.179634996 | 0.267116388 | -0.672497098 |
| Bzw1 | 2353.558553 | -0.149600459 | 0.094503418 | -1.583016386 |
| Clk1 | 1464.941916 | 0.118608295 | 0.088847192 | 1.334969547 |
| Ppil3 | 142.411199 | -0.104682362 | 0.095845668 | -1.092197119 |
| Nif3l1 | 211.5522702 | -0.049135007 | 0.091497147 | -0.537011353 |
| Orc2 | 434.4142678 | -0.114665919 | 0.093901715 | -1.221126997 |
| Gm23966 | 4.595236341 | 0.914918122 | 0.63624291 | 1.438001285 |
| Gm15834 | 26.01032292 | -0.177955105 | 0.258416809 | -0.688635952 |
| Fam126b | 3027.102502 | -0.157544697 | 0.086029209 | -1.831293104 |
| Ndufb3 | 313.2837583 | 0.167864522 | 0.111769787 | 1.501877443 |
| Gm29389 | 0 | NA | NA | NA |

| | | | | |
|---|---|---|---|---|
| Gm18802 | 0 | NA | NA | NA |
| Gm10068 | 3.8142007 | -0.321029532 | 0.517479901 | -0.620371015 |
| Als2cr12 | 0.590977814 | -1.351913465 | 1.169126244 | -1.156345152 |
| Gm20257 | 14.75086306 | -0.12923065 | 0.339128878 | -0.381066487 |
| Gm28803 | 0 | NA | NA | NA |
| Cflar | 951.4956611 | -0.78989962 | 0.12883747 | -6.130977423 |
| Gm3841 | 0 | NA | NA | NA |
| Casp8 | 157.9166945 | -1.371391407 | 0.20113672 | -6.818205098 |
| Gm37760 | 18.59560002 | -0.690155127 | 0.315481686 | -2.187623431 |
| Gm28802 | 1.046132847 | -2.394067367 | 1.079657328 | -2.217432611 |
| Trak2 | 2825.777811 | -0.007261437 | 0.069838242 | -0.10397509 |
| Stradb | 332.0964291 | 0.07099005 | 0.144522851 | 0.491202945 |
| Gm33589 | 0.118371171 | 0.037073798 | 3.724945049 | 0.009952844 |
| Gm29018 | 0.22549826 | -0.882933078 | 2.128349392 | -0.414844048 |
| Als2cr11 | 0.178692882 | 0.037075318 | 2.707499386 | 0.013693565 |
| Gm25502 | 0 | NA | NA | NA |
| Tmem237 | 159.7002594 | 0.041561146 | 0.10694073 | 0.388637205 |
| G730003C15Rik | 13.69276046 | -0.155947138 | 0.306567132 | -0.508688381 |
| Mpp4 | 26.50791038 | -0.013104178 | 0.274300578 | -0.04777306 |
| Gm29017 | 0.060174537 | 0.517957688 | 3.724945049 | 0.139051095 |
| Gm19125 | 0 | NA | NA | NA |
| Als2 | 1234.116172 | -0.000436342 | 0.107681966 | -0.004052141 |
| Cdk15 | 4.87154909 | -1.223054857 | 0.573407779 | -2.132958256 |
| Gm29016 | 0 | NA | NA | NA |
| Gm37737 | 0.176324247 | 0.037071534 | 2.764794017 | 0.013408425 |
| Fzd7 | 304.579202 | -0.466166847 | 0.155356635 | -3.000624008 |
| Gm26813 | 0.044191549 | 0.037062677 | 3.724945049 | 0.009949859 |
| Gm973 | 103.013939 | 0.206459907 | 0.154252212 | 1.3384567 |
| Gm28411 | 0.118371171 | 0.037073798 | 3.724945049 | 0.009952844 |
| 1700122D07Rik | 0.136120942 | -0.876404342 | 3.595683781 | -0.24373788 |
| Sumo1 | 630.9605461 | 0.05914767 | 0.128112762 | 0.461684446 |
| Gm24545 | 0.226675736 | 0.517974316 | 2.143799153 | 0.241615133 |
| D430013B06Rik | 32.24759346 | 0.524502718 | 0.308164322 | 1.702022852 |
| Gm29596 | 1.33134391 | -0.204820405 | 0.990387554 | -0.206808339 |
| Nop58 | 820.0043818 | 0.009402423 | 0.106289606 | 0.088460419 |
| Snord70 | 4.644160981 | 0.707060945 | 0.491838565 | 1.437587442 |
| Gm26293 | 0 | NA | NA | NA |
| Gm26287 | 2.814965633 | 1.085455109 | 0.648650011 | 1.673406444 |
| Snord11 | 1.304845187 | -0.548601707 | 1.00508334 | -0.545827082 |
| Bmpr2 | 5482.687723 | 0.117171164 | 0.055968542 | 2.093518224 |
| Gm38125 | 29.65508468 | -0.001462287 | 0.268268347 | -0.005450836 |
| Gm37977 | 0.18514425 | 0.037065053 | 3.305081369 | 0.011214566 |
| Fam117b | 1173.743163 | 0.270996877 | 0.069857414 | 3.879285845 |
| Gm37531 | 63.74214748 | 0.598479553 | 0.278422453 | 2.149537673 |
| Gm18829 | 0 | NA | NA | NA |
| Ica1l | 384.5618481 | 0.528856516 | 0.07820091 | 6.762792361 |
| Wdr12 | 444.9492766 | -0.09921955 | 0.077973112 | -1.27248416 |

| | | | | |
|---|---|---|---|---|
| Carf | 373.7108772 | 0.552347551 | 0.1065855 | 5.182201598 |
| Gm15464 | 5.596760458 | 0.505367436 | 0.475306142 | 1.063246174 |
| Nbeal1 | 1755.709698 | -0.229866408 | 0.083659024 | -2.747658272 |
| Gm29332 | 0 | NA | NA | NA |
| Cyp20a1 | 130.1781194 | -0.540680356 | 0.155200198 | -3.48376073 |
| 6030460B20Rik | 3.025787973 | 0.170876838 | 0.636206709 | 0.268586979 |
| Abi2 | 3457.462493 | 0.507059135 | 0.076140667 | 6.659504787 |
| Raph1 | 1729.282084 | 0.050756008 | 0.104768012 | 0.484460928 |
| Gm11575 | 9.518548027 | 0.513749236 | 0.396154108 | 1.296841874 |
| Gm11576 | 0.448283844 | -0.443684669 | 1.476781946 | -0.300440204 |
| Gm11578 | 18.19495742 | 0.167874395 | 0.422077116 | 0.397733942 |
| Gm38388 | 0.262256756 | 0.517974805 | 2.055088451 | 0.252045018 |
| Gm23762 | 0 | NA | NA | NA |
| Gm11577 | 0 | NA | NA | NA |
| Eif4a-ps4 | 428.6153442 | -0.489879611 | 0.115439736 | -4.243596071 |
| Cd28 | 53.32452365 | -0.585696163 | 0.229603793 | -2.550899338 |
| Gm11579 | 0.085327104 | -0.905414734 | 3.723500899 | -0.243162217 |
| 2310016D23Rik | 0.256420178 | -0.900571506 | 2.450467766 | -0.36751004 |
| Gm37198 | 0.546045768 | -1.681577669 | 1.294533501 | -1.298983509 |
| Gm11609 | 0.066050875 | -0.443818988 | 3.724945049 | -0.119147795 |
| Gm11581 | 0.146451277 | -0.443820591 | 2.882358847 | -0.153978257 |
| Gm38137 | 1.155240215 | 0.556294806 | 1.13772399 | 0.488954096 |
| Ctla4 | 27.96364358 | -0.194167921 | 0.381981637 | -0.508317421 |
| Rpl18-ps1 | 12.04120041 | 0.010633164 | 0.354316238 | 0.030010378 |
| Gm11582 | 0.048949532 | 0.037064902 | 3.724945049 | 0.009950456 |
| Icos | 90.9935698 | -1.188212206 | 0.481091457 | -2.469826037 |
| Gm37039 | 2.679819182 | -1.130583109 | 0.771156121 | -1.466088484 |
| Rpl17-ps1 | 1.010478028 | 0.147324282 | 1.036355012 | 0.142156192 |
| Gm11590 | 0 | NA | NA | NA |
| Gm11591 | 0 | NA | NA | NA |
| Gm28083 | 0 | NA | NA | NA |
| Gm11587 | 0.127971488 | -0.95589744 | 3.6855341 | -0.259364698 |
| 9530026F06Rik | 0 | NA | NA | NA |
| Gm11588 | 0 | NA | NA | NA |
| Gm25839 | 0 | NA | NA | NA |
| Gm28449 | 0.11147366 | -0.955981391 | 3.722845977 | -0.25678779 |
| Gm36943 | 0 | NA | NA | NA |
| Gm28094 | 0.046345709 | -0.443818988 | 3.724945049 | -0.119147795 |
| 4930587A21Rik | 0 | NA | NA | NA |
| Pard3b | 166.061435 | -0.230039589 | 0.131463244 | -1.749839593 |
| Gm37205 | 11.98801516 | -0.198144308 | 0.340359845 | -0.582161235 |
| Gm38012 | 7.816280804 | -0.208624717 | 0.407704222 | -0.511706049 |
| Gm38278 | 3.411970267 | -1.281603697 | 0.643169316 | -1.99263812 |
| Pard3bos1 | 0 | NA | NA | NA |
| Gm38075 | 5.595486472 | -0.599178705 | 0.45839616 | -1.307119817 |
| Gm29640 | 0 | NA | NA | NA |
| Gm29641 | 0.044191549 | 0.037062677 | 3.724945049 | 0.009949859 |

| | | | | |
|---|---|---|---|---|
| Pard3bos2 | 0.453587956 | -1.250855111 | 1.355769012 | -0.922616685 |
| Gm38351 | 3.154252535 | 0.859902035 | 0.750790613 | 1.145328698 |
| Pard3bos3 | 0.304629397 | -0.12664272 | 2.010387665 | -0.062994179 |
| Gm29084 | 2.918337729 | -0.745820593 | 0.682573391 | -1.092659929 |
| Gm11600 | 0 | NA | NA | NA |
| Gm11599 | 0.770699074 | 0.950841991 | 1.188036916 | 0.800347176 |
| Nrp2 | 974.5911931 | -0.452947567 | 0.090641577 | -4.997128037 |
| Gm29083 | 5.840962313 | 0.02742328 | 0.435128623 | 0.063023389 |
| Gm37121 | 13.64577828 | 0.40151001 | 0.398563068 | 1.007393917 |
| Gm4208 | 0.515886961 | -0.864329943 | 1.428042482 | -0.605255063 |
| Ino80d | 1352.80268 | -0.028306752 | 0.066213407 | -0.427507858 |
| AL645685.1 | 0 | NA | NA | NA |
| Gm24942 | 0 | NA | NA | NA |
| Rps13-ps7 | 0 | NA | NA | NA |
| Rps27-ps1 | 0 | NA | NA | NA |
| Ino80dos | 70.0990274 | 0.196520137 | 0.146164446 | 1.344513959 |
| Gm20342 | 231.0873436 | 0.180686435 | 0.229852146 | 0.786098534 |
| Ndufs1 | 1507.181369 | 0.176197755 | 0.053654559 | 3.283928868 |
| Gm24557 | 0.222111871 | 1.705571485 | 3.112888666 | 0.547906356 |
| Eef1b2 | 871.8357536 | -0.23409539 | 0.133305466 | -1.756082455 |
| Gm27390 | 0 | NA | NA | NA |
| Gm26457 | 0.157314084 | 0.037075025 | 3.435173482 | 0.010792766 |
| snoZ196 | 0 | NA | NA | NA |
| Snora41 | 4.156744695 | -0.481720293 | 0.513961686 | -0.937268877 |
| Gpr1 | 4.89974049 | 0.181754854 | 0.501556329 | 0.362381738 |
| Gm27512 | 0.049553047 | -0.443818988 | 3.724945049 | -0.119147795 |
| Platr12 | 0.049553047 | -0.443818988 | 3.724945049 | -0.119147795 |
| 1700039I01Rik | 0.097899064 | 0.037062677 | 3.724945049 | 0.009949859 |
| Gm11608 | 0.112029109 | 0.037060453 | 3.724945049 | 0.009949262 |
| Gm24513 | 0.109124069 | 0.517955464 | 3.724945049 | 0.139050498 |
| Zdbf2 | 969.9858277 | -0.017852049 | 0.15270693 | -0.116903984 |
| Gm23448 | 0.180187022 | 0.998861527 | 2.700353499 | 0.369900284 |
| Gm11606 | 0 | NA | NA | NA |
| Gm11605 | 3.581982469 | 0.036576639 | 0.528747093 | 0.069176057 |
| Adam23 | 3251.808784 | 0.51092059 | 0.059472922 | 8.590810243 |
| Gm22281 | 0 | NA | NA | NA |
| Dytn | 1.029483559 | 0.513099868 | 1.010327658 | 0.507854916 |
| Gm12748 | 0 | NA | NA | NA |
| Mdh1b | 8.408068065 | 0.189628738 | 0.39266256 | 0.482930529 |
| Fastkd2 | 291.2035401 | -0.283231644 | 0.127161953 | -2.227330082 |
| 4933402D24Rik | 1.548786521 | -0.12645704 | 0.782389109 | -0.161629347 |
| Gm13751 | 0.449839025 | -1.789404325 | 1.479553093 | -1.209422179 |
| Gm13754 | 0.863190071 | 2.246022413 | 1.221648251 | 1.838518093 |
| Gm13749 | 1.958865323 | 0.58595187 | 0.994020272 | 0.589476781 |
| Klf7 | 1205.650868 | 0.065142354 | 0.103357931 | 0.630259853 |
| Gm26649 | 0.52188035 | -0.92766875 | 1.442789128 | -0.642969046 |
| Mir6899 | 0.842388825 | 0.332313045 | 1.167878291 | 0.284544243 |

| | | | | |
|---|---|---|---|---|
| Gm13748 | 0.622058045 | -0.58410949 | 1.426777921 | -0.409390614 |
| Gm13750 | 1.235703043 | -0.359268152 | 0.894401861 | -0.401685381 |
| Gm28981 | 0.770914299 | -1.225990864 | 1.254383626 | -0.977365168 |
| Gm25748 | 0 | NA | NA | NA |
| Gm28980 | 0 | NA | NA | NA |
| Creb1 | 1118.909764 | -0.169862661 | 0.085434205 | -1.988227799 |
| Mettl21a | 95.46113856 | -0.035977889 | 0.14367035 | -0.250419718 |
| Gm28982 | 1.624436933 | -0.585324169 | 0.808271407 | -0.724167853 |
| Gm29152 | 0 | NA | NA | NA |
| 2810408I11Rik | 4.24223269 | -1.222632787 | 0.495593903 | -2.467005306 |
| Ccnyl1 | 180.7599231 | -0.357895195 | 0.135396387 | -2.643314213 |
| Fzd5 | 105.8313593 | -0.085853995 | 0.185443181 | -0.462966581 |
| Gm38058 | 2.181430594 | 0.673070899 | 0.701054796 | 0.960083152 |
| Plekhm3 | 907.4639202 | -0.141781669 | 0.057986538 | -2.445079041 |
| Rpl31-ps24 | 0 | NA | NA | NA |
| Rpl10a-ps1 | 39.84853431 | -0.189791995 | 0.194773038 | -0.974426422 |
| Gm24788 | 0 | NA | NA | NA |
| Akr1cl | 0.229212118 | 1.46143958 | 2.536135019 | 0.576246757 |
| Gm8550 | 0 | NA | NA | NA |
| Cryge | 0 | NA | NA | NA |
| Crygd | 0 | NA | NA | NA |
| Crygc | 0 | NA | NA | NA |
| Crygb | 0.150086409 | 1.027495679 | 3.723011807 | 0.275985071 |
| Gm7329 | 4.40840556 | -0.297726284 | 0.479930711 | -0.62035264 |
| Cryga | 0.97358459 | 2.489104925 | 1.251602927 | 1.988733704 |
| Gm28845 | 0.23654427 | 1.568164077 | 2.541886388 | 0.616929256 |
| D630023F18Rik | 33.29031821 | 0.740038061 | 0.200667349 | 3.687884779 |
| Idh1 | 461.6797512 | -0.002489373 | 0.127116218 | -0.019583438 |
| Pikfyve | 2046.792619 | 0.007292864 | 0.071097549 | 0.10257546 |
| Pth2r | 15.63521041 | -0.445167297 | 0.342463383 | -1.299897506 |
| Gm28927 | 0 | NA | NA | NA |
| Gm28926 | 0 | NA | NA | NA |
| Gm28925 | 0 | NA | NA | NA |
| Gm28923 | 0 | NA | NA | NA |
| Gm28922 | 0 | NA | NA | NA |
| Gm28728 | 0 | NA | NA | NA |
| Gm5255 | 0 | NA | NA | NA |
| Gm8805 | 0 | NA | NA | NA |
| Crygf | 0 | NA | NA | NA |
| Gm8809 | 0 | NA | NA | NA |
| Gm15659 | 0 | NA | NA | NA |
| Gm8812 | 0 | NA | NA | NA |
| Map2 | 21973.53413 | 0.43582612 | 0.096477866 | 4.517368969 |
| Gm24729 | 8.150369374 | 0.843569712 | 0.481492525 | 1.751989217 |
| Gm10558 | 1.106847758 | -0.125327554 | 0.98659295 | -0.12703066 |
| Gm25411 | 10.74259572 | 0.310823167 | 0.367087204 | 0.846728417 |
| Unc80 | 7728.076732 | 0.598159958 | 0.066056369 | 9.055295719 |

| | | | | |
|---|---|---|---|---|
| Gm37181 | 0.701668669 | 1.367047197 | 1.447902028 | 0.94415725 |
| Rpe | 507.539684 | -0.306269876 | 0.08334511 | -3.674719196 |
| Kansl1l | 470.4693895 | -0.051153991 | 0.081834817 | -0.625088344 |
| Gm37570 | 61.3256198 | 0.395902644 | 0.248728694 | 1.591704748 |
| Gm38387 | 64.00509308 | 0.614684777 | 0.195218225 | 3.148705907 |
| Gm38162 | 96.75562574 | 0.286255493 | 0.190445369 | 1.503084552 |
| Gm15789 | 30.55906383 | 0.377542387 | 0.214941573 | 1.756488433 |
| 6820402A03Rik | 78.68479837 | 0.427286911 | 0.188800876 | 2.263161697 |
| Gm37285 | 31.02313678 | 0.051858108 | 0.229493914 | 0.225967249 |
| Gm37755 | 15.79173624 | 0.088397972 | 0.345618717 | 0.255767317 |
| Gm38218 | 8.540597391 | 0.509542003 | 0.343455511 | 1.483574981 |
| Acadl | 460.9663652 | -0.473358666 | 0.126310989 | -3.74756518 |
| Gm15793 | 6.435362979 | -0.473926196 | 0.370609467 | -1.278775203 |
| Gm27934 | 0.075043205 | 0.517957688 | 3.724945049 | 0.139051095 |
| Gm25832 | 0.12092566 | 0.998850474 | 3.724945049 | 0.268151734 |
| Myl1 | 8.248283288 | 1.197321956 | 0.399929146 | 2.993835202 |
| Rpl31-ps14 | 4.920430513 | -0.151991921 | 0.474169178 | -0.320543655 |
| Gm15826 | 0.731126313 | 1.936336618 | 1.36779101 | 1.415667017 |
| Lancl1 | 1237.017238 | 0.514945488 | 0.047930704 | 10.74354105 |
| Cps1 | 8.332986231 | 0.935224369 | 0.363610264 | 2.572051621 |
| Gm15668 | 0.088534442 | -0.905414734 | 3.723500899 | -0.243162217 |
| Gm26342 | 0 | NA | NA | NA |
| Kif22-ps | 0 | NA | NA | NA |
| Gm22548 | 0 | NA | NA | NA |
| Gm15669 | 0 | NA | NA | NA |
| Gm37957 | 0 | NA | NA | NA |
| Gm23189 | 0 | NA | NA | NA |
| Gm15671 | 0 | NA | NA | NA |
| Erbb4 | 1732.331576 | 0.661406487 | 0.067006307 | 9.870809392 |
| Gm37100 | 8.451353278 | 0.703759916 | 0.424698134 | 1.657082666 |
| Gm15670 | 8.648476067 | 1.171185213 | 0.483535696 | 2.422127722 |
| Gm25542 | 1.041370757 | 1.601751564 | 1.095094865 | 1.462660099 |
| Gm19335 | 10.47773709 | 0.077463032 | 0.392769183 | 0.197222784 |
| Gm37735 | 8.967735426 | 0.061683163 | 0.387869691 | 0.159030633 |
| Gm37769 | 40.95679204 | 0.229647092 | 0.239228536 | 0.959948575 |
| Gm37061 | 10.97852267 | 0.547571398 | 0.467713667 | 1.170740641 |
| Gm16076 | 36.27563093 | 0.29085316 | 0.283010284 | 1.027712337 |
| Gm8840 | 0 | NA | NA | NA |
| Pced1c-ps | 0 | NA | NA | NA |
| Gm28497 | 0.047948698 | 0.037064902 | 3.724945049 | 0.009950456 |
| Gm29113 | 9.644466874 | 0.812915086 | 0.33683649 | 2.413381893 |
| Gm29114 | 0 | NA | NA | NA |
| Ikzf2 | 269.8286907 | -0.201350853 | 0.105804233 | -1.903051009 |
| Gm29112 | 12.57536217 | -0.87973122 | 0.364762961 | -2.411788787 |
| Gm28112 | 19.23386575 | 0.068711943 | 0.313171395 | 0.219406831 |
| Gm37930 | 0.044191549 | 0.037062677 | 3.724945049 | 0.009949859 |
| Spag16 | 19.54478478 | 0.089100781 | 0.241210637 | 0.36938993 |

| | | | | |
|---|---|---|---|---|
| Gm22665 | 0 | NA | NA | NA |
| Gm23879 | 0 | NA | NA | NA |
| Gm28580 | 0 | NA | NA | NA |
| Gm38272 | 0.042716945 | 0.037062677 | 3.724945049 | 0.009949859 |
| Gm37456 | 0.272865106 | 2.116370698 | 2.408694967 | 0.878637904 |
| Gm23422 | 0 | NA | NA | NA |
| Vwc2l | 194.8593581 | 0.186820778 | 0.093841978 | 1.99080179 |
| Bard1 | 62.05484921 | -0.426717296 | 0.21282232 | -2.005040149 |
| Gm16236 | 1.002654625 | -0.919010134 | 1.07687952 | -0.853401069 |
| Gm8870 | 0 | NA | NA | NA |
| Abca12 | 5.342748304 | 0.707016157 | 0.546703163 | 1.293235902 |
| Gm6947 | 0 | NA | NA | NA |
| Atic | 353.5973081 | -0.569855918 | 0.110463427 | -5.158774553 |
| Fn1 | 1063.525137 | -1.023664299 | 0.224426169 | -4.561251941 |
| Apol7d | 1.07675232 | -1.405350599 | 1.091165611 | -1.287935199 |
| Gm5829 | 1.23306525 | -0.234227266 | 0.832025925 | -0.281514384 |
| Gm5256 | 3.462762875 | -0.202171568 | 0.563479049 | -0.358791633 |
| Gm37217 | 0 | NA | NA | NA |
| Gm8883 | 0 | NA | NA | NA |
| Gm28818 | 0.213332449 | 0.476297135 | 2.16768839 | 0.219725832 |
| 4933417E11Rik | 0.198105924 | -0.44382372 | 2.645781003 | -0.167747716 |
| 4930556G22Rik | 0 | NA | NA | NA |
| Gm5528 | 0 | NA | NA | NA |
| Gm8885 | 0.131209559 | -0.409992761 | 2.972496229 | -0.137928774 |
| Gm4319 | 0.520182433 | -0.00446856 | 1.291119689 | -0.003460996 |
| Mreg | 141.6531737 | -0.13089836 | 0.133165397 | -0.98297578 |
| Gm25360 | 67.27497613 | -0.819806424 | 0.214504676 | -3.821858062 |
| D230017M19Rik | 46.21119991 | 0.390232736 | 0.171923893 | 2.269799317 |
| Gm23472 | 572.9451334 | 0.734842725 | 0.804476308 | 0.913442345 |
| Gm24497 | 54.97402124 | 0.505997734 | 0.374498222 | 1.351135212 |
| Gm25939 | 1027.212574 | -0.245536819 | 0.171843348 | -1.428840981 |
| Pecr | 29.32302085 | 0.505771551 | 0.201420636 | 2.511021524 |
| Tmem169 | 181.7981465 | 0.221740368 | 0.114719663 | 1.932888947 |
| Xrcc5 | 253.261693 | -0.144639427 | 0.078968014 | -1.83162042 |
| 4-Mar | 295.8236201 | 0.013928312 | 0.126261626 | 0.110313105 |
| Gm15843 | 0.209261529 | 0.034789973 | 2.615696522 | 0.013300462 |
| Smarcal1 | 689.8282747 | 0.10252203 | 0.055659013 | 1.841966352 |
| Ankar | 0.251200408 | -0.993104798 | 2.090477133 | -0.475061306 |
| Rpl37a | 999.4688314 | -0.068916079 | 0.127300766 | -0.54136421 |
| Igfbp2 | 490.8609578 | -0.606711259 | 0.190701506 | -3.181470735 |
| Igfbp5 | 3859.554287 | 0.435377142 | 0.140174802 | 3.105958668 |
| 1700027A15Rik | 1.147127115 | -2.337443986 | 1.081772739 | -2.160753274 |
| Tnp1 | 0 | NA | NA | NA |
| Gm38106 | 0 | NA | NA | NA |
| Gm29185 | 0.044544648 | 0.517957688 | 3.724945049 | 0.139051095 |
| Gm29184 | 0 | NA | NA | NA |
| Pinc | 0.243249674 | 0.517969664 | 2.557831642 | 0.202503423 |

| | | | | |
|---|---|---|---|---|
| Gm27315 | 0 | NA | NA | NA |
| D530049I02Rik | 0 | NA | NA | NA |
| Gm9553 | 0.046345709 | -0.443818988 | 3.724945049 | -0.119147795 |
| C530043A13Rik | 1.12921268 | -1.685824846 | 0.952908102 | -1.769136858 |
| Gm29186 | 0 | NA | NA | NA |
| 6030407O03Rik | 6.071023221 | 0.400228123 | 0.474243514 | 0.843929567 |
| Gm29183 | 1.411578533 | 0.722188417 | 1.078608922 | 0.66955539 |
| Gm25329 | 0 | NA | NA | NA |
| Mir6351 | 0.439338454 | 1.421662818 | 1.473260538 | 0.964977192 |
| Tns1 | 929.0676011 | 0.030402531 | 0.115815105 | 0.262509207 |
| Rufy4 | 2.528566672 | -1.905461322 | 0.709500392 | -2.685638151 |
| Cxcr2 | 2.628643724 | -0.335383585 | 0.554563984 | -0.604769863 |
| Cxcr1 | 0.182828849 | -1.328698802 | 2.253829453 | -0.589529434 |
| Mir1928 | 0 | NA | NA | NA |
| Gm25524 | 0 | NA | NA | NA |
| Arpc2 | 1724.327588 | -0.198781579 | 0.085643732 | -2.321028915 |
| Gpbar1 | 0.083526043 | 0.037073798 | 3.724945049 | 0.009952844 |
| Aamp | 1126.602311 | -0.051292452 | 0.081628678 | -0.628363138 |
| Gm28364 | 0.299075359 | -1.624059866 | 1.72316471 | -0.942486726 |
| Pnkd | 1381.963466 | 0.358541355 | 0.092337772 | 3.882932699 |
| Gm37503 | 2.04875246 | 0.535737712 | 0.721600806 | 0.742429481 |
| Tmbim1 | 824.4000618 | -0.84115261 | 0.152072 | -5.531278667 |
| Gm29358 | 41.10390762 | 0.648304175 | 0.348736674 | 1.859007737 |
| Gm37931 | 25.45680512 | 1.04700466 | 0.459547873 | 2.278336429 |
| Catip | 16.47013026 | -0.408612346 | 0.263137545 | -1.552846998 |
| Slc11a1 | 156.1085199 | -1.046028821 | 0.209238787 | -4.999210881 |
| Ctdsp1 | 480.9035942 | -0.364240761 | 0.120744059 | -3.016635062 |
| Mir26b | 0.460912027 | 0.025255588 | 1.444402505 | 0.017485146 |
| Vil1 | 4.671085706 | -0.766226105 | 0.465824017 | -1.64488321 |
| Usp37 | 682.5650065 | -0.195751425 | 0.072987515 | -2.681985055 |
| Gm37733 | 34.22973971 | 0.011888119 | 0.300689111 | 0.039536248 |
| Rqcd1 | 401.5594137 | -0.335263111 | 0.063614836 | -5.270203187 |
| Plcd4 | 276.691242 | -0.347428753 | 0.094098113 | -3.692196799 |
| Zfp142 | 808.2705847 | -0.120063372 | 0.081215884 | -1.478323772 |
| Gm25035 | 2.10081791 | 1.054987739 | 0.753077705 | 1.400901568 |
| Bcs1l | 123.2340065 | -0.198184245 | 0.130487106 | -1.518803285 |
| Rnf25 | 329.113652 | -0.126581445 | 0.092908289 | -1.362434362 |
| Stk36 | 70.93351887 | -0.732431996 | 0.172281526 | -4.251367004 |
| Ttll4 | 457.8466587 | -0.131896972 | 0.111602609 | -1.181844884 |
| Cyp27a1 | 60.63009196 | -0.11085031 | 0.170609667 | -0.649730537 |
| Prkag3 | 1.196546384 | -2.215622117 | 1.019616881 | -2.172994738 |
| Gm27507 | 0 | NA | NA | NA |
| Wnt6 | 20.88210294 | 0.235939184 | 0.316397485 | 0.745704994 |
| Gm29539 | 0.08169834 | -0.443821212 | 3.724945049 | -0.119148392 |
| Wnt10a | 51.88315531 | -0.447563434 | 0.239313995 | -1.870193316 |
| Gm16582 | 1.058309834 | -1.044186089 | 1.213146211 | -0.860725673 |
| Cdk5r2 | 2117.926955 | 0.597883472 | 0.114335993 | 5.229179839 |

| | | | | |
|---|---|---|---|---|
| Fev | 2.243593674 | -1.105096885 | 0.944988285 | -1.169429189 |
| Cryba2 | 3.578913291 | -2.585824012 | 0.718819848 | -3.597318602 |
| Mir375 | 0 | NA | NA | NA |
| Ccdc108 | 54.38524299 | 0.538451098 | 0.1949892 | 2.761440622 |
| Gm15841 | 0.048949532 | 0.037064902 | 3.724945049 | 0.009950456 |
| Ihh | 2.70016216 | -0.36907945 | 0.67151796 | -0.549619627 |
| Gm37744 | 0.044544648 | 0.517957688 | 3.724945049 | 0.139051095 |
| Nhej1 | 18.60744933 | -0.002045669 | 0.271017217 | -0.007548113 |
| Slc23a3 | 6.004310424 | -0.923516903 | 0.484102258 | -1.907689725 |
| Cnppd1 | 349.0250262 | -0.31839694 | 0.094818443 | -3.357964235 |
| Fam134a | 1705.406746 | 0.064272767 | 0.048856957 | 1.315529482 |
| Zfand2b | 100.1118125 | -0.05747293 | 0.140318531 | -0.409589024 |
| Abcb6 | 14.09945832 | 0.68887175 | 0.327981432 | 2.100337652 |
| Gm29253 | 0.323844909 | -0.506665605 | 2.034980216 | -0.248978148 |
| Atg9a | 756.6561104 | 0.336753523 | 0.057202778 | 5.887013431 |
| Ankzf1 | 160.4436551 | 0.090263314 | 0.148134737 | 0.60933253 |
| Glb1l | 98.7643794 | -0.043091259 | 0.130044591 | -0.331357563 |
| Gm27943 | 0 | NA | NA | NA |
| Stk16 | 338.2768 | -0.320276367 | 0.068125684 | -4.701257257 |
| Tuba4a | 2899.871314 | 0.499113664 | 0.071380436 | 6.992303435 |
| A630095N17Rik | 0.826064167 | -0.87171846 | 1.09555592 | -0.795685957 |
| Dnajb2 | 784.3513935 | -0.316333048 | 0.082087288 | -3.853617984 |
| Ptprn | 4437.838423 | -0.076467932 | 0.069530394 | -1.099777057 |
| Gm28294 | 1.007042841 | 1.558233529 | 1.121275225 | 1.389697636 |
| Resp18 | 711.8094129 | -0.05782681 | 0.154233462 | -0.374930378 |
| Gm28902 | 0.705697236 | -2.102387638 | 1.264729878 | -1.662321476 |
| Dnpep | 296.9380932 | -0.228204823 | 0.111303199 | -2.050298873 |
| Des | 14.87844252 | -1.060443628 | 0.27664501 | -3.833228825 |
| Gm15179 | 4.05260616 | -0.120303724 | 0.580199651 | -0.207348838 |
| Speg | 1539.141981 | 0.447356232 | 0.141999149 | 3.15041488 |
| Gm15178 | 0 | NA | NA | NA |
| Gmppa | 265.8934165 | -0.264338501 | 0.083891517 | -3.150956286 |
| Asic4 | 155.5102748 | 0.578859006 | 0.137704688 | 4.203625996 |
| Gm37264 | 4.269864839 | -0.14482693 | 0.491417707 | -0.294712478 |
| Chpf | 816.6335401 | -0.094871801 | 0.087594992 | -1.083073343 |
| Obsl1 | 195.5669334 | -0.157921021 | 0.094826782 | -1.665363078 |
| Tmem198 | 376.0167297 | -0.310781117 | 0.129309297 | -2.403393455 |
| Inha | 132.6233029 | 0.732404665 | 0.188107573 | 3.893541613 |
| Gm15181 | 0 | NA | NA | NA |
| Stk11ip | 336.2431366 | 0.005593722 | 0.082331582 | 0.067941394 |
| Gm15183 | 0.113999573 | -0.443821212 | 3.724945049 | -0.119148392 |
| Slc4a3 | 1347.377089 | 0.507734078 | 0.083446194 | 6.084568397 |
| Gm15180 | 0.245985165 | 0.197082229 | 2.48650074 | 0.079260877 |
| Gm15182 | 0.115603922 | -0.93550548 | 3.724188147 | -0.251197158 |
| Gm5257 | 0.104637558 | -0.443821212 | 3.724945049 | -0.119148392 |
| Gm29065 | 0 | NA | NA | NA |
| Gm29068 | 0 | NA | NA | NA |

| | | | | |
|---|---|---|---|---|
| Gm29069 | 0.135858026 | 0.037065849 | 2.948997537 | 0.012568966 |
| Gm17751 | 4.194728469 | 0.007113344 | 0.472550786 | 0.015053077 |
| Mir6343 | 0 | NA | NA | NA |
| Gm816 | 0.297786538 | 0.517973814 | 1.616984636 | 0.32033317 |
| Gm38265 | 0 | NA | NA | NA |
| Gm25974 | 0 | NA | NA | NA |
| Epha4 | 2103.014444 | 1.110632107 | 0.129019329 | 8.608261405 |
| Mir6352 | 3.045583208 | 1.411495689 | 0.819916159 | 1.721512221 |
| Gm28386 | 2.146862605 | 1.799949343 | 0.864035668 | 2.083188704 |
| Gm28385 | 0.301967822 | 0.03704835 | 3.724945049 | 0.009946012 |
| Gm26263 | 0 | NA | NA | NA |
| Gm28387 | 0.044544648 | 0.517957688 | 3.724945049 | 0.139051095 |
| Pax3 | 26.95793292 | -0.258078102 | 0.231938345 | -1.112701317 |
| Sgpp2 | 367.8063322 | -0.119022249 | 0.111235529 | -1.070002097 |
| Gm28410 | 0.413627519 | 2.506901039 | 1.431057275 | 1.751782464 |
| Gm37884 | 3.006274965 | 0.313727592 | 0.702261879 | 0.446738747 |
| Farsb | 522.3359989 | -0.26979066 | 0.0815946 | -3.306476911 |
| Gm28995 | 0 | NA | NA | NA |
| 4833412K13Rik | 33.12266117 | 0.386919541 | 0.279830015 | 1.382694921 |
| BC035947 | 16.4961278 | -0.147412137 | 0.311860173 | -0.472686638 |
| Mogat1 | 1.455068488 | 0.611707503 | 0.914540539 | 0.668868657 |
| 5730419F03Rik | 0.372487069 | -1.966570667 | 1.703287144 | -1.154573774 |
| Gm29611 | 0 | NA | NA | NA |
| Acsl3 | 1660.961095 | 0.201246909 | 0.077816848 | 2.586161138 |
| Utp14b | 706.5540839 | -0.080876482 | 0.089549694 | -0.903146388 |
| Gm6159 | 2.778261606 | -0.702581327 | 0.603921095 | -1.163366097 |
| Gm29536 | 0.417935612 | -1.274670192 | 1.689108507 | -0.754640798 |
| Kcne4 | 23.89478986 | -0.193819119 | 0.283777414 | -0.682996989 |
| Gm29187 | 0.218130886 | 0.539436107 | 2.170555645 | 0.248524431 |
| Gm5830 | 3.384143917 | -0.072863662 | 0.578149595 | -0.126029081 |
| Scg2 | 2880.675548 | -0.51966782 | 0.146286557 | -3.552396261 |
| Gm19027 | 0.061920613 | -0.443816764 | 3.724945049 | -0.119147198 |
| Gm24160 | 0 | NA | NA | NA |
| Ap1s3 | 62.26140866 | -0.289425287 | 0.190237914 | -1.521385936 |
| Gm37886 | 2.695407072 | 0.511344594 | 0.706005523 | 0.724278461 |
| Wdfy1 | 946.0370216 | -0.429429431 | 0.064679119 | -6.639382776 |
| Gm10555 | 0.378505399 | -1.837504122 | 1.596593791 | -1.150890184 |
| Mrpl44 | 178.4373768 | -0.133688349 | 0.093079137 | -1.436286939 |
| Serpine2 | 1567.911986 | -0.12083401 | 0.124503351 | -0.970528172 |
| Gm5529 | 18.26108814 | -0.062897814 | 0.26753998 | -0.235096878 |
| Gm28071 | 2.944880665 | 0.158084302 | 0.646737358 | 0.24443354 |
| Gm28058 | 0 | NA | NA | NA |
| Gm23934 | 0.310678561 | -0.409103266 | 1.609402526 | -0.25419574 |
| Gm38339 | 0.456870417 | -1.605281012 | 3.386811124 | -0.473980081 |
| 9830004L10Rik | 0 | NA | NA | NA |
| Fam124b | 3.258857179 | -1.630278431 | 0.553935494 | -2.943083535 |
| 2310015K22Rik | 0 | NA | NA | NA |

| | | | | |
|---|---|---|---|---|
| Gm37932 | 4.562962678 | -1.495552396 | 0.559617077 | -2.67245668 |
| Cul3 | 2324.447608 | 0.010834087 | 0.080223852 | 0.135048205 |
| Gm37902 | 63.67772732 | 0.106905323 | 0.166670564 | 0.641416939 |
| Gm37262 | 28.6341495 | -0.336927208 | 0.237773791 | -1.417007343 |
| Gm29125 | 1.604780936 | -0.417673801 | 0.786440121 | -0.531094217 |
| Gm6189 | 0.828249129 | -2.130996153 | 1.256885231 | -1.695458026 |
| 1700016L21Rik | 0.083172944 | -0.443821212 | 3.724945049 | -0.119148392 |
| Dock10 | 1627.450511 | -0.333123711 | 0.093072911 | -3.579169352 |
| Gm37645 | 36.84305094 | -0.210258865 | 0.241623633 | -0.870191639 |
| Gm38062 | 55.87989744 | -0.054622677 | 0.337309776 | -0.161936239 |
| A630081D01Rik | 65.65902623 | -0.309951787 | 0.227656075 | -1.361491398 |
| Gm29124 | 0 | NA | NA | NA |
| Gm28066 | 0 | NA | NA | NA |
| Nyap2 | 954.1496905 | 0.626293404 | 0.094432638 | 6.632171019 |
| Gm37210 | 0 | NA | NA | NA |
| Gm6198 | 0 | NA | NA | NA |
| Gm5530 | 3.003564229 | -1.280438574 | 0.574422642 | -2.229087922 |
| Gm7516 | 0 | NA | NA | NA |
| Gm37114 | 0 | NA | NA | NA |
| Mir6344 | 0 | NA | NA | NA |
| Irs1 | 784.393355 | -0.103907262 | 0.079456338 | -1.307727797 |
| Gm9747 | 0.131672813 | -1.283459693 | 3.660707039 | -0.350604317 |
| Rhbdd1 | 192.9071811 | -0.733264221 | 0.162711836 | -4.506520469 |
| Gm19552 | 8.274522055 | -0.171908503 | 0.374008405 | -0.459638074 |
| Gm28940 | 0.217873223 | 0.527468522 | 2.548329196 | 0.206986022 |
| Gm22112 | 0.107803068 | 0.036092186 | 3.724874622 | 0.009689503 |
| Col4a4 | 19.40116672 | 0.139702063 | 0.270334255 | 0.516775288 |
| Col4a3 | 9.983396163 | 0.721078111 | 0.350159848 | 2.059282685 |
| n-R5s213 | 0 | NA | NA | NA |
| C430014B12Rik | 2.511728143 | -0.671864155 | 0.613271952 | -1.095540327 |
| Mff | 1852.754622 | 0.142266787 | 0.045305048 | 3.140197218 |
| Gm28941 | 40.41534512 | 0.442187525 | 0.266709014 | 1.657939932 |
| Tm4sf20 | 0.089062654 | -0.443823436 | 3.724945049 | -0.119148989 |
| Gm19257 | 0 | NA | NA | NA |
| Gm36999 | 0.100902008 | -0.876201588 | 3.72093227 | -0.235479048 |
| Gm28942 | 0.253304709 | -0.387734367 | 1.810964926 | -0.214103742 |
| Gm17764 | 0 | NA | NA | NA |
| Gm22396 | 0.22174029 | -0.975012761 | 3.152579238 | -0.309274625 |
| Agfg1 | 1873.69111 | -0.172968039 | 0.060602177 | -2.85415553 |
| Gm7553 | 0.943831312 | -0.128675379 | 1.064713504 | -0.120854463 |
| Slc19a3 | 12.62212474 | 0.433145726 | 0.33910673 | 1.277313857 |
| Gm7539 | 0 | NA | NA | NA |
| Krtap28-10 | 0 | NA | NA | NA |
| A030005K14Rik | 0 | NA | NA | NA |
| Krtap28-13 | 0.044191549 | 0.037062677 | 3.724945049 | 0.009949859 |
| Gm29518 | 0.066050875 | -0.443818988 | 3.724945049 | -0.119147795 |
| Ccl20 | 0.166497232 | -1.455331858 | 3.388676174 | -0.429469145 |

| | | | | |
|---|---|---|---|---|
| Gm29519 | 0 | NA | NA | NA |
| Daw1 | 3.764882787 | 0.20784488 | 0.604429495 | 0.34386952 |
| Gm6244 | 0 | NA | NA | NA |
| n-R5s214 | 0 | NA | NA | NA |
| Sphkap | 3563.73903 | 0.420418688 | 0.101792448 | 4.130155987 |
| Gm6374 | 0.068097359 | 0.037064902 | 3.724945049 | 0.009950456 |
| Gm37517 | 48.29894096 | 0.786300604 | 0.369886349 | 2.125789738 |
| Gm37989 | 0 | NA | NA | NA |
| 4933436I20Rik | 1.025359323 | -1.056103378 | 1.2080745 | -0.874203849 |
| Gm25754 | 0.150086409 | 1.027495679 | 3.723011807 | 0.275985071 |
| Gm24555 | 0 | NA | NA | NA |
| Pid1 | 538.7136745 | -0.178513657 | 0.081992929 | -2.177183557 |
| Gm37058 | 25.37561462 | -0.457592331 | 0.332653201 | -1.375583729 |
| Gm18304 | 0 | NA | NA | NA |
| Mir6353 | 3.815514299 | -0.664788213 | 0.720238833 | -0.923010788 |
| Dner | 2450.532823 | 0.129726221 | 0.108801367 | 1.192321609 |
| Gm37959 | 19.69556321 | 0.574633085 | 0.29568222 | 1.943414406 |
| Gm37925 | 43.42924116 | 0.554477361 | 0.280736802 | 1.975078993 |
| Mir5126 | 0.291915594 | 1.054952088 | 2.008064771 | 0.5253576 |
| Trip12 | 5110.806766 | -0.139616137 | 0.052446514 | -2.662067049 |
| Gm38374 | 71.20713703 | -0.103524747 | 0.195325697 | -0.530010893 |
| Gm37738 | 93.58139767 | 0.081834503 | 0.250671761 | 0.326460799 |
| Fbxo36 | 28.2317926 | 0.148381483 | 0.255470674 | 0.580816109 |
| Slc16a14 | 112.1372938 | 0.33258733 | 0.149831808 | 2.219737818 |
| Gm15433 | 3.38607774 | -2.805872359 | 0.661552378 | -4.24134574 |
| Gm29284 | 0.509711911 | 1.984592879 | 1.432560704 | 1.385346445 |
| C130026I21Rik | 27.88048967 | -1.200569734 | 0.323063836 | -3.716199714 |
| Rpl19-ps1 | 2.447118844 | 0.182977736 | 0.674624616 | 0.271228965 |
| Gm27907 | 0 | NA | NA | NA |
| Gm7582 | 5.427132103 | -1.370812398 | 0.522877024 | -2.621672659 |
| Gm2389 | 0 | NA | NA | NA |
| Gm18342 | 8.221653517 | -1.983779805 | 0.405978763 | -4.886412746 |
| A530032D15Rik | 14.11902256 | -1.647641234 | 0.318555344 | -5.172229148 |
| Gm10553 | 0.099106094 | -0.93550548 | 3.724188147 | -0.251197158 |
| A530040E14Rik | 19.7285084 | -1.582640979 | 0.326934138 | -4.840855675 |
| Gm16038 | 0.038981395 | -0.443816764 | 3.724945049 | -0.119147198 |
| Gm2635 | 0.088534442 | -0.905414734 | 3.723500899 | -0.243162217 |
| Gm6264 | 1.445224006 | -0.870148847 | 1.06536899 | -0.816758189 |
| Gm7609 | 9.276052769 | -1.708683714 | 0.35175407 | -4.857608933 |
| Gm16028 | 0.132101749 | -0.93550548 | 3.724188147 | -0.251197158 |
| Gm16026 | 7.688870986 | -1.153244269 | 0.369031802 | -3.125053892 |
| Gm2619 | 4.165531338 | -0.789204572 | 0.458088127 | -1.722822588 |
| Gm16025 | 3.376741903 | -0.785613942 | 0.537261278 | -1.462256771 |
| Gm7281 | 0.301876039 | -0.722594432 | 1.728139822 | -0.41813424 |
| Gm2666 | 0.742190307 | -2.302955951 | 1.188987715 | -1.936904748 |
| Gm2427 | 1.636398745 | -2.111766773 | 0.965979825 | -2.186139625 |
| Gm16092 | 29.37301029 | 0.357925413 | 0.27764758 | 1.289135721 |

| | | | | |
|---|---|---|---|---|
| Gm7592 | 2.43285919 | -3.08406805 | 0.941578708 | -3.275422462 |
| G530012D18Rik | 0 | NA | NA | NA |
| Sp110 | 246.5019786 | -0.928151982 | 0.155439422 | -5.9711492 |
| Gm16094 | 4.539762321 | -1.398912611 | 0.554642417 | -2.522188292 |
| Sp140 | 215.8469477 | -1.354031029 | 0.178537164 | -7.584028981 |
| Gm10552 | 1.59243937 | -2.253318819 | 0.863535455 | -2.60941089 |
| Gm17017 | 6.466014695 | 0.000620358 | 0.430193386 | 0.001442045 |
| Sp100 | 678.1539777 | -1.371584441 | 0.24973468 | -5.492166494 |
| n-R5s215 | 0 | NA | NA | NA |
| A630001G21Rik | 22.19689395 | -0.650992965 | 0.247874625 | -2.626299339 |
| Mir8096 | 0 | NA | NA | NA |
| Gm25472 | 0 | NA | NA | NA |
| 9930111H07Rik | 0.389848623 | -2.274231277 | 1.651352731 | -1.377192913 |
| Gm22906 | 0 | NA | NA | NA |
| Cab39 | 2445.880552 | 0.028203669 | 0.088974074 | 0.316987494 |
| Itm2c | 5468.611581 | 0.438852134 | 0.073723634 | 5.952665508 |
| 4933407L21Rik | 23.38036317 | 0.498639027 | 0.283565675 | 1.758460458 |
| Gm28884 | 0.751763609 | 2.277904003 | 1.23572901 | 1.843368557 |
| Gpr55 | 7.561829528 | -0.511991586 | 0.403459604 | -1.269003342 |
| Gm28100 | 0.240280111 | 0.037077443 | 2.463083609 | 0.015053262 |
| Gm18180 | 25.54930956 | -0.497248668 | 0.244120698 | -2.036896802 |
| Spata3 | 2.778035728 | -0.334923106 | 0.660369444 | -0.507175354 |
| 2810459M11Rik | 146.6634441 | -1.203090344 | 0.158387624 | -7.595860799 |
| Psmd1 | 1536.480455 | -0.159652542 | 0.063727671 | -2.505231093 |
| Gm22786 | 4.654196711 | 0.710023182 | 0.495681478 | 1.432418223 |
| Htr2b | 18.41870533 | -2.468337031 | 0.405114063 | -6.092943338 |
| Gm37914 | 41.69621495 | -0.241215501 | 0.271128457 | -0.889672383 |
| Gm21972 | 0.320475213 | 0.03706974 | 2.355682608 | 0.015736305 |
| Armc9 | 417.345602 | -0.053867956 | 0.177844692 | -0.302893243 |
| Gm16341 | 0.537632031 | -1.689359321 | 1.527289032 | -1.106116318 |
| B3gnt7 | 21.27627369 | -0.949765371 | 0.274512675 | -3.459823376 |
| Gm28626 | 1.81028421 | 0.809904675 | 0.838980432 | 0.965343939 |
| Gm5258 | 0 | NA | NA | NA |
| Gm28627 | 0 | NA | NA | NA |
| Ncl | 3170.074209 | -0.047944362 | 0.101009868 | -0.474650283 |
| Snora75 | 0 | NA | NA | NA |
| mmu-mir-3535 | 0.451658936 | 0.859516477 | 1.295399957 | 0.663514363 |
| Gm24148 | 0 | NA | NA | NA |
| Snord82 | 0.870629472 | 1.610590296 | 1.112475995 | 1.447752853 |
| C130036L24Rik | 32.9831117 | 0.090418628 | 0.245786917 | 0.367874045 |
| Nmur1 | 2.498311769 | 0.200298586 | 0.584526408 | 0.342668155 |
| 1700019O17Rik | 0.04393175 | 0.037062677 | 3.724945049 | 0.009949859 |
| Rpl30-ps6 | 0 | NA | NA | NA |
| Ptma | 1852.709718 | -0.143936755 | 0.136903917 | -1.051370611 |
| Pde6d | 314.3105157 | -0.221533511 | 0.109946058 | -2.014929093 |
| Gm37152 | 0.163519603 | 0.966857846 | 2.774974014 | 0.348420504 |
| Cops7b | 225.9968041 | -0.005986886 | 0.083134265 | -0.07201466 |

| | | | | |
|---|---|---|---|---|
| Gm22955 | 0 | NA | NA | NA |
| Gm37022 | 4.10661082 | 0.707404955 | 0.581936184 | 1.215605723 |
| Nppc | 53.44822697 | 0.069058269 | 0.240249242 | 0.287444275 |
| Dis3l2 | 456.7534953 | 0.300111842 | 0.080612971 | 3.722872864 |
| Gm29097 | 0.04393175 | 0.037062677 | 3.724945049 | 0.009949859 |
| Gm29055 | 12.11508896 | 0.513708106 | 0.428799189 | 1.198015575 |
| Gm37485 | 23.25905464 | 0.583355712 | 0.319296977 | 1.827000424 |
| Gm38142 | 24.33645793 | 0.250225187 | 0.323831598 | 0.772701578 |
| Gm38010 | 20.30950254 | -0.170110586 | 0.293445573 | -0.579700637 |
| Gm37541 | 10.5940186 | 0.624227112 | 0.440030986 | 1.418598079 |
| Gm38021 | 12.72797742 | 0.875814466 | 0.343098357 | 2.552662955 |
| Gm6136 | 179.3671153 | -0.187527737 | 0.111399315 | -1.683383218 |
| Gm37967 | 11.04668228 | 0.377862164 | 0.373160323 | 1.012600056 |
| Gm28375 | 0.045882455 | 0.517959912 | 3.724945049 | 0.139051692 |
| Gm37017 | 0.873848832 | 1.686539984 | 1.260750086 | 1.337727439 |
| Gm37447 | 21.46227591 | 0.638365055 | 0.316653049 | 2.015976344 |
| Akp-ps1 | 0.752438969 | 0.122582809 | 1.17421578 | 0.10439547 |
| Gm29371 | 0 | NA | NA | NA |
| Alppl2 | 0 | NA | NA | NA |
| Alpi | 0 | NA | NA | NA |
| Gm29372 | 0 | NA | NA | NA |
| Akp3 | 0.332159689 | -1.613390556 | 1.924908899 | -0.83816463 |
| Gm29374 | 5.217580992 | -0.846466828 | 0.425128666 | -1.991083867 |
| Ecel1 | 298.3676729 | -0.812760667 | 0.141408389 | -5.747612803 |
| Prss56 | 20.4462505 | -1.696656973 | 0.771895434 | -2.198039914 |
| Chrnd | 0 | NA | NA | NA |
| Chrng | 0.171131556 | -0.444767681 | 3.383981516 | -0.131433248 |
| Eif4e2 | 578.568997 | -0.194264276 | 0.062971098 | -3.084975199 |
| Gm26187 | 0.955307282 | 2.175983854 | 1.083669139 | 2.007978059 |
| Efhd1 | 197.7653228 | 0.258885032 | 0.15441872 | 1.676513269 |
| Efhd1os | 0.337798032 | 0.992985591 | 1.923179117 | 0.516325069 |
| Gigyf2 | 1430.757424 | -0.076584361 | 0.082931539 | -0.923464839 |
| Gm22549 | 0 | NA | NA | NA |
| Kcnj13 | 41.11332588 | 0.891217338 | 0.270886393 | 3.290004072 |
| Gm23504 | 0 | NA | NA | NA |
| 3110079O15Rik | 0.366579243 | 0.564899943 | 1.596962291 | 0.353734052 |
| Ngef | 1590.707428 | 0.34128175 | 0.12954538 | 2.634457138 |
| Neu2 | 18.78655172 | -0.301118638 | 0.288533288 | -1.043618363 |
| Gm17181 | 0.432611478 | 0.043197763 | 1.48546466 | 0.029080304 |
| Gm6153 | 1.379575143 | 0.201435537 | 0.809276755 | 0.248908096 |
| Inpp5d | 416.1094285 | -0.427686466 | 0.12608164 | -3.39213913 |
| Atg16l1 | 673.2924239 | 0.246284473 | 0.123644147 | 1.991881377 |
| Gm25395 | 742.8621918 | -0.003690415 | 0.185294873 | -0.019916442 |
| Scarna6 | 154.5847017 | 0.002766705 | 0.156744824 | 0.017651013 |
| Sag | 17.39870792 | -0.218683302 | 0.280561397 | -0.779449007 |
| Dgkd | 1519.408834 | 0.005039881 | 0.072527779 | 0.069488974 |
| Gm19582 | 6.959264896 | -0.122970188 | 0.468980708 | -0.26220735 |

| | | | | |
|---|---|---|---|---|
| 4833421G17Rik | 11.9590053 | 0.422253634 | 0.344785158 | 1.224686229 |
| Gm38365 | 57.83653722 | 0.647059493 | 0.319297209 | 2.026511583 |
| Usp40 | 453.472211 | -0.226270982 | 0.115418805 | -1.96043428 |
| Gm25198 | 0 | NA | NA | NA |
| Gm5832 | 0.105852363 | -0.443821212 | 3.724945049 | -0.119148392 |
| Gm15375 | 0.049553047 | -0.443818988 | 3.724945049 | -0.119147795 |
| Ugt1a10 | 0.161949631 | -1.309098506 | 2.76260123 | -0.473864448 |
| Gm20528 | 0.038981395 | -0.443816764 | 3.724945049 | -0.119147198 |
| Ugt1a9 | 0.038981395 | -0.443816764 | 3.724945049 | -0.119147198 |
| Ugt1a8 | 0 | NA | NA | NA |
| Ugt1a7c | 12.7903233 | -0.797521882 | 0.311546609 | -2.559879839 |
| Gm15373 | 0.042716945 | 0.037062677 | 3.724945049 | 0.009949859 |
| Ugt1a6b | 1.167440347 | -0.724138525 | 0.924134091 | -0.783585988 |
| Gm15372 | 1.333048035 | -1.760267179 | 1.10239046 | -1.59677287 |
| Gm15370 | 0.204364325 | 0.036222055 | 2.645665836 | 0.013691092 |
| Ugt1a6a | 1.585662245 | -0.545673028 | 0.774655149 | -0.70440767 |
| Gm15369 | 0.593490712 | -1.146411526 | 1.201631977 | -0.954045455 |
| Gm15374 | 1.464328245 | 0.538685109 | 1.12280041 | 0.479769249 |
| Gm15371 | 0.512448221 | 1.07513399 | 1.224286856 | 0.878171635 |
| Ugt1a5 | 1.213945837 | 1.437222073 | 0.971559341 | 1.479294173 |
| Gm15368 | 0.624217637 | -0.317958841 | 1.364958173 | -0.232944018 |
| Gm15376 | 0.474048072 | -0.151926022 | 1.448466687 | -0.104887481 |
| Ugt1a2 | 0.875486306 | -1.287588924 | 1.248537799 | -1.031277487 |
| Dnajb3 | 7.809597006 | -0.719080449 | 0.380073078 | -1.891953126 |
| Ugt1a1 | 0.279519214 | 0.532039066 | 1.798254808 | 0.295864114 |
| Mroh2a | 3.442503238 | 0.013171176 | 0.621347332 | 0.021197768 |
| Hjurp | 549.4911097 | 0.08162956 | 0.084170387 | 0.969813295 |
| Trpm8 | 6.249743756 | 0.865073755 | 0.410877471 | 2.105430002 |
| 4930453O03Rik | 0.121388914 | 0.037071574 | 3.724945049 | 0.009952247 |
| Spp2 | 1.584204494 | -0.443724662 | 0.775621349 | -0.572089284 |
| Gm29538 | 0.330861516 | -0.924715759 | 1.794449783 | -0.515319943 |
| Gm28888 | 0 | NA | NA | NA |
| Glrp1 | 2.778605519 | -1.154085548 | 0.553870243 | -2.083674943 |
| Gm19589 | 0.038981395 | -0.443816764 | 3.724945049 | -0.119147198 |
| Gm38130 | 0.038981395 | -0.443816764 | 3.724945049 | -0.119147198 |
| Gm29336 | 0.085433889 | 0.037058229 | 3.724945049 | 0.009948665 |
| Arl4c | 850.2294767 | 0.033533166 | 0.101071312 | 0.331777288 |
| Gm29335 | 0 | NA | NA | NA |
| Platr5 | 0.048949532 | 0.037064902 | 3.724945049 | 0.009950456 |
| Gm29337 | 0 | NA | NA | NA |
| Gm4753 | 0 | NA | NA | NA |
| 1700067G17Rik | 0 | NA | NA | NA |
| Gm18699 | 0 | NA | NA | NA |
| Sh3bp4 | 304.2424635 | 0.16723936 | 0.140409229 | 1.19108524 |
| Gm38312 | 5.824867419 | 0.250195475 | 0.515076736 | 0.48574408 |
| Gm5259 | 0.045882455 | 0.517959912 | 3.724945049 | 0.139051692 |
| Gm28342 | 0.049553047 | -0.443818988 | 3.724945049 | -0.119147795 |

| | | | | |
|---|---|---|---|---|
| C030007H22Rik | 0.466539735 | -0.093218223 | 3.676981926 | -0.025351831 |
| Agap1 | 3469.23624 | 0.107806184 | 0.065543609 | 1.644800868 |
| Gm37297 | 49.83626593 | 0.291624828 | 0.344007348 | 0.847728484 |
| Gm25180 | 3.378117529 | 0.7887562 | 0.789448058 | 0.999123617 |
| Gm28499 | 9.617358175 | 0.59853589 | 0.485130248 | 1.233763289 |
| 4933400F21Rik | 0.445470297 | 0.930036081 | 1.604446252 | 0.579661725 |
| Gm26037 | 7.202871343 | 1.230992433 | 0.533876294 | 2.305763429 |
| Gm37521 | 23.73742709 | 0.640663895 | 0.359369491 | 1.782744252 |
| Gbx2 | 22.87581777 | -0.46411324 | 0.280935384 | -1.652028424 |
| D130058E05Rik | 7.849306383 | 0.205027569 | 0.462361384 | 0.443435754 |
| Asb18 | 37.68452797 | 0.754183297 | 0.195425974 | 3.859176352 |
| Iqca | 7.094191665 | 0.015809303 | 0.434841191 | 0.036356499 |
| 4930434B07Rik | 0.106465261 | 0.037073798 | 3.724945049 | 0.009952844 |
| Ackr3 | 127.6655458 | -0.301638287 | 0.238614315 | -1.264124857 |
| Gm38277 | 0.04393175 | 0.037062677 | 3.724945049 | 0.009949859 |
| Gm28721 | 0.047948698 | 0.037064902 | 3.724945049 | 0.009950456 |
| Gm28723 | 1.104344843 | 0.54680729 | 1.035605218 | 0.528007469 |
| Gm27648 | 0 | NA | NA | NA |
| Gm28722 | 0.261620403 | -1.724337405 | 2.050257562 | -0.84103453 |
| Gm35048 | 0.834783721 | 0.593667445 | 1.038717914 | 0.571538661 |
| Cops8 | 733.5162779 | -0.073731272 | 0.075620973 | -0.975010892 |
| Gm9991 | 0.061920613 | -0.443816764 | 3.724945049 | -0.119147198 |
| Col6a3 | 94.47715798 | 0.059179624 | 0.156865501 | 0.377263473 |
| Gm36970 | 0 | NA | NA | NA |
| 4930474B08Rik | 0 | NA | NA | NA |
| Mlph | 41.52328994 | 0.75662746 | 0.213754877 | 3.539696821 |
| Prlh | 0.989340548 | -1.658358585 | 1.274168241 | -1.301522462 |
| Rab17 | 0.340708601 | 0.630181762 | 1.744892814 | 0.361157864 |
| Gm28199 | 0 | NA | NA | NA |
| Lrrfip1 | 605.1451411 | -0.237158443 | 0.118861216 | -1.995255054 |
| Rbm44 | 1.28530837 | -2.248543296 | 1.113449765 | -2.019438476 |
| Gm7776 | 0.372084114 | 0.155997964 | 1.939054785 | 0.080450519 |
| Ramp1 | 339.6276143 | 0.219963747 | 0.093378872 | 2.355605099 |
| Ube2f | 531.3694709 | 0.116823392 | 0.122797488 | 0.951350012 |
| Gm7785 | 19.25179799 | 0.096794467 | 0.314314602 | 0.307954088 |
| Gm26683 | 0.0885994 | 0.517955464 | 3.724945049 | 0.139050498 |
| Scly | 285.0654636 | -0.294830466 | 0.092896045 | -3.173767692 |
| Gm17090 | 0.087930927 | -0.443164241 | 3.724898049 | -0.118973522 |
| Espnl | 4.028718013 | -0.648311476 | 0.534639691 | -1.212613818 |
| Klhl30 | 7.180205943 | -1.355158052 | 0.601660898 | -2.252361848 |
| Fam132b | 9.846500243 | -1.365244896 | 0.500596064 | -2.727238577 |
| Ilkap | 381.4142211 | 0.05184279 | 0.10641834 | 0.487160293 |
| 1700020N18Rik | 1.184503545 | 1.575996907 | 0.977641643 | 1.61203946 |
| Hes6 | 86.06443169 | 0.4274973 | 0.139836388 | 3.057124868 |
| Per2 | 585.5330981 | 0.638330634 | 0.197149834 | 3.237794419 |
| Gm28381 | 0 | NA | NA | NA |
| Gm28382 | 0 | NA | NA | NA |

| | | | | |
|---|---|---|---|---|
| Gm19085 | 0.292057199 | 0.006954299 | 2.431316602 | 0.002860302 |
| Traf3ip1 | 223.3191922 | -0.073394429 | 0.109655059 | -0.669320963 |
| Asb1 | 423.9897668 | 0.144813611 | 0.083536754 | 1.73353169 |
| Gm28380 | 0.180091664 | 0.037072677 | 2.750740793 | 0.013477343 |
| Gm29100 | 8.346887551 | 2.054645516 | 0.500726827 | 4.103326218 |
| Twist2 | 5.328893884 | -0.050563869 | 0.448084535 | -0.112844487 |
| Gm29101 | 0.685283607 | 0.46291765 | 1.354019094 | 0.341884137 |
| Gm37600 | 0 | NA | NA | NA |
| Hdac4 | 1369.404079 | 0.089521831 | 0.081082179 | 1.104087636 |
| Gm37048 | 0.281888464 | 1.904064318 | 1.75382793 | 1.085661988 |
| Gm26720 | 0 | NA | NA | NA |
| Gm37036 | 19.75259723 | 0.357577756 | 0.340506794 | 1.050133985 |
| Gm29099 | 0 | NA | NA | NA |
| Gm7837 | 0.143650742 | -0.429650722 | 3.581766334 | -0.119954984 |
| Ndufa10 | 1126.271048 | 0.149746884 | 0.055681579 | 2.689343362 |
| Mir6900 | 0.79482971 | 0.058585774 | 1.280335073 | 0.045758158 |
| Olfr1416 | 0 | NA | NA | NA |
| Olfr1415 | 0 | NA | NA | NA |
| Olfr1414 | 0 | NA | NA | NA |
| 1700054K02Rik | 0 | NA | NA | NA |
| Olfr1413 | 0 | NA | NA | NA |
| Gm28478 | 0 | NA | NA | NA |
| Gm28479 | 0 | NA | NA | NA |
| Olfr1412 | 0 | NA | NA | NA |
| Olfr1411 | 0 | NA | NA | NA |
| Olfr1410 | 0 | NA | NA | NA |
| Olfr12 | 0 | NA | NA | NA |
| Myeov2 | 377.7320121 | 0.05445671 | 0.113883068 | 0.478180921 |
| Otos | 0.964645659 | 0.036012045 | 1.157505228 | 0.031111778 |
| Gm29483 | 1.804989916 | 0.584372218 | 0.767241785 | 0.761653275 |
| Gm29482 | 0 | NA | NA | NA |
| Gm29481 | 0.153668717 | 0.998859713 | 2.839403574 | 0.351785045 |
| Gpc1 | 1142.72074 | -0.014806602 | 0.079912469 | -0.185285257 |
| Gm29480 | 6.109138398 | 0.780261308 | 0.431441192 | 1.808499797 |
| Mir149 | 1.829161676 | 0.909713386 | 0.814201058 | 1.117308037 |
| Ankmy1 | 5.039844727 | -0.058584772 | 0.451608967 | -0.129724555 |
| Dusp28 | 267.5008543 | 0.079741731 | 0.164317274 | 0.485291227 |
| 5033417F24Rik | 38.62813476 | 0.12331943 | 0.188833505 | 0.653059055 |
| Rnpepl1 | 468.7456855 | -0.222973948 | 0.092580322 | -2.408437817 |
| 9430060I03Rik | 3.610602274 | 0.307536579 | 0.626264432 | 0.491065057 |
| Capn10 | 256.0341623 | -0.148073429 | 0.081244978 | -1.822554849 |
| Gpr35 | 139.3197398 | -1.645176205 | 0.336774451 | -4.885098029 |
| Gm18697 | 0.51839282 | -1.754214168 | 1.597612591 | -1.098022247 |
| Aqp12 | 0.175102337 | -1.16604509 | 3.306428121 | -0.352660045 |
| Kif1a | 18866.61453 | 0.261921108 | 0.054125157 | 4.839175007 |
| Gm28086 | 0.753737496 | 1.402483659 | 1.117380493 | 1.255153162 |
| Agxt | 0.11197992 | 0.037073798 | 3.724945049 | 0.009952844 |

| | | | | |
|---|---|---|---|---|
| 2310007B03Rik | 0.800599448 | 0.235639203 | 1.04779481 | 0.2248906 |
| Crocc2 | 17.59480686 | -0.233631068 | 0.284367629 | -0.821581096 |
| Sned1 | 257.5400578 | 0.205626399 | 0.127698164 | 1.610253374 |
| Mir6901 | 0 | NA | NA | NA |
| Mterf4 | 142.5738671 | 0.150568506 | 0.108552951 | 1.387051243 |
| Gm28535 | 1.935100933 | 0.85071784 | 0.719219496 | 1.182834789 |
| Gm7867 | 1.11968618 | 0.028711195 | 0.950340268 | 0.030211489 |
| Pask | 68.98058163 | -0.335029045 | 0.215613795 | -1.553838639 |
| Ppp1r7 | 1320.763185 | 0.286980415 | 0.06252619 | 4.589763309 |
| mmu-mir-6902 | 0.22467814 | 0.027764878 | 2.16939739 | 0.012798429 |
| Mir6345 | 0.94748184 | 0.799398206 | 1.130458114 | 0.707145356 |
| Gm26446 | 2.680156834 | 1.279015416 | 0.757540504 | 1.688378917 |
| Ano7 | 2.740637547 | 0.888747715 | 0.66264413 | 1.341214197 |
| Hdlbp | 3229.635038 | -0.213150179 | 0.048493838 | -4.395407539 |
| Gm17415 | 4.04170504 | -1.459716363 | 0.644563176 | -2.264659878 |
| 2-Sep | 1408.822543 | -0.341563373 | 0.084099029 | -4.061442511 |
| Farp2 | 211.4310669 | 0.100580397 | 0.117719686 | 0.854405925 |
| Gm37250 | 18.96035902 | 0.292490153 | 0.366944047 | 0.797097418 |
| Stk25 | 1506.774651 | 0.282654789 | 0.061425196 | 4.601609853 |
| Gm28536 | 1.8285836 | -0.754358769 | 0.898315666 | -0.839747983 |
| Bok | 205.8186137 | 0.212852874 | 0.094772905 | 2.245925398 |
| Thap4 | 221.3880966 | -0.055213834 | 0.08205429 | -0.672893928 |
| Gm10550 | 0.177745132 | 0.037630679 | 2.300135837 | 0.016360198 |
| Atg4b | 616.7307566 | -0.052187446 | 0.049352452 | -1.057443831 |
| Dtymk | 176.2881088 | -0.242821559 | 0.114120766 | -2.12775963 |
| Ing5 | 203.0410209 | -0.166556265 | 0.082794191 | -2.011690238 |
| Gm22089 | 0 | NA | NA | NA |
| D2hgdh | 49.47669206 | -0.174604714 | 0.224002283 | -0.779477387 |
| Gal3st2 | 0.259214904 | 0.969513794 | 1.812226079 | 0.534985014 |
| Gm18666 | 2.840417843 | 0.427770874 | 0.770207779 | 0.555396721 |
| Gm9994 | 1.809180368 | -0.254303376 | 0.781983574 | -0.325202964 |
| Gm7889 | 3.175371776 | -0.741517831 | 0.627958338 | -1.180839215 |
| Gm6086 | 0.846372671 | -0.290668111 | 1.136542705 | -0.255747637 |
| Neu4 | 89.65460407 | 0.247066434 | 0.200026249 | 1.23517006 |
| Pdcd1 | 42.57420289 | -0.599903079 | 0.585874827 | -1.023944111 |
| Gm7891 | 0.255524584 | 0.005723045 | 2.485195634 | 0.002302855 |
| Gm7895 | 67.17479994 | 0.020433575 | 0.197852311 | 0.103276909 |
| Gm23389 | 0 | NA | NA | NA |
| 4930440C22Rik | 0.220980774 | -1.406009479 | 3.104094945 | -0.452953116 |
| Gm23181 | 0 | NA | NA | NA |
| Gm5264 | 0.086908494 | 0.037058229 | 3.724945049 | 0.009948665 |
| Fam174a | 321.8174294 | 0.067080597 | 0.141124705 | 0.47532852 |
| BC055308 | 22.09994013 | 0.216953685 | 0.272497773 | 0.796166819 |
| Gm15427 | 268.0268351 | -0.182436335 | 0.127252078 | -1.433660951 |
| St8sia4 | 341.5027806 | -0.171749314 | 0.158068882 | -1.086547285 |
| Gm37642 | 12.46481921 | 0.151031899 | 0.288977206 | 0.522642949 |
| 4930598F16Rik | 0 | NA | NA | NA |

| | | | | |
|---|---|---|---|---|
| Gm37761 | 0 | NA | NA | NA |
| Gm22360 | 1.558628739 | -0.841714396 | 0.93638626 | -0.898896569 |
| 1700063A18Rik | 0 | NA | NA | NA |
| Gm24586 | 0 | NA | NA | NA |
| Gm22917 | 0 | NA | NA | NA |
| Gm37076 | 0 | NA | NA | NA |
| Gm29601 | 0.044544648 | 0.517957688 | 3.724945049 | 0.139051095 |
| 4930533P14Rik | 0 | NA | NA | NA |
| Gm26245 | 0 | NA | NA | NA |
| Gm1833 | 0.161949631 | -1.309098506 | 2.76260123 | -0.473864448 |
| Gm26078 | 0 | NA | NA | NA |
| Slco4c1 | 49.5126318 | 0.079621305 | 0.198187168 | 0.401748031 |
| Panct2 | 37.25735819 | 0.024002042 | 0.233890887 | 0.102620682 |
| Slco6b1 | 0.161026707 | -1.437459239 | 3.451444183 | -0.416480512 |
| Gm9499 | 0 | NA | NA | NA |
| Gm28901 | 0 | NA | NA | NA |
| Gm6430 | 14.31094521 | -0.411084076 | 0.272792573 | -1.50694746 |
| Slco6c1 | 4.084609871 | 0.699940454 | 0.523503884 | 1.337030107 |
| Gm7133 | 0.216806889 | 0.738466102 | 2.595246904 | 0.284545606 |
| Gm7135 | 0.141094079 | 0.037072457 | 3.563850528 | 0.01040236 |
| Gm37171 | 2.848103185 | 0.927563672 | 0.593222493 | 1.563601657 |
| D1Ertd622e | 880.9560972 | 0.078700482 | 0.062737884 | 1.254433149 |
| Ppip5k2 | 911.1116556 | 0.485010419 | 0.096213153 | 5.040999126 |
| Gin1 | 94.83341611 | -0.043227687 | 0.125322904 | -0.344930457 |
| Pam | 2285.737651 | 0.309365111 | 0.062641915 | 4.938627962 |
| Gm37140 | 76.97524024 | 0.77792109 | 0.242361479 | 3.209755503 |
| Gm25413 | 0 | NA | NA | NA |
| Gm3531 | 31.35057007 | -0.093915771 | 0.197662859 | -0.475131096 |
| B230216N24Rik | 130.1992663 | 0.835786565 | 0.234959099 | 3.557157687 |
| Gm24465 | 0.049553047 | -0.443818988 | 3.724945049 | -0.119147795 |
| 1810006J02Rik | 0.496068496 | -2.555794991 | 1.303370371 | -1.960912299 |
| Gm29461 | 0.196785726 | 1.413728005 | 2.209521759 | 0.639834389 |
| Gm7952 | 0 | NA | NA | NA |
| Gm29462 | 0.477857232 | -1.281656415 | 1.501671855 | -0.85348634 |
| Slco6d1 | 0.368527898 | 0.41572942 | 1.569854707 | 0.264820316 |
| Gm3555 | 28.33472675 | -0.513587183 | 0.23403778 | -2.19446272 |
| Gm26134 | 0 | NA | NA | NA |
| Gm7967 | 10.59292213 | 0.569447119 | 0.349653492 | 1.628604123 |
| Gm29460 | 1.272524668 | 0.623949491 | 0.94328988 | 0.661461025 |
| Gm23434 | 0.075043205 | 0.517957688 | 3.724945049 | 0.139051095 |
| Gm18248 | 0.075043205 | 0.517957688 | 3.724945049 | 0.139051095 |
| Cntnap5b | 377.3845069 | 0.509825567 | 0.095859495 | 5.318467051 |
| Tdpx-ps1 | 13.77002096 | 0.110538177 | 0.271001747 | 0.407887321 |
| Gm37877 | 23.5485973 | 0.253762507 | 0.296916583 | 0.85465926 |
| Gm19029 | 3.564592169 | 1.050989171 | 0.692839769 | 1.516929625 |
| Gm23602 | 0 | NA | NA | NA |
| Gm29668 | 2.625920046 | 0.534597014 | 0.657796794 | 0.812708451 |

| | | | | |
|---|---|---|---|---|
| Gm29667 | 14.43481637 | 0.850314288 | 0.337605932 | 2.518659208 |
| Gm18700 | 0 | NA | NA | NA |
| Gm18185 | 0 | NA | NA | NA |
| Gm29334 | 0 | NA | NA | NA |
| Gm25537 | 0 | NA | NA | NA |
| Gm20268 | 0 | NA | NA | NA |
| Gm28801 | 0 | NA | NA | NA |
| Psmb6-ps2 | 0.576768135 | -0.201645668 | 1.311120529 | -0.153796439 |
| Gm18969 | 0 | NA | NA | NA |
| Gm28188 | 0.072025462 | 0.517959912 | 3.724945049 | 0.139051692 |
| Gm28187 | 1.345496586 | 0.015873527 | 0.884042107 | 0.017955623 |
| Gm24122 | 0 | NA | NA | NA |
| Gm28623 | 0 | NA | NA | NA |
| Gm23036 | 0 | NA | NA | NA |
| Gm3116 | 0 | NA | NA | NA |
| Gm20281 | 0 | NA | NA | NA |
| Gm22034 | 0 | NA | NA | NA |
| Gm18728 | 0 | NA | NA | NA |
| Gm23965 | 0 | NA | NA | NA |
| Gm29141 | 1.908747754 | -0.432557713 | 0.694008652 | -0.623274236 |
| Gm37789 | 0.149576918 | -0.443202057 | 2.863264737 | -0.15478906 |
| Gm18447 | 0 | NA | NA | NA |
| Gm29140 | 0.04393175 | 0.037062677 | 3.724945049 | 0.009949859 |
| Gm23537 | 0 | NA | NA | NA |
| Gm5260 | 0 | NA | NA | NA |
| Gm29592 | 0 | NA | NA | NA |
| Gm29011 | 0.220980774 | -1.406009479 | 3.104094945 | -0.452953116 |
| Cdh20 | 327.0013875 | -0.169915293 | 0.108313132 | -1.568741385 |
| Gm38031 | 85.93314083 | -0.440703301 | 0.199277307 | -2.211507707 |
| n-R5s216 | 0 | NA | NA | NA |
| Gm36969 | 21.4981108 | 0.357111603 | 0.356510687 | 1.001685547 |
| Gm15699 | 0 | NA | NA | NA |
| Gm29012 | 0.089089296 | 0.957155977 | 3.723939561 | 0.257027796 |
| Gm29013 | 0.046345709 | -0.443818988 | 3.724945049 | -0.119147795 |
| Rnf152 | 731.5992459 | 0.065579832 | 0.090245119 | 0.726685642 |
| Gm29014 | 0 | NA | NA | NA |
| Gm17634 | 9.160375891 | 0.425390515 | 0.362366583 | 1.173923133 |
| Gm28403 | 1.125166341 | 1.184161547 | 1.00462383 | 1.178711386 |
| Gm3608 | 115.3994121 | -0.842928226 | 0.228246373 | -3.693062957 |
| Pign | 477.5995311 | -0.19847745 | 0.113294513 | -1.751871682 |
| Gm28404 | 2.874152792 | -0.323382882 | 0.660878485 | -0.489322757 |
| Gm20302 | 1.560084669 | -1.827017269 | 0.924839806 | -1.975495926 |
| Gm22426 | 0 | NA | NA | NA |
| Gm8004 | 0 | NA | NA | NA |
| 2310035C23Rik | 1589.964662 | 0.208964429 | 0.079510947 | 2.628121512 |
| Gm37566 | 68.8298285 | 0.460949568 | 0.190441472 | 2.420426405 |
| Gm10193 | 1.224645236 | 0.223590397 | 0.978576221 | 0.228485418 |

| | | | | |
|---|---|---|---|---|
| Tnfrsf11a | 174.3773544 | -0.482375549 | 0.154314076 | -3.125933551 |
| Gm37779 | 0.176451472 | 0.539299713 | 2.721126888 | 0.198189844 |
| Gm26214 | 0 | NA | NA | NA |
| Gm18801 | 2.226781727 | -2.439219287 | 0.786496337 | -3.101373996 |
| Gm7160 | 28.83074711 | -0.917833673 | 0.25987069 | -3.531886087 |
| Zcchc2 | 822.9552979 | -0.459867944 | 0.10974387 | -4.190374787 |
| Gm37372 | 0.112396584 | -0.917692504 | 3.724434845 | -0.246397787 |
| Gm26509 | 3.937004826 | 0.091166764 | 0.538250599 | 0.169376057 |
| Gm20753 | 4.418263327 | 1.199586614 | 0.557849326 | 2.150377453 |
| Phlpp1 | 1704.681168 | 0.040494744 | 0.07648368 | 0.529456009 |
| Gm38235 | 158.3680684 | 0.231872633 | 0.246883759 | 0.939197596 |
| A530053M12Rik | 0.224329914 | -0.965962928 | 2.570323892 | -0.375813698 |
| Gm6651 | 0.568438163 | -1.16206601 | 1.270502652 | -0.914650598 |
| Bcl2 | 493.9864849 | -0.300201488 | 0.114409719 | -2.623915953 |
| D630008O14Rik | 0.626527047 | 0.910732719 | 1.552128401 | 0.586763774 |
| Mir3473f | 0.393472992 | -0.616825852 | 1.784312239 | -0.345693898 |
| Gm37053 | 17.84686388 | -0.134176094 | 0.387103685 | -0.346615387 |
| Kdsr | 626.2639973 | -0.032701862 | 0.062108531 | -0.526527693 |
| Vps4b | 768.3375885 | 0.074697258 | 0.076653703 | 0.974476832 |
| Serpinb5 | 4.060285521 | -2.213112224 | 0.854933166 | -2.5886377 |
| Pou2f3-rs1 | 0 | NA | NA | NA |
| Serpinb12 | 0.340899075 | 0.133745241 | 1.965980986 | 0.068029773 |
| Serpinb13 | 0 | NA | NA | NA |
| Gm8080 | 0.048949532 | 0.037064902 | 3.724945049 | 0.009950456 |
| Serpinb3a | 0.112642008 | 0.517955464 | 3.724945049 | 0.139050498 |
| Serpinb3d | 0.044544648 | 0.517957688 | 3.724945049 | 0.139051095 |
| Gm15388 | 0 | NA | NA | NA |
| Serpinb3b | 0.36088792 | 2.328389491 | 3.684451251 | 0.631950142 |
| Gm8084 | 0.045882455 | 0.517959912 | 3.724945049 | 0.139051692 |
| Gm15390 | 0 | NA | NA | NA |
| Serpinb3c | 0.256447508 | 0.010978678 | 2.469852793 | 0.004445074 |
| Gm8089 | 0.10503227 | -0.905414734 | 3.723500899 | -0.243162217 |
| Serpinb11 | 0.313959609 | -0.893873489 | 1.590601442 | -0.56197201 |
| Gm15391 | 0 | NA | NA | NA |
| Serpinb7 | 0.394543937 | -1.407876223 | 1.661166386 | -0.847522701 |
| Gm24553 | 0 | NA | NA | NA |
| Serpinb2 | 8.118383424 | -2.395985887 | 0.611925802 | -3.915484327 |
| Serpinb10 | 0.745572437 | -1.694732999 | 1.202632749 | -1.409185806 |
| Serpinb8 | 106.9109276 | -0.516123235 | 0.178492597 | -2.891566624 |
| Gm15389 | 0 | NA | NA | NA |
| Gm24267 | 0 | NA | NA | NA |
| Gm22438 | 0.044544648 | 0.517957688 | 3.724945049 | 0.139051095 |
| D830032E09Rik | 0 | NA | NA | NA |
| Gm18448 | 0.11147366 | -0.955981391 | 3.722845977 | -0.25678779 |
| Gm28282 | 0 | NA | NA | NA |
| Gm5702 | 0 | NA | NA | NA |
| Gm38212 | 0.044191549 | 0.037062677 | 3.724945049 | 0.009949859 |

| | | | | |
|---|---|---|---|---|
| Gm28283 | 0 | NA | NA | NA |
| Gm6679 | 0 | NA | NA | NA |
| Gm17847 | 0 | NA | NA | NA |
| Gm28281 | 0 | NA | NA | NA |
| Gm22159 | 0.068097359 | 0.037064902 | 3.724945049 | 0.009950456 |
| Gm25293 | 0 | NA | NA | NA |
| Gm8141 | 0.177100366 | 0.037058322 | 3.38847586 | 0.010936576 |
| Cdh7 | 599.3121521 | 0.26882058 | 0.120612781 | 2.22879016 |
| Gm18552 | 0 | NA | NA | NA |
| Gm3332 | 0 | NA | NA | NA |
| Gm6693 | 0 | NA | NA | NA |
| Cdh19 | 348.6925766 | -0.044968685 | 0.131323443 | -0.342426938 |
| Gm38255 | 57.38585182 | 0.368307226 | 0.211673346 | 1.739979237 |
| Gm36944 | 56.47739328 | 0.502997828 | 0.249141097 | 2.018927567 |
| Gm29088 | 1.097863377 | -0.443720316 | 0.982793781 | -0.45148873 |
| Gm19927 | 0.068097359 | 0.037064902 | 3.724945049 | 0.009950456 |
| Gm8173 | 15.82173517 | -0.417256905 | 0.392453917 | -1.063199747 |
| Gm37934 | 0 | NA | NA | NA |
| Dsel | 373.0186971 | 0.047960184 | 0.082084493 | 0.584278259 |
| Gm6028 | 0.150520013 | 0.037075864 | 2.857840531 | 0.012973384 |
| Gm17752 | 0 | NA | NA | NA |
| Gm6717 | 0 | NA | NA | NA |
| Gm19086 | 0 | NA | NA | NA |
| Gm18406 | 9.163048501 | -0.20290179 | 0.355363211 | -0.570970163 |
| Gm22331 | 0 | NA | NA | NA |
| Gm8204 | 0 | NA | NA | NA |
| Gm28190 | 0 | NA | NA | NA |
| Gm28189 | 0 | NA | NA | NA |
| Gm17946 | 0 | NA | NA | NA |
| Gm18181 | 0 | NA | NA | NA |
| Gm24937 | 0 | NA | NA | NA |
| 9330185C12Rik | 1.904432073 | 0.031624398 | 0.706543394 | 0.044759314 |
| Gm4854 | 0 | NA | NA | NA |
| Gm17916 | 16.58061403 | 0.641740497 | 0.286404807 | 2.240676421 |
| Cntnap5a | 1043.802266 | 0.83524713 | 0.085060977 | 9.819392611 |
| Gm23393 | 0 | NA | NA | NA |
| Gm28263 | 0.417031551 | 1.847792271 | 1.518634845 | 1.216745603 |
| Gm19965 | 1.062057355 | 2.04962331 | 0.988618946 | 2.073218724 |
| Gm17777 | 0 | NA | NA | NA |
| Gm24030 | 0 | NA | NA | NA |
| Gm27785 | 0 | NA | NA | NA |
| Gm28361 | 0 | NA | NA | NA |
| Zfp813-ps | 0 | NA | NA | NA |
| Gm28362 | 0 | NA | NA | NA |
| Gm28363 | 0 | NA | NA | NA |
| Gm28359 | 0 | NA | NA | NA |
| Gm18182 | 0 | NA | NA | NA |

| | | | | |
|---|---|---|---|---|
| B020011L13Rik | 0 | NA | NA | NA |
| Gm28360 | 0 | NA | NA | NA |
| Gm28169 | 0.218500166 | 0.064659972 | 2.167165066 | 0.029836201 |
| Gm7195 | 0 | NA | NA | NA |
| Gm28168 | 0 | NA | NA | NA |
| Gm7145 | 0 | NA | NA | NA |
| Gm28358 | 0 | NA | NA | NA |
| Gm29489 | 0 | NA | NA | NA |
| Gm29490 | 0.044191549 | 0.037062677 | 3.724945049 | 0.009949859 |
| Gm29104 | 0 | NA | NA | NA |
| Gm29106 | 0.046345709 | -0.443818988 | 3.724945049 | -0.119147795 |
| Gm7200 | 0 | NA | NA | NA |
| Tsn | 868.3708414 | 0.025458175 | 0.069686122 | 0.365326328 |
| Nifk | 240.6864645 | -0.164149993 | 0.119228484 | -1.376768266 |
| Gm28867 | 0.180050447 | -0.925657836 | 3.327036994 | -0.278222886 |
| Clasp1 | 2304.76087 | 0.131847118 | 0.064102329 | 2.056822573 |
| Gm26080 | 0 | NA | NA | NA |
| Gm22710 | 2.569247428 | 0.725832297 | 1.032640027 | 0.70288995 |
| Gm28466 | 0 | NA | NA | NA |
| Tfcp2l1 | 115.5571692 | 0.243781825 | 0.152008323 | 1.603739977 |
| Gm22094 | 0 | NA | NA | NA |
| Gm28882 | 0 | NA | NA | NA |
| Gm23497 | 0 | NA | NA | NA |
| Gm28467 | 0 | NA | NA | NA |
| Gli2 | 130.8498884 | -0.046247911 | 0.17169164 | -0.269366117 |
| Mir6346 | 0 | NA | NA | NA |
| Gm26831 | 0.564941401 | 0.528521699 | 1.36074838 | 0.388405165 |
| Gm8321 | 0 | NA | NA | NA |
| Gm3508 | 0 | NA | NA | NA |
| Gm29456 | 0 | NA | NA | NA |
| Gm29455 | 2.475683411 | 0.261524131 | 0.744213555 | 0.351410061 |
| Inhbb | 109.178341 | -0.026240713 | 0.190589523 | -0.137681824 |
| Ralb | 386.9036816 | -0.490181198 | 0.095533802 | -5.130971332 |
| Gm29454 | 0 | NA | NA | NA |
| Gm27184 | 4.248472554 | 0.626068325 | 0.542396053 | 1.15426416 |
| Tmem185b | 22.02441361 | -0.38632697 | 0.253417214 | -1.524470113 |
| AC132393.1 | 0 | NA | NA | NA |
| 3830432H09Rik | 2.926677728 | -0.309548892 | 0.589607425 | -0.52500847 |
| Gm25322 | 0.15596086 | 0.037059316 | 3.490591407 | 0.010616916 |
| Epb4.1l5 | 218.5589968 | -0.072369909 | 0.091800456 | -0.788339321 |
| Gm37219 | 2.565042467 | 1.077415683 | 0.69936794 | 1.540556296 |
| Ptpn4 | 3441.616975 | 0.195060903 | 0.089631841 | 2.17624564 |
| Gm3851 | 0.583829782 | -0.013816454 | 1.328081339 | -0.010403319 |
| Gm37745 | 125.9235229 | 0.522349216 | 0.244168021 | 2.13930233 |
| Gm29657 | 0 | NA | NA | NA |
| Gm8338 | 0 | NA | NA | NA |
| Gm29658 | 0.060174537 | 0.517957688 | 3.724945049 | 0.139051095 |

| | | | | |
|---|---|---|---|---|
| Gm3551 | 0.190043611 | 0.037075574 | 2.655002333 | 0.013964423 |
| Gm28209 | 0 | NA | NA | NA |
| Tmem177 | 83.02492162 | -0.393678237 | 0.136338674 | -2.887502315 |
| Cfap221 | 5.416510947 | 0.235628161 | 0.475516185 | 0.495520802 |
| Gm17973 | 0 | NA | NA | NA |
| Sctr | 5.377310654 | 0.850002037 | 0.464863664 | 1.828497478 |
| Tmem37 | 24.45334209 | -0.888353721 | 0.297786404 | -2.983191002 |
| Dbi | 1017.882369 | 0.018629018 | 0.144243403 | 0.129149881 |
| 3110009E18Rik | 15.78515723 | -0.038643995 | 0.28854965 | -0.133924943 |
| Steap3 | 67.01145874 | -1.036044036 | 0.252252659 | -4.107167956 |
| C1ql2 | 92.61537039 | -0.222365723 | 0.16760143 | -1.326753133 |
| 1700012E03Rik | 0.060174537 | 0.517957688 | 3.724945049 | 0.139051095 |
| Marco | 0.34718572 | -1.014238494 | 1.739723668 | -0.582988272 |
| En1 | 26.69343009 | -0.428618998 | 0.222303148 | -1.928083347 |
| RP24-358J4.2 | 0.343046485 | 1.35457638 | 1.527802436 | 0.886617502 |
| 2610027F03Rik | 0 | NA | NA | NA |
| Gm29346 | 0.099155932 | 0.037073798 | 3.724945049 | 0.009952844 |
| Gm29345 | 0.060174537 | 0.517957688 | 3.724945049 | 0.139051095 |
| Gm29344 | 0 | NA | NA | NA |
| Gm29348 | 0.333474911 | -1.813384597 | 1.724949145 | -1.05126844 |
| Gm29347 | 0.045882455 | 0.517959912 | 3.724945049 | 0.139051692 |
| Gm37043 | 0 | NA | NA | NA |
| Celrr | 13.48815634 | 0.961589051 | 0.471445243 | 2.03966222 |
| Gm29359 | 0 | NA | NA | NA |
| Gm38283 | 4.643660642 | 0.098434673 | 0.563189275 | 0.174780802 |
| Insig2 | 471.7823211 | -0.233018078 | 0.084404063 | -2.760744803 |
| Gm37174 | 21.08112134 | 0.426918149 | 0.262981809 | 1.623375208 |
| 2210011K15Rik | 0.099106094 | -0.93550548 | 3.724188147 | -0.251197158 |
| Ccdc93 | 456.8414215 | -0.221339777 | 0.10529889 | -2.102014339 |
| Gm15392 | 0 | NA | NA | NA |
| Gm15393 | 0.061920613 | -0.443816764 | 3.724945049 | -0.119147198 |
| Htr5b | 20.85714652 | 0.515832734 | 0.323891019 | 1.592612031 |
| Gm8217 | 0 | NA | NA | NA |
| Ddx18 | 253.9984606 | -0.395491323 | 0.080621248 | -4.905547052 |
| Gm28591 | 0 | NA | NA | NA |
| Gm37920 | 0 | NA | NA | NA |
| Gm15394 | 0 | NA | NA | NA |
| Gm8392 | 0 | NA | NA | NA |
| Gm26472 | 0 | NA | NA | NA |
| Gm28590 | 0 | NA | NA | NA |
| Gm24547 | 0 | NA | NA | NA |
| Gm36982 | 0 | NA | NA | NA |
| Gm25771 | 0 | NA | NA | NA |
| Gm29192 | 0 | NA | NA | NA |
| Gm37970 | 0.148650935 | 0.966592573 | 2.859657381 | 0.338009924 |
| Gm8403 | 0.085327104 | -0.905414734 | 3.723500899 | -0.243162217 |
| Gm28911 | 0.104106287 | 0.51795324 | 3.724945049 | 0.139049901 |

| | | | | |
|---|---|---|---|---|
| Gm20058 | 0.094097695 | 0.037071574 | 3.724945049 | 0.009952247 |
| Gm28912 | 0 | NA | NA | NA |
| Dpp10 | 1832.665455 | 0.634491757 | 0.08754699 | 7.247442252 |
| Gm37478 | 62.41200189 | 0.946263328 | 0.27880432 | 3.394005259 |
| Gm37221 | 43.7843784 | 0.786509621 | 0.286819344 | 2.742177745 |
| Gm24791 | 5.806203063 | 0.537036037 | 0.600873329 | 0.893759151 |
| Gm37066 | 38.05353759 | 0.55125171 | 0.30621735 | 1.800197505 |
| Gm37781 | 35.98403262 | 0.68690268 | 0.331298978 | 2.073361905 |
| Gm37396 | 18.40949471 | 0.233922926 | 0.480941786 | 0.486385114 |
| Gm38263 | 58.17025342 | 0.742027958 | 0.310898287 | 2.386722565 |
| Gm28929 | 0 | NA | NA | NA |
| Gm18875 | 20.78588443 | 0.615292604 | 0.406173369 | 1.514852158 |
| Gm37551 | 88.74313094 | 0.444121172 | 0.301143724 | 1.474781431 |
| Gm38073 | 157.9840511 | 1.251942474 | 0.446385996 | 2.804618618 |
| Gm25578 | 0 | NA | NA | NA |
| Gm28928 | 91.8867847 | 1.047875273 | 0.422322783 | 2.481218904 |
| Gm28299 | 17.41287975 | 0.677918743 | 0.401867533 | 1.686920906 |
| Gm37717 | 149.0700901 | 1.002717674 | 0.461915063 | 2.170783667 |
| 1700121L03Rik | 0.097139547 | -0.443816764 | 3.724945049 | -0.119147198 |
| Actr3 | 2282.827765 | -0.254440016 | 0.086987226 | -2.925027368 |
| Gm28706 | 0 | NA | NA | NA |
| Slc35f5 | 263.8309605 | -0.366101 | 0.09387406 | -3.899916548 |
| Gm18444 | 0 | NA | NA | NA |
| Gpr39 | 2.657777857 | 0.584962182 | 0.633629186 | 0.923193241 |
| Lypd1 | 535.5413066 | 1.012803073 | 0.087129469 | 11.62411625 |
| Nckap5 | 192.1177816 | 0.127173067 | 0.147920923 | 0.859736835 |
| Gm37161 | 5.830223875 | 0.384031564 | 0.4805856 | 0.799090867 |
| Gm37936 | 6.414345313 | 0.735758067 | 0.493497719 | 1.490904696 |
| Gm38177 | 4.318687175 | 0.029601376 | 0.555581862 | 0.053279953 |
| 4930599A14Rik | 0 | NA | NA | NA |
| Gm5261 | 0 | NA | NA | NA |
| Mgat5 | 2217.417936 | 0.145531119 | 0.061321395 | 2.373251921 |
| Gm37772 | 11.00123342 | 0.523601665 | 0.40695039 | 1.286647408 |
| Gm37510 | 48.12361392 | 0.11049508 | 0.328642798 | 0.336216344 |
| Gm8451 | 18.99290832 | -0.466942718 | 0.321015749 | -1.454578847 |
| SCARNA4 | 0.095435502 | 0.037073798 | 3.724945049 | 0.009952844 |
| SCARNA15 | 0 | NA | NA | NA |
| Gm23734 | 4.197363489 | 1.293528957 | 0.585928086 | 2.207658224 |
| Gm38301 | 6.512391526 | 0.375175533 | 0.495113856 | 0.757756077 |
| Tmem163 | 261.7619097 | 0.178325496 | 0.080590577 | 2.212733819 |
| Acmsd | 0.917997538 | -0.134245769 | 1.038824001 | -0.129228597 |
| 2900009J06Rik | 4.736596173 | -0.357316632 | 0.510482733 | -0.69995831 |
| Ccnt2 | 1690.484171 | 0.272014662 | 0.13248641 | 2.053151432 |
| Map3k19 | 75.93146868 | -0.183771337 | 0.193982473 | -0.947360521 |
| Rab3gap1 | 1284.44217 | -0.012324848 | 0.047557742 | -0.259155457 |
| Zranb3 | 101.8848498 | 0.170014561 | 0.138957588 | 1.22349966 |
| Gm28800 | 35.95127173 | 0.767949054 | 0.204174624 | 3.761236523 |

| | | | | |
|---|---|---|---|---|
| Gm38248 | 15.05177351 | 0.274154818 | 0.415530293 | 0.659770955 |
| Rpl28-ps1 | 66.91960972 | -0.505189031 | 0.163935487 | -3.08163315 |
| Gm37625 | 7.67879382 | 0.30868946 | 0.412026146 | 0.749198717 |
| Gm38026 | 3.696876593 | -1.315784794 | 0.706489644 | -1.862426158 |
| R3hdm1 | 4415.135964 | 0.318123429 | 0.148345598 | 2.14447501 |
| Gm29383 | 0 | NA | NA | NA |
| Gm37588 | 55.83216539 | 0.372873923 | 0.238942861 | 1.560515018 |
| Gm37868 | 81.33316158 | 0.762349348 | 0.282977634 | 2.694026867 |
| Gm37407 | 115.1470443 | 0.73073729 | 0.205096636 | 3.562892608 |
| Mir128-1 | 17.81336244 | 1.262563389 | 0.323370658 | 3.904384518 |
| Ubxn4 | 978.701728 | -0.341988596 | 0.09326938 | -3.666675968 |
| Lct | 31.88336858 | 0.660633583 | 0.327508462 | 2.01714966 |
| Mcm6 | 72.35290303 | -1.03685488 | 0.243859963 | -4.251845468 |
| Dars | 621.5308942 | -0.337501858 | 0.1001296 | -3.370650212 |
| Gm23902 | 0 | NA | NA | NA |
| Gm29598 | 0 | NA | NA | NA |
| Gm29597 | 0 | NA | NA | NA |
| Gm25384 | 0 | NA | NA | NA |
| Gm6170 | 19.62597736 | -0.315882446 | 0.273840315 | -1.153527908 |
| Cxcr4 | 127.9107672 | -1.272964915 | 0.264683746 | -4.809380766 |
| Gm29599 | 0.045882455 | 0.517959912 | 3.724945049 | 0.139051692 |
| Gm26686 | 0.389802664 | 0.006900418 | 1.417770612 | 0.00486709 |
| Gm23056 | 0 | NA | NA | NA |
| Thsd7b | 229.954111 | 0.502432553 | 0.125283799 | 4.010355359 |
| Gm38240 | 12.58215362 | 0.078622007 | 0.340239402 | 0.231078489 |
| Gm38330 | 4.986765232 | 0.099361941 | 0.536648227 | 0.185152836 |
| Gm36951 | 10.68103066 | 0.5737416 | 0.361247318 | 1.588223831 |
| Gm15790 | 8.257872952 | -0.668682866 | 0.388515702 | -1.721121853 |
| Rpl31-ps9 | 4.393169526 | 0.187585502 | 0.503687404 | 0.372424445 |
| Gm37479 | 19.45303257 | 0.851356345 | 0.270985205 | 3.141707851 |
| Gm37064 | 10.99610456 | 0.850707473 | 0.322412519 | 2.638568367 |
| Gm37278 | 4.9996304 | 0.945396793 | 0.472919246 | 1.999066016 |
| Gm29428 | 1.653651842 | 1.988783193 | 0.993905072 | 2.000979015 |
| Gm16081 | 0 | NA | NA | NA |
| Daf2 | 0.977993353 | -0.163587668 | 1.019143398 | -0.160514868 |
| Gm15674 | 0.068097359 | 0.037064902 | 3.724945049 | 0.009950456 |
| Gm15675 | 15.76810382 | 0.782175365 | 0.364565621 | 2.14549952 |
| Cd55 | 67.64943717 | 0.222182754 | 0.146701936 | 1.514518215 |
| Gm16083 | 14.70625333 | 0.523541765 | 0.300396092 | 1.742838137 |
| Gm37109 | 7.99913161 | 0.730366679 | 0.516876501 | 1.413039048 |
| Gm37283 | 5.739234123 | 1.050071064 | 0.542792358 | 1.934572305 |
| Zp3r | 2.754922463 | 0.57327005 | 0.654299542 | 0.876158415 |
| Gm23623 | 0.044544648 | 0.517957688 | 3.724945049 | 0.139051095 |
| C4bp | 2.027159789 | -1.307127982 | 0.67425011 | -1.938639628 |
| Gm29427 | 1.964410769 | -0.358658592 | 0.803345403 | -0.44645627 |
| C4bp-ps1 | 4.021019734 | 0.269450984 | 0.508129438 | 0.530280208 |
| Pfkfb2 | 323.8491531 | 0.31748437 | 0.121423148 | 2.614693957 |

| | | | | |
|---|---|---|---|---|
| Yod1 | 565.3277476 | 0.121081668 | 0.110334137 | 1.097408937 |
| Gm37163 | 0.049553047 | -0.443818988 | 3.724945049 | -0.119147795 |
| AA986860 | 42.90179849 | -0.143205324 | 0.18880492 | -0.758483012 |
| Gm28857 | 0.066050875 | -0.443818988 | 3.724945049 | -0.119147795 |
| Gm28856 | 0.17254495 | 0.037075168 | 2.737704633 | 0.013542428 |
| Fcamr | 0.566848388 | -3.112001692 | 1.401952313 | -2.219762872 |
| Gm15848 | 0.042716945 | 0.037062677 | 3.724945049 | 0.009949859 |
| Pigr | 0.796384302 | -0.746540833 | 1.052198964 | -0.709505387 |
| Fcmr | 1.860507999 | 0.263011896 | 0.693639398 | 0.379176697 |
| Il24 | 0.119234754 | 0.51795324 | 3.724945049 | 0.139049901 |
| Il20 | 0.112396584 | -0.917692504 | 3.724434845 | -0.246397787 |
| Il19 | 1.495078259 | 0.785128147 | 0.970406841 | 0.809071117 |
| Il10 | 32.26081669 | -1.047064395 | 0.619524776 | -1.690108992 |
| Mapkapk2 | 535.196025 | -0.429642472 | 0.137301737 | -3.129184526 |
| Gm25549 | 2.426982663 | 0.558456535 | 0.643114839 | 0.868362074 |
| Gm37084 | 67.84258375 | -0.158382527 | 0.276060468 | -0.573724039 |
| Gm25717 | 0.038981395 | -0.443816764 | 3.724945049 | -0.119147198 |
| Dyrk3 | 126.7275624 | 0.03436215 | 0.179261816 | 0.191686945 |
| Eif2d | 340.1033352 | -0.051572754 | 0.131484338 | -0.392234957 |
| Rassf5 | 359.522077 | -0.293443266 | 0.087858634 | -3.339947981 |
| Gm17796 | 0.165470308 | 1.408943363 | 2.753318612 | 0.511725507 |
| Ikbke | 98.54742827 | -1.172027546 | 0.219005019 | -5.351601314 |
| Gm28914 | 1.174162655 | 0.642400303 | 1.100416984 | 0.583778979 |
| Gm28913 | 1.688430464 | 0.21634661 | 0.792469892 | 0.273002939 |
| Srgap2 | 1739.750525 | 0.149011959 | 0.080195421 | 1.858110578 |
| Gm38262 | 21.98513846 | 0.218419447 | 0.318837877 | 0.68504862 |
| Gm37954 | 37.75257877 | 0.509471101 | 0.302460322 | 1.684422928 |
| Gm29488 | 96.98125143 | 0.151208543 | 0.234839288 | 0.643880948 |
| Gm29487 | 5.722531327 | -0.134874278 | 0.602506167 | -0.223855431 |
| Gm24273 | 9.213624824 | -0.044105409 | 0.417405454 | -0.105665628 |
| Fam72a | 24.35166078 | -0.350917884 | 0.226144538 | -1.551741588 |
| Avpr1b | 0.995114343 | -0.147574433 | 1.276530922 | -0.115605843 |
| Gm8532 | 1.429318806 | -0.930307491 | 0.814293348 | -1.142472174 |
| Ctse | 45.61320862 | -0.699049311 | 0.264040404 | -2.647508873 |
| Rab7b | 58.99444633 | -1.372743285 | 0.305996078 | -4.486146665 |
| Slc26a9 | 1.868919118 | 0.516257153 | 0.664244111 | 0.77720998 |
| Pm20d1 | 45.2287229 | 0.217661721 | 0.266915391 | 0.815470849 |
| Slc41a1 | 1235.334159 | 0.006281699 | 0.1103374 | 0.056931731 |
| Rab29 | 79.93970694 | -0.604894438 | 0.205996195 | -2.936435007 |
| Gm29105 | 0 | NA | NA | NA |
| Nucks1 | 3423.866732 | 0.268526292 | 0.076522739 | 3.509104565 |
| Gm38067 | 40.11111897 | 0.554726995 | 0.279464329 | 1.984965295 |
| Gm37676 | 0.045882455 | 0.517959912 | 3.724945049 | 0.139051692 |
| Gm29103 | 0 | NA | NA | NA |
| Slc45a3 | 23.9080149 | -0.367215702 | 0.252457685 | -1.454563374 |
| Elk4 | 760.1203314 | -0.152289261 | 0.071549651 | -2.128441683 |
| Mfsd4 | 533.3513531 | 0.722036725 | 0.096069361 | 7.515785647 |

| | | | | |
|---|---|---|---|---|
| Gm26892 | 3.660297346 | 0.04514652 | 0.619695275 | 0.072852775 |
| Gm26034 | 0.106465261 | 0.037073798 | 3.724945049 | 0.009952844 |
| Gm37120 | 6.573103098 | 0.674857862 | 0.476823227 | 1.415320865 |
| Cdk18 | 350.945801 | -0.238029825 | 0.128772509 | -1.848452173 |
| Gm29629 | 1.14102893 | 0.010253295 | 0.898292904 | 0.011414201 |
| Lemd1 | 70.07703057 | -0.567643076 | 0.231625975 | -2.450688335 |
| Mir135b | 0.432115038 | -1.259792593 | 1.647699051 | -0.764576876 |
| 1700037F24Rik | 0 | NA | NA | NA |
| Gm10188 | 0.044544648 | 0.517957688 | 3.724945049 | 0.139051095 |
| Gm29630 | 0 | NA | NA | NA |
| Gm7241 | 0.177688696 | 1.324991508 | 2.718670365 | 0.487367474 |
| Klhdc8a | 280.7923049 | -0.405764296 | 0.150979707 | -2.687541945 |
| Nuak2 | 109.3043444 | -0.698653359 | 0.234387553 | -2.98076135 |
| F730311O21Rik | 4.129176112 | -1.458569745 | 0.525284472 | -2.776723514 |
| Tmcc2 | 2513.213478 | 0.438166154 | 0.097394263 | 4.49889082 |
| Gm15849 | 0.048949532 | 0.037064902 | 3.724945049 | 0.009950456 |
| Dstyk | 977.2630414 | -0.059107941 | 0.051760201 | -1.141957338 |
| 6030442K20Rik | 15.5449471 | -0.164251787 | 0.35235255 | -0.466157509 |
| Rbbp5 | 612.646602 | -0.120757406 | 0.058986849 | -2.047191999 |
| Tmem81 | 45.06443145 | 0.271064853 | 0.169848624 | 1.595920215 |
| Cntn2 | 3239.191646 | -0.048270084 | 0.08432557 | -0.572425228 |
| Nfasc | 5772.548432 | 0.040046211 | 0.057495771 | 0.69650707 |
| Gm10538 | 0.168483864 | 1.432078318 | 2.741982648 | 0.522278403 |
| Gm22609 | 0.089089296 | 0.957155977 | 3.723939561 | 0.257027796 |
| Lrrn2 | 1233.035506 | 0.811195181 | 0.095705729 | 8.475931235 |
| AC124095.1 | 0.988133447 | 3.043349122 | 1.33017254 | 2.28793561 |
| Gm38308 | 6.071337244 | 1.492406633 | 0.613335002 | 2.433265065 |
| Mdm4 | 1609.398272 | 0.043632219 | 0.117891531 | 0.370104778 |
| Gm29257 | 3.367574844 | -0.466107111 | 0.521105759 | -0.8944578 |
| Pik3c2b | 418.6707706 | 0.055734506 | 0.133330209 | 0.418018588 |
| Gm28609 | 1.26125615 | -1.160939782 | 0.865101388 | -1.341969621 |
| Gm26616 | 7.818633818 | -0.166240359 | 0.39295702 | -0.423049723 |
| Ppp1r15b | 1586.837848 | -0.44659647 | 0.121892523 | -3.663854516 |
| Plekha6 | 1798.908558 | 0.164191514 | 0.087728901 | 1.871578378 |
| Gm19461 | 22.93070562 | 0.068749657 | 0.240499823 | 0.285861572 |
| Golt1a | 0.589467498 | -0.113956646 | 1.595143313 | -0.071439754 |
| Gm28040 | 0.066050875 | -0.443818988 | 3.724945049 | -0.119147795 |
| Kiss1 | 0.282483437 | 0.955922622 | 1.770585618 | 0.539890651 |
| Ren1 | 0.441022512 | 0.410436162 | 1.542113325 | 0.266151751 |
| Etnk2 | 45.75414376 | 0.70442271 | 0.216568337 | 3.25265789 |
| Sox13 | 89.13343506 | -0.062947603 | 0.144408193 | -0.4359005 |
| Gm8596 | 0 | NA | NA | NA |
| Gm23939 | 0 | NA | NA | NA |
| Snrpe | 146.4574117 | -0.189571336 | 0.145146948 | -1.306064911 |
| Gm26706 | 0.926160546 | 0.902550607 | 1.097416892 | 0.822431852 |
| Zc3h11a | 237.5869313 | 0.034447835 | 0.122217894 | 0.281855904 |
| Gm38394 | 135.9246581 | 0.168301244 | 0.162512308 | 1.035621525 |

| | | | | |
|---|---|---|---|---|
| Zbed6 | 0 | NA | NA | NA |
| Lax1 | 23.20725604 | -0.541798004 | 0.223964592 | -2.419123486 |
| Atp2b4 | 2855.857049 | 0.391553737 | 0.100532182 | 3.894809908 |
| Mir6903 | 0.148659141 | -1.239386499 | 3.716315506 | -0.333498729 |
| Gm37368 | 12.07769997 | -0.12066208 | 0.421371911 | -0.2863553 |
| Gm10537 | 0.578745683 | -1.311949635 | 1.208350645 | -1.085735867 |
| Gm28441 | 0.061920613 | -0.443816764 | 3.724945049 | -0.119147198 |
| Optc | 6.651424553 | -0.148419623 | 0.393355668 | -0.377316598 |
| Gm15851 | 0.236932206 | 1.045543731 | 2.533571309 | 0.412675865 |
| Prelp | 162.3998482 | -0.205139827 | 0.216339432 | -0.948231328 |
| Gm1627 | 0.141089779 | 0.037067668 | 2.906109796 | 0.012755082 |
| Gm18970 | 0.109124069 | 0.517955464 | 3.724945049 | 0.139050498 |
| Fmod | 152.7762662 | 1.538970159 | 0.368528155 | 4.17599073 |
| Btg2 | 283.4370204 | -0.488921758 | 0.291617409 | -1.676586317 |
| Gm38140 | 0.628069627 | -0.398381588 | 1.284227556 | -0.310211058 |
| Gm8618 | 0.104637558 | -0.443821212 | 3.724945049 | -0.119148392 |
| Chit1 | 1.035276703 | 1.406728604 | 1.019462196 | 1.379873241 |
| Chil1 | 168.3470407 | 0.145974578 | 0.248482504 | 0.587464211 |
| Mybph | 3.909958598 | 0.071281659 | 0.498239712 | 0.143066996 |
| Adora1 | 2485.958102 | 0.186506199 | 0.073462884 | 2.538781328 |
| Gm24517 | 0.075043205 | 0.517957688 | 3.724945049 | 0.139051095 |
| Myog | 0.249503106 | -0.866803305 | 2.112394465 | -0.41034159 |
| Ppfia4 | 1225.389552 | 0.557775308 | 0.11381115 | 4.900884572 |
| Tmem183a | 645.1063955 | -0.047702412 | 0.085082186 | -0.560662744 |
| 4933406M09Rik | 0 | NA | NA | NA |
| Cyb5r1 | 155.8595938 | -0.934013864 | 0.113416782 | -8.235235136 |
| Adipor1 | 881.11371 | -0.347172044 | 0.074389065 | -4.666976818 |
| Platr1 | 1.832214876 | -1.920076133 | 1.058072274 | -1.814692796 |
| Gm37935 | 2.337573936 | 1.227250152 | 0.762613852 | 1.609268108 |
| Klhl12 | 493.9899022 | 0.284666354 | 0.08099036 | 3.514817735 |
| Rabif | 301.4519563 | -0.259127498 | 0.084393553 | -3.070465572 |
| Gm25612 | 0.720424393 | 0.843310408 | 1.163455701 | 0.724832417 |
| Gm15454 | 0.589661845 | -1.194134964 | 1.322513701 | -0.902928236 |
| Gm29376 | 0.461892809 | 1.002382224 | 1.552979017 | 0.645457674 |
| Mgat4e | 0.04393175 | 0.037062677 | 3.724945049 | 0.009949859 |
| Kdm5b | 1415.896103 | -0.058219968 | 0.082235568 | -0.707965771 |
| Gm26783 | 4.085067263 | 1.178486509 | 0.55243427 | 2.133261047 |
| Syt2 | 2912.71677 | 0.142156588 | 0.134449256 | 1.057325209 |
| Ppp1r12b | 1746.350506 | 0.272469685 | 0.095789009 | 2.844477535 |
| Gm37949 | 36.84559847 | 0.275415472 | 0.338704674 | 0.813143405 |
| Gm22232 | 0.044191549 | 0.037062677 | 3.724945049 | 0.009949859 |
| Gm37677 | 160.3400663 | 0.253316026 | 0.244722537 | 1.035115233 |
| Gm28892 | 1.029532974 | -0.367786326 | 1.442974649 | -0.254880657 |
| Ube2t | 14.99856726 | -0.11101559 | 0.271959231 | -0.408206735 |
| Lgr6 | 136.8876649 | -0.125071403 | 0.27822254 | -0.449537278 |
| Ptprv | 31.89205934 | 1.823103287 | 0.356178526 | 5.118509826 |
| Gm10535 | 0.133946075 | 0.036100024 | 3.626570338 | 0.009954315 |

| | | | | |
|---|---|---|---|---|
| Ptpn7 | 31.7926034 | -0.792837688 | 0.280709417 | -2.82440716 |
| Arl8a | 2134.195625 | -0.126270762 | 0.061167741 | -2.064335878 |
| Gm26280 | 0.068097359 | 0.037064902 | 3.724945049 | 0.009950456 |
| Gm15445 | 11.41385022 | 0.267747545 | 0.438930692 | 0.609999596 |
| Gpr37l1 | 2282.91187 | -0.228802005 | 0.148735401 | -1.538315714 |
| Gm4204 | 333.7656317 | 0.100072143 | 0.10655905 | 0.939123829 |
| Gm26642 | 0.168999367 | -0.868028675 | 2.739468898 | -0.316860204 |
| Elf3 | 3.18304154 | -0.645432763 | 0.597084175 | -1.080974492 |
| Rnpep | 506.4393182 | -0.462748659 | 0.077881981 | -5.941665223 |
| Timm17a | 833.5513504 | 0.048474832 | 0.066326078 | 0.730856305 |
| Lmod1 | 6.027761192 | 0.244842615 | 0.476963841 | 0.513335799 |
| Gm22796 | 0.04393175 | 0.037062677 | 3.724945049 | 0.009949859 |
| Shisa4 | 891.8069629 | -0.003160699 | 0.08746784 | -0.036135551 |
| Ipo9 | 1823.820265 | 0.1373238 | 0.047797859 | 2.873011516 |
| Gm28277 | 10.85246338 | 0.581822828 | 0.463180184 | 1.256147926 |
| Nav1 | 1957.583854 | 0.266788704 | 0.101428322 | 2.630317641 |
| Mir1231 | 0 | NA | NA | NA |
| 2610012C04Rik | 6.062032631 | 1.059494417 | 0.48610174 | 2.17957339 |
| Gm37333 | 28.22584607 | 0.215856533 | 0.41735325 | 0.517203432 |
| Gm4793 | 13.11457074 | 0.336298811 | 0.340419092 | 0.987896446 |
| Gm38399 | 26.61474392 | -0.494500861 | 0.207141549 | -2.387260611 |
| Csrp1 | 2908.739873 | -0.248427497 | 0.152938686 | -1.624360084 |
| Phlda3 | 300.6871226 | -0.617158415 | 0.156093595 | -3.953771553 |
| Tnni1 | 43.38678691 | -0.366104079 | 0.217514713 | -1.683123292 |
| Lad1 | 13.14689316 | 0.880672241 | 0.397254428 | 2.21689723 |
| Tnnt2 | 21.42389219 | 1.270190889 | 0.309082019 | 4.109559311 |
| Gm28883 | 1.230445023 | 0.367271651 | 0.922837951 | 0.397980654 |
| Pkp1 | 4.016384854 | -0.213992548 | 0.684673749 | -0.312546739 |
| Gm36938 | 0.07796279 | -0.798301147 | 3.721829328 | -0.214491605 |
| Igfn1 | 106.9666211 | 0.491159352 | 0.300490675 | 1.634524438 |
| Tmem9 | 526.3167638 | 0.024122464 | 0.110274069 | 0.218750101 |
| Ascl5 | 1.900208836 | -0.494249455 | 0.73161459 | -0.675559867 |
| Cacna1s | 20.1854872 | 0.319271952 | 0.263104148 | 1.213481259 |
| Gm37014 | 0 | NA | NA | NA |
| Gm37759 | 0.707021654 | -1.273390307 | 1.437336081 | -0.885937759 |
| Gm15850 | 4.717623853 | -1.064326298 | 0.601329989 | -1.769953798 |
| Kif21b | 1190.983809 | 0.07092221 | 0.09604796 | 0.738404121 |
| Mroh3 | 7.987918312 | -0.717028654 | 0.378424946 | -1.894771106 |
| 5730559C18Rik | 56.42049381 | -0.720540628 | 0.176093646 | -4.091803684 |
| Gm26568 | 0.061920613 | -0.443816764 | 3.724945049 | -0.119147198 |
| A430034D21Rik | 0 | NA | NA | NA |
| Gm37552 | 0.476395801 | -1.303677442 | 1.353194458 | -0.963407317 |
| Gm26781 | 4.326125698 | -0.242396371 | 0.48136624 | -0.503559143 |
| Gpr25 | 16.1131376 | 0.338101532 | 0.285610861 | 1.183783875 |
| Camsap2 | 3490.88954 | 0.184053901 | 0.051822682 | 3.551608931 |
| 9230116N13Rik | 6.573994958 | 0.24091095 | 0.504493863 | 0.47752999 |
| Ddx59 | 80.65548895 | -0.048250802 | 0.135192873 | -0.356903447 |

| | | | | |
|---|---|---|---|---|
| Kif14 | 11.46902475 | -0.408566707 | 0.347100434 | -1.17708498 |
| Gm29486 | 0.097139547 | -0.443816764 | 3.724945049 | -0.119147198 |
| Gm29485 | 0.285872948 | 0.037074884 | 2.115041849 | 0.017529149 |
| Gm37799 | 0.092691418 | -0.908687078 | 3.724560878 | -0.2439716 |
| Gm33994 | 0.860407559 | -0.935850919 | 0.981499211 | -0.95349126 |
| 4933409D19Rik | 0.218406194 | -1.118221202 | 2.130880312 | -0.524769597 |
| Platr23 | 2.85951047 | -0.343070209 | 0.582476483 | -0.588985512 |
| Zfp281 | 1236.560802 | -0.353430078 | 0.10902497 | -3.241735155 |
| Gm19705 | 24.95245937 | -0.322758577 | 0.216369215 | -1.491702859 |
| Platr22 | 0.971167388 | 1.080539836 | 0.936165345 | 1.154219008 |
| Gm24086 | 0.066050875 | -0.443818988 | 3.724945049 | -0.119147795 |
| Gm22132 | 0 | NA | NA | NA |
| Gm8762 | 0 | NA | NA | NA |
| Gm29718 | 0.856212859 | -0.739334869 | 1.262502668 | -0.58561054 |
| Gm17781 | 0 | NA | NA | NA |
| Nr5a2 | 2.338360347 | -0.967405869 | 0.719676958 | -1.344222374 |
| Gm23763 | 0 | NA | NA | NA |
| Gm25609 | 0.046345709 | -0.443818988 | 3.724945049 | -0.119147795 |
| Gm37057 | 0.333318405 | 1.781324172 | 1.697104687 | 1.049625392 |
| 1700006P03Rik | 0 | NA | NA | NA |
| Gm23534 | 0 | NA | NA | NA |
| Gm37903 | 0.499362493 | 0.998869046 | 1.687622538 | 0.591879418 |
| Gm22727 | 0 | NA | NA | NA |
| Gm37099 | 0.251787852 | 1.910899889 | 2.039326176 | 0.937025137 |
| Gm37293 | 0 | NA | NA | NA |
| Gm4258 | 79.79539825 | 0.042982612 | 0.254587202 | 0.16883257 |
| Gm37298 | 205.364216 | 0.096194116 | 0.200575455 | 0.479590668 |
| Gm26979 | 0 | NA | NA | NA |
| Gm37124 | 62.91602665 | 0.114106464 | 0.240774462 | 0.473914314 |
| A130050O07Rik | 109.5267668 | -0.040344701 | 0.264293577 | -0.152651084 |
| Gm26936 | 0 | NA | NA | NA |
| Gm26930 | 92.36817408 | 0.07464695 | 0.279220256 | 0.267340742 |
| D130019J16Rik | 127.5229106 | -0.239858342 | 0.217033567 | -1.10516703 |
| Gm37534 | 242.1756396 | -0.262294499 | 0.201739719 | -1.300162906 |
| Mir181a-1 | 0.399769715 | -1.430166476 | 1.575311133 | -0.907862864 |
| A430106G13Rik | 56.85494254 | -0.376527741 | 0.283203596 | -1.329530225 |
| Mir181b-1 | 0 | NA | NA | NA |
| Gm37632 | 17.06609017 | -0.691140236 | 0.29298313 | -2.358976216 |
| Gm23782 | 0 | NA | NA | NA |
| Ptprc | 1013.820159 | -0.938031616 | 0.180418528 | -5.199197801 |
| A130071D04Rik | 74.31188421 | -0.744974877 | 0.287758217 | -2.588891759 |
| Gm18667 | 0 | NA | NA | NA |
| Gm28556 | 0 | NA | NA | NA |
| 1700113B19Rik | 0 | NA | NA | NA |
| Atp6v1g3 | 0.18576184 | -1.327298715 | 3.713354821 | -0.357439237 |
| Gm28500 | 0.232206954 | 1.298057402 | 2.545374422 | 0.509967174 |
| Nek7 | 865.5502947 | 0.059278022 | 0.121910598 | 0.486241744 |

| | | | | |
|---|---|---|---|---|
| Gm28501 | 0.215166026 | 1.019404318 | 2.554583188 | 0.399049177 |
| Gm37080 | 8.379318968 | 0.445623566 | 0.435175868 | 1.02400799 |
| Gm37101 | 16.00512593 | -0.101579775 | 0.351352267 | -0.289110914 |
| Gm5833 | 0.04393175 | 0.037062677 | 3.724945049 | 0.009949859 |
| Gm8790 | 0 | NA | NA | NA |
| Lhx9 | 105.8300051 | 0.28228468 | 0.105669886 | 2.671382459 |
| 2310009B15Rik | 35.43042701 | 0.20989882 | 0.188529372 | 1.113348109 |
| 1700019P21Rik | 0 | NA | NA | NA |
| n-R5s217 | 0.075043205 | 0.517957688 | 3.724945049 | 0.139051095 |
| Gm3933 | 0 | NA | NA | NA |
| Gm37045 | 1.708528675 | -0.641900986 | 0.761074387 | -0.843414254 |
| Dennd1b | 1082.456846 | -0.028427736 | 0.075541511 | -0.3763194 |
| Gm37423 | 58.57700844 | -0.078121798 | 0.257304338 | -0.303616326 |
| AC117623.1 | 17.28746858 | -0.156247153 | 0.361225279 | -0.43254767 |
| Gm18553 | 0.072025462 | 0.517959912 | 3.724945049 | 0.139051692 |
| Gm16305 | 0 | NA | NA | NA |
| Gm37470 | 63.54316749 | 0.107543484 | 0.196590146 | 0.547044124 |
| RP23-114L12.4 | 43.15415538 | 0.311644621 | 0.249759842 | 1.247777135 |
| Gm37655 | 0.107078754 | -0.443164241 | 3.724898049 | -0.118973522 |
| Gm16304 | 0 | NA | NA | NA |
| Gm37718 | 31.22466683 | 0.309564681 | 0.253384305 | 1.22172003 |
| Crb1 | 30.89787103 | -0.084249062 | 0.269151886 | -0.3130168 |
| 4933436E23Rik | 0.112288909 | 0.037060453 | 3.724945049 | 0.009949262 |
| Gm34816 | 0 | NA | NA | NA |
| Zbtb41 | 682.7308389 | -0.173020919 | 0.073346334 | -2.358957962 |
| Aspm | 33.81106085 | -0.636242341 | 0.341618736 | -1.862433978 |
| F13b | 0.541494639 | -2.625429136 | 1.419468068 | -1.849586613 |
| Cfhr1 | 0.094294407 | -0.443821212 | 3.724945049 | -0.119148392 |
| Cfhr3 | 0 | NA | NA | NA |
| Gm26271 | 0 | NA | NA | NA |
| Gm4788 | 0.298414574 | -0.461261685 | 1.816931621 | -0.253868489 |
| Gm25619 | 0 | NA | NA | NA |
| Cfhr2 | 0.451902873 | 0.086737827 | 1.350426616 | 0.064229945 |
| Gm16332 | 4.277969054 | -0.414938343 | 0.451880483 | -0.918247986 |
| Gm5837 | 9.46935442 | -0.621540865 | 0.362321256 | -1.715441354 |
| Gm5834 | 1.243360777 | -0.105624177 | 1.053019599 | -0.100305994 |
| Gm38266 | 0 | NA | NA | NA |
| Gm26048 | 0 | NA | NA | NA |
| Gm38019 | 0.981599412 | 0.306193349 | 1.034087696 | 0.296099983 |
| Cfh | 496.518771 | -0.415380739 | 0.130651015 | -3.179315055 |
| Kcnt2 | 523.8644755 | 0.856221808 | 0.119223863 | 7.181631166 |
| Gm37777 | 49.81740225 | 1.30535994 | 0.253305542 | 5.153301935 |
| 4930590L20Rik | 0 | NA | NA | NA |
| Gm37371 | 0.255648062 | -0.895944232 | 1.84256953 | -0.486247177 |
| Gm4845 | 0 | NA | NA | NA |
| Gm25457 | 0 | NA | NA | NA |
| Gm22016 | 0 | NA | NA | NA |

| | | | | |
|---|---|---|---|---|
| Gm37599 | 0.182775946 | 1.568991847 | 3.32402584 | 0.472015539 |
| Gm37142 | 0 | NA | NA | NA |
| Gm37604 | 0 | NA | NA | NA |
| Gm18183 | 0.075043205 | 0.517957688 | 3.724945049 | 0.139051095 |
| Gm28724 | 0 | NA | NA | NA |
| Gm37796 | 0 | NA | NA | NA |
| Gm5835 | 7.977887189 | -0.499685468 | 0.399776105 | -1.249913293 |
| Cdc73 | 1577.910083 | 0.128415017 | 0.054991982 | 2.33515891 |
| B3galt2 | 609.7109376 | 0.948300377 | 0.087797185 | 10.80103392 |
| Gm6397 | 3.497495822 | 0.37105651 | 0.548579761 | 0.676394822 |
| Glrx2 | 603.4570244 | 0.253353836 | 0.064549637 | 3.924945945 |
| Trove2 | 1714.422575 | -0.121989595 | 0.098457871 | -1.239002971 |
| Uchl5 | 265.6573748 | 0.302310041 | 0.141618736 | 2.134675454 |
| Gm29170 | 125.043805 | 0.202888157 | 0.121215585 | 1.673779467 |
| Gm25663 | 0 | NA | NA | NA |
| Rgs2 | 585.4803157 | -0.593447649 | 0.203571414 | -2.915181643 |
| Rgs13 | 2.523201682 | 0.03391014 | 0.623216446 | 0.054411498 |
| Rbm6-ps2 | 0.220340683 | -0.049041869 | 2.186381828 | -0.022430606 |
| Rgs1 | 170.7604979 | -0.731897806 | 0.435156039 | -1.681920367 |
| Gm7266 | 0.253428164 | 0.383984973 | 2.081456452 | 0.184478985 |
| Rgs21 | 0 | NA | NA | NA |
| Gm23672 | 0 | NA | NA | NA |
| Rgs18 | 3.121267796 | -0.584705591 | 0.587773621 | -0.994780252 |
| Gm8856 | 0 | NA | NA | NA |
| Gm22681 | 0 | NA | NA | NA |
| Gm25708 | 0 | NA | NA | NA |
| Gm6550 | 0 | NA | NA | NA |
| Gm5262 | 0 | NA | NA | NA |
| Gm25460 | 0 | NA | NA | NA |
| Gm28392 | 0 | NA | NA | NA |
| Gm38269 | 0.279407529 | 0.535161812 | 2.073798664 | 0.258058712 |
| Gm29514 | 42.66995015 | 0.050338031 | 0.229656876 | 0.219187999 |
| Gm5263 | 0.047948698 | 0.037064902 | 3.724945049 | 0.009950456 |
| Gm38187 | 45.19927449 | 0.004125264 | 0.299574135 | 0.013770427 |
| Brinp3 | 455.9540705 | 0.285924976 | 0.089731924 | 3.186435362 |
| Gm15583 | 12.60536537 | 0.049200547 | 0.406017661 | 0.121178342 |
| A230059L01Rik | 0.205212924 | 1.000555958 | 2.635865574 | 0.379592938 |
| Gm15584 | 0.127515837 | -1.208141086 | 3.685333689 | -0.327824069 |
| Gm37749 | 0.118371171 | 0.037073798 | 3.724945049 | 0.009952844 |
| Gm9931 | 0 | NA | NA | NA |
| Gm17867 | 0.136963818 | 0.037074342 | 3.599544405 | 0.010299732 |
| Gm38284 | 0.044544648 | 0.517957688 | 3.724945049 | 0.139051095 |
| Gm29516 | 0 | NA | NA | NA |
| Gm29515 | 0 | NA | NA | NA |
| Gm22966 | 0 | NA | NA | NA |
| Gm19891 | 0 | NA | NA | NA |
| Gm18505 | 0.044191549 | 0.037062677 | 3.724945049 | 0.009949859 |

| | | | | |
|---|---|---|---|---|
| Gm4034 | 0 | NA | NA | NA |
| Gm29022 | 0 | NA | NA | NA |
| Gm5703 | 0 | NA | NA | NA |
| Gm15443 | 3.976562986 | -0.023145671 | 0.491436877 | -0.047097952 |
| Gm18751 | 0 | NA | NA | NA |
| Gm29020 | 0 | NA | NA | NA |
| Gm18701 | 0 | NA | NA | NA |
| Gm37630 | 0.062426873 | 0.517959912 | 3.724945049 | 0.139051692 |
| Gm38189 | 0.200297358 | 1.01969918 | 2.613669874 | 0.390140771 |
| Gm29398 | 0.199606257 | 0.490665623 | 2.214366477 | 0.221582844 |
| Gm37096 | 0.086648695 | 0.037058229 | 3.724945049 | 0.009948665 |
| Gm37775 | 0.044544648 | 0.517957688 | 3.724945049 | 0.139051095 |
| Gm23535 | 0 | NA | NA | NA |
| n-R5s218 | 0 | NA | NA | NA |
| Pla2g4a | 121.1500737 | -0.929177713 | 0.208947044 | -4.446953131 |
| n-R5s219 | 0 | NA | NA | NA |
| Gm19136 | 0.350167968 | -0.865540424 | 1.59737164 | -0.54185288 |
| AC114655.1 | 0.088736197 | 0.51795324 | 3.724945049 | 0.139049901 |
| Ptgs2os | 10.82272229 | -0.997670431 | 0.372575215 | -2.677769189 |
| Ptgs2 | 389.8698901 | -1.149255144 | 0.351276192 | -3.271656806 |
| Gm9687 | 0.529068976 | -0.888177554 | 1.410838192 | -0.629538922 |
| Ptgs2os2 | 1.911122854 | 0.27825363 | 0.706587922 | 0.393799017 |
| Gm15453 | 44.68224246 | 0.362659183 | 0.215866162 | 1.680018678 |
| Gm4322 | 0.586444394 | -0.32808145 | 1.375001007 | -0.238604516 |
| Pdc | 0.117760149 | 0.51795324 | 3.724945049 | 0.139049901 |
| 2310030A07Rik | 0.378962075 | 1.88926034 | 1.421649993 | 1.328920866 |
| Gm37764 | 0 | NA | NA | NA |
| BC003331 | 544.608875 | 0.041252019 | 0.064030995 | 0.644250784 |
| Tpr | 2374.975998 | -0.011621078 | 0.081621591 | -0.142377496 |
| Prg4 | 160.3573172 | -0.561556253 | 0.381824604 | -1.470717828 |
| Ska2l-ps | 0.636218932 | 0.500787775 | 1.33199626 | 0.375967854 |
| Hmcn1 | 64.89396023 | 0.436858232 | 0.176561926 | 2.474249363 |
| Gm20631 | 0.097139547 | -0.443816764 | 3.724945049 | -0.119147198 |
| Gm29188 | 0 | NA | NA | NA |
| Gm22756 | 0 | NA | NA | NA |
| Gm37671 | 0.106358623 | 0.517955464 | 3.724945049 | 0.139050498 |
| Gm19087 | 0 | NA | NA | NA |
| Gm8941 | 0.044544648 | 0.517957688 | 3.724945049 | 0.139051095 |
| C730036E19Rik | 0.469314954 | 1.565777341 | 1.434505666 | 1.091510043 |
| Gm24402 | 0 | NA | NA | NA |
| Gm10138 | 35.28446059 | 0.325347085 | 0.218552933 | 1.488642043 |
| Ivns1abp | 1648.737111 | 0.151726312 | 0.075254985 | 2.016162927 |
| Swt1 | 331.9812924 | -0.276233102 | 0.11600301 | -2.381258057 |
| Trmt1l | 460.2408708 | -0.62034756 | 0.19500012 | -3.181267585 |
| mmu-mir-7682 | 1.074470705 | -1.198364998 | 1.042197015 | -1.149844972 |
| Gm37893 | 50.46302794 | -0.308684662 | 0.312447701 | -0.987956258 |
| Rnf2 | 494.0673276 | 0.024996645 | 0.096908009 | 0.257941994 |

| | | | | |
|---|---|---|---|---|
| Gm36527 | 43.64518431 | 0.29867992 | 0.350466814 | 0.852234527 |
| Gm29072 | 0 | NA | NA | NA |
| Gm19503 | 0.173853943 | -0.499202846 | 2.738479247 | -0.182291995 |
| Gm28181 | 0.049553047 | -0.443818988 | 3.724945049 | -0.119147795 |
| Fam129a | 103.521438 | -0.542294591 | 0.12810708 | -4.233135193 |
| 2810414N06Rik | 13.51786206 | 0.040864167 | 0.311485034 | 0.131191429 |
| Edem3 | 1232.170216 | -0.212437179 | 0.063673518 | -3.336350594 |
| 1700025G04Rik | 1157.852342 | 0.177356324 | 0.063059437 | 2.812526289 |
| Gm25279 | 0.423681387 | -0.495580429 | 1.720177146 | -0.288098485 |
| Gm28610 | 0.118371171 | 0.037073798 | 3.724945049 | 0.009952844 |
| Gm8964 | 0 | NA | NA | NA |
| Gm38238 | 0.047948698 | 0.037064902 | 3.724945049 | 0.009950456 |
| Tsen15 | 205.9704903 | -0.08894458 | 0.092245318 | -0.964217826 |
| Colgalt2 | 66.2940545 | 0.155650032 | 0.150775374 | 1.032330602 |
| Gm15618 | 0.134880151 | -1.305960502 | 2.927087508 | -0.446163806 |
| Rgl1 | 1379.502362 | -0.001793125 | 0.09001994 | -0.019919198 |
| Gm15479 | 6.574668662 | 0.36987326 | 0.523828155 | 0.706096564 |
| Gm7278 | 19.06484076 | 0.776605099 | 0.332172671 | 2.337956031 |
| Gm28792 | 7.043371214 | 1.59247073 | 0.5055626 | 3.149898211 |
| Apobec4 | 0.396489138 | 1.136633716 | 1.478462002 | 0.768794676 |
| Arpc5 | 1009.34046 | -0.388548599 | 0.119602493 | -3.248666396 |
| Ncf2 | 128.1024071 | -1.05084814 | 0.182562977 | -5.756085699 |
| Smg7 | 2499.293702 | -0.199409227 | 0.085879836 | -2.32195632 |
| Gm28791 | 48.57147184 | 0.234913362 | 0.25665665 | 0.915282584 |
| 1700040K01Rik | 0.217189781 | -1.229375415 | 2.165401332 | -0.567735596 |
| Gm8976 | 0.43436012 | 1.071769991 | 1.672966054 | 0.640640608 |
| Nmnat2 | 3031.207894 | 0.275060516 | 0.076267374 | 3.606529256 |
| Gm28960 | 0 | NA | NA | NA |
| Lamc2 | 154.9877848 | -0.716701497 | 0.139472789 | -5.138647483 |
| Lamc1 | 1030.31094 | -0.263605589 | 0.121456206 | -2.170375626 |
| Gm38009 | 27.75065565 | 0.463891748 | 0.292551329 | 1.585676433 |
| Gm8818 | 0.931703337 | -2.163280929 | 1.142026134 | -1.894248183 |
| E330020D12Rik | 5.955571085 | -0.342723547 | 0.387430875 | -0.884605666 |
| Shcbp1l | 22.81337923 | 0.732852587 | 0.295489168 | 2.480133511 |
| Gm25588 | 0.946605489 | 0.047914776 | 1.443249402 | 0.033199235 |
| Gm37669 | 3.285918002 | 0.708779435 | 0.692990566 | 1.022783671 |
| Dhx9 | 2116.613299 | 0.005868262 | 0.077468784 | 0.07575002 |
| Npl | 38.83715233 | 0.129413156 | 0.212758901 | 0.608262007 |
| Gm6652 | 5.116277746 | 0.281078015 | 0.553513007 | 0.507807425 |
| Rgs8 | 2498.85835 | 0.187407479 | 0.160204019 | 1.169805103 |
| Gm29529 | 0.238359071 | -0.947557347 | 2.115672711 | -0.447875204 |
| 5530400K19Rik | 2.012029913 | -1.901657931 | 0.842315181 | -2.25765601 |
| C230024C17Rik | 0.83423333 | -0.247703708 | 1.150564777 | -0.215288798 |
| Rgs16 | 197.3246427 | -0.029101735 | 0.133244608 | -0.218408351 |
| Gm28513 | 0.329560409 | 0.979138742 | 1.718234682 | 0.569851576 |
| Rnasel | 213.55156 | -0.1179435 | 0.141481375 | -0.833632695 |
| Gm28512 | 4.354440161 | 0.657251719 | 0.552843686 | 1.188856335 |

| | | | | |
|---|---|---|---|---|
| Rgsl1 | 23.12760851 | 1.247234272 | 0.328313811 | 3.798908943 |
| Teddm2 | 33.79161557 | 1.068791718 | 0.303326502 | 3.523568536 |
| Teddm1b | 0.047948698 | 0.037064902 | 3.724945049 | 0.009950456 |
| Gm38352 | 5.642899575 | 1.098650911 | 0.586368227 | 1.87365355 |
| Teddm1a | 0.061920613 | -0.443816764 | 3.724945049 | -0.119147198 |
| Mir8114 | 0.170693195 | -0.49936863 | 2.754366169 | -0.181300742 |
| Glul | 15318.94443 | 0.427763231 | 0.158674498 | 2.695853692 |
| Gm29290 | 0.209920398 | -0.934053945 | 2.624885147 | -0.355845644 |
| Gm29291 | 0.447162105 | -1.73210344 | 1.409803269 | -1.228613579 |
| Gm28286 | 0.825983375 | 1.785113672 | 1.268440388 | 1.407329575 |
| A830008E24Rik | 2.805075798 | 1.019360832 | 0.644126932 | 1.582546517 |
| Gm25336 | 0 | NA | NA | NA |
| Zfp648 | 4.036848141 | 0.368905167 | 0.538489349 | 0.685074214 |
| 4930452N14Rik | 0.785070532 | -1.097832955 | 1.235627315 | -0.888482264 |
| 4930532M18Rik | 0.278668908 | 0.517975066 | 1.798586127 | 0.287990137 |
| n-R5s220 | 0.221251082 | 0.054611765 | 2.575702265 | 0.02120267 |
| Gm18727 | 0.332553061 | -0.396703152 | 1.951391719 | -0.203292424 |
| Cacna1e | 5959.244093 | 0.689866449 | 0.157843858 | 4.370562506 |
| Gm29282 | 1.600249739 | -0.321301539 | 0.805671401 | -0.398799732 |
| Gm29441 | 0.044544648 | 0.517957688 | 3.724945049 | 0.139051095 |
| Gm9530 | 4.308798137 | 0.255955904 | 0.520019084 | 0.492204829 |
| Ier5 | 750.1768822 | -0.76250639 | 0.146782945 | -5.194788745 |
| Mr1 | 88.50137408 | -0.123698709 | 0.151977032 | -0.813930284 |
| Stx6 | 782.513869 | 0.109056688 | 0.053677026 | 2.03172 |
| Gm8850 | 14.83397756 | 0.887225972 | 0.381983245 | 2.322682957 |
| BC034090 | 156.7399653 | -0.918346257 | 0.106830735 | -8.596273863 |
| Gm37622 | 0.256551671 | 0.979241194 | 2.455008643 | 0.398874846 |
| Xpr1 | 2184.107984 | -0.052883803 | 0.059297426 | -0.89183976 |
| Gm22569 | 0 | NA | NA | NA |
| Gm5532 | 8.533642962 | -1.253779974 | 0.479809093 | -2.613080898 |
| Acbd6 | 499.0889261 | 0.114014192 | 0.082449773 | 1.382832097 |
| Lhx4 | 8.533691663 | -0.168870451 | 0.875011609 | -0.192992241 |
| Gm37089 | 0 | NA | NA | NA |
| Qsox1 | 389.4008561 | -0.228239402 | 0.087464819 | -2.60949949 |
| Gm37539 | 0 | NA | NA | NA |
| Gm37571 | 0.346370079 | -0.998773977 | 2.004837073 | -0.498182117 |
| Cep350 | 1964.651825 | -0.086787972 | 0.075633572 | -1.147479486 |
| Gm38038 | 0 | NA | NA | NA |
| Gm36934 | 0.108309328 | 0.967119648 | 3.722483401 | 0.259804959 |
| Gm37383 | 6.027087886 | -0.225471963 | 0.491236492 | -0.458988627 |
| Gm23744 | 0.061920613 | -0.443816764 | 3.724945049 | -0.119147198 |
| Gm9694 | 3.918626087 | 1.303527329 | 0.612142348 | 2.129451316 |
| Tor1aip1 | 1314.956962 | -0.561748147 | 0.140927791 | -3.986070757 |
| 9430034N14Rik | 86.42678797 | 0.200480689 | 0.275233679 | 0.728401733 |
| Tor1aip2 | 1490.887513 | -0.567901195 | 0.130627861 | -4.347473733 |
| Fam163a | 99.41997446 | -0.009227268 | 0.146668776 | -0.062912287 |
| Gm38113 | 0 | NA | NA | NA |

| | | | | |
|---|---|---|---|---|
| Tdrd5 | 36.39050195 | 0.84226396 | 0.240413851 | 3.503391993 |
| A430050A11Rik | 1.164548605 | 1.178096963 | 1.062716318 | 1.108571444 |
| 4930518J20Rik | 0.382980858 | -0.449573353 | 1.573934691 | -0.2856366 |
| Nphs2 | 3.238706364 | -0.628394309 | 0.590891696 | -1.063467829 |
| Axdnd1 | 8.537287892 | 0.756517265 | 0.382512219 | 1.977759737 |
| Gm2000 | 140.1670121 | -0.079749866 | 0.14449454 | -0.551923039 |
| Soat1 | 740.7127753 | -0.953387713 | 0.148630984 | -6.414461423 |
| Gm37648 | 0.139856492 | -0.443820827 | 3.574411099 | -0.124166139 |
| Gm10031 | 2.652532074 | 0.732999199 | 0.616204746 | 1.189538387 |
| AC114588.1 | 0 | NA | NA | NA |
| Abl2 | 1607.186212 | -0.011117059 | 0.107697842 | -0.103224531 |
| Gm38043 | 100.360574 | 0.32318821 | 0.302416598 | 1.068685425 |
| Tor3a | 2329.182894 | -0.870837739 | 0.244013783 | -3.568805533 |
| Fam20b | 1176.797779 | 0.030739255 | 0.051050659 | 0.602132393 |
| Gm19918 | 0 | NA | NA | NA |
| Gm15428 | 0.152810971 | -0.444701676 | 2.84449009 | -0.156337924 |
| Ralgps2 | 727.6286885 | 0.115984544 | 0.089625015 | 1.294109065 |
| Gm28693 | 10.3212494 | 0.120972699 | 0.436695439 | 0.277018463 |
| Angptl1 | 9.659502383 | 0.405099239 | 0.336612294 | 1.203459431 |
| Gm37802 | 27.9942778 | 0.557574128 | 0.295075828 | 1.889596079 |
| Gm37943 | 22.83945072 | 0.699490236 | 0.40765811 | 1.715874696 |
| 4930439D14Rik | 2.895518095 | 0.263690531 | 0.68413791 | 0.38543476 |
| Gm28694 | 0 | NA | NA | NA |
| Gm10531 | 0.066050875 | -0.443818988 | 3.724945049 | -0.119147795 |
| Tex35 | 6.225377881 | 2.160719371 | 0.824763484 | 2.619804845 |
| Rasal2 | 2052.323456 | 0.250487883 | 0.064605544 | 3.877188687 |
| Gm37788 | 31.13971489 | 0.014484767 | 0.250759154 | 0.057763661 |
| Gm37578 | 17.57312196 | -0.061292109 | 0.360951753 | -0.169806931 |
| Gm6058 | 12.1668676 | 0.60941395 | 0.38876864 | 1.567549146 |
| Gm37060 | 72.32838032 | 0.227936314 | 0.231765853 | 0.983476691 |
| 2810025M15Rik | 74.04682442 | -0.323230855 | 0.152409175 | -2.120809688 |
| BC026585 | 13.64787826 | -0.511167461 | 0.38627351 | -1.323330355 |
| Gm15486 | 0.137416547 | -0.443827336 | 3.639393827 | -0.121950896 |
| Sec16b | 13.26555679 | -0.093215994 | 0.287696857 | -0.32400769 |
| Gm37682 | 0 | NA | NA | NA |
| Gm7412 | 0 | NA | NA | NA |
| Gm37713 | 0 | NA | NA | NA |
| Gm38213 | 0 | NA | NA | NA |
| Gm38256 | 0.119234754 | 0.51795324 | 3.724945049 | 0.139049901 |
| Brinp2 | 1002.677425 | 0.628811335 | 0.125220315 | 5.021639928 |
| Astn1 | 3735.831764 | 0.669467805 | 0.063249295 | 10.5845892 |
| Gm37861 | 45.70823912 | 0.755495336 | 0.349725371 | 2.160253157 |
| Gm37679 | 1.033175386 | 2.532009662 | 1.13953922 | 2.221959207 |
| Mir488 | 2.924473085 | 1.193247514 | 0.780120549 | 1.52956811 |
| Gm37367 | 35.2279183 | 0.812391153 | 0.36509273 | 2.225163872 |
| Gm37848 | 2.466912353 | 0.774828692 | 1.06113107 | 0.730191315 |
| Pappa2 | 44.61674889 | -0.130966464 | 0.25587589 | -0.51183589 |

| | | | | |
|---|---|---|---|---|
| 1600012P17Rik | 0.046345709 | -0.443818988 | 3.724945049 | -0.119147795 |
| 4930562F17Rik | 0 | NA | NA | NA |
| Gm4953 | 0 | NA | NA | NA |
| Gm7430 | 0.038981395 | -0.443816764 | 3.724945049 | -0.119147198 |
| Rfwd2 | 802.379122 | 0.151465505 | 0.104039411 | 1.455847388 |
| Gm37115 | 30.16231669 | 0.278656071 | 0.283338445 | 0.983474273 |
| Gm37088 | 0.075043205 | 0.517957688 | 3.724945049 | 0.139051095 |
| Scarna3a | 3.950984661 | 1.534664054 | 0.53686817 | 2.858549156 |
| mmu-mir-1843b | 0 | NA | NA | NA |
| Gm31256 | 1.235093012 | 2.069030576 | 0.997384936 | 2.07445541 |
| Gm37757 | 0.066050875 | -0.443818988 | 3.724945049 | -0.119147795 |
| Tnr | 2666.999354 | 0.427981087 | 0.080127514 | 5.341250068 |
| Gm37731 | 39.71725266 | 0.520089768 | 0.303165366 | 1.715531608 |
| Gm10530 | 19.32941874 | 0.698393649 | 0.425721603 | 1.640493798 |
| Gm38039 | 0.270352934 | -0.898141164 | 1.788475188 | -0.502182625 |
| Gm24719 | 0 | NA | NA | NA |
| Gm37294 | 0 | NA | NA | NA |
| 4930523C07Rik | 65.68144704 | 0.031864616 | 0.185916229 | 0.17139233 |
| 4930562F07Rik | 0.26443487 | -0.822121535 | 1.783335421 | -0.46100219 |
| Tnn | 13.34929327 | -2.140608348 | 0.370420472 | -5.778860805 |
| Mrps14 | 388.8406474 | -0.104213541 | 0.092953506 | -1.121136206 |
| Cacybp | 851.1120869 | -0.282521462 | 0.131941598 | -2.141261492 |
| 4933417C20Rik | 5.899375495 | -0.192118313 | 0.466070689 | -0.412208529 |
| Rabgap1l | 3165.401473 | 0.316829312 | 0.064446675 | 4.916146753 |
| Gm36975 | 0.049553047 | -0.443818988 | 3.724945049 | -0.119147795 |
| Mir1927 | 6.385949591 | 0.1727268 | 0.449616822 | 0.384164451 |
| 9430087J23Rik | 22.80069575 | 0.13740431 | 0.282717769 | 0.48601229 |
| Gm37529 | 68.10051105 | 0.180995371 | 0.307905113 | 0.587828403 |
| Gm37212 | 65.71378451 | 0.23235958 | 0.255673241 | 0.908814625 |
| Gm37215 | 27.68718647 | -0.248290784 | 0.30384772 | -0.81715533 |
| Gm38302 | 37.68841078 | -0.211952512 | 0.366299874 | -0.578631134 |
| 6720464F23Rik | 73.41064915 | -0.044462476 | 0.257938473 | -0.172376287 |
| Gm23528 | 0 | NA | NA | NA |
| Gm37873 | 11.00141775 | 0.095739969 | 0.386360848 | 0.247799356 |
| Gm37328 | 9.255702839 | -0.554160661 | 0.399575534 | -1.386873355 |
| Gm38329 | 12.82265238 | 0.155627686 | 0.288887121 | 0.538714516 |
| Gm38126 | 0.106465261 | 0.037073798 | 3.724945049 | 0.009952844 |
| Gm37083 | 0.417055923 | -1.16253884 | 1.545125355 | -0.752391278 |
| Gm38060 | 0.123841226 | -1.017823543 | 3.719440177 | -0.273649661 |
| Gm24988 | 0 | NA | NA | NA |
| Gm6177 | 0.068097359 | 0.037064902 | 3.724945049 | 0.009950456 |
| Rc3h1 | 1747.242785 | 0.004149127 | 0.047459982 | 0.087423691 |
| Gm37052 | 30.04310741 | -0.088865588 | 0.258443512 | -0.343849174 |
| Gm37809 | 11.17045412 | -0.100067217 | 0.396457033 | -0.252403686 |
| Gm37653 | 63.56315506 | 0.083752026 | 0.227689999 | 0.367833571 |
| Serpinc1 | 4.738076569 | -0.874475141 | 0.527885249 | -1.656562944 |
| Gm37729 | 0.047948698 | 0.037064902 | 3.724945049 | 0.009950456 |

| | | | | |
|---|---|---|---|---|
| Gm37072 | 1.390520144 | 0.460749901 | 0.919443201 | 0.501118395 |
| Zbtb37 | 638.7082528 | 0.094625167 | 0.097479172 | 0.970721894 |
| Gm37362 | 0.22549826 | -0.882933078 | 2.128349392 | -0.414844048 |
| Gas5 | 565.6631599 | 0.008763178 | 0.116225527 | 0.07539805 |
| Gm26224 | 0 | NA | NA | NA |
| Gm22489 | 0 | NA | NA | NA |
| Gm22357 | 0 | NA | NA | NA |
| Gm23212 | 0 | NA | NA | NA |
| Snord78 | 0 | NA | NA | NA |
| Mir5117 | 0 | NA | NA | NA |
| Snord80 | 0 | NA | NA | NA |
| SNORD77 | 0 | NA | NA | NA |
| Snord47 | 0 | NA | NA | NA |
| Gm25789 | 0 | NA | NA | NA |
| Dars2 | 123.0133515 | -0.217690005 | 0.135107818 | -1.61123174 |
| Gm37634 | 2.14016008 | 1.520421981 | 0.784178944 | 1.938871214 |
| Cenpl | 31.81213049 | -0.258410643 | 0.262411691 | -0.984752782 |
| Klhl20 | 300.3613963 | 0.160280119 | 0.115251412 | 1.390699835 |
| Ankrd45 | 441.8861501 | 0.151061213 | 0.122701696 | 1.231125711 |
| 4930469G21Rik | 0 | NA | NA | NA |
| Gm6185 | 1.432525857 | -1.119249722 | 0.848826792 | -1.318584347 |
| Prdx6 | 2507.892268 | -0.943102898 | 0.222486808 | -4.238916032 |
| Gm19057 | 0 | NA | NA | NA |
| Tnfsf4 | 0.470075119 | -1.992395049 | 1.474090903 | -1.351609351 |
| Tnfsf18 | 2.177622544 | -1.201137254 | 0.736742099 | -1.630336119 |
| Gm25488 | 0 | NA | NA | NA |
| Gm37638 | 0 | NA | NA | NA |
| Gm15429 | 9.641298891 | 0.077849464 | 0.323950046 | 0.240313174 |
| Gm5049 | 0 | NA | NA | NA |
| Fasl | 19.90409191 | -0.97236505 | 0.264048575 | -3.682523376 |
| Gm37357 | 16.51548777 | 0.759330831 | 0.328034639 | 2.314788563 |
| Suco | 1095.050559 | 0.018057594 | 0.093222742 | 0.193703743 |
| Gm7496 | 0 | NA | NA | NA |
| Gm31925 | 0.049553047 | -0.443818988 | 3.724945049 | -0.119147795 |
| 4930558K02Rik | 1.743960802 | 0.163370611 | 0.843327797 | 0.193721364 |
| Pigc | 130.768612 | -0.207886338 | 0.132908486 | -1.564131415 |
| Gm38304 | 0.16282664 | -0.51179507 | 3.411119175 | -0.150037288 |
| Dnm3 | 4093.372687 | 0.371718849 | 0.059323919 | 6.265918604 |
| 2810442N19Rik | 0.321884861 | 0.517974668 | 1.782148614 | 0.290646169 |
| Gm37767 | 5.981304473 | 1.283747631 | 0.537259781 | 2.389435568 |
| Dnm3os | 3.891078412 | -1.154569186 | 0.54726851 | -2.109694173 |
| Mir199a-2 | 0 | NA | NA | NA |
| Mir214 | 0 | NA | NA | NA |
| Gm16101 | 0 | NA | NA | NA |
| Gm10176 | 1.890308503 | 1.080617395 | 0.970200154 | 1.113808724 |
| Mettl13 | 182.4821947 | -0.217011125 | 0.094542781 | -2.295374873 |
| Vamp4 | 850.2108809 | -0.018659106 | 0.056888954 | -0.327991727 |

| | | | | |
|---|---|---|---|---|
| Gm5705 | 0.038981395 | -0.443816764 | 3.724945049 | -0.119147198 |
| Myoc | 87.94067266 | 0.53099204 | 0.456133563 | 1.164115256 |
| 7420461P10Rik | 0.321637403 | -1.035830241 | 1.993003485 | -0.519733281 |
| Prrc2c | 5776.579411 | 0.041927218 | 0.0734581 | 0.570763709 |
| Gm27616 | 4.734205595 | -0.596532691 | 0.468731256 | -1.272653964 |
| Fmo4 | 5.221071454 | -0.787036563 | 0.601597365 | -1.308244698 |
| Fmo1 | 94.46220945 | 0.029804693 | 0.178704405 | 0.166782084 |
| Fmo2 | 41.24708577 | 0.033082505 | 0.273936318 | 0.120767137 |
| Fmo6 | 0.536071372 | -1.274300434 | 1.452113883 | -0.877548551 |
| Fmo3 | 0.266623809 | -0.41727176 | 1.80678292 | -0.230947368 |
| Gm37273 | 0.04393175 | 0.037062677 | 3.724945049 | 0.009949859 |
| Mroh9 | 0.04393175 | 0.037062677 | 3.724945049 | 0.009949859 |
| Gm22671 | 0 | NA | NA | NA |
| Gm37049 | 0.128271896 | 0.517956073 | 3.679089552 | 0.140783763 |
| Gm22434 | 0 | NA | NA | NA |
| Prrx1 | 291.7242276 | -1.096672564 | 0.184467434 | -5.945074085 |
| Gm20471 | 0.226778023 | 0.517974318 | 2.143518348 | 0.241646785 |
| Gm26523 | 7.941957124 | -0.511563552 | 0.4148761 | -1.23305139 |
| Gm37644 | 11.16339375 | -1.55865041 | 0.385674067 | -4.041366902 |
| Gorab | 151.8566572 | -0.089435229 | 0.119203159 | -0.750275667 |
| Gm24940 | 0 | NA | NA | NA |
| Gm29500 | 0.061920613 | -0.443816764 | 3.724945049 | -0.119147198 |
| Rpsa-ps1 | 2.716655785 | -0.061194934 | 0.571903803 | -0.107002145 |
| Mettl11b | 4.382455216 | 3.057311735 | 0.751972893 | 4.065720673 |
| Gm2343 | 0 | NA | NA | NA |
| Kifap3 | 2725.982408 | 0.421927432 | 0.052375542 | 8.055810241 |
| Scyl3 | 425.2181155 | 0.109486265 | 0.103062836 | 1.062325367 |
| BC055324 | 25.44071188 | -0.269733015 | 0.232770634 | -1.158793144 |
| Mettl18 | 30.62632286 | -0.359804226 | 0.207739592 | -1.731996406 |
| Gm37119 | 0 | NA | NA | NA |
| Sele | 15.49838402 | -0.573506291 | 0.815628424 | -0.703146524 |
| Sell | 313.8249993 | -1.959138427 | 0.497534653 | -3.937692407 |
| Selp | 27.22107536 | -1.236782069 | 0.403770465 | -3.063082059 |
| Gm16587 | 0.246813481 | -1.236567058 | 1.785372264 | -0.692610209 |
| Gm16548 | 0.29488824 | 0.503878895 | 1.833142123 | 0.274871702 |
| F5 | 60.57846656 | 0.034237607 | 0.24080964 | 0.142177061 |
| Gm16588 | 0.11197992 | 0.037073798 | 3.724945049 | 0.009952844 |
| Slc19a2 | 224.9421596 | -0.030310552 | 0.111153596 | -0.272690698 |
| Ccdc181 | 152.8998847 | -0.045830452 | 0.153127933 | -0.299295179 |
| Blzf1 | 283.9203218 | -0.063579618 | 0.083698162 | -0.759629799 |
| Nme7 | 157.2285977 | 0.270232658 | 0.108922286 | 2.480967562 |
| Gm37411 | 15.43722212 | 0.323774982 | 0.426350394 | 0.759410539 |
| Atp1b1 | 14543.60903 | 0.215546642 | 0.058738253 | 3.669612769 |
| Gm32391 | 15.06111228 | 0.248093601 | 0.285493604 | 0.868998804 |
| Gm37732 | 0 | NA | NA | NA |
| Gm26685 | 0.956865499 | 0.204620866 | 0.942538843 | 0.21709542 |
| D630023O14Rik | 0.652111363 | 0.42043819 | 1.164245056 | 0.361125167 |

| | | | | |
|---|---|---|---|---|
| Gm37754 | 0.289849351 | 1.433637773 | 1.751103892 | 0.818705149 |
| Gm37853 | 0 | NA | NA | NA |
| Gm37184 | 2.721089389 | 1.228148629 | 0.604655531 | 2.031154213 |
| Gm36945 | 0 | NA | NA | NA |
| Dpt | 2.400487258 | -0.499510903 | 0.704365361 | -0.709164491 |
| Gm32569 | 2.899492925 | 0.088617572 | 0.596113786 | 0.148658819 |
| Gm20743 | 18.05075087 | -0.340863624 | 0.294463056 | -1.157576875 |
| Gm37403 | 2.741653471 | -0.744241699 | 0.687493899 | -1.082542987 |
| Xcl1 | 1.167394941 | -0.990076999 | 0.840686023 | -1.177701273 |
| 1700029M03Rik | 0.062426873 | 0.517959912 | 3.724945049 | 0.139051692 |
| Gm26042 | 0 | NA | NA | NA |
| 4930568G15Rik | 2.644077932 | -0.12879192 | 0.612671482 | -0.210213669 |
| Tbx19 | 0.755699022 | -1.687980204 | 1.388331868 | -1.215833364 |
| Gm16550 | 1.768574416 | -0.415787547 | 0.743396798 | -0.559307691 |
| Sft2d2 | 402.8459024 | -0.066427062 | 0.126611402 | -0.524653079 |
| Tiprl | 413.2944695 | 0.052172808 | 0.092483564 | 0.564130595 |
| Gm38190 | 112.7353609 | 0.40975092 | 0.240704751 | 1.702296768 |
| Gpr161 | 332.1613004 | 0.648721567 | 0.105650025 | 6.140287892 |
| Dcaf6 | 876.2034936 | 0.272770173 | 0.065380318 | 4.172053354 |
| Gm38368 | 0.038981395 | -0.443816764 | 3.724945049 | -0.119147198 |
| Gm24637 | 0.087930927 | -0.443164241 | 3.724898049 | -0.118973522 |
| Gm37864 | 59.27064257 | 0.182474305 | 0.280907842 | 0.649587792 |
| Mpc2 | 532.4916413 | 0.003135261 | 0.058740559 | 0.053374724 |
| Gm37860 | 11.90237977 | 1.233556306 | 0.431969519 | 2.855655901 |
| Adcy10 | 12.42222246 | 0.103867664 | 0.313985993 | 0.330803494 |
| Gm18186 | 0.674049174 | -0.270209496 | 1.417126567 | -0.190674215 |
| Gm38325 | 1.250018007 | 1.63572747 | 1.119462539 | 1.461172137 |
| Gm23599 | 0.39032937 | 1.378282953 | 3.278820199 | 0.420359419 |
| Gm23208 | 0.712615371 | -0.00566051 | 1.177848356 | -0.004805805 |
| Gm38210 | 0.496695103 | -1.020840925 | 1.331396064 | -0.766744737 |
| Mpzl1 | 254.7787603 | -0.06469823 | 0.111503355 | -0.580235726 |
| Gm37559 | 11.16705096 | 1.376069736 | 0.580611377 | 2.370035775 |
| Gm18407 | 0.233595724 | 0.89637001 | 2.539297648 | 0.352999189 |
| Rcsd1 | 77.36566827 | 0.141227699 | 0.131806814 | 1.071474947 |
| AC124587.1 | 0 | NA | NA | NA |
| Gm37073 | 0.097139547 | -0.443816764 | 3.724945049 | -0.119147198 |
| Gm34767 | 0.288888109 | 1.069216709 | 2.062365076 | 0.518442017 |
| Creg1 | 384.0157734 | -0.383598009 | 0.156446008 | -2.451951396 |
| Gm36972 | 0 | NA | NA | NA |
| Gm16565 | 4.973903582 | -0.642239181 | 0.438961207 | -1.463088697 |
| Gm24808 | 0 | NA | NA | NA |
| Cd247 | 27.67774213 | -0.242547013 | 0.266928174 | -0.908660221 |
| Gm23402 | 0.088736197 | 0.51795324 | 3.724945049 | 0.139049901 |
| Gm32999 | 0.098502579 | -0.443821212 | 3.724945049 | -0.119148392 |
| Gm37856 | 0.167459142 | -0.410658575 | 2.753748719 | -0.149127105 |
| Gm37469 | 0.04393175 | 0.037062677 | 3.724945049 | 0.009949859 |
| Pou2f1 | 796.3877359 | 0.274957708 | 0.110127561 | 2.496720217 |

| | | | | |
|---|---|---|---|---|
| Gm17976 | 0 | NA | NA | NA |
| Gm26665 | 0 | NA | NA | NA |
| Gm7626 | 0 | NA | NA | NA |
| Gm37923 | 0.047948698 | 0.037064902 | 3.724945049 | 0.009950456 |
| Dusp27 | 9.458378533 | -0.121964802 | 0.392168329 | -0.311001151 |
| Gpa33 | 0.260737493 | 0.326681388 | 2.044176515 | 0.159810753 |
| Mael | 2.369035115 | 0.467850856 | 0.934558857 | 0.500611441 |
| Ildr2 | 1753.501267 | 0.154133117 | 0.097810112 | 1.57584032 |
| 5330438I03Rik | 1.619090465 | -1.376973462 | 0.94718771 | -1.453749291 |
| Gm15853 | 26.8966923 | 0.016933978 | 0.221896112 | 0.076314892 |
| Tada1 | 290.2844388 | -0.185981181 | 0.085676111 | -2.170747243 |
| Pogk | 1000.722577 | -0.006988444 | 0.070526168 | -0.099090085 |
| Gm16418 | 40.55579905 | 0.074799174 | 0.210941753 | 0.354596339 |
| Gm33354 | 0 | NA | NA | NA |
| Gm4846 | 0 | NA | NA | NA |
| Gm22992 | 0.042716945 | 0.037062677 | 3.724945049 | 0.009949859 |
| Gm4847 | 0.370767229 | 1.326470416 | 1.661248102 | 0.798478213 |
| Fmo9 | 0.364366871 | -1.351365876 | 1.521052183 | -0.888441495 |
| Gm4954 | 0 | NA | NA | NA |
| Gm37904 | 0.072025462 | 0.517959912 | 3.724945049 | 0.139051692 |
| Gm6286 | 20.94810405 | -0.205669317 | 0.25005002 | -0.822512696 |
| Gm16701 | 90.39573473 | 0.103149508 | 0.133580627 | 0.772189128 |
| Fam78b | 369.0147211 | -0.09673223 | 0.087778943 | -1.101998111 |
| Gm17868 | 0.454564407 | -1.248686342 | 1.705198605 | -0.732282057 |
| Gm37994 | 0.075043205 | 0.517957688 | 3.724945049 | 0.139051095 |
| Uck2 | 476.1615417 | -0.42962246 | 0.113759051 | -3.776600258 |
| Gm37982 | 0.042716945 | 0.037062677 | 3.724945049 | 0.009949859 |
| Gm2453 | 1.234537134 | -0.124422971 | 0.963574895 | -0.129126415 |
| Mir6347 | 0 | NA | NA | NA |
| Gm37232 | 0.061920613 | -0.443816764 | 3.724945049 | -0.119147198 |
| Tmco1 | 304.4300019 | -0.453875784 | 0.111881608 | -4.056750645 |
| Gm25911 | 23.94632856 | -0.312351336 | 1.056607168 | -0.295617279 |
| Gm24182 | 0.992449309 | 0.384670222 | 1.095090248 | 0.351268056 |
| Aldh9a1 | 305.1290703 | 0.092277643 | 0.116410627 | 0.792690883 |
| Mgst3 | 487.8742445 | 0.698965977 | 0.117612899 | 5.94293636 |
| Lrrc52 | 0.467496845 | -0.366389205 | 1.378918762 | -0.265707607 |
| Rxrg | 90.84925895 | 0.338860205 | 0.148914125 | 2.275541051 |
| Lmx1a | 12.68353485 | -0.039369597 | 0.343693782 | -0.11454847 |
| Gm20711 | 0.175176114 | -0.423385941 | 3.359108055 | -0.126041179 |
| Pbx1 | 3030.795283 | 0.227346207 | 0.082803483 | 2.745611634 |
| Gm37524 | 93.77681393 | 0.304190694 | 0.270227734 | 1.125682731 |
| Gm38381 | 118.8604427 | 0.279205108 | 0.255592462 | 1.092383968 |
| Mir6348 | 0.044544648 | 0.517957688 | 3.724945049 | 0.139051095 |
| 1700063I16Rik | 0.141317957 | 0.482895681 | 3.595358959 | 0.13431084 |
| Mir6354 | 0 | NA | NA | NA |
| Gm37839 | 0 | NA | NA | NA |
| Gm5265 | 0 | NA | NA | NA |

| | | | | |
|---|---|---|---|---|
| Nuf2 | 31.87590655 | 0.063254488 | 0.345661167 | 0.182995645 |
| Gm18900 | 0.075043205 | 0.517957688 | 3.724945049 | 0.139051095 |
| Gm36980 | 0 | NA | NA | NA |
| Rgs5 | 507.967864 | 0.303534832 | 0.172499751 | 1.759624753 |
| Rgs4 | 2513.987794 | 0.023386585 | 0.151298462 | 0.154572525 |
| Gm22114 | 0 | NA | NA | NA |
| Gm37586 | 0 | NA | NA | NA |
| 1700084C01Rik | 205.568575 | -0.016996824 | 0.177596814 | -0.095704557 |
| Gm15852 | 1.264098847 | -0.117944629 | 0.913422739 | -0.129123815 |
| Hsd17b7 | 333.9167156 | -0.198035333 | 0.128605006 | -1.539872658 |
| 3110045C21Rik | 6.064642518 | -0.68659588 | 0.391470998 | -1.753886967 |
| Ddr2 | 402.5981112 | -0.839707501 | 0.135667401 | -6.18945669 |
| Uap1 | 505.8478976 | -0.052230771 | 0.069718305 | -0.749168682 |
| Gm37748 | 2.515876782 | -0.672836037 | 0.720662934 | -0.933634859 |
| Gm37502 | 0.166709681 | 0.517964155 | 2.76763122 | 0.187150713 |
| Uhmk1 | 4307.233663 | 0.146450135 | 0.065862872 | 2.223561326 |
| 4930500M09Rik | 0.668413162 | -1.324944507 | 1.240880071 | -1.067745818 |
| Sh2d1b2 | 3.139971286 | -2.32659306 | 0.706663451 | -3.292363652 |
| Gm6345 | 2.814645699 | 0.619719081 | 0.605161429 | 1.024055816 |
| Sh2d1b1 | 8.798894628 | -1.392654227 | 0.398599864 | -3.493865285 |
| Gm7694 | 71.28747091 | -0.165657681 | 0.146527537 | -1.130556647 |
| Nos1ap | 949.0976046 | 0.55352901 | 0.171136035 | 3.23443867 |
| 1700015E13Rik | 0 | NA | NA | NA |
| Gm7299 | 0.686918141 | -2.002102851 | 1.234146383 | -1.622257197 |
| Olfml2b | 107.0665219 | 0.583512396 | 0.120648022 | 4.836485378 |
| Atf6 | 1426.693892 | 0.134895168 | 0.057425934 | 2.349028707 |
| Gm23523 | 2.532308763 | 0.860878825 | 0.772898399 | 1.113831812 |
| Gm25235 | 4.986884554 | 1.05844381 | 0.69727747 | 1.517966455 |
| Gm9929 | 5.148652476 | 1.284743494 | 0.543722442 | 2.362866408 |
| Dusp12 | 141.0307956 | -0.047534117 | 0.126505741 | -0.375746717 |
| Gm26620 | 3.756679203 | 0.86801777 | 0.528218383 | 1.643293378 |
| Fcrlb | 20.36960655 | -1.294105474 | 0.400970545 | -3.227432762 |
| Fcrla | 4.0864204 | -1.345259698 | 0.794940093 | -1.692278085 |
| Gm2962 | 27.33055226 | -0.336619357 | 0.248044896 | -1.357090439 |
| Gm37030 | 0.072025462 | 0.517959912 | 3.724945049 | 0.139051692 |
| Fcgr2b | 292.4194751 | -0.329390133 | 0.188738628 | -1.745218435 |
| Gm38204 | 0.062426873 | 0.517959912 | 3.724945049 | 0.139051692 |
| Gm10522 | 0.106465261 | 0.037073798 | 3.724945049 | 0.009952844 |
| Fcgr4 | 282.2570289 | -1.177613015 | 0.323677954 | -3.638224357 |
| Gm26110 | 0.545665175 | -2.992176275 | 1.267709306 | -2.360301578 |
| Fcgr3 | 124.3411756 | -0.505862282 | 0.139724202 | -3.620434223 |
| Mir6546 | 0.741170898 | -1.718113022 | 1.144404327 | -1.501316433 |
| Gm27552 | 0.046345709 | -0.443818988 | 3.724945049 | -0.119147795 |
| Gm27844 | 0 | NA | NA | NA |
| Cfap126 | 16.16273733 | -0.140962665 | 0.350026493 | -0.402719988 |
| Sdhc | 664.894161 | -0.171245009 | 0.107934849 | -1.586559021 |
| Mpz | 3.16266539 | -0.12112975 | 0.59616374 | -0.203182015 |

| | | | | |
|---|---|---|---|---|
| Pcp4l1 | 1008.326566 | 0.150734156 | 0.109793817 | 1.372883832 |
| Nr1i3 | 2.346028351 | 1.088925122 | 0.703705084 | 1.547416876 |
| Tomm40l | 561.1166494 | 0.045102272 | 0.092859232 | 0.485705853 |
| Apoa2 | 0.57971966 | 0.952231632 | 1.303773434 | 0.73036588 |
| Fcer1g | 255.0199519 | -0.929734111 | 0.223769535 | -4.154873496 |
| Ndufs2 | 1322.178398 | -0.004449434 | 0.074190553 | -0.059973055 |
| Adamts4 | 381.9510903 | -0.50132109 | 0.123925702 | -4.045335877 |
| B4galt3 | 352.3072303 | 0.001377218 | 0.086711916 | 0.01588268 |
| Ppox | 128.0565181 | 0.104268879 | 0.137314144 | 0.759345511 |
| Usp21 | 286.6155179 | -0.076974101 | 0.100608799 | -0.765083193 |
| Ufc1 | 373.6742995 | 0.095283715 | 0.093548712 | 1.018546515 |
| Rpl27-ps1 | 1.899360943 | -0.604454975 | 0.723664742 | -0.835269346 |
| Gm25151 | 0 | NA | NA | NA |
| Gm20045 | 91.15361111 | -0.13771032 | 0.229025831 | -0.601287286 |
| Dedd | 384.4686255 | -0.093867384 | 0.08613114 | -1.089819371 |
| Nit1 | 175.9259044 | -0.109036555 | 0.10223802 | -1.06649713 |
| Pfdn2 | 372.1457708 | 0.232150888 | 0.170868218 | 1.358654585 |
| Gm37154 | 10.37578366 | 0.892613107 | 0.40001713 | 2.231437208 |
| Klhdc9 | 28.71586709 | 1.220380544 | 0.233645777 | 5.223208224 |
| Gm37097 | 0.580816694 | 2.054385608 | 1.364552418 | 1.505538065 |
| Pvrl4 | 89.21914686 | -0.238164892 | 0.124217486 | -1.917321779 |
| Arhgap30 | 261.1304874 | -0.844736477 | 0.165047602 | -5.118138446 |
| Usf1 | 598.6805458 | 0.216465107 | 0.079841242 | 2.711194136 |
| Tstd1 | 8.585507017 | -0.566412603 | 0.369123717 | -1.534479029 |
| Gm26641 | 0.661565226 | 0.555314065 | 1.244815694 | 0.446101433 |
| F11r | 61.91267919 | -0.631166342 | 0.15584314 | -4.050010432 |
| B930036N10Rik | 1.495946937 | 0.711549413 | 0.85025198 | 0.836868868 |
| Gm24871 | 0 | NA | NA | NA |
| Alyref2 | 25.03101443 | -0.383114333 | 0.328040464 | -1.167887426 |
| Gm9765 | 0.094831987 | 0.517957688 | 3.724945049 | 0.139051095 |
| Itln1 | 0.674729533 | 1.854366042 | 1.415761975 | 1.309800712 |
| Gm37450 | 0 | NA | NA | NA |
| Cd244 | 24.4635172 | -0.606505726 | 0.291238144 | -2.082507869 |
| Ly9 | 63.06084566 | -1.069095695 | 0.234136622 | -4.566119065 |
| Slamf7 | 242.896543 | -1.864798601 | 0.432694822 | -4.309731723 |
| Gm37787 | 0.62421388 | 1.138880527 | 1.328680979 | 0.857151224 |
| Mir7683 | 0.465770878 | -1.229591699 | 1.642385578 | -0.748662017 |
| A630035G10Rik | 0.725223345 | -0.694719962 | 1.212507424 | -0.572961409 |
| Gm37981 | 0.097139547 | -0.443816764 | 3.724945049 | -0.119147198 |
| Cd48 | 54.8175684 | -1.091662998 | 0.26616303 | -4.101482448 |
| Gm37010 | 3.46582388 | -0.788032164 | 0.631110724 | -1.24864328 |
| Tma7-ps | 109.4366652 | -0.06848486 | 0.185543937 | -0.369103195 |
| Gm37693 | 0 | NA | NA | NA |
| Gm19030 | 0.512648943 | -2.173220435 | 3.239288849 | -0.670894303 |
| Gm26412 | 0 | NA | NA | NA |
| Gm22950 | 0 | NA | NA | NA |
| Slamf1 | 7.891299907 | -0.205478462 | 0.340941509 | -0.602679511 |

| | | | | |
|---|---|---|---|---|
| Cd84 | 247.5967198 | -1.404264453 | 0.229108335 | -6.129259574 |
| Gm10521 | 6.960180967 | -1.444064104 | 0.548152494 | -2.634420384 |
| Gm37065 | 0.847774459 | 1.048597938 | 1.375350189 | 0.762422506 |
| Slamf6 | 33.56933231 | -0.795275415 | 0.247041879 | -3.219192701 |
| Vangl2 | 264.8998109 | 0.416134869 | 0.100549575 | 4.138603948 |
| Nhlh1 | 18.19327503 | 1.671161488 | 0.456860926 | 3.657921687 |
| Ncstn | 716.1302082 | -0.139600041 | 0.073571114 | -1.897484391 |
| Copa | 2652.049841 | -0.199551083 | 0.066806142 | -2.987017 |
| Gm38164 | 3.312973942 | 0.731930528 | 0.577773042 | 1.266813221 |
| Gm37756 | 0 | NA | NA | NA |
| Pex19 | 729.4998742 | 0.089713991 | 0.078798371 | 1.138525964 |
| Gm10171 | 8.347475085 | -0.233958076 | 0.362336493 | -0.645692832 |
| Dcaf8 | 1703.591968 | 0.301783095 | 0.051809739 | 5.824833312 |
| Pea15a | 5072.520127 | -0.273492228 | 0.129743013 | -2.107953415 |
| Casq1 | 10.6483708 | 0.515562293 | 0.333317179 | 1.546761839 |
| Gm17224 | 5.574451365 | 1.777037136 | 0.869419688 | 2.043934776 |
| Atp1a4 | 4.799918733 | 1.638290598 | 0.797621168 | 2.053970811 |
| Gm37950 | 1.138245853 | 2.794194731 | 1.37318983 | 2.034820437 |
| Igsf8 | 2030.429245 | 0.402930282 | 0.092933734 | 4.335672999 |
| Atp1a2 | 9906.868086 | 0.243983767 | 0.137433621 | 1.77528442 |
| Kcnj9 | 920.7093185 | 0.455344711 | 0.167970329 | 2.710863952 |
| Gm36937 | 1.071485715 | 0.259990816 | 0.980640365 | 0.26512351 |
| Kcnj10 | 4831.592137 | -0.116685496 | 0.155409562 | -0.750825718 |
| Pigm | 357.594723 | -0.079354199 | 0.126671089 | -0.626458646 |
| Gm37145 | 0.222528853 | 0.012312771 | 2.572758162 | 0.004785825 |
| Gm7804 | 0.35063021 | -1.35322823 | 1.90975902 | -0.708585856 |
| Slamf9 | 59.21569036 | -0.68007562 | 0.21120948 | -3.219910487 |
| Igsf9 | 57.64957815 | 0.005579492 | 0.193960634 | 0.028766105 |
| Tagln2 | 422.7865102 | -1.122807511 | 0.206151541 | -5.446515237 |
| Gm37125 | 0.235978602 | -0.466730289 | 2.12183998 | -0.219964886 |
| Cfap45 | 24.01669692 | -0.361101995 | 0.317759785 | -1.136399294 |
| 4933439K11Rik | 2.636750292 | -0.199251591 | 0.764216605 | -0.260726592 |
| Vsig8 | 4.194719996 | -0.402917617 | 0.482609587 | -0.834872799 |
| Slamf8 | 27.5563302 | 0.109476488 | 0.260103041 | 0.420896609 |
| Fcrl6 | 0.659227985 | -2.286644411 | 1.373973817 | -1.664256176 |
| Dusp23 | 20.89498152 | -0.089892959 | 0.280796847 | -0.320135215 |
| Gm2710 | 3.560626721 | 0.524692081 | 0.53295045 | 0.984504435 |
| Gm22383 | 0 | NA | NA | NA |
| Crp | 0.134618652 | 0.998858087 | 2.958498178 | 0.337623357 |
| Gm38051 | 0 | NA | NA | NA |
| Gm24817 | 0 | NA | NA | NA |
| 9430083B18Rik | 0 | NA | NA | NA |
| Apcs | 0.049553047 | -0.443818988 | 3.724945049 | -0.119147795 |
| Gm26077 | 0.04393175 | 0.037062677 | 3.724945049 | 0.009949859 |
| Olfr16 | 0 | NA | NA | NA |
| RP23-127L7.9 | 0.046345709 | -0.443818988 | 3.724945049 | -0.119147795 |
| RP23-127L7.8 | 0 | NA | NA | NA |

| | | | | |
|---|---|---|---|---|
| Olfr1409-ps1 | 0 | NA | NA | NA |
| Olfr1408 | 0.042716945 | 0.037062677 | 3.724945049 | 0.009949859 |
| Olfr1407-ps1 | 0 | NA | NA | NA |
| Olfr1406 | 0.153801061 | -0.443820463 | 2.838638591 | -0.15634976 |
| Olfr218 | 0 | NA | NA | NA |
| Olfr1404 | 0 | NA | NA | NA |
| Fcer1a | 0.138573873 | -0.444788617 | 3.62882948 | -0.122570823 |
| Olfr418 | 0 | NA | NA | NA |
| Mptx2 | 0.116944185 | -1.028246102 | 3.719337068 | -0.27645951 |
| Ackr1 | 337.5743771 | 0.151859105 | 0.114554095 | 1.325654096 |
| Cadm3 | 5210.19823 | 0.454954746 | 0.102527246 | 4.437403348 |
| Aim2 | 59.97870837 | -0.227385043 | 0.181859205 | -1.250335628 |
| Gm4955 | 118.0231463 | -1.416092829 | 0.395005322 | -3.584996831 |
| BC094916 | 233.0659656 | -1.82971691 | 0.460369963 | -3.974448938 |
| AC084073.1 | 0 | NA | NA | NA |
| Pydc4 | 70.39675328 | -1.259397863 | 0.338124959 | -3.72465217 |
| Pyhin1 | 372.8526333 | -1.464244449 | 0.34644708 | -4.226459204 |
| AC117682.1 | 0 | NA | NA | NA |
| Pydc3 | 163.3167361 | -1.371153433 | 0.360553908 | -3.80290826 |
| Gm18752 | 7.837557856 | -1.673104182 | 0.44634494 | -3.748455579 |
| Gm7871 | 0.290529814 | 0.506947466 | 1.986293731 | 0.255222809 |
| AI607873 | 591.904346 | -1.636707574 | 0.327933001 | -4.990981593 |
| Ifi204 | 992.1935414 | -1.703771808 | 0.360470516 | -4.726521958 |
| Gm16340 | 27.30976778 | -1.894540965 | 0.281400687 | -6.732538518 |
| Gm7897 | 1.063697798 | -3.502800827 | 1.100991804 | -3.181495826 |
| Gm38050 | 43.49727825 | -0.591462971 | 0.299430878 | -1.975290508 |
| Mndal | 289.3519551 | -1.263519754 | 0.280016895 | -4.512298287 |
| Mnda | 254.7490093 | -1.512844734 | 0.466796899 | -3.240905709 |
| Ifi203 | 735.2718426 | -1.405161578 | 0.305601502 | -4.598019213 |
| Ifi202b | 1532.515879 | -1.728354433 | 0.391768242 | -4.411675704 |
| Gm18445 | 3.120173769 | -1.08145288 | 0.759473142 | -1.423951448 |
| Ifi205 | 186.9498831 | -1.797956448 | 0.557399506 | -3.225615428 |
| Olfr433 | 1.889349206 | -1.746730563 | 0.812435889 | -2.149991879 |
| Olfr432 | 0.498029041 | -2.156156527 | 1.590913763 | -1.355294408 |
| Olfr431-ps1 | 0.481158361 | -2.133405736 | 1.372631754 | -1.554244778 |
| Olfr430 | 0 | NA | NA | NA |
| Olfr429 | 0.127515837 | -1.208141086 | 3.685333689 | -0.327824069 |
| RP23-343E12.13 | 0.066050875 | -0.443818988 | 3.724945049 | -0.119147795 |
| Olfr427 | 0 | NA | NA | NA |
| Gm37417 | 0 | NA | NA | NA |
| Olfr231 | 0 | NA | NA | NA |
| Olfr424 | 0 | NA | NA | NA |
| Olfr258-ps1 | 0 | NA | NA | NA |
| Olfr421-ps1 | 0.097139547 | -0.443816764 | 3.724945049 | -0.119147198 |
| Olfr420 | 0 | NA | NA | NA |
| RP23-343E12.14 | 0 | NA | NA | NA |
| Spta1 | 4.811948516 | 0.584433278 | 0.513910747 | 1.137227197 |

| | | | | |
|---|---|---|---|---|
| Olfr419 | 0 | NA | NA | NA |
| Mptx1 | 0.066050875 | -0.443818988 | 3.724945049 | -0.119147795 |
| Olfr417 | 0 | NA | NA | NA |
| Olfr416-ps1 | 0 | NA | NA | NA |
| Olfr248 | 0 | NA | NA | NA |
| Olfr414 | 0.117046891 | 0.037062677 | 3.724945049 | 0.009949859 |
| Olfr220 | 0.244461041 | -1.004375557 | 2.513992492 | -0.399514143 |
| Fmn2 | 1375.015257 | 0.191641273 | 0.057753079 | 3.318286665 |
| Grem2 | 314.3592299 | -0.40417165 | 0.170734085 | -2.367258127 |
| Rgs7 | 979.1331399 | 0.120679759 | 0.094392635 | 1.278487025 |
| 5730408A14Rik | 66.38004792 | 0.217714025 | 0.242384362 | 0.898218114 |
| Gm36307 | 0.200008102 | -1.706133596 | 2.610702492 | -0.653515137 |
| Fh1 | 588.3356071 | -0.090604844 | 0.078985374 | -1.147109134 |
| Kmo | 33.53845944 | -0.499553454 | 0.2687352 | -1.85890592 |
| Opn3 | 46.18549766 | -0.528704751 | 0.212081752 | -2.492928999 |
| Chml | 277.8742486 | -0.120260425 | 0.082385466 | -1.459728657 |
| Wdr64 | 0.404309506 | 0.037077296 | 1.593046192 | 0.023274464 |
| Gm25560 | 0.044191549 | 0.037062677 | 3.724945049 | 0.009949859 |
| Exo1 | 11.02950793 | -0.267023086 | 0.318237025 | -0.839069829 |
| Gm23805 | 0 | NA | NA | NA |
| Gm38100 | 0.094097695 | 0.037071574 | 3.724945049 | 0.009952247 |
| Gm37668 | 9.723507462 | 0.151222372 | 0.412763322 | 0.366365819 |
| Pld5 | 273.9533392 | -0.00513891 | 0.10157425 | -0.050592645 |
| B020018G12Rik | 7.849527246 | 0.331243287 | 0.471130612 | 0.703081647 |
| Gm38081 | 26.68633732 | 0.607329675 | 0.337637262 | 1.798763774 |
| Gm37916 | 0.048949532 | 0.037064902 | 3.724945049 | 0.009950456 |
| Gm38101 | 8.572675195 | -0.308098207 | 0.386270835 | -0.797622236 |
| Gm17965 | 0.042716945 | 0.037062677 | 3.724945049 | 0.009949859 |
| Gm38152 | 0.194279094 | -1.017823543 | 3.719440177 | -0.273649661 |
| Gm36536 | 0.394395078 | -0.068072052 | 1.616733835 | -0.042104675 |
| Gm17966 | 0.161076545 | -0.410543693 | 2.787893798 | -0.147259445 |
| Gm26104 | 0 | NA | NA | NA |
| Cep170 | 2565.856711 | 0.047374858 | 0.060991506 | 0.77674518 |
| 3110062G12Rik | 32.76611244 | 0.215017988 | 0.225737636 | 0.952512803 |
| Gm38104 | 128.4296084 | 0.057465922 | 0.188839041 | 0.304311658 |
| Gm36992 | 0.232229192 | 0.037061516 | 3.136228082 | 0.011817226 |
| Mir350 | 3.5612593 | 0.11015291 | 0.57192931 | 0.19259882 |
| Gm25288 | 0 | NA | NA | NA |
| Sdccag8 | 235.1494786 | 0.039860672 | 0.169476831 | 0.235198355 |
| Gm38158 | 1.079846962 | 0.302596135 | 1.001050869 | 0.302278479 |
| Gm38053 | 0.047948698 | 0.037064902 | 3.724945049 | 0.009950456 |
| Gm25993 | 2.439810687 | -0.545647819 | 0.750324905 | -0.72721539 |
| Gm37246 | 8.305594911 | 0.259999983 | 0.501368076 | 0.51858105 |
| Gm37490 | 6.173150445 | -0.278515991 | 0.476260282 | -0.584797855 |
| Gm15423 | 5.069600121 | 0.992966493 | 0.498168474 | 1.993234306 |
| Gm37519 | 17.14460392 | -0.128979425 | 0.461956994 | -0.279202234 |
| Akt3 | 2779.8936 | 0.137911564 | 0.061625965 | 2.237880815 |

| | | | | |
|---|---|---|---|---|
| Gm37960 | 79.23742208 | 0.211456533 | 0.219700613 | 0.962475842 |
| Gm37106 | 32.57544906 | 0.265319469 | 0.2673449 | 0.992423901 |
| Gm37598 | 54.69327838 | 0.529406474 | 0.294904041 | 1.795182161 |
| Gm38146 | 68.83238682 | 0.512739124 | 0.202266832 | 2.534963929 |
| Gm37706 | 171.6006066 | 0.158803573 | 0.242948251 | 0.653651846 |
| Gm37463 | 61.45825268 | 0.229472398 | 0.278356614 | 0.824382777 |
| Gm38328 | 3.391924355 | -0.336713518 | 0.611243104 | -0.550866776 |
| Gm26801 | 7.020008498 | -0.467191851 | 0.392190095 | -1.191238272 |
| Zbtb18 | 3593.895032 | 0.436332193 | 0.103792836 | 4.203875818 |
| Gm29856 | 8.835626795 | 2.3439389 | 0.528754993 | 4.432939513 |
| Gm37306 | 1.917068816 | 2.391963973 | 0.951295638 | 2.514427563 |
| Gm37280 | 0.17486102 | -0.738756738 | 2.713408578 | -0.272261518 |
| Gm37858 | 0.209701617 | -0.800773287 | 2.585494854 | -0.30971761 |
| 2310043L19Rik | 0.252619382 | 0.555176813 | 2.082463083 | 0.266596233 |
| 1700016C15Rik | 0.143022002 | 0.03610674 | 3.547735707 | 0.010177404 |
| Adss | 882.0855769 | 0.110466676 | 0.071236622 | 1.550700653 |
| Gm16585 | 0 | NA | NA | NA |
| Gm7068 | 2.009888605 | 1.882176094 | 0.934812947 | 2.01342536 |
| Gm16432 | 157.8944835 | 0.817643343 | 0.139448868 | 5.863391749 |
| Gm37313 | 0.130017973 | -0.444798045 | 3.662714201 | -0.121439463 |
| Gm25312 | 0 | NA | NA | NA |
| Gm8010 | 62.50190762 | 0.313183472 | 0.14265161 | 2.195442948 |
| Desi2 | 449.7295651 | -0.157379031 | 0.083505077 | -1.884664217 |
| Gm8023 | 0.25372768 | -1.447354027 | 2.070088829 | -0.699174841 |
| 4930527J03Rik | 0.49872596 | -0.913308002 | 1.600175024 | -0.570755067 |
| Gm37486 | 42.26792344 | -0.08846292 | 0.182373771 | -0.485063829 |
| B230369F24Rik | 55.62345512 | -0.328096149 | 0.161044665 | -2.037299094 |
| Cox20 | 107.0345127 | -0.230647034 | 0.192272428 | -1.199584547 |
| Gm16586 | 574.5462637 | 0.274640203 | 0.105165626 | 2.611501616 |
| Gm28062 | 3.424742753 | 1.132105919 | 0.621978077 | 1.820170134 |
| Hnrnpu | 2062.938983 | -0.08894923 | 0.065641229 | -1.355081723 |
| Gm38300 | 25.20762262 | -0.014653137 | 0.248784497 | -0.058898915 |
| Gm27458 | 0.042716945 | 0.037062677 | 3.724945049 | 0.009949859 |
| Gm24919 | 0.075043205 | 0.517957688 | 3.724945049 | 0.139051095 |
| Efcab2 | 203.061846 | 0.374026046 | 0.095872577 | 3.901282927 |
| Kif26b | 843.0367765 | 0.297769164 | 0.110847536 | 2.686294839 |
| Gm24405 | 1.624132796 | 0.545280441 | 0.922645468 | 0.590996715 |
| Gm34176 | 4.619527546 | 1.561863369 | 0.639865798 | 2.440923355 |
| Gm16564 | 2.140983095 | 1.739613779 | 0.8569811 | 2.02993249 |
| Smyd3 | 427.0445891 | 0.455194021 | 0.079729248 | 5.709247639 |
| Gm5561 | 0.317629712 | 0.551551936 | 1.798381837 | 0.306693453 |
| Tfb2m | 166.5859778 | 0.201821717 | 0.095229696 | 2.119314937 |
| Gm15480 | 0 | NA | NA | NA |
| Cnst | 707.9421141 | 0.155565327 | 0.057337376 | 2.713157428 |
| Sccpdh | 630.1321158 | -0.381339828 | 0.096555773 | -3.949425461 |
| Gm1305 | 0.735920534 | -1.102824255 | 1.225016775 | -0.90025237 |
| Ahctf1 | 1340.102284 | 0.084802666 | 0.081946363 | 1.034855753 |

| | | | | |
|---|---|---|---|---|
| Gm10518 | 1.39994924 | 0.995174971 | 0.918705499 | 1.083236111 |
| Gm37070 | 7.2364301 | -0.113540348 | 0.402225126 | -0.282280595 |
| Cdc42bpa | 5990.889057 | 0.366849126 | 0.059723301 | 6.14247907 |
| Gm38169 | 41.0227843 | 0.675292084 | 0.283676444 | 2.380501088 |
| Gm38331 | 0.311669542 | -0.444361441 | 1.750282711 | -0.25387981 |
| Adck3 | 280.2963435 | 0.64165096 | 0.12830489 | 5.000986009 |
| Gm37336 | 1.602826546 | 1.034137824 | 0.939175759 | 1.101112133 |
| Psen2 | 248.7304506 | -0.346263124 | 0.125510884 | -2.75882946 |
| Gm31728 | 3.493469674 | 0.008641728 | 0.648973898 | 0.013315986 |
| Gm5069 | 31.42240588 | 0.23312992 | 0.223977885 | 1.040861336 |
| Itpkb | 1738.483568 | -0.506762875 | 0.134770293 | -3.760197166 |
| Gm37033 | 72.29879847 | -0.431898779 | 0.231113992 | -1.868769503 |
| Gm37390 | 9.932302052 | -0.121697004 | 0.384015579 | -0.316906424 |
| Gm36933 | 22.96547666 | -0.690423672 | 0.318987677 | -2.164421144 |
| 6330403A02Rik | 2006.529945 | 1.641922455 | 0.230341027 | 7.128224086 |
| Gm37984 | 25.25468169 | 1.642235287 | 0.394062418 | 4.16744965 |
| Gm37267 | 0.197212531 | 1.332580241 | 2.626830053 | 0.507295948 |
| Parp1 | 872.0659159 | -0.05118298 | 0.087575236 | -0.584445809 |
| Mir6904 | 1.534744651 | 0.816170597 | 1.088205998 | 0.750014793 |
| Lin9 | 91.26341556 | -0.127673829 | 0.156629056 | -0.815135019 |
| Mixl1 | 0.441773728 | -1.250630585 | 1.358665622 | -0.920484456 |
| Gm8101 | 0 | NA | NA | NA |
| Acbd3 | 598.1054057 | -0.410037153 | 0.080824574 | -5.073174335 |
| Gm37768 | 0.122991902 | 0.517955464 | 3.724945049 | 0.139050498 |
| Gm38293 | 0.815026891 | 0.715973802 | 1.179067773 | 0.60723719 |
| H3f3a | 906.3881747 | -0.05672577 | 0.113708939 | -0.49886817 |
| Gm17275 | 0.368617932 | -0.590429608 | 1.728628142 | -0.341559641 |
| H3f3aos | 16.19441088 | -0.100275186 | 0.300481638 | -0.333714854 |
| Gm24836 | 0 | NA | NA | NA |
| Sde2 | 325.6542778 | -0.389466909 | 0.101747074 | -3.827794695 |
| Gm20305 | 1.024514732 | -0.101537472 | 0.895811535 | -0.113346912 |
| Gm18036 | 0 | NA | NA | NA |
| Lefty2 | 3.909599014 | 3.439210598 | 0.903879311 | 3.804944485 |
| Pycr2 | 201.3612007 | -0.282753633 | 0.11029171 | -2.563688896 |
| Lefty1 | 9.891254432 | 0.724717795 | 0.442956718 | 1.636091668 |
| Tmem63a | 404.5987899 | -0.278575132 | 0.124185202 | -2.243223252 |
| 2210411M09Rik | 0.048949532 | 0.037064902 | 3.724945049 | 0.009950456 |
| Gm36961 | 0 | NA | NA | NA |
| Ephx1 | 210.2340185 | 0.274479348 | 0.132756665 | 2.067537223 |
| Gm37431 | 0.049553047 | -0.443818988 | 3.724945049 | -0.119147795 |
| Gm37086 | 0 | NA | NA | NA |
| 9130409I23Rik | 0.044544648 | 0.517957688 | 3.724945049 | 0.139051095 |
| Gm17967 | 0.193566712 | -1.675392576 | 2.630757503 | -0.63684797 |
| Gm37406 | 0 | NA | NA | NA |
| Gm6606 | 0.061920613 | -0.443816764 | 3.724945049 | -0.119147198 |
| Nvl | 960.5660003 | 0.067858656 | 0.120168788 | 0.564694519 |
| Fgfr3-ps | 0 | NA | NA | NA |

| | | | | |
|---|---|---|---|---|
| Cnih4 | 409.2563578 | -0.280903705 | 0.072722892 | -3.862658609 |
| Wdr26 | 2480.374956 | -0.017821862 | 0.077069547 | -0.231243895 |
| A430110L20Rik | 18.50512536 | 0.161527041 | 0.351806057 | 0.459136611 |
| Rpl35a-ps2 | 28.95887429 | -0.59509574 | 0.220608877 | -2.697514936 |
| Gm8146 | 0.858591824 | -0.268580361 | 1.13392767 | -0.236858459 |
| Cnih3 | 290.3343059 | 0.500036679 | 0.122480333 | 4.082587513 |
| 2900069G24Rik | 4.593070302 | 0.660531155 | 0.625122588 | 1.056642597 |
| Gm16547 | 0 | NA | NA | NA |
| Gm16539 | 0 | NA | NA | NA |
| Ccdc121 | 5.062754143 | -0.446221265 | 0.529222818 | -0.843163314 |
| Dnah14 | 10.40201644 | 1.134710379 | 0.336476318 | 3.372333559 |
| Gm37664 | 0.09176491 | 0.921395572 | 3.723061103 | 0.247483333 |
| Gm23690 | 1.324611736 | 0.211106632 | 1.058666751 | 0.199408012 |
| Lbr | 304.5066716 | -0.194673188 | 0.105710696 | -1.841565676 |
| Gm26004 | 0.608946563 | -2.273964976 | 1.171929899 | -1.940359212 |
| Gm5533 | 0.046345709 | -0.443818988 | 3.724945049 | -0.119147795 |
| Enah | 3997.842318 | 0.21020174 | 0.064558107 | 3.25600844 |
| 2700078F05Rik | 123.0838922 | 0.395688353 | 0.210655215 | 1.878369609 |
| Gm38359 | 39.36611411 | 0.45509415 | 0.256808309 | 1.772116144 |
| Gm37018 | 80.20486681 | 0.353197167 | 0.283296531 | 1.246740176 |
| Gm37258 | 42.3525549 | 0.253416138 | 0.243263412 | 1.041735527 |
| Gm37177 | 0.238685809 | 1.596407295 | 2.530977483 | 0.630747332 |
| Srp9 | 514.3250057 | -0.022568328 | 0.133301814 | -0.169302485 |
| Gm17968 | 0.075043205 | 0.517957688 | 3.724945049 | 0.139051095 |
| Vmn1r1 | 0 | NA | NA | NA |
| Degs1 | 871.076032 | -0.127117066 | 0.092185461 | -1.378927484 |
| 1700047M11Rik | 71.18991338 | 0.796273963 | 0.250969394 | 3.172793107 |
| Fbxo28 | 708.639217 | -0.292845575 | 0.102451727 | -2.858376164 |
| Gm5706 | 0.042716945 | 0.037062677 | 3.724945049 | 0.009949859 |
| Gm10517 | 2.305351242 | -0.060785718 | 0.750143584 | -0.081032111 |
| Trp53bp2 | 449.2976652 | 0.116946265 | 0.082471113 | 1.418027005 |
| Capn2 | 1213.867559 | -0.870114123 | 0.105180634 | -8.272569696 |
| Gm37885 | 64.78973926 | 1.396239694 | 0.285394327 | 4.892317616 |
| Gm37069 | 19.58732916 | 1.592844931 | 0.485740295 | 3.279211027 |
| Gm8197 | 0 | NA | NA | NA |
| Capn8 | 0.352854448 | 1.506068274 | 1.558269985 | 0.966500214 |
| Gm37516 | 0.044544648 | 0.517957688 | 3.724945049 | 0.139051095 |
| Gm37218 | 0 | NA | NA | NA |
| Ccdc185 | 0.590978791 | -0.766591725 | 1.205135558 | -0.636104146 |
| Susd4 | 1048.966416 | 0.39146724 | 0.061217012 | 6.394745977 |
| Gm37542 | 0.158320866 | -0.93971554 | 2.813533518 | -0.333998346 |
| Tlr5 | 15.84910574 | -1.03392214 | 0.471622913 | -2.192264437 |
| Gm38079 | 0.535911413 | -1.142094422 | 1.330288261 | -0.858531535 |
| Gm24975 | 0.165856615 | 0.955583004 | 2.759407178 | 0.346300108 |
| Disp1 | 84.25718148 | -0.286282294 | 0.125112426 | -2.288200338 |
| BC085271 | 7.295759161 | -0.144712927 | 0.435335663 | -0.33241689 |
| Brox | 846.8803349 | -0.211922495 | 0.057799006 | -3.666542199 |

| | | | | |
|---|---|---|---|---|
| Aida | 1155.608412 | -0.060809642 | 0.147391393 | -0.412572546 |
| Mia3 | 1173.748108 | 0.057238879 | 0.063489378 | 0.901550477 |
| Gm37339 | 0.824550338 | 0.332005115 | 1.101656244 | 0.301369068 |
| Taf1a | 110.8777169 | -0.344770043 | 0.149960408 | -2.299073784 |
| Gm37986 | 3.238027032 | -2.399609879 | 0.878395899 | -2.731809063 |
| Hhipl2 | 10.87507933 | -1.586126695 | 0.416402319 | -3.809120703 |
| Gm37214 | 0.761058687 | -1.897000513 | 1.231247984 | -1.540713599 |
| Gm33973 | 0.152667883 | 0.998859635 | 2.845214387 | 0.351066563 |
| Gm8214 | 0.087930927 | -0.443164241 | 3.724898049 | -0.118973522 |
| Gm34068 | 0.293108411 | 1.003668245 | 3.717558017 | 0.26998052 |
| Gm38332 | 0.075043205 | 0.517957688 | 3.724945049 | 0.139051095 |
| Gm38108 | 0.089089296 | 0.957155977 | 3.723939561 | 0.257027796 |
| Dusp10 | 189.900222 | -0.310698505 | 0.160867205 | -1.931397426 |
| 1700056E22Rik | 3.04764041 | -1.246599432 | 0.568129673 | -2.194216375 |
| Gm37223 | 0.046345709 | -0.443818988 | 3.724945049 | -0.119147795 |
| Gm37712 | 0 | NA | NA | NA |
| 2900092O11Rik | 0.068097359 | 0.037064902 | 3.724945049 | 0.009950456 |
| 1700112H15Rik | 0.11193333 | 0.037073798 | 3.724945049 | 0.009952844 |
| Gm23349 | 0 | NA | NA | NA |
| Gm37800 | 0.214735602 | -0.274798115 | 2.168920121 | -0.126698126 |
| Rpl21-ps1 | 1.040400475 | 1.588512924 | 0.995361442 | 1.595915671 |
| Gm38358 | 0.24875112 | 0.517974759 | 1.849989491 | 0.279987947 |
| 4930488B22Rik | 0 | NA | NA | NA |
| Hlx | 28.3948279 | -0.693762926 | 0.202647411 | -3.423497613 |
| Gm34342 | 0.33098618 | -1.529167838 | 3.693033812 | -0.414068193 |
| 1-Mar | 1.626573545 | -0.292430396 | 0.76618907 | -0.381668712 |
| 2-Mar | 436.1785284 | -0.083986854 | 0.108953756 | -0.770848641 |
| Mir1981 | 0 | NA | NA | NA |
| Gm23780 | 0 | NA | NA | NA |
| C130074G19Rik | 294.2439593 | 0.341282093 | 0.174962203 | 1.950604682 |
| Mark1 | 1064.898702 | 0.105686308 | 0.079157625 | 1.335137435 |
| Gm38251 | 0.231207844 | -1.640880984 | 3.039940608 | -0.539774027 |
| Gm34732 | 0.152865955 | -0.389869207 | 3.499663388 | -0.111401916 |
| 4930433J02Rik | 0 | NA | NA | NA |
| Gm37662 | 0 | NA | NA | NA |
| Rab3gap2 | 1548.083402 | -0.162768997 | 0.057238995 | -2.843673199 |
| Snora36b | 0.393059373 | -0.445563502 | 1.596207927 | -0.27913876 |
| mmu-mir-664 | 0 | NA | NA | NA |
| Gm34882 | 0 | NA | NA | NA |
| Iars2 | 594.1626988 | 0.059165329 | 0.055930651 | 1.057833717 |
| 2010103J01Rik | 0 | NA | NA | NA |
| Mir194-1 | 0 | NA | NA | NA |
| Mir215 | 0 | NA | NA | NA |
| Gm37785 | 5.812857154 | 0.536602685 | 0.42707745 | 1.256452866 |
| Bpnt1 | 456.4610138 | -0.139866058 | 0.082751927 | -1.690184921 |
| 4930532G15Rik | 1.624021655 | 1.291856202 | 1.020443394 | 1.265975369 |
| Eprs | 2033.228255 | -0.183324851 | 0.087996145 | -2.083328222 |

| | | | | |
|---|---|---|---|---|
| 9630028B13Rik | 4.198644665 | 1.135998708 | 0.596742562 | 1.903666307 |
| Gm2061 | 20.60515688 | 1.280687385 | 0.267729804 | 4.783506975 |
| Slc30a10 | 131.5005642 | 0.759815734 | 0.186830305 | 4.066876279 |
| 5033404E19Rik | 1.621213957 | 1.93002446 | 0.991679017 | 1.946218914 |
| Gm38093 | 0 | NA | NA | NA |
| Gm9722 | 0.233998759 | 0.560262676 | 2.129076758 | 0.263148181 |
| Lyplal1 | 31.22453688 | -0.17401349 | 0.202667374 | -0.858616198 |
| Gm38268 | 1.323601962 | -0.420972851 | 1.239857868 | -0.339533152 |
| Gm23821 | 0 | NA | NA | NA |
| Gm37134 | 0 | NA | NA | NA |
| Gm37272 | 0 | NA | NA | NA |
| Gm37491 | 0 | NA | NA | NA |
| A730004F24Rik | 0.094097695 | 0.037071574 | 3.724945049 | 0.009952247 |
| Tgfb2 | 267.8778118 | -0.066733274 | 0.11890649 | -0.561224823 |
| Gm26169 | 0.061920613 | -0.443816764 | 3.724945049 | -0.119147198 |
| Gm29491 | 2.076177089 | -0.541124642 | 0.751032299 | -0.720507817 |
| Gm37849 | 4.193292652 | -0.35549893 | 0.608532374 | -0.584190661 |
| Rrp15 | 96.16436869 | 0.173160256 | 0.177460396 | 0.975768454 |
| A430105J06Rik | 9.038545814 | -0.032286782 | 0.421524778 | -0.076595218 |
| 1700023G09Rik | 0 | NA | NA | NA |
| Gm3809 | 0.939714841 | -0.826261335 | 1.067809848 | -0.773790705 |
| Gm37912 | 0.240869569 | -1.419319044 | 3.089662856 | -0.459376673 |
| D1Pas1 | 0.137647365 | 1.167618838 | 3.719469251 | 0.31392082 |
| Gm19058 | 0.109869311 | -0.444802825 | 3.724874622 | -0.119414174 |
| Spata17 | 10.75281961 | -0.965500011 | 0.336119913 | -2.872486794 |
| Gm15508 | 0.503467097 | -0.020187464 | 1.370792747 | -0.014726854 |
| RP23-438D3.4 | 1.299896474 | -0.028086915 | 1.047804986 | -0.02680548 |
| RP23-438D3.3 | 1.714015964 | -2.314757176 | 1.037706884 | -2.230646451 |
| Gm15509 | 0.816619922 | -1.492263723 | 1.163355166 | -1.282724113 |
| 1700007P06Rik | 4.898989025 | 0.648481923 | 0.558805323 | 1.160479144 |
| Gpatch2 | 371.7543322 | 0.062607556 | 0.084514909 | 0.740787118 |
| Gm21710 | 76.9518533 | 0.12782339 | 0.207607551 | 0.615697209 |
| Gm38155 | 0.312806312 | -0.414257038 | 1.747309888 | -0.237082753 |
| Gm37896 | 5.45108928 | 2.070737736 | 0.554144625 | 3.736818229 |
| Esrrg | 699.5845546 | 0.184533838 | 0.099988365 | 1.845553109 |
| Gm36388 | 5.428564864 | -0.374284448 | 0.551022627 | -0.679254226 |
| Gm37929 | 0 | NA | NA | NA |
| 9330162B11Rik | 0.045882455 | 0.517959912 | 3.724945049 | 0.139051692 |
| Gm38315 | 9.475868802 | 1.237206424 | 0.493846005 | 2.505247408 |
| Gm38064 | 0.045882455 | 0.517959912 | 3.724945049 | 0.139051692 |
| Ush2a | 7.7862644 | 1.14958666 | 0.412465701 | 2.787108496 |
| Gm25095 | 0 | NA | NA | NA |
| Gm25269 | 0 | NA | NA | NA |
| Kctd3 | 958.3543934 | 0.056069473 | 0.062924858 | 0.891054425 |
| Gm37178 | 0.897014366 | 1.007841132 | 1.074838878 | 0.937667173 |
| Gm18698 | 0.26837777 | 0.505737452 | 2.09021956 | 0.241954224 |
| Kcnk2 | 433.3427616 | 0.42404846 | 0.10406337 | 4.074906078 |

| | | | | |
|---|---|---|---|---|
| 2900042K21Rik | 2.348091134 | 3.019897632 | 1.049926189 | 2.876295175 |
| Gm3837 | 46.20088099 | -0.166780073 | 0.284766322 | -0.585673445 |
| A430027H14Rik | 10.33216563 | 0.586922919 | 0.354483594 | 1.655712502 |
| Gm24092 | 0.042716945 | 0.037062677 | 3.724945049 | 0.009949859 |
| Gm38122 | 0.072025462 | 0.517959912 | 3.724945049 | 0.139051692 |
| Cenpf | 46.80573964 | -0.273840797 | 0.375691988 | -0.728897091 |
| Gm37239 | 0.092881282 | 0.037060453 | 3.724945049 | 0.009949262 |
| Ptpn14 | 196.8365016 | 0.0998852 | 0.148156063 | 0.674189083 |
| Gm30280 | 0.99636849 | -1.262909773 | 1.175496161 | -1.074363163 |
| Gm26574 | 0.255031674 | -1.987100823 | 2.981746952 | -0.666421683 |
| Gm37422 | 1.509855825 | -0.025399994 | 0.866603182 | -0.029309832 |
| Smyd2 | 468.8475306 | 0.101480372 | 0.075886503 | 1.337265097 |
| Gm38245 | 23.42935459 | 0.661571853 | 0.396340567 | 1.669200451 |
| Prox1 | 753.5982863 | -0.053110436 | 0.072472653 | -0.732834155 |
| Gm28172 | 0.53361457 | 1.210736897 | 1.427941913 | 0.84788946 |
| Prox1os | 9.870830895 | 0.002954643 | 0.321291294 | 0.00919615 |
| Gm29678 | 0.175900428 | 0.0093455 | 2.724668585 | 0.003429958 |
| Gm30725 | 0.194279094 | -1.017823543 | 3.719440177 | -0.273649661 |
| Gm23153 | 0.062426873 | 0.517959912 | 3.724945049 | 0.139051692 |
| AC124613.1 | 0 | NA | NA | NA |
| Gm30932 | 0 | NA | NA | NA |
| Rps6kc1 | 776.7345912 | 0.07374459 | 0.06229727 | 1.183753144 |
| Angel2 | 626.1745066 | 0.016446207 | 0.055179896 | 0.298047078 |
| Vash2 | 94.15710728 | 0.043949474 | 0.172581124 | 0.254659797 |
| Gm31373 | 0.061920613 | -0.443816764 | 3.724945049 | -0.119147198 |
| Gm37095 | 4.062122998 | 0.691828042 | 0.603211847 | 1.146907253 |
| Gm8407 | 6.023452583 | 1.412439574 | 0.513022174 | 2.753174514 |
| Mfsd7b | 308.2483388 | -0.095444132 | 0.101993739 | -0.935784225 |
| A230020J21Rik | 0.378086136 | 1.916747241 | 1.719761541 | 1.11454245 |
| Spata45 | 0.216916081 | 0.539470301 | 2.174076706 | 0.248137657 |
| Tatdn3 | 79.86263131 | 0.242388851 | 0.143868099 | 1.684799145 |
| Nsl1 | 10.37462125 | 0.146256834 | 0.301834504 | 0.484559693 |
| Gm38188 | 0 | NA | NA | NA |
| Batf3 | 9.56190983 | 0.485452904 | 0.308901873 | 1.571544064 |
| Gm38161 | 0 | NA | NA | NA |
| Fam71a | 0.398364559 | 0.253378392 | 1.655554305 | 0.153047466 |
| Atf3 | 253.8533324 | -1.814239679 | 0.428424055 | -4.2346821 |
| Gm24748 | 0 | NA | NA | NA |
| D730003I15Rik | 30.70567579 | -0.493529824 | 0.369898186 | -1.334231534 |
| Gm37074 | 9.818226126 | -0.658479045 | 0.584920351 | -1.12575848 |
| Gm37168 | 4.648304818 | -0.634309376 | 0.494105259 | -1.283753543 |
| Nenf | 240.7507074 | 0.212740099 | 0.107112667 | 1.986133911 |
| Tmem206 | 104.9785531 | -0.224701979 | 0.123772094 | -1.815449442 |
| Ppp2r5a | 813.5882951 | -0.280209218 | 0.080849761 | -3.465801435 |
| Gm2272 | 0 | NA | NA | NA |
| Gm32200 | 9.706590373 | 0.963029944 | 0.389675704 | 2.471362554 |
| Gm37688 | 0.11193333 | 0.037073798 | 3.724945049 | 0.009952844 |

| | | | | |
|---|---|---|---|---|
| Gm37432 | 0.52136618 | -0.571782975 | 1.496130037 | -0.382174651 |
| 9430037O13Rik | 2.32288591 | -0.02970348 | 0.739222938 | -0.040182033 |
| Dtl | 77.14293698 | -0.570270723 | 0.176946587 | -3.222841041 |
| Ints7 | 338.6046142 | 0.046053608 | 0.073592791 | 0.625789657 |
| Gm37349 | 5.323050413 | 0.59916366 | 0.519288197 | 1.153817213 |
| Lpgat1 | 3755.069215 | 0.126103427 | 0.058391282 | 2.159627645 |
| Gm22446 | 0 | NA | NA | NA |
| Gm26203 | 0 | NA | NA | NA |
| Gm20203 | 0 | NA | NA | NA |
| Gm38037 | 1.052686423 | 1.207434758 | 1.134168098 | 1.064599471 |
| Nek2 | 70.81536854 | 0.051848044 | 0.315543336 | 0.164313544 |
| 1700034H15Rik | 43.54077039 | 0.585147263 | 0.19502674 | 3.00034376 |
| Slc30a1 | 668.252428 | -0.000710265 | 0.115149565 | -0.006168193 |
| Rd3 | 12.96523121 | 0.270426873 | 0.274006856 | 0.986934692 |
| Gm26670 | 1.595190335 | -1.139800274 | 0.886990633 | -1.285019516 |
| Gm32460 | 0.281516365 | 1.432598892 | 1.772004532 | 0.808462318 |
| Traf5 | 35.39043388 | -0.429865943 | 0.20944721 | -2.052383235 |
| Mir3473c | 0 | NA | NA | NA |
| Gm3934 | 136.9920633 | 0.121632796 | 0.121094596 | 1.004444452 |
| Gm26879 | 0 | NA | NA | NA |
| Rcor3 | 532.0108606 | 0.086052635 | 0.11200719 | 0.76827778 |
| Gm37055 | 0.109124069 | 0.517955464 | 3.724945049 | 0.139050498 |
| Gm10516 | 32.23956246 | 0.440174527 | 0.194487469 | 2.263253924 |
| Kcnh1 | 1278.299755 | 0.410695922 | 0.14768256 | 2.780937174 |
| Hhat | 36.35158037 | 0.337359156 | 0.23423831 | 1.440239027 |
| Gm22265 | 0.377885927 | -1.551262146 | 1.648582944 | -0.940966999 |
| Gm38360 | 0 | NA | NA | NA |
| A730013G03Rik | 1.204108136 | 1.774469292 | 0.940294833 | 1.887141384 |
| 1700065J18Rik | 0.258344737 | -0.992252453 | 2.12023067 | -0.467992689 |
| Sertad4 | 214.5306265 | 0.449937734 | 0.121531967 | 3.702217175 |
| Gm15867 | 2.355781424 | -0.305500039 | 0.680946672 | -0.448640182 |
| Gm2367 | 0.23837135 | -0.363235373 | 2.545605237 | -0.142691164 |
| Syt14 | 711.6526152 | 0.822233875 | 0.11069909 | 7.42764801 |
| Gm23583 | 5.178783273 | 1.319361596 | 0.530648902 | 2.486317396 |
| Gm31406 | 1.718171857 | -0.153080037 | 0.776689753 | -0.197092901 |
| Gm37103 | 0 | NA | NA | NA |
| Diexf | 261.3112737 | 0.188641482 | 0.096183208 | 1.961272511 |
| Irf6 | 4.847835626 | -0.042550625 | 0.477765654 | -0.089061708 |
| A130010J15Rik | 56.29500924 | 0.054380452 | 0.158181164 | 0.343785889 |
| Traf3ip3 | 31.36915684 | -0.064411127 | 0.192306404 | -0.334940107 |
| Gm15872 | 0.108123235 | 0.517955464 | 3.724945049 | 0.139050498 |
| 4930570N18Rik | 0.062426873 | 0.517959912 | 3.724945049 | 0.139051692 |
| Lamb3 | 38.67870685 | -0.654398275 | 0.226917006 | -2.883866166 |
| Hsd11b1 | 94.82910089 | 0.311434526 | 0.190068641 | 1.638537131 |
| Gm38118 | 1.886096619 | 0.782108587 | 0.737469834 | 1.060529598 |
| G0s2 | 30.35106119 | -0.01473218 | 0.202000679 | -0.07293134 |
| Gm37691 | 3.727288002 | 0.049367412 | 0.566095685 | 0.087206832 |

| | | | | |
|---|---|---|---|---|
| Gm37650 | 0 | NA | NA | NA |
| Camk1g | 290.0946269 | 0.706079573 | 0.151021712 | 4.675351397 |
| Gm37942 | 0 | NA | NA | NA |
| Gm38022 | 0 | NA | NA | NA |
| Gm38322 | 0.094097695 | 0.037071574 | 3.724945049 | 0.009952247 |
| 4631405K08Rik | 0 | NA | NA | NA |
| Mir205 | 0 | NA | NA | NA |
| Gm21362 | 0 | NA | NA | NA |
| 4930503O07Rik | 0.33841178 | 1.438330407 | 3.706323023 | 0.388074757 |
| Plxna2 | 2388.321215 | 0.492100743 | 0.10106491 | 4.869155292 |
| Gm37783 | 41.75688494 | 0.680841219 | 0.299309028 | 2.274709934 |
| Gm37674 | 39.82514733 | 0.817032509 | 0.320737624 | 2.547354748 |
| Gm38036 | 64.68369131 | 0.720915789 | 0.337839676 | 2.133899125 |
| 2900035J10Rik | 6.043773575 | 1.004766195 | 0.499123771 | 2.013060193 |
| Gm19777 | 0.047948698 | 0.037064902 | 3.724945049 | 0.009950456 |
| Mir3962 | 0.624784964 | 0.066488232 | 2.763401071 | 0.02406029 |
| Gm23885 | 0.198152624 | -1.482926123 | 3.70739678 | -0.39999121 |
| Cd34 | 275.1221153 | 0.901642379 | 0.136427619 | 6.608943152 |
| Gm16897 | 7.560911453 | -0.32427048 | 0.394682081 | -0.821599194 |
| A330023F24Rik | 1658.07212 | 0.269964701 | 0.287408185 | 0.93930763 |
| Gm32250 | 0.045882455 | 0.517959912 | 3.724945049 | 0.139051692 |
| Gm37334 | 228.9475402 | 0.607031088 | 0.237410933 | 2.556879247 |
| 9630010A21Rik | 156.1316392 | 0.665275434 | 0.277423523 | 2.39804984 |
| Gm37027 | 0.870290022 | 0.152492178 | 1.122167483 | 0.135890748 |
| Cd46 | 147.5920642 | 0.657150266 | 0.202498922 | 3.245203774 |
| Mir29b-2 | 0 | NA | NA | NA |
| Mir29c | 0 | NA | NA | NA |
| Gm37366 | 35.61469171 | 0.797365441 | 0.251513487 | 3.170269123 |
| Gm37132 | 36.12461116 | 1.236546505 | 0.321159407 | 3.850257774 |
| Cr1l | 392.2221467 | -0.06996775 | 0.084304974 | -0.829936207 |
| Cr2 | 5.163701346 | -0.222551867 | 0.475878704 | -0.46766511 |
| Gm37887 | 0.066050875 | -0.443818988 | 3.724945049 | -0.119147795 |
| Gm24802 | 0.044191549 | 0.037062677 | 3.724945049 | 0.009949859 |
| Gm27940 | 0 | NA | NA | NA |
| Gm38046 | 0 | NA | NA | NA |
| Gm38194 | 0.659436564 | 0.377560839 | 1.251624128 | 0.301656728 |
| 9230019H11Rik | 0 | NA | NA | NA |
| H60c | 0 | NA | NA | NA |
| Gm26752 | 1.987374958 | 0.953892406 | 0.90848784 | 1.049978177 |
| Ppp1r14c | 150.8264208 | 0.054551455 | 0.10256464 | 0.531873893 |
| Gm24482 | 0.106358623 | 0.517955464 | 3.724945049 | 0.139050498 |
| Iyd | 3.589799332 | 1.218551476 | 0.733355266 | 1.661611408 |
| Plekhg1 | 783.4816878 | 0.300539768 | 0.078803552 | 3.813784551 |
| Gm16149 | 0.045882455 | 0.517959912 | 3.724945049 | 0.139051692 |
| Gm23023 | 0 | NA | NA | NA |
| Gm16148 | 0.049553047 | -0.443818988 | 3.724945049 | -0.119147795 |
| Mthfd1l | 264.3319854 | 0.260378832 | 0.099377311 | 2.620103419 |

| | | | | |
|---|---|---|---|---|
| Gm16074 | 3.696842725 | 0.778505138 | 0.794843345 | 0.979444746 |
| Gm25515 | 0.67351111 | 1.24860312 | 1.381727276 | 0.903653812 |
| Gm25369 | 0.211762239 | 0.998861477 | 2.61665017 | 0.381732907 |
| Akap12 | 1387.377891 | -0.541815923 | 0.272194913 | -1.990543899 |
| Zbtb2 | 206.2791963 | -0.04898148 | 0.165134683 | -0.296615337 |
| Gm21781 | 207.0306887 | 0.441634679 | 0.248160921 | 1.779630236 |
| Rmnd1 | 211.8870746 | -0.006214401 | 0.149484422 | -0.041572228 |
| Gm16153 | 6.393807032 | -1.687130347 | 0.541256244 | -3.117063986 |
| Armt1 | 185.8306212 | 0.110362793 | 0.09797013 | 1.126494296 |
| Ccdc170 | 10.84234273 | -0.337406985 | 0.337715272 | -0.999087138 |
| Gm22739 | 0.087261593 | 0.51795324 | 3.724945049 | 0.139049901 |
| Esr1 | 111.3424344 | -0.176090648 | 0.156236278 | -1.127079124 |
| Syne1 | 6019.024116 | 0.650114652 | 0.122993445 | 5.285766668 |
| Gm25694 | 9.729705334 | 0.442952618 | 0.472576911 | 0.937313288 |
| Gm10097 | 0.451409952 | -2.218975188 | 1.360288298 | -1.631253603 |
| Gm23573 | 0 | NA | NA | NA |
| Myct1 | 1.954060925 | -1.484413186 | 0.783277679 | -1.895130203 |
| Vip | 156.4683625 | 0.954224218 | 0.163228244 | 5.845950416 |
| Fbxo5 | 28.97769736 | -0.367208906 | 0.189484533 | -1.937936045 |
| Mtrf1l | 83.2976707 | -0.180146038 | 0.137143139 | -1.313562164 |
| Rgs17 | 1666.499129 | 0.001425948 | 0.084126438 | 0.016950056 |
| Gm10945 | 0.044544648 | 0.517957688 | 3.724945049 | 0.139051095 |
| Oprm1 | 120.7739671 | 0.108452094 | 0.140671564 | 0.770959609 |
| Ipcef1 | 966.4889659 | 0.223220642 | 0.153265834 | 1.456427938 |
| Gm15536 | 9.233390511 | 0.210294117 | 0.335104832 | 0.627547254 |
| Cnksr3 | 268.184003 | 0.121073767 | 0.149864804 | 0.807886605 |
| Ulbp1 | 7.246076538 | -0.290720698 | 0.532371193 | -0.546086456 |
| Lrp11 | 1467.900515 | 0.218698514 | 0.052996616 | 4.126650524 |
| Gm16254 | 0.342362695 | 0.363546991 | 3.693717558 | 0.098423062 |
| Pcmt1 | 1204.251679 | 0.20750196 | 0.051897981 | 3.998266558 |
| Nup43 | 42.06222489 | -0.813516934 | 0.270848255 | -3.003589347 |
| Lats1 | 1258.401443 | -0.189180646 | 0.079363769 | -2.383715515 |
| Katna1 | 226.4998142 | -0.550718648 | 0.111352963 | -4.945702696 |
| Gm20470 | 0.150550172 | 0.517963085 | 2.857661753 | 0.181254161 |
| Ginm1 | 172.1070177 | -0.472103992 | 0.128563628 | -3.672142736 |
| B430219N15Rik | 0.972892922 | -0.361433007 | 0.934102126 | -0.386930933 |
| Ppil4 | 480.6494858 | 0.176751135 | 0.106055173 | 1.666596072 |
| Gm15560 | 0.152667883 | 0.998859635 | 2.845214387 | 0.351066563 |
| Zc3h12d | 64.81458113 | -1.669361152 | 0.346578093 | -4.816695534 |
| Mir5104 | 0.256698001 | -0.869795783 | 2.443903287 | -0.355904338 |
| Gm24726 | 0 | NA | NA | NA |
| Tab2 | 1189.099294 | -0.001577924 | 0.076518127 | -0.020621569 |
| Ust | 495.4936296 | 0.089373481 | 0.095216322 | 0.938636141 |
| Gm24374 | 0 | NA | NA | NA |
| Sash1 | 1877.282141 | -0.187018582 | 0.11597288 | -1.612606174 |
| Gm25410 | 1.528398176 | -0.655056158 | 0.88007607 | -0.744317657 |
| Gm26674 | 0.49731986 | -0.839156029 | 1.349475055 | -0.621838859 |

| | | | | |
|---|---|---|---|---|
| Gm28904 | 9.097189116 | 0.773297078 | 0.40345312 | 1.916696238 |
| Gm9930 | 39.07755842 | 0.838540564 | 0.391671642 | 2.140927436 |
| Samd5 | 168.1922456 | -0.021644665 | 0.122370584 | -0.176878009 |
| Stxbp5 | 2460.054131 | 0.162282074 | 0.08168274 | 1.986736413 |
| Gm25635 | 3.189004039 | -0.257346411 | 0.755066551 | -0.34082613 |
| Gm28905 | 7.03479442 | 1.785510916 | 0.479664597 | 3.722415469 |
| Gm25682 | 1.985393048 | 1.111774407 | 0.983546578 | 1.130372909 |
| Adgb | 11.41173491 | -0.435123271 | 0.355076992 | -1.225433585 |
| Gm15568 | 0.044544648 | 0.517957688 | 3.724945049 | 0.139051095 |
| Rab32 | 75.72339442 | -1.430019704 | 0.206032526 | -6.940747309 |
| Gm5177 | 0.397592075 | 0.294570304 | 1.71387284 | 0.171874072 |
| Gm23601 | 0 | NA | NA | NA |
| Gm10944 | 0 | NA | NA | NA |
| Grm1 | 1592.406977 | 0.284455952 | 0.092571165 | 3.072835395 |
| Shprh | 1486.660042 | 0.185971549 | 0.088499397 | 2.101387755 |
| Gm16577 | 1.038701827 | -1.163894425 | 1.021628494 | -1.139254075 |
| 4930432B10Rik | 21.06911396 | 0.133689001 | 0.397463527 | 0.336355394 |
| Fbxo30 | 323.9844007 | -0.563195944 | 0.095762345 | -5.881183736 |
| Gm25979 | 0.044191549 | 0.037062677 | 3.724945049 | 0.009949859 |
| Epm2a | 429.8223933 | 0.550379112 | 0.104999753 | 5.24171814 |
| Rpl17-ps11 | 0 | NA | NA | NA |
| Gm9797 | 3.625096175 | -0.008168956 | 0.56857854 | -0.01436733 |
| Gm22546 | 0.038981395 | -0.443816764 | 3.724945049 | -0.119147198 |
| Gm22194 | 0 | NA | NA | NA |
| Gm25011 | 0 | NA | NA | NA |
| Gm23680 | 0 | NA | NA | NA |
| Gm25139 | 0 | NA | NA | NA |
| Utrn | 1645.064442 | -0.308113292 | 0.056691112 | -5.434948846 |
| Gm23044 | 1.167901279 | -0.323456641 | 0.875152842 | -0.369600172 |
| B230208H11Rik | 2.934711433 | -1.060037877 | 0.696855224 | -1.521173755 |
| Stx11 | 71.27238607 | -1.207621836 | 0.435398514 | -2.773601192 |
| Sf3b5 | 177.9606754 | -0.219903268 | 0.136410568 | -1.612069145 |
| Plagl1 | 355.2404399 | -0.401275109 | 0.106126335 | -3.781107778 |
| Gm27741 | 0.062426873 | 0.517959912 | 3.724945049 | 0.139051692 |
| Zc2hc1b | 0.135956459 | 0.963714371 | 3.641541882 | 0.264644593 |
| Ltv1 | 143.8273288 | -0.343007904 | 0.134549238 | -2.549311385 |
| Phactr2 | 555.9939026 | 0.233815849 | 0.082166928 | 2.845619957 |
| Fuca2 | 561.995271 | -0.354250095 | 0.097717026 | -3.625264816 |
| Pex3 | 290.4366472 | 0.088683758 | 0.077593999 | 1.142920315 |
| Adat2 | 56.9225531 | 0.09657313 | 0.149264982 | 0.646991204 |
| Aig1 | 695.0458195 | -0.145993969 | 0.062371263 | -2.340724922 |
| Gm22995 | 0 | NA | NA | NA |
| Gm8330 | 0.285805375 | -0.817648585 | 1.791770044 | -0.456335671 |
| Hivep2 | 5288.46622 | 0.309818679 | 0.092992267 | 3.331660685 |
| Gm26835 | 0.195722939 | -0.401381472 | 2.666087843 | -0.15055073 |
| Gm28289 | 0.0878635 | 0.037058229 | 3.724945049 | 0.009948665 |
| Gm24343 | 0.447929572 | 0.921856665 | 1.420662443 | 0.648892121 |

| | | | | |
|---|---|---|---|---|
| Gm37299 | 0.353350569 | -0.372449201 | 1.686771732 | -0.22080593 |
| Adgrg6 | 79.60354276 | -0.432919644 | 0.176265569 | -2.456064706 |
| Gm10335 | 39.8430328 | -0.238766379 | 0.262559039 | -0.909381675 |
| Vta1 | 390.7247282 | -0.128056988 | 0.065785167 | -1.946593648 |
| Gm16234 | 0.304277014 | 0.06753762 | 2.026771559 | 0.033322759 |
| Nmbr | 18.58630088 | 0.515391077 | 0.278927484 | 1.847760104 |
| Gje1 | 0.044191549 | 0.037062677 | 3.724945049 | 0.009949859 |
| Gm8355 | 90.21878376 | -0.563241578 | 0.149263226 | -3.773478534 |
| B230364G03Rik | 0.277885724 | 0.963342539 | 1.795273201 | 0.536599409 |
| Gm23892 | 0 | NA | NA | NA |
| Gm20125 | 6.549262826 | -0.665940981 | 0.478428054 | -1.391935477 |
| Gm25382 | 0 | NA | NA | NA |
| Cited2 | 577.6609852 | -0.313090297 | 0.148059958 | -2.1146183 |
| Gm20655 | 8.589832509 | 0.458777121 | 0.401676281 | 1.142156366 |
| Txlnb | 15.59385469 | 0.402040309 | 0.332718558 | 1.208349518 |
| Gm7584 | 0.106056992 | 0.998850474 | 3.724945049 | 0.268151734 |
| Heca | 505.4761868 | 0.217119655 | 0.092535467 | 2.346339857 |
| Abracl | 74.37190115 | -0.077236287 | 0.180151316 | -0.428730072 |
| Gm17711 | 0.109982625 | -0.443823436 | 3.724945049 | -0.119148989 |
| Reps1 | 606.0807053 | 0.167421306 | 0.093639561 | 1.78793348 |
| Ect2l | 5.746393814 | -0.203684478 | 0.435039609 | -0.468197547 |
| Ccdc28a | 182.412726 | -0.027673976 | 0.145133243 | -0.190679788 |
| Gm10827 | 7.130661091 | -1.434361757 | 0.562541279 | -2.549789341 |
| Nhsl1 | 1061.864252 | -0.299614417 | 0.09799937 | -3.057309631 |
| Hebp2 | 41.95025653 | -0.986375142 | 0.236470666 | -4.171236787 |
| Arfgef3 | 2592.376606 | 0.373332461 | 0.091674638 | 4.072363609 |
| Gm4922 | 0 | NA | NA | NA |
| Perp | 62.70509499 | -0.125973241 | 0.205426834 | -0.613226803 |
| Gm38175 | 0 | NA | NA | NA |
| Tnfaip3 | 387.4893776 | -0.402687911 | 0.369399546 | -1.090114797 |
| Olig3 | 0.674822466 | -1.029598249 | 1.257097617 | -0.819028081 |
| Ifngr1 | 246.1757685 | -0.22155023 | 0.122961219 | -1.801789468 |
| Il22ra2 | 0.165580463 | 0.037073754 | 2.773571312 | 0.013366793 |
| 9230106D20Rik | 0 | NA | NA | NA |
| Il20ra | 10.38114758 | 0.052722596 | 0.352767169 | 0.149454372 |
| Gm25256 | 0.617475372 | 0.391034971 | 1.245075113 | 0.314065366 |
| Slc35d3 | 65.99433708 | 0.475211827 | 0.170243856 | 2.791359626 |
| Pex7 | 193.8965027 | 0.315292076 | 0.085189423 | 3.701070692 |
| Map3k5 | 659.4939879 | 0.299708801 | 0.076008191 | 3.943111894 |
| Map7 | 996.5614196 | 0.41707817 | 0.086564997 | 4.81809259 |
| Mir6413 | 0 | NA | NA | NA |
| 4933406P04Rik | 7.134388926 | -0.495372297 | 0.41253015 | -1.200814768 |
| Bclaf1 | 2858.72397 | 0.242364009 | 0.069547716 | 3.484859367 |
| Mtfr2 | 15.4157451 | -1.034857519 | 0.356798099 | -2.900400874 |
| Pde7b | 825.652375 | 0.212240557 | 0.111610664 | 1.901615387 |
| 4930403O15Rik | 14.60270139 | -0.04298765 | 0.308562486 | -0.139315865 |
| Gm17229 | 23.00707965 | 0.068007677 | 0.351596819 | 0.193425178 |

| | | | | |
|---|---|---|---|---|
| Gm17230 | 1.594890904 | -3.352476124 | 0.994238108 | -3.371904676 |
| Ahi1 | 2789.699633 | 0.355382346 | 0.059265765 | 5.996418739 |
| Myb | 21.21849641 | -0.553135935 | 0.24950148 | -2.216964546 |
| Gm26577 | 5.818997596 | -0.142594144 | 0.452709929 | -0.314979051 |
| Gm2467 | 0 | NA | NA | NA |
| Hbs1l | 606.1812586 | -0.132882469 | 0.089694398 | -1.481502424 |
| Aldh8a1 | 4.539391149 | -1.437953272 | 0.50029503 | -2.874210585 |
| 1700021A07Rik | 0.178864798 | -1.319733861 | 3.264222506 | -0.404302666 |
| 1700020N01Rik | 0.490798134 | -1.495092902 | 1.459949148 | -1.024071903 |
| Gm5420 | 0.09349418 | 0.517955464 | 3.724945049 | 0.139050498 |
| Gm29323 | 0 | NA | NA | NA |
| Platr6 | 2.028201447 | 0.940608598 | 0.767656829 | 1.225298288 |
| Gm23839 | 0.264066224 | 1.48041419 | 2.437241944 | 0.607413717 |
| Sgk1 | 2023.552573 | 0.25918412 | 0.155696985 | 1.664670128 |
| Gm8540 | 0.066050875 | -0.443818988 | 3.724945049 | -0.119147795 |
| Gm7672 | 1.404341276 | 0.894085181 | 0.974354868 | 0.917617605 |
| Gm34267 | 0 | NA | NA | NA |
| 4930444G20Rik | 0 | NA | NA | NA |
| Gm2539 | 0.516867762 | -0.256551047 | 1.585983658 | -0.16176147 |
| Gm19791 | 0.170186936 | -1.44442222 | 3.399009296 | -0.424953889 |
| E030030I06Rik | 0.513032606 | 0.944726966 | 1.352231233 | 0.698643059 |
| Gm7678 | 0 | NA | NA | NA |
| Gm4895 | 29.59563694 | 0.077666098 | 0.345423244 | 0.224843287 |
| Raet1e | 4.132199462 | -3.213810342 | 0.785590374 | -4.090949241 |
| H60b | 4.341397877 | 0.394714905 | 0.518596106 | 0.761121999 |
| Gm26740 | 3.074037044 | -0.051175403 | 0.560367464 | -0.091324723 |
| Gm26581 | 0.35312086 | -2.423278183 | 3.679225572 | -0.658638112 |
| Gm1972 | 0.209168331 | -1.642358585 | 2.554678236 | -0.642882756 |
| Hmgb1-ps8 | 0.251652017 | 0.84076032 | 2.082118399 | 0.403800437 |
| Gm26528 | 1.473604354 | -1.158467101 | 0.865418666 | -1.338620424 |
| C920009B18Rik | 12.28007475 | -0.955224356 | 0.33577574 | -2.844828385 |
| Gm5421 | 64.80517792 | 0.098170708 | 0.160169409 | 0.612917963 |
| Raet1d | 2.579584798 | -3.027385607 | 0.829914074 | -3.647830181 |
| Gm10825 | 0.745135914 | -2.822034689 | 1.248648236 | -2.260071818 |
| Gm16531 | 0.100902008 | -0.876201588 | 3.72093227 | -0.235479048 |
| Gm26585 | 0.066050875 | -0.443818988 | 3.724945049 | -0.119147795 |
| Slc2a12 | 91.64299392 | 0.158129135 | 0.16937279 | 0.933615928 |
| Tbpl1 | 649.6904834 | 0.065513708 | 0.050671455 | 1.292911531 |
| Mir7663 | 9.275244636 | 0.566659293 | 0.415051943 | 1.365273196 |
| Tcf21 | 0.038981395 | -0.443816764 | 3.724945049 | -0.119147198 |
| Eya4 | 83.33865471 | 0.224193338 | 0.120415837 | 1.861826017 |
| Gm24655 | 0.526083665 | -0.752958763 | 1.361212201 | -0.55315311 |
| Rps12 | 490.9795335 | -0.313343861 | 0.08633119 | -3.629555679 |
| Snora33 | 5.090834635 | -0.629230714 | 0.533629091 | -1.179153695 |
| Snord100 | 3.200779369 | 0.423808962 | 0.567093733 | 0.747334942 |
| Gm23130 | 6.373645778 | 0.779318792 | 0.536995622 | 1.451257255 |
| Slc18b1 | 157.4498406 | 0.028357738 | 0.122055237 | 0.232335284 |

| | | | | |
|---|---|---|---|---|
| Vnn3 | 3.620663213 | -0.542211949 | 0.509930777 | -1.063305009 |
| Vnn1 | 16.91961705 | -0.096844614 | 0.268817163 | -0.360262021 |
| Taar1 | 0.366161499 | -0.640981276 | 1.71299513 | -0.374187448 |
| Taar2 | 0 | NA | NA | NA |
| Taar3 | 0.061920613 | -0.443816764 | 3.724945049 | -0.119147198 |
| Taar4 | 0.07796279 | -0.798301147 | 3.721829328 | -0.214491605 |
| Taar5 | 0 | NA | NA | NA |
| Taar6 | 0 | NA | NA | NA |
| Taar7a | 0 | NA | NA | NA |
| Taar7b | 0.046345709 | -0.443818988 | 3.724945049 | -0.119147795 |
| Taar7c-ps | 0 | NA | NA | NA |
| Gm15136 | 0 | NA | NA | NA |
| Taar7d | 0 | NA | NA | NA |
| Taar7e | 0 | NA | NA | NA |
| Gm6884 | 0 | NA | NA | NA |
| Taar7f | 0 | NA | NA | NA |
| Taar8a | 0 | NA | NA | NA |
| Taar8b | 0 | NA | NA | NA |
| Taar8c | 0 | NA | NA | NA |
| Taar9 | 0 | NA | NA | NA |
| Gm15137 | 0 | NA | NA | NA |
| Stx7 | 1125.137819 | 0.049529725 | 0.061473326 | 0.80571084 |
| Moxd1 | 76.92434119 | -0.093874962 | 0.135635477 | -0.692112154 |
| Gm15271 | 0.245390278 | 0.037076276 | 2.153456308 | 0.017217102 |
| Gm15272 | 0.315650473 | 0.037076998 | 1.767501089 | 0.020977072 |
| Gm15270 | 6.965063545 | -0.773120034 | 0.462367236 | -1.672090872 |
| Gm6893 | 0.385058506 | -0.931243407 | 1.553312913 | -0.599520804 |
| Ctgf | 300.6738772 | -0.036052679 | 0.205103386 | -0.175778079 |
| Enpp1 | 114.2531846 | -0.560545665 | 0.131894887 | -4.249942341 |
| Enpp3 | 5.598650687 | -0.252898493 | 0.443519771 | -0.570207935 |
| Med23 | 495.4221233 | 0.082377907 | 0.08862243 | 0.929537893 |
| Gm23410 | 0 | NA | NA | NA |
| Mir6905 | 0.793572527 | 0.397827771 | 1.227019429 | 0.324222878 |
| Arg1 | 19.82112316 | -2.721777095 | 0.364705198 | -7.462951202 |
| Gm22566 | 0.337983087 | 1.495839963 | 1.584970973 | 0.943764894 |
| Akap7 | 458.6354203 | 0.446693823 | 0.073955072 | 6.040070118 |
| Gm25016 | 0.821005853 | 1.723914471 | 1.261348242 | 1.366723648 |
| Gm26167 | 1.164232618 | 1.234783425 | 1.045194195 | 1.181391392 |
| Epb4.1l2 | 1228.23948 | -0.27076862 | 0.084123657 | -3.218697669 |
| Smlr1 | 0.237992309 | 0.037077212 | 2.113697599 | 0.017541399 |
| Gm29571 | 1.193153915 | 0.510040198 | 1.011196284 | 0.504392872 |
| Tmem200a | 188.3161123 | -0.194155792 | 0.148035633 | -1.311547685 |
| Gm9767 | 4.750744946 | 0.561552169 | 0.551849485 | 1.017582121 |
| Samd3 | 18.30979764 | 0.305314495 | 0.289235507 | 1.055591333 |
| L3mbtl3 | 344.0952442 | -0.102193722 | 0.085819161 | -1.190803083 |
| Gm27754 | 1.24360625 | -0.324355024 | 1.069878 | -0.303170104 |
| Gm23757 | 0 | NA | NA | NA |

| | | | | |
|---|---|---|---|---|
| Gm8709 | 0 | NA | NA | NA |
| Arhgap18 | 194.7190015 | -0.113815239 | 0.11198267 | -1.016364751 |
| Lama2 | 360.511942 | 0.217566772 | 0.156075883 | 1.393980721 |
| 4930579H20Rik | 0.409009542 | -0.034906196 | 1.695364717 | -0.020589196 |
| Gm23353 | 0.042716945 | 0.037062677 | 3.724945049 | 0.009949859 |
| Gm36962 | 9.987815949 | 0.021396285 | 0.374803505 | 0.057086672 |
| Gm10145 | 22.9043597 | -0.273905625 | 0.256532213 | -1.067724095 |
| Ptprk | 857.1284641 | 0.462218692 | 0.066839943 | 6.915306533 |
| Gm22370 | 0 | NA | NA | NA |
| Themis | 33.46669582 | -0.199324468 | 0.246801893 | -0.807629411 |
| 4930519F09Rik | 0.493807599 | 0.312033302 | 1.344157991 | 0.232140346 |
| Gm22811 | 0 | NA | NA | NA |
| 2310057J18Rik | 0.049553047 | -0.443818988 | 3.724945049 | -0.119147795 |
| Gm9824 | 52.77535247 | 0.310088429 | 0.194323309 | 1.595734606 |
| Gm9996 | 16.71528001 | 0.592784532 | 0.24584411 | 2.411221205 |
| Soga3 | 1394.424857 | -0.197515517 | 0.071895882 | -2.747243806 |
| 9330159F19Rik | 1284.714598 | -0.052979811 | 0.076489445 | -0.692642109 |
| Echdc1 | 288.0513829 | -0.139756747 | 0.11409023 | -1.224966828 |
| Rnf146 | 973.2977547 | 0.303668723 | 0.112607093 | 2.696710427 |
| Gm25596 | 5.415374794 | 0.647466621 | 0.46169299 | 1.402374813 |
| Gm24286 | 0 | NA | NA | NA |
| Rspo3 | 164.9902427 | -0.326600844 | 0.147358885 | -2.216363433 |
| Gm10275 | 252.6440296 | -0.163320975 | 0.141673406 | -1.152799104 |
| Cenpw | 14.39510704 | 0.718209982 | 0.374005339 | 1.920320129 |
| Gm22623 | 0.372763955 | -0.804291909 | 1.41687807 | -0.567650757 |
| Trmt11 | 140.2023999 | -0.139713095 | 0.099122026 | -1.409506047 |
| Hint3 | 162.1893017 | 0.101340168 | 0.116239746 | 0.871820279 |
| Ncoa7 | 1617.001063 | 0.125160346 | 0.097282807 | 1.286561827 |
| Gm24189 | 3.823607117 | 0.413764625 | 0.556569103 | 0.743420039 |
| Hey2 | 73.53388155 | 0.011885498 | 0.256712069 | 0.046298946 |
| Gm24168 | 0 | NA | NA | NA |
| Gm23790 | 0 | NA | NA | NA |
| Gm5422 | 32.64956114 | -0.619429071 | 0.213495797 | -2.901364237 |
| Hddc2 | 80.54346223 | 0.300030503 | 0.156322278 | 1.919307388 |
| n-R5s76 | 0 | NA | NA | NA |
| Tpd52l1 | 150.8148291 | 0.318224614 | 0.107485361 | 2.960632131 |
| Rnf217 | 787.7636301 | 0.064375537 | 0.087729263 | 0.733797767 |
| Gm7909 | 13.66660191 | 0.190236763 | 0.317310044 | 0.599529597 |
| Gm8793 | 0.447136588 | -0.688015688 | 1.377590217 | -0.499434215 |
| Nkain2 | 470.4928905 | 0.81470246 | 0.078908349 | 10.32466748 |
| Trdn | 7.453187596 | -0.01370969 | 0.754619767 | -0.01816768 |
| Clvs2 | 440.1642126 | 0.400891369 | 0.097512246 | 4.111190004 |
| Gm15939 | 3.414634292 | -0.062107564 | 0.551498358 | -0.112616045 |
| Gm4794 | 0 | NA | NA | NA |
| Gm10327 | 7.81528087 | -0.713933703 | 0.402788051 | -1.772479848 |
| Sult3a1 | 0.245625509 | 0.950765884 | 1.834325944 | 0.518318943 |
| Rsph4a | 69.28623393 | 0.291241355 | 0.173264782 | 1.680903363 |

| | | | | |
|---|---|---|---|---|
| Zufsp | 477.2138068 | -0.045432225 | 0.131559563 | -0.345335788 |
| Gm17567 | 17.61824852 | 0.855979392 | 0.292638659 | 2.925038661 |
| Rwdd1 | 291.9026565 | -0.036187468 | 0.099241432 | -0.364640722 |
| Trappc3l | 1.780324605 | 1.018572958 | 0.781782385 | 1.302885531 |
| Fam26d | 0.532386167 | -2.85509988 | 1.496994108 | -1.907221855 |
| Fam26e | 35.94992537 | 0.193078043 | 0.195618077 | 0.987015344 |
| Fam26f | 40.02314751 | -1.198971967 | 0.353050269 | -3.396037539 |
| Dse | 148.6253122 | -0.645998627 | 0.150131235 | -4.302892904 |
| Tspyl1 | 2089.883738 | 0.002773431 | 0.066745234 | 0.041552502 |
| Nt5dc1 | 71.24363843 | 0.033168313 | 0.154618828 | 0.21451665 |
| Gm26564 | 26.65952218 | -0.241459295 | 0.271067335 | -0.89077238 |
| Tspyl4 | 5615.800342 | 0.25232521 | 0.070641662 | 3.571903667 |
| Col10a1 | 2.713943931 | -0.441268694 | 0.649185984 | -0.679726156 |
| Frk | 10.56880692 | -1.037151181 | 0.34908836 | -2.971027686 |
| Gm26341 | 0 | NA | NA | NA |
| Amd2 | 40.7342373 | -0.326466596 | 0.173886269 | -1.877471962 |
| Hs3st5 | 126.4096297 | -0.104737971 | 0.157572795 | -0.664695774 |
| Gm22554 | 0 | NA | NA | NA |
| Hdac2 | 735.3892852 | -0.115366466 | 0.060741867 | -1.899290764 |
| Gm38231 | 0 | NA | NA | NA |
| Marcks | 2411.516162 | -0.110075387 | 0.115854941 | -0.950113872 |
| 5930403N24Rik | 7.127291511 | 0.323298963 | 0.448101576 | 0.721485887 |
| Gm26535 | 0.348734193 | -1.963817006 | 1.659865274 | -1.183118315 |
| Gm24710 | 0 | NA | NA | NA |
| Gm24889 | 0 | NA | NA | NA |
| Gm24297 | 0 | NA | NA | NA |
| Gm22911 | 0 | NA | NA | NA |
| Rfpl4b | 0.174199136 | 0.517971765 | 2.729447667 | 0.189771642 |
| Lama4 | 161.5765428 | -0.257900836 | 0.107832768 | -2.39167407 |
| Fam229b | 68.30184625 | 0.278890648 | 0.197588648 | 1.411471009 |
| Tube1 | 37.79269275 | 0.060946481 | 0.192545046 | 0.316531026 |
| Wisp3 | 1.507103499 | 0.457464127 | 0.855047983 | 0.535015738 |
| Fyn | 1361.385442 | 0.042934738 | 0.062628039 | 0.685551369 |
| Gm6963 | 0 | NA | NA | NA |
| Gm16364 | 1.605310233 | 0.569201956 | 0.702341302 | 0.810434976 |
| Gm16365 | 6.794706781 | -0.612094135 | 0.448854226 | -1.363681345 |
| Traf3ip2 | 50.94917886 | -0.407347351 | 0.229902768 | -1.771824473 |
| E130307A14Rik | 318.1438882 | 0.354057203 | 0.122013975 | 2.901775813 |
| Rev3l | 1908.960953 | 0.160450999 | 0.096462966 | 1.663342998 |
| Gm22105 | 5.849095356 | 0.439380547 | 0.513324698 | 0.855950528 |
| Gm16210 | 0 | NA | NA | NA |
| Gm26824 | 0.048949532 | 0.037064902 | 3.724945049 | 0.009950456 |
| AA474331 | 80.17615184 | 0.019528122 | 0.176854229 | 0.110419311 |
| 4930547M16Rik | 7.251366017 | 0.646019866 | 0.452544073 | 1.427529174 |
| 2010001E11Rik | 0.234609624 | 0.538139408 | 2.484844901 | 0.216568611 |
| G630090E17Rik | 3.289605183 | -3.204535935 | 0.827189356 | -3.874005281 |
| BC021785 | 2.350275864 | -0.26983782 | 0.620177301 | -0.435097866 |

| | | | | |
|---|---|---|---|---|
| AI317395 | 5.519524843 | -0.184667357 | 0.433199917 | -0.426286686 |
| Gm8055 | 1.673936437 | 0.747732171 | 0.806786971 | 0.926802486 |
| Slc16a10 | 102.3548944 | -0.602857189 | 0.176221523 | -3.421019057 |
| Gm25613 | 0 | NA | NA | NA |
| Gm37786 | 11.97329313 | -0.217747235 | 0.344053227 | -0.632888222 |
| Rpf2 | 140.9629893 | -0.319189129 | 0.125019451 | -2.553115746 |
| Gtf3c6 | 109.1455228 | -0.138786128 | 0.147933145 | -0.938167902 |
| U3 | 0.099155932 | 0.037073798 | 3.724945049 | 0.009952844 |
| Rnu3a | 0 | NA | NA | NA |
| Amd1 | 757.3382757 | -0.170913017 | 0.070178912 | -2.435389949 |
| Cdk19os | 1.037245751 | -0.044312092 | 0.871272593 | -0.050859045 |
| Cdk19 | 1184.871889 | 0.035603002 | 0.096021551 | 0.370781366 |
| Slc22a16 | 0 | NA | NA | NA |
| Ddo | 72.37407717 | -0.293988663 | 0.180486815 | -1.628865034 |
| Mettl24 | 13.24789061 | 0.753216627 | 0.319446266 | 2.357882081 |
| Cdc40 | 528.5442356 | 0.342506867 | 0.067325076 | 5.08735954 |
| Gm22948 | 2.318641632 | 0.992200146 | 0.869543917 | 1.141058119 |
| Wasf1 | 1683.342798 | 0.765208588 | 0.101048905 | 7.572655916 |
| Gpr6 | 59.52113351 | 0.90378926 | 0.269249198 | 3.356701772 |
| Fig4 | 411.9453483 | 0.161210206 | 0.079311719 | 2.032615209 |
| Gm25526 | 0 | NA | NA | NA |
| Ak9 | 11.82815708 | -0.006770531 | 0.448743215 | -0.015087762 |
| Gm25626 | 0.159956389 | 0.055131606 | 2.79840974 | 0.019701048 |
| Gm27783 | 0 | NA | NA | NA |
| Zbtb24 | 171.9864541 | -0.195382527 | 0.090017013 | -2.170506679 |
| Mical1 | 186.6857392 | -0.329345913 | 0.12383886 | -2.659471458 |
| Smpd2 | 91.56976239 | 0.041961871 | 0.148327803 | 0.282899565 |
| Ppil6 | 32.28546686 | 0.299034215 | 0.226006948 | 1.323119565 |
| Cd164 | 1390.831552 | -0.529195195 | 0.128529658 | -4.117300265 |
| Ccdc162 | 21.51099621 | -0.806968594 | 0.246949485 | -3.267747625 |
| Gm17196 | 0.189991899 | -0.805636447 | 2.666886348 | -0.302088781 |
| 5730435O14Rik | 0.634867864 | -2.543539999 | 1.220596898 | -2.083849307 |
| Gm29095 | 0.661465078 | -0.723938771 | 1.1373684 | -0.636503327 |
| Gm29096 | 5.25281502 | -0.433601822 | 0.442015873 | -0.980964371 |
| Gm6976 | 9.172692315 | -0.09968165 | 0.395153494 | -0.252260582 |
| Mir3473b | 0 | NA | NA | NA |
| Cep57l1 | 78.55169042 | 0.070968356 | 0.196611394 | 0.360957496 |
| BC048559 | 0 | NA | NA | NA |
| Sesn1 | 535.6668866 | 0.496961182 | 0.08080239 | 6.150327753 |
| Armc2 | 66.24966393 | 0.495407476 | 0.290334463 | 1.706333691 |
| Gm9855 | 93.44015898 | 2.93E-05 | 0.147814536 | 0.000198536 |
| Foxo3 | 2059.889726 | 0.319667954 | 0.066659819 | 4.795511912 |
| Gm23533 | 0 | NA | NA | NA |
| Lace1 | 140.8858273 | -0.263537988 | 0.114895184 | -2.293725274 |
| Gm15197 | 3.036843357 | 0.104760463 | 0.736738541 | 0.142194899 |
| Gm15198 | 0 | NA | NA | NA |
| Gm25277 | 0 | NA | NA | NA |

| | | | | |
|---|---|---|---|---|
| Snx3 | 1059.401976 | -0.377653512 | 0.072149237 | -5.2343383 |
| Nr2e1 | 94.72567818 | -0.051205035 | 0.21836594 | -0.234491859 |
| Ostm1 | 722.7865687 | -0.12537542 | 0.081559479 | -1.537226835 |
| Gm15200 | 0.23382787 | -0.749962889 | 2.121198477 | -0.353556207 |
| Gm15199 | 0.066050875 | -0.443818988 | 3.724945049 | -0.119147795 |
| Sec63 | 1231.43905 | 0.105195937 | 0.065044526 | 1.617291171 |
| Gm26860 | 0.331010926 | -0.167116806 | 1.736877965 | -0.096216781 |
| Gm26464 | 0.049553047 | -0.443818988 | 3.724945049 | -0.119147795 |
| Scml4 | 87.95950352 | 0.327089098 | 0.156051938 | 2.096027147 |
| Sobp | 1030.323232 | 0.450032309 | 0.085476167 | 5.265003391 |
| Gm29245 | 0 | NA | NA | NA |
| Gm29246 | 2.290242257 | 0.355689122 | 0.842573962 | 0.422145875 |
| 9030612E09Rik | 15.14098368 | 0.094625151 | 0.284523566 | 0.332574038 |
| Pdss2 | 173.4444041 | 0.093109915 | 0.096643705 | 0.96343487 |
| Gm16546 | 3.059482485 | 1.071656607 | 0.692947095 | 1.546520095 |
| Rps19-ps11 | 0 | NA | NA | NA |
| Gm3699 | 19.13216094 | -0.67595067 | 0.284548001 | -2.375524228 |
| Bend3 | 116.2620847 | 0.226710467 | 0.176004463 | 1.288094993 |
| Gm9803 | 11.92917661 | 0.164241797 | 0.334283782 | 0.491324456 |
| 1700021F05Rik | 133.4447187 | 0.034522179 | 0.131272918 | 0.26298021 |
| Cd24a | 328.2481463 | -1.206478295 | 0.214392667 | -5.627423325 |
| F930017D23Rik | 0.149263833 | 1.27732144 | 3.486034464 | 0.366411019 |
| Gm9034 | 0.045882455 | 0.517959912 | 3.724945049 | 0.139051692 |
| Qrsl1 | 139.8918007 | -0.046394237 | 0.097245726 | -0.477082532 |
| Rtn4ip1 | 176.0176104 | -0.334398581 | 0.088818731 | -3.764955638 |
| Aim1 | 288.648516 | -1.417867214 | 0.293942017 | -4.823628922 |
| Gm10812 | 1.879239285 | -0.806552248 | 0.742491484 | -1.086278113 |
| Speer5-ps1 | 0.049553047 | -0.443818988 | 3.724945049 | -0.119147795 |
| Atg5 | 276.1062705 | -0.329304239 | 0.098405815 | -3.346390023 |
| Mir1929 | 2.194932386 | -0.609602104 | 0.830159642 | -0.734319128 |
| Prdm1 | 54.05971938 | -0.846772792 | 0.321171228 | -2.636515102 |
| Gm22087 | 0 | NA | NA | NA |
| Gm27977 | 0 | NA | NA | NA |
| Gm22503 | 0 | NA | NA | NA |
| Gm24271 | 0 | NA | NA | NA |
| Gm4795 | 0.55021313 | -0.988300798 | 1.283272872 | -0.770140801 |
| Prep | 326.8201584 | -0.199776523 | 0.068803864 | -2.903565473 |
| Gm23766 | 0 | NA | NA | NA |
| Popdc3 | 11.87580344 | -0.471654269 | 0.306367425 | -1.539505282 |
| Gm27723 | 0 | NA | NA | NA |
| Gm26016 | 0 | NA | NA | NA |
| Gm15934 | 0.066050875 | -0.443818988 | 3.724945049 | -0.119147795 |
| Bves | 10.58168931 | -0.309658243 | 0.375698533 | -0.824219993 |
| Lin28b | 106.2737535 | 0.497081159 | 0.161766677 | 3.072827905 |
| Gm27731 | 0 | NA | NA | NA |
| Hace1 | 690.7874818 | 0.570931643 | 0.104028523 | 5.48822212 |
| Gm27410 | 0 | NA | NA | NA |

| | | | | |
|---|---|---|---|---|
| Gm25740 | 0 | NA | NA | NA |
| Gm23481 | 0 | NA | NA | NA |
| Gm25650 | 0 | NA | NA | NA |
| Gm23584 | 0 | NA | NA | NA |
| Gm24256 | 0 | NA | NA | NA |
| Grik2 | 2090.882152 | 0.269505194 | 0.096690191 | 2.787306451 |
| Gm26543 | 111.823433 | 0.334345156 | 0.19514717 | 1.713297488 |
| Ascc3 | 1896.390272 | -0.564336846 | 0.092826047 | -6.079509607 |
| Gm23883 | 0 | NA | NA | NA |
| Sim1 | 14.16261422 | 0.133898983 | 0.322172941 | 0.415612133 |
| Gm26257 | 0 | NA | NA | NA |
| Gp49a | 332.5075168 | -1.247238254 | 0.46049609 | -2.708466544 |
| Lilrb4 | 459.2260594 | -1.153719148 | 0.399910355 | -2.884944423 |
| Fam162b | 0 | NA | NA | NA |
| Gprc6a | 0.291554621 | 1.341026407 | 1.768019389 | 0.758490781 |
| Rfx6 | 0.232111299 | 0.889564523 | 2.5035866 | 0.355316059 |
| 4933411G06Rik | 0 | NA | NA | NA |
| Gm23906 | 0 | NA | NA | NA |
| Vgll2 | 1.144635215 | 0.299711797 | 0.965148237 | 0.310534471 |
| Gm24084 | 0.042716945 | 0.037062677 | 3.724945049 | 0.009949859 |
| Ros1 | 0 | NA | NA | NA |
| Gm20630 | 0 | NA | NA | NA |
| Gm18513 | 0 | NA | NA | NA |
| Dcbld1 | 120.8113824 | 0.392484757 | 0.128774445 | 3.047846627 |
| Gopc | 773.3070799 | 0.166216627 | 0.068892933 | 2.412680363 |
| Gm26741 | 0 | NA | NA | NA |
| Rps19-ps10 | 0 | NA | NA | NA |
| Nepn | 0.397314471 | -0.473581249 | 1.600722629 | -0.29585466 |
| Nus1 | 1151.136521 | -0.110021976 | 0.084212658 | -1.306477887 |
| Zfa-ps | 7.158863588 | 0.060839352 | 0.480050101 | 0.126735423 |
| Slc35f1 | 918.4030169 | 0.404841243 | 0.065114071 | 6.217415601 |
| Gm25664 | 1.948438448 | 1.355046678 | 0.919160446 | 1.474222139 |
| Cep85l | 359.2181035 | 0.324562492 | 0.124006959 | 2.617292562 |
| Pln | 12.74356708 | 0.39787796 | 0.337658601 | 1.178343921 |
| Gm5423 | 0.736769121 | -0.160791332 | 1.366480956 | -0.117668183 |
| Mcm9 | 170.9008201 | -0.083444629 | 0.103753883 | -0.804255473 |
| Asf1a | 203.4365805 | -0.358347134 | 0.11023832 | -3.250658518 |
| Fam184a | 493.2642014 | 0.131331642 | 0.093016788 | 1.411913318 |
| Man1a | 671.9857947 | -0.212992155 | 0.078940922 | -2.698120931 |
| Gm16998 | 3.462133719 | 1.529386906 | 0.586321973 | 2.608442077 |
| Gm26177 | 0 | NA | NA | NA |
| Gm25602 | 0 | NA | NA | NA |
| Tbc1d32 | 271.3183319 | 0.158663753 | 0.108764127 | 1.458787538 |
| Msl3l2 | 168.4746344 | -0.341103489 | 0.101756745 | -3.35214622 |
| Rps8-ps1 | 3.747707297 | 0.464370903 | 0.501538068 | 0.925893632 |
| Gja1 | 2426.848436 | 0.232706028 | 0.175597701 | 1.325222523 |
| Gm9795 | 5.035917182 | -0.156614799 | 0.46351714 | -0.337883511 |

| | | | | |
|---|---|---|---|---|
| Gm9956 | 0.322113463 | -0.495325571 | 1.775621403 | -0.278959 |
| Gm24285 | 0 | NA | NA | NA |
| 4930467K11Rik | 0.534824616 | 0.725903462 | 1.337044054 | 0.542916638 |
| Hsf2 | 367.3738673 | -0.147981497 | 0.081968895 | -1.805337214 |
| Serinc1 | 9098.13279 | 0.316219162 | 0.07006691 | 4.513102712 |
| Rpl48-ps1 | 0 | NA | NA | NA |
| Pkib | 225.7964083 | 0.124781393 | 0.159001194 | 0.784782742 |
| Gm5777 | 0.168032776 | -0.985228429 | 3.411705311 | -0.288778877 |
| Fabp7 | 502.0611919 | -0.919612646 | 0.334359671 | -2.750369512 |
| Smpdl3a | 299.0070279 | -0.79028592 | 0.173692861 | -4.549904452 |
| AW822073 | 0.119587853 | 0.981063299 | 3.723618439 | 0.263470416 |
| Gm10807 | 0.089089296 | 0.957155977 | 3.723939561 | 0.257027796 |
| Duxf3 | 0.097139547 | -0.443816764 | 3.724945049 | -0.119147198 |
| Gm4981 | 0.424461334 | 1.260114766 | 3.090336206 | 0.407759765 |
| Gm19459 | 0.218736334 | 0.517972312 | 2.218267419 | 0.233503097 |
| Gcc2 | 1409.09042 | 0.183642939 | 0.063808168 | 2.8780475 |
| Lims1 | 608.1636776 | -0.506645692 | 0.105359721 | -4.808722793 |
| Gm23058 | 0 | NA | NA | NA |
| Ranbp2 | 4662.349267 | -0.161582124 | 0.075470044 | -2.14101006 |
| Ccdc138 | 45.03437437 | -0.045418215 | 0.206751753 | -0.219675115 |
| Edar | 4.213958645 | 0.026101348 | 0.507764698 | 0.051404416 |
| BB019430 | 0 | NA | NA | NA |
| Gm25055 | 0 | NA | NA | NA |
| Gm17542 | 0 | NA | NA | NA |
| Gm9987 | 7.178240949 | 0.970429538 | 0.545402281 | 1.779291308 |
| Sh3rf3 | 355.2242412 | 0.2115554 | 0.176984572 | 1.195332434 |
| Gm27672 | 1.813029451 | 1.151523442 | 0.921498133 | 1.249621026 |
| 10-Sep | 84.1994719 | -0.664151159 | 0.183074197 | -3.627770427 |
| Gm23223 | 0.038981395 | -0.443816764 | 3.724945049 | -0.119147198 |
| AC153941.1 | 0 | NA | NA | NA |
| Sowahc | 0 | NA | NA | NA |
| P4ha1 | 725.4105145 | -0.514431241 | 0.101519863 | -5.067296424 |
| Gm10273 | 13.1323639 | -0.12473893 | 0.324891047 | -0.383940805 |
| Pla2g12b | 0.130524232 | 0.517958497 | 3.657871415 | 0.141601067 |
| Oit3 | 3.202472028 | -0.627288834 | 0.567604311 | -1.105151638 |
| Mcu | 514.9080926 | 0.03505284 | 0.065464748 | 0.535446034 |
| Gm10322 | 0.281194119 | 0.036404207 | 2.01426096 | 0.018073232 |
| Micu1 | 809.3638985 | -0.076327831 | 0.06005431 | -1.270980079 |
| Gm17148 | 2.409158253 | 0.421953167 | 0.721916157 | 0.584490544 |
| Gm17059 | 0.044544648 | 0.517957688 | 3.724945049 | 0.139051095 |
| Gm7399 | 5.948605025 | -0.265598916 | 0.524259233 | -0.506617526 |
| Gm23803 | 0 | NA | NA | NA |
| Dnajb12 | 642.9386256 | 0.152702793 | 0.065325014 | 2.337585315 |
| Ddit4 | 632.8851748 | 0.76499295 | 0.174365015 | 4.387307571 |
| Anapc16 | 407.1600245 | 0.059820439 | 0.084116666 | 0.711160369 |
| Ascc1 | 125.3215762 | 0.119624195 | 0.106655342 | 1.121595908 |
| Spock2 | 9371.899727 | 0.438506501 | 0.061973113 | 7.075753954 |

| | | | | |
|---|---|---|---|---|
| mmu-mir-6906 | 1.418442629 | 0.531443387 | 0.946933641 | 0.561225586 |
| Chst3 | 65.59010548 | -0.989717097 | 0.180132824 | -5.494373952 |
| Gm22117 | 0.042716945 | 0.037062677 | 3.724945049 | 0.009949859 |
| Psap | 17356.86525 | -0.306562078 | 0.103836775 | -2.9523459 |
| Cdh23 | 85.39716704 | -0.284489825 | 0.22723524 | -1.251961733 |
| 4632428N05Rik | 287.6489372 | -0.563509758 | 0.179350213 | -3.141951981 |
| Gm17455 | 4.08792871 | -1.617585305 | 0.543378591 | -2.976902902 |
| Slc29a3 | 540.1404158 | -0.836484975 | 0.130229118 | -6.423179288 |
| Unc5b | 525.6811087 | -0.204006215 | 0.100202429 | -2.035940816 |
| Mir6408 | 0 | NA | NA | NA |
| Pldi | 0.297702773 | -2.187227153 | 1.882745695 | -1.161722031 |
| Gm26947 | 9.224316412 | -0.38826841 | 0.382868435 | -1.014103994 |
| Mir466j | 0.085433889 | 0.037058229 | 3.724945049 | 0.009948665 |
| Gm20611 | 0.272151178 | 0.882392488 | 2.476534869 | 0.356301257 |
| Gm20625 | 0.049553047 | -0.443818988 | 3.724945049 | -0.119147795 |
| Gm20610 | 0 | NA | NA | NA |
| Pcbd1 | 47.59642594 | 0.078725994 | 0.186786076 | 0.421476779 |
| Sgpl1 | 896.1568652 | -0.761497054 | 0.132040956 | -5.767127673 |
| AC123530.1 | 1.014434794 | 1.564696719 | 1.132128247 | 1.382084338 |
| Tbata | 34.89983757 | 0.83865037 | 0.384503794 | 2.181123783 |
| Adamts14 | 36.59661118 | -0.834016558 | 0.275882361 | -3.023087653 |
| Prf1 | 187.0444178 | -1.669391128 | 0.593495171 | -2.812813328 |
| Rpl27a-ps1 | 0.285326127 | 0.018678305 | 1.800941828 | 0.010371409 |
| Pald1 | 146.0309485 | 0.094029529 | 0.140770809 | 0.667961838 |
| Nodal | 1.922318947 | -0.507437952 | 0.819303174 | -0.61935309 |
| Eif4ebp2 | 324.7426737 | -0.152161927 | 0.097143382 | -1.566364308 |
| D830039M14Rik | 3.979968375 | 0.309672329 | 0.578214514 | 0.53556651 |
| Lrrc20 | 191.831978 | 0.300070879 | 0.100346637 | 2.990343152 |
| Gm28447 | 0.497289099 | 1.374795489 | 1.352497017 | 1.016486891 |
| Npffr1 | 23.64047625 | -0.276978048 | 0.234438681 | -1.181452 |
| Ppa1 | 944.9657682 | -0.251474624 | 0.137240098 | -1.832369903 |
| Sar1a | 1213.944438 | -0.157134662 | 0.103553565 | -1.517423968 |
| Tysnd1 | 151.8466627 | -0.203894545 | 0.100554766 | -2.027696476 |
| Aifm2 | 86.05177613 | -0.432761584 | 0.156044826 | -2.773315823 |
| H2afy2 | 209.0800445 | 0.175869679 | 0.114297695 | 1.538698385 |
| Mir5108 | 0 | NA | NA | NA |
| Col13a1 | 11.90106664 | -0.19868368 | 0.368781597 | -0.538757036 |
| Gm5424 | 120.3562083 | -0.232243305 | 0.158232362 | -1.467735823 |
| 2010107G23Rik | 182.7260212 | 0.176827042 | 0.110205261 | 1.604524497 |
| Neurog3 | 0.108123235 | 0.517955464 | 3.724945049 | 0.139050498 |
| Tspan15 | 244.3145109 | -0.512063795 | 0.150395422 | -3.404783115 |
| Mir7662 | 1.416932329 | -0.144561124 | 0.923646472 | -0.156511315 |
| Tacr2 | 2.342138215 | -1.236314572 | 0.709459675 | -1.742614295 |
| Hk1 | 4224.69477 | 0.06473311 | 0.056037998 | 1.155164575 |
| Hk1os | 8.888663959 | -0.656994245 | 0.351450547 | -1.869378922 |
| Hkdc1 | 74.93408703 | 0.764550929 | 0.185969342 | 4.111166513 |
| Supv3l1 | 235.6531214 | 0.129285245 | 0.102194968 | 1.265084251 |

| | | | | |
|---|---|---|---|---|
| 4930507D05Rik | 5.319148665 | -0.75339118 | 0.47277117 | -1.593564132 |
| Vps26a | 797.6808432 | -0.162560525 | 0.081002573 | -2.006856302 |
| Srgn | 508.9808035 | -0.948067127 | 0.37658248 | -2.517555055 |
| Kif1bp | 1369.8405 | 0.181978144 | 0.0625822 | 2.907825948 |
| RP24-255E8.1 | 2.889734049 | 1.073993618 | 0.583012044 | 1.842146536 |
| Ddx21 | 587.0568038 | -0.436290618 | 0.11655319 | -3.743274795 |
| Ddx50 | 804.8841463 | -0.047206159 | 0.07554542 | -0.624871228 |
| Stox1 | 37.04183788 | 0.721652593 | 0.245939367 | 2.934270357 |
| mmu-mir-7215 | 0 | NA | NA | NA |
| Ccar1 | 875.9317859 | -0.162710811 | 0.09085026 | -1.790977935 |
| Snord98 | 8.868178155 | 0.187452931 | 0.424059479 | 0.442043958 |
| Tet1 | 553.5602397 | -0.015305646 | 0.121755018 | -0.125708547 |
| Gm26479 | 0 | NA | NA | NA |
| Gm22803 | 0.09349418 | 0.517955464 | 3.724945049 | 0.139050498 |
| Slc25a16 | 401.0961555 | 0.439117524 | 0.092664508 | 4.738788714 |
| Gm25862 | 0 | NA | NA | NA |
| Gm16128 | 5.788109619 | 1.815340372 | 0.575514395 | 3.154291863 |
| Dna2 | 37.65220541 | 0.302118652 | 0.240640796 | 1.255475616 |
| Rufy2 | 990.6841044 | 0.381753516 | 0.110737315 | 3.447379191 |
| Gm16135 | 8.108709463 | 0.80808161 | 0.502416279 | 1.608390579 |
| Hnrnph3 | 390.3370152 | 0.15730043 | 0.136643423 | 1.151174543 |
| Pbld2 | 14.50480443 | 0.24736552 | 0.288813807 | 0.85648786 |
| Gm16143 | 0.144050924 | 1.05357957 | 3.721447096 | 0.283110183 |
| Gm38063 | 0.592709862 | -0.848485861 | 1.284529732 | -0.660542018 |
| Atoh7 | 10.98456152 | 0.067085421 | 0.422563906 | 0.158758049 |
| Mypn | 24.12272005 | 0.498593129 | 0.248899809 | 2.003188075 |
| 1700120B22Rik | 2.541048356 | -0.779099547 | 0.659210578 | -1.181867484 |
| Herc4 | 532.4503219 | -0.061157802 | 0.065832367 | -0.928992907 |
| Gm26789 | 0.091666476 | 0.037060453 | 3.724945049 | 0.009949262 |
| Sirt1 | 314.9020831 | -0.135797086 | 0.084208086 | -1.61263714 |
| Gm16145 | 0.37202309 | 1.021820079 | 1.589705658 | 0.642773128 |
| Dnajc12 | 114.8880949 | -0.257865201 | 0.135253881 | -1.906527187 |
| Gm7075 | 0.075043205 | 0.517957688 | 3.724945049 | 0.139051095 |
| Ctnna3 | 57.98869911 | 0.470809524 | 0.184984978 | 2.545123012 |
| Gm10118 | 118.090575 | 1.068625354 | 0.219883954 | 4.859951513 |
| Lrrtm3 | 650.2187372 | 0.603798769 | 0.082730747 | 7.298359963 |
| AC116138.1 | 1.265617311 | 0.077700513 | 1.093093016 | 0.071083167 |
| Gm23854 | 0 | NA | NA | NA |
| Gm28881 | 0 | NA | NA | NA |
| 1700023F02Rik | 0.746473191 | 0.779634714 | 1.301177286 | 0.599176394 |
| Gm22829 | 0 | NA | NA | NA |
| Gm22594 | 0 | NA | NA | NA |
| Gm28139 | 0 | NA | NA | NA |
| 1110002J07Rik | 9.655946214 | -0.558985153 | 0.408252541 | -1.369214143 |
| Gm26576 | 2.265170197 | 0.084796107 | 0.629680019 | 0.134665393 |
| Reep3 | 781.8290574 | -0.38786493 | 0.086110899 | -4.504249012 |
| Jmjd1c | 3049.812434 | 0.059501393 | 0.093104676 | 0.639080609 |

| | | | | |
|---|---|---|---|---|
| Gm26359 | 0 | NA | NA | NA |
| Nrbf2 | 127.8010227 | 0.032328984 | 0.098295663 | 0.328895326 |
| Egr2 | 185.0575354 | -1.604028289 | 0.529491367 | -3.029375712 |
| Ado | 961.6363525 | -0.045420245 | 0.082807016 | -0.548507201 |
| Gm10797 | 2.84885825 | 1.045755511 | 0.649222432 | 1.610781542 |
| Zfp365 | 3160.688214 | 0.532188337 | 0.126804847 | 4.19690846 |
| Rtkn2 | 179.8255072 | -0.613244766 | 0.147251916 | -4.164596182 |
| Gm16212 | 0.075043205 | 0.517957688 | 3.724945049 | 0.139051095 |
| Arid5b | 664.009088 | -0.398808028 | 0.106126062 | -3.757870772 |
| 1700040L02Rik | 9.415632204 | 1.066792979 | 0.412949802 | 2.583347842 |
| Tmem26 | 24.00413336 | -2.047210483 | 0.280991085 | -7.285677701 |
| Gm24172 | 0 | NA | NA | NA |
| Rhobtb1 | 46.1805193 | 0.289178493 | 0.191236225 | 1.512153327 |
| A930033H14Rik | 22.60880403 | -0.071626426 | 0.249405401 | -0.287188751 |
| Cdk1 | 20.93667416 | -0.649131293 | 0.439724766 | -1.476221816 |
| Ank3 | 10782.92156 | 0.280939576 | 0.090999666 | 3.087259425 |
| Ccdc6 | 1500.046561 | -0.070898567 | 0.088500712 | -0.801107311 |
| Mrln | 8.103814522 | -0.038047781 | 0.411122108 | -0.092546182 |
| Slc16a9 | 87.74711258 | -0.794959775 | 0.181837962 | -4.371803148 |
| Gm7097 | 0.061920613 | -0.443816764 | 3.724945049 | -0.119147198 |
| Gm9877 | 0.095435502 | 0.037073798 | 3.724945049 | 0.009952844 |
| Fam13c | 629.0128233 | -0.205027105 | 0.097913808 | -2.093954966 |
| Gm22320 | 0.300192116 | 0.294317683 | 3.721680097 | 0.07908194 |
| Phyhipl | 1745.320192 | -0.101668061 | 0.057627834 | -1.764217993 |
| 4930533K18Rik | 0.42462922 | -0.477114639 | 1.569333224 | -0.304023793 |
| Gm24486 | 0.340478385 | 0.037727123 | 1.509368908 | 0.024995296 |
| Bicc1 | 210.8407525 | -0.470327952 | 0.130514206 | -3.603653317 |
| Gm15647 | 0 | NA | NA | NA |
| Tfam | 397.4176102 | -0.18771706 | 0.104906148 | -1.789380929 |
| Ube2d1 | 468.1548561 | 0.036893799 | 0.059314858 | 0.621999277 |
| Gm20609 | 0.553453779 | 1.653631683 | 1.228644481 | 1.345899249 |
| Cisd1 | 406.1124699 | 0.074777736 | 0.074532486 | 1.003290504 |
| Ipmk | 440.7198033 | 0.019503259 | 0.073113407 | 0.266753526 |
| Gm22937 | 0 | NA | NA | NA |
| 1700049L16Rik | 0 | NA | NA | NA |
| Gm9923 | 40.21032448 | -0.017539093 | 0.174740614 | -0.10037216 |
| Gm24984 | 0 | NA | NA | NA |
| Zwint | 7079.662798 | 0.096447772 | 0.074678238 | 1.291511092 |
| Pcdh15 | 611.2248178 | -0.30110118 | 0.189846672 | -1.586022954 |
| AC159477.1 | 0 | NA | NA | NA |
| Gm37438 | 0 | NA | NA | NA |
| Gm15398 | 0.158818843 | -0.47083079 | 2.809622806 | -0.167577936 |
| Gm5778 | 18.43137054 | -0.344313819 | 0.524360034 | -0.656636274 |
| Gm15397 | 18.38539889 | -0.377445717 | 0.336261092 | -1.122478116 |
| Gm6407 | 0 | NA | NA | NA |
| Gm6419 | 9.498614017 | 0.233359066 | 0.346135652 | 0.674183849 |
| Rsph14 | 2.362931495 | -0.272740832 | 0.767587788 | -0.355322006 |

| | | | | |
|---|---|---|---|---|
| Gnaz | 847.2906744 | 0.228099217 | 0.095042265 | 2.399976647 |
| Rab36 | 225.5149465 | 0.440972789 | 0.129571356 | 3.403320016 |
| Bcr | 2079.456346 | 0.145177097 | 0.072350732 | 2.006573987 |
| Specc1l | 984.3676726 | -0.058880206 | 0.050182565 | -1.173319971 |
| Adora2a | 245.9031525 | 1.332651191 | 0.17905631 | 7.442637406 |
| Gucd1 | 173.5614678 | -0.092824248 | 0.097061362 | -0.956346021 |
| Upb1 | 7.757773806 | 0.301442807 | 0.399651689 | 0.754263814 |
| Gm25545 | 0.075043205 | 0.517957688 | 3.724945049 | 0.139051095 |
| Snrpd3 | 401.6260286 | 0.109859195 | 0.09010743 | 1.219202402 |
| Lrrc75b | 242.2640389 | 0.231252926 | 0.105221888 | 2.197764465 |
| Ggt1 | 11.84378073 | -0.209137941 | 0.336571965 | -0.621376592 |
| Ggt5 | 50.67091169 | -0.55323067 | 0.249523033 | -2.217152714 |
| Gm9985 | 0.329446336 | -1.904043931 | 1.718739576 | -1.107814097 |
| Gm26402 | 0 | NA | NA | NA |
| Susd2 | 127.9307858 | 0.237275241 | 0.122610271 | 1.935198736 |
| Cabin1 | 1380.689238 | -0.007218136 | 0.071674209 | -0.100707573 |
| Ddt | 134.0537612 | -0.08553441 | 0.112832726 | -0.758063846 |
| Gm20441 | 0 | NA | NA | NA |
| Gstt3 | 21.78233339 | -0.207907103 | 0.267931453 | -0.775971245 |
| Gstt1 | 101.288934 | -0.126501054 | 0.186208403 | -0.679352018 |
| Gstt4 | 0.243832141 | -1.654175408 | 3.023649585 | -0.547079071 |
| Gstt2 | 19.40726938 | -0.215990103 | 0.233212299 | -0.926152286 |
| Mif | 1051.951317 | 0.190703484 | 0.102818344 | 1.85476128 |
| Derl3 | 13.80484017 | -0.315861567 | 0.316457505 | -0.998116848 |
| Smarcb1 | 629.9954016 | -0.1523453 | 0.063384639 | -2.40350503 |
| Mmp11 | 15.83720611 | -1.380363579 | 0.281247711 | -4.907999339 |
| Chchd10 | 1094.33119 | -0.10786804 | 0.09231963 | -1.168419333 |
| Gm867 | 15.27197981 | -1.324079315 | 0.272424519 | -4.860352957 |
| Vpreb3 | 1.181927862 | -1.711807861 | 1.014020713 | -1.688138949 |
| Gm16221 | 0.106112162 | -0.443821212 | 3.724945049 | -0.119148392 |
| Gm5134 | 1.323702316 | -0.242063431 | 0.909180469 | -0.266243545 |
| Gm16219 | 0 | NA | NA | NA |
| Gm16220 | 0 | NA | NA | NA |
| Mir6409 | 0 | NA | NA | NA |
| Zfp280b | 318.8465618 | 0.150195118 | 0.097802452 | 1.535698901 |
| Slc5a4b | 0.19103638 | 0.03707656 | 2.252891159 | 0.016457324 |
| Gm16222 | 0.688368266 | 0.682013476 | 1.401219453 | 0.486728524 |
| Slc5a4a | 0.337090955 | -1.487470349 | 1.704525055 | -0.872659715 |
| Prmt2 | 399.9587037 | -0.29664052 | 0.109719739 | -2.703620358 |
| Mir678 | 0 | NA | NA | NA |
| S100b | 1036.502615 | 0.166724096 | 0.147846615 | 1.127682876 |
| Dip2a | 1191.452903 | 0.013292818 | 0.087185712 | 0.152465555 |
| Gm16240 | 0.04393175 | 0.037062677 | 3.724945049 | 0.009949859 |
| Pcnt | 701.5738582 | 0.447211181 | 0.089589032 | 4.991807244 |
| 1700094J05Rik | 0.456503428 | 1.680727737 | 1.510314925 | 1.112832635 |
| 2610028H24Rik | 0.44299472 | -1.021080156 | 1.431994393 | -0.713047594 |
| Ybey | 77.35572085 | -0.379360588 | 0.132263422 | -2.868219964 |

| | | | | |
|---|---|---|---|---|
| Mcm3ap | 786.2211096 | 0.007826508 | 0.089956603 | 0.087003147 |
| Lss | 497.5698186 | 0.09708296 | 0.07329733 | 1.324508821 |
| 4930483K19Rik | 0 | NA | NA | NA |
| Spatc1l | 0.383664054 | -1.397487722 | 1.5855043 | -0.881415284 |
| Gm15343 | 0 | NA | NA | NA |
| Ftcd | 0.710593519 | -1.884449862 | 1.289283364 | -1.461625826 |
| Col6a2 | 230.4261296 | -0.499874426 | 0.140135147 | -3.567088185 |
| Col6a1 | 323.7855034 | -0.326099804 | 0.186605468 | -1.747536159 |
| Pcbp3 | 627.0954025 | -0.216678286 | 0.116462944 | -1.860491229 |
| Gm37782 | 0.137011352 | -0.443822158 | 2.942485906 | -0.150832382 |
| Gm10787 | 0.044544648 | 0.517957688 | 3.724945049 | 0.139051095 |
| Slc19a1 | 146.9527464 | 0.199550161 | 0.102899991 | 1.939263148 |
| Col18a1 | 325.5916588 | -0.461539954 | 0.165466428 | -2.78932687 |
| Gm10941 | 3.197615884 | -0.011624193 | 0.671825255 | -0.017302405 |
| Pofut2 | 316.8806264 | -0.431618975 | 0.085321378 | -5.058743589 |
| Adarb1 | 3284.804805 | 0.164831332 | 0.073951922 | 2.228898535 |
| Fam207a | 347.3627384 | -0.371598453 | 0.099435965 | -3.73706289 |
| 2810425M01Rik | 0.856869739 | 0.754289314 | 1.182634265 | 0.63780438 |
| Itgb2 | 410.5844244 | -0.957283122 | 0.162904366 | -5.87635031 |
| Pttg1ip | 760.5419193 | -0.595314628 | 0.124215518 | -4.792594651 |
| Sumo3 | 1158.6635 | -0.003629412 | 0.088763811 | -0.040888421 |
| Ube2g2 | 330.4207087 | -0.23765692 | 0.088347158 | -2.690034682 |
| Mir1930 | 0 | NA | NA | NA |
| Gm19402 | 0 | NA | NA | NA |
| Gm10272 | 0.197138355 | 0.517972414 | 2.624157036 | 0.197386211 |
| Gm10024 | 0.182127695 | -0.443828229 | 3.361117991 | -0.132047798 |
| Gm10142 | 0 | NA | NA | NA |
| Krtap12-1 | 0.19999695 | -0.497864915 | 2.619555419 | -0.190057027 |
| Gm10100 | 0.235931372 | 1.045582562 | 2.53687691 | 0.412153446 |
| Gm3233 | 0 | NA | NA | NA |
| Gm3238 | 0 | NA | NA | NA |
| Gm7138 | 0 | NA | NA | NA |
| Gm3250 | 0.062426873 | 0.517959912 | 3.724945049 | 0.139051692 |
| Gm7137 | 0 | NA | NA | NA |
| Gm19668 | 0 | NA | NA | NA |
| Gm9507 | 0.060174537 | 0.517957688 | 3.724945049 | 0.139051095 |
| Gm2696 | 0 | NA | NA | NA |
| Krtap10-4 | 0 | NA | NA | NA |
| AC153874.1 | 0 | NA | NA | NA |
| Tspear | 5.130515031 | -0.17254596 | 0.458805864 | -0.376076188 |
| Gm10318 | 0.127104694 | -0.443822432 | 3.01099858 | -0.147400412 |
| 1700009J07Rik | 0.532715572 | 1.538605782 | 1.449813203 | 1.061244151 |
| Lrrc3 | 123.618813 | 0.917403024 | 0.206442207 | 4.443873357 |
| Trpm2 | 446.088107 | 0.702973732 | 0.140428539 | 5.005917848 |
| Gm22394 | 0.066050875 | -0.443818988 | 3.724945049 | -0.119147795 |
| Gm22871 | 0.044544648 | 0.517957688 | 3.724945049 | 0.139051095 |
| 1810043G02Rik | 263.2113337 | 0.338544951 | 0.085376528 | 3.965316454 |

| | | | | |
|---|---|---|---|---|
| Pfkl | 1143.605665 | 0.188883472 | 0.082847255 | 2.279900196 |
| Aire | 5.516484393 | 3.156038958 | 0.64061804 | 4.926553364 |
| Dnmt3l | 4.167386937 | -2.201105383 | 0.547327765 | -4.021548922 |
| Icosl | 151.2157367 | 0.002186348 | 0.154134098 | 0.014184716 |
| D10Jhu81e | 415.5710078 | 0.302841635 | 0.088691298 | 3.414558619 |
| Pwp2 | 271.1694499 | -0.149583135 | 0.070451509 | -2.123206968 |
| Trappc10 | 1524.4217 | 0.004003718 | 0.042867537 | 0.093397426 |
| Agpat3 | 3193.730264 | 0.147293905 | 0.063927812 | 2.304066095 |
| n-R5s77 | 0 | NA | NA | NA |
| Mir7659 | 1.249419009 | -1.620138477 | 1.109665183 | -1.460024612 |
| Gm22514 | 0 | NA | NA | NA |
| Gm10146 | 20.48723405 | 0.020681716 | 0.254164364 | 0.081371424 |
| Rrp1 | 2027.885291 | -0.020282539 | 0.089195064 | -0.227395304 |
| Mir6907 | 2.924483123 | -0.069748909 | 0.762168125 | -0.09151381 |
| Cstb | 362.581673 | -1.199959652 | 0.19808448 | -6.057817605 |
| Mir6410 | 0 | NA | NA | NA |
| Pdxk | 2399.16992 | 0.334568395 | 0.082066696 | 4.076786483 |
| Mir6908 | 0.341053521 | 0.401168487 | 1.720920835 | 0.233112691 |
| Gm9978 | 0.176220594 | 0.517965899 | 2.719488438 | 0.190464461 |
| Gm25164 | 0 | NA | NA | NA |
| Gm27541 | 0.044191549 | 0.037062677 | 3.724945049 | 0.009949859 |
| Ilvbl | 215.6243573 | 0.056995386 | 0.121702699 | 0.468316531 |
| Syde1 | 248.2945122 | -0.597984602 | 0.157975303 | -3.785304343 |
| Gm37514 | 0.458997541 | -0.389546078 | 1.468578027 | -0.26525392 |
| Olfr1357 | 0 | NA | NA | NA |
| Casp14 | 0.09176491 | 0.921395572 | 3.723061103 | 0.247483333 |
| Ccdc105 | 3.747387982 | 0.140905538 | 0.507001587 | 0.277919323 |
| Slc1a6 | 295.8032966 | 0.27106008 | 0.167599659 | 1.61730687 |
| Olfr1356 | 0 | NA | NA | NA |
| Olfr1355 | 0.089814205 | 0.517955464 | 3.724945049 | 0.139050498 |
| Olfr1354 | 0.11147366 | -0.955981391 | 3.722845977 | -0.25678779 |
| Olfr8 | 0 | NA | NA | NA |
| Olfr1353 | 0.097139547 | -0.443816764 | 3.724945049 | -0.119147198 |
| Olfr1352 | 0 | NA | NA | NA |
| Olfr1351 | 0 | NA | NA | NA |
| Olfr57 | 0.139856492 | -0.443820827 | 3.574411099 | -0.124166139 |
| 2610008E11Rik | 487.1302078 | 0.034567352 | 0.080002006 | 0.432081068 |
| Vmn2r80 | 0 | NA | NA | NA |
| Vmn2r81 | 0 | NA | NA | NA |
| Vmn2r82 | 0.049553047 | -0.443818988 | 3.724945049 | -0.119147795 |
| Vmn2r83 | 0 | NA | NA | NA |
| Ppap2c | 40.71318606 | -0.75733288 | 0.231215707 | -3.275438729 |
| Mir6909 | 0.048949532 | 0.037064902 | 3.724945049 | 0.009950456 |
| Mier2 | 291.2295781 | -0.09331887 | 0.092986241 | -1.003577185 |
| Theg | 1.989322217 | -0.769827287 | 0.814899631 | -0.944689699 |
| C2cd4c | 123.155375 | 0.260256293 | 0.172219941 | 1.511185588 |
| Shc2 | 358.7182919 | 0.02037372 | 0.164435474 | 0.123901003 |

| | | | | |
|---|---|---|---|---|
| Odf3l2 | 1.872511713 | 1.310928093 | 0.963112051 | 1.36113767 |
| Madcam1 | 2.384226188 | -0.517327346 | 0.784809483 | -0.659175707 |
| Tpgs1 | 315.7362571 | -0.086475287 | 0.099015784 | -0.873348504 |
| Cdc34 | 422.4012987 | -0.297635825 | 0.099928655 | -2.978483239 |
| Gzmm | 21.34813532 | -0.424278579 | 0.333166459 | -1.273473264 |
| Gm25794 | 0 | NA | NA | NA |
| Bsg | 2599.444032 | -0.047244718 | 0.082143738 | -0.575146929 |
| Hcn2 | 1794.794068 | 0.195381825 | 0.095058709 | 2.05538059 |
| Polrmt | 300.0243024 | -0.019865213 | 0.107741702 | -0.184378122 |
| Fgf22 | 7.710007984 | 0.963620579 | 0.417073842 | 2.310431591 |
| Rnf126 | 503.0038443 | 0.080465656 | 0.076230016 | 1.055563936 |
| Fstl3 | 34.70897125 | -1.69576383 | 0.309998546 | -5.470231558 |
| Prss57 | 5.311167766 | -0.485257538 | 0.449759158 | -1.07892753 |
| Palm | 1414.659549 | 0.385908422 | 0.082473974 | 4.679153945 |
| Misp | 27.35574496 | -1.050406024 | 0.287435854 | -3.654401535 |
| E130317F20Rik | 72.27215724 | 0.442904405 | 0.150043241 | 2.951845089 |
| Ptbp1 | 656.5387669 | -0.595964903 | 0.118554781 | -5.0269158 |
| Lppr3 | 374.0295548 | 0.700904157 | 0.115105595 | 6.089227527 |
| Mir6910 | 0 | NA | NA | NA |
| Gm17134 | 0.135598227 | 0.037065847 | 2.950266134 | 0.01256356 |
| Prtn3 | 7.905640509 | -0.415058005 | 0.475965195 | -0.872034361 |
| Elane | 0.826815689 | -1.477854003 | 1.120583103 | -1.318825885 |
| Cfd | 0.08486385 | 0.037730769 | 3.724898049 | 0.010129343 |
| Med16 | 415.3832138 | 0.398024742 | 0.070848165 | 5.617996526 |
| Gm24019 | 2.36422992 | -1.949240785 | 0.817175011 | -2.385340665 |
| R3hdm4 | 1784.00259 | 0.532435371 | 0.080289616 | 6.631435027 |
| Gm26602 | 0.222471884 | 0.924750855 | 2.601894483 | 0.355414434 |
| Kiss1r | 45.63901687 | 0.400307216 | 0.176650872 | 2.266092497 |
| Arid3a | 140.4446512 | -0.163871982 | 0.128869983 | -1.271607078 |
| Wdr18 | 517.0323779 | -0.120207945 | 0.06889495 | -1.744800522 |
| Grin3b | 6.451345326 | -0.265240564 | 0.484324939 | -0.547650024 |
| Tmem259 | 829.796043 | -0.018573186 | 0.064900567 | -0.2861791 |
| Gm26243 | 1.926543226 | -2.042919282 | 0.942382646 | -2.167823538 |
| Cnn2 | 130.197291 | -0.708318016 | 0.176358949 | -4.016342917 |
| Abca7 | 370.3957148 | -0.214490593 | 0.100612109 | -2.131856655 |
| Hmha1 | 218.2921494 | -0.653953646 | 0.147539467 | -4.432398051 |
| Polr2e | 305.7771488 | 0.039232649 | 0.077880622 | 0.503753673 |
| Gpx4 | 1189.407151 | -0.160674613 | 0.116877262 | -1.374729443 |
| Sbno2 | 850.3215173 | -1.018955066 | 0.235062128 | -4.334832989 |
| Stk11 | 787.6366792 | -0.205492621 | 0.067951727 | -3.02409709 |
| Dos | 2722.989964 | 0.16633563 | 0.098722703 | 1.684877181 |
| Atp5d | 1392.897457 | -0.078145963 | 0.080010737 | -0.97669345 |
| Midn | 1117.501778 | -0.620710416 | 0.128071065 | -4.846609321 |
| Cirbp | 669.8346141 | 1.003688861 | 0.137047122 | 7.323677051 |
| 1600002K03Rik | 19.50566266 | -0.372216756 | 0.273401872 | -1.361427241 |
| Efna2 | 60.39084732 | -0.216918445 | 0.173568633 | -1.249756024 |
| Mum1 | 291.2372399 | 0.335930155 | 0.082607618 | 4.066575959 |

| | | | | |
|---|---|---|---|---|
| Ndufs7 | 538.4769027 | -0.053687694 | 0.076790211 | -0.699147624 |
| Gamt | 97.19118593 | 0.580492181 | 0.146938067 | 3.950590846 |
| Dazap1 | 600.5946169 | -0.106115252 | 0.092962265 | -1.141487372 |
| Mir6911 | 0.644009716 | 1.032567109 | 1.388157692 | 0.743839922 |
| Gm15122 | 0 | NA | NA | NA |
| Rps15 | 2187.489196 | -0.013400646 | 0.109634602 | -0.122230075 |
| Apc2 | 2618.790862 | 0.515247558 | 0.050477169 | 10.20753673 |
| 2310011J03Rik | 240.725416 | -0.45929093 | 0.095943145 | -4.787115646 |
| Pcsk4 | 31.63242481 | 1.027605192 | 0.264549807 | 3.884354347 |
| Reep6 | 48.59301236 | 0.60225645 | 0.1707444 | 3.527239828 |
| Adamtsl5 | 57.94529312 | -0.399914456 | 0.195087391 | -2.049924678 |
| Plk5 | 149.4888237 | 0.449999251 | 0.222947113 | 2.018412551 |
| Gm15123 | 0 | NA | NA | NA |
| Gm15124 | 0.780980462 | -1.090863967 | 1.209989763 | -0.901548096 |
| Mex3d | 199.4244648 | -0.18010268 | 0.110606101 | -1.628325005 |
| Gm22721 | 0 | NA | NA | NA |
| Mbd3 | 699.1099546 | -0.030787696 | 0.083579933 | -0.3683623 |
| Uqcr11 | 557.7849959 | 0.164920591 | 0.125969452 | 1.309210993 |
| Tcf3 | 302.852696 | -0.292764554 | 0.109054749 | -2.684564917 |
| Gm25044 | 0.104637558 | -0.443821212 | 3.724945049 | -0.119148392 |
| Onecut3 | 5.071639938 | -0.773787237 | 0.523584459 | -1.477865174 |
| 2610034O05Rik | 0.287784626 | 0.037076308 | 1.809550247 | 0.020489239 |
| Atp8b3 | 7.756145753 | -0.763902207 | 0.376656288 | -2.028114839 |
| Rexo1 | 665.8855935 | 0.251607011 | 0.094978236 | 2.649101751 |
| Klf16 | 435.0080941 | 0.168476585 | 0.123836443 | 1.360476621 |
| Gm26710 | 2.145254238 | 0.670292471 | 0.753815854 | 0.889199222 |
| Abhd17a | 822.7097596 | 0.2045187 | 0.080341256 | 2.545624885 |
| Gm29093 | 14.5563475 | 0.215384604 | 0.369678416 | 0.58262694 |
| Adat3 | 37.40970625 | 0.063860685 | 0.2871144 | 0.222422439 |
| Scamp4 | 2.483253496 | -0.942192732 | 0.755081806 | -1.247802191 |
| Mir6912 | 0.975114657 | -0.649747611 | 1.195319623 | -0.543576461 |
| Csnk1g2 | 1099.862112 | 0.170830068 | 0.057609255 | 2.965323321 |
| Btbd2 | 650.9398297 | 0.265681763 | 0.089628729 | 2.964247806 |
| Mknk2 | 631.5413471 | -0.286271953 | 0.100018061 | -2.862202572 |
| Gm26541 | 0.467763849 | -1.269840993 | 1.497720738 | -0.847848976 |
| Mob3a | 177.9043315 | -0.773681772 | 0.167247604 | -4.625966242 |
| Izumo4 | 105.0430274 | 0.934528619 | 0.190693117 | 4.900694023 |
| Ap3d1 | 2190.778951 | -0.170950888 | 0.050909678 | -3.357925112 |
| Dot1l | 1006.566257 | -0.388083997 | 0.154281832 | -2.515422534 |
| Gm17151 | 0.089089296 | 0.957155977 | 3.723939561 | 0.257027796 |
| Plekhj1 | 280.4915415 | -0.348962895 | 0.086694649 | -4.0251953 |
| Sf3a2 | 462.5313837 | -0.118884157 | 0.108791147 | -1.092774181 |
| Amh | 11.52291177 | 0.181960323 | 0.431244754 | 0.421942113 |
| Jsrp1 | 14.91585505 | -1.367176016 | 0.431398304 | -3.16917337 |
| Oaz1 | 1026.280487 | -0.063794734 | 0.106716934 | -0.597793922 |
| Mir1982 | 0 | NA | NA | NA |
| Lingo3 | 479.2193401 | 0.801197032 | 0.126123754 | 6.352467364 |

| | | | | |
|---|---|---|---|---|
| Lsm7 | 91.89504733 | -0.007541283 | 0.143793165 | -0.052445353 |
| Sppl2b | 350.2345351 | 0.030891204 | 0.077816987 | 0.396972494 |
| Tmprss9 | 6.250140971 | -0.752387491 | 0.452384085 | -1.663160833 |
| Timm13 | 280.1181752 | -0.219430698 | 0.112516906 | -1.95020203 |
| Lmnb2 | 221.9066422 | -0.325942785 | 0.099871463 | -3.263622821 |
| Gadd45b | 364.2798403 | -0.847504468 | 0.308050948 | -2.751182792 |
| Gng7 | 1229.082418 | 0.995204295 | 0.241620556 | 4.118872634 |
| Gm3828 | 3.291353755 | 0.736492077 | 0.89490758 | 0.822981159 |
| Diras1 | 1155.444641 | 0.155519598 | 0.068449698 | 2.272027522 |
| Gm16099 | 2.516192357 | 0.480939308 | 0.66862467 | 0.719296386 |
| Slc39a3 | 692.1380104 | 0.110406553 | 0.074791543 | 1.476190331 |
| Sgta | 1016.513989 | -0.068911204 | 0.05562527 | -1.238847083 |
| Thop1 | 406.0563204 | -0.247258764 | 0.066674215 | -3.708461577 |
| Creb3l3 | 2.644382858 | -0.176521819 | 0.66265041 | -0.26638755 |
| mmu-mir-5615-1 | 0.100902008 | -0.876201588 | 3.72093227 | -0.235479048 |
| Mir5615-2 | 0 | NA | NA | NA |
| Map2k2 | 675.7442005 | -0.365819443 | 0.087025966 | -4.203566602 |
| Zbtb7a | 2167.929022 | -0.015177386 | 0.070448909 | -0.215438191 |
| Pias4 | 256.0514789 | -0.221079032 | 0.102114509 | -2.165010979 |
| Eef2 | 12379.31089 | -0.15532559 | 0.074267534 | -2.091433261 |
| Snord37 | 3.244100523 | -1.183941875 | 0.645002466 | -1.835561781 |
| 4930442H23Rik | 0.124853745 | 1.05357957 | 3.721447096 | 0.283110183 |
| Dapk3 | 292.8022491 | -0.177231101 | 0.092674194 | -1.912410489 |
| Atcayos | 2.229235169 | -1.654234621 | 0.837869206 | -1.974335145 |
| Nmrk2 | 2.519283505 | -1.486024551 | 0.684263996 | -2.171712322 |
| Atcay | 1736.526661 | 0.314803927 | 0.074739641 | 4.212007482 |
| Zfr2 | 517.1457258 | 0.497842105 | 0.139929291 | 3.557811959 |
| Matk | 696.4187115 | 0.701373932 | 0.14879126 | 4.713811351 |
| Mrpl54 | 179.3062319 | -0.24815758 | 0.121165414 | -2.048089231 |
| Apba3 | 193.0437716 | -0.790333216 | 0.134495116 | -5.876296781 |
| Mir3057 | 0.237113133 | -0.416570844 | 1.860946973 | -0.223848852 |
| Tjp3 | 12.76057694 | -0.408309192 | 0.28311175 | -1.442219168 |
| Pip5k1c | 3939.793252 | 0.211806387 | 0.090330024 | 2.344806052 |
| Gm16315 | 2.415294132 | -0.506508543 | 0.661664043 | -0.765507131 |
| Cactin | 253.0737681 | -0.197285477 | 0.104777062 | -1.882907129 |
| Tbxa2r | 4.861979082 | -1.890638302 | 0.521096694 | -3.628190938 |
| Gipc3 | 3.688286577 | -1.216339168 | 0.547672004 | -2.220926318 |
| Hmg20b | 157.6554839 | -0.512892883 | 0.105077631 | -4.881085353 |
| Mfsd12 | 595.8008876 | -0.563749138 | 0.111254872 | -5.067186086 |
| 4930404N11Rik | 5.09738268 | 0.798857866 | 0.628683338 | 1.270684011 |
| Fzr1 | 541.0904011 | 0.008810938 | 0.061184707 | 0.144005562 |
| Dohh | 443.5414654 | -0.017095563 | 0.076227935 | -0.224269009 |
| Mir6913 | 0.75335248 | -0.351765749 | 1.141918041 | -0.308048158 |
| Smim24 | 5.77545002 | -0.843395434 | 0.517420213 | -1.630000941 |
| Nfic | 2277.670415 | -0.090578401 | 0.064972689 | -1.394099635 |
| Gm16104 | 0.970068437 | -1.841948578 | 1.038320436 | -1.773969301 |
| mmu-mir-1191b | 3.498764646 | 0.858068677 | 0.705317931 | 1.216570059 |

| | | | | |
|---|---|---|---|---|
| Celf5 | 1725.007547 | 0.652988475 | 0.172931059 | 3.776004592 |
| Gm16105 | 0.342577373 | 0.621156905 | 1.74310967 | 0.356349871 |
| Ncln | 601.4543055 | -0.311225973 | 0.068443808 | -4.54717501 |
| S1pr4 | 11.27433504 | -0.618195544 | 0.299923335 | -2.061178548 |
| Gna15 | 25.30164811 | -0.025311031 | 0.294522625 | -0.085939174 |
| Gm16106 | 0.086908494 | 0.037058229 | 3.724945049 | 0.009948665 |
| Gm25595 | 0 | NA | NA | NA |
| Gna11 | 1114.577472 | 0.244085006 | 0.064725244 | 3.771094418 |
| Aes | 3534.40744 | 0.207834811 | 0.07363202 | 2.822614548 |
| Gm15917 | 0.222899254 | -0.445120444 | 2.163630285 | -0.205728514 |
| Tle2 | 226.3530908 | 0.862458707 | 0.221958642 | 3.885673019 |
| Tle6 | 20.29999438 | 0.258024821 | 0.245585449 | 1.050651909 |
| BC025920 | 34.61278247 | 0.360906879 | 0.219862803 | 1.641509499 |
| Gm26896 | 0.097501745 | -0.443821212 | 3.724945049 | -0.119148392 |
| Sirt6 | 169.5015077 | -0.095550967 | 0.09636942 | -0.991507124 |
| Ankrd24 | 225.4576987 | 0.565458966 | 0.169348401 | 3.339027479 |
| Gm10778 | 8.434511479 | -0.51408294 | 0.392057098 | -1.311245078 |
| Gm8112 | 0.038981395 | -0.443816764 | 3.724945049 | -0.119147198 |
| Zfp433 | 19.00695932 | 0.234160904 | 0.236441066 | 0.990356321 |
| Gm3055 | 1.030732002 | -1.118699135 | 1.110725441 | -1.007178816 |
| Gm4767 | 1.089198015 | -0.145096769 | 1.312950653 | -0.110511975 |
| Zfp781 | 1.228365715 | -0.563789848 | 1.021814973 | -0.551753363 |
| Zfp873 | 74.79504174 | -0.230680549 | 0.166014382 | -1.389521476 |
| AU041133 | 155.8068101 | -0.161122743 | 0.111970121 | -1.438979805 |
| Gm15608 | 10.40952602 | -0.607095337 | 0.401590048 | -1.511729038 |
| Zfp938 | 255.4037447 | 0.074984201 | 0.104731861 | 0.715963608 |
| 4932415D10Rik | 0.046345709 | -0.443818988 | 3.724945049 | -0.119147795 |
| Gm4924 | 47.95002397 | -0.098461156 | 0.177005221 | -0.556261308 |
| Gm1553 | 0 | NA | NA | NA |
| Ap3m1-ps | 10.60277313 | 0.239989986 | 0.334054403 | 0.718415874 |
| 1190007I07Rik | 27.31729701 | 0.121609208 | 0.302355796 | 0.402205645 |
| Tdg | 118.6799162 | 0.024700096 | 0.148959649 | 0.165817359 |
| Glt8d2 | 72.76759065 | 0.792179162 | 0.283484053 | 2.794439952 |
| Hcfc2 | 483.7093496 | -0.134790928 | 0.071793399 | -1.877483583 |
| Nfyb | 378.4988816 | 0.121440329 | 0.076293522 | 1.591751509 |
| 1700028I16Rik | 0.19184735 | -0.799014536 | 2.659049215 | -0.300488811 |
| Txnrd1 | 1189.720619 | -0.850042659 | 0.133032149 | -6.389753624 |
| Gm4799 | 4.084400745 | -1.088246827 | 0.556050768 | -1.957099763 |
| Chst11 | 1110.164625 | -0.845983769 | 0.1065897 | -7.936824716 |
| 1700025N21Rik | 0.075043205 | 0.517957688 | 3.724945049 | 0.139051095 |
| Gm23122 | 0 | NA | NA | NA |
| Slc41a2 | 340.536907 | -0.176865965 | 0.066387552 | -2.664143486 |
| D10Wsu102e | 409.3325013 | -0.012000468 | 0.080633225 | -0.148827826 |
| Aldh1l2 | 203.3414108 | -0.671043979 | 0.13146882 | -5.104206313 |
| A230046K03Rik | 1351.145369 | -0.211522356 | 0.078974055 | -2.678377815 |
| Appl2 | 956.0063235 | 0.235082942 | 0.184103197 | 1.27690853 |
| 1500009L16Rik | 86.40011601 | 0.260370153 | 0.137081211 | 1.899386145 |

| | | | | |
|---|---|---|---|---|
| Gm37908 | 0 | NA | NA | NA |
| Nuak1 | 1233.243768 | 0.309336201 | 0.106494487 | 2.904715622 |
| 4930463O16Rik | 0.038981395 | -0.443816764 | 3.724945049 | -0.119147198 |
| Ckap4 | 392.0257556 | -0.4776313 | 0.115939291 | -4.119667236 |
| Platr7 | 0.182269687 | 0.517967699 | 3.31332368 | 0.156328735 |
| Tcp11l2 | 249.9597353 | 0.080480903 | 0.115403554 | 0.697386691 |
| Gm25657 | 0.602022574 | -0.840170712 | 1.330250379 | -0.631588402 |
| Gm17249 | 4.368992961 | -1.427770425 | 0.637993399 | -2.237907834 |
| Polr3b | 386.4218719 | -0.364373465 | 0.070558686 | -5.164119183 |
| Rfx4 | 218.129252 | 0.108718174 | 0.149976148 | 0.724903099 |
| Gm24226 | 0.075043205 | 0.517957688 | 3.724945049 | 0.139051095 |
| Ric8b | 669.9951763 | 0.127781982 | 0.053636985 | 2.382348339 |
| Fhl4 | 12.95936291 | -0.434464052 | 0.33164355 | -1.310033175 |
| Tmem263 | 569.4293162 | -0.173697575 | 0.0825473 | -2.104218731 |
| Mterf2 | 117.1077398 | 0.256418344 | 0.153854008 | 1.666634152 |
| Cry1 | 350.2670687 | 0.129607355 | 0.152938524 | 0.847447401 |
| Gm27392 | 0 | NA | NA | NA |
| Gm8394 | 52.17199378 | -0.099679239 | 0.200492024 | -0.497173086 |
| Gm25281 | 0 | NA | NA | NA |
| Btbd11 | 507.0576605 | 0.336636034 | 0.099430623 | 3.385637376 |
| Pwp1 | 247.3605164 | -0.270300561 | 0.090216277 | -2.996139601 |
| Prdm4 | 408.5061188 | -0.284856295 | 0.071579264 | -3.979592401 |
| Ascl4 | 2.415104612 | 1.463799385 | 0.740860932 | 1.975808579 |
| Rtcb | 1117.008586 | -0.102851996 | 0.082307624 | -1.249604729 |
| Bpifc | 3.577274519 | -0.239122557 | 0.54095249 | -0.442039848 |
| A230060F14Rik | 5.145365424 | -0.117758237 | 0.455026206 | -0.258794406 |
| Fbxo7 | 420.5312043 | 0.021443491 | 0.085495236 | 0.25081504 |
| Gm24430 | 0.537185458 | -1.247529307 | 1.402347807 | -0.889600497 |
| Syn3 | 338.3405326 | 0.432121643 | 0.17059488 | 2.533028205 |
| 4930486F22Rik | 0.066050875 | -0.443818988 | 3.724945049 | -0.119147795 |
| Gm15990 | 3.87042643 | -1.959514078 | 0.539780979 | -3.630202162 |
| Timp3 | 2834.917203 | -0.904627062 | 0.13712899 | -6.596906017 |
| Gm6729 | 0 | NA | NA | NA |
| Gm5174 | 1.40433107 | 1.854074818 | 0.923300608 | 2.008094441 |
| 1810014B01Rik | 16.79727865 | 0.31322512 | 0.281873872 | 1.111224385 |
| Hsp90b1 | 4072.803122 | -0.378302584 | 0.10615014 | -3.563844404 |
| Gm15344 | 13.86957958 | 0.981943193 | 0.351074231 | 2.796967443 |
| Gm25745 | 0 | NA | NA | NA |
| Ttc41 | 22.08483189 | -0.332977106 | 0.296279238 | -1.123862437 |
| Fabp3-ps1 | 8.356713702 | -0.038345369 | 0.446010328 | -0.085974173 |
| Nt5dc3 | 1819.324601 | 0.13414111 | 0.085488628 | 1.569110569 |
| Gm16268 | 9.292864951 | 0.447110662 | 0.366353607 | 1.220434721 |
| Stab2 | 7.172886002 | 0.704932635 | 0.457572698 | 1.540591556 |
| Gm16280 | 0 | NA | NA | NA |
| Gm16270 | 0.987182834 | -1.205439225 | 0.980194512 | -1.229795933 |
| Gm16271 | 0.570993603 | -0.307279156 | 1.526650953 | -0.201276628 |
| Gm16269 | 0 | NA | NA | NA |

| | | | | |
|---|---|---|---|---|
| 1700113H08Rik | 0.362813327 | -0.419245184 | 1.605633436 | -0.261108902 |
| Gm10764 | 0.093831153 | 0.517957688 | 3.724945049 | 0.139051095 |
| Gm23191 | 0 | NA | NA | NA |
| Ascl1 | 36.60879247 | 0.395134523 | 0.231927928 | 1.703695309 |
| Pah | 2.972707099 | 0.785140312 | 0.651776659 | 1.20461557 |
| Igf1os | 0.184770788 | 0.57041104 | 2.68592176 | 0.212370683 |
| Igf1 | 193.7112454 | 0.011734126 | 0.117889839 | 0.099534667 |
| Pmch | 109.1827471 | 0.578935535 | 0.171564633 | 3.374445678 |
| Parpbp | 16.45101383 | 0.502502161 | 0.329589613 | 1.524629845 |
| Nup37 | 44.56539038 | -0.729397724 | 0.196522031 | -3.711531584 |
| Gm26357 | 0 | NA | NA | NA |
| Ccdc53 | 129.2902162 | -0.068176334 | 0.142714758 | -0.477710464 |
| Dram1 | 46.36865287 | -0.84272526 | 0.240318896 | -3.506695782 |
| Gnptab | 1221.700717 | -0.347710131 | 0.054602903 | -6.367978846 |
| Gm26546 | 4.34695503 | -1.191818308 | 0.504310358 | -2.36326359 |
| Gm16235 | 0.918591783 | -0.677976509 | 1.036855728 | -0.653877382 |
| Chpt1 | 1036.030604 | 0.11786178 | 0.104130489 | 1.131866189 |
| Sycp3 | 7.536531091 | 0.424253465 | 0.439656208 | 0.964966393 |
| Mybpc1 | 139.6857782 | 0.113136561 | 0.230938211 | 0.4898997 |
| Spic | 2.940423805 | -1.403653376 | 0.573682635 | -2.446741962 |
| n-R5s79 | 0 | NA | NA | NA |
| Gm4925 | 6.36087602 | -0.009839345 | 0.406320243 | -0.02421574 |
| Arl1 | 1132.198676 | -0.065152666 | 0.083089096 | -0.784130153 |
| Utp20 | 565.1990458 | -0.303565625 | 0.073534877 | -4.128185657 |
| Slc5a8 | 3.475576094 | 0.981170518 | 0.576085683 | 1.703167682 |
| Ano4 | 411.0563822 | 0.035555035 | 0.103293372 | 0.3442141 |
| Gas2l3 | 106.5758252 | -0.766070224 | 0.248729932 | -3.079927762 |
| Nr1h4 | 10.81026118 | 1.213734482 | 0.559434003 | 2.169575814 |
| Slc17a8 | 21.16662692 | -0.022415565 | 0.327376615 | -0.068470269 |
| Scyl2 | 725.9700778 | -0.053766982 | 0.07783745 | -0.690759802 |
| 1500026H17Rik | 20.55184199 | -0.252253097 | 0.255293896 | -0.988089026 |
| Actr6 | 166.1310719 | -0.220842417 | 0.113807419 | -1.940492275 |
| Uhrf1bp1l | 1814.744137 | -0.142178067 | 0.065904195 | -2.157344722 |
| Gm26180 | 0.045882455 | 0.517959912 | 3.724945049 | 0.139051692 |
| Anks1b | 4400.438249 | 0.424566319 | 0.093579263 | 4.536970121 |
| Gm22602 | 2.708289425 | 0.747340321 | 0.73744367 | 1.013420213 |
| Apaf1 | 344.5692301 | -0.474994976 | 0.114539731 | -4.146988747 |
| Gm25390 | 0 | NA | NA | NA |
| Ikbip | 137.6747652 | -0.295794448 | 0.139421492 | -2.121584298 |
| Slc25a3 | 3200.487187 | -0.091199357 | 0.056009088 | -1.628295713 |
| Gm24119 | 26.8554828 | -0.349303564 | 0.302313331 | -1.155435532 |
| Tmpo | 631.4065621 | -0.372497558 | 0.086710523 | -4.295874883 |
| Gm26765 | 0.307480012 | -1.017953445 | 2.009644638 | -0.506534054 |
| Rmst | 270.1421029 | 0.167305199 | 0.121669719 | 1.375076726 |
| Gm27544 | 10.08854525 | -0.413813911 | 0.324757495 | -1.27422436 |
| Mir135a-2 | 4.853900475 | 0.031338103 | 0.47908486 | 0.065412426 |
| Gm27999 | 0 | NA | NA | NA |

| | | | | |
|---|---|---|---|---|
| Gm27608 | 0 | NA | NA | NA |
| Gm27588 | 11.46356049 | 0.223759526 | 0.329353463 | 0.679390232 |
| Gm27753 | 0 | NA | NA | NA |
| Gm27582 | 0.044191549 | 0.037062677 | 3.724945049 | 0.009949859 |
| Gm27366 | 0.132101749 | -0.93550548 | 3.724188147 | -0.251197158 |
| Gm27419 | 0 | NA | NA | NA |
| Mir1251 | 5.125747786 | 0.739931526 | 0.452803406 | 1.634112105 |
| Gm26700 | 0.516149922 | -0.302511061 | 1.354021638 | -0.223416711 |
| Gm20757 | 0.625729796 | 0.831153476 | 1.250321095 | 0.664752022 |
| Gm25884 | 0 | NA | NA | NA |
| Gm22582 | 0 | NA | NA | NA |
| Nedd1 | 87.64473989 | -0.932501093 | 0.171374264 | -5.441313488 |
| Gm24241 | 0.242925194 | -0.44382119 | 2.136946501 | -0.207689425 |
| Cfap54 | 546.4457364 | 0.584113763 | 0.181607719 | 3.216348763 |
| Cdk17 | 2255.31025 | 0.307238623 | 0.069490078 | 4.42133078 |
| Mir1931 | 9.166638643 | -0.024594644 | 0.40832063 | -0.060233655 |
| Elk3 | 176.0696239 | -1.129657644 | 0.181993704 | -6.207124847 |
| Mir7688 | 0.168606834 | -0.443823518 | 2.785398186 | -0.159339343 |
| Lta4h | 347.4203626 | -0.053887329 | 0.085794894 | -0.628094825 |
| Hal | 122.7623701 | -2.46761726 | 0.525722798 | -4.693761181 |
| Amdhd1 | 3.584129096 | -1.673740368 | 0.577457549 | -2.898464779 |
| Ccdc38 | 1.217260431 | -0.548802286 | 0.875769544 | -0.626651486 |
| Snrpf | 77.19710455 | -0.250083486 | 0.179600153 | -1.392445843 |
| 4933408J17Rik | 0.132790949 | 0.517961689 | 2.970953037 | 0.174341931 |
| Ntn4 | 142.6461492 | -0.100398854 | 0.161284835 | -0.622494075 |
| Gm8580 | 0.167271369 | 0.517971541 | 2.764694888 | 0.187352154 |
| Gm15915 | 0.796817125 | -0.190522199 | 0.983123202 | -0.193792801 |
| Gm15963 | 1.049255278 | 0.403156569 | 1.045049351 | 0.385777542 |
| Gm3571 | 24.10692972 | -0.652870535 | 0.23668554 | -2.758387923 |
| Gm26239 | 0 | NA | NA | NA |
| Usp44 | 0.703834089 | 0.149257467 | 1.100457998 | 0.135632134 |
| Metap2 | 993.8475694 | -0.126236213 | 0.097630695 | -1.292997188 |
| 4930471D02Rik | 0.044191549 | 0.037062677 | 3.724945049 | 0.009949859 |
| Gm25962 | 0 | NA | NA | NA |
| Vezt | 1354.242168 | 0.05287604 | 0.121664107 | 0.434606732 |
| Mir331 | 0 | NA | NA | NA |
| Fgd6 | 264.4343943 | -0.53705608 | 0.118691388 | -4.524810838 |
| Nr2c1 | 165.6055157 | -0.155923163 | 0.116021418 | -1.343917061 |
| Gm26122 | 0 | NA | NA | NA |
| Ndufa12 | 388.6068376 | 0.113260457 | 0.093874706 | 1.206506646 |
| Gm4792 | 0.047948698 | 0.037064902 | 3.724945049 | 0.009950456 |
| Tmcc3 | 859.2376155 | 0.081836674 | 0.115251443 | 0.710070713 |
| Gm16155 | 3.245903696 | -0.064162198 | 0.675271668 | -0.095016867 |
| Tmcc3os | 2.419038749 | 1.16658401 | 0.632053986 | 1.845703114 |
| Mir7211 | 0 | NA | NA | NA |
| Cep83os | 296.969172 | 0.454736526 | 0.085378433 | 5.326128725 |
| Cep83 | 388.069701 | -0.077241707 | 0.086561942 | -0.89232873 |

| | | | | |
|---|---|---|---|---|
| Plxnc1 | 869.6228163 | 0.158591148 | 0.080475488 | 1.970676445 |
| Gm24186 | 1.18741419 | 0.284378263 | 0.853648833 | 0.333132609 |
| Gm25266 | 0.715554978 | 0.667246795 | 1.298096688 | 0.514019334 |
| Cradd | 45.57421372 | -0.094389032 | 0.162295312 | -0.581588162 |
| 2310039L15Rik | 0 | NA | NA | NA |
| Socs2 | 273.8451504 | 0.055237957 | 0.130078384 | 0.424651318 |
| 5730420D15Rik | 1.317555436 | 0.196941168 | 0.872342282 | 0.225761346 |
| Mrpl42 | 211.5063124 | -0.059206005 | 0.114682242 | -0.516261313 |
| Ube2n | 1238.439884 | 0.095262813 | 0.057057003 | 1.66960773 |
| Nudt4 | 1713.099017 | 0.112646771 | 0.119507747 | 0.942589691 |
| Mir3058 | 2.797676517 | 0.257055476 | 0.596990935 | 0.430585224 |
| Gm23776 | 0 | NA | NA | NA |
| Anapc15-ps | 1.031766027 | 0.739535419 | 0.952563249 | 0.776363584 |
| 4732465J04Rik | 0.435690537 | -0.776813191 | 1.358865109 | -0.571663211 |
| Gm24433 | 0 | NA | NA | NA |
| Eea1 | 1239.903993 | -0.194930608 | 0.060896326 | -3.201024132 |
| Gm5426 | 0 | NA | NA | NA |
| Gm22913 | 0 | NA | NA | NA |
| Gm20091 | 0.131209559 | -0.409992761 | 2.972496229 | -0.137928774 |
| Gm8601 | 0 | NA | NA | NA |
| Btg1 | 1120.140172 | -0.350806831 | 0.126472972 | -2.773769179 |
| Gm24587 | 0 | NA | NA | NA |
| mmu-mir-3966 | 0 | NA | NA | NA |
| Dcn | 434.2514662 | -0.927981952 | 0.174037572 | -5.332078247 |
| Lum | 32.29644331 | -1.064667503 | 0.242709868 | -4.386585146 |
| Kera | 0.075043205 | 0.517957688 | 3.724945049 | 0.139051095 |
| Epyc | 1.611010591 | -0.170783656 | 0.706931301 | -0.241584516 |
| Gm10754 | 7.111199995 | 1.737414515 | 0.519555317 | 3.34404145 |
| Ccer1 | 0 | NA | NA | NA |
| C030005K15Rik | 6.350231153 | 0.702663809 | 0.567728548 | 1.237675667 |
| Gm25658 | 1.646274761 | 0.703183352 | 1.094937675 | 0.64221313 |
| Gm37631 | 0.066050875 | -0.443818988 | 3.724945049 | -0.119147795 |
| Atp2b1 | 8738.965953 | 0.572118292 | 0.115472787 | 4.954572456 |
| Poc1b | 99.61505393 | -0.403357781 | 0.121377832 | -3.323158562 |
| Galnt4 | 92.75288412 | -0.289548733 | 0.150541325 | -1.923383715 |
| Phxr2 | 0 | NA | NA | NA |
| Gm16239 | 0.066050875 | -0.443818988 | 3.724945049 | -0.119147795 |
| Dusp6 | 640.3874158 | -0.17037062 | 0.188132245 | -0.905589682 |
| B530045E10Rik | 1.636950402 | 0.634759577 | 0.8173714 | 0.776586478 |
| Gad1-ps | 4.900567814 | -1.013456788 | 0.560244581 | -1.808954198 |
| Csl | 0.893710704 | -1.226408673 | 1.206792522 | -1.016254783 |
| Kitl | 948.0407053 | -0.197749622 | 0.131492577 | -1.503884294 |
| Gm22918 | 0 | NA | NA | NA |
| Gm25287 | 0 | NA | NA | NA |
| Gm9476 | 0 | NA | NA | NA |
| Gm8701 | 0 | NA | NA | NA |
| Gm4301 | 0 | NA | NA | NA |

| | | | | |
|---|---|---|---|---|
| Gm4302 | 0 | NA | NA | NA |
| Gm4303 | 0 | NA | NA | NA |
| Gm4305 | 0 | NA | NA | NA |
| Gm8711 | 0 | NA | NA | NA |
| Gm4307 | 0 | NA | NA | NA |
| Gm4308 | 0 | NA | NA | NA |
| Gm8723 | 0 | NA | NA | NA |
| Gm4312 | 0 | NA | NA | NA |
| Gm4781 | 0 | NA | NA | NA |
| Tmtc3 | 401.2372202 | -0.267554731 | 0.077660053 | -3.445204081 |
| Cep290 | 566.5526866 | -0.077800811 | 0.097509268 | -0.797881193 |
| 4930430F08Rik | 178.6427657 | -0.060893996 | 0.156822266 | -0.388299428 |
| 1700017N19Rik | 0.256989293 | 0.463557294 | 2.086824834 | 0.22213522 |
| Gm38219 | 0 | NA | NA | NA |
| Gm26278 | 0.136194719 | 0.037062677 | 3.724945049 | 0.009949859 |
| Mgat4c | 312.1695497 | 0.247670526 | 0.095339902 | 2.597763584 |
| Mir3059 | 1.49611593 | -1.107777635 | 0.983484455 | -1.126380421 |
| Gm26988 | 0.108767819 | -0.443823436 | 3.724945049 | -0.119148989 |
| Nts | 88.60440578 | -0.364870128 | 0.234857537 | -1.553580664 |
| Rassf9 | 16.58629026 | -0.619921306 | 0.287162159 | -2.158784805 |
| Alx1 | 1.374376221 | 0.486731237 | 0.953403388 | 0.510519727 |
| Gm17028 | 0 | NA | NA | NA |
| Lrriq1 | 58.42473187 | 0.167735901 | 0.188513366 | 0.889782536 |
| Gm25376 | 0 | NA | NA | NA |
| Slc6a15 | 616.6613696 | 0.510327502 | 0.085521986 | 5.967208274 |
| Gm26923 | 0 | NA | NA | NA |
| Gm25143 | 0 | NA | NA | NA |
| Gm6763 | 0 | NA | NA | NA |
| Gm21293 | 0 | NA | NA | NA |
| Gm8764 | 0 | NA | NA | NA |
| Gm21304 | 0 | NA | NA | NA |
| Gm21312 | 0 | NA | NA | NA |
| Gm20765 | 0 | NA | NA | NA |
| Gm4340 | 0 | NA | NA | NA |
| Gm22339 | 0 | NA | NA | NA |
| Mir6411 | 0 | NA | NA | NA |
| Gm22945 | 0 | NA | NA | NA |
| Gm25522 | 0 | NA | NA | NA |
| Gm25587 | 0 | NA | NA | NA |
| Gm25422 | 0.254945911 | 1.78416951 | 2.060262565 | 0.865991326 |
| Gm36997 | 0.361186056 | 0.023676733 | 1.572115408 | 0.01506043 |
| Tmtc2 | 369.308983 | -0.014917296 | 0.10425722 | -0.143081663 |
| Gm15662 | 0 | NA | NA | NA |
| Gm15666 | 12.30954061 | 0.397051833 | 0.420802373 | 0.943558921 |
| Gm15664 | 0 | NA | NA | NA |
| Gm15663 | 93.64214882 | -0.012723246 | 0.139286321 | -0.091345981 |
| Mettl25 | 107.6127449 | 0.313864193 | 0.144573662 | 2.170963848 |

| | | | | |
|---|---|---|---|---|
| n-R5s80 | 1.30631546 | 1.437422239 | 0.951498166 | 1.510693651 |
| Ccdc59 | 282.5902914 | -0.0471104 | 0.112974014 | -0.417002087 |
| Gm24237 | 0 | NA | NA | NA |
| Gm25342 | 0 | NA | NA | NA |
| Rps15a-ps1 | 0 | NA | NA | NA |
| C230021G24Rik | 14.38138754 | -0.470432184 | 0.310098086 | -1.517043171 |
| Ppfia2 | 1751.263262 | 0.322615437 | 0.099691351 | 3.236142695 |
| Acss3 | 21.10431072 | 0.101744449 | 0.268331432 | 0.379174548 |
| Gm24745 | 0.047948698 | 0.037064902 | 3.724945049 | 0.009950456 |
| Lin7a | 1127.089259 | 0.235778125 | 0.106012305 | 2.224063763 |
| Gm38117 | 0 | NA | NA | NA |
| AC155168.1 | 0.358057361 | 1.665328836 | 3.702420844 | 0.449794582 |
| Myf5 | 0.154585317 | -1.377635992 | 2.821196527 | -0.488316209 |
| Myf6 | 0.075043205 | 0.517957688 | 3.724945049 | 0.139051095 |
| Ptprq | 5.567795988 | 0.228544392 | 0.440679629 | 0.51861801 |
| Otogl | 7.088242965 | 0.421852302 | 0.417885334 | 1.009492959 |
| Ppp1r12a | 1620.397248 | -0.105986154 | 0.07636605 | -1.387870042 |
| Pawr | 30.19438579 | -0.648569023 | 0.250435661 | -2.589763059 |
| Gm23105 | 0.203286941 | -0.771106728 | 2.603555075 | -0.296174541 |
| Gm25591 | 0 | NA | NA | NA |
| Syt1 | 12677.56377 | 0.721654469 | 0.114091135 | 6.325245759 |
| Gm22097 | 0.164132501 | 1.299746508 | 3.390982409 | 0.383294972 |
| AC125531.1 | 0.108123235 | 0.517955464 | 3.724945049 | 0.139050498 |
| Gm23041 | 0.2701572 | 0.532245637 | 1.813461505 | 0.29349707 |
| Nav3 | 1988.147118 | 0.572880024 | 0.116241328 | 4.928367851 |
| Gm36935 | 26.04041665 | 0.244104256 | 0.420498166 | 0.58051206 |
| E2f7 | 15.83057167 | 0.313374973 | 0.294228219 | 1.0650745 |
| 1700020G17Rik | 0 | NA | NA | NA |
| Gm23698 | 0 | NA | NA | NA |
| Gm22186 | 0 | NA | NA | NA |
| Gm22236 | 0 | NA | NA | NA |
| Csrp2 | 26.62848996 | -0.798435449 | 0.230740804 | -3.460313191 |
| Zdhhc17 | 1450.600249 | 0.200500217 | 0.053052927 | 3.779248897 |
| Gm5428 | 96.95467252 | -0.236129828 | 0.165826473 | -1.423957366 |
| Gm22101 | 0 | NA | NA | NA |
| Osbpl8 | 2307.979124 | 0.064750244 | 0.051701116 | 1.25239547 |
| Bbs10 | 50.75359481 | -0.151356012 | 0.167278663 | -0.904813614 |
| Nap1l1 | 1669.332245 | 0.210084343 | 0.071762199 | 2.927507048 |
| Gm25117 | 163.1343521 | 0.237368448 | 0.470756049 | 0.504228143 |
| Phlda1 | 275.8361962 | -0.701597179 | 0.180918385 | -3.87797613 |
| Gm28592 | 7.265044534 | -0.319477133 | 0.478762892 | -0.667297191 |
| 4933440J02Rik | 0 | NA | NA | NA |
| Gm37189 | 0.109727584 | 0.037071574 | 3.724945049 | 0.009952247 |
| Krr1 | 496.8011956 | 0.055152383 | 0.073218097 | 0.753261631 |
| Glipr1 | 31.42829398 | -0.404537402 | 0.258518558 | -1.564829252 |
| Glipr1l1 | 0.138226157 | -0.443822126 | 2.934480201 | -0.151243865 |
| Glipr1l2 | 0.144013665 | -1.209023387 | 3.538333554 | -0.341692881 |

| | | | | |
|---|---|---|---|---|
| Gm6808 | 0 | NA | NA | NA |
| Gm15914 | 0 | NA | NA | NA |
| Caps2 | 5.498926737 | -0.548927201 | 0.467856629 | -1.173280803 |
| Kcnc2 | 1574.614437 | 0.636795114 | 0.085819604 | 7.42015909 |
| 1700010J16Rik | 0.193455382 | 1.245696396 | 2.621157943 | 0.475246598 |
| Atxn7l3b | 2117.555783 | 0.153052215 | 0.049958689 | 3.063575475 |
| Gm26596 | 1.459047308 | -0.501742034 | 0.896695168 | -0.559545821 |
| Gm22507 | 0.128477747 | 0.037074008 | 3.677132019 | 0.010082316 |
| Gm38333 | 0 | NA | NA | NA |
| Gm38281 | 0 | NA | NA | NA |
| Trhde | 612.5170607 | 0.554624006 | 0.094224099 | 5.886222466 |
| 4930473D10Rik | 0 | NA | NA | NA |
| Gm15723 | 23.33627371 | 0.214953013 | 0.266745026 | 0.805837005 |
| Gm25319 | 0 | NA | NA | NA |
| Gm22070 | 0 | NA | NA | NA |
| Tph2 | 55.63792861 | -0.484096564 | 0.659709703 | -0.733802401 |
| Tbc1d15 | 653.7365103 | -0.006899631 | 0.096997153 | -0.071132303 |
| Gm22794 | 0.044544648 | 0.517957688 | 3.724945049 | 0.139051095 |
| Rab21 | 1138.336811 | -0.195294175 | 0.094281105 | -2.071403107 |
| Tmem19 | 316.2960843 | -0.038729644 | 0.079274582 | -0.488550589 |
| Thap2 | 234.4484823 | -0.645228292 | 0.240867745 | -2.67876586 |
| Zfc3h1 | 1151.166255 | 0.038100933 | 0.101897772 | 0.37391331 |
| Lgr5 | 129.3954019 | -0.35987979 | 0.280901143 | -1.281161714 |
| A930009A15Rik | 4.998511455 | -0.530314746 | 0.591865951 | -0.896004821 |
| Tspan8 | 5.527738138 | -0.21600488 | 0.450234453 | -0.479760886 |
| Ptprr | 373.9766246 | 0.407477571 | 0.105979537 | 3.844870275 |
| 4933416C03Rik | 0.062426873 | 0.517959912 | 3.724945049 | 0.139051692 |
| Ptprb | 234.5662353 | -0.194960641 | 0.197852875 | -0.985381895 |
| Gm26495 | 0.042716945 | 0.037062677 | 3.724945049 | 0.009949859 |
| Kcnmb4 | 239.9415874 | 0.55608845 | 0.131871916 | 4.216883072 |
| Kcnmb4os1 | 0.534709659 | 0.235121264 | 1.602343423 | 0.146735875 |
| Kcnmb4os2 | 19.7696678 | 0.562365101 | 0.249259916 | 2.256139333 |
| Cnot2 | 821.4440981 | 0.021775225 | 0.057078607 | 0.38149539 |
| Gm25190 | 6.442501395 | 0.655635781 | 0.438175747 | 1.496284961 |
| 5330438D12Rik | 61.66108682 | 0.177500588 | 0.168865762 | 1.051134262 |
| Gm26579 | 0.044191549 | 0.037062677 | 3.724945049 | 0.009949859 |
| 1700030O20Rik | 0.066050875 | -0.443818988 | 3.724945049 | -0.119147795 |
| Myrfl | 6.514367323 | 0.313692004 | 0.488490887 | 0.642165519 |
| Rab3ip | 355.5556215 | -0.016028048 | 0.082626983 | -0.193980798 |
| Gm10271 | 0 | NA | NA | NA |
| D630029K05Rik | 3.06831776 | 0.427759082 | 0.505258332 | 0.846614603 |
| Best3 | 10.8055888 | 0.349177807 | 0.310720759 | 1.123767232 |
| Lrrc10 | 0.159795471 | -0.939694907 | 2.805348586 | -0.33496547 |
| Cct2 | 1508.454392 | -0.245563412 | 0.072936557 | -3.366808383 |
| Frs2 | 1041.72447 | 0.185700039 | 0.064049088 | 2.899339331 |
| Gm26311 | 0 | NA | NA | NA |
| Yeats4 | 230.0577702 | -0.081430976 | 0.115672515 | -0.703978607 |

| | | | | |
|---|---|---|---|---|
| 9530003J23Rik | 0.138076337 | 0.037074383 | 3.589802603 | 0.010327694 |
| Gm10936 | 0 | NA | NA | NA |
| Lyz2 | 917.3809082 | -2.565874935 | 0.448557859 | -5.720276405 |
| Lyz1 | 2.092478253 | -0.368123526 | 0.769947122 | -0.478115335 |
| Cpsf6 | 2549.871472 | 0.116620939 | 0.069841558 | 1.669792918 |
| Gm9002 | 0 | NA | NA | NA |
| Kifc5c-ps | 0 | NA | NA | NA |
| Cpm | 200.0726009 | 0.046203559 | 0.138633358 | 0.333278799 |
| Gm25709 | 2.906247223 | -0.425895988 | 0.693122066 | -0.614460294 |
| Mdm2 | 840.5416714 | -0.365574606 | 0.095985932 | -3.808626951 |
| Slc35e3 | 296.2709981 | -0.081718237 | 0.082864326 | -0.986169085 |
| Nup107 | 192.1877859 | 0.097692864 | 0.10373193 | 0.941781996 |
| Rap1b | 1149.527098 | -0.565016063 | 0.125114215 | -4.516002127 |
| Gm37852 | 0 | NA | NA | NA |
| Mdm1 | 91.75534052 | 0.271111756 | 0.130893658 | 2.071236762 |
| Il22 | 14.70729829 | -0.397211246 | 0.403284118 | -0.984941454 |
| Iltifb | 0.488214936 | 0.790032777 | 1.377027629 | 0.573723258 |
| Gm9040 | 0 | NA | NA | NA |
| Gm9044 | 0 | NA | NA | NA |
| Gm9045 | 0.068097359 | 0.037064902 | 3.724945049 | 0.009950456 |
| Gm9046 | 0 | NA | NA | NA |
| Gm9048 | 0 | NA | NA | NA |
| Gm9049 | 0.219658751 | 0.037050927 | 3.724945049 | 0.009946704 |
| Ifng | 22.72435143 | -1.165071792 | 0.61957413 | -1.880439702 |
| Dyrk2 | 260.899164 | 0.138999948 | 0.111707293 | 1.244322947 |
| 4932442E05Rik | 0.27373663 | 0.459218743 | 1.78187239 | 0.257716964 |
| Gm26976 | 0 | NA | NA | NA |
| 1700064J06Rik | 0 | NA | NA | NA |
| Cand1 | 2796.110183 | 0.010095883 | 0.043498687 | 0.232096269 |
| 1700025F24Rik | 0.942170168 | 0.876031191 | 1.019522304 | 0.859256524 |
| Grip1os3 | 1.395336821 | -0.164020877 | 1.079916203 | -0.151882967 |
| Gm16321 | 1.40953533 | 0.37115315 | 1.066864678 | 0.347891497 |
| Grip1 | 365.1462489 | 0.088159556 | 0.124770473 | 0.706573869 |
| Grip1os2 | 0 | NA | NA | NA |
| Grip1os1 | 0.072025462 | 0.517959912 | 3.724945049 | 0.139051692 |
| Helb | 127.7033369 | -0.676509922 | 0.114425009 | -5.912255814 |
| Irak3 | 30.23155639 | -0.527042541 | 0.230018553 | -2.291304483 |
| Tmbim4 | 275.0985624 | -0.553459145 | 0.123429694 | -4.48400321 |
| Llph | 97.51439187 | -0.220686815 | 0.168355422 | -1.31083877 |
| Gm38141 | 0 | NA | NA | NA |
| Hmga2 | 52.04420527 | -2.38609927 | 0.563155641 | -4.237015664 |
| 1700006J14Rik | 0.548916986 | -0.247627773 | 1.328067757 | -0.186457183 |
| 9230105E05Rik | 0 | NA | NA | NA |
| Mir763 | 0.331210058 | -0.971525162 | 1.982932424 | -0.489943656 |
| Gm24298 | 0 | NA | NA | NA |
| Gm38289 | 0.370157025 | -1.762458028 | 1.614528413 | -1.091624039 |
| Gm37505 | 0.097139547 | -0.443816764 | 3.724945049 | -0.119147198 |

| | | | | |
|---|---|---|---|---|
| 4921513I03Rik | 0 | NA | NA | NA |
| Msrb3 | 98.13132724 | -0.32524814 | 0.124643543 | -2.609426308 |
| Gm15961 | 1.450891998 | 0.245563494 | 0.86623863 | 0.283482502 |
| Gm15910 | 3.363554134 | -0.807499862 | 0.549188569 | -1.470350819 |
| Gm4473 | 0.74372259 | -0.462578951 | 1.360344774 | -0.340045376 |
| Lemd3 | 732.3495551 | 0.059862337 | 0.068234488 | 0.87730323 |
| Gm16166 | 0.157815785 | 0.483090695 | 3.465799425 | 0.139387955 |
| Gm23632 | 0.389039412 | -0.430427426 | 1.595432183 | -0.269787353 |
| Wif1 | 23.54109142 | -0.934606533 | 0.313975579 | -2.976685432 |
| Tbc1d30 | 832.5656543 | 0.460738656 | 0.101092225 | 4.557607236 |
| Gns | 1953.99692 | -0.334982413 | 0.104674453 | -3.200230851 |
| Rassf3 | 242.4931995 | 0.147546697 | 0.126121279 | 1.169879484 |
| Tbk1 | 841.1038649 | 0.004762916 | 0.091324153 | 0.052153957 |
| Xpot | 1103.074494 | -0.033008289 | 0.052253382 | -0.631696695 |
| Gm24330 | 0.750750929 | 0.490060367 | 1.283670283 | 0.381764985 |
| D930020B18Rik | 4.785986039 | 0.180584375 | 0.513854033 | 0.35143127 |
| BC048403 | 125.9765068 | -0.103767364 | 0.104484421 | -0.993137187 |
| C030027H14Rik | 0.046345709 | -0.443818988 | 3.724945049 | -0.119147795 |
| Srgap1 | 808.6398788 | -0.468949779 | 0.079602 | -5.891180867 |
| Tmem5 | 317.9777111 | 0.026399697 | 0.094745776 | 0.278637194 |
| Gm9081 | 0.90419666 | -0.025423809 | 1.060331026 | -0.023977238 |
| Avpr1a | 7.942965041 | -0.342126795 | 0.365007397 | -0.93731469 |
| Mir6412 | 0 | NA | NA | NA |
| Ppm1h | 1340.411078 | -0.166471101 | 0.079613612 | -2.09098793 |
| Mir8104 | 0 | NA | NA | NA |
| Mirlet7i | 1.909998494 | 0.856531905 | 0.671714603 | 1.275142601 |
| Mon2 | 1822.582098 | -0.085315213 | 0.045982019 | -1.85540379 |
| Usp15 | 1257.977968 | -0.071162052 | 0.057212074 | -1.243829277 |
| Gm27920 | 0 | NA | NA | NA |
| Fam19a2 | 696.2404132 | 0.508325768 | 0.08717248 | 5.83126425 |
| Gm22958 | 7.051847066 | 1.039283532 | 0.535621374 | 1.940332449 |
| Gm23777 | 0 | NA | NA | NA |
| Slc16a7 | 304.2321303 | 0.282855336 | 0.155127327 | 1.823375298 |
| Lrig3 | 67.70650913 | 0.160672367 | 0.148663404 | 1.080779548 |
| Gm22183 | 0 | NA | NA | NA |
| Mir378d | 0.20330765 | 0.517969191 | 2.214952998 | 0.233851098 |
| Gm24179 | 0.30098968 | 1.893077056 | 1.992832781 | 0.949942752 |
| Xrcc6bp1 | 46.49479728 | 0.246292289 | 0.187030914 | 1.316853369 |
| Ctdsp2 | 1189.150957 | -0.140629506 | 0.089309075 | -1.574638471 |
| Mir26a-2 | 0.925632266 | 0.298900283 | 1.209639361 | 0.247098674 |
| Mir546 | 0 | NA | NA | NA |
| Avil | 16.18070083 | -1.172632406 | 0.307904397 | -3.80843021 |
| Tsfm | 123.6160451 | -0.344598778 | 0.113594297 | -3.033592233 |
| Mettl21b | 2.262034746 | 0.444844343 | 0.702281239 | 0.633427633 |
| Mettl1 | 114.9215301 | -0.719354866 | 0.1390999 | -5.171498072 |
| Cyp27b1 | 5.379193204 | 0.18639502 | 0.432987392 | 0.430486023 |
| 9-Mar | 204.9397292 | 0.238739924 | 0.113686057 | 2.099992994 |

| | | | | |
|---|---|---|---|---|
| Cdk4 | 393.057798 | -0.634617243 | 0.130099565 | -4.877935157 |
| Tspan31 | 608.5951114 | -0.468011851 | 0.107633592 | -4.348195034 |
| Agap2 | 8823.335821 | 0.91253985 | 0.139544044 | 6.539439614 |
| Os9 | 1552.713354 | 0.063772606 | 0.078884176 | 0.808433438 |
| Gm23664 | 0.045882455 | 0.517959912 | 3.724945049 | 0.139051692 |
| B4galnt1 | 858.1814942 | 0.269034487 | 0.071942946 | 3.739553381 |
| Slc26a10 | 18.10165712 | 0.784116057 | 0.461848677 | 1.69777699 |
| Arhgef25 | 709.5156223 | 0.850672535 | 0.10630122 | 8.002471952 |
| Dtx3 | 1335.531959 | 0.251643683 | 0.051657882 | 4.871351148 |
| F420014N23Rik | 11.7378327 | 0.069067902 | 0.412651446 | 0.167375887 |
| Pip4k2c | 1116.600625 | 0.041390296 | 0.062315309 | 0.664207503 |
| Kif5a | 22627.74193 | 0.353991679 | 0.105148331 | 3.366593408 |
| Dctn2 | 1693.964189 | 0.035983312 | 0.0495153 | 0.726710976 |
| Mbd6 | 504.0837198 | -0.157174069 | 0.140231604 | -1.120817733 |
| Gm20492 | 1.795058199 | 0.186157067 | 0.780983055 | 0.238362492 |
| Gm4189 | 3.902576099 | 1.294711028 | 0.620461333 | 2.086690919 |
| Ddit3 | 291.5705915 | -0.959736004 | 0.195035066 | -4.920838206 |
| Mars | 495.381841 | -0.436223389 | 0.099300917 | -4.392944224 |
| Gm6283 | 1.594842726 | 1.435894491 | 1.047781588 | 1.370413936 |
| Arhgap9 | 51.31007067 | -1.052386913 | 0.240515918 | -4.375539555 |
| Gli1 | 50.51559619 | 0.543362094 | 0.262297727 | 2.071547094 |
| Inhbe | 0.872322242 | -0.452114537 | 1.328937784 | -0.340207452 |
| Inhbc | 0.072025462 | 0.517959912 | 3.724945049 | 0.139051692 |
| R3hdm2 | 2616.368291 | 0.214799878 | 0.073881061 | 2.907374027 |
| Gm23819 | 0 | NA | NA | NA |
| Stac3 | 24.43958761 | -0.774246223 | 0.433283602 | -1.78692713 |
| Ndufa4l2 | 7.948266496 | 0.084683202 | 0.373145475 | 0.226944202 |
| Shmt2 | 276.9083029 | -0.70864349 | 0.094803771 | -7.474844945 |
| Nxph4 | 143.5244743 | -0.122900639 | 0.227943896 | -0.539170565 |
| Lrp1 | 13094.28162 | -0.091689976 | 0.084248342 | -1.088329739 |
| Gm16217 | 0.962459693 | -1.93718353 | 1.094923356 | -1.769241216 |
| Stat6 | 224.5914028 | -0.837653454 | 0.167155437 | -5.011224709 |
| Nab2 | 546.0189877 | -0.487773974 | 0.143297901 | -3.403915689 |
| Gm16229 | 0.705361133 | -1.633729791 | 1.179131968 | -1.385536001 |
| Tmem194 | 106.2750348 | -0.305273434 | 0.171721827 | -1.77772063 |
| Gm16230 | 1.3230785 | -0.344632147 | 0.857642234 | -0.401836726 |
| Myo1a | 8.599191626 | 0.053078997 | 0.365988385 | 0.145029183 |
| Tac2 | 49.86492223 | 0.1069604 | 0.194093251 | 0.551077382 |
| Zbtb39 | 235.0509989 | 0.010987726 | 0.130426319 | 0.084244701 |
| Gpr182 | 35.51224453 | -0.437577376 | 0.224881517 | -1.945812982 |
| Rdh1 | 7.125358113 | 0.812360581 | 0.516625116 | 1.572437257 |
| Gm28046 | 0 | NA | NA | NA |
| Rdh9 | 1.759257097 | 2.189087436 | 0.931925217 | 2.348994742 |
| Rdh16 | 4.45419811 | 0.952077198 | 0.504652698 | 1.886598848 |
| Rdh18-ps | 0.308514759 | 1.066550242 | 1.753477658 | 0.608248549 |
| Rdh19 | 0.061920613 | -0.443816764 | 3.724945049 | -0.119147198 |
| BC089597 | 0.047948698 | 0.037064902 | 3.724945049 | 0.009950456 |

| | | | | |
|---|---|---|---|---|
| Rdh7 | 0 | NA | NA | NA |
| Sdr9c7 | 0.145038387 | 0.50369485 | 2.88682069 | 0.174480823 |
| Gm15905 | 0 | NA | NA | NA |
| Gm15900 | 1.613670225 | 0.40708292 | 1.189675181 | 0.342179888 |
| Gm23996 | 0 | NA | NA | NA |
| Hsd17b6 | 0.048949532 | 0.037064902 | 3.724945049 | 0.009950456 |
| Prim1 | 70.56909489 | -0.38627663 | 0.142458977 | -2.711493783 |
| Naca | 1262.918694 | -0.109018915 | 0.064335878 | -1.694527503 |
| Ptges3 | 1465.432757 | -0.434643771 | 0.095078882 | -4.571401806 |
| Atp5b | 9103.422248 | -0.040593226 | 0.072687638 | -0.558461207 |
| Snord59a | 1.560301263 | 1.163114425 | 0.899937225 | 1.29243951 |
| mmu-mir-677 | 0.075043205 | 0.517957688 | 3.724945049 | 0.139051095 |
| Mir677 | 0 | NA | NA | NA |
| Baz2a | 1409.129041 | -0.139923483 | 0.086025059 | -1.626543293 |
| Rbms2 | 301.1430177 | -0.523688814 | 0.140895551 | -3.716858413 |
| Gls2 | 183.4405106 | 0.215971477 | 0.131625789 | 1.64079911 |
| Spryd4 | 66.56019931 | -0.248557111 | 0.165988756 | -1.497433424 |
| Mip | 0 | NA | NA | NA |
| Gm26847 | 5.379666823 | 1.29312432 | 0.550498139 | 2.349007613 |
| Timeless | 115.0545271 | -0.579309503 | 0.141567939 | -4.092095346 |
| Apon | 0.725473741 | -2.20609192 | 1.198890862 | -1.840110714 |
| Apof | 3.028410459 | -0.666849133 | 0.566047451 | -1.178079914 |
| Stat2 | 2927.543515 | -0.377260809 | 0.102796805 | -3.669966294 |
| Il23a | 1.801413292 | -1.767306511 | 0.982907375 | -1.79803973 |
| Gm23241 | 0 | NA | NA | NA |
| Pan2 | 211.7421926 | 0.039930978 | 0.131746992 | 0.303088344 |
| Cnpy2 | 199.2616733 | -0.462999011 | 0.12837911 | -3.606498052 |
| Cs | 2382.585503 | -0.076891902 | 0.066521276 | -1.155899396 |
| Gm23182 | 1.59776671 | 0.10051217 | 0.788365504 | 0.127494378 |
| Coq10a | 213.594349 | 0.001916288 | 0.085475416 | 0.022419171 |
| Ankrd52 | 1433.771392 | -0.280760983 | 0.079339479 | -3.538729869 |
| Gm26308 | 0 | NA | NA | NA |
| Mir6914 | 0.577876106 | -0.553212284 | 1.46892268 | -0.376610894 |
| mmu-mir-6915 | 0.129647038 | -0.443820923 | 2.992845889 | -0.148293945 |
| Gm17201 | 0.247536319 | 0.507002133 | 1.847383112 | 0.274443417 |
| Slc39a5 | 1.555265169 | 2.026804364 | 1.043643957 | 1.942045801 |
| Nabp2 | 369.4297074 | -0.14226407 | 0.073849876 | -1.926395525 |
| Gm25361 | 0 | NA | NA | NA |
| Rnf41 | 702.2743574 | 0.097187003 | 0.064635473 | 1.503617116 |
| Gm26347 | 3.130125749 | 0.06210577 | 0.741350428 | 0.083773837 |
| Mir8105 | 0.208009692 | -1.017917153 | 2.593857713 | -0.39243369 |
| Smarcc2 | 3642.728046 | 0.159467155 | 0.07142389 | 2.232686493 |
| Myl6 | 1464.017352 | -0.339275853 | 0.111101825 | -3.053737885 |
| Myl6b | 108.9152618 | 0.121259486 | 0.13410267 | 0.904228718 |
| A430046D13Rik | 90.5350024 | 0.261842169 | 0.171700908 | 1.524990007 |
| Esyt1 | 580.33528 | -0.701567143 | 0.184549944 | -3.801502868 |
| Mir6916 | 0 | NA | NA | NA |

| | | | | |
|---|---|---|---|---|
| Gm29585 | 0.520597108 | 0.268959964 | 1.405045792 | 0.191424341 |
| Zc3h10 | 205.4911794 | -0.086314107 | 0.088858193 | -0.971369146 |
| Rpl41 | 1706.429966 | -0.325888322 | 0.16043061 | -2.031335055 |
| Pa2g4 | 643.314172 | -0.217302878 | 0.090729772 | -2.395055908 |
| AC117232.1 | 0 | NA | NA | NA |
| Erbb3 | 358.7695039 | -1.001462949 | 0.166980061 | -5.997500195 |
| Mir6917 | 0.044191549 | 0.037062677 | 3.724945049 | 0.009949859 |
| Rps26 | 1049.123509 | -0.231769503 | 0.102067072 | -2.270756841 |
| Ikzf4 | 257.6485544 | 0.014131991 | 0.138988593 | 0.101677341 |
| Gm26876 | 8.215402088 | -0.06672823 | 0.36871165 | -0.180976733 |
| Suox | 127.1525145 | 0.111780265 | 0.127990337 | 0.873349256 |
| Rab5b | 1465.571473 | 0.018745812 | 0.063165457 | 0.296773159 |
| Cdk2 | 38.99389103 | -0.532736472 | 0.217461992 | -2.449791196 |
| Pmel | 8.731260839 | 0.93872882 | 0.375203087 | 2.501921899 |
| Dgka | 134.8647648 | -0.425772295 | 0.127667273 | -3.335015205 |
| Wibg | 245.4724351 | -0.099432188 | 0.127189943 | -0.781761399 |
| Rpsa-ps2 | 30.10672328 | -0.065357428 | 0.232245929 | -0.28141474 |
| Mmp19 | 189.6992867 | -1.916479309 | 0.498917176 | -3.841277475 |
| Tmem198b | 157.6632237 | -0.514349231 | 0.150771978 | -3.411437831 |
| Dnajc14 | 675.6783784 | -0.189650637 | 0.055717164 | -3.403809944 |
| Ormdl2 | 72.79417923 | -0.707612279 | 0.212028642 | -3.337342885 |
| Sarnp | 164.5829855 | 0.087952113 | 0.128212792 | 0.685985479 |
| Gdf11 | 478.8249761 | -0.365693044 | 0.13291185 | -2.751395336 |
| Cd63 | 793.3225511 | -1.216245864 | 0.233908069 | -5.199674677 |
| Rdh5 | 84.87448236 | -1.47235573 | 0.177612273 | -8.289718397 |
| Bloc1s1 | 41.20713241 | -0.081982555 | 0.267472962 | -0.306507823 |
| Itga7 | 368.0914646 | -0.890823572 | 0.134246487 | -6.635730956 |
| Gm25494 | 0.763054649 | -0.421599063 | 1.184379286 | -0.355966258 |
| Mettl7b | 0.038981395 | -0.443816764 | 3.724945049 | -0.119147198 |
| 9030616G12Rik | 0 | NA | NA | NA |
| Olfr9 | 0.042716945 | 0.037062677 | 3.724945049 | 0.009949859 |
| Olfr763 | 0 | NA | NA | NA |
| Olfr765 | 0 | NA | NA | NA |
| Olfr767 | 0 | NA | NA | NA |
| Olfr768 | 0 | NA | NA | NA |
| Olfr769 | 0 | NA | NA | NA |
| Olfr770 | 0 | NA | NA | NA |
| Olfr771 | 0 | NA | NA | NA |
| Gm25807 | 0 | NA | NA | NA |
| Olfr772 | 0 | NA | NA | NA |
| Olfr773 | 0 | NA | NA | NA |
| Olfr774 | 0.045882455 | 0.517959912 | 3.724945049 | 0.139051692 |
| Olfr775 | 0 | NA | NA | NA |
| Olfr776 | 0 | NA | NA | NA |
| Olfr777 | 0 | NA | NA | NA |
| Gm25764 | 0 | NA | NA | NA |
| Olfr779 | 0 | NA | NA | NA |

| | | | | |
|---|---|---|---|---|
| Olfr780 | 0 | NA | NA | NA |
| Olfr781 | 0 | NA | NA | NA |
| Olfr782 | 0.044544648 | 0.517957688 | 3.724945049 | 0.139051095 |
| Olfr784 | 0 | NA | NA | NA |
| Olfr786 | 0 | NA | NA | NA |
| Olfr787 | 0 | NA | NA | NA |
| Olfr788 | 0 | NA | NA | NA |
| Olfr790 | 0.08169834 | -0.443821212 | 3.724945049 | -0.119148392 |
| Olfr791 | 0 | NA | NA | NA |
| Olfr792 | 0 | NA | NA | NA |
| Olfr794 | 0 | NA | NA | NA |
| Gm23252 | 0 | NA | NA | NA |
| Olfr796 | 0 | NA | NA | NA |
| Olfr798 | 0 | NA | NA | NA |
| Olfr799 | 0.046345709 | -0.443818988 | 3.724945049 | -0.119147795 |
| Olfr800 | 0 | NA | NA | NA |
| Olfr801 | 0 | NA | NA | NA |
| Olfr802 | 0 | NA | NA | NA |
| Olfr803 | 0 | NA | NA | NA |
| Gm24717 | 0 | NA | NA | NA |
| Olfr804 | 0 | NA | NA | NA |
| Olfr805 | 0 | NA | NA | NA |
| Gm24485 | 0 | NA | NA | NA |
| Olfr806 | 0 | NA | NA | NA |
| Olfr807 | 0 | NA | NA | NA |
| Olfr808 | 0 | NA | NA | NA |
| Olfr809 | 0.046345709 | -0.443818988 | 3.724945049 | -0.119147795 |
| Olfr810 | 0.066050875 | -0.443818988 | 3.724945049 | -0.119147795 |
| Olfr811 | 0 | NA | NA | NA |
| Olfr812 | 0 | NA | NA | NA |
| Olfr813 | 0 | NA | NA | NA |
| Olfr814 | 0 | NA | NA | NA |
| Olfr815 | 0 | NA | NA | NA |
| Olfr816 | 0.045882455 | 0.517959912 | 3.724945049 | 0.139051692 |
| Olfr818 | 0 | NA | NA | NA |
| Olfr819 | 0 | NA | NA | NA |
| Olfr247 | 0 | NA | NA | NA |
| Gm10310 | 0 | NA | NA | NA |
| Olfr820 | 0 | NA | NA | NA |
| Olfr821 | 0 | NA | NA | NA |
| Olfr822 | 0 | NA | NA | NA |
| Olfr823 | 0.045882455 | 0.517959912 | 3.724945049 | 0.139051692 |
| Gm5181 | 0 | NA | NA | NA |
| Olfr824 | 0 | NA | NA | NA |
| Olfr825 | 0.062426873 | 0.517959912 | 3.724945049 | 0.139051692 |
| Olfr826 | 0 | NA | NA | NA |
| Olfr827 | 0.062426873 | 0.517959912 | 3.724945049 | 0.139051692 |

| | | | | |
|---|---|---|---|---|
| Neurod4 | 9.261960518 | 0.504938974 | 0.38003086 | 1.328678872 |
| Tespa1 | 3.163043473 | -0.011275382 | 0.505291065 | -0.022314628 |
| Vmn2r84 | 31.40616345 | 0.838034236 | 0.291254525 | 2.877326059 |
| Gm9770 | 48.13481903 | -0.226563366 | 0.158272871 | -1.431473154 |
| Vmn2r85 | 40.51229935 | 0.833380519 | 0.283036021 | 2.944432713 |
| Vmn2r86 | 19.28575388 | 0.546422484 | 0.368609097 | 1.482390121 |
| Vmn2r87 | 47.38212013 | 0.766955763 | 0.303121592 | 2.530191791 |
| Gm23793 | 0 | NA | NA | NA |
| Gm26445 | 0.178837887 | -0.891757593 | 2.696671676 | -0.330688233 |
| Gm20746 | 0.242679635 | 0.279246689 | 2.490377907 | 0.112130247 |
| Pisd-ps1 | 253.7945557 | 0.166081628 | 0.126424297 | 1.313684403 |
| Sfi1 | 373.2312864 | -0.074375051 | 0.150932184 | -0.492771318 |
| Gm11399 | 1.590475537 | -0.088856948 | 0.807236083 | -0.11007554 |
| Gm11400 | 21.45170634 | -0.204243344 | 0.307168073 | -0.664923742 |
| Drg1 | 586.4306311 | -0.209276146 | 0.07461737 | -2.804657218 |
| Gm12735 | 2.937817048 | -1.563307379 | 0.693799638 | -2.25325482 |
| Fau-ps2 | 1.291849242 | 0.999433235 | 1.088002075 | 0.918594971 |
| Eif4enif1 | 683.4614762 | 0.068854268 | 0.074773481 | 0.920838068 |
| Patz1 | 528.6289566 | 0.149814288 | 0.083467845 | 1.794874289 |
| Gm12592 | 46.92358016 | 0.553474153 | 0.222310604 | 2.489643513 |
| Gm11944 | 38.0840153 | -0.029454785 | 0.239601751 | -0.122932263 |
| Pik3ip1 | 504.3401145 | 0.318804201 | 0.090853482 | 3.508992652 |
| Limk2 | 962.7434137 | -0.150870106 | 0.050836585 | -2.967746658 |
| Gm11948 | 5.24095402 | 1.235221322 | 0.569378313 | 2.169421091 |
| Rnf185 | 379.3246065 | -0.12232762 | 0.082768409 | -1.477950601 |
| 8430429K09Rik | 61.46987694 | -0.004793217 | 0.161512519 | -0.029677062 |
| Gm11945 | 1.421692497 | 0.11435719 | 0.895458142 | 0.127708025 |
| Gm24439 | 0.228704471 | 0.998557651 | 2.201211124 | 0.453640107 |
| Pla2g3 | 97.55501564 | 0.489010676 | 0.230007396 | 2.126065005 |
| Inpp5j | 911.3473185 | 0.139499037 | 0.08253595 | 1.690160923 |
| Selm | 738.0868402 | 0.165362774 | 0.141108178 | 1.171886532 |
| Smtn | 69.98786474 | -0.273483581 | 0.184065218 | -1.485797178 |
| Gm11946 | 0.933952761 | -0.114973946 | 1.071049492 | -0.107346997 |
| Gm11947 | 0 | NA | NA | NA |
| Gm11949 | 0 | NA | NA | NA |
| Gm11950 | 0 | NA | NA | NA |
| Gm11951 | 0.453565911 | 0.091785812 | 1.484900804 | 0.061812757 |
| Tug1 | 1777.476593 | -0.115776965 | 0.071466325 | -1.620021252 |
| Gm27624 | 0 | NA | NA | NA |
| Gm27517 | 0 | NA | NA | NA |
| Gm27937 | 0 | NA | NA | NA |
| Gm27640 | 0 | NA | NA | NA |
| Morc2a | 721.9875827 | -0.286298902 | 0.075106681 | -3.811896628 |
| Gm26393 | 0.141094079 | 0.037072457 | 3.563850528 | 0.01040236 |
| Gm16518 | 10.42668312 | 0.826706688 | 0.397460175 | 2.07997364 |
| Osbp2 | 906.3048646 | 0.491218187 | 0.101443032 | 4.842305846 |
| Gm24013 | 0.11193333 | 0.037073798 | 3.724945049 | 0.009952844 |

| | | | | |
|---|---|---|---|---|
| Gm11952 | 0 | NA | NA | NA |
| 4921536K21Rik | 11.5351327 | -0.920594542 | 0.336896186 | -2.732576326 |
| Dusp18 | 270.8170531 | -0.396252386 | 0.099127364 | -3.997406683 |
| Slc35e4 | 326.5983649 | -0.193348923 | 0.131086272 | -1.474974608 |
| Tcn2 | 256.4564119 | -0.488448965 | 0.142835 | -3.419672802 |
| 4930556J24Rik | 2.805439532 | 0.739087698 | 0.637913575 | 1.158601614 |
| Gm11953 | 2.130460637 | 1.375392188 | 0.801617265 | 1.715771664 |
| Pes1 | 430.3711271 | -0.437158445 | 0.092716591 | -4.714996945 |
| Gal3st1 | 117.3315211 | -0.310571461 | 0.148986315 | -2.08456368 |
| Gm11954 | 0.875788929 | 2.376588624 | 1.185877389 | 2.004076178 |
| Sec14l4 | 4.148966471 | 0.879815195 | 0.57864345 | 1.520478967 |
| Sec14l3 | 15.8893622 | 1.011992212 | 0.511908135 | 1.976901992 |
| Mtfp1 | 343.8323402 | 0.481740757 | 0.115221391 | 4.181001046 |
| Sec14l2 | 346.4809089 | 0.34923599 | 0.112578159 | 3.102164684 |
| Gm11955 | 0 | NA | NA | NA |
| Gm11957 | 1.061319949 | 0.393548498 | 1.000233714 | 0.393456541 |
| Rnf215 | 199.4153862 | -0.397333245 | 0.095215977 | -4.172968233 |
| Mir3060 | 6.092409896 | 0.299559076 | 0.486340846 | 0.615944718 |
| Ccdc157 | 315.6146368 | -0.07106574 | 0.08625925 | -0.823862259 |
| Sf3a1 | 976.4741568 | -0.111146473 | 0.060566238 | -1.835122607 |
| Tbc1d10a | 286.7296832 | -0.538525509 | 0.10984862 | -4.90243306 |
| Gatsl3 | 28.49626455 | -0.615347068 | 0.281471894 | -2.18617589 |
| Gm11956 | 0.045882455 | 0.517959912 | 3.724945049 | 0.139051692 |
| Osm | 17.76436873 | -1.351117664 | 0.329852722 | -4.096124044 |
| Lif | 163.2509148 | -2.181040697 | 0.473518237 | -4.606033153 |
| Gm11959 | 0 | NA | NA | NA |
| Gm11958 | 0 | NA | NA | NA |
| Gm24803 | 1.135110426 | 0.866921525 | 1.040156164 | 0.83345324 |
| Hormad2 | 2.594629735 | -0.495202984 | 0.61610454 | -0.803764542 |
| Mtmr3 | 1608.347275 | 0.050327833 | 0.069173084 | 0.727563816 |
| Gm11960 | 2.472144487 | 0.492590172 | 0.660193624 | 0.746129853 |
| Gm11032 | 0 | NA | NA | NA |
| Gm11961 | 0.364460012 | -0.059372744 | 1.532040925 | -0.03875402 |
| Ascc2 | 346.5615331 | -0.615229435 | 0.07100629 | -8.664435757 |
| Uqcr10 | 496.5398793 | -0.127066724 | 0.109531255 | -1.160095573 |
| Zmat5 | 89.65500592 | -0.347151813 | 0.131332377 | -2.64330717 |
| Cabp7 | 502.185691 | -0.016925921 | 0.130116486 | -0.13008283 |
| Nf2 | 1204.431154 | -0.057155563 | 0.054384284 | -1.050957356 |
| Nipsnap1 | 497.6595648 | 0.313218104 | 0.082684113 | 3.788129219 |
| Mir6918 | 0.256523886 | 0.998869203 | 1.83437832 | 0.54452737 |
| Thoc5 | 333.4021351 | -0.138500326 | 0.069068135 | -2.005270953 |
| AA413626 | 4.943749683 | 1.052575149 | 0.540737381 | 1.946555178 |
| Nefh | 1154.799543 | -0.16074075 | 0.232757156 | -0.690594232 |
| Ap1b1 | 2045.497515 | 0.112231062 | 0.045995231 | 2.440058673 |
| AL645522.1 | 0.147068667 | 0.998853574 | 3.51441668 | 0.284216035 |
| Gm24958 | 0 | NA | NA | NA |
| Gas2l1 | 586.1682792 | -0.235259353 | 0.067918098 | -3.463868416 |

| | | | | |
|---|---|---|---|---|
| Rasl10a | 112.452118 | -0.260685543 | 0.267769416 | -0.973544878 |
| Ewsr1 | 2343.895997 | 0.151990336 | 0.09592787 | 1.58442313 |
| Rhbdd3 | 147.1104437 | -0.013431175 | 0.125010933 | -0.107440002 |
| Emid1 | 46.41330166 | 0.13985254 | 0.185335521 | 0.754591125 |
| Kremen1 | 346.9264262 | -0.417859319 | 0.115107105 | -3.630178327 |
| Gm25142 | 0 | NA | NA | NA |
| Znrf3 | 372.0960024 | -0.0287901 | 0.09391553 | -0.306553134 |
| Gm11962 | 0 | NA | NA | NA |
| Gm11963 | 0 | NA | NA | NA |
| Xbp1 | 893.6785691 | -0.802202124 | 0.101408202 | -7.910623662 |
| Ccdc117 | 216.8499613 | -0.224024773 | 0.105062347 | -2.132303153 |
| Gm22471 | 0 | NA | NA | NA |
| Ankrd36 | 0.048949532 | 0.037064902 | 3.724945049 | 0.009950456 |
| Gm11964 | 12.76498303 | -0.237882087 | 0.317844522 | -0.7484228 |
| Gm11965 | 0.126225412 | 0.03707172 | 3.698755067 | 0.010022756 |
| Mrps24 | 181.2008852 | -0.40062096 | 0.114245609 | -3.506663953 |
| Urgcp | 807.8473661 | 0.176591623 | 0.068716902 | 2.569842624 |
| 2210015D19Rik | 182.3089558 | 0.232577737 | 0.085774219 | 2.711510984 |
| Dbnl | 1162.267136 | -0.368789226 | 0.12490206 | -2.952627241 |
| Pgam2 | 10.44641268 | 1.334396922 | 0.394532169 | 3.382225904 |
| Gm11966 | 36.99000587 | -0.05884654 | 0.204793826 | -0.287345284 |
| Polm | 78.87651427 | -0.030542382 | 0.135712846 | -0.225051518 |
| Aebp1 | 394.2398871 | -0.737740996 | 0.177947369 | -4.145838175 |
| Pold2 | 110.4180631 | -0.306649601 | 0.145936147 | -2.101258711 |
| Myl7 | 0.376384159 | 1.406066523 | 1.720062268 | 0.817450943 |
| Gck | 15.14292826 | -0.062599985 | 0.32500398 | -0.192612978 |
| Gm11967 | 0.249409806 | -1.346410794 | 2.518891846 | -0.534525052 |
| Ykt6 | 770.8424576 | -0.212113543 | 0.06244082 | -3.397033284 |
| Camk2b | 7644.2209 | 0.498705784 | 0.132927692 | 3.751707225 |
| Nudcd3 | 1857.880843 | 0.172992483 | 0.036821081 | 4.698191298 |
| Rps15a-ps6 | 9.077851534 | 0.524951839 | 0.380558537 | 1.379424684 |
| Gm11969 | 6.242235417 | 0.281067646 | 0.487554308 | 0.576484796 |
| A730071L15Rik | 1.959824955 | -0.190502717 | 0.676556216 | -0.281577071 |
| Npc1l1 | 17.38551341 | -0.267245568 | 0.398890698 | -0.669971923 |
| Gm11401 | 0.198115332 | -0.476840274 | 3.232275238 | -0.14752465 |
| Ddx56 | 347.1312207 | -0.224182 | 0.08235772 | -2.722052032 |
| Tmed4 | 598.9634342 | -0.063277964 | 0.086219651 | -0.733915791 |
| n-R5s67 | 0 | NA | NA | NA |
| Ogdh | 3550.241053 | -0.049584141 | 0.063730472 | -0.77802877 |
| Zmiz2 | 2387.144131 | 0.456758147 | 0.068122743 | 6.704928902 |
| Mir7651 | 2.015150011 | 0.938848204 | 0.855029064 | 1.09803075 |
| Ppia | 2699.446977 | 0.175047483 | 0.092856787 | 1.885133968 |
| H2afv | 161.7851873 | -0.040335684 | 0.143186673 | -0.28169999 |
| Gm11970 | 0.194008847 | -0.73756515 | 2.638008782 | -0.27959162 |
| Gm11971 | 0.305478853 | 0.396568976 | 1.756724924 | 0.225743354 |
| Gm11972 | 0.155921052 | 1.020628401 | 2.829228019 | 0.360744483 |
| Purb | 8286.574662 | 0.278752761 | 0.08438269 | 3.303435332 |

| | | | | |
|---|---|---|---|---|
| Gm22123 | 0 | NA | NA | NA |
| Gm11973 | 8.076220047 | -0.035675526 | 0.356091699 | -0.100186347 |
| Myo1g | 77.4741113 | -0.941280896 | 0.232324421 | -4.051579643 |
| Gm11974 | 44.13713644 | -0.710762945 | 0.24587991 | -2.890691419 |
| Gm22457 | 0 | NA | NA | NA |
| Gm24000 | 0.045882455 | 0.517959912 | 3.724945049 | 0.139051692 |
| Ccm2 | 367.7041191 | -0.341027523 | 0.115791509 | -2.945185931 |
| Gm11975 | 0 | NA | NA | NA |
| Nacad | 676.8149995 | -0.004917285 | 0.076196069 | -0.064534629 |
| Tbrg4 | 350.0488541 | -0.198822996 | 0.090861677 | -2.188194231 |
| Gm24313 | 0.992467499 | -0.273036105 | 1.116375667 | -0.24457368 |
| Snora5c | 0.582740348 | 1.377337543 | 1.258775635 | 1.094188277 |
| Wap | 0.066050875 | -0.443818988 | 3.724945049 | -0.119147795 |
| Gm11978 | 0.38240426 | 0.8775935 | 1.563708675 | 0.561225703 |
| Ramp3 | 89.90826615 | 0.337082756 | 0.150932918 | 2.233328284 |
| Gm11980 | 0.066050875 | -0.443818988 | 3.724945049 | -0.119147795 |
| Gm11981 | 0 | NA | NA | NA |
| Gm11976 | 0 | NA | NA | NA |
| Gm11977 | 0.369792422 | -0.843842843 | 1.566970918 | -0.538518509 |
| Gm11979 | 0 | NA | NA | NA |
| Gm11982 | 0 | NA | NA | NA |
| Gm11983 | 0.178864798 | -1.319733861 | 3.264222506 | -0.404302666 |
| Gm11984 | 0 | NA | NA | NA |
| Adcy1 | 9022.462185 | 0.744356314 | 0.153070863 | 4.862821705 |
| Gm11985 | 0 | NA | NA | NA |
| Igfbp1 | 2.637679718 | 0.896556865 | 0.648659289 | 1.382169162 |
| Igfbp3 | 602.9842508 | -1.725247691 | 0.315380712 | -5.470365262 |
| Gm11986 | 0.276998953 | 0.586106077 | 1.786237716 | 0.328123223 |
| Gm27393 | 0 | NA | NA | NA |
| Gm11987 | 0 | NA | NA | NA |
| Gm24051 | 0 | NA | NA | NA |
| Gm11988 | 0 | NA | NA | NA |
| Gm11989 | 0.796528503 | 1.160244881 | 1.12220262 | 1.033899636 |
| Tns3 | 1722.397234 | -0.456074788 | 0.131595816 | -3.465724084 |
| Gm11990 | 0 | NA | NA | NA |
| Pkd1l1 | 3.97064273 | 0.300411768 | 0.508081711 | 0.591266643 |
| Gm11991 | 0 | NA | NA | NA |
| Hus1 | 255.0186718 | -0.357668016 | 0.118557163 | -3.016840217 |
| Sun3 | 0.968963158 | -0.840436925 | 1.060555626 | -0.792449641 |
| Gm11992 | 17.30127652 | 0.008288255 | 0.299551205 | 0.02766891 |
| Upp1 | 71.12326318 | -1.22390914 | 0.301451253 | -4.060056575 |
| Gm11993 | 27.56449686 | -0.307703385 | 0.22422603 | -1.372291099 |
| Abca13 | 10.03132759 | -0.574804326 | 0.398762928 | -1.441468816 |
| Gm36954 | 0 | NA | NA | NA |
| Eif3s6-ps1 | 4.535504711 | 0.059994135 | 0.497277244 | 0.120645245 |
| Gm11996 | 0 | NA | NA | NA |
| Gm11997 | 0 | NA | NA | NA |

| | | | | |
|---|---|---|---|---|
| Vwc2 | 143.5725646 | 0.016919647 | 0.107952845 | 0.156731827 |
| Gm23065 | 0.966893809 | 1.873331976 | 1.064991827 | 1.759010659 |
| Zpbp | 24.98626369 | 0.440227686 | 0.225681421 | 1.950659846 |
| Gm12598 | 0.344633608 | 2.231891011 | 1.890394648 | 1.180648185 |
| Gm26458 | 0.27289602 | 0.967555054 | 1.848376103 | 0.523462218 |
| Gm12599 | 2.05315699 | 0.416336858 | 0.708637734 | 0.587517202 |
| 4930415F15Rik | 5.574285272 | -0.123358911 | 0.435809852 | -0.283056729 |
| Gm11998 | 0.161486377 | -0.444693834 | 2.795640213 | -0.159066904 |
| 4930512M02Rik | 0.106358623 | 0.517955464 | 3.724945049 | 0.139050498 |
| Gm11999 | 0.204942615 | -0.537224224 | 2.606107066 | -0.206140504 |
| Ikzf1 | 243.3512954 | -0.636563817 | 0.158542001 | -4.015111524 |
| Gm12000 | 0.060174537 | 0.517957688 | 3.724945049 | 0.139051095 |
| Fignl1 | 18.28682388 | -1.491574366 | 0.33715303 | -4.42402777 |
| Ddc | 130.1558318 | 0.44054225 | 0.224123245 | 1.965624981 |
| 1700042O10Rik | 0.263601517 | -0.412503951 | 2.075096713 | -0.198787819 |
| Gm25058 | 0 | NA | NA | NA |
| Grb10 | 839.2894725 | 0.123567817 | 0.071371191 | 1.731340282 |
| Gm24036 | 0 | NA | NA | NA |
| Gm12001 | 0.097139547 | -0.443816764 | 3.724945049 | -0.119147198 |
| Cobl | 1095.669369 | 0.202253537 | 0.128668301 | 1.571898718 |
| Gm12002 | 1.538964163 | -1.716751519 | 0.902396863 | -1.90243516 |
| Gm22156 | 0 | NA | NA | NA |
| Gm22789 | 0 | NA | NA | NA |
| Gm12003 | 0.812725683 | -0.880744255 | 1.128278356 | -0.780609014 |
| Gm12004 | 0 | NA | NA | NA |
| 4930554G24Rik | 0.349921031 | -2.194914359 | 1.927423665 | -1.138781472 |
| Gm12005 | 0 | NA | NA | NA |
| Gm12006 | 0 | NA | NA | NA |
| 1700046C09Rik | 0 | NA | NA | NA |
| Pom121l12 | 0 | NA | NA | NA |
| Gm12007 | 0 | NA | NA | NA |
| Gm12008 | 0 | NA | NA | NA |
| Gm12009 | 7.023367752 | -0.694860326 | 0.401392908 | -1.731122579 |
| Gm24707 | 0 | NA | NA | NA |
| Gm25490 | 0 | NA | NA | NA |
| Gm12010 | 0 | NA | NA | NA |
| Gm37319 | 0 | NA | NA | NA |
| Gm12011 | 0 | NA | NA | NA |
| Vstm2a | 820.6459809 | 0.510439963 | 0.086680765 | 5.888733943 |
| Gm12012 | 0.950406788 | 2.512482592 | 1.323771721 | 1.897972703 |
| Sec61g | 266.2425794 | -0.453375509 | 0.150838991 | -3.005691729 |
| Akt2-ps | 59.61674162 | -0.334508574 | 0.143283114 | -2.334598719 |
| Gm12663 | 89.45054939 | 0.09018237 | 0.116772382 | 0.772291942 |
| Gm12664 | 0 | NA | NA | NA |
| Gm24714 | 0 | NA | NA | NA |
| Gm25698 | 0 | NA | NA | NA |
| Egfr | 584.8862284 | -0.02158185 | 0.124494194 | -0.173356276 |

| | | | | |
|---|---|---|---|---|
| Egfros | 2.163206028 | -1.964740138 | 0.759762339 | -2.585993064 |
| Eldr | 6.358422579 | -1.854653405 | 0.487126006 | -3.807338104 |
| Fbxo48 | 10.03527172 | -0.217651739 | 0.332255311 | -0.655073769 |
| Plek | 727.3535995 | -0.514782791 | 0.118184372 | -4.355760272 |
| Gm12014 | 0 | NA | NA | NA |
| Cnrip1 | 500.2213394 | 0.285836921 | 0.077860285 | 3.67115174 |
| Gm12013 | 1.807586739 | -0.359873391 | 0.744045747 | -0.483671055 |
| Ppp3r1 | 5062.853256 | 0.615586196 | 0.083457651 | 7.376030694 |
| Wdr92 | 87.67747582 | -0.476939996 | 0.158761811 | -3.004122932 |
| Pno1 | 283.54597 | -0.470729991 | 0.121671118 | -3.868872077 |
| C1d | 398.4995549 | 0.015919355 | 0.12572481 | 0.126620631 |
| Gm12015 | 0 | NA | NA | NA |
| Gm23959 | 0 | NA | NA | NA |
| Gm12016 | 0 | NA | NA | NA |
| Etaa1 | 152.4498586 | -0.052527499 | 0.107582447 | -0.488253434 |
| Etaa1os | 0.910758945 | -0.398226123 | 1.004456678 | -0.396459232 |
| Gm12017 | 1.313957777 | -0.137266056 | 0.924253309 | -0.148515623 |
| Gm12018 | 0.106112162 | -0.443821212 | 3.724945049 | -0.119148392 |
| Gm12019 | 0 | NA | NA | NA |
| Gm12020 | 0.546458135 | -0.853491249 | 1.32371477 | -0.644769755 |
| Gm37869 | 0 | NA | NA | NA |
| Gm12022 | 0.803435494 | 0.5073563 | 1.07955963 | 0.46996598 |
| Gm12021 | 0.841174947 | 0.943238076 | 1.146830107 | 0.822474114 |
| Gm12024 | 0.313606012 | 1.699826049 | 3.704357424 | 0.45887204 |
| Gm12023 | 2.23920711 | 1.934219481 | 1.033634074 | 1.871280687 |
| Gm23093 | 0 | NA | NA | NA |
| Gm28401 | 1.961032719 | 1.946043758 | 0.933188371 | 2.085370777 |
| Gm23021 | 0 | NA | NA | NA |
| 8430419K02Rik | 12.33011639 | 1.795202208 | 0.469386199 | 3.824573907 |
| Meis1 | 395.3527126 | 0.026649992 | 0.106271035 | 0.250773807 |
| Gm37818 | 22.45192434 | 0.324781278 | 0.315021774 | 1.03098041 |
| 2900018N21Rik | 18.03296757 | 0.281697894 | 0.367375011 | 0.766785672 |
| 2610001A08Rik | 15.58260202 | -0.231921697 | 0.372873281 | -0.621985293 |
| Gm16140 | 0.421709722 | -0.058929521 | 1.389834052 | -0.042400401 |
| Gm16141 | 1.235942255 | -2.296274654 | 0.983464385 | -2.334883388 |
| 4933406G16Rik | 0.427771506 | -0.029186778 | 1.577305452 | -0.018504202 |
| Gm37439 | 0.296404033 | 1.591266391 | 1.774224361 | 0.896880026 |
| Gm12025 | 0.044191549 | 0.037062677 | 3.724945049 | 0.009949859 |
| Gm12026 | 0.088534442 | -0.905414734 | 3.723500899 | -0.243162217 |
| Gm12027 | 0.123841226 | -1.017823543 | 3.719440177 | -0.273649661 |
| Gm23386 | 0.075043205 | 0.517957688 | 3.724945049 | 0.139051095 |
| Glns-ps1 | 73.63675014 | 0.458279054 | 0.171623222 | 2.670262503 |
| Gm12028 | 0 | NA | NA | NA |
| Gm12029 | 0 | NA | NA | NA |
| Spred2 | 1270.960265 | -0.204791464 | 0.082448682 | -2.483865825 |
| Actr2 | 3727.967054 | -0.005580269 | 0.053982127 | -0.103372529 |
| Gm12030 | 0 | NA | NA | NA |

| | | | | |
|---|---|---|---|---|
| Gm12031 | 1.129362261 | 2.136970275 | 1.077900958 | 1.98252934 |
| Gm26253 | 0.30153761 | 1.076328364 | 1.767889926 | 0.608820916 |
| Rab1 | 1943.110902 | -0.147814759 | 0.080913449 | -1.8268256 |
| Cep68 | 468.5192915 | -0.037882325 | 0.105067505 | -0.360552248 |
| Rpl12-ps2 | 0 | NA | NA | NA |
| Slc1a4 | 846.6499244 | -0.021096876 | 0.083163717 | -0.253678853 |
| Gm12033 | 100.5140019 | 0.109418081 | 0.145968521 | 0.749600533 |
| Gm12034 | 2.661321104 | 1.22680044 | 0.698664903 | 1.755921093 |
| Sertad2 | 498.1303247 | -0.066419385 | 0.092706654 | -0.716446796 |
| Gm12035 | 0 | NA | NA | NA |
| Aftph | 1713.653678 | -0.219100926 | 0.064453762 | -3.399350478 |
| Gm12036 | 0 | NA | NA | NA |
| Lgalsl | 1035.137919 | 0.02684585 | 0.079577313 | 0.337355571 |
| Gm12037 | 2.870345996 | 1.221767508 | 0.583096172 | 2.095310457 |
| Gm12038 | 0.044191549 | 0.037062677 | 3.724945049 | 0.009949859 |
| Gm22807 | 0 | NA | NA | NA |
| Gm23681 | 0 | NA | NA | NA |
| Peli1 | 911.878915 | -0.208861179 | 0.099653516 | -2.095873661 |
| 2610024D14Rik | 21.08894881 | -0.064147999 | 0.312051792 | -0.205568437 |
| Gm22600 | 0 | NA | NA | NA |
| Gm12040 | 0.133633945 | 1.204093506 | 3.720899422 | 0.323602809 |
| Gm12041 | 0.223001177 | -0.121575297 | 2.537599379 | -0.047909571 |
| Vps54 | 967.0089129 | -0.222697737 | 0.087080983 | -2.557363602 |
| Gm23582 | 0 | NA | NA | NA |
| Gm12039 | 0 | NA | NA | NA |
| Gm12042 | 9.247728932 | -0.101463321 | 0.37947176 | -0.267380427 |
| Ugp2 | 931.9428543 | -0.282410832 | 0.113704187 | -2.483732912 |
| Mir1933 | 1.993695893 | -0.452744786 | 0.87211743 | -0.519132826 |
| Gm12043 | 3.417719091 | 0.168846927 | 0.5448134 | 0.309916986 |
| Gm12044 | 0 | NA | NA | NA |
| Gm12045 | 0 | NA | NA | NA |
| Gm12046 | 0 | NA | NA | NA |
| 4932414J04Rik | 0 | NA | NA | NA |
| Mdh1 | 4531.387594 | 0.297676475 | 0.097403335 | 3.056121994 |
| Gm22990 | 2.17550669 | 1.067034116 | 0.820496918 | 1.300473033 |
| Wdpcp | 154.1019773 | 0.131734419 | 0.115248873 | 1.143043014 |
| Pgam1-ps1 | 0.548058809 | -1.230601942 | 1.280419001 | -0.961093159 |
| Gm12047 | 0.149223819 | -0.924098271 | 2.865376951 | -0.322504957 |
| Rpl15-ps1 | 0.048949532 | 0.037064902 | 3.724945049 | 0.009950456 |
| Otx1 | 56.99576563 | 0.16804319 | 0.155126515 | 1.083265423 |
| Ehbp1 | 1086.104915 | 0.320074217 | 0.06487265 | 4.93388534 |
| Rpsa-ps3 | 0 | NA | NA | NA |
| Gm12050 | 0.136201742 | -0.443823494 | 2.946222509 | -0.150641539 |
| Gm12051 | 0 | NA | NA | NA |
| Tmem17 | 31.45379496 | -0.463909476 | 0.215575029 | -2.151962952 |
| Gm26829 | 0.106971521 | 0.998850474 | 3.724945049 | 0.268151734 |
| Gm12052 | 0.878931054 | 0.091511446 | 1.236618491 | 0.074001357 |

| | | | | |
|---|---|---|---|---|
| Gm12053 | 0.158674157 | 0.517963644 | 2.811010914 | 0.184262409 |
| Gm12055 | 0.424314013 | 1.314581964 | 1.548343863 | 0.849024558 |
| Gm12056 | 0 | NA | NA | NA |
| Gm12057 | 7.744039244 | 0.460190749 | 0.368043934 | 1.250369062 |
| Gm12058 | 1.980218008 | 0.421751968 | 0.834220144 | 0.505564354 |
| Gm28048 | 0.044544648 | 0.517957688 | 3.724945049 | 0.139051095 |
| B3gnt2 | 14.36761034 | -0.13600243 | 0.329598452 | -0.412630671 |
| Gm20456 | 0.099106094 | -0.93550548 | 3.724188147 | -0.251197158 |
| Gm23772 | 0 | NA | NA | NA |
| 9130230N09Rik | 6.400237851 | -0.731909724 | 0.412568645 | -1.774031384 |
| Gm24917 | 0 | NA | NA | NA |
| Commd1 | 92.82422243 | -0.019437399 | 0.169813928 | -0.114462923 |
| Zrsr1 | 521.7180302 | 0.370297565 | 0.102095608 | 3.626968599 |
| Cct4 | 1159.74844 | -0.577172411 | 0.09746536 | -5.921820925 |
| Fam161a | 45.44904334 | -0.410567883 | 0.225227053 | -1.822906612 |
| Gm23740 | 0.228592689 | 0.037076365 | 2.138563796 | 0.01733704 |
| Gm12059 | 13.03468439 | -1.105034217 | 0.325858092 | -3.391151683 |
| 1700061J23Rik | 0.046345709 | -0.443818988 | 3.724945049 | -0.119147795 |
| Gm24398 | 0 | NA | NA | NA |
| Xpo1 | 1829.109104 | -0.435312907 | 0.088254323 | -4.932482535 |
| Usp34 | 4434.045314 | 0.185710144 | 0.063655198 | 2.917438811 |
| Gm22753 | 2.309867597 | -0.372982052 | 0.787506711 | -0.473623966 |
| Ahsa2 | 295.3081191 | -0.267941454 | 0.116155958 | -2.306738789 |
| AL672049.1 | 0.184646192 | 0.925953388 | 2.705003907 | 0.342311294 |
| 1700030C12Rik | 0.940659763 | -0.265577563 | 1.062428275 | -0.249972228 |
| 1700093K21Rik | 3.270068486 | 1.304613105 | 0.606052708 | 2.152639676 |
| Gm12060 | 7.122622078 | 0.817208623 | 0.472362318 | 1.730046178 |
| 0610010F05Rik | 731.0037471 | 0.161057116 | 0.067112079 | 2.399823095 |
| Gm23114 | 5.978366796 | 0.571016671 | 0.696252188 | 0.820129087 |
| Pex13 | 566.0191946 | -0.145088958 | 0.086836402 | -1.670831062 |
| Pus10 | 234.206064 | 0.118776239 | 0.102291387 | 1.161155816 |
| Rel | 254.6607661 | -0.467738457 | 0.16704959 | -2.799997628 |
| Gm23827 | 0.061920613 | -0.443816764 | 3.724945049 | -0.119147198 |
| Papolg | 392.4489342 | -0.09431746 | 0.091053061 | -1.035851605 |
| Gm12061 | 0.546436044 | 0.034198786 | 1.600348956 | 0.021369581 |
| Gm12062 | 0 | NA | NA | NA |
| AL662810.1 | 0 | NA | NA | NA |
| A830031A19Rik | 4.718729471 | 0.778318718 | 0.493773981 | 1.57626515 |
| Gm12063 | 6.626652096 | 1.42270857 | 0.475392183 | 2.992705017 |
| Bcl11a | 1084.849886 | 0.768933962 | 0.154491478 | 4.977193392 |
| Gm12064 | 0 | NA | NA | NA |
| 4930538E20Rik | 0.409305803 | 1.63420276 | 1.523564668 | 1.072617916 |
| Gm12065 | 0 | NA | NA | NA |
| Gm12066 | 0.089089296 | 0.957155977 | 3.723939561 | 0.257027796 |
| Gm12068 | 0.5449234 | 0.084005261 | 1.226691097 | 0.068481186 |
| 4933430M04Rik | 0.038981395 | -0.443816764 | 3.724945049 | -0.119147198 |
| Gm12067 | 0 | NA | NA | NA |

| | | | | |
|---|---|---|---|---|
| Gm22787 | 0 | NA | NA | NA |
| Gm25507 | 0 | NA | NA | NA |
| Gm10466 | 0 | NA | NA | NA |
| Gm12665 | 0 | NA | NA | NA |
| 4933427E13Rik | 0.566718995 | 1.458927666 | 1.258903891 | 1.158887249 |
| Gm24255 | 0 | NA | NA | NA |
| Gm12069 | 0.137762903 | 0.51796349 | 2.937523392 | 0.176326592 |
| 5730522E02Rik | 70.17051206 | 1.490556704 | 0.258355509 | 5.769401659 |
| Gm23514 | 0 | NA | NA | NA |
| Mir6372 | 0 | NA | NA | NA |
| Gm22868 | 0 | NA | NA | NA |
| Fancl | 76.3059331 | -0.153226108 | 0.165203689 | -0.927498099 |
| Gm12713 | 5.450982021 | 0.290336924 | 0.526037676 | 0.551931805 |
| Vrk2 | 67.55354626 | -0.537719743 | 0.185526923 | -2.898338066 |
| Gm6899 | 0.12209515 | 0.037073798 | 3.724945049 | 0.009952844 |
| Gm12071 | 0 | NA | NA | NA |
| Gm12070 | 46.42488409 | 0.274466178 | 0.167102485 | 1.642502073 |
| Gm12072 | 0.172942268 | 0.517958134 | 3.372003067 | 0.153605475 |
| Gm12073 | 0 | NA | NA | NA |
| Gm12074 | 0 | NA | NA | NA |
| n-R5s68 | 0 | NA | NA | NA |
| Gm24850 | 0 | NA | NA | NA |
| Gm25387 | 0 | NA | NA | NA |
| Gm12075 | 0 | NA | NA | NA |
| Gm12077 | 0 | NA | NA | NA |
| Gm3810 | 0 | NA | NA | NA |
| Gm12079 | 2.754754232 | -0.413914213 | 0.606450181 | -0.682519728 |
| Gm12080 | 0 | NA | NA | NA |
| Gm6685 | 30.4764154 | -0.401983514 | 0.199217111 | -2.017816197 |
| Gm12081 | 0 | NA | NA | NA |
| Ccdc85a | 925.7833576 | 0.748767585 | 0.076016064 | 9.850123118 |
| 2810471M01Rik | 0.525803784 | -0.968682427 | 1.446460057 | -0.669691791 |
| Mir216b | 0 | NA | NA | NA |
| mmu-mir-216c | 0 | NA | NA | NA |
| Gm12082 | 0.832920528 | -0.546397876 | 1.082239369 | -0.504877102 |
| Mir216a | 0.066050875 | -0.443818988 | 3.724945049 | -0.119147795 |
| Mir217 | 0 | NA | NA | NA |
| Efemp1 | 96.95456267 | -0.550970583 | 0.194425707 | -2.833836072 |
| Rpsa-ps5 | 0.463552209 | 0.71762823 | 1.378214634 | 0.520694101 |
| Pnpt1 | 855.1922142 | -0.309911965 | 0.112813732 | -2.747112066 |
| A630052C17Rik | 26.90946008 | -0.260680978 | 0.267503337 | -0.974496175 |
| Smek2 | 1087.669138 | -0.183698545 | 0.055632618 | -3.301993534 |
| Cfap36 | 532.1389956 | 0.10622093 | 0.084922936 | 1.250792006 |
| Gm12085 | 0 | NA | NA | NA |
| Gm23109 | 0.11197992 | 0.037073798 | 3.724945049 | 0.009952844 |
| Gm12086 | 0 | NA | NA | NA |
| Gm12087 | 0.124596252 | 0.037071632 | 3.714681146 | 0.009979761 |

| | | | | |
|---|---|---|---|---|
| Ccdc88a | 2613.854868 | -0.05040781 | 0.058478994 | -0.861981491 |
| Prorsd1 | 38.98209458 | -0.626647979 | 0.233775094 | -2.680559197 |
| Gm12089 | 10.3364996 | 0.585688588 | 0.447191567 | 1.309704007 |
| Mtif2 | 334.856477 | 0.107541386 | 0.077585968 | 1.386093236 |
| Rps27a | 597.2006918 | -0.133964685 | 0.251625975 | -0.532396088 |
| Clhc1 | 9.730852973 | -0.026690604 | 0.342568497 | -0.07791319 |
| Gm12090 | 0 | NA | NA | NA |
| Gm12088 | 0.547473135 | 0.077624814 | 1.330130195 | 0.058358809 |
| Gm12091 | 2.208204681 | 0.020700654 | 0.686918987 | 0.030135511 |
| Gm12092 | 0.662335159 | -1.310141372 | 1.248557776 | -1.049323785 |
| Gm12093 | 0.303276099 | 0.048971695 | 1.620490821 | 0.030220285 |
| Rtn4 | 7648.216462 | -0.013840408 | 0.071766841 | -0.192852408 |
| Eml6 | 546.8534648 | 0.451841152 | 0.106235116 | 4.253218409 |
| 4931440F15Rik | 25.51154845 | 0.759299549 | 0.310191758 | 2.447839211 |
| Gm12094 | 0.298205325 | -1.19788285 | 2.024921553 | -0.591570003 |
| Gm22588 | 0 | NA | NA | NA |
| Gm12095 | 0 | NA | NA | NA |
| Sptbn1 | 16443.0819 | 0.315192714 | 0.052831031 | 5.966052672 |
| Gm29237 | 0.044544648 | 0.517957688 | 3.724945049 | 0.139051095 |
| Gm8098 | 47.87148983 | 0.518218387 | 0.244059245 | 2.123330285 |
| 4930505A04Rik | 3.929212317 | 1.170123578 | 0.613436206 | 1.907490244 |
| Acyp2 | 181.980473 | 0.064251023 | 0.138008115 | 0.465559745 |
| Gm12096 | 0.263964426 | -1.461898583 | 2.097717825 | -0.696899538 |
| Gm12098 | 9.953545217 | 0.441119634 | 0.496043798 | 0.889275576 |
| Gm12097 | 33.54705527 | 0.392055548 | 0.309792078 | 1.265544138 |
| Gm12099 | 0 | NA | NA | NA |
| Gm12100 | 0.45607993 | 2.053638194 | 1.72033887 | 1.193740507 |
| Gm12101 | 0.170229941 | 0.448751302 | 2.740258101 | 0.163762421 |
| Psme4 | 1642.607637 | 0.222711462 | 0.113948837 | 1.954486495 |
| Gm12103 | 0.045882455 | 0.517959912 | 3.724945049 | 0.139051692 |
| Gpr75 | 147.2383283 | 0.000871872 | 0.117085384 | 0.00744646 |
| Gm12104 | 4.359450978 | 0.711858611 | 0.522819355 | 1.361576623 |
| Erlec1 | 677.4619536 | 0.255681384 | 0.060830705 | 4.203163234 |
| Asb3 | 233.2081696 | 0.083153675 | 0.100678271 | 0.825934674 |
| Chac2 | 85.03464214 | -0.259761241 | 0.170894111 | -1.520012833 |
| Gm12102 | 0 | NA | NA | NA |
| Gm12105 | 4.176201957 | 0.298855822 | 0.599557015 | 0.498461056 |
| Gm12106 | 0 | NA | NA | NA |
| Gm17332 | 0 | NA | NA | NA |
| Stc2 | 26.77438353 | -1.244537179 | 0.323237085 | -3.850230182 |
| Bod1 | 387.4332913 | -0.239085698 | 0.082698205 | -2.891062733 |
| Gm38061 | 0.047948698 | 0.037064902 | 3.724945049 | 0.009950456 |
| D630024D03Rik | 1.773596836 | -0.193093121 | 0.812481539 | -0.237658472 |
| Gm12107 | 0 | NA | NA | NA |
| Cpeb4 | 3230.776281 | 0.053706634 | 0.070265576 | 0.764337776 |
| 4930524B15Rik | 1.33642055 | 0.035528337 | 0.832537581 | 0.042674755 |
| Nsg2 | 3948.965464 | 0.661355893 | 0.068503914 | 9.654278937 |

| | | | | |
|---|---|---|---|---|
| Gm12108 | 0.431234269 | -0.461086654 | 1.54940896 | -0.297588736 |
| Gm12109 | 0.060174537 | 0.517957688 | 3.724945049 | 0.139051095 |
| Il9r | 0.166808115 | 1.33436098 | 3.380229911 | 0.394754504 |
| Snrnp25 | 64.99433264 | 0.107578641 | 0.221714372 | 0.485212755 |
| Rhbdf1 | 175.977258 | -0.698812213 | 0.166206078 | -4.204492516 |
| Nprl3 | 290.7020933 | 0.212000758 | 0.093518274 | 2.26694472 |
| Mpg | 101.0618394 | -0.019599282 | 0.126066513 | -0.155467789 |
| Hba-x | 0.066050875 | -0.443818988 | 3.724945049 | -0.119147795 |
| Hba-a1 | 2.403392355 | 2.181725131 | 1.144936397 | 1.905542646 |
| Hbq1b | 0.787662625 | -0.03600209 | 1.282814298 | -0.028064927 |
| Hba-a2 | 2.268919436 | 2.82925589 | 1.995356965 | 1.417919671 |
| Hbq1a | 1.119917166 | 2.424489966 | 1.213676756 | 1.997640602 |
| Sh3pxd2b | 817.112173 | -0.971480788 | 0.135274045 | -7.181575639 |
| Ubtd2 | 162.9170653 | 0.302382169 | 0.121476753 | 2.489218411 |
| Efcab9 | 6.264832892 | -0.122383186 | 0.41768975 | -0.293000214 |
| Stk10 | 237.0838909 | -0.368194716 | 0.132827594 | -2.771974596 |
| Fbxw11 | 1952.482481 | -0.033026589 | 0.068658967 | -0.481023687 |
| AL669951.2 | 0 | NA | NA | NA |
| AL669951.1 | 0 | NA | NA | NA |
| Gm8192 | 0.049553047 | -0.443818988 | 3.724945049 | -0.119147795 |
| 4930555O08Rik | 0 | NA | NA | NA |
| Smim23 | 0 | NA | NA | NA |
| Gm12110 | 0 | NA | NA | NA |
| Gm12111 | 0.20222093 | 0.502994613 | 2.21040094 | 0.227558089 |
| Gm12112 | 32.64724094 | 0.139859207 | 0.19641152 | 0.712072319 |
| Gm26052 | 0 | NA | NA | NA |
| Rpsa-ps4 | 1.265922642 | -0.304567597 | 1.037365551 | -0.293597177 |
| Gm12114 | 0.044191549 | 0.037062677 | 3.724945049 | 0.009949859 |
| Fgf18 | 14.860062 | -0.07177096 | 0.295649522 | -0.242756896 |
| Npm1 | 1433.536217 | -0.136835578 | 0.082432564 | -1.659969933 |
| Gm12115 | 4.703682786 | 0.185023673 | 0.444207105 | 0.416525694 |
| Tlx3 | 9.956390658 | -0.585985142 | 0.412750784 | -1.419706916 |
| Gm12116 | 10.20532615 | 0.587405225 | 0.570296164 | 1.03000031 |
| Ranbp17 | 205.143616 | 0.598806789 | 0.153656681 | 3.897043625 |
| Gm12117 | 3.560310751 | -0.0333887 | 0.581020581 | -0.057465606 |
| Anp32-ps | 6.77190338 | 1.135340807 | 0.453553079 | 2.503214857 |
| Gm12118 | 0.07796279 | -0.798301147 | 3.721829328 | -0.214491605 |
| Gabrp | 0.357753142 | 0.023604355 | 1.574930797 | 0.01498755 |
| D130052B06Rik | 0 | NA | NA | NA |
| Kcnip1 | 277.5432354 | 0.390608522 | 0.083305489 | 4.688868991 |
| Gm12119 | 0.042716945 | 0.037062677 | 3.724945049 | 0.009949859 |
| Gm12120 | 0.062426873 | 0.517959912 | 3.724945049 | 0.139051692 |
| Kcnmb1 | 5.253596414 | -0.714825215 | 0.488733243 | -1.462608132 |
| 4930469K13Rik | 1.763134642 | 1.278602724 | 1.061096319 | 1.204982715 |
| Lcp2 | 190.2229504 | -0.387786737 | 0.152598429 | -2.541223654 |
| Gm12121 | 0 | NA | NA | NA |
| Foxi1 | 0.275123771 | -0.06807324 | 1.796023502 | -0.037902199 |

| | | | | |
|---|---|---|---|---|
| Dock2 | 455.1128129 | -0.754548176 | 0.150963308 | -4.998222321 |
| 4930403D09Rik | 0.062426873 | 0.517959912 | 3.724945049 | 0.139051692 |
| Fam196b | 116.3261388 | -0.511034376 | 0.161900076 | -3.156480156 |
| Spdl1 | 20.02293662 | -1.059578863 | 0.242921516 | -4.361815627 |
| Gm22022 | 0 | NA | NA | NA |
| Gm12711 | 0 | NA | NA | NA |
| Gm25799 | 0 | NA | NA | NA |
| Slit3 | 590.7003565 | 0.200911874 | 0.146677054 | 1.369756671 |
| Gm26070 | 1.06710716 | 2.844507023 | 1.299740208 | 2.18851968 |
| Gm12123 | 0 | NA | NA | NA |
| Gm12122 | 14.36718598 | 0.044545327 | 0.464847229 | 0.095827886 |
| Mir218-2 | 2.110899703 | 0.913040195 | 0.783136559 | 1.165876097 |
| Pank3 | 1420.981158 | -0.020412166 | 0.0673433 | -0.303106113 |
| Mir103-1 | 0.791513252 | -0.788836372 | 1.126753393 | -0.700096735 |
| Gm12124 | 1.289573466 | -0.130306255 | 0.867544989 | -0.15020115 |
| Fbll1 | 180.3183456 | 0.129291157 | 0.123855024 | 1.043891096 |
| Rars | 540.4176717 | -0.323848198 | 0.100380568 | -3.226204093 |
| Wwc1 | 973.1767011 | -0.015757777 | 0.069650618 | -0.2262403 |
| Gm12125 | 0 | NA | NA | NA |
| Tenm2 | 1701.648525 | 0.745832071 | 0.109992379 | 6.780761313 |
| Gm12126 | 0 | NA | NA | NA |
| Gm12127 | 0 | NA | NA | NA |
| Gm22127 | 0 | NA | NA | NA |
| Gm12128 | 0.048949532 | 0.037064902 | 3.724945049 | 0.009950456 |
| Gm25693 | 0.047948698 | 0.037064902 | 3.724945049 | 0.009950456 |
| Gm12129 | 0.066050875 | -0.443818988 | 3.724945049 | -0.119147795 |
| Gm12130 | 1.911894038 | 1.150188201 | 0.881336688 | 1.305049724 |
| Gm23520 | 0.097501745 | -0.443821212 | 3.724945049 | -0.119148392 |
| Gm12131 | 1.727409906 | -0.400929778 | 1.049272797 | -0.382102518 |
| Gm24192 | 0 | NA | NA | NA |
| Gm12132 | 0 | NA | NA | NA |
| Gm12133 | 0 | NA | NA | NA |
| Gm12134 | 0 | NA | NA | NA |
| 3110004A20Rik | 0.20299191 | -0.97861201 | 2.607194669 | -0.375350572 |
| Gm12135 | 0 | NA | NA | NA |
| Gm12136 | 0 | NA | NA | NA |
| Gm12137 | 0 | NA | NA | NA |
| Gm12138 | 2.171605266 | 0.801480557 | 0.685788632 | 1.1686991 |
| Mat2b | 871.6522152 | 0.383392296 | 0.075082325 | 5.106292278 |
| Hmmr | 18.26964906 | -0.372098044 | 0.355854244 | -1.045647343 |
| Nudcd2 | 124.2665417 | -0.177151979 | 0.12513983 | -1.415632248 |
| Ccng1 | 1174.108608 | -0.247453354 | 0.109747953 | -2.254742321 |
| Gm24446 | 0 | NA | NA | NA |
| Hmgb1-ps1 | 6.658866634 | 0.772995493 | 0.387868762 | 1.992930518 |
| Platr13 | 0 | NA | NA | NA |
| Gm12140 | 0 | NA | NA | NA |
| Hspd1-ps3 | 127.1742578 | -0.363109076 | 0.189182532 | -1.91935837 |

| | | | | |
|---|---|---|---|---|
| Gm12142 | 0 | NA | NA | NA |
| A230004M16Rik | 12.26251621 | -1.789726819 | 0.446017657 | -4.012681541 |
| Gm12143 | 0 | NA | NA | NA |
| Gabrg2 | 1919.016568 | 0.39359977 | 0.075026768 | 5.246124567 |
| Gabra1 | 5062.268846 | 0.463393968 | 0.087837104 | 5.275606175 |
| Gabra6 | 1128.6417 | -0.152547208 | 0.37551742 | -0.406232042 |
| Gabrb2 | 2861.342192 | 0.458192129 | 0.091253116 | 5.021112139 |
| Gm25543 | 0 | NA | NA | NA |
| Gm9972 | 0.444159033 | 0.037075307 | 1.651966785 | 0.022443131 |
| Gm12145 | 0 | NA | NA | NA |
| Atp10b | 25.10547728 | 0.235425258 | 0.236486343 | 0.995513125 |
| Gm12146 | 1.289012822 | -0.946431701 | 0.885706149 | -1.068561737 |
| Gm12144 | 0.970234074 | 0.649010507 | 1.037920996 | 0.625298563 |
| Gm12147 | 0 | NA | NA | NA |
| Mir146 | 0.061920613 | -0.443816764 | 3.724945049 | -0.119147198 |
| Gm12148 | 0 | NA | NA | NA |
| Pttg1 | 36.67629004 | -0.508789787 | 0.228192208 | -2.229654522 |
| Slu7 | 647.400309 | 0.016382155 | 0.070135068 | 0.233580088 |
| Gm12151 | 0 | NA | NA | NA |
| C1qtnf2 | 6.55721845 | 0.502555516 | 0.409917574 | 1.225991633 |
| Ccnjl | 56.86863016 | 0.597897328 | 0.1761993 | 3.393301386 |
| Gm12150 | 0 | NA | NA | NA |
| 4933415A04Rik | 9.074042005 | 0.960849237 | 0.346568432 | 2.772466124 |
| Fabp6 | 0 | NA | NA | NA |
| Gm12149 | 0 | NA | NA | NA |
| Gm12152 | 0 | NA | NA | NA |
| Pwwp2a | 556.3087863 | 0.023821976 | 0.085292511 | 0.279297396 |
| Ttc1 | 454.7934483 | -0.391511505 | 0.105757018 | -3.701990762 |
| Adra1b | 167.6708051 | 0.356871504 | 0.127146613 | 2.806771614 |
| Gm12153 | 0.799763445 | -0.984132172 | 1.139189321 | -0.863888164 |
| Gm12154 | 0 | NA | NA | NA |
| Gm12155 | 0.038981395 | -0.443816764 | 3.724945049 | -0.119147198 |
| Il12b | 80.53366365 | 0.92091466 | 0.959556668 | 0.959729312 |
| Ublcp1 | 282.0163601 | 0.029972911 | 0.101992818 | 0.293872758 |
| Gm22121 | 0 | NA | NA | NA |
| Gm12156 | 0.547961811 | 1.815475045 | 1.420263632 | 1.278266235 |
| Rnf145 | 1154.947274 | 0.091274024 | 0.050381644 | 1.811652352 |
| 4930597A21Rik | 0.434349921 | 0.834346506 | 1.453125026 | 0.57417393 |
| Gm37695 | 1.050344328 | 1.771275606 | 1.180842498 | 1.500010043 |
| AL603913.1 | 0 | NA | NA | NA |
| Ebf1 | 264.2278728 | 0.136955102 | 0.125761253 | 1.089008728 |
| Gm12158 | 0 | NA | NA | NA |
| Gm12159 | 0 | NA | NA | NA |
| Gm12160 | 0 | NA | NA | NA |
| Gm38087 | 0 | NA | NA | NA |
| Gm37312 | 0 | NA | NA | NA |
| Gm12161 | 1.855823375 | -1.555693752 | 1.004651891 | -1.548490343 |

| | | | | |
|---|---|---|---|---|
| Gm12162 | 0 | NA | NA | NA |
| Gm22284 | 0 | NA | NA | NA |
| F630206G17Rik | 0.072025462 | 0.517959912 | 3.724945049 | 0.139051692 |
| Gm12163 | 0.042716945 | 0.037062677 | 3.724945049 | 0.009949859 |
| Gm12164 | 1.520812657 | -0.764394854 | 0.785457613 | -0.973184092 |
| Gm22751 | 0 | NA | NA | NA |
| Clint1 | 800.9757842 | -0.213489093 | 0.063704106 | -3.351261107 |
| Lsm11 | 391.468034 | -0.139453244 | 0.113094973 | -1.233063147 |
| Thg1l | 79.70507417 | -0.141824281 | 0.127561244 | -1.111813252 |
| Sox30 | 1.344747471 | -1.869167621 | 0.945335244 | -1.977253711 |
| Gm12165 | 0 | NA | NA | NA |
| Gm12166 | 16.95436423 | -0.334479113 | 0.249035187 | -1.343099809 |
| Adam19 | 340.8717861 | -0.524750811 | 0.144866957 | -3.62229469 |
| Mir8100 | 0.635485296 | -1.232022718 | 1.560993793 | -0.78925536 |
| Gm16033 | 0.647121192 | -0.997558332 | 1.163777506 | -0.857172721 |
| Gm16034 | 0.200316428 | 0.037073269 | 2.661959355 | 0.01392706 |
| Nipal4 | 54.85333669 | -0.406729221 | 0.203414546 | -1.999509032 |
| Mir7239 | 0.223868884 | -1.005693403 | 2.582883473 | -0.389368477 |
| Cyfip2 | 10488.44967 | 0.627526651 | 0.086530703 | 7.252069212 |
| Fndc9 | 118.2743247 | -2.17281213 | 0.397796664 | -5.462117522 |
| Gm12568 | 12.41784929 | 0.68473252 | 0.415690368 | 1.647217671 |
| Itk | 74.98247743 | -0.498209311 | 0.216214003 | -2.304241651 |
| Gm12167 | 0.124347486 | 0.036092774 | 3.717245565 | 0.009709548 |
| Fam71b | 0.55691522 | 0.512602303 | 1.474722396 | 0.347592404 |
| Med7 | 299.1592794 | -0.58858328 | 0.105641278 | -5.571527448 |
| 2310031A07Rik | 13.18712137 | -1.431426319 | 0.424781636 | -3.369793323 |
| Gm12168 | 5.395875324 | -0.420672059 | 0.472126213 | -0.891016105 |
| Havcr2 | 109.3330086 | -0.433719065 | 0.189584709 | -2.287732319 |
| Gm12169 | 0.124308499 | -1.207977564 | 3.71482282 | -0.325177706 |
| Gm12170 | 0 | NA | NA | NA |
| Gm12171 | 0.288794137 | -0.33319807 | 1.787706598 | -0.186382973 |
| Rpl9-ps2 | 0 | NA | NA | NA |
| BC053393 | 0.265064467 | 0.020424883 | 1.803417537 | 0.011325654 |
| Dppa1 | 0.143022002 | 0.03610674 | 3.547735707 | 0.010177404 |
| Gm4926 | 0.476246927 | 1.424159147 | 1.423783197 | 1.00026405 |
| Gm12173 | 0 | NA | NA | NA |
| Timd2 | 0.095435502 | 0.037073798 | 3.724945049 | 0.009952844 |
| Gm12174 | 30.95181581 | -0.140460472 | 0.257242133 | -0.546024363 |
| Havcr1 | 1.532951352 | -1.65293261 | 0.965374361 | -1.712219297 |
| Timd4 | 7.437687587 | -0.735868359 | 0.376915214 | -1.952344536 |
| Gm12175 | 0 | NA | NA | NA |
| Sgcd | 134.9314582 | 0.728929878 | 0.135216322 | 5.390842373 |
| Gm12176 | 0.97553329 | 1.540657737 | 1.0607839 | 1.452376622 |
| Gm23039 | 4.120196972 | 1.128460787 | 0.648685006 | 1.739612873 |
| Gm12178 | 2.707360892 | 0.286736428 | 0.612072138 | 0.468468355 |
| Gm12179 | 0.104637558 | -0.443821212 | 3.724945049 | -0.119148392 |
| Gm12180 | 0 | NA | NA | NA |

| | | | | |
|---|---|---|---|---|
| Gm12181 | 0.114443069 | -0.443821212 | 3.724945049 | -0.119148392 |
| Gm24422 | 0 | NA | NA | NA |
| Gm12182 | 0.476797581 | -1.743737594 | 1.316062924 | -1.324965215 |
| Gm12183 | 218.1818688 | 0.0478051 | 0.112919932 | 0.423353958 |
| Gnb2l1 | 1496.591963 | -0.598087567 | 0.117756984 | -5.078998682 |
| Gm25296 | 0 | NA | NA | NA |
| Snord96a | 0 | NA | NA | NA |
| Snord95 | 0 | NA | NA | NA |
| Trim41 | 551.7216766 | 0.061388773 | 0.064519634 | 0.9514743 |
| Gm12184 | 52.49590673 | -0.719956266 | 0.167915325 | -4.287615003 |
| Trim7 | 26.45757724 | 0.355729509 | 0.2383798 | 1.492280422 |
| Gm16170 | 11.86762837 | 1.545211332 | 0.459332403 | 3.364037287 |
| Irgm1 | 5470.912599 | -0.721191087 | 0.176582817 | -4.08415212 |
| Gm5431 | 192.2399859 | -1.451682355 | 0.229866735 | -6.315321591 |
| Gm12185 | 296.7910131 | -2.022893487 | 0.397974091 | -5.082977842 |
| Psme2b | 176.4926986 | -0.74540025 | 0.170541287 | -4.37079057 |
| 9930111J21Rik1 | 96.17708593 | -0.967185596 | 0.229132246 | -4.221080248 |
| Olfr56 | 27.72815539 | -0.896956163 | 0.572321114 | -1.567225358 |
| Tgtp1 | 654.3535308 | -1.600545932 | 0.351428983 | -4.554393662 |
| Gm12186 | 0.311786641 | -1.984785226 | 1.9263494 | -1.030335009 |
| 9930111J21Rik2 | 219.1877081 | -0.653455045 | 0.175430234 | -3.724871312 |
| Tgtp2 | 962.1752517 | -1.544212518 | 0.308904953 | -4.99898918 |
| Gm12187 | 0.675604184 | -2.061864505 | 1.324562105 | -1.556638603 |
| Ifi47 | 1143.870919 | -1.120281076 | 0.323332322 | -3.464797674 |
| Gm12188 | 0.333018454 | -0.798975248 | 1.699853714 | -0.470025886 |
| Olfr1396 | 0 | NA | NA | NA |
| Olfr1395 | 0.571466717 | 2.099390376 | 1.616112928 | 1.29903693 |
| Olfr1394 | 0.066050875 | -0.443818988 | 3.724945049 | -0.119147795 |
| Btnl9 | 1.653010361 | -0.370344059 | 0.775642308 | -0.477467584 |
| Zfp62 | 822.4603642 | 0.012965152 | 0.052310005 | 0.247852249 |
| Mgat1 | 662.2687355 | -0.306728012 | 0.076813124 | -3.993171937 |
| Olfr1393 | 2.448580025 | 0.692183824 | 0.789331088 | 0.876924568 |
| Olfr1392 | 1.186358578 | 0.318453693 | 1.009453975 | 0.315471236 |
| Olfr10 | 0.306560087 | 1.428084495 | 1.598929143 | 0.893150582 |
| Olfr1391 | 0.124596252 | 0.037071632 | 3.714681146 | 0.009979761 |
| Olfr1390 | 0.188595572 | 1.569860932 | 3.285103736 | 0.47787256 |
| Gm12189 | 0 | NA | NA | NA |
| Olfr1389 | 0.061920613 | -0.443816764 | 3.724945049 | -0.119147198 |
| Olfr1388 | 0 | NA | NA | NA |
| Olfr1387 | 0.046345709 | -0.443818988 | 3.724945049 | -0.119147795 |
| Olfr1386 | 0 | NA | NA | NA |
| Olfr1385 | 0 | NA | NA | NA |
| Olfr1384 | 0.045882455 | 0.517959912 | 3.724945049 | 0.139051692 |
| Olfr1383 | 0 | NA | NA | NA |
| Olfr1382 | 0 | NA | NA | NA |
| Olfr1381 | 0 | NA | NA | NA |
| Olfr1380 | 0 | NA | NA | NA |

| | | | | |
|---|---|---|---|---|
| 1700024J04Rik | 0.046345709 | -0.443818988 | 3.724945049 | -0.119147795 |
| Gm12190 | 0 | NA | NA | NA |
| Flt4 | 66.9739403 | -0.758565405 | 0.200664709 | -3.780263153 |
| Scgb3a1 | 5.818159514 | 0.074680698 | 0.456281769 | 0.163672326 |
| Cnot6 | 1091.733663 | -0.109473072 | 0.078004782 | -1.403414886 |
| Gm12191 | 162.606518 | -0.149157372 | 0.142098942 | -1.049672637 |
| Gm12192 | 0 | NA | NA | NA |
| Gm23813 | 0 | NA | NA | NA |
| Gm12193 | 0.060174537 | 0.517957688 | 3.724945049 | 0.139051095 |
| Gfpt2 | 101.9890799 | -0.431768128 | 0.13193369 | -3.272614662 |
| Mapk9 | 2585.547055 | 0.219381969 | 0.049310178 | 4.44901999 |
| Gm23492 | 0.042716945 | 0.037062677 | 3.724945049 | 0.009949859 |
| Rasgef1c | 167.346068 | 0.123547765 | 0.138043083 | 0.894994244 |
| Rnf130 | 704.3708933 | -0.171542797 | 0.057695089 | -2.97326512 |
| Mir340 | 5.005408619 | 0.343560526 | 0.533608574 | 0.643843714 |
| Gm12194 | 0.07796279 | -0.798301147 | 3.721829328 | -0.214491605 |
| Tbc1d9b | 2240.773314 | 0.275367755 | 0.079689458 | 3.455510474 |
| Gm12195 | 0 | NA | NA | NA |
| 3010026O09Rik | 75.01584386 | 0.480998307 | 0.145863333 | 3.297595743 |
| Sqstm1 | 5575.809125 | -0.408871533 | 0.098835115 | -4.136905538 |
| Mgat4b | 410.2977067 | 0.217302672 | 0.096612557 | 2.249217678 |
| Mir6919 | 0.119234754 | 0.51795324 | 3.724945049 | 0.139049901 |
| Ltc4s | 8.403515007 | 2.006361168 | 0.53023362 | 3.783919187 |
| Gm26542 | 2.8898106 | -0.003715372 | 0.644835 | -0.005761741 |
| Maml1 | 333.2785951 | -0.207714865 | 0.127160778 | -1.633482184 |
| Canx | 5951.612133 | -0.220759585 | 0.081937509 | -2.694243309 |
| Gm12196 | 0.048949532 | 0.037064902 | 3.724945049 | 0.009950456 |
| Cby3 | 1.08873821 | -0.23070149 | 1.051479237 | -0.219406606 |
| Mir804 | 0.587766037 | -0.375645244 | 1.229456372 | -0.305537677 |
| Hnrnph1 | 2118.885193 | -0.198221178 | 0.08585717 | -2.308731779 |
| Rufy1 | 315.9868514 | -0.026279937 | 0.085306465 | -0.308065007 |
| Gm23824 | 0 | NA | NA | NA |
| Gm12198 | 7.905317881 | 0.832179201 | 0.38627642 | 2.154361897 |
| Adamts2 | 218.5703262 | 0.613432189 | 0.101410711 | 6.048988123 |
| Gm12199 | 0 | NA | NA | NA |
| Gm12200 | 0.062426873 | 0.517959912 | 3.724945049 | 0.139051692 |
| Gm23392 | 0 | NA | NA | NA |
| Zfp354c | 737.6052082 | 0.291028128 | 0.077729672 | 3.744105937 |
| Zfp879 | 31.60590681 | 0.277177618 | 0.222471849 | 1.245899736 |
| Grm6 | 0 | NA | NA | NA |
| Gm12660 | 0 | NA | NA | NA |
| Zfp454 | 55.6120256 | -0.204474594 | 0.170911114 | -1.196379744 |
| Zfp2 | 299.1331036 | 0.414143663 | 0.08340252 | 4.965601286 |
| Zfp354b | 46.21141538 | -0.045523201 | 0.195806215 | -0.232491091 |
| 4933414I15Rik | 0.060174537 | 0.517957688 | 3.724945049 | 0.139051095 |
| Prop1 | 0.410687608 | 0.159031685 | 1.668644195 | 0.095305929 |
| Olfr1379-ps1 | 0.047948698 | 0.037064902 | 3.724945049 | 0.009950456 |

| | | | | |
|---|---|---|---|---|
| Olfr1378 | 0.11037557 | 0.517957688 | 3.724945049 | 0.139051095 |
| Gm12661 | 0 | NA | NA | NA |
| Olfr1377 | 0.049553047 | -0.443818988 | 3.724945049 | -0.119147795 |
| Olfr1376-ps1 | 0 | NA | NA | NA |
| Olfr51 | 0 | NA | NA | NA |
| Olfr54 | 0 | NA | NA | NA |
| Olfr1375 | 0 | NA | NA | NA |
| Zfp354a | 104.6133705 | 0.20145451 | 0.155437245 | 1.296050439 |
| 9230009I02Rik | 1.062461154 | 1.127168418 | 1.152123712 | 0.978339745 |
| Gm12202 | 0 | NA | NA | NA |
| Gm12201 | 0.113999573 | -0.443821212 | 3.724945049 | -0.119148392 |
| Olfr1374-ps1 | 0 | NA | NA | NA |
| Gm12203 | 0 | NA | NA | NA |
| Platr20 | 0.066050875 | -0.443818988 | 3.724945049 | -0.119147795 |
| Gm12569 | 0.212513282 | 1.689178857 | 3.159662247 | 0.534607412 |
| BC049762 | 1.440851686 | -0.643884222 | 0.717474079 | -0.897432033 |
| Clk4 | 881.9365492 | -0.008994949 | 0.090225742 | -0.099693816 |
| Gm25082 | 22.05723362 | 0.203940827 | 0.299102212 | 0.681843258 |
| Col23a1 | 236.6432467 | 0.011122749 | 0.145376336 | 0.07651004 |
| Phykpl | 90.42005935 | -0.026260914 | 0.188921127 | -0.139004646 |
| Hnrnpab | 1099.124842 | -0.206257805 | 0.123931595 | -1.664287508 |
| Nhp2 | 191.8689779 | 0.01572079 | 0.109206477 | 0.143954738 |
| Rmnd5b | 328.920745 | 0.279047346 | 0.07286806 | 3.829487817 |
| N4bp3 | 118.9340086 | 0.186195667 | 0.10784104 | 1.726575216 |
| D930048N14Rik | 22.3543129 | 0.55956529 | 0.39248132 | 1.425711904 |
| 0610009B22Rik | 109.4926551 | -0.136127447 | 0.156334201 | -0.870746431 |
| Sec24a | 739.2067926 | -0.232774394 | 0.096779242 | -2.405209945 |
| Gm25291 | 176.1642292 | 0.266774473 | 0.307812432 | 0.86667868 |
| Sar1b | 418.9258972 | -0.048123966 | 0.089726498 | -0.536340627 |
| Jade2 | 1658.936098 | 0.109121676 | 0.095548272 | 1.142058083 |
| Gm26551 | 2.891936054 | -0.27212762 | 0.791171293 | -0.343955376 |
| Gm26061 | 0 | NA | NA | NA |
| Cdkn2aipnl | 319.6958028 | -0.001382187 | 0.088766203 | -0.01557109 |
| Ube2b | 1147.001563 | -0.080389768 | 0.061806445 | -1.300669665 |
| Gm12204 | 0.487092162 | -1.192669768 | 1.503761799 | -0.79312413 |
| Cdkl3 | 149.4357486 | 0.153493422 | 0.095779282 | 1.602574368 |
| Gm12205 | 0 | NA | NA | NA |
| Ppp2ca | 3811.689111 | -0.015212056 | 0.080832535 | -0.188192247 |
| Gm24157 | 3.140872897 | 0.125771077 | 0.566179 | 0.222140131 |
| 9430098F02Rik | 0.363523191 | 1.716340844 | 1.875788402 | 0.914997044 |
| Mir3061 | 0 | NA | NA | NA |
| Olfr1373 | 0 | NA | NA | NA |
| Olfr1372-ps1 | 1.830548353 | -0.256476699 | 0.859196122 | -0.298507748 |
| Gm12206 | 0.049553047 | -0.443818988 | 3.724945049 | -0.119147795 |
| Olfr1371 | 0 | NA | NA | NA |
| Skp1a | 2595.731114 | 0.137000324 | 0.077521531 | 1.767255125 |
| Gm12207 | 10.46687507 | 0.174727733 | 0.430354789 | 0.40600857 |

| | | | | |
|---|---|---|---|---|
| Tcf7 | 44.21850987 | -0.387016878 | 0.200158783 | -1.93354931 |
| A630014C17Rik | 2.221200599 | -1.427848711 | 0.752073054 | -1.89855055 |
| Vdac1 | 3622.033645 | 0.150434203 | 0.05455743 | 2.757354991 |
| 9530068E07Rik | 1167.821488 | -0.189978002 | 0.093808071 | -2.025177585 |
| Gm12208 | 2.679228202 | 0.337831645 | 0.690768906 | 0.489066086 |
| Gm12209 | 0 | NA | NA | NA |
| Gm12210 | 0 | NA | NA | NA |
| Fstl4 | 404.2876281 | 0.232245322 | 0.149627461 | 1.552157077 |
| Gm11186 | 0.223616349 | -0.828518003 | 1.862306615 | -0.444888074 |
| Gm11189 | 2.178677355 | -0.587888518 | 0.685859571 | -0.857155812 |
| Hspa4 | 2862.524315 | -0.352330608 | 0.080141307 | -4.396367112 |
| Zcchc10 | 74.48748572 | -0.201374803 | 0.132925208 | -1.514948189 |
| Gm11188 | 0.045882455 | 0.517959912 | 3.724945049 | 0.139051692 |
| Aff4 | 5179.750789 | 0.057997284 | 0.04890589 | 1.185895686 |
| Gm25930 | 3.800172548 | 1.279524103 | 0.650909398 | 1.965748394 |
| mmu-mir-6920 | 1.519428967 | 0.126088876 | 0.806062687 | 0.156425646 |
| Leap2 | 0.397769051 | 0.549578524 | 1.538665759 | 0.357178628 |
| Uqcrq | 757.3825943 | -0.233358018 | 0.133139523 | -1.752732875 |
| Gdf9 | 9.225389834 | -0.665691625 | 0.384672846 | -1.730539685 |
| Gm10447 | 1.155224236 | 3.002953511 | 1.094955761 | 2.742534098 |
| Shroom1 | 58.79842685 | 0.336688504 | 0.156225273 | 2.155147489 |
| Gm12212 | 3.290890966 | 1.500230697 | 0.638642538 | 2.349092972 |
| Gm9837 | 0.267041986 | -0.443849189 | 1.997170985 | -0.222238953 |
| Sowaha | 702.0530503 | 1.385895342 | 0.140454662 | 9.86720786 |
| Gm9945 | 10.46328265 | 0.465437328 | 0.38286075 | 1.215683061 |
| Rpl9-ps3 | 1.040046165 | -0.228275705 | 1.10896554 | -0.205845625 |
| 8-Sep | 2499.725176 | 0.06081362 | 0.066532059 | 0.914049877 |
| A430108G06Rik | 9.283107748 | 0.418953542 | 0.377200274 | 1.110692571 |
| Kif3a | 1223.698615 | 0.303577319 | 0.078264699 | 3.878853708 |
| Il4 | 0.170190954 | -0.70542992 | 2.744856668 | -0.25700064 |
| Il13 | 0 | NA | NA | NA |
| Rps2-ps4 | 0.4561058 | -1.358880619 | 1.56814466 | -0.866553101 |
| Rad50 | 251.7638459 | -0.024373019 | 0.129390732 | -0.188367578 |
| Gm22275 | 0 | NA | NA | NA |
| Il5 | 1.025767513 | 1.730920193 | 1.119022557 | 1.546814389 |
| Gm12214 | 0.972493339 | 1.608179817 | 1.277272903 | 1.259072993 |
| Gm12213 | 0 | NA | NA | NA |
| Mir7671 | 0.940546779 | 0.716202674 | 1.029982377 | 0.695354298 |
| Irf1 | 1452.897363 | -0.62933638 | 0.264434553 | -2.379932474 |
| Gm17334 | 5.090110731 | 0.535944487 | 0.427802734 | 1.252784157 |
| Gm12216 | 33.70519998 | 0.178293028 | 0.298490494 | 0.597315599 |
| Slc22a5 | 121.7288286 | 0.018989793 | 0.120942319 | 0.157015286 |
| Gm12217 | 0 | NA | NA | NA |
| Gm12218 | 0 | NA | NA | NA |
| Gm12220 | 0 | NA | NA | NA |
| Slc22a21 | 25.5234154 | 0.141726917 | 0.298960823 | 0.474065182 |
| Gm12219 | 2.781362659 | 1.218366791 | 0.75306833 | 1.617870175 |

| | | | | |
|---|---|---|---|---|
| Slc22a4 | 168.0969617 | -0.08355498 | 0.12159735 | -0.687144741 |
| Pdlim4 | 206.5108102 | -1.103906605 | 0.158164222 | -6.979496304 |
| P4ha2 | 113.5551428 | -0.055582264 | 0.159307457 | -0.348899327 |
| Gm12221 | 0.192293863 | -0.452147617 | 2.249617968 | -0.200988623 |
| 4933405E24Rik | 0 | NA | NA | NA |
| Gm12222 | 0 | NA | NA | NA |
| Csf2 | 0.705254363 | -2.202556612 | 1.161421303 | -1.896432076 |
| Gm12223 | 0.986197153 | 0.139966616 | 0.996292828 | 0.140487427 |
| Il3 | 0.048949532 | 0.037064902 | 3.724945049 | 0.009950456 |
| Acsl6 | 2735.937845 | -0.047476751 | 0.064655875 | -0.734299104 |
| Gm12224 | 0.727497348 | -1.124196683 | 1.223322894 | -0.918969709 |
| Meikin | 5.386405023 | -0.256301843 | 0.549479636 | -0.466444661 |
| Gm12226 | 3.723226009 | -0.542701219 | 0.543048592 | -0.999360327 |
| Gm12225 | 0 | NA | NA | NA |
| Fnip1 | 1691.944238 | -0.161331718 | 0.068011624 | -2.372119751 |
| Gm24198 | 6.506667764 | -0.181878242 | 0.537888433 | -0.338133767 |
| Rapgef6 | 1160.993809 | 0.26888448 | 0.068177593 | 3.943883428 |
| Gm26116 | 0 | NA | NA | NA |
| Cdc42se2 | 1261.495251 | 0.130753885 | 0.078170994 | 1.672664989 |
| Gm12227 | 0.36191905 | -0.871482587 | 1.580438731 | -0.551418141 |
| Gm12229 | 9.801174999 | 0.446340093 | 0.385043387 | 1.159194284 |
| Lyrm7os | 0.089089296 | 0.957155977 | 3.723939561 | 0.257027796 |
| Lyrm7 | 91.39166592 | 0.226335794 | 0.135839654 | 1.66619825 |
| Gm25426 | 0 | NA | NA | NA |
| Hint1 | 1051.934254 | -0.241331282 | 0.135294983 | -1.78374154 |
| Gpx3 | 335.6017487 | -0.774240432 | 0.276802337 | -2.797087769 |
| Tnip1 | 367.6412068 | -0.048485175 | 0.094243299 | -0.51446814 |
| Gm12230 | 0.092228164 | 0.037073798 | 3.724945049 | 0.009952844 |
| Anxa6 | 1191.581291 | -0.174736408 | 0.073185691 | -2.38757612 |
| Ccdc69 | 10.64187061 | -0.006197233 | 0.30971468 | -0.020009491 |
| Gm2a | 793.8364917 | 0.043971479 | 0.099765239 | 0.440749497 |
| Gm12231 | 50.21335086 | -0.033540423 | 0.201822559 | -0.166187682 |
| Mup-ps22 | 0 | NA | NA | NA |
| Slc36a3 | 0.361016523 | 0.028446313 | 3.152303285 | 0.009023977 |
| Slc36a3os | 0.686935554 | 0.153355388 | 1.094200465 | 0.140152918 |
| Slc36a2 | 7.128692757 | 0.992871678 | 0.494116701 | 2.009387003 |
| Slc36a1os | 0.119234754 | 0.51795324 | 3.724945049 | 0.139049901 |
| Slc36a1 | 1018.826301 | 0.01175742 | 0.053480159 | 0.219846388 |
| Fat2 | 2711.320312 | -0.073682755 | 0.274561766 | -0.268364953 |
| Sparc | 5655.720685 | -0.37555428 | 0.164906146 | -2.277381942 |
| Atox1 | 160.8036094 | -0.318820138 | 0.136826316 | -2.33010832 |
| Gm12235 | 0 | NA | NA | NA |
| G3bp1 | 943.5798911 | -0.465214816 | 0.074997001 | -6.203112227 |
| Gm22884 | 0.510616014 | 0.369300194 | 1.383556966 | 0.266920845 |
| Gm12238 | 0 | NA | NA | NA |
| Mir7652 | 0 | NA | NA | NA |
| Glra1 | 265.7926779 | -0.333378784 | 0.386784072 | -0.8619248 |

| | | | | |
|---|---|---|---|---|
| Gm12236 | 0.101408268 | 0.037730769 | 3.724898049 | 0.010129343 |
| Gm12237 | 0.066050875 | -0.443818988 | 3.724945049 | -0.119147795 |
| Gm12239 | 4.674577041 | 0.282147231 | 0.520195871 | 0.542386525 |
| Nmur2 | 2.399470382 | -0.329940854 | 0.649234793 | -0.508199587 |
| Gm12240 | 0.124308499 | -1.207977564 | 3.71482282 | -0.325177706 |
| Gm24253 | 0 | NA | NA | NA |
| Gm12241 | 0.724742635 | 1.002971097 | 1.188322639 | 0.844022544 |
| Gm22603 | 0 | NA | NA | NA |
| Gria1 | 3990.825874 | 0.608777434 | 0.110336158 | 5.517479002 |
| Gm12242 | 0.047948698 | 0.037064902 | 3.724945049 | 0.009950456 |
| Fam114a2 | 280.8245509 | -0.43150077 | 0.078729162 | -5.480825142 |
| Mfap3 | 364.5954244 | -0.354058707 | 0.079856034 | -4.433712637 |
| Gm12243 | 0.840033521 | 1.587763901 | 1.388020431 | 1.143905281 |
| 4933424L21Rik | 0 | NA | NA | NA |
| Galnt10 | 170.7549866 | -0.355228428 | 0.129704641 | -2.738748792 |
| 4933426K07Rik | 0 | NA | NA | NA |
| Gm24582 | 0.170988841 | 0.517964413 | 2.745561902 | 0.188655157 |
| n-R5s69 | 0.143140564 | 0.517957789 | 3.546633296 | 0.146042104 |
| 2010001A14Rik | 26.6509321 | 0.003215743 | 0.215972843 | 0.014889569 |
| Sap30l | 239.499411 | -0.112205571 | 0.111573492 | -1.005665138 |
| Hand1 | 0.104366086 | 0.51795324 | 3.724945049 | 0.139049901 |
| Gm12244 | 0 | NA | NA | NA |
| Gm12245 | 0.101408268 | 0.037730769 | 3.724898049 | 0.010129343 |
| Gm26025 | 0 | NA | NA | NA |
| Gm12249 | 0 | NA | NA | NA |
| Larp1 | 2687.565109 | -0.156759738 | 0.066084513 | -2.372110073 |
| Gm12248 | 0.378899548 | 0.300160976 | 1.715099412 | 0.175010833 |
| Gm12247 | 0.394465599 | 0.078286404 | 1.721716666 | 0.045469969 |
| Cnot8 | 374.7702201 | -0.010723962 | 0.061827518 | -0.173449666 |
| Gemin5 | 376.1235968 | -0.368092864 | 0.071712122 | -5.132923891 |
| Gm12246 | 2.686282868 | 0.360930823 | 0.838645319 | 0.430373622 |
| Mrpl22 | 119.8290135 | -0.003871728 | 0.124490615 | -0.031100561 |
| Gm12250 | 1005.570989 | -1.107325051 | 0.304173977 | -3.640433222 |
| Igtp | 4765.845661 | -1.051186274 | 0.235050188 | -4.472177975 |
| Irgm2 | 1836.176375 | -1.106381108 | 0.178817886 | -6.187194876 |
| 4930438A08Rik | 0.296006519 | 0.123193424 | 2.027288081 | 0.060767596 |
| Zfp692 | 130.8218891 | 0.16932914 | 0.20574716 | 0.822996249 |
| Zfp672 | 340.7570754 | -0.117804788 | 0.08136042 | -1.447937317 |
| 1700047K16Rik | 2.233748514 | 0.76945573 | 0.61138483 | 1.25854567 |
| Sh3bp5l | 520.8892961 | -0.287564084 | 0.084899467 | -3.387112932 |
| Lypd8 | 0.598341843 | -1.381076034 | 1.381861774 | -0.999431391 |
| Gm12251 | 32.14483831 | 0.016816187 | 0.218726385 | 0.076882295 |
| Gm12252 | 0 | NA | NA | NA |
| 1810065E05Rik | 0.095898756 | -0.917692504 | 3.724434845 | -0.246397787 |
| Gm12253 | 8.666853529 | -1.356906253 | 0.404606339 | -3.35364556 |
| 4930504O13Rik | 0 | NA | NA | NA |
| Olfr30 | 0 | NA | NA | NA |

| | | | | |
|---|---|---|---|---|
| Olfr333-ps1 | 0 | NA | NA | NA |
| Olfr332 | 0.045882455 | 0.517959912 | 3.724945049 | 0.139051692 |
| Olfr331 | 0 | NA | NA | NA |
| Gm12254 | 27.93282506 | -0.232753649 | 0.211125545 | -1.102441909 |
| Olfr330 | 0.072025462 | 0.517959912 | 3.724945049 | 0.139051692 |
| Gm12255 | 0 | NA | NA | NA |
| Olfr329-ps | 0.0878635 | 0.037058229 | 3.724945049 | 0.009948665 |
| Olfr328 | 0.068097359 | 0.037064902 | 3.724945049 | 0.009950456 |
| Olfr224 | 0 | NA | NA | NA |
| Olfr325 | 0 | NA | NA | NA |
| Olfr324 | 0.389730154 | 1.133446487 | 1.618976851 | 0.700100491 |
| 2210407C18Rik | 0.08169834 | -0.443821212 | 3.724945049 | -0.119148392 |
| Olfr323 | 0.122991902 | 0.517955464 | 3.724945049 | 0.139050498 |
| Trim58 | 1.195637209 | 2.096171348 | 1.062695265 | 1.972504647 |
| Gm22599 | 0 | NA | NA | NA |
| Olfr322 | 0 | NA | NA | NA |
| Olfr320 | 0 | NA | NA | NA |
| Gm12570 | 0 | NA | NA | NA |
| Olfr319 | 0 | NA | NA | NA |
| Olfr318 | 0 | NA | NA | NA |
| Olfr317 | 0 | NA | NA | NA |
| Olfr316 | 0 | NA | NA | NA |
| Olfr315 | 0.049553047 | -0.443818988 | 3.724945049 | -0.119147795 |
| Olfr314 | 0 | NA | NA | NA |
| Fam183b | 11.51123108 | -0.16650045 | 0.458770632 | -0.362927438 |
| Olfr313 | 0 | NA | NA | NA |
| Olfr312 | 0 | NA | NA | NA |
| Olfr311 | 0 | NA | NA | NA |
| Gm12258 | 234.0799246 | 0.117657155 | 0.098840371 | 1.19037548 |
| Gm12259 | 3.878080432 | 0.691388955 | 0.703230542 | 0.98316116 |
| 2810021J22Rik | 159.8131489 | -0.004756676 | 0.092829149 | -0.051241193 |
| Zfp39 | 105.0647483 | 0.202659089 | 0.152685672 | 1.327296049 |
| Gm12256 | 0.146451277 | -0.443820591 | 2.882358847 | -0.153978257 |
| Gm12257 | 0.048949532 | 0.037064902 | 3.724945049 | 0.009950456 |
| Btnl10 | 1.324943677 | -0.106446901 | 1.076279911 | -0.098902618 |
| Rnf187 | 2464.147731 | 0.071131626 | 0.059439241 | 1.196711543 |
| Hist3h2ba | 174.4290419 | 0.45219812 | 0.138300159 | 3.269686205 |
| Hist3h2bb-ps | 13.70298843 | 0.988626528 | 0.297989054 | 3.317660553 |
| Trim17 | 59.87795607 | -0.460601674 | 0.204897937 | -2.24795662 |
| Hist3h2a | 106.0988195 | -0.092248453 | 0.155305532 | -0.593980467 |
| AL662809.1 | 1.743011301 | 1.578050639 | 0.943624259 | 1.672329451 |
| Gm12260 | 5.27939244 | 0.577573223 | 0.458129954 | 1.260719185 |
| 2310058D17Rik | 14.57587431 | -0.578200401 | 0.356019581 | -1.6240691 |
| Trim11 | 219.4000615 | 0.064754151 | 0.120073193 | 0.539288994 |
| Obscn | 127.4374054 | 1.312712099 | 0.288647282 | 4.547806887 |
| Gm10435 | 0.809248464 | -0.369269281 | 1.225641063 | -0.301286643 |
| Iba57 | 77.73534886 | -0.090762163 | 0.14438607 | -0.628607479 |

| | | | | |
|---|---|---|---|---|
| Gjc2 | 99.21955442 | -0.332150265 | 0.182920926 | -1.815813374 |
| Guk1 | 493.4726625 | 0.147970708 | 0.115466022 | 1.281508673 |
| 2610507I01Rik | 3.133265115 | 0.139330871 | 0.638211569 | 0.218314549 |
| Mrpl55 | 89.18665174 | -0.044978108 | 0.178600133 | -0.251836926 |
| Gm10434 | 1.747669216 | -1.50226839 | 0.815998176 | -1.8410193 |
| 2310033P09Rik | 92.90886109 | -0.156298798 | 0.144363971 | -1.082671779 |
| Arf1 | 2962.793734 | -0.110559397 | 0.069505152 | -1.590664778 |
| Wnt3a | 0.23148996 | -0.445500109 | 2.554735304 | -0.174382101 |
| Wnt9a | 76.15545421 | 0.283966333 | 0.174535158 | 1.626986429 |
| Gm24756 | 0 | NA | NA | NA |
| Prss38 | 1.918719673 | 0.117398728 | 0.874326849 | 0.134273274 |
| Gm15755 | 0 | NA | NA | NA |
| Snap47 | 2340.344067 | 0.195988134 | 0.053130185 | 3.688828338 |
| Jmjd4 | 428.2804086 | 0.015402342 | 0.065393878 | 0.235531862 |
| Zfp867 | 99.85436412 | -0.13893949 | 0.17596481 | -0.789586797 |
| Gm12263 | 27.6794227 | -0.274132121 | 0.220262608 | -1.244569486 |
| Gm12262 | 0 | NA | NA | NA |
| Zkscan17 | 266.3947386 | 0.082384581 | 0.123319475 | 0.668058158 |
| 4933439C10Rik | 96.1816591 | 0.349727815 | 0.190646092 | 1.834434747 |
| Nlrp3 | 44.3755037 | -0.602466041 | 0.16529327 | -3.644831037 |
| Olfr222 | 0.26446025 | -0.511098862 | 2.059683529 | -0.248144365 |
| Gm12261 | 0.418705261 | -0.749754871 | 1.379701864 | -0.543418032 |
| Olfr223 | 0 | NA | NA | NA |
| Olfr225 | 0 | NA | NA | NA |
| Mprip | 6005.10193 | -0.024122865 | 0.060161595 | -0.400967851 |
| 1700007J10Rik | 1.636732283 | 0.022627699 | 0.967457873 | 0.023388821 |
| Gm12264 | 0.290485957 | 0.037077033 | 1.816818675 | 0.020407669 |
| Pld6 | 8.31364424 | 0.106930519 | 0.406539603 | 0.263026082 |
| Flcn | 628.2729707 | -0.163419222 | 0.063904481 | -2.557241986 |
| Gm16062 | 8.661166123 | -0.777233043 | 0.377992867 | -2.056210875 |
| Gm24939 | 0.352988078 | 0.534720708 | 1.485369491 | 0.359991713 |
| Cops3 | 618.5280119 | 0.033168331 | 0.081898928 | 0.404991032 |
| Nt5m | 258.4297601 | 0.073102401 | 0.079176713 | 0.923281583 |
| Gm12714 | 0.04393175 | 0.037062677 | 3.724945049 | 0.009949859 |
| 1810063I02Rik | 0.161948271 | -0.443822416 | 3.446645162 | -0.128769396 |
| Med9os | 2.791248013 | -0.633396034 | 0.668016047 | -0.948174878 |
| Med9 | 292.2435254 | -0.073719837 | 0.093835923 | -0.785624891 |
| Rasd1 | 132.7603455 | 0.625852759 | 0.198624823 | 3.150929222 |
| Pemt | 14.2244874 | -0.085043907 | 0.288967451 | -0.294302719 |
| Gm27511 | 0 | NA | NA | NA |
| Rai1 | 1366.993626 | 0.098630932 | 0.080607252 | 1.223598746 |
| 4930412M03Rik | 2.074250591 | -0.452464833 | 0.62379877 | -0.72533781 |
| Srebf1 | 894.7642396 | 0.058676447 | 0.126006932 | 0.465660468 |
| Mir6921 | 0 | NA | NA | NA |
| Mir6922 | 0 | NA | NA | NA |
| Tom1l2 | 3217.981596 | 0.195902023 | 0.0690516 | 2.837038126 |
| Gm12265 | 0.343978253 | -1.47553883 | 1.486690269 | -0.992499152 |

| | | | | |
|---|---|---|---|---|
| Gm27711 | 2.937723515 | 0.554059742 | 0.778756548 | 0.711467202 |
| Gm12266 | 0 | NA | NA | NA |
| Lrrc48 | 90.36745919 | 0.169405825 | 0.122248287 | 1.385752142 |
| Atpaf2 | 218.7768415 | -0.126985081 | 0.075406826 | -1.683999818 |
| Gid4 | 732.8917463 | 0.235375914 | 0.091042312 | 2.585346413 |
| Gm12268 | 1.686071008 | -0.39497417 | 1.004548194 | -0.393185884 |
| Gm12267 | 0.287879534 | 0.344103652 | 1.712304925 | 0.200959331 |
| Drg2 | 463.374575 | -0.165946824 | 0.065388524 | -2.537858546 |
| Myo15 | 13.54234393 | -0.154075498 | 0.447319615 | -0.344441632 |
| Alkbh5 | 1342.8845 | -0.191537485 | 0.068568798 | -2.793362145 |
| Gm12622 | 0 | NA | NA | NA |
| Gm12628 | 0 | NA | NA | NA |
| Gm26837 | 0 | NA | NA | NA |
| Gm12623 | 0.044191549 | 0.037062677 | 3.724945049 | 0.009949859 |
| Gm12621 | 0 | NA | NA | NA |
| Gm12627 | 0 | NA | NA | NA |
| Gm12613 | 0 | NA | NA | NA |
| Gm12614 | 0 | NA | NA | NA |
| Gm12615 | 0 | NA | NA | NA |
| Gm12626 | 0 | NA | NA | NA |
| Gm12625 | 0 | NA | NA | NA |
| Gm12619 | 0 | NA | NA | NA |
| Gm12624 | 0 | NA | NA | NA |
| Gm12612 | 0 | NA | NA | NA |
| Gm12616 | 0 | NA | NA | NA |
| Gm12620 | 0 | NA | NA | NA |
| Gm12617 | 0 | NA | NA | NA |
| Llgl1 | 814.2720307 | -0.234055154 | 0.087584798 | -2.672326251 |
| Flii | 919.6804464 | -0.609538006 | 0.096689518 | -6.304075352 |
| Mief2 | 152.1320646 | -0.097826553 | 0.110737032 | -0.883413175 |
| Mir5100 | 0.418037274 | -0.925697691 | 3.339398695 | -0.277204903 |
| Gm23341 | 0.048949532 | 0.037064902 | 3.724945049 | 0.009950456 |
| Top3a | 117.1519205 | -0.101362536 | 0.112769334 | -0.898848408 |
| Smcr8 | 1137.612935 | -0.292391274 | 0.077577586 | -3.769017412 |
| Shmt1 | 28.85367593 | 0.458965195 | 0.202388546 | 2.26774293 |
| Gm24945 | 0.047948698 | 0.037064902 | 3.724945049 | 0.009950456 |
| Gm12611 | 0.097139547 | -0.443816764 | 3.724945049 | -0.119147198 |
| Gm12618 | 0.132101749 | -0.93550548 | 3.724188147 | -0.251197158 |
| Dhrs7b | 155.5720981 | -0.246484738 | 0.143151681 | -1.721843128 |
| Tmem11 | 223.7050137 | -0.065733482 | 0.10557877 | -0.622601324 |
| Gm26534 | 14.74527245 | -0.131125908 | 0.298961686 | -0.43860439 |
| Natd1 | 186.5605054 | -0.184735006 | 0.127277217 | -1.451438129 |
| Map2k3os | 10.95972877 | -1.670660031 | 0.391762919 | -4.264466973 |
| Map2k3 | 195.4668182 | -0.64309333 | 0.151592966 | -4.24223727 |
| Kcnj12 | 243.7727264 | 0.14273218 | 0.144546787 | 0.987446229 |
| Tnfrsf13b | 25.743611 | -0.735955355 | 0.310150446 | -2.372897942 |
| Gm12269 | 0 | NA | NA | NA |

| | | | | |
|---|---|---|---|---|
| Usp22 | 3131.14818 | 0.210727778 | 0.053409024 | 3.945546298 |
| Rps13-ps5 | 0.882161971 | 0.32698676 | 1.04763742 | 0.312118252 |
| Aldh3a1 | 6.417984165 | 0.384191992 | 0.43802989 | 0.877090812 |
| Aldh3a2 | 629.8388114 | -0.196847101 | 0.097908375 | -2.010523625 |
| Slc47a2 | 0.919894618 | -1.960232333 | 1.52420708 | -1.286066939 |
| Gm25929 | 0 | NA | NA | NA |
| Slc47a1 | 39.2732027 | -0.544773857 | 0.25743325 | -2.116175193 |
| Gm12271 | 0.275857165 | 1.871886634 | 2.431909914 | 0.76971874 |
| Rnf112 | 1605.26594 | 0.768803388 | 0.135491943 | 5.674163132 |
| Mfap4 | 9.173601336 | -0.148636973 | 0.380835374 | -0.390291931 |
| Mapk7 | 210.9337897 | -0.623120129 | 0.106699791 | -5.839937664 |
| B9d1os | 5.30594644 | 0.089733771 | 0.508532216 | 0.176456414 |
| B9d1 | 90.68224864 | -0.146270078 | 0.130175592 | -1.123636747 |
| Epn2 | 1283.27487 | 0.175429509 | 0.06308274 | 2.780943068 |
| Gm12272 | 1.010954397 | 1.303684952 | 1.172270784 | 1.11210223 |
| Grap | 28.95643677 | -0.098512515 | 0.21228267 | -0.464062916 |
| Slc5a10 | 1.756770038 | 1.21393614 | 0.802805996 | 1.512116435 |
| Fam83g | 11.17234595 | -1.945886358 | 0.395480992 | -4.920303121 |
| Prpsap2 | 311.9887896 | -0.17816765 | 0.108873408 | -1.636466177 |
| Ulk2 | 1353.293992 | 0.281554915 | 0.091280185 | 3.084512989 |
| A530017D24Rik | 232.7074356 | 0.204248559 | 0.082768011 | 2.467723432 |
| Gm26964 | 0.259490603 | 0.517967068 | 2.469022377 | 0.2097863 |
| Akap10 | 671.4108366 | -0.042039079 | 0.061380072 | -0.684897847 |
| Gm23721 | 0 | NA | NA | NA |
| Gm12273 | 0.316938377 | -1.269142517 | 2.030432497 | -0.625060187 |
| Specc1 | 1898.855375 | -0.031385163 | 0.065333947 | -0.480380643 |
| Gm12274 | 0.049553047 | -0.443818988 | 3.724945049 | -0.119147795 |
| Adora2b | 44.79440823 | 0.416557332 | 0.212106674 | 1.963904882 |
| Zswim7 | 50.11354397 | 0.342959634 | 0.154737616 | 2.216394712 |
| Ttc19 | 1111.563203 | 0.268992445 | 0.086611806 | 3.105724932 |
| Gm12275 | 2.211543673 | 1.090365498 | 0.767450688 | 1.420762943 |
| Ncor1 | 3691.866875 | 0.265397753 | 0.072660908 | 3.652552117 |
| Gm12276 | 22.41048438 | -0.480548753 | 0.251178266 | -1.913178081 |
| Pigl | 79.7396089 | -0.169613358 | 0.14707714 | -1.1532272 |
| Gm12278 | 0.637771319 | -1.298355495 | 1.247022161 | -1.041164733 |
| Gm12949 | 0 | NA | NA | NA |
| Cenpv | 151.2123072 | 0.087151476 | 0.113627984 | 0.766989543 |
| Gm12279 | 7.228218235 | 0.397054691 | 0.414875615 | 0.957045138 |
| Ubb | 6377.498152 | -0.112689474 | 0.10457163 | -1.077629504 |
| Gm12280 | 0 | NA | NA | NA |
| Trpv2 | 211.9525486 | -0.488380272 | 0.124112306 | -3.934986675 |
| 2410006H16Rik | 257.1498311 | -0.439505179 | 0.179800309 | -2.444407254 |
| Snord49b | 2.860297155 | 0.378655783 | 0.590451651 | 0.641298543 |
| Snord49a | 6.687144773 | 0.212667529 | 0.441929394 | 0.481225128 |
| Snord65 | 1.151518338 | -0.390745699 | 0.987322402 | -0.395763023 |
| Lrrc75a | 93.92612434 | -0.302783425 | 0.17085893 | -1.772125251 |
| Lrrc75aos2 | 8.114142353 | 0.331188743 | 0.444757588 | 0.744650012 |

| | | | | |
|---|---|---|---|---|
| Lrrc75aos1 | 0 | NA | NA | NA |
| Mmgt2 | 82.28640359 | -0.657375419 | 0.12629816 | -5.204948507 |
| Gm12281 | 0.339850366 | -0.519206541 | 1.74538235 | -0.297474385 |
| Zfp287 | 282.6640166 | 0.007374724 | 0.103527116 | 0.071234714 |
| Gm27194 | 0.484088067 | -0.147325558 | 1.444815464 | -0.101968426 |
| Wsb2-ps | 33.81592792 | 0.194042653 | 0.196942189 | 0.985277223 |
| Gm10428 | 0.042716945 | 0.037062677 | 3.724945049 | 0.009949859 |
| Zfp286 | 136.9541373 | 0.180419699 | 0.114811111 | 1.571448071 |
| Zfp286it | 0.804776955 | 1.235219819 | 1.152927255 | 1.071377066 |
| Zfp286os | 0 | NA | NA | NA |
| Gm24151 | 0 | NA | NA | NA |
| Trim16 | 38.08170184 | -0.415590075 | 0.212041966 | -1.959942565 |
| Fbxw10 | 9.908817 | -0.189235886 | 0.347406978 | -0.544709514 |
| Tvp23b | 188.0397332 | -0.365634421 | 0.156822039 | -2.331524464 |
| Tvp23bos | 2.044779979 | -0.288051857 | 0.683024857 | -0.421729684 |
| Cdrt4os2 | 0 | NA | NA | NA |
| Cdrt4 | 2.232638995 | -0.840704905 | 0.674099591 | -1.247152373 |
| Cdrt4os1 | 0 | NA | NA | NA |
| Tekt3 | 0.583929304 | 0.418204801 | 1.309534448 | 0.319353799 |
| Pmp22 | 317.1591218 | -0.213142882 | 0.159261502 | -1.33832018 |
| B430319H21Rik | 0.78030421 | 0.656613299 | 1.107846617 | 0.592693329 |
| Gm12285 | 0 | NA | NA | NA |
| Gm12286 | 16.66800335 | 0.503667297 | 0.253922119 | 1.983550311 |
| B130011K05Rik | 1.258106493 | 1.262651935 | 0.951705378 | 1.326725649 |
| Gm12287 | 0.044544648 | 0.517957688 | 3.724945049 | 0.139051095 |
| Gm12288 | 0 | NA | NA | NA |
| Hmgb1-ps3 | 8.21809902 | -0.092037549 | 0.404711689 | -0.2274151 |
| Hs3st3b1 | 91.21008015 | 0.003462559 | 0.186721593 | 0.018543969 |
| Cox10 | 247.0159409 | -0.05236644 | 0.086090433 | -0.608272466 |
| 9630013K17Rik | 0.09689823 | 0.037062677 | 3.724945049 | 0.009949859 |
| Gm12289 | 0 | NA | NA | NA |
| 2810001G20Rik | 35.83958181 | 0.051862236 | 0.196544652 | 0.263869995 |
| Gm12290 | 1.421699425 | -0.398427011 | 0.79311506 | -0.502357137 |
| F930015N05Rik | 5.181464156 | 1.820388979 | 0.588314215 | 3.094246123 |
| Hs3st3a1 | 37.68402565 | 0.287795245 | 0.253629628 | 1.134706724 |
| Gm24275 | 0 | NA | NA | NA |
| Gm12291 | 0.332139225 | -1.16122603 | 1.73928523 | -0.667645542 |
| Gm12292 | 0.254955332 | -0.113834189 | 2.456176447 | -0.046346096 |
| Elac2 | 339.1600695 | -0.0818601 | 0.087507391 | -0.935464984 |
| Arhgap44 | 1922.5473 | 0.374554102 | 0.074652691 | 5.017288693 |
| Gm12294 | 19.04726638 | 0.974778261 | 0.470116163 | 2.073483827 |
| Gm25991 | 1.580643696 | 0.217381622 | 0.914879574 | 0.237606815 |
| 1700086D15Rik | 40.20348781 | 1.101996213 | 0.399251143 | 2.760157939 |
| Gm12293 | 0.046345709 | -0.443818988 | 3.724945049 | -0.119147795 |
| Myocd | 4.652030534 | 0.078851823 | 0.50278374 | 0.156830496 |
| Gm12295 | 2.004716341 | -3.145208281 | 0.979842016 | -3.20991367 |
| Gm26128 | 0 | NA | NA | NA |

| | | | | |
|---|---|---|---|---|
| Map2k4 | 1997.328949 | 0.205499355 | 0.046283467 | 4.440016482 |
| Mir744 | 16.92311974 | 0.12846346 | 0.315714933 | 0.406897003 |
| Zkscan6 | 133.418627 | -0.152016223 | 0.111289624 | -1.365951454 |
| Dnah9 | 161.9890621 | 0.37588158 | 0.128686389 | 2.920911699 |
| Gm12296 | 7.015874426 | 1.038572772 | 0.509448074 | 2.038623414 |
| Shisa6 | 795.9631443 | 0.30726292 | 0.132830296 | 2.313199096 |
| Gm12297 | 1.996095088 | 0.514056642 | 0.776235865 | 0.662242837 |
| Gm12298 | 0 | NA | NA | NA |
| Pirt | 6.080719795 | 0.52677413 | 0.493288208 | 1.067883077 |
| 9130409J20Rik | 0 | NA | NA | NA |
| Gm12299 | 0.127580795 | 0.037075204 | 3.00756786 | 0.012327304 |
| Tmem220 | 8.211970812 | -0.620141613 | 0.353568822 | -1.753948805 |
| Adprm | 137.339951 | -0.449408085 | 0.123751613 | -3.631533132 |
| Sco1 | 243.6408562 | -0.183571676 | 0.1516181 | -1.210750406 |
| Myh3 | 10.50794026 | 1.251801728 | 0.402598644 | 3.10930438 |
| Mir6923 | 0 | NA | NA | NA |
| 2310065F04Rik | 0.176426445 | 0.037076448 | 2.718482223 | 0.013638658 |
| Myh2 | 5.858603139 | 0.770001766 | 0.55202257 | 1.394873704 |
| 9430073C21Rik | 0.072025462 | 0.517959912 | 3.724945049 | 0.139051692 |
| Myh1 | 7.368640863 | 1.095888398 | 0.455285187 | 2.407037235 |
| Gm12300 | 1.913586201 | 3.751222192 | 0.995204276 | 3.769298707 |
| Myh4 | 5.497083594 | 1.092257374 | 0.512776397 | 2.130085121 |
| Myh8 | 7.157414437 | 1.916716617 | 0.489069569 | 3.919108322 |
| Myh13 | 1.635466931 | 0.249178051 | 0.831245181 | 0.299764807 |
| Gm26476 | 0 | NA | NA | NA |
| Gas7 | 7031.634939 | 0.536832786 | 0.106394862 | 5.045664591 |
| Gm12301 | 0.159545055 | 0.037074822 | 2.806181499 | 0.01321184 |
| Glp2r | 34.73267854 | 0.439643691 | 0.237720493 | 1.849414349 |
| Rcvrn | 8.311551091 | 0.487660866 | 0.491433267 | 0.992323676 |
| Gm12302 | 0.259617586 | -2.019675445 | 2.408645394 | -0.838510911 |
| Dhrs7c | 0.646105857 | -2.912752979 | 1.400730296 | -2.079453116 |
| Usp43 | 27.84987753 | 0.264505313 | 0.216983371 | 1.219011909 |
| 9130017K11Rik | 3.804522088 | 0.181013692 | 0.578776973 | 0.312752062 |
| Cfap52 | 13.68582642 | -0.695611752 | 0.302081222 | -2.302730863 |
| Stx8 | 213.3769252 | -0.193138623 | 0.091909966 | -2.101389336 |
| Gm12303 | 0.044544648 | 0.517957688 | 3.724945049 | 0.139051095 |
| Gm26157 | 1.267602396 | 0.946140993 | 1.009703993 | 0.937047887 |
| Gm12304 | 0 | NA | NA | NA |
| C78197 | 2.497471105 | 0.294570484 | 0.637824896 | 0.461835978 |
| Ntn1 | 541.1663475 | -0.420410859 | 0.153457832 | -2.739585546 |
| Mir6406 | 0.083526043 | 0.037073798 | 3.724945049 | 0.009952844 |
| Gm12305 | 1.238804443 | -1.105614305 | 1.097591473 | -1.007309488 |
| Pik3r5 | 167.1648793 | -1.521729969 | 0.248068286 | -6.134318876 |
| Pik3r6 | 32.88554754 | -0.338841996 | 0.265505009 | -1.276216962 |
| Mfsd6l | 0.872192936 | -2.221524798 | 1.224306369 | -1.814517064 |
| Ccdc42os | 0 | NA | NA | NA |
| Ccdc42 | 2.972495032 | 0.94165068 | 0.692378708 | 1.360022584 |

| | | | | |
|---|---|---|---|---|
| Myh10 | 3525.532232 | 0.368682733 | 0.091613685 | 4.024319419 |
| Ndel1 | 867.2338058 | -0.182903788 | 0.070531527 | -2.593220286 |
| Rnf222 | 1.039519431 | -0.098484544 | 1.080018328 | -0.091187845 |
| Rpl26 | 246.1278693 | -0.217693536 | 0.159986458 | -1.360699762 |
| Gm22608 | 0.141071297 | -0.443820801 | 3.564043909 | -0.12452731 |
| Odf4 | 1.2405255 | 0.782659237 | 1.096517947 | 0.713767831 |
| Arhgef15 | 134.1597504 | -0.225431198 | 0.158513223 | -1.422160201 |
| Slc25a35 | 55.2247238 | 0.067153289 | 0.22556918 | 0.297705959 |
| Rangrf | 38.50784585 | -1.071467633 | 0.225486874 | -4.751796029 |
| Pfas | 417.0734811 | -0.2258321 | 0.072798854 | -3.102138103 |
| Mir3062 | 0 | NA | NA | NA |
| Gm25819 | 0 | NA | NA | NA |
| Ctc1 | 356.1546317 | 0.237470719 | 0.10027134 | 2.36828109 |
| Aurkb | 11.54692778 | -0.038226907 | 0.422703013 | -0.090434432 |
| 9330160F10Rik | 16.80478976 | 1.175058348 | 0.299723216 | 3.920478243 |
| 2310047M10Rik | 219.3535084 | 0.105426048 | 0.105610248 | 0.998255844 |
| Tmem107 | 35.40669845 | -0.617609986 | 0.235243389 | -2.625408468 |
| Snord118 | 16.03095282 | -0.501239344 | 0.363270535 | -1.37979631 |
| Gm12306 | 0.293428113 | -0.458762653 | 1.638117849 | -0.280054731 |
| Gm25371 | 0.515366098 | -0.024296297 | 1.369501554 | -0.017740978 |
| Vamp2 | 8741.82778 | 0.485874449 | 0.078537792 | 6.186505062 |
| Per1 | 1912.220295 | 0.515259954 | 0.16435507 | 3.135041438 |
| 9130213A22Rik | 7.871844516 | 0.618317323 | 0.387160259 | 1.597057831 |
| Hes7 | 8.964305686 | 0.769023395 | 0.391456094 | 1.964520177 |
| Aloxe3 | 69.09659255 | 0.286013679 | 0.183081717 | 1.562218691 |
| Alox12b | 38.03970433 | 2.40163176 | 0.315052218 | 7.622964148 |
| Alox8 | 33.51618146 | 0.280461991 | 0.211953003 | 1.323227255 |
| Gucy2e | 40.20224287 | -0.156159479 | 0.243084619 | -0.642407896 |
| Gm23194 | 0 | NA | NA | NA |
| Cntrobos | 0 | NA | NA | NA |
| Cntrob | 149.0597755 | 0.113536313 | 0.128919545 | 0.880675719 |
| Trappc1 | 165.2164685 | -0.081038467 | 0.126476974 | -0.640736926 |
| Kcnab3 | 622.5251017 | 0.370749843 | 0.105698723 | 3.507609491 |
| Kcnab3os | 3.132524194 | -0.14581029 | 0.595318234 | -0.244928312 |
| Chd3os | 937.6597823 | 0.160662827 | 0.077553725 | 2.071632613 |
| Chd3 | 5496.775121 | 0.804078946 | 0.128300552 | 6.267151087 |
| Gm22442 | 482.6561501 | 0.158241623 | 0.123702172 | 1.279214584 |
| Gm17305 | 8.479929393 | 0.983264017 | 0.425762299 | 2.309420115 |
| Cyb5d1 | 166.3102935 | -0.063368256 | 0.094382815 | -0.671396124 |
| Naa38 | 194.4625342 | 0.178228677 | 0.14865597 | 1.198933865 |
| Tmem88 | 12.86887999 | -0.678769155 | 0.348702783 | -1.946555029 |
| Kdm6b | 1048.172561 | -0.515272298 | 0.163401897 | -3.153404631 |
| Kdm6bos | 0.21690863 | 0.037075247 | 2.219567793 | 0.016703814 |
| Dnah2 | 63.85404554 | 0.188538721 | 0.159080743 | 1.185176267 |
| Dnah2os | 1.999759116 | -0.292374825 | 0.679679223 | -0.4301659 |
| Gm27493 | 0.038981395 | -0.443816764 | 3.724945049 | -0.119147198 |
| Efnb3 | 1089.578534 | 0.187127783 | 0.080897124 | 2.31315742 |

| | | | | |
|---|---|---|---|---|
| Wrap53 | 98.47005554 | -0.162726467 | 0.118591793 | -1.372156229 |
| Trp53 | 281.3127673 | -0.626098976 | 0.126725693 | -4.94058437 |
| Atp1b2 | 7719.462683 | 0.072855565 | 0.113719463 | 0.64066048 |
| Shbg | 1.970655031 | -0.559565004 | 0.689011418 | -0.812127331 |
| Sat2 | 69.39285511 | 0.534203553 | 0.145719205 | 3.665979058 |
| Gm12308 | 0 | NA | NA | NA |
| Gm12309 | 0 | NA | NA | NA |
| Fxr2 | 1268.108725 | -0.025097199 | 0.056479551 | -0.444359035 |
| Mir467f | 0 | NA | NA | NA |
| Sox15 | 1.197362358 | 1.260519164 | 1.074788745 | 1.172806442 |
| Mpdu1 | 282.855519 | -0.063063162 | 0.094297746 | -0.668766377 |
| Mir1934 | 0.521325481 | 0.927354571 | 1.589027488 | 0.583598823 |
| Cd68 | 498.24161 | -1.883401154 | 0.333176024 | -5.652871208 |
| Eif4a1 | 2251.416923 | -0.390097609 | 0.113501178 | -3.436947679 |
| Gm24029 | 0 | NA | NA | NA |
| Gm25835 | 93.03237133 | -0.747895868 | 0.202360045 | -3.695867267 |
| Gm22988 | 0 | NA | NA | NA |
| Senp3 | 708.1528605 | -0.224643274 | 0.058248829 | -3.856614394 |
| Tnfsf13os | 8.601013049 | -0.357637505 | 0.348175888 | -1.02717482 |
| Tnfsfm13 | 5.612199682 | -0.867257637 | 0.496028571 | -1.748402588 |
| Tnfsf13 | 5.550421209 | -0.546326131 | 0.438428579 | -1.246100635 |
| Tnfsf12 | 55.83410333 | -0.789113158 | 0.176472414 | -4.471594954 |
| Gm12307 | 0.075043205 | 0.517957688 | 3.724945049 | 0.139051095 |
| Polr2a | 2028.090813 | -0.199831411 | 0.077426514 | -2.580917065 |
| Slc35g3 | 0.474471242 | -0.467057239 | 1.500636358 | -0.311239453 |
| Zbtb4 | 2349.635844 | 0.235556713 | 0.056030465 | 4.204082779 |
| Chrnb1 | 9.482089685 | -0.698502304 | 0.406748609 | -1.717282587 |
| Fgf11 | 217.0904447 | -5.42E-05 | 0.111002396 | -0.000488375 |
| Tmem102 | 2.228356223 | -0.687108758 | 0.801634455 | -0.857134762 |
| 4933402P03Rik | 0 | NA | NA | NA |
| Spem1 | 3.349245703 | -0.867774764 | 0.575867834 | -1.506899175 |
| Nlgn2 | 4433.469855 | 0.345528743 | 0.092892182 | 3.719675185 |
| Tmem256 | 162.3407789 | -0.434327083 | 0.106907006 | -4.062662487 |
| Plscr3 | 159.9402606 | -0.508349191 | 0.113606066 | -4.474665899 |
| Tnk1 | 0.927088552 | -0.018491432 | 1.10788563 | -0.016690741 |
| Tmem95 | 1.006289673 | -3.538409634 | 1.108006604 | -3.193491465 |
| Kctd11 | 45.25365547 | -1.716289873 | 0.280901444 | -6.109936109 |
| Acap1 | 25.63139517 | -0.856541207 | 0.285104439 | -3.004306804 |
| mmu-mir-6924 | 0 | NA | NA | NA |
| 2810408A11Rik | 1.623343953 | -0.928991426 | 0.910786651 | -1.019987969 |
| Neurl4 | 1197.492728 | 0.090912318 | 0.055927769 | 1.625530927 |
| Gps2 | 171.2820456 | 0.027339645 | 0.154217169 | 0.177280167 |
| Eif5a | 3116.497447 | -0.188552391 | 0.096793001 | -1.947996134 |
| Ybx2 | 14.27143314 | 0.806856389 | 0.390771863 | 2.064776063 |
| Slc2a4 | 20.6317304 | 0.853289024 | 0.355872007 | 2.397741345 |
| Gm12310 | 0 | NA | NA | NA |
| Cldn7 | 0.279349812 | 0.037077994 | 1.802372587 | 0.02057177 |

| | | | | |
|---|---|---|---|---|
| Elp5 | 407.5792987 | -0.164265781 | 0.100914286 | -1.62777529 |
| AL596185.1 | 14.58297401 | -0.162010051 | 0.309914348 | -0.522757503 |
| Ctdnep1 | 632.1150543 | -0.382472081 | 0.07139394 | -5.357206532 |
| Gabarap | 1356.650847 | -0.593723412 | 0.127011249 | -4.674573436 |
| Phf23 | 440.621247 | -0.662627544 | 0.09234936 | -7.175226197 |
| Dvl2 | 217.8185947 | -0.192423434 | 0.135644014 | -1.418591413 |
| Acadvl | 388.2749648 | -0.430633609 | 0.09311001 | -4.624998011 |
| Mir324 | 0 | NA | NA | NA |
| Dlg4 | 6142.381317 | 0.644904282 | 0.09972763 | 6.466656069 |
| Asgr1 | 14.73418541 | 0.090861953 | 0.311092495 | 0.292073754 |
| Asgr2 | 0.670311827 | -1.941979003 | 1.400734556 | -1.386400439 |
| Gm12311 | 0 | NA | NA | NA |
| Mgl2 | 3.864736009 | -1.30876655 | 0.593938772 | -2.203537826 |
| Clec10a | 7.893594261 | -0.576810921 | 0.370439968 | -1.557096887 |
| Slc16a11 | 154.4850972 | 0.749144327 | 0.179829462 | 4.165859805 |
| Slc16a13 | 67.2589845 | 0.082564446 | 0.177336692 | 0.465580165 |
| Bcl6b | 16.16416338 | -0.495339781 | 0.27174146 | -1.822834765 |
| Mir497b | 0.072025462 | 0.517959912 | 3.724945049 | 0.139051692 |
| mmu-mir-497a | 0.695279289 | 0.683054127 | 1.327317653 | 0.514612403 |
| Mir195a | 1.190084389 | -1.272292371 | 1.134991024 | -1.120971306 |
| 0610010K14Rik | 9.73724523 | 0.290805003 | 0.390570046 | 0.744565554 |
| Gm21988 | 3.655964835 | 0.053511279 | 0.5420182 | 0.098725983 |
| Rnasek | 9.847312144 | 0.369456462 | 0.342715877 | 1.07802552 |
| Alox12 | 4.024321177 | -1.51305359 | 0.645893866 | -2.342573092 |
| Alox12e | 0.602894331 | 2.166485349 | 1.377439066 | 1.572835709 |
| Alox15 | 1.707423372 | -0.655484042 | 0.746725375 | -0.877811394 |
| Gm12312 | 2.105566461 | -0.984053323 | 0.726045274 | -1.355360828 |
| Pelp1 | 421.1587153 | -0.015280039 | 0.056698911 | -0.269494399 |
| Gm12313 | 0 | NA | NA | NA |
| Gm12314 | 0.049553047 | -0.443818988 | 3.724945049 | -0.119147795 |
| Arrb2 | 535.4570107 | -0.164222618 | 0.075964291 | -2.161839682 |
| Mir7115 | 0 | NA | NA | NA |
| Med11 | 69.16924485 | -0.626401279 | 0.206628977 | -3.031526781 |
| Cxcl16 | 296.0926 | -0.446299321 | 0.17569159 | -2.540242941 |
| Zmynd15 | 30.5256007 | -0.717446587 | 0.230952014 | -3.106474694 |
| Tm4sf5 | 2.723137057 | -0.277972855 | 0.620327274 | -0.448106776 |
| Vmo1 | 1.167525075 | 1.12193307 | 1.086258777 | 1.032841431 |
| Gm12317 | 0.120679735 | -0.837634339 | 3.722822199 | -0.224999824 |
| Gltpd2 | 0.276014283 | 0.068478278 | 2.082254203 | 0.032886608 |
| Psmb6 | 866.9721086 | -0.299884227 | 0.090001816 | -3.331979738 |
| Gm10418 | 0.929389619 | -0.932719272 | 1.138648978 | -0.819145576 |
| Pld2 | 260.2865872 | -0.444092318 | 0.130613321 | -3.400053808 |
| Mink1 | 3061.414012 | 0.31461442 | 0.091772302 | 3.428206698 |
| Gm12316 | 0.706900582 | -0.762204444 | 1.335539744 | -0.570708919 |
| Chrne | 2.081055214 | -0.663982724 | 0.871853126 | -0.761576353 |
| 4930544D05Rik | 1.506657277 | 0.961378612 | 1.004917652 | 0.956674022 |
| Gm12315 | 0.418629604 | 0.075103216 | 1.377061098 | 0.054538768 |

| | | | | |
|---|---|---|---|---|
| Gp1ba | 0.170190954 | -0.70542992 | 2.744856668 | -0.25700064 |
| Slc25a11 | 758.6093433 | 0.214603345 | 0.073728418 | 2.910727643 |
| Rnf167 | 591.5226223 | 0.273257037 | 0.062144873 | 4.397097071 |
| Pfn1 | 1454.63144 | -0.700480204 | 0.130047004 | -5.386361725 |
| Gm12319 | 3.626546672 | -0.135546501 | 0.58824662 | -0.230424614 |
| Eno3 | 42.27077307 | 0.578732105 | 0.208139481 | 2.780501334 |
| Spag7 | 215.7681227 | 0.116297242 | 0.130185742 | 0.893317814 |
| Camta2 | 2839.805525 | 0.745482742 | 0.095064461 | 7.841865727 |
| Inca1 | 17.29699935 | 0.535030252 | 0.264769053 | 2.020743162 |
| Kif1c | 1453.545255 | -0.223553396 | 0.081442949 | -2.744907921 |
| mmu-mir-6925 | 0 | NA | NA | NA |
| Gm12320 | 0.135956459 | 0.963714371 | 3.641541882 | 0.264644593 |
| Gm12318 | 0 | NA | NA | NA |
| Gm22760 | 0 | NA | NA | NA |
| Zfp3 | 139.4854402 | -0.011849977 | 0.108105105 | -0.109615328 |
| Scimp | 44.06491336 | -1.261400378 | 0.297457841 | -4.240602216 |
| Rabep1 | 1978.442608 | -0.047854176 | 0.062791561 | -0.762111579 |
| Gm19967 | 38.13002371 | 0.025774465 | 0.243733352 | 0.105748619 |
| Rpl23a-ps2 | 9.688741338 | 0.405431862 | 0.438865189 | 0.923818686 |
| Gm12322 | 0.177934686 | 1.051638768 | 2.718161642 | 0.386893388 |
| Nup88 | 578.9786411 | 0.155746156 | 0.080142884 | 1.943356025 |
| Rpain | 118.6108747 | -0.262462261 | 0.119799574 | -2.190844688 |
| C1qbp | 471.9490247 | -0.147311867 | 0.124522637 | -1.183012745 |
| Dhx33 | 545.9733439 | 0.355055152 | 0.087558275 | 4.055072485 |
| Gm22297 | 1.938621463 | 0.366345735 | 0.804754171 | 0.455226885 |
| Derl2 | 390.884449 | -0.471391477 | 0.091129345 | -5.172773693 |
| Mis12 | 261.7075808 | -0.596824163 | 0.134360486 | -4.441961928 |
| 6330403K07Rik | 1104.505225 | -0.047049309 | 0.105072205 | -0.447780737 |
| Gm12321 | 0 | NA | NA | NA |
| Gm15377 | 0 | NA | NA | NA |
| Nlrp1a | 0.174317197 | -1.489079052 | 2.733201959 | -0.544811205 |
| Nlrp1b | 15.20331131 | -1.393703434 | 0.2686486 | -5.187830629 |
| Gm23266 | 0 | NA | NA | NA |
| Nlrp1c-ps | 0 | NA | NA | NA |
| Gm23311 | 0 | NA | NA | NA |
| Gm16013 | 0.447195792 | -1.869076789 | 1.509589413 | -1.238135862 |
| 9230020A06Rik | 0.161949631 | -1.309098506 | 2.76260123 | -0.473864448 |
| Gm12324 | 1.538502375 | -1.565739231 | 1.056648101 | -1.481798178 |
| Gm12325 | 0 | NA | NA | NA |
| Gm12326 | 2.668994113 | -0.154552203 | 0.672037644 | -0.229975515 |
| Wscd1 | 774.0552541 | -0.233973901 | 0.083799998 | -2.792051384 |
| Gm22927 | 0 | NA | NA | NA |
| 4930401O10Rik | 0.10503227 | -0.905414734 | 3.723500899 | -0.243162217 |
| Gm23226 | 0 | NA | NA | NA |
| Aipl1 | 1.181230752 | -0.006074142 | 0.858589772 | -0.007074556 |
| Fam64a | 7.10051421 | -1.02025937 | 0.577613453 | -1.766335885 |
| Pitpnm3 | 1123.234306 | 0.255722792 | 0.114988173 | 2.22390517 |

| | | | | |
|---|---|---|---|---|
| 4933427D14Rik | 402.8306143 | 0.034113371 | 0.155166914 | 0.219849516 |
| Txndc17 | 137.0373374 | -0.283076557 | 0.184357471 | -1.535476458 |
| Med31 | 91.72905903 | 0.020060384 | 0.149623046 | 0.134072824 |
| 4930563E22Rik | 22.70421901 | 0.587254853 | 0.269894788 | 2.175865854 |
| Slc13a5 | 115.5056273 | 0.77168777 | 0.185506847 | 4.159888348 |
| 1700051A21Rik | 0.48812072 | 1.822458294 | 1.646710613 | 1.106726513 |
| n-R5s70 | 0 | NA | NA | NA |
| Xaf1 | 848.1965907 | -0.497819032 | 0.123085539 | -4.044496503 |
| Fbxo39 | 1.50133588 | -0.383308484 | 0.728507765 | -0.52615566 |
| Gm23224 | 0 | NA | NA | NA |
| Tekt1 | 14.19528596 | -0.346180097 | 0.29434892 | -1.176087541 |
| Mir6338 | 0 | NA | NA | NA |
| Smtnl2 | 23.5679464 | 0.163383115 | 0.361389569 | 0.452096931 |
| Ggt6 | 0.550474514 | 0.872326 | 1.354809198 | 0.643873692 |
| Mybbp1a | 930.9934562 | -0.583170771 | 0.085511975 | -6.819755612 |
| Spns2 | 584.9244595 | 0.355989443 | 0.158133022 | 2.251202424 |
| Spns3 | 6.967261194 | -0.37798405 | 0.457704747 | -0.825825061 |
| Gm12712 | 1.554484234 | 1.093520953 | 0.839767309 | 1.302171377 |
| Ube2g1 | 638.5559752 | 0.109292111 | 0.078265324 | 1.39643082 |
| Gm24143 | 0 | NA | NA | NA |
| Gm25678 | 1.584377093 | 0.9826633 | 0.854009471 | 1.15064684 |
| Ankfy1 | 1808.623818 | -0.438587579 | 0.077613622 | -5.650909826 |
| Cyb5d2 | 155.8716509 | -0.033334975 | 0.129815625 | -0.256787081 |
| Zzef1 | 1733.026892 | 0.021063624 | 0.073976203 | 0.284735137 |
| Gm22075 | 0 | NA | NA | NA |
| Gm6733 | 29.33660425 | -0.114405677 | 0.272003235 | -0.420604103 |
| Gm26121 | 4.21018102 | 0.917062547 | 0.605424209 | 1.514743767 |
| AL663082.1 | 0 | NA | NA | NA |
| Atp2a3 | 446.9867994 | -0.003706297 | 0.264494632 | -0.014012749 |
| P2rx1 | 3.999946171 | 0.843370186 | 0.52654151 | 1.601716428 |
| 4732414G09Rik | 8.889423667 | 0.661705602 | 0.402865031 | 1.642499478 |
| Camkk1 | 1051.478754 | 0.676685596 | 0.132142072 | 5.12089438 |
| 1200014J11Rik | 480.4026274 | -0.254217594 | 0.088205834 | -2.882095024 |
| Itgae | 5.448280777 | 0.276662716 | 0.660135327 | 0.4191 |
| Gsg2 | 7.452195508 | -0.530118381 | 0.404761193 | -1.309706538 |
| P2rx5 | 25.59531151 | -0.767141518 | 0.249499035 | -3.074727399 |
| Emc6 | 323.5210552 | -0.29646478 | 0.107888291 | -2.747886512 |
| Tax1bp3 | 46.14509597 | -0.877283478 | 0.228701536 | -3.835931723 |
| Ctns | 210.1024968 | -0.53946976 | 0.116563056 | -4.62813672 |
| Shpk | 35.71037531 | -0.161406057 | 0.216975229 | -0.743891633 |
| Gm16021 | 0 | NA | NA | NA |
| Trpv1 | 1.771085664 | -0.181408872 | 0.744743108 | -0.243585836 |
| Trpv3 | 13.35021764 | -0.128089846 | 0.332230552 | -0.385545054 |
| Aspa | 279.3669201 | 0.022410639 | 0.190983981 | 0.117343029 |
| Spata22 | 0.720179912 | 0.004441211 | 1.252134358 | 0.003546913 |
| Olfr20 | 0 | NA | NA | NA |
| Olfr376 | 0 | NA | NA | NA |

| | | | | |
|---|---|---|---|---|
| Olfr377-ps1 | 0 | NA | NA | NA |
| Olfr1 | 0.108309328 | 0.967119648 | 3.722483401 | 0.259804959 |
| Olfr75-ps1 | 0 | NA | NA | NA |
| Olfr378 | 0 | NA | NA | NA |
| Olfr379-ps1 | 0 | NA | NA | NA |
| Olfr380 | 0 | NA | NA | NA |
| Olfr381 | 0 | NA | NA | NA |
| Olfr382 | 0 | NA | NA | NA |
| Olfr383-ps1 | 0 | NA | NA | NA |
| Olfr385 | 0 | NA | NA | NA |
| Olfr384 | 0 | NA | NA | NA |
| Gm12327 | 0 | NA | NA | NA |
| Olfr387-ps1 | 0 | NA | NA | NA |
| Zfp735 | 0 | NA | NA | NA |
| Olfr388-ps1 | 0 | NA | NA | NA |
| Gm22381 | 0 | NA | NA | NA |
| Gm12328 | 0.100902008 | -0.876201588 | 3.72093227 | -0.235479048 |
| Olfr389 | 0 | NA | NA | NA |
| Olfr390 | 0.097139547 | -0.443816764 | 3.724945049 | -0.119147198 |
| Olfr391-ps | 0 | NA | NA | NA |
| Olfr392 | 0.046345709 | -0.443818988 | 3.724945049 | -0.119147795 |
| Gm12329 | 0 | NA | NA | NA |
| Olfr393 | 0 | NA | NA | NA |
| Olfr394 | 0 | NA | NA | NA |
| Olfr395 | 0 | NA | NA | NA |
| Olfr396-ps1 | 0 | NA | NA | NA |
| Olfr23 | 0 | NA | NA | NA |
| Olfr22-ps1 | 0 | NA | NA | NA |
| Olfr397 | 0 | NA | NA | NA |
| Olfr398 | 0.042716945 | 0.037062677 | 3.724945049 | 0.009949859 |
| Olfr139 | 0 | NA | NA | NA |
| Olfr399 | 0 | NA | NA | NA |
| Zfp616 | 0 | NA | NA | NA |
| Gm12331 | 0.304339188 | -1.16948761 | 1.729474313 | -0.676209876 |
| Olfr400-ps1 | 0 | NA | NA | NA |
| Olfr401 | 0 | NA | NA | NA |
| Olfr402 | 0.044191549 | 0.037062677 | 3.724945049 | 0.009949859 |
| Olfr403 | 0 | NA | NA | NA |
| Olfr43 | 0 | NA | NA | NA |
| Olfr404-ps1 | 0 | NA | NA | NA |
| Olfr405-ps1 | 0 | NA | NA | NA |
| Olfr406 | 0 | NA | NA | NA |
| Olfr407-ps1 | 0.066050875 | -0.443818988 | 3.724945049 | -0.119147795 |
| Olfr59 | 0.062426873 | 0.517959912 | 3.724945049 | 0.139051692 |
| Olfr408-ps1 | 0 | NA | NA | NA |
| Olfr409-ps1 | 0 | NA | NA | NA |
| n-R5s71 | 0 | NA | NA | NA |

| | | | | |
|---|---|---|---|---|
| Gm12332 | 0 | NA | NA | NA |
| Olfr410 | 0.049553047 | -0.443818988 | 3.724945049 | -0.119147795 |
| Olfr411 | 0.046345709 | -0.443818988 | 3.724945049 | -0.119147795 |
| Olfr412 | 0.048949532 | 0.037064902 | 3.724945049 | 0.009950456 |
| Rap1gap2 | 2227.61315 | 0.210954551 | 0.104035911 | 2.027708989 |
| Gm22993 | 0.552195079 | 0.355975971 | 1.189782782 | 0.299194085 |
| E130309D14Rik | 443.6514258 | 0.297772508 | 0.070593647 | 4.218120465 |
| Cluh | 1336.662119 | -0.065362117 | 0.049091466 | -1.331435432 |
| Pafah1b1 | 8824.289057 | 0.097222496 | 0.057860822 | 1.680281983 |
| Gm24211 | 0 | NA | NA | NA |
| Gm16032 | 0.075043205 | 0.517957688 | 3.724945049 | 0.139051095 |
| Mettl16 | 528.4778054 | -0.368745824 | 0.076088013 | -4.846306412 |
| Mnt | 707.5314908 | 0.187808609 | 0.097912579 | 1.918125445 |
| Sgsm2 | 1130.427896 | 0.401072948 | 0.081926132 | 4.895543562 |
| Mir6926 | 0 | NA | NA | NA |
| Tsr1 | 408.3790304 | -0.392431981 | 0.119474243 | -3.284657615 |
| Snord91a | 2.110678478 | 0.776047024 | 0.733641581 | 1.057801308 |
| Gm22771 | 1.808802678 | -1.095591892 | 0.885294098 | -1.237545687 |
| Srr | 740.5480199 | 0.376763024 | 0.114334714 | 3.295263637 |
| Smg6 | 855.9187194 | 0.228980303 | 0.076382087 | 2.997827243 |
| Gm22733 | 0 | NA | NA | NA |
| Gm28180 | 0 | NA | NA | NA |
| Gm12333 | 0 | NA | NA | NA |
| Gm12335 | 2.169066299 | 0.739121643 | 0.7508292 | 0.984407163 |
| Rps12-ps19 | 2.189863395 | -0.346563594 | 0.61884175 | -0.560019736 |
| Hic1 | 51.99884426 | -0.997593992 | 0.230217378 | -4.33326971 |
| 1700016P03Rik | 48.88953987 | -0.713857359 | 0.499810809 | -1.428255145 |
| Mir212 | 22.07890698 | -0.523543076 | 0.710448041 | -0.736919585 |
| Mir132 | 4.115963635 | -0.803657506 | 0.65020542 | -1.236005547 |
| Ovca2 | 77.25714499 | -0.357658342 | 0.163810766 | -2.183362855 |
| Dph1 | 85.00803555 | -0.130823119 | 0.136343171 | -0.959513544 |
| Rtn4rl1 | 700.6189518 | 0.325870613 | 0.154542115 | 2.108620127 |
| Rpa1 | 558.9030021 | -0.182493577 | 0.083786729 | -2.178072575 |
| AL603834.1 | 0.047948698 | 0.037064902 | 3.724945049 | 0.009950456 |
| Smyd4 | 164.5240954 | -0.278420913 | 0.09844425 | -2.828209004 |
| Serpinf1 | 126.7245348 | -0.510210922 | 0.201604563 | -2.530750863 |
| Gm12336 | 0.089062654 | -0.443823436 | 3.724945049 | -0.119148989 |
| Gm12337 | 0.219222703 | 0.85696078 | 2.169844874 | 0.394941035 |
| Serpinf2 | 5.648606599 | -0.45636202 | 0.448536774 | -1.017446163 |
| Wdr81 | 600.0628896 | -0.404360757 | 0.098162331 | -4.119306794 |
| Mir22hg | 200.1675278 | -0.019643528 | 0.170798051 | -0.11501026 |
| Tlcd2 | 9.434003272 | -0.53602398 | 0.339299079 | -1.579797923 |
| Mir22 | 0 | NA | NA | NA |
| Prpf8 | 4246.36712 | -0.030278189 | 0.058572037 | -0.516939316 |
| Rilp | 13.6021336 | -0.111683014 | 0.282479291 | -0.395367086 |
| Scarf1 | 30.35591607 | -1.335360738 | 0.251470845 | -5.310201023 |
| Slc43a2 | 1420.800711 | -0.396123502 | 0.077306959 | -5.124034236 |

| | | | | |
|---|---|---|---|---|
| Mir3971 | 2.406714335 | 0.908009389 | 0.899863031 | 1.009052886 |
| Pitpna | 3114.811598 | 0.049383908 | 0.062613314 | 0.788712569 |
| 4931413K12Rik | 0 | NA | NA | NA |
| Gm12338 | 94.1563446 | -0.159436206 | 0.13609024 | -1.171547693 |
| Inpp5k | 474.3717951 | 0.436901078 | 0.113652271 | 3.844191367 |
| Myo1c | 416.9620297 | -0.624974729 | 0.144816627 | -4.315628256 |
| Crk | 1895.988957 | -0.359581074 | 0.091158567 | -3.944566979 |
| Ywhae | 6464.299146 | -0.002136275 | 0.063068948 | -0.03387206 |
| Gm26836 | 0.047948698 | 0.037064902 | 3.724945049 | 0.009950456 |
| Doc2b | 509.5744778 | 0.261371259 | 0.136024031 | 1.92150796 |
| Gm12339 | 4.361932539 | 0.940306978 | 0.594595783 | 1.581422212 |
| Rph3al | 23.25949346 | -0.102991594 | 0.257598771 | -0.399813994 |
| Gm16075 | 3.522107247 | 1.521310647 | 0.705341509 | 2.156842647 |
| 1700016K19Rik | 11.13294416 | -0.727522118 | 0.367384981 | -1.980271798 |
| Fam101b | 120.2091784 | 0.166903636 | 0.157477615 | 1.059856256 |
| Vps53 | 790.8122356 | 0.046709282 | 0.066709911 | 0.700185046 |
| Glod4 | 354.7645622 | 0.344258864 | 0.076319951 | 4.510732252 |
| Gm12340 | 0 | NA | NA | NA |
| Fam57a | 179.7700705 | -0.002140113 | 0.103230301 | -0.020731444 |
| Gemin4 | 5.229107669 | -0.112958578 | 0.529032811 | -0.213519041 |
| Dbil5 | 3.336932514 | 0.071707826 | 0.591776451 | 0.121173842 |
| Rnmtl1 | 118.9831452 | -0.336245893 | 0.113734134 | -2.956420222 |
| Nxn | 198.759646 | -0.21437931 | 0.129152664 | -1.659890729 |
| Timm22 | 253.9457515 | -0.29555215 | 0.118167026 | -2.501138938 |
| Abr | 5618.476176 | 0.335421426 | 0.068752972 | 4.878646205 |
| Gm12341 | 0.351219581 | 1.020148796 | 1.604513352 | 0.635799505 |
| Rpl36-ps2 | 4.05888357 | -0.034636602 | 0.582003502 | -0.059512704 |
| Gm26780 | 0.189892101 | -1.47044595 | 3.278134527 | -0.448561808 |
| Bhlha9 | 8.304293226 | 0.64379142 | 0.397438049 | 1.619853512 |
| Tusc5 | 10.29048676 | -0.318986457 | 0.37587571 | -0.848648764 |
| Gosr1 | 767.1062309 | -0.083388371 | 0.065079496 | -1.28133092 |
| Cpd | 2040.908043 | -0.057290972 | 0.04756826 | -1.204394946 |
| Tmigd1 | 0.164253626 | 0.503873089 | 2.776247997 | 0.181494265 |
| Blmh | 568.2821817 | 0.040460977 | 0.069623616 | 0.58113869 |
| Gm22611 | 3.32088423 | 0.449526542 | 0.587815732 | 0.764740577 |
| Slc6a4 | 46.2681185 | -0.341223309 | 0.794093989 | -0.429701413 |
| Gm12343 | 0 | NA | NA | NA |
| Ccdc55 | 161.7852798 | -0.023965295 | 0.19956819 | -0.120085745 |
| Gm22772 | 0 | NA | NA | NA |
| Mir423 | 0.516177223 | -0.434471259 | 1.384802907 | -0.313742307 |
| Efcab5 | 66.97098547 | 0.416305029 | 0.180689129 | 2.303984921 |
| Gm12344 | 1.005780181 | 1.478474894 | 1.232746687 | 1.199333902 |
| Gm12345 | 29.19047675 | 0.049173671 | 0.186489858 | 0.263680138 |
| Ssh2 | 1266.750364 | 0.149946225 | 0.057764997 | 2.595797328 |
| Gm24989 | 0 | NA | NA | NA |
| Gm12346 | 262.7722694 | -0.336603403 | 0.106129103 | -3.171640904 |
| Coro6 | 239.6645014 | 1.057838879 | 0.204657215 | 5.168832564 |

| | | | | |
|---|---|---|---|---|
| Ankrd13b | 701.4661961 | 0.059541283 | 0.094547228 | 0.629751761 |
| 2210008F06Rik | 16.34395686 | -0.009840796 | 0.293420847 | -0.033538162 |
| Git1 | 3031.997312 | -0.056233588 | 0.066617179 | -0.844130432 |
| Trp53i13 | 48.23483047 | -0.341579835 | 0.179251332 | -1.905591619 |
| Abhd15 | 33.33330842 | 0.317810089 | 0.212014794 | 1.498999587 |
| Gm10392 | 0.289147236 | 0.135538409 | 2.04387025 | 0.066314586 |
| Taok1 | 6408.541416 | 0.18571583 | 0.047991733 | 3.869746292 |
| Gm12734 | 2.565340877 | -0.405602961 | 0.614077997 | -0.660507237 |
| Gm12732 | 0 | NA | NA | NA |
| Gm12733 | 0 | NA | NA | NA |
| Gm23132 | 0.137483974 | -0.891006646 | 2.929862383 | -0.304112115 |
| Nufip2 | 1887.858992 | -0.218150884 | 0.061012483 | -3.575512278 |
| Cryba1 | 0.167183452 | 0.517964184 | 2.765153692 | 0.187318407 |
| Gm11191 | 0.293994645 | 1.449892997 | 2.013913678 | 0.719938006 |
| Myo18a | 2948.916186 | -0.157516581 | 0.039776405 | -3.960050736 |
| Gm10277 | 0.30796268 | -0.389087627 | 1.777278053 | -0.218923329 |
| Pipox | 16.89892386 | -0.534840349 | 0.29216483 | -1.830611678 |
| Gm11190 | 4.356108356 | -0.763682675 | 0.522780232 | -1.46081016 |
| Sez6 | 1855.373666 | 0.399057146 | 0.136302674 | 2.927727925 |
| Phf12 | 804.2323745 | 0.187347977 | 0.101955566 | 1.837545361 |
| Dhrs13os | 5.524772491 | -0.312075952 | 0.486855644 | -0.641003049 |
| Dhrs13 | 73.22834646 | -0.318556816 | 0.144101337 | -2.210644415 |
| Flot2 | 917.7646301 | 0.152707174 | 0.080325942 | 1.901094109 |
| Mir144 | 0 | NA | NA | NA |
| Mir451a | 0 | NA | NA | NA |
| mmu-mir-451a | 0.164553454 | -0.925587584 | 2.779126865 | -0.333049777 |
| Eral1 | 174.3755178 | -0.25502541 | 0.085199267 | -2.993281727 |
| Fam222b | 474.7107881 | -0.170678114 | 0.071810208 | -2.376794603 |
| Gm12571 | 0.999428373 | -0.604885288 | 1.150370401 | -0.525817848 |
| Traf4 | 100.814411 | -0.261751072 | 0.150809143 | -1.735644583 |
| Nek8 | 29.99850749 | -0.384707039 | 0.233711627 | -1.646075741 |
| Tlcd1 | 129.4641411 | 0.698918695 | 0.168391814 | 4.150550304 |
| Mir7653 | 0 | NA | NA | NA |
| Rpl23a | 798.9907631 | -0.017012295 | 0.125195559 | -0.13588577 |
| Snord42a | 0.955875529 | 0.295319497 | 1.062881041 | 0.277848118 |
| Snord4a | 1.972325891 | 0.701565439 | 0.77419286 | 0.906189497 |
| Snord42b | 3.467268438 | -0.928180078 | 0.622177057 | -1.49182627 |
| Rab34 | 131.9099265 | 0.083647355 | 0.148673299 | 0.562625266 |
| Proca1 | 20.46478121 | 0.353084169 | 0.290125904 | 1.217003255 |
| Supt6 | 2745.970221 | 0.116472544 | 0.090794047 | 1.28282137 |
| Sdf2 | 269.515758 | -0.149125452 | 0.095855599 | -1.555730216 |
| 2610507B11Rik | 4624.687977 | -0.088123257 | 0.056019463 | -1.573082865 |
| Gm11192 | 0.159819677 | -0.473500379 | 3.463168831 | -0.136724602 |
| BC030499 | 82.88575948 | 1.689896621 | 0.4672038 | 3.617043828 |
| Gm11193 | 0.044191549 | 0.037062677 | 3.724945049 | 0.009949859 |
| Spag5 | 131.8137158 | 0.155603717 | 0.176228559 | 0.88296538 |
| Aldoc | 6307.64853 | 0.54153101 | 0.148724994 | 3.641156708 |

| | | | | |
|---|---|---|---|---|
| Pigs | 527.1172141 | -0.116053765 | 0.090334783 | -1.284707416 |
| Unc119 | 149.7412516 | 0.156777034 | 0.131176757 | 1.195158636 |
| Foxn1 | 0.172802443 | 1.046239481 | 3.382542082 | 0.309305681 |
| Slc13a2os | 0 | NA | NA | NA |
| Slc13a2 | 0 | NA | NA | NA |
| Gm11195 | 0.045882455 | 0.517959912 | 3.724945049 | 0.139051692 |
| Slc46a1 | 58.13216195 | -0.133342214 | 0.169356931 | -0.7873443 |
| Sarm1 | 378.1139612 | 0.23256799 | 0.083013728 | 2.801560605 |
| Vtn | 396.8074538 | 0.113649702 | 0.158197858 | 0.718402279 |
| Sebox | 24.6969001 | 1.100966661 | 0.259602988 | 4.240962973 |
| Tmem199 | 242.7545441 | -0.087239543 | 0.090110011 | -0.968144848 |
| Poldip2 | 775.1605727 | 0.046373786 | 0.082444631 | 0.562484 |
| Tnfaip1 | 559.6315322 | -0.347355883 | 0.077970572 | -4.454961349 |
| Ift20 | 265.7227452 | 0.257939294 | 0.07717946 | 3.342071777 |
| Tmem97 | 90.94084935 | -0.040761206 | 0.149878558 | -0.271961554 |
| D11Bhm181e | 1.727533833 | 1.23078578 | 1.155264517 | 1.065371404 |
| Gm22947 | 0.742175153 | 0.517973678 | 1.374255725 | 0.376912149 |
| Nlk | 1172.726608 | 0.283688741 | 0.098050619 | 2.893288616 |
| Gm17235 | 0 | NA | NA | NA |
| Gm11196 | 6.623448048 | 0.551850005 | 0.513054049 | 1.075617679 |
| Gm11197 | 0 | NA | NA | NA |
| Fam58b | 92.53061276 | -0.118211583 | 0.13093782 | -0.902807018 |
| Gm23840 | 0 | NA | NA | NA |
| Gm23219 | 0 | NA | NA | NA |
| Lyrm9 | 495.4572617 | 0.443286921 | 0.084458953 | 5.248548637 |
| Nos2 | 95.34328637 | -2.260886551 | 0.673659805 | -3.356125057 |
| Lgals9 | 1210.742748 | -0.67393735 | 0.242170616 | -2.782903071 |
| Ksr1 | 731.8760987 | 0.335846283 | 0.088856643 | 3.779641812 |
| Gm11201 | 0.406995362 | -2.24505081 | 1.671879223 | -1.342830738 |
| Gm11200 | 0 | NA | NA | NA |
| Wsb1 | 1011.052735 | -0.269699885 | 0.070357791 | -3.833262543 |
| Gm9964 | 0 | NA | NA | NA |
| Nf1 | 4025.801747 | 0.043475605 | 0.037301271 | 1.165526113 |
| Gm11198 | 0 | NA | NA | NA |
| Gm11199 | 0 | NA | NA | NA |
| AU040972 | 4.210711181 | 1.724024686 | 0.555208965 | 3.105181642 |
| Omg | 735.6960469 | 0.642131489 | 0.083002366 | 7.736303426 |
| Gm21975 | 2.443568159 | -1.52497376 | 0.62272813 | -2.448859601 |
| Evi2b | 0 | NA | NA | NA |
| Evi2a | 264.6184616 | -0.378419412 | 0.122437044 | -3.09072646 |
| Rab11fip4 | 3582.291547 | 0.448472376 | 0.096154573 | 4.664077448 |
| Gm23293 | 1.882325847 | 1.539225701 | 0.986642842 | 1.560063719 |
| Rab11fip4os1 | 0.112642008 | 0.517955464 | 3.724945049 | 0.139050498 |
| Gm25867 | 0 | NA | NA | NA |
| Gm24887 | 0 | NA | NA | NA |
| Rab11fip4os2 | 0.045882455 | 0.517959912 | 3.724945049 | 0.139051692 |
| Gm11205 | 0.304750214 | 0.880296241 | 1.974039209 | 0.445936553 |

| | | | | |
|---|---|---|---|---|
| Mir193a | 0.044191549 | 0.037062677 | 3.724945049 | 0.009949859 |
| Mir365-2 | 0.211497531 | -0.915969573 | 2.173517569 | -0.421422668 |
| 9130204K15Rik | 0.414175607 | -1.1973581 | 1.646103509 | -0.727389312 |
| Gm24454 | 0.133633945 | 1.204093506 | 3.720899422 | 0.323602809 |
| Gm11203 | 0 | NA | NA | NA |
| Gm11204 | 0.206494471 | 0.037076023 | 2.202635634 | 0.016832572 |
| Utp6 | 497.722905 | -0.150972131 | 0.076589362 | -1.971189291 |
| Suz12 | 822.7645743 | -0.039180853 | 0.070621825 | -0.55479807 |
| Gm26563 | 1.315499399 | 1.099039396 | 0.888915673 | 1.236382065 |
| Crlf3 | 219.9408582 | -0.332194916 | 0.098759234 | -3.36368461 |
| Atad5 | 155.6883303 | 0.406299842 | 0.183592652 | 2.213050677 |
| Tefm | 62.86592296 | -0.33230001 | 0.152972822 | -2.172281354 |
| Adap2 | 173.6780421 | -0.732385688 | 0.169869485 | -4.311461159 |
| Adap2os | 0.240869569 | -1.419319044 | 3.089662856 | -0.459376673 |
| Rnf135 | 133.3790234 | -1.322967104 | 0.185107787 | -7.147009455 |
| Rhot1 | 976.0750427 | 0.007495055 | 0.046259417 | 0.162022251 |
| Rhbdl3 | 250.4651554 | 0.099412278 | 0.14366541 | 0.691970867 |
| 5730455P16Rik | 801.0455395 | 0.313748414 | 0.070880697 | 4.426429558 |
| Zfp207 | 1918.849716 | -0.021395684 | 0.077544149 | -0.27591616 |
| Psmd11 | 1260.672455 | -0.35430301 | 0.087743393 | -4.037945161 |
| Cdk5r1 | 2328.335231 | 0.346574576 | 0.098733195 | 3.510213331 |
| Myo1d | 327.8254873 | -0.264856617 | 0.093995385 | -2.817761944 |
| C030013C21Rik | 2.678984608 | 0.110817373 | 0.791627961 | 0.139986683 |
| Tmem98 | 68.95410002 | -0.520635368 | 0.219913784 | -2.367452188 |
| Gm11207 | 0 | NA | NA | NA |
| 4930507D10Rik | 0.549547734 | -1.311190055 | 1.369175912 | -0.957649082 |
| Spaca3 | 0.045882455 | 0.517959912 | 3.724945049 | 0.139051692 |
| Asic2 | 625.5974346 | 0.527185885 | 0.072557175 | 7.265799472 |
| Gm11416 | 0 | NA | NA | NA |
| Gm11417 | 0.04393175 | 0.037062677 | 3.724945049 | 0.009949859 |
| 4930527B05Rik | 0 | NA | NA | NA |
| 1700071K01Rik | 9.922463048 | 1.129127802 | 0.508570519 | 2.220199089 |
| Gm11418 | 0 | NA | NA | NA |
| 5530401A14Rik | 0.732425175 | -1.430397341 | 1.229479795 | -1.163416713 |
| Gm11419 | 2.867374077 | 0.438761026 | 0.712060195 | 0.616185302 |
| Gm17268 | 0.061920613 | -0.443816764 | 3.724945049 | -0.119147198 |
| Ccl2 | 1280.074291 | -0.916073662 | 0.658185107 | -1.391817669 |
| Ccl7 | 524.7321755 | -1.756955642 | 0.608257961 | -2.888504145 |
| Ccl11 | 7.798742698 | -0.724331807 | 0.429626803 | -1.685955815 |
| Ccl12 | 226.9053113 | -0.092396723 | 0.362143694 | -0.25513829 |
| Ccl8 | 52.35493119 | -2.332653807 | 0.601922043 | -3.875342052 |
| Ccl1 | 0.343821097 | -0.365271692 | 1.992799746 | -0.183295734 |
| Tmem132e | 318.4462841 | 0.071263104 | 0.130171034 | 0.547457466 |
| Gm24612 | 0 | NA | NA | NA |
| Gm11426 | 0 | NA | NA | NA |
| Cct6b | 3.620020008 | 1.49868431 | 0.617310334 | 2.427764816 |
| Gm11421 | 0 | NA | NA | NA |

| | | | | |
|---|---|---|---|---|
| Gm11422 | 0 | NA | NA | NA |
| Gm11420 | 0.621419653 | 2.044045091 | 1.219718247 | 1.675833821 |
| Zfp830 | 109.1762647 | -0.125571956 | 0.117084308 | -1.072491763 |
| Gm11423 | 5.437272543 | -0.30858573 | 0.491522509 | -0.627816071 |
| Lig3 | 405.8341476 | -0.287530209 | 0.068038908 | -4.225967421 |
| Rffl | 192.7097897 | -0.205436511 | 0.129858316 | -1.582005038 |
| Rad51d | 202.7225488 | 0.072523034 | 0.084267354 | 0.86063025 |
| Fndc8 | 0.20460011 | 0.517955348 | 3.243234116 | 0.159703348 |
| Nle1 | 99.69246992 | -0.166307019 | 0.118077865 | -1.408452118 |
| Unc45b | 11.03143117 | -0.193344953 | 0.345304923 | -0.559925273 |
| Unc45bos | 2.092676817 | -0.26120754 | 0.719281563 | -0.363150612 |
| Slfn5os | 0.252317246 | 0.005653035 | 2.494815902 | 0.002265913 |
| Slfn5 | 854.2001971 | -1.502836699 | 0.276405185 | -5.437078528 |
| Gm11424 | 0.243199696 | -1.167642884 | 2.442366572 | -0.478078474 |
| Slfn9 | 576.8930042 | -1.081535785 | 0.246597682 | -4.38583111 |
| Mir7679 | 0.98936666 | -1.606263354 | 0.979572088 | -1.639760233 |
| Slfn8 | 1181.313578 | -1.140750251 | 0.223946881 | -5.093843004 |
| Gm11425 | 15.12308696 | -1.860802308 | 0.333220863 | -5.584291135 |
| Slfn10-ps | 120.7759095 | -1.737402633 | 0.379286263 | -4.58071595 |
| Gm27598 | 0.379788221 | -1.835444103 | 1.843085272 | -0.995854142 |
| Slfn2 | 559.7972036 | -1.153686184 | 0.307926451 | -3.746629034 |
| Slfn1 | 299.5776399 | -1.400925722 | 0.395427421 | -3.542813795 |
| Slfn4 | 2011.837015 | -2.056494352 | 0.434966436 | -4.727938022 |
| Slfn3 | 17.21960857 | -0.290940625 | 0.283244453 | -1.027171483 |
| AI662270 | 42.49283015 | -1.124299765 | 0.30939501 | -3.633865219 |
| Gm11427 | 4.559114128 | -1.143486723 | 0.505829625 | -2.260616357 |
| Gm23444 | 10.89494943 | -0.806612128 | 0.40535015 | -1.989914466 |
| Slfn14 | 0.407088326 | 0.539288534 | 1.564258333 | 0.344756695 |
| AA465934 | 34.77868168 | 0.363082193 | 0.196926754 | 1.843742337 |
| Snord7 | 2.473802751 | -0.114330012 | 0.744701916 | -0.15352453 |
| Pex12 | 83.19986102 | -0.363073653 | 0.158807345 | -2.286252266 |
| Ap2b1 | 4452.733103 | 0.084048806 | 0.053796306 | 1.562352756 |
| Gm23951 | 0 | NA | NA | NA |
| Rasl10b | 2095.394555 | 0.63292718 | 0.120517508 | 5.251744684 |
| Gas2l2 | 4.963609251 | -0.505073145 | 0.517069534 | -0.976799273 |
| 1700020L24Rik | 3.62909452 | -0.214471748 | 0.529082576 | -0.405365358 |
| Mmp28 | 45.16190099 | -0.736725262 | 0.206394047 | -3.569508298 |
| Taf15 | 585.2888291 | -0.164748263 | 0.128887563 | -1.278232429 |
| Heatr9 | 16.53926368 | -1.639145688 | 0.472215344 | -3.471182609 |
| Gm23247 | 1.077303893 | 0.245839936 | 0.908847295 | 0.270496417 |
| Ccl5 | 670.2920451 | -0.570025704 | 0.439784648 | -1.296147344 |
| Ccl9 | 101.7341889 | -1.645271561 | 0.46789602 | -3.516318777 |
| E230016K23Rik | 2.428642307 | -1.046383141 | 0.63015831 | -1.660508358 |
| Ccl6 | 71.52556139 | -1.693465952 | 0.333115325 | -5.083722732 |
| Rpl9-ps1 | 0 | NA | NA | NA |
| Ccl3 | 250.7978955 | -0.161171854 | 0.816558576 | -0.197379415 |
| Ccl4 | 181.900904 | -0.98857421 | 0.572340141 | -1.727249479 |

| | | | | |
|---|---|---|---|---|
| Gm11433 | 0.066050875 | -0.443818988 | 3.724945049 | -0.119147795 |
| Gm11438 | 0 | NA | NA | NA |
| Gm11432 | 0.038981395 | -0.443816764 | 3.724945049 | -0.119147198 |
| Gm11430 | 0 | NA | NA | NA |
| Gm11429 | 0 | NA | NA | NA |
| Wfdc17 | 58.81490581 | -2.139200032 | 0.506560766 | -4.222987993 |
| Wfdc18 | 2.91100996 | -0.453290404 | 0.610306717 | -0.742725569 |
| Wfdc21 | 0.36705964 | 1.999929755 | 3.69587791 | 0.541124411 |
| Heatr6 | 295.8490656 | -0.198332067 | 0.087665104 | -2.262383299 |
| Gm22311 | 0 | NA | NA | NA |
| Gm11434 | 0.042716945 | 0.037062677 | 3.724945049 | 0.009949859 |
| Gm12576 | 1.282016488 | 0.696048745 | 1.041414338 | 0.668368698 |
| Gm23211 | 0 | NA | NA | NA |
| Hnf1b | 4.558351918 | -0.140118544 | 0.536568178 | -0.261138378 |
| Ddx52 | 333.6973482 | -0.222078536 | 0.107825757 | -2.059605622 |
| Synrg | 1329.598341 | -0.038581978 | 0.05675538 | -0.679794195 |
| Dusp14 | 294.0225346 | -0.229455329 | 0.18027934 | -1.272776617 |
| Tada2a | 253.1517061 | 0.334410757 | 0.154268056 | 2.167725231 |
| Acaca | 1627.488358 | -0.021786053 | 0.060153978 | -0.362171449 |
| Gm11437 | 0.12092566 | 0.998850474 | 3.724945049 | 0.268151734 |
| Gm37391 | 10.91891655 | 0.343491669 | 0.345238093 | 0.994941392 |
| Aatf | 235.0359788 | 0.050489852 | 0.14113469 | 0.357742321 |
| Lhx1 | 140.9707341 | -0.178754108 | 0.234019671 | -0.763842231 |
| Lhx1os | 31.23601852 | 0.14639928 | 0.236284821 | 0.619588172 |
| 1700109G15Rik | 0 | NA | NA | NA |
| Gm11431 | 0 | NA | NA | NA |
| Mrm1 | 101.0285373 | -0.488833473 | 0.123938819 | -3.944151454 |
| Dhrs11 | 57.07916713 | -0.121902995 | 0.17169992 | -0.709977006 |
| 4930502E09Rik | 0.377064288 | -1.536000541 | 3.707562836 | -0.414288472 |
| Ggnbp2 | 1167.177578 | 0.10895029 | 0.084373548 | 1.291284914 |
| Pigw | 46.32962589 | -0.165013961 | 0.16472182 | -1.001773543 |
| Myo19 | 116.4020599 | -0.086756443 | 0.201390355 | -0.430787477 |
| Gm23564 | 1.319357828 | -0.083782704 | 0.947097023 | -0.088462642 |
| Znhit3 | 108.0352272 | 0.344895764 | 0.119653196 | 2.882461784 |
| Car4 | 225.7880006 | 0.449884953 | 0.205686509 | 2.187236076 |
| Usp32 | 2485.609366 | 0.067227305 | 0.04950918 | 1.357875548 |
| Gm11440 | 7.169804879 | 0.824287251 | 0.420262597 | 1.96136239 |
| Gm11439 | 9.641612845 | 0.409638232 | 0.503725053 | 0.813217905 |
| Rpl13-ps1 | 6.979134821 | 1.683643906 | 0.522219303 | 3.224016991 |
| Gm23506 | 0 | NA | NA | NA |
| 1700125H20Rik | 0.788910745 | -1.681840934 | 1.130792983 | -1.487311082 |
| Appbp2 | 1916.277606 | 0.060594576 | 0.061743892 | 0.981385747 |
| Akirin1-ps | 0.295845421 | 1.404022212 | 2.064726812 | 0.680003865 |
| Appbp2os | 6.2436769 | -0.387988093 | 0.52244735 | -0.742635776 |
| Gm22587 | 0 | NA | NA | NA |
| Ppm1d | 315.9501097 | 0.005893391 | 0.110175194 | 0.05349109 |
| Bcas3 | 671.5506855 | 0.512303118 | 0.070755025 | 7.24051922 |

| | | | | |
|---|---|---|---|---|
| Gm23765 | 0.716442652 | 0.926789798 | 1.236215352 | 0.749699312 |
| Bcas3os1 | 0.047948698 | 0.037064902 | 3.724945049 | 0.009950456 |
| Mir5110 | 0 | NA | NA | NA |
| Bcas3os2 | 0 | NA | NA | NA |
| Gm23473 | 1.868429801 | 1.817966844 | 0.96721822 | 1.879582918 |
| 2610027K06Rik | 0.597123318 | -1.908465676 | 1.391019962 | -1.371990143 |
| Tbx2 | 23.31208816 | 0.084004922 | 0.231050065 | 0.363578871 |
| Gm11444 | 0 | NA | NA | NA |
| Tbx4 | 0.57584922 | -0.226490037 | 1.469440254 | -0.154133546 |
| Gm11443 | 0 | NA | NA | NA |
| Brip1 | 34.57829007 | -0.24287917 | 0.227777521 | -1.066299998 |
| Atp5l-ps2 | 2.273527578 | 1.448049843 | 0.860437692 | 1.682922373 |
| Brip1os | 315.6828442 | -0.680610938 | 0.144395861 | -4.713507261 |
| Ints2 | 398.3495836 | 0.110768957 | 0.129222256 | 0.857197205 |
| Med13 | 3263.787859 | -0.106518242 | 0.050453362 | -2.111221882 |
| Gm25427 | 0 | NA | NA | NA |
| Gm23667 | 0 | NA | NA | NA |
| Gm26439 | 0 | NA | NA | NA |
| Rnft1 | 257.0685552 | -0.219770027 | 0.101750338 | -2.159894818 |
| Rps6kb1 | 899.7579894 | 0.077932521 | 0.05856842 | 1.33062359 |
| Tubd1 | 49.92708075 | -0.339096257 | 0.196350072 | -1.726998384 |
| Vmp1 | 1340.608191 | -0.626165269 | 0.105974431 | -5.908644803 |
| Mir21a | 0 | NA | NA | NA |
| Gm11478 | 49.42339314 | -0.206276167 | 0.170699424 | -1.208417474 |
| Ptrh2 | 123.2383261 | -0.255062633 | 0.120168412 | -2.122543087 |
| Cltc | 12039.16266 | -0.088843747 | 0.05690848 | -1.561168854 |
| Dhx40 | 393.4698973 | -0.202649387 | 0.084683738 | -2.393014195 |
| Ypel2 | 824.3745057 | 0.091921577 | 0.111638118 | 0.823388812 |
| Gm22387 | 0 | NA | NA | NA |
| Gdpd1 | 414.5698482 | -0.063987541 | 0.075497324 | -0.847547145 |
| Gm11479 | 0.152667883 | 0.998859635 | 2.845214387 | 0.351066563 |
| Smg8 | 236.0766196 | -0.019633919 | 0.121759631 | -0.16125147 |
| Gm9378 | 0.044191549 | 0.037062677 | 3.724945049 | 0.009949859 |
| Prr11 | 19.51012886 | 0.002999782 | 0.264973254 | 0.011321076 |
| Ska2 | 149.134786 | -0.198723157 | 0.127550458 | -1.557996421 |
| Mir301 | 1.204075239 | 0.33455915 | 0.907214634 | 0.368776183 |
| Gm11491 | 0.332371241 | -0.443830769 | 3.724945049 | -0.119150957 |
| Trim37 | 3977.710216 | 0.535882184 | 0.077589961 | 6.906591756 |
| Ppm1e | 3157.400212 | 0.236292452 | 0.099508377 | 2.374598601 |
| Gm11490 | 0 | NA | NA | NA |
| Gm22883 | 12.43333056 | 0.873514729 | 0.501966223 | 1.740186268 |
| Rad51c | 45.74186524 | -0.797592907 | 0.209436729 | -3.80827619 |
| Tex14 | 23.09397913 | -0.97151209 | 0.238118664 | -4.079949356 |
| Rnu1a1 | 44.69699138 | 0.329392756 | 0.50467797 | 0.652679086 |
| Gm22068 | 772.7219214 | -0.058019312 | 0.166622437 | -0.34820828 |
| Rnu3b1 | 0.136120942 | -0.876404342 | 3.595683781 | -0.24373788 |
| Gm23137 | 0 | NA | NA | NA |

| | | | | |
|---|---|---|---|---|
| Rnu3b3 | 0.045882455 | 0.517959912 | 3.724945049 | 0.139051692 |
| Rnu3b4 | 0 | NA | NA | NA |
| Gm11489 | 0.231904818 | 0.51796906 | 2.173137077 | 0.238350846 |
| Rnu3b2 | 0 | NA | NA | NA |
| Gm11492 | 0.278403928 | -0.443820842 | 2.057010628 | -0.215760111 |
| 4-Sep | 1581.659409 | 0.164122381 | 0.089230891 | 1.839300029 |
| Mtmr4 | 1614.493203 | -0.115124969 | 0.062476286 | -1.842698652 |
| Hsf5 | 20.47148481 | 0.434562113 | 0.338321006 | 1.284466837 |
| Rnf43 | 59.17937029 | -0.004991202 | 0.175759243 | -0.028397947 |
| Gm25968 | 0 | NA | NA | NA |
| 1110028F11Rik | 2.71851656 | 0.121896003 | 0.710722453 | 0.171509993 |
| Supt4a | 181.1208407 | -0.021467104 | 0.084464431 | -0.254155548 |
| Mir142hg | 12.62452238 | -1.042563347 | 0.303618565 | -3.433793141 |
| Mir142b | 0.614441445 | 0.080740176 | 1.28707583 | 0.062731483 |
| mmu-mir-142a | 0.12092566 | 0.998850474 | 3.724945049 | 0.268151734 |
| Bzrap1 | 2131.147747 | 0.621823104 | 0.155419934 | 4.000922457 |
| Mpo | 4.50923424 | -2.310274474 | 0.574997556 | -4.017885729 |
| Gm11505 | 0.14164095 | -0.925681839 | 3.603106436 | -0.256912155 |
| Lpo | 20.44324603 | -0.522253498 | 0.247790013 | -2.107645468 |
| Mks1 | 36.78772966 | -0.437301635 | 0.195069999 | -2.241767758 |
| Epx | 0.366894703 | -1.686377055 | 1.589295319 | -1.061084769 |
| Olfr462 | 0.183788242 | -0.443821179 | 2.683430479 | -0.16539321 |
| Olfr463 | 0.038981395 | -0.443816764 | 3.724945049 | -0.119147198 |
| Olfr464 | 0.707581183 | -1.058169201 | 1.171526578 | -0.903239603 |
| Gm11504 | 0 | NA | NA | NA |
| Gm11507 | 0.066050875 | -0.443818988 | 3.724945049 | -0.119147795 |
| Dynll2 | 5064.124974 | 0.185136495 | 0.076754542 | 2.412059134 |
| Gm11508 | 7.373820318 | -0.578652809 | 0.427505408 | -1.353556699 |
| 2010015M23Rik | 13.47151829 | 0.358702229 | 0.304766978 | 1.176972098 |
| Srsf1 | 2301.674563 | -0.214220943 | 0.049815577 | -4.300280312 |
| Gm15892 | 24.85441836 | 0.469653763 | 0.293209132 | 1.601770582 |
| Vezf1 | 819.1262531 | -0.338789295 | 0.088133443 | -3.844049262 |
| Cuedc1 | 496.2529045 | -0.015267208 | 0.072250987 | -0.211307944 |
| Mrps23 | 225.4695307 | -0.263571642 | 0.101581127 | -2.594691063 |
| 1700106J16Rik | 0.47539415 | 0.027223824 | 1.469691093 | 0.018523501 |
| Gm11509 | 0 | NA | NA | NA |
| Msi2 | 2918.689029 | 0.12901434 | 0.07283644 | 1.771288382 |
| Mir378b | 0 | NA | NA | NA |
| Gm15893 | 0.758254207 | -1.06842292 | 1.272192678 | -0.839827911 |
| Gm11510 | 2.984155966 | -0.092785903 | 0.631798749 | -0.146859902 |
| C030037D09Rik | 74.7814881 | -0.289407652 | 0.164580738 | -1.758453967 |
| Gm11493 | 0 | NA | NA | NA |
| Akap1 | 422.0526709 | 0.161653136 | 0.100259129 | 1.612353286 |
| 4930556N13Rik | 0.0878635 | 0.037058229 | 3.724945049 | 0.009948665 |
| Gm23507 | 0 | NA | NA | NA |
| Scpep1os | 0 | NA | NA | NA |
| Scpep1 | 257.6478549 | -0.991813519 | 0.185059632 | -5.359426625 |

| | | | | |
|---|---|---|---|---|
| Gm15698 | 0.068097359 | 0.037064902 | 3.724945049 | 0.009950456 |
| Coil | 170.2213894 | 0.270841877 | 0.091318195 | 2.965913608 |
| 2210409E12Rik | 0.042716945 | 0.037062677 | 3.724945049 | 0.009949859 |
| Gm11496 | 0.352842203 | -0.475896242 | 1.956110647 | -0.243286975 |
| Trim25 | 943.4408156 | -1.180419875 | 0.182404396 | -6.471444217 |
| Dgke | 1158.574667 | 0.28706081 | 0.070378831 | 4.078794783 |
| Dgkeos | 4.476524561 | 0.165508847 | 0.473197552 | 0.349766913 |
| Gm525 | 0 | NA | NA | NA |
| Gm22534 | 0.038981395 | -0.443816764 | 3.724945049 | -0.119147198 |
| Gm11497 | 0 | NA | NA | NA |
| Gm24734 | 0 | NA | NA | NA |
| Nog | 67.55626535 | 0.292775753 | 0.183271416 | 1.597498173 |
| Gm2018 | 0.338611762 | 1.038275063 | 2.081548477 | 0.498799367 |
| 4932411E22Rik | 35.54587762 | 0.766048892 | 0.232720288 | 3.291715132 |
| Ankfn1 | 45.04732695 | 0.523482309 | 0.183394619 | 2.85440387 |
| Pctp | 14.03537449 | -0.634998572 | 0.294430131 | -2.156703765 |
| Tmem100 | 84.40838063 | 0.042872456 | 0.149987721 | 0.285839769 |
| Gm11494 | 0 | NA | NA | NA |
| n-R5s72 | 0 | NA | NA | NA |
| Mmd | 1332.487735 | 0.676896305 | 0.100462598 | 6.737794138 |
| Hlf | 2065.341962 | 1.394012782 | 0.132306962 | 10.53620125 |
| Stxbp4 | 295.2899188 | 0.133969694 | 0.100392109 | 1.334464387 |
| Gm11511 | 3.871387691 | 0.864498517 | 0.631616656 | 1.368707598 |
| Gm11512 | 2.280412116 | 0.247454188 | 0.757561693 | 0.326645593 |
| Cox11 | 152.2450534 | -0.066298497 | 0.123849368 | -0.535315591 |
| Tom1l1 | 226.4499419 | -0.083389758 | 0.11612827 | -0.718083186 |
| Gm22762 | 0 | NA | NA | NA |
| 4930405D11Rik | 0.230025872 | 0.505750912 | 2.549675566 | 0.198358928 |
| Gm11516 | 0 | NA | NA | NA |
| Kif2b | 0 | NA | NA | NA |
| Gm11498 | 0 | NA | NA | NA |
| Gm22702 | 0 | NA | NA | NA |
| Gm11500 | 0 | NA | NA | NA |
| Gm11501 | 0 | NA | NA | NA |
| Gm25113 | 0 | NA | NA | NA |
| Car10 | 1148.248468 | 0.265371132 | 0.089603409 | 2.961618708 |
| Gm24856 | 0 | NA | NA | NA |
| Gm11502 | 0.044544648 | 0.517957688 | 3.724945049 | 0.139051095 |
| Gm11506 | 0 | NA | NA | NA |
| Utp18 | 272.3077803 | -0.204331924 | 0.085730168 | -2.383430801 |
| Mbtd1 | 630.0243432 | 0.230427204 | 0.090212831 | 2.554261967 |
| Nme2 | 16.60911457 | -0.453112464 | 0.344108074 | -1.316773706 |
| Gm20390 | 0.049553047 | -0.443818988 | 3.724945049 | -0.119147795 |
| Nme1 | 441.0854408 | -0.242189125 | 0.079245653 | -3.056181843 |
| Spag9 | 5268.156952 | 0.323079932 | 0.049916969 | 6.472346684 |
| Gm11518 | 36.08732431 | -0.529982507 | 0.222514762 | -2.381785827 |
| B230206L02Rik | 10.54869608 | 0.833758861 | 0.374499725 | 2.226327032 |

| | | | | |
|---|---|---|---|---|
| Tob1 | 560.157712 | 0.18537228 | 0.084569325 | 2.191956475 |
| Wfikkn2 | 33.87170838 | 0.266264995 | 0.219316472 | 1.214067476 |
| Gm21885 | 0.047948698 | 0.037064902 | 3.724945049 | 0.009950456 |
| Luc7l3 | 1814.337726 | 0.243868908 | 0.097631089 | 2.497861183 |
| Ankrd40 | 1549.21346 | -0.145138023 | 0.090548405 | -1.602877744 |
| Abcc3 | 101.8740177 | -1.676091417 | 0.304082273 | -5.511966872 |
| Cacna1g | 1923.935302 | -0.114220654 | 0.137399503 | -0.831303253 |
| Spata20 | 1.880027439 | -0.156232809 | 0.794856397 | -0.196554761 |
| Epn3 | 81.88700119 | 0.295327141 | 0.16224524 | 1.820251498 |
| Mycbpap | 22.79270077 | -0.504425062 | 0.261051087 | -1.932284856 |
| Rsad1 | 129.9462632 | -0.234171099 | 0.149986597 | -1.561280173 |
| Acsf2 | 203.4382502 | 0.24567437 | 0.166878114 | 1.472178491 |
| Chad | 2.13663281 | 0.204087634 | 0.712508321 | 0.28643544 |
| Gm11540 | 0 | NA | NA | NA |
| Lrrc59 | 1158.226995 | -0.455269512 | 0.108936976 | -4.179200953 |
| Eme1 | 6.299531046 | -1.386196247 | 0.562003522 | -2.466525907 |
| Mrpl27 | 283.0008542 | -0.095675302 | 0.110193221 | -0.86825035 |
| Xylt2 | 348.7120337 | 0.225725285 | 0.091715674 | 2.461141857 |
| Gm11542 | 2.690072256 | 0.370159187 | 0.641198352 | 0.577292791 |
| Gm11541 | 0.211252453 | 0.037076869 | 2.188118072 | 0.016944638 |
| Cdc34-ps | 2.829894638 | 0.223186759 | 0.612223573 | 0.36455107 |
| Gm11545 | 0.145451803 | -1.224261167 | 3.529361758 | -0.346878912 |
| Tmem92 | 0.060174537 | 0.517957688 | 3.724945049 | 0.139051095 |
| Gm11546 | 0 | NA | NA | NA |
| AA623943 | 0 | NA | NA | NA |
| Gm11543 | 0 | NA | NA | NA |
| Gm11544 | 0 | NA | NA | NA |
| A430060F13Rik | 0 | NA | NA | NA |
| Gm22456 | 0 | NA | NA | NA |
| Col1a1 | 391.7651446 | -0.744491011 | 0.160572086 | -4.63649088 |
| Gm24877 | 0 | NA | NA | NA |
| Sgca | 2.248524686 | 0.406284176 | 0.738384122 | 0.550234172 |
| Hils1 | 0 | NA | NA | NA |
| Ppp1r9b | 8227.147287 | 0.580820119 | 0.100845642 | 5.759496481 |
| Samd14 | 673.600847 | 0.264444744 | 0.117179452 | 2.25675013 |
| Pdk2 | 946.573687 | 0.507433018 | 0.100608448 | 5.043642242 |
| Gm11513 | 1.036123199 | 1.456956544 | 1.189385384 | 1.224965906 |
| Itga3 | 513.8169415 | -0.203746983 | 0.090414654 | -2.253473015 |
| Dlx3 | 0.948391139 | 1.23066667 | 1.127670013 | 1.091335813 |
| Dlx4 | 3.24303925 | 0.674587442 | 0.588703452 | 1.145886675 |
| A730090H04Rik | 3.582880236 | 0.974736516 | 0.580976042 | 1.67775682 |
| Gm11514 | 1.617110251 | -0.827860346 | 0.825525902 | -1.002827827 |
| Gm11515 | 0.488879413 | -2.270787499 | 1.481160584 | -1.533113644 |
| Tac4 | 0.201378205 | 0.959998295 | 2.643723124 | 0.363123614 |
| Kat7 | 1091.977074 | 0.011145499 | 0.077431379 | 0.143940334 |
| Gm11520 | 2.683034273 | 0.2230138 | 0.633894931 | 0.351815087 |
| Gm11521 | 0.240698147 | 1.662157314 | 3.707812846 | 0.448285117 |

| | | | | |
|---|---|---|---|---|
| Fam117a | 84.85769561 | 0.477755399 | 0.158562665 | 3.013038407 |
| Gm29477 | 0.19690755 | -0.934164037 | 2.67725883 | -0.34892556 |
| Slc35b1 | 393.4153695 | -0.190277919 | 0.106953116 | -1.779077846 |
| Spop | 1831.86833 | 0.086405323 | 0.072714509 | 1.188281736 |
| Gm11522 | 0 | NA | NA | NA |
| Nxph3 | 195.7791226 | 0.683830699 | 0.125919732 | 5.430687376 |
| Ngfr | 42.86765523 | -0.557675241 | 0.271478296 | -2.054216669 |
| Gm11528 | 0 | NA | NA | NA |
| Phb | 513.3109458 | -0.361140597 | 0.117479888 | -3.074063184 |
| Gm26830 | 2.519878091 | 0.64362488 | 0.791861231 | 0.81280009 |
| Gm9796 | 6.527712466 | -0.524357527 | 0.463342499 | -1.131684505 |
| Zfp652os | 6.322034818 | 0.550610442 | 0.541467028 | 1.016886374 |
| Zfp652 | 1286.698194 | 0.141506251 | 0.059491972 | 2.378577228 |
| B130006D01Rik | 36.80483133 | 0.414446711 | 0.272221872 | 1.522459268 |
| Polr2k-ps | 4.087166043 | 0.579811323 | 0.51212173 | 1.13217481 |
| Phospho1 | 32.25678491 | -0.612283303 | 0.208935071 | -2.930495577 |
| Abi3 | 109.2635649 | -0.448781857 | 0.171793384 | -2.61233493 |
| Gngt2 | 37.40057289 | -1.031814232 | 0.257111768 | -4.013096101 |
| Gm11527 | 0.940895551 | -2.098853254 | 1.09880563 | -1.910122406 |
| 4833417C18Rik | 16.33142681 | 0.410696112 | 0.312971756 | 1.312246565 |
| B4galnt2 | 0.576309813 | -2.278912138 | 1.412217107 | -1.613712316 |
| Gm11534 | 0 | NA | NA | NA |
| Gm24725 | 0 | NA | NA | NA |
| Igf2bp1 | 1.321944539 | 0.488098433 | 0.840741053 | 0.580557392 |
| Mir3063 | 0 | NA | NA | NA |
| Gm29202 | 0 | NA | NA | NA |
| Gm37179 | 0 | NA | NA | NA |
| Gip | 0 | NA | NA | NA |
| Gm24949 | 0 | NA | NA | NA |
| Snf8 | 318.1547224 | -0.074217286 | 0.101769647 | -0.729267406 |
| Ube2z | 1093.216647 | -0.209874128 | 0.063084474 | -3.326874525 |
| Atp5g1 | 503.7452782 | 0.036116716 | 0.089309419 | 0.404399855 |
| Gm22423 | 0 | NA | NA | NA |
| Calcoco2 | 0.681926656 | 1.419833502 | 1.114719084 | 1.273714179 |
| Ttll6 | 7.585280821 | -0.736221979 | 0.424537158 | -1.734175596 |
| Gm11535 | 0 | NA | NA | NA |
| Hoxb13 | 0 | NA | NA | NA |
| Gm27898 | 0 | NA | NA | NA |
| Gm27605 | 0 | NA | NA | NA |
| Gm11539 | 1.7546716 | -0.349425056 | 0.771504446 | -0.45291386 |
| Gm53 | 0.110242424 | -0.443823436 | 3.724945049 | -0.119148989 |
| Mir196a-1 | 0 | NA | NA | NA |
| Hoxb9 | 0.072025462 | 0.517959912 | 3.724945049 | 0.139051692 |
| Hoxb8 | 30.1408604 | -1.526297304 | 1.731721369 | -0.881375798 |
| Hoxb7 | 3.606077423 | -1.320379478 | 1.442120491 | -0.915581941 |
| Hoxb5os | 0.345588754 | -0.49884404 | 3.207331033 | -0.155532446 |
| Hoxb6 | 10.70947983 | -1.139353658 | 1.243465711 | -0.916272679 |

| | | | | |
|---|---|---|---|---|
| Hoxb5 | 50.63370746 | -0.849181668 | 1.019693101 | -0.832781616 |
| Mir10a | 0.182096932 | -0.443820047 | 2.691326182 | -0.164907565 |
| Hoxb4 | 17.2026822 | -1.523425068 | 0.791895914 | -1.923769325 |
| Hoxb3 | 35.54127179 | -0.959223349 | 0.747972614 | -1.282431108 |
| Gm11536 | 0.751775467 | 0.210418111 | 1.130912951 | 0.186060396 |
| Hoxb3os | 4.383958145 | -0.489279461 | 0.547226373 | -0.894107969 |
| Hoxb2 | 10.86433349 | -1.135378845 | 0.872316637 | -1.30156734 |
| Hoxb1 | 0.038981395 | -0.443816764 | 3.724945049 | -0.119147198 |
| Gm11531 | 0.072025462 | 0.517959912 | 3.724945049 | 0.139051692 |
| Gm11529 | 0.471635551 | 0.451625904 | 1.393149489 | 0.324176197 |
| Skap1 | 41.01891481 | 0.007671236 | 0.23493811 | 0.032652157 |
| 2010300F17Rik | 0.313173757 | -1.398312045 | 2.047254153 | -0.683018297 |
| Gm11537 | 0.487078227 | 0.336697407 | 1.232233251 | 0.273241618 |
| Snx11 | 313.2983275 | -0.274082338 | 0.080716505 | -3.395617011 |
| Cbx1 | 382.9098145 | -0.235249033 | 0.097341006 | -2.416751619 |
| Gm11517 | 5.580449154 | 0.017451802 | 0.434518708 | 0.040163522 |
| Nfe2l1 | 4454.295115 | -0.006768869 | 0.046099577 | -0.146831483 |
| Copz2 | 26.60350595 | 0.290403249 | 0.245896268 | 1.180999007 |
| Mir152 | 0.090665643 | 0.037060453 | 3.724945049 | 0.009949262 |
| Gm11523 | 0.271332793 | 0.603789201 | 2.06270173 | 0.292717649 |
| Gm11524 | 0 | NA | NA | NA |
| Cdk5rap3 | 188.9212374 | 0.050477216 | 0.10605256 | 0.475964141 |
| D030028A08Rik | 76.08496205 | -0.008378885 | 0.145775916 | -0.057477843 |
| Gm11525 | 3.501911378 | 0.676238229 | 0.570849946 | 1.184616436 |
| Prr15l | 1.00014491 | 0.076131689 | 1.014753502 | 0.075024811 |
| Pnpo | 446.5624225 | 0.194211749 | 0.073885275 | 2.628558252 |
| Sp2 | 265.9732879 | 0.178811796 | 0.131892505 | 1.355738867 |
| Sp6 | 5.037085609 | 0.120437391 | 0.469329291 | 0.256615969 |
| Gm11532 | 1.602391015 | -0.832024071 | 0.833010095 | -0.998816313 |
| Scrn2 | 30.51638522 | -0.307919467 | 0.256110174 | -1.202292988 |
| Lrrc46 | 5.763699656 | -1.221802892 | 0.477636257 | -2.558019568 |
| Gm11533 | 0.306021358 | 0.05082147 | 1.799454184 | 0.028242714 |
| Mrpl10 | 403.0972378 | -0.198847419 | 0.08388807 | -2.370389725 |
| Gm11574 | 2.609456233 | 0.021916211 | 0.655256913 | 0.033446745 |
| Osbpl7 | 231.9002821 | 0.019511143 | 0.085637175 | 0.227834963 |
| Mir8103 | 1.700203044 | 0.131910609 | 1.021165066 | 0.129176578 |
| Gm11573 | 0 | NA | NA | NA |
| Tbx21 | 34.3699015 | -0.843395267 | 0.418195041 | -2.016751002 |
| Gm11572 | 0.075043205 | 0.517957688 | 3.724945049 | 0.139051095 |
| Tbkbp1 | 732.199412 | 0.000447975 | 0.071699593 | 0.006247938 |
| Mir7235 | 0 | NA | NA | NA |
| Kpnb1 | 2804.956967 | -0.247500414 | 0.061277518 | -4.039008436 |
| Gm11583 | 2.829785719 | 0.022317872 | 0.657772431 | 0.033929473 |
| Npepps | 2128.553621 | -0.024879439 | 0.057558933 | -0.432242878 |
| Gm11592 | 3.097463869 | 0.271664438 | 0.596445747 | 0.455472168 |
| Mrpl45 | 265.8735872 | -0.380910294 | 0.10702371 | -3.559120641 |
| Gm11594 | 0.046345709 | -0.443818988 | 3.724945049 | -0.119147795 |

| | | | | |
|---|---|---|---|---|
| Gpr179 | 53.52322391 | 0.369960301 | 0.259374345 | 1.426356567 |
| Socs7 | 1787.58141 | 0.210824789 | 0.0719322 | 2.930881988 |
| Arhgap23 | 1777.390419 | -0.26448561 | 0.064364399 | -4.109191036 |
| 4933428G20Rik | 1.863773006 | -0.128992484 | 0.913338184 | -0.141231896 |
| Srcin1 | 3857.360928 | 0.578648312 | 0.090808763 | 6.372163781 |
| Gm11611 | 105.1460556 | 0.19960081 | 0.222143942 | 0.898520158 |
| Gm11612 | 0.519932157 | 1.351970251 | 1.464275436 | 0.923303238 |
| 2410003L11Rik | 0.232084656 | -0.466845891 | 2.132105974 | -0.218959984 |
| E130012A19Rik | 333.7269305 | 1.224354567 | 0.158738317 | 7.713037333 |
| Gm11613 | 75.52422125 | 0.626228958 | 0.178138458 | 3.515405751 |
| Gm24158 | 0 | NA | NA | NA |
| Mllt6 | 2810.779345 | 0.37993651 | 0.061800732 | 6.147767166 |
| Mir6927 | 0 | NA | NA | NA |
| Cisd3 | 84.26726889 | -0.070836698 | 0.168465614 | -0.420481641 |
| Pcgf2 | 208.1768514 | 0.304986813 | 0.148081708 | 2.059584646 |
| Psmb3 | 579.8326605 | -0.299076756 | 0.143198145 | -2.088551884 |
| Pip4k2b | 1705.574018 | 0.275840592 | 0.064516399 | 4.275511289 |
| Gm11614 | 0 | NA | NA | NA |
| Mir8102 | 0.581519725 | 0.633594171 | 1.278554606 | 0.495555034 |
| Cwc25 | 283.5958626 | -0.05887863 | 0.112944081 | -0.5213078 |
| Atp5l2-ps | 17.66472391 | 0.187871578 | 0.329718409 | 0.569794021 |
| 1700001P01Rik | 6.295635024 | -1.152754436 | 0.464355646 | -2.48248179 |
| Rpl23 | 782.0632473 | -0.162042423 | 0.125210116 | -1.294163987 |
| Gm27326 | 2.560534601 | -1.071443218 | 0.746690726 | -1.434922359 |
| Snora21 | 15.9140621 | -0.043633266 | 0.287533433 | -0.151750234 |
| Lasp1 | 1454.162824 | -0.299514326 | 0.081849589 | -3.659325952 |
| B230217C12Rik | 275.8448458 | 0.552872159 | 0.136468698 | 4.05127452 |
| Fbxo47 | 12.48405167 | -0.155819253 | 0.305651345 | -0.509794103 |
| Gm11630 | 0.282072957 | 0.037078026 | 1.796565559 | 0.020638281 |
| Plxdc1 | 370.3130502 | 0.358033967 | 0.129427501 | 2.766289726 |
| Gm22461 | 0.138038828 | 0.998847613 | 3.63333328 | 0.274912191 |
| Gm11633 | 0.1140246 | 0.037073798 | 3.724945049 | 0.009952844 |
| Arl5c | 23.21351478 | -0.154499413 | 0.284754736 | -0.542570126 |
| Cacnb1 | 1086.374005 | 0.228129353 | 0.142874306 | 1.596713645 |
| Gm11629 | 3.893533808 | 0.619978887 | 0.553532695 | 1.12004023 |
| Rpl19 | 1377.971411 | -0.279371483 | 0.143907668 | -1.941324506 |
| Gm27597 | 0 | NA | NA | NA |
| Stac2 | 733.8978338 | 0.068347307 | 0.097645175 | 0.699955804 |
| Gm11631 | 0.045882455 | 0.517959912 | 3.724945049 | 0.139051692 |
| Fbxl20 | 783.1636922 | 0.333908351 | 0.098272924 | 3.397765487 |
| Gm11632 | 0 | NA | NA | NA |
| Med1 | 1253.203097 | -0.162626629 | 0.052252984 | -3.11229363 |
| Gm11604 | 1.649565372 | 1.009729378 | 0.961784035 | 1.049850425 |
| Cdk12 | 1154.320364 | -0.084693856 | 0.094336217 | -0.897787288 |
| Gm27399 | 1.487186809 | -0.101378343 | 0.841887952 | -0.120417857 |
| Mir5119 | 5.066509227 | -0.115358846 | 0.492596371 | -0.234185334 |
| n-R5s73 | 0 | NA | NA | NA |

| | | | | |
|---|---|---|---|---|
| Gm20644 | 4.541882812 | 1.415195621 | 0.734015916 | 1.928017623 |
| Neurod2 | 846.9061386 | 0.620305876 | 0.135858887 | 4.56581008 |
| Ppp1r1b | 2292.267063 | 0.934834013 | 0.258087791 | 3.62215512 |
| 1700003D09Rik | 3.143667968 | 0.000251529 | 0.79627739 | 0.000315881 |
| Stard3 | 334.9821919 | -0.521994063 | 0.105990348 | -4.924920728 |
| Tcap | 15.2625128 | 1.224597374 | 0.333544197 | 3.671469581 |
| Pnmt | 1.554873959 | -0.095337392 | 0.881020138 | -0.1082125 |
| Pgap3 | 82.83766852 | 0.221849588 | 0.164750596 | 1.346578368 |
| Erbb2 | 73.62724615 | -0.721981779 | 0.185731833 | -3.887226905 |
| Gm25048 | 0 | NA | NA | NA |
| Mien1 | 288.359261 | -0.355788506 | 0.135350307 | -2.62864942 |
| Gm12352 | 1.915443829 | -0.106780363 | 0.731599225 | -0.145954724 |
| Grb7 | 32.20120666 | -0.017261148 | 0.212938639 | -0.081061605 |
| Ikzf3 | 75.08787135 | -0.534023558 | 0.283442078 | -1.884065912 |
| Gm25106 | 0.044544648 | 0.517957688 | 3.724945049 | 0.139051095 |
| Zpbp2 | 0.628980615 | 0.105425801 | 1.153375142 | 0.09140634 |
| Ormdl3 | 555.4362379 | -0.225793515 | 0.074723131 | -3.021735188 |
| Gm12 | 0 | NA | NA | NA |
| Gm12355 | 8.225470329 | -0.474610359 | 0.352781963 | -1.34533624 |
| Gsdma3 | 0.182775946 | 1.568991847 | 3.32402584 | 0.472015539 |
| Gsdma2 | 0 | NA | NA | NA |
| Gsdma | 1.768266679 | -1.018565807 | 0.807503138 | -1.261376902 |
| Psmd3 | 1360.534359 | -0.238848961 | 0.091981038 | -2.596719577 |
| Gm12356 | 1.14577023 | -0.337723351 | 0.923963698 | -0.365515822 |
| Csf3 | 0.907385681 | 3.742855702 | 1.177548625 | 3.178514773 |
| Med24 | 900.5465924 | 0.337371828 | 0.072589998 | 4.647635161 |
| Gm22059 | 0.481050399 | 0.505370546 | 1.662194253 | 0.304038198 |
| Thra | 4319.830123 | 0.689279015 | 0.076632173 | 8.994642655 |
| Nr1d1 | 1275.640339 | 0.445437209 | 0.090765337 | 4.907569585 |
| Msl1 | 1734.681355 | 0.100046101 | 0.068099435 | 1.469117923 |
| Gm12359 | 6.105237519 | 0.138032871 | 0.444005542 | 0.310880964 |
| Casc3 | 869.1967834 | -0.05470282 | 0.055776993 | -0.980741639 |
| Rapgefl1 | 1200.68795 | 0.889895065 | 0.144954275 | 6.139143297 |
| Wipf2 | 1511.917858 | 0.210027803 | 0.049148485 | 4.273332203 |
| Gm23640 | 0 | NA | NA | NA |
| Cdc6 | 14.16964528 | -1.161826069 | 0.292972578 | -3.965647829 |
| Rara | 285.8551299 | -0.986463965 | 0.162464506 | -6.071873711 |
| Gm22061 | 0.570559668 | 0.395590508 | 1.246182211 | 0.317441947 |
| Gjd3 | 5.273691302 | 0.22285883 | 0.57464 | 0.387823386 |
| Top2a | 129.6343859 | -0.725278308 | 0.382342956 | -1.896931267 |
| Gm26394 | 0 | NA | NA | NA |
| Gm23451 | 0 | NA | NA | NA |
| Igfbp4 | 440.1552601 | -0.323232365 | 0.122494952 | -2.638740291 |
| Tns4 | 6.644394917 | 0.261540007 | 0.377544559 | 0.692739441 |
| Ccr7 | 12.21422624 | -0.845944661 | 0.349153919 | -2.42284166 |
| Smarce1 | 477.9356527 | -0.084916004 | 0.07276504 | -1.166989027 |
| Krt222 | 403.0542806 | 0.236052792 | 0.07088933 | 3.329877581 |

| | | | | |
|---|---|---|---|---|
| Krt24 | 0.665888905 | 2.470516279 | 1.489345484 | 1.658793279 |
| Gm12358 | 0 | NA | NA | NA |
| Krt25 | 4.739584563 | -0.114666909 | 0.505848138 | -0.226682476 |
| Krt26 | 0.63227042 | -1.750244426 | 1.383217429 | -1.265342952 |
| Krt27 | 0.305696448 | 1.391673568 | 1.983502629 | 0.701624262 |
| Krt28 | 1.347390254 | 1.002416291 | 0.865018605 | 1.158837839 |
| Krt10 | 43.27398083 | 0.835366757 | 0.407719817 | 2.048874551 |
| Gm11940 | 7.488360699 | -0.901054608 | 0.39388172 | -2.28762738 |
| Krt12 | 43.87784049 | 1.15569202 | 0.223410501 | 5.172953002 |
| A830036E02Rik | 268.3212654 | 1.201167694 | 0.36064294 | 3.330628608 |
| Krt20 | 0.572048112 | 1.064773809 | 1.453686326 | 0.732464625 |
| Krt23 | 0.336553411 | 0.951843466 | 2.007941266 | 0.474039496 |
| Krt39 | 0 | NA | NA | NA |
| Krt40 | 0 | NA | NA | NA |
| Krtap3-3 | 0 | NA | NA | NA |
| Krtap3-2 | 0 | NA | NA | NA |
| Gm11939 | 0 | NA | NA | NA |
| Krtap3-1 | 0 | NA | NA | NA |
| Krtap1-5 | 0 | NA | NA | NA |
| Krtap1-4 | 0 | NA | NA | NA |
| Krtap1-3 | 0 | NA | NA | NA |
| Krtap9-3 | 0 | NA | NA | NA |
| Gm11938 | 0 | NA | NA | NA |
| Gm11937 | 0 | NA | NA | NA |
| Krtap2-4 | 0 | NA | NA | NA |
| Gm11562 | 0 | NA | NA | NA |
| Krtap4-1 | 0 | NA | NA | NA |
| Krtap4-2 | 0 | NA | NA | NA |
| Krtap4-7 | 0 | NA | NA | NA |
| Gm11555 | 0 | NA | NA | NA |
| Gm11563 | 0.072025462 | 0.517959912 | 3.724945049 | 0.139051692 |
| Gm14182 | 0 | NA | NA | NA |
| Krtap4-6 | 0 | NA | NA | NA |
| Gm11557 | 0.202171817 | -1.352528503 | 2.5871409 | -0.522788884 |
| 4930415H17Rik | 0 | NA | NA | NA |
| Gm14190 | 0 | NA | NA | NA |
| Gm14186 | 0 | NA | NA | NA |
| Gm14193 | 0 | NA | NA | NA |
| Gm14180 | 0 | NA | NA | NA |
| Gm14188 | 0 | NA | NA | NA |
| Gm11595 | 0 | NA | NA | NA |
| Krtap4-8 | 0.046345709 | -0.443818988 | 3.724945049 | -0.119147795 |
| Krtap4-9 | 0 | NA | NA | NA |
| Gm11596 | 0 | NA | NA | NA |
| Gm11569 | 0 | NA | NA | NA |
| Gm11554 | 0 | NA | NA | NA |
| Krtap4-13 | 0 | NA | NA | NA |

| | | | | |
|---|---|---|---|---|
| Gm11564 | 0 | NA | NA | NA |
| 2300003K06Rik | 0 | NA | NA | NA |
| Gm11556 | 0 | NA | NA | NA |
| Krtap4-16 | 0.075043205 | 0.517957688 | 3.724945049 | 0.139051095 |
| Gm26324 | 0 | NA | NA | NA |
| Gm11568 | 0 | NA | NA | NA |
| Gm11559 | 0 | NA | NA | NA |
| Krtap9-1 | 0 | NA | NA | NA |
| Gm11567 | 0 | NA | NA | NA |
| Gm11566 | 0 | NA | NA | NA |
| Krtap31-1 | 0.038981395 | -0.443816764 | 3.724945049 | -0.119147198 |
| Gm11565 | 0 | NA | NA | NA |
| Gm11560 | 17.37014342 | -0.505007246 | 0.353368494 | -1.429123573 |
| Krtap31-2 | 0 | NA | NA | NA |
| Krtap9-5 | 0 | NA | NA | NA |
| Gm11561 | 0 | NA | NA | NA |
| Gm11553 | 0 | NA | NA | NA |
| Krtap29-1 | 0.072025462 | 0.517959912 | 3.724945049 | 0.139051692 |
| Krtap16-1 | 0.141781211 | -0.444711846 | 2.911624983 | -0.15273665 |
| Gm11558 | 0 | NA | NA | NA |
| Krtap17-1 | 0.608261712 | -1.156961958 | 1.306033021 | -0.885859652 |
| Krt33a | 0 | NA | NA | NA |
| Krt33b | 0 | NA | NA | NA |
| Krt34 | 0 | NA | NA | NA |
| Krt31 | 0 | NA | NA | NA |
| Gm11571 | 0 | NA | NA | NA |
| Krt32 | 0 | NA | NA | NA |
| Krt35 | 0 | NA | NA | NA |
| Krt36 | 0.15956642 | 0.036118649 | 3.418731032 | 0.010564929 |
| Gm12347 | 0 | NA | NA | NA |
| Krt13 | 0.652975239 | 2.771203822 | 1.612779856 | 1.718277799 |
| Krt15 | 1.82600251 | -1.286696265 | 0.972429006 | -1.323177586 |
| Krt19 | 0.608478225 | -1.760329154 | 1.558221345 | -1.129704172 |
| Krt9 | 26.56616351 | -0.485831661 | 0.297357875 | -1.633828128 |
| Krt14 | 1.447260382 | 0.140162007 | 1.266442104 | 0.110673837 |
| Gm11597 | 0 | NA | NA | NA |
| Gm11598 | 0.706088228 | 0.586180354 | 1.212759936 | 0.483344096 |
| Krt16 | 2.108251394 | -3.299321763 | 1.195700754 | -2.759320635 |
| Krt17 | 3.233276867 | -1.753309998 | 0.817066479 | -2.145859661 |
| Krt42 | 0 | NA | NA | NA |
| Gm14206 | 0.67116138 | 0.119215361 | 1.193762575 | 0.099865219 |
| Gm12349 | 7.416241275 | -1.313426146 | 0.924637353 | -1.420477056 |
| Eif1 | 1981.137352 | -0.244668111 | 0.095872136 | -2.552025249 |
| Gast | 0.589408234 | -1.533715173 | 1.314762357 | -1.166534138 |
| Hap1 | 3690.110612 | 0.184949433 | 0.142134608 | 1.301227305 |
| Gm20394 | 9.564810652 | -0.139915247 | 0.491545024 | -0.284643806 |
| Gm10039 | 471.8830643 | -0.031416291 | 0.09032929 | -0.34779739 |

| | | | | |
|---|---|---|---|---|
| Jup | 876.4788789 | -0.548779674 | 0.098861396 | -5.551000635 |
| Gm12348 | 0.360464465 | -0.924715651 | 1.479814766 | -0.624886082 |
| P3h4 | 107.8090644 | -0.060191292 | 0.116350528 | -0.517327196 |
| Fkbp10 | 103.2641343 | -0.672668905 | 0.159066209 | -4.22886111 |
| Nt5c3b | 220.2515258 | 0.042010133 | 0.089408921 | 0.469865109 |
| Klhl10 | 2.707144304 | -0.446392698 | 0.650424204 | -0.686310096 |
| Gm27374 | 1.132676578 | 1.065929722 | 1.021004125 | 1.044001386 |
| Klhl11 | 985.9766124 | 0.087005994 | 0.063397395 | 1.372390684 |
| Acly | 2346.506125 | -0.172785946 | 0.084405452 | -2.047094605 |
| mmu-mir-7116 | 0.285551684 | 0.111600237 | 2.048526276 | 0.054478304 |
| Ttc25 | 14.3848452 | -0.68532847 | 0.278192926 | -2.463500705 |
| Cnp | 4776.985164 | 0.062965571 | 0.115826016 | 0.543622 |
| Dnajc7 | 1151.346391 | 0.029255403 | 0.071938724 | 0.406671142 |
| Nkiras2 | 157.3130789 | -0.605630347 | 0.105170748 | -5.758543713 |
| Zfp385c | 87.71940538 | 0.2269015 | 0.228495701 | 0.99302306 |
| Dhx58os | 2.117912561 | 0.463332175 | 0.666779912 | 0.694880225 |
| Dhx58 | 526.9953047 | -0.70623364 | 0.203013415 | -3.478753557 |
| Kat2a | 785.4970274 | 0.092779338 | 0.057384654 | 1.616797025 |
| Hspb9 | 0.468943176 | -2.697744138 | 3.221979915 | -0.837293903 |
| Rab5c | 1897.413452 | -0.414550033 | 0.127869101 | -3.24198755 |
| Kcnh4 | 95.02347477 | 0.873041955 | 0.268531363 | 3.251173142 |
| Hcrt | 38.8486744 | 0.208302215 | 0.210358821 | 0.990223342 |
| Ghdc | 106.0321 | -0.974024054 | 0.177921388 | -5.474463006 |
| Gm24358 | 0.048949532 | 0.037064902 | 3.724945049 | 0.009950456 |
| Stat5b | 602.2188295 | -0.345560037 | 0.106117116 | -3.256402458 |
| Stat5a | 70.6584695 | -0.779699765 | 0.177996405 | -4.380424216 |
| Stat3 | 2401.024981 | -0.448173415 | 0.164863265 | -2.718455283 |
| Ptrf | 400.0738531 | -1.099641274 | 0.178008957 | -6.177449114 |
| Atp6v0a1 | 5069.409655 | 0.242282182 | 0.063293774 | 3.827899101 |
| Mir6928 | 2.18332649 | 0.857206335 | 0.834899418 | 1.026718088 |
| Naglu | 145.2410652 | -0.59411864 | 0.149680397 | -3.969248168 |
| Hsd17b1 | 3.534966268 | -1.094781737 | 0.764623187 | -1.431792491 |
| Coasy | 224.3589791 | -0.278172989 | 0.093531117 | -2.974122398 |
| Mlx | 272.5349141 | -0.52553957 | 0.104845934 | -5.012493584 |
| Psmc3ip | 15.59208128 | -0.30159387 | 0.294282197 | -1.024845785 |
| Fam134c | 741.7829735 | -0.324275532 | 0.066980071 | -4.841373347 |
| Tubg1 | 286.1670221 | -0.100747266 | 0.088766565 | -1.134968615 |
| Gm27626 | 0 | NA | NA | NA |
| Tubg2 | 274.8310085 | 0.694723293 | 0.08635886 | 8.044609396 |
| Plekhh3 | 197.5858469 | -0.261341209 | 0.118788403 | -2.200056597 |
| Cntnap1 | 2524.742535 | 0.676442925 | 0.085377458 | 7.922968678 |
| Ccr10 | 4.101636231 | -0.256656748 | 0.502574745 | -0.510683734 |
| Ezh1 | 709.9648608 | 0.15647662 | 0.080348321 | 1.947478395 |
| Ramp2 | 134.6263258 | -0.029861793 | 0.152070691 | -0.196367837 |
| Vps25 | 18.38762957 | 0.327596287 | 0.277252397 | 1.181581441 |
| Wnk4 | 77.9757232 | -0.044291197 | 0.174825493 | -0.253345184 |
| Gm11615 | 0.280085221 | 0.801887587 | 2.084990492 | 0.384600117 |

| | | | | |
|---|---|---|---|---|
| Coa3 | 343.5032495 | -0.223575477 | 0.142569914 | -1.568181325 |
| Cntd1 | 4.658726077 | -0.602553568 | 0.526463342 | -1.144530909 |
| Becn1 | 989.2074398 | -0.049850813 | 0.068902707 | -0.723495702 |
| Gm11617 | 0 | NA | NA | NA |
| Psme3 | 1184.471621 | -0.01039685 | 0.065924039 | -0.157709548 |
| Aoc2 | 39.91492154 | -0.651248111 | 0.304930851 | -2.135723914 |
| Aoc3 | 5.919817619 | -2.14346783 | 0.517860297 | -4.139085087 |
| Gm28156 | 0 | NA | NA | NA |
| Gm11618 | 0 | NA | NA | NA |
| Gm11619 | 0 | NA | NA | NA |
| Gm25593 | 0 | NA | NA | NA |
| G6pc | 0.591033593 | 0.649561125 | 1.344436841 | 0.483147371 |
| Gm25475 | 0 | NA | NA | NA |
| Gm27029 | 8.009665686 | -0.223324654 | 0.484129686 | -0.461290974 |
| Aarsd1 | 19.81807774 | 0.135397105 | 0.282894402 | 0.47861359 |
| Ptges3l | 9.728067973 | -0.434173043 | 0.346307243 | -1.253722099 |
| Mir6929 | 0.072025462 | 0.517959912 | 3.724945049 | 0.139051692 |
| Rundc1 | 403.5552599 | -0.115136832 | 0.084113533 | -1.3688265 |
| Rpl27 | 526.9415286 | -0.316004982 | 0.127606412 | -2.476403626 |
| Ifi35 | 471.7219268 | -0.721272988 | 0.203290913 | -3.547984404 |
| Gm11626 | 1.005912905 | -1.496689785 | 1.10590072 | -1.353367222 |
| Vat1 | 770.8644348 | -0.874475955 | 0.163185873 | -5.358772425 |
| Rnd2 | 698.8704723 | 0.307093028 | 0.150065745 | 2.046389923 |
| Gm11625 | 0.085327104 | -0.905414734 | 3.723500899 | -0.243162217 |
| Brca1 | 51.12254382 | -0.147213884 | 0.257991918 | -0.570614323 |
| Nbr1 | 2218.562145 | -0.066996923 | 0.038947511 | -1.720184948 |
| Gm27456 | 0 | NA | NA | NA |
| Tmem106a | 300.3975511 | -1.728728258 | 0.303066235 | -5.704126886 |
| Gm11634 | 0 | NA | NA | NA |
| Gm26316 | 17.08146439 | -0.277980653 | 0.350626287 | -0.792811786 |
| Gm11635 | 0.049553047 | -0.443818988 | 3.724945049 | -0.119147795 |
| Rdm1 | 27.87786844 | -0.36151504 | 0.235982888 | -1.531954471 |
| Gm23849 | 1.056059865 | -0.141393951 | 1.041537624 | -0.13575501 |
| Gm24950 | 3.757983618 | 0.147431845 | 0.561108006 | 0.262751277 |
| Gm23971 | 333.7774647 | -0.39213952 | 0.195167617 | -2.0092448 |
| Rnu2-10 | 3.530281436 | -0.319772978 | 0.593942611 | -0.538390364 |
| Arl4d | 211.6319964 | -0.291179753 | 0.335685186 | -0.867419132 |
| Dhx8 | 571.733905 | -0.180028347 | 0.067652572 | -2.661071716 |
| Etv4 | 83.85799785 | -0.102216636 | 0.149355305 | -0.684385703 |
| Meox1 | 5.667363994 | -0.494034966 | 0.428916564 | -1.151820675 |
| Gm11551 | 0.198035107 | 0.998854246 | 3.22244265 | 0.309968044 |
| Rpl27-ps2 | 0.195392278 | 1.462541818 | 3.230774363 | 0.452690796 |
| 4930417O22Rik | 0.123841226 | -1.017823543 | 3.719440177 | -0.273649661 |
| Sost | 2.176193535 | -2.275213702 | 0.979519891 | -2.322784582 |
| Gm20659 | 0.323889686 | 2.130655568 | 3.69255725 | 0.57701355 |
| Dusp3 | 1716.840804 | 0.088677307 | 0.081592175 | 1.08683593 |
| 1700006E09Rik | 0.21733455 | 0.037076198 | 2.170157969 | 0.017084562 |

| | | | | |
|---|---|---|---|---|
| Mpp3 | 656.0571513 | 0.359635932 | 0.112888819 | 3.185753327 |
| E130111B04Rik | 2.471955472 | 0.246949675 | 0.670063096 | 0.368546898 |
| Cd300lg | 0.955408868 | -0.718523459 | 0.920529424 | -0.780554581 |
| Mpp2 | 1306.923634 | 0.553985827 | 0.101913825 | 5.435826047 |
| Ppy | 0.061920613 | -0.443816764 | 3.724945049 | -0.119147198 |
| Pyy | 0.755118542 | -0.091501459 | 1.437058012 | -0.063672766 |
| Gm11584 | 0.066050875 | -0.443818988 | 3.724945049 | -0.119147795 |
| Gm11585 | 0.714893038 | 0.297693562 | 1.181699816 | 0.251919784 |
| Nags | 0.527376817 | -1.56324329 | 1.265042464 | -1.235723965 |
| Gm11586 | 0 | NA | NA | NA |
| Tmem101 | 154.1546923 | -0.205995178 | 0.111488968 | -1.847673197 |
| Lsm12 | 620.6014926 | -0.34377013 | 0.09138382 | -3.761827108 |
| G6pc3 | 331.7313334 | -0.059289929 | 0.101212884 | -0.585794286 |
| Hdac5 | 1648.304379 | 0.07028454 | 0.104525101 | 0.672417814 |
| Mir8101 | 0 | NA | NA | NA |
| BC030867 | 13.51292698 | -0.292197375 | 0.360714698 | -0.810051204 |
| Asb16 | 47.65270219 | 0.707763006 | 0.182937488 | 3.868879009 |
| Tmub2 | 469.4932909 | 0.122299788 | 0.063662857 | 1.92105402 |
| Atxn7l3 | 1951.581005 | 0.247474167 | 0.054171937 | 4.568309355 |
| Ubtf | 1192.509439 | -0.097540665 | 0.070108943 | -1.39127279 |
| Slc4a1 | 10.15205109 | 0.740239999 | 0.62107192 | 1.191874846 |
| Bloodlinc | 0.118974955 | 0.51795324 | 3.724945049 | 0.139049901 |
| Rundc3a | 2248.259499 | 0.255332422 | 0.059694209 | 4.277339895 |
| Slc25a39 | 589.2097923 | -0.522095215 | 0.101524143 | -5.142571981 |
| Mir6930 | 0 | NA | NA | NA |
| Grn | 3148.773445 | -0.872556194 | 0.159026502 | -5.48686027 |
| Fam171a2 | 531.5743851 | 0.229057076 | 0.092166275 | 2.485259113 |
| Itga2b | 26.87028233 | -0.778478804 | 0.223808911 | -3.478319069 |
| Gm23563 | 0.090427103 | 0.998850474 | 3.724945049 | 0.268151734 |
| Gpatch8 | 1536.270915 | 0.176295172 | 0.098616033 | 1.787692797 |
| Gm11628 | 8.708300683 | 0.799675978 | 0.517576488 | 1.54503923 |
| Mdk-ps1 | 0 | NA | NA | NA |
| Gm25337 | 4.232730408 | 0.990883507 | 0.563712507 | 1.757781661 |
| Gm11627 | 10.86367426 | 2.101334178 | 0.607162623 | 3.460908326 |
| Fzd2 | 124.2975267 | -0.312422534 | 0.272694984 | -1.14568493 |
| Gm11636 | 0.35856548 | 0.5179731 | 1.830183679 | 0.283017003 |
| 2810433D01Rik | 2.836445391 | -1.48851131 | 0.654410305 | -2.274584154 |
| Psmb5-ps | 0 | NA | NA | NA |
| Gm22920 | 0 | NA | NA | NA |
| Gm1564 | 0.341943343 | 1.439120933 | 1.720667532 | 0.83637362 |
| Ccdc43 | 214.8599409 | -0.070241848 | 0.087017046 | -0.807219407 |
| Gm11637 | 5.095273339 | 0.716886582 | 0.458202289 | 1.564563511 |
| Gm16342 | 0.648681289 | 1.412965073 | 1.486056759 | 0.950815011 |
| Adam11 | 1411.899054 | 0.55826743 | 0.121900254 | 4.579706863 |
| Gjc1 | 65.81803581 | 0.06522015 | 0.174006319 | 0.374814835 |
| Gm22683 | 0 | NA | NA | NA |
| Higd1b | 5.836578858 | -0.519235155 | 0.470660516 | -1.103205257 |

| | | | | |
|---|---|---|---|---|
| Eftud2 | 907.904142 | 0.050070616 | 0.072359456 | 0.69197059 |
| Gm26668 | 0.400913792 | 0.154589092 | 1.756398596 | 0.088014812 |
| Ccdc103 | 10.14206194 | -0.162724569 | 0.33139901 | -0.491023098 |
| Fam187a | 0.4546328 | -0.446474247 | 1.501989706 | -0.297255197 |
| Gfap | 17257.76253 | -0.807982421 | 0.171158358 | -4.720671726 |
| Kif18b | 11.6304259 | -0.902621269 | 0.489883781 | -1.842521236 |
| C1ql1 | 74.68787927 | -0.296951863 | 0.195145796 | -1.521692341 |
| 2410004I01Rik | 2.828652251 | -0.339614061 | 0.649072458 | -0.523229813 |
| Dcakd | 326.4044415 | -0.366054913 | 0.103692511 | -3.530196256 |
| Mir6931 | 0.170867779 | 0.037052907 | 3.724945049 | 0.009947236 |
| Nmt1 | 1910.096591 | 0.039372337 | 0.04641476 | 0.848271905 |
| Plcd3 | 130.7093725 | 0.159676485 | 0.104253904 | 1.531611554 |
| Acbd4 | 248.1541703 | 0.146290187 | 0.097852027 | 1.495014375 |
| Hexim1 | 470.7862717 | -0.110486439 | 0.105505671 | -1.047208537 |
| Hexim2 | 107.5740266 | 0.611455207 | 0.147567658 | 4.143558385 |
| Fmnl1 | 978.2301766 | -0.315853745 | 0.110550278 | -2.857104925 |
| Gm20511 | 5.853403199 | 0.007893582 | 0.508500605 | 0.01552325 |
| 1700023F06Rik | 21.3606864 | 0.336084579 | 0.267561722 | 1.256101121 |
| Spata32 | 11.37911894 | 1.553243063 | 0.401611974 | 3.867521793 |
| Map3k14 | 131.8227025 | -0.482524977 | 0.185572931 | -2.600190521 |
| 1700028N14Rik | 3.893030208 | 0.204293337 | 0.542042521 | 0.376895409 |
| Arhgap27 | 221.5069098 | -0.241751275 | 0.098810001 | -2.446627604 |
| Arhgap27os3 | 8.082714316 | -0.286936203 | 0.392620665 | -0.730822977 |
| Arhgap27os2 | 8.179740595 | 0.285646406 | 0.373168259 | 0.765462763 |
| Arhgap27os1 | 2.613037336 | -0.196154671 | 0.637372966 | -0.307754927 |
| Plekhm1 | 513.8582558 | -0.798865952 | 0.087279179 | -9.152995768 |
| Lrrc37a | 0.598753221 | -2.50031094 | 1.332263551 | -1.876738981 |
| Gm884 | 1.391014422 | -0.359884425 | 0.842199826 | -0.427314771 |
| Gm11643 | 0.038981395 | -0.443816764 | 3.724945049 | -0.119147198 |
| Lyzl6 | 0.124596252 | 0.037071632 | 3.714681146 | 0.009979761 |
| Rprml | 232.2274179 | -0.120070311 | 0.199943478 | -0.60052127 |
| Gosr2 | 931.0627658 | -0.424127075 | 0.094887006 | -4.469811974 |
| C130046K22Rik | 12.71673352 | -0.250666781 | 0.290705707 | -0.8622699 |
| Gm11642 | 9.924921471 | -0.225327338 | 0.358820883 | -0.627966065 |
| Wnt9b | 31.73347887 | -1.260147108 | 0.296811079 | -4.245620186 |
| Wnt3 | 86.98556824 | 0.549194548 | 0.216242666 | 2.539714095 |
| Nsf | 6788.932109 | 0.390345216 | 0.061162571 | 6.382092947 |
| Gm8738 | 9.128726587 | 0.695185198 | 0.532541813 | 1.305409605 |
| Arf2 | 510.9708085 | -0.292437892 | 0.106796001 | -2.73828503 |
| Crhr1 | 208.0710271 | 0.022459184 | 0.117487208 | 0.191162801 |
| Sppl2c | 0.098502579 | -0.443821212 | 3.724945049 | -0.119148392 |
| Mapt | 4636.496193 | 0.286527725 | 0.068128585 | 4.205690269 |
| Gm11659 | 0.104366086 | 0.51795324 | 3.724945049 | 0.139049901 |
| Gm22743 | 0 | NA | NA | NA |
| AL596383.1 | 0 | NA | NA | NA |
| Kansl1 | 1511.632219 | 0.015682986 | 0.036967765 | 0.424234099 |
| AL593843.1 | 0.534702693 | -0.636221366 | 1.499042471 | -0.424418506 |

| | | | | |
|---|---|---|---|---|
| Gm11660 | 0 | NA | NA | NA |
| Cdc27 | 1303.554971 | -0.062857569 | 0.05394026 | -1.165318245 |
| Myl4 | 65.04662385 | 0.80263275 | 0.22912214 | 3.503078094 |
| Gm11665 | 0 | NA | NA | NA |
| Gm22000 | 0.267335191 | -0.87962182 | 1.79215852 | -0.490816973 |
| Gm11661 | 0 | NA | NA | NA |
| Gm11663 | 0.542039732 | -0.540354428 | 1.475017019 | -0.366337758 |
| Itgb3 | 53.54022327 | -0.604177034 | 0.191143329 | -3.160858594 |
| Gm11662 | 0.044191549 | 0.037062677 | 3.724945049 | 0.009949859 |
| Gm11639 | 8.031358769 | 0.710485953 | 0.382539865 | 1.857286044 |
| Gm11623 | 0.110595523 | 0.037071574 | 3.724945049 | 0.009952247 |
| Gm11621 | 0.116046057 | 0.037062677 | 3.724945049 | 0.009949859 |
| Gm11622 | 0.090074004 | 0.517955464 | 3.724945049 | 0.139050498 |
| Gm11624 | 0.048949532 | 0.037064902 | 3.724945049 | 0.009950456 |
| Efcab3 | 4.279745736 | 0.769188205 | 0.603857504 | 1.273790919 |
| Gm25362 | 0 | NA | NA | NA |
| Mettl2 | 133.1323383 | -0.061337017 | 0.128134996 | -0.478690589 |
| Gm24290 | 0.451210867 | 0.400696576 | 1.469037432 | 0.272761311 |
| Gm10842 | 0.310932149 | -0.914698086 | 1.778566541 | -0.514289494 |
| Gm11620 | 9.032629779 | 0.12840921 | 0.365647172 | 0.351183381 |
| 1700052K11Rik | 167.7122578 | -0.164579696 | 0.099114926 | -1.660493553 |
| Tlk2 | 1033.874952 | -0.219827847 | 0.101463758 | -2.166565193 |
| Mrc2 | 195.2873509 | -0.596650814 | 0.154800421 | -3.854322948 |
| 10-Mar | 11.52093314 | 0.278372052 | 0.357505602 | 0.77865088 |
| Gm11638 | 3.337525259 | -0.757524626 | 0.819797347 | -0.924038885 |
| Gm11640 | 0.456339188 | -0.67636846 | 1.339923594 | -0.504781365 |
| Tanc2 | 3818.814185 | 0.364939002 | 0.105517417 | 3.45856649 |
| Gm23990 | 3.257021581 | -0.548433564 | 0.660192832 | -0.830717235 |
| Gm11652 | 25.69936743 | 0.274039607 | 0.294498996 | 0.930528151 |
| Gm11653 | 35.85399802 | 0.236051537 | 0.305438271 | 0.772828946 |
| Cyb561 | 385.0963259 | 0.181824297 | 0.069959575 | 2.598990888 |
| Gm9910 | 2.885346812 | -0.372247424 | 0.594695233 | -0.62594654 |
| Gm11651 | 0.18135768 | 0.517962113 | 2.746620309 | 0.188581622 |
| Ace | 389.3701917 | 0.460591564 | 0.111384706 | 4.135141882 |
| Ace3 | 0.089089296 | 0.957155977 | 3.723939561 | 0.257027796 |
| Kcnh6 | 35.00512634 | 0.178848365 | 0.197682589 | 0.904724919 |
| Dcaf7 | 2563.358122 | 0.361972484 | 0.06220783 | 5.818760856 |
| Gm11646 | 0.25779132 | -0.172539517 | 1.99993844 | -0.086272414 |
| Taco1os | 4.692172118 | -0.711920894 | 0.459312613 | -1.549970267 |
| Taco1 | 111.1047001 | -0.382657788 | 0.164790852 | -2.322081497 |
| Map3k3 | 378.7212756 | -0.002404456 | 0.119363994 | -0.020143893 |
| Gm11672 | 0.164893722 | 1.344872572 | 3.387385485 | 0.397023775 |
| Limd2 | 471.2960561 | 0.556854222 | 0.108517808 | 5.131454771 |
| Gm10840 | 0.825556236 | 0.998366476 | 1.084970084 | 0.920178806 |
| Strada | 263.1762033 | 0.106665346 | 0.12607566 | 0.846042336 |
| Gm25556 | 0 | NA | NA | NA |
| Ccdc47 | 1290.145382 | -0.044216863 | 0.067045603 | -0.659504291 |

| | | | | |
|---|---|---|---|---|
| Ddx42 | 1058.979334 | 0.084578022 | 0.085366052 | 0.990768812 |
| Ftsj3 | 311.2842433 | -0.510557082 | 0.130476632 | -3.913015505 |
| Psmc5 | 1264.146288 | -0.23891633 | 0.097900201 | -2.440406948 |
| Gm23645 | 2.33541923 | 1.531252783 | 0.747128201 | 2.049518117 |
| Smarcd2 | 283.2968771 | -0.099569004 | 0.115350339 | -0.863187793 |
| Tcam1 | 4.654232259 | -0.022487083 | 0.447547626 | -0.050245118 |
| Gh | 110.6277303 | -4.980781894 | 2.433151312 | -2.047049795 |
| Cd79b | 1.812060818 | -0.172490541 | 0.658404361 | -0.261982683 |
| Scn4a | 1.34423325 | -0.164205214 | 0.842676753 | -0.19486145 |
| 2310007L24Rik | 2.639496584 | -0.790867691 | 0.625147661 | -1.265089418 |
| Icam2 | 17.72490718 | -1.041050826 | 0.272728686 | -3.817166589 |
| Ern1 | 301.964099 | -0.715930976 | 0.105588268 | -6.780402709 |
| Snord104 | 16.93328137 | -0.43700662 | 0.274073595 | -1.594486402 |
| SNORA50 | 0.197850883 | 0.037075115 | 2.231290102 | 0.016615999 |
| Gm22711 | 0 | NA | NA | NA |
| Tex2 | 1392.018177 | -0.013881672 | 0.051373117 | -0.270212767 |
| Pecam1 | 139.8937637 | -0.926742584 | 0.140814438 | -6.581303705 |
| Milr1 | 37.7867777 | -1.348226878 | 0.289055889 | -4.664242896 |
| Gm25889 | 0.53783135 | -0.86981491 | 1.423636082 | -0.61098122 |
| Polg2 | 47.1941341 | -0.322568617 | 0.262059106 | -1.230900245 |
| Ddx5 | 9164.25171 | 0.134640212 | 0.07733594 | 1.740978548 |
| Mir3064 | 0 | NA | NA | NA |
| Gm25994 | 0 | NA | NA | NA |
| Cep95 | 165.3431975 | -0.088133705 | 0.159365555 | -0.553028572 |
| Gm11706 | 1.894273405 | 0.435343094 | 0.875763017 | 0.497101483 |
| Smurf2 | 621.2522263 | -0.160558038 | 0.061360743 | -2.616624748 |
| Gm27322 | 0.075043205 | 0.517957688 | 3.724945049 | 0.139051095 |
| Gm11707 | 0.436105163 | -0.236084511 | 1.618241657 | -0.145889528 |
| Kpna2 | 179.0519059 | -0.31019303 | 0.130039653 | -2.385372644 |
| Gm11708 | 0 | NA | NA | NA |
| Gm11705 | 2.194360983 | 0.363063269 | 0.725213113 | 0.500629763 |
| 1810010H24Rik | 17.13476079 | 0.776081353 | 0.282006864 | 2.751994547 |
| Bptf | 2737.946275 | 0.053865262 | 0.06349967 | 0.848276247 |
| Gm11715 | 0.139856492 | -0.443820827 | 3.574411099 | -0.124166139 |
| Nol11 | 402.0846801 | -0.201963118 | 0.089269013 | -2.262410114 |
| Rps12-ps26 | 3.470784895 | 0.996235068 | 0.623107922 | 1.598816244 |
| Gm11718 | 0.038981395 | -0.443816764 | 3.724945049 | -0.119147198 |
| Pitpnc1 | 980.2405448 | 0.177658192 | 0.066753343 | 2.661412666 |
| Gm11719 | 0 | NA | NA | NA |
| Gm11716 | 0.710007532 | -1.267927911 | 1.14857214 | -1.103916651 |
| Gm11712 | 0.072025462 | 0.517959912 | 3.724945049 | 0.139051692 |
| Gm11713 | 0.068097359 | 0.037064902 | 3.724945049 | 0.009950456 |
| Gm11714 | 1.466365607 | -0.327421354 | 0.761223121 | -0.43012534 |
| Psmd12 | 782.060693 | -0.261957738 | 0.096216892 | -2.722575343 |
| Gm11722 | 4.228535416 | 0.567118011 | 0.520121132 | 1.090357564 |
| C330019F10Rik | 0.3156755 | 0.53130023 | 1.774865837 | 0.299346699 |
| A830035A12Rik | 0.580962004 | 0.431702292 | 1.273641944 | 0.338951064 |

| | | | | |
|---|---|---|---|---|
| Gm11649 | 0 | NA | NA | NA |
| Helz | 1652.750466 | 0.064868063 | 0.090405116 | 0.717526463 |
| Mir6932 | 0.570554915 | 0.024578707 | 1.551816439 | 0.015838669 |
| Cacng1 | 1.900742604 | 1.940518046 | 0.816763239 | 2.375863596 |
| Cacng4 | 438.0588544 | -0.013262459 | 0.110911744 | -0.119576684 |
| Gm11650 | 3.119570997 | 0.484486053 | 0.68449345 | 0.707802321 |
| Gm23120 | 0 | NA | NA | NA |
| Gm27595 | 0 | NA | NA | NA |
| Cacng5 | 366.3860086 | -0.260657927 | 0.154911115 | -1.682628947 |
| Prkca | 1907.502606 | 0.443280338 | 0.0948127 | 4.675326575 |
| Gm11657 | 0 | NA | NA | NA |
| Mir7223 | 0 | NA | NA | NA |
| Gm11655 | 0.048949532 | 0.037064902 | 3.724945049 | 0.009950456 |
| Apoh | 2.789102959 | 0.116074865 | 0.648233719 | 0.179063293 |
| Gm11656 | 0.072025462 | 0.517959912 | 3.724945049 | 0.139051692 |
| Gm11667 | 0 | NA | NA | NA |
| Cep112 | 85.63833006 | 0.407642061 | 0.181957265 | 2.240317591 |
| Cep112os1 | 0 | NA | NA | NA |
| Olfr1397-ps1 | 0.047948698 | 0.037064902 | 3.724945049 | 0.009950456 |
| Rpl23a-ps14 | 5.868082076 | 0.108979603 | 0.450042716 | 0.24215391 |
| Cep112os2 | 13.20886531 | 0.529920615 | 0.468211898 | 1.131796559 |
| Axin2 | 366.2212925 | 0.35410839 | 0.111001408 | 3.190125212 |
| Gm11670 | 1.328031477 | -0.700420382 | 0.889862745 | -0.787110581 |
| Gm11668 | 0.544841187 | -1.555350603 | 1.411282296 | -1.102083266 |
| Gm24149 | 0.044544648 | 0.517957688 | 3.724945049 | 0.139051095 |
| E030025P04Rik | 0.237045706 | 0.037075228 | 2.144038011 | 0.017292244 |
| Gm11658 | 0.044544648 | 0.517957688 | 3.724945049 | 0.139051095 |
| Rgs9 | 522.9399079 | 0.365676204 | 0.246757757 | 1.481923846 |
| 1700096J18Rik | 0 | NA | NA | NA |
| Gm11696 | 30.54970298 | -0.174781282 | 0.204013888 | -0.856712668 |
| Gna13 | 2195.672482 | -0.518433116 | 0.131078154 | -3.955145082 |
| 9930022D16Rik | 1.783663422 | -0.14959131 | 0.741608874 | -0.201711866 |
| Amz2 | 356.9352975 | -0.270061082 | 0.119737369 | -2.255445259 |
| Gm15642 | 0 | NA | NA | NA |
| Slc16a6 | 246.9242258 | -0.108776921 | 0.170777722 | -0.636950297 |
| Gm25540 | 3.251877087 | 0.389359669 | 0.714412206 | 0.545007022 |
| Arsg | 200.3840775 | 0.209026816 | 0.133904093 | 1.56101887 |
| Gm22378 | 0 | NA | NA | NA |
| Wipi1 | 268.8479832 | -0.307432044 | 0.090294192 | -3.404782067 |
| Gm11685 | 0.075043205 | 0.517957688 | 3.724945049 | 0.139051095 |
| Prkar1a | 7255.166353 | -0.001281861 | 0.058133948 | -0.022050132 |
| Fam20a | 97.31167799 | 0.416156415 | 0.137915998 | 3.017462947 |
| Gm11684 | 0 | NA | NA | NA |
| 1700012B07Rik | 0.10503227 | -0.905414734 | 3.723500899 | -0.243162217 |
| Gm11683 | 0 | NA | NA | NA |
| 1700023C21Rik | 0 | NA | NA | NA |
| Abca8b | 635.0819168 | 0.262213069 | 0.110691674 | 2.368859905 |

| | | | | |
|---|---|---|---|---|
| Gm11697 | 0 | NA | NA | NA |
| Abca8a | 98.13962934 | 0.6589919 | 0.184442816 | 3.572879198 |
| Abca9 | 172.9692926 | 0.202538836 | 0.222277217 | 0.911199262 |
| Abca6 | 37.25579249 | -0.142879787 | 0.23480031 | -0.608516178 |
| Abca5 | 1374.997336 | 0.310834746 | 0.058569457 | 5.30711336 |
| Map2k6 | 151.8243697 | 0.7848768 | 0.145933108 | 5.378332638 |
| Gm11682 | 0.63216095 | 0.410185393 | 1.257630407 | 0.326157343 |
| Kcnj16 | 546.3597821 | 0.199314043 | 0.157083979 | 1.268837494 |
| Kcnj2 | 326.6825132 | 0.426582998 | 0.102631762 | 4.156442316 |
| Gm11675 | 0.837406104 | 0.245497676 | 1.025150109 | 0.239474857 |
| Gm11676 | 0 | NA | NA | NA |
| Gm11677 | 0.110870145 | -0.444802825 | 3.724874622 | -0.119414174 |
| Gm11674 | 0.062426873 | 0.517959912 | 3.724945049 | 0.139051692 |
| Gm11679 | 0 | NA | NA | NA |
| Gm11680 | 0 | NA | NA | NA |
| BC006965 | 12.81790452 | 0.916856733 | 0.361265259 | 2.537904521 |
| Sox9 | 1126.317198 | -0.620634849 | 0.133049014 | -4.664708359 |
| Gm11681 | 4.401306684 | 0.392893099 | 0.475953008 | 0.825487166 |
| 4933434M16Rik | 0 | NA | NA | NA |
| 2610035D17Rik | 77.49257512 | -0.255477263 | 0.144227451 | -1.771349779 |
| 4732490B19Rik | 0.42613393 | -2.07399274 | 1.535846499 | -1.350390642 |
| Slc39a11 | 151.2282737 | -0.289132569 | 0.102581538 | -2.818563419 |
| Gm16487 | 1.032826873 | -2.226321354 | 1.128360991 | -1.97305771 |
| Sstr2 | 160.0653886 | -1.4714633 | 0.328051635 | -4.485462484 |
| Cog1 | 598.811076 | -0.079188752 | 0.067651846 | -1.170533501 |
| Fam104a | 378.1237319 | -0.16534594 | 0.07276883 | -2.27220831 |
| Gm11693 | 0.456148335 | -1.419781399 | 1.632493784 | -0.869700952 |
| D11Wsu47e | 95.6758204 | -0.384897331 | 0.161515911 | -2.383030439 |
| Cpsf4l | 2.642413763 | -0.61899069 | 0.642378968 | -0.963591153 |
| Cdc42ep4 | 690.6837532 | -0.397573561 | 0.141684432 | -2.806049724 |
| Gm11736 | 0.197525616 | -0.427474636 | 2.225104434 | -0.192114415 |
| Sdk2 | 727.2965037 | 0.248831451 | 0.119552712 | 2.081353461 |
| 1700025D23Rik | 0.122991902 | 0.517955464 | 3.724945049 | 0.139050498 |
| 1700001J04Rik | 0.288935699 | -1.547730241 | 2.061811502 | -0.750665247 |
| 1700092K14Rik | 0.758667428 | 0.431913951 | 1.268701332 | 0.340437848 |
| Gm11690 | 0.042716945 | 0.037062677 | 3.724945049 | 0.009949859 |
| Gm11691 | 0.047948698 | 0.037064902 | 3.724945049 | 0.009950456 |
| Gm11692 | 0 | NA | NA | NA |
| 4932435O22Rik | 2.584340838 | 0.179049761 | 0.750462558 | 0.238585868 |
| Gm11689 | 0 | NA | NA | NA |
| Rpl38 | 617.2880742 | -0.046461246 | 0.198425311 | -0.234149794 |
| Ttyh2 | 758.4082506 | 0.135005254 | 0.097511086 | 1.384511849 |
| Dnaic2 | 33.61650857 | -0.835639229 | 0.231157973 | -3.615013652 |
| Kif19a | 46.14809591 | 0.367618313 | 0.180409851 | 2.037684258 |
| Btbd17 | 82.92747472 | 0.839921972 | 0.278397481 | 3.016988409 |
| Gpr142 | 0.138152447 | 0.037074147 | 2.93496362 | 0.012631893 |
| Gprc5c | 168.3894624 | -0.161552633 | 0.196572189 | -0.821848877 |

| | | | | |
|---|---|---|---|---|
| Cd300a | 55.24117626 | -0.514573475 | 0.260491731 | -1.975392748 |
| Gm11698 | 0.946192865 | 0.873544161 | 0.968560388 | 0.901899533 |
| Cd300lb | 7.999125163 | -3.407551522 | 0.546470353 | -6.235565208 |
| Gm11699 | 0 | NA | NA | NA |
| Cd300c | 0.985558492 | 0.604763 | 2.099846967 | 0.288003369 |
| Cd300ld | 9.206469351 | -2.19288817 | 0.432562748 | -5.069526167 |
| AF251705 | 73.50869836 | -1.018909542 | 0.186598189 | -5.460447116 |
| Gm11709 | 0.932213635 | -1.563602159 | 1.267413076 | -1.233695776 |
| Gm11710 | 0.854414486 | -1.233993137 | 1.156891844 | -1.066645205 |
| Gm11711 | 0.991446059 | -2.745021856 | 1.262781967 | -2.173789243 |
| Cd300lh | 2.74015677 | -1.822795844 | 0.704959586 | -2.585674244 |
| Cd300e | 5.001744579 | -1.642051129 | 0.480740911 | -3.41566755 |
| Rab37 | 152.6386923 | 0.319004908 | 0.116677831 | 2.734066157 |
| Cd300lf | 291.2418054 | -1.149050937 | 0.316415922 | -3.631457389 |
| Slc9a3r1 | 582.5242563 | -0.310003435 | 0.139822536 | -2.21712067 |
| Nat9 | 95.23938015 | -0.228032863 | 0.134179576 | -1.699460299 |
| Tmem104 | 386.5952224 | -0.471348905 | 0.106377602 | -4.430903645 |
| Grin2c | 638.1417609 | 0.559157523 | 0.163956893 | 3.410393513 |
| Gm25837 | 0.089089296 | 0.957155977 | 3.723939561 | 0.257027796 |
| Fdxr | 101.2810521 | 0.002906692 | 0.121860115 | 0.023852692 |
| Fads6 | 408.9834072 | 0.072768587 | 0.103732281 | 0.701503778 |
| Otop2 | 12.03184821 | -0.194021597 | 0.509608884 | -0.380726481 |
| Ush1g | 18.68407142 | -0.678002344 | 0.340560354 | -1.990843431 |
| Otop3 | 2.248585396 | -1.682355072 | 0.809222103 | -2.078978151 |
| Hid1 | 846.0541876 | 0.187062617 | 0.079556743 | 2.351310649 |
| Cdr2l | 554.9718017 | -0.005610934 | 0.083268598 | -0.067383556 |
| Ict1 | 293.2357144 | 0.041190093 | 0.074190726 | 0.55519194 |
| Ict1os | 0.802129109 | -0.238171541 | 1.279670919 | -0.186119367 |
| Atp5h | 982.123738 | 0.015924323 | 0.088626837 | 0.17967834 |
| Kctd2 | 366.4040735 | 0.027887824 | 0.070562599 | 0.395221038 |
| Trim80 | 1.020577158 | 0.255354862 | 1.219823605 | 0.209337531 |
| Mir3968 | 0 | NA | NA | NA |
| Gm11695 | 0.907486357 | 1.063446896 | 1.115004334 | 0.953760325 |
| Slc16a5 | 0.45393394 | -1.149491591 | 3.33259115 | -0.344924276 |
| Armc7 | 103.9306984 | -0.344621547 | 0.136751648 | -2.520054061 |
| Nt5c | 193.1442848 | -0.097758511 | 0.086718451 | -1.127309233 |
| Gm11694 | 1.069943461 | -2.457761175 | 1.076955589 | -2.282137908 |
| Hn1 | 386.0221868 | -0.366533358 | 0.112504252 | -3.257951146 |
| Sumo2 | 322.8270685 | 0.12493494 | 0.11271811 | 1.108383908 |
| Nup85 | 257.3817318 | -0.086866856 | 0.080546565 | -1.078467537 |
| Gga3 | 719.1270991 | -0.137731344 | 0.078078389 | -1.764013642 |
| Gm25364 | 0 | NA | NA | NA |
| Mrps7 | 278.078163 | -0.38596502 | 0.103243616 | -3.738391164 |
| Mif4gd | 118.7681913 | -0.327526784 | 0.140161671 | -2.33677853 |
| Slc25a19 | 241.7616655 | -0.075249062 | 0.082165673 | -0.915821166 |
| Gm26613 | 0.042716945 | 0.037062677 | 3.724945049 | 0.009949859 |
| Grb2 | 1605.895681 | -0.113618531 | 0.059285682 | -1.916458193 |

| | | | | |
|---|---|---|---|---|
| Gm11702 | 38.96983336 | 0.293772368 | 0.191594846 | 1.533299952 |
| Gm11703 | 12.20134384 | 0.255324984 | 0.343176223 | 0.74400546 |
| Gm23699 | 0 | NA | NA | NA |
| Gm11704 | 0 | NA | NA | NA |
| 2310067B10Rik | 722.2047825 | -0.099586526 | 0.09718828 | -1.02467629 |
| Mir5621 | 0.180265876 | 0.517963268 | 2.748190349 | 0.188474306 |
| Caskin2 | 336.692886 | -0.299881552 | 0.112690753 | -2.661101682 |
| Tsen54 | 70.37095567 | -0.388566819 | 0.132087253 | -2.941743502 |
| Llgl2 | 24.64085341 | -0.095457001 | 0.219586584 | -0.434712356 |
| Myo15b | 2.674434539 | -0.919786896 | 0.887776243 | -1.036057119 |
| Recql5 | 210.8391631 | -0.190123917 | 0.154928716 | -1.227170291 |
| Smim5 | 3.739367707 | -0.228459479 | 0.497363895 | -0.459340698 |
| Smim6 | 0.320213338 | 0.037077015 | 1.762949024 | 0.021031246 |
| Recql5os1 | 0.062426873 | 0.517959912 | 3.724945049 | 0.139051692 |
| Sap30bp | 450.623263 | -0.112372605 | 0.076828246 | -1.462647022 |
| Sap30bpos | 16.7515477 | -0.825108887 | 0.292115039 | -2.824602562 |
| Itgb4 | 300.6627537 | -0.208354313 | 0.139769412 | -1.490700362 |
| B230344G16Rik | 10.95198732 | -0.162753668 | 0.328872382 | -0.494883964 |
| Galk1 | 66.2841132 | 0.132478622 | 0.175493746 | 0.754890843 |
| H3f3b | 3094.200091 | -0.209462231 | 0.170895877 | -1.225671649 |
| Unk | 347.7817187 | -0.008066145 | 0.063771649 | -0.126484817 |
| Unc13d | 61.4735399 | -1.132722751 | 0.238534418 | -4.748676363 |
| Wbp2 | 3196.52038 | 0.355596353 | 0.072671296 | 4.893216027 |
| Trim47 | 116.7135195 | -1.564273642 | 0.210522282 | -7.430442155 |
| Trim65 | 57.48636768 | -0.142531776 | 0.155587661 | -0.916086631 |
| Mrpl38 | 328.9494862 | -0.382804653 | 0.074116122 | -5.164930939 |
| Fbf1 | 655.435408 | 0.378673538 | 0.100976441 | 3.750117677 |
| Acox1 | 2125.616724 | -0.036249496 | 0.051925307 | -0.698108461 |
| Ten1 | 127.8018245 | -0.018274663 | 0.113723144 | -0.160694315 |
| Cdk3-ps | 6.063886428 | 0.471740316 | 0.440179849 | 1.071699028 |
| Evpl | 18.36159365 | -0.252415443 | 0.246134946 | -1.025516475 |
| Srp68 | 664.4255585 | -0.331637086 | 0.070562933 | -4.699876744 |
| Galr2 | 1.093534925 | 1.187062364 | 1.096643771 | 1.082450286 |
| Exoc7 | 686.1507893 | 0.04967481 | 0.050396164 | 0.985686319 |
| Gm26730 | 1.17710504 | -0.927330739 | 1.052539042 | -0.881041654 |
| n-R5s74 | 0 | NA | NA | NA |
| Foxj1 | 234.7792888 | -0.921571149 | 0.162176609 | -5.682515851 |
| Rnf157 | 2678.61245 | 0.507700687 | 0.081842343 | 6.20339877 |
| Rps11-ps2 | 3.484535615 | -0.604286709 | 0.608141093 | -0.993662023 |
| Ubald2 | 148.6109574 | 0.03456706 | 0.130781281 | 0.264311987 |
| Qrich2 | 0.384783943 | 0.070291165 | 1.545951946 | 0.045467885 |
| Gm11739 | 0 | NA | NA | NA |
| Prpsap1 | 399.2746756 | -0.256588105 | 0.074233966 | -3.456478482 |
| Rpl36-ps1 | 0 | NA | NA | NA |
| Gm29292 | 10.23524976 | -0.738270793 | 0.462796234 | -1.595239413 |
| Sphk1 | 151.140251 | -1.034958133 | 0.358353261 | -2.888094645 |
| Ube2o | 1590.155959 | 0.192521455 | 0.074642751 | 2.579238486 |

| | | | | |
|---|---|---|---|---|
| Gm11742 | 0 | NA | NA | NA |
| Aanat | 3.437531617 | 1.402648777 | 0.686549541 | 2.043040879 |
| Rhbdf2 | 162.7293829 | -1.217407415 | 0.254154772 | -4.790023834 |
| Cygb | 534.7165052 | 0.36675209 | 0.08776287 | 4.1788981 |
| Gm11744 | 17.7695872 | 0.585844707 | 0.314883069 | 1.860515108 |
| 1810032O08Rik | 43.33906545 | -0.080949008 | 0.182125191 | -0.444469035 |
| Snord1c | 0 | NA | NA | NA |
| Snord1b | 0 | NA | NA | NA |
| Snord1a | 0 | NA | NA | NA |
| St6galnac2 | 21.17771716 | 0.74146527 | 0.241359362 | 3.072038569 |
| Gm11735 | 0.061920613 | -0.443816764 | 3.724945049 | -0.119147198 |
| BC018473 | 0 | NA | NA | NA |
| St6galnac1 | 0.21290388 | -1.471117409 | 2.620256331 | -0.561440265 |
| Mxra7 | 93.40309728 | -0.19857296 | 0.137095254 | -1.448430589 |
| Jmjd6 | 264.2573602 | -0.211909579 | 0.120584735 | -1.75734995 |
| Gm27401 | 0 | NA | NA | NA |
| Mettl23 | 60.97083361 | -0.23020563 | 0.159969165 | -1.439062524 |
| Srsf2 | 1239.606384 | -0.183943742 | 0.072298464 | -2.544227523 |
| Mfsd11 | 294.976682 | -0.355435018 | 0.080251125 | -4.429034737 |
| Mgat5b | 569.5109985 | 0.492862065 | 0.11668586 | 4.223837105 |
| Gm11728 | 0.813341144 | -0.154533516 | 1.182968153 | -0.130632017 |
| Gm11731 | 0.068097359 | 0.037064902 | 3.724945049 | 0.009950456 |
| Snhg20 | 102.2173952 | 0.496325815 | 0.154328609 | 3.216032464 |
| Gm26083 | 0 | NA | NA | NA |
| mmu-mir-6516 | 0 | NA | NA | NA |
| Gm24323 | 0.154433973 | -0.443826563 | 3.50312282 | -0.126694548 |
| Gm11730 | 1.74719428 | -0.541962906 | 0.768352916 | -0.705356738 |
| Sec14l1 | 2162.074218 | 0.161398798 | 0.100837266 | 1.600586803 |
| Gm22667 | 7.017353583 | -0.457621431 | 0.572606782 | -0.79918968 |
| Gm11745 | 7.216095472 | 0.445122252 | 0.639380615 | 0.696177271 |
| 9-Sep | 1015.394361 | -0.672431113 | 0.114132955 | -5.891647274 |
| Gm16045 | 1.18260799 | 1.062544415 | 1.08426277 | 0.979969473 |
| Gm11729 | 0.295494355 | -0.941144508 | 1.811229342 | -0.519616421 |
| Gm8624 | 0.24265998 | 0.036251342 | 2.508899134 | 0.014449103 |
| Gm11732 | 4.722481653 | -0.064089793 | 0.500575306 | -0.12803227 |
| Gm11733 | 2.291093954 | -0.138963483 | 0.610980131 | -0.227443538 |
| Gm11734 | 3.524723029 | -0.679863133 | 0.562457441 | -1.208737022 |
| 2900041M22Rik | 4.279435866 | 0.391951162 | 0.533067329 | 0.735275152 |
| Tnrc6c | 1645.798208 | 0.385913006 | 0.12000879 | 3.215706172 |
| Tmc6 | 62.04352302 | -0.46655135 | 0.185779387 | -2.511319244 |
| Tmc8 | 13.88221168 | -0.567457201 | 0.283793414 | -1.999543238 |
| Gm11723 | 0.109124069 | 0.517955464 | 3.724945049 | 0.139050498 |
| Gm11724 | 0.044544648 | 0.517957688 | 3.724945049 | 0.139051095 |
| 6030468B19Rik | 0.072025462 | 0.517959912 | 3.724945049 | 0.139051692 |
| Gm20708 | 0.075043205 | 0.517957688 | 3.724945049 | 0.139051095 |
| Syngr2 | 214.0619962 | -0.69293955 | 0.143612781 | -4.825054876 |
| Tk1 | 11.63018472 | -0.416949829 | 0.362724496 | -1.149494545 |

| | | | | |
|---|---|---|---|---|
| Afmid | 13.88472388 | 0.476363427 | 0.304006725 | 1.566950294 |
| Birc5 | 15.65536197 | 0.052950378 | 0.300712378 | 0.176083133 |
| Tmem235 | 0.809771811 | -0.874949148 | 1.199653152 | -0.729335097 |
| Tha1 | 35.97253463 | -0.036757349 | 0.205176621 | -0.17914979 |
| Gm11725 | 4.41486742 | 0.330344504 | 0.543857382 | 0.607410168 |
| Gm11726 | 0.061920613 | -0.443816764 | 3.724945049 | -0.119147198 |
| Socs3 | 775.4504325 | -0.865738899 | 0.402652276 | -2.150090663 |
| Pgs1 | 896.2403674 | -0.643511663 | 0.089220987 | -7.212559343 |
| Mir6933 | 0.186399948 | 0.575238522 | 3.295513719 | 0.174552003 |
| Dnah17 | 7.417164814 | -0.514236068 | 0.368878079 | -1.394054289 |
| Gm11738 | 6.391237767 | 1.750288999 | 0.522363637 | 3.350709877 |
| Cyth1 | 1001.744073 | 0.132561894 | 0.059727864 | 2.21943135 |
| Gm24060 | 17.80235189 | 0.877361183 | 0.294549303 | 2.978656457 |
| Gm11737 | 8.112751975 | -0.151489347 | 0.48662499 | -0.31130614 |
| Usp36 | 634.1888172 | 0.045016668 | 0.151913906 | 0.296330134 |
| Timp2 | 2901.777889 | 0.05589899 | 0.066046825 | 0.846353933 |
| BC100451 | 15.86145379 | 0.041068727 | 0.354098223 | 0.115981174 |
| Lgals3bp | 4154.63823 | -0.371554291 | 0.167840835 | -2.213730002 |
| Cant1 | 623.271791 | -0.205720623 | 0.060542589 | -3.397948865 |
| C1qtnf1 | 107.3234082 | -0.291264629 | 0.191574637 | -1.520371553 |
| Gm11747 | 0.291849543 | 0.037516573 | 1.778128177 | 0.021098914 |
| Gm11748 | 1.545561197 | -0.801146315 | 0.856446775 | -0.93543036 |
| Engase | 86.50548769 | -0.010129091 | 0.147146536 | -0.068836765 |
| Rbfox3 | 2325.205722 | 0.531014274 | 0.104691142 | 5.072198709 |
| Gm11750 | 0.497147205 | -1.613518596 | 1.565737767 | -1.030516495 |
| Gm25040 | 1.127514739 | 1.749645021 | 1.187826957 | 1.472979722 |
| Gm11749 | 17.63452542 | 0.122081743 | 0.371824747 | 0.32833141 |
| Enpp7 | 0.038981395 | -0.443816764 | 3.724945049 | -0.119147198 |
| Cbx2 | 46.53041809 | 0.616793143 | 0.253494789 | 2.433159064 |
| Cbx8 | 139.4173976 | 0.064090167 | 0.12029356 | 0.532781365 |
| Gm26508 | 4.027668366 | -0.381436591 | 0.562177308 | -0.678498733 |
| Cbx4 | 968.7534903 | 0.367010356 | 0.10249648 | 3.580711809 |
| Gm11754 | 0.08486385 | 0.037730769 | 3.724898049 | 0.010129343 |
| Gm11755 | 2.180156761 | 0.872947981 | 0.86156929 | 1.013206937 |
| Tbc1d16 | 764.1409362 | 0.500502252 | 0.121858913 | 4.107227297 |
| Gm26888 | 0.416129427 | -0.860330983 | 1.622998797 | -0.530087259 |
| Mir6934 | 0 | NA | NA | NA |
| Gm11753 | 1.914943955 | -0.588588286 | 0.831165892 | -0.708147786 |
| Gm11752 | 0.044191549 | 0.037062677 | 3.724945049 | 0.009949859 |
| Ccdc40 | 32.62461688 | -0.754986821 | 0.222083363 | -3.399564963 |
| Gaa | 3712.581302 | 0.181586263 | 0.04620598 | 3.92992994 |
| Eif4a3 | 482.0859025 | -0.245558966 | 0.085982486 | -2.855918423 |
| Card14 | 10.20988058 | -0.419606633 | 0.361027989 | -1.162255131 |
| Sgsh | 132.7659391 | -0.418903275 | 0.135217267 | -3.098001337 |
| Slc26a11 | 53.44579997 | -0.405075776 | 0.169110614 | -2.395330287 |
| Mir1932 | 5.939787235 | -0.285153499 | 0.433360838 | -0.658004772 |
| Rnf213 | 13237.76603 | -0.738856651 | 0.161639906 | -4.571003966 |

| | | | | |
|---|---|---|---|---|
| Endov | 287.30136 | -0.196617744 | 0.130327367 | -1.508645103 |
| Nptx1 | 4410.742837 | 0.217503312 | 0.106373838 | 2.044706818 |
| Gm11762 | 16.8669638 | -0.645257444 | 0.294259976 | -2.192814162 |
| Rptor | 1323.078836 | -0.045275751 | 0.040767116 | -1.110594897 |
| Gm11761 | 0 | NA | NA | NA |
| Gm23663 | 1.256631142 | -1.511379105 | 1.016909442 | -1.486247489 |
| Rptoros | 0.455959994 | -0.533129073 | 1.51578562 | -0.351717991 |
| Chmp6 | 176.3066057 | -0.037032257 | 0.109174273 | -0.339203151 |
| Gm11767 | 18.81677128 | -0.036304882 | 0.339931727 | -0.106800511 |
| Gm11766 | 0.898050108 | 0.011822015 | 1.168230728 | 0.010119589 |
| Baiap2 | 2183.791868 | 0.299695488 | 0.134609133 | 2.226412736 |
| Aatk | 3564.404415 | 0.251995863 | 0.046471822 | 5.422551868 |
| Mir338 | 4.047384828 | 0.803334931 | 0.605161542 | 1.327471883 |
| mmu-mir-3065 | 0.136882968 | -0.87044322 | 3.630339257 | -0.239769112 |
| Cep131 | 180.3487011 | 0.48106964 | 0.177371474 | 2.712215378 |
| Gm25948 | 0.585001534 | 0.86441412 | 1.430977571 | 0.604072445 |
| Enthd2 | 135.2998032 | -0.27172254 | 0.125944243 | -2.15748282 |
| 1810043H04Rik | 75.99610051 | -0.167477235 | 0.191803248 | -0.873172044 |
| Slc38a10 | 947.089944 | -0.272628108 | 0.055630165 | -4.900724432 |
| Gm11769 | 0.670691275 | -0.638621539 | 1.180808643 | -0.540834066 |
| 2810410L24Rik | 42.09187015 | 0.51632084 | 0.211650937 | 2.439492337 |
| Gm11770 | 0.567693313 | 0.966007418 | 1.240660538 | 0.778623474 |
| 2900052L18Rik | 38.54445378 | 0.693907976 | 0.28663499 | 2.420876723 |
| Bahcc1 | 551.7191568 | 0.305451899 | 0.097332048 | 3.138245872 |
| Gm11772 | 31.98353754 | 0.829447385 | 0.315125942 | 2.632113943 |
| AL928567.1 | 0 | NA | NA | NA |
| Actg1 | 4258.481376 | -0.307178121 | 0.116535379 | -2.635921597 |
| Gm23419 | 0 | NA | NA | NA |
| Mir6935 | 0 | NA | NA | NA |
| 0610009L18Rik | 11.33403993 | 0.191512711 | 0.333586295 | 0.574102455 |
| Fscn2 | 1.822354792 | 2.057573957 | 0.935411646 | 2.199645435 |
| 2310003H01Rik | 147.9010322 | -0.175130654 | 0.113808351 | -1.53882077 |
| Nploc4 | 1106.700852 | -0.367286298 | 0.076379831 | -4.808681727 |
| Tspan10 | 1.198353109 | -0.213129636 | 0.89990827 | -0.236834846 |
| Pde6g | 3.767627917 | -1.034253839 | 0.585476016 | -1.766517861 |
| Oxld1 | 37.49257177 | 0.172692514 | 0.235272447 | 0.734010788 |
| Ccdc137 | 228.8428678 | -0.545342517 | 0.125901623 | -4.331497115 |
| Arl16 | 155.9283109 | 0.151345971 | 0.110325168 | 1.371817271 |
| Hgs | 966.1986603 | -0.316599227 | 0.062108589 | -5.097511195 |
| Mrpl12 | 411.1473201 | -0.193385792 | 0.122282478 | -1.58146773 |
| Gm16755 | 0.616632713 | 0.914523748 | 1.275455065 | 0.717017614 |
| Gm11788 | 0.154696864 | 0.037074681 | 2.833481357 | 0.013084498 |
| Slc25a10 | 142.1098057 | 0.098300356 | 0.124507149 | 0.78951576 |
| Gm11789 | 0.038981395 | -0.443816764 | 3.724945049 | -0.119147198 |
| Gcgr | 1.708960799 | -0.589024639 | 0.790721615 | -0.744920371 |
| Fam195b | 400.1399544 | -0.198122148 | 0.100319319 | -1.974915199 |
| Ppp1r27 | 0.150086409 | 1.027495679 | 3.723011807 | 0.275985071 |

| | | | | |
|---|---|---|---|---|
| P4hb | 1498.417556 | -0.571298511 | 0.103363079 | -5.527104219 |
| Arhgdia | 2780.717337 | -0.369746074 | 0.095204281 | -3.883712699 |
| Gm3807 | 3.342078322 | -0.672831107 | 0.606150654 | -1.110006403 |
| Alyref | 326.2916262 | 0.015519772 | 0.142155353 | 0.109174729 |
| Anapc11 | 300.5463482 | -0.011730213 | 0.111313052 | -0.105380395 |
| Npb | 4.91649906 | 0.266862387 | 0.458845282 | 0.581595579 |
| Pcyt2 | 527.1846474 | 0.023554595 | 0.076875962 | 0.3063974 |
| Sirt7 | 232.8371362 | -0.352287674 | 0.114235824 | -3.083863378 |
| Mir6936 | 0 | NA | NA | NA |
| Mafg | 947.7400382 | -0.311937382 | 0.087653944 | -3.558737539 |
| AL663030.1 | 0 | NA | NA | NA |
| Gm17586 | 36.02489538 | 0.368197205 | 0.201116352 | 1.830767127 |
| Pycr1 | 25.50189758 | -0.478135892 | 0.226293626 | -2.112900395 |
| Myadml2 | 26.50837914 | 0.347998465 | 0.222651337 | 1.562974961 |
| Myadml2os | 1.355090691 | 2.21038291 | 1.168593283 | 1.891490343 |
| Notum | 9.73076196 | -0.575396103 | 0.390462688 | -1.473626344 |
| Notumos | 2.072861717 | 0.480639682 | 0.742840564 | 0.647029397 |
| Aspscr1 | 368.7567786 | -0.068974773 | 0.080343055 | -0.85850325 |
| Gm17178 | 0.221007548 | 0.973136931 | 2.177307151 | 0.446945177 |
| Stra13 | 57.79830129 | -0.067079103 | 0.178495613 | -0.375802529 |
| Lrrc45 | 209.0769499 | 0.166144239 | 0.208372905 | 0.797340898 |
| Rac3 | 88.08912298 | 0.070731841 | 0.144691062 | 0.488847344 |
| Dcxr | 57.27746725 | -0.014548876 | 0.194798773 | -0.07468669 |
| Cbr2 | 7.287723491 | -0.529035489 | 0.679913312 | -0.778092559 |
| Hmga1-rs1 | 524.4408412 | -1.298975792 | 0.259920664 | -4.997585702 |
| Gm11771 | 0 | NA | NA | NA |
| Rfng | 389.6471229 | 0.0945075 | 0.070550459 | 1.339573131 |
| Gps1 | 549.026855 | -0.174929162 | 0.085944693 | -2.035368972 |
| Dus1l | 255.7077501 | -0.149515774 | 0.090144086 | -1.658630981 |
| Fasn | 4709.556278 | 0.077517 | 0.085654949 | 0.904991493 |
| Ccdc57 | 96.87917321 | 0.394432298 | 0.21607031 | 1.82548124 |
| Gm11773 | 0.216076386 | 1.49879282 | 3.709588964 | 0.404032046 |
| Slc16a3 | 72.70316503 | -1.039418846 | 0.226821254 | -4.582546074 |
| Csnk1d | 1494.74923 | -0.127709397 | 0.057217422 | -2.232001946 |
| Gm28192 | 0 | NA | NA | NA |
| Gm11774 | 0.066050875 | -0.443818988 | 3.724945049 | -0.119147795 |
| Gm11775 | 0.090277459 | -0.443823436 | 3.724945049 | -0.119148989 |
| Gm22152 | 0.044191549 | 0.037062677 | 3.724945049 | 0.009949859 |
| Cd7 | 13.98101131 | -0.014191815 | 0.297975975 | -0.04762738 |
| Sectm1b | 0.607868506 | -2.046048621 | 1.37075202 | -1.492646803 |
| Sectm1a | 17.65934462 | -2.018507332 | 0.402768835 | -5.011577752 |
| Gm11792 | 0 | NA | NA | NA |
| Gm11791 | 0 | NA | NA | NA |
| Gm11793 | 1.517674278 | -0.315399716 | 0.741505057 | -0.425350729 |
| Tex19.2 | 0 | NA | NA | NA |
| Tex19.1 | 0.038981395 | -0.443816764 | 3.724945049 | -0.119147198 |
| Uts2r | 2.195192171 | 0.271483993 | 0.620247902 | 0.437702396 |

| | | | | |
|---|---|---|---|---|
| Gm11790 | 0.802756328 | 3.491327211 | 1.22932969 | 2.840025129 |
| Ogfod3 | 69.70424457 | -0.033204162 | 0.158571603 | -0.209395384 |
| Hexdc | 139.714264 | 0.322057342 | 0.113751212 | 2.831243168 |
| BC017643 | 297.1303026 | -0.586472869 | 0.124232631 | -4.7207635 |
| Narf | 577.2846721 | 0.329129055 | 0.096030515 | 3.427338217 |
| Foxk2 | 1306.171416 | -0.11800883 | 0.049072153 | -2.404802369 |
| Wdr45b | 650.5471699 | -0.201039557 | 0.077048842 | -2.609248273 |
| Rab40b | 176.4499084 | 0.447204012 | 0.161648442 | 2.766522258 |
| Mir7236 | 2.550224353 | 0.452414782 | 0.745093922 | 0.607191616 |
| Gm11866 | 0 | NA | NA | NA |
| Fn3krp | 434.4560457 | 0.005971872 | 0.105168466 | 0.056783865 |
| Fn3k | 227.4287037 | 0.424779052 | 0.133684619 | 3.177471397 |
| Tbcd | 616.9440393 | -0.015427462 | 0.063236323 | -0.24396519 |
| Zfp750 | 17.50032145 | -0.078624266 | 0.285442785 | -0.275446674 |
| B3gntl1 | 72.02965567 | -0.035428868 | 0.180481783 | -0.196301629 |
| Metrnl | 107.2840231 | -1.298709616 | 0.204593382 | -6.347759642 |
| Gm12589 | 0.398050276 | -1.099727112 | 1.590329138 | -0.691509126 |
| Gm29513 | 0 | NA | NA | NA |
| Gm15643 | 0 | NA | NA | NA |
| Ptchd3 | 0 | NA | NA | NA |
| Gm12586 | 0 | NA | NA | NA |
| Gm12590 | 0 | NA | NA | NA |
| Gm12587 | 0 | NA | NA | NA |
| Gm12591 | 0 | NA | NA | NA |
| Gm9071 | 0 | NA | NA | NA |
| Gm9075 | 0 | NA | NA | NA |
| Rab10os | 387.6915989 | 0.193005841 | 0.100446067 | 1.921487286 |
| Rab10 | 2113.636369 | -0.21541095 | 0.078424735 | -2.74672208 |
| Gm26520 | 3.045321195 | 1.217494214 | 0.643296873 | 1.892585312 |
| Kif3c | 2815.319838 | 0.250625926 | 0.060244557 | 4.160142202 |
| 1110002L01Rik | 127.452233 | -0.581934221 | 0.117548579 | -4.950584903 |
| Asxl2 | 1098.710745 | 0.074081629 | 0.076418918 | 0.969414791 |
| Dtnb | 785.8508435 | 0.457800462 | 0.086679014 | 5.281560582 |
| Gm26050 | 2.458093412 | 0.04572811 | 0.842497018 | 0.05427688 |
| Dtnbos | 0.849323088 | 1.048250846 | 1.112819265 | 0.941977622 |
| Dnmt3a | 1911.391238 | 0.005039977 | 0.092383426 | 0.054554986 |
| Dnmt3aos | 1.674671121 | -1.515281464 | 0.807477811 | -1.876561118 |
| Pomc | 33.59521228 | -1.782705751 | 0.91551336 | -1.947219811 |
| Efr3b | 2749.842645 | 0.045397351 | 0.054893499 | 0.82700779 |
| Dnajc27 | 1474.757923 | 0.394037997 | 0.065311978 | 6.033165914 |
| Adcy3 | 597.0799168 | -0.058099201 | 0.081721095 | -0.710944967 |
| Cenpo | 63.78631368 | -0.302515243 | 0.138487223 | -2.184427094 |
| Ptrhd1 | 115.3757405 | 0.335818618 | 0.114211473 | 2.940322974 |
| Ncoa1 | 2595.887549 | 0.216126244 | 0.053246818 | 4.058951355 |
| Gm23756 | 0 | NA | NA | NA |
| Gm22343 | 0 | NA | NA | NA |
| Gm3625 | 2.933921127 | 0.442262075 | 0.585728037 | 0.755063864 |

| | | | | |
|---|---|---|---|---|
| Itsn2 | 887.6567088 | -0.138702963 | 0.06518949 | -2.127689026 |
| Gm17541 | 0.106465261 | 0.037073798 | 3.724945049 | 0.009952844 |
| Fam228a | 28.71218686 | 0.631267999 | 0.357741648 | 1.764591855 |
| Fam228b | 74.82727959 | 0.532686064 | 0.195639349 | 2.722796132 |
| Gm23676 | 0 | NA | NA | NA |
| Gm22003 | 0 | NA | NA | NA |
| Pfn4 | 43.32535029 | 0.895808705 | 0.184450019 | 4.856647396 |
| Gm6682 | 21.88487653 | -0.466652901 | 0.223651505 | -2.086518043 |
| Sf3b6 | 380.7093426 | -0.21825184 | 0.146169798 | -1.493139101 |
| Fkbp1b | 121.655647 | 0.430030651 | 0.128901306 | 3.33612331 |
| BC068281 | 110.397684 | -0.42052642 | 0.138950192 | -3.026454398 |
| Mfsd2b | 23.20891546 | 0.505931951 | 0.41029089 | 1.233105496 |
| Ubxn2a | 760.0083217 | 0.042939721 | 0.079338975 | 0.5412185 |
| Atad2b | 541.029005 | 0.157018358 | 0.093785573 | 1.674227203 |
| Gm23167 | 0 | NA | NA | NA |
| Klhl29 | 743.3499042 | 0.084346982 | 0.121348298 | 0.695081706 |
| 2810032G03Rik | 93.60943587 | 0.248079103 | 0.119422477 | 2.077323374 |
| Gm5652 | 0.048949532 | 0.037064902 | 3.724945049 | 0.009950456 |
| Gm23752 | 0.431854921 | 1.540583579 | 1.529594603 | 1.007184241 |
| 1700101O22Rik | 0.245087052 | 0.517974584 | 2.095767594 | 0.247152683 |
| Gm9144 | 9.249347624 | -0.180178113 | 0.349210727 | -0.515958127 |
| Apob | 11.59973813 | 1.395491919 | 0.452836505 | 3.081668339 |
| Ldah | 329.5450486 | -0.45111005 | 0.112512404 | -4.009425062 |
| Gdf7 | 3.829800073 | 1.161328235 | 0.597778241 | 1.942740896 |
| Hs1bp3 | 177.2730511 | -1.022641968 | 0.115168196 | -8.879551892 |
| Rhob | 3017.567704 | 0.129245137 | 0.115004285 | 1.123828879 |
| Slc7a15 | 0.061920613 | -0.443816764 | 3.724945049 | -0.119147198 |
| Pum2 | 3606.55936 | -0.115175671 | 0.066450834 | -1.733246414 |
| Gm4755 | 0.187019432 | 0.037072891 | 2.718962484 | 0.01363494 |
| Sdc1 | 95.2175769 | -1.820638713 | 0.228805298 | -7.957152764 |
| 9930038B18Rik | 0.162099466 | 0.984742474 | 2.787845923 | 0.353227008 |
| Laptm4a | 2359.716549 | -0.355781181 | 0.118961274 | -2.990731093 |
| Matn3 | 0.908179219 | 0.225035691 | 1.055715221 | 0.213159464 |
| Wdr35 | 500.8882983 | 0.068179234 | 0.072047051 | 0.946315399 |
| Ttc32 | 25.69466073 | -0.451300841 | 0.229384075 | -1.967446259 |
| Gm7099 | 8.593693136 | -0.548302099 | 0.435122705 | -1.260109144 |
| 1700022H16Rik | 0 | NA | NA | NA |
| Osr1 | 21.77471978 | 0.124024173 | 0.290346601 | 0.427159033 |
| Gm23979 | 0 | NA | NA | NA |
| Gm22845 | 0 | NA | NA | NA |
| Nt5c1b | 1.964636517 | -0.035234478 | 0.914009043 | -0.038549376 |
| Rdh14 | 214.0479375 | 0.04127882 | 0.107446793 | 0.384179172 |
| Pgk1-rs7 | 21.9025282 | 0.027935385 | 0.234707193 | 0.119022279 |
| Kcns3 | 56.27639958 | 0.136459437 | 0.175898857 | 0.775783533 |
| Msgn1 | 0 | NA | NA | NA |
| 1700034J04Rik | 0.087261593 | 0.51795324 | 3.724945049 | 0.139049901 |
| Gen1 | 22.65896892 | -0.176907532 | 0.284929862 | -0.620880978 |

| | | | | |
|---|---|---|---|---|
| Smc6 | 783.5087835 | -0.149410435 | 0.065668333 | -2.275228075 |
| Mir7675 | 1.906635116 | 0.348393793 | 0.744332949 | 0.468061763 |
| Vsnl1 | 3374.486255 | 0.671201702 | 0.084339103 | 7.958369032 |
| 4930511A02Rik | 1.096741448 | -1.086850524 | 1.030152977 | -1.055037988 |
| Rad51ap2 | 31.8005175 | 0.952009543 | 0.274002721 | 3.474452878 |
| Gm23781 | 0 | NA | NA | NA |
| Gm25610 | 0.211102819 | -0.470949454 | 2.622609133 | -0.179572872 |
| Gm4294 | 93.43490713 | -0.41419389 | 0.181671366 | -2.279907392 |
| Gm26171 | 0 | NA | NA | NA |
| Fam49a | 2242.155381 | 0.334084467 | 0.073561433 | 4.541570921 |
| Gm37414 | 0.066050875 | -0.443818988 | 3.724945049 | -0.119147795 |
| Gm27952 | 0 | NA | NA | NA |
| Gm25538 | 0 | NA | NA | NA |
| Platr19 | 0 | NA | NA | NA |
| 4930519A11Rik | 0.060174537 | 0.517957688 | 3.724945049 | 0.139051095 |
| Rpl36-ps3 | 0.314164845 | 0.03819394 | 1.637608449 | 0.023322999 |
| Mycn | 47.66468018 | 0.252434855 | 0.209853147 | 1.202911935 |
| Gm24208 | 0.042716945 | 0.037062677 | 3.724945049 | 0.009949859 |
| Ddx1 | 1202.104352 | -0.022888 | 0.059132549 | -0.387062642 |
| Nbas | 1083.615517 | -0.143843752 | 0.052307953 | -2.749940366 |
| Gm6736 | 3.145663366 | 0.207140283 | 0.519802714 | 0.398497886 |
| Gm24142 | 0 | NA | NA | NA |
| Gm22170 | 0.191089262 | 0.517966625 | 2.650365354 | 0.19543216 |
| Gm4929 | 0.355586071 | -0.982119019 | 1.990769217 | -0.49333645 |
| Fam84a | 632.1102194 | 0.573245211 | 0.118443292 | 4.839828412 |
| Gm9847 | 0.673365293 | -1.265324735 | 1.199647261 | -1.054747322 |
| Gm25414 | 0.044191549 | 0.037062677 | 3.724945049 | 0.009949859 |
| Gm26449 | 0 | NA | NA | NA |
| Gm27682 | 0.049553047 | -0.443818988 | 3.724945049 | -0.119147795 |
| Gm4804 | 2.483041552 | 0.659091213 | 0.739728841 | 0.890990287 |
| Trib2 | 460.7235091 | -0.123353761 | 0.151154128 | -0.816079345 |
| Mir6387 | 0.891236725 | 0.05299838 | 1.292239841 | 0.041012805 |
| Gm5432 | 2.197748528 | -0.927895271 | 0.729505189 | -1.271951571 |
| Lpin1 | 478.9431543 | 0.328250846 | 0.110355069 | 2.974497214 |
| Ntsr2 | 425.5275216 | 1.047734454 | 0.168759144 | 6.208460351 |
| Greb1 | 74.66542959 | -0.386078835 | 0.15272569 | -2.527923335 |
| AC122228.1 | 0 | NA | NA | NA |
| E2f6 | 314.0510662 | -0.367198021 | 0.093955648 | -3.908205945 |
| Rock2 | 3956.78452 | 0.110668508 | 0.076879419 | 1.439507604 |
| Gm22303 | 1.492607829 | 0.324442363 | 0.867783922 | 0.373874595 |
| Pqlc3 | 54.75605864 | -0.814700823 | 0.205283363 | -3.96866464 |
| 2410004P03Rik | 23.25089268 | -0.725087641 | 0.280992979 | -2.580447543 |
| Kcnf1 | 695.0270978 | 0.501750879 | 0.159378269 | 3.148176237 |
| Pdia6 | 1198.63775 | -0.765233673 | 0.118676012 | -6.44809056 |
| Atp6v1c2 | 10.10846665 | 1.57987504 | 0.409502826 | 3.858032079 |
| Nol10 | 367.5541281 | 0.159964566 | 0.125770658 | 1.271875084 |
| Mir3066 | 1.524462984 | -0.758469998 | 0.863522461 | -0.878344262 |

| | | | | |
|---|---|---|---|---|
| Odc1 | 613.484201 | -0.822074597 | 0.16843345 | -4.880708657 |
| Gm22748 | 7.258283802 | -0.60662081 | 0.432821437 | -1.401549825 |
| Gm25925 | 0 | NA | NA | NA |
| Hpcal1 | 649.1744098 | 0.056051456 | 0.142989586 | 0.391996769 |
| B430203G13Rik | 10.4805672 | 0.200297767 | 0.390178799 | 0.513348669 |
| Gm10480 | 62.44942269 | -0.014235145 | 0.147537055 | -0.096485217 |
| 5730507C01Rik | 3.246809683 | 0.291825619 | 0.653243251 | 0.446733462 |
| Gm3944 | 0.500379391 | -1.545768515 | 1.469483928 | -1.051912502 |
| Gm5784 | 1.293338167 | 0.501012363 | 1.07409231 | 0.466451867 |
| Gm6802 | 0 | NA | NA | NA |
| Gm9257 | 0.060174537 | 0.517957688 | 3.724945049 | 0.139051095 |
| Gm22189 | 0 | NA | NA | NA |
| Gm21863 | 0.047948698 | 0.037064902 | 3.724945049 | 0.009950456 |
| Gm3993 | 0.060174537 | 0.517957688 | 3.724945049 | 0.139051095 |
| Gm10479 | 0.270569244 | -0.90939633 | 2.073291382 | -0.438624468 |
| 1700030C10Rik | 18.72189217 | 0.647049526 | 0.271471693 | 2.383488015 |
| Gm28503 | 0 | NA | NA | NA |
| 3110053B16Rik | 90.02746546 | 0.283806669 | 0.162830641 | 1.742956165 |
| Gm10478 | 5.792289935 | -0.654736374 | 0.520426548 | -1.258076429 |
| Zfp125 | 38.28518153 | -0.258005648 | 0.195071266 | -1.322622511 |
| Gm21989 | 3.535470757 | 0.545574326 | 0.624715349 | 0.873316665 |
| Gm5977 | 1.165717276 | 1.248176966 | 1.203895635 | 1.036781702 |
| Asap2 | 769.9183214 | -0.014546534 | 0.086944591 | -0.167308094 |
| Itgb1bp1 | 249.9318875 | 0.001881192 | 0.094392858 | 0.019929394 |
| Gm20535 | 1.863766393 | -1.116611188 | 0.975781645 | -1.144324854 |
| Cpsf3 | 408.1523475 | -0.115616347 | 0.069537466 | -1.662648264 |
| Gm25821 | 2.089596347 | 0.020090874 | 0.690289335 | 0.029105005 |
| Iah1 | 73.05919511 | 0.085470948 | 0.156021408 | 0.547815514 |
| Adam17 | 298.4801547 | -0.528413372 | 0.117335756 | -4.503430063 |
| Gm22766 | 0 | NA | NA | NA |
| Ywhaq | 1407.12046 | 0.134023763 | 0.0760068 | 1.763312788 |
| Gm4419 | 4.781372743 | 0.134071567 | 0.501745118 | 0.267210507 |
| Gm5633 | 5.363150736 | -0.141465728 | 0.485453338 | -0.291409528 |
| Gm22908 | 0 | NA | NA | NA |
| Gm4425 | 1.228836612 | 0.385932403 | 0.952213693 | 0.405300203 |
| Gm20486 | 0 | NA | NA | NA |
| Gm6838 | 0.045882455 | 0.517959912 | 3.724945049 | 0.139051692 |
| Gm10476 | 0.988361678 | 0.256439636 | 1.209012287 | 0.212106724 |
| Gm17391 | 0.141071297 | -0.443820801 | 3.564043909 | -0.12452731 |
| 2410018L13Rik | 0.412847196 | -1.266313588 | 1.586241459 | -0.798310737 |
| Gm26140 | 0 | NA | NA | NA |
| Gm20472 | 0.391014225 | -0.745444776 | 1.433138159 | -0.520148578 |
| Gm20473 | 0 | NA | NA | NA |
| Gm9285 | 0.258012988 | 1.539579558 | 2.115178261 | 0.727872249 |
| Gm9286 | 0 | NA | NA | NA |
| Gm20474 | 0.284703419 | -0.454650177 | 2.010683822 | -0.22611719 |
| Gm10330 | 0.04393175 | 0.037062677 | 3.724945049 | 0.009949859 |

| | | | | |
|---|---|---|---|---|
| Gm17330 | 0 | NA | NA | NA |
| 9030624G23Rik | 0.15672128 | 0.037076017 | 2.821957958 | 0.013138402 |
| Gm9312 | 0.046345709 | -0.443818988 | 3.724945049 | -0.119147795 |
| Gm16372 | 0.480740522 | -0.560416476 | 1.518913374 | -0.368958813 |
| Taf1b | 209.6766852 | 0.004694212 | 0.087050861 | 0.05392493 |
| Grhl1 | 110.9007615 | 0.450626406 | 0.172330616 | 2.614894644 |
| Gm6969 | 32.27026886 | 0.067701445 | 0.303068611 | 0.223386528 |
| Klf11 | 102.5281956 | -0.059798274 | 0.154765458 | -0.386379979 |
| Cys1 | 49.24944908 | 0.697329134 | 0.251122725 | 2.776845996 |
| Rrm2 | 38.30661462 | -1.019000233 | 0.400853507 | -2.542076384 |
| Gm26211 | 0 | NA | NA | NA |
| Mboat2 | 425.7860528 | 0.344081781 | 0.105725492 | 3.254482683 |
| Kidins220 | 4110.912429 | 0.315596508 | 0.057302938 | 5.50751009 |
| Id2 | 1035.883774 | 0.112648462 | 0.148386059 | 0.759157984 |
| Gm38052 | 0 | NA | NA | NA |
| Gm22527 | 0 | NA | NA | NA |
| Rnf144a | 371.6551165 | 0.112826076 | 0.091772761 | 1.229407013 |
| Mir6538 | 1.774752459 | 1.17623688 | 0.999963976 | 1.176279253 |
| Rsad2 | 3584.881863 | -1.087107917 | 0.397346757 | -2.735917426 |
| Cmpk2 | 2770.91634 | -0.557737027 | 0.264532823 | -2.108384963 |
| Gm9866 | 90.23738068 | 0.46753537 | 0.145299876 | 3.217727243 |
| Sox11 | 612.1246314 | -0.347200987 | 0.126457006 | -2.745604997 |
| Gm24326 | 0 | NA | NA | NA |
| Gm25923 | 0 | NA | NA | NA |
| Gm25564 | 0 | NA | NA | NA |
| Dcdc2c | 0.829077587 | 0.873836052 | 1.379188181 | 0.633587254 |
| 1700022F17Rik | 0.238210844 | -0.982796769 | 3.037335205 | -0.323572047 |
| Allc | 2.669291208 | 1.641902552 | 0.839352815 | 1.956153028 |
| Colec11 | 3.087887419 | 0.007326179 | 0.526365373 | 0.013918428 |
| Rps7 | 1086.152334 | -0.179455662 | 0.124659605 | -1.439565465 |
| Rnaseh1 | 69.48037499 | -0.092650865 | 0.162783694 | -0.569165516 |
| Adi1 | 240.4881119 | -0.002357193 | 0.11228932 | -0.020992142 |
| Mir6937 | 0 | NA | NA | NA |
| Trappc12 | 622.1510625 | 0.142499741 | 0.081595056 | 1.746426166 |
| Tssc1 | 236.185517 | 0.051357966 | 0.083223484 | 0.617109062 |
| Gm23342 | 0 | NA | NA | NA |
| Myt1l | 2705.158939 | 0.687023969 | 0.127119668 | 5.404545017 |
| Pxdn | 585.0750425 | 0.116934217 | 0.184144546 | 0.635013196 |
| Tpo | 6.12441587 | 0.293341529 | 0.428253143 | 0.684972275 |
| Gm15691 | 3.03177422 | 2.178907999 | 0.742561819 | 2.934311925 |
| Sntg2 | 50.41524398 | 0.471475878 | 0.20257702 | 2.327390729 |
| 2310016D03Rik | 0.251964224 | -0.466265384 | 2.081917787 | -0.223959556 |
| Gm25865 | 0 | NA | NA | NA |
| Gm28806 | 0 | NA | NA | NA |
| Gm25243 | 0.038981395 | -0.443816764 | 3.724945049 | -0.119147198 |
| Tmem18 | 291.1926203 | -0.029415008 | 0.090276764 | -0.325831445 |
| Fam150b | 13.83564331 | -1.360099539 | 0.312528508 | -4.351921522 |

| | | | | |
|---|---|---|---|---|
| Acp1 | 179.9360824 | 0.017055585 | 0.10356866 | 0.164679019 |
| Sh3yl1 | 69.38233294 | 0.027853383 | 0.14519519 | 0.191834059 |
| Gm25408 | 0 | NA | NA | NA |
| Fam110c | 3.004281714 | -1.49463952 | 0.60658733 | -2.464013747 |
| 6030469F06Rik | 0.048949532 | 0.037064902 | 3.724945049 | 0.009950456 |
| Gm24613 | 0.553521388 | 1.013928019 | 1.333209655 | 0.760516559 |
| Lamb1 | 296.7992853 | 0.166963342 | 0.114233861 | 1.4615924 |
| Dld | 1183.816382 | -0.042712465 | 0.084670441 | -0.504455443 |
| Slc26a3 | 2.221929775 | 1.2339091 | 0.746267381 | 1.653441021 |
| Gm9359 | 0 | NA | NA | NA |
| Cbll1 | 375.0362158 | -0.09992978 | 0.07383918 | -1.353343579 |
| Gm29542 | 0 | NA | NA | NA |
| Gm29541 | 0 | NA | NA | NA |
| Slc26a4 | 32.11330933 | -0.415329728 | 0.239597143 | -1.733450253 |
| Bcap29 | 377.0266182 | 0.134934165 | 0.089642101 | 1.505254372 |
| Dus4l | 47.02044193 | -0.651552586 | 0.179902521 | -3.621697922 |
| Cog5 | 397.0078528 | 0.086236212 | 0.078374809 | 1.10030522 |
| Gpr22 | 1006.691692 | 0.235523547 | 0.101437809 | 2.321851673 |
| Gm25913 | 1.857927479 | 0.558443693 | 0.800466023 | 0.697648216 |
| Hbp1 | 434.1401043 | 0.18103074 | 0.104152733 | 1.738127601 |
| Prkar2b | 606.5567347 | 0.296156515 | 0.100106065 | 2.9584273 |
| Pik3cg | 109.1690308 | 0.064713987 | 0.114631421 | 0.564539689 |
| Gm23875 | 0 | NA | NA | NA |
| Ccdc71l | 284.7411437 | -0.497965792 | 0.122723062 | -4.057638256 |
| Nampt | 2161.206624 | -0.176140091 | 0.122513941 | -1.43771467 |
| Gdap10 | 195.362713 | -0.02515797 | 0.191042346 | -0.131687925 |
| 4933406C10Rik | 19.31754882 | 1.126203447 | 0.312601581 | 3.602679941 |
| Sypl | 610.6402324 | -0.168099977 | 0.087370508 | -1.923989916 |
| Gm24922 | 0 | NA | NA | NA |
| Cdhr3 | 9.236620993 | -0.093593502 | 0.349512076 | -0.267783314 |
| Gm22904 | 0.044544648 | 0.517957688 | 3.724945049 | 0.139051095 |
| F730043M19Rik | 202.5005712 | 0.073281028 | 0.141324162 | 0.518531487 |
| Gm16267 | 0 | NA | NA | NA |
| Atxn7l1 | 715.5692364 | 0.121826622 | 0.087854163 | 1.386691507 |
| Atxn7l1os1 | 0.109124069 | 0.517955464 | 3.724945049 | 0.139050498 |
| Atxn7l1os2 | 0 | NA | NA | NA |
| Efcab10 | 8.606127557 | 0.507970949 | 0.373839861 | 1.3587929 |
| Twistnb | 221.7759397 | -0.187255086 | 0.095203897 | -1.966884685 |
| Gm23529 | 0 | NA | NA | NA |
| Ferd3l | 0.671239363 | -1.629881106 | 1.390221869 | -1.1723892 |
| Twist1 | 6.585054892 | -1.035649258 | 0.413710181 | -2.503320697 |
| Gm18025 | 5.060744911 | -0.701124654 | 0.516879028 | -1.356457927 |
| Hdac9 | 424.2109994 | 0.381038145 | 0.131067368 | 2.907193069 |
| 4921508M14Rik | 2.520821878 | 0.275774789 | 0.705554088 | 0.390862718 |
| Gm24429 | 0 | NA | NA | NA |
| Prps1l1 | 0 | NA | NA | NA |
| AC133459.1 | 0 | NA | NA | NA |

| | | | | |
|---|---|---|---|---|
| Gm24359 | 0.075043205 | 0.517957688 | 3.724945049 | 0.139051095 |
| Snx13 | 1558.690179 | 0.154921885 | 0.052097127 | 2.973712665 |
| Mir680-3 | 0 | NA | NA | NA |
| AC132434.1 | 0 | NA | NA | NA |
| Gm26071 | 0 | NA | NA | NA |
| Ahr | 138.7406453 | -0.334036443 | 0.101816736 | -3.280761652 |
| Agr3 | 0.095295241 | -0.443821212 | 3.724945049 | -0.119148392 |
| Agr2 | 0.23039375 | -0.919033024 | 2.131410771 | -0.431185315 |
| Tspan13 | 1131.407557 | 0.156768138 | 0.070239655 | 2.231903581 |
| Gm5434 | 2.449668862 | -0.490500202 | 0.604996901 | -0.810748289 |
| Bzw2 | 293.0080855 | 0.274610263 | 0.095918061 | 2.862967207 |
| Ankmy2 | 458.9611624 | 0.099697604 | 0.066019144 | 1.51013173 |
| Lrrc72 | 0.323815284 | -0.007702512 | 1.731849594 | -0.004447564 |
| Sostdc1 | 41.87300568 | 1.575132995 | 0.28096635 | 5.60612683 |
| Gm25438 | 0.046345709 | -0.443818988 | 3.724945049 | -0.119147795 |
| D630036H23Rik | 7.277711592 | 0.982882094 | 0.386776143 | 2.541216961 |
| Ispd | 147.0734498 | 0.107498226 | 0.095939637 | 1.120477718 |
| Mir5099 | 0 | NA | NA | NA |
| CT010456.1 | 0 | NA | NA | NA |
| Gm29007 | 0 | NA | NA | NA |
| Meox2 | 0.449709139 | 0.03707677 | 1.580464494 | 0.023459414 |
| Agmo | 60.54347211 | 0.62247425 | 0.192304342 | 3.236922487 |
| Dgkb | 1454.497695 | 0.797035142 | 0.150504348 | 5.295761558 |
| Etv1 | 1246.89087 | 0.285207194 | 0.145281344 | 1.96313708 |
| Gm24069 | 0 | NA | NA | NA |
| Gm24434 | 0 | NA | NA | NA |
| Arl4a | 499.8111158 | 0.395732026 | 0.099044012 | 3.995516914 |
| Arl4aos | 1.421581723 | 2.18142405 | 1.00774187 | 2.164665491 |
| Scin | 6.368096109 | -0.23410276 | 0.383620092 | -0.610246348 |
| Lsmem1 | 17.83132388 | -0.012460435 | 0.273634486 | -0.045536787 |
| Gm17056 | 0.249945515 | -0.44441496 | 1.847817605 | -0.240508024 |
| Ifrd1 | 942.5158603 | -0.659839469 | 0.14326221 | -4.605816636 |
| Gm17024 | 29.97646992 | 0.148810674 | 0.319007799 | 0.466479735 |
| Mir1938 | 6.231293264 | 0.308179801 | 0.475966835 | 0.64748167 |
| Gm7008 | 11.68594954 | -0.134866311 | 0.316139939 | -0.426603205 |
| Zfp277 | 160.924537 | 0.03046156 | 0.133416095 | 0.228319981 |
| Dock4 | 2536.949238 | 0.392652092 | 0.101438702 | 3.870831208 |
| Immp2l | 31.50920012 | 0.087752062 | 0.198237249 | 0.442661822 |
| Gm25497 | 0.147068667 | 0.998853574 | 3.51441668 | 0.284216035 |
| Lrrn3 | 669.4920803 | 0.350874016 | 0.074641637 | 4.700781357 |
| Gm25248 | 0 | NA | NA | NA |
| Gm26326 | 0 | NA | NA | NA |
| Dnajb9 | 682.3485262 | -0.320715538 | 0.109451897 | -2.930196237 |
| Mir5627 | 0.044191549 | 0.037062677 | 3.724945049 | 0.009949859 |
| Pnpla8 | 1553.715183 | 0.201744417 | 0.065402561 | 3.084656205 |
| Nrcam | 5819.560715 | -0.039004484 | 0.07917699 | -0.492623985 |
| Stxbp6 | 733.0217604 | 0.37134208 | 0.10719081 | 3.464308942 |

| | | | | |
|---|---|---|---|---|
| Gm15901 | 0.132101749 | -0.93550548 | 3.724188147 | -0.251197158 |
| Gm15902 | 13.4453144 | 0.54064503 | 0.335225499 | 1.612780147 |
| Gm24731 | 0.429113465 | -0.816631494 | 1.417701872 | -0.57602484 |
| Gm22076 | 0 | NA | NA | NA |
| Gm9921 | 0.047948698 | 0.037064902 | 3.724945049 | 0.009950456 |
| Gm24948 | 0 | NA | NA | NA |
| Nova1 | 1267.849016 | 0.132270178 | 0.072940517 | 1.813397867 |
| Gm25051 | 0 | NA | NA | NA |
| Gm36971 | 0 | NA | NA | NA |
| Gm1818 | 26.89079701 | 0.2430884 | 0.236436774 | 1.02813279 |
| Gm26454 | 0 | NA | NA | NA |
| Gm7456 | 0 | NA | NA | NA |
| Foxg1 | 626.865741 | 0.336543601 | 0.136909114 | 2.458153375 |
| 3110039M20Rik | 111.2821837 | 1.060838568 | 0.250528859 | 4.234396682 |
| RP24-490H13.4 | 48.60029554 | 0.906635737 | 0.244193243 | 3.712779785 |
| Gm9804 | 13.1091654 | 0.435963799 | 0.43134168 | 1.01071568 |
| 1810007C17Rik | 0.183032319 | 0.647812728 | 3.311598859 | 0.195619323 |
| Gm25015 | 0 | NA | NA | NA |
| Gm7481 | 0.044191549 | 0.037062677 | 3.724945049 | 0.009949859 |
| Prkd1 | 131.3087976 | -0.244446052 | 0.113887088 | -2.146389526 |
| Gm22088 | 0 | NA | NA | NA |
| Rps11-ps4 | 0.060174537 | 0.517957688 | 3.724945049 | 0.139051095 |
| G2e3 | 315.7949451 | -0.008377887 | 0.105631923 | -0.079312079 |
| Scfd1 | 573.8494792 | -0.240546972 | 0.096562273 | -2.491107194 |
| Coch | 246.9813543 | 0.690348613 | 0.154348787 | 4.472653318 |
| Strn3 | 1743.354882 | 0.005894887 | 0.055107681 | 0.10697033 |
| Ap4s1 | 296.6051223 | 0.211744883 | 0.081287676 | 2.604882979 |
| Gm22663 | 0.270436937 | -0.4438227 | 2.401776452 | -0.184789346 |
| Hectd1 | 2718.891165 | -0.028879766 | 0.040620759 | -0.71096076 |
| Heatr5a | 367.9070692 | -0.524556641 | 0.129721426 | -4.043716283 |
| Gm26517 | 0.276231724 | -0.372945106 | 2.059360791 | -0.181097507 |
| Dtd2 | 62.41787465 | -0.010965698 | 0.162848839 | -0.067336666 |
| Gpr33 | 0.154148777 | -0.500251271 | 2.843849781 | -0.175906363 |
| Nubpl | 61.33971456 | 0.20999038 | 0.153571025 | 1.367382815 |
| Arhgap5 | 3781.700427 | 0.002346456 | 0.104503397 | 0.022453397 |
| 1700031P21Rik | 0.115603922 | -0.93550548 | 3.724188147 | -0.251197158 |
| Gm24859 | 1.957101456 | -0.093574354 | 0.733493669 | -0.127573499 |
| Gm23296 | 1.92086539 | -0.312357581 | 0.783269993 | -0.398786605 |
| Akap6 | 5339.749417 | 0.351570673 | 0.063193502 | 5.563399114 |
| n-R5s58 | 0 | NA | NA | NA |
| Npas3 | 426.0647591 | -0.025524304 | 0.082218575 | -0.3104445 |
| 1700060O08Rik | 0.07796279 | -0.798301147 | 3.721829328 | -0.214491605 |
| Egln3 | 112.0375452 | -0.372504438 | 0.125426933 | -2.969891947 |
| Gm22220 | 0 | NA | NA | NA |
| Gm24692 | 0 | NA | NA | NA |
| Sptssa | 171.7818652 | -0.41292746 | 0.143924787 | -2.869050358 |
| mmu-mir-1892 | 0 | NA | NA | NA |

| | | | | |
|---|---|---|---|---|
| Gm24971 | 0.26923737 | -0.447805413 | 2.126779793 | -0.210555609 |
| Eapp | 505.3675571 | -0.059627502 | 0.075089272 | -0.794088163 |
| Gm22513 | 8.835653036 | -0.229291525 | 0.414847207 | -0.552713195 |
| Rpl31-ps17 | 1.178692824 | 0.072228321 | 0.839734105 | 0.086013324 |
| Gm22634 | 17.48739027 | -0.084915872 | 0.542570896 | -0.156506499 |
| Gm23804 | 1.13792396 | -0.008283782 | 0.954851489 | -0.008675466 |
| Gm25679 | 43.52464816 | 0.140533349 | 0.254109234 | 0.553043061 |
| Gm26444 | 126.6439584 | 0.010786178 | 0.154028597 | 0.070027112 |
| Gm24305 | 91.6640994 | -0.253664185 | 0.203907877 | -1.244013664 |
| Gm22317 | 529.3228771 | -0.099263998 | 0.240463827 | -0.412802205 |
| Snx6 | 482.8085486 | -0.459519239 | 0.087393572 | -5.258043883 |
| Cfl2 | 953.9436151 | -0.261159992 | 0.095546308 | -2.733334217 |
| Baz1a | 191.7919684 | -1.260490317 | 0.178120261 | -7.07662514 |
| Gm27014 | 0 | NA | NA | NA |
| Gm20403 | 0 | NA | NA | NA |
| Gm24296 | 0 | NA | NA | NA |
| 2700097O09Rik | 100.6248846 | -0.000483542 | 0.176210099 | -0.002744124 |
| Srp54a | 0.941593011 | -0.700528416 | 1.143619612 | -0.612553692 |
| Fam177a | 2.46611504 | 0.206696809 | 0.656138747 | 0.315019971 |
| Srp54b | 195.0106852 | -0.004533347 | 0.143625243 | -0.031563717 |
| 1700047I17Rik2 | 3.457287012 | -0.257083951 | 0.644560812 | -0.398851352 |
| Ppp2r3c | 188.8955074 | -0.251270372 | 0.113234636 | -2.219023967 |
| 1110008L16Rik | 73.54196958 | 0.00354457 | 0.122873918 | 0.02884721 |
| Psma6 | 718.3364319 | -0.418917693 | 0.129331515 | -3.239099877 |
| Rpl18a-ps1 | 0.963303439 | -0.486663126 | 1.016331396 | -0.478842952 |
| Nfkbia | 657.3079934 | -0.020177352 | 0.283253918 | -0.071234151 |
| Aldoart2 | 0.61825299 | -0.414559749 | 1.227381954 | -0.337759365 |
| Insm2 | 3.842056669 | -0.367322557 | 0.526356952 | -0.697858279 |
| Ralgapa1 | 2973.912485 | 0.192945081 | 0.065627158 | 2.940018856 |
| Gm9812 | 2.978437241 | 0.617278423 | 0.590839219 | 1.044748559 |
| Gm23394 | 0 | NA | NA | NA |
| Brms1l | 538.8067484 | 0.013253207 | 0.082343083 | 0.160951065 |
| Gm22296 | 0 | NA | NA | NA |
| Gm23799 | 0 | NA | NA | NA |
| Gm25917 | 0.075043205 | 0.517957688 | 3.724945049 | 0.139051095 |
| Gm24685 | 0.291934259 | 0.5524322 | 1.782490249 | 0.309921583 |
| Mbip | 262.0904343 | 0.458112956 | 0.149266856 | 3.069086921 |
| Gm6265 | 9.407196128 | 0.270268202 | 0.571274604 | 0.473096826 |
| Nkx2-1 | 18.9524978 | -0.053056842 | 0.303470015 | -0.174833886 |
| Gm26973 | 0.859979859 | 2.005015113 | 1.367182975 | 1.466530194 |
| Gm25760 | 0 | NA | NA | NA |
| C87198 | 0.421608343 | 0.037076167 | 1.69725876 | 0.021844734 |
| Gm38103 | 1.370820674 | -1.117561722 | 1.015332252 | -1.100685731 |
| Nkx2-9 | 4.229279226 | -0.964165086 | 0.636324739 | -1.515209179 |
| Gm15524 | 0 | NA | NA | NA |
| Pax9 | 0.231631338 | -0.43871099 | 2.122910229 | -0.20665546 |
| Slc25a21 | 5.399330309 | 0.384160274 | 0.450164978 | 0.853376635 |

| | | | | |
|---|---|---|---|---|
| Prps1l3 | 132.0803981 | -0.269412581 | 0.099855307 | -2.698029674 |
| Mipol1 | 126.8360654 | 0.34019824 | 0.11261825 | 3.020809138 |
| Gm11025 | 0 | NA | NA | NA |
| Gm16246 | 0.044544648 | 0.517957688 | 3.724945049 | 0.139051095 |
| Gm2568 | 0 | NA | NA | NA |
| Gm37291 | 0 | NA | NA | NA |
| Foxa1 | 3.942515582 | 0.1520493 | 0.6155799 | 0.247001729 |
| 4921506M07Rik | 10.13732351 | 0.88619118 | 0.395779717 | 2.239102061 |
| 4921518K17Rik | 0.038981395 | -0.443816764 | 3.724945049 | -0.119147198 |
| Rpl26-ps2 | 0 | NA | NA | NA |
| Sstr1 | 99.2471505 | -0.211669363 | 0.172454622 | -1.227391655 |
| Clec14a | 49.14502572 | -0.697097946 | 0.19258572 | -3.619676188 |
| Gm17529 | 0.072025462 | 0.517959912 | 3.724945049 | 0.139051692 |
| Sec23a | 1353.977473 | 0.149056191 | 0.052819207 | 2.822007361 |
| Gemin2 | 59.82869919 | -0.486596646 | 0.144416152 | -3.369405976 |
| Gm22973 | 1.521163403 | 0.396972054 | 0.859698784 | 0.461757143 |
| Trappc6b | 482.938189 | 0.052207266 | 0.071048209 | 0.734814657 |
| Pnn | 591.1787483 | 0.147635646 | 0.154995116 | 0.952518052 |
| Gm5786 | 32.33765274 | 0.261289419 | 0.276754187 | 0.944120924 |
| Rpl21-ps3 | 1.284833043 | 0.796866593 | 0.852611125 | 0.934619042 |
| Mia2 | 5.838823621 | 0.351436773 | 0.54616275 | 0.643465292 |
| Ctage5 | 709.7395958 | 0.015759012 | 0.076603313 | 0.205722332 |
| Gm24233 | 4.255553331 | 1.210468438 | 0.574154957 | 2.108260884 |
| Fbxo33 | 714.5346483 | 0.062356214 | 0.117317559 | 0.531516459 |
| Gm27284 | 0 | NA | NA | NA |
| Gm7719 | 0.04393175 | 0.037062677 | 3.724945049 | 0.009949859 |
| Gm24377 | 0 | NA | NA | NA |
| Gm22673 | 0 | NA | NA | NA |
| Gm23059 | 0 | NA | NA | NA |
| Gm38172 | 0 | NA | NA | NA |
| Gm7745 | 2.081729379 | 0.021883465 | 0.757570357 | 0.028886379 |
| Nanog-ps2 | 0 | NA | NA | NA |
| Lrfn5 | 1144.044751 | 0.579763608 | 0.080163673 | 7.232248506 |
| Gm23213 | 11.15017964 | 0.72194188 | 0.39320772 | 1.836031804 |
| Gm24750 | 0 | NA | NA | NA |
| Gm24590 | 0 | NA | NA | NA |
| Spanxn4 | 0 | NA | NA | NA |
| Gm7763 | 0 | NA | NA | NA |
| Gm5185 | 0 | NA | NA | NA |
| Gm24921 | 0 | NA | NA | NA |
| Fscb | 0.143022002 | 0.03610674 | 3.547735707 | 0.010177404 |
| Gm24066 | 0 | NA | NA | NA |
| Gm6395 | 108.7166347 | 0.173190309 | 0.155419278 | 1.114342515 |
| Gm527 | 45.68336777 | 0.358388553 | 0.162006809 | 2.212182034 |
| Klhl28 | 276.6149016 | -0.312374356 | 0.087696884 | -3.561977839 |
| Fam179b | 1164.381407 | -0.045006601 | 0.073592464 | -0.61156536 |
| Prpf39 | 881.5198811 | 0.117998914 | 0.103575924 | 1.139250411 |

| | | | | |
|---|---|---|---|---|
| Gm25970 | 3.058502859 | 1.047235186 | 0.753593607 | 1.389655082 |
| Fkbp3 | 818.3394673 | 0.38994284 | 0.101706345 | 3.834007018 |
| Fancm | 325.0406225 | 0.187790448 | 0.124029526 | 1.514078576 |
| Mis18bp1 | 20.94898307 | -0.110717808 | 0.317758321 | -0.348434015 |
| Wdr20rt | 0.136963818 | 0.037074342 | 3.599544405 | 0.010299732 |
| Gm26015 | 0 | NA | NA | NA |
| Gm25599 | 0 | NA | NA | NA |
| Gm23948 | 0 | NA | NA | NA |
| Gm22782 | 0 | NA | NA | NA |
| Rpl10l | 0.061920613 | -0.443816764 | 3.724945049 | -0.119147198 |
| Mdga2 | 1092.640628 | 0.488587902 | 0.107484503 | 4.545659036 |
| Gm23826 | 1.550082411 | 1.079180087 | 0.891680777 | 1.210276272 |
| n-R5s60 | 0 | NA | NA | NA |
| Gm26256 | 0 | NA | NA | NA |
| n-R5s61 | 0 | NA | NA | NA |
| Gm15571 | 0.136963818 | 0.037074342 | 3.599544405 | 0.010299732 |
| Gm24499 | 0.13681464 | 0.037072711 | 2.943790034 | 0.012593531 |
| Rps29 | 1758.697591 | -0.063315415 | 0.105088878 | -0.602493961 |
| Rn7s1 | 42650.12024 | -0.457868814 | 0.309749182 | -1.478192167 |
| AC099934.2 | 0 | NA | NA | NA |
| AC099934.1 | 0 | NA | NA | NA |
| Lrr1 | 2.041322758 | -1.878702178 | 0.958059116 | -1.960945986 |
| Rpl36al | 266.3361645 | -0.376506158 | 0.1590962 | -2.366531438 |
| Mgat2 | 450.2752201 | -0.396616537 | 0.111342969 | -3.562115702 |
| Dnaaf2 | 60.5402635 | -0.280332228 | 0.169854945 | -1.65042135 |
| 9330151L19Rik | 197.4609258 | 0.311490012 | 0.100503889 | 3.099283164 |
| Pole2 | 9.423356029 | -0.593630727 | 0.335762276 | -1.768009003 |
| Klhdc1 | 176.7937862 | 0.191602658 | 0.149896915 | 1.278229493 |
| Gm15561 | 0 | NA | NA | NA |
| Klhdc2 | 1197.243616 | 0.091616389 | 0.060793906 | 1.506999561 |
| Nemf | 637.5580402 | -0.031343546 | 0.079578264 | -0.393870692 |
| AC157822.1 | 0 | NA | NA | NA |
| Rn7s2 | 34914.20044 | 0.002169831 | 0.206979487 | 0.010483313 |
| AC157822.2 | 0 | NA | NA | NA |
| Gm9887 | 2.974284523 | -0.020530992 | 0.588592263 | -0.034881518 |
| Arf6 | 1024.329735 | -0.531716361 | 0.093532933 | -5.684803651 |
| Gm23018 | 0.038981395 | -0.443816764 | 3.724945049 | -0.119147198 |
| Vcpkmt | 182.6319952 | -0.170894153 | 0.099861167 | -1.711317406 |
| Sos2 | 821.7158266 | -0.128812005 | 0.058960145 | -2.184730145 |
| Gm24449 | 1.413161897 | -0.605455086 | 0.928311537 | -0.652211097 |
| L2hgdh | 176.5242931 | -0.003783026 | 0.123622757 | -0.030601374 |
| Atp5s | 115.1168851 | 0.378129984 | 0.127361336 | 2.968954291 |
| Cdkl1 | 209.0876337 | 0.513533018 | 0.108822288 | 4.719005871 |
| Mir681 | 2.100525097 | -0.447443225 | 0.805236904 | -0.555666565 |
| 4930512B01Rik | 3.315021913 | 0.622832013 | 0.573924553 | 1.085215834 |
| Map4k5 | 890.2594534 | 0.126793146 | 0.115569334 | 1.097117561 |
| 4931403G20Rik | 0.07796279 | -0.798301147 | 3.721829328 | -0.214491605 |

| | | | | |
|---|---|---|---|---|
| Atl1 | 920.937231 | 0.545193262 | 0.060349814 | 9.033884712 |
| Gm6316 | 3.637700379 | 0.608553903 | 0.582872161 | 1.044060677 |
| Gm22058 | 5.202496254 | 0.704715763 | 0.50341939 | 1.399858204 |
| Gm3086 | 7.899535637 | 0.156719456 | 0.510005313 | 0.30728985 |
| Sav1 | 344.851752 | -0.131067013 | 0.130017615 | -1.008071198 |
| Gm23579 | 0 | NA | NA | NA |
| Nin | 902.9475348 | 0.044907664 | 0.138496199 | 0.324251961 |
| Abhd12b | 8.085658348 | 0.156637704 | 0.416936378 | 0.375687303 |
| Pygl | 47.59505423 | -0.612827311 | 0.173954361 | -3.522920076 |
| F730035M05Rik | 0.381387581 | 0.119566692 | 1.535441288 | 0.077871224 |
| Trim9 | 3338.081355 | 0.266056999 | 0.104136509 | 2.55488686 |
| Tmx1 | 634.4816424 | -0.256185907 | 0.096478561 | -2.655366158 |
| Gm24474 | 0.044544648 | 0.517957688 | 3.724945049 | 0.139051095 |
| Frmd6 | 461.8109092 | -0.075183809 | 0.139895148 | -0.537429713 |
| Gm24857 | 0 | NA | NA | NA |
| Actr10 | 1289.027537 | 0.04654562 | 0.066027396 | 0.704944054 |
| Psma3 | 949.5019891 | -0.164095856 | 0.075826705 | -2.16409054 |
| 3110056K07Rik | 60.20099725 | 0.133230768 | 0.188288482 | 0.707588517 |
| Arid4a | 748.6510582 | 0.150435695 | 0.108047403 | 1.392311999 |
| Tomm20l | 0.146692594 | -0.955495625 | 3.521109961 | -0.271362052 |
| Timm9 | 217.381226 | 0.125678473 | 0.074016291 | 1.697983924 |
| 2700049A03Rik | 196.0398049 | 0.092593649 | 0.127065945 | 0.728705468 |
| Dact1 | 166.182804 | -0.056891136 | 0.159160413 | -0.357445268 |
| 4930474H06Rik | 0.409321926 | -0.779662913 | 1.454521164 | -0.536027204 |
| 1700083H02Rik | 0 | NA | NA | NA |
| Gm22959 | 0 | NA | NA | NA |
| Daam1 | 1425.073008 | 0.049360381 | 0.048634521 | 1.014924781 |
| Gm7985 | 3.193668984 | 0.884626642 | 0.685529175 | 1.290428875 |
| Gpr135 | 77.04711869 | 0.4137643 | 0.139329432 | 2.969683378 |
| L3hypdh | 19.58973956 | -0.28181715 | 0.274803006 | -1.025524262 |
| Jkamp | 337.9089615 | -0.001813383 | 0.101674552 | -0.017835172 |
| Ccdc175 | 2.241016658 | -1.040849659 | 0.733575706 | -1.418871496 |
| Gm24466 | 0 | NA | NA | NA |
| Rtn1 | 7105.395095 | 0.584180641 | 0.074231988 | 7.869661829 |
| Lrrc9 | 75.53725959 | -0.282614852 | 0.150527103 | -1.877501433 |
| AC103353.1 | 0 | NA | NA | NA |
| Pcnxl4 | 483.304744 | 0.036339694 | 0.069910406 | 0.519803788 |
| Dhrs7 | 149.9058334 | -0.131291854 | 0.117504617 | -1.117333571 |
| Ppm1a | 1736.427688 | 0.203354876 | 0.062068446 | 3.276300405 |
| 4930447C04Rik | 136.3889103 | 0.620020275 | 0.195020728 | 3.179253205 |
| Six6 | 2.974111705 | -0.423833704 | 0.873902115 | -0.484989905 |
| Gm26709 | 0 | NA | NA | NA |
| Gm38170 | 0 | NA | NA | NA |
| Six1 | 9.772493715 | -0.650343777 | 0.376668667 | -1.726567232 |
| Six4 | 52.62636027 | 0.134477128 | 0.196716677 | 0.683608169 |
| Mnat1 | 196.833954 | 0.030319823 | 0.086801243 | 0.349301718 |
| Gm23910 | 0 | NA | NA | NA |

| | | | | |
|---|---|---|---|---|
| Trmt5 | 87.0705171 | -0.273023195 | 0.188114476 | -1.451367273 |
| Slc38a6 | 110.6102002 | -0.026531484 | 0.179247657 | -0.148015791 |
| D830013O20Rik | 0 | NA | NA | NA |
| Tmem30b | 3.209381466 | -0.341406829 | 0.570585322 | -0.598344921 |
| Prkch | 135.4372229 | -0.248643803 | 0.142278826 | -1.747581217 |
| Hif1a | 2116.265102 | -0.375346418 | 0.129286991 | -2.903203284 |
| Gm15283 | 3.904436585 | 1.052129533 | 0.599908222 | 1.753817492 |
| Gm5068 | 0 | NA | NA | NA |
| Snapc1 | 313.850414 | -0.046208515 | 0.092173592 | -0.501320543 |
| Syt16 | 456.7941473 | 0.939531375 | 0.147193064 | 6.382986742 |
| Gm29587 | 0 | NA | NA | NA |
| 1700086L19Rik | 251.9530011 | 0.174107054 | 0.176159387 | 0.988349567 |
| Dbpht2 | 497.2679655 | 0.259026797 | 0.157070654 | 1.649110061 |
| Gm11042 | 0.897295135 | 0.444187197 | 1.297355019 | 0.342379064 |
| Kcnh5 | 625.0782806 | 0.835350307 | 0.146994631 | 5.682862707 |
| Rhoj | 239.1716956 | -0.557785363 | 0.318980341 | -1.74865122 |
| Gphb5 | 1.844461247 | -1.663266348 | 0.873363491 | -1.904437689 |
| Ppp2r5e | 1376.244815 | -0.089971707 | 0.046754271 | -1.924352694 |
| Gm29236 | 0.285890061 | -0.802727966 | 1.761238266 | -0.45577477 |
| Wdr89 | 187.0451121 | -0.350021677 | 0.108112829 | -3.237559133 |
| Sgpp1 | 600.9576085 | 0.210149076 | 0.089658318 | 2.343888238 |
| Syne2 | 912.2047069 | -0.394804144 | 0.076166651 | -5.183425308 |
| Mir5101 | 0.042716945 | 0.037062677 | 3.724945049 | 0.009949859 |
| Esr2 | 7.599516194 | -0.020029285 | 0.358527778 | -0.055865364 |
| Gm7862 | 4.308722595 | -0.517404905 | 0.532211931 | -0.972178328 |
| Tex21 | 0.318741927 | -0.382104548 | 1.719574842 | -0.222208734 |
| Mthfd1 | 536.2563463 | -0.232593026 | 0.071451295 | -3.255266763 |
| Akap5 | 895.5134347 | 1.601225554 | 0.187497039 | 8.540004489 |
| Zbtb25 | 191.2371808 | 0.298770969 | 0.090767995 | 3.291589392 |
| Zbtb1 | 407.9327547 | -0.337047062 | 0.08691697 | -3.877805001 |
| 4930426I24Rik | 3.588158775 | 0.628143648 | 0.593796842 | 1.05784269 |
| Hspa2 | 433.2408539 | -0.160080385 | 0.133113214 | -1.202588237 |
| Ppp1r36 | 10.56443581 | -0.77772154 | 0.31634112 | -2.458490191 |
| Gm25563 | 0 | NA | NA | NA |
| Gm10451 | 3.439471239 | 0.544002286 | 0.559278282 | 0.972686235 |
| Mir5135 | 0.047948698 | 0.037064902 | 3.724945049 | 0.009950456 |
| Plekhg3 | 262.9259336 | 0.152498642 | 0.108252605 | 1.408729536 |
| Sptb | 3404.075389 | 0.309454219 | 0.093368901 | 3.31431789 |
| Gm24010 | 0 | NA | NA | NA |
| Churc1 | 76.27497148 | -0.070199905 | 0.148809976 | -0.471741926 |
| Fntb | 334.4021306 | -0.111850529 | 0.082940563 | -1.34856245 |
| Gpx2 | 1.498903582 | -0.513491773 | 0.841020353 | -0.610558082 |
| Rab15 | 1748.719287 | 0.03755249 | 0.124734481 | 0.301059418 |
| Max | 974.4805696 | -0.366723954 | 0.089287677 | -4.107218008 |
| Gm22696 | 0 | NA | NA | NA |
| Fut8 | 862.4398163 | 0.077497703 | 0.073454366 | 1.055045561 |
| Gm24775 | 2.254287688 | 1.024476465 | 0.860536396 | 1.190509164 |

| | | | | |
|---|---|---|---|---|
| Gm25926 | 5.521549126 | 0.259526832 | 0.511090164 | 0.507790699 |
| Gm24070 | 0 | NA | NA | NA |
| Gm24994 | 0 | NA | NA | NA |
| Gm6657 | 0 | NA | NA | NA |
| Gphn | 1083.325677 | 0.210395105 | 0.06516047 | 3.228876412 |
| Fam71d | 7.845798074 | -0.149228324 | 0.471791121 | -0.31630168 |
| Mpp5 | 811.6373095 | -0.255952381 | 0.097503493 | -2.62505858 |
| Gm25225 | 0.097139547 | -0.443816764 | 3.724945049 | -0.119147198 |
| Atp6v1d | 1557.396393 | 0.201991409 | 0.057775721 | 3.496129595 |
| Eif2s1 | 534.2845462 | -0.576087717 | 0.105199381 | -5.476151182 |
| Plek2 | 8.61393615 | -0.917002959 | 0.359335016 | -2.551944336 |
| Tmem229b | 976.9241392 | -0.270584849 | 0.115184451 | -2.349143887 |
| Plekhh1 | 575.4863948 | -0.16522402 | 0.104631445 | -1.579104826 |
| Pigh | 108.6090619 | -0.487072838 | 0.134837655 | -3.612290919 |
| Gm27297 | 0 | NA | NA | NA |
| Arg2 | 76.92116969 | -1.096005477 | 0.283350097 | -3.86802577 |
| Vti1b | 794.0198813 | -0.125538654 | 0.076194348 | -1.647611115 |
| Gm17195 | 12.61275266 | -0.276044708 | 0.366106275 | -0.754001574 |
| Gm17194 | 0 | NA | NA | NA |
| Rdh11 | 314.5614071 | -0.231406549 | 0.072582627 | -3.188180965 |
| Rdh12 | 16.14501922 | 0.10740113 | 0.277954551 | 0.386398172 |
| Zfyve26 | 591.7739187 | -0.200771879 | 0.094729556 | -2.119421721 |
| Rad51b | 1.767988423 | -0.254079884 | 0.771526475 | -0.329321018 |
| Gm28056 | 0 | NA | NA | NA |
| Zfp36l1 | 660.9038404 | -0.479426939 | 0.119077697 | -4.026169071 |
| Gm26669 | 0.480640422 | -0.858016688 | 1.303094123 | -0.658445674 |
| 2310015A10Rik | 112.2249857 | 0.031669191 | 0.107460866 | 0.29470441 |
| Actn1 | 1100.046826 | -0.846848943 | 0.092112757 | -9.19361198 |
| Dcaf5 | 1241.49422 | 0.221387945 | 0.070116511 | 3.157429564 |
| mmu-mir-1843a | 0 | NA | NA | NA |
| Scarna3b | 2.198311352 | -0.599925938 | 0.89312745 | -0.671713694 |
| Exd2 | 554.3707735 | 0.408088147 | 0.102834991 | 3.968378306 |
| Gm26777 | 28.6766377 | 0.961906011 | 0.2425385 | 3.96599307 |
| Galnt16 | 1559.333316 | 0.493409274 | 0.064689439 | 7.627354315 |
| Gm23817 | 0.07796279 | -0.798301147 | 3.721829328 | -0.214491605 |
| Erh | 198.2980328 | -0.327824908 | 0.103332608 | -3.172521395 |
| Slc39a9 | 697.3314638 | -0.137876913 | 0.066118321 | -2.085305724 |
| Plekhd1os | 1.509627474 | 0.460257259 | 0.880909564 | 0.52247958 |
| Plekhd1 | 169.5676397 | 0.271110358 | 0.154341993 | 1.756556022 |
| Ccdc177 | 193.1722448 | 0.37790188 | 0.127063246 | 2.974124234 |
| Gm26796 | 1.873447992 | 0.636768871 | 0.713588765 | 0.892347108 |
| Susd6 | 1200.089806 | -0.165090033 | 0.108842925 | -1.516773209 |
| Gm26545 | 0 | NA | NA | NA |
| Gm23298 | 0.127971488 | -0.95589744 | 3.6855341 | -0.259364698 |
| 1700052I22Rik | 0 | NA | NA | NA |
| Srsf5 | 1958.310891 | 0.351366502 | 0.085993627 | 4.085959774 |
| Slc10a1 | 0.066050875 | -0.443818988 | 3.724945049 | -0.119147795 |

| | | | | |
|---|---|---|---|---|
| Smoc1 | 160.7710477 | -0.312769468 | 0.16693074 | -1.87364812 |
| Mir3067 | 0.784624355 | 1.23950893 | 1.140858503 | 1.086470344 |
| Slc8a3 | 615.5791632 | 0.145660993 | 0.102562379 | 1.420218544 |
| Gm3693 | 13.26815873 | 0.520379524 | 0.501787288 | 1.037052027 |
| Gm20498 | 37.16701504 | 0.256080095 | 0.194504432 | 1.316577173 |
| Cox16 | 98.00354183 | 0.017597286 | 0.126461363 | 0.139151478 |
| Gm3695 | 17.50620631 | 0.375347255 | 0.413412594 | 0.907924095 |
| Gm4787 | 60.98925875 | 0.214115646 | 0.201966055 | 1.060156598 |
| Hspe1-ps2 | 0 | NA | NA | NA |
| Adam4 | 66.87698665 | 0.307571288 | 0.25847271 | 1.189956529 |
| Gm28370 | 1.129831383 | 0.597105349 | 1.076243003 | 0.554805325 |
| Synj2bp | 590.4251372 | -0.195281205 | 0.106679047 | -1.830548833 |
| Gm24853 | 0.038981395 | -0.443816764 | 3.724945049 | -0.119147198 |
| Gm17191 | 9.020757225 | 0.223964014 | 0.391239764 | 0.572446962 |
| 6530401F13Rik | 4.716167235 | -0.058840721 | 0.633279528 | -0.092914295 |
| Adam21 | 15.49915397 | 0.752499549 | 0.266274318 | 2.82603127 |
| Med6 | 165.5733909 | -0.048323625 | 0.121510947 | -0.397689475 |
| Gm16572 | 0.53222664 | -1.905673019 | 1.665430458 | -1.144252532 |
| Gm16570 | 4.946459833 | 0.895494876 | 0.570342638 | 1.570099825 |
| Ttc9 | 455.2096388 | -0.306475075 | 0.088435561 | -3.46551851 |
| Map3k9 | 1178.392681 | 0.587730709 | 0.101232521 | 5.805750016 |
| Pcnx | 2449.47711 | 0.135693732 | 0.070044315 | 1.937255454 |
| n-R5s62 | 0 | NA | NA | NA |
| Sipa1l1 | 4759.581652 | 0.682127659 | 0.127672359 | 5.342798287 |
| Gm5435 | 1.384865981 | 1.008584147 | 0.827018951 | 1.21954176 |
| Gm8358 | 0 | NA | NA | NA |
| n-R5s63 | 0 | NA | NA | NA |
| Rgs6 | 645.1774152 | 0.15048489 | 0.106284766 | 1.415865088 |
| SNORA43 | 0 | NA | NA | NA |
| Gm22149 | 0 | NA | NA | NA |
| 1700085C21Rik | 0.21789665 | 0.517974176 | 2.168530754 | 0.238859502 |
| 4930423C22Rik | 0.847184061 | 2.001685195 | 1.200861965 | 1.666873673 |
| Gm29530 | 0.044544648 | 0.517957688 | 3.724945049 | 0.139051095 |
| 4933436F18Rik | 0 | NA | NA | NA |
| 1700030I03Rik | 2.414365681 | 0.382974275 | 0.745505002 | 0.513711207 |
| Dpf3 | 155.1908164 | 0.061605521 | 0.146741825 | 0.419822512 |
| Dcaf4 | 167.3521746 | 0.166987467 | 0.106284045 | 1.571143312 |
| Zfyve1 | 610.9257158 | -0.098857202 | 0.0584195 | -1.692195272 |
| Gm26623 | 0.048949532 | 0.037064902 | 3.724945049 | 0.009950456 |
| Rbm25 | 2301.160026 | 0.273664348 | 0.110574086 | 2.474941083 |
| Gm29361 | 5.081951061 | -0.481365403 | 0.537818051 | -0.895033929 |
| Psen1 | 837.0310577 | -0.17537893 | 0.065163787 | -2.69135569 |
| Papln | 26.40192091 | 0.909813281 | 0.255809572 | 3.55660375 |
| Numb | 544.7112012 | -0.231375542 | 0.089486751 | -2.585584342 |
| 2410016O06Rik | 126.434428 | -0.039107834 | 0.133413526 | -0.293132451 |
| Gm26571 | 3.09834508 | 0.175285236 | 0.544938339 | 0.321660679 |
| Gm17673 | 1.031598502 | 2.597004192 | 1.201784036 | 2.160957473 |

| | | | | |
|---|---|---|---|---|
| Acot2 | 62.86110075 | 0.091158407 | 0.1530776 | 0.595504548 |
| Acot1 | 68.7388213 | -0.160980008 | 0.244645004 | -0.658014696 |
| Acot4 | 13.42834362 | 0.384605888 | 0.296397783 | 1.29760042 |
| 4732463B04Rik | 11.2321051 | 0.892673878 | 0.368461181 | 2.422708074 |
| Acot3 | 12.40614317 | 0.844128292 | 0.362782557 | 2.326816097 |
| Acot5 | 14.23358867 | 1.09124844 | 0.337415427 | 3.234139141 |
| Gm8385 | 0 | NA | NA | NA |
| Acot6 | 73.66342787 | -0.189673339 | 0.192805806 | -0.983753256 |
| Dnal1 | 1112.279329 | 0.46416867 | 0.095591861 | 4.855734216 |
| Pnma1 | 28.80769789 | 0.775999497 | 0.239525592 | 3.239735225 |
| Elmsan1 | 799.7188921 | 0.290195851 | 0.17086362 | 1.698406312 |
| Gm5436 | 39.89268335 | -0.224023115 | 0.217403902 | -1.030446618 |
| Ptgr2 | 431.9967278 | -0.159101008 | 0.098864269 | -1.609287257 |
| Zfp410 | 288.7236925 | -0.073926806 | 0.089244941 | -0.828358515 |
| Fam161b | 141.580014 | -0.284649621 | 0.186020313 | -1.530207194 |
| Coq6 | 126.6661858 | -0.200390081 | 0.113354926 | -1.767810968 |
| Entpd5 | 391.9413108 | -0.029991336 | 0.073469074 | -0.408217154 |
| Ccdc176 | 64.5472327 | 0.272130716 | 0.197622707 | 1.377021496 |
| AC120402.1 | 0 | NA | NA | NA |
| Rnf113a2 | 0 | NA | NA | NA |
| Aldh6a1 | 576.5073902 | 0.637268778 | 0.129517514 | 4.920328992 |
| Lin52 | 260.6951333 | 0.098268562 | 0.089828741 | 1.093954575 |
| Vsx2 | 9.446677945 | -0.180215461 | 0.485277412 | -0.371365855 |
| Abcd4 | 114.3789344 | -0.05708506 | 0.129007171 | -0.442495244 |
| Vrtn | 1.178380111 | 0.355137005 | 0.963860495 | 0.368452704 |
| Syndig1l | 316.8534421 | 0.208653472 | 0.114987734 | 1.814571571 |
| Npc2 | 1928.697367 | -1.128422773 | 0.158155865 | -7.134877833 |
| Isca2 | 199.1382904 | -0.191352478 | 0.09016158 | -2.122328366 |
| Ltbp2 | 65.48633923 | 0.320716896 | 0.215954862 | 1.485110789 |
| D030025P21Rik | 2.762732937 | 0.416746207 | 0.618530434 | 0.673768313 |
| Gm17139 | 0.226559374 | 1.806164991 | 2.511711466 | 0.719097323 |
| Arel1 | 2266.135036 | 0.04021724 | 0.071057945 | 0.56597809 |
| Gm17193 | 24.019554 | -0.634684536 | 0.334822075 | -1.895587491 |
| Fcf1 | 140.4858461 | -0.470327628 | 0.133864322 | -3.51346512 |
| Ylpm1 | 2692.241911 | 0.246850051 | 0.105313935 | 2.343944811 |
| Prox2os | 42.00672927 | 0.044938759 | 0.310771002 | 0.144604094 |
| Prox2 | 14.36306314 | -0.375953428 | 0.274899011 | -1.367605602 |
| Gm38355 | 1.352228905 | -0.163408738 | 0.913374631 | -0.178906587 |
| Dlst | 1232.377248 | -0.12682305 | 0.05669438 | -2.236959803 |
| Rps6kl1 | 284.8098805 | 0.136944397 | 0.118367457 | 1.156942972 |
| Pgf | 33.44763411 | 0.069237031 | 0.246537022 | 0.280838273 |
| Eif2b2 | 302.5668293 | -0.272967183 | 0.109349582 | -2.496280069 |
| Mlh3 | 324.2556209 | 0.001365002 | 0.108305636 | 0.012603243 |
| Mir6938 | 0.348080155 | -1.500890419 | 1.96078343 | -0.765454459 |
| Acyp1 | 106.7121191 | 0.172753097 | 0.135243576 | 1.277347888 |
| Zc2hc1c | 44.15230839 | -0.403483888 | 0.171112117 | -2.35800886 |
| Nek9 | 985.3989254 | -0.281245035 | 0.069468708 | -4.048513963 |

| | | | | |
|---|---|---|---|---|
| Tmed10 | 1862.85624 | -0.198420717 | 0.097634438 | -2.03228205 |
| Fos | 757.1228931 | -1.018800969 | 0.462841918 | -2.201185607 |
| Jdp2 | 344.9332217 | 0.091451084 | 0.092624183 | 0.987334846 |
| Batf | 16.13625052 | -0.657554375 | 0.31007621 | -2.120621815 |
| n-R5s64 | 0 | NA | NA | NA |
| AC117240.1 | 0 | NA | NA | NA |
| Mfsd7c | 17.4725598 | 0.087345424 | 0.286976805 | 0.304364055 |
| 0610007P14Rik | 325.4729982 | -0.189511131 | 0.065702897 | -2.884364911 |
| Ttll5 | 894.6595886 | 0.094156311 | 0.072151571 | 1.304979354 |
| Gm26531 | 0.135217741 | 1.027273954 | 3.618332993 | 0.283908075 |
| Tgfb3 | 281.890318 | -0.630506475 | 0.127764796 | -4.934899885 |
| Gm37804 | 1.282550361 | 0.732737809 | 0.918709628 | 0.797572798 |
| Ift43 | 63.1959967 | -0.341089872 | 0.150650898 | -2.264107791 |
| Gm805 | 0.516561414 | 0.767324671 | 1.330443956 | 0.576743325 |
| Gpatch2l | 583.809052 | -0.075742897 | 0.067747762 | -1.118013273 |
| 4732487G21Rik | 0.11657011 | 0.998850474 | 3.724945049 | 0.268151734 |
| Esrrb | 65.56798319 | 0.078982231 | 0.157756399 | 0.500659444 |
| Gm27993 | 0 | NA | NA | NA |
| Gm22004 | 0 | NA | NA | NA |
| Vash1 | 371.0212885 | 0.143830678 | 0.073345814 | 1.960993689 |
| Angel1 | 126.5566759 | -0.14968542 | 0.12510683 | -1.196460825 |
| Gm17138 | 0 | NA | NA | NA |
| Lrrc74a | 0.176451472 | 0.539299713 | 2.721126888 | 0.198189844 |
| Gm25888 | 0 | NA | NA | NA |
| Irf2bpl | 1475.011036 | -0.570736146 | 0.111531293 | -5.117273631 |
| Gm26698 | 1.907481327 | -0.472570661 | 0.719873627 | -0.656463361 |
| Cipc | 1257.563404 | -0.028855942 | 0.04338609 | -0.665096623 |
| Zdhhc22 | 154.2953736 | -0.102787723 | 0.117534793 | -0.874530169 |
| Tmem63c | 504.9714176 | 0.096594269 | 0.114467934 | 0.843854388 |
| Gm25847 | 6.00824128 | -0.015461601 | 0.569329328 | -0.027157569 |
| Gm15745 | 1.796415912 | 2.284829771 | 1.01405193 | 2.253168406 |
| Ngb | 44.68760207 | 0.353033788 | 0.230404767 | 1.532233004 |
| Pomt2 | 325.0464884 | -0.070364659 | 0.071021045 | -0.990757862 |
| Gstz1 | 241.2031799 | 0.266959546 | 0.087601623 | 3.04742693 |
| Tmed8 | 929.2418369 | 0.140494037 | 0.0754894 | 1.861109472 |
| Samd15 | 21.85261841 | 0.006379588 | 0.265890746 | 0.023993267 |
| Noxred1 | 12.44760366 | -1.339280886 | 0.397963599 | -3.365335145 |
| Vipas39 | 515.9231004 | -0.204761918 | 0.056847879 | -3.601927127 |
| Ahsa1 | 741.40917 | -0.321629066 | 0.089536784 | -3.592144487 |
| AC151967.1 | 0.038981395 | -0.443816764 | 3.724945049 | -0.119147198 |
| Ism2 | 2.941330987 | -0.310183587 | 0.625397419 | -0.495978361 |
| Sptlc2 | 820.5122325 | -0.416972372 | 0.130760217 | -3.18883206 |
| Gm26764 | 0.618129248 | -1.444211795 | 1.340489017 | -1.07737682 |
| Alkbh1 | 295.3910853 | 0.083065007 | 0.152366943 | 0.54516423 |
| mmu-mir-3068 | 0.086930093 | -0.443164241 | 3.724898049 | -0.118973522 |
| Mir3068 | 0.060174537 | 0.517957688 | 3.724945049 | 0.139051095 |
| Gm29362 | 0 | NA | NA | NA |

| | | | | |
|---|---|---|---|---|
| Slirp | 288.8250465 | -0.323721858 | 0.125859937 | -2.572080238 |
| Snw1 | 789.2603872 | -0.105765135 | 0.081000999 | -1.305726312 |
| Gm8300 | 0 | NA | NA | NA |
| Oog1 | 0 | NA | NA | NA |
| Gm4027 | 0 | NA | NA | NA |
| Gm16381 | 0 | NA | NA | NA |
| Gm21319 | 0 | NA | NA | NA |
| Gm21936 | 0 | NA | NA | NA |
| BB287469 | 0 | NA | NA | NA |
| Gm2001 | 0 | NA | NA | NA |
| Gm2016 | 0 | NA | NA | NA |
| Gm2022 | 0 | NA | NA | NA |
| Gm2035 | 0 | NA | NA | NA |
| Gm2042 | 0 | NA | NA | NA |
| Gm2046 | 0 | NA | NA | NA |
| Gm2075 | 0 | NA | NA | NA |
| Gm6803 | 0 | NA | NA | NA |
| Gm2056 | 0 | NA | NA | NA |
| Gm16368 | 0 | NA | NA | NA |
| RP23-174O6.3 | 0 | NA | NA | NA |
| Gm8587 | 0.789916185 | -0.157169265 | 1.17750441 | -0.133476583 |
| Gm10436 | 0 | NA | NA | NA |
| Gm8332 | 0 | NA | NA | NA |
| Gm5662 | 0 | NA | NA | NA |
| Gm7104 | 0 | NA | NA | NA |
| Gm5039 | 0.128271896 | 0.517956073 | 3.679089552 | 0.140783763 |
| Gm10264 | 0 | NA | NA | NA |
| Adck1 | 270.5190383 | -0.146032802 | 0.082809069 | -1.76348804 |
| Nrxn3 | 5644.729856 | 0.282061031 | 0.105619595 | 2.670536939 |
| Gm37980 | 0.244108046 | 0.517964358 | 2.509695299 | 0.206385356 |
| Gm27313 | 0 | NA | NA | NA |
| Gm23989 | 9.194225744 | 0.772684136 | 0.482157784 | 1.602554521 |
| Dio2 | 725.4291824 | 1.010241566 | 0.188261751 | 5.366154088 |
| Gm26512 | 30.26736896 | 0.958901092 | 0.228114501 | 4.203595512 |
| 4930544I03Rik | 0.534296218 | 2.688257946 | 1.379513838 | 1.948699514 |
| n-R5s65 | 0 | NA | NA | NA |
| Gm23195 | 0 | NA | NA | NA |
| Gm26477 | 0 | NA | NA | NA |
| Cep128 | 104.2602885 | 0.251676916 | 0.12630288 | 1.9926459 |
| 5430427M07Rik | 0 | NA | NA | NA |
| CR974487.1 | 0 | NA | NA | NA |
| Tshr | 7.627433336 | 0.320385995 | 0.398469409 | 0.804041634 |
| Gm18827 | 0.108266322 | -0.955981391 | 3.722845977 | -0.25678779 |
| Gtf2a1 | 1089.805438 | -0.265639176 | 0.062700583 | -4.236630069 |
| Gm25776 | 271.3150006 | -0.624429394 | 0.224907039 | -2.776388846 |
| Ston2 | 295.1567159 | -0.294920874 | 0.122085825 | -2.415684825 |
| Gm8378 | 10.50589885 | 0.325318028 | 0.371144075 | 0.876527609 |

| | | | | |
|---|---|---|---|---|
| Gm16876 | 0.085327104 | -0.905414734 | 3.723500899 | -0.243162217 |
| Gm22015 | 0 | NA | NA | NA |
| Sel1l | 2575.426926 | 0.007546459 | 0.039204412 | 0.192490058 |
| Rpl31-ps1 | 0.104719185 | 0.981063299 | 3.723618439 | 0.263470416 |
| Gm21614 | 0 | NA | NA | NA |
| Gm23249 | 0 | NA | NA | NA |
| Gm9726 | 14.69313774 | -0.468054552 | 0.293365131 | -1.595467565 |
| Mir8099-1 | 0.066050875 | -0.443818988 | 3.724945049 | -0.119147795 |
| Gm26055 | 0.060174537 | 0.517957688 | 3.724945049 | 0.139051095 |
| Gm22246 | 0.682665822 | 0.112827726 | 1.149363127 | 0.098165431 |
| Flrt2 | 1031.326902 | 0.311701854 | 0.074567448 | 4.180133054 |
| 1700019M22Rik | 0 | NA | NA | NA |
| Gm26480 | 0 | NA | NA | NA |
| Gm6863 | 145.6421742 | -0.146800706 | 0.137772971 | -1.065526164 |
| Gm23558 | 0 | NA | NA | NA |
| Galc | 301.800896 | 0.169786314 | 0.098913481 | 1.716513384 |
| Gpr65 | 55.39552026 | -0.361062229 | 0.22089147 | -1.634568452 |
| Kcnk10 | 201.901865 | -0.474078109 | 0.132189455 | -3.586353451 |
| Gm25486 | 0 | NA | NA | NA |
| Spata7 | 282.5135196 | 0.524369352 | 0.143055693 | 3.66549097 |
| Ptpn21 | 244.2151489 | -0.654079498 | 0.090426904 | -7.233239995 |
| Zc3h14 | 1282.947868 | 0.094444471 | 0.059025168 | 1.600071192 |
| Eml5 | 1567.483707 | 0.447869246 | 0.151040794 | 2.965220417 |
| Ttc8 | 210.9206023 | 0.040709268 | 0.15690983 | 0.259443704 |
| Rpl30-ps8 | 10.08383252 | -0.344058923 | 0.329743688 | -1.043413216 |
| Gm28933 | 0.301731016 | -1.759016082 | 2.02810192 | -0.867321344 |
| Foxn3 | 762.8771155 | 0.050786392 | 0.054377881 | 0.933953127 |
| Gm19898 | 0.254453631 | -0.50137576 | 2.971545213 | -0.168725604 |
| Gm20682 | 16.54715445 | 0.632488218 | 0.395929775 | 1.597475759 |
| 3300002A11Rik | 4.616394162 | 0.190407866 | 0.57862544 | 0.329069295 |
| Gm26839 | 2.688780499 | 0.583464609 | 0.659799323 | 0.884306165 |
| 1700064M15Rik | 1.043907264 | 0.814654491 | 0.983220859 | 0.828556964 |
| Efcab11 | 4.011499476 | 0.153392624 | 0.524340647 | 0.292543835 |
| Tdp1 | 173.9666548 | -0.145003547 | 0.163014029 | -0.889515755 |
| Kcnk13 | 123.3766849 | -0.985982527 | 0.148677706 | -6.631677038 |
| Gm23291 | 0.044544648 | 0.517957688 | 3.724945049 | 0.139051095 |
| Psmc1 | 1000.106951 | -0.2014043 | 0.112126889 | -1.796217682 |
| Nrde2 | 211.3513749 | 0.248061989 | 0.103721587 | 2.391613895 |
| Gm10433 | 3.50734834 | 0.712602953 | 0.565762952 | 1.259543332 |
| Calm1 | 27013.61596 | 0.544973062 | 0.082370223 | 6.616141642 |
| Gm26723 | 0.970285468 | 0.481668929 | 0.933696923 | 0.515872889 |
| Ttc7b | 1662.607921 | 0.34643734 | 0.069962147 | 4.951782569 |
| Rps6ka5 | 469.9143002 | 0.658611757 | 0.091674758 | 7.184221359 |
| Gm25801 | 0.225129614 | 1.580880368 | 3.710727143 | 0.426029807 |
| 9030617O03Rik | 99.5661288 | -0.336766822 | 0.168654275 | -1.996787934 |
| Gm8482 | 2.942155206 | 1.41653509 | 0.840096998 | 1.686156591 |
| Gpr68 | 165.6196338 | -0.20127482 | 0.120384352 | -1.671935077 |

| | | | | |
|---|---|---|---|---|
| Ccdc88c | 393.2724972 | 0.261232325 | 0.149278151 | 1.749970258 |
| A630072L19Rik | 0.314467885 | 1.092836081 | 2.014475605 | 0.542491593 |
| Mir1190 | 0.80558184 | 1.005296897 | 1.343722286 | 0.748143354 |
| Smek1 | 938.2289803 | -0.313275199 | 0.053282252 | -5.879541228 |
| D130020L05Rik | 15.63426209 | 0.273776903 | 0.285673163 | 0.958357097 |
| Kif4-ps | 0 | NA | NA | NA |
| Gm18848 | 0 | NA | NA | NA |
| Gm22749 | 0 | NA | NA | NA |
| Catsperb | 0.068097359 | 0.037064902 | 3.724945049 | 0.009950456 |
| Gm24542 | 0 | NA | NA | NA |
| Tc2n | 7.321160109 | 1.163732886 | 0.480166157 | 2.423604557 |
| Gm16285 | 0 | NA | NA | NA |
| Fbln5 | 104.6927819 | 0.084434427 | 0.196303474 | 0.43012192 |
| Trip11 | 1573.263012 | -0.220108216 | 0.048818888 | -4.508669192 |
| Gm20701 | 0 | NA | NA | NA |
| Gm20702 | 0 | NA | NA | NA |
| Gm20036 | 4.340709641 | -0.445411413 | 0.553169463 | -0.805198845 |
| Atxn3 | 451.4130845 | -0.08206701 | 0.073647867 | -1.114316187 |
| Cpsf2 | 797.9273037 | -0.039871538 | 0.076818945 | -0.519032615 |
| Gm16339 | 11.18950982 | 1.757842402 | 0.467007293 | 3.764057709 |
| Slc24a4 | 531.3783664 | 0.725549417 | 0.185349287 | 3.914498028 |
| Gm25203 | 0 | NA | NA | NA |
| Rin3 | 105.1190873 | -0.603836244 | 0.17157954 | -3.519278831 |
| Lgmn | 2861.87464 | -0.805034734 | 0.132035961 | -6.097086959 |
| Golga5 | 336.8030054 | -0.054327993 | 0.089247917 | -0.608731222 |
| Chga | 1338.904208 | -0.57182245 | 0.177386157 | -3.22360245 |
| Itpk1 | 872.6122047 | -0.008926991 | 0.122241763 | -0.073027341 |
| Gm37019 | 35.26459873 | -0.075940592 | 0.318475967 | -0.238449994 |
| Gm20069 | 0.551997393 | 0.278849236 | 1.283838444 | 0.217199631 |
| Gm28373 | 39.99975009 | -0.000560385 | 0.298112487 | -0.001879776 |
| Mir1936 | 2.100202168 | -0.156763294 | 0.766474413 | -0.20452515 |
| Gm28051 | 0.060174537 | 0.517957688 | 3.724945049 | 0.139051095 |
| Gm21971 | 0 | NA | NA | NA |
| Gm20604 | 131.2845531 | -0.087012268 | 0.116719886 | -0.745479376 |
| Moap1 | 4.795281827 | -0.699057478 | 0.538425703 | -1.298336008 |
| Tmem251 | 141.6121926 | -0.500176973 | 0.15352268 | -3.258000523 |
| AK010878 | 58.12541358 | -0.383066877 | 0.154615445 | -2.477545996 |
| Ubr7 | 418.4523783 | -0.055731343 | 0.072054597 | -0.773459921 |
| Btbd7 | 452.2462838 | -0.241527883 | 0.07677773 | -3.145806518 |
| Cox8c | 0 | NA | NA | NA |
| Unc79 | 1662.520299 | 0.293657482 | 0.115483486 | 2.542852596 |
| AC122367.1 | 17.42179249 | 0.780853273 | 0.399261725 | 1.955742871 |
| Gm27976 | 0.999255461 | -0.667779085 | 1.114610237 | -0.599114438 |
| Prima1 | 64.20214066 | -0.153571994 | 0.168461779 | -0.911613274 |
| Fam181a | 11.59340835 | 0.871905485 | 0.358676933 | 2.430893671 |
| Gm29508 | 2.89487079 | 0.386521067 | 0.776152282 | 0.497996432 |
| Asb2 | 49.37814463 | -0.164574958 | 0.214877258 | -0.76590217 |

| | | | | |
|---|---|---|---|---|
| Gm15523 | 0.643207634 | -0.383421302 | 1.382484742 | -0.277342158 |
| Otub2 | 164.6469116 | -0.330610122 | 0.170467263 | -1.939434676 |
| 9330161L09Rik | 14.42659375 | 0.067671767 | 0.281127917 | 0.240715214 |
| Ddx24 | 2634.287795 | 0.120811983 | 0.097550386 | 1.238457247 |
| Ifi27 | 153.7870265 | -0.174677304 | 0.24227225 | -0.720995923 |
| Ifi27l2a | 1103.207706 | -0.915236486 | 0.261688916 | -3.49742167 |
| Ifi27l2b | 0.044191549 | 0.037062677 | 3.724945049 | 0.009949859 |
| Ppp4r4 | 572.2723124 | 0.173432612 | 0.096320046 | 1.800586884 |
| Serpina10 | 1.913476902 | -1.368097647 | 0.836815064 | -1.634886496 |
| Gm18501 | 0 | NA | NA | NA |
| Serpina6 | 0.146451277 | -0.443820591 | 2.882358847 | -0.153978257 |
| Serpina16 | 0 | NA | NA | NA |
| Serpina1f | 0.066050875 | -0.443818988 | 3.724945049 | -0.119147795 |
| Gm5656 | 0 | NA | NA | NA |
| Gm17142 | 0 | NA | NA | NA |
| Serpina1b | 7.892462027 | -0.323049599 | 0.390633591 | -0.826988784 |
| Gm17042 | 0 | NA | NA | NA |
| Serpina1d | 0 | NA | NA | NA |
| Gm17043 | 0 | NA | NA | NA |
| Gm8893 | 0.487995331 | -0.431172119 | 1.457877426 | -0.29575334 |
| Gm6918 | 0 | NA | NA | NA |
| Serpina1a | 0.839309821 | 0.309094247 | 1.00121627 | 0.308718762 |
| Gm8895 | 0.092691418 | -0.908687078 | 3.724560878 | -0.2439716 |
| Serpina1c | 0.721092599 | 1.044563217 | 1.155608089 | 0.903907844 |
| B430119L08Rik | 0.437661561 | -1.450224413 | 1.371259905 | -1.05758537 |
| Gm17198 | 0 | NA | NA | NA |
| Serpina1e | 0.347991053 | 0.047789315 | 1.736479313 | 0.027520809 |
| Serpina11 | 2.314302159 | -1.070753284 | 0.641712139 | -1.668588172 |
| Gm28577 | 0 | NA | NA | NA |
| Serpina9 | 42.26228332 | 0.925081681 | 0.251697277 | 3.675374204 |
| Gm37496 | 0.085433889 | 0.037058229 | 3.724945049 | 0.009948665 |
| Serpina12 | 0.359936507 | 1.737246044 | 3.700439905 | 0.469470141 |
| Serpina4-ps1 | 0.088534442 | -0.905414734 | 3.723500899 | -0.243162217 |
| Serpina5 | 0.066050875 | -0.443818988 | 3.724945049 | -0.119147795 |
| Gm23289 | 0 | NA | NA | NA |
| Serpina3a | 0.228000506 | -1.632571791 | 2.476811158 | -0.659142618 |
| Serpina3b | 0.099106094 | -0.93550548 | 3.724188147 | -0.251197158 |
| Serpina3c | 2.76240905 | -0.0853284 | 0.735693916 | -0.115983561 |
| Serpina3d-ps | 0 | NA | NA | NA |
| Serpina3e-ps | 7.226489367 | -0.530206815 | 0.442057475 | -1.199406966 |
| Serpina3f | 27.21369777 | -0.67261131 | 0.451038989 | -1.491248711 |
| Gm8918 | 0 | NA | NA | NA |
| Serpina3g | 192.2982361 | -1.715036419 | 0.469516953 | -3.652767825 |
| Serpina3h | 81.26495241 | -0.916004801 | 0.42241353 | -2.168502514 |
| Serpina3i | 29.39647476 | -0.464204165 | 0.445571719 | -1.041816942 |
| Serpina3j | 0 | NA | NA | NA |
| Serpina3m | 31.36521844 | -0.833867738 | 0.352035604 | -2.368702846 |

| | | | | |
|---|---|---|---|---|
| Serpina3k | 1.059265352 | -0.5888668 | 1.040129877 | -0.566147375 |
| Serpina3l-ps | 0 | NA | NA | NA |
| Serpina3n | 2233.484408 | -0.534389398 | 0.307242241 | -1.739309662 |
| Gm24334 | 0 | NA | NA | NA |
| Gsc | 0.456948362 | 0.249898685 | 1.478451243 | 0.169027343 |
| Gm10000 | 0.122601409 | 0.998850474 | 3.724945049 | 0.268151734 |
| Dicer1 | 1104.172179 | 0.185659973 | 0.098158569 | 1.89142909 |
| Clmn | 1628.39178 | 0.712188727 | 0.104088725 | 6.842131318 |
| 4930408O17Rik | 1.425419836 | -0.970662927 | 0.98957676 | -0.980886947 |
| Gm28875 | 11.6389744 | -0.396666238 | 0.388894983 | -1.019982916 |
| Syne3 | 100.3522481 | -0.513342989 | 0.147554469 | -3.479006714 |
| Scarna13 | 339.6868988 | 0.272561538 | 0.290203111 | 0.939209565 |
| mmu-mir-3069 | 0 | NA | NA | NA |
| Glrx5 | 236.9685444 | -0.0104862 | 0.103514867 | -0.101301388 |
| Tcl1b2 | 0.273490436 | 0.743541448 | 2.416262625 | 0.307723772 |
| Tcl1b1 | 0.12092566 | 0.998850474 | 3.724945049 | 0.268151734 |
| Tcl1b5 | 0 | NA | NA | NA |
| Tcl1b3 | 0 | NA | NA | NA |
| Tcl1b4 | 0 | NA | NA | NA |
| Tcl1 | 0 | NA | NA | NA |
| Tunar | 293.4716965 | -0.150773504 | 0.104515957 | -1.442588369 |
| D430019H16Rik | 1275.143419 | -0.08928217 | 0.140039778 | -0.637548641 |
| Bdkrb2 | 21.21549069 | -1.110365828 | 0.319010349 | -3.480657704 |
| Bdkrb1 | 6.675953194 | -0.384951866 | 0.561168758 | -0.685982355 |
| Atg2b | 1471.256828 | 0.066373129 | 0.056391596 | 1.177003912 |
| Gm22790 | 0 | NA | NA | NA |
| Gskip | 216.9176705 | -0.438962844 | 0.133561566 | -3.286595506 |
| Ak7 | 31.57741441 | -0.105111311 | 0.204362726 | -0.514336998 |
| Papola | 1915.550768 | -0.352917452 | 0.081782933 | -4.31529463 |
| Gm22120 | 0 | NA | NA | NA |
| Vrk1 | 161.0238951 | 0.123223801 | 0.114788847 | 1.073482349 |
| Gm24986 | 0 | NA | NA | NA |
| Gm17032 | 0.342848402 | -1.505607016 | 1.967430463 | -0.765265682 |
| 1700121N20Rik | 0 | NA | NA | NA |
| Gm3191 | 0.406963084 | -1.895212471 | 3.680822654 | -0.514888287 |
| Gm17031 | 0.100902008 | -0.876201588 | 3.72093227 | -0.235479048 |
| Gm17053 | 0.121578509 | 0.037073798 | 3.724945049 | 0.009952844 |
| Gm17033 | 0 | NA | NA | NA |
| 4933406K04Rik | 0 | NA | NA | NA |
| 1700013N06Rik | 0 | NA | NA | NA |
| Gm9599 | 0 | NA | NA | NA |
| Gm16087 | 0.365883897 | 0.517974004 | 1.567381877 | 0.330470839 |
| Gm16086 | 0.197846125 | -0.443827572 | 3.270942157 | -0.135687992 |
| Gm16085 | 0 | NA | NA | NA |
| Gm16084 | 0 | NA | NA | NA |
| Gm2800 | 0.20562365 | 0.517963344 | 2.612685484 | 0.198249406 |
| Gm5187 | 0 | NA | NA | NA |

| | | | | |
|---|---|---|---|---|
| Gm17052 | 0.203140137 | -0.948632908 | 2.216123492 | -0.428059588 |
| 3110018I06Rik | 0 | NA | NA | NA |
| Gm9006 | 0 | NA | NA | NA |
| 4930465M20Rik | 0.148569308 | -0.443821873 | 2.869495721 | -0.154668944 |
| Gm15208 | 0.15596086 | 0.037059316 | 3.490591407 | 0.010616916 |
| Bcl11b | 1306.155659 | 0.982247606 | 0.152597584 | 6.436849003 |
| Setd3 | 1675.172341 | 0.159581636 | 0.054134875 | 2.94785266 |
| Ccnk | 411.8626785 | -0.208361735 | 0.10718804 | -1.943889767 |
| Ccdc85c | 333.5260322 | 0.111747973 | 0.085275546 | 1.310433978 |
| Hhipl1 | 69.09591632 | -0.273758595 | 0.168608519 | -1.623634418 |
| 4930478K11Rik | 1.119709845 | -1.313276484 | 1.176597155 | -1.116164933 |
| Cyp46a1 | 1040.049922 | 0.977573441 | 0.15375132 | 6.358146641 |
| Eml1 | 662.3640817 | -0.346035914 | 0.099294531 | -3.484944341 |
| Gm15636 | 0 | NA | NA | NA |
| Gm16596 | 7.416129088 | 0.343896694 | 0.382653955 | 0.898714596 |
| Evl | 1049.201664 | 0.241520891 | 0.056419854 | 4.28077841 |
| Mir342 | 1.676187767 | 0.716337191 | 1.078237155 | 0.664359587 |
| Degs2 | 4.353830809 | 0.02111867 | 0.751459571 | 0.028103535 |
| Gm23872 | 0 | NA | NA | NA |
| Yy1 | 536.7063009 | -0.291205643 | 0.081993197 | -3.551582989 |
| Slc25a29 | 103.2788333 | 0.120297669 | 0.13568038 | 0.886625385 |
| Mir345 | 1.700378639 | -0.002471693 | 0.872502387 | -0.002832878 |
| Slc25a47 | 16.0241689 | 0.05915796 | 0.317201261 | 0.186499763 |
| Wars | 1271.197342 | -0.695880602 | 0.116002589 | -5.998836819 |
| Gm22079 | 2.165001819 | 0.019556935 | 0.810678812 | 0.024124147 |
| Wdr25 | 96.15949304 | -0.032356111 | 0.111528921 | -0.290114089 |
| Begain | 792.2631617 | 0.599386802 | 0.109397697 | 5.478970935 |
| Gm28498 | 1.537157928 | 0.943940679 | 1.012576865 | 0.93221632 |
| Gm22234 | 0 | NA | NA | NA |
| Gm25297 | 0 | NA | NA | NA |
| Dlk1 | 383.0032136 | -0.113385381 | 0.175880452 | -0.644673015 |
| Gm27528 | 0 | NA | NA | NA |
| Meg3 | 23754.92684 | 0.954255799 | 0.443936982 | 2.149529861 |
| Gm27596 | 7.046539481 | 2.039221333 | 0.598204537 | 3.408903156 |
| Gm27300 | 0 | NA | NA | NA |
| Mir1906-1 | 0 | NA | NA | NA |
| Mir770 | 0 | NA | NA | NA |
| Gm26906 | 0.900331428 | 0.211439557 | 1.363800922 | 0.155036966 |
| Mir673 | 3.122023216 | 1.496700246 | 0.88539147 | 1.690438972 |
| Mir493 | 16.71408191 | 1.025983888 | 0.49650606 | 2.066407585 |
| Mir337 | 13.9759078 | 0.779869789 | 0.383875095 | 2.031571727 |
| mmu-mir-3544 | 0 | NA | NA | NA |
| Mir540 | 13.29036576 | 1.013479697 | 0.448229761 | 2.261071853 |
| Mir665 | 11.41153131 | 1.165383679 | 0.47075039 | 2.475587283 |
| Mir3070a | 19.33383777 | 0.742615528 | 0.454455949 | 1.634075933 |
| Mir3070b | 15.52429422 | 0.150826384 | 0.418154509 | 0.360695342 |
| Rtl1 | 49.79748458 | -0.617415586 | 0.269680019 | -2.289437643 |

| | | | | |
|---|---|---|---|---|
| Mir431 | 8.591852434 | 0.519944569 | 0.465152403 | 1.117794009 |
| Mir433 | 30.4253769 | 0.827331491 | 0.383131891 | 2.159390826 |
| Mir127 | 8.254821574 | 0.843100094 | 0.465092942 | 1.812756159 |
| Mir434 | 6.186359681 | 0.190100993 | 0.471826287 | 0.402904625 |
| Mir432 | 5.368193358 | 0.286280382 | 0.503436723 | 0.568652164 |
| Mir136 | 0 | NA | NA | NA |
| mmu-mir-136 | 4.439960643 | 1.169207323 | 0.592729761 | 1.972580761 |
| Gm27350 | 1.495241998 | 1.200499603 | 1.147968744 | 1.045759834 |
| Rian | 7362.023447 | 0.440235338 | 0.112785526 | 3.903296409 |
| Mir341 | 46.98560136 | 0.491035916 | 0.321979837 | 1.525051758 |
| Mir1188 | 40.87071727 | 0.564858887 | 0.405661285 | 1.392439722 |
| Gm23912 | 5.84972083 | 1.204043658 | 0.6870729 | 1.752424899 |
| Mir370 | 7.676564023 | 0.597527161 | 0.625209055 | 0.95572378 |
| Gm24564 | 32.85825865 | -0.082203701 | 0.349670974 | -0.235088717 |
| Gm23474 | 0 | NA | NA | NA |
| AF357425 | 0 | NA | NA | NA |
| Gm26945 | 332.9786148 | 0.0867621 | 0.208567339 | 0.415990826 |
| Gm24899 | 0 | NA | NA | NA |
| Gm23600 | 67.41334034 | 0.928642832 | 0.182089824 | 5.099916135 |
| Gm24895 | 0 | NA | NA | NA |
| DQ267100 | 0 | NA | NA | NA |
| DQ267101 | 11.01360181 | 0.53829092 | 0.345008728 | 1.560224064 |
| Gm23508 | 0 | NA | NA | NA |
| Gm26922 | 0 | NA | NA | NA |
| DQ267102 | 24.0816795 | 0.253523909 | 0.283432114 | 0.894478419 |
| Gm25357 | 8.037931983 | 0.640128001 | 0.416769553 | 1.535927941 |
| Gm27604 | 0 | NA | NA | NA |
| Gm37899 | 685.7979278 | 0.777450825 | 0.29348105 | 2.649066524 |
| Gm23347 | 0 | NA | NA | NA |
| AF357428 | 5.570026761 | 0.663377495 | 0.595420344 | 1.11413307 |
| Gm23736 | 2.72795131 | 0.621966364 | 0.625308475 | 0.99465526 |
| AF357341 | 4.666739356 | 0.753824107 | 0.640803386 | 1.176373476 |
| Mir882 | 2.263396373 | -0.063345689 | 0.816258842 | -0.077604904 |
| Gm22981 | 1.907638496 | 1.887031307 | 0.853218487 | 2.211662469 |
| Gm25856 | 14.93485102 | 0.978649505 | 0.369227488 | 2.650532628 |
| Gm23787 | 13.95901734 | 0.723472518 | 0.393884816 | 1.836761634 |
| Gm25854 | 10.14198828 | 0.720770168 | 0.394615595 | 1.82651212 |
| Gm22205 | 21.52538062 | 0.472749507 | 0.295940856 | 1.597445901 |
| Gm25224 | 10.64312297 | 0.62303878 | 0.382926251 | 1.627046405 |
| B830012L14Rik | 195.3433677 | 0.784789305 | 0.357530544 | 2.195027297 |
| Mir379 | 4.424652029 | 0.898997893 | 0.684704155 | 1.312972743 |
| Mir411 | 6.496099058 | 0.654638342 | 0.491721284 | 1.331319922 |
| mmu-mir-299a | 2.459243483 | 0.311014464 | 0.807832137 | 0.384998875 |
| Mir299b | 0 | NA | NA | NA |
| Gm25750 | 0 | NA | NA | NA |
| mmu-mir-380 | 4.04787115 | 0.359968224 | 0.601285237 | 0.598664663 |
| Mir1197 | 3.390250948 | 1.026498022 | 0.662397024 | 1.549671851 |

| | | | | |
|---|---|---|---|---|
| Mir323 | 3.255378368 | 0.943166518 | 0.862901005 | 1.093018217 |
| Mir758 | 6.502832914 | 0.477520291 | 0.499709087 | 0.955596575 |
| Mir329 | 5.909624248 | 0.623388254 | 0.582569828 | 1.070066151 |
| Mir494 | 6.573050005 | 0.505081919 | 0.488806056 | 1.033297179 |
| Mir679 | 11.72323245 | 1.386886011 | 0.407323177 | 3.404878707 |
| Mir1193 | 10.23592663 | 0.677805542 | 0.518133803 | 1.308166998 |
| Mir666 | 14.05147067 | 0.516228476 | 0.412260187 | 1.252190953 |
| Mir543 | 4.380404366 | 0.875406654 | 0.73225899 | 1.195487752 |
| Mir495 | 2.202405632 | -0.039060313 | 0.840179171 | -0.046490456 |
| Mir667 | 9.470118255 | 0.771510348 | 0.406441277 | 1.898208649 |
| Mir376b | 10.35390795 | 0.636022674 | 0.443975795 | 1.432561597 |
| Mir654 | 8.477009777 | 0.247624956 | 0.444344313 | 0.557281703 |
| mmu-mir-376b | 2.209145549 | 0.389212604 | 0.720820338 | 0.539957855 |
| Gm25252 | 0.047948698 | 0.037064902 | 3.724945049 | 0.009950456 |
| mmu-mir-376a | 5.585566604 | -0.327957776 | 0.582728933 | -0.562796453 |
| Mir300 | 6.166566501 | -0.053581772 | 0.551219364 | -0.097205895 |
| Mir381 | 6.601995524 | 0.755482674 | 0.565488981 | 1.335981248 |
| Mir487b | 4.545755139 | 1.953744997 | 0.70688101 | 2.763895151 |
| AC121784.1 | 0 | NA | NA | NA |
| Mir539 | 4.506246945 | -0.328603434 | 0.559499087 | -0.587317194 |
| Gm27379 | 8.227541344 | 0.912209918 | 0.553994023 | 1.646606064 |
| Mir544 | 5.245180705 | 0.712988212 | 0.593013224 | 1.202314187 |
| Mirg | 537.7305624 | 0.212748711 | 0.271224557 | 0.784400624 |
| Mir382 | 2.007426773 | 0.691013701 | 0.771714281 | 0.895426866 |
| Mir134 | 0 | NA | NA | NA |
| Mir668 | 0 | NA | NA | NA |
| Mir485 | 0 | NA | NA | NA |
| Mir453 | 14.69478301 | 0.684324624 | 0.44155191 | 1.549816929 |
| Mir154 | 6.301721876 | -0.279698755 | 0.485918148 | -0.575608785 |
| Mir496a | 7.410274305 | 0.335477912 | 0.45532834 | 0.736782411 |
| Mir377 | 0 | NA | NA | NA |
| Mir541 | 8.228173234 | 0.484085439 | 0.454627873 | 1.064794898 |
| mmu-mir-409 | 3.297680665 | 0.015770506 | 0.600507881 | 0.026261946 |
| Gm24321 | 0 | NA | NA | NA |
| Mir412 | 2.651183465 | 1.322322935 | 0.829796826 | 1.593550246 |
| Mir369 | 1.213203356 | 0.745006403 | 1.045806729 | 0.712374842 |
| Mir410 | 0 | NA | NA | NA |
| Mir3072 | 1.399651712 | -0.056306017 | 1.253787288 | -0.044908748 |
| Mir1247 | 0.141331096 | -0.443820795 | 3.561840904 | -0.124604329 |
| Dio3 | 10.88659069 | 1.432738082 | 0.380563071 | 3.764784846 |
| Ppp2r5c | 1467.571291 | 0.177880344 | 0.054854666 | 3.242756877 |
| Dync1h1 | 16373.17254 | -0.109054771 | 0.045925216 | -2.374616375 |
| Gm17111 | 0.611020101 | 1.919938146 | 1.347672028 | 1.424633075 |
| 1700001K19Rik | 0.241252684 | -1.786647812 | 3.028958552 | -0.589855484 |
| Gm27299 | 0.462939631 | 0.883852479 | 1.461979552 | 0.604558715 |
| Hsp90aa1 | 8408.664402 | -0.227347713 | 0.118853367 | -1.912842015 |
| Wdr20 | 372.1537339 | 0.065435024 | 0.098841389 | 0.662020482 |

| | | | | |
|---|---|---|---|---|
| Mok | 93.83344369 | -0.190648769 | 0.133829288 | -1.424566861 |
| 4921507G05Rik | 0.309276477 | 0.517971476 | 1.790548415 | 0.28928091 |
| Gm26912 | 1.084888343 | -0.435485056 | 1.030335045 | -0.422663539 |
| Zfp839 | 119.0822902 | 0.05260908 | 0.140200552 | 0.375241608 |
| Mpc1-ps | 240.9835416 | 0.125157865 | 0.113973421 | 1.09813204 |
| Cinp | 250.286756 | -0.155910116 | 0.089519937 | -1.741624506 |
| Tecpr2 | 926.178597 | 0.435698513 | 0.080026166 | 5.444450674 |
| Rps19-ps6 | 18.4757316 | -0.204790764 | 0.261716299 | -0.782491442 |
| Ankrd9 | 131.61924 | 0.11024674 | 0.102307376 | 1.077603046 |
| Rcor1 | 394.3624715 | -0.439680708 | 0.090907799 | -4.836556524 |
| 4930595D18Rik | 0.920001838 | 0.428581886 | 1.035287907 | 0.413973623 |
| Traf3 | 916.582985 | -0.051205946 | 0.062462482 | -0.819787253 |
| Amn | 31.33882718 | 1.079927369 | 0.23132425 | 4.66845723 |
| A230087F16Rik | 11.27729028 | 0.344474206 | 0.399424307 | 0.862426748 |
| Cdc42bpb | 2200.324733 | 0.183471043 | 0.064086483 | 2.862866453 |
| A230065H16Rik | 14.31749866 | -0.319301326 | 0.358999995 | -0.889418747 |
| Exoc3l4 | 26.39184743 | -0.51057917 | 0.258198178 | -1.977470071 |
| Gm10425 | 4.124014502 | -1.538682578 | 0.645770272 | -2.382708905 |
| Tnfaip2 | 430.9565934 | -0.777039287 | 0.536658744 | -1.447920668 |
| Gm266 | 27.84625617 | -0.196767176 | 0.239920928 | -0.820133442 |
| Eif5 | 3419.805663 | -0.1760462 | 0.107027759 | -1.644864867 |
| Snora28 | 29.94917886 | 0.5235521 | 0.240966716 | 2.172715422 |
| 2810029C07Rik | 61.53833547 | 0.66652604 | 0.192986787 | 3.45373924 |
| Mark3 | 1177.712359 | -0.101099294 | 0.071497938 | -1.414016922 |
| Ckb | 9080.847841 | 0.020805488 | 0.093063807 | 0.223561543 |
| Trmt61a | 154.0841474 | -0.400573028 | 0.105273266 | -3.805078367 |
| Bag5 | 421.2921291 | 0.005962724 | 0.074002046 | 0.080575123 |
| Apopt1 | 168.4724147 | 0.061156411 | 0.105557063 | 0.579368247 |
| Gm15995 | 0.042716945 | 0.037062677 | 3.724945049 | 0.009949859 |
| Gm15996 | 0.04393175 | 0.037062677 | 3.724945049 | 0.009949859 |
| Klc1 | 4978.643089 | 0.082897427 | 0.075804607 | 1.093567136 |
| Xrcc3 | 62.34707954 | 0.266403694 | 0.176596325 | 1.508546083 |
| Zfyve21 | 80.73079552 | 0.005631117 | 0.159828467 | 0.035232252 |
| Ppp1r13b | 928.8327789 | 0.400548927 | 0.075978294 | 5.271886306 |
| 2010107E04Rik | 287.253024 | -0.016780601 | 0.165336575 | -0.101493579 |
| Tdrd9 | 6.653230718 | 0.2535957 | 0.462189527 | 0.548683354 |
| Rd3l | 8.12228703 | -0.390787017 | 0.455131553 | -0.858624314 |
| Aspg | 140.2133012 | -0.753120836 | 0.187853451 | -4.009087036 |
| Mir3073a | 0.372180926 | -0.034238338 | 1.568912456 | -0.021822975 |
| mmu-mir-3073b | 0 | NA | NA | NA |
| Mir203 | 0 | NA | NA | NA |
| Kif26a | 86.52596024 | 0.34928797 | 0.166390178 | 2.099210259 |
| Gm38123 | 0.492632247 | -1.54920805 | 1.427356065 | -1.085369017 |
| A730018C14Rik | 2.250253578 | -0.223601173 | 0.764041008 | -0.292655984 |
| A530016L24Rik | 0 | NA | NA | NA |
| Tmem179 | 794.5266479 | 0.238397853 | 0.062420723 | 3.819210049 |
| Inf2 | 1281.719595 | 0.055904881 | 0.086121177 | 0.649142095 |

| | | | | |
|---|---|---|---|---|
| Adssl1 | 268.3142132 | -0.282963836 | 0.111505706 | -2.537662382 |
| Siva1 | 44.47611933 | -0.195225252 | 0.200893714 | -0.971783778 |
| Akt1 | 1539.989558 | 0.009584048 | 0.07293188 | 0.13141096 |
| Mir6939 | 0.990373052 | 0.229273982 | 0.951147151 | 0.241049959 |
| Zbtb42 | 29.83589809 | -0.519676408 | 0.246724109 | -2.106305746 |
| Gm30238 | 24.73544283 | 0.080390244 | 0.36214331 | 0.221984616 |
| Cep170b | 3968.143558 | 0.308411099 | 0.076562215 | 4.028241623 |
| Mir6940 | 0.393711421 | 0.600863589 | 1.727786089 | 0.347765034 |
| Pld4 | 304.2279336 | -0.440415 | 0.141580109 | -3.110712395 |
| Ahnak2 | 140.5418072 | -0.964106172 | 0.248933504 | -3.872946616 |
| BC022687 | 106.0209119 | -0.446828045 | 0.127449976 | -3.505909226 |
| Cdca4 | 92.32272779 | -0.318820805 | 0.153602737 | -2.075619295 |
| Gpr132 | 12.16430624 | -0.616964675 | 0.346681197 | -1.779631202 |
| Jag2 | 271.294481 | 0.34511443 | 0.118231123 | 2.918981253 |
| Mir6941 | 0.653922745 | -0.796129835 | 1.621756836 | -0.490905799 |
| Nudt14 | 37.15232537 | -0.091965637 | 0.197405439 | -0.465871848 |
| Gm26583 | 4.273896503 | 1.600218195 | 0.740766149 | 2.160220466 |
| Brf1 | 375.6428438 | 0.199489648 | 0.078131344 | 2.553260163 |
| Btbd6 | 332.183834 | 0.258868936 | 0.071325853 | 3.629384386 |
| 9230104M06Rik | 1.025204639 | 1.002088622 | 1.39245541 | 0.719655807 |
| Pacs2 | 2506.266713 | -0.212317494 | 0.092616954 | -2.292425788 |
| AC073562.1 | 0.303558719 | -1.27811143 | 2.024687248 | -0.631263634 |
| Tex22 | 3.833641248 | 0.515598455 | 0.595928181 | 0.865202337 |
| Mta1 | 765.704057 | -0.140153067 | 0.059460863 | -2.357064135 |
| Crip2 | 531.6473017 | -0.263027728 | 0.1019546 | -2.579851508 |
| Crip1 | 42.16258074 | -1.050779636 | 0.248422055 | -4.22981621 |
| 4930427A07Rik | 20.00319577 | -0.119619669 | 0.264492007 | -0.452261942 |
| RP24-468O24.5 | 0.060174537 | 0.517957688 | 3.724945049 | 0.139051095 |
| Tmem121 | 107.667371 | 0.064788152 | 0.11929998 | 0.543069259 |
| Gm25622 | 0 | NA | NA | NA |
| Igha | 7.387870444 | -2.866689867 | 1.68855013 | -1.697722689 |
| Ighe | 0.136120942 | -0.876404342 | 3.595683781 | -0.24373788 |
| Ighg2c | 6.374334057 | -2.092256136 | 1.087496995 | -1.923919005 |
| Ighg2b | 4.144711669 | -1.237041469 | 1.300788436 | -0.950993593 |
| Gm37944 | 0.048949532 | 0.037064902 | 3.724945049 | 0.009950456 |
| Ighg1 | 1.048355144 | -1.088931214 | 2.200588211 | -0.49483643 |
| Ighg3 | 2.594242155 | 0.099128155 | 1.652467118 | 0.059987974 |
| Gm17421 | 0.038981395 | -0.443816764 | 3.724945049 | -0.119147198 |
| Gm9134 | 2.929093634 | 0.254532613 | 0.606721275 | 0.419521489 |
| Gm9136 | 0 | NA | NA | NA |
| Gm5190 | 0.038981395 | -0.443816764 | 3.724945049 | -0.119147198 |
| Ighd | 2.086187063 | 1.157278761 | 0.805270456 | 1.437130534 |
| Ighm | 86.67814311 | -0.488414583 | 0.445657566 | -1.095941415 |
| Gm25747 | 0 | NA | NA | NA |
| Gm24022 | 0 | NA | NA | NA |
| AC073553.1 | 0 | NA | NA | NA |
| Gm22175 | 0 | NA | NA | NA |

| | | | | |
|---|---|---|---|---|
| Ighj4 | 0.495312865 | 0.299341738 | 1.678734778 | 0.178313896 |
| Ighj3 | 0.07796279 | -0.798301147 | 3.721829328 | -0.214491605 |
| Ighj2 | 0.566709933 | -0.332790824 | 1.317265983 | -0.25263753 |
| Ighj1 | 0.937655582 | 0.987719821 | 1.170237792 | 0.844033433 |
| Ighd4-1 | 0 | NA | NA | NA |
| Ighd3-2 | 0 | NA | NA | NA |
| Ighd5-6 | 0 | NA | NA | NA |
| Ighd2-8 | 0 | NA | NA | NA |
| Ighd5-5 | 0 | NA | NA | NA |
| Ighd2-7 | 0 | NA | NA | NA |
| Ighd5-8 | 0 | NA | NA | NA |
| Ighd5-4 | 0 | NA | NA | NA |
| Ighd2-6 | 0 | NA | NA | NA |
| Ighd5-7 | 0 | NA | NA | NA |
| Ighd5-3 | 0 | NA | NA | NA |
| Ighd2-5 | 0 | NA | NA | NA |
| Gm37327 | 0 | NA | NA | NA |
| Ighd5-2 | 0 | NA | NA | NA |
| Ighd2-4 | 0 | NA | NA | NA |
| Ighd6-2 | 0 | NA | NA | NA |
| Gm37227 | 0 | NA | NA | NA |
| Ighd2-3 | 0 | NA | NA | NA |
| Ighd6-1 | 0 | NA | NA | NA |
| Ighd1-1 | 0 | NA | NA | NA |
| Adam6b | 0.203125505 | 0.517963919 | 2.630882593 | 0.196878386 |
| Ighd3-1 | 0 | NA | NA | NA |
| Ighd5-1 | 0 | NA | NA | NA |
| Gm16968 | 0 | NA | NA | NA |
| Adam6a | 0 | NA | NA | NA |
| Ighv5-1 | 0 | NA | NA | NA |
| Ighv2-1 | 0 | NA | NA | NA |
| Ighv5-2 | 0 | NA | NA | NA |
| Ighv2-2 | 0 | NA | NA | NA |
| Ighv5-3 | 0 | NA | NA | NA |
| Ighv5-4 | 0.345657073 | -2.280878757 | 1.908297619 | -1.195242678 |
| Ighv6-1 | 0 | NA | NA | NA |
| Ighv2-3 | 0.072025462 | 0.517959912 | 3.724945049 | 0.139051692 |
| Ighv5-5 | 0 | NA | NA | NA |
| Ighv5-6 | 0.066050875 | -0.443818988 | 3.724945049 | -0.119147795 |
| Gm9517 | 109.6899113 | 0.234062765 | 0.13685144 | 1.710341998 |
| Ighv5-7 | 0 | NA | NA | NA |
| Ighv2-4 | 0 | NA | NA | NA |
| Ighv5-8 | 0 | NA | NA | NA |
| Ighv5-9 | 0.097139547 | -0.443816764 | 3.724945049 | -0.119147198 |
| Ighv5-10 | 0 | NA | NA | NA |
| Gm37187 | 0 | NA | NA | NA |
| Ighv2-5 | 0 | NA | NA | NA |

| | | | | |
|---|---|---|---|---|
| Ighv5-11 | 0 | NA | NA | NA |
| Ighv5-12 | 0 | NA | NA | NA |
| Ighv2-6 | 0.060174537 | 0.517957688 | 3.724945049 | 0.139051095 |
| Gm37976 | 0 | NA | NA | NA |
| Gm37418 | 0 | NA | NA | NA |
| Ighv5-9-1 | 0 | NA | NA | NA |
| Gm38203 | 0 | NA | NA | NA |
| Ighv5-12-4 | 0 | NA | NA | NA |
| Gm38205 | 0 | NA | NA | NA |
| Ighv2-9-1 | 0 | NA | NA | NA |
| Gm37722 | 0 | NA | NA | NA |
| Ighv5-13 | 0 | NA | NA | NA |
| Ighv2-6-8 | 0.046345709 | -0.443818988 | 3.724945049 | -0.119147795 |
| Gm7003 | 0 | NA | NA | NA |
| Ighv2-7 | 0 | NA | NA | NA |
| Rpl26-ps3 | 0 | NA | NA | NA |
| Ighv5-15 | 0 | NA | NA | NA |
| Ighv5-16 | 0 | NA | NA | NA |
| Ighv5-17 | 0.427539584 | -2.031269592 | 3.695652097 | -0.549637666 |
| Ighv5-18 | 0 | NA | NA | NA |
| Ighv2-8 | 0 | NA | NA | NA |
| Ighv2-9 | 0.631149539 | -3.117018567 | 3.661083925 | -0.851392274 |
| Ighv5-19 | 0 | NA | NA | NA |
| Ighv7-1 | 0 | NA | NA | NA |
| Ighv7-2 | 0 | NA | NA | NA |
| Ighv14-1 | 0.480014214 | -0.058866977 | 3.681263755 | -0.01599097 |
| Ighv4-1 | 0.072025462 | 0.517959912 | 3.724945049 | 0.139051692 |
| Ighv3-1 | 0 | NA | NA | NA |
| Gm37961 | 0 | NA | NA | NA |
| Ighv11-1 | 0 | NA | NA | NA |
| Ighv14-2 | 0.20773507 | -0.444654215 | 2.58074016 | -0.172297166 |
| Ighv4-2 | 0 | NA | NA | NA |
| Ighv3-2 | 0 | NA | NA | NA |
| Ighv11-2 | 0 | NA | NA | NA |
| Ighv14-3 | 0 | NA | NA | NA |
| Ighv16-1 | 0 | NA | NA | NA |
| Ighv6-2 | 0 | NA | NA | NA |
| Ighv9-1 | 0 | NA | NA | NA |
| Ighv12-1 | 0 | NA | NA | NA |
| Ighv9-2 | 0 | NA | NA | NA |
| Ighv12-2 | 0 | NA | NA | NA |
| Ighv9-3 | 0 | NA | NA | NA |
| Ighv7-3 | 0 | NA | NA | NA |
| Ighv15-1 | 0 | NA | NA | NA |
| Ighv14-4 | 0 | NA | NA | NA |
| Gm37838 | 0 | NA | NA | NA |
| Ighv3-3 | 0 | NA | NA | NA |

| | | | | |
|---|---|---|---|---|
| Ighv7-4 | 0 | NA | NA | NA |
| Ighv3-4 | 0 | NA | NA | NA |
| Ighv3-5 | 0 | NA | NA | NA |
| Gm18551 | 0 | NA | NA | NA |
| Ighv13-1 | 0 | NA | NA | NA |
| Ighv3-6 | 0.625300531 | -1.492387601 | 3.667459234 | -0.406926841 |
| Ighv9-4 | 0 | NA | NA | NA |
| Ighv3-7 | 0 | NA | NA | NA |
| Ighv5-21 | 0 | NA | NA | NA |
| Ighv3-8 | 0.038981395 | -0.443816764 | 3.724945049 | -0.119147198 |
| Ighv8-1 | 0 | NA | NA | NA |
| Ighv1-1 | 0 | NA | NA | NA |
| Ighv13-2 | 0 | NA | NA | NA |
| Ighv12-3 | 0 | NA | NA | NA |
| Ighv6-3 | 0 | NA | NA | NA |
| Ighv6-4 | 0 | NA | NA | NA |
| Ighv6-5 | 0 | NA | NA | NA |
| Ighv6-6 | 0.211582616 | -0.978186923 | 2.572998811 | -0.380173873 |
| Ighv6-7 | 0 | NA | NA | NA |
| Ighv8-2 | 0 | NA | NA | NA |
| Ighv1-2 | 0 | NA | NA | NA |
| mmu-mir-7094-1 | 0 | NA | NA | NA |
| Ighv10-1 | 0.049553047 | -0.443818988 | 3.724945049 | -0.119147795 |
| Ighv1-3 | 0 | NA | NA | NA |
| Ighv1-4 | 0 | NA | NA | NA |
| Ighv10-2 | 0 | NA | NA | NA |
| Ighv1-5 | 0 | NA | NA | NA |
| Mir7094-2 | 0 | NA | NA | NA |
| Ighv10-3 | 0 | NA | NA | NA |
| Ighv1-6 | 0 | NA | NA | NA |
| Ighv1-7 | 0 | NA | NA | NA |
| Ighv10-4 | 0 | NA | NA | NA |
| Ighv1-8 | 0 | NA | NA | NA |
| Ighv15-2 | 0 | NA | NA | NA |
| Ighv1-9 | 0.287786674 | -0.982516562 | 1.999459422 | -0.491391099 |
| Ighv1-10 | 0 | NA | NA | NA |
| Ighv1-11 | 0 | NA | NA | NA |
| Ighv1-12 | 0 | NA | NA | NA |
| Ighv1-13 | 0 | NA | NA | NA |
| Ighv1-14 | 0 | NA | NA | NA |
| Ighv1-15 | 0 | NA | NA | NA |
| Ighv1-16 | 0 | NA | NA | NA |
| Ighv1-17-1 | 0 | NA | NA | NA |
| Ighv1-17 | 0 | NA | NA | NA |
| Ighv1-18 | 0.194279094 | -1.017823543 | 3.719440177 | -0.273649661 |
| Ighv1-19-1 | 0 | NA | NA | NA |
| Ighv1-19 | 0 | NA | NA | NA |

| | | | | |
|---|---|---|---|---|
| Ighv1-20 | 0 | NA | NA | NA |
| Ighv1-21-1 | 0 | NA | NA | NA |
| Ighv1-21 | 0 | NA | NA | NA |
| Ighv1-22 | 0 | NA | NA | NA |
| Ighv1-23 | 0.097139547 | -0.443816764 | 3.724945049 | -0.119147198 |
| Ighv1-24 | 0 | NA | NA | NA |
| Ighv1-25 | 0 | NA | NA | NA |
| Ighv1-26 | 0 | NA | NA | NA |
| Ighv1-27 | 0 | NA | NA | NA |
| Ighv1-28 | 0 | NA | NA | NA |
| Ighv1-29 | 0 | NA | NA | NA |
| Ighv1-30 | 0 | NA | NA | NA |
| Ighv1-31 | 0 | NA | NA | NA |
| Ighv1-32 | 0 | NA | NA | NA |
| Ighv1-33 | 0.097139547 | -0.443816764 | 3.724945049 | -0.119147198 |
| Ighv1-34 | 0 | NA | NA | NA |
| Ighv1-35 | 0 | NA | NA | NA |
| Ighv1-36 | 0 | NA | NA | NA |
| Ighv1-37 | 0 | NA | NA | NA |
| Ighv1-38 | 0 | NA | NA | NA |
| Ighv1-39 | 0 | NA | NA | NA |
| Ighv1-40 | 0 | NA | NA | NA |
| Ighv1-41 | 0 | NA | NA | NA |
| Ighv1-42 | 0 | NA | NA | NA |
| Ighv1-43 | 0 | NA | NA | NA |
| Ighv1-44 | 0 | NA | NA | NA |
| Ighv1-45 | 0 | NA | NA | NA |
| Ighv1-46 | 0 | NA | NA | NA |
| Ighv1-47 | 0 | NA | NA | NA |
| Ighv8-3 | 0 | NA | NA | NA |
| Ighv8-4 | 0 | NA | NA | NA |
| Ighv1-48 | 0 | NA | NA | NA |
| Ighv1-49 | 0 | NA | NA | NA |
| Ighv8-5 | 0 | NA | NA | NA |
| Ighv1-50 | 0 | NA | NA | NA |
| Ighv1-51 | 0 | NA | NA | NA |
| Ighv1-52 | 0.072025462 | 0.517959912 | 3.724945049 | 0.139051692 |
| Ighv1-53 | 0 | NA | NA | NA |
| Ighv8-6 | 0 | NA | NA | NA |
| Gm9227 | 0 | NA | NA | NA |
| Ighv1-54 | 0 | NA | NA | NA |
| Ighv1-55 | 0 | NA | NA | NA |
| Gm7142 | 0 | NA | NA | NA |
| Ighv1-56 | 0 | NA | NA | NA |
| Gm7019 | 1.468020652 | 0.426370869 | 0.810283596 | 0.526199557 |
| Ighv8-7 | 0 | NA | NA | NA |
| Gm30948 | 0 | NA | NA | NA |

| | | | | |
|---|---|---|---|---|
| Ighv1-57 | 0 | NA | NA | NA |
| Ighv8-8 | 0.144050924 | 1.05357957 | 3.721447096 | 0.283110183 |
| Ighv1-58 | 0 | NA | NA | NA |
| Ighv1-59 | 0 | NA | NA | NA |
| Ighv1-60 | 0 | NA | NA | NA |
| Ighv1-61 | 0 | NA | NA | NA |
| Ighv1-62 | 0 | NA | NA | NA |
| Ighv1-62-1 | 0 | NA | NA | NA |
| Gm19331 | 0 | NA | NA | NA |
| Gm37511 | 0 | NA | NA | NA |
| Gm38184 | 0 | NA | NA | NA |
| Ighv1-62-2 | 0 | NA | NA | NA |
| Ighv1-62-3 | 0 | NA | NA | NA |
| Ighv8-9 | 0 | NA | NA | NA |
| RP24-350F17.8 | 0 | NA | NA | NA |
| Ighv1-63 | 0.046345709 | -0.443818988 | 3.724945049 | -0.119147795 |
| Gm27722 | 0 | NA | NA | NA |
| Ighv1-64 | 0 | NA | NA | NA |
| Ighv8-10 | 0 | NA | NA | NA |
| RP23-218B2.1 | 0 | NA | NA | NA |
| Ighv1-65 | 0 | NA | NA | NA |
| RP23-218B2.3 | 0.230454321 | -0.444554157 | 2.18616566 | -0.203348797 |
| Ighv8-11 | 0.066050875 | -0.443818988 | 3.724945049 | -0.119147795 |
| RP23-218B2.5 | 0 | NA | NA | NA |
| Ighv1-66 | 0 | NA | NA | NA |
| Ighv1-67 | 0.060174537 | 0.517957688 | 3.724945049 | 0.139051095 |
| Ighv1-68 | 0 | NA | NA | NA |
| RP23-218B2.9 | 0 | NA | NA | NA |
| RP23-218B2.10 | 0 | NA | NA | NA |
| Ighv1-69 | 0 | NA | NA | NA |
| Ighv8-12 | 0 | NA | NA | NA |
| RP23-218B2.13 | 0 | NA | NA | NA |
| Ighv1-70 | 0 | NA | NA | NA |
| Mir6388 | 0 | NA | NA | NA |
| Ighv1-71 | 0 | NA | NA | NA |
| Ighv1-72 | 0 | NA | NA | NA |
| Ighv8-13 | 0 | NA | NA | NA |
| RP23-123N21.1 | 0 | NA | NA | NA |
| Ighv1-73 | 0 | NA | NA | NA |
| Ighv1-74 | 0.136120942 | -0.876404342 | 3.595683781 | -0.24373788 |
| Ighv8-14 | 0 | NA | NA | NA |
| RP23-123N21.5 | 0 | NA | NA | NA |
| RP23-123N21.6 | 0 | NA | NA | NA |
| Ighv1-75 | 0 | NA | NA | NA |
| Ighv8-15 | 0 | NA | NA | NA |
| Ighv1-76 | 0 | NA | NA | NA |
| Ighv8-16 | 0 | NA | NA | NA |

| | | | | |
|---|---|---|---|---|
| Ighv1-77 | 0.046345709 | -0.443818988 | 3.724945049 | -0.119147795 |
| Ighv1-78 | 0 | NA | NA | NA |
| RP23-172K2.2 | 0 | NA | NA | NA |
| RP23-172K2.1 | 0 | NA | NA | NA |
| Ighv1-79 | 0 | NA | NA | NA |
| RP23-172K2.4 | 2.956099666 | 0.760376121 | 0.729940716 | 1.041695723 |
| Ighv1-80 | 0 | NA | NA | NA |
| Ighv1-81 | 0 | NA | NA | NA |
| RP23-172K2.8 | 0 | NA | NA | NA |
| Olfr241-ps1 | 0 | NA | NA | NA |
| Ighv1-82 | 0 | NA | NA | NA |
| Ighv1-83 | 0 | NA | NA | NA |
| Ighv1-84 | 0 | NA | NA | NA |
| RP23-172K2.12 | 5.084652238 | -0.979174399 | 0.500017766 | -1.958279215 |
| Ighv1-85 | 0.072025462 | 0.517959912 | 3.724945049 | 0.139051692 |
| Ighv1-86 | 0 | NA | NA | NA |
| Zfp386 | 523.1915969 | 0.147400902 | 0.082586458 | 1.784807161 |
| Vipr2 | 33.917483 | -0.202550585 | 0.24903056 | -0.813356342 |
| Gm20658 | 0.517478627 | -0.546010192 | 1.229280125 | -0.444170682 |
| AC105950.1 | 0 | NA | NA | NA |
| Gm23732 | 0 | NA | NA | NA |
| Wdr60 | 178.5716657 | 0.030845158 | 0.113045215 | 0.272856822 |
| Gm23190 | 0.086648695 | 0.037058229 | 3.724945049 | 0.009948665 |
| Gm11027 | 0 | NA | NA | NA |
| Esyt2 | 1140.517075 | 0.174209695 | 0.055791326 | 3.122522913 |
| Ncapg2 | 136.9875684 | -0.564545936 | 0.16925955 | -3.335386018 |
| Gm25112 | 0.475792389 | 0.846530332 | 1.435986058 | 0.589511526 |
| Gm27649 | 0.04393175 | 0.037062677 | 3.724945049 | 0.009949859 |
| Ptprn2 | 2936.458257 | 0.844604385 | 0.099377399 | 8.498958478 |
| Gm6750 | 0.877833841 | 0.019885255 | 1.183826996 | 0.016797433 |
| Gm17807 | 4.466807706 | 0.781552492 | 0.642779359 | 1.215895441 |
| Gm10421 | 10.74823256 | 0.973610336 | 0.360447868 | 2.701112763 |
| Gm5441 | 1.251482101 | 0.289989283 | 0.94031192 | 0.308396903 |
| Mir153 | 1.154465371 | 2.961563135 | 1.14419434 | 2.588339263 |
| Rapgef5 | 853.7827578 | 0.176093982 | 0.097604511 | 1.804158237 |
| Gm24741 | 1.59128742 | 1.003813665 | 1.107662113 | 0.906245373 |
| Cdca7l | 18.95984674 | -0.56067478 | 0.30593279 | -1.832673054 |
| Gm20613 | 0.046345709 | -0.443818988 | 3.724945049 | -0.119147795 |
| Dnah11 | 127.3083906 | -0.193327735 | 0.153770117 | -1.257251659 |
| Sp4 | 671.9126227 | 0.155180671 | 0.097064862 | 1.598731697 |
| Gm25577 | 0.046345709 | -0.443818988 | 3.724945049 | -0.119147795 |
| Gm22537 | 0 | NA | NA | NA |
| Sp8 | 103.1593826 | -0.695185454 | 0.575742146 | -1.207459725 |
| Abcb5 | 0 | NA | NA | NA |
| Itgb8 | 2067.75423 | -0.476718568 | 0.147116921 | -3.2404061 |
| Gm6768 | 116.6792349 | -0.186579639 | 0.107782018 | -1.731083184 |
| Macc1 | 2.369395584 | -0.550304676 | 0.692521061 | -0.794639625 |

| | | | | |
|---|---|---|---|---|
| Gm25675 | 0 | NA | NA | NA |
| Tmem196 | 135.2367115 | 0.306983202 | 0.108719064 | 2.823637286 |
| Gm27236 | 2.533080183 | -0.37635818 | 0.693661234 | -0.5425677 |
| Gm17177 | 0 | NA | NA | NA |
| Speer6-ps1 | 0 | NA | NA | NA |
| Gm16505 | 1.0973389 | 2.4400402 | 1.220763326 | 1.998782359 |
| 2810429I04Rik | 28.4214383 | 0.559025862 | 0.223815886 | 2.497704131 |
| Gdi2 | 2568.627456 | -0.156714391 | 0.081927196 | -1.912849442 |
| Fam208b | 1293.848084 | -0.024263048 | 0.092246657 | -0.263023602 |
| Asb13 | 641.9525158 | -0.025613913 | 0.11506764 | -0.222598752 |
| Gm23084 | 0.981575226 | -1.712099238 | 1.065269014 | -1.607198948 |
| Calml3 | 0.554260052 | 1.891714997 | 3.682330046 | 0.513727714 |
| Calm4 | 0.780001735 | 2.799193108 | 3.676122419 | 0.761452636 |
| Calm5 | 0.045882455 | 0.517959912 | 3.724945049 | 0.139051692 |
| AC132239.1 | 0 | NA | NA | NA |
| Net1 | 365.646222 | 0.341114602 | 0.091502096 | 3.727943039 |
| Tubal3 | 0.038981395 | -0.443816764 | 3.724945049 | -0.119147198 |
| Ucn3 | 2.74335279 | 0.79128265 | 0.696195246 | 1.136581518 |
| Gm22460 | 0 | NA | NA | NA |
| Akr1c14 | 23.64608385 | -0.896674749 | 0.375001657 | -2.391122097 |
| Akr1c18 | 8.892648978 | 0.144684908 | 0.366684403 | 0.394576118 |
| Akr1c13 | 6.039600961 | -1.575443985 | 0.423724696 | -3.718083939 |
| Akr1c19 | 0.060174537 | 0.517957688 | 3.724945049 | 0.139051095 |
| Marcksl1-ps4 | 0 | NA | NA | NA |
| Akr1c12 | 0.642786378 | 0.406326472 | 1.328226685 | 0.30591651 |
| Akr1c6 | 0.085327104 | -0.905414734 | 3.723500899 | -0.243162217 |
| Akr1c20 | 0.875592853 | -1.333790022 | 1.391759508 | -0.958348058 |
| Akr1c21 | 0 | NA | NA | NA |
| Akr1e1 | 194.9019369 | -0.229831313 | 0.119311233 | -1.92631747 |
| Rpl29-ps2 | 3.103436511 | 0.036843711 | 0.625897388 | 0.058865417 |
| Gm5444 | 0.134148234 | -0.443820953 | 3.624632483 | -0.122445781 |
| Gm24790 | 0 | NA | NA | NA |
| Gm26043 | 0 | NA | NA | NA |
| 1700016G22Rik | 0.413724333 | -1.920785493 | 1.586475015 | -1.210725334 |
| Klf6 | 1264.230787 | -0.640043821 | 0.169698166 | -3.77166022 |
| Gm26259 | 0 | NA | NA | NA |
| Gm28465 | 0 | NA | NA | NA |
| Pitrm1 | 461.3651909 | -0.298854169 | 0.074747977 | -3.998157283 |
| Pfkp | 2522.968723 | -0.140925411 | 0.074540356 | -1.890592128 |
| Gm10029 | 6.775184329 | -0.139829757 | 0.378471887 | -0.369458767 |
| Gm22689 | 0 | NA | NA | NA |
| Gm9742 | 0.283836568 | -1.107085401 | 2.075593713 | -0.533382518 |
| Adarb2 | 565.2457729 | 0.437687527 | 0.074264208 | 5.893653766 |
| Gm10357 | 4.418419051 | 0.540336248 | 0.464313792 | 1.163730773 |
| Wdr37 | 1368.014662 | 0.195432103 | 0.072160267 | 2.708306266 |
| Idi1 | 182.2969244 | -0.512539794 | 0.17985167 | -2.849791678 |
| Gm22750 | 0.171083144 | -1.487610616 | 3.383974877 | -0.43960451 |

| | | | | |
|---|---|---|---|---|
| Rpl10a-ps2 | 0.785454149 | -1.105998911 | 1.068519034 | -1.035076471 |
| Gm9745 | 0.104637558 | -0.443821212 | 3.724945049 | -0.119148392 |
| Gm25274 | 0.066050875 | -0.443818988 | 3.724945049 | -0.119147795 |
| Gm22115 | 0.115603922 | -0.93550548 | 3.724188147 | -0.251197158 |
| Idi2 | 0.326649557 | -0.884474504 | 1.491204233 | -0.593127678 |
| Gm23923 | 0.066050875 | -0.443818988 | 3.724945049 | -0.119147795 |
| Gtpbp4 | 492.4961443 | -0.244064334 | 0.114259837 | -2.136046591 |
| Larp4b | 1451.804564 | -0.066690348 | 0.053245799 | -1.252499707 |
| Gm23553 | 10.68005917 | -0.032781535 | 0.382387943 | -0.085728475 |
| Gm28155 | 0 | NA | NA | NA |
| Dip2c | 2087.5135 | -0.034843756 | 0.071691446 | -0.486023904 |
| Gm26601 | 155.8395443 | 0.232078363 | 0.389484943 | 0.595859653 |
| Gm23496 | 0.048949532 | 0.037064902 | 3.724945049 | 0.009950456 |
| Zmynd11 | 1907.336184 | 0.246412883 | 0.075871218 | 3.247778151 |
| Gm24187 | 251.7405501 | -0.413177174 | 0.979798004 | -0.42169628 |
| Chrm3 | 417.2410317 | 0.889779209 | 0.110044334 | 8.085643075 |
| Gm26861 | 0.849695755 | 0.086197053 | 1.12792922 | 0.076420623 |
| Gm23415 | 0 | NA | NA | NA |
| Gm25496 | 0 | NA | NA | NA |
| Ryr2 | 4975.297084 | 0.708045752 | 0.152972494 | 4.628582142 |
| Gm26956 | 0.116944185 | -1.028246102 | 3.719337068 | -0.27645951 |
| Mtr | 314.5275728 | -0.27993021 | 0.117013956 | -2.39228054 |
| Actn2 | 215.0662038 | 0.679099917 | 0.206051018 | 3.295785296 |
| Gm5445 | 1.336133173 | 0.583600316 | 0.976084293 | 0.597899506 |
| Heatr1 | 496.059456 | -0.606653707 | 0.091046664 | -6.663107471 |
| Lgals8 | 902.3953984 | -0.513605655 | 0.140458162 | -3.656645134 |
| Edaradd | 6.858033054 | 0.932361387 | 0.440627758 | 2.115984229 |
| Ero1lb | 286.3774935 | -0.103956467 | 0.088261048 | -1.177829516 |
| Gm25483 | 3.19383198 | 1.206601402 | 0.651499392 | 1.852037647 |
| Gpr137b-ps | 100.0559785 | -0.214982357 | 0.154614366 | -1.390442318 |
| Gm24065 | 0 | NA | NA | NA |
| Gm2399 | 0.304316145 | -0.004209266 | 1.601144046 | -0.002628912 |
| Prl2c3 | 1.422091401 | 0.703663608 | 1.018113337 | 0.691144672 |
| Prl2c2 | 0.494725653 | -0.646690374 | 1.5611948 | -0.414227855 |
| Prl2c5 | 0.049553047 | -0.443818988 | 3.724945049 | -0.119147795 |
| Gm2423 | 35.43526429 | -0.157966713 | 0.252763818 | -0.624957773 |
| Gm26133 | 0 | NA | NA | NA |
| Gm25804 | 0 | NA | NA | NA |
| Gpr137b | 160.9115362 | -0.311287717 | 0.116862411 | -2.663711232 |
| Gm23081 | 0 | NA | NA | NA |
| Gm7426 | 0 | NA | NA | NA |
| Nid1 | 422.1371149 | -0.643213154 | 0.14492309 | -4.438306919 |
| Lyst | 2594.837581 | 0.00921121 | 0.101007417 | 0.091193404 |
| Gng4 | 505.6373642 | -0.36483264 | 0.25452127 | -1.433407275 |
| Gm25271 | 3.698276596 | 0.750369329 | 0.715997642 | 1.048005308 |
| B3galnt2 | 227.2140593 | 0.082836806 | 0.111366127 | 0.743824077 |
| Gm18856 | 1.228493641 | 0.890364636 | 0.89465581 | 0.995203548 |

| | | | | |
|---|---|---|---|---|
| Tbce | 525.6659255 | 0.181094713 | 0.104134498 | 1.739046297 |
| Ggps1 | 552.9079473 | -0.083900385 | 0.081200649 | -1.033247709 |
| Arid4b | 1466.024357 | 0.12503207 | 0.080980641 | 1.543974808 |
| Gm26129 | 7.167244977 | 0.210972554 | 0.408792537 | 0.516087099 |
| n-R5s52 | 0 | NA | NA | NA |
| Hecw1 | 1632.790211 | 0.447748646 | 0.09777131 | 4.579550452 |
| Mrpl32 | 170.5037384 | -0.310150117 | 0.113322003 | -2.736892292 |
| Psma2 | 925.2712897 | -0.324202341 | 0.113253721 | -2.862619768 |
| Gm25444 | 0.046345709 | -0.443818988 | 3.724945049 | -0.119147795 |
| AW209491 | 188.160879 | -0.248326295 | 0.148694474 | -1.670043874 |
| Gm6556 | 34.49633554 | 1.337692716 | 0.234449145 | 5.705683927 |
| Gm26645 | 23.14899875 | 1.781540369 | 0.299429622 | 5.949779982 |
| Gli3 | 305.6293553 | -0.152206362 | 0.116210834 | -1.309743306 |
| Inhba | 124.8662631 | -2.61113172 | 0.407716679 | -6.404279865 |
| Gm22628 | 0.250684468 | 0.037075444 | 2.148577679 | 0.017255808 |
| Sugct | 24.65360541 | 0.24098023 | 0.242765144 | 0.992647569 |
| Gm26098 | 0 | NA | NA | NA |
| Gm27043 | 0 | NA | NA | NA |
| Gm25020 | 0 | NA | NA | NA |
| Gm24873 | 1.427842546 | -0.65859319 | 0.931207564 | -0.707246393 |
| Mplkip | 150.6828418 | -0.623543481 | 0.215454618 | -2.894082689 |
| Cdk13 | 1371.430534 | -0.008642749 | 0.059721687 | -0.144717092 |
| Mir466i | 0.78113724 | -1.046763781 | 1.066343955 | -0.981638031 |
| Gm24018 | 0.047948698 | 0.037064902 | 3.724945049 | 0.009950456 |
| Gm18859 | 0.219031842 | -1.401822469 | 2.578180455 | -0.543725505 |
| Rala | 653.2611828 | -0.299145974 | 0.089147934 | -3.355613089 |
| Yae1d1 | 237.983152 | -0.298649866 | 0.094523293 | -3.159537258 |
| Gm37888 | 0.089089296 | 0.957155977 | 3.723939561 | 0.257027796 |
| Vdac3-ps1 | 215.80711 | -0.113450178 | 0.117871578 | -0.962489679 |
| Pou6f2 | 51.8533924 | 0.344123464 | 0.177832133 | 1.935102836 |
| Vps41 | 1555.895699 | 0.194601715 | 0.059255887 | 3.284090814 |
| Gm26172 | 0.049553047 | -0.443818988 | 3.724945049 | -0.119147795 |
| Amph | 1915.694854 | 0.401462273 | 0.073243679 | 5.481186605 |
| RP23-167F18.7 | 0 | NA | NA | NA |
| Tcrg-V7 | 0.107803068 | 0.036092186 | 3.724874622 | 0.009689503 |
| Tcrg-V4 | 0.038981395 | -0.443816764 | 3.724945049 | -0.119147198 |
| Tcrg-V6 | 0.075043205 | 0.517957688 | 3.724945049 | 0.139051095 |
| Tcrg-V5 | 0.192276286 | -0.525344508 | 3.26041873 | -0.161127926 |
| Trgj1 | 0 | NA | NA | NA |
| Tcrg-C1 | 0.641965332 | -1.293930794 | 1.322787072 | -0.978185244 |
| Tcrg-V3 | 0 | NA | NA | NA |
| Trgj3 | 0 | NA | NA | NA |
| Tcrg-C3 | 0.092228164 | 0.037073798 | 3.724945049 | 0.009952844 |
| Tcrg-C2 | 0.588064751 | 1.282987408 | 1.338220116 | 0.958726739 |
| Trgj2 | 0 | NA | NA | NA |
| Trgv2 | 0.097139547 | -0.443816764 | 3.724945049 | -0.119147198 |
| Tcrg-V1 | 0 | NA | NA | NA |

| | | | | |
|---|---|---|---|---|
| Trgj4 | 0 | NA | NA | NA |
| Tcrg-C4 | 0.312934076 | -0.067673133 | 2.024606331 | -0.033425329 |
| Stard3nl | 346.4093214 | -0.465603447 | 0.092352716 | -5.041578298 |
| RP23-306G1.11 | 25.52656834 | -0.161741456 | 0.295544298 | -0.547266374 |
| RP23-306G1.12 | 0.525666135 | -0.461395697 | 1.489758262 | -0.309711789 |
| Epdr1 | 994.4722949 | -0.089788284 | 0.073594515 | -1.220040444 |
| Sfrp4 | 29.48509951 | -0.688458679 | 0.219448464 | -3.137222595 |
| Nme8 | 0.465692922 | 0.444250474 | 1.305488445 | 0.340294451 |
| A530099J19Rik | 0.780625029 | -2.268850147 | 1.185930856 | -1.913138642 |
| Gpr141 | 71.74378744 | -1.602117341 | 0.302079861 | -5.303621812 |
| Gm5446 | 4.31637155 | -0.049268374 | 0.453950411 | -0.108532502 |
| Elmo1 | 1541.907887 | 0.138398361 | 0.056502084 | 2.449438187 |
| Gm7611 | 0 | NA | NA | NA |
| Aoah | 31.82302678 | -0.2281168 | 0.3151967 | -0.723728391 |
| Gm25605 | 0.583760085 | 0.179666974 | 1.210734667 | 0.148395003 |
| Olfr1370 | 0 | NA | NA | NA |
| Olfr1369-ps1 | 0 | NA | NA | NA |
| Olfr263 | 0.213549163 | -1.3011867 | 2.569305668 | -0.50643515 |
| Olfr1368 | 0 | NA | NA | NA |
| Trim27 | 275.0576437 | -0.390197311 | 0.080407543 | -4.852745094 |
| Gpx5 | 0 | NA | NA | NA |
| Gm11270 | 0 | NA | NA | NA |
| Gpx6 | 2.180372029 | -0.932874731 | 0.760495689 | -1.226666693 |
| Olfr1367 | 0.046345709 | -0.443818988 | 3.724945049 | -0.119147795 |
| Zscan12 | 64.42265566 | 0.196041287 | 0.149414741 | 1.312061215 |
| Gm11271 | 0.280341511 | 0.037078014 | 1.799282482 | 0.020607111 |
| Zkscan3 | 573.1062161 | -0.239284127 | 0.133419756 | -1.793468474 |
| Mir6942 | 2.650403497 | -0.825757215 | 0.746517413 | -1.106145953 |
| Pgbd1 | 43.29911835 | -0.470171026 | 0.198009747 | -2.374484255 |
| 4930470G03Rik | 1.381930285 | -0.137953574 | 0.863149844 | -0.159825753 |
| Zscan26 | 980.2046494 | 0.02064201 | 0.058633609 | 0.352050821 |
| Mir1896 | 0 | NA | NA | NA |
| Nkapl | 5.734391589 | -0.715851901 | 0.495429318 | -1.44491227 |
| Zkscan4 | 37.6093419 | 0.068743362 | 0.191681304 | 0.358633632 |
| Gm10065 | 0.298142035 | 0.113263718 | 3.717879548 | 0.030464601 |
| Gm23779 | 0 | NA | NA | NA |
| AK157302 | 180.0808189 | 0.187808208 | 0.092587584 | 2.028438361 |
| AL589742.1 | 0 | NA | NA | NA |
| AL589742.2 | 0 | NA | NA | NA |
| Gm11273 | 0 | NA | NA | NA |
| Zfp389 | 4.935521334 | -0.333322348 | 0.482691069 | -0.690550062 |
| Zkscan8 | 581.7328674 | -0.057623544 | 0.102337079 | -0.563075914 |
| Olfr1366 | 0.049553047 | -0.443818988 | 3.724945049 | -0.119147795 |
| Olfr1535 | 0 | NA | NA | NA |
| Olfr1364 | 0.097899064 | 0.037062677 | 3.724945049 | 0.009949859 |
| Olfr1362 | 0 | NA | NA | NA |
| Olfr11 | 0.091666476 | 0.037060453 | 3.724945049 | 0.009949262 |

| | | | | |
|---|---|---|---|---|
| Olfr1361 | 0.061920613 | -0.443816764 | 3.724945049 | -0.119147198 |
| Olfr1360 | 0 | NA | NA | NA |
| Olfr1359 | 0.062426873 | 0.517959912 | 3.724945049 | 0.139051692 |
| Hist1h2bl | 17.04762427 | -0.646721213 | 0.361223506 | -1.790363034 |
| Hist1h2ai | 8.571363186 | -0.307425985 | 0.377358033 | -0.814679847 |
| AL589651.1 | 0.12092566 | 0.998850474 | 3.724945049 | 0.268151734 |
| Hist1h3h | 40.08083796 | -0.366984028 | 0.229038927 | -1.602277977 |
| AL589651.2 | 0 | NA | NA | NA |
| Hist1h2aj | 0.635935163 | -1.523775468 | 1.22433229 | -1.244576722 |
| Hist1h2bm | 22.0061962 | -0.031179312 | 0.297096012 | -0.104946922 |
| Hist1h4j | 57.33528979 | -0.263663123 | 0.21922845 | -1.202686615 |
| Hist1h4k | 181.1049781 | -0.040624342 | 0.148934699 | -0.272766136 |
| Hist1h2ak | 39.92311535 | -0.105313095 | 0.274696144 | -0.383380316 |
| AL589651.3 | 0.07796279 | -0.798301147 | 3.721829328 | -0.214491605 |
| Hist1h2bn | 10.00898834 | -0.556509992 | 0.371084465 | -1.499685501 |
| Gm11274 | 1.966252851 | -1.851773343 | 0.800872593 | -2.312194672 |
| Hist1h1b | 62.13502671 | 0.022825541 | 0.218732206 | 0.10435382 |
| Hist1h3i | 77.67695009 | -0.218934572 | 0.237251128 | -0.922796759 |
| Hist1h2an | 2.07791248 | -0.517093446 | 0.678369376 | -0.762259419 |
| AL589651.4 | 0 | NA | NA | NA |
| Hist1h2bp | 6.144754156 | -0.210420645 | 0.405760285 | -0.518583639 |
| Hist1h2bq | 12.73633966 | -0.576597457 | 0.303847666 | -1.897653072 |
| Hist1h2ao | 3.983646474 | -0.623153684 | 0.537984234 | -1.158312168 |
| AL589879.1 | 0.072025462 | 0.517959912 | 3.724945049 | 0.139051692 |
| Hist1h4m | 15.4295438 | -0.637349129 | 0.316077878 | -2.016430677 |
| Hist1h4n | 166.7008044 | -0.525976166 | 0.166490789 | -3.159190784 |
| Hist1h2ap | 1.700876845 | 0.147447073 | 0.909537218 | 0.162112193 |
| AL589879.3 | 0.060174537 | 0.517957688 | 3.724945049 | 0.139051095 |
| Hist1h2br | 3.568372849 | -0.607517891 | 0.594821207 | -1.021345378 |
| Gm11278 | 0 | NA | NA | NA |
| Mir1983 | 1.672508104 | -1.512323911 | 0.946563454 | -1.597699452 |
| Gm11279 | 0.061920613 | -0.443816764 | 3.724945049 | -0.119147198 |
| AL589879.2 | 0.514605876 | -2.027999939 | 1.428256654 | -1.419912824 |
| Gm11280 | 0.088534442 | -0.905414734 | 3.723500899 | -0.243162217 |
| Zfp184 | 35.438407 | 0.069152136 | 0.189695135 | 0.36454354 |
| Gm11281 | 0 | NA | NA | NA |
| Gm11282 | 0.068097359 | 0.037064902 | 3.724945049 | 0.009950456 |
| Pom121l2 | 0.476324185 | 1.620092834 | 1.505619671 | 1.076030598 |
| Prss16 | 7.136397789 | 0.746818132 | 0.492635417 | 1.515965165 |
| Gm11290 | 0 | NA | NA | NA |
| Gm11291 | 0 | NA | NA | NA |
| Gm11292 | 0.121388914 | 0.037071574 | 3.724945049 | 0.009952247 |
| AL590614.1 | 0.110870145 | -0.444802825 | 3.724874622 | -0.119414174 |
| Hist1h2ah | 3.414149285 | -1.418836619 | 0.670548278 | -2.115935072 |
| Hist1h2bk | 6.592695331 | -0.747535747 | 0.413191419 | -1.809175393 |
| Hist1h4i | 181.0810821 | -0.64281705 | 0.179913331 | -3.57292618 |
| Hist1h2ag | 7.85075097 | -0.115417316 | 0.376172319 | -0.306820333 |

| | | | | |
|---|---|---|---|---|
| AL590614.2 | 0.038981395 | -0.443816764 | 3.724945049 | -0.119147198 |
| Hist1h2bj | 11.38635759 | -0.262058973 | 0.355205506 | -0.737767205 |
| Vmn1r-ps92 | 0 | NA | NA | NA |
| Vmn1r-ps93 | 0 | NA | NA | NA |
| Gm11285 | 0 | NA | NA | NA |
| Gm26300 | 0 | NA | NA | NA |
| Vmn1r188 | 0 | NA | NA | NA |
| Vmn1r-ps94 | 0.341912768 | -0.909700248 | 1.968296601 | -0.462176406 |
| Vmn1r189 | 0 | NA | NA | NA |
| Vmn1r-ps95 | 0 | NA | NA | NA |
| Vmn1r-ps97 | 0 | NA | NA | NA |
| Vmn1r190-ps | 0 | NA | NA | NA |
| Gm11284 | 0.374772292 | 0.806942681 | 1.510978963 | 0.534052889 |
| Vmn1r-ps99 | 0 | NA | NA | NA |
| Vmn1r191 | 0 | NA | NA | NA |
| Vmn1r192 | 0 | NA | NA | NA |
| Gm25084 | 0 | NA | NA | NA |
| Gm23282 | 0.060174537 | 0.517957688 | 3.724945049 | 0.139051095 |
| Vmn1r193 | 0 | NA | NA | NA |
| Vmn1r194 | 0.049553047 | -0.443818988 | 3.724945049 | -0.119147795 |
| Vmn1r195 | 0 | NA | NA | NA |
| Gm11295 | 0.521690028 | 0.133289382 | 1.311307169 | 0.101646193 |
| Vmn1r196 | 0 | NA | NA | NA |
| Vmn1r-ps100 | 0 | NA | NA | NA |
| Vmn1r197 | 0.066050875 | -0.443818988 | 3.724945049 | -0.119147795 |
| Vmn1r198 | 0 | NA | NA | NA |
| Vmn1r199 | 0.038981395 | -0.443816764 | 3.724945049 | -0.119147198 |
| Vmn1r-ps102 | 0 | NA | NA | NA |
| Vmn1r200 | 0.302366008 | -1.493738912 | 1.988402753 | -0.75122553 |
| Vmn1r-ps103 | 0 | NA | NA | NA |
| Vmn1r-ps104 | 0 | NA | NA | NA |
| Gm11298 | 0.060174537 | 0.517957688 | 3.724945049 | 0.139051095 |
| Vmn1r201 | 0 | NA | NA | NA |
| Vmn1r-ps106 | 0 | NA | NA | NA |
| Vmn1r-ps107 | 0 | NA | NA | NA |
| Vmn1r-ps108 | 0 | NA | NA | NA |
| Vmn1r202 | 0 | NA | NA | NA |
| Gm11300 | 0.038981395 | -0.443816764 | 3.724945049 | -0.119147198 |
| Vmn1r203 | 0 | NA | NA | NA |
| Vmn1r204 | 0 | NA | NA | NA |
| Vmn1r205 | 0.044191549 | 0.037062677 | 3.724945049 | 0.009949859 |
| Vmn1r206 | 1.041252144 | 0.472767582 | 1.230388146 | 0.384242634 |
| Vmn1r-ps109 | 0.733882059 | -0.384066455 | 1.281936724 | -0.299598606 |
| Vmn1r-ps110 | 1.137431431 | 0.067317781 | 1.086015145 | 0.061986042 |
| Gm11307 | 0 | NA | NA | NA |
| Vmn1r-ps111 | 0.097139547 | -0.443816764 | 3.724945049 | -0.119147198 |
| Gm11309 | 0 | NA | NA | NA |

| | | | | |
|---|---|---|---|---|
| Vmn1r207-ps | 0.447844902 | -0.260514294 | 1.506320501 | -0.172947453 |
| Vmn1r-ps112 | 0 | NA | NA | NA |
| Vmn1r-ps113 | 1.019683961 | -1.087355074 | 1.177799189 | -0.923209222 |
| Gm11313 | 0 | NA | NA | NA |
| Vmn1r208 | 0 | NA | NA | NA |
| Gm11314 | 0 | NA | NA | NA |
| Vmn1r209 | 0.344645118 | -0.847115452 | 1.588970782 | -0.533122108 |
| Vmn1r210 | 0.134880151 | -1.305960502 | 2.927087508 | -0.446163806 |
| Vmn1r-ps114 | 0 | NA | NA | NA |
| Vmn1r211 | 0 | NA | NA | NA |
| Vmn1r-ps116 | 0 | NA | NA | NA |
| Gm23774 | 0 | NA | NA | NA |
| Vmn1r212 | 0.038981395 | -0.443816764 | 3.724945049 | -0.119147198 |
| Gm11320 | 0 | NA | NA | NA |
| Gm11321 | 0 | NA | NA | NA |
| Vmn1r-ps120 | 0 | NA | NA | NA |
| Gm11322 | 0 | NA | NA | NA |
| Gm11318 | 0.044544648 | 0.517957688 | 3.724945049 | 0.139051095 |
| Vmn1r-ps123 | 0 | NA | NA | NA |
| Vmn1r213 | 0.109982625 | -0.443823436 | 3.724945049 | -0.119148989 |
| Vmn1r-ps125 | 0 | NA | NA | NA |
| Vmn1r214 | 0 | NA | NA | NA |
| Vmn1r-ps126 | 0 | NA | NA | NA |
| Vmn1r215 | 0 | NA | NA | NA |
| Vmn1r216 | 0 | NA | NA | NA |
| Gm11325 | 0 | NA | NA | NA |
| Vmn1r-ps127 | 0 | NA | NA | NA |
| Vmn1r217 | 0 | NA | NA | NA |
| Vmn1r-ps128 | 0 | NA | NA | NA |
| Vmn1r218 | 0 | NA | NA | NA |
| Vmn1r219 | 0 | NA | NA | NA |
| Vmn1r220 | 0.038981395 | -0.443816764 | 3.724945049 | -0.119147198 |
| Vmn1r-ps132 | 0 | NA | NA | NA |
| Vmn1r221 | 0 | NA | NA | NA |
| Vmn1r222 | 0 | NA | NA | NA |
| Vmn1r223 | 0 | NA | NA | NA |
| Gm11331 | 0 | NA | NA | NA |
| Vmn1r-ps135 | 0 | NA | NA | NA |
| 4930557F10Rik | 0.136120942 | -0.876404342 | 3.595683781 | -0.24373788 |
| 4933404K08Rik | 0.044544648 | 0.517957688 | 3.724945049 | 0.139051095 |
| Gm11333 | 0 | NA | NA | NA |
| 4930586N03Rik | 0.124596252 | 0.037071632 | 3.714681146 | 0.009979761 |
| Zfp322a | 495.523674 | -0.123726349 | 0.114051249 | -1.084831162 |
| Gm11334 | 0 | NA | NA | NA |
| Abt1 | 128.8824264 | -0.125876889 | 0.12941854 | -0.972634135 |
| C230035I16Rik | 12.59447301 | 0.649271527 | 0.391312627 | 1.659214352 |
| Btn1a1 | 18.16397336 | 1.059529817 | 0.495581716 | 2.137951789 |

| | | | | |
|---|---|---|---|---|
| Btn2a2 | 7.935836238 | 0.635722708 | 0.451371884 | 1.408423367 |
| AC034285.1 | 0 | NA | NA | NA |
| Gm11335 | 0.44640837 | -0.007585666 | 1.357188921 | -0.005589249 |
| Hist1h4h | 362.5692348 | -0.144609049 | 0.18039546 | -0.801622438 |
| Hist1h2af | 13.35131425 | -0.293104029 | 0.450524143 | -0.650584511 |
| AC034285.3 | 0.10503227 | -0.905414734 | 3.723500899 | -0.243162217 |
| Hist1h3g | 12.55213016 | -0.118985657 | 0.372737217 | -0.319221295 |
| Hist1h2bh | 13.53059552 | 0.050046842 | 0.356661906 | 0.140320122 |
| Hist1h3f | 117.9595612 | 0.09733439 | 0.176190568 | 0.552438143 |
| Hist1h4f | 133.4200694 | 0.01946659 | 0.186040205 | 0.10463647 |
| Hist1h1d | 190.8763881 | 0.357684945 | 0.191915157 | 1.863766001 |
| Hist1h3e | 180.3700906 | 0.415519444 | 0.135109876 | 3.075418731 |
| Hist1h2ae | 13.93887753 | -0.18836477 | 0.384186974 | -0.490294525 |
| AC034285.2 | 0.480269614 | 1.117656853 | 1.315493747 | 0.84961016 |
| Hist1h2bg | 107.7133796 | 0.012150529 | 0.108731868 | 0.111747629 |
| Hist1h2bf | 17.61300926 | -0.147774907 | 0.344544983 | -0.428898733 |
| Hist1h2ad | 7.472743362 | -0.90642685 | 0.417230682 | -2.172483685 |
| AC034285.4 | 0.068097359 | 0.037064902 | 3.724945049 | 0.009950456 |
| Hist1h3d | 38.7205137 | -0.214760014 | 0.238196367 | -0.901609111 |
| Hist1h4d | 957.8385019 | 0.008518928 | 0.126145959 | 0.06753231 |
| Hist1h2be | 153.5786973 | 0.052272193 | 0.145515731 | 0.359220223 |
| Gm11398 | 0 | NA | NA | NA |
| Hist1h1e | 485.7404665 | 0.39680002 | 0.179899902 | 2.205671135 |
| Gm22452 | 0 | NA | NA | NA |
| Gm24991 | 0.1140246 | 0.037073798 | 3.724945049 | 0.009952844 |
| Hist1h2ac | 9.474268641 | -0.30060995 | 0.388342675 | -0.774084254 |
| AL592149.1 | 0.048949532 | 0.037064902 | 3.724945049 | 0.009950456 |
| Hist1h2bc | 147.070612 | -0.715399487 | 0.141640268 | -5.050819902 |
| Hist1h1t | 0.61465116 | -1.480941814 | 1.212878363 | -1.22101429 |
| Hist1h4c | 207.5103062 | -0.117254362 | 0.167693017 | -0.699220299 |
| Hfe | 68.88156831 | -0.267662978 | 0.19561364 | -1.368324712 |
| Gm11338 | 0 | NA | NA | NA |
| Hist1h1c | 665.5210639 | 0.147195957 | 0.164154043 | 0.896694069 |
| Hist1h3c | 20.18455272 | -0.178458568 | 0.302948467 | -0.58907236 |
| Gm11336 | 0.090890357 | 0.037071574 | 3.724945049 | 0.009952247 |
| AL590388.2 | 0.068097359 | 0.037064902 | 3.724945049 | 0.009950456 |
| Hist1h2bb | 38.92068366 | 0.013502278 | 0.286170788 | 0.047182586 |
| Hist1h2ab | 19.95108352 | -0.526621353 | 0.236557972 | -2.226183073 |
| AL590388.1 | 0.038981395 | -0.443816764 | 3.724945049 | -0.119147198 |
| Hist1h3b | 12.81895394 | -0.258403547 | 0.331113859 | -0.780406923 |
| 4930558J22Rik | 0.487944759 | 0.058312406 | 1.461941578 | 0.039886961 |
| Hist1h4b | 52.56315827 | -0.529995443 | 0.243012565 | -2.180938429 |
| Hist1h4a | 71.08590372 | -0.400787577 | 0.152647368 | -2.62557804 |
| Hist1h3a | 21.90840992 | -0.651893623 | 0.262156492 | -2.486658327 |
| Hist1h1a | 77.50845308 | -0.044418411 | 0.195714629 | -0.226954985 |
| Trim38 | 1.17402693 | 0.165727849 | 0.866048683 | 0.191360892 |
| Gm11337 | 0 | NA | NA | NA |

| | | | | |
|---|---|---|---|---|
| Slc17a2 | 0.366957926 | -1.570059614 | 1.38336165 | -1.134959622 |
| Slc17a3 | 0.879126543 | 1.336938996 | 1.22215129 | 1.093922664 |
| Slc17a1 | 0.066050875 | -0.443818988 | 3.724945049 | -0.119147795 |
| Slc17a4 | 0 | NA | NA | NA |
| Snrp1c-ps2 | 0 | NA | NA | NA |
| Hist1h2ba | 0.304971633 | -1.410124194 | 1.718719925 | -0.820450251 |
| Hist1h2aa | 0 | NA | NA | NA |
| AL606464.1 | 0 | NA | NA | NA |
| Scgn | 17.90384641 | -1.232630479 | 0.564363883 | -2.184105887 |
| Gm11339 | 0 | NA | NA | NA |
| Lrrc16a | 976.9628538 | 0.188785549 | 0.090936142 | 2.076023282 |
| Gm11343 | 0 | NA | NA | NA |
| Gm11340 | 0.044191549 | 0.037062677 | 3.724945049 | 0.009949859 |
| Gm23340 | 0 | NA | NA | NA |
| Gm11341 | 0 | NA | NA | NA |
| Cmah | 7.485215113 | -0.906082946 | 0.457993569 | -1.978374823 |
| Gm11342 | 0.062426873 | 0.517959912 | 3.724945049 | 0.139051692 |
| Gm11345 | 0.130840244 | 0.037064923 | 2.960883856 | 0.012518195 |
| Gm11344 | 0 | NA | NA | NA |
| Gm22013 | 0 | NA | NA | NA |
| Fam65b | 791.2498213 | 0.188188464 | 0.116321804 | 1.617826214 |
| Gm11346 | 3.822611367 | -0.259055638 | 0.509948867 | -0.508003163 |
| C530050E15Rik | 25.2429927 | 1.018739776 | 0.346312347 | 2.941679047 |
| 1700016G14Rik | 1.09380521 | -0.188022674 | 0.927852185 | -0.202642918 |
| Gmnn | 28.95427006 | -0.154041571 | 0.253879764 | -0.606750096 |
| BC005537 | 3337.74108 | -0.345104101 | 0.051345891 | -6.72116301 |
| Gm22358 | 0.227031315 | 0.517969832 | 2.148964399 | 0.2410323 |
| Acot13 | 187.4166705 | 0.303935335 | 0.158826163 | 1.913635198 |
| Tdp2 | 199.2904976 | 0.216255371 | 0.093239756 | 2.319347253 |
| D130043K22Rik | 395.2440717 | 0.423722075 | 0.124223326 | 3.410970283 |
| 4932702P03Rik | 1.792617364 | 0.866420463 | 0.793794442 | 1.091492227 |
| Aldh5a1 | 1047.224437 | 0.40165762 | 0.06744007 | 5.955771131 |
| 9330162012Rik | 49.19052179 | 0.730488021 | 0.249705549 | 2.925397636 |
| Gpld1 | 489.1648644 | 0.673502604 | 0.135029101 | 4.987833001 |
| Mrs2 | 387.1535434 | 0.069272431 | 0.087278897 | 0.793690491 |
| Dcdc2a | 119.9340571 | -0.032556787 | 0.161189244 | -0.201978656 |
| Gm11348 | 0 | NA | NA | NA |
| Nrsn1 | 2860.65622 | 0.40392863 | 0.088797748 | 4.548861211 |
| Tpi-rs8 | 0.046345709 | -0.443818988 | 3.724945049 | -0.119147795 |
| Gm11349 | 2.028713949 | -0.693826271 | 0.746246652 | -0.929754618 |
| Gm11350 | 0 | NA | NA | NA |
| Gm11351 | 0.19500485 | -1.493677758 | 3.21608855 | -0.464439251 |
| Gm11352 | 0.044544648 | 0.517957688 | 3.724945049 | 0.139051095 |
| 1700092E19Rik | 0 | NA | NA | NA |
| Gm11353 | 25.4493118 | -0.228201902 | 0.241460359 | -0.945090543 |
| Hdgfl1 | 0 | NA | NA | NA |
| Gm11354 | 0 | NA | NA | NA |

| | | | | |
|---|---|---|---|---|
| Gm11355 | 0 | NA | NA | NA |
| Prl | 159.9440114 | -4.336261213 | 2.886392088 | -1.502311911 |
| Prl3d1 | 0.046345709 | -0.443818988 | 3.724945049 | -0.119147795 |
| Prl3d2 | 0 | NA | NA | NA |
| Prl3d3 | 0 | NA | NA | NA |
| Prl3c1 | 0 | NA | NA | NA |
| Prl3b1 | 0 | NA | NA | NA |
| Prl3a1 | 0 | NA | NA | NA |
| 4930511O05Rik | 0 | NA | NA | NA |
| Gm11356 | 0 | NA | NA | NA |
| Prl6a1 | 0.046345709 | -0.443818988 | 3.724945049 | -0.119147795 |
| Gm11357 | 0 | NA | NA | NA |
| Gm11358 | 0 | NA | NA | NA |
| Prl8a2 | 0 | NA | NA | NA |
| Prl2b1 | 0 | NA | NA | NA |
| Prl8a6 | 0 | NA | NA | NA |
| Prl8a8 | 0 | NA | NA | NA |
| Prl8a9 | 0 | NA | NA | NA |
| Prl8a1 | 0 | NA | NA | NA |
| Prl7b1 | 0 | NA | NA | NA |
| Prl7a1 | 0 | NA | NA | NA |
| Prl7a2 | 0 | NA | NA | NA |
| Gm11359 | 0.049553047 | -0.443818988 | 3.724945049 | -0.119147795 |
| Prl7d1 | 0.045882455 | 0.517959912 | 3.724945049 | 0.139051692 |
| Prl7c1 | 0.062426873 | 0.517959912 | 3.724945049 | 0.139051692 |
| Prl2a1 | 0 | NA | NA | NA |
| Prl2c1 | 0 | NA | NA | NA |
| Gm22476 | 0 | NA | NA | NA |
| Gm11360 | 0 | NA | NA | NA |
| Prl4a1 | 0 | NA | NA | NA |
| Prl5a1 | 0 | NA | NA | NA |
| Gm11361 | 310.4479338 | -0.548688446 | 0.151437323 | -3.623204873 |
| 2610307P16Rik | 11.882246 | 0.241320779 | 0.331485739 | 0.72799747 |
| Gm6081 | 0.66818121 | -0.804691686 | 1.276480327 | -0.630398815 |
| Mir6368 | 0.123992736 | 0.517955521 | 3.720643114 | 0.139211288 |
| Gm17528 | 0 | NA | NA | NA |
| Gm11362 | 0.126225412 | 0.03707172 | 3.698755067 | 0.010022756 |
| Gm26735 | 7.2496826 | 0.296553336 | 0.407922246 | 0.726984957 |
| Sox4 | 422.4467281 | -0.330732576 | 0.151241484 | -2.186784782 |
| A330102I10Rik | 83.11002475 | 0.255352922 | 0.246218272 | 1.037099806 |
| Gm11363 | 0.134452335 | 1.053217666 | 3.628238184 | 0.290283496 |
| Cdkal1 | 296.7585439 | -0.053553361 | 0.110949417 | -0.482682671 |
| Gm11364 | 0 | NA | NA | NA |
| E2f3 | 366.9818336 | -0.036326741 | 0.072507721 | -0.501005143 |
| Gm11365 | 0.042716945 | 0.037062677 | 3.724945049 | 0.009949859 |
| Gm11368 | 0.105852363 | -0.443821212 | 3.724945049 | -0.119148392 |
| Gm11367 | 0 | NA | NA | NA |

| | | | | |
|---|---|---|---|---|
| Mboat1 | 33.58227926 | -1.25437968 | 0.24058233 | -5.213931039 |
| Gm23206 | 0 | NA | NA | NA |
| Gm11366 | 0.322434302 | -0.722568952 | 1.736154582 | -0.416189295 |
| Agtr1a | 5.923613152 | -0.272962974 | 0.483344299 | -0.564738169 |
| Gm11369 | 0.085327104 | -0.905414734 | 3.723500899 | -0.243162217 |
| Uqcrfs1 | 840.2137481 | -0.035128566 | 0.113260278 | -0.310157864 |
| 4930519D14Rik | 0.04393175 | 0.037062677 | 3.724945049 | 0.009949859 |
| Dusp22 | 130.6997309 | -0.013036732 | 0.119398761 | -0.109186493 |
| Gm11370 | 0 | NA | NA | NA |
| Irf4 | 27.40671112 | -0.98556358 | 0.268909243 | -3.66504167 |
| Exoc2 | 757.5578618 | 0.019392336 | 0.045345292 | 0.427659301 |
| Hus1b | 7.907650439 | 0.185983242 | 0.423698539 | 0.438951815 |
| Gm5447 | 0.689547818 | -1.015614157 | 1.267381988 | -0.801348107 |
| Gm11371 | 0.046345709 | -0.443818988 | 3.724945049 | -0.119147795 |
| Gm11372 | 0 | NA | NA | NA |
| 4930401O12Rik | 0.147083031 | 0.517961574 | 2.877346641 | 0.180013616 |
| Gm11376 | 0 | NA | NA | NA |
| Gm11373 | 0.313638404 | 0.517974583 | 1.794586695 | 0.288631686 |
| Gm11374 | 0.506145433 | 2.272433483 | 1.384422399 | 1.64143074 |
| G630018N14Rik | 1.020023321 | 2.00840501 | 1.205780067 | 1.665647878 |
| Gm22330 | 0 | NA | NA | NA |
| Gm11375 | 1.368265809 | 2.638436201 | 1.060869888 | 2.487049761 |
| Gm23351 | 0 | NA | NA | NA |
| A530084C06Rik | 4.406782827 | -0.68988209 | 0.473157227 | -1.458039844 |
| Foxq1 | 10.28519308 | 0.143901883 | 0.364322088 | 0.394985339 |
| Gm11377 | 0 | NA | NA | NA |
| 1700018A04Rik | 0.705754361 | -0.870045335 | 1.158892881 | -0.750755612 |
| Gm11378 | 0 | NA | NA | NA |
| Foxf2 | 22.59941583 | -0.456787196 | 0.316007083 | -1.445496702 |
| Gm27516 | 0.075043205 | 0.517957688 | 3.724945049 | 0.139051095 |
| Gm25186 | 0.382576472 | -0.074384297 | 1.725472997 | -0.043109511 |
| Gm11379 | 0.403183719 | 0.507731363 | 1.427398449 | 0.355704017 |
| Foxc1 | 125.7748194 | -0.358685145 | 0.193226935 | -1.856289574 |
| Gmds | 102.4660629 | -0.057917401 | 0.124515884 | -0.465140667 |
| Gm11380 | 0.044544648 | 0.517957688 | 3.724945049 | 0.139051095 |
| Gm11381 | 0 | NA | NA | NA |
| Mylk4 | 2.177335128 | 0.79769753 | 0.707220516 | 1.127933244 |
| Gm27342 | 0 | NA | NA | NA |
| Wrnip1 | 563.9693568 | -0.166691453 | 0.070914682 | -2.35059156 |
| D930007J09Rik | 3.931524232 | -0.171180134 | 0.525054533 | -0.326023534 |
| Serpinb1a | 58.83320495 | -0.054304513 | 0.235834542 | -0.230265306 |
| Serpinb1-ps1 | 0.259499533 | 0.516092277 | 2.494191193 | 0.206917689 |
| Serpinb1c | 2.097314789 | -2.180143253 | 0.782891458 | -2.784732457 |
| Gm11382 | 0.61025173 | -3.044941305 | 1.324503166 | -2.298930937 |
| Serpinb6b | 64.34938334 | -1.13729556 | 0.307068376 | -3.703720895 |
| Serpinb9 | 320.5775695 | -0.836412238 | 0.219841327 | -3.804617849 |
| Serpinb9b | 16.14145208 | -1.834091505 | 0.50034781 | -3.66563312 |

| | | | | |
|---|---|---|---|---|
| Serpinb1b | 57.17042438 | -0.596353726 | 0.334158561 | -1.784642968 |
| Gm11384 | 0 | NA | NA | NA |
| Gm11383 | 0 | NA | NA | NA |
| Serpinb9c | 0.260452197 | -0.436625895 | 1.843891755 | -0.236795839 |
| Serpinb9d | 0 | NA | NA | NA |
| Gm11385 | 0 | NA | NA | NA |
| Serpinb9e | 0.061920613 | -0.443816764 | 3.724945049 | -0.119147198 |
| Serpinb9f | 0 | NA | NA | NA |
| Gm11397 | 0 | NA | NA | NA |
| Serpinb9g | 0 | NA | NA | NA |
| Gm11387 | 0 | NA | NA | NA |
| Gm11386 | 0 | NA | NA | NA |
| Gm11393 | 0 | NA | NA | NA |
| Gm25043 | 0 | NA | NA | NA |
| Gm11388 | 0 | NA | NA | NA |
| Serpinb6d | 0 | NA | NA | NA |
| Gm11389 | 0 | NA | NA | NA |
| Gm26701 | 0 | NA | NA | NA |
| Gm11392 | 0 | NA | NA | NA |
| Gm11394 | 0 | NA | NA | NA |
| Gm11395 | 0 | NA | NA | NA |
| Vmn2r-ps94 | 0 | NA | NA | NA |
| Gm16566 | 0 | NA | NA | NA |
| Serpinb6e | 0.733278584 | -0.112970512 | 1.373275884 | -0.082263523 |
| Serpinb6c | 3.256068972 | 0.019163635 | 0.550554091 | 0.034807905 |
| Serpinb6a | 261.2870453 | -0.578593346 | 0.108809487 | -5.317489899 |
| 1110046J04Rik | 30.45720878 | -0.080717593 | 0.229797734 | -0.351254955 |
| Nqo2 | 104.6663154 | -0.492656501 | 0.155874052 | -3.160606241 |
| Ripk1 | 212.9648014 | -0.635537881 | 0.159630368 | -3.981309367 |
| Gm17083 | 0.214446769 | 0.828506796 | 2.554573618 | 0.324322928 |
| Gm17084 | 5.185998495 | 0.169861759 | 0.462440736 | 0.367315736 |
| Bphl | 141.617793 | 0.207267216 | 0.103143139 | 2.009510452 |
| Tubb2a | 3268.123541 | -0.400653003 | 0.114370906 | -3.503102479 |
| Tubb2b | 1731.528415 | -1.135628946 | 0.172918894 | -6.567408117 |
| Psmg4 | 78.61758626 | -0.477131849 | 0.178534718 | -2.672487753 |
| Slc22a23 | 1092.32252 | -0.173209567 | 0.050293558 | -3.443971207 |
| Gm24530 | 6.119618873 | 0.58420847 | 0.491891501 | 1.187677503 |
| Pxdc1 | 58.73051673 | -0.657146401 | 0.207484852 | -3.167201825 |
| Gm15908 | 0.16390591 | 0.517965222 | 2.782471991 | 0.186152897 |
| Fam50b | 0.097139547 | -0.443816764 | 3.724945049 | -0.119147198 |
| Prpf4b | 962.083236 | 0.027062198 | 0.09862448 | 0.27439636 |
| Fam217a | 2.489500889 | -0.585916896 | 0.726399433 | -0.806604286 |
| 4933417A18Rik | 0.093141081 | 0.037060453 | 3.724945049 | 0.009949262 |
| Gm8245 | 0.066050875 | -0.443818988 | 3.724945049 | -0.119147795 |
| Eci3 | 1.298423698 | -1.391604165 | 0.908608188 | -1.531577838 |
| Gm16984 | 0.868294842 | -0.79256981 | 1.184606264 | -0.669057588 |
| Eci2 | 255.3897712 | -0.114928879 | 0.130988989 | -0.877393438 |

| | | | | |
|---|---|---|---|---|
| 1700011B04Rik | 0.157356115 | -0.500078175 | 2.825615622 | -0.176980256 |
| Gm22674 | 0 | NA | NA | NA |
| Cdyl | 458.3848708 | -0.050918246 | 0.113290636 | -0.449447962 |
| Gm22126 | 0 | NA | NA | NA |
| Rpp40 | 47.71410153 | -0.563579911 | 0.210726251 | -2.674464661 |
| Ppp1r3g | 58.16805527 | 0.702349459 | 0.302129421 | 2.324664234 |
| Lyrm4 | 177.4291997 | 0.013815706 | 0.113146541 | 0.122104538 |
| Fars2 | 206.9435174 | 0.100449572 | 0.11646174 | 0.862511342 |
| Gm26395 | 0.075043205 | 0.517957688 | 3.724945049 | 0.139051095 |
| Nrn1 | 1029.09811 | 0.142281728 | 0.160061947 | 0.888916639 |
| F13a1 | 144.5358196 | -1.903073228 | 0.416606099 | -4.568039769 |
| Ly86 | 116.6552724 | -0.156017221 | 0.270761511 | -0.576216393 |
| Gm4035 | 0.308888673 | -0.443835108 | 3.724945049 | -0.119152122 |
| Gm29459 | 0 | NA | NA | NA |
| Gm29458 | 0.240304698 | 0.555737012 | 2.112815245 | 0.263031523 |
| Gm29590 | 0 | NA | NA | NA |
| Gm37149 | 0 | NA | NA | NA |
| Rreb1 | 710.6869941 | 0.350821135 | 0.174613962 | 2.009124187 |
| Ssr1 | 2094.214592 | -0.051173433 | 0.045968345 | -1.113232007 |
| Cage1 | 29.07551637 | 0.033908178 | 0.238242733 | 0.14232618 |
| Riok1 | 219.0593976 | -0.062345757 | 0.1224569 | -0.509124086 |
| Gm27387 | 0 | NA | NA | NA |
| Gm10129 | 0.169723681 | -0.52143512 | 3.40726837 | -0.153036117 |
| Dsp | 96.30191792 | 0.785621496 | 0.169882197 | 4.624507512 |
| Snrnp48 | 159.0698413 | -0.31364204 | 0.169945938 | -1.845540081 |
| Bmp6 | 104.867172 | 0.041919954 | 0.19827171 | 0.211426805 |
| Txndc5 | 655.6323935 | -0.337803225 | 0.082786217 | -4.080428349 |
| Bloc1s5 | 130.6301522 | -0.237076644 | 0.108685986 | -2.181299104 |
| Eef1e1 | 223.262343 | -0.457714578 | 0.124973243 | -3.662500609 |
| Gm22035 | 0.046345709 | -0.443818988 | 3.724945049 | -0.119147795 |
| Slc35b3 | 116.4993211 | 0.151591851 | 0.121109722 | 1.251690186 |
| Gm26514 | 39.29548882 | 0.317891825 | 0.232910171 | 1.364868798 |
| 5033403F01Rik | 3.803373425 | 0.640747513 | 0.529140085 | 1.210922272 |
| Gm38206 | 0 | NA | NA | NA |
| Gm23972 | 0 | NA | NA | NA |
| Ofcc1 | 0.044191549 | 0.037062677 | 3.724945049 | 0.009949859 |
| Gm9979 | 0.471418172 | -1.198090503 | 1.467138387 | -0.816617242 |
| Tfap2a | 38.58151316 | -0.028206241 | 0.240414354 | -0.117323447 |
| Gm26688 | 1.413787916 | 0.569827745 | 0.836734437 | 0.681013855 |
| Gm37823 | 0 | NA | NA | NA |
| Gm6245 | 0.59099902 | 0.430183623 | 1.265213213 | 0.340008797 |
| Gcnt2 | 477.1222298 | -0.612383463 | 0.140874314 | -4.347020023 |
| Mir5124a | 0.74668639 | -0.772675425 | 1.198438565 | -0.644735114 |
| A730081D07Rik | 27.84660377 | 0.246980458 | 0.295164631 | 0.836754923 |
| Pak1ip1 | 411.5182475 | -0.362123568 | 0.116456295 | -3.109523344 |
| Tmem14c | 229.1318417 | -0.391047805 | 0.102605091 | -3.811192998 |
| Mak | 31.22020358 | -0.260529536 | 0.209276358 | -1.24490668 |

| | | | | |
|---|---|---|---|---|
| Gcm2 | 0.462408164 | -0.083002367 | 1.36247069 | -0.060920479 |
| Sycp2l | 3.696698795 | 1.09908664 | 0.576080766 | 1.90786901 |
| Elovl2 | 197.6053173 | 0.28670078 | 0.124411695 | 2.304452003 |
| Smim13 | 2327.736401 | 0.306242964 | 0.086114982 | 3.556210038 |
| Gm17364 | 0.17692102 | -0.979939382 | 2.723401576 | -0.359821846 |
| Nedd9 | 334.1506425 | -0.488160105 | 0.182674126 | -2.672300204 |
| Gm26877 | 0.927545067 | -1.907200393 | 1.207521935 | -1.579433331 |
| Gm24819 | 0.43165004 | 0.052092507 | 1.40219551 | 0.037150673 |
| Tmem170b | 1737.683987 | 0.19445172 | 0.04934333 | 3.940790421 |
| Gm5082 | 0.47229425 | -0.428221253 | 3.697230667 | -0.115822163 |
| Adtrp | 1.749374567 | -0.128435306 | 0.900217132 | -0.142671474 |
| 1700061E18Rik | 0.094294407 | -0.443821212 | 3.724945049 | -0.119148392 |
| Gm28564 | 0.066050875 | -0.443818988 | 3.724945049 | -0.119147795 |
| Gm37698 | 0 | NA | NA | NA |
| Gm22157 | 0.123841226 | -1.017823543 | 3.719440177 | -0.273649661 |
| Gm28707 | 0.850789295 | -1.559876422 | 1.284110452 | -1.214752531 |
| Hivep1 | 1475.754828 | -0.11501857 | 0.061713137 | -1.86376153 |
| Edn1 | 31.52376167 | -0.114492306 | 0.301068285 | -0.380286838 |
| Gm23086 | 0 | NA | NA | NA |
| Phactr1 | 3991.264433 | 0.654824109 | 0.117675639 | 5.564653093 |
| Gm15810 | 0.418886444 | 0.529600887 | 1.396594057 | 0.379208893 |
| Gm15812 | 0 | NA | NA | NA |
| Gm15809 | 21.27740935 | 0.673812323 | 0.336711194 | 2.00115807 |
| Gm26086 | 0 | NA | NA | NA |
| Gm15813 | 0.11193333 | 0.037073798 | 3.724945049 | 0.009952844 |
| Tbc1d7 | 105.3200195 | 0.028411689 | 0.118963936 | 0.238826072 |
| Gfod1 | 1583.085298 | -0.004129326 | 0.090213745 | -0.045772696 |
| Sirt5 | 74.72223409 | 0.221618711 | 0.148748529 | 1.489888421 |
| Nol7 | 275.008259 | -0.135738536 | 0.138705641 | -0.978608622 |
| Ranbp9 | 1368.855223 | 0.038484131 | 0.075962089 | 0.506622862 |
| Gm26064 | 7.226595754 | 1.609125958 | 0.499192542 | 3.223457527 |
| Mcur1 | 233.0888101 | 0.001738401 | 0.105828207 | 0.016426634 |
| Rnf182 | 236.6714893 | -0.257314405 | 0.110327926 | -2.332269018 |
| Cd83 | 231.474017 | 0.246266779 | 0.12905372 | 1.908250131 |
| Gm23117 | 0 | NA | NA | NA |
| Gm5083 | 5.023184258 | -0.831422544 | 0.504411175 | -1.64830318 |
| Gm27007 | 0 | NA | NA | NA |
| Gm27036 | 0.066050875 | -0.443818988 | 3.724945049 | -0.119147795 |
| 1700029N11Rik | 0 | NA | NA | NA |
| Jarid2 | 797.878151 | -0.159564425 | 0.088510235 | -1.802779358 |
| Gm22213 | 0.047948698 | 0.037064902 | 3.724945049 | 0.009950456 |
| Dtnbp1 | 383.1011531 | 0.04146677 | 0.111211218 | 0.37286499 |
| Gm23104 | 0 | NA | NA | NA |
| Hsp25-ps1 | 4.708514708 | -1.406532708 | 0.684490941 | -2.054859495 |
| Gm9817 | 0.309768565 | -0.373071314 | 1.771795054 | -0.210561212 |
| Gm8513 | 0.148910734 | 0.966597255 | 2.858091675 | 0.338196729 |
| Gm23755 | 0 | NA | NA | NA |

| | | | | |
|---|---|---|---|---|
| Mylip | 196.9662467 | -0.320417907 | 0.103167669 | -3.105797684 |
| Gm23387 | 0 | NA | NA | NA |
| Gmpr | 269.5668325 | 0.379169845 | 0.090013479 | 4.212367429 |
| Atxn1 | 2899.31005 | 0.338101371 | 0.082111591 | 4.117583951 |
| 5033430I15Rik | 31.65815391 | -0.012951229 | 0.179320015 | -0.072224112 |
| Gm10113 | 0.097139547 | -0.443816764 | 3.724945049 | -0.119147198 |
| Stmnd1 | 2.201457459 | 0.627226497 | 0.694266144 | 0.903438117 |
| Rbm24 | 161.1267455 | 0.510513443 | 0.142064895 | 3.593522831 |
| Cap2 | 1272.593817 | 0.020754677 | 0.135365969 | 0.153322706 |
| Gm24915 | 2.015269402 | 0.514537185 | 0.935848421 | 0.549808252 |
| C78339 | 1105.225121 | -0.104693794 | 0.081564624 | -1.283568647 |
| Nup153 | 1192.694187 | -0.379337961 | 0.059831472 | -6.340107414 |
| Kif13a | 663.6183412 | -0.065017777 | 0.086650028 | -0.750349169 |
| Gm24620 | 4.394585356 | 0.386207979 | 0.525392391 | 0.735084835 |
| Gm22960 | 0 | NA | NA | NA |
| Gm24769 | 0 | NA | NA | NA |
| 2010001K21Rik | 1.984554007 | -1.201972386 | 0.743999096 | -1.615556245 |
| Nhlrc1 | 147.1966146 | -0.016388876 | 0.101884417 | -0.160857535 |
| Tpmt | 47.2986511 | 0.05008588 | 0.180537747 | 0.277426082 |
| Kdm1b | 181.3782971 | -0.276482441 | 0.121752159 | -2.270862745 |
| Dek | 884.7800832 | 0.024775267 | 0.120667888 | 0.205317816 |
| Rnf144b | 226.1613996 | -0.275710161 | 0.117023016 | -2.356033629 |
| 4931429P17Rik | 0.349470464 | 0.546310364 | 1.810019352 | 0.301825704 |
| A330033J07Rik | 16.48077188 | 2.05293866 | 0.381072236 | 5.387269042 |
| Id4 | 466.0183473 | 0.159215413 | 0.163304067 | 0.974962942 |
| A330048O09Rik | 5.209108024 | 0.273112613 | 0.529743113 | 0.515556703 |
| Zfp169 | 346.1617771 | 0.190419753 | 0.09835318 | 1.936081303 |
| A830005F24Rik | 4.046338601 | -0.620858924 | 0.625453489 | -0.992654026 |
| 6720427I07Rik | 280.6546708 | -0.102466462 | 0.102224957 | -1.00236249 |
| Mirlet7d | 0 | NA | NA | NA |
| Gm24111 | 0 | NA | NA | NA |
| mmu-let-7f-1 | 12.7179762 | -0.248367334 | 0.309535716 | -0.802386675 |
| Mirlet7a-1 | 9.148397703 | 0.02458348 | 0.438452997 | 0.056068678 |
| Gm37238 | 83.05504175 | 0.252692689 | 0.237890416 | 1.062223073 |
| Gm25232 | 0 | NA | NA | NA |
| Ptpdc1 | 475.0012653 | 0.156735716 | 0.078643104 | 1.993000133 |
| Barx1 | 0 | NA | NA | NA |
| Phf2 | 871.8668907 | -0.039749307 | 0.0645108 | -0.616165151 |
| Phf2os1 | 14.57948276 | 0.364347411 | 0.361023501 | 1.00920691 |
| Fam120a | 2396.718561 | -0.243488205 | 0.058994157 | -4.127327497 |
| Fam120aos | 129.3350566 | 0.471311701 | 0.150157406 | 3.138784249 |
| Wnk2 | 2363.287208 | 0.576787708 | 0.112327682 | 5.134866985 |
| Ninj1 | 228.5158888 | -0.51933094 | 0.169808333 | -3.058336018 |
| 1110007C09Rik | 111.7269501 | -0.781357775 | 0.179702131 | -4.348071839 |
| Susd3 | 9.113448631 | -0.184537685 | 0.34025857 | -0.542345445 |
| Fgd3 | 292.1672564 | -1.11037076 | 0.159705641 | -6.952608267 |
| Bicd2 | 1037.207963 | -0.166571567 | 0.058959655 | -2.82517881 |

| | | | | |
|---|---|---|---|---|
| Ippk | 334.3509724 | -0.090675381 | 0.092379163 | -0.981556634 |
| Cenpp | 10.13615864 | 0.242900101 | 0.409041714 | 0.593827212 |
| Tes3-ps | 44.94843152 | 0.174247845 | 0.295943202 | 0.58878813 |
| Ecm2 | 93.28506817 | -0.36439175 | 0.175321038 | -2.078425694 |
| Aspn | 20.67132123 | -0.458534301 | 0.311447613 | -1.472267832 |
| Omd | 21.95627783 | 0.832043958 | 0.316929912 | 2.625324799 |
| Ogn | 102.8893265 | 0.303237591 | 0.221417957 | 1.369525737 |
| Nol8 | 394.8682349 | -0.092550838 | 0.064718319 | -1.430056269 |
| Iars | 963.3246765 | -0.564292651 | 0.074313114 | -7.593446446 |
| Gm25394 | 8.321309178 | -0.26373509 | 0.451716201 | -0.583851297 |
| Gm23482 | 2.859218534 | 0.176302782 | 0.707963633 | 0.249028021 |
| Gm7251 | 0.539950272 | 0.651881958 | 1.373693573 | 0.47454685 |
| Gm8674 | 0 | NA | NA | NA |
| Gm10784 | 0 | NA | NA | NA |
| Gm17611 | 0 | NA | NA | NA |
| Gm24300 | 0 | NA | NA | NA |
| Gm906 | 0 | NA | NA | NA |
| Gm24017 | 0 | NA | NA | NA |
| Gm8739 | 0 | NA | NA | NA |
| Fbxw17 | 179.6250207 | -0.548380406 | 0.157352478 | -3.485044611 |
| Nutm2 | 0.276311821 | 1.001655484 | 2.407135238 | 0.416119322 |
| Mir683-1 | 0 | NA | NA | NA |
| Mir683-2 | 0.088123299 | 0.037058229 | 3.724945049 | 0.009948665 |
| Gm904 | 0.038981395 | -0.443816764 | 3.724945049 | -0.119147198 |
| Gm23952 | 0 | NA | NA | NA |
| Gm8765 | 0 | NA | NA | NA |
| Gm22199 | 0 | NA | NA | NA |
| Spin1 | 2716.345326 | 0.236233953 | 0.065440962 | 3.609878974 |
| Nxnl2 | 1.249332652 | 0.489458016 | 0.893656825 | 0.547702432 |
| Hist1h2al | 110.2101923 | -0.03858169 | 0.162433512 | -0.237522969 |
| AC160122.1 | 0.260264958 | -0.295937411 | 2.081708207 | -0.142160852 |
| S1pr3 | 554.2109028 | -1.486061713 | 0.290910891 | -5.108305536 |
| Shc3 | 425.7683225 | 0.457585055 | 0.152290089 | 3.004693604 |
| Cks2 | 12.99324003 | -0.456622748 | 0.375829352 | -1.214973619 |
| Gm22806 | 0.426919779 | 0.183478687 | 1.723603957 | 0.106450607 |
| Secisbp2 | 291.1100637 | -0.204526456 | 0.088629095 | -2.307667216 |
| Gm15440 | 205.1826132 | 0.289743688 | 0.1198935 | 2.416675543 |
| Sema4d | 1108.340496 | -0.3612357 | 0.091066988 | -3.96670307 |
| Gm23566 | 0 | NA | NA | NA |
| Gadd45g | 275.2348046 | -1.013450934 | 0.320942474 | -3.157733911 |
| Gm26651 | 0.10503227 | -0.905414734 | 3.723500899 | -0.243162217 |
| Gm37872 | 0 | NA | NA | NA |
| Diras2 | 2681.114 | 0.125187509 | 0.107604926 | 1.163399422 |
| Syk | 204.4789825 | -1.044309854 | 0.219968709 | -4.74753822 |
| Auh | 575.8122667 | 0.064916778 | 0.057578618 | 1.127445916 |
| n-R5s54 | 0.832946694 | -0.862307684 | 1.246996874 | -0.691507494 |
| Nfil3 | 206.9757982 | -0.671530441 | 0.342341602 | -1.961580002 |

| | | | | |
|---|---|---|---|---|
| Ror2 | 25.4305805 | -0.081137753 | 0.226615494 | -0.358041507 |
| Sptlc1 | 374.6291411 | -0.454960417 | 0.093164889 | -4.883389248 |
| RP23-340E6.2 | 12.21145091 | 0.284077573 | 0.336282031 | 0.844759893 |
| RP23-340E6.3 | 0 | NA | NA | NA |
| Msx2 | 19.08535284 | -0.752204596 | 0.341009395 | -2.205817811 |
| Gm5449 | 0.111006857 | 0.037730769 | 3.724898049 | 0.010129343 |
| Drd1 | 333.3471759 | 0.295773708 | 0.222720633 | 1.328003177 |
| Sfxn1 | 561.8273373 | 0.101661677 | 0.067918845 | 1.496811044 |
| Hrh2 | 63.56667738 | -0.001904016 | 0.178755648 | -0.010651501 |
| Gm16578 | 0.107803068 | 0.036092186 | 3.724874622 | 0.009689503 |
| Cplx2 | 9264.934547 | 0.877446792 | 0.12126734 | 7.235639789 |
| Gm16248 | 0 | NA | NA | NA |
| Gm22668 | 0 | NA | NA | NA |
| Thoc3 | 293.4655856 | -0.043094629 | 0.095356262 | -0.451932874 |
| Gm2830 | 71.59267597 | 0.357483706 | 0.158215035 | 2.259479995 |
| Simc1 | 267.6907852 | -0.004572261 | 0.09107847 | -0.050201334 |
| 4833439L19Rik | 1366.025163 | -0.054529454 | 0.061779829 | -0.882641719 |
| Arl10 | 135.7640415 | -0.243393725 | 0.132976031 | -1.830357868 |
| Gm24195 | 1.207241537 | 2.585330238 | 1.148423247 | 2.251199847 |
| Nop16 | 117.0143181 | -0.435128035 | 0.141089927 | -3.084047497 |
| Higd2a | 413.1629045 | 0.107281056 | 0.095966134 | 1.117905368 |
| Cltb | 2228.333823 | -0.532717829 | 0.09362626 | -5.689833496 |
| Faf2 | 759.1273305 | -0.289310696 | 0.071535978 | -4.044268398 |
| Rnf44 | 1513.683017 | -0.014089821 | 0.064481351 | -0.218510019 |
| Cdhr2 | 4.783323946 | 0.370079375 | 0.480693939 | 0.769885669 |
| Gprin1 | 1231.562756 | 0.261337618 | 0.097608547 | 2.677405068 |
| Sncb | 3503.722137 | 0.710568662 | 0.101828933 | 6.978062569 |
| Gm16249 | 0 | NA | NA | NA |
| Eif4e1b | 0 | NA | NA | NA |
| Tspan17 | 212.3703016 | 0.08357832 | 0.132144325 | 0.632477555 |
| Gm16250 | 0 | NA | NA | NA |
| Gm29431 | 0.504101427 | -1.820684775 | 1.440908805 | -1.263566972 |
| 5330429C05Rik | 1.674398058 | -0.618311134 | 0.762504196 | -0.810895386 |
| 4930526F13Rik | 0.237545104 | 1.56811443 | 2.538594424 | 0.617709712 |
| Unc5a | 875.420532 | 0.763408646 | 0.126424813 | 6.038439999 |
| Hk3 | 302.6549373 | -1.868067327 | 0.335419866 | -5.569340159 |
| Uimc1 | 267.0504781 | -0.015104197 | 0.117819646 | -0.128197611 |
| Gm6344 | 1.345796852 | -0.93743195 | 0.82540411 | -1.13572484 |
| Zfp346 | 338.7158424 | -0.043682508 | 0.101292113 | -0.431252807 |
| Gm17617 | 0 | NA | NA | NA |
| Fgfr4 | 13.86132062 | -1.82315717 | 0.401884228 | -4.53652332 |
| Nsd1 | 4468.214251 | 0.060409715 | 0.069347719 | 0.871113222 |
| Rab24 | 365.9432809 | 0.320285525 | 0.070705702 | 4.52984011 |
| Prelid1 | 846.0882492 | -0.121107281 | 0.091322319 | -1.326152059 |
| Mxd3 | 8.517488434 | 0.401566013 | 0.533654952 | 0.752482502 |
| Lman2 | 1016.200266 | -0.46464361 | 0.086099443 | -5.396592497 |
| Rgs14 | 301.6653793 | 1.036333489 | 0.118561585 | 8.740887637 |

| | | | | |
|---|---|---|---|---|
| Slc34a1 | 0.764795395 | 0.397696718 | 1.132107033 | 0.351288974 |
| Pfn3 | 0.525579064 | 1.888574622 | 1.444960271 | 1.307007992 |
| F12 | 5.68547348 | -1.62670346 | 0.651740896 | -2.495935837 |
| Grk6 | 508.6179697 | -0.052247122 | 0.056980438 | -0.916930866 |
| Mir6943 | 0 | NA | NA | NA |
| Prr7 | 196.075479 | 0.101120943 | 0.127269946 | 0.794539052 |
| Dbn1 | 1526.862046 | 0.538096377 | 0.102421016 | 5.253769152 |
| Mir6944 | 0 | NA | NA | NA |
| Pdlim7 | 426.9912292 | -0.575508387 | 0.088076331 | -6.534200305 |
| Mir6945 | 0 | NA | NA | NA |
| Dok3 | 92.02924505 | -1.172223977 | 0.225231559 | -5.204528089 |
| Ddx41 | 338.5120362 | -0.346149941 | 0.109150672 | -3.171303799 |
| Fam193b | 537.9735023 | 0.137655047 | 0.181625586 | 0.757905589 |
| Tmed9 | 877.8577826 | -0.219366059 | 0.07789739 | -2.81608998 |
| B4galt7 | 189.7572043 | 0.028015584 | 0.08750534 | 0.320158558 |
| Gm15911 | 0.088476398 | 0.51795324 | 3.724945049 | 0.139049901 |
| Caml | 264.5844916 | -0.242590939 | 0.080083055 | -3.029241819 |
| Ddx46 | 1095.580132 | 0.179750196 | 0.080261173 | 2.239566037 |
| B230219D22Rik | 1459.612451 | 0.033518431 | 0.074456238 | 0.450176263 |
| Txndc15 | 462.6097524 | -0.175274837 | 0.10027401 | -1.747958789 |
| Pcbd2 | 43.31228221 | -0.017168996 | 0.19655204 | -0.087350892 |
| 4930451E10Rik | 0 | NA | NA | NA |
| Catsper3 | 9.093479106 | 0.378686505 | 0.394337987 | 0.960309475 |
| Pitx1 | 1.253999855 | 0.244305393 | 0.932639799 | 0.261950427 |
| Gm28760 | 0.04393175 | 0.037062677 | 3.724945049 | 0.009949859 |
| H2afy | 903.7078744 | -0.166488877 | 0.072218055 | -2.305363631 |
| Gm25148 | 0 | NA | NA | NA |
| Tifab | 116.9424394 | -1.055736587 | 0.285301418 | -3.700425307 |
| 4930550C17Rik | 0 | NA | NA | NA |
| Neurog1 | 0.267034963 | 0.03707203 | 2.110058379 | 0.017569196 |
| Cxcl14 | 505.7540529 | 0.45118382 | 0.124014005 | 3.638168289 |
| Gm24716 | 0 | NA | NA | NA |
| Gm10782 | 0 | NA | NA | NA |
| Slc25a48 | 3.125560887 | -1.145316069 | 0.798404727 | -1.434505621 |
| Il9 | 0 | NA | NA | NA |
| Fbxl21 | 83.03568542 | 0.562936174 | 0.134811279 | 4.175734976 |
| Lect2 | 0.045882455 | 0.517959912 | 3.724945049 | 0.139051692 |
| Tgfbi | 1350.337662 | -1.342817733 | 0.309635465 | -4.336769792 |
| Gm27698 | 0 | NA | NA | NA |
| Smad5 | 771.3450184 | -0.268026273 | 0.079668646 | -3.364262932 |
| Gm27301 | 0.049553047 | -0.443818988 | 3.724945049 | -0.119147795 |
| Gm27834 | 0.26282868 | -0.926044447 | 1.82374036 | -0.507772086 |
| Gm27271 | 2.902142703 | 0.41755103 | 0.684386929 | 0.610109591 |
| Trpc7 | 38.7170757 | -0.26187831 | 0.181999529 | -1.438895538 |
| Gm22777 | 0 | NA | NA | NA |
| Spock1 | 3106.932593 | 0.497370851 | 0.072732131 | 6.838392383 |
| Klhl3 | 316.2281292 | 0.328135026 | 0.162401899 | 2.020512243 |

| | | | | |
|---|---|---|---|---|
| Mir874 | 4.532224978 | -0.463995383 | 0.663957934 | -0.6988325 |
| Hnrnpa0 | 1502.786418 | -0.006815575 | 0.100310914 | -0.067944499 |
| 1700066J03Rik | 0.225300421 | 1.220222262 | 2.56572005 | 0.475586673 |
| Idnk | 305.8515994 | -0.134844169 | 0.155286881 | -0.868355187 |
| Ubqln1 | 2320.123517 | -0.233727893 | 0.06289292 | -3.716283028 |
| Gkap1 | 124.6432152 | 0.094206106 | 0.141576111 | 0.665409615 |
| Gm26555 | 4.731241973 | 0.73297551 | 0.604341919 | 1.212849029 |
| Kif27 | 62.59338873 | 0.258057979 | 0.1772952 | 1.455527157 |
| Mir6369 | 0.042716945 | 0.037062677 | 3.724945049 | 0.009949859 |
| Gm24284 | 0 | NA | NA | NA |
| 2210016F16Rik | 128.8048172 | -0.814536645 | 0.162502244 | -5.012463992 |
| Hnrnpk | 4038.157108 | -0.047839897 | 0.076778739 | -0.623087811 |
| Mir7-1 | 15.77224037 | 0.601812627 | 0.270949974 | 2.221120812 |
| Rmi1 | 123.5749391 | -0.075324876 | 0.111162803 | -0.677608644 |
| Slc28a3 | 2.52226614 | 0.334905437 | 0.837303218 | 0.39998107 |
| Ntrk2 | 6695.738885 | 0.487739838 | 0.10582475 | 4.608939207 |
| Agtpbp1 | 2886.668731 | 0.420519213 | 0.069169271 | 6.079566913 |
| Naa35 | 495.813822 | -0.106845015 | 0.057115507 | -1.870683116 |
| Golm1 | 357.7061881 | 0.240745469 | 0.117366322 | 2.051231267 |
| 4921517D22Rik | 0.150248541 | 0.998859444 | 2.859451703 | 0.349318522 |
| Spata31d1a | 2.564266786 | -0.239071735 | 0.665401246 | -0.359289581 |
| Spata31d1b | 1.567374066 | -0.187195238 | 0.803687611 | -0.232920397 |
| Spata31d1d | 3.578253317 | 1.19182549 | 0.639330819 | 1.864176502 |
| 4930528D03Rik | 0.109124069 | 0.517955464 | 3.724945049 | 0.139050498 |
| 1700014D04Rik | 13.37826059 | 0.758324348 | 0.310812035 | 2.439816552 |
| Isca1 | 1058.772678 | 0.309153575 | 0.055952854 | 5.525251278 |
| Etohd2 | 12.41969927 | -0.47791487 | 0.357001077 | -1.338693074 |
| Zcchc6 | 1650.158349 | -0.127217494 | 0.097708545 | -1.302009909 |
| Gas1 | 102.0490903 | -0.597933432 | 0.189027061 | -3.16321604 |
| Gm5084 | 0.588182734 | -1.743386979 | 1.557394943 | -1.119425093 |
| Gm24999 | 0 | NA | NA | NA |
| Gm25000 | 0 | NA | NA | NA |
| Dapk1 | 1289.496349 | 0.557423087 | 0.121135419 | 4.601652347 |
| Gm10312 | 0 | NA | NA | NA |
| BC051665 | 0.048949532 | 0.037064902 | 3.724945049 | 0.009950456 |
| Ctsll3 | 0 | NA | NA | NA |
| 4930486L24Rik | 7.005733943 | -0.667407094 | 0.415917265 | -1.604663114 |
| Ctla2b | 52.79599323 | -0.693990545 | 0.274895959 | -2.5245571 |
| Tpbpb | 0 | NA | NA | NA |
| Gm24594 | 0 | NA | NA | NA |
| Ctla2a | 216.8941751 | 0.215624162 | 0.244841587 | 0.880668048 |
| Tpbpa | 0 | NA | NA | NA |
| Ctsj | 0 | NA | NA | NA |
| Ctsq | 0 | NA | NA | NA |
| Ctsr | 0 | NA | NA | NA |
| Cts6 | 0 | NA | NA | NA |
| Cts8 | 0 | NA | NA | NA |

| | | | | |
|---|---|---|---|---|
| Gm27704 | 0 | NA | NA | NA |
| Cts7 | 0 | NA | NA | NA |
| Ctsm | 0 | NA | NA | NA |
| Cts3 | 0 | NA | NA | NA |
| Gm19961 | 1.330930655 | -1.346211194 | 0.882876213 | -1.52480175 |
| Gm23739 | 0 | NA | NA | NA |
| Gm24193 | 0 | NA | NA | NA |
| Gm20563 | 3.002659297 | 0.903439726 | 0.725589511 | 1.245111338 |
| Gm25649 | 0.044544648 | 0.517957688 | 3.724945049 | 0.139051095 |
| Gm6478 | 2.109479346 | 0.279009383 | 0.663890175 | 0.420264364 |
| Zfp808 | 54.99002886 | -0.172877367 | 0.196821669 | -0.878345195 |
| Gm23713 | 0.185738947 | 0.037076627 | 2.674436242 | 0.013863343 |
| Gm24539 | 0.901339752 | -0.496184264 | 1.184139321 | -0.419025241 |
| Gm3604 | 26.94184347 | 0.126589872 | 0.231515164 | 0.546788685 |
| Gm25933 | 0.061920613 | -0.443816764 | 3.724945049 | -0.119147198 |
| Gm23855 | 0 | NA | NA | NA |
| Zfp935 | 99.08513468 | -0.177396557 | 0.137645355 | -1.288794357 |
| Gm22256 | 0.093484797 | -0.443823436 | 3.724945049 | -0.119148989 |
| Zfp934 | 37.404617 | -0.139891631 | 0.219916923 | -0.636111259 |
| Gm7664 | 14.88251027 | -0.179687708 | 0.277604227 | -0.647280156 |
| Gm24095 | 0.244748102 | -0.843965679 | 2.506620305 | -0.336694663 |
| Gm22615 | 1.125305835 | -1.570544679 | 1.087722837 | -1.443883152 |
| 6720489N17Rik | 53.55286888 | 0.101327606 | 0.194595978 | 0.520707609 |
| Gm24020 | 0.042716945 | 0.037062677 | 3.724945049 | 0.009949859 |
| Platr25 | 30.64885467 | 0.286427178 | 0.319898658 | 0.89536849 |
| Mir713 | 0 | NA | NA | NA |
| Gm5141 | 99.08605519 | 0.039737066 | 0.16784802 | 0.236744322 |
| Gm21963 | 0 | NA | NA | NA |
| Gm24588 | 0 | NA | NA | NA |
| Gm25448 | 0 | NA | NA | NA |
| Fbp2 | 0.981774026 | 0.499835276 | 1.251573224 | 0.399365587 |
| Fbp1 | 2.587088777 | -0.951318763 | 0.672618143 | -1.41435192 |
| 2010111I01Rik | 498.5900927 | -0.075952429 | 0.091129813 | -0.833453146 |
| Gm16907 | 5.155791298 | 0.653439897 | 0.438319629 | 1.490784017 |
| Mir23b | 8.607625547 | 0.023271952 | 0.428738983 | 0.054280001 |
| Mir27b | 11.22797161 | 0.385883081 | 0.366629778 | 1.052514291 |
| Mir3074-1 | 0.352569346 | -0.383294368 | 1.585308368 | -0.241779061 |
| mmu-mir-24-1 | 9.164853348 | 0.119306622 | 0.357611891 | 0.3336204 |
| Fancc | 74.63926839 | 0.111809269 | 0.140605237 | 0.79519989 |
| Gm16133 | 0.546112947 | 0.624928515 | 1.257561083 | 0.496936907 |
| Gm16132 | 1.982187692 | 1.116187159 | 0.68636194 | 1.626236966 |
| Gm16134 | 0 | NA | NA | NA |
| Ptch1 | 1765.336376 | 0.162172493 | 0.103237572 | 1.57086698 |
| A930032L01Rik | 2.69933516 | -1.072009946 | 0.711902431 | -1.505838298 |
| Ercc6l2 | 388.7591555 | -0.1093188 | 0.084094087 | -1.299958226 |
| Hsd17b3 | 0.044191549 | 0.037062677 | 3.724945049 | 0.009949859 |
| Slc35d2 | 16.4056275 | -1.401007021 | 0.305374474 | -4.587832777 |

| | | | | |
|---|---|---|---|---|
| Zfp367 | 307.36151 | -0.099949417 | 0.125666105 | -0.795357005 |
| Habp4 | 1131.608262 | 0.268449954 | 0.080235376 | 3.345780467 |
| Cdc14b | 233.8312342 | 0.342946252 | 0.101162497 | 3.39005325 |
| 1810034E14Rik | 1.766063332 | -1.042497939 | 0.704663891 | -1.479425796 |
| Aaed1 | 90.98287385 | -0.015503072 | 0.146911675 | -0.10552648 |
| Gm25654 | 2.492378667 | 0.339019548 | 0.681457096 | 0.497492139 |
| Ctsl | 1539.723944 | -0.944379967 | 0.17951951 | -5.260597966 |
| Cdk20 | 89.79521824 | -0.225237503 | 0.123018483 | -1.830924087 |
| 1190003K10Rik | 0.204782255 | 0.517969162 | 2.213323883 | 0.234023211 |
| Gm4810 | 1.123243924 | 3.140189933 | 1.075384214 | 2.920063259 |
| Cntnap3 | 108.0176306 | 0.657670782 | 0.137853602 | 4.770791436 |
| Spata31 | 0 | NA | NA | NA |
| Spata31d1c | 0 | NA | NA | NA |
| Prss47 | 0.08486385 | 0.037730769 | 3.724898049 | 0.010129343 |
| Hiatl1 | 800.205226 | -0.230000947 | 0.08228434 | -2.79519707 |
| Olfr466 | 0 | NA | NA | NA |
| Gm24130 | 0 | NA | NA | NA |
| Nlrp4f | 0.716239967 | -1.659747969 | 1.184245232 | -1.401523877 |
| Gm10775 | 2.001003253 | -0.140757212 | 0.774981762 | -0.181626483 |
| Zfp369 | 347.6968427 | -0.062500119 | 0.108667738 | -0.575148799 |
| Gm5792 | 0 | NA | NA | NA |
| Gm10139 | 0 | NA | NA | NA |
| Gm6901 | 0.223727156 | -0.451806035 | 2.152942534 | -0.20985513 |
| Gm26639 | 0 | NA | NA | NA |
| Platr2 | 0 | NA | NA | NA |
| Gm6917 | 0 | NA | NA | NA |
| Gm6910 | 5.181120878 | 0.672443763 | 0.450560499 | 1.492460535 |
| Gm10324 | 0 | NA | NA | NA |
| Gm17514 | 0 | NA | NA | NA |
| Gm10772 | 0 | NA | NA | NA |
| Gm26754 | 0 | NA | NA | NA |
| Gm26715 | 0 | NA | NA | NA |
| Vmn2r-ps104 | 0 | NA | NA | NA |
| 2410141K09Rik | 0 | NA | NA | NA |
| Gm26806 | 0 | NA | NA | NA |
| Gm27501 | 0 | NA | NA | NA |
| Zfp640 | 0 | NA | NA | NA |
| Gm17404 | 0 | NA | NA | NA |
| Gm17449 | 0 | NA | NA | NA |
| Gm10323 | 0 | NA | NA | NA |
| Gm5196 | 0.205773706 | -0.4438211 | 2.20735399 | -0.20106476 |
| Gm5451 | 26.41010186 | -0.576468609 | 0.243951548 | -2.363045505 |
| Uqcrb | 583.1177258 | 0.085578486 | 0.100399512 | 0.852379504 |
| Gm10767 | 12.81661586 | 0.18206084 | 0.293882946 | 0.619501206 |
| Mterf3 | 229.6188801 | 0.006477137 | 0.082013066 | 0.078976893 |
| Ptdss1 | 668.8494159 | -0.079802092 | 0.070136013 | -1.137819054 |
| Zfp712 | 93.19383338 | 0.0097958 | 0.176467541 | 0.055510489 |

| | | | | |
|---|---|---|---|---|
| Zfp708 | 0.458268618 | -0.443818503 | 1.484240643 | -0.299020584 |
| Gm28557 | 4.418616242 | -0.980977066 | 0.594517918 | -1.650037848 |
| Rslcan18 | 4.299918267 | -0.971126909 | 0.583846972 | -1.663324391 |
| Zfp759 | 131.3437174 | -0.062211166 | 0.159328615 | -0.390458213 |
| Rsl1 | 30.30590387 | -0.521975729 | 0.277806867 | -1.878915859 |
| Zfp455 | 45.95318099 | -0.595114712 | 0.272535541 | -2.1836224 |
| Zfp458 | 240.4665658 | -0.262810811 | 0.135680604 | -1.936981433 |
| Zfp457 | 21.39111743 | 0.128517664 | 0.29022407 | 0.442822209 |
| Zfp595 | 65.32548619 | -0.567780411 | 0.20133105 | -2.820133366 |
| Gm28044 | 0.23851432 | 0.037076503 | 2.11235448 | 0.017552216 |
| Zfp953 | 36.77585063 | -0.541939948 | 0.208353916 | -2.60105477 |
| Gm28041 | 0.153723802 | 0.037075944 | 2.839085074 | 0.013059117 |
| Gm17039 | 2.363953117 | -0.390841439 | 0.747361678 | -0.522961574 |
| Zfp456 | 19.86136891 | -1.07073107 | 0.264379689 | -4.049974778 |
| Zfp429 | 33.84957497 | -0.753598039 | 0.209899299 | -3.590283728 |
| Zfp459 | 66.34388881 | 0.166544365 | 0.158796768 | 1.048789389 |
| Zfp874a | 90.85288293 | -0.509290841 | 0.157601509 | -3.231509927 |
| Gm38307 | 0.282611191 | -0.458439376 | 1.649525598 | -0.277921953 |
| Gm7928 | 0 | NA | NA | NA |
| Zfp874b | 171.5607145 | -0.307881663 | 0.160049836 | -1.92366122 |
| Zfp58 | 50.28513376 | -0.372524689 | 0.177302499 | -2.101068463 |
| Gm26965 | 0 | NA | NA | NA |
| Zfp87 | 184.3921584 | -0.250282598 | 0.106302337 | -2.354441153 |
| Zfp748 | 247.3472852 | -0.295283127 | 0.135610032 | -2.177443093 |
| 9430065F17Rik | 2.796353279 | 0.48773759 | 0.653431471 | 0.746425005 |
| Zfp729b | 275.2153557 | -0.149737307 | 0.118072854 | -1.268177238 |
| Gm37276 | 23.42523615 | 0.178877063 | 0.24501491 | 0.730066032 |
| Zfp729a | 293.0100012 | -0.108851836 | 0.087462921 | -1.244548362 |
| Gm26587 | 0 | NA | NA | NA |
| Zfp738 | 303.073871 | -0.230420024 | 0.133464475 | -1.726452113 |
| Gm26875 | 0 | NA | NA | NA |
| Zfp65 | 240.5209492 | -0.296139679 | 0.11455479 | -2.585135709 |
| Zfp85os | 44.02786501 | 0.085043382 | 0.191802251 | 0.443390951 |
| Zfp85 | 74.6551353 | -0.100920332 | 0.161782039 | -0.623804303 |
| Gm9894 | 0 | NA | NA | NA |
| Zfp493 | 42.58231206 | -0.423747804 | 0.284848592 | -1.48762471 |
| 4930525G20Rik | 7.15404337 | 0.100567409 | 0.456823459 | 0.220145019 |
| Gm9625 | 125.9289179 | -0.423708008 | 0.15446584 | -2.743053147 |
| Zfp273 | 60.68720866 | 0.020296362 | 0.164775394 | 0.123175928 |
| Gm10037 | 3.305635379 | -0.259179571 | 0.561279123 | -0.461765921 |
| BC048507 | 0.806474979 | 0.966741043 | 1.153280923 | 0.838252869 |
| Mtrr | 182.8372363 | -0.427671087 | 0.102113608 | -4.188188984 |
| Fastkd3 | 102.6350786 | -0.373707473 | 0.126709586 | -2.949322819 |
| 1700001L19Rik | 42.40692636 | 0.701632124 | 0.180501274 | 3.887131152 |
| Adcy2 | 1089.553893 | 0.530304603 | 0.091637406 | 5.786988371 |
| Gm26929 | 0 | NA | NA | NA |
| Gm37122 | 2.65229942 | 1.164419503 | 0.8054182 | 1.445732792 |

| | | | | |
|---|---|---|---|---|
| Gm26844 | 0 | NA | NA | NA |
| Gm4812 | 0 | NA | NA | NA |
| AC161592.1 | 0 | NA | NA | NA |
| Papd7 | 947.3406657 | -0.225607715 | 0.080904409 | -2.788571308 |
| Nsun2 | 820.5404197 | -0.172335588 | 0.063725867 | -2.704327085 |
| A530095I07Rik | 0.197913552 | -0.908377233 | 2.22383349 | -0.408473583 |
| Srd5a1 | 47.64261977 | 0.217642866 | 0.178594035 | 1.218645775 |
| Gm17108 | 3.053155272 | 0.682992141 | 0.629372697 | 1.085195059 |
| Ube2ql1 | 1424.867787 | -0.024647909 | 0.085766891 | -0.287382568 |
| Med10 | 183.4117316 | -0.416335349 | 0.11078946 | -3.757896728 |
| Gm26819 | 0.108772582 | 0.037073798 | 3.724945049 | 0.009952844 |
| 1700084F23Rik | 0.092691418 | -0.908687078 | 3.724560878 | -0.2439716 |
| AC123920.1 | 0.239188194 | 0.462050438 | 1.844853859 | 0.25045368 |
| Gm26018 | 0 | NA | NA | NA |
| 1700100L14Rik | 0 | NA | NA | NA |
| Ice1 | 1670.349411 | 0.177443041 | 0.111121226 | 1.596842007 |
| Adamts16 | 57.86550676 | 0.73596986 | 0.245332514 | 2.999887165 |
| 8030423J24Rik | 1.260089781 | -0.166584085 | 0.984289563 | -0.169242965 |
| Gm38191 | 0.247225956 | 0.517970033 | 2.99204733 | 0.173115588 |
| Mir466f-4 | 0 | NA | NA | NA |
| Irx1 | 50.49064827 | -0.0473716 | 0.212205455 | -0.2232346 |
| Rpl9-ps4 | 43.5655805 | -0.443898784 | 0.189937007 | -2.337084226 |
| Gm20554 | 3.453678518 | 0.120010304 | 0.525690318 | 0.228290878 |
| Irx2 | 81.72096566 | -0.055376641 | 0.211673633 | -0.261613315 |
| Irx4 | 3.045905106 | -0.162638162 | 0.569603836 | -0.285528558 |
| Gm10263 | 1.909878088 | 0.843717155 | 0.761811037 | 1.107515006 |
| Ndufs6 | 294.0100797 | 0.057694595 | 0.078959904 | 0.730682181 |
| Mrpl36 | 129.8851055 | -0.438989219 | 0.179289429 | -2.448494707 |
| Lpcat1 | 574.2185807 | -0.308290431 | 0.088080009 | -3.500118087 |
| Slc6a3 | 74.17620595 | 1.228052616 | 0.297161616 | 4.132608489 |
| Clptm1l | 878.6178926 | -0.278363867 | 0.064304615 | -4.328831888 |
| Tert | 17.88891691 | 0.484601839 | 0.314091532 | 1.542868205 |
| Slc6a18 | 0.178490723 | 0.539256917 | 2.711241605 | 0.198896666 |
| Slc6a19 | 6.934680317 | -0.469961482 | 0.374069963 | -1.25634648 |
| Slc6a19os | 0.042716945 | 0.037062677 | 3.724945049 | 0.009949859 |
| Slc12a7 | 159.1782715 | -0.487952188 | 0.165651022 | -2.945663608 |
| Nkd2 | 44.99299481 | 0.362319716 | 0.200234336 | 1.809478447 |
| Gm8062 | 0 | NA | NA | NA |
| Trip13 | 13.34211523 | -0.219952981 | 0.268865758 | -0.81807733 |
| Gm25197 | 0.235280811 | 0.517970028 | 2.126140285 | 0.243619874 |
| Brd9 | 453.7906305 | 0.491065803 | 0.092959768 | 5.282562713 |
| Zdhhc11 | 0.149223819 | -0.924098271 | 2.865376951 | -0.322504957 |
| Gm10126 | 0.987133562 | 1.857731991 | 1.124749407 | 1.651685237 |
| Gm15912 | 22.38223563 | 0.478232535 | 0.260456205 | 1.836134159 |
| Tppp | 4530.77903 | 0.34426455 | 0.080947221 | 4.252950801 |
| Gm6263 | 0.459143553 | 0.106275493 | 1.30908277 | 0.081183173 |
| Cep72 | 28.65246828 | -0.150766451 | 0.231729368 | -0.650614343 |

| | | | | |
|---|---|---|---|---|
| Slc9a3 | 15.35249687 | 0.445592581 | 0.350954837 | 1.2696579 |
| Exoc3 | 1219.916005 | 0.039648866 | 0.047951122 | 0.82686003 |
| Ahrr | 8.460366272 | -0.11451314 | 0.367998777 | -0.311178046 |
| Pdcd6 | 440.5530706 | -0.236062194 | 0.083808061 | -2.81670033 |
| Sdha | 3363.70453 | 0.124411582 | 0.061399174 | 2.026274504 |
| Gm16125 | 0 | NA | NA | NA |
| Ccdc127 | 908.4665307 | -0.033374997 | 0.063277478 | -0.5274388 |
| Lrrc14b | 12.94872141 | -1.397330823 | 0.419489855 | -3.331024112 |
| Zfp72 | 58.03635434 | -0.347962256 | 0.218171469 | -1.59490266 |
| Gm10116 | 125.931969 | -1.290898355 | 0.209051525 | -6.175024815 |
| Mir692-3 | 0 | NA | NA | NA |
| Zfp825 | 69.54643972 | -0.356187912 | 0.18836721 | -1.890923117 |
| Erap1 | 747.9723258 | -0.432960034 | 0.115103782 | -3.761475306 |
| Cast | 225.7625238 | -0.526119279 | 0.144147398 | -3.649870099 |
| Hnrnpa1l2-ps | 0.737427016 | -0.674108476 | 1.195058927 | -0.564079696 |
| Pcsk1 | 401.8836238 | -0.222367638 | 0.165555873 | -1.343157658 |
| Gm4149 | 23.46034018 | -0.908756389 | 0.656996586 | -1.383198038 |
| Mir682 | 0 | NA | NA | NA |
| Gm23012 | 0 | NA | NA | NA |
| Ell2 | 509.0942641 | -0.412807086 | 0.099455108 | -4.150687609 |
| Gm23127 | 1.778293916 | 0.394891354 | 0.77418573 | 0.51007315 |
| Glrx | 434.8245854 | -0.568691555 | 0.17283815 | -3.290312675 |
| Rhobtb3 | 441.2117491 | -0.012681446 | 0.130544705 | -0.097142557 |
| Gm24163 | 0 | NA | NA | NA |
| Gm15528 | 4.932600309 | 0.397399775 | 0.552671904 | 0.719051886 |
| Gm6311 | 20.93266756 | 0.178717272 | 0.294912777 | 0.606000438 |
| Gm23319 | 0.214976432 | 0.037076921 | 2.177043914 | 0.017030856 |
| Spata9 | 2.596037066 | -0.419464423 | 0.64265473 | -0.652705727 |
| Rfesd | 170.1520903 | 0.029241496 | 0.102117969 | 0.286350153 |
| Gpr150 | 6.580668916 | 0.920041851 | 0.435926047 | 2.110545714 |
| Arsk | 95.44909683 | -0.230945113 | 0.144683892 | -1.596204725 |
| Ttc37 | 526.0109267 | -0.204895933 | 0.068220961 | -3.003416117 |
| Gm24025 | 0 | NA | NA | NA |
| Mctp1 | 1090.396185 | 0.91382782 | 0.097100948 | 9.411111235 |
| Gm24519 | 0 | NA | NA | NA |
| Gm21959 | 0 | NA | NA | NA |
| Gm15740 | 8.875000737 | 0.823333065 | 0.393250568 | 2.093660204 |
| Gm8185 | 13.50337661 | 0.681226868 | 0.401379594 | 1.697213504 |
| Ankrd32 | 492.724753 | 0.327197042 | 0.083809733 | 3.904045863 |
| 2210408I21Rik | 307.5143814 | 0.399025393 | 0.082455034 | 4.839309083 |
| Gm26088 | 0.042716945 | 0.037062677 | 3.724945049 | 0.009949859 |
| Fam172a | 517.357941 | 0.142358852 | 0.067990139 | 2.093816163 |
| Pou5f2 | 26.05560058 | 0.336190251 | 0.371175656 | 0.905744344 |
| Nr2f1 | 507.987299 | 0.30330032 | 0.107310459 | 2.826381715 |
| A830082K12Rik | 207.7537088 | 0.638346436 | 0.162169812 | 3.936283992 |
| Gm6317 | 0 | NA | NA | NA |
| Gm25383 | 0 | NA | NA | NA |

| | | | | |
|---|---|---|---|---|
| Gm29318 | 0.046345709 | -0.443818988 | 3.724945049 | -0.119147795 |
| Gm38349 | 0 | NA | NA | NA |
| Gm24863 | 0.296104695 | 0.51797522 | 1.77208077 | 0.292297749 |
| Gm28526 | 0.110242424 | -0.443823436 | 3.724945049 | -0.119148989 |
| Gm24635 | 0 | NA | NA | NA |
| Arrdc3 | 1031.103706 | 0.16720033 | 0.098296042 | 1.700987422 |
| 1700023H06Rik | 0.473445911 | -1.896187606 | 1.319582497 | -1.436960259 |
| Gm4211 | 15.12852727 | 0.471544559 | 0.30639633 | 1.539001983 |
| 9330111N05Rik | 6.889709052 | 1.679373692 | 0.503305541 | 3.336688267 |
| Gm8399 | 71.44903874 | -0.093539533 | 0.151152288 | -0.61884298 |
| Gm25241 | 0 | NA | NA | NA |
| Adgrv1 | 98.32611678 | -0.327934269 | 0.161571733 | -2.029651242 |
| Lysmd3 | 266.3986882 | -0.139215629 | 0.105675245 | -1.317391117 |
| Polr3g | 69.8321072 | 0.337746904 | 0.135972871 | 2.483928605 |
| Mblac2 | 357.5721259 | 0.443487021 | 0.103742293 | 4.274891266 |
| Gm17259 | 17.35136828 | -0.512899473 | 0.298097007 | -1.720579078 |
| Cetn3 | 274.7140815 | 0.22761246 | 0.139472113 | 1.631956775 |
| Gm24295 | 0 | NA | NA | NA |
| mmu-mir-3961 | 0 | NA | NA | NA |
| Mef2c | 4656.118685 | 1.216621387 | 0.158719469 | 7.665230975 |
| C130071C03Rik | 839.3578359 | 0.233406555 | 0.127821476 | 1.826035521 |
| Gm26803 | 22.51607708 | 0.617640172 | 0.248058558 | 2.489896643 |
| Gm33366 | 4.40666377 | 0.301610553 | 0.464156069 | 0.649804179 |
| Mir9-2 | 0 | NA | NA | NA |
| 2810049E08Rik | 34.32347081 | 0.632347113 | 0.258672509 | 2.444585688 |
| Gm17750 | 9.902264217 | 0.130357889 | 0.368539854 | 0.353714497 |
| Gm24331 | 0 | NA | NA | NA |
| A230107N01Rik | 12.62433136 | -0.611276745 | 0.416215892 | -1.468653064 |
| Tmem161b | 374.8315931 | 0.144173757 | 0.098714933 | 1.460506044 |
| Gm26927 | 0.174715781 | 0.517965821 | 2.726888692 | 0.189947548 |
| Gm26913 | 0.044191549 | 0.037062677 | 3.724945049 | 0.009949859 |
| Gm4076 | 2.353414094 | -0.260131953 | 0.720667072 | -0.360959954 |
| Ccnh | 371.5577082 | -0.097675198 | 0.099537403 | -0.981291404 |
| Rasa1 | 1059.457239 | -0.12671415 | 0.077990899 | -1.624729952 |
| Gm37641 | 0 | NA | NA | NA |
| Cox7c | 960.0717164 | 0.049994401 | 0.11157696 | 0.448071014 |
| Gm22574 | 1.539931973 | -0.845213462 | 0.886092186 | -0.953866285 |
| Gm27025 | 0 | NA | NA | NA |
| Gm25700 | 0 | NA | NA | NA |
| Gm24935 | 0 | NA | NA | NA |
| Gm21232 | 0 | NA | NA | NA |
| Gm27044 | 0 | NA | NA | NA |
| Gm8526 | 0 | NA | NA | NA |
| Edil3 | 2220.587599 | 0.218548303 | 0.091471216 | 2.389257644 |
| Hapln1 | 201.6502422 | 0.799637276 | 0.218323223 | 3.662630407 |
| Gm8546 | 11.32043437 | 0.16230122 | 0.350178264 | 0.463481709 |
| Vcan | 1385.185601 | -0.3079609 | 0.122927978 | -2.505214069 |

| | | | | |
|---|---|---|---|---|
| Gm16318 | 0.048949532 | 0.037064902 | 3.724945049 | 0.009950456 |
| Gm4117 | 32.71244237 | -0.069186879 | 0.224501115 | -0.308180557 |
| Xrcc4 | 102.7407697 | -0.212185032 | 0.135900416 | -1.561327315 |
| Gm24498 | 3.65796296 | 0.157168968 | 0.624177212 | 0.251801836 |
| Tmem167 | 992.695371 | -0.046172773 | 0.057012939 | -0.809864807 |
| Gm24507 | 5.663741534 | 0.217993648 | 0.424291523 | 0.51378271 |
| Gm37727 | 1.381073991 | 0.151457961 | 0.938657597 | 0.161355921 |
| Gm37054 | 0.576775271 | 1.013255834 | 1.310163785 | 0.773381043 |
| Gm37708 | 0 | NA | NA | NA |
| Gm21726 | 0.255118679 | 0.051282787 | 2.474427558 | 0.020725111 |
| Gm36966 | 0 | NA | NA | NA |
| AC147633.1 | 0 | NA | NA | NA |
| Atp6ap1l | 69.06133185 | 0.554318697 | 0.205354637 | 2.699323984 |
| Rps23 | 1420.870961 | -0.014777009 | 0.091222027 | -0.161989482 |
| Atg10 | 83.32890499 | 0.168563297 | 0.12208234 | 1.380734486 |
| Gm17450 | 0 | NA | NA | NA |
| A830009L08Rik | 17.31106247 | 1.150925481 | 0.348894818 | 3.298774936 |
| Gm29540 | 0.823808628 | 2.105783215 | 1.441237341 | 1.461093989 |
| Ssbp2 | 1111.715029 | 0.450701373 | 0.084658299 | 5.323770714 |
| 4833422C13Rik | 42.21647888 | -0.707785066 | 0.188866816 | -3.74753534 |
| Gm22778 | 1.426922496 | 1.045949517 | 0.99620905 | 1.049929748 |
| Acot12 | 6.478410357 | -0.434801797 | 0.439839799 | -0.988545824 |
| Gm24597 | 0 | NA | NA | NA |
| Gm27656 | 0 | NA | NA | NA |
| Zcchc9 | 113.9382792 | -0.19809219 | 0.116165779 | -1.705254263 |
| Ckmt2 | 1.022167862 | -0.426815772 | 1.009843253 | -0.422655467 |
| 1700119I11Rik | 3.377993944 | 0.135906272 | 0.6323026 | 0.214938658 |
| Rasgrf2 | 1981.271647 | 0.574254916 | 0.095219305 | 6.030866511 |
| Msh3 | 384.3197434 | 0.016004511 | 0.103032872 | 0.155334028 |
| Gm20379 | 0 | NA | NA | NA |
| AC161588.1 | 0 | NA | NA | NA |
| Dhfr | 139.8089927 | -1.162097073 | 0.190465475 | -6.101352872 |
| C030017D09Rik | 1.139049109 | -1.704494723 | 1.098262591 | -1.55199197 |
| Ankrd34b | 374.0828976 | 0.270784887 | 0.103477653 | 2.616844114 |
| Fam151b | 60.4619812 | -0.169187562 | 0.149335007 | -1.132939725 |
| Zfyve16 | 462.5719769 | -0.235965452 | 0.098219719 | -2.402424429 |
| Spz1 | 0 | NA | NA | NA |
| Serinc5 | 1004.030055 | -0.051149609 | 0.083035958 | -0.615993487 |
| Thbs4 | 44.88367789 | 0.480577549 | 0.352516044 | 1.363278515 |
| Mtx3 | 682.3862295 | 0.288658841 | 0.076857302 | 3.75577638 |
| Cmya5 | 96.50354882 | 0.510653323 | 0.153323984 | 3.330550823 |
| Papd4 | 358.1413963 | -0.018589414 | 0.119363963 | -0.155737241 |
| Gm2076 | 0.653050501 | 0.683785561 | 1.200507387 | 0.56958047 |
| Homer1 | 3096.738791 | -0.337360464 | 0.24294223 | -1.388644796 |
| Jmy | 1417.927992 | 0.211374978 | 0.053732133 | 3.933865366 |
| Bhmt | 2.668832678 | 1.623635211 | 0.730491643 | 2.222660899 |
| Gm15622 | 0.30042186 | 0.542532674 | 1.612105307 | 0.336536746 |

| | | | | |
|---|---|---|---|---|
| Bhmt2 | 0.855497761 | 1.03497928 | 1.155603065 | 0.895618324 |
| Dmgdh | 2.316670105 | 0.598765766 | 0.819086798 | 0.731016258 |
| AW495222 | 7.0482006 | 1.031501279 | 0.461414542 | 2.235519656 |
| Gm15620 | 0 | NA | NA | NA |
| Arsb | 966.5384739 | 0.629820028 | 0.128736618 | 4.892314549 |
| Mir5624 | 0.311503589 | -0.529129692 | 1.805909926 | -0.292998939 |
| Gm25534 | 0.045882455 | 0.517959912 | 3.724945049 | 0.139051692 |
| Gm26527 | 0.345934107 | -0.427257767 | 1.725732131 | -0.247580583 |
| Gm24737 | 0.579409632 | 2.805433293 | 1.286397095 | 2.180845482 |
| Lhfpl2 | 534.543113 | -1.197672337 | 0.156456888 | -7.654967158 |
| Gm15907 | 0.284494328 | 0.507488516 | 2.051211533 | 0.247409157 |
| Scamp1 | 2084.156384 | 0.351729517 | 0.053784076 | 6.53965897 |
| Gm9776 | 34.06058694 | 0.491320957 | 0.180310172 | 2.72486545 |
| Ap3b1 | 770.6625569 | -0.201098 | 0.064920994 | -3.097580412 |
| Gm25109 | 1.803894262 | -0.461234895 | 0.934737016 | -0.49343814 |
| Gm36991 | 0.478351411 | 2.353208172 | 3.687373693 | 0.638180008 |
| Tbca | 287.401201 | -0.399196454 | 0.123948806 | -3.220655908 |
| Otp | 27.26391552 | -0.036501443 | 0.332924941 | -0.109638657 |
| Wdr41 | 404.2473427 | 0.021375189 | 0.069849002 | 0.30601996 |
| Pde8b | 739.5660274 | 0.235900606 | 0.140000317 | 1.685000515 |
| Gm16243 | 0.075043205 | 0.517957688 | 3.724945049 | 0.139051095 |
| Zbed3 | 136.959666 | 0.094174715 | 0.154950319 | 0.607773613 |
| Snora47 | 6.619789565 | 0.357337245 | 0.395756716 | 0.902921494 |
| Gm22661 | 5.694343269 | 0.434589051 | 0.429488809 | 1.011875145 |
| Aggf1 | 620.7099307 | -0.117846974 | 0.076246934 | -1.545596231 |
| Crhbp | 115.8235207 | -0.475982271 | 0.154040041 | -3.089990559 |
| S100z | 0 | NA | NA | NA |
| F2rl1 | 1.572027216 | 0.132698921 | 0.870906905 | 0.152368663 |
| F2r | 208.5978153 | -0.101833165 | 0.140072496 | -0.727003287 |
| Iqgap2 | 782.9040417 | -0.108739044 | 0.104232536 | -1.043235142 |
| F2rl2 | 16.42119601 | -0.719626468 | 0.303311597 | -2.372564965 |
| Gm24032 | 0.090427103 | 0.998850474 | 3.724945049 | 0.268151734 |
| Sv2c | 1728.872108 | -0.850832767 | 0.136608389 | -6.228261493 |
| Gm17190 | 0.114742407 | 0.517955464 | 3.724945049 | 0.139050498 |
| Gm23955 | 0.962110563 | -0.456467907 | 1.14485435 | -0.398712646 |
| Gm29543 | 3.164183009 | -0.544337387 | 0.597272048 | -0.911372614 |
| Gm25213 | 0 | NA | NA | NA |
| Poc5 | 406.0730432 | 0.067202228 | 0.079603469 | 0.844212296 |
| Ankdd1b | 3.542632829 | 0.208707449 | 0.584191476 | 0.357258635 |
| Polk | 176.7680094 | -0.587623926 | 0.123535917 | -4.756705112 |
| Gm17622 | 0 | NA | NA | NA |
| Col4a3bp | 1491.199334 | -0.009819702 | 0.051535586 | -0.190542171 |
| Hmgcr | 1361.035643 | -0.194810646 | 0.094209752 | -2.067839482 |
| Gm27792 | 0 | NA | NA | NA |
| Gcnt4 | 101.3429506 | -0.319607889 | 0.153835484 | -2.077595363 |
| 1700029F12Rik | 0 | NA | NA | NA |
| Fam169a | 842.6174546 | 0.255836193 | 0.075802207 | 3.375049413 |

| | | | | |
|---|---|---|---|---|
| Gm6169 | 6.279356964 | -1.306381527 | 0.422332377 | -3.093254503 |
| Nsa2 | 317.8801227 | -0.143622949 | 0.126424411 | -1.136038111 |
| Gfm2 | 387.8031722 | -0.060280218 | 0.123313999 | -0.488835159 |
| Hexb | 1048.089142 | 0.286804047 | 0.171075711 | 1.676474379 |
| Enc1 | 4686.802237 | 0.668458168 | 0.131772947 | 5.072802735 |
| C430039J16Rik | 0.150086409 | 1.027495679 | 3.723011807 | 0.275985071 |
| Lincenc1 | 22.69480111 | -0.013218676 | 0.24968334 | -0.052941762 |
| Gm8756 | 1.313089905 | 0.601150025 | 0.895557599 | 0.671257801 |
| 5330416C01Rik | 28.18081706 | -1.255594522 | 0.327198092 | -3.83741395 |
| Gm26619 | 4.459207448 | -0.382116498 | 0.512458474 | -0.745653584 |
| Gm10260 | 388.9520094 | -0.454090553 | 0.139186012 | -3.262472614 |
| Arhgef28 | 438.5791712 | 0.544076292 | 0.092388865 | 5.888981215 |
| Utp15 | 405.8469762 | 0.047867468 | 0.105737608 | 0.452700502 |
| Ankra2 | 370.3092848 | 0.21246339 | 0.092579871 | 2.294919916 |
| Gm21976 | 0.640962652 | -2.115648289 | 1.165746184 | -1.814844705 |
| Btf3 | 823.2235028 | -0.195904947 | 0.088426616 | -2.215452266 |
| Gm9828 | 25.61986482 | 0.127428772 | 0.250245662 | 0.509214711 |
| Foxd1 | 28.76092711 | -0.74191068 | 0.293047748 | -2.531705789 |
| Gm2445 | 3.175554283 | -0.277287004 | 0.575552671 | -0.481775201 |
| 2310005E17Rik | 0 | NA | NA | NA |
| Gm29501 | 0 | NA | NA | NA |
| Gm10320 | 0.789918879 | 0.067004152 | 1.039024846 | 0.064487536 |
| Gm37425 | 0 | NA | NA | NA |
| Gm22698 | 0 | NA | NA | NA |
| Tmem174 | 0.124347486 | 0.036092774 | 3.717245565 | 0.009709548 |
| Tmem171 | 14.34229216 | -1.373043875 | 0.324262189 | -4.234363177 |
| Fcho2 | 753.4025212 | 0.073796784 | 0.067710275 | 1.089890476 |
| Tnpo1 | 2400.224482 | -0.396076311 | 0.087362548 | -4.533708314 |
| Gm5453 | 2.269219555 | -0.575858793 | 0.630396759 | -0.913486284 |
| 1700024P04Rik | 0 | NA | NA | NA |
| Gm24733 | 0.085433889 | 0.037058229 | 3.724945049 | 0.009948665 |
| 2310020H05Rik | 0 | NA | NA | NA |
| Gm807 | 7.497632429 | -0.909095557 | 0.464696206 | -1.956322316 |
| Zfp366 | 15.07922646 | -1.071784528 | 0.319487105 | -3.354703558 |
| Ptcd2 | 245.1837604 | -0.078458955 | 0.076193584 | -1.029731788 |
| Mrps27 | 194.2719257 | -0.062254342 | 0.085526097 | -0.727898787 |
| 6430562O15Rik | 0.528537426 | -1.204734565 | 1.380152829 | -0.872899392 |
| Map1b | 25160.03304 | 0.278394271 | 0.096042616 | 2.898653574 |
| Gm26559 | 14.25506631 | 1.281556748 | 0.365492982 | 3.506378536 |
| 1700012J22Rik | 0.18753903 | 0.037072906 | 2.716639151 | 0.013646607 |
| Gm24471 | 0 | NA | NA | NA |
| Cartpt | 99.17765322 | -0.293703838 | 0.235346052 | -1.247965863 |
| Mccc2 | 207.1112753 | 0.511646642 | 0.091562788 | 5.587932112 |
| Bdp1 | 1242.860492 | -0.053066759 | 0.08114425 | -0.653980518 |
| BC001981 | 0.782345574 | 0.006475159 | 1.059090382 | 0.006113887 |
| Serf1 | 47.59636433 | 0.097638599 | 0.186080031 | 0.524712932 |
| Smn1 | 181.1517505 | -0.158146022 | 0.092249639 | -1.714326724 |

| | | | | |
|---|---|---|---|---|
| Naip2 | 165.0036826 | -0.998105852 | 0.184915692 | -5.397626556 |
| Naip5 | 40.97982787 | -0.848954508 | 0.243945919 | -3.480093092 |
| Naip6 | 36.81044465 | -0.742556102 | 0.243654272 | -3.047580889 |
| Naip3 | 1.709236637 | -0.665755263 | 0.670952774 | -0.992253537 |
| Naip1 | 6.360266967 | -0.031016108 | 0.448916657 | -0.069091016 |
| Rpl31-ps13 | 5.091748836 | -0.624312893 | 0.53437361 | -1.16830787 |
| Gtf2h2 | 239.663237 | 0.21625504 | 0.127249754 | 1.699453509 |
| Ocln | 88.83136681 | -0.211757129 | 0.159501068 | -1.32762201 |
| Marveld2 | 42.42039179 | 0.048742003 | 0.201164934 | 0.242298707 |
| Gm24261 | 0 | NA | NA | NA |
| Rad17 | 254.2430578 | -0.141182641 | 0.100240202 | -1.4084433 |
| Ak6 | 71.85688038 | -0.596771598 | 0.146456031 | -4.074749233 |
| Taf9 | 6.955395866 | 0.051002812 | 0.425733447 | 0.119799871 |
| Ccdc125 | 6.403646332 | -0.67254192 | 0.362620048 | -1.854673849 |
| Cdk7 | 820.0282203 | -0.022733581 | 0.056937773 | -0.399270634 |
| Mrps36 | 129.039131 | -0.323688855 | 0.131663738 | -2.458451046 |
| Cenph | 6.451376144 | -0.279429903 | 0.583388522 | -0.478977374 |
| Gm16335 | 0.346789506 | -0.772145031 | 1.435891343 | -0.537746143 |
| Ccnb1 | 20.0589336 | -0.439467548 | 0.383762829 | -1.145154025 |
| Slc30a5 | 515.9001532 | -0.12499001 | 0.076787745 | -1.62773382 |
| Gm29502 | 6.014679714 | 0.100383339 | 0.432072625 | 0.232329783 |
| Gm37830 | 0.152961209 | 0.055211374 | 2.838062297 | 0.019453898 |
| Gm10257 | 10.11133075 | 0.238979109 | 0.3166875 | 0.754621224 |
| Gm6114 | 0 | NA | NA | NA |
| Hmgb1-ps9 | 1.368962209 | 1.206835491 | 0.972476147 | 1.240992383 |
| 4932411K12Rik | 0.386457151 | -2.307898953 | 1.660738651 | -1.389682207 |
| Gm36994 | 0 | NA | NA | NA |
| Gm29341 | 0 | NA | NA | NA |
| Pik3r1 | 3272.596283 | 0.081610118 | 0.06907 | 1.181556653 |
| Gm36638 | 65.83543421 | 0.241614224 | 0.245035037 | 0.986039493 |
| Gm38133 | 0 | NA | NA | NA |
| Gm25808 | 0 | NA | NA | NA |
| Cd180 | 333.6593441 | -1.32946083 | 0.209408442 | -6.34864965 |
| Mast4 | 1713.710742 | 0.192915198 | 0.081819764 | 2.357806842 |
| 1700099I09Rik | 0.905219458 | -0.13565136 | 1.049185766 | -0.129292032 |
| Gm17160 | 0.372057435 | -1.057609112 | 1.480643031 | -0.714290406 |
| Gm6211 | 10.83872605 | 1.716867641 | 0.473254132 | 3.627792183 |
| Gm5454 | 142.1539982 | 0.239360708 | 0.188194062 | 1.271882366 |
| n-R5s56 | 0 | NA | NA | NA |
| Gm16416 | 0.30254151 | 1.442576143 | 1.776619124 | 0.811978281 |
| Gm24441 | 0 | NA | NA | NA |
| Gm24870 | 0 | NA | NA | NA |
| Srek1 | 793.2803317 | 0.215402627 | 0.1525759 | 1.411773589 |
| Erbb2ip | 3344.991394 | -0.340823641 | 0.132230332 | -2.577499691 |
| Nln | 399.6247544 | -0.059525256 | 0.080676989 | -0.737821982 |
| Sgtb | 1145.826265 | 0.504540737 | 0.089492967 | 5.637769689 |
| Trappc13 | 722.8966951 | -0.069983484 | 0.072045381 | -0.971380576 |

| | | | | |
|---|---|---|---|---|
| Trim23 | 1039.038422 | 0.295904194 | 0.091376115 | 3.238310066 |
| Ppwd1 | 182.8657831 | -0.071371319 | 0.099867598 | -0.714659414 |
| Cenpk | 9.293126097 | -0.185880095 | 0.370794214 | -0.501302577 |
| Adamts6 | 345.3432011 | 0.009963415 | 0.146879261 | 0.067834051 |
| Cwc27 | 204.665214 | 0.312877958 | 0.163831309 | 1.90975681 |
| 2610204G07Rik | 0.047948698 | 0.037064902 | 3.724945049 | 0.009950456 |
| Srek1ip1 | 192.1915529 | -0.33319742 | 0.128248415 | -2.598062673 |
| Fam159b | 3.808355539 | -0.300957022 | 0.568791001 | -0.529117059 |
| Gm4938 | 0.726615386 | 1.086345843 | 1.114646895 | 0.974609849 |
| Rgs7bp | 3653.625861 | 0.436293793 | 0.099214156 | 4.397495386 |
| 4933425L06Rik | 0.717517214 | -1.194474167 | 1.093095011 | -1.092745054 |
| Rnf180 | 161.367705 | 0.167086008 | 0.098137411 | 1.702571994 |
| Gm25631 | 2.548041448 | 0.033016694 | 0.67471582 | 0.048934222 |
| Htr1a | 192.2048649 | 0.332915676 | 0.111868969 | 2.975943 |
| Dph3b-ps | 0 | NA | NA | NA |
| Gm24527 | 0.388660859 | 1.058873633 | 1.590854252 | 0.665600656 |
| Gm28989 | 0.486723951 | 0.4568716 | 1.539453638 | 0.29677516 |
| Ipo11 | 842.5268281 | 0.156844635 | 0.06150446 | 2.55013432 |
| Gm28988 | 0.166709681 | 0.517964155 | 2.76763122 | 0.187150713 |
| Dimt1 | 147.6538309 | -0.212417235 | 0.098573236 | -2.154917933 |
| Kif2a | 1440.730209 | -0.041895564 | 0.065937671 | -0.635381304 |
| Gm24431 | 2.158765044 | 0.215664639 | 0.891328236 | 0.24195872 |
| 3830408C21Rik | 12.03259843 | 0.308430257 | 0.365226108 | 0.844491263 |
| Apoo-ps | 41.16296874 | -0.57836079 | 0.281993513 | -2.050971968 |
| 1700006H21Rik | 0.426601203 | -2.109322949 | 3.210286541 | -0.657051301 |
| Zswim6 | 635.8544846 | -0.159945959 | 0.096166504 | -1.663219032 |
| Gm22118 | 4.848883265 | 0.26792811 | 0.568628304 | 0.471183211 |
| Smim15 | 306.3096422 | -0.236916587 | 0.106678787 | -2.220840665 |
| Ndufaf2 | 61.73980177 | -0.064420717 | 0.258827548 | -0.24889436 |
| Ercc8 | 184.4501039 | 0.250149687 | 0.113687091 | 2.200335008 |
| Elovl7 | 210.6233394 | -0.232324707 | 0.146527062 | -1.585541288 |
| Gm2822 | 0.316450815 | 0.496241232 | 1.770936295 | 0.28021405 |
| Depdc1b | 9.399548456 | -0.710508705 | 0.378905532 | -1.875160549 |
| Pde4d | 1367.797614 | 0.366917218 | 0.086947987 | 4.219962191 |
| Gm27572 | 0 | NA | NA | NA |
| Gm28016 | 0 | NA | NA | NA |
| Rps3a3 | 38.33507003 | -0.178926388 | 0.1887455 | -0.947976977 |
| Mir582 | 2.019407445 | 0.643354898 | 0.716584698 | 0.897807197 |
| AC161585.1 | 9.306437351 | 0.888009376 | 0.427882742 | 2.07535684 |
| Mir1904 | 1.272510498 | 1.203286724 | 0.917678074 | 1.311229676 |
| Rab3c | 4667.9155 | 0.308751891 | 0.092796991 | 3.327175657 |
| Gm5455 | 0.386702458 | -0.729436827 | 1.70099353 | -0.428829865 |
| Gapt | 3.214970131 | -1.658860554 | 0.686917015 | -2.414935892 |
| Plk2 | 1844.679949 | 0.000112385 | 0.140240954 | 0.000801369 |
| Actbl2 | 1.958193336 | -0.968480863 | 0.965077009 | -1.003527029 |
| Gm15285 | 0.507331407 | -1.052760145 | 1.357644637 | -0.775431299 |
| Gm15289 | 0 | NA | NA | NA |

| | | | | |
|---|---|---|---|---|
| Gm6270 | 0 | NA | NA | NA |
| Gpbp1 | 1669.061245 | -0.073607472 | 0.071358191 | -1.031520995 |
| Gm15290 | 0 | NA | NA | NA |
| Gm15288 | 0 | NA | NA | NA |
| Gm15286 | 0.075043205 | 0.517957688 | 3.724945049 | 0.139051095 |
| Gm10198 | 0 | NA | NA | NA |
| Gm10736 | 67.77195831 | -0.40672123 | 0.176298967 | -2.30699724 |
| Mier3 | 595.0131209 | -0.105019653 | 0.091438771 | -1.148524323 |
| Gm15488 | 0.182466651 | -1.346561657 | 2.672339661 | -0.503888662 |
| Gm15287 | 0.237498462 | -1.691621985 | 2.099246106 | -0.805823567 |
| Map3k1 | 549.7994848 | -0.752717247 | 0.15660985 | -4.806321223 |
| Gm15327 | 2.461737376 | 0.132013592 | 0.724290762 | 0.182266016 |
| Gm15325 | 0 | NA | NA | NA |
| Gm15324 | 0 | NA | NA | NA |
| Gm15326 | 3.336705128 | -1.203975526 | 0.521173586 | -2.31012384 |
| Gm9025 | 2.20249356 | -0.544881723 | 0.659569426 | -0.826117314 |
| Gm15322 | 0.293377407 | -0.443835017 | 3.724945049 | -0.119152098 |
| Gm15323 | 0.143781519 | 0.517957382 | 3.582979241 | 0.144560531 |
| Gm37427 | 0 | NA | NA | NA |
| Ankrd55 | 97.25075632 | -0.273901233 | 0.175886062 | -1.557265138 |
| Gm22768 | 0.696415487 | 0.034118763 | 1.372089812 | 0.024866275 |
| Il6st | 2388.948021 | -0.43556813 | 0.070926091 | -6.141155165 |
| Il31ra | 12.44555901 | -0.43250587 | 0.352202148 | -1.228004633 |
| Ddx4 | 13.81354613 | -1.14499268 | 0.32438068 | -3.529780752 |
| Slc38a9 | 402.4228987 | -0.238367093 | 0.127884254 | -1.863928407 |
| Ppap2a | 177.5947723 | 0.026010798 | 0.09587021 | 0.27131262 |
| Skiv2l2 | 776.3773442 | -0.165549125 | 0.059191498 | -2.796839571 |
| Gm26943 | 0.235215884 | -0.49615023 | 2.487520751 | -0.199455715 |
| Dhx29 | 365.0039571 | -0.404457268 | 0.069612304 | -5.810140491 |
| Ccno | 13.15772638 | 0.923666714 | 0.401961208 | 2.297900133 |
| Mcidas | 3.060161834 | 0.556942335 | 0.638418804 | 0.872377711 |
| Cdc20b | 1.498584608 | -0.836548492 | 0.933052705 | -0.89657153 |
| Mir449c | 0.04393175 | 0.037062677 | 3.724945049 | 0.009949859 |
| Mir449b | 0.038981395 | -0.443816764 | 3.724945049 | -0.119147198 |
| Mir449a | 0 | NA | NA | NA |
| Gm10735 | 1.512916718 | 0.988979941 | 0.787764046 | 1.255426604 |
| Gpx8 | 87.13244342 | -1.103484256 | 0.199967116 | -5.518328616 |
| Gzma | 199.9537804 | -1.520948958 | 0.455266774 | -3.340786201 |
| Gm9848 | 0 | NA | NA | NA |
| Gzmk | 39.86666651 | -0.130665683 | 0.308974901 | -0.422900639 |
| Esm1 | 8.383027344 | 0.416017836 | 0.399126476 | 1.042320822 |
| BC067074 | 57.61241419 | 0.263978766 | 0.178106307 | 1.482141597 |
| Gm22385 | 0 | NA | NA | NA |
| Snx18 | 579.4915124 | -0.283235798 | 0.104037018 | -2.722452088 |
| Hspb3 | 12.0017422 | 0.065167674 | 0.294822169 | 0.221040618 |
| Arl15 | 416.4383343 | 0.524170863 | 0.094282681 | 5.559566828 |
| Gm37907 | 0.135812198 | 0.037057153 | 3.653259119 | 0.010143587 |

| | | | | |
|---|---|---|---|---|
| Ndufs4 | 556.7189796 | 0.037531266 | 0.101893031 | 0.368339877 |
| Fst | 27.88029129 | -0.882992882 | 0.291754706 | -3.026490622 |
| Gm25950 | 0 | NA | NA | NA |
| 4930544M13Rik | 0 | NA | NA | NA |
| Mocs2 | 459.8435434 | -0.0461055 | 0.072064889 | -0.63977757 |
| Itga2 | 10.13867756 | -1.220273617 | 0.396715718 | -3.075939674 |
| Itga1 | 103.8467935 | -0.801492429 | 0.180395628 | -4.442970367 |
| Pelo | 139.7216595 | -0.412088321 | 0.113780996 | -3.621767575 |
| Gm25985 | 0 | NA | NA | NA |
| Gm6404 | 0 | NA | NA | NA |
| Isl1 | 36.11947267 | 0.29911358 | 0.189760464 | 1.576269224 |
| Gm38193 | 6.0191764 | -0.25933865 | 0.522708981 | -0.496143475 |
| Gm23198 | 0 | NA | NA | NA |
| Parp8 | 540.5637389 | 0.11121482 | 0.113221015 | 0.982280718 |
| Gm17509 | 11.37141038 | 0.24694296 | 0.391091663 | 0.631419648 |
| Emb | 394.0332575 | 0.077026609 | 0.123481636 | 0.623789997 |
| Gm6421 | 12.74783509 | -0.177407666 | 0.341377132 | -0.519682337 |
| Hcn1 | 981.9976838 | 0.013737354 | 0.068782332 | 0.199722129 |
| 4933413L06Rik | 0 | NA | NA | NA |
| Gm26349 | 0.072025462 | 0.517959912 | 3.724945049 | 0.139051692 |
| Mrps30 | 237.5140506 | -0.204146062 | 0.095026206 | -2.148313305 |
| B430218F22Rik | 1.974484148 | 0.104383554 | 0.665797977 | 0.15677962 |
| 1700003P14Rik | 0 | NA | NA | NA |
| Fgf10 | 52.0629943 | 0.480147488 | 0.166905008 | 2.876771016 |
| Gm16263 | 1.381163524 | 1.447308916 | 1.073030036 | 1.348805594 |
| AC154241.1 | 0 | NA | NA | NA |
| Nnt | 359.3284767 | -0.17974544 | 0.114229827 | -1.57354209 |
| Paip1 | 990.7888142 | -0.091772812 | 0.073333614 | -1.251442649 |
| 4833420G17Rik | 380.1582038 | 0.261079782 | 0.104778405 | 2.491732727 |
| Gm9752 | 0.269061364 | 1.539303632 | 3.70868876 | 0.415053333 |
| 3110070M22Rik | 13.70697502 | 0.772726352 | 0.293398336 | 2.633710751 |
| Gm7120 | 31.73718637 | -0.230870565 | 0.21049015 | -1.096823608 |
| Gm21967 | 80.16892809 | -0.217589441 | 0.154574534 | -1.407666811 |
| Ccl28 | 18.99182376 | 1.058832845 | 0.295153499 | 3.587397231 |
| 1700074H08Rik | 0 | NA | NA | NA |
| Hmgcs1 | 2950.403097 | -0.248951728 | 0.130270752 | -1.911033172 |
| Nim1k | 238.1724293 | 0.409881733 | 0.135868118 | 3.016761701 |
| Zfp131 | 574.5633546 | -0.121678842 | 0.087022575 | -1.398244567 |
| Gm23308 | 0 | NA | NA | NA |
| AF067063 | 0 | NA | NA | NA |
| D13Ertd608e | 1.077945179 | 0.18318437 | 1.225932309 | 0.149424539 |
| Gm21358 | 0 | NA | NA | NA |
| Gm37619 | 0.171181394 | 0.539411076 | 2.747344444 | 0.196339078 |
| Tcstv1 | 0 | NA | NA | NA |
| Gm21761 | 0 | NA | NA | NA |
| Gm21181 | 0 | NA | NA | NA |
| B020031M17Rik | 0 | NA | NA | NA |

| | | | | |
|---|---|---|---|---|
| Gm20784 | 1.793435801 | -0.095240921 | 0.768612483 | -0.123912795 |
| Gm22108 | 0 | NA | NA | NA |
| Gm21370 | 0 | NA | NA | NA |
| Gm21188 | 0.521673782 | -1.228682254 | 1.329782894 | -0.92397207 |
| Gm24798 | 0 | NA | NA | NA |
| Gm21378 | 13.07532112 | -0.08935682 | 0.305635373 | -0.292364129 |
| Gm20767 | 0 | NA | NA | NA |
| Gm21818 | 0.095898756 | -0.917692504 | 3.724434845 | -0.246397787 |
| Gm21762 | 1.958730402 | 0.054667758 | 0.790814262 | 0.069128443 |
| Gm21731 | 0 | NA | NA | NA |
| AF067061 | 0 | NA | NA | NA |
| Gm20900 | 10.70701964 | 0.119708604 | 0.360609496 | 0.331961874 |
| BC147527 | 33.1074984 | -1.74911728 | 0.423400789 | -4.131114836 |
| Tcstv3 | 0 | NA | NA | NA |
| Gm2888 | 0 | NA | NA | NA |
| Gm2897 | 10.21470906 | -0.617644694 | 0.348281868 | -1.773404676 |
| Gm2904 | 0.731111494 | -2.326545293 | 1.313313306 | -1.771508202 |
| Gm17051 | 0 | NA | NA | NA |
| Gm10340 | 0.412722675 | 0.70397032 | 1.610486791 | 0.437116482 |
| Gm17221 | 0 | NA | NA | NA |
| Gm17188 | 0 | NA | NA | NA |
| Gm5795 | 0.0878635 | 0.037058229 | 3.724945049 | 0.009948665 |
| Gm17222 | 1.483555279 | -0.958417537 | 0.742277994 | -1.291184091 |
| D830030K20Rik | 12.49083266 | 0.098675556 | 0.324458264 | 0.304124035 |
| Gm4904 | 0 | NA | NA | NA |
| 1700110I01Rik | 0.924198543 | -1.25073783 | 1.201616748 | -1.040879159 |
| Gm2956 | 0.84966665 | 0.703804504 | 1.374235607 | 0.512142532 |
| Gm17223 | 0 | NA | NA | NA |
| Gm10413 | 0 | NA | NA | NA |
| Gm10409 | 0.801287066 | -0.210111951 | 1.460963745 | -0.143817361 |
| Gm3005 | 2.629406015 | 0.188977175 | 0.836231096 | 0.225986783 |
| Gm26817 | 0 | NA | NA | NA |
| Gm10408 | 0.044191549 | 0.037062677 | 3.724945049 | 0.009949859 |
| Gm8323 | 0 | NA | NA | NA |
| Gm8324 | 0 | NA | NA | NA |
| Gm11109 | 0 | NA | NA | NA |
| Gm17107 | 0 | NA | NA | NA |
| Gm3012 | 0.072025462 | 0.517959912 | 3.724945049 | 0.139051692 |
| Gm3020 | 0.88261066 | -0.426962236 | 1.259712983 | -0.338936124 |
| Gm2866 | 0 | NA | NA | NA |
| Gm3043 | 0 | NA | NA | NA |
| Gm2871 | 0 | NA | NA | NA |
| Gm2877 | 0 | NA | NA | NA |
| Gm21965 | 0.038981395 | -0.443816764 | 3.724945049 | -0.119147198 |
| Gm3002 | 2.349117343 | -0.13433554 | 0.638154936 | -0.210506153 |
| Gm26886 | 0 | NA | NA | NA |
| Gm26570 | 0 | NA | NA | NA |

| | | | | |
|---|---|---|---|---|
| Gm3033 | 0.403254479 | -0.150216865 | 1.615892057 | -0.09296219 |
| Gm2907 | 0 | NA | NA | NA |
| Gm2916 | 0.044191549 | 0.037062677 | 3.724945049 | 0.009949859 |
| Gm3029 | 0 | NA | NA | NA |
| Gm3095 | 0.864161856 | -0.056763822 | 1.154227261 | -0.049179069 |
| Gm17129 | 0 | NA | NA | NA |
| Gm3099 | 0.095898756 | -0.917692504 | 3.724434845 | -0.246397787 |
| Gm5796 | 5.645051647 | 0.003005292 | 0.404696963 | 0.007426031 |
| Gm11107 | 0 | NA | NA | NA |
| Gm3115 | 0.045882455 | 0.517959912 | 3.724945049 | 0.139051692 |
| Gm8108 | 0.608376789 | 0.809316213 | 1.368200848 | 0.591518573 |
| Gm17152 | 0 | NA | NA | NA |
| Gm3127 | 0.0878635 | 0.037058229 | 3.724945049 | 0.009948665 |
| Gm2974 | 9.121692315 | 0.505573086 | 0.327489497 | 1.543784126 |
| Gm11105 | 0 | NA | NA | NA |
| Gm3138 | 0.101408268 | 0.037730769 | 3.724898049 | 0.010129343 |
| Gm17085 | 0 | NA | NA | NA |
| Gm3149 | 0.060174537 | 0.517957688 | 3.724945049 | 0.139051095 |
| 2610042L04Rik | 9.926510475 | -0.001412452 | 0.312547693 | -0.004519155 |
| Gm17086 | 0 | NA | NA | NA |
| Gm3159 | 0.075043205 | 0.517957688 | 3.724945049 | 0.139051095 |
| Gm3164 | 5.22863904 | 0.043116538 | 0.521100511 | 0.082741308 |
| Gm3173 | 3.269157551 | 0.954520861 | 0.54002375 | 1.767553486 |
| Gm17170 | 0 | NA | NA | NA |
| Gm3182 | 0.048949532 | 0.037064902 | 3.724945049 | 0.009950456 |
| Gm17074 | 0 | NA | NA | NA |
| Gm3030 | 0.11997416 | 0.517957688 | 3.724945049 | 0.139051095 |
| Gm26655 | 0 | NA | NA | NA |
| Gm3047 | 0.107078754 | -0.443164241 | 3.724898049 | -0.118973522 |
| Gm8164 | 0 | NA | NA | NA |
| Gm26898 | 0.093141081 | 0.037060453 | 3.724945049 | 0.009949262 |
| Gm8159 | 0 | NA | NA | NA |
| Gm3239 | 1.60056093 | 1.022251679 | 0.862982685 | 1.184556419 |
| Gm17073 | 0 | NA | NA | NA |
| Gm7876 | 0.046345709 | -0.443818988 | 3.724945049 | -0.119147795 |
| Gm3252 | 10.87374526 | 0.47934438 | 0.390714655 | 1.226840031 |
| Gm3084 | 0 | NA | NA | NA |
| Gm9602 | 0.3514278 | -0.462145159 | 1.7562912 | -0.263136978 |
| Gm3091 | 0 | NA | NA | NA |
| Gm3269 | 0 | NA | NA | NA |
| Gm3264 | 2.10462685 | 0.471764408 | 0.656854805 | 0.718217183 |
| Gm26650 | 0 | NA | NA | NA |
| Gm3278 | 0.274437282 | 0.037077402 | 1.662026427 | 0.022308551 |
| Gm8305 | 0 | NA | NA | NA |
| Gm10410 | 0 | NA | NA | NA |
| Gm8297 | 0 | NA | NA | NA |
| Gm3298 | 0.938975218 | -0.327541086 | 0.947782436 | -0.345586786 |

| | | | | |
|---|---|---|---|---|
| 4930555G01Rik | 0.088534442 | -0.905414734 | 3.723500899 | -0.243162217 |
| Gm8281 | 11.41715526 | 0.636677086 | 0.340956063 | 1.86732883 |
| Gm17069 | 0 | NA | NA | NA |
| Gm8279 | 0.163190422 | -0.955151543 | 3.396300892 | -0.281232898 |
| Gm26552 | 0 | NA | NA | NA |
| Gm3317 | 1.057189525 | 2.28364524 | 1.102285974 | 2.071735732 |
| Gm3140 | 0 | NA | NA | NA |
| Gm8271 | 0.068097359 | 0.037064902 | 3.724945049 | 0.009950456 |
| Gm3326 | 0 | NA | NA | NA |
| Gm3526 | 0.137762903 | 0.51796349 | 2.937523392 | 0.176326592 |
| Gm3339 | 0 | NA | NA | NA |
| Gm3542 | 0 | NA | NA | NA |
| Gm17126 | 0 | NA | NA | NA |
| Gm3500 | 0.712486263 | -1.350141521 | 1.183550472 | -1.14075534 |
| Gm3348 | 0 | NA | NA | NA |
| Gm3194 | 0.375475346 | 0.876653742 | 1.667558779 | 0.525710849 |
| Gm8246 | 0 | NA | NA | NA |
| Gm3187 | 0.044191549 | 0.037062677 | 3.724945049 | 0.009949859 |
| Gm3488 | 1.283947918 | 0.092599035 | 0.874451827 | 0.105893809 |
| Gm26893 | 0 | NA | NA | NA |
| Gm26746 | 0 | NA | NA | NA |
| Gm3532 | 0 | NA | NA | NA |
| Gm8266 | 0 | NA | NA | NA |
| Gm26731 | 0 | NA | NA | NA |
| Gm3383 | 0.828062179 | 0.706178945 | 1.163379658 | 0.607006441 |
| Gm3344 | 0 | NA | NA | NA |
| Gm8265 | 0 | NA | NA | NA |
| Gm9603 | 0.111006857 | 0.037730769 | 3.724898049 | 0.010129343 |
| Gm3373 | 0.817666292 | -0.443062666 | 1.099991343 | -0.402787411 |
| Gm3170 | 0 | NA | NA | NA |
| Gm26557 | 0 | NA | NA | NA |
| Gm3424 | 0.342449771 | -1.637870898 | 1.546079738 | -1.059370263 |
| Gm17158 | 0 | NA | NA | NA |
| Gm8237 | 12.34732519 | -0.585458524 | 0.380837534 | -1.53729208 |
| Gm17046 | 0.139037128 | -1.190058937 | 3.717743703 | -0.320102469 |
| Gm3242 | 0 | NA | NA | NA |
| Gm17047 | 0 | NA | NA | NA |
| Gm3248 | 20.30228851 | 0.120548559 | 0.228436932 | 0.527710461 |
| Gm17048 | 0 | NA | NA | NA |
| Gm3453 | 0.066050875 | -0.443818988 | 3.724945049 | -0.119147795 |
| Gm3261 | 0 | NA | NA | NA |
| Gm8206 | 2.632186321 | -0.131229856 | 0.587126302 | -0.223512139 |
| Gm3468 | 14.54750696 | 0.067416265 | 0.355265902 | 0.189762836 |
| Gm17045 | 0.385383671 | 0.466854375 | 1.611708883 | 0.289664207 |
| Gm6337 | 0 | NA | NA | NA |
| Gm3275 | 0 | NA | NA | NA |
| Gm3476 | 0.611787247 | 0.338094557 | 1.399484129 | 0.241585132 |

| | | | | |
|---|---|---|---|---|
| Gm26991 | 0 | NA | NA | NA |
| Gm3208 | 0 | NA | NA | NA |
| Gm21560 | 0 | NA | NA | NA |
| Gm21772 | 0 | NA | NA | NA |
| Gm3411 | 2.644118259 | 0.989879366 | 0.637374348 | 1.553058057 |
| Gm21103 | 0 | NA | NA | NA |
| Gm3618 | 0 | NA | NA | NA |
| Gm26630 | 0 | NA | NA | NA |
| Gm8340 | 0 | NA | NA | NA |
| Gm16440 | 0.365173462 | 0.882474762 | 1.450256051 | 0.608495831 |
| Gm3591 | 2.764354947 | -0.469663564 | 0.591848079 | -0.793554261 |
| Gm3594 | 0 | NA | NA | NA |
| Gm8356 | 0.045882455 | 0.517959912 | 3.724945049 | 0.139051692 |
| Gm3629 | 1.032577368 | 1.880692514 | 1.121478101 | 1.676976584 |
| Gm3460 | 0 | NA | NA | NA |
| Gm10251 | 0.095435502 | 0.037073798 | 3.724945049 | 0.009952844 |
| Gm26680 | 0 | NA | NA | NA |
| Gm3466 | 0 | NA | NA | NA |
| Gm8050 | 0.427185557 | 0.044797605 | 1.582811027 | 0.028302561 |
| Gm3636 | 17.74053186 | -0.217325993 | 0.250481617 | -0.867632505 |
| Gm8362 | 0 | NA | NA | NA |
| Gm3642 | 0.049553047 | -0.443818988 | 3.724945049 | -0.119147795 |
| Gm3667 | 13.76281871 | 0.441102741 | 0.303196488 | 1.454841192 |
| Gm6356 | 0.072025462 | 0.517959912 | 3.724945049 | 0.139051692 |
| Gm10406 | 10.70611474 | 0.249236209 | 0.324948344 | 0.76700255 |
| Gm3685 | 0.098502579 | -0.443821212 | 3.724945049 | -0.119148392 |
| Gm3696 | 16.41020923 | 0.306700323 | 0.257642865 | 1.190408756 |
| Gm6676 | 0.092269992 | -0.443823436 | 3.724945049 | -0.119148989 |
| Gm3512 | 4.130618934 | 0.104887066 | 0.493477909 | 0.212546628 |
| Gm16434 | 0 | NA | NA | NA |
| Gm3715 | 1.793529573 | 0.076969433 | 0.693525443 | 0.110982855 |
| Gm5456 | 0.437842326 | -0.408159744 | 1.387628495 | -0.294141945 |
| Gm3727 | 0.094294407 | -0.443821212 | 3.724945049 | -0.119148392 |
| Gm3739 | 19.15242329 | 0.063797401 | 0.289908239 | 0.22006067 |
| Gm5797 | 0.049553047 | -0.443818988 | 3.724945049 | -0.119147795 |
| Gm8374 | 0.038981395 | -0.443816764 | 3.724945049 | -0.119147198 |
| Gm10339 | 0 | NA | NA | NA |
| 4933406F09Rik | 0.083526043 | 0.037073798 | 3.724945049 | 0.009952844 |
| Gm8379 | 0.046345709 | -0.443818988 | 3.724945049 | -0.119147795 |
| Gm3755 | 0.852403115 | 0.66968155 | 1.076300284 | 0.622206981 |
| Gm3752 | 3.599181491 | 0.050222888 | 0.680850848 | 0.07376489 |
| Gm3760 | 0.150455055 | -1.336248778 | 2.838957719 | -0.470682874 |
| Gm3558 | 7.230339464 | 0.097791176 | 0.38218056 | 0.255876898 |
| Gm17121 | 0.092140247 | 0.037060453 | 3.724945049 | 0.009949262 |
| Gm10338 | 0 | NA | NA | NA |
| Gm10128 | 0.356838182 | 0.972002342 | 1.708103934 | 0.569053394 |
| Gm10044 | 5.427300065 | 0.068684558 | 0.487086946 | 0.141010879 |

| | | | | |
|---|---|---|---|---|
| Gm16741 | 0.543837322 | -0.899357645 | 1.411856952 | -0.637003376 |
| Gm26661 | 0.047948698 | 0.037064902 | 3.724945049 | 0.009950456 |
| Gm8396 | 0.090277459 | -0.443823436 | 3.724945049 | -0.119148989 |
| Flnb | 1267.997399 | -0.157721993 | 0.150881565 | -1.045336406 |
| Dnase1l3 | 17.4478056 | 0.502687279 | 0.293899658 | 1.710404433 |
| Abhd6 | 413.5173793 | 0.092943476 | 0.07971876 | 1.165892145 |
| Rpp14 | 225.7839547 | -0.196430585 | 0.082986339 | -2.367023145 |
| Pxk | 710.6699321 | -0.50285661 | 0.08671886 | -5.79869948 |
| Pdhb | 939.7685917 | 0.110373603 | 0.08695606 | 1.269303168 |
| Kctd6 | 337.2401318 | 0.214469776 | 0.144800648 | 1.481138233 |
| Acox2 | 3.533179683 | -0.096403107 | 0.589397705 | -0.163562068 |
| Fam107a | 4664.005083 | 0.537804987 | 0.182630847 | 2.944765338 |
| Oit1 | 0 | NA | NA | NA |
| 4930452B06Rik | 90.32014768 | 1.112545078 | 0.149898174 | 7.422005551 |
| 4930455B14Rik | 0 | NA | NA | NA |
| Gm9800 | 154.1573682 | -0.183198201 | 0.171823071 | -1.066202573 |
| Fhit | 18.766523 | 0.42163408 | 0.261300151 | 1.6136006 |
| Hmgb3-ps | 0 | NA | NA | NA |
| Mnd1-ps | 0.293028276 | 0.51794538 | 3.724945049 | 0.139047791 |
| Gm22053 | 0 | NA | NA | NA |
| Fhitos | 0 | NA | NA | NA |
| Slc25a5-ps | 0.541865536 | 0.97036735 | 1.417120594 | 0.684745782 |
| Rpl21-ps4 | 0.428504882 | -1.29856364 | 1.765153602 | -0.735666085 |
| Gm3839 | 0.806310386 | 1.543605189 | 1.167208179 | 1.32247633 |
| AC158396.1 | 0 | NA | NA | NA |
| Mir5124b | 0 | NA | NA | NA |
| Gm25466 | 0 | NA | NA | NA |
| Ptprg | 1480.140492 | 0.2911338 | 0.061081995 | 4.766278472 |
| Gm3848 | 0.118974955 | 0.51795324 | 3.724945049 | 0.139049901 |
| 3830406C13Rik | 205.9760077 | 0.091324438 | 0.110251815 | 0.828325941 |
| Gm24578 | 0.075043205 | 0.517957688 | 3.724945049 | 0.139051095 |
| Fezf2 | 140.4500694 | 0.525418601 | 0.131975781 | 3.981174404 |
| Cadps | 3883.453482 | 0.410665001 | 0.053170121 | 7.723604765 |
| n-R5s45 | 5.980340237 | 1.95947956 | 0.566894016 | 3.456518333 |
| Gm5457 | 6.294359381 | 0.70029673 | 0.461314487 | 1.518046257 |
| Gm5087 | 8.752821209 | 0.856288825 | 0.373387777 | 2.293296348 |
| Synpr | 975.6483675 | 0.592749442 | 0.167796164 | 3.532556584 |
| Gm15913 | 0 | NA | NA | NA |
| Sntn | 3.353049333 | -0.804969633 | 0.635506806 | -1.266657769 |
| Rpl19-ps4 | 1.071806319 | -0.245058022 | 1.125773495 | -0.217679687 |
| Gm27506 | 0 | NA | NA | NA |
| Gm281 | 0.82274574 | -1.848594232 | 1.113629346 | -1.659972629 |
| Gm25964 | 0 | NA | NA | NA |
| Gm11100 | 1.662603624 | -1.779785521 | 0.839963035 | -2.118885532 |
| Thoc7 | 350.5995921 | -0.152930805 | 0.097733508 | -1.564773512 |
| Atxn7 | 575.3815957 | 0.063127516 | 0.093259861 | 0.676899093 |
| Psmd6 | 699.5893116 | -0.209967089 | 0.087669303 | -2.394989829 |

| | | | | |
|---|---|---|---|---|
| Olfr720 | 0 | NA | NA | NA |
| Olfr31 | 0.115957212 | 0.517955464 | 3.724945049 | 0.139050498 |
| Il3ra | 52.25554781 | -1.111196326 | 0.291068194 | -3.817649431 |
| Olfr721-ps1 | 0 | NA | NA | NA |
| Slc4a7 | 745.5514613 | -0.042835031 | 0.088379091 | -0.484673816 |
| Nek10 | 32.25918065 | 1.638535277 | 0.232295964 | 7.053653677 |
| Lrrc3b | 129.9205039 | 0.335317356 | 0.103471294 | 3.240680018 |
| B230110C06Rik | 12.80781388 | 0.764968993 | 0.367085157 | 2.083900638 |
| Oxsm | 84.03786033 | -0.359557474 | 0.157132882 | -2.288238273 |
| Ngly1 | 371.6785756 | 0.118233407 | 0.070388178 | 1.679733867 |
| Top2b | 2165.3358 | 0.141472683 | 0.067580679 | 2.09338948 |
| Rarb | 169.0811627 | 0.498459177 | 0.117234269 | 4.251821417 |
| Thrb | 548.3553106 | 0.412140121 | 0.10002815 | 4.12024138 |
| Gm24140 | 0 | NA | NA | NA |
| Gm27851 | 0 | NA | NA | NA |
| Gm6773 | 0.143022002 | 0.03610674 | 3.547735707 | 0.010177404 |
| CU024892.1 | 0 | NA | NA | NA |
| Nr1d2 | 2201.828554 | 0.259651271 | 0.063124926 | 4.113292274 |
| Rpl15 | 1078.184859 | -0.258481379 | 0.120750315 | -2.140626961 |
| Nkiras1 | 524.1011176 | 0.333186004 | 0.083666 | 3.982334572 |
| Ube2e1 | 611.3267082 | -0.120832879 | 0.100040032 | -1.207845264 |
| Gm22350 | 0.766686841 | 0.765750035 | 1.278042337 | 0.599158583 |
| Gm6781 | 3.825407769 | 0.90085841 | 0.541184262 | 1.664605705 |
| Ube2e2 | 649.6422344 | 0.418500799 | 0.084313778 | 4.963611067 |
| Gm20678 | 11.63441953 | 0.090228251 | 0.461494636 | 0.195513109 |
| Gm20699 | 63.74099188 | 0.566644869 | 0.320586024 | 1.767528297 |
| Gm22499 | 0 | NA | NA | NA |
| Gm15916 | 0 | NA | NA | NA |
| Gm20677 | 0 | NA | NA | NA |
| Gm8582 | 0.356401535 | 1.139872628 | 1.719286223 | 0.662991777 |
| Gm8584 | 0.374155823 | 1.064104173 | 1.722565805 | 0.6177437 |
| Gm9637 | 0 | NA | NA | NA |
| Gm21738 | 1.310113234 | -1.005511578 | 0.855064201 | -1.175948632 |
| Gm7352 | 0 | NA | NA | NA |
| Gm2244 | 0.202043434 | 0.010039337 | 2.6073024 | 0.003850469 |
| Gm2237 | 15.97977484 | 0.29333862 | 0.274894823 | 1.067094011 |
| Gm5458 | 0.372874528 | 0.517975087 | 1.596311107 | 0.324482543 |
| 4930469B13Rik | 0 | NA | NA | NA |
| Gm17030 | 0.08169834 | -0.443821212 | 3.724945049 | -0.119148392 |
| Nid2 | 107.7830705 | -0.631633852 | 0.166881438 | -3.784925751 |
| 2700060E02Rik | 556.5044864 | 0.022193536 | 0.099271316 | 0.223564442 |
| Gng2 | 1773.216353 | 0.39599099 | 0.065246907 | 6.06911515 |
| Saysd1 | 59.98590933 | -0.570899831 | 0.157334838 | -3.628565922 |
| Kcnk5 | 24.84306935 | -0.467082176 | 0.25157229 | -1.856651925 |
| Gm24912 | 0 | NA | NA | NA |
| Kcnk16 | 0.038981395 | -0.443816764 | 3.724945049 | -0.119147198 |
| Nudt13 | 124.3202546 | -0.144479678 | 0.126059893 | -1.146119315 |

| | | | | |
|---|---|---|---|---|
| Ecd | 421.6024906 | -0.200254538 | 0.072719012 | -2.753812708 |
| Fam149b | 307.5778511 | 0.152322775 | 0.087551124 | 1.739815174 |
| Dnajc9 | 241.2584147 | -0.15365871 | 0.120073052 | -1.279710204 |
| Mrps16 | 150.6596876 | -0.260249684 | 0.106989795 | -2.432472031 |
| Cfap70 | 17.61224759 | 0.177778021 | 0.262064194 | 0.678375853 |
| 7330404K18Rik | 0 | NA | NA | NA |
| Anxa7 | 995.7442115 | -0.231589404 | 0.083872363 | -2.761212336 |
| Mss51 | 28.46984622 | -0.395488522 | 0.279680056 | -1.41407481 |
| Ppp3cb | 3014.123295 | 0.389618875 | 0.060511319 | 6.438776809 |
| 1810062O18Rik | 11.1715319 | 0.280990474 | 0.33562463 | 0.837216488 |
| Usp54 | 1368.558277 | 0.011658395 | 0.072546737 | 0.16070185 |
| 2810402E24Rik | 26.39841974 | 0.248104705 | 0.215551942 | 1.151020505 |
| Myoz1 | 2.494795275 | -0.317006437 | 0.800412408 | -0.396053876 |
| Synpo2l | 4.120754503 | 0.736722829 | 0.552960206 | 1.332325222 |
| Sec24c | 1144.691689 | -0.439292197 | 0.073189124 | -6.002151347 |
| 6230400D17Rik | 14.90222924 | 0.156003185 | 0.280910094 | 0.555349161 |
| Mir6946 | 0.27962944 | -0.895710052 | 2.49662091 | -0.358768946 |
| Fut11 | 188.4790295 | -0.160457789 | 0.098299427 | -1.632336972 |
| Gm25328 | 3.501233001 | -0.054498202 | 0.618175944 | -0.088159693 |
| Chchd1 | 206.0508832 | -0.375643422 | 0.112350554 | -3.343494163 |
| Zswim8 | 1959.999352 | -0.057368585 | 0.072907886 | -0.786863926 |
| Ndst2 | 264.5758868 | -0.394421815 | 0.110526929 | -3.568558526 |
| Camk2g | 2341.047828 | 0.595688808 | 0.056706623 | 10.50474842 |
| Plau | 140.9153563 | -1.474143216 | 0.285949584 | -5.155255675 |
| Vcl | 823.9348543 | -0.752150833 | 0.113571533 | -6.622705653 |
| Ap3m1 | 731.0242943 | -0.096141839 | 0.068816746 | -1.397070408 |
| Adk | 509.0668211 | 0.30357285 | 0.130749111 | 2.321796673 |
| Gm26372 | 4.248258162 | 0.29647261 | 0.572186641 | 0.518139691 |
| Gm25864 | 0.26743452 | -0.926119628 | 2.097871979 | -0.441456694 |
| Kat6b | 1504.25356 | 0.222191762 | 0.079813153 | 2.78389907 |
| Dupd1 | 0.592353126 | 0.753317917 | 1.230369805 | 0.61226951 |
| Dusp13 | 5.809763333 | -0.846151334 | 0.468339747 | -1.806704084 |
| Gm15935 | 0.68377221 | -1.384504267 | 1.342712825 | -1.031124631 |
| Samd8 | 1631.759024 | 0.011799289 | 0.075397085 | 0.156495299 |
| Gm23502 | 0.371001759 | 0.517953172 | 3.234181938 | 0.160149671 |
| Vdac2 | 1632.626953 | -0.029182943 | 0.068240611 | -0.427647747 |
| Comtd1 | 88.38321911 | -0.063759994 | 0.121875575 | -0.523156457 |
| A430057M04Rik | 12.38941375 | -0.187864429 | 0.343643319 | -0.546684363 |
| Gm37590 | 0 | NA | NA | NA |
| Zfp503 | 121.7214339 | 0.237077088 | 0.157904026 | 1.501399896 |
| C130012C08Rik | 2.040813782 | 0.830265222 | 0.765123186 | 1.085139278 |
| 1700112E06Rik | 23.04170766 | -0.110468793 | 0.299886773 | -0.368368342 |
| Gm27532 | 0 | NA | NA | NA |
| Gm5670 | 0 | NA | NA | NA |
| Gm10248 | 1.061926442 | 0.584589473 | 0.917589594 | 0.637092527 |
| Kcnma1 | 2520.351437 | 0.414329233 | 0.120826799 | 3.429117007 |
| A830039N20Rik | 615.447544 | 0.530490226 | 0.114997753 | 4.613048626 |

| | | | | |
|---|---|---|---|---|
| Gm29626 | 0 | NA | NA | NA |
| Gm6158 | 5.934383314 | -0.081220824 | 0.425864174 | -0.190720022 |
| mmu-mir-7210 | 0 | NA | NA | NA |
| Dlg5 | 927.9578943 | -0.135071606 | 0.083272736 | -1.62203877 |
| Gm17105 | 11.22543593 | 0.020962755 | 0.414324887 | 0.050594969 |
| E330034G19Rik | 0.520501943 | 1.032807123 | 1.472339014 | 0.701473718 |
| Polr3a | 517.7839581 | 0.035441235 | 0.067697488 | 0.523523637 |
| Rps24 | 2358.715317 | -0.009204003 | 0.106376478 | -0.086522913 |
| 4930542C16Rik | 0.048949532 | 0.037064902 | 3.724945049 | 0.009950456 |
| 4931403M11Rik | 0.541236733 | -1.165430049 | 1.618155925 | -0.720221106 |
| 4930428N03Rik | 0.347568383 | -1.397573561 | 1.72534598 | -0.810025107 |
| Gm10398 | 0 | NA | NA | NA |
| Gm26660 | 26.92887363 | 0.150021665 | 0.301212236 | 0.498059663 |
| Zmiz1os1 | 24.20908764 | -0.101944387 | 0.224811127 | -0.45346682 |
| Zmiz1 | 3086.44194 | -0.24326834 | 0.081582782 | -2.981858849 |
| Mir3075 | 1.765392608 | 0.006323026 | 0.888716607 | 0.007114784 |
| Gm26772 | 0.433939115 | -1.341118189 | 1.42954441 | -0.938143775 |
| Ppif | 256.0875963 | -0.029766582 | 0.081001159 | -0.367483414 |
| 1700054O19Rik | 0.226439421 | 0.556382932 | 2.149532657 | 0.258839022 |
| Zcchc24 | 1699.51765 | -0.199967927 | 0.127319712 | -1.570596762 |
| Gm27185 | 0 | NA | NA | NA |
| Gm23701 | 0 | NA | NA | NA |
| Anxa11 | 152.4094582 | -0.061447549 | 0.09745623 | -0.630514327 |
| Anxa11os | 1.923336751 | -1.62699223 | 0.734498079 | -2.215107537 |
| Plac9a | 0 | NA | NA | NA |
| Tmem254a | 0 | NA | NA | NA |
| Cphx1 | 0 | NA | NA | NA |
| Gm19013 | 0 | NA | NA | NA |
| Duxbl1 | 0.094294407 | -0.443821212 | 3.724945049 | -0.119148392 |
| Gm2260 | 1.263942296 | -2.139352123 | 1.363082737 | -1.569495428 |
| Gm9780 | 0.038981395 | -0.443816764 | 3.724945049 | -0.119147198 |
| Tmem254c | 0.63148455 | 1.243083454 | 1.392792347 | 0.892511692 |
| Gm19014 | 0 | NA | NA | NA |
| Cphx2 | 0 | NA | NA | NA |
| Cphx3 | 0 | NA | NA | NA |
| Duxbl2 | 0 | NA | NA | NA |
| Gm2274 | 0.075043205 | 0.517957688 | 3.724945049 | 0.139051095 |
| Plac9b | 0 | NA | NA | NA |
| Tmem254b | 0 | NA | NA | NA |
| Gm19015 | 0 | NA | NA | NA |
| Duxbl3 | 0 | NA | NA | NA |
| 4933413J09Rik | 0 | NA | NA | NA |
| Gm16245 | 0.158569248 | 0.51797124 | 2.811594837 | 0.184226843 |
| Slmapos2 | 0 | NA | NA | NA |
| Slmap | 1697.427193 | 0.180883811 | 0.046027059 | 3.929945009 |
| Gm24031 | 0 | NA | NA | NA |
| Gm2178 | 0 | NA | NA | NA |

| | | | | |
|---|---|---|---|---|
| Dennd6a | 497.1097104 | -0.117986865 | 0.083318252 | -1.416098668 |
| Arf4os | 0.854956099 | -0.404187994 | 1.083149627 | -0.373159889 |
| Arf4 | 1525.733304 | -0.317445505 | 0.129207096 | -2.456873612 |
| Pde12 | 370.8916435 | 0.01464 | 0.139076299 | 0.105265956 |
| Mir7672 | 0 | NA | NA | NA |
| 9930004E17Rik | 4.759985783 | 0.891838393 | 0.478095418 | 1.865398326 |
| Dnah12 | 39.20255926 | -0.270764132 | 0.231430352 | -1.169959469 |
| Asb14 | 0.828364824 | -0.690937725 | 1.286744499 | -0.53696575 |
| Appl1 | 1598.16168 | 0.186201448 | 0.050470014 | 3.689348054 |
| Hesx1 | 0 | NA | NA | NA |
| Il17rd | 75.9257241 | 0.143982974 | 0.18251853 | 0.788867708 |
| Arhgef3 | 850.6410935 | 0.347500309 | 0.061100246 | 5.687379891 |
| Gm23633 | 0 | NA | NA | NA |
| Fam208a | 1486.670546 | -0.258482961 | 0.120384703 | -2.147141235 |
| Ccdc66 | 229.1061877 | -0.272919879 | 0.127873928 | -2.134288697 |
| Gm22485 | 2.784173388 | 0.423941211 | 0.730930923 | 0.580001745 |
| Gm25697 | 0 | NA | NA | NA |
| Erc2 | 1776.688752 | 0.525362783 | 0.112258414 | 4.679941277 |
| Wnt5a | 214.1936815 | -0.122267685 | 0.092021255 | -1.328689613 |
| Gm2670 | 2.740285017 | -0.912030646 | 0.596235624 | -1.529648027 |
| Cacna2d3 | 929.5729355 | 0.662793276 | 0.116968957 | 5.666403218 |
| Gm23999 | 4.885903765 | 1.580151149 | 0.682633273 | 2.314787766 |
| Lrtm1 | 0.20890592 | 0.08591469 | 2.612155806 | 0.032890339 |
| Gm6043 | 0.823931708 | -0.729705813 | 1.212754645 | -0.601692862 |
| Selk | 760.4563827 | -0.133510146 | 0.108858914 | -1.226451208 |
| Actr8 | 526.6961103 | 0.1447647 | 0.069205606 | 2.091805973 |
| mmu-mir-6947 | 1.07422636 | 1.616170662 | 1.105953055 | 1.46133749 |
| Il17rb | 16.19919013 | -0.405266898 | 0.260940991 | -1.553097873 |
| Chdh | 62.42317444 | -0.371651168 | 0.245548766 | -1.51355339 |
| Cacna1d | 1000.382172 | 0.325480247 | 0.148844116 | 2.186718938 |
| Dcp1a | 375.2765008 | -0.229581101 | 0.069180634 | -3.318574686 |
| Tkt | 1261.009206 | -0.055808056 | 0.075611283 | -0.73809164 |
| Mir3076 | 0.281790215 | -0.045249818 | 2.059803956 | -0.021968022 |
| Prkcd | 783.4000836 | 0.136996385 | 0.099185687 | 1.381211229 |
| Rft1 | 140.5032926 | 0.269143441 | 0.114587388 | 2.348805091 |
| Sfmbt1 | 877.7713785 | 0.051907046 | 0.061780031 | 0.840191331 |
| 1700087M22Rik | 0.048949532 | 0.037064902 | 3.724945049 | 0.009950456 |
| Gm15667 | 10.02380282 | 0.670048897 | 0.371222351 | 1.804979943 |
| Tmem110 | 229.3938144 | 0.207707869 | 0.094794002 | 2.191149912 |
| Mustn1 | 28.82764176 | -1.347922135 | 0.312524923 | -4.313006848 |
| Itih4 | 5.953922204 | -0.350052177 | 0.492713607 | -0.710457702 |
| Itih3 | 692.5237228 | 0.492186511 | 0.165498506 | 2.973963473 |
| Itih1 | 0 | NA | NA | NA |
| Nek4 | 254.004222 | -0.31438582 | 0.093211406 | -3.372825638 |
| Spcs1 | 557.2353825 | -0.301485478 | 0.146778853 | -2.054011683 |
| Glt8d1 | 210.2471507 | -0.149147029 | 0.104757546 | -1.423735433 |
| Gnl3 | 342.5965878 | -0.076865929 | 0.167946729 | -0.457680418 |

| | | | | |
|---|---|---|---|---|
| Snord69 | 6.499084164 | 0.265113029 | 0.440687718 | 0.60158933 |
| Gm24916 | 0.901079044 | -1.415132803 | 1.136888891 | -1.244741517 |
| Snord19 | 6.156095358 | 0.973553616 | 0.500151003 | 1.946519369 |
| Pbrm1 | 1674.501212 | -0.038900602 | 0.045322819 | -0.858300596 |
| Gm5459 | 0.740733539 | -0.424502014 | 1.448166533 | -0.293130661 |
| Smim4 | 39.16029245 | -0.04593727 | 0.180270669 | -0.254823872 |
| Nt5dc2 | 56.55209549 | -0.234923125 | 0.176920098 | -1.327848712 |
| Stab1 | 251.2128195 | -0.643234249 | 0.140386013 | -4.581896965 |
| Nisch | 5669.263495 | 0.215230057 | 0.043338178 | 4.966292285 |
| Tnnc1 | 26.93316509 | 0.589674525 | 0.264156685 | 2.232290751 |
| Sema3g | 57.06689575 | -0.976878488 | 0.228930359 | -4.267142601 |
| Phf7 | 44.60374799 | 0.181045871 | 0.189122158 | 0.957295926 |
| Bap1 | 1295.484164 | 0.04713552 | 0.06649237 | 0.708886152 |
| Dnah1 | 99.51919862 | 0.07958099 | 0.128688154 | 0.618401825 |
| Capn7 | 711.6756629 | -0.035825147 | 0.068993992 | -0.519250241 |
| Sh3bp5 | 520.8229502 | 0.07503534 | 0.091536744 | 0.819729181 |
| Mettl6 | 226.22855 | -0.05242299 | 0.104875733 | -0.499858148 |
| Eaf1 | 382.8875853 | -0.62156602 | 0.12198776 | -5.095314633 |
| Colq | 11.36104049 | -0.440987963 | 0.398955075 | -1.105357449 |
| Gm24057 | 0.166923653 | 0.517964169 | 2.766511212 | 0.187226484 |
| Hacl1 | 56.72216686 | -0.078915966 | 0.234408594 | -0.336659867 |
| Btd | 112.7781148 | -0.369041541 | 0.140998821 | -2.61733778 |
| Ankrd28 | 1173.696225 | -0.155935715 | 0.072102659 | -2.162690211 |
| Gm22876 | 0.047948698 | 0.037064902 | 3.724945049 | 0.009950456 |
| D830044D21Rik | 0.364803694 | 1.897845802 | 1.690398179 | 1.122721158 |
| Gm26045 | 0 | NA | NA | NA |
| Galnt15 | 194.1661234 | -0.454583215 | 0.237896188 | -1.910846991 |
| Dph3 | 331.9002246 | -0.088018681 | 0.09755795 | -0.902219467 |
| Oxnad1 | 151.3315786 | -0.577463569 | 0.130018748 | -4.441386927 |
| Msmb | 0 | NA | NA | NA |
| Ncoa4 | 556.7801885 | -0.150323937 | 0.075998229 | -1.97799264 |
| Timm23 | 256.1091139 | -0.167871731 | 0.118846452 | -1.412509404 |
| Gm23946 | 1.072725949 | 0.671078988 | 1.025063922 | 0.654670381 |
| Parg | 736.7172274 | -0.021373073 | 0.054584137 | -0.391561984 |
| Gm7707 | 0.203659793 | -0.444657567 | 2.597136576 | -0.171210698 |
| Gm17210 | 0.136194719 | 0.037062677 | 3.724945049 | 0.009949859 |
| Ogdhl | 1425.208207 | 0.17470074 | 0.069502137 | 2.513602409 |
| 1700024G13Rik | 0.700875826 | 0.915136828 | 1.24733358 | 0.73367449 |
| Chat | 45.18468771 | -0.022014179 | 0.204689644 | -0.10754906 |
| Slc18a3 | 49.59830663 | -0.441218328 | 0.339972487 | -1.297805985 |
| Gm2990 | 1.893473903 | 0.651128798 | 0.880034836 | 0.739889799 |
| Ercc6 | 588.9859623 | -0.136317849 | 0.114515776 | -1.190384886 |
| Prrxl1 | 7.584775115 | -0.396857201 | 0.614602036 | -0.645714102 |
| 3425401B19Rik | 4.622495198 | 0.514906334 | 0.513511308 | 1.002716642 |
| Gm28651 | 0.300197857 | -1.227865471 | 1.753572081 | -0.700208154 |
| 1810011H11Rik | 3.566868347 | -0.557714406 | 0.597538186 | -0.933353582 |
| Fam170b | 0.045882455 | 0.517959912 | 3.724945049 | 0.139051692 |

| | | | | |
|---|---|---|---|---|
| Vstm4 | 53.0402276 | -0.374260683 | 0.258794513 | -1.446169313 |
| Gm7734 | 0.640869663 | 0.516444742 | 1.287505194 | 0.401120512 |
| Wdfy4 | 177.0804989 | -0.442102357 | 0.173084388 | -2.554259 |
| Lrrc18 | 16.544867 | -0.085463617 | 0.294949618 | -0.289756663 |
| Gm25498 | 0 | NA | NA | NA |
| Arhgap22 | 86.59583876 | -0.399887959 | 0.139970849 | -2.856937434 |
| Gm15512 | 6.43024678 | -0.084407903 | 0.537941146 | -0.156909179 |
| Mapk8 | 1298.322725 | 0.222518957 | 0.064291099 | 3.461116073 |
| Ptpn20 | 4.379735502 | -0.310706838 | 0.505277707 | -0.614922909 |
| Gm26228 | 0 | NA | NA | NA |
| Gdf10 | 323.3389736 | 0.050836359 | 0.354680204 | 0.143330128 |
| Gdf2 | 0.41755988 | -0.476479598 | 1.446528173 | -0.329395311 |
| Rbp3 | 3.927501353 | 0.259740406 | 0.686653525 | 0.378269967 |
| Zfp488 | 48.54553077 | -0.259539795 | 0.188581489 | -1.376273971 |
| Gm5460 | 0 | NA | NA | NA |
| A630023A22Rik | 0.092691418 | -0.908687078 | 3.724560878 | -0.2439716 |
| Antxrl | 0 | NA | NA | NA |
| Anxa8 | 7.103537313 | -0.414617158 | 0.483016277 | -0.858391689 |
| Gm24998 | 0 | NA | NA | NA |
| Npy4r | 1.230904645 | 0.195534654 | 1.035213196 | 0.188883463 |
| Rpl23a-ps3 | 207.5274121 | -0.15793341 | 0.138497154 | -1.140336856 |
| Gprin2 | 18.00709799 | 0.328285689 | 0.298052233 | 1.101436771 |
| Syt15 | 3.163297818 | -0.905452191 | 0.599574325 | -1.51015838 |
| Fam35a | 121.8552786 | 0.479606267 | 0.132405399 | 3.622256117 |
| Glud1 | 2362.144562 | 0.160871233 | 0.105075436 | 1.531007045 |
| Fam25c | 0.591489534 | 0.000649568 | 1.267818706 | 0.000512351 |
| Sncg | 85.14162449 | -0.130723608 | 0.328423204 | -0.39803402 |
| Mmrn2 | 26.86649225 | -1.176175635 | 0.281742659 | -4.174645188 |
| Bmpr1a | 1071.470205 | -0.01581133 | 0.094014583 | -0.168179551 |
| Gm17110 | 1.151183522 | 0.598824367 | 1.019643687 | 0.587287868 |
| 9230112D13Rik | 0.075043205 | 0.517957688 | 3.724945049 | 0.139051095 |
| Ldb3 | 91.38560832 | 0.463018234 | 0.157589327 | 2.938131942 |
| Opn4 | 1.339915753 | -1.894522017 | 1.011521671 | -1.872942589 |
| Wapal | 1670.7941 | -0.071374149 | 0.070551998 | -1.011653116 |
| A930038B10Rik | 0.504415982 | -0.449494883 | 1.473888318 | -0.304972146 |
| Grid1 | 726.7856293 | 0.310776405 | 0.06723245 | 4.622416767 |
| Mir346 | 1.214291556 | 0.878757992 | 1.102080786 | 0.797362592 |
| Gm27617 | 0.732444829 | 0.324754697 | 1.079033685 | 0.300968081 |
| 4930596D02Rik | 0 | NA | NA | NA |
| Gm7853 | 0 | NA | NA | NA |
| 4930474N05Rik | 0 | NA | NA | NA |
| Gm24394 | 0 | NA | NA | NA |
| Gm25204 | 0 | NA | NA | NA |
| n-R5s46 | 0.182014725 | 1.476593388 | 2.685235118 | 0.549893519 |
| Ccser2 | 2137.802388 | 0.095143876 | 0.038306524 | 2.483751243 |
| Rgr | 0.246474914 | 0.037077308 | 2.092340239 | 0.017720497 |
| Lrit1 | 0.184428051 | -0.705365233 | 2.6619757 | -0.264978089 |

| | | | | |
|---|---|---|---|---|
| Lrit2 | 0.338204618 | -2.185401137 | 3.687884436 | -0.592589376 |
| Cdhr1 | 304.1165299 | -0.934715422 | 0.588919185 | -1.587170948 |
| 2610528A11Rik | 2.326728918 | -1.896598548 | 0.748117722 | -2.535160566 |
| Ghitm | 4045.339356 | -0.11951108 | 0.090227675 | -1.324550151 |
| Gm22469 | 0 | NA | NA | NA |
| Gm23909 | 0 | NA | NA | NA |
| Nrg3 | 1077.56296 | 0.431333165 | 0.136164839 | 3.167727947 |
| Gm20641 | 0 | NA | NA | NA |
| Nrg3os | 0 | NA | NA | NA |
| Gm27332 | 2.333541239 | -0.329331859 | 0.696656793 | -0.472731857 |
| Gm20642 | 0.139856492 | -0.443820827 | 3.574411099 | -0.124166139 |
| 1700109I08Rik | 0 | NA | NA | NA |
| Eif1-ps1 | 3.471245013 | -0.323217263 | 0.583799457 | -0.553644336 |
| Gm25012 | 0 | NA | NA | NA |
| Gm22191 | 0.04393175 | 0.037062677 | 3.724945049 | 0.009949859 |
| Gm22726 | 0.094831987 | 0.517957688 | 3.724945049 | 0.139051095 |
| Gm24388 | 0 | NA | NA | NA |
| Gm5205 | 0.0885994 | 0.517955464 | 3.724945049 | 0.139050498 |
| Sh2d4b | 40.5303677 | 0.325677179 | 0.254326031 | 1.280549923 |
| Tspan14 | 417.092454 | -0.217267431 | 0.0896289 | -2.424077851 |
| Fam213a | 601.8361932 | 0.288750954 | 0.114688203 | 2.517704067 |
| Dydc2 | 4.886726711 | -0.09544145 | 0.477131542 | -0.200031734 |
| Dydc1 | 0.704807757 | 0.354548684 | 1.161130663 | 0.305347792 |
| Mat1a | 4.404352952 | -0.718487832 | 0.60072321 | -1.196038077 |
| Sftpa1 | 0.045882455 | 0.517959912 | 3.724945049 | 0.139051692 |
| Mbl1 | 0 | NA | NA | NA |
| Sftpd | 0 | NA | NA | NA |
| Gm7925 | 0 | NA | NA | NA |
| Gm22823 | 0.476740939 | -0.005182751 | 1.592163911 | -0.003255161 |
| Gm2832 | 0.567915064 | -0.083196767 | 1.231969332 | -0.067531525 |
| Gm9671 | 0 | NA | NA | NA |
| Gm5798 | 0 | NA | NA | NA |
| Gm2855 | 0 | NA | NA | NA |
| Gm7945 | 0.044544648 | 0.517957688 | 3.724945049 | 0.139051095 |
| Gm6482 | 0.047948698 | 0.037064902 | 3.724945049 | 0.009950456 |
| Gm7953 | 0 | NA | NA | NA |
| Gm7954 | 0 | NA | NA | NA |
| Gm2878 | 0 | NA | NA | NA |
| Gm3486 | 0 | NA | NA | NA |
| Gm6490 | 0 | NA | NA | NA |
| Gm17061 | 0 | NA | NA | NA |
| Gm7970 | 0 | NA | NA | NA |
| 1700049E17Rik2 | 0 | NA | NA | NA |
| Gm17159 | 0 | NA | NA | NA |
| Gm3072 | 0 | NA | NA | NA |
| Gm3077 | 0 | NA | NA | NA |
| Gm3676 | 0 | NA | NA | NA |

| | | | | |
|---|---|---|---|---|
| Gm8065 | 0 | NA | NA | NA |
| Gm8068 | 0 | NA | NA | NA |
| Gm7929 | 0 | NA | NA | NA |
| Gm17062 | 0 | NA | NA | NA |
| Gm7980 | 0 | NA | NA | NA |
| 1700049E17Rik1 | 0 | NA | NA | NA |
| 1700024B05Rik | 0 | NA | NA | NA |
| Gm9890 | 0 | NA | NA | NA |
| Gm6401 | 0 | NA | NA | NA |
| Gm3543 | 0 | NA | NA | NA |
| Gm17063 | 0 | NA | NA | NA |
| Gm2930 | 0 | NA | NA | NA |
| Gm7991 | 0 | NA | NA | NA |
| 1700091H14Rik | 0 | NA | NA | NA |
| Gm7951 | 0 | NA | NA | NA |
| Gm17064 | 0 | NA | NA | NA |
| Gm17027 | 0 | NA | NA | NA |
| Gm17166 | 0 | NA | NA | NA |
| Gm3573 | 0.045882455 | 0.517959912 | 3.724945049 | 0.139051692 |
| Gm17165 | 0 | NA | NA | NA |
| Gm17038 | 0 | NA | NA | NA |
| Gm17026 | 0 | NA | NA | NA |
| Gm25208 | 0 | NA | NA | NA |
| Gm25892 | 0 | NA | NA | NA |
| Gm9611 | 0 | NA | NA | NA |
| Gm7995 | 0 | NA | NA | NA |
| Gm17164 | 0 | NA | NA | NA |
| Gm2951 | 0 | NA | NA | NA |
| Gm2959 | 0 | NA | NA | NA |
| Gm8005 | 0 | NA | NA | NA |
| Gm8011 | 0 | NA | NA | NA |
| Gm2966 | 0 | NA | NA | NA |
| Gm8020 | 0 | NA | NA | NA |
| Gm8024 | 0 | NA | NA | NA |
| Gm8032 | 0.048949532 | 0.037064902 | 3.724945049 | 0.009950456 |
| Gm17204 | 0 | NA | NA | NA |
| Gm17124 | 0 | NA | NA | NA |
| Gm25228 | 0 | NA | NA | NA |
| Gm23620 | 0.048949532 | 0.037064902 | 3.724945049 | 0.009950456 |
| Gm3633 | 0 | NA | NA | NA |
| Gm10378 | 0 | NA | NA | NA |
| Gm17162 | 0 | NA | NA | NA |
| Gm5929 | 0 | NA | NA | NA |
| Gm3008 | 0 | NA | NA | NA |
| Gm17161 | 0 | NA | NA | NA |
| Gm3015 | 0 | NA | NA | NA |
| Gm3025 | 0 | NA | NA | NA |

| | | | | |
|---|---|---|---|---|
| Gm10377 | 0 | NA | NA | NA |
| Gm7152 | 0 | NA | NA | NA |
| Gm21977 | 0 | NA | NA | NA |
| Gm3102 | 0 | NA | NA | NA |
| Gm8082 | 0 | NA | NA | NA |
| Gm3111 | 0 | NA | NA | NA |
| Gm10376 | 0 | NA | NA | NA |
| Gm6536 | 0 | NA | NA | NA |
| Gm3123 | 0 | NA | NA | NA |
| Gm8094 | 0.086930093 | -0.443164241 | 3.724898049 | -0.118973522 |
| Gm8104 | 2.686914903 | 0.104634066 | 0.746678776 | 0.140132638 |
| Gm7236 | 14.77965937 | -0.357454693 | 0.262917826 | -1.359568116 |
| Gm8114 | 0 | NA | NA | NA |
| Gm3141 | 0.276783799 | -0.411518441 | 2.502828957 | -0.16442132 |
| Gm7233 | 2.888901366 | 1.347468706 | 0.707228846 | 1.905279619 |
| Gm7316 | 0 | NA | NA | NA |
| Gm8122 | 0 | NA | NA | NA |
| Gm8126 | 0.513218277 | 0.36094857 | 1.270908649 | 0.284008273 |
| Gm17153 | 0 | NA | NA | NA |
| Gm8127 | 0 | NA | NA | NA |
| Gm9732 | 2.036409272 | -0.664754434 | 0.719363628 | -0.924086802 |
| Gm17155 | 0 | NA | NA | NA |
| Gm17154 | 0 | NA | NA | NA |
| 1700001F09Rik | 0.08486385 | 0.037730769 | 3.724898049 | 0.010129343 |
| Gm9597 | 0.358995975 | -1.560660071 | 3.694581539 | -0.42241863 |
| Gm8138 | 0 | NA | NA | NA |
| Gm5799 | 0.049553047 | -0.443818988 | 3.724945049 | -0.119147795 |
| Gm3216 | 0 | NA | NA | NA |
| Gm17654 | 0 | NA | NA | NA |
| Gm3221 | 0 | NA | NA | NA |
| Gm10375 | 0 | NA | NA | NA |
| Gm8046 | 0.277407222 | -0.394620399 | 1.756603746 | -0.22464964 |
| Gm8161 | 0 | NA | NA | NA |
| Gm8165 | 0 | NA | NA | NA |
| Gm9672 | 0 | NA | NA | NA |
| Gm17208 | 0.132101749 | -0.93550548 | 3.724188147 | -0.251197158 |
| Gm7324 | 15.17033637 | -0.674152913 | 0.280259157 | -2.405462576 |
| Gm16506 | 0 | NA | NA | NA |
| Gm6526 | 0.404621698 | -1.198496057 | 1.645587319 | -0.728309002 |
| Gm9587 | 0 | NA | NA | NA |
| Gm8180 | 0 | NA | NA | NA |
| Ear1 | 0.104637558 | -0.443821212 | 3.724945049 | -0.119148392 |
| Ear-ps12 | 0 | NA | NA | NA |
| Ang-ps2 | 0 | NA | NA | NA |
| Ear-ps9 | 0 | NA | NA | NA |
| Ear-ps8 | 0.045882455 | 0.517959912 | 3.724945049 | 0.139051692 |
| Ang-ps3 | 0 | NA | NA | NA |

| | | | | |
|---|---|---|---|---|
| Ear10 | 0 | NA | NA | NA |
| Gm8113 | 3.368233017 | 1.045564683 | 0.615089006 | 1.699859163 |
| Gm9550 | 0 | NA | NA | NA |
| Ang5 | 0 | NA | NA | NA |
| Ear-ps7 | 0 | NA | NA | NA |
| Ear-ps10 | 0 | NA | NA | NA |
| Ang6 | 0 | NA | NA | NA |
| Gm8194 | 0 | NA | NA | NA |
| Gm3287 | 0 | NA | NA | NA |
| Ear-ps2 | 0 | NA | NA | NA |
| Ang-ps1 | 0.046345709 | -0.443818988 | 3.724945049 | -0.119147795 |
| Gm16545 | 0 | NA | NA | NA |
| Ear2 | 0.663448651 | -1.524581954 | 1.21574407 | -1.254031989 |
| Gm3327 | 0 | NA | NA | NA |
| Gm8207 | 0 | NA | NA | NA |
| 4930503E14Rik | 0.57934571 | -1.652134557 | 1.289045578 | -1.281672723 |
| Gm8212 | 0.068097359 | 0.037064902 | 3.724945049 | 0.009950456 |
| Gm8220 | 0.072025462 | 0.517959912 | 3.724945049 | 0.139051692 |
| Gm8224 | 0 | NA | NA | NA |
| Gm5930 | 0.049553047 | -0.443818988 | 3.724945049 | -0.119147795 |
| Gm8229 | 0 | NA | NA | NA |
| Gm3370 | 0 | NA | NA | NA |
| Gm3371 | 0 | NA | NA | NA |
| Gm8232 | 0.045882455 | 0.517959912 | 3.724945049 | 0.139051692 |
| Gm21991 | 0 | NA | NA | NA |
| Gm17184 | 0.465126201 | 0.957021326 | 1.662419609 | 0.575679762 |
| BC061237 | 0.120349074 | 1.027495679 | 3.723011807 | 0.275985071 |
| Gm17093 | 0 | NA | NA | NA |
| Gm17183 | 0 | NA | NA | NA |
| Gm5624 | 0.172257197 | -1.186514766 | 2.734354793 | -0.433928607 |
| Gm8247 | 0.04393175 | 0.037062677 | 3.724945049 | 0.009949859 |
| Gm16439 | 6.205147171 | 0.705199158 | 0.417483611 | 1.689166086 |
| Gm8256 | 0 | NA | NA | NA |
| Gm9593 | 0 | NA | NA | NA |
| Gm8257 | 0 | NA | NA | NA |
| Gm21754 | 0 | NA | NA | NA |
| Gm6541 | 0 | NA | NA | NA |
| Gm8267 | 0.060174537 | 0.517957688 | 3.724945049 | 0.139051095 |
| Gm17187 | 0 | NA | NA | NA |
| Gm17070 | 0 | NA | NA | NA |
| Ptgdr | 10.70749769 | -1.364955282 | 0.478153682 | -2.85463719 |
| Gm16976 | 1.330389483 | -1.222671453 | 1.012284439 | -1.207833892 |
| mmu-mir-327 | 0 | NA | NA | NA |
| 4930431P03Rik | 1.669097723 | 0.070006304 | 0.757740167 | 0.092388271 |
| Gm17173 | 1.687842655 | -1.188200763 | 0.776243532 | -1.530706168 |
| Ptger2 | 14.92504073 | -1.422479001 | 0.331902948 | -4.285828163 |
| Txndc16 | 1225.114205 | 0.309478237 | 0.068976649 | 4.486710265 |

| | | | | |
|---|---|---|---|---|
| Gm15765 | 0 | NA | NA | NA |
| Gpr137c | 408.4638035 | 0.123690379 | 0.084695271 | 1.460416586 |
| Ero1l | 547.1939839 | -0.159137195 | 0.055933165 | -2.845131256 |
| Psmc6 | 1042.749598 | -0.307221895 | 0.100323246 | -3.062320123 |
| Styx | 68.64544997 | 0.102035222 | 0.216420591 | 0.471467254 |
| Gnpnat1 | 150.1275212 | -0.288891272 | 0.152122311 | -1.899072328 |
| Gm25353 | 0 | NA | NA | NA |
| Fermt2 | 877.773119 | -0.351454805 | 0.074344888 | -4.727356733 |
| Gm22637 | 3.849995044 | 0.749831673 | 0.590610078 | 1.269588348 |
| Gm15601 | 0.688336999 | -2.000188563 | 1.205616272 | -1.659059031 |
| Ddhd1 | 1001.935963 | -0.044824736 | 0.081670859 | -0.548846146 |
| Mir5131 | 0 | NA | NA | NA |
| A530076I17Rik | 4.579416614 | 0.309419496 | 0.451122245 | 0.685888358 |
| Gm15516 | 0.119234754 | 0.51795324 | 3.724945049 | 0.139049901 |
| Gm7206 | 18.53655595 | -0.152958831 | 0.251395416 | -0.608439221 |
| Gm15218 | 0.089089296 | 0.957155977 | 3.723939561 | 0.257027796 |
| Gm8317 | 0.802853338 | 0.56067172 | 1.209152337 | 0.463689895 |
| Gm15220 | 0 | NA | NA | NA |
| Gm6580 | 0.146060149 | -0.762452964 | 3.524367124 | -0.216337554 |
| Gm37257 | 0 | NA | NA | NA |
| Gm1821 | 160.7341621 | -0.205008073 | 0.163450434 | -1.254252241 |
| Gm15217 | 0.045882455 | 0.517959912 | 3.724945049 | 0.139051692 |
| Bmp4 | 27.37682956 | -0.290192398 | 0.252991343 | -1.147044774 |
| Gm15222 | 0.402965827 | -0.034139755 | 1.589099455 | -0.021483712 |
| Gm15221 | 0.062426873 | 0.517959912 | 3.724945049 | 0.139051692 |
| E130120K24Rik | 0.066050875 | -0.443818988 | 3.724945049 | -0.119147795 |
| Rubie | 0.049553047 | -0.443818988 | 3.724945049 | -0.119147795 |
| Cdkn3 | 4.411638001 | -0.71784477 | 0.470554204 | -1.525530456 |
| Cnih1 | 494.7336958 | -0.208020921 | 0.077242692 | -2.693082224 |
| Gmfb | 1861.18559 | -0.168311333 | 0.091392161 | -1.841638607 |
| Gm10101 | 2.636333195 | 0.51140198 | 0.633242802 | 0.807592251 |
| Cgrrf1 | 236.9675109 | -0.007251751 | 0.094863733 | -0.076443871 |
| Samd4 | 517.3838205 | 0.139047967 | 0.078425263 | 1.772999702 |
| Mir378c | 0 | NA | NA | NA |
| Gm15562 | 5.089503423 | -0.468575844 | 0.472555235 | -0.991578993 |
| Gm10371 | 0.119587853 | 0.981063299 | 3.723618439 | 0.263470416 |
| Gch1 | 83.87125827 | -1.5014887 | 0.288324668 | -5.207631766 |
| Wdhd1 | 97.88412802 | -0.040855012 | 0.168499116 | -0.242464249 |
| Gm24378 | 1.312748754 | -0.21427605 | 0.973291033 | -0.220156195 |
| Socs4 | 478.1944072 | -0.356904757 | 0.07363933 | -4.846659459 |
| Mapk1ip1l | 616.9594476 | -0.239046704 | 0.072591502 | -3.293039776 |
| Lgals3 | 777.9121779 | -2.25229358 | 0.431449759 | -5.220291664 |
| Dlgap5 | 14.65831982 | -0.842902979 | 0.403312006 | -2.089952608 |
| Fbxo34 | 544.4943191 | 0.329090079 | 0.07236604 | 4.547576159 |
| Atg14 | 247.0243887 | -0.058737343 | 0.075626049 | -0.776681366 |
| Ktn1 | 1793.170985 | -0.094416358 | 0.079613212 | -1.185938303 |
| Gm37874 | 52.01025927 | -0.005448208 | 0.228699126 | -0.023822599 |

| | | | | |
|---|---|---|---|---|
| Gm6055 | 178.1871439 | -0.008515982 | 0.10848013 | -0.078502688 |
| Peli2 | 376.3843311 | 0.193461767 | 0.09417771 | 2.054220338 |
| Gm6498 | 16.31733287 | 0.701857914 | 0.355176065 | 1.97608449 |
| Gm23881 | 0 | NA | NA | NA |
| Gm3534 | 2.621659813 | -0.331704636 | 0.571266726 | -0.58064757 |
| Tmem260 | 200.6788583 | -0.192327426 | 0.17054061 | -1.127751487 |
| Gm23469 | 0 | NA | NA | NA |
| Otx2 | 54.10925475 | -0.000916032 | 0.144726505 | -0.006329402 |
| Otx2os1 | 2.535154825 | -0.621095318 | 0.752427886 | -0.825454943 |
| Gm25380 | 0 | NA | NA | NA |
| Gm22699 | 0 | NA | NA | NA |
| Gm23470 | 0.573140788 | 1.20810981 | 1.302918911 | 0.927233306 |
| Exoc5 | 1321.293313 | -0.065943257 | 0.081679309 | -0.807343469 |
| Ap5m1 | 326.8043512 | -0.211295491 | 0.079266245 | -2.665642764 |
| Naa30 | 880.0118064 | 0.104753822 | 0.104170557 | 1.005599141 |
| 1700011H14Rik | 0 | NA | NA | NA |
| Slc35f4 | 116.8448588 | 0.620917524 | 0.129028926 | 4.812235095 |
| Gm22116 | 5.008872367 | 1.498848022 | 0.61473809 | 2.438189604 |
| Gm22949 | 0 | NA | NA | NA |
| 3632451O06Rik | 470.0630739 | 0.027185713 | 0.107434812 | 0.253043801 |
| Gm16082 | 1.838486464 | 0.271171149 | 0.950691314 | 0.285235749 |
| Gm22989 | 0 | NA | NA | NA |
| Olfr722 | 0 | NA | NA | NA |
| Olfr723 | 0 | NA | NA | NA |
| Olfr724 | 0 | NA | NA | NA |
| Olfr725 | 0 | NA | NA | NA |
| Gm37194 | 0.41257556 | -0.400002837 | 1.567230859 | -0.255229046 |
| Olfr726 | 0 | NA | NA | NA |
| Olfr727 | 0 | NA | NA | NA |
| Olfr728 | 0.068097359 | 0.037064902 | 3.724945049 | 0.009950456 |
| Olfr729 | 0 | NA | NA | NA |
| Olfr730 | 0 | NA | NA | NA |
| Olfr731 | 0.046345709 | -0.443818988 | 3.724945049 | -0.119147795 |
| Olfr732 | 0 | NA | NA | NA |
| Olfr733 | 0.092881282 | 0.037060453 | 3.724945049 | 0.009949262 |
| Olfr734 | 0.109869311 | -0.444802825 | 3.724874622 | -0.119414174 |
| Olfr735 | 0.066050875 | -0.443818988 | 3.724945049 | -0.119147795 |
| Tlr11 | 0.527131164 | 0.253447983 | 1.586953314 | 0.159707271 |
| Olfr736 | 0 | NA | NA | NA |
| Olfr738 | 0 | NA | NA | NA |
| Olfr739 | 0 | NA | NA | NA |
| Olfr740 | 0 | NA | NA | NA |
| Olfr741 | 0.048949532 | 0.037064902 | 3.724945049 | 0.009950456 |
| Olfr742 | 0 | NA | NA | NA |
| Olfr743 | 0 | NA | NA | NA |
| Olfr744 | 0 | NA | NA | NA |
| Olfr745 | 0 | NA | NA | NA |

| | | | | |
|---|---|---|---|---|
| Olfr746 | 0 | NA | NA | NA |
| Olfr747 | 0 | NA | NA | NA |
| Olfr748 | 0 | NA | NA | NA |
| Gm24316 | 0 | NA | NA | NA |
| Olfr749 | 0.172571121 | -0.433506555 | 2.733273128 | -0.158603453 |
| Ttc5 | 245.1817073 | -0.099437981 | 0.093908152 | -1.058885508 |
| Gm10916 | 0.925454891 | 0.253741141 | 0.910592133 | 0.278655099 |
| Ccnb1ip1 | 5.389104856 | -0.619947264 | 0.526180052 | -1.178203662 |
| Rpph1 | 38912.15563 | 0.00221283 | 0.218596392 | 0.010122901 |
| Parp2 | 165.0646803 | -0.096956265 | 0.119029477 | -0.814556762 |
| Tep1 | 233.8406273 | -0.991549551 | 0.115431914 | -8.589908277 |
| Gm26782 | 38.7123841 | -0.101319254 | 0.237705943 | -0.426237783 |
| Klhl33 | 73.15949532 | 0.310263383 | 0.183348437 | 1.69220631 |
| A930018M24Rik | 44.75620921 | 0.137521604 | 0.18490279 | 0.743750833 |
| Osgep | 153.0925057 | 0.252942876 | 0.102006066 | 2.47968465 |
| Gm24689 | 0 | NA | NA | NA |
| Apex1 | 280.3220763 | -0.041938948 | 0.099838146 | -0.420069379 |
| Tmem55b | 636.4352874 | 0.01124927 | 0.058478441 | 0.192366107 |
| Pnp | 408.8476884 | -0.998283127 | 0.270573549 | -3.68950745 |
| Gm38316 | 0.038981395 | -0.443816764 | 3.724945049 | -0.119147198 |
| Pnp2 | 4.854385997 | -0.760668712 | 0.420525015 | -1.808854847 |
| Rnase10 | 0.957491927 | -2.634550431 | 1.114610592 | -2.36365099 |
| Rnase9 | 0.066050875 | -0.443818988 | 3.724945049 | -0.119147795 |
| Rnase11 | 0.038981395 | -0.443816764 | 3.724945049 | -0.119147198 |
| Rnase12 | 0.062426873 | 0.517959912 | 3.724945049 | 0.139051692 |
| Olfr750 | 0.092493346 | 0.517955464 | 3.724945049 | 0.139050498 |
| Rnase4 | 86.11690773 | -0.119266885 | 0.22247395 | -0.536093707 |
| Ang | 3.984262514 | -1.037909045 | 0.505896706 | -2.051622463 |
| Eddm3b | 0 | NA | NA | NA |
| Rnase6 | 15.68530113 | -1.140726945 | 0.381347394 | -2.991306518 |
| Rnase1 | 3.783653538 | 0.074906167 | 0.543791983 | 0.137747832 |
| Rnase2b | 0.112642008 | 0.517955464 | 3.724945049 | 0.139050498 |
| Ang2 | 0.049553047 | -0.443818988 | 3.724945049 | -0.119147795 |
| Ear14 | 0.049553047 | -0.443818988 | 3.724945049 | -0.119147795 |
| Rnase2a | 0.216167348 | -0.915520905 | 1.888434231 | -0.484804231 |
| Gm21718 | 0 | NA | NA | NA |
| Gm3719 | 0 | NA | NA | NA |
| Gm16289 | 0 | NA | NA | NA |
| Gm7247 | 0.038981395 | -0.443816764 | 3.724945049 | -0.119147198 |
| Vmn2r-ps111 | 0 | NA | NA | NA |
| Vmn2r88 | 0 | NA | NA | NA |
| Vmn2r89 | 0 | NA | NA | NA |
| Gm5622 | 0 | NA | NA | NA |
| Gm16290 | 0 | NA | NA | NA |
| Gm17175 | 0.075043205 | 0.517957688 | 3.724945049 | 0.139051095 |
| Gm17174 | 0 | NA | NA | NA |
| Gm17078 | 0 | NA | NA | NA |

| | | | | |
|---|---|---|---|---|
| Gm4181 | 0 | NA | NA | NA |
| Gm17186 | 0 | NA | NA | NA |
| Gm17079 | 0 | NA | NA | NA |
| Gm5800 | 0.136813032 | 0.037057272 | 3.644480825 | 0.010168052 |
| Ear-ps3 | 0 | NA | NA | NA |
| Ang4 | 0.095435502 | 0.037073798 | 3.724945049 | 0.009952844 |
| Ear-ps4 | 0 | NA | NA | NA |
| AY358078 | 0 | NA | NA | NA |
| Gm6047 | 0.182466651 | -1.346561657 | 2.672339661 | -0.503888662 |
| Gm27375 | 0 | NA | NA | NA |
| Ear-ps5 | 0 | NA | NA | NA |
| Ear6 | 0 | NA | NA | NA |
| Mettl17 | 160.3336998 | 0.153366938 | 0.159977141 | 0.958680326 |
| Slc39a2 | 25.14959081 | -0.855361001 | 0.293448352 | -2.914860465 |
| Ndrg2 | 5788.517411 | 0.391167573 | 0.150639525 | 2.596712743 |
| Tppp2 | 0.205880021 | -0.443822128 | 2.637684363 | -0.168262031 |
| Rnase13 | 0.16050186 | 0.998860224 | 2.800912447 | 0.356619581 |
| Arhgef40 | 469.4959225 | -0.505271616 | 0.086419345 | -5.846741995 |
| Gm16617 | 1.237774572 | 3.263278991 | 1.194090118 | 2.732858217 |
| Zfp219 | 473.6521586 | -0.31031796 | 0.090899814 | -3.413845936 |
| Tmem253 | 3.25900046 | -0.602237127 | 0.585060477 | -1.02935876 |
| Olfr221 | 0.038981395 | -0.443816764 | 3.724945049 | -0.119147198 |
| Gm22354 | 0.087261593 | 0.51795324 | 3.724945049 | 0.139049901 |
| Hnrnpc | 1799.363866 | -0.295493664 | 0.07376053 | -4.006121777 |
| Rpgrip1 | 13.58329738 | -0.167768433 | 0.304200467 | -0.551506164 |
| Supt16 | 1326.937056 | -0.059973037 | 0.080130495 | -0.748442119 |
| Gm26590 | 3.018699778 | -1.655408108 | 0.630108811 | -2.627178163 |
| Chd8 | 2275.929444 | 0.07978236 | 0.076571083 | 1.041938504 |
| Snord8 | 7.001247873 | -0.183903227 | 0.442600806 | -0.415505857 |
| RP24-223I19.6 | 3.473627573 | 0.669833556 | 0.583617567 | 1.147726857 |
| Rab2b | 331.1686235 | 0.083191251 | 0.074069756 | 1.123147368 |
| Gm23758 | 0 | NA | NA | NA |
| Tox4 | 931.9067073 | -0.285127645 | 0.06744711 | -4.227425688 |
| Mettl3 | 190.3858702 | -0.556477203 | 0.140856065 | -3.95067974 |
| Sall2 | 992.2939578 | 0.022549787 | 0.065826777 | 0.342562519 |
| Gm26328 | 0.151841013 | 0.517963177 | 2.85004976 | 0.181738292 |
| Olfr1513 | 0 | NA | NA | NA |
| RP24-461L18.3 | 26.78966518 | -0.172224373 | 0.207241441 | -0.831032499 |
| Olfr1512 | 0 | NA | NA | NA |
| Gm37504 | 0 | NA | NA | NA |
| Olfr1511 | 0 | NA | NA | NA |
| Olfr1510 | 0.062426873 | 0.517959912 | 3.724945049 | 0.139051692 |
| RP24-461L18.7 | 0.061920613 | -0.443816764 | 3.724945049 | -0.119147198 |
| Trav1 | 0.280850934 | 1.785115884 | 2.004912203 | 0.8903711 |
| Olfr1509 | 0.473637868 | 0.517973938 | 1.63631638 | 0.31654877 |
| Olfr1508 | 0.186324499 | 0.969221451 | 2.713146912 | 0.357231467 |
| RP23-117I19.7 | 0 | NA | NA | NA |

| | | | | |
|---|---|---|---|---|
| Olfr1507 | 1.123152174 | 0.017369496 | 0.996748534 | 0.017426157 |
| RP23-117I19.2 | 0.060174537 | 0.517957688 | 3.724945049 | 0.139051095 |
| RP23-117I19.3 | 0.172571121 | -0.433506555 | 2.733273128 | -0.158603453 |
| Trav2 | 2.527889093 | 1.763481997 | 0.667110504 | 2.643463093 |
| Trav3-1 | 0.371273269 | 0.940752053 | 1.684932402 | 0.558332222 |
| Trav4-1 | 0 | NA | NA | NA |
| Trav5-1 | 0.4844674 | -1.668776969 | 1.371435199 | -1.216810659 |
| Trav6-1 | 0.59120296 | -1.400934607 | 1.411526077 | -0.99249644 |
| RP24-447O5.2 | 0 | NA | NA | NA |
| Trav7-1 | 0.214323314 | 0.01351181 | 2.181940532 | 0.006192565 |
| Trav6-2 | 0.503770797 | -0.31271481 | 1.429931294 | -0.218692193 |
| Trav7d-2 | 0 | NA | NA | NA |
| Trav4d-2 | 0 | NA | NA | NA |
| Trav6d-3 | 0 | NA | NA | NA |
| Trav7d-3 | 0.231728546 | -1.654126091 | 3.701790695 | -0.446844846 |
| Trav6d-4 | 0.360686062 | 0.359070041 | 1.517557068 | 0.236610569 |
| Trav7d-4 | 0.038981395 | -0.443816764 | 3.724945049 | -0.119147198 |
| RP23-452K12.17 | 0 | NA | NA | NA |
| Trav8d-1 | 0.480868668 | -1.554699996 | 1.559744411 | -0.996765871 |
| Trav9d-1 | 0.075043205 | 0.517957688 | 3.724945049 | 0.139051095 |
| Trav6d-5 | 0.045882455 | 0.517959912 | 3.724945049 | 0.139051692 |
| Trav10d | 0.08486385 | 0.037730769 | 3.724898049 | 0.010129343 |
| Trav6d-6 | 0.038981395 | -0.443816764 | 3.724945049 | -0.119147198 |
| Trav11d | 0 | NA | NA | NA |
| Trav7d-5 | 0 | NA | NA | NA |
| Trav12d-1 | 0 | NA | NA | NA |
| Trav13d-1 | 0.061920613 | -0.443816764 | 3.724945049 | -0.119147198 |
| Trav14d-1 | 0 | NA | NA | NA |
| RP23-452K12.18 | 0.160559904 | -0.410534351 | 2.790729611 | -0.147106459 |
| Trav15d-1-dv6d-1 | 0 | NA | NA | NA |
| Trav3d-2 | 0 | NA | NA | NA |
| Trav9d-2 | 0 | NA | NA | NA |
| Trav4d-3 | 0 | NA | NA | NA |
| Rps19-ps2 | 0.546944188 | -0.763519971 | 1.438998496 | -0.530591222 |
| Trav5d-2 | 0 | NA | NA | NA |
| Trav12d-2 | 0.075043205 | 0.517957688 | 3.724945049 | 0.139051095 |
| Trav9d-3 | 0.216532645 | -1.775198458 | 3.14528588 | -0.564399716 |
| Trav5d-3 | 0 | NA | NA | NA |
| Trav12d-3 | 0 | NA | NA | NA |
| Trav13d-2 | 0 | NA | NA | NA |
| Trav14d-2 | 0 | NA | NA | NA |
| Trav15d-2-dv6d-2 | 0 | NA | NA | NA |
| RP24-548I6.12 | 0 | NA | NA | NA |
| Trav3d-3 | 0 | NA | NA | NA |
| Trav9d-4 | 0.11147366 | -0.955981391 | 3.722845977 | -0.25678779 |
| Trav4d-4 | 0 | NA | NA | NA |
| RP24-334B8.4 | 0 | NA | NA | NA |

| | | | | |
|---|---|---|---|---|
| Trav5d-4 | 0 | NA | NA | NA |
| Trav6d-7 | 0.046345709 | -0.443818988 | 3.724945049 | -0.119147795 |
| Trav7d-6 | 0 | NA | NA | NA |
| RP24-334B8.8 | 0.414202207 | 1.253984347 | 1.487696466 | 0.842903358 |
| Trav13d-3 | 0 | NA | NA | NA |
| Trav8d-2 | 0.305990011 | -2.247381407 | 1.924156795 | -1.167982471 |
| Trav16d-dv11 | 0.583486461 | -1.328407972 | 3.358700284 | -0.395512508 |
| Trav13d-4 | 0.066050875 | -0.443818988 | 3.724945049 | -0.119147795 |
| Trav14d-3-dv8 | 0.35026241 | -1.617619492 | 1.64490821 | -0.983410188 |
| Trav15d-3 | 0 | NA | NA | NA |
| Trav7n-4 | 0 | NA | NA | NA |
| Trav9n-1 | 0.141094079 | 0.037072457 | 3.563850528 | 0.01040236 |
| Trav6n-5 | 0 | NA | NA | NA |
| Trav10n | 0 | NA | NA | NA |
| Trav6n-6 | 0.230315653 | -1.159588941 | 2.106304418 | -0.550532454 |
| Trav11n | 0 | NA | NA | NA |
| Trav7n-5 | 0 | NA | NA | NA |
| Trav12n-1 | 0 | NA | NA | NA |
| Trav13n-1 | 0.049553047 | -0.443818988 | 3.724945049 | -0.119147795 |
| Trav14n-1 | 0 | NA | NA | NA |
| RP23-371K2.5 | 0.606297752 | -0.230253883 | 1.216323537 | -0.189303155 |
| Trav15n-1 | 0 | NA | NA | NA |
| Trav3n-2 | 0 | NA | NA | NA |
| Trav9n-2 | 0 | NA | NA | NA |
| Trav4n-3 | 0 | NA | NA | NA |
| Rps19-ps1 | 0 | NA | NA | NA |
| Trav5n-2 | 0 | NA | NA | NA |
| Trav12n-2 | 0 | NA | NA | NA |
| Trav9n-3 | 0 | NA | NA | NA |
| Trav5n-3 | 0 | NA | NA | NA |
| Trav12n-3 | 0 | NA | NA | NA |
| Trav13n-2 | 0 | NA | NA | NA |
| Trav14n-2 | 0 | NA | NA | NA |
| Trav15n-2 | 0 | NA | NA | NA |
| RP23-125A3.6 | 0 | NA | NA | NA |
| Trav3n-3 | 0 | NA | NA | NA |
| Trav9n-4 | 0.08486385 | 0.037730769 | 3.724898049 | 0.010129343 |
| Trav4n-4 | 0.12092566 | 0.998850474 | 3.724945049 | 0.268151734 |
| Trav5n-4 | 0 | NA | NA | NA |
| Trav6n-7 | 0 | NA | NA | NA |
| Trav7n-6 | 0 | NA | NA | NA |
| RP24-334B8.20 | 0 | NA | NA | NA |
| Trav13n-3 | 0 | NA | NA | NA |
| Trav8n-2 | 0.072025462 | 0.517959912 | 3.724945049 | 0.139051692 |
| Trav16n | 0.219234697 | -0.496916407 | 2.544014286 | -0.195327679 |
| Trav13n-4 | 0.049553047 | -0.443818988 | 3.724945049 | -0.119147795 |
| Trav14n-3 | 0.163190422 | -0.955151543 | 3.396300892 | -0.281232898 |

| | | | | |
|---|---|---|---|---|
| Trav15n-3 | 0 | NA | NA | NA |
| Trav7-2 | 0 | NA | NA | NA |
| Trav4-2 | 0.045882455 | 0.517959912 | 3.724945049 | 0.139051692 |
| Trav6-3 | 0.15956642 | 0.036118649 | 3.418731032 | 0.010564929 |
| Trav7-3 | 0.350945957 | -0.53357162 | 1.691134946 | -0.315510966 |
| Trav6-4 | 0.163577647 | -0.443208269 | 2.784038015 | -0.159196199 |
| Trav7-4 | 0.394584743 | -0.447667136 | 3.687206643 | -0.121410916 |
| RP23-230L15.11 | 0.04393175 | 0.037062677 | 3.724945049 | 0.009949859 |
| Trav8-1 | 0 | NA | NA | NA |
| Trav9-1 | 0.21728767 | -0.698126703 | 2.530005581 | -0.2759388 |
| Trav6-5 | 0.124347486 | 0.036092774 | 3.717245565 | 0.009709548 |
| Trav10 | 0.044544648 | 0.517957688 | 3.724945049 | 0.139051095 |
| Trav6-6 | 0.141781211 | -0.444711846 | 2.911624983 | -0.15273665 |
| Trav11 | 0 | NA | NA | NA |
| Trav7-5 | 0 | NA | NA | NA |
| Trav12-1 | 0.061920613 | -0.443816764 | 3.724945049 | -0.119147198 |
| Trav13-1 | 0.353543486 | 0.464439115 | 1.641205264 | 0.282986611 |
| Trav14-1 | 0 | NA | NA | NA |
| Trav15-1-dv6-1 | 0.099106094 | -0.93550548 | 3.724188147 | -0.251197158 |
| Trav3-2 | 0 | NA | NA | NA |
| Trav9-2 | 0.219744001 | -1.121555118 | 2.125982586 | -0.527546709 |
| Trav4-3 | 0 | NA | NA | NA |
| Trav5-2 | 0 | NA | NA | NA |
| Trav12-2 | 0 | NA | NA | NA |
| Trav12-3 | 0.264301748 | 0.051180706 | 2.447660897 | 0.020910048 |
| Trav13-2 | 0.124853745 | 1.05357957 | 3.721447096 | 0.283110183 |
| Trav14-2 | 0 | NA | NA | NA |
| Trav15-2-dv6-2 | 0 | NA | NA | NA |
| RP24-100P9.4 | 0 | NA | NA | NA |
| Trav3-3 | 2.07315737 | 0.433055575 | 0.679787517 | 0.637045495 |
| Trav9-4 | 0.183309527 | -0.444674696 | 2.685766745 | -0.165567132 |
| Trav4-4-dv10 | 0.315232318 | 1.48314361 | 3.69734468 | 0.401137502 |
| RP24-100P9.5 | 0 | NA | NA | NA |
| Trav5-4 | 0 | NA | NA | NA |
| Trav6-7-dv9 | 0.09176491 | 0.921395572 | 3.723061103 | 0.247483333 |
| Trav7-6 | 0 | NA | NA | NA |
| Trav13-3 | 0 | NA | NA | NA |
| Trav8-2 | 0 | NA | NA | NA |
| Trav16 | 0 | NA | NA | NA |
| Trav13-4-dv7 | 0.143485256 | -0.955563601 | 3.547405725 | -0.269369696 |
| Trav14-3 | 0 | NA | NA | NA |
| Trav15-3 | 0 | NA | NA | NA |
| Trav3-4 | 0.305831651 | 1.662524113 | 2.017130992 | 0.824202354 |
| Trav12-4 | 0.062426873 | 0.517959912 | 3.724945049 | 0.139051692 |
| B230359F08Rik | 0.108266322 | -0.955981391 | 3.722845977 | -0.25678779 |
| Trav17 | 0.511617548 | 0.2476695 | 1.294434958 | 0.191334063 |
| Trav18 | 0 | NA | NA | NA |

| | | | | |
|---|---|---|---|---|
| Trav19 | 0 | NA | NA | NA |
| Trav20 | 0 | NA | NA | NA |
| Trav21-dv12 | 0.038981395 | -0.443816764 | 3.724945049 | -0.119147198 |
| Trdv1 | 0.294928389 | -1.485728356 | 3.697061147 | -0.401867401 |
| RP23-339J4.3 | 0.166286699 | 0.037075006 | 2.769850463 | 0.013385201 |
| Trav22 | 0 | NA | NA | NA |
| Trdv2-1 | 0 | NA | NA | NA |
| Trdv2-2 | 0.066050875 | -0.443818988 | 3.724945049 | -0.119147795 |
| Trav23 | 0.0878635 | 0.037058229 | 3.724945049 | 0.009948665 |
| Trdv3 | 0 | NA | NA | NA |
| RP23-6A14.5 | 0.391491935 | 0.465841247 | 1.404520315 | 0.331672844 |
| Trdv4 | 0.800063227 | -0.265133412 | 1.005893967 | -0.263579881 |
| Trdd1 | 0 | NA | NA | NA |
| Gm23704 | 0 | NA | NA | NA |
| Trdd2 | 0.215047464 | 0.45069739 | 2.542337718 | 0.177276759 |
| Trdj1 | 0.138110619 | 0.482857281 | 3.622723928 | 0.133285696 |
| RP23-6A14.10 | 1.151756246 | -0.856284937 | 0.977857144 | -0.87567488 |
| Trdj2 | 0.506381433 | -1.552008345 | 1.674612937 | -0.92678631 |
| Trdc | 6.909450248 | -1.078810564 | 0.422246204 | -2.554932537 |
| Trdv5 | 0.261680116 | 0.037076478 | 2.115320361 | 0.017527595 |
| Traj61 | 0.355005915 | 1.014255792 | 1.565405648 | 0.647918827 |
| Traj60 | 0 | NA | NA | NA |
| Traj59 | 0.384053089 | -0.834991828 | 1.98402139 | -0.420858279 |
| Traj58 | 0.060174537 | 0.517957688 | 3.724945049 | 0.139051095 |
| Traj57 | 0.066050875 | -0.443818988 | 3.724945049 | -0.119147795 |
| Traj56 | 0.049553047 | -0.443818988 | 3.724945049 | -0.119147795 |
| Traj55 | 0.038981395 | -0.443816764 | 3.724945049 | -0.119147198 |
| Traj54 | 0 | NA | NA | NA |
| Traj53 | 0.235191775 | -0.019972378 | 2.103668607 | -0.00949407 |
| Traj52 | 0.049553047 | -0.443818988 | 3.724945049 | -0.119147795 |
| Traj51 | 0.143140564 | 0.517957789 | 3.546633296 | 0.146042104 |
| Traj50 | 0.066050875 | -0.443818988 | 3.724945049 | -0.119147795 |
| Traj49 | 0.094097695 | 0.037071574 | 3.724945049 | 0.009952247 |
| Traj48 | 0.268211629 | 0.505735922 | 2.090580679 | 0.241911698 |
| Traj47 | 0 | NA | NA | NA |
| Traj46 | 0.173853943 | -0.499202846 | 2.738479247 | -0.182291995 |
| Traj45 | 0.089089296 | 0.957155977 | 3.723939561 | 0.257027796 |
| Traj44 | 0.378197461 | -0.722764524 | 1.784616934 | -0.404997011 |
| Traj43 | 0.372057435 | -1.057609112 | 1.480643031 | -0.714290406 |
| Traj42 | 0.270744093 | -0.511621865 | 2.407044487 | -0.212551894 |
| Traj41 | 0.062426873 | 0.517959912 | 3.724945049 | 0.139051692 |
| Traj40 | 0.373373093 | -0.153886851 | 1.524662528 | -0.100931746 |
| Traj39 | 0.520484338 | -2.21618237 | 1.547202515 | -1.432380278 |
| Traj38 | 0.304511964 | -2.196338631 | 1.948639819 | -1.127113697 |
| Traj37 | 0.321009791 | -2.236945537 | 1.669021082 | -1.340273985 |
| Traj36 | 1.007510561 | -2.562238831 | 1.241357751 | -2.064061571 |
| Traj35 | 0.11147366 | -0.955981391 | 3.722845977 | -0.25678779 |

| | | | | |
|---|---|---|---|---|
| Traj34 | 0.174360202 | 0.503964903 | 2.724220683 | 0.184994155 |
| Traj33 | 0.478372902 | -0.65903567 | 3.462983831 | -0.190308619 |
| Traj32 | 0.292144397 | -2.081180284 | 3.690602698 | -0.563913391 |
| Traj31 | 0.534577455 | -0.087914341 | 1.288647275 | -0.068222191 |
| Traj30 | 0.493327417 | -0.04058939 | 1.300854715 | -0.031202093 |
| Traj29 | 0.56458922 | -2.481555568 | 1.305521185 | -1.900816008 |
| Traj28 | 0.229902237 | 0.476673 | 2.51110688 | 0.189825851 |
| Traj27 | 0.668611641 | -0.001898834 | 1.145003087 | -0.001658366 |
| Traj26 | 0.066050875 | -0.443818988 | 3.724945049 | -0.119147795 |
| Traj25 | 0.632730142 | -1.27052139 | 1.148974285 | -1.10578749 |
| Traj24 | 0.739637387 | 0.705478793 | 1.256722694 | 0.561363932 |
| Traj23 | 0.722755445 | -0.519887053 | 1.213481195 | -0.428426131 |
| Traj22 | 0.569236379 | -0.293965935 | 1.189713654 | -0.247089654 |
| Traj21 | 0.962101734 | 0.932871778 | 0.962243682 | 0.969475608 |
| Traj20 | 0.661355215 | 0.254924603 | 1.097442604 | 0.23228969 |
| Traj19 | 0.223817141 | 1.521216541 | 2.152523449 | 0.70671311 |
| Traj18 | 0.206905613 | -0.848785521 | 2.172787516 | -0.390643592 |
| Traj17 | 0.414345469 | -1.149038292 | 1.647518141 | -0.697435897 |
| Traj16 | 0.11193333 | 0.037073798 | 3.724945049 | 0.009952844 |
| Traj15 | 0.43445422 | -0.453243413 | 1.671454287 | -0.271167101 |
| Traj14 | 0.377056126 | -0.15011369 | 3.28217214 | -0.045736081 |
| Traj13 | 0.208613207 | -1.518073047 | 2.589972439 | -0.586134827 |
| Traj12 | 0.486308198 | -0.272903288 | 1.446379673 | -0.188680257 |
| Traj11 | 0.728352697 | -0.047149621 | 1.064516287 | -0.044292062 |
| Traj9 | 0.154585317 | -1.377635992 | 2.821196527 | -0.488316209 |
| Traj8 | 0.203604809 | -0.497684777 | 2.604693872 | -0.191072272 |
| Traj7 | 0.161486377 | -0.444693834 | 2.795640213 | -0.159066904 |
| Traj6 | 0.490533913 | -1.096720046 | 1.407035125 | -0.779454632 |
| Traj5 | 0.517714159 | -1.386659189 | 1.372747632 | -1.010134096 |
| Traj4 | 0.260182639 | -1.248638299 | 2.054475287 | -0.607765061 |
| Traj3 | 0 | NA | NA | NA |
| Traj2 | 0.364308502 | -1.177404889 | 1.683107773 | -0.699542185 |
| Traj1 | 0.178447459 | -1.389338828 | 3.327416506 | -0.417542807 |
| Trac | 42.85427748 | -0.282197887 | 0.282851887 | -0.997687835 |
| Dad1 | 331.339338 | -0.184074143 | 0.096813276 | -1.901331621 |
| Abhd4 | 923.6765976 | -0.292310468 | 0.136705069 | -2.13825625 |
| Olfr49 | 0.04393175 | 0.037062677 | 3.724945049 | 0.009949859 |
| RP24-378G4.3 | 0.161488833 | -0.59752174 | 3.431296725 | -0.174138755 |
| RP24-378G4.4 | 0 | NA | NA | NA |
| RP24-378G4.5 | 0 | NA | NA | NA |
| RP24-378G4.6 | 0.093484797 | -0.443823436 | 3.724945049 | -0.119148989 |
| Oxa1l | 446.5440451 | 0.059763788 | 0.062290574 | 0.959435512 |
| Slc7a7 | 63.95572232 | -0.858158745 | 0.169262958 | -5.069973709 |
| Mrpl52 | 179.7423914 | -0.45913742 | 0.115827094 | -3.963989796 |
| Mmp14 | 516.3861542 | -0.836550124 | 0.187646116 | -4.458126508 |
| Gm24424 | 0 | NA | NA | NA |
| Lrp10 | 683.6561126 | -0.711681027 | 0.136982587 | -5.195412357 |

| | | | | |
|---|---|---|---|---|
| Rem2 | 153.9580932 | -1.027602629 | 0.336031716 | -3.058052496 |
| Prmt5 | 476.4345395 | -0.354696055 | 0.061458079 | -5.771349537 |
| Haus4 | 37.03329486 | 0.019563073 | 0.196515877 | 0.099549578 |
| Ajuba | 48.90169317 | -1.131689489 | 0.259128185 | -4.367296007 |
| Gm24705 | 0 | NA | NA | NA |
| 4931414P19Rik | 61.32594599 | -0.06201941 | 0.140481027 | -0.441478904 |
| Psmb5 | 352.4206726 | -0.176780433 | 0.100073339 | -1.766508796 |
| Mir686 | 0 | NA | NA | NA |
| Psmb11 | 6.261784242 | 1.633386207 | 0.495019881 | 3.299637593 |
| Cdh24 | 130.6564516 | 0.488492551 | 0.129501705 | 3.772093581 |
| Gm20726 | 0.104106287 | 0.51795324 | 3.724945049 | 0.139049901 |
| Gm17606 | 1.502658278 | -1.91355462 | 1.107419341 | -1.727940401 |
| Acin1 | 1506.43145 | -0.154589452 | 0.09819898 | -1.574247028 |
| mmu-mir-6948 | 0.536873645 | 0.003842565 | 1.371486879 | 0.002801751 |
| 4930579G18Rik | 0.898085247 | -1.958907057 | 1.149997779 | -1.703400731 |
| 1700123O20Rik | 137.487668 | -0.439315425 | 0.133538223 | -3.289810329 |
| Cebpe | 1.473260831 | -2.056205196 | 0.941508926 | -2.183946577 |
| Slc7a8 | 825.5824292 | -0.036897642 | 0.085033888 | -0.433916911 |
| Gm8865 | 0 | NA | NA | NA |
| Homez | 148.0548322 | -9.23E-05 | 0.157010367 | -0.000588139 |
| Ppp1r3e | 55.21987148 | 0.184969091 | 0.183749109 | 1.006639392 |
| Gm37271 | 5.557962796 | 0.061391013 | 0.506425152 | 0.121224257 |
| Bcl2l2 | 1139.670049 | 0.256214412 | 0.06549061 | 3.912231259 |
| Gm20521 | 0.375216023 | 2.079291887 | 3.695373159 | 0.562674403 |
| Pabpn1 | 264.758843 | 0.148053729 | 0.216866751 | 0.682694455 |
| Slc22a17 | 4291.308412 | 0.50622133 | 0.05521312 | 9.168497049 |
| Efs | 132.9613014 | -0.278908094 | 0.182590913 | -1.527502594 |
| Il25 | 0.855745269 | -1.418471287 | 1.075718057 | -1.318627383 |
| Cmtm5 | 177.6437715 | 0.002710226 | 0.192516519 | 0.014077888 |
| Myh6 | 15.5941189 | 0.443792576 | 0.355106794 | 1.249743977 |
| Mir208a | 0 | NA | NA | NA |
| Mhrt | 3.992004508 | 2.070193203 | 0.702954326 | 2.944989631 |
| Myh7 | 58.29767794 | 0.265718505 | 0.274037644 | 0.969642349 |
| Gm29015 | 0.163519603 | 0.966857846 | 2.774974014 | 0.348420504 |
| Mir208b | 0 | NA | NA | NA |
| Gm17428 | 0.038981395 | -0.443816764 | 3.724945049 | -0.119147198 |
| Ngdn | 176.4773903 | -0.401998208 | 0.134408867 | -2.990860773 |
| Gm10364 | 2.051660529 | -0.377983817 | 0.739775614 | -0.510943872 |
| Gm20687 | 0.094831987 | 0.517957688 | 3.724945049 | 0.139051095 |
| Thtpa | 328.5777116 | 0.17281427 | 0.084439328 | 2.046608787 |
| Zfhx2 | 1084.53136 | 0.045664172 | 0.173335456 | 0.263443921 |
| Zfhx2os | 47.4886808 | 0.503751475 | 0.391280096 | 1.287444671 |
| Gm37034 | 35.49877453 | 0.473599649 | 0.365042003 | 1.297383987 |
| Gm38129 | 19.96999586 | 0.123571922 | 0.389633254 | 0.31714932 |
| Ap1g2 | 74.95724327 | 0.343188597 | 0.210922273 | 1.627085618 |
| Jph4 | 3324.850563 | 1.239423508 | 0.129253505 | 9.589090125 |
| Gm23596 | 0 | NA | NA | NA |

| | | | | |
|---|---|---|---|---|
| Dhrs2 | 1.791652276 | -1.466527607 | 1.027267342 | -1.427600729 |
| Gm8894 | 12.48756017 | -0.439341582 | 0.325292077 | -1.350606462 |
| Dhrs4 | 75.51960051 | -0.213423645 | 0.158968519 | -1.342552889 |
| Lrrc16b | 392.3729988 | 0.523829488 | 0.139224232 | 3.762487887 |
| Cpne6 | 868.2829338 | 0.839475482 | 0.12820049 | 6.548145664 |
| Nrl | 2.663415179 | -0.155501017 | 0.701957138 | -0.221524946 |
| Pck2 | 235.7861946 | -0.038394561 | 0.10249563 | -0.374597054 |
| A730061H03Rik | 1.918506749 | 0.989220165 | 0.882911102 | 1.120407438 |
| Dcaf11 | 703.1883374 | 0.046291037 | 0.053070293 | 0.872258934 |
| Gm15932 | 0.28189265 | 1.602181017 | 3.706919401 | 0.43221361 |
| Fitm1 | 0.170186936 | -1.44442222 | 3.399009296 | -0.424953889 |
| Psme1 | 1237.910375 | -0.258288881 | 0.15293118 | -1.688922307 |
| Emc9 | 130.4834388 | -0.021129706 | 0.14280487 | -0.14796208 |
| Psme2 | 385.2943989 | -0.425567965 | 0.155536708 | -2.736125578 |
| Rnf31 | 653.5334504 | -0.123458681 | 0.107961699 | -1.143541479 |
| Irf9 | 958.7618358 | -0.425327557 | 0.131520789 | -3.233918835 |
| Rec8 | 6.735313399 | 0.289034064 | 0.408619192 | 0.707343341 |
| Ipo4 | 859.5313815 | 0.038383302 | 0.047956943 | 0.800370067 |
| Tm9sf1 | 264.973998 | -0.399295536 | 0.086041469 | -4.640733577 |
| Tssk4 | 3.653146442 | 0.233884473 | 0.567847539 | 0.411878994 |
| Mdp1 | 234.6429327 | 0.148260113 | 0.09765226 | 1.518245586 |
| Nedd8 | 570.9432889 | -0.114500768 | 0.110457137 | -1.036608146 |
| Gmpr2 | 130.1104825 | -0.104320007 | 0.135463 | -0.770099637 |
| Tinf2 | 129.962745 | -0.135325132 | 0.182203733 | -0.742713283 |
| Gm25887 | 0.119587853 | 0.981063299 | 3.723618439 | 0.263470416 |
| Tgm1 | 450.7877214 | -1.959873079 | 0.401940119 | -4.876032486 |
| Rabggta | 331.6413799 | -0.129384827 | 0.066727423 | -1.939005319 |
| Dhrs1 | 1107.416416 | -1.08376752 | 0.130431242 | -8.309109846 |
| Nop9 | 182.2591248 | -0.726424697 | 0.142623302 | -5.093310036 |
| Cideb | 12.60069031 | 0.328208037 | 0.376564774 | 0.871584544 |
| Ltb4r2 | 1.834185785 | -1.365753321 | 0.80228932 | -1.702320208 |
| Ltb4r1 | 13.33054419 | -1.971469085 | 0.502723411 | -3.921578035 |
| Adcy4 | 30.55724621 | -0.79247719 | 0.300971278 | -2.633065833 |
| Ripk3 | 29.79384862 | -1.043531833 | 0.322319109 | -3.237573587 |
| Nfatc4 | 25.04774085 | -0.688828262 | 0.231829839 | -2.971266619 |
| Nynrin | 300.1840239 | -0.054562961 | 0.107353329 | -0.508255884 |
| Gm26751 | 4.016269843 | 1.894019202 | 0.670907287 | 2.823071439 |
| Cbln3 | 2048.884425 | 0.139163377 | 0.317432542 | 0.438402994 |
| Khnyn | 336.7112361 | -0.169624882 | 0.091147527 | -1.860992702 |
| Sdr39u1 | 401.4242335 | 0.108308769 | 0.09564754 | 1.132373803 |
| Cma1 | 1.421829526 | -3.153101369 | 1.091614865 | -2.888474197 |
| Cma2 | 0.212113984 | 1.775136953 | 3.153803221 | 0.562855964 |
| Gm5801 | 19.00002645 | -0.327129555 | 0.250597567 | -1.305397971 |
| Mcpt1 | 0.342615221 | -1.622833662 | 1.4864665 | -1.091739142 |
| Mcpt9 | 0 | NA | NA | NA |
| Mcpt2 | 0.74687904 | -1.428951802 | 3.553668042 | -0.402106158 |
| Mcpt4 | 0.384964691 | -0.896981534 | 1.455094269 | -0.616442215 |

| | | | | |
|---|---|---|---|---|
| Mcpt8 | 0 | NA | NA | NA |
| Mir6949 | 0 | NA | NA | NA |
| Ctsg | 0.191334258 | -0.81959484 | 2.66041249 | -0.308070588 |
| Gzme | 0.045882455 | 0.517959912 | 3.724945049 | 0.139051692 |
| Gzmd | 0.066050875 | -0.443818988 | 3.724945049 | -0.119147795 |
| Gzmg | 0 | NA | NA | NA |
| Gzmn | 0 | NA | NA | NA |
| Gzmf | 1.629358998 | -1.964563142 | 1.087000741 | -1.807324566 |
| Gzmc | 1.857191415 | -0.963719669 | 0.842368632 | -1.144059302 |
| Gm24118 | 0 | NA | NA | NA |
| Gm4819 | 0 | NA | NA | NA |
| Gzmb | 1148.613576 | -0.965642912 | 0.573365925 | -1.684165155 |
| Atp12a | 1.447939701 | -0.406668432 | 0.904469493 | -0.449620949 |
| Rnf17 | 3.730754265 | -0.462083706 | 0.632694554 | -0.730342475 |
| Gm24392 | 0.066050875 | -0.443818988 | 3.724945049 | -0.119147795 |
| Cenpj | 130.4607887 | -0.903287405 | 0.200272446 | -4.510292976 |
| Gm22431 | 0 | NA | NA | NA |
| Parp4 | 406.4978698 | -0.318303688 | 0.106298761 | -2.994425182 |
| Gm16573 | 0 | NA | NA | NA |
| Mphosph8 | 456.601835 | -0.082062124 | 0.123306059 | -0.665515743 |
| Gm16973 | 101.7464204 | -0.295090426 | 0.148125691 | -1.992162355 |
| Pspc1 | 679.2798492 | 0.148245468 | 0.07516232 | 1.972337578 |
| C030013D06Rik | 71.86212202 | 0.822197006 | 0.305609394 | 2.690352532 |
| Gm4479 | 0.082913145 | -0.443821212 | 3.724945049 | -0.119148392 |
| Zmym5 | 534.3723691 | 0.096051607 | 0.080872159 | 1.18769684 |
| 2410022M11Rik | 46.59279127 | 0.144219951 | 0.164588112 | 0.876247675 |
| Gm22498 | 0 | NA | NA | NA |
| Gm27635 | 0 | NA | NA | NA |
| Gm22218 | 0 | NA | NA | NA |
| Zmym2 | 1925.381415 | 0.181620695 | 0.073185884 | 2.481635602 |
| B020004C17Rik | 0.385592413 | -1.343904708 | 3.693583715 | -0.363848449 |
| Gja3 | 0.839270164 | -3.247717601 | 1.365445572 | -2.378503888 |
| Gjb2 | 207.1465136 | -0.96161783 | 0.186765726 | -5.148791743 |
| Gm4491 | 3.32060927 | 1.069297018 | 0.712071733 | 1.501670364 |
| Gjb6 | 476.1042231 | 0.746419805 | 0.166011959 | 4.496180937 |
| Cryl1 | 92.72897768 | 0.556219001 | 0.156962241 | 3.543648453 |
| Gm23593 | 0.852011583 | -1.469399124 | 1.235683827 | -1.189138428 |
| Ift88 | 288.14491 | 0.2283011 | 0.101603957 | 2.246970559 |
| Gm15836 | 0.068097359 | 0.037064902 | 3.724945049 | 0.009950456 |
| Ift88os | 3.791653849 | 2.368461216 | 0.688908194 | 3.437992515 |
| Il17d | 56.83289221 | 0.39804771 | 0.181792364 | 2.18957332 |
| N6amt2 | 87.43391111 | -0.331316784 | 0.155672991 | -2.128286876 |
| 1700039M10Rik | 1.307453931 | 1.896041741 | 0.933876455 | 2.030291833 |
| Xpo4 | 372.4125732 | 0.145204414 | 0.110059982 | 1.319320714 |
| Lats2 | 654.4618862 | -0.333861417 | 0.113548197 | -2.940261717 |
| Gm26440 | 0.104366086 | 0.51795324 | 3.724945049 | 0.139049901 |
| Gm20430 | 0.779952531 | 0.427745899 | 1.185708152 | 0.360751419 |

| | | | | |
|---|---|---|---|---|
| Sap18 | 492.5086436 | 0.065006038 | 0.094550163 | 0.68752963 |
| Mir3077 | 0 | NA | NA | NA |
| Ska3 | 10.9912364 | -0.373879918 | 0.467229721 | -0.800205768 |
| Mrpl57 | 188.5806294 | -0.054937169 | 0.123032867 | -0.446524337 |
| Gm22738 | 2.721618137 | 0.582227393 | 0.675172482 | 0.862338748 |
| Zdhhc20 | 1037.770143 | 0.154242162 | 0.099257431 | 1.553960854 |
| Micu2 | 298.7097166 | -0.393975223 | 0.112313509 | -3.507816872 |
| 3110083C13Rik | 6.409078304 | -0.252498542 | 0.562984309 | -0.448500142 |
| Fgf9 | 261.7121968 | 0.027109036 | 0.086173462 | 0.314586823 |
| Gm17109 | 0 | NA | NA | NA |
| Gm25614 | 0 | NA | NA | NA |
| Gm9022 | 0.171181394 | 0.539411076 | 2.747344444 | 0.196339078 |
| Rpl13-ps3 | 2.686924039 | 0.156074917 | 0.630889829 | 0.247388546 |
| 1700129C05Rik | 0.097139547 | -0.443816764 | 3.724945049 | -0.119147198 |
| Gm5142 | 0 | NA | NA | NA |
| Rcbtb1 | 723.0799953 | 0.117087154 | 0.084111163 | 1.392052495 |
| Gm6904 | 9.706043522 | -1.867905371 | 0.440297453 | -4.24237151 |
| Phf11a | 116.1058508 | -1.279451493 | 0.37909625 | -3.375004345 |
| Phf11b | 380.8323055 | -1.040041193 | 0.337285603 | -3.08356237 |
| Phf11d | 768.3922217 | -1.080823343 | 0.368722211 | -2.931267256 |
| Phf11c | 63.97794148 | -0.623461763 | 0.21930924 | -2.84284312 |
| Setdb2 | 303.1628553 | -0.040790634 | 0.199513687 | -0.204450306 |
| Gm16301 | 4.198859835 | -0.142741631 | 0.620790148 | -0.2299354 |
| Cab39l | 377.7726523 | -0.120866711 | 0.095382768 | -1.267175554 |
| Cdadc1 | 398.4046235 | 0.195446216 | 0.077078967 | 2.535662126 |
| Shisa2 | 57.22810011 | -0.683528343 | 0.269704366 | -2.534361427 |
| Atp8a2 | 620.7074487 | 0.127689334 | 0.095593936 | 1.335747214 |
| Gm9013 | 6.16443421 | 0.553923035 | 0.524555322 | 1.055985922 |
| 4930563I02Rik | 0.208097268 | -0.451972809 | 2.198688536 | -0.205564727 |
| Gm6913 | 0 | NA | NA | NA |
| Gm16261 | 0 | NA | NA | NA |
| Nupl1 | 647.3223968 | -0.116365821 | 0.071085553 | -1.636982703 |
| Mir719 | 0.1633926 | -0.44382031 | 2.785222646 | -0.159348234 |
| Mtmr6 | 1302.815769 | -0.061899059 | 0.0528256 | -1.171762536 |
| Amer2 | 525.7164864 | 0.200570654 | 0.108077903 | 1.855797057 |
| Gm29266 | 1.126110934 | 2.414158772 | 1.159910686 | 2.081331607 |
| Spata13 | 889.6645646 | -0.04676689 | 0.129777115 | -0.360363152 |
| C1qtnf9 | 1.516716675 | -0.150228794 | 0.824457305 | -0.182215371 |
| Mipepos | 0.63216027 | -0.316168382 | 1.266623399 | -0.249615144 |
| Mipep | 261.2781114 | 0.074535233 | 0.085810127 | 0.868606484 |
| Tnfrsf19 | 737.4688333 | -0.473645278 | 0.126869703 | -3.733320617 |
| Sacs | 3375.54935 | 0.060330373 | 0.069793652 | 0.864410608 |
| Sgcg | 2.524458954 | -0.513803149 | 0.724967886 | -0.708725392 |
| 1700109G14Rik | 1.219573449 | 1.186474162 | 0.992724773 | 1.195169291 |
| Arl11 | 45.06214385 | -1.241463492 | 0.245356066 | -5.059844296 |
| Ebpl | 74.2502746 | -0.149370797 | 0.156990713 | -0.951462629 |
| Gm25716 | 0.646183563 | -0.69884564 | 1.255747044 | -0.556517846 |

| | | | | |
|---|---|---|---|---|
| Kpna3 | 1362.917462 | 0.140238342 | 0.072035937 | 1.946783054 |
| Spryd7 | 646.9547196 | 0.065664077 | 0.109727441 | 0.598428953 |
| Trim13 | 93.97118903 | 0.047540936 | 0.146090561 | 0.325420999 |
| Dleu2 | 509.193176 | -0.29964096 | 0.138591239 | -2.162048342 |
| Kcnrg | 4.493413694 | 0.55302132 | 0.494213511 | 1.118992717 |
| Gm27010 | 39.15760298 | -0.190538693 | 0.26779111 | -0.711519859 |
| Gm27845 | 0.122991902 | 0.517955464 | 3.724945049 | 0.139050498 |
| Gm26916 | 0.226559374 | 1.806164991 | 2.511711466 | 0.719097323 |
| Mir16-1 | 0.351804329 | -0.392230773 | 1.472391129 | -0.266390339 |
| Mir15a | 2.378988063 | -1.108260094 | 0.793618099 | -1.396465246 |
| Gm27747 | 0.12092566 | 0.998850474 | 3.724945049 | 0.268151734 |
| Gm27972 | 0 | NA | NA | NA |
| Gm27017 | 1.269792884 | -1.138306639 | 0.875674363 | -1.299920024 |
| Gm27928 | 0.047948698 | 0.037064902 | 3.724945049 | 0.009950456 |
| Gm37420 | 92.02312766 | 0.133449764 | 0.232282721 | 0.574514385 |
| Gm37472 | 129.5225664 | 0.295700174 | 0.24238208 | 1.219975394 |
| Gm37820 | 1.452044762 | 0.702445583 | 0.920536311 | 0.763082971 |
| Gm26969 | 0 | NA | NA | NA |
| Dleu7 | 46.2767513 | 0.18959751 | 0.183932842 | 1.030797479 |
| Rnaseh2b | 84.37813906 | 0.235357333 | 0.13279144 | 1.772383316 |
| Gucy1b2 | 4.464211597 | -0.113309296 | 0.46001566 | -0.246316171 |
| Gm4131 | 0.940148893 | -2.587625734 | 1.066645931 | -2.425946286 |
| Gm24625 | 0 | NA | NA | NA |
| Fam124a | 214.0573691 | -0.112662502 | 0.098768919 | -1.140667563 |
| Gm23816 | 0 | NA | NA | NA |
| Serpine3 | 0.736171263 | 0.199692051 | 1.178617576 | 0.169429046 |
| Gm25162 | 0 | NA | NA | NA |
| Ints6 | 590.4517194 | -0.37844314 | 0.119732552 | -3.160737283 |
| 4931440J10Rik | 4.963742587 | 0.760033257 | 0.574415217 | 1.323142623 |
| Gm26536 | 1.065495214 | 0.107654226 | 0.930806858 | 0.115656889 |
| Wdfy2 | 169.1506073 | -0.260245828 | 0.101488943 | -2.564277659 |
| Mir8098 | 0 | NA | NA | NA |
| Mir6541 | 1.551969969 | 0.148631362 | 0.809814837 | 0.183537465 |
| n-R5s47 | 0.9747039 | 0.860069799 | 1.159624377 | 0.741679647 |
| Defb48 | 0 | NA | NA | NA |
| Defb47 | 0.450831908 | -0.363888639 | 1.346732778 | -0.270201071 |
| Defb43 | 0 | NA | NA | NA |
| Defb30 | 0.101408268 | 0.037730769 | 3.724898049 | 0.010129343 |
| Defb42 | 4.17522043 | 1.225373472 | 0.759525636 | 1.613340504 |
| Gm23629 | 0 | NA | NA | NA |
| Ctsb | 9370.229512 | -0.78236627 | 0.145443592 | -5.379173171 |
| Fdft1 | 819.3075922 | -0.106833476 | 0.117404299 | -0.909962211 |
| Neil2 | 16.96188054 | -0.227931894 | 0.313046936 | -0.728107729 |
| Gata4 | 0.136623169 | -0.870437938 | 3.632616033 | -0.23961738 |
| Blk | 0.439667587 | 0.468561285 | 1.36888619 | 0.342293821 |
| Gm21430 | 0.110595523 | 0.037071574 | 3.724945049 | 0.009952247 |
| Fam167a | 87.58736384 | -0.873829558 | 0.195181971 | -4.476999346 |

| | | | | |
|---|---|---|---|---|
| Tdh | 0.432707378 | -0.476616457 | 1.562356237 | -0.305062601 |
| Gm17232 | 0.288435958 | 0.899959046 | 2.023116196 | 0.444838041 |
| Mtmr9 | 857.9831177 | -0.052795783 | 0.052916737 | -0.997714253 |
| Gm15918 | 7.496844332 | -0.372544434 | 0.400219164 | -0.930851064 |
| Xkr6 | 411.7970283 | 0.071230671 | 0.160485028 | 0.44384621 |
| Mir598 | 1.860426698 | 0.642478805 | 0.873135668 | 0.735829297 |
| Pinx1 | 73.25514319 | 0.157866693 | 0.142759783 | 1.10582049 |
| Sox7 | 14.05727425 | -0.154325736 | 0.308416812 | -0.500380426 |
| 4930578I06Rik | 0.15100991 | 0.51796601 | 3.527597456 | 0.146832516 |
| Rp1l1 | 0.46004984 | 0.969908134 | 1.495206875 | 0.64867822 |
| Prss55 | 1.808854001 | 0.363148418 | 0.80539643 | 0.450894 |
| Prss51 | 0 | NA | NA | NA |
| 4930471C04Rik | 6.910637616 | 0.7528959 | 0.570734242 | 1.319170719 |
| Prss52 | 0.240888117 | -0.969127062 | 2.529674579 | -0.383103451 |
| Gm17116 | 16.40399705 | 0.248078502 | 0.403664335 | 0.614566313 |
| Msra | 295.1015591 | 0.246692094 | 0.09076876 | 2.71780834 |
| Mir124a-1hg | 1670.155338 | 0.347877335 | 0.36040861 | 0.965230366 |
| Mir124a-1 | 0 | NA | NA | NA |
| Mir3078 | 0 | NA | NA | NA |
| Kif13b | 733.5243985 | -0.068580217 | 0.107536258 | -0.637740411 |
| Gm26225 | 1.137833813 | 0.859484741 | 1.038598362 | 0.827542939 |
| Hmbox1 | 1181.12912 | 0.10831535 | 0.05463914 | 1.982376575 |
| Gm37183 | 36.08914756 | 0.218250756 | 0.33120506 | 0.658959606 |
| Gm20111 | 0 | NA | NA | NA |
| Ints9 | 171.8987803 | -0.414280042 | 0.104973034 | -3.946537759 |
| Extl3 | 2015.132037 | -0.132700834 | 0.063166772 | -2.100801272 |
| Fzd3 | 1027.303391 | 0.432315326 | 0.076831725 | 5.626781477 |
| Gm27263 | 1.700332359 | -0.062062082 | 0.8304459 | -0.074733444 |
| Fbxo16 | 53.77017717 | 0.030176495 | 0.152575868 | 0.197780264 |
| Zfp395 | 203.6050118 | 0.144114116 | 0.124854024 | 1.154260883 |
| Pnoc | 87.06028916 | -0.487423956 | 0.145767373 | -3.343848115 |
| Elp3 | 772.4740043 | -0.084593518 | 0.061744525 | -1.370056985 |
| Nuggc | 2.07464996 | -3.3567996 | 1.015174662 | -3.306622719 |
| Scara5 | 27.21228639 | -0.485205322 | 0.344583661 | -1.408091494 |
| Pbk | 12.44226385 | -1.393450788 | 0.579238622 | -2.405659319 |
| Esco2 | 8.165872688 | -0.730715735 | 0.485773455 | -1.504231504 |
| Ccdc25 | 465.8537132 | -0.137289949 | 0.127080783 | -1.080336031 |
| Scara3 | 275.1443948 | 0.230117456 | 0.099092259 | 2.322254605 |
| Clu | 7658.013756 | -0.016401383 | 0.108716467 | -0.150863838 |
| Gulo | 1.585533927 | -0.541993063 | 1.110415951 | -0.488099133 |
| Adam2 | 0.443182709 | -0.228055228 | 1.703673066 | -0.133860911 |
| Gm10233 | 0.231561134 | -0.451724624 | 2.131538859 | -0.21192418 |
| Ephx2 | 103.3385784 | 0.50446086 | 0.1450107 | 3.478783709 |
| Chrna2 | 9.44470098 | 0.102511132 | 0.362684547 | 0.282645436 |
| Ptk2b | 3860.082922 | 0.647038677 | 0.15102385 | 4.284347639 |
| 1700001G11Rik | 4.137181521 | -1.317348928 | 0.582346361 | -2.26213988 |
| Trim35 | 1878.620081 | -0.090088665 | 0.069126821 | -1.303237498 |

| | | | | |
|---|---|---|---|---|
| Stmn4 | 1213.698151 | 0.330278286 | 0.098918324 | 3.338898935 |
| Gm23899 | 0 | NA | NA | NA |
| Adra1a | 340.4045469 | -0.35170465 | 0.116093051 | -3.029506481 |
| Gm10032 | 0.180493943 | -0.925573715 | 2.699005356 | -0.342931411 |
| Dpysl2 | 11326.80766 | 0.172379048 | 0.054521241 | 3.161686043 |
| Gm5464 | 7.318891932 | -0.374113322 | 0.412504508 | -0.906931477 |
| Pnma2 | 428.5622982 | 0.35783167 | 0.12232368 | 2.925285362 |
| 4930578I07Rik | 0 | NA | NA | NA |
| Gm24258 | 0 | NA | NA | NA |
| Bnip3l | 1147.218713 | 0.154973004 | 0.098683501 | 1.5704044 |
| Ppp2r2a | 1306.363493 | -0.154143519 | 0.072110628 | -2.137597781 |
| Gm27647 | 0 | NA | NA | NA |
| Gm20675 | 0.146692594 | -0.955495625 | 3.521109961 | -0.271362052 |
| Ebf2 | 68.02274029 | 0.054854907 | 0.18086406 | 0.303293571 |
| Gm23178 | 0 | NA | NA | NA |
| Gm6878 | 0.454723033 | 1.080394056 | 1.776334695 | 0.608215366 |
| 1700120O09Rik | 0 | NA | NA | NA |
| 4930438E09Rik | 0 | NA | NA | NA |
| Gm10860 | 0 | NA | NA | NA |
| Gm24981 | 0 | NA | NA | NA |
| Gm20667 | 1.359754244 | -0.121690109 | 0.953692088 | -0.12759895 |
| Cdca2 | 13.24071265 | -0.153568854 | 0.375163897 | -0.40933804 |
| Kctd9 | 293.1358824 | -0.061696389 | 0.139249977 | -0.443062114 |
| Gnrh1 | 6.140945712 | 0.341952254 | 0.509541329 | 0.671098171 |
| Dock5 | 320.7362463 | -0.600304796 | 0.119424237 | -5.026657976 |
| Mir6539 | 0.673941775 | 1.845606997 | 1.348761379 | 1.368371771 |
| Nefm | 2120.125399 | -0.021606907 | 0.220863468 | -0.09782925 |
| Nefl | 4244.358884 | 0.263216179 | 0.134380215 | 1.958742061 |
| Gm22319 | 0 | NA | NA | NA |
| Adam7 | 0.28902587 | -0.162958674 | 2.072842497 | -0.078616042 |
| Adamdec1 | 0.646331757 | -1.73984251 | 1.323818311 | -1.314260798 |
| Adam28 | 1.311804504 | -0.536739833 | 1.184419149 | -0.453167135 |
| Stc1 | 48.95481958 | -0.402022083 | 0.20253562 | -1.98494508 |
| Rps2-ps5 | 0.934890173 | -1.529406958 | 1.184730275 | -1.29093262 |
| 1700092C10Rik | 0 | NA | NA | NA |
| Nkx2-6 | 0 | NA | NA | NA |
| Nkx3-1 | 12.50312542 | 0.633678056 | 0.313097791 | 2.0238982 |
| Gm37762 | 0.256447508 | 0.010978678 | 2.469852793 | 0.004445074 |
| Slc25a37 | 364.4713757 | -0.356309932 | 0.120359984 | -2.96036872 |
| Gm27222 | 0 | NA | NA | NA |
| Gm20236 | 0.061920613 | -0.443816764 | 3.724945049 | -0.119147198 |
| Gm31748 | 0 | NA | NA | NA |
| Gm27223 | 0 | NA | NA | NA |
| Synb | 0 | NA | NA | NA |
| Gm16677 | 0 | NA | NA | NA |
| Entpd4 | 0.136880458 | -0.425048655 | 3.63688887 | -0.116871499 |
| Gm27221 | 0 | NA | NA | NA |

| | | | | |
|---|---|---|---|---|
| Gm28049 | 0.042716945 | 0.037062677 | 3.724945049 | 0.009949859 |
| Gm21451 | 0 | NA | NA | NA |
| Gm38378 | 0 | NA | NA | NA |
| Gm37094 | 0 | NA | NA | NA |
| Gm37847 | 0.111006857 | 0.037730769 | 3.724898049 | 0.010129343 |
| Gm27175 | 0.061920613 | -0.443816764 | 3.724945049 | -0.119147198 |
| Gm27179 | 0 | NA | NA | NA |
| Gm27174 | 0 | NA | NA | NA |
| Gm21685 | 17.74312811 | -0.197848714 | 0.32658989 | -0.60580171 |
| Gm27176 | 0 | NA | NA | NA |
| Gm27177 | 0.075043205 | 0.517957688 | 3.724945049 | 0.139051095 |
| Loxl2 | 21.89650722 | -1.106310902 | 0.309499867 | -3.574511724 |
| Gm37513 | 0 | NA | NA | NA |
| Mir6950 | 0 | NA | NA | NA |
| R3hcc1 | 176.6082625 | 0.078897335 | 0.098764041 | 0.79884677 |
| Chmp7 | 739.9914465 | 0.00639583 | 0.059554246 | 0.10739503 |
| 4930480K23Rik | 35.48654686 | 0.387411859 | 0.19436085 | 1.993260777 |
| Tnfrsf10b | 17.05474737 | -1.472135825 | 0.303150984 | -4.85611429 |
| Rhobtb2 | 1292.582811 | 0.285782741 | 0.099995146 | 2.857966137 |
| Pebp4 | 0.585213848 | 0.477269128 | 1.277006649 | 0.373740519 |
| Egr3 | 501.7132685 | -0.197333404 | 0.321706344 | -0.613396061 |
| Bin3 | 135.9347836 | -0.5045793 | 0.105882514 | -4.765463925 |
| Ccar2 | 987.7091471 | 0.233016344 | 0.052334239 | 4.452464536 |
| 9930012K11Rik | 11.13584856 | 0.1540513 | 0.341487091 | 0.451118956 |
| Pdlim2 | 142.1719157 | 0.068348633 | 0.139536085 | 0.489827653 |
| Sorbs3 | 390.0743824 | -0.18545243 | 0.117393385 | -1.579751963 |
| Gm26908 | 10.16694102 | 0.246723352 | 0.356037768 | 0.692969607 |
| Ppp3cc | 243.8924496 | 0.034216146 | 0.089614566 | 0.381814557 |
| AC139294.1 | 1.827339133 | -0.500593023 | 0.803049353 | -0.623365203 |
| Slc39a14 | 953.2277302 | -0.433841426 | 0.177623844 | -2.442472905 |
| Gm9174 | 1.024625068 | -1.433978468 | 1.039656709 | -1.379280733 |
| Piwil2 | 21.10693623 | -0.815220205 | 0.225209565 | -3.619829394 |
| Gm22725 | 0.106465261 | 0.037073798 | 3.724945049 | 0.009952844 |
| Gm24890 | 0.088476398 | 0.51795324 | 3.724945049 | 0.139049901 |
| Polr3d | 217.1158271 | -0.237684393 | 0.115659074 | -2.055043196 |
| Mir320 | 1.741823475 | 0.998114332 | 0.936644703 | 1.065627477 |
| Phyhip | 4018.25641 | 0.915780274 | 0.11322352 | 8.088251191 |
| Bmp1 | 483.1780716 | -0.43211067 | 0.100216176 | -4.311785651 |
| Sftpc | 3.468929289 | 1.133338772 | 0.57703174 | 1.964083939 |
| Lgi3 | 1208.980655 | 0.421667057 | 0.085873197 | 4.910345405 |
| Reep4 | 37.19800003 | -0.461004893 | 0.1969678 | -2.340508929 |
| Hr | 708.6183582 | -0.132483437 | 0.116130838 | -1.140811863 |
| Nudt18 | 347.76934 | -0.03290503 | 0.077191071 | -0.426280266 |
| Fam160b2 | 415.787422 | -0.012820232 | 0.094980567 | -0.134977416 |
| Dmtn | 2327.269897 | 0.441426187 | 0.085474895 | 5.164395763 |
| Fgf17 | 6.855534712 | -0.450782176 | 0.485410796 | -0.928661207 |
| Npm2 | 6.446150388 | -0.160904915 | 0.427666615 | -0.37623913 |

| | | | | |
|---|---|---|---|---|
| Xpo7 | 1222.808954 | 0.066427038 | 0.054866965 | 1.210692779 |
| Dok2 | 40.29440054 | -1.306320036 | 0.212917124 | -6.135345114 |
| 2410012E07Rik | 0.066050875 | -0.443818988 | 3.724945049 | -0.119147795 |
| Gfra2 | 372.4700337 | -0.226136975 | 0.098712102 | -2.290873872 |
| Gm37455 | 0 | NA | NA | NA |
| Gm9192 | 0 | NA | NA | NA |
| Gm23735 | 0 | NA | NA | NA |
| Fndc3a | 2663.159039 | -0.66551221 | 0.099710853 | -6.674420987 |
| Gm16549 | 0.514076732 | -0.611041652 | 1.502141239 | -0.406780425 |
| Cysltr2 | 20.03900042 | -0.877538167 | 0.332337651 | -2.640501803 |
| Rcbtb2 | 290.6187941 | -0.0114369 | 0.090585088 | -0.126255883 |
| Gm17233 | 4.303707252 | 0.452349539 | 0.606676646 | 0.745618843 |
| Rb1 | 791.2567668 | -0.295120624 | 0.070939992 | -4.160144615 |
| Mir687 | 2.668100091 | -0.623731645 | 0.729266361 | -0.85528646 |
| Lpar6 | 59.93468125 | -0.70922517 | 0.183187903 | -3.871572079 |
| Itm2b | 7614.81607 | -0.039176622 | 0.102629006 | -0.381730506 |
| Med4 | 170.1772693 | 0.076765058 | 0.134061797 | 0.572609491 |
| Nudt15 | 22.3670934 | 0.424584361 | 0.242948625 | 1.747630228 |
| Sucla2 | 1247.988688 | 0.159025435 | 0.087214855 | 1.823375563 |
| Gm21750 | 0.774729477 | -0.943161656 | 1.051086574 | -0.89732062 |
| Gm24540 | 2.261342538 | -1.093701315 | 0.689699106 | -1.585765889 |
| Gm16409 | 3.92403765 | -0.842541844 | 0.496001097 | -1.698669315 |
| Gm22164 | 0 | NA | NA | NA |
| Gm6984 | 34.159564 | 0.246935924 | 0.187094101 | 1.319848797 |
| Htr2a | 177.8320817 | 0.219461519 | 0.18078982 | 1.213904184 |
| Esd | 684.0533462 | -1.094329219 | 0.138029178 | -7.928245544 |
| Lrch1 | 437.4414308 | -0.110484355 | 0.078482495 | -1.407757926 |
| 5031414D18Rik | 62.98482591 | -0.475358966 | 0.200121558 | -2.375351121 |
| Lrrc63 | 0.699855268 | -0.251320255 | 1.359004466 | -0.184929676 |
| Lcp1 | 670.0106953 | -0.707283749 | 0.157788854 | -4.482469659 |
| Gm15628 | 0 | NA | NA | NA |
| Gm15629 | 0.174360202 | 0.503964903 | 2.724220683 | 0.184994155 |
| Cpb2 | 0 | NA | NA | NA |
| AC161877.1 | 0 | NA | NA | NA |
| Zc3h13 | 935.6372034 | 0.217940294 | 0.128147938 | 1.700692945 |
| Siah3 | 24.32705589 | 0.057881786 | 0.236532115 | 0.244710049 |
| Spert | 0.0885994 | 0.517955464 | 3.724945049 | 0.139050498 |
| 4930564B18Rik | 0.087261593 | 0.51795324 | 3.724945049 | 0.139049901 |
| Gm23622 | 0 | NA | NA | NA |
| Cog3 | 540.4936263 | -0.024676614 | 0.073991908 | -0.333504225 |
| Gm24876 | 5.352175477 | 0.345957734 | 0.530834805 | 0.651723909 |
| Slc25a30 | 95.05997151 | -0.126935332 | 0.15041734 | -0.84388763 |
| Gm4285 | 34.11371507 | 0.568519578 | 0.216264753 | 2.628812926 |
| Tpt1 | 4005.741273 | -0.572075276 | 0.098167413 | -5.827547651 |
| Snora31 | 5.414963088 | -0.554112616 | 0.472953319 | -1.171601074 |
| Gtf2f2 | 148.747947 | -0.419566591 | 0.106318475 | -3.946318745 |
| Kctd4 | 189.0479674 | 0.84556942 | 0.146168602 | 5.784890941 |

| | | | | |
|---|---|---|---|---|
| Gm25517 | 1.093745531 | -2.009627816 | 1.130586979 | -1.777508368 |
| Gpalpp1 | 612.613148 | 0.151474712 | 0.057949548 | 2.613906718 |
| Nufip1 | 200.4613363 | -0.145737406 | 0.108944255 | -1.337724568 |
| Rps2-ps6 | 4.539665628 | 0.283020302 | 0.439683724 | 0.643690649 |
| Tsc22d1 | 3622.389764 | 0.141969603 | 0.065148013 | 2.179185475 |
| 4930444M15Rik | 1.224655951 | 0.648129215 | 0.951836868 | 0.68092468 |
| Serp2 | 345.0104461 | 0.196658506 | 0.10722002 | 1.834158452 |
| 1700108F19Rik | 0.956737924 | 0.428359807 | 1.161546674 | 0.368783981 |
| Lacc1 | 58.05462338 | 0.032289751 | 0.174350447 | 0.185200278 |
| Ccdc122 | 14.09176844 | -0.269928474 | 0.275389542 | -0.980169661 |
| Enox1 | 345.9653688 | -0.019056946 | 0.12581467 | -0.151468396 |
| Gm10132 | 18.18438502 | -0.067775461 | 0.280881356 | -0.241295693 |
| Gm6994 | 0 | NA | NA | NA |
| n-R5s48 | 4.967990864 | 0.7041717 | 0.653146019 | 1.07812293 |
| Gm23823 | 0 | NA | NA | NA |
| Gm1587 | 0.104719185 | 0.981063299 | 3.723618439 | 0.263470416 |
| Dnajc15 | 183.7589994 | -0.237549914 | 0.097476188 | -2.437004557 |
| Epsti1 | 187.7081649 | -1.041762102 | 0.220859111 | -4.716862695 |
| Gm19301 | 0 | NA | NA | NA |
| Fam216b | 12.69086337 | -0.896215512 | 0.324868567 | -2.758701836 |
| Mir1971 | 0 | NA | NA | NA |
| Gm23652 | 0 | NA | NA | NA |
| Tnfsf11 | 1.022937004 | -0.571781403 | 0.959531232 | -0.595896605 |
| Gm26197 | 0 | NA | NA | NA |
| Akap11 | 5712.428926 | 0.294105949 | 0.060046791 | 4.897946123 |
| Dgkh | 929.9045385 | 0.25356782 | 0.118579297 | 2.138381876 |
| Gm26251 | 0.072025462 | 0.517959912 | 3.724945049 | 0.139051692 |
| Vwa8 | 1020.041174 | 0.164350287 | 0.066060457 | 2.487876921 |
| Zfp957 | 0 | NA | NA | NA |
| Rgcc | 194.1349241 | -0.375353823 | 0.146456178 | -2.562908774 |
| Naa16 | 214.9779865 | -0.20005765 | 0.135253027 | -1.479136212 |
| Mtrf1 | 71.56468441 | 0.26599869 | 0.133683153 | 1.989769722 |
| Kbtbd7 | 573.16381 | 0.024453658 | 0.078762049 | 0.310475133 |
| Kbtbd6 | 5.414111675 | 0.220620086 | 0.451791538 | 0.488322749 |
| AC124745.1 | 0 | NA | NA | NA |
| Wbp4 | 689.3854996 | -0.000254109 | 0.088843756 | -0.00286018 |
| Elf1 | 298.3952992 | -0.895098909 | 0.155272328 | -5.764703351 |
| Gm27857 | 0 | NA | NA | NA |
| Sugt1 | 598.6834534 | -0.146000911 | 0.084543134 | -1.726939895 |
| Lect1 | 10.35988659 | -0.159363521 | 0.345627698 | -0.461084346 |
| Mir759 | 0 | NA | NA | NA |
| Pcdh8 | 429.8832926 | -0.096274759 | 0.159151671 | -0.60492459 |
| Gm9748 | 3.167342963 | -0.22918446 | 0.532855605 | -0.430106126 |
| Gm6999 | 2.919730593 | -1.256963969 | 0.643275346 | -1.954006129 |
| Gm10845 | 0.072025462 | 0.517959912 | 3.724945049 | 0.139051692 |
| Olfm4 | 11.07339153 | -0.344864841 | 0.346802035 | -0.994414121 |
| Gm9578 | 0 | NA | NA | NA |

| | | | | |
|---|---|---|---|---|
| Gm22891 | 0 | NA | NA | NA |
| Gm24774 | 0 | NA | NA | NA |
| Pcdh17 | 1873.471252 | -0.021100823 | 0.064931278 | -0.324971631 |
| Gm23926 | 0 | NA | NA | NA |
| Gm25115 | 0.075043205 | 0.517957688 | 3.724945049 | 0.139051095 |
| Diap3 | 34.4707947 | -0.394000373 | 0.235214937 | -1.675065277 |
| Gm23438 | 0 | NA | NA | NA |
| Gm37002 | 0 | NA | NA | NA |
| Gm23278 | 0 | NA | NA | NA |
| Tdrd3 | 499.7196116 | 0.11219197 | 0.057710824 | 1.944036888 |
| Rps3a2 | 226.2208805 | -0.195658082 | 0.145680476 | -1.343063165 |
| Gm26405 | 0.047948698 | 0.037064902 | 3.724945049 | 0.009950456 |
| Gm9274 | 0.048949532 | 0.037064902 | 3.724945049 | 0.009950456 |
| Pcdh20 | 350.8125221 | 0.358606593 | 0.159842798 | 2.24349547 |
| Gm36946 | 0 | NA | NA | NA |
| Gm25886 | 0 | NA | NA | NA |
| Gm25415 | 0.048949532 | 0.037064902 | 3.724945049 | 0.009950456 |
| Gm10110 | 26.30868621 | -0.590999567 | 0.222843626 | -2.652081991 |
| Gm25495 | 0 | NA | NA | NA |
| Gm23225 | 0 | NA | NA | NA |
| Gm24127 | 0 | NA | NA | NA |
| Gm37341 | 0 | NA | NA | NA |
| Pcdh9 | 3659.38219 | 0.305933542 | 0.073915142 | 4.138983353 |
| Gm23509 | 13.87461055 | 0.310293255 | 0.417455686 | 0.743296273 |
| Gm23324 | 0 | NA | NA | NA |
| Gm24680 | 0 | NA | NA | NA |
| 4921530L21Rik | 0.197024265 | 0.517969061 | 2.231719505 | 0.232094159 |
| Klhl1 | 133.1425135 | -0.067405127 | 0.192957431 | -0.349326414 |
| Gm15515 | 0.04393175 | 0.037062677 | 3.724945049 | 0.009949859 |
| Gm24043 | 0 | NA | NA | NA |
| Gm25133 | 0 | NA | NA | NA |
| Gm24770 | 0 | NA | NA | NA |
| Gm27441 | 0 | NA | NA | NA |
| Gm23276 | 0.217428854 | -1.735499013 | 2.547916594 | -0.681144358 |
| Dach1 | 251.06346 | 0.347687184 | 0.11000558 | 3.160632256 |
| Gm25831 | 2.466058065 | -0.03826947 | 0.700087353 | -0.05466385 |
| Gm16331 | 0 | NA | NA | NA |
| Gm26006 | 0 | NA | NA | NA |
| Gm26139 | 0 | NA | NA | NA |
| Gm27034 | 0 | NA | NA | NA |
| Rpl36a-ps1 | 11.21814357 | -0.02096381 | 0.30761562 | -0.068149367 |
| Mzt1 | 677.8415215 | 0.149592766 | 0.073425616 | 2.037337575 |
| Bora | 35.47661271 | 0.143257618 | 0.233652139 | 0.613123502 |
| Dis3 | 309.1793791 | -0.068823685 | 0.131929162 | -0.521671512 |
| Pibf1 | 250.2980066 | 0.068240197 | 0.069445125 | 0.982649206 |
| Gm4412 | 0 | NA | NA | NA |
| Klf5 | 148.4473388 | -0.605180663 | 0.169932488 | -3.561300543 |

| | | | | |
|---|---|---|---|---|
| Gm22984 | 0 | NA | NA | NA |
| Gm25130 | 0 | NA | NA | NA |
| Gm37093 | 0.13998015 | 0.517970503 | 2.923064931 | 0.177201162 |
| Gm22970 | 0 | NA | NA | NA |
| Gm5958 | 0.163532861 | 0.037076175 | 2.784469987 | 0.013315344 |
| Klf12 | 318.1322375 | 0.350919183 | 0.091700708 | 3.826788134 |
| 4930517O19Rik | 0.663801701 | 2.144587759 | 1.511542178 | 1.418807752 |
| Gm16260 | 0 | NA | NA | NA |
| Gm26367 | 0 | NA | NA | NA |
| Gm24345 | 0 | NA | NA | NA |
| Gm22401 | 0 | NA | NA | NA |
| Gm23538 | 0 | NA | NA | NA |
| Prr30 | 0.448613728 | -0.050533398 | 1.3559704 | -0.037267332 |
| Gm26173 | 0 | NA | NA | NA |
| Tbc1d4 | 374.3985729 | -0.2170388 | 0.089371588 | -2.428498851 |
| Commd6 | 158.7771116 | -0.142867284 | 0.116543166 | -1.225874409 |
| Uchl3 | 201.0806668 | -0.143700552 | 0.100849186 | -1.424905427 |
| Gm9922 | 0.794139519 | -1.088866081 | 1.099756285 | -0.99009762 |
| Lmo7 | 977.0007035 | 1.229596601 | 0.184335079 | 6.6704428 |
| Gm22347 | 0 | NA | NA | NA |
| Gm23764 | 0 | NA | NA | NA |
| Kctd12 | 489.5908408 | 0.066794576 | 0.141202593 | 0.473040721 |
| AC102815.1 | 0 | NA | NA | NA |
| Gm26778 | 8.453551642 | 0.808470729 | 0.389034752 | 2.078145267 |
| Mir5130 | 0 | NA | NA | NA |
| Irg1 | 221.3397588 | -1.494392429 | 0.490113483 | -3.049074307 |
| Cln5 | 245.9919061 | -0.710396776 | 0.134439867 | -5.284122883 |
| Fbxl3 | 1187.753289 | 0.052836665 | 0.065371539 | 0.808251815 |
| Mycbp2 | 9169.971231 | 0.322013117 | 0.083851735 | 3.84026777 |
| Gm4654 | 6.877366695 | 0.151838258 | 0.404298374 | 0.375559904 |
| Scel | 26.78325797 | -0.465007219 | 0.229092067 | -2.029783159 |
| Slain1 | 561.6860723 | 0.021403047 | 0.082281705 | 0.26011915 |
| Gm22877 | 0.140122821 | 0.517959509 | 3.572128867 | 0.145000231 |
| Slain1os | 0.09349418 | 0.517955464 | 3.724945049 | 0.139050498 |
| Ednrb | 961.6585733 | -0.703971054 | 0.177321979 | -3.970015766 |
| Gm37910 | 0.066050875 | -0.443818988 | 3.724945049 | -0.119147795 |
| Pou4f1 | 42.92036986 | -0.159099607 | 0.336626558 | -0.472629399 |
| Rnf219 | 139.6039557 | 0.003751884 | 0.098356842 | 0.038145636 |
| Gm22021 | 0 | NA | NA | NA |
| Gm25724 | 0 | NA | NA | NA |
| Gm24280 | 0 | NA | NA | NA |
| Rbm26 | 1630.5591 | 0.21988647 | 0.095638174 | 2.299149597 |
| Gm17066 | 187.5789837 | 0.228626577 | 0.125276112 | 1.824981421 |
| Gm22290 | 0 | NA | NA | NA |
| Ndfip2 | 1106.337769 | -0.034792673 | 0.064008525 | -0.543563111 |
| Gm24207 | 0 | NA | NA | NA |
| Gm16259 | 0.115603922 | -0.93550548 | 3.724188147 | -0.251197158 |

| | | | | |
|---|---|---|---|---|
| 4930449E01Rik | 0.297771212 | -0.438660228 | 1.645420783 | -0.266594559 |
| Mir6390 | 7.744732698 | 1.464432312 | 0.47461785 | 3.085497756 |
| Gm10076 | 151.4983943 | -0.520015361 | 0.168460402 | -3.086869993 |
| Gm22406 | 0 | NA | NA | NA |
| Spry2 | 675.0338336 | -0.694356764 | 0.131426602 | -5.283228455 |
| Gm4775 | 0 | NA | NA | NA |
| Trim52 | 0 | NA | NA | NA |
| Gm23459 | 0 | NA | NA | NA |
| 1700128A07Rik | 0 | NA | NA | NA |
| Gm24634 | 0 | NA | NA | NA |
| Gm17366 | 0 | NA | NA | NA |
| Gm23437 | 0 | NA | NA | NA |
| Gm24818 | 0 | NA | NA | NA |
| Gm24229 | 0 | NA | NA | NA |
| Rpl27a-ps2 | 2.255766892 | 1.621304901 | 0.801701681 | 2.022329427 |
| Slitrk1 | 1192.574154 | 0.509082836 | 0.094853704 | 5.367031696 |
| Gm25901 | 0 | NA | NA | NA |
| Gm24369 | 0 | NA | NA | NA |
| n-R5s50 | 0 | NA | NA | NA |
| Gm23242 | 0 | NA | NA | NA |
| Gm25670 | 0 | NA | NA | NA |
| Gm26255 | 0 | NA | NA | NA |
| Slitrk6 | 94.50009073 | 0.452177071 | 0.135346328 | 3.340889102 |
| Gm22802 | 0 | NA | NA | NA |
| Slitrk5 | 1071.418396 | 0.436965587 | 0.099783671 | 4.3791292 |
| Gm22764 | 0 | NA | NA | NA |
| Gm4822 | 0 | NA | NA | NA |
| Tpm3-rs7 | 867.6432377 | -0.488835284 | 0.111224735 | -4.395023157 |
| Gm4487 | 0 | NA | NA | NA |
| Gm37495 | 0 | NA | NA | NA |
| Gm24073 | 0 | NA | NA | NA |
| Mir17hg | 18.52657897 | -0.582498049 | 0.275281325 | -2.116010045 |
| Mir17 | 0 | NA | NA | NA |
| Mir18 | 0 | NA | NA | NA |
| Mir19a | 0 | NA | NA | NA |
| Mir20a | 0 | NA | NA | NA |
| Mir19b-1 | 0 | NA | NA | NA |
| Mir92-1 | 0 | NA | NA | NA |
| Gpc5 | 280.2541914 | 0.638439765 | 0.137441004 | 4.645191378 |
| 4930505G20Rik | 0 | NA | NA | NA |
| Gm18367 | 0 | NA | NA | NA |
| Gm20713 | 0.048949532 | 0.037064902 | 3.724945049 | 0.009950456 |
| Gm38045 | 0 | NA | NA | NA |
| Gpc6 | 374.2491254 | 0.143865499 | 0.083883314 | 1.715066952 |
| Mir6239 | 0 | NA | NA | NA |
| Dct | 6.057780134 | -0.171018607 | 0.448473116 | -0.381335248 |
| Gm26773 | 0.075043205 | 0.517957688 | 3.724945049 | 0.139051095 |

| | | | | |
|---|---|---|---|---|
| Tgds | 62.65483873 | -0.683082022 | 0.201202459 | -3.394998381 |
| Gpr180 | 179.3319749 | 0.07034495 | 0.090045215 | 0.78121808 |
| Sox21 | 98.70143058 | -0.200998645 | 0.239558688 | -0.839037176 |
| Gm4675 | 4.582536418 | 0.567527481 | 0.484634651 | 1.171041896 |
| Mir6241 | 0.075043205 | 0.517957688 | 3.724945049 | 0.139051095 |
| Gm9376 | 0 | NA | NA | NA |
| Gm23669 | 0 | NA | NA | NA |
| Gm26791 | 0.061920613 | -0.443816764 | 3.724945049 | -0.119147198 |
| 1700044C05Rik | 0 | NA | NA | NA |
| Abcc4 | 177.0567019 | -0.362074903 | 0.141996116 | -2.549893012 |
| Gm23302 | 0 | NA | NA | NA |
| Mir6391 | 0.804474276 | 0.646915715 | 1.06772428 | 0.605882743 |
| Gm27198 | 0 | NA | NA | NA |
| Gm22379 | 0.042716945 | 0.037062677 | 3.724945049 | 0.009949859 |
| Cldn10 | 133.9096642 | 0.776320122 | 0.194006668 | 4.001512569 |
| n-R5s51 | 0 | NA | NA | NA |
| Dzip1 | 1031.570717 | 0.471692958 | 0.075704027 | 6.230751233 |
| Dnajc3 | 1082.666445 | -0.352578955 | 0.07548979 | -4.670551529 |
| Uggt2 | 618.685887 | 0.326447931 | 0.090621655 | 3.602317024 |
| Gm16835 | 39.44922461 | 0.426753661 | 0.27926242 | 1.528145677 |
| Hs6st3 | 282.2429297 | 0.722865151 | 0.161643106 | 4.47198255 |
| 1700006F04Rik | 17.63169857 | 0.620550823 | 0.359193185 | 1.727624156 |
| Oxgr1 | 3.717291957 | 0.106652772 | 0.590943496 | 0.180478799 |
| Mbnl2 | 3584.706982 | -0.079556887 | 0.068223102 | -1.166128251 |
| Gm26679 | 5.318848262 | 1.753521864 | 0.601559675 | 2.914959124 |
| Rap2a | 997.0783507 | 0.259610552 | 0.070779538 | 3.667875755 |
| Gm17613 | 0.138137327 | -0.357090338 | 3.608654548 | -0.098953871 |
| Gm25781 | 0 | NA | NA | NA |
| Ipo5 | 1921.943571 | -0.279531459 | 0.062792652 | -4.451658744 |
| Farp1 | 948.5700461 | 0.163973778 | 0.074025914 | 2.215086164 |
| B930095G15Rik | 416.2277983 | 0.780197688 | 0.116084875 | 6.720924568 |
| Stk24 | 646.4051468 | 0.13022418 | 0.084279814 | 1.545140819 |
| Slc15a1 | 1.066426709 | -0.94335326 | 0.975483183 | -0.967062555 |
| Dock9 | 2161.334577 | 0.442621402 | 0.091755383 | 4.823928418 |
| 1810041H14Rik | 1.973916092 | 0.760565867 | 0.762881691 | 0.996964373 |
| Ubac2 | 353.8238038 | 0.01074934 | 0.093612941 | 0.114827502 |
| Gpr18 | 9.840232673 | -0.13776646 | 0.45214908 | -0.30469256 |
| Gpr183 | 14.82188085 | -0.177622992 | 0.303698455 | -0.584866301 |
| Timm8a2 | 1.438846906 | 1.310819088 | 0.839017307 | 1.562326637 |
| A330035P11Rik | 10.89891607 | -0.182503542 | 0.30469227 | -0.598976609 |
| Tm9sf2 | 1465.863616 | -0.152262447 | 0.069762666 | -2.182577809 |
| Gm6254 | 34.64912996 | 0.042548865 | 0.323527359 | 0.131515507 |
| Gm17082 | 26.87597815 | 0.440045594 | 0.258622749 | 1.701496082 |
| Clybl | 80.6719109 | 0.179004129 | 0.150990903 | 1.185529232 |
| 1700108J01Rik | 0.0878635 | 0.037058229 | 3.724945049 | 0.009948665 |
| Gm25464 | 0.76305848 | -1.231782448 | 1.290897433 | -0.954206289 |
| Gm5089 | 8.639524202 | 0.487728252 | 0.517157431 | 0.943094351 |

| | | | | |
|---|---|---|---|---|
| Zic5 | 171.9911989 | -0.30234943 | 0.146788334 | -2.059764707 |
| 2610035F20Rik | 11.57667783 | 0.403726601 | 0.337111356 | 1.197606054 |
| Zic2 | 736.6056247 | 0.103753106 | 0.192581618 | 0.538748749 |
| Gm10837 | 14.11353788 | 0.942138172 | 0.454626011 | 2.072336709 |
| Gm9399 | 0.044544648 | 0.517957688 | 3.724945049 | 0.139051095 |
| Pcca | 484.4393937 | 0.375744654 | 0.11195232 | 3.356291794 |
| Ggact | 197.5556307 | 0.040168295 | 0.125455625 | 0.320179307 |
| 4930594M22Rik | 104.9172262 | 0.458910033 | 0.202918336 | 2.261550346 |
| Tmtc4 | 235.3928922 | 0.003090035 | 0.108288029 | 0.028535334 |
| Gm15735 | 9.699810692 | 0.589889532 | 0.445498093 | 1.324112361 |
| Nalcn | 1752.798546 | 0.639997779 | 0.059550403 | 10.74716122 |
| Itgbl1 | 26.65978906 | 0.770926548 | 0.326360311 | 2.36219455 |
| Gm9308 | 0 | NA | NA | NA |
| Gm25272 | 0.047948698 | 0.037064902 | 3.724945049 | 0.009950456 |
| Fgf14 | 940.1745018 | 0.643936389 | 0.091708972 | 7.02152005 |
| Gm28932 | 1.468722268 | 2.197861182 | 0.989482165 | 2.221223646 |
| Gm15422 | 16.80940044 | 0.775380201 | 0.370073182 | 2.095207755 |
| 1700024B18Rik | 0.075043205 | 0.517957688 | 3.724945049 | 0.139051095 |
| Gm17615 | 0.320086762 | 1.502599667 | 1.588177612 | 0.946115633 |
| Sepp1 | 2912.542392 | 0.31031432 | 0.136257888 | 2.277404441 |
| Ccdc152 | 0.495781838 | -0.768218251 | 1.432214066 | -0.536385076 |
| Ghr | 164.1599138 | -0.125098993 | 0.104550216 | -1.196544567 |
| Gm22031 | 0 | NA | NA | NA |
| Gm4823 | 0.066050875 | -0.443818988 | 3.724945049 | -0.119147795 |
| Gm23228 | 0 | NA | NA | NA |
| Fbxo4 | 73.81450778 | -0.55198449 | 0.13859473 | -3.982723508 |
| AW549877 | 1232.520806 | -0.102933878 | 0.072934584 | -1.411317804 |
| Gm2093 | 0.919878397 | 0.203908655 | 0.966179906 | 0.21104626 |
| BC037032 | 34.71017491 | 0.707240185 | 0.265107319 | 2.667750509 |
| Oxct1 | 2597.175127 | 0.130581625 | 0.088253449 | 1.479620637 |
| Gm15632 | 0 | NA | NA | NA |
| Plcxd3 | 320.4411957 | 0.339129837 | 0.111096933 | 3.052558045 |
| C6 | 2.125003216 | 0.619208542 | 0.76002033 | 0.814726288 |
| Mroh2b | 0.19444009 | 1.024066476 | 2.689656809 | 0.380742433 |
| C7 | 8.962534464 | 0.719265686 | 0.384971032 | 1.86836314 |
| Gm25652 | 0 | NA | NA | NA |
| Gm16077 | 0.04393175 | 0.037062677 | 3.724945049 | 0.009949859 |
| Card6 | 43.77270322 | -0.170942665 | 0.160283502 | -1.066501938 |
| Rpl37 | 814.5795284 | -0.070552051 | 0.109708567 | -0.643086063 |
| Snord72 | 5.885017498 | -0.336353447 | 0.45533764 | -0.738690189 |
| Gm10250 | 114.0624678 | 0.065203411 | 0.114840986 | 0.567771258 |
| Prkaa1 | 690.6189185 | -0.286713241 | 0.104478924 | -2.744220845 |
| Gm15938 | 2.619102942 | 0.013147284 | 0.757545412 | 0.01735511 |
| Ttc33 | 464.303568 | 0.159057733 | 0.070731233 | 2.248762346 |
| Ptger4 | 41.58960014 | -0.444767714 | 0.225478824 | -1.97254761 |
| Gm15937 | 14.4175352 | 0.354827323 | 0.337305862 | 1.051945319 |
| Gm27760 | 0.173816988 | 0.037058464 | 3.34140675 | 0.011090677 |

| | | | | |
|---|---|---|---|---|
| Gm23139 | 0 | NA | NA | NA |
| Dab2 | 418.7976734 | -0.770294938 | 0.182661465 | -4.217063182 |
| Gm16311 | 0 | NA | NA | NA |
| C9 | 0.135879625 | -0.426108425 | 2.940622282 | -0.144904168 |
| Gm23218 | 0 | NA | NA | NA |
| Fyb | 220.4543545 | -0.642459634 | 0.171552847 | -3.744966315 |
| Gm2245 | 0.436872751 | -0.78408584 | 1.635302115 | -0.479474608 |
| Gm7666 | 27.5160756 | -0.458899293 | 0.196388755 | -2.336688236 |
| Rictor | 1871.956267 | 0.098916937 | 0.066094011 | 1.49660968 |
| Osmr | 628.1045177 | -0.659101392 | 0.283749491 | -2.322828453 |
| Gm38282 | 0.159330469 | 0.517966511 | 3.465376739 | 0.149469033 |
| Lifr | 1758.792625 | -0.144014905 | 0.071646654 | -2.010071588 |
| Egflam | 43.8423358 | -0.325804086 | 0.174397236 | -1.868172305 |
| Gm16030 | 0 | NA | NA | NA |
| Gm16029 | 0.048949532 | 0.037064902 | 3.724945049 | 0.009950456 |
| Gm37743 | 0.061920613 | -0.443816764 | 3.724945049 | -0.119147198 |
| Gdnf | 23.58641988 | -0.988559274 | 0.241474094 | -4.093852295 |
| Wdr70 | 191.8981038 | 0.010436551 | 0.107164984 | 0.097387695 |
| Gm27529 | 0.871987541 | 0.497583748 | 1.193062004 | 0.41706445 |
| Nup155 | 488.0096314 | 0.180582488 | 0.115263722 | 1.566689712 |
| 2410089E03Rik | 1512.066955 | 0.360883253 | 0.16211123 | 2.226145913 |
| Gm24144 | 14.79135611 | 1.073601473 | 0.467663782 | 2.295669481 |
| Nipbl | 1929.6716 | -0.095427541 | 0.064215937 | -1.486041396 |
| Gm2310 | 0 | NA | NA | NA |
| Gm2348 | 0.104106287 | 0.51795324 | 3.724945049 | 0.139049901 |
| Slc1a3 | 7685.648105 | 0.059577206 | 0.116770464 | 0.510207836 |
| Gm37310 | 119.6237137 | 0.955451105 | 0.298481706 | 3.201037398 |
| RP24-341L14.2 | 0.280881662 | -0.125787571 | 2.065493055 | -0.060899537 |
| Gm5210 | 0 | NA | NA | NA |
| Ranbp3l | 56.33245746 | 0.162462608 | 0.259204199 | 0.626774598 |
| Nadk2 | 494.6776195 | -0.041564058 | 0.111527665 | -0.372679351 |
| Skp2 | 125.2851504 | -0.012436714 | 0.137509331 | -0.090442691 |
| Lmbrd2 | 1052.783199 | 0.225798847 | 0.05494731 | 4.109370365 |
| Ugt3a1 | 0.150086409 | 1.027495679 | 3.723011807 | 0.275985071 |
| Ugt3a2 | 0.171181394 | 0.539411076 | 2.747344444 | 0.196339078 |
| Capsl | 15.44470158 | -0.070072433 | 0.28564224 | -0.245315375 |
| Il7r | 194.8832465 | -2.140812193 | 0.425780893 | -5.027966803 |
| Spef2 | 48.63543957 | -0.396589338 | 0.188771036 | -2.100901424 |
| Gm26350 | 0 | NA | NA | NA |
| Prlr | 148.8562054 | 0.537467148 | 0.182770575 | 2.940665626 |
| Gm21973 | 0 | NA | NA | NA |
| Agxt2 | 0.124596252 | 0.037071632 | 3.714681146 | 0.009979761 |
| Dnajc21 | 254.7197003 | 0.117238309 | 0.110046793 | 1.065349627 |
| Brix1 | 263.1432847 | -0.349635561 | 0.109726497 | -3.186427795 |
| Rad1 | 85.02316572 | -0.532225159 | 0.143534854 | -3.70798552 |
| Ttc23l | 0.267080418 | 0.517974954 | 1.817541951 | 0.284986519 |
| Gm17044 | 0.038981395 | -0.443816764 | 3.724945049 | -0.119147198 |

| | | | | |
|---|---|---|---|---|
| Gm8174 | 0.386177252 | -0.904087391 | 1.585250209 | -0.57031211 |
| Rai14 | 244.3821753 | -0.297150355 | 0.130572546 | -2.275749105 |
| 4930556M19Rik | 7.969462537 | -0.174353599 | 0.369895278 | -0.471359354 |
| C1qtnf3 | 3.38967256 | 0.014430141 | 0.542554391 | 0.026596672 |
| Amacr | 74.0218532 | -0.291030212 | 0.157620269 | -1.846400932 |
| Slc45a2 | 0.883742299 | -1.260927322 | 1.20998722 | -1.042099702 |
| Rxfp3 | 34.39250656 | -0.522810399 | 0.208176003 | -2.511386479 |
| Adamts12 | 17.29016175 | -0.157564365 | 0.279967325 | -0.562795552 |
| Gm24844 | 0 | NA | NA | NA |
| Tars | 425.9557127 | -0.562912674 | 0.119706302 | -4.702448114 |
| Npr3 | 146.1473597 | 1.388566791 | 0.140439773 | 9.88727595 |
| 1700047G03Rik | 0.146692594 | -0.955495625 | 3.521109961 | -0.271362052 |
| Gm24074 | 0 | NA | NA | NA |
| Sub1 | 1754.400611 | 0.570097398 | 0.079279073 | 7.191019972 |
| RP24-105K16.2 | 0.060174537 | 0.517957688 | 3.724945049 | 0.139051095 |
| Zfr | 4001.152899 | 0.019529962 | 0.057441833 | 0.339995451 |
| Gm25713 | 0 | NA | NA | NA |
| Mir1898 | 18.01255643 | -0.021306009 | 0.262152118 | -0.081273457 |
| Mtmr12 | 542.6306912 | 0.493631707 | 0.094693882 | 5.212920785 |
| 1810049J17Rik | 2.812275527 | 0.863970056 | 0.641625741 | 1.346532722 |
| Golph3 | 1317.912574 | -0.274354145 | 0.091707483 | -2.991622242 |
| Pdzd2 | 1799.644411 | 0.346135466 | 0.166749655 | 2.075779198 |
| Gm24135 | 1.998901016 | 1.183425212 | 0.867887552 | 1.36356975 |
| Gm2606 | 25.26788695 | 0.42591988 | 0.24989596 | 1.704388823 |
| F830212C03Rik | 14.03902647 | 0.959982066 | 0.438897695 | 2.187257025 |
| Gm24302 | 0 | NA | NA | NA |
| 6030458C11Rik | 453.2360569 | 0.082732977 | 0.102996813 | 0.803257643 |
| Drosha | 1331.381267 | 0.043073608 | 0.050903029 | 0.846189489 |
| AC133189.1 | 0 | NA | NA | NA |
| Gm24664 | 0.315718849 | 2.100241129 | 1.673673474 | 1.254869102 |
| Cdh6 | 210.1792556 | 0.222112479 | 0.119134874 | 1.864378341 |
| Gm9630 | 0 | NA | NA | NA |
| Gm24080 | 0 | NA | NA | NA |
| Gm25550 | 0 | NA | NA | NA |
| Cdh9 | 185.9667162 | 0.869592177 | 0.137250243 | 6.335815231 |
| Gm22032 | 0 | NA | NA | NA |
| Gm22895 | 0 | NA | NA | NA |
| Gm24713 | 0.044544648 | 0.517957688 | 3.724945049 | 0.139051095 |
| 4921515E04Rik | 0 | NA | NA | NA |
| Gm8318 | 1.912323087 | 0.501027069 | 0.69426395 | 0.721666549 |
| C030047K22Rik | 5.198307323 | 0.577441882 | 0.564044912 | 1.023751601 |
| Cdh10 | 687.055273 | 0.527843817 | 0.120624032 | 4.375942408 |
| Gm25976 | 0 | NA | NA | NA |
| Gm37815 | 0 | NA | NA | NA |
| Acot10 | 1.155605027 | -0.235211605 | 0.98926808 | -0.237763262 |
| Gm25711 | 4.168298335 | 1.940292098 | 0.688631595 | 2.817605396 |
| Gm27397 | 3.423615971 | 0.851632681 | 0.683023166 | 1.246857681 |

| | | | | |
|---|---|---|---|---|
| Cdh12 | 473.520525 | 0.565069111 | 0.110767111 | 5.101415985 |
| Gm25101 | 4.033556537 | 1.515057385 | 0.740164191 | 2.046920676 |
| Gm38234 | 6.221313409 | -0.459903785 | 0.45542049 | -1.009844298 |
| AC166159.1 | 0 | NA | NA | NA |
| Gm27525 | 0 | NA | NA | NA |
| Cdh18 | 540.5763889 | 0.241810469 | 0.104438995 | 2.315327427 |
| Gm5803 | 0.68701006 | -1.801486763 | 1.235597828 | -1.457987964 |
| Gm19840 | 13.48907376 | 0.362770814 | 0.328459673 | 1.104460742 |
| Gm24656 | 2.60150024 | 0.929680141 | 0.707843903 | 1.313397115 |
| Gm23980 | 0 | NA | NA | NA |
| Gm22810 | 0 | NA | NA | NA |
| Gm2824 | 69.97913716 | 1.315132772 | 0.220228773 | 5.97166644 |
| 9230109A22Rik | 0.817550754 | 0.080976062 | 1.008748095 | 0.080273819 |
| Gm38341 | 0.308655657 | 1.353223238 | 3.704213799 | 0.36531996 |
| Basp1 | 2855.029311 | 0.244702295 | 0.107590222 | 2.274391586 |
| Gm5468 | 5.212139954 | -0.275305061 | 0.573138411 | -0.480346554 |
| Myo10 | 988.6576999 | -0.660689906 | 0.1202377 | -5.494864816 |
| Gm2862 | 0.999928428 | 0.809248487 | 1.068381638 | 0.757452635 |
| Fam134b | 559.6848899 | -0.007294584 | 0.083140924 | -0.087737585 |
| Mir7212 | 1.890715168 | -0.194355213 | 0.975307314 | -0.199275869 |
| Zfp622 | 385.825252 | -0.013496839 | 0.097488881 | -0.13844491 |
| 11-Mar | 93.58712996 | -0.142849335 | 0.132227714 | -1.08032825 |
| Fbxl7 | 13.24742036 | 0.276458208 | 0.346261622 | 0.798408459 |
| Gm6576 | 4.198207553 | -0.07492004 | 0.464919283 | -0.161146338 |
| Gm23903 | 0 | NA | NA | NA |
| Ank | 1809.490276 | 0.346453814 | 0.051417974 | 6.737990362 |
| Mir7117 | 0 | NA | NA | NA |
| Otulin | 178.6870836 | -0.018936974 | 0.101909594 | -0.185821312 |
| Fam105a | 144.2450621 | -0.15926438 | 0.122284547 | -1.302408065 |
| Trio | 4183.182376 | 0.082062039 | 0.085726544 | 0.957253553 |
| Dnah5 | 99.5374478 | -0.036328311 | 0.135962436 | -0.267193736 |
| Mir3964 | 0 | NA | NA | NA |
| Gm23033 | 0 | NA | NA | NA |
| Gm24554 | 0 | NA | NA | NA |
| Gm23327 | 0 | NA | NA | NA |
| Gm26163 | 0 | NA | NA | NA |
| Ctnnd2 | 3344.046455 | 0.132905411 | 0.061920562 | 2.146385725 |
| Gm23956 | 8.765840549 | 0.352009141 | 0.465876815 | 0.755584158 |
| Gm24487 | 2.253680491 | 1.040120425 | 0.731266927 | 1.422353982 |
| Gm26416 | 0.503286714 | 2.028873281 | 3.679457773 | 0.551405508 |
| Dap | 102.7869284 | -0.813890934 | 0.193248896 | -4.211620094 |
| Fam136b-ps | 0.047948698 | 0.037064902 | 3.724945049 | 0.009950456 |
| Ankrd33b | 1536.741971 | 0.546439255 | 0.161618492 | 3.381044144 |
| Ropn1l | 5.055244084 | -0.495231855 | 0.429187673 | -1.153881823 |
| Gm6361 | 5.200498561 | 0.476911887 | 0.48218051 | 0.989073339 |
| 6-Mar | 3828.438788 | 0.262118324 | 0.048317477 | 5.424917493 |
| Cmbl | 49.64000691 | 0.659600784 | 0.198887652 | 3.316449148 |

| | | | | |
|---|---|---|---|---|
| Cct5 | 1440.817807 | -0.362024903 | 0.077427278 | -4.675676477 |
| Fam173b | 140.20352 | -0.201990826 | 0.236293011 | -0.854831997 |
| Tas2r119 | 0 | NA | NA | NA |
| Snhg18 | 13.92677096 | -0.568357589 | 0.301452937 | -1.885394099 |
| Snord123 | 0.145446656 | -0.410258174 | 2.878908473 | -0.142504764 |
| Sema5a | 1219.347837 | 0.148100551 | 0.081032644 | 1.827665301 |
| Sdc2 | 591.0431138 | -0.098655101 | 0.104295399 | -0.945919975 |
| Cpq | 169.9525733 | -0.197230515 | 0.170163354 | -1.159065752 |
| 1700084J12Rik | 0.496918936 | -0.777533515 | 3.594798464 | -0.216294049 |
| Gm24476 | 0.306418947 | -2.150783454 | 3.688184606 | -0.583155044 |
| Tspyl5 | 513.6623244 | 0.048393531 | 0.096056423 | 0.503803179 |
| Mtdh | 1529.028053 | -0.003124337 | 0.061860935 | -0.050505823 |
| Gm24500 | 0 | NA | NA | NA |
| Laptm4b | 633.1682788 | 0.017035295 | 0.104164346 | 0.163542476 |
| Gm25809 | 0 | NA | NA | NA |
| Matn2 | 218.3605345 | 0.312842783 | 0.167634356 | 1.866221169 |
| 9430069I07Rik | 0.203825279 | 0.032006124 | 2.61982179 | 0.012216909 |
| Rpl30 | 585.149464 | -0.114809287 | 0.080983948 | -1.417679546 |
| Gm24523 | 0 | NA | NA | NA |
| Erich5 | 14.29563529 | 1.710573295 | 0.407356404 | 4.199205601 |
| Hrsp12 | 208.6478451 | 0.191073484 | 0.141720216 | 1.34824437 |
| Pop1 | 119.2294828 | -0.137334595 | 0.162328152 | -0.846030671 |
| Gm22208 | 2.694090619 | 0.121994764 | 0.623475415 | 0.195668924 |
| Nipal2 | 159.5133862 | -0.00447911 | 0.098998353 | -0.04524429 |
| Kcns2 | 254.6488734 | 0.222597324 | 0.101064833 | 2.202520076 |
| Stk3 | 354.1359327 | 0.079159154 | 0.089918805 | 0.880340368 |
| Osr2 | 0.114742407 | 0.517955464 | 3.724945049 | 0.139050498 |
| Vps13b | 2358.901886 | 0.273223126 | 0.070602951 | 3.869854209 |
| Gm24771 | 4.664523441 | 0.505110046 | 0.579880606 | 0.871058698 |
| Mir599 | 0 | NA | NA | NA |
| Mir875 | 0 | NA | NA | NA |
| Cox6c | 1187.10824 | 0.35961565 | 0.149553126 | 2.404601351 |
| Rgs22 | 32.16090155 | 0.417963434 | 0.195485346 | 2.138080638 |
| Gm25372 | 0 | NA | NA | NA |
| Gm18889 | 2.742579127 | -0.393440428 | 0.596107215 | -0.660016214 |
| Fbxo43 | 1.809171643 | -1.730243886 | 0.864189267 | -2.002158499 |
| Gm22979 | 0 | NA | NA | NA |
| Polr2k | 87.53921573 | -0.12201793 | 0.13871538 | -0.879627988 |
| Spag1 | 177.5961547 | -0.221762704 | 0.136759404 | -1.621553602 |
| Gm23754 | 0.706791778 | -1.069621198 | 1.254319173 | -0.852750417 |
| Rnf19a | 663.4722742 | 0.013269548 | 0.100596318 | 0.131908886 |
| Mir8097 | 0 | NA | NA | NA |
| Rpl7a-ps3 | 1.338063731 | -0.07331634 | 0.863561803 | -0.084899934 |
| Ankrd46 | 1500.096336 | 0.218712412 | 0.059145568 | 3.697866457 |
| Snx31 | 4.226100588 | 0.276757172 | 0.500303658 | 0.55317839 |
| AC137871.1 | 0 | NA | NA | NA |
| Pabpc1 | 2806.56132 | -0.323074385 | 0.075817377 | -4.261218154 |

| | | | | |
|---|---|---|---|---|
| Ywhaz | 10312.43491 | 0.18616818 | 0.066104028 | 2.816291018 |
| Gm25303 | 0 | NA | NA | NA |
| Gm10384 | 1.188161708 | -0.104272477 | 1.143340429 | -0.091199851 |
| Zfp706 | 1437.63576 | -0.040000632 | 0.057139907 | -0.700047193 |
| Gm26766 | 0.392978793 | -0.976590993 | 1.710578995 | -0.570912537 |
| Gm16136 | 0.154527273 | 0.037073388 | 2.83445495 | 0.013079548 |
| Grhl2 | 2.049607807 | -0.310310923 | 0.922603402 | -0.336342704 |
| Gm16137 | 0 | NA | NA | NA |
| Gm16138 | 0.695937234 | 0.721769956 | 2.357050904 | 0.306217382 |
| Ncald | 3437.612725 | 0.562987549 | 0.107079257 | 5.257671421 |
| Gm28221 | 0.522018477 | 0.034653276 | 1.284515521 | 0.026977701 |
| Gm23654 | 2.696581576 | 0.649460791 | 0.711948398 | 0.912230146 |
| Gm15941 | 47.58132438 | 0.800804348 | 0.382229013 | 2.09509043 |
| Gm8668 | 0 | NA | NA | NA |
| 4930447A16Rik | 0.047948698 | 0.037064902 | 3.724945049 | 0.009950456 |
| Gm3267 | 0 | NA | NA | NA |
| Gm24833 | 0.580994137 | 0.491689416 | 1.435274118 | 0.342575268 |
| Gm15942 | 0.537603422 | 1.759978601 | 1.313724867 | 1.33968584 |
| Gm15940 | 9.81108231 | 0.883253303 | 0.436127358 | 2.02521875 |
| Rps19-ps5 | 1.865250533 | 1.317254415 | 0.917929361 | 1.435028088 |
| Rrm2b | 427.3554432 | 0.053698822 | 0.070158637 | 0.765391455 |
| Gm3362 | 13.70112511 | -0.172622063 | 0.294720765 | -0.585713949 |
| Ubr5 | 4115.687366 | -0.021285387 | 0.060138495 | -0.353939471 |
| Gm22353 | 0 | NA | NA | NA |
| Odf1 | 0.128271896 | 0.517956073 | 3.679089552 | 0.140783763 |
| Gm24098 | 0 | NA | NA | NA |
| Gm8705 | 0.498759394 | 0.29973434 | 1.63768561 | 0.183023126 |
| Klf10 | 430.8187853 | -1.116670233 | 0.251948976 | -4.432128489 |
| Gm26863 | 0.301278804 | -0.44823364 | 1.63027145 | -0.274944176 |
| Azin1 | 2466.052833 | -0.132472128 | 0.082181223 | -1.611951282 |
| Mir6951 | 0.490407978 | 1.615747819 | 1.306351183 | 1.236840323 |
| 2310043O21Rik | 0.088534442 | -0.905414734 | 3.723500899 | -0.243162217 |
| Atp6v1c1 | 1712.938684 | 0.075817449 | 0.044383916 | 1.708218985 |
| Gm26704 | 1.33281409 | -1.011653856 | 0.893008301 | -1.132860528 |
| Baalc | 276.5131432 | -0.180686946 | 0.093579852 | -1.930831714 |
| Fzd6 | 96.39112974 | -0.634796604 | 0.134431361 | -4.722087171 |
| Gm24789 | 0.09176491 | 0.921395572 | 3.723061103 | 0.247483333 |
| Cthrc1 | 75.42071906 | -0.426540705 | 0.1603106 | -2.660714296 |
| Slc25a32 | 170.269925 | -0.321885524 | 0.092449549 | -3.481742504 |
| Dcaf13 | 241.4209333 | -0.249412097 | 0.089894749 | -2.774490173 |
| n-R5s39 | 0.07796279 | -0.798301147 | 3.721829328 | -0.214491605 |
| Gm9522 | 11.31287606 | 0.29628673 | 0.338153545 | 0.876189927 |
| Rims2 | 2518.228734 | 0.423328714 | 0.089264547 | 4.742405909 |
| Gm25381 | 0 | NA | NA | NA |
| Dcstamp | 2.546063767 | -0.770663294 | 0.61811871 | -1.246788492 |
| Dpys | 1.614744503 | 0.760122849 | 0.807312474 | 0.941547261 |
| Gm16291 | 9.213264714 | -0.203901422 | 0.437010107 | -0.466582853 |

| | | | | |
|---|---|---|---|---|
| Lrp12 | 640.2551409 | 0.087236196 | 0.080409525 | 1.084898783 |
| 9330182O14Rik | 0 | NA | NA | NA |
| Gm25958 | 0 | NA | NA | NA |
| Gm16294 | 6.25597695 | 0.383033611 | 0.449968806 | 0.851244811 |
| Zfpm2 | 484.4239322 | 0.373065592 | 0.115273812 | 3.236342989 |
| 4930555K19Rik | 0 | NA | NA | NA |
| Oxr1 | 3719.638992 | 0.477731625 | 0.051174122 | 9.33541428 |
| Gm26854 | 0.181780715 | -0.003164606 | 3.307972889 | -0.00095666 |
| Abra | 0.899802893 | 1.356221218 | 0.996719091 | 1.360685504 |
| Gm24751 | 0.127971488 | -0.95589744 | 3.6855341 | -0.259364698 |
| Angpt1 | 263.5197957 | 0.19171018 | 0.161164788 | 1.189528942 |
| Gm17473 | 35.3169963 | 0.367136958 | 0.27002785 | 1.359626269 |
| Rspo2 | 100.8944359 | 0.155845559 | 0.158534792 | 0.983036954 |
| Gm9658 | 1.11165528 | 0.453584981 | 0.884657886 | 0.512723606 |
| Gm22037 | 0.385685737 | -1.01281945 | 1.709989764 | -0.592295622 |
| Eif3e | 970.451907 | -0.079912965 | 0.103624691 | -0.771176874 |
| Gm10373 | 1.964982618 | 1.084186456 | 0.770325864 | 1.40743873 |
| Emc2 | 503.3907252 | 0.014015626 | 0.0990983 | 0.141431548 |
| Tmem74 | 82.76716195 | -0.084389587 | 0.145014148 | -0.581940371 |
| Trhr | 65.62486328 | 0.208310891 | 0.140193099 | 1.485885488 |
| Gm24903 | 0 | NA | NA | NA |
| Gm25093 | 0 | NA | NA | NA |
| Nudcd1 | 289.802079 | -0.224582685 | 0.128592366 | -1.74646981 |
| Eny2 | 530.2164271 | 0.33237659 | 0.076374126 | 4.351952798 |
| Gm22392 | 8.528292636 | 0.162090538 | 0.353690971 | 0.45828294 |
| Pkhd1l1 | 13.16766861 | -0.122414639 | 0.29666855 | -0.412630995 |
| Ebag9 | 211.2230902 | -0.004522147 | 0.088307055 | -0.051209347 |
| Sybu | 684.3366891 | 0.265009726 | 0.067138344 | 3.947218671 |
| 2310069G16Rik | 44.1689834 | 0.38821041 | 0.176760772 | 2.196247537 |
| A930017M01Rik | 111.592861 | 0.376624306 | 0.147716396 | 2.549644564 |
| Gm24768 | 0.351351704 | -0.296213087 | 1.568717913 | -0.188824954 |
| Gm23343 | 0.299746944 | 0.477714831 | 1.59980181 | 0.298608758 |
| Gm5471 | 0.105143817 | 0.517955464 | 3.724945049 | 0.139050498 |
| Kcnv1 | 528.0842402 | 1.100085296 | 0.153414605 | 7.170668658 |
| Gm5472 | 0.382197862 | 0.958843097 | 1.572277649 | 0.609843368 |
| Gm26063 | 0 | NA | NA | NA |
| 1700022A22Rik | 0 | NA | NA | NA |
| Gm23562 | 0 | NA | NA | NA |
| 4930548G14Rik | 0.066050875 | -0.443818988 | 3.724945049 | -0.119147795 |
| Gm23665 | 0 | NA | NA | NA |
| Csmd3 | 1405.512621 | 0.736634377 | 0.115477718 | 6.379017439 |
| Gm16300 | 8.844286263 | 1.214859792 | 0.40633992 | 2.989762347 |
| Gm23530 | 0.060174537 | 0.517957688 | 3.724945049 | 0.139051095 |
| Gm25007 | 0 | NA | NA | NA |
| Gm23185 | 0 | NA | NA | NA |
| Gm23848 | 0 | NA | NA | NA |
| Trps1 | 1156.791181 | -0.013138674 | 0.063743869 | -0.206116672 |

| | | | | |
|---|---|---|---|---|
| Mir1907 | 3.164287563 | 0.370947765 | 0.68091581 | 0.544777724 |
| Eif3h | 1020.195418 | -0.061195495 | 0.08369685 | -0.731156494 |
| Utp23 | 94.48504293 | -0.523987667 | 0.14420059 | -3.633741486 |
| Rad21 | 1630.277696 | 0.073258015 | 0.089145479 | 0.821780484 |
| Aard | 24.65755326 | 0.008436205 | 0.251133882 | 0.033592459 |
| Slc30a8 | 0.5812948 | -1.579994251 | 1.314849698 | -1.201653888 |
| Gm10020 | 9.983419005 | -0.223458093 | 0.430319702 | -0.5192839 |
| Gm23545 | 0 | NA | NA | NA |
| Gm23267 | 0 | NA | NA | NA |
| Med30 | 115.6898845 | -0.225104948 | 0.140968066 | -1.596850644 |
| Ext1 | 627.9625793 | -0.016839245 | 0.065072449 | -0.258776872 |
| Gm27926 | 0.066050875 | -0.443818988 | 3.724945049 | -0.119147795 |
| Samd12 | 154.297153 | 0.509066315 | 0.119582258 | 4.257038834 |
| Gm23200 | 7.643354964 | 0.478253532 | 0.498441805 | 0.959497232 |
| mmu-mir-28c | 7.291715215 | 0.650377608 | 0.652021134 | 0.997479338 |
| Gm7489 | 0 | NA | NA | NA |
| Gm26933 | 0 | NA | NA | NA |
| Tnfrsf11b | 55.23446549 | -0.855315674 | 0.236872406 | -3.610870889 |
| Gm5215 | 0 | NA | NA | NA |
| Colec10 | 0.252184553 | 0.051169697 | 1.80930506 | 0.02828141 |
| Mal2 | 1117.686592 | 0.393072667 | 0.10240712 | 3.83833336 |
| Nov | 659.4404828 | 1.102847761 | 0.14101845 | 7.820592018 |
| Gm7577 | 0 | NA | NA | NA |
| Enpp2 | 3338.021252 | 0.569716203 | 0.137903046 | 4.131280769 |
| Gm26684 | 4.138921123 | 3.806583016 | 0.898432632 | 4.236915357 |
| Taf2 | 560.7641421 | -0.091701669 | 0.073852292 | -1.241690221 |
| Dscc1 | 3.002576494 | -0.042752314 | 0.566391718 | -0.075481885 |
| Gm9920 | 2.459937312 | 0.021385334 | 0.715395505 | 0.029893022 |
| Deptor | 493.9836769 | -0.053104176 | 0.08549622 | -0.621128931 |
| Gm26296 | 0 | NA | NA | NA |
| Col14a1 | 17.16314811 | -0.037892149 | 0.273200045 | -0.138697448 |
| 1700040F17Rik | 0.075043205 | 0.517957688 | 3.724945049 | 0.139051095 |
| Mrpl13 | 304.6669654 | -0.295131759 | 0.083103717 | -3.551366532 |
| Mtbp | 67.98096681 | -0.341122973 | 0.172749722 | -1.97466583 |
| Sntb1 | 66.74833637 | -0.017905571 | 0.163708803 | -0.109374515 |
| Has2 | 80.3213104 | -1.630555302 | 0.478817623 | -3.405378633 |
| Has2os | 3.183229149 | -0.737476361 | 0.662749053 | -1.11275355 |
| Gm26178 | 0 | NA | NA | NA |
| Slc22a22 | 0.334826625 | -0.013228981 | 1.575117108 | -0.008398728 |
| Zhx2 | 102.0661113 | -0.009343587 | 0.15426249 | -0.060569403 |
| Gm16006 | 1.944759997 | -0.018206644 | 0.700348437 | -0.025996551 |
| 9330154K18Rik | 0.788164698 | -0.697089744 | 1.186344823 | -0.587594543 |
| Derl1 | 846.7041548 | -0.369550061 | 0.089912412 | -4.110111757 |
| Tbc1d31 | 114.9980324 | -0.268565697 | 0.200776121 | -1.337637644 |
| Fam83a | 0.697414605 | -3.019341478 | 1.295659051 | -2.330351858 |
| 9130401M01Rik | 105.150685 | -0.155240569 | 0.118617336 | -1.308751103 |
| Gm29394 | 4.265622792 | 0.684811637 | 0.678747983 | 1.008933587 |

| | | | | |
|---|---|---|---|---|
| Gm22299 | 0.817774247 | 0.326441705 | 1.299519855 | 0.251201784 |
| Zhx1 | 1069.677375 | 0.129426111 | 0.068238724 | 1.896666623 |
| Atad2 | 218.8401466 | -0.255270627 | 0.163752452 | -1.558881251 |
| Wdyhv1 | 104.6557226 | -0.423445997 | 0.11564052 | -3.66174414 |
| Gm23475 | 0 | NA | NA | NA |
| Gm22488 | 1.584432859 | 0.850766779 | 0.959167694 | 0.886984397 |
| Fbxo32 | 391.5402532 | -0.095918688 | 0.104717054 | -0.915979625 |
| Gm15943 | 6.572403966 | 0.237094469 | 0.448473141 | 0.528670386 |
| Klhl38 | 13.89074863 | -0.86592006 | 0.337912939 | -2.562553723 |
| Anxa13 | 0 | NA | NA | NA |
| D15Ertd621e | 1138.565379 | -0.091059486 | 0.05549148 | -1.640963377 |
| Fer1l6 | 7.081223484 | -0.541030081 | 0.426348007 | -1.268987005 |
| Gm23789 | 0 | NA | NA | NA |
| Gm20712 | 48.91645952 | 0.608210021 | 0.188001629 | 3.235131655 |
| Tmem65 | 1185.359275 | -0.184100428 | 0.062399765 | -2.950338461 |
| Gm24041 | 11.06577238 | 0.274068648 | 0.355936721 | 0.769992617 |
| Ube2d4 | 0 | NA | NA | NA |
| Trmt12 | 90.91210585 | -0.361267231 | 0.12275673 | -2.942952559 |
| Rnf139 | 607.7861192 | -0.23128319 | 0.107330383 | -2.154871565 |
| Tatdn1 | 116.8025155 | -0.237796421 | 0.133679177 | -1.7788591 |
| Ndufb9 | 863.2536908 | -0.040418621 | 0.12340803 | -0.327520185 |
| Mtss1 | 982.1708213 | -0.082419155 | 0.099891952 | -0.825083041 |
| Gm5045 | 4.498534705 | 0.235957033 | 0.587263182 | 0.401790952 |
| Zfp572 | 0.238889773 | 0.998865751 | 2.150309179 | 0.464521921 |
| Sqle | 640.1137635 | -0.430589439 | 0.11449403 | -3.760802546 |
| E430025E21Rik | 722.4473385 | -0.399413978 | 0.087232707 | -4.578718132 |
| Nsmce2 | 214.2833592 | -0.083491553 | 0.079688335 | -1.04772615 |
| Trib1 | 585.2550097 | -0.919654 | 0.257197185 | -3.575676773 |
| Gm7083 | 0.061920613 | -0.443816764 | 3.724945049 | -0.119147198 |
| Gm7713 | 0 | NA | NA | NA |
| 4933412E24Rik | 0 | NA | NA | NA |
| Gm25875 | 0.044544648 | 0.517957688 | 3.724945049 | 0.139051095 |
| Gm23835 | 0.042716945 | 0.037062677 | 3.724945049 | 0.009949859 |
| Fam84b | 212.0412458 | -0.094095518 | 0.153204493 | -0.614182498 |
| 9930014A18Rik | 33.67317411 | 0.879886497 | 0.196555305 | 4.476533953 |
| A1bg | 0.161114758 | 1.389564196 | 3.426332673 | 0.405554372 |
| Gm24696 | 0 | NA | NA | NA |
| Gm37947 | 0.038981395 | -0.443816764 | 3.724945049 | -0.119147198 |
| Myc | 144.727581 | -0.707186621 | 0.267345184 | -2.645219229 |
| Pvt1 | 146.0044478 | -0.15211096 | 0.144610227 | -1.051868619 |
| Gm27913 | 0 | NA | NA | NA |
| Gm27782 | 1.314786873 | -1.43009438 | 1.076143526 | -1.328906735 |
| Gm27718 | 0 | NA | NA | NA |
| Gm27957 | 0.583672784 | -0.40621738 | 1.343898794 | -0.302267836 |
| H2afy3 | 0.122507438 | -0.356739328 | 3.722822199 | -0.095824971 |
| Gm24810 | 0 | NA | NA | NA |
| Gm22521 | 0.240278643 | 0.051171979 | 1.831981643 | 0.027932583 |

| | | | | |
|---|---|---|---|---|
| RP23-55P19.1 | 7.30667397 | 0.123838349 | 0.397805911 | 0.31130344 |
| Gm25702 | 0.042716945 | 0.037062677 | 3.724945049 | 0.009949859 |
| Gm37615 | 0 | NA | NA | NA |
| Gm5473 | 0.91428204 | 1.685516688 | 1.147320532 | 1.469089624 |
| Gsdmc | 0.046345709 | -0.443818988 | 3.724945049 | -0.119147795 |
| 1700065I16Rik | 0 | NA | NA | NA |
| Gsdmc2 | 0.112396584 | -0.917692504 | 3.724434845 | -0.246397787 |
| Gsdmcl1 | 0 | NA | NA | NA |
| Gsdmc3 | 0.258721378 | -2.000869561 | 1.99997042 | -1.000449577 |
| Gsdmcl2 | 0 | NA | NA | NA |
| Gsdmc4 | 1.216796211 | -2.021084045 | 1.094975197 | -1.845780663 |
| Gsdmcl-ps | 0 | NA | NA | NA |
| Fam49b | 1750.706656 | 0.145086335 | 0.049335855 | 2.940788875 |
| Gm25628 | 0 | NA | NA | NA |
| Asap1 | 1466.62248 | 0.023180487 | 0.061662391 | 0.375925855 |
| Gm20717 | 0.527708665 | 0.371899433 | 1.396408222 | 0.266325726 |
| Adcy8 | 473.5293802 | 0.065494939 | 0.075179446 | 0.871181454 |
| Gm21798 | 0 | NA | NA | NA |
| Gm21961 | 0 | NA | NA | NA |
| Gm5046 | 0.097139547 | -0.443816764 | 3.724945049 | -0.119147198 |
| Gm23907 | 0 | NA | NA | NA |
| Efr3a | 2113.727247 | 0.083867821 | 0.065422699 | 1.281937643 |
| Gm20405 | 22.08023906 | 1.287156642 | 0.394421846 | 3.263400991 |
| Oc90 | 0.439163765 | 0.440244023 | 1.411367889 | 0.311927193 |
| Hhla1 | 1.7811158 | 2.112278381 | 1.038615243 | 2.03374483 |
| Gm2991 | 7.931298069 | -0.061574191 | 0.46380043 | -0.132760099 |
| Kcnq3 | 2910.472199 | 0.892772509 | 0.092585811 | 9.642649334 |
| Gm27153 | 0.044544648 | 0.517957688 | 3.724945049 | 0.139051095 |
| Gm27242 | 2.088118819 | 2.298452665 | 0.822979183 | 2.792844234 |
| Lrrc6 | 31.55937458 | 0.629723567 | 0.223585405 | 2.816478862 |
| Tmem71 | 8.010071188 | -0.15988699 | 0.330381344 | -0.483946788 |
| Phf20l1 | 1418.308876 | 0.116978945 | 0.071151817 | 1.644075286 |
| Tg | 5.923287473 | -1.028452486 | 0.428997933 | -2.397336696 |
| Gm17140 | 0 | NA | NA | NA |
| Sla | 260.5870413 | 0.244063623 | 0.107415467 | 2.272145981 |
| Wisp1 | 38.31189845 | -1.751371904 | 0.332817931 | -5.262252239 |
| Ndrg1 | 857.5715411 | -0.363736352 | 0.133175179 | -2.731262345 |
| Gm17035 | 0.30380643 | -0.966347364 | 2.021969672 | -0.477923768 |
| Gm2895 | 1.340196514 | 1.39566343 | 1.259966384 | 1.107698941 |
| St3gal1 | 321.0877622 | -0.388442553 | 0.109870344 | -3.53546314 |
| 1700012I11Rik | 0.522753486 | 0.992540905 | 1.416670179 | 0.700615373 |
| Zfat | 173.6083377 | -0.033892325 | 0.132990448 | -0.254847816 |
| Gm10240 | 8.242000484 | -0.355075716 | 0.454026378 | -0.782059662 |
| Gm37320 | 57.92546311 | -0.940353153 | 0.493899276 | -1.903937094 |
| Gm26621 | 67.44066586 | -0.476329022 | 0.315524902 | -1.50964003 |
| Mir30b | 2.936662866 | 0.005618283 | 0.690588398 | 0.008135501 |
| Gm37296 | 72.21542741 | -0.376954388 | 0.2112865 | -1.784091215 |

| | | | | |
|---|---|---|---|---|
| Gm20732 | 11.82845525 | -1.125215779 | 0.375579393 | -2.995946527 |
| Mir30d | 0 | NA | NA | NA |
| Gm25987 | 0 | NA | NA | NA |
| Gm7125 | 2.234104145 | -0.257693767 | 0.69477243 | -0.370903847 |
| Khdrbs3 | 814.1907786 | 0.071707007 | 0.117065195 | 0.612539081 |
| Gm24232 | 0 | NA | NA | NA |
| Gm23987 | 0 | NA | NA | NA |
| Fam135b | 709.6683967 | -0.270586121 | 0.118776819 | -2.278105474 |
| AC157602.1 | 0 | NA | NA | NA |
| Gm22519 | 0.044191549 | 0.037062677 | 3.724945049 | 0.009949859 |
| Col22a1 | 42.46963243 | -0.51726314 | 0.203873995 | -2.537170767 |
| Gm16308 | 0.108309328 | 0.967119648 | 3.722483401 | 0.259804959 |
| Gm23217 | 4.535027026 | 0.606997281 | 0.552472928 | 1.098691448 |
| Gm24390 | 0 | NA | NA | NA |
| Kcnk9 | 245.3309406 | 0.394933482 | 0.097338435 | 4.05732313 |
| Trappc9 | 814.4156458 | 0.277439576 | 0.045027843 | 6.161511584 |
| Gm28020 | 1.075827988 | 1.889744797 | 1.127787331 | 1.675621586 |
| Peg13 | 3399.486185 | 0.29234938 | 0.052502807 | 5.568261944 |
| Gm3150 | 0.456099668 | -1.009376749 | 1.424622745 | -0.708522135 |
| Chrac1 | 76.46489437 | 0.173970541 | 0.161056067 | 1.080186199 |
| Ago2 | 2822.203106 | -0.307698849 | 0.072227101 | -4.260157822 |
| Ptk2 | 1504.870551 | 0.175462525 | 0.06895835 | 2.544471053 |
| Mir151 | 5.981125562 | 1.202483416 | 0.607688817 | 1.978781544 |
| Gm24787 | 0 | NA | NA | NA |
| Dennd3 | 115.7213391 | 0.246149238 | 0.219376759 | 1.122038813 |
| Slc45a4 | 787.9133075 | 0.306803687 | 0.072942167 | 4.206122452 |
| 1700010B13Rik | 0.609775176 | 1.291405462 | 1.226772358 | 1.052685491 |
| Gpr20 | 1.015418215 | 1.724894998 | 1.063002323 | 1.622663432 |
| Ptp4a3 | 340.214372 | -0.094503545 | 0.119380396 | -0.79161695 |
| Gm22106 | 2.287044039 | 0.058536498 | 0.775089696 | 0.075522224 |
| Mroh5 | 17.84213286 | 0.215350964 | 0.259632758 | 0.829444502 |
| Gm3244 | 3.864046566 | 0.380006335 | 0.559505967 | 0.679181916 |
| Gm6569 | 1.915143953 | -0.635969058 | 0.707118107 | -0.899381662 |
| Gm7935 | 7.955384209 | -0.334440998 | 0.358012703 | -0.934159585 |
| Gm15387 | 13.22477098 | -0.000390187 | 0.488586623 | -0.000798603 |
| Adgrb1 | 3078.93178 | 0.209731588 | 0.146009444 | 1.436424809 |
| Gm26117 | 0 | NA | NA | NA |
| Mroh4 | 0.178490723 | 0.539256917 | 2.711241605 | 0.198896666 |
| Arc | 2012.660561 | -1.3923741 | 0.498484282 | -2.793215652 |
| Jrk | 151.5292579 | -0.223198706 | 0.107760079 | -2.071255972 |
| 4933427E11Rik | 4.797225747 | -0.606122016 | 0.566066161 | -1.070761791 |
| Psca | 1.592565788 | -1.116256746 | 0.781535616 | -1.428286469 |
| Gm28976 | 0 | NA | NA | NA |
| Them6 | 413.7288862 | 0.194774275 | 0.079519519 | 2.449389505 |
| Slurp1 | 1.036256916 | 1.37336833 | 1.025415187 | 1.339329032 |
| Lypd2 | 2.552977391 | 0.035279699 | 0.622408494 | 0.056682547 |
| 2300005B03Rik | 0.58378306 | 0.474541162 | 1.528344398 | 0.310493605 |

| | | | | |
|---|---|---|---|---|
| Lynx1 | 4130.871836 | 0.008650222 | 0.070247261 | 0.123139638 |
| Ly6d | 10.57966156 | -0.531569898 | 0.488240434 | -1.088746161 |
| D730001G18Rik | 0.049553047 | -0.443818988 | 3.724945049 | -0.119147795 |
| Gm17189 | 0.097139547 | -0.443816764 | 3.724945049 | -0.119147198 |
| Ly6k | 0.732543458 | -0.869330406 | 1.135919012 | -0.7653102 |
| Gml | 0 | NA | NA | NA |
| Hemt1 | 0 | NA | NA | NA |
| Cyp11b1 | 0.755541319 | 0.12899713 | 1.078429677 | 0.119615709 |
| Cyp11b2 | 1.063164669 | -1.191546734 | 1.123520726 | -1.060547177 |
| 2010109I03Rik | 3.337759678 | -1.894158163 | 0.660761444 | -2.866629372 |
| Gm6610 | 1.171300456 | -2.121974175 | 1.025941585 | -2.06831871 |
| Gm29126 | 0.728772687 | -1.47048464 | 1.105479436 | -1.330178195 |
| Ly6e | 7038.794082 | -0.842719211 | 0.167197171 | -5.040271952 |
| Ly6i | 26.75927167 | -1.390311841 | 0.578726754 | -2.402363172 |
| Gm8000 | 1.59659966 | -0.850108002 | 0.719924622 | -1.180829182 |
| Gm37042 | 1.174124634 | -0.992733274 | 0.945432712 | -1.050030596 |
| Ly6a | 4998.068703 | -1.305720251 | 0.310357001 | -4.207155787 |
| Gm28068 | 1.632511506 | -0.321997117 | 0.711462609 | -0.452584735 |
| Gm28069 | 0.045882455 | 0.517959912 | 3.724945049 | 0.139051692 |
| Ly6c1 | 1373.160618 | -1.62290697 | 0.360007258 | -4.50798403 |
| Gm5960 | 192.4448412 | -2.253737578 | 0.66772266 | -3.375259989 |
| Gm28502 | 0.175945213 | -0.410809826 | 2.710746807 | -0.151548579 |
| Ly6c2 | 840.6845555 | -1.795805146 | 0.452547876 | -3.96821031 |
| I830127L07Rik | 12.15111785 | -2.066092215 | 0.491578324 | -4.202976648 |
| Ly6g | 0.512981325 | -2.047944559 | 1.341004848 | -1.52717163 |
| Gm28117 | 0 | NA | NA | NA |
| BC025446 | 0.044544648 | 0.517957688 | 3.724945049 | 0.139051095 |
| Gm10238 | 0 | NA | NA | NA |
| Gm28115 | 0 | NA | NA | NA |
| Ly6f | 0 | NA | NA | NA |
| Gm28116 | 0 | NA | NA | NA |
| Gm3454 | 0.07796279 | -0.798301147 | 3.721829328 | -0.214491605 |
| Gm28113 | 0.038981395 | -0.443816764 | 3.724945049 | -0.119147198 |
| Gm29517 | 0 | NA | NA | NA |
| Gm28114 | 0.149988252 | -0.281740967 | 3.492946197 | -0.080659979 |
| Gm3142 | 0 | NA | NA | NA |
| 9030619P08Rik | 0.161532967 | -0.444773734 | 3.449701211 | -0.128931089 |
| Gm20654 | 0 | NA | NA | NA |
| Ly6h | 944.0802929 | 0.32573139 | 0.096402718 | 3.378861054 |
| Gpihbp1 | 0.278632317 | -0.33247557 | 1.803623687 | -0.184337549 |
| Zfp41 | 98.87672504 | -0.267234096 | 0.130511073 | -2.047597109 |
| 2810039B14Rik | 19.1597757 | -0.380748383 | 0.268171293 | -1.419795457 |
| Top1mt | 124.7567236 | -0.255358585 | 0.118837884 | -2.148797816 |
| Gm15945 | 0.585634519 | -0.033608414 | 1.401065969 | -0.023987746 |
| Rhpn1 | 84.34622614 | 0.19644512 | 0.147857144 | 1.328614322 |
| Mafa | 7.826514485 | -0.045099008 | 0.42134753 | -0.107035178 |
| Zc3h3 | 159.3577311 | -0.058580623 | 0.154184637 | -0.379938136 |

| | | | | |
|---|---|---|---|---|
| Gm23747 | 0 | NA | NA | NA |
| Gm9568 | 0.083526043 | 0.037073798 | 3.724945049 | 0.009952844 |
| Gsdmd | 138.4714216 | -0.500378702 | 0.175745226 | -2.847182331 |
| Mroh6 | 1.934735662 | -0.565554179 | 0.785395715 | -0.720088191 |
| Naprt | 43.80295543 | 0.023747908 | 0.23911065 | 0.09931765 |
| Eef1d | 688.7842089 | -0.111224094 | 0.066179697 | -1.680637698 |
| Tigd5 | 16.71972297 | -0.16982605 | 0.271619154 | -0.625235914 |
| AC116520.1 | 0 | NA | NA | NA |
| Pycrl | 118.6008582 | -0.554846647 | 0.172769096 | -3.211492444 |
| Tsta3 | 168.0293222 | -0.24250395 | 0.12668303 | -1.914257577 |
| Zfp623 | 159.2088236 | 0.018183755 | 0.134533012 | 0.13516203 |
| Zfp707 | 75.27886443 | 0.24351116 | 0.144293939 | 1.687604909 |
| Ccdc166 | 61.81052859 | 0.270311668 | 0.148046599 | 1.825855299 |
| Mapk15 | 15.79109506 | 0.243578194 | 0.340596669 | 0.715151427 |
| Fam83h | 77.85627554 | -0.115267484 | 0.135264844 | -0.852161442 |
| K230010J24Rik | 12.00267003 | 0.324316136 | 0.328668742 | 0.986756861 |
| Gm23027 | 0.654384519 | -1.99722433 | 1.446912032 | -1.380335699 |
| Scrib | 592.0251915 | -0.182938108 | 0.076911988 | -2.378538281 |
| Mir6952 | 0 | NA | NA | NA |
| Puf60 | 1138.944899 | -0.162320975 | 0.086778327 | -1.870524373 |
| Nrbp2 | 715.0247032 | 0.602144949 | 0.117243876 | 5.135832859 |
| BC024139 | 9.779656096 | 0.254445603 | 0.329292048 | 0.772704974 |
| Plec | 5010.938922 | -0.347891218 | 0.07851886 | -4.430670766 |
| Gm25720 | 3.45116213 | -0.899573641 | 0.733568297 | -1.226298415 |
| Mir1942 | 3.221922816 | 0.24469736 | 0.711217648 | 0.344054117 |
| Parp10 | 866.3362482 | -0.545087918 | 0.168515996 | -3.234636061 |
| Grina | 4908.811971 | 0.029594743 | 0.085402539 | 0.34653236 |
| Mir6953 | 0 | NA | NA | NA |
| Gm10872 | 0.972832988 | -2.593898287 | 1.159491597 | -2.237099685 |
| Spatc1 | 0.136963818 | 0.037074342 | 3.599544405 | 0.010299732 |
| Mir3079 | 0 | NA | NA | NA |
| Smpd5 | 4.917982605 | 0.006951053 | 0.457208563 | 0.015203243 |
| Oplah | 309.3643565 | 0.431245522 | 0.11786094 | 3.65893502 |
| Exosc4 | 178.5313927 | -0.026894491 | 0.11240337 | -0.239267652 |
| Gpaa1 | 246.0551654 | 0.061778558 | 0.090036796 | 0.6861479 |
| Rps6-ps1 | 8.203160127 | -0.064566504 | 0.35033716 | -0.184298189 |
| Cyc1 | 1071.038313 | -0.164862554 | 0.09310124 | -1.770787962 |
| Sharpin | 316.269203 | -0.131953439 | 0.083891996 | -1.572896648 |
| Maf1 | 531.081438 | 0.111492629 | 0.080956814 | 1.377186473 |
| Hgh1 | 82.07062805 | -0.465007878 | 0.187857034 | -2.475328538 |
| Tssk5 | 6.475749014 | 0.59661601 | 0.45316133 | 1.316564258 |
| Mroh1 | 1129.595412 | 0.070446746 | 0.087306428 | 0.80689071 |
| Mir6954 | 0 | NA | NA | NA |
| Bop1 | 410.1160048 | -0.418810054 | 0.064575029 | -6.48563479 |
| Scx | 29.69111678 | -0.100066317 | 0.224431686 | -0.445865372 |
| Hsf1 | 313.4887952 | -0.17096894 | 0.099688376 | -1.715033861 |
| Dgat1 | 157.725041 | -0.24624645 | 0.123418643 | -1.99521275 |

| | | | | |
|---|---|---|---|---|
| Scrt1 | 1878.856055 | 0.61139458 | 0.079298454 | 7.710044151 |
| Fbxl6 | 146.2430035 | -0.090992784 | 0.096943972 | -0.938612094 |
| Slc52a2 | 132.5881942 | -0.240756185 | 0.107521237 | -2.239150058 |
| Adck5 | 90.29773825 | -0.177939538 | 0.125580197 | -1.416939469 |
| Cpsf1 | 643.6551908 | -0.086480959 | 0.078492902 | -1.101767895 |
| Slc39a4 | 6.380351065 | -0.002312352 | 0.417058281 | -0.005544433 |
| Tonsl | 636.9176432 | 0.004235223 | 0.085204176 | 0.049706756 |
| Cyhr1 | 972.3595972 | 0.002738253 | 0.057770798 | 0.047398568 |
| Kifc2 | 2493.40518 | 1.331832281 | 0.139853734 | 9.523036979 |
| Foxh1 | 0.909626782 | 1.470207077 | 1.085162667 | 1.354826444 |
| Ppp1r16a | 515.0001235 | 0.170610663 | 0.070858247 | 2.407774266 |
| Gpt | 84.05780739 | 0.130993071 | 0.20232141 | 0.647450367 |
| Mfsd3 | 34.06178562 | -0.189320148 | 0.223647773 | -0.846510321 |
| Recql4 | 21.17447754 | -0.245908176 | 0.301431883 | -0.815800151 |
| Lrrc14 | 184.1576737 | -0.077105411 | 0.086153548 | -0.89497661 |
| Lrrc24 | 405.4748953 | 0.191808377 | 0.080530004 | 2.381825009 |
| Arhgap39 | 936.4403093 | 0.327674186 | 0.094218772 | 3.477801497 |
| Gm17271 | 0.060174537 | 0.517957688 | 3.724945049 | 0.139051095 |
| Zfp251 | 456.6486897 | 0.147346825 | 0.068466651 | 2.152096273 |
| Zfp7 | 92.43301678 | -0.258150079 | 0.124089408 | -2.080355463 |
| Commd5 | 140.0820505 | -0.132146431 | 0.129731601 | -1.018614045 |
| Rpl8 | 2132.795094 | -0.314000571 | 0.101755524 | -3.085833179 |
| Zfp647 | 28.94240596 | 0.136464131 | 0.206731449 | 0.660103394 |
| 1110038F14Rik | 133.089957 | -0.093597431 | 0.105427793 | -0.887787066 |
| Mb | 1.797034208 | 0.705413822 | 0.825148091 | 0.8548936 |
| Apol6 | 78.23562678 | -0.445511969 | 0.275362521 | -1.6179107 |
| Gm24753 | 2.783881763 | 0.030592313 | 0.695840058 | 0.043964575 |
| Rbfox2 | 3954.719732 | 0.10266038 | 0.080602525 | 1.27366209 |
| 1700109K24Rik | 0.04393175 | 0.037062677 | 3.724945049 | 0.009949859 |
| Gm24056 | 9.581094532 | 0.173885665 | 0.455919633 | 0.381395432 |
| Apol7a | 18.60379838 | 0.803866392 | 0.424813064 | 1.892282655 |
| Gm20688 | 0.311586316 | 0.387786511 | 1.961379695 | 0.197711087 |
| Apol9a | 239.6974814 | -1.104075758 | 0.391872508 | -2.817436121 |
| Apol7b | 0.961738709 | -0.527399322 | 1.097188238 | -0.480682625 |
| 4930571N24Rik | 0.157815785 | 0.483090695 | 3.465799425 | 0.139387955 |
| Gm6506 | 0.409269261 | -0.033577393 | 1.431917035 | -0.023449259 |
| Apol10a | 0.15100991 | 0.51796601 | 3.527597456 | 0.146832516 |
| Apol11a | 0 | NA | NA | NA |
| Apol7c | 0.292232717 | -1.416258942 | 1.949435343 | -0.726497007 |
| 1700025B11Rik | 0.199129965 | -0.912181976 | 2.212011822 | -0.412376628 |
| Apol10b | 22.02299047 | -1.48437025 | 0.375476088 | -3.953301678 |
| Gm8221 | 0.475668934 | -0.038621146 | 1.293538253 | -0.02985698 |
| Apol11b | 3.059439512 | -0.201165468 | 0.564722191 | -0.356220229 |
| Apol7e | 1.633903828 | -0.179665689 | 0.760491817 | -0.236249339 |
| Apol9b | 163.6719493 | -1.121726585 | 0.360033513 | -3.115617141 |
| Apol8 | 54.12186927 | 0.144250763 | 0.160465008 | 0.89895464 |
| Myh9 | 2978.335968 | -0.737761921 | 0.130197955 | -5.666463218 |

| | | | | |
|---|---|---|---|---|
| Gm22107 | 5.021758635 | -0.67056278 | 0.43826929 | -1.530024563 |
| Txn2 | 707.7197206 | -0.082626384 | 0.072532496 | -1.139163661 |
| Foxred2 | 319.3024116 | 0.298058083 | 0.095300149 | 3.127572056 |
| Eif3d | 737.7459319 | -0.550349641 | 0.095725862 | -5.749226245 |
| Gm17638 | 11.65945967 | 0.408807899 | 0.372727113 | 1.096802152 |
| Cacng2 | 970.2458113 | 0.29440023 | 0.089878734 | 3.275527096 |
| A730060N03Rik | 73.25342862 | 0.255888669 | 0.178730379 | 1.431702154 |
| Ift27 | 107.8713168 | 0.256613067 | 0.133797472 | 1.917921636 |
| Pvalb | 1103.933019 | 0.002431418 | 0.152292785 | 0.015965415 |
| Ncf4 | 100.2792551 | -1.133311717 | 0.225480998 | -5.026196109 |
| Csf2rb2 | 172.9194105 | -0.976116018 | 0.21801519 | -4.47728444 |
| Mir7676-1 | 0 | NA | NA | NA |
| Csf2rb | 284.283327 | -1.15297423 | 0.211255125 | -5.457733773 |
| Mir7676-2 | 0 | NA | NA | NA |
| Gm26329 | 0 | NA | NA | NA |
| Tex33 | 0.613719218 | 0.005220339 | 1.141924089 | 0.004571529 |
| Tst | 291.9109073 | -0.788107415 | 0.156072597 | -5.0496207 |
| Mpst | 172.564482 | -0.002754835 | 0.137925563 | -0.01997335 |
| Kctd17 | 1094.573162 | 0.256465193 | 0.059976309 | 4.276108273 |
| Tmprss6 | 5.777471174 | 0.766796478 | 0.394550797 | 1.943467062 |
| Il2rb | 153.2540098 | -0.943893531 | 0.365264395 | -2.584137802 |
| C1qtnf6 | 29.05525177 | -0.618812657 | 0.203362096 | -3.042910501 |
| Sstr3 | 170.7394951 | -0.555608986 | 0.16774366 | -3.312250281 |
| Gm6723 | 0.648304949 | -0.730497446 | 1.210096745 | -0.603668631 |
| Rac2 | 192.5000097 | -0.729123738 | 0.19531966 | -3.732976693 |
| Cyth4 | 257.1621641 | -0.870396468 | 0.136477572 | -6.377578796 |
| Elfn2 | 1215.602818 | 0.473006244 | 0.122174643 | 3.871558228 |
| Mfng | 23.1312152 | -0.130646412 | 0.269923998 | -0.484011842 |
| Card10 | 133.9315844 | -0.259031687 | 0.178959098 | -1.447435143 |
| Cdc42ep1 | 428.9155534 | -0.733375408 | 0.165553236 | -4.4298464 |
| Lgals2 | 1.456018128 | -3.091621946 | 0.985041636 | -3.138569817 |
| Gm26634 | 0.085327104 | -0.905414734 | 3.723500899 | -0.243162217 |
| Gga1 | 712.034101 | -0.181862749 | 0.069437917 | -2.619069773 |
| Mir6955 | 1.342239839 | -0.221484446 | 1.054725023 | -0.209992596 |
| Sh3bp1 | 1228.874401 | 0.45686056 | 0.057914311 | 7.888560674 |
| 1700027A07Rik | 1.879021917 | 0.638343589 | 0.807632303 | 0.790388877 |
| Lgals1 | 413.7512805 | -1.85681904 | 0.328452451 | -5.653235446 |
| Nol12 | 155.5073409 | -0.465121573 | 0.111833845 | -4.159041216 |
| Triobp | 275.7571062 | -0.922148182 | 0.167759956 | -5.496831331 |
| Mir6956 | 0 | NA | NA | NA |
| H1f0 | 1010.011615 | 0.196140131 | 0.102892962 | 1.90625409 |
| Gcat | 70.63897891 | -0.272880087 | 0.152499607 | -1.789382239 |
| Ankrd54 | 161.076979 | -0.102253931 | 0.144823265 | -0.70606011 |
| Eif3l | 629.3207194 | -0.252042428 | 0.076702996 | -3.285952832 |
| Gm25072 | 0 | NA | NA | NA |
| Micall1 | 863.8231058 | -0.415406829 | 0.081631346 | -5.088815162 |
| 1700088E04Rik | 67.10927695 | -0.365877843 | 0.145010777 | -2.523107937 |

| | | | | |
|---|---|---|---|---|
| Polr2f | 144.8684693 | -0.403748187 | 0.192455781 | -2.097875081 |
| Sox10 | 508.9917057 | -0.588691784 | 0.108106655 | -5.445472175 |
| Gm10863 | 17.85705234 | -1.920293943 | 0.308479922 | -6.225020848 |
| Pick1 | 359.5554522 | 0.169236254 | 0.09183314 | 1.842866891 |
| Slc16a8 | 3.295224942 | 3.62352439 | 0.954371579 | 3.796764771 |
| Lgals1-ps2 | 0 | NA | NA | NA |
| Baiap2l2 | 8.450121399 | 0.029132469 | 0.463135994 | 0.062902622 |
| Pla2g6 | 246.7582411 | -0.24128355 | 0.10790197 | -2.236136659 |
| Gm20420 | 0 | NA | NA | NA |
| Maff | 106.5092356 | -0.440132771 | 0.455288227 | -0.966712392 |
| Tmem184b | 1712.559106 | -0.05416638 | 0.12727922 | -0.425571279 |
| Mir1943 | 2.514149661 | 0.440012563 | 0.693104079 | 0.634843418 |
| Csnk1e | 892.3901685 | 0.23936829 | 0.079453678 | 3.012677277 |
| Gm16059 | 0.387099777 | -0.248545523 | 1.888108157 | -0.131637333 |
| AC165278.1 | 0 | NA | NA | NA |
| Kcnj4 | 701.1998511 | 0.472052039 | 0.153441642 | 3.076427196 |
| Kdelr3 | 10.19668411 | -1.305201543 | 0.408436365 | -3.195605617 |
| Ddx17 | 3680.454381 | -0.152169614 | 0.06341734 | -2.399495383 |
| Dmc1 | 9.014701878 | -0.417279421 | 0.411167078 | -1.014865839 |
| Gm27903 | 0.358948382 | 0.931675471 | 1.468536525 | 0.634424446 |
| Fam227a | 83.10424768 | 0.065891544 | 0.254865633 | 0.25853444 |
| Cby1 | 177.468213 | 0.250184044 | 0.109702502 | 2.280568265 |
| Tomm22 | 344.3752573 | -0.241375483 | 0.098206868 | -2.457826914 |
| Josd1 | 713.1554615 | -0.0426024 | 0.08640454 | -0.493057429 |
| Gm26884 | 1.046429311 | -0.717443399 | 1.043692278 | -0.687408937 |
| Gtpbp1 | 638.7878775 | -0.07905282 | 0.062979577 | -1.255213576 |
| n-R5s40 | 1.378629347 | -0.266258438 | 0.955625951 | -0.278622025 |
| Sun2 | 786.962264 | 0.121318367 | 0.071379077 | 1.699634858 |
| Gm16575 | 0.364940181 | -1.473661379 | 1.722384289 | -0.855593835 |
| Gm16576 | 14.37465387 | -0.656089351 | 0.333182504 | -1.969159075 |
| Dnal4 | 230.0228909 | 0.038499474 | 0.086045289 | 0.447432683 |
| Nptxr | 4627.776829 | 0.787187339 | 0.152898783 | 5.148421207 |
| Npcd | 0.498288516 | 2.505826628 | 1.338640699 | 1.871918752 |
| Cbx6 | 4151.173169 | 0.348786767 | 0.088771203 | 3.929053062 |
| Apobec3 | 148.4452752 | -0.796246506 | 0.162254941 | -4.907379098 |
| Cbx7 | 316.6582599 | 0.334337641 | 0.088085652 | 3.795597064 |
| Pdgfb | 148.5994786 | -0.146764645 | 0.160035199 | -0.917077283 |
| Mir7213 | 0 | NA | NA | NA |
| Rpl3 | 1750.858766 | -0.056236021 | 0.095424704 | -0.589323502 |
| Snord83b | 1.863983961 | 0.165099707 | 0.746179256 | 0.221260113 |
| Gm22077 | 0 | NA | NA | NA |
| AC161199.1 | 2.341657818 | -0.628912969 | 0.713534342 | -0.881405326 |
| Gm24204 | 0.553939527 | 0.069464212 | 1.411165536 | 0.049224708 |
| Syngr1 | 4833.272579 | 0.452271597 | 0.060446373 | 7.482195766 |
| Tab1 | 241.9134514 | 0.026631694 | 0.086475234 | 0.30796903 |
| Mgat3 | 1733.281968 | 0.186070606 | 0.071463433 | 2.603717711 |
| Mief1 | 547.7891883 | -0.225653443 | 0.080368416 | -2.807737837 |

| | | | | |
|---|---|---|---|---|
| Atf4 | 1610.207181 | -0.64567136 | 0.157830396 | -4.090918964 |
| Rps19bp1 | 105.055002 | -0.621761051 | 0.15189309 | -4.093412344 |
| Cacna1i | 1385.507454 | 0.205372034 | 0.121335088 | 1.692602176 |
| Enthd1 | 1.187168766 | -0.082760205 | 0.817268506 | -0.101264401 |
| Gm23220 | 0 | NA | NA | NA |
| Grap2 | 30.59568962 | -0.638239481 | 0.268752654 | -2.37482113 |
| Mir6957 | 0 | NA | NA | NA |
| Fam83f | 18.37852399 | -0.890418827 | 0.357012526 | -2.494082872 |
| Tnrc6b | 3449.044664 | 0.073994878 | 0.077950282 | 0.949257358 |
| Mir5113 | 0 | NA | NA | NA |
| Adsl | 284.5366327 | -0.494723146 | 0.077729829 | -6.364649836 |
| Gm6612 | 1.14679352 | -1.145740902 | 1.194994332 | -0.958783545 |
| Sgsm3 | 461.0312359 | -0.112001012 | 0.069150194 | -1.619677491 |
| Mkl1 | 795.5391381 | 0.387122482 | 0.084770174 | 4.56672982 |
| Gm25131 | 0 | NA | NA | NA |
| 4930483J18Rik | 0.130580445 | 0.037055688 | 3.67861741 | 0.010073265 |
| Mchr1 | 166.8774582 | 0.093344272 | 0.187318509 | 0.498318467 |
| Slc25a17 | 303.5824259 | -0.023540913 | 0.097503787 | -0.241435883 |
| St13 | 1980.343047 | -0.164957723 | 0.064822815 | -2.544747904 |
| Xpnpep3 | 220.8414576 | -0.306105206 | 0.110525035 | -2.769555393 |
| Dnajb7 | 0.134148234 | -0.443820953 | 3.624632483 | -0.122445781 |
| Gm17025 | 0.441788749 | 0.522844291 | 1.366993348 | 0.382477568 |
| Rbx1 | 792.4993545 | -0.134488065 | 0.08201528 | -1.639792791 |
| Gm5218 | 0.038981395 | -0.443816764 | 3.724945049 | -0.119147198 |
| 1110025M09Rik | 12.66128553 | -0.296116861 | 0.312701442 | -0.94696353 |
| Ep300 | 2415.831866 | -0.108524901 | 0.063888125 | -1.698670929 |
| Gm23880 | 0 | NA | NA | NA |
| Gm22067 | 12.43328101 | -0.265865885 | 0.317187587 | -0.838197635 |
| L3mbtl2 | 382.5292509 | 0.102759125 | 0.065462922 | 1.569730189 |
| Gm18284 | 12.73109718 | 0.052192768 | 0.295769338 | 0.176464431 |
| Chadl | 117.5180505 | 0.13386753 | 0.135079954 | 0.9910244 |
| Rangap1 | 1965.883829 | -0.176289284 | 0.066257783 | -2.66065774 |
| Zc3h7b | 3000.898726 | 0.213516975 | 0.079749626 | 2.677341386 |
| Gm17597 | 7.602086333 | 0.692729617 | 0.409394819 | 1.692082032 |
| Tef | 3374.752106 | 0.963974998 | 0.107473022 | 8.969460255 |
| Gm8444 | 1.234579709 | 0.606352693 | 0.934269146 | 0.649012862 |
| Tob2 | 667.3737093 | -0.008331636 | 0.111167266 | -0.074946843 |
| Phf5a | 193.4131225 | -0.337439126 | 0.134360571 | -2.51144457 |
| Aco2 | 4367.876781 | 0.235187535 | 0.068204341 | 3.448278099 |
| Polr3h | 303.152543 | -0.084761759 | 0.095944189 | -0.883448592 |
| Csdc2 | 982.7828558 | -0.097859782 | 0.113003535 | -0.865988677 |
| Pmm1 | 652.8758253 | 0.193264306 | 0.058884463 | 3.282093392 |
| Gm5805 | 68.93469726 | -0.115653987 | 0.209968392 | -0.550816178 |
| 1700029P11Rik | 0.075043205 | 0.517957688 | 3.724945049 | 0.139051095 |
| Xrcc6 | 153.2673552 | -0.336738086 | 0.120634179 | -2.791398658 |
| Desi1 | 1377.267412 | -0.221154436 | 0.078846154 | -2.804885536 |
| Nhp2l1 | 547.7572345 | -0.206284621 | 0.105729658 | -1.951057301 |

| | | | | |
|---|---|---|---|---|
| 4930407I10Rik | 0.062426873 | 0.517959912 | 3.724945049 | 0.139051692 |
| Mei1 | 19.77925728 | -0.643999911 | 0.293353542 | -2.195302998 |
| Ccdc134 | 79.33616336 | -0.57200255 | 0.148334157 | -3.85617555 |
| Srebf2 | 2183.707136 | -0.088227972 | 0.079553549 | -1.109038799 |
| Mir33 | 0.155921052 | 1.020628401 | 2.829228019 | 0.360744483 |
| Shisa8 | 125.8398763 | -0.016835643 | 0.204021245 | -0.08251907 |
| Tnfrsf13c | 6.674742414 | -0.12870557 | 0.414224632 | -0.31071443 |
| Cenpm | 13.06399243 | -0.098319962 | 0.315241342 | -0.311887907 |
| 1500009C09Rik | 1247.839228 | 0.093631171 | 0.136554069 | 0.685671045 |
| 3-Sep | 4869.520885 | 0.617379354 | 0.069619379 | 8.867923858 |
| Wbp2nl | 0.428891288 | 0.401490505 | 1.51503468 | 0.265004168 |
| Naga | 263.3133539 | -0.315962916 | 0.119622435 | -2.641334938 |
| Fam109b | 4.886605523 | -0.133820794 | 0.470710883 | -0.284295092 |
| Smdt1 | 561.4413452 | -0.087972369 | 0.109441452 | -0.803830432 |
| n-R5s41 | 0 | NA | NA | NA |
| Ndufa6 | 517.8924798 | -0.043870929 | 0.108145193 | -0.405666932 |
| Cyp2d22 | 207.7231852 | -0.270476499 | 0.134152504 | -2.016186735 |
| Cyp2d11 | 0 | NA | NA | NA |
| Mir7684 | 0 | NA | NA | NA |
| Cyp2d10 | 0 | NA | NA | NA |
| Gm27331 | 0 | NA | NA | NA |
| Gm25990 | 0 | NA | NA | NA |
| Cyp2d9 | 0.049553047 | -0.443818988 | 3.724945049 | -0.119147795 |
| Gm27622 | 0 | NA | NA | NA |
| Gm27825 | 0 | NA | NA | NA |
| Gm5806 | 3.903672454 | -0.054551253 | 0.541154785 | -0.100805268 |
| Gm28023 | 0 | NA | NA | NA |
| Cyp2d12 | 0.066050875 | -0.443818988 | 3.724945049 | -0.119147795 |
| Gm27924 | 0 | NA | NA | NA |
| Gm27742 | 0 | NA | NA | NA |
| Cyp2d34 | 0.18989612 | -0.707205106 | 2.65062973 | -0.266806449 |
| Gm27463 | 0.194279094 | -1.017823543 | 3.719440177 | -0.273649661 |
| Cyp2d37-ps | 0.066050875 | -0.443818988 | 3.724945049 | -0.119147795 |
| Cyp2d40 | 0.493912024 | -0.380945015 | 1.435097491 | -0.265448875 |
| Cyp2d41-ps | 0.567195248 | -0.58455307 | 1.317878194 | -0.44355622 |
| Gm27618 | 0 | NA | NA | NA |
| Cyp2d26 | 2.149324495 | -0.231857935 | 0.636714964 | -0.364147143 |
| Tcf20 | 4578.253049 | 0.398251088 | 0.097279573 | 4.093881938 |
| Gm20324 | 2.615112804 | -0.087332503 | 0.65103483 | -0.134144133 |
| Mir3080 | 2.487744732 | 2.229399348 | 0.859739303 | 2.593110889 |
| Gm29019 | 5.11266864 | 0.900993124 | 0.625151244 | 1.441240231 |
| AW121686 | 10.56313198 | -0.009729945 | 0.358794338 | -0.027118447 |
| Nfam1 | 67.01581601 | -1.033193123 | 0.217749873 | -4.744862107 |
| Serhl | 61.58247766 | -0.195427889 | 0.176726816 | -1.105819104 |
| Gm17206 | 0.066050875 | -0.443818988 | 3.724945049 | -0.119147795 |
| Rrp7a | 918.7979819 | -0.219930504 | 0.075363306 | -2.918270408 |
| Poldip3 | 1119.919797 | -0.241761496 | 0.077244194 | -3.129833908 |

| | | | | |
|---|---|---|---|---|
| Rnu12 | 255.543187 | 0.156988066 | 0.166483255 | 0.9429661 |
| Cyb5r3 | 666.3127489 | -0.452473605 | 0.117123455 | -3.863219419 |
| A4galt | 36.02363458 | -0.861497663 | 0.24575386 | -3.505530552 |
| Arfgap3 | 242.8767015 | -0.411941159 | 0.076969025 | -5.352038172 |
| 1700001L05Rik | 36.6738785 | 0.382745655 | 0.211553911 | 1.809210959 |
| Pacsin2 | 643.9682478 | -0.400125487 | 0.089275144 | -4.481936079 |
| Gm5417 | 28.18046227 | -0.004866365 | 0.252613276 | -0.01926409 |
| Ttll1 | 251.422686 | -0.010041475 | 0.114358291 | -0.087807143 |
| Gm24575 | 0 | NA | NA | NA |
| Bik | 0.708533877 | -1.14152793 | 1.146329889 | -0.995811014 |
| Mcat | 123.3968876 | 0.06901566 | 0.117407951 | 0.587827822 |
| Tspo | 459.8898812 | -1.056283263 | 0.310432738 | -3.402615551 |
| Ttll12 | 215.0437835 | -0.664237025 | 0.113337256 | -5.860712086 |
| Scube1 | 480.4921202 | 0.5706579 | 0.17475496 | 3.265474696 |
| Mpped1 | 824.048876 | 1.084554757 | 0.144734145 | 7.493427043 |
| Efcab6 | 85.58806344 | -0.439168751 | 0.295117442 | -1.488115196 |
| Rpl31-ps22 | 0 | NA | NA | NA |
| Sult4a1 | 2771.319555 | 0.191345947 | 0.086442958 | 2.213551584 |
| Pnpla5 | 0 | NA | NA | NA |
| Pnpla3 | 102.6316791 | 0.52720255 | 0.195778291 | 2.692854995 |
| Mir6392 | 3.244976357 | 0.772190957 | 0.621561438 | 1.242340515 |
| Samm50 | 528.8198655 | -0.014991884 | 0.071116282 | -0.210808043 |
| Parvb | 380.2549727 | -0.618070092 | 0.10133721 | -6.099142598 |
| Gm22890 | 0.046345709 | -0.443818988 | 3.724945049 | -0.119147795 |
| Parvg | 82.64255846 | -0.089495576 | 0.170106201 | -0.526115896 |
| 1810041L15Rik | 649.3280647 | -0.180716465 | 0.12751015 | -1.417271209 |
| Ldoc1l | 614.5341085 | 0.129984669 | 0.083965581 | 1.548070859 |
| Prr5 | 124.3176796 | -0.003156193 | 0.133631454 | -0.02361864 |
| Arhgap8 | 19.60055845 | 0.217879178 | 0.292983527 | 0.743656754 |
| Phf21b | 132.294507 | 0.297274897 | 0.179997323 | 1.651551771 |
| Gm4217 | 13.23963404 | -0.300419379 | 0.360269093 | -0.83387497 |
| Gm29666 | 48.12944454 | 0.408459515 | 0.223950117 | 1.823886141 |
| Nup50 | 602.2802685 | -0.342392435 | 0.082374318 | -4.156543502 |
| 5031439G07Rik | 1832.693521 | -0.289116265 | 0.057203448 | -5.054175448 |
| Mir1249 | 0 | NA | NA | NA |
| Upk3a | 0.636985128 | 0.068360884 | 1.432426234 | 0.047723842 |
| Fam118a | 232.4075255 | -0.220592016 | 0.085040682 | -2.593958694 |
| Smc1b | 5.078404989 | 0.081177455 | 0.47499162 | 0.170902919 |
| Gm10923 | 2.764924986 | -1.014264114 | 0.612344087 | -1.656363041 |
| Ribc2 | 3.994063061 | -0.128686624 | 0.475781387 | -0.270474272 |
| Fbln1 | 204.5288485 | -0.501619821 | 0.176377461 | -2.844013159 |
| Atxn10 | 2123.828845 | -0.097831748 | 0.062681093 | -1.56078561 |
| Gm23517 | 0 | NA | NA | NA |
| 7530416G11Rik | 0 | NA | NA | NA |
| Gm4825 | 0.947696482 | -0.147640379 | 0.914213594 | -0.161494403 |
| Wnt7b | 268.9427193 | -0.221565248 | 0.134537334 | -1.646868128 |
| AU022754 | 9.118481155 | -0.152418259 | 0.368899642 | -0.413169984 |

| | | | | |
|---|---|---|---|---|
| D130051D11Rik | 5.759629403 | 0.354113844 | 0.46531932 | 0.761012555 |
| Gm37699 | 45.08239792 | 0.051780404 | 0.261324169 | 0.198146251 |
| Mirlet7c-2 | 5.369208344 | 1.114439869 | 0.471326618 | 2.364474711 |
| Mirlet7b | 13.61717214 | 0.534151999 | 0.364971113 | 1.463545965 |
| Ppara | 39.64814777 | 0.427137294 | 0.219883486 | 1.942561952 |
| Cdpf1 | 48.27012333 | -0.357142634 | 0.191895184 | -1.86113391 |
| Pkdrej | 11.56756557 | 0.832372027 | 0.321732342 | 2.587156829 |
| Ttc38 | 59.42181862 | 0.816161132 | 0.156746502 | 5.206885769 |
| Gtse1 | 17.82322006 | -0.502189815 | 0.301973721 | -1.663024892 |
| Trmu | 131.3669997 | 0.146511656 | 0.126839911 | 1.155091132 |
| Celsr1 | 124.8124239 | 0.190757252 | 0.159441144 | 1.196411707 |
| Gramd4 | 450.6965781 | -0.317911405 | 0.071975423 | -4.416943895 |
| Gm15722 | 0 | NA | NA | NA |
| Cerk | 1311.25289 | -0.208550466 | 0.08369558 | -2.491773944 |
| Gm24011 | 0 | NA | NA | NA |
| Gm15569 | 1.568615446 | -0.263555833 | 0.894931697 | -0.294498266 |
| Gm22818 | 0.772793883 | 0.929619757 | 1.288612545 | 0.721411382 |
| Tbc1d22a | 375.15338 | 0.135809932 | 0.070709777 | 1.920666941 |
| Gm23416 | 0.044544648 | 0.517957688 | 3.724945049 | 0.139051095 |
| n-R5s42 | 0 | NA | NA | NA |
| Fam19a5 | 1108.298912 | 0.227119818 | 0.076702157 | 2.961061683 |
| Mir6393 | 0 | NA | NA | NA |
| Zdhhc25 | 0 | NA | NA | NA |
| Brd1 | 988.6755636 | 0.121290005 | 0.115351143 | 1.051485074 |
| Zbed4 | 103.903103 | -0.243393228 | 0.1147895 | -2.120344006 |
| Alg12 | 119.1075428 | -0.362001671 | 0.113910522 | -3.17794761 |
| Creld2 | 178.5176635 | -1.109700204 | 0.161433654 | -6.874032604 |
| Gm23144 | 0 | NA | NA | NA |
| Pim3 | 592.2908598 | -0.240520196 | 0.171297561 | -1.404107534 |
| Gm20621 | 2.301900616 | -0.103363896 | 0.71555435 | -0.144452893 |
| Ttll8 | 19.07562425 | 0.634449445 | 0.383271002 | 1.655354676 |
| Mlc1 | 1260.237927 | -0.041798802 | 0.129982633 | -0.321572204 |
| Mov10l1 | 0.232306593 | -0.924714546 | 2.191977153 | -0.421863223 |
| Gm23737 | 0 | NA | NA | NA |
| Panx2 | 872.8564831 | 0.441461963 | 0.100681443 | 4.384740141 |
| 1810021B22Rik | 9.773541415 | 0.145261878 | 0.341282548 | 0.425635235 |
| Trabd | 275.7218993 | -0.312629745 | 0.091517433 | -3.416067678 |
| Selo | 248.0954824 | 0.265191411 | 0.092138643 | 2.878177968 |
| Tubgcp6 | 650.1368911 | 0.058581671 | 0.084512226 | 0.693173924 |
| Hdac10 | 82.94525066 | 0.102841963 | 0.161356938 | 0.637356931 |
| Mapk12 | 96.05029267 | 0.438981016 | 0.1734049 | 2.531537559 |
| Mapk11 | 187.7470408 | 0.028862849 | 0.156927355 | 0.183924904 |
| Plxnb2 | 1637.125997 | -0.738445481 | 0.125799814 | -5.87000457 |
| mmu-mir-7118 | 0 | NA | NA | NA |
| Dennd6b | 661.1196248 | -0.058492912 | 0.18238003 | -0.320719936 |
| Mir6958 | 0 | NA | NA | NA |
| Gm26798 | 0.47411084 | 1.34495199 | 1.349314033 | 0.996767214 |

| | | | | |
|---|---|---|---|---|
| Ppp6r2 | 1298.582558 | 0.36300835 | 0.054719213 | 6.634019915 |
| Sbf1 | 2446.17373 | 0.102380111 | 0.053088447 | 1.928481937 |
| Mir6959 | 1.302949929 | 1.821999938 | 0.969730667 | 1.878872144 |
| Adm2 | 0.810290415 | -1.91240039 | 1.174953629 | -1.627639034 |
| Miox | 0.11193333 | 0.037073798 | 3.724945049 | 0.009952844 |
| Lmf2 | 236.2729926 | -0.567767897 | 0.115621872 | -4.910557896 |
| Ncaph2 | 806.3681184 | -0.041741287 | 0.063906425 | -0.653162607 |
| Sco2 | 38.3484847 | 0.109609944 | 0.181120232 | 0.605177801 |
| Tymp | 10.59392881 | -0.670556501 | 0.334744616 | -2.00318831 |
| Odf3b | 13.0856276 | 0.224976044 | 0.310582258 | 0.72436863 |
| Klhdc7b | 7.376514687 | -0.211403913 | 0.434559713 | -0.486478398 |
| Syce3 | 0 | NA | NA | NA |
| Chkb | 231.7816191 | 0.077122964 | 0.096861554 | 0.796218533 |
| Cpt1b | 2.011645178 | -0.082829504 | 0.737422659 | -0.112322972 |
| C730034F03Rik | 13.21654071 | -0.930877868 | 0.294956986 | -3.155978368 |
| Mapk8ip2 | 3763.788627 | 0.299917257 | 0.055084609 | 5.444665293 |
| Arsa | 324.9924385 | -0.000210549 | 0.099089798 | -0.002124831 |
| C230037L18Rik | 26.54613617 | 0.887938624 | 0.302342139 | 2.936866909 |
| Shank3 | 2814.903766 | 0.682009534 | 0.133465174 | 5.110018705 |
| Gm15609 | 24.97572873 | 0.491015962 | 0.365930637 | 1.341827966 |
| Acr | 23.80281401 | 1.930529478 | 0.438507156 | 4.402503932 |
| Rabl2 | 143.4438333 | 0.016123151 | 0.137751015 | 0.117045609 |
| Syt10 | 129.1730868 | 0.14113889 | 0.181117659 | 0.779266314 |
| Gm24643 | 0 | NA | NA | NA |
| Gm24415 | 0 | NA | NA | NA |
| Alg10b | 588.5402971 | 0.103507673 | 0.081279643 | 1.27347598 |
| Mir7648 | 0 | NA | NA | NA |
| Cpne8 | 261.9200795 | 0.102221472 | 0.107273696 | 0.952903418 |
| Gm6746 | 0 | NA | NA | NA |
| Rpl31-ps8 | 113.0537037 | -0.056406529 | 0.241329175 | -0.23373274 |
| Tcea1-ps1 | 127.5401109 | 0.280119506 | 0.110167946 | 2.542658884 |
| Kif21a | 3097.142762 | 0.106150611 | 0.060562868 | 1.752734214 |
| Abcd2 | 509.0775856 | 0.700311128 | 0.099835689 | 7.01463712 |
| CN725425 | 2.738760097 | 0.330288056 | 0.736306453 | 0.44857417 |
| Slc2a13 | 1660.411825 | 0.405156759 | 0.101232475 | 4.002240963 |
| Gm15383 | 16.38285574 | 0.689361075 | 0.418292442 | 1.64803617 |
| Gm15381 | 25.44622861 | 1.094435249 | 0.32787559 | 3.337958919 |
| Gm24647 | 4.156244294 | 1.505539831 | 0.694448333 | 2.167965218 |
| 4933438A12Rik | 0.060174537 | 0.517957688 | 3.724945049 | 0.139051095 |
| Gm25389 | 0 | NA | NA | NA |
| Lrrk2 | 1067.387671 | -0.18412529 | 0.139900813 | -1.316113081 |
| Gm15382 | 4.663180715 | -0.816520624 | 0.53901827 | -1.514829216 |
| Smgc | 0.890372413 | -2.191295666 | 1.220924081 | -1.794784541 |
| Muc19 | 0.988759962 | -2.304973281 | 1.226096598 | -1.879927964 |
| Cntn1 | 4874.212289 | 0.61283426 | 0.067063503 | 9.138118785 |
| Gm26760 | 2.85183316 | 0.866768782 | 0.672031645 | 1.289773761 |
| Pdzrn4 | 94.36282265 | 0.261429611 | 0.121237073 | 2.1563504 |

| | | | | |
|---|---|---|---|---|
| Gm37304 | 37.54844209 | 0.857497941 | 0.327061516 | 2.621824635 |
| Gm4335 | 6.712093293 | 0.132767918 | 0.445480988 | 0.298032736 |
| Gm24447 | 11246.06972 | 0.038377164 | 0.147229513 | 0.26066217 |
| Gxylt1 | 308.8367025 | 0.182882087 | 0.099429955 | 1.839305741 |
| Yaf2 | 648.3196561 | 0.273115028 | 0.076664807 | 3.562456371 |
| Gm26371 | 0.04393175 | 0.037062677 | 3.724945049 | 0.009949859 |
| Zcrb1 | 380.1198483 | 0.083210414 | 0.124754996 | 0.666990631 |
| Pphln1 | 620.7309094 | -0.070699106 | 0.056174725 | -1.258557233 |
| Prickle1 | 831.5948449 | 0.260772851 | 0.108066229 | 2.413083659 |
| Adamts20 | 204.9752347 | 0.866506258 | 0.203117721 | 4.266029843 |
| Pus7l | 33.70876001 | -0.184352977 | 0.191105388 | -0.964666557 |
| Irak4 | 54.36204711 | -0.910128402 | 0.233422043 | -3.899067931 |
| Twf1 | 752.74737 | -0.425461906 | 0.082853014 | -5.135140952 |
| Tmem117 | 131.7499303 | -0.213781682 | 0.106921879 | -1.999419437 |
| Gm25546 | 0 | NA | NA | NA |
| Gm23129 | 0 | NA | NA | NA |
| 1700129L04Rik | 4.754384054 | 0.377967545 | 0.650868091 | 0.580712974 |
| Nell2 | 3266.316947 | 0.900635534 | 0.123401983 | 7.29838787 |
| Gm24668 | 0 | NA | NA | NA |
| Dbx2 | 163.3891668 | -0.517538343 | 0.150101156 | -3.447930426 |
| Gm8849 | 1.166878852 | 0.660236563 | 0.921984198 | 0.716103991 |
| A130051J06Rik | 4.097955438 | -1.024810546 | 0.506486497 | -2.023371901 |
| Ano6 | 622.2228295 | -0.635950598 | 0.100296354 | -6.34071504 |
| Gm17546 | 0.627185525 | -0.335675529 | 1.211162171 | -0.277151596 |
| Gm25070 | 0.044544648 | 0.517957688 | 3.724945049 | 0.139051095 |
| D030018L15Rik | 0.718519848 | -1.721429368 | 1.258460995 | -1.367884562 |
| 2610037D02Rik | 115.5236633 | -0.091047144 | 0.229785447 | -0.396226766 |
| 4833422M21Rik | 10.72830272 | -0.691828869 | 0.369010837 | -1.874819922 |
| Arid2 | 997.3079313 | -0.070066095 | 0.070491443 | -0.99396597 |
| Gm25397 | 0 | NA | NA | NA |
| Scaf11 | 1520.248971 | -0.030135187 | 0.067434237 | -0.446882605 |
| Slc38a1 | 3775.966176 | -0.003930445 | 0.048410719 | -0.081189553 |
| Gm22045 | 0 | NA | NA | NA |
| Slc38a2 | 2617.581037 | -0.175752498 | 0.076821551 | -2.287802002 |
| Gm38144 | 0.240186774 | -0.466606409 | 2.110992137 | -0.221036545 |
| Gm8888 | 0 | NA | NA | NA |
| Slc38a4 | 9.499651179 | 0.491247078 | 0.368399252 | 1.333463829 |
| Amigo2 | 212.2260951 | 0.478929352 | 0.167380824 | 2.861315528 |
| Pced1b | 100.5813245 | 0.367538287 | 0.129732456 | 2.833048087 |
| Rpap3 | 291.5755603 | 0.15342089 | 0.078300631 | 1.959382545 |
| Endou | 46.06882177 | 0.272060774 | 0.199634544 | 1.362794078 |
| Rapgef3 | 390.02648 | 0.276349875 | 0.114613381 | 2.411148442 |
| Rapgef3os1 | 0.072025462 | 0.517959912 | 3.724945049 | 0.139051692 |
| Rapgef3os2 | 3.594151187 | 1.262457283 | 0.597376575 | 2.11333577 |
| Slc48a1 | 1148.386551 | -0.100728106 | 0.077579696 | -1.298382334 |
| Hdac7 | 379.135267 | -0.020717601 | 0.164660511 | -0.125820093 |
| Vdr | 23.71495927 | -0.796203123 | 0.289017572 | -2.754860605 |

| | | | | |
|---|---|---|---|---|
| Gm22202 | 0 | NA | NA | NA |
| Tmem106c | 169.9404391 | -0.158861343 | 0.145343715 | -1.093004554 |
| Col2a1 | 22.89251041 | -0.346951373 | 0.298713872 | -1.161483968 |
| Senp1 | 454.6842185 | -0.215197064 | 0.078583021 | -2.738467709 |
| Gm25668 | 0.044544648 | 0.517957688 | 3.724945049 | 0.139051095 |
| Pfkm | 3358.190513 | 0.138987029 | 0.06507177 | 2.135903607 |
| Gm26513 | 7.485808625 | 0.380026749 | 0.408510277 | 0.930274636 |
| Asb8 | 417.0210594 | 0.023287245 | 0.095755473 | 0.243194928 |
| Ccdc184 | 256.2935464 | -0.194364354 | 0.127874679 | -1.519959665 |
| Olfr288 | 1.13985991 | 1.336329844 | 1.011841334 | 1.3206911 |
| Olfr287 | 28.52921329 | 0.622096436 | 0.242452268 | 2.565851179 |
| Olfr286 | 0.16123844 | 0.037068524 | 2.802306608 | 0.013227862 |
| H1fnt | 0 | NA | NA | NA |
| Zfp641 | 111.9084074 | 0.101400167 | 0.130378495 | 0.777736903 |
| Olfr285 | 0 | NA | NA | NA |
| Olfr284 | 0 | NA | NA | NA |
| Olfr283 | 0.082913145 | -0.443821212 | 3.724945049 | -0.119148392 |
| 1700031M16Rik | 0.045882455 | 0.517959912 | 3.724945049 | 0.139051692 |
| Olfr282 | 0 | NA | NA | NA |
| Olfr281 | 0.061920613 | -0.443816764 | 3.724945049 | -0.119147198 |
| Lalba | 0.845223607 | -3.464989721 | 1.150817669 | -3.010893744 |
| Olfr279 | 0.133280846 | 0.915483429 | 3.652577767 | 0.250640366 |
| Kansl2 | 403.2058431 | -0.195454499 | 0.075337928 | -2.594370496 |
| Snora34 | 1.248930875 | 0.206282187 | 0.918177975 | 0.224664708 |
| mmu-mir-1291 | 1.156326783 | -0.390762251 | 1.141070856 | -0.342452223 |
| Snora2b | 4.632530903 | -0.020491393 | 0.546370209 | -0.037504594 |
| Ccnt1 | 886.6569998 | -0.09859517 | 0.046825327 | -2.105594899 |
| 9330020H09Rik | 18.2002576 | 0.391137477 | 0.293471613 | 1.332794927 |
| 4930415O20Rik | 0.192434788 | -0.97914261 | 2.651844871 | -0.369230727 |
| Gm29331 | 0.089062654 | -0.443823436 | 3.724945049 | -0.119148989 |
| Adcy6 | 518.3618143 | -0.040820291 | 0.098299831 | -0.415263085 |
| Cacnb3 | 1158.604023 | 0.749311941 | 0.097824524 | 7.659755561 |
| Ddx23 | 691.4598963 | -0.038067251 | 0.093468084 | -0.407275392 |
| Rnd1 | 252.5003933 | -0.294327735 | 0.218523352 | -1.346893741 |
| Ccdc65 | 49.36712147 | 0.137088044 | 0.159470568 | 0.859644795 |
| Fkbp11 | 8.263769739 | -0.461362189 | 0.400215402 | -1.152784691 |
| Arf3 | 8081.452134 | 0.475365683 | 0.068193542 | 6.970831364 |
| Wnt10b | 8.994095922 | 0.938556005 | 0.489874243 | 1.915912133 |
| Wnt1 | 0.424046041 | -0.762726955 | 1.583228387 | -0.481754219 |
| Ddn | 6153.970663 | 1.397935881 | 0.183576734 | 7.614994823 |
| B130046B21Rik | 36.69080057 | 0.576295476 | 0.252943213 | 2.278359123 |
| Prkag1 | 322.4183075 | -0.140075129 | 0.085120092 | -1.64561769 |
| Kmt2d | 5538.759192 | 0.076440213 | 0.082276408 | 0.929066003 |
| Rhebl1 | 33.80626636 | 0.243870472 | 0.189973055 | 1.283710853 |
| Dhh | 5.749732932 | -0.922036454 | 0.419015894 | -2.200480855 |
| Lmbr1l | 116.8431874 | -0.071274605 | 0.156328914 | -0.455927209 |
| Tuba1b | 2389.31285 | -0.231371268 | 0.133655529 | -1.731101355 |

| | | | | |
|---|---|---|---|---|
| Tuba1a | 4753.134624 | 0.001329497 | 0.116007935 | 0.011460395 |
| 4930578M01Rik | 10.95583967 | 0.443093694 | 0.393622085 | 1.125683011 |
| Gm8973 | 0.060174537 | 0.517957688 | 3.724945049 | 0.139051095 |
| Tuba1c | 280.0639814 | -0.847831154 | 0.188619676 | -4.494924248 |
| Prph | 45.07156546 | -0.532862539 | 0.331669285 | -1.606608036 |
| Troap | 4.659496152 | -0.189197806 | 0.592672456 | -0.319228275 |
| Mir6960 | 0 | NA | NA | NA |
| C1ql4 | 6.463082896 | -0.1798816 | 0.417441229 | -0.430914791 |
| Dnajc22 | 1.159556269 | 2.396727621 | 1.056501112 | 2.268551916 |
| Spats2 | 344.7749557 | 0.091030982 | 0.064937223 | 1.401830537 |
| Gm25183 | 0 | NA | NA | NA |
| Gm25057 | 0 | NA | NA | NA |
| Kcnh3 | 619.2177828 | 1.450947849 | 0.167217779 | 8.676995119 |
| Mcrs1 | 447.4221575 | 0.117097404 | 0.083889462 | 1.395853571 |
| 1700120C14Rik | 12.4281098 | 0.018038669 | 0.328792166 | 0.05486344 |
| Gm25999 | 0 | NA | NA | NA |
| Fam186b | 17.25303227 | 0.514756673 | 0.288719037 | 1.782898278 |
| Prpf40b | 472.5579373 | 0.078556301 | 0.133155254 | 0.589960204 |
| Fmnl3 | 135.4055067 | -0.134863001 | 0.145958574 | -0.923981353 |
| Gm22951 | 0.744072464 | -0.810796968 | 1.400346767 | -0.578997279 |
| Tmbim6 | 4439.906879 | -0.360762615 | 0.122709611 | -2.93997033 |
| Nckap5l | 263.9144233 | 0.403933153 | 0.108473955 | 3.723780072 |
| Nckap5los | 0.092140247 | 0.037060453 | 3.724945049 | 0.009949262 |
| Bcdin3d | 22.52486403 | -0.454408984 | 0.222589503 | -2.041466365 |
| Faim2 | 4926.385511 | 0.507874338 | 0.068501878 | 7.414020687 |
| Gm5808 | 0.066050875 | -0.443818988 | 3.724945049 | -0.119147795 |
| Aqp2 | 0.235388486 | -0.438161674 | 2.118161588 | -0.206859418 |
| Aqp5 | 2.102526643 | -0.4660099 | 0.782960922 | -0.59518922 |
| Aqp6 | 109.2123799 | -0.162419552 | 0.22718369 | -0.714926111 |
| Racgap1 | 122.2464084 | -0.132727436 | 0.12815829 | -1.035652367 |
| Asic1 | 756.9424049 | -0.030743077 | 0.072074625 | -0.426545083 |
| Gm17349 | 9.498973506 | -0.172149692 | 0.353542672 | -0.486927621 |
| Smarcd1 | 1391.542852 | 0.418631513 | 0.08186967 | 5.113389508 |
| Gm16537 | 2.123900911 | 0.282284615 | 0.805910284 | 0.35026804 |
| Gpd1 | 1414.234084 | -0.480042032 | 0.104083806 | -4.612072246 |
| Cox14 | 343.5443005 | -0.117438734 | 0.093656401 | -1.253931742 |
| Cers5 | 400.0721546 | -0.153489665 | 0.07431376 | -2.06542725 |
| Gm17058 | 0.41865154 | 0.088111867 | 1.527365219 | 0.0576888 |
| Lima1 | 217.8937591 | -0.53065429 | 0.110725453 | -4.792523086 |
| Gm25897 | 0.60093359 | 1.230410823 | 1.236147746 | 0.995359031 |
| Gm4468 | 0.964311207 | 0.151420453 | 1.045245029 | 0.144865987 |
| Gm17057 | 0.099106094 | -0.93550548 | 3.724188147 | -0.251197158 |
| Gm9763 | 0.448689577 | 0.448917597 | 1.349378847 | 0.332684626 |
| Fam186a | 4.175781228 | 0.330296069 | 0.5855334 | 0.564094326 |
| Gm21917 | 0.34674037 | 0.517975826 | 1.587424757 | 0.326299451 |
| Gm23133 | 0 | NA | NA | NA |
| Larp4 | 1360.21083 | -0.130530202 | 0.084978907 | -1.536030602 |

| | | | | |
|---|---|---|---|---|
| 2310068J16Rik | 5.082496618 | 1.178851253 | 0.491172531 | 2.400075692 |
| Dip2b | 2171.940491 | 0.020607704 | 0.045643446 | 0.451493162 |
| Atf1 | 263.8766334 | -0.373140249 | 0.127012804 | -2.93781602 |
| Tmprss12 | 0.281502737 | -0.926775555 | 2.063470175 | -0.449134456 |
| Mettl7a1 | 199.3407439 | -0.202897684 | 0.218275845 | -0.929547126 |
| Mettl7a3 | 1.559777635 | 1.669965895 | 0.918214011 | 1.81871097 |
| Methig1 | 0 | NA | NA | NA |
| Mettl7a2 | 0.046345709 | -0.443818988 | 3.724945049 | -0.119147795 |
| Higd1c | 0 | NA | NA | NA |
| Slc11a2 | 350.6928657 | -0.153528817 | 0.097183758 | -1.579778547 |
| Gm5475 | 0.572323712 | 0.323528054 | 1.412308208 | 0.229077515 |
| n-R5s43 | 0 | NA | NA | NA |
| 5330439K02Rik | 13.54861615 | -0.194434964 | 0.293239931 | -0.663057599 |
| Letmd1 | 357.2468629 | -0.016701964 | 0.081327229 | -0.205367429 |
| Csrnp2 | 483.3671844 | 0.282120472 | 0.129137319 | 2.184654857 |
| Tfcp2 | 253.237872 | -0.205029759 | 0.111878675 | -1.832608044 |
| Pou6f1 | 1003.020766 | 0.244108968 | 0.081680867 | 2.988569736 |
| Gm20680 | 0.523790777 | 0.226798438 | 1.333836788 | 0.170034625 |
| Gm27209 | 0.085433889 | 0.037058229 | 3.724945049 | 0.009948665 |
| C330013E15Rik | 16.45784469 | 0.564571818 | 0.27192399 | 2.076211877 |
| Dazap2 | 1246.964278 | -0.105662144 | 0.107559896 | -0.982356323 |
| Smagp | 10.67246666 | -0.864515365 | 0.317075566 | -2.726527859 |
| Bin2 | 77.96195219 | -0.406138941 | 0.202872828 | -2.001938582 |
| Mir6961 | 0 | NA | NA | NA |
| Cela1 | 27.94372578 | -1.210712208 | 0.24726585 | -4.896398782 |
| Galnt6 | 328.7283006 | -0.835281057 | 0.188967767 | -4.420230338 |
| Galnt6os | 0.119234754 | 0.51795324 | 3.724945049 | 0.139049901 |
| Slc4a8 | 2144.399781 | 0.266063771 | 0.083105669 | 3.201511675 |
| Scn8a | 4962.265033 | 0.674899937 | 0.113685578 | 5.936548417 |
| Fignl2 | 63.27470895 | -0.255191257 | 0.165711673 | -1.539971525 |
| Ankrd33 | 0.864742418 | 0.05357805 | 1.143734266 | 0.046844841 |
| Acvrl1 | 107.1769413 | -0.802865803 | 0.138043488 | -5.816035325 |
| Gm16031 | 0.097139547 | -0.443816764 | 3.724945049 | -0.119147198 |
| Acvr1b | 1299.302991 | 0.366722942 | 0.066369155 | 5.52550269 |
| mmu-mir-6962 | 0.93249843 | -1.635829965 | 1.172289522 | -1.395414643 |
| A330009N23Rik | 60.10913543 | -0.21183751 | 0.184384974 | -1.148887059 |
| Grasp | 286.1441832 | -0.264923232 | 0.216973243 | -1.220994943 |
| Nr4a1 | 763.9124175 | -0.182378456 | 0.422952847 | -0.431202809 |
| Atg101 | 182.2922892 | -0.182530356 | 0.108101345 | -1.688511422 |
| 6030408B16Rik | 2.433334847 | -0.326376457 | 0.683527353 | -0.477488509 |
| Krt80 | 28.28862397 | -0.776890901 | 0.219111057 | -3.545649001 |
| Mir1941 | 0.066050875 | -0.443818988 | 3.724945049 | -0.119147795 |
| Krt7 | 2.609690864 | -0.868181932 | 0.666842014 | -1.301930463 |
| Krt83 | 0.375360954 | 1.603652888 | 1.542600511 | 1.039577568 |
| 1700011A15Rik | 0.04393175 | 0.037062677 | 3.724945049 | 0.009949859 |
| Krt81 | 0 | NA | NA | NA |
| Krt86 | 0.614518174 | 0.320485753 | 1.300204376 | 0.246488752 |

| | | | | |
|---|---|---|---|---|
| 5430421N21Rik | 0.095897396 | 0.037062677 | 3.724945049 | 0.009949859 |
| Krt84 | 0 | NA | NA | NA |
| Krt82 | 0 | NA | NA | NA |
| 4732456N10Rik | 0.738905749 | -0.513428614 | 1.561940182 | -0.328712085 |
| AC103674.1 | 0 | NA | NA | NA |
| Krt75 | 1.991379766 | -4.457392745 | 1.021278255 | -4.364523307 |
| AC103674.2 | 0 | NA | NA | NA |
| Gm5414 | 0 | NA | NA | NA |
| Gm5478 | 0 | NA | NA | NA |
| AC120787.3 | 0 | NA | NA | NA |
| Krt6b | 0.519670895 | -0.711870126 | 1.697709574 | -0.419312076 |
| AC120787.2 | 0 | NA | NA | NA |
| Krt6a | 4.257533425 | -5.007299129 | 1.060915128 | -4.719792372 |
| AC120787.4 | 0 | NA | NA | NA |
| Krt5 | 1.350984256 | 0.849351575 | 0.957487841 | 0.887062518 |
| AC120787.1 | 0 | NA | NA | NA |
| Krt71 | 0.51940401 | 0.452339299 | 1.469289811 | 0.307862544 |
| Krt74 | 0 | NA | NA | NA |
| Krt72 | 0.233626327 | 0.942025819 | 3.105378667 | 0.303352963 |
| Krt73 | 9.672351053 | -0.706602456 | 0.374098983 | -1.888811486 |
| Krt2 | 16.20338829 | 1.830456236 | 0.858575563 | 2.131968711 |
| Krt1 | 24.4159562 | 0.471763383 | 0.312137559 | 1.511395761 |
| Krt77 | 35.13814104 | 1.188089674 | 0.301691996 | 3.938088152 |
| Krt76 | 0.099106094 | -0.93550548 | 3.724188147 | -0.251197158 |
| AC139844.1 | 0 | NA | NA | NA |
| Krt4 | 0.254192696 | 1.482809323 | 2.470011019 | 0.600324983 |
| Krt79 | 0.300336086 | -0.088359302 | 3.716827973 | -0.023772772 |
| Krt78 | 0.207810967 | 0.853672974 | 2.593681201 | 0.32913566 |
| Krt8 | 17.9984126 | -0.969127523 | 0.351801495 | -2.754756697 |
| Krt18 | 21.72902479 | -0.313563443 | 0.33490767 | -0.936268325 |
| Eif4b | 2876.650182 | 0.209216363 | 0.058478933 | 3.577636456 |
| Tns2 | 268.4246283 | 0.347272232 | 0.121932144 | 2.848077797 |
| Spryd3 | 1083.158277 | 0.174704586 | 0.068933356 | 2.534398381 |
| Igfbp6 | 171.4122214 | -0.16483552 | 0.115459687 | -1.427645657 |
| Soat2 | 21.31164465 | -1.21433355 | 0.29333377 | -4.139767304 |
| Csad | 172.5508944 | 0.519990738 | 0.107753654 | 4.825736469 |
| Zfp740 | 877.8255617 | 0.153656818 | 0.067149645 | 2.288274474 |
| Itgb7 | 116.7476434 | -1.997860193 | 0.376130847 | -5.31160954 |
| Rarg | 131.1736488 | -0.351635402 | 0.116905711 | -3.007854769 |
| Gm9918 | 0.827590884 | 0.474490803 | 1.260475585 | 0.376437916 |
| Mfsd5 | 351.832301 | -0.334394045 | 0.102356457 | -3.266956029 |
| Espl1 | 27.92326781 | -0.683011946 | 0.30041763 | -2.273541489 |
| Pfdn5 | 660.9445207 | 0.054731739 | 0.094496108 | 0.579195694 |
| Myg1 | 214.9838279 | -0.235102955 | 0.090945506 | -2.585097001 |
| Aaas | 128.6515988 | -0.105981172 | 0.170522658 | -0.621507859 |
| Sp7 | 27.79878053 | -1.067761769 | 0.358280074 | -2.98024324 |
| Gm23590 | 0 | NA | NA | NA |

| | | | | |
|---|---|---|---|---|
| Sp1 | 861.7402076 | -0.284564258 | 0.097851957 | -2.908110021 |
| Amhr2 | 7.396618114 | -1.168980171 | 0.428475389 | -2.728231776 |
| Gm26518 | 0.703641397 | -0.782696528 | 1.284997924 | -0.609103341 |
| Gm24128 | 0.061920613 | -0.443816764 | 3.724945049 | -0.119147198 |
| Prr13 | 456.4408238 | -0.458118359 | 0.113858094 | -4.023590621 |
| Pcbp2 | 2462.12792 | -0.074656129 | 0.071026637 | -1.051100443 |
| Gm27406 | 2.84356551 | 0.552890252 | 0.557623627 | 0.991511523 |
| Map3k12 | 1206.534751 | 0.335249272 | 0.068267408 | 4.910824668 |
| Gm10337 | 2.690945547 | 0.367041656 | 0.751737709 | 0.488257609 |
| Tarbp2 | 121.0001225 | 0.087800382 | 0.11836823 | 0.741756309 |
| Npff | 8.501785748 | -0.10814631 | 0.596830777 | -0.181200961 |
| Gm28047 | 0 | NA | NA | NA |
| Atf7 | 596.7832338 | -0.180968554 | 0.069958695 | -2.586791435 |
| Gm25141 | 0 | NA | NA | NA |
| Rpl39-ps | 40.47241404 | -0.588270342 | 0.24150187 | -2.435883179 |
| Atp5g2 | 389.3075053 | -0.059144332 | 0.113658967 | -0.520366611 |
| Mir688 | 0 | NA | NA | NA |
| Calcoco1 | 660.9432622 | 0.322076444 | 0.096785319 | 3.327740675 |
| Hoxc13 | 0.157352566 | -0.41047621 | 2.808585592 | -0.146150508 |
| Hoxc12 | 0 | NA | NA | NA |
| Hotair | 0 | NA | NA | NA |
| Gm27337 | 0 | NA | NA | NA |
| Gm27443 | 0 | NA | NA | NA |
| Gm28011 | 0 | NA | NA | NA |
| Gm27535 | 0 | NA | NA | NA |
| Hoxc11 | 0 | NA | NA | NA |
| Hoxc10 | 0 | NA | NA | NA |
| Hoxc5 | 4.717838401 | -1.275381716 | 1.249587897 | -1.02064186 |
| Mir196a-2 | 0 | NA | NA | NA |
| Hoxc9 | 0.095435502 | 0.037073798 | 3.724945049 | 0.009952844 |
| Hoxc8 | 0.100902008 | -0.876201588 | 3.72093227 | -0.235479048 |
| Hoxc6 | 1.521766428 | -2.231108866 | 2.082402515 | -1.071410954 |
| Mir615 | 0 | NA | NA | NA |
| Hoxc4 | 16.48418288 | -0.570771478 | 1.027744881 | -0.555362998 |
| 5730585A16Rik | 0.314534502 | -0.443831296 | 3.724945049 | -0.119151099 |
| Gm28876 | 4.308624721 | -0.536419028 | 0.539959887 | -0.993442368 |
| Gm10830 | 0.40206694 | -0.903772392 | 1.57328677 | -0.574448606 |
| Gm28265 | 0.496950623 | 1.386148872 | 1.74729282 | 0.793312292 |
| D930007P13Rik | 1.440959972 | 1.164751709 | 1.131816473 | 1.02909945 |
| Smug1 | 140.5464891 | -0.282605581 | 0.120838432 | -2.338706123 |
| Cbx5 | 3979.323211 | 0.043983402 | 0.065460697 | 0.671905498 |
| Hnrnpa1 | 1468.269564 | 0.054102481 | 0.093301132 | 0.5798695 |
| Nfe2 | 3.16484666 | -2.544399769 | 0.696739924 | -3.651864461 |
| Copz1 | 570.9806916 | -0.294007222 | 0.092454829 | -3.180009348 |
| Mir148b | 1.892187285 | 1.287397697 | 0.80378727 | 1.6016647 |
| Gpr84 | 34.27634392 | 1.291989578 | 0.3072645 | 4.204812399 |
| Zfp385a | 1024.49315 | -0.044065792 | 0.082485687 | -0.534223491 |

| | | | | |
|---|---|---|---|---|
| Itga5 | 497.2109644 | -1.568541026 | 0.335828754 | -4.670657309 |
| Mir6963 | 0.095435502 | 0.037073798 | 3.724945049 | 0.009952844 |
| Gm37322 | 0.248806291 | -0.926001524 | 2.141483241 | -0.432411287 |
| Gtsf1 | 0.666252615 | -1.180334058 | 1.262400056 | -0.934992083 |
| BC048502 | 0.169045294 | 1.022313391 | 3.40049023 | 0.300637062 |
| Nckap1l | 484.3215996 | -0.854310061 | 0.141943989 | -6.01864206 |
| Pde1b | 1849.40123 | 0.688373001 | 0.124863637 | 5.512998183 |
| Ppp1r1a | 376.4709638 | 0.265563341 | 0.137433301 | 1.932307088 |
| Glycam1 | 0.54398204 | 1.667186 | 1.31310664 | 1.269650118 |
| Mucl1 | 0.04393175 | 0.037062677 | 3.724945049 | 0.009949859 |
| Gm22852 | 0 | NA | NA | NA |
| Spt1 | 0.136548098 | 0.517963397 | 2.945560596 | 0.175845439 |
| Gm23215 | 0 | NA | NA | NA |
| Gm22862 | 0 | NA | NA | NA |
| Gm22598 | 0 | NA | NA | NA |
| Olfr161 | 0 | NA | NA | NA |
| Mefv | 48.54285063 | -1.57617814 | 0.519577153 | -3.033578616 |
| Tdgf1-ps1 | 0.267098211 | -0.393443719 | 2.459809065 | -0.159948886 |
| Zfp263 | 304.9461567 | -0.125212318 | 0.105829567 | -1.183150628 |
| Olfr15 | 0.061920613 | -0.443816764 | 3.724945049 | -0.119147198 |
| Zfp174 | 93.47116396 | 0.140082373 | 0.177175863 | 0.790640278 |
| Zfp597 | 424.0273003 | -0.120158822 | 0.106522728 | -1.128011117 |
| Gm26675 | 8.550105792 | -0.51794061 | 0.449091959 | -1.153306355 |
| Naa60 | 734.2670764 | 0.127448552 | 0.065495369 | 1.945916995 |
| Gm20695 | 1.863767 | 1.608940099 | 0.901742482 | 1.784256738 |
| 1700037C18Rik | 31.49051414 | -0.583214307 | 0.229387567 | -2.542484384 |
| Cluap1 | 161.5850996 | 0.087614348 | 0.10461989 | 0.837454031 |
| 1700016D08Rik | 0 | NA | NA | NA |
| Gm15538 | 10.93859038 | 0.013987039 | 0.462346773 | 0.030252269 |
| Gm23493 | 2.266218546 | -0.01382041 | 0.691877354 | -0.01997523 |
| Gm15539 | 10.65583297 | -0.025335406 | 0.439004423 | -0.05771105 |
| Gm15537 | 0.604736192 | -1.359994322 | 1.232001775 | -1.103889905 |
| Nlrc3 | 40.04733098 | -0.55443645 | 0.204640461 | -2.70931979 |
| Slx4 | 453.7919637 | -0.334659579 | 0.093912629 | -3.563520494 |
| Gm15879 | 36.75047168 | -0.284244658 | 0.212985193 | -1.334574736 |
| Dnase1 | 0.317440118 | 0.037077927 | 1.619936442 | 0.022888508 |
| Trap1 | 710.530572 | -0.126576146 | 0.055859967 | -2.265954536 |
| Gm23413 | 3.363014587 | 0.43302541 | 0.621182022 | 0.69709907 |
| Crebbp | 3534.609602 | -0.005444944 | 0.053234634 | -0.102281976 |
| Gm37186 | 92.75499785 | 0.0004067 | 0.318352751 | 0.001277515 |
| Gm5766 | 0.297064053 | 0.036524311 | 1.770719164 | 0.020626823 |
| Adcy9 | 953.9955112 | 0.898930938 | 0.129789 | 6.926094935 |
| Gm6142 | 13.94176219 | 1.128553872 | 0.338051229 | 3.338410791 |
| Srl | 63.68776743 | -0.038655791 | 0.165795874 | -0.23315291 |
| Gm15885 | 0 | NA | NA | NA |
| Tfap4 | 61.90644834 | 0.204621464 | 0.173682136 | 1.178137651 |
| Glis2 | 152.6195355 | -0.315904517 | 0.104955935 | -3.009877591 |

| | | | | |
|---|---|---|---|---|
| Pam16 | 156.721799 | -0.263377315 | 0.151977379 | -1.733003398 |
| Coro7 | 608.4531973 | 0.443310232 | 0.109029133 | 4.065979626 |
| Vasn | 199.7417839 | -1.082084781 | 0.213858982 | -5.059805159 |
| Dnaja3 | 623.6357018 | -0.394147182 | 0.086432441 | -4.56017646 |
| Nmral1 | 147.3588111 | -0.023333834 | 0.137253731 | -0.1700051 |
| Hmox2 | 693.4042527 | -0.060273527 | 0.081141718 | -0.74281798 |
| Gm15859 | 12.58934561 | -0.318591485 | 0.396141238 | -0.804237112 |
| Cdip1 | 1931.022938 | 0.129250162 | 0.05465102 | 2.365009133 |
| Gm15835 | 24.15715004 | 0.492574125 | 0.322663879 | 1.526585892 |
| 4930562C15Rik | 0.450624642 | 0.136438587 | 1.459026005 | 0.093513472 |
| Ubald1 | 776.8048839 | 0.202263848 | 0.087291299 | 2.317113504 |
| Mgrn1 | 1884.371733 | 0.068517392 | 0.051375898 | 1.33364854 |
| Gm16861 | 17.52762606 | 0.499662698 | 0.255550187 | 1.95524294 |
| Nudt16l1 | 313.4024066 | 0.063154139 | 0.086780947 | 0.727741986 |
| Anks3 | 314.2732224 | 0.063838439 | 0.099502038 | 0.641579212 |
| 4930451G09Rik | 27.70571581 | 0.033160725 | 0.217746728 | 0.152290347 |
| 12-Sep | 0.042716945 | 0.037062677 | 3.724945049 | 0.009949859 |
| Smim22 | 0.651416456 | 0.661763041 | 1.228160762 | 0.538824445 |
| RP23-450O11.3 | 0.373087082 | -0.672822212 | 1.683379206 | -0.399685472 |
| Rogdi | 663.6085146 | 0.715382392 | 0.07925224 | 9.02665206 |
| Glyr1 | 1981.897547 | 0.149486913 | 0.054998519 | 2.718017032 |
| Ubn1 | 976.2780589 | -0.147735652 | 0.08065543 | -1.831688849 |
| Gm23483 | 0 | NA | NA | NA |
| Ppl | 31.86021987 | -1.343003412 | 0.260473595 | -5.156005974 |
| Sec14l5 | 164.8073848 | -0.344083052 | 0.13424208 | -2.563153472 |
| Nagpa | 106.0130434 | -0.447660646 | 0.120926335 | -3.701928505 |
| AU021092 | 9.404176926 | -1.015991159 | 0.406650783 | -2.498436497 |
| Mir6364 | 0.049553047 | -0.443818988 | 3.724945049 | -0.119147795 |
| Gm23689 | 0.15591167 | 0.037074717 | 2.826544765 | 0.013116621 |
| Gm26095 | 0.09689823 | 0.037062677 | 3.724945049 | 0.009949859 |
| Alg1 | 199.3302287 | -0.224069305 | 0.109278981 | -2.0504337 |
| Eef2kmt | 50.93323876 | -0.944046274 | 0.215868632 | -4.373244337 |
| Gm15983 | 5.210713431 | 0.339908275 | 0.444600657 | 0.764524904 |
| n-R5s30 | 0 | NA | NA | NA |
| 1700123O21Rik | 37.36951086 | 0.722768563 | 0.390876404 | 1.849097451 |
| Rbfox1 | 3132.04572 | 0.799897137 | 0.08104056 | 9.870330869 |
| Gm23476 | 0 | NA | NA | NA |
| Gm26159 | 0 | NA | NA | NA |
| Gm24579 | 0 | NA | NA | NA |
| Tmem114 | 8.994908763 | 0.777434018 | 0.416738359 | 1.865520659 |
| Gm25782 | 0 | NA | NA | NA |
| Mettl22 | 149.3545907 | 0.161510112 | 0.091237063 | 1.770224805 |
| Gm23706 | 0 | NA | NA | NA |
| Abat | 3761.014197 | 0.440415639 | 0.085640266 | 5.142623416 |
| Gm22137 | 0 | NA | NA | NA |
| Gm7541 | 1.437230806 | 0.047905081 | 0.941504572 | 0.050881411 |
| Tmem186 | 191.5203021 | -0.034400389 | 0.10109688 | -0.34027152 |

| | | | | |
|---|---|---|---|---|
| Pmm2 | 257.7422973 | -0.082136305 | 0.100939423 | -0.813718787 |
| Carhsp1 | 736.470531 | -0.417392501 | 0.154794213 | -2.696434796 |
| RP24-183G19.6 | 0.790581521 | 2.146023158 | 1.410719905 | 1.521225546 |
| Usp7 | 2331.28947 | 0.099965952 | 0.068799098 | 1.453012536 |
| Gm25805 | 0 | NA | NA | NA |
| 1810013L24Rik | 1107.486804 | -0.316226058 | 0.067249855 | -4.702256318 |
| Gm22224 | 0 | NA | NA | NA |
| Gm10247 | 1.54124956 | -0.021379091 | 0.848623004 | -0.025192684 |
| Grin2a | 4615.108378 | 0.859422094 | 0.161719625 | 5.314272109 |
| Gm7572 | 0 | NA | NA | NA |
| Gm3890 | 0 | NA | NA | NA |
| Rpl39l | 0.823456907 | 0.539414883 | 0.992202677 | 0.543653929 |
| Atf7ip2 | 0.994634002 | 1.476283868 | 1.019687914 | 1.447780099 |
| Gm24667 | 0 | NA | NA | NA |
| Gm22932 | 0 | NA | NA | NA |
| Emp2 | 291.2677059 | -0.794326449 | 0.133376264 | -5.955530788 |
| Tekt5 | 64.82616325 | 0.143878947 | 0.186808685 | 0.7701941 |
| Nubp1 | 155.2730509 | -0.455630809 | 0.136153289 | -3.346454664 |
| Tvp23a | 456.5008628 | -0.58272071 | 0.107271387 | -5.432210062 |
| Ciita | 53.45720072 | -0.656612227 | 0.28845128 | -2.276336676 |
| Dexi | 379.7312674 | -0.005439395 | 0.08068634 | -0.067414073 |
| Clec16a | 1806.749408 | 0.173639683 | 0.067809015 | 2.560716789 |
| Gm15558 | 0.339464816 | 0.489879927 | 1.745380012 | 0.28067236 |
| Socs1 | 257.5468929 | -1.074535463 | 0.351878878 | -3.053708337 |
| Tnp2 | 0.356459053 | -0.901345019 | 1.592080164 | -0.566142987 |
| Prm3 | 0 | NA | NA | NA |
| Prm2 | 0.134416897 | -0.410053006 | 2.950415516 | -0.138981443 |
| Prm1 | 0.978261292 | -3.915048487 | 1.163730679 | -3.364222117 |
| Gm10343 | 5.069691734 | 0.660772424 | 0.455155621 | 1.451750551 |
| Gm11172 | 0.044544648 | 0.517957688 | 3.724945049 | 0.139051095 |
| Gm26822 | 0.840740177 | 2.434851934 | 1.234060544 | 1.973040907 |
| Rmi2 | 18.46361415 | -0.508830086 | 0.257085424 | -1.97922573 |
| Gm21859 | 0.468571393 | -2.467090484 | 1.41506106 | -1.743451611 |
| Litaf | 694.90215 | -1.008799403 | 0.255385789 | -3.950099992 |
| Gm26268 | 0 | NA | NA | NA |
| Gm24961 | 0 | NA | NA | NA |
| Gm4262 | 0.72304365 | -2.733647468 | 1.177189242 | -2.322181829 |
| Snn | 1735.197761 | 0.580774795 | 0.080308387 | 7.23180754 |
| Txndc11 | 466.704329 | -0.031271966 | 0.093806477 | -0.333366815 |
| Zc3h7a | 853.6995849 | -0.084424966 | 0.126383069 | -0.668008509 |
| Gm23935 | 15251.03865 | 0.965405776 | 0.922164585 | 1.046890969 |
| Gm24145 | 5.507590669 | 0.237113606 | 0.548180402 | 0.432546667 |
| 4930509G22Rik | 5.036420442 | 0.337065852 | 0.444019704 | 0.759123636 |
| Gm7638 | 2.798124148 | 0.529750658 | 0.6580383 | 0.805045326 |
| Gm6305 | 2.348678771 | -0.973306686 | 0.710868617 | -1.369179428 |
| Rsl1d1 | 606.0058677 | -0.186534124 | 0.100857639 | -1.84947938 |
| 2610020C07Rik | 17.10628321 | 1.081183435 | 0.277507315 | 3.896053819 |

| | | | | |
|---|---|---|---|---|
| Gspt1 | 1482.382663 | -0.16257483 | 0.052564096 | -3.092887404 |
| Mir1945 | 0.407802109 | -1.565497201 | 1.37961064 | -1.134738422 |
| Tnfrsf17 | 1.11445967 | -1.263561609 | 1.096638774 | -1.152213143 |
| Snx29 | 234.388902 | 0.202537674 | 0.123571091 | 1.639037677 |
| Gm15897 | 0.089089296 | 0.957155977 | 3.723939561 | 0.257027796 |
| Gm4279 | 0.739397817 | -0.918002359 | 1.218648508 | -0.753295436 |
| Cpped1 | 428.5321132 | -0.007406125 | 0.081649356 | -0.090706473 |
| Gm9961 | 0.136963818 | 0.037074342 | 3.599544405 | 0.010299732 |
| Shisa9 | 635.7404322 | 0.297740885 | 0.102387341 | 2.90798532 |
| Gm6327 | 0.086930093 | -0.443164241 | 3.724898049 | -0.118973522 |
| 1700003L19Rik | 0.364102347 | -0.443820616 | 1.740461278 | -0.255001718 |
| Ercc4 | 327.5978949 | 0.121408212 | 0.117074711 | 1.037014831 |
| Mkl2 | 3224.420774 | 0.674295539 | 0.122585521 | 5.500613254 |
| Gm15738 | 0.13861434 | 0.037059817 | 3.618284367 | 0.010242373 |
| Mir193b | 0.600656357 | -0.472859196 | 1.504199293 | -0.314359406 |
| Mir365-1 | 1.735414367 | 0.417629302 | 1.112507023 | 0.375394756 |
| Mir6365 | 0 | NA | NA | NA |
| 2310015D24Rik | 1.269356065 | 0.879670818 | 0.961323028 | 0.915062672 |
| Gm25276 | 0.11037557 | 0.517957688 | 3.724945049 | 0.139051095 |
| Parn | 231.9657214 | 0.201250637 | 0.087754054 | 2.293348607 |
| Bfar | 419.0822541 | -0.008722913 | 0.096217624 | -0.090658159 |
| 3110001I22Rik | 41.39457374 | 0.04979549 | 0.177897155 | 0.279911671 |
| Pla2g10 | 0 | NA | NA | NA |
| Pla2g10os | 0.090890357 | 0.037071574 | 3.724945049 | 0.009952247 |
| Rrn3 | 908.1970389 | -0.176965877 | 0.065024297 | -2.721534634 |
| Gm27734 | 0.941805692 | 1.740738324 | 1.115768799 | 1.560124575 |
| Ntan1 | 347.9210283 | 0.065890333 | 0.090026734 | 0.731897405 |
| Pdxdc1 | 751.5294376 | -0.198124192 | 0.061022534 | -3.24673822 |
| Mpv17l | 545.4884099 | 0.516466132 | 0.08311952 | 6.213535995 |
| Gm15950 | 0.843513109 | -1.264380059 | 1.368275893 | -0.924068067 |
| Gm26213 | 0.252317246 | 0.005653035 | 2.494815902 | 0.002265913 |
| Ifitm7 | 16.50735806 | -0.298945018 | 0.29788964 | -1.003542849 |
| 2900011O08Rik | 1845.656536 | 0.325004455 | 0.061485389 | 5.285881077 |
| Gm15806 | 0.061920613 | -0.443816764 | 3.724945049 | -0.119147198 |
| Gm15807 | 0 | NA | NA | NA |
| Gm15808 | 7.400018289 | 0.04745718 | 0.639976864 | 0.074154524 |
| Marf1 | 2957.81909 | 0.241356548 | 0.075409518 | 3.200611182 |
| Gm27671 | 0 | NA | NA | NA |
| Mir484 | 0.262798536 | 1.016751513 | 1.818294286 | 0.559178743 |
| Nde1 | 330.4944427 | -0.27361129 | 0.144153228 | -1.898058714 |
| Myh11 | 25.31456723 | 0.530893903 | 0.288794934 | 1.838307532 |
| Fopnl | 265.3373334 | -0.067592293 | 0.094188236 | -0.717629885 |
| Gm15868 | 0 | NA | NA | NA |
| Gm15869 | 19.70131526 | 0.373272732 | 0.289956437 | 1.287340729 |
| Abcc1 | 333.839765 | -0.113613791 | 0.114176106 | -0.995075017 |
| Gm23116 | 0.060174537 | 0.517957688 | 3.724945049 | 0.139051095 |
| Gm1758 | 0.227501076 | -0.438120676 | 2.139256666 | -0.204800426 |

| | | | | |
|---|---|---|---|---|
| A630010A05Rik | 0 | NA | NA | NA |
| Snai2 | 6.714012416 | 0.161962786 | 0.38236679 | 0.423579637 |
| Gm25547 | 1.090696383 | -0.882756221 | 0.926197653 | -0.953097018 |
| Efcab1 | 65.3173956 | 0.478446574 | 0.16468823 | 2.905165559 |
| Gm7731 | 0 | NA | NA | NA |
| Gm21897 | 0.148910734 | 0.966597255 | 2.858091675 | 0.338196729 |
| Gm6031 | 0 | NA | NA | NA |
| Ube2v2 | 393.0830429 | 0.248603425 | 0.099197691 | 2.506141239 |
| Mcm4 | 207.5248428 | -0.392148657 | 0.161511293 | -2.427995275 |
| Prkdc | 719.6145477 | 0.311330117 | 0.124141817 | 2.507858553 |
| Mzt2 | 126.2984278 | 0.278027754 | 0.132118708 | 2.104378384 |
| Cebpd | 477.928814 | -1.035100585 | 0.33862984 | -3.056731753 |
| Spidr | 147.239883 | -1.555960054 | 0.257797254 | -6.035595909 |
| n-R5s31 | 0.50758071 | -1.323563449 | 1.267479074 | -1.044248758 |
| Pkp2 | 124.1564146 | 0.669055061 | 0.137731659 | 4.857670825 |
| Yars2 | 199.1181058 | -0.380812959 | 0.110418466 | -3.448815893 |
| Dnm1l | 3082.769836 | 0.218970213 | 0.043992187 | 4.977479548 |
| Gm7765 | 0 | NA | NA | NA |
| Fgd4 | 1158.129079 | -0.151817832 | 0.085536277 | -1.774894088 |
| Gm16139 | 0 | NA | NA | NA |
| Olfr19 | 0.447268164 | -0.758020118 | 1.417162743 | -0.534885723 |
| Spag6 | 66.08212979 | 0.3305787 | 0.162385454 | 2.035765472 |
| Rpl31-ps12 | 2.913194718 | 1.053893038 | 0.604932111 | 1.742167458 |
| Gm15714 | 0.095898756 | -0.917692504 | 3.724434845 | -0.246397787 |
| Igll1 | 0.075043205 | 0.517957688 | 3.724945049 | 0.139051095 |
| Vpreb1 | 0.913331526 | -0.384829423 | 1.251823004 | -0.307415203 |
| Top3b | 435.3366088 | 0.053599316 | 0.120365101 | 0.445306117 |
| Ppm1f | 789.0102064 | -0.094195842 | 0.090514981 | -1.040665764 |
| Mapk1 | 4968.897064 | 0.362489061 | 0.070440405 | 5.146038842 |
| Gm22558 | 0 | NA | NA | NA |
| 1700056N10Rik | 40.39149775 | 0.805730592 | 0.320607326 | 2.513138431 |
| Ypel1 | 118.1725456 | 1.038436314 | 0.214234613 | 4.847192062 |
| Gm9974 | 6.459321561 | 0.719033521 | 0.44199793 | 1.62678029 |
| Ppil2 | 680.5363805 | -0.169736407 | 0.054681389 | -3.104098285 |
| Gm15585 | 2.078701808 | 1.093274163 | 0.84640251 | 1.291671693 |
| 2610318N02Rik | 4.21806971 | 0.624261921 | 0.493918657 | 1.263896213 |
| Mir130b | 0.075043205 | 0.517957688 | 3.724945049 | 0.139051095 |
| Mir301b | 0.118974955 | 0.51795324 | 3.724945049 | 0.139049901 |
| Sdf2l1 | 144.324589 | -0.927996087 | 0.157388552 | -5.896210856 |
| Ccdc116 | 27.44048453 | 0.811179854 | 0.240387326 | 3.374470146 |
| Gm15646 | 0 | NA | NA | NA |
| Ydjc | 88.53278356 | 0.266115798 | 0.136787149 | 1.945473682 |
| Ube2l3 | 1433.809399 | -0.114008296 | 0.08112782 | -1.405292237 |
| Gm15648 | 2.296535557 | -0.014617978 | 0.730745189 | -0.020004207 |
| Gm22129 | 0.093744596 | -0.443823436 | 3.724945049 | -0.119148989 |
| Rimbp3 | 28.96339831 | 0.465559839 | 0.236380652 | 1.969534455 |
| Hic2 | 126.3818273 | 0.786748225 | 0.182030372 | 4.322071194 |

| | | | | |
|---|---|---|---|---|
| Gm26635 | 0.934835317 | 0.577694676 | 1.137938693 | 0.507667663 |
| Tmem191c | 275.0346229 | 1.063210849 | 0.161404655 | 6.587237811 |
| Pi4ka | 4405.809757 | 0.048105879 | 0.080825248 | 0.595183812 |
| Serpind1 | 61.15174061 | 0.098047415 | 0.195596886 | 0.501272882 |
| Gm24927 | 3.890661782 | 0.042368893 | 0.55118493 | 0.076868744 |
| Snap29 | 384.8317826 | -0.706973684 | 0.144812149 | -4.88200533 |
| Crkl | 918.3467102 | -0.297680238 | 0.082871476 | -3.592071154 |
| Gm22770 | 0 | NA | NA | NA |
| Aifm3 | 673.3284071 | 0.937090077 | 0.175858504 | 5.328659429 |
| Lztr1 | 1143.994835 | 0.046073519 | 0.052960934 | 0.869952921 |
| Thap7 | 186.8558046 | -0.455872928 | 0.105084927 | -4.33813813 |
| Lrrc74b | 11.80310187 | -0.502396944 | 0.328183801 | -1.530840163 |
| P2rx6 | 28.77111211 | -1.117823861 | 0.248743614 | -4.493879626 |
| Slc7a4 | 422.6595786 | 0.135980693 | 0.107342335 | 1.266794622 |
| Smpd4 | 474.152001 | 0.256082026 | 0.143390869 | 1.785901902 |
| Ccdc74a | 148.3892086 | 0.687369813 | 0.195953905 | 3.507813812 |
| Med15 | 779.2021573 | -0.241533738 | 0.0669138 | -3.60962517 |
| Rpl26-ps4 | 0 | NA | NA | NA |
| Klhl22 | 742.0166947 | 0.258833097 | 0.047206777 | 5.482964805 |
| Scarf2 | 44.76671024 | 0.614352853 | 0.184744939 | 3.32541101 |
| Car15 | 16.28727946 | -0.163952873 | 0.322781822 | -0.507937133 |
| Gm20518 | 10.60745708 | 0.618950879 | 0.43838974 | 1.411873551 |
| Dgcr2 | 954.3027326 | 0.005940729 | 0.071346061 | 0.083266387 |
| Tssk1 | 0.674642588 | -0.157006294 | 1.132304671 | -0.138660819 |
| Tssk2 | 0.04393175 | 0.037062677 | 3.724945049 | 0.009949859 |
| Dgcr14 | 286.5719584 | 0.023093606 | 0.06995443 | 0.330123569 |
| Gsc2 | 0.267511063 | 0.517964829 | 2.522289024 | 0.205355066 |
| AA914427 | 7.57215109 | -0.185769865 | 0.415227458 | -0.447393016 |
| Slc25a1 | 386.8663658 | -0.327123622 | 0.117332643 | -2.788001813 |
| Vpreb2 | 0.042716945 | 0.037062677 | 3.724945049 | 0.009949859 |
| Dgcr6 | 244.4688473 | -0.002958485 | 0.110732638 | -0.026717372 |
| Prodh | 187.7866812 | 1.128715837 | 0.167095223 | 6.754925824 |
| Gm15954 | 5.737854463 | 0.45001396 | 0.519231731 | 0.866691946 |
| Rtn4r | 444.6635611 | 0.530571723 | 0.125693582 | 4.221152054 |
| Mir6366 | 0 | NA | NA | NA |
| Gm7873 | 0.093141081 | 0.037060453 | 3.724945049 | 0.009949262 |
| Zdhhc8 | 854.0005545 | 0.249990918 | 0.08771604 | 2.85000232 |
| Ranbp1 | 621.7089953 | -0.085408985 | 0.1491361 | -0.572691553 |
| Gm25777 | 2.388976987 | 0.187691693 | 0.798948361 | 0.234923435 |
| Trmt2a | 346.4940552 | -0.221644548 | 0.076380788 | -2.90183637 |
| Dgcr8 | 527.8985729 | 0.095280672 | 0.106147339 | 0.897626573 |
| Mir1306 | 0 | NA | NA | NA |
| Gm24572 | 0 | NA | NA | NA |
| Tango2 | 629.321088 | 0.194630171 | 0.095471298 | 2.038624957 |
| Mir185 | 2.706681505 | -0.912642843 | 0.724450079 | -1.259773267 |
| Arvcf | 417.1584827 | 0.015941389 | 0.067885638 | 0.234827126 |
| Gm26170 | 1.640007924 | -0.606868063 | 0.884382723 | -0.686205245 |

| | | | | |
|---|---|---|---|---|
| Gm26219 | 0 | NA | NA | NA |
| Comt | 659.6866933 | -0.246832076 | 0.058652544 | -4.208377999 |
| Gm15764 | 1.82265688 | -1.505147194 | 0.778880405 | -1.932449685 |
| Txnrd2 | 113.8682495 | -0.03841135 | 0.137785133 | -0.278777176 |
| Gnb1l | 33.08347225 | 0.119467894 | 0.190587301 | 0.626840786 |
| Gm16314 | 20.96206684 | 0.030776477 | 0.243955203 | 0.126156265 |
| Tbx1 | 2.659687333 | -0.529347159 | 0.591958461 | -0.894230244 |
| 4930588K23Rik | 0.132314849 | 0.037055789 | 3.665114298 | 0.010110405 |
| Gp1bb | 61.84750879 | 0.204244347 | 0.194977847 | 1.047525909 |
| 5-Sep | 2743.444644 | 0.615927242 | 0.110826181 | 5.557596902 |
| Cldn5 | 120.886244 | -0.456897443 | 0.169550978 | -2.694749672 |
| Cdc45 | 22.31723845 | -0.509305409 | 0.276223151 | -1.843818693 |
| Ufd1l | 707.9940102 | 0.107131793 | 0.055199496 | 1.940811067 |
| 2510002D24Rik | 76.81866928 | 0.127391967 | 0.131491391 | 0.968823635 |
| Gm28539 | 0.048949532 | 0.037064902 | 3.724945049 | 0.009950456 |
| Mrpl40 | 175.5484898 | -0.098874406 | 0.103368447 | -0.956524056 |
| Hira | 202.5271098 | 0.107427391 | 0.12203509 | 0.880299196 |
| Gm15799 | 5.251177882 | 0.503404023 | 0.626997146 | 0.802880883 |
| Gm15798 | 0 | NA | NA | NA |
| RP23-374P12.1 | 0 | NA | NA | NA |
| Gm10088 | 0.415109666 | -0.373913796 | 1.547994858 | -0.241547182 |
| RP23-247I11.8 | 0.329535318 | -0.378832269 | 1.704168349 | -0.222297444 |
| Iglc1 | 0.667277359 | -0.706990141 | 2.4108366 | -0.293255105 |
| Iglj1 | 0 | NA | NA | NA |
| Iglc3 | 1.77022427 | -1.361615806 | 0.744619575 | -1.828605979 |
| Iglj3p | 0 | NA | NA | NA |
| Iglj3 | 0.072025462 | 0.517959912 | 3.724945049 | 0.139051692 |
| Iglv1 | 0.611248024 | -0.932861337 | 2.493400203 | -0.374132213 |
| RP23-247I11.7 | 0 | NA | NA | NA |
| Iglc4 | 0 | NA | NA | NA |
| Iglj4 | 0 | NA | NA | NA |
| Iglc2 | 0.938589659 | -2.559523765 | 1.401000912 | -1.826925124 |
| Iglj2 | 0 | NA | NA | NA |
| RP23-272P19.8 | 0.046345709 | -0.443818988 | 3.724945049 | -0.119147795 |
| RP23-272P19.7 | 0.343041433 | 0.039265659 | 1.717560081 | 0.022861301 |
| Iglv3 | 0.097139547 | -0.443816764 | 3.724945049 | -0.119147198 |
| RP23-272P19.10 | 1.305037701 | -0.316317272 | 1.102360696 | -0.286945346 |
| Iglv2 | 0.097139547 | -0.443816764 | 3.724945049 | -0.119147198 |
| Olfr164 | 0.207953851 | -0.443819746 | 2.579760025 | -0.172039159 |
| Gm22158 | 0.391389582 | -1.458129711 | 3.360791054 | -0.43386503 |
| Gm23612 | 0.049553047 | -0.443818988 | 3.724945049 | -0.119147795 |
| Olfr165 | 0 | NA | NA | NA |
| Gm25762 | 0.144848288 | -0.92560533 | 2.892350527 | -0.32001838 |
| Olfr166 | 0.296045541 | 0.900222559 | 2.00819374 | 0.448274756 |
| Olfr167 | 0 | NA | NA | NA |
| Olfr168 | 0 | NA | NA | NA |
| Olfr169 | 0.0878635 | 0.037058229 | 3.724945049 | 0.009948665 |

| | | | | |
|---|---|---|---|---|
| Olfr170 | 0 | NA | NA | NA |
| Olfr171 | 0 | NA | NA | NA |
| Lamp3 | 0.505840948 | 1.07696412 | 1.25967817 | 0.8549518 |
| Gm17799 | 0.062426873 | 0.517959912 | 3.724945049 | 0.139051692 |
| B3gnt5 | 145.4656175 | -0.346746255 | 0.145018238 | -2.391052741 |
| A930003A15Rik | 2.53052731 | -0.43675733 | 0.568773042 | -0.767893866 |
| Gm27883 | 0 | NA | NA | NA |
| Klhl6 | 70.61673169 | 0.054062689 | 0.261047881 | 0.207098747 |
| Klhl24 | 1378.234747 | 0.203691729 | 0.079453203 | 2.563669201 |
| Yeats2 | 481.9769426 | 0.041742807 | 0.131542062 | 0.317334292 |
| Gm10241 | 24.31474395 | 0.153099168 | 0.265955261 | 0.575657602 |
| Map6d1 | 369.8382378 | 0.451284681 | 0.107109847 | 4.213288459 |
| Parl | 312.4995396 | -0.044810242 | 0.072199807 | -0.620642122 |
| Cyp2ab1 | 1.187519348 | -0.210067588 | 0.849169492 | -0.247380046 |
| Abcc5 | 1948.385061 | 0.079289267 | 0.138097422 | 0.574154579 |
| Gm20614 | 0.26923737 | -0.447805413 | 2.126779793 | -0.210555609 |
| Gm23332 | 0 | NA | NA | NA |
| Gm20615 | 0.04393175 | 0.037062677 | 3.724945049 | 0.009949859 |
| Eif2b5 | 489.2421831 | -0.193382987 | 0.063921983 | -3.02529705 |
| Gm16618 | 0.359481004 | -0.472231527 | 1.800413619 | -0.262290577 |
| Dvl3 | 617.5796496 | -0.153840291 | 0.074654497 | -2.060696895 |
| Ap2m1 | 2464.108253 | -0.046739779 | 0.041156331 | -1.13566436 |
| Gm15760 | 0.195053328 | 0.037673265 | 2.683734851 | 0.014037626 |
| Abcf3 | 658.1946899 | -0.163776542 | 0.065372356 | -2.505287412 |
| Vwa5b2 | 365.3797502 | 0.827580434 | 0.204395142 | 4.048924187 |
| Mir1224 | 0 | NA | NA | NA |
| Alg3 | 92.27766346 | -0.403579515 | 0.12791278 | -3.155114884 |
| Ece2 | 434.908798 | -0.051884262 | 0.102106074 | -0.508140797 |
| Camk2n2 | 831.1974061 | 0.249547843 | 0.09927462 | 2.513712401 |
| Psmd2 | 2330.871484 | -0.362492154 | 0.082237074 | -4.407892171 |
| Gm4462 | 1.62367863 | 0.503790829 | 0.847451707 | 0.594477332 |
| Eif4g1 | 4413.780657 | -0.280751388 | 0.045409201 | -6.182698197 |
| Snord66 | 1.086592418 | 0.037487371 | 0.987747627 | 0.037952378 |
| Fam131a | 1339.029144 | 0.947333869 | 0.112445187 | 8.424850312 |
| Clcn2 | 427.9328786 | 0.565580821 | 0.110945821 | 5.097810957 |
| Polr2h | 104.4427863 | 0.142002074 | 0.117576037 | 1.207746734 |
| Thpo | 27.14520041 | 0.56310831 | 0.403124418 | 1.396859839 |
| Chrd | 234.488815 | 0.72667817 | 0.211132093 | 3.441817669 |
| Gm22741 | 0.075043205 | 0.517957688 | 3.724945049 | 0.139051095 |
| Rps10-ps2 | 203.4377443 | -0.393317789 | 0.133916487 | -2.937037833 |
| Gm9697 | 0 | NA | NA | NA |
| Teddm3 | 0.044544648 | 0.517957688 | 3.724945049 | 0.139051095 |
| AC168061.1 | 0 | NA | NA | NA |
| Ephb3 | 170.6997336 | -0.041443502 | 0.136922124 | -0.302679365 |
| Gm16863 | 0.106358623 | 0.517955464 | 3.724945049 | 0.139050498 |
| Mir7680 | 0 | NA | NA | NA |
| Vps8 | 810.4907143 | 0.079293872 | 0.057901942 | 1.369450998 |

| | | | | |
|---|---|---|---|---|
| 2510009E07Rik | 1017.889882 | -0.007399976 | 0.088929296 | -0.0832119 |
| Ehhadh | 44.10308811 | 0.056153686 | 0.219616513 | 0.255689725 |
| Gm24898 | 0 | NA | NA | NA |
| 1300002E11Rik | 152.2564682 | 0.152040345 | 0.18489948 | 0.822286491 |
| Gm26744 | 2.147345651 | 0.492784301 | 0.6384515 | 0.771842969 |
| Map3k13 | 489.4826492 | -0.031189343 | 0.083627544 | -0.372955392 |
| Gm6467 | 24.4487381 | -0.311268027 | 0.220031896 | -1.414649569 |
| 2610020F03Rik | 0.233260489 | -1.395414586 | 3.03991854 | -0.459030256 |
| Tmem41a | 97.70389257 | 0.07850745 | 0.134577543 | 0.583362192 |
| Liph | 10.28461881 | -0.90104241 | 0.330082866 | -2.729746085 |
| Senp2 | 1234.002812 | 0.079232177 | 0.073090754 | 1.084024617 |
| Gm4943 | 1.131366322 | 0.213221905 | 1.013940851 | 0.21029028 |
| Igf2bp2 | 94.65434489 | -2.314245273 | 0.278457318 | -8.310951538 |
| Gm25810 | 0 | NA | NA | NA |
| Gm5809 | 0.299370719 | -0.751562118 | 1.750048626 | -0.429452135 |
| Gm26980 | 0 | NA | NA | NA |
| Tra2b | 1042.494832 | -0.156859649 | 0.083085046 | -1.887940806 |
| Gm15776 | 0.868382071 | -1.431982047 | 1.055788781 | -1.356314892 |
| Etv5 | 804.3619116 | -0.054382529 | 0.119805093 | -0.453925019 |
| Dgkg | 917.1223497 | 0.229415372 | 0.102023505 | 2.248652131 |
| 9230117E06Rik | 0.105852363 | -0.443821212 | 3.724945049 | -0.119148392 |
| Mir8095 | 6.066797584 | 0.080719305 | 0.540623455 | 0.149307811 |
| Gm15651 | 2.236386351 | -0.284404784 | 0.718752259 | -0.395692369 |
| Gm22481 | 0 | NA | NA | NA |
| Mir6362 | 0 | NA | NA | NA |
| Crygs | 0.564059227 | -1.906702959 | 1.355616604 | -1.406520806 |
| Tbccd1 | 144.3603587 | -0.222037061 | 0.120072123 | -1.849197427 |
| Dnajb11 | 661.7688257 | -0.493377745 | 0.134436938 | -3.669956727 |
| Ahsg | 1.513507497 | -0.119290908 | 1.075785933 | -0.110887217 |
| Fetub | 1.838490909 | -1.325630132 | 0.783957169 | -1.690947138 |
| Hrg | 0.529344572 | 0.075289862 | 1.342418253 | 0.056085249 |
| Kng2 | 1.792037434 | -2.365884506 | 0.869533928 | -2.720865088 |
| Gm6493 | 33.39922536 | -0.704613715 | 0.196868954 | -3.579100208 |
| Gm23715 | 0 | NA | NA | NA |
| Kng1 | 0.385993897 | -1.5604941 | 1.701170361 | -0.917306189 |
| Eif4a2 | 5039.606027 | 0.311957554 | 0.06338957 | 4.921275782 |
| Snord2 | 0 | NA | NA | NA |
| Gm24201 | 0 | NA | NA | NA |
| AC154540.1 | 0 | NA | NA | NA |
| Snora81 | 0 | NA | NA | NA |
| Gm26447 | 0 | NA | NA | NA |
| Gm24616 | 7.869682849 | -0.229551301 | 0.369375603 | -0.621457667 |
| Rfc4 | 32.64420203 | -0.47472419 | 0.201600639 | -2.354775229 |
| Adipoq | 0.218718758 | 0.503206125 | 2.160441116 | 0.232918232 |
| B630019A10Rik | 0.40993559 | 0.051049426 | 1.596498313 | 0.031975872 |
| Gm6630 | 0.062426873 | 0.517959912 | 3.724945049 | 0.139051692 |
| Gm15746 | 0 | NA | NA | NA |

| | | | | |
|---|---|---|---|---|
| BC106179 | 11.069689 | 0.461892715 | 0.360329718 | 1.281861284 |
| St6gal1 | 316.7496297 | -0.335684357 | 0.101531699 | -3.306202503 |
| Gm23430 | 0 | NA | NA | NA |
| Gm23066 | 0 | NA | NA | NA |
| Rtp1 | 5.214837505 | -0.423313683 | 0.640333209 | -0.661083443 |
| Masp1 | 117.7213065 | 0.011367012 | 0.109841518 | 0.103485569 |
| Rtp4 | 910.9235272 | -0.619517616 | 0.152440445 | -4.063997698 |
| Gm6640 | 0.597217552 | 1.511756109 | 1.262070606 | 1.197837982 |
| Sst | 654.4591436 | 0.240260616 | 0.151367661 | 1.587265175 |
| Rtp2 | 3.004889707 | 0.02159905 | 0.77271581 | 0.027952126 |
| Bcl6 | 818.2508927 | -0.412773442 | 0.133948984 | -3.081572016 |
| Gm37419 | 0.567687411 | -0.101032618 | 1.322871301 | -0.076373732 |
| Lppos | 20.64171431 | -0.154708337 | 0.267141314 | -0.579125461 |
| Lpp | 689.3135003 | -0.125144747 | 0.131419635 | -0.952253039 |
| Gm24440 | 0 | NA | NA | NA |
| Mir28a | 2.122337162 | 0.260270806 | 0.921541787 | 0.282429739 |
| Gm22672 | 0.704522364 | 0.290471504 | 1.132183924 | 0.256558584 |
| A230028O05Rik | 10.09423595 | -0.478657709 | 0.43217611 | -1.107552449 |
| Tprg | 0.366915913 | -0.361759439 | 1.503539534 | -0.240605206 |
| Trp63 | 10.83564004 | -0.066018494 | 0.369046246 | -0.178889488 |
| Gm24928 | 0 | NA | NA | NA |
| Gm27005 | 0.212482393 | 0.036341181 | 2.18452096 | 0.016635766 |
| Gm4524 | 1.767199681 | -1.207701577 | 0.962242761 | -1.255090322 |
| P3h2 | 56.59938758 | -0.059878558 | 0.161569349 | -0.370605928 |
| AU015336 | 6.800499267 | 0.473434847 | 0.446564379 | 1.060171544 |
| Cldn1 | 56.29871389 | 0.119723513 | 0.172352576 | 0.694643018 |
| Cldn16 | 0 | NA | NA | NA |
| Tmem207 | 0.473315337 | 1.428550649 | 1.360701159 | 1.049863623 |
| Il1rap | 1004.716365 | 0.015485548 | 0.069399167 | 0.223137378 |
| Gm20319 | 0.326503072 | -0.89646882 | 1.584760166 | -0.565681066 |
| Gmnc | 8.309823615 | 0.113672695 | 0.468864732 | 0.242442408 |
| Ostn | 1.447793589 | -1.757912657 | 0.879833921 | -1.998005094 |
| Gm15970 | 0 | NA | NA | NA |
| Uts2b | 1.222794617 | -0.654157452 | 1.215189213 | -0.538317363 |
| Ccdc50 | 1385.300587 | 0.050348618 | 0.067094387 | 0.750414756 |
| Fgf12 | 2033.20002 | 0.559322221 | 0.082458686 | 6.783060087 |
| Mir690 | 11.63860447 | 0.063528363 | 0.434165822 | 0.146322809 |
| Mb21d2 | 459.0744955 | 0.270470138 | 0.084402706 | 3.204519727 |
| Hrasls | 202.7297218 | 0.775621079 | 0.088760507 | 8.738357947 |
| Atp13a5 | 116.1853686 | 0.561933417 | 0.161245097 | 3.484964371 |
| Gm15752 | 0.667341687 | -0.840295451 | 1.342576305 | -0.625882826 |
| Atp13a4 | 513.9834346 | -0.641540029 | 0.151436708 | -4.236357459 |
| Opa1 | 2535.740764 | 0.227424358 | 0.055712689 | 4.082092675 |
| Gm26834 | 2.849524632 | -0.044219462 | 0.646708347 | -0.068376204 |
| Gm26569 | 0.312910946 | -1.501209532 | 1.657524967 | -0.905693466 |
| Gm1968 | 0.303184894 | -2.078967951 | 3.690206733 | -0.563374386 |
| 4632428C04Rik | 13.7946314 | 0.220703157 | 0.321813616 | 0.685810502 |

| | | | | |
|---|---|---|---|---|
| Hes1 | 94.25849431 | 0.080544648 | 0.127990793 | 0.629300326 |
| n-R5s32 | 0 | NA | NA | NA |
| 1700025H01Rik | 0 | NA | NA | NA |
| Gm26562 | 0.090074004 | 0.517955464 | 3.724945049 | 0.139050498 |
| Cpn2 | 0.207381971 | -0.925551248 | 2.582144409 | -0.358442868 |
| Lrrc15 | 1.556998443 | -0.672375632 | 0.778903384 | -0.863233678 |
| Gp5 | 14.94000079 | 0.167763226 | 0.297711504 | 0.563509383 |
| Atp13a3 | 1577.269787 | -0.048078481 | 0.054963969 | -0.874727245 |
| Tmem44 | 273.2629563 | 0.847155146 | 0.131216164 | 6.45617978 |
| Lsg1 | 245.3569069 | -0.342584404 | 0.102376695 | -3.346312386 |
| Fam43a | 229.047573 | -0.385057048 | 0.096016111 | -4.010337891 |
| Gm26295 | 0.327620686 | -0.932530965 | 2.009364099 | -0.464092578 |
| Gm25375 | 0 | NA | NA | NA |
| Xxylt1 | 202.0539679 | -0.140639469 | 0.093047354 | -1.511482725 |
| Gm15742 | 2.112839908 | -0.210716013 | 0.89205379 | -0.23621447 |
| Acap2 | 1866.515927 | 0.277026162 | 0.081006567 | 3.419798795 |
| Ppp1r2 | 1509.036798 | 0.275277949 | 0.079129974 | 3.478807514 |
| Gm27415 | 0 | NA | NA | NA |
| Apod | 8482.021723 | -0.598789866 | 0.259180372 | -2.310321041 |
| 1700007E05Rik | 2.68870073 | -0.304033307 | 0.705319561 | -0.431057528 |
| Bdh1 | 477.7842783 | 0.199100664 | 0.066643266 | 2.987558642 |
| Gm15743 | 22.43508857 | 0.657544283 | 0.318955424 | 2.061555423 |
| Dlg1 | 2632.440439 | 0.046834631 | 0.046633474 | 1.004313574 |
| Gm24879 | 0 | NA | NA | NA |
| Mfi2 | 8.341406001 | -0.827305607 | 0.415492028 | -1.991146762 |
| Gm15703 | 0.299449808 | -1.332610219 | 2.008679919 | -0.663425868 |
| Pigz | 89.07597162 | 0.528391335 | 0.159639479 | 3.309903903 |
| NCBP2-AS2 | 84.09966569 | 0.100964067 | 0.136407276 | 0.740166282 |
| 0610012G03Rik | 0 | NA | NA | NA |
| Ncbp2 | 349.8016824 | -0.457986374 | 0.127236134 | -3.599499277 |
| Senp5 | 762.9688528 | -0.158424007 | 0.069567219 | -2.277279579 |
| Gm15696 | 12.05015047 | 0.806029604 | 0.366792625 | 2.197507663 |
| Gm26269 | 0.072025462 | 0.517959912 | 3.724945049 | 0.139051692 |
| Gm15694 | 6.232026559 | -0.675173977 | 0.434588359 | -1.553594253 |
| Hmgb1-ps6 | 11.46318819 | 0.156291242 | 0.365883499 | 0.427161221 |
| Pak2 | 1045.01706 | -0.525950415 | 0.082738336 | -6.35679227 |
| Gm26419 | 0 | NA | NA | NA |
| Pigx | 116.9372743 | -0.180507532 | 0.137497132 | -1.312809437 |
| Cep19 | 336.7429659 | 0.186473524 | 0.093090387 | 2.003144786 |
| Gm15729 | 0.971776759 | -0.696504927 | 1.140305725 | -0.610805429 |
| Nrros | 147.122329 | -0.371508218 | 0.127600397 | -2.911497342 |
| Bex6 | 0.377709955 | 0.086588808 | 1.46658967 | 0.059040923 |
| Fbxo45 | 678.3823491 | 0.14314514 | 0.074406364 | 1.92382927 |
| Gm25848 | 45.86024405 | -0.78182228 | 0.32601861 | -2.398090956 |
| Wdr53 | 71.48528253 | -0.044136977 | 0.147777069 | -0.298672706 |
| Mir1946a | 0 | NA | NA | NA |
| Smco1 | 1.487580774 | 0.823393869 | 0.765307533 | 1.075899339 |

| | | | | |
|---|---|---|---|---|
| Rnf168 | 482.2284669 | -0.072371036 | 0.060016303 | -1.205856278 |
| Ubxn7 | 1865.438353 | 0.349325648 | 0.07513864 | 4.649081294 |
| Tm4sf19 | 8.203405253 | -2.327343669 | 0.577910905 | -4.027166902 |
| Gm20056 | 14.90719541 | -1.017645125 | 0.331324508 | -3.071445365 |
| Tctex1d2 | 113.226245 | -0.226610905 | 0.108593903 | -2.086773733 |
| Pcyt1a | 752.3083677 | -0.425980226 | 0.092784739 | -4.591059147 |
| Slc51a | 0 | NA | NA | NA |
| Zdhhc19 | 0.768268477 | -1.813419592 | 1.221331743 | -1.484788718 |
| Gm5405 | 0 | NA | NA | NA |
| Tfrc | 1390.85657 | 0.011984172 | 0.113974313 | 0.105148009 |
| Tnk2 | 2566.202347 | 0.327457187 | 0.048508187 | 6.750555053 |
| Tnk2os | 0.364228253 | -0.919430053 | 1.583977465 | -0.580456524 |
| Gm9556 | 0 | NA | NA | NA |
| Muc4 | 0.409426173 | -1.096189151 | 1.621032117 | -0.676229138 |
| Muc20 | 0.330893424 | 0.868620078 | 1.538730759 | 0.564504266 |
| Gm933 | 0.097139547 | -0.443816764 | 3.724945049 | -0.119147198 |
| Gm26769 | 0 | NA | NA | NA |
| 1700021K19Rik | 991.8281154 | -0.490399569 | 0.104523455 | -4.691765782 |
| Fyttd1 | 1180.673463 | 0.065986503 | 0.094357324 | 0.699325721 |
| Lrch3 | 977.5154375 | -0.228748271 | 0.065501056 | -3.492283689 |
| Gm6611 | 33.95866455 | 0.04980503 | 0.353262075 | 0.140986065 |
| Gm17106 | 0.097139547 | -0.443816764 | 3.724945049 | -0.119147198 |
| n-R5s33 | 0.130017973 | -0.444798045 | 3.662714201 | -0.121439463 |
| Iqcg | 37.65510366 | -0.438340251 | 0.197521753 | -2.21919988 |
| Rpl35a | 394.3677081 | -0.138402516 | 0.092717894 | -1.492727129 |
| Lmln | 221.22393 | 0.040447999 | 0.092579078 | 0.436902157 |
| Gm22231 | 0.549303492 | -0.323979109 | 1.372335575 | -0.236078635 |
| Mir1947 | 0.769571883 | 1.231202807 | 1.113492492 | 1.105712716 |
| Gm23916 | 0 | NA | NA | NA |
| Osbpl11 | 369.2866785 | -0.351773936 | 0.094210663 | -3.733907882 |
| Snx4 | 448.4376452 | -0.011944513 | 0.082439439 | -0.144888334 |
| 1700007L15Rik | 6.048360371 | -0.033323252 | 0.41264439 | -0.080755373 |
| Zfp148 | 2196.281864 | 0.151765217 | 0.048159012 | 3.151335749 |
| Slc12a8 | 28.22738804 | -0.251302646 | 0.220496265 | -1.139713847 |
| Heg1 | 831.6435367 | -0.375375198 | 0.077872442 | -4.820385594 |
| Gm15658 | 0 | NA | NA | NA |
| Gm15657 | 0.194279094 | -1.017823543 | 3.719440177 | -0.273649661 |
| Muc13 | 4.915050087 | -0.424150487 | 0.466014809 | -0.910165255 |
| Itgb5 | 478.5994184 | -0.277685274 | 0.198576696 | -1.398377953 |
| Umps | 206.1452455 | -0.591175624 | 0.090331303 | -6.5445267 |
| Gm15829 | 0.247806867 | 0.877305135 | 2.094906202 | 0.41878015 |
| Kalrn | 4921.446429 | 0.869026613 | 0.183196514 | 4.743685311 |
| Ropn1 | 0.318826904 | -1.18343438 | 1.769863929 | -0.668658399 |
| Ccdc14 | 47.35523537 | 0.063550791 | 0.176219384 | 0.360634509 |
| Mylk | 164.66998 | -0.189982132 | 0.128989615 | -1.472848278 |
| 1700119H24Rik | 0.164132501 | 1.299746508 | 3.390982409 | 0.383294972 |
| Hacd2 | 451.6459504 | 0.175546174 | 0.094451405 | 1.85858723 |

| | | | | |
|---|---|---|---|---|
| Adcy5 | 2609.480313 | 0.822829408 | 0.132011228 | 6.233025935 |
| Gm25179 | 0 | NA | NA | NA |
| Sec22a | 183.1160017 | 0.040097219 | 0.114953288 | 0.348813153 |
| Gm25079 | 0 | NA | NA | NA |
| Gm23518 | 0.038981395 | -0.443816764 | 3.724945049 | -0.119147198 |
| Pdia5 | 30.59877438 | -0.620619813 | 0.232557524 | -2.668672262 |
| Gm25548 | 0.246683951 | -0.444418058 | 1.853897639 | -0.239720926 |
| 1600019K03Rik | 0.049553047 | -0.443818988 | 3.724945049 | -0.119147795 |
| Sema5b | 130.570813 | 0.112977554 | 0.12331672 | 0.916157634 |
| Gm5963 | 36.49740693 | -0.039125554 | 0.293229795 | -0.133429668 |
| Dirc2 | 652.6475954 | 0.296388484 | 0.079248792 | 3.739974795 |
| Gm25967 | 2.125672366 | 1.325852162 | 0.767758473 | 1.726913096 |
| Hspbap1 | 70.57271391 | -0.758127821 | 0.206181868 | -3.676985907 |
| Gm26838 | 0 | NA | NA | NA |
| Parp14 | 2858.919116 | -0.848113989 | 0.164706723 | -5.149237236 |
| Gm10237 | 0.042716945 | 0.037062677 | 3.724945049 | 0.009949859 |
| Dtx3l | 1910.456235 | -0.715015869 | 0.15364999 | -4.653536708 |
| Parp9 | 1102.792659 | -0.696331694 | 0.143174655 | -4.863512273 |
| Gm15564 | 359.9778991 | -0.31311964 | 0.972316337 | -0.322034741 |
| Kpna1 | 1948.470927 | 0.096075321 | 0.058082823 | 1.654109012 |
| Wdr5b | 49.21408818 | 0.070372488 | 0.169843345 | 0.414337624 |
| Fam162a | 149.3833853 | -0.397898427 | 0.122809438 | -3.239966182 |
| Ccdc58 | 55.1879007 | -0.035083302 | 0.166108287 | -0.211207416 |
| Csta1 | 0 | NA | NA | NA |
| Gm10092 | 0.045882455 | 0.517959912 | 3.724945049 | 0.139051692 |
| Gm23080 | 0 | NA | NA | NA |
| Stfa2l1 | 0.423264953 | -0.578576647 | 1.348531119 | -0.429042118 |
| Gm22885 | 0 | NA | NA | NA |
| Gm5483 | 0.535697641 | -0.095226098 | 1.606848431 | -0.059262651 |
| Gm25024 | 0 | NA | NA | NA |
| Gm10913 | 0.156524568 | 0.517971166 | 2.823069785 | 0.183477989 |
| Gm5416 | 0.092228164 | 0.037073798 | 3.724945049 | 0.009952844 |
| Gm24801 | 0 | NA | NA | NA |
| 2010005H15Rik | 0.699696385 | -0.049329308 | 1.063348364 | -0.046390543 |
| Gm10813 | 0 | NA | NA | NA |
| Gm24569 | 0 | NA | NA | NA |
| Stfa1 | 0 | NA | NA | NA |
| Gm23306 | 0 | NA | NA | NA |
| Gm4758 | 0 | NA | NA | NA |
| Gm25254 | 0 | NA | NA | NA |
| Gm24437 | 0 | NA | NA | NA |
| BC117090 | 0 | NA | NA | NA |
| Gm25465 | 0.060174537 | 0.517957688 | 3.724945049 | 0.139051095 |
| BC100530 | 0.115000406 | -0.443821212 | 3.724945049 | -0.119148392 |
| Gm15845 | 0.075043205 | 0.517957688 | 3.724945049 | 0.139051095 |
| Gm24218 | 0 | NA | NA | NA |
| Gm22349 | 0 | NA | NA | NA |

| | | | | |
|---|---|---|---|---|
| Stfa2 | 0.143022002 | 0.03610674 | 3.547735707 | 0.010177404 |
| Gm22263 | 0 | NA | NA | NA |
| Gm25903 | 0 | NA | NA | NA |
| Stfa3 | 0.131209559 | -0.409992761 | 2.972496229 | -0.137928774 |
| Casr | 17.16410259 | -0.118172287 | 0.278707829 | -0.424000602 |
| Gm25902 | 0 | NA | NA | NA |
| Cd86 | 81.75402155 | -0.017836331 | 0.160654178 | -0.111023137 |
| Ildr1 | 14.35142951 | 0.389004901 | 0.323508389 | 1.202456918 |
| Slc15a2 | 296.0330333 | 0.592582685 | 0.221543332 | 2.674793595 |
| Gm8425 | 0.186025035 | 0.037076632 | 2.67312714 | 0.013870134 |
| Gm22617 | 0 | NA | NA | NA |
| Eaf2 | 14.00843118 | -0.110126928 | 0.273111346 | -0.40323088 |
| Iqcb1 | 366.8902262 | 0.18734367 | 0.085139184 | 2.200440041 |
| Golgb1 | 1930.593704 | 0.015833715 | 0.108171676 | 0.14637579 |
| 4930565N06Rik | 0.233785498 | -0.77237807 | 2.48979613 | -0.310217395 |
| Hcls1 | 216.1493986 | -0.569754071 | 0.167371121 | -3.404136076 |
| Fbxo40 | 25.85214164 | -0.286571906 | 0.311312538 | -0.920527992 |
| Polq | 42.09947402 | 0.14235225 | 0.293183154 | 0.485540346 |
| Stxbp5l | 2108.032249 | 0.335094034 | 0.095432178 | 3.511331725 |
| Gm15665 | 42.97331767 | 0.716412995 | 0.308972568 | 2.3186945 |
| Gtf2e1 | 231.0323693 | -0.581981187 | 0.133535015 | -4.358266524 |
| Rabl3 | 128.7194147 | -0.173200103 | 0.119685626 | -1.44712535 |
| Hgd | 1.882553937 | -0.031908702 | 0.77010244 | -0.04143436 |
| Ndufb4 | 405.4328191 | 0.237225009 | 0.136588807 | 1.736782199 |
| Fstl1 | 650.539503 | -0.363340729 | 0.174470091 | -2.082538777 |
| n-R5s34 | 0 | NA | NA | NA |
| Lrrc58 | 3637.314835 | 0.192457143 | 0.1002852 | 1.919098169 |
| Gm15725 | 0 | NA | NA | NA |
| Gpr156 | 12.90207318 | 0.776434107 | 0.465716205 | 1.66718293 |
| Gm25140 | 0.106056992 | 0.998850474 | 3.724945049 | 0.268151734 |
| Gsk3b | 4829.972571 | 0.150641778 | 0.045765739 | 3.291584094 |
| SNORA17 | 0 | NA | NA | NA |
| Gm25617 | 0 | NA | NA | NA |
| Gm27931 | 0 | NA | NA | NA |
| Nr1i2 | 0.200534954 | -0.443820861 | 2.645328197 | -0.167775349 |
| Maats1 | 18.76607067 | -0.245354084 | 0.247733016 | -0.990397194 |
| Maats1os | 0.072025462 | 0.517959912 | 3.724945049 | 0.139051692 |
| Cox17 | 102.4271693 | -0.106790876 | 0.210293608 | -0.507817986 |
| Gm21987 | 0.2386254 | 0.036366338 | 2.112154822 | 0.017217648 |
| Popdc2 | 3.459091227 | 0.232002957 | 0.530382824 | 0.43742547 |
| Pla1a | 56.89416345 | -0.710305869 | 0.289642008 | -2.452357912 |
| Adprh | 679.8642621 | -0.132861034 | 0.084243919 | -1.577099402 |
| D930030I03Rik | 2.891043216 | -0.08928386 | 0.642917497 | -0.138872966 |
| Cd80 | 14.42150522 | -0.863102127 | 0.37477318 | -2.302998651 |
| Timmdc1 | 207.9193946 | -0.57985361 | 0.134705286 | -4.304609185 |
| Gm15953 | 0 | NA | NA | NA |
| Poglut1 | 522.8527119 | -0.064787242 | 0.062646341 | -1.034174395 |

| | | | | |
|---|---|---|---|---|
| Tmem39a | 165.5822775 | -0.47048678 | 0.131972552 | -3.56503509 |
| Gm17103 | 0.159330469 | 0.517966511 | 3.465376739 | 0.149469033 |
| Arhgap31 | 855.9405782 | -0.381801885 | 0.093903479 | -4.065897102 |
| Gm15530 | 5.747139663 | -0.299924125 | 0.46313136 | -0.647600554 |
| B4galt4 | 168.6399155 | 0.034989549 | 0.100764795 | 0.34723982 |
| Gm15575 | 0 | NA | NA | NA |
| Upk1b | 9.705317616 | 0.418137559 | 0.395925654 | 1.056101201 |
| 4930435E12Rik | 0 | NA | NA | NA |
| Gm15803 | 0 | NA | NA | NA |
| Gm15802 | 0 | NA | NA | NA |
| Igsf11 | 559.2207678 | -0.436376973 | 0.138368381 | -3.153733314 |
| Gm22500 | 7.621091703 | -0.161772116 | 0.539241752 | -0.29999924 |
| Gm24795 | 0 | NA | NA | NA |
| Gm8609 | 0.167459142 | -0.410658575 | 2.753748719 | -0.149127105 |
| Lsamp | 5421.197146 | 0.603536533 | 0.06342092 | 9.516363632 |
| Gm37508 | 3.928946461 | 0.60447728 | 0.62245127 | 0.97112386 |
| Gm22543 | 16.99085758 | 0.708811248 | 0.381609427 | 1.85742594 |
| Gm36939 | 48.56639706 | 0.812456713 | 0.245179406 | 3.313723304 |
| Gm27887 | 0 | NA | NA | NA |
| Gm27717 | 0 | NA | NA | NA |
| Gm26381 | 1.397116197 | 1.746246749 | 0.992433767 | 1.75955999 |
| AC124193.1 | 0 | NA | NA | NA |
| Gm27587 | 0 | NA | NA | NA |
| Gm28750 | 0 | NA | NA | NA |
| Gap43 | 1479.213936 | 0.13207815 | 0.09647131 | 1.369092524 |
| Mir6540 | 2.543989632 | -0.115400368 | 0.761560084 | -0.151531534 |
| 4932412D23Rik | 1.816140692 | -0.509182315 | 0.83806925 | -0.60756592 |
| Zbtb20 | 6323.085067 | -0.066830758 | 0.083965074 | -0.795935208 |
| BC002163 | 27.11636185 | 0.079599569 | 0.226225852 | 0.351858852 |
| Gm25873 | 5.983328485 | 0.403056921 | 0.442054631 | 0.91178079 |
| Gm15712 | 29.69002061 | 0.584377403 | 0.364308737 | 1.604071887 |
| Gm37946 | 0 | NA | NA | NA |
| Gm15711 | 21.32018645 | 0.42231416 | 0.325780524 | 1.296314938 |
| Gm9968 | 1.290099097 | 0.279503719 | 0.892318605 | 0.313233096 |
| Gm15713 | 0.087930927 | -0.443164241 | 3.724898049 | -0.118973522 |
| Mir568 | 0 | NA | NA | NA |
| Tigit | 22.00729436 | -0.97475739 | 0.407371385 | -2.392797886 |
| Drd3 | 8.262393275 | 0.54921484 | 0.467034479 | 1.175962085 |
| Gm25996 | 0.1580736 | 0.517956762 | 3.474509474 | 0.149073349 |
| Gm26074 | 0 | NA | NA | NA |
| Qtrtd1 | 200.210793 | -0.267250995 | 0.112284472 | -2.380124251 |
| Gm25989 | 9.070232471 | -0.516726539 | 0.435793644 | -1.185713803 |
| 2610015P09Rik | 57.77977805 | -0.084316947 | 0.227176378 | -0.371151913 |
| Zdhhc23 | 82.33084375 | 0.202172651 | 0.160413221 | 1.260324114 |
| Gramd1c | 24.07085944 | 0.236382259 | 0.219243037 | 1.078174532 |
| Mir3470b | 0 | NA | NA | NA |
| Atp6v1a | 5242.393468 | 0.029268261 | 0.060084188 | 0.487120849 |

| | | | | |
|---|---|---|---|---|
| Naa50 | 1580.032754 | -0.403942827 | 0.090187685 | -4.478913345 |
| Gm608 | 1399.505967 | 0.233934913 | 0.073549919 | 3.180627753 |
| Sidt1 | 676.9943462 | 0.92016602 | 0.141861662 | 6.486361479 |
| Gm26732 | 6.046421774 | 0.504861731 | 0.491154606 | 1.027907965 |
| Spice1 | 128.0135962 | -0.242317407 | 0.172506484 | -1.404685789 |
| Cfap44 | 93.24835868 | -0.003196022 | 0.179941402 | -0.017761458 |
| Boc | 445.6221661 | -0.252562895 | 0.16788965 | -1.504338687 |
| Mir3081 | 1.772274333 | -0.784605411 | 0.924478694 | -0.848700372 |
| BC027231 | 116.0115484 | -0.256141643 | 0.165722268 | -1.545607876 |
| Gtpbp8 | 357.6076025 | 0.078080306 | 0.08487242 | 0.919972662 |
| Cd200r1 | 51.42872498 | -0.888325531 | 0.285670797 | -3.109612678 |
| Cd200r4 | 69.15376312 | -2.238894473 | 0.349949411 | -6.397766076 |
| Cd200r2 | 1.386349714 | 0.278453739 | 0.780902717 | 0.356579294 |
| Cd200r3 | 0.534659868 | -0.445381421 | 1.322317367 | -0.336818855 |
| Ccdc80 | 217.6073964 | -1.312188934 | 0.152039975 | -8.630552153 |
| Slc35a5 | 550.0251731 | -0.201435883 | 0.10213832 | -1.97218717 |
| Atg3 | 539.1722443 | -0.226216129 | 0.070139767 | -3.225219285 |
| Gm6030 | 17.33716352 | -0.172155627 | 0.297483558 | -0.578706359 |
| Btla | 10.07865719 | -0.882459458 | 0.314662235 | -2.804465738 |
| Cd200 | 1252.555305 | -0.012263563 | 0.103861832 | -0.118075741 |
| Gm5406 | 0.966602524 | -1.111363587 | 1.087638417 | -1.021813472 |
| Gm609 | 0.707603298 | 0.502822584 | 1.374445307 | 0.365836735 |
| Gm16011 | 0.581873452 | -0.749806547 | 1.240945314 | -0.60422207 |
| Gm6903 | 0.049553047 | -0.443818988 | 3.724945049 | -0.119147795 |
| Slc9c1 | 0.038981395 | -0.443816764 | 3.724945049 | -0.119147198 |
| Gcsam | 0.711975465 | 0.057637944 | 1.196841579 | 0.048158373 |
| BC016579 | 3.578730896 | -0.669908839 | 0.712406665 | -0.940346113 |
| Tmprss7 | 24.10678365 | -0.391980354 | 0.320105895 | -1.224533383 |
| Tagln3 | 1506.536885 | 0.086224999 | 0.096138842 | 0.896879938 |
| Abhd10 | 203.4530973 | -0.143577849 | 0.095702472 | -1.500252247 |
| Gm15591 | 0.060174537 | 0.517957688 | 3.724945049 | 0.139051095 |
| Phldb2 | 129.9886621 | -0.466736823 | 0.127669235 | -3.655828446 |
| Gm15640 | 0.229952095 | -0.399236177 | 2.111088744 | -0.189113877 |
| Gm15638 | 0.108266322 | -0.955981391 | 3.722845977 | -0.25678779 |
| Plcxd2 | 1552.118851 | 0.097235153 | 0.079600853 | 1.221534064 |
| Gm24754 | 0 | NA | NA | NA |
| Cd96 | 3.576396818 | 0.315731502 | 0.479101368 | 0.659007723 |
| Gm4737 | 112.4227191 | -0.091799101 | 0.165059066 | -0.556159094 |
| Gm26297 | 0 | NA | NA | NA |
| Pvrl3 | 517.9500622 | -0.238302483 | 0.088707361 | -2.686389055 |
| Gm6912 | 0 | NA | NA | NA |
| Gm24651 | 0 | NA | NA | NA |
| Gm25318 | 0 | NA | NA | NA |
| Gm25339 | 0 | NA | NA | NA |
| Gm7275 | 0 | NA | NA | NA |
| Gm7204 | 1.57829385 | -3.885116093 | 1.024095445 | -3.793705081 |
| Dppa4 | 0 | NA | NA | NA |

| | | | | |
|---|---|---|---|---|
| Dppa2 | 0.379773675 | -1.81877464 | 1.586810101 | -1.146182923 |
| n-R5s36 | 0 | NA | NA | NA |
| Morc1 | 1.278798984 | -1.597659014 | 0.943823226 | -1.692752381 |
| Trat1 | 6.136963414 | 0.126597461 | 0.398388133 | 0.317774176 |
| Retnlb | 0 | NA | NA | NA |
| Retnla | 0.060174537 | 0.517957688 | 3.724945049 | 0.139051095 |
| Retnlg | 1.932622373 | -2.330565252 | 0.914757382 | -2.547741398 |
| Dzip3 | 1926.33653 | -0.122644421 | 0.095264915 | -1.287403883 |
| Gm22576 | 7.017922756 | -0.130012062 | 0.443504097 | -0.293147375 |
| C330027C09Rik | 62.11505808 | 0.046876974 | 0.16338288 | 0.286914848 |
| Myh15 | 9.416055846 | -0.002013512 | 0.349952246 | -0.005753677 |
| 1700026J12Rik | 0.047948698 | 0.037064902 | 3.724945049 | 0.009950456 |
| Gm5407 | 0.075043205 | 0.517957688 | 3.724945049 | 0.139051095 |
| Gm6931 | 0.04393175 | 0.037062677 | 3.724945049 | 0.009949859 |
| Ift57 | 340.5122929 | 0.205835052 | 0.087778712 | 2.34493134 |
| Gm15518 | 0.295614739 | 0.052756612 | 2.085469147 | 0.025297239 |
| Gm16619 | 0.436453079 | 0.665366672 | 1.356410112 | 0.490535028 |
| Cd47 | 3446.906187 | 0.090465393 | 0.098558409 | 0.917886093 |
| AC107830.1 | 0 | NA | NA | NA |
| Gm8824 | 0.047948698 | 0.037064902 | 3.724945049 | 0.009950456 |
| Bbx | 1297.513501 | 0.042908244 | 0.11172375 | 0.384056604 |
| G730013B05Rik | 0.417980132 | -0.754943482 | 1.471319109 | -0.513106557 |
| Ccdc54 | 0.042716945 | 0.037062677 | 3.724945049 | 0.009949859 |
| Gm9575 | 16.04186629 | -0.072747122 | 0.277327423 | -0.262314925 |
| 5330426P16Rik | 100.6912749 | -0.597702912 | 0.18114705 | -3.299545381 |
| Mir6363 | 0 | NA | NA | NA |
| 1700116B05Rik | 0 | NA | NA | NA |
| Gm24925 | 0 | NA | NA | NA |
| Gm6767 | 3.585344364 | -0.043750196 | 0.530956617 | -0.082398815 |
| Cblb | 789.0152622 | 0.091752674 | 0.092245824 | 0.994653968 |
| Gm23513 | 0 | NA | NA | NA |
| Alcam | 2206.12951 | 0.507208414 | 0.071938066 | 7.050626206 |
| Gm25723 | 0 | NA | NA | NA |
| Gm22459 | 0 | NA | NA | NA |
| Gm25041 | 0 | NA | NA | NA |
| Gm25853 | 0 | NA | NA | NA |
| Gm23180 | 0 | NA | NA | NA |
| Gm22977 | 0 | NA | NA | NA |
| Zpld1 | 1.833497833 | 0.672676307 | 0.928385554 | 0.724565676 |
| Mir5118 | 0 | NA | NA | NA |
| Gm23003 | 0 | NA | NA | NA |
| Gm26369 | 0 | NA | NA | NA |
| Nfkbiz | 257.9280466 | -0.464155864 | 0.160724546 | -2.88789656 |
| Nxpe3 | 312.8495634 | 0.312041804 | 0.096583228 | 3.230807367 |
| Cep97 | 386.1470673 | -0.02074383 | 0.149122044 | -0.139106396 |
| Rpl24 | 16.95551249 | -0.040920895 | 0.302944367 | -0.135077259 |
| Gm28037 | 0 | NA | NA | NA |

| | | | | |
|---|---|---|---|---|
| Zbtb11os1 | 37.78932254 | 0.325292107 | 0.200403779 | 1.6231835 |
| Zbtb11 | 1088.837139 | 0.036723633 | 0.069529818 | 0.52817099 |
| Pcnp | 1051.54738 | 0.083012897 | 0.078211093 | 1.061395434 |
| Trmt10c | 191.2919896 | -0.253218963 | 0.099971308 | -2.532916384 |
| Gm22422 | 89.64062021 | -0.025980183 | 0.389787536 | -0.066652164 |
| Senp7 | 795.4829463 | -0.025124919 | 0.120005288 | -0.209365098 |
| Gm24047 | 0.105143817 | 0.517955464 | 3.724945049 | 0.139050498 |
| Impg2 | 63.00063838 | -0.624701984 | 0.147605955 | -4.232227516 |
| Abi3bp | 266.089939 | -0.001178056 | 0.206760046 | -0.005697698 |
| Tfg | 756.0033368 | -0.28966426 | 0.057546678 | -5.033553124 |
| Adgrg7 | 0.562361829 | 0.376227033 | 1.362022069 | 0.276226826 |
| Gm15693 | 0.044191549 | 0.037062677 | 3.724945049 | 0.009949859 |
| Tmem45a | 10.44425536 | -0.672025416 | 0.313438838 | -2.144040029 |
| Gm15839 | 0.181290818 | -0.428757036 | 2.739792868 | -0.1564925 |
| Gm15840 | 0 | NA | NA | NA |
| Tmem45a2 | 3.654246218 | 0.408142325 | 0.517898519 | 0.788073937 |
| Tomm70a | 2858.851143 | 0.196665663 | 0.081292306 | 2.419240806 |
| Nit2 | 67.55208235 | -0.092562771 | 0.145011394 | -0.638313777 |
| Gm16892 | 2.540529182 | 0.494569883 | 0.634178835 | 0.779858702 |
| Tbc1d23 | 546.2969052 | -0.1381682 | 0.058056152 | -2.379906264 |
| Tmem30c | 8.630201669 | 0.36059243 | 0.419234034 | 0.860122035 |
| Gm26800 | 0.820767868 | -0.788422274 | 1.22690995 | -0.642608102 |
| Cmss1 | 65.09916204 | 0.216600184 | 0.133090244 | 1.627468537 |
| Filip1l | 99.78162097 | -0.433652198 | 0.284254326 | -1.525578182 |
| Col8a1 | 91.62796141 | 0.421734031 | 0.16964301 | 2.486008886 |
| Gm24039 | 0 | NA | NA | NA |
| Gm37458 | 0.072025462 | 0.517959912 | 3.724945049 | 0.139051692 |
| Gm37825 | 0 | NA | NA | NA |
| 4930461C15Rik | 1.662681963 | 0.956479084 | 0.933374689 | 1.024753613 |
| Dcbld2 | 658.0902064 | 0.318155253 | 0.094416797 | 3.369689102 |
| St3gal6 | 198.8642845 | -0.146118361 | 0.10054626 | -1.453245108 |
| Gm813 | 1.602843074 | -0.704241615 | 0.778522618 | -0.904587225 |
| E330017A01Rik | 0 | NA | NA | NA |
| Cpox | 461.6201025 | 0.11022396 | 0.063061526 | 1.747879679 |
| Gpr15 | 6.100871677 | -2.411909552 | 0.591471999 | -4.077808514 |
| Cldnd1 | 811.4291424 | -0.052642572 | 0.088198994 | -0.596861367 |
| Gm27564 | 0 | NA | NA | NA |
| Gm26019 | 0 | NA | NA | NA |
| Olfr172 | 0 | NA | NA | NA |
| Olfr173 | 0 | NA | NA | NA |
| Olfr175-ps1 | 0.049553047 | -0.443818988 | 3.724945049 | -0.119147795 |
| Olfr177 | 0 | NA | NA | NA |
| Olfr178 | 0 | NA | NA | NA |
| Olfr180 | 0 | NA | NA | NA |
| Olfr181 | 0 | NA | NA | NA |
| Olfr183 | 0 | NA | NA | NA |
| Olfr186 | 0.172571121 | -0.433506555 | 2.733273128 | -0.158603453 |

| | | | | |
|---|---|---|---|---|
| Olfr187 | 0.106520246 | 0.037071574 | 3.724945049 | 0.009952247 |
| Olfr190 | 0 | NA | NA | NA |
| Olfr191 | 0 | NA | NA | NA |
| Olfr192 | 0.046345709 | -0.443818988 | 3.724945049 | -0.119147795 |
| Olfr193 | 0 | NA | NA | NA |
| Olfr194 | 0 | NA | NA | NA |
| Olfr195 | 0.097139547 | -0.443816764 | 3.724945049 | -0.119147198 |
| Olfr196 | 0 | NA | NA | NA |
| Olfr197 | 0 | NA | NA | NA |
| Olfr198 | 0.194201709 | -0.443827523 | 3.285796589 | -0.135074558 |
| Olfr199 | 0 | NA | NA | NA |
| Olfr201 | 0 | NA | NA | NA |
| Olfr202 | 0 | NA | NA | NA |
| Olfr203 | 0 | NA | NA | NA |
| Olfr204 | 0.068097359 | 0.037064902 | 3.724945049 | 0.009950456 |
| Olfr205 | 0.046345709 | -0.443818988 | 3.724945049 | -0.119147795 |
| Olfr206 | 0.075043205 | 0.517957688 | 3.724945049 | 0.139051095 |
| Olfr209 | 0.048949532 | 0.037064902 | 3.724945049 | 0.009950456 |
| Gabrr3 | 1.001170369 | 1.062782743 | 0.917762057 | 1.158015561 |
| 1700022E09Rik | 0.097139547 | -0.443816764 | 3.724945049 | -0.119147198 |
| Mina | 105.9311312 | -0.475390319 | 0.136746712 | -3.476429623 |
| Crybg3 | 125.2170434 | -0.089317689 | 0.111633542 | -0.800097241 |
| Gm9581 | 73.34347339 | -0.169196383 | 0.127969106 | -1.322165852 |
| Arl6 | 230.3950377 | -0.071517889 | 0.104219688 | -0.686222444 |
| Epha6 | 866.3468396 | 0.779212755 | 0.125548319 | 6.206477001 |
| Gm24755 | 0.366471122 | 0.037079586 | 1.614349086 | 0.022968753 |
| Gm4660 | 10.78697284 | 0.421199675 | 0.456454127 | 0.922764522 |
| Gm9017 | 5.516248819 | 0.069038557 | 0.439587265 | 0.157053133 |
| Gm27001 | 0.095897396 | 0.037062677 | 3.724945049 | 0.009949859 |
| Gm27784 | 0 | NA | NA | NA |
| Gm17065 | 0 | NA | NA | NA |
| Gm23852 | 0 | NA | NA | NA |
| Gm17054 | 0 | NA | NA | NA |
| n-R5s37 | 0 | NA | NA | NA |
| Gm9027 | 1.878461702 | -0.269852826 | 0.718023249 | -0.375827421 |
| Gm17192 | 0 | NA | NA | NA |
| Gm24164 | 0 | NA | NA | NA |
| Gm9816 | 144.6420699 | -0.294247449 | 0.113649602 | -2.589075933 |
| Nsun3 | 164.1841752 | -0.036358746 | 0.095297826 | -0.38152755 |
| Arl13b | 185.8996067 | -0.024251232 | 0.093503355 | -0.259362163 |
| Gm25986 | 0 | NA | NA | NA |
| Stx19 | 0.062426873 | 0.517959912 | 3.724945049 | 0.139051692 |
| Pros1 | 274.6408184 | -0.722517292 | 0.160911319 | -4.490158273 |
| Epha3 | 257.2072295 | 0.187291461 | 0.158634883 | 1.180644871 |
| Gm22769 | 4.369109362 | 0.821797514 | 0.53518983 | 1.535525281 |
| Csnka2ip | 2.872518017 | 0.150730891 | 0.628528204 | 0.239815636 |
| 4930453N24Rik | 641.6073447 | -0.136122958 | 0.113817878 | -1.195971674 |

| | | | | |
|---|---|---|---|---|
| Zfp654 | 590.4534688 | 0.165055281 | 0.120623487 | 1.3683511 |
| Cggbp1 | 1135.996861 | -0.355809398 | 0.094927118 | -3.748237658 |
| Gm9061 | 0.350048956 | -0.823105823 | 1.555628181 | -0.529114754 |
| Gm23374 | 2.380614889 | 0.206527929 | 0.746553634 | 0.276641784 |
| Htr1f | 54.47759132 | 0.487244432 | 0.170152546 | 2.86357415 |
| Gm22429 | 0 | NA | NA | NA |
| Gm23659 | 0 | NA | NA | NA |
| Pou1f1 | 0.726361874 | -3.015288495 | 3.664956597 | -0.822735117 |
| Chmp2b | 560.9048391 | 0.04542455 | 0.085910454 | 0.528742991 |
| Vgll3 | 46.32731703 | 0.560888824 | 0.163511368 | 3.430274184 |
| Gm22217 | 0.045882455 | 0.517959912 | 3.724945049 | 0.139051692 |
| Gm22448 | 0 | NA | NA | NA |
| Cadm2 | 4980.463628 | 0.806374349 | 0.091952401 | 8.769475691 |
| 1700010K23Rik | 0.914990437 | 0.834180744 | 1.094563705 | 0.762112557 |
| 4933411O13Rik | 0 | NA | NA | NA |
| Gm15828 | 81.5457743 | 0.268246769 | 0.301543891 | 0.88957786 |
| Gm24681 | 0 | NA | NA | NA |
| Gm23591 | 0 | NA | NA | NA |
| RP24-377L12.1 | 0.046345709 | -0.443818988 | 3.724945049 | -0.119147795 |
| Speer2 | 0.175920186 | -0.979991501 | 2.728285742 | -0.359196798 |
| Gbe1 | 229.1812539 | -0.062355542 | 0.091956611 | -0.678097436 |
| Gm24968 | 0.168853147 | -0.75832428 | 2.756733419 | -0.275080744 |
| Gm22797 | 0 | NA | NA | NA |
| Robo1 | 968.3837098 | 0.666675019 | 0.092857226 | 7.179570738 |
| Gm25699 | 0 | NA | NA | NA |
| Robo2 | 1762.408431 | 0.637975849 | 0.108930995 | 5.856697169 |
| Gm24272 | 0 | NA | NA | NA |
| Gm22163 | 1.874863541 | 1.877930421 | 0.980114229 | 1.916032199 |
| mmu-mir-691 | 8.12823547 | 0.193338897 | 0.468869653 | 0.412351057 |
| Gm26307 | 0 | NA | NA | NA |
| Gm26937 | 0.185762564 | 0.037060026 | 3.327558188 | 0.011137304 |
| Lipi | 0.34836924 | -0.391932899 | 1.553649965 | -0.252265895 |
| Rbm11 | 76.42304198 | 0.938350401 | 0.182628991 | 5.138014497 |
| Gm15553 | 0 | NA | NA | NA |
| Hspa13 | 738.3975908 | -0.292995159 | 0.076568858 | -3.826557777 |
| Gm22923 | 0 | NA | NA | NA |
| Gm15554 | 0.120974994 | 0.517957688 | 3.724945049 | 0.139051095 |
| Samsn1 | 95.65148686 | -0.469322472 | 0.245475184 | -1.911893755 |
| Gm15555 | 0 | NA | NA | NA |
| Gm37705 | 0 | NA | NA | NA |
| Gm22268 | 0 | NA | NA | NA |
| Gm26915 | 0 | NA | NA | NA |
| Nrip1 | 1272.517009 | 0.346551818 | 0.090274258 | 3.83887748 |
| Gm9843 | 41.81605505 | -0.105783748 | 0.197484235 | -0.535656673 |
| 1700063K16Rik | 0.044544648 | 0.517957688 | 3.724945049 | 0.139051095 |
| 1700041M19Rik | 0.061920613 | -0.443816764 | 3.724945049 | -0.119147198 |
| Usp25 | 1277.058717 | -0.466753091 | 0.110049164 | -4.241314266 |

| | | | | |
|---|---|---|---|---|
| Mir99ahg | 442.0252824 | -0.056616748 | 0.130671636 | -0.43327496 |
| Gm31258 | 0.728787387 | 1.778579007 | 1.252796633 | 1.41968693 |
| Gm37466 | 10.46537345 | -0.193957703 | 0.389661967 | -0.497758877 |
| Gm21816 | 50.06920957 | -0.185637336 | 0.239838493 | -0.774009765 |
| 9430092D12Rik | 19.231994 | 0.249047156 | 0.284713669 | 0.874728485 |
| Gm38071 | 38.74659047 | 0.066894757 | 0.249834354 | 0.26775644 |
| 9430053O09Rik | 25.41324745 | 0.678601842 | 0.213867744 | 3.172997619 |
| Gm37063 | 47.20854318 | 0.167319707 | 0.209584651 | 0.798339506 |
| Gm37241 | 6.880611024 | 0.586369793 | 0.516251383 | 1.135822224 |
| Gm36963 | 34.35472376 | -0.153597258 | 0.231579166 | -0.663260262 |
| Gm37606 | 95.09549254 | 0.379613557 | 0.219662758 | 1.728165306 |
| Gm11146 | 0.04393175 | 0.037062677 | 3.724945049 | 0.009949859 |
| Mir99a | 7.589586068 | -0.049252689 | 0.397196627 | -0.124000773 |
| Mirlet7c-1 | 8.265456032 | 0.101761913 | 0.377131805 | 0.269831161 |
| Gm37311 | 115.4089426 | -0.023612783 | 0.206346442 | -0.114432714 |
| Mir125b-2 | 0.487613095 | -1.479076551 | 1.529418504 | -0.967084253 |
| C130023A14Rik | 179.7393386 | 0.00924967 | 0.149368005 | 0.061925374 |
| 9530003O04Rik | 61.28675002 | 0.157454578 | 0.168573446 | 0.934041404 |
| Gm37694 | 93.40348466 | -0.178266605 | 0.213539817 | -0.834816697 |
| Gm17333 | 0.423282238 | 0.079208034 | 3.681436415 | 0.021515524 |
| Gm25025 | 0.066050875 | -0.443818988 | 3.724945049 | -0.119147795 |
| Gm25916 | 0 | NA | NA | NA |
| Cxadr | 476.9468567 | -0.041820935 | 0.145057604 | -0.288305709 |
| Btg3 | 111.9560118 | -0.152928123 | 0.235277136 | -0.649991434 |
| Gm25038 | 0 | NA | NA | NA |
| D16Ertd472e | 110.9694825 | -0.163312135 | 0.12718586 | -1.284043166 |
| Gm24861 | 0 | NA | NA | NA |
| Chodl | 16.16523681 | 0.013317809 | 0.311889666 | 0.042700387 |
| Tmprss15 | 0.287833757 | -0.146118617 | 2.028698232 | -0.072025802 |
| Gm15692 | 0 | NA | NA | NA |
| Gm37709 | 0 | NA | NA | NA |
| Rps19-ps12 | 0 | NA | NA | NA |
| 1700066C05Rik | 0 | NA | NA | NA |
| Gm23083 | 0 | NA | NA | NA |
| Gm24773 | 0 | NA | NA | NA |
| Ncam2 | 1411.136076 | 0.311749532 | 0.078062432 | 3.993592344 |
| Gm26023 | 0.228677828 | -0.443821575 | 2.181307319 | -0.203465862 |
| Gm23355 | 0 | NA | NA | NA |
| Gm21833 | 0.117046891 | 0.037062677 | 3.724945049 | 0.009949859 |
| Rpl21-ps5 | 0 | NA | NA | NA |
| Gm24508 | 0 | NA | NA | NA |
| Gm25292 | 0.193566712 | -1.675392576 | 2.630757503 | -0.63684797 |
| Gm26667 | 11.04978208 | -0.959705815 | 0.328263552 | -2.923583229 |
| Mir155 | 0 | NA | NA | NA |
| Mrpl39 | 254.7822395 | 0.003775277 | 0.102999634 | 0.036653302 |
| Gm26153 | 0.072025462 | 0.517959912 | 3.724945049 | 0.139051692 |
| Jam2 | 457.3571526 | -0.214321535 | 0.157230502 | -1.36310406 |

| | | | | |
|---|---|---|---|---|
| Gm25908 | 0.500634304 | -0.406656216 | 1.425326431 | -0.285307427 |
| Atp5j | 806.9086276 | 0.15874657 | 0.131070326 | 1.211155686 |
| Gabpa | 471.5751915 | -0.287301468 | 0.060770609 | -4.72763849 |
| App | 15888.22377 | 0.399075625 | 0.05867347 | 6.801636691 |
| Gm27295 | 0 | NA | NA | NA |
| Cyyr1 | 66.70143812 | -0.738829903 | 0.16106414 | -4.587178155 |
| Gm2541 | 17.77359248 | 0.141132441 | 0.277987835 | 0.507692868 |
| Adamts1 | 613.1894008 | -0.232305443 | 0.301611628 | -0.770213817 |
| Adamts5 | 105.3116989 | -0.240380757 | 0.19853756 | -1.210757083 |
| Gm25715 | 0 | NA | NA | NA |
| N6amt1 | 193.0747588 | 0.210610528 | 0.114706341 | 1.836084432 |
| Ltn1 | 1170.966149 | -0.152642855 | 0.042208694 | -3.61638421 |
| Gm25896 | 1.990209358 | -0.643534265 | 0.915346075 | -0.703050226 |
| Rwdd2b | 73.7279705 | 0.10272228 | 0.13176634 | 0.779579062 |
| Usp16 | 561.8276137 | 0.163615456 | 0.123059183 | 1.329567221 |
| Cct8 | 1716.316278 | -0.209073804 | 0.087241402 | -2.39649752 |
| B130034C11Rik | 16.77205567 | 0.581376235 | 0.346943071 | 1.67571075 |
| Gm24891 | 0 | NA | NA | NA |
| Rpl31-ps4 | 0.092228164 | 0.037073798 | 3.724945049 | 0.009952844 |
| Map3k7cl | 1.664220238 | 0.488478666 | 0.800078145 | 0.610538695 |
| Gm22808 | 0 | NA | NA | NA |
| Bach1 | 550.4479081 | -0.656651382 | 0.123075994 | -5.335332748 |
| Gm23994 | 0 | NA | NA | NA |
| 2810407A14Rik | 5.016221076 | 2.4008516 | 0.797301319 | 3.011222411 |
| Grik1 | 248.488386 | 0.373939738 | 0.099295622 | 3.765923721 |
| Cldn17 | 0.086930093 | -0.443164241 | 3.724898049 | -0.118973522 |
| Cldn8 | 0.133745955 | 0.517951857 | 3.671135057 | 0.141087661 |
| Krtap24-1 | 0 | NA | NA | NA |
| Gm23587 | 0 | NA | NA | NA |
| 2310079G19Rik | 0 | NA | NA | NA |
| Krtap26-1 | 0 | NA | NA | NA |
| Krtap27-1 | 0 | NA | NA | NA |
| 2310061N02Rik | 0 | NA | NA | NA |
| Krtap13-1 | 0.115603922 | -0.93550548 | 3.724188147 | -0.251197158 |
| Krtap13 | 0 | NA | NA | NA |
| 2310034C09Rik | 0 | NA | NA | NA |
| AC125199.1 | 0 | NA | NA | NA |
| 2310057N15Rik | 0 | NA | NA | NA |
| Gm5965 | 0 | NA | NA | NA |
| Krtap14 | 0.061920613 | -0.443816764 | 3.724945049 | -0.119147198 |
| Krtap15 | 0 | NA | NA | NA |
| Krtap19-1 | 0 | NA | NA | NA |
| Krtap19-2 | 0 | NA | NA | NA |
| Krtap19-3 | 0 | NA | NA | NA |
| Krtap19-4 | 0 | NA | NA | NA |
| Krtap19-5 | 0 | NA | NA | NA |
| Krtap19-9a | 0 | NA | NA | NA |

| | | | | |
|---|---|---|---|---|
| Krtap19-9b | 0 | NA | NA | NA |
| Krtap16-3 | 0 | NA | NA | NA |
| AC140194.1 | 0 | NA | NA | NA |
| Krtap22-2 | 0 | NA | NA | NA |
| Gm10229 | 0 | NA | NA | NA |
| Krtap6-1 | 0 | NA | NA | NA |
| Gm10228 | 0 | NA | NA | NA |
| Krtap6-5 | 0 | NA | NA | NA |
| 1110025L11Rik | 0 | NA | NA | NA |
| Krtap6-3 | 0 | NA | NA | NA |
| Gm6358 | 0 | NA | NA | NA |
| Gm10061 | 0 | NA | NA | NA |
| Gm9789 | 0 | NA | NA | NA |
| Gm7735 | 0 | NA | NA | NA |
| 1110057P08Rik | 0 | NA | NA | NA |
| Krtap20-2 | 0 | NA | NA | NA |
| Krtap21-1 | 0 | NA | NA | NA |
| Krtap6-2 | 0 | NA | NA | NA |
| AC121818.1 | 0 | NA | NA | NA |
| Gm23655 | 0.049553047 | -0.443818988 | 3.724945049 | -0.119147795 |
| Krtap8-1 | 0 | NA | NA | NA |
| Krtap7-1 | 0.106618422 | 0.517955464 | 3.724945049 | 0.139050498 |
| Krtap11-1 | 0.045882455 | 0.517959912 | 3.724945049 | 0.139051692 |
| Tiam1 | 2404.733508 | 0.304189754 | 0.130559078 | 2.329901215 |
| Sod1 | 1216.13554 | 0.009570296 | 0.095127907 | 0.100604508 |
| Scaf4 | 656.9093113 | -0.042449344 | 0.108023372 | -0.392964437 |
| Hunk | 320.5353761 | -0.093964886 | 0.070220301 | -1.338144159 |
| 1110008E08Rik | 0.670543963 | -2.682723197 | 1.310010803 | -2.047863416 |
| Gm23406 | 0.090074004 | 0.517955464 | 3.724945049 | 0.139050498 |
| Mis18a | 41.5770124 | -0.034516793 | 0.20928755 | -0.164925211 |
| Gm17518 | 6.352540116 | 0.261693777 | 0.433921498 | 0.603090141 |
| Mrap | 10.74916131 | -0.513318808 | 0.524479843 | -0.978719802 |
| Urb1 | 305.5173951 | -0.071531144 | 0.08820987 | -0.810919961 |
| Eva1c | 41.11539447 | 0.301377748 | 0.202412891 | 1.488925661 |
| Gm7831 | 1.072190385 | 2.511733145 | 1.052569704 | 2.386286756 |
| 1110004E09Rik | 190.7191731 | -0.247614794 | 0.116478636 | -2.125838715 |
| Synj1 | 5333.078176 | 0.2807233 | 0.061931924 | 4.532772147 |
| Gm22933 | 0 | NA | NA | NA |
| Gm15965 | 0.045882455 | 0.517959912 | 3.724945049 | 0.139051692 |
| 4930404I05Rik | 0.517542306 | -0.617532581 | 1.290556746 | -0.478500913 |
| Paxbp1 | 967.3107701 | 0.188822944 | 0.135783073 | 1.390622119 |
| 4931406G06Rik | 3.965964068 | -0.168092213 | 0.51219844 | -0.328177908 |
| 4932438H23Rik | 22.83793654 | 0.079075636 | 0.217542936 | 0.363494386 |
| H3f3a-ps2 | 2.549826182 | 0.550386734 | 0.673603126 | 0.817078652 |
| Gm9881 | 5.470790701 | 1.292121667 | 0.558021718 | 2.315540105 |
| Olig2 | 347.143824 | -0.514410325 | 0.161189498 | -3.191338962 |
| Gm37016 | 0.089089296 | 0.957155977 | 3.723939561 | 0.257027796 |

| | | | | |
|---|---|---|---|---|
| Olig1 | 1119.055513 | -0.293394446 | 0.148021689 | -1.982104434 |
| Gm15966 | 0 | NA | NA | NA |
| Gm24695 | 0 | NA | NA | NA |
| Ifnar2 | 243.5630765 | -0.675198955 | 0.115800416 | -5.830712704 |
| Gm21970 | 0 | NA | NA | NA |
| Il10rb | 126.5063742 | -1.458058904 | 0.185472684 | -7.861313425 |
| A930006K02Rik | 5.618572222 | 0.570073918 | 0.467094823 | 1.220467215 |
| Ifnar1 | 638.5446908 | -0.182557512 | 0.068506295 | -2.664828267 |
| Gm7856 | 0.270435483 | 1.765898977 | 3.704343652 | 0.476710355 |
| Mir6367 | 0 | NA | NA | NA |
| Gm7860 | 1.38062972 | -0.621552762 | 0.994750816 | -0.624832623 |
| Ifngr2 | 476.6547008 | -0.23220108 | 0.066846021 | -3.473670917 |
| Tmem50b | 796.4973955 | 0.040909888 | 0.060289832 | 0.678553696 |
| Gm15964 | 4.772112467 | -0.819270701 | 0.453824576 | -1.805258562 |
| Dnajc28 | 95.3526294 | -0.191840499 | 0.181034109 | -1.059692568 |
| Gart | 409.6463015 | -0.403921208 | 0.09910311 | -4.075767222 |
| Son | 6238.367876 | 0.045589269 | 0.060547252 | 0.75295357 |
| Donson | 181.5744247 | -0.163734061 | 0.100480965 | -1.629503261 |
| Atp5o | 1115.945799 | 0.001721568 | 0.076666216 | 0.022455363 |
| Gm10785 | 5.027936496 | 0.185640164 | 0.495587785 | 0.374585835 |
| Cryzl1 | 541.6613726 | 0.000517199 | 0.080817676 | 0.006399581 |
| Itsn1 | 2861.680614 | 0.139065726 | 0.05839851 | 2.381323198 |
| Gm25245 | 0.133144048 | 0.998857947 | 2.968531487 | 0.33648218 |
| Gm15976 | 0.272598396 | 0.593766382 | 1.806920579 | 0.328606796 |
| D430001F17Rik | 1.721321887 | -0.065763499 | 0.753399914 | -0.087288966 |
| Gm27773 | 0 | NA | NA | NA |
| Gm23210 | 0 | NA | NA | NA |
| Mrps6 | 345.9199876 | -1.361358729 | 0.168840554 | -8.062984266 |
| Slc5a3 | 2337.043536 | -1.002544606 | 0.131663117 | -7.614468092 |
| Gm16310 | 32.27711759 | -0.584764417 | 0.333238753 | -1.754791159 |
| Kcne2 | 13.78272213 | 2.543954263 | 0.606414276 | 4.195076474 |
| Smim11 | 43.31674533 | -0.245178908 | 0.228051372 | -1.075103852 |
| 1700048M11Rik | 0 | NA | NA | NA |
| 4930563D23Rik | 0.422874972 | -1.342444611 | 1.684517667 | -0.796931156 |
| Kcne1 | 0.561001133 | 0.69403471 | 1.313266295 | 0.528479801 |
| Rcan1 | 683.9052406 | -0.534644375 | 0.15689027 | -3.407759929 |
| 2410124H12Rik | 1.475924581 | 0.370636398 | 0.989803126 | 0.374454665 |
| Clic6 | 121.7859387 | 0.849089197 | 0.173326832 | 4.898775254 |
| Runx1 | 156.9186044 | -1.17406134 | 0.168838588 | -6.953750043 |
| Gm26626 | 3.202063597 | -2.544831277 | 0.631406791 | -4.030414803 |
| Gm28003 | 0 | NA | NA | NA |
| 1810053B23Rik | 0 | NA | NA | NA |
| Mir802 | 0 | NA | NA | NA |
| 1700029J03Rik | 0.049553047 | -0.443818988 | 3.724945049 | -0.119147795 |
| Setd4 | 64.56323875 | 0.391622999 | 0.17022019 | 2.300684778 |
| Cbr1 | 336.8011914 | 0.193403037 | 0.085109608 | 2.272399569 |
| Gm5678 | 9.467024167 | 0.556282149 | 0.380389536 | 1.462401294 |

| | | | | |
|---|---|---|---|---|
| Gm22344 | 0 | NA | NA | NA |
| Cbr3 | 89.79952323 | 0.380293769 | 0.159972817 | 2.377239936 |
| Dopey2 | 1643.428677 | 0.116910985 | 0.065041882 | 1.797472353 |
| 2310043M15Rik | 4.325146265 | -1.340806645 | 0.48817445 | -2.746572754 |
| Morc3 | 612.5180941 | -0.386038684 | 0.089216126 | -4.327005702 |
| Chaf1b | 23.6865518 | -0.598285853 | 0.28159653 | -2.124620828 |
| Cldn14 | 11.98292911 | 0.373992659 | 0.382028229 | 0.978966031 |
| Sim2 | 1.471461872 | -0.080335278 | 0.800268905 | -0.100385354 |
| Hlcs | 243.6874178 | 0.282199471 | 0.08443609 | 3.342166508 |
| Gm23062 | 1.571274675 | 0.314729001 | 1.068451189 | 0.294565633 |
| Ripply3 | 0.878604591 | -1.625973858 | 1.243367772 | -1.307717552 |
| Pigp | 188.369765 | -0.236370242 | 0.160998795 | -1.468149139 |
| Gm15310 | 0.048949532 | 0.037064902 | 3.724945049 | 0.009950456 |
| Ttc3 | 12670.90535 | 0.491506608 | 0.057806897 | 8.502559989 |
| Gm23692 | 8.647424846 | 0.290253857 | 0.420627953 | 0.690048904 |
| Dscr3 | 356.437628 | -0.827872668 | 0.166536931 | -4.971105583 |
| Dyrk1a | 1886.330775 | 0.014564027 | 0.085773333 | 0.16979668 |
| Gm15971 | 19.22130654 | 0.301080553 | 0.359574629 | 0.83732424 |
| Kcnj6 | 616.7000442 | 0.243764451 | 0.11955664 | 2.038903494 |
| Gm22745 | 0 | NA | NA | NA |
| Gm7976 | 0.168942356 | 0.037073858 | 2.756030596 | 0.013451904 |
| Kcnj15 | 5.024696343 | -2.523699717 | 0.546667994 | -4.616512661 |
| Erg | 22.49114717 | -0.319261498 | 0.237153808 | -1.34622126 |
| Ets2 | 1062.543237 | -0.02243089 | 0.106452195 | -0.210713268 |
| Gm37259 | 0.066050875 | -0.443818988 | 3.724945049 | -0.119147795 |
| 2810404F17Rik | 0.08169834 | -0.443821212 | 3.724945049 | -0.119148392 |
| 1600002D24Rik | 0.782188385 | -1.825089861 | 1.167838184 | -1.562793446 |
| Gm15340 | 0 | NA | NA | NA |
| Psmg1 | 92.11413496 | -0.486337626 | 0.128328678 | -3.789781327 |
| Brwd1 | 2327.039944 | 0.316685955 | 0.115391265 | 2.744453456 |
| Gm15342 | 0 | NA | NA | NA |
| Gm27833 | 5.512382878 | 0.722352817 | 0.529381504 | 1.364522204 |
| 1700093J21Rik | 4.649104205 | 0.566833149 | 0.572406032 | 0.990264109 |
| Gm15341 | 0.325655381 | 2.02089862 | 1.900201575 | 1.063518022 |
| Hmgn1 | 496.6434735 | -0.524114657 | 0.110650258 | -4.736678138 |
| Wrb | 502.1431527 | 0.046211294 | 0.090829547 | 0.508769399 |
| Gm15317 | 0.04393175 | 0.037062677 | 3.724945049 | 0.009949859 |
| Lca5l | 22.78202919 | 0.204136036 | 0.293191967 | 0.69625385 |
| Sh3bgr | 25.544519 | 0.120933573 | 0.24703062 | 0.489548918 |
| B3galt5 | 665.4843436 | 0.220795648 | 0.136141948 | 1.621804681 |
| Igsf5 | 4.211675288 | -0.42346198 | 0.50592051 | -0.837012874 |
| Itgb2l | 0.181155367 | 0.037075376 | 2.695761678 | 0.01375321 |
| Pcp4 | 324.4405224 | 0.528480928 | 0.173101462 | 3.053012507 |
| Dscam | 1116.965166 | 0.498244911 | 0.091477191 | 5.446657329 |
| Bace2 | 46.32598539 | -0.174413472 | 0.174641274 | -0.998695595 |
| Mx1 | 1975.925507 | -0.495781257 | 0.347856662 | -1.425245829 |
| Fam3b | 2.301576675 | -0.866013429 | 0.573782612 | -1.509305809 |

| | | | | |
|---|---|---|---|---|
| Gm9242 | 1.910909441 | -0.686741633 | 0.742725854 | -0.924623304 |
| Gm24777 | 0 | NA | NA | NA |
| Mx2 | 1286.777433 | -0.427350447 | 0.227251722 | -1.880515772 |
| Tmprss2 | 0.798754509 | -0.014818861 | 1.355412079 | -0.010933104 |
| Ripk4 | 12.75005239 | -0.036376797 | 0.293895601 | -0.123774553 |
| Prdm15 | 265.211607 | -0.34998468 | 0.152396116 | -2.296545929 |
| C2cd2 | 212.6215641 | -0.386212905 | 0.09951344 | -3.881012519 |
| mmu-mir-6964 | 0.047948698 | 0.037064902 | 3.724945049 | 0.009950456 |
| Zbtb21 | 472.0135521 | -0.335421321 | 0.092842047 | -3.612816962 |
| C030010L15Rik | 1.895179359 | 1.019353072 | 0.773445165 | 1.317938386 |
| B230307C23Rik | 102.4124531 | -0.274885996 | 0.132577422 | -2.073399778 |
| Gm10232 | 17.35894481 | -0.03251636 | 0.29743386 | -0.109322994 |
| Gm16569 | 0 | NA | NA | NA |
| Gm5479 | 0 | NA | NA | NA |
| Gm25909 | 0.276643538 | -0.881191996 | 2.118355232 | -0.415979333 |
| Pisd-ps2 | 107.0691569 | 0.155752138 | 0.130115222 | 1.197032412 |
| Scaf8 | 1244.518299 | 0.162622407 | 0.095478654 | 1.70323314 |
| Tiam2 | 517.9732827 | 0.021278708 | 0.121365978 | 0.175326798 |
| Tfb1m | 51.82495722 | -0.187051893 | 0.181837466 | -1.028676303 |
| Cldn20 | 1.178313167 | 0.665544372 | 0.950676455 | 0.700074529 |
| Nox3 | 0.272415186 | 1.035992877 | 1.80161399 | 0.575035986 |
| Gm23186 | 0 | NA | NA | NA |
| Arid1b | 2127.270207 | 0.284869595 | 0.063841899 | 4.462110293 |
| Gm15599 | 0.896855871 | -0.731205839 | 1.268505829 | -0.576430807 |
| Gm29050 | 0.223558305 | 0.378998553 | 2.162675168 | 0.175245251 |
| Tmem242 | 341.4840666 | -0.064364841 | 0.09795848 | -0.657062472 |
| Ldhal6b | 8.737325657 | 1.091359955 | 0.483911613 | 2.255287797 |
| Gm22475 | 0.038981395 | -0.443816764 | 3.724945049 | -0.119147198 |
| Zdhhc14 | 490.2302194 | -0.047910974 | 0.114124094 | -0.419814715 |
| 3300005D01Rik | 0.549721405 | 1.162705804 | 1.404785454 | 0.827675002 |
| Gm10231 | 1.299158781 | 0.269062177 | 1.129175902 | 0.238281898 |
| Gm26595 | 0.095897396 | 0.037062677 | 3.724945049 | 0.009949859 |
| Gm26622 | 0 | NA | NA | NA |
| Snx9 | 182.1926714 | -0.642671457 | 0.126646206 | -5.074541727 |
| Synj2 | 857.4048739 | -0.041557416 | 0.092066224 | -0.451386119 |
| Serac1 | 260.8621082 | 0.047731204 | 0.124201755 | 0.384303779 |
| Gtf2h5 | 444.1448567 | 0.182383385 | 0.083871268 | 2.174563355 |
| Tulp4 | 5049.784843 | -0.122689007 | 0.066907028 | -1.833723751 |
| Gm5812 | 12.25279831 | -0.021574081 | 0.309517191 | -0.069702367 |
| Gm22774 | 1.94531079 | -2.684542078 | 0.887101876 | -3.026193665 |
| Gm15590 | 16.93295172 | -1.196181318 | 0.307362264 | -3.891763757 |
| n-R5s26 | 7.164646703 | -0.505010957 | 0.445697756 | -1.133079425 |
| Ppp1r2-ps6 | 4.838948672 | -0.304448423 | 0.461545912 | -0.659627602 |
| Tmem181a | 366.4664287 | 0.339930843 | 0.128428362 | 2.646851825 |
| Dynlt1a | 11.06668227 | 0.038975654 | 0.34533569 | 0.112863092 |
| Gm26848 | 0.173022395 | 0.517971728 | 2.735311408 | 0.189364811 |
| Ppp1r2-ps1 | 3.995967374 | 0.640140477 | 0.578203865 | 1.107118988 |

| | | | | |
|---|---|---|---|---|
| Dynlt1b | 39.38951723 | -0.04977551 | 0.205375942 | -0.242362907 |
| Tmem181b-ps | 797.3566581 | 0.667930789 | 0.259091717 | 2.577970448 |
| Dynlt1c | 50.2770515 | -0.414151269 | 0.21627382 | -1.914939448 |
| Tmem181c-ps | 423.9445639 | 0.6381026 | 0.269809125 | 2.365014901 |
| Dynlt1f | 15.90420978 | -0.635194049 | 0.280481696 | -2.264654191 |
| Sytl3 | 19.32529137 | 0.338332846 | 0.36446392 | 0.928302712 |
| Ezr | 1074.367285 | -0.733143753 | 0.129279271 | -5.671007801 |
| Gm2885 | 11.26381867 | 0.63355907 | 0.326420764 | 1.940927599 |
| Mir692-1 | 0.048949532 | 0.037064902 | 3.724945049 | 0.009950456 |
| Gm29205 | 0 | NA | NA | NA |
| Rsph3b | 25.73551809 | -0.078222602 | 0.241104921 | -0.324433868 |
| Tagap1 | 187.261273 | 0.129820953 | 0.086698032 | 1.497392154 |
| Rnaset2b | 43.79098034 | -1.151580757 | 0.493870397 | -2.331746878 |
| Gm1604b | 6.225126559 | 0.895874322 | 0.630840733 | 1.420127578 |
| Gm25119 | 0 | NA | NA | NA |
| Rps6ka2 | 480.6686781 | 0.313475581 | 0.093870636 | 3.339442398 |
| Tcp10a | 0.109727584 | 0.037071574 | 3.724945049 | 0.009952247 |
| Ttll2 | 0.198340327 | -0.467888748 | 2.231534997 | -0.209671257 |
| Gm9992 | 2.241760946 | -0.409328646 | 0.641231885 | -0.638347306 |
| Gm26057 | 0 | NA | NA | NA |
| Gm25872 | 0 | NA | NA | NA |
| Fndc1 | 124.9694906 | 0.257943536 | 0.174665072 | 1.476789449 |
| E430024P14Rik | 37.33337678 | 0.485404693 | 0.234703192 | 2.068164002 |
| Tagap | 96.63924738 | -0.421414087 | 0.14640116 | -2.878488718 |
| Rsph3a | 115.5642771 | -0.439342461 | 0.104155282 | -4.218148657 |
| Gm1604a | 4.428521402 | 1.516653608 | 0.613839722 | 2.47076485 |
| Gm22416 | 0 | NA | NA | NA |
| Rnaset2a | 127.2262442 | -0.292621779 | 0.438665318 | -0.667072975 |
| Fgfr1op | 286.1447221 | -0.067325952 | 0.102813977 | -0.654832676 |
| Ccr6 | 7.182727363 | 1.582889745 | 0.74915295 | 2.112905977 |
| Mpc1 | 409.8670687 | 0.3717191 | 0.067755638 | 5.486172269 |
| 4930506C21Rik | 16.13295338 | -0.397960073 | 0.265272995 | -1.500190676 |
| Sft2d1 | 202.5780256 | -0.385684277 | 0.136903816 | -2.81719157 |
| Prr18 | 521.3498503 | 0.173261889 | 0.162934923 | 1.063380922 |
| T2 | 5.357486873 | 0.718928656 | 0.461957333 | 1.556266356 |
| Gm16702 | 11.86087499 | 0.622163515 | 0.300636293 | 2.069489046 |
| T | 0.764042974 | -3.221439023 | 1.130567235 | -2.849400656 |
| Pde10a | 2524.291417 | 0.842133898 | 0.183740831 | 4.583270334 |
| Gm15425 | 0 | NA | NA | NA |
| Gm17087 | 0.0878635 | 0.037058229 | 3.724945049 | 0.009948665 |
| Gm15426 | 0.048949532 | 0.037064902 | 3.724945049 | 0.009950456 |
| 1700010I14Rik | 16.46611213 | -0.773226327 | 0.26875978 | -2.877016517 |
| 6530411M01Rik | 2.907305987 | 0.391886933 | 0.705943906 | 0.555124748 |
| Gm8492 | 0.83813809 | 0.953133821 | 1.056682681 | 0.902005719 |
| Gm17748 | 3.088466909 | -0.172461333 | 0.604591074 | -0.28525286 |
| Gm17728 | 0 | NA | NA | NA |
| Pabpc6 | 0.088534442 | -0.905414734 | 3.723500899 | -0.243162217 |

| | | | | |
|---|---|---|---|---|
| Qk | 8468.123674 | -0.143919707 | 0.120904429 | -1.19035927 |
| 1700110C19Rik | 10.49805674 | -0.491868611 | 0.491831604 | -1.000075243 |
| Pacrg | 86.69560762 | -0.094899353 | 0.137180975 | -0.691782173 |
| A230009B12Rik | 11.05179156 | -1.139516218 | 0.406224574 | -2.805138565 |
| Gm16168 | 29.62534574 | 0.087086587 | 0.331321766 | 0.262845958 |
| Gm16169 | 3.038906809 | -0.211524902 | 0.598739915 | -0.353283449 |
| Park2 | 207.7195687 | 1.008977996 | 0.23305721 | 4.329314659 |
| Gm3145 | 3.839486989 | 0.580047307 | 0.584668772 | 0.992095586 |
| Gm28505 | 0.112396584 | -0.917692504 | 3.724434845 | -0.246397787 |
| Gm37402 | 18.41567043 | 1.113833896 | 0.318772875 | 3.494130096 |
| Gm37332 | 0.321071396 | -1.485728356 | 3.697061147 | -0.401867401 |
| Gm28118 | 0 | NA | NA | NA |
| Gm10513 | 0.405560335 | -0.601872005 | 1.479620361 | -0.406774617 |
| Agpat4 | 470.2599501 | -0.367540808 | 0.110728997 | -3.319282388 |
| Tdgf1-ps2 | 2.738925537 | 0.472294694 | 0.668943369 | 0.706030907 |
| Map3k4 | 824.0897494 | 0.16641273 | 0.060644678 | 2.744061577 |
| Gm24794 | 0 | NA | NA | NA |
| 4732491K20Rik | 48.6473752 | 0.193317055 | 0.182236994 | 1.060800281 |
| Plg | 1.917939655 | 0.221524549 | 0.673863006 | 0.328738255 |
| Slc22a3 | 26.84679186 | 1.212076576 | 0.237741926 | 5.098287031 |
| Slc22a2 | 15.39931173 | 0.392797798 | 0.312377506 | 1.25744585 |
| Slc22a1 | 2.772496349 | -0.64241427 | 0.673470807 | -0.953885846 |
| Igf2r | 1261.515538 | -0.388983626 | 0.065951317 | -5.898041829 |
| Airn | 54.16339736 | 0.138710305 | 0.246918401 | 0.56176577 |
| Gm23833 | 6.456263911 | 0.516424705 | 0.544481044 | 0.948471412 |
| Gm16574 | 6.618925518 | 0.040911016 | 0.550797856 | 0.074275917 |
| Gm7162 | 0.709135383 | 2.21055454 | 1.374181152 | 1.608634013 |
| Mas1 | 77.7093898 | 0.506486646 | 0.214611046 | 2.360021334 |
| Gm24085 | 0 | NA | NA | NA |
| Mrgprh | 1.689186202 | -2.898280096 | 1.016333211 | -2.851702635 |
| Pnldc1 | 15.6955681 | -0.010034113 | 0.38437657 | -0.026104904 |
| Mrpl18 | 276.4221612 | -0.272280282 | 0.09379795 | -2.902838302 |
| Tcp1 | 1522.478764 | -0.197579167 | 0.065734278 | -3.005725078 |
| Gm26130 | 0 | NA | NA | NA |
| Snora20 | 1.9528199 | 1.19926404 | 0.85539614 | 1.401998424 |
| Acat3 | 48.71270492 | -0.060589947 | 0.196938374 | -0.307659427 |
| Acat2 | 268.88389 | 0.038920148 | 0.088596047 | 0.439298924 |
| Gm15946 | 1.917568322 | -0.748880342 | 0.78711446 | -0.951424958 |
| Wtap | 592.5866412 | -0.026379698 | 0.074296863 | -0.355058026 |
| Sod2 | 2289.929684 | 0.28748 | 0.067135326 | 4.282097312 |
| Gpr31b | 8.443869178 | -0.614263705 | 0.393976807 | -1.559136714 |
| Tcp10b | 7.010379163 | 0.801881216 | 0.601927924 | 1.332188099 |
| Unc93a | 6.630768804 | -1.300587601 | 0.38400122 | -3.386936115 |
| Gm23050 | 0 | NA | NA | NA |
| Smok2a | 0.046345709 | -0.443818988 | 3.724945049 | -0.119147795 |
| Smok2b | 0 | NA | NA | NA |
| Gm26500 | 0.250452015 | -0.451534 | 2.083611478 | -0.216707388 |

| | | | | |
|---|---|---|---|---|
| Tcp10c | 0.224267024 | -0.451800379 | 2.151436484 | -0.209999404 |
| Gm8597 | 4.592470227 | -0.24862237 | 0.53130481 | -0.467946771 |
| Gm25730 | 0.075043205 | 0.517957688 | 3.724945049 | 0.139051095 |
| Rnu6 | 0.503427435 | 0.023262824 | 1.562361543 | 0.014889527 |
| Tcte2 | 120.1207726 | 0.534356453 | 0.211660851 | 2.524588045 |
| Gm16050 | 0.048949532 | 0.037064902 | 3.724945049 | 0.009950456 |
| Gm8603 | 0 | NA | NA | NA |
| Smok4a | 0 | NA | NA | NA |
| Gm26375 | 1.918309794 | 0.436413336 | 0.802451799 | 0.543849906 |
| Gm24867 | 0 | NA | NA | NA |
| Gm16046 | 10.12097898 | 0.13893779 | 0.513587042 | 0.27052433 |
| Gm16052 | 12.43848109 | 0.171745671 | 0.325977468 | 0.526863628 |
| Gm16049 | 10.2923578 | -0.337037807 | 0.383391716 | -0.879095174 |
| Mllt4 | 2109.491735 | -0.017599911 | 0.082452653 | -0.213454758 |
| Gm3355 | 0.441163179 | -0.387523538 | 1.466646439 | -0.264224238 |
| Gm7168 | 0.355971994 | -2.31531152 | 1.858096305 | -1.246066479 |
| Gm7356 | 0 | NA | NA | NA |
| Gm7358 | 0 | NA | NA | NA |
| 4930488N24Rik | 0.072025462 | 0.517959912 | 3.724945049 | 0.139051692 |
| Dact2 | 350.5464997 | 0.419768263 | 0.140571348 | 2.986158055 |
| Smoc2 | 139.6069329 | -0.353139762 | 0.210667079 | -1.676293056 |
| Thbs2 | 224.1080342 | -0.682955651 | 0.180680327 | -3.779911536 |
| Gm3222 | 8.619157282 | 0.46162838 | 0.353279387 | 1.30669492 |
| Gm23352 | 0 | NA | NA | NA |
| Wdr27 | 15.49212616 | 0.309970339 | 0.339351384 | 0.913419993 |
| Gm7393 | 0 | NA | NA | NA |
| 1600012H06Rik | 204.8958293 | -0.119750113 | 0.09506329 | -1.25968828 |
| Phf10 | 695.9833845 | -0.150592549 | 0.077942602 | -1.932095487 |
| Gm3448 | 0.554247143 | -1.437714084 | 1.35061545 | -1.064488107 |
| Ermard | 167.2744434 | -0.42515898 | 0.147044937 | -2.891354091 |
| Tcte3 | 0 | NA | NA | NA |
| Gm28869 | 9.40787293 | -0.412311182 | 0.329371855 | -1.251810608 |
| Gm28873 | 3.819986632 | -0.805929251 | 0.560265415 | -1.438477603 |
| Gm5091 | 0 | NA | NA | NA |
| Dll1 | 136.4076956 | 0.08418389 | 0.144070661 | 0.584323622 |
| A930024N18Rik | 109.5960546 | 0.359227394 | 0.153631719 | 2.338237158 |
| Fam120b | 782.860338 | 0.047271913 | 0.091694701 | 0.515535933 |
| Psmb1 | 1128.534319 | -0.079942001 | 0.118915034 | -0.672261518 |
| Tbp | 458.9414412 | 0.308922057 | 0.131467136 | 2.349804421 |
| Pdcd2 | 107.0321996 | -0.35063259 | 0.138100324 | -2.538970087 |
| Prdm9 | 37.41141876 | 0.176549987 | 0.187019021 | 0.944021554 |
| Gm6686 | 0 | NA | NA | NA |
| Chd1 | 779.4457597 | -0.177238917 | 0.091542913 | -1.936129314 |
| Rgmb | 695.6204429 | 0.165503216 | 0.076928545 | 2.151388874 |
| Gm25598 | 0 | NA | NA | NA |
| BC002059 | 100.3680048 | -0.221738771 | 0.15107423 | -1.467747157 |
| Zfp960 | 12.66666622 | 0.11708537 | 0.45713605 | 0.25612806 |

| | | | | |
|---|---|---|---|---|
| Zfp97 | 21.19205506 | -0.522911828 | 0.269730042 | -1.938648819 |
| Gm6712 | 39.55212394 | -0.572033622 | 0.203646907 | -2.80894825 |
| Oaz1-ps | 23.5739542 | 0.087586241 | 0.245966663 | 0.356089887 |
| Riok2 | 310.94823 | -0.273401606 | 0.076796502 | -3.560078875 |
| Gm26873 | 0.801515891 | 1.853217816 | 1.138708931 | 1.627472804 |
| Lix1 | 332.7540407 | 0.847939255 | 0.164952105 | 5.140517946 |
| Lnpep | 2780.673188 | -0.096292115 | 0.056297464 | -1.710416553 |
| Vmn2r90 | 0.038981395 | -0.443816764 | 3.724945049 | -0.119147198 |
| Gm23837 | 0 | NA | NA | NA |
| Mir99b | 6.303641128 | 0.407343824 | 0.506659818 | 0.803978941 |
| Gm25927 | 0.066050875 | -0.443818988 | 3.724945049 | -0.119147795 |
| mmu-let-7e | 6.339667104 | 0.51679354 | 0.456150852 | 1.13294437 |
| Spaca6 | 577.2041235 | 0.087580775 | 0.29194683 | 0.299988785 |
| Mir125a | 0 | NA | NA | NA |
| Has1 | 11.28386909 | -0.938515731 | 0.356199269 | -2.634805328 |
| Fpr1 | 8.843961322 | -1.834495248 | 0.456266443 | -4.0206666 |
| Fpr2 | 37.5235979 | -1.100084721 | 0.402969988 | -2.729942066 |
| Fpr3 | 0.406865304 | -2.691602397 | 1.50072701 | -1.793532321 |
| Gm7535 | 0.668988238 | -1.467372715 | 1.262921331 | -1.161887664 |
| Fpr-rs4 | 0.097139547 | -0.443816764 | 3.724945049 | -0.119147198 |
| Vmn2r124 | 0.527106987 | -0.361327243 | 1.335131792 | -0.270630394 |
| Vmn2r91 | 0.129871752 | -0.443193064 | 2.991078735 | -0.148171647 |
| Vmn2r92 | 0.150961315 | -0.410359602 | 2.845512226 | -0.144212911 |
| Vmn2r94 | 0 | NA | NA | NA |
| Vmn2r93 | 0 | NA | NA | NA |
| Vmn2r95 | 0.066050875 | -0.443818988 | 3.724945049 | -0.119147795 |
| Vmn2r-ps114 | 0 | NA | NA | NA |
| Vmn2r96 | 0 | NA | NA | NA |
| Vmn2r-ps115 | 0 | NA | NA | NA |
| Vmn2r-ps116 | 0 | NA | NA | NA |
| Vmn2r-ps117 | 0 | NA | NA | NA |
| Vmn2r97 | 0 | NA | NA | NA |
| Vmn2r-ps118 | 0 | NA | NA | NA |
| Vmn2r98 | 0 | NA | NA | NA |
| Vmn2r-ps119 | 0 | NA | NA | NA |
| Vmn2r-ps120 | 0 | NA | NA | NA |
| Vmn2r99 | 0.062426873 | 0.517959912 | 3.724945049 | 0.139051692 |
| Vmn2r-ps121 | 0 | NA | NA | NA |
| Vmn2r100 | 0 | NA | NA | NA |
| Vmn2r101 | 0 | NA | NA | NA |
| Vmn2r102 | 0 | NA | NA | NA |
| Vmn2r-ps122 | 0 | NA | NA | NA |
| Vmn2r103 | 0 | NA | NA | NA |
| Vmn2r-ps123 | 0 | NA | NA | NA |
| Vmn2r-ps124 | 0 | NA | NA | NA |
| Vmn2r-ps125 | 0 | NA | NA | NA |
| Vmn2r104 | 0 | NA | NA | NA |

| | | | | |
|---|---|---|---|---|
| Vmn2r-ps126 | 0 | NA | NA | NA |
| Fpr-rs7 | 0 | NA | NA | NA |
| Fpr-rs6 | 0 | NA | NA | NA |
| Vmn2r105 | 0 | NA | NA | NA |
| Vmn2r106 | 0 | NA | NA | NA |
| Vmn2r-ps127 | 0 | NA | NA | NA |
| Vmn2r107 | 0 | NA | NA | NA |
| Vmn1r-ps141 | 0 | NA | NA | NA |
| Vmn1r224 | 0 | NA | NA | NA |
| Vmn1r-ps142 | 0 | NA | NA | NA |
| Vmn1r-ps143 | 0 | NA | NA | NA |
| Vmn2r108 | 0 | NA | NA | NA |
| Vmn1r225 | 0 | NA | NA | NA |
| Vmn2r109 | 0.225617294 | -0.496608103 | 2.520825427 | -0.197002179 |
| Gm5145 | 2.222558862 | -0.946170579 | 0.72271492 | -1.309189216 |
| Vmn2r110 | 0.133946075 | 0.036100024 | 3.626570338 | 0.009954315 |
| Fpr-rs3 | 0 | NA | NA | NA |
| Vmn1r226 | 0 | NA | NA | NA |
| Vmn1r227 | 0 | NA | NA | NA |
| Vmn1r228 | 0.088534442 | -0.905414734 | 3.723500899 | -0.243162217 |
| Vmn1r229 | 0 | NA | NA | NA |
| Vmn1r230 | 0 | NA | NA | NA |
| Vmn1r231 | 0.108767819 | -0.443823436 | 3.724945049 | -0.119148989 |
| Vmn1r232 | 0.192963196 | -1.188374014 | 2.637066134 | -0.450642477 |
| Vmn1r-ps144 | 0 | NA | NA | NA |
| Ppp2r1a | 4099.317594 | 0.186685607 | 0.060675883 | 3.076767873 |
| Vmn1r-ps145 | 0 | NA | NA | NA |
| Vmn1r233 | 0 | NA | NA | NA |
| Zfp160 | 354.561668 | -0.102290108 | 0.084080599 | -1.216572065 |
| Gm5681 | 0 | NA | NA | NA |
| Vmn1r-ps146 | 0 | NA | NA | NA |
| Gm5092 | 0.092140247 | 0.037060453 | 3.724945049 | 0.009949262 |
| Gm6811 | 0.490429956 | -0.551807754 | 1.290119766 | -0.427718239 |
| Gm7732 | 0.049553047 | -0.443818988 | 3.724945049 | -0.119147795 |
| Gm7736 | 0 | NA | NA | NA |
| Gm7740 | 0 | NA | NA | NA |
| Gm7742 | 0 | NA | NA | NA |
| Vmn1r-ps147 | 0 | NA | NA | NA |
| Vmn1r234 | 0 | NA | NA | NA |
| Vmn1r-ps148 | 0 | NA | NA | NA |
| Vmn1r235 | 0.11193333 | 0.037073798 | 3.724945049 | 0.009952844 |
| Vmn1r-ps149 | 0 | NA | NA | NA |
| Gm26384 | 1.87298797 | -2.405038345 | 0.866544291 | -2.775436142 |
| Vmn1r236 | 0 | NA | NA | NA |
| Vmn1r237 | 0.097139547 | -0.443816764 | 3.724945049 | -0.119147198 |
| Vmn1r-ps150 | 0 | NA | NA | NA |
| Zfp677 | 55.30162814 | -0.668982562 | 0.16762505 | -3.990946232 |

| | | | | |
|---|---|---|---|---|
| Zfp54 | 27.41881594 | -1.243996696 | 0.273308276 | -4.551624684 |
| Zfp51 | 105.8080167 | -0.507374067 | 0.130680184 | -3.882563148 |
| Gm26753 | 1.899580325 | -0.584047253 | 0.77469281 | -0.753908189 |
| Zfp53 | 100.822271 | -0.355640503 | 0.14140114 | -2.515117643 |
| 9330136K24Rik | 11.4954026 | -0.036541682 | 0.46295824 | -0.078930839 |
| Zfp52 | 67.11583658 | -0.413375769 | 0.176960559 | -2.335976845 |
| Zfp948 | 235.9906384 | -0.155455633 | 0.122726324 | -1.266685316 |
| 3110052M02Rik | 172.7372577 | -0.250872408 | 0.108276873 | -2.316952824 |
| Gm10509 | 18.52972673 | -0.021451111 | 0.245270887 | -0.087458855 |
| Gm10226 | 26.0094674 | -0.082923785 | 0.219026281 | -0.378601985 |
| Zfp760 | 279.5658373 | -0.206586646 | 0.150160046 | -1.375776395 |
| Zfp229 | 105.1619278 | -0.36729065 | 0.181327902 | -2.025560582 |
| 4930515G13Rik | 0 | NA | NA | NA |
| Gm9818 | 0.049553047 | -0.443818988 | 3.724945049 | -0.119147795 |
| Zfp820 | 16.99717544 | -0.440913266 | 0.276276203 | -1.595914743 |
| 2210404O09Rik | 62.10937004 | -0.11968683 | 0.146531594 | -0.816798801 |
| Zfp942 | 313.3301051 | 0.00963302 | 0.098421764 | 0.097874896 |
| Zfp943 | 306.7466701 | 0.130433615 | 0.101365687 | 1.286762997 |
| Gm7805 | 0 | NA | NA | NA |
| Gm7809 | 5.578788543 | -1.762200976 | 0.498751104 | -3.533227216 |
| Gm9772 | 0 | NA | NA | NA |
| Gm7052 | 0.038981395 | -0.443816764 | 3.724945049 | -0.119147198 |
| Zfp947 | 19.48152888 | -0.399673643 | 0.253083495 | -1.57921655 |
| Gm4944 | 183.4875709 | -0.188957745 | 0.115991654 | -1.629063289 |
| Zfp944 | 179.0922888 | 0.077203587 | 0.102770379 | 0.751224113 |
| Zfp758 | 147.1029375 | -0.433590765 | 0.106113675 | -4.086096978 |
| Zfp946 | 267.7372296 | 0.015494909 | 0.094695959 | 0.163627983 |
| Vmn2r111 | 0.571153248 | -2.5561859 | 1.490375343 | -1.715128952 |
| Vmn2r112 | 0 | NA | NA | NA |
| Gm9805 | 0.259702268 | -0.466045761 | 2.063779569 | -0.225821482 |
| Gm5224 | 0 | NA | NA | NA |
| Gm5493 | 1.253500652 | -0.122477323 | 0.867582911 | -0.141170741 |
| Gm16386 | 36.63223039 | -0.313177578 | 0.223543352 | -1.400970215 |
| Zfp945 | 660.5015324 | -0.127494855 | 0.1193288 | -1.068433224 |
| Vmn2r-ps128 | 0 | NA | NA | NA |
| Vmn2r113 | 0.044544648 | 0.517957688 | 3.724945049 | 0.139051095 |
| Vmn2r-ps129 | 0 | NA | NA | NA |
| Gm18415 | 0 | NA | NA | NA |
| Gm18396 | 0 | NA | NA | NA |
| Vmn2r-ps130 | 0 | NA | NA | NA |
| Gm18397 | 0 | NA | NA | NA |
| Gm18416 | 0 | NA | NA | NA |
| Vmn2r-ps131 | 0 | NA | NA | NA |
| Vmn2r-ps132 | 0.415262948 | -0.430961352 | 1.392939392 | -0.30938988 |
| Zfp40 | 220.4414794 | 0.045213331 | 0.131665939 | 0.343394285 |
| Vmn2r-ps133 | 0 | NA | NA | NA |
| Vmn2r114 | 0 | NA | NA | NA |

| | | | | |
|---|---|---|---|---|
| Vmn2r115 | 0 | NA | NA | NA |
| Vmn2r116 | 0 | NA | NA | NA |
| Vmn2r-ps134 | 0 | NA | NA | NA |
| Vmn2r117 | 0.156477978 | 0.517971164 | 2.82333336 | 0.183460859 |
| Casp16 | 0.468340293 | -0.340941924 | 1.439663442 | -0.236820575 |
| Gm26695 | 3.060190583 | 0.704035982 | 0.680799918 | 1.034130533 |
| Zfp213 | 78.04727618 | -0.366434715 | 0.139946129 | -2.618398357 |
| Zfp13 | 172.7206419 | 0.406711439 | 0.1051936 | 3.866313526 |
| Zscan10 | 0.304370145 | 2.269092941 | 3.68669199 | 0.615482103 |
| Mmp25 | 15.55909309 | -0.985063253 | 0.33758662 | -2.91795704 |
| Ccdc64b | 0.263919366 | 1.28687532 | 2.035124248 | 0.632332558 |
| Thoc6 | 57.82215382 | -0.624679307 | 0.175680069 | -3.555777893 |
| Hcfc1r1 | 508.7659468 | 0.118965484 | 0.097757516 | 1.216944632 |
| Tnfrsf12a | 409.2070675 | -1.662431184 | 0.38994582 | -4.263236328 |
| Cldn6 | 0.270371171 | 0.532240858 | 1.813105727 | 0.293552025 |
| Cldn9 | 1.372057613 | -0.509722534 | 0.833822015 | -0.611308558 |
| 1520401A03Rik | 0.641047896 | 0.560900243 | 1.445437037 | 0.388048894 |
| Pkmyt1 | 17.10398538 | -0.112187558 | 0.405292365 | -0.276806493 |
| Paqr4 | 569.8115097 | 0.082384492 | 0.100718593 | 0.817967066 |
| Kremen2 | 6.737691786 | 0.031048527 | 0.431142326 | 0.072014564 |
| 9530082P21Rik | 164.7586404 | -0.619717497 | 0.209795754 | -2.953908668 |
| Flywch1 | 2434.68631 | 0.046537651 | 0.059433961 | 0.783014459 |
| Gm16275 | 0.151377979 | -1.354346162 | 2.835615389 | -0.477619838 |
| Flywch2 | 102.393296 | 0.321621354 | 0.144940966 | 2.21898171 |
| D930048G16Rik | 12.89624846 | 0.438933922 | 0.376554007 | 1.165659941 |
| Srrm2 | 8917.247412 | 0.182143809 | 0.0979935 | 1.858733583 |
| Mir5125 | 32.86136734 | 0.833609897 | 0.361402344 | 2.306597924 |
| Tceb2 | 859.0037201 | -0.069356819 | 0.0987077 | -0.702648516 |
| Prss33 | 1.317762752 | 2.430610917 | 1.180509996 | 2.058949882 |
| Prss41 | 25.3916789 | 0.10283623 | 0.219819581 | 0.467821062 |
| Prss32 | 1.380959146 | -0.898079412 | 1.063961838 | -0.844089873 |
| Gm15947 | 0.088534442 | -0.905414734 | 3.723500899 | -0.243162217 |
| Prss21 | 0 | NA | NA | NA |
| Dcpp1 | 0 | NA | NA | NA |
| Dcpp2 | 0 | NA | NA | NA |
| Dcpp3 | 0 | NA | NA | NA |
| Sbp | 0 | NA | NA | NA |
| Sbpl | 0 | NA | NA | NA |
| Prss30 | 3.148853658 | -2.165802775 | 0.664230677 | -3.260618412 |
| Prss22 | 4.912676513 | -2.211784706 | 0.549132025 | -4.027783128 |
| Gm27700 | 0 | NA | NA | NA |
| Gm25092 | 0 | NA | NA | NA |
| Gm5225 | 0 | NA | NA | NA |
| Prss27 | 0.22024148 | -1.469686779 | 2.161641247 | -0.679893937 |
| Kctd5 | 261.9043685 | -0.464840171 | 0.110964035 | -4.189106581 |
| Pdpk1 | 2152.703641 | -0.067152318 | 0.073319047 | -0.91589186 |
| Amdhd2 | 151.6535398 | -0.908711355 | 0.16031934 | -5.668133089 |

| | | | | |
|---|---|---|---|---|
| RP23-27H19.7 | 0 | NA | NA | NA |
| Atp6v0c | 323.8959897 | 0.214040136 | 0.128975076 | 1.659546493 |
| Tbc1d24 | 880.7733529 | 0.258818483 | 0.111216775 | 2.327153286 |
| BC028777 | 3.029716819 | 0.26299512 | 0.58402884 | 0.450311871 |
| 1600002H07Rik | 75.97695628 | -0.163066104 | 0.164701978 | -0.990067668 |
| Ccnf | 89.28741565 | -0.942817695 | 0.180022954 | -5.237208211 |
| Mir5134 | 0.102891481 | 0.51795324 | 3.724945049 | 0.139049901 |
| Mir6965 | 0.049553047 | -0.443818988 | 3.724945049 | -0.119147795 |
| Abca17 | 4.624081096 | 0.20557042 | 0.481284021 | 0.427129119 |
| Gm24427 | 0 | NA | NA | NA |
| Abca3 | 1345.901901 | -0.052377707 | 0.066782757 | -0.784299849 |
| Gm25618 | 0 | NA | NA | NA |
| D330041H03Rik | 54.50038671 | 0.111295573 | 0.225157211 | 0.494301614 |
| Rnps1 | 386.2518119 | -0.060510698 | 0.065816449 | -0.919385635 |
| Eci1 | 99.86578218 | -0.432747696 | 0.176107987 | -2.457286033 |
| Dnase1l2 | 15.71919709 | -0.597582697 | 0.586802208 | -1.018371589 |
| E4f1 | 179.8701777 | -0.006291655 | 0.104581903 | -0.06016007 |
| Pgp | 366.4948428 | -0.208097653 | 0.121680771 | -1.710193407 |
| Mlst8 | 286.6817983 | -0.056193426 | 0.070770672 | -0.794021368 |
| Bricd5 | 7.409630445 | 0.630707708 | 0.412619741 | 1.528544677 |
| Caskin1 | 1583.967537 | 0.360205753 | 0.071532635 | 5.035544281 |
| Traf7 | 402.3587103 | -0.166315004 | 0.067873228 | -2.450377102 |
| Rab26 | 113.4819396 | 0.929823021 | 0.171522003 | 5.42101307 |
| Rab26os | 18.9116278 | -0.095809878 | 0.282569889 | -0.339066127 |
| Snord60 | 0 | NA | NA | NA |
| Pkd1 | 2223.361591 | 0.270505041 | 0.139833359 | 1.93448146 |
| Gm24589 | 0 | NA | NA | NA |
| Tsc2 | 1269.544961 | 0.046699069 | 0.04910979 | 0.950911598 |
| Nthl1 | 19.09256242 | -0.750139141 | 0.295949246 | -2.534688465 |
| Slc9a3r2 | 379.4006015 | -0.286231703 | 0.080346223 | -3.562478648 |
| Npw | 2.00579695 | -3.708245716 | 1.010764824 | -3.668752242 |
| Zfp598 | 484.0735371 | -0.312523067 | 0.113389107 | -2.756200077 |
| Syngr3 | 1091.758285 | 0.226496349 | 0.064094413 | 3.533792422 |
| Gfer | 118.1883123 | -0.535976024 | 0.13298043 | -4.030487968 |
| Noxo1 | 16.22553485 | 0.044269696 | 0.29433718 | 0.150404703 |
| Tbl3 | 217.5084557 | -0.210059212 | 0.104952032 | -2.00147828 |
| Rnf151 | 7.038343901 | -0.943530828 | 0.423747713 | -2.226633441 |
| Rps2 | 374.06437 | -0.029234051 | 0.134535223 | -0.217296636 |
| Snhg9 | 0.044191549 | 0.037062677 | 3.724945049 | 0.009949859 |
| Snora78 | 10.19340988 | -1.013113522 | 0.44204462 | -2.291880673 |
| Snora64 | 0 | NA | NA | NA |
| Ndufb10 | 578.8159123 | -0.008462288 | 0.101729109 | -0.083184525 |
| Rpl3l | 1.739503778 | 0.337697506 | 0.830360326 | 0.406687911 |
| Msrb1 | 303.5260477 | -1.125314626 | 0.194601956 | -5.78264807 |
| Hs3st6 | 1.483788266 | 0.462961329 | 1.069592238 | 0.432839088 |
| Meiob | 0.700302977 | -0.538983042 | 1.089965592 | -0.494495465 |
| Hagh | 454.8144511 | -0.193399428 | 0.082500289 | -2.344227294 |

| | | | | |
|---|---|---|---|---|
| Fahd1 | 88.20798843 | -0.043112842 | 0.144363387 | -0.298641106 |
| Igfals | 0.769314682 | 2.381861831 | 3.623184436 | 0.657394586 |
| Nubp2 | 229.2468458 | -0.483905444 | 0.130970209 | -3.694774924 |
| Spsb3 | 316.2208862 | -0.146699167 | 0.112101264 | -1.308630802 |
| Eme2 | 125.0526026 | -0.359980543 | 0.135548451 | -2.655733366 |
| Mapk8ip3 | 5120.798928 | 0.136164664 | 0.050075479 | 2.719188414 |
| Mrps34 | 292.3664307 | 0.108407365 | 0.102199018 | 1.060747618 |
| Nme3 | 0.049553047 | -0.443818988 | 3.724945049 | -0.119147795 |
| Hn1l | 155.596658 | -1.041842482 | 0.193376334 | -5.387642111 |
| Cramp1l | 709.8548994 | 0.146664613 | 0.100632863 | 1.45742264 |
| Ift140 | 319.2473791 | 0.165604544 | 0.129931315 | 1.274554516 |
| Tmem204 | 25.86076114 | 0.315310854 | 0.229345062 | 1.374831667 |
| Telo2 | 173.173594 | -0.004423928 | 0.123499703 | -0.035821366 |
| Ptx4 | 2.281795086 | 1.874300304 | 0.738697689 | 2.5373036 |
| Clcn7 | 760.3513654 | -0.541115296 | 0.100431653 | -5.387895957 |
| Ccdc154 | 1.912657768 | -1.712624293 | 0.800020093 | -2.140726597 |
| BC003965 | 211.743186 | 0.024803288 | 0.109877334 | 0.225736169 |
| Unkl | 277.8387526 | 0.270111666 | 0.127609424 | 2.116706256 |
| Gnptg | 438.2214292 | -0.019734264 | 0.073714347 | -0.267712656 |
| Tsr3 | 147.8211637 | -0.067430281 | 0.130838887 | -0.515368806 |
| Baiap3 | 567.9275415 | 0.178545316 | 0.160065466 | 1.115451825 |
| Mir3547 | 0 | NA | NA | NA |
| Ube2i | 715.8321189 | 0.234092782 | 0.106720106 | 2.19352089 |
| Prss34 | 0.061920613 | -0.443816764 | 3.724945049 | -0.119147198 |
| Gm25921 | 0 | NA | NA | NA |
| Prss28 | 0 | NA | NA | NA |
| Prss29 | 0 | NA | NA | NA |
| Gm5430 | 0.892004414 | 0.624822129 | 1.063233864 | 0.587661991 |
| Tpsab1 | 3.207720559 | -0.5533049 | 0.54163864 | -1.021538826 |
| Tpsb2 | 1.275377428 | 0.270802091 | 0.918161459 | 0.294939511 |
| Tpsg1 | 0.042716945 | 0.037062677 | 3.724945049 | 0.009949859 |
| Cacna1h | 727.8784213 | 0.265933405 | 0.129926318 | 2.046801676 |
| Tekt4 | 20.49047743 | 0.535683909 | 0.373768443 | 1.43319726 |
| Sstr5 | 11.97499174 | 0.348361684 | 0.332964345 | 1.046243207 |
| Sox8 | 497.4054169 | 0.543326149 | 0.117924973 | 4.607388372 |
| 2810468N07Rik | 100.8375544 | 0.760575847 | 0.196645797 | 3.867745248 |
| Lmf1 | 167.6280357 | 0.254716035 | 0.138622091 | 1.837485163 |
| Gm22645 | 0 | NA | NA | NA |
| Gng13 | 254.443737 | 0.207735662 | 0.20376269 | 1.019498033 |
| Chtf18 | 22.59824114 | -0.581566539 | 0.335427583 | -1.733806546 |
| Rpusd1 | 275.4402358 | 0.343978908 | 0.089540239 | 3.841612583 |
| Mslnl | 3.752562866 | -1.242630457 | 0.598566873 | -2.076009403 |
| Gm25083 | 0 | NA | NA | NA |
| Msln | 26.85201184 | -0.711250259 | 0.313612577 | -2.26792645 |
| Narfl | 208.0663513 | -0.126343244 | 0.078205461 | -1.615529681 |
| Haghl | 395.0534574 | 0.521709036 | 0.101069172 | 5.161900739 |
| Mir6966 | 0 | NA | NA | NA |

| | | | | |
|---|---|---|---|---|
| Ccdc78 | 1.842483344 | -0.239153133 | 0.79524195 | -0.300730027 |
| Fam173a | 248.2616466 | -0.062119143 | 0.123661044 | -0.502333971 |
| Metrn | 475.4136771 | -0.534446922 | 0.260607506 | -2.050773329 |
| Gm26694 | 2.65610448 | -0.524753791 | 0.59781613 | -0.877784598 |
| Fbxl16 | 5262.479825 | 1.119750147 | 0.127985945 | 8.7490087 |
| Wdr24 | 242.1513502 | -0.130230835 | 0.080455437 | -1.618670417 |
| Jmjd8 | 285.0629585 | 0.050945271 | 0.090673836 | 0.56185195 |
| Stub1 | 735.6538842 | -0.083098316 | 0.09050339 | -0.918179039 |
| Mir3082 | 1.578690045 | 0.457899456 | 0.866975651 | 0.528157228 |
| Rhbdl1 | 30.55610411 | 0.425836113 | 0.222930573 | 1.910173676 |
| Gm20683 | 0 | NA | NA | NA |
| Rhot2 | 307.1541338 | 0.150769418 | 0.08947022 | 1.685135212 |
| Wdr90 | 125.7964112 | 0.349020809 | 0.198581496 | 1.757569638 |
| Fam195a | 55.95550801 | -0.121541241 | 0.160369465 | -0.757882686 |
| 0610011F06Rik | 119.2220752 | 0.131561076 | 0.126015853 | 1.044004172 |
| Wfikkn1 | 12.97584509 | 1.750313624 | 0.612459751 | 2.857842691 |
| Rab40c | 356.9302055 | 0.264431956 | 0.108548572 | 2.436070332 |
| Gm10012 | 55.0896093 | 0.245118226 | 0.205837332 | 1.190834645 |
| Gm16278 | 14.17840066 | 0.078506985 | 0.325151176 | 0.241447644 |
| Pigq | 882.6485576 | -0.126215189 | 0.101468703 | -1.243882944 |
| Nhlrc4 | 4.440385187 | 0.382032386 | 0.513165094 | 0.744462923 |
| A930017K11Rik | 39.98571024 | 1.05059532 | 0.21850978 | 4.80800136 |
| D630044L22Rik | 2.092098959 | 0.486566971 | 0.867220602 | 0.56106482 |
| Capn15 | 519.3883034 | -0.040558873 | 0.102035061 | -0.397499374 |
| 1700022N22Rik | 1.65831888 | -1.755995915 | 0.86419326 | -2.031948172 |
| Rab11fip3 | 1324.716031 | 0.309582733 | 0.083115607 | 3.724724439 |
| Decr2 | 128.9572076 | 0.29959164 | 0.12235013 | 2.448641782 |
| Nme4 | 11.32066076 | 0.17438963 | 0.331994357 | 0.525278898 |
| Gm8186 | 9.968334961 | -0.655500402 | 0.37895635 | -1.72975173 |
| Tmem8 | 155.1022499 | -0.140616886 | 0.108289562 | -1.298526689 |
| Mrpl28 | 258.4561109 | -0.27180376 | 0.101627127 | -2.674519768 |
| Axin1 | 402.0340246 | -0.324602834 | 0.089906749 | -3.610439022 |
| Pdia2 | 0.090277459 | -0.443823436 | 3.724945049 | -0.119148989 |
| Arhgdig | 330.2245481 | 0.300479427 | 0.100054797 | 3.003148621 |
| Rgs11 | 126.1618791 | 0.69740472 | 0.292924375 | 2.380835397 |
| Itfg3 | 218.9037408 | -0.463415275 | 0.140188592 | -3.305656112 |
| Gm23123 | 4.679423458 | -0.308352588 | 0.57714847 | -0.534269091 |
| Luc7l | 1216.152592 | -0.032903926 | 0.071439522 | -0.46058435 |
| Gm16580 | 14.51804168 | 0.153103794 | 0.263095267 | 0.58193291 |
| Hba-ps4 | 0.121388914 | 0.037071574 | 3.724945049 | 0.009952247 |
| Neurl1b | 522.8334351 | 1.205979915 | 0.177362409 | 6.799523754 |
| Dusp1 | 584.3483468 | -0.420459819 | 0.314023219 | -1.338945 |
| Gm8225 | 0.51166646 | -1.518772536 | 1.333841062 | -1.13864581 |
| 1700049J03Rik | 1.147468153 | 1.689083958 | 1.115231196 | 1.51455946 |
| Ergic1 | 1341.076534 | 0.013705934 | 0.054994574 | 0.249223391 |
| Gm22420 | 0 | NA | NA | NA |
| Gm17218 | 0 | NA | NA | NA |

| | | | | |
|---|---|---|---|---|
| Atp6v0e | 321.5954019 | -0.482582538 | 0.1286776 | -3.750322799 |
| Crebrf | 2489.558306 | 0.176961344 | 0.091375064 | 1.936648112 |
| Bnip1 | 161.7726201 | 0.144276746 | 0.158335933 | 0.911206592 |
| Nkx2-5 | 0 | NA | NA | NA |
| Gm17382 | 7.241949452 | 0.970284986 | 0.383296564 | 2.531421036 |
| Gm20468 | 0.607877193 | 0.707829264 | 1.236877663 | 0.572271038 |
| Kifc5b | 20.39272261 | 0.097849159 | 0.269422886 | 0.363180577 |
| Gm27775 | 0 | NA | NA | NA |
| Phf1 | 420.9294313 | 0.012759651 | 0.085358702 | 0.149482725 |
| Mir6968 | 0.061920613 | -0.443816764 | 3.724945049 | -0.119147198 |
| Cuta | 316.9454618 | -0.204832942 | 0.107858105 | -1.899096425 |
| Syngap1 | 4379.193043 | 1.078815088 | 0.135752526 | 7.946924586 |
| Mir3083 | 0 | NA | NA | NA |
| Zbtb9 | 0 | NA | NA | NA |
| Ggnbp1 | 24.84235746 | 0.81395381 | 0.271116536 | 3.00222857 |
| Bak1 | 376.0418335 | -0.876955859 | 0.159614108 | -5.494225218 |
| Itpr3 | 158.6747885 | -0.529268871 | 0.162508871 | -3.256861415 |
| Mir7677 | 0 | NA | NA | NA |
| Itpr3os | 0 | NA | NA | NA |
| Uqcc2 | 404.6628392 | -0.087805316 | 0.106772951 | -0.822355427 |
| Ip6k3 | 19.80013244 | 1.347790435 | 0.287675518 | 4.685106487 |
| Lemd2 | 337.854379 | -0.431920893 | 0.086244421 | -5.008102426 |
| Gm26724 | 0.046345709 | -0.443818988 | 3.724945049 | -0.119147795 |
| mmu-mir-7214 | 0.072025462 | 0.517959912 | 3.724945049 | 0.139051692 |
| Mir7216 | 0 | NA | NA | NA |
| Gm10505 | 1.106753324 | -0.803658171 | 0.88897826 | -0.904024549 |
| mmu-mir-7217 | 0 | NA | NA | NA |
| mmu-mir-7218 | 0 | NA | NA | NA |
| Grm4 | 1362.166969 | -0.200794483 | 0.169726815 | -1.183045135 |
| Gm16005 | 0 | NA | NA | NA |
| Gm9703 | 3.905880329 | -0.751542959 | 0.533057235 | -1.409872917 |
| Hmga1 | 483.005091 | -1.23556089 | 0.261993464 | -4.715998914 |
| AI413582 | 165.7897589 | -0.034671076 | 0.128452159 | -0.269914309 |
| Nudt3 | 2683.122826 | 0.080369491 | 0.065119335 | 1.234187836 |
| Rps10 | 731.2072688 | -0.391734488 | 0.115129543 | -3.402554017 |
| Gm15420 | 1.006594715 | 0.133711614 | 1.042611543 | 0.128246819 |
| Pacsin1 | 4703.755317 | 0.242325651 | 0.075202217 | 3.222320566 |
| Spdef | 9.032787089 | 1.528046554 | 0.562233435 | 2.717815159 |
| Gm15458 | 0.243671637 | -0.369045282 | 2.123491625 | -0.173791729 |
| D17Wsu92e | 3051.297663 | 0.097833483 | 0.078029759 | 1.253797066 |
| Rps2-ps9 | 0.733389001 | 0.650738496 | 1.406917774 | 0.462527738 |
| Snrpc | 219.5654359 | -0.264092468 | 0.08739204 | -3.021928154 |
| Uhrf1bp1 | 565.7822905 | -0.156987016 | 0.10004288 | -1.569197285 |
| Taf11 | 229.7296066 | 0.175722482 | 0.085305421 | 2.059921628 |
| Anks1 | 569.0372512 | -0.392968032 | 0.066455912 | -5.913214087 |
| Gm15597 | 0 | NA | NA | NA |
| Gm15598 | 0.962678808 | 0.670072494 | 1.108409735 | 0.604535013 |

| | | | | |
|---|---|---|---|---|
| Tcp11 | 4.048470876 | -0.14935845 | 0.542001172 | -0.275568501 |
| 4930526A20Rik | 2.494837672 | 0.222834021 | 0.719206518 | 0.30983315 |
| Scube3 | 58.8908513 | 0.61539506 | 0.245289088 | 2.50885624 |
| Zfp523 | 600.1388987 | 0.220233702 | 0.107029713 | 2.057687503 |
| Def6 | 48.95250913 | -0.20940745 | 0.189347475 | -1.10594266 |
| Ppard | 357.9241621 | -0.231462091 | 0.070929075 | -3.263289289 |
| Fance | 139.6977314 | 0.129401637 | 0.111052595 | 1.165228397 |
| Rpl10a | 628.5301939 | -0.184675791 | 0.091733447 | -2.013178355 |
| Tead3 | 55.35341921 | -0.449065394 | 0.248895588 | -1.804232035 |
| Tulp1 | 1.121467505 | 1.493903908 | 1.203874037 | 1.240913802 |
| Fkbp5 | 903.5710338 | 0.521096589 | 0.171252867 | 3.042848842 |
| Gm22146 | 0 | NA | NA | NA |
| Gm20109 | 2.702534539 | 0.682354386 | 0.622511166 | 1.096131962 |
| E230001N04Rik | 14.53897155 | -0.084892979 | 0.284052969 | -0.2988632 |
| Gm25195 | 0 | NA | NA | NA |
| Armc12 | 0.464894546 | -0.9717641 | 1.553823612 | -0.625401811 |
| Clpsl2 | 0 | NA | NA | NA |
| Clps | 0.10503227 | -0.905414734 | 3.723500899 | -0.243162217 |
| Mir6969 | 0 | NA | NA | NA |
| Lhfpl5 | 32.4049498 | -0.339404271 | 0.25471052 | -1.332509828 |
| Srpk1 | 762.4953308 | -0.184389052 | 0.084147192 | -2.19126802 |
| Slc26a8 | 26.07395248 | 0.956731614 | 0.251350608 | 3.806362847 |
| Mapk14 | 951.4501698 | -0.016888234 | 0.063187614 | -0.267271266 |
| Mapk13 | 6.079458248 | 0.078952229 | 0.456992064 | 0.172764989 |
| Brpf3 | 576.6212084 | -0.011976477 | 0.081298268 | -0.147315274 |
| Pnpla1os | 0.18989612 | -0.707205106 | 2.65062973 | -0.266806449 |
| Pnpla1 | 2.646101737 | 0.481598013 | 0.600617697 | 0.801837868 |
| 4930539E08Rik | 83.46186464 | 0.576729349 | 0.161970755 | 3.560700503 |
| 1700030A11Rik | 0.307834097 | -0.008731097 | 1.753240997 | -0.004979975 |
| Gm16191 | 0.066050875 | -0.443818988 | 3.724945049 | -0.119147795 |
| Pxt1 | 0.042716945 | 0.037062677 | 3.724945049 | 0.009949859 |
| Kctd20 | 733.6283113 | -0.211843108 | 0.057073813 | -3.711739191 |
| Stk38 | 479.5171983 | -0.246090462 | 0.116934693 | -2.104511977 |
| Gm23887 | 7.551778246 | 1.177536259 | 0.524171036 | 2.246473341 |
| Gm16196 | 0.628440879 | -1.264572724 | 1.405008799 | -0.900046124 |
| Gm16195 | 0.038981395 | -0.443816764 | 3.724945049 | -0.119147198 |
| Srsf3 | 1460.401065 | -0.179149538 | 0.086673046 | -2.06695791 |
| Trp53cor1 | 8.461310733 | -0.533669143 | 0.394726906 | -1.351995862 |
| Gm27599 | 0.04393175 | 0.037062677 | 3.724945049 | 0.009949859 |
| Cdkn1a | 962.23884 | -0.427517189 | 0.356243389 | -1.200070522 |
| Gm16194 | 0.143847454 | -0.92560625 | 2.898586737 | -0.319330189 |
| Rab44 | 2.105186647 | -1.368375962 | 0.664040741 | -2.060680733 |
| Rpl35a-ps3 | 1.427311429 | 0.411433954 | 0.853672468 | 0.481957623 |
| Cpne5 | 630.1572259 | 0.611425428 | 0.247530487 | 2.470101506 |
| Ppil1 | 83.28828306 | -0.672919459 | 0.155649646 | -4.323295782 |
| BC004004 | 500.5361452 | -0.036257306 | 0.062020222 | -0.584604575 |
| Gm10045 | 2.961333848 | 0.437435688 | 0.671304779 | 0.651620101 |

| | | | | |
|---|---|---|---|---|
| Pi16 | 15.76972868 | -0.742144206 | 0.272052658 | -2.727943231 |
| Mtch1 | 2170.561962 | 0.118623332 | 0.06606513 | 1.795551321 |
| Fgd2 | 41.20869392 | 0.726241171 | 0.309583062 | 2.345868559 |
| Gm26885 | 6.791476013 | -0.021370561 | 0.411028552 | -0.051992888 |
| Pim1 | 294.2735208 | -0.991817095 | 0.300374506 | -3.301935005 |
| Gm17657 | 0 | NA | NA | NA |
| Tmem217 | 0.817974253 | 1.212099925 | 1.494677003 | 0.810944387 |
| Tbc1d22b | 597.8773199 | 0.218793899 | 0.064490454 | 3.392655579 |
| Gm28052 | 0 | NA | NA | NA |
| Tbc1d22bos | 0 | NA | NA | NA |
| Mir7667 | 0.28873988 | 0.986038588 | 1.829129957 | 0.539075195 |
| Rnf8 | 73.9322441 | 0.276028056 | 0.167042051 | 1.652446524 |
| Gm28043 | 40.56409608 | 0.373415144 | 0.338239569 | 1.103996039 |
| Cmtr1 | 326.8605501 | 0.345916913 | 0.118238208 | 2.925593329 |
| Ccdc167 | 100.0491434 | -0.175165803 | 0.141533982 | -1.237623649 |
| Mdga1 | 427.208838 | 0.047484276 | 0.144732132 | 0.328083855 |
| Gm16758 | 7.249476259 | 0.178312721 | 0.444657995 | 0.40101094 |
| Gm22327 | 0 | NA | NA | NA |
| Zfand3 | 884.6551979 | -0.373580008 | 0.083104923 | -4.495281307 |
| Btbd9 | 1023.020761 | 0.070378934 | 0.059653816 | 1.179789306 |
| Gm25049 | 0 | NA | NA | NA |
| Gm9874 | 0.097139547 | -0.443816764 | 3.724945049 | -0.119147198 |
| Glo1 | 537.9219683 | -0.053778021 | 0.093411329 | -0.575711981 |
| 1700097N02Rik | 0.406736653 | -0.073639273 | 1.532619442 | -0.048047983 |
| Dnah8 | 41.46003794 | -0.847652938 | 0.251767973 | -3.366802086 |
| Gm24661 | 0 | NA | NA | NA |
| Glp1r | 10.69878445 | -0.387623667 | 0.308769755 | -1.255380944 |
| Umodl1 | 10.985597 | 0.139775703 | 0.603138519 | 0.231747267 |
| Gm24026 | 0 | NA | NA | NA |
| Abcg1 | 1340.501293 | -0.319437437 | 0.0760084 | -4.202659636 |
| Gm25447 | 2.050495351 | 0.192104173 | 0.758579527 | 0.253241969 |
| Tff3 | 1.029781442 | 1.072672878 | 1.114185071 | 0.962742103 |
| Tff2 | 0 | NA | NA | NA |
| Gm15318 | 0.237699072 | -0.987095762 | 2.113983205 | -0.466936426 |
| Tff1 | 0.320114016 | -0.242143695 | 1.97148245 | -0.122823156 |
| Tmprss3 | 1.885701583 | -0.144876857 | 0.913245936 | -0.158639476 |
| Ubash3a | 6.422161135 | -0.332524745 | 0.390858874 | -0.850753986 |
| Rsph1 | 24.15190014 | -0.221945427 | 0.261036977 | -0.850245162 |
| Slc37a1 | 77.30229299 | -0.160367049 | 0.134711222 | -1.190450555 |
| Pde9a | 243.102937 | 0.176961773 | 0.140161581 | 1.26255549 |
| Wdr4 | 235.3192353 | -0.370182379 | 0.104166474 | -3.553757422 |
| Ndufv3 | 462.2896515 | 0.216585009 | 0.089208069 | 2.427863424 |
| Gm38099 | 1.2404959 | 0.424822773 | 0.995460926 | 0.426759868 |
| 4833413E03Rik | 0.117046891 | 0.037062677 | 3.724945049 | 0.009949859 |
| Pknox1 | 214.0257755 | -0.488509294 | 0.092872988 | -5.25997177 |
| Gm24970 | 0.09349418 | 0.517955464 | 3.724945049 | 0.139050498 |
| Cbs | 185.6562434 | 0.613315175 | 0.155743495 | 3.937982618 |

| | | | | |
|---|---|---|---|---|
| U2af1 | 321.058565 | -0.350226935 | 0.101452938 | -3.452112299 |
| Cryaa | 0.372981075 | -1.470805837 | 1.558088982 | -0.943980642 |
| Sik1 | 432.2003957 | -0.144425408 | 0.188392183 | -0.766621019 |
| Hsf2bp | 3.787275169 | -1.036274998 | 0.614804959 | -1.685534547 |
| Gm17572 | 0 | NA | NA | NA |
| Gm25883 | 0 | NA | NA | NA |
| Rrp1b | 198.546731 | -0.172787369 | 0.147377012 | -1.172417373 |
| Pdxk-ps | 1.082653786 | -0.220163153 | 1.092462236 | -0.201529303 |
| Gm17276 | 31.48193691 | 0.934828967 | 0.254931436 | 3.666981918 |
| Notch3 | 246.8071195 | 0.17088616 | 0.201729518 | 0.84710538 |
| Ephx3 | 2.938467936 | -0.441730517 | 0.6055432 | -0.729478123 |
| Brd4 | 1978.347052 | -0.004687795 | 0.074699915 | -0.062755025 |
| Gm26549 | 15.20514642 | -0.314718397 | 0.41440529 | -0.759445897 |
| Akap8 | 845.3496397 | 0.329337895 | 0.132832619 | 2.479345037 |
| Akap8l | 594.3929537 | 0.253868776 | 0.139459266 | 1.820379403 |
| Wiz | 755.6427441 | 0.084146891 | 0.063579921 | 1.323482148 |
| Gm26858 | 0.274350536 | 0.587499254 | 2.081008455 | 0.282314689 |
| Gm22201 | 0.044191549 | 0.037062677 | 3.724945049 | 0.009949859 |
| Rasal3 | 63.09606561 | 0.048248128 | 0.241007926 | 0.200193118 |
| Pglyrp2 | 9.032459669 | -0.959693286 | 0.408339958 | -2.350231139 |
| Cyp4f39 | 0 | NA | NA | NA |
| Cyp4f17 | 39.89199084 | 0.281458553 | 0.191434343 | 1.470261546 |
| Cyp4f16 | 69.22286846 | -0.910353209 | 0.183930019 | -4.949454225 |
| Cyp4f37 | 0.44264929 | -2.070809577 | 1.367975845 | -1.51377642 |
| Cyp4f40 | 0.108772582 | 0.037073798 | 3.724945049 | 0.009952844 |
| Cyp4f15 | 57.06117773 | 2.07756629 | 0.241532973 | 8.601584568 |
| Zfp871 | 1423.833684 | 0.163068715 | 0.060922614 | 2.676653272 |
| Gm17115 | 5.126403706 | 0.981530146 | 0.479131701 | 2.048560225 |
| Zfp811 | 109.0107342 | 0.08556576 | 0.170351006 | 0.502290897 |
| Gm23294 | 1.960852514 | 0.875131885 | 0.733984729 | 1.192302579 |
| Gm25555 | 1.60505521 | 2.284636345 | 1.079264839 | 2.11684497 |
| Gm26693 | 0.210884633 | -1.396893933 | 2.175374804 | -0.642139428 |
| Zfp799 | 476.0807173 | -0.03049673 | 0.09673697 | -0.315254138 |
| Zfp870 | 141.8172216 | -0.067410961 | 0.097433977 | -0.691862973 |
| Cyp4f14 | 50.57166942 | 0.75620763 | 0.202516223 | 3.734059519 |
| Cyp4f13 | 93.77020389 | -0.235302706 | 0.154062659 | -1.527318224 |
| Cyp4f41-ps | 1.046059986 | 0.041853401 | 1.029705487 | 0.040645992 |
| Gm25667 | 0.310541583 | 1.067907484 | 3.71555192 | 0.287415573 |
| Zfp472 | 71.14659937 | 0.003305515 | 0.147711556 | 0.022378173 |
| Zfp952 | 219.1287848 | -0.048763367 | 0.090493814 | -0.538858571 |
| Zfp763 | 281.1564641 | 0.300524882 | 0.099537121 | 3.019224173 |
| 4921501E09Rik | 0.687686063 | -0.231591596 | 1.215857917 | -0.190475871 |
| Zfp563 | 131.0379676 | -0.013642181 | 0.128096051 | -0.106499625 |
| Morc2b | 49.27469275 | 0.051319728 | 0.369067853 | 0.139052284 |
| Olfr55 | 0.072025462 | 0.517959912 | 3.724945049 | 0.139051692 |
| Olfr239 | 0.04393175 | 0.037062677 | 3.724945049 | 0.009949859 |
| Gm4461 | 9.889894451 | 1.271439526 | 0.449787874 | 2.826753675 |

| | | | | |
|---|---|---|---|---|
| Zfp955a | 185.3926737 | 0.103054691 | 0.132908067 | 0.775383265 |
| Olfr63 | 0.330364319 | -1.257220782 | 1.706788135 | -0.736600376 |
| Zfp955b | 228.1232927 | 0.014762929 | 0.089761855 | 0.164467739 |
| Zfp422-rs1 | 1.757111958 | 1.049947029 | 0.772144111 | 1.359781178 |
| Zfp81 | 498.1008011 | 0.254253795 | 0.08556662 | 2.971413327 |
| Zfp101 | 191.0430263 | 0.337179432 | 0.103965345 | 3.243190614 |
| CT030734.1 | 0 | NA | NA | NA |
| Actl9 | 0 | NA | NA | NA |
| Adamts10 | 190.1840441 | 0.11830143 | 0.149446351 | 0.791597983 |
| Myo1f | 224.7996339 | -1.424490177 | 0.209250554 | -6.80758139 |
| Zfp414 | 163.0997787 | 0.073618076 | 0.104673269 | 0.703313049 |
| Gm20507 | 3.129648303 | 0.869305495 | 0.704525638 | 1.233887665 |
| Pram1 | 0.821047175 | -1.459385557 | 1.203951045 | -1.212163537 |
| Hnrnpm | 845.7194937 | -0.08473099 | 0.145317476 | -0.583075017 |
| 2-Mar | 559.2133353 | 0.121354696 | 0.0640129 | 1.895784998 |
| Gm22943 | 0 | NA | NA | NA |
| Rab11b | 1217.267332 | 0.188864218 | 0.076741104 | 2.461056825 |
| Gm17251 | 14.95472732 | 0.126614814 | 0.265000128 | 0.477791521 |
| Angptl4 | 147.9950521 | -0.86415732 | 0.426244301 | -2.027375656 |
| 4931413I07Rik | 0.260232477 | -0.249057025 | 2.450694468 | -0.101627122 |
| Kank3 | 84.23068527 | -0.453896659 | 0.202507742 | -2.241379286 |
| Rps28 | 433.7597313 | 0.047215799 | 0.168800313 | 0.279713929 |
| Ndufa7 | 437.6705464 | -0.144335072 | 0.097173782 | -1.485329367 |
| Cd320 | 102.4821458 | 0.04389366 | 0.132603912 | 0.331013313 |
| Gm4134 | 0.061920613 | -0.443816764 | 3.724945049 | -0.119147198 |
| Gm7029 | 1.064012846 | -0.502837474 | 1.02591472 | -0.490135743 |
| Kifc1 | 18.16257989 | -0.324810804 | 0.2691202 | -1.206935801 |
| Gm27740 | 0 | NA | NA | NA |
| Platr17 | 2.945475634 | -0.945487538 | 0.683494088 | -1.383314874 |
| BC051226 | 10.35737011 | -0.421012551 | 0.393465428 | -1.070011547 |
| Daxx | 959.7967857 | -0.604894063 | 0.172693326 | -3.502706666 |
| Tapbp | 4153.144377 | -0.253978789 | 0.158791859 | -1.599444642 |
| Zbtb22 | 294.4051857 | 0.252973579 | 0.079323763 | 3.189127294 |
| Gm19412 | 30.80291434 | 0.514410218 | 0.208581204 | 2.466234774 |
| Rgl2 | 188.6076046 | -0.706052724 | 0.137509077 | -5.134589938 |
| Pfdn6 | 222.2854681 | 0.02528416 | 0.14147423 | 0.17871919 |
| Wdr46 | 114.1398418 | -0.688506157 | 0.145424268 | -4.734465333 |
| B3galt4 | 35.09101575 | 0.250883471 | 0.173609559 | 1.445101714 |
| Rps18 | 253.3875219 | -0.291852697 | 0.200964934 | -1.452256825 |
| Vps52 | 738.2973603 | 0.076201782 | 0.06586688 | 1.156905899 |
| Gm23111 | 0.99397625 | 2.435474673 | 1.054416038 | 2.309785309 |
| H2-K2 | 47.37552283 | -0.841400524 | 0.251585346 | -3.344394005 |
| Gm4152 | 0 | NA | NA | NA |
| H2-K1 | 2006.445401 | -0.847785083 | 0.154993872 | -5.469797437 |
| Ring1 | 223.6236612 | 0.066284457 | 0.082923733 | 0.799342419 |
| Gm26940 | 0.048949532 | 0.037064902 | 3.724945049 | 0.009950456 |
| Mir219a-1 | 0.047948698 | 0.037064902 | 3.724945049 | 0.009950456 |

| | | | | |
|---|---|---|---|---|
| mmu-mir-219c | 1.228891946 | -0.419590137 | 0.942193027 | -0.44533352 |
| H2-Ke6 | 1.732078677 | -0.250241852 | 0.797926226 | -0.313615273 |
| Gm20427 | 1.185090727 | -1.753388503 | 0.981755506 | -1.78597267 |
| Slc39a7 | 92.16732074 | -0.163428773 | 0.152416918 | -1.072248246 |
| Rxrb | 612.8230062 | -0.007002045 | 0.057440991 | -0.121899797 |
| Col11a2 | 1625.713743 | 0.279605493 | 0.243938873 | 1.146211302 |
| H2-Pb | 0.204910391 | 0.037067724 | 2.659251144 | 0.013939159 |
| BC051537 | 0.820567722 | 3.279770897 | 1.175555236 | 2.789976 |
| H2-Pa | 0 | NA | NA | NA |
| H2-Oa | 2.48498024 | 1.711547466 | 0.645685311 | 2.650745553 |
| Brd2 | 3845.362694 | -0.219451445 | 0.080018789 | -2.742498933 |
| Gm20493 | 0 | NA | NA | NA |
| H2-DMa | 136.095749 | -0.435550586 | 0.158344655 | -2.750649116 |
| H2-DMb2 | 12.8997608 | -0.575114744 | 0.397428716 | -1.447089053 |
| H2-DMb1 | 15.23037961 | -0.872779648 | 0.325969895 | -2.677485444 |
| Psmb9 | 510.8582445 | -0.826554073 | 0.194311422 | -4.253759589 |
| Tap1 | 2068.383747 | -0.370293424 | 0.159142461 | -2.326804691 |
| Psmb8 | 1292.910119 | -0.621260063 | 0.224692263 | -2.76493749 |
| Gm20496 | 0.676590931 | -0.514266855 | 1.351443805 | -0.380531439 |
| Tap2 | 1370.634535 | -0.76799947 | 0.159913698 | -4.802587127 |
| Gm15821 | 0.89598866 | -0.533973179 | 1.228808501 | -0.434545479 |
| H2-Ob | 4.820269621 | -0.505249981 | 0.477365037 | -1.0584143 |
| Gm20506 | 0.159280631 | -0.429728558 | 3.461051974 | -0.124161255 |
| H2-Ab1 | 139.2632816 | -0.467603449 | 0.246916269 | -1.893773341 |
| H2-Aa | 90.90688896 | -0.236216116 | 0.274065018 | -0.861898093 |
| Gm20513 | 0.289806465 | -0.460732806 | 1.956555846 | -0.235481551 |
| H2-Eb1 | 122.6882474 | -0.563775574 | 0.454884622 | -1.239381475 |
| H2-Eb2 | 1.375804068 | -0.906069003 | 0.899917799 | -1.006835295 |
| H2-Ea-ps | 95.63606511 | -0.382845821 | 0.306812742 | -1.247815909 |
| Btnl2 | 0.44006443 | -1.076716598 | 1.471290179 | -0.731817974 |
| Btnl1 | 1.318555613 | -0.088017879 | 0.866963322 | -0.10152434 |
| Gm15320 | 0 | NA | NA | NA |
| Gm15321 | 0 | NA | NA | NA |
| BC051142 | 7.899042157 | 0.914772488 | 0.371132057 | 2.46481669 |
| Btnl4 | 0.635806793 | -1.460618175 | 1.300760071 | -1.122895919 |
| Gm15332 | 0 | NA | NA | NA |
| Btnl5-ps | 0 | NA | NA | NA |
| Gm15331 | 0 | NA | NA | NA |
| Btnl6 | 0 | NA | NA | NA |
| Gm15329 | 0 | NA | NA | NA |
| Btnl7-ps | 0.643414179 | -0.726607802 | 1.109588288 | -0.654844513 |
| Notch4 | 57.84156358 | -0.87842466 | 0.180514339 | -4.866232043 |
| Gm20462 | 0 | NA | NA | NA |
| Gpsm3 | 60.20938841 | -0.772950601 | 0.239251397 | -3.230704644 |
| Pbx2 | 551.6966075 | 0.014191557 | 0.070550464 | 0.201154698 |
| Ager | 5.903964042 | 1.307108149 | 0.530228997 | 2.465176663 |
| Rnf5 | 318.1261628 | 0.184539346 | 0.076940959 | 2.398453925 |

| | | | | |
|---|---|---|---|---|
| Gm20463 | 0.104106287 | 0.51795324 | 3.724945049 | 0.139049901 |
| Agpat1 | 1285.566767 | -0.090537291 | 0.062008943 | -1.460068291 |
| Egfl8 | 5.399883896 | 0.311671422 | 0.544847258 | 0.572034487 |
| Gm20460 | 0 | NA | NA | NA |
| Ppt2 | 144.3949312 | -0.247590846 | 0.100965509 | -2.452231946 |
| Prrt1 | 1088.589987 | 0.552877158 | 0.119329679 | 4.633190701 |
| Gm20461 | 0.042716945 | 0.037062677 | 3.724945049 | 0.009949859 |
| Fkbpl | 63.97067545 | 0.049479912 | 0.140761905 | 0.351514936 |
| Atf6b | 618.9161125 | -0.341140296 | 0.080568554 | -4.234161831 |
| Tnxb | 123.7697874 | -0.268487165 | 0.200164143 | -1.341334969 |
| Gm24389 | 0.068097359 | 0.037064902 | 3.724945049 | 0.009950456 |
| Cyp21a2-ps | 0.84670602 | -0.191854994 | 1.171426246 | -0.163778979 |
| C4b | 4516.244341 | -0.485167647 | 0.140501625 | -3.453110571 |
| Stk19-ps1 | 0.301345325 | -1.024116546 | 2.002124482 | -0.511514921 |
| Skiv2l-ps1 | 0.092269992 | -0.443823436 | 3.724945049 | -0.119148989 |
| Gm27471 | 0 | NA | NA | NA |
| Tnxa | 0.049553047 | -0.443818988 | 3.724945049 | -0.119147795 |
| Cyp21a1 | 0.116944185 | -1.028246102 | 3.719337068 | -0.27645951 |
| C4a | 141.065081 | -0.593495718 | 0.196495617 | -3.020401815 |
| Stk19 | 195.7826537 | -0.087127329 | 0.166626833 | -0.522888948 |
| Dxo | 129.6180202 | -0.31680397 | 0.12862225 | -2.463057295 |
| Skiv2l | 714.6748977 | 0.054671196 | 0.056133149 | 0.973955631 |
| Mir6971 | 0 | NA | NA | NA |
| Mir6970 | 0 | NA | NA | NA |
| Nelfe | 167.1258369 | -0.242638748 | 0.089076071 | -2.723949818 |
| Cfb | 85.69436929 | -1.165577553 | 0.30221566 | -3.856774171 |
| Gm20547 | 0 | NA | NA | NA |
| Mir6972 | 0 | NA | NA | NA |
| C2 | 78.60544563 | -0.474333191 | 0.204045228 | -2.324647312 |
| Zbtb12 | 59.55277683 | -0.398114327 | 0.244230931 | -1.630073332 |
| Ehmt2 | 1573.434019 | 0.159782689 | 0.073177735 | 2.183487778 |
| Slc44a4 | 2.383600578 | 0.687414378 | 0.707845302 | 0.971136456 |
| Neu1 | 530.836773 | -0.135533061 | 0.079702307 | -1.700491071 |
| 1110038B12Rik | 128.359516 | -0.788754689 | 0.167503607 | -4.708881804 |
| Snord52 | 0 | NA | NA | NA |
| Gm25744 | 0 | NA | NA | NA |
| Gm8696 | 0.535596869 | -1.559735151 | 1.449204407 | -1.076269948 |
| Hspa1b | 205.5992703 | -2.941488867 | 0.621275603 | -4.734595813 |
| Hspa1a | 168.1854328 | -2.881733526 | 0.575211003 | -5.009872049 |
| Gm20481 | 0.568897959 | -2.178107613 | 1.482841004 | -1.468874685 |
| Hspa1l | 8.258336928 | -0.979607541 | 0.371079391 | -2.639886677 |
| Gm10501 | 57.48590907 | 0.304561138 | 0.181677417 | 1.676384124 |
| Lsm2 | 72.82678909 | -0.377727405 | 0.13249011 | -2.850985666 |
| Vars | 655.1030319 | -0.033128897 | 0.066866741 | -0.495446565 |
| Vwa7 | 91.21471042 | 0.142683266 | 0.199593708 | 0.714868557 |
| Sapcd1 | 2.507914302 | 1.768906888 | 0.678351073 | 2.607656946 |
| Msh5 | 15.17733262 | 0.248635297 | 0.381802028 | 0.651215235 |

| | | | | |
|---|---|---|---|---|
| Gm23421 | 0 | NA | NA | NA |
| Clic1 | 397.1204603 | -1.273606348 | 0.202512937 | -6.289012278 |
| Ddah2 | 89.30682263 | -0.476081879 | 0.164338126 | -2.896965489 |
| G6b | 1.225313711 | 0.566857356 | 1.071629868 | 0.528967485 |
| Ly6g6c | 0.500444145 | -2.052655818 | 1.449199283 | -1.416406868 |
| Ly6g6d | 2.936279196 | 0.47270726 | 0.681031449 | 0.694104892 |
| Ly6g6e | 18.02936065 | -1.224959128 | 0.453766581 | -2.699535796 |
| Ly6g6f | 3.98262746 | -0.417352468 | 0.584849705 | -0.713606358 |
| Abhd16a | 1009.474395 | 0.105181241 | 0.103550171 | 1.015751496 |
| Ly6g5c | 0.083526043 | 0.037073798 | 3.724945049 | 0.009952844 |
| Ly6g5b | 3.278244484 | -0.222972036 | 0.563524524 | -0.395674059 |
| Csnk2b | 658.9798502 | -0.246471877 | 0.068244632 | -3.611593616 |
| Gpank1 | 122.9329401 | 0.001029656 | 0.127813465 | 0.008055931 |
| Gm20522 | 3.138335889 | -1.069861602 | 0.633722993 | -1.688216481 |
| D17H6S53E | 194.6956857 | -0.298944884 | 0.09500807 | -3.146520952 |
| Apom | 0.669558094 | -0.032818565 | 1.105728907 | -0.02968048 |
| Bag6 | 2358.238324 | 0.084800832 | 0.046742501 | 1.814212535 |
| Prrc2a | 6282.854268 | -0.072680949 | 0.076439219 | -0.950833229 |
| Gm23442 | 7.065932701 | 0.216421437 | 0.452724248 | 0.478042512 |
| Gm17705 | 6.475434284 | 0.523533 | 0.451558459 | 1.159391416 |
| Aif1 | 114.5567538 | -0.221096953 | 0.175474538 | -1.259994501 |
| Ncr3-ps | 0.383232264 | 1.694249069 | 1.661725601 | 1.019572106 |
| Lst1 | 34.76546749 | -0.939854865 | 0.238421309 | -3.941991891 |
| Ltb | 23.43129296 | -0.653291425 | 0.281973256 | -2.316855982 |
| Mir6973a | 0.196800764 | -1.695339541 | 2.170504729 | -0.781080786 |
| Tnf | 84.48636142 | -0.567473527 | 0.442215015 | -1.283252508 |
| Lta | 7.270911819 | 1.633821468 | 0.51045286 | 3.200729383 |
| Mir6974 | 0 | NA | NA | NA |
| Nfkbil1 | 124.1921778 | -0.101960836 | 0.153584731 | -0.663873518 |
| Gm16181 | 0.333516142 | -0.507186323 | 1.733129227 | -0.292641954 |
| Atp6v1g2 | 2556.244658 | 0.692251554 | 0.094705451 | 7.30952178 |
| Ddx39b | 1299.932362 | -0.066366233 | 0.071232777 | -0.931681124 |
| Gm22589 | 0 | NA | NA | NA |
| Mir6975 | 0.574701389 | -0.815875481 | 1.343133492 | -0.607441841 |
| Gm25128 | 2.512166706 | 0.155091081 | 0.741690595 | 0.209104823 |
| Mir8094 | 0.904930538 | 0.193683681 | 1.314053336 | 0.147394079 |
| H2-D1 | 7783.618613 | -0.882026857 | 0.185233794 | -4.761695148 |
| Gm18733 | 1.69076615 | -1.66030601 | 0.841128355 | -1.97390327 |
| Gm18734 | 0.938757343 | 0.717630427 | 0.99441945 | 0.721657673 |
| H2-Q1 | 113.5987717 | -0.910524585 | 0.230311345 | -3.953450867 |
| Gm8741 | 9.465940354 | 0.014145553 | 0.37519262 | 0.037702108 |
| H2-Q2 | 32.27579733 | -0.858255766 | 0.23096233 | -3.715998905 |
| H2-Q3 | 0.044544648 | 0.517957688 | 3.724945049 | 0.139051095 |
| Gm11131 | 30.17597747 | -1.344472605 | 0.247639518 | -5.429152093 |
| H2-Q4 | 2741.587666 | -0.60130444 | 0.22373082 | -2.687624537 |
| H2-Q5 | 203.0857095 | -0.919328491 | 0.180253831 | -5.100188368 |
| Gm8750 | 0.56888582 | -0.772768425 | 1.241048063 | -0.622674052 |

| | | | | |
|---|---|---|---|---|
| H2-Q6 | 1065.781777 | -0.972960846 | 0.261081774 | -3.726651737 |
| H2-Q7 | 469.1606159 | -1.103035577 | 0.21273303 | -5.185069652 |
| Gm8752 | 0.72059279 | -0.611359912 | 1.175439139 | -0.520111924 |
| H2-Q10 | 15.54474821 | -0.589775679 | 0.294892534 | -1.999968162 |
| Pou5f1 | 0 | NA | NA | NA |
| Gm19553 | 0 | NA | NA | NA |
| Tcf19 | 44.18076177 | 0.102604041 | 0.317251189 | 0.323415781 |
| Cchcr1 | 54.80302612 | -0.129094995 | 0.185233344 | -0.69693173 |
| Psors1c2 | 0.771204239 | -0.307892692 | 1.361424157 | -0.226154862 |
| Cdsn | 7.227748083 | -1.679393448 | 0.413276886 | -4.063603617 |
| Gm9613 | 0.0885994 | 0.517955464 | 3.724945049 | 0.139050498 |
| 2300002M23Rik | 0 | NA | NA | NA |
| Sfta2 | 0.116217011 | 0.517955464 | 3.724945049 | 0.139050498 |
| Gm9573 | 0.122991902 | 0.517955464 | 3.724945049 | 0.139050498 |
| Dpcr1 | 0.220032676 | -1.401785252 | 2.574515855 | -0.544484995 |
| Gm20483 | 0.193987248 | -0.443820568 | 2.24898814 | -0.197342334 |
| Vars2 | 218.6187569 | 0.232478296 | 0.096612105 | 2.406306078 |
| Gtf2h4 | 76.61225677 | -0.609247522 | 0.148772193 | -4.095170672 |
| Ddr1 | 1349.901542 | -0.683919095 | 0.113319138 | -6.03533621 |
| Gm23864 | 3.332665857 | 0.04450244 | 0.570284808 | 0.078035464 |
| Gm4577 | 0.622983114 | 2.37180967 | 1.191770989 | 1.990155569 |
| Gm20443 | 0.385495486 | -0.451906464 | 1.537762771 | -0.293872678 |
| Gm20442 | 0.201858732 | 0.517972533 | 2.604406155 | 0.198883163 |
| 4833427F10Rik | 0.08486385 | 0.037730769 | 3.724898049 | 0.010129343 |
| Ier3 | 124.9554162 | -0.294253788 | 0.247688335 | -1.188000183 |
| Flot1 | 628.8586158 | 0.106092376 | 0.089821493 | 1.181146888 |
| Tubb5 | 3619.132628 | 0.053158778 | 0.093311597 | 0.569691007 |
| Mdc1 | 303.644182 | -0.489604535 | 0.115601726 | -4.23527012 |
| Ppp1r18os | 5.57648041 | -0.731684825 | 0.451768321 | -1.619601887 |
| Nrm | 18.32685686 | 0.136534587 | 0.245006318 | 0.557269658 |
| Ppp1r18 | 331.6778556 | -0.699744535 | 0.201190896 | -3.478012912 |
| Gm28033 | 0.098502579 | -0.443821212 | 3.724945049 | -0.119148392 |
| Dhx16 | 338.5688034 | -0.445794227 | 0.099613375 | -4.475244695 |
| 2310061I04Rik | 406.7426439 | -0.07711181 | 0.084465537 | -0.912938142 |
| Gm16279 | 3.489032565 | 0.355198608 | 0.676918204 | 0.524728995 |
| Atat1 | 349.1959485 | 0.200624226 | 0.081262198 | 2.468850605 |
| Mrps18b | 109.9134655 | -0.343922776 | 0.121213693 | -2.837326105 |
| Ppp1r10 | 838.888173 | -0.271770022 | 0.081757823 | -3.32408585 |
| Mir1894 | 0 | NA | NA | NA |
| Gm9840 | 3.211017017 | -0.385061221 | 0.574043675 | -0.670787324 |
| Gm8801 | 0.654072159 | 0.109548351 | 1.298651638 | 0.084355456 |
| Abcf1 | 700.9733728 | -0.218069773 | 0.085706083 | -2.544390848 |
| Mir877 | 0 | NA | NA | NA |
| Prr3 | 210.8648339 | -0.096164898 | 0.127358583 | -0.755071989 |
| Gnl1 | 1616.148979 | 0.173964046 | 0.068473292 | 2.540611694 |
| Gm20508 | 2.081650308 | -0.952636087 | 0.798719077 | -1.192704813 |
| A930015D03Rik | 113.1905163 | -0.597362642 | 0.159769706 | -3.73889805 |

| | | | | |
|---|---|---|---|---|
| H2-T24 | 905.0134114 | -0.918564875 | 0.225756248 | -4.068834783 |
| H2-T23 | 3145.49009 | -0.560486078 | 0.181126539 | -3.094444815 |
| H2-T22 | 1287.107806 | -0.589602312 | 0.15371552 | -3.835671967 |
| Gm6034 | 25.06496248 | -0.336895708 | 0.243882737 | -1.381383991 |
| Gm11127 | 246.9603189 | -0.862621315 | 0.18345107 | -4.702187437 |
| Gm8810 | 46.0581403 | -0.651524986 | 0.241240451 | -2.700728604 |
| H2-Bl | 5.276086801 | -1.003506064 | 0.489245528 | -2.051129762 |
| Gm20530 | 0 | NA | NA | NA |
| Gm8815 | 1.211501154 | -0.976676702 | 0.858217417 | -1.138029458 |
| C920025E04Rik | 19.33663716 | -0.84622105 | 0.324395778 | -2.608606857 |
| Gm7030 | 5.681141433 | -0.592652219 | 0.509889974 | -1.16231393 |
| BC023719 | 0.374687972 | -0.65729467 | 1.519665306 | -0.432525943 |
| H2-T10 | 170.5753034 | -0.593513604 | 0.178115386 | -3.332186038 |
| Gm19684 | 10.16906943 | -0.184334546 | 0.365930319 | -0.503742205 |
| Gm9574 | 0.115603922 | -0.93550548 | 3.724188147 | -0.251197158 |
| Gm10499 | 19.27648584 | -0.597651001 | 0.36121708 | -1.654548009 |
| 2410017I17Rik | 0.142780685 | 0.517965309 | 2.900967263 | 0.178549174 |
| Gm17782 | 0.038981395 | -0.443816764 | 3.724945049 | -0.119147198 |
| Gm6659 | 0.124347486 | 0.036092774 | 3.717245565 | 0.009709548 |
| Gm8909 | 5.891852665 | -0.329436457 | 0.387333838 | -0.850523307 |
| Gm20478 | 1.20453066 | -1.555611685 | 1.260290929 | -1.234327446 |
| Gm6623 | 0.15621502 | -0.981036059 | 2.83205109 | -0.346404788 |
| Gm20495 | 0 | NA | NA | NA |
| H2-T3 | 0.239012627 | -0.434983463 | 2.092274933 | -0.207899763 |
| Gm4246 | 0.068097359 | 0.037064902 | 3.724945049 | 0.009950456 |
| Gm20447 | 0 | NA | NA | NA |
| Gm8835 | 0.696575882 | -3.337896837 | 1.220385052 | -2.735117767 |
| Gm5682 | 0.219744001 | -1.121555118 | 2.125982586 | -0.527546709 |
| Gm20392 | 0.238833577 | 1.768112476 | 2.461453536 | 0.718320476 |
| Gm10074 | 1.752125518 | 0.098434227 | 0.715340638 | 0.13760469 |
| Gm20545 | 0.204225461 | 1.631603513 | 3.2048902 | 0.5090981 |
| Gm29402 | 0 | NA | NA | NA |
| Gm18604 | 0 | NA | NA | NA |
| Gm20391 | 0 | NA | NA | NA |
| Gm22453 | 0 | NA | NA | NA |
| Gm4252 | 0 | NA | NA | NA |
| Gm20546 | 0.114742407 | 0.517955464 | 3.724945049 | 0.139050498 |
| Gm6633 | 0.13733263 | 0.037057285 | 3.640803958 | 0.010178325 |
| Rpp21 | 97.94756067 | -0.116555742 | 0.126001246 | -0.925036424 |
| Trim39 | 333.3226075 | -0.007282568 | 0.132282878 | -0.05505299 |
| H2-M10.2 | 0.048949532 | 0.037064902 | 3.724945049 | 0.009950456 |
| H2-M10.1 | 0 | NA | NA | NA |
| H2-M10.3 | 0 | NA | NA | NA |
| Stk-ps1 | 0 | NA | NA | NA |
| H2-M10.5-ps1 | 0 | NA | NA | NA |
| Lgals1-ps1 | 0 | NA | NA | NA |
| Rpl9-ps9 | 0 | NA | NA | NA |

| | | | | |
|---|---|---|---|---|
| Obox3-ps8 | 0.062426873 | 0.517959912 | 3.724945049 | 0.139051692 |
| H2-M10.4 | 0 | NA | NA | NA |
| Gm20452 | 0.139037128 | -1.190058937 | 3.717743703 | -0.320102469 |
| Gm20451 | 0.072025462 | 0.517959912 | 3.724945049 | 0.139051692 |
| Gm20454 | 0 | NA | NA | NA |
| Gm20453 | 0 | NA | NA | NA |
| H2-M11 | 0.066050875 | -0.443818988 | 3.724945049 | -0.119147795 |
| Gm8868 | 0 | NA | NA | NA |
| Gm20450 | 0 | NA | NA | NA |
| H2-M9 | 0.541282201 | -2.522245538 | 1.294476656 | -1.94846738 |
| Rpl9-ps5 | 0.163190422 | -0.955151543 | 3.396300892 | -0.281232898 |
| H2-M1 | 0.163289894 | -1.433956951 | 3.406547043 | -0.420941479 |
| Gm8877 | 0 | NA | NA | NA |
| Gm20549 | 0 | NA | NA | NA |
| Gm20523 | 0 | NA | NA | NA |
| Gm20525 | 0 | NA | NA | NA |
| Gm20526 | 0 | NA | NA | NA |
| Gm4271 | 0 | NA | NA | NA |
| Gm20401 | 0 | NA | NA | NA |
| Gm8878 | 0.22323753 | 0.479505308 | 2.560144731 | 0.187296172 |
| H2-M10.5 | 0.066050875 | -0.443818988 | 3.724945049 | -0.119147795 |
| H2-M10.6 | 0 | NA | NA | NA |
| Trim26 | 648.1818355 | -0.389694832 | 0.099104156 | -3.932174464 |
| Gm25973 | 0 | NA | NA | NA |
| Trim15 | 1.517773654 | -1.58770029 | 0.873134273 | -1.818391899 |
| Trim10 | 2.069903214 | 0.620056141 | 0.744509324 | 0.832838652 |
| Trim40 | 1.721525315 | 0.833528206 | 0.892837647 | 0.933571975 |
| Trim31 | 0.590394638 | 0.073980433 | 1.143379603 | 0.0647033 |
| 1700031A10Rik | 0.066050875 | -0.443818988 | 3.724945049 | -0.119147795 |
| Rnf39 | 43.61237388 | 0.021900824 | 0.249234433 | 0.087872384 |
| Ppp1r11 | 632.5865216 | -0.240341881 | 0.096983909 | -2.478162446 |
| Znrd1 | 116.024339 | -0.243478605 | 0.136454341 | -1.784322898 |
| Znrd1as | 20.09275304 | -0.499023585 | 0.257523711 | -1.93777724 |
| Gm27735 | 0 | NA | NA | NA |
| H2-M6-ps | 4.48790112 | 1.631362726 | 0.59456395 | 2.743796904 |
| 2410137M14Rik | 2.456754703 | 2.059604597 | 0.815489681 | 2.525604732 |
| H2-M5 | 10.19594007 | 1.208884346 | 0.44743009 | 2.701839623 |
| Zfp57 | 63.14501023 | 0.29514979 | 0.230082565 | 1.282799458 |
| Mog | 603.8852982 | 0.079133709 | 0.156715202 | 0.504952347 |
| Gm22388 | 0.655055538 | 1.739672327 | 1.230607896 | 1.413669076 |
| Gabbr1 | 8020.403664 | 0.400701193 | 0.070844923 | 5.656032615 |
| Olfr90 | 0.046345709 | -0.443818988 | 3.724945049 | -0.119147795 |
| Olfr91 | 0.106520246 | 0.037071574 | 3.724945049 | 0.009952247 |
| Olfr92 | 0.119587853 | 0.981063299 | 3.723618439 | 0.263470416 |
| Gm4831 | 0.046345709 | -0.443818988 | 3.724945049 | -0.119147795 |
| Gm19807 | 2.281184893 | -1.445941505 | 0.724998667 | -1.994405742 |
| Olfr93 | 0.291730981 | -0.402646272 | 1.75253492 | -0.229750784 |

| | | | | |
|---|---|---|---|---|
| Vhl-ps1 | 3.944458201 | -0.662891605 | 0.529678738 | -1.251497478 |
| Olfr753-ps1 | 11.84344878 | -1.283780891 | 0.458210901 | -2.8017249 |
| Gm20497 | 17.12714754 | -0.748893492 | 0.452012273 | -1.656799022 |
| Gm7031 | 0.038981395 | -0.443816764 | 3.724945049 | -0.119147198 |
| Ubd | 60.38255723 | -1.811135935 | 0.629948079 | -2.875055888 |
| Olfr94 | 0.247779669 | 0.495277106 | 1.818914916 | 0.272292619 |
| Olfr95 | 0.082913145 | -0.443821212 | 3.724945049 | -0.119148392 |
| Olfr96 | 0.046345709 | -0.443818988 | 3.724945049 | -0.119147795 |
| Olfr97 | 0 | NA | NA | NA |
| Gm20429 | 1.971464702 | -0.415998118 | 0.722902124 | -0.575455658 |
| Olfr754-ps1 | 0.99187381 | -0.176211394 | 0.958249315 | -0.183888881 |
| Olfr98 | 0 | NA | NA | NA |
| Gm25104 | 0.045882455 | 0.517959912 | 3.724945049 | 0.139051692 |
| H2-M3 | 192.7482628 | -0.702986463 | 0.182128474 | -3.859838311 |
| Olfr755-ps1 | 6.424303794 | -0.60820473 | 0.490040106 | -1.241132557 |
| Olfr99 | 0 | NA | NA | NA |
| Olfr756-ps1 | 0 | NA | NA | NA |
| Olfr101 | 0.110870145 | -0.444802825 | 3.724874622 | -0.119414174 |
| Olfr102 | 0.068097359 | 0.037064902 | 3.724945049 | 0.009950456 |
| Olfr103 | 0 | NA | NA | NA |
| Olfr104-ps | 0 | NA | NA | NA |
| Olfr757-ps1 | 0 | NA | NA | NA |
| Olfr105-ps | 0 | NA | NA | NA |
| Olfr758-ps1 | 0 | NA | NA | NA |
| Olfr106-ps | 0 | NA | NA | NA |
| Olfr107 | 0 | NA | NA | NA |
| Olfr759-ps1 | 0 | NA | NA | NA |
| Gm18808 | 0 | NA | NA | NA |
| Olfr108 | 0 | NA | NA | NA |
| Olfr760-ps1 | 0 | NA | NA | NA |
| Olfr109 | 0.108772582 | 0.037073798 | 3.724945049 | 0.009952844 |
| H2-M2 | 1.126195538 | 0.452966244 | 1.070918823 | 0.422969729 |
| Olfr110 | 0.784115673 | 1.329784074 | 1.171203593 | 1.135399585 |
| Olfr111 | 1.850398219 | 0.263598067 | 0.73401579 | 0.359117706 |
| Olfr112 | 0.062426873 | 0.517959912 | 3.724945049 | 0.139051692 |
| Olfr113 | 0 | NA | NA | NA |
| CR974439.2 | 0 | NA | NA | NA |
| Olfr114 | 0 | NA | NA | NA |
| Olfr115 | 0.046345709 | -0.443818988 | 3.724945049 | -0.119147795 |
| CR974439.1 | 0 | NA | NA | NA |
| Olfr116 | 0.046345709 | -0.443818988 | 3.724945049 | -0.119147795 |
| Olfr117 | 0 | NA | NA | NA |
| Olfr118 | 0.061920613 | -0.443816764 | 3.724945049 | -0.119147198 |
| Olfr119 | 0 | NA | NA | NA |
| Gm7335 | 43.44352658 | 0.268953343 | 0.203970846 | 1.318587186 |
| Olfr120 | 0.155118291 | -0.444777837 | 3.496459604 | -0.127208058 |
| Olfr121 | 0 | NA | NA | NA |

| | | | | |
|---|---|---|---|---|
| Olfr122 | 0.08486385 | 0.037730769 | 3.724898049 | 0.010129343 |
| Olfr123 | 0 | NA | NA | NA |
| Gm20524 | 0.042716945 | 0.037062677 | 3.724945049 | 0.009949859 |
| Olfr124 | 0 | NA | NA | NA |
| Olfr125 | 0 | NA | NA | NA |
| Gm9577 | 0.148659141 | -1.239386499 | 3.716315506 | -0.333498729 |
| Olfr126 | 0 | NA | NA | NA |
| Gm18737 | 0.194588561 | 0.037075813 | 2.240900734 | 0.016545049 |
| Gm18520 | 0 | NA | NA | NA |
| Olfr127 | 0 | NA | NA | NA |
| Olfr128 | 0 | NA | NA | NA |
| Gm20543 | 0 | NA | NA | NA |
| Gm20542 | 0 | NA | NA | NA |
| Gm5967 | 0 | NA | NA | NA |
| Olfr761 | 0 | NA | NA | NA |
| Gm18522 | 0 | NA | NA | NA |
| Gm20541 | 1.213329596 | 0.336128781 | 1.011575071 | 0.332282587 |
| Olfr129 | 0 | NA | NA | NA |
| Olfr130 | 0.068097359 | 0.037064902 | 3.724945049 | 0.009950456 |
| Gm18833 | 0 | NA | NA | NA |
| Olfr131 | 0.048949532 | 0.037064902 | 3.724945049 | 0.009950456 |
| Olfr132 | 0.042716945 | 0.037062677 | 3.724945049 | 0.009949859 |
| Olfr133 | 0.061920613 | -0.443816764 | 3.724945049 | -0.119147198 |
| Olfr134 | 0 | NA | NA | NA |
| Olfr135 | 0 | NA | NA | NA |
| Olfr138 | 0 | NA | NA | NA |
| Olfr137 | 0 | NA | NA | NA |
| Olfr762-ps1 | 0 | NA | NA | NA |
| Olfr136 | 0.089062654 | -0.443823436 | 3.724945049 | -0.119148989 |
| Esp36 | 0 | NA | NA | NA |
| Gm20409 | 0 | NA | NA | NA |
| Gm6726 | 0 | NA | NA | NA |
| Esp34 | 0 | NA | NA | NA |
| Esp31 | 0.04393175 | 0.037062677 | 3.724945049 | 0.009949859 |
| Esp24 | 0 | NA | NA | NA |
| Gm21903 | 0 | NA | NA | NA |
| Esp23 | 0 | NA | NA | NA |
| Esp18 | 0 | NA | NA | NA |
| Gm26429 | 0 | NA | NA | NA |
| Esp16 | 0 | NA | NA | NA |
| Esp15 | 0 | NA | NA | NA |
| Gm26917 | 335746.0181 | -0.30197303 | 0.104827496 | -2.880666259 |
| Rn18s-rs5 | 102.7885986 | 0.579116995 | 0.818877357 | 0.70720846 |
| Yam1 | 128743.8113 | 0.134324955 | 0.635051632 | 0.211518164 |
| Esp38 | 0 | NA | NA | NA |
| Gm26376 | 0 | NA | NA | NA |
| Tgif2-ps2 | 0.524319023 | -0.015344536 | 1.352944692 | -0.011341584 |

| | | | | |
|---|---|---|---|---|
| Pgk2 | 0 | NA | NA | NA |
| Crisp3 | 0 | NA | NA | NA |
| Crisp1 | 0.331267484 | -0.329858051 | 1.726177403 | -0.191091629 |
| Esp8 | 0 | NA | NA | NA |
| Esp6Esp5 | 0 | NA | NA | NA |
| Esp6 | 0 | NA | NA | NA |
| Esp4 | 0 | NA | NA | NA |
| Esp3 | 0 | NA | NA | NA |
| Esp1 | 0 | NA | NA | NA |
| Crisp2 | 0.293737151 | 0.499159306 | 3.702560026 | 0.134814643 |
| Rhag | 0.351655166 | 0.875829155 | 3.698748799 | 0.236790656 |
| Gm20574 | 0 | NA | NA | NA |
| 9130008F23Rik | 2.649375936 | -2.299636 | 0.740821219 | -3.104171345 |
| Glyatl3 | 0.199876958 | -0.443823165 | 2.661322747 | -0.166767885 |
| Gm23078 | 0.852119224 | 0.974449457 | 1.13522718 | 0.858373966 |
| Cenpq | 63.31617002 | -0.307564175 | 0.159770966 | -1.925031702 |
| Mut | 485.5759982 | 0.112902973 | 0.101534021 | 1.111971847 |
| Gm6771 | 0.062426873 | 0.517959912 | 3.724945049 | 0.139051692 |
| Ptchd4 | 384.0058745 | 0.047503131 | 0.081019453 | 0.586317603 |
| Opn5 | 0.714665336 | 0.06927709 | 1.141214554 | 0.060704702 |
| Adgrf4 | 19.83835518 | -0.886747234 | 0.265517874 | -3.339689419 |
| Adgrf2 | 18.64465502 | 0.205438212 | 0.30915761 | 0.664509641 |
| Cd2ap | 541.0115779 | 0.083977947 | 0.141437129 | 0.593747536 |
| Tnfrsf21 | 965.9785103 | 0.26607248 | 0.067811067 | 3.923732407 |
| Adgrf1 | 1.021366362 | 0.870257254 | 0.991741136 | 0.877504444 |
| Adgrf5 | 194.5698432 | -0.567285649 | 0.180451122 | -3.143708067 |
| Gm25135 | 0.066050875 | -0.443818988 | 3.724945049 | -0.119147795 |
| Mep1a | 1.39389272 | -2.01789934 | 1.018832383 | -1.980599923 |
| Ankrd66 | 10.82149774 | -2.235751837 | 0.439131932 | -5.091298704 |
| Pla2g7 | 1523.964145 | -0.429278678 | 0.129858156 | -3.305750613 |
| 1700071M16Rik | 0.275776889 | 0.018608714 | 2.068079672 | 0.008998065 |
| Tdrd6 | 25.59334993 | 0.750968582 | 0.295066897 | 2.545079069 |
| Slc25a27 | 391.4686422 | 0.398236018 | 0.13588595 | 2.930663674 |
| Cyp39a1 | 17.12697954 | -0.33092856 | 0.307153546 | -1.077404328 |
| Gm25783 | 0 | NA | NA | NA |
| Rcan2 | 1857.55946 | 0.242162592 | 0.101672292 | 2.381795356 |
| Gm24532 | 0.062426873 | 0.517959912 | 3.724945049 | 0.139051692 |
| Enpp5 | 2499.132028 | 0.474498043 | 0.051109473 | 9.283954929 |
| Enpp4 | 696.8122402 | -0.449206019 | 0.165392338 | -2.716002591 |
| Clic5 | 79.94705142 | -0.797003754 | 0.195156387 | -4.083923489 |
| Runx2 | 205.0751855 | -0.366519801 | 0.167726353 | -2.185224895 |
| Runx2os3 | 0 | NA | NA | NA |
| n-R5s27 | 0.766229633 | -0.47406196 | 1.12595863 | -0.421029643 |
| Runx2os2 | 0.092269992 | -0.443823436 | 3.724945049 | -0.119148989 |
| Runx2os1 | 7.863194607 | -1.03595425 | 0.460554573 | -2.249362641 |
| Supt3 | 134.9145071 | -0.435073143 | 0.126529661 | -3.438507139 |
| Gm24979 | 0 | NA | NA | NA |

| | | | | |
|---|---|---|---|---|
| Cdc5l | 544.4221431 | -0.227323747 | 0.066462038 | -3.42035476 |
| B230354K17Rik | 184.8456956 | -0.428978096 | 0.132856034 | -3.228894349 |
| Spats1 | 16.79904519 | 0.035499465 | 0.285779503 | 0.124219774 |
| Gm9104 | 1.30365031 | 0.397052695 | 0.96478971 | 0.411543252 |
| Aars2 | 207.7781551 | -0.036896743 | 0.093461964 | -0.39477817 |
| Tcte1 | 13.23124629 | 0.316395516 | 0.323477929 | 0.978105422 |
| Tmem151b | 1924.460039 | 0.383754554 | 0.086962396 | 4.41287928 |
| Nfkbie | 174.6593782 | -0.168211405 | 0.26011897 | -0.646671041 |
| Slc35b2 | 231.0133132 | -0.321254498 | 0.147633862 | -2.176021772 |
| Hsp90ab1 | 12795.80252 | -0.138602869 | 0.083479178 | -1.660328628 |
| Slc29a1 | 163.1308914 | 0.185595267 | 0.102969556 | 1.802428543 |
| Gm17080 | 1.296318896 | -0.573532279 | 0.977358978 | -0.586818448 |
| Gm7325 | 0 | NA | NA | NA |
| Gm25008 | 0 | NA | NA | NA |
| Capn11 | 4.929061397 | -0.418362285 | 0.566948704 | -0.737919113 |
| Tmem63b | 1144.513863 | 0.289158716 | 0.074347536 | 3.889284441 |
| Gm16172 | 0.25133133 | 0.707014142 | 2.414259629 | 0.292849258 |
| Mrpl14 | 161.5517627 | -0.406005623 | 0.135990853 | -2.985536259 |
| 1600014C23Rik | 0.431841237 | -0.003024126 | 1.448649873 | -0.002087548 |
| F630040K05Rik | 2.147486072 | 3.110594581 | 0.985305924 | 3.156983538 |
| Vegfa | 569.830768 | -0.275460888 | 0.102710443 | -2.681917059 |
| Mrps18a | 383.939429 | -0.30099028 | 0.080299873 | -3.748328222 |
| Rsph9 | 56.95556391 | 0.295348489 | 0.14412408 | 2.049265388 |
| Mad2l1bp | 60.28485607 | -0.543338873 | 0.161643085 | -3.361349317 |
| Gtpbp2 | 597.9990834 | -0.16685071 | 0.083145546 | -2.006730585 |
| Polh | 84.99680479 | -0.540521511 | 0.154925224 | -3.488918703 |
| Xpo5 | 708.0881202 | -0.140776683 | 0.05936333 | -2.371441827 |
| Mir693 | 0.626433637 | -0.418911036 | 1.370742609 | -0.305608823 |
| Gm22474 | 0 | NA | NA | NA |
| Polr1c | 186.1516209 | -0.30824578 | 0.111523589 | -2.763951402 |
| Yipf3 | 497.8450709 | -0.180272886 | 0.072432059 | -2.488854933 |
| Lrrc73 | 196.1398283 | 0.766745091 | 0.120091878 | 6.384653988 |
| Tjap1 | 262.6446001 | -0.533881129 | 0.108919504 | -4.901611821 |
| Gm26785 | 0.440626437 | 0.847222907 | 1.618755834 | 0.52337906 |
| Dlk2 | 119.3259875 | 0.543492214 | 0.185321094 | 2.932705616 |
| Abcc10 | 137.5955981 | 0.001998113 | 0.172535235 | 0.011580899 |
| Zfp318 | 978.2373152 | 0.304276288 | 0.094693121 | 3.213288199 |
| Gm23037 | 4.835503307 | 1.332631615 | 0.565089836 | 2.358265056 |
| Crip3 | 13.66803183 | -0.543441537 | 0.436567405 | -1.24480557 |
| Slc22a7 | 3.396474255 | -0.176425679 | 0.51671411 | -0.341437702 |
| Gm5093 | 4.877282298 | -0.284187429 | 0.572018493 | -0.496815108 |
| Ttbk1 | 1698.649847 | 0.124392989 | 0.08985386 | 1.384392266 |
| Gm26904 | 0.300318634 | -0.441313512 | 1.792996333 | -0.246131854 |
| Dnph1 | 20.31270367 | -0.2896203 | 0.256187704 | -1.130500392 |
| Cul9 | 1048.399803 | 0.46784245 | 0.076897737 | 6.083956046 |
| Srf | 574.4040805 | -0.050155376 | 0.13605067 | -0.36865218 |
| mmu-mir-6976 | 0.656458977 | 1.838539512 | 1.162724274 | 1.581234307 |

| | | | | |
|---|---|---|---|---|
| Ptk7 | 194.7380974 | -0.132687399 | 0.090268327 | -1.469921985 |
| Klc4 | 177.2125805 | -0.135471142 | 0.09938428 | -1.363104327 |
| Mrpl2 | 228.4860238 | 0.110223646 | 0.115487104 | 0.954423849 |
| Cul7 | 465.2672236 | -0.063542203 | 0.095531179 | -0.665146225 |
| Rrp36 | 135.9417813 | -0.625003609 | 0.100982719 | -6.189213487 |
| Klhdc3 | 1295.864183 | 0.178999786 | 0.050673607 | 3.532406673 |
| Mea1 | 401.4088604 | -0.182433816 | 0.123867962 | -1.472808724 |
| Ppp2r5d | 1040.331458 | 0.176711531 | 0.066801016 | 2.645341979 |
| Pex6 | 410.5596741 | -0.31398084 | 0.132948939 | -2.361664883 |
| Gnmt | 15.57555665 | 0.730090438 | 0.307385539 | 2.375161957 |
| Cnpy3 | 552.2456984 | 0.041477711 | 0.079113504 | 0.524281048 |
| Ptcra | 0.771865818 | 0.025680206 | 1.101757468 | 0.023308402 |
| 2310039H08Rik | 29.97597163 | -0.350242535 | 0.219485519 | -1.595743247 |
| Rpl7l1 | 408.9885597 | -0.618761737 | 0.104348491 | -5.92976221 |
| Gltscr1l | 1089.761893 | 0.12032968 | 0.082938285 | 1.45083395 |
| A330017A19Rik | 35.0263174 | 0.05721716 | 0.285294908 | 0.20055444 |
| Tbcc | 165.35841 | -0.272559322 | 0.130273604 | -2.092206821 |
| Prph2 | 0.359258722 | -0.344717678 | 1.453923345 | -0.237094809 |
| Ubr2 | 1461.655788 | -0.138954386 | 0.055889817 | -2.486220099 |
| Gm23657 | 4.719513391 | 0.740014347 | 0.509179932 | 1.453345469 |
| Gm16494 | 2.052894931 | -0.689240734 | 0.896099375 | -0.769156584 |
| Gm4945 | 0 | NA | NA | NA |
| Trerf1 | 592.2320734 | 0.81373904 | 0.156058441 | 5.214322484 |
| Gm26216 | 1.145909063 | 1.05775622 | 1.074001288 | 0.984874257 |
| Mrps10 | 197.4832179 | -0.129749745 | 0.101735196 | -1.275367333 |
| Guca1b | 0.605175792 | -1.337144967 | 1.272962323 | -1.050419909 |
| Guca1a | 2.149392939 | -0.006962518 | 0.824632128 | -0.008443181 |
| Gm5814 | 2.49590342 | -0.126288264 | 0.580213932 | -0.217658103 |
| 1700001C19Rik | 0.29699235 | 0.531667079 | 1.771717418 | 0.30008571 |
| AI661453 | 6.189948269 | 0.228277795 | 0.383057361 | 0.595936322 |
| Taf8 | 186.7542863 | -0.068332448 | 0.091967265 | -0.743008373 |
| Ccnd3 | 416.7992574 | -0.484799229 | 0.127343976 | -3.807005604 |
| Bysl | 253.0965139 | -0.356999304 | 0.091137722 | -3.917140957 |
| Med20 | 240.4788844 | -0.156709529 | 0.121280048 | -1.292129512 |
| Gm20517 | 21.19396069 | 1.250132673 | 0.340239221 | 3.674275617 |
| Kpna2-ps | 1.554960773 | -0.588824243 | 0.846600977 | -0.695515667 |
| Usp49 | 129.8291219 | -0.009877812 | 0.130146279 | -0.07589777 |
| Tomm6 | 2.522522481 | 0.727803307 | 0.877529095 | 0.829377978 |
| Gm21981 | 0.57942775 | 2.336110028 | 1.210656964 | 1.929621765 |
| Tomm6os | 39.84955966 | 0.39395023 | 0.181783603 | 2.16713842 |
| Prickle4 | 1.072312625 | 2.104055939 | 1.128692133 | 1.864153987 |
| Frs3 | 230.2853897 | 0.339835046 | 0.111176196 | 3.056724908 |
| Frs3os | 0.363326102 | 0.036133905 | 1.712113419 | 0.021104855 |
| Pgc | 0.329750641 | 1.080143106 | 1.730054774 | 0.62434041 |
| Tfeb | 165.6362251 | -0.060734873 | 0.130683675 | -0.464747202 |
| Mdfi | 11.42660599 | -0.595420403 | 0.335131932 | -1.776674638 |
| Gm25201 | 0.048949532 | 0.037064902 | 3.724945049 | 0.009950456 |

| | | | | |
|---|---|---|---|---|
| n-R5s28 | 0 | NA | NA | NA |
| Foxp4 | 579.0805347 | 0.435396108 | 0.105849426 | 4.113353523 |
| Gm15556 | 0.196245641 | -1.436545123 | 3.230301694 | -0.444709275 |
| Gm5228 | 0 | NA | NA | NA |
| Gm14871 | 0 | NA | NA | NA |
| 1700122O11Rik | 0 | NA | NA | NA |
| 1700067P10Rik | 0 | NA | NA | NA |
| 9830107B12Rik | 0 | NA | NA | NA |
| A530064D06Rik | 30.08457509 | -2.09968485 | 0.442393609 | -4.746191648 |
| Trem1 | 13.84162536 | -1.64455692 | 0.546347248 | -3.010094633 |
| Trem3 | 3.961576269 | -2.141024237 | 0.577984274 | -3.704294967 |
| Treml4 | 2.160387598 | -0.812007184 | 0.877359738 | -0.925512249 |
| Treml2 | 120.9667195 | -1.177142788 | 0.219727629 | -5.357281629 |
| B430306N03Rik | 110.3507723 | -1.937902853 | 0.39690039 | -4.882592469 |
| Trem2 | 83.40596837 | -0.746821152 | 0.490099732 | -1.523814651 |
| Treml1 | 1.368647145 | 0.092304816 | 0.906248842 | 0.10185372 |
| Nfya | 266.8544357 | -0.814113186 | 0.129454741 | -6.288786193 |
| Oard1 | 77.30484208 | -0.184897316 | 0.147301897 | -1.255226984 |
| Apobec2 | 1.336996185 | -0.596305833 | 0.948897898 | -0.628419384 |
| Tspo2 | 0 | NA | NA | NA |
| Unc5cl | 6.179264061 | -0.752159587 | 0.45663671 | -1.64717284 |
| Gm24071 | 0 | NA | NA | NA |
| Gm23642 | 0 | NA | NA | NA |
| Lrfn2 | 299.7160112 | 0.30280861 | 0.137893677 | 2.195957172 |
| AC154247.1 | 0.135598227 | 0.037065847 | 2.950266134 | 0.01256356 |
| 1700008K24Rik | 0 | NA | NA | NA |
| Gm22978 | 0.046345709 | -0.443818988 | 3.724945049 | -0.119147795 |
| Gm20540 | 0.410699371 | -0.354785412 | 1.461080262 | -0.24282404 |
| Mocs1 | 175.5053655 | -0.055654117 | 0.114670134 | -0.485340992 |
| Daam2 | 1383.444506 | 0.374987638 | 0.100003381 | 3.749749588 |
| Kif6 | 28.85323937 | 0.171050211 | 0.245883901 | 0.695654373 |
| Gm16555 | 0 | NA | NA | NA |
| Gm16554 | 0.075043205 | 0.517957688 | 3.724945049 | 0.139051095 |
| Rftn1 | 212.415448 | -0.833817773 | 0.139177955 | -5.991019001 |
| Gm37173 | 0 | NA | NA | NA |
| AY702103 | 0 | NA | NA | NA |
| Dazl | 26.34726813 | 0.717440139 | 0.282616066 | 2.538568133 |
| Plcl2 | 1538.397032 | 0.293459057 | 0.083189245 | 3.527608149 |
| Gm7334 | 7.373441669 | -0.363845732 | 0.403433095 | -0.901873783 |
| Tbc1d5 | 596.1325065 | 0.096380544 | 0.062956306 | 1.530911689 |
| Gm25177 | 2.526566463 | -0.103057543 | 0.70816009 | -0.145528595 |
| Satb1 | 1280.976831 | 0.354878738 | 0.10978106 | 3.232604407 |
| E430014B02Rik | 30.56272651 | -0.178171944 | 0.334762632 | -0.53223367 |
| C230085N15Rik | 40.73930876 | 0.282234056 | 0.342771752 | 0.823387732 |
| Gm36931 | 44.82250833 | 0.431358486 | 0.23990018 | 1.798074872 |
| 5830444F18Rik | 19.30851287 | 0.479839773 | 0.397450535 | 1.207294319 |
| Gm37248 | 39.69177475 | 0.494321333 | 0.341204107 | 1.448755519 |

| | | | | |
|---|---|---|---|---|
| Gm6934 | 12.03475686 | 0.454025588 | 0.472333503 | 0.961239432 |
| Gm37593 | 18.0879042 | 0.431869959 | 0.44427044 | 0.97208799 |
| Gm37266 | 10.80255973 | -0.388074977 | 0.478742928 | -0.810612449 |
| Gm37176 | 9.616469211 | 0.235288819 | 0.562012019 | 0.418654426 |
| Gm19585 | 5.697516218 | -0.500259209 | 0.467300926 | -1.070529034 |
| Gm27217 | 0.396216368 | 0.506050529 | 1.74365448 | 0.290224087 |
| Kcnh8 | 23.78866619 | 0.217654497 | 0.260392681 | 0.835870257 |
| Efhb | 28.27811259 | -0.303368225 | 0.232935894 | -1.302367873 |
| Rab5a | 1148.748334 | -0.075780835 | 0.085660756 | -0.884662224 |
| Pp2d1 | 0.305447682 | 1.434748657 | 1.71895512 | 0.834663244 |
| Kat2b | 803.7724207 | -0.212954939 | 0.099761204 | -2.134646846 |
| Gm6919 | 0.108772582 | 0.037073798 | 3.724945049 | 0.009952844 |
| Sgol1 | 6.962193728 | -0.113039074 | 0.587237973 | -0.192492787 |
| Gm19518 | 0.186767869 | 1.022050669 | 3.289032417 | 0.310745088 |
| 4932415M13Rik | 0 | NA | NA | NA |
| Ck-ps1 | 0.277663158 | 0.989600091 | 3.716337573 | 0.266283692 |
| RP24-173D2.6 | 0 | NA | NA | NA |
| RP24-173D2.8 | 0 | NA | NA | NA |
| RP24-173D2.9 | 0 | NA | NA | NA |
| Sult1c2 | 0.362371572 | 1.780657795 | 1.71642964 | 1.037419626 |
| Mrps36-ps1 | 2.335139537 | -0.478160837 | 0.760786323 | -0.628508719 |
| RP24-173D2.10 | 0 | NA | NA | NA |
| RP24-173D2.11 | 0 | NA | NA | NA |
| RP23-416K11.2 | 0 | NA | NA | NA |
| RP23-416K11.3 | 0 | NA | NA | NA |
| RP23-416K11.4 | 0 | NA | NA | NA |
| Sult1c1 | 1.161929334 | 0.460905613 | 1.181568488 | 0.390079473 |
| RP23-416K11.5 | 0 | NA | NA | NA |
| Gm37966 | 0 | NA | NA | NA |
| Slc5a7 | 146.1741277 | -0.071040577 | 0.248247452 | -0.286168403 |
| 1700025K24Rik | 0.046345709 | -0.443818988 | 3.724945049 | -0.119147795 |
| Trp53-ps | 0 | NA | NA | NA |
| Gm26291 | 0 | NA | NA | NA |
| Gm17377 | 2.328145098 | -0.189484717 | 0.758634837 | -0.249770651 |
| St6gal2 | 374.6218881 | 0.644983492 | 0.163298308 | 3.949725513 |
| Gm26547 | 0.778497894 | -1.666262865 | 1.444654715 | -1.153398696 |
| Vmn2r118 | 1.673678531 | -0.915979232 | 0.780090588 | -1.17419598 |
| Pot1b | 63.94887526 | -0.021967205 | 0.156683026 | -0.140201562 |
| Gm22654 | 0 | NA | NA | NA |
| Adgre4 | 3.084562812 | -1.39838236 | 0.66280827 | -2.109784115 |
| Gm18417 | 1.232924928 | 1.10739495 | 0.896559449 | 1.235160648 |
| Zfp119a | 72.0705982 | -0.032719794 | 0.13613785 | -0.24034311 |
| Zfp959 | 70.36865212 | -0.393750209 | 0.141585268 | -2.781011148 |
| Rpl21-ps6 | 2.92891158 | 1.07766816 | 0.56753496 | 1.898857754 |
| Zfp119b | 52.43127221 | -0.114492967 | 0.186620275 | -0.61350765 |
| Ebi3 | 12.63508742 | -0.225235291 | 0.300473637 | -0.749600839 |
| Gm16712 | 2.434106523 | 0.085550237 | 0.642259816 | 0.133201915 |

| | | | | |
|---|---|---|---|---|
| Ccdc94 | 63.31143036 | -0.159043974 | 0.161287478 | -0.986090034 |
| Shd | 180.526558 | 0.553633909 | 0.091326548 | 6.062135511 |
| Fsd1 | 267.0610619 | 0.098831285 | 0.087844139 | 1.125075453 |
| Stap2 | 23.62948936 | -0.021033885 | 0.231690007 | -0.0907846 |
| Mpnd | 574.8071085 | 0.20851603 | 0.060155704 | 3.466271951 |
| Gm16276 | 0.401552702 | 0.451833856 | 1.578837575 | 0.286181342 |
| Sh3gl1 | 328.504642 | -0.162313602 | 0.085719325 | -1.89354736 |
| Chaf1a | 59.75683498 | 0.028858052 | 0.214087772 | 0.134795424 |
| Ubxn6 | 593.1748788 | 0.140254248 | 0.053058382 | 2.643394714 |
| Hdgfrp2 | 630.2550697 | 0.189142231 | 0.07745133 | 2.442078561 |
| Plin4 | 580.4887041 | 0.88394578 | 0.279131114 | 3.166776238 |
| Plin5 | 6.445794394 | 0.508180766 | 0.407911068 | 1.245812644 |
| Lrg1 | 47.22509705 | -0.508581338 | 0.312267095 | -1.628674127 |
| Sema6b | 1185.955188 | 0.731936176 | 0.095230096 | 7.685975394 |
| Tnfaip8l1 | 28.12050717 | 0.312465709 | 0.236981856 | 1.318521654 |
| AC026385.1 | 0.119587853 | 0.981063299 | 3.723618439 | 0.263470416 |
| Mydgf | 321.3040017 | -0.427233535 | 0.121252304 | -3.523508604 |
| Dpp9 | 687.0448644 | -0.263489654 | 0.057343103 | -4.594966794 |
| Mir7b | 2.668141924 | -1.339642731 | 0.817747895 | -1.63820994 |
| Fem1a | 839.6826125 | 0.154046186 | 0.07782763 | 1.979325156 |
| Ticam1 | 133.3080346 | -0.756623924 | 0.142354854 | -5.315055332 |
| Plin3 | 209.6287355 | -0.582397733 | 0.196118235 | -2.969625611 |
| Arrdc5 | 0 | NA | NA | NA |
| Uhrf1 | 83.37304998 | -1.258139528 | 0.310276419 | -4.054898954 |
| Kdm4b | 809.6209241 | 0.260107562 | 0.08169701 | 3.183807626 |
| Ptprs | 5796.573917 | 0.429559219 | 0.087713554 | 4.897295819 |
| Mir6977 | 0 | NA | NA | NA |
| Znrf4 | 0.092691418 | -0.908687078 | 3.724560878 | -0.2439716 |
| Safb2 | 645.9551208 | 0.121279042 | 0.179225599 | 0.6766837 |
| Safb | 970.0585309 | 0.185084505 | 0.160115053 | 1.15594693 |
| 2410015M20Rik | 237.0366201 | 0.047714449 | 0.142822066 | 0.334083171 |
| Rpl36 | 557.6166036 | -0.276835978 | 0.151066798 | -1.832540185 |
| Lonp1 | 634.6361425 | -0.31009802 | 0.075550205 | -4.104529146 |
| Catsperd | 6.596655121 | 0.234366902 | 0.445622777 | 0.525931156 |
| Ranbp3 | 724.4876957 | -0.122900438 | 0.059046289 | -2.081425281 |
| Vmac | 91.17240515 | 0.4806793 | 0.125377994 | 3.833841059 |
| Ndufa11 | 437.8651622 | -0.009862435 | 0.107883418 | -0.091417528 |
| Nrtn | 25.07553935 | 0.350797236 | 0.231723561 | 1.513860887 |
| Dus3l | 477.2858719 | 0.337308728 | 0.091897874 | 3.670473692 |
| Prr22 | 6.373319375 | 0.371963309 | 0.412998522 | 0.900640775 |
| Rfx2 | 53.12569894 | -0.771859504 | 0.171349802 | -4.504583568 |
| Acsbg2 | 0.382360723 | -1.577248473 | 1.651869552 | -0.954826288 |
| 1700061G19Rik | 1.170313971 | -0.396372711 | 1.112648587 | -0.356242497 |
| Mllt1 | 783.4489317 | -0.071123916 | 0.070558394 | -1.008014951 |
| Acer1 | 0.046345709 | -0.443818988 | 3.724945049 | -0.119147795 |
| Clpp | 295.2646874 | -0.224374001 | 0.088365377 | -2.539161926 |
| Alkbh7 | 82.81251037 | -0.125245492 | 0.186634206 | -0.671074693 |

| | | | | |
|---|---|---|---|---|
| Pspn | 0.342416647 | 0.53418333 | 1.997715296 | 0.267397127 |
| Gtf2f1 | 546.3864564 | -0.110218324 | 0.130745341 | -0.843000014 |
| Khsrp | 1672.094708 | -0.196066024 | 0.041329322 | -4.743993268 |
| Slc25a41 | 1.408439352 | -1.952341076 | 1.073417659 | -1.81880842 |
| Gm17168 | 0.17329739 | 0.037058443 | 3.344182904 | 0.011081464 |
| Slc25a23 | 2717.008405 | 0.666069664 | 0.090043192 | 7.397223991 |
| Crb3 | 6.90059268 | 0.347684704 | 0.421582901 | 0.824712537 |
| Dennd1c | 54.0186804 | -0.987003598 | 0.228230028 | -4.324600075 |
| Tubb4a | 6699.351768 | 0.434894234 | 0.095391385 | 4.559051469 |
| Gm11110 | 11.24476628 | 0.550265911 | 0.336047147 | 1.637466397 |
| Tnfsf9 | 20.88192311 | -0.286598791 | 0.319370352 | -0.897386963 |
| Cd70 | 3.132506103 | -0.634046448 | 0.562326395 | -1.127541679 |
| Tnfsf14 | 6.752466611 | -0.810727745 | 0.607739827 | -1.334004632 |
| C3 | 1371.649863 | -1.001075501 | 0.461533505 | -2.169020213 |
| Mir6978 | 0 | NA | NA | NA |
| Gpr108 | 284.67282 | -0.317875235 | 0.097518739 | -3.259632328 |
| Trip10 | 80.25548548 | -0.499163298 | 0.169323388 | -2.947987905 |
| Vav1 | 298.6647182 | -0.945217099 | 0.169183764 | -5.586925586 |
| Adgre1 | 147.8605273 | -0.975700962 | 0.237399924 | -4.109946394 |
| Vmn2r120 | 0 | NA | NA | NA |
| Gm21834 | 0.060174537 | 0.517957688 | 3.724945049 | 0.139051095 |
| Gm25102 | 0 | NA | NA | NA |
| Cntnap5c | 204.371656 | 0.163641026 | 0.150266671 | 1.089004131 |
| Rpl7a-ps5 | 12.52156343 | -0.030519985 | 0.444848457 | -0.0686076 |
| Gm5815 | 49.22368419 | 0.294725972 | 0.193658865 | 1.521882163 |
| Pdzph1 | 37.42674745 | 0.916538464 | 0.230489878 | 3.976480314 |
| Nudt12os | 0.099106094 | -0.93550548 | 3.724188147 | -0.251197158 |
| Nudt12 | 83.70631604 | -0.643228874 | 0.177700633 | -3.61973316 |
| Gm23769 | 0 | NA | NA | NA |
| Gm23203 | 0 | NA | NA | NA |
| Gm29051 | 0 | NA | NA | NA |
| Gm23447 | 0 | NA | NA | NA |
| Gm23773 | 0 | NA | NA | NA |
| Obox6-ps1 | 0 | NA | NA | NA |
| Gm4518 | 4.468190482 | 0.717408192 | 0.506689535 | 1.415873317 |
| Gm27023 | 0.131193343 | 0.517960218 | 2.980368697 | 0.173790652 |
| Gm25800 | 0 | NA | NA | NA |
| Efna5 | 380.5177091 | 0.079006225 | 0.088928082 | 0.888428308 |
| Fbxl17 | 855.8478773 | 0.302699624 | 0.061822506 | 4.896269086 |
| Gm25348 | 2.594574602 | 0.469709438 | 0.672396903 | 0.698559788 |
| Gm24730 | 0 | NA | NA | NA |
| 4930405O22Rik | 44.37612019 | 0.479678375 | 0.285791809 | 1.678418902 |
| Gm17228 | 21.45301415 | 0.245332319 | 0.317322998 | 0.773131227 |
| Gm24813 | 0 | NA | NA | NA |
| A930002H24Rik | 1.337378336 | 0.1763858 | 0.990129612 | 0.178144152 |
| Fer | 419.2667044 | 0.050471675 | 0.063937395 | 0.789392116 |
| Pja2 | 6703.040193 | 0.345785408 | 0.059422408 | 5.819107943 |

| | | | | |
|---|---|---|---|---|
| Gm24194 | 3.223858552 | 0.769502019 | 0.734315922 | 1.047916838 |
| Mir6420 | 0 | NA | NA | NA |
| AU016765 | 0 | NA | NA | NA |
| Man2a1 | 771.0017978 | -0.624345994 | 0.109983341 | -5.676732383 |
| Gm17133 | 0 | NA | NA | NA |
| 4930583I09Rik | 0 | NA | NA | NA |
| Gm25344 | 0 | NA | NA | NA |
| Tmem232 | 14.97100193 | -0.017378045 | 0.271976885 | -0.0638953 |
| Gm25731 | 0 | NA | NA | NA |
| Vapa | 1462.977574 | -0.233952386 | 0.090634472 | -2.581273768 |
| Txndc2 | 4.780619236 | 0.231987778 | 0.472279355 | 0.491208805 |
| Rab31 | 1383.126595 | -0.453418508 | 0.128708426 | -3.522834686 |
| Ppp4r1 | 548.0723031 | -0.535647799 | 0.087090681 | -6.150460565 |
| Ralbp1 | 849.2774884 | -0.113477779 | 0.084440693 | -1.343875507 |
| Twsg1 | 633.6219113 | -0.318362718 | 0.105392152 | -3.02074408 |
| Ankrd12 | 2379.847845 | 0.291881125 | 0.072605347 | 4.020105096 |
| Ndufv2 | 1210.518907 | 0.023791146 | 0.096167023 | 0.247394015 |
| Gm23264 | 0.127717592 | 0.037073759 | 3.006584803 | 0.012330854 |
| Wash1 | 277.8590539 | -0.150136557 | 0.077306085 | -1.942105292 |
| Ddx11 | 26.93519898 | -0.742047714 | 0.239092828 | -3.103596708 |
| Mtcl1 | 1748.393845 | 0.216635517 | 0.141686146 | 1.528981645 |
| Gm4705 | 22.01963504 | 0.525811455 | 0.282041311 | 1.864306517 |
| Rab12 | 884.2363581 | 0.157956555 | 0.085872029 | 1.839441285 |
| Themis3 | 0.652038697 | -0.514559649 | 1.260690892 | -0.408156871 |
| Ptprm | 817.855665 | 0.225380554 | 0.067760841 | 3.326118009 |
| Mir5709 | 2.098507799 | 0.405244334 | 0.746974825 | 0.542514045 |
| Gm22277 | 0 | NA | NA | NA |
| Lrrc30 | 0.047948698 | 0.037064902 | 3.724945049 | 0.009950456 |
| Lama1 | 126.6409752 | -0.044298202 | 0.180000653 | -0.24610023 |
| Arhgap28 | 21.82346039 | -0.31644818 | 0.224778958 | -1.407819412 |
| Gm17097 | 0.284087786 | 0.535064749 | 1.783398313 | 0.300025375 |
| L3mbtl4 | 19.07932733 | 0.379541532 | 0.259163307 | 1.464487919 |
| Gm15974 | 0.045882455 | 0.517959912 | 3.724945049 | 0.139051692 |
| Tmem200c | 195.2513106 | -0.296474984 | 0.098174257 | -3.019885166 |
| Epb4.1l3 | 2609.206492 | 0.303220234 | 0.071058912 | 4.267166862 |
| 2410021H03Rik | 0.229803803 | -0.368395176 | 2.159245057 | -0.170612953 |
| Zbtb14 | 266.5595265 | 0.052386943 | 0.086737754 | 0.603969333 |
| A930029G22Rik | 17.70809633 | -0.219511213 | 0.289023577 | -0.759492409 |
| C030034I22Rik | 43.35133415 | -0.01966381 | 0.18972933 | -0.103641384 |
| A330050F15Rik | 100.5892871 | -0.027784617 | 0.129621791 | -0.214351436 |
| Gm22967 | 0 | NA | NA | NA |
| Dlgap1 | 5541.017591 | 0.53066077 | 0.10431994 | 5.086858457 |
| 5031415H12Rik | 2.165235617 | -1.693268638 | 0.708127288 | -2.391192468 |
| Gm20703 | 2.008244415 | 2.815738476 | 0.844228923 | 3.335278379 |
| Tgif1 | 173.2492349 | -1.44150836 | 0.257515715 | -5.597749087 |
| Gm28727 | 2.554427972 | -0.397539111 | 0.590501333 | -0.673223055 |
| Mir1195 | 0 | NA | NA | NA |

| | | | | |
|---|---|---|---|---|
| Gm26510 | 1.053857694 | 0.367586834 | 0.884401262 | 0.415633548 |
| Gm9320 | 7.10377163 | -0.170103187 | 0.415240591 | -0.40964971 |
| Gm26561 | 0.274682661 | 1.058421007 | 3.708688692 | 0.285389553 |
| Gm16519 | 0 | NA | NA | NA |
| Myl12b | 1231.826391 | 0.190464068 | 0.101198888 | 1.882076701 |
| Myl12a | 318.06527 | -0.742937762 | 0.184093918 | -4.035645341 |
| Myom1 | 11.02360487 | -0.159547413 | 0.341950189 | -0.466580859 |
| Lpin2 | 1181.411907 | -0.019076817 | 0.060322861 | -0.316245236 |
| Emilin2 | 338.4431336 | -1.252254895 | 0.286951429 | -4.363996023 |
| Smchd1 | 1270.998503 | -0.377700852 | 0.071819576 | -5.259023698 |
| Gm37639 | 37.94753101 | -0.005433549 | 0.262285697 | -0.020716147 |
| Gm38220 | 18.1195503 | 0.620210947 | 0.408077863 | 1.519834824 |
| Gm4707 | 30.94223545 | -0.03228008 | 0.332276229 | -0.097148329 |
| Ndc80 | 7.673135682 | 0.344438406 | 0.411484068 | 0.837063773 |
| Gm23115 | 0.060174537 | 0.517957688 | 3.724945049 | 0.139051095 |
| Mettl4-ps1 | 0.604171475 | -1.825609296 | 1.20322718 | -1.517260686 |
| Spdya | 9.137166137 | 1.165461511 | 0.373109627 | 3.123643632 |
| Trmt61b | 64.18131499 | -0.207696295 | 0.153930622 | -1.349285104 |
| Wdr43 | 540.8743764 | -0.240664549 | 0.105238402 | -2.286851033 |
| Snord92 | 1.968787839 | -0.000375508 | 0.9432077 | -0.000398118 |
| Snord53 | 3.604104618 | 1.109726771 | 0.690797808 | 1.606442229 |
| Gm22858 | 0.576946659 | 2.026988942 | 3.575595887 | 0.566895423 |
| Fam179a | 77.43913187 | -0.261011015 | 0.175211872 | -1.489687955 |
| Gm15641 | 0 | NA | NA | NA |
| BC027072 | 1.833725576 | -0.280622789 | 0.729512895 | -0.384671458 |
| Clip4 | 625.3662385 | -0.033421757 | 0.055896481 | -0.59792238 |
| Alk | 126.1837875 | 0.092603305 | 0.112149771 | 0.825711041 |
| Gm24736 | 0.908322817 | 0.613590442 | 1.236858141 | 0.496087968 |
| Gm26963 | 0 | NA | NA | NA |
| Ypel5 | 905.935313 | -0.070160516 | 0.0794601 | -0.882965363 |
| Lbh | 1015.797448 | -0.844495781 | 0.135354291 | -6.23915042 |
| Gm23649 | 0.075043205 | 0.517957688 | 3.724945049 | 0.139051095 |
| Lclat1 | 432.1555874 | -0.192113863 | 0.082177803 | -2.337782917 |
| Gm9311 | 0.089089296 | 0.957155977 | 3.723939561 | 0.257027796 |
| Gm25406 | 3.791534426 | -1.210093291 | 0.684374741 | -1.768173513 |
| Capn13 | 0.17782045 | -0.427193696 | 2.294087073 | -0.186215119 |
| Galnt14 | 193.3014098 | -0.060926845 | 0.153103851 | -0.397944562 |
| D630014O11Rik | 13.27744998 | 0.480383668 | 0.445982753 | 1.077135079 |
| Ehd3 | 1261.372702 | -0.287696323 | 0.085453864 | -3.366685964 |
| Xdh | 746.6583068 | -1.260285723 | 0.2471394 | -5.099493329 |
| Gm24475 | 0 | NA | NA | NA |
| Srd5a2 | 1.0712781 | -0.055351881 | 1.035178685 | -0.053470847 |
| Memo1 | 287.5100735 | -0.12041288 | 0.085684948 | -1.405297931 |
| Dpy30 | 145.3608656 | 0.005847813 | 0.122026406 | 0.047922521 |
| Spast | 787.6234006 | -0.003529954 | 0.055607717 | -0.06347957 |
| Slc30a6 | 112.2934621 | -0.21554471 | 0.133455689 | -1.615103191 |
| Nlrc4 | 11.8016311 | -0.756867615 | 0.299885335 | -2.523856714 |

| | | | | |
|---|---|---|---|---|
| Yipf4 | 489.5110183 | -0.280274266 | 0.098403237 | -2.848222001 |
| Birc6 | 5732.824248 | -0.063205276 | 0.065011725 | -0.972213491 |
| Gm26749 | 0.249552943 | 0.037538806 | 1.84829201 | 0.020309998 |
| Ttc27 | 207.7543264 | -0.06953651 | 0.080078601 | -0.868353207 |
| Ltbp1 | 508.8403771 | -0.66409706 | 0.178807008 | -3.714043803 |
| Gm6276 | 0.110870145 | -0.444802825 | 3.724874622 | -0.119414174 |
| Rasgrp3 | 150.64243 | -0.089932313 | 0.189690604 | -0.474099989 |
| Fam98a | 451.495364 | -0.343481227 | 0.089071989 | -3.856220466 |
| Gm24126 | 0 | NA | NA | NA |
| Gm26124 | 0 | NA | NA | NA |
| Gm16391 | 0 | NA | NA | NA |
| Gm17538 | 12.6024622 | -0.0132283 | 0.324857173 | -0.040720358 |
| Gm5229 | 0.072025462 | 0.517959912 | 3.724945049 | 0.139051692 |
| Crim1 | 1289.41108 | 0.4766043 | 0.077541793 | 6.14641839 |
| Gm25069 | 0.582565216 | -0.810358655 | 1.47041927 | -0.551107206 |
| Fez2 | 418.1889862 | -0.180872637 | 0.092928542 | -1.94636258 |
| Gm10093 | 94.11779407 | -0.324850866 | 0.156510997 | -2.075578532 |
| Vit | 19.4303546 | 0.157333233 | 0.251006747 | 0.62680878 |
| Gm22562 | 0.092493346 | 0.517955464 | 3.724945049 | 0.139050498 |
| Strn | 1780.921145 | 0.288212418 | 0.060870866 | 4.734817091 |
| Heatr5b | 1002.913214 | 0.043129077 | 0.061224361 | 0.70444308 |
| Gpatch11 | 308.9091273 | -0.050696656 | 0.126452857 | -0.40091349 |
| Gm6548 | 198.422454 | -0.554801069 | 0.16341774 | -3.394986792 |
| Eif2ak2 | 2234.053102 | -0.151082509 | 0.15774436 | -0.957768062 |
| n-R5s29 | 0.136963818 | 0.037074342 | 3.599544405 | 0.010299732 |
| Gm26637 | 5.907155527 | 0.01473861 | 0.410287477 | 0.035922642 |
| Sult6b1 | 1.803778753 | -1.446108554 | 0.956600272 | -1.511716645 |
| Cebpzos | 71.15640056 | 0.31491915 | 0.155316478 | 2.02759652 |
| Cebpz | 422.2918863 | -0.206538022 | 0.08954846 | -2.306438577 |
| Ndufaf7 | 198.2430765 | 0.021382671 | 0.085850917 | 0.249067473 |
| Prkd3 | 324.5446214 | -0.466148638 | 0.117624084 | -3.963037349 |
| Gm26957 | 0 | NA | NA | NA |
| Qpct | 101.3108535 | -0.080379915 | 0.163023132 | -0.493058342 |
| Cdc42ep3 | 151.9232374 | 0.044200575 | 0.123217707 | 0.35871934 |
| Gm17315 | 8.227418531 | 0.428733526 | 0.378415639 | 1.132969894 |
| Rmdn2 | 100.9181395 | 0.139480203 | 0.114794724 | 1.215040187 |
| 4921513D11Rik | 17.43713135 | -0.268396332 | 0.286645658 | -0.936334895 |
| Cyp1b1 | 282.3238744 | -1.060413143 | 0.221603303 | -4.785186538 |
| Gm37555 | 0.136963818 | 0.037074342 | 3.599544405 | 0.010299732 |
| Atl2 | 973.0030504 | 0.169654819 | 0.069924474 | 2.426258055 |
| Hnrnpll | 974.8453895 | 0.053576893 | 0.085816489 | 0.624319332 |
| Gm22215 | 0 | NA | NA | NA |
| Gm23125 | 0 | NA | NA | NA |
| Galm | 27.71090595 | -0.835487256 | 0.313246477 | -2.667188034 |
| Gm25706 | 3.582960898 | -0.775700942 | 0.639751471 | -1.212503571 |
| Srsf7 | 905.8057965 | -0.201952411 | 0.072640691 | -2.780155426 |
| Ttc39d | 1.912265302 | -1.840471995 | 0.865024491 | -2.127653048 |

| | | | | |
|---|---|---|---|---|
| Gemin6 | 32.80368467 | -0.268101519 | 0.208492354 | -1.285905761 |
| Dhx57 | 544.389297 | 0.059326442 | 0.105748156 | 0.561016322 |
| Morn2 | 84.79043694 | -0.033428739 | 0.167268553 | -0.199850707 |
| Arhgef33 | 272.4084982 | 0.065317911 | 0.303500173 | 0.2152154 |
| Gm10190 | 0.322675615 | 0.951990748 | 1.493223773 | 0.637540578 |
| Sos1 | 1511.796088 | -0.036965781 | 0.053215234 | -0.694646586 |
| Cdkl4 | 159.4655411 | 1.62446765 | 0.163297186 | 9.947921882 |
| Map4k3 | 1163.46542 | -0.008757531 | 0.051334133 | -0.170598596 |
| Gm9959 | 0.271824581 | -0.934515538 | 1.81171471 | -0.515818265 |
| Gm9386 | 0.459282249 | 1.07919565 | 1.432336113 | 0.7534514 |
| Tmem178 | 593.6338004 | 0.4711783 | 0.091420885 | 5.153945947 |
| Thumpd2 | 64.74403033 | -0.128646568 | 0.150562678 | -0.854438623 |
| Slc8a1 | 2319.537813 | 0.387323955 | 0.087732959 | 4.414805529 |
| Gm11096 | 0.461313532 | 1.633817822 | 1.310488819 | 1.24672397 |
| Rpl31-ps25 | 0 | NA | NA | NA |
| Gm27359 | 0 | NA | NA | NA |
| Gm22386 | 0 | NA | NA | NA |
| Gm23400 | 0 | NA | NA | NA |
| Gm6594 | 14.12482822 | -0.504238767 | 0.313027385 | -1.610845539 |
| Gm29052 | 0 | NA | NA | NA |
| Pkdcc | 80.91185254 | -0.177724409 | 0.150057834 | -1.184372741 |
| Gm24240 | 0 | NA | NA | NA |
| Rpl21-ps7 | 2.037653348 | 0.495281641 | 0.69344577 | 0.714232694 |
| Eml4 | 575.2161808 | -0.024543543 | 0.080802543 | -0.303747161 |
| Cox7a2l | 853.8877004 | 0.104592333 | 0.079282244 | 1.319240319 |
| Kcng3 | 81.41433254 | 0.676431713 | 0.193606045 | 3.493856372 |
| Mta3 | 891.3237956 | 0.163689384 | 0.058915797 | 2.77836154 |
| Haao | 4.556506885 | 0.671587076 | 0.54408833 | 1.234334647 |
| C430042M11Rik | 1.362936493 | -1.172399925 | 0.914758205 | -1.281650078 |
| Gm23905 | 0.061920613 | -0.443816764 | 3.724945049 | -0.119147198 |
| Gm37549 | 0.452867449 | 1.275436997 | 1.771641219 | 0.719918335 |
| Zfp36l2 | 928.29596 | -0.745697293 | 0.225133366 | -3.312246899 |
| Thada | 445.9157641 | -0.145084501 | 0.0930925 | -1.558498276 |
| Gm24492 | 0.916014489 | -0.539043645 | 1.061602018 | -0.507764337 |
| Gm24722 | 0.550570057 | 0.169191868 | 1.295436651 | 0.130606053 |
| Plekhh2 | 185.6708911 | 0.030483158 | 0.11369424 | 0.268115233 |
| Gm26066 | 0.13205505 | 0.037055774 | 3.667120883 | 0.010104868 |
| Dync2li1 | 99.93135035 | 0.084263129 | 0.146084498 | 0.576810884 |
| Abcg5 | 0.732414602 | -0.237401302 | 1.331166699 | -0.178340776 |
| Abcg8 | 0.118974955 | 0.51795324 | 3.724945049 | 0.139049901 |
| Lrpprc | 1213.010164 | 0.096651574 | 0.073352601 | 1.317629816 |
| 1110020A21Rik | 4.802739388 | -0.071513264 | 0.471741846 | -0.151594064 |
| Ppm1b | 1207.845512 | 0.171402619 | 0.063487532 | 2.69978396 |
| Slc3a1 | 11.21614055 | 0.083205266 | 0.396069836 | 0.210077261 |
| Prepl | 4734.893635 | 0.389291159 | 0.054153338 | 7.188682618 |
| Camkmt | 72.64996259 | 0.077881591 | 0.145035286 | 0.536983742 |
| Rpl31-ps16 | 2.862600439 | 0.365205497 | 0.65199582 | 0.560134721 |

| | | | | |
|---|---|---|---|---|
| Gm27178 | 0 | NA | NA | NA |
| Six3os1 | 52.19109397 | 0.01441354 | 0.232757583 | 0.061925116 |
| Gm27435 | 0.089062654 | -0.443823436 | 3.724945049 | -0.119148989 |
| Gm27801 | 0.250814404 | 0.51797199 | 1.827301929 | 0.283462728 |
| Six3 | 109.2161158 | 0.101058901 | 0.155377662 | 0.650408169 |
| Gm27610 | 0.226876767 | 0.517968201 | 2.186165849 | 0.236929966 |
| Gm27283 | 0.388809915 | 1.465058723 | 1.731501968 | 0.846120161 |
| Gm27275 | 0.04393175 | 0.037062677 | 3.724945049 | 0.009949859 |
| Gm27502 | 0.090665643 | 0.037060453 | 3.724945049 | 0.009949262 |
| Gm27427 | 0.359940881 | -0.952500877 | 1.784888635 | -0.53364723 |
| Gm28528 | 1.48446612 | 0.507476599 | 0.928705565 | 0.546434326 |
| Six2 | 7.803849726 | -0.312400482 | 0.349325031 | -0.894297444 |
| Gm29418 | 0.434220697 | 1.372766122 | 1.354047486 | 1.013824209 |
| Srbd1 | 196.7237653 | -0.450107944 | 0.119793177 | -3.757375465 |
| Prkce | 4692.067451 | 0.539124902 | 0.093442863 | 5.769567453 |
| 2010106C02Rik | 0 | NA | NA | NA |
| Gm10309 | 10.12416963 | 0.612649866 | 0.468824713 | 1.306778096 |
| Gm22235 | 0 | NA | NA | NA |
| Epas1 | 1364.101885 | -0.08996291 | 0.114201485 | -0.787756047 |
| Gm29168 | 0.150086409 | 1.027495679 | 3.723011807 | 0.275985071 |
| Tmem247 | 0 | NA | NA | NA |
| Atp6v1e2 | 0.434133818 | -0.467334675 | 1.562340515 | -0.299124724 |
| Rhoq | 718.01171 | -0.654127069 | 0.106178601 | -6.160629921 |
| Pigf | 100.958299 | -0.441826476 | 0.15556383 | -2.84016199 |
| Cript | 263.4740334 | -0.374161181 | 0.142197592 | -2.631276482 |
| Gm24648 | 0 | NA | NA | NA |
| Gm5499 | 18.0195093 | -0.170510773 | 0.26652012 | -0.639766983 |
| Socs5 | 1211.318638 | 0.083380057 | 0.093454224 | 0.892202129 |
| Gm22346 | 0.044191549 | 0.037062677 | 3.724945049 | 0.009949859 |
| Mcfd2 | 543.0470171 | -0.457316101 | 0.085869461 | -5.325712984 |
| 4833418N02Rik | 13.1105544 | 0.234085184 | 0.296873732 | 0.788500829 |
| Ttc7 | 196.4810743 | -0.749006117 | 0.147084181 | -5.092363512 |
| Gm15978 | 0.6240965 | 0.971385139 | 1.283929473 | 0.756572039 |
| 0610012D04Rik | 0.332525094 | -1.741549379 | 1.700877371 | -1.023912369 |
| 1700011E24Rik | 3.15672782 | -0.238770214 | 0.548827448 | -0.435055161 |
| Calm2 | 8582.912101 | 0.417192901 | 0.104431949 | 3.994878053 |
| Gm26005 | 0.04393175 | 0.037062677 | 3.724945049 | 0.009949859 |
| Gm28676 | 0.161026707 | -1.437459239 | 3.451444183 | -0.416480512 |
| Epcam | 12.12751263 | -1.977347273 | 0.437994674 | -4.514546388 |
| Msh2 | 333.7442204 | 0.218162385 | 0.099206892 | 2.199064797 |
| Kcnk12 | 130.455902 | -0.45338533 | 0.142027694 | -3.192231854 |
| Rpl36-ps4 | 0.224455361 | 1.369258445 | 3.133385076 | 0.43699016 |
| Msh6 | 331.2527847 | -0.215904075 | 0.078681196 | -2.744036511 |
| Fbxo11 | 1810.721455 | 0.108099346 | 0.06670361 | 1.620592149 |
| Gm4832 | 0.217012189 | -0.882850138 | 2.152598169 | -0.410132346 |
| Gm26612 | 0.40569442 | 1.783915662 | 1.414618112 | 1.261058123 |
| Gm22542 | 0.253603277 | 0.532830245 | 2.124113839 | 0.250848253 |

| | | | | |
|---|---|---|---|---|
| Gm23947 | 0 | NA | NA | NA |
| Foxn2 | 537.5500919 | -0.263224289 | 0.093972503 | -2.801077772 |
| Ppp1r21 | 678.815562 | 0.195810576 | 0.058690403 | 3.336330383 |
| Gm38109 | 8.730948186 | -0.168650376 | 0.452396744 | -0.372793082 |
| Ston1 | 93.35916316 | 0.16228874 | 0.173343766 | 0.936224843 |
| Gtf2a1l | 2.669826932 | 0.072615045 | 0.615668846 | 0.117944973 |
| Lhcgr | 3.114871368 | 1.104629222 | 0.7133732 | 1.548459097 |
| Fshr | 0.045882455 | 0.517959912 | 3.724945049 | 0.139051692 |
| Gm10184 | 3.494817459 | -0.413512635 | 0.548025538 | -0.754549937 |
| Nrxn1 | 7760.329357 | 0.479721028 | 0.098301632 | 4.880092214 |
| Gm24510 | 0 | NA | NA | NA |
| Gm10493 | 3.600385196 | -0.015365975 | 0.578906076 | -0.026543123 |
| Gm19815 | 7.90218533 | 0.04240786 | 0.568967128 | 0.074534816 |
| Gm20491 | 88.08114114 | 0.403167351 | 0.317579816 | 1.269499289 |
| 4930480K15Rik | 0.097899064 | 0.037062677 | 3.724945049 | 0.009949859 |
| AC151286.1 | 0.277228681 | 0.037052164 | 3.724945049 | 0.009947036 |
| Gm10308 | 1.537590905 | 1.212706696 | 0.861712476 | 1.407321735 |
| Gm27475 | 0 | NA | NA | NA |
| Gm6741 | 17.12533191 | 0.183414723 | 0.365483652 | 0.501841114 |
| Gm15404 | 0.089089296 | 0.957155977 | 3.723939561 | 0.257027796 |
| Gm15403 | 0 | NA | NA | NA |
| Gm24244 | 0 | NA | NA | NA |
| AC131325.1 | 0 | NA | NA | NA |
| Gm27039 | 0 | NA | NA | NA |
| Adcyap1 | 181.5271029 | -0.409634082 | 0.183901623 | -2.227463107 |
| Gm25935 | 0 | NA | NA | NA |
| Gm22593 | 0.196742721 | -0.437955803 | 2.231534635 | -0.196257677 |
| Mettl4 | 150.9550503 | -0.260169075 | 0.164901256 | -1.577726459 |
| 2700099C18Rik | 6.495084589 | -0.327115052 | 0.424159728 | -0.771207236 |
| Gm27038 | 3.867668655 | 2.127897787 | 0.63321711 | 3.360455289 |
| Gm1976 | 94.15986379 | 0.763164993 | 0.124868973 | 6.111726336 |
| Gm23964 | 0.139856492 | -0.443820827 | 3.574411099 | -0.124166139 |
| Gm26734 | 6.887014156 | 0.260342043 | 0.435981298 | 0.597140392 |
| Gm23102 | 0 | NA | NA | NA |
| Gm20939 | 23.37788906 | -0.888513267 | 0.36196926 | -2.454664984 |
| Vmn1r-ps151 | 0 | NA | NA | NA |
| Vmn1r-ps152 | 0.092691418 | -0.908687078 | 3.724560878 | -0.2439716 |
| Vmn1r238 | 0.090890357 | 0.037071574 | 3.724945049 | 0.009952247 |
| Crem | 361.0165932 | -0.675605474 | 0.161746515 | -4.176939913 |
| Gm6225 | 15.32231194 | 0.095998804 | 0.361148842 | 0.265815069 |
| Cul2 | 847.8838813 | -0.042825688 | 0.080778891 | -0.530159397 |
| G430049J08Rik | 0.148708979 | -0.443201667 | 2.868467393 | -0.154508177 |
| Bambi | 60.79279935 | -0.373227236 | 0.176946407 | -2.109267102 |
| Gm23643 | 0 | NA | NA | NA |
| Gm24632 | 0 | NA | NA | NA |
| Lyzl1 | 0 | NA | NA | NA |
| Gm10557 | 201.5185117 | -0.152367983 | 0.146832413 | -1.03769992 |

| | | | | |
|---|---|---|---|---|
| Map3k8 | 99.88469274 | -0.519144855 | 0.1675511 | -3.098427015 |
| Gm26865 | 0.332096027 | -0.434263518 | 1.736708917 | -0.250049686 |
| Mtpap | 376.5909295 | -0.039085688 | 0.096080923 | -0.406799669 |
| 9430020K01Rik | 1287.14579 | -0.125268298 | 0.112143468 | -1.117036065 |
| Gm10556 | 0.381271471 | -1.253318124 | 1.649782392 | -0.759686932 |
| Svil | 295.6710269 | -0.548620984 | 0.105596518 | -5.195445778 |
| Gm26682 | 7.326093239 | -0.00248888 | 0.448902668 | -0.005544365 |
| Zfp438 | 70.78775138 | -0.047233582 | 0.152984758 | -0.308746978 |
| Gm10125 | 71.18097263 | 0.578522976 | 0.190062499 | 3.043856513 |
| Gm26575 | 19.11664666 | 0.53016956 | 0.337671118 | 1.570076713 |
| Gm2238 | 59.71255148 | 0.370425602 | 0.202367111 | 1.830463459 |
| Zeb1 | 1461.053986 | -0.068700422 | 0.05241583 | -1.310680808 |
| Gm16147 | 0 | NA | NA | NA |
| Gm22709 | 4.178956302 | -1.029436758 | 0.613931886 | -1.676793114 |
| Arhgap12 | 782.7285597 | -0.113771709 | 0.076363613 | -1.489868069 |
| Kif5b | 4777.179878 | -0.170699169 | 0.115665457 | -1.47580076 |
| Gm17036 | 1.215361072 | -1.240369969 | 1.040912432 | -1.19161798 |
| Rpl27-ps3 | 9.972102879 | -0.725678591 | 0.371468319 | -1.953540998 |
| Gm6291 | 0 | NA | NA | NA |
| Epc1 | 763.0696826 | 0.024905935 | 0.074977894 | 0.332177039 |
| Gm28529 | 13.22339979 | 0.12486619 | 0.354972842 | 0.351762658 |
| Mir1893 | 0 | NA | NA | NA |
| 4921524L21Rik | 0.072025462 | 0.517959912 | 3.724945049 | 0.139051692 |
| Rab18 | 1241.658341 | -0.108152837 | 0.080771992 | -1.338989358 |
| Mkx | 204.0449348 | 0.61211878 | 0.123863872 | 4.941866995 |
| Armc4 | 20.81306997 | 0.60184336 | 0.236785779 | 2.541720882 |
| Mpp7 | 246.8020213 | 0.325183127 | 0.104511772 | 3.111449748 |
| Fabp5l2 | 0.240849226 | -0.443353875 | 1.86479929 | -0.237748844 |
| Gm24292 | 0 | NA | NA | NA |
| Wac | 1913.852717 | 0.055289317 | 0.06321699 | 0.874595851 |
| Gm7502 | 0.768676318 | 1.742511411 | 1.254556587 | 1.388946046 |
| Gm5819 | 9.13781552 | -0.283771381 | 0.370442891 | -0.766032736 |
| Gm26119 | 0 | NA | NA | NA |
| Gm37148 | 0 | NA | NA | NA |
| Fzd8 | 66.68569282 | -0.453783432 | 0.15872369 | -2.85895213 |
| Gjd4 | 0.815409053 | -0.576409684 | 1.061191452 | -0.543172189 |
| Ccny | 1305.979739 | 0.254575371 | 0.051744611 | 4.919843187 |
| Gm22765 | 0 | NA | NA | NA |
| Cetn1 | 0.038981395 | -0.443816764 | 3.724945049 | -0.119147198 |
| Colec12 | 279.2173052 | -0.435963106 | 0.14271145 | -3.054857232 |
| Gm17430 | 0 | NA | NA | NA |
| Gm7497 | 4.858631947 | -0.365350664 | 0.597247312 | -0.61172425 |
| Gm23350 | 0 | NA | NA | NA |
| Gm23637 | 0 | NA | NA | NA |
| Gm5501 | 0 | NA | NA | NA |
| Thoc1 | 432.2063309 | 0.169325502 | 0.080229909 | 2.11050347 |
| Usp14 | 1395.726703 | 0.122652799 | 0.050876838 | 2.410778727 |

| | | | | |
|---|---|---|---|---|
| Rock1 | 1035.477942 | -0.204630246 | 0.056992875 | -3.590453113 |
| Gm2531 | 25.92851566 | 0.17168197 | 0.323870099 | 0.530095152 |
| Gm26244 | 55.4930898 | 0.031580124 | 0.190147269 | 0.16608245 |
| Greb1l | 215.658039 | 1.040431345 | 0.145580865 | 7.146758935 |
| AC124740.1 | 0 | NA | NA | NA |
| Gm10349 | 1.210176943 | 1.00708444 | 0.912299756 | 1.103896426 |
| Gm24894 | 2.156355716 | -0.230505269 | 0.746721737 | -0.308689647 |
| Esco1 | 664.6422751 | -0.08725893 | 0.081573936 | -1.069691298 |
| Snrpd1 | 236.8850658 | -0.193708716 | 0.131609485 | -1.471844645 |
| Abhd3 | 409.5174636 | 0.320614446 | 0.146448505 | 2.189264048 |
| 4930563E18Rik | 1.299310135 | 0.319734654 | 0.965907628 | 0.331019908 |
| Mib1 | 3835.229447 | 0.086471855 | 0.060626918 | 1.426294762 |
| Gm25116 | 4.118368001 | 0.83098227 | 0.73212511 | 1.135027687 |
| Gm21747 | 0.049553047 | -0.443818988 | 3.724945049 | -0.119147795 |
| Mir133a-1 | 0 | NA | NA | NA |
| Mir1b | 7.73100226 | 0.739241157 | 0.395857353 | 1.867443286 |
| mmu-mir-1a-2 | 0 | NA | NA | NA |
| 1010001N08Rik | 0.099155932 | 0.037073798 | 3.724945049 | 0.009952844 |
| Gata6 | 1.608823659 | -0.134656864 | 0.817335574 | -0.164751014 |
| Rbbp8 | 116.5073145 | -0.450596722 | 0.116513036 | -3.86735028 |
| Gm6277 | 35.32926637 | 0.380494949 | 0.246179145 | 1.545601879 |
| Cables1 | 273.0530706 | 0.425898801 | 0.122187206 | 3.485625181 |
| Mir1901 | 6.131714793 | 0.249692734 | 0.407384548 | 0.612916557 |
| Tmem241 | 88.72054137 | 0.155338719 | 0.164414185 | 0.944801199 |
| Riok3 | 978.9154087 | -0.26878094 | 0.084109054 | -3.195624319 |
| Gm25289 | 0.089062654 | -0.443823436 | 3.724945049 | -0.119148989 |
| 3110002H16Rik | 281.027959 | -0.166961987 | 0.092826022 | -1.79865498 |
| Npc1 | 1148.204684 | -0.328167116 | 0.087382278 | -3.755534012 |
| Gm15956 | 8.077703934 | -0.834539786 | 0.420259619 | -1.985772005 |
| Ankrd29 | 171.8859524 | 0.041228254 | 0.094558004 | 0.436010195 |
| Gm16072 | 0.87006192 | -0.322047448 | 1.276613528 | -0.252266987 |
| Lama3 | 110.321609 | 0.000778761 | 0.170956012 | 0.004555327 |
| Gm29201 | 0.436251183 | -0.441596739 | 1.82337032 | -0.242187083 |
| Gm29199 | 0 | NA | NA | NA |
| Gm29200 | 0.582012116 | -1.127561928 | 1.232235192 | -0.915054151 |
| Gm22340 | 0.038981395 | -0.443816764 | 3.724945049 | -0.119147198 |
| Ttc39c | 360.5212832 | 0.01280731 | 0.104967595 | 0.122012034 |
| Mir1948 | 2.898584899 | 0.603427761 | 0.696042059 | 0.86694152 |
| Cabyr | 18.12948923 | 0.598699656 | 0.423984233 | 1.41208 |
| Gm7599 | 1.737466599 | 1.867176181 | 0.912745424 | 2.045670274 |
| Osbpl1a | 2161.262522 | 0.790642761 | 0.070149354 | 11.27084881 |
| Gm5687 | 0.269295688 | -0.700747777 | 2.064664678 | -0.339400284 |
| Gm26407 | 27.03744251 | 0.255397784 | 0.304986112 | 0.837407913 |
| Impact | 3627.068778 | 0.075599273 | 0.083528822 | 0.905068106 |
| Hrh4 | 0.228099782 | -1.390403623 | 3.115459228 | -0.446291709 |
| Gm22251 | 0 | NA | NA | NA |
| Zfp521 | 803.5141072 | -0.184256418 | 0.103804985 | -1.775024754 |

| | | | | |
|---|---|---|---|---|
| Gm5160 | 3.237902636 | 0.003669336 | 0.597290868 | 0.006143299 |
| Gm6457 | 44.70599757 | -0.066356617 | 0.154407908 | -0.429748827 |
| Ss18 | 416.8945723 | -0.380334649 | 0.097750851 | -3.890857673 |
| Gm24924 | 13.08384193 | 0.186377906 | 0.294094731 | 0.633734257 |
| Psma8 | 5.559946366 | -0.528287822 | 0.465705638 | -1.134381418 |
| Taf4b | 120.7625759 | 0.092473072 | 0.13558329 | 0.682038857 |
| Kctd1 | 603.8212261 | 0.705950671 | 0.10049658 | 7.024623799 |
| Gm6304 | 6.355730337 | 0.790753159 | 0.453996075 | 1.741762103 |
| Aqp4 | 4262.513783 | -0.146743859 | 0.140752942 | -1.042563351 |
| Chst9 | 8.958957809 | 1.187546885 | 0.380620977 | 3.12002479 |
| Gm10036 | 42.03144848 | -0.629471913 | 0.260971262 | -2.412035361 |
| Gm24228 | 0.072025462 | 0.517959912 | 3.724945049 | 0.139051692 |
| Gm4835 | 1.43143147 | 0.070088922 | 0.851037476 | 0.082357034 |
| Gm22389 | 1.98025533 | 0.485187575 | 0.814599452 | 0.595614904 |
| Gm24385 | 0 | NA | NA | NA |
| Gm7665 | 11.35479647 | -1.790457271 | 0.499676687 | -3.583231552 |
| Cdh2 | 2013.677982 | 0.074910008 | 0.062279427 | 1.202805026 |
| Gm15485 | 0 | NA | NA | NA |
| Gm15328 | 30.8296316 | 0.177335386 | 0.306025927 | 0.579478306 |
| Gm7670 | 0 | NA | NA | NA |
| 1700001G01Rik | 0.046345709 | -0.443818988 | 3.724945049 | -0.119147795 |
| Gm22470 | 0 | NA | NA | NA |
| Gm24015 | 0 | NA | NA | NA |
| Gm26235 | 0.048949532 | 0.037064902 | 3.724945049 | 0.009950456 |
| Gm25392 | 0 | NA | NA | NA |
| Gm24238 | 0 | NA | NA | NA |
| Dsc3 | 13.88762389 | 0.041278541 | 0.298055094 | 0.138492989 |
| Dsc2 | 11.39730211 | 0.128335357 | 0.376327781 | 0.341020152 |
| Dsc1 | 2.506558332 | 0.097139719 | 0.984912551 | 0.09862776 |
| Dsg1c | 4.094471938 | 2.202196345 | 0.790500323 | 2.78582599 |
| Dsg1a | 1.903639553 | 4.310367192 | 2.969750173 | 1.451424174 |
| Dsg1b | 0.835693444 | 1.687117484 | 1.260402407 | 1.338554635 |
| Dsg4 | 0 | NA | NA | NA |
| Dsg3 | 1.159328825 | -0.158475676 | 0.898390323 | -0.176399581 |
| Dsg2 | 36.65644163 | -0.666107479 | 0.207883348 | -3.204236829 |
| Gm16090 | 0.646850989 | 0.395913 | 1.348724787 | 0.293546173 |
| Ttr | 2265.392118 | 2.156549556 | 0.335565527 | 6.426612336 |
| Gm10269 | 7.797321979 | 0.141149616 | 0.413319374 | 0.341502541 |
| B4galt6 | 2045.964961 | 0.184899703 | 0.063159981 | 2.927481936 |
| Trappc8 | 1142.654653 | -0.308845826 | 0.053996203 | -5.719769376 |
| Gm24959 | 4.935114692 | -0.099992802 | 0.48954532 | -0.204256476 |
| Rnf125 | 36.46586033 | -0.352121877 | 0.33383021 | -1.054793325 |
| Rnf138 | 267.4053105 | -0.146072256 | 0.088367147 | -1.653015409 |
| Mep1b | 0 | NA | NA | NA |
| Gm6378 | 0.155469778 | 0.037074704 | 2.829060425 | 0.013104953 |
| Garem | 494.2667245 | 0.246615898 | 0.100719903 | 2.448531932 |
| Gm24372 | 0.044191549 | 0.037062677 | 3.724945049 | 0.009949859 |

| | | | | |
|---|---|---|---|---|
| Gm22886 | 0 | NA | NA | NA |
| Klhl14 | 23.8882674 | 0.066839522 | 0.261219868 | 0.255874572 |
| 4930426D05Rik | 6.412194933 | 0.724372836 | 0.447769354 | 1.617736519 |
| Ccdc178 | 1.26832265 | 0.032818475 | 0.917455481 | 0.035771191 |
| Asxl3 | 371.9529472 | 0.384913526 | 0.117974804 | 3.26267571 |
| Nol4 | 882.8164278 | 0.564819052 | 0.096176727 | 5.87272064 |
| Gm22495 | 0 | NA | NA | NA |
| Dtna | 1930.500953 | -0.202142287 | 0.08500527 | -2.377997103 |
| Gm15972 | 1.928713869 | 1.546202425 | 0.814468247 | 1.898419529 |
| Mapre2 | 3079.479868 | 0.238130569 | 0.060753895 | 3.919593443 |
| Mir6360 | 0 | NA | NA | NA |
| Gm23207 | 0 | NA | NA | NA |
| Zfp397 | 704.3356492 | -0.325405976 | 0.084396909 | -3.855662243 |
| Zscan30 | 8.824857606 | -0.443954978 | 0.364605855 | -1.217629865 |
| Zfp35 | 243.5789265 | -0.392688851 | 0.095741639 | -4.101547183 |
| Zfp191 | 825.6271123 | -0.159603637 | 0.097152176 | -1.642821014 |
| Ino80c | 294.4170576 | 0.001229524 | 0.108125838 | 0.011371233 |
| Galnt1 | 801.5304355 | -0.020246749 | 0.088961299 | -0.227590532 |
| Gm7701 | 14.47583842 | 0.306710519 | 0.30361857 | 1.010183661 |
| Mir187 | 0.214710016 | -1.576433344 | 3.129503726 | -0.503732694 |
| 2700062C07Rik | 106.228182 | 0.112760489 | 0.11430699 | 0.986470631 |
| Rprd1a | 820.2179591 | 0.095638788 | 0.069736689 | 1.371427144 |
| Gm23606 | 0.061920613 | -0.443816764 | 3.724945049 | -0.119147198 |
| Slc39a6 | 544.4852359 | -0.210198983 | 0.094089437 | -2.234033804 |
| Elp2 | 877.5310991 | 0.197132135 | 0.050492946 | 3.904151961 |
| Mocos | 101.3743352 | -1.199677008 | 0.179663903 | -6.677340236 |
| n-R5s23 | 0.045882455 | 0.517959912 | 3.724945049 | 0.139051692 |
| Gm9955 | 0.957795399 | 0.845871707 | 1.022931761 | 0.826909222 |
| Fhod3 | 695.6586359 | 0.104383576 | 0.131321297 | 0.794871648 |
| Gm24294 | 1.176033076 | -0.981761238 | 1.144386223 | -0.857893269 |
| Tpgs2 | 457.1009357 | 0.221593972 | 0.07734205 | 2.865116369 |
| AW554918 | 270.5192402 | 0.346868789 | 0.080110541 | 4.329876981 |
| Gm16558 | 5.405763959 | 1.21131515 | 0.521839841 | 2.321239307 |
| Celf4 | 4385.110905 | 0.379802246 | 0.090979728 | 4.174581032 |
| Gm25845 | 9.017131998 | 0.79265419 | 0.420308104 | 1.885888428 |
| Gm7847 | 0 | NA | NA | NA |
| Gm23674 | 0 | NA | NA | NA |
| Gm26106 | 0 | NA | NA | NA |
| Gm7729 | 7.595035811 | 0.199448107 | 0.415428778 | 0.480101808 |
| Gm24246 | 0 | NA | NA | NA |
| Pik3c3 | 522.6692759 | 0.206915129 | 0.059105192 | 3.500794489 |
| Rpl29-ps5 | 0.839167439 | -0.556418313 | 1.128387503 | -0.493109248 |
| Gm22870 | 0 | NA | NA | NA |
| Gm7936 | 0.203976984 | -0.416833431 | 2.177355301 | -0.191440245 |
| Rit2 | 621.7940845 | 0.184424657 | 0.093424765 | 1.974044629 |
| Gm23283 | 0 | NA | NA | NA |
| Syt4 | 3383.452968 | 0.001157391 | 0.077730494 | 0.014889795 |

| | | | | |
|---|---|---|---|---|
| Gm26658 | 4.064855943 | -0.135625764 | 0.511901765 | -0.264944904 |
| Gm25396 | 0 | NA | NA | NA |
| Slc25a46 | 1147.053562 | -0.057409137 | 0.081210023 | -0.706921812 |
| B930094E09Rik | 4.156831401 | -0.254026002 | 0.50845481 | -0.499603893 |
| Sap130 | 827.4019423 | -0.041398918 | 0.056296797 | -0.73536898 |
| Ammecr1l | 554.7126231 | -0.525195042 | 0.07924696 | -6.627320976 |
| Gm26533 | 2.98239301 | -0.030279141 | 0.588515006 | -0.051450075 |
| Polr2d | 122.288999 | -0.258800385 | 0.118298525 | -2.187689016 |
| Wdr33 | 970.9585418 | -0.00924492 | 0.05411394 | -0.170841741 |
| Gm6665 | 1.827239475 | -0.442620486 | 0.701938761 | -0.63056852 |
| Gm26823 | 0.062426873 | 0.517959912 | 3.724945049 | 0.139051692 |
| Sft2d3 | 54.7958328 | -0.157116614 | 0.156497663 | -1.003955018 |
| Lims2 | 61.79027259 | -0.191344765 | 0.165263258 | -1.157817938 |
| Gpr17 | 643.1742382 | -0.530938211 | 0.285525845 | -1.859510166 |
| Myo7b | 0.405683672 | -0.933758628 | 1.666697366 | -0.560244857 |
| Gm16344 | 0.401548357 | 0.110854396 | 1.564226182 | 0.070868521 |
| Gm16345 | 0.075043205 | 0.517957688 | 3.724945049 | 0.139051095 |
| Iws1 | 844.5236566 | 0.016096135 | 0.078017279 | 0.206315008 |
| Proc | 0.048949532 | 0.037064902 | 3.724945049 | 0.009950456 |
| Gm26717 | 0.046345709 | -0.443818988 | 3.724945049 | -0.119147795 |
| Map3k2 | 1196.837555 | 0.059931977 | 0.088917462 | 0.674018079 |
| Ercc3 | 374.8626789 | -0.174132165 | 0.066947644 | -2.601020041 |
| Gm22325 | 0 | NA | NA | NA |
| A830052D11Rik | 8.708922263 | 1.514825354 | 0.42319389 | 3.579506673 |
| Bin1 | 1973.364051 | 0.272578987 | 0.041651781 | 6.5442336 |
| Gypc | 57.79268309 | -0.263802683 | 0.2660266 | -0.991640247 |
| Gm25826 | 0 | NA | NA | NA |
| 4930455D15Rik | 0.382692223 | -0.888564296 | 1.545913253 | -0.574782767 |
| Tslp | 1.429800265 | -0.944899489 | 0.831364303 | -1.136564903 |
| Gm26378 | 0.072025462 | 0.517959912 | 3.724945049 | 0.139051692 |
| Wdr36 | 397.0988872 | -0.181834304 | 0.063803041 | -2.849931625 |
| Mir6361 | 0 | NA | NA | NA |
| 2310026I22Rik | 3.939202308 | 0.604583592 | 0.531301508 | 1.137929373 |
| Camk4 | 6160.809613 | 0.890412367 | 0.157909496 | 5.638751237 |
| Gm22814 | 0 | NA | NA | NA |
| Stard4 | 406.2855849 | -0.24194845 | 0.102023983 | -2.37148602 |
| Nrep | 1304.95571 | 0.040702634 | 0.157293002 | 0.258769516 |
| Gm10549 | 1.094754251 | -0.65386658 | 1.172416164 | -0.557708602 |
| Epb4.1l4aos | 32.18986031 | -0.222584801 | 0.239238311 | -0.930389451 |
| Gm23639 | 3.703464063 | -0.174576689 | 0.482744802 | -0.361633494 |
| Epb4.1l4a | 142.1446161 | -0.50966881 | 0.132867575 | -3.835915648 |
| Gm22788 | 0 | NA | NA | NA |
| Gm24432 | 0 | NA | NA | NA |
| Gm23411 | 0 | NA | NA | NA |
| Gm10548 | 0.669147434 | 0.427849388 | 1.104053651 | 0.387525903 |
| Apc | 7569.545602 | 0.251285671 | 0.075099946 | 3.346016661 |
| Srp19 | 313.7947171 | -0.038795919 | 0.091161218 | -0.42557482 |

| | | | | |
|---|---|---|---|---|
| Reep5 | 3174.602423 | 0.163708569 | 0.064637871 | 2.53270361 |
| Pkd2l2 | 2.995780043 | -2.021121448 | 0.737918979 | -2.738947642 |
| Fam13b | 1185.027783 | -0.192507281 | 0.0651479 | -2.95492687 |
| n-R5s24 | 0.131453142 | 0.517960224 | 2.979065385 | 0.173866685 |
| Wnt8a | 0.202010919 | -1.362830177 | 2.194321454 | -0.621071345 |
| Nme5 | 50.66745939 | 0.102390478 | 0.171249215 | 0.597903345 |
| 4933408B17Rik | 16.70085405 | 0.568358344 | 0.328265942 | 1.731396014 |
| Brd8 | 925.5748569 | 0.296109473 | 0.07475749 | 3.960933841 |
| Kif20a | 22.72681451 | -0.644374946 | 0.38430404 | -1.676732168 |
| Cdc23 | 555.3324902 | 0.009156909 | 0.074073592 | 0.12361908 |
| n-R5s25 | 0 | NA | NA | NA |
| Gfra3 | 0 | NA | NA | NA |
| Cdc25c | 5.025978301 | -0.116858259 | 0.433044883 | -0.269852534 |
| Gm3550 | 0.207873913 | -1.281610677 | 2.5926923 | -0.494316536 |
| 2010110K18Rik | 0.294784161 | 1.095959467 | 1.997959082 | 0.548539495 |
| Fam53c | 1037.28919 | -0.020066217 | 0.055474299 | -0.36172097 |
| Kdm3b | 1100.794087 | -0.020282368 | 0.051520785 | -0.393673511 |
| Gm26538 | 0.390497984 | -0.425044494 | 1.541720009 | -0.275694998 |
| Reep2 | 1197.302419 | 0.209500934 | 0.068675021 | 3.050613329 |
| Egr1 | 3179.097212 | -0.485541637 | 0.353981116 | -1.371659713 |
| Etf1 | 1390.357339 | -0.445533718 | 0.099453694 | -4.479810651 |
| Hspa9 | 2869.344892 | -0.32034054 | 0.10676076 | -3.000545703 |
| Gm22200 | 4.566833944 | -0.269593138 | 0.540204452 | -0.4990576 |
| Gm26109 | 3.869151513 | 1.088906417 | 0.58376562 | 1.865314399 |
| Ctnna1 | 941.2579512 | -0.567807288 | 0.102052182 | -5.563891688 |
| Lrrtm2 | 1096.201602 | 0.080383231 | 0.07949019 | 1.011234604 |
| Sil1 | 202.6507413 | 0.071485006 | 0.096570937 | 0.740233119 |
| Gm5239 | 0.164165709 | 0.517965237 | 2.781083822 | 0.18624582 |
| Snhg4 | 73.54980766 | 0.032882314 | 0.153477619 | 0.214248267 |
| Gm28285 | 6.985251871 | -0.22711201 | 0.369851727 | -0.614062321 |
| Mir1949 | 0 | NA | NA | NA |
| mmu-mir-1949 | 0 | NA | NA | NA |
| Snora74a | 0 | NA | NA | NA |
| AC121821.1 | 0.048949532 | 0.037064902 | 3.724945049 | 0.009950456 |
| Matr3 | 4954.108798 | 0.225181273 | 0.068755593 | 3.275097523 |
| Gm36989 | 219.4274089 | 0.068124083 | 0.138929242 | 0.490350926 |
| Paip2 | 1058.0764 | 0.009652317 | 0.096098668 | 0.10044174 |
| Slc23a1 | 0.196018198 | 1.019785506 | 2.631724125 | 0.387497115 |
| Mzb1 | 4.18320868 | -0.944768435 | 0.609303516 | -1.550571121 |
| Prob1 | 99.4461201 | 0.126371999 | 0.155575273 | 0.812288458 |
| Spata24 | 30.05154039 | 0.039018223 | 0.21384786 | 0.182457861 |
| Dnajc18 | 1508.902544 | 0.113987373 | 0.045600382 | 2.499702169 |
| 1700066B19Rik | 1.640932998 | -0.353750065 | 0.793614994 | -0.445745189 |
| Ecscr | 35.18133015 | -0.680730987 | 0.226918176 | -2.999896258 |
| Tmem173 | 105.9553805 | -0.828432254 | 0.18654262 | -4.440981113 |
| Gm16490 | 0 | NA | NA | NA |
| Ube2d2a | 1153.052153 | -0.187421596 | 0.076969371 | -2.435015284 |

| | | | | |
|---|---|---|---|---|
| Cxxc5 | 830.8617964 | -0.281438956 | 0.063981511 | -4.398754456 |
| Gm29417 | 3.083648494 | -0.473760746 | 0.626560975 | -0.756128716 |
| Gm23882 | 0 | NA | NA | NA |
| Psd2 | 681.0294117 | 0.186046248 | 0.144245688 | 1.289787238 |
| Nrg2 | 144.6574193 | 0.190350772 | 0.128999126 | 1.475597374 |
| Pura | 8433.573605 | 0.118168155 | 0.077471399 | 1.525313284 |
| Cystm1 | 401.2118129 | -0.180502546 | 0.100594903 | -1.794350815 |
| Pfdn1 | 311.4709643 | -0.384304488 | 0.134329398 | -2.860911256 |
| Hbegf | 582.7227112 | -1.972551833 | 0.314223169 | -6.277550571 |
| Slc4a9 | 2.54483004 | 1.217653208 | 0.776147004 | 1.568843533 |
| Ankhd1 | 1581.43401 | -0.104389807 | 0.084898299 | -1.229586547 |
| Gm6542 | 84.43238 | -0.006916839 | 0.138104402 | -0.050084131 |
| Eif4ebp3 | 7.191909191 | 0.141493007 | 0.426131171 | 0.332040968 |
| Sra1 | 541.6420646 | 0.018659323 | 0.069200262 | 0.269642372 |
| Slc35a4 | 487.0228943 | -0.281056875 | 0.075743577 | -3.710636425 |
| E230025N22Rik | 1.365445969 | -1.047835525 | 0.950318098 | -1.102615564 |
| Cd14 | 75.49450281 | -0.99184387 | 0.310219078 | -3.197236858 |
| Tmco6 | 58.12891111 | -0.317330112 | 0.179089216 | -1.771910775 |
| Ndufa2 | 320.0471109 | -0.100685431 | 0.117320481 | -0.858208475 |
| Ik | 860.0388227 | 0.022160969 | 0.117832373 | 0.188071988 |
| Wdr55 | 90.11495044 | -0.367630303 | 0.140930538 | -2.608592208 |
| Dnd1 | 14.61022796 | -1.253125487 | 0.332545059 | -3.768287801 |
| Hars | 700.2311977 | -0.11952811 | 0.125153626 | -0.955051116 |
| Hars2 | 250.6053811 | 0.049857039 | 0.108923129 | 0.457726834 |
| Zmat2 | 1268.32983 | 0.312003557 | 0.062599818 | 4.984096897 |
| Vaultrc5 | 18.77737098 | -1.126408321 | 0.405671652 | -2.776650317 |
| Gm23424 | 0.049553047 | -0.443818988 | 3.724945049 | -0.119147795 |
| Gm6756 | 1.105122771 | -0.849537832 | 1.057548872 | -0.803308343 |
| Pcdha1 | 15.4569917 | 0.761344551 | 0.29868206 | 2.549013328 |
| Gm37751 | 3.355297506 | 0.517739787 | 0.59413668 | 0.871415289 |
| Pcdha2 | 32.39827103 | 0.075668674 | 0.216318925 | 0.349801451 |
| Pcdha3 | 47.11146249 | 0.22083781 | 0.198404506 | 1.113068522 |
| Pcdha4 | 76.22283242 | 0.301818196 | 0.165096595 | 1.82813096 |
| Gm38225 | 2.632135046 | -0.058843479 | 0.794442415 | -0.074068904 |
| Pcdha5 | 55.26370248 | 0.360922844 | 0.199796449 | 1.806452751 |
| Gm37753 | 3.33123091 | 0.207128905 | 0.690146874 | 0.300122934 |
| Pcdha6 | 52.26043541 | 0.430672129 | 0.179821082 | 2.395003541 |
| Gm37013 | 0 | NA | NA | NA |
| Pcdha7 | 45.39741467 | 0.565801681 | 0.215734441 | 2.622676652 |
| Gm37388 | 0 | NA | NA | NA |
| Gm36858 | 26.49116316 | 0.431176116 | 0.30510932 | 1.413185659 |
| Pcdha8 | 34.08858227 | 0.22388754 | 0.327437381 | 0.683756813 |
| Pcdha9 | 61.46070133 | 0.139156785 | 0.183810643 | 0.75706598 |
| Pcdha11 | 154.8993467 | 0.33988469 | 0.178726219 | 1.901705814 |
| Gm10545 | 8.703390065 | 0.674048051 | 0.378104248 | 1.782704253 |
| Gm18150 | 13.89624275 | 0.126879589 | 0.453882679 | 0.2795427 |
| Pcdha12 | 117.7268123 | 0.217513342 | 0.145692401 | 1.49296285 |

| | | | | |
|---|---|---|---|---|
| Gm19035 | 45.21062936 | 0.472805969 | 0.296220971 | 1.596125916 |
| Pcdhac1 | 92.62921019 | 0.275354881 | 0.132912352 | 2.07170272 |
| Pcdhac2 | 545.5711865 | 0.398728269 | 0.098345709 | 4.054353487 |
| Gm10544 | 2.182918281 | -1.091783766 | 0.78100443 | -1.397922629 |
| Gm38097 | 0.07796279 | -0.798301147 | 3.721829328 | -0.214491605 |
| Pcdhb1 | 5.985971091 | 0.655003584 | 0.456583781 | 1.434574796 |
| Pcdhb2 | 60.01309998 | 0.425419359 | 0.187281529 | 2.271550006 |
| Pcdhb3 | 83.43922558 | 0.43678319 | 0.137937625 | 3.16652684 |
| Gm18529 | 8.138222198 | -0.415256167 | 0.388889119 | -1.067800942 |
| Pcdhb4 | 60.73868439 | 0.024269006 | 0.177077704 | 0.137052858 |
| Pcdhb5 | 51.69657475 | 0.576398947 | 0.185224407 | 3.111895229 |
| Pcdhb6 | 68.6643995 | 0.707473738 | 0.168862642 | 4.189640352 |
| Pcdhb7 | 114.8261978 | 0.417203158 | 0.151210687 | 2.75908514 |
| Pcdhb8 | 89.35877465 | 0.445519213 | 0.165622131 | 2.689973918 |
| Pcdhb9 | 170.4389826 | 0.441155134 | 0.108848742 | 4.052918992 |
| Pcdhb10 | 105.3601154 | 0.32795619 | 0.132162703 | 2.481457955 |
| Pcdhb11 | 148.8037279 | 0.540409469 | 0.125417016 | 4.308900694 |
| Gm37446 | 0.821410149 | -0.590875947 | 1.391571859 | -0.424610445 |
| Gm20162 | 21.72530242 | 0.777863981 | 0.329150575 | 2.36324661 |
| Pcdhb12 | 117.9366013 | 0.756597708 | 0.162018024 | 4.66983665 |
| Pcdhb13 | 90.40139946 | 0.746934008 | 0.175848559 | 4.247598122 |
| Pcdhb14 | 293.0642983 | 0.488102936 | 0.120129869 | 4.06312719 |
| Pcdhb15 | 108.4402084 | 0.44258876 | 0.161091415 | 2.747438537 |
| Pcdhb16 | 347.3551605 | 0.27188635 | 0.082408041 | 3.299269653 |
| Pcdhb17 | 319.6024478 | 0.438742764 | 0.112458389 | 3.901378692 |
| Pcdhb18 | 229.3400325 | 0.375949845 | 0.093931945 | 4.002364102 |
| Pcdhb19 | 338.4815422 | 0.542788996 | 0.141561678 | 3.834293316 |
| Pcdhb20 | 206.2975924 | 0.298517318 | 0.127870526 | 2.334527953 |
| Pcdhb21 | 135.2304553 | 0.347031073 | 0.133272565 | 2.603919827 |
| Pcdhb22 | 238.2686593 | 0.366551904 | 0.084124126 | 4.357274462 |
| 4930517L18Rik | 0.044544648 | 0.517957688 | 3.724945049 | 0.139051095 |
| Slc25a2 | 10.03606488 | -0.652298201 | 0.388844307 | -1.677530544 |
| Taf7 | 183.7126313 | -0.42146291 | 0.116074091 | -3.630981771 |
| Gm8242 | 6.947208237 | 0.252920209 | 0.432132419 | 0.585284042 |
| 3222401L13Rik | 32.8963234 | -0.308468931 | 0.201893747 | -1.527877588 |
| Pcdhga1 | 89.5175488 | 0.098421203 | 0.131518013 | 0.748347704 |
| Gm37118 | 0.066050875 | -0.443818988 | 3.724945049 | -0.119147795 |
| Pcdhga2 | 78.94781358 | 0.523757744 | 0.151289244 | 3.461962862 |
| Pcdhga3 | 73.82529401 | 0.254349503 | 0.155946322 | 1.63100675 |
| Pcdhgb1 | 82.71764577 | 0.127984466 | 0.158525856 | 0.807341271 |
| Pcdhga4 | 165.6903328 | 0.232904965 | 0.099355797 | 2.344150734 |
| Pcdhgb2 | 147.8807373 | 0.298392029 | 0.138225116 | 2.158739578 |
| Pcdhga5 | 172.6956286 | 0.116977749 | 0.116764264 | 1.001828341 |
| Gm24401 | 7.516116141 | 1.03431787 | 0.571065524 | 1.811206991 |
| Pcdhga6 | 188.1446477 | 0.168836875 | 0.114906691 | 1.469338935 |
| Gm37165 | 0.949659951 | -0.328232191 | 1.066846876 | -0.3076657 |
| Pcdhga7 | 242.4294219 | 0.196747155 | 0.107105222 | 1.836952023 |

| | | | | |
|---|---|---|---|---|
| Pcdhgb4 | 169.0788982 | 0.084613816 | 0.160708758 | 0.526504073 |
| Pcdhga8 | 424.9442205 | 0.299823933 | 0.095065735 | 3.153859112 |
| Pcdhgb5 | 111.8894427 | -0.021502922 | 0.137808713 | -0.156034558 |
| Pcdhga9 | 152.843188 | 0.174414662 | 0.107888924 | 1.616613234 |
| Pcdhgb6 | 97.63560197 | 0.1357526 | 0.197721678 | 0.686584301 |
| Gm26672 | 13.40694889 | 0.096104436 | 0.309171921 | 0.310844645 |
| Pcdhga10 | 152.3565402 | 0.100572059 | 0.14090204 | 0.713772909 |
| Pcdhgb7 | 116.7201272 | 0.199972916 | 0.144358183 | 1.385255146 |
| Pcdhga11 | 97.79522649 | -0.011955441 | 0.188883704 | -0.063295247 |
| Pcdhgb8 | 105.1432484 | 0.256648201 | 0.17232623 | 1.489315936 |
| Pcdhga12 | 164.3031905 | 0.369077184 | 0.167719458 | 2.200562705 |
| BC037039 | 16.91683977 | -0.369071619 | 0.27104943 | -1.361639532 |
| Pcdhgc3 | 1154.35759 | 0.019245932 | 0.107574245 | 0.178908363 |
| Pcdhgc5 | 972.9292026 | 0.234105064 | 0.105092325 | 2.227613322 |
| Gm38182 | 0 | NA | NA | NA |
| Gm29994 | 1.935845755 | 0.483353104 | 0.877822429 | 0.550627426 |
| Diap1 | 993.6614968 | -0.427296671 | 0.095021024 | -4.496864503 |
| Mir6979 | 0.451714327 | -1.099049651 | 1.467424597 | -0.748964992 |
| Hdac3 | 567.0461211 | 0.076597917 | 0.071941165 | 1.06473 |
| Rell2 | 1076.046952 | 0.31452043 | 0.087136913 | 3.609497063 |
| Fchsd1 | 97.53749871 | -0.595164861 | 0.120152489 | -4.953412679 |
| Arap3 | 55.62706146 | -0.212641502 | 0.304320012 | -0.69874308 |
| Mir6981 | 0 | NA | NA | NA |
| Mir6980 | 0 | NA | NA | NA |
| Pcdh1 | 2873.63663 | 0.178983522 | 0.111156037 | 1.610200636 |
| 2010320O07Rik | 1.189163173 | 0.696562457 | 1.020531874 | 0.682548458 |
| 1700086O06Rik | 44.53713014 | 0.151194845 | 0.186804599 | 0.809374314 |
| Gm10268 | 1.309431784 | 0.206225662 | 0.840707595 | 0.245300105 |
| 0610009O20Rik | 346.6488783 | 0.129075257 | 0.078776749 | 1.63849435 |
| Pcdh12 | 4.608338074 | -0.748008843 | 0.476876114 | -1.568560095 |
| Rnf14 | 3142.218856 | 0.178946146 | 0.071894614 | 2.489006293 |
| Gm23205 | 10.03026786 | 0.878503539 | 0.523769722 | 1.677270567 |
| Gnpda1 | 269.096148 | -1.156226172 | 0.111581702 | -10.36214856 |
| Gm24690 | 0 | NA | NA | NA |
| Ndfip1 | 2572.872965 | -0.016332652 | 0.070814045 | -0.230641424 |
| Spry4 | 260.3722342 | -0.433849933 | 0.154662514 | -2.805139535 |
| Gm27369 | 0 | NA | NA | NA |
| Gm27804 | 0 | NA | NA | NA |
| Arhgap26 | 2536.520982 | 0.311051554 | 0.0958922 | 3.243762823 |
| Gm19774 | 0.900419082 | 0.21325796 | 1.16989454 | 0.182288191 |
| Gm15335 | 0 | NA | NA | NA |
| Gm8302 | 0.044544648 | 0.517957688 | 3.724945049 | 0.139051095 |
| Fgf1 | 1013.624655 | 0.139562516 | 0.123365638 | 1.131291646 |
| Gm15334 | 1.714908477 | 3.469696161 | 0.978687163 | 3.545255615 |
| Gm15336 | 0.09349418 | 0.517955464 | 3.724945049 | 0.139050498 |
| Gm15333 | 0 | NA | NA | NA |
| Gm15337 | 14.68375219 | 0.921410528 | 0.326251565 | 2.824233278 |

| | | | | |
|---|---|---|---|---|
| Nr3c1 | 1778.771107 | 0.037517359 | 0.053665517 | 0.699096188 |
| Pabpc2 | 0.066050875 | -0.443818988 | 3.724945049 | -0.119147795 |
| Yipf5 | 514.5136962 | -0.333987084 | 0.084975216 | -3.930405846 |
| 2900055J20Rik | 110.5064999 | 0.83105095 | 0.206799314 | 4.018634946 |
| Kctd16 | 294.0495108 | 0.624521354 | 0.1930659 | 3.234757429 |
| Rps19-ps13 | 0.270136144 | 0.561265622 | 2.098434325 | 0.267468758 |
| Gm22515 | 0 | NA | NA | NA |
| Gm22483 | 0.046345709 | -0.443818988 | 3.724945049 | -0.119147795 |
| AC149064.1 | 0 | NA | NA | NA |
| Prelid2 | 7.690273632 | -0.814698233 | 0.381200353 | -2.137191705 |
| Grxcr2 | 0.311588305 | 0.025136989 | 2.036622473 | 0.012342488 |
| Sh3rf2 | 92.9302299 | 0.116991609 | 0.282150525 | 0.414642537 |
| Gm5689 | 0.669351884 | -1.222984286 | 1.210117297 | -1.010632844 |
| Plac8l1 | 0.74423012 | -2.196251102 | 1.463896234 | -1.500277855 |
| Lars | 747.0783727 | -0.502894622 | 0.113645437 | -4.425119334 |
| Rps19-ps4 | 0.931580594 | -0.760560384 | 1.126942893 | -0.674888133 |
| Rbm27 | 1071.37103 | -0.026208437 | 0.063455471 | -0.413020925 |
| Gm22697 | 0 | NA | NA | NA |
| Pou4f3 | 1.201894167 | -1.92805417 | 1.069476603 | -1.802801637 |
| Gm16415 | 0.834345451 | -0.536266181 | 1.173290291 | -0.457061807 |
| Tcerg1 | 1111.337889 | 0.138887812 | 0.087346633 | 1.590076314 |
| Gm27457 | 2.628954575 | 0.795225902 | 0.681951541 | 1.166103241 |
| Gpr151 | 33.37780463 | -1.646663516 | 0.275064768 | -5.986457401 |
| Ppp2r2b | 978.0231163 | 0.149994092 | 0.068010145 | 2.20546643 |
| 4933407I08Rik | 0.075043205 | 0.517957688 | 3.724945049 | 0.139051095 |
| Gm22264 | 0 | NA | NA | NA |
| Gm8181 | 9.04075607 | -0.098100217 | 0.473214762 | -0.20730591 |
| Stk32a | 162.2987704 | -0.256340375 | 0.142482194 | -1.799104633 |
| Dpysl3 | 999.9963044 | -0.195727294 | 0.069224319 | -2.827435453 |
| Gm3650 | 17.99992091 | -0.147114493 | 0.35761121 | -0.411381101 |
| Eif3j2 | 165.1918275 | -0.435517016 | 0.119409178 | -3.647265834 |
| Gm27325 | 0 | NA | NA | NA |
| Jakmip2 | 846.4346633 | 0.343821423 | 0.078089565 | 4.402911239 |
| Spink1 | 0 | NA | NA | NA |
| Gm23894 | 0 | NA | NA | NA |
| Scgb3a2 | 0.047948698 | 0.037064902 | 3.724945049 | 0.009950456 |
| Gm94 | 0.18352982 | 1.388044403 | 3.713044316 | 0.373829205 |
| Spink5 | 0.975055063 | 2.563595101 | 3.524349361 | 0.727395283 |
| Spink14 | 0 | NA | NA | NA |
| Spink6 | 0 | NA | NA | NA |
| Spink12 | 0 | NA | NA | NA |
| Gm37797 | 0 | NA | NA | NA |
| Gm10267 | 0 | NA | NA | NA |
| Spinkl | 0 | NA | NA | NA |
| Spink11 | 0.047948698 | 0.037064902 | 3.724945049 | 0.009950456 |
| Gm10542 | 0 | NA | NA | NA |
| Npy6r | 0.261451683 | -0.966418703 | 2.092973515 | -0.461744354 |

| | | | | |
|---|---|---|---|---|
| Myot | 0.836501452 | -0.789097519 | 1.194554245 | -0.66057906 |
| Dcp2 | 1781.048542 | -0.038553183 | 0.094519107 | -0.407887715 |
| Mcc | 632.7057759 | -0.097057418 | 0.092995878 | -1.043674415 |
| A930012L18Rik | 13.61962428 | 0.633801464 | 0.313331521 | 2.02278233 |
| RP23-342A15.2 | 0 | NA | NA | NA |
| Ythdc2 | 676.2351253 | 0.175041327 | 0.084867642 | 2.062521383 |
| Gm26928 | 0 | NA | NA | NA |
| Kcnn2 | 557.7293241 | 0.379553412 | 0.111469469 | 3.40499884 |
| Gm10540 | 0.30732292 | 1.336595121 | 1.558251153 | 0.857753334 |
| A330093E20Rik | 2.019082719 | 0.196274599 | 0.981656065 | 0.199942328 |
| Gm26262 | 0 | NA | NA | NA |
| Gm38337 | 0.335887269 | 1.292097681 | 1.52224509 | 0.848810542 |
| Trim36 | 95.57263784 | -0.100346263 | 0.178451147 | -0.562317837 |
| 1700018A14Rik | 0.75323169 | -2.639113365 | 1.34044869 | -1.968828337 |
| Pggt1b | 266.2078042 | -0.248563661 | 0.089725673 | -2.77026243 |
| Ccdc112 | 108.5123134 | 0.389722089 | 0.157162087 | 2.479746211 |
| Gm24617 | 0.282138413 | -0.368999018 | 3.716196214 | -0.099294816 |
| Mospd4 | 3.756273683 | 0.29713556 | 0.542389278 | 0.547827128 |
| Gm24076 | 0.776630353 | 2.052576998 | 1.194683271 | 1.718093028 |
| Fem1c | 864.4434414 | -0.19687428 | 0.083935112 | -2.345553309 |
| Ticam2 | 12.34676471 | -0.539951124 | 0.371903911 | -1.451856539 |
| Tmed7 | 1507.489644 | -0.201265502 | 0.107354519 | -1.87477437 |
| Eif1a | 755.0916563 | -0.19785854 | 0.148509941 | -1.332291555 |
| Cdo1 | 339.5541898 | -0.312694811 | 0.148603631 | -2.104220536 |
| Atg12 | 650.7230075 | -0.353031269 | 0.087102089 | -4.053074645 |
| Ap3s1 | 461.3279815 | -0.324700307 | 0.087796315 | -3.698336383 |
| Lvrn | 10.74167226 | -0.565527149 | 0.400308412 | -1.412728616 |
| Gm23610 | 0 | NA | NA | NA |
| Arl14epl | 1.575196263 | -0.248560557 | 0.768955922 | -0.32324422 |
| Commd10 | 179.0565603 | 0.027223034 | 0.091249991 | 0.298334653 |
| Gm22791 | 0 | NA | NA | NA |
| Gm25036 | 0 | NA | NA | NA |
| Sema6a | 831.0817601 | 0.027802351 | 0.060363027 | 0.460585764 |
| Eno1b | 521.4844324 | -0.339994316 | 0.123584511 | -2.751107845 |
| Gm5839 | 0 | NA | NA | NA |
| 1700044K03Rik | 0 | NA | NA | NA |
| Gm25440 | 0 | NA | NA | NA |
| Dtwd2 | 52.5847649 | 0.249176503 | 0.154881477 | 1.608820551 |
| Gm16283 | 2.090780166 | -0.245292899 | 0.726717234 | -0.337535548 |
| 1700065O20Rik | 0 | NA | NA | NA |
| Dmxl1 | 3104.946172 | -0.036612653 | 0.050053244 | -0.731474113 |
| Tnfaip8 | 137.5382232 | -0.668866545 | 0.142956665 | -4.678806311 |
| C030005K06Rik | 2.787674368 | 0.737796114 | 1.080533326 | 0.682807365 |
| Hsd17b4 | 542.6081984 | -0.116129027 | 0.08744321 | -1.32805083 |
| Cd63-ps | 108.0511914 | -1.349065053 | 0.200811791 | -6.71805699 |
| Gm8529 | 0 | NA | NA | NA |
| Fam170a | 0 | NA | NA | NA |

| | | | | |
|---|---|---|---|---|
| Hdhd1a | 0.113999573 | -0.443821212 | 3.724945049 | -0.119148392 |
| Prr16 | 138.718701 | 0.488882519 | 0.142683094 | 3.426352098 |
| Gm25739 | 0 | NA | NA | NA |
| Gm25785 | 0 | NA | NA | NA |
| Gm4950 | 20.25727402 | -0.175514284 | 0.247699234 | -0.708578229 |
| Gm22259 | 0 | NA | NA | NA |
| Ftmt | 0 | NA | NA | NA |
| Srfbp1 | 140.6912451 | -0.138713775 | 0.114142613 | -1.215267208 |
| Gm27026 | 0 | NA | NA | NA |
| Lox | 47.85031176 | -0.486849863 | 0.317877232 | -1.531565692 |
| Zfp474 | 3.585501729 | -1.244340912 | 0.581768868 | -2.138892231 |
| 1700034E13Rik | 0.108266322 | -0.955981391 | 3.722845977 | -0.25678779 |
| Gykl1 | 0 | NA | NA | NA |
| Sncaip | 113.2002606 | 0.226057614 | 0.111387838 | 2.029464068 |
| Snx2 | 781.8559442 | -0.362407584 | 0.084041097 | -4.312266237 |
| Snx24 | 137.1094418 | 0.073521027 | 0.101428973 | 0.724852329 |
| Ppic | 20.57672658 | -0.786016377 | 0.235882869 | -3.332231712 |
| Prdm6 | 12.21164467 | -0.717543644 | 0.33944507 | -2.113872632 |
| Cep120 | 755.1587019 | -0.166399926 | 0.061598944 | -2.701343821 |
| Csnk1g3 | 932.5335226 | 0.04549788 | 0.061227552 | 0.74309488 |
| Gm5507 | 5.787175505 | 0.482147492 | 0.440606914 | 1.094280358 |
| Redrum | 0.747472414 | -0.657680632 | 1.027234542 | -0.640243883 |
| Zfp608 | 685.0116038 | 0.068901153 | 0.073418238 | 0.938474629 |
| Gm26742 | 0.583001974 | -1.792693772 | 1.249266045 | -1.434997597 |
| Gm4221 | 12.78354649 | 0.707504742 | 0.39362953 | 1.797387362 |
| Gm24183 | 0 | NA | NA | NA |
| Gm34073 | 0.428599116 | 1.903959762 | 3.319240644 | 0.573613054 |
| Gm22597 | 0 | NA | NA | NA |
| Gm37828 | 0 | NA | NA | NA |
| Gm37337 | 0.587544172 | -0.513948668 | 1.251110797 | -0.410793887 |
| Gm24514 | 0.493679557 | 0.037076958 | 1.568141242 | 0.02364389 |
| Gm26959 | 0 | NA | NA | NA |
| Gm8614 | 0.29391207 | -0.443820661 | 2.066089338 | -0.214811941 |
| Gm23335 | 0 | NA | NA | NA |
| Gm6944 | 0 | NA | NA | NA |
| Gm22484 | 0.068097359 | 0.037064902 | 3.724945049 | 0.009950456 |
| Gm23555 | 0 | NA | NA | NA |
| Gm25476 | 0 | NA | NA | NA |
| Gramd3 | 170.2177708 | -0.298513441 | 0.110655419 | -2.697684803 |
| Aldh7a1 | 399.3017201 | -0.152029767 | 0.078601202 | -1.934191357 |
| Mir1258 | 0 | NA | NA | NA |
| Phax | 381.2748711 | 0.037682881 | 0.101977142 | 0.369522823 |
| Tex43 | 2.674966445 | -0.781730981 | 0.668410677 | -1.169536944 |
| Lmnb1 | 214.1019998 | -0.491437964 | 0.144839724 | -3.392977776 |
| 3-Mar | 67.75458388 | -0.157908183 | 0.176069357 | -0.896852159 |
| Gm15345 | 0.999978248 | -1.164691851 | 1.047476397 | -1.111902716 |
| C330018D20Rik | 87.14994236 | -0.323692609 | 0.14612828 | -2.215126388 |

| | | | | |
|---|---|---|---|---|
| Megf10 | 704.5314188 | -0.363376962 | 0.126571745 | -2.870916909 |
| Prrc1 | 265.116747 | -0.241387761 | 0.082826999 | -2.914360832 |
| 4930511M06Rik | 2.334218344 | -0.437028572 | 0.598678149 | -0.729989182 |
| Ctxn3 | 52.35061762 | -0.002829064 | 0.191435736 | -0.014778141 |
| 1700066O22Rik | 0.436302845 | -0.530559183 | 1.356966458 | -0.390989166 |
| 1700011I03Rik | 2.706585767 | -0.511277056 | 0.6709918 | -0.761972138 |
| Gm26038 | 0 | NA | NA | NA |
| AC102232.1 | 0.04393175 | 0.037062677 | 3.724945049 | 0.009949859 |
| Slc12a2 | 1824.003985 | 0.008142296 | 0.088957892 | 0.091529778 |
| Fbn2 | 140.4520598 | 0.118832123 | 0.173422652 | 0.685216845 |
| Gm26507 | 13.7398474 | 0.542022309 | 0.333176546 | 1.626832129 |
| Gm25660 | 0 | NA | NA | NA |
| Slc27a6 | 3.148605881 | -1.004102563 | 0.610480818 | -1.644773323 |
| Gm23270 | 0 | NA | NA | NA |
| Isoc1 | 459.7670982 | -0.005514744 | 0.123930263 | -0.044498771 |
| Adamts19 | 19.03947336 | 0.373717816 | 0.354209624 | 1.055075272 |
| A730017C20Rik | 501.0808718 | 0.320942287 | 0.108621241 | 2.954691779 |
| Chsy3 | 230.6243843 | 0.192442338 | 0.158694712 | 1.212657529 |
| Mir6355 | 0 | NA | NA | NA |
| Gm26028 | 0 | NA | NA | NA |
| Gm23576 | 0 | NA | NA | NA |
| Gm4951 | 925.2780923 | -0.949869382 | 0.230149649 | -4.127181539 |
| Gm5970 | 122.1217523 | -0.417479023 | 0.240951932 | -1.732623677 |
| Gm4841 | 260.7014249 | -1.162935131 | 0.393851358 | -2.952725964 |
| F830016B08Rik | 944.1534938 | -0.942505061 | 0.220863097 | -4.267372284 |
| Iigp1 | 5189.009943 | -0.808569325 | 0.194561413 | -4.155856554 |
| Gm6960 | 0.926812825 | 0.359325845 | 0.954546614 | 0.376436142 |
| BC023105 | 164.7922582 | -1.602724173 | 0.347716218 | -4.609287953 |
| Smim3 | 173.6181185 | -0.249347567 | 0.222223207 | -1.122059082 |
| Dctn4 | 1434.033031 | -0.130236295 | 0.059137914 | -2.20224701 |
| Rbm22 | 390.7352864 | 0.08231601 | 0.1424476 | 0.577868703 |
| Myoz3 | 0.060174537 | 0.517957688 | 3.724945049 | 0.139051095 |
| Synpo | 1700.826908 | 0.058810364 | 0.129562575 | 0.453914751 |
| Ndst1 | 2081.012663 | -0.241578765 | 0.090692113 | -2.663724071 |
| Rps14 | 1385.960924 | -0.152218542 | 0.137234893 | -1.109182499 |
| Gm8731 | 0.109869311 | -0.444802825 | 3.724874622 | -0.119414174 |
| Cd74 | 447.6522266 | -0.379396494 | 0.317950413 | -1.193256804 |
| Mir5107 | 0.060174537 | 0.517957688 | 3.724945049 | 0.139051095 |
| Tcof1 | 457.500707 | -0.451022029 | 0.094614101 | -4.766964153 |
| Arsi | 22.1297461 | -0.500455902 | 0.267869578 | -1.868281965 |
| Camk2a | 16056.95831 | 1.454847043 | 0.175338451 | 8.297364541 |
| Mir7220 | 4.383661021 | 0.905550606 | 0.679832137 | 1.332020886 |
| Mir6982 | 3.753094389 | 1.382311727 | 0.75232344 | 1.837390216 |
| Slc6a7 | 440.8262577 | 0.513265259 | 0.144641275 | 3.548539378 |
| Cdx1 | 0.329049156 | 0.037076754 | 1.775088275 | 0.020887273 |
| Pdgfrb | 307.8251967 | -0.409587607 | 0.091723622 | -4.465453927 |
| Csf1r | 863.5867635 | 0.848923997 | 0.301364499 | 2.816934308 |

| | | | | |
|---|---|---|---|---|
| Mir6983 | 0 | NA | NA | NA |
| Hmgxb3 | 688.8088613 | 0.020603905 | 0.060299712 | 0.341691595 |
| Gm26403 | 2.513420025 | 0.813673841 | 0.750838892 | 1.083686326 |
| Slc26a2 | 293.7967227 | -0.269132324 | 0.101775412 | -2.644374698 |
| Pde6a | 3.821803709 | -0.398248592 | 0.53419901 | -0.745506047 |
| Rps2-ps10 | 1236.213371 | -0.051888219 | 0.129582114 | -0.400427324 |
| Ppargc1b | 407.0618076 | 0.336388355 | 0.129529163 | 2.597008644 |
| Gm25301 | 0 | NA | NA | NA |
| mmu-mir-378a | 3.818950962 | 0.161267513 | 0.60835361 | 0.265088446 |
| Gm25834 | 0.038981395 | -0.443816764 | 3.724945049 | -0.119147198 |
| Arhgef37 | 140.4268379 | -0.287398318 | 0.140499719 | -2.045543716 |
| Csnk1a1 | 4557.884024 | -0.142129513 | 0.06697027 | -2.122277727 |
| Bvht | 5.444855971 | -0.80998933 | 0.524213274 | -1.545152269 |
| Mir145a | 0.194631057 | 1.456047977 | 3.23213671 | 0.450490839 |
| Mir143 | 0.147068667 | 0.998853574 | 3.51441668 | 0.284216035 |
| Mir143hg | 8.471314973 | 0.771550582 | 0.477065405 | 1.617284704 |
| Il17b | 2.212656516 | -0.341533931 | 0.806490828 | -0.423481482 |
| Gm38165 | 0.662591016 | 0.343262536 | 1.275861005 | 0.269043834 |
| Pcyox1l | 207.9956365 | -0.060299301 | 0.139488479 | -0.432288759 |
| Grpel2 | 247.9051696 | -0.046562217 | 0.098072416 | -0.474773833 |
| 1500015A07Rik | 35.15775905 | 0.236948258 | 0.197549056 | 1.199440092 |
| Afap1l1 | 138.1820362 | 0.290174901 | 0.097703738 | 2.969946773 |
| Ablim3 | 445.2700515 | 0.157920146 | 0.118552934 | 1.33206442 |
| Gm37774 | 0 | NA | NA | NA |
| Sh3tc2 | 44.0296305 | -0.466940635 | 0.225865675 | -2.067337747 |
| Adrb2 | 16.87312987 | 0.380336614 | 0.275575899 | 1.380151949 |
| Gm9949 | 1.519386227 | 0.765355975 | 0.787579704 | 0.971782247 |
| Htr4 | 92.71409538 | 0.346874806 | 0.146643768 | 2.365424803 |
| Fbxo38 | 754.8506995 | -0.108732476 | 0.057510088 | -1.890667891 |
| Spink10 | 11.52016113 | -0.1946504 | 0.329613816 | -0.59054078 |
| Spink7 | 0.099106094 | -0.93550548 | 3.724188147 | -0.251197158 |
| Spink13 | 0.546984718 | -1.243117363 | 1.319034496 | -0.942444922 |
| Gm17732 | 5.889985807 | 0.512031477 | 0.456894916 | 1.120676677 |
| 2700046A07Rik | 9.548929167 | 0.33730587 | 0.462681384 | 0.72902408 |
| Apcdd1 | 206.8351242 | 0.360268859 | 0.166230302 | 2.167287518 |
| Napg | 1597.555114 | 0.204768002 | 0.056628234 | 3.616005446 |
| Piezo2 | 47.03296206 | -0.432310035 | 0.200585225 | -2.155243661 |
| Gm26972 | 0 | NA | NA | NA |
| Gm23581 | 0 | NA | NA | NA |
| Gm22560 | 0 | NA | NA | NA |
| Gm22210 | 0 | NA | NA | NA |
| Gm24848 | 0 | NA | NA | NA |
| Txnl1 | 825.0032602 | -0.291503551 | 0.096355117 | -3.025304312 |
| Wdr7 | 3498.348225 | 0.318179774 | 0.045284264 | 7.026276778 |
| Gm24384 | 0 | NA | NA | NA |
| St8sia3os | 1.428069919 | -0.593803584 | 0.899289993 | -0.66030267 |
| St8sia3 | 1740.780446 | 0.375404042 | 0.081444484 | 4.609324344 |

| | | | | |
|---|---|---|---|---|
| Onecut2 | 194.3572374 | 0.179140345 | 0.156718858 | 1.143068212 |
| Amd-ps3 | 30.2089109 | -0.199622784 | 0.194832206 | -1.024588224 |
| Fech | 621.1316025 | 0.142032669 | 0.068198418 | 2.08263875 |
| Nars | 1964.183135 | -0.285592352 | 0.088279184 | -3.235104126 |
| Atp8b1 | 51.70323717 | -0.833146017 | 0.20486877 | -4.066730206 |
| Gm23016 | 0 | NA | NA | NA |
| Gm24504 | 0 | NA | NA | NA |
| Nedd4l | 2403.215903 | 0.196468921 | 0.105252807 | 1.866638297 |
| Mir122 | 0.049553047 | -0.443818988 | 3.724945049 | -0.119147795 |
| Alpk2 | 11.61777028 | 0.098089319 | 0.365806466 | 0.268145395 |
| Gm22567 | 0.046345709 | -0.443818988 | 3.724945049 | -0.119147795 |
| Malt1 | 178.3264262 | -0.15587314 | 0.177522311 | -0.878048172 |
| Zfp532 | 1029.175328 | 0.049418669 | 0.061202981 | 0.807455255 |
| Gm37643 | 15.88324691 | -0.154160683 | 0.382220238 | -0.403329463 |
| Oacyl | 19.6943626 | 1.489252435 | 0.379724852 | 3.921925121 |
| Sec11c | 480.2888366 | -0.051788772 | 0.087523217 | -0.591714675 |
| Grp | 19.80146899 | 0.240571429 | 0.272281318 | 0.883539975 |
| Gm25150 | 0 | NA | NA | NA |
| Rax | 0.423125362 | -1.150340254 | 1.547204003 | -0.743496172 |
| Raxos1 | 0 | NA | NA | NA |
| Cplx4 | 0.598788582 | -0.46330466 | 1.272967253 | -0.363956464 |
| Lman1 | 670.4253144 | -0.299919917 | 0.063748074 | -4.704768311 |
| Ccbe1 | 53.63741432 | 1.210949843 | 0.222368169 | 5.445697768 |
| Mir694 | 0.217748738 | 0.037067974 | 2.49659883 | 0.014847389 |
| Gm15958 | 0.904555577 | 1.889635124 | 1.160464551 | 1.628343686 |
| Pmaip1 | 46.65352456 | -0.772999034 | 0.167375498 | -4.618352415 |
| Gm24679 | 0 | NA | NA | NA |
| Gm22219 | 0 | NA | NA | NA |
| Mc4r | 36.60476633 | 0.406147736 | 0.223357025 | 1.818379053 |
| Gnal | 2244.602178 | 0.25445691 | 0.095544342 | 2.663233693 |
| Gm23512 | 0.291502873 | 0.517970935 | 2.047224824 | 0.253011262 |
| Mppe1 | 130.6858995 | -0.081832073 | 0.108355025 | -0.755221765 |
| Impa2 | 34.6996934 | 0.297605305 | 0.230710098 | 1.289953525 |
| B430212C06Rik | 2.198202818 | 1.589760662 | 0.915051352 | 1.737345843 |
| Cidea | 13.25459769 | 1.070674686 | 0.318599493 | 3.360566192 |
| Tubb6 | 218.1172078 | -1.623768094 | 0.312260703 | -5.200039838 |
| Afg3l2 | 820.858718 | 0.073996846 | 0.078228575 | 0.945905579 |
| Gm24900 | 0 | NA | NA | NA |
| Slmo1 | 256.90097 | 0.775881047 | 0.102790955 | 7.548145153 |
| Spire1 | 2405.877136 | 0.317557129 | 0.072063182 | 4.406648747 |
| Gm17669 | 0.12092566 | 0.998850474 | 3.724945049 | 0.268151734 |
| Cep76 | 301.4880058 | -0.063633662 | 0.160834525 | -0.395646783 |
| Psmg2 | 96.05648597 | -0.25635911 | 0.161344735 | -1.588890461 |
| Ptpn2 | 330.6143394 | -0.163543272 | 0.09595638 | -1.704350163 |
| Gm26910 | 20.45356711 | 0.445308763 | 0.319040123 | 1.395776677 |
| Seh1l | 391.9333236 | -0.091738361 | 0.064193075 | -1.429100589 |
| Cep192 | 279.5064016 | -0.413257786 | 0.128471528 | -3.216726636 |

| | | | | |
|---|---|---|---|---|
| Ldlrad4 | 528.0018019 | -0.037989627 | 0.098598787 | -0.385295074 |
| Fam210a | 679.3179832 | 0.284693127 | 0.073236581 | 3.887307752 |
| Mir7219 | 0 | NA | NA | NA |
| Rnmt | 878.6655392 | 0.20425192 | 0.05128765 | 3.982477691 |
| Mc5r | 18.63850153 | 0.443380414 | 0.316371411 | 1.401455377 |
| Mc2r | 1.528578462 | -0.122439841 | 0.836084465 | -0.146444344 |
| Mir6356 | 0 | NA | NA | NA |
| 4930546C10Rik | 0.160559904 | -0.410534351 | 2.790729611 | -0.147106459 |
| Mir145b | 0 | NA | NA | NA |
| Gm23536 | 0.335787014 | 0.540624325 | 1.722958932 | 0.313776675 |
| Gm4842 | 0 | NA | NA | NA |
| Tcf4 | 6112.811251 | 0.175326003 | 0.08806668 | 1.990832427 |
| Gm24845 | 5.630675305 | 1.073485128 | 0.516152801 | 2.079781655 |
| Gm22508 | 0 | NA | NA | NA |
| Ccdc68 | 4.327574142 | -0.534234567 | 0.529880628 | -1.008216831 |
| 1700061H18Rik | 0.072025462 | 0.517959912 | 3.724945049 | 0.139051692 |
| 4930448D08Rik | 0.605479419 | 0.068867153 | 1.200945174 | 0.057344128 |
| Rab27b | 340.9708684 | 0.205431217 | 0.103120636 | 1.992144583 |
| Dynap | 0.072025462 | 0.517959912 | 3.724945049 | 0.139051692 |
| 4930503L19Rik | 92.03394658 | -0.453014342 | 0.132956381 | -3.407240309 |
| Stard6 | 8.232924272 | -0.13963366 | 0.359403224 | -0.388515323 |
| Poli | 92.01233665 | -0.140554615 | 0.123812481 | -1.1352217 |
| Mir6357 | 0.407891471 | 1.016095402 | 1.453607766 | 0.699016217 |
| Mbd2 | 568.7990903 | -0.518227122 | 0.124273621 | -4.170049284 |
| Dcc | 395.337002 | 0.637025623 | 0.124786601 | 5.104920042 |
| Gm25509 | 0 | NA | NA | NA |
| n-R5-8s1 | 0 | NA | NA | NA |
| Mex3c | 685.3858209 | -0.613416473 | 0.094982375 | -6.458213639 |
| Gm26438 | 0 | NA | NA | NA |
| Gm22065 | 0.042716945 | 0.037062677 | 3.724945049 | 0.009949859 |
| Smad4 | 842.5159027 | -0.154207815 | 0.063521412 | -2.42765094 |
| Elac1 | 466.4413671 | 0.349694012 | 0.086426978 | 4.046121033 |
| Me2 | 344.6597954 | -0.206037377 | 0.072946727 | -2.824491051 |
| Mro | 101.9368696 | 0.204377144 | 0.127495229 | 1.603017971 |
| Mapk4 | 1333.921859 | -0.170892146 | 0.098208433 | -1.740096458 |
| D730045A05Rik | 2.135082683 | -3.781909478 | 0.988368239 | -3.826417453 |
| Gm9925 | 7.46780898 | 0.828577168 | 0.473956326 | 1.748214177 |
| Ska1 | 2.02584455 | 0.23220832 | 0.677396405 | 0.342795324 |
| Cxxc1 | 352.5600067 | -0.355722638 | 0.091154388 | -3.902419252 |
| Mbd1 | 382.0312851 | 0.121228825 | 0.138614722 | 0.874573951 |
| Cfap53 | 11.98097365 | 0.579305977 | 0.464778329 | 1.246413486 |
| Gm14328 | 4.493268881 | 0.292216726 | 0.468519665 | 0.623702158 |
| Myo5b | 428.355463 | 0.419161602 | 0.109805827 | 3.81729836 |
| 1700120E14Rik | 0 | NA | NA | NA |
| Scarna17 | 10.66151498 | 0.039246293 | 0.401493082 | 0.097750856 |
| Gm23119 | 43.96335493 | -0.199108011 | 0.177412126 | -1.12229088 |
| Acaa2 | 104.6941341 | 0.056442993 | 0.180473944 | 0.312748709 |

| | | | | |
|---|---|---|---|---|
| Gm24559 | 0 | NA | NA | NA |
| Gm25594 | 0 | NA | NA | NA |
| Lipg | 453.85713 | -1.915180411 | 0.421340078 | -4.545450365 |
| Rpl17 | 766.1645459 | -0.152213532 | 0.076676475 | -1.985139927 |
| Snord58b | 1.376506768 | 0.183910722 | 0.952709349 | 0.193039695 |
| Gm23301 | 2.714581473 | 0.505602768 | 0.641550762 | 0.78809472 |
| Gm26202 | 2.364995654 | 1.004774756 | 0.68488448 | 1.467071873 |
| BC031181 | 600.9122168 | -0.098710754 | 0.094853249 | -1.040668138 |
| Dym | 677.9260573 | 0.179223676 | 0.060130592 | 2.98057395 |
| Gm27781 | 0.317560373 | -0.942168405 | 3.720459044 | -0.253239827 |
| Gm8807 | 0 | NA | NA | NA |
| 2010010A06Rik | 0 | NA | NA | NA |
| Gm20544 | 9.279688315 | 0.029360924 | 0.338716882 | 0.086682788 |
| Smad7 | 616.9288546 | -0.465674327 | 0.081548018 | -5.710430994 |
| Ctif | 1554.441727 | 0.34744507 | 0.066840262 | 5.198140419 |
| Gm10532 | 5.803090283 | -0.292849044 | 0.494499272 | -0.592213297 |
| 1700034B16Rik | 0 | NA | NA | NA |
| Zbtb7c | 113.4256275 | 0.739292355 | 0.135029434 | 5.475045933 |
| Mir6358 | 0 | NA | NA | NA |
| Smad2 | 378.1534108 | -0.093142382 | 0.080768168 | -1.153206576 |
| Skor2 | 31.77545133 | -0.11567893 | 0.30977915 | -0.373423872 |
| Ier3ip1 | 232.6003113 | -0.145092273 | 0.0936573 | -1.549182749 |
| Hdhd2 | 753.876256 | -0.019994727 | 0.060793007 | -0.328898469 |
| Katnal2 | 40.6261297 | -0.178801656 | 0.219896776 | -0.813116317 |
| Pias2 | 1238.28436 | 0.089122115 | 0.053619714 | 1.66211469 |
| Gm7276 | 15.00176566 | 0.248842101 | 0.38187739 | 0.651628266 |
| St8sia5 | 623.4113925 | 0.208456646 | 0.108588711 | 1.919689852 |
| Gm15957 | 0 | NA | NA | NA |
| Loxhd1 | 23.89937881 | -0.794532168 | 0.622730923 | -1.275883594 |
| Rnf165 | 750.5636098 | 0.503162766 | 0.161525719 | 3.115062838 |
| 8030462N17Rik | 630.218415 | -0.047623163 | 0.068375181 | -0.696497807 |
| Gm26014 | 0 | NA | NA | NA |
| 4930465K10Rik | 2.048990233 | 0.029355287 | 0.762033289 | 0.038522316 |
| A330094K24Rik | 6.372749315 | 0.376071023 | 0.414789191 | 0.906655794 |
| Haus1 | 42.89733926 | 0.08956601 | 0.169801308 | 0.527475383 |
| Atp5a1 | 6848.041245 | 0.105692527 | 0.069166336 | 1.52809203 |
| F830208F22Rik | 1.770308639 | -0.569624483 | 0.76914768 | -0.740591823 |
| Pstpip2 | 100.7389169 | 0.391584082 | 0.136102864 | 2.877118602 |
| Gm23954 | 0.113999573 | -0.443821212 | 3.724945049 | -0.119148392 |
| Epg5 | 1099.858229 | -0.146307065 | 0.08930995 | -1.638194452 |
| Siglec15 | 0.39439978 | 0.308867819 | 1.55193573 | 0.199021011 |
| Slc14a1 | 350.1029007 | -0.208104103 | 0.122997285 | -1.691940631 |
| Slc14a2 | 4.084512964 | -0.594479671 | 0.588659113 | -1.009887826 |
| Gm6133 | 2.464583914 | 1.222067785 | 0.721825179 | 1.6930246 |
| Gm26430 | 0 | NA | NA | NA |
| Gm23895 | 0 | NA | NA | NA |
| Setbp1 | 1103.372977 | 0.103991551 | 0.088650349 | 1.173052919 |

| | | | | |
|---|---|---|---|---|
| Gm25824 | 0 | NA | NA | NA |
| Gm26977 | 0.099106094 | -0.93550548 | 3.724188147 | -0.251197158 |
| Pard6g | 127.4897943 | 0.038183573 | 0.145689791 | 0.262088187 |
| Gm21886 | 0.269331125 | 0.603778214 | 2.063939841 | 0.292536731 |
| Adnp2 | 487.4784465 | -0.304533344 | 0.097408071 | -3.126366634 |
| Rbfa | 151.797274 | 0.118290886 | 0.093696738 | 1.2624867 |
| Gm16286 | 543.6190761 | -0.021370213 | 0.097766854 | -0.218583414 |
| Txnl4a | 164.8076413 | -0.605011869 | 0.115592697 | -5.233997327 |
| Hsbp1l1 | 8.924943958 | 0.477334362 | 0.365458057 | 1.30612625 |
| Pqlc1 | 221.7266294 | -0.239460656 | 0.102166879 | -2.34381884 |
| Kcng2 | 77.54725198 | 1.0001101 | 0.192968165 | 5.18277251 |
| Gm37015 | 4.004740963 | 1.739806048 | 0.651429124 | 2.670752634 |
| Gm26676 | 5.505563926 | 0.04457758 | 0.448208969 | 0.099457135 |
| D330025C20Rik | 2.748297996 | 1.121396828 | 0.58669807 | 1.911369554 |
| Ctdp1 | 381.7265928 | -0.024980561 | 0.099107272 | -0.252055778 |
| Nfatc1 | 215.3281497 | -0.483567552 | 0.129134281 | -3.74468769 |
| Atp9b | 951.6403139 | 0.017591307 | 0.07356524 | 0.239125259 |
| Gm25071 | 1.130492263 | -1.050907586 | 1.032722071 | -1.017609302 |
| Sall3 | 188.0210551 | -0.390585615 | 0.179056834 | -2.181349941 |
| Gm27238 | 1.264061948 | -2.175978819 | 1.196281852 | -1.818951625 |
| Gm27239 | 2.200041218 | 1.007018341 | 0.918807244 | 1.096006096 |
| Gm25718 | 0 | NA | NA | NA |
| 4930594M17Rik | 0.045882455 | 0.517959912 | 3.724945049 | 0.139051692 |
| Gm26012 | 0 | NA | NA | NA |
| Mir5127 | 0 | NA | NA | NA |
| Gm17383 | 86.5616375 | -0.234065942 | 0.175105322 | -1.336715184 |
| Gm25060 | 0 | NA | NA | NA |
| Galr1 | 70.21229872 | -0.645963806 | 0.261828457 | -2.467126041 |
| Mbp | 30062.83706 | 0.365923108 | 0.176858354 | 2.069017941 |
| Rpl21-ps8 | 1.385867572 | -1.610118562 | 0.900823495 | -1.787385177 |
| Zfp236 | 860.9371065 | -0.194623893 | 0.110477293 | -1.761664201 |
| Gm10524 | 101.1240389 | 0.371666379 | 0.12902837 | 2.880501229 |
| Mir5112 | 0.320492979 | -0.394032629 | 1.601741781 | -0.246002591 |
| 2210420H20Rik | 0.04393175 | 0.037062677 | 3.724945049 | 0.009949859 |
| 4930445N18Rik | 0.552122516 | 0.165276968 | 1.470975104 | 0.11235878 |
| Zfp516 | 555.826877 | -0.127029623 | 0.091669118 | -1.385740651 |
| 1700095A13Rik | 0 | NA | NA | NA |
| Gm24720 | 0 | NA | NA | NA |
| Tshz1 | 687.2218015 | -0.186911526 | 0.141275699 | -1.323026724 |
| Zadh2 | 324.0628898 | -0.126678104 | 0.111316928 | -1.137994971 |
| Zfp407 | 728.9401691 | 0.062368933 | 0.099473551 | 0.626990114 |
| Gm37216 | 25.25091243 | 0.494466596 | 0.300221375 | 1.647006632 |
| Cndp1 | 9.704862909 | 1.02438034 | 0.342167242 | 2.993800144 |
| Gm7289 | 1.584876195 | -1.171191496 | 0.873859348 | -1.340251723 |
| Cndp2 | 787.0114979 | -0.689667474 | 0.160705205 | -4.291506754 |
| D030046N08Rik | 4.544185544 | 0.318013304 | 0.508338018 | 0.625594177 |
| Fam69c | 47.9173264 | 0.507964447 | 0.192282641 | 2.641759258 |

| | | | | |
|---|---|---|---|---|
| Gm22014 | 0 | NA | NA | NA |
| Gm25005 | 0 | NA | NA | NA |
| Gm22819 | 0 | NA | NA | NA |
| Cyb5a | 343.7727805 | -0.032221957 | 0.077994847 | -0.41312931 |
| Gm16146 | 0.241190471 | 0.594215967 | 3.027989488 | 0.196241093 |
| Fbxo15 | 2.684576295 | -0.53720466 | 0.614265384 | -0.874548159 |
| Timm21 | 78.28501918 | -0.142453977 | 0.206763884 | -0.68896934 |
| Gm25453 | 0 | NA | NA | NA |
| Neto1 | 920.988223 | 0.430855104 | 0.113378503 | 3.800148109 |
| Cbln2 | 282.7539475 | 0.157792155 | 0.113584484 | 1.389205195 |
| Gm5096 | 0.150520013 | 0.037075864 | 2.857840531 | 0.012973384 |
| Gm24987 | 0 | NA | NA | NA |
| Socs6 | 448.8133279 | 0.035404925 | 0.096942243 | 0.365216687 |
| Rttn | 192.02834 | 0.025083657 | 0.099216276 | 0.252817965 |
| Mir6359 | 0.075043205 | 0.517957688 | 3.724945049 | 0.139051095 |
| Cd226 | 7.086717468 | -0.514515281 | 0.477655583 | -1.077167942 |
| 9330132A10Rik | 749.6793612 | 0.36963464 | 0.098706579 | 3.744782183 |
| Dok6 | 388.457898 | 0.448219652 | 0.106585856 | 4.205245112 |
| Gm16296 | 5.46580188 | 0.240856517 | 0.458164804 | 0.525698428 |
| Gm16297 | 0 | NA | NA | NA |
| Gm22694 | 0 | NA | NA | NA |
| Gm27037 | 0 | NA | NA | NA |
| Gm23859 | 0 | NA | NA | NA |
| Eif3s6-ps2 | 6.153886045 | -0.460043661 | 0.504872099 | -0.911208329 |
| Gm25418 | 0 | NA | NA | NA |
| Tmx3 | 1329.967682 | -0.105668198 | 0.089773383 | -1.177054872 |
| 1700030N03Rik | 0.163190422 | -0.955151543 | 3.396300892 | -0.281232898 |
| Gm25431 | 0 | NA | NA | NA |
| Ighmbp2 | 158.9101558 | 0.167002058 | 0.11417272 | 1.462714198 |
| Mrpl21 | 165.3835846 | -0.298210401 | 0.115787526 | -2.575496786 |
| Mir6984 | 0.950361064 | 1.691603433 | 1.204367901 | 1.404557055 |
| Cpt1a | 573.796439 | -0.513321724 | 0.142515709 | -3.601860642 |
| Mtl5 | 3.167783942 | -0.236330688 | 0.689185841 | -0.34291286 |
| Gal | 59.25410188 | -1.555479337 | 0.364469527 | -4.267789822 |
| Ppp6r3 | 1233.355573 | -0.059755319 | 0.061614385 | -0.969827406 |
| Gm23940 | 0 | NA | NA | NA |
| Lrp5 | 186.6160727 | -0.303943584 | 0.14997788 | -2.02658942 |
| 1810055G02Rik | 195.6912383 | -0.293092803 | 0.099707756 | -2.939518593 |
| Suv420h1 | 939.8295677 | -0.034693583 | 0.074239798 | -0.467317854 |
| Gm16066 | 1.927026077 | 1.119106687 | 0.774848488 | 1.444290986 |
| Chka | 1064.49316 | 0.341029707 | 0.169049085 | 2.017341339 |
| Tcirg1 | 611.8661545 | -1.106663033 | 0.165366566 | -6.692181238 |
| Ndufs8 | 433.8570073 | 0.267705756 | 0.111856316 | 2.393300309 |
| Aldh3b1 | 78.94818307 | -0.790639986 | 0.226190016 | -3.4954681 |
| Unc93b1 | 424.5399235 | -0.860526882 | 0.151399675 | -5.683809326 |
| Aldh3b3 | 1.060607173 | -0.183601117 | 0.913513253 | -0.200983528 |
| Aldh3b2 | 8.359127856 | 0.255996946 | 0.397052861 | 0.644742731 |

| | | | | |
|---|---|---|---|---|
| Acy3 | 20.90646341 | 0.161305112 | 0.240544517 | 0.6705832 |
| Tbx10 | 0.479449896 | -0.732194543 | 1.425568715 | -0.513615749 |
| Nudt8 | 43.46713457 | 0.014267995 | 0.18604305 | 0.076691902 |
| Gm16312 | 0.283672814 | -0.871029422 | 1.622485371 | -0.53684886 |
| Doc2g | 55.67602762 | -0.642096366 | 0.619768744 | -1.03602573 |
| Ndufv1 | 1083.51582 | 0.018101115 | 0.065780766 | 0.275173374 |
| Gstp1 | 541.1525819 | 0.049639374 | 0.139650557 | 0.355454176 |
| Gstp2 | 1.939495377 | -1.604470804 | 0.981579418 | -1.634580732 |
| BC021614 | 1.429393014 | 0.845392052 | 0.831163686 | 1.017118608 |
| Gm23927 | 0 | NA | NA | NA |
| Cabp2 | 3.994848174 | -0.63616191 | 0.737697567 | -0.862361405 |
| Cdk2ap2 | 150.3217526 | -0.454018944 | 0.133841092 | -3.392223855 |
| Pitpnm1 | 2202.207616 | 0.553455382 | 0.070206775 | 7.883218988 |
| Aip | 570.1830399 | 0.099602771 | 0.082751381 | 1.20363877 |
| Tmem134 | 248.6189701 | 0.026219197 | 0.097262045 | 0.269572751 |
| Cabp4 | 3.477720489 | -0.181115314 | 0.526854693 | -0.343767107 |
| Gpr152 | 2.789673889 | -2.142569084 | 0.797899411 | -2.685262146 |
| Coro1b | 812.1320772 | -0.579055231 | 0.091981275 | -6.295359936 |
| Ptprcap | 24.66290501 | -0.504243344 | 0.330492372 | -1.525733682 |
| Rps6kb2 | 211.501265 | 0.219467246 | 0.150459649 | 1.458645204 |
| Carns1 | 117.9629228 | 0.225898033 | 0.129223511 | 1.748118678 |
| Gm17552 | 4.682108124 | 0.19642206 | 0.474338304 | 0.414096981 |
| Tbc1d10c | 30.14122667 | 0.216026927 | 0.20536086 | 1.051938168 |
| Ppp1ca | 1432.944828 | -0.202553803 | 0.084214613 | -2.40520969 |
| Rad9a | 127.978386 | -0.725908344 | 0.120525437 | -6.022864254 |
| Clcf1 | 180.1899159 | -2.317607569 | 0.351630026 | -6.591040008 |
| Gm26115 | 0.127364494 | -0.44382566 | 3.724945049 | -0.119149586 |
| Pold4 | 108.3674615 | -0.580238545 | 0.137225108 | -4.228370116 |
| Ssh3 | 269.6560211 | 0.145679544 | 0.151924176 | 0.958896391 |
| Mir6985 | 0 | NA | NA | NA |
| Ankrd13d | 332.9753029 | 0.552737094 | 0.113709308 | 4.860966105 |
| Adrbk1 | 2117.90304 | 0.156388106 | 0.059852729 | 2.61288178 |
| Kdm2a | 1799.114924 | -0.122120873 | 0.062384154 | -1.957562381 |
| Rhod | 32.92487569 | -1.819698145 | 0.302870618 | -6.008169947 |
| A930001C03Rik | 2.701377055 | -1.501176763 | 0.703582104 | -2.133619878 |
| Syt12 | 590.78774 | -0.152890924 | 0.116849222 | -1.308446236 |
| 2010003K11Rik | 0.302393035 | -1.258104782 | 1.867480761 | -0.673690893 |
| Pcx | 931.2379665 | 0.465736705 | 0.105188944 | 4.427620322 |
| Lrfn4 | 397.610308 | -0.080347713 | 0.093494224 | -0.859386922 |
| Rce1 | 129.1447744 | -0.509703733 | 0.147982263 | -3.444356924 |
| Mir6986 | 0.044191549 | 0.037062677 | 3.724945049 | 0.009949859 |
| Gm960 | 25.50013179 | 0.779010705 | 0.231314035 | 3.367762382 |
| Sptbn2 | 9930.18535 | 0.758493128 | 0.103401134 | 7.335443002 |
| Gm21992 | 41.25034136 | 0.363018572 | 0.191343587 | 1.897207937 |
| Rbm4b | 188.1038022 | -0.366538742 | 0.092264986 | -3.972674315 |
| Gm37206 | 190.853055 | 0.391227273 | 0.202244147 | 1.934430627 |
| Rbm4 | 178.0962253 | 0.064817085 | 0.17652058 | 0.367192795 |

| | | | | |
|---|---|---|---|---|
| Rbm14 | 411.7655184 | -0.099111054 | 0.086916488 | -1.140302099 |
| Gm21844 | 0.416305032 | -2.453185822 | 1.432209959 | -1.712867451 |
| Ccs | 117.7739688 | -0.125450817 | 0.138113117 | -0.908319349 |
| Ccdc87 | 18.82604018 | 0.037455809 | 0.281582408 | 0.133018995 |
| Ctsf | 447.0332469 | 0.548619492 | 0.086643087 | 6.331947674 |
| Actn3 | 10.89038947 | 0.237714906 | 0.335138588 | 0.709303299 |
| Zdhhc24 | 191.9093747 | -0.244818687 | 0.094140606 | -2.600564186 |
| Bbs1 | 425.8384116 | 0.054794687 | 0.079853124 | 0.686193397 |
| Gm25334 | 0 | NA | NA | NA |
| Dpp3 | 394.3142871 | -0.214367137 | 0.081580295 | -2.627682789 |
| Peli3 | 250.3912601 | 0.209142181 | 0.093728859 | 2.23135312 |
| Mrpl11 | 282.4075369 | 0.067036404 | 0.088295989 | 0.759223661 |
| Npas4 | 211.216604 | -0.795739838 | 0.600981561 | -1.324066976 |
| Slc29a2 | 100.667391 | -0.261407697 | 0.126812589 | -2.061370241 |
| B4gat1 | 647.79448 | 0.542991174 | 0.063986288 | 8.486055239 |
| Brms1 | 180.8653628 | -0.241455053 | 0.103724642 | -2.32784659 |
| Rin1 | 659.0041585 | 0.142734486 | 0.180923391 | 0.788922236 |
| Cd248 | 53.27825101 | -0.738570159 | 0.201062644 | -3.673333564 |
| Tmem151a | 1868.678076 | 0.28786912 | 0.069537427 | 4.13977237 |
| Yif1a | 156.5502458 | -0.416247295 | 0.147169374 | -2.828355416 |
| Cnih2 | 897.9476727 | 0.533394537 | 0.099341512 | 5.369301598 |
| Rab1b | 1115.051491 | -0.003236235 | 0.097617782 | -0.033152108 |
| Klc2 | 2105.216524 | 0.36641008 | 0.070258876 | 5.215142926 |
| Pacs1 | 821.0770873 | 0.383235428 | 0.07194796 | 5.326564189 |
| Gm27468 | 0.909851095 | 0.489303358 | 1.242209747 | 0.393897535 |
| Sf3b2 | 2394.762425 | -0.04876722 | 0.061664375 | -0.790849166 |
| Gal3st3 | 390.9376513 | 0.873861027 | 0.112176466 | 7.790056677 |
| Catsper1 | 0 | NA | NA | NA |
| Cst6 | 23.01852267 | 0.78334262 | 0.258801788 | 3.02680529 |
| Banf1 | 258.7929532 | -0.251489581 | 0.09545084 | -2.634755037 |
| Eif1ad | 254.8369414 | -0.221484564 | 0.107589269 | -2.058612032 |
| Sart1 | 490.457391 | 0.009198107 | 0.148017677 | 0.06214195 |
| Tsga10ip | 0.362047852 | 1.301885436 | 1.537570634 | 0.846715856 |
| 4930481A15Rik | 2.199134214 | 2.65032614 | 0.851354398 | 3.113070358 |
| Gm6293 | 0.045882455 | 0.517959912 | 3.724945049 | 0.139051692 |
| Drap1 | 882.2497186 | 0.105288595 | 0.090890352 | 1.158413326 |
| AI837181 | 578.5093112 | 0.198688396 | 0.114406694 | 1.736685056 |
| Gm25432 | 0 | NA | NA | NA |
| Fosl1 | 112.2972541 | -2.074980185 | 0.588677462 | -3.524816758 |
| Ccdc85b | 470.1620099 | 0.060721669 | 0.106439631 | 0.570479896 |
| Fibp | 216.1359764 | -0.086497467 | 0.078476566 | -1.102207599 |
| Ctsw | 32.65133826 | -0.257025893 | 0.248601836 | -1.033885741 |
| Efemp2 | 186.5996474 | -0.97240881 | 0.153530897 | -6.333635969 |
| Mus81 | 195.7583366 | -0.090871544 | 0.117990056 | -0.770162731 |
| Cfl1 | 6985.640524 | -0.046834151 | 0.069333657 | -0.675489405 |
| Snx32 | 463.4435586 | 0.366066124 | 0.08536035 | 4.288479668 |
| 1700020D05Rik | 15.41874029 | 0.338010842 | 0.433795552 | 0.779193886 |

| | | | | |
|---|---|---|---|---|
| Ovol1 | 1.251933375 | -1.057403785 | 1.192990918 | -0.886346886 |
| Ap5b1 | 86.41281867 | -0.617040448 | 0.137202155 | -4.497308734 |
| Rnaseh2c | 82.70313133 | -0.026184893 | 0.180953277 | -0.144705273 |
| Kat5 | 511.7440207 | 0.015412909 | 0.064965525 | 0.237247515 |
| Rela | 372.9586236 | -0.303439919 | 0.16152093 | -1.878641479 |
| Sipa1 | 217.0465711 | -0.482431688 | 0.148139576 | -3.256602316 |
| Pcnxl3 | 996.3058461 | -0.016748854 | 0.072669363 | -0.230480264 |
| Mir6987 | 0 | NA | NA | NA |
| Map3k11 | 329.8028705 | -0.288797221 | 0.108884365 | -2.652329569 |
| Kcnk7 | 0.811426433 | -1.32281129 | 1.169049074 | -1.131527598 |
| Ehbp1l1 | 284.648246 | -0.473520064 | 0.109464671 | -4.325779795 |
| Gm16538 | 0.233003758 | -1.874069331 | 2.523043414 | -0.742781246 |
| Fam89b | 0.325597809 | 1.009232137 | 1.757215222 | 0.5743361 |
| Sssca1 | 172.9191299 | -0.026502422 | 0.103320432 | -0.256507074 |
| Ltbp3 | 514.8847917 | -0.012074263 | 0.076027302 | -0.158814825 |
| Scyl1 | 711.1667025 | -0.097642238 | 0.073062959 | -1.336412325 |
| Malat1 | 158003.9038 | 0.087453325 | 0.226883796 | 0.385454256 |
| Gm20417 | 3.344837808 | 0.156411131 | 0.925319627 | 0.169034706 |
| Gm27694 | 0 | NA | NA | NA |
| Gm37376 | 17.03796044 | 0.704846826 | 0.96758917 | 0.728456713 |
| Gm27702 | 0 | NA | NA | NA |
| Neat1 | 2624.857955 | -0.120163695 | 0.288988532 | -0.415807834 |
| Gm27533 | 0 | NA | NA | NA |
| Gm27505 | 0 | NA | NA | NA |
| Gm28006 | 0 | NA | NA | NA |
| Frmd8os | 2.120407615 | -1.192603267 | 0.826358334 | -1.443203532 |
| Frmd8 | 356.4512706 | -1.192873221 | 0.186180627 | -6.407074899 |
| Slc25a45 | 48.19350137 | -0.793793361 | 0.225989562 | -3.512522241 |
| Tigd3 | 29.86989943 | 0.31613847 | 0.214274017 | 1.475393398 |
| Dpf2 | 712.9918355 | -0.308412905 | 0.067417919 | -4.574642935 |
| Cdc42ep2 | 113.290494 | -0.34464606 | 0.165347677 | -2.084371949 |
| Pola2 | 84.75758802 | -0.246652513 | 0.144170872 | -1.71083457 |
| Slc22a20 | 0.141071297 | -0.443820801 | 3.564043909 | -0.12452731 |
| Capn1 | 318.3485814 | -0.397093567 | 0.10001265 | -3.970433398 |
| Gm10814 | 14.06531991 | 0.407642435 | 0.307606256 | 1.325208534 |
| Gm8034 | 7.330356508 | -0.249919498 | 0.423295104 | -0.590414335 |
| Syvn1 | 680.7779014 | -0.186937891 | 0.082568324 | -2.264038836 |
| Mir6988 | 0 | NA | NA | NA |
| Mrpl49 | 285.2862106 | -0.204477464 | 0.092828961 | -2.202733513 |
| Fau | 1011.116782 | -0.145485308 | 0.099953779 | -1.455525834 |
| Znhit2 | 165.2069863 | 0.071447051 | 0.098178267 | 0.72772777 |
| Tm7sf2 | 103.8839401 | 0.653794505 | 0.152557556 | 4.285559632 |
| Vps51 | 374.0248763 | 0.059178358 | 0.071929042 | 0.822732459 |
| Gm28374 | 0.132725991 | -0.92409061 | 2.971216227 | -0.311014258 |
| BC048609 | 0.046345709 | -0.443818988 | 3.724945049 | -0.119147795 |
| Zfpl1 | 181.431277 | 0.115173475 | 0.11131095 | 1.034700318 |
| Cdca5 | 6.165523022 | -0.710898582 | 0.478284617 | -1.486350505 |

| | | | | |
|---|---|---|---|---|
| Naaladl1 | 0.666154845 | 0.038466192 | 1.36584345 | 0.028162959 |
| Sac3d1 | 83.16707754 | -0.242292271 | 0.144162248 | -1.680691536 |
| Snx15 | 211.7518732 | 0.125811501 | 0.084003973 | 1.497685136 |
| Arl2 | 383.7572647 | 0.015186635 | 0.098976841 | 0.153436242 |
| Batf2 | 180.1949883 | -1.216678163 | 0.370375299 | -3.284987329 |
| n-R5s19 | 0 | NA | NA | NA |
| Gm5510 | 2.351247315 | -0.591247081 | 0.646367942 | -0.914722162 |
| 1700123I01Rik | 7.996023092 | -0.347112527 | 0.323402839 | -1.07331317 |
| Gpha2 | 0.377319533 | 1.350881666 | 1.531744327 | 0.881923727 |
| Ppp2r5b | 1407.304023 | 0.239864015 | 0.053710825 | 4.465841191 |
| Atg2a | 650.2802182 | 0.068171414 | 0.061700336 | 1.104879144 |
| Mir194-2 | 0.099106094 | -0.93550548 | 3.724188147 | -0.251197158 |
| Mir192 | 0.254293348 | 0.517972323 | 2.47280135 | 0.209467826 |
| Gm14963 | 2.026491629 | -0.353058805 | 0.789314516 | -0.447298 |
| Ehd1 | 703.3125589 | -0.281135723 | 0.118136528 | -2.379752713 |
| Cdc42bpg | 176.7180393 | 0.127760975 | 0.154938814 | 0.824589859 |
| Men1 | 585.1800558 | -0.062043385 | 0.054558679 | -1.137186341 |
| Map4k2 | 638.3859077 | 0.427197225 | 0.099348144 | 4.300002075 |
| Mir6989 | 0 | NA | NA | NA |
| Gm14966 | 12.5179983 | 0.19772699 | 0.330786719 | 0.597747667 |
| Sf1 | 2104.915733 | -0.017091799 | 0.082516632 | -0.207131567 |
| Pygm | 267.9090404 | 0.389343646 | 0.172688239 | 2.254604296 |
| Rasgrp2 | 263.4585251 | 0.75877699 | 0.200249935 | 3.789149742 |
| Gm14965 | 2.017609801 | -1.851246856 | 0.803656244 | -2.303530731 |
| Nrxn2 | 6243.867174 | 0.293399695 | 0.076490323 | 3.83577533 |
| Gm14964 | 23.8586541 | 0.267145135 | 0.277587796 | 0.96238069 |
| Slc22a12 | 5.133294896 | -0.122537445 | 0.545406525 | -0.224671762 |
| Gm14967 | 0 | NA | NA | NA |
| Gm26460 | 0.404356784 | -0.015771186 | 1.501706587 | -0.010502175 |
| Hmgb1-ps4 | 0.663061751 | -1.52622636 | 1.270528876 | -1.201252792 |
| 2410152P15Rik | 0 | NA | NA | NA |
| Gm14968 | 0.11037557 | 0.517957688 | 3.724945049 | 0.139051095 |
| Rps6ka4 | 374.0853682 | -0.091279792 | 0.081136989 | -1.125008364 |
| mmu-mir-5046 | 0 | NA | NA | NA |
| Ccdc88b | 121.8708714 | -0.71180411 | 0.159333115 | -4.46739592 |
| Prdx5 | 1538.036727 | -0.176714928 | 0.0861099 | -2.052202223 |
| Trmt112 | 108.60505 | -0.252505351 | 0.162325536 | -1.55554916 |
| Esrra | 457.3522009 | -0.134044281 | 0.085992793 | -1.55878507 |
| Mir6990 | 0 | NA | NA | NA |
| Tex40 | 21.48008408 | 0.805184211 | 0.325739184 | 2.471867836 |
| Kcnk4 | 51.05303909 | 1.040339919 | 0.225766885 | 4.608027067 |
| Gpr137 | 725.8468719 | 0.020237339 | 0.061037332 | 0.331556738 |
| Bad | 198.4137653 | -0.187302159 | 0.121866707 | -1.536942807 |
| Plcb3 | 323.134626 | -0.339608934 | 0.12067676 | -2.814203283 |
| Ppp1r14b | 149.5878577 | -0.949824877 | 0.167431235 | -5.672925229 |
| Fkbp2 | 300.3276835 | 0.263790364 | 0.283700889 | 0.929818602 |
| Vegfb | 398.1291403 | 0.054864161 | 0.107883215 | 0.50855141 |

| | | | | |
|---|---|---|---|---|
| Dnajc4 | 95.01009773 | -0.057004478 | 0.137152512 | -0.415628392 |
| Nudt22 | 67.37727446 | -0.302363783 | 0.146302602 | -2.06670133 |
| Trpt1 | 56.60693119 | 0.076122099 | 0.155921916 | 0.488206538 |
| Fermt3 | 175.2517023 | -0.513809748 | 0.141750232 | -3.624754194 |
| Stip1 | 1604.279865 | -0.382167083 | 0.096616051 | -3.955523747 |
| Macrod1 | 69.5949187 | 0.353191291 | 0.147743221 | 2.390575271 |
| Flrt1 | 608.6636325 | 0.210944537 | 0.127792898 | 1.650674958 |
| Otub1 | 1632.950069 | 0.265177187 | 0.065981849 | 4.018941407 |
| Cox8a | 2092.571017 | 0.174852765 | 0.074970684 | 2.33228184 |
| Naa40 | 276.9429562 | -0.067512943 | 0.123249934 | -0.547772653 |
| Gm22113 | 0 | NA | NA | NA |
| Gm23997 | 0 | NA | NA | NA |
| Rcor2 | 69.31615873 | 1.047067477 | 0.194213178 | 5.39133073 |
| Mark2 | 1468.643762 | 0.04290317 | 0.078979911 | 0.543216239 |
| Gm17227 | 0.066050875 | -0.443818988 | 3.724945049 | -0.119147795 |
| AI846148 | 273.2784663 | -0.172338547 | 0.10254039 | -1.68068941 |
| 1700105P06Rik | 6.754696947 | 0.046611397 | 0.434378445 | 0.107305963 |
| Rab11b-ps2 | 191.3767199 | 0.207023162 | 0.125922638 | 1.644050389 |
| 2700081O15Rik | 415.5788108 | -0.018946839 | 0.105726319 | -0.179206452 |
| Mir6991 | 0.662460401 | 1.274495168 | 1.456696231 | 0.874921717 |
| Rtn3 | 9090.002618 | 0.442900848 | 0.05151687 | 8.597200332 |
| Atl3 | 817.3787042 | -0.182617856 | 0.062750651 | -2.910214511 |
| Pla2g16 | 615.7304892 | -0.157921088 | 0.130466364 | -1.210435264 |
| Lgals12 | 1.476747611 | 0.81447524 | 1.045508249 | 0.779023256 |
| Hrasls5 | 1.687958028 | 2.045244882 | 0.891546657 | 2.294041334 |
| Gm16437 | 26.12384685 | 0.013875758 | 0.261470786 | 0.053068101 |
| Slc22a19 | 0 | NA | NA | NA |
| Gm15967 | 0 | NA | NA | NA |
| Gm15968 | 0 | NA | NA | NA |
| Slc22a26 | 0.04393175 | 0.037062677 | 3.724945049 | 0.009949859 |
| Slc22a27 | 0 | NA | NA | NA |
| Gm24951 | 0 | NA | NA | NA |
| Slc22a28 | 0 | NA | NA | NA |
| Slc22a29 | 1.523828694 | -1.34166227 | 0.894822856 | -1.499360751 |
| Slc22a30 | 0.083526043 | 0.037073798 | 3.724945049 | 0.009952844 |
| Gm6425 | 1.341472408 | -1.804865412 | 1.02006738 | -1.769359013 |
| Slc22a8 | 69.67438683 | 0.641929753 | 0.29680352 | 2.162810446 |
| Slc22a6 | 62.43237234 | 0.409778868 | 0.276253775 | 1.483342148 |
| Chrm1 | 1319.762724 | 0.9826616 | 0.177396069 | 5.539365131 |
| Gm22820 | 0.118974955 | 0.51795324 | 3.724945049 | 0.139049901 |
| Slc3a2 | 1681.539151 | -0.771540023 | 0.142031745 | -5.432166035 |
| Gm25855 | 7.213846394 | 0.181765401 | 0.392619939 | 0.462955094 |
| Gm25894 | 7.96667369 | 0.225018379 | 0.41853529 | 0.53763299 |
| Gm24452 | 7.467891793 | 0.866822378 | 0.397158708 | 2.182559164 |
| Gm24453 | 6.941141185 | -0.246908081 | 0.360470117 | -0.684961303 |
| Gm22744 | 7.168190812 | -0.478528381 | 0.411273338 | -1.163528817 |
| Gm22680 | 5.248670073 | 0.193775557 | 0.443499159 | 0.436924294 |

| | | | | |
|---|---|---|---|---|
| Gm23246 | 6.637239202 | 0.3168515 | 0.409728046 | 0.773321481 |
| Snord22 | 166.9869455 | -0.491733969 | 0.242861036 | -2.024754475 |
| Wdr74 | 196.0682104 | -0.23960381 | 0.114296053 | -2.096343689 |
| 1700092M07Rik | 1.245725704 | 0.844688918 | 1.100483295 | 0.767561781 |
| Stx5a | 534.5973207 | -0.122188971 | 0.081969842 | -1.490657637 |
| Mir6992 | 0 | NA | NA | NA |
| Nxf1 | 945.8212657 | -0.01346449 | 0.068642895 | -0.196152707 |
| Taf6l | 170.5846256 | -0.478495348 | 0.111521179 | -4.290623115 |
| Polr2g | 319.7014694 | -0.053229012 | 0.078381323 | -0.67910326 |
| Zbtb3 | 40.55560979 | -0.743989949 | 0.199077681 | -3.737184127 |
| Ttc9c | 517.0751682 | -0.146567505 | 0.078028384 | -1.878387041 |
| Hnrnpul2 | 2234.773551 | 0.010897275 | 0.111304789 | 0.097904813 |
| Gng3 | 1184.138002 | 0.313076081 | 0.071874816 | 4.355852306 |
| Bscl2 | 767.6870937 | -0.083023904 | 0.069346548 | -1.197231953 |
| Lrrn4cl | 17.94000658 | -0.775642972 | 0.253517798 | -3.059520779 |
| Ubxn1 | 553.7697328 | -0.179689079 | 0.091878476 | -1.955725496 |
| Uqcc3 | 121.6763236 | -0.140570831 | 0.141539631 | -0.993155273 |
| Lbhd1 | 49.53080462 | 0.187620292 | 0.180485448 | 1.03953141 |
| Snora57 | 16.08879982 | -0.554680532 | 0.320067257 | -1.733012421 |
| Mir5136 | 0.568319789 | 1.309411772 | 1.374140872 | 0.952894858 |
| 1810009A15Rik | 2.938865308 | 0.760406636 | 0.626279256 | 1.214165452 |
| Ints5 | 306.8553324 | -0.325522659 | 0.102238667 | -3.183948581 |
| Ganab | 1308.618353 | -0.270557655 | 0.062292236 | -4.343360815 |
| B3gat3 | 910.6532437 | 0.086607962 | 0.091833008 | 0.943102743 |
| Rom1 | 80.3855847 | -0.328200236 | 0.163555241 | -2.006662907 |
| Eml3 | 209.4625949 | -0.242699871 | 0.108986481 | -2.226880511 |
| Mta2 | 687.4573254 | -0.339760137 | 0.080212609 | -4.235744734 |
| Tut1 | 264.2884532 | -0.262488256 | 0.095300717 | -2.754315659 |
| Eef1g | 1677.123862 | -0.378836937 | 0.083676108 | -4.527420649 |
| Ahnak | 4057.702084 | -1.209676156 | 0.172884683 | -6.997011751 |
| Scgb1a1 | 0 | NA | NA | NA |
| Asrgl1 | 1213.125555 | 0.207741278 | 0.119971694 | 1.731585766 |
| Gm9750 | 3.69099669 | -0.145992946 | 0.537386246 | -0.271672279 |
| Gm24543 | 0 | NA | NA | NA |
| Pcna-ps2 | 9.410251089 | -0.46298291 | 0.335772713 | -1.378858056 |
| Gm2617 | 0.087261593 | 0.51795324 | 3.724945049 | 0.139049901 |
| Stxbp3-ps | 23.11005125 | -0.54309959 | 0.231022308 | -2.350853451 |
| Gm29231 | 0.2218918 | 0.51795986 | 3.143193936 | 0.164787751 |
| Gm6445 | 1.943633249 | 0.843155448 | 0.723655487 | 1.165133773 |
| Scgb2a2 | 0.152314784 | 0.51796321 | 2.84727539 | 0.181915389 |
| Gm22369 | 0 | NA | NA | NA |
| Incenp | 144.1045022 | 0.212654428 | 0.128046607 | 1.660758016 |
| Gm24484 | 0.061920613 | -0.443816764 | 3.724945049 | -0.119147198 |
| Fth1 | 13460.26008 | -0.285585321 | 0.167192796 | -1.708119771 |
| Best1 | 1.289165993 | 0.380746085 | 0.786154991 | 0.484314276 |
| Rab3il1 | 131.0561169 | -0.465948737 | 0.170592445 | -2.731356227 |
| Fads3 | 258.367544 | -0.241970416 | 0.089489085 | -2.703909834 |

| | | | | |
|---|---|---|---|---|
| Fads2 | 1025.070797 | 0.071434036 | 0.119574468 | 0.597402078 |
| Fads1 | 1910.999182 | 0.169867041 | 0.098547317 | 1.72371046 |
| Mir6993 | 1.831497397 | -0.411671051 | 0.800734791 | -0.514116603 |
| Gm10143 | 2.039768525 | 0.431923354 | 0.83241125 | 0.518882168 |
| Fen1 | 177.2425644 | 0.02556908 | 0.143741675 | 0.177882161 |
| Tmem258 | 152.1698681 | -0.359890476 | 0.166034229 | -2.16756797 |
| Myrf | 1402.777588 | -0.276386374 | 0.136521875 | -2.024484157 |
| Dagla | 1427.846811 | 0.531388367 | 0.113668481 | 4.674896346 |
| Syt7 | 3169.726564 | 0.61217105 | 0.104129229 | 5.878955009 |
| Gm26859 | 2.155457834 | -0.086012626 | 0.759478704 | -0.11325219 |
| Lrrc10b | 214.7262662 | 1.127813433 | 0.21734342 | 5.189084774 |
| Ppp1r32 | 18.15485456 | -0.445320958 | 0.229866578 | -1.937301896 |
| Sdhaf2 | 369.9013947 | -0.352947977 | 0.072896292 | -4.841782325 |
| Cpsf7 | 982.7429653 | 0.041374935 | 0.080052038 | 0.516850489 |
| Tmem216 | 43.25074293 | -0.671266872 | 0.187634118 | -3.577530986 |
| Tmem138 | 84.89562639 | -0.235669081 | 0.122543602 | -1.923144719 |
| Cyb561a3 | 70.26552882 | -0.632636897 | 0.192721327 | -3.282651208 |
| Tkfc | 98.20098442 | 0.138944131 | 0.13452422 | 1.032855872 |
| Ddb1 | 2897.624334 | -0.145568107 | 0.0553439 | -2.630246639 |
| Vwce | 6.549817523 | -0.535241205 | 0.461433414 | -1.159953286 |
| Pga5 | 0.066050875 | -0.443818988 | 3.724945049 | -0.119147795 |
| Vps37c | 286.9571528 | -0.168585563 | 0.09950935 | -1.694168077 |
| Cd5 | 21.9583943 | -0.54840315 | 0.269503785 | -2.034862514 |
| A430093F15Rik | 5.018136537 | -0.781097655 | 0.4875154 | -1.602200986 |
| Cd6 | 55.03509925 | -0.057357687 | 0.199428875 | -0.28760974 |
| Slc15a3 | 354.5772757 | -0.944004646 | 0.277485322 | -3.401998485 |
| Tmem132a | 2470.423893 | 0.520984806 | 0.093474607 | 5.573543686 |
| Tmem109 | 292.8556633 | -0.248053102 | 0.122795837 | -2.020044878 |
| Prpf19 | 1647.79328 | -0.010689316 | 0.065770453 | -0.162524591 |
| Zp1 | 0.369197529 | 0.457484564 | 1.432489053 | 0.319363393 |
| Ptgdr2 | 2.273986841 | -0.677597219 | 0.691255413 | -0.980241465 |
| Ccdc86 | 181.1409562 | -0.334716314 | 0.133159428 | -2.513650882 |
| Ms4a10 | 0.044544648 | 0.517957688 | 3.724945049 | 0.139051095 |
| Ms4a15 | 28.11515156 | -1.737163121 | 1.140695603 | -1.52289806 |
| Ms4a18 | 0.263774563 | -1.952630542 | 2.376524226 | -0.821632921 |
| Gm28347 | 2.460283742 | -1.6557509 | 0.652786861 | -2.53643417 |
| AW112010 | 268.3498837 | -0.967135981 | 0.362489899 | -2.668035676 |
| Ms4a8a | 62.44774592 | -2.133971888 | 0.664959257 | -3.209176901 |
| Gm336 | 28.43566127 | -0.127877079 | 0.199662556 | -0.640466002 |
| 1700017D01Rik | 0.314055832 | -0.021049594 | 1.715556695 | -0.012269833 |
| Gm6501 | 23.72387051 | -0.364292162 | 0.266242986 | -1.368269518 |
| 1700025F22Rik | 0.715113835 | -1.009683138 | 1.325663137 | -0.761643822 |
| Ms4a13 | 0 | NA | NA | NA |
| 4930526L06Rik | 0.358247177 | 0.044541699 | 1.574073781 | 0.028297085 |
| Ms4a12 | 0.470940457 | -2.184819206 | 1.53191496 | -1.426201365 |
| Ms4a1 | 1.60259925 | -1.208084533 | 0.735724572 | -1.642033689 |
| Ms4a5 | 0 | NA | NA | NA |

| | | | | |
|---|---|---|---|---|
| Ms4a14 | 14.20065461 | -1.706251827 | 0.411550169 | -4.145914533 |
| Ms4a7 | 116.0378029 | -2.113789034 | 0.295501881 | -7.153216858 |
| Gm28935 | 0.610760527 | -0.035206733 | 1.343513342 | -0.026204975 |
| Ms4a4a | 77.63966184 | -1.152836413 | 0.342361476 | -3.367307633 |
| Gm5692 | 0.269752699 | -1.344850337 | 1.76912721 | -0.760177295 |
| Ms4a4c | 622.4266349 | -1.701657834 | 0.388085047 | -4.38475496 |
| Gm5511 | 2.864211081 | 0.55023294 | 0.55258126 | 0.995750272 |
| Ms4a4b | 229.0039815 | -0.928258615 | 0.348099175 | -2.666649853 |
| Gm37387 | 7.980888526 | -1.252677357 | 0.504354007 | -2.483726388 |
| Ms4a6c | 638.4433901 | -1.278879241 | 0.280789303 | -4.554586763 |
| BE692007 | 14.17246462 | -0.486371724 | 0.369149108 | -1.317548149 |
| Gm8369 | 26.70344983 | -0.350266866 | 0.2813711 | -1.244857294 |
| Gm19261 | 0.679959982 | -1.071241319 | 1.107869058 | -0.966938566 |
| Ms4a6b | 404.189003 | -1.053299611 | 0.267885229 | -3.931906263 |
| Ms4a4d | 12.73072494 | -1.396501287 | 0.390551452 | -3.575716543 |
| Gm10212 | 1.028441291 | -1.948960818 | 2.079159714 | -0.937379079 |
| Gm38275 | 1.398451413 | -3.187196249 | 1.030765808 | -3.092066329 |
| Ms4a6d | 780.7598453 | -1.451895265 | 0.345805404 | -4.198590448 |
| Gm24599 | 0 | NA | NA | NA |
| Ms4a2 | 1.450910466 | -0.796239553 | 0.790505362 | -1.00725383 |
| Ms4a3 | 0.922047842 | -0.352900725 | 0.974385711 | -0.362177648 |
| Oosp2 | 0.454235466 | -0.046571654 | 1.308074399 | -0.035603215 |
| Oosp1 | 0.420049603 | -1.970620276 | 1.596107823 | -1.234641074 |
| Gm26183 | 0 | NA | NA | NA |
| Gm25443 | 0 | NA | NA | NA |
| Oosp3 | 0 | NA | NA | NA |
| Gm19224 | 0.308730509 | -0.177816597 | 3.712624185 | -0.047895124 |
| Gif | 0.046345709 | -0.443818988 | 3.724945049 | -0.119147795 |
| Mrpl16 | 136.9670042 | 0.064076116 | 0.118317165 | 0.5415623 |
| Stx3 | 268.7199862 | -0.106625715 | 0.114437039 | -0.931741295 |
| Olfr1417 | 0.045882455 | 0.517959912 | 3.724945049 | 0.139051692 |
| Olfr1418 | 0 | NA | NA | NA |
| Olfr1419 | 0.385885129 | -1.262218509 | 1.54432789 | -0.817325464 |
| Gm6365 | 6.926587766 | 0.064418371 | 0.360760068 | 0.178562919 |
| Olfr1420 | 0.092228164 | 0.037073798 | 3.724945049 | 0.009952844 |
| Patl1 | 512.1726233 | -0.389818773 | 0.08333889 | -4.67751339 |
| mmu-mir-6994 | 0.26492599 | 0.037530493 | 1.820983064 | 0.020610018 |
| Gm22272 | 0 | NA | NA | NA |
| Osbp | 1002.556198 | -0.083167559 | 0.047971194 | -1.733697922 |
| Olfr1423 | 0 | NA | NA | NA |
| Olfr1424 | 0.049553047 | -0.443818988 | 3.724945049 | -0.119147795 |
| Olfr1425 | 0.045882455 | 0.517959912 | 3.724945049 | 0.139051692 |
| Olfr1426 | 0 | NA | NA | NA |
| Olfr1427 | 0 | NA | NA | NA |
| Olfr1428 | 0.090277459 | -0.443823436 | 3.724945049 | -0.119148989 |
| Olfr76 | 0.045882455 | 0.517959912 | 3.724945049 | 0.139051692 |
| Olfr1431 | 0 | NA | NA | NA |

| | | | | |
|---|---|---|---|---|
| Olfr1432 | 0 | NA | NA | NA |
| Olfr262 | 0 | NA | NA | NA |
| Gm26864 | 0 | NA | NA | NA |
| Olfr235 | 0.119234754 | 0.51795324 | 3.724945049 | 0.139049901 |
| Olfr1434 | 0 | NA | NA | NA |
| Olfr1436 | 0 | NA | NA | NA |
| Olfr1437 | 0.435432306 | -2.459845484 | 3.679085321 | -0.668602457 |
| Olfr1440 | 0.060174537 | 0.517957688 | 3.724945049 | 0.139051095 |
| Olfr1441 | 0.105852363 | -0.443821212 | 3.724945049 | -0.119148392 |
| Pfpl | 0 | NA | NA | NA |
| Mpeg1 | 2850.264192 | -1.076144199 | 0.182908422 | -5.883513669 |
| Dtx4 | 733.6416179 | -0.309141359 | 0.095065623 | -3.25187328 |
| Fam111a | 54.9166158 | -1.742726949 | 0.332378297 | -5.243203198 |
| Gm16274 | 2.331142904 | 0.883019123 | 0.685577499 | 1.28799315 |
| A330040F15Rik | 6.392297317 | 0.065321285 | 0.420721371 | 0.155260201 |
| Gm4952 | 2.655904937 | -2.223075434 | 0.670870762 | -3.313716382 |
| Glyat | 2.115680473 | -1.997679961 | 0.938911898 | -2.127654326 |
| Gm24521 | 0.066050875 | -0.443818988 | 3.724945049 | -0.119147795 |
| Olfr1442 | 0.259089178 | -1.840490482 | 2.017824654 | -0.912116164 |
| Olfr1443 | 0.092691418 | -0.908687078 | 3.724560878 | -0.2439716 |
| Keg1 | 0.403815225 | -2.51422446 | 3.676495668 | -0.683864388 |
| Gm15962 | 0 | NA | NA | NA |
| Cntf | 4.709263031 | 0.220331715 | 0.493241297 | 0.446701677 |
| Zfp91 | 2396.758906 | 0.12720001 | 0.061451996 | 2.069908521 |
| Lpxn | 113.3029764 | -1.44034376 | 0.251902308 | -5.717866465 |
| Gm5244 | 0.635603962 | -0.133583192 | 1.214122399 | -0.110024486 |
| Olfr1444 | 0.544199285 | -2.086886003 | 1.23909025 | -1.684208235 |
| Olfr1445 | 0.066050875 | -0.443818988 | 3.724945049 | -0.119147795 |
| Olfr1446 | 0 | NA | NA | NA |
| Olfr1447 | 0 | NA | NA | NA |
| Olfr1448 | 0.092228164 | 0.037073798 | 3.724945049 | 0.009952844 |
| Olfr1449 | 0.066050875 | -0.443818988 | 3.724945049 | -0.119147795 |
| Olfr1450 | 0 | NA | NA | NA |
| Olfr1451 | 0 | NA | NA | NA |
| Olfr1452-ps1 | 0 | NA | NA | NA |
| Olfr1453 | 0 | NA | NA | NA |
| Olfr1454 | 0 | NA | NA | NA |
| Olfr1456-ps1 | 0 | NA | NA | NA |
| Olfr1457 | 0 | NA | NA | NA |
| Olfr1458 | 0.049553047 | -0.443818988 | 3.724945049 | -0.119147795 |
| Olfr1459 | 0 | NA | NA | NA |
| Olfr1461 | 0 | NA | NA | NA |
| Olfr1462 | 0 | NA | NA | NA |
| Olfr1463 | 0 | NA | NA | NA |
| Gm22443 | 0 | NA | NA | NA |
| Olfr1465 | 0 | NA | NA | NA |
| Olfr1466 | 0.062426873 | 0.517959912 | 3.724945049 | 0.139051692 |

| | | | | |
|---|---|---|---|---|
| Olfr1467 | 0 | NA | NA | NA |
| Olfr1469 | 0 | NA | NA | NA |
| Olfr1471 | 0 | NA | NA | NA |
| Olfr1472 | 0 | NA | NA | NA |
| Olfr1474 | 0 | NA | NA | NA |
| Olfr1475 | 0 | NA | NA | NA |
| Olfr1477 | 0 | NA | NA | NA |
| Olfr1480 | 0 | NA | NA | NA |
| Olfr1484 | 0.068097359 | 0.037064902 | 3.724945049 | 0.009950456 |
| Gm22323 | 0 | NA | NA | NA |
| Gm23931 | 0 | NA | NA | NA |
| Olfr1487 | 0 | NA | NA | NA |
| Olfr1489 | 0 | NA | NA | NA |
| Gm24124 | 0 | NA | NA | NA |
| Olfr1490 | 0 | NA | NA | NA |
| Olfr1491 | 0 | NA | NA | NA |
| Olfr1494 | 0.061920613 | -0.443816764 | 3.724945049 | -0.119147198 |
| Olfr1495 | 0 | NA | NA | NA |
| Olfr1496 | 0 | NA | NA | NA |
| Olfr1497 | 0.04393175 | 0.037062677 | 3.724945049 | 0.009949859 |
| Olfr1499 | 0 | NA | NA | NA |
| Olfr1500 | 0 | NA | NA | NA |
| Olfr1501 | 0.044191549 | 0.037062677 | 3.724945049 | 0.009949859 |
| Olfr1502 | 0 | NA | NA | NA |
| Olfr1503-ps1 | 0 | NA | NA | NA |
| Olfr1504 | 0 | NA | NA | NA |
| Olfr1505 | 0 | NA | NA | NA |
| Gm26993 | 0 | NA | NA | NA |
| Tle4 | 1024.20378 | 0.15571499 | 0.069297417 | 2.247053299 |
| Gm37997 | 0 | NA | NA | NA |
| Gm26026 | 0 | NA | NA | NA |
| Gm5513 | 1.595119001 | -0.180455452 | 0.750850503 | -0.240334729 |
| Gm24319 | 0 | NA | NA | NA |
| Psat1 | 926.2091933 | -0.227945026 | 0.126364712 | -1.803866143 |
| Gm3329 | 0 | NA | NA | NA |
| Cep78 | 106.2535858 | 0.288360395 | 0.129051673 | 2.234456843 |
| C130060C02Rik | 7.941562867 | 0.197539094 | 0.510401744 | 0.387026682 |
| Gnaq | 3032.47275 | 0.259941833 | 0.038289882 | 6.788786443 |
| E030024N20Rik | 20.01620722 | 0.197365574 | 0.234678983 | 0.841002341 |
| Gm10819 | 108.4716668 | -0.005689909 | 0.131276943 | -0.043342793 |
| Gm17068 | 25.77906067 | 0.252221509 | 0.38006098 | 0.663634317 |
| Mir496b | 0 | NA | NA | NA |
| Gna14 | 16.24861454 | -0.8584074 | 0.443567893 | -1.935233398 |
| Vps13a | 1858.188761 | 0.359964847 | 0.073742693 | 4.881362942 |
| Gm9806 | 0.46212231 | -0.957674416 | 1.534757723 | -0.623990615 |
| Foxb2 | 1.423250569 | -0.228356825 | 0.950485919 | -0.240252717 |
| Prune2 | 1550.090265 | 0.033329773 | 0.114059253 | 0.292214548 |

| | | | | |
|---|---|---|---|---|
| Gm27970 | 0 | NA | NA | NA |
| Gm27796 | 0 | NA | NA | NA |
| Gcnt1 | 110.511412 | -0.502079983 | 0.192995848 | -2.601506649 |
| Rfk | 726.378061 | -0.218516849 | 0.109116823 | -2.00259541 |
| Pcsk5 | 144.4692259 | 0.089449071 | 0.191402569 | 0.467334744 |
| Gm22880 | 0 | NA | NA | NA |
| Gm23063 | 0 | NA | NA | NA |
| Gm23061 | 0 | NA | NA | NA |
| Ostf1 | 237.4482545 | -0.792392543 | 0.141580876 | -5.596748414 |
| Gm8250 | 0.075043205 | 0.517957688 | 3.724945049 | 0.139051095 |
| Nmrk1 | 21.71250768 | 0.13047452 | 0.243812432 | 0.535143016 |
| 2410127L17Rik | 57.66373482 | 0.119023788 | 0.157110804 | 0.757578633 |
| D030056L22Rik | 161.6691014 | -0.371950363 | 0.107409005 | -3.462934638 |
| Gm23238 | 1902.741476 | -0.122316389 | 0.398662424 | -0.30681695 |
| Trpm6 | 41.27246436 | -0.613305836 | 0.205106548 | -2.99018165 |
| Gm24190 | 0 | NA | NA | NA |
| Rorb | 1457.656088 | -0.204048221 | 0.083960431 | -2.430290305 |
| Gm25437 | 0 | NA | NA | NA |
| Gm22684 | 0 | NA | NA | NA |
| Anxa1 | 128.5436956 | -1.739101695 | 0.291688869 | -5.96218053 |
| 1500015L24Rik | 2.063333161 | 0.179802177 | 0.812523018 | 0.221288718 |
| Gm6684 | 0 | NA | NA | NA |
| Aldh1a1 | 547.0244569 | -0.125995297 | 0.187030949 | -0.673660147 |
| Aldh1a7 | 11.46744072 | 0.386327373 | 0.508191643 | 0.760200169 |
| Tmc1 | 1.743158711 | 0.316746672 | 0.866219038 | 0.365665794 |
| Zfand5 | 1899.320038 | -0.127825356 | 0.074500184 | -1.715772369 |
| Gda | 1770.327303 | 0.280276215 | 0.105085512 | 2.667125186 |
| Gm3443 | 0.062426873 | 0.517959912 | 3.724945049 | 0.139051692 |
| 1110059E24Rik | 203.1103973 | -0.148710236 | 0.134089102 | -1.109040433 |
| Abhd17b | 456.3664533 | 0.058368154 | 0.077066811 | 0.757370826 |
| Tmem2 | 509.6691455 | -0.409455602 | 0.105744638 | -3.872116941 |
| Gm5514 | 10.26669978 | 0.454760165 | 0.373835904 | 1.216470008 |
| Trpm3 | 1663.018852 | 0.420511942 | 0.082075308 | 5.12348905 |
| Gm22506 | 0 | NA | NA | NA |
| Gm27151 | 4.77538895 | 0.787006841 | 0.505551749 | 1.556728551 |
| Mir204 | 3.450472164 | 1.306112465 | 0.717472504 | 1.820435567 |
| C330002G04Rik | 1.481948713 | -0.752863814 | 0.798734242 | -0.942571102 |
| 2410080I02Rik | 2.836846521 | 0.728592732 | 0.662113485 | 1.100404612 |
| Klf9 | 2856.577684 | 0.398958114 | 0.121841646 | 3.274398605 |
| Mir1192 | 10.09841093 | 0.626044349 | 0.41146148 | 1.521513872 |
| Smc5 | 682.7510276 | 0.155818348 | 0.089294817 | 1.744987608 |
| Gm24263 | 0.146692594 | -0.955495625 | 3.521109961 | -0.271362052 |
| Mamdc2 | 41.82610674 | -0.080616127 | 0.23093839 | -0.349080667 |
| 1700028P14Rik | 7.326566073 | -0.866881192 | 0.405000591 | -2.140444264 |
| Gm9493 | 43.04589625 | -0.209239666 | 0.195335507 | -1.071180909 |
| Gm22144 | 0 | NA | NA | NA |
| Gm21542 | 0 | NA | NA | NA |

| | | | | |
|---|---|---|---|---|
| Gm6563 | 117.5227222 | 0.014564616 | 0.144043312 | 0.101112751 |
| Ptar1 | 312.4373579 | -0.022102304 | 0.085864151 | -0.257410146 |
| Gm9938 | 199.0723875 | -0.014347709 | 0.09671154 | -0.14835571 |
| Apba1 | 2979.940385 | 0.186168462 | 0.06848776 | 2.718273473 |
| Fam189a2 | 83.27374125 | 0.084044508 | 0.150413695 | 0.558755693 |
| 1700021P04Rik | 0.123841226 | -1.017823543 | 3.719440177 | -0.273649661 |
| Tjp2 | 802.6063976 | -0.49906949 | 0.108703578 | -4.591104524 |
| Fxn | 57.39510889 | -0.150508851 | 0.158565581 | -0.949189913 |
| Gm16216 | 0.110870145 | -0.444802825 | 3.724874622 | -0.119414174 |
| Pip5k1b | 239.2249923 | 0.440336904 | 0.112102995 | 3.927967355 |
| Pip5k1bos | 0.196777805 | -0.443820458 | 2.233055396 | -0.198750313 |
| Fam122a | 94.19792559 | 0.060894186 | 0.160799502 | 0.378696357 |
| Tmem252 | 56.70084035 | -0.541597248 | 0.288687493 | -1.876067588 |
| Pgm5 | 91.2960709 | -1.506353778 | 0.214273378 | -7.030055667 |
| Gm17800 | 0.452161616 | -0.370080995 | 1.364174751 | -0.271285621 |
| Gm10053 | 118.3751616 | -0.35008386 | 0.160851218 | -2.176445196 |
| Foxd4 | 0.431881386 | 1.302305531 | 1.63048029 | 0.798725099 |
| Cbwd1 | 142.8925874 | -0.042623713 | 0.103654296 | -0.411210288 |
| mmu-mir-3084-1 | 0 | NA | NA | NA |
| Gm23658 | 0.086648695 | 0.037058229 | 3.724945049 | 0.009948665 |
| Dock8 | 246.4602774 | -0.792344242 | 0.165304663 | -4.793235882 |
| Gm24252 | 0 | NA | NA | NA |
| Kank1 | 435.2904436 | -0.272677667 | 0.132500431 | -2.05793797 |
| Gm26207 | 0 | NA | NA | NA |
| Dmrt1 | 0 | NA | NA | NA |
| Dmrt3 | 3.281054849 | -0.107449407 | 0.514635253 | -0.208787498 |
| 2610016A17Rik | 0.172734917 | -0.443820173 | 2.736751429 | -0.162170436 |
| Dmrt2 | 7.322241276 | 0.885690441 | 0.424332271 | 2.087256853 |
| Gm8825 | 0 | NA | NA | NA |
| Gm25176 | 0.061920613 | -0.443816764 | 3.724945049 | -0.119147198 |
| Smarca2 | 2885.991631 | 0.528420915 | 0.070015818 | 7.547164777 |
| Gm20616 | 0.038981395 | -0.443816764 | 3.724945049 | -0.119147198 |
| 4931403E22Rik | 1.142959485 | -0.617825839 | 0.98651296 | -0.6262724 |
| Gm815 | 0.098502579 | -0.443821212 | 3.724945049 | -0.119148392 |
| Vldlr | 1529.815537 | -0.137637695 | 0.063335362 | -2.173157165 |
| Kcnv2 | 1.799299276 | -0.31977169 | 0.880837034 | -0.363031614 |
| D19Bwg1357e | 318.8741132 | -0.346493264 | 0.098441498 | -3.519788611 |
| Gm24782 | 0 | NA | NA | NA |
| Rfx3 | 1251.313148 | 0.210206472 | 0.11782103 | 1.784116734 |
| Gm8412 | 0.124853745 | 1.05357957 | 3.721447096 | 0.283110183 |
| Gm20407 | 0 | NA | NA | NA |
| Gm28228 | 0.047948698 | 0.037064902 | 3.724945049 | 0.009950456 |
| Glis3 | 334.0083546 | -0.823809177 | 0.115990263 | -7.10239944 |
| 4933413C19Rik | 10.33626457 | -0.509937622 | 0.382081531 | -1.334630388 |
| D930032P07Rik | 0.131806241 | 0.915443182 | 3.666020831 | 0.249710305 |
| Rps15a-ps2 | 0.107078754 | -0.443164241 | 3.724898049 | -0.118973522 |
| Gm6788 | 2.966347917 | -0.561242381 | 0.591509209 | -0.94883118 |

| | | | | |
|---|---|---|---|---|
| Gm26942 | 0.038981395 | -0.443816764 | 3.724945049 | -0.119147198 |
| Slc1a1 | 1257.835077 | 0.287561861 | 0.080915554 | 3.553851483 |
| Gm5822 | 3.84996234 | 0.128380609 | 0.531277416 | 0.241645146 |
| 4430402I18Rik | 33.49194021 | 1.14291583 | 0.211906555 | 5.393489734 |
| Ppapdc2 | 422.5883542 | 0.328692625 | 0.084736132 | 3.879013809 |
| Cdc37l1 | 944.3468794 | 0.1242612 | 0.060357685 | 2.058746944 |
| Gm10136 | 20.53681307 | -0.32354737 | 0.236993006 | -1.365219066 |
| Ak3 | 872.3625232 | -0.219810644 | 0.118115606 | -1.860978843 |
| 1700018L02Rik | 10.4047421 | 0.82986111 | 0.378898468 | 2.190193893 |
| Rcl1 | 158.5573071 | -0.080634174 | 0.090564926 | -0.890346597 |
| Mir101b | 0.39937024 | 0.037077173 | 1.55349072 | 0.023867007 |
| Gm5518 | 30.36473408 | -0.02444861 | 0.27619199 | -0.088520344 |
| Jak2 | 1349.483123 | -0.456338352 | 0.096905877 | -4.70908853 |
| Insl6 | 6.983240532 | -0.201877728 | 0.536761528 | -0.3761032 |
| Rln1 | 4.858786034 | -0.167706716 | 0.518114971 | -0.323686297 |
| Plgrkt | 208.1764301 | -0.218624225 | 0.103706261 | -2.108110174 |
| Cd274 | 1796.888883 | -0.658164727 | 0.246482349 | -2.670230666 |
| Pdcd1lg2 | 5.725591314 | -1.292939828 | 0.445812325 | -2.900188615 |
| A930007I19Rik | 28.01513506 | -0.782622647 | 0.213732858 | -3.661686159 |
| Ric1 | 1103.079595 | -0.084033429 | 0.085159753 | -0.986773984 |
| Ermp1 | 862.3818722 | 0.143714412 | 0.082109705 | 1.750273158 |
| Mlana | 1.722373549 | 0.734020119 | 0.71411373 | 1.027875656 |
| 9930021J03Rik | 1199.788204 | 0.011573083 | 0.050256589 | 0.230279912 |
| Ranbp6 | 784.8181587 | 0.017936808 | 0.092444906 | 0.194027 |
| Il33 | 391.409633 | -1.395031643 | 0.288357504 | -4.837854487 |
| Gm26046 | 0 | NA | NA | NA |
| Trpd52l3 | 0 | NA | NA | NA |
| Uhrf2 | 805.07859 | -0.370217123 | 0.068252148 | -5.424255979 |
| Gldc | 269.8156168 | -0.35523137 | 0.172375822 | -2.060795795 |
| Rpl31-ps20 | 0 | NA | NA | NA |
| Mbl2 | 0.229843471 | 0.00615129 | 2.549000163 | 0.002413217 |
| Gm24108 | 0 | NA | NA | NA |
| Gm24883 | 0 | NA | NA | NA |
| Ppp1r2-ps3 | 1.44902218 | -0.256819922 | 0.83033237 | -0.309297736 |
| Dkk1 | 0.239409371 | 0.998865747 | 2.149976935 | 0.464593704 |
| Prkg1 | 254.0169383 | 0.472298953 | 0.128609269 | 3.672355486 |
| Gm23426 | 0 | NA | NA | NA |
| Cstf2t | 1157.615467 | -0.012612085 | 0.05504318 | -0.229130747 |
| A1cf | 0.581447236 | -0.400261424 | 1.250477548 | -0.320086854 |
| Asah2 | 367.16758 | -0.033634747 | 0.14491603 | -0.232098185 |
| Sgms1 | 487.606945 | 0.069692982 | 0.094579869 | 0.736869088 |
| 2700046G09Rik | 25.94331831 | 0.236273046 | 0.241457916 | 0.978526821 |
| Rpl9-ps6 | 47.62432018 | -0.089615165 | 0.175759261 | -0.509874499 |
| Minpp1 | 627.6333895 | 0.234315177 | 0.080939152 | 2.894954659 |
| Papss2 | 133.0147862 | -0.52553121 | 0.181289403 | -2.898852339 |
| Atad1 | 1547.98136 | 0.098812384 | 0.101450795 | 0.973993196 |
| Pten | 3804.391786 | 0.408620143 | 0.070050306 | 5.833238488 |

| | | | | |
|---|---|---|---|---|
| Gm22482 | 0.072025462 | 0.517959912 | 3.724945049 | 0.139051692 |
| Rnls | 22.89813653 | -0.405691876 | 0.235516187 | -1.722564726 |
| mmu-mir-3970 | 0 | NA | NA | NA |
| Gm7237 | 0.127971488 | -0.95589744 | 3.6855341 | -0.259364698 |
| Gm8975 | 0 | NA | NA | NA |
| Lipo4 | 0.195256443 | 0.475842931 | 2.223427336 | 0.214013259 |
| Lipo1 | 94.81852167 | -0.518665236 | 0.119110764 | -4.354478304 |
| Gm27440 | 0 | NA | NA | NA |
| Gm8978 | 0 | NA | NA | NA |
| Gm8980 | 0.107078754 | -0.443164241 | 3.724898049 | -0.118973522 |
| Lipo2 | 1.691184024 | 0.141009126 | 0.886924177 | 0.158986675 |
| Gm5097 | 0.090537259 | -0.443823436 | 3.724945049 | -0.119148989 |
| Gm5519 | 0.296824366 | 0.229369004 | 2.364571914 | 0.097002338 |
| Gm8988 | 0 | NA | NA | NA |
| Gm23095 | 0.106971521 | 0.998850474 | 3.724945049 | 0.268151734 |
| Lipf | 0.066050875 | -0.443818988 | 3.724945049 | -0.119147795 |
| Lipk | 0.091880448 | 0.037060453 | 3.724945049 | 0.009949262 |
| Lipn | 0.251288325 | -1.017542178 | 2.091452386 | -0.48652419 |
| Gm23334 | 0 | NA | NA | NA |
| Lipm | 1.074639982 | 1.348389062 | 1.052869719 | 1.280679877 |
| Ankrd22 | 0.527224786 | 0.85622582 | 1.610111452 | 0.531780467 |
| Stambpl1 | 217.0575449 | 0.242430351 | 0.087455872 | 2.772030572 |
| Acta2 | 30.64504911 | -0.981289216 | 0.289219752 | -3.392884507 |
| Fas | 57.72959901 | -0.562571814 | 0.283919452 | -1.98144865 |
| Gm27818 | 0.914190395 | 0.664428149 | 1.179894774 | 0.563124919 |
| Gm23060 | 0 | NA | NA | NA |
| Ch25h | 163.957365 | -1.111432078 | 0.355254445 | -3.128552205 |
| Gm26902 | 14.57275529 | 0.270855638 | 0.373401358 | 0.725374002 |
| Lipa | 315.4234948 | -0.90292032 | 0.153073225 | -5.898616969 |
| Ifit2 | 10029.24711 | -0.529107263 | 0.277676866 | -1.90547837 |
| Ifit3 | 3311.123546 | -0.946156405 | 0.227368383 | -4.161336739 |
| Gm14446 | 387.1233819 | -1.982023763 | 0.450656678 | -4.3980792 |
| Ifit3b | 2018.601927 | -1.023482538 | 0.238699127 | -4.287751499 |
| 2010002M12Rik | 153.7706214 | -0.665736578 | 0.216702605 | -3.072120789 |
| Ifit1 | 4083.025541 | -0.542156009 | 0.239506848 | -2.263634692 |
| Slc16a12 | 39.40020055 | 0.532744734 | 0.228562683 | 2.330847394 |
| Gm27476 | 0 | NA | NA | NA |
| Gm22082 | 0 | NA | NA | NA |
| Pank1 | 222.2087571 | -0.014067077 | 0.109407443 | -0.128575138 |
| Mir107 | 0.848910726 | 2.947012274 | 1.338018213 | 2.202520299 |
| Kif20b | 26.85031262 | -0.208757219 | 0.244664374 | -0.853239135 |
| Mir1950 | 0 | NA | NA | NA |
| Gm5248 | 0 | NA | NA | NA |
| Gm24006 | 0 | NA | NA | NA |
| Gm23075 | 0 | NA | NA | NA |
| Gm22853 | 0 | NA | NA | NA |
| Gm25611 | 0.204668643 | 0.517963338 | 2.614970123 | 0.198076197 |

| | | | | |
|---|---|---|---|---|
| A830019P07Rik | 16.97608067 | -0.279182024 | 0.313293434 | -0.891119934 |
| Htr7 | 177.6287066 | -0.12138019 | 0.110475338 | -1.098708468 |
| Rpp30 | 118.3865456 | -0.29048339 | 0.119555611 | -2.429692649 |
| Ankrd1 | 0.384174469 | 0.769096734 | 1.636296574 | 0.470022823 |
| Pcgf5 | 290.0103938 | -0.525067467 | 0.107014205 | -4.906521234 |
| Gm23170 | 0 | NA | NA | NA |
| Hectd2 | 578.0733075 | -0.335853338 | 0.106098709 | -3.165479977 |
| 1500017E21Rik | 5.192905494 | 0.192434546 | 0.509609568 | 0.377611721 |
| Ppp1r3c | 367.6091572 | -0.14755958 | 0.180144461 | -0.819118051 |
| Tnks2 | 2533.779749 | -0.08083399 | 0.059255704 | -1.364155427 |
| Fgfbp3 | 45.89989236 | -0.333539913 | 0.207072168 | -1.610742363 |
| Btaf1 | 981.4939569 | -0.178378316 | 0.090316142 | -1.97504357 |
| Cpeb3 | 1630.611145 | 0.11410962 | 0.051367297 | 2.221444876 |
| Gm22714 | 7.127945666 | 0.386661777 | 0.482819913 | 0.800840574 |
| A330032B11Rik | 4.137844924 | 0.675379416 | 0.660526659 | 1.022486233 |
| Gm23918 | 1.204149766 | 0.168661747 | 1.002943835 | 0.168166692 |
| 5-Mar | 1036.41543 | -0.202671202 | 0.120038188 | -1.688389384 |
| Gm25268 | 0 | NA | NA | NA |
| Ide | 826.1150129 | -0.289442577 | 0.068114367 | -4.249361624 |
| Rpl10-ps6 | 9.176167568 | 0.291949448 | 0.371523479 | 0.785816951 |
| Kif11 | 42.94126884 | -0.415153602 | 0.266252109 | -1.559250006 |
| Gm38345 | 1.850005102 | -1.042223935 | 0.731812798 | -1.424167408 |
| Hhex | 20.51235511 | -0.593945764 | 0.31575795 | -1.881016022 |
| Gm23026 | 0 | NA | NA | NA |
| Exoc6 | 443.4655257 | 0.155789683 | 0.078198674 | 1.992229202 |
| Cyp26c1 | 3.181018617 | -0.344160109 | 0.714058546 | -0.481977438 |
| Cyp26a1 | 1.890572446 | 0.435330938 | 0.69663958 | 0.624901241 |
| Myof | 324.4012162 | -1.668060435 | 0.273075188 | -6.108429145 |
| I830134H01Rik | 0.540537234 | 0.037077929 | 1.426876203 | 0.025985386 |
| Cep55 | 11.79181242 | -0.776617881 | 0.344183217 | -2.25640834 |
| Gm23300 | 0 | NA | NA | NA |
| Ffar4 | 0.430805093 | 0.933681469 | 1.577985062 | 0.59169221 |
| Rbp4 | 74.47465664 | 0.067263644 | 0.186578917 | 0.36051042 |
| Pde6c | 1.452976049 | -0.485736545 | 0.845086288 | -0.574777454 |
| Fra10ac1 | 83.34982365 | -0.006582292 | 0.177333132 | -0.037118229 |
| Lgi1 | 1139.244008 | 0.558730684 | 0.093397549 | 5.982284203 |
| RP24-417P12.1 | 0.567844938 | -0.082549919 | 1.301709307 | -0.063416554 |
| Slc35g1 | 45.27465032 | -0.108478074 | 0.18753376 | -0.578445579 |
| Gm24930 | 0 | NA | NA | NA |
| Plce1 | 890.1324761 | -0.168416389 | 0.134741992 | -1.249917609 |
| Gm28991 | 0 | NA | NA | NA |
| Noc3l | 300.5445623 | -0.320481147 | 0.079814255 | -4.015337218 |
| Tbc1d12 | 321.3981898 | -0.106447173 | 0.091403076 | -1.164590709 |
| Gm25838 | 2.422369998 | 0.115752477 | 0.740516114 | 0.156313246 |
| Hells | 136.3095865 | -0.364782174 | 0.144581178 | -2.523026708 |
| Cyp2c55 | 1.660109277 | -0.844543408 | 0.770421493 | -1.096209563 |
| Gm23800 | 0 | NA | NA | NA |

| | | | | |
|---|---|---|---|---|
| Cyp2c65 | 0.792025016 | -0.260819145 | 1.068025967 | -0.244206745 |
| Cyp2c66 | 0.061920613 | -0.443816764 | 3.724945049 | -0.119147198 |
| Cyp2c53-ps | 0.038981395 | -0.443816764 | 3.724945049 | -0.119147198 |
| Cyp2c29 | 0.220032676 | -1.401785252 | 2.574515855 | -0.544484995 |
| Cyp2c38 | 0.045882455 | 0.517959912 | 3.724945049 | 0.139051692 |
| Cyp2c39 | 0.206165386 | -1.006862472 | 2.649120745 | -0.380074209 |
| Cyp2c67 | 0 | NA | NA | NA |
| Cyp2c68 | 0.045882455 | 0.517959912 | 3.724945049 | 0.139051692 |
| Cyp2c40 | 0.138763737 | 0.517963565 | 2.930963636 | 0.176721252 |
| Cyp2c69 | 0.075043205 | 0.517957688 | 3.724945049 | 0.139051095 |
| Cyp2c37 | 0.045882455 | 0.517959912 | 3.724945049 | 0.139051692 |
| Cyp2c54 | 0.045882455 | 0.517959912 | 3.724945049 | 0.139051692 |
| Cyp2c50 | 0.153424464 | -0.891301076 | 2.83106242 | -0.314829185 |
| Cyp2c70 | 0.309994989 | 0.022695528 | 1.740415166 | 0.013040296 |
| Gm16470 | 3.026502306 | -0.043567788 | 0.61782333 | -0.070518198 |
| Pdlim1 | 63.17543641 | -0.756453931 | 0.183882091 | -4.113798826 |
| Sorbs1 | 1628.853159 | -0.115977493 | 0.093015323 | -1.246864378 |
| Aldh18a1 | 344.8097266 | -0.738186435 | 0.116513484 | -6.335630925 |
| Gm15801 | 18.90733818 | -0.448162569 | 0.245768528 | -1.823514883 |
| Gm27042 | 1.18341529 | -1.405123308 | 0.953361076 | -1.473862677 |
| Tctn3 | 216.4391839 | 0.278096963 | 0.104696466 | 2.656221116 |
| Entpd1 | 233.1810674 | -0.396430647 | 0.181089009 | -2.189148022 |
| Gm16027 | 0.848235736 | 1.790489689 | 1.228860846 | 1.457032091 |
| Gm8783 | 30.04226012 | -0.474172391 | 0.187303367 | -2.531574306 |
| Ccnj | 128.7530246 | -0.03535935 | 0.13016862 | -0.271642658 |
| E030044B06Rik | 1.397451337 | 0.316524911 | 0.861271952 | 0.367508671 |
| Mir8092 | 0 | NA | NA | NA |
| Zfp518a | 366.4925213 | -0.086785495 | 0.113709209 | -0.763223102 |
| Blnk | 124.657933 | 0.012545411 | 0.186114264 | 0.067407039 |
| Dntt | 1.408550033 | -2.179487014 | 1.079983801 | -2.0180738 |
| Opalin | 167.6099803 | 0.026937119 | 0.138409921 | 0.19461841 |
| Tll2 | 11.98457804 | -0.108473623 | 0.307761957 | -0.352459491 |
| Tm9sf3 | 2559.714795 | 0.071497465 | 0.058612573 | 1.219831533 |
| Pik3ap1 | 444.6212228 | -0.958546808 | 0.16671677 | -5.749552404 |
| Gm9788 | 0.072025462 | 0.517959912 | 3.724945049 | 0.139051692 |
| Lcor | 579.5255258 | 0.270696674 | 0.11458033 | 2.362505612 |
| Gm340 | 323.3061025 | 0.298587757 | 0.113757264 | 2.624779692 |
| Mir8091 | 15.23441784 | 0.084643327 | 0.292618358 | 0.289261848 |
| AI606181 | 120.0613128 | 0.244874371 | 0.127098692 | 1.926647452 |
| Slit1 | 868.1752054 | 0.849146307 | 0.100082446 | 8.484467995 |
| Gm22601 | 0.086648695 | 0.037058229 | 3.724945049 | 0.009948665 |
| Arhgap19 | 33.0013039 | 0.168722912 | 0.190229555 | 0.886943735 |
| Frat1 | 29.17957322 | -0.15572726 | 0.208945193 | -0.745301949 |
| Frat2 | 52.5338819 | 0.46662114 | 0.161685586 | 2.885978587 |
| Rrp12 | 552.962244 | -0.38637943 | 0.08548491 | -4.51985539 |
| Pgam1 | 1666.384237 | -0.153942873 | 0.061615359 | -2.498449674 |
| Exosc1 | 140.0755093 | -0.16445165 | 0.09854275 | -1.668835602 |

| | | | | |
|---|---|---|---|---|
| Zdhhc16 | 217.4843809 | -0.113336916 | 0.087582873 | -1.294053422 |
| Mms19 | 424.2345625 | -0.021252884 | 0.116414333 | -0.182562431 |
| Gm23113 | 0 | NA | NA | NA |
| Ubtd1 | 125.2316124 | -0.890422704 | 0.151147636 | -5.891079263 |
| Ankrd2 | 4.166437747 | 0.355845969 | 0.510033843 | 0.697690897 |
| Hoga1 | 9.277047816 | -0.443613318 | 0.399238511 | -1.111148616 |
| 4933411K16Rik | 1.522479514 | 2.055194778 | 1.050487679 | 1.956419688 |
| Morn4 | 316.6990762 | 0.262165009 | 0.102957625 | 2.546338934 |
| Pi4k2a | 1103.365836 | -0.241728986 | 0.106140075 | -2.277452559 |
| Avpi1 | 112.560822 | -0.261115486 | 0.132708323 | -1.967589373 |
| Marveld1 | 84.73861597 | -0.020825519 | 0.127098839 | -0.163852947 |
| Zfyve27 | 786.8275757 | 0.079454802 | 0.074944153 | 1.060186798 |
| Sfrp5 | 34.79157235 | 0.835004804 | 0.361280239 | 2.311238516 |
| Golga7b | 962.3374335 | -0.057543189 | 0.070760456 | -0.813211115 |
| Gm26339 | 1.445692662 | 0.41238955 | 0.844854426 | 0.488119061 |
| Mir3085 | 0.135858026 | 0.037065849 | 2.948997537 | 0.012568966 |
| Crtac1 | 458.2694328 | -0.488446608 | 0.104314627 | -4.682436429 |
| R3hcc1l | 325.8532248 | -0.145647591 | 0.093905909 | -1.550994951 |
| Gm16541 | 0.261817563 | 1.001732022 | 2.443998749 | 0.409874196 |
| Loxl4 | 7.914160501 | -0.581243231 | 0.339582999 | -1.711638194 |
| Gm25216 | 0 | NA | NA | NA |
| Pyroxd2 | 33.56925608 | 0.578925174 | 0.254724846 | 2.272747173 |
| Hps1 | 90.94188656 | 0.038512957 | 0.149172632 | 0.258177098 |
| Gm16244 | 3.388720562 | 1.864481556 | 0.703002846 | 2.652167865 |
| Hpse2 | 1.819823472 | 1.592795887 | 0.754519859 | 2.111005917 |
| Gm22135 | 0 | NA | NA | NA |
| Gm6776 | 3.379576263 | -0.933058198 | 0.606521347 | -1.538376517 |
| Cnnm1 | 1191.472548 | 0.298604545 | 0.08168606 | 3.655514105 |
| Got1 | 2463.067521 | 0.185208632 | 0.0633021 | 2.925789693 |
| Gm20467 | 0.07796279 | -0.798301147 | 3.721829328 | -0.214491605 |
| Nkx2-3 | 0.04393175 | 0.037062677 | 3.724945049 | 0.009949859 |
| Gm22646 | 0 | NA | NA | NA |
| Slc25a28 | 398.2739226 | 0.215101482 | 0.133198262 | 1.614897062 |
| Entpd7 | 424.4745962 | -0.180328731 | 0.09598664 | -1.878685733 |
| Cox15 | 306.6258307 | -0.138948294 | 0.116130677 | -1.196482257 |
| Cutc | 59.7964683 | 0.095247574 | 0.180658791 | 0.527223575 |
| Abcc2 | 1.945998912 | 0.431276935 | 0.679473511 | 0.63472222 |
| Dnmbp | 248.0196757 | -0.055278754 | 0.110058435 | -0.502267312 |
| Gm24400 | 2.333191932 | 1.195265713 | 0.850098845 | 1.406031452 |
| n-R5s21 | 0 | NA | NA | NA |
| Cpn1 | 2.984315308 | -1.04997337 | 0.614841441 | -1.707714055 |
| Cyp2c44 | 1.49985572 | -3.001360445 | 1.020415001 | -2.941313527 |
| Erlin1 | 312.6811813 | -0.235248132 | 0.078883387 | -2.982226573 |
| Chuk | 597.7287007 | -0.333239062 | 0.086366158 | -3.85844491 |
| Cwf19l1 | 148.8001304 | 0.146252885 | 0.101278724 | 1.444063265 |
| Gm24336 | 70.39117647 | -0.058959907 | 0.144293188 | -0.408611856 |
| Bloc1s2 | 164.8635678 | -0.318999841 | 0.120306771 | -2.651553518 |

| | | | | |
|---|---|---|---|---|
| Pkd2l1 | 8.957326903 | -1.646748906 | 0.400849833 | -4.108144174 |
| Scd3 | 92.32162226 | -0.048283017 | 0.157229283 | -0.307086669 |
| Scd2 | 13180.9979 | 0.443085219 | 0.125950328 | 3.517936205 |
| Mir5114 | 0 | NA | NA | NA |
| Scd4 | 17.24753904 | -0.508488278 | 0.274943458 | -1.849428541 |
| Scd1 | 2469.814438 | 0.256714442 | 0.142203333 | 1.805263191 |
| Wnt8b | 3.609855578 | 0.977804848 | 0.590316063 | 1.656409014 |
| Sec31b | 4.76996336 | -0.611849083 | 0.511766171 | -1.195563752 |
| Gm20538 | 0.061920613 | -0.443816764 | 3.724945049 | -0.119147198 |
| Ndufb8 | 791.3026735 | 0.008892355 | 0.119944815 | 0.074137053 |
| Hif1an | 511.8430662 | -0.329237452 | 0.073432696 | -4.483526716 |
| Gm26644 | 1.498020843 | 0.020732549 | 0.905462983 | 0.02289718 |
| Pax2 | 56.35874315 | -0.160132304 | 0.303967724 | -0.526806932 |
| Gm20395 | 0 | NA | NA | NA |
| Mir6405 | 0.042716945 | 0.037062677 | 3.724945049 | 0.009949859 |
| 1700039E22Rik | 0.097899064 | 0.037062677 | 3.724945049 | 0.009949859 |
| Fam178a | 1155.011791 | -0.049857062 | 0.091823081 | -0.542968736 |
| Gm25482 | 3.723338729 | 0.575062128 | 0.656668774 | 0.875726319 |
| Sema4g | 448.4563521 | 0.506187493 | 0.108477223 | 4.666302078 |
| Mrpl43 | 278.1537674 | -0.161756223 | 0.095729011 | -1.68973042 |
| Peo1 | 143.5811533 | -0.314920154 | 0.116186756 | -2.710465164 |
| Lzts2 | 639.0048677 | -0.24843528 | 0.090593725 | -2.742301193 |
| Pdzd7 | 38.25076259 | 0.235360901 | 0.330655662 | 0.711800609 |
| Sfxn3 | 897.777185 | 0.133931557 | 0.044123908 | 3.035351198 |
| Kazald1 | 26.8175025 | -0.113022633 | 0.215749006 | -0.523861661 |
| Tlx1 | 0.675592865 | 0.367328961 | 1.177381592 | 0.311988028 |
| Tlx1os | 0 | NA | NA | NA |
| Lbx1 | 26.55161603 | -0.234046506 | 0.333650055 | -0.701473002 |
| Gm29595 | 9.837264003 | 0.037170503 | 0.32836733 | 0.113197934 |
| Gm29593 | 4.239070863 | -1.4422527 | 0.589388018 | -2.447034308 |
| Gm29594 | 0.57177649 | 0.413740733 | 1.347999259 | 0.306929496 |
| Gm28578 | 19.28251988 | -0.738092219 | 0.379388261 | -1.945479852 |
| Btrc | 1150.587465 | 0.219562517 | 0.06570994 | 3.341389721 |
| Gm6807 | 12.07198287 | -0.099169614 | 0.301754032 | -0.328643873 |
| Poll | 98.22017834 | -0.131837172 | 0.139523596 | -0.944909503 |
| Dpcd | 221.260172 | 0.150320022 | 0.141264787 | 1.064101152 |
| Fbxw4 | 248.8152368 | -0.191032436 | 0.10244265 | -1.864774452 |
| Fgf8 | 0.275408967 | 0.037076002 | 2.08174708 | 0.017810041 |
| Npm3 | 55.30170471 | -1.17146449 | 0.200512664 | -5.842346653 |
| Gm15491 | 0.865681968 | -1.386166209 | 1.08473956 | -1.277879281 |
| Mgea5 | 3029.551855 | -0.061822687 | 0.106232304 | -0.581957509 |
| Kcnip2 | 682.9003887 | 1.259187947 | 0.161325139 | 7.805280389 |
| 9130011E15Rik | 279.4855959 | 0.066009706 | 0.078440122 | 0.84152987 |
| 4930505N22Rik | 0.892093866 | -2.276544697 | 1.100542648 | -2.068565631 |
| Hps6 | 95.93381243 | -0.429070661 | 0.155859051 | -2.752940293 |
| Ldb1 | 957.9423255 | 0.282991094 | 0.077817413 | 3.636603734 |
| Pprc1 | 454.9862607 | -0.508259551 | 0.089772024 | -5.66166973 |

| | | | | |
|---|---|---|---|---|
| Nolc1 | 518.4413524 | -0.115438098 | 0.11099683 | -1.040012563 |
| Elovl3 | 0.899697409 | -3.059060948 | 1.177343338 | -2.59827431 |
| Pitx3 | 0.694683682 | 0.302493334 | 1.225978569 | 0.246736233 |
| Gbf1 | 1560.460353 | -0.237785304 | 0.051392345 | -4.626862342 |
| Gm8974 | 0.178004931 | 0.03763054 | 2.299166841 | 0.016367033 |
| Gm16726 | 0.21650593 | -1.791584846 | 2.13065922 | -0.840859406 |
| Nfkb2 | 234.2002576 | -0.752375056 | 0.229537199 | -3.277791399 |
| Psd | 2382.095847 | 1.080138972 | 0.120510814 | 8.963004495 |
| Fbxl15 | 156.8858204 | 0.080172586 | 0.115186464 | 0.696024364 |
| Cuedc2 | 723.9696748 | 0.055837491 | 0.082915085 | 0.673429823 |
| Gm26792 | 1.450243399 | -1.303731254 | 0.996582963 | -1.308201427 |
| Tmem180 | 197.5786801 | 0.400091505 | 0.108757358 | 3.678753435 |
| Mir146b | 0.495786544 | -1.578754678 | 1.367684822 | -1.154326386 |
| 2310034G01Rik | 6.1931368 | -0.32566901 | 0.541922381 | -0.600951395 |
| Gm24610 | 1.876595299 | 0.741455593 | 0.898441405 | 0.825268725 |
| Actr1a | 1762.40465 | -0.070190182 | 0.051305325 | -1.36808766 |
| Sufu | 283.8186178 | -0.152399667 | 0.09519274 | -1.600958932 |
| Trim8 | 1307.724394 | 0.029511508 | 0.071525975 | 0.41259847 |
| Arl3 | 347.6175142 | 0.025151715 | 0.119823064 | 0.209907124 |
| Sfxn2 | 104.3476033 | 0.271572609 | 0.151046831 | 1.797936485 |
| Wbp1l | 711.1586369 | -0.343913036 | 0.097127735 | -3.540832456 |
| Cyp17a1 | 0.280011045 | 0.036509843 | 1.796497865 | 0.020322786 |
| Gm6967 | 0.062426873 | 0.517959912 | 3.724945049 | 0.139051692 |
| 2010012O05Rik | 189.8383209 | 0.424816039 | 0.09400116 | 4.519263802 |
| As3mt | 170.4653533 | -0.565928904 | 0.136155228 | -4.15649778 |
| Cnnm2 | 423.4727026 | 0.14065663 | 0.071119506 | 1.977750396 |
| Nt5c2 | 724.1452932 | 0.190479895 | 0.079248328 | 2.403582508 |
| Ina | 2102.910469 | -0.021295783 | 0.113453235 | -0.187705382 |
| Pcgf6 | 132.4800137 | -0.093310539 | 0.154089175 | -0.605561937 |
| Gm23140 | 0.042716945 | 0.037062677 | 3.724945049 | 0.009949859 |
| Taf5 | 105.5145098 | -0.329558803 | 0.167418834 | -1.9684691 |
| Usmg5 | 854.0068898 | -0.000932607 | 0.141039429 | -0.006612386 |
| Pdcd11 | 426.8069572 | -0.535602475 | 0.08917098 | -6.006466164 |
| Calhm2 | 55.28273003 | 0.185933951 | 0.250061085 | 0.743554126 |
| Calhm1 | 0 | NA | NA | NA |
| Calhm3 | 0.222639008 | -1.336773467 | 2.55171024 | -0.523873536 |
| Gm6970 | 2.783210482 | 1.011953564 | 0.671307792 | 1.507436048 |
| Neurl1a | 1174.052599 | 0.564161625 | 0.1045367 | 5.396780509 |
| Sh3pxd2a | 901.8376151 | 0.308796723 | 0.080717671 | 3.825639651 |
| Mir6995 | 1.143944469 | 1.10697524 | 1.154403625 | 0.958915249 |
| Obfc1 | 85.46752745 | -0.359794585 | 0.136789 | -2.630288865 |
| Gm19557 | 1.777092953 | -0.775991926 | 0.788025355 | -0.984729643 |
| Slk | 1543.68933 | 0.158400636 | 0.074122718 | 2.137005225 |
| Gm16068 | 0.225284295 | 0.998868557 | 1.89559601 | 0.526941686 |
| Col17a1 | 13.5201549 | 0.566634546 | 0.302842938 | 1.871050881 |
| Gm26085 | 0 | NA | NA | NA |
| Gm24805 | 0 | NA | NA | NA |

| | | | | |
|---|---|---|---|---|
| Sfr1 | 586.6289326 | -0.092909275 | 0.099734137 | -0.931569441 |
| Cfap43 | 93.17411037 | -0.74860938 | 0.180072392 | -4.157269046 |
| Mir8090 | 0.046345709 | -0.443818988 | 3.724945049 | -0.119147795 |
| Gsto1 | 187.9833352 | 0.171733099 | 0.111319563 | 1.542703675 |
| Gsto2 | 5.405473811 | -0.63446878 | 0.469888375 | -1.350254259 |
| Gm22240 | 0 | NA | NA | NA |
| Cfap58 | 6.680071795 | 0.538628862 | 0.370227485 | 1.4548592 |
| Sorcs3 | 648.2072618 | -0.341743667 | 0.14255117 | -2.397340318 |
| Gm23857 | 0 | NA | NA | NA |
| Gm27935 | 0 | NA | NA | NA |
| Rpl13a-ps1 | 0.376112881 | 0.578113996 | 3.288570459 | 0.175794925 |
| Sorcs1 | 756.9127055 | -0.053925972 | 0.155383996 | -0.347049719 |
| Gm27749 | 0 | NA | NA | NA |
| Gm26629 | 0.136194719 | 0.037062677 | 3.724945049 | 0.009949859 |
| Gm25659 | 0 | NA | NA | NA |
| Gm25644 | 0 | NA | NA | NA |
| Rpl31-ps18 | 0 | NA | NA | NA |
| Ins1 | 0 | NA | NA | NA |
| Xpnpep1 | 575.104864 | -0.202286028 | 0.084012592 | -2.407806065 |
| Mir6407 | 1.567823505 | 1.352915144 | 0.911972595 | 1.483504167 |
| 1700054A03Rik | 0.092691418 | -0.908687078 | 3.724560878 | -0.2439716 |
| Add3 | 1815.758105 | -0.439888575 | 0.094808616 | -4.639753144 |
| Gm24724 | 0 | NA | NA | NA |
| Mxi1 | 685.6681876 | 0.282958435 | 0.071935203 | 3.933518284 |
| Gm10197 | 0.330485341 | 0.425954912 | 1.754581483 | 0.242767244 |
| Smndc1 | 390.3324799 | -0.163326922 | 0.122039373 | -1.338313353 |
| 5830416P10Rik | 35.48233613 | -0.43134786 | 0.249208407 | -1.730872022 |
| 4833407H14Rik | 1.343421229 | 0.341508642 | 0.83081794 | 0.411051117 |
| Dusp5 | 436.4752899 | -0.902503585 | 0.26462422 | -3.410510134 |
| Nutf2-ps1 | 25.95248433 | -0.850865993 | 0.273889023 | -3.106608596 |
| Smc3 | 928.9007826 | 0.113271276 | 0.088412354 | 1.281170235 |
| Rbm20 | 75.31453989 | -0.022125216 | 0.139540267 | -0.15855793 |
| Gm16298 | 0.11037557 | 0.517957688 | 3.724945049 | 0.139051095 |
| Gm16299 | 1.831283359 | 0.786717341 | 0.834880536 | 0.942311273 |
| Pdcd4 | 830.0440107 | 0.521651754 | 0.077837939 | 6.701767282 |
| Bbip1 | 292.1551775 | -0.160109778 | 0.105567112 | -1.516663433 |
| Shoc2 | 875.8097201 | -0.119364349 | 0.057074142 | -2.091391038 |
| Adra2a | 223.1903651 | 0.715812692 | 0.158737288 | 4.509417435 |
| Gpam | 844.7644142 | -0.016354405 | 0.087584797 | -0.186726528 |
| Tectb | 0.98697613 | -0.026483592 | 1.059041439 | -0.025007135 |
| Mir6715 | 0 | NA | NA | NA |
| Gucy2g | 23.72729764 | 0.206262434 | 0.296560708 | 0.695515045 |
| Acsl5 | 698.4865369 | -0.387936196 | 0.068666614 | -5.649560604 |
| Zdhhc6 | 305.077262 | -0.405897746 | 0.084877355 | -4.782167709 |
| Vti1a | 524.7365872 | 0.071697674 | 0.064485793 | 1.111836741 |
| 4930552P12Rik | 3.0018596 | -0.910483852 | 0.538386331 | -1.691134785 |
| Tcf7l2 | 729.2267912 | 0.072894343 | 0.084531285 | 0.862335684 |

| | | | | |
|---|---|---|---|---|
| Ppnr | 24.84417134 | -0.179791879 | 0.403348067 | -0.445748706 |
| Gm22271 | 0 | NA | NA | NA |
| Habp2 | 0.044544648 | 0.517957688 | 3.724945049 | 0.139051095 |
| Nrap | 37.22294834 | 0.014022407 | 0.25073317 | 0.055925617 |
| Gm17197 | 0.54074031 | -0.2740338 | 1.290396372 | -0.21236405 |
| Casp7 | 137.0862569 | -0.852548086 | 0.155921093 | -5.467817533 |
| Plekhs1 | 3.487526041 | 0.027601319 | 0.525594841 | 0.052514441 |
| Dclre1a | 154.6814328 | -0.1686445 | 0.103344558 | -1.631866295 |
| Nhlrc2 | 312.5790796 | -0.289020466 | 0.129945532 | -2.224166241 |
| Adrb1 | 210.0560459 | 0.285047292 | 0.106852981 | 2.667658779 |
| Gm6990 | 0.88041345 | -0.387460597 | 1.207716269 | -0.32082088 |
| Ccdc186 | 1033.921743 | 0.02062867 | 0.061250258 | 0.336793193 |
| Tdrd1 | 1.956606073 | 0.53757683 | 0.736695211 | 0.729714028 |
| Vwa2 | 0.389464624 | -0.32684835 | 1.523038894 | -0.21460276 |
| Afap1l2 | 288.9123761 | -0.504635145 | 0.13933733 | -3.621679456 |
| Ablim1 | 1533.129369 | 0.074920238 | 0.064666856 | 1.158556982 |
| B230217O12Rik | 56.18862831 | 0.312614105 | 0.162402969 | 1.924928509 |
| Fam160b1 | 577.2204722 | 0.128542771 | 0.074467864 | 1.726150894 |
| Trub1 | 145.5468936 | 0.334916537 | 0.150509395 | 2.225220144 |
| Gm26874 | 0 | NA | NA | NA |
| Atrnl1 | 1851.542762 | 0.292091543 | 0.085952813 | 3.39827788 |
| Mir5623 | 3.511196682 | -0.165645928 | 0.78725098 | -0.21041057 |
| Gm16277 | 0.097501745 | -0.443821212 | 3.724945049 | -0.119148392 |
| Gfra1 | 699.0851246 | -0.219016948 | 0.092480003 | -2.368262771 |
| Ccdc172 | 1.84083352 | -0.768628689 | 0.76259853 | -1.007907383 |
| 1810007D17Rik | 0.526835603 | -0.7944455 | 1.568931754 | -0.506360776 |
| Pnlip | 3.471747287 | 0.354409287 | 0.534543504 | 0.663012991 |
| 1810018F18Rik | 0 | NA | NA | NA |
| Pnliprp1 | 0.093484797 | -0.443823436 | 3.724945049 | -0.119148989 |
| Pnliprp2 | 0.200944149 | 0.036329567 | 2.220206532 | 0.016363148 |
| 1700019N19Rik | 1.010445843 | -0.57968807 | 0.901969264 | -0.6426916 |
| Gm22561 | 0 | NA | NA | NA |
| Gm17150 | 0 | NA | NA | NA |
| Hspa12a | 2209.460681 | 0.512723421 | 0.080758555 | 6.348843389 |
| Mir3086 | 1.922482983 | 2.072668071 | 0.983973593 | 2.10642652 |
| Eno4 | 32.39598207 | 0.062447757 | 0.185427887 | 0.33677651 |
| Shtn1 | 427.0917043 | 0.470304475 | 0.107511547 | 4.374455454 |
| Vax1 | 9.031124418 | -0.075702449 | 0.40141712 | -0.188587994 |
| Kcnk18 | 0.044191549 | 0.037062677 | 3.724945049 | 0.009949859 |
| Gm29261 | 1.407425821 | 0.662807735 | 0.966307987 | 0.685917682 |
| Slc18a2 | 109.7658993 | 0.214046119 | 0.302230832 | 0.708220658 |
| Pdzd8 | 1685.906693 | -0.006224313 | 0.063817595 | -0.097532865 |
| Rps12-ps3 | 13.1319493 | -0.423910641 | 0.313705133 | -1.351302851 |
| Emx2os | 17.8875887 | 0.865261113 | 0.345738936 | 2.502642955 |
| Emx2 | 54.25553675 | 0.234932682 | 0.17735848 | 1.32462052 |
| 2700089I24Rik | 0.185374628 | 0.517968595 | 2.272545264 | 0.227924435 |
| Gm17203 | 36.89967017 | 0.271281407 | 0.246378854 | 1.101074231 |

| | | | | |
|---|---|---|---|---|
| Rab11fip2 | 973.2053457 | 0.349649339 | 0.069967835 | 4.997286842 |
| Gm4219 | 3.097868212 | -0.908378662 | 0.60039822 | -1.512960284 |
| E330013P04Rik | 2.49875752 | -1.566269473 | 0.763692844 | -2.050915476 |
| Fam204a | 247.5678764 | -0.03258545 | 0.106612622 | -0.305643452 |
| 4933412A08Rik | 0.53212218 | 0.483891758 | 1.267449404 | 0.381783885 |
| Gm28351 | 0 | NA | NA | NA |
| Prlhr | 4.925174813 | -0.041699385 | 0.466395155 | -0.089407843 |
| Cacul1 | 885.5461787 | -0.224138139 | 0.078980557 | -2.837890061 |
| Gm25238 | 3.512009492 | 0.699767287 | 0.590125278 | 1.185794463 |
| Nanos1 | 296.0425935 | 0.329831388 | 0.10697915 | 3.08313712 |
| Eif3a | 2000.929003 | -0.28341025 | 0.118271698 | -2.396264326 |
| Snora19 | 2.074201168 | 0.006267333 | 0.69791882 | 0.008980031 |
| Gm26158 | 3.094791288 | 5.31E-06 | 0.541040711 | 9.81E-06 |
| Fam45a | 537.0189958 | 0.089026449 | 0.106610287 | 0.835064344 |
| Gm15959 | 0 | NA | NA | NA |
| Sfxn4 | 143.5137547 | 0.147247534 | 0.148080964 | 0.994371797 |
| Prdx3 | 403.3254737 | -0.161396531 | 0.123995489 | -1.301632278 |
| Grk5 | 237.0149773 | -0.016565303 | 0.093960983 | -0.176299803 |
| Gm9529 | 0.17796667 | -0.471322379 | 2.715209716 | -0.173585995 |
| Gm22520 | 0 | NA | NA | NA |
| 5830428H23Rik | 346.0755543 | -0.338976264 | 0.224879295 | -1.507369832 |
| Gm22365 | 0 | NA | NA | NA |
| Gm7102 | 0.828685883 | -0.423984141 | 1.234290036 | -0.343504467 |
| Gm6020 | 0 | NA | NA | NA |
| Csf2ra | 189.2145181 | -0.791803687 | 0.150032759 | -5.27753867 |
| Gm21060 | 0 | NA | NA | NA |
| Gm37392 | 0 | NA | NA | NA |
| Gm27306 | 0 | NA | NA | NA |
| Fam171a1 | 789.1479404 | -0.0564935 | 0.077591004 | -0.728093428 |
| Nmt2 | 1499.866317 | 0.340752411 | 0.070643958 | 4.823518113 |
| Gm22005 | 0.075043205 | 0.517957688 | 3.724945049 | 0.139051095 |
| Rpp38 | 43.57699851 | -0.727244916 | 0.214280999 | -3.393884297 |
| Acbd7 | 3.006698544 | 0.235623236 | 0.730587107 | 0.322512174 |
| Olah | 0.742270129 | 1.0805141 | 1.033651331 | 1.045337115 |
| Gm37525 | 0.038981395 | -0.443816764 | 3.724945049 | -0.119147198 |
| Meig1 | 9.115918498 | -0.613133507 | 0.342397774 | -1.790705293 |
| Dclre1c | 259.4107841 | -0.163827473 | 0.152080249 | -1.077243584 |
| Suv39h2 | 54.79723866 | -0.046668025 | 0.167269624 | -0.278998802 |
| Gm13184 | 2.921456852 | 0.325532302 | 0.66693339 | 0.48810317 |
| Hspa14 | 484.0326706 | -0.009889437 | 0.114556984 | -0.086327669 |
| Cdnf | 15.49217047 | -0.021397263 | 0.272987175 | -0.078381935 |
| Gm13186 | 0.075043205 | 0.517957688 | 3.724945049 | 0.139051095 |
| Gm13182 | 0.275763028 | -1.290014649 | 1.968841301 | -0.65521515 |
| Fam107b | 448.5992859 | -0.604069656 | 0.175366551 | -3.4446116 |
| Gm13183 | 0 | NA | NA | NA |
| Gm13181 | 0.648853987 | -1.491734447 | 1.342080391 | -1.111509011 |
| Gm13185 | 0.04393175 | 0.037062677 | 3.724945049 | 0.009949859 |

| | | | | |
|---|---|---|---|---|
| Gm13180 | 0.088736197 | 0.51795324 | 3.724945049 | 0.139049901 |
| Gm13191 | 0.045882455 | 0.517959912 | 3.724945049 | 0.139051692 |
| Frmd4a | 1510.085395 | -0.361120077 | 0.062703841 | -5.759138019 |
| Gm2639 | 0.385855529 | 0.51795975 | 3.279807079 | 0.157923847 |
| Gm13188 | 0 | NA | NA | NA |
| Gm10862 | 12.42465393 | 1.701163535 | 0.35286402 | 4.821017268 |
| Gm38085 | 10.53103825 | 1.542735779 | 0.460048467 | 3.353420103 |
| 1700080N15Rik | 0 | NA | NA | NA |
| Gm13189 | 2.288528955 | 2.264947043 | 0.921179214 | 2.458747449 |
| Gm37814 | 13.6296294 | 1.825346438 | 0.468114628 | 3.89935783 |
| Gm37107 | 25.13242011 | 0.978506602 | 0.381974945 | 2.561703628 |
| Gm38348 | 18.53952862 | 0.98795889 | 0.444226081 | 2.224000193 |
| Gm13187 | 1.991338835 | -1.601013842 | 0.783806728 | -2.042613038 |
| Gm36988 | 22.65632828 | -0.080564232 | 0.359784518 | -0.223923564 |
| Gm23691 | 0 | NA | NA | NA |
| Gm13175 | 0 | NA | NA | NA |
| Gm13179 | 0.804355203 | -0.436943224 | 1.023700389 | -0.426827252 |
| Prpf18 | 625.4248675 | -0.054951993 | 0.065319901 | -0.841274904 |
| Gm13196 | 0.049553047 | -0.443818988 | 3.724945049 | -0.119147795 |
| Bend7 | 58.21736025 | -0.353468622 | 0.148390529 | -2.382016057 |
| Gm13176 | 0 | NA | NA | NA |
| Sephs1 | 350.7542985 | 0.162703508 | 0.081216475 | 2.003331323 |
| Phyh | 939.6459375 | 0.128827618 | 0.083143202 | 1.549466634 |
| Gm13194 | 2.056906148 | 0.763839331 | 0.874185514 | 0.873772579 |
| Gm13193 | 0 | NA | NA | NA |
| Gm13192 | 14.54002108 | 0.239403266 | 0.271107158 | 0.883057709 |
| Ucma | 8.601202928 | -0.164163268 | 0.366961244 | -0.4473586 |
| Mcm10 | 26.1128149 | -0.588503799 | 0.383681293 | -1.533835008 |
| Optn | 314.9999699 | 0.097583896 | 0.09098563 | 1.072519875 |
| 4930551O13Rik | 0.150455055 | -1.336248778 | 2.838957719 | -0.470682874 |
| Gm26776 | 1.257166043 | 0.694099821 | 1.077748199 | 0.644027818 |
| Ccdc3 | 115.3722469 | -0.366293471 | 0.11235481 | -3.260149452 |
| Gm13198 | 1.380091768 | -0.110233664 | 0.943759429 | -0.116802715 |
| Gm23118 | 0 | NA | NA | NA |
| Gm13197 | 0.068097359 | 0.037064902 | 3.724945049 | 0.009950456 |
| Camk1d | 1938.800003 | 0.386861144 | 0.091864793 | 4.211201367 |
| Gm13216 | 0.097501745 | -0.443821212 | 3.724945049 | -0.119148392 |
| Cdc123 | 504.6193172 | 0.034782782 | 0.071266312 | 0.488067657 |
| Nudt5 | 187.3923638 | -0.360964311 | 0.131464104 | -2.74572525 |
| Gm13199 | 8.870008164 | -0.508370628 | 0.403701405 | -1.259273863 |
| Sec61a2 | 789.7387573 | 0.044013646 | 0.113516204 | 0.387730073 |
| Gm13267 | 0.3753498 | -1.404487968 | 1.607188348 | -0.873878889 |
| Dhtkd1 | 85.77264844 | 0.537040708 | 0.155304267 | 3.457990692 |
| Upf2 | 873.5434251 | -0.1522413 | 0.079773864 | -1.908410763 |
| Proser2 | 48.48109559 | -0.068763915 | 0.171690361 | -0.400511216 |
| Gm13384 | 0 | NA | NA | NA |
| Gm10857 | 0.297167372 | -0.24930374 | 1.710440953 | -0.145754076 |

| | | | | |
|---|---|---|---|---|
| Echdc3 | 8.401856911 | -0.767777948 | 0.371682089 | -2.065684552 |
| A230108P19Rik | 0.266247585 | 0.998861067 | 2.508393958 | 0.398207412 |
| Gm13383 | 0.729279448 | -1.238332203 | 1.232623858 | -1.004631052 |
| Usp6nl | 655.1216898 | 0.008077205 | 0.071906795 | 0.112328812 |
| Gm13388 | 0.223035371 | 0.980643623 | 2.557900868 | 0.383378275 |
| Gm13391 | 5.144468292 | -0.029875804 | 0.449150335 | -0.066516268 |
| Gm38386 | 2.604362067 | -2.346777727 | 0.791249495 | -2.965913712 |
| Celf2 | 6175.024994 | 0.475486686 | 0.120601687 | 3.942620514 |
| Gm13389 | 0.654952557 | 0.114020428 | 1.385321893 | 0.08230609 |
| Gm10855 | 169.6995702 | 0.773638566 | 0.499906593 | 1.54756624 |
| Gm37340 | 0.066050875 | -0.443818988 | 3.724945049 | -0.119147795 |
| Gm10115 | 101.9444587 | 1.056784583 | 0.474039163 | 2.229319146 |
| Gm26315 | 191.8007175 | -0.054165317 | 0.253145905 | -0.213968768 |
| Gm26565 | 0.531771505 | 1.050189009 | 1.453504364 | 0.722522089 |
| Gm24340 | 0 | NA | NA | NA |
| Gm28641 | 0.190780345 | 0.03706236 | 3.299406433 | 0.011233039 |
| Gm13210 | 0.681349771 | -2.664460476 | 1.511952656 | -1.762264489 |
| Gm13211 | 0.138642343 | 0.434745388 | 3.605171816 | 0.120589367 |
| Gm13254 | 0.062426873 | 0.517959912 | 3.724945049 | 0.139051692 |
| Gm24534 | 0 | NA | NA | NA |
| Gm13255 | 0 | NA | NA | NA |
| Gm13217 | 27.36624143 | 0.310440091 | 0.197890719 | 1.568745076 |
| 1700061F12Rik | 0 | NA | NA | NA |
| Gm23963 | 0 | NA | NA | NA |
| Gm13221 | 0 | NA | NA | NA |
| Gm13219 | 0 | NA | NA | NA |
| Gm13220 | 0 | NA | NA | NA |
| Gm13218 | 0 | NA | NA | NA |
| Gm37866 | 0 | NA | NA | NA |
| Gata3 | 51.8974266 | -0.198183331 | 0.160828289 | -1.232266615 |
| Gm13256 | 0.066050875 | -0.443818988 | 3.724945049 | -0.119147795 |
| 4930412O13Rik | 0.725537781 | 0.089093494 | 1.477607579 | 0.060295775 |
| 9230102O04Rik | 0.685158741 | 0.479804663 | 1.290691983 | 0.371742189 |
| Gm13262 | 0 | NA | NA | NA |
| Taf3 | 355.5580828 | 0.019331279 | 0.066197165 | 0.292025779 |
| C630004M23Rik | 3.262821873 | -0.20792256 | 0.706486978 | -0.294304872 |
| Atp5c1 | 1526.152544 | 0.077086081 | 0.079582519 | 0.96863082 |
| Gm23608 | 6.323166952 | 0.723425583 | 0.45764227 | 1.580766529 |
| Kin | 91.12058367 | -0.21714817 | 0.131993426 | -1.645143824 |
| Itih2 | 90.97624014 | -0.618859598 | 0.222904412 | -2.77634522 |
| Itih5 | 570.7931782 | 0.366209459 | 0.154735421 | 2.366681497 |
| Gm13261 | 0.038981395 | -0.443816764 | 3.724945049 | -0.119147198 |
| Sfmbt2 | 174.706509 | -0.223850113 | 0.122523541 | -1.826996765 |
| Gm22072 | 0.909783392 | 0.831085519 | 1.131336878 | 0.734604816 |
| Gm22677 | 0.580675992 | -0.880789537 | 1.63816194 | -0.537669393 |
| Mir466m | 0.940874839 | 1.117818943 | 1.117568259 | 1.000224312 |
| Mir466f-1 | 0.953625643 | 0.257543771 | 1.151290255 | 0.223700122 |

| | | | | |
|---|---|---|---|---|
| Mir669f | 1.177175292 | -1.653410211 | 0.959692173 | -1.722854741 |
| Mir669e | 0.868322942 | 0.768494286 | 1.020959548 | 0.752717664 |
| Mir669b | 1.131281954 | -0.193214367 | 1.058191229 | -0.182589273 |
| Gm23578 | 1.362203202 | -0.298608283 | 0.852448097 | -0.350294973 |
| Mir669d | 0.802169344 | 0.499536606 | 1.118138328 | 0.44675743 |
| Mir466f-2 | 0.667839854 | -0.891988344 | 1.324668591 | -0.673367172 |
| Mir669l | 0.610120006 | 1.02486457 | 1.331389243 | 0.769770805 |
| Mir669d-2 | 0.709755772 | 0.221251038 | 1.223029335 | 0.180904114 |
| Mir466f-3 | 0.856346278 | 1.429756905 | 1.109180878 | 1.289020514 |
| Mir297a-2 | 0.78950392 | -1.615228979 | 1.208384612 | -1.336684498 |
| Mir466o | 1.891729189 | 0.241589962 | 0.720953274 | 0.335097947 |
| Gm26073 | 1.013961535 | 0.26458522 | 0.947966179 | 0.279108291 |
| Mir467c | 0.194094924 | -1.37604891 | 2.224736657 | -0.618522155 |
| mmu-mir-466b-1 | 0.068097359 | 0.037064902 | 3.724945049 | 0.009950456 |
| Mir669a-3 | 0.890972926 | 1.095835338 | 1.313612962 | 0.834214772 |
| Mir669k | 1.409213138 | 0.187207884 | 0.939466505 | 0.199270419 |
| Gm26491 | 1.682866984 | 0.88820337 | 0.846630201 | 1.049104282 |
| Mir467a-1 | 0.921542032 | 0.111333765 | 1.105327988 | 0.100724642 |
| Mir466b-8 | 0.330565531 | -0.413968905 | 1.696012797 | -0.244083598 |
| Mir669a-1 | 1.495866474 | 0.310040119 | 0.80130858 | 0.386917259 |
| Mir669g | 0.495109938 | 0.471441226 | 1.330531683 | 0.354325441 |
| Gm25701 | 0.561285994 | -0.650964464 | 1.208808433 | -0.538517474 |
| Mir669j | 1.067120961 | 0.245900012 | 1.159944674 | 0.21199288 |
| Mir467a-2 | 0.612405649 | -0.325788879 | 1.223425327 | -0.26629241 |
| mmu-mir-466e | 0.75911489 | -0.917595953 | 1.313818263 | -0.69841924 |
| mmu-mir-669a-4 | 2.460216864 | 0.341458965 | 0.637199421 | 0.535874568 |
| Gm24315 | 0.616248996 | -0.122667884 | 1.256656652 | -0.097614479 |
| Gm25451 | 0.626970746 | 1.322367316 | 1.440309661 | 0.918113203 |
| Mir467b | 1.221212496 | -0.327468878 | 0.920385545 | -0.35579533 |
| mmu-mir-466c-1 | 1.101702777 | -1.205177293 | 1.066964883 | -1.129537919 |
| Gm24761 | 0.825397108 | -1.143238811 | 1.010397446 | -1.131474368 |
| Gm23651 | 1.10380611 | 0.99878716 | 1.050526886 | 0.950748784 |
| Mir467a-3 | 0 | NA | NA | NA |
| Mir466c-2 | 0 | NA | NA | NA |
| Gm23108 | 0.643541246 | 0.356074269 | 1.483547798 | 0.240015367 |
| Gm25680 | 0 | NA | NA | NA |
| Gm22775 | 0.32589515 | -0.480303127 | 1.763008662 | -0.272433787 |
| Mir467a-4 | 0.173861546 | -1.573537539 | 2.700165011 | -0.582756066 |
| Mir466b-4 | 0 | NA | NA | NA |
| Gm26060 | 0 | NA | NA | NA |
| Gm26431 | 0.062426873 | 0.517959912 | 3.724945049 | 0.139051692 |
| Gm26229 | 0.045882455 | 0.517959912 | 3.724945049 | 0.139051692 |
| Mir467a-5 | 0 | NA | NA | NA |
| Mir466b-5 | 0 | NA | NA | NA |
| Mir669p-1 | 0 | NA | NA | NA |
| Gm24167 | 0.038981395 | -0.443816764 | 3.724945049 | -0.119147198 |
| Gm24640 | 0 | NA | NA | NA |

| | | | | |
|---|---|---|---|---|
| Gm23288 | 0.060174537 | 0.517957688 | 3.724945049 | 0.139051095 |
| Mir467a-6 | 0 | NA | NA | NA |
| Gm25285 | 0.094831987 | 0.517957688 | 3.724945049 | 0.139051095 |
| Gm24673 | 0.458955183 | 1.026440579 | 1.423382849 | 0.721127545 |
| Gm22635 | 0.309435895 | 0.517948876 | 3.724945049 | 0.139048729 |
| Gm23569 | 0.231961655 | -0.438143924 | 2.127211574 | -0.205971014 |
| Mir467a-7 | 0 | NA | NA | NA |
| Mir466b-6 | 0 | NA | NA | NA |
| Gm26426 | 0 | NA | NA | NA |
| Gm24488 | 0.108266322 | -0.955981391 | 3.722845977 | -0.25678779 |
| Gm25158 | 0.038981395 | -0.443816764 | 3.724945049 | -0.119147198 |
| Mir467a-8 | 0 | NA | NA | NA |
| Mir466b-7 | 0.158320866 | -0.93971554 | 2.813533518 | -0.333998346 |
| Mir669p-2 | 0 | NA | NA | NA |
| Gm24028 | 0.085433889 | 0.037058229 | 3.724945049 | 0.009948665 |
| Gm22626 | 0.305016863 | -0.546171093 | 2.026948286 | -0.269454873 |
| Gm22166 | 0.060174537 | 0.517957688 | 3.724945049 | 0.139051095 |
| Mir467a-9 | 0.048949532 | 0.037064902 | 3.724945049 | 0.009950456 |
| Mir466b-2 | 0.150451996 | 0.037073243 | 2.858243881 | 0.012970637 |
| Gm26156 | 1.082398122 | -1.489062732 | 1.10729516 | -1.344774894 |
| Gm23571 | 0.95850512 | 0.004427061 | 1.135794827 | 0.003897765 |
| Gm25316 | 0.639171633 | 0.029503051 | 1.29501797 | 0.022781962 |
| Gm25022 | 0.159191956 | -0.443821639 | 2.808135686 | -0.158048502 |
| Gm25943 | 0.342780899 | -0.391303352 | 1.579368702 | -0.247759343 |
| Gm23988 | 0.869956765 | -0.721051398 | 1.026407992 | -0.702499789 |
| Gm25673 | 0.68128338 | -0.258695019 | 1.180448759 | -0.219149724 |
| Gm25753 | 0.533146218 | -1.366518859 | 1.39310676 | -0.980914671 |
| Mir467a-10 | 0 | NA | NA | NA |
| Mir466b-3 | 0.655224956 | 0.983653405 | 1.338968136 | 0.73463541 |
| Gm26398 | 0.832395869 | -1.846236417 | 1.196016667 | -1.543654421 |
| Mir467e | 0.572335319 | 0.660063971 | 1.250275483 | 0.527934827 |
| Mir466p | 1.677881506 | -0.589235337 | 0.837243343 | -0.70378026 |
| Gm22051 | 0.979291814 | -0.765428092 | 1.194599112 | -0.64074055 |
| Gm22678 | 0.741352231 | 0.818197847 | 1.18720022 | 0.689182695 |
| Mir467d | 0.189382193 | -0.443822435 | 2.706644442 | -0.163975152 |
| Mir466 | 0.816872591 | -0.360164287 | 1.131534837 | -0.318297126 |
| Gm22091 | 0.829130969 | 0.039377343 | 1.258323231 | 0.031293504 |
| Mir297c | 0.615674597 | -1.128194744 | 1.341633514 | -0.840911272 |
| Mir669c | 0.542133415 | 2.092050106 | 1.642140136 | 1.273977817 |
| Gm24342 | 0.492872509 | 0.036561438 | 1.394256981 | 0.026222884 |
| Gm25263 | 0.34438493 | 0.496892061 | 1.473338827 | 0.337255798 |
| Mir669a-2 | 0.768366922 | -0.502944212 | 1.153778337 | -0.435910605 |
| Gm22239 | 0.500671887 | 0.437711753 | 1.514869043 | 0.288943625 |
| Gm22925 | 0.21460851 | 0.037072591 | 2.607481384 | 0.014217778 |
| Gm23542 | 0.931192987 | 0.385298584 | 0.956614357 | 0.402773156 |
| Mir297b | 0.635701853 | -0.034927392 | 1.258052911 | -0.027763055 |
| Mir466d | 0.422902843 | 0.539838167 | 1.54298817 | 0.349865396 |

| | | | | |
|---|---|---|---|---|
| Gm26092 | 0.669827454 | 2.342391041 | 1.323153172 | 1.77030981 |
| Mir669m-1 | 0.636524932 | -2.593832896 | 1.247198995 | -2.079726576 |
| Mir669m-2 | 0.70834526 | 0.278307981 | 1.20290217 | 0.23136377 |
| Mir466n | 0.60136891 | -1.192893815 | 1.256993896 | -0.949005257 |
| Mir669o | 1.139979212 | -0.275368647 | 0.97173901 | -0.283377166 |
| Mir466g | 0.512204714 | -0.416899326 | 1.333648425 | -0.312600621 |
| Mir466h | 0.440637317 | 0.756499098 | 3.385008554 | 0.223485136 |
| Mir297a-3 | 0.589191525 | 0.203814201 | 1.478146342 | 0.137884995 |
| Mir466l | 0.747968401 | 1.117495098 | 1.149343048 | 0.972290301 |
| Mir297a-4 | 0.343551435 | -1.429247173 | 1.736202032 | -0.82320326 |
| Gm24806 | 0.654401101 | 0.310961848 | 1.156636855 | 0.268850026 |
| Mir669i | 0.597540007 | -0.761064852 | 1.288901857 | -0.59047541 |
| Mir669h | 0.264963494 | 0.018395373 | 2.091368767 | 0.008795853 |
| Gm18547 | 0 | NA | NA | NA |
| Gm13265 | 0.236919184 | 1.852419947 | 2.480798692 | 0.746703049 |
| Gm13263 | 0.565796537 | 2.413280202 | 1.405508206 | 1.717016088 |
| Gm37343 | 0 | NA | NA | NA |
| Gm23877 | 0 | NA | NA | NA |
| Gm13264 | 0.423046101 | -0.369557857 | 1.652586033 | -0.223623974 |
| Gm26478 | 0 | NA | NA | NA |
| Gm13297 | 0 | NA | NA | NA |
| Gm13294 | 1.544067245 | 0.92774369 | 0.829797016 | 1.118036908 |
| Prkcq | 193.2675698 | -0.182481859 | 0.120969279 | -1.508497541 |
| Gm36932 | 3.868039749 | -0.627851356 | 0.535221549 | -1.173068156 |
| Gm37520 | 0.177524535 | -1.515341479 | 2.72178371 | -0.556745737 |
| Gm37766 | 3.664845353 | -0.766430458 | 0.694538015 | -1.103511171 |
| Gm38171 | 1.860369092 | -1.809049128 | 0.847816497 | -2.133774389 |
| Gm37851 | 0 | NA | NA | NA |
| 8030442B05Rik | 2.076510791 | -1.243872185 | 0.811690956 | -1.532445539 |
| Gm13293 | 46.58195302 | -0.422860207 | 0.176821911 | -2.39144687 |
| Gm37730 | 0.669761723 | 0.521526942 | 1.426660851 | 0.365557758 |
| 4933403L11Rik | 0.631690809 | -0.450318127 | 1.318879637 | -0.341439897 |
| Gm13291 | 0.045882455 | 0.517959912 | 3.724945049 | 0.139051692 |
| Gm13292 | 11.7837924 | 0.403049373 | 0.296069801 | 1.361332268 |
| Gm13296 | 0 | NA | NA | NA |
| Pfkfb3 | 910.8026235 | -0.746244638 | 0.130373918 | -5.723879808 |
| Gm10851 | 1.168248907 | -2.511061671 | 1.020445795 | -2.460749687 |
| Gm37975 | 7.09582318 | 0.40220645 | 0.56761492 | 0.708590342 |
| Rbm17 | 378.7991564 | -0.153983965 | 0.130337809 | -1.18142208 |
| Gm13260 | 0.138087489 | -1.320984914 | 3.611429328 | -0.365778974 |
| Gm17490 | 0 | NA | NA | NA |
| Il2ra | 71.84653869 | -0.81584649 | 0.293042014 | -2.784059799 |
| Gm37881 | 0.909342915 | -1.861992616 | 2.500052234 | -0.744781485 |
| Il15ra | 192.4286151 | -0.425368771 | 0.212616888 | -2.000634918 |
| Fbxo18 | 817.0147377 | -0.05450946 | 0.136319655 | -0.399865006 |
| Ankrd16 | 279.6460949 | 0.486090879 | 0.17741375 | 2.739871505 |
| Gm34768 | 0.194631057 | 1.456047977 | 3.23213671 | 0.450490839 |

| | | | | |
|---|---|---|---|---|
| Gm13310 | 0.075043205 | 0.517957688 | 3.724945049 | 0.139051095 |
| Itga8 | 97.06132719 | 1.322822398 | 0.227281338 | 5.820198031 |
| Gm37701 | 1.141257353 | 1.97895153 | 1.121361086 | 1.764776355 |
| E030013I19Rik | 6.292898879 | 1.085882413 | 0.515212166 | 2.107641248 |
| Fam188a | 576.4656474 | 0.02357192 | 0.088325971 | 0.266874162 |
| Gm37289 | 14.48869372 | 0.624917525 | 0.342026837 | 1.827100852 |
| Gm37255 | 40.61645279 | 0.383018453 | 0.312177794 | 1.226924081 |
| Gm38014 | 40.65078467 | 0.517282273 | 0.328526633 | 1.574552018 |
| Gm13321 | 0 | NA | NA | NA |
| Gm37565 | 0.527338585 | -3.009285922 | 3.145289142 | -0.956759708 |
| Pter | 24.4696158 | -0.53276423 | 0.232659889 | -2.28988431 |
| C1ql3 | 535.3698455 | 0.649803851 | 0.083791427 | 7.755015898 |
| Gm37824 | 3.481665934 | 0.338304623 | 0.605324183 | 0.558881724 |
| Gm37811 | 9.289570475 | 0.793008353 | 0.3691089 | 2.148440077 |
| Gm37356 | 2.692269002 | 1.610998325 | 0.724601199 | 2.22328962 |
| Gm37742 | 0.85289683 | 1.798868963 | 1.203021494 | 1.495292454 |
| Rsu1 | 237.1157615 | -0.441896147 | 0.11834577 | -3.733941218 |
| Gm38156 | 11.10230631 | -0.761181624 | 0.439991912 | -1.729990037 |
| Gm37780 | 17.42481316 | -0.608014497 | 0.304336463 | -1.99783651 |
| Gm27540 | 0 | NA | NA | NA |
| Gm37160 | 10.15377686 | -0.387539522 | 0.359318183 | -1.078541362 |
| Gm13270 | 8.36952636 | -0.546437557 | 0.403748354 | -1.353411232 |
| Cubn | 24.44027687 | -0.661151279 | 0.265910059 | -2.48637183 |
| Trdmt1 | 114.6086781 | -0.161505106 | 0.12410518 | -1.301356689 |
| Vim | 4040.904579 | -2.082350501 | 0.282261014 | -7.377393245 |
| St8sia6 | 68.51491012 | 0.130821045 | 0.163915172 | 0.798102109 |
| Gm37126 | 0 | NA | NA | NA |
| Hacd1 | 81.82130967 | -0.070089259 | 0.143617165 | -0.488028427 |
| D930036K23Rik | 55.51995555 | 0.169131692 | 0.216172702 | 0.782391533 |
| Gm37894 | 18.6674481 | 0.758231186 | 0.321517981 | 2.358285479 |
| Stamos | 3.400319483 | -0.916617947 | 0.587813044 | -1.559369867 |
| Stam | 1143.237138 | -0.058612456 | 0.058872273 | -0.995586754 |
| Gm37499 | 111.4820659 | 0.151856886 | 0.133050579 | 1.141347052 |
| Tmem236 | 7.691974964 | -1.355226822 | 0.410250789 | -3.30341064 |
| Gm13320 | 0.278513079 | 1.096415486 | 2.083018116 | 0.526359074 |
| Mrc1 | 62.46851479 | 0.385430811 | 0.283388295 | 1.36008021 |
| Mir511 | 0.109727584 | 0.037071574 | 3.724945049 | 0.009952247 |
| Gm37697 | 0.16123844 | 0.037068524 | 2.802306608 | 0.013227862 |
| Slc39a12 | 432.812937 | 0.680897114 | 0.177150599 | 3.843606051 |
| Gm13266 | 1.979745462 | 0.968326386 | 0.756515141 | 1.279982825 |
| Cacnb2 | 1007.083002 | 0.693425443 | 0.151511302 | 4.576724222 |
| Gm10849 | 2.270217088 | 1.042283181 | 0.794720968 | 1.311508345 |
| Gm38105 | 15.2328292 | 1.07999641 | 0.32230806 | 3.350820356 |
| Gm13268 | 1.66036403 | 0.074928987 | 0.713492083 | 0.105017265 |
| Gm13312 | 24.04166377 | 0.966793554 | 0.448191885 | 2.157097408 |
| 2900092N22Rik | 49.08839729 | 1.020010901 | 0.321245661 | 3.175174096 |
| Gm13314 | 0.346172528 | 0.706442623 | 1.716688929 | 0.411514638 |

| | | | | |
|---|---|---|---|---|
| Gm13311 | 0.601079571 | 1.711950685 | 1.290092016 | 1.326998899 |
| Gm10848 | 3.555860714 | 0.794709082 | 0.617640609 | 1.286685283 |
| Gm37006 | 66.77607684 | 0.910043205 | 0.352006313 | 2.585303649 |
| Gm13315 | 24.30272673 | 1.211499686 | 0.438425461 | 2.763296832 |
| Gm38273 | 9.78169311 | -0.098992163 | 0.38185042 | -0.2592433 |
| Gm13269 | 0.198547815 | -0.387702284 | 2.601214258 | -0.149046655 |
| Gm13316 | 0.175141309 | -0.427154801 | 2.304192649 | -0.185381548 |
| Gm37437 | 53.34605896 | 1.080348139 | 0.380117321 | 2.842143939 |
| Gm37891 | 58.0044897 | 1.07504438 | 0.261010049 | 4.118785407 |
| Nsun6 | 110.3532385 | -0.06204921 | 0.158797662 | -0.390743854 |
| Gm36930 | 6.501557468 | 0.966443244 | 0.415495291 | 2.326002884 |
| Arl5b | 609.464288 | -0.086730416 | 0.102494162 | -0.846198595 |
| Gm38257 | 38.93688588 | 0.235394661 | 0.301722725 | 0.780168814 |
| Gm13313 | 0.521845371 | -0.329777891 | 1.320813455 | -0.24967787 |
| Gm13367 | 0 | NA | NA | NA |
| Gm13366 | 0 | NA | NA | NA |
| Gm13365 | 0.11997416 | 0.517957688 | 3.724945049 | 0.139051095 |
| Gm13368 | 0 | NA | NA | NA |
| Malrd1 | 1.059665568 | 0.65288928 | 0.906846537 | 0.719955641 |
| Gm37595 | 0 | NA | NA | NA |
| Gm6587 | 0 | NA | NA | NA |
| Gm37416 | 2.356264561 | 0.448820239 | 0.672903689 | 0.666990308 |
| Plxdc2 | 861.0341044 | 0.41833126 | 0.074973843 | 5.579696128 |
| Gm24933 | 0.072025462 | 0.517959912 | 3.724945049 | 0.139051692 |
| Gm13326 | 1.471932881 | 0.836388323 | 1.074790838 | 0.77818706 |
| Gm13325 | 0 | NA | NA | NA |
| Gm13322 | 3.121057985 | 1.143112526 | 0.747006428 | 1.530257952 |
| Nebl | 1622.729757 | -0.113421067 | 0.06386663 | -1.775904975 |
| Gm13324 | 0.045882455 | 0.517959912 | 3.724945049 | 0.139051692 |
| Mir6419 | 0 | NA | NA | NA |
| Gm13323 | 0 | NA | NA | NA |
| H2afb1 | 0.165156969 | -1.323927776 | 3.401541256 | -0.389214087 |
| Gm13329 | 0.121578509 | 0.037073798 | 3.724945049 | 0.009952844 |
| A930004D18Rik | 38.49094849 | -0.537501738 | 0.181735776 | -2.957600042 |
| Gm13330 | 0.172363982 | -0.443822677 | 2.784585521 | -0.159385543 |
| Skida1 | 111.6125948 | 0.139096575 | 0.121787847 | 1.142121958 |
| 1810059C17Rik | 0.960935127 | -1.310218461 | 1.029977541 | -1.272084495 |
| Mllt10 | 761.3719482 | 0.017437375 | 0.09588794 | 0.181851599 |
| Mir7655 | 0.321829426 | 0.517971997 | 1.764580388 | 0.29353834 |
| Gm23638 | 0 | NA | NA | NA |
| Gm25147 | 6.022279029 | -0.15116416 | 0.521729321 | -0.289736755 |
| Gm13351 | 0.9210764 | -0.078834265 | 2.063906559 | -0.038196625 |
| Dnajc1 | 546.8891029 | -0.220293777 | 0.070162652 | -3.139758404 |
| Gm13350 | 4.932017945 | 1.375360594 | 0.624532044 | 2.202225822 |
| Gm13349 | 8.865425504 | -0.262562411 | 0.44003397 | -0.596686687 |
| Gm13353 | 0 | NA | NA | NA |
| Gm13355 | 0.097139547 | -0.443816764 | 3.724945049 | -0.119147198 |

| | | | | |
|---|---|---|---|---|
| Gm13352 | 0.195196415 | -1.316741042 | 2.208871262 | -0.596114887 |
| Commd3 | 408.9733762 | -0.053023271 | 0.105991959 | -0.500257487 |
| Bmi1 | 684.4399484 | -0.238162808 | 0.096136 | -2.477353005 |
| Spag6l | 18.02665997 | 1.02549045 | 0.307575065 | 3.334114394 |
| Gm20539 | 2.239606267 | -1.137088756 | 0.75928588 | -1.497576587 |
| Gm13333 | 0.068097359 | 0.037064902 | 3.724945049 | 0.009950456 |
| Carlr | 12.90806688 | 0.135728671 | 0.275983259 | 0.491800376 |
| Pip4k2a | 1173.833449 | -0.091116071 | 0.112210848 | -0.812007682 |
| 4930426L09Rik | 9.715138516 | -0.116457723 | 0.323462463 | -0.360034737 |
| Gm13332 | 0 | NA | NA | NA |
| Armc3 | 8.367447745 | -0.105658109 | 0.389269708 | -0.271426486 |
| 4930447M23Rik | 25.2336097 | 1.125073903 | 0.235638661 | 4.774572643 |
| Msrb2 | 108.7981829 | -0.006951224 | 0.128706443 | -0.054008361 |
| Gm13344 | 0 | NA | NA | NA |
| Ptf1aos | 0 | NA | NA | NA |
| Ptf1a | 0.167536551 | 0.998860717 | 2.763312802 | 0.361472185 |
| 1810010K12Rik | 2.426680169 | 0.628717288 | 0.795440142 | 0.79040176 |
| 4921504E06Rik | 0.234904022 | -1.194726108 | 2.102427536 | -0.568260303 |
| Gm13345 | 0 | NA | NA | NA |
| Gm13348 | 0.146089079 | -0.444786384 | 3.522464394 | -0.126271364 |
| Gm13343 | 0.231405101 | 0.037075053 | 2.545874441 | 0.014562797 |
| Otud1 | 292.6909858 | -0.720623331 | 0.162969339 | -4.421833781 |
| Gm13346 | 0.525489152 | -1.632221106 | 1.55121521 | -1.052220927 |
| Gm13347 | 0.240250011 | 0.049552149 | 2.111286354 | 0.023470122 |
| Gm24670 | 0 | NA | NA | NA |
| BX571892.1 | 0.123992736 | 0.517955521 | 3.720643114 | 0.139211288 |
| Etl4 | 1406.655128 | 0.481353596 | 0.096631053 | 4.981355171 |
| Gm25859 | 5.580134033 | 1.091787866 | 0.5519012 | 1.978230643 |
| Gm13361 | 0.132101749 | -0.93550548 | 3.724188147 | -0.251197158 |
| Gm13363 | 9.874902262 | 0.083217701 | 0.413376911 | 0.201311922 |
| Gm13360 | 0.049553047 | -0.443818988 | 3.724945049 | -0.119147795 |
| Gm13328 | 2.887214733 | 0.883737963 | 0.583291011 | 1.51508929 |
| Gm23970 | 0.864953071 | 0.658711027 | 1.192859022 | 0.552211967 |
| Gm17171 | 1.518325143 | -0.203100698 | 0.865933247 | -0.234545445 |
| Gm13335 | 0.728721824 | 2.032868681 | 1.299335041 | 1.564545415 |
| Gm27446 | 0.851828701 | -0.442079682 | 1.290102635 | -0.342670164 |
| Gm13362 | 0.31548088 | -1.285457183 | 1.746574006 | -0.735987813 |
| Arhgap21 | 2959.631058 | 0.370065895 | 0.083673624 | 4.422730583 |
| Gm13380 | 18.50560142 | 0.50391981 | 0.391319507 | 1.287745182 |
| Gm13375 | 61.63895035 | 0.598974245 | 0.171283325 | 3.496979325 |
| Gm13377 | 0.2958732 | 0.956502823 | 1.754558718 | 0.545152929 |
| Gm13376 | 0.075043205 | 0.517957688 | 3.724945049 | 0.139051095 |
| Gm25368 | 0 | NA | NA | NA |
| Enkur | 19.33843805 | -0.610640661 | 0.273093092 | -2.236016505 |
| Gm23007 | 0.130861843 | -0.443820897 | 2.984315003 | -0.148717845 |
| Thnsl1 | 325.2700769 | -0.045267365 | 0.117382631 | -0.385639382 |
| Gm13378 | 1.494280535 | -0.365088019 | 0.873932099 | -0.417753301 |

| | | | | |
|---|---|---|---|---|
| Gm13327 | 3.973417911 | -0.084061173 | 0.533001179 | -0.157712922 |
| Gpr158 | 2965.115641 | 0.547740891 | 0.096959577 | 5.649167482 |
| Gm13379 | 0 | NA | NA | NA |
| Gm24886 | 0 | NA | NA | NA |
| Gm13337 | 5.556964523 | 0.050938622 | 0.482018934 | 0.105677638 |
| Myo3a | 0.925384949 | -0.270398999 | 1.067755288 | -0.253240608 |
| Gm13342 | 2.124844546 | -0.2475437 | 0.644352778 | -0.38417418 |
| Gm13341 | 22.54937325 | 0.485212511 | 0.339241186 | 1.430287748 |
| Gm13340 | 244.019484 | 0.288979436 | 0.143620259 | 2.012107749 |
| Gm13339 | 4.846303826 | 0.694964425 | 0.872807047 | 0.79624062 |
| Gad2 | 3413.524295 | -0.104390385 | 0.088284454 | -1.182432242 |
| mmu-mir-3967 | 2.839739068 | 0.140017721 | 0.644077793 | 0.217392562 |
| Gm23636 | 1.903429752 | -0.043691563 | 0.735482189 | -0.059405332 |
| Gm13338 | 0.254763098 | 0.492206883 | 1.829084814 | 0.269100087 |
| 1700092C17Rik | 0.045882455 | 0.517959912 | 3.724945049 | 0.139051692 |
| Gm13356 | 0.169165009 | 0.036125373 | 3.351346561 | 0.01077936 |
| Apbb1ip | 159.2402503 | -0.285029145 | 0.109297065 | -2.607838959 |
| A130006I12Rik | 3.281667885 | -0.316546998 | 0.606606326 | -0.521832669 |
| Pdss1 | 223.7617087 | 0.207317965 | 0.131922134 | 1.571517674 |
| Gm25544 | 3.664064784 | 0.430863626 | 0.671488118 | 0.641654878 |
| Gm14441 | 0 | NA | NA | NA |
| Abi1 | 1539.731744 | 0.033906053 | 0.049946716 | 0.678844504 |
| Gm13357 | 12.25840608 | 0.836529129 | 0.389306147 | 2.148769382 |
| Acbd5 | 986.3592934 | 0.152622337 | 0.076031169 | 2.007365389 |
| Mastl | 28.61317683 | -0.413955351 | 0.303869676 | -1.362279238 |
| Yme1l1 | 1605.641626 | 0.018124129 | 0.064707964 | 0.280091161 |
| Gm13423 | 14.64943441 | 0.3560951 | 0.298066613 | 1.194682951 |
| 4931423N10Rik | 14.857103 | 0.36055556 | 0.306476493 | 1.17645421 |
| Gm13422 | 10.18410332 | 0.207526228 | 0.320609055 | 0.647287484 |
| Nxph2 | 18.63269461 | 0.446991012 | 0.253559804 | 1.762862268 |
| Gm38309 | 0.149576918 | -0.443202057 | 2.863264737 | -0.15478906 |
| Spopl | 699.9338885 | 0.167285421 | 0.093948079 | 1.78061565 |
| Gm13331 | 22.66616647 | 0.143622961 | 0.326791212 | 0.439494563 |
| Sh3d2c-ps1 | 0.219384069 | 0.03707698 | 1.908351561 | 0.0194288 |
| Gm26323 | 0 | NA | NA | NA |
| Gm13392 | 0.616639959 | -0.03954951 | 1.369768179 | -0.028873141 |
| Hnmt | 147.4068612 | 0.463044612 | 0.127199208 | 3.640310504 |
| Tbpl2 | 0 | NA | NA | NA |
| Il1f8 | 0 | NA | NA | NA |
| Gm13410 | 44.62557519 | -0.04549371 | 0.164245243 | -0.276986468 |
| Il1f9 | 2.482099328 | -1.360771473 | 0.712570388 | -1.909666043 |
| Il1f6 | 0 | NA | NA | NA |
| Gm37703 | 0 | NA | NA | NA |
| Gm13409 | 0.143485256 | -0.955563601 | 3.547405725 | -0.269369696 |
| Il1f5 | 0.291332888 | 0.988804016 | 3.700633271 | 0.267198596 |
| Il1f10 | 0 | NA | NA | NA |
| Gm13411 | 0.061920613 | -0.443816764 | 3.724945049 | -0.119147198 |

| | | | | |
|---|---|---|---|---|
| Il1rn | 704.4699326 | -1.159424019 | 0.661241456 | -1.753404914 |
| Psd4 | 122.4570999 | -1.524124051 | 0.231057274 | -6.596304136 |
| Pax8 | 14.32991156 | 0.021791647 | 0.487151314 | 0.044732811 |
| Gm13415 | 0.371620636 | -0.509591414 | 1.690971074 | -0.301360219 |
| Gm27193 | 0 | NA | NA | NA |
| Gm13418 | 0 | NA | NA | NA |
| Gm13417 | 0.292450921 | -0.432105816 | 2.095743266 | -0.20618261 |
| Gm38383 | 0 | NA | NA | NA |
| Gm13416 | 20.11579464 | -0.255073309 | 0.25184852 | -1.012804476 |
| Gm25455 | 0 | NA | NA | NA |
| Cacna1b | 2031.063943 | 0.54679458 | 0.130834741 | 4.179276663 |
| Gm13459 | 0 | NA | NA | NA |
| Ehmt1 | 682.9022921 | 0.140449147 | 0.105707344 | 1.328660266 |
| Gm38147 | 36.95672247 | 0.367274504 | 0.315316546 | 1.164780311 |
| Gm37139 | 53.01011326 | 0.574774049 | 0.306054203 | 1.878013908 |
| Arrdc1 | 89.73200459 | 0.055117522 | 0.123339663 | 0.446875894 |
| Gm22875 | 0.044544648 | 0.517957688 | 3.724945049 | 0.139051095 |
| Zmynd19 | 269.41162 | 0.159064458 | 0.121560636 | 1.308519461 |
| Dph7 | 184.0627944 | 0.053048081 | 0.088915889 | 0.596609684 |
| Mrpl41 | 329.4748938 | 0.15225115 | 0.119842869 | 1.270423106 |
| Pnpla7 | 332.8704237 | -0.07913501 | 0.108886697 | -0.726764727 |
| Gm13370 | 5.147326582 | 0.433509032 | 0.443143 | 0.978259913 |
| Gm13369 | 0.218114541 | 0.010450048 | 2.544355953 | 0.004107148 |
| Gm25518 | 0 | NA | NA | NA |
| Gm24875 | 0 | NA | NA | NA |
| Nsmf | 2802.591237 | 0.499263072 | 0.092260789 | 5.411432903 |
| Gm24328 | 0 | NA | NA | NA |
| Mir7664 | 1.286742338 | 1.755035133 | 1.155366764 | 1.519028578 |
| Entpd8 | 0.593920364 | -1.751273542 | 1.269845043 | -1.379123816 |
| Noxa1 | 0 | NA | NA | NA |
| Gm13387 | 1.538924929 | 0.364082364 | 0.962844673 | 0.378131982 |
| Nrarp | 119.8106717 | 0.904424078 | 0.266933921 | 3.388194636 |
| Gm38287 | 0.509714849 | 0.373519204 | 1.554409208 | 0.240296572 |
| Tor4a | 84.46660317 | -1.098904773 | 0.204538188 | -5.372614188 |
| Nelfb | 451.1973938 | -0.183337747 | 0.066505405 | -2.756734525 |
| Gm22572 | 0 | NA | NA | NA |
| 4933433C11Rik | 0.117650406 | -0.443821212 | 3.724945049 | -0.119148392 |
| Fam166a | 0.395870588 | 1.396566329 | 1.42093149 | 0.982852684 |
| Tubb4b | 1827.495376 | -0.213933882 | 0.098989763 | -2.161171768 |
| Slc34a3 | 1.160340464 | 0.027704564 | 0.896216006 | 0.030912819 |
| Rnf224 | 0.060174537 | 0.517957688 | 3.724945049 | 0.139051095 |
| Cysrt1 | 0.419553954 | 0.324632226 | 1.610592305 | 0.20156077 |
| Rnf208 | 937.0051333 | 0.486443817 | 0.067347563 | 7.222886739 |
| Ndor1 | 276.8690761 | -0.109666129 | 0.147265238 | -0.744684424 |
| Tmem203 | 61.6019059 | 0.135780736 | 0.17340221 | 0.783039244 |
| Tprn | 478.8306303 | 0.08586068 | 0.144769887 | 0.593083838 |
| Ssna1 | 171.2042094 | -0.329444039 | 0.123168204 | -2.674749059 |

| | | | | |
|---|---|---|---|---|
| Anapc2 | 780.9388462 | -0.074710578 | 0.074151397 | -1.007541074 |
| Tmem210 | 0.270178086 | -1.068497054 | 3.707630273 | -0.288188675 |
| Lrrc26 | 0.469296467 | -1.329234658 | 3.262428113 | -0.407437225 |
| Grin1 | 5090.601218 | 0.654764089 | 0.120259252 | 5.444604729 |
| Grin1os | 5.022424966 | 0.630494905 | 0.517746583 | 1.217767389 |
| AA543186 | 4.244450911 | -0.144419275 | 0.585831966 | -0.246519965 |
| Man1b1 | 806.4183462 | -0.064247307 | 0.061451723 | -1.04549236 |
| Gm25220 | 2.58962508 | 0.107653605 | 0.750173502 | 0.143504943 |
| Dpp7 | 187.4441202 | -0.431885524 | 0.108039431 | -3.997480559 |
| Uap1l1 | 215.8935137 | -0.640872503 | 0.140663705 | -4.556061582 |
| Sapcd2 | 9.831372002 | 0.339009943 | 0.413974841 | 0.818914362 |
| Entpd2 | 57.81792083 | -0.148088553 | 0.206889458 | -0.715785881 |
| Npdc1 | 1413.945948 | 0.10693743 | 0.0503557 | 2.123641005 |
| Fut7 | 5.453509397 | 0.453292809 | 0.591120331 | 0.766836776 |
| Abca2 | 5084.226177 | 0.279921133 | 0.070188435 | 3.988137523 |
| mmu-mir-3087 | 0 | NA | NA | NA |
| BC029214 | 267.2170635 | 0.377460969 | 0.085654571 | 4.406781392 |
| Clic3 | 1.915678769 | 0.188641619 | 0.766141681 | 0.24622289 |
| Ptgds | 4437.782382 | 0.704804929 | 0.160028516 | 4.404245871 |
| Lcn12 | 0.07796279 | -0.798301147 | 3.721829328 | -0.214491605 |
| C8g | 3.739276094 | 0.308926726 | 0.560545173 | 0.551118341 |
| Fbxw5 | 594.4106511 | 0.201264895 | 0.059768353 | 3.367415769 |
| Traf2 | 170.0261398 | -0.600148525 | 0.130285424 | -4.606413408 |
| Gm13433 | 0 | NA | NA | NA |
| Gm13432 | 0.086930093 | -0.443164241 | 3.724898049 | -0.118973522 |
| Edf1 | 525.3660091 | -0.161503049 | 0.200973495 | -0.803603722 |
| Mamdc4 | 22.73292264 | -0.277744871 | 0.323810792 | -0.857738155 |
| Phpt1 | 63.01003723 | 0.595239159 | 0.167759002 | 3.548180132 |
| Gm996 | 1271.384592 | 1.037521187 | 0.129842088 | 7.990638473 |
| Gm26702 | 5.898309265 | 1.168494443 | 0.606717819 | 1.925927352 |
| Rabl6 | 1439.377497 | 0.182895415 | 0.057014229 | 3.207890723 |
| Fcnaos | 8.104967773 | 0.358751294 | 0.377607911 | 0.950062972 |
| Ccdc183 | 0.262568881 | -0.926098635 | 1.823734561 | -0.507803413 |
| Tmem141 | 79.20590439 | 0.260886553 | 0.178743445 | 1.459558717 |
| Fcna | 2.11389961 | 0.691369312 | 0.838600687 | 0.824432083 |
| Lcn8 | 0.091666476 | 0.037060453 | 3.724945049 | 0.009949262 |
| Lcn5 | 0.13733263 | 0.037057285 | 3.640803958 | 0.010178325 |
| Lcn6 | 0.208472373 | 1.037039053 | 2.638636856 | 0.393020756 |
| Lcn10 | 0.112396584 | -0.917692504 | 3.724434845 | -0.246397787 |
| Obp2a | 0.229116515 | -0.396900707 | 2.177291514 | -0.182291027 |
| A230005M16Rik | 4.640067633 | 0.841597713 | 0.48248661 | 1.74429237 |
| Bmyc | 286.5842141 | 0.259739268 | 0.099114393 | 2.620600896 |
| Obp2b | 1.10611924 | -2.450890968 | 1.248640524 | -1.96284753 |
| Lcn3 | 0.068097359 | 0.037064902 | 3.724945049 | 0.009950456 |
| Lcn11 | 0 | NA | NA | NA |
| Gm13538 | 0.049553047 | -0.443818988 | 3.724945049 | -0.119147795 |
| Gm13539 | 0.066050875 | -0.443818988 | 3.724945049 | -0.119147795 |

| | | | | |
|---|---|---|---|---|
| Glt6d1 | 0 | NA | NA | NA |
| Lcn9 | 0.042716945 | 0.037062677 | 3.724945049 | 0.009949859 |
| Sohlh1 | 0.142434329 | -0.44382202 | 2.907370995 | -0.152654072 |
| Kcnt1 | 1559.190496 | 0.579358293 | 0.182475906 | 3.174985149 |
| Camsap1 | 1832.116154 | 0.015641823 | 0.069469756 | 0.225160182 |
| Gm13542 | 4.306872891 | 0.654445835 | 0.638170866 | 1.025502527 |
| Ubac1 | 350.8632011 | -0.062888721 | 0.067449875 | -0.93237713 |
| Nacc2 | 2614.976181 | -0.09860072 | 0.10185816 | -0.968019847 |
| Gm13554 | 0.124596252 | 0.037071632 | 3.714681146 | 0.009979761 |
| C330006A16Rik | 755.4666773 | 0.023178131 | 0.075849641 | 0.305579968 |
| 1810012K08Rik | 1.703298555 | -0.145669073 | 1.016527385 | -0.143300688 |
| Gm13553 | 0.097139547 | -0.443816764 | 3.724945049 | -0.119147198 |
| Lhx3 | 3.930402932 | -0.013090884 | 0.503533404 | -0.025998045 |
| Gm27196 | 0.679199126 | 0.106725551 | 1.18576852 | 0.090005384 |
| Qsox2 | 316.2840522 | -0.239896014 | 0.081791819 | -2.933007435 |
| C030048H21Rik | 9.452175138 | 0.509817377 | 0.408232581 | 1.248840491 |
| 4932418E24Rik | 35.63244048 | 1.390586691 | 0.258098754 | 5.387808623 |
| Gpsm1 | 740.3008245 | 0.207702356 | 0.091738309 | 2.264074388 |
| Dnlz | 309.3938586 | -0.341016089 | 0.110371679 | -3.089706447 |
| Card9 | 28.78353682 | -0.251860805 | 0.234888145 | -1.072258478 |
| Gm13562 | 6.216571396 | 0.175210739 | 0.487692174 | 0.35926502 |
| Snapc4 | 357.5941975 | 0.196095711 | 0.173509559 | 1.130172379 |
| Gm13563 | 26.0969972 | 0.147487115 | 0.285746379 | 0.516146923 |
| Sdccag3 | 585.8077495 | -0.045027374 | 0.074009286 | -0.608401687 |
| Pmpca | 864.0585262 | -0.044678419 | 0.066328563 | -0.673592452 |
| Inpp5e | 354.6391807 | -0.041932069 | 0.091526344 | -0.458142077 |
| Sec16a | 1481.474974 | -0.238783426 | 0.045620802 | -5.23409089 |
| 0610009E02Rik | 13.44346893 | 0.08061059 | 0.35183704 | 0.229113428 |
| Gm13568 | 2.19974275 | -0.283394874 | 0.68924071 | -0.411169669 |
| Notch1 | 1250.567747 | -0.763120266 | 0.117945097 | -6.47013134 |
| Mir6996 | 0.216707685 | -0.451880205 | 2.172962412 | -0.207955831 |
| Gm13358 | 0 | NA | NA | NA |
| Egfl7 | 168.1436807 | -0.704303117 | 0.122123437 | -5.76714129 |
| Gm20532 | 0 | NA | NA | NA |
| Mir126a | 0 | NA | NA | NA |
| mmu-mir-126b | 0 | NA | NA | NA |
| Agpat2 | 19.91019668 | -1.098205872 | 0.282225889 | -3.891230095 |
| Fam69b | 271.5122921 | 0.22872609 | 0.074800863 | 3.057800155 |
| Snhg7 | 0.151994617 | 0.037075901 | 2.849149129 | 0.013012973 |
| Snora43 | 0 | NA | NA | NA |
| Snora17 | 8.121164419 | -0.891801088 | 0.404000756 | -2.207424307 |
| Snhg7os | 0.566020539 | -0.038904955 | 1.374912225 | -0.028296319 |
| Lcn4 | 0.485200684 | -0.93561506 | 1.607581633 | -0.582001586 |
| Gm13359 | 0 | NA | NA | NA |
| Abo | 0.44885175 | -0.281521213 | 1.337834689 | -0.210430493 |
| Surf6 | 237.2764312 | -0.227545582 | 0.09165243 | -2.482701014 |
| Med22 | 318.481323 | -0.263284461 | 0.104866931 | -2.510652873 |

| | | | | |
|---|---|---|---|---|
| Rpl7a | 773.724984 | -0.21896077 | 0.131878132 | -1.6603266 |
| Gm23969 | 2.942191425 | 0.149519585 | 0.641925804 | 0.232923468 |
| Gm22879 | 0 | NA | NA | NA |
| Gm24134 | 0 | NA | NA | NA |
| Surf1 | 388.1526091 | -0.048420211 | 0.083386532 | -0.580671834 |
| Surf2 | 198.1272666 | -0.051121432 | 0.085944429 | -0.594819613 |
| Surf4 | 907.7278515 | -0.481468424 | 0.104617039 | -4.602198921 |
| Stkld1 | 2.796069476 | -0.398725529 | 0.7068743 | -0.564068503 |
| Rexo4 | 357.7792058 | -0.259390657 | 0.107982333 | -2.402158287 |
| Adamts13 | 7.574428338 | 0.419682471 | 0.40285221 | 1.041777754 |
| Cacfd1 | 416.4720264 | -0.145307352 | 0.072903721 | -1.9931404 |
| Slc2a6 | 213.3264522 | 0.386215521 | 0.103323395 | 3.737929075 |
| Gm13397 | 0.26409243 | -1.827653419 | 2.011062787 | -0.90879978 |
| Gm13398 | 0.795847279 | -0.401311071 | 1.117609772 | -0.35907978 |
| Tmem8c | 3.290439168 | 0.52265057 | 0.621459052 | 0.841005644 |
| Adamtsl2 | 40.03111391 | -0.013263852 | 0.218476324 | -0.060710705 |
| Fam163b | 723.0454923 | 0.777543993 | 0.149438275 | 5.203111411 |
| Dbhos | 1.810426623 | -0.589899129 | 0.81741481 | -0.721664352 |
| Dbh | 21.92637974 | -0.067517316 | 0.351349869 | -0.192165481 |
| Sardh | 177.0436753 | -0.623186446 | 0.166002764 | -3.754072712 |
| Sardhos | 0 | NA | NA | NA |
| Vav2 | 234.323641 | -0.037096181 | 0.095489849 | -0.388482974 |
| AA645442 | 0 | NA | NA | NA |
| Gm24049 | 0 | NA | NA | NA |
| Brd3 | 790.2310845 | 0.055093642 | 0.068908471 | 0.799519152 |
| Mir7578 | 0.414979285 | 0.538373862 | 1.474494407 | 0.365124384 |
| Wdr5 | 327.3142271 | -0.243058379 | 0.101225772 | -2.401151149 |
| Gm25541 | 8.286915501 | 0.619750399 | 0.38368172 | 1.615272157 |
| Gm13421 | 4.157031413 | -0.115686332 | 0.561077871 | -0.206185875 |
| Rxra | 493.0791904 | 0.038771849 | 0.080292772 | 0.482880932 |
| 2810430I11Rik | 3.721301919 | -0.550187083 | 0.618642866 | -0.889345232 |
| Col5a1 | 208.2168104 | -0.243519721 | 0.146369571 | -1.663731883 |
| Gm13372 | 0 | NA | NA | NA |
| Fcnb | 0.737107465 | -0.877573973 | 1.080380138 | -0.812282587 |
| Gm13371 | 0.299853729 | 1.434474065 | 1.790892285 | 0.800982883 |
| F730016J06Rik | 3.372121501 | 0.891821936 | 0.584700745 | 1.525262184 |
| Olfm1 | 4803.122293 | 0.377238134 | 0.095254547 | 3.960316269 |
| Gm13373 | 2.464667239 | 0.442462238 | 0.656750383 | 0.673714473 |
| Ppp1r26 | 345.0378359 | 0.391441649 | 0.069846307 | 5.604328476 |
| 1700007K13Rik | 15.71640074 | -0.309451485 | 0.317090656 | -0.975908556 |
| Mrps2 | 321.9683543 | -0.269904451 | 0.092686341 | -2.912019707 |
| Rpsa-ps9 | 16.7483217 | -0.069589632 | 0.288733148 | -0.241017122 |
| Gbgt1 | 30.60507794 | -0.205330467 | 0.222880288 | -0.92125898 |
| Gm10134 | 5.669552113 | 0.179801813 | 0.455326236 | 0.394885686 |
| Ralgds | 1434.688426 | -0.036394501 | 0.103620896 | -0.351227426 |
| Cel | 0.371662472 | -0.443819695 | 1.70749775 | -0.259924029 |
| Gm13381 | 0.066050875 | -0.443818988 | 3.724945049 | -0.119147795 |

| | | | | |
|---|---|---|---|---|
| Gtf3c5 | 232.8597514 | -0.33216728 | 0.08688137 | -3.823227927 |
| Gm22675 | 0 | NA | NA | NA |
| Gm22824 | 0.072025462 | 0.517959912 | 3.724945049 | 0.139051692 |
| Gfi1b | 0.585889909 | 0.94130614 | 1.412368139 | 0.666473644 |
| Tsc1 | 1195.112856 | 0.072580454 | 0.086419615 | 0.839860882 |
| 1700026L06Rik | 2.561385301 | -0.094500396 | 0.67162687 | -0.140703715 |
| Ak8 | 10.0543632 | 0.223014256 | 0.353157349 | 0.631486945 |
| Mir3088 | 0 | NA | NA | NA |
| Gm13385 | 0 | NA | NA | NA |
| Gm13386 | 0.066050875 | -0.443818988 | 3.724945049 | -0.119147795 |
| Gtf3c4 | 803.8265441 | -0.181911866 | 0.056995889 | -3.191666414 |
| Ddx31 | 110.419689 | -0.280358889 | 0.114485921 | -2.448850354 |
| Barhl1 | 48.18788692 | -0.00536112 | 0.255710071 | -0.020965618 |
| 1700101E01Rik | 8.146087367 | -0.537783106 | 0.344001749 | -1.563315033 |
| Gm13394 | 2432.869465 | 0.226735592 | 0.07509969 | 3.019128183 |
| Ttf1 | 162.3829832 | -0.502181487 | 0.141871242 | -3.539698961 |
| Gm13393 | 0.138076337 | 0.037074383 | 3.589802603 | 0.010327694 |
| Gm13396 | 0.120349074 | 1.027495679 | 3.723011807 | 0.275985071 |
| Setx | 1922.416912 | -0.020363171 | 0.067910175 | -0.299854486 |
| Ntng2 | 269.5695528 | 0.152962025 | 0.128110109 | 1.193988717 |
| 6530402F18Rik | 198.2560696 | -0.052994845 | 0.104876364 | -0.505307802 |
| Med27 | 188.5821441 | -0.109395726 | 0.104085634 | -1.051016569 |
| Gm13402 | 0.13742593 | 0.517962079 | 2.939744525 | 0.176192888 |
| Gm24976 | 1.183229494 | 1.73782199 | 1.100456614 | 1.579182648 |
| Mir133c | 0 | NA | NA | NA |
| Gm13401 | 1.918047756 | -0.19791181 | 0.920481408 | -0.215009025 |
| Rapgef1 | 1745.697021 | 0.363378998 | 0.059249091 | 6.133072984 |
| Gm9823 | 0.449135467 | -1.341176737 | 1.308495312 | -1.024976341 |
| Gm13420 | 1.634298146 | -0.750776897 | 0.992589663 | -0.756381942 |
| Gm13547 | 0.049553047 | -0.443818988 | 3.724945049 | -0.119147795 |
| Trub2 | 328.3662552 | 0.029768756 | 0.082997039 | 0.358672506 |
| Coq4 | 113.9900504 | -0.229858938 | 0.108080946 | -2.126729512 |
| Slc27a4 | 871.144694 | -0.084634247 | 0.053610234 | -1.578695704 |
| Urm1 | 399.0492176 | -0.129725618 | 0.087158472 | -1.488387931 |
| Mir219a-2 | 4.701385445 | 0.011365612 | 0.655155912 | 0.01734795 |
| mmu-mir-219b | 2.33080191 | 0.761591463 | 0.723263327 | 1.052993335 |
| 2600006K01Rik | 2.48771953 | -1.60831025 | 0.802987725 | -2.00290764 |
| Cercam | 101.0078076 | -0.095942813 | 0.146649721 | -0.654231134 |
| Odf2 | 917.3723799 | 0.336640677 | 0.148660905 | 2.264486937 |
| Gm23865 | 0 | NA | NA | NA |
| Gle1 | 489.083346 | 0.235986691 | 0.070001498 | 3.371166287 |
| Sptan1 | 16083.58909 | 0.230001541 | 0.070529813 | 3.26105416 |
| Mir6997 | 1.080983412 | 1.66056484 | 1.230299612 | 1.34972394 |
| Wdr34 | 186.8631446 | 0.204760629 | 0.122498614 | 1.671534246 |
| Set | 1857.674102 | -0.15417726 | 0.079678879 | -1.934982806 |
| Pkn3 | 10.32177187 | -1.131987364 | 0.324872002 | -3.484410339 |
| Zdhhc12 | 49.78225683 | -0.221056648 | 0.146939357 | -1.504407346 |

| | | | | |
|---|---|---|---|---|
| Zer1 | 1722.618374 | 0.222838778 | 0.055511809 | 4.014259014 |
| Tbc1d13 | 421.3439125 | -0.212828158 | 0.088161108 | -2.414082155 |
| Slc39a1-ps | 5.938480901 | -0.931124846 | 0.458574847 | -2.030475182 |
| Endog | 111.3222115 | -0.042453681 | 0.137678307 | -0.30835418 |
| D2Wsu81e | 251.2755507 | -0.010784543 | 0.092717663 | -0.116315955 |
| Ccbl1 | 92.26571604 | 0.267492521 | 0.153923035 | 1.737832937 |
| 1700084E18Rik | 9.307150498 | 0.41047177 | 0.337886546 | 1.214821291 |
| Lrrc8a | 523.7153376 | 0.105422216 | 0.101946838 | 1.0340901 |
| Gm28035 | 4.247830997 | 0.759278931 | 0.530110041 | 1.432304373 |
| Phyhd1 | 42.27143342 | 0.118621046 | 0.250588595 | 0.473369691 |
| Gm28038 | 0 | NA | NA | NA |
| Dolk | 191.5944489 | -0.072362753 | 0.10789413 | -0.670682943 |
| Nup188 | 388.5180733 | -0.372764585 | 0.091064295 | -4.093421949 |
| Gm14471 | 0 | NA | NA | NA |
| Sh3glb2 | 1245.303918 | 0.446175162 | 0.067773866 | 6.583292194 |
| Fam73b | 803.1893246 | 0.068725725 | 0.064543083 | 1.064803866 |
| AL954388.1 | 0 | NA | NA | NA |
| Dolpp1 | 175.3732175 | -0.136539483 | 0.09154142 | -1.49155959 |
| Crat | 430.9212061 | 0.079167799 | 0.081747161 | 0.968447076 |
| Gm16323 | 10.6181011 | 0.311012122 | 0.41764765 | 0.744675859 |
| Ppp2r4 | 1830.296797 | 0.055636822 | 0.074040742 | 0.751435233 |
| Ier5l | 217.6479524 | -0.326223644 | 0.168263195 | -1.938770057 |
| Gm14487 | 1.256366316 | 1.994079246 | 1.290242077 | 1.54550784 |
| Cstad | 12.63152331 | 0.978283423 | 0.326103903 | 2.999913264 |
| 9330198N18Rik | 2.48591285 | 0.56074755 | 0.764376929 | 0.73360083 |
| Gm14486 | 1.55717633 | 0.869157484 | 0.722280471 | 1.203351771 |
| Gm14488 | 4.29828934 | -1.139121761 | 0.471454163 | -2.416187723 |
| Mir3089 | 0.367473112 | 0.037077569 | 1.598317177 | 0.023197879 |
| 1700001O22Rik | 34.98746724 | 0.162085341 | 0.211268549 | 0.767200523 |
| 4930527E20Rik | 0.124415284 | -0.44382566 | 3.724945049 | -0.119149586 |
| Ntmt1 | 195.850967 | -0.034017193 | 0.120726949 | -0.281769672 |
| Asb6 | 292.993291 | -0.162739105 | 0.083093998 | -1.958494096 |
| Prrx2 | 13.36258994 | -0.752900713 | 0.301504707 | -2.497144139 |
| Ptges | 99.72996419 | 0.013796286 | 0.274573395 | 0.050246261 |
| Gm13400 | 1.094212761 | -0.491465939 | 0.993791345 | -0.494536344 |
| Tor1b | 368.7015473 | -0.175668004 | 0.077001282 | -2.281364655 |
| Tor1a | 257.103606 | -0.031705503 | 0.085249569 | -0.371913947 |
| BC005624 | 537.4115427 | 0.113102589 | 0.100096757 | 1.129932605 |
| Usp20 | 792.5786421 | 0.285467132 | 0.071027566 | 4.019103385 |
| Fnbp1 | 2552.528974 | -0.069482245 | 0.071972059 | -0.965405826 |
| D330023K18Rik | 31.26496909 | -0.17697428 | 0.226872364 | -0.780060987 |
| Gpr107 | 862.5963975 | -0.203030462 | 0.048778234 | -4.162316792 |
| Gm22516 | 4.36673596 | -0.088903635 | 0.550823182 | -0.161401404 |
| Gm13405 | 0.116046057 | 0.037062677 | 3.724945049 | 0.009949859 |
| Ncs1 | 3353.155418 | -0.002032714 | 0.105799159 | -0.019212951 |
| Hmcn2 | 32.42021316 | -1.785392105 | 0.291278372 | -6.129504547 |
| Ass1 | 313.9716636 | -0.118379109 | 0.131427813 | -0.900715814 |

| | | | | |
|---|---|---|---|---|
| Gm13426 | 0 | NA | NA | NA |
| Fubp3 | 562.5375217 | -0.022852695 | 0.080717431 | -0.283119701 |
| Mir6998 | 1.273153199 | 0.714091495 | 1.096764579 | 0.651089129 |
| Gm13425 | 3.449187758 | 1.039159908 | 0.561077052 | 1.852080571 |
| Prdm12 | 5.973420824 | -0.108442154 | 0.471905082 | -0.229796537 |
| Gm13424 | 0 | NA | NA | NA |
| Exosc2 | 146.3618478 | -0.206014576 | 0.101776243 | -2.02419121 |
| Gm13427 | 1.245568681 | -0.021310734 | 0.908162514 | -0.023465772 |
| Abl1 | 1015.182759 | 0.072929512 | 0.053317429 | 1.367836247 |
| Qrfp | 1.096023364 | -0.681781276 | 0.8934958 | -0.763049223 |
| Fibcd1 | 171.8501216 | 0.852188821 | 0.146779446 | 5.805913853 |
| Gm13441 | 0.122601409 | 0.998850474 | 3.724945049 | 0.268151734 |
| Lamc3 | 85.86094398 | -0.050824629 | 0.187162873 | -0.271552942 |
| Aif1l | 75.90635778 | -0.695953495 | 0.221796513 | -3.13780178 |
| Gm13442 | 0.192796986 | -0.948964596 | 2.249928049 | -0.42177553 |
| Nup214 | 817.8687593 | 0.043464515 | 0.073734643 | 0.589472103 |
| Gm13608 | 10.74285294 | 0.933610877 | 0.482065709 | 1.936688005 |
| Gm13609 | 0.288698636 | -0.020002212 | 2.053651327 | -0.009739829 |
| Fam78a | 86.39529847 | 0.329686006 | 0.145337372 | 2.268418658 |
| Gm16534 | 0.731853201 | 2.215794517 | 1.205586726 | 1.837938714 |
| Mir7674 | 0.20460011 | 0.517955348 | 3.243234116 | 0.159703348 |
| Ppapdc3 | 209.3998458 | 0.055034951 | 0.095336578 | 0.57727005 |
| Prrc2b | 8454.239442 | 0.255005417 | 0.069884211 | 3.648970378 |
| Gm27805 | 3.918894907 | 0.948383144 | 0.613522119 | 1.545801064 |
| Gm22192 | 0.800855801 | 0.514357959 | 1.264763116 | 0.406683238 |
| Gm25632 | 3.196964654 | 0.45075407 | 0.775115713 | 0.581531328 |
| Gm13610 | 1.740926027 | -0.861145996 | 0.839108341 | -1.026263181 |
| Pomt1 | 154.4461912 | -0.336305942 | 0.171130927 | -1.965196755 |
| Uck1 | 264.0364267 | -0.127490197 | 0.096596925 | -1.319816317 |
| Gm13611 | 4.282612097 | -0.034308654 | 0.508151536 | -0.06751658 |
| Swi5 | 679.3934042 | -0.14815772 | 0.075849596 | -1.95330929 |
| Golga2 | 1066.72792 | 0.088060413 | 0.083034992 | 1.060521729 |
| Dnm1 | 10321.66057 | 0.605702691 | 0.084510791 | 7.167163906 |
| Gm23363 | 0.044544648 | 0.517957688 | 3.724945049 | 0.139051095 |
| Mir199b | 0.042716945 | 0.037062677 | 3.724945049 | 0.009949859 |
| Ciz1 | 515.2909118 | -0.137091867 | 0.077075277 | -1.778674983 |
| 1110008P14Rik | 542.4132862 | 0.149528192 | 0.099245706 | 1.506646468 |
| Lcn2 | 343.4844549 | 0.258736255 | 0.757297332 | 0.341657424 |
| Ptges2 | 252.589667 | -0.200755443 | 0.090940522 | -2.207546636 |
| Gm13413 | 0.473410537 | -1.072499441 | 1.401029387 | -0.765508169 |
| Slc25a25 | 955.9872403 | -0.286361029 | 0.131795905 | -2.172761192 |
| Naif1 | 76.9182076 | -0.504486048 | 0.152233766 | -3.313890619 |
| Gm37169 | 0 | NA | NA | NA |
| Gm13412 | 0 | NA | NA | NA |
| Fam102a | 1133.257023 | -0.146179818 | 0.08893019 | -1.643759193 |
| Dpm2 | 258.7392704 | -0.321217259 | 0.084596238 | -3.79706317 |
| 9430097D07Rik | 4.085120842 | -0.351839888 | 0.561348592 | -0.626776112 |

| | | | | |
|---|---|---|---|---|
| Pip5kl1 | 31.38216028 | -0.390686875 | 0.309677443 | -1.261592938 |
| St6galnac4 | 520.5645504 | -0.633016501 | 0.089371748 | -7.082959838 |
| Gm23369 | 0.044544648 | 0.517957688 | 3.724945049 | 0.139051095 |
| St6galnac6 | 748.2103617 | 0.656695076 | 0.090019205 | 7.295055242 |
| Ak1 | 1088.578713 | 0.00679279 | 0.072051909 | 0.094276342 |
| Eng | 245.6138286 | -0.646027841 | 0.164966435 | -3.916116882 |
| Mir1954 | 0.415365334 | -2.507781702 | 3.225004956 | -0.777605534 |
| AL772271.1 | 0 | NA | NA | NA |
| Gm26236 | 0.112642008 | 0.517955464 | 3.724945049 | 0.139050498 |
| Fpgs | 67.89073458 | -0.410335463 | 0.177395861 | -2.313106188 |
| Mir8093 | 0 | NA | NA | NA |
| Cdk9 | 551.2940997 | -0.10982785 | 0.071627831 | -1.53331253 |
| Mir2861 | 0 | NA | NA | NA |
| mmu-mir-3960 | 0 | NA | NA | NA |
| Sh2d3c | 241.4715273 | -0.207745095 | 0.117698639 | -1.765059457 |
| 6330409D20Rik | 20.79074414 | 0.264006864 | 0.246717016 | 1.070079675 |
| Ttc16 | 12.6100887 | -0.713865941 | 0.356419322 | -2.002882271 |
| Tor2a | 191.8598868 | -0.280903684 | 0.098889221 | -2.840589515 |
| Ptrh1 | 37.42700904 | -1.153228706 | 0.21595001 | -5.340257715 |
| 1700019L03Rik | 4.625372364 | -1.158263928 | 0.544384934 | -2.127656105 |
| Stxbp1 | 9943.087999 | 0.375136922 | 0.062130579 | 6.037879037 |
| Gm24165 | 0 | NA | NA | NA |
| Gm13524 | 5.575737892 | -0.335909625 | 0.478263107 | -0.702353202 |
| Gm13523 | 5.508606278 | 0.44108519 | 0.452519408 | 0.974732094 |
| Fam129b | 816.2277192 | -1.183687564 | 0.171856806 | -6.887638554 |
| Lrsam1 | 401.4005097 | 0.120748189 | 0.076474678 | 1.578930325 |
| Rpl12 | 1113.211754 | -0.234528964 | 0.098688471 | -2.376457568 |
| Snora65 | 0 | NA | NA | NA |
| Slc2a8 | 138.0519622 | 0.434809002 | 0.131442648 | 3.307975054 |
| Garnl3 | 1194.21391 | 0.704285644 | 0.128772058 | 5.469242746 |
| Gm38299 | 48.15575876 | 1.275135482 | 0.416108821 | 3.064427903 |
| Ralgps1 | 1094.760979 | 0.329618274 | 0.098594367 | 3.343175531 |
| Gm13528 | 0.134209209 | -0.443827508 | 3.667652278 | -0.121011338 |
| Gm23546 | 4.82094354 | -0.267193454 | 0.567300644 | -0.470990923 |
| Angptl2 | 62.62522016 | -0.676714701 | 0.170100496 | -3.97832292 |
| Gm13537 | 0.097139547 | -0.443816764 | 3.724945049 | -0.119147198 |
| Gm25998 | 3.056188022 | 1.344627186 | 0.645312844 | 2.083682664 |
| Zbtb34 | 281.4807306 | 0.034210002 | 0.100962643 | 0.338838216 |
| Gm13536 | 1.068960359 | -2.045161404 | 1.199877398 | -1.704475314 |
| Zbtb43 | 400.5586369 | -0.121939027 | 0.105472884 | -1.156117317 |
| Gm13530 | 1.493521525 | 1.062869773 | 0.994812838 | 1.068411798 |
| Lmx1b | 29.823212 | -0.415352692 | 0.379420745 | -1.094702114 |
| Gm38011 | 0 | NA | NA | NA |
| C130021I20Rik | 27.33416534 | -0.470287021 | 0.385104136 | -1.221194417 |
| Gm13529 | 0.108772582 | 0.037073798 | 3.724945049 | 0.009952844 |
| 9430024E24Rik | 0.066050875 | -0.443818988 | 3.724945049 | -0.119147795 |
| Mvb12b | 1603.937987 | 0.04775629 | 0.058941322 | 0.810234452 |

| | | | | |
|---|---|---|---|---|
| Nron | 0.158932157 | -0.443821645 | 2.809576876 | -0.157967432 |
| Gm13403 | 3.213919713 | -0.134006811 | 0.607912295 | -0.220437737 |
| Gm13404 | 2.447136968 | 0.195362663 | 0.612866093 | 0.31876892 |
| C230014O12Rik | 11.75502483 | 0.097628846 | 0.298821454 | 0.326712972 |
| Gm38389 | 0.847399432 | 0.511072959 | 1.365003506 | 0.374411462 |
| Pbx3 | 410.290469 | -0.190050032 | 0.159706919 | -1.189992475 |
| C79798 | 0.252257838 | -0.975990755 | 2.475517975 | -0.394257188 |
| Mapkap1 | 523.1598314 | -0.182004371 | 0.094774143 | -1.920401123 |
| Gm13407 | 0 | NA | NA | NA |
| Gm13406 | 0 | NA | NA | NA |
| Gm13408 | 58.61577562 | -0.184513114 | 0.194610883 | -0.94811303 |
| Gm13414 | 23.60150067 | 0.032935501 | 0.273572947 | 0.120390198 |
| Gapvd1 | 1564.103693 | -0.117009631 | 0.043420457 | -2.694804188 |
| Hspa5 | 4684.153117 | -0.76583038 | 0.123289433 | -6.211646564 |
| Rabepk | 186.7520506 | -0.122268049 | 0.10030079 | -1.219013815 |
| Fbxw2 | 731.5136731 | 0.141151769 | 0.077478431 | 1.821820182 |
| Gm13419 | 0.060174537 | 0.517957688 | 3.724945049 | 0.139051095 |
| RP23-446N16.6 | 0.320849018 | 0.838088582 | 1.73508733 | 0.483023861 |
| Psmd5 | 541.5249382 | -0.147763813 | 0.067616999 | -2.185305705 |
| Cutal | 4.992381895 | -0.791331554 | 0.472840794 | -1.673568703 |
| Phf19 | 22.86602922 | -0.417472737 | 0.379612277 | -1.09973455 |
| Gm13448 | 0 | NA | NA | NA |
| Gm24905 | 0 | NA | NA | NA |
| Traf1 | 68.5058879 | -0.641334768 | 0.214611272 | -2.988355473 |
| Gm13450 | 0.044544648 | 0.517957688 | 3.724945049 | 0.139051095 |
| Hc | 5.496526028 | -0.643523437 | 0.426943691 | -1.507279417 |
| Gm13449 | 0.579463306 | -1.109805386 | 1.538549417 | -0.721332298 |
| AI182371 | 3.278154351 | -0.138117166 | 0.56025873 | -0.246523897 |
| Cntrl | 338.5002484 | 0.005076775 | 0.152696963 | 0.033247388 |
| Rab14 | 2869.966926 | -0.003582533 | 0.038631346 | -0.092736438 |
| Gm13605 | 0 | NA | NA | NA |
| Gsn | 990.1649303 | -0.367541724 | 0.135152507 | -2.719459167 |
| RP23-186B18.8 | 6.833187758 | -0.570573375 | 0.466301046 | -1.223615902 |
| Stom | 265.4412871 | -0.601378688 | 0.142286767 | -4.226525763 |
| Gm13443 | 73.68671835 | -0.310546007 | 0.144800593 | -2.144645957 |
| 4930568D16Rik | 0 | NA | NA | NA |
| 4930402F06Rik | 0.092691418 | -0.908687078 | 3.724560878 | -0.2439716 |
| Ggta1 | 165.7903113 | -1.240866347 | 0.180583469 | -6.871428231 |
| RP23-458A10.4 | 2.488423977 | -2.465570028 | 0.908105744 | -2.715069302 |
| Gm13444 | 0.047948698 | 0.037064902 | 3.724945049 | 0.009950456 |
| Gm13445 | 0 | NA | NA | NA |
| Gm13447 | 0.202171817 | -1.352528503 | 2.5871409 | -0.522788884 |
| Gm13446 | 33.55988347 | 0.830812827 | 0.233169861 | 3.563122714 |
| Dab2ip | 1883.686994 | 0.211904282 | 0.077521382 | 2.733494628 |
| Ttll11 | 316.1783459 | 0.002501016 | 0.115586358 | 0.021637638 |
| RP23-366N13.3 | 20.18499025 | 0.203630787 | 0.431061685 | 0.472393612 |
| 9030204H09Rik | 0.047948698 | 0.037064902 | 3.724945049 | 0.009950456 |

| | | | | |
|---|---|---|---|---|
| RP23-366N13.2 | 0.655197851 | -2.21767085 | 1.300391627 | -1.705386903 |
| Ndufa8 | 520.9048915 | -0.073385998 | 0.104878274 | -0.699725454 |
| Morn5 | 2.567728017 | -1.484774791 | 0.750216271 | -1.979129016 |
| RP23-169L4.6 | 0 | NA | NA | NA |
| Lhx6 | 117.6126763 | 0.270382349 | 0.133802018 | 2.020764355 |
| Gm27953 | 0 | NA | NA | NA |
| Rbm18 | 518.3893294 | -0.111599105 | 0.077019375 | -1.448974432 |
| Mrrf | 178.907885 | -0.306198978 | 0.101937651 | -3.00378686 |
| Gm13428 | 0 | NA | NA | NA |
| Gm13429 | 0.948980887 | -0.393090048 | 1.118489337 | -0.351447292 |
| Gm13431 | 0.366933406 | 0.679577294 | 1.69889659 | 0.400010983 |
| Ptgs1 | 131.6150916 | -0.677777535 | 0.22576227 | -3.00217364 |
| RP23-454A14.10 | 0.048949532 | 0.037064902 | 3.724945049 | 0.009950456 |
| Olfr334-ps1 | 0.427301431 | 0.026619769 | 3.345755114 | 0.007956281 |
| Olfr335-ps | 0.092140247 | 0.037060453 | 3.724945049 | 0.009949262 |
| Olfr336-ps1 | 0.11657011 | 0.998850474 | 3.724945049 | 0.268151734 |
| Olfr337-ps1 | 0.194279094 | -1.017823543 | 3.719440177 | -0.273649661 |
| Olfr338 | 0.195866688 | -0.520814814 | 3.248878229 | -0.160306043 |
| Gm13430 | 9.735155595 | 0.195329697 | 0.398502112 | 0.490159754 |
| Olfr339 | 0 | NA | NA | NA |
| Olfr340 | 0 | NA | NA | NA |
| Olfr341 | 0 | NA | NA | NA |
| Olfr342 | 0.194279094 | -1.017823543 | 3.719440177 | -0.273649661 |
| Olfr343-ps1 | 0 | NA | NA | NA |
| Olfr344 | 0.046345709 | -0.443818988 | 3.724945049 | -0.119147795 |
| Olfr345 | 0 | NA | NA | NA |
| Olfr346 | 0 | NA | NA | NA |
| Olfr347 | 0 | NA | NA | NA |
| Gm13438 | 0 | NA | NA | NA |
| Olfr348 | 0 | NA | NA | NA |
| Olfr50 | 0 | NA | NA | NA |
| Gm13436 | 42.25227907 | -0.554398471 | 0.211311458 | -2.623608187 |
| Olfr3 | 0 | NA | NA | NA |
| Olfr349-ps1 | 0 | NA | NA | NA |
| Olfr350 | 0 | NA | NA | NA |
| Olfr351 | 0.097139547 | -0.443816764 | 3.724945049 | -0.119147198 |
| Olfr352 | 0 | NA | NA | NA |
| Olfr353 | 0 | NA | NA | NA |
| Olfr354 | 0 | NA | NA | NA |
| Gm13440 | 0 | NA | NA | NA |
| Olfr355 | 0 | NA | NA | NA |
| Olfr356 | 0 | NA | NA | NA |
| Olfr357 | 0 | NA | NA | NA |
| Olfr358 | 0 | NA | NA | NA |
| Olfr359-ps1 | 0 | NA | NA | NA |
| Gm13439 | 0 | NA | NA | NA |
| Olfr360 | 0 | NA | NA | NA |

| | | | | |
|---|---|---|---|---|
| Gm13437 | 0 | NA | NA | NA |
| Olfr361 | 0 | NA | NA | NA |
| Gm13434 | 0 | NA | NA | NA |
| Olfr362 | 0 | NA | NA | NA |
| Olfr363-ps | 0 | NA | NA | NA |
| Olfr364-ps1 | 0 | NA | NA | NA |
| Olfr365 | 0 | NA | NA | NA |
| Gm13435 | 0 | NA | NA | NA |
| Olfr366 | 0.068097359 | 0.037064902 | 3.724945049 | 0.009950456 |
| Olfr367-ps | 0 | NA | NA | NA |
| Gm22306 | 0 | NA | NA | NA |
| Olfr368 | 0 | NA | NA | NA |
| Pdcl | 328.2885349 | -0.306214654 | 0.12341611 | -2.481156272 |
| Rc3h2 | 3083.559831 | 0.016138252 | 0.037694704 | 0.428130482 |
| Snord90 | 1.013447778 | 0.246795694 | 1.110287688 | 0.222280853 |
| Zbtb6 | 434.5484348 | -0.288948941 | 0.076059145 | -3.799003287 |
| Zbtb26 | 221.5652181 | -0.483305225 | 0.131535624 | -3.674329505 |
| Rabgap1 | 1677.895686 | 0.066011562 | 0.043679247 | 1.51127977 |
| Strbp | 4912.750144 | 0.580239249 | 0.13206654 | 4.393537164 |
| Gpr21 | 129.8264781 | 0.214043312 | 0.155756933 | 1.374213708 |
| Gm13543 | 0 | NA | NA | NA |
| Mir5128 | 8.038112018 | 0.58516787 | 0.487679438 | 1.199902691 |
| Crb2 | 102.1842241 | 0.043008194 | 0.161370925 | 0.266517612 |
| Dennd1a | 974.3781944 | 0.037640664 | 0.060832681 | 0.618757275 |
| AL929186.1 | 4.630423038 | -0.107102894 | 0.557750046 | -0.192026687 |
| Gm13556 | 0.04393175 | 0.037062677 | 3.724945049 | 0.009949859 |
| Lhx2 | 320.8917483 | 0.379407659 | 0.126433811 | 3.000840179 |
| Gm27197 | 1.480046296 | 1.630110468 | 1.002622284 | 1.625847035 |
| Gm13584 | 0 | NA | NA | NA |
| Gm13589 | 3.470889967 | -0.242739837 | 0.548163052 | -0.442824149 |
| Gm25081 | 0.13733263 | 0.037057285 | 3.640803958 | 0.010178325 |
| Gm13588 | 0.045882455 | 0.517959912 | 3.724945049 | 0.139051692 |
| Nek6 | 512.9966335 | -0.852185422 | 0.136570387 | -6.239899016 |
| Psmb7 | 966.8015143 | -0.220840819 | 0.085546131 | -2.581540688 |
| Gm13586 | 0.536876103 | -1.719155631 | 1.473054778 | -1.167068365 |
| Gm27634 | 1.100154067 | 0.165703253 | 1.163506875 | 0.142417081 |
| Adgrd2-ps | 0 | NA | NA | NA |
| Gm13587 | 0 | NA | NA | NA |
| Nr5a1 | 5.753592802 | 0.383961364 | 0.504834955 | 0.760568102 |
| Nr5a1os | 0.104719185 | 0.981063299 | 3.723618439 | 0.263470416 |
| Nr6a1 | 153.9253641 | 0.140571663 | 0.145841502 | 0.963865986 |
| Rpl15-ps4 | 0 | NA | NA | NA |
| Nr6a1os | 2.499784555 | 0.720749727 | 0.844404517 | 0.853559772 |
| Mir181a-2 | 0.139226991 | -0.443822101 | 2.927946011 | -0.151581381 |
| Mir181b-2 | 0.377368388 | 0.020275465 | 1.720871306 | 0.011782093 |
| Olfml2a | 59.91228772 | -0.07008918 | 0.214174776 | -0.327252262 |
| Gm13473 | 0 | NA | NA | NA |

| | | | | |
|---|---|---|---|---|
| Gm13474 | 0.110595523 | 0.037071574 | 3.724945049 | 0.009952247 |
| Gm13496 | 0.873180464 | -1.331103188 | 1.136590972 | -1.171136513 |
| Wdr38 | 12.8811223 | -0.283088381 | 0.308818111 | -0.916683224 |
| Rpl35 | 950.799867 | -0.220409209 | 0.102778983 | -2.144496892 |
| Arpc5l | 615.8797353 | -0.02394602 | 0.098645341 | -0.242748612 |
| Golga1 | 507.3639855 | -0.146206086 | 0.087280699 | -1.675125059 |
| Gm26127 | 0 | NA | NA | NA |
| Scai | 1810.878198 | 0.507033854 | 0.074266289 | 6.827240987 |
| Gm16523 | 10.92727314 | -0.422448559 | 0.451402076 | -0.935858699 |
| Ppp6c | 892.2623954 | -0.235882647 | 0.083000187 | -2.841953216 |
| Gm13511 | 6.921421994 | 0.735906907 | 0.492326132 | 1.494754918 |
| Gm13504 | 0 | NA | NA | NA |
| Vmn2r-ps2 | 0.066050875 | -0.443818988 | 3.724945049 | -0.119147795 |
| Gm13506 | 0 | NA | NA | NA |
| Gm13454 | 0 | NA | NA | NA |
| Gm13451 | 0 | NA | NA | NA |
| Gm13452 | 0.066050875 | -0.443818988 | 3.724945049 | -0.119147795 |
| Gm13453 | 7.860772947 | -0.485326189 | 0.369417827 | -1.313759522 |
| Gm13457 | 0 | NA | NA | NA |
| Gm13458 | 0.061920613 | -0.443816764 | 3.724945049 | -0.119147198 |
| Gm13455 | 0 | NA | NA | NA |
| Gm24467 | 0 | NA | NA | NA |
| Gm13456 | 339.2160491 | -0.287920887 | 0.114012875 | -2.525336607 |
| Lrp1b | 2840.147083 | 0.514060492 | 0.123104368 | 4.175810337 |
| Gm27348 | 0.290738398 | 0.425409681 | 1.797850735 | 0.236621246 |
| Gm13462 | 0 | NA | NA | NA |
| Gm13460 | 0 | NA | NA | NA |
| Gm13461 | 25.32005373 | -0.304252436 | 0.220297309 | -1.381099195 |
| Gm13463 | 0 | NA | NA | NA |
| Mir6336 | 9.1407384 | 0.684647093 | 0.399252853 | 1.714820792 |
| Gm24350 | 0 | NA | NA | NA |
| Gm13464 | 14.15448784 | 0.192630783 | 0.307689402 | 0.626055957 |
| Kynu | 6.170416941 | -0.977654224 | 0.392643725 | -2.489927033 |
| Arhgap15 | 136.0233683 | -0.09458769 | 0.158579821 | -0.596467379 |
| Arhgap15os | 0.169875191 | 0.984813711 | 2.746838256 | 0.358526283 |
| Gm22867 | 0.042716945 | 0.037062677 | 3.724945049 | 0.009949859 |
| Gtdc1 | 447.6665794 | 0.60016623 | 0.098362478 | 6.10157699 |
| Gm13477 | 0.271375346 | 1.863367467 | 2.046214235 | 0.910641435 |
| Zeb2 | 3739.770028 | -0.225069321 | 0.097168187 | -2.316286106 |
| Mir5129 | 1.526907099 | -1.083962028 | 0.966430537 | -1.121614008 |
| Zeb2os | 56.92489478 | -0.476839328 | 0.190391072 | -2.504525672 |
| Gm27761 | 0 | NA | NA | NA |
| Gm27676 | 0.743251506 | -0.174563846 | 1.046118817 | -0.166868087 |
| Gm27404 | 0 | NA | NA | NA |
| Gm27645 | 0 | NA | NA | NA |
| Gm28643 | 0.172790758 | -1.002257145 | 3.387088684 | -0.295905197 |
| Vis1 | 2.225936112 | -1.640241137 | 0.68292066 | -2.401803363 |

| | | | | |
|---|---|---|---|---|
| Gm13475 | 0.044191549 | 0.037062677 | 3.724945049 | 0.009949859 |
| Gm13479 | 1.673299741 | 0.1320744 | 0.788638738 | 0.167471357 |
| Gm13478 | 0 | NA | NA | NA |
| 1700019E08Rik | 0 | NA | NA | NA |
| Gm13465 | 0.044191549 | 0.037062677 | 3.724945049 | 0.009949859 |
| Gm13467 | 0 | NA | NA | NA |
| Gm13466 | 0 | NA | NA | NA |
| Gm23072 | 0.060174537 | 0.517957688 | 3.724945049 | 0.139051095 |
| Gm13469 | 0 | NA | NA | NA |
| Gm13470 | 0.459694823 | 0.494901594 | 1.451453031 | 0.340969762 |
| Gm25264 | 0 | NA | NA | NA |
| Gm25959 | 0.869297867 | -0.910785338 | 0.947956527 | -0.960788086 |
| Gm13481 | 2.425955829 | -0.127418362 | 0.673715504 | -0.189127846 |
| Gm13471 | 0 | NA | NA | NA |
| Gm13482 | 0 | NA | NA | NA |
| Gm25338 | 0.121578509 | 0.037073798 | 3.724945049 | 0.009952844 |
| Gm13472 | 0.062426873 | 0.517959912 | 3.724945049 | 0.139051692 |
| Gm13489 | 0.524554913 | 2.163661411 | 1.372014174 | 1.576996399 |
| Acvr2a | 849.6809722 | 0.587329393 | 0.064009751 | 9.175623744 |
| Orc4 | 1017.373273 | 0.013056672 | 0.089800135 | 0.145397019 |
| Mbd5 | 1674.173319 | 0.309448998 | 0.096965762 | 3.191322295 |
| Gm13510 | 17.67851426 | 0.256018982 | 0.313571971 | 0.816460033 |
| Gm13509 | 582.147492 | -0.010119917 | 0.098109244 | -0.103149477 |
| Gm13508 | 0 | NA | NA | NA |
| Gm25215 | 0.07796279 | -0.798301147 | 3.721829328 | -0.214491605 |
| Epc2 | 915.7051038 | 0.291178009 | 0.098520473 | 2.955507615 |
| Kif5c | 11473.79799 | 0.30665278 | 0.086574838 | 3.542054345 |
| Gm13525 | 21.45150064 | 0.467345424 | 0.45511201 | 1.026880008 |
| Lypd6b | 138.4529101 | 0.668125284 | 0.159295154 | 4.194259939 |
| Gm13480 | 0.55522199 | 0.762616894 | 1.255793388 | 0.607278953 |
| Gm25223 | 0 | NA | NA | NA |
| Lypd6 | 201.0019211 | 0.2947405 | 0.116172302 | 2.537097869 |
| Mmadhc | 325.9114616 | -0.313871516 | 0.094928775 | -3.306389597 |
| Gm13483 | 20.39339472 | 0.191181694 | 0.278628738 | 0.6861521 |
| Gm13486 | 0.761951221 | 1.093100401 | 1.191426093 | 0.917472269 |
| Gm13488 | 0.134148234 | -0.443820953 | 3.624632483 | -0.122445781 |
| Gm13487 | 0.658756006 | -0.558350698 | 1.290974152 | -0.432503391 |
| Gm13485 | 1.138498193 | 1.193487941 | 0.990050439 | 1.205481958 |
| Gm13484 | 0 | NA | NA | NA |
| Gm13498 | 2.899184114 | 0.706314991 | 0.547806008 | 1.289352399 |
| Rnd3 | 459.0834406 | -0.599010608 | 0.165128566 | -3.627540787 |
| Gm28639 | 0 | NA | NA | NA |
| Gm13500 | 0 | NA | NA | NA |
| Gm13497 | 0.338153501 | -0.416551409 | 1.485160149 | -0.280475751 |
| Gm13499 | 0.047948698 | 0.037064902 | 3.724945049 | 0.009950456 |
| 1700057H21Rik | 0.253674139 | -1.321443172 | 2.123438654 | -0.622312855 |
| Gm23505 | 0.049553047 | -0.443818988 | 3.724945049 | -0.119147795 |

| | | | | |
|---|---|---|---|---|
| Gm13494 | 0 | NA | NA | NA |
| Gm13491 | 0.045882455 | 0.517959912 | 3.724945049 | 0.139051692 |
| Tas2r134 | 0 | NA | NA | NA |
| Gm13490 | 0.088383099 | 0.037058229 | 3.724945049 | 0.009948665 |
| Gm13492 | 0 | NA | NA | NA |
| Gm13493 | 88.56531709 | -0.020150881 | 0.134310522 | -0.150032035 |
| Rbm43 | 483.6855949 | -0.174094724 | 0.150900366 | -1.153706439 |
| Gm13495 | 1.701798027 | 0.762153407 | 0.928420802 | 0.820913755 |
| Nmi | 215.7031363 | -0.375480128 | 0.232248395 | -1.616717859 |
| Tnfaip6 | 134.9876113 | -0.33198924 | 0.17925211 | -1.852079961 |
| Rif1 | 1558.481454 | -0.112676097 | 0.070661091 | -1.594598888 |
| Neb | 246.6161145 | 0.392737526 | 0.161765044 | 2.42782691 |
| Gm22095 | 0 | NA | NA | NA |
| Gm13521 | 0 | NA | NA | NA |
| Gm13522 | 0.21580698 | 0.957735505 | 2.580683689 | 0.371116968 |
| Arl5a | 1365.664992 | 0.048575517 | 0.065151566 | 0.745577111 |
| A430018G15Rik | 10.01027238 | -0.1574607 | 0.373262666 | -0.421849582 |
| Cacnb4 | 3963.510753 | 0.325712999 | 0.068154146 | 4.779063589 |
| Gm13541 | 0 | NA | NA | NA |
| Bloc1s2-ps | 5.732035973 | 0.321441587 | 0.429512383 | 0.748387242 |
| Stam2 | 454.7228158 | -0.558099732 | 0.077424909 | -7.20827105 |
| Gm13548 | 1.65657422 | 0.220781449 | 0.883356604 | 0.249934679 |
| Fmnl2 | 1786.717828 | -0.19092163 | 0.115788738 | -1.6488791 |
| Gm26603 | 0.266304556 | -0.602115226 | 3.715438719 | -0.162057639 |
| Prpf40a | 1253.218375 | 0.002370274 | 0.094837727 | 0.024992944 |
| Arl6ip6 | 155.8469731 | -0.301863329 | 0.104350266 | -2.89278926 |
| Gm25898 | 0 | NA | NA | NA |
| Gm13501 | 0 | NA | NA | NA |
| Gm13503 | 0 | NA | NA | NA |
| Gm13502 | 9.659148329 | -0.89445165 | 0.351053591 | -2.547906285 |
| Gm23315 | 0 | NA | NA | NA |
| Rprm | 258.4970102 | 0.347239134 | 0.097283375 | 3.569357406 |
| Gm22298 | 0.222122807 | 0.037074875 | 2.578296755 | 0.0143796 |
| Gm13505 | 0 | NA | NA | NA |
| Gm14035 | 0 | NA | NA | NA |
| Gm14034 | 0.195609542 | 0.764980429 | 2.611791971 | 0.292894854 |
| Galnt13 | 988.7175182 | 0.421751741 | 0.104723473 | 4.027289499 |
| Gm24549 | 0 | NA | NA | NA |
| Gm25489 | 4.670143882 | 0.964865888 | 0.589841804 | 1.635804518 |
| Gm14032 | 0.804512061 | 0.628677948 | 1.129197509 | 0.556747551 |
| Gm14033 | 39.47582969 | -0.011440336 | 0.245483384 | -0.046603301 |
| Gm26794 | 31.41234798 | 0.899305276 | 0.212382984 | 4.234356539 |
| Kcnj3 | 1639.12147 | 0.420221648 | 0.094945059 | 4.425945415 |
| Gm13517 | 0.255816663 | -0.46615561 | 2.072796085 | -0.22489217 |
| Gm13615 | 0 | NA | NA | NA |
| Gm13518 | 0.061920613 | -0.443816764 | 3.724945049 | -0.119147198 |
| Mir195b | 0 | NA | NA | NA |

| | | | | |
|---|---|---|---|---|
| Gm13527 | 0 | NA | NA | NA |
| Nr4a2 | 294.6002765 | 0.490991338 | 0.19569422 | 2.508972105 |
| BB557941 | 2.213086149 | 1.796521572 | 0.844280122 | 2.127873824 |
| 4930555B11Rik | 0.042716945 | 0.037062677 | 3.724945049 | 0.009949859 |
| Gm13526 | 0 | NA | NA | NA |
| A930012O16Rik | 7.994888912 | 0.145819722 | 0.37036353 | 0.393720521 |
| Gpd2 | 1202.471657 | 0.212852661 | 0.088443378 | 2.406654589 |
| Gm13535 | 0 | NA | NA | NA |
| Gm13531 | 0 | NA | NA | NA |
| Gm13532 | 2.612576842 | -0.904286328 | 0.584444302 | -1.547258354 |
| Gm13533 | 0 | NA | NA | NA |
| Gm25388 | 0 | NA | NA | NA |
| Galnt5 | 0.79124893 | 1.06553135 | 1.207489921 | 0.882434985 |
| Ermn | 1375.435044 | -0.50805025 | 0.176490229 | -2.878631038 |
| Gm13534 | 0.908099955 | -1.616716762 | 1.137908026 | -1.420779821 |
| Cytip | 92.50638567 | -0.662528326 | 0.268439088 | -2.468076955 |
| Gm13546 | 1.506085986 | -0.382227556 | 0.746441495 | -0.512066329 |
| Acvr1c | 451.2463121 | -0.288806267 | 0.171804755 | -1.6810144 |
| Gm13544 | 2.162167769 | 0.943422259 | 0.65330515 | 1.444075955 |
| Gm13545 | 0.038981395 | -0.443816764 | 3.724945049 | -0.119147198 |
| Acvr1 | 465.7691716 | -0.409687585 | 0.095111914 | -4.307426589 |
| Gm17409 | 0 | NA | NA | NA |
| Upp2 | 286.6269545 | 0.592468336 | 0.104519689 | 5.668485431 |
| Gm13558 | 10.32835312 | 0.826906625 | 0.363717622 | 2.273485185 |
| Gm13559 | 1.310073478 | 1.100722981 | 0.979791388 | 1.123425858 |
| Ccdc148 | 215.227496 | 0.754984734 | 0.11255544 | 6.707669858 |
| Gm11099 | 0.095897396 | 0.037062677 | 3.724945049 | 0.009949859 |
| Gm13560 | 0.045882455 | 0.517959912 | 3.724945049 | 0.139051692 |
| Gm13557 | 1.02920101 | 1.015833512 | 1.532995447 | 0.662646137 |
| Gm13550 | 2.191863138 | 0.880372155 | 0.774401474 | 1.136842045 |
| Pkp4 | 2887.099339 | 0.176609108 | 0.058907984 | 2.998050426 |
| 5330411J11Rik | 0.17957102 | -0.979801808 | 2.710633689 | -0.36146596 |
| Gm13551 | 0 | NA | NA | NA |
| Gm13549 | 0 | NA | NA | NA |
| Dapl1 | 3.189629895 | 0.906934125 | 0.701786002 | 1.292322907 |
| Gm13552 | 6.37558536 | -0.392107928 | 0.40725303 | -0.962811566 |
| Tanc1 | 1294.003133 | -0.031788934 | 0.08705777 | -0.365147585 |
| Wdsub1 | 122.8229018 | -0.016020005 | 0.126339337 | -0.126801404 |
| Baz2b | 1280.473641 | 0.15257996 | 0.100209168 | 1.522614771 |
| Gm13574 | 8.429661362 | 0.681815732 | 0.393411026 | 1.7330875 |
| Gm13572 | 1.149049355 | -0.342805002 | 0.939141918 | -0.365019382 |
| Gm13570 | 9.56573459 | 0.012955764 | 0.374904929 | 0.034557464 |
| Gm13620 | 2.37993816 | -1.430477387 | 0.76959067 | -1.858750946 |
| Ldha-ps | 0.208487022 | -0.24635703 | 2.577735318 | -0.095571112 |
| 7-Mar | 1192.427979 | 0.038506215 | 0.095896589 | 0.401538945 |
| Cd302 | 130.1829803 | 0.245822731 | 0.159381671 | 1.54235258 |
| Gm13573 | 2.148767599 | 0.201037719 | 0.677085373 | 0.296916353 |

| | | | | |
|---|---|---|---|---|
| Ly75 | 45.62619954 | -0.770197979 | 0.214028647 | -3.598574251 |
| Gm13571 | 1.148968944 | -1.237012486 | 1.00195965 | -1.234593115 |
| Gm13580 | 0.165156969 | -1.323927776 | 3.401541256 | -0.389214087 |
| Pla2r1 | 18.48570985 | -1.014936547 | 0.289152711 | -3.510036417 |
| Gm13583 | 0 | NA | NA | NA |
| Gm13566 | 0 | NA | NA | NA |
| Gm13567 | 0 | NA | NA | NA |
| Itgb6 | 5.350769152 | -0.289168015 | 0.426468467 | -0.67805251 |
| Rbms1 | 693.4012186 | -0.650789355 | 0.123326911 | -5.276945249 |
| Gm13582 | 0.21728767 | -0.698126703 | 2.530005581 | -0.2759388 |
| Gm13579 | 0 | NA | NA | NA |
| Gm13581 | 0 | NA | NA | NA |
| Gm26422 | 0 | NA | NA | NA |
| Gm22338 | 0 | NA | NA | NA |
| Tank | 472.8096782 | -0.310232211 | 0.084419534 | -3.674886561 |
| Gm13626 | 0.359719716 | 0.020984381 | 1.571536015 | 0.013352784 |
| Psmd14 | 942.4841296 | -0.20497108 | 0.074397139 | -2.755093568 |
| Gm22403 | 0 | NA | NA | NA |
| Gm25966 | 3.561922538 | 0.615753991 | 0.541131283 | 1.1379013 |
| Tbr1 | 776.6407633 | 1.268840975 | 0.18199567 | 6.971819568 |
| Gm13555 | 0 | NA | NA | NA |
| Slc4a10 | 3386.482564 | 0.402587659 | 0.083200518 | 4.83876384 |
| Dpp4 | 80.82233867 | -0.139718355 | 0.157050956 | -0.889637088 |
| Gm13561 | 3.915321939 | -0.395643362 | 0.602210821 | -0.656984809 |
| Gm13565 | 0 | NA | NA | NA |
| Gm13564 | 0.340218975 | 1.439943106 | 1.978769402 | 0.727696267 |
| Gcg | 1.314123019 | -1.434358898 | 1.048368616 | -1.368181835 |
| Fap | 70.46597968 | 0.232870023 | 0.311856503 | 0.746721715 |
| Ifih1 | 2560.45954 | -0.385246051 | 0.229416504 | -1.679242972 |
| Gca | 214.646939 | -0.224937019 | 0.120883037 | -1.860782321 |
| Gm13590 | 0.316156321 | -0.4444863 | 2.002529216 | -0.221962454 |
| Kcnh7 | 1930.150615 | 0.780508019 | 0.15675814 | 4.979058938 |
| Gm22020 | 4.453022388 | 2.069982942 | 0.668195891 | 3.09786841 |
| Gm13569 | 0.048949532 | 0.037064902 | 3.724945049 | 0.009950456 |
| Gm23503 | 0 | NA | NA | NA |
| Fign | 177.2882869 | 0.120760403 | 0.096406059 | 1.25262255 |
| Gm13575 | 0 | NA | NA | NA |
| Gm13577 | 10.38401885 | -0.08815043 | 0.328361319 | -0.268455584 |
| Gm13576 | 0 | NA | NA | NA |
| Gm13578 | 0.248954575 | -0.417434765 | 1.841272786 | -0.226709898 |
| Grb14 | 287.0919978 | 0.283072794 | 0.097174813 | 2.91302638 |
| Cobll1 | 254.6797965 | -0.418527818 | 0.131464585 | -3.183578429 |
| Gm13594 | 1.366457952 | -0.237137884 | 0.948369727 | -0.250047927 |
| Gm13592 | 0 | NA | NA | NA |
| Gm13593 | 0 | NA | NA | NA |
| Slc38a11 | 11.09653691 | 1.036061218 | 0.394472374 | 2.626448104 |
| Mir6337 | 0 | NA | NA | NA |

| | | | | |
|---|---|---|---|---|
| Gm24138 | 0.150086409 | 1.027495679 | 3.723011807 | 0.275985071 |
| Scn3a | 1030.005389 | 0.248068015 | 0.075462376 | 3.287307259 |
| Gm13595 | 0.072025462 | 0.517959912 | 3.724945049 | 0.139051692 |
| Scn2a1 | 6263.540254 | 0.69470801 | 0.110778706 | 6.27113311 |
| Csrnp3 | 2276.632116 | 0.418092489 | 0.080513406 | 5.192830714 |
| Gm13619 | 2.502007469 | 0.037510653 | 0.705163458 | 0.053194267 |
| Gm13617 | 8.016718732 | -0.457717745 | 0.437041669 | -1.047309164 |
| Galnt3 | 26.28681886 | 0.162556795 | 0.243183742 | 0.668452561 |
| Gm28638 | 0.099106094 | -0.93550548 | 3.724188147 | -0.251197158 |
| Gm13616 | 0 | NA | NA | NA |
| Ttc21b | 309.885117 | -0.124696282 | 0.171349174 | -0.727732028 |
| Gm13618 | 0.367269401 | 0.540371405 | 1.569389702 | 0.344319454 |
| Scn1a | 3205.479907 | 0.480770851 | 0.066448542 | 7.235235498 |
| Gm13630 | 13.73615189 | 0.255176634 | 0.322042009 | 0.792370644 |
| Gm13629 | 78.07883946 | 0.346814536 | 0.167047072 | 2.076148544 |
| Rpl35-ps1 | 0.152586256 | -0.443820484 | 2.845690328 | -0.155962326 |
| Scn9a | 113.7734046 | -0.187073918 | 0.176181456 | -1.061825249 |
| Gm13627 | 0.809136868 | 1.788609327 | 1.236534467 | 1.446469448 |
| Gm23046 | 0.7960769 | 0.799098 | 1.132479206 | 0.705618254 |
| Scn7a | 68.38610524 | 0.167300317 | 0.183459171 | 0.911921253 |
| Gm13600 | 0.068097359 | 0.037064902 | 3.724945049 | 0.009950456 |
| Gm13602 | 0.090277459 | -0.443823436 | 3.724945049 | -0.119148989 |
| Gm13598 | 0 | NA | NA | NA |
| Gm1322 | 0 | NA | NA | NA |
| Gm13599 | 0 | NA | NA | NA |
| Gm21830 | 0 | NA | NA | NA |
| Gm23328 | 0 | NA | NA | NA |
| Gm26727 | 0 | NA | NA | NA |
| Xirp2 | 1.65068616 | -0.329212801 | 0.800823778 | -0.41109269 |
| Gm13601 | 10.77684248 | 0.81166229 | 0.415954753 | 1.951323514 |
| B3galt1 | 1121.981432 | 0.105937245 | 0.085564962 | 1.238091427 |
| Mir7224 | 5.441005428 | 0.197954045 | 0.427356435 | 0.463205954 |
| Gm26743 | 0.083526043 | 0.037073798 | 3.724945049 | 0.009952844 |
| Gm13604 | 0.106618422 | 0.517955464 | 3.724945049 | 0.139050498 |
| Gm13603 | 9.345943703 | -0.214737249 | 0.441646857 | -0.486219352 |
| Mir6335 | 8.512614594 | -0.294695199 | 0.472966712 | -0.623078097 |
| Gm37964 | 115.6859084 | -0.034438388 | 0.235717533 | -0.146100239 |
| Gm13607 | 9.156164614 | 0.15241153 | 0.488116265 | 0.312244317 |
| Stk39 | 1177.70026 | 0.16301237 | 0.059246887 | 2.751408215 |
| Gm13596 | 0.577259679 | 0.02793043 | 1.42404424 | 0.019613457 |
| Gm26321 | 0.044191549 | 0.037062677 | 3.724945049 | 0.009949859 |
| Gm13597 | 10.00332192 | -1.137851597 | 0.356616239 | -3.190689237 |
| 4933409G03Rik | 0.150086409 | 1.027495679 | 3.723011807 | 0.275985071 |
| 4932414N04Rik | 0.339353417 | -0.715773315 | 1.547699381 | -0.462475674 |
| Gm13612 | 0.421611163 | 0.528765148 | 1.465834163 | 0.360726446 |
| Cers6 | 1364.374441 | -0.283804661 | 0.049273495 | -5.759783471 |
| Gm37159 | 76.06150798 | -0.472352988 | 0.241482717 | -1.956052977 |

| | | | | |
|---|---|---|---|---|
| Gm38236 | 46.00811634 | 0.00296004 | 0.314747282 | 0.009404499 |
| Gm37651 | 29.9890416 | -0.152273926 | 0.34657589 | -0.43936676 |
| Gm37747 | 50.62951173 | -0.262677269 | 0.256970752 | -1.022206873 |
| Gm37666 | 22.07432209 | 0.043758483 | 0.326071207 | 0.134199161 |
| Gm38377 | 75.00233876 | -0.058405605 | 0.200657901 | -0.291070547 |
| Nostrin | 27.43524618 | 0.124839419 | 0.234835875 | 0.531602845 |
| Spc25 | 14.006815 | -0.761162715 | 0.37224377 | -2.044796383 |
| G6pc2 | 0.378093966 | -1.731854282 | 3.681950294 | -0.470363298 |
| Abcb11 | 1.943768998 | -1.529998512 | 0.821805177 | -1.861753315 |
| Gm13613 | 0 | NA | NA | NA |
| Gm13614 | 0 | NA | NA | NA |
| Dhrs9 | 8.333878881 | -2.569887276 | 0.542336287 | -4.738549376 |
| Lrp2 | 28.81441089 | -0.800334779 | 0.300402425 | -2.664208779 |
| Bbs5 | 149.8333186 | 0.078291821 | 0.121063647 | 0.646699672 |
| Klhl41 | 8.40861694 | 0.278537761 | 0.392037249 | 0.710488001 |
| Fastkd1 | 167.3175697 | 0.128955667 | 0.095315805 | 1.35293058 |
| Ppig | 1069.962896 | 0.098002078 | 0.080925093 | 1.211022127 |
| Gm13622 | 1.277078827 | 0.759623274 | 0.960132793 | 0.791164805 |
| Ccdc173 | 33.00873947 | -0.141724593 | 0.213352322 | -0.664274906 |
| Gm13623 | 0.146451277 | -0.443820591 | 2.882358847 | -0.153978257 |
| Phospho2 | 316.3809453 | -0.38736541 | 0.124138726 | -3.120423597 |
| Gm13624 | 0.441815039 | 1.880158759 | 3.361288971 | 0.559356478 |
| Klhl23 | 607.9612626 | 0.366699715 | 0.09428691 | 3.889190095 |
| Ssb | 1044.348789 | 0.058112045 | 0.092211781 | 0.630201956 |
| Mettl5 | 79.04134513 | -0.361989233 | 0.133566833 | -2.710173067 |
| Gm13625 | 2.609485436 | 0.148836373 | 0.665015612 | 0.223808841 |
| Mettl5os | 3.090821495 | 1.051105033 | 0.618554123 | 1.699293553 |
| Ubr3 | 4154.724246 | 0.453103375 | 0.069032426 | 6.563631015 |
| Gm13631 | 0.295881152 | -0.092214325 | 2.034805595 | -0.045318494 |
| Myo3b | 31.90249792 | -0.872986984 | 0.275727324 | -3.16612431 |
| Rpl9-ps7 | 51.73231683 | -0.605144745 | 0.20674002 | -2.927080806 |
| Sp5 | 11.17023444 | -0.695540834 | 0.31606677 | -2.200613603 |
| Gad1os | 2.192682401 | -1.360846166 | 0.675444803 | -2.014740746 |
| Erich2os | 3.966806659 | -1.947075072 | 0.541501404 | -3.595697184 |
| Erich2 | 9.894614024 | -1.807034643 | 0.383197004 | -4.715680509 |
| Gm37026 | 0.230844305 | 0.010068291 | 2.130602731 | 0.00472556 |
| Gad1 | 4252.286034 | -0.110748625 | 0.092388527 | -1.198727041 |
| Gm16510 | 0.089814205 | 0.517955464 | 3.724945049 | 0.139050498 |
| Gm26558 | 1.60906251 | -0.035661756 | 0.822266877 | -0.04337005 |
| Gorasp2 | 1047.261539 | -0.122938543 | 0.068695705 | -1.789610311 |
| Tlk1 | 1317.27295 | 0.181622329 | 0.057351402 | 3.166833289 |
| Gm13633 | 0.219273179 | -0.836486873 | 2.133770486 | -0.39202289 |
| Gm13632 | 0.086908494 | 0.037058229 | 3.724945049 | 0.009948665 |
| Mettl8 | 114.1898919 | -0.071381144 | 0.164466825 | -0.434015454 |
| Dcaf17 | 286.7511595 | 0.039768829 | 0.132543525 | 0.300043548 |
| Cybrd1 | 58.06687379 | -0.092725425 | 0.281260144 | -0.329678511 |
| Dync1i2 | 1658.410193 | -0.049085249 | 0.043594096 | -1.125960921 |

| | | | | |
|---|---|---|---|---|
| Slc25a12 | 1679.553867 | 0.249071685 | 0.046404794 | 5.367369685 |
| Gm13641 | 0.124853745 | 1.05357957 | 3.721447096 | 0.283110183 |
| Hat1 | 224.6790855 | -0.350772322 | 0.12984728 | -2.701422168 |
| Gm13642 | 0.843921636 | 0.513550154 | 1.019578684 | 0.503688594 |
| Gm23166 | 0 | NA | NA | NA |
| Metap1d | 226.2128202 | 0.287747352 | 0.096464855 | 2.982924219 |
| Gm23253 | 0.412462835 | -0.219643923 | 1.508748106 | -0.145580247 |
| Dlx1as | 32.54297881 | 0.352645055 | 0.314726979 | 1.120479265 |
| Dlx1 | 196.3712922 | 0.012903704 | 0.300769273 | 0.042902334 |
| Dlx2 | 66.09861157 | 0.031041295 | 0.270648415 | 0.114692322 |
| Platr26 | 0.08486385 | 0.037730769 | 3.724898049 | 0.010129343 |
| Gm13647 | 0.342086031 | 2.215526155 | 3.689232929 | 0.600538431 |
| Itga6 | 388.4782796 | -0.411412309 | 0.140059362 | -2.937413847 |
| Gm13663 | 0.038981395 | -0.443816764 | 3.724945049 | -0.119147198 |
| Gm17250 | 1.158557209 | -0.271842156 | 1.002019526 | -0.27129427 |
| Gm13662 | 0 | NA | NA | NA |
| Gm13739 | 0 | NA | NA | NA |
| Pdk1 | 468.9053974 | 0.726486279 | 0.152521888 | 4.763160815 |
| Gm13746 | 0 | NA | NA | NA |
| Rapgef4 | 3912.521623 | 0.515798771 | 0.112585433 | 4.581398826 |
| Gm26408 | 0 | NA | NA | NA |
| Rapgef4os1 | 0.258731086 | 0.036384675 | 2.063229975 | 0.017634813 |
| Rapgef4os2 | 0.479549184 | 0.45715143 | 1.461286169 | 0.312841824 |
| Rapgef4os3 | 1.315840983 | 2.004984441 | 0.941017 | 2.130656983 |
| Zak | 235.4261635 | -0.325402665 | 0.089359201 | -3.641512673 |
| Zakit | 0.427144302 | 0.077993194 | 1.434597262 | 0.054365915 |
| Rpl36a-ps4 | 0.431106946 | -0.407938823 | 1.391605687 | -0.293142538 |
| Cdca7 | 32.93658422 | -0.597592144 | 0.208591128 | -2.864897222 |
| Rps13-ps8 | 0 | NA | NA | NA |
| Eif1-ps3 | 0 | NA | NA | NA |
| Pex13-ps | 0.061920613 | -0.443816764 | 3.724945049 | -0.119147198 |
| Ak3l2-ps | 18.80802623 | -0.033910785 | 0.25853844 | -0.131163415 |
| 8430437L04Rik | 0.222986776 | 0.119371009 | 2.143889557 | 0.055679645 |
| 6430710C18Rik | 1.790214168 | -1.350041173 | 0.869745333 | -1.552225831 |
| Sp3 | 1150.979175 | -0.165204298 | 0.087100008 | -1.896719668 |
| Gm25960 | 2.212404147 | 0.01501666 | 0.776624194 | 0.019335813 |
| Hspe1-ps5 | 0 | NA | NA | NA |
| Gm11084 | 3.542390089 | -0.543092433 | 0.548355444 | -0.990402191 |
| Sp3os | 52.06250527 | 0.549735024 | 0.180222143 | 3.05031899 |
| Ola1 | 736.2602298 | 0.146822906 | 0.070296362 | 2.088627363 |
| Gm13665 | 0 | NA | NA | NA |
| Gm13666 | 0.293512495 | 0.003511811 | 2.003138436 | 0.001753154 |
| Gm13664 | 0 | NA | NA | NA |
| Sp9 | 90.07522838 | 0.291615296 | 0.249725969 | 1.167741173 |
| Cir1 | 188.1924414 | 0.371201886 | 0.110352185 | 3.363792791 |
| Scrn3 | 297.9077961 | -0.09337656 | 0.111932047 | -0.834225431 |
| Gpr155 | 760.0828086 | 0.381045385 | 0.072005617 | 5.291884166 |

| | | | | |
|---|---|---|---|---|
| Gm13703 | 0.773921687 | 0.10782776 | 1.131873889 | 0.095264818 |
| Gm13709 | 0.142443712 | 0.51795508 | 3.596298777 | 0.144024485 |
| D230022J07Rik | 3.614518035 | -0.064719503 | 0.558414462 | -0.115898688 |
| Gm13707 | 1.060348673 | -0.309068265 | 0.985806034 | -0.313518333 |
| Wipf1 | 548.8928039 | -0.580071112 | 0.141701942 | -4.093600306 |
| Gm13708 | 0 | NA | NA | NA |
| Chrna1 | 3.856870735 | -0.799814985 | 0.629494839 | -1.27056639 |
| Chrna1os | 4.004686001 | 0.113157187 | 0.518747781 | 0.218135269 |
| Chn1os3 | 3.866965461 | -0.627300985 | 0.533640764 | -1.175511745 |
| Chn1 | 4895.40377 | 0.936045655 | 0.117681675 | 7.95404772 |
| Chn1os1 | 6.863703603 | 0.833808554 | 0.420053306 | 1.985006528 |
| Gm13640 | 0 | NA | NA | NA |
| Atf2 | 1720.92591 | 0.09664557 | 0.057491235 | 1.681048767 |
| Gm27421 | 0 | NA | NA | NA |
| Gm10822 | 3.167618981 | -0.966529412 | 0.655811662 | -1.473791133 |
| Atp5g3 | 1417.808151 | -0.077662196 | 0.094916048 | -0.818219863 |
| Gm13668 | 0 | NA | NA | NA |
| A630050E04Rik | 0.102891481 | 0.51795324 | 3.724945049 | 0.139049901 |
| Gm28640 | 0 | NA | NA | NA |
| Gm13667 | 0 | NA | NA | NA |
| 4930441J16Rik | 0.04393175 | 0.037062677 | 3.724945049 | 0.009949859 |
| Lnp | 1061.691288 | 0.158338595 | 0.07297074 | 2.169891588 |
| 4930445N08Rik | 0 | NA | NA | NA |
| Mrpl23-ps1 | 10.74443234 | 0.180383606 | 0.438175032 | 0.411670208 |
| Gm22975 | 0 | NA | NA | NA |
| Gm13672 | 0 | NA | NA | NA |
| Evx2 | 6.495853999 | -0.463089161 | 0.412512806 | -1.12260554 |
| Hoxd13 | 0.292684482 | 0.479841686 | 1.613521571 | 0.297387835 |
| Hoxd12 | 0 | NA | NA | NA |
| Hoxd11 | 0 | NA | NA | NA |
| Gm28309 | 0 | NA | NA | NA |
| Hoxd10 | 0 | NA | NA | NA |
| Gm14396 | 0 | NA | NA | NA |
| Gm28793 | 0 | NA | NA | NA |
| Hoxd9 | 0 | NA | NA | NA |
| AL928644.1 | 0 | NA | NA | NA |
| Hoxd3os1 | 0.132812548 | 0.055445511 | 2.965536047 | 0.018696624 |
| Hoxd8 | 0.583508756 | -2.432199142 | 3.418768007 | -0.711425618 |
| Hoxd3 | 4.727112919 | -0.739367855 | 0.69780494 | -1.059562369 |
| Hoxd4 | 1.678255395 | 0.071850804 | 0.983065642 | 0.073088511 |
| Mir10b | 0.042716945 | 0.037062677 | 3.724945049 | 0.009949859 |
| Gm28230 | 0.047948698 | 0.037064902 | 3.724945049 | 0.009950456 |
| Haglr | 0.983322447 | -1.796712455 | 1.356076014 | -1.324934913 |
| Gm14424 | 0.086930093 | -0.443164241 | 3.724898049 | -0.118973522 |
| Gm14426 | 0 | NA | NA | NA |
| Hoxd1 | 1.509597968 | -1.363859809 | 1.009100238 | -1.351560289 |
| Mtx2 | 366.8248721 | -0.046408612 | 0.081775197 | -0.567514529 |

| | | | | |
|---|---|---|---|---|
| Gm14425 | 13.47600813 | 0.510122282 | 0.341165148 | 1.495235622 |
| Gm14427 | 0 | NA | NA | NA |
| n-R5s198 | 0 | NA | NA | NA |
| Gm13652 | 1.412897596 | -0.625274647 | 0.873151585 | -0.716112365 |
| 2600014E21Rik | 28.30992819 | 0.629996116 | 0.235137287 | 2.679269308 |
| A330043C09Rik | 1.006727404 | 0.615449342 | 0.900094692 | 0.683760662 |
| Rps6-ps4 | 71.2506606 | -0.280576449 | 0.14153424 | -1.98239274 |
| Gm13653 | 0.886214715 | -0.327362142 | 1.075608315 | -0.304350698 |
| Gm37243 | 0 | NA | NA | NA |
| Gm13658 | 0.044544648 | 0.517957688 | 3.724945049 | 0.139051095 |
| Gm13660 | 0 | NA | NA | NA |
| Gm13655 | 0.042716945 | 0.037062677 | 3.724945049 | 0.009949859 |
| Gm24574 | 0.04393175 | 0.037062677 | 3.724945049 | 0.009949859 |
| Hnrnpa3 | 910.2237922 | 0.215021198 | 0.077206561 | 2.785011987 |
| Nfe2l2 | 396.3091985 | -0.898943835 | 0.154495704 | -5.818568502 |
| Gm24996 | 0.882521852 | -0.895310531 | 1.136485614 | -0.787788706 |
| E030042O20Rik | 0.923975021 | -2.257845193 | 1.083878839 | -2.083115853 |
| Gm13656 | 0 | NA | NA | NA |
| Gm13657 | 0.629646617 | -1.784092596 | 1.439630588 | -1.239271109 |
| Gm13669 | 1.634157898 | -0.457005821 | 0.760087406 | -0.6012543 |
| Gm13670 | 1.659798175 | -0.925440308 | 1.019420828 | -0.907809888 |
| Agps | 976.4418691 | -0.106142967 | 0.059051463 | -1.797465491 |
| Ttc30b | 125.5045713 | -0.031625996 | 0.120952009 | -0.261475576 |
| Ttc30a2 | 2.514664855 | 0.669347051 | 0.618052597 | 1.082993672 |
| Ttc30a1 | 46.83376903 | -0.443984322 | 0.194022086 | -2.288318469 |
| Pde11a | 85.82156519 | 0.380629958 | 0.127316883 | 2.989626737 |
| AL845373.1 | 0 | NA | NA | NA |
| Cyct | 0 | NA | NA | NA |
| Rbm45 | 248.2191173 | -0.278665137 | 0.081361656 | -3.425018024 |
| Osbpl6 | 1518.253489 | 0.129751434 | 0.089831729 | 1.444383125 |
| Gm23164 | 1.915350769 | 0.649979201 | 0.893315794 | 0.727602943 |
| Prkra | 162.1113476 | -0.193511103 | 0.096254028 | -2.010420834 |
| Dfnb59 | 0.622664233 | 1.927930803 | 1.344953769 | 1.433455073 |
| Fkbp7 | 23.5112496 | 0.220857375 | 0.212139317 | 1.04109591 |
| Plekha3 | 466.6346452 | -0.240890572 | 0.077979425 | -3.089155546 |
| Ttn | 182.6205874 | 0.102830678 | 0.223938908 | 0.459190764 |
| Gm13938 | 1.375616079 | 0.840311884 | 1.059134283 | 0.793395037 |
| Ccdc141 | 278.830304 | -0.033054157 | 0.117432866 | -0.281472798 |
| AL928681.1 | 1.008651653 | 1.716798126 | 1.149334342 | 1.493732557 |
| Sestd1 | 1270.998036 | 0.254008355 | 0.076172045 | 3.334666358 |
| Gm13650 | 19.02386222 | -0.422349919 | 0.414754124 | -1.01831397 |
| Gm13651 | 3.760012592 | 0.112444442 | 0.520535221 | 0.216016971 |
| Gm13943 | 0 | NA | NA | NA |
| Gm13649 | 0.072025462 | 0.517959912 | 3.724945049 | 0.139051692 |
| Zfp385b | 633.2440097 | 0.340917747 | 0.101448187 | 3.360510993 |
| Gm13944 | 5.255555274 | -1.095974397 | 0.650993028 | -1.683542451 |
| Gm13661 | 1.194926524 | 2.022448226 | 0.979769162 | 2.064208901 |

| | | | | |
|---|---|---|---|---|
| Gm13727 | 0.386897144 | 0.497443336 | 1.458827695 | 0.34098841 |
| Cwc22 | 371.0030043 | 0.273992857 | 0.105948711 | 2.58608958 |
| Gm13726 | 0 | NA | NA | NA |
| 4930440I19Rik | 0.335959249 | -0.746316241 | 2.01267548 | -0.370808035 |
| Gm14461 | 0.674729951 | -1.792061405 | 1.215446457 | -1.474405882 |
| Gm14460 | 0 | NA | NA | NA |
| Gm23149 | 0 | NA | NA | NA |
| Gm14464 | 0.757266105 | -0.507005056 | 1.108628942 | -0.457326195 |
| Gm14463 | 0.136194719 | 0.037062677 | 3.724945049 | 0.009949859 |
| Gm17014 | 0 | NA | NA | NA |
| Ube2e3 | 827.2373707 | -0.049423694 | 0.084564204 | -0.584451717 |
| Gm37004 | 1.348672834 | -1.951400026 | 0.915984098 | -2.130386357 |
| Gm14466 | 0.233003758 | -1.874069331 | 2.523043414 | -0.742781246 |
| Gm14469 | 0.066050875 | -0.443818988 | 3.724945049 | -0.119147795 |
| Gm14465 | 0.075043205 | 0.517957688 | 3.724945049 | 0.139051095 |
| Itga4 | 452.6472453 | -0.173936554 | 0.138212333 | -1.258473463 |
| Cerkl | 55.52887644 | 0.577357527 | 0.2732218 | 2.113145901 |
| Gm22606 | 0 | NA | NA | NA |
| Gm13673 | 0.097139547 | -0.443816764 | 3.724945049 | -0.119147198 |
| Neurod1 | 741.6134161 | 0.035371416 | 0.28517926 | 0.124032219 |
| Gm26755 | 3.778983902 | 0.23191261 | 0.508275872 | 0.456273105 |
| Ssfa2 | 1354.956421 | -0.360588 | 0.137649351 | -2.619612784 |
| Ppp1r1c | 7.321389695 | 0.30783709 | 0.365479838 | 0.842282001 |
| Pde1a | 1335.258322 | 0.481525453 | 0.13975284 | 3.445550403 |
| Gm13674 | 0 | NA | NA | NA |
| Prdx6b | 0 | NA | NA | NA |
| Dnajc10 | 1132.405827 | 0.032610221 | 0.050691857 | 0.643302946 |
| Gm13752 | 0 | NA | NA | NA |
| Frzb | 132.7433967 | 0.833452358 | 0.179256798 | 4.649488155 |
| Gm13687 | 0 | NA | NA | NA |
| Nckap1 | 5774.673254 | 0.181267474 | 0.04689471 | 3.865414118 |
| Gm13689 | 37.9418997 | 0.4983576 | 0.310515276 | 1.604937468 |
| Gm24461 | 0 | NA | NA | NA |
| Gm13688 | 12.81351646 | -0.324110156 | 0.319515246 | -1.014380877 |
| Dusp19 | 119.9719043 | -0.223397562 | 0.121145154 | -1.844048686 |
| Nup35 | 181.7556722 | -0.694934283 | 0.130776301 | -5.313916024 |
| mmu-mir-684-1 | 0.866431109 | -0.664605932 | 1.097161845 | -0.605750131 |
| Gm13690 | 0.627304132 | 0.948512905 | 1.284617598 | 0.738362067 |
| Gm25420 | 0.262453343 | -0.039886507 | 2.101965409 | -0.018975815 |
| Gm4735 | 399.2680756 | -0.28947069 | 0.115350517 | -2.509487587 |
| Gm22652 | 0 | NA | NA | NA |
| Gm13700 | 0 | NA | NA | NA |
| Gm13755 | 0.332110845 | 0.578091932 | 1.56862409 | 0.368534396 |
| Gm13756 | 0 | NA | NA | NA |
| Gm23452 | 0 | NA | NA | NA |
| Gm23900 | 0 | NA | NA | NA |
| Zfp804a | 500.2371726 | 0.07496909 | 0.126705663 | 0.591679078 |

| | | | | |
|---|---|---|---|---|
| Gm13677 | 0 | NA | NA | NA |
| A130030D18Rik | 0.048949532 | 0.037064902 | 3.724945049 | 0.009950456 |
| Gm13675 | 0.141830814 | 0.037060062 | 3.593924434 | 0.010311865 |
| Gm13676 | 0 | NA | NA | NA |
| Gm13678 | 0 | NA | NA | NA |
| Fsip2 | 0.617250809 | 2.216926141 | 1.193270756 | 1.857856761 |
| Gm28675 | 0.121388914 | 0.037071574 | 3.724945049 | 0.009952247 |
| Gm13680 | 10.62432231 | -0.284293384 | 0.314933038 | -0.902710574 |
| Gm13679 | 0 | NA | NA | NA |
| Gm13682 | 0 | NA | NA | NA |
| Gm13681 | 0 | NA | NA | NA |
| Gm13683 | 0 | NA | NA | NA |
| Gm13685 | 0 | NA | NA | NA |
| Zc3h15 | 953.3117506 | -0.038918402 | 0.104750844 | -0.371533059 |
| Gm13686 | 0 | NA | NA | NA |
| Itgav | 2001.084212 | -0.149265535 | 0.07760135 | -1.923491466 |
| Gm13684 | 0.113999573 | -0.443821212 | 3.724945049 | -0.119148392 |
| Dnmt3l-ps1 | 0.097139547 | -0.443816764 | 3.724945049 | -0.119147198 |
| Fam171b | 2724.632699 | 0.55398548 | 0.086124575 | 6.432374035 |
| Zswim2 | 0.405375818 | 0.893099525 | 1.555548678 | 0.574137947 |
| Gm13698 | 0 | NA | NA | NA |
| Gm13693 | 0 | NA | NA | NA |
| Gm13695 | 0 | NA | NA | NA |
| Gm13696 | 0 | NA | NA | NA |
| Gm13694 | 0.047948698 | 0.037064902 | 3.724945049 | 0.009950456 |
| Gm13697 | 0.061920613 | -0.443816764 | 3.724945049 | -0.119147198 |
| Gm13691 | 0 | NA | NA | NA |
| Gm13692 | 0 | NA | NA | NA |
| Gm13702 | 0 | NA | NA | NA |
| Gm25972 | 0.044191549 | 0.037062677 | 3.724945049 | 0.009949859 |
| Gm13711 | 0.180501107 | 1.347540021 | 2.704438636 | 0.498269771 |
| Calcrl | 175.101438 | -0.42941875 | 0.123526483 | -3.476329433 |
| Tfpi | 63.47998308 | -0.382464225 | 0.141729301 | -2.698554384 |
| Gm13710 | 7.944981639 | 0.599852427 | 0.516663708 | 1.161011346 |
| Ctnnd1 | 284.3298198 | -0.050298814 | 0.093679517 | -0.536924354 |
| Gm28635 | 0 | NA | NA | NA |
| Btbd18 | 0.061920613 | -0.443816764 | 3.724945049 | -0.119147198 |
| 2700094K13Rik | 93.3688872 | -0.033115436 | 0.113781926 | -0.291043024 |
| Tmx2 | 1256.227034 | -0.044873815 | 0.057963995 | -0.774167048 |
| Med19 | 347.2590899 | -0.248837107 | 0.089432401 | -2.782404422 |
| Zdhhc5 | 1049.31998 | -0.401349893 | 0.063975783 | -6.273465858 |
| n-R5s200 | 2.639998601 | -0.658734555 | 0.789570913 | -0.834294355 |
| Gm27942 | 0.072025462 | 0.517959912 | 3.724945049 | 0.139051692 |
| Clp1 | 87.12695073 | -0.531053837 | 0.163593629 | -3.246176757 |
| Ypel4 | 577.9281882 | 0.349602164 | 0.116231999 | 3.007796196 |
| Mir130a | 0.283042562 | -1.199661825 | 3.714760153 | -0.322944625 |
| Gm19426 | 6.293410423 | -0.260902782 | 0.470077033 | -0.555021334 |

| | | | | |
|---|---|---|---|---|
| Serping1 | 1376.102164 | -0.870131853 | 0.272372362 | -3.194640771 |
| Ube2l6 | 958.9461895 | -0.466585988 | 0.264711648 | -1.762619786 |
| Smtnl1 | 0.1140246 | 0.037073798 | 3.724945049 | 0.009952844 |
| Timm10 | 121.5002785 | -0.060613753 | 0.121358178 | -0.499461629 |
| Slc43a1 | 4.475148185 | -0.523416006 | 0.494612437 | -1.058234623 |
| Rtn4rl2 | 239.9515241 | 0.0565988 | 0.190546203 | 0.297034522 |
| Gm24332 | 0.173129629 | -0.891656386 | 2.724725303 | -0.327246341 |
| Slc43a3 | 68.69465978 | -0.332555465 | 0.30074899 | -1.105757544 |
| Prg2 | 0.484678492 | -2.204844908 | 3.689460308 | -0.59760635 |
| Prg3 | 0.205912129 | -0.497570127 | 2.595365188 | -0.191714881 |
| P2rx3 | 14.94560384 | 0.629695134 | 0.285829443 | 2.203045029 |
| Ssrp1 | 920.1403945 | -0.201815379 | 0.068181911 | -2.959954839 |
| Tnks1bp1 | 1348.991537 | 0.040971266 | 0.083319423 | 0.491737279 |
| 4930443O20Rik | 0.247670153 | 0.973498534 | 2.108404709 | 0.461722804 |
| Aplnr | 18.49192457 | -1.069575729 | 0.268167743 | -3.988457808 |
| Gm13712 | 0.447666499 | 0.070197922 | 1.673586875 | 0.041944594 |
| Gm13713 | 6.805315018 | -0.631676765 | 0.372631846 | -1.695176544 |
| Lrrc55 | 258.0702094 | -0.333666362 | 0.128815781 | -2.590259985 |
| Gm13716 | 0.932468966 | 1.003570295 | 1.240486569 | 0.809013431 |
| Gm13715 | 0.832268342 | -0.499183885 | 0.969003544 | -0.515151764 |
| Gm13714 | 0.047948698 | 0.037064902 | 3.724945049 | 0.009950456 |
| Olfr987 | 0.046345709 | -0.443818988 | 3.724945049 | -0.119147795 |
| Olfr988 | 0.042716945 | 0.037062677 | 3.724945049 | 0.009949859 |
| Olfr990-ps1 | 0 | NA | NA | NA |
| Olfr992 | 0 | NA | NA | NA |
| Olfr993 | 0 | NA | NA | NA |
| Gm23716 | 0 | NA | NA | NA |
| Olfr994 | 0.042716945 | 0.037062677 | 3.724945049 | 0.009949859 |
| Olfr995 | 0 | NA | NA | NA |
| 4833423E24Rik | 3.225753529 | -0.42324246 | 0.590513902 | -0.716735811 |
| Olfr996 | 0 | NA | NA | NA |
| Olfr997-ps1 | 0 | NA | NA | NA |
| Olfr998 | 0.038981395 | -0.443816764 | 3.724945049 | -0.119147198 |
| Olfr999-ps1 | 0 | NA | NA | NA |
| Olfr1000 | 0 | NA | NA | NA |
| Olfr1001-ps1 | 0.046345709 | -0.443818988 | 3.724945049 | -0.119147795 |
| Olfr1002 | 0 | NA | NA | NA |
| Olfr1003-ps1 | 0 | NA | NA | NA |
| Olfr1004-ps1 | 0 | NA | NA | NA |
| Olfr154 | 0 | NA | NA | NA |
| Olfr1006 | 0.048949532 | 0.037064902 | 3.724945049 | 0.009950456 |
| Olfr1008 | 0 | NA | NA | NA |
| Olfr1009 | 0 | NA | NA | NA |
| Olfr1010 | 0.038981395 | -0.443816764 | 3.724945049 | -0.119147198 |
| Olfr1012 | 0.089062654 | -0.443823436 | 3.724945049 | -0.119148989 |
| Olfr1013 | 0 | NA | NA | NA |
| Olfr1014 | 0 | NA | NA | NA |

| | | | | |
|---|---|---|---|---|
| Olfr1015 | 0 | NA | NA | NA |
| Olfr1016 | 0.04393175 | 0.037062677 | 3.724945049 | 0.009949859 |
| Olfr1017-ps1 | 0 | NA | NA | NA |
| Gm13720 | 0 | NA | NA | NA |
| Olfr1018 | 0 | NA | NA | NA |
| Olfr1019 | 0.062426873 | 0.517959912 | 3.724945049 | 0.139051692 |
| Olfr1020 | 0 | NA | NA | NA |
| Olfr1021-ps1 | 0 | NA | NA | NA |
| Olfr1022 | 0 | NA | NA | NA |
| Gm24037 | 0 | NA | NA | NA |
| Olfr1023 | 0 | NA | NA | NA |
| Olfr1024 | 0 | NA | NA | NA |
| Olfr1025-ps1 | 0 | NA | NA | NA |
| Olfr1026 | 0 | NA | NA | NA |
| Olfr1027-ps1 | 0 | NA | NA | NA |
| Olfr1028 | 0 | NA | NA | NA |
| Olfr1029 | 0.108772582 | 0.037073798 | 3.724945049 | 0.009952844 |
| Olfr1030 | 0 | NA | NA | NA |
| Olfr1031 | 0 | NA | NA | NA |
| Olfr1032 | 0 | NA | NA | NA |
| Olfr1033 | 0 | NA | NA | NA |
| Olfr1034 | 0 | NA | NA | NA |
| Olfr1035-ps1 | 0 | NA | NA | NA |
| Olfr1036 | 0 | NA | NA | NA |
| Olfr1037 | 0 | NA | NA | NA |
| Gm13722 | 0.13742593 | 0.517962079 | 2.939744525 | 0.176192888 |
| Olfr1038-ps | 0 | NA | NA | NA |
| Olfr1039 | 0 | NA | NA | NA |
| Olfr1040 | 0 | NA | NA | NA |
| Olfr1042 | 0 | NA | NA | NA |
| Olfr1043 | 0 | NA | NA | NA |
| Olfr1044 | 0 | NA | NA | NA |
| Olfr52 | 0.108772582 | 0.037073798 | 3.724945049 | 0.009952844 |
| Olfr1045 | 0 | NA | NA | NA |
| Olfr1046 | 0 | NA | NA | NA |
| Olfr1047 | 0 | NA | NA | NA |
| Olfr1048 | 0 | NA | NA | NA |
| Olfr1049 | 0 | NA | NA | NA |
| Olfr1050-ps1 | 0 | NA | NA | NA |
| Olfr1051 | 0 | NA | NA | NA |
| Olfr1052 | 0.044544648 | 0.517957688 | 3.724945049 | 0.139051095 |
| Olfr1053 | 0 | NA | NA | NA |
| Gm13719 | 2.60552406 | 0.552674133 | 0.628217528 | 0.879749623 |
| Olfr1054 | 0 | NA | NA | NA |
| Olfr1055 | 0 | NA | NA | NA |
| Olfr1056 | 0 | NA | NA | NA |
| Olfr1057 | 0 | NA | NA | NA |

| | | | | |
|---|---|---|---|---|
| Olfr1058 | 0 | NA | NA | NA |
| Olfr1060 | 0 | NA | NA | NA |
| Olfr1061 | 0 | NA | NA | NA |
| Olfr1062 | 0 | NA | NA | NA |
| Olfr1063-ps1 | 0 | NA | NA | NA |
| Olfr1065 | 0 | NA | NA | NA |
| Olfr1066 | 0 | NA | NA | NA |
| Olfr1069-ps1 | 0 | NA | NA | NA |
| Olfr228 | 0 | NA | NA | NA |
| Olfr1073-ps1 | 0 | NA | NA | NA |
| Olfr1076 | 0 | NA | NA | NA |
| Olfr1077-ps1 | 0 | NA | NA | NA |
| Olfr1078-ps1 | 0 | NA | NA | NA |
| Olfr1079 | 0 | NA | NA | NA |
| Olfr1080 | 0 | NA | NA | NA |
| Olfr1081-ps1 | 0 | NA | NA | NA |
| Olfr1082 | 0 | NA | NA | NA |
| Olfr1083-ps | 0 | NA | NA | NA |
| Olfr1084 | 0 | NA | NA | NA |
| Olfr1085 | 0 | NA | NA | NA |
| Olfr1086 | 0 | NA | NA | NA |
| Olfr1087 | 0 | NA | NA | NA |
| Olfr1089 | 0 | NA | NA | NA |
| Olfr1090 | 0 | NA | NA | NA |
| Olfr1093 | 0 | NA | NA | NA |
| BX284685.1 | 0 | NA | NA | NA |
| Olfr141 | 0 | NA | NA | NA |
| Olfr1094 | 0 | NA | NA | NA |
| Olfr1095 | 0 | NA | NA | NA |
| Olfr1096-ps1 | 0 | NA | NA | NA |
| Gm13723 | 0 | NA | NA | NA |
| Olfr1097 | 0 | NA | NA | NA |
| Olfr1098 | 0 | NA | NA | NA |
| Olfr1099 | 0.048949532 | 0.037064902 | 3.724945049 | 0.009950456 |
| Olfr1100 | 0 | NA | NA | NA |
| Olfr1101 | 0 | NA | NA | NA |
| Olfr1102 | 0 | NA | NA | NA |
| Olfr1103-ps1 | 0 | NA | NA | NA |
| Olfr1104 | 0 | NA | NA | NA |
| Olfr1105 | 0 | NA | NA | NA |
| Olfr1106 | 0 | NA | NA | NA |
| Olfr1107 | 0 | NA | NA | NA |
| Olfr1109 | 0.046345709 | -0.443818988 | 3.724945049 | -0.119147795 |
| Olfr259 | 0 | NA | NA | NA |
| Gm13733 | 0 | NA | NA | NA |
| Olfr1110 | 0 | NA | NA | NA |
| Gm13734 | 0 | NA | NA | NA |

| | | | | |
|---|---|---|---|---|
| Olfr1111 | 0 | NA | NA | NA |
| Nlk-ps1 | 0.21574249 | 0.998866801 | 2.174796342 | 0.45929211 |
| Olfr1112 | 0 | NA | NA | NA |
| Gm13730 | 0 | NA | NA | NA |
| Olfr1113 | 0 | NA | NA | NA |
| Gm13735 | 0 | NA | NA | NA |
| Olfr1115 | 0 | NA | NA | NA |
| Gm27438 | 0 | NA | NA | NA |
| Gm13732 | 0 | NA | NA | NA |
| Olfr1116 | 0 | NA | NA | NA |
| Gm13728 | 0.042716945 | 0.037062677 | 3.724945049 | 0.009949859 |
| Olfr1117-ps1 | 0 | NA | NA | NA |
| Gm13729 | 0 | NA | NA | NA |
| Olfr1118 | 0 | NA | NA | NA |
| Gm13737 | 0 | NA | NA | NA |
| Gm13731 | 0 | NA | NA | NA |
| Olfr1120 | 0 | NA | NA | NA |
| Olfr1121 | 0 | NA | NA | NA |
| Olfr1122 | 0 | NA | NA | NA |
| Gm1826 | 8.12848589 | -0.862317141 | 0.38680871 | -2.229311593 |
| Olfr1123 | 0 | NA | NA | NA |
| Olfr1124 | 0 | NA | NA | NA |
| Olfr1126 | 0 | NA | NA | NA |
| Pramel7 | 0 | NA | NA | NA |
| Pramel6 | 0 | NA | NA | NA |
| Gm13738 | 0 | NA | NA | NA |
| Olfr1127-ps1 | 0 | NA | NA | NA |
| Gm13736 | 22.02889899 | 0.030022767 | 0.27020699 | 0.111110253 |
| Olfr153 | 0 | NA | NA | NA |
| Olfr1128 | 0 | NA | NA | NA |
| Gm13740 | 0.18866314 | 0.855473539 | 2.660683079 | 0.321524027 |
| Gm13744 | 0 | NA | NA | NA |
| Gm13743 | 3.080495725 | 0.394998627 | 0.565874645 | 0.69803203 |
| Olfr1129 | 0 | NA | NA | NA |
| Olfr1130 | 0 | NA | NA | NA |
| Olfr1131 | 0 | NA | NA | NA |
| Olfr1132 | 0 | NA | NA | NA |
| Gm13742 | 0 | NA | NA | NA |
| Olfr1133 | 0 | NA | NA | NA |
| Gm13741 | 0 | NA | NA | NA |
| Olfr1134 | 0 | NA | NA | NA |
| Olfr1135 | 0 | NA | NA | NA |
| Olfr1136 | 0 | NA | NA | NA |
| Olfr1137 | 0 | NA | NA | NA |
| Olfr1138 | 0 | NA | NA | NA |
| Olfr1140 | 0 | NA | NA | NA |
| Olfr1141 | 0 | NA | NA | NA |

| | | | | |
|---|---|---|---|---|
| Olfr152 | 0 | NA | NA | NA |
| Olfr1143 | 0 | NA | NA | NA |
| Olfr1144-ps1 | 0 | NA | NA | NA |
| Olfr1145 | 0 | NA | NA | NA |
| Olfr1146-ps1 | 0 | NA | NA | NA |
| Olfr1147-ps1 | 0 | NA | NA | NA |
| Olfr1148 | 0 | NA | NA | NA |
| Olfr1149-ps1 | 0 | NA | NA | NA |
| Olfr1150-ps1 | 0 | NA | NA | NA |
| Olfr1151 | 0 | NA | NA | NA |
| Olfr1152 | 0 | NA | NA | NA |
| Gm13745 | 0 | NA | NA | NA |
| Olfr1153 | 0 | NA | NA | NA |
| Olfr1154 | 0 | NA | NA | NA |
| Olfr1155 | 0 | NA | NA | NA |
| Olfr1156 | 0 | NA | NA | NA |
| Olfr1157 | 0 | NA | NA | NA |
| Olfr74 | 0 | NA | NA | NA |
| Olfr1158 | 0 | NA | NA | NA |
| Olfr1159-ps1 | 0 | NA | NA | NA |
| Olfr1160 | 0 | NA | NA | NA |
| Olfr1161 | 0 | NA | NA | NA |
| Olfr73 | 0 | NA | NA | NA |
| Gm13747 | 0 | NA | NA | NA |
| Olfr1162 | 0 | NA | NA | NA |
| Olfr1163 | 0 | NA | NA | NA |
| Olfr1164 | 0 | NA | NA | NA |
| Olfr1165-ps | 0.046345709 | -0.443818988 | 3.724945049 | -0.119147795 |
| Olfr1166 | 0 | NA | NA | NA |
| Olfr1167 | 0 | NA | NA | NA |
| Olfr1168 | 0.046345709 | -0.443818988 | 3.724945049 | -0.119147795 |
| Olfr1169-ps1 | 0 | NA | NA | NA |
| Olfr1170 | 0 | NA | NA | NA |
| Olfr1171-ps1 | 0 | NA | NA | NA |
| Gm13753 | 0 | NA | NA | NA |
| Olfr1172-ps1 | 0 | NA | NA | NA |
| Olfr1173 | 0 | NA | NA | NA |
| Olfr1174-ps | 0 | NA | NA | NA |
| Olfr1175-ps | 0 | NA | NA | NA |
| Olfr1176 | 0.061920613 | -0.443816764 | 3.724945049 | -0.119147198 |
| Olfr1177-ps | 0 | NA | NA | NA |
| Olfr1178 | 0 | NA | NA | NA |
| Olfr1179 | 0 | NA | NA | NA |
| Olfr1180 | 0 | NA | NA | NA |
| Olfr1181 | 0 | NA | NA | NA |
| Olfr1182 | 0 | NA | NA | NA |
| Gm13757 | 0 | NA | NA | NA |

| | | | | |
|---|---|---|---|---|
| Olfr1183 | 0 | NA | NA | NA |
| Olfr1184 | 0 | NA | NA | NA |
| Olfr1185-ps1 | 0 | NA | NA | NA |
| Olfr1186 | 0 | NA | NA | NA |
| Olfr1187-ps1 | 0 | NA | NA | NA |
| Olfr1188 | 0 | NA | NA | NA |
| Gm13758 | 0 | NA | NA | NA |
| Olfr1189 | 0 | NA | NA | NA |
| Olfr475-ps1 | 0 | NA | NA | NA |
| Olfr1191-ps1 | 0.062426873 | 0.517959912 | 3.724945049 | 0.139051692 |
| Gm13759 | 0 | NA | NA | NA |
| Olfr1192-ps1 | 0 | NA | NA | NA |
| Olfr1193 | 0.042716945 | 0.037062677 | 3.724945049 | 0.009949859 |
| Olfr1195 | 0.131862677 | -0.443820876 | 2.977354665 | -0.149065505 |
| Olfr1196 | 0 | NA | NA | NA |
| Olfr1197 | 0 | NA | NA | NA |
| Olfr1198 | 0 | NA | NA | NA |
| Olfr1199 | 0 | NA | NA | NA |
| Gm13760 | 0 | NA | NA | NA |
| Olfr1200 | 0 | NA | NA | NA |
| Olfr1201 | 0 | NA | NA | NA |
| Olfr1202 | 0 | NA | NA | NA |
| Olfr1205 | 0 | NA | NA | NA |
| Olfr1204 | 0 | NA | NA | NA |
| Olfr1206 | 0 | NA | NA | NA |
| Olfr1208 | 0 | NA | NA | NA |
| Olfr1209 | 0 | NA | NA | NA |
| Olfr1211 | 0.068097359 | 0.037064902 | 3.724945049 | 0.009950456 |
| Gm13763 | 0 | NA | NA | NA |
| Olfr1212 | 0 | NA | NA | NA |
| Olfr1213 | 0 | NA | NA | NA |
| Gm13762 | 0 | NA | NA | NA |
| Olfr1214 | 0 | NA | NA | NA |
| Olfr1215 | 0 | NA | NA | NA |
| Olfr1216 | 0 | NA | NA | NA |
| Olfr1217 | 0 | NA | NA | NA |
| Olfr1218 | 0 | NA | NA | NA |
| Olfr1219 | 0.045882455 | 0.517959912 | 3.724945049 | 0.139051692 |
| Olfr1220 | 0.095435502 | 0.037073798 | 3.724945049 | 0.009952844 |
| Olfr1221 | 0 | NA | NA | NA |
| Gm13764 | 0.048949532 | 0.037064902 | 3.724945049 | 0.009950456 |
| Olfr1222 | 0.048949532 | 0.037064902 | 3.724945049 | 0.009950456 |
| Olfr1223 | 0 | NA | NA | NA |
| Olfr1224-ps1 | 0 | NA | NA | NA |
| Olfr1225 | 0 | NA | NA | NA |
| Gm13761 | 0 | NA | NA | NA |
| Olfr1226 | 0 | NA | NA | NA |

| | | | | |
|---|---|---|---|---|
| Olfr1228 | 0.062426873 | 0.517959912 | 3.724945049 | 0.139051692 |
| Olfr1229 | 0 | NA | NA | NA |
| Olfr1230 | 0 | NA | NA | NA |
| Olfr1231 | 0.046345709 | -0.443818988 | 3.724945049 | -0.119147795 |
| Olfr1232 | 0 | NA | NA | NA |
| Olfr1233 | 0.048949532 | 0.037064902 | 3.724945049 | 0.009950456 |
| Olfr1234 | 0.048949532 | 0.037064902 | 3.724945049 | 0.009950456 |
| Olfr1235-ps1 | 0 | NA | NA | NA |
| Olfr1238 | 0 | NA | NA | NA |
| Olfr1239 | 0 | NA | NA | NA |
| Olfr1240 | 0 | NA | NA | NA |
| Olfr1241 | 0 | NA | NA | NA |
| Olfr1242 | 0 | NA | NA | NA |
| Olfr1243 | 0.106618422 | 0.517955464 | 3.724945049 | 0.139050498 |
| Olfr1245 | 0 | NA | NA | NA |
| Olfr1246 | 0.045882455 | 0.517959912 | 3.724945049 | 0.139051692 |
| Olfr1247 | 0 | NA | NA | NA |
| Olfr1248 | 0 | NA | NA | NA |
| Olfr1249 | 0 | NA | NA | NA |
| Olfr1250 | 0.042716945 | 0.037062677 | 3.724945049 | 0.009949859 |
| Olfr1251 | 0 | NA | NA | NA |
| Olfr1252 | 0 | NA | NA | NA |
| Olfr1253 | 0 | NA | NA | NA |
| Olfr1254 | 0.04393175 | 0.037062677 | 3.724945049 | 0.009949859 |
| Olfr1255 | 0 | NA | NA | NA |
| Olfr1256 | 0 | NA | NA | NA |
| Olfr48 | 0.156220659 | 0.03706712 | 2.825874445 | 0.013117044 |
| Olfr1257 | 0 | NA | NA | NA |
| Olfr1258 | 0.254495662 | -1.017189418 | 2.083804207 | -0.488140591 |
| Olfr1259 | 0 | NA | NA | NA |
| Olfr1260 | 0.042716945 | 0.037062677 | 3.724945049 | 0.009949859 |
| Olfr1261 | 0.198041555 | -1.296529483 | 2.587933318 | -0.500990297 |
| Olfr1262 | 0.31782415 | -1.420235472 | 3.698238508 | -0.384030253 |
| Olfr1263 | 0 | NA | NA | NA |
| Olfr1264 | 0.068097359 | 0.037064902 | 3.724945049 | 0.009950456 |
| Olfr1265 | 0.097139547 | -0.443816764 | 3.724945049 | -0.119147198 |
| Olfr140 | 0.229124067 | -1.47395843 | 2.560995672 | -0.575541164 |
| Olfr1267-ps1 | 0.049553047 | -0.443818988 | 3.724945049 | -0.119147795 |
| Olfr1268-ps1 | 0.202674043 | -1.2519774 | 2.606063873 | -0.48040933 |
| Olfr1269 | 0.045882455 | 0.517959912 | 3.724945049 | 0.139051692 |
| Olfr32 | 0.631605561 | -0.8254762 | 1.461544638 | -0.564797118 |
| Gm4222 | 0 | NA | NA | NA |
| Olfr1270 | 0.066050875 | -0.443818988 | 3.724945049 | -0.119147795 |
| Gm13768 | 4.397907185 | 0.286984698 | 0.585770695 | 0.489926691 |
| Olfr1506 | 0 | NA | NA | NA |
| Gm18953 | 1.627769934 | -1.781308427 | 0.908158549 | -1.961450927 |
| Olfr142 | 0.123841226 | -1.017823543 | 3.719440177 | -0.273649661 |

| | | | | |
|---|---|---|---|---|
| Olfr1271 | 0 | NA | NA | NA |
| Olfr1272 | 0 | NA | NA | NA |
| Olfr1273-ps | 0.044544648 | 0.517957688 | 3.724945049 | 0.139051095 |
| Gm13769 | 0 | NA | NA | NA |
| Gm13767 | 0.326846603 | -0.40549398 | 1.579320082 | -0.256752247 |
| Olfr1274-ps | 0.163996116 | -0.925588077 | 2.782099537 | -0.332694091 |
| Ptprj | 2106.140895 | 0.023410092 | 0.077417413 | 0.30238794 |
| 4933423P22Rik | 0.57303442 | 0.262164211 | 1.280362794 | 0.204757755 |
| Nup160 | 358.4498607 | -0.151164563 | 0.108351432 | -1.395132118 |
| Fnbp4 | 883.9475744 | 0.380901642 | 0.154821498 | 2.460263251 |
| Rpl30-ps3 | 4.003505758 | 0.213520273 | 0.536793036 | 0.397770199 |
| Agbl2 | 38.0546618 | -0.303726384 | 0.207089091 | -1.466645986 |
| Gm13772 | 2.379780159 | 0.490512222 | 0.808724929 | 0.606525414 |
| Mtch2 | 1065.574097 | -0.054006066 | 0.054962617 | -0.982596334 |
| C1qtnf4 | 729.131802 | 0.53427838 | 0.09699862 | 5.508102925 |
| Ndufs3 | 556.9779417 | 0.043909709 | 0.054952274 | 0.799051708 |
| Kbtbd4 | 295.8165819 | -0.045798966 | 0.079454225 | -0.576419523 |
| Ptpmt1 | 185.7476871 | 0.023319687 | 0.091664979 | 0.25440127 |
| Gm17661 | 14.75789179 | -0.08970422 | 0.366760908 | -0.244585008 |
| Celf1 | 3282.919782 | 0.096594101 | 0.069554323 | 1.388757698 |
| Gm13777 | 0.066050875 | -0.443818988 | 3.724945049 | -0.119147795 |
| Rapsn | 7.482500097 | 0.439152182 | 0.412189157 | 1.065414202 |
| Gm13778 | 1.349943917 | -0.426922074 | 1.008967247 | -0.423127783 |
| Psmc3 | 1077.312818 | -0.207967507 | 0.091961076 | -2.261473172 |
| Slc39a13 | 259.1457734 | -0.151570219 | 0.082921103 | -1.827884737 |
| Spi1 | 202.8166777 | -0.835481247 | 0.201448189 | -4.147375321 |
| Mybpc3 | 140.415035 | 0.375433674 | 0.246839347 | 1.520963647 |
| Madd | 2828.091406 | 0.143096459 | 0.077770006 | 1.839995459 |
| Nr1h3 | 80.38358335 | -1.388694408 | 0.214741998 | -6.466803976 |
| Gm13783 | 1.952547309 | -0.122923201 | 0.856444798 | -0.14352729 |
| Gm23390 | 0 | NA | NA | NA |
| Acp2 | 985.571155 | -0.014500039 | 0.044772946 | -0.323857159 |
| Ddb2 | 90.31751625 | 0.553670067 | 0.213882661 | 2.588662704 |
| A330069E16Rik | 10.08996547 | -1.135968434 | 0.359790877 | -3.157301938 |
| Gm17281 | 10.85400332 | 0.410690964 | 0.399682768 | 1.027542334 |
| Pacsin3 | 255.622615 | 0.020869986 | 0.125324837 | 0.166527133 |
| Arfgap2 | 730.4068622 | -0.206358758 | 0.074606405 | -2.765965697 |
| 1110051M20Rik | 549.9156413 | 0.260751595 | 0.058817361 | 4.43324197 |
| Gm13787 | 4.652905194 | -0.218570923 | 0.585514988 | -0.373296888 |
| Gm22071 | 0.442430462 | -1.980361646 | 3.352718207 | -0.590673455 |
| Lrp4 | 768.9458671 | 0.054484364 | 0.142560609 | 0.382183859 |
| Gm13770 | 1.121024803 | 0.45451667 | 1.032621548 | 0.440158033 |
| Ckap5 | 3392.946938 | 0.062949522 | 0.056616282 | 1.11186252 |
| Snord67 | 0.174512195 | 0.037058139 | 3.362865018 | 0.011019812 |
| F2 | 4.096713608 | -0.977785741 | 0.55776447 | -1.75304415 |
| Gm13771 | 3.006370356 | -0.302039626 | 0.665169601 | -0.454079118 |
| Zfp408 | 162.7132082 | -0.14849921 | 0.17107611 | -0.868030085 |

| | | | | |
|---|---|---|---|---|
| Arhgap1 | 635.3928673 | -0.129008726 | 0.063091841 | -2.044776682 |
| Atg13 | 1068.978477 | 0.249581387 | 0.051398623 | 4.855799067 |
| Harbi1 | 67.29079701 | -0.266443863 | 0.163790317 | -1.626737577 |
| Gm13776 | 0 | NA | NA | NA |
| Ambra1 | 1182.397324 | 0.145068541 | 0.069436932 | 2.089213001 |
| Gm10805 | 2.718864041 | 0.207304965 | 0.758135184 | 0.273440633 |
| Chrm4 | 179.7457586 | 0.688811619 | 0.16664501 | 4.133406804 |
| Mdk | 159.9674338 | -0.682538776 | 0.184078333 | -3.707871351 |
| Dgkz | 3070.276697 | 0.356519476 | 0.092044353 | 3.873344357 |
| Mir6999 | 2.840504449 | 0.223211523 | 0.796817795 | 0.280128688 |
| Gm9821 | 0.506210706 | -0.953840539 | 1.649985733 | -0.578090174 |
| Creb3l1 | 136.2000294 | -0.910317652 | 0.220182341 | -4.13438084 |
| Gm13780 | 0 | NA | NA | NA |
| 2900072N19Rik | 2.504782524 | 0.644562466 | 0.846778946 | 0.761193306 |
| Phf21a | 888.8724875 | -0.065259298 | 0.083247503 | -0.783918982 |
| Mir1955 | 9.919958007 | -0.322130148 | 0.36948452 | -0.871836656 |
| Gm13786 | 17.47248729 | -0.136180172 | 0.319723585 | -0.425930954 |
| Gm13785 | 11.39462224 | 0.641538926 | 0.359416669 | 1.784944833 |
| Gyltl1b | 4.391866736 | -0.198348399 | 0.542544049 | -0.365589485 |
| Pex16 | 172.030486 | -0.110077945 | 0.100953879 | -1.09037856 |
| 1700029I15Rik | 16.98714461 | 0.245992372 | 0.292851978 | 0.839988766 |
| Mapk8ip1 | 3383.018323 | 0.072660137 | 0.083951908 | 0.865497141 |
| Mir7000 | 0.999544483 | 0.401943243 | 1.191878308 | 0.337235136 |
| Cry2 | 1477.051092 | 0.629528945 | 0.124275712 | 5.065583107 |
| D930015M05Rik | 1.112373992 | 0.374887529 | 1.014061648 | 0.369689091 |
| Slc35c1 | 209.4076668 | -0.120949107 | 0.098557283 | -1.227196033 |
| Gm13814 | 0.592656722 | -1.447712587 | 1.325126287 | -1.092509146 |
| Gm13817 | 0 | NA | NA | NA |
| Gm13815 | 0.046345709 | -0.443818988 | 3.724945049 | -0.119147795 |
| Mir7221 | 0 | NA | NA | NA |
| Mir7222 | 0 | NA | NA | NA |
| Chst1 | 1513.4659 | 0.543112235 | 0.104719049 | 5.186374779 |
| Gm13816 | 0.371244445 | -2.000933495 | 1.626148822 | -1.230473784 |
| Gm13791 | 2.292177245 | -0.131114868 | 0.720593288 | -0.181954052 |
| Gm13792 | 0.47588132 | 1.974587733 | 1.721500892 | 1.147015225 |
| Syt13 | 2064.734027 | 0.218748269 | 0.077931442 | 2.806932157 |
| Prdm11 | 417.3168277 | 0.26101846 | 0.114778563 | 2.274104606 |
| 4631405J19Rik | 0.851482706 | 0.52019573 | 1.32706912 | 0.391988422 |
| Gm13802 | 0 | NA | NA | NA |
| Trp53i11 | 736.2819127 | 0.123034633 | 0.113756359 | 1.081562683 |
| Tspan18 | 97.99565433 | 0.527576627 | 0.145923615 | 3.615430081 |
| Gm23380 | 0 | NA | NA | NA |
| Gm13807 | 0 | NA | NA | NA |
| Cd82 | 303.0056944 | -0.369202172 | 0.140845603 | -2.621325512 |
| Mir7001 | 0.756442295 | 0.96183057 | 1.301735784 | 0.738883099 |
| Gm10804 | 0.060174537 | 0.517957688 | 3.724945049 | 0.139051095 |
| Gm10803 | 0 | NA | NA | NA |

| | | | | |
|---|---|---|---|---|
| Alx4 | 48.87403417 | -0.651506276 | 0.35918149 | -1.81386373 |
| Ext2 | 641.1248268 | -0.038709435 | 0.076318307 | -0.50721034 |
| Gm13818 | 0 | NA | NA | NA |
| Accs | 46.96754752 | 0.098582506 | 0.197575081 | 0.498962242 |
| Accsl | 3.620414628 | 0.662471487 | 0.59198698 | 1.119064285 |
| Gm1335 | 0.929738683 | 0.798817018 | 1.06663225 | 0.748915119 |
| Gm13889 | 181.9860749 | -0.326879526 | 0.341739948 | -0.956515407 |
| AL772372.1 | 0 | NA | NA | NA |
| Gm27027 | 1.373972637 | 0.579270871 | 0.953881885 | 0.607277359 |
| 4921507L20Rik | 0.891453265 | -1.503796712 | 1.063847071 | -1.413545944 |
| Alkbh3 | 168.1665116 | 0.170010536 | 0.139269687 | 1.22072893 |
| Hsd17b12 | 914.0604724 | -0.4021178 | 0.126678193 | -3.174325352 |
| Gm23630 | 0.694735342 | -1.245473978 | 1.270557601 | -0.980257783 |
| Mir129-2 | 0.549517402 | 0.279105918 | 1.255805252 | 0.222252548 |
| mmu-mir-129b | 0 | NA | NA | NA |
| E530001K10Rik | 26.89108908 | -0.510837087 | 0.307565311 | -1.660906054 |
| Mir670 | 0.304542382 | 1.046896366 | 2.037664264 | 0.513772747 |
| Ttc17 | 942.8773192 | 0.162440381 | 0.095899912 | 1.693853282 |
| Itpa-ps1 | 137.8875215 | -0.086288877 | 0.120823279 | -0.71417427 |
| 2810002D19Rik | 26.44289622 | -0.148477663 | 0.231778186 | -0.640602404 |
| Api5 | 1200.933468 | -0.16227494 | 0.064756965 | -2.505907144 |
| AL672251.1 | 0 | NA | NA | NA |
| Gm13788 | 0 | NA | NA | NA |
| Gm24318 | 0 | NA | NA | NA |
| Gm13793 | 0 | NA | NA | NA |
| Gm26396 | 0 | NA | NA | NA |
| Gm13794 | 0 | NA | NA | NA |
| Gm13798 | 0 | NA | NA | NA |
| Gm13797 | 0 | NA | NA | NA |
| Gm13799 | 0 | NA | NA | NA |
| Gm37175 | 0 | NA | NA | NA |
| Gm13801 | 0 | NA | NA | NA |
| Lrrc4c | 1313.773726 | 0.642994141 | 0.084569653 | 7.603130884 |
| 4930445B16Rik | 0 | NA | NA | NA |
| Gm17075 | 11.93369344 | 0.163550323 | 0.329351777 | 0.496582482 |
| Gm17076 | 16.91552809 | 1.452146276 | 0.364634153 | 3.98247466 |
| Gm22957 | 5.852693499 | 1.322093428 | 0.583256964 | 2.266742636 |
| Gm22313 | 8.145373654 | 1.436812712 | 0.629363408 | 2.282961949 |
| Gm23317 | 0 | NA | NA | NA |
| BX813336.1 | 0 | NA | NA | NA |
| Gm13803 | 11.5628337 | 0.794415394 | 0.34776172 | 2.284366993 |
| Gm13804 | 0 | NA | NA | NA |
| Gm13806 | 0 | NA | NA | NA |
| Gm10801 | 0.412640929 | 0.974705879 | 1.59144105 | 0.612467473 |
| Gm10800 | 8.185264119 | 0.586431906 | 0.492490624 | 1.190747353 |
| Gm29235 | 0 | NA | NA | NA |
| Gm13809 | 0 | NA | NA | NA |

| | | | | |
|---|---|---|---|---|
| Gm13808 | 0 | NA | NA | NA |
| Gm13811 | 0.109727584 | 0.037071574 | 3.724945049 | 0.009952247 |
| Gm13813 | 0 | NA | NA | NA |
| Gm13812 | 0.047948698 | 0.037064902 | 3.724945049 | 0.009950456 |
| Gm13810 | 0 | NA | NA | NA |
| B230118H07Rik | 178.272976 | 0.062114013 | 0.150688092 | 0.412202529 |
| Gm20693 | 0 | NA | NA | NA |
| Rag2 | 0.139883403 | -1.137320593 | 3.551352803 | -0.32024996 |
| Rag1 | 5.95628265 | 0.052493717 | 0.416052133 | 0.126171008 |
| Traf6 | 437.7135397 | -0.073961199 | 0.107233246 | -0.689722657 |
| Prr5l | 253.5858244 | -0.562918468 | 0.13816888 | -4.074133535 |
| Gm13905 | 0.083526043 | 0.037073798 | 3.724945049 | 0.009952844 |
| Commd9 | 191.447391 | -0.333413372 | 0.117358584 | -2.840979851 |
| Ldlrad3 | 167.5338321 | -0.36122367 | 0.128501667 | -2.811042682 |
| Gm13919 | 2.169032929 | -0.188799165 | 0.773634639 | -0.244041768 |
| Gm13920 | 4.969440295 | -1.940304578 | 0.600848264 | -3.229275503 |
| Gm13921 | 0.311841693 | 1.267968538 | 1.718579694 | 0.737800256 |
| Gm13866 | 1.228132247 | 0.436107709 | 1.048941498 | 0.415759802 |
| Trim44 | 3706.544855 | 0.200660396 | 0.046988279 | 4.270435113 |
| Gm13868 | 13.87276866 | 0.528502607 | 0.411957193 | 1.282906613 |
| Fjx1 | 429.7281807 | 0.180921276 | 0.09477139 | 1.909028411 |
| Pamr1 | 172.126028 | 0.080405938 | 0.15505773 | 0.518554851 |
| Slc1a2 | 36923.0076 | 1.123357786 | 0.16063204 | 6.99336064 |
| Gm13872 | 1.821369377 | -0.121472115 | 0.9020623 | -0.134660449 |
| Cd44 | 1546.027959 | -1.764588912 | 0.329215604 | -5.359979574 |
| Gm23731 | 0 | NA | NA | NA |
| Pdhx | 517.8134996 | 0.07420726 | 0.069304657 | 1.070739885 |
| Apip | 121.8280582 | -0.174468969 | 0.108660738 | -1.605630258 |
| Gm10038 | 4.822154624 | 0.293411175 | 0.487951555 | 0.601312101 |
| Gm13874 | 0.503556157 | -0.640513873 | 1.65885489 | -0.386118085 |
| Gm13869 | 0.217279565 | 0.037076952 | 2.170317435 | 0.017083654 |
| Gm13873 | 0 | NA | NA | NA |
| Ehf | 2.553683234 | -0.963753679 | 0.627123544 | -1.536784398 |
| A930006I01Rik | 0 | NA | NA | NA |
| Gm37220 | 0 | NA | NA | NA |
| BC016548 | 0.652016589 | -0.839912367 | 1.294867225 | -0.648647484 |
| Elf5 | 2.42823633 | 1.780889836 | 0.892854562 | 1.994602383 |
| Cat | 444.9583296 | -0.185172489 | 0.116433247 | -1.590374697 |
| Gm13996 | 0 | NA | NA | NA |
| Abtb2 | 322.9374325 | -0.428570384 | 0.140300137 | -3.05466833 |
| Nat10 | 403.567961 | 0.236435 | 0.170583402 | 1.386037552 |
| Caprin1 | 3734.751293 | -0.351823999 | 0.05242419 | -6.711100321 |
| 4930547E08Rik | 0.171199454 | 0.541396852 | 3.386170951 | 0.159884678 |
| Gm13878 | 0 | NA | NA | NA |
| Gm13880 | 2.206276467 | -0.607715855 | 0.666957805 | -0.911175864 |
| Gm13877 | 0 | NA | NA | NA |
| Gm13875 | 0 | NA | NA | NA |

| | | | | |
|---|---|---|---|---|
| Gm22757 | 0 | NA | NA | NA |
| Gm13879 | 0 | NA | NA | NA |
| Gm22761 | 0 | NA | NA | NA |
| Gm13876 | 0 | NA | NA | NA |
| Gm13881 | 0.097899064 | 0.037062677 | 3.724945049 | 0.009949859 |
| Lmo2 | 263.3668701 | 0.332490625 | 0.106885578 | 3.110715509 |
| 4931422A03Rik | 0.485755142 | 1.254690455 | 1.313672783 | 0.955101203 |
| Fbxo3 | 1097.464659 | 0.146797885 | 0.055130977 | 2.66271146 |
| Gm24644 | 12.13107012 | 1.107020233 | 0.383837213 | 2.884087825 |
| Gm10912 | 3.939700132 | -1.424175869 | 0.588746308 | -2.418997538 |
| Gm10799 | 0.09689823 | 0.037062677 | 3.724945049 | 0.009949859 |
| Cd59b | 12.1589774 | -0.351729107 | 0.320366755 | -1.097895152 |
| Cd59a | 104.2603775 | -0.540660057 | 0.216002363 | -2.503028441 |
| A930018P22Rik | 1.374894827 | -1.612244115 | 0.951825144 | -1.693844847 |
| D430041D05Rik | 2766.772851 | 0.579687371 | 0.150819116 | 3.843593472 |
| Gm22501 | 3.530580087 | 1.657097583 | 0.694219661 | 2.386993161 |
| Gm13883 | 6.270791699 | 0.017748704 | 0.531103681 | 0.03341853 |
| Gm25554 | 0 | NA | NA | NA |
| Gm13882 | 5.232096263 | 0.254002489 | 0.432775242 | 0.586915481 |
| Hipk3 | 2248.948924 | -0.060507661 | 0.062492983 | -0.968231289 |
| Mir1902 | 0 | NA | NA | NA |
| Gm13870 | 0 | NA | NA | NA |
| Gm13884 | 0.326587101 | -1.800073387 | 1.98566249 | -0.906535424 |
| Platr14 | 2.284861264 | 0.873120863 | 0.736408196 | 1.185647942 |
| Gm22439 | 0 | NA | NA | NA |
| Gm13885 | 0.266822755 | 0.540036961 | 2.422294538 | 0.222944383 |
| Cstf3 | 425.9326561 | 0.004489398 | 0.075587022 | 0.05939377 |
| Tcp11l1 | 978.2370623 | -0.030085624 | 0.072485572 | -0.41505672 |
| Pin1rt1 | 0.178918687 | -0.443824103 | 2.727252327 | -0.162736722 |
| Depdc7 | 17.9411153 | -0.338818989 | 0.29798484 | -1.137034317 |
| Qser1 | 622.2252455 | -0.093435301 | 0.066413028 | -1.406882107 |
| Gm13886 | 11.02501732 | -0.290306011 | 0.348530261 | -0.832943487 |
| Prrg4 | 12.12341216 | -0.702778573 | 0.316364583 | -2.221419875 |
| Ccdc73 | 1.808963643 | 0.569404333 | 0.774001113 | 0.735663454 |
| Gm13887 | 0.209356743 | 0.503086776 | 2.188175767 | 0.229911501 |
| Eif3m | 484.4513724 | 0.016562557 | 0.081045093 | 0.204362241 |
| Gm13888 | 0 | NA | NA | NA |
| Wt1os | 0.172571121 | -0.433506555 | 2.733273128 | -0.158603453 |
| Gm11060 | 0 | NA | NA | NA |
| Gm27715 | 0 | NA | NA | NA |
| Gm27538 | 0 | NA | NA | NA |
| Gm27636 | 0 | NA | NA | NA |
| Gm27308 | 0 | NA | NA | NA |
| Wt1 | 11.92317521 | -3.496889246 | 0.532728578 | -6.564110488 |
| Them7 | 0.493638219 | -0.398266164 | 1.424282224 | -0.279625876 |
| Rcn1 | 399.3853322 | -0.262769615 | 0.126932941 | -2.070145173 |
| 4930527A07Rik | 0 | NA | NA | NA |

| | | | | |
|---|---|---|---|---|
| Pax6os1 | 4.443791399 | 0.94957356 | 0.574352762 | 1.653293278 |
| Gm13890 | 0.122991902 | 0.517955464 | 3.724945049 | 0.139050498 |
| Rpl10-ps1 | 5.581713243 | -0.898555385 | 0.428028538 | -2.099288495 |
| Paupar | 4.745398909 | 0.579112748 | 0.463776501 | 1.248689286 |
| Pax6 | 311.7227136 | 0.006132302 | 0.162928892 | 0.037637907 |
| Elp4 | 237.3869113 | 0.040879713 | 0.086484983 | 0.472679904 |
| Gm13954 | 0.303403639 | 1.412060586 | 1.988292925 | 0.710187402 |
| Immp1l | 117.1741609 | -0.628827582 | 0.195257815 | -3.220498914 |
| Dnajc24 | 115.5254544 | -0.187689195 | 0.103388847 | -1.815371773 |
| Gm22002 | 0.104637558 | -0.443821212 | 3.724945049 | -0.119148392 |
| n-R5s201 | 0 | NA | NA | NA |
| Dcdc5 | 20.38028449 | 0.060996643 | 0.290294107 | 0.210120157 |
| Gm13992 | 0.047948698 | 0.037064902 | 3.724945049 | 0.009950456 |
| Gm29053 | 0 | NA | NA | NA |
| Gm14013 | 0.459558249 | 0.525023259 | 1.434800192 | 0.365920818 |
| Gm14015 | 1.812101644 | 1.791841662 | 0.887996121 | 2.017848524 |
| Gm14014 | 0.072025462 | 0.517959912 | 3.724945049 | 0.139051692 |
| BB218582 | 7.281320362 | 1.424157226 | 0.465792647 | 3.057491858 |
| Mpped2 | 264.5495323 | 0.412291236 | 0.103249888 | 3.993139776 |
| Gm13901 | 0.066050875 | -0.443818988 | 3.724945049 | -0.119147795 |
| Gm13902 | 0.292336004 | 0.507651731 | 1.633450362 | 0.310784915 |
| Arl14ep | 637.6380912 | -0.166804642 | 0.079369606 | -2.101618635 |
| Gm13900 | 0.111006857 | 0.037730769 | 3.724898049 | 0.010129343 |
| Gm13904 | 0.527697161 | 1.371532504 | 1.417610497 | 0.967496013 |
| Fshb | 0.248616998 | -1.950849138 | 3.001053554 | -0.650054757 |
| Gm24885 | 0 | NA | NA | NA |
| Gm13903 | 0 | NA | NA | NA |
| Kcna4 | 692.1728847 | 0.224868033 | 0.11645764 | 1.93089979 |
| Gm9864 | 1.048421568 | -0.630364003 | 0.992233424 | -0.635298094 |
| Gm23439 | 0 | NA | NA | NA |
| Gm23749 | 0.105852363 | -0.443821212 | 3.724945049 | -0.119148392 |
| Gm13912 | 1.945135077 | -1.788299614 | 0.74056283 | -2.414784462 |
| Gm13916 | 0.123845245 | -0.280519638 | 3.720871772 | -0.075390837 |
| Gm13915 | 0.038981395 | -0.443816764 | 3.724945049 | -0.119147198 |
| Gm13913 | 0 | NA | NA | NA |
| Gm13914 | 0 | NA | NA | NA |
| Rpl35a-ps4 | 1.664460293 | -0.694664002 | 0.810961879 | -0.856592671 |
| Gm13910 | 140.7366961 | -0.283612035 | 0.157170439 | -1.804487134 |
| Gm13923 | 0 | NA | NA | NA |
| Mettl15 | 54.80204305 | -0.217421059 | 0.174852518 | -1.24345398 |
| Gm24115 | 0.713863726 | -0.960713738 | 1.155334814 | -0.831545736 |
| Kif18a | 32.64308567 | -0.179476199 | 0.213901148 | -0.839061409 |
| Gm13924 | 0 | NA | NA | NA |
| Gm13925 | 0 | NA | NA | NA |
| Gm20638 | 0.165580463 | 0.037073754 | 2.773571312 | 0.013366793 |
| Bdnf | 331.5846942 | -1.001542031 | 0.21836641 | -4.586520576 |
| Gm13932 | 0.182620985 | 0.517972018 | 2.688869904 | 0.192635582 |

| | | | | |
|---|---|---|---|---|
| Lin7c | 1365.827837 | 0.206159054 | 0.063395883 | 3.251931269 |
| Platr8 | 0 | NA | NA | NA |
| Lgr4 | 521.05221 | 0.020925492 | 0.10022602 | 0.208783033 |
| Ccdc34 | 111.7137007 | 0.083007659 | 0.151820925 | 0.546747156 |
| Ccdc34os | 0 | NA | NA | NA |
| Gm13936 | 1.575920251 | 1.247219426 | 0.911168038 | 1.368813846 |
| Gm13937 | 0.157911564 | 0.517964864 | 2.815266196 | 0.18398433 |
| Gm22997 | 0.40088837 | 1.383787025 | 1.415056824 | 0.977902089 |
| Bbox1 | 28.0529557 | 0.246053392 | 0.288689454 | 0.852311675 |
| Fibin | 60.99905906 | 0.264789272 | 0.212698016 | 1.244907109 |
| Gm13935 | 0.04393175 | 0.037062677 | 3.724945049 | 0.009949859 |
| Gm10794 | 0.046345709 | -0.443818988 | 3.724945049 | -0.119147795 |
| Slc5a12 | 0.290078826 | -0.40909945 | 1.817895213 | -0.225040171 |
| Ano3 | 1378.79017 | 1.056680095 | 0.166797322 | 6.335114263 |
| Muc15 | 15.98576495 | -0.9997141 | 0.272541467 | -3.668117406 |
| Gm13942 | 0 | NA | NA | NA |
| Gm13941 | 0 | NA | NA | NA |
| Gm15130 | 0 | NA | NA | NA |
| 4930430A15Rik | 0.123841226 | -1.017823543 | 3.719440177 | -0.273649661 |
| Gm13961 | 21.20792997 | -0.405469662 | 0.235181248 | -1.724073092 |
| Olfr1275 | 0 | NA | NA | NA |
| Gm13962 | 0 | NA | NA | NA |
| Olfr1276 | 0.075043205 | 0.517957688 | 3.724945049 | 0.139051095 |
| Olfr1277 | 0 | NA | NA | NA |
| Olfr1278 | 0 | NA | NA | NA |
| Olfr1279 | 0 | NA | NA | NA |
| Olfr1280 | 0 | NA | NA | NA |
| Olfr1281 | 0 | NA | NA | NA |
| Olfr1282 | 0 | NA | NA | NA |
| Olfr1283 | 0.038981395 | -0.443816764 | 3.724945049 | -0.119147198 |
| Olfr1284 | 0 | NA | NA | NA |
| Olfr1285 | 0 | NA | NA | NA |
| Olfr1286 | 0 | NA | NA | NA |
| Olfr1287 | 0 | NA | NA | NA |
| Olfr1288 | 0.11197992 | 0.037073798 | 3.724945049 | 0.009952844 |
| Olfr1289 | 0 | NA | NA | NA |
| Olfr1290 | 0.062426873 | 0.517959912 | 3.724945049 | 0.139051692 |
| Olfr1291-ps1 | 0 | NA | NA | NA |
| Olfr1292-ps1 | 0 | NA | NA | NA |
| Olfr1293-ps | 0 | NA | NA | NA |
| Olfr1294 | 0.134986936 | -0.443827263 | 3.656698115 | -0.121373777 |
| Olfr1295 | 0.08486385 | 0.037730769 | 3.724898049 | 0.010129343 |
| Olfr1296-ps1 | 0 | NA | NA | NA |
| Olfr1297 | 0 | NA | NA | NA |
| Olfr1298 | 0 | NA | NA | NA |
| Olfr1299 | 0 | NA | NA | NA |
| Olfr1300-ps1 | 0 | NA | NA | NA |

| | | | | |
|---|---|---|---|---|
| Gm13922 | 0 | NA | NA | NA |
| Gm13930 | 0 | NA | NA | NA |
| Olfr1301 | 0.046345709 | -0.443818988 | 3.724945049 | -0.119147795 |
| Olfr1302 | 0 | NA | NA | NA |
| Olfr1303 | 0.046345709 | -0.443818988 | 3.724945049 | -0.119147795 |
| Gm26421 | 0 | NA | NA | NA |
| Olfr268-ps1 | 0.068097359 | 0.037064902 | 3.724945049 | 0.009950456 |
| Olfr1305 | 0 | NA | NA | NA |
| Olfr1306 | 0 | NA | NA | NA |
| Gm13928 | 0 | NA | NA | NA |
| Olfr1307 | 0 | NA | NA | NA |
| Olfr1308 | 0 | NA | NA | NA |
| Olfr1309 | 0 | NA | NA | NA |
| Gm13929 | 0 | NA | NA | NA |
| Olfr1310 | 0.048949532 | 0.037064902 | 3.724945049 | 0.009950456 |
| Olfr1311 | 0 | NA | NA | NA |
| Olfr1312 | 0 | NA | NA | NA |
| Olfr1313 | 0.066050875 | -0.443818988 | 3.724945049 | -0.119147795 |
| Olfr1314 | 0 | NA | NA | NA |
| Olfr1315-ps1 | 0 | NA | NA | NA |
| Olfr1316 | 0.049553047 | -0.443818988 | 3.724945049 | -0.119147795 |
| Olfr1317 | 0 | NA | NA | NA |
| Olfr1318 | 0 | NA | NA | NA |
| Lpcat4 | 780.3235412 | 0.400807274 | 0.095939185 | 4.177722315 |
| Nutm1 | 0.463200932 | -0.741262645 | 1.499770642 | -0.49425067 |
| Nop10 | 173.0751968 | 0.122710518 | 0.119701685 | 1.025136098 |
| Slc12a6 | 925.7716682 | 0.346702737 | 0.084762951 | 4.090262704 |
| Gm21985 | 0.288125237 | 1.723641176 | 3.707988908 | 0.464845289 |
| Emc4 | 460.0520838 | 0.108584958 | 0.074594892 | 1.45566211 |
| Katnbl1 | 227.6757466 | -0.186667 | 0.105935013 | -1.762089745 |
| Gm13940 | 0 | NA | NA | NA |
| Emc7 | 1191.037724 | -0.105364371 | 0.066230351 | -1.590877429 |
| n-R5s202 | 0 | NA | NA | NA |
| Chrm5 | 22.04456264 | 0.72813301 | 0.244248399 | 2.981116812 |
| Aven | 92.20401975 | -0.405595045 | 0.116068553 | -3.49444388 |
| Gm22676 | 0.183828322 | -0.417220156 | 2.675955811 | -0.155914442 |
| AL691423.1 | 0.566015516 | 1.912754289 | 1.526256628 | 1.253232421 |
| Ryr3 | 1324.13524 | 0.64948588 | 0.270713246 | 2.399165502 |
| Tmco5b | 1.585639959 | 0.89959959 | 0.744577533 | 1.208201363 |
| Gm26505 | 0.194120612 | 0.538934232 | 2.639891497 | 0.204150145 |
| Fmn1 | 837.5690741 | 0.362361001 | 0.076364355 | 4.745158959 |
| Gm13967 | 4.347705094 | 0.916777686 | 0.57256748 | 1.601169675 |
| Gm13963 | 0.697976008 | -2.719219621 | 1.302534697 | -2.087636995 |
| Gm13965 | 7.79987822 | 0.642053961 | 0.426966511 | 1.503757189 |
| Gm13966 | 0.044191549 | 0.037062677 | 3.724945049 | 0.009949859 |
| AL691437.1 | 0 | NA | NA | NA |
| Gm13964 | 0.060174537 | 0.517957688 | 3.724945049 | 0.139051095 |

| | | | | |
|---|---|---|---|---|
| 4930533B01Rik | 0.045882455 | 0.517959912 | 3.724945049 | 0.139051692 |
| Grem1 | 36.57688901 | 0.118914836 | 0.226245002 | 0.525602047 |
| Scg5 | 2876.760524 | 0.466286459 | 0.095716358 | 4.871544121 |
| Arhgap11a | 55.95224419 | 0.104363443 | 0.238137154 | 0.438249306 |
| Gm21596 | 8.155034204 | 0.219013528 | 0.391088379 | 0.560010319 |
| Gjd2 | 128.1457705 | 0.065907566 | 0.136162468 | 0.484036223 |
| A530058N18Rik | 102.4745166 | 0.443665056 | 0.242514035 | 1.829440743 |
| Actc1 | 0.709411339 | -2.155368105 | 1.20251611 | -1.792381897 |
| C130080G10Rik | 3.984159091 | 0.789758252 | 0.515134386 | 1.533111114 |
| Aqr | 833.2625079 | -0.338081747 | 0.047315792 | -7.145220084 |
| Mir7002 | 0.230049058 | 0.517969349 | 2.134872547 | 0.242623078 |
| Zfp770 | 474.7123103 | -0.100114028 | 0.089117471 | -1.123393947 |
| Gm25103 | 0.044544648 | 0.517957688 | 3.724945049 | 0.139051095 |
| Gm29234 | 0.986742042 | -0.084649845 | 1.162379063 | -0.072824647 |
| Dph6 | 238.7373716 | 0.100679819 | 0.117837602 | 0.85439467 |
| Gm13972 | 0.263371022 | 0.464172407 | 1.812290192 | 0.256124769 |
| Gm13975 | 0.044191549 | 0.037062677 | 3.724945049 | 0.009949859 |
| Gm13974 | 2.444458607 | 0.636101484 | 0.717898982 | 0.886059878 |
| Gm28494 | 0 | NA | NA | NA |
| Gm28492 | 0.508786284 | 0.038561068 | 1.397921249 | 0.027584578 |
| Gm13973 | 0 | NA | NA | NA |
| Gm28493 | 0.257385133 | -1.162345646 | 2.027400099 | -0.573318333 |
| 4930528P14Rik | 0.479723347 | -0.804014941 | 1.352017374 | -0.594677965 |
| Gm13976 | 0.060174537 | 0.517957688 | 3.724945049 | 0.139051095 |
| 3110099E03Rik | 2.086914652 | -0.636368967 | 0.673761176 | -0.94450228 |
| Gm13977 | 0.219276728 | -0.94542478 | 2.590889308 | -0.364903579 |
| BC052040 | 104.7146314 | -0.288909154 | 0.1462082 | -1.976011979 |
| Mir1951 | 0.245748936 | -0.846257543 | 2.066281856 | -0.409555715 |
| Meis2 | 1352.514286 | 0.269842643 | 0.141905362 | 1.901567626 |
| 2700033N17Rik | 6.560169035 | -0.04817225 | 0.395262823 | -0.121873972 |
| G630016G05Rik | 19.18698179 | 0.984842554 | 0.339294795 | 2.902616155 |
| 2810405F15Rik | 1.233431567 | 0.399169654 | 1.111243093 | 0.35921002 |
| Gm13991 | 44.848571 | -0.096113525 | 0.167725445 | -0.573040812 |
| Gm13990 | 0.730890257 | -1.035561632 | 1.357840649 | -0.762653285 |
| Tmco5 | 9.517001854 | -0.078620455 | 0.359011184 | -0.218991661 |
| D330050G23Rik | 15.28741942 | -0.73891092 | 0.296670092 | -2.490682208 |
| Gm24914 | 0 | NA | NA | NA |
| Gm25514 | 0.127971488 | -0.95589744 | 3.6855341 | -0.259364698 |
| Gm29340 | 5.225310162 | -0.73981305 | 0.462542765 | -1.599447891 |
| Gm13981 | 0.117907917 | 0.967119648 | 3.722483401 | 0.259804959 |
| Spred1 | 3571.650489 | -0.192436591 | 0.088346422 | -2.178204702 |
| Mir674 | 5.596050554 | 0.167143135 | 0.447010937 | 0.373912852 |
| Rap1a-ps2 | 0.847725382 | -1.789227302 | 1.151847054 | -1.553354931 |
| Gm25189 | 0.284781514 | -0.444509644 | 2.060972187 | -0.215679594 |
| Fam98b | 385.3078931 | -0.113983087 | 0.087505099 | -1.302587937 |
| Rasgrp1 | 4351.652765 | 0.661694188 | 0.13929922 | 4.750164349 |
| Gm13982 | 0.169723681 | -0.52143512 | 3.40726837 | -0.153036117 |

| | | | | |
|---|---|---|---|---|
| Gm28183 | 0.166952883 | -1.336542448 | 2.746539428 | -0.486627803 |
| Gm13985 | 0.090890357 | 0.037071574 | 3.724945049 | 0.009952247 |
| 4930412B13Rik | 0 | NA | NA | NA |
| Gm13986 | 9.239123008 | -1.173286577 | 0.406468088 | -2.886540451 |
| Gm36979 | 0 | NA | NA | NA |
| Gm25308 | 0.048949532 | 0.037064902 | 3.724945049 | 0.009950456 |
| Thbs1 | 1597.215543 | -1.938672034 | 0.453885395 | -4.271280939 |
| Gm29233 | 0.136201742 | -0.443823494 | 2.946222509 | -0.150641539 |
| Fsip1 | 6.681599266 | 0.230948144 | 0.384360786 | 0.600862919 |
| Gm22800 | 0 | NA | NA | NA |
| Gm29232 | 0.049553047 | -0.443818988 | 3.724945049 | -0.119147795 |
| A930104D05Rik | 0.722394163 | -0.218527418 | 1.341916909 | -0.162847205 |
| Gpr176 | 129.1472356 | -0.155184656 | 0.124542392 | -1.246038822 |
| 1700054M17Rik | 0.459216712 | 1.629051344 | 1.500430316 | 1.08572276 |
| Gm22651 | 0 | NA | NA | NA |
| Eif2ak4 | 429.1232767 | 0.305252006 | 0.139030349 | 2.195578224 |
| Srp14 | 522.9120673 | -0.242314237 | 0.105339431 | -2.300318458 |
| Gm13983 | 0 | NA | NA | NA |
| Bmf | 64.79741328 | 0.397156689 | 0.149914738 | 2.649217103 |
| Gm14016 | 0 | NA | NA | NA |
| Bub1b | 14.01725363 | 0.099955912 | 0.330595031 | 0.302351525 |
| Pak6 | 596.5307493 | 0.026087513 | 0.151296236 | 0.172426713 |
| Ankrd63 | 286.9989274 | 0.959815464 | 0.334742685 | 2.86732319 |
| Plcb2 | 54.89159527 | 0.027749697 | 0.24132172 | 0.114990467 |
| Inafm2 | 508.6639642 | -0.071366624 | 0.088749039 | -0.804139683 |
| A430105I19Rik | 127.3481097 | -0.28954175 | 0.139744076 | -2.071942929 |
| Phgr1 | 0.264111009 | 1.16916734 | 3.710686547 | 0.315081138 |
| Disp2 | 4917.700072 | 0.131941417 | 0.059249863 | 2.226864505 |
| Knstrn | 21.59877154 | -0.026677338 | 0.373972871 | -0.071334955 |
| Gm14091 | 1.226992653 | -0.260596521 | 1.214263368 | -0.214612849 |
| Ivd | 475.053727 | 0.389330588 | 0.083495271 | 4.662905867 |
| Bahd1 | 560.8711464 | -0.193348906 | 0.090641846 | -2.13310866 |
| Chst14 | 36.12922769 | -0.489342318 | 0.199264993 | -2.455736507 |
| Gm14089 | 0.273591019 | 0.11571315 | 2.019156627 | 0.057307664 |
| Gm14088 | 0.141331096 | -0.443820795 | 3.561840904 | -0.124604329 |
| Ccdc32 | 448.3274695 | -0.119580889 | 0.117760907 | -1.01545489 |
| Rpusd2 | 91.45199017 | 0.054073674 | 0.145552849 | 0.371505429 |
| Casc5 | 31.52527001 | -0.199654668 | 0.325832793 | -0.612751915 |
| Rad51 | 19.29236894 | -0.592317586 | 0.33610039 | -1.762323409 |
| Rmdn3 | 789.494809 | -0.191113799 | 0.102201872 | -1.869963786 |
| Gchfr | 7.98091292 | 0.095239456 | 0.443252524 | 0.214865004 |
| Dnajc17 | 78.13003113 | 0.189550494 | 0.14082386 | 1.346011204 |
| Gm14137 | 2.178995086 | -2.414888183 | 0.777633602 | -3.105431885 |
| Gm14138 | 1.456796327 | 0.059082 | 0.875516409 | 0.067482459 |
| Zfyve19 | 182.5094088 | 0.022748483 | 0.088306981 | 0.257606845 |
| Ppp1r14d | 0.089814205 | 0.517955464 | 3.724945049 | 0.139050498 |
| Spint1 | 12.20675764 | -0.215580364 | 0.284405625 | -0.758003164 |

| | | | | |
|---|---|---|---|---|
| Rhov | 153.8616767 | -0.977821386 | 0.164951559 | -5.927930561 |
| Gm14175 | 0 | NA | NA | NA |
| Vps18 | 675.7131133 | -0.177043466 | 0.055130011 | -3.211380911 |
| Gm14207 | 0.327809541 | 1.540100242 | 1.896473823 | 0.812086211 |
| Dll4 | 18.4801647 | 0.6294885 | 0.391979171 | 1.605923342 |
| Chac1 | 175.096179 | -1.395649177 | 0.257992954 | -5.409640666 |
| Ino80 | 813.323168 | -0.085137312 | 0.063802308 | -1.334392354 |
| Gm14208 | 0.083526043 | 0.037073798 | 3.724945049 | 0.009952844 |
| Exd1 | 16.27157243 | -0.773227692 | 0.293681495 | -2.632878496 |
| Chp1 | 1163.742042 | 0.038777584 | 0.055974065 | 0.692777687 |
| 1700020I14Rik | 6679.70898 | 0.463380633 | 0.088496848 | 5.236125827 |
| Oip5 | 7.956782189 | 0.742543897 | 0.404731959 | 1.834655949 |
| Nusap1 | 22.39643515 | -0.097095381 | 0.376825637 | -0.257666602 |
| n-R5s204 | 0 | NA | NA | NA |
| Ndufaf1 | 192.2611336 | -0.187189052 | 0.078458138 | -2.385846219 |
| H3f3c | 22.18271704 | -0.026456148 | 0.250407752 | -0.105652272 |
| Gm14383 | 0.267580615 | 0.532303409 | 1.817776819 | 0.292832103 |
| Rtf1 | 860.2757225 | 0.270254866 | 0.080670938 | 3.350089529 |
| Gm25857 | 1.36269325 | 3.095477079 | 1.045062406 | 2.962002135 |
| Itpka | 639.0705542 | 1.210692417 | 0.134836104 | 8.978992873 |
| Ltk | 62.0682925 | 1.128525237 | 0.161780714 | 6.975647512 |
| Rpap1 | 285.1701272 | -0.022408507 | 0.096227626 | -0.232869787 |
| Gm23814 | 0 | NA | NA | NA |
| Tyro3 | 1444.38765 | 0.389114604 | 0.087399116 | 4.452157194 |
| Gm13998 | 0 | NA | NA | NA |
| Mga | 2323.037544 | -0.15196147 | 0.066231397 | -2.294402315 |
| Mapkbp1 | 844.8660816 | 0.24420378 | 0.112973672 | 2.161599034 |
| Mir7665 | 0.088383099 | 0.037058229 | 3.724945049 | 0.009948665 |
| Gm13999 | 0 | NA | NA | NA |
| Jmjd7 | 2.162814427 | -0.367345061 | 0.727399807 | -0.505011217 |
| Gm28042 | 0.10503227 | -0.905414734 | 3.723500899 | -0.243162217 |
| Pla2g4b | 5.643927168 | -0.247475906 | 0.446600248 | -0.554132934 |
| Sptbn5 | 13.1047588 | 0.592656673 | 0.358196725 | 1.65455637 |
| Ehd4 | 458.2061878 | -0.727612606 | 0.153752765 | -4.732354594 |
| Pla2g4e | 246.4736302 | 0.707162186 | 0.141755992 | 4.988587623 |
| Gm13997 | 1.380678606 | 0.026615227 | 0.910567535 | 0.029229273 |
| Pla2g4d | 1.305860104 | 0.421937822 | 0.889833434 | 0.474176184 |
| Pla2g4f | 1.649860538 | -0.64523648 | 0.805875558 | -0.800665157 |
| Vps39 | 1015.694296 | 0.119985256 | 0.055961759 | 2.144057971 |
| Tmem87a | 799.4634507 | -0.014231886 | 0.065155699 | -0.218428877 |
| Ganc | 158.6786428 | 0.01238062 | 0.109957264 | 0.112594833 |
| Capn3 | 66.72612438 | 1.731855847 | 0.242919106 | 7.129352143 |
| Zfp106 | 4291.41446 | 0.165768327 | 0.048418281 | 3.423672331 |
| Gm26899 | 5.004844821 | -0.086192231 | 0.507007856 | -0.170001766 |
| Snap23 | 220.6488251 | -0.449720437 | 0.141791708 | -3.171697721 |
| Lrrc57 | 243.5709634 | 0.162512321 | 0.088726525 | 1.831609209 |
| Haus2 | 232.8508983 | -0.14295281 | 0.118159962 | -1.20982445 |

| | | | | |
|---|---|---|---|---|
| Stard9 | 314.5610356 | 0.224589634 | 0.174813542 | 1.284738192 |
| n-R5s205 | 0 | NA | NA | NA |
| Cdan1 | 293.1784353 | 0.274116835 | 0.157937884 | 1.735599003 |
| Ttbk2 | 1910.365418 | 0.029370391 | 0.065804085 | 0.446330813 |
| AV039307 | 6.926568459 | -0.140616662 | 0.446219175 | -0.315129133 |
| Ubr1 | 1554.190815 | 0.111823407 | 0.047260123 | 2.366126004 |
| Tmem62 | 383.5196607 | -0.072722401 | 0.073686599 | -0.986914873 |
| AL844548.1 | 1.862757 | 0.900055155 | 0.757015561 | 1.188951986 |
| Gm22865 | 3.966952568 | 0.793582706 | 0.652499058 | 1.216220464 |
| Ccndbp1 | 483.2632691 | 0.035136173 | 0.075114716 | 0.467766837 |
| Epb4.2 | 1.132377036 | -0.105571407 | 0.967354319 | -0.109134166 |
| Tgm5 | 0.516597926 | -1.02950211 | 1.444263008 | -0.712821767 |
| Gm14213 | 0 | NA | NA | NA |
| Tgm7 | 0.64993548 | -1.583460212 | 1.213560235 | -1.304805617 |
| Lcmt2 | 75.2947259 | -0.229866655 | 0.158908354 | -1.446536002 |
| Gm23858 | 0.195161473 | 0.037072169 | 2.679514636 | 0.013835404 |
| Adal | 331.7277646 | -0.060164383 | 0.065629079 | -0.916733611 |
| Zscan29 | 154.5557525 | -0.822449957 | 0.129924985 | -6.330190898 |
| Tubgcp4 | 399.137152 | -0.331961974 | 0.090782916 | -3.656656861 |
| Trp53bp1 | 1866.10298 | 0.013982756 | 0.07607728 | 0.183796741 |
| Map1a | 23447.45946 | 0.475330578 | 0.068407991 | 6.948465679 |
| Gm14978 | 2.354129177 | 0.324733552 | 0.787929007 | 0.412135547 |
| Ppip5k1 | 1660.739469 | -0.015789209 | 0.084968894 | -0.1858234 |
| Ckmt1 | 1627.233016 | 0.420922306 | 0.067443416 | 6.241117832 |
| Strc | 2.337199581 | 0.461103667 | 0.659014304 | 0.699686887 |
| Catsper2 | 52.60120326 | 0.24153575 | 0.200997028 | 1.201688167 |
| Gm14001 | 3.014229995 | -0.511264854 | 0.721203269 | -0.708905348 |
| Gm14000 | 0 | NA | NA | NA |
| Pdia3 | 2693.142216 | -0.538107387 | 0.102914544 | -5.228681647 |
| Ell3 | 0.594313001 | 1.197896887 | 1.271202255 | 0.942333828 |
| Serinc4 | 3.508801613 | 0.286568317 | 0.514946224 | 0.556501443 |
| Serf2 | 753.224674 | -0.144090397 | 0.070359683 | -2.047911403 |
| Gm22953 | 0.938485331 | -0.68834077 | 1.057371084 | -0.650992617 |
| Hypk | 31.77497691 | 0.825003722 | 0.304537194 | 2.709040929 |
| Mfap1b | 106.0742756 | -0.272716141 | 0.132786001 | -2.053801896 |
| Gm14017 | 2.307785566 | 0.277992268 | 0.738284044 | 0.376538366 |
| Gm14018 | 0.899881821 | -0.679028012 | 1.132417824 | -0.599626743 |
| Mfap1a | 281.266986 | 0.111343417 | 0.064742547 | 1.719787398 |
| Wdr76 | 70.42691052 | -0.231765743 | 0.139742762 | -1.658516975 |
| Frmd5 | 605.5814555 | 0.036961383 | 0.074392511 | 0.496842787 |
| Gm14021 | 44.18792118 | 0.578970422 | 0.270318395 | 2.141809183 |
| Gm14020 | 0.04393175 | 0.037062677 | 3.724945049 | 0.009949859 |
| Gm14019 | 0.838491377 | -1.671463602 | 1.161678924 | -1.438834404 |
| Casc4 | 1001.124057 | 0.141893187 | 0.065368535 | 2.170664928 |
| Gm23983 | 0 | NA | NA | NA |
| Mageb3 | 1.086684801 | -0.653168006 | 1.001346067 | -0.65228998 |
| Ctdspl2 | 624.7685435 | 0.045314786 | 0.08655131 | 0.523559796 |

| | | | | |
|---|---|---|---|---|
| Eif3j1 | 235.1963187 | -0.022529061 | 0.130663409 | -0.172420584 |
| AL845457.1 | 0 | NA | NA | NA |
| Spg11 | 412.0809888 | -0.347846272 | 0.095233267 | -3.652571007 |
| Patl2 | 2.452851275 | 0.544431332 | 0.672443624 | 0.80963119 |
| B2m | 9604.418411 | -0.700333122 | 0.158559717 | -4.416841396 |
| Trim69 | 8.404189488 | -2.655012619 | 0.458622016 | -5.789108518 |
| 4933406J08Rik | 0.139037128 | -1.190058937 | 3.717743703 | -0.320102469 |
| Gm14050 | 0 | NA | NA | NA |
| Gm14058 | 0.04393175 | 0.037062677 | 3.724945049 | 0.009949859 |
| Rps12-ps10 | 0.709725355 | 0.299516196 | 1.279946948 | 0.234006727 |
| Sord | 185.688388 | 0.30224586 | 0.088536858 | 3.41378572 |
| Duox2 | 2.808703517 | -1.14268782 | 0.706392668 | -1.617638281 |
| Duoxa2 | 0.276302848 | 0.877247705 | 2.029676132 | 0.432210682 |
| Duoxa1 | 3.074242624 | -0.108333622 | 0.891961959 | -0.121455429 |
| Duox1 | 2.917555992 | 1.229414319 | 0.666367667 | 1.844948936 |
| Shf | 467.0710296 | 0.177969584 | 0.113331118 | 1.570350551 |
| Gm24409 | 0 | NA | NA | NA |
| Slc28a2 | 188.3325738 | -2.715948975 | 0.655060864 | -4.146101721 |
| Gm14086 | 0 | NA | NA | NA |
| Bambi-ps1 | 12.14750959 | 0.252245237 | 0.352703223 | 0.71517701 |
| Gm14085 | 41.28799092 | -2.275369085 | 0.399348944 | -5.697696504 |
| Gatm | 1118.00039 | -0.118150798 | 0.154661278 | -0.763932636 |
| Spata5l1 | 53.6862279 | -0.878780742 | 0.198052308 | -4.437114365 |
| AA467197 | 88.13508126 | -1.562184113 | 0.430847595 | -3.625839231 |
| Mir147 | 0 | NA | NA | NA |
| Slc30a4 | 1022.722383 | 0.2201287 | 0.083349657 | 2.641027061 |
| 4930417H01Rik | 4.957477421 | 0.97862534 | 0.546527519 | 1.790624088 |
| Bloc1s6os | 0.578202636 | -0.935349893 | 1.324582275 | -0.706147071 |
| Bloc1s6 | 473.7227365 | -0.207661757 | 0.102199354 | -2.031928265 |
| Sqrdl | 62.88662421 | -0.811080539 | 0.211423457 | -3.836284532 |
| Gm13988 | 0.046345709 | -0.443818988 | 3.724945049 | -0.119147795 |
| Sema6d | 1571.33555 | -0.120592992 | 0.077970766 | -1.546643667 |
| 4930517E11Rik | 0.136309558 | 1.320344047 | 3.610467153 | 0.36569895 |
| 4930583P06Rik | 0.286786342 | -1.86457592 | 1.989387664 | -0.937261225 |
| Gm13994 | 0 | NA | NA | NA |
| Slc24a5 | 5.913197406 | 1.313500717 | 0.480131275 | 2.73571164 |
| Myef2 | 1104.874051 | 0.209548894 | 0.088681851 | 2.362928735 |
| Gm22352 | 0 | NA | NA | NA |
| Gm14004 | 11.63604071 | 0.680810055 | 0.333812787 | 2.039496631 |
| Ctxn2 | 55.00778503 | 0.2660818 | 0.184883674 | 1.439184944 |
| Slc12a1 | 4.418703675 | 0.211209609 | 0.490405589 | 0.430683528 |
| Gm14002 | 3.728554256 | 0.311008885 | 0.583595827 | 0.532918281 |
| Dut | 174.499205 | -0.05683757 | 0.105764748 | -0.53739616 |
| A530010F05Rik | 0 | NA | NA | NA |
| Gm14003 | 0.116944185 | -1.028246102 | 3.719337068 | -0.27645951 |
| Fbn1 | 344.1427708 | -0.137736584 | 0.128682814 | -1.070357256 |
| Gm9913 | 2.575840985 | -0.295223322 | 0.591956189 | -0.498724953 |

| | | | | |
|---|---|---|---|---|
| Cep152 | 85.54108715 | -0.119951216 | 0.156365469 | -0.767120877 |
| Shc4 | 125.3363377 | -0.442993844 | 0.144297286 | -3.070008149 |
| Eid1 | 1534.982027 | 0.466286816 | 0.072175763 | 6.460434843 |
| Secisbp2l | 3169.337731 | 0.231835762 | 0.043320749 | 5.351610184 |
| Cops2 | 1452.800378 | 0.053807384 | 0.108258577 | 0.497026518 |
| Galk2 | 221.865611 | -0.008563492 | 0.090863993 | -0.09424516 |
| Fam227b | 26.91957889 | 0.944666088 | 0.333291318 | 2.834355527 |
| Fgf7 | 42.37332393 | -0.129623455 | 0.246017875 | -0.526886327 |
| Gm26496 | 0.514662899 | -1.61649116 | 1.326191764 | -1.218896998 |
| Dtwd1 | 64.96816098 | -0.110870763 | 0.138267686 | -0.801855925 |
| Gm17555 | 0 | NA | NA | NA |
| Gm24739 | 0 | NA | NA | NA |
| Atp8b4 | 80.28755012 | -1.193167958 | 0.254254106 | -4.692816876 |
| Gm26697 | 1.020000067 | -1.28773737 | 1.11148478 | -1.158574003 |
| Slc27a2 | 66.76116308 | 0.242919037 | 0.155891253 | 1.558259573 |
| Hdc | 76.84591766 | -0.669713297 | 0.254908514 | -2.627269237 |
| Gabpb1 | 280.3852881 | -0.048605124 | 0.116859112 | -0.415929263 |
| Gm23753 | 1.578568981 | 1.826485354 | 1.010550928 | 1.807415444 |
| Gm27003 | 13.76199491 | 0.153949736 | 0.375140353 | 0.410379034 |
| Usp8 | 1065.714832 | -0.148353069 | 0.054464635 | -2.72384216 |
| Usp50 | 5.50178608 | -0.180747362 | 0.474743101 | -0.380726675 |
| Gm10774 | 0 | NA | NA | NA |
| Trpm7 | 1310.402208 | 0.131937124 | 0.10071526 | 1.310001325 |
| Gm14212 | 0 | NA | NA | NA |
| Sppl2a | 1346.162496 | -0.342515866 | 0.100511595 | -3.407724913 |
| Mir3473g | 5.710082525 | 0.466776348 | 0.594526591 | 0.785122743 |
| Ap4e1 | 265.198369 | 0.133751876 | 0.127920132 | 1.04558894 |
| Gm24081 | 0 | NA | NA | NA |
| Blvra | 139.0631896 | -0.571552384 | 0.135203528 | -4.227348159 |
| Ncaph | 24.30005576 | -0.176885003 | 0.25998781 | -0.680358833 |
| Itpripl1 | 51.57497973 | -0.425652497 | 0.1813735 | -2.346828485 |
| 1810024B03Rik | 6.848270579 | 1.00420735 | 0.426767082 | 2.35305719 |
| Snrnp200 | 2169.627635 | -0.053707391 | 0.054956251 | -0.977275382 |
| Ciao1 | 447.7927703 | -0.411617913 | 0.099131548 | -4.152239318 |
| Tmem127 | 1631.67063 | -0.127907802 | 0.062206369 | -2.056184975 |
| Gm10766 | 3.235620746 | -0.233728382 | 0.675676958 | -0.345917349 |
| Stard7 | 1005.190084 | 0.229056143 | 0.053615817 | 4.272174842 |
| Dusp2 | 53.83102264 | -0.52364084 | 0.333424719 | -1.570491957 |
| Astl | 1.38321721 | -0.439172711 | 0.894035895 | -0.49122492 |
| Adra2b | 12.20916545 | 0.874977842 | 0.43395956 | 2.016265852 |
| Gpat2 | 10.37835042 | 0.866799291 | 0.37191224 | 2.330655454 |
| Fahd2a | 142.0280488 | 0.505224583 | 0.112266508 | 4.50022533 |
| Kcnip3 | 821.4776332 | -0.185109502 | 0.079137899 | -2.339075251 |
| Gm22613 | 5.411512881 | 0.200188847 | 0.583427983 | 0.343125206 |
| Prom2 | 9.157968272 | -0.194521208 | 0.380107945 | -0.511752545 |
| Gm14229 | 0 | NA | NA | NA |
| Zfp661 | 72.87159097 | -0.272835803 | 0.141009828 | -1.934870824 |

| | | | | |
|---|---|---|---|---|
| Mrps5 | 469.1048793 | 0.055783441 | 0.067998545 | 0.820362273 |
| Mal | 1616.595918 | 0.470950933 | 0.095666757 | 4.922827426 |
| Gm22889 | 0 | NA | NA | NA |
| Gm14245 | 0 | NA | NA | NA |
| Gm14244 | 0.04393175 | 0.037062677 | 3.724945049 | 0.009949859 |
| Mall | 5.752169691 | -0.350787584 | 0.449432295 | -0.780512633 |
| Nphp1 | 102.8958306 | 0.192960006 | 0.110994725 | 1.738461043 |
| 1500011K16Rik | 147.7564642 | -0.039046918 | 0.180223238 | -0.216658617 |
| Bub1 | 15.64735813 | -0.55805688 | 0.392931004 | -1.420241403 |
| Acoxl | 0 | NA | NA | NA |
| Gm22411 | 0.044191549 | 0.037062677 | 3.724945049 | 0.009949859 |
| Gm23101 | 0 | NA | NA | NA |
| Bcl2l11 | 158.1317849 | -0.589117698 | 0.175826999 | -3.350553105 |
| Gm14009 | 0.120974994 | 0.517957688 | 3.724945049 | 0.139051095 |
| Gm14005 | 19.18807827 | -1.698078918 | 0.40863037 | -4.155537725 |
| Gm14012 | 0.247225956 | 0.517970033 | 2.99204733 | 0.173115588 |
| Gm14010 | 0.594047536 | -0.941545853 | 1.306939926 | -0.720420147 |
| Gm29010 | 0 | NA | NA | NA |
| Gm14006 | 0.075043205 | 0.517957688 | 3.724945049 | 0.139051095 |
| Gm14008 | 0.04393175 | 0.037062677 | 3.724945049 | 0.009949859 |
| Gm355 | 0 | NA | NA | NA |
| Gm14007 | 0 | NA | NA | NA |
| Anapc1 | 1911.795479 | -0.073306283 | 0.058989219 | -1.242706457 |
| Mertk | 498.8880327 | -0.424486258 | 0.13423828 | -3.162184873 |
| Gm14011 | 0 | NA | NA | NA |
| Gm23172 | 3.964087094 | 0.849965602 | 0.767768448 | 1.107059823 |
| Tmem87b | 666.3521244 | 0.124069513 | 0.075254837 | 1.648658315 |
| Fbln7 | 48.08116317 | -0.430688896 | 0.209252185 | -2.058228908 |
| Gm22859 | 0 | NA | NA | NA |
| Zc3h8 | 36.68443589 | -0.238940513 | 0.256738546 | -0.930676427 |
| Gm14028 | 0 | NA | NA | NA |
| 4933427J07Rik | 0.358075244 | 1.397864417 | 1.746132524 | 0.800548869 |
| Gm10762 | 30.64394088 | 0.147665059 | 0.2082713 | 0.709003396 |
| Zc3h6 | 404.6004255 | 0.479686606 | 0.110298965 | 4.348967445 |
| Gm14027 | 11.52035073 | 0.382485665 | 0.336133826 | 1.13789698 |
| Gm14025 | 1.764317597 | -1.028286416 | 0.811365977 | -1.267352151 |
| Gm14022 | 5.545450415 | 0.053562415 | 0.505229819 | 0.106015942 |
| Ttl | 570.9256818 | -0.052752571 | 0.097617254 | -0.540402118 |
| Polr1b | 151.5190305 | -0.228715566 | 0.151818028 | -1.506511241 |
| Chchd5 | 87.19880141 | -0.136721724 | 0.154742115 | -0.883545664 |
| AI847159 | 3.26828567 | -1.597109857 | 0.601390926 | -2.655693307 |
| Gm14026 | 27.00074528 | 0.263044481 | 0.238250821 | 1.104065373 |
| Gm14029 | 5.503856369 | -1.152181064 | 0.438633682 | -2.626750089 |
| Gm4430 | 83.58514794 | -0.102531061 | 0.133330299 | -0.76900046 |
| Slc20a1 | 1626.484812 | -0.120078956 | 0.079944943 | -1.502020656 |
| 9830144P21Rik | 0.631002787 | -0.536341219 | 1.202174329 | -0.446142632 |
| A730036I17Rik | 9.097055571 | 0.31584942 | 0.416475575 | 0.75838642 |

| | | | | |
|---|---|---|---|---|
| Gm14024 | 0.180493943 | -0.925573715 | 2.699005356 | -0.342931411 |
| Ckap2l | 17.79315673 | -0.671897335 | 0.463981237 | -1.448113159 |
| Gm14023 | 2.610220779 | -0.196228746 | 0.683104604 | -0.287260172 |
| Il1a | 25.73372477 | 1.393761236 | 0.300913659 | 4.631764611 |
| Gm25703 | 0 | NA | NA | NA |
| Gm14039 | 0 | NA | NA | NA |
| Il1b | 17.42109151 | -1.158904644 | 0.275622547 | -4.204680112 |
| Il1bos | 0.58991373 | -0.321459627 | 1.33185196 | -0.241362882 |
| Spcs2-ps | 12.01699771 | -0.379706142 | 0.282478377 | -1.344195427 |
| F830045P16Rik | 7.997923877 | 1.478132103 | 0.556832756 | 2.654535115 |
| Gm14040 | 0.25468147 | -0.927769677 | 2.112214077 | -0.439240363 |
| Gm14041 | 2.468877162 | 0.119561465 | 0.795944943 | 0.150213235 |
| Sirpa | 3103.836334 | 0.210163377 | 0.0842589 | 2.494257308 |
| Pdyn | 260.8278238 | 0.428907825 | 0.152416781 | 2.814045955 |
| Gm14046 | 0.049553047 | -0.443818988 | 3.724945049 | -0.119147795 |
| Stk35 | 654.0656122 | 0.107398636 | 0.064994147 | 1.652435505 |
| Gm28196 | 0.664320874 | 0.185254754 | 1.61608659 | 0.1146317 |
| Tgm3 | 28.71390692 | 0.972210453 | 0.271143445 | 3.585594528 |
| Mir6339 | 0 | NA | NA | NA |
| Gm22424 | 0.046345709 | -0.443818988 | 3.724945049 | -0.119147795 |
| Gm24665 | 0.038981395 | -0.443816764 | 3.724945049 | -0.119147198 |
| AU015228 | 0.066050875 | -0.443818988 | 3.724945049 | -0.119147795 |
| Tgm6 | 0.648101363 | 0.884250712 | 1.225057044 | 0.721803704 |
| Snrpb | 546.91693 | -0.08566735 | 0.106043013 | -0.80785473 |
| Tmc2 | 0.617199072 | 1.913072147 | 1.362951287 | 1.403624741 |
| Nop56 | 546.9667433 | -0.062114518 | 0.102290345 | -0.607237354 |
| Snord110 | 0.459163063 | 0.520456678 | 1.402866524 | 0.370995151 |
| Gm24451 | 0 | NA | NA | NA |
| Gm23187 | 0 | NA | NA | NA |
| Gm23650 | 0 | NA | NA | NA |
| Snord57 | 0 | NA | NA | NA |
| Idh3b | 1254.967299 | 0.11056482 | 0.057885118 | 1.910073324 |
| Ebf4 | 86.18724845 | 0.432357911 | 0.13644777 | 3.168669668 |
| Gm14044 | 0.90691343 | -1.06023282 | 1.062099301 | -0.99824265 |
| Cpxm1 | 57.79122351 | 0.54568573 | 0.277467994 | 1.96666189 |
| Gm14042 | 0.049553047 | -0.443818988 | 3.724945049 | -0.119147795 |
| Gm28372 | 0 | NA | NA | NA |
| 1700020A23Rik | 0.230076657 | 0.8885161 | 3.108237376 | 0.285858508 |
| Tmem239 | 0.066050875 | -0.443818988 | 3.724945049 | -0.119147795 |
| Pced1a | 228.252696 | 0.119550479 | 0.100959977 | 1.184137343 |
| Vps16 | 502.9920582 | 0.069469011 | 0.064407195 | 1.078590842 |
| Ptpra | 1842.435841 | 0.0542841 | 0.062007386 | 0.875445708 |
| Gm23876 | 0 | NA | NA | NA |
| 4930473A02Rik | 0.931693893 | -0.225110411 | 0.984930004 | -0.22855473 |
| Mrps26 | 372.6054324 | 0.01096671 | 0.12669137 | 0.086562404 |
| Oxt | 55.53220185 | 0.15038664 | 0.530861119 | 0.283288104 |
| Avp | 105.9017596 | 0.166400177 | 0.312175439 | 0.533034171 |

| | | | | |
|---|---|---|---|---|
| Ubox5 | 144.7706665 | -0.069019921 | 0.111296241 | -0.620146018 |
| Fastkd5 | 53.00231664 | -0.617892187 | 0.176718414 | -3.496478789 |
| Lzts3 | 1886.840697 | 1.091596621 | 0.136660725 | 7.987639633 |
| Ddrgk1 | 422.8448348 | 0.07095056 | 0.088299873 | 0.803518253 |
| Itpa | 134.0024633 | 0.109320727 | 0.117532178 | 0.930134444 |
| Slc4a11 | 82.95192658 | -0.268089106 | 0.216508085 | -1.238240622 |
| Gm24893 | 1.856422655 | 0.967510455 | 0.767460014 | 1.26066562 |
| 4930402H24Rik | 1406.23195 | -0.198017367 | 0.112495119 | -1.760230748 |
| Gm14057 | 0.232395771 | 0.052946473 | 2.12208108 | 0.024950259 |
| A730017L22Rik | 0.834338752 | 0.342820977 | 1.053489121 | 0.325414824 |
| Gm14048 | 32.5064405 | -0.575466248 | 0.205923299 | -2.794565995 |
| Atrn | 2962.760329 | 0.355028723 | 0.072181431 | 4.918560305 |
| Gm14049 | 10.85113572 | 0.670642676 | 0.374169303 | 1.792350872 |
| Gfra4 | 152.1835605 | 0.481949222 | 0.138138163 | 3.488892644 |
| Mir6973b | 0 | NA | NA | NA |
| Adam33 | 45.18950367 | 0.249134697 | 0.22022738 | 1.131261231 |
| Siglec1 | 680.3250049 | -0.957988848 | 0.160944393 | -5.952297146 |
| Hspa12b | 36.83173324 | -0.411139195 | 0.197889797 | -2.077616948 |
| Gm11037 | 0.127971488 | -0.95589744 | 3.6855341 | -0.259364698 |
| 1700037H04Rik | 296.061265 | 0.15383334 | 0.090677665 | 1.696485445 |
| Gm14232 | 0.292476601 | 1.916739985 | 1.73816959 | 1.102734736 |
| Spef1 | 54.9579452 | -0.120710966 | 0.162235375 | -0.744048368 |
| Cenpb | 1691.510661 | 0.016162078 | 0.087277399 | 0.185180562 |
| Cdc25b | 96.01317364 | 0.442751201 | 0.161418864 | 2.742871496 |
| Ap5s1 | 93.61776372 | -0.177826362 | 0.120402019 | -1.476938367 |
| Mavs | 109.308156 | -0.979345418 | 0.194256657 | -5.041502473 |
| Pank2 | 601.5473895 | -0.130642928 | 0.068069181 | -1.919266918 |
| Mir103-2 | 2.477700388 | -0.33669461 | 0.699700335 | -0.481198298 |
| Rnf24 | 607.3774196 | 0.255052962 | 0.064720194 | 3.940855954 |
| Gm14233 | 1.848697502 | 0.143179016 | 0.889524164 | 0.160961356 |
| Gm14284 | 25.84299431 | -0.571695973 | 0.206858996 | -2.763698866 |
| Gm22902 | 0 | NA | NA | NA |
| Gm14283 | 0 | NA | NA | NA |
| Gm14304 | 0.048949532 | 0.037064902 | 3.724945049 | 0.009950456 |
| Smox | 612.36022 | -0.440346575 | 0.115099939 | -3.825775917 |
| Adra1d | 160.5552724 | 0.408915303 | 0.128373553 | 3.185354724 |
| Gm14285 | 9.998759481 | 0.9585125 | 0.418247505 | 2.291735127 |
| Gm14280 | 0.350616066 | 1.507567939 | 1.641532226 | 0.918390705 |
| 5330413P13Rik | 17.0164514 | 0.062436886 | 0.319290352 | 0.195548929 |
| Gm14282 | 0 | NA | NA | NA |
| Gm14281 | 0.072025462 | 0.517959912 | 3.724945049 | 0.139051692 |
| Erv3 | 13.96015691 | -2.223173477 | 0.426827266 | -5.2086023 |
| Lamr1-ps1 | 0 | NA | NA | NA |
| Prnp | 7268.086947 | 0.130761621 | 0.091763018 | 1.424992598 |
| Gm28045 | 0.311063295 | -0.806801333 | 1.735779873 | -0.464806249 |
| Prnd | 0.302678705 | 1.471468935 | 1.73250642 | 0.849329571 |
| Rassf2 | 1100.292024 | -0.228473877 | 0.113676984 | -2.009851682 |

| | | | | |
|---|---|---|---|---|
| Slc23a2 | 2945.124527 | 0.286530577 | 0.071293258 | 4.019041718 |
| Gm14052 | 0.072025462 | 0.517959912 | 3.724945049 | 0.139051692 |
| Gm14051 | 0.790290772 | 1.146123852 | 1.235189318 | 0.927893268 |
| Tmem230 | 195.6462545 | -0.377716548 | 0.102720887 | -3.677115325 |
| Pcna | 440.6784249 | -0.513784779 | 0.156064847 | -3.292123675 |
| AV099323 | 3.007737845 | -0.690819394 | 0.739010276 | -0.934789971 |
| Cds2 | 6744.842328 | 0.263466369 | 0.060959652 | 4.321979582 |
| Prokr2 | 44.48347877 | -0.766448251 | 0.219027537 | -3.499323697 |
| 4921508D12Rik | 3.197768018 | -0.173617768 | 0.571654089 | -0.303711232 |
| Gm14095 | 0 | NA | NA | NA |
| 1700026D11Rik | 0.269634638 | 0.535593608 | 1.806416715 | 0.296495047 |
| Gpcpd1 | 1339.978017 | 0.124149664 | 0.093690164 | 1.325108829 |
| AU019990 | 0.221462189 | -0.45661298 | 2.211490944 | -0.206472914 |
| Rpl23a-ps4 | 0 | NA | NA | NA |
| Gm14099 | 0 | NA | NA | NA |
| Gm14098 | 0.075043205 | 0.517957688 | 3.724945049 | 0.139051095 |
| 1110034G24Rik | 59.26919426 | -0.017976828 | 0.190516499 | -0.094358381 |
| Gm14097 | 0.654170826 | -0.588812147 | 1.260271296 | -0.467210631 |
| Gm22245 | 1.105592398 | 0.174999591 | 1.158005532 | 0.15112155 |
| Chgb | 2627.906697 | 0.40441965 | 0.252955871 | 1.598775505 |
| Trmt6 | 322.2692483 | -0.0281191 | 0.082288943 | -0.341711761 |
| Mcm8 | 58.14385559 | 0.173138226 | 0.180225914 | 0.960673313 |
| Gm14104 | 1.507066517 | -0.296452458 | 0.873491765 | -0.339387811 |
| Crls1 | 253.7650353 | -0.155046698 | 0.107284688 | -1.445189436 |
| Lrrn4 | 5.522564292 | 0.733873283 | 0.58117474 | 1.262741191 |
| Fermt1 | 3.449481863 | 0.526260497 | 0.718121745 | 0.732829078 |
| Gm14102 | 0.088736197 | 0.51795324 | 3.724945049 | 0.139049901 |
| Gm14101 | 0 | NA | NA | NA |
| Gm25704 | 0 | NA | NA | NA |
| Gm14103 | 0 | NA | NA | NA |
| Gm14094 | 0.049553047 | -0.443818988 | 3.724945049 | -0.119147795 |
| Gm27449 | 0 | NA | NA | NA |
| A430048G15Rik | 0.11037557 | 0.517957688 | 3.724945049 | 0.139051095 |
| Gm14100 | 0 | NA | NA | NA |
| Bmp2 | 72.0830543 | -0.309434267 | 0.175985366 | -1.758295447 |
| Mir3090 | 0 | NA | NA | NA |
| Gm25258 | 0 | NA | NA | NA |
| Hao1 | 8.182416264 | 1.330419772 | 0.644479034 | 2.064333676 |
| Tmx4 | 4150.07463 | 0.159926309 | 0.066724963 | 2.396798761 |
| Gm14037 | 0.044191549 | 0.037062677 | 3.724945049 | 0.009949859 |
| Gm14038 | 8.079079625 | 0.662832629 | 0.379994285 | 1.744322625 |
| Plcb1 | 2558.946918 | 0.577236444 | 0.097806526 | 5.901819302 |
| Gm14036 | 21.28777894 | 0.343702187 | 0.243398407 | 1.412097109 |
| 4930545L23Rik | 0.686786262 | 0.157143662 | 1.191435221 | 0.131894424 |
| 9630028H03Rik | 3.015362371 | 1.06348514 | 0.657179282 | 1.618257254 |
| Platr3 | 0.132101749 | -0.93550548 | 3.724188147 | -0.251197158 |
| Plcb4 | 1658.213635 | 0.165960504 | 0.06049725 | 2.743273523 |

| | | | | |
|---|---|---|---|---|
| Gm14209 | 16.51388486 | 0.458983428 | 0.363728006 | 1.261886411 |
| Gm14211 | 0.046345709 | -0.443818988 | 3.724945049 | -0.119147795 |
| Gm14210 | 0 | NA | NA | NA |
| Gm26227 | 0.166709681 | 0.517964155 | 2.76763122 | 0.187150713 |
| Lamp5 | 675.5566326 | 1.073601045 | 0.154573678 | 6.945561898 |
| Pak7 | 618.9095195 | 0.182557269 | 0.121255755 | 1.505555501 |
| Gm14218 | 0 | NA | NA | NA |
| Gm14056 | 0.369423471 | -1.627513519 | 1.503554272 | -1.082444146 |
| Gm14054 | 0.515606452 | 1.001615483 | 1.667004642 | 0.600847447 |
| Ankef1 | 28.51623535 | 0.320002304 | 0.211261625 | 1.514720447 |
| Gm36967 | 0.044544648 | 0.517957688 | 3.724945049 | 0.139051095 |
| Snap25 | 16400.01604 | 0.437942969 | 0.076374955 | 5.734117571 |
| Gm14053 | 0.15906016 | -1.015882374 | 3.433492635 | -0.295874342 |
| Mkks | 278.0206276 | -0.434876574 | 0.150855366 | -2.882738513 |
| Slx4ip | 166.0135647 | -0.464351507 | 0.121924246 | -3.808524746 |
| Jag1 | 129.0559854 | -1.206779926 | 0.139453889 | -8.653612583 |
| Gm28214 | 0.110595523 | 0.037071574 | 3.724945049 | 0.009952247 |
| Gm14055 | 0.284116082 | 0.015522488 | 2.011004169 | 0.007718775 |
| Gm14064 | 0 | NA | NA | NA |
| Gm14062 | 10.87638177 | 1.993623867 | 0.441297209 | 4.517644399 |
| Btbd3 | 2584.985912 | 0.216090133 | 0.108296576 | 1.99535518 |
| Gm14061 | 4.211247599 | 0.782267787 | 0.540447474 | 1.447444616 |
| Mir1952 | 0 | NA | NA | NA |
| Gm14065 | 0.143485256 | -0.955563601 | 3.547405725 | -0.269369696 |
| Gm14067 | 0 | NA | NA | NA |
| Gm14069 | 0 | NA | NA | NA |
| Gm14066 | 25.91994233 | 0.907222252 | 0.239490712 | 3.788131253 |
| Gm14071 | 0 | NA | NA | NA |
| Sptlc3 | 1.944739822 | 0.083156525 | 0.851333873 | 0.097677923 |
| Gm27385 | 0 | NA | NA | NA |
| Ism1 | 46.24466388 | 0.833934618 | 0.179461654 | 4.64686801 |
| Tasp1 | 470.1320558 | 0.115748606 | 0.090780596 | 1.275036855 |
| Gm14070 | 0 | NA | NA | NA |
| Rps19-ps7 | 1.19142324 | -1.069279852 | 0.999407185 | -1.069914113 |
| Gm14073 | 0.066050875 | -0.443818988 | 3.724945049 | -0.119147795 |
| Esf1 | 372.9243209 | 0.014300441 | 0.08957028 | 0.159656088 |
| Gm17374 | 0 | NA | NA | NA |
| Ndufaf5 | 256.6486657 | -0.030648709 | 0.081945511 | -0.374013271 |
| Sel1l2 | 0 | NA | NA | NA |
| Macrod2 | 727.1380872 | 0.684368451 | 0.091502138 | 7.479261895 |
| Gm23846 | 0 | NA | NA | NA |
| Flrt3 | 365.9243577 | 0.610181521 | 0.079953496 | 7.631705358 |
| Calr-ps | 0.108772582 | 0.037073798 | 3.724945049 | 0.009952844 |
| Zcchc9-ps | 30.12202849 | 0.791589028 | 0.391774454 | 2.020522322 |
| Macrod2os2 | 0 | NA | NA | NA |
| Macrod2os1 | 0 | NA | NA | NA |
| Gm25583 | 2.242176843 | 2.094472405 | 0.86817074 | 2.412512089 |

| | | | | |
|---|---|---|---|---|
| Fau-ps1 | 8.271865471 | 1.12328762 | 0.55087248 | 2.039106438 |
| Gm23179 | 0.743559953 | 1.499330547 | 1.380083786 | 1.08640545 |
| Rpl26-ps5 | 0 | NA | NA | NA |
| Rbmx2-ps | 0.144013665 | -1.209023387 | 3.538333554 | -0.341692881 |
| Kif16b | 346.7941092 | -0.04461035 | 0.104630759 | -0.426359806 |
| Gm22310 | 0.089089296 | 0.957155977 | 3.723939561 | 0.257027796 |
| Kif16bos | 0.612922179 | 0.039724374 | 1.232203692 | 0.03223848 |
| Snrpb2 | 243.6495243 | 0.116483129 | 0.132983861 | 0.875919289 |
| Otor | 0.348607679 | -2.375931785 | 1.892362917 | -1.255537066 |
| Gm14083 | 0 | NA | NA | NA |
| Gm14082 | 0 | NA | NA | NA |
| Gm11644 | 0 | NA | NA | NA |
| Gm11645 | 3.276776717 | 1.278044163 | 0.766208077 | 1.668011864 |
| Pcsk2os1 | 65.67909268 | 0.716586013 | 0.17916884 | 3.999501312 |
| Pcsk2 | 3501.594654 | 0.90847582 | 0.105358164 | 8.622737728 |
| Pcsk2os2 | 0.594248913 | 0.420345432 | 1.412728294 | 0.297541597 |
| Bfsp1 | 5.877618629 | -0.038547072 | 0.418322191 | -0.09214685 |
| Rps13-ps6 | 0.132101749 | -0.93550548 | 3.724188147 | -0.251197158 |
| Gm11673 | 0 | NA | NA | NA |
| Gm11686 | 0.0878635 | 0.037058229 | 3.724945049 | 0.009948665 |
| Dstn | 1131.896076 | -0.156048172 | 0.091249409 | -1.710128034 |
| Rrbp1 | 1060.324141 | -0.542013584 | 0.170459682 | -3.179717207 |
| Banf2 | 0 | NA | NA | NA |
| Banf2os | 0.710948324 | 0.748850659 | 1.200568128 | 0.623746909 |
| Gm11687 | 1.624027788 | -0.784449353 | 0.79035983 | -0.992521789 |
| Gm5535 | 4.621059661 | -0.244631973 | 0.448072547 | -0.5459651 |
| 4930444E06Rik | 0.044191549 | 0.037062677 | 3.724945049 | 0.009949859 |
| Snx5 | 1040.74114 | -0.603709444 | 0.096900614 | -6.230192118 |
| Gm11688 | 3.796974204 | -0.063235567 | 0.503026315 | -0.125710257 |
| Gm26165 | 17.87795363 | -0.445503461 | 0.293045383 | -1.520254155 |
| Snord17 | 426.2071553 | -0.162637008 | 0.14485819 | -1.122732574 |
| Mgme1 | 58.73843915 | -0.896383622 | 0.163900959 | -5.469056601 |
| Ovol2 | 23.1269052 | 0.333912806 | 0.300274879 | 1.11202378 |
| 1700108N11Rik | 1.600434186 | 0.436070369 | 0.722428883 | 0.60361702 |
| Gm11700 | 0 | NA | NA | NA |
| Gm11701 | 1.140052013 | 0.401976716 | 0.993995686 | 0.40440489 |
| Csrp2bp | 346.2940874 | -0.012232354 | 0.066993653 | -0.182589751 |
| Pet117 | 0 | NA | NA | NA |
| Zfp133-ps | 17.17923113 | -0.984827394 | 0.305320441 | -3.225553424 |
| Dzank1 | 3640.794771 | 0.529283982 | 0.06248735 | 8.470258085 |
| Polr3f | 438.0816624 | 0.049785775 | 0.116462717 | 0.42748251 |
| Rbbp9 | 270.2515903 | -0.170692623 | 0.137120258 | -1.24483884 |
| Sec23b | 663.5913093 | -0.391009133 | 0.090166577 | -4.336519641 |
| Gm561 | 84.95980064 | -0.235465539 | 0.154567067 | -1.523387504 |
| Dtd1 | 350.4218442 | 0.209286225 | 0.074746334 | 2.799953018 |
| Gm24823 | 0 | NA | NA | NA |
| Gm11743 | 12.81133691 | 0.538557871 | 0.424528657 | 1.268601922 |

| | | | | |
|---|---|---|---|---|
| Scp2d1 | 0.431157666 | 0.714674039 | 1.479226953 | 0.483140222 |
| Gm11746 | 0 | NA | NA | NA |
| Slc24a3 | 1355.879502 | 0.563996238 | 0.078110351 | 7.220505775 |
| Gm14093 | 14.45402812 | 0.474333237 | 0.347505176 | 1.364967402 |
| Gm22731 | 3.697861772 | 0.994664873 | 0.678325353 | 1.466353673 |
| Gm22390 | 0 | NA | NA | NA |
| Gm11759 | 0.254720735 | 0.037535981 | 1.838858546 | 0.020412653 |
| Gm14092 | 0.048949532 | 0.037064902 | 3.724945049 | 0.009950456 |
| BC039771 | 5.098733496 | -0.452419316 | 0.514244444 | -0.879774826 |
| Gm11763 | 0.066050875 | -0.443818988 | 3.724945049 | -0.119147795 |
| Rin2 | 761.9091662 | -0.622321906 | 0.122883571 | -5.064321486 |
| Gm25026 | 1.82714634 | -0.833758281 | 0.855651225 | -0.974413705 |
| Gm11764 | 0 | NA | NA | NA |
| Naa20 | 524.4015188 | -0.292247233 | 0.098703779 | -2.9608515 |
| Crnkl1 | 313.9592317 | -0.137173864 | 0.074321123 | -1.845691489 |
| Cfap61 | 40.89814708 | 0.313161048 | 0.199827698 | 1.567155362 |
| Insm1 | 43.73815878 | 0.061563617 | 0.202783618 | 0.303592654 |
| Ralgapa2 | 804.7755793 | 0.280973571 | 0.152032403 | 1.848116354 |
| A930019D19Rik | 0.11147366 | -0.955981391 | 3.722845977 | -0.25678779 |
| Gm14117 | 89.00856018 | 0.278865493 | 0.134914746 | 2.066975641 |
| 4933406D12Rik | 0 | NA | NA | NA |
| Gm14111 | 0.398068774 | 1.322643017 | 3.687638593 | 0.358669372 |
| Gm14114 | 1.234305601 | 0.244843848 | 0.928389975 | 0.263729526 |
| Kiz | 194.5668815 | -0.004504093 | 0.102159064 | -0.044089024 |
| Xrn2 | 1167.834356 | 0.068223367 | 0.08716366 | 0.782704249 |
| Nkx2-4 | 1.324777524 | -1.556089126 | 1.244421145 | -1.250452174 |
| Gm14116 | 0.119587853 | 0.981063299 | 3.723618439 | 0.263470416 |
| Nkx2-2 | 88.16107763 | -0.46767959 | 0.142782976 | -3.275457648 |
| Nkx2-2os | 6.403733378 | -0.092315925 | 0.419812605 | -0.219897935 |
| 6430503K07Rik | 6.531065735 | 0.605897001 | 0.455942796 | 1.328888201 |
| Gm22261 | 0 | NA | NA | NA |
| Gm14115 | 0.048949532 | 0.037064902 | 3.724945049 | 0.009950456 |
| Gm14113 | 3.843754305 | -0.158429929 | 0.491257222 | -0.322498931 |
| Gm14110 | 0 | NA | NA | NA |
| AI646519 | 0 | NA | NA | NA |
| Pax1 | 0.59047672 | -0.769007094 | 1.318448203 | -0.583266822 |
| Gm14109 | 0 | NA | NA | NA |
| Gm14107 | 0 | NA | NA | NA |
| Gm14106 | 0.066050875 | -0.443818988 | 3.724945049 | -0.119147795 |
| A530006G24Rik | 0 | NA | NA | NA |
| Gm25516 | 0 | NA | NA | NA |
| 9030622O22Rik | 0.80144685 | 0.032180273 | 1.353829049 | 0.02376982 |
| Foxa2 | 2.785346695 | 0.373214638 | 0.812748487 | 0.459200656 |
| Gm27420 | 0 | NA | NA | NA |
| Gm14108 | 0 | NA | NA | NA |
| Gm24221 | 0 | NA | NA | NA |
| Sstr4 | 86.59783816 | 0.768090486 | 0.159918626 | 4.803008282 |

| | | | | |
|---|---|---|---|---|
| Thbd | 190.3799444 | -0.770789063 | 0.172684639 | -4.46356471 |
| Gm14119 | 0.109727584 | 0.037071574 | 3.724945049 | 0.009952247 |
| Cd93 | 290.705625 | -1.134643998 | 0.191380355 | -5.928738068 |
| Gm14120 | 2.718509683 | 2.741887186 | 0.945031235 | 2.901372021 |
| Gm25866 | 0 | NA | NA | NA |
| Nxt1 | 62.52770177 | -0.575626169 | 0.192457529 | -2.990925695 |
| Gzf1 | 464.5606654 | -0.13252822 | 0.063775804 | -2.078032909 |
| Napb | 4683.537772 | 0.321219149 | 0.049806712 | 6.449314458 |
| Cstl1 | 7.743612027 | -0.255543886 | 0.377022205 | -0.677795321 |
| Cst11 | 0.068097359 | 0.037064902 | 3.724945049 | 0.009950456 |
| 8030411F24Rik | 0.116944185 | -1.028246102 | 3.719337068 | -0.27645951 |
| Cst12 | 0.585343961 | 0.438188711 | 1.473784566 | 0.297322093 |
| Cst8 | 0.132101749 | -0.93550548 | 3.724188147 | -0.251197158 |
| Cst13 | 0.138076337 | 0.037074383 | 3.589802603 | 0.010327694 |
| Cst9 | 0.038981395 | -0.443816764 | 3.724945049 | -0.119147198 |
| 9230104L09Rik | 0.693909001 | -1.022886949 | 1.177658831 | -0.868576639 |
| Cst3 | 6207.776889 | 0.241689024 | 0.12516445 | 1.930971801 |
| Gm38221 | 0.060174537 | 0.517957688 | 3.724945049 | 0.139051095 |
| Gm14129 | 0 | NA | NA | NA |
| Gm1330 | 0 | NA | NA | NA |
| Gm10750 | 0 | NA | NA | NA |
| Gm14135 | 0 | NA | NA | NA |
| Gm14132 | 0.875345121 | 1.901128085 | 1.100108405 | 1.728127952 |
| Gm14131 | 0.1140246 | 0.037073798 | 3.724945049 | 0.009952844 |
| Cst10 | 0 | NA | NA | NA |
| Gm24551 | 0.086648695 | 0.037058229 | 3.724945049 | 0.009948665 |
| Gm14130 | 2.711335961 | 0.031822073 | 0.744655131 | 0.042733974 |
| Gm14133 | 2.358317775 | 0.372873828 | 0.743775926 | 0.501325486 |
| Syndig1 | 375.012134 | 0.089266176 | 0.0886357 | 1.007113109 |
| C530025M09Rik | 3.362260024 | -0.922891929 | 0.673650758 | -1.369985735 |
| Gm14140 | 1.613620997 | -0.70854418 | 0.775924356 | -0.913161411 |
| Zfp120 | 248.9582029 | -0.153214092 | 0.133150116 | -1.150686878 |
| Gm10770 | 0 | NA | NA | NA |
| Gm14139 | 0.236020217 | 0.531476836 | 2.508763135 | 0.211848153 |
| Zfp937 | 71.56539432 | -0.39375509 | 0.167120821 | -2.356110306 |
| Gm21994 | 0.580893185 | 0.204996644 | 1.487730984 | 0.137791473 |
| Gm14124 | 2.427470934 | 1.37862039 | 0.786802363 | 1.752181303 |
| Gm14125 | 0 | NA | NA | NA |
| 3300002I08Rik | 53.22250888 | -0.197061483 | 0.205101834 | -0.960798246 |
| Gm10130 | 6.196226755 | 0.117961774 | 0.389737182 | 0.302670053 |
| Gm14126 | 0.606327527 | 1.944050895 | 1.280300551 | 1.518433226 |
| Gm14122 | 12.30982907 | 0.203595617 | 0.323391049 | 0.629564788 |
| Zfp442 | 70.96873342 | -0.443425108 | 0.18076654 | -2.453026475 |
| Gm14127 | 0.190396633 | 0.541112798 | 3.26794622 | 0.165581917 |
| Zfp345 | 0.580755766 | -0.020076409 | 1.224393866 | -0.016397019 |
| Gm14121 | 87.55734381 | 0.024977277 | 0.173285809 | 0.144139193 |
| Gm14128 | 0 | NA | NA | NA |

| | | | | |
|---|---|---|---|---|
| Cst7 | 47.58322733 | 0.493987488 | 0.205616828 | 2.402466241 |
| Apmap | 559.647641 | 0.22997989 | 0.090932316 | 2.529132658 |
| Gm14123 | 1.105496479 | -0.240188343 | 0.85371442 | -0.28134507 |
| Acss1 | 208.0155599 | 0.310859346 | 0.123147309 | 2.524288576 |
| E130215H24Rik | 2.738071508 | 1.099897421 | 0.691419826 | 1.590780854 |
| Vsx1 | 2.657225507 | 0.007681246 | 0.741554666 | 0.010358301 |
| Platr15 | 0.232351398 | 1.0602475 | 2.542298253 | 0.417042925 |
| Gm28450 | 0 | NA | NA | NA |
| Gm26681 | 0.300525961 | -1.559557319 | 3.691933976 | -0.422422863 |
| Entpd6 | 438.5268923 | 0.200302687 | 0.074549929 | 2.686825976 |
| Pygb | 1906.36878 | 0.084321719 | 0.10596598 | 0.795743304 |
| Abhd12 | 2275.162201 | 0.154586844 | 0.048776573 | 3.169284623 |
| Gm23598 | 0 | NA | NA | NA |
| Gins1 | 16.63953435 | -0.142590781 | 0.29260588 | -0.487313449 |
| Gm14143 | 0.083526043 | 0.037073798 | 3.724945049 | 0.009952844 |
| Ninl | 67.5515232 | 0.351948991 | 0.166944467 | 2.10818003 |
| Gm14142 | 0.423021739 | 1.362574032 | 1.666825039 | 0.817466741 |
| Gm14141 | 0 | NA | NA | NA |
| Nanp | 8.071585247 | 0.475605594 | 0.483048785 | 0.984591223 |
| Gm22187 | 0 | NA | NA | NA |
| Gm24199 | 0 | NA | NA | NA |
| Gm14151 | 0 | NA | NA | NA |
| Gm14147 | 0 | NA | NA | NA |
| Gm14145 | 0 | NA | NA | NA |
| Gm22517 | 0 | NA | NA | NA |
| Gm14149 | 0.579212071 | 0.590147007 | 1.398656627 | 0.421938448 |
| Gm14144 | 0 | NA | NA | NA |
| Gm23005 | 0 | NA | NA | NA |
| Gm24077 | 0 | NA | NA | NA |
| 4930442J19Rik | 0 | NA | NA | NA |
| Gm23261 | 0 | NA | NA | NA |
| Gm14148 | 0.134276636 | -0.890946624 | 2.951368388 | -0.301875777 |
| Gm14152 | 0 | NA | NA | NA |
| Gm24868 | 0 | NA | NA | NA |
| Gm14150 | 75.86947207 | -0.145948564 | 0.217876053 | -0.669869691 |
| Gm14146 | 0 | NA | NA | NA |
| 4921509C19Rik | 0.322748876 | -1.14211116 | 3.700038777 | -0.308675457 |
| Nsfl1c | 781.3049052 | -0.125552278 | 0.059717081 | -2.102451708 |
| Gm14167 | 3.663554111 | 0.817075354 | 0.529064885 | 1.544376461 |
| Fkbp1a | 3786.330142 | 0.299526951 | 0.067495814 | 4.437711496 |
| Sdcbp2 | 21.61712664 | -0.988292417 | 0.351744646 | -2.809687162 |
| Snph | 3369.871354 | 0.618758202 | 0.092363057 | 6.699195808 |
| Rad21l | 5.277545605 | 0.497893887 | 0.466294398 | 1.067767249 |
| Tmem74bos | 12.83654197 | 0.111496201 | 0.340198032 | 0.327739113 |
| Gm14159 | 1.598224206 | 1.844134507 | 0.892709387 | 2.065772504 |
| Tmem74b | 29.74572494 | 0.228301519 | 0.239271222 | 0.95415369 |
| Psmf1 | 502.2961145 | 0.040450123 | 0.098568747 | 0.410374729 |

| | | | | |
|---|---|---|---|---|
| 4930556L07Rik | 0.192951122 | 1.461101854 | 2.632329567 | 0.555060382 |
| Gm14153 | 0 | NA | NA | NA |
| Gm14155 | 0 | NA | NA | NA |
| Rspo4 | 1.536911596 | -0.622238987 | 0.944867687 | -0.65854616 |
| Rspo4os | 0.417636614 | 0.531361641 | 1.449630627 | 0.366549679 |
| Angpt4 | 4.455728532 | -1.774945376 | 0.566569163 | -3.132795589 |
| Gm14154 | 0.934869149 | 1.713986834 | 1.093587008 | 1.567307238 |
| Mir1953 | 0.11193333 | 0.037073798 | 3.724945049 | 0.009952844 |
| Fam110a | 57.87769406 | -0.183028478 | 0.153810723 | -1.189959156 |
| Slc52a3 | 22.60185597 | -0.886017549 | 0.227378048 | -3.896671445 |
| AA387200 | 2.676545693 | -1.688757769 | 0.748461712 | -2.256304819 |
| Scrt2 | 297.4913887 | -0.338286399 | 0.143925255 | -2.350431121 |
| Srxn1 | 1496.550893 | -1.181060502 | 0.111333121 | -10.60834809 |
| Tcf15 | 3.917975245 | -0.722458325 | 0.555160149 | -1.301351197 |
| Rps15a-ps7 | 16.65743553 | -0.355138254 | 0.256169243 | -1.386342289 |
| Csnk2a1 | 1499.289359 | -0.019039584 | 0.038962426 | -0.488665247 |
| Tbc1d20 | 711.5116405 | -0.139337894 | 0.05534922 | -2.517431961 |
| Rbck1 | 775.7363636 | 0.028813644 | 0.06564663 | 0.43892038 |
| Trib3 | 48.90459884 | -1.548806268 | 0.338659369 | -4.573345408 |
| Gm14164 | 0.458177261 | -0.875441971 | 1.576118732 | -0.555441639 |
| Gm14165 | 3.039451885 | -0.528486481 | 0.585220305 | -0.903055613 |
| Nrsn2 | 894.8007339 | 0.29624205 | 0.087225921 | 3.396261641 |
| Sox12 | 175.956736 | 0.274846176 | 0.145103153 | 1.894143371 |
| Zcchc3 | 147.4086379 | -0.015394259 | 0.120041715 | -0.128240915 |
| 6820408C15Rik | 27.31515061 | -0.024539464 | 0.19942231 | -0.123052753 |
| Gm27343 | 3.046712631 | -0.069538047 | 0.626162598 | -0.111054297 |
| Gm14158 | 0.634981691 | -0.030741484 | 1.31493803 | -0.023378656 |
| Defb23 | 0.097139547 | -0.443816764 | 3.724945049 | -0.119147198 |
| Gm14156 | 0 | NA | NA | NA |
| Defb20 | 0.545971167 | 0.10186619 | 1.406291716 | 0.07243603 |
| Defb22 | 0.061920613 | -0.443816764 | 3.724945049 | -0.119147198 |
| Defb26 | 0 | NA | NA | NA |
| Defb28 | 0 | NA | NA | NA |
| Defb29 | 0 | NA | NA | NA |
| Gm14161 | 0.118371171 | 0.037073798 | 3.724945049 | 0.009952844 |
| Gm17416 | 0 | NA | NA | NA |
| Defb21 | 0.04393175 | 0.037062677 | 3.724945049 | 0.009949859 |
| Defb19 | 0 | NA | NA | NA |
| Defb45 | 0.07796279 | -0.798301147 | 3.721829328 | -0.214491605 |
| Defb36 | 0 | NA | NA | NA |
| Gm14163 | 0 | NA | NA | NA |
| Defb25 | 0.234671876 | 0.505418163 | 2.170256769 | 0.23288404 |
| Rem1 | 3.041196478 | -0.233204316 | 0.692416944 | -0.336797529 |
| 1700030C14Rik | 0.290525896 | 1.558530218 | 1.784208722 | 0.873513395 |
| H13 | 832.2285858 | -0.324012543 | 0.07285345 | -4.447456436 |
| Mcts2 | 80.0283442 | -0.054726076 | 0.125209099 | -0.437077467 |
| Gm26841 | 0.794703889 | -0.666704652 | 1.260343082 | -0.52898664 |

| | | | | |
|---|---|---|---|---|
| Id1 | 158.7587211 | -0.623269251 | 0.199806178 | -3.119369267 |
| Gm14162 | 0.044191549 | 0.037062677 | 3.724945049 | 0.009949859 |
| Cox4i2 | 12.07102929 | 0.489088888 | 0.321264667 | 1.522386178 |
| Gm23802 | 0.93732221 | 1.476786907 | 1.151823299 | 1.28212974 |
| Bcl2l1 | 1656.202641 | -0.274615111 | 0.111984269 | -2.452265062 |
| Tpx2 | 97.61010764 | -0.531481927 | 0.221928181 | -2.394837485 |
| Mir5622 | 0 | NA | NA | NA |
| Mylk2 | 13.81209412 | -2.337398102 | 0.539256801 | -4.334480526 |
| Foxs1 | 7.663944057 | -1.262614862 | 0.419626642 | -3.008900614 |
| Dusp15 | 161.0227663 | 0.247648585 | 0.126198486 | 1.96237366 |
| Ttll9 | 15.9748206 | 0.090893858 | 0.332174665 | 0.273632724 |
| Gm14199 | 0.260717194 | -0.883269996 | 2.039123787 | -0.433161538 |
| Pdrg1 | 314.6393574 | -0.570521456 | 0.089731803 | -6.358074163 |
| Xkr7 | 77.04392738 | -0.193663216 | 0.170799394 | -1.133863601 |
| Ccm2l | 14.10545208 | -0.113707233 | 0.272914194 | -0.416640965 |
| Hck | 217.2183995 | -0.634522075 | 0.169727033 | -3.738485644 |
| Gm25643 | 0 | NA | NA | NA |
| Tm9sf4 | 988.450612 | -0.099889378 | 0.043253725 | -2.309382118 |
| Tspyl3 | 233.7002 | 0.463925506 | 0.096294372 | 4.817784201 |
| Plagl2 | 263.3156821 | -0.325222916 | 0.111111891 | -2.926985688 |
| Pofut1 | 393.8680983 | 0.042291797 | 0.08508609 | 0.497047136 |
| Kif3b | 984.8683119 | -0.077802671 | 0.046261939 | -1.681785773 |
| 2500004C02Rik | 39.85926624 | 0.092931045 | 0.23565235 | 0.394356536 |
| Asxl1 | 918.9589658 | 0.21500036 | 0.083266456 | 2.582076506 |
| Nol4l | 699.5239319 | 0.282396907 | 0.126354048 | 2.234965251 |
| Gm14472 | 0.227745227 | 1.572768169 | 2.573036673 | 0.611249807 |
| 4930404H24Rik | 0.825833058 | -0.801406077 | 1.157783912 | -0.692189682 |
| Commd7 | 204.4885226 | -0.337390915 | 0.093877643 | -3.593943174 |
| Dnmt3bos | 0 | NA | NA | NA |
| Dnmt3b | 18.90429361 | 0.220612922 | 0.287858309 | 0.766394142 |
| Gm14490 | 1.779788623 | -1.427238201 | 0.766360602 | -1.862358527 |
| Mapre1 | 1211.391518 | -0.104141679 | 0.071170233 | -1.463275777 |
| Efcab8 | 3.424510338 | 0.57012212 | 0.729860199 | 0.781138801 |
| Gm14494 | 80.10444224 | 0.142870007 | 0.131694249 | 1.084861395 |
| Sun5 | 0.124596252 | 0.037071632 | 3.714681146 | 0.009979761 |
| Bpifb2 | 0.955311604 | -1.020372863 | 1.123872017 | -0.907908416 |
| Bpifb6 | 1.501918467 | -0.806661898 | 0.844224631 | -0.955506234 |
| Bpifb3 | 1.988936712 | -1.691590387 | 1.580371097 | -1.070375426 |
| Bpifb4 | 3.358904097 | -0.469635298 | 1.21553589 | -0.386360701 |
| Bpifa6 | 0.496470236 | -1.210407037 | 3.706396085 | -0.3265725 |
| Bpifa2 | 0 | NA | NA | NA |
| Gm21850 | 0 | NA | NA | NA |
| Gm14497 | 0.15596086 | 0.037059316 | 3.490591407 | 0.010616916 |
| Bpifa3 | 0 | NA | NA | NA |
| Bpifa1 | 1.270782233 | -3.059064749 | 1.30789234 | -2.338927032 |
| Bpifa5 | 0.243228626 | -1.073592254 | 3.022466967 | -0.355203966 |
| Bpifb1 | 0.700162942 | -1.148505507 | 1.147675955 | -1.00072281 |

| | | | | |
|---|---|---|---|---|
| Bpifb5 | 0.765485319 | 0.10182525 | 1.253043535 | 0.08126234 |
| Bpifb9a | 1.362283058 | 0.409458399 | 0.969403321 | 0.422381882 |
| Gm14197 | 1.834732021 | -0.33155854 | 0.752044801 | -0.440876049 |
| Bpifb9b | 3.934499224 | -0.268721489 | 1.531286206 | -0.175487435 |
| Cdk5rap1 | 105.428526 | -0.017153932 | 0.13811213 | -0.124202936 |
| Snta1 | 269.5638173 | -0.132016786 | 0.115155196 | -1.146424914 |
| Cbfa2t2 | 875.8606125 | 0.045530878 | 0.051499477 | 0.884103692 |
| Necab3 | 257.0570188 | 0.808263135 | 0.176384271 | 4.582399157 |
| 1700003F12Rik | 3.150202034 | 0.972772159 | 0.701165272 | 1.387365003 |
| Actl10 | 0.266983968 | -0.488349435 | 1.818389719 | -0.268561481 |
| E2f1 | 73.04566475 | 0.312750953 | 0.231997327 | 1.348079984 |
| Pxmp4 | 135.3858191 | 0.404857724 | 0.117992622 | 3.431212189 |
| 4930519P11Rik | 1.196740622 | 1.018013838 | 0.930609131 | 1.09392204 |
| Zfp341 | 104.6788915 | -0.007639409 | 0.136931134 | -0.055790153 |
| Gm14198 | 0.162949104 | -0.443820316 | 2.787607702 | -0.159211899 |
| Chmp4b | 2004.387491 | -0.268786761 | 0.136301991 | -1.971994389 |
| Rpl5-ps2 | 5.302412506 | -0.472813831 | 0.412130346 | -1.147243426 |
| Gm14215 | 0 | NA | NA | NA |
| Gm14216 | 0.099155932 | 0.037073798 | 3.724945049 | 0.009952844 |
| Gm14214 | 0 | NA | NA | NA |
| Raly | 850.5641207 | -0.216786 | 0.086457195 | -2.50743735 |
| a | 0.140640116 | -0.925682484 | 3.611524051 | -0.256313532 |
| Eif2s2 | 723.9443759 | -0.233199182 | 0.114656645 | -2.033891556 |
| Gm14226 | 0.71216146 | -0.3063215 | 1.243711429 | -0.246296281 |
| Ahcy | 112.8096952 | -0.198994617 | 0.147222346 | -1.351660417 |
| 2310005A03Rik | 0.04393175 | 0.037062677 | 3.724945049 | 0.009949859 |
| Itch | 1259.365133 | -0.309512609 | 0.062920693 | -4.919090895 |
| Gm14227 | 0.141071297 | -0.443820801 | 3.564043909 | -0.12452731 |
| Dynlrb1 | 713.8695448 | 0.173166237 | 0.088668296 | 1.952966777 |
| Map1lc3a | 781.5540749 | -0.068035194 | 0.122327647 | -0.556171847 |
| Pigu | 261.7083534 | -0.378019761 | 0.11159589 | -3.387398597 |
| Mir695 | 0 | NA | NA | NA |
| Trp53inp2 | 2971.079894 | -0.095148655 | 0.106995369 | -0.88927826 |
| Ncoa6 | 1310.679018 | 0.038697813 | 0.06940554 | 0.557560861 |
| Gm14239 | 28.68396613 | 0.175840865 | 0.3166055 | 0.555394221 |
| Gm14238 | 0.886238366 | 1.920362743 | 1.050281918 | 1.828425979 |
| Ggt7 | 845.8338809 | 0.432158148 | 0.073696778 | 5.864003253 |
| Acss2 | 635.0895078 | 0.404879434 | 0.066992678 | 6.043637089 |
| Mipep-ps | 1.837594399 | -0.549856325 | 0.856812577 | -0.641746329 |
| Gm20637 | 2.163212633 | 1.081067282 | 0.725437668 | 1.490227665 |
| Acss2os | 0.277114854 | -0.996289447 | 2.090542526 | -0.476569806 |
| Gss | 143.0865772 | -0.255326745 | 0.126532904 | -2.017868371 |
| Gssos2 | 1.802868249 | 0.561575258 | 0.912320453 | 0.615546058 |
| Gssos1 | 1.992642508 | 2.529735738 | 0.984244398 | 2.57023128 |
| Gm14257 | 0.038981395 | -0.443816764 | 3.724945049 | -0.119147198 |
| Myh7b | 88.10465837 | 0.667537915 | 0.272332757 | 2.451184799 |
| Mir499 | 0 | NA | NA | NA |

| | | | | |
|---|---|---|---|---|
| Trpc4ap | 1703.710218 | -0.033837296 | 0.054989109 | -0.615345417 |
| Gm22334 | 1.228515858 | -1.216696 | 1.17542646 | -1.03511027 |
| Edem2 | 274.0937995 | -0.241396138 | 0.090078455 | -2.679843221 |
| Gm17581 | 0 | NA | NA | NA |
| Procr | 21.11771009 | -1.209407357 | 0.310173445 | -3.899132486 |
| Mmp24 | 793.763571 | 0.013410036 | 0.133664189 | 0.100326321 |
| BC029722 | 543.0851477 | 0.164234655 | 0.093028182 | 1.765429046 |
| Eif6 | 475.8961878 | -0.633570543 | 0.117736774 | -5.38124599 |
| Gm16098 | 1.335672075 | -0.719245066 | 0.830972053 | -0.865546637 |
| Fam83c | 0.240161549 | -0.594704831 | 3.035026595 | -0.195947156 |
| Uqcc1 | 374.2692931 | -0.066226999 | 0.083997195 | -0.788442985 |
| Gm15557 | 0.53525542 | -0.108895394 | 1.430668027 | -0.076115068 |
| Gdf5 | 4.286069218 | 1.126151576 | 0.519501783 | 2.167753054 |
| Cep250 | 337.5416032 | 0.232158568 | 0.159403509 | 1.456420685 |
| 6430550D23Rik | 7.007522312 | -0.822298345 | 0.482607322 | -1.703866283 |
| Ergic3 | 697.3852742 | -0.004660614 | 0.089473682 | -0.052089218 |
| Fer1l4 | 0.699011511 | -1.07118319 | 2.530400501 | -0.423325552 |
| Gm14240 | 0 | NA | NA | NA |
| Spag4 | 20.11396154 | 0.159039294 | 0.327236669 | 0.486006945 |
| Cpne1 | 7.154347536 | 0.400163119 | 0.485325544 | 0.824525153 |
| Gm28036 | 0.689591474 | -0.662303001 | 1.166856964 | -0.567595705 |
| Gm37274 | 40.70937969 | -0.30485078 | 0.338687301 | -0.900095099 |
| Rbm12 | 333.4415097 | -0.003650762 | 0.084857179 | -0.043022429 |
| Nfs1 | 286.4122944 | 0.032656199 | 0.091005866 | 0.358836201 |
| Romo1 | 128.8625568 | -0.087822243 | 0.181971267 | -0.482615988 |
| Rbm39 | 3275.570639 | 0.128448402 | 0.088186244 | 1.456558265 |
| Phf20 | 1427.32006 | 0.291628134 | 0.052078187 | 5.599813461 |
| Scand1 | 367.1199098 | -0.369170463 | 0.17586287 | -2.09919503 |
| Cnbd2 | 68.49456786 | -0.247770869 | 0.137276513 | -1.804903578 |
| Gm14251 | 0.489024281 | -0.908694455 | 1.591986008 | -0.570792991 |
| Gm22759 | 0 | NA | NA | NA |
| Gm14252 | 0.106056992 | 0.998850474 | 3.724945049 | 0.268151734 |
| Gm14253 | 61.94935748 | -0.087777368 | 0.194284117 | -0.451798989 |
| Gm14170 | 0.179599912 | -0.847875541 | 2.728846284 | -0.310708429 |
| Gm22316 | 0.234323835 | 0.14026979 | 2.52962103 | 0.055450911 |
| Rpl21-ps9 | 0.666172099 | 0.416389617 | 1.330629662 | 0.312926751 |
| 2900097C17Rik | 2086.303339 | 0.479168609 | 0.092553878 | 5.177185656 |
| Gm14173 | 0.760931553 | -1.55042181 | 1.141822137 | -1.357848792 |
| Gm22758 | 0 | NA | NA | NA |
| Epb4.1l1 | 5881.508901 | 0.094482927 | 0.078305223 | 1.206598018 |
| Gm14225 | 0 | NA | NA | NA |
| Gm14224 | 0.038981395 | -0.443816764 | 3.724945049 | -0.119147198 |
| Aar2 | 389.6193864 | -0.029917221 | 0.071300159 | -0.419595441 |
| Gm14168 | 20.52960382 | 0.045270015 | 0.301559892 | 0.150119484 |
| Gm14169 | 37.69901258 | 0.192316333 | 0.218772371 | 0.879070481 |
| Dlgap4 | 2838.730653 | 0.144306378 | 0.101239919 | 1.425390091 |
| Gm14174 | 0.748568137 | 1.552111256 | 1.23396235 | 1.257827077 |

| | | | | |
|---|---|---|---|---|
| Gm14172 | 0 | NA | NA | NA |
| 4930405A21Rik | 7.408513515 | -0.105516894 | 0.375270149 | -0.281175826 |
| Myl9 | 37.34594635 | -0.158872012 | 0.271688127 | -0.584758758 |
| Gm14231 | 0.108266322 | -0.955981391 | 3.722845977 | -0.25678779 |
| Gm14230 | 5.976257995 | -0.734526618 | 0.425302001 | -1.727070685 |
| Tgif2 | 38.14849271 | -0.628362441 | 0.294636475 | -2.132670236 |
| 5430405H02Rik | 85.38075778 | -0.058152299 | 0.174128369 | -0.333962231 |
| Gm37790 | 6.47799121 | 0.206705796 | 0.441620612 | 0.46806193 |
| 4930518I15Rik | 26.40534196 | -0.249676335 | 0.264939462 | -0.942390136 |
| Gm27651 | 0 | NA | NA | NA |
| 1110008F13Rik | 336.8881828 | -0.423254792 | 0.1112646 | -3.804038239 |
| Sla2 | 15.40209609 | -0.889857891 | 0.34728071 | -2.562359111 |
| Gm14278 | 0.888132063 | -0.6501677 | 1.267103395 | -0.513113375 |
| Gm14248 | 0 | NA | NA | NA |
| Gm14247 | 0.534422126 | 1.039416014 | 1.753684618 | 0.592704072 |
| Ndrg3 | 3682.906368 | 0.440861431 | 0.078371872 | 5.625250732 |
| Gm27653 | 9.457828482 | 0.962749064 | 0.469107949 | 2.052297485 |
| Gm14276 | 8.337030673 | -0.072605681 | 0.380328281 | -0.19090266 |
| Dsn1 | 45.4075692 | -0.064580688 | 0.204057361 | -0.316483011 |
| Soga1 | 1303.081876 | 0.194951787 | 0.120396933 | 1.619242134 |
| Tldc2 | 4.0710617 | 0.431540877 | 0.537817452 | 0.802392849 |
| Samhd1 | 1799.230623 | -1.087559527 | 0.181734546 | -5.984330179 |
| Rbl1 | 152.3466098 | -0.631198693 | 0.150345086 | -4.198332715 |
| Gm14277 | 3.35648137 | -0.640196705 | 0.530072523 | -1.207753046 |
| Mroh8 | 16.34303778 | 0.167374262 | 0.301165269 | 0.555755526 |
| Rpn2 | 1339.45037 | -0.30986006 | 0.076828579 | -4.033135375 |
| Ghrh | 3.638464397 | -0.380758239 | 0.590713302 | -0.644573666 |
| Manbal | 220.2483533 | 0.098848234 | 0.128106093 | 0.771612271 |
| Src | 1335.75724 | 0.041451889 | 0.062131825 | 0.66716034 |
| Gm14279 | 2.044822302 | 0.141599085 | 0.611667495 | 0.231496829 |
| Gm14287 | 40.93809312 | 0.099747193 | 0.259059871 | 0.385035292 |
| Gm14286 | 22.25133751 | 0.976347641 | 0.361907178 | 2.697784679 |
| Blcap | 862.3755201 | -0.137637931 | 0.063474142 | -2.168409462 |
| Nnat | 1274.354139 | -0.029768937 | 0.16777516 | -0.17743352 |
| Gm23134 | 19.82322296 | 0.382097068 | 0.455248161 | 0.839316006 |
| Platr27 | 0.11147366 | -0.955981391 | 3.722845977 | -0.25678779 |
| Gm25407 | 0 | NA | NA | NA |
| Gm23463 | 0 | NA | NA | NA |
| Ctnnbl1 | 279.0876754 | -0.087139109 | 0.081900329 | -1.063965311 |
| Gm14176 | 7.679034499 | -0.120326503 | 0.35809781 | -0.336015747 |
| Vstm2l | 466.6791405 | 0.276360203 | 0.099720241 | 2.771355141 |
| Tti1 | 247.1897018 | -0.412095388 | 0.1215099 | -3.391455259 |
| Rprd1b | 501.6807887 | -0.089399828 | 0.066474113 | -1.344881837 |
| 2010009K17Rik | 0.075043205 | 0.517957688 | 3.724945049 | 0.139051095 |
| Tgm2 | 2012.035572 | -1.287681594 | 0.326325986 | -3.945997715 |
| Gm20412 | 1.717954688 | -0.276719091 | 1.158431567 | -0.238873921 |
| D630003M21Rik | 59.49929145 | -0.110763195 | 0.18848011 | -0.587665168 |

| | | | | |
|---|---|---|---|---|
| 1700060C20Rik | 0.29895891 | 0.391684908 | 2.018578617 | 0.194039957 |
| Mir7685 | 0 | NA | NA | NA |
| Bpi | 0.044544648 | 0.517957688 | 3.724945049 | 0.139051095 |
| Lbp | 183.630573 | -0.825811853 | 0.245085899 | -3.369479256 |
| Snhg17 | 148.0843631 | -0.134983456 | 0.186541172 | -0.723612138 |
| Gm24411 | 0 | NA | NA | NA |
| Gm26003 | 0 | NA | NA | NA |
| Gm23925 | 0 | NA | NA | NA |
| Snhg11 | 9158.007078 | 0.913684789 | 0.334622901 | 2.730490908 |
| Gm25187 | 0 | NA | NA | NA |
| Gm25129 | 29.82494218 | 0.776213456 | 0.30374526 | 2.555475124 |
| Gm14201 | 0 | NA | NA | NA |
| Gm14202 | 0 | NA | NA | NA |
| Ralgapb | 2672.82956 | 0.125294775 | 0.052913269 | 2.367927329 |
| Gm14200 | 0.166639798 | 0.51797152 | 2.767997395 | 0.187128616 |
| Gm24379 | 4.859614805 | 0.139666812 | 0.486930291 | 0.286831225 |
| Adig | 1.688935393 | -1.779319319 | 0.967811231 | -1.838498317 |
| Arhgap40 | 1.707670524 | -1.350315406 | 0.933177791 | -1.447007653 |
| Gm14204 | 129.1982071 | -0.236672913 | 0.151205559 | -1.565239499 |
| Slc32a1 | 1229.442361 | 0.108681585 | 0.062959196 | 1.72622256 |
| Gm14205 | 1.579080267 | -0.61236964 | 0.801093358 | -0.764417323 |
| Actr5 | 113.0741844 | -0.074784663 | 0.171009667 | -0.437312485 |
| Mir3474 | 0.201505633 | 0.03707581 | 2.605863092 | 0.014227843 |
| Ppp1r16b | 2425.103286 | 0.167554106 | 0.077642871 | 2.158010197 |
| Gm26141 | 1.537427175 | -1.005106032 | 0.939575204 | -1.069745166 |
| Fam83d | 51.99511303 | -0.038122001 | 0.218657054 | -0.174346083 |
| Dhx35 | 216.8272346 | -0.129519438 | 0.111575897 | -1.160819148 |
| AL669913.1 | 0.255320825 | 1.936461352 | 2.379358218 | 0.813858686 |
| Gm25090 | 0 | NA | NA | NA |
| Gm14219 | 0.112288909 | 0.037060453 | 3.724945049 | 0.009949262 |
| Gm11445 | 0 | NA | NA | NA |
| Gm11446 | 0 | NA | NA | NA |
| Gm14223 | 0 | NA | NA | NA |
| 4933416E03Rik | 0 | NA | NA | NA |
| Gm14220 | 0 | NA | NA | NA |
| 9130015L21Rik | 0 | NA | NA | NA |
| Gm826 | 0.8433133 | 1.276295359 | 1.049110045 | 1.216550509 |
| Mafb | 493.6009209 | -1.044914158 | 0.169009661 | -6.182570595 |
| Gm14221 | 0.304607851 | -0.931635881 | 1.976727818 | -0.471302054 |
| Gm14222 | 0 | NA | NA | NA |
| Gm23968 | 0.062426873 | 0.517959912 | 3.724945049 | 0.139051692 |
| Top1 | 1394.007593 | -0.191033808 | 0.063703728 | -2.99878539 |
| Plcg1 | 938.5103693 | 0.187954458 | 0.06889255 | 2.728226177 |
| Gm27206 | 3.426664286 | 0.679076448 | 0.533521782 | 1.272818602 |
| Zhx3 | 1591.941502 | 0.167629814 | 0.053349934 | 3.142080987 |
| Gm16751 | 10.07322116 | -0.484208106 | 0.338626166 | -1.429919349 |
| Gm11447 | 0.089814205 | 0.517955464 | 3.724945049 | 0.139050498 |

| | | | | |
|---|---|---|---|---|
| Lpin3 | 58.0580359 | -1.140671085 | 0.194585331 | -5.862061036 |
| Emilin3 | 11.35137183 | 0.237296196 | 0.336994184 | 0.704155165 |
| Gm14228 | 0.449436652 | 2.281081716 | 3.689435046 | 0.618273987 |
| Chd6 | 2195.755601 | 0.207762925 | 0.069341754 | 2.99621674 |
| Gm25561 | 0 | NA | NA | NA |
| Gm14243 | 0.04393175 | 0.037062677 | 3.724945049 | 0.009949859 |
| Gm14242 | 0 | NA | NA | NA |
| Gm5270 | 0.106618422 | 0.517955464 | 3.724945049 | 0.139050498 |
| Gm11448 | 0.467761363 | -0.345537017 | 1.428928178 | -0.241815525 |
| Gm14241 | 0 | NA | NA | NA |
| Gm11449 | 0.644287204 | 0.938453353 | 1.191523256 | 0.787608087 |
| Gm29229 | 0.291544834 | -1.327439066 | 1.597590527 | -0.830900687 |
| Ptprt | 2587.731476 | 0.428710663 | 0.077069468 | 5.562652432 |
| Ptprtos | 0.352863859 | -1.060668871 | 1.86454058 | -0.568863388 |
| 9430021M05Rik | 23.91964892 | 1.019625086 | 0.272436578 | 3.742614501 |
| Gm14246 | 8.592019774 | 1.032955685 | 0.435170278 | 2.373681609 |
| Gm22936 | 0 | NA | NA | NA |
| Gm11450 | 0 | NA | NA | NA |
| Gm24478 | 0 | NA | NA | NA |
| Gm14255 | 0.044191549 | 0.037062677 | 3.724945049 | 0.009949859 |
| Gm11451 | 16.98984007 | -0.165252971 | 0.310504064 | -0.532208723 |
| Gm11452 | 0.150711795 | 0.037073253 | 2.856704404 | 0.01297763 |
| Srsf6 | 1346.747418 | -0.082367543 | 0.082129192 | -1.002902146 |
| L3mbtl1 | 55.09161921 | 0.598286904 | 0.378988451 | 1.578641522 |
| Gm11453 | 0.038981395 | -0.443816764 | 3.724945049 | -0.119147198 |
| Sgk2 | 21.73993536 | 0.825829699 | 0.304239816 | 2.714403751 |
| Ift52 | 322.3681409 | -0.153887776 | 0.07694805 | -1.999891804 |
| Mybl2 | 17.83251159 | -0.62264245 | 0.292197723 | -2.130894258 |
| Mir7003 | 0.049553047 | -0.443818988 | 3.724945049 | -0.119147795 |
| Gtsf1l | 0.489956056 | 0.074715674 | 1.333626614 | 0.056024432 |
| Gm22174 | 0 | NA | NA | NA |
| Gm11454 | 2.386488514 | 0.093041209 | 0.690868376 | 0.134672844 |
| Gm14254 | 0 | NA | NA | NA |
| Tox2 | 401.2220022 | 0.166594513 | 0.105137725 | 1.584536026 |
| Jph2 | 9.352170419 | -0.310705575 | 0.363989505 | -0.853611356 |
| Oser1 | 111.6600522 | -0.422873318 | 0.133774777 | -3.161084067 |
| 2900093K20Rik | 14.33683216 | -0.555749245 | 0.294261469 | -1.888623904 |
| Gdap1l1 | 668.5251384 | 0.078656899 | 0.090632775 | 0.86786374 |
| Fitm2 | 424.7360197 | -0.270317879 | 0.067564593 | -4.000880721 |
| 2310001K24Rik | 0.198365723 | -0.443823723 | 2.645106319 | -0.167790504 |
| R3hdml | 0.349069061 | 0.074337802 | 1.77477479 | 0.041885766 |
| Hnf4aos | 0 | NA | NA | NA |
| Hnf4a | 0.497304619 | -0.942938992 | 1.500798316 | -0.628291611 |
| Gm25881 | 0 | NA | NA | NA |
| Ttpal | 655.7681734 | -0.115771827 | 0.089968228 | -1.286807911 |
| Serinc3 | 3221.802251 | -0.207111715 | 0.071924673 | -2.879564215 |
| 0610039K10Rik | 0.721809532 | -0.15848364 | 1.063119411 | -0.149074167 |

| | | | | |
|---|---|---|---|---|
| Pkig | 413.2230987 | -0.208361122 | 0.078336952 | -2.659806331 |
| Gm16316 | 3.804176175 | -1.199398636 | 0.50880187 | -2.357299977 |
| Ada | 48.39031174 | -0.700205701 | 0.230913174 | -3.032333276 |
| Wisp2 | 23.40200403 | -1.905693763 | 0.430048628 | -4.431344832 |
| Kcnk15 | 8.160985404 | -1.079621288 | 0.533825329 | -2.022424245 |
| Rims4 | 1059.645651 | 0.012551739 | 0.083566514 | 0.150200578 |
| Ywhab | 4440.302886 | 0.156763373 | 0.059130943 | 2.651122499 |
| Pabpc1l | 10.92999753 | -0.55066357 | 0.448039757 | -1.229050686 |
| Tomm34 | 543.6160348 | 0.269875688 | 0.070655696 | 3.819588574 |
| Stk4 | 685.9737304 | -0.294357423 | 0.076878941 | -3.828843382 |
| Gm11455 | 4.303662858 | 0.365480336 | 0.632627286 | 0.577718262 |
| n-R5s207 | 0 | NA | NA | NA |
| Kcns1 | 75.78496726 | 0.33607156 | 0.205788201 | 1.633094412 |
| Wfdc5 | 0.042716945 | 0.037062677 | 3.724945049 | 0.009949859 |
| A730032A03Rik | 0.162822621 | -1.238016119 | 3.396660554 | -0.364480377 |
| Wfdc12 | 0.231631338 | -0.43871099 | 2.122910229 | -0.20665546 |
| Wfdc15a | 0.066050875 | -0.443818988 | 3.724945049 | -0.119147795 |
| Wfdc15b | 0 | NA | NA | NA |
| Svs2 | 0 | NA | NA | NA |
| Svs3b | 0.141331096 | -0.443820795 | 3.561840904 | -0.124604329 |
| Svs4 | 0 | NA | NA | NA |
| Svs3a | 0 | NA | NA | NA |
| Svs6 | 0.038981395 | -0.443816764 | 3.724945049 | -0.119147198 |
| Svs5 | 0.282943117 | -1.583212474 | 2.020630507 | -0.783523989 |
| Slpi | 106.5214453 | -2.321195203 | 0.508446679 | -4.565267706 |
| Mir7678 | 0.061920613 | -0.443816764 | 3.724945049 | -0.119147198 |
| Matn4 | 90.53204317 | -0.307498759 | 0.18135671 | -1.695546638 |
| Rbpjl | 13.41464962 | -0.022608899 | 0.330331317 | -0.068443097 |
| Sdc4 | 2785.760239 | -0.778765535 | 0.251536973 | -3.0960281 |
| Sys1 | 225.5397579 | -0.196429259 | 0.081901614 | -2.398356396 |
| Gm14302 | 1.474490841 | 1.044358187 | 0.881657376 | 1.184539727 |
| Gm20458 | 0.478777135 | 0.768889654 | 1.294530684 | 0.593952437 |
| Trp53tg5 | 0.109982625 | -0.443823436 | 3.724945049 | -0.119148989 |
| Dbndd2 | 527.4245514 | 0.078205973 | 0.093498344 | 0.836442331 |
| Pigt | 917.1375041 | 0.015599096 | 0.083170138 | 0.187556458 |
| Gm14317 | 0.567797178 | 1.187530343 | 1.263188247 | 0.940105599 |
| Wfdc2 | 6.468348977 | -0.055717371 | 0.419476477 | -0.132825972 |
| Spint3 | 0 | NA | NA | NA |
| Wfdc6a | 0.062426873 | 0.517959912 | 3.724945049 | 0.139051692 |
| Eppin | 0.044544648 | 0.517957688 | 3.724945049 | 0.139051095 |
| Wfdc8 | 0.123845245 | -0.280519638 | 3.720871772 | -0.075390837 |
| Gm25512 | 0 | NA | NA | NA |
| Wfdc6b | 0 | NA | NA | NA |
| Gm24473 | 0 | NA | NA | NA |
| Wfdc16 | 0.038981395 | -0.443816764 | 3.724945049 | -0.119147198 |
| Gm11456 | 0 | NA | NA | NA |
| Wfdc9 | 0.046345709 | -0.443818988 | 3.724945049 | -0.119147795 |

| | | | | |
|---|---|---|---|---|
| Wfdc10 | 0.171131556 | -0.444767681 | 3.383981516 | -0.131433248 |
| Wfdc11 | 0 | NA | NA | NA |
| Wfdc13 | 0 | NA | NA | NA |
| Spint4 | 0 | NA | NA | NA |
| Spint5 | 0 | NA | NA | NA |
| Wfdc3 | 9.036228213 | -0.767911108 | 0.364124217 | -2.108926219 |
| Dnttip1 | 245.2841238 | -0.348361742 | 0.107807628 | -3.231327398 |
| Gm11457 | 0.366597908 | 0.034095102 | 1.585493799 | 0.021504406 |
| Gm22596 | 5.545541213 | 0.169695328 | 0.49960396 | 0.339659693 |
| Ube2c | 12.17535567 | -0.588767206 | 0.423861244 | -1.389056476 |
| Tnnc2 | 0.984947528 | -0.387599105 | 1.164051017 | -0.332974328 |
| Snx21 | 316.0422822 | 0.173987808 | 0.073034898 | 2.382255786 |
| Acot8 | 121.1509639 | -0.574952665 | 0.112258623 | -5.121679304 |
| Zswim3 | 75.20523749 | -0.463462457 | 0.183421335 | -2.526764166 |
| Zswim1 | 246.2299447 | 0.144001401 | 0.075033584 | 1.919159313 |
| Spata25 | 2.304449883 | -0.171212178 | 0.807834651 | -0.211939631 |
| Neurl2 | 13.0448324 | 0.350336436 | 0.3024964 | 1.158150762 |
| Ctsa | 1093.099412 | -0.399541615 | 0.092183302 | -4.33420811 |
| Pltp | 377.8729552 | -0.451799331 | 0.180094535 | -2.508678743 |
| Pcif1 | 806.4495592 | 0.188051896 | 0.092490988 | 2.033191549 |
| Zfp335 | 457.5970439 | 0.044282249 | 0.10212408 | 0.43361222 |
| Zfp335os | 35.33703191 | 0.09833144 | 0.199910332 | 0.491877728 |
| Mmp9 | 18.18569822 | 0.231802563 | 0.254622775 | 0.910376392 |
| Slc12a5 | 6336.594774 | 0.579519736 | 0.073660082 | 7.867487035 |
| Ncoa5 | 554.329722 | -0.391063276 | 0.102183408 | -3.827072154 |
| Cd40 | 64.99215734 | -0.931842192 | 0.26943414 | -3.458515658 |
| 1700025C18Rik | 0 | NA | NA | NA |
| Cdh22 | 506.5456601 | 0.076887127 | 0.106682318 | 0.720711065 |
| Gm14397 | 0 | NA | NA | NA |
| Gm25569 | 0 | NA | NA | NA |
| Slc35c2 | 388.3991378 | 0.079749235 | 0.063264158 | 1.26057531 |
| Elmo2 | 1659.397843 | 0.254412008 | 0.071790933 | 3.543790244 |
| Zfp663 | 0.749070597 | -1.65201183 | 1.416678556 | -1.166116211 |
| Zfp334 | 573.5096397 | -0.006450672 | 0.062041735 | -0.103973111 |
| Ocstamp | 4.715999993 | -1.070858326 | 0.715056798 | -1.497584987 |
| Slc13a3 | 348.6971527 | 0.063980647 | 0.130242473 | 0.491242566 |
| Gm14437 | 0 | NA | NA | NA |
| Trp53rka | 121.9068042 | -0.203407042 | 0.141581079 | -1.43668238 |
| Slc2a10 | 29.04143457 | -0.558966534 | 0.26749095 | -2.08966522 |
| Eya2 | 64.95457155 | -0.178329405 | 0.171105762 | -1.042217418 |
| Gm28163 | 0 | NA | NA | NA |
| Gm11460 | 0.325497449 | 1.019139761 | 1.593812955 | 0.639434984 |
| Zmynd8 | 1768.666945 | 0.026456246 | 0.0947975 | 0.279081687 |
| BC046401 | 0.509709817 | -0.136739459 | 1.286833928 | -0.106260378 |
| Gm11465 | 2.089642419 | 0.546730374 | 0.706209865 | 0.774175499 |
| Gm22692 | 0.117907917 | 0.967119648 | 3.722483401 | 0.259804959 |
| Gm11461 | 0 | NA | NA | NA |

| | | | | |
|---|---|---|---|---|
| Gm11462 | 0 | NA | NA | NA |
| Platr29 | 0 | NA | NA | NA |
| Gm11464 | 0.663210203 | 0.030893234 | 1.135413767 | 0.027208789 |
| Ncoa3 | 801.9318103 | -0.40989985 | 0.093040769 | -4.405593965 |
| Sulf2 | 1316.311321 | -0.209685999 | 0.063249309 | -3.3152299 |
| Gm22687 | 0 | NA | NA | NA |
| Gm11466 | 0 | NA | NA | NA |
| Gm11467 | 0 | NA | NA | NA |
| Gm11468 | 0.423089777 | 0.56105284 | 1.451385614 | 0.386563595 |
| Gm14268 | 0.409889595 | -1.70563229 | 1.603372283 | -1.063778081 |
| 5031425F14Rik | 0.951684498 | -1.10609071 | 1.025662733 | -1.078415618 |
| Gm11469 | 0.922271222 | -1.633478646 | 1.103300466 | -1.480538345 |
| Gm14267 | 0.809026523 | 1.313955607 | 1.172500447 | 1.120644013 |
| Prex1 | 2399.621753 | -0.419047808 | 0.071275292 | -5.879285772 |
| Gm23152 | 1.245355593 | -0.949423605 | 1.104292833 | -0.859757101 |
| Gm25265 | 0 | NA | NA | NA |
| Gm24477 | 0 | NA | NA | NA |
| Trp53rkb | 54.71816161 | 0.067303214 | 0.153485829 | 0.438497897 |
| Arfgef2 | 1429.215814 | -0.141756037 | 0.044295465 | -3.200238152 |
| Cse1l | 859.0610123 | -0.076661118 | 0.069882234 | -1.097004397 |
| Gm17096 | 7.764297659 | 0.789773296 | 0.536990935 | 1.4707386 |
| Stau1 | 911.3828237 | -0.101092234 | 0.069820749 | -1.447882408 |
| Gm11470 | 0.342301797 | 1.532513677 | 3.697442554 | 0.414479375 |
| Ddx27 | 291.9835414 | -0.16176701 | 0.121838604 | -1.327715557 |
| Znfx1 | 5280.077714 | -0.58813787 | 0.166286754 | -3.536889472 |
| 1500012F01Rik | 269.2484598 | -0.079326333 | 0.142319701 | -0.557381247 |
| Gm27494 | 0 | NA | NA | NA |
| Gm25878 | 0 | NA | NA | NA |
| Gm23201 | 0 | NA | NA | NA |
| Snord12 | 0 | NA | NA | NA |
| Gm11472 | 0.906216104 | 1.696410254 | 1.096256605 | 1.54745727 |
| Gm11471 | 0.062426873 | 0.517959912 | 3.724945049 | 0.139051692 |
| Gm14290 | 607.9128014 | 0.466032281 | 0.176951568 | 2.633671384 |
| Kcnb1 | 1138.507184 | 0.427539448 | 0.149202969 | 2.865488866 |
| 1110018N20Rik | 50.53414942 | 0.216842775 | 0.219419202 | 0.988257971 |
| Ptgis | 12.61769256 | -0.106098246 | 0.28222616 | -0.375933421 |
| Gm14291 | 10.76029111 | 0.64575427 | 0.387643808 | 1.665844409 |
| Gm11473 | 0.150086409 | 1.027495679 | 3.723011807 | 0.275985071 |
| B4galt5 | 1227.37508 | -0.870161227 | 0.106781863 | -8.148960916 |
| Slc9a8 | 501.9560524 | 0.001262883 | 0.075596111 | 0.016705666 |
| Spata2 | 691.9053702 | 0.330027452 | 0.088903645 | 3.712192585 |
| Rnf114 | 1225.538309 | -0.247845649 | 0.121045744 | -2.047537085 |
| Gm11474 | 0.049553047 | -0.443818988 | 3.724945049 | -0.119147795 |
| Snai1 | 13.66662012 | -0.847644007 | 0.318052539 | -2.665106869 |
| Gm11476 | 0.445249545 | -0.037599733 | 1.439507989 | -0.02611985 |
| Gm11475 | 0 | NA | NA | NA |
| Ube2v1 | 21.15999022 | -0.149883853 | 0.243853393 | -0.614647396 |

| | | | | |
|---|---|---|---|---|
| Gm20431 | 0 | NA | NA | NA |
| Tmem189 | 432.2688695 | -0.277156905 | 0.093223578 | -2.973034398 |
| Gm22355 | 0.959597608 | 0.520389842 | 1.056794075 | 0.492423126 |
| Gm14320 | 1.589793752 | -0.736209008 | 0.920411025 | -0.799869828 |
| Cebpb | 250.3011784 | -1.15523455 | 0.278244346 | -4.151870706 |
| A530013C23Rik | 6.464581192 | -0.616590289 | 0.495979684 | -1.243176502 |
| Gm14321 | 4.54434231 | -0.260146786 | 0.578117032 | -0.449989832 |
| 9230111E07Rik | 2.522272708 | 0.754743341 | 0.672637237 | 1.122065951 |
| 1200007C13Rik | 1.134624916 | 0.963199466 | 1.139941707 | 0.844955019 |
| Gm14319 | 0.048949532 | 0.037064902 | 3.724945049 | 0.009950456 |
| Ptpn1 | 822.4208813 | -0.302751462 | 0.105538363 | -2.868638981 |
| Fam65c | 5.824423608 | 0.757804012 | 0.530491661 | 1.428493731 |
| Gm14236 | 0.314827273 | 1.355072703 | 1.55504208 | 0.871405809 |
| Gm24327 | 0 | NA | NA | NA |
| Pard6b | 76.59336974 | -0.1979268 | 0.206460003 | -0.95866898 |
| Gm14235 | 0.229025006 | -0.427905877 | 2.131143139 | -0.200787019 |
| E130018N17Rik | 1.228602664 | 0.477389403 | 1.118525796 | 0.426802319 |
| Gm24561 | 2.66823802 | 1.231948224 | 0.675994514 | 1.822423405 |
| Adnp | 84.6346395 | -0.140460567 | 0.183053692 | -0.767318948 |
| Gm20716 | 2.018522609 | 0.262782279 | 0.702974916 | 0.373814589 |
| Dpm1 | 240.6952631 | 0.473035137 | 0.19246774 | 2.457737267 |
| Mocs3 | 110.2099467 | -0.439475072 | 0.152860619 | -2.87500518 |
| Gm14237 | 0.075043205 | 0.517957688 | 3.724945049 | 0.139051095 |
| Kcng1 | 163.4779742 | 0.206564158 | 0.20336811 | 1.015715583 |
| Gm14262 | 0 | NA | NA | NA |
| Gm22704 | 0.118974955 | 0.51795324 | 3.724945049 | 0.139049901 |
| Gm14234 | 0.075043205 | 0.517957688 | 3.724945049 | 0.139051095 |
| Nfatc2 | 337.8978929 | -0.09629353 | 0.086766806 | -1.109796872 |
| Atp9a | 3839.633382 | 0.303324413 | 0.039043269 | 7.768929654 |
| mmu-mir-7004 | 0.834414618 | -0.109568523 | 1.246835044 | -0.08787732 |
| Sall4 | 2.654068103 | 0.25553253 | 0.743764642 | 0.343566386 |
| Gm14261 | 0.095897396 | 0.037062677 | 3.724945049 | 0.009949859 |
| Gm25214 | 0.150086409 | 1.027495679 | 3.723011807 | 0.275985071 |
| Zfp64 | 184.749999 | 0.070856796 | 0.110508894 | 0.641186363 |
| 1700017J07Rik | 0.04393175 | 0.037062677 | 3.724945049 | 0.009949859 |
| Gm14260 | 0 | NA | NA | NA |
| Gm9873 | 2.521204246 | 0.377607442 | 0.655103486 | 0.576408842 |
| Gm26469 | 0.075043205 | 0.517957688 | 3.724945049 | 0.139051095 |
| Gm14258 | 0.182681975 | 0.079779833 | 2.723875079 | 0.029289094 |
| 9430093N23Rik | 0 | NA | NA | NA |
| Gm14250 | 0 | NA | NA | NA |
| 4930529I22Rik | 0 | NA | NA | NA |
| Gm14249 | 0.152853976 | 1.404519388 | 2.820813914 | 0.497912812 |
| Gm11011 | 0 | NA | NA | NA |
| Gm34294 | 5.209546091 | 0.051140888 | 0.541277525 | 0.094481825 |
| Tshz2 | 720.7148521 | 0.493364993 | 0.107403889 | 4.593548679 |
| Gm26883 | 2.358426798 | -0.364283968 | 0.652463782 | -0.558320597 |

| | | | | |
|---|---|---|---|---|
| AY702102 | 0 | NA | NA | NA |
| A630075F10Rik | 0 | NA | NA | NA |
| Zfp217 | 142.5419381 | -0.4876218 | 0.14967336 | -3.257906426 |
| Gm17619 | 0.100902008 | -0.876201588 | 3.72093227 | -0.235479048 |
| Gm14270 | 11.75654133 | -0.473429165 | 0.292624535 | -1.617872424 |
| Bcas1 | 1958.212372 | 0.093968674 | 0.142068228 | 0.661433424 |
| Bcas1os2 | 1.937199938 | -0.193323365 | 0.832978209 | -0.232086941 |
| Bcas1os1 | 3.671955489 | -0.511447191 | 0.559528246 | -0.914068583 |
| Gm14269 | 52.69193393 | 0.079173083 | 0.167112922 | 0.473769963 |
| Cyp24a1 | 1.170018213 | -0.301844416 | 1.052434608 | -0.286805863 |
| Pfdn4 | 118.4952192 | 0.016004091 | 0.192863513 | 0.082981435 |
| Gm16796 | 10.39617679 | -0.198319236 | 0.357604574 | -0.554576899 |
| Gm14637 | 2.883279791 | 0.716961244 | 0.5739519 | 1.249166078 |
| 4930470P17Rik | 0.106465261 | 0.037073798 | 3.724945049 | 0.009952844 |
| Gm14264 | 0.597452826 | -0.695876571 | 1.353836487 | -0.51400341 |
| Dok5 | 201.9773433 | -0.996108529 | 0.15020989 | -6.631444367 |
| Gm14263 | 0 | NA | NA | NA |
| Gm14266 | 0.20369824 | 0.735459297 | 3.19191532 | 0.230413161 |
| Gm14640 | 0 | NA | NA | NA |
| 1700007M16Rik | 0.088534442 | -0.905414734 | 3.723500899 | -0.243162217 |
| Gm26489 | 0 | NA | NA | NA |
| 1700028P15Rik | 0 | NA | NA | NA |
| Gm14272 | 0 | NA | NA | NA |
| Gm14641 | 0 | NA | NA | NA |
| Cbln4 | 171.4002324 | 0.212108902 | 0.139918466 | 1.515946455 |
| Gm14271 | 0.25568908 | -0.965554221 | 1.825923532 | -0.528803208 |
| Gm14274 | 0 | NA | NA | NA |
| AL844846.1 | 0 | NA | NA | NA |
| Mc3r | 7.192941036 | -0.173155984 | 0.432289098 | -0.400555983 |
| Gm14275 | 0 | NA | NA | NA |
| Gm14273 | 0.454387946 | -1.827745216 | 1.479020834 | -1.235780574 |
| Fam210b | 297.4704023 | 0.154722907 | 0.098878459 | 1.564778711 |
| Aurka | 15.8043844 | -0.476335099 | 0.30272295 | -1.573501773 |
| Cstf1 | 247.1233043 | -0.119867378 | 0.080796277 | -1.483575505 |
| Cass4 | 5.704398591 | -0.926071524 | 0.495517297 | -1.868898479 |
| Gm14455 | 6.260709098 | -0.062596995 | 0.454689806 | -0.137669669 |
| Rtfdc1 | 1259.401244 | -0.209677958 | 0.064163885 | -3.267850128 |
| Gcnt7 | 0.721712508 | -1.858200463 | 1.247117306 | -1.489996534 |
| Fam209 | 5.869341775 | 0.564740578 | 0.542342315 | 1.041299123 |
| Gm14303 | 72.23225202 | -0.253890535 | 0.192769027 | -1.317071206 |
| Tfap2c | 3.754962934 | -0.114294819 | 0.720202355 | -0.158698202 |
| Gm22773 | 0 | NA | NA | NA |
| Bmp7 | 187.5782872 | -0.355554385 | 0.19580776 | -1.815833985 |
| Spo11 | 3.986507545 | 0.47093848 | 0.528305443 | 0.891413265 |
| Rae1 | 377.0248253 | 0.153966916 | 0.081604546 | 1.886744343 |
| Rbm38 | 50.55568089 | -0.477973766 | 0.188923077 | -2.529991431 |
| Gm20490 | 0 | NA | NA | NA |

| | | | | |
|---|---|---|---|---|
| Gm14453 | 0 | NA | NA | NA |
| Ctcfl | 0.389334344 | -0.722809375 | 1.426188853 | -0.506811825 |
| Ctcflos | 0.803203798 | 0.822062577 | 1.101436019 | 0.746355269 |
| Pck1 | 0.852421781 | -1.084943028 | 1.336794299 | -0.81160058 |
| Zbp1 | 1311.11427 | -1.574799307 | 0.349257144 | -4.508996692 |
| Pmepa1 | 867.4379167 | -0.965038187 | 0.158579794 | -6.085505365 |
| Pmepa1os | 1.650675276 | -0.572772317 | 0.73026873 | -0.784330882 |
| Gm14642 | 0 | NA | NA | NA |
| 1700021F07Rik | 0 | NA | NA | NA |
| Ankrd60 | 0.203084296 | -0.443823099 | 2.648251873 | -0.167590969 |
| Ppp4r1l-ps | 195.0349838 | 0.022670466 | 0.157762794 | 0.143699702 |
| Rab22a | 871.4199496 | -0.121003712 | 0.074155408 | -1.631758433 |
| Mir6340 | 0 | NA | NA | NA |
| 1700010B08Rik | 0.15795635 | 0.037074776 | 2.815015606 | 0.013170363 |
| Vapb | 1573.7862 | 0.049514903 | 0.062245044 | 0.795483468 |
| Gm14371 | 0 | NA | NA | NA |
| Gm14336 | 0 | NA | NA | NA |
| Atp5k-ps2 | 1.191345417 | -0.554839342 | 0.967364386 | -0.573557751 |
| Vamp7-ps | 9.408148121 | 0.287336724 | 0.406438284 | 0.706962744 |
| Stx16 | 905.3860616 | 0.125660674 | 0.071818228 | 1.749704477 |
| Npepl1 | 158.3376538 | -0.445327139 | 0.135892554 | -3.277053263 |
| Gm10714 | 1.585474913 | 1.051383582 | 1.06149619 | 0.990473251 |
| Gm27591 | 0 | NA | NA | NA |
| Mir296 | 0.229181457 | 0.517969277 | 2.603735265 | 0.198933157 |
| Mir298 | 0.18819539 | -0.044930485 | 3.278252393 | -0.013705621 |
| Nespas | 11.25411321 | 0.202994085 | 0.376240345 | 0.539533008 |
| Gnas | 10595.54034 | 0.052253699 | 0.079693601 | 0.655682494 |
| Gm27899 | 0.248714005 | 0.517971423 | 1.863371772 | 0.277975351 |
| Gm27869 | 0.247908139 | 0.535784769 | 2.143973186 | 0.249902738 |
| Gm27849 | 0.638233968 | 2.328508525 | 2.447989937 | 0.951192033 |
| Gm20721 | 37.27970981 | -0.290214543 | 0.271607443 | -1.068507326 |
| Nelfcd | 290.7180538 | 0.027366562 | 0.067083496 | 0.407947768 |
| Ctsz | 1102.778525 | -0.76135821 | 0.146432396 | -5.199383683 |
| Rpl30-ps5 | 0 | NA | NA | NA |
| Tubb1 | 0.687228199 | -1.57700301 | 1.167527992 | -1.350719658 |
| Atp5e | 358.6752708 | -0.352243656 | 0.138163795 | -2.549464247 |
| Slmo2 | 386.0940015 | -0.290597143 | 0.099853145 | -2.910245261 |
| Gm14385 | 0 | NA | NA | NA |
| Vmn1r-ps1 | 0.042716945 | 0.037062677 | 3.724945049 | 0.009949859 |
| Zfp831 | 392.3466882 | 0.375886617 | 0.191364531 | 1.964243923 |
| Edn3 | 25.09549224 | 1.273629469 | 0.445276187 | 2.860313456 |
| Gm14617 | 0.43206172 | -2.205008336 | 3.290709812 | -0.670070733 |
| Gm14618 | 0.490955376 | -1.181406121 | 3.691205594 | -0.320059691 |
| Gm14616 | 0 | NA | NA | NA |
| Gm14444 | 1.336470638 | -0.566037462 | 1.046702881 | -0.540781412 |
| Gm16354 | 2.08236629 | 1.69790991 | 0.967169895 | 1.755544624 |
| Gm14445 | 0.131100043 | 0.037055718 | 3.674543085 | 0.010084442 |

| | | | | |
|---|---|---|---|---|
| Gm14393 | 21.09049984 | 0.467911757 | 0.329663553 | 1.419361506 |
| Gm14439 | 0.167183452 | 0.517964184 | 2.765153692 | 0.187318407 |
| Gm14387 | 3.01954408 | -0.610180067 | 0.661420484 | -0.922529739 |
| Gm14392 | 2.964599863 | 0.254526677 | 0.630271611 | 0.403836493 |
| Gm14399 | 171.4815682 | 0.090650644 | 0.145616927 | 0.622528203 |
| Gm16355 | 0 | NA | NA | NA |
| Gm14443 | 181.6545351 | 0.088509944 | 0.202484247 | 0.437120147 |
| Gm14391 | 24.59774181 | 0.18637895 | 0.268865546 | 0.693205034 |
| Gm14442 | 0.132475575 | 0.037075359 | 2.973122152 | 0.012470177 |
| Gm14438 | 2.078804666 | 0.256901172 | 0.662798838 | 0.387600516 |
| Gm14440 | 1.419637689 | -1.137540585 | 0.891992387 | -1.275280598 |
| Gm14298 | 0 | NA | NA | NA |
| Gm4631 | 1.996170162 | -0.254665962 | 0.751405247 | -0.338919595 |
| Gm4723 | 1.755952801 | 2.988762654 | 0.970915066 | 3.07829465 |
| Gm14297 | 0 | NA | NA | NA |
| Gm4724 | 1.988766949 | -0.641018895 | 0.817540582 | -0.78408205 |
| Gm8923 | 0.572800032 | 0.606996253 | 1.347060046 | 0.450608164 |
| Gm11007 | 1.812554613 | 0.290494327 | 0.784557191 | 0.370265329 |
| Gm16359 | 0 | NA | NA | NA |
| Gm4245 | 2.26497711 | -0.600698239 | 0.765933091 | -0.784269862 |
| Gm2007 | 0.319719969 | 0.824128371 | 2.03565787 | 0.404846209 |
| Gm16358 | 0 | NA | NA | NA |
| Gm14288 | 19.90464905 | -0.124169228 | 0.331117004 | -0.375001061 |
| Rps8-ps2 | 2.50324314 | 0.291117419 | 0.70317297 | 0.414005418 |
| Gm2020 | 0.047948698 | 0.037064902 | 3.724945049 | 0.009950456 |
| Gm6710 | 28.49017795 | -0.350213199 | 0.249598895 | -1.403103962 |
| Gm16356 | 0 | NA | NA | NA |
| Gm8898 | 2.984855315 | -1.429910048 | 0.762701256 | -1.874797029 |
| Gm2026 | 2.773167693 | -1.472492019 | 0.873941618 | -1.684886025 |
| Gm11008 | 0.226297694 | 0.945557518 | 2.123090057 | 0.44536854 |
| Gm16357 | 0.13861434 | 0.037059817 | 3.618284367 | 0.010242373 |
| Gm2004 | 6.050452602 | 0.879408267 | 0.481080563 | 1.827985446 |
| Gm6182 | 0.19897497 | 0.971544339 | 2.216505307 | 0.438322586 |
| 2210418O10Rik | 24.2680911 | -0.243916207 | 0.337990227 | -0.721666448 |
| Gm17258 | 4.979667379 | -0.452303066 | 0.674511334 | -0.670564071 |
| Gm1980 | 0.288549458 | 0.020997735 | 1.637060886 | 0.012826484 |
| Gm11009 | 0.13733263 | 0.037057285 | 3.640803958 | 0.010178325 |
| Gm14435 | 0.646335998 | -0.340204685 | 1.289330815 | -0.26386144 |
| Gm16362 | 0 | NA | NA | NA |
| Gm14434 | 1.597696658 | -0.34058316 | 0.964079313 | -0.353272968 |
| Gm14431 | 1.363863261 | 2.084864356 | 1.008989671 | 2.066289096 |
| Gm14307 | 0 | NA | NA | NA |
| Gm14308 | 2.466484677 | -0.772418276 | 0.853695102 | -0.904794082 |
| Gm14306 | 4.680219553 | 0.064928724 | 0.525315779 | 0.123599417 |
| Gm14305 | 173.9135921 | 0.188369058 | 0.129327167 | 1.456531228 |
| Gm14301 | 0.675521499 | -0.651915013 | 1.483289359 | -0.439506297 |
| Gm14390 | 0.066050875 | -0.443818988 | 3.724945049 | -0.119147795 |

| | | | | |
|---|---|---|---|---|
| Gm14295 | 67.51412393 | -0.01764499 | 0.260197154 | -0.067813925 |
| Gm14408 | 0.284066144 | -1.300588538 | 2.090846072 | -0.622039353 |
| Gm14405 | 1.448471955 | 1.110850517 | 0.91719343 | 1.211140945 |
| Gm14296 | 30.11651545 | 0.508202788 | 0.308130578 | 1.649309821 |
| Mup-ps25 | 0 | NA | NA | NA |
| Gm14417 | 0.204225461 | 1.631603513 | 3.2048902 | 0.5090981 |
| Gm14419 | 4.87970569 | 0.320868501 | 0.479264793 | 0.669501507 |
| Gm14402 | 1.138899903 | -1.340975189 | 1.103805884 | -1.214865049 |
| Gm14421 | 5.323853793 | -0.073070118 | 0.48716625 | -0.149990107 |
| Gm14401 | 0.702992681 | 0.4671194 | 1.172624667 | 0.398353722 |
| Gm14407 | 3.984100659 | 0.547484525 | 0.573824308 | 0.954097827 |
| Gm14415 | 18.81040368 | -0.039064313 | 0.342677882 | -0.113997182 |
| Gm14411 | 16.94595019 | 0.276241001 | 0.331664132 | 0.832893807 |
| Gm14416 | 3.389752464 | 0.318276503 | 0.586506655 | 0.542664777 |
| Gm14413 | 14.03696361 | 0.328180755 | 0.327011824 | 1.003574583 |
| Gm14410 | 143.7066193 | 0.213664983 | 0.153821043 | 1.389049108 |
| Gm14400 | 0.439370512 | -0.78061378 | 1.309930441 | -0.595920024 |
| Gm14409 | 0.902861002 | 2.545945959 | 1.088205635 | 2.339581671 |
| Gm14412 | 64.87765343 | 0.71115906 | 0.241818182 | 2.940883331 |
| Gm14418 | 19.13716893 | -0.184225946 | 0.271920689 | -0.67749882 |
| Gm14404 | 0.1140246 | 0.037073798 | 3.724945049 | 0.009952844 |
| Gm14414 | 1.844525372 | -0.653741565 | 0.703747636 | -0.928943177 |
| Gm14420 | 72.73691108 | 0.067190107 | 0.194804465 | 0.344910505 |
| Gm14403 | 33.59942204 | -0.006634924 | 0.251726525 | -0.026357667 |
| Gm16363 | 0.136194719 | 0.037062677 | 3.724945049 | 0.009949859 |
| Gm14406 | 1.816762255 | 1.030297478 | 0.87595552 | 1.176198396 |
| Gm14324 | 0.130017973 | -0.444798045 | 3.662714201 | -0.121439463 |
| Gm14322 | 38.40290494 | 0.133475456 | 0.209061799 | 0.638449765 |
| Gm14325 | 156.7524774 | -0.204019162 | 0.172801451 | -1.180656536 |
| Gm14429 | 5.50850624 | 0.411482412 | 0.472717966 | 0.8704607 |
| Gm14323 | 1.346967727 | 1.347152697 | 1.074994267 | 1.253171983 |
| Gm14327 | 9.478232069 | 0.598756049 | 0.351081619 | 1.705461112 |
| C330013J21Rik | 6.088359734 | 0.555019333 | 0.557219653 | 0.996051253 |
| Gm14326 | 207.3808432 | 0.074647655 | 0.116274173 | 0.641996866 |
| Gm14428 | 0.751182044 | 0.106341477 | 1.119087688 | 0.095025152 |
| Etohi1 | 168.5832989 | 0.029305902 | 0.126690415 | 0.231319016 |
| Gm26869 | 1.499946283 | -1.780115929 | 0.999287751 | -1.781384719 |
| Zfp931 | 152.8081414 | -0.312508411 | 0.165072178 | -1.893162218 |
| Phactr3 | 1341.248435 | 0.916258225 | 0.100767942 | 9.092755182 |
| Sycp2 | 51.90291932 | 0.884162855 | 0.189106096 | 4.675485739 |
| Ppp1r3d | 90.63451093 | 0.492231227 | 0.169895118 | 2.897265275 |
| Fam217b | 760.5922785 | 0.558089774 | 0.087153241 | 6.403545847 |
| Cdh26 | 2.606367607 | -2.302743813 | 0.853364506 | -2.698429331 |
| Gm14314 | 0 | NA | NA | NA |
| Gm14292 | 70.00844022 | -0.068651001 | 0.128071075 | -0.536038299 |
| Gm14293 | 0 | NA | NA | NA |
| Gm14294 | 0 | NA | NA | NA |

| | | | | |
|---|---|---|---|---|
| 4930591A17Rik | 0 | NA | NA | NA |
| Cdh4 | 374.0015759 | 0.261522585 | 0.091620583 | 2.854408639 |
| Gm14300 | 0.042716945 | 0.037062677 | 3.724945049 | 0.009949859 |
| Taf4a | 305.5847678 | -0.13477376 | 0.100472814 | -1.341395288 |
| 4921531C22Rik | 29.89290028 | -0.355671973 | 0.287184712 | -1.238478088 |
| Gm14299 | 0.10503227 | -0.905414734 | 3.723500899 | -0.243162217 |
| Lsm14b | 804.4044537 | 0.201248031 | 0.07398427 | 2.720146183 |
| Psma7 | 1015.88698 | -0.212739467 | 0.115932775 | -1.835024368 |
| Ss18l1 | 608.8536376 | 0.327488967 | 0.067830039 | 4.828081661 |
| Mtg2 | 112.4777664 | -0.579000204 | 0.119340838 | -4.851651883 |
| Hrh3 | 541.8584486 | 1.290657367 | 0.147792039 | 8.732928914 |
| Gm17180 | 0.40273248 | 0.854661105 | 3.384235278 | 0.25254187 |
| Osbpl2 | 658.7052841 | 0.054376233 | 0.053935952 | 1.008163029 |
| Adrm1 | 690.902229 | -0.222588305 | 0.083357459 | -2.670286575 |
| Lama5 | 192.3097236 | 0.134820423 | 0.192970206 | 0.698659268 |
| Mir7005 | 0 | NA | NA | NA |
| Rps21 | 796.5226521 | -0.093582545 | 0.14734802 | -0.635112334 |
| Mir3091 | 3.756198497 | 0.474263065 | 0.613134947 | 0.773505193 |
| Cables2 | 296.6979083 | -0.050686844 | 0.086582308 | -0.585418025 |
| Rbbp8nl | 0.042716945 | 0.037062677 | 3.724945049 | 0.009949859 |
| Gata5 | 0.107803068 | 0.036092186 | 3.724874622 | 0.009689503 |
| Gata5os | 0 | NA | NA | NA |
| B230312C02Rik | 9.425892428 | 0.493066666 | 0.414176181 | 1.190475668 |
| Mir1a-1 | 0.044544648 | 0.517957688 | 3.724945049 | 0.139051095 |
| Mir133a-2 | 0.230754588 | 0.037068127 | 2.584843641 | 0.014340568 |
| Slco4a1 | 65.52497335 | 0.297761769 | 0.15184459 | 1.960963958 |
| Ntsr1 | 81.28326963 | 0.681587626 | 0.147597467 | 4.617881585 |
| Mrgbp | 164.3286957 | -0.23933302 | 0.089975331 | -2.659984872 |
| Ogfr | 1008.536141 | -0.543551007 | 0.130119066 | -4.177335619 |
| Col9a3 | 196.0069601 | 0.196342889 | 0.119749849 | 1.639608663 |
| Tcfl5 | 5.628819471 | -0.237178245 | 0.458538341 | -0.517248447 |
| Dido1 | 1582.103518 | 0.094325974 | 0.089995689 | 1.048116585 |
| Gm22502 | 0 | NA | NA | NA |
| Gid8 | 550.3352372 | 0.017723664 | 0.058552056 | 0.30269926 |
| Slc17a9 | 7.930149013 | 0.937660631 | 0.530394781 | 1.767854181 |
| Gm14343 | 0 | NA | NA | NA |
| Bhlhe23 | 1.454032994 | 1.096558935 | 0.903892209 | 1.213152325 |
| Gm14340 | 2.077246534 | -0.577368015 | 0.676376093 | -0.853619785 |
| Gm14342 | 4.321485189 | 0.934948883 | 0.507970805 | 1.840556334 |
| Gm14344 | 0.429573641 | -0.500113805 | 1.465057488 | -0.341361216 |
| Gm22842 | 0.112288909 | 0.037060453 | 3.724945049 | 0.009949262 |
| Gm14339 | 0.202674043 | -1.2519774 | 2.606063873 | -0.48040933 |
| Mir124a-3 | 0.458109599 | -0.147335147 | 1.727877932 | -0.085269419 |
| Gm27032 | 102.6108423 | 0.317270094 | 0.196918776 | 1.611172386 |
| Ythdf1 | 817.9793704 | -0.113565991 | 0.072535043 | -1.565670688 |
| Birc7 | 1.578816077 | 0.052480654 | 1.183007686 | 0.044362057 |
| Nkain4 | 83.76982434 | 1.165910882 | 0.259887035 | 4.486221801 |

| | | | | |
|---|---|---|---|---|
| Gm14341 | 1.152403953 | 0.866793687 | 1.044028154 | 0.830239763 |
| Arfgap1 | 898.465573 | 0.217444636 | 0.095421118 | 2.278789433 |
| 9230112E08Rik | 145.4154449 | 0.165724382 | 0.198399295 | 0.835307315 |
| Col20a1 | 78.79971225 | -0.309210038 | 0.22501324 | -1.374185974 |
| Chrna4 | 653.8085059 | 0.852858098 | 0.124440567 | 6.853537539 |
| Kcnq2 | 4581.893317 | 0.296783453 | 0.128007621 | 2.318482681 |
| Eef1a2 | 7678.32573 | 0.120513006 | 0.074313872 | 1.621675784 |
| Gm24892 | 0 | NA | NA | NA |
| Ppdpf | 209.3303033 | -0.237135877 | 0.097765647 | -2.425554221 |
| Ptk6 | 1.572767014 | -3.306273668 | 0.982659554 | -3.36461764 |
| Srms | 1.881316667 | 0.016787983 | 0.716484664 | 0.023431043 |
| Gm14398 | 0.631396267 | 0.698437661 | 1.192297833 | 0.585791269 |
| BC051628 | 0.900425091 | -0.159322271 | 1.01540598 | -0.156904996 |
| Helz2 | 1488.152321 | -1.457229837 | 0.267356684 | -5.450508355 |
| Gmeb2 | 338.4791566 | 0.003162333 | 0.117558503 | 0.026900081 |
| Gm7645 | 4.486974988 | -0.410267502 | 0.593320819 | -0.691476666 |
| Stmn3 | 2848.78216 | 0.202601844 | 0.106514442 | 1.902106803 |
| Rtel1 | 189.1322798 | 0.115989662 | 0.153495421 | 0.755655514 |
| Arfrp1 | 263.798925 | -0.052389987 | 0.099474415 | -0.526667957 |
| Zgpat | 398.0436084 | -0.124060373 | 0.123689651 | -1.002997191 |
| Lime1 | 3.160497059 | 0.848947752 | 0.734339259 | 1.156070224 |
| Slc2a4rg-ps | 70.84153722 | -0.253739942 | 0.357646594 | -0.709471155 |
| Zbtb46 | 247.6912476 | 0.198013746 | 0.137418749 | 1.440951447 |
| Gm22708 | 0.139376635 | 0.984529594 | 2.922770152 | 0.336848107 |
| Abhd16b | 0 | NA | NA | NA |
| Tpd52l2 | 603.1097156 | -0.129503565 | 0.062740729 | -2.06410678 |
| Gm25184 | 2.339668925 | 1.546796768 | 0.84548783 | 1.82947254 |
| Dnajc5 | 4139.797779 | 0.103093503 | 0.042891316 | 2.403598527 |
| Uckl1 | 187.8382027 | -0.311740108 | 0.129757748 | -2.402477787 |
| Uckl1os | 3.735935155 | -0.465190874 | 0.603877514 | -0.770339785 |
| Znf512b | 722.5500756 | 0.364782187 | 0.114945626 | 3.17351953 |
| Mir7006 | 0 | NA | NA | NA |
| Prpf6 | 758.9138183 | -0.320060415 | 0.082392847 | -3.884565564 |
| Samd10 | 162.7504342 | -0.031205961 | 0.104196801 | -0.299490587 |
| Gm25879 | 0 | NA | NA | NA |
| Sox18 | 27.0834614 | -0.728469863 | 0.222761563 | -3.270177562 |
| Tcea2 | 231.4654535 | 0.815507907 | 0.150111058 | 5.432697087 |
| Rgs19 | 143.057256 | -0.646154193 | 0.117674227 | -5.491042607 |
| Lkaaear1 | 0 | NA | NA | NA |
| Oprl1 | 406.903062 | 0.463292408 | 0.087241727 | 5.310445167 |
| Gm14489 | 0 | NA | NA | NA |
| Myt1 | 485.0253362 | 0.2131191 | 0.136744232 | 1.558523512 |
| Pcmtd2 | 953.1475173 | 0.077469765 | 0.089486607 | 0.865713511 |
| Polr3k | 406.1158035 | -0.022297927 | 0.06338452 | -0.351788207 |
| Gm5466 | 17.7294434 | -0.05870484 | 0.282966993 | -0.207461795 |
| Gm14496 | 0.046345709 | -0.443818988 | 3.724945049 | -0.119147795 |
| Gm28002 | 0 | NA | NA | NA |

| | | | | |
|---|---|---|---|---|
| Gm27460 | 0 | NA | NA | NA |
| Gm37338 | 0 | NA | NA | NA |
| Gm8747 | 0 | NA | NA | NA |
| Hnf4g | 1.172273398 | 0.447931147 | 0.87994741 | 0.509043089 |
| Gm37000 | 0.047948698 | 0.037064902 | 3.724945049 | 0.009950456 |
| Gm23237 | 0 | NA | NA | NA |
| Rps24-ps2 | 0.229688899 | 0.517966448 | 2.561073377 | 0.202245844 |
| Gm2071 | 0 | NA | NA | NA |
| Gm2077 | 0.293065915 | -0.479643756 | 1.779570664 | -0.269527794 |
| Gm8775 | 6.987551743 | -0.249851312 | 0.380846995 | -0.65604118 |
| Gm23976 | 0.841305426 | -0.521296624 | 1.340389079 | -0.388914407 |
| n-R5s194 | 0 | NA | NA | NA |
| Gm34023 | 0 | NA | NA | NA |
| Gm38131 | 0 | NA | NA | NA |
| Gm37029 | 0 | NA | NA | NA |
| Gm22944 | 0 | NA | NA | NA |
| Gm37763 | 0 | NA | NA | NA |
| Gm37801 | 0 | NA | NA | NA |
| 1110015O18Rik | 13.63005333 | 0.174337711 | 0.330501906 | 0.527493814 |
| Gm6140 | 0 | NA | NA | NA |
| Gm17308 | 1.32255022 | -0.058958753 | 1.004973418 | -0.058666978 |
| 2700069I18Rik | 10.07409941 | 0.556266939 | 0.358994228 | 1.549514994 |
| Gm10748 | 0.529004033 | -0.370818499 | 1.419786837 | -0.261178995 |
| Zfhx4 | 931.4280682 | 0.093662695 | 0.073744604 | 1.270095569 |
| 4930555M17Rik | 0 | NA | NA | NA |
| Pex2 | 255.6278662 | -0.151462767 | 0.105148409 | -1.440466554 |
| Gm8797 | 20.70001869 | -0.160194808 | 0.323306 | -0.495489746 |
| Gm37350 | 0.062426873 | 0.517959912 | 3.724945049 | 0.139051692 |
| Gm25352 | 0 | NA | NA | NA |
| Gm25144 | 0 | NA | NA | NA |
| Gm17934 | 0.363515313 | 1.9251287 | 1.938902465 | 0.992896102 |
| Gm6162 | 0.35319577 | 0.550072276 | 1.54035041 | 0.357108533 |
| Gm38298 | 0 | NA | NA | NA |
| Gm23147 | 0.044544648 | 0.517957688 | 3.724945049 | 0.139051095 |
| Gm18658 | 0 | NA | NA | NA |
| Gm5272 | 0 | NA | NA | NA |
| Gm38291 | 0 | NA | NA | NA |
| 1700008P02Rik | 0.048949532 | 0.037064902 | 3.724945049 | 0.009950456 |
| Olfr289-ps1 | 0.152964131 | 0.037074629 | 2.843489623 | 0.013038426 |
| Gm22074 | 0 | NA | NA | NA |
| Gm8823 | 0 | NA | NA | NA |
| Pkia | 2228.089358 | 0.292535362 | 0.08312155 | 3.519368446 |
| Gm37918 | 0 | NA | NA | NA |
| Gm8826 | 0.880107612 | 0.297538914 | 1.069920036 | 0.278094534 |
| Gm23592 | 0 | NA | NA | NA |
| Zc2hc1a | 623.5639214 | 0.10172674 | 0.07247507 | 1.403610102 |
| Il7 | 17.75567427 | -0.656493455 | 0.275333794 | -2.38435481 |

| | | | | |
|---|---|---|---|---|
| Gm16685 | 14.13537915 | 0.02841037 | 0.386274815 | 0.073549631 |
| Gm29747 | 0.364508673 | -1.707469632 | 1.571390727 | -1.086597752 |
| 1700010I02Rik | 0.136120942 | -0.876404342 | 3.595683781 | -0.24373788 |
| Gm7103 | 0.4787828 | 0.965230281 | 1.395108777 | 0.691867399 |
| Gm5841 | 5.450194582 | -0.522582955 | 0.432666238 | -1.207820044 |
| Gm24614 | 0 | NA | NA | NA |
| 1700003N22Rik | 1.26814241 | 0.29637516 | 1.067248791 | 0.27770016 |
| Stmn2 | 2333.139224 | 0.210323646 | 0.068716413 | 3.060748323 |
| Gm18822 | 4.048703541 | 0.701450911 | 0.495770035 | 1.414871536 |
| Gm32496 | 1.115586983 | 3.380999684 | 1.22250576 | 2.765630883 |
| Hey1 | 408.0829732 | 0.025628826 | 0.0707733 | 0.362125633 |
| Gm38007 | 0.33971137 | 0.518281253 | 1.715233225 | 0.30216372 |
| Gm23670 | 0 | NA | NA | NA |
| C230057A21Rik | 0.383304402 | -1.804162942 | 1.665923466 | -1.082980688 |
| Gm37647 | 0.199458489 | 0.037073246 | 2.665489594 | 0.013908606 |
| Mrps28 | 80.2242516 | -0.207048112 | 0.155305284 | -1.333168501 |
| Gm15467 | 0 | NA | NA | NA |
| Gm15466 | 6.37553239 | 0.586048575 | 0.475329929 | 1.232930096 |
| Tpd52 | 690.3315485 | -0.174706289 | 0.071209545 | -2.453411098 |
| Gm37616 | 0 | NA | NA | NA |
| 4930539M17Rik | 0.310536081 | 0.488004808 | 1.652861368 | 0.295248481 |
| Gm8860 | 0.282898957 | 0.977976546 | 2.051177959 | 0.476787761 |
| Zbtb10 | 267.8821127 | -0.018058815 | 0.101527814 | -0.177870616 |
| Gm22604 | 0.061920613 | -0.443816764 | 3.724945049 | -0.119147198 |
| C030034L19Rik | 1.859233056 | 0.471618219 | 0.760897929 | 0.619817982 |
| Zfp704 | 1430.007501 | 0.298521903 | 0.092452936 | 3.228906667 |
| Gm38001 | 0 | NA | NA | NA |
| Gm17806 | 0 | NA | NA | NA |
| Pag1 | 670.6925854 | 0.113402667 | 0.113895762 | 0.995670641 |
| Gm16337 | 0 | NA | NA | NA |
| Gm24786 | 3.953076191 | 0.537937547 | 0.606050365 | 0.887611952 |
| Gm6236 | 0 | NA | NA | NA |
| Gm17877 | 0.154612032 | -1.394968402 | 3.490355212 | -0.399663735 |
| Fabp5 | 368.7894691 | -0.145508765 | 0.129352625 | -1.124899979 |
| Gm38335 | 0.498917054 | -1.209525307 | 1.380441741 | -0.876187145 |
| Gm9833 | 92.22302499 | -0.154423891 | 0.169293164 | -0.9121685 |
| Gm37308 | 0 | NA | NA | NA |
| Pmp2 | 0 | NA | NA | NA |
| Gm37389 | 0 | NA | NA | NA |
| Fabp9 | 0 | NA | NA | NA |
| Fabp4 | 3.144700407 | -1.380252708 | 0.588189417 | -2.346612616 |
| Gm38310 | 0.547572158 | -3.036878737 | 1.490786503 | -2.037098358 |
| Gm37421 | 0 | NA | NA | NA |
| Gm37384 | 0 | NA | NA | NA |
| Fabp12 | 1.277907819 | -1.372588208 | 0.949779033 | -1.445165833 |
| Gm37316 | 0.044544648 | 0.517957688 | 3.724945049 | 0.139051095 |
| Gm38364 | 0 | NA | NA | NA |

| | | | | |
|---|---|---|---|---|
| Impa1 | 639.4989438 | 0.009669414 | 0.056818684 | 0.170180176 |
| Gm38303 | 0 | NA | NA | NA |
| Slc10a5 | 0.840861136 | -1.68681831 | 1.175417199 | -1.435080507 |
| Zfand1 | 152.3432722 | 0.405828839 | 0.139445903 | 2.910295892 |
| Chmp4c | 3.570328421 | -0.02241253 | 0.537832466 | -0.041671955 |
| Snx16 | 391.7704708 | -0.400202633 | 0.108561363 | -3.686418641 |
| Gm37831 | 0.049553047 | -0.443818988 | 3.724945049 | -0.119147795 |
| Gm18692 | 0 | NA | NA | NA |
| Gm37244 | 0.092140247 | 0.037060453 | 3.724945049 | 0.009949262 |
| Gm22795 | 0 | NA | NA | NA |
| Gm21631 | 0 | NA | NA | NA |
| Gm37832 | 0 | NA | NA | NA |
| Gm22547 | 0.047948698 | 0.037064902 | 3.724945049 | 0.009950456 |
| Gm8919 | 0 | NA | NA | NA |
| Gm10745 | 0 | NA | NA | NA |
| Gm37996 | 0 | NA | NA | NA |
| Gm2429 | 0 | NA | NA | NA |
| Gm8935 | 0.243796007 | -0.416733313 | 1.847899854 | -0.225517261 |
| Gm8939 | 0 | NA | NA | NA |
| Gm38317 | 9.626198242 | -0.416775511 | 0.364949302 | -1.142009338 |
| Gm37813 | 0.0885994 | 0.517955464 | 3.724945049 | 0.139050498 |
| Gm37197 | 0.044544648 | 0.517957688 | 3.724945049 | 0.139051095 |
| Gm2464 | 10.69073118 | -0.538091261 | 0.353079506 | -1.523994601 |
| Gm38112 | 362.4382461 | 0.221110067 | 0.1449108 | 1.525835664 |
| Gm37164 | 1.862656823 | -0.216335776 | 0.75520335 | -0.286460297 |
| Gm2474 | 11.24932943 | 0.883483389 | 0.384578876 | 2.297274875 |
| Gm38400 | 64.08375772 | 0.571517809 | 0.17859569 | 3.200064964 |
| Gm19114 | 0.044544648 | 0.517957688 | 3.724945049 | 0.139051095 |
| Gm25981 | 24.11327504 | 0.157415536 | 0.306962792 | 0.512816341 |
| Ralyl | 450.4153493 | 0.406611985 | 0.078475833 | 5.181365651 |
| Gm22026 | 0.132101749 | -0.93550548 | 3.724188147 | -0.251197158 |
| Gm23112 | 0 | NA | NA | NA |
| Gm38311 | 0.286827043 | 1.009439072 | 2.03675267 | 0.495612004 |
| Gm6300 | 1.130984576 | 0.858533998 | 0.917390343 | 0.935843727 |
| Gm37358 | 0 | NA | NA | NA |
| Gm8955 | 0.038981395 | -0.443816764 | 3.724945049 | -0.119147198 |
| Slc7a12 | 0 | NA | NA | NA |
| Gm8957 | 0.226376398 | -0.467017071 | 2.147568223 | -0.217463206 |
| Lrrcc1 | 419.00625 | 0.361997959 | 0.092537238 | 3.91191662 |
| Gm17119 | 0 | NA | NA | NA |
| E2f5 | 109.6456114 | -0.363371788 | 0.124012134 | -2.930130926 |
| 1810022K09Rik | 264.8506355 | -0.033339797 | 0.117187254 | -0.284500195 |
| Gm38151 | 0.061920613 | -0.443816764 | 3.724945049 | -0.119147198 |
| Car13 | 144.2580462 | -0.987677191 | 0.231889851 | -4.259251477 |
| Gm5843 | 0 | NA | NA | NA |
| Car1 | 0.04393175 | 0.037062677 | 3.724945049 | 0.009949859 |
| Gm16399 | 0.953016895 | 1.214144678 | 1.139709008 | 1.065311119 |

| | | | | |
|---|---|---|---|---|
| Car3 | 3.829701666 | 0.315141131 | 0.596936781 | 0.527930496 |
| Car2 | 1336.217829 | -0.496697811 | 0.182696853 | -2.718699327 |
| A930001A20Rik | 0.572612294 | -1.877165624 | 1.328494102 | -1.413002602 |
| Gm7132 | 0 | NA | NA | NA |
| Gm33819 | 0.263469446 | 0.036370832 | 2.105945695 | 0.017270546 |
| Gm9733 | 1.473887978 | -3.341866272 | 1.11606623 | -2.994326127 |
| Sirpb1a | 36.66926605 | -2.058658809 | 0.384272332 | -5.357291272 |
| Sirpb1b | 7.832067487 | -1.558935938 | 0.519805452 | -2.999075771 |
| Gm37468 | 0 | NA | NA | NA |
| Sirpb1c | 25.99497885 | -1.889687648 | 0.610653214 | -3.094534842 |
| Gm6321 | 0.066050875 | -0.443818988 | 3.724945049 | -0.119147795 |
| Gm7343 | 0.430172323 | -2.629761164 | 1.546369882 | -1.700602937 |
| Gm5150 | 10.40898798 | -2.321064907 | 0.477299807 | -4.862907698 |
| Gm24694 | 0 | NA | NA | NA |
| Gm37836 | 0 | NA | NA | NA |
| Gm25863 | 0 | NA | NA | NA |
| Ythdf3 | 1623.468186 | -0.09387219 | 0.049158571 | -1.909579287 |
| Gm26485 | 0 | NA | NA | NA |
| Gm5283 | 0.46900631 | -0.316390621 | 1.287664726 | -0.245708852 |
| Gm24963 | 0.253979914 | 0.98798747 | 1.838957507 | 0.537254105 |
| Gm6350 | 0.222549579 | 0.506695678 | 1.897660875 | 0.267010658 |
| Gm30340 | 13.55485612 | 0.104883179 | 0.325024793 | 0.322692856 |
| Gm19694 | 0 | NA | NA | NA |
| Gm38154 | 0 | NA | NA | NA |
| Mir124-2hg | 730.7247633 | 0.513311849 | 0.093360712 | 5.498156972 |
| Mir124a-2 | 2.558881719 | 0.326036203 | 0.650238446 | 0.501410221 |
| Gm23441 | 0 | NA | NA | NA |
| RP24-84E6.5 | 0.192405858 | -1.188324619 | 2.639510532 | -0.450206432 |
| Cypt12 | 0.109982625 | -0.443823436 | 3.724945049 | -0.119148989 |
| Gm5844 | 33.81437118 | -0.273559089 | 0.198679146 | -1.376888787 |
| Bhlhe22 | 439.9718104 | 0.89586527 | 0.159276479 | 5.624592371 |
| Gm23726 | 0 | NA | NA | NA |
| Cyp7b1 | 183.0659675 | 0.259406214 | 0.133205032 | 1.947420522 |
| Gm23686 | 3.150048424 | 1.24131861 | 0.609256031 | 2.037433439 |
| RP23-77D22.2 | 2.298046648 | 0.162925761 | 0.659597069 | 0.247008012 |
| Gm30667 | 0 | NA | NA | NA |
| 4930433B08Rik | 0 | NA | NA | NA |
| Gm6369 | 0 | NA | NA | NA |
| RP23-427B4.3 | 0 | NA | NA | NA |
| Gm37317 | 2.158753835 | 2.088689667 | 0.833458936 | 2.50604988 |
| RP23-427B4.2 | 0.137548931 | 0.517963473 | 2.938933035 | 0.176242013 |
| Gm30341 | 0 | NA | NA | NA |
| Armc1 | 876.5532839 | -0.002224249 | 0.080146438 | -0.027752317 |
| Gm37979 | 14.15384062 | 0.305642027 | 0.332692232 | 0.918693007 |
| Mtfr1 | 148.0536149 | -0.49575881 | 0.134761579 | -3.678784525 |
| Pde7a | 422.8626837 | 0.197596179 | 0.108680491 | 1.818138448 |
| Gm16093 | 2.00382768 | 0.954481803 | 0.820560764 | 1.16320673 |

| | | | | |
|---|---|---|---|---|
| Gm22282 | 0.538570099 | 1.352348048 | 1.363533858 | 0.991796456 |
| Dnajc5b | 1.101603561 | -2.912675694 | 1.020972184 | -2.852845298 |
| Gm23330 | 0.225375977 | 0.036353782 | 2.147468566 | 0.016928668 |
| Gm6141 | 0.044191549 | 0.037062677 | 3.724945049 | 0.009949859 |
| 1700064H15Rik | 1.062875418 | 3.410405688 | 1.061210822 | 3.213692903 |
| Trim55 | 5.56456899 | 1.001200748 | 0.595394459 | 1.681575521 |
| BC002189 | 0 | NA | NA | NA |
| Crh | 42.16080477 | -0.031417029 | 0.281045447 | -0.111786292 |
| Gm18864 | 0.120349074 | 1.027495679 | 3.723011807 | 0.275985071 |
| RP23-105K2.2 | 0 | NA | NA | NA |
| 4632415L05Rik | 336.8514353 | -0.08258574 | 0.074086253 | -1.114724214 |
| Gm37523 | 0.176667124 | 0.037076309 | 2.30438001 | 0.016089494 |
| Gm17771 | 0.368202395 | -0.970716378 | 1.46272496 | -0.663635615 |
| Cp | 857.3323739 | 0.096398347 | 0.24914971 | 0.386909328 |
| Hps3 | 339.45544 | 0.087217367 | 0.085428787 | 1.020936511 |
| Hltf | 534.1407693 | 0.038023246 | 0.07251931 | 0.524318919 |
| Gyg | 294.265161 | -0.481171154 | 0.12778013 | -3.765617975 |
| Cpa3 | 1.364377633 | -1.456066141 | 0.959033168 | -1.518264633 |
| Mir7007 | 0 | NA | NA | NA |
| Cpb1 | 1.12517466 | -1.539087668 | 0.976725698 | -1.575762438 |
| RP24-190F20.3 | 0.04393175 | 0.037062677 | 3.724945049 | 0.009949859 |
| Agtr1b | 1.030384379 | 1.014428578 | 0.960093168 | 1.056593893 |
| Gm18426 | 0.046345709 | -0.443818988 | 3.724945049 | -0.119147795 |
| RP24-190F20.4 | 0 | NA | NA | NA |
| Gm31466 | 0 | NA | NA | NA |
| Gm18491 | 0 | NA | NA | NA |
| Gm38323 | 0 | NA | NA | NA |
| Gm18493 | 0 | NA | NA | NA |
| Gm37480 | 0.122601409 | 0.998850474 | 3.724945049 | 0.268151734 |
| Gm7488 | 13.25965715 | 0.328990714 | 0.306842399 | 1.072181402 |
| RP23-105J10.1 | 0.180386268 | 0.037062359 | 3.342495489 | 0.01108823 |
| Gm29137 | 0.309067702 | -0.71669826 | 3.714525094 | -0.192944789 |
| RP23-458I4.1 | 0 | NA | NA | NA |
| RP23-458I4.2 | 0 | NA | NA | NA |
| RP23-458I4.3 | 0 | NA | NA | NA |
| 7530428D23Rik | 1.976191175 | 0.900306612 | 0.745174787 | 1.208181795 |
| RP23-105M18.2 | 4.557716346 | 0.868755269 | 0.551951239 | 1.573971045 |
| RP23-105M18.1 | 0 | NA | NA | NA |
| RP24-487F9.4 | 40.33606232 | 0.174666914 | 0.178401565 | 0.979066042 |
| Tbl1xr1 | 1839.527346 | 0.386502185 | 0.071985061 | 5.369199972 |
| Gm5842 | 2.255157929 | -0.074729294 | 0.628070587 | -0.118982317 |
| Rprl2 | 0.403845505 | -0.972775416 | 1.571613659 | -0.618965997 |
| Gm38324 | 0 | NA | NA | NA |
| Gm37636 | 0 | NA | NA | NA |
| Gm26354 | 0 | NA | NA | NA |
| Spin3-ps | 2.893792528 | -0.456698218 | 0.725820522 | -0.629216459 |
| Gm37889 | 0.097139547 | -0.443816764 | 3.724945049 | -0.119147198 |

| | | | | |
|---|---|---|---|---|
| Gm36983 | 1.90140223 | 0.149231765 | 0.728983153 | 0.204712228 |
| Gm37021 | 1.244068253 | -0.878127779 | 0.85064789 | -1.032304657 |
| Gm37492 | 0 | NA | NA | NA |
| 4930412E21Rik | 0.700626182 | 0.545259703 | 1.062508204 | 0.51318164 |
| Naaladl2 | 18.35136831 | 0.391647186 | 0.283561715 | 1.381170888 |
| Gm7536 | 51.1862964 | -0.260434171 | 0.192106338 | -1.35567714 |
| Gm24704 | 0 | NA | NA | NA |
| Gm24062 | 0 | NA | NA | NA |
| RP23-152K14.2 | 3.653026081 | -0.235842461 | 0.719236767 | -0.327906569 |
| Gm37136 | 0.341775432 | 0.75266308 | 2.042743217 | 0.368457021 |
| Gm38002 | 0 | NA | NA | NA |
| Gm10259 | 3.08375672 | 0.491154257 | 0.545092447 | 0.901047629 |
| Nlgn1 | 2154.133087 | 0.508075475 | 0.094515848 | 5.375558556 |
| Gm24250 | 0 | NA | NA | NA |
| Gm37573 | 125.0600198 | 0.221561359 | 0.196123684 | 1.129702212 |
| Gm6197 | 0.660619988 | 1.323655398 | 1.192283904 | 1.110184741 |
| A830092H15Rik | 35.52709241 | -0.292268585 | 0.28010411 | -1.043428406 |
| Gm4855 | 1.087037337 | -0.2400289 | 1.2310035 | -0.194986367 |
| Gm25773 | 0.04393175 | 0.037062677 | 3.724945049 | 0.009949859 |
| Spata16 | 0.312179078 | 1.728796917 | 1.710592531 | 1.010642152 |
| Gm37659 | 0 | NA | NA | NA |
| AC108829.1 | 0 | NA | NA | NA |
| Gm7558 | 0 | NA | NA | NA |
| Ect2 | 26.23418059 | -0.845785483 | 0.395694958 | -2.137468438 |
| Gm25152 | 0 | NA | NA | NA |
| 1700125G22Rik | 0.384795782 | 1.48536028 | 1.474339947 | 1.007474757 |
| Nceh1 | 1766.692954 | 0.063572005 | 0.075113472 | 0.846346247 |
| Gm37191 | 0.357797561 | 1.68321042 | 1.938905944 | 0.868123812 |
| RP23-131B7.3 | 0.060174537 | 0.517957688 | 3.724945049 | 0.139051095 |
| Tnfsf10 | 460.2181555 | -1.16196874 | 0.315520406 | -3.682705519 |
| Ghsr | 15.57801817 | -0.126074395 | 0.322072681 | -0.391447031 |
| Fndc3b | 605.684463 | -0.717305557 | 0.079399955 | -9.034080095 |
| RP24-62O16.3 | 0 | NA | NA | NA |
| Mir3092 | 1.750169107 | -0.562690294 | 0.774332069 | -0.726678278 |
| Tmem212 | 11.59725055 | 0.145398468 | 0.349203406 | 0.416371851 |
| Gm26040 | 0 | NA | NA | NA |
| Pld1 | 212.9629629 | -0.048715463 | 0.120215281 | -0.405235195 |
| Gm37098 | 0 | NA | NA | NA |
| Tnik | 3513.493683 | 0.546768262 | 0.130240153 | 4.198154312 |
| Gm37537 | 40.89549279 | 0.788038889 | 0.318700202 | 2.472665172 |
| Mir466q | 6.638222057 | 0.853344481 | 0.468546993 | 1.821256979 |
| Slc2a2 | 4.994744966 | -1.620957438 | 0.480012539 | -3.37690645 |
| 1700112D23Rik | 5.161670123 | 0.413065852 | 0.467124822 | 0.884272966 |
| Gm6505 | 17.67248122 | -0.009276506 | 0.25594734 | -0.036243805 |
| Eif5a2 | 1024.993172 | 0.378431596 | 0.122608973 | 3.086491854 |
| Gm32950 | 1.23849881 | 0.546474694 | 0.989006504 | 0.552549141 |
| Rpl22l1 | 357.7709375 | -0.245939725 | 0.145583865 | -1.689333669 |

| | | | | |
|---|---|---|---|---|
| Gm33051 | 0.1140246 | 0.037073798 | 3.724945049 | 0.009952844 |
| Gm37867 | 0 | NA | NA | NA |
| Gm37924 | 0 | NA | NA | NA |
| Gm1527 | 0.240368072 | -1.875670652 | 2.497392314 | -0.751051663 |
| Egfem1 | 145.805512 | 1.331065922 | 0.148327152 | 8.973852104 |
| Gm38029 | 61.7718118 | 1.630878409 | 0.378970297 | 4.303446528 |
| Gm37965 | 30.75954022 | 1.086628455 | 0.277648855 | 3.913678867 |
| Mir551b | 1.726387458 | 2.73908883 | 1.033501727 | 2.650299228 |
| Mir21b | 0.755943838 | 0.887076866 | 1.270998939 | 0.697936748 |
| Gm37557 | 0.247197661 | 1.078801341 | 2.09622298 | 0.514640547 |
| Gm33206 | 0.045882455 | 0.517959912 | 3.724945049 | 0.139051692 |
| Mannr | 0.312486075 | 1.393003837 | 1.7490085 | 0.796453441 |
| Gm37281 | 0.306751849 | 0.530471059 | 1.622435939 | 0.326959633 |
| Mecom | 32.25235124 | -0.284627033 | 0.253184355 | -1.124188865 |
| Mecomos | 0.404625785 | -0.590889524 | 1.776526753 | -0.332609415 |
| Gm38066 | 1.468378907 | -0.570215651 | 1.053209181 | -0.541407786 |
| Gm38197 | 1.573498709 | -0.547486131 | 0.870116098 | -0.629210437 |
| Gm10258 | 0.786098488 | 0.321165307 | 1.276258061 | 0.251646056 |
| Gm38362 | 0.449676329 | -1.292827523 | 1.43716634 | -0.899567076 |
| Gm37024 | 0 | NA | NA | NA |
| Gm37652 | 2.839259381 | -0.185864198 | 0.739065939 | -0.251485271 |
| Gm38258 | 0.044191549 | 0.037062677 | 3.724945049 | 0.009949859 |
| Gm38153 | 0.060174537 | 0.517957688 | 3.724945049 | 0.139051095 |
| 1600017P15Rik | 2.396959378 | 0.603037022 | 0.655816919 | 0.919520379 |
| Actrt3 | 12.26637337 | 0.500682481 | 0.316030756 | 1.584284031 |
| Mynn | 359.7065454 | -0.049927195 | 0.071314369 | -0.700100078 |
| Lrrc34 | 2.945986073 | 0.516471531 | 0.685424017 | 0.753506615 |
| Lrriq4 | 0.139037128 | -1.190058937 | 3.717743703 | -0.320102469 |
| Lrrc31 | 0.163642604 | 0.998860448 | 2.783881642 | 0.358801334 |
| Gm15462 | 0.843690133 | 0.444627278 | 1.20347869 | 0.369451725 |
| Samd7 | 0.989999884 | -1.059738755 | 0.970752263 | -1.091667561 |
| 4933429H19Rik | 2.000591198 | 0.908056445 | 0.687175505 | 1.321433082 |
| Sec62 | 1867.564255 | 0.014470701 | 0.05906935 | 0.244978169 |
| Gm38080 | 0.150086409 | 1.027495679 | 3.723011807 | 0.275985071 |
| Gpr160 | 12.95102122 | -0.190934474 | 0.339167714 | -0.562950026 |
| Phc3 | 2283.871593 | -0.021434203 | 0.072910846 | -0.293978249 |
| 9530022L04Rik | 9.381402519 | 0.348572082 | 0.372020001 | 0.93697135 |
| Gm2979 | 0 | NA | NA | NA |
| Prkci | 784.3797283 | -0.096136108 | 0.057263098 | -1.678849246 |
| Mir7008 | 0.429612315 | -1.423109527 | 1.479595033 | -0.961823671 |
| Skil | 1820.158226 | -0.391898965 | 0.095053723 | -4.122920734 |
| Cldn11 | 3469.821532 | -0.297799649 | 0.142153453 | -2.094916753 |
| Slc7a14 | 2555.251066 | 0.099109244 | 0.061925478 | 1.600459888 |
| Gm15496 | 15.67699621 | 0.545357961 | 0.257487008 | 2.118001861 |
| Gm38025 | 1.47797824 | 0.671285344 | 0.925515593 | 0.725309599 |
| Gm6558 | 0 | NA | NA | NA |
| Kcnmb2 | 45.73165487 | 0.148312494 | 0.18652387 | 0.795139483 |

| | | | | |
|---|---|---|---|---|
| Gm37834 | 14.36667755 | 0.640946538 | 0.335587043 | 1.909926356 |
| Gm37770 | 0 | NA | NA | NA |
| Zmat3 | 1732.179252 | 0.113794597 | 0.062498178 | 1.820766637 |
| Gm36990 | 0.044544648 | 0.517957688 | 3.724945049 | 0.139051095 |
| Pik3ca | 1418.733806 | -0.068811858 | 0.042160455 | -1.632142187 |
| Gm37123 | 26.09608814 | 0.450999208 | 0.337522192 | 1.336206087 |
| Kcnmb3 | 0.414197093 | -0.88620477 | 1.767697477 | -0.501332825 |
| Zfp639 | 349.0890839 | 0.118891864 | 0.090262821 | 1.317174263 |
| Gm37592 | 0 | NA | NA | NA |
| Mfn1 | 1164.365532 | -0.369129995 | 0.069695111 | -5.296354187 |
| Gnb4 | 421.880452 | 0.119039061 | 0.107230581 | 1.110122315 |
| Gm37640 | 0.609267175 | 0.006171477 | 1.30709009 | 0.004721539 |
| Actl6a | 159.1766196 | -0.013358863 | 0.151900388 | -0.087944892 |
| Gm37353 | 0.579969052 | 0.046078787 | 1.403907719 | 0.032821806 |
| Mrpl47 | 199.0986341 | -0.334852809 | 0.095827284 | -3.49433685 |
| Ndufb5 | 763.5212508 | -0.010532421 | 0.077539308 | -0.135833307 |
| Usp13 | 553.4368463 | -0.237785287 | 0.090997447 | -2.613098438 |
| Gm15574 | 0 | NA | NA | NA |
| Gm37459 | 3.866617621 | -0.248523485 | 0.618610533 | -0.401744671 |
| Pex5l | 2341.167297 | 0.175515113 | 0.08126167 | 2.159875784 |
| Gm23214 | 0 | NA | NA | NA |
| 4930419G24Rik | 1.068758569 | -0.643231639 | 0.999218062 | -0.643735 |
| Gm37443 | 87.11784037 | 0.609815547 | 0.317421782 | 1.921152173 |
| Gm19445 | 130.8739924 | 0.478650241 | 0.297234365 | 1.610346239 |
| Gm37128 | 0.066050875 | -0.443818988 | 3.724945049 | -0.119147795 |
| Gm27974 | 0 | NA | NA | NA |
| Gm37031 | 0.110870145 | -0.444802825 | 3.724874622 | -0.119414174 |
| Gm24780 | 0 | NA | NA | NA |
| Gm37398 | 0.221866436 | 1.052488932 | 2.162979903 | 0.486592099 |
| Gm5845 | 1.420437351 | -0.502278326 | 0.808616571 | -0.621157597 |
| RP23-373G12.3 | 0.234927675 | 0.037076959 | 1.888353269 | 0.019634546 |
| RP23-373G12.1 | 1.69071356 | -0.225277638 | 0.992055667 | -0.22708165 |
| RP23-373G12.4 | 0.110242424 | -0.443823436 | 3.724945049 | -0.119148989 |
| Ttc14 | 2035.534113 | 0.211944332 | 0.132059383 | 1.604916878 |
| Ccdc39 | 244.5153087 | 0.306637504 | 0.187911002 | 1.631823043 |
| RP23-373G12.5 | 10.01486383 | 0.573915736 | 0.392091348 | 1.463729662 |
| RP23-373G12.2 | 0 | NA | NA | NA |
| RP24-457B22.2 | 0 | NA | NA | NA |
| Gm23175 | 0 | NA | NA | NA |
| Gm9791 | 24.4382261 | -0.423569936 | 0.251088197 | -1.686936863 |
| RP24-457B22.3 | 1.01049493 | 2.865726222 | 1.081506156 | 2.649754887 |
| Fxr1 | 1205.298799 | -0.069232544 | 0.053836274 | -1.285983204 |
| RP23-418O21.3 | 27.80606419 | -0.560947734 | 0.251205318 | -2.233024911 |
| Dnajc19 | 148.9300614 | 0.114528135 | 0.141685802 | 0.808324709 |
| RP23-418O21.5 | 200.8974819 | 0.524766401 | 0.166401629 | 3.153613359 |
| Sox2ot | 356.2706365 | -0.306531256 | 0.105308381 | -2.910796388 |
| RP23-418O21.1 | 0.658880446 | -1.221130351 | 1.371791817 | -0.890171771 |

| | | | | |
|---|---|---|---|---|
| Gm20446 | 0 | NA | NA | NA |
| RP24-396D16.3 | 2.524695083 | 1.27726493 | 0.916636811 | 1.393425308 |
| Gm20515 | 12.14791881 | -0.385591964 | 0.387175648 | -0.99590965 |
| Gm29135 | 0 | NA | NA | NA |
| Gm27403 | 0 | NA | NA | NA |
| RP24-277H8.2 | 0.177597096 | -1.302077284 | 2.263528882 | -0.575242178 |
| RP24-277H8.3 | 0.133275802 | -0.89092784 | 2.958201129 | -0.301172166 |
| Mir1897 | 0 | NA | NA | NA |
| mmu-mir-1897 | 0 | NA | NA | NA |
| RP24-277H8.1 | 54.03438621 | -0.319601532 | 0.243270753 | -1.313768827 |
| Sox2 | 458.0693419 | -0.546727442 | 0.127277919 | -4.295540394 |
| Gm27611 | 0 | NA | NA | NA |
| RP23-423J10.6 | 3.912046527 | 0.136031955 | 0.535939715 | 0.253819508 |
| RP23-423J10.7 | 0.80598721 | -0.801497486 | 1.244050203 | -0.644264584 |
| RP23-423J10.4 | 0 | NA | NA | NA |
| RP23-423J10.2 | 0.090665643 | 0.037060453 | 3.724945049 | 0.009949262 |
| Mir6378 | 0 | NA | NA | NA |
| RP23-477L20.1 | 1.570552818 | -0.835822328 | 0.777276705 | -1.075321468 |
| Gm25442 | 0.241676147 | -0.443819749 | 1.875130429 | -0.236687402 |
| RP24-141D20.1 | 2.674854522 | 0.586530912 | 0.664633863 | 0.882487252 |
| RP23-112A11.3 | 2.926346155 | -0.155675855 | 0.631593324 | -0.24648116 |
| Ube2l3-ps1 | 0 | NA | NA | NA |
| RP23-112A11.1 | 0.159330469 | 0.517966511 | 3.465376739 | 0.149469033 |
| Gm25696 | 0 | NA | NA | NA |
| Atp11b | 1725.242478 | 0.241097833 | 0.106947676 | 2.254353174 |
| RP23-331L5.1 | 12.60277648 | 0.1299092 | 0.423496718 | 0.306753736 |
| Gm7741 | 2.653371563 | 0.396242152 | 0.787546731 | 0.50313478 |
| Dcun1d1 | 831.5482228 | 0.10647424 | 0.064127538 | 1.660351291 |
| Gm15952 | 9.495619415 | 0.686353375 | 0.406035972 | 1.690375784 |
| RP23-111M12.3 | 21.60654816 | 0.391696843 | 0.411509546 | 0.951853601 |
| Mccc1 | 363.2980275 | 0.315320667 | 0.148728178 | 2.120113835 |
| RP23-111M12.2 | 53.110291 | 0.540952396 | 0.293433439 | 1.843526759 |
| AC116677.4 | 0.320207964 | 0.444608627 | 1.726225056 | 0.257561217 |
| Mccc1os | 32.77909455 | 0.321316315 | 0.200145557 | 1.605413175 |
| Ccdc144b | 0.1140246 | 0.037073798 | 3.724945049 | 0.009952844 |
| Acad9 | 366.3759439 | 0.096808513 | 0.066875304 | 1.44759736 |
| RP23-218L17.3 | 0 | NA | NA | NA |
| RP23-218L17.2 | 12.80704659 | -0.197180257 | 0.296265613 | -0.665552291 |
| D3Ertd254e | 555.8188512 | 0.279059003 | 0.093799018 | 2.975073817 |
| Qrfpr | 20.7155208 | -0.133378454 | 0.251831193 | -0.529634366 |
| RP24-389H4.1 | 1.921125148 | 1.361489248 | 0.756881761 | 1.798813656 |
| RP24-389H4.2 | 1.29860622 | 2.444036805 | 1.101726924 | 2.218368955 |
| RP24-363L5.1 | 0.700674929 | -0.919421928 | 1.134230172 | -0.810613181 |
| RP23-165D1.3 | 0.038981395 | -0.443816764 | 3.724945049 | -0.119147198 |
| RP23-165D1.2 | 0 | NA | NA | NA |
| Anxa5 | 799.2156265 | -0.860859227 | 0.144518205 | -5.956752837 |
| Mir7009 | 0 | NA | NA | NA |

| | | | | |
|---|---|---|---|---|
| 1810062G17Rik | 1.725215873 | 0.477997913 | 0.802860457 | 0.595368608 |
| Gm11548 | 0.199639456 | 0.574790039 | 3.22081537 | 0.178461033 |
| Gm11549 | 318.3834355 | 0.8979078 | 0.28237968 | 3.179789003 |
| RP23-151D16.2 | 14.17505885 | -0.03998873 | 0.354043757 | -0.112948553 |
| Gm11550 | 0 | NA | NA | NA |
| Exosc9 | 304.3832438 | 0.07512647 | 0.101240651 | 0.742058341 |
| Ccna2 | 45.411053 | -0.072834469 | 0.278497906 | -0.261526092 |
| Bbs7 | 206.6797326 | 0.257075457 | 0.137055395 | 1.87570477 |
| Trpc3 | 351.384491 | 0.065327878 | 0.115036359 | 0.567888962 |
| Gm12377 | 0 | NA | NA | NA |
| Gm12376 | 0 | NA | NA | NA |
| 4932438A13Rik | 5270.27695 | 0.202811628 | 0.067667646 | 2.997172807 |
| Gm12531 | 0.264018088 | -0.92132221 | 2.097564717 | -0.439234224 |
| Adad1 | 0.098502579 | -0.443821212 | 3.724945049 | -0.119148392 |
| Gm12582 | 0 | NA | NA | NA |
| Gm12583 | 0.046345709 | -0.443818988 | 3.724945049 | -0.119147795 |
| Gm12584 | 0 | NA | NA | NA |
| Il2 | 0.52475134 | -0.385796102 | 1.270981044 | -0.30354198 |
| Gm12532 | 1.446123796 | 0.815442501 | 0.88733263 | 0.918981758 |
| Il21 | 4.872996577 | -0.849764458 | 0.452362377 | -1.878503831 |
| Gm12534 | 11.60045621 | -0.518287342 | 0.525197978 | -0.986841844 |
| RP23-243E17.13 | 4.833786608 | 0.297388898 | 0.63857593 | 0.4657064 |
| Gm24169 | 0.075043205 | 0.517957688 | 3.724945049 | 0.139051095 |
| Cetn4 | 69.0187925 | -0.160528649 | 0.157085589 | -1.021918372 |
| RP23-328P19.3 | 0 | NA | NA | NA |
| Bbs12 | 67.61927239 | -0.578690684 | 0.206247183 | -2.80581134 |
| Gm12540 | 6.941958254 | 0.092170046 | 0.459243676 | 0.200699653 |
| Fgf2os | 0.170190954 | -0.70542992 | 2.744856668 | -0.25700064 |
| RP23-137F6.5 | 13.65974216 | -0.23046995 | 0.362329551 | -0.63607826 |
| Fgf2 | 254.8186985 | -0.527191482 | 0.221509014 | -2.380000135 |
| RP23-328P19.4 | 0.559063306 | 0.434261246 | 1.246939845 | 0.348261585 |
| Nudt6 | 28.00862341 | 0.218722483 | 0.227072638 | 0.963226943 |
| Spata5 | 377.9700528 | -0.101363777 | 0.090258357 | -1.123040357 |
| RP23-465A17.7 | 107.3750371 | 0.323396689 | 0.253262672 | 1.276922045 |
| Gm12564 | 0 | NA | NA | NA |
| Gm12565 | 0.230175392 | -1.625458588 | 2.102973631 | -0.772933414 |
| 4930594O21Rik | 0.042716945 | 0.037062677 | 3.724945049 | 0.009949859 |
| RP23-382L1.6 | 4.839488669 | 1.292093606 | 0.559929672 | 2.307599812 |
| Spry1 | 135.2416231 | -0.153279352 | 0.196495466 | -0.780065595 |
| RP23-382L1.4 | 8.157939626 | 0.721076445 | 0.415520851 | 1.735355621 |
| RP23-382L1.3 | 1.313092963 | 1.055706838 | 1.025974417 | 1.02897969 |
| Gm5148 | 59.44844248 | 0.852710265 | 0.222383977 | 3.834405149 |
| Rps23-ps1 | 35.46711007 | 0.079933842 | 0.178971755 | 0.44662825 |
| Rpl31-ps15 | 0.453368568 | 0.883543406 | 1.505807366 | 0.586757262 |
| RP23-382L1.7 | 0.0878635 | 0.037058229 | 3.724945049 | 0.009948665 |
| Gm26404 | 0 | NA | NA | NA |
| RP23-175A22.1 | 0.138375801 | 0.984520051 | 2.929303049 | 0.336093615 |

| | | | | |
|---|---|---|---|---|
| RP23-175A22.2 | 0.167536551 | 0.998860717 | 2.763312802 | 0.361472185 |
| RP23-112C14.1 | 0.320788412 | 0.137959259 | 1.720285035 | 0.080195582 |
| Gm7799 | 0 | NA | NA | NA |
| RP23-112C14.2 | 0 | NA | NA | NA |
| Gm22899 | 0 | NA | NA | NA |
| RP24-537K9.3 | 0.194322771 | 0.037075665 | 2.636227018 | 0.014063912 |
| RP24-537K9.1 | 6.837594151 | -0.620360996 | 0.417112152 | -1.487276245 |
| RP24-537K9.2 | 0 | NA | NA | NA |
| RP24-392E13.1 | 0 | NA | NA | NA |
| Gm25574 | 0 | NA | NA | NA |
| RP23-334N1.5 | 0 | NA | NA | NA |
| RP23-334N1.3 | 0.614372065 | 1.805717604 | 1.403234754 | 1.28682503 |
| RP23-334N1.1 | 2.007620227 | 3.222102535 | 1.093841729 | 2.945675275 |
| RP23-334N1.2 | 0.353731249 | 0.517972636 | 1.738376039 | 0.297963516 |
| Ankrd50 | 1197.125447 | -0.249305137 | 0.066862971 | -3.728597961 |
| Gm24462 | 0 | NA | NA | NA |
| RP23-33I24.2 | 0 | NA | NA | NA |
| RP23-33I24.1 | 0 | NA | NA | NA |
| C230034O21Rik | 45.21372464 | 0.509349863 | 0.188697017 | 2.69930003 |
| Fat4 | 722.5535298 | 0.580903063 | 0.101900039 | 5.700714822 |
| RP23-194E5.1 | 0 | NA | NA | NA |
| RP23-382L1.5 | 0.075043205 | 0.517957688 | 3.724945049 | 0.139051095 |
| RP23-49O22.2 | 0.068097359 | 0.037064902 | 3.724945049 | 0.009950456 |
| Gm9845 | 9.009532912 | -0.728357028 | 0.335432169 | -2.17139886 |
| n-R5s195 | 0 | NA | NA | NA |
| RP24-85O24.1 | 0.162691105 | 0.517965151 | 2.788998835 | 0.185717235 |
| Gm23796 | 0 | NA | NA | NA |
| Gm23831 | 0 | NA | NA | NA |
| RP23-256D7.1 | 0 | NA | NA | NA |
| RP23-256D7.2 | 0.608405602 | 1.550258049 | 1.593636649 | 0.972780119 |
| 1700017G19Rik | 0.863538605 | -1.129185089 | 1.076177587 | -1.049255348 |
| Gm23810 | 0 | NA | NA | NA |
| Intu | 226.3559068 | -0.13340272 | 0.090971934 | -1.466416229 |
| Gm16114 | 0 | NA | NA | NA |
| Slc25a31 | 2.917089469 | 0.417091868 | 0.766304902 | 0.544289704 |
| Hspa4l | 1914.656164 | -0.008806556 | 0.073283534 | -0.120171005 |
| Plk4 | 7.648973505 | -0.248275863 | 0.367796953 | -0.675035128 |
| AC146980.1 | 0 | NA | NA | NA |
| Mfsd8 | 158.5183234 | 0.01487294 | 0.107628469 | 0.138187789 |
| 3110057O12Rik | 161.8450865 | 0.537678226 | 0.194848758 | 2.759464482 |
| Gm10731 | 0.47056225 | 1.847196245 | 1.500453591 | 1.231091889 |
| 6430590A07Rik | 35.09747937 | -0.050364622 | 0.334773354 | -0.150443937 |
| 1700034I23Rik | 0.704088048 | 0.674777819 | 1.233266363 | 0.547146861 |
| Larp1b | 264.3748089 | 0.393943339 | 0.161643464 | 2.437112694 |
| Gm22043 | 4.896223194 | 0.632301134 | 0.502067624 | 1.25939436 |
| RP23-235G13.3 | 37.44091296 | 0.677525588 | 0.345249024 | 1.96242579 |
| RP23-235G13.4 | 1.041424966 | 0.761350199 | 0.967939052 | 0.786568325 |

| | | | | |
|---|---|---|---|---|
| Pgrmc2 | 739.9745203 | 0.010316013 | 0.08033955 | 0.128405167 |
| RP23-144N19.1 | 0.207481443 | -1.482962728 | 2.601748577 | -0.569986947 |
| RP23-149I5.1 | 0 | NA | NA | NA |
| RP23-325M18.3 | 0 | NA | NA | NA |
| Gm25487 | 0 | NA | NA | NA |
| RP23-325M18.5 | 0.106618422 | 0.517955464 | 3.724945049 | 0.139050498 |
| Platr4 | 0.110814304 | 0.037060453 | 3.724945049 | 0.009949262 |
| RP23-325M18.2 | 0.374922842 | 0.982232796 | 1.684535814 | 0.583088105 |
| RP23-325M18.1 | 0.284631292 | 0.03706806 | 2.413321104 | 0.015359771 |
| Jade1 | 693.3829749 | 0.141148013 | 0.068789093 | 2.051895259 |
| Gm31266 | 2.246458349 | 1.083173122 | 0.755034003 | 1.434601775 |
| Sclt1 | 316.1906754 | 0.335949109 | 0.125376824 | 2.67951524 |
| Gm38047 | 3.02071253 | -0.237091733 | 0.626788511 | -0.378264325 |
| D3Ertd751e | 153.9838688 | 0.48216628 | 0.158804712 | 3.036221496 |
| Gm37268 | 18.97111997 | 0.3595229 | 0.341936987 | 1.051430273 |
| Gm7899 | 12.35707777 | -0.628021578 | 0.342376997 | -1.834298398 |
| Gm37348 | 0.897478403 | 0.435139492 | 1.179528128 | 0.368909805 |
| Gm37846 | 1.825474869 | 0.277650286 | 0.894490729 | 0.310400406 |
| Gm38274 | 0 | NA | NA | NA |
| Gm37533 | 0.066050875 | -0.443818988 | 3.724945049 | -0.119147795 |
| Gm38044 | 0 | NA | NA | NA |
| Gm9723 | 0 | NA | NA | NA |
| Gm18865 | 0 | NA | NA | NA |
| Gm36976 | 0.179149492 | 1.051578331 | 2.712320496 | 0.387704304 |
| Gm29739 | 0 | NA | NA | NA |
| Gm25714 | 0 | NA | NA | NA |
| Gm32175 | 0 | NA | NA | NA |
| Gm37746 | 0 | NA | NA | NA |
| Gm37365 | 0 | NA | NA | NA |
| Gm38128 | 0 | NA | NA | NA |
| Gm37597 | 0 | NA | NA | NA |
| Gm24997 | 0 | NA | NA | NA |
| Gm27989 | 0 | NA | NA | NA |
| Gm38124 | 15.15835603 | 0.224153895 | 0.322469784 | 0.695115965 |
| 5930409G06Rik | 29.06761066 | 0.217786188 | 0.24834157 | 0.876962274 |
| 2610316D01Rik | 48.48921872 | -0.099072635 | 0.20474016 | -0.483894491 |
| Gm36996 | 0.744898608 | 0.011515749 | 1.279594496 | 0.00899953 |
| Pcdh10 | 3146.497889 | 0.393613796 | 0.080829507 | 4.869679521 |
| 1700027H10Rik | 0.202816535 | 0.262125258 | 3.198091588 | 0.081963024 |
| Gm2136 | 79.20150034 | 0.646765144 | 0.136889562 | 4.724722146 |
| Gm37345 | 48.63821402 | 0.990697259 | 0.215361876 | 4.600151499 |
| Gm37835 | 0 | NA | NA | NA |
| Gm38166 | 0.402076344 | 1.49630196 | 1.711413622 | 0.874307614 |
| Gm32340 | 0.086930093 | -0.443164241 | 3.724898049 | -0.118973522 |
| Gm25796 | 0 | NA | NA | NA |
| Gm23337 | 0 | NA | NA | NA |
| Gm26081 | 0 | NA | NA | NA |

| | | | | |
|---|---|---|---|---|
| Gm18843 | 0 | NA | NA | NA |
| Pabpc4l | 7.77852774 | -0.702805241 | 0.469391924 | -1.497267432 |
| Gm37445 | 0 | NA | NA | NA |
| Mir6379 | 0 | NA | NA | NA |
| Gm10356 | 11.08110355 | 0.170321867 | 0.313165757 | 0.5438713 |
| Gm5274 | 0 | NA | NA | NA |
| Gm23891 | 0 | NA | NA | NA |
| Rpl23a-ps5 | 0.044544648 | 0.517957688 | 3.724945049 | 0.139051095 |
| Gm10730 | 0 | NA | NA | NA |
| Gm2229 | 0 | NA | NA | NA |
| Gm7977 | 1.207653455 | -0.150506922 | 0.931252387 | -0.161617757 |
| Gm37295 | 0 | NA | NA | NA |
| 1700018B24Rik | 0 | NA | NA | NA |
| Gm37190 | 0.042716945 | 0.037062677 | 3.724945049 | 0.009949859 |
| Gm23511 | 0 | NA | NA | NA |
| Gm31415 | 0 | NA | NA | NA |
| Gm37819 | 0.126225412 | 0.03707172 | 3.698755067 | 0.010022756 |
| Gm37684 | 0 | NA | NA | NA |
| Gm20557 | 0.243142889 | -0.411081944 | 2.122743798 | -0.193655939 |
| Gm5846 | 20.89105308 | -0.302652731 | 0.250587761 | -1.207771398 |
| Gm22480 | 0 | NA | NA | NA |
| Gm37424 | 0 | NA | NA | NA |
| Gm38261 | 0.543563254 | -0.37300495 | 1.376379275 | -0.27100448 |
| Gm38088 | 0 | NA | NA | NA |
| Gm2328 | 0 | NA | NA | NA |
| Gm29741 | 0 | NA | NA | NA |
| Gm23096 | 0 | NA | NA | NA |
| Pcdh18 | 119.7553779 | -0.003143511 | 0.167239993 | -0.018796408 |
| Gm37550 | 0.066050875 | -0.443818988 | 3.724945049 | -0.119147795 |
| Slc7a11 | 1346.770682 | -0.58097661 | 0.147210666 | -3.946566016 |
| Gm37854 | 0 | NA | NA | NA |
| Gm37826 | 0 | NA | NA | NA |
| Gm37692 | 0 | NA | NA | NA |
| Gm2345 | 0 | NA | NA | NA |
| Gm37498 | 31.49812525 | -0.730273162 | 0.303375595 | -2.407158565 |
| Gm6209 | 0.762494802 | 0.035603084 | 1.094624116 | 0.032525397 |
| 5830415G21Rik | 0.198884431 | -0.50322743 | 3.225284697 | -0.156025739 |
| Gm38237 | 0.062426873 | 0.517959912 | 3.724945049 | 0.139051692 |
| Gm37199 | 2.566445083 | -1.80598473 | 0.641333383 | -2.815984289 |
| Gm37461 | 0.509388139 | -0.6142155 | 1.439589089 | -0.426660291 |
| Gm24503 | 0.088534442 | -0.905414734 | 3.723500899 | -0.243162217 |
| Gm37548 | 1.788766466 | -0.929762368 | 0.860133852 | -1.080950791 |
| Gm37843 | 13.03457715 | -1.033943589 | 0.384987442 | -2.685655367 |
| Gm37209 | 0.2407585 | -1.317939465 | 2.075179277 | -0.635096678 |
| Gm29230 | 0.044191549 | 0.037062677 | 3.724945049 | 0.009949859 |
| Gm38246 | 0 | NA | NA | NA |
| Ccrn4l | 783.7619654 | -0.948264065 | 0.12735404 | -7.445889134 |

| | | | | |
|---|---|---|---|---|
| Gm38357 | 107.4530042 | -1.204360732 | 0.235393766 | -5.116366306 |
| Elf2 | 758.9211253 | -0.175872298 | 0.078352262 | -2.244635878 |
| 4930577N17Rik | 6.225444594 | 0.366343418 | 0.471858277 | 0.776384427 |
| Gm24282 | 0 | NA | NA | NA |
| Gm37399 | 7.483535934 | 0.440228016 | 0.385783564 | 1.141126933 |
| Gm37203 | 0.04393175 | 0.037062677 | 3.724945049 | 0.009949859 |
| Gm5103 | 0.703642298 | 0.220757115 | 1.344272622 | 0.164220495 |
| Mgarp | 3.540390614 | -0.428638446 | 0.602769291 | -0.711115267 |
| Gm9442 | 4.775535441 | -0.523815037 | 0.605222486 | -0.865491698 |
| Ndufc1 | 324.928662 | -0.197481484 | 0.10152976 | -1.945060087 |
| Naa15 | 1426.402046 | -0.073951242 | 0.07025906 | -1.052550978 |
| Gm37646 | 5.007836009 | -0.172840316 | 0.585348032 | -0.295277862 |
| Gm38160 | 2.044538346 | -0.918945008 | 0.799890048 | -1.148839157 |
| Rab33b | 363.4884665 | 0.008548084 | 0.095509045 | 0.089500254 |
| Gm37968 | 16.21162866 | 0.223010721 | 0.352788479 | 0.632137199 |
| Setd7 | 3529.863564 | 0.247324579 | 0.086211929 | 2.868797637 |
| Gm38200 | 38.03688965 | 0.46782673 | 0.303348203 | 1.542210323 |
| 5031434O11Rik | 65.52831371 | 0.643573994 | 0.172393581 | 3.733166814 |
| Gm38334 | 0 | NA | NA | NA |
| 5430433H01Rik | 0.38231627 | 2.002815626 | 3.698171442 | 0.541569167 |
| Gm38247 | 0 | NA | NA | NA |
| Mgst2 | 4.6105502 | -0.149317826 | 0.426421573 | -0.350164804 |
| Gm37261 | 0.485780372 | -0.643876074 | 1.236173794 | -0.520862097 |
| Maml3 | 325.2409999 | -0.005602795 | 0.095791394 | -0.058489549 |
| Gm10729 | 0.120349074 | 1.027495679 | 3.723011807 | 0.275985071 |
| Gm37342 | 22.56178576 | -0.345630237 | 0.286234373 | -1.207507796 |
| Gm37229 | 0.104366086 | 0.51795324 | 3.724945049 | 0.139049901 |
| n-R5s196 | 0.213343419 | 0.830199536 | 2.562041708 | 0.32403826 |
| 3110080O07Rik | 13.48253032 | 0.004618097 | 0.465679581 | 0.009916899 |
| Gm38000 | 2.827666343 | -1.462547279 | 0.848425185 | -1.723837653 |
| Gm37675 | 6.636235977 | 0.276467109 | 0.516434349 | 0.535338344 |
| Gm38252 | 9.035061384 | 0.540067934 | 0.478537354 | 1.128580516 |
| Gm10728 | 9.423739219 | 1.416799475 | 0.505538534 | 2.802554861 |
| Gm20089 | 0.225870551 | -1.005563361 | 2.575825953 | -0.390384824 |
| Gm38055 | 26.67308699 | 0.467671802 | 0.329274025 | 1.420311857 |
| Gm37526 | 8.63944635 | 0.324037212 | 0.413145994 | 0.784316482 |
| Gm37465 | 35.32081551 | 0.261350179 | 0.334128536 | 0.782184553 |
| C130089K02Rik | 104.1019996 | -0.026141675 | 0.318682908 | -0.082030365 |
| 5430420F09Rik | 30.46786594 | 0.068836586 | 0.304528599 | 0.226043091 |
| Gm37202 | 0 | NA | NA | NA |
| Gm30074 | 0.368444795 | -1.27109476 | 1.42526424 | -0.891830949 |
| Gm38132 | 0 | NA | NA | NA |
| Gm37170 | 1.831335546 | -0.57000859 | 0.82764298 | -0.688713134 |
| Olfr1398-ps1 | 0 | NA | NA | NA |
| Foxo1 | 708.0426262 | 0.339408934 | 0.106001335 | 3.20193074 |
| Gm20402 | 0.491186624 | 2.724374485 | 3.296379507 | 0.826474767 |
| Gm38034 | 38.7415979 | 0.324612376 | 0.324214283 | 1.00122787 |

| | | | | |
|---|---|---|---|---|
| Gm38098 | 1.419478831 | -0.023626802 | 0.870146765 | -0.027152663 |
| Gm30173 | 1.249461252 | -0.822887936 | 1.030684546 | -0.798389711 |
| Gm10293 | 11.59886104 | 0.186842432 | 0.308529194 | 0.605590771 |
| Gm22798 | 7.790214668 | 0.338396364 | 0.447430755 | 0.75631002 |
| Gm2447 | 1.231911519 | 2.264804019 | 1.150516171 | 1.968511245 |
| Gm30292 | 0.646295659 | -0.86862595 | 1.27713698 | -0.680135305 |
| Gm20750 | 0.229084632 | -0.393775147 | 2.576071145 | -0.152858801 |
| Gm37007 | 0 | NA | NA | NA |
| Cog6 | 435.3182281 | -0.144470721 | 0.070906402 | -2.037484862 |
| RP23-256O22.3 | 10.07518621 | 0.300817772 | 0.334479325 | 0.899361335 |
| Lhfp | 547.2260578 | -0.336959908 | 0.101739248 | -3.311995266 |
| RP23-372C7.5 | 0.218578798 | -0.777486553 | 2.579319704 | -0.301430859 |
| RP23-372C7.4 | 86.73692889 | 0.215435674 | 0.295361451 | 0.729396721 |
| RP23-372C7.2 | 134.5349933 | 0.177609996 | 0.364130813 | 0.487764259 |
| RP23-372C7.7 | 21.66084825 | 0.137526888 | 0.342020839 | 0.402100904 |
| RP23-372C7.6 | 0 | NA | NA | NA |
| RP23-372C7.8 | 30.07124974 | -0.003569463 | 0.385341369 | -0.00926312 |
| RP23-335E20.1 | 0 | NA | NA | NA |
| RP23-335E20.2 | 0.902201473 | 0.294232197 | 1.11147535 | 0.264722197 |
| Nhlrc3 | 292.4384838 | -0.241276668 | 0.105530677 | -2.286317822 |
| Gm16206 | 0.175945213 | -0.410809826 | 2.710746807 | -0.151548579 |
| Proser1 | 377.6447841 | 0.175210965 | 0.115843023 | 1.512486134 |
| Stoml3 | 10.96884363 | 0.01235277 | 0.817109844 | 0.015117637 |
| RP23-335E20.3 | 0.440404983 | 0.510370286 | 1.574080235 | 0.324233972 |
| Gm23675 | 0.061920613 | -0.443816764 | 3.724945049 | -0.119147198 |
| Frem2 | 82.40792598 | 0.453063409 | 0.143177762 | 3.164342021 |
| RP24-291E8.1 | 0 | NA | NA | NA |
| RP24-291E8.2 | 16.32404086 | 0.045117226 | 0.268034069 | 0.168326459 |
| RP24-291E8.4 | 0.325009107 | 1.061518688 | 1.77894636 | 0.596712027 |
| RP24-291E8.3 | 0.344979474 | 0.631488266 | 1.757183387 | 0.359375277 |
| RP24-211P15.1 | 0.713939428 | 0.38691734 | 1.252125037 | 0.309008549 |
| Gm24851 | 0 | NA | NA | NA |
| Gm10985 | 0 | NA | NA | NA |
| Ufm1 | 702.309027 | -0.167656271 | 0.061874293 | -2.709627283 |
| RP23-391P1.1 | 0 | NA | NA | NA |
| Gm8109 | 0.115603922 | -0.93550548 | 3.724188147 | -0.251197158 |
| RP23-234G15.2 | 0 | NA | NA | NA |
| Trpc4 | 278.9395309 | 0.277329679 | 0.102987623 | 2.692844744 |
| Postn | 86.52036441 | -0.383436892 | 0.273594172 | -1.40148048 |
| Gm5641 | 12.39303311 | 0.381401885 | 0.448816606 | 0.849794504 |
| Gm26678 | 0 | NA | NA | NA |
| RP23-382K4.1 | 0 | NA | NA | NA |
| Supt20 | 511.0808974 | -0.144603169 | 0.100968666 | -1.432158854 |
| Gm21958 | 0.229069447 | 1.094997994 | 2.533285998 | 0.432244127 |
| Exosc8 | 123.3892681 | 0.047108059 | 0.118022141 | 0.399145948 |
| Alg5 | 141.422727 | -0.143632322 | 0.128942551 | -1.113924927 |
| Smad9 | 371.890028 | 0.262226627 | 0.103375144 | 2.536650649 |

| | | | | |
|---|---|---|---|---|
| Rfxap | 213.8530842 | 0.171192102 | 0.115337666 | 1.484268825 |
| Gm10254 | 54.85742217 | -0.475580912 | 0.187133153 | -2.541403829 |
| Sertm1 | 270.0278678 | -0.363981515 | 0.167744774 | -2.169853081 |
| Ccna1 | 5.128124108 | 0.565892221 | 0.554922164 | 1.019768641 |
| RP23-137P16.2 | 0.532719758 | 1.895237283 | 1.477341149 | 1.282870436 |
| RP23-137P16.1 | 8.16035312 | 0.615558646 | 0.380906403 | 1.616036488 |
| Spg20 | 505.3869147 | -0.07903442 | 0.080601308 | -0.980560009 |
| Ccdc169 | 8.621555654 | -0.175590815 | 0.394133618 | -0.445510881 |
| Gm22029 | 0.119587853 | 0.981063299 | 3.723618439 | 0.263470416 |
| Sohlh2 | 0.439952846 | -2.023382742 | 3.230544269 | -0.626328746 |
| Dclk1 | 10037.27044 | 0.137080397 | 0.119440226 | 1.147690373 |
| Gm25132 | 4.787591675 | 1.253354793 | 0.614614946 | 2.039252057 |
| RP24-372J8.2 | 44.9034577 | 0.307805182 | 0.327203331 | 0.940715308 |
| RP24-372J8.3 | 0.907267504 | -1.630433555 | 1.053804369 | -1.547188076 |
| Gm21954 | 0.194279094 | -1.017823543 | 3.719440177 | -0.273649661 |
| RP24-555P13.2 | 122.1621789 | 0.536316862 | 0.245638803 | 2.183355627 |
| RP24-555P13.4 | 139.4117742 | 0.492301976 | 0.265179703 | 1.85648438 |
| Gm25404 | 0.125889889 | -0.443822468 | 3.019818834 | -0.146969898 |
| WI1-49P9.1 | 0.657052355 | -0.756521248 | 1.123490369 | -0.673366918 |
| WI1-49P9.2 | 2.567514962 | -0.379526689 | 0.578255122 | -0.656330873 |
| Nbea | 6573.173349 | 0.441979211 | 0.087817716 | 5.032916254 |
| Mab21l1 | 155.2590693 | -0.027645686 | 0.141819643 | -0.194935524 |
| RP24-549I13.1 | 0.075043205 | 0.517957688 | 3.724945049 | 0.139051095 |
| RP24-549I13.2 | 0.126225412 | 0.03707172 | 3.698755067 | 0.010022756 |
| Gm22784 | 14.91209993 | 0.438935833 | 0.31111706 | 1.4108382 |
| RP24-293C5.5 | 80.47609759 | 0.420844968 | 0.280057746 | 1.502707829 |
| RP24-80L24.3 | 126.7577974 | 0.828638472 | 0.391098283 | 2.118747401 |
| RP24-80L24.2 | 91.10912563 | 0.702632396 | 0.422949612 | 1.661267386 |
| RP24-80L24.1 | 244.6441849 | 0.720942916 | 0.298387757 | 2.41612767 |
| Gm25727 | 0 | NA | NA | NA |
| RP24-111L5.2 | 0.108266322 | -0.955981391 | 3.722845977 | -0.25678779 |
| Gm22269 | 0.061920613 | -0.443816764 | 3.724945049 | -0.119147198 |
| Gm8177 | 1.764273044 | 0.096656613 | 0.727197895 | 0.132916519 |
| RP24-218M19.4 | 0.066050875 | -0.443818988 | 3.724945049 | -0.119147795 |
| RP24-218M19.5 | 0 | NA | NA | NA |
| Tm4sf1 | 103.6301723 | -1.362278015 | 0.230789728 | -5.902680439 |
| RP23-234F9.2 | 0.925167547 | 0.048321512 | 1.040610583 | 0.04643573 |
| Tm4sf4 | 0.133813382 | 0.055433725 | 2.958683727 | 0.018735941 |
| Wwtr1 | 465.1289583 | -0.956603551 | 0.144237243 | -6.632153621 |
| Gm16016 | 0 | NA | NA | NA |
| Gm26671 | 2.020290962 | 0.208553949 | 0.691149788 | 0.301749278 |
| Mir6377 | 0 | NA | NA | NA |
| RP23-132D7.2 | 73.00516792 | 0.224710755 | 0.175255187 | 1.282191755 |
| Commd2 | 168.5798793 | -0.386812298 | 0.110114952 | -3.512804481 |
| RP23-132D7.4 | 6.836736274 | 1.122994212 | 0.478562695 | 2.346597892 |
| Ankub1 | 37.54081804 | -0.183860578 | 0.268795279 | -0.68401714 |
| RP23-132D7.5 | 0 | NA | NA | NA |

| | | | | |
|---|---|---|---|---|
| RP24-352B14.1 | 0.588070563 | 0.699448398 | 1.348731122 | 0.518597359 |
| Rnf13 | 1345.661625 | 0.067719532 | 0.108581416 | 0.623675156 |
| RP24-352B14.2 | 0.154696864 | 0.037074681 | 2.833481357 | 0.013084498 |
| RP24-352B14.3 | 0.210855403 | 0.51796373 | 2.621995755 | 0.197545602 |
| Pfn2 | 3248.658378 | 0.272555957 | 0.080101428 | 3.402635447 |
| Gm24531 | 0.132790949 | 0.517961689 | 2.970953037 | 0.174341931 |
| RP23-1B20.1 | 0.115957212 | 0.517955464 | 3.724945049 | 0.139050498 |
| RP23-1B20.2 | 6.805132405 | 1.298488305 | 0.49979131 | 2.59806099 |
| RP23-1B20.3 | 0.497476243 | -1.122520618 | 1.518048875 | -0.73944959 |
| RP23-1B20.4 | 0.383276703 | 0.460273322 | 1.700499767 | 0.270669441 |
| Gm5537 | 1.183520696 | 1.244718381 | 1.005585931 | 1.237804093 |
| RP23-375H15.2 | 20.27877935 | 1.490072421 | 0.336373464 | 4.429815614 |
| 1700007F19Rik | 6.275202345 | 0.112423699 | 0.519228728 | 0.216520567 |
| Gm26166 | 0 | NA | NA | NA |
| RP24-376N4.1 | 0 | NA | NA | NA |
| RP23-413C3.1 | 0 | NA | NA | NA |
| Tsc22d2 | 1435.051753 | -0.227668503 | 0.070318415 | -3.237679686 |
| Serp1 | 878.7409205 | -0.430771819 | 0.13046554 | -3.301805372 |
| Eif2a | 501.9842971 | 0.187809363 | 0.071276464 | 2.634942197 |
| RP23-413C3.4 | 13.02020907 | -0.170408432 | 0.329120217 | -0.517769568 |
| Selt | 2800.757624 | -0.160137984 | 0.094998674 | -1.685686525 |
| Erich6 | 23.0190923 | 0.033045769 | 0.252141845 | 0.131060232 |
| RP24-275F18.1 | 4.381802891 | 1.029074708 | 0.558555219 | 1.842386702 |
| RP24-275F18.2 | 3.555894033 | -1.127195291 | 0.74130816 | -1.520548878 |
| Siah2 | 196.4969752 | -0.105589333 | 0.137256014 | -0.769287473 |
| RP24-275F18.4 | 3.656147857 | 0.149659426 | 0.562164333 | 0.266220065 |
| 4930593A02Rik | 1.120771396 | 0.976910799 | 1.028917883 | 0.949454583 |
| Fam188b2 | 11.97822246 | 0.68636571 | 0.352888111 | 1.944995277 |
| Clrn1 | 6.849902671 | 0.269106525 | 0.394779002 | 0.681663725 |
| Gm22491 | 0 | NA | NA | NA |
| Gm25572 | 0.089089296 | 0.957155977 | 3.723939561 | 0.257027796 |
| Gm10071 | 7.465476509 | -0.231650057 | 0.388723314 | -0.595925298 |
| Gm27793 | 0.08169834 | -0.443821212 | 3.724945049 | -0.119148392 |
| RP24-493L18.4 | 21.6695156 | 0.540309365 | 0.276844976 | 1.951667582 |
| Med12l | 1187.74355 | 0.457756562 | 0.094877717 | 4.824700407 |
| Gpr171 | 49.42350281 | -0.988211937 | 0.365694428 | -2.702288748 |
| P2ry14 | 82.16983007 | -0.946843719 | 0.231071994 | -4.097613494 |
| Gpr87 | 0.049553047 | -0.443818988 | 3.724945049 | -0.119147795 |
| P2ry13 | 94.23721791 | -0.026322492 | 0.186731858 | -0.14096412 |
| P2ry12 | 174.5045895 | 1.157400454 | 0.234260881 | 4.940647586 |
| RP24-427G24.5 | 16.25700884 | 1.083979007 | 0.423382834 | 2.560280958 |
| Igsf10 | 258.1667321 | -0.272487413 | 0.181759501 | -1.499164621 |
| Gm38186 | 1.889693139 | -0.606941842 | 0.825062474 | -0.735631374 |
| Gm37715 | 0.038981395 | -0.443816764 | 3.724945049 | -0.119147198 |
| Gm8276 | 1.405995652 | 0.960648568 | 0.914084427 | 1.050940744 |
| Gm38288 | 0 | NA | NA | NA |
| Gm37771 | 0 | NA | NA | NA |

| | | | | |
|---|---|---|---|---|
| Gm4856 | 0.075043205 | 0.517957688 | 3.724945049 | 0.139051095 |
| Gm5709 | 0 | NA | NA | NA |
| Gm37926 | 0 | NA | NA | NA |
| Gm5538 | 0 | NA | NA | NA |
| Gm38208 | 0 | NA | NA | NA |
| Gm37579 | 0 | NA | NA | NA |
| Gm5539 | 2.248417857 | -0.066315895 | 0.68878893 | -0.096278979 |
| 4930449A18Rik | 0.066050875 | -0.443818988 | 3.724945049 | -0.119147795 |
| Gm2756 | 0 | NA | NA | NA |
| Gm8298 | 0.066050875 | -0.443818988 | 3.724945049 | -0.119147795 |
| C130079G13Rik | 0.049553047 | -0.443818988 | 3.724945049 | -0.119147795 |
| Gm9696 | 0.049553047 | -0.443818988 | 3.724945049 | -0.119147795 |
| Aadacl2 | 0 | NA | NA | NA |
| Gm18427 | 0 | NA | NA | NA |
| Aadac | 0.048949532 | 0.037064902 | 3.724945049 | 0.009950456 |
| Sucnr1 | 0.697362497 | -0.615195622 | 1.36459904 | -0.450825191 |
| Gm24382 | 0 | NA | NA | NA |
| RP23-254F12.1 | 1.05607488 | 1.87968821 | 1.111249804 | 1.691508248 |
| Mbnl1 | 2752.745771 | 0.017588253 | 0.049907345 | 0.352418134 |
| Gm37589 | 184.5056286 | -0.150834097 | 0.200899607 | -0.750793389 |
| Gm37488 | 59.18839802 | 0.164893739 | 0.211676644 | 0.778988823 |
| Gm38280 | 0.072025462 | 0.517959912 | 3.724945049 | 0.139051692 |
| Gm33707 | 0.263895486 | 0.037058996 | 3.724945049 | 0.009948871 |
| Gm38326 | 0 | NA | NA | NA |
| Gm36985 | 0.325677113 | 0.520691186 | 1.583336457 | 0.328856943 |
| Gm8325 | 112.612681 | 0.143262733 | 0.290558502 | 0.493059857 |
| Gm37035 | 1.517989225 | 1.116045858 | 1.056307623 | 1.056553824 |
| P2ry1 | 94.52974778 | 0.206614131 | 0.141784613 | 1.457239449 |
| Gm37719 | 0.066050875 | -0.443818988 | 3.724945049 | -0.119147795 |
| Gm19901 | 0.038981395 | -0.443816764 | 3.724945049 | -0.119147198 |
| Gm37696 | 26.28877038 | 0.537514744 | 0.230944245 | 2.327465423 |
| Rap2b | 869.3150327 | 0.061336657 | 0.081756429 | 0.750236505 |
| B430305J03Rik | 35.36276701 | 0.039581456 | 0.197403799 | 0.200510102 |
| Gm8349 | 2.822290259 | 0.072916063 | 0.628284332 | 0.116055836 |
| Gm9700 | 0.470561891 | 1.912027072 | 1.539195175 | 1.242225225 |
| Gm37404 | 0.15100991 | 0.51796601 | 3.527597456 | 0.146832516 |
| 9330121J05Rik | 22.77484088 | 0.473833038 | 0.2477395 | 1.912626114 |
| Arhgef26 | 647.6477621 | -0.373521065 | 0.110393876 | -3.383530672 |
| Dhx36 | 1564.510516 | -0.049104563 | 0.063606775 | -0.772002097 |
| Gpr149 | 48.93600836 | 0.574967083 | 0.184395761 | 3.118114432 |
| Gm18382 | 0.246177344 | -0.443821758 | 2.50940469 | -0.176863365 |
| Gm9701 | 0.309747539 | -0.224968724 | 3.720736698 | -0.060463489 |
| Gm22433 | 0 | NA | NA | NA |
| Gm8388 | 2.905967886 | -1.251680823 | 0.604728712 | -2.069822052 |
| Mme | 252.6871563 | 0.667794376 | 0.093592492 | 7.135127593 |
| Gm34240 | 0 | NA | NA | NA |
| Gm34302 | 0.608629602 | 0.361575678 | 1.349696728 | 0.267894017 |

| | | | | |
|---|---|---|---|---|
| Gm37279 | 0 | NA | NA | NA |
| Gm24946 | 0 | NA | NA | NA |
| Plch1 | 346.9457733 | 0.072984934 | 0.092698562 | 0.7873362 |
| Gm23500 | 0 | NA | NA | NA |
| Gm16162 | 0 | NA | NA | NA |
| Gm16129 | 0 | NA | NA | NA |
| Gm34379 | 0.085327104 | -0.905414734 | 3.723500899 | -0.243162217 |
| E130311K13Rik | 62.02924702 | 0.357979771 | 0.193401058 | 1.85097111 |
| Slc33a1 | 460.1328625 | -0.249553937 | 0.084892102 | -2.939660246 |
| Gm26850 | 5.346930916 | 0.464086708 | 0.43509664 | 1.066629031 |
| Gmps | 2065.915186 | 0.131529324 | 0.054638716 | 2.407255013 |
| Vmn2r-ps3 | 0.09176491 | 0.921395572 | 3.723061103 | 0.247483333 |
| Vmn2r1 | 7.920741964 | 0.335248854 | 0.360725443 | 0.929374017 |
| Gm22165 | 0.115603922 | -0.93550548 | 3.724188147 | -0.251197158 |
| Vmn2r2 | 2.02639444 | -0.15772848 | 0.72951678 | -0.216209529 |
| Gm5139 | 0.217472118 | 0.119789578 | 2.159708166 | 0.055465632 |
| Gm18294 | 0.049553047 | -0.443818988 | 3.724945049 | -0.119147795 |
| Vmn2r3 | 0.204914957 | 0.027628924 | 2.19975699 | 0.012559989 |
| Gm23897 | 0.049553047 | -0.443818988 | 3.724945049 | -0.119147795 |
| Vmn2r-ps4 | 0.124853745 | 1.05357957 | 3.721447096 | 0.283110183 |
| Vmn2r-ps5 | 0.174149299 | -0.429800891 | 3.359371215 | -0.127940874 |
| Vmn2r4 | 0.145501641 | -0.443200222 | 2.888011192 | -0.153462086 |
| Vmn2r-ps6 | 0 | NA | NA | NA |
| Vmn2r-ps7 | 0 | NA | NA | NA |
| Gm20656 | 0.439965521 | 1.41770379 | 1.536321247 | 0.922791241 |
| Vmn2r-ps8 | 0 | NA | NA | NA |
| Vmn2r5 | 0.344244685 | -0.543095608 | 1.831209261 | -0.296577578 |
| Vmn2r6 | 0.549943695 | -0.251061782 | 1.456480148 | -0.172375698 |
| Vmn2r-ps9 | 0.898522788 | -0.034767571 | 1.074490555 | -0.03235726 |
| Gm20657 | 0.48442959 | 0.364860431 | 1.362649188 | 0.267758154 |
| Vmn2r-ps10 | 0 | NA | NA | NA |
| Vmn2r-ps11 | 0.791552024 | 0.021192311 | 1.089486979 | 0.019451642 |
| Gm26939 | 0 | NA | NA | NA |
| Vmn2r7 | 3.934381616 | 0.647490541 | 0.580243086 | 1.115895314 |
| Vmn2r-ps12 | 0.044544648 | 0.517957688 | 3.724945049 | 0.139051095 |
| Gm20698 | 0 | NA | NA | NA |
| Gm25222 | 0.066050875 | -0.443818988 | 3.724945049 | -0.119147795 |
| Vmn2r-ps13 | 0 | NA | NA | NA |
| Gm38048 | 50.77510117 | 0.412290343 | 0.225406786 | 1.829094633 |
| Kcnab1 | 1328.585016 | 0.394594621 | 0.094081814 | 4.194164684 |
| A330015K06Rik | 13.82607597 | 0.895065435 | 0.370139737 | 2.418182498 |
| A730090N16Rik | 2.24970321 | 0.080082812 | 0.716576822 | 0.111757469 |
| Ssr3 | 1935.252712 | -0.175155282 | 0.092327047 | -1.897117779 |
| Gm5540 | 6.757178092 | -0.790749372 | 0.41157527 | -1.92127523 |
| Gm34780 | 0.183309527 | -0.444674696 | 2.685766745 | -0.165567132 |
| Gm37009 | 0.786503428 | 2.041272412 | 1.216284113 | 1.678285846 |
| 4931440P22Rik | 18.83552398 | -0.812656003 | 0.287785983 | -2.823820653 |

| | | | | |
|---|---|---|---|---|
| Gm22279 | 0 | NA | NA | NA |
| Tiparp | 657.7234625 | -0.732649617 | 0.23343035 | -3.138621939 |
| Gm5847 | 0.982102764 | -0.790548563 | 1.14818963 | -0.688517421 |
| Gm38233 | 0 | NA | NA | NA |
| Mir8120 | 0.138110619 | 0.482857281 | 3.622723928 | 0.133285696 |
| Lekr1 | 106.5364298 | -0.132723667 | 0.198106903 | -0.669959828 |
| Gm37776 | 35.53555177 | 0.232448212 | 0.328297263 | 0.708041884 |
| Gm25755 | 0 | NA | NA | NA |
| Gm37359 | 0.166952883 | -1.336542448 | 2.746539428 | -0.486627803 |
| Gm37131 | 0.161076545 | -0.410543693 | 2.787893798 | -0.147259445 |
| Gm37037 | 0.659599365 | -1.461331024 | 1.291122663 | -1.131829736 |
| Ccnl1 | 947.3859406 | -0.035196238 | 0.135781482 | -0.259212353 |
| Gm37305 | 112.0234715 | -0.114138483 | 0.10829785 | -1.053931202 |
| Gm35106 | 0.709932405 | -1.225572391 | 1.260520287 | -0.972275023 |
| Gm37078 | 4.562323805 | -0.617038728 | 0.545681952 | -1.130766239 |
| Gm19024 | 0 | NA | NA | NA |
| Gm6546 | 0 | NA | NA | NA |
| Veph1 | 5.036914537 | -0.863814121 | 0.606829057 | -1.423488396 |
| Gm37822 | 0 | NA | NA | NA |
| Gm26442 | 0 | NA | NA | NA |
| Gm36973 | 0.066050875 | -0.443818988 | 3.724945049 | -0.119147795 |
| Tpi-rs2 | 0 | NA | NA | NA |
| Gm37330 | 0 | NA | NA | NA |
| Ptx3 | 208.6062493 | -1.165704839 | 0.575332831 | -2.026139959 |
| Gm17952 | 0.276493613 | 1.436715596 | 1.780142965 | 0.807078771 |
| Gm6555 | 0 | NA | NA | NA |
| Gm38356 | 0.174174109 | 0.037075208 | 2.729571866 | 0.013582793 |
| RP23-103D17.4 | 0 | NA | NA | NA |
| Shox2 | 134.0978986 | 0.144124138 | 0.14878183 | 0.968694479 |
| Rsrc1 | 525.7525617 | 0.280255035 | 0.064330744 | 4.356471214 |
| Gm22295 | 1.86515265 | 0.299346683 | 0.771581666 | 0.387965003 |
| Gm17402 | 0 | NA | NA | NA |
| Mlf1 | 15.4417281 | -0.536765396 | 0.308139049 | -1.741958371 |
| Gfm1 | 616.4629507 | -0.073895005 | 0.050989795 | -1.44921164 |
| Lxn | 276.0893367 | -0.069864241 | 0.160422301 | -0.435502051 |
| Gm37453 | 18.12819328 | 0.15956648 | 0.293244784 | 0.544140898 |
| Rarres1 | 12.46569881 | 0.164270516 | 0.338849146 | 0.484789523 |
| Gm35299 | 0.137416547 | -0.443827336 | 3.639393827 | -0.121950896 |
| 4930402C01Rik | 0 | NA | NA | NA |
| Gm25919 | 0 | NA | NA | NA |
| Mfsd1 | 388.2294903 | -0.565563812 | 0.111953805 | -5.051760502 |
| Gm8515 | 6.537093261 | -1.196805628 | 0.417298313 | -2.867985783 |
| Gm412 | 0 | NA | NA | NA |
| Gm37601 | 0.128477747 | 0.037074008 | 3.677132019 | 0.010082316 |
| Gm18717 | 0.045882455 | 0.517959912 | 3.724945049 | 0.139051692 |
| Iqcj | 18.38164937 | 0.987958614 | 0.387384359 | 2.550331708 |
| Schip1 | 1612.457602 | 0.293516318 | 0.088207478 | 3.327567274 |

| | | | | |
|---|---|---|---|---|
| Gm10292 | 11.84136725 | 0.998300219 | 0.492758101 | 2.0259438 |
| Gm10723 | 64.31874068 | 0.882026889 | 0.354339123 | 2.489216772 |
| Gm22073 | 0 | NA | NA | NA |
| Il12a | 10.96521039 | -0.312508777 | 0.36297901 | -0.860955505 |
| Gm10040 | 0.437154627 | 0.020527864 | 1.394834734 | 0.014717058 |
| 4930535E02Rik | 0 | NA | NA | NA |
| Gm35584 | 0.044191549 | 0.037062677 | 3.724945049 | 0.009949859 |
| Gm7270 | 0 | NA | NA | NA |
| Gm17641 | 6.199809403 | 1.657365809 | 0.555951633 | 2.98113309 |
| 1110032F04Rik | 89.81938694 | 0.393578129 | 0.188468376 | 2.088297984 |
| Ift80 | 422.7464304 | -0.070173186 | 0.126541848 | -0.554545292 |
| Gm20689 | 16.19484603 | -0.233219885 | 0.275704817 | -0.84590428 |
| Gm20690 | 0.088383099 | 0.037058229 | 3.724945049 | 0.009948665 |
| Gm18588 | 78.97133165 | -0.334862557 | 0.165363227 | -2.025012229 |
| Gm20691 | 11.60463875 | -0.816595817 | 0.317309114 | -2.573502561 |
| Smc4 | 201.8629177 | -0.317121761 | 0.136948544 | -2.315627109 |
| Mir15b | 0.132199999 | 0.998851712 | 3.642365922 | 0.274231566 |
| Mir16-2 | 0.358973896 | -0.852515457 | 1.460508761 | -0.583711293 |
| Trim59 | 149.282414 | -0.145354661 | 0.14948553 | -0.972366095 |
| Kpna4 | 1744.547512 | -0.356011361 | 0.057907242 | -6.147959157 |
| Gm22009 | 385.5335551 | 0.215076854 | 0.278502241 | 0.772262562 |
| Gm37558 | 126.3325027 | 0.190971986 | 0.216916774 | 0.880392893 |
| Gm1647 | 0.255219096 | -0.924714571 | 1.847884026 | -0.500418077 |
| Arl14 | 0.146692594 | -0.955495625 | 3.521109961 | -0.271362052 |
| Ppm1l | 3121.106828 | 0.275450287 | 0.076907647 | 3.581572138 |
| Gm17214 | 0 | NA | NA | NA |
| Gm17212 | 0 | NA | NA | NA |
| Gm17213 | 0 | NA | NA | NA |
| Gm8565 | 1.479948724 | 0.874733031 | 1.062483836 | 0.823290672 |
| B3galnt1 | 333.6927738 | 0.150525177 | 0.09151035 | 1.644897833 |
| Gm37380 | 0.038981395 | -0.443816764 | 3.724945049 | -0.119147198 |
| Gm25621 | 0 | NA | NA | NA |
| Rpl32-ps | 0.038981395 | -0.443816764 | 3.724945049 | -0.119147198 |
| Nmd3 | 566.0817292 | 0.006206182 | 0.101210767 | 0.061319382 |
| Sptssb | 37.86867121 | -0.11116405 | 0.199122751 | -0.558268954 |
| Gm17895 | 0 | NA | NA | NA |
| Gm23484 | 0 | NA | NA | NA |
| Gm37784 | 14.73966199 | -0.08214659 | 0.293035117 | -0.280330191 |
| Otol1 | 0.139883403 | -1.137320593 | 3.551352803 | -0.32024996 |
| Gm37681 | 0.143485256 | -0.955563601 | 3.547405725 | -0.269369696 |
| Gm37585 | 0 | NA | NA | NA |
| Gm23477 | 0 | NA | NA | NA |
| Gm6631 | 0 | NA | NA | NA |
| Gm37213 | 0 | NA | NA | NA |
| Gm10780 | 0 | NA | NA | NA |
| Gm6634 | 0.207012398 | 0.037073445 | 2.635059324 | 0.014069302 |
| Gm8599 | 0 | NA | NA | NA |

| | | | | |
|---|---|---|---|---|
| Gm38305 | 0 | NA | NA | NA |
| Gm19194 | 0 | NA | NA | NA |
| Gm8604 | 0 | NA | NA | NA |
| Gm37617 | 0 | NA | NA | NA |
| Gm3351 | 0 | NA | NA | NA |
| Gm37603 | 0 | NA | NA | NA |
| Gm27465 | 0 | NA | NA | NA |
| Gm37901 | 0 | NA | NA | NA |
| Gm22451 | 0 | NA | NA | NA |
| Sis | 3.661596079 | 2.433664759 | 0.688194932 | 3.536301482 |
| Gm37056 | 46.70151825 | 0.769772253 | 0.167821829 | 4.586842236 |
| Slitrk3 | 736.758703 | 0.645255731 | 0.085727624 | 7.526812222 |
| Gm20754 | 35.42307591 | 1.295189076 | 0.22354006 | 5.793990901 |
| Gm24544 | 0.953051642 | 1.82323842 | 1.286101379 | 1.417647512 |
| Gm37725 | 9.175969031 | 1.146090863 | 0.384713043 | 2.979079819 |
| Gm22538 | 0 | NA | NA | NA |
| Gm38353 | 21.39725853 | 1.134298465 | 0.42540672 | 2.666385866 |
| 4930509J09Rik | 0 | NA | NA | NA |
| Bche | 53.21720014 | 0.042519705 | 0.177350792 | 0.239749167 |
| Gm8625 | 0 | NA | NA | NA |
| Gm20356 | 0 | NA | NA | NA |
| Gm6098 | 0 | NA | NA | NA |
| Gm37050 | 0 | NA | NA | NA |
| Gm8643 | 0 | NA | NA | NA |
| Gm37464 | 5.385716294 | -0.682886577 | 0.478735973 | -1.426436733 |
| Zbbx | 20.8812694 | 0.031767414 | 0.279210095 | 0.113776021 |
| Serpini2 | 1.184533449 | 0.080746385 | 0.797386597 | 0.101263785 |
| Wdr49 | 44.34530745 | -0.828973782 | 0.283661055 | -2.922409573 |
| Gm17836 | 0 | NA | NA | NA |
| Pdcd10 | 402.9200995 | -0.250769181 | 0.129346271 | -1.938743033 |
| Serpini1 | 2225.756587 | 0.823940738 | 0.095477508 | 8.629684177 |
| Gm37672 | 32.21841053 | 0.975700062 | 0.273363696 | 3.569237891 |
| Platr10 | 0.223571151 | 1.630765231 | 2.536259989 | 0.642980309 |
| Gm37193 | 0 | NA | NA | NA |
| Gm25846 | 0.062426873 | 0.517959912 | 3.724945049 | 0.139051692 |
| Gm29133 | 0 | NA | NA | NA |
| Golim4 | 319.1997903 | -0.160166914 | 0.095609549 | -1.675218805 |
| Gm37685 | 0 | NA | NA | NA |
| Gm37256 | 0.048949532 | 0.037064902 | 3.724945049 | 0.009950456 |
| Gm18428 | 0 | NA | NA | NA |
| Fstl5 | 496.3711575 | 0.275080097 | 0.101599841 | 2.707485499 |
| Gm37897 | 17.02018816 | 0.498329982 | 0.337137171 | 1.478122336 |
| Gm37290 | 0.044544648 | 0.517957688 | 3.724945049 | 0.139051095 |
| Gm19231 | 0.236516182 | 0.491413771 | 2.107367791 | 0.233188423 |
| Gm23644 | 0 | NA | NA | NA |
| Dnmt3a-ps1 | 0.044544648 | 0.517957688 | 3.724945049 | 0.139051095 |
| Gm37806 | 0.072025462 | 0.517959912 | 3.724945049 | 0.139051692 |

| | | | | |
|---|---|---|---|---|
| Gm38195 | 0 | NA | NA | NA |
| Gm37576 | 0 | NA | NA | NA |
| RP23-257I21.2 | 0 | NA | NA | NA |
| Gm6677 | 0 | NA | NA | NA |
| Gm8684 | 0.072025462 | 0.517959912 | 3.724945049 | 0.139051692 |
| Gm6680 | 0 | NA | NA | NA |
| RP23-399J17.1 | 4.954402907 | -0.204776249 | 0.514800944 | -0.397777533 |
| Gm15442 | 3.380166461 | 0.088850695 | 0.534952529 | 0.166090803 |
| Gm18952 | 0 | NA | NA | NA |
| Gm24749 | 0 | NA | NA | NA |
| Gm17953 | 0.338292759 | -2.167381122 | 3.68934737 | -0.58747006 |
| Gm5277 | 0.066050875 | -0.443818988 | 3.724945049 | -0.119147795 |
| Gm10291 | 12.97480729 | 0.041242829 | 0.345518939 | 0.119364887 |
| Gm9762 | 88.84803059 | -0.017013386 | 0.178991623 | -0.095051297 |
| Gm23440 | 0 | NA | NA | NA |
| Rapgef2 | 2966.610995 | 0.521244888 | 0.076989813 | 6.770309871 |
| Gm24150 | 0 | NA | NA | NA |
| RP24-253I2.4 | 0.238666214 | 0.222990047 | 2.527364247 | 0.088230277 |
| 6430573P05Rik | 8.415433566 | -0.107818822 | 0.369624975 | -0.29169788 |
| Gm17359 | 0.372745468 | 0.893659942 | 1.431829979 | 0.624138309 |
| Fnip2 | 1106.231662 | -0.670774286 | 0.146664064 | -4.57354219 |
| Gm3513 | 13.83725982 | -0.301585003 | 0.334390702 | -0.901894106 |
| 4930589L23Rik | 0.532739632 | -0.839772036 | 1.243275169 | -0.675451466 |
| Gm35067 | 0.044544648 | 0.517957688 | 3.724945049 | 0.139051095 |
| Ppid | 668.4459743 | -0.136707119 | 0.075561845 | -1.809208326 |
| Etfdh | 350.8915493 | -0.281340373 | 0.105186154 | -2.674690179 |
| Gm37973 | 5.608718336 | -0.376204924 | 0.470526076 | -0.799541074 |
| 4930579G24Rik | 15.15182425 | -0.268113967 | 0.281665408 | -0.951888159 |
| Rxfp1 | 93.03486081 | 1.288811703 | 0.257537996 | 5.004355567 |
| Gm26984 | 4.172577245 | 0.646600059 | 0.672511887 | 0.961470082 |
| Tmem144 | 92.24781446 | 0.435922723 | 0.184908338 | 2.357507118 |
| Gm26420 | 0 | NA | NA | NA |
| Gm36569 | 0.140714442 | 0.882076007 | 3.579135506 | 0.246449458 |
| Fam198b | 79.79527804 | 0.265626295 | 0.182017794 | 1.459342464 |
| A330069K06Rik | 1.631297931 | 1.682607549 | 0.969848211 | 1.734918444 |
| Gm19066 | 0.045882455 | 0.517959912 | 3.724945049 | 0.139051692 |
| RP23-121N19.4 | 0.091880448 | 0.037060453 | 3.724945049 | 0.009949262 |
| Gm38295 | 0.117760149 | 0.51795324 | 3.724945049 | 0.139049901 |
| Gm37892 | 0.469035539 | 2.275419884 | 1.481764803 | 1.535614748 |
| Gm37971 | 2.078197752 | -0.925225321 | 0.702571217 | -1.316913216 |
| Gm6706 | 0.213298592 | -1.775259985 | 2.136310083 | -0.83099359 |
| Gria2 | 8463.13547 | 0.867116677 | 0.114254763 | 7.589326282 |
| Gm25749 | 0 | NA | NA | NA |
| Glrb | 2044.481699 | 0.179397418 | 0.067148608 | 2.671647595 |
| RP24-445D5.2 | 0 | NA | NA | NA |
| Gm29808 | 0.687463866 | 0.099900177 | 1.283763544 | 0.077818207 |
| RP23-187I18.1 | 0.148659141 | -1.239386499 | 3.716315506 | -0.333498729 |

| | | | | |
|---|---|---|---|---|
| Pdgfc | 119.0878968 | 0.007778731 | 0.117126935 | 0.06641283 |
| Gm16000 | 0.137483974 | -0.891006646 | 2.929862383 | -0.304112115 |
| Gm37300 | 0 | NA | NA | NA |
| Gm4857 | 0 | NA | NA | NA |
| RP23-144D4.1 | 0 | NA | NA | NA |
| A830029E22Rik | 4.685490163 | -0.035762662 | 0.498016468 | -0.0718102 |
| Gm22531 | 0 | NA | NA | NA |
| Gm9989 | 1.473237501 | 0.687158306 | 0.916234887 | 0.749980508 |
| Ctso | 423.7109783 | 0.058826522 | 0.112635218 | 0.522274672 |
| Tdo2 | 9.666145708 | 0.102960347 | 0.371849056 | 0.276887476 |
| Asic5 | 0.518949687 | 1.713203829 | 1.350534455 | 1.268537669 |
| Gucy1b3 | 1608.78633 | 0.42109879 | 0.091347975 | 4.609831698 |
| Gucy1a3 | 860.8261742 | 0.890867817 | 0.110358887 | 8.072461073 |
| Mir7010 | 0.40751274 | 1.251608295 | 1.406562996 | 0.88983451 |
| Map9 | 1296.963444 | 0.477277518 | 0.07118175 | 6.705054617 |
| Npy2r | 123.6004455 | -0.410278683 | 0.206140703 | -1.990284676 |
| Gm35323 | 0 | NA | NA | NA |
| Gm29999 | 0.156071933 | 0.517956562 | 3.489245341 | 0.148443721 |
| RP23-469J3.5 | 0.062426873 | 0.517959912 | 3.724945049 | 0.139051692 |
| RP23-223I2.3 | 0 | NA | NA | NA |
| Gm22038 | 0 | NA | NA | NA |
| RP23-223I2.2 | 0 | NA | NA | NA |
| Rbm46 | 23.88977253 | -0.182894492 | 0.24460151 | -0.747724298 |
| Rbm46os | 0.56520525 | -0.377817359 | 1.476361584 | -0.255911128 |
| Lrat | 10.45023781 | -0.21037304 | 0.499100571 | -0.421504306 |
| Gm30097 | 0.374039128 | 0.409121325 | 1.59952185 | 0.255777266 |
| Fgg | 0.874079576 | -0.637639116 | 1.103090505 | -0.578047869 |
| Fga | 10.18503337 | -0.878012753 | 0.561946291 | -1.562449592 |
| Fgb | 1.174237097 | -1.009236706 | 1.305603325 | -0.773004087 |
| Plrg1 | 372.6327839 | -0.290330778 | 0.096413376 | -3.011312238 |
| Gm10710 | 19.45997954 | 1.15264603 | 0.294130724 | 3.918822271 |
| Dchs2 | 240.9884021 | 0.639687384 | 0.1169864 | 5.468049148 |
| Gm38096 | 1.042898341 | 1.201182402 | 1.097940943 | 1.094031887 |
| 1700028M03Rik | 0.205907367 | -0.925552451 | 2.588042188 | -0.357626493 |
| Gm22342 | 0.083172944 | -0.443821212 | 3.724945049 | -0.119148392 |
| Rpl28-ps2 | 0 | NA | NA | NA |
| Gm24584 | 0 | NA | NA | NA |
| Sfrp2 | 19.78329196 | 0.192357339 | 0.283258737 | 0.679087047 |
| Gm26771 | 5.06408741 | 0.408860419 | 0.46702553 | 0.875456251 |
| Gm7115 | 0 | NA | NA | NA |
| Tlr2 | 267.1921433 | 0.053173547 | 0.210179797 | 0.252990761 |
| D930015E06Rik | 251.151783 | 0.238188566 | 0.152114572 | 1.565849761 |
| Gm36981 | 2.682934602 | -0.609318289 | 0.656572823 | -0.928028494 |
| Mnd1 | 6.972706414 | -0.371970059 | 0.587351156 | -0.633300974 |
| Trim2 | 5153.10411 | 0.235850188 | 0.061473714 | 3.836602261 |
| Gm6525 | 0.265820157 | 0.037068004 | 2.500340898 | 0.01482518 |
| 4930565D16Rik | 0.075043205 | 0.517957688 | 3.724945049 | 0.139051095 |

| | | | | |
|---|---|---|---|---|
| Fhdc1 | 188.3832 | -0.583123797 | 0.105636305 | -5.520107835 |
| Arfip1 | 76.98979055 | -0.117390779 | 0.126063751 | -0.931201697 |
| Gm37240 | 3.253759892 | 0.814126354 | 0.618482014 | 1.316329879 |
| Gm8813 | 4.188906716 | -0.208696949 | 0.563152343 | -0.370587022 |
| Gm38241 | 5.062387905 | 0.67585533 | 0.50023372 | 1.35107911 |
| Tigd4 | 1.127018203 | 0.517961015 | 0.996647668 | 0.519703232 |
| Tmem154 | 14.23682184 | -0.818875316 | 0.394726163 | -2.074540255 |
| Fbxw7 | 2169.618475 | 0.228000665 | 0.067601095 | 3.372736256 |
| Gm37726 | 37.63354527 | 0.133608908 | 0.285110267 | 0.468621874 |
| AC124496.1 | 0 | NA | NA | NA |
| Gm38041 | 0 | NA | NA | NA |
| Gm38313 | 0 | NA | NA | NA |
| 1700036G14Rik | 0.202694752 | 0.03707203 | 2.227489178 | 0.016642968 |
| RP23-94M23.3 | 2.853198453 | 0.60462624 | 0.649786892 | 0.930499287 |
| Mir466k | 0 | NA | NA | NA |
| RP24-389G6.1 | 1.736188672 | -1.835876502 | 1.00251055 | -1.831278985 |
| Gm15535 | 15.003366 | -0.152223585 | 0.522542228 | -0.291313461 |
| Gatb | 159.5508096 | -0.00014629 | 0.109958266 | -0.001330416 |
| Fam160a1 | 54.14219974 | 0.72274187 | 0.150427501 | 4.804586035 |
| Glt28d2 | 6.204872336 | 0.85515569 | 0.442731471 | 1.931544845 |
| RP23-329L9.3 | 0.488911873 | 0.504262491 | 1.476171236 | 0.341601624 |
| Gm26204 | 0 | NA | NA | NA |
| Gm38030 | 0 | NA | NA | NA |
| Gm9790 | 12.80588666 | -0.337779661 | 0.351398493 | -0.961243907 |
| Gm25876 | 0 | NA | NA | NA |
| Gm24501 | 0 | NA | NA | NA |
| Gm27811 | 0 | NA | NA | NA |
| Sh3d19 | 475.5047945 | -0.290025909 | 0.113997424 | -2.544144411 |
| Prss48 | 0.087261593 | 0.51795324 | 3.724945049 | 0.139049901 |
| Gm25188 | 1.524225208 | 0.675239722 | 0.999217612 | 0.675768436 |
| Rps3a1 | 1477.132456 | -0.129138719 | 0.095796772 | -1.34804876 |
| Snord73a | 1.006316758 | 0.067199229 | 0.924199579 | 0.072710733 |
| Rnu73b | 4.371881307 | -0.18933786 | 0.552011535 | -0.342996202 |
| Gm37933 | 3.2042328 | 0.256778283 | 0.611905662 | 0.419637043 |
| Gm37493 | 0 | NA | NA | NA |
| Gm23025 | 0 | NA | NA | NA |
| Lrba | 1713.427863 | 0.273757544 | 0.072650507 | 3.768143613 |
| Gm37288 | 0 | NA | NA | NA |
| Gm26343 | 0 | NA | NA | NA |
| Gm23850 | 3.443253566 | 0.462081358 | 0.724594605 | 0.637710182 |
| Gm25039 | 5.355852112 | 0.552281906 | 0.432160644 | 1.277955114 |
| Mab21l2 | 76.13297546 | -0.499404546 | 0.433064391 | -1.153187739 |
| Gm3788 | 95.68068835 | -0.454557426 | 0.281258457 | -1.616155585 |
| Gm37876 | 2.813292308 | 0.350566746 | 0.764939663 | 0.458293331 |
| Dclk2 | 1006.44227 | 0.073880554 | 0.07705504 | 0.958802358 |
| Gm37025 | 0.066050875 | -0.443818988 | 3.724945049 | -0.119147795 |
| Gm23376 | 0 | NA | NA | NA |

| | | | | |
|---|---|---|---|---|
| Gm37207 | 0.045882455 | 0.517959912 | 3.724945049 | 0.139051692 |
| Cd1d2 | 0.436249931 | -2.192035327 | 1.35851551 | -1.613551933 |
| Cd1d1 | 24.21988694 | -0.575250281 | 0.205968811 | -2.792899946 |
| Gm8869 | 0 | NA | NA | NA |
| Gm18403 | 0 | NA | NA | NA |
| Gm37575 | 0 | NA | NA | NA |
| Gm24472 | 0 | NA | NA | NA |
| Kirrel | 258.042677 | -0.71723194 | 0.148158972 | -4.840961901 |
| Gm37855 | 0.445528452 | -0.430065531 | 1.381694507 | -0.311259493 |
| Gm10705 | 72.15716491 | 0.221451 | 0.147862444 | 1.497682541 |
| Fcrls | 15.92947883 | 1.293622443 | 0.574046578 | 2.253514771 |
| Cd5l | 3.379428938 | -2.02505106 | 0.745453065 | -2.716537305 |
| Fcrl1 | 1.476596612 | -1.786009585 | 0.824653666 | -2.165769292 |
| Fcrl5 | 1.398885825 | -1.628964782 | 0.984527132 | -1.654565658 |
| Etv3 | 674.3690235 | -0.317773148 | 0.11422828 | -2.781913085 |
| Etv3l | 0.263217861 | -0.07228098 | 2.103755092 | -0.034358077 |
| Gm25167 | 0 | NA | NA | NA |
| Gm24109 | 0 | NA | NA | NA |
| Arhgef11 | 1362.693994 | 0.078608255 | 0.040949853 | 1.919622385 |
| Lrrc71 | 7.798021523 | 0.105512721 | 0.434522352 | 0.242824612 |
| Pear1 | 78.16134709 | -0.455541805 | 0.140678887 | -3.238167527 |
| Ntrk1 | 25.65422587 | 0.297516104 | 0.261933947 | 1.135844009 |
| Insrr | 23.94499803 | -0.503345429 | 0.222282717 | -2.264437989 |
| Sh2d2a | 48.27836269 | -0.309116055 | 0.322628088 | -0.958118856 |
| Prcc | 536.6201964 | -0.07759717 | 0.062646417 | -1.238652968 |
| Hdgf | 1352.852318 | -0.236923263 | 0.058613677 | -4.042115654 |
| Mrpl24 | 277.5276893 | -0.434969834 | 0.092617605 | -4.69640554 |
| Rrnad1 | 232.3898889 | 0.077329873 | 0.126085783 | 0.613311601 |
| Isg20l2 | 217.9624788 | -0.401598048 | 0.092686317 | -4.332873098 |
| Crabp2 | 45.98201362 | -1.631523861 | 0.270113954 | -6.040131709 |
| Gm3745 | 0 | NA | NA | NA |
| Nes | 844.5112463 | -2.548901116 | 0.404308582 | -6.304345818 |
| Bcan | 1604.188113 | 1.141723218 | 0.148914748 | 7.666958687 |
| Hapln2 | 312.1547389 | 0.154293218 | 0.168533713 | 0.915503583 |
| RP23-240E15.3 | 115.0424175 | 0.325534977 | 0.196442449 | 1.657151897 |
| Gpatch4 | 115.8497267 | -0.181188899 | 0.183765394 | -0.985979431 |
| Apoa1bp | 265.3851978 | -0.082968434 | 0.10350268 | -0.801606625 |
| Ttc24 | 0.042716945 | 0.037062677 | 3.724945049 | 0.009949859 |
| Iqgap3 | 18.40657348 | -0.430714225 | 0.436465773 | -0.986822454 |
| Mef2d | 2901.836846 | 0.306799888 | 0.077485433 | 3.959452444 |
| 1700113A16Rik | 8.541957373 | -0.447609137 | 0.405773378 | -1.103101291 |
| AI849053 | 5.915945368 | 0.852856723 | 0.485086518 | 1.758153836 |
| AW047730 | 113.9278857 | 0.164806243 | 0.206051869 | 0.799828916 |
| Gm3764 | 811.9285128 | -0.241403893 | 0.181405605 | -1.330741088 |
| Mir3093 | 13.97251529 | -1.180534098 | 0.343678552 | -3.434994974 |
| Gm25641 | 0.574683629 | -0.92511113 | 1.382511061 | -0.669152787 |
| mmu-mir-9-1 | 11.97633615 | -0.509875853 | 0.364977155 | -1.397007583 |

| | | | | |
|---|---|---|---|---|
| Rhbg | 2.720079777 | -1.041756651 | 0.675921165 | -1.541239873 |
| Gm38392 | 2.080370332 | 0.45315263 | 0.795939971 | 0.569330158 |
| Tsacc | 7.84413125 | -0.626896123 | 0.407857379 | -1.537047399 |
| Gm37116 | 2.788796254 | -0.409766445 | 0.727074935 | -0.563582136 |
| Cct3 | 1136.589819 | -0.525355544 | 0.083611799 | -6.283270452 |
| Glmp | 457.6296344 | -0.516072696 | 0.125698794 | -4.105629663 |
| Tmem79 | 9.024460536 | -0.238565766 | 0.328056344 | -0.727209734 |
| Smg5 | 1241.608425 | -0.045147877 | 0.069761172 | -0.647177725 |
| Paqr6 | 85.91657556 | 0.339954386 | 0.176895875 | 1.921776789 |
| Bglap3 | 2.927464552 | -1.765688256 | 0.727095964 | -2.428411577 |
| Gm21956 | 0.097139547 | -0.443816764 | 3.724945049 | -0.119147198 |
| Bglap2 | 0.555158567 | -1.341117528 | 1.220090144 | -1.099195444 |
| Gm6821 | 0.068097359 | 0.037064902 | 3.724945049 | 0.009950456 |
| Bglap | 0.801792645 | -1.261620663 | 1.201366322 | -1.050154844 |
| Pmf1 | 65.8586388 | -0.148376643 | 0.179293153 | -0.827564473 |
| Slc25a44 | 1273.147583 | 0.080731362 | 0.06430976 | 1.255351643 |
| RP24-298F19.6 | 2.520287764 | -0.908240662 | 0.599144229 | -1.515896537 |
| Sema4a | 442.3120329 | -0.486839082 | 0.101336578 | -4.804179198 |
| Mir7011 | 0 | NA | NA | NA |
| Gm5979 | 0 | NA | NA | NA |
| Lmna | 660.5220702 | -0.780626169 | 0.168404227 | -4.635430958 |
| Gm37584 | 6.065917723 | -0.019387573 | 0.485694764 | -0.039917196 |
| Mex3a | 75.1280634 | -0.448210632 | 0.166089481 | -2.69860939 |
| Mir1905 | 0.318342637 | -0.443835465 | 3.724945049 | -0.119152218 |
| Rab25 | 1.64905476 | 0.332695617 | 0.7318109 | 0.454619653 |
| Lamtor2 | 189.4900279 | 0.000893615 | 0.114976109 | 0.007772177 |
| Ubqln4 | 1064.387744 | 0.079729292 | 0.063750615 | 1.250643486 |
| Ssr2 | 493.374432 | -0.223848295 | 0.086131379 | -2.598916888 |
| Gm10704 | 39.754764 | 0.160283186 | 0.225515298 | 0.710741966 |
| Arhgef2 | 1402.732759 | -0.13121309 | 0.064988437 | -2.019022102 |
| Gm20652 | 1.311025493 | 1.073871149 | 0.914600482 | 1.174142339 |
| Rxfp4 | 0 | NA | NA | NA |
| RP23-199B2.8 | 2.30646632 | 0.68259994 | 0.687507606 | 0.992861656 |
| 2810403A07Rik | 753.6655769 | -0.083098715 | 0.097418544 | -0.853007156 |
| Gm25945 | 6.725632374 | 0.330640974 | 0.4213511 | 0.784716059 |
| Rit1 | 282.0700927 | -0.412457655 | 0.10888061 | -3.788164451 |
| Gm17146 | 5.547323716 | 0.11889062 | 0.541433333 | 0.219584966 |
| Gm10253 | 5.431051021 | 1.35733075 | 0.595602651 | 2.278919928 |
| Syt11 | 7534.20102 | 0.386917299 | 0.051433902 | 7.522612236 |
| 5830417I10Rik | 63.92099027 | -0.376050126 | 0.157553344 | -2.386811458 |
| 1500004A13Rik | 4.959186419 | 0.287651478 | 0.515891799 | 0.557581026 |
| Gon4l | 808.0428841 | -0.002646576 | 0.073913431 | -0.035806432 |
| RP23-51N3.3 | 0.176220594 | 0.517965899 | 2.719488438 | 0.190464461 |
| Gm23054 | 2.17116889 | -0.724099504 | 0.824417438 | -0.87831658 |
| Msto1 | 187.4319594 | -0.085531875 | 0.085952698 | -0.995104018 |
| n-R5s197 | 0.163066018 | 1.056936228 | 3.445567766 | 0.306752414 |
| Dap3 | 627.5926466 | 0.061841966 | 0.056311751 | 1.098207121 |

| | | | | |
|---|---|---|---|---|
| Ash1l | 4532.149695 | 0.158272582 | 0.061691342 | 2.565555823 |
| Gm26465 | 0 | NA | NA | NA |
| Gm24403 | 0 | NA | NA | NA |
| RP24-394C15.7 | 0 | NA | NA | NA |
| Rusc1 | 959.1680519 | 0.426631352 | 0.065279061 | 6.535500729 |
| Fdps | 34.50620784 | -0.215435713 | 0.208117757 | -1.035162571 |
| Gm22935 | 1.101484168 | 2.01250506 | 1.194318637 | 1.68506544 |
| RP24-394C15.4 | 0 | NA | NA | NA |
| Pklr | 0.541363579 | 1.010765188 | 1.34547287 | 0.751234165 |
| Hcn3 | 139.9590133 | -0.381490582 | 0.13300774 | -2.868183336 |
| Clk2 | 311.1271416 | 0.018826182 | 0.146303315 | 0.128679123 |
| Scamp3 | 397.6184147 | -0.295245756 | 0.093876601 | -3.145040976 |
| Gm16069 | 1.160335885 | 0.888536928 | 1.083302452 | 0.820211314 |
| Fam189b | 352.3484391 | 0.263030997 | 0.070883994 | 3.710724824 |
| Gm23269 | 0 | NA | NA | NA |
| Gba | 494.4063209 | -0.725041731 | 0.115905876 | -6.255435484 |
| Mtx1 | 203.9215083 | 0.071923684 | 0.086405833 | 0.83239385 |
| RP23-350K3.2 | 104.7257883 | -0.35695165 | 0.112702705 | -3.167196818 |
| Thbs3 | 91.591488 | 0.207310666 | 0.193847509 | 1.069452307 |
| Mir92b | 0.280267168 | -0.819747739 | 1.796298784 | -0.45635378 |
| Muc1 | 17.55319324 | -0.066873472 | 0.293965813 | -0.227487242 |
| Trim46 | 576.4645691 | 0.218024955 | 0.128155419 | 1.701254277 |
| Krtcap2 | 231.132102 | -0.594636219 | 0.106544558 | -5.581103606 |
| Dpm3 | 102.4945791 | -0.606148939 | 0.136029224 | -4.456019961 |
| Slc50a1 | 178.718743 | 0.156311193 | 0.10102594 | 1.547238196 |
| Efna1 | 37.54655563 | -0.127039042 | 0.199390199 | -0.637137844 |
| Gm22642 | 0 | NA | NA | NA |
| Efna3 | 366.6839132 | 0.289005345 | 0.148320726 | 1.948516251 |
| Gm15998 | 1.818279517 | 0.369025227 | 0.871453015 | 0.423459694 |
| Efna4 | 3.626077548 | -1.051443717 | 0.548507624 | -1.916917233 |
| 4731419I09Rik | 0.772156528 | -0.547219792 | 1.136109965 | -0.481660938 |
| Adam15 | 928.1765902 | 0.060900406 | 0.075611619 | 0.805437134 |
| Dcst1 | 1.444845127 | 0.307482528 | 0.766335261 | 0.401237609 |
| Gm10702 | 17.15096642 | 1.048925777 | 0.309694419 | 3.386970222 |
| Zbtb7b | 489.1333668 | -0.404301751 | 0.133064436 | -3.038390746 |
| Gm15417 | 31.01182027 | 0.160191698 | 0.191248769 | 0.83760904 |
| Lenep | 0.241023442 | 0.04890402 | 1.845135142 | 0.026504302 |
| Flad1 | 214.0339016 | -0.205534347 | 0.089382404 | -2.299494506 |
| Cks1b | 21.556343 | -0.481096628 | 0.358945525 | -1.340305407 |
| Shc1 | 291.0704285 | -0.606068773 | 0.121305877 | -4.996202893 |
| Pygo2 | 447.2134043 | -0.260053036 | 0.097742497 | -2.660593339 |
| Pbxip1 | 1167.4111 | -0.568985285 | 0.127854446 | -4.450258122 |
| Pmvk | 242.2372548 | -0.05668311 | 0.122136457 | -0.464096567 |
| Kcnn3 | 548.2658225 | 0.345806194 | 0.096550114 | 3.581623878 |
| Adar | 3619.123941 | -0.178086126 | 0.090730989 | -1.962792731 |
| Chrnb2 | 737.8398015 | 0.307953927 | 0.071388351 | 4.313784035 |
| 4632404H12Rik | 215.8736812 | -0.123115809 | 0.136728362 | -0.900440896 |

| | | | | |
|---|---|---|---|---|
| Ube2q1 | 886.7801256 | -0.08326847 | 0.05526764 | -1.506640582 |
| RP24-122I22.4 | 0 | NA | NA | NA |
| She | 25.99300754 | -0.82605122 | 0.279864046 | -2.951616089 |
| Il6ra | 238.4815568 | -0.897923741 | 0.123216755 | -7.287350995 |
| RP24-122I22.6 | 1.56556854 | -0.588783071 | 0.888363718 | -0.662772532 |
| Atp8b2 | 1163.874269 | 0.19540316 | 0.098341316 | 1.986989472 |
| Aqp10-ps | 0.173853943 | -0.499202846 | 2.738479247 | -0.182291995 |
| Gm24046 | 0.066050875 | -0.443818988 | 3.724945049 | -0.119147795 |
| Hax1 | 320.365027 | -0.840410047 | 0.116764731 | -7.197464866 |
| RP23-114P22.5 | 1.940516017 | -0.169382 | 0.804589013 | -0.210519902 |
| Ubap2l | 1956.985286 | -0.035327916 | 0.068593098 | -0.515036008 |
| Gm24608 | 5.024933871 | 1.62732147 | 0.616608391 | 2.639149085 |
| Mir7669 | 4.045153519 | 0.91729386 | 0.63845541 | 1.436739115 |
| 4933434E20Rik | 645.1823872 | -0.165780145 | 0.089326208 | -1.855895917 |
| Gm16540 | 6.216095886 | 0.663549505 | 0.52811509 | 1.256448675 |
| Mir190b | 0.194279094 | -1.017823543 | 3.719440177 | -0.273649661 |
| Tpm3 | 824.8814748 | -0.3627145 | 0.105671955 | -3.43245755 |
| Nup210l | 14.08624278 | -0.331004863 | 0.359338333 | -0.921150994 |
| Gm23723 | 0.35561486 | -0.362386205 | 1.574835096 | -0.230110572 |
| RP24-257C17.2 | 1.986149933 | -0.582737662 | 0.725350101 | -0.803388132 |
| Gm24185 | 0 | NA | NA | NA |
| Rps27 | 1242.923509 | -0.149215806 | 0.143080315 | -1.042881451 |
| Rab13 | 74.74244885 | -0.437529857 | 0.192886371 | -2.26832956 |
| Jtb | 232.6321241 | -0.260505628 | 0.098286416 | -2.650474368 |
| Creb3l4 | 5.167911745 | -0.466329373 | 0.491706203 | -0.948390258 |
| Slc39a1 | 919.2059152 | -0.645663231 | 0.133273348 | -4.84465378 |
| Crtc2 | 364.8494084 | -0.177571713 | 0.115238571 | -1.540905199 |
| Dennd4b | 682.1714222 | 0.178625467 | 0.065420824 | 2.730406843 |
| Mir7012 | 0 | NA | NA | NA |
| RP23-358O22.1 | 2.468095164 | -0.35878619 | 0.656605539 | -0.546425775 |
| Gatad2b | 2548.376701 | 0.175455058 | 0.063382204 | 2.768206961 |
| Slc27a3 | 56.99060069 | 0.148784179 | 0.169059367 | 0.880070603 |
| Ints3 | 947.6918643 | -0.287870927 | 0.08382956 | -3.434002584 |
| Npr1 | 184.6399887 | 0.061021217 | 0.205593545 | 0.296805123 |
| Gm16048 | 1.106761898 | -1.375425881 | 1.089098552 | -1.262903047 |
| RP24-333B11.3 | 0.066050875 | -0.443818988 | 3.724945049 | -0.119147795 |
| Ilf2 | 670.5056944 | 0.222812794 | 0.08378847 | 2.659229779 |
| Snapin | 374.92744 | 0.052337297 | 0.078356953 | 0.667934311 |
| Chtop | 1458.724535 | -0.162990234 | 0.075921221 | -2.146833666 |
| S100a1 | 401.8878366 | -0.030982633 | 0.151871105 | -0.204006106 |
| S100a13 | 63.775755 | -0.573683646 | 0.163565036 | -3.50737333 |
| S100a14 | 1.045271382 | 2.029735663 | 1.173051313 | 1.730304242 |
| S100a16 | 533.8051986 | -0.747745151 | 0.168381291 | -4.440785234 |
| Gm24593 | 0 | NA | NA | NA |
| Gm23464 | 0 | NA | NA | NA |
| S100a3 | 9.213372739 | -0.176968306 | 0.343127623 | -0.515750685 |
| S100a2 | 0.475453923 | 1.251255839 | 1.461109422 | 0.856373808 |

| | | | | |
|---|---|---|---|---|
| RP23-326D6.5 | 0 | NA | NA | NA |
| S100a4 | 14.92865454 | -1.573343486 | 0.400162365 | -3.931762767 |
| S100a5 | 23.65689645 | -1.578240918 | 1.467001536 | -1.07582772 |
| S100a6 | 362.9019992 | -1.576608296 | 0.264889138 | -5.951955256 |
| RP24-174D10.3 | 0 | NA | NA | NA |
| S100a7a | 5.924278019 | -1.387395971 | 0.445191109 | -3.116405391 |
| S100a8 | 13.16848317 | -2.217079823 | 0.532514124 | -4.163419753 |
| S100a9 | 10.76590429 | -1.169819419 | 0.407022332 | -2.874091488 |
| RP24-174D10.4 | 0 | NA | NA | NA |
| RP24-174D10.5 | 0 | NA | NA | NA |
| Pglyrp4 | 0.060174537 | 0.517957688 | 3.724945049 | 0.139051095 |
| RP24-174D10.6 | 2.240082864 | -0.497113234 | 0.75084831 | -0.662068793 |
| Gm5849 | 0 | NA | NA | NA |
| RP23-111A11.1 | 0 | NA | NA | NA |
| RP23-309N14.1 | 0 | NA | NA | NA |
| RP24-296O9.1 | 0 | NA | NA | NA |
| 4930529C04Rik | 0.183030605 | 0.03707559 | 2.280962899 | 0.016254359 |
| 9130204L05Rik | 0 | NA | NA | NA |
| Gm23697 | 0 | NA | NA | NA |
| Pglyrp3 | 2.336438979 | -0.829065472 | 0.743918574 | -1.114457281 |
| Mif-ps3 | 0.176744298 | 0.037631212 | 2.303885 | 0.016333807 |
| Gm25145 | 0 | NA | NA | NA |
| Lor | 15.81042869 | 0.332663237 | 0.318781305 | 1.043546882 |
| RP23-381I16.5 | 0 | NA | NA | NA |
| RP23-381I16.3 | 0 | NA | NA | NA |
| RP23-381I16.4 | 0 | NA | NA | NA |
| Prr9 | 0 | NA | NA | NA |
| Lelp1 | 0 | NA | NA | NA |
| RP23-381I16.6 | 0.291617713 | 1.693047594 | 1.731084925 | 0.978026884 |
| Sprr2a1 | 0 | NA | NA | NA |
| Sprr2a2 | 0 | NA | NA | NA |
| Gm5850 | 0.099106094 | -0.93550548 | 3.724188147 | -0.251197158 |
| Gm37087 | 0.047948698 | 0.037064902 | 3.724945049 | 0.009950456 |
| Gm5851 | 0.36081128 | -2.20441996 | 1.665879719 | -1.323276786 |
| Gm37752 | 0 | NA | NA | NA |
| Sprr2a3 | 0 | NA | NA | NA |
| Sprr2b | 0.207368832 | 0.009858303 | 2.637090101 | 0.003738326 |
| Gm37428 | 0 | NA | NA | NA |
| Sprr2c-ps | 0 | NA | NA | NA |
| Sprr2d | 0 | NA | NA | NA |
| Sprr2e | 0.186228844 | 0.51796788 | 3.289587573 | 0.157456784 |
| Sprr2f | 0 | NA | NA | NA |
| Sprr2g | 0.097139547 | -0.443816764 | 3.724945049 | -0.119147198 |
| Sprr2h | 0 | NA | NA | NA |
| Sprr2i | 0 | NA | NA | NA |
| Sprr2j-ps | 0 | NA | NA | NA |
| Gm9774 | 28.33359312 | -0.791760783 | 0.233478781 | -3.391146635 |

| | | | | |
|---|---|---|---|---|
| Sprr2k | 0 | NA | NA | NA |
| Sprr1b | 0.278361807 | 1.539667399 | 3.70512207 | 0.415551059 |
| Sprr3 | 0 | NA | NA | NA |
| Gm37377 | 0 | NA | NA | NA |
| Sprr1a | 21.48993395 | -3.014995794 | 0.744945746 | -4.04726896 |
| Sprr4 | 0 | NA | NA | NA |
| 2310046K23Rik | 0.227721369 | 1.433869842 | 3.108233563 | 0.461313416 |
| Ivl | 0 | NA | NA | NA |
| Smcp | 0 | NA | NA | NA |
| 2210017I01Rik | 0 | NA | NA | NA |
| Lce6a | 0.09176491 | 0.921395572 | 3.723061103 | 0.247483333 |
| Lce1a1 | 0.475368968 | 2.245094412 | 3.690247583 | 0.60838585 |
| Lce1b | 0.18352982 | 1.388044403 | 3.713044316 | 0.373829205 |
| Rpl13-ps4 | 0 | NA | NA | NA |
| Lce1a2 | 0 | NA | NA | NA |
| Lce1c | 0.27529473 | 1.72897849 | 3.703391444 | 0.466863554 |
| Lce1d | 0.45492314 | 2.196644884 | 3.690964336 | 0.595141184 |
| Gm4202 | 299.4188111 | -0.282686033 | 0.173413438 | -1.63012761 |
| Lce1e | 0 | NA | NA | NA |
| Lce1f | 0.373862045 | -1.879916649 | 1.612559379 | -1.165796853 |
| Gm38119 | 0 | NA | NA | NA |
| Lce1g | 0.655362467 | -1.542265114 | 2.398000398 | -0.643146313 |
| Lce1h | 0.27529473 | 1.72897849 | 3.703391444 | 0.466863554 |
| Lce1i | 0.17962841 | 0.998850067 | 3.374012501 | 0.296042195 |
| Lce1j | 0.09176491 | 0.921395572 | 3.723061103 | 0.247483333 |
| Lce1k | 0 | NA | NA | NA |
| Kprp | 0.596471915 | 2.484999492 | 3.684311218 | 0.674481428 |
| Gm38201 | 0 | NA | NA | NA |
| Lce1l | 0.09176491 | 0.921395572 | 3.723061103 | 0.247483333 |
| 2310050C09Rik | 0.31922648 | 1.79924161 | 3.70283485 | 0.485909224 |
| Lce3a | 0 | NA | NA | NA |
| Lce3b | 0 | NA | NA | NA |
| Gm38078 | 0 | NA | NA | NA |
| Lce3c | 0.342841614 | 0.577010955 | 2.008268103 | 0.287317691 |
| Gm24481 | 0 | NA | NA | NA |
| Lce3d | 0 | NA | NA | NA |
| Gm22577 | 0 | NA | NA | NA |
| Lce3e | 0 | NA | NA | NA |
| Lce3f | 0.038981395 | -0.443816764 | 3.724945049 | -0.119147198 |
| Crct1 | 1.039606943 | -0.761331188 | 0.953526488 | -0.798437377 |
| Lce1m | 0.045882455 | 0.517959912 | 3.724945049 | 0.139051692 |
| Gm38040 | 0.139839889 | 0.037074206 | 2.923971603 | 0.0126794 |
| Gm4858 | 0 | NA | NA | NA |
| Gm37361 | 0 | NA | NA | NA |
| Crnn | 0 | NA | NA | NA |
| Gm37501 | 0 | NA | NA | NA |
| Flg2 | 0.73411928 | 2.717324032 | 3.67867585 | 0.73866906 |

| | | | | |
|---|---|---|---|---|
| Gm18432 | 0.092691418 | -0.908687078 | 3.724560878 | -0.2439716 |
| Flg | 1.695700131 | 4.053838292 | 3.648662627 | 1.11104772 |
| Rplp0-ps1 | 2.415407705 | -1.277638516 | 0.681673574 | -1.874267341 |
| Gm37922 | 0 | NA | NA | NA |
| Gm37245 | 0.045882455 | 0.517959912 | 3.724945049 | 0.139051692 |
| Gm37913 | 0.038981395 | -0.443816764 | 3.724945049 | -0.119147198 |
| Hrnr | 2.939544198 | 4.306555469 | 3.644903919 | 1.181527844 |
| AC123859.1 | 0 | NA | NA | NA |
| Gm37857 | 0 | NA | NA | NA |
| Rptn | 0.124596252 | 0.037071632 | 3.714681146 | 0.009979761 |
| Gm18296 | 0.119587853 | 0.981063299 | 3.723618439 | 0.263470416 |
| Gm22752 | 0.140442044 | 0.517954843 | 3.61309026 | 0.14335508 |
| Gm5278 | 0.680250709 | 0.803005801 | 1.561215862 | 0.514346428 |
| Tchh | 32.61220021 | 0.028836969 | 0.247050221 | 0.116725129 |
| Tchhl1 | 0.282572244 | 0.559651036 | 2.066711947 | 0.270792955 |
| Gm36953 | 0 | NA | NA | NA |
| S100a11 | 216.4239304 | -1.48839018 | 0.372050373 | -4.000507153 |
| Gm29950 | 0 | NA | NA | NA |
| S100a10 | 413.4047504 | -1.199177775 | 0.191739325 | -6.254208807 |
| Gm5541 | 0 | NA | NA | NA |
| Gm38207 | 0 | NA | NA | NA |
| Tdpoz2 | 0 | NA | NA | NA |
| Tdpoz1 | 0 | NA | NA | NA |
| Gm18433 | 0 | NA | NA | NA |
| Gm37596 | 0 | NA | NA | NA |
| Gm38086 | 0 | NA | NA | NA |
| Gm5852 | 0 | NA | NA | NA |
| Gm5773 | 0 | NA | NA | NA |
| Tdpoz4 | 0 | NA | NA | NA |
| Tdpoz3 | 0 | NA | NA | NA |
| Gm37430 | 0 | NA | NA | NA |
| Gm9113 | 0.044544648 | 0.517957688 | 3.724945049 | 0.139051095 |
| Gm9116 | 0 | NA | NA | NA |
| Gm37185 | 0 | NA | NA | NA |
| Gm9117 | 0 | NA | NA | NA |
| Gm37104 | 0 | NA | NA | NA |
| Tdpoz5 | 0 | NA | NA | NA |
| Gm37807 | 0 | NA | NA | NA |
| Gm37481 | 0 | NA | NA | NA |
| Gm9121 | 0.045882455 | 0.517959912 | 3.724945049 | 0.139051692 |
| Gm9124 | 0 | NA | NA | NA |
| Gm37085 | 0 | NA | NA | NA |
| Gm9125 | 0 | NA | NA | NA |
| Gm37081 | 0 | NA | NA | NA |
| Gm10697 | 0 | NA | NA | NA |
| Gm37778 | 0 | NA | NA | NA |
| Gm37155 | 0 | NA | NA | NA |

| | | | | |
|---|---|---|---|---|
| Gm5542 | 0 | NA | NA | NA |
| Gm9131 | 0 | NA | NA | NA |
| Gm37882 | 0 | NA | NA | NA |
| Gm5543 | 0 | NA | NA | NA |
| Gm37001 | 0 | NA | NA | NA |
| Gm16560 | 0 | NA | NA | NA |
| Gm10696 | 0.045882455 | 0.517959912 | 3.724945049 | 0.139051692 |
| Gm5286 | 0 | NA | NA | NA |
| Gm16584 | 0 | NA | NA | NA |
| Gm4982 | 0.038981395 | -0.443816764 | 3.724945049 | -0.119147198 |
| Gm9141 | 0 | NA | NA | NA |
| Gm4778 | 0 | NA | NA | NA |
| Gm16561 | 0 | NA | NA | NA |
| Them4 | 202.6068946 | 0.206883611 | 0.091888596 | 2.251461236 |
| RP23-148G12.2 | 0 | NA | NA | NA |
| Them5 | 0.249936274 | 1.18768375 | 2.489289 | 0.477117663 |
| RP23-148G12.3 | 4.416068108 | -0.050589928 | 0.550507707 | -0.091896857 |
| C2cd4d | 4.518953556 | -0.781510669 | 0.509944641 | -1.532540213 |
| RP23-148G12.5 | 0.049553047 | -0.443818988 | 3.724945049 | -0.119147795 |
| Rorc | 43.04669428 | 0.637262279 | 0.243754788 | 2.614357996 |
| RP23-298M23.3 | 0.086648695 | 0.037058229 | 3.724945049 | 0.009948665 |
| Lingo4 | 68.20621523 | 0.116674595 | 0.249647753 | 0.467356879 |
| 1700040D17Rik | 7.265577632 | -0.540596719 | 0.434288071 | -1.244788321 |
| Tdrkh | 464.6804853 | -0.099244824 | 0.089305072 | -1.111301099 |
| RP23-298M23.7 | 44.92890297 | -0.134854747 | 0.273930875 | -0.492294805 |
| RP23-298M23.8 | 0.072025462 | 0.517959912 | 3.724945049 | 0.139051692 |
| Oaz3 | 2.190110068 | 1.188877903 | 0.983129916 | 1.209278534 |
| Mrpl9 | 334.7621041 | -0.238238518 | 0.086836912 | -2.743516687 |
| RP23-298M23.9 | 0.961922688 | -0.570954537 | 1.031015017 | -0.55377907 |
| Riiad1 | 12.40794134 | 0.534340738 | 0.332924566 | 1.604990415 |
| Celf3 | 927.0754436 | 0.55068954 | 0.143159936 | 3.846673563 |
| Snx27 | 1992.373795 | -0.092212121 | 0.057705893 | -1.597967144 |
| RP23-298M23.11 | 19.53939973 | 0.325221509 | 0.359404674 | 0.90488948 |
| RP23-298M23.12 | 7.754593511 | 0.386913179 | 0.506626889 | 0.76370439 |
| RP24-276A6.1 | 23.82234211 | 0.773600017 | 0.334856632 | 2.310242481 |
| AC127252.1 | 0.108123235 | 0.517955464 | 3.724945049 | 0.139050498 |
| Rpl31-ps11 | 1.676330775 | 0.209449941 | 0.900437374 | 0.232609115 |
| Tuft1 | 38.81743166 | -0.327690274 | 0.21712064 | -1.509254364 |
| Gm10972 | 0 | NA | NA | NA |
| Gm15234 | 0.403742549 | 0.901063514 | 1.431529691 | 0.629441024 |
| BC021767 | 1.492922606 | 0.970644006 | 0.833497572 | 1.164543291 |
| Selenbp2 | 2.718408089 | 0.032905475 | 0.695293793 | 0.047326001 |
| Gm15264 | 3.338245402 | 0.714685948 | 0.664121602 | 1.076137181 |
| RP23-449N19.1 | 0 | NA | NA | NA |
| Cgn | 50.96569556 | 0.533414048 | 0.192794768 | 2.766745457 |
| Gm15263 | 0.664861716 | 2.282312735 | 1.419857771 | 1.607423491 |
| Pogz | 1033.996817 | 0.041218462 | 0.062980478 | 0.654464107 |

| | | | | |
|---|---|---|---|---|
| Psmb4 | 959.4736378 | -0.451422196 | 0.114747782 | -3.934038523 |
| Gm26279 | 0.212513282 | 1.689178857 | 3.159662247 | 0.534607412 |
| Selenbp1 | 39.74177677 | 0.210279388 | 0.260266617 | 0.807938377 |
| Rfx5 | 386.701994 | 0.183367429 | 0.074914772 | 2.447680542 |
| B230398E01Rik | 27.22757332 | 1.210630611 | 0.390244883 | 3.102233142 |
| Pi4kb | 609.4035235 | -0.166239183 | 0.065166206 | -2.551002938 |
| A730011C13Rik | 177.9491982 | -0.077532542 | 0.180351303 | -0.429897322 |
| Gm15265 | 34.26348499 | 0.75039544 | 0.270009304 | 2.779146604 |
| Mir7013 | 0.881607526 | 1.496673004 | 1.189444115 | 1.258296195 |
| Zfp687 | 719.9016467 | 0.238506868 | 0.088579282 | 2.692580723 |
| 4930481B07Rik | 9.171066605 | -0.040839908 | 0.344141892 | -0.118671714 |
| Psmd4 | 779.7769061 | -0.277219476 | 0.097159462 | -2.853242184 |
| Pip5k1a | 758.4226424 | 0.161079708 | 0.086352561 | 1.865372681 |
| Gm24920 | 5.105149463 | 0.649502527 | 0.617529651 | 1.051775451 |
| Gm22301 | 0 | NA | NA | NA |
| Vps72 | 306.4355309 | -0.158153126 | 0.06674231 | -2.369608201 |
| Tmod4 | 6.657288501 | -0.403963602 | 0.516497379 | -0.782121301 |
| Scnm1 | 114.2649021 | -0.046975208 | 0.112353272 | -0.418102712 |
| Lysmd1 | 47.85101064 | 0.059125345 | 0.161877634 | 0.365247153 |
| Mir8099-2 | 0.049553047 | -0.443818988 | 3.724945049 | -0.119147795 |
| Tnfaip8l2 | 38.08952461 | -0.968125373 | 0.209836098 | -4.613721772 |
| Gm9169 | 2.545959807 | -1.079801771 | 0.679035692 | -1.590198842 |
| Sema6c | 68.04043718 | 0.518750703 | 0.199058182 | 2.606025527 |
| AC140190.1 | 0 | NA | NA | NA |
| Gabpb2 | 547.3167726 | -0.014332968 | 0.110983153 | -0.129145436 |
| Gm16740 | 8.225987562 | -0.082415624 | 0.37742429 | -0.218363329 |
| Mllt11 | 2083.533929 | 0.069149406 | 0.072918859 | 0.948306197 |
| Cdc42se1 | 925.5629768 | -0.597592362 | 0.110330518 | -5.416383172 |
| Gm128 | 1.475208495 | -0.568623289 | 1.041286412 | -0.5460777 |
| Bnipl | 3.731006451 | -0.454625065 | 0.729502369 | -0.623198887 |
| Prune | 303.723384 | -0.041819298 | 0.077625533 | -0.538731214 |
| Fam63a | 346.9866243 | -0.296790135 | 0.100863432 | -2.942494921 |
| Anxa9 | 8.904347822 | -0.942435588 | 0.412989476 | -2.281984515 |
| 6330562C20Rik | 7.360490999 | -0.42171805 | 0.382212441 | -1.103360344 |
| Cers2 | 766.0749113 | -0.46259719 | 0.106838458 | -4.32987521 |
| Setdb1 | 649.1102505 | -0.05476103 | 0.093102726 | -0.588178587 |
| RP23-407B16.2 | 0 | NA | NA | NA |
| Gm37500 | 1.278257442 | 1.514442192 | 1.039969499 | 1.456237124 |
| RP23-407B16.3 | 0 | NA | NA | NA |
| Gm5070 | 0.368600158 | 0.434225133 | 1.521117645 | 0.285464529 |
| RP23-407B16.4 | 0.048949532 | 0.037064902 | 3.724945049 | 0.009950456 |
| Gm4349 | 3.988126414 | 0.080122026 | 0.588566655 | 0.13613076 |
| Arnt | 674.0580543 | 0.030131837 | 0.110828933 | 0.271876993 |
| RP23-422N18.1 | 29.38944538 | 0.721561082 | 0.279703042 | 2.579739845 |
| Ctsk | 101.6290436 | -0.689288116 | 0.186713028 | -3.691698031 |
| Ctss | 2705.512219 | -0.955785255 | 0.181119165 | -5.277107235 |
| Hmgb1-ps5 | 4.681085585 | -0.995927946 | 0.51474431 | -1.93480127 |

| | | | | |
|---|---|---|---|---|
| Hormad1 | 0.108266322 | -0.955981391 | 3.722845977 | -0.25678779 |
| Golph3l | 291.0130091 | -0.429320837 | 0.109857458 | -3.907980812 |
| RP23-422N18.3 | 2.848305637 | 0.189123932 | 0.735994919 | 0.256963639 |
| Rps10-ps1 | 80.90969664 | -0.492784498 | 0.186616827 | -2.640621995 |
| Ensa | 2238.110929 | -0.171925524 | 0.073619655 | -2.335320989 |
| RP23-231L15.4 | 47.75963671 | 0.127766767 | 0.269752375 | 0.473644642 |
| Mcl1 | 1783.072468 | -0.730838477 | 0.141639595 | -5.159845863 |
| Adamtsl4 | 317.4111096 | -0.731456766 | 0.141272108 | -5.177644581 |
| Ecm1 | 388.3138544 | -2.116725536 | 0.336137956 | -6.297192861 |
| Mir7014 | 0 | NA | NA | NA |
| Tars2 | 297.5330298 | -0.288128572 | 0.090348704 | -3.189072552 |
| Rprd2 | 1419.095113 | 0.091373024 | 0.055724183 | 1.63973734 |
| RP23-218K6.1 | 0.165206806 | -0.443208982 | 2.775355575 | -0.159694486 |
| Gm26361 | 3.436253382 | 0.571184655 | 0.695173839 | 0.821642909 |
| RP23-218K6.4 | 0.306564387 | 1.337935898 | 1.964695212 | 0.680989036 |
| Prpf3 | 300.0577214 | -0.268536328 | 0.11561596 | -2.322657947 |
| Gm26594 | 0.250472921 | -0.96481344 | 2.484005718 | -0.388410314 |
| Mrps21 | 159.8335192 | -0.226660605 | 0.123776224 | -1.831212796 |
| C920021L13Rik | 8.391545128 | 0.734833788 | 0.399632386 | 1.838774366 |
| Ciart | 209.0128952 | 1.544273863 | 0.198261336 | 7.789082282 |
| BC028528 | 36.09890605 | -0.695937273 | 0.173626023 | -4.008254408 |
| Aph1a | 300.5160635 | -0.236349779 | 0.102091862 | -2.315069721 |
| Car14 | 134.9063813 | 0.534101464 | 0.139124859 | 3.839008122 |
| Anp32e | 879.4154652 | -0.037015125 | 0.062781435 | -0.58958711 |
| RP23-49O17.4 | 9.354061232 | 0.600502028 | 0.443484569 | 1.354053939 |
| RP23-49O17.5 | 0 | NA | NA | NA |
| Plekho1 | 424.3660083 | -0.532769335 | 0.121991456 | -4.367267623 |
| Vps45 | 380.4768023 | -0.098019953 | 0.096492325 | -1.015831605 |
| RP23-261M19.2 | 0.092691418 | -0.908687078 | 3.724560878 | -0.2439716 |
| Otud7b | 1265.281876 | -0.014535788 | 0.061998843 | -0.234452572 |
| Mtmr11 | 145.3552089 | -0.653223161 | 0.154570303 | -4.226058627 |
| Gm17690 | 21.29907499 | -0.800274488 | 0.284820863 | -2.809746727 |
| Sf3b4 | 445.2184839 | -0.370220063 | 0.11273212 | -3.28406902 |
| Gm22027 | 1.636315489 | 0.331754423 | 0.809265988 | 0.409944849 |
| Sv2a | 5873.135983 | 0.285648889 | 0.041339912 | 6.909760475 |
| Bola1 | 50.41452498 | 0.045938178 | 0.18468816 | 0.248733746 |
| Gm15444 | 1.046383212 | -0.588480668 | 1.039524227 | -0.566105774 |
| Hist2h2ab | 34.08697731 | 0.311273245 | 0.303204963 | 1.026609996 |
| Hist2h2ac | 86.71087739 | -0.032237049 | 0.169718699 | -0.189944005 |
| AC125099.2 | 1.307878832 | 0.543193612 | 0.857280551 | 0.633624094 |
| Hist2h2be | 54.11456325 | -0.119552359 | 0.221605979 | -0.539481651 |
| Hist2h3c2 | 19.58774007 | -0.352950197 | 0.234148004 | -1.507380763 |
| Gm20632 | 14.19037399 | 0.940598142 | 0.295739474 | 3.180495756 |
| Hist2h2aa2 | 13.3101306 | -0.318967854 | 0.284381196 | -1.121620765 |
| AC125099.3 | 0 | NA | NA | NA |
| Gm20633 | 28.70123197 | 0.041285045 | 0.227923688 | 0.181135384 |
| Hist2h2aa1 | 0.135217741 | 1.027273954 | 3.618332993 | 0.283908075 |

| | | | | |
|---|---|---|---|---|
| Gm20634 | 1.134122097 | -1.58237754 | 0.945574705 | -1.673455868 |
| AC125099.1 | 0 | NA | NA | NA |
| Hist2h3c1 | 0 | NA | NA | NA |
| Gm20627 | 9.369992915 | -1.410870591 | 0.461841286 | -3.054881909 |
| Hist2h4 | 368.633472 | -0.318136097 | 0.15490063 | -2.053807631 |
| Gm38227 | 4.279540226 | -0.077891295 | 0.486867026 | -0.159984742 |
| Gm20628 | 12.09656833 | 0.07116562 | 0.332906995 | 0.213770277 |
| Hist2h3b | 50.50959981 | 0.037648849 | 0.284826857 | 0.132181526 |
| Hist2h2bb | 201.6384358 | -0.232011684 | 0.154894425 | -1.497869816 |
| Fcgr1 | 583.6067119 | -1.64284778 | 0.264471977 | -6.211802842 |
| Gm24830 | 0 | NA | NA | NA |
| Platr30 | 0 | NA | NA | NA |
| Gm26553 | 0 | NA | NA | NA |
| Rnu1b1 | 6.056136525 | 0.485267939 | 0.500642321 | 0.969290688 |
| Rnu1b2 | 0.421929236 | 0.879914745 | 1.429868943 | 0.615381395 |
| Gm17575 | 0 | NA | NA | NA |
| Gm26689 | 0 | NA | NA | NA |
| 4930477E14Rik | 0 | NA | NA | NA |
| Gm26654 | 2.969550979 | -4.763082229 | 0.930757222 | -5.117427099 |
| Pafah1b1-ps1 | 0.705160482 | 0.472512008 | 1.17235523 | 0.403045081 |
| Gm21960 | 0.551586351 | -1.116534917 | 1.305703403 | -0.855121396 |
| Terc | 7.890901767 | -1.188965915 | 0.453825795 | -2.61987293 |
| Gm24136 | 0 | NA | NA | NA |
| BC107364 | 0.047948698 | 0.037064902 | 3.724945049 | 0.009950456 |
| Gm26232 | 0 | NA | NA | NA |
| RP23-404J7.8 | 2.243072942 | 0.749058374 | 0.721431139 | 1.038295041 |
| RP23-404J7.10 | 0 | NA | NA | NA |
| Rnu1b6 | 0.046345709 | -0.443818988 | 3.724945049 | -0.119147795 |
| RP23-404J7.9 | 0 | NA | NA | NA |
| Gm28001 | 0 | NA | NA | NA |
| Gm25890 | 0 | NA | NA | NA |
| Gm22614 | 1.309217219 | 0.402781515 | 0.814844313 | 0.494304873 |
| Hfe2 | 3.500549162 | -0.129873734 | 0.600108584 | -0.216417057 |
| Gm15441 | 0.831325084 | 0.575613527 | 1.029176486 | 0.559295256 |
| Txnip | 698.8955186 | -0.16413561 | 0.182980723 | -0.897010396 |
| Gm16253 | 0.327232518 | -0.796751911 | 2.217046622 | -0.359375352 |
| Polr3gl | 127.7419174 | 0.087628941 | 0.128186382 | 0.683605692 |
| Ankrd34a | 497.5908006 | 0.178649319 | 0.109691515 | 1.628652124 |
| Lix1l | 406.2282066 | -0.59140092 | 0.150870946 | -3.919912597 |
| 6330549D23Rik | 6.560446392 | 0.130054097 | 0.471633651 | 0.275752371 |
| Rbm8a | 404.2810953 | -0.225586056 | 0.12164435 | -1.854472122 |
| Pex11b | 270.344613 | 0.178309233 | 0.078298868 | 2.277290046 |
| Itga10 | 23.71082201 | -0.347510748 | 0.289242556 | -1.201450965 |
| RP23-204O17.2 | 0.060174537 | 0.517957688 | 3.724945049 | 0.139051095 |
| Ankrd35 | 72.85459675 | 0.414079409 | 0.1771027 | 2.338075081 |
| Pias3 | 236.294852 | 0.235811074 | 0.100109484 | 2.3555318 |
| Nudt17 | 21.37927619 | 0.938973811 | 0.250795109 | 3.743987724 |

| | | | | |
|---|---|---|---|---|
| Polr3c | 240.883843 | -0.141010524 | 0.094768307 | -1.487950227 |
| Gm22581 | 3.657530093 | 0.835620614 | 0.6276734 | 1.331298432 |
| RP23-204O17.4 | 9.840503469 | 0.734913689 | 0.376046509 | 1.954315948 |
| Rnf115 | 868.8846253 | 0.08311916 | 0.096368335 | 0.862515263 |
| RP23-395B13.2 | 10.83277785 | 0.498894918 | 0.43719657 | 1.141122671 |
| Cd160 | 2.898004674 | -1.07570404 | 0.57692996 | -1.864531423 |
| Pdzk1 | 2.857891658 | -1.601243213 | 0.686006077 | -2.334153102 |
| 4930442L01Rik | 0.150086409 | 1.027495679 | 3.723011807 | 0.275985071 |
| Gpr89 | 501.075764 | -0.0306326 | 0.062359135 | -0.491228755 |
| Gja5 | 5.28099507 | -1.634420991 | 0.489281201 | -3.340453279 |
| Gja8 | 0 | NA | NA | NA |
| RP24-86K23.1 | 0 | NA | NA | NA |
| RP23-97A19.1 | 0.188929682 | 0.517973644 | 2.260134588 | 0.22917823 |
| RP23-97A19.2 | 0.09349418 | 0.517955464 | 3.724945049 | 0.139050498 |
| Gm24435 | 0 | NA | NA | NA |
| Acp6 | 115.9459266 | -0.023308004 | 0.186610447 | -0.124901927 |
| Bcl9 | 929.9899206 | 0.08740565 | 0.060895927 | 1.435328341 |
| RP23-394K8.5 | 0.094831987 | 0.517957688 | 3.724945049 | 0.139051095 |
| RP23-394K8.3 | 12.87106279 | 1.419377939 | 0.418765264 | 3.389435708 |
| Olfr1402 | 0 | NA | NA | NA |
| RP23-290C4.1 | 0 | NA | NA | NA |
| Chd1l | 117.2581915 | -0.082741938 | 0.134911526 | -0.613305183 |
| RP24-316D12.2 | 2.928805847 | -1.11947275 | 0.757479072 | -1.477892647 |
| Fmo5 | 26.93201772 | -0.070467628 | 0.290955306 | -0.242193996 |
| Prkab2 | 567.66263 | -0.329223196 | 0.126724648 | -2.597941296 |
| Gm17651 | 0 | NA | NA | NA |
| Pde4dip | 3395.191701 | -0.040772201 | 0.093752243 | -0.43489307 |
| Mir7225 | 0 | NA | NA | NA |
| RP24-124P9.3 | 0.671139028 | 0.385583794 | 1.246416321 | 0.309353935 |
| Gm15999 | 0.08486385 | 0.037730769 | 3.724898049 | 0.010129343 |
| Sec22b | 837.8239496 | -0.05608853 | 0.063041565 | -0.88970714 |
| Gm5544 | 1.267397472 | -3.593799939 | 1.035641558 | -3.470119475 |
| Notch2 | 1217.583416 | -0.781063808 | 0.133332159 | -5.858030147 |
| RP23-97M20.1 | 25.75372241 | -0.224119921 | 0.278053102 | -0.806032799 |
| RP23-3C4.2 | 11.28696547 | 0.297477611 | 0.358329952 | 0.83017791 |
| Adam30 | 0.456191127 | 1.015924542 | 3.239790793 | 0.313577205 |
| RP23-3C4.5 | 0.046345709 | -0.443818988 | 3.724945049 | -0.119147795 |
| Reg4 | 0.197005158 | -0.954793305 | 3.277479893 | -0.291319348 |
| RP23-348H21.1 | 0 | NA | NA | NA |
| Hmgcs2 | 85.88327543 | 0.801706462 | 0.286668132 | 2.796636151 |
| RP23-348H21.2 | 2.607862967 | 1.20596553 | 0.793754557 | 1.51931793 |
| Phgdh | 330.7849704 | -0.523247419 | 0.183072146 | -2.858148709 |
| Zfp697 | 358.3939029 | 0.318215061 | 0.137537596 | 2.313658734 |
| Gm4450 | 0 | NA | NA | NA |
| RP23-381G13.3 | 0.131100043 | 0.037055718 | 3.674543085 | 0.010084442 |
| RP23-381G13.4 | 0 | NA | NA | NA |
| Hsd3b4 | 0 | NA | NA | NA |

| | | | | |
|---|---|---|---|---|
| RP23-381G13.6 | 0 | NA | NA | NA |
| Gm10681 | 0 | NA | NA | NA |
| RP24-285G1.2 | 0.109727584 | 0.037071574 | 3.724945049 | 0.009952247 |
| RP24-285G1.1 | 0 | NA | NA | NA |
| Hsd3b5 | 0 | NA | NA | NA |
| Gm12399 | 0 | NA | NA | NA |
| Gm12400 | 0 | NA | NA | NA |
| Hsd3b2 | 0.319559307 | -0.927952116 | 1.619949743 | -0.572827719 |
| Hsd3b3 | 0.048949532 | 0.037064902 | 3.724945049 | 0.009950456 |
| Hsd3b6 | 0 | NA | NA | NA |
| Gm12427 | 0 | NA | NA | NA |
| Gm12428 | 10.43905456 | 0.459449779 | 0.37562959 | 1.223145863 |
| Gm12431 | 0 | NA | NA | NA |
| Gm12432 | 0.046345709 | -0.443818988 | 3.724945049 | -0.119147795 |
| Gm12433 | 0.907786306 | 1.748980235 | 1.243692894 | 1.406279832 |
| Hsd3b1 | 0.244145947 | -1.302897789 | 1.817780426 | -0.716752018 |
| Hao2 | 0.110595523 | 0.037071574 | 3.724945049 | 0.009952247 |
| 5730437C11Rik | 0 | NA | NA | NA |
| Gm12440 | 0.266043621 | 1.648449889 | 2.452600323 | 0.672123327 |
| Gm12441 | 0 | NA | NA | NA |
| Gm22341 | 0 | NA | NA | NA |
| Gm24216 | 0 | NA | NA | NA |
| Wars2 | 117.2102137 | -0.183231785 | 0.157800988 | -1.161157405 |
| RP23-200E11.1 | 19.63971965 | -0.129557439 | 0.326413805 | -0.396911642 |
| RP23-200E11.3 | 0 | NA | NA | NA |
| Mir6481 | 0 | NA | NA | NA |
| RP23-200E11.2 | 15.98219274 | 0.12924466 | 0.354012127 | 0.365085403 |
| Tbx15 | 21.3154785 | 0.010769199 | 0.29061963 | 0.037055994 |
| RP23-200E11.4 | 0 | NA | NA | NA |
| Gm12449 | 0 | NA | NA | NA |
| M6pr-ps | 7.110781671 | -0.749982949 | 0.483990279 | -1.549582671 |
| Gm12448 | 0 | NA | NA | NA |
| Gm22505 | 0 | NA | NA | NA |
| Spag17 | 14.25668977 | 0.048624466 | 0.303398884 | 0.160265803 |
| Rpl36-ps7 | 0 | NA | NA | NA |
| Tra2b-ps | 0 | NA | NA | NA |
| Spag17os | 4.312461576 | 0.854339121 | 0.584011974 | 1.46287946 |
| Wdr3 | 327.3352847 | -0.192626165 | 0.08549785 | -2.252994256 |
| Gdap2 | 404.7187721 | 0.017953857 | 0.059947991 | 0.299490558 |
| Gm25802 | 0 | NA | NA | NA |
| Gm12464 | 0 | NA | NA | NA |
| 4930406D18Rik | 0 | NA | NA | NA |
| RP23-335A19.3 | 0 | NA | NA | NA |
| Fam46c | 258.8118225 | -0.819436868 | 0.165554403 | -4.949653133 |
| Gm12474 | 1.5464145 | -0.654128728 | 0.886314617 | -0.738032202 |
| RP23-356K2.5 | 3.883316289 | -1.018438272 | 0.639908049 | -1.591538462 |
| Man1a2 | 2251.936784 | 0.32509119 | 0.067136196 | 4.842264056 |

| | | | | |
|---|---|---|---|---|
| RP23-356K2.6 | 73.44238526 | 0.474147409 | 0.218444986 | 2.170557529 |
| RP23-267O21.4 | 93.44924604 | 0.673550511 | 0.214618801 | 3.138357441 |
| RP23-267O21.3 | 152.3385727 | 0.649123636 | 0.244914229 | 2.650412101 |
| RP23-267O21.2 | 27.15683575 | 0.498913064 | 0.276800208 | 1.802430232 |
| Gm12485 | 0 | NA | NA | NA |
| RP23-267O21.5 | 2.75295346 | -1.852139525 | 0.635474822 | -2.914575781 |
| Gm23465 | 0 | NA | NA | NA |
| Vtcn1 | 9.162540417 | -1.324917875 | 0.560726102 | -2.362861066 |
| Trim45 | 89.1891917 | -0.096320494 | 0.132198688 | -0.728604007 |
| Ttf2 | 41.18065889 | -0.859749142 | 0.224195291 | -3.83482248 |
| Gm12488 | 0 | NA | NA | NA |
| Cd101 | 11.43243995 | -0.274247819 | 0.345173451 | -0.794521764 |
| Ptgfrn | 610.398054 | -0.230068225 | 0.090737652 | -2.535532055 |
| RP23-434L7.5 | 6.750937273 | 1.256691075 | 0.493904482 | 2.544401034 |
| Gm12486 | 0 | NA | NA | NA |
| Tpt1-ps1 | 0.068097359 | 0.037064902 | 3.724945049 | 0.009950456 |
| Gm12490 | 0.045882455 | 0.517959912 | 3.724945049 | 0.139051692 |
| Cd2 | 30.67976892 | -0.404788611 | 0.265552892 | -1.524323863 |
| Gm10355 | 2.030855555 | 1.041320557 | 0.766271182 | 1.358945215 |
| RP23-347G20.3 | 0 | NA | NA | NA |
| Igsf3 | 593.9020174 | -0.226506667 | 0.103456396 | -2.189392594 |
| RP23-347G20.4 | 4.3771345 | -0.524287142 | 0.584493874 | -0.896993391 |
| RP23-347G20.5 | 2.646385505 | 0.017278777 | 0.799566783 | 0.021610174 |
| RP23-347G20.6 | 0 | NA | NA | NA |
| Atp1a1 | 7569.239803 | 0.139063463 | 0.076672334 | 1.813737188 |
| RP23-103P11.2 | 13.65833158 | 0.08148926 | 0.377114248 | 0.2160864 |
| RP23-103P11.1 | 32.33019698 | 0.595673448 | 0.338307448 | 1.76074589 |
| RP24-115G23.1 | 0.049553047 | -0.443818988 | 3.724945049 | -0.119147795 |
| RP24-115G23.2 | 0.117046891 | 0.037062677 | 3.724945049 | 0.009949859 |
| Mab21l3 | 2.984303933 | -0.95951336 | 0.585812675 | -1.6379184 |
| Slc22a15 | 163.9002955 | 0.02959529 | 0.119421079 | 0.247823001 |
| Nhlh2 | 53.60793837 | -0.103845208 | 0.185386945 | -0.560153837 |
| RP23-226J22.2 | 0 | NA | NA | NA |
| RP23-226J22.3 | 0 | NA | NA | NA |
| Casq2 | 54.88658604 | -0.240082958 | 0.219444127 | -1.094050507 |
| RP24-89C11.1 | 0.931900496 | -0.542695152 | 1.094989299 | -0.495616854 |
| RP24-89C11.2 | 3.004992861 | 0.41523853 | 0.696252618 | 0.59639062 |
| Vangl1 | 127.7913766 | -0.453216996 | 0.117693346 | -3.850829401 |
| Gm16160 | 0.666323006 | -2.393688693 | 1.220269847 | -1.961606032 |
| RP23-191N23.1 | 0.144846928 | 0.037063327 | 3.535641013 | 0.010482774 |
| RP23-63A19.1 | 24.83821061 | -0.256606698 | 0.249572241 | -1.028186055 |
| RP24-409P22.1 | 3.653697379 | 0.700426481 | 0.619920845 | 1.129864379 |
| RP24-409P22.2 | 0 | NA | NA | NA |
| Ngf | 23.55789567 | -1.203264013 | 0.219089387 | -5.492114572 |
| RP23-63A19.3 | 5.480294069 | -1.25684802 | 0.460122121 | -2.731553131 |
| RP24-89D17.1 | 0.141094079 | 0.037072457 | 3.563850528 | 0.01040236 |
| RP24-89D17.3 | 0.352483205 | 0.995705011 | 1.732647513 | 0.574672577 |

| | | | | |
|---|---|---|---|---|
| RP24-89D17.2 | 0 | NA | NA | NA |
| RP23-183L19.1 | 0.184015763 | -0.503753035 | 3.310013942 | -0.152190608 |
| Tspan2os | 4.207560025 | 0.598067481 | 0.499442825 | 1.197469361 |
| Tspan2 | 1566.869316 | 0.608280434 | 0.098231815 | 6.192295584 |
| Tshb | 0.532135551 | -1.194728309 | 1.357716765 | -0.879954008 |
| RP23-183L19.3 | 0.133946075 | 0.036100024 | 3.626570338 | 0.009954315 |
| Sycp1 | 43.46231598 | -0.019836098 | 0.317598502 | -0.062456523 |
| RP23-183L19.5 | 0 | NA | NA | NA |
| RP23-183L19.6 | 0 | NA | NA | NA |
| RP23-119F18.1 | 0 | NA | NA | NA |
| Nr1h5 | 0.37220402 | 1.168537535 | 1.479455393 | 0.789843033 |
| Gm22826 | 0.042716945 | 0.037062677 | 3.724945049 | 0.009949859 |
| Sike1 | 543.965291 | -0.249307942 | 0.076309615 | -3.26705804 |
| Csde1 | 5133.717668 | 0.059260385 | 0.076513765 | 0.774506195 |
| Nras | 746.5356171 | -0.486076051 | 0.091221155 | -5.328545232 |
| RP23-106G24.2 | 13.15425346 | 0.321073682 | 0.335568013 | 0.956806578 |
| Ampd1 | 2.035201819 | -1.119525748 | 0.79967374 | -1.39997813 |
| Gm23820 | 0 | NA | NA | NA |
| RP24-409D4.3 | 0.140443404 | -0.411242499 | 2.909819489 | -0.141329213 |
| Dennd2c | 11.7142143 | -0.851893884 | 0.314042841 | -2.712667738 |
| Bcas2 | 429.1205886 | -0.031374091 | 0.065972457 | -0.475563481 |
| Gm27024 | 0 | NA | NA | NA |
| 2410024N13Rik | 0 | NA | NA | NA |
| RP23-70G12.2 | 0 | NA | NA | NA |
| Trim33 | 1476.330058 | 0.005080868 | 0.062914079 | 0.080758844 |
| RP23-70G12.3 | 47.1675988 | 0.061583098 | 0.356024719 | 0.17297422 |
| Gm25009 | 0 | NA | NA | NA |
| Gm25869 | 2.797214178 | -0.263880089 | 0.546523872 | -0.482833601 |
| Gm25199 | 0 | NA | NA | NA |
| Syt6 | 477.8102627 | -0.658219844 | 0.174945352 | -3.762431164 |
| RP23-184N6.1 | 10.82643969 | -0.871820128 | 0.478634856 | -1.821472294 |
| Atg4a-ps | 7.430379725 | -0.040452066 | 0.34889608 | -0.115943024 |
| Gm25325 | 0 | NA | NA | NA |
| Gm15986 | 0 | NA | NA | NA |
| Olfml3 | 289.0413337 | -0.544011256 | 0.159102151 | -3.419257703 |
| Gm10964 | 0 | NA | NA | NA |
| Hipk1 | 3690.970568 | -0.136365298 | 0.064137745 | -2.126131784 |
| RP24-559G13.2 | 18.01004 | 0.343662799 | 0.266090575 | 1.291525634 |
| Gm15886 | 0.872687401 | -0.261515793 | 1.056359567 | -0.247563236 |
| Gm16520 | 20.13453085 | 0.110048964 | 0.331550449 | 0.331922228 |
| Dclre1b | 139.879516 | 0.153322551 | 0.101865058 | 1.505153526 |
| Gm15471 | 0.37827039 | -1.216017947 | 1.536300873 | -0.791523307 |
| Ap4b1 | 32.06009994 | -0.070849675 | 0.208024234 | -0.340583758 |
| RP24-180F7.1 | 1.345434513 | 1.25950812 | 1.132977942 | 1.111679294 |
| Bcl2l15 | 59.73746658 | 0.147068608 | 0.214617245 | 0.685259976 |
| Ptpn22 | 140.1073571 | -0.105141046 | 0.338647293 | -0.310473604 |
| Gm15472 | 7.402758977 | -0.040935172 | 0.591839682 | -0.069165981 |

| | | | | |
|---|---|---|---|---|
| RP24-180F7.2 | 12.50366051 | -0.414213863 | 0.383141955 | -1.08109764 |
| Gm15602 | 0.044544648 | 0.517957688 | 3.724945049 | 0.139051095 |
| Rsbn1 | 982.5522449 | 0.047234182 | 0.055861701 | 0.845555743 |
| Phtf1os | 24.52540072 | -0.084713659 | 0.220378847 | -0.384400138 |
| Phtf1 | 538.6159203 | -0.051786986 | 0.099183609 | -0.522132503 |
| Magi3 | 898.9838019 | 0.135876699 | 0.087900944 | 1.545793394 |
| RP24-535M14.2 | 5.341278601 | 1.288752954 | 0.553479483 | 2.328456598 |
| RP24-535M14.1 | 11.88755228 | 0.594834745 | 0.341507241 | 1.741792484 |
| Gm23830 | 0 | NA | NA | NA |
| RP23-314M6.1 | 7.461541511 | 0.450142701 | 0.452319694 | 0.995187047 |
| RP23-333I5.9 | 6.272582582 | 0.413931099 | 0.597745055 | 0.692487701 |
| RP23-333I5.8 | 0.141094079 | 0.037072457 | 3.563850528 | 0.01040236 |
| RP23-333I5.7 | 34.16158652 | 0.669050789 | 0.326506921 | 2.049116715 |
| RP23-333I5.6 | 17.9477799 | 0.226457074 | 0.293609131 | 0.771287571 |
| RP23-333I5.5 | 22.39591524 | 0.577102474 | 0.374877083 | 1.539444528 |
| RP23-333I5.4 | 10.78090146 | 1.258827527 | 0.381812864 | 3.296975161 |
| RP23-333I5.3 | 57.87337 | 0.33343572 | 0.263925054 | 1.263372748 |
| RP23-333I5.2 | 44.8221039 | 0.298582061 | 0.300078992 | 0.995011541 |
| RP23-333I5.1 | 53.3220374 | 0.337428698 | 0.273315187 | 1.234577198 |
| Gm23077 | 0 | NA | NA | NA |
| Gm5546 | 0.706390846 | 1.350314149 | 1.286660667 | 1.049471849 |
| Lrig2 | 847.580948 | 0.065594833 | 0.065434528 | 1.002449865 |
| RP23-357D17.5 | 0.044191549 | 0.037062677 | 3.724945049 | 0.009949859 |
| RP23-357D17.6 | 0.315700266 | 1.025564089 | 1.784311159 | 0.574767514 |
| RP23-357D17.4 | 84.69226509 | 0.293674678 | 0.273029613 | 1.075614747 |
| RP23-357D17.3 | 51.88363519 | 0.84843626 | 0.264561148 | 3.206957131 |
| RP23-38L11.4 | 32.89618772 | 0.344669084 | 0.221737704 | 1.554399984 |
| Gm26091 | 0.643663771 | 0.148761341 | 1.246452227 | 0.119347808 |
| RP23-38L11.3 | 7.967952249 | -0.117328409 | 0.445477239 | -0.263376889 |
| Slc16a1 | 692.2362281 | -0.2402264 | 0.112847272 | -2.128774546 |
| RP23-38L11.1 | 26.81542658 | 0.249682824 | 0.290048453 | 0.860831427 |
| RP23-38L11.2 | 8.709900708 | 0.667539852 | 0.456221879 | 1.463191231 |
| Gm29561 | 0 | NA | NA | NA |
| Gm29560 | 0.214789954 | -0.978029719 | 2.560687419 | -0.381940299 |
| Gm24223 | 0 | NA | NA | NA |
| Gm29572 | 0 | NA | NA | NA |
| Gm6485 | 49.19972077 | -0.063564775 | 0.199679478 | -0.318334042 |
| RP24-503L4.2 | 0.134880151 | -1.305960502 | 2.927087508 | -0.446163806 |
| Fam19a3 | 6.79823111 | -0.90463192 | 0.509853438 | -1.774297968 |
| Ppm1j | 22.89084466 | -0.847755017 | 0.361419905 | -2.345623487 |
| Rhoc | 431.5423868 | -0.923910685 | 0.184172074 | -5.016562313 |
| Mov10 | 458.979696 | -0.75565231 | 0.143114768 | -5.280044282 |
| Capza1 | 545.1140594 | -0.547589592 | 0.091744598 | -5.968630319 |
| RP24-503L4.4 | 28.08267805 | 0.69084027 | 0.33510936 | 2.061536776 |
| RP24-503L4.3 | 42.80149299 | 0.838626958 | 0.342005772 | 2.452084224 |
| St7l | 574.3995767 | -0.187934226 | 0.067992438 | -2.764046008 |
| RP24-274H19.1 | 0.085327104 | -0.905414734 | 3.723500899 | -0.243162217 |

| | | | | |
|---|---|---|---|---|
| Wnt2b | 46.92012226 | -0.843659965 | 0.248383769 | -3.39659861 |
| RP24-274H19.3 | 0.479138389 | 2.133584261 | 1.381742854 | 1.544125418 |
| RP24-274H19.5 | 3.753741655 | -0.125080272 | 0.589150816 | -0.212306031 |
| Cttnbp2nl | 873.2904401 | -0.643702245 | 0.080424604 | -8.00379754 |
| 4930564D02Rik | 0.045882455 | 0.517959912 | 3.724945049 | 0.139051692 |
| Kcnd3os | 14.28524435 | 0.042579406 | 0.412025272 | 0.103341734 |
| Kcnd3 | 1457.767906 | 0.232693567 | 0.069098279 | 3.367574004 |
| RP23-443O1.1 | 42.61810582 | 0.550106233 | 0.268482538 | 2.048946039 |
| Ddx20 | 175.0365538 | 0.076799706 | 0.108194254 | 0.709831652 |
| Fam212b | 966.0315667 | 0.290391859 | 0.173929574 | 1.669594493 |
| RP24-500I24.6 | 43.0223173 | 0.371881569 | 0.208586858 | 1.78286193 |
| Rap1a | 552.0763281 | -0.139052753 | 0.107371854 | -1.295057763 |
| RP24-500I24.4 | 37.31668493 | 0.455632275 | 0.28016571 | 1.626295651 |
| RP24-500I24.3 | 48.64596038 | 0.065606361 | 0.259002083 | 0.253304375 |
| RP24-500I24.2 | 32.60373694 | 0.09620651 | 0.26864606 | 0.358116214 |
| RP24-500I24.5 | 33.83971767 | -0.172886644 | 0.273453025 | -0.63223526 |
| RP24-500I24.7 | 2.098595915 | -0.280037728 | 0.687196307 | -0.407507615 |
| Adora3 | 9.886871762 | 0.262144747 | 0.351122862 | 0.7465898 |
| Tmigd3 | 0.650450437 | -0.283969902 | 1.28268221 | -0.221387574 |
| I830077J02Rik | 45.37358814 | -0.685203952 | 0.267784534 | -2.558788367 |
| Atp5f1 | 2177.75384 | 0.005230703 | 0.084513231 | 0.061892125 |
| Wdr77 | 315.9051542 | -0.093767333 | 0.076292321 | -1.229053347 |
| RP23-169H9.3 | 8.021508243 | 0.850836924 | 0.437744404 | 1.943684295 |
| Ovgp1 | 35.01689641 | 0.583645068 | 0.304217399 | 1.918513109 |
| Pifo | 4.561976979 | 0.241913243 | 0.502888414 | 0.481047557 |
| RP23-169H9.6 | 8.449170836 | 1.496633909 | 0.585611343 | 2.555677801 |
| RP23-169H9.5 | 0 | NA | NA | NA |
| Chil5 | 6.28406316 | -1.015337164 | 0.408938136 | -2.482862502 |
| Gm4540 | 10.64549809 | -0.328169462 | 0.318896627 | -1.029077869 |
| RP23-169H9.8 | 0 | NA | NA | NA |
| RP23-62K10.1 | 0.969048608 | 2.607754988 | 1.134869799 | 2.297845084 |
| Chia1 | 8.605265352 | 1.481382309 | 0.472919105 | 3.132422208 |
| Chil3 | 390.2692537 | -3.044364176 | 1.026127506 | -2.966847843 |
| RP23-169H9.9 | 0 | NA | NA | NA |
| Chil4 | 0.092691418 | -0.908687078 | 3.724560878 | -0.2439716 |
| RP23-191E1.1 | 0.325695176 | -2.306998722 | 1.869789257 | -1.233828205 |
| RP23-191E1.2 | 0 | NA | NA | NA |
| Chil6 | 1.21043771 | -0.379979839 | 0.97496513 | -0.389736851 |
| Gm38244 | 0 | NA | NA | NA |
| 2010016I18Rik | 10.3751689 | -0.025452381 | 0.345624431 | -0.073641729 |
| Dennd2d | 26.9763545 | -0.198634767 | 0.226877783 | -0.875514404 |
| Cept1 | 592.7431419 | -0.504400722 | 0.115894129 | -4.352254312 |
| Dram2 | 391.8607469 | -0.268382724 | 0.111586227 | -2.405159948 |
| RP23-164N7.2 | 0 | NA | NA | NA |
| RP23-164N7.3 | 46.01383439 | 0.284077252 | 0.289947403 | 0.979754428 |
| Gm38271 | 1.062454588 | 0.455325889 | 0.879680081 | 0.517603954 |
| Lrif1 | 343.0611023 | -0.224743317 | 0.100541219 | -2.23533511 |

| | | | | |
|---|---|---|---|---|
| Gm38253 | 70.59424867 | -0.115577964 | 0.232993227 | -0.496057186 |
| Cd53 | 595.4541458 | -1.320895756 | 0.28365708 | -4.656664159 |
| Olfr266 | 0.317967701 | 1.008454904 | 1.782395635 | 0.565786229 |
| RP24-481E4.4 | 1.491118976 | 1.510274688 | 0.824124294 | 1.8325812 |
| Gm27008 | 4.962650854 | 0.417747113 | 0.526398664 | 0.793594554 |
| Kcna3 | 229.4345153 | -0.276651099 | 0.09819297 | -2.817422672 |
| AI504432 | 766.1594696 | -0.171435799 | 0.093799961 | -1.827674526 |
| A930002I21Rik | 0.066050875 | -0.443818988 | 3.724945049 | -0.119147795 |
| Kcna2 | 5020.598395 | 0.365889454 | 0.074178068 | 4.932582679 |
| Kcna10 | 0.10503227 | -0.905414734 | 3.723500899 | -0.243162217 |
| Cym | 1.145893073 | -0.522825964 | 0.881142669 | -0.593349956 |
| A630076J17Rik | 0 | NA | NA | NA |
| Prok1 | 4.595448637 | -0.164215846 | 0.639336869 | -0.256853396 |
| Lamtor5 | 258.6284024 | -0.250116527 | 0.136851504 | -1.827649099 |
| Slc16a4 | 15.80845376 | -0.186969032 | 0.344861185 | -0.542157368 |
| Rbm15 | 884.0175483 | -0.181098373 | 0.067058634 | -2.700597416 |
| RP23-332I1.7 | 0.048949532 | 0.037064902 | 3.724945049 | 0.009950456 |
| Gm24383 | 0.089089296 | 0.957155977 | 3.723939561 | 0.257027796 |
| Kcnc4 | 618.4655694 | 0.277454181 | 0.082539373 | 3.361476738 |
| Slc6a17 | 6021.526622 | -0.126208665 | 0.110287136 | -1.144364331 |
| Ubl4b | 0.42233865 | -0.884646451 | 1.518095684 | -0.582734316 |
| Alx3 | 13.73531081 | -1.047516961 | 0.457140108 | -2.291457133 |
| Strip1 | 672.0896654 | -0.098391416 | 0.0739748 | -1.330066681 |
| Gm10961 | 0.150451996 | 0.037073243 | 2.858243881 | 0.012970637 |
| Ahcyl1 | 4251.746979 | 0.254257982 | 0.077013837 | 3.301458415 |
| Csf1 | 1357.448556 | -0.709784824 | 0.227543445 | -3.119337598 |
| RP23-455E11.2 | 0.180126469 | 0.03706234 | 3.344092834 | 0.011082928 |
| RP23-455E11.1 | 0.049553047 | -0.443818988 | 3.724945049 | -0.119147795 |
| RP23-313F7.2 | 1.373329198 | 0.290998295 | 0.874862548 | 0.33262173 |
| RP23-313F7.8 | 0 | NA | NA | NA |
| Eps8l3 | 0.290796633 | 0.037078033 | 1.633135155 | 0.022703591 |
| RP23-313F7.4 | 0 | NA | NA | NA |
| 4933431E20Rik | 826.6688943 | 0.743494788 | 0.120651869 | 6.162314713 |
| Gstm5 | 640.7472393 | 0.518590361 | 0.131798484 | 3.934721745 |
| RP23-313F7.6 | 0.038981395 | -0.443816764 | 3.724945049 | -0.119147198 |
| Gm12489 | 0 | NA | NA | NA |
| Gstm7 | 114.4264593 | 0.289052607 | 0.188283738 | 1.53519688 |
| Gstm6 | 23.14962032 | 0.089414908 | 0.265427767 | 0.336870963 |
| Gm12494 | 1.001132476 | -2.432982143 | 1.206157378 | -2.017134901 |
| Gstm3 | 0.425236713 | -1.080813352 | 1.683551081 | -0.641984294 |
| Gm12497 | 0 | NA | NA | NA |
| Gstm2 | 20.7015664 | -0.07164296 | 0.32038711 | -0.22361374 |
| RP23-313F7.7 | 0.044544648 | 0.517957688 | 3.724945049 | 0.139051095 |
| Gstm1 | 1044.038813 | 0.526024466 | 0.181517739 | 2.897923189 |
| Gm12498 | 0.194528622 | -0.444753359 | 3.242027121 | -0.137183726 |
| Gm12502 | 2.704774846 | 0.339931384 | 0.609573145 | 0.557654789 |
| Gm12499 | 0 | NA | NA | NA |

| | | | | |
|---|---|---|---|---|
| Gm25592 | 0 | NA | NA | NA |
| Gstm4 | 56.39273402 | 0.146011222 | 0.178145426 | 0.819618135 |
| Ampd2 | 616.2955637 | -0.026898488 | 0.073253687 | -0.367196374 |
| Gm12500 | 6.670440749 | -0.259086928 | 0.402682032 | -0.64340325 |
| Gnat2 | 1.625657436 | -1.31584176 | 0.85919172 | -1.531487943 |
| Gnai3 | 600.7292191 | -0.608639882 | 0.132215565 | -4.603390543 |
| Gpr61 | 149.8257791 | 0.353150248 | 0.132250983 | 2.670303385 |
| Gm12524 | 0.213105667 | -1.381832878 | 2.596471244 | -0.532196488 |
| Amigo1 | 1089.407811 | 0.19840354 | 0.068988844 | 2.875878581 |
| Cyb561d1 | 232.812123 | -0.103152922 | 0.081771845 | -1.261472352 |
| Atxn7l2 | 171.8889948 | 0.502344983 | 0.169090717 | 2.970860792 |
| Sypl2 | 13.67443026 | 0.957433885 | 0.391867979 | 2.443256244 |
| RP23-89M15.15 | 1.384376546 | 2.781601804 | 1.160230981 | 2.397455205 |
| Gm12501 | 0.378789762 | 0.037077 | 1.703498107 | 0.021765213 |
| 1700010K24Rik | 1.755101273 | -1.788540966 | 0.976597483 | -1.831400343 |
| Psma5 | 407.5269465 | -0.429600994 | 0.120026166 | -3.579227817 |
| Gm12523 | 1.090677807 | -0.203244308 | 0.956046462 | -0.212588317 |
| Sort1 | 4425.028415 | 0.856600159 | 0.104226689 | 8.218625834 |
| RP23-27B23.9 | 51.99739385 | 0.769564081 | 0.389863155 | 1.973933854 |
| Gm12525 | 7.569909558 | 0.456252988 | 0.503502619 | 0.906158122 |
| Mybphl | 0.67227865 | -0.430122094 | 1.379273308 | -0.311846892 |
| Gm12522 | 10.72785558 | 1.533259308 | 0.422576368 | 3.62836028 |
| Psrc1 | 41.89133445 | 0.502404094 | 0.228174698 | 2.201839637 |
| Celsr2 | 4197.124515 | 0.718942479 | 0.121980555 | 5.893910524 |
| Sars | 1527.302422 | -0.274932574 | 0.082816675 | -3.319773148 |
| Gm22942 | 0 | NA | NA | NA |
| RP23-37G19.3 | 0.129625439 | 0.037055631 | 3.686164085 | 0.010052627 |
| 5330417C22Rik | 475.2428445 | 0.272427809 | 0.131275343 | 2.075239749 |
| Gm23336 | 0 | NA | NA | NA |
| Gm37859 | 19.39367035 | -0.177846823 | 0.496823784 | -0.357967611 |
| 1700013F07Rik | 7.727506971 | -0.037563236 | 0.398064504 | -0.094364696 |
| Scarna2 | 566.7431568 | -0.237775446 | 0.235938882 | -1.007784069 |
| RP23-128G10.7 | 0 | NA | NA | NA |
| Tmem167b | 537.5116706 | -0.183953353 | 0.066752291 | -2.755760887 |
| Gm22860 | 0.047948698 | 0.037064902 | 3.724945049 | 0.009950456 |
| Taf13 | 217.8213017 | -0.261450371 | 0.118984338 | -2.197351145 |
| Wdr47 | 1240.073746 | 0.218168423 | 0.074229962 | 2.939088458 |
| Gm12517 | 0 | NA | NA | NA |
| Clcc1 | 305.3641183 | 0.011263495 | 0.080887616 | 0.139248698 |
| Gpsm2 | 151.0149954 | -0.534589554 | 0.163896753 | -3.261745847 |
| Gm27244 | 0 | NA | NA | NA |
| Aknad1 | 0.537146556 | -0.46895007 | 1.25897061 | -0.372486908 |
| Stxbp3 | 425.8909719 | -0.457560789 | 0.108565014 | -4.214624705 |
| Fndc7 | 3.245667884 | -1.38942071 | 0.558465287 | -2.487926722 |
| RP23-316M11.8 | 0.10503227 | -0.905414734 | 3.723500899 | -0.243162217 |
| Prpf38b | 771.447708 | -0.13394935 | 0.115760907 | -1.157120777 |
| Henmt1 | 3.246767699 | 0.144929128 | 0.629486719 | 0.230233814 |

| | | | | |
|---|---|---|---|---|
| Gm9857 | 0.597180582 | 0.958004221 | 1.249089318 | 0.766962144 |
| Fam102b | 647.6814876 | -0.076427583 | 0.112759998 | -0.677789853 |
| RP23-316M11.10 | 0 | NA | NA | NA |
| AL671921.1 | 0 | NA | NA | NA |
| RP23-288B16.2 | 0 | NA | NA | NA |
| Slc25a54 | 0 | NA | NA | NA |
| RP23-288B16.4 | 8.294058351 | 0.306179251 | 0.422903619 | 0.723992981 |
| Slc25a24 | 72.40842471 | -1.390776244 | 0.219514262 | -6.335698778 |
| Gm13865 | 0 | NA | NA | NA |
| Vav3 | 309.6866449 | -0.517379076 | 0.078293384 | -6.608209436 |
| Ntng1 | 634.8071448 | 0.495257188 | 0.095509037 | 5.185448442 |
| Gm25681 | 0 | NA | NA | NA |
| Gm12535 | 0 | NA | NA | NA |
| RP23-327M1.2 | 2.031725428 | 0.174065923 | 0.640338277 | 0.271834324 |
| Prmt6 | 217.8360219 | -0.109340483 | 0.10312584 | -1.060262714 |
| RP23-327M1.1 | 30.13062775 | -0.325525835 | 0.214803701 | -1.515457289 |
| RP23-440G4.1 | 0 | NA | NA | NA |
| RP23-440G4.2 | 0.155610039 | 0.517971132 | 2.828260995 | 0.183141207 |
| RP23-43A4.1 | 0 | NA | NA | NA |
| Gm26076 | 0 | NA | NA | NA |
| RP24-366N1.1 | 0.538110248 | -0.526343552 | 1.55600148 | -0.338266743 |
| RP24-223F21.1 | 0.075043205 | 0.517957688 | 3.724945049 | 0.139051095 |
| RP24-230D18.1 | 0 | NA | NA | NA |
| RP23-14C8.1 | 0 | NA | NA | NA |
| RP23-14C8.2 | 0 | NA | NA | NA |
| RP24-93G19.1 | 0.12092566 | 0.998850474 | 3.724945049 | 0.268151734 |
| Gm25519 | 0 | NA | NA | NA |
| RP24-325G15.1 | 0 | NA | NA | NA |
| RP24-325G15.2 | 0.39879887 | -0.697014949 | 1.734434214 | -0.401868773 |
| RP23-117K10.1 | 0 | NA | NA | NA |
| RP24-221P14.1 | 0 | NA | NA | NA |
| RP23-430D14.1 | 5.464530712 | -0.011113749 | 0.495761291 | -0.02241754 |
| RP24-141M5.1 | 0 | NA | NA | NA |
| Gm5548 | 0.097139547 | -0.443816764 | 3.724945049 | -0.119147198 |
| RP23-470O13.1 | 0 | NA | NA | NA |
| Amy2b | 0.038981395 | -0.443816764 | 3.724945049 | -0.119147198 |
| Amy2-ps1 | 0 | NA | NA | NA |
| Amy2a5 | 0.143022002 | 0.03610674 | 3.547735707 | 0.010177404 |
| Amy2a4 | 0.089089296 | 0.957155977 | 3.723939561 | 0.257027796 |
| Amy2a3 | 0.060174537 | 0.517957688 | 3.724945049 | 0.139051095 |
| Amy2a2 | 0 | NA | NA | NA |
| Amy2a1 | 0.175778459 | -0.458577645 | 2.722156135 | -0.168461184 |
| Amy1 | 192.4654876 | 0.664067119 | 0.187742883 | 3.537109413 |
| Rnpc3 | 532.6768979 | 0.427562477 | 0.083588421 | 5.115092168 |
| RP24-237D17.3 | 0 | NA | NA | NA |
| RP24-182K13.1 | 0.088383099 | 0.037058229 | 3.724945049 | 0.009948665 |
| RP23-256J12.1 | 0.075043205 | 0.517957688 | 3.724945049 | 0.139051095 |

| | | | | |
|---|---|---|---|---|
| Col11a1 | 184.655142 | -0.71170843 | 0.151381303 | -4.701428883 |
| RP23-281B17.2 | 0 | NA | NA | NA |
| Gm23750 | 0 | NA | NA | NA |
| RP23-292P14.1 | 0 | NA | NA | NA |
| Olfm3 | 941.997379 | 0.23198307 | 0.14775147 | 1.570089758 |
| RP23-231J6.1 | 0.046345709 | -0.443818988 | 3.724945049 | -0.119147795 |
| Gm22337 | 0 | NA | NA | NA |
| Gm25347 | 0 | NA | NA | NA |
| RP23-238A12.1 | 0.12092566 | 0.998850474 | 3.724945049 | 0.268151734 |
| S1pr1 | 1328.879257 | -0.332499798 | 0.098855113 | -3.363506329 |
| Gm9889 | 3.625155807 | 0.86047683 | 0.517686717 | 1.662157443 |
| A930005H10Rik | 72.61233888 | -0.046970275 | 0.140917313 | -0.333317989 |
| Dph5 | 126.6011858 | -0.218422663 | 0.121322721 | -1.800344253 |
| RP24-155C13.8 | 0.199959441 | 0.517971038 | 2.663425918 | 0.194475481 |
| RP24-155C13.6 | 0.510666263 | 0.413422764 | 1.355217218 | 0.305060147 |
| RP24-155C13.7 | 0.595761752 | 0.954035497 | 1.244414083 | 0.766654372 |
| Slc30a7 | 561.2209546 | -0.059314524 | 0.088464003 | -0.670493331 |
| RP24-245I16.1 | 1.76977757 | 1.67700335 | 0.997911416 | 1.680513243 |
| RP24-155C13.3 | 4.122231334 | 0.30287713 | 0.505763551 | 0.59885124 |
| Mir669n | 0 | NA | NA | NA |
| Extl2 | 975.1969821 | 0.235777712 | 0.079833217 | 2.953378582 |
| Gm26344 | 0 | NA | NA | NA |
| Vcam1 | 1522.902751 | -0.889189794 | 0.213811581 | -4.158754123 |
| RP24-245I16.3 | 1.059433203 | -0.831613326 | 1.087879832 | -0.764434915 |
| RP23-93F11.2 | 1.275102148 | 0.273574214 | 0.831288734 | 0.329096501 |
| Gm26544 | 0.239854152 | 0.561426601 | 2.523748012 | 0.222457471 |
| RP23-93F11.4 | 0.265934997 | 1.410148674 | 2.10568339 | 0.669686944 |
| Gpr88 | 1159.493482 | 1.263391908 | 0.155339255 | 8.133114242 |
| Cdc14a | 108.7934714 | -0.552357193 | 0.12856765 | -4.29623775 |
| Gm29151 | 0 | NA | NA | NA |
| Gm22361 | 0 | NA | NA | NA |
| Gm26358 | 0 | NA | NA | NA |
| Rtca | 414.5859796 | 0.110164593 | 0.110650227 | 0.995611088 |
| Gm26530 | 0.207691825 | -1.343202105 | 2.608991152 | -0.514835822 |
| Dbt | 216.6118277 | 0.013538186 | 0.102920131 | 0.131540698 |
| Gm4596 | 16.93547575 | -0.199431878 | 0.268911078 | -0.741627603 |
| RP23-453M11.3 | 3.08271666 | 3.12991997 | 0.897068789 | 3.489052353 |
| Gm9761 | 55.52035785 | -0.197863533 | 0.157514433 | -1.256161289 |
| Lrrc39 | 5.281818806 | -0.677718408 | 0.544384509 | -1.244925961 |
| RP23-453M11.4 | 10.63896767 | 0.263086388 | 0.430651819 | 0.610902767 |
| Trmt13 | 134.1213703 | 0.02628322 | 0.13502376 | 0.194656257 |
| Sass6 | 170.9486267 | -0.868606932 | 0.114005094 | -7.619018601 |
| RP24-565B3.4 | 8.173995324 | 0.397388598 | 0.372382028 | 1.067152999 |
| Hiat1 | 617.2974484 | -0.031878271 | 0.067077276 | -0.47524695 |
| RP24-565B3.2 | 6.147691891 | -0.86728217 | 0.490929999 | -1.766610659 |
| RP24-565B3.5 | 0 | NA | NA | NA |
| RP24-565B3.3 | 0.186953752 | 0.037076649 | 2.668894834 | 0.013892136 |

| | | | | |
|---|---|---|---|---|
| Slc35a3 | 453.1222322 | -0.058573431 | 0.084345057 | -0.694450073 |
| Agl | 702.330067 | -0.145955837 | 0.076875309 | -1.898604888 |
| RP24-423B3.1 | 0.424532791 | 0.480585905 | 1.404440891 | 0.342190197 |
| RP24-423B3.2 | 0.346647033 | 0.037053993 | 1.629566311 | 0.022738561 |
| RP24-539I8.1 | 0 | NA | NA | NA |
| CH25-626J19.1 | 0.097139547 | -0.443816764 | 3.724945049 | -0.119147198 |
| Frrs1 | 66.10065566 | -1.045226972 | 0.203407411 | -5.138588433 |
| Palmd | 159.5277226 | -0.076896247 | 0.164184994 | -0.468351249 |
| RP24-370B22.3 | 26.73605718 | 0.981238705 | 0.379955916 | 2.582506718 |
| 4930455H04Rik | 0.11657011 | 0.998850474 | 3.724945049 | 0.268151734 |
| RP24-341P14.2 | 0 | NA | NA | NA |
| 1700061I17Rik | 0 | NA | NA | NA |
| RP24-83G17.1 | 0.038981395 | -0.443816764 | 3.724945049 | -0.119147198 |
| RP24-83G17.2 | 0 | NA | NA | NA |
| RP23-173L24.3 | 0.766169159 | 0.736930437 | 1.363346098 | 0.540530712 |
| RP23-173L24.2 | 1.114955615 | -2.862957435 | 1.183529021 | -2.41900062 |
| Lppr4 | 2571.916126 | 0.127428596 | 0.136613043 | 0.932770352 |
| Gm22408 | 0.596872081 | -0.387711876 | 1.452835245 | -0.266865687 |
| Lppr5 | 515.7492512 | 0.085704247 | 0.113908261 | 0.752397114 |
| Snx7 | 152.0630282 | -0.085013227 | 0.113459361 | -0.749283498 |
| Gm24811 | 0.346782673 | 0.85391985 | 1.569068926 | 0.544220739 |
| CH25-626J19.2 | 0 | NA | NA | NA |
| RP24-294N3.1 | 0 | NA | NA | NA |
| Gm26871 | 573.8481393 | 0.696331394 | 0.258189635 | 2.696976564 |
| Mir137 | 0 | NA | NA | NA |
| Gm24155 | 3.962088627 | 1.746459311 | 0.659692039 | 2.647385761 |
| Gm9916 | 100.2258943 | 0.919801688 | 0.391987079 | 2.346510221 |
| Gm37773 | 129.487562 | 0.635799523 | 0.17386759 | 3.65680299 |
| Gm37928 | 218.6973413 | 0.916944717 | 0.246734132 | 3.716327001 |
| Gm32444 | 100.5808754 | 0.3001274 | 0.194239898 | 1.545137758 |
| Dpyd | 164.1135791 | 0.516254713 | 0.117378025 | 4.398222866 |
| RP24-173A7.2 | 9.88657072 | 0.995478412 | 0.342728846 | 2.904565587 |
| Gm23432 | 0 | NA | NA | NA |
| RP23-235E15.1 | 152.0784296 | -0.271006192 | 0.144820627 | -1.87132316 |
| Ptbp2 | 2043.522665 | 0.48000605 | 0.117907299 | 4.071046103 |
| RP23-235E15.5 | 196.3079525 | 0.995880141 | 0.260023917 | 3.829955921 |
| RP23-235E15.6 | 474.5268742 | 0.818875484 | 0.211183157 | 3.877560573 |
| RP23-235E15.7 | 0 | NA | NA | NA |
| Gm24744 | 0 | NA | NA | NA |
| Gm26137 | 0 | NA | NA | NA |
| RP23-427K12.1 | 0 | NA | NA | NA |
| Gm23733 | 0 | NA | NA | NA |
| 6530403H02Rik | 55.37544644 | 1.018099996 | 0.208371547 | 4.885983762 |
| Gm25013 | 0.075043205 | 0.517957688 | 3.724945049 | 0.139051095 |
| RP23-324P1.1 | 0.575937436 | 1.719234294 | 1.253569148 | 1.371471448 |
| RP24-295P14.2 | 0.374268822 | -1.220874763 | 3.717386859 | -0.328422843 |
| RP24-295P14.1 | 0.066050875 | -0.443818988 | 3.724945049 | -0.119147795 |

| | | | | |
|---|---|---|---|---|
| Rwdd3 | 95.46200367 | 0.273798503 | 0.19685516 | 1.390862723 |
| RP24-210J6.4 | 0.060174537 | 0.517957688 | 3.724945049 | 0.139051095 |
| Gm5710 | 0 | NA | NA | NA |
| Tmem56 | 921.2771509 | 0.368124404 | 0.073740423 | 4.992165631 |
| RP24-291L17.1 | 8.847422939 | 0.356023872 | 0.488860735 | 0.728272587 |
| RP24-291L17.2 | 0.233612452 | 0.979519378 | 2.527723168 | 0.387510543 |
| Alg14 | 122.1469618 | -0.164617137 | 0.126801231 | -1.298229798 |
| Gm26027 | 1.314748965 | -0.140319196 | 1.04406815 | -0.134396587 |
| RP24-277J22.6 | 0 | NA | NA | NA |
| RP24-277J22.4 | 33.98944694 | -0.86393867 | 0.420502587 | -2.054538301 |
| Cnn3 | 2221.060378 | -0.966222672 | 0.16944763 | -5.702190547 |
| RP24-277J22.3 | 26.71658222 | -0.307197455 | 0.326324931 | -0.941385184 |
| Slc44a3 | 22.72182752 | -0.76873767 | 0.399614006 | -1.923700519 |
| A530020G20Rik | 0.979355264 | -1.414735462 | 0.991935795 | -1.42623693 |
| A730020M07Rik | 12.26765879 | -0.222776515 | 0.310791056 | -0.716804781 |
| 4930432M17Rik | 0 | NA | NA | NA |
| RP24-404P10.3 | 0.136120942 | -0.876404342 | 3.595683781 | -0.24373788 |
| RP24-404P10.4 | 0 | NA | NA | NA |
| RP24-404P10.2 | 5.861481922 | 0.404099063 | 0.403687936 | 1.001018428 |
| F3 | 702.2491587 | -0.290109945 | 0.149953619 | -1.934664511 |
| RP24-404P10.8 | 9.75452295 | 0.149603002 | 0.399251151 | 0.374709006 |
| Abcd3 | 888.7244778 | -0.076341132 | 0.085711996 | -0.890670335 |
| RP24-404P10.7 | 19.74428529 | 0.121115211 | 0.273407724 | 0.442983868 |
| RP24-404P10.6 | 21.98336171 | 0.095385109 | 0.3191125 | 0.298907467 |
| RP24-404P10.9 | 0.234491734 | -0.968032212 | 2.118441136 | -0.456954973 |
| RP23-54L16.1 | 0 | NA | NA | NA |
| Arhgap29 | 378.6264762 | -0.45480312 | 0.081730212 | -5.564687852 |
| RP23-54L16.2 | 5.712404952 | 0.945349082 | 0.473715115 | 1.995606753 |
| Abca4 | 76.95354951 | 0.495645236 | 0.191697857 | 2.585554393 |
| RP24-471B15.3 | 0.334549462 | 1.084265695 | 1.63888328 | 0.661588112 |
| RP24-471B15.2 | 0 | NA | NA | NA |
| RP24-471B15.1 | 0.04393175 | 0.037062677 | 3.724945049 | 0.009949859 |
| Gm4609 | 10.08219113 | 0.699873534 | 0.334001677 | 2.09541922 |
| Gm22903 | 0.085433889 | 0.037058229 | 3.724945049 | 0.009948665 |
| Gclm | 1174.677346 | -0.22204942 | 0.105083322 | -2.113079556 |
| Gm17494 | 67.68281203 | 0.71941755 | 0.284573969 | 2.528051148 |
| Gm6061 | 18.81787436 | 0.68755682 | 0.244962468 | 2.806784337 |
| Dnttip2 | 377.7972389 | -0.204128521 | 0.118157835 | -1.727591923 |
| RP23-71G16.7 | 1.463879734 | -0.688385592 | 1.021825136 | -0.673682383 |
| Mir760 | 0 | NA | NA | NA |
| Bcar3 | 197.0241314 | -0.060155874 | 0.121555682 | -0.494883277 |
| RP23-71G16.6 | 1.51534681 | -0.564872395 | 0.740507632 | -0.76281779 |
| RP23-192O12.4 | 0.11657011 | 0.998850474 | 3.724945049 | 0.268151734 |
| Gm25153 | 0.892441177 | 0.95024497 | 1.25910616 | 0.754698055 |
| Fnbp1l | 1523.630982 | 0.190322165 | 0.10199158 | 1.866057625 |
| mmu-mir-7657 | 0 | NA | NA | NA |
| RP23-1A12.4 | 0.048949532 | 0.037064902 | 3.724945049 | 0.009950456 |

| | | | | |
|---|---|---|---|---|
| RP23-1A12.2 | 29.85261811 | 0.509878781 | 0.294878825 | 1.729112905 |
| RP23-1A12.5 | 0.187986659 | -1.264069549 | 3.306965886 | -0.382244508 |
| RP23-1A12.1 | 0.044544648 | 0.517957688 | 3.724945049 | 0.139051095 |
| Pde5a | 353.2225523 | -0.114625578 | 0.166825143 | -0.68710013 |
| RP24-182A8.1 | 4.735351793 | 0.105889629 | 0.50016149 | 0.21171088 |
| RP24-182A8.2 | 0.332478248 | 0.893910591 | 1.993864527 | 0.448330656 |
| RP24-70L3.3 | 0.663651821 | -0.304003843 | 1.16022761 | -0.262020866 |
| 4933405D12Rik | 1.378174623 | -0.761914729 | 0.913812595 | -0.833775692 |
| Fabp2 | 0 | NA | NA | NA |
| Gm10959 | 0.205306308 | 1.332944249 | 2.603665266 | 0.511949161 |
| 1810037I17Rik | 393.3493054 | -0.472147372 | 0.134682276 | -3.505638506 |
| Usp53 | 810.7985606 | -0.046826163 | 0.113854611 | -0.411280339 |
| RP23-48L9.1 | 1.574470888 | -0.174387289 | 0.814827234 | -0.214017501 |
| Myoz2 | 0.617569467 | 0.692063064 | 1.23421002 | 0.560733629 |
| Gm22066 | 0.060174537 | 0.517957688 | 3.724945049 | 0.139051095 |
| RP23-48L9.2 | 0 | NA | NA | NA |
| Synpo2 | 278.5160077 | 0.398588757 | 0.133835092 | 2.978208125 |
| RP23-48L9.4 | 0.075043205 | 0.517957688 | 3.724945049 | 0.139051095 |
| RP23-48L9.3 | 2.800041622 | 0.476530825 | 0.654019752 | 0.728618399 |
| RP24-488I7.1 | 9.542608144 | 0.898676744 | 0.431100128 | 2.084612565 |
| RP24-488I7.2 | 12.44003729 | 0.2301113 | 0.381629441 | 0.602970512 |
| Sec24d | 210.6853567 | -0.749264927 | 0.128115788 | -5.848341879 |
| Mettl14 | 164.8105688 | -0.049510013 | 0.097399837 | -0.508317202 |
| Gm22713 | 0.066050875 | -0.443818988 | 3.724945049 | -0.119147795 |
| RP23-113O9.4 | 1.285708346 | 1.424433431 | 0.888042473 | 1.604014981 |
| Prss12 | 146.351687 | 0.421252708 | 0.162549751 | 2.591530935 |
| RP23-113O9.3 | 0.319644482 | 0.956469469 | 1.581176878 | 0.60490985 |
| Snhg8 | 70.39141616 | 0.006941639 | 0.340283682 | 0.020399566 |
| Snora24 | 0 | NA | NA | NA |
| RP24-149B7.2 | 0.066050875 | -0.443818988 | 3.724945049 | -0.119147795 |
| Ndst3 | 624.8561942 | 0.071523835 | 0.167308059 | 0.427497851 |
| RP24-374F22.1 | 20.67476562 | 0.136037191 | 0.314317427 | 0.432801936 |
| Gm24456 | 0 | NA | NA | NA |
| Gm15400 | 0.119587853 | 0.981063299 | 3.723618439 | 0.263470416 |
| Gm15399 | 0.060174537 | 0.517957688 | 3.724945049 | 0.139051095 |
| RP24-107N14.2 | 0 | NA | NA | NA |
| Gm9372 | 0.194279094 | -1.017823543 | 3.719440177 | -0.273649661 |
| RP24-107N14.4 | 0 | NA | NA | NA |
| Tram1l1 | 242.7416911 | 0.135024772 | 0.092150453 | 1.465264343 |
| Gm4617 | 19.97291991 | -0.210476065 | 0.33288065 | -0.632286871 |
| RP23-412L13.3 | 0 | NA | NA | NA |
| RP23-412L13.5 | 0 | NA | NA | NA |
| 1700006A11Rik | 0.255320825 | 1.936461352 | 2.379358218 | 0.813858686 |
| Gm22575 | 0.066050875 | -0.443818988 | 3.724945049 | -0.119147795 |
| RP23-412L13.4 | 0 | NA | NA | NA |
| Gm22900 | 0 | NA | NA | NA |
| 1700003H04Rik | 0 | NA | NA | NA |

| | | | | |
|---|---|---|---|---|
| RP24-332L11.2 | 0 | NA | NA | NA |
| RP23-24O21.1 | 0.192951104 | 1.045927396 | 3.260557335 | 0.320781783 |
| Gm23673 | 0.219094128 | 1.630590776 | 3.130988095 | 0.520791113 |
| Gm9381 | 0.088383099 | 0.037058229 | 3.724945049 | 0.009948665 |
| RP23-446O11.1 | 0 | NA | NA | NA |
| Ndst4 | 306.8135137 | 0.217332532 | 0.139689586 | 1.555824872 |
| Ugt8a | 1013.869147 | -0.343815775 | 0.225302622 | -1.526017636 |
| Gm26275 | 0.551648795 | -0.517260978 | 1.716609575 | -0.301327096 |
| RP24-297G3.1 | 0 | NA | NA | NA |
| RP23-301O12.1 | 0.159882028 | 0.037076092 | 2.804321448 | 0.013221056 |
| Arsj | 51.42050542 | -0.025388824 | 0.198935297 | -0.127623525 |
| Gm17225 | 0.384085174 | 0.083316743 | 3.275794943 | 0.025434053 |
| RP23-60D3.4 | 6.256257677 | 0.105970671 | 0.551725036 | 0.192071528 |
| RP23-60D3.3 | 21.83940584 | 0.530845856 | 0.332260722 | 1.597678634 |
| Camk2d | 2946.282952 | -0.104887374 | 0.049453495 | -2.120929454 |
| RP23-293K21.1 | 122.6972469 | -0.167377544 | 0.238159698 | -0.702795419 |
| RP23-293K21.2 | 86.00755271 | -0.300467824 | 0.215081621 | -1.396994418 |
| RP23-293K21.3 | 29.22029167 | -0.359272239 | 0.275632581 | -1.303446197 |
| RP23-293K21.4 | 16.36398547 | -0.356313972 | 0.399170102 | -0.892636923 |
| RP23-293K21.5 | 4.108293731 | -0.893898478 | 0.626713863 | -1.426326319 |
| Gm15551 | 0 | NA | NA | NA |
| RP23-293K21.7 | 52.63659607 | -0.421187157 | 0.275876754 | -1.526722173 |
| RP23-293K21.8 | 19.23585701 | -0.086563277 | 0.296824873 | -0.291630808 |
| Ank2 | 12996.91815 | 0.159507298 | 0.054770054 | 2.912308587 |
| RP24-423L4.5 | 54.25629816 | 0.786551622 | 0.295159978 | 2.66483155 |
| RP24-423L4.4 | 84.96617208 | 0.326314071 | 0.250856718 | 1.300798611 |
| RP24-423L4.3 | 17.18518708 | 0.569724497 | 0.324803293 | 1.7540601 |
| Gm25501 | 0 | NA | NA | NA |
| RP24-423L4.2 | 66.13758907 | 0.701724156 | 0.336373273 | 2.086147184 |
| RP24-423L4.1 | 97.87426476 | 0.306446799 | 0.282443401 | 1.084984805 |
| RP23-50E10.7 | 69.57910019 | -0.002280547 | 0.233966501 | -0.009747323 |
| Gm16958 | 3.547171809 | -0.212173446 | 0.508761364 | -0.417039226 |
| RP23-50E10.5 | 105.9269057 | -0.106526249 | 0.262850002 | -0.405273914 |
| RP23-50E10.4 | 141.5973348 | -0.022960092 | 0.214651184 | -0.106964666 |
| RP23-50E10.3 | 170.8359331 | 0.255918792 | 0.373727855 | 0.684773127 |
| RP23-50E10.2 | 42.82266199 | 0.153997602 | 0.352258244 | 0.437172457 |
| RP23-50E10.1 | 29.57400322 | 0.370061514 | 0.426461082 | 0.867749788 |
| RP24-163F11.1 | 44.61355707 | 0.604859173 | 0.321145458 | 1.883443024 |
| RP24-260H22.2 | 2.710738714 | -0.350792451 | 0.703807582 | -0.498420961 |
| RP24-260H22.1 | 2.472653692 | 0.787505986 | 0.868626931 | 0.906610143 |
| AC157817.1 | 0 | NA | NA | NA |
| RP24-166J20.2 | 0 | NA | NA | NA |
| Larp7 | 118.9071558 | -0.092328791 | 0.14973115 | -0.616630482 |
| Mir302b | 0 | NA | NA | NA |
| Mir302c | 0 | NA | NA | NA |
| Mir302a | 0 | NA | NA | NA |
| Mir302d | 0 | NA | NA | NA |

| | | | | |
|---|---|---|---|---|
| Mir367 | 0 | NA | NA | NA |
| Zgrf1 | 113.3311546 | 0.293037367 | 0.147334202 | 1.98892968 |
| RP23-182M12.2 | 0 | NA | NA | NA |
| Neurog2 | 4.696444918 | 0.338895267 | 0.490771785 | 0.690535351 |
| Alpk1 | 333.345234 | 0.045652638 | 0.160326921 | 0.284747174 |
| Gm16238 | 2.994207206 | -0.242087576 | 0.582879218 | -0.415330601 |
| RP23-182M12.4 | 9.869931754 | 0.000430949 | 0.433907191 | 0.000993183 |
| RP23-182M12.3 | 4.2109437 | -0.079176896 | 0.629909368 | -0.125695695 |
| Gm23279 | 0 | NA | NA | NA |
| Tifa | 352.3487698 | 0.399400352 | 0.158018156 | 2.527559883 |
| Ap1ar | 641.0884169 | 0.333785768 | 0.080994333 | 4.121100273 |
| RP23-173N16.4 | 70.08836606 | 0.795191604 | 0.354182271 | 2.245147964 |
| RP23-173N16.3 | 2.767049531 | 0.15778098 | 0.605078384 | 0.260761224 |
| RP23-173N16.5 | 0.803713496 | 1.127448645 | 1.381078733 | 0.816353636 |
| RP23-173N16.8 | 0.269164771 | 0.321300173 | 2.0471671 | 0.156948679 |
| 5730508B09Rik | 54.45665569 | -0.842583257 | 0.213356274 | -3.949184349 |
| RP24-285A14.1 | 0.588650639 | 1.769510187 | 1.332198438 | 1.328263221 |
| RP24-285A14.2 | 0.048949532 | 0.037064902 | 3.724945049 | 0.009950456 |
| Gm10650 | 0 | NA | NA | NA |
| RP24-380G1.1 | 1.298106709 | -0.714092508 | 0.872049051 | -0.818867364 |
| RP24-380G1.2 | 0 | NA | NA | NA |
| Gm22293 | 0 | NA | NA | NA |
| RP24-196G4.1 | 0 | NA | NA | NA |
| RP23-61G16.1 | 0.062426873 | 0.517959912 | 3.724945049 | 0.139051692 |
| RP24-261K17.1 | 0 | NA | NA | NA |
| Gm9387 | 8.054043807 | -0.476700095 | 0.390997688 | -1.219189038 |
| Gm28543 | 0 | NA | NA | NA |
| Pitx2 | 11.30362075 | 0.230570603 | 0.346144372 | 0.666111085 |
| Gm27195 | 0.265852463 | 0.063284409 | 1.819377419 | 0.034783552 |
| RP23-446B17.3 | 0.066050875 | -0.443818988 | 3.724945049 | -0.119147795 |
| Enpep | 24.04966124 | 0.110012338 | 0.27827873 | 0.395331465 |
| RP24-140K12.2 | 0 | NA | NA | NA |
| RP24-140K12.4 | 0 | NA | NA | NA |
| RP23-257P11.1 | 0 | NA | NA | NA |
| RP23-257P11.2 | 0.095897396 | 0.037062677 | 3.724945049 | 0.009949859 |
| Rpl7a-ps7 | 2.979957284 | 0.422205546 | 0.611238892 | 0.690737372 |
| RP23-257P11.4 | 0.174406792 | 0.541348768 | 3.365276146 | 0.160863104 |
| Elovl6 | 655.176359 | -0.256303947 | 0.078991334 | -3.244709688 |
| RP23-3L20.3 | 30.90056436 | 0.686476426 | 0.268538762 | 2.556340179 |
| Egf | 35.62468515 | 0.583697567 | 0.236192923 | 2.471274587 |
| Gm24515 | 0.098502579 | -0.443821212 | 3.724945049 | -0.119148392 |
| RP23-3L20.6 | 0.263299413 | -0.443820983 | 2.148013626 | -0.206619259 |
| RP23-3L20.8 | 0 | NA | NA | NA |
| RP23-3L20.5 | 1.405849262 | 0.55620683 | 0.937401928 | 0.593349356 |
| Lrit3 | 3.350603553 | 0.299412017 | 0.755471287 | 0.396324813 |
| Rrh | 25.69789624 | 1.034639428 | 0.46468202 | 2.22655361 |
| Gar1 | 114.8672286 | 0.002503023 | 0.204206727 | 0.012257299 |

| | | | | |
|---|---|---|---|---|
| Cfi | 0.976809389 | -0.401144561 | 1.192910071 | -0.336273933 |
| Pla2g12a | 138.6047462 | 0.070848354 | 0.114574245 | 0.618361958 |
| Casp6 | 39.32840354 | -0.109934622 | 0.191113591 | -0.575231837 |
| Ccdc109b | 33.80637658 | -1.088171481 | 0.24957815 | -4.36004307 |
| Sec24b | 993.3623414 | -0.106739669 | 0.049603182 | -2.151871396 |
| Gm9396 | 8.082634732 | -0.38004265 | 0.418776933 | -0.90750617 |
| RP23-123C15.1 | 0 | NA | NA | NA |
| Col25a1 | 767.761363 | 0.698896538 | 0.107025352 | 6.53019614 |
| RP23-123C15.2 | 0.147068667 | 0.998853574 | 3.51441668 | 0.284216035 |
| Gm22682 | 2.066582291 | 0.088676356 | 0.77226378 | 0.114826511 |
| RP23-105B8.2 | 34.01273389 | 1.140756541 | 0.32064603 | 3.557681789 |
| RP23-105B8.3 | 9.17532523 | 1.460586284 | 0.394947278 | 3.698180405 |
| RP23-105B8.4 | 6.910068968 | 1.260430725 | 0.451888955 | 2.789248801 |
| Etnppl | 214.5413129 | 1.081993392 | 0.190199902 | 5.688716885 |
| Ostc | 293.8716669 | -0.595331847 | 0.148153444 | -4.018346336 |
| RP23-246L14.2 | 8.0379673 | -0.380575655 | 0.482662386 | -0.788492467 |
| Rpl34 | 837.1533585 | -0.356860752 | 0.131139166 | -2.721237007 |
| RP23-246L14.3 | 2.763783036 | 0.65117366 | 0.640795526 | 1.016195703 |
| RP23-246L14.4 | 0.231616962 | 0.984260761 | 2.124643584 | 0.463259235 |
| RP23-246L14.5 | 0.678557026 | 0.496723214 | 1.297106782 | 0.382947049 |
| RP23-349J3.1 | 0 | NA | NA | NA |
| RP23-349J3.2 | 0.048949532 | 0.037064902 | 3.724945049 | 0.009950456 |
| RP23-349J3.3 | 0.086908494 | 0.037058229 | 3.724945049 | 0.009948665 |
| Gm26090 | 0 | NA | NA | NA |
| RP23-349J3.4 | 6.268689147 | -1.951905831 | 0.4951381 | -3.942144289 |
| RP23-349J3.5 | 0.049553047 | -0.443818988 | 3.724945049 | -0.119147795 |
| RP23-349J3.6 | 0 | NA | NA | NA |
| RP23-349J3.7 | 0.075043205 | 0.517957688 | 3.724945049 | 0.139051095 |
| RP23-349J3.8 | 18.61381697 | -0.117869362 | 0.242357776 | -0.486344462 |
| RP24-176P19.1 | 0.066050875 | -0.443818988 | 3.724945049 | -0.119147795 |
| RP24-176P19.3 | 3.678638758 | -2.054192769 | 0.681685291 | -3.013403394 |
| Lef1 | 120.0536891 | 0.381383123 | 0.149640337 | 2.548665227 |
| RP24-176P19.6 | 5.471034591 | 0.617029305 | 0.496854813 | 1.241870439 |
| Hadh | 222.3007882 | -0.039674355 | 0.146608304 | -0.270614648 |
| RP24-434E24.5 | 0.259720851 | -1.185320894 | 2.026354886 | -0.584952272 |
| Cyp2u1 | 190.2662416 | 0.231068131 | 0.102060657 | 2.264027467 |
| 4930534D22Rik | 5.431134866 | -0.16713794 | 0.461489159 | -0.362170891 |
| Sgms2 | 102.4694722 | -0.388383866 | 0.192439058 | -2.018217455 |
| RP23-86I19.3 | 0 | NA | NA | NA |
| Papss1 | 709.3146802 | 0.053490827 | 0.084689846 | 0.631608499 |
| RP24-188G23.3 | 0.068097359 | 0.037064902 | 3.724945049 | 0.009950456 |
| RP23-292C19.1 | 0.108772582 | 0.037073798 | 3.724945049 | 0.009952844 |
| Olfr375-ps1 | 0 | NA | NA | NA |
| Gm26103 | 0 | NA | NA | NA |
| Gm22421 | 0.047948698 | 0.037064902 | 3.724945049 | 0.009950456 |
| Dkk2 | 4.638032695 | -0.778256137 | 0.52814845 | -1.47355566 |
| RP23-435E17.1 | 0 | NA | NA | NA |

| | | | | |
|---|---|---|---|---|
| Gm25307 | 0 | NA | NA | NA |
| Gm5549 | 0 | NA | NA | NA |
| RP24-450N10.1 | 0.627611986 | 1.027764024 | 1.171972644 | 0.87695223 |
| Aimp1 | 305.8677058 | -0.106053653 | 0.118901199 | -0.891947718 |
| Tbck | 868.7732137 | 0.1171079 | 0.064734711 | 1.809043363 |
| RP24-450N10.3 | 11.22849639 | 0.682803256 | 0.353079115 | 1.933853426 |
| Gm22322 | 0 | NA | NA | NA |
| RP23-447N11.5 | 18.07808909 | 0.286371073 | 0.29815663 | 0.960471926 |
| RP23-447N11.2 | 0.788050755 | 0.828272469 | 1.196173489 | 0.692435066 |
| RP23-447N11.3 | 31.71702897 | 0.177744605 | 0.255198889 | 0.696494429 |
| Npnt | 229.9057745 | 0.529432366 | 0.137784413 | 3.842469226 |
| Gm23196 | 0 | NA | NA | NA |
| Gstcd | 129.9010644 | -0.411594519 | 0.137378286 | -2.996066777 |
| RP23-10D15.4 | 0 | NA | NA | NA |
| RP23-10D15.3 | 6.257617145 | 0.017748884 | 0.582944272 | 0.030446966 |
| RP23-10D15.2 | 9.313879906 | -0.107232057 | 0.439133206 | -0.244190273 |
| Ints12 | 209.4893335 | -0.408307874 | 0.101818982 | -4.01013511 |
| Arhgef38 | 3.684370567 | -1.195093959 | 0.576294781 | -2.073754608 |
| RP23-335C9.1 | 28.10162786 | 0.178212556 | 0.220480159 | 0.80829294 |
| Ppa2 | 314.4988618 | 0.132476638 | 0.078426181 | 1.689188959 |
| Gm25042 | 0.060174537 | 0.517957688 | 3.724945049 | 0.139051095 |
| Tet2 | 1445.087248 | 0.325669788 | 0.150106685 | 2.169588833 |
| RP24-333H9.2 | 2.368693662 | -0.633179178 | 0.708572021 | -0.8935989 |
| RP24-333H9.1 | 0 | NA | NA | NA |
| Gm23011 | 0 | NA | NA | NA |
| Gm37228 | 0 | NA | NA | NA |
| RP24-289J19.2 | 0 | NA | NA | NA |
| RP23-437C24.1 | 0.044544648 | 0.517957688 | 3.724945049 | 0.139051095 |
| RP23-437C24.2 | 0.116046057 | 0.037062677 | 3.724945049 | 0.009949859 |
| Gm26691 | 49.78358579 | 0.754770911 | 0.241961912 | 3.119379016 |
| AC123608.1 | 1.334504357 | 3.063567239 | 1.039830955 | 2.946216619 |
| RP23-304A10.3 | 0 | NA | NA | NA |
| RP23-304A10.1 | 0 | NA | NA | NA |
| RP23-304A10.2 | 4.293971641 | 0.597561798 | 0.537664974 | 1.111401759 |
| Cxxc4 | 699.6168967 | 0.181919172 | 0.082072454 | 2.216567967 |
| Mir1895 | 0 | NA | NA | NA |
| RP23-384A11.6 | 0.213180531 | 0.079167746 | 2.596008885 | 0.030495946 |
| RP23-384A11.5 | 0.844415878 | 1.033457955 | 1.16629396 | 0.886104181 |
| RP23-384A11.4 | 0.116217011 | 0.517955464 | 3.724945049 | 0.139050498 |
| RP23-424L10.1 | 0 | NA | NA | NA |
| Gm26820 | 0.046345709 | -0.443818988 | 3.724945049 | -0.119147795 |
| Tacr3 | 61.25689593 | 0.0011832 | 0.177737761 | 0.006657 |
| RP23-465M13.2 | 0.165933562 | 0.486011036 | 2.762106923 | 0.175956634 |
| RP23-465M13.3 | 0.044544648 | 0.517957688 | 3.724945049 | 0.139051095 |
| RP23-465M13.4 | 0 | NA | NA | NA |
| RP23-93G15.1 | 0 | NA | NA | NA |
| Gm24048 | 0 | NA | NA | NA |

| | | | | |
|---|---|---|---|---|
| RP23-93G15.2 | 0 | NA | NA | NA |
| RP23-453I14.1 | 0 | NA | NA | NA |
| Cenpe | 78.90502966 | 0.452356259 | 0.177124389 | 2.553890299 |
| Bdh2 | 27.37528258 | -0.176507465 | 0.320358314 | -0.55096889 |
| Slc9b2 | 28.12143682 | -0.091722732 | 0.238127372 | -0.385183491 |
| Slc9b1 | 1.329624306 | 0.324563238 | 0.966727714 | 0.335733872 |
| Cisd2 | 673.1771875 | 0.103710025 | 0.059415982 | 1.74549039 |
| RP24-372J15.5 | 30.69028919 | 0.545818962 | 0.262769204 | 2.077180104 |
| RP24-372J15.4 | 100.7051577 | 0.660551216 | 0.19943176 | 3.312166608 |
| 4930539J05Rik | 36.18476307 | 0.48797252 | 0.266995797 | 1.827641205 |
| Ube2d3 | 3995.493788 | -0.109286912 | 0.079804478 | -1.369433338 |
| RP23-167C5.2 | 0 | NA | NA | NA |
| Manba | 173.3995577 | -0.240186584 | 0.105624439 | -2.273967904 |
| Gm16100 | 2.944243352 | -0.478952997 | 0.696051148 | -0.68810029 |
| Oaz2-ps | 15.79173318 | -0.202470846 | 0.264789355 | -0.764648737 |
| Nfkb1 | 567.2400552 | -0.256256556 | 0.123389054 | -2.076817581 |
| RP23-167C5.1 | 62.27141224 | 0.174605142 | 0.364890751 | 0.478513477 |
| Slc39a8 | 39.47806482 | -0.400945389 | 0.19973934 | -2.00734311 |
| RP24-349N15.3 | 0.185615945 | 0.037075477 | 2.674999861 | 0.013859992 |
| RP24-349N15.4 | 1.794573941 | -0.075929046 | 0.715432654 | -0.10613025 |
| RP24-349N15.5 | 1.687301136 | -0.015976802 | 0.876456429 | -0.018228861 |
| RP24-349N15.6 | 0 | NA | NA | NA |
| RP24-349N15.2 | 0.623481245 | 0.968527236 | 1.293576964 | 0.748720225 |
| RP24-349N15.1 | 0.362626786 | -0.530113366 | 1.740600433 | -0.304557758 |
| RP24-326F21.2 | 0.246385664 | 0.917129815 | 2.060318213 | 0.445139886 |
| Bank1 | 11.35624835 | 0.12464032 | 0.329881619 | 0.37783348 |
| RP24-335I21.1 | 0.047948698 | 0.037064902 | 3.724945049 | 0.009950456 |
| RP24-335I21.2 | 0.075043205 | 0.517957688 | 3.724945049 | 0.139051095 |
| 1700030L20Rik | 0.443470786 | -1.388288401 | 1.528708971 | -0.908144341 |
| Ppp3ca | 9465.004104 | 0.735941585 | 0.112399078 | 6.547576716 |
| RP24-164P24.2 | 107.1091577 | 0.320720349 | 0.384002954 | 0.835202817 |
| RP24-164P24.5 | 0.044191549 | 0.037062677 | 3.724945049 | 0.009949859 |
| AC124177.1 | 0 | NA | NA | NA |
| RP23-174N20.1 | 0.068097359 | 0.037064902 | 3.724945049 | 0.009950456 |
| RP23-174N20.3 | 0.099155932 | 0.037073798 | 3.724945049 | 0.009952844 |
| RP23-174N20.2 | 0 | NA | NA | NA |
| Gm26107 | 0 | NA | NA | NA |
| RP23-77E5.1 | 0.132101749 | -0.93550548 | 3.724188147 | -0.251197158 |
| Emcn | 39.45898547 | -0.497027363 | 0.2494421 | -1.99255604 |
| Gm4861 | 0.22850592 | 0.517965306 | 2.564978803 | 0.201937461 |
| Gm22468 | 0.075043205 | 0.517957688 | 3.724945049 | 0.139051095 |
| Ddit4l | 50.60904352 | 0.599147814 | 0.202112043 | 2.964434011 |
| RP24-134B9.2 | 0.132574648 | 0.037054914 | 3.724945049 | 0.009947775 |
| Gm21962 | 0.087261593 | 0.51795324 | 3.724945049 | 0.139049901 |
| RP24-134B9.3 | 0 | NA | NA | NA |
| Gm25080 | 0 | NA | NA | NA |
| Mir6380 | 0 | NA | NA | NA |

| | | | | |
|---|---|---|---|---|
| RP24-380C2.1 | 0 | NA | NA | NA |
| RP24-380C2.2 | 1.550909849 | 0.023610056 | 0.96428538 | 0.024484511 |
| H2afz | 499.3132004 | 0.060003907 | 0.097004909 | 0.618565678 |
| Dnajb14 | 1867.871122 | 0.198713062 | 0.049993914 | 3.974745049 |
| RP23-209N19.2 | 0 | NA | NA | NA |
| RP23-209N19.1 | 20.11453365 | 0.380164755 | 0.358183089 | 1.061369914 |
| Gm15460 | 2.001670347 | 0.20997596 | 0.766400769 | 0.273976709 |
| Gm25920 | 0 | NA | NA | NA |
| RP23-209N19.4 | 0.841319663 | 0.372227693 | 1.057370533 | 0.352031461 |
| Lamtor3 | 343.7921853 | -0.184938259 | 0.083876394 | -2.204890441 |
| Dapp1 | 62.05077585 | -0.82625226 | 0.207884437 | -3.974574878 |
| RP23-209N19.6 | 0.453752633 | 1.218232967 | 1.309100507 | 0.930587805 |
| Gm5105 | 1.256408573 | -0.099652759 | 0.87817159 | -0.113477548 |
| 1110002E22Rik | 61.32006928 | -0.286235123 | 0.180944836 | -1.581891639 |
| Mttp | 190.0120384 | 0.123955984 | 0.109416593 | 1.132881043 |
| RP24-553B2.3 | 0.388301457 | -2.5051218 | 3.676631089 | -0.681363384 |
| Trmt10a | 78.50456046 | -0.22331708 | 0.153645366 | -1.453457952 |
| 4930579F01Rik | 0.038981395 | -0.443816764 | 3.724945049 | -0.119147198 |
| 0610031O16Rik | 0 | NA | NA | NA |
| Adh7 | 0.650173001 | 0.665540534 | 1.399840098 | 0.475440399 |
| Adh1 | 2.324828864 | -0.752620106 | 0.769934234 | -0.977512198 |
| Gm16559 | 0.066050875 | -0.443818988 | 3.724945049 | -0.119147795 |
| Adh6a | 0.234786912 | 0.616844331 | 3.046705743 | 0.20246272 |
| Adh6b | 0.808183924 | -0.79990534 | 1.150470587 | -0.695285346 |
| Gm23467 | 0 | NA | NA | NA |
| Adh6-ps1 | 0 | NA | NA | NA |
| Adh4 | 0.195798127 | 0.03706017 | 3.724945049 | 0.009949186 |
| Adh5 | 412.48678 | -0.121011325 | 0.097690993 | -1.238715267 |
| Metap1 | 515.9695598 | -0.238834672 | 0.053173833 | -4.491582784 |
| RP24-129H9.2 | 0.047948698 | 0.037064902 | 3.724945049 | 0.009950456 |
| RP24-129H9.1 | 0.20323857 | -0.118973025 | 2.636289133 | -0.045128975 |
| Eif4e | 1261.786848 | -0.111404169 | 0.065319768 | -1.705519977 |
| Mir1956 | 0 | NA | NA | NA |
| RP24-129H9.7 | 0 | NA | NA | NA |
| RP24-129H9.6 | 0.597553932 | 0.037077385 | 1.511068084 | 0.024537203 |
| RP24-129H9.8 | 0 | NA | NA | NA |
| RP24-93M7.1 | 0 | NA | NA | NA |
| RP24-93M7.2 | 0.061920613 | -0.443816764 | 3.724945049 | -0.119147198 |
| Tspan5 | 1750.479894 | -0.067269623 | 0.080368157 | -0.83701836 |
| Gm16060 | 0.740177848 | -0.568797239 | 1.186939658 | -0.479213274 |
| Rap1gds1 | 3054.008212 | 0.152882946 | 0.049768604 | 3.071875321 |
| RP24-428N8.5 | 39.16231688 | -0.028738534 | 0.301228921 | -0.0954043 |
| Gm4862 | 0 | NA | NA | NA |
| Stpg2 | 4.136924382 | 0.566543842 | 0.511852934 | 1.106848871 |
| RP23-276I11.1 | 0.091666476 | 0.037060453 | 3.724945049 | 0.009949262 |
| Gm24260 | 1.280600893 | 0.838604241 | 0.854787337 | 0.981067694 |
| RP23-4F17.1 | 21.90285516 | 0.674025859 | 0.262212472 | 2.570533177 |

| | | | | |
|---|---|---|---|---|
| RP23-434M21.1 | 0.112396584 | -0.917692504 | 3.724434845 | -0.246397787 |
| RP23-434M21.2 | 0.115957212 | 0.517955464 | 3.724945049 | 0.139050498 |
| Mir7j | 0.266304556 | -0.602115226 | 3.715438719 | -0.162057639 |
| RP24-221D2.1 | 0 | NA | NA | NA |
| RP24-63C3.1 | 0.099155932 | 0.037073798 | 3.724945049 | 0.009952844 |
| RP23-201A9.1 | 0 | NA | NA | NA |
| RP23-460F15.1 | 0 | NA | NA | NA |
| RP23-205G9.1 | 0 | NA | NA | NA |
| Gm6214 | 0 | NA | NA | NA |
| RP23-148J20.2 | 0 | NA | NA | NA |
| RP23-148J20.3 | 0.226297694 | 0.945557518 | 2.123090057 | 0.44536854 |
| RP23-148J20.4 | 0.157356115 | -0.500078175 | 2.825615622 | -0.176980256 |
| Pdha2 | 0 | NA | NA | NA |
| Gm15688 | 15.75159018 | 0.618155586 | 0.305583294 | 2.022871005 |
| Unc5c | 1697.764115 | 0.088332806 | 0.089881791 | 0.982766424 |
| Gm15689 | 1.903171754 | 0.477238311 | 0.668247305 | 0.714164215 |
| RP23-97D22.3 | 43.75796597 | 0.466499314 | 0.286151016 | 1.630255662 |
| Bmpr1b | 318.2842301 | 0.032377151 | 0.156665916 | 0.206663654 |
| RP23-100B7.1 | 0.094097695 | 0.037071574 | 3.724945049 | 0.009952247 |
| Pdlim5 | 665.5298056 | -0.860838159 | 0.107757976 | -7.988625899 |
| Gbp5 | 1952.644448 | -0.317434093 | 0.388446892 | -0.817187883 |
| Gbp7 | 3441.7084 | -0.788435625 | 0.229342736 | -3.437805085 |
| RP23-100J23.5 | 0 | NA | NA | NA |
| Gbp3 | 3129.531274 | -0.332461243 | 0.22599178 | -1.471120954 |
| Gbp2b | 4052.460821 | -1.243236279 | 0.315436504 | -3.941320246 |
| Gbp2 | 2406.349028 | -0.865153545 | 0.261872361 | -3.303722253 |
| Mir6381 | 0.56675494 | -2.226482559 | 1.251975895 | -1.77837494 |
| RP24-314I8.2 | 0 | NA | NA | NA |
| RP24-314I8.3 | 0.091666476 | 0.037060453 | 3.724945049 | 0.009949262 |
| Ccbl2 | 63.67129629 | 0.721701474 | 0.146134661 | 4.938605722 |
| Gm15540 | 0.047948698 | 0.037064902 | 3.724945049 | 0.009950456 |
| Gtf2b | 311.3568098 | -0.084764099 | 0.076729865 | -1.104708044 |
| Gm24457 | 0 | NA | NA | NA |
| Pkn2 | 955.7411507 | -0.282541355 | 0.080535675 | -3.50827572 |
| Gm25965 | 0 | NA | NA | NA |
| RP24-445M20.2 | 0.255858694 | -1.01722863 | 2.132293333 | -0.477058487 |
| 9530052C20Rik | 8.046682377 | 0.326832378 | 0.395705224 | 0.82594911 |
| RP24-445M20.3 | 0 | NA | NA | NA |
| Gm2574 | 15.8626213 | 0.09274087 | 0.288634229 | 0.321309329 |
| RP23-49B23.1 | 0.306809375 | -1.856334454 | 1.716692804 | -1.081343412 |
| RP23-49B23.2 | 0.297888126 | 0.808609961 | 1.743347855 | 0.463825942 |
| RP24-377E5.1 | 10.93206221 | -0.161497419 | 0.351679401 | -0.459217738 |
| RP24-337A2.1 | 0.793655395 | -1.714705674 | 1.432954955 | -1.196622175 |
| RP24-337A2.2 | 1.407833646 | -1.100066652 | 0.91735334 | -1.199174411 |
| RP24-337A2.3 | 0.046345709 | -0.443818988 | 3.724945049 | -0.119147795 |
| RP23-402L6.2 | 0.617903777 | 1.695354334 | 1.222751945 | 1.386507166 |
| RP23-402L6.1 | 1.629102473 | 1.036543979 | 0.897390862 | 1.155064112 |

| | | | | |
|---|---|---|---|---|
| RP24-273H20.2 | 0.222190708 | 0.517970413 | 2.165824457 | 0.23915623 |
| Gm38006 | 0 | NA | NA | NA |
| RP24-273H20.3 | 0.068097359 | 0.037064902 | 3.724945049 | 0.009950456 |
| RP24-273H20.4 | 3.820552698 | 1.549040476 | 0.742392066 | 2.086553114 |
| Gm24728 | 0 | NA | NA | NA |
| RP23-229C5.1 | 0.29112196 | 0.507671076 | 1.634483334 | 0.310600338 |
| RP23-229C5.2 | 0.131251387 | -0.924089913 | 2.981416013 | -0.309950007 |
| RP24-224P21.1 | 0.177934686 | 1.051638768 | 2.718161642 | 0.386893388 |
| RP23-327I19.1 | 2.34560619 | 0.473586119 | 0.7262234 | 0.652121811 |
| RP24-224P21.2 | 0.044544648 | 0.517957688 | 3.724945049 | 0.139051095 |
| RP23-327I19.2 | 0.038981395 | -0.443816764 | 3.724945049 | -0.119147198 |
| Lmo4 | 1293.515492 | 0.069230652 | 0.075482919 | 0.917169778 |
| RP23-461P7.1 | 43.89163664 | 0.502919425 | 0.253652078 | 1.982713599 |
| RP23-461P7.2 | 0 | NA | NA | NA |
| RP24-232N19.1 | 0.421951652 | 2.545255327 | 1.66894423 | 1.525069131 |
| RP24-224M8.1 | 22.99821006 | 1.255992086 | 0.29196397 | 4.301873567 |
| Hs2st1 | 927.2177343 | 0.075114104 | 0.055075926 | 1.363828255 |
| RP23-225M12.2 | 46.28994543 | 0.348273438 | 0.260826601 | 1.335268093 |
| RP23-225M12.1 | 14.69975535 | 0.36122292 | 0.389428588 | 0.927571655 |
| 15-Sep | 1591.561157 | -0.297068777 | 0.123999366 | -2.395728197 |
| Gm24406 | 0 | NA | NA | NA |
| Rpl9-ps8 | 0.045882455 | 0.517959912 | 3.724945049 | 0.139051692 |
| Sh3glb1 | 1627.178665 | -0.221928789 | 0.083723518 | -2.65073415 |
| RP24-321M14.4 | 0.899213277 | -0.398961357 | 1.172641124 | -0.340224599 |
| Clca3a1 | 41.15834729 | -0.528797153 | 0.546816971 | -0.96704598 |
| RP24-321M14.5 | 0.600994565 | -0.734814067 | 1.282811909 | -0.572815127 |
| RP24-271G16.1 | 1.394083107 | 1.520820465 | 1.040789883 | 1.461217572 |
| Clca3a2 | 1.360869654 | 0.552360443 | 0.819999481 | 0.673610723 |
| RP24-271G16.2 | 0.93582689 | -0.605947107 | 1.088129421 | -0.556870438 |
| Clca3b | 0.093141081 | 0.037060453 | 3.724945049 | 0.009949262 |
| Gm16215 | 0.201808035 | -0.71779893 | 3.188241939 | -0.225139417 |
| Clca4c-ps | 0.221289295 | 1.766416798 | 2.520812829 | 0.700733024 |
| Clca4b | 0 | NA | NA | NA |
| Gm16233 | 0.233058144 | 0.037077731 | 1.871568562 | 0.019811046 |
| Clca4a | 0 | NA | NA | NA |
| RP24-277E17.1 | 0.140122821 | 0.517959509 | 3.572128867 | 0.145000231 |
| Clca1 | 0.274296751 | -1.109907705 | 3.710575206 | -0.299120121 |
| Clca2 | 0.311801716 | 1.827047175 | 3.701979814 | 0.493532452 |
| Odf2l | 262.3245237 | -0.073448887 | 0.108899065 | -0.674467567 |
| Col24a1 | 47.56753531 | 0.389256722 | 0.255257329 | 1.524958061 |
| RP23-307M9.1 | 0 | NA | NA | NA |
| RP23-307M9.2 | 0.046345709 | -0.443818988 | 3.724945049 | -0.119147795 |
| Znhit6 | 139.6637941 | 0.062094585 | 0.139380863 | 0.445502949 |
| Cyr61 | 545.855505 | -0.543634859 | 0.584107505 | -0.930710279 |
| Gm17501 | 9.319672861 | -1.220310709 | 0.375018624 | -3.254000284 |
| RP24-267M22.1 | 0 | NA | NA | NA |
| Ddah1 | 1181.961696 | -0.562589318 | 0.105993753 | -5.307759209 |

| | | | | |
|---|---|---|---|---|
| Rpl36a-ps2 | 93.94039235 | 0.045266555 | 0.162735112 | 0.278160961 |
| Bcl10 | 200.6606098 | -0.453487767 | 0.112502011 | -4.030930311 |
| 2410004B18Rik | 209.7691853 | -0.241988581 | 0.12287244 | -1.969429283 |
| Syde2 | 65.37702174 | 0.066356368 | 0.157495917 | 0.421321196 |
| RP24-267M11.2 | 0 | NA | NA | NA |
| Wdr63 | 17.44671922 | -0.319899171 | 0.314285721 | -1.017860974 |
| Gm25627 | 0 | NA | NA | NA |
| Mcoln3 | 1.139337436 | -0.655692198 | 0.861320334 | -0.761264041 |
| Mcoln2 | 13.29155609 | -1.413647053 | 0.348148478 | -4.060471732 |
| Lpar3 | 6.729774449 | -0.747587252 | 0.411456534 | -1.81692886 |
| RP24-455A5.3 | 0 | NA | NA | NA |
| RP24-364E17.1 | 0.399538249 | -0.998039469 | 1.624078933 | -0.614526455 |
| Gm22078 | 0 | NA | NA | NA |
| Gm22901 | 0 | NA | NA | NA |
| Ssx2ip | 795.7799406 | 0.320843903 | 0.080705076 | 3.975510806 |
| Ctbs | 130.2632904 | -0.132015968 | 0.144072423 | -0.916316705 |
| Spata1 | 56.80384134 | -0.220059604 | 0.187293055 | -1.174948022 |
| Gng5 | 243.1666755 | -0.917480903 | 0.154058737 | -5.955396768 |
| Rpf1 | 164.2090017 | 0.171754165 | 0.113238781 | 1.516743315 |
| RP24-100G24.2 | 0 | NA | NA | NA |
| Uox | 7.769118165 | -2.499839414 | 0.512207185 | -4.880523915 |
| Dnase2b | 0.061920613 | -0.443816764 | 3.724945049 | -0.119147198 |
| Gm16325 | 1.444443621 | -0.870678749 | 0.783154493 | -1.111758607 |
| 4930503B20Rik | 2.396082498 | -1.96189049 | 0.743241405 | -2.639641005 |
| Samd13 | 1.10620882 | -1.105548135 | 0.970528361 | -1.139119865 |
| RP24-86C15.1 | 0.038981395 | -0.443816764 | 3.724945049 | -0.119147198 |
| Prkacb | 5477.753168 | 0.280015448 | 0.047540257 | 5.890070148 |
| RP24-409N2.3 | 0.044544648 | 0.517957688 | 3.724945049 | 0.139051095 |
| Gm10288 | 30.25828668 | -0.489620338 | 0.249063095 | -1.965848607 |
| Ttll7 | 4265.777113 | 0.26837408 | 0.072211989 | 3.716475373 |
| Gm29771 | 2.774619275 | 1.031218439 | 0.733602376 | 1.405691246 |
| Gm6074 | 0 | NA | NA | NA |
| RP24-293H7.2 | 0 | NA | NA | NA |
| Gm37803 | 0 | NA | NA | NA |
| Gm37041 | 0.332175987 | 0.526664084 | 3.709664405 | 0.141970816 |
| Gm36888 | 0 | NA | NA | NA |
| RP24-184C18.1 | 0.044191549 | 0.037062677 | 3.724945049 | 0.009949859 |
| Gm23131 | 0 | NA | NA | NA |
| RP23-290J2.1 | 0 | NA | NA | NA |
| Gm38068 | 0 | NA | NA | NA |
| RP24-196A21.2 | 0 | NA | NA | NA |
| Adgrl2 | 1106.274129 | 0.596217792 | 0.12773009 | 4.667794335 |
| RP24-224A19.1 | 3.670148074 | 0.539622205 | 0.603132648 | 0.894699045 |
| RP24-224A19.2 | 2.247096996 | 0.420979054 | 0.813888167 | 0.517244348 |
| Gm25127 | 1.189027437 | -0.869113339 | 0.932912631 | -0.93161279 |
| RP23-7K18.3 | 0.083172944 | -0.443821212 | 3.724945049 | -0.119148392 |
| Gm10287 | 19.5831566 | 0.790396374 | 0.389523571 | 2.029136187 |

| | | | | |
|---|---|---|---|---|
| RP23-473G18.1 | 38.52654686 | 1.030448467 | 0.250068636 | 4.120662562 |
| RP24-512E12.1 | 0.471094982 | 0.009963856 | 1.432363538 | 0.006956234 |
| Gm26468 | 0.506521265 | -0.38683925 | 1.38269405 | -0.279772123 |
| RP24-248L23.1 | 0 | NA | NA | NA |
| Rpsa-ps10 | 128.4715849 | -0.318328949 | 0.129719092 | -2.453986869 |
| RP24-134A5.1 | 4.381038091 | -0.775877234 | 0.490223638 | -1.582700574 |
| RP24-257N6.1 | 0 | NA | NA | NA |
| Mir3963 | 0 | NA | NA | NA |
| RP24-232H5.1 | 0 | NA | NA | NA |
| RP24-232H5.2 | 0.216927137 | 0.491021149 | 2.16135484 | 0.227182108 |
| RP24-232H5.3 | 0.510381651 | 0.660444695 | 1.384204642 | 0.477129374 |
| RP23-239A13.1 | 0.433950198 | -2.354893118 | 1.336027973 | -1.762607644 |
| RP23-239A13.2 | 0.049553047 | -0.443818988 | 3.724945049 | -0.119147795 |
| Adgrl4 | 62.50733234 | 0.155216341 | 0.226580844 | 0.685037351 |
| Gm24458 | 0.493466105 | -1.743818494 | 1.543510401 | -1.129774372 |
| Ifi44 | 3535.414574 | -0.790199771 | 0.266469236 | -2.96544465 |
| Ifi44l | 509.0915789 | -1.230052689 | 0.342127218 | -3.595307897 |
| Ptgfr | 13.2398592 | -0.62784399 | 0.397498441 | -1.579487931 |
| RP24-267D17.1 | 0.162302921 | 0.037074899 | 2.791096991 | 0.013283271 |
| RP24-267D17.2 | 0 | NA | NA | NA |
| Gm25056 | 0 | NA | NA | NA |
| RP23-245O13.3 | 0.112396584 | -0.917692504 | 3.724434845 | -0.246397787 |
| RP23-245O13.1 | 0.13307909 | -0.443194549 | 2.968793206 | -0.149284412 |
| RP23-245O13.2 | 0.060174537 | 0.517957688 | 3.724945049 | 0.139051095 |
| RP24-499N1.1 | 0.129258854 | -0.924088969 | 2.995409137 | -0.308501753 |
| Gipc2 | 25.12877053 | -0.076327231 | 0.21289034 | -0.358528389 |
| RP24-499N1.2 | 2.888459128 | 0.083176586 | 0.690299438 | 0.120493487 |
| Gm16198 | 0.109869311 | -0.444802825 | 3.724874622 | -0.119414174 |
| D630002J18Rik | 0.719682426 | 0.509166459 | 1.102039926 | 0.462021789 |
| Dnajb4 | 820.1226806 | 0.025941924 | 0.079493352 | 0.3263408 |
| Fubp1 | 2019.104171 | -0.017527528 | 0.092462734 | -0.189563155 |
| Nexn | 60.15238185 | 0.051279515 | 0.196359339 | 0.261151394 |
| RP23-181C16.4 | 5.705262924 | -1.034833578 | 0.572578802 | -1.807320798 |
| Fam73a | 1902.314384 | 0.289767993 | 0.055895692 | 5.184084542 |
| Usp33 | 1820.049344 | 0.159437687 | 0.066135197 | 2.410784184 |
| RP24-511F2.1 | 7.968776044 | 0.581675254 | 0.369164673 | 1.575652539 |
| RP24-511F2.2 | 7.38870447 | -0.163605543 | 0.435986836 | -0.375253402 |
| RP24-511F2.3 | 58.00030583 | 0.854284689 | 0.381721413 | 2.237979478 |
| RP24-511F2.4 | 45.38790308 | 0.05056982 | 0.225043447 | 0.224711367 |
| Zzz3 | 889.1435612 | 0.07556086 | 0.068857048 | 1.097358405 |
| Ak5 | 942.1680984 | 0.545795257 | 0.130186119 | 4.192422835 |
| AI115009 | 13.76623626 | 0.233642113 | 0.299974022 | 0.77887449 |
| Pigk | 806.3801702 | 0.158728467 | 0.04667399 | 3.400790615 |
| Rpl29-ps1 | 0.333447913 | 1.661917932 | 1.657765223 | 1.002505005 |
| St6galnac5 | 598.912246 | 0.520429865 | 0.126434973 | 4.116185998 |
| Gm16232 | 3.327150153 | 1.501258488 | 0.743876465 | 2.018155646 |
| Gm16213 | 1.998664546 | -2.15455039 | 0.839243652 | -2.56725253 |

| | | | | |
|---|---|---|---|---|
| Gm16231 | 0.342796299 | -0.544990503 | 1.763419731 | -0.309053196 |
| RP23-394P9.1 | 0.038981395 | -0.443816764 | 3.724945049 | -0.119147198 |
| RP24-117O8.1 | 6.160904613 | 1.519541295 | 0.559011125 | 2.718266647 |
| St6galnac3 | 220.6741086 | 0.20605552 | 0.114864287 | 1.793904135 |
| RP24-303G10.2 | 9.238644217 | 0.48945842 | 0.559803925 | 0.874339027 |
| RP24-303G10.1 | 48.22872844 | 0.472228326 | 0.315568909 | 1.496434891 |
| Gm27405 | 0.046345709 | -0.443818988 | 3.724945049 | -0.119147795 |
| RP24-104D12.1 | 4.330890979 | 0.346084122 | 0.627779447 | 0.551282975 |
| RP24-104D12.2 | 7.42840581 | -0.225343286 | 0.525760678 | -0.428604297 |
| Gm22136 | 0 | NA | NA | NA |
| Gm22206 | 0 | NA | NA | NA |
| RP24-186A10.2 | 0 | NA | NA | NA |
| Gm10042 | 0 | NA | NA | NA |
| 5730460C07Rik | 0 | NA | NA | NA |
| Gm25757 | 0 | NA | NA | NA |
| Asb17os | 1.419315775 | 1.069226901 | 0.961593904 | 1.111931862 |
| Asb17 | 0.04393175 | 0.037062677 | 3.724945049 | 0.009949859 |
| Msh4 | 11.18809076 | 0.112865362 | 0.334553014 | 0.337361665 |
| Gm24395 | 0 | NA | NA | NA |
| Rabggtb | 426.99162 | -0.369463223 | 0.099975282 | -3.695545681 |
| Snord45b | 0 | NA | NA | NA |
| Gm24494 | 6.926426429 | 0.242707163 | 0.425987696 | 0.56975158 |
| Snord45c | 0 | NA | NA | NA |
| RP23-403C5.3 | 0.748230394 | 0.048296646 | 1.168083245 | 0.041346921 |
| Acadm | 231.7967069 | -0.308564576 | 0.131584188 | -2.344997383 |
| Slc44a5 | 191.2716275 | 0.880757156 | 0.154464282 | 5.702011794 |
| Gm3076 | 3.513132756 | 0.384028477 | 0.677629012 | 0.566723783 |
| RP24-334O8.1 | 5.951295928 | 1.855341721 | 0.630216482 | 2.943975245 |
| RP24-527P22.4 | 0 | NA | NA | NA |
| Lhx8 | 17.86896063 | -0.056018761 | 0.27509706 | -0.203632713 |
| AI606473 | 4.337481582 | 0.813996426 | 0.511930622 | 1.590052228 |
| Tyw3 | 99.42373595 | 0.403268103 | 0.134144363 | 3.006224737 |
| Cryz | 179.0070392 | 0.230006193 | 0.092515505 | 2.486136719 |
| Gm38139 | 1.950077631 | -0.259318841 | 0.913478229 | -0.283880702 |
| Erich3 | 339.8935089 | 0.127418253 | 0.153046174 | 0.832547783 |
| Gm20389 | 0.04393175 | 0.037062677 | 3.724945049 | 0.009949859 |
| Tnni3k | 1.221396067 | 0.306193138 | 1.006962277 | 0.304076076 |
| Gm37890 | 0 | NA | NA | NA |
| Fpgt | 339.8878321 | -0.336325541 | 0.090666629 | -3.709474412 |
| Gm26456 | 0 | NA | NA | NA |
| Lrriq3 | 12.459711 | -0.228088937 | 0.300058031 | -0.760149416 |
| Gm37482 | 1.621103067 | -0.845076399 | 0.840822635 | -1.005059051 |
| Gm5285 | 0 | NA | NA | NA |
| Scp2-ps1 | 0.068097359 | 0.037064902 | 3.724945049 | 0.009950456 |
| Gm37449 | 0.554117557 | 1.962352922 | 1.276002118 | 1.537891586 |
| Gm37324 | 0.156124879 | 0.037074723 | 2.825334053 | 0.013122244 |
| Gm23038 | 0 | NA | NA | NA |

| | | | | |
|---|---|---|---|---|
| Gm37102 | 0.042716945 | 0.037062677 | 3.724945049 | 0.009949859 |
| Gm37515 | 0.408926061 | 0.044308376 | 1.48283719 | 0.02988081 |
| RP23-322I5.1 | 0.044191549 | 0.037062677 | 3.724945049 | 0.009949859 |
| RP23-322I5.2 | 0 | NA | NA | NA |
| RP23-322I5.3 | 0 | NA | NA | NA |
| RP23-187E3.1 | 0 | NA | NA | NA |
| 4930570G19Rik | 328.3713293 | 0.805086055 | 0.169651399 | 4.745531472 |
| Negr1 | 3447.761138 | 0.707286022 | 0.091644128 | 7.717745097 |
| RP23-354G14.1 | 240.3499704 | 0.584230946 | 0.315500794 | 1.851757449 |
| RP23-354G14.2 | 93.63872821 | 0.493801352 | 0.263630285 | 1.873082796 |
| RP23-354G14.3 | 137.6961956 | 0.455388132 | 0.272682482 | 1.670030758 |
| RP23-354G14.4 | 101.0414782 | 0.565766042 | 0.287213777 | 1.969842979 |
| Gm15578 | 21.23400752 | 0.884770856 | 0.308827451 | 2.864935919 |
| Gm15577 | 12.27858454 | -0.964614638 | 0.409880888 | -2.353402333 |
| RP24-176M3.1 | 45.69791764 | 0.72797559 | 0.342443811 | 2.125824929 |
| RP23-103L12.1 | 51.84091573 | 1.02511492 | 0.270067577 | 3.795771897 |
| 4930566N20Rik | 0.044544648 | 0.517957688 | 3.724945049 | 0.139051095 |
| Gm24373 | 15.34325161 | 0.850024804 | 0.493562667 | 1.72222265 |
| Gm23725 | 0 | NA | NA | NA |
| Gm22458 | 0.088383099 | 0.037058229 | 3.724945049 | 0.009948665 |
| Gm23379 | 0 | NA | NA | NA |
| RP23-64P1.1 | 0.244387265 | -1.745341046 | 2.046368895 | -0.852896587 |
| RP23-210I6.1 | 1.128024331 | -0.030027225 | 0.989685276 | -0.030340175 |
| Zranb2 | 1982.94177 | 0.156385265 | 0.081780222 | 1.91226266 |
| Mir186 | 29.35376171 | 0.385759176 | 0.249314479 | 1.547279478 |
| Ptger3 | 53.06310274 | -0.211416865 | 0.189260117 | -1.117070346 |
| RP24-411A22.2 | 0.062426873 | 0.517959912 | 3.724945049 | 0.139051692 |
| RP23-287M3.1 | 0.332723394 | -0.287897037 | 1.529226393 | -0.188263189 |
| RP23-287M3.3 | 0 | NA | NA | NA |
| Cth | 54.84234108 | -1.03490301 | 0.189486024 | -5.461632402 |
| RP23-287M3.2 | 3.057191251 | -0.127621931 | 0.641194292 | -0.199037846 |
| 1810013D15Rik | 0.328075509 | 1.393049985 | 1.728074516 | 0.806128423 |
| Ankrd13c | 1568.866954 | -0.234239534 | 0.065267559 | -3.588912118 |
| RP24-68G17.2 | 78.17925785 | 0.345227368 | 0.304278108 | 1.134578397 |
| Srsf11 | 1647.601996 | 0.207088259 | 0.121149262 | 1.709364586 |
| RP24-68G17.3 | 48.08364435 | 0.564792976 | 0.267949279 | 2.107835404 |
| Lrrc40 | 326.5754133 | 0.196983258 | 0.073931766 | 2.664392715 |
| RP24-68G17.4 | 1897.008325 | 0.644525683 | 0.229668895 | 2.806325529 |
| Lrrc7 | 2770.36321 | 0.52889996 | 0.219795076 | 2.406332162 |
| RP24-213M16.2 | 0.56553622 | 0.604983795 | 3.432782858 | 0.176237129 |
| RP24-213M16.1 | 0.354487963 | 0.023534802 | 1.577648786 | 0.014917644 |
| RP24-481I21.3 | 0.26582355 | 0.517965402 | 2.495627865 | 0.207549134 |
| RP24-481I21.2 | 0 | NA | NA | NA |
| Gm9423 | 0 | NA | NA | NA |
| RP23-324D2.1 | 2.321564718 | 3.114966487 | 1.106811488 | 2.814360459 |
| RP24-209L5.1 | 0.316660866 | -0.943964459 | 1.928334505 | -0.489523191 |
| Depdc1a | 5.945976878 | -0.102981283 | 0.543043963 | -0.189637101 |

| | | | | |
|---|---|---|---|---|
| Rpe65 | 5.700803298 | 1.62480666 | 0.813330589 | 1.997719848 |
| Wls | 574.2412302 | -0.382909526 | 0.103937052 | -3.684052198 |
| Wl1-1749A12.1 | 12.79191494 | -0.090079003 | 0.424774141 | -0.212063293 |
| Gm23941 | 0 | NA | NA | NA |
| Gm27878 | 0 | NA | NA | NA |
| Vmn1r-ps2 | 0 | NA | NA | NA |
| Gm11812 | 0 | NA | NA | NA |
| Vmn1r2 | 0 | NA | NA | NA |
| Vmn1r3 | 0 | NA | NA | NA |
| AI838599 | 1.421121277 | -0.165810193 | 0.976578032 | -0.169786937 |
| Gm11783 | 0.04393175 | 0.037062677 | 3.724945049 | 0.009949859 |
| Gm11786 | 0 | NA | NA | NA |
| Gm11784 | 0 | NA | NA | NA |
| Gm11785 | 0.044544648 | 0.517957688 | 3.724945049 | 0.139051095 |
| Gm11787 | 2.521183062 | -0.476561855 | 0.58938567 | -0.808573875 |
| Mir3471-2 | 0 | NA | NA | NA |
| Tmem68 | 334.4267812 | 0.041849497 | 0.071335152 | 0.586660236 |
| Tgs1 | 548.3289277 | -0.069908445 | 0.090447177 | -0.772920143 |
| Gm22541 | 0.302788994 | -0.460496095 | 1.810744524 | -0.254313123 |
| 2210414B05Rik | 0.54041206 | 0.647588356 | 1.40313339 | 0.461530144 |
| Lyn | 538.8172436 | -0.80929239 | 0.132552671 | -6.105440091 |
| Gm26857 | 5.106476976 | -1.496380675 | 0.48141612 | -3.10828951 |
| Gm11805 | 0.72046455 | 2.22965562 | 1.286341352 | 1.733331216 |
| Gm22781 | 2.17130535 | 0.555953391 | 0.73922003 | 0.75208107 |
| 9430025C20Rik | 38.03548056 | -0.005728713 | 0.292406307 | -0.01959162 |
| Rps20 | 938.4544041 | -0.113303453 | 0.0973917 | -1.163378935 |
| Gm24016 | 0 | NA | NA | NA |
| Gm24125 | 0 | NA | NA | NA |
| Mos | 0.721375344 | 0.48431555 | 1.27718488 | 0.379205515 |
| Plag1 | 101.48303 | 0.619834149 | 0.159161146 | 3.894381042 |
| Chchd7 | 221.4013722 | 0.412535025 | 0.108336625 | 3.807899911 |
| Gm11808 | 144.4071686 | 0.043847258 | 0.211190443 | 0.207619517 |
| Rps10-ps3 | 0 | NA | NA | NA |
| Sdr16c5 | 0.092228164 | 0.037073798 | 3.724945049 | 0.009952844 |
| Sdr16c6 | 0.09176491 | 0.921395572 | 3.723061103 | 0.247483333 |
| Penk | 1553.820092 | -0.338288879 | 0.222064833 | -1.523378887 |
| A830012C17Rik | 4.915491126 | 0.343705238 | 0.495205336 | 0.694066104 |
| Gm11780 | 0.085433889 | 0.037058229 | 3.724945049 | 0.009948665 |
| Gm26436 | 0 | NA | NA | NA |
| Impad1 | 2842.589061 | 0.174441486 | 0.08500331 | 2.052172878 |
| Gm11779 | 0 | NA | NA | NA |
| Gm11782 | 0 | NA | NA | NA |
| Gm11781 | 0 | NA | NA | NA |
| 4930423M02Rik | 0 | NA | NA | NA |
| Fam110b | 526.7316921 | 0.289585283 | 0.079589339 | 3.638493394 |
| Gm24337 | 0 | NA | NA | NA |
| Gm11796 | 0 | NA | NA | NA |

| | | | | |
|---|---|---|---|---|
| Gm11797 | 0.624599851 | 1.492390098 | 3.614112904 | 0.412934 |
| Ubxn2b | 545.8183538 | -0.015373022 | 0.125612662 | -0.122384333 |
| Gm11798 | 0 | NA | NA | NA |
| Cyp7a1 | 0.264608745 | 0.517965434 | 2.496817873 | 0.207450227 |
| 4930430E12Rik | 36.79472905 | -1.659176044 | 0.418455158 | -3.965003211 |
| Sdcbp | 2354.562233 | -0.638344888 | 0.126501565 | -5.046142206 |
| Gm37689 | 8.169802865 | -0.372606956 | 0.459935617 | -0.810128511 |
| Nsmaf | 844.8221621 | 0.006720514 | 0.112068171 | 0.059968092 |
| BX510318.1 | 42.81714402 | 0.209510349 | 0.282577846 | 0.741425245 |
| Gm11803 | 0 | NA | NA | NA |
| Gm11801 | 0 | NA | NA | NA |
| Gm22473 | 0 | NA | NA | NA |
| Tox | 323.9376703 | 0.46765148 | 0.124766668 | 3.748208448 |
| Gm11802 | 0.913351404 | -1.440209104 | 1.041262648 | -1.3831372 |
| Gm11804 | 0 | NA | NA | NA |
| 8430436N08Rik | 0.23059471 | 1.038092715 | 2.129271528 | 0.487534211 |
| Gm11795 | 0.166709681 | 0.517964155 | 2.76763122 | 0.187150713 |
| Gm11794 | 0.481267046 | -0.60018703 | 1.316562442 | -0.455874337 |
| Gm23860 | 0 | NA | NA | NA |
| Gm11800 | 0 | NA | NA | NA |
| Car8 | 1324.707954 | -0.108608584 | 0.304879242 | -0.356234762 |
| Gm11799 | 5.921281838 | 0.673918977 | 0.496813045 | 1.356484061 |
| Gm25355 | 0 | NA | NA | NA |
| Gm37386 | 0.066050875 | -0.443818988 | 3.724945049 | -0.119147795 |
| Gm11810 | 0.196054161 | 0.037621059 | 2.235824779 | 0.016826479 |
| Rab2a | 2207.111245 | 0.125493913 | 0.067822701 | 1.850323138 |
| Gm11809 | 0 | NA | NA | NA |
| Chd7 | 1522.223342 | 0.049339172 | 0.136119241 | 0.362470223 |
| Rps18-ps2 | 0.644285774 | 0.939989347 | 1.276374415 | 0.736452671 |
| Gm23423 | 0.075043205 | 0.517957688 | 3.724945049 | 0.139051095 |
| Gm26548 | 1.161740484 | -0.440012505 | 0.889388013 | -0.494736267 |
| Clvs1 | 941.1106066 | 0.20822838 | 0.079593512 | 2.616147656 |
| Gm11816 | 3.30561101 | -0.536373986 | 0.581957558 | -0.921672 |
| Gm11817 | 0.206997213 | 1.755076924 | 2.574827492 | 0.681628936 |
| Asph | 1754.934329 | 0.212947671 | 0.104720062 | 2.033494515 |
| Gm25677 | 0.044191549 | 0.037062677 | 3.724945049 | 0.009949859 |
| Gm24152 | 0 | NA | NA | NA |
| 4930412C18Rik | 9.306510091 | 0.195241723 | 0.367423686 | 0.531380339 |
| Gdf6 | 2.455729561 | 2.086365808 | 0.990475871 | 2.106427697 |
| 4930448K20Rik | 0 | NA | NA | NA |
| Gm11834 | 0 | NA | NA | NA |
| Gm11835 | 0 | NA | NA | NA |
| Gm11814 | 0.850521408 | -0.759650074 | 1.048373328 | -0.724598818 |
| 1700123O12Rik | 0 | NA | NA | NA |
| Gm11815 | 0.853137171 | 1.555275367 | 1.211631688 | 1.283620577 |
| Gm11813 | 0 | NA | NA | NA |
| Gm12920 | 0.044191549 | 0.037062677 | 3.724945049 | 0.009949859 |

| | | | | |
|---|---|---|---|---|
| Mir3471-1 | 0 | NA | NA | NA |
| Gm12918 | 39.37875012 | -0.346934633 | 0.209139472 | -1.658867307 |
| Gm12919 | 1.164100891 | -1.548387558 | 1.112637945 | -1.391636484 |
| AL672160.1 | 0.087930927 | -0.443164241 | 3.724898049 | -0.118973522 |
| 2610301B20Rik | 374.8592291 | -0.210825374 | 0.108475287 | -1.943533693 |
| Rps11-ps3 | 0.170186936 | -1.44442222 | 3.399009296 | -0.424953889 |
| Plekhf2 | 240.1964193 | -0.800282431 | 0.139895964 | -5.720554102 |
| Ndufaf6 | 73.7669564 | 0.059539579 | 0.168609225 | 0.35312172 |
| Gm11827 | 1.479390131 | 1.885585349 | 0.929507922 | 2.028584484 |
| Gm11829 | 0 | NA | NA | NA |
| mmu-mir-684-2 | 0.974719181 | -1.642793018 | 1.3521702 | -1.214930649 |
| Gm11830 | 0.15906016 | -1.015882374 | 3.433492635 | -0.295874342 |
| Trp53inp1 | 416.2178088 | 0.303765706 | 0.125091048 | 2.428356863 |
| Ccne2 | 114.245643 | 0.150666399 | 0.149444194 | 1.008178338 |
| Ints8 | 627.572806 | -0.244148901 | 0.090901586 | -2.685859642 |
| Dpy19l4 | 659.7126594 | 0.198845042 | 0.071459015 | 2.782644582 |
| Gm22940 | 3.154500895 | 0.584228304 | 0.78254226 | 0.746577321 |
| Gm26663 | 0.326609485 | 0.022911483 | 1.602572843 | 0.014296687 |
| Esrp1 | 1.320956406 | -0.375071981 | 1.057475756 | -0.354686128 |
| Gm11821 | 0.370194986 | 0.934522532 | 1.546707291 | 0.604201285 |
| Gm11822 | 0 | NA | NA | NA |
| 1110037F02Rik | 957.3336285 | -0.184726738 | 0.053906943 | -3.426770796 |
| Gm11831 | 0 | NA | NA | NA |
| Rad54b | 15.9562118 | -0.363921879 | 0.315504908 | -1.153458694 |
| Fsbp | 18.80141561 | 0.651299892 | 0.251229538 | 2.592449508 |
| Gm11832 | 0 | NA | NA | NA |
| Gm11833 | 0 | NA | NA | NA |
| Gem | 164.20295 | -1.159310448 | 0.351934694 | -3.294106739 |
| Cdh17 | 2.087534489 | -1.967226724 | 0.754398256 | -2.60767666 |
| Gm22477 | 0.061920613 | -0.443816764 | 3.724945049 | -0.119147198 |
| Gm25002 | 0 | NA | NA | NA |
| Pdp1 | 1783.616816 | 0.308630743 | 0.126544518 | 2.43891041 |
| 1700123M08Rik | 21.10087427 | 0.136574563 | 0.255880658 | 0.533743205 |
| Gm10604 | 0.068097359 | 0.037064902 | 3.724945049 | 0.009950456 |
| Gm26895 | 2.384116323 | -0.332445862 | 0.813755614 | -0.408532803 |
| Gm11840 | 0.048949532 | 0.037064902 | 3.724945049 | 0.009950456 |
| Gm11842 | 0.297840172 | 0.037078118 | 1.774381382 | 0.020896363 |
| Tmem67 | 313.6983145 | -0.065287641 | 0.10385638 | -0.628633899 |
| Rbm12b2 | 274.5941422 | -0.30213959 | 0.086750924 | -3.482840008 |
| Gm11839 | 15.15531981 | 0.558820906 | 0.385258 | 1.450510843 |
| Gm11841 | 7.772074058 | 0.309791594 | 0.46735043 | 0.662867891 |
| Gm11843 | 1.645711418 | -1.670946078 | 0.831733309 | -2.008992618 |
| Rbm12b1 | 103.5381567 | -0.182975432 | 0.175423939 | -1.043047108 |
| Fam92a | 431.8026252 | 0.151386878 | 0.073441875 | 2.061315539 |
| Gm11847 | 135.4062284 | -0.282999568 | 0.12600931 | -2.245862373 |
| Gm11846 | 0.044191549 | 0.037062677 | 3.724945049 | 0.009949859 |
| Gm37985 | 1.26802963 | -1.085505929 | 0.897177857 | -1.209911636 |

| | | | | |
|---|---|---|---|---|
| n-R5s182 | 0 | NA | NA | NA |
| Triqk | 145.0715167 | 0.600588796 | 0.139382786 | 4.308916554 |
| Gm11820 | 0.208450558 | 0.998866561 | 2.196616105 | 0.454729691 |
| Gm26250 | 0 | NA | NA | NA |
| Gm11819 | 0 | NA | NA | NA |
| Gm11826 | 3.374863659 | -0.206330094 | 0.568534019 | -0.362916004 |
| Gm11825 | 0.044544648 | 0.517957688 | 3.724945049 | 0.139051095 |
| Gm23162 | 0 | NA | NA | NA |
| Runx1t1 | 591.5908782 | 0.244150822 | 0.099370578 | 2.456972938 |
| Gm11823 | 0.253691411 | -1.917269603 | 2.430601013 | -0.788804741 |
| Gm24068 | 0 | NA | NA | NA |
| Gm11824 | 3.085586268 | -1.392066434 | 0.802753285 | -1.734114901 |
| 1700120G11Rik | 0 | NA | NA | NA |
| Gm24908 | 0 | NA | NA | NA |
| Gm11836 | 2.257326888 | 0.069311004 | 0.74400938 | 0.093158777 |
| Slc26a7 | 38.39735321 | -0.005856147 | 0.233217305 | -0.025110259 |
| Lrrc69 | 6.895850206 | -0.571178252 | 0.498423584 | -1.145969553 |
| Gm11838 | 0 | NA | NA | NA |
| Otud6b | 619.2074641 | 0.091696312 | 0.063664759 | 1.440299364 |
| Tmem55a | 534.2645095 | -0.317045782 | 0.092334419 | -3.433668458 |
| Gm11837 | 8.354147893 | 0.108559638 | 0.35444734 | 0.306278608 |
| Necab1 | 1188.342868 | 0.693115937 | 0.113099922 | 6.128350288 |
| Gm11844 | 0 | NA | NA | NA |
| Gm11845 | 0 | NA | NA | NA |
| Tmem64 | 564.4500706 | -0.057006374 | 0.087447771 | -0.651890532 |
| Gm24317 | 0 | NA | NA | NA |
| Gm11856 | 0 | NA | NA | NA |
| Gm11859 | 0.314457495 | 0.035824252 | 1.78355326 | 0.020085888 |
| Gm11857 | 0 | NA | NA | NA |
| Gm11858 | 0.914915251 | 0.802838689 | 1.206960447 | 0.665173984 |
| Gm11855 | 11.73213907 | 0.488008535 | 0.355818887 | 1.371508237 |
| Gm24567 | 0 | NA | NA | NA |
| Calb1 | 2166.896649 | 0.453244874 | 0.156391343 | 2.89814555 |
| Gm26148 | 0.061920613 | -0.443816764 | 3.724945049 | -0.119147198 |
| Decr1 | 158.7523432 | -0.418046704 | 0.134691624 | -3.103732002 |
| Mir6400 | 0.947067764 | -2.8326981 | 1.05984027 | -2.672759452 |
| Nbn | 245.1479976 | -0.123048252 | 0.080264742 | -1.533029934 |
| Gm11869 | 0.066050875 | -0.443818988 | 3.724945049 | -0.119147795 |
| Osgin2 | 252.2581438 | -0.360796352 | 0.099943869 | -3.60998985 |
| Gm11870 | 26.42157704 | 0.09475546 | 0.27753338 | 0.341420048 |
| Gm11874 | 4.610084374 | 0.814951191 | 0.480129782 | 1.697356051 |
| Ripk2 | 184.8884919 | -0.047256571 | 0.175580578 | -0.269144637 |
| Gm22690 | 1.641288957 | -0.06876313 | 0.84272451 | -0.081596214 |
| A530072M11Rik | 6.322029942 | -1.180096953 | 0.437821092 | -2.695386253 |
| Gm11865 | 0.577397264 | -0.337123675 | 1.32853753 | -0.253755477 |
| Gm37355 | 0.136963818 | 0.037074342 | 3.599544405 | 0.010299732 |
| Gm11864 | 0.066050875 | -0.443818988 | 3.724945049 | -0.119147795 |

| | | | | |
|---|---|---|---|---|
| Gm11861 | 0.117650406 | -0.443821212 | 3.724945049 | -0.119148392 |
| Gm24978 | 0.911472295 | 1.536112077 | 1.059309226 | 1.450107334 |
| Gm11862 | 0 | NA | NA | NA |
| Gm23975 | 0 | NA | NA | NA |
| Gm11848 | 0.044191549 | 0.037062677 | 3.724945049 | 0.009949859 |
| Gm11850 | 0 | NA | NA | NA |
| Gm11849 | 0.314045685 | -1.341164447 | 1.766709786 | -0.759131159 |
| Gm23384 | 1.375344072 | -1.827270491 | 1.102703887 | -1.657081753 |
| Gm11851 | 0.112396584 | -0.917692504 | 3.724434845 | -0.246397787 |
| Mmp16 | 803.4952648 | 0.283595349 | 0.094806487 | 2.991307441 |
| Gm11860 | 0 | NA | NA | NA |
| Gm11852 | 0.075043205 | 0.517957688 | 3.724945049 | 0.139051095 |
| Gm11867 | 0.108266322 | -0.955981391 | 3.722845977 | -0.25678779 |
| Gm11854 | 0 | NA | NA | NA |
| Gm11853 | 0 | NA | NA | NA |
| Gm25971 | 0 | NA | NA | NA |
| Gm24419 | 0 | NA | NA | NA |
| Gm11868 | 68.16477173 | -0.596759889 | 0.170757782 | -3.494774177 |
| Cnbd1 | 0 | NA | NA | NA |
| Gm12392 | 0 | NA | NA | NA |
| Gm12393 | 0 | NA | NA | NA |
| Cngb3 | 8.051515703 | -0.324659747 | 0.514839294 | -0.630604057 |
| Gm11877 | 0 | NA | NA | NA |
| Gm25655 | 0.263406493 | -0.528947964 | 2.066862864 | -0.255918268 |
| Cpne3 | 505.9295095 | -0.623040847 | 0.114273759 | -5.452177743 |
| Rmdn1 | 159.5744036 | -0.032968435 | 0.105098176 | -0.313691794 |
| Gm12353 | 18.78038428 | 0.046779302 | 0.28717315 | 0.162895806 |
| Wwp1 | 1103.425899 | -0.057026395 | 0.064214175 | -0.888065525 |
| Gm12354 | 0 | NA | NA | NA |
| Slc7a13 | 0.367515839 | -1.190705773 | 1.682649677 | -0.707637359 |
| Atp6v0d2 | 1.127083422 | -2.142784675 | 1.057724154 | -2.025844514 |
| 4930480G23Rik | 0.075043205 | 0.517957688 | 3.724945049 | 0.139051095 |
| Ttpa | 39.70977185 | -0.119871111 | 0.266030193 | -0.450592129 |
| 4930598A11Rik | 0.729378188 | -1.128321861 | 1.247017907 | -0.904816086 |
| Ggh | 127.1892972 | -0.581637558 | 0.134286719 | -4.331311109 |
| Nkain3 | 142.3883735 | 0.370616541 | 0.120022393 | 3.087894957 |
| Gm11872 | 3.989504448 | 0.244959966 | 0.49977171 | 0.490143722 |
| Gm11873 | 0.255186079 | 0.974143334 | 1.829844025 | 0.532364137 |
| Gm11871 | 0 | NA | NA | NA |
| Gm22817 | 0 | NA | NA | NA |
| Gm25977 | 0 | NA | NA | NA |
| Gm11875 | 0 | NA | NA | NA |
| Gm11876 | 0 | NA | NA | NA |
| Prdm13 | 1.264085207 | -0.807301624 | 1.147970888 | -0.703242244 |
| Ccnc | 534.707253 | -0.079805543 | 0.083474283 | -0.956049466 |
| Tstd3 | 155.9862153 | -0.073780554 | 0.10158805 | -0.726271972 |
| Usp45 | 1038.908871 | -0.210984045 | 0.05585202 | -3.777554424 |

| | | | | |
|---|---|---|---|---|
| Pnisr | 1930.889929 | 0.353623441 | 0.141781587 | 2.49414221 |
| Coq3 | 168.8413342 | -0.023067721 | 0.114880524 | -0.200797489 |
| Faxc | 940.4861022 | 0.303762212 | 0.091035982 | 3.336726924 |
| Gm11885 | 0.049553047 | -0.443818988 | 3.724945049 | -0.119147795 |
| Gm11880 | 0 | NA | NA | NA |
| Gm11882 | 0 | NA | NA | NA |
| Fbxl4 | 229.5017738 | 0.273179535 | 0.09763892 | 2.797854947 |
| Gm24607 | 2.356080661 | 1.308618466 | 0.776343735 | 1.685617345 |
| Gm37900 | 25.72877726 | 0.740235167 | 0.359552498 | 2.058767973 |
| Pou3f2 | 455.224007 | 0.40770854 | 0.130805034 | 3.116917816 |
| Gm11878 | 0.318934723 | -0.862388713 | 1.581236473 | -0.545388832 |
| Gm11881 | 0 | NA | NA | NA |
| Gm11879 | 0 | NA | NA | NA |
| Gm24078 | 0 | NA | NA | NA |
| 1700025O08Rik | 0 | NA | NA | NA |
| Gm11884 | 0.044544648 | 0.517957688 | 3.724945049 | 0.139051095 |
| Gm11888 | 0 | NA | NA | NA |
| Gm11889 | 2.706150538 | 0.165747202 | 0.712608703 | 0.232592166 |
| Gm25978 | 0.086648695 | 0.037058229 | 3.724945049 | 0.009948665 |
| Gm11890 | 1.145011917 | 1.208160798 | 1.156776006 | 1.044420693 |
| Gm28448 | 1.698407129 | 2.025326522 | 1.015224976 | 1.994953404 |
| Gm11892 | 0.26738533 | -0.945058378 | 2.089224358 | -0.452348918 |
| Gm11891 | 0 | NA | NA | NA |
| Gm27243 | 8.985200294 | 1.26235283 | 0.462260327 | 2.73082667 |
| Mms22l | 56.96292425 | -0.152000653 | 0.263843168 | -0.57610229 |
| Klhl32 | 321.673524 | -0.036126781 | 0.093830915 | -0.385020021 |
| Gm11899 | 0.512549389 | 0.74064986 | 1.380708629 | 0.536427342 |
| Ndufaf4 | 345.0672365 | -0.261811156 | 0.10420835 | -2.512381739 |
| C230012O17Rik | 12.79852409 | 0.135647819 | 0.315797375 | 0.429540679 |
| Gpr63 | 190.3210986 | 0.122079381 | 0.156439845 | 0.780359894 |
| Gm11906 | 2.251356483 | -0.377187158 | 0.756145535 | -0.498828784 |
| Fhl5 | 2.616435038 | -0.497706708 | 0.705224258 | -0.705742468 |
| Ufl1 | 454.4895708 | -0.0572056 | 0.075778725 | -0.754903175 |
| Prdx6-ps2 | 0.044191549 | 0.037062677 | 3.724945049 | 0.009949859 |
| Gm11897 | 0 | NA | NA | NA |
| Gm11894 | 0.044544648 | 0.517957688 | 3.724945049 | 0.139051095 |
| Gm25975 | 0.109124069 | 0.517955464 | 3.724945049 | 0.139050498 |
| Gm11896 | 0 | NA | NA | NA |
| Gm11893 | 3.313083698 | -0.900887162 | 0.563678201 | -1.598229558 |
| Gm24977 | 0 | NA | NA | NA |
| Fut9 | 4215.601891 | 0.510082883 | 0.102315059 | 4.985413547 |
| Gm24042 | 0 | NA | NA | NA |
| Gm11898 | 0 | NA | NA | NA |
| Gm26067 | 0 | NA | NA | NA |
| Gm11895 | 0 | NA | NA | NA |
| Gm24965 | 0 | NA | NA | NA |
| Manea | 627.7666335 | -0.004023122 | 0.112350173 | -0.035808774 |

| | | | | |
|---|---|---|---|---|
| Bhmt-ps1 | 0 | NA | NA | NA |
| Gm11903 | 0 | NA | NA | NA |
| Gng2-ps1 | 0.3346585 | 0.504133533 | 1.781586464 | 0.282968884 |
| Gm11905 | 0 | NA | NA | NA |
| 4930548K13Rik | 0 | NA | NA | NA |
| Gm11904 | 0 | NA | NA | NA |
| Gm11902 | 0 | NA | NA | NA |
| Gm11901 | 0.045882455 | 0.517959912 | 3.724945049 | 0.139051692 |
| Gm25173 | 0 | NA | NA | NA |
| Gm11914 | 3.923996896 | 0.614649453 | 0.53394107 | 1.151155976 |
| Tpm3-rs2 | 0 | NA | NA | NA |
| Gm11911 | 0 | NA | NA | NA |
| Gm11907 | 0.192582685 | 0.954468024 | 2.681495147 | 0.355946206 |
| Gm25378 | 0 | NA | NA | NA |
| Gm11908 | 0 | NA | NA | NA |
| Gm11909 | 0 | NA | NA | NA |
| Gm11910 | 0 | NA | NA | NA |
| Gm11912 | 0 | NA | NA | NA |
| Gm11913 | 0 | NA | NA | NA |
| Epha7 | 823.6749401 | 0.253104923 | 0.142085398 | 1.781357731 |
| Gm11915 | 4.082662799 | 0.503980992 | 0.506589166 | 0.9948515 |
| Gm11917 | 0.126225412 | 0.03707172 | 3.698755067 | 0.010022756 |
| Gm11916 | 0 | NA | NA | NA |
| Gm11918 | 0 | NA | NA | NA |
| Gm11919 | 0 | NA | NA | NA |
| Gm11923 | 0 | NA | NA | NA |
| Gm11925 | 0 | NA | NA | NA |
| Gm11924 | 0.272022412 | 0.006078806 | 2.438162277 | 0.002493192 |
| 4930556G01Rik | 0.046345709 | -0.443818988 | 3.724945049 | -0.119147795 |
| Gm22831 | 0 | NA | NA | NA |
| Gm11920 | 0.174512195 | 0.037058139 | 3.362865018 | 0.011019812 |
| Gm25705 | 0.044544648 | 0.517957688 | 3.724945049 | 0.139051095 |
| Gm11922 | 0.132199999 | 0.998851712 | 3.642365922 | 0.274231566 |
| Map3k7 | 821.1190469 | -0.000269469 | 0.055000752 | -0.004899369 |
| Gm11926 | 0 | NA | NA | NA |
| Gm26254 | 0 | NA | NA | NA |
| Gm11927 | 0.564158409 | -0.804109238 | 1.541273505 | -0.521717421 |
| Gm11928 | 0 | NA | NA | NA |
| Gm11929 | 0 | NA | NA | NA |
| Bach2 | 252.756904 | -0.083551103 | 0.084453878 | -0.98931044 |
| Bach2it1 | 12.10687002 | 0.339955672 | 0.426229319 | 0.797588662 |
| Gm24371 | 4.271398782 | -0.398133569 | 0.687155637 | -0.579393588 |
| Bach2os | 18.90185043 | -2.014610161 | 0.310013203 | -6.498465686 |
| Gja10 | 0.072025462 | 0.517959912 | 3.724945049 | 0.139051692 |
| Gm11933 | 0.233612452 | 0.979519378 | 2.527723168 | 0.387510543 |
| Casp8ap2 | 404.3248676 | -0.126935973 | 0.096676868 | -1.312992194 |
| Gm11936 | 0 | NA | NA | NA |

| | | | | |
|---|---|---|---|---|
| Mdn1 | 1768.721666 | -0.166422425 | 0.118362894 | -1.406035448 |
| Lyrm2 | 76.45699897 | -0.029560686 | 0.1826605 | -0.161834038 |
| Ankrd6 | 449.0953625 | -0.022213284 | 0.096177023 | -0.230962482 |
| Gm11941 | 0 | NA | NA | NA |
| Gm11942 | 24.81150096 | -0.463706952 | 0.2700669 | -1.717007718 |
| Rragd | 1716.34318 | 0.182627504 | 0.098652258 | 1.85122478 |
| Gm11943 | 0.062426873 | 0.517959912 | 3.724945049 | 0.139051692 |
| 4933421O10Rik | 104.7690149 | 0.083673937 | 0.132092113 | 0.633451427 |
| Ube2j1 | 1400.842074 | -0.004563741 | 0.065437965 | -0.069741488 |
| Gm11931 | 0 | NA | NA | NA |
| Gabrr2 | 12.63021313 | 0.450806327 | 0.311379294 | 1.447772333 |
| Gabrr1 | 1.88379093 | -1.039050076 | 0.787150576 | -1.320014375 |
| Pm20d2 | 35.06690641 | 0.849332698 | 0.200994413 | 4.225653277 |
| Gm11930 | 0 | NA | NA | NA |
| Srsf12 | 183.1024447 | 0.432168525 | 0.105250214 | 4.106105905 |
| Pnrc1 | 961.8622753 | 0.494247803 | 0.097700401 | 5.058810404 |
| Gm11934 | 0.906232762 | 0.360964667 | 1.037942962 | 0.347769271 |
| Rngtt | 426.6224941 | -0.15399032 | 0.063513503 | -2.424528839 |
| Mir8118 | 1.705950494 | -0.41389202 | 0.804461182 | -0.51449595 |
| Gm11935 | 0 | NA | NA | NA |
| Gm23304 | 0.144050924 | 1.05357957 | 3.721447096 | 0.283110183 |
| Gm24341 | 0 | NA | NA | NA |
| Cnr1 | 3191.885636 | 0.258845943 | 0.103377187 | 2.503898095 |
| Gm22010 | 0 | NA | NA | NA |
| Spaca1 | 0.461160426 | 0.052397774 | 1.327121437 | 0.039482275 |
| Gm25623 | 0 | NA | NA | NA |
| Gm12753 | 0.179340113 | -0.847972862 | 2.729607159 | -0.310657473 |
| Gm12751 | 94.08612612 | -0.114289121 | 0.127815371 | -0.894173521 |
| Gm12752 | 0.088534442 | -0.905414734 | 3.723500899 | -0.243162217 |
| Gm12754 | 0.046345709 | -0.443818988 | 3.724945049 | -0.119147795 |
| Gm23466 | 0 | NA | NA | NA |
| Gm12351 | 0 | NA | NA | NA |
| Gm12350 | 35.03170047 | -0.332144373 | 0.209721472 | -1.583740429 |
| Akirin2 | 531.2879269 | -0.260668831 | 0.071964387 | -3.622192049 |
| Orc3 | 556.144257 | 0.074809386 | 0.098935604 | 0.756142209 |
| Rars2 | 246.197839 | 0.037198641 | 0.10015389 | 0.371414837 |
| Slc35a1 | 169.2461004 | -0.059808192 | 0.109918772 | -0.544112631 |
| Gm22849 | 0.748256776 | 0.686598502 | 1.09267186 | 0.628366601 |
| 1700003M02Rik | 17.88018956 | -0.148676753 | 0.29802335 | -0.498876187 |
| Gm136 | 1.409557441 | 1.459343103 | 1.211510251 | 1.204565212 |
| Smim8 | 84.47012772 | -0.197987393 | 0.130761266 | -1.514113463 |
| Zfp292 | 1650.746143 | 0.248636053 | 0.094738436 | 2.624447526 |
| Gm12363 | 0 | NA | NA | NA |
| Cga | 2.158593695 | -4.311198294 | 1.623051133 | -2.656230728 |
| Platr9 | 0 | NA | NA | NA |
| Gm12361 | 0.042716945 | 0.037062677 | 3.724945049 | 0.009949859 |
| Gm12362 | 0.090537259 | -0.443823436 | 3.724945049 | -0.119148989 |

| | | | | |
|---|---|---|---|---|
| Mob3b | 188.1401066 | -0.178492654 | 0.162227756 | -1.100259647 |
| Gm12364 | 0 | NA | NA | NA |
| Ifnk | 0.038981395 | -0.443816764 | 3.724945049 | -0.119147198 |
| 3110043O21Rik | 349.8244095 | -0.144836665 | 0.076726206 | -1.887707901 |
| Gm12366 | 24.68870437 | -0.034646881 | 0.230588633 | -0.150254073 |
| Gm12367 | 4.278993874 | -0.190017293 | 0.574560507 | -0.330717636 |
| Gm12368 | 0 | NA | NA | NA |
| Gm12365 | 0 | NA | NA | NA |
| Lingo2 | 668.3004972 | 0.713083399 | 0.123961658 | 5.752451289 |
| Gm12369 | 25.01436732 | 1.14547659 | 0.280903276 | 4.077832795 |
| Gm12370 | 0 | NA | NA | NA |
| Mir876 | 9.32634141 | 0.301203047 | 0.358918595 | 0.839195994 |
| Mir873a | 8.772029568 | 0.3955234 | 0.386889912 | 1.022315102 |
| mmu-mir-873a | 0 | NA | NA | NA |
| Gm23314 | 0 | NA | NA | NA |
| AL844160.1 | 0 | NA | NA | NA |
| Gm12371 | 46.23436975 | 1.294174161 | 0.187453284 | 6.903982334 |
| Gm12372 | 1.336268985 | -0.575186064 | 0.816664598 | -0.70431125 |
| Rpl23a-ps6 | 0 | NA | NA | NA |
| Gm12374 | 2.339895364 | 0.327002267 | 0.699756328 | 0.467308768 |
| Gm22732 | 0.04393175 | 0.037062677 | 3.724945049 | 0.009949859 |
| Gm12375 | 0 | NA | NA | NA |
| Gm12378 | 0 | NA | NA | NA |
| Gm12379 | 0 | NA | NA | NA |
| Gm12380 | 0.504525611 | 0.692135442 | 1.295921487 | 0.534087481 |
| Gm12381 | 0 | NA | NA | NA |
| Gm12382 | 0 | NA | NA | NA |
| Gm25581 | 0 | NA | NA | NA |
| Gm12383 | 0 | NA | NA | NA |
| Gm23607 | 0 | NA | NA | NA |
| Gm17094 | 0.175174943 | 0.037075233 | 2.724622297 | 0.013607476 |
| 1700009N14Rik | 0 | NA | NA | NA |
| Gm12385 | 0.045882455 | 0.517959912 | 3.724945049 | 0.139051692 |
| Gm12384 | 0.119234754 | 0.51795324 | 3.724945049 | 0.139049901 |
| Gm12386 | 0 | NA | NA | NA |
| Gm22850 | 0 | NA | NA | NA |
| Gm12387 | 0.097139547 | -0.443816764 | 3.724945049 | -0.119147198 |
| Gm26087 | 0 | NA | NA | NA |
| Gm12388 | 0.062426873 | 0.517959912 | 3.724945049 | 0.139051692 |
| Gm12389 | 0 | NA | NA | NA |
| Aco1 | 373.8664707 | -0.165692328 | 0.097517478 | -1.69910391 |
| Ddx58 | 2204.242023 | -0.407820517 | 0.214238601 | -1.903580939 |
| Topors | 450.6972402 | -0.119290076 | 0.072609059 | -1.642908991 |
| Toporsos | 15.19562275 | -0.387044084 | 0.32033866 | -1.208234072 |
| Ndufb6 | 365.4765144 | -0.130526654 | 0.106856894 | -1.221508965 |
| 4930509K18Rik | 0 | NA | NA | NA |
| Gm23338 | 0.239936926 | 1.671895677 | 2.427468652 | 0.688740378 |

| | | | | |
|---|---|---|---|---|
| Tmem215 | 126.9200701 | 1.103581737 | 0.206466261 | 5.345094794 |
| 4933428C19Rik | 0.268438559 | 0.077210075 | 1.804838436 | 0.042779494 |
| Gm12391 | 0 | NA | NA | NA |
| Aptx | 267.7610802 | 0.154783936 | 0.093142343 | 1.661799895 |
| Gm12390 | 0.717871974 | 1.186566228 | 1.293015456 | 0.917673662 |
| Gm6297 | 15.05970898 | 0.151629611 | 0.299474119 | 0.506319582 |
| Dnaja1 | 2433.261954 | -0.205853209 | 0.102537388 | -2.00759169 |
| Mir207 | 0.128150834 | 0.037055157 | 3.724945049 | 0.00994784 |
| Smu1 | 643.1185477 | -0.095308121 | 0.078545673 | -1.213410204 |
| Gm12396 | 8.729157278 | -0.654466184 | 0.346428817 | -1.889179399 |
| B4galt1 | 382.463159 | -0.764246871 | 0.189204557 | -4.039262486 |
| Mir5123 | 0.217530652 | 0.013610755 | 2.172555185 | 0.00626486 |
| Gm24112 | 0.24398126 | 0.517971605 | 1.852930388 | 0.279541859 |
| Gm25931 | 0.653402772 | -0.175477332 | 1.305874577 | -0.134375334 |
| Gm12397 | 7.076756442 | -0.77279123 | 0.430599795 | -1.794685552 |
| Spink4 | 0.044544648 | 0.517957688 | 3.724945049 | 0.139051095 |
| Bag1 | 715.7527563 | -0.038218116 | 0.090554658 | -0.422044731 |
| Chmp5 | 658.9006136 | -0.099141377 | 0.114599279 | -0.865113447 |
| Nfx1 | 1354.303035 | 0.161334994 | 0.091852339 | 1.756460381 |
| Mir3094 | 0.798730366 | -0.007448915 | 1.064470749 | -0.006997764 |
| Aqp7 | 0.174839421 | -0.443820766 | 2.312507876 | -0.19192184 |
| Gm25637 | 0 | NA | NA | NA |
| Aqp3 | 1.154822103 | -0.378939762 | 0.914643926 | -0.414303043 |
| Nol6 | 1475.373276 | 0.1945913 | 0.046603315 | 4.175481921 |
| Gm12402 | 0.230106605 | -0.443823796 | 2.593199487 | -0.171149114 |
| Ube2r2 | 789.6006133 | -0.094878887 | 0.064591956 | -1.468896331 |
| Gm12403 | 11.82053707 | -0.272181357 | 0.439671232 | -0.619056551 |
| Gm24650 | 0 | NA | NA | NA |
| Ubap2 | 693.4431244 | -0.390507851 | 0.085580468 | -4.563048761 |
| Gm22888 | 2.630926952 | 0.681636232 | 0.726270997 | 0.938542548 |
| Gm24837 | 2.20124 | 0.537332858 | 0.783302215 | 0.685984092 |
| Gm23090 | 0.341439828 | 0.020427559 | 1.587616133 | 0.012866813 |
| Gm12401 | 0.164132501 | 1.299746508 | 3.390982409 | 0.383294972 |
| Dcaf12 | 631.7207173 | -0.412010487 | 0.086278344 | -4.775363897 |
| n-R5s183 | 0 | NA | NA | NA |
| Gm26049 | 4.518367824 | 0.267903274 | 0.638241791 | 0.419752009 |
| Gm26084 | 0.449030389 | 0.037074986 | 1.601438816 | 0.023151048 |
| Ubap1 | 662.4155343 | -0.377781563 | 0.071125186 | -5.311501938 |
| Kif24 | 28.6589064 | -0.182019998 | 0.192389874 | -0.946099676 |
| Nudt2 | 96.39035252 | 0.033099567 | 0.117678861 | 0.281270288 |
| AI464131 | 299.7303309 | 0.123528931 | 0.202599581 | 0.609719576 |
| 1110017D15Rik | 25.98366852 | -0.336565897 | 0.209712123 | -1.604894808 |
| Fam219aos | 0.808497328 | 0.497787871 | 1.146333764 | 0.4342434 |
| Fam219a | 1771.430822 | 0.150345081 | 0.052052023 | 2.888361931 |
| Gm12404 | 1.315597923 | 0.087719088 | 0.841322519 | 0.104263331 |
| Dnaic1 | 23.7871792 | 0.143704037 | 0.285760349 | 0.502883056 |
| Enho | 315.6348979 | 0.128445092 | 0.132095795 | 0.972363216 |

| | | | | |
|---|---|---|---|---|
| Gm12406 | 1.108658207 | 0.030271305 | 1.116893415 | 0.027103128 |
| Gm12405 | 0.625648376 | 0.998868123 | 1.679102925 | 0.59488201 |
| Cntfr | 660.2301978 | 0.28582886 | 0.105841428 | 2.700538571 |
| Il11ra1 | 183.2536711 | -0.110616167 | 0.105958902 | -1.043953506 |
| Rpp25l | 90.6293161 | -0.414645661 | 0.126731468 | -3.271844545 |
| Dctn3 | 483.0857412 | 0.08836336 | 0.070041515 | 1.261585503 |
| Arid3c | 1.591794818 | -0.527434112 | 1.046334249 | -0.504078035 |
| Sigmar1 | 271.0497585 | -0.49522532 | 0.088520093 | -5.594496139 |
| Galt | 238.390805 | 0.35068485 | 0.088386902 | 3.967611067 |
| Ccl27a | 57.35970255 | 0.542560547 | 0.180135076 | 3.011965026 |
| Gm12407 | 0.693596057 | -0.555098229 | 1.146027597 | -0.484367244 |
| Gm10601 | 0.106056992 | 0.998850474 | 3.724945049 | 0.268151734 |
| Gm13307 | 0 | NA | NA | NA |
| Gm13302 | 0 | NA | NA | NA |
| Gm21541 | 0 | NA | NA | NA |
| Ccl19-ps1 | 0 | NA | NA | NA |
| Gm20878 | 0 | NA | NA | NA |
| Gm20938 | 0 | NA | NA | NA |
| Gm13303 | 0 | NA | NA | NA |
| Gm21093 | 0.097139547 | -0.443816764 | 3.724945049 | -0.119147198 |
| Gm37530 | 0.382550573 | 0.497380338 | 1.461164132 | 0.340400047 |
| Gm3892 | 0 | NA | NA | NA |
| Gm3893 | 0 | NA | NA | NA |
| Gm13304 | 0.049553047 | -0.443818988 | 3.724945049 | -0.119147795 |
| Gm13299 | 0.097501745 | -0.443821212 | 3.724945049 | -0.119148392 |
| Ccl19-ps3 | 0.089089296 | 0.957155977 | 3.723939561 | 0.257027796 |
| Gm13306 | 1.067961214 | -0.020338377 | 1.002684227 | -0.020283931 |
| Gm13305 | 0.062426873 | 0.517959912 | 3.724945049 | 0.139051692 |
| 1700045I11Rik | 0 | NA | NA | NA |
| Gm17167 | 0.21074972 | -0.951854522 | 2.623964242 | -0.36275438 |
| Gm13301 | 0 | NA | NA | NA |
| Gm5859 | 0 | NA | NA | NA |
| Gm17081 | 0 | NA | NA | NA |
| Gm10600 | 0 | NA | NA | NA |
| Ccl21b | 0 | NA | NA | NA |
| Gm21980 | 0 | NA | NA | NA |
| Gm21953 | 8.239491219 | 0.472745901 | 0.41949555 | 1.126939012 |
| Gm21598 | 0 | NA | NA | NA |
| Gm21586 | 0.193603971 | 0.541066362 | 3.24974582 | 0.166494979 |
| Gm3883 | 0 | NA | NA | NA |
| Gm2163 | 0 | NA | NA | NA |
| Gm13298 | 0.089814205 | 0.517955464 | 3.724945049 | 0.139050498 |
| Gm10593 | 0.444623086 | 2.138264757 | 3.202684215 | 0.667647702 |
| Gm10591 | 0 | NA | NA | NA |
| Gm2564 | 0 | NA | NA | NA |
| Ccl27b | 1.785252223 | 0.619648586 | 0.709931575 | 0.872828604 |
| Il11ra2 | 0.424152111 | 0.12884561 | 1.685560439 | 0.076440813 |

| | | | | |
|---|---|---|---|---|
| Gm21955 | 83.35882431 | 0.466363434 | 0.22054064 | 2.114637167 |
| 4930578G10Rik | 26.07152981 | 2.07177757 | 0.392787921 | 5.274545017 |
| Gm12395 | 6.47297608 | 1.807539874 | 0.658885455 | 2.74332945 |
| Ccl19 | 26.89383427 | -0.019364747 | 0.327827365 | -0.059069952 |
| Ccl21a | 1.739286695 | -1.026410225 | 0.706609805 | -1.452584181 |
| Gm12394 | 0.61514822 | 0.835976344 | 1.301511634 | 0.642311849 |
| Gm12429 | 2.41835347 | 1.388914553 | 0.864339427 | 1.60690871 |
| BC049635 | 0.963622687 | -1.219015092 | 1.080448786 | -1.128248843 |
| N28178 | 2446.431538 | 0.791978414 | 0.151016339 | 5.244322709 |
| Dnajb5 | 1123.231011 | -0.302112221 | 0.166008447 | -1.81986054 |
| 1700022I11Rik | 5.07266936 | -2.77198606 | 0.584136299 | -4.745443935 |
| Vcp | 2874.0526 | -0.25581785 | 0.065484793 | -3.906523017 |
| Fancg | 91.45811176 | -0.14129039 | 0.173969346 | -0.812156816 |
| Pigo | 262.034385 | 0.040627947 | 0.0841784 | 0.482640999 |
| Stoml2 | 348.4296267 | -0.151112815 | 0.095680447 | -1.579348974 |
| Gm26643 | 0.357756446 | -0.443819083 | 1.475315764 | -0.300829893 |
| Fam214b | 535.3240349 | -0.478115997 | 0.081079897 | -5.896850098 |
| Gm26881 | 11.80662906 | 0.218492772 | 0.362328142 | 0.60302457 |
| Unc13b | 466.7629952 | 0.012291728 | 0.186658729 | 0.065851341 |
| Gm23709 | 0 | NA | NA | NA |
| Gm25010 | 0 | NA | NA | NA |
| Atp8b5 | 7.649160145 | -1.263250403 | 0.398591102 | -3.169289023 |
| Rusc2 | 1981.730262 | 0.229600663 | 0.071049006 | 3.231581656 |
| Fam166b | 3.554918289 | 1.091109242 | 0.645416288 | 1.69055114 |
| Tesk1 | 834.3782588 | 0.102634881 | 0.064747353 | 1.58515948 |
| Cd72 | 162.2518301 | -0.244687143 | 0.22493767 | -1.087799759 |
| Sit1 | 2.004396688 | 0.345387214 | 0.669037109 | 0.516245227 |
| Rmrp | 17396.1871 | -0.392197311 | 0.169908438 | -2.308286251 |
| Ccdc107 | 211.9356722 | -0.112821999 | 0.151590134 | -0.744256876 |
| Arhgef39 | 4.559760072 | 0.225757751 | 0.501524863 | 0.450142689 |
| Gm12454 | 3.511636557 | 0.204049197 | 0.593209125 | 0.343975148 |
| Car9 | 11.98985167 | 0.11624605 | 0.363844846 | 0.319493464 |
| Tpm2 | 58.51175456 | -0.114481615 | 0.206062124 | -0.555568448 |
| Tln1 | 1972.671443 | -0.776139114 | 0.129890922 | -5.975314512 |
| Creb3 | 390.8704704 | -0.305492193 | 0.126114527 | -2.422339445 |
| Gba2 | 788.9852365 | 0.252654663 | 0.054658702 | 4.622405114 |
| Rgp1 | 609.9554589 | -0.141082864 | 0.056011734 | -2.518809073 |
| Msmp | 0.198131738 | 0.037076674 | 2.229210881 | 0.016632197 |
| Gm12472 | 0.580871419 | 0.406226498 | 1.294109493 | 0.313904272 |
| Gm25262 | 9.682546495 | -0.22213949 | 0.442291256 | -0.502247076 |
| Gm12473 | 1.944300039 | -0.146081867 | 0.743060439 | -0.196594865 |
| Npr2 | 447.7307936 | 0.081422392 | 0.097985315 | 0.830965259 |
| Spag8 | 14.00717955 | -0.068856781 | 0.318365175 | -0.216282392 |
| Hint2 | 86.66528165 | 0.066568032 | 0.153557909 | 0.433504416 |
| Fam221b | 17.22801957 | -0.248349121 | 0.29837699 | -0.832333356 |
| Gm12481 | 74.19163858 | -0.340693301 | 0.17761279 | -1.91818 |
| Tmem8b | 757.5108737 | 0.626085566 | 0.08196854 | 7.638120285 |

| | | | | |
|---|---|---|---|---|
| Gm23257 | 4.230344492 | -0.553524737 | 0.582329401 | -0.950535446 |
| Olfr70 | 0 | NA | NA | NA |
| Olfr71 | 0 | NA | NA | NA |
| Hrct1 | 4.20546811 | -0.694474301 | 0.536997487 | -1.293254285 |
| 5430416O09Rik | 3.015746376 | -0.134509103 | 0.5655657 | -0.237831082 |
| A630077J23Rik | 0.321964759 | 0.228289911 | 1.975833074 | 0.115541092 |
| Olfr269-ps1 | 0 | NA | NA | NA |
| Olfr159 | 0 | NA | NA | NA |
| Olfr29-ps1 | 0 | NA | NA | NA |
| Gm12492 | 0 | NA | NA | NA |
| Olfr156 | 0.048949532 | 0.037064902 | 3.724945049 | 0.009950456 |
| Olfr157 | 0 | NA | NA | NA |
| Olfr155 | 0.08169834 | -0.443821212 | 3.724945049 | -0.119148392 |
| Reck | 121.3301511 | 0.331505436 | 0.187895162 | 1.764310653 |
| Gm27265 | 2.022028428 | 0.491474722 | 0.808129553 | 0.608163282 |
| Glipr2 | 555.7186848 | -1.105748234 | 0.338376002 | -3.267809268 |
| Ccin | 0.952506814 | -0.358922253 | 1.019898172 | -0.351919694 |
| Clta | 1302.037201 | -0.112139485 | 0.080817975 | -1.387556239 |
| Gne | 417.5634539 | -0.142417841 | 0.079379056 | -1.794148823 |
| Gm22706 | 0.185638728 | 0.486352713 | 2.665023679 | 0.182494706 |
| Gm12503 | 0.506185493 | -1.360642182 | 1.342178628 | -1.013756406 |
| Gm12504 | 0.323274704 | 1.046671224 | 1.598393872 | 0.65482685 |
| Rnf38 | 803.2801551 | 0.116202082 | 0.069617257 | 1.669156287 |
| Mir5106 | 2.834065591 | 0.163295225 | 0.61912528 | 0.263751507 |
| Melk | 12.45686811 | -0.514441343 | 0.318551638 | -1.614938622 |
| Gm12677 | 0 | NA | NA | NA |
| Gm12455 | 0 | NA | NA | NA |
| Gm24376 | 0 | NA | NA | NA |
| Pax5 | 62.51624943 | -0.405740798 | 0.182623081 | -2.221738866 |
| Mir5120 | 0.090665643 | 0.037060453 | 3.724945049 | 0.009949262 |
| Gm12462 | 5.015296868 | -0.28977512 | 0.478401964 | -0.605714737 |
| 4930517J16Rik | 1.109399417 | -0.396644312 | 1.120295644 | -0.354053248 |
| 5730488B01Rik | 1.238426347 | 0.453792482 | 0.907073691 | 0.500281825 |
| Gm12463 | 0.758386275 | -1.309713387 | 1.193597268 | -1.097282494 |
| Zcchc7 | 970.4223863 | 0.003594578 | 0.148107639 | 0.024270041 |
| Gm22639 | 3.577120001 | -0.93700246 | 0.617576742 | -1.517224332 |
| Gm12678 | 0 | NA | NA | NA |
| Gm12493 | 0 | NA | NA | NA |
| Gm12679 | 2.474621302 | 0.937683418 | 0.613952916 | 1.527288809 |
| Grhpr | 156.0778864 | 0.294222376 | 0.130028084 | 2.262760213 |
| Zbtb5 | 270.4928444 | -0.136250704 | 0.109235273 | -1.247314179 |
| 1700055D18Rik | 14.55774235 | 0.13575481 | 0.268462066 | 0.505675949 |
| Polr1e | 68.48577099 | -0.725879014 | 0.154456678 | -4.699563799 |
| Fbxo10 | 581.7003137 | 0.394629795 | 0.071656119 | 5.507272756 |
| Tomm5 | 134.9204341 | -0.16285353 | 0.147147945 | -1.106733298 |
| Gm12566 | 0 | NA | NA | NA |
| Frmpd1 | 273.0835449 | 0.285266274 | 0.138080107 | 2.065947664 |

| | | | | |
|---|---|---|---|---|
| Frmpd1os | 0.15456712 | 0.517964654 | 2.834226082 | 0.182753471 |
| Trmt10b | 109.9286799 | -0.006098155 | 0.107414946 | -0.056771937 |
| Exosc3 | 78.17585261 | -0.472921679 | 0.163985871 | -2.883917227 |
| Gm12585 | 0 | NA | NA | NA |
| Dcaf10 | 447.5914563 | -0.082946525 | 0.106982591 | -0.775327314 |
| Slc25a51 | 825.9827367 | 0.086916815 | 0.046852742 | 1.855106254 |
| Gm25692 | 0 | NA | NA | NA |
| Shb | 131.2113877 | -0.315661016 | 0.122273063 | -2.581607172 |
| Gm22518 | 0.171044682 | 0.037076338 | 2.745278419 | 0.013505493 |
| Gm22055 | 0 | NA | NA | NA |
| Gm12411 | 5.818705562 | 0.032437795 | 0.499059481 | 0.064997854 |
| Gm12408 | 2.792582105 | 0.552975807 | 0.616884983 | 0.896400175 |
| Gm829 | 0.154612032 | -1.394968402 | 3.490355212 | -0.399663735 |
| Gm12410 | 0.044191549 | 0.037062677 | 3.724945049 | 0.009949859 |
| Gm12409 | 0.36880298 | 0.120310418 | 1.537715548 | 0.078239709 |
| Gm12412 | 0.044544648 | 0.517957688 | 3.724945049 | 0.139051095 |
| Aldh1b1 | 125.3782894 | 0.025597795 | 0.172879588 | 0.148067189 |
| Igfbpl1 | 40.81830737 | -1.249091068 | 0.26577552 | -4.699797285 |
| Stra6l | 52.92012026 | -1.994840101 | 0.310828445 | -6.417817075 |
| E230008N13Rik | 12.72585994 | -0.547887508 | 0.353558777 | -1.549636281 |
| Tdrd7 | 1191.587 | -0.513673858 | 0.123998383 | -4.142585129 |
| Tmod1 | 407.2062115 | 0.038553552 | 0.131206889 | 0.293837866 |
| Tstd2 | 221.2458736 | -0.579591424 | 0.107887639 | -5.372176377 |
| Gm23250 | 0.397571503 | 0.002673679 | 3.262307451 | 0.000819567 |
| Ncbp1 | 647.6804023 | -0.118793595 | 0.071940663 | -1.65127189 |
| Xpa | 75.64072436 | 0.445560199 | 0.176504265 | 2.52435939 |
| Gm12444 | 2.312647122 | 0.490933958 | 0.706422148 | 0.694958332 |
| Gm12445 | 0 | NA | NA | NA |
| Gm12446 | 0.091666476 | 0.037060453 | 3.724945049 | 0.009949262 |
| Gm12447 | 2.955086367 | 1.073284801 | 0.659585189 | 1.627211798 |
| Foxe1 | 0.36004552 | 0.965087452 | 1.70581481 | 0.565763321 |
| 5830415F09Rik | 17.35131692 | -0.712641649 | 0.280100597 | -2.544234666 |
| Hemgn | 1.129829066 | -0.17147864 | 1.189791384 | -0.144124964 |
| Gm12443 | 0.104366086 | 0.51795324 | 3.724945049 | 0.139049901 |
| Anp32b | 407.7977659 | -0.422667858 | 0.121252391 | -3.485851749 |
| Nans | 132.4373756 | -0.491799966 | 0.146680302 | -3.352869876 |
| Trim14 | 69.63745902 | -1.138688344 | 0.214701975 | -5.303576473 |
| Gm22247 | 0.26079097 | 0.037075547 | 2.45364588 | 0.01511039 |
| Gm16731 | 0.468050163 | -0.593773513 | 1.340544229 | -0.442934668 |
| Coro2a | 311.7321339 | 0.076240104 | 0.082461657 | 0.924552172 |
| Tbc1d2 | 54.86226986 | -0.496634534 | 0.199674358 | -2.487222392 |
| Gabbr2 | 3104.900051 | 0.553360001 | 0.118118997 | 4.68476719 |
| Gm568 | 0.609243813 | 0.071368883 | 1.49464371 | 0.047749763 |
| Anks6 | 149.640635 | 0.26148264 | 0.199056939 | 1.313607256 |
| Gm12425 | 0 | NA | NA | NA |
| Gm12422 | 0.506756615 | 2.81889164 | 1.346177164 | 2.093997518 |
| Galnt12 | 14.31060596 | 0.730471795 | 0.328842255 | 2.221344073 |

| | | | | |
|---|---|---|---|---|
| Gm12424 | 0.961043033 | 0.482097967 | 1.214660101 | 0.396899484 |
| Gm12423 | 0 | NA | NA | NA |
| Gm12426 | 1.368259282 | 0.158166994 | 0.90570864 | 0.174633417 |
| Col15a1 | 61.55752535 | -0.000820952 | 0.187865241 | -0.004369898 |
| Tgfbr1 | 481.364332 | 0.16317564 | 0.145371497 | 1.122473407 |
| Gm12430 | 5.138410126 | 0.059514362 | 0.416650259 | 0.142840094 |
| Alg2 | 1382.70197 | 0.114595365 | 0.06424898 | 1.783613772 |
| Sec61b | 158.1254426 | -0.789551706 | 0.156874301 | -5.03302135 |
| Gm27601 | 0 | NA | NA | NA |
| Gm27668 | 0 | NA | NA | NA |
| Gm38343 | 0.066050875 | -0.443818988 | 3.724945049 | -0.119147795 |
| Gm22670 | 0 | NA | NA | NA |
| Gm12434 | 0 | NA | NA | NA |
| Nr4a3 | 542.7975049 | -0.212708549 | 0.310192739 | -0.685730264 |
| Stx17 | 355.1670537 | 0.119618117 | 0.074977741 | 1.595381705 |
| Erp44 | 524.436621 | -0.445292861 | 0.092932788 | -4.791558197 |
| Mir1958 | 0.694929783 | 0.512606995 | 1.420402756 | 0.360888482 |
| Invs | 182.2944049 | -0.423818789 | 0.116798101 | -3.628644519 |
| Gm12435 | 13.93294907 | -0.674291159 | 0.270963016 | -2.488498869 |
| Tex10 | 345.3321695 | -0.186317767 | 0.078168224 | -2.383548671 |
| Msantd3 | 126.8937499 | -0.012102305 | 0.115679473 | -0.104619293 |
| Tmeff1 | 650.557545 | 0.199181829 | 0.115119947 | 1.730211273 |
| Murc | 22.05245246 | 0.530217496 | 0.254226726 | 2.085608793 |
| Gm12439 | 2.996720251 | -0.776371336 | 0.755302319 | -1.027894813 |
| Gm12438 | 0 | NA | NA | NA |
| Gm24573 | 0 | NA | NA | NA |
| Gm23746 | 0 | NA | NA | NA |
| Gm12436 | 0.044544648 | 0.517957688 | 3.724945049 | 0.139051095 |
| Gm12442 | 0 | NA | NA | NA |
| Lppr1 | 119.1937067 | 0.94993402 | 0.154584404 | 6.14508316 |
| Gm22685 | 0.467201629 | 1.695005316 | 1.504345829 | 1.126739134 |
| Gm12453 | 0.624449418 | 0.428302627 | 1.402694111 | 0.305342857 |
| Rbpsuh-ps1 | 0.090074004 | 0.517955464 | 3.724945049 | 0.139050498 |
| Acnat2 | 0.218471152 | 0.037556477 | 1.91119973 | 0.019650734 |
| Gm12451 | 0.314054052 | 1.593882629 | 3.703755135 | 0.430342334 |
| Gm12452 | 0.075043205 | 0.517957688 | 3.724945049 | 0.139051095 |
| Acnat1 | 2.07728567 | -0.35006546 | 0.789866952 | -0.443195477 |
| Gm26424 | 0 | NA | NA | NA |
| Baat | 6.000090479 | -0.697167594 | 0.49913876 | -1.396741047 |
| Mrpl50 | 361.7674449 | 0.08903755 | 0.090488043 | 0.983970341 |
| Zfp189 | 205.1948466 | 0.494605112 | 0.099743739 | 4.958758463 |
| Aldob | 16.11872095 | 1.603033855 | 0.328944907 | 4.87325938 |
| Tmem246 | 593.6062876 | 0.109032894 | 0.075613275 | 1.441980838 |
| Rnf20 | 1042.085228 | 0.026112578 | 0.071563517 | 0.364886737 |
| Grin3a | 866.8310369 | 0.509978064 | 0.076359988 | 6.678603237 |
| Ppp3r2 | 15.47209461 | 0.496973746 | 0.354970227 | 1.400043462 |
| Gm12459 | 3.099509663 | 1.568920139 | 0.740301218 | 2.119299687 |

| | | | | |
|---|---|---|---|---|
| Gm12458 | 0.048949532 | 0.037064902 | 3.724945049 | 0.009950456 |
| Gm25485 | 0 | NA | NA | NA |
| Gm12460 | 0.083526043 | 0.037073798 | 3.724945049 | 0.009952844 |
| Cylc2 | 0 | NA | NA | NA |
| Amd-ps4 | 32.8894957 | 0.200804535 | 0.221477351 | 0.906659456 |
| Gm22563 | 0 | NA | NA | NA |
| Gm23043 | 0 | NA | NA | NA |
| Gm24943 | 0 | NA | NA | NA |
| Gm22042 | 0.482432794 | -0.499294953 | 1.385504012 | -0.360370629 |
| Gm12461 | 0 | NA | NA | NA |
| C630028M04Rik | 0.154174174 | -0.443823039 | 2.837762037 | -0.156398963 |
| Gm12465 | 0 | NA | NA | NA |
| Gm12466 | 0.144050924 | 1.05357957 | 3.721447096 | 0.283110183 |
| Gm12467 | 0 | NA | NA | NA |
| Gm12475 | 1.233559488 | 0.141643817 | 0.920381977 | 0.153896774 |
| Smc2os | 3.507361272 | 0.395283504 | 0.639337872 | 0.618270122 |
| Smc2 | 184.4916281 | -0.206611126 | 0.172379995 | -1.198579483 |
| Tmc4-ps | 0 | NA | NA | NA |
| Vma21-ps | 0.038981395 | -0.443816764 | 3.724945049 | -0.119147198 |
| Toporsl | 0 | NA | NA | NA |
| 4930552N02Rik | 0 | NA | NA | NA |
| Gm12482 | 0.399060758 | -1.897099024 | 1.671357559 | -1.135064734 |
| Olfr275 | 0 | NA | NA | NA |
| Olfr273 | 0.062426873 | 0.517959912 | 3.724945049 | 0.139051692 |
| Cct3-ps1 | 0.33824293 | -1.82968546 | 1.885292431 | -0.970504856 |
| Olfr272 | 0.046345709 | -0.443818988 | 3.724945049 | -0.119147795 |
| Olfr271-ps1 | 0.118974955 | 0.51795324 | 3.724945049 | 0.139049901 |
| Olfr270 | 0 | NA | NA | NA |
| Nipsnap3a | 0 | NA | NA | NA |
| Nipsnap3b | 127.01825 | -0.48967215 | 0.185406103 | -2.641078916 |
| Abca1 | 2154.257843 | -0.399396064 | 0.145458264 | -2.745777744 |
| 4930412L05Rik | 1.32543013 | 0.427367034 | 0.922858713 | 0.463090425 |
| 4930522O17Rik | 0.038981395 | -0.443816764 | 3.724945049 | -0.119147198 |
| AI427809 | 3.06421064 | -0.805219141 | 0.639897556 | -1.25835633 |
| Gm12495 | 0 | NA | NA | NA |
| Gm12496 | 0.503001268 | -0.917578134 | 1.540221107 | -0.595744423 |
| Slc44a1 | 1434.826822 | -0.160262307 | 0.086481499 | -1.853139792 |
| 1700060J05Rik | 0.251076704 | -0.558941193 | 2.492945678 | -0.224209135 |
| Fsd1l | 701.9750228 | 0.052189754 | 0.063194776 | 0.825855509 |
| Gm12470 | 15.34197034 | 0.655400842 | 0.379398153 | 1.727475044 |
| Fktn | 705.1094383 | 0.013368201 | 0.064117082 | 0.208496711 |
| Gm25445 | 0 | NA | NA | NA |
| Tal2 | 3.478949006 | -0.125815695 | 0.515738922 | -0.243952297 |
| Rps15a-ps8 | 0 | NA | NA | NA |
| Tmem38b | 167.1596389 | -0.249604921 | 0.112027352 | -2.228071243 |
| n-R5s185 | 0 | NA | NA | NA |
| Gm12468 | 0.068097359 | 0.037064902 | 3.724945049 | 0.009950456 |

| | | | | |
|---|---|---|---|---|
| Gm23745 | 0.047948698 | 0.037064902 | 3.724945049 | 0.009950456 |
| Gm24177 | 0 | NA | NA | NA |
| Gm12484 | 0 | NA | NA | NA |
| Gm12469 | 0.094831987 | 0.517957688 | 3.724945049 | 0.139051095 |
| Gm25687 | 0 | NA | NA | NA |
| Tpt1-ps2 | 0 | NA | NA | NA |
| Gm29067 | 0 | NA | NA | NA |
| Gm12478 | 0 | NA | NA | NA |
| Gm12480 | 0.086930093 | -0.443164241 | 3.724898049 | -0.118973522 |
| Gm12477 | 0 | NA | NA | NA |
| Gm24628 | 0 | NA | NA | NA |
| Gm12479 | 0 | NA | NA | NA |
| Zfp462 | 1321.864971 | 0.064904702 | 0.077251417 | 0.840174905 |
| Gm12515 | 0.097139547 | -0.443816764 | 3.724945049 | -0.119147198 |
| Gm12514 | 0.145451803 | -1.224261167 | 3.529361758 | -0.346878912 |
| Gm12509 | 0 | NA | NA | NA |
| Gm12508 | 0.167320703 | 0.037076259 | 2.764437563 | 0.013411863 |
| Gm12512 | 0 | NA | NA | NA |
| Gm25419 | 0 | NA | NA | NA |
| Rad23b | 2280.331224 | -0.168036069 | 0.073776331 | -2.277641991 |
| Gm12516 | 0 | NA | NA | NA |
| Gm12513 | 0 | NA | NA | NA |
| Gm12505 | 0.048949532 | 0.037064902 | 3.724945049 | 0.009950456 |
| Gm12510 | 0 | NA | NA | NA |
| Gm22150 | 0 | NA | NA | NA |
| Gm12507 | 0 | NA | NA | NA |
| Klf4 | 85.90030529 | -0.103022177 | 0.258154859 | -0.39907123 |
| Gm12511 | 0.772413307 | 0.02762949 | 1.191793795 | 0.023183113 |
| Gm12506 | 1.081462396 | -0.931084407 | 1.00811206 | -0.923592172 |
| Gm12519 | 0.384295504 | -0.97446815 | 1.579077484 | -0.617112307 |
| Gm12520 | 0 | NA | NA | NA |
| 2310081O03Rik | 1.122999083 | -1.713318628 | 0.979595547 | -1.749006141 |
| Gm12518 | 0 | NA | NA | NA |
| Gm26144 | 0.110595523 | 0.037071574 | 3.724945049 | 0.009952247 |
| Actl7b | 1.554701746 | 0.380831508 | 0.892875509 | 0.426522516 |
| Gm26657 | 0 | NA | NA | NA |
| Actl7a | 0.04393175 | 0.037062677 | 3.724945049 | 0.009949859 |
| Ikbkap | 924.710914 | 0.271512302 | 0.089227023 | 3.04293804 |
| Gm22952 | 4.197905476 | 1.388804218 | 0.559160291 | 2.483731838 |
| Fam206a | 278.4213948 | -0.181432293 | 0.092476745 | -1.961923426 |
| Ctnnal1 | 338.824844 | -0.022307191 | 0.132444792 | -0.168426337 |
| Tmem245 | 2131.110561 | 0.478114432 | 0.092095315 | 5.191517443 |
| Mir32 | 0.646815127 | 0.539982164 | 1.548758513 | 0.348654848 |
| Gm25053 | 0.048949532 | 0.037064902 | 3.724945049 | 0.009950456 |
| Gm22300 | 0.061920613 | -0.443816764 | 3.724945049 | -0.119147198 |
| Frrs1l | 3263.550938 | 0.542400022 | 0.103949192 | 5.217933987 |
| Epb4.1l4b | 724.2344794 | 0.513186448 | 0.085900915 | 5.974167456 |

| | | | | |
|---|---|---|---|---|
| Gm12530 | 6.592824524 | -1.437813962 | 0.525596286 | -2.735586229 |
| Ptpn3 | 752.3310739 | 0.420686842 | 0.117991057 | 3.565412954 |
| Gm12536 | 3.13007401 | 0.301271435 | 0.626936075 | 0.480545701 |
| 1700042G15Rik | 0 | NA | NA | NA |
| Gm12537 | 23.16944423 | 0.209192847 | 0.232368454 | 0.900263538 |
| Gm12538 | 1.143639422 | -0.947297458 | 1.29545561 | -0.731246559 |
| Palm2 | 230.289783 | 0.301208853 | 0.119018831 | 2.530766358 |
| Gm12539 | 0 | NA | NA | NA |
| Pakap | 9.557701205 | 1.395555881 | 0.359612008 | 3.880726593 |
| Akap2 | 22.84245767 | 0.045907369 | 0.252766431 | 0.181619722 |
| Gm12526 | 0.201955748 | 0.065235438 | 2.217323108 | 0.029420808 |
| Gm24277 | 1.91044629 | -0.147661568 | 0.866000455 | -0.170509805 |
| Gm26138 | 0 | NA | NA | NA |
| D630039A03Rik | 3.846205949 | -0.258750088 | 0.583600961 | -0.443368166 |
| Txn1 | 650.0859016 | -0.684069597 | 0.163208007 | -4.191397286 |
| Txndc8 | 0 | NA | NA | NA |
| Svep1 | 270.025923 | 0.144845969 | 0.219713257 | 0.659250019 |
| Gm26889 | 1.802555565 | -1.103406746 | 0.908246624 | -1.214875692 |
| Musk | 19.66503011 | 0.365519997 | 0.22981695 | 1.590483196 |
| Lpar1 | 579.1101207 | -0.047967407 | 0.138085884 | -0.347373721 |
| Mir3095 | 0.494865952 | -0.023359404 | 1.487288926 | -0.01570603 |
| Gm12577 | 4.644629704 | -1.182650573 | 0.639917152 | -1.848130761 |
| Gm12578 | 0 | NA | NA | NA |
| Gm12579 | 0.044191549 | 0.037062677 | 3.724945049 | 0.009949859 |
| Gm12580 | 0.526179982 | -2.168575648 | 1.262902685 | -1.717135987 |
| Olfr267 | 2.961215905 | -1.593061362 | 0.594075479 | -2.681580739 |
| AI314180 | 2945.683157 | 0.094361134 | 0.074393752 | 1.268401336 |
| Gm26400 | 0 | NA | NA | NA |
| Zkscan16 | 430.9535622 | 0.842712931 | 0.099955065 | 8.430917723 |
| Ptgr1 | 21.48653641 | -1.443918782 | 0.377566089 | -3.824280898 |
| Dnajc25 | 87.02722299 | 0.000921325 | 0.163670647 | 0.005629142 |
| Gm20503 | 0 | NA | NA | NA |
| Gng10 | 14.26907646 | -0.260417325 | 0.302708157 | -0.860291734 |
| AI481877 | 0.167924218 | -0.444688103 | 2.761407197 | -0.161036773 |
| Gm12593 | 0.308655657 | 1.353223238 | 3.704213799 | 0.36531996 |
| Gm12594 | 0 | NA | NA | NA |
| Rpsa-ps11 | 1.26617739 | -0.991178473 | 0.897649008 | -1.104193804 |
| Ugcg | 1300.167576 | -0.20045093 | 0.057895008 | -3.462317999 |
| Gm12596 | 0.346136167 | -0.85564061 | 1.643016208 | -0.520774296 |
| Susd1 | 107.3121581 | 0.396056672 | 0.167908682 | 2.358762326 |
| Gm12527 | 0.551374303 | 0.965882803 | 1.370086039 | 0.70497967 |
| Ptbp3 | 1030.313491 | -0.649754446 | 0.116152503 | -5.593977128 |
| Gm12529 | 0 | NA | NA | NA |
| 1700018M17Rik | 1.139841548 | 0.802152533 | 0.912237994 | 0.879323749 |
| Hsdl2 | 568.5576942 | -0.144684678 | 0.117345721 | -1.232977872 |
| E130308A19Rik | 348.4769703 | 0.072649892 | 0.07066392 | 1.028104465 |
| Inip | 231.4780127 | -0.171057635 | 0.087310402 | -1.959189636 |

| | | | | |
|---|---|---|---|---|
| Snx30 | 979.4902318 | 0.058982065 | 0.067572992 | 0.872864486 |
| Gm12543 | 7.392584082 | 0.713791156 | 0.450644024 | 1.583935696 |
| Gm12542 | 0.038981395 | -0.443816764 | 3.724945049 | -0.119147198 |
| Slc46a2 | 0 | NA | NA | NA |
| Mup4 | 0.187777844 | 0.037062266 | 3.313226725 | 0.011186154 |
| Mup6 | 0.095435502 | 0.037073798 | 3.724945049 | 0.009952844 |
| Mup7 | 0.045882455 | 0.517959912 | 3.724945049 | 0.139051692 |
| Mup-ps1 | 0 | NA | NA | NA |
| Mup2 | 0 | NA | NA | NA |
| Mup-ps2 | 0 | NA | NA | NA |
| Mup8 | 0 | NA | NA | NA |
| Mup-ps3 | 0 | NA | NA | NA |
| Mup-ps29 | 0 | NA | NA | NA |
| Mup-ps28 | 0 | NA | NA | NA |
| Mup9 | 0 | NA | NA | NA |
| Mup-ps4 | 0 | NA | NA | NA |
| Mup-ps26 | 0 | NA | NA | NA |
| Mup1 | 0 | NA | NA | NA |
| Gm13773 | 0 | NA | NA | NA |
| Mup-ps6 | 0 | NA | NA | NA |
| Mup-ps24 | 0 | NA | NA | NA |
| Mup10 | 0 | NA | NA | NA |
| Mup-ps5 | 0 | NA | NA | NA |
| Mup11 | 0 | NA | NA | NA |
| Mup-ps7 | 0 | NA | NA | NA |
| Mup12 | 0 | NA | NA | NA |
| Mup-ps8 | 0 | NA | NA | NA |
| Gm21286 | 0 | NA | NA | NA |
| Mup-ps9 | 0 | NA | NA | NA |
| Mup13 | 0 | NA | NA | NA |
| Mup-ps10 | 0 | NA | NA | NA |
| Mup14 | 0 | NA | NA | NA |
| Mup-ps11 | 0 | NA | NA | NA |
| Mup15 | 0 | NA | NA | NA |
| Mup-ps12 | 0 | NA | NA | NA |
| Mup16 | 0 | NA | NA | NA |
| Mup-ps13 | 0 | NA | NA | NA |
| Mup17 | 0 | NA | NA | NA |
| Mup-ps14 | 0 | NA | NA | NA |
| Mup18 | 0 | NA | NA | NA |
| Mup-ps15 | 0 | NA | NA | NA |
| Gm12573 | 0 | NA | NA | NA |
| Mup19 | 0 | NA | NA | NA |
| Gm13775 | 0 | NA | NA | NA |
| Mup-ps16 | 0 | NA | NA | NA |
| Mup5 | 0.135858026 | 0.037065849 | 2.948997537 | 0.012568966 |
| Mup-ps23 | 0 | NA | NA | NA |

| | | | | |
|---|---|---|---|---|
| Mup-ps17 | 0 | NA | NA | NA |
| Mup-ps18 | 0 | NA | NA | NA |
| Mup-ps19 | 0 | NA | NA | NA |
| Mup-ps20 | 0 | NA | NA | NA |
| Mup-ps21 | 0 | NA | NA | NA |
| Mup20 | 0 | NA | NA | NA |
| Mup3 | 1.273710888 | 0.396137796 | 0.956369252 | 0.414210092 |
| Gm12909 | 0 | NA | NA | NA |
| Gm12910 | 0 | NA | NA | NA |
| Mup21 | 0.088476398 | 0.51795324 | 3.724945049 | 0.139049901 |
| Zfp37 | 301.4106421 | 0.093189911 | 0.078118094 | 1.192936312 |
| Slc31a2 | 164.8023296 | -0.884408716 | 0.20805401 | -4.250861194 |
| Gm23148 | 0 | NA | NA | NA |
| Fkbp15 | 744.3892095 | 0.014761991 | 0.074159779 | 0.199056572 |
| Slc31a1 | 348.3933879 | -0.340226432 | 0.112785908 | -3.016568642 |
| Cdc26 | 169.15404 | -0.041543607 | 0.106721608 | -0.389270809 |
| Prpf4 | 294.0494031 | -0.256765639 | 0.076448913 | -3.358656503 |
| Rnf183 | 1.151952751 | 0.020856251 | 0.992389202 | 0.021016201 |
| Wdr31 | 53.77822627 | 0.297318216 | 0.162416107 | 1.830595632 |
| Bspry | 7.199256456 | -1.006177197 | 0.407771 | -2.46750553 |
| Hdhd3 | 119.7174087 | -0.153445407 | 0.146228915 | -1.049350652 |
| Alad | 452.599456 | -0.004518398 | 0.085198703 | -0.053033644 |
| Pole3 | 241.0396122 | -0.105711949 | 0.097680551 | -1.082221059 |
| 4933430I17Rik | 8.456414667 | -0.399014874 | 0.351287102 | -1.135865428 |
| Rgs3 | 280.2490358 | -0.022576111 | 0.080212115 | -0.281455128 |
| Gm11209 | 0.195038611 | -0.472858308 | 3.240402907 | -0.145925776 |
| Gm11210 | 0.707441185 | -0.352116985 | 1.123851219 | -0.313312811 |
| Gm11211 | 0.066050875 | -0.443818988 | 3.724945049 | -0.119147795 |
| Gm24117 | 0 | NA | NA | NA |
| Gm25391 | 0 | NA | NA | NA |
| Gm11481 | 0 | NA | NA | NA |
| Gm11480 | 0 | NA | NA | NA |
| Zfp618 | 184.4983496 | 0.164767799 | 0.109354546 | 1.50673022 |
| Ambp | 0.720892307 | 0.204446569 | 1.39322762 | 0.146743121 |
| Kif12 | 0.229693479 | 0.539115056 | 2.138260958 | 0.252127812 |
| Col27a1 | 179.4777456 | -0.143545645 | 0.202639288 | -0.708380129 |
| Mir455 | 0.26766946 | 0.052505024 | 2.099918455 | 0.025003363 |
| Orm1 | 0.137647365 | 1.167618838 | 3.719469251 | 0.31392082 |
| Gm11212 | 0 | NA | NA | NA |
| Orm3 | 0.157314084 | 0.037075025 | 3.435173482 | 0.010792766 |
| Orm2 | 0.751984476 | -0.191867866 | 1.029791276 | -0.186317238 |
| Akna | 219.0544516 | -0.519429306 | 0.10880644 | -4.77388386 |
| Aknaos | 0.425011668 | 1.207801145 | 1.474504726 | 0.819123278 |
| Whrn | 563.7172746 | 0.303263622 | 0.0846399 | 3.582986524 |
| Atp6v1g1 | 617.2868754 | -0.198627783 | 0.113193164 | -1.754768361 |
| 6330416G13Rik | 190.6195337 | 0.228764047 | 0.11550186 | 1.980609213 |
| Gm11483 | 0 | NA | NA | NA |

| | | | | |
|---|---|---|---|---|
| Gm24448 | 0 | NA | NA | NA |
| Gm11213 | 0 | NA | NA | NA |
| 1700018C11Rik | 0 | NA | NA | NA |
| Gm24234 | 0 | NA | NA | NA |
| Gm11482 | 1.060662479 | -0.080970604 | 1.007574919 | -0.080361869 |
| Gm11214 | 0 | NA | NA | NA |
| Tnfsf15 | 4.678869644 | -1.448466158 | 0.525476138 | -2.756483224 |
| Tnfsf8 | 49.08655836 | -1.064845974 | 0.277846934 | -3.832491352 |
| Rpl17-ps4 | 3.619380876 | -0.950537775 | 0.521268615 | -1.823508548 |
| Tnc | 854.476176 | -1.825668812 | 0.319059636 | -5.722030008 |
| 8030451A03Rik | 1.099630792 | -0.162476191 | 0.902620671 | -0.180004952 |
| Gm11216 | 0.038981395 | -0.443816764 | 3.724945049 | -0.119147198 |
| Gm11217 | 0 | NA | NA | NA |
| Gm23950 | 0 | NA | NA | NA |
| Gm11218 | 0.060174537 | 0.517957688 | 3.724945049 | 0.139051095 |
| Gm11219 | 0 | NA | NA | NA |
| Pappa | 141.7302013 | -1.139753204 | 0.216391283 | -5.267093896 |
| Astn2 | 1043.79549 | 0.294102463 | 0.087369337 | 3.366197713 |
| Trim32 | 1762.038928 | 0.315561867 | 0.060165694 | 5.244880348 |
| Gm11484 | 0 | NA | NA | NA |
| Gm25480 | 0 | NA | NA | NA |
| Gm11220 | 0 | NA | NA | NA |
| Tlr4 | 178.7502272 | -1.301358065 | 0.232804917 | -5.589907992 |
| Gm11403 | 0 | NA | NA | NA |
| Hmgb1-rs18 | 0 | NA | NA | NA |
| AL589845.1 | 0 | NA | NA | NA |
| Gm11249 | 12.40046496 | -0.318078052 | 0.367514756 | -0.865483759 |
| Gm11751 | 0.126225412 | 0.03707172 | 3.698755067 | 0.010022756 |
| Gm23738 | 0 | NA | NA | NA |
| Tcp1-ps1 | 0.143022002 | 0.03610674 | 3.547735707 | 0.010177404 |
| Gm12911 | 0 | NA | NA | NA |
| Brinp1 | 2163.011251 | 0.240676515 | 0.116935636 | 2.058196492 |
| Gm24697 | 0 | NA | NA | NA |
| Gm11404 | 1.959659191 | -2.297651606 | 1.258477826 | -1.825738649 |
| Gm11221 | 11.46002724 | 0.09758741 | 0.324185446 | 0.301023415 |
| Gm11222 | 0.623048784 | -0.755978088 | 1.289235553 | -0.586377009 |
| Gm11223 | 93.55249628 | 0.636726908 | 0.162411098 | 3.920464284 |
| Gm11224 | 5.294292594 | -0.407879601 | 0.452672113 | -0.90104866 |
| Gm11225 | 0.044191549 | 0.037062677 | 3.724945049 | 0.009949859 |
| Cdk5rap2 | 668.7817866 | -0.500177546 | 0.098644532 | -5.070504527 |
| Gm11226 | 21.32553314 | 0.514915153 | 0.261600691 | 1.968324898 |
| Gm26320 | 0.207279166 | 0.517971242 | 2.634010951 | 0.196647338 |
| Megf9 | 2380.976153 | 0.581143427 | 0.083162088 | 6.988081257 |
| Gm11227 | 0 | NA | NA | NA |
| Gm11228 | 0 | NA | NA | NA |
| Gm11229 | 0 | NA | NA | NA |
| Rps18-ps1 | 0.478648982 | 0.114237403 | 1.330279344 | 0.085874747 |

| | | | | |
|---|---|---|---|---|
| Gm11231 | 0.345321076 | 0.517974812 | 1.757208973 | 0.294771322 |
| Gm11232 | 0.115000406 | -0.443821212 | 3.724945049 | -0.119148392 |
| Gm11233 | 0 | NA | NA | NA |
| Gm11234 | 0 | NA | NA | NA |
| Tle1 | 623.9206616 | 0.278763496 | 0.098067672 | 2.842562586 |
| Gm11250 | 0 | NA | NA | NA |
| C630043F03Rik | 39.86748014 | 0.643359803 | 0.204367838 | 3.148048185 |
| Gm11235 | 0.263040051 | -0.034869245 | 1.790965744 | -0.019469521 |
| Aldoart1 | 2.66134452 | 0.418671569 | 0.657381941 | 0.636877198 |
| Gm5287 | 0 | NA | NA | NA |
| Gm25653 | 0 | NA | NA | NA |
| Gm11251 | 0 | NA | NA | NA |
| Gm25769 | 0 | NA | NA | NA |
| Gm11487 | 0 | NA | NA | NA |
| Gm11486 | 0 | NA | NA | NA |
| Gm11485 | 0.097139547 | -0.443816764 | 3.724945049 | -0.119147198 |
| Gm11488 | 14.4078922 | 0.676471107 | 0.334632466 | 2.021534598 |
| Gm24170 | 0 | NA | NA | NA |
| Gm11238 | 0 | NA | NA | NA |
| Gm11237 | 0 | NA | NA | NA |
| Gm11236 | 0 | NA | NA | NA |
| Gm11239 | 0 | NA | NA | NA |
| Gm428 | 0 | NA | NA | NA |
| Rasef | 2.980195284 | 0.45611984 | 0.631933166 | 0.721784936 |
| Gm11240 | 0.068097359 | 0.037064902 | 3.724945049 | 0.009950456 |
| Gm13867 | 0.251494677 | 1.116785124 | 2.494820455 | 0.447641481 |
| Gm11758 | 0 | NA | NA | NA |
| Gm11757 | 0 | NA | NA | NA |
| Gm13871 | 0 | NA | NA | NA |
| Gm11756 | 0 | NA | NA | NA |
| 2310002L09Rik | 0.092493346 | 0.517955464 | 3.724945049 | 0.139050498 |
| Frmd3 | 144.1270821 | -0.049655327 | 0.129644671 | -0.383010935 |
| Gm11760 | 4.795815343 | -0.12237464 | 0.595615248 | -0.205459212 |
| Gm11405 | 10.60243355 | -0.654979191 | 0.380289603 | -1.722316849 |
| Kdm4c | 648.3243287 | 0.111490012 | 0.086624936 | 1.287042928 |
| A230083N12Rik | 2.600875038 | 1.288677135 | 0.715168111 | 1.801921975 |
| Gm26379 | 0 | NA | NA | NA |
| Gm11254 | 0.066050875 | -0.443818988 | 3.724945049 | -0.119147795 |
| Gm11258 | 0 | NA | NA | NA |
| Gm11253 | 0 | NA | NA | NA |
| Gm11257 | 0 | NA | NA | NA |
| Gm12912 | 0.235437891 | -0.913164016 | 2.582676892 | -0.35357269 |
| Tmem261 | 148.8567893 | -0.09900947 | 0.116454113 | -0.850201574 |
| Gm22864 | 0 | NA | NA | NA |
| Gm11255 | 0 | NA | NA | NA |
| Gm11259 | 0 | NA | NA | NA |
| Gm11256 | 0 | NA | NA | NA |

| | | | | |
|---|---|---|---|---|
| Gm11241 | 0.044544648 | 0.517957688 | 3.724945049 | 0.139051095 |
| Ptprd | 4811.876378 | 0.607766016 | 0.099943808 | 6.081077252 |
| Gm11252 | 0 | NA | NA | NA |
| Gm11242 | 0.10503227 | -0.905414734 | 3.723500899 | -0.243162217 |
| Gm11243 | 0 | NA | NA | NA |
| Gm11244 | 7.743757494 | 1.05902439 | 0.431813433 | 2.452504505 |
| Gm11245 | 0 | NA | NA | NA |
| Gm11246 | 0.158325629 | -0.444696675 | 2.813062183 | -0.158082775 |
| Gm23159 | 10.5795326 | 1.061326419 | 0.444455627 | 2.387924359 |
| B230208B08Rik | 26.21483297 | 0.83337594 | 0.281013982 | 2.965603113 |
| Gm11260 | 0.431719086 | 0.516191754 | 1.466844327 | 0.351906296 |
| Gm11261 | 0.068097359 | 0.037064902 | 3.724945049 | 0.009950456 |
| Gm22262 | 0 | NA | NA | NA |
| Gm11262 | 0.136963818 | 0.037074342 | 3.599544405 | 0.010299732 |
| Gm25836 | 0 | NA | NA | NA |
| Gm11263 | 32.27155379 | 0.079640368 | 0.227885745 | 0.349474986 |
| Gm11409 | 0 | NA | NA | NA |
| Gm11407 | 1.154096829 | 1.692727546 | 1.149739098 | 1.472271013 |
| Gm11408 | 0.245813264 | 0.88965116 | 2.05284656 | 0.433374407 |
| Gm11410 | 0.40441216 | -0.535559295 | 1.674770059 | -0.319780792 |
| Gm11406 | 0 | NA | NA | NA |
| Tyrp1 | 7.568395809 | -0.502550807 | 0.432847943 | -1.161033141 |
| Lurap1l | 131.167045 | 0.587539469 | 0.117081861 | 5.018193788 |
| Gm27452 | 1.818514136 | 1.43406114 | 0.898616463 | 1.595854515 |
| Mpdz | 900.4834607 | -0.093453591 | 0.073963583 | -1.263508169 |
| Gm25803 | 3.471866242 | -0.041303856 | 0.599164355 | -0.06893577 |
| Gm11765 | 0 | NA | NA | NA |
| Gm11411 | 0.049553047 | -0.443818988 | 3.724945049 | -0.119147795 |
| Gm11412 | 0.477185738 | -0.41455269 | 1.4234814 | -0.291224522 |
| n-R5s187 | 0 | NA | NA | NA |
| Gm12914 | 0 | NA | NA | NA |
| Gm11264 | 0.922800452 | 0.495955273 | 1.131743194 | 0.438222448 |
| Gm22402 | 0 | NA | NA | NA |
| Gm5860 | 3.25720724 | 0.688974644 | 0.579137531 | 1.189656354 |
| Nfib | 2628.924338 | 0.17076911 | 0.098200022 | 1.738992587 |
| n-R5s188 | 0 | NA | NA | NA |
| Gm11266 | 18.26086893 | 0.242992711 | 0.296836567 | 0.81860774 |
| Gm11267 | 0.708875568 | -1.298372011 | 1.236187953 | -1.050303077 |
| Gm11268 | 0.189675868 | 0.037076698 | 2.656640771 | 0.013956233 |
| Gm11269 | 0.044544648 | 0.517957688 | 3.724945049 | 0.139051095 |
| Zdhhc21 | 1208.957435 | 0.037822073 | 0.053538848 | 0.706441661 |
| Cer1 | 0.481690985 | 1.87834375 | 1.425527533 | 1.317648173 |
| Frem1 | 91.57009446 | 0.615486176 | 0.180204414 | 3.415488913 |
| Gm24884 | 0 | NA | NA | NA |
| Gm11248 | 0 | NA | NA | NA |
| Gm11185 | 2.941272167 | -0.177365853 | 0.700199694 | -0.253307527 |
| Ttc39b | 1497.137622 | -0.021060224 | 0.099337609 | -0.212006552 |

| | | | | |
|---|---|---|---|---|
| Gm23412 | 0.392867355 | -0.808446477 | 1.921807376 | -0.420669879 |
| Gm11413 | 1.891350131 | 0.435558915 | 0.874376736 | 0.498136441 |
| Gm11184 | 0 | NA | NA | NA |
| Snapc3 | 534.9796752 | -0.049163861 | 0.099142445 | -0.495891152 |
| Psip1 | 2060.877521 | 0.132074053 | 0.076795885 | 1.719806379 |
| Gm11414 | 0.092493346 | 0.517955464 | 3.724945049 | 0.139050498 |
| Gm10154 | 11.65614726 | -0.243093544 | 0.491065549 | -0.495032779 |
| Ccdc171 | 44.96192503 | 0.087828478 | 0.178009221 | 0.493392859 |
| Gm25899 | 0 | NA | NA | NA |
| Gm27046 | 0.323557773 | -1.469759417 | 1.730896193 | -0.84913204 |
| Cbx3-ps3 | 0.066050875 | -0.443818988 | 3.724945049 | -0.119147795 |
| Gm26968 | 0.264809544 | 0.03707587 | 2.10454422 | 0.017617054 |
| Gm11415 | 0 | NA | NA | NA |
| 6030471H07Rik | 0 | NA | NA | NA |
| Gm12416 | 6.021123113 | -0.770049025 | 0.425898844 | -1.808056153 |
| Gm12414 | 0.092269992 | -0.443823436 | 3.724945049 | -0.119148989 |
| Bnc2 | 94.00517019 | -0.034118019 | 0.16405184 | -0.20797096 |
| Gm12421 | 0.473868817 | 1.424859388 | 1.417856502 | 1.004939065 |
| Gm12420 | 0 | NA | NA | NA |
| Gm12415 | 0.047948698 | 0.037064902 | 3.724945049 | 0.009950456 |
| Cntln | 269.0535505 | 0.322433505 | 0.093800505 | 3.437438905 |
| Gm12418 | 9.82225973 | -0.37004573 | 0.338278612 | -1.093908147 |
| Gm12419 | 0.575593679 | -1.648391902 | 1.273925668 | -1.29394669 |
| Gm12417 | 0 | NA | NA | NA |
| Sh3gl2 | 2324.421738 | 0.344450948 | 0.060111663 | 5.730184992 |
| Gm12413 | 0 | NA | NA | NA |
| Gm12680 | 8.074609009 | 0.633687739 | 0.434043482 | 1.459963727 |
| Gm26069 | 1.027412341 | 0.030275852 | 1.04719472 | 0.028911388 |
| Gm25811 | 0 | NA | NA | NA |
| Adamtsl1 | 125.2776745 | -0.210417231 | 0.135722989 | -1.550343336 |
| Gm12550 | 0.075043205 | 0.517957688 | 3.724945049 | 0.139051095 |
| Saxo1 | 0.416337256 | 1.91337199 | 1.689616794 | 1.132429553 |
| Gm26017 | 0 | NA | NA | NA |
| Saxo1os | 0 | NA | NA | NA |
| Rraga | 797.0060883 | -0.059190217 | 0.086590328 | -0.683566145 |
| Haus6 | 193.2480707 | 0.103903336 | 0.088712336 | 1.171238869 |
| Scarna8 | 0.537445765 | 0.913691609 | 1.364516896 | 0.669608131 |
| Gm12551 | 0.925310778 | -2.174208569 | 1.228148868 | -1.770313539 |
| Plin2 | 668.7166373 | -2.136070972 | 0.349147715 | -6.117957765 |
| Dennd4c | 657.3655427 | -0.360482042 | 0.072587669 | -4.966160902 |
| Gm12630 | 2.006602685 | 1.457885415 | 0.938488203 | 1.553440321 |
| Rps6 | 1228.330895 | -0.164046163 | 0.068777506 | -2.385171725 |
| Acer2 | 61.2692064 | -0.170809179 | 0.195869448 | -0.872056262 |
| Slc24a2 | 8155.610505 | 0.762304448 | 0.103736868 | 7.348442899 |
| Gm12600 | 0.561663198 | -0.439727043 | 1.247095547 | -0.352600925 |
| Gm23154 | 0 | NA | NA | NA |
| Gm12604 | 0.075043205 | 0.517957688 | 3.724945049 | 0.139051095 |

| | | | | |
|---|---|---|---|---|
| Mllt3 | 907.8146563 | 0.005153162 | 0.07196951 | 0.071602021 |
| Gm12631 | 3.345683444 | 0.427372052 | 0.532629244 | 0.802381875 |
| Gm12646 | 0.591515762 | -0.523393369 | 1.375078097 | -0.380628104 |
| Focad | 1053.950382 | 0.393411811 | 0.109362816 | 3.597308721 |
| Mir491 | 10.7430229 | 0.784960444 | 0.350982454 | 2.236466342 |
| Gm12645 | 3.761156174 | 0.315557121 | 0.570633178 | 0.552994697 |
| Hacd4 | 63.03965297 | -1.2418338 | 0.2244243 | -5.533419511 |
| Ifnb1 | 44.66037375 | -0.690113141 | 0.887216644 | -0.777840616 |
| Ifna15 | 0.599468885 | 0.797397084 | 1.42282307 | 0.560433058 |
| Ifna14 | 0.481692262 | -0.310026918 | 1.589923868 | -0.194994819 |
| Ifna-ps1 | 0 | NA | NA | NA |
| Ifna9 | 0.316561663 | -2.210012743 | 1.886262212 | -1.171636016 |
| Mrpl48-ps | 38.64384611 | 0.08693124 | 0.245508472 | 0.354086519 |
| Ifna12 | 0.176391548 | 0.998861293 | 2.7186527 | 0.367410406 |
| Gm12601 | 0.215725693 | -0.299472843 | 2.176745926 | -0.137578226 |
| C87499 | 0.779996687 | -0.425915966 | 1.069751945 | -0.398144605 |
| Ifna13 | 0.075043205 | 0.517957688 | 3.724945049 | 0.139051095 |
| Gm13281 | 0.046345709 | -0.443818988 | 3.724945049 | -0.119147795 |
| Ifna16 | 0.145446656 | -0.410258174 | 2.878908473 | -0.142504764 |
| Ifna2 | 8.251847137 | 0.407368608 | 0.68865101 | 0.591545794 |
| Ifnab | 1.397532748 | -0.195348612 | 0.784669077 | -0.248956685 |
| Gm13273 | 0.313664056 | 0.599258659 | 3.70621055 | 0.161690398 |
| Gm13282 | 2.459004477 | 1.031062588 | 0.755006277 | 1.365634459 |
| Klhl9 | 1871.695579 | -0.018845975 | 0.087612696 | -0.215105522 |
| Gm26566 | 4.232091902 | -0.237183317 | 0.516908216 | -0.458849965 |
| Gm13274 | 0.044191549 | 0.037062677 | 3.724945049 | 0.009949859 |
| Gm13271 | 0.08486385 | 0.037730769 | 3.724898049 | 0.010129343 |
| Gm16686 | 0 | NA | NA | NA |
| Gm13286 | 0 | NA | NA | NA |
| Gm10583 | 0.144244773 | -0.4438229 | 3.581436381 | -0.123923156 |
| Gm13283 | 0 | NA | NA | NA |
| Gm10582 | 0 | NA | NA | NA |
| Gm13284 | 0 | NA | NA | NA |
| Gm10581 | 0 | NA | NA | NA |
| Gm13290 | 0 | NA | NA | NA |
| Gm10580 | 0 | NA | NA | NA |
| Gm13289 | 0 | NA | NA | NA |
| Gm13272 | 0 | NA | NA | NA |
| Ifnz | 0 | NA | NA | NA |
| Gm26525 | 0.092269992 | -0.443823436 | 3.724945049 | -0.119148989 |
| Gm13276 | 0 | NA | NA | NA |
| Gm13277 | 0 | NA | NA | NA |
| Gm13278 | 0 | NA | NA | NA |
| Gm13275 | 0 | NA | NA | NA |
| Gm26867 | 0.045882455 | 0.517959912 | 3.724945049 | 0.139051692 |
| Gm13279 | 0 | NA | NA | NA |
| Gm13285 | 0 | NA | NA | NA |

| | | | | |
|---|---|---|---|---|
| Gm13287 | 0 | NA | NA | NA |
| Gm13288 | 0 | NA | NA | NA |
| Ifna7 | 0.305381988 | 0.829485332 | 2.415839697 | 0.343352803 |
| Ifna11 | 0.275757984 | 1.135955144 | 3.707986664 | 0.306353622 |
| Ifna6 | 0.418480079 | 2.065958022 | 3.678030225 | 0.561702296 |
| Ifna5 | 11.73825278 | -0.311380295 | 0.981359488 | -0.317294834 |
| Ifna4 | 14.65900129 | 0.311649629 | 1.135123379 | 0.274551326 |
| Ifna1 | 2.156732396 | 0.201290851 | 1.067923384 | 0.188488101 |
| 4930553M12Rik | 0.266535354 | -1.456331022 | 2.105556087 | -0.691660997 |
| Ifne | 0.083526043 | 0.037073798 | 3.724945049 | 0.009952844 |
| Mir31 | 1.252737907 | 1.467821997 | 0.907916406 | 1.616692888 |
| Gm12603 | 0.048949532 | 0.037064902 | 3.724945049 | 0.009950456 |
| Gm12602 | 0 | NA | NA | NA |
| Mtap | 151.2844411 | -0.452188104 | 0.151034557 | -2.993938039 |
| Gm12607 | 0 | NA | NA | NA |
| Gm26490 | 0 | NA | NA | NA |
| Gm12606 | 0.046345709 | -0.443818988 | 3.724945049 | -0.119147795 |
| Cdkn2a | 2.014914513 | -0.198279607 | 0.690817123 | -0.287021848 |
| Cdkn2b | 6.964193097 | -0.094306293 | 0.443544863 | -0.212619513 |
| Gm12609 | 0.942397274 | -0.144603986 | 1.111853863 | -0.130056648 |
| Gm12610 | 0.811625864 | 2.037377239 | 1.115479388 | 1.826458885 |
| Gm12608 | 0 | NA | NA | NA |
| Gm27580 | 0.090427103 | 0.998850474 | 3.724945049 | 0.268151734 |
| Dmrta1 | 4.432480129 | 0.672953307 | 0.510412987 | 1.318448638 |
| Gm12629 | 0 | NA | NA | NA |
| Zfp352 | 0 | NA | NA | NA |
| Gm12633 | 0.176993581 | -0.908078785 | 2.297186023 | -0.395300501 |
| Gm12634 | 0 | NA | NA | NA |
| Gm12635 | 0 | NA | NA | NA |
| Gm12636 | 0 | NA | NA | NA |
| Gm12632 | 0 | NA | NA | NA |
| Gm25244 | 0.236157963 | 1.841742413 | 2.481497191 | 0.742190005 |
| Gm12643 | 0 | NA | NA | NA |
| Gm12644 | 0.116217011 | 0.517955464 | 3.724945049 | 0.139050498 |
| Elavl2 | 1479.215199 | -0.000928943 | 0.122641989 | -0.007574426 |
| Gm12653 | 0 | NA | NA | NA |
| Mir6402 | 6.56355345 | -0.164189239 | 0.479807907 | -0.342197859 |
| Gm12668 | 0.048949532 | 0.037064902 | 3.724945049 | 0.009950456 |
| Gm12670 | 0.219222703 | 0.85696078 | 2.169844874 | 0.394941035 |
| Gm12667 | 0.171083144 | -1.487610616 | 3.383974877 | -0.43960451 |
| Gm12669 | 20.03569431 | -0.262504764 | 0.231394214 | -1.134448264 |
| Gm12671 | 13.22935111 | -0.199409478 | 0.287166806 | -0.694402951 |
| 4930577H14Rik | 0 | NA | NA | NA |
| Izumo3 | 0 | NA | NA | NA |
| Gm12666 | 1.628724105 | 0.964993235 | 0.941403473 | 1.025058078 |
| Gm12638 | 0 | NA | NA | NA |
| Gm12637 | 0 | NA | NA | NA |

| | | | | |
|---|---|---|---|---|
| Gm22605 | 0 | NA | NA | NA |
| Gm12639 | 0 | NA | NA | NA |
| Gm12641 | 0 | NA | NA | NA |
| Gm12642 | 0 | NA | NA | NA |
| Tusc1 | 50.66388116 | 0.040502601 | 0.170288957 | 0.237846314 |
| Gm12649 | 0.203196539 | 0.504219968 | 2.59432468 | 0.194354998 |
| Gm23443 | 0 | NA | NA | NA |
| Gm12640 | 0 | NA | NA | NA |
| Gm24796 | 0 | NA | NA | NA |
| Gm22619 | 0 | NA | NA | NA |
| Gm12650 | 0 | NA | NA | NA |
| Gm12647 | 0 | NA | NA | NA |
| Gm12648 | 0.255640523 | -1.390516243 | 2.106863521 | -0.659993507 |
| Gm23836 | 0 | NA | NA | NA |
| Gm12654 | 0 | NA | NA | NA |
| Gm26410 | 0.038981395 | -0.443816764 | 3.724945049 | -0.119147198 |
| Gm12655 | 23.30350443 | -0.13396855 | 0.231225331 | -0.579385267 |
| Gm22804 | 0 | NA | NA | NA |
| Gm12656 | 0 | NA | NA | NA |
| Caap1 | 152.6588818 | -0.334417637 | 0.119418746 | -2.800378075 |
| Gm10306 | 0.49643671 | 0.517974464 | 1.556629178 | 0.332753922 |
| Plaa | 900.5907788 | -0.236212975 | 0.084314822 | -2.801559305 |
| Gm12657 | 14.22801902 | -0.103027439 | 0.293209589 | -0.351378137 |
| Ift74 | 232.3827732 | -0.110127086 | 0.091184082 | -1.207744636 |
| Lrrc19 | 0.551940858 | -0.76068192 | 1.416206881 | -0.53712627 |
| Mir872 | 1.137124893 | 0.601860608 | 0.981226951 | 0.613375537 |
| Tek | 47.11216226 | -0.676463175 | 0.204814725 | -3.302805377 |
| Eqtn | 11.24146221 | -0.931929801 | 0.396940221 | -2.347783748 |
| Gm12693 | 29.98890503 | -0.056517941 | 0.197106798 | -0.286737653 |
| Mysm1 | 851.8413644 | -0.015470617 | 0.085157656 | -0.181670302 |
| Gm12694 | 0.203088869 | 0.479139288 | 2.636766596 | 0.181714714 |
| Gm12703 | 1.47600346 | -1.35212094 | 0.932797964 | -1.449532473 |
| Gm12704 | 0.126912322 | -0.76154727 | 3.693829068 | -0.206167437 |
| Jun | 1515.464005 | -0.910123885 | 0.231105039 | -3.938139516 |
| Junos | 29.43953063 | -0.164759428 | 0.229609865 | -0.717562498 |
| n-R5s189 | 0.792393139 | 1.84114185 | 1.121443179 | 1.641761157 |
| 4930551L18Rik | 0.046345709 | -0.443818988 | 3.724945049 | -0.119147795 |
| Gm28096 | 1.499901305 | 0.817156293 | 0.969993013 | 0.842435236 |
| Gm12708 | 1.926663799 | -1.731333761 | 0.776489044 | -2.229695028 |
| Gm29064 | 0 | NA | NA | NA |
| Gm830 | 0.19500349 | -0.954828506 | 3.288425201 | -0.290360415 |
| Gm12707 | 0 | NA | NA | NA |
| Gm25004 | 0 | NA | NA | NA |
| Fggy | 21.91263025 | 0.723685749 | 0.275197291 | 2.629697942 |
| Gm20651 | 0 | NA | NA | NA |
| 9530080O11Rik | 18.68130564 | 0.486342233 | 0.240343619 | 2.023528795 |
| Hook1 | 1088.16238 | 0.352224607 | 0.11160082 | 3.156111289 |

| | | | | |
|---|---|---|---|---|
| Cyp2j13 | 7.229154167 | 1.057153072 | 0.406341124 | 2.601639386 |
| Hspe1-ps6 | 0.042716945 | 0.037062677 | 3.724945049 | 0.009949859 |
| Cyp2j12 | 25.97268065 | -0.199510041 | 0.224237385 | -0.889726931 |
| Cyp2j7-ps3 | 0 | NA | NA | NA |
| Cyp2j7-ps2 | 0 | NA | NA | NA |
| Cyp2j7-ps1 | 0 | NA | NA | NA |
| Cyp2j7 | 0.068097359 | 0.037064902 | 3.724945049 | 0.009950456 |
| Cyp2j15-ps | 0 | NA | NA | NA |
| Cyp2j11 | 0.27158854 | -1.382834899 | 2.073996395 | -0.66674894 |
| Cyp2j14-ps | 0 | NA | NA | NA |
| Cyp2j8 | 2.219411473 | 1.200170077 | 0.670484282 | 1.790004792 |
| Cyp2j6 | 245.6472783 | -0.11383717 | 0.131898403 | -0.863067085 |
| Cyp2j9 | 418.0043846 | 0.376285869 | 0.149365733 | 2.519224874 |
| Cyp2j5 | 0.490888186 | -1.322455749 | 1.669351427 | -0.792197333 |
| Gm12695 | 0.642811059 | 0.895151307 | 1.260107889 | 0.710376719 |
| Gm27579 | 0 | NA | NA | NA |
| Gm27521 | 0 | NA | NA | NA |
| Gm10192 | 0 | NA | NA | NA |
| Gm12696 | 28.72011168 | -0.348576967 | 0.20988234 | -1.66082085 |
| Gm12697 | 3.534004104 | 0.607044545 | 0.810613482 | 0.748870527 |
| E130114P18Rik | 46.0958998 | 0.072655786 | 0.178226229 | 0.407660456 |
| Gm12676 | 1.650824257 | -0.130241103 | 1.223396878 | -0.106458587 |
| Nfia | 1906.233348 | 0.039984286 | 0.082392618 | 0.485289668 |
| Gm12691 | 1.556603733 | 0.106831532 | 0.870170624 | 0.122770787 |
| Gm12692 | 0.373707575 | 0.998280524 | 1.601256566 | 0.62343571 |
| Gm12787 | 0 | NA | NA | NA |
| Gm24203 | 0 | NA | NA | NA |
| 0610025J13Rik | 0.422542347 | 0.025429437 | 1.458898275 | 0.017430576 |
| Gm12788 | 0 | NA | NA | NA |
| Tm2d1 | 205.9564824 | -0.14220589 | 0.107824008 | -1.318870375 |
| Gm24960 | 0 | NA | NA | NA |
| Inadl | 702.0612203 | 0.035399793 | 0.129076753 | 0.274253823 |
| Gm22625 | 0.042716945 | 0.037062677 | 3.724945049 | 0.009949859 |
| Gm26154 | 4.026424441 | 0.156763626 | 0.623013881 | 0.251621401 |
| Gm12846 | 13.02039719 | 0.81024095 | 0.398393106 | 2.033772518 |
| L1td1 | 1.260641324 | -0.317595573 | 0.960458745 | -0.330670708 |
| Gm17376 | 0.049553047 | -0.443818988 | 3.724945049 | -0.119147795 |
| Kank4 | 103.4907107 | -0.042981729 | 0.166544921 | -0.258078895 |
| Kank4os | 0.479560471 | 0.744371207 | 1.64858503 | 0.45152127 |
| I0C0044D17Rik | 0.072025462 | 0.517959912 | 3.724945049 | 0.139051692 |
| Gm12851 | 0 | NA | NA | NA |
| Usp1 | 392.4990393 | -0.125578474 | 0.102871174 | -1.220735304 |
| Dock7 | 964.8693113 | 0.010627276 | 0.083215989 | 0.127707137 |
| Angptl3 | 2.293039322 | -0.018322353 | 0.618001677 | -0.02964774 |
| Gm12852 | 6.635996975 | 0.762664545 | 0.572006902 | 1.333313536 |
| Gm12705 | 0.112642008 | 0.517955464 | 3.724945049 | 0.139050498 |
| Gm12681 | 0.364815146 | -1.291729409 | 1.57844957 | -0.818353297 |

| | | | | |
|---|---|---|---|---|
| Atg4c | 539.9595983 | 0.212189573 | 0.089900264 | 2.360277529 |
| Gm10305 | 27.16950442 | 0.491717554 | 0.292792192 | 1.679408014 |
| Gm12689 | 9.012307495 | -0.050786792 | 0.456692994 | -0.111205541 |
| Gm37556 | 0 | NA | NA | NA |
| Gm12685 | 0 | NA | NA | NA |
| Gm12686 | 0 | NA | NA | NA |
| Gm12684 | 0 | NA | NA | NA |
| Gm12683 | 0 | NA | NA | NA |
| Gm37117 | 0 | NA | NA | NA |
| Gm12690 | 0 | NA | NA | NA |
| Gm12687 | 0 | NA | NA | NA |
| Gm12688 | 2.558789337 | -0.43888648 | 0.863243576 | -0.508415576 |
| Foxd3 | 16.0637295 | -1.704697508 | 1.064953538 | -1.600724771 |
| Gm23366 | 0 | NA | NA | NA |
| Gm12682 | 0 | NA | NA | NA |
| Alg6 | 154.9068683 | 0.140417185 | 0.162279983 | 0.865277295 |
| Itgb3bp | 53.12261853 | -0.273486035 | 0.16388203 | -1.668798194 |
| Efcab7 | 42.69675528 | -0.562642056 | 0.185473024 | -3.03355196 |
| Gm26425 | 0.198326984 | 0.540004917 | 2.231313137 | 0.242012162 |
| Pgm2 | 636.1507098 | 0.143215969 | 0.083004597 | 1.725398043 |
| Gm12702 | 0 | NA | NA | NA |
| Ror1 | 118.724752 | 0.30657408 | 0.154417612 | 1.985356954 |
| Gm12701 | 11.03703998 | 0.494793203 | 0.465155276 | 1.063716199 |
| Gm12706 | 0.200296414 | 0.490678553 | 2.212362422 | 0.221789409 |
| Ube2u | 0.310183902 | 0.455205916 | 1.746621787 | 0.260620771 |
| Ube2uos | 0 | NA | NA | NA |
| Ccdc50-ps | 0.574153598 | 1.399525404 | 1.349145711 | 1.037341921 |
| Gm12700 | 0 | NA | NA | NA |
| Cachd1 | 422.3160165 | -0.100229523 | 0.075898255 | -1.32057743 |
| Gm10577 | 0 | NA | NA | NA |
| Gm10576 | 0 | NA | NA | NA |
| Raver2 | 272.6356933 | 0.369766038 | 0.190494364 | 1.941086498 |
| Gm22100 | 0.061920613 | -0.443816764 | 3.724945049 | -0.119147198 |
| Jak1 | 2519.077659 | -0.178999614 | 0.062951217 | -2.843465495 |
| Gm24468 | 0.165470308 | 1.408943363 | 2.753318612 | 0.511725507 |
| Gm12785 | 2.331474832 | -0.424240899 | 0.776238242 | -0.546534396 |
| Gm25124 | 0 | NA | NA | NA |
| Gm12801 | 0 | NA | NA | NA |
| Gm12796 | 0.250633908 | 0.053548174 | 1.830387588 | 0.029255101 |
| Gm12793 | 0.066050875 | -0.443818988 | 3.724945049 | -0.119147795 |
| E130102H24Rik | 106.6384263 | 0.220209424 | 0.21105764 | 1.04336154 |
| Mir101a | 0 | NA | NA | NA |
| 0610043K17Rik | 25.29227341 | -0.140211824 | 0.272625978 | -0.514301039 |
| Gm12795 | 0 | NA | NA | NA |
| Gm12798 | 2.114628594 | 0.705235771 | 0.800484041 | 0.881011656 |
| Ak4 | 432.3296171 | 0.039423759 | 0.149205477 | 0.264224611 |
| Dnajc6 | 4085.768198 | 0.37949101 | 0.05796583 | 6.546805392 |

| | | | | |
|---|---|---|---|---|
| Gm22533 | 11.8805187 | 0.857227442 | 0.41148854 | 2.083235277 |
| Leprot | 268.8974141 | -0.464059978 | 0.11662006 | -3.979246607 |
| Lepr | 95.56977508 | -0.262527285 | 0.14246369 | -1.842766292 |
| B020004J07Rik | 0.102891481 | 0.51795324 | 3.724945049 | 0.139049901 |
| Gm12791 | 3.064345978 | -0.38450904 | 0.575269185 | -0.668398464 |
| Gm12797 | 0.048949532 | 0.037064902 | 3.724945049 | 0.009950456 |
| C130073F10Rik | 0 | NA | NA | NA |
| Gm12792 | 0 | NA | NA | NA |
| Gm12800 | 0.092691418 | -0.908687078 | 3.724560878 | -0.2439716 |
| Gm12794 | 0.223364959 | -0.444641595 | 2.521503375 | -0.176339877 |
| Gm12790 | 0.047948698 | 0.037064902 | 3.724945049 | 0.009950456 |
| Gm12789 | 0.194279094 | -1.017823543 | 3.719440177 | -0.273649661 |
| Gm12799 | 4.949076095 | -0.620999963 | 0.487175719 | -1.274693993 |
| Pde4b | 1759.695653 | 0.185163556 | 0.094118627 | 1.967342302 |
| Gm24869 | 0 | NA | NA | NA |
| Sgip1 | 2873.427705 | 0.429540951 | 0.07619039 | 5.63773138 |
| Gm12709 | 0.447471404 | 1.065871195 | 1.615125133 | 0.659931031 |
| Tctex1d1 | 5.63377873 | -0.668498134 | 0.51256621 | -1.304218112 |
| BB031773 | 3.502581246 | 3.237175856 | 0.919415693 | 3.520905592 |
| Insl5 | 5.684128039 | -0.564097078 | 0.509061216 | -1.108112463 |
| Wdr78 | 212.7570655 | -0.453795806 | 0.163386619 | -2.777435556 |
| Gm12710 | 0.045882455 | 0.517959912 | 3.724945049 | 0.139051692 |
| Mier1 | 957.8111048 | -0.080627464 | 0.059949241 | -1.344928853 |
| Slc35d1 | 196.800066 | -0.124739444 | 0.106982148 | -1.16598373 |
| 4921539E11Rik | 0.165933562 | 0.486011036 | 2.762106923 | 0.175956634 |
| 4930456L15Rik | 0.038981395 | -0.443816764 | 3.724945049 | -0.119147198 |
| Oma1 | 95.40654656 | -0.587656833 | 0.174775933 | -3.362344126 |
| Gm12718 | 0.250798881 | -0.451530573 | 2.082776455 | -0.216792624 |
| Gm23064 | 0 | NA | NA | NA |
| Gm12720 | 0.060174537 | 0.517957688 | 3.724945049 | 0.139051095 |
| Gm12716 | 0 | NA | NA | NA |
| Gm12715 | 393.1600641 | -0.264888842 | 0.107143259 | -2.472286593 |
| Gm22801 | 0 | NA | NA | NA |
| Gm25877 | 0 | NA | NA | NA |
| Dab1 | 1235.718576 | 0.563639366 | 0.07838835 | 7.190346096 |
| Gm12719 | 0 | NA | NA | NA |
| Gm12717 | 3.781002861 | 2.691297451 | 0.76956049 | 3.497187663 |
| C8b | 6.711931076 | 2.35470402 | 0.978167435 | 2.407260696 |
| C8a | 0.626920446 | -0.917712891 | 1.164970009 | -0.787756667 |
| Gm17662 | 0.225129614 | 1.580880368 | 3.710727143 | 0.426029807 |
| 1700024P16Rik | 18.7794585 | 1.719906454 | 0.342606464 | 5.020064231 |
| Gm12721 | 0.110242424 | -0.443823436 | 3.724945049 | -0.119148989 |
| Prkaa2 | 803.30275 | 0.446654592 | 0.052346814 | 8.53260318 |
| Ppap2b | 1987.101825 | 0.266275456 | 0.131864714 | 2.019307882 |
| Gm12722 | 2.944400151 | 0.243887186 | 0.582318462 | 0.418820975 |
| Gm12723 | 0 | NA | NA | NA |
| Gm12726 | 0.106112162 | -0.443821212 | 3.724945049 | -0.119148392 |

| | | | | |
|---|---|---|---|---|
| Gm12728 | 0.174579622 | -0.443820762 | 2.312978349 | -0.1918828 |
| Gm24363 | 0 | NA | NA | NA |
| Gm12729 | 0.086930093 | -0.443164241 | 3.724898049 | -0.118973522 |
| Gm12727 | 0.640223289 | -0.034739012 | 1.285436721 | -0.027025066 |
| Gm26047 | 0 | NA | NA | NA |
| Hspd1-ps4 | 0.60453213 | -0.395105347 | 1.322456374 | -0.298766262 |
| Gm12724 | 0.146823232 | 0.517961571 | 2.878417312 | 0.179946656 |
| Gm12731 | 0 | NA | NA | NA |
| Gm12730 | 0.072025462 | 0.517959912 | 3.724945049 | 0.139051692 |
| Gm37253 | 0 | NA | NA | NA |
| Usp24 | 2364.206895 | -0.088702634 | 0.07546638 | -1.175392713 |
| Pcsk9 | 22.35786243 | 0.588898826 | 0.299180091 | 1.968375716 |
| Bsnd | 0.044191549 | 0.037062677 | 3.724945049 | 0.009949859 |
| Tmem61 | 0 | NA | NA | NA |
| 2700068H02Rik | 1.22619721 | 0.881629054 | 1.140390362 | 0.773094095 |
| Dhcr24 | 1131.153985 | -0.065467681 | 0.113997698 | -0.574289504 |
| Gm12744 | 0.280446515 | -0.920255159 | 1.846438175 | -0.498394786 |
| BC055111 | 2.227745575 | 0.218720108 | 0.650447611 | 0.336260914 |
| Ttc22 | 5.229776616 | 0.127910131 | 0.463173609 | 0.276160232 |
| Pars2 | 65.0940951 | 0.296502739 | 0.147044426 | 2.016416036 |
| Ttc4 | 363.5749445 | -0.011377112 | 0.062497571 | -0.182040872 |
| Mroh7 | 34.09442817 | 0.867489734 | 0.321984667 | 2.694195788 |
| Fam151a | 2.097884303 | -1.874437465 | 0.818404558 | -2.290355602 |
| Acot11 | 491.6282229 | 0.422133186 | 0.147434118 | 2.863198777 |
| Gm12745 | 0 | NA | NA | NA |
| Gm12746 | 0.048949532 | 0.037064902 | 3.724945049 | 0.009950456 |
| Ssbp3 | 1314.52194 | 0.074178127 | 0.100028325 | 0.741571218 |
| Gm12786 | 0.08486385 | 0.037730769 | 3.724898049 | 0.010129343 |
| Mrpl37 | 409.2304581 | 0.043383125 | 0.079693344 | 0.544375761 |
| Cyb5rl | 41.42110935 | 0.055305816 | 0.189734284 | 0.291490895 |
| Cdcp2 | 0.295634611 | -1.010197124 | 2.04297351 | -0.494473922 |
| Gm12802 | 0.337683083 | -1.826191643 | 1.723126409 | -1.059812927 |
| Tceanc2 | 245.3997247 | -0.3818684 | 0.09770361 | -3.908436966 |
| Tmem59 | 1671.351481 | -0.13565889 | 0.073650369 | -1.84193089 |
| Ldlrad1 | 1.155427534 | -0.298171921 | 1.060540282 | -0.281150962 |
| Lrrc42 | 182.0305453 | -0.566916049 | 0.113214548 | -5.007448774 |
| Hspb11 | 48.05584592 | -0.462637096 | 0.157525196 | -2.936908554 |
| Gm12850 | 2.64217102 | 0.467301993 | 0.753562589 | 0.620123663 |
| Dio1 | 0.200957741 | -1.699401354 | 3.22964838 | -0.526187731 |
| Yipf1 | 215.0974917 | -0.117816669 | 0.084098176 | -1.40094203 |
| Gm12870 | 0.418005171 | 0.029272995 | 1.485225957 | 0.019709456 |
| Gm12869 | 0 | NA | NA | NA |
| Ndc1 | 286.6300775 | -0.027974711 | 0.085764568 | -0.326180286 |
| Gm24804 | 0 | NA | NA | NA |
| Glis1 | 17.53901492 | -0.215256331 | 0.284195771 | -0.75742271 |
| Dmrtb1 | 26.37534565 | -0.04994691 | 0.218241035 | -0.22886122 |
| Gm12899 | 0 | NA | NA | NA |

| | | | | |
|---|---|---|---|---|
| 1700047F07Rik | 0.364322736 | -0.443820742 | 1.758062946 | -0.252448721 |
| Gm25123 | 0 | NA | NA | NA |
| Gm12906 | 0.718440912 | -1.233903934 | 1.312903157 | -0.939828598 |
| Gm12907 | 0 | NA | NA | NA |
| 4933424M12Rik | 0.383698231 | -1.443781707 | 1.694098613 | -0.852241833 |
| Lrp8os3 | 15.16761459 | 0.343838117 | 0.331281406 | 1.037903458 |
| Lrp8 | 1123.019012 | -0.211664379 | 0.098568086 | -2.147392597 |
| Lrp8os2 | 23.3868331 | 1.377874667 | 0.242889014 | 5.672857108 |
| Lrp8os1 | 0.049553047 | -0.443818988 | 3.724945049 | -0.119147795 |
| Magoh | 160.5650247 | -0.424651557 | 0.143281096 | -2.963765411 |
| 0610037L13Rik | 286.4495153 | 0.100748823 | 0.088384239 | 1.139895799 |
| Cpt2 | 89.15985385 | -0.342743106 | 0.191530287 | -1.78949821 |
| 4930407G08Rik | 0.217432726 | 0.517969565 | 2.175998451 | 0.238037653 |
| Slc1a7 | 1.139422845 | -1.376439796 | 1.087031102 | -1.26623773 |
| Podn | 28.84068978 | 0.318161195 | 0.296153291 | 1.07431254 |
| Gm23354 | 0.487093236 | 1.493874549 | 1.438289605 | 1.038646559 |
| Scp2 | 720.4369207 | 0.03029982 | 0.084462898 | 0.358735261 |
| Mir6397 | 0 | NA | NA | NA |
| Echdc2 | 50.39875141 | 0.194712753 | 0.2129218 | 0.914480115 |
| Zyg11a | 0.650364221 | 0.955485201 | 1.216442597 | 0.785474961 |
| Zyg11b | 3195.301796 | 0.251107157 | 0.047626963 | 5.272373888 |
| Gm12742 | 0 | NA | NA | NA |
| Gm12740 | 0 | NA | NA | NA |
| Coa7 | 154.9104418 | -0.16930821 | 0.119152375 | -1.420938604 |
| Fam159a | 1.112155548 | 1.093990238 | 1.051417222 | 1.040491078 |
| Gpx7 | 31.9655395 | -0.556451403 | 0.207918155 | -2.676300213 |
| Gm12738 | 0.049553047 | -0.443818988 | 3.724945049 | -0.119147795 |
| Zcchc11 | 1174.50692 | 0.16719162 | 0.095019726 | 1.759546429 |
| Gm24967 | 6.345655539 | 0.070255908 | 0.442915911 | 0.158621323 |
| Gm12739 | 0.090277459 | -0.443823436 | 3.724945049 | -0.119148989 |
| Gm23138 | 4.286477608 | 1.03679441 | 0.624575849 | 1.659997599 |
| Gm12737 | 23.64047355 | 0.180448496 | 0.375486258 | 0.480572837 |
| Gm25827 | 0.189729909 | -0.468167298 | 2.260391221 | -0.207117818 |
| Gm22345 | 0.183160349 | -0.388000196 | 2.259866259 | -0.171691663 |
| Prpf38a | 397.5696076 | -0.095188442 | 0.093571955 | -1.017275334 |
| Orc1 | 6.089138634 | -0.299607914 | 0.378754382 | -0.791034847 |
| Gm12743 | 18.25140362 | 0.147024866 | 0.302877637 | 0.485426614 |
| Cc2d1b | 307.3753207 | -0.196397874 | 0.105890798 | -1.854720874 |
| Gm20731 | 6.967362313 | 0.465522763 | 0.441728776 | 1.053865601 |
| Gm17354 | 78.27363649 | 0.189886082 | 0.197478017 | 0.961555546 |
| Zfyve9 | 2056.889909 | 0.130612122 | 0.068846791 | 1.897141759 |
| 3110021N24Rik | 19.00627441 | 0.634228297 | 0.278849974 | 2.274442732 |
| Gm12741 | 0 | NA | NA | NA |
| Btf3l4 | 356.4128181 | 0.396742043 | 0.086910058 | 4.564972736 |
| Txndc12 | 312.2245455 | -0.43313903 | 0.120537121 | -3.593407801 |
| A730015C16Rik | 2.046195117 | -0.013355607 | 0.71202911 | -0.018757108 |
| Kti12 | 133.8561621 | -0.173228615 | 0.104551649 | -1.656871186 |

| | | | | |
|---|---|---|---|---|
| Rab3b | 288.1114516 | 0.436589625 | 0.144364545 | 3.024216403 |
| Gm12736 | 0 | NA | NA | NA |
| 8030443G20Rik | 0.272094973 | 0.08291632 | 2.077868097 | 0.039904516 |
| Gm23589 | 0.08486385 | 0.037730769 | 3.724898049 | 0.010129343 |
| Nrd1 | 1613.530245 | -0.124419854 | 0.053528912 | -2.32434865 |
| Mir761 | 0.40746677 | -0.443820229 | 1.537091689 | -0.288740244 |
| Osbpl9 | 801.6924287 | -0.217222844 | 0.069148218 | -3.141409134 |
| Gm23743 | 1.374639076 | 0.313000027 | 0.972922721 | 0.321711088 |
| Calr4 | 3.300543358 | -0.692026333 | 0.640646669 | -1.080199688 |
| Eps15 | 2866.84323 | 0.126486217 | 0.09232467 | 1.370015368 |
| Gm12749 | 6.563129853 | -0.361942287 | 0.44188899 | -0.819079669 |
| Ttc39aos1 | 29.58635856 | 0.750856839 | 0.243955933 | 3.077837991 |
| Ttc39a | 61.40979687 | 0.423682593 | 0.190237233 | 2.2271276 |
| Rnf11 | 2222.627844 | 0.072924928 | 0.084473395 | 0.863288709 |
| 4930522H14Rik | 0 | NA | NA | NA |
| Gm22497 | 0 | NA | NA | NA |
| Gm12811 | 0.075043205 | 0.517957688 | 3.724945049 | 0.139051095 |
| Gm12803 | 0 | NA | NA | NA |
| 9630013D21Rik | 4.08627299 | 0.630578286 | 0.617188352 | 1.021695053 |
| Cdkn2c | 32.18597743 | -0.323688664 | 0.281180304 | -1.151178301 |
| Faf1 | 667.4892566 | -0.153667129 | 0.071081519 | -2.161843644 |
| Gm12808 | 0 | NA | NA | NA |
| Dmrta2os | 0.681178858 | 1.23202609 | 1.262446366 | 0.975903708 |
| Dmrta2 | 37.20715882 | -0.405505807 | 0.201694103 | -2.010499075 |
| Elavl4 | 1395.565762 | 0.156671401 | 0.079599112 | 1.968255636 |
| Agbl4 | 173.4577221 | 0.578862073 | 0.173135721 | 3.343400593 |
| Gm12806 | 4.671720804 | 1.132523409 | 0.525311355 | 2.155908868 |
| Gm12807 | 1.537250626 | 0.332817551 | 0.798659804 | 0.416720046 |
| Gm12804 | 7.93214406 | 0.231884683 | 0.430585157 | 0.538533851 |
| Gm12805 | 2.823428998 | 0.405809623 | 0.608338488 | 0.667078659 |
| Bend5 | 101.6608683 | 0.604687368 | 0.171317954 | 3.529620543 |
| n-R5s191 | 0 | NA | NA | NA |
| Spata6 | 105.6804718 | -0.509780237 | 0.134237149 | -3.797609235 |
| Gm12961 | 0.049553047 | -0.443818988 | 3.724945049 | -0.119147795 |
| Gm12960 | 11.3450459 | 0.135299336 | 0.386516152 | 0.350048337 |
| Slc5a9 | 0.45131048 | -2.047475847 | 1.626101382 | -1.259131731 |
| Skint8 | 0.165933562 | 0.486011036 | 2.762106923 | 0.175956634 |
| Skint7 | 0.891009806 | 0.201784215 | 1.106064519 | 0.182434399 |
| Gm25844 | 0 | NA | NA | NA |
| Skint1 | 0.404673549 | -1.211290731 | 1.522427466 | -0.79563116 |
| Skint4 | 0 | NA | NA | NA |
| Gm12820 | 0 | NA | NA | NA |
| Gm12815 | 0 | NA | NA | NA |
| Skint3 | 0 | NA | NA | NA |
| Skint9 | 0 | NA | NA | NA |
| Gm12819 | 0 | NA | NA | NA |
| Gm12821 | 0.143485256 | -0.955563601 | 3.547405725 | -0.269369696 |

| | | | | |
|---|---|---|---|---|
| Skint2 | 0.106112162 | -0.443821212 | 3.724945049 | -0.119148392 |
| Gm12814 | 0 | NA | NA | NA |
| Gm24099 | 0 | NA | NA | NA |
| Gm12817 | 0 | NA | NA | NA |
| Skint10 | 0 | NA | NA | NA |
| Skint6 | 0 | NA | NA | NA |
| Gm12823 | 0.15269957 | -0.443823005 | 2.84335149 | -0.156091502 |
| Gm12816 | 17.15975745 | -0.264471624 | 0.270283091 | -0.978498594 |
| Gm12818 | 0.762181831 | -0.170191874 | 1.198283646 | -0.142029706 |
| Gm12813 | 0 | NA | NA | NA |
| Skint5 | 0 | NA | NA | NA |
| Gm12825 | 0 | NA | NA | NA |
| Skint11 | 0 | NA | NA | NA |
| Gm15741 | 0 | NA | NA | NA |
| Trabd2b | 136.8658561 | 0.182957024 | 0.137796125 | 1.327737077 |
| Gm28864 | 0.044544648 | 0.517957688 | 3.724945049 | 0.139051095 |
| Gm12829 | 0.792430516 | 1.334998301 | 1.194900126 | 1.117246766 |
| 9130410C08Rik | 0.223616349 | -0.828518003 | 1.862306615 | -0.444888074 |
| Gm12830 | 0.072025462 | 0.517959912 | 3.724945049 | 0.139051692 |
| Gm23230 | 0 | NA | NA | NA |
| Foxd2 | 5.895850479 | -0.669991031 | 0.448267173 | -1.494624348 |
| Foxd2os | 3.321918732 | -0.582055122 | 0.533293553 | -1.09143476 |
| Foxe3 | 0.279606198 | -1.881106017 | 3.70069412 | -0.508311672 |
| Cmpk1 | 803.5483932 | -0.056516596 | 0.099587856 | -0.567504902 |
| Gm24045 | 1.058755838 | 0.619437184 | 1.24272328 | 0.498451421 |
| Stil | 19.48329208 | 0.310135682 | 0.298317491 | 1.039616149 |
| Tal1 | 51.3179016 | -0.070933435 | 0.166258538 | -0.426645368 |
| Pdzk1ip1 | 3.920075909 | -0.829703702 | 0.541832144 | -1.531292875 |
| Cyp4x1 | 182.705986 | 0.375656067 | 0.105236546 | 3.569635097 |
| Cyp4x1os | 7.5998123 | 0.494410786 | 0.415780539 | 1.189114786 |
| Cyp4a28-ps | 0 | NA | NA | NA |
| Cyp4a29-ps1 | 0 | NA | NA | NA |
| Gm12834 | 0.117650406 | -0.443821212 | 3.724945049 | -0.119148392 |
| Cyp4a29 | 0.128150834 | 0.037055157 | 3.724945049 | 0.00994784 |
| Cyp4a12a | 0.066050875 | -0.443818988 | 3.724945049 | -0.119147795 |
| Cyp4a30-ps | 0 | NA | NA | NA |
| Cyp4a12b | 2.349144555 | -0.399541873 | 0.64305392 | -0.621319396 |
| Gm12837 | 0 | NA | NA | NA |
| Cyp4a30b | 0 | NA | NA | NA |
| Gm12838 | 0 | NA | NA | NA |
| Gm12833 | 0 | NA | NA | NA |
| Cyp4a14 | 0.301400513 | -0.901765317 | 2.357561887 | -0.382499107 |
| Cyp4a10 | 0.11147366 | -0.955981391 | 3.722845977 | -0.25678779 |
| Cyp4b1-ps1 | 0 | NA | NA | NA |
| Cyp4a31 | 0 | NA | NA | NA |
| Cyp4b1-ps2 | 0 | NA | NA | NA |
| Cyp4a32 | 0 | NA | NA | NA |

| | | | | |
|---|---|---|---|---|
| Cyp4b1 | 3.175114547 | 0.389140554 | 0.60676503 | 0.641336488 |
| Gm12848 | 0 | NA | NA | NA |
| Gm12849 | 0 | NA | NA | NA |
| Efcab14 | 1217.01994 | -0.294629335 | 0.095529385 | -3.084174937 |
| Tex38 | 1.456205946 | -0.168011052 | 0.898530274 | -0.186984298 |
| Atpaf1 | 375.0164648 | 0.487401373 | 0.073448739 | 6.635939242 |
| Gm12847 | 4.394170468 | 0.883858999 | 0.640682573 | 1.379558359 |
| Mob3c | 80.03911941 | -0.292125372 | 0.162348901 | -1.799367728 |
| Mknk1 | 268.8412447 | 0.047956713 | 0.077302104 | 0.620380435 |
| Gm29434 | 0.203831703 | 1.332845623 | 2.609621517 | 0.510742885 |
| Kncn | 2.503085161 | 0.079960293 | 0.629846415 | 0.126952048 |
| 6430628N08Rik | 3.312938502 | -1.358906884 | 0.637528419 | -2.13152362 |
| Dmbx1 | 11.05645536 | -0.310217383 | 0.34353998 | -0.903002272 |
| Faah | 605.5629304 | 0.905675375 | 0.103286735 | 8.768554568 |
| Nsun4 | 130.3320412 | -0.287794872 | 0.130975128 | -2.197324616 |
| Uqcrh | 768.3040327 | -0.061874515 | 0.150157 | -0.41206547 |
| Gm12854 | 9.36533067 | -1.240425878 | 0.415106173 | -2.988213522 |
| Lrrc41 | 643.5531935 | -0.185845161 | 0.089957662 | -2.065918087 |
| Rad54l | 35.66809566 | 0.045262769 | 0.292998731 | 0.15448111 |
| Pomgnt1 | 564.1232995 | -0.081143198 | 0.062722944 | -1.293676478 |
| Lurap1 | 633.8593911 | 0.147962129 | 0.155381304 | 0.952251818 |
| Tspan1 | 1.564275958 | -1.601349406 | 0.91195854 | -1.755945403 |
| 1700042G07Rik | 0.107078754 | -0.443164241 | 3.724898049 | -0.118973522 |
| Gm12951 | 1.402628926 | -1.359222594 | 1.051773605 | -1.292314798 |
| Llph-ps1 | 0.397229082 | 0.037076889 | 1.751476676 | 0.021168931 |
| Pik3r3 | 1226.032443 | 0.137034154 | 0.07105284 | 1.928623178 |
| Mast2 | 1537.82531 | -0.139103338 | 0.055030237 | -2.527761943 |
| Gm12950 | 0 | NA | NA | NA |
| Ipp | 171.4951214 | 0.187945821 | 0.0997042 | 1.88503415 |
| Tmem69 | 92.62743766 | -0.410786238 | 0.152282662 | -2.697524675 |
| Gm12953 | 2.041941669 | 0.659230985 | 0.723645942 | 0.910985534 |
| Gpbp1l1 | 655.1959422 | -0.01101172 | 0.068970052 | -0.159659445 |
| C530005A16Rik | 44.26883829 | -0.008428864 | 0.20553245 | -0.041009896 |
| Ccdc17 | 25.14108292 | 0.493935533 | 0.235624553 | 2.096282101 |
| Nasp | 221.6735445 | -0.230058934 | 0.130011726 | -1.769524495 |
| Akr1a1 | 2348.648558 | -0.278886494 | 0.097774096 | -2.852355643 |
| Gm25137 | 0 | NA | NA | NA |
| Prdx1 | 1494.062489 | -0.766527289 | 0.17245287 | -4.444850878 |
| Mmachc | 150.583502 | -0.216858573 | 0.109196045 | -1.985956296 |
| Ccdc163 | 28.56972285 | -0.479150907 | 0.262253756 | -1.827050697 |
| Tesk2 | 80.19330176 | -0.209752287 | 0.123568803 | -1.697453423 |
| Gm27688 | 0.669412454 | 0.471623026 | 1.24232167 | 0.379630362 |
| Toe1 | 130.6486763 | -0.286690864 | 0.10392067 | -2.758747275 |
| Mutyh | 16.21670338 | 0.21625695 | 0.273706537 | 0.790105171 |
| Hpdl | 10.50466602 | -0.321367884 | 0.358478158 | -0.896478284 |
| Gm12996 | 2.848165926 | 0.8329643 | 0.652679391 | 1.27622277 |
| Rpl36-ps8 | 0 | NA | NA | NA |

| | | | | |
|---|---|---|---|---|
| Gm12994 | 0.047948698 | 0.037064902 | 3.724945049 | 0.009950456 |
| Gm12993 | 8.790287878 | 0.585550076 | 0.37733584 | 1.551800847 |
| Zswim5 | 280.6732534 | -0.010499109 | 0.105740093 | -0.099291659 |
| Gm26355 | 2.145613104 | 0.822863288 | 0.807395141 | 1.019158088 |
| Gm25435 | 4.493034422 | -0.0541741 | 0.519328703 | -0.104315629 |
| Gm22092 | 0.088123299 | 0.037058229 | 3.724945049 | 0.009948665 |
| Gm12998 | 0.333118728 | 1.73464427 | 3.689232798 | 0.470191057 |
| Urod | 271.8023691 | -0.145421706 | 0.093497796 | -1.555349017 |
| Hectd3 | 678.4462804 | 0.109261591 | 0.056065872 | 1.94880747 |
| 1700021J08Rik | 0.083172944 | -0.443821212 | 3.724945049 | -0.119148392 |
| Gm12997 | 0 | NA | NA | NA |
| Eif2b3 | 158.1507724 | -0.012837188 | 0.087064818 | -0.147444031 |
| Gm22335 | 0.400019583 | -0.434562261 | 1.575391284 | -0.275844018 |
| Ptch2 | 34.03195546 | 0.875449277 | 0.238998587 | 3.662989337 |
| Btbd19 | 68.25176527 | 0.012127881 | 0.128351439 | 0.094489634 |
| Tctex1d4 | 5.254348095 | 0.342677129 | 0.466803652 | 0.734092648 |
| Plk3 | 168.7442851 | -0.524206904 | 0.133327822 | -3.931714304 |
| Best4-ps | 0.256651301 | 0.049328349 | 2.09727511 | 0.023520209 |
| Gm13015 | 0.494552988 | -0.323312639 | 1.354336703 | -0.238723973 |
| Rps8 | 1469.97404 | -0.246596247 | 0.112075118 | -2.200276491 |
| Gm22980 | 0.6528699 | 0.723040075 | 1.231574332 | 0.587086022 |
| Snord38a | 0 | NA | NA | NA |
| Gm26330 | 0 | NA | NA | NA |
| Snord55 | 5.259902018 | -0.000104649 | 0.426237649 | -0.000245518 |
| Gm22398 | 0 | NA | NA | NA |
| Kif2c | 10.61551923 | -0.230463247 | 0.400897497 | -0.574868262 |
| Gm25099 | 10.85160868 | -0.017405976 | 0.371596627 | -0.04684105 |
| Gm23143 | 1.628265577 | 0.800602598 | 0.734303603 | 1.09028826 |
| Gm23287 | 0.580091356 | 0.715935207 | 1.428539928 | 0.501165696 |
| Gm1661 | 2.47324152 | -0.103211302 | 0.705948431 | -0.146202325 |
| 1700012C08Rik | 0 | NA | NA | NA |
| Tmem53 | 15.28288558 | -0.248568643 | 0.262471165 | -0.947032194 |
| Rnf220 | 1260.662527 | 0.193633876 | 0.061255582 | 3.161081338 |
| Gm12843 | 2.662738786 | -0.040231064 | 0.551133297 | -0.072996977 |
| Gm12828 | 0.106520246 | 0.037071574 | 3.724945049 | 0.009952247 |
| Gm12827 | 1.000155967 | 1.04955862 | 1.147834641 | 0.914381377 |
| Gm12826 | 0.267394713 | 0.01492753 | 1.82090162 | 0.008197878 |
| Eri3 | 969.1178252 | 0.202298558 | 0.053789853 | 3.760905568 |
| Gm25559 | 3.182023478 | 2.019873009 | 0.689861677 | 2.927939145 |
| Dmap1 | 212.1899085 | -0.024765732 | 0.096365288 | -0.256998472 |
| Gm12840 | 0.644938119 | 0.662969525 | 1.134980431 | 0.58412419 |
| Gm12845 | 0.176646398 | -0.526386109 | 3.35399333 | -0.1569431 |
| Gm12844 | 4.659505038 | -0.440588736 | 0.480526736 | -0.916887038 |
| Klf17 | 0.667883529 | 0.572859904 | 1.364993289 | 0.419679649 |
| Gm12842 | 0.226488584 | 0.037076514 | 1.896341764 | 0.019551599 |
| Slc6a9 | 973.6643676 | -0.180571862 | 0.139633553 | -1.293183895 |
| Gm17114 | 0.046345709 | -0.443818988 | 3.724945049 | -0.119147795 |

| | | | | |
|---|---|---|---|---|
| Ccdc24 | 3.944649576 | -0.576243353 | 0.557138844 | -1.034290392 |
| B4galt2 | 391.215399 | 0.462578888 | 0.088746998 | 5.212332779 |
| Gm12841 | 2.986054906 | -0.148774197 | 0.67560295 | -0.220209514 |
| Atp6v0b | 1543.475499 | -0.238854776 | 0.067035912 | -3.563086828 |
| Dph2 | 90.95128011 | -0.586531449 | 0.117032248 | -5.011707974 |
| Ipo13 | 1183.676242 | 0.095969598 | 0.056121144 | 1.710043512 |
| Artn | 7.776272826 | -0.548959599 | 0.498020212 | -1.102283773 |
| St3gal3 | 444.5428053 | -0.077902077 | 0.063402342 | -1.228693989 |
| 9530034E10Rik | 0.241524581 | -0.513032745 | 2.114341686 | -0.242644199 |
| Kdm4a | 879.391759 | -0.125448333 | 0.067202287 | -1.86672714 |
| Gm23382 | 0 | NA | NA | NA |
| Ptprf | 1597.948165 | -0.270388294 | 0.04705744 | -5.745920121 |
| Mir7226 | 0 | NA | NA | NA |
| Hyi | 25.72014861 | 0.055477975 | 0.215865288 | 0.25700276 |
| Szt2 | 768.797515 | 0.12637914 | 0.123896503 | 1.020037993 |
| Med8 | 175.0860242 | -0.405038049 | 0.138379474 | -2.927009608 |
| Elovl1 | 262.3958902 | -0.762325576 | 0.136541529 | -5.583104149 |
| Cdc20 | 20.13486822 | -0.344653602 | 0.298726993 | -1.153741077 |
| Gm12858 | 0.124415284 | -0.44382566 | 3.724945049 | -0.119149586 |
| Mpl | 4.268644651 | 0.761376332 | 0.615957693 | 1.236085433 |
| Tie1 | 38.04113095 | -0.603813852 | 0.196941664 | -3.065952837 |
| Gm12859 | 0 | NA | NA | NA |
| Gm12857 | 0 | NA | NA | NA |
| 2610528J11Rik | 0.045882455 | 0.517959912 | 3.724945049 | 0.139051692 |
| Tmem125 | 85.91567833 | 0.04944599 | 0.194284451 | 0.254503072 |
| Cfap57 | 12.09672597 | -0.218626048 | 0.315125936 | -0.693773578 |
| Ebna1bp2 | 374.5410988 | -0.041773321 | 0.089547492 | -0.466493478 |
| Gm12853 | 0 | NA | NA | NA |
| Olfr62 | 0.260073237 | -1.968162266 | 3.694038657 | -0.532794171 |
| Olfr1343-ps1 | 0 | NA | NA | NA |
| Olfr1342 | 0.045882455 | 0.517959912 | 3.724945049 | 0.139051692 |
| Olfr1341 | 0.138076337 | 0.037074383 | 3.589802603 | 0.010327694 |
| Olfr1340 | 0.163066018 | 1.056936228 | 3.445567766 | 0.306752414 |
| Olfr1339 | 0.060174537 | 0.517957688 | 3.724945049 | 0.139051095 |
| Olfr1338 | 0 | NA | NA | NA |
| Olfr1337 | 0 | NA | NA | NA |
| Gm12855 | 0 | NA | NA | NA |
| Olfr1335 | 0 | NA | NA | NA |
| Olfr1333 | 0 | NA | NA | NA |
| Olfr1332-ps1 | 0 | NA | NA | NA |
| Olfr1331 | 0 | NA | NA | NA |
| Olfr1330 | 0 | NA | NA | NA |
| Olfr1329 | 0 | NA | NA | NA |
| Olfr1328 | 0.212007022 | 0.517968917 | 3.149720668 | 0.16444916 |
| Gm12856 | 0.270533821 | -1.860085064 | 2.409117009 | -0.772102416 |
| Gm12863 | 0.232395771 | 0.052946473 | 2.12208108 | 0.024950259 |
| Lao1 | 0.148650935 | 0.966592573 | 2.859657381 | 0.338009924 |

| | | | | |
|---|---|---|---|---|
| Gm12861 | 0 | NA | NA | NA |
| Gm24352 | 0 | NA | NA | NA |
| Gm12865 | 0.488817361 | -0.773355984 | 1.501735831 | -0.514974716 |
| Gm12864 | 0 | NA | NA | NA |
| Gm12862 | 0.177984204 | -0.444763424 | 3.339898761 | -0.133166738 |
| Gm12866 | 0.218085611 | -0.247095795 | 2.540840353 | -0.097249634 |
| Slc2a1 | 824.8283364 | -0.275794965 | 0.109356046 | -2.52199102 |
| Mir1957a | 1.374216805 | -0.215589213 | 0.980433612 | -0.219891699 |
| Gm12867 | 0.088534442 | -0.905414734 | 3.723500899 | -0.243162217 |
| Gm12868 | 0.130524232 | 0.517958497 | 3.657871415 | 0.141601067 |
| Zfp691 | 52.30840471 | -0.430889737 | 0.149989693 | -2.872795642 |
| Ermap | 7.770417513 | -0.81505584 | 0.391664196 | -2.081006759 |
| Ccdc23 | 99.45512681 | -0.304317273 | 0.175896889 | -1.730088998 |
| AU022252 | 374.3096476 | -0.115618252 | 0.110084441 | -1.050268785 |
| 4930538K18Rik | 9.22375745 | 0.109828862 | 0.369086633 | 0.297569331 |
| Gm12898 | 0.061920613 | -0.443816764 | 3.724945049 | -0.119147198 |
| P3h1 | 112.7927286 | -0.47403909 | 0.138570276 | -3.420929104 |
| Gm12927 | 0.131862677 | -0.443820876 | 2.977354665 | -0.149065505 |
| Cldn19 | 10.33246628 | -0.187196608 | 0.384159212 | -0.487289129 |
| Ybx1 | 865.8775829 | -0.282703969 | 0.127635087 | -2.2149393 |
| Gm37817 | 0.046345709 | -0.443818988 | 3.724945049 | -0.119147795 |
| Ppih | 73.62859532 | 0.446535994 | 0.169574094 | 2.633279548 |
| Ccdc30 | 150.4461269 | 0.222165702 | 0.202235281 | 1.098550665 |
| Ldha-ps2 | 11.09854899 | -0.29553333 | 0.341738685 | -0.864793314 |
| Ppcs | 48.56475404 | -0.292972869 | 0.15911801 | -1.841230094 |
| Zmynd12 | 4.300457922 | -0.575729236 | 0.515051632 | -1.117808778 |
| Rimkla | 301.1007889 | -0.062364354 | 0.13008862 | -0.479399 |
| Gm12954 | 0.144050924 | 1.05357957 | 3.721447096 | 0.283110183 |
| Gm12956 | 0.301349932 | 0.971853661 | 1.605923204 | 0.605168204 |
| Gm12955 | 1.414308396 | 0.750538585 | 0.920295057 | 0.815541254 |
| Gm12957 | 0.066050875 | -0.443818988 | 3.724945049 | -0.119147795 |
| Rnu6-ps2 | 0 | NA | NA | NA |
| AA415398 | 75.1633106 | 0.416995828 | 0.147886305 | 2.819705509 |
| Foxj3 | 1047.552468 | 0.055075752 | 0.076480647 | 0.720126645 |
| Gm12959 | 0 | NA | NA | NA |
| Guca2a | 0.577807171 | 1.424572162 | 1.236663782 | 1.151947832 |
| Guca2b | 0.198152624 | -1.482926123 | 3.70739678 | -0.39999121 |
| Hivep3 | 1525.201888 | -0.263328761 | 0.109154698 | -2.412436344 |
| Edn2 | 0.478833344 | 0.746501079 | 1.316727362 | 0.56693671 |
| Foxo6 | 102.9223404 | 0.272119683 | 0.147044755 | 1.850590883 |
| Foxo6os | 0.889815719 | -1.513390134 | 1.120780255 | -1.350300496 |
| Scmh1 | 665.7514625 | 0.169199078 | 0.094077839 | 1.798500893 |
| Slfnl1 | 5.927427543 | 0.718810336 | 0.540767091 | 1.329242012 |
| Ctps | 504.9997313 | -0.242513273 | 0.108809219 | -2.228793436 |
| Gm8439 | 0.147068667 | 0.998853574 | 3.51441668 | 0.284216035 |
| Gm12860 | 1.253221006 | -0.531584924 | 0.923579427 | -0.575570339 |
| Cited4 | 44.06063388 | 0.80167522 | 0.189616411 | 4.227878874 |

| | | | | |
|---|---|---|---|---|
| Kcnq4 | 121.7819068 | 0.240690815 | 0.159590512 | 1.508177467 |
| Nfyc | 366.6066488 | 0.036004497 | 0.089090546 | 0.404133754 |
| Mir30c-1 | 1.678046897 | 0.527935083 | 0.882701634 | 0.598090071 |
| mmu-mir-30e | 0 | NA | NA | NA |
| Mir30e | 2.389660156 | -0.955924826 | 0.755402344 | -1.265451231 |
| Gm23936 | 0.479339789 | 0.505499657 | 1.396895798 | 0.361873561 |
| Rims3 | 2593.580875 | 0.423146946 | 0.086628216 | 4.88463187 |
| Gm12871 | 0.804633503 | 0.625734213 | 1.05044241 | 0.595686358 |
| Gm24678 | 0.394656821 | 1.644554756 | 3.312264434 | 0.496504669 |
| Exo5 | 86.65934631 | -0.308506368 | 0.148113944 | -2.082898883 |
| Zfp69 | 17.8840471 | -0.506433179 | 0.279614093 | -1.811186177 |
| Smap2 | 1633.072005 | 0.109949702 | 0.077503855 | 1.418635271 |
| mmu-mir-7015 | 1.055430728 | -1.082401705 | 1.199786565 | -0.902161882 |
| Col9a2 | 128.0701482 | 0.393818223 | 0.140257399 | 2.807824945 |
| Zmpste24 | 545.8456237 | -0.292253709 | 0.077764102 | -3.758208509 |
| Gm26186 | 0 | NA | NA | NA |
| Gm23540 | 0.845517282 | 0.25429353 | 1.055908707 | 0.240829087 |
| Tmco2 | 0.224241997 | -0.933936084 | 2.571903794 | -0.363130256 |
| Gm12882 | 0 | NA | NA | NA |
| Gm12881 | 0.075043205 | 0.517957688 | 3.724945049 | 0.139051095 |
| Gm12879 | 0 | NA | NA | NA |
| Gm22982 | 0 | NA | NA | NA |
| Rlf | 853.6771386 | 0.200516241 | 0.116154653 | 1.726286766 |
| Gm12890 | 0 | NA | NA | NA |
| Gm12888 | 0 | NA | NA | NA |
| Gm12889 | 0 | NA | NA | NA |
| Gm12885 | 0 | NA | NA | NA |
| Gm12884 | 0 | NA | NA | NA |
| Gm12883 | 0.060174537 | 0.517957688 | 3.724945049 | 0.139051095 |
| Gm12886 | 0 | NA | NA | NA |
| Gm12880 | 0 | NA | NA | NA |
| Gm26162 | 0 | NA | NA | NA |
| Gm12887 | 0 | NA | NA | NA |
| Gm12878 | 0 | NA | NA | NA |
| Gm12892 | 61.92315877 | -0.067995311 | 0.177044938 | -0.384056794 |
| Gm12896 | 0 | NA | NA | NA |
| Gm12897 | 0 | NA | NA | NA |
| Gm12893 | 0 | NA | NA | NA |
| Gm12895 | 0 | NA | NA | NA |
| Gm12877 | 0.066050875 | -0.443818988 | 3.724945049 | -0.119147795 |
| Gm12874 | 0 | NA | NA | NA |
| Gm12875 | 0 | NA | NA | NA |
| Gm12876 | 0.038981395 | -0.443816764 | 3.724945049 | -0.119147198 |
| 9530002B09Rik | 0 | NA | NA | NA |
| Gm12872 | 0 | NA | NA | NA |
| Gm12873 | 0.136623169 | -0.870437938 | 3.632616033 | -0.23961738 |
| Ppt1 | 1536.560467 | 0.006423943 | 0.054245252 | 0.11842406 |

| | | | | |
|---|---|---|---|---|
| Cap1 | 2420.151979 | -0.456623501 | 0.065650291 | -6.955391868 |
| Gm12891 | 0 | NA | NA | NA |
| Mfsd2a | 324.2124731 | -0.358766902 | 0.151646392 | -2.365812323 |
| 4933421A08Rik | 2.139806348 | -0.411550262 | 0.658018839 | -0.625438419 |
| Mycl | 128.6415578 | 0.84123679 | 0.192226533 | 4.376278214 |
| Trit1 | 181.2275785 | -0.051761237 | 0.170851248 | -0.302960835 |
| Gm12923 | 0.164553454 | -0.925587584 | 2.779126865 | -0.333049777 |
| Bmp8b | 1.325872221 | -0.119130243 | 0.783198969 | -0.152107252 |
| Gm17244 | 11.29059836 | 0.771254339 | 0.314503072 | 2.452295091 |
| Oxct2b | 0.96273357 | 1.792722081 | 1.161910038 | 1.542909539 |
| Ppie | 115.1280594 | -0.558589179 | 0.134727422 | -4.146068947 |
| Hpcal4 | 5044.886741 | 0.355048534 | 0.163811555 | 2.167420571 |
| Nt5c1a | 53.31890087 | -0.143955094 | 0.169658705 | -0.848498133 |
| Heyl | 92.72271586 | 0.237951323 | 0.189649039 | 1.254693003 |
| Gm12901 | 0.083526043 | 0.037073798 | 3.724945049 | 0.009952844 |
| Pabpc4 | 351.2920336 | -0.432717561 | 0.104442686 | -4.143110239 |
| Gm12900 | 0 | NA | NA | NA |
| Gm25788 | 10.10737276 | 0.186940935 | 0.326650488 | 0.572296511 |
| Gm22154 | 9.014346252 | -0.472294355 | 0.364924597 | -1.294224504 |
| Bmp8a | 2.710969876 | -0.342306796 | 0.645215151 | -0.530531242 |
| Oxct2a | 0.793872245 | 1.445487166 | 1.227772851 | 1.177324588 |
| Macf1 | 11900.70528 | 0.003711414 | 0.06148728 | 0.06036068 |
| Gm12924 | 0.060174537 | 0.517957688 | 3.724945049 | 0.139051095 |
| Gm23548 | 0 | NA | NA | NA |
| Mir6398 | 0 | NA | NA | NA |
| Gm12926 | 0 | NA | NA | NA |
| Gm12925 | 2.717467516 | -1.046564688 | 0.647103255 | -1.617307098 |
| Gm12922 | 4.003874815 | 0.4059216 | 0.543475235 | 0.746899903 |
| Ndufs5 | 423.3196819 | -0.010752342 | 0.097604335 | -0.11016255 |
| Akirin1 | 729.4963505 | -0.28810952 | 0.097685377 | -2.949361796 |
| Rhbdl2 | 5.725445938 | 0.136698354 | 0.408001518 | 0.33504374 |
| 1700121C08Rik | 6.402408373 | 1.23529359 | 0.514344624 | 2.401684651 |
| Gm12904 | 0.192693611 | 0.037075631 | 2.643312858 | 0.014026199 |
| 4933427I04Rik | 0.332356096 | 1.357617741 | 3.70407012 | 0.366520529 |
| Gm26606 | 53.85320551 | 0.089141326 | 0.245406573 | 0.363239357 |
| D130007C19Rik | 26.17483582 | 0.065322446 | 0.310700285 | 0.210242632 |
| Gm12903 | 0 | NA | NA | NA |
| Mycbp | 64.1662213 | 0.062627424 | 0.138265333 | 0.452951019 |
| Gm12905 | 20.80298438 | -0.435284149 | 0.306137414 | -1.421858713 |
| Rragc | 863.0862859 | -0.218588039 | 0.087945109 | -2.485505358 |
| 4930535I16Rik | 1.505311134 | 0.722853296 | 0.830838247 | 0.870028912 |
| Gm12902 | 7.717556106 | 0.852296552 | 0.417833631 | 2.03979883 |
| Gm37667 | 0.274851136 | -0.439969788 | 3.713816915 | -0.118468357 |
| Gm12916 | 0.134481855 | 0.963696503 | 3.654633248 | 0.263691713 |
| 1700021L23Rik | 0.106971521 | 0.998850474 | 3.724945049 | 0.268151734 |
| 4933435F18Rik | 0.562550613 | -0.068614406 | 1.467980783 | -0.046740671 |
| Gm22983 | 0 | NA | NA | NA |

| | | | | |
|---|---|---|---|---|
| 1700057H15Rik | 1.747775439 | 0.617932427 | 0.838113581 | 0.7372896 |
| Gm27247 | 15.84162281 | 0.569306592 | 0.333690115 | 1.706093669 |
| 4933407E24Rik | 0 | NA | NA | NA |
| 3100002H09Rik | 4.323581818 | 0.528458208 | 0.672707514 | 0.785569059 |
| Pou3f1 | 370.5457929 | -0.098716236 | 0.129436624 | -0.76266077 |
| Utp11l | 181.9343619 | -0.546429276 | 0.086223663 | -6.337347042 |
| Gm24480 | 0 | NA | NA | NA |
| Fhl3 | 24.63805473 | -0.652229457 | 0.229664144 | -2.839927235 |
| Sf3a3 | 314.4333351 | -0.185332335 | 0.092292244 | -2.008103036 |
| Gm12917 | 6.099625106 | 0.111390671 | 0.621834449 | 0.179132357 |
| Mir697 | 1.448093177 | 0.545314128 | 0.912373899 | 0.59768712 |
| Gm12915 | 2.377012089 | 0.631040326 | 0.658292871 | 0.958601186 |
| Gm24721 | 0.117760149 | 0.51795324 | 3.724945049 | 0.139049901 |
| Inpp5b | 535.9436542 | -0.089598424 | 0.07372463 | -1.215311962 |
| Mir698 | 0.286901049 | 0.908031192 | 2.022259381 | 0.449018163 |
| Mtf1 | 507.9230992 | -0.213678158 | 0.093458222 | -2.286349495 |
| n-R5s192 | 1.205568873 | 1.219346546 | 1.105918178 | 1.102564883 |
| 1110065P20Rik | 188.1940773 | 0.055673404 | 0.113053081 | 0.492453667 |
| Yrdc | 152.5953946 | -0.228360716 | 0.194128322 | -1.176338998 |
| Maneal | 554.488513 | 0.262781898 | 0.083447783 | 3.149057865 |
| Epha10 | 353.1436211 | 0.682842046 | 0.173571143 | 3.934075876 |
| 1700125G02Rik | 0.77578221 | -0.442247493 | 1.205002353 | -0.367009651 |
| Cdca8 | 18.02129889 | -0.736209689 | 0.336287288 | -2.18922842 |
| 9930104L06Rik | 211.7189511 | -0.067183755 | 0.081483939 | -0.824503042 |
| Gm12930 | 0.650770932 | -0.348074037 | 1.321927163 | -0.26330803 |
| Rspo1 | 34.80671183 | 0.391450559 | 0.274327178 | 1.426947787 |
| Gnl2 | 487.0670719 | -0.049775766 | 0.103065307 | -0.482953647 |
| Gm12931 | 0 | NA | NA | NA |
| Dnali1 | 15.85509031 | -0.919527668 | 0.299808553 | -3.067049486 |
| Snip1 | 158.6932189 | -0.413716312 | 0.105351609 | -3.927005176 |
| Gm12932 | 3.658838888 | -0.248370477 | 0.550167591 | -0.451445124 |
| Meaf6 | 803.9924443 | 0.443529993 | 0.104185645 | 4.257112336 |
| Zc3h12a | 71.74715942 | -0.74610759 | 0.20285852 | -3.677970195 |
| 1700041M05Rik | 0 | NA | NA | NA |
| Grik3 | 1453.049993 | 0.152079959 | 0.082030135 | 1.85395233 |
| Ftl2 | 1.736617851 | -1.48827829 | 0.886102053 | -1.679578876 |
| Mir692-2 | 0.541671891 | -0.609898717 | 1.42702865 | -0.427390661 |
| 2610028E06Rik | 11.39828975 | 0.552450844 | 0.318959545 | 1.732040483 |
| Csf3r | 194.1912691 | -1.103266407 | 0.183044119 | -6.027325072 |
| Mrps15 | 201.7560872 | 0.095282543 | 0.106472316 | 0.894904394 |
| Oscp1 | 268.9501632 | 0.387682615 | 0.097591765 | 3.972493127 |
| Rpl28-ps3 | 19.78160788 | 0.133732957 | 0.253016393 | 0.528554513 |
| Lsm10 | 59.05863203 | -0.962401818 | 0.17736825 | -5.426009542 |
| Stk40 | 956.9610872 | -1.01719586 | 0.18900627 | -5.381810138 |
| Gm12946 | 0.363304151 | -1.382858393 | 1.659574137 | -0.833260993 |
| Eva1b | 26.65916579 | -0.926431666 | 0.241489548 | -3.836322004 |
| Sh3d21 | 14.96427342 | 0.062089104 | 0.280716154 | 0.221181085 |

| | | | | |
|---|---|---|---|---|
| Thrap3 | 1545.929057 | -0.019503685 | 0.059537032 | -0.327589143 |
| Gm25323 | 1.633037017 | 0.328790416 | 1.016398012 | 0.32348589 |
| Map7d1 | 3375.866482 | -0.231903929 | 0.076217678 | -3.042652766 |
| Trappc3 | 541.5073919 | -0.089269901 | 0.073696303 | -1.211321292 |
| Col8a2 | 71.84796312 | 0.493444621 | 0.223527178 | 2.207537467 |
| Adprhl2 | 208.9240687 | -0.084584599 | 0.105805994 | -0.79943107 |
| Tekt2 | 9.162665626 | 0.277498058 | 0.384975374 | 0.720820284 |
| Gm12945 | 0.40938346 | -0.074876892 | 1.415066792 | -0.052914034 |
| Ago3 | 1555.337107 | -0.065151916 | 0.104936529 | -0.620869741 |
| Ago1 | 1116.199882 | 0.055402438 | 0.07379077 | 0.750804444 |
| Gm12929 | 1.033962832 | -1.328167972 | 0.993887109 | -1.336336853 |
| Gm12928 | 17.68241393 | 0.107425097 | 0.269593345 | 0.398470878 |
| Ago4 | 190.9073592 | 0.193712931 | 0.128293521 | 1.509919824 |
| Gm12935 | 0.038981395 | -0.443816764 | 3.724945049 | -0.119147198 |
| Gm12934 | 4.064846038 | 0.585727539 | 0.667311224 | 0.877742675 |
| Clspn | 29.12900793 | -0.617086486 | 0.401811476 | -1.535761228 |
| Mir7119 | 0 | NA | NA | NA |
| Gm12936 | 0.066050875 | -0.443818988 | 3.724945049 | -0.119147795 |
| 5730409E04Rik | 892.9733379 | 0.515701585 | 0.06454433 | 7.989882077 |
| Gm12933 | 1.508740896 | -0.167293541 | 0.871251429 | -0.192015227 |
| Psmb2 | 628.3734742 | -0.214155841 | 0.108224361 | -1.978813639 |
| Tfap2e | 0.181508466 | 0.517971985 | 2.694094937 | 0.192261965 |
| Ncdn | 11330.12844 | 0.497840129 | 0.078574134 | 6.335928924 |
| AU040320 | 846.8326345 | -0.046288874 | 0.050420257 | -0.91806105 |
| Gm12937 | 0 | NA | NA | NA |
| Gm24161 | 0 | NA | NA | NA |
| Gm25604 | 3.572392357 | 0.892312471 | 0.539920183 | 1.65267478 |
| Zmym4 | 1413.511916 | -0.08353099 | 0.0458279 | -1.822710413 |
| Gm12939 | 0 | NA | NA | NA |
| Sfpq | 2483.364451 | -0.241787958 | 0.094243727 | -2.565560225 |
| Gm12940 | 393.2352481 | -0.231213499 | 0.256063163 | -0.902954943 |
| Gm12941 | 12.76876565 | -1.08473075 | 0.583052241 | -1.860434922 |
| Zmym1 | 160.873064 | -0.191633039 | 0.142783741 | -1.34212087 |
| Gm12944 | 1.541842 | -0.428863045 | 0.829679829 | -0.516901858 |
| Zmym6 | 362.4092715 | -0.084575513 | 0.119576515 | -0.707292001 |
| Gm25600 | 0 | NA | NA | NA |
| Gm12942 | 27.99737021 | -0.48067272 | 0.262045581 | -1.834309584 |
| Gm12943 | 0.338503588 | 0.037077617 | 1.601161525 | 0.0231567 |
| Dlgap3 | 2650.38369 | 0.730535495 | 0.123517982 | 5.914406029 |
| Smim12 | 221.697071 | -0.066510654 | 0.090153251 | -0.73775103 |
| Gm22221 | 0 | NA | NA | NA |
| Gm12948 | 0 | NA | NA | NA |
| Gja4 | 31.84417143 | -0.149095608 | 0.234585663 | -0.635569992 |
| Gjb3 | 2.20869273 | -1.329753423 | 0.692676606 | -1.919731968 |
| Gjb4 | 1.851903907 | -2.109260038 | 0.84484203 | -2.496632463 |
| Gjb5 | 0.85820172 | -2.018795151 | 1.295331456 | -1.558516271 |
| Gm12947 | 0 | NA | NA | NA |

| | | | | |
|---|---|---|---|---|
| 4930471C06Rik | 0.072025462 | 0.517959912 | 3.724945049 | 0.139051692 |
| 1700080G11Rik | 0.674986302 | -0.302162314 | 1.216059258 | -0.248476637 |
| A630031M04Rik | 0.919688069 | -0.659864142 | 0.960084737 | -0.687297815 |
| 1700112K13Rik | 0.099155932 | 0.037073798 | 3.724945049 | 0.009952844 |
| CK137956 | 8.83914943 | 0.042643205 | 0.357076436 | 0.119423183 |
| Csmd2 | 1037.327866 | -0.011543167 | 0.119607222 | -0.096508948 |
| Csmd2os | 0.11657011 | 0.998850474 | 3.724945049 | 0.268151734 |
| Hmgb4 | 0.309883016 | -2.192905912 | 3.687548901 | -0.594678463 |
| Hmgb4os | 3.44465028 | 0.176283182 | 0.725707475 | 0.242912176 |
| Zscan20 | 123.2244835 | -0.251340826 | 0.16132188 | -1.558008292 |
| Tlr12 | 8.189155563 | 0.943686304 | 0.534103902 | 1.766859035 |
| Gm12958 | 0 | NA | NA | NA |
| Phc2 | 1259.457039 | -0.34228857 | 0.093731484 | -3.651799331 |
| A3galt2 | 16.37679573 | -0.715781292 | 0.319632254 | -2.239390058 |
| Zfp362 | 336.5354883 | 0.111832717 | 0.093018556 | 1.202262448 |
| Gm12968 | 0.072025462 | 0.517959912 | 3.724945049 | 0.139051692 |
| Gm12969 | 0.192749185 | 0.517953186 | 3.29964367 | 0.15697246 |
| Trim62 | 671.0414608 | 0.4087465 | 0.156000507 | 2.620161362 |
| Azin2 | 295.6170531 | 0.114110127 | 0.088634923 | 1.287417234 |
| Gm15904 | 0.046345709 | -0.443818988 | 3.724945049 | -0.119147795 |
| Ak2 | 283.4084168 | -0.627255338 | 0.106198562 | -5.906439091 |
| 1700086P04Rik | 1.876204568 | 0.380335961 | 0.677418098 | 0.561449364 |
| Rnf19b | 1117.961967 | -0.527238755 | 0.161243507 | -3.26982937 |
| Tmem54 | 5.394435422 | 1.18230007 | 0.48324732 | 2.446573463 |
| Hpca | 2746.890777 | 1.391393262 | 0.156998584 | 8.862457417 |
| Gm15906 | 1.230936373 | -0.486290983 | 0.997612799 | -0.487454635 |
| Fndc5 | 973.7975932 | 0.206409542 | 0.09652084 | 2.138497153 |
| S100pbp | 678.3393157 | 0.151421344 | 0.070162028 | 2.158166588 |
| Yars | 969.2323716 | -0.288867558 | 0.084024209 | -3.437908692 |
| C77080 | 318.9680605 | 0.358663107 | 0.112379918 | 3.191523116 |
| Gm12976 | 2.076702053 | 0.423937711 | 0.796916442 | 0.5319726 |
| Gm22994 | 0 | NA | NA | NA |
| Sync | 16.23627725 | -0.266482304 | 0.271586126 | -0.981207352 |
| Rbbp4 | 1422.281671 | -0.049650265 | 0.071793586 | -0.691569645 |
| Gm26722 | 3.033581745 | -0.726571485 | 0.562506385 | -1.291667977 |
| Zbtb8os | 112.0160394 | 0.115150542 | 0.115470071 | 0.997232797 |
| Zbtb8a | 48.71610654 | 0.404314059 | 0.1753586 | 2.305641461 |
| Gm12980 | 0.075043205 | 0.517957688 | 3.724945049 | 0.139051095 |
| Zbtb8b | 38.62991341 | 0.527618675 | 0.250193402 | 2.108843283 |
| Gm24621 | 0 | NA | NA | NA |
| Bsdc1 | 654.7636716 | 0.057271518 | 0.06623533 | 0.864667211 |
| Tssk3 | 1.78535976 | 1.996636655 | 0.941863327 | 2.11987939 |
| Fam229a | 1.511769237 | -0.811338651 | 0.945582483 | -0.858030543 |
| Gm12979 | 0 | NA | NA | NA |
| Marcksl1 | 1179.090896 | -0.208577061 | 0.128214173 | -1.626786302 |
| Hdac1 | 211.8204638 | -0.448679938 | 0.113616502 | -3.949073681 |
| Lck | 123.7661852 | -0.412014648 | 0.221994732 | -1.855965875 |

| | | | | |
|---|---|---|---|---|
| Mir8119 | 0.156477978 | 0.517971164 | 2.82333336 | 0.183460859 |
| Fam167b | 2.785268692 | -1.380374413 | 0.723296838 | -1.908448012 |
| Eif3i | 596.7175796 | -0.181725496 | 0.086105641 | -2.110494674 |
| Tmem234 | 475.3810905 | -0.105112549 | 0.056858088 | -1.848682452 |
| Dcdc2b | 6.370853605 | 0.126222591 | 0.502957398 | 0.250960799 |
| Iqcc | 93.52464096 | 0.107184869 | 0.153542015 | 0.698081686 |
| Ccdc28b | 159.6210152 | 0.159586442 | 0.130707824 | 1.220940238 |
| Txlna | 633.3282999 | -0.217658004 | 0.067466145 | -3.226181119 |
| Kpna6 | 1687.26505 | -0.001528834 | 0.050985487 | -0.029985662 |
| Tmem39b | 67.54451293 | -0.175031102 | 0.174231727 | -1.004587998 |
| Mir7016 | 0.152586256 | -0.443820484 | 2.845690328 | -0.155962326 |
| Khdrbs1 | 1365.931483 | -0.063364579 | 0.062504688 | -1.013757227 |
| Gm12967 | 0 | NA | NA | NA |
| Gm28017 | 0 | NA | NA | NA |
| Ptp4a2 | 1712.805092 | -0.171300206 | 0.085277104 | -2.008747928 |
| Gm10571 | 11.18052223 | 0.251147525 | 0.377295589 | 0.665651898 |
| Gm23845 | 0.075043205 | 0.517957688 | 3.724945049 | 0.139051095 |
| Gm12966 | 92.3427904 | 0.461433179 | 0.18923278 | 2.438442111 |
| E330017L17Rik | 0.177020223 | 0.517970312 | 2.765325987 | 0.187308952 |
| Spocd1 | 2.098579804 | 1.052551511 | 0.816823447 | 1.288591206 |
| 1700003M07Rik | 24.71805353 | 0.33647651 | 0.289457841 | 1.162437018 |
| Adgrb2 | 3754.218321 | 0.455226955 | 0.129199195 | 3.523450409 |
| Gm12963 | 2.392410068 | 1.41872636 | 0.782389773 | 1.813324264 |
| Col16a1 | 419.3408002 | -0.782965171 | 0.150044158 | -5.218231628 |
| Pef1 | 447.0267585 | -0.352842467 | 0.095975876 | -3.676366218 |
| Hcrtr1 | 16.529312 | -0.956688917 | 0.32876721 | -2.909928024 |
| Tinagl1 | 47.82351247 | -0.70457571 | 0.230603753 | -3.055352316 |
| Gm853 | 1.50545653 | 1.153411425 | 0.970189148 | 1.18885212 |
| Serinc2 | 80.11973323 | -1.119492627 | 0.1756933 | -6.371857258 |
| Gm10570 | 4.998772346 | -0.486553087 | 0.46391365 | -1.048800972 |
| Fabp3 | 388.7351392 | 0.535714348 | 0.105805552 | 5.063196946 |
| Zcchc17 | 445.3923863 | 0.00953903 | 0.07614573 | 0.125273341 |
| Snrnp40 | 199.2041572 | -0.005348468 | 0.081166153 | -0.065895299 |
| Nkain1 | 233.0882025 | -0.452509733 | 0.126790787 | -3.568948047 |
| Pum1 | 1545.221209 | 0.064457723 | 0.056935018 | 1.132127905 |
| Snord85 | 3.381283184 | 0.310098054 | 0.587842701 | 0.527518762 |
| Gm12971 | 8.510907694 | 0.031152063 | 0.443519355 | 0.070238339 |
| Gm24091 | 2.986937453 | -0.85385319 | 0.66747739 | -1.279224141 |
| Gm26516 | 0.225317292 | -1.21975365 | 2.559353818 | -0.476586567 |
| Gm23711 | 1.081483968 | -1.70948198 | 1.137722521 | -1.502547368 |
| Gm12972 | 0.415245445 | 0.028936726 | 1.446605799 | 0.020003187 |
| Sdc3 | 6090.612101 | -0.207312306 | 0.12217497 | -1.696847618 |
| Gm12970 | 0.131862677 | -0.443820876 | 2.977354665 | -0.149065505 |
| Gm26716 | 0.436796958 | -1.278151478 | 1.583668496 | -0.80708272 |
| Gm22660 | 0 | NA | NA | NA |
| Laptm5 | 869.0779485 | -0.532723372 | 0.162798144 | -3.272293891 |
| Matn1 | 10.36744075 | -0.492205107 | 0.361498647 | -1.361568323 |

| | | | | |
|---|---|---|---|---|
| Gm12973 | 0.042716945 | 0.037062677 | 3.724945049 | 0.009949859 |
| A930031H19Rik | 0.159888962 | 0.51795502 | 3.461347823 | 0.149639691 |
| Gm25261 | 0 | NA | NA | NA |
| Gm831 | 0.963234904 | 0.068130996 | 0.913943549 | 0.074546175 |
| Gm12962 | 0.066050875 | -0.443818988 | 3.724945049 | -0.119147795 |
| Gm16080 | 0.066050875 | -0.443818988 | 3.724945049 | -0.119147795 |
| Ptpru | 445.2308485 | 0.482473963 | 0.091854652 | 5.252580612 |
| Mecr | 135.1425178 | -0.158081749 | 0.125551049 | -1.259103372 |
| Srsf4 | 621.5934249 | -0.0787998 | 0.075809472 | -1.039445298 |
| A930004J17Rik | 191.8201737 | 0.370273601 | 0.240869666 | 1.537236332 |
| Gm12992 | 46.74477129 | 0.102704768 | 0.20763911 | 0.494631131 |
| Gm12978 | 0 | NA | NA | NA |
| Tmem200b | 45.76488705 | -0.131848219 | 0.236267739 | -0.5580458 |
| Epb4.1 | 1119.732651 | 0.238755437 | 0.137463988 | 1.736858073 |
| Gm27445 | 1.407831039 | 0.91256732 | 0.901142663 | 1.012677966 |
| Gm13063 | 0.62973509 | -0.947635401 | 1.167374817 | -0.811766185 |
| Gm10300 | 1.165032807 | 0.517022676 | 0.960181827 | 0.538463301 |
| Gm13214 | 0 | NA | NA | NA |
| Gm37711 | 81.49722938 | 0.01963292 | 0.221869797 | 0.088488476 |
| Oprd1 | 249.9746901 | 0.16402175 | 0.155533819 | 1.054572895 |
| Gm13215 | 7.792582945 | 0.354051081 | 0.394174067 | 0.898209982 |
| Ythdf2 | 645.7985944 | -0.126992896 | 0.104645392 | -1.21355459 |
| Gm22033 | 0.373327567 | -1.277989815 | 3.698704696 | -0.345523614 |
| Rps15a-ps4 | 8.387413765 | -0.184403009 | 0.359010979 | -0.5136417 |
| Gmeb1 | 299.6588852 | -0.018652655 | 0.111189164 | -0.167756048 |
| Gm13252 | 3.851277915 | 0.002228076 | 0.56714196 | 0.003928603 |
| Gm28874 | 1.719071184 | 0.389772954 | 0.963958678 | 0.404346122 |
| Gm28872 | 0.044544648 | 0.517957688 | 3.724945049 | 0.139051095 |
| Rnu11 | 24.6768285 | -0.160431871 | 0.259351362 | -0.61858889 |
| Taf12 | 193.1512937 | -0.57648498 | 0.088413695 | -6.520313171 |
| Rab42 | 6.121133301 | -0.819769959 | 0.47347631 | -1.731385375 |
| Snhg12 | 177.2717645 | -0.250384584 | 0.158309327 | -1.581616123 |
| snopsi28S-3327 | 0 | NA | NA | NA |
| Snora16a | 0 | NA | NA | NA |
| Snora44 | 0 | NA | NA | NA |
| Snora61 | 0 | NA | NA | NA |
| Snord99 | 0 | NA | NA | NA |
| Trnau1ap | 131.2572649 | -0.200396282 | 0.108878537 | -1.840549 |
| Rcc1 | 103.1344041 | -0.422720793 | 0.131950525 | -3.203631007 |
| Snhg3 | 63.93668591 | -0.22762199 | 0.144372629 | -1.576628413 |
| Gm17300 | 0.711576293 | 0.81930442 | 1.293417781 | 0.633441438 |
| Snora73b | 425.1513821 | -0.530165716 | 0.212034347 | -2.500376578 |
| Snora73a | 0.690105543 | -0.568569808 | 1.352558923 | -0.420366017 |
| Phactr4 | 351.1978573 | -0.123097913 | 0.140809615 | -0.874215255 |
| Gm25707 | 0 | NA | NA | NA |
| Gm23055 | 0 | NA | NA | NA |
| Med18 | 61.32883924 | -0.132136195 | 0.144292553 | -0.915752008 |

| | | | | |
|---|---|---|---|---|
| Sesn2 | 307.2550336 | -0.837811019 | 0.173215553 | -4.836811746 |
| Gm12981 | 4.426758316 | -0.867499451 | 0.514733855 | -1.68533591 |
| Atpif1 | 785.2790718 | 0.389151223 | 0.132392009 | 2.939386049 |
| Gm12999 | 2.573520391 | 0.827616189 | 0.666819227 | 1.241140261 |
| Dnajc8 | 648.8436146 | 0.107656103 | 0.08831472 | 1.219005201 |
| Gm24762 | 0 | NA | NA | NA |
| Ptafr | 215.886901 | -1.419582867 | 0.350197561 | -4.053662914 |
| Eya3 | 516.9045441 | -0.440211703 | 0.073199143 | -6.013891509 |
| Xkr8 | 74.57135612 | -0.287694328 | 0.142871334 | -2.013660266 |
| Smpdl3b | 33.7182008 | 0.51844574 | 0.210888435 | 2.458388669 |
| Rpa2 | 85.55301726 | -0.087759157 | 0.158334214 | -0.554265276 |
| Themis2 | 329.3702901 | -1.292876193 | 0.301981045 | -4.281315712 |
| Gm13022 | 0 | NA | NA | NA |
| Ppp1r8 | 368.0675459 | -0.154468272 | 0.080530083 | -1.918143709 |
| Gm22767 | 11.91181268 | 0.121059845 | 0.347217395 | 0.348657201 |
| Stx12 | 1297.871004 | 0.000734796 | 0.062996047 | 0.011664159 |
| Gm13033 | 2.338964691 | 0.51980897 | 0.690685921 | 0.752598185 |
| Fam76a | 504.2948798 | 0.161891588 | 0.075085235 | 2.156104184 |
| Gm24913 | 0 | NA | NA | NA |
| Fgr | 92.45807742 | -1.548465496 | 0.317549029 | -4.876303663 |
| Ahdc1 | 1531.214203 | 0.231235157 | 0.097367547 | 2.374868867 |
| Gm26615 | 5.273376998 | 0.698869888 | 0.598860663 | 1.166999157 |
| Wasf2 | 693.9247449 | -0.327247404 | 0.11834274 | -2.765251188 |
| Gm24636 | 0 | NA | NA | NA |
| Mir7017 | 0.269321163 | 1.539303632 | 3.70868876 | 0.415053333 |
| Gpr3 | 98.5239047 | -0.678667865 | 0.258044313 | -2.63004387 |
| Cd164l2 | 13.0383265 | -0.628446251 | 0.315547837 | -1.991603738 |
| Map3k6 | 67.01031991 | 0.730271104 | 0.360876297 | 2.023605071 |
| Sytl1 | 19.69897721 | -0.061905459 | 0.29752347 | -0.208069161 |
| Tmem222 | 442.6011969 | -0.040133109 | 0.089171608 | -0.450066005 |
| Gm13259 | 0.441858849 | 0.524120781 | 1.46115245 | 0.358703693 |
| Wdtc1 | 1171.200647 | -0.288326101 | 0.089580934 | -3.21861013 |
| Slc9a1 | 963.2912325 | -0.044828398 | 0.070042864 | -0.640013782 |
| Mir5122 | 0 | NA | NA | NA |
| Gm13257 | 0.719481786 | -0.400963815 | 1.190791366 | -0.336720459 |
| Fam46b | 4.767060263 | -0.278558207 | 0.537881086 | -0.517880651 |
| Trnp1 | 753.4664742 | 0.013638873 | 0.116628167 | 0.116943219 |
| Kdf1 | 1.792464502 | -0.254999972 | 0.828547191 | -0.307767589 |
| Nudc | 578.9021612 | -0.043492257 | 0.169358489 | -0.256805886 |
| Nr0b2 | 0.783927492 | 1.708324615 | 1.181918241 | 1.44538307 |
| Gm23158 | 0.049553047 | -0.443818988 | 3.724945049 | -0.119147795 |
| Gpatch3 | 56.4655971 | -0.060941784 | 0.168091312 | -0.362551661 |
| Gpn2 | 58.10242144 | -0.541125796 | 0.192906089 | -2.805125533 |
| Sfn | 14.04179457 | -1.000873996 | 0.404910609 | -2.471839395 |
| Zdhhc18 | 588.2128307 | -0.680583643 | 0.08937044 | -7.615310457 |
| Gm13213 | 0.097899064 | 0.037062677 | 3.724945049 | 0.009949859 |
| Gm22662 | 0 | NA | NA | NA |

| | | | | |
|---|---|---|---|---|
| Pigv | 97.35604423 | -0.314502796 | 0.171259623 | -1.836409483 |
| Arid1a | 2860.653318 | 0.028889985 | 0.064605529 | 0.447175111 |
| mmu-mir-7227 | 2.041220482 | -0.115927277 | 0.883339261 | -0.131237546 |
| Gm25270 | 0 | NA | NA | NA |
| Gm12974 | 0.15596086 | 0.037059316 | 3.490591407 | 0.010616916 |
| Rps6ka1 | 359.3082904 | -0.120616806 | 0.115465341 | -1.044614823 |
| Gm12977 | 0 | NA | NA | NA |
| Gm12985 | 2.189170689 | -1.350817037 | 0.890919844 | -1.516204904 |
| Gm22523 | 2.911568235 | 0.938539878 | 0.821839898 | 1.141998435 |
| Hmgn2 | 174.4388056 | 0.018705624 | 0.168848159 | 0.110783701 |
| Dhdds | 602.5039709 | 0.1262068 | 0.061482152 | 2.052738809 |
| Lin28a | 0.305648149 | -0.885774609 | 1.734082725 | -0.510802971 |
| Gm13061 | 0 | NA | NA | NA |
| Zfp683 | 0.393365814 | -2.382334682 | 1.430731855 | -1.665116125 |
| Aim1l | 11.08135394 | -0.705072346 | 0.308927032 | -2.282326481 |
| Cd52 | 180.6648992 | -0.680104595 | 0.305684811 | -2.2248557 |
| Ubxn11 | 97.44407537 | 0.322524153 | 0.206580281 | 1.561253339 |
| Sh3bgrl3 | 569.2295487 | -0.324576157 | 0.088975406 | -3.647931165 |
| Cep85 | 164.0264212 | -0.496499738 | 0.099615686 | -4.984152178 |
| Gm27619 | 0.047948698 | 0.037064902 | 3.724945049 | 0.009950456 |
| Gm7534 | 0 | NA | NA | NA |
| Gm13131 | 1.316625395 | -0.725798665 | 0.928316965 | -0.781843586 |
| Catsper4 | 0.735041978 | 0.699477817 | 1.117664406 | 0.625838859 |
| Cnksr1 | 18.73766402 | -0.533210607 | 0.351986763 | -1.514859826 |
| Zfp593 | 102.2427118 | -0.425124229 | 0.167653699 | -2.535728304 |
| E130218I03Rik | 0.926475344 | -0.880417734 | 1.100768895 | -0.799820688 |
| Grrp1 | 25.39142091 | -1.19439776 | 0.238662354 | -5.004550315 |
| Pdik1l | 224.1146395 | 0.004185521 | 0.087353759 | 0.047914602 |
| Trim63 | 3.249610149 | -0.294713995 | 0.619087328 | -0.476045918 |
| Gm13195 | 0 | NA | NA | NA |
| Slc30a2 | 1.413987658 | -0.522289719 | 0.80974584 | -0.64500451 |
| Extl1 | 547.7075812 | 1.163286211 | 0.187505997 | 6.20399469 |
| Pafah2 | 65.33746165 | -0.62370936 | 0.210186305 | -2.967411987 |
| 1700021N21Rik | 2.462361163 | 0.314631234 | 0.647444995 | 0.485958246 |
| Stmn1 | 1361.893308 | 0.77948536 | 0.093320119 | 8.352811506 |
| Gm23634 | 0.446933609 | 1.178332556 | 1.71171642 | 0.688392389 |
| Gm13250 | 0 | NA | NA | NA |
| Paqr7 | 388.1743221 | 0.261927941 | 0.104220409 | 2.513211595 |
| Aunip | 1.169101948 | -0.051950393 | 1.043147177 | -0.049801595 |
| Mtfr1l | 639.4674001 | -0.159159414 | 0.069271552 | -2.297615815 |
| Sepn1 | 472.582935 | -0.439161279 | 0.082134939 | -5.346826601 |
| Man1c1 | 640.4685701 | -0.316229111 | 0.115105639 | -2.7472947 |
| Mir6403 | 1.273845189 | 1.21034656 | 1.015971411 | 1.191319507 |
| Ldlrap1 | 252.7861691 | -0.475880586 | 0.161584505 | -2.945087994 |
| Tmem57 | 886.6149259 | 0.273153278 | 0.07621732 | 3.583874097 |
| Rhd | 1.579581474 | -0.423132464 | 0.904266383 | -0.467929 |
| Tmem50a | 553.5327818 | 0.047454896 | 0.105065045 | 0.451671592 |

| | | | | |
|---|---|---|---|---|
| Rsrp1 | 2711.383038 | 0.015082252 | 0.091831088 | 0.16423907 |
| Syf2 | 379.4524818 | -0.151880672 | 0.134395668 | -1.130100953 |
| Gm16225 | 0 | NA | NA | NA |
| Runx3 | 97.20798678 | -0.62882078 | 0.303588409 | -2.071293766 |
| Gm16224 | 0 | NA | NA | NA |
| Clic4 | 3776.355182 | -0.840760892 | 0.232662323 | -3.613652958 |
| Gm12983 | 0.072025462 | 0.517959912 | 3.724945049 | 0.139051692 |
| Gm12984 | 0 | NA | NA | NA |
| Gm12982 | 0 | NA | NA | NA |
| Srrm1 | 1767.897914 | 0.233823711 | 0.095237827 | 2.455155884 |
| Ncmap | 11.39922601 | 1.261514105 | 0.338722942 | 3.724324366 |
| Gm12991 | 0 | NA | NA | NA |
| Gm12990 | 3.613492829 | -0.165103775 | 0.632669164 | -0.260963842 |
| Rcan3 | 591.8608699 | -0.130810504 | 0.095978965 | -1.362908055 |
| Mir700 | 0.366383937 | 1.469362674 | 1.742672109 | 0.84316646 |
| Gm15979 | 4.479156939 | 0.353505448 | 0.59399373 | 0.595133298 |
| Nipal3 | 1375.805614 | 0.309571681 | 0.065355202 | 4.736756533 |
| Gm37270 | 0.048949532 | 0.037064902 | 3.724945049 | 0.009950456 |
| Stpg1 | 13.32983798 | 0.828043865 | 0.315757205 | 2.622406876 |
| Gm25317 | 0 | NA | NA | NA |
| Grhl3 | 12.09498146 | -0.036007343 | 0.309577925 | -0.116311082 |
| Gm23106 | 0.133280846 | 0.915483429 | 3.652577767 | 0.250640366 |
| 1700029M20Rik | 0.369889 | -0.864491245 | 1.566978442 | -0.551693133 |
| Ifnlr1 | 9.250335689 | -0.412307102 | 0.32114371 | -1.283871019 |
| Gm12989 | 39.77014755 | -0.279372349 | 0.235502381 | -1.186282483 |
| Il22ra1 | 3.4240698 | 0.262181674 | 0.537873812 | 0.48744086 |
| Gm12988 | 0.337511725 | -0.483066975 | 1.583905613 | -0.304984698 |
| Myom3 | 10.58892132 | 1.61385532 | 0.415975593 | 3.879687532 |
| Gm13000 | 0 | NA | NA | NA |
| Srsf10 | 1164.791877 | -0.025230629 | 0.078020221 | -0.32338577 |
| Pnrc2 | 595.9273528 | -0.132185691 | 0.102200494 | -1.293395807 |
| Gm13006 | 0 | NA | NA | NA |
| Cnr2 | 15.77841223 | -0.624559807 | 0.390078518 | -1.601113053 |
| Fuca1 | 282.912463 | -0.206103052 | 0.093466575 | -2.205099016 |
| Hmgcl | 258.9557325 | -0.455369552 | 0.095309702 | -4.777788026 |
| Gale | 89.88904888 | -0.344347473 | 0.127645388 | -2.697688326 |
| Lypla2 | 893.641742 | -0.545122411 | 0.129826198 | -4.198862949 |
| Pithd1 | 571.8837247 | -0.139731785 | 0.080496427 | -1.73587561 |
| Gm13007 | 0.235668049 | 1.045592789 | 2.537749714 | 0.412015725 |
| Tceb3 | 686.4047522 | -0.230461612 | 0.077739167 | -2.964549535 |
| Gm13008 | 0.394976806 | 1.467266948 | 1.535887293 | 0.955322018 |
| Gm24362 | 2.378997451 | -0.288907597 | 0.899764424 | -0.321092487 |
| Rpl11 | 1068.809872 | -0.241167701 | 0.137370233 | -1.755603788 |
| Gm13009 | 0 | NA | NA | NA |
| Gm26001 | 0 | NA | NA | NA |
| Id3 | 323.3694017 | -0.528799287 | 0.160623127 | -3.292174029 |
| E2f2 | 28.44497551 | -0.358690251 | 0.271486001 | -1.32121085 |

| | | | | |
|---|---|---|---|---|
| Asap3 | 274.1575355 | -0.510478156 | 0.183658595 | -2.779495055 |
| Tcea3 | 4.626838346 | 0.192024847 | 0.489194523 | 0.3925327 |
| Zfp46 | 472.2796557 | 0.4132901 | 0.126438268 | 3.268710545 |
| Gm13013 | 0 | NA | NA | NA |
| Hnrnpr | 1542.097552 | 0.115882698 | 0.078936575 | 1.468048227 |
| Gm27022 | 0 | NA | NA | NA |
| Htr1d | 41.88636825 | -0.232852141 | 0.204164618 | -1.140511726 |
| 6030445D17Rik | 4.632584323 | -1.4581174 | 0.496722243 | -2.935478371 |
| Luzp1 | 1615.824644 | -0.030798561 | 0.061994961 | -0.496791359 |
| Kdm1a | 902.2554757 | -0.184455765 | 0.071761314 | -2.57040675 |
| Gm12987 | 8.499369334 | 0.024968454 | 0.529260427 | 0.047176122 |
| Gm12986 | 0 | NA | NA | NA |
| Gm22180 | 0 | NA | NA | NA |
| 4930549C01Rik | 0 | NA | NA | NA |
| Lactbl1 | 0.276289708 | 1.419714192 | 2.410215448 | 0.589040367 |
| Gm37583 | 0.251203948 | 0.03707547 | 2.107362209 | 0.017593307 |
| Ephb2 | 367.915314 | -0.035690202 | 0.103397496 | -0.345174722 |
| C1qb | 684.4735841 | 0.63994543 | 0.117931376 | 5.426422132 |
| C1qc | 767.5830084 | 0.599700876 | 0.150686012 | 3.979804557 |
| C1qa | 854.8487578 | 0.639734469 | 0.168195392 | 3.803519595 |
| Epha8 | 193.1729168 | 0.430808002 | 0.109146071 | 3.947077514 |
| Zbtb40 | 195.6296531 | -0.662585381 | 0.174912117 | -3.78810451 |
| Gm23834 | 0.094097695 | 0.037071574 | 3.724945049 | 0.009952247 |
| 1700037C06Rik | 0 | NA | NA | NA |
| Gm13001 | 0.262972053 | 0.940738934 | 1.799994086 | 0.522634458 |
| Gm13003 | 0.049553047 | -0.443818988 | 3.724945049 | -0.119147795 |
| Gm25399 | 0 | NA | NA | NA |
| Gm23263 | 0 | NA | NA | NA |
| Gm13005 | 0.48771898 | 0.037076702 | 1.556139257 | 0.023826082 |
| Gm13002 | 0.904140992 | -1.852467304 | 1.201661341 | -1.5415885 |
| Wnt4 | 161.8183686 | -0.242166423 | 0.141306699 | -1.713764628 |
| Cdc42 | 2845.278538 | -0.268048467 | 0.088199913 | -3.039101282 |
| Gm25772 | 0 | NA | NA | NA |
| 2810405F17Rik | 18.73872419 | 0.183656148 | 0.357456938 | 0.513785378 |
| Rpl31-ps10 | 1.205425868 | 1.111036254 | 0.896103461 | 1.239852654 |
| Cela3a | 0 | NA | NA | NA |
| Cela3b | 0 | NA | NA | NA |
| Gm13010 | 0.090890357 | 0.037071574 | 3.724945049 | 0.009952247 |
| 1700013G24Rik | 0 | NA | NA | NA |
| Hspg2 | 519.9526523 | -0.949391118 | 0.141347548 | -6.716714451 |
| Mir7018 | 0 | NA | NA | NA |
| Ldlrad2 | 0.178810541 | -0.443821193 | 2.741380009 | -0.161896998 |
| Usp48 | 1402.55254 | 0.300994841 | 0.125110392 | 2.405834056 |
| Rap1gap | 2225.616142 | 0.576160068 | 0.094739044 | 6.081548284 |
| Rap1gapos | 0.214365142 | -0.451905219 | 2.179830129 | -0.207312127 |
| Alpl | 62.50808463 | -0.367108678 | 0.173040689 | -2.121516507 |
| Ece1 | 1063.140229 | -0.51048853 | 0.089344416 | -5.713714999 |

| | | | | |
|---|---|---|---|---|
| Gm13012 | 0.199351704 | -0.760524772 | 2.615059374 | -0.290825049 |
| Eif4g3 | 3297.877854 | 0.022716218 | 0.072222202 | 0.314532337 |
| 1700095J12Rik | 0 | NA | NA | NA |
| Mir6399 | 2.27686558 | 0.697392167 | 0.768032676 | 0.908024085 |
| 2310026L22Rik | 1.951947418 | -0.839322905 | 0.776455132 | -1.080967683 |
| Hp1bp3 | 1903.997062 | 0.051072068 | 0.067662158 | 0.754809921 |
| Sh2d5 | 591.0024521 | -0.107641101 | 0.134740924 | -0.798874597 |
| Kif17 | 150.6944098 | 0.076805558 | 0.207791474 | 0.369628051 |
| Ddost | 796.5474421 | -0.423845471 | 0.086566269 | -4.896196606 |
| Pink1 | 2455.55194 | 0.259015752 | 0.095185069 | 2.721180485 |
| Mir7019 | 0 | NA | NA | NA |
| Gm27646 | 0 | NA | NA | NA |
| Cda | 12.16163641 | -0.117290642 | 0.345170371 | -0.339805071 |
| Fam43b | 205.4390261 | -0.089307829 | 0.145247453 | -0.614866749 |
| AB041806 | 16.25226339 | 0.883991105 | 0.30968376 | 2.854496161 |
| Rpl38-ps1 | 1.247502082 | 0.082185428 | 0.904402863 | 0.090872587 |
| Mul1 | 449.0477894 | -0.096247712 | 0.11535802 | -0.834339149 |
| Camk2n1 | 9074.818375 | 0.95318486 | 0.123189095 | 7.737575019 |
| Vwa5b1 | 72.56742202 | 0.480510636 | 0.193774992 | 2.479735039 |
| Gm22961 | 0.044191549 | 0.037062677 | 3.724945049 | 0.009949859 |
| Ubxn10 | 30.26165497 | -0.015908146 | 0.192653114 | -0.082574039 |
| Pla2g2c | 2.097114995 | -1.866332918 | 0.83386503 | -2.238171468 |
| Pla2g2f | 0.721011721 | 2.416460793 | 1.246795045 | 1.938137951 |
| Pla2g2d | 2.315173619 | 0.244464754 | 0.638906848 | 0.382629728 |
| Pla2g5 | 22.3998674 | 0.434876883 | 0.265772304 | 1.636276156 |
| Gm25280 | 0.126225412 | 0.03707172 | 3.698755067 | 0.010022756 |
| Pla2g2a | 0 | NA | NA | NA |
| Gm13030 | 0.123845245 | -0.280519638 | 3.720871772 | -0.075390837 |
| Pla2g2e | 6.503439099 | -1.417827057 | 0.48090542 | -2.948245117 |
| Otud3 | 118.1387416 | -0.216361435 | 0.138282551 | -1.564632936 |
| Rnf186 | 0.191999622 | 1.109115749 | 3.264119849 | 0.339790143 |
| Tmco4 | 88.13299696 | -0.811702246 | 0.200686682 | -4.044624372 |
| Htr6 | 39.32244043 | 0.563856541 | 0.20718787 | 2.721474675 |
| Nbl1 | 578.5377914 | -0.073895636 | 0.12873104 | -0.574031217 |
| Minos1 | 533.822241 | 0.010262171 | 0.101785823 | 0.100821224 |
| Hspe1-ps4 | 0 | NA | NA | NA |
| Gm16287 | 5.315121335 | 0.017528682 | 0.454865037 | 0.038536006 |
| Capzb | 1556.163037 | -0.118983242 | 0.064588326 | -1.842178744 |
| Pqlc2 | 134.0855149 | -0.130451619 | 0.12086622 | -1.079305858 |
| Akr7a5 | 131.3216444 | -0.152504274 | 0.129628268 | -1.176473897 |
| Gm25732 | 0 | NA | NA | NA |
| Gm24245 | 1.005478281 | 2.453964777 | 2.172215227 | 1.129706093 |
| Gm13019 | 0 | NA | NA | NA |
| Ccnd3-ps | 1.016942265 | -1.723023507 | 1.128065176 | -1.527414855 |
| Mrto4 | 197.6742609 | -0.121028207 | 0.106528289 | -1.136113315 |
| Emc1 | 1429.8396 | 0.001058443 | 0.071554878 | 0.01479204 |
| Ubr4 | 6917.676742 | -0.252120006 | 0.05073972 | -4.968888356 |

| | | | | |
|---|---|---|---|---|
| 2700016F22Rik | 3.508143679 | -0.612750366 | 0.597611966 | -1.025331487 |
| Iffo2 | 505.8284133 | -0.07161657 | 0.155546947 | -0.460417712 |
| Gm25571 | 0 | NA | NA | NA |
| Gm21969 | 1.469723789 | 0.533709916 | 0.967532892 | 0.551619403 |
| Aldh4a1 | 91.67888502 | 0.038321948 | 0.128890355 | 0.297322076 |
| Mir7020 | 0.163190422 | -0.955151543 | 3.396300892 | -0.281232898 |
| Tas1r2 | 0.187591875 | 0.47885132 | 2.702695179 | 0.177175481 |
| Pax7 | 25.98741359 | 0.214717447 | 0.242569017 | 0.885180844 |
| Gm1667 | 0.227050272 | 0.054033371 | 2.168712488 | 0.024914954 |
| Gm13028 | 0.115000406 | -0.443821212 | 3.724945049 | -0.119148392 |
| 4933427I22Rik | 0.61103801 | -1.04587193 | 1.458033436 | -0.717316836 |
| Klhdc7a | 224.24311 | 0.136561917 | 0.194956023 | 0.700475496 |
| Mir2139 | 0 | NA | NA | NA |
| Igsf21 | 330.4037649 | 0.507889785 | 0.087021919 | 5.836343212 |
| Gm13027 | 0.336994846 | 0.036555634 | 1.717995587 | 0.021278072 |
| Gm13029 | 0.895881185 | 0.592276585 | 1.13930679 | 0.519856978 |
| Gm9867 | 0 | NA | NA | NA |
| Gm13016 | 1.130346611 | -0.447498661 | 1.147084699 | -0.390118238 |
| Gm13017 | 0.248149725 | -1.813863362 | 2.39039049 | -0.758814666 |
| Gm13021 | 0 | NA | NA | NA |
| Gm13026 | 0.049553047 | -0.443818988 | 3.724945049 | -0.119147795 |
| Arhgef10l | 696.9675906 | 0.093466014 | 0.081813991 | 1.142420907 |
| Gm13025 | 0.470217137 | -0.085213525 | 1.367277853 | -0.062323488 |
| Gm23045 | 0 | NA | NA | NA |
| Rcc2 | 1056.188668 | -0.317278775 | 0.060483755 | -5.245685788 |
| Gm25951 | 0 | NA | NA | NA |
| Padi6 | 0.601311955 | 0.475623795 | 1.286825467 | 0.369610182 |
| Padi4 | 0.880251114 | -0.427149479 | 1.055363106 | -0.40474172 |
| Padi3 | 0.34999831 | -1.653010873 | 1.531458876 | -1.07937007 |
| Gm13032 | 0.111006857 | 0.037730769 | 3.724898049 | 0.010129343 |
| Padi1 | 3.269897503 | -0.066464032 | 0.610099167 | -0.10893972 |
| 4930515B02Rik | 0.062426873 | 0.517959912 | 3.724945049 | 0.139051692 |
| Padi2 | 199.5704136 | 0.245668075 | 0.236552763 | 1.038533948 |
| Gm26226 | 0.275809893 | 0.03707393 | 2.034285109 | 0.01822455 |
| Gm13031 | 7.163063276 | -0.031655954 | 0.484054404 | -0.065397512 |
| Sdhb | 803.5181823 | -0.083432315 | 0.072882936 | -1.144744159 |
| Atp13a2 | 1396.203774 | 0.282068751 | 0.066427475 | 4.246266362 |
| Mfap2 | 17.61408699 | 0.226413364 | 0.268769822 | 0.842406199 |
| Crocc | 384.0657555 | 0.894200002 | 0.179241191 | 4.988808648 |
| Necap2 | 429.1644651 | -0.500376908 | 0.120777191 | -4.142975227 |
| Gm13047 | 0 | NA | NA | NA |
| Spata21 | 3.445363304 | 1.202244955 | 0.656979149 | 1.829959073 |
| Szrd1 | 732.8062499 | -0.48819261 | 0.116401989 | -4.19402292 |
| 4921514A10Rik | 0.424480717 | -0.924738711 | 1.504821104 | -0.614517373 |
| Fbxo42 | 663.8042497 | -0.055871426 | 0.079793478 | -0.700200411 |
| Rsg1 | 15.89625249 | 1.052433847 | 0.317852171 | 3.311079622 |
| C630004L07Rik | 0.566706395 | -1.309375328 | 1.45420574 | -0.900405831 |

| | | | | |
|---|---|---|---|---|
| Arhgef19 | 124.7706795 | -0.109448302 | 0.204334665 | -0.535632571 |
| Gm13055 | 0.062426873 | 0.517959912 | 3.724945049 | 0.139051692 |
| Gm13056 | 0.112396584 | -0.917692504 | 3.724434845 | -0.246397787 |
| Epha2 | 56.26488158 | -0.456201267 | 0.244223722 | -1.867964603 |
| Gm25690 | 0 | NA | NA | NA |
| Gm13074 | 0 | NA | NA | NA |
| Gm13076 | 0 | NA | NA | NA |
| Fam131c | 33.32296123 | 0.192420413 | 0.219684549 | 0.875894158 |
| Clcnka | 0.814794334 | 0.586243873 | 1.061892079 | 0.552074815 |
| Gm13075 | 0.049553047 | -0.443818988 | 3.724945049 | -0.119147795 |
| Clcnkb | 0.117046891 | 0.037062677 | 3.724945049 | 0.009949859 |
| Hspb7 | 5.320519917 | -1.664648486 | 0.653215586 | -2.54839064 |
| Gm694 | 13.84823866 | -0.894625989 | 0.350672236 | -2.551174283 |
| Zbtb17 | 276.1721208 | -0.256339566 | 0.108154569 | -2.370122404 |
| Spen | 2269.995177 | 0.024570681 | 0.070718299 | 0.347444461 |
| B330016D10Rik | 28.20952649 | 0.113978075 | 0.252784954 | 0.450889474 |
| Fblim1 | 31.34586486 | -1.393789232 | 0.270125349 | -5.159786885 |
| Tmem82 | 7.023389443 | -0.22032427 | 0.379291215 | -0.580884188 |
| AI507597 | 0.761152721 | -0.148532201 | 1.273855325 | -0.116600526 |
| Slc25a34 | 67.64740712 | 0.555702649 | 0.290188286 | 1.914972715 |
| Plekhm2 | 1488.934527 | 0.010811172 | 0.0972748 | 0.111140526 |
| Ddi2 | 701.2059972 | -0.228699862 | 0.076515813 | -2.988922866 |
| Rsc1a1 | 0 | NA | NA | NA |
| Agmat | 14.27242461 | -0.474343557 | 0.353698096 | -1.341097286 |
| Dnajc16 | 741.3636435 | 0.184940375 | 0.060351054 | 3.064410042 |
| Casp9 | 369.2534085 | 0.372457035 | 0.070687797 | 5.269042941 |
| Cela2a | 0.706300188 | 0.330233387 | 1.539428706 | 0.214516844 |
| Ctrc | 0 | NA | NA | NA |
| Ctrcos | 0 | NA | NA | NA |
| Efhd2 | 2399.369846 | -0.608081299 | 0.09565883 | -6.356771263 |
| Gm29367 | 15.45718579 | 0.11749726 | 0.285769831 | 0.411160475 |
| Fhad1os2 | 0.207873529 | 0.120528095 | 2.188505336 | 0.055073247 |
| Fhad1 | 63.64227438 | -0.294403887 | 0.215767426 | -1.364450101 |
| Gm13059 | 0 | NA | NA | NA |
| Fhad1os1 | 0 | NA | NA | NA |
| 4930455G09Rik | 2.445960431 | -1.837339953 | 0.711215613 | -2.583379666 |
| Tmem51 | 80.10815602 | -0.708295174 | 0.194926784 | -3.633647261 |
| Gm13053 | 1.791442492 | -0.464400037 | 0.76968734 | -0.603361928 |
| Tmem51os1 | 3.504632306 | -0.074784793 | 0.548978702 | -0.136225309 |
| Kazn | 1222.293754 | 0.090908481 | 0.067743262 | 1.341956055 |
| Gm13062 | 0.107803068 | 0.036092186 | 3.724874622 | 0.009689503 |
| Gm13052 | 20.91063599 | 0.679863837 | 0.488302022 | 1.392301907 |
| Gm37624 | 0 | NA | NA | NA |
| Prdm2 | 1439.09533 | 0.023673003 | 0.052472971 | 0.451146605 |
| mmu-mir-7021 | 0.613445446 | -0.218770987 | 1.340935989 | -0.163147972 |
| Gm13038 | 0 | NA | NA | NA |
| Gm13039 | 0.205676178 | 0.068904158 | 2.641342059 | 0.026086798 |

| | | | | |
|---|---|---|---|---|
| Gm22039 | 0 | NA | NA | NA |
| Gm13050 | 32.91936983 | -0.276977159 | 0.19700826 | -1.405916481 |
| Pdpn | 594.7265377 | -1.397712098 | 0.252710241 | -5.530888226 |
| Gm26313 | 0 | NA | NA | NA |
| Lrrc38 | 24.86342066 | -0.463982054 | 0.220074671 | -2.108293748 |
| Anp32b-ps1 | 18.32970893 | -0.47192381 | 0.328912509 | -1.434800434 |
| Pramel1 | 0.631666884 | -0.27689681 | 1.187768541 | -0.233123542 |
| Gm13045 | 0.09176491 | 0.921395572 | 3.723061103 | 0.247483333 |
| Pramef8 | 229.9222564 | -0.301836206 | 0.102268708 | -2.95140333 |
| Oog4 | 0 | NA | NA | NA |
| Gm13042 | 1.564646527 | -1.225888508 | 0.811514181 | -1.510618713 |
| Gm13046 | 0 | NA | NA | NA |
| Gm13041 | 0 | NA | NA | NA |
| Gm13043 | 0 | NA | NA | NA |
| Gm13040 | 0 | NA | NA | NA |
| BC080695 | 0 | NA | NA | NA |
| Gm13057 | 0 | NA | NA | NA |
| Gm13058 | 0 | NA | NA | NA |
| Gm13080 | 5.317189742 | -0.511591155 | 0.498242617 | -1.026791241 |
| Gm13083 | 0 | NA | NA | NA |
| Gm13088 | 0 | NA | NA | NA |
| Gm13085 | 0 | NA | NA | NA |
| Gm13082 | 2.119392185 | 0.095489681 | 0.776656444 | 0.122949706 |
| Gm13089 | 0 | NA | NA | NA |
| Gm13078 | 0 | NA | NA | NA |
| Gm13087 | 0 | NA | NA | NA |
| Gm13081 | 0 | NA | NA | NA |
| Gm13023 | 0 | NA | NA | NA |
| Gm13084 | 0 | NA | NA | NA |
| Gm13103 | 0 | NA | NA | NA |
| Pramef6 | 0 | NA | NA | NA |
| Gm13100 | 0 | NA | NA | NA |
| Gm13106 | 0 | NA | NA | NA |
| Pramef25 | 0 | NA | NA | NA |
| Gm13101 | 0 | NA | NA | NA |
| Pramef17 | 0 | NA | NA | NA |
| Gm13108 | 0 | NA | NA | NA |
| Gm13104 | 0 | NA | NA | NA |
| Pramel4 | 0 | NA | NA | NA |
| Gm13102 | 0 | NA | NA | NA |
| Oog3 | 0 | NA | NA | NA |
| Oog2 | 0 | NA | NA | NA |
| C87977 | 0 | NA | NA | NA |
| Gm13123 | 21.87672395 | 0.131034533 | 0.23254898 | 0.563470684 |
| Pramel5 | 0 | NA | NA | NA |
| Gm13120 | 0 | NA | NA | NA |
| Gm13130 | 0 | NA | NA | NA |

| | | | | |
|---|---|---|---|---|
| Gm13128 | 0 | NA | NA | NA |
| Gm13122 | 0 | NA | NA | NA |
| Gm13119 | 0 | NA | NA | NA |
| Gm13125 | 0 | NA | NA | NA |
| Pramef12 | 0 | NA | NA | NA |
| Pramef12os | 0 | NA | NA | NA |
| 1700012P22Rik | 0.273864066 | -0.500684205 | 2.42630673 | -0.206356517 |
| Gm9944 | 0.048949532 | 0.037064902 | 3.724945049 | 0.009950456 |
| Aadacl3 | 0 | NA | NA | NA |
| Gm13118 | 0 | NA | NA | NA |
| 9430007A20Rik | 0.068097359 | 0.037064902 | 3.724945049 | 0.009950456 |
| Gm13121 | 0.042716945 | 0.037062677 | 3.724945049 | 0.009949859 |
| Gm13126 | 0.046345709 | -0.443818988 | 3.724945049 | -0.119147795 |
| Gm13124 | 0 | NA | NA | NA |
| Gm13127 | 28.94697736 | 0.010462788 | 0.220363874 | 0.047479598 |
| Gm13177 | 0 | NA | NA | NA |
| Gm436 | 0 | NA | NA | NA |
| Gm13178 | 0.072025462 | 0.517959912 | 3.724945049 | 0.139051692 |
| Gm25938 | 0 | NA | NA | NA |
| Gm438 | 0.169902121 | 0.575810967 | 3.398864629 | 0.169412739 |
| Prdx2-rs3 | 0.044191549 | 0.037062677 | 3.724945049 | 0.009949859 |
| Dhrs3 | 254.6119046 | -1.161721384 | 0.126351679 | -9.194348593 |
| Gm38074 | 1.747383901 | -0.285679063 | 0.894208837 | -0.319476896 |
| Vps13d | 3201.77473 | 0.179471062 | 0.072040347 | 2.491257607 |
| Tnfrsf1b | 457.2834194 | -1.233933511 | 0.28338853 | -4.354211196 |
| Tnfrsf8 | 21.18745374 | -0.628564588 | 0.234320879 | -2.682495014 |
| Gm13024 | 0.288426457 | -0.082776801 | 2.06538327 | -0.040078179 |
| Gm13224 | 0 | NA | NA | NA |
| Platr16 | 0 | NA | NA | NA |
| Gm13230 | 0.097899064 | 0.037062677 | 3.724945049 | 0.009949859 |
| Gm13223 | 0.093141081 | 0.037060453 | 3.724945049 | 0.009949262 |
| Smarca5-ps | 7.147813683 | 0.230253322 | 0.385989221 | 0.596527855 |
| Gm13229 | 0 | NA | NA | NA |
| Gm13225 | 0.120679735 | -0.837634339 | 3.722822199 | -0.224999824 |
| Gm21978 | 0.060174537 | 0.517957688 | 3.724945049 | 0.139051095 |
| Gm13228 | 0 | NA | NA | NA |
| Gm13226 | 9.748555663 | -0.047043514 | 0.347164789 | -0.135507734 |
| Gm13222 | 0 | NA | NA | NA |
| Gm13212 | 8.306006014 | -1.135022546 | 0.388084068 | -2.924682152 |
| Gm26763 | 0 | NA | NA | NA |
| Gm13234 | 0.712482738 | -0.688745158 | 1.180235234 | -0.583566003 |
| Gm13231 | 1.074501331 | 0.748985078 | 1.104460314 | 0.678145759 |
| Gm13249 | 0 | NA | NA | NA |
| Gm13242 | 0.150086409 | 1.027495679 | 3.723011807 | 0.275985071 |
| Gm13239 | 0 | NA | NA | NA |
| Gm13232 | 0 | NA | NA | NA |
| Gm13244 | 0.205478627 | -1.648305888 | 3.178545695 | -0.518572343 |

| | | | | |
|---|---|---|---|---|
| Gm13236 | 0 | NA | NA | NA |
| Gm13243 | 1.675672734 | -0.488300807 | 0.806771582 | -0.60525286 |
| Vmn2r-ps15 | 0.062426873 | 0.517959912 | 3.724945049 | 0.139051692 |
| Znf41-ps | 66.78666031 | -0.23848058 | 0.208490686 | -1.143842848 |
| Gm13238 | 8.42982011 | 0.030341049 | 0.392591907 | 0.077283939 |
| Gm13246 | 0 | NA | NA | NA |
| Gm13235 | 2.295964185 | 0.12882384 | 0.629827746 | 0.20453821 |
| Gm13036 | 0 | NA | NA | NA |
| Gm13034 | 8.179144896 | -0.223135088 | 0.360257709 | -0.619376302 |
| Gm13035 | 0 | NA | NA | NA |
| Gm13051 | 0 | NA | NA | NA |
| Gm26573 | 0 | NA | NA | NA |
| Zfp600 | 0.463671963 | 1.83421967 | 1.35243568 | 1.356234309 |
| Gm13165 | 0 | NA | NA | NA |
| Gm13170 | 0 | NA | NA | NA |
| Gm13167 | 0 | NA | NA | NA |
| Gm13169 | 0.687288638 | 0.516691923 | 1.337785461 | 0.386229286 |
| Gm13166 | 0.061920613 | -0.443816764 | 3.724945049 | -0.119147198 |
| Gm13168 | 0.090665643 | 0.037060453 | 3.724945049 | 0.009949262 |
| Vmn2r-ps17 | 0 | NA | NA | NA |
| Gm13251 | 42.78458666 | -0.22785764 | 0.179949874 | -1.266228392 |
| Gm13241 | 5.328295998 | -0.666933908 | 0.452768646 | -1.473012572 |
| Gm13245 | 1.259358607 | 0.250169901 | 0.91274113 | 0.274086368 |
| Gm13247 | 3.086345102 | 1.654794085 | 0.755871914 | 2.189251982 |
| Gm26880 | 0 | NA | NA | NA |
| 1700095A21Rik | 0 | NA | NA | NA |
| Gm13237 | 0.2312517 | 0.487687317 | 2.17205337 | 0.224528238 |
| Gm13233 | 0.066050875 | -0.443818988 | 3.724945049 | -0.119147795 |
| Gm13240 | 0.060174537 | 0.517957688 | 3.724945049 | 0.139051095 |
| Vmn2r-ps16 | 0.194279094 | -1.017823543 | 3.719440177 | -0.273649661 |
| Gm13248 | 6.591240675 | 0.790750804 | 0.456240672 | 1.73318788 |
| Gm21411 | 0.046345709 | -0.443818988 | 3.724945049 | -0.119147795 |
| Gm20760 | 0.333659237 | -0.265537762 | 1.717921572 | -0.1545692 |
| Gm13140 | 10.11168674 | -0.27801314 | 0.354115833 | -0.785090962 |
| Gm13149 | 0 | NA | NA | NA |
| Gm13150 | 0 | NA | NA | NA |
| C230088H06Rik | 0 | NA | NA | NA |
| Rex2 | 0.066050875 | -0.443818988 | 3.724945049 | -0.119147795 |
| Gm13153 | 0 | NA | NA | NA |
| 4933438K21Rik | 0 | NA | NA | NA |
| Gm13160 | 0.210629592 | -0.882787293 | 2.171652341 | -0.406504889 |
| Gm13142 | 2.102244023 | -0.836038965 | 0.666480957 | -1.254407881 |
| Gm13162 | 1.043892331 | -0.439120402 | 0.911855649 | -0.481567891 |
| Vmn2r-ps18 | 0.642632734 | 0.356042638 | 1.235307644 | 0.288221837 |
| Gm13139 | 85.76215223 | -0.012722794 | 0.130465552 | -0.097518416 |
| Gm26624 | 9.940525084 | -0.395639748 | 0.357464891 | -1.106793304 |
| Gm13146 | 6.463804552 | -0.290849432 | 0.427314102 | -0.680645526 |

| | | | | |
|---|---|---|---|---|
| Gm13163 | 0 | NA | NA | NA |
| Gm13147 | 1.544988629 | -0.939461665 | 0.830929458 | -1.130615428 |
| Gm13158 | 0.256692543 | 0.037077365 | 1.834077987 | 0.020215806 |
| Gm13135 | 13.09013645 | -0.144408867 | 0.362639661 | -0.398215867 |
| Gm13144 | 0 | NA | NA | NA |
| Gm13151 | 2.603243443 | -0.370612363 | 0.703616753 | -0.526724757 |
| Gm13145 | 0.352653913 | -1.548143977 | 1.954024143 | -0.792284979 |
| Gm8178 | 0 | NA | NA | NA |
| Gm13161 | 3.441294106 | 0.115924809 | 0.637292195 | 0.181902131 |
| Vmn2r-ps19 | 2.482748844 | -1.245124594 | 0.757808922 | -1.643058768 |
| Gm13152 | 33.96153438 | -0.004575777 | 0.237530469 | -0.01926396 |
| Gm15951 | 0 | NA | NA | NA |
| Vmn2r-ps20 | 4.516610571 | -0.462107467 | 0.53844202 | -0.85823069 |
| Gm13141 | 0 | NA | NA | NA |
| Gm16503 | 25.68570846 | 0.152181373 | 0.348111447 | 0.437162794 |
| Gm13154 | 0.973397315 | -0.423976048 | 0.99259655 | -0.427138345 |
| 2610305D13Rik | 21.91887679 | -0.751286858 | 0.268311395 | -2.800055726 |
| Gm13155 | 0.092140247 | 0.037060453 | 3.724945049 | 0.009949262 |
| Gm16211 | 0.547750696 | 1.072237483 | 1.466254293 | 0.731276619 |
| Zfp534 | 0.518969261 | -0.11823091 | 1.275557481 | -0.092689598 |
| Gm13137 | 0 | NA | NA | NA |
| Gm13136 | 14.06422963 | -0.586657515 | 0.272287424 | -2.154552371 |
| Gm13148 | 0.33580603 | -1.737511543 | 1.645956616 | -1.055624144 |
| Gm13157 | 104.6671989 | -0.432051261 | 0.134370006 | -3.215384699 |
| Gm20707 | 14.6133478 | 0.105601215 | 0.335442089 | 0.314812061 |
| Rps19-ps3 | 30.20379719 | 0.267025692 | 0.212500567 | 1.256588141 |
| Zfp933 | 375.230941 | -0.113898655 | 0.084415534 | -1.34926179 |
| Miip | 134.4730676 | 0.482779939 | 0.166324571 | 2.902637502 |
| Fv1 | 52.36557907 | -0.28138748 | 0.179764029 | -1.565315824 |
| Mfn2 | 2014.021368 | 0.083703207 | 0.038131625 | 2.195112535 |
| Gm13156 | 0.836555448 | -0.663689842 | 1.161267742 | -0.57152181 |
| Plod1 | 436.6411955 | -0.621495531 | 0.117699692 | -5.280349685 |
| 2510039O18Rik | 318.750318 | -0.359359363 | 0.101252965 | -3.549124332 |
| Nppb | 0.91071486 | -0.824139764 | 1.041205693 | -0.791524451 |
| Nppa | 2.986404308 | -0.512541416 | 0.776446087 | -0.660112048 |
| Gm13054 | 12.47324529 | -0.402084344 | 0.380080986 | -1.057891237 |
| Clcn6 | 1341.943323 | 0.032939264 | 0.047610689 | 0.691845985 |
| Gm13207 | 0.045882455 | 0.517959912 | 3.724945049 | 0.139051692 |
| Mthfr | 731.1208707 | -0.796936737 | 0.114045566 | -6.987880067 |
| Gm13202 | 5.417309824 | 0.188291775 | 0.441784183 | 0.426207596 |
| Gm13201 | 0.498362359 | 1.403881637 | 1.562611352 | 0.898420221 |
| Agtrap | 214.8520272 | -0.294678851 | 0.142117969 | -2.073480596 |
| Draxin | 11.61009598 | 0.064101293 | 0.335968191 | 0.19079572 |
| Mad2l2 | 105.7912503 | -0.450565767 | 0.1550522 | -2.905897284 |
| Fbxo6 | 181.4057765 | -0.632351147 | 0.113160627 | -5.588084504 |
| Mir7022 | 0.141872876 | 0.037074276 | 2.910933192 | 0.012736217 |
| Fbxo44 | 836.5623358 | 0.227174093 | 0.054875938 | 4.139776013 |

| | | | | |
|---|---|---|---|---|
| Fbxo2 | 551.7280945 | 0.198901428 | 0.116228476 | 1.711296886 |
| Ptchd2 | 379.3452522 | 0.328990202 | 0.128247954 | 2.565266669 |
| Gm13206 | 0.047948698 | 0.037064902 | 3.724945049 | 0.009950456 |
| Gm13200 | 2.555034347 | 1.750007636 | 0.759408873 | 2.304434013 |
| Ubiad1 | 175.5010009 | -0.449317093 | 0.120569935 | -3.726609733 |
| Gm23303 | 1.89929672 | -0.999246594 | 0.853775227 | -1.170386025 |
| Mtor | 2020.048309 | 0.152403032 | 0.036545693 | 4.170204984 |
| Angptl7 | 17.72480738 | -0.361113777 | 0.309107661 | -1.168245961 |
| Gm24002 | 0.589151384 | -0.94447352 | 1.374200706 | -0.687289357 |
| Gm23318 | 1.238646735 | -0.674703124 | 0.964562357 | -0.699491452 |
| Exosc10 | 484.0039437 | -0.096926266 | 0.080567645 | -1.203042063 |
| Gm13204 | 0 | NA | NA | NA |
| Gm13209 | 0.331673312 | -1.676160953 | 1.68476255 | -0.994894476 |
| Srm | 303.9037999 | -0.370726585 | 0.09114236 | -4.067555235 |
| Masp2 | 31.81998135 | -0.953174828 | 0.306312268 | -3.111774908 |
| Tardbp | 2054.279046 | -0.018375094 | 0.07461859 | -0.246253565 |
| Gm572 | 0.786597823 | 0.435423711 | 1.06387683 | 0.40928019 |
| Gm13203 | 0 | NA | NA | NA |
| Casz1 | 60.05755845 | -0.15901657 | 0.177425991 | -0.896241691 |
| Gm15969 | 0 | NA | NA | NA |
| Gm13205 | 27.32691937 | 0.530139874 | 0.24144334 | 2.195711314 |
| Gm17029 | 1.365296087 | -0.795263979 | 0.882720938 | -0.900923435 |
| Pex14 | 521.3504447 | 0.089962244 | 0.087909414 | 1.023351647 |
| Gm9506 | 0 | NA | NA | NA |
| Dffa | 268.8330888 | -0.091225696 | 0.076792225 | -1.18795485 |
| Cort | 13.59908539 | 1.079651949 | 0.333583262 | 3.236529143 |
| Apitd1 | 26.0334167 | 0.17909406 | 0.221379742 | 0.808990282 |
| Rpsa-ps12 | 20.84511033 | -0.249516049 | 0.340875617 | -0.731985616 |
| Pgd | 591.4610587 | -0.568047418 | 0.098008723 | -5.795886307 |
| Kif1b | 15537.60889 | 0.244747004 | 0.050691539 | 4.828162865 |
| Ube4b | 1942.055481 | 0.101053361 | 0.046011673 | 2.19625486 |
| Ube4bos3 | 0.106056992 | 0.998850474 | 3.724945049 | 0.268151734 |
| Ube4bos2 | 0.038981395 | -0.443816764 | 3.724945049 | -0.119147198 |
| Ube4bos1 | 0.415914162 | -1.085803747 | 3.337904263 | -0.325295054 |
| Rbp7 | 1.09427696 | 0.000216481 | 0.937895229 | 0.000230816 |
| Trmt112-ps2 | 31.55425196 | -0.417766047 | 0.231357348 | -1.805717655 |
| Nmnat1 | 53.5119218 | -0.300435873 | 0.182025817 | -1.65051243 |
| Gm13097 | 0.04393175 | 0.037062677 | 3.724945049 | 0.009949859 |
| Lzic | 141.3885089 | -0.495361421 | 0.115585683 | -4.28566417 |
| Gm13071 | 9.973492174 | 0.539959634 | 0.327091194 | 1.650792332 |
| Gm24381 | 0 | NA | NA | NA |
| Ctnnbip1 | 373.0545822 | 0.13751966 | 0.100449839 | 1.369038132 |
| Mir5616 | 0.908927132 | 0.017169745 | 1.038470479 | 0.016533686 |
| Gm13066 | 0.3242121 | -0.343248644 | 1.78314124 | -0.19249661 |
| Clstn1 | 10198.28505 | 0.643840132 | 0.081977665 | 7.853848149 |
| Pik3cd | 275.4671803 | -0.655559587 | 0.11698432 | -5.603824377 |
| Mir7023 | 0 | NA | NA | NA |

| | | | | |
|---|---|---|---|---|
| Tmem201 | 968.9539549 | 0.185721932 | 0.09050013 | 2.052173088 |
| Slc25a33 | 270.7812472 | -0.017412454 | 0.120867622 | -0.14406219 |
| Gm13068 | 0.185281328 | 0.037067357 | 2.71551244 | 0.013650226 |
| Gm16188 | 0.247276735 | -1.908410802 | 1.996545594 | -0.955856359 |
| Gm13073 | 0.234101041 | 0.037547443 | 1.87816433 | 0.019991564 |
| Gm13065 | 0.044544648 | 0.517957688 | 3.724945049 | 0.139051095 |
| Gm13064 | 0.150086409 | 1.027495679 | 3.723011807 | 0.275985071 |
| n-R5s193 | 0 | NA | NA | NA |
| AL626808.1 | 0 | NA | NA | NA |
| Spsb1 | 673.0771038 | -0.25526458 | 0.184036697 | -1.387030871 |
| Gm13070 | 0.062426873 | 0.517959912 | 3.724945049 | 0.139051692 |
| H6pd | 247.2321755 | -0.508921908 | 0.137699717 | -3.695882027 |
| Gm13067 | 8.956120355 | -0.196238929 | 0.414671651 | -0.473239318 |
| Mir34a | 0.164553454 | -0.925587584 | 2.779126865 | -0.333049777 |
| Gm37795 | 24.19206811 | 0.101120569 | 0.341438806 | 0.296160153 |
| Gpr157 | 52.57716929 | -0.059457235 | 0.184137996 | -0.322894982 |
| Gm13093 | 6.364908752 | 0.694018895 | 0.509404499 | 1.362412181 |
| Slc2a5 | 12.42258672 | 0.202324214 | 0.306222781 | 0.660709218 |
| Slc2a7 | 0 | NA | NA | NA |
| Car6 | 1.391048348 | -3.743002226 | 1.184193744 | -3.16080223 |
| Gm13094 | 0.173900533 | -0.499200409 | 2.73824775 | -0.182306517 |
| Eno1 | 2763.278328 | -0.304372786 | 0.066768133 | -4.558653556 |
| Rere | 3154.94565 | 0.246069015 | 0.061171746 | 4.022592644 |
| Gm13092 | 5.881560967 | 0.071425149 | 0.617773106 | 0.115617123 |
| Gm23209 | 6.850924464 | 0.463053625 | 0.584897933 | 0.791682785 |
| Gm13091 | 33.11693011 | 1.121879217 | 0.239715294 | 4.680048559 |
| Slc45a1 | 569.9271968 | 0.123051878 | 0.064637348 | 1.903727209 |
| Gm16079 | 0 | NA | NA | NA |
| Gm13049 | 0 | NA | NA | NA |
| 1700045H11Rik | 0.109982625 | -0.443823436 | 3.724945049 | -0.119148989 |
| Rps15a-ps3 | 6.113901122 | -0.25932872 | 0.396370786 | -0.654257905 |
| Errfi1 | 790.4060845 | 0.038648937 | 0.235654705 | 0.164006643 |
| Park7 | 548.339284 | 0.007382731 | 0.102704381 | 0.071883308 |
| Tnfrsf9 | 42.91515327 | -1.310852772 | 0.520110492 | -2.520335184 |
| Camta1 | 4644.702699 | 0.329351371 | 0.053871788 | 6.113614983 |
| Gm23405 | 0 | NA | NA | NA |
| Uts2 | 1.009670171 | -0.634413566 | 1.00636575 | -0.630400593 |
| Per3 | 705.3281863 | 1.381827581 | 0.238508237 | 5.793626235 |
| Vamp3 | 420.1066725 | -0.427386009 | 0.119002213 | -3.591412291 |
| Gm13090 | 0.329009797 | -1.062093277 | 1.519302054 | -0.699066571 |
| 4930589P08Rik | 0.062426873 | 0.517959912 | 3.724945049 | 0.139051692 |
| Gm37141 | 0.038981395 | -0.443816764 | 3.724945049 | -0.119147198 |
| Dnajc11 | 693.2058939 | -0.086614238 | 0.062944935 | -1.376031896 |
| Thap3 | 106.1287797 | -0.033228182 | 0.129258659 | -0.257067359 |
| Phf13 | 200.4894639 | -0.467649749 | 0.165536977 | -2.82504705 |
| Klhl21 | 246.8014045 | -0.511448983 | 0.112958505 | -4.527759838 |
| Zbtb48 | 85.35332904 | -0.17498154 | 0.161772809 | -1.081649884 |

| | | | | |
|---|---|---|---|---|
| Tas1r1 | 4.276683327 | -0.44443654 | 0.589290828 | -0.754188795 |
| Nol9 | 273.9583489 | -0.368349765 | 0.083107613 | -4.432202458 |
| Plekhg5 | 606.8905356 | 0.492460514 | 0.166444509 | 2.958706889 |
| Tnfrsf25 | 29.08316862 | -1.232277565 | 0.629658221 | -1.957057851 |
| Espn | 20.79395242 | 0.961181072 | 0.225283618 | 4.266537793 |
| Gm13077 | 2.185424001 | -0.139737562 | 0.629229815 | -0.222077147 |
| Hes2 | 0.293168959 | -0.496848055 | 2.04437427 | -0.243031847 |
| Acot7 | 1614.713023 | -0.019119511 | 0.073412553 | -0.260439267 |
| Gm27500 | 0 | NA | NA | NA |
| Gm13096 | 2.546102757 | -0.002632456 | 0.852514021 | -0.003087875 |
| Gpr153 | 338.048994 | -0.044059736 | 0.09556131 | -0.461062493 |
| Hes3 | 57.0044881 | -0.073767726 | 0.26568696 | -0.277649025 |
| Icmt | 1103.865666 | 0.034330614 | 0.118840738 | 0.288879171 |
| Rnf207 | 32.84951904 | 1.150560588 | 0.273438083 | 4.207755465 |
| Rpl22 | 661.306753 | -0.19935176 | 0.117822893 | -1.691961165 |
| Chd5 | 2858.157954 | 0.449951139 | 0.095890016 | 4.692366929 |
| Gm13645 | 1.241205481 | 1.694242271 | 1.197122128 | 1.41526268 |
| Kcnab2 | 3430.815311 | 0.684969823 | 0.080184809 | 8.542388958 |
| Gm16333 | 0.211164147 | -0.411420197 | 2.556588203 | -0.160925485 |
| Gm16334 | 14.13596111 | 0.646204081 | 0.368596965 | 1.753145421 |
| Nphp4 | 205.6488101 | 0.293155001 | 0.194706663 | 1.505623877 |
| Gm13172 | 0 | NA | NA | NA |
| Gm833 | 0.678447637 | -0.344334907 | 1.250455182 | -0.275367652 |
| Gm25779 | 0 | NA | NA | NA |
| Gm13173 | 0.184770788 | 0.57041104 | 2.68592176 | 0.212370683 |
| Gm13174 | 0.2360389 | -0.805269119 | 2.132224089 | -0.37766627 |
| Ajap1 | 418.755463 | 0.329125956 | 0.116756676 | 2.8189048 |
| Gm25880 | 0 | NA | NA | NA |
| Gm13115 | 0.044191549 | 0.037062677 | 3.724945049 | 0.009949859 |
| BC039966 | 5.183547717 | 0.781967103 | 0.522400644 | 1.496872395 |
| A430005L14Rik | 129.8028141 | -0.428514402 | 0.122342169 | -3.502589552 |
| Dffb | 64.42929542 | 0.685660807 | 0.203875561 | 3.363133879 |
| Cep104 | 594.1324898 | -0.232617691 | 0.070980677 | -3.277197426 |
| Lrrc47 | 577.7083269 | 0.0238386 | 0.096094306 | 0.248075057 |
| Smim1 | 121.5585543 | -0.006326243 | 0.147439207 | -0.042907466 |
| Ccdc27 | 0.478716937 | 0.04333089 | 1.551366911 | 0.027930781 |
| Gm13117 | 0 | NA | NA | NA |
| Gm13114 | 0 | NA | NA | NA |
| Trp73 | 20.9206457 | 0.692373873 | 0.313195161 | 2.210678703 |
| Trp73os | 0.141071297 | -0.443820801 | 3.564043909 | -0.12452731 |
| Gm26036 | 0 | NA | NA | NA |
| Wrap73 | 158.2566248 | -0.037251087 | 0.101067922 | -0.368574781 |
| Tprgl | 931.6745304 | 0.272124984 | 0.090447859 | 3.008639298 |
| Megf6 | 28.54359773 | -0.195599981 | 0.212372583 | -0.921022754 |
| Gm13132 | 0 | NA | NA | NA |
| Arhgef16 | 13.58180626 | -0.37515462 | 0.314024104 | -1.194668229 |
| Prdm16 | 164.5019658 | -0.294128834 | 0.127852716 | -2.300528638 |

| | | | | |
|---|---|---|---|---|
| B230104I21Rik | 0.989733885 | 1.849541446 | 1.295069356 | 1.428140846 |
| Gm13133 | 0.742042093 | -2.607107566 | 1.271848756 | -2.049856599 |
| Gm13134 | 0.593269415 | -1.372545736 | 1.224147576 | -1.121225711 |
| Gm13111 | 52.71521894 | 0.362509823 | 0.287954412 | 1.258913937 |
| Gm27202 | 0 | NA | NA | NA |
| Gm13110 | 1.937464243 | -0.242923497 | 0.818416834 | -0.296821237 |
| Actrt2 | 0 | NA | NA | NA |
| Gm13113 | 1.711017467 | 0.349213895 | 0.736090847 | 0.474416842 |
| Ttc34 | 32.00110063 | 0.355880933 | 0.192688642 | 1.846922216 |
| Gm13112 | 3.984283232 | 0.732697933 | 0.570154007 | 1.285087755 |
| Mmel1 | 12.70547386 | 0.091469416 | 0.325799899 | 0.280753361 |
| Fam213b | 258.5981968 | 0.431384579 | 0.136869058 | 3.151804996 |
| Tnfrsf14 | 53.00596033 | -0.67787196 | 0.205210652 | -3.303298124 |
| Gm20421 | 6.460871859 | 0.146357994 | 0.479859857 | 0.305001536 |
| Hes5 | 56.4435291 | 0.495571221 | 0.359237867 | 1.379507191 |
| Pank4 | 318.1852538 | 0.038408496 | 0.074675921 | 0.514335758 |
| Plch2 | 799.0502262 | 0.508221603 | 0.149583932 | 3.397568153 |
| Gm10564 | 2.918309484 | 2.706367967 | 0.821147819 | 3.295835296 |
| Pex10 | 129.4440069 | 0.086529992 | 0.121359094 | 0.713007893 |
| Rer1 | 559.9984354 | -0.471477514 | 0.089756824 | -5.252831961 |
| Morn1 | 92.30872075 | 0.523284358 | 0.235452773 | 2.222459952 |
| Ski | 2357.896731 | 0.018185886 | 0.090433265 | 0.201097302 |
| 2610002J02Rik | 142.9826346 | -0.553199147 | 0.118468633 | -4.669583257 |
| Prkcz | 1482.735323 | 0.336609856 | 0.107495051 | 3.131398623 |
| Gm27200 | 1.590510877 | 1.254333227 | 0.824270844 | 1.521748873 |
| Gabrd | 409.0842723 | 0.654804188 | 0.16413964 | 3.989311724 |
| 5830444B04Rik | 0.822034208 | -1.24493214 | 1.168415787 | -1.065487264 |
| Cfap74 | 161.3175935 | 0.36087629 | 0.188080165 | 1.918736562 |
| Tmem52 | 14.34551441 | 0.31438588 | 0.304982344 | 1.03083305 |
| Gnb1 | 6791.840275 | 0.167158546 | 0.056790295 | 2.94343505 |
| Gm13171 | 0.045882455 | 0.517959912 | 3.724945049 | 0.139051692 |
| Nadk | 615.6859532 | -0.377635484 | 0.093127535 | -4.055035764 |
| Slc35e2 | 942.3227332 | 0.059718418 | 0.060974129 | 0.979405833 |
| Gm16023 | 7.870775316 | -1.403801028 | 0.493577955 | -2.844132347 |
| Gm10563 | 0.110242424 | -0.443823436 | 3.724945049 | -0.119148989 |
| Gm16024 | 3.554695965 | 0.921510049 | 0.679448197 | 1.35626241 |
| Cdk11b | 500.6534106 | -0.234404687 | 0.131506198 | -1.782461143 |
| Mmp23 | 18.56308676 | 0.06422193 | 0.272947648 | 0.235290285 |
| Mib2 | 880.1110826 | 0.011752668 | 0.06795055 | 0.172959136 |
| AL670236.1 | 0 | NA | NA | NA |
| B930041F14Rik | 0 | NA | NA | NA |
| Ssu72 | 463.6570633 | -0.243395579 | 0.079709356 | -3.053538398 |
| Gm22573 | 0.048949532 | 0.037064902 | 3.724945049 | 0.009950456 |
| 1500002C15Rik | 18.92769097 | 0.06777206 | 0.266907136 | 0.253916254 |
| Tmem240 | 359.2263579 | 0.184329352 | 0.110904088 | 1.66206093 |
| Atad3a | 323.5868663 | -0.462824265 | 0.105578744 | -4.383687927 |
| Atad3aos | 2.955322402 | -0.970230836 | 0.723518987 | -1.340988769 |

| | | | | |
|---|---|---|---|---|
| Vwa1 | 254.228511 | -0.592074139 | 0.110350817 | -5.365380656 |
| Gm26840 | 23.01521584 | -0.556707909 | 0.232908497 | -2.39024302 |
| Tmem88b | 1074.722227 | -0.321489703 | 0.138017696 | -2.329336832 |
| Gm27743 | 0 | NA | NA | NA |
| Ankrd65 | 0.81397646 | 0.372177768 | 1.2104837 | 0.307462023 |
| Mrpl20 | 295.0369231 | -0.163688835 | 0.138286495 | -1.183693576 |
| Ccnl2 | 1143.506995 | 0.072367009 | 0.132844533 | 0.54474962 |
| Aurkaip1 | 435.2140168 | -0.176044013 | 0.112566546 | -1.563910582 |
| Mxra8os | 0.102891481 | 0.51795324 | 3.724945049 | 0.139049901 |
| Mxra8 | 155.5730168 | -0.065894456 | 0.177900801 | -0.370399994 |
| Dvl1 | 770.5694415 | -0.134270333 | 0.076471426 | -1.755823568 |
| Tas1r3 | 8.894659495 | 0.06280413 | 0.364890525 | 0.172117733 |
| Cptp | 215.1678806 | -0.149648244 | 0.101287539 | -1.477459575 |
| Cpsf3l | 232.6024494 | -0.52754832 | 0.075874467 | -6.952909718 |
| Pusl1 | 121.0895384 | 0.309011544 | 0.137085478 | 2.254152292 |
| Acap3 | 1132.056167 | 0.537502635 | 0.107782555 | 4.986916789 |
| Gm22991 | 0 | NA | NA | NA |
| Ube2j2 | 313.5792266 | -0.182641082 | 0.097099714 | -1.880964154 |
| Fam132a | 40.80633962 | 0.81125035 | 0.256160805 | 3.166957373 |
| B3galt6 | 79.13773638 | 0.077403994 | 0.167009677 | 0.463470118 |
| Sdf4 | 2638.964062 | 0.139936597 | 0.055329111 | 2.529167624 |
| Tnfrsf4 | 12.95165872 | 0.294722319 | 0.433170296 | 0.680384418 |
| Gm37090 | 21.13329923 | 0.204154311 | 0.319490679 | 0.638999271 |
| Gm16008 | 35.8777959 | 0.060351511 | 0.226017794 | 0.267021061 |
| Gm25982 | 1.203677616 | -1.051911582 | 1.095493753 | -0.960216869 |
| Tnfrsf18 | 8.73841548 | -0.759122183 | 0.521992133 | -1.454278972 |
| Ttll10 | 1.729677859 | 0.15767516 | 0.797756629 | 0.197648197 |
| Gm13648 | 0.840567365 | -0.307300972 | 1.309594041 | -0.234653612 |
| Mir429 | 0 | NA | NA | NA |
| Mir200a | 0.075043205 | 0.517957688 | 3.724945049 | 0.139051095 |
| Mir200b | 0.044191549 | 0.037062677 | 3.724945049 | 0.009949859 |
| 9430015G10Rik | 166.5177331 | 0.209214261 | 0.14914373 | 1.402769399 |
| Agrn | 2086.461589 | 0.14888737 | 0.094166865 | 1.581101472 |
| Isg15 | 2298.739811 | -0.909899313 | 0.349950218 | -2.600082144 |
| AW011738 | 181.0853581 | -0.45342138 | 0.20410498 | -2.221510619 |
| Perm1 | 7.861063657 | -0.233334405 | 0.365814008 | -0.637849837 |
| Plekhn1 | 36.96362493 | -0.821859791 | 0.19630957 | -4.186549808 |
| Klhl17 | 254.2446718 | 0.217726052 | 0.122816901 | 1.772769469 |
| Mir7658 | 0 | NA | NA | NA |
| Noc2l | 659.9224954 | -0.122384868 | 0.073986299 | -1.654155845 |
| Samd11 | 37.37872384 | 0.042661227 | 0.219595322 | 0.194272023 |
| Vmn2r-ps159 | 0.208220181 | 0.517962109 | 3.175804864 | 0.163096327 |
| Gm20782 | 0 | NA | NA | NA |
| Vmn1r-ps5 | 0 | NA | NA | NA |
| Gm3027 | 0 | NA | NA | NA |
| RP24-481N17.5 | 0 | NA | NA | NA |
| RP24-481N17.7 | 0 | NA | NA | NA |

| | | | | |
|---|---|---|---|---|
| RP24-481N17.8 | 0 | NA | NA | NA |
| Gm6486 | 0 | NA | NA | NA |
| Gm3053 | 0 | NA | NA | NA |
| RP23-128P4.1 | 0 | NA | NA | NA |
| Gm8715 | 0 | NA | NA | NA |
| RP23-135F23.1 | 0 | NA | NA | NA |
| Gm15772 | 876.5883103 | -0.230650136 | 0.165147859 | -1.396628076 |
| RP23-135F23.2 | 0 | NA | NA | NA |
| Cdk6 | 185.3456827 | -0.575069658 | 0.168100408 | -3.420989072 |
| RP23-454P8.1 | 0.082913145 | -0.443821212 | 3.724945049 | -0.119148392 |
| RP23-454P8.3 | 0.295949093 | -0.508527699 | 2.040368845 | -0.249233221 |
| Fam133b | 414.4885661 | 0.124703893 | 0.091322675 | 1.365530443 |
| RP23-434H17.4 | 7.609880518 | 0.369643565 | 0.429055313 | 0.861528932 |
| n-R5s169 | 0 | NA | NA | NA |
| 1700109H08Rik | 123.3221518 | 1.013193325 | 0.378351777 | 2.677913482 |
| Rbm48 | 131.5051051 | 0.204667948 | 0.136108787 | 1.503708553 |
| RP23-434H17.3 | 18.64070939 | 0.55053375 | 0.323123041 | 1.703789828 |
| Pex1 | 339.3891102 | 0.043559484 | 0.133335512 | 0.32669079 |
| RP23-434H17.5 | 1.254627161 | 0.474141535 | 0.845495661 | 0.560785296 |
| RP23-434H17.6 | 46.52130311 | 0.259537894 | 0.16559448 | 1.567310053 |
| Gatad1 | 720.1271276 | -0.132198802 | 0.086274546 | -1.532303658 |
| Tmbim7 | 0.149576918 | -0.443202057 | 2.863264737 | -0.15478906 |
| Gm26238 | 0.120349074 | 1.027495679 | 3.723011807 | 0.275985071 |
| Ankib1 | 1655.656652 | -0.034398878 | 0.059833163 | -0.574913243 |
| Gm22926 | 0 | NA | NA | NA |
| RP23-425I8.2 | 106.4878435 | 0.355580021 | 0.248855993 | 1.428858581 |
| Krit1 | 1289.563298 | -0.0038463 | 0.107480592 | -0.035785991 |
| Lrrd1 | 7.878715828 | -0.624846458 | 0.361301639 | -1.729431561 |
| Rps4x-ps | 7.918913376 | -0.458146559 | 0.358970015 | -1.276280859 |
| Mterf1a | 31.30236157 | -0.03644366 | 0.244988872 | -0.148756388 |
| Akap9 | 2881.578872 | 0.106910639 | 0.100144691 | 1.067561721 |
| Wdr46-ps | 30.12730886 | 0.793993407 | 0.282561467 | 2.809984727 |
| RP23-369O21.1 | 30.72591236 | 0.129829414 | 0.273355196 | 0.474947672 |
| Cyp51 | 835.8806558 | -0.222698826 | 0.13791935 | -1.61470328 |
| Mterf1b | 11.84616759 | 0.134605712 | 0.359594664 | 0.374326221 |
| Fzd1 | 479.4036875 | -0.061601342 | 0.140046622 | -0.439863105 |
| Gm15750 | 1.582330525 | 0.554701605 | 0.898970852 | 0.617040701 |
| RP23-114F4.1 | 8.620768254 | 1.186285194 | 0.469249279 | 2.528049048 |
| AC026231.1 | 0 | NA | NA | NA |
| Cdk14 | 2684.227082 | 0.800692963 | 0.08249482 | 9.705978649 |
| RP23-114F4.2 | 25.16791745 | 1.121023672 | 0.367720658 | 3.048574091 |
| RP23-114F4.3 | 17.31984568 | 0.738662023 | 0.319958479 | 2.308618375 |
| Gm25037 | 1.520499152 | -0.220392431 | 0.857518327 | -0.25701192 |
| RP23-114F4.4 | 36.06826517 | 0.820178227 | 0.418029187 | 1.962011871 |
| RP23-282L6.2 | 14.6407167 | 0.914702367 | 0.349458236 | 2.617486934 |
| RP23-282L6.1 | 20.90691181 | 0.620088428 | 0.325910179 | 1.902635964 |
| RP23-407F12.1 | 44.77564562 | 0.37600182 | 0.252817037 | 1.487248743 |

| | | | | |
|---|---|---|---|---|
| RP23-396K2.1 | 12.0291979 | -0.658930966 | 0.356098408 | -1.850418175 |
| RP23-396K2.2 | 12.35285954 | -0.376534176 | 0.365745027 | -1.029499099 |
| 1700015F17Rik | 1.819296732 | 0.06091102 | 0.712985071 | 0.08543099 |
| Cldn12 | 558.7465676 | -0.239970667 | 0.077892308 | -3.080800557 |
| Gtpbp10 | 131.1217465 | 0.129714126 | 0.160770581 | 0.8068275 |
| RP23-119M19.2 | 0 | NA | NA | NA |
| Gm8773 | 3.281282651 | -0.312500904 | 0.512919348 | -0.60925934 |
| Cfap69 | 276.8459991 | -0.055141436 | 0.119109345 | -0.462948022 |
| Steap2 | 544.5451271 | -0.181335548 | 0.112963351 | -1.605259998 |
| RP23-261D18.1 | 0.350305046 | 1.799501296 | 1.735587647 | 1.036825366 |
| Steap1 | 8.981601178 | 0.113461034 | 0.353049796 | 0.321374024 |
| Gm26825 | 3.449447184 | -0.331506411 | 0.607266303 | -0.545899566 |
| Gm15459 | 4780.466862 | -0.544633259 | 0.104257046 | -5.223946782 |
| RP23-261D18.2 | 0 | NA | NA | NA |
| Gm23838 | 0 | NA | NA | NA |
| RP24-323L24.1 | 0.269679299 | 0.037077302 | 1.645340052 | 0.022534735 |
| RP24-323L24.2 | 0 | NA | NA | NA |
| Gm24686 | 0 | NA | NA | NA |
| RP24-338C23.1 | 0.161949631 | -1.309098506 | 2.76260123 | -0.473864448 |
| RP24-338C23.2 | 0.322473324 | 0.437357251 | 1.754330445 | 0.249301523 |
| RP23-189N11.1 | 0.187072314 | 0.517965296 | 2.668356435 | 0.194113983 |
| RP23-74J16.1 | 1.533852754 | 2.078264662 | 1.031135186 | 2.015511342 |
| Zfp804b | 64.00592618 | 0.600413171 | 0.150109167 | 3.999843472 |
| Gm24525 | 5.266732454 | -0.085855194 | 0.559401285 | -0.153476934 |
| Tubb4b-ps1 | 53.69871365 | -0.251156902 | 0.186553508 | -1.34629954 |
| RP24-72E12.3 | 27.34862605 | 0.834553305 | 0.34985933 | 2.385396743 |
| RP24-72E12.2 | 24.54249589 | 0.193563779 | 0.308336029 | 0.627768932 |
| RP24-72E12.4 | 0 | NA | NA | NA |
| 4921511H03Rik | 0 | NA | NA | NA |
| Gm4960 | 0.287827876 | 1.009397469 | 2.034751743 | 0.496078931 |
| RP23-465J5.1 | 0.130017973 | -0.444798045 | 3.662714201 | -0.121439463 |
| Gm8802 | 0.060174537 | 0.517957688 | 3.724945049 | 0.139051095 |
| Gm22196 | 0 | NA | NA | NA |
| RP23-212F6.1 | 0 | NA | NA | NA |
| Steap4 | 41.56055496 | 0.163741732 | 0.624563627 | 0.262169817 |
| Gm15731 | 0.761917678 | 0.166656951 | 1.064243181 | 0.156596682 |
| Sri | 562.266319 | -0.209292477 | 0.089390998 | -2.341314909 |
| Gm15730 | 0.062426873 | 0.517959912 | 3.724945049 | 0.139051692 |
| Adam22 | 4541.002357 | 0.640743126 | 0.092367262 | 6.936907188 |
| Gm21759 | 0.083172944 | -0.443821212 | 3.724945049 | -0.119148392 |
| Gm24196 | 4.833961456 | 0.892398644 | 0.58109097 | 1.53572967 |
| Gm23993 | 0 | NA | NA | NA |
| RP23-201E13.1 | 54.61168029 | 0.547619961 | 0.290825734 | 1.882983163 |
| Dbf4 | 47.80200343 | -0.310982686 | 0.252613946 | -1.231059056 |
| Slc25a40 | 206.1233218 | -0.05902624 | 0.101143042 | -0.583591703 |
| Rundc3b | 435.3356137 | 0.321228561 | 0.096721983 | 3.321153601 |
| RP24-159A20.4 | 29.56068002 | 0.619572169 | 0.299929656 | 2.065724937 |

| | | | | |
|---|---|---|---|---|
| Abcb1a | 172.8364366 | -0.286237234 | 0.159761665 | -1.791651552 |
| RP24-223N5.1 | 5.526269997 | -0.225478472 | 0.529771617 | -0.425614481 |
| RP23-162P5.1 | 1.143206505 | 0.749163076 | 0.958460865 | 0.781631367 |
| Gm5106 | 1.599390384 | 0.403525066 | 0.800175117 | 0.504295944 |
| Abcb1b | 71.1266548 | -0.117719656 | 0.156488842 | -0.75225591 |
| RP23-338P12.2 | 0.602373977 | 0.182282222 | 1.285612038 | 0.141786337 |
| Abcb4 | 48.46738024 | 1.052940415 | 0.194469617 | 5.414421179 |
| RP23-338P12.3 | 11.71418499 | -1.111679687 | 0.333744014 | -3.330935208 |
| Crot | 573.9541497 | -0.129453211 | 0.121448728 | -1.06590833 |
| Gm15610 | 11.65246704 | -0.331881963 | 0.452776845 | -0.732992348 |
| Gm37153 | 0.09689823 | 0.037062677 | 3.724945049 | 0.009949859 |
| Gm27941 | 0 | NA | NA | NA |
| Gm15611 | 0.120349074 | 1.027495679 | 3.723011807 | 0.275985071 |
| Tmem243 | 183.1811946 | -0.226393125 | 0.124306182 | -1.821253958 |
| Dmtf1 | 881.7559076 | -0.215978939 | 0.10035505 | -2.152148189 |
| Gm24666 | 1.171812973 | -0.739110815 | 0.988412623 | -0.747775572 |
| Gm7332 | 10.24110484 | -0.587815495 | 0.361016644 | -1.6282227 |
| Gm15734 | 0.410151891 | 0.457454159 | 1.562174814 | 0.292831606 |
| 9330182L06Rik | 795.1077193 | 0.272546563 | 0.05818951 | 4.683774805 |
| Gm15733 | 3.241155436 | -0.017003311 | 0.693390256 | -0.024521994 |
| Mir879 | 0.599062483 | 1.098363972 | 1.285284656 | 0.854568649 |
| Grm3 | 803.8407714 | 1.013282446 | 0.130423047 | 7.769197767 |
| RP24-118C7.3 | 12.97903413 | 1.100995846 | 0.403169809 | 2.730848939 |
| Gm23312 | 0 | NA | NA | NA |
| RP23-193N17.1 | 0.49396662 | 1.194971263 | 1.378365152 | 0.866948255 |
| RP23-193C5.2 | 0.302344664 | -0.8744243 | 3.702087462 | -0.236197634 |
| Gm6586 | 0.313880683 | -0.069463139 | 1.767752006 | -0.039294618 |
| RP23-193C5.3 | 0.217113707 | 0.010424806 | 2.548098894 | 0.004091209 |
| Gm10482 | 0.386285301 | 0.257293099 | 1.534445274 | 0.167678251 |
| Gm28685 | 0.419836731 | -1.487709781 | 1.366928689 | -1.088359468 |
| Gm5152 | 0.268081508 | -0.882098281 | 2.017854759 | -0.437146567 |
| RP23-412L2.1 | 0.048949532 | 0.037064902 | 3.724945049 | 0.009950456 |
| Gm17091 | 0.90048758 | 2.2167482 | 1.114726093 | 1.98860349 |
| RP24-383C4.1 | 0.834920584 | 2.272767627 | 1.316474786 | 1.726404221 |
| RP23-281K10.3 | 0 | NA | NA | NA |
| RP23-281K10.1 | 0 | NA | NA | NA |
| RP23-281K10.2 | 0 | NA | NA | NA |
| Gm6455 | 0 | NA | NA | NA |
| RP24-113C21.2 | 0 | NA | NA | NA |
| RP23-307D8.3 | 0 | NA | NA | NA |
| Gm8871 | 0 | NA | NA | NA |
| Gm8857 | 0 | NA | NA | NA |
| RP24-79B1.2 | 0 | NA | NA | NA |
| RP23-307D8.4 | 0 | NA | NA | NA |
| RP24-79B1.3 | 0 | NA | NA | NA |
| RP23-307D8.5 | 0 | NA | NA | NA |
| Gm8879 | 0 | NA | NA | NA |

| | | | | |
|---|---|---|---|---|
| RP23-92D22.3 | 0 | NA | NA | NA |
| RP23-307D8.7 | 0 | NA | NA | NA |
| Gm5861 | 0 | NA | NA | NA |
| RP23-307D8.9 | 0 | NA | NA | NA |
| Gm8890 | 0 | NA | NA | NA |
| RP24-79B1.4 | 0 | NA | NA | NA |
| RP23-307D8.8 | 0 | NA | NA | NA |
| Speer1 | 0 | NA | NA | NA |
| RP24-390F5.2 | 0 | NA | NA | NA |
| RP23-307D8.10 | 0 | NA | NA | NA |
| Gm8897 | 0 | NA | NA | NA |
| RP23-299O17.3 | 0 | NA | NA | NA |
| RP23-299O17.4 | 0.739845808 | -0.065146527 | 1.3789238 | -0.047244472 |
| RP23-307D8.11 | 0 | NA | NA | NA |
| Gm8906 | 0 | NA | NA | NA |
| RP23-299O17.5 | 0 | NA | NA | NA |
| RP23-307D8.12 | 0 | NA | NA | NA |
| Gm6460 | 0 | NA | NA | NA |
| RP23-395E24.4 | 0 | NA | NA | NA |
| RP23-307D8.6 | 0 | NA | NA | NA |
| Gm8922 | 0 | NA | NA | NA |
| 1700003C15Rik | 0 | NA | NA | NA |
| RP23-395E24.3 | 0 | NA | NA | NA |
| Gm8926 | 0 | NA | NA | NA |
| RP23-408K8.3 | 0 | NA | NA | NA |
| RP23-307D8.13 | 0 | NA | NA | NA |
| Gm6465 | 0 | NA | NA | NA |
| RP24-341O14.4 | 0 | NA | NA | NA |
| RP23-307D8.14 | 0 | NA | NA | NA |
| 4933402N22Rik | 0 | NA | NA | NA |
| RP24-341O14.3 | 0.201351563 | -0.443821124 | 2.219042163 | -0.20000572 |
| Gm29066 | 0 | NA | NA | NA |
| Gm38242 | 0 | NA | NA | NA |
| RP24-235L14.3 | 0.152964131 | 0.037074629 | 2.843489623 | 0.013038426 |
| RP23-364E18.2 | 0.046345709 | -0.443818988 | 3.724945049 | -0.119147795 |
| Sema3d | 247.8894151 | -0.189683819 | 0.152323187 | -1.245272122 |
| RP23-237H9.3 | 0 | NA | NA | NA |
| RP23-237H9.2 | 4.98564735 | -0.028093801 | 0.465716216 | -0.060323862 |
| RP24-346C14.1 | 0 | NA | NA | NA |
| Gm23243 | 0 | NA | NA | NA |
| Gm10481 | 11.05583305 | -0.010743175 | 0.318629361 | -0.03371684 |
| RP23-30B3.2 | 0 | NA | NA | NA |
| Gm10108 | 1.450945253 | -0.801361423 | 0.753967951 | -1.062858735 |
| Sema3a | 401.2948366 | 0.380105664 | 0.092666534 | 4.101865546 |
| RP23-30B3.4 | 0 | NA | NA | NA |
| RP23-10F17.2 | 15.6478416 | 0.526125366 | 0.378780527 | 1.388997924 |
| RP23-10F17.3 | 7.940230631 | -0.071555017 | 0.400160699 | -0.178815704 |

| | | | | |
|---|---|---|---|---|
| RP23-10F17.4 | 8.41488845 | 0.45731668 | 0.481502274 | 0.94977055 |
| RP23-192D21.1 | 16.18937152 | -0.042598063 | 0.272909267 | -0.156088737 |
| RP23-192D21.2 | 0 | NA | NA | NA |
| RP24-405C17.3 | 0 | NA | NA | NA |
| RP24-405C17.4 | 0.101408268 | 0.037730769 | 3.724898049 | 0.010129343 |
| Gm6647 | 0.217530652 | 0.013610755 | 2.172555185 | 0.00626486 |
| Speer3 | 0 | NA | NA | NA |
| Gm6650 | 0 | NA | NA | NA |
| RP24-405C17.5 | 0 | NA | NA | NA |
| RP24-405C17.6 | 0 | NA | NA | NA |
| RP24-405C17.7 | 0 | NA | NA | NA |
| Gm24054 | 0 | NA | NA | NA |
| Sema3e | 550.5002642 | -0.411418959 | 0.116446339 | -3.533120593 |
| RP24-194J1.1 | 17.56400965 | -0.374168525 | 0.314870892 | -1.188323643 |
| Gm4128 | 0 | NA | NA | NA |
| RP24-481P22.1 | 12.20307049 | 0.092710236 | 0.317878886 | 0.291652703 |
| Pclo | 8266.158298 | 0.460678776 | 0.108217178 | 4.256983818 |
| Gm26918 | 1.104625153 | 0.546750814 | 1.152138176 | 0.474553162 |
| RP23-342M4.2 | 153.5905569 | 0.349302392 | 0.239439031 | 1.458836473 |
| RP23-342M4.3 | 100.8300785 | 0.374960901 | 0.238746112 | 1.57054244 |
| RP23-342M4.4 | 73.09923858 | 0.132546991 | 0.204331114 | 0.64868726 |
| RP23-342M4.5 | 112.0484156 | 0.331529826 | 0.263220935 | 1.25951162 |
| RP23-342M4.6 | 56.2336579 | 0.476904553 | 0.291231889 | 1.637542355 |
| RP23-342M4.7 | 8.648288359 | 0.512721088 | 0.398404019 | 1.28693754 |
| Gm26954 | 0.454340154 | -0.418934777 | 1.382664373 | -0.302990939 |
| RP23-451C1.2 | 88.75190297 | 0.527398529 | 0.21904104 | 2.407761257 |
| RP23-451C1.1 | 51.00169416 | 0.335479165 | 0.272978244 | 1.22895935 |
| Gm9758 | 0.061920613 | -0.443816764 | 3.724945049 | -0.119147198 |
| Speer4e | 0.072025462 | 0.517959912 | 3.724945049 | 0.139051692 |
| RP23-239L21.4 | 0.368800866 | 1.455864774 | 1.593854885 | 0.913423667 |
| Gm10354 | 0.066050875 | -0.443818988 | 3.724945049 | -0.119147795 |
| Gm17019 | 0.062426873 | 0.517959912 | 3.724945049 | 0.139051692 |
| RP24-152F23.2 | 0.163328881 | -0.387020395 | 2.769442679 | -0.139746671 |
| Gm8579 | 0.295956696 | -1.483381249 | 1.677164301 | -0.88445792 |
| Gm21847 | 0.231556444 | -0.882505508 | 2.111331605 | -0.417985269 |
| RP24-152F23.3 | 0.112396584 | -0.917692504 | 3.724434845 | -0.246397787 |
| RP23-465P18.1 | 0.409112724 | -1.00977862 | 1.687251656 | -0.598475406 |
| Gm21655 | 0 | NA | NA | NA |
| Speer4d | 0 | NA | NA | NA |
| 4930572O03Rik | 0 | NA | NA | NA |
| Speer4cos | 0.174149299 | -0.429800891 | 3.359371215 | -0.127940874 |
| Speer4c | 0.220272213 | -0.817476051 | 2.175258299 | -0.375806428 |
| RP23-77E18.1 | 0 | NA | NA | NA |
| RP23-77E18.2 | 0 | NA | NA | NA |
| RP23-77E18.3 | 0 | NA | NA | NA |
| RP23-77E18.5 | 0.109124069 | 0.517955464 | 3.724945049 | 0.139050498 |
| RP23-77E18.6 | 0 | NA | NA | NA |

| | | | | |
|---|---|---|---|---|
| Gm3510 | 30.33973269 | 0.618094937 | 0.236226286 | 2.616537503 |
| Cacna2d1 | 2942.12003 | 0.573721403 | 0.098772419 | 5.808518293 |
| RP23-123P10.5 | 0.04393175 | 0.037062677 | 3.724945049 | 0.009949859 |
| RP24-220P21.4 | 77.60211899 | 0.240813507 | 0.26178379 | 0.919894646 |
| RP24-372A2.2 | 0 | NA | NA | NA |
| Gm8984 | 0 | NA | NA | NA |
| Hgf | 116.1833121 | 0.32789548 | 0.158495832 | 2.068795594 |
| RP24-295A5.2 | 4.076536194 | 0.15196435 | 0.550568774 | 0.276013383 |
| RP24-295A5.3 | 3.112827446 | -0.104386478 | 0.666708413 | -0.156569913 |
| Gm8991 | 9.192051872 | -0.330219316 | 0.353414029 | -0.934369573 |
| Gm28710 | 0.299775621 | -0.369111044 | 1.753317299 | -0.210521532 |
| RP24-363D4.1 | 1.258311994 | -0.280682511 | 0.891529317 | -0.31483262 |
| Gm6673 | 0 | NA | NA | NA |
| Gm3495 | 0 | NA | NA | NA |
| Gm21999 | 0 | NA | NA | NA |
| Speer4f | 0 | NA | NA | NA |
| RP23-76L18.2 | 0 | NA | NA | NA |
| Sema3c | 248.8209663 | 0.351074946 | 0.128561837 | 2.730786645 |
| Cd36 | 46.54322315 | -0.732825242 | 0.536426485 | -1.366124274 |
| Gnat3 | 0.097139547 | -0.443816764 | 3.724945049 | -0.119147198 |
| Gnai1 | 1646.477261 | 0.375440867 | 0.075662088 | 4.96207385 |
| RP24-302O2.1 | 0 | NA | NA | NA |
| RP24-459D3.1 | 0.038981395 | -0.443816764 | 3.724945049 | -0.119147198 |
| RP24-332E15.1 | 0 | NA | NA | NA |
| Gm23822 | 0.060174537 | 0.517957688 | 3.724945049 | 0.139051095 |
| RP23-350E1.2 | 51.82168262 | 0.260656955 | 0.160242052 | 1.626645146 |
| 4921504A21Rik | 48.57949567 | 0.224982714 | 0.19872595 | 1.132125495 |
| RP23-350E1.3 | 0 | NA | NA | NA |
| Magi2 | 1985.599496 | 0.482638876 | 0.107055226 | 4.508316826 |
| RP24-388K6.6 | 0.090665643 | 0.037060453 | 3.724945049 | 0.009949262 |
| Gm25335 | 0 | NA | NA | NA |
| Gm23570 | 4.591749267 | 1.020504053 | 0.615177279 | 1.658877997 |
| Gm25761 | 0 | NA | NA | NA |
| RP24-531D1.4 | 0.048949532 | 0.037064902 | 3.724945049 | 0.009950456 |
| Gm22755 | 8.57913668 | 0.220839058 | 0.389955716 | 0.5663183 |
| RP23-281M20.1 | 6.048354155 | 1.701247001 | 0.534141183 | 3.185013728 |
| Gm29254 | 8.172606336 | 0.811566251 | 0.448852742 | 1.808090213 |
| RP23-343F16.2 | 0.689035189 | -0.298124058 | 1.224701217 | -0.243425951 |
| Phtf2 | 298.2117683 | 0.011802745 | 0.108044325 | 0.109239838 |
| Rpl31-ps5 | 0.299266739 | 0.036526131 | 1.767544483 | 0.0206649 |
| Tmem60 | 129.5434812 | 0.011229798 | 0.140171619 | 0.080114638 |
| Rsbn1l | 755.8552143 | 0.039055664 | 0.07024897 | 0.55596067 |
| A630072M18Rik | 116.1077269 | 0.259275852 | 0.132740994 | 1.953246274 |
| Ptpn12 | 569.2757863 | -0.637314496 | 0.09155817 | -6.960760546 |
| RP23-234P16.4 | 0.061920613 | -0.443816764 | 3.724945049 | -0.119147198 |
| Gm22490 | 0 | NA | NA | NA |
| Rpl31-ps21 | 0 | NA | NA | NA |

| | | | | |
|---|---|---|---|---|
| RP23-234P16.2 | 0.792202851 | -0.41310818 | 1.316292864 | -0.313842148 |
| RP23-234P16.3 | 0.272741579 | 0.990107606 | 1.795751344 | 0.551361194 |
| Gsap | 232.3497055 | -0.676715568 | 0.144655535 | -4.678117343 |
| RP23-250D22.2 | 3.212737209 | -0.446851906 | 0.557422287 | -0.801639827 |
| RP23-250D22.3 | 0 | NA | NA | NA |
| Ccdc146 | 10.59368012 | -0.504014697 | 0.368625793 | -1.367280061 |
| Fgl2 | 737.5148379 | -1.240724496 | 0.259106911 | -4.788465469 |
| Fam185a | 169.3008059 | 0.093413192 | 0.132970242 | 0.702512009 |
| Fbxl13 | 5.258800805 | 0.493105108 | 0.513717734 | 0.959875581 |
| Lrrc17 | 1.924082696 | 0.548376786 | 0.772014498 | 0.710319284 |
| RP23-6C10.3 | 0.285998089 | 0.075386796 | 2.095067783 | 0.035982987 |
| RP23-6C10.4 | 1.597009952 | -0.102969483 | 0.780455226 | -0.131935157 |
| Armc10 | 162.4313593 | 0.070457259 | 0.097582822 | 0.722025221 |
| Napepld | 190.2960083 | 0.358636102 | 0.115183292 | 3.11361219 |
| Gm15719 | 0.801310233 | 0.784204367 | 1.132665099 | 0.692353254 |
| Pmpcb | 597.1185698 | -0.087690418 | 0.076071316 | -1.1527396 |
| Gm15715 | 0.120974994 | 0.517957688 | 3.724945049 | 0.139051095 |
| Dnajc2 | 320.2722277 | -0.214150504 | 0.124372223 | -1.721851541 |
| Psmc2 | 881.0786388 | -0.280251853 | 0.084447182 | -3.318664377 |
| Slc26a5 | 5.142851575 | 0.96639114 | 0.504017667 | 1.917375528 |
| Reln | 2935.282946 | 0.229728637 | 0.128263719 | 1.791064841 |
| Gm16111 | 0 | NA | NA | NA |
| Gm10475 | 1.275826341 | 0.120784179 | 0.948204227 | 0.12738203 |
| Dnaja1-ps | 7.533972655 | 0.824666762 | 0.503038624 | 1.639370663 |
| RP23-205D3.1 | 11.79089672 | 0.423560532 | 0.435836206 | 0.971834205 |
| Gm16113 | 6.656687351 | -0.327664383 | 0.495104981 | -0.661807892 |
| Gm16112 | 0 | NA | NA | NA |
| Gm16110 | 0 | NA | NA | NA |
| Gm25459 | 0 | NA | NA | NA |
| RP23-143M9.2 | 0.121578509 | 0.037073798 | 3.724945049 | 0.009952844 |
| n-R5s170 | 0 | NA | NA | NA |
| RP24-86E4.1 | 1.834562507 | -0.584304896 | 0.865671466 | -0.674973034 |
| Orc5 | 252.3162393 | -0.024688066 | 0.099769076 | -0.247452088 |
| Gm15421 | 12.14808016 | -0.42614143 | 0.316625406 | -1.345885145 |
| Gm22887 | 3.183676125 | -0.251299319 | 0.680161014 | -0.369470337 |
| 6030443J06Rik | 71.11363277 | 0.087126591 | 0.232332498 | 0.375008197 |
| RP23-258C13.1 | 3.322405445 | -0.009082366 | 0.601318433 | -0.015104088 |
| Rpl17-ps5 | 55.4808862 | -0.047249917 | 0.164322015 | -0.287544656 |
| Gm24512 | 0 | NA | NA | NA |
| RP24-324L19.2 | 13.10249877 | -0.303165711 | 0.379214592 | -0.799456871 |
| RP24-324L19.3 | 161.9541949 | -0.16172362 | 0.116657342 | -1.386313263 |
| RP24-324L19.4 | 19.8545456 | -0.497774778 | 0.228682026 | -2.176711422 |
| AC164538.1 | 0.163066018 | 1.056936228 | 3.445567766 | 0.306752414 |
| Lhfpl3 | 375.6236598 | 0.008829942 | 0.099963829 | 0.088331367 |
| Gm21846 | 71.52268729 | 0.278280289 | 0.321930165 | 0.864411971 |
| RP23-211E12.6 | 0.049553047 | -0.443818988 | 3.724945049 | -0.119147795 |
| Gm6564 | 0 | NA | NA | NA |

| | | | | |
|---|---|---|---|---|
| RP23-240A11.2 | 0.172588697 | -0.43782042 | 2.317759933 | -0.188898088 |
| Gm28022 | 0 | NA | NA | NA |
| RP23-443N7.4 | 0.181654796 | -1.405197014 | 3.311020957 | -0.424399915 |
| RP23-443N7.5 | 0 | NA | NA | NA |
| 5031425E22Rik | 328.5648936 | -0.410590308 | 0.167025241 | -2.458253041 |
| RP23-443N7.6 | 0 | NA | NA | NA |
| RP23-443N7.10 | 6.257156534 | -0.424028202 | 0.577171716 | -0.734665595 |
| RP23-443N7.7 | 0.066050875 | -0.443818988 | 3.724945049 | -0.119147795 |
| Kmt2e | 2575.672994 | 0.159028908 | 0.078669382 | 2.021484149 |
| Gm25219 | 2.780058051 | 1.914543545 | 0.698426814 | 2.741222856 |
| Srpk2 | 2903.573116 | 0.332934531 | 0.047326857 | 7.034790596 |
| Gm24009 | 0.699061148 | -0.318318296 | 1.436722862 | -0.221558593 |
| RP23-110C17.1 | 13.01527202 | -0.000210602 | 0.303285712 | -0.000694401 |
| RP23-110C17.2 | 20.4004732 | -0.708434165 | 0.274767546 | -2.578303648 |
| Mir3096 | 0.548185132 | -0.358333395 | 1.4051332 | -0.255017386 |
| Pus7 | 222.5424342 | 0.015118438 | 0.141975641 | 0.106486143 |
| Rint1 | 243.1730144 | -0.134221629 | 0.099721667 | -1.345962546 |
| RP23-354E8.1 | 9.469473219 | -0.113656487 | 0.383161279 | -0.296628322 |
| 4933427G23Rik | 0.868720842 | 0.882254141 | 1.331380694 | 0.662661059 |
| Gm22054 | 0.593565611 | 0.851713475 | 1.273779679 | 0.668650544 |
| Tomm7 | 416.4948252 | 0.029908987 | 0.109268899 | 0.273719119 |
| 2700038G22Rik | 16.39405462 | -0.160389989 | 0.3155186 | -0.508337668 |
| Snord93 | 2.473235781 | -0.606365177 | 0.720772417 | -0.841271339 |
| Fam126a | 374.7933423 | -0.090816493 | 0.094140311 | -0.964692934 |
| RP23-170E15.2 | 3.709199715 | -0.003367725 | 0.581254647 | -0.005793889 |
| RP23-451E8.3 | 0 | NA | NA | NA |
| Klhl7 | 1038.582517 | 0.093490193 | 0.063676565 | 1.468204096 |
| RP23-451E8.2 | 13.50599551 | -0.24655637 | 0.285068817 | -0.864901227 |
| RP23-451E8.7 | 63.10412136 | 0.964216246 | 0.268066069 | 3.596935068 |
| Nupl2 | 116.6198412 | 0.202403395 | 0.134793208 | 1.501584527 |
| Gm5575 | 2.4388784 | -0.254423122 | 0.690325482 | -0.36855531 |
| RP23-451E8.8 | 0 | NA | NA | NA |
| RP23-451E8.9 | 0.083526043 | 0.037073798 | 3.724945049 | 0.009952844 |
| Gm9836 | 12.95280587 | 0.066939939 | 0.281789514 | 0.237552982 |
| Kcnh2 | 620.4579014 | -0.001555075 | 0.110011241 | -0.0141356 |
| Gm15589 | 0 | NA | NA | NA |
| Nos3 | 66.34634559 | -0.481868855 | 0.176239441 | -2.734171498 |
| Gm15587 | 5.162842253 | 0.413331156 | 0.725969377 | 0.569350677 |
| Atg9b | 48.1694312 | 0.283779192 | 0.184892437 | 1.534833966 |
| Abcb8 | 361.5327256 | 0.240608067 | 0.08735162 | 2.754477453 |
| Asic3 | 3.0483297 | 1.87382736 | 0.792417117 | 2.36469824 |
| Cdk5 | 656.490936 | 0.263421633 | 0.055874792 | 4.714498722 |
| Slc4a2 | 638.8880438 | -0.274690305 | 0.116756681 | -2.352673126 |
| Fastk | 605.3488211 | 0.06601045 | 0.062486588 | 1.056393888 |
| Gm22333 | 0 | NA | NA | NA |
| Tmub1 | 167.7441116 | -0.03138283 | 0.112554379 | -0.278823718 |
| RP24-239E2.2 | 2.781790008 | -0.774322102 | 0.673120025 | -1.150347744 |

| | | | | |
|---|---|---|---|---|
| Gm10472 | 2.635270342 | 0.148971005 | 0.670666677 | 0.222123761 |
| Agap3 | 2182.413688 | 0.371116779 | 0.065224535 | 5.689834053 |
| Gbx1 | 12.22572069 | 0.032804649 | 0.310793268 | 0.10555135 |
| Asb10 | 0.545953117 | 0.033283001 | 1.406301404 | 0.023667047 |
| 4931409K22Rik | 0.666655512 | -0.281881401 | 1.304961072 | -0.216007517 |
| Abcf2 | 862.9471066 | -0.038577238 | 0.069476461 | -0.555256223 |
| Gm10221 | 170.6058012 | -0.173145091 | 0.151949377 | -1.139491942 |
| Chpf2 | 822.9892482 | 0.015004172 | 0.0812686 | 0.184624469 |
| Smarcd3 | 501.6330187 | 0.277431466 | 0.084981357 | 3.264615634 |
| Mir671 | 0 | NA | NA | NA |
| 1700022A21Rik | 0 | NA | NA | NA |
| Nub1 | 1661.721015 | 0.128976143 | 0.081605411 | 1.580485183 |
| Gm26648 | 1.867746029 | -0.851205151 | 0.805879719 | -1.056243421 |
| Wdr86 | 10.62884738 | 0.879105679 | 0.452901892 | 1.941051021 |
| Crygn | 1.666884929 | 0.681390456 | 0.833246988 | 0.817753277 |
| Rheb | 967.3691609 | -0.444151375 | 0.103874876 | -4.275830629 |
| Gm25363 | 7.580824313 | -0.170960835 | 0.435677729 | -0.392402053 |
| Prkag2 | 719.0261586 | -0.003201729 | 0.113455284 | -0.028220186 |
| Prkag2os2 | 1.088852924 | 0.263244114 | 0.992562839 | 0.265216572 |
| Prkag2os1 | 0 | NA | NA | NA |
| 1500035N22Rik | 3.98331357 | 0.705736088 | 0.562940243 | 1.253660751 |
| 2900005J15Rik | 14.08399665 | 0.529808508 | 0.347436625 | 1.524906902 |
| RP24-84G9.2 | 1.333500337 | 0.542397914 | 1.064863808 | 0.509358953 |
| Gm26051 | 0 | NA | NA | NA |
| Galntl5 | 0.110595523 | 0.037071574 | 3.724945049 | 0.009952247 |
| Gm26356 | 0 | NA | NA | NA |
| Galnt11 | 598.673327 | 0.136505386 | 0.073189317 | 1.865099877 |
| E130116L18Rik | 0 | NA | NA | NA |
| Kmt2c | 3673.91328 | 0.030002574 | 0.079190845 | 0.378864168 |
| RP23-171F16.5 | 0.112029109 | 0.037060453 | 3.724945049 | 0.009949262 |
| Hspe1-ps3 | 0.038981395 | -0.443816764 | 3.724945049 | -0.119147198 |
| Gm20432 | 5.780307253 | 0.210000319 | 0.547854698 | 0.383313897 |
| Gm18343 | 0 | NA | NA | NA |
| 4831440E17Rik | 12.8027681 | 1.00495475 | 0.364592089 | 2.756381116 |
| Gm22656 | 0.372618932 | -1.83241919 | 1.547128507 | -1.184400121 |
| Cct8l1 | 0.82877443 | -0.183009963 | 1.156047461 | -0.158306617 |
| 1700096K18Rik | 15.86316258 | 0.2629219 | 0.280311849 | 0.937962135 |
| RP23-405O11.3 | 0.04393175 | 0.037062677 | 3.724945049 | 0.009949859 |
| Gm10062 | 3.869150979 | 1.555303049 | 0.642364807 | 2.421214601 |
| RP24-121I11.1 | 0 | NA | NA | NA |
| RP24-121I11.4 | 17.85167491 | 0.765635862 | 0.31492215 | 2.431190893 |
| Xrcc2 | 97.35748713 | 0.083072723 | 0.151629178 | 0.547867657 |
| RP24-121I11.5 | 0.245367999 | -0.443356401 | 1.856129566 | -0.238860697 |
| Actr3b | 678.1794346 | 0.692038894 | 0.07556181 | 9.158580192 |
| RP23-56M18.4 | 0 | NA | NA | NA |
| RP23-56M18.5 | 0.232282403 | 0.9953969 | 2.537884712 | 0.392215176 |
| Gm21671 | 0.275686258 | 0.113390441 | 1.803869522 | 0.062859558 |

| | | | | |
|---|---|---|---|---|
| RP23-56M18.8 | 24.28531059 | 0.569528993 | 0.345984897 | 1.646109403 |
| RP23-56M18.6 | 0.144833758 | -0.891143198 | 2.882712686 | -0.309133547 |
| Gm21698 | 0 | NA | NA | NA |
| Gm1979 | 0.35614329 | 0.394433221 | 1.702875876 | 0.231627699 |
| Gm5067 | 39.82603458 | 0.080126023 | 0.25473315 | 0.314548864 |
| Gm5862 | 0 | NA | NA | NA |
| Speer4a | 0.739238851 | 0.471501437 | 1.118887095 | 0.421402159 |
| Gm7347 | 0.182828849 | -1.328698802 | 2.253829453 | -0.589529434 |
| RP24-162F23.9 | 8.999765542 | 0.077453939 | 0.45392867 | 0.170630198 |
| Gm10471 | 0.138573873 | -0.444788617 | 3.62882948 | -0.122570823 |
| 5031410I06Rik | 0 | NA | NA | NA |
| Gm10220 | 0.108266322 | -0.955981391 | 3.722845977 | -0.25678779 |
| Gm7361 | 6.292791096 | 0.326123083 | 0.414666544 | 0.78647069 |
| RP24-362M8.2 | 0.097139547 | -0.443816764 | 3.724945049 | -0.119147198 |
| 4930584F24Rik | 0 | NA | NA | NA |
| RP24-165O21.1 | 0 | NA | NA | NA |
| Dpp6 | 3300.38036 | 0.408624475 | 0.063538231 | 6.431159114 |
| RP23-474A22.1 | 0 | NA | NA | NA |
| Gm16056 | 0 | NA | NA | NA |
| Gm16057 | 0 | NA | NA | NA |
| RP23-478E6.1 | 84.06891195 | 0.433241839 | 0.314766683 | 1.376390392 |
| RP23-478E6.3 | 79.08630011 | 0.859673435 | 0.314026797 | 2.737579857 |
| Speer4b | 0.378318939 | 0.975719328 | 1.698155794 | 0.574575861 |
| Gm16058 | 0.092228164 | 0.037073798 | 3.724945049 | 0.009952844 |
| Paxip1 | 503.8218817 | 0.321952663 | 0.080634355 | 3.992748045 |
| RP23-397C5.2 | 0.110870145 | -0.444802825 | 3.724874622 | -0.119414174 |
| RP23-397C5.1 | 0 | NA | NA | NA |
| Htr5a | 258.8451686 | 0.167230836 | 0.12415778 | 1.346921924 |
| RP23-397C5.5 | 4.181737927 | -0.145994858 | 0.528710809 | -0.276133673 |
| RP23-397C5.4 | 1.516892051 | -0.310033672 | 0.879474978 | -0.35252131 |
| RP23-256A5.2 | 3.457092485 | -0.124043947 | 0.644606996 | -0.192433447 |
| Gm26608 | 2.045375739 | 0.443295626 | 0.671043009 | 0.660606876 |
| RP23-256A5.4 | 1.504629277 | 0.4978106 | 1.013926194 | 0.490973212 |
| Insig1 | 653.7555455 | 0.095759349 | 0.138541091 | 0.691198173 |
| En2 | 252.7745475 | -0.25342308 | 0.283257815 | -0.894672864 |
| RP23-358F24.3 | 0.075043205 | 0.517957688 | 3.724945049 | 0.139051095 |
| Cnpy1 | 429.1835849 | 0.353498474 | 0.297168326 | 1.189556366 |
| Rbm33 | 1626.312609 | 0.417204842 | 0.125516774 | 3.323897103 |
| Shh | 57.43406997 | -0.089548398 | 0.156598026 | -0.57183606 |
| 9530036O11Rik | 6.185525992 | 1.24582933 | 0.492708005 | 2.528534789 |
| Gm26894 | 16.72981752 | 0.839209736 | 0.301546743 | 2.783017079 |
| RP23-94F17.1 | 1.563650339 | -0.690776581 | 0.808831645 | -0.854042476 |
| RP23-94F17.2 | 0.559314221 | 0.23874732 | 1.566767261 | 0.152382122 |
| Rnf32 | 99.31368546 | 0.327078275 | 0.161002374 | 2.031512124 |
| Gm7420 | 6.011270768 | 1.380446027 | 0.700835302 | 1.969715313 |
| Lmbr1 | 450.4254583 | 0.393084382 | 0.080827576 | 4.863245979 |
| RP23-396H16.2 | 45.39463601 | 0.615753005 | 0.264916475 | 2.324328844 |

| | | | | |
|---|---|---|---|---|
| Gm23801 | 1.101786575 | -1.528969665 | 1.117754735 | -1.367893704 |
| 4632411P08Rik | 8.069396325 | 0.422565973 | 0.394023029 | 1.072439787 |
| Nom1 | 466.9739484 | -0.103988254 | 0.073727932 | -1.410432258 |
| Mnx1 | 0.180050447 | -0.925657836 | 3.327036994 | -0.278222886 |
| Gm24538 | 0 | NA | NA | NA |
| Gm6083 | 0 | NA | NA | NA |
| Ube3c | 1443.177322 | 0.027859457 | 0.047515058 | 0.586329026 |
| RP24-224N3.5 | 0.401137426 | 1.43292208 | 1.531850439 | 0.935419048 |
| Gm26346 | 0.288530209 | 0.866718157 | 1.731817688 | 0.50046732 |
| RP24-224N3.4 | 0 | NA | NA | NA |
| Gm5129 | 0.619575296 | -0.36884852 | 1.329600594 | -0.277413023 |
| Dnajb6 | 825.5467343 | 0.166384072 | 0.057459976 | 2.895651625 |
| RP24-224N3.8 | 0.381425259 | -0.791815383 | 1.534508731 | -0.516005785 |
| RP24-224N3.9 | 0 | NA | NA | NA |
| RP24-176O2.1 | 0 | NA | NA | NA |
| RP24-176O2.2 | 0.121388914 | 0.037071574 | 3.724945049 | 0.009952247 |
| RP24-176O2.4 | 0.21891683 | -0.771763925 | 2.542683888 | -0.303523347 |
| RP24-418J2.4 | 0.04393175 | 0.037062677 | 3.724945049 | 0.009949859 |
| Il6 | 43.66316143 | -0.7732886 | 0.806794672 | -0.958470138 |
| Tyms | 45.1820026 | -0.183134179 | 0.205612968 | -0.890674265 |
| RP24-418J2.5 | 0.550684032 | 1.2565541 | 1.263934534 | 0.994160746 |
| 3110082J24Rik | 7.185718025 | 0.389453352 | 0.511500211 | 0.761394314 |
| Gareml | 196.41537 | 0.299499663 | 0.161828129 | 1.85072685 |
| Hadha | 925.8447827 | -0.448227504 | 0.087898536 | -5.099373955 |
| Hadhb | 496.8674058 | -0.070337672 | 0.085814176 | -0.819650961 |
| Mir1960 | 2.414716579 | -0.653499316 | 0.86767618 | -0.753160373 |
| Adgrf3 | 4.779796961 | 0.958073981 | 0.475717444 | 2.01395596 |
| Ept1 | 1152.189136 | -0.036062485 | 0.051510473 | -0.70010005 |
| Gm16288 | 0.621982117 | -2.10528256 | 1.373618318 | -1.532654692 |
| Drc1 | 103.3566621 | -0.726450022 | 0.190412811 | -3.815132077 |
| WI1-2080J1.2 | 1.385170032 | 0.810434484 | 1.207671898 | 0.671071742 |
| Otof | 183.3860885 | 2.036254823 | 0.218027752 | 9.339429538 |
| 1700001C02Rik | 2.869726659 | 0.409881431 | 0.67034232 | 0.611450925 |
| Cib4 | 0.038981395 | -0.443816764 | 3.724945049 | -0.119147198 |
| Gm25666 | 0 | NA | NA | NA |
| RP23-446J15.1 | 0.136419714 | 0.037074086 | 2.946414848 | 0.012582779 |
| Gm9899 | 195.7601427 | 0.020961844 | 0.17822453 | 0.117614808 |
| Kcnk3 | 462.3649798 | -0.152690458 | 0.120975398 | -1.262161237 |
| Slc35f6 | 587.4667661 | -0.579135572 | 0.095158759 | -6.085993309 |
| Mir5625 | 0 | NA | NA | NA |
| Cenpa | 31.10520093 | 0.161613613 | 0.256805438 | 0.62932317 |
| Dpysl5 | 464.621931 | -0.129087567 | 0.092751068 | -1.391763664 |
| RP23-342N8.1 | 3.407592442 | 1.493769143 | 0.749273508 | 1.993623326 |
| Mapre3 | 1596.309188 | 0.287581669 | 0.065333298 | 4.401762612 |
| RP23-342N8.2 | 49.78286253 | 0.097944537 | 0.335405989 | 0.292017855 |
| Gm15461 | 0 | NA | NA | NA |
| Tmem214 | 593.0310336 | -0.271738476 | 0.061951377 | -4.386318577 |

| | | | | |
|---|---|---|---|---|
| RP23-113M20.5 | 2.578624624 | 1.207813546 | 0.816665297 | 1.478957843 |
| Agbl5 | 130.4690619 | 0.106111114 | 0.146059429 | 0.726492737 |
| RP23-113M20.4 | 16.2135406 | -0.428547148 | 0.296413336 | -1.44577553 |
| Ost4 | 231.3485647 | -0.38544973 | 0.148950939 | -2.587763006 |
| Emilin1 | 69.27241042 | -1.049517507 | 0.177395927 | -5.916243545 |
| Khk | 59.85891304 | 0.261698846 | 0.170199661 | 1.537599097 |
| Cgref1 | 143.7238615 | -0.550160198 | 0.194781959 | -2.824492581 |
| Abhd1 | 97.11033058 | -0.278138484 | 0.144416474 | -1.925947056 |
| Preb | 614.623067 | -0.150727425 | 0.066682087 | -2.260388519 |
| Tcf23 | 3.89979361 | 0.008625042 | 0.498708539 | 0.017294755 |
| Slc5a6 | 215.3588507 | 0.061322619 | 0.094107737 | 0.651621432 |
| Atraid | 300.310359 | 0.049475591 | 0.109254621 | 0.452846664 |
| Cad | 344.3964736 | -0.499622861 | 0.079254983 | -6.303993054 |
| Slc30a3 | 365.9223408 | 0.937037679 | 0.151174832 | 6.198370886 |
| Gm9924 | 4.140852306 | 0.234021533 | 0.508003864 | 0.460668806 |
| Dnajc5g | 0.288733551 | -1.791745799 | 2.009258318 | -0.891744871 |
| Trim54 | 0.834511584 | 1.185069825 | 1.215096643 | 0.975288535 |
| Gm15469 | 0 | NA | NA | NA |
| Ucn | 1.659798835 | 0.781117794 | 1.290586498 | 0.605242496 |
| Mpv17 | 189.2554659 | -0.080768788 | 0.105683304 | -0.764253051 |
| Gm29609 | 14.60460443 | 0.401948816 | 0.332255465 | 1.209758328 |
| Gtf3c2 | 322.248693 | 0.251100082 | 0.100418275 | 2.50054168 |
| Eif2b4 | 322.6252131 | -0.271125974 | 0.078963759 | -3.433549486 |
| Snx17 | 719.475255 | -0.17998474 | 0.072102139 | -2.496246871 |
| Zfp513 | 310.9180716 | 0.059484856 | 0.097594055 | 0.609513111 |
| Ppm1g | 655.6864786 | -0.197711362 | 0.07773399 | -2.543435143 |
| RP24-86I23.4 | 34.84001459 | 0.673109809 | 0.370310857 | 1.817688561 |
| Gm9970 | 1.14535829 | -0.658252953 | 0.992134403 | -0.663471553 |
| Nrbp1 | 774.3138472 | -0.017943279 | 0.06220736 | -0.288443021 |
| Krtcap3 | 6.967827923 | -1.129058996 | 0.438539577 | -2.574588602 |
| Ift172 | 621.7871443 | -0.068391288 | 0.083904941 | -0.815104413 |
| Fndc4 | 773.732385 | -0.122345972 | 0.088639991 | -1.38025705 |
| Gckr | 6.742714648 | 0.520959941 | 0.412674839 | 1.262398122 |
| RP23-383E21.3 | 0 | NA | NA | NA |
| RP23-383E21.5 | 0 | NA | NA | NA |
| RP23-383E21.4 | 0.38059087 | -0.422399543 | 1.710344102 | -0.24696758 |
| Zfp512 | 587.0033922 | -0.14604428 | 0.066099424 | -2.209463749 |
| RP23-383E21.1 | 18.84380616 | -0.092508675 | 0.343734274 | -0.269128457 |
| 4930548H24Rik | 1.368337452 | -0.038174846 | 0.935590523 | -0.040802942 |
| 4930566F21Rik | 0.183097766 | -0.443820034 | 2.686642852 | -0.165195025 |
| Gpn1 | 250.7486316 | -0.064098456 | 0.078742251 | -0.814028755 |
| Supt7l | 184.2270805 | 0.021433308 | 0.099360811 | 0.21571189 |
| Slc4a1ap | 405.404987 | -0.08109874 | 0.071180769 | -1.139334987 |
| RP24-80F7.3 | 2.627266017 | -2.867782509 | 0.762354746 | -3.761742841 |
| RP24-80F7.5 | 6.412813486 | -2.547336357 | 0.497297104 | -5.122363143 |
| RP24-80F7.6 | 3.22349573 | 1.947080051 | 0.652597125 | 2.983586623 |
| Mrpl33 | 188.1679828 | -0.439300157 | 0.11837561 | -3.711069843 |

| | | | | |
|---|---|---|---|---|
| RP24-80F7.7 | 0 | NA | NA | NA |
| Rbks | 30.53545577 | -0.409120963 | 0.20987486 | -1.949356693 |
| Mir3473e | 1.131450813 | -0.33743149 | 1.003645152 | -0.336205969 |
| Bre | 293.3718634 | 0.0136658 | 0.070820014 | 0.19296522 |
| RP24-80F7.8 | 3.022895243 | -0.261511189 | 0.67921616 | -0.385019091 |
| Gm24436 | 2.112795873 | 0.046926188 | 0.844279803 | 0.055581323 |
| RP23-181I1.4 | 6.442610278 | -0.213418245 | 0.397089474 | -0.537456315 |
| Gm17130 | 0.230862969 | -0.451731822 | 2.133407855 | -0.211741895 |
| Gm10463 | 2.064454091 | -0.667859988 | 0.703732062 | -0.949025949 |
| Fosl2 | 1087.482308 | -0.883399907 | 0.309965319 | -2.849995956 |
| Gm26335 | 0 | NA | NA | NA |
| RP23-415E17.3 | 0 | NA | NA | NA |
| Plb1 | 16.55472255 | 0.004684139 | 0.280362144 | 0.016707459 |
| Gm15615 | 0 | NA | NA | NA |
| RP23-262J12.5 | 0.060174537 | 0.517957688 | 3.724945049 | 0.139051095 |
| RP23-262J12.4 | 0 | NA | NA | NA |
| Gm5864 | 1.456934103 | -0.020751917 | 0.804808672 | -0.025784907 |
| Ppp1cb | 4028.577039 | 0.050704203 | 0.087533734 | 0.579253283 |
| RP23-262J12.3 | 12.26687181 | 0.544831864 | 0.340749303 | 1.598922901 |
| RP23-262J12.2 | 50.45321847 | 0.642215389 | 0.288993229 | 2.222250644 |
| Gm15613 | 0 | NA | NA | NA |
| Gm15614 | 0.985784616 | 3.063195392 | 1.274252276 | 2.403915967 |
| RP23-149A1.7 | 0.075043205 | 0.517957688 | 3.724945049 | 0.139051095 |
| RP23-149A1.5 | 0.156588133 | -0.914325396 | 2.818547681 | -0.32439593 |
| RP23-149A1.6 | 21.7617041 | 0.475103112 | 0.262553388 | 1.809548584 |
| Yes1 | 334.4835077 | -0.213940987 | 0.095987024 | -2.228853212 |
| RP23-149A1.2 | 1.316494506 | -0.217026633 | 0.862072617 | -0.251749828 |
| RP23-353M5.2 | 7.746880428 | 0.138344064 | 0.467050481 | 0.296207945 |
| Gm2420 | 4.384803307 | 0.577644514 | 0.619112379 | 0.933020456 |
| Pisd | 289.4875556 | -0.14565668 | 0.07728014 | -1.884787995 |
| Gm20671 | 0 | NA | NA | NA |
| RP23-353M5.1 | 0 | NA | NA | NA |
| Prr14l | 1555.512628 | -0.040383642 | 0.055592997 | -0.726415989 |
| RP23-118L1.4 | 11.68546513 | 1.232631031 | 0.441106589 | 2.794406297 |
| RP23-118L1.5 | 0 | NA | NA | NA |
| Depdc5 | 821.0906113 | 0.152281317 | 0.076037143 | 2.002722763 |
| Gm24003 | 1.689516722 | 1.105180679 | 1.127641163 | 0.980081887 |
| RP23-69J7.1 | 15.99339509 | 0.749682904 | 0.4170219 | 1.79770632 |
| Ywhah | 4942.475359 | 0.229468223 | 0.074857416 | 3.065404014 |
| RP23-69J7.4 | 0 | NA | NA | NA |
| Rpl35a-ps5 | 4.14321256 | 0.557796066 | 0.496940678 | 1.122460066 |
| Slc5a1 | 30.58515113 | -0.179392003 | 0.299478521 | -0.599014589 |
| Gm15936 | 0.038981395 | -0.443816764 | 3.724945049 | -0.119147198 |
| 5033423K11Rik | 0.310320078 | -0.266052208 | 1.95702531 | -0.13594725 |
| Spon2 | 4.327007151 | 0.141103736 | 0.619387296 | 0.227811803 |
| Ctbp1 | 1818.873768 | -0.041427535 | 0.074411317 | -0.556737019 |
| RP24-406N3.3 | 0 | NA | NA | NA |

| | | | | |
|---|---|---|---|---|
| Gm20465 | 0 | NA | NA | NA |
| Maea | 757.2802516 | -0.258576305 | 0.077138006 | -3.352125879 |
| Uvssa | 220.4651291 | 0.505002157 | 0.131660059 | 3.835651917 |
| Nkx1-1 | 0.498903533 | -2.306381789 | 1.517999008 | -1.519356585 |
| RP24-94L5.3 | 0.038981395 | -0.443816764 | 3.724945049 | -0.119147198 |
| Fam53a | 291.3243653 | -0.014667193 | 0.078004122 | -0.188030995 |
| Gm9903 | 6.764630232 | -0.331467613 | 0.42393717 | -0.781879099 |
| Slbp | 223.0429402 | -0.150249309 | 0.094624484 | -1.587848113 |
| RP23-391P21.2 | 0.046345709 | -0.443818988 | 3.724945049 | -0.119147795 |
| Tmem129 | 242.2710187 | -0.511473195 | 0.099671839 | -5.131571776 |
| Tacc3 | 66.98387769 | 0.063789778 | 0.206924046 | 0.308276291 |
| RP23-391P21.5 | 0 | NA | NA | NA |
| RP23-391P21.4 | 17.89361774 | 0.883108193 | 0.2879238 | 3.067159415 |
| Fgfr3 | 1284.08441 | 0.027227122 | 0.13480252 | 0.201977838 |
| Letm1 | 945.73928 | 0.016389705 | 0.049787115 | 0.329195717 |
| Gm10059 | 2.485764843 | 0.137542402 | 0.716095169 | 0.192072797 |
| RP23-391P21.3 | 0 | NA | NA | NA |
| Whsc1 | 2442.311656 | -0.004631547 | 0.045718704 | -0.101305303 |
| Nelfa | 291.9774525 | -0.023976588 | 0.090363225 | -0.265335679 |
| Mir7024 | 0 | NA | NA | NA |
| RP24-294K15.5 | 2.887331315 | 0.024310222 | 0.572867368 | 0.042436039 |
| RP24-294K15.4 | 13.77895346 | 0.264610563 | 0.332996494 | 0.794634681 |
| Gm1673 | 134.0090932 | -0.467359862 | 0.165848009 | -2.81800104 |
| RP24-294K15.7 | 56.16396208 | 0.58610262 | 0.165768181 | 3.53567624 |
| Nat8l | 3474.475194 | 0.466059746 | 0.070583983 | 6.602910872 |
| Poln | 11.42523575 | -0.28699548 | 0.313986102 | -0.914038799 |
| RP23-60E7.2 | 0.517722191 | 0.67237862 | 1.240462759 | 0.542038538 |
| RP23-60E7.3 | 0.042716945 | 0.037062677 | 3.724945049 | 0.009949859 |
| RP23-60E7.1 | 0.122601409 | 0.998850474 | 3.724945049 | 0.268151734 |
| Haus3 | 129.8840804 | -0.186024819 | 0.109652788 | -1.696489643 |
| Gm25918 | 0 | NA | NA | NA |
| Mxd4 | 934.5782134 | 0.498513401 | 0.11738247 | 4.246915236 |
| RP24-230P10.6 | 0.165470308 | 1.408943363 | 2.753318612 | 0.511725507 |
| Zfyve28 | 544.0876334 | 0.389542842 | 0.130671234 | 2.981091005 |
| Gm15513 | 73.95229317 | 0.62997191 | 0.313019992 | 2.012561259 |
| RP23-201D22.6 | 0 | NA | NA | NA |
| RP23-201D22.4 | 0.535802326 | -0.203249027 | 1.312987778 | -0.154798872 |
| RP23-201D22.5 | 0.419184927 | 1.644660558 | 3.327918175 | 0.49420102 |
| Rnf4 | 1197.186287 | -0.297327104 | 0.083128359 | -3.576722866 |
| Fam193a | 1067.198729 | -0.048972633 | 0.070212377 | -0.697492879 |
| RP23-429E19.4 | 44.14393845 | -0.030278702 | 0.292104795 | -0.103656983 |
| Gm26931 | 0.110870145 | -0.444802825 | 3.724874622 | -0.119414174 |
| RP23-429E19.5 | 37.84079439 | -0.165808368 | 0.254678878 | -0.651048764 |
| Tnip2 | 144.594261 | -0.07915473 | 0.162070723 | -0.48839623 |
| Sh3bp2 | 240.9989662 | -1.042226167 | 0.162749911 | -6.403850896 |
| Gm22847 | 1.623325927 | -1.203583169 | 0.833254296 | -1.4444368 |
| Add1 | 3389.415062 | 0.175601447 | 0.051626095 | 3.401408663 |

| | | | | |
|---|---|---|---|---|
| Mir7036b | 0 | NA | NA | NA |
| RP24-229D18.2 | 13.74406399 | 0.521822625 | 0.457973852 | 1.139415762 |
| Mfsd10 | 128.2944897 | 0.07184574 | 0.156700218 | 0.458491638 |
| Nop14 | 276.7616417 | -0.241225361 | 0.139384495 | -1.730647027 |
| Gm15522 | 0.123992736 | 0.517955521 | 3.720643114 | 0.139211288 |
| Grk4 | 52.31774275 | -0.197665427 | 0.200194106 | -0.987368863 |
| RP23-44L13.1 | 0.042716945 | 0.037062677 | 3.724945049 | 0.009949859 |
| Mir467g | 0 | NA | NA | NA |
| Htt | 3899.683035 | -0.051001229 | 0.065797533 | -0.775123731 |
| Msantd1 | 17.05411229 | 0.465952667 | 0.258609484 | 1.801761713 |
| Rgs12 | 414.5610989 | 0.381698091 | 0.094175236 | 4.053062208 |
| Mir3097 | 0.476896902 | 0.827698112 | 1.44866161 | 0.571353659 |
| RP23-352G19.2 | 0.30060319 | 0.477740494 | 1.598879143 | 0.298797127 |
| Hgfac | 2.316699699 | -0.121800087 | 0.681516791 | -0.178719128 |
| Dok7 | 162.3460107 | 0.565019869 | 0.140360333 | 4.025495374 |
| Lrpap1 | 1373.379439 | -0.008939161 | 0.101747679 | -0.08785617 |
| Adra2c | 169.5646234 | 0.822808236 | 0.124384147 | 6.61505712 |
| RP24-276N7.5 | 0.097139547 | -0.443816764 | 3.724945049 | -0.119147198 |
| E130018O15Rik | 0 | NA | NA | NA |
| Hmx1 | 0.038981395 | -0.443816764 | 3.724945049 | -0.119147198 |
| 4930442P19Rik | 0 | NA | NA | NA |
| RP24-276N7.7 | 0 | NA | NA | NA |
| Cpz | 3.532213938 | -2.035224669 | 0.687287311 | -2.961242898 |
| RP23-26P10.1 | 28.70925171 | -0.460037101 | 0.317291883 | -1.449886133 |
| Plk-ps1 | 0 | NA | NA | NA |
| RP23-26P10.2 | 15.37615298 | -0.235457955 | 0.331034378 | -0.711279464 |
| Trmt44 | 83.81309991 | -0.234895932 | 0.167846152 | -1.399471653 |
| 4931431C16Rik | 1.224874833 | -0.03479277 | 0.877911811 | -0.039631281 |
| Acox3 | 452.6412082 | 0.228439046 | 0.084392355 | 2.706868964 |
| Htra3 | 58.84703746 | -0.595993743 | 0.204055189 | -2.920747798 |
| Sh3tc1 | 32.64252681 | 0.10856437 | 0.214339457 | 0.506506698 |
| Gm15650 | 0.369202626 | -0.377712091 | 1.742868217 | -0.216718675 |
| Ablim2 | 1153.537665 | 0.690730262 | 0.130954329 | 5.274588999 |
| Afap1 | 482.3908662 | -0.173796549 | 0.07005812 | -2.48074809 |
| H2af-ps | 0.473803596 | 0.511049363 | 1.466624932 | 0.348452663 |
| Sorcs2 | 714.7892678 | -0.13696394 | 0.078186374 | -1.751762279 |
| Psapl1 | 0.670061835 | -0.942625645 | 1.2254281 | -0.769221503 |
| Mir7689 | 0.235888945 | -1.377681506 | 2.154632347 | -0.639404448 |
| 2210406O10Rik | 10.33515927 | -0.342973941 | 0.329512265 | -1.040853337 |
| Grpel1 | 361.9003875 | -0.242205482 | 0.112793333 | -2.14733863 |
| Tada2b | 496.4167252 | 0.182327597 | 0.107399442 | 1.6976587 |
| Ccdc96 | 19.96485888 | -0.682963541 | 0.255696164 | -2.670996425 |
| Tbc1d14 | 875.8556229 | 0.008183069 | 0.053558016 | 0.152788873 |
| RP23-463H10.2 | 0 | NA | NA | NA |
| D5Ertd579e | 1861.527523 | 0.094914557 | 0.056137478 | 1.69075205 |
| RP23-463H10.1 | 0 | NA | NA | NA |
| Gm24878 | 6.521421862 | 0.493197073 | 0.460151767 | 1.071813927 |

| | | | | |
|---|---|---|---|---|
| Gm25767 | 0 | NA | NA | NA |
| Bloc1s4 | 95.95579344 | -0.557528441 | 0.159155845 | -3.503034656 |
| RP23-382O15.2 | 0.348252234 | -0.867020132 | 1.474404055 | -0.588047848 |
| Mrfap1 | 2250.019853 | -0.066299836 | 0.077310583 | -0.85757775 |
| Man2b2 | 514.8886766 | -0.215529193 | 0.108388892 | -1.988480456 |
| Ppp2r2c | 4810.614684 | 0.207555165 | 0.099573544 | 2.084440869 |
| Ppp2r2cos | 0.150455055 | -1.336248778 | 2.838957719 | -0.470682874 |
| RP23-16N23.2 | 5.207155389 | 0.131914908 | 0.57742404 | 0.228454133 |
| Wfs1 | 1615.855355 | -0.346892327 | 0.134354612 | -2.581916043 |
| Jakmip1 | 619.5416328 | 0.591411041 | 0.187841765 | 3.148453381 |
| Gm26647 | 0.176363682 | -0.979968395 | 2.726117188 | -0.359474053 |
| Gm25551 | 0 | NA | NA | NA |
| RP23-106D12.1 | 30.55419668 | 0.323095346 | 0.301450422 | 1.071802601 |
| Gm1043 | 54.30076238 | 0.176582773 | 0.232214431 | 0.760429797 |
| Crmp1 | 1273.345192 | 0.409277815 | 0.074599641 | 5.486324182 |
| Evc | 77.51767331 | -0.284883513 | 0.2066725 | -1.3784297 |
| RP23-339H11.3 | 0 | NA | NA | NA |
| Evc2 | 92.04870115 | -0.088440852 | 0.138681457 | -0.637726586 |
| Gm15732 | 0 | NA | NA | NA |
| Stk32b | 51.08340483 | -0.369519354 | 0.184523738 | -2.002557279 |
| Cytl1 | 1.87157638 | 0.104146374 | 0.821131555 | 0.126832751 |
| RP24-199D21.1 | 0 | NA | NA | NA |
| Msx1 | 35.29830261 | -0.20899391 | 0.266865626 | -0.783142861 |
| RP24-292O13.1 | 0.159287055 | 0.998860136 | 2.807608854 | 0.355768979 |
| RP23-453O15.1 | 0.961470315 | -0.87799976 | 1.182992625 | -0.742185321 |
| RP23-453O15.4 | 0.066050875 | -0.443818988 | 3.724945049 | -0.119147795 |
| Stx18 | 243.7354772 | -0.146367399 | 0.085628699 | -1.709326433 |
| RP24-535K22.2 | 24.02896761 | -0.149089448 | 0.238998028 | -0.623810369 |
| Nsg1 | 2866.052638 | 0.208592175 | 0.08560965 | 2.436549775 |
| Zbtb49 | 139.0035971 | 0.393922669 | 0.139773372 | 2.81829553 |
| Lyar | 86.46303146 | -0.165643441 | 0.216470108 | -0.765202378 |
| Tmem128 | 186.1267044 | -0.397514103 | 0.126896633 | -3.132581953 |
| Otop1 | 1.049274302 | -0.36495855 | 0.927363589 | -0.393544187 |
| Gm26560 | 1.795106397 | -1.7319956 | 0.785431576 | -2.205151478 |
| Drd5 | 35.97084852 | -0.325797418 | 0.19219029 | -1.695181467 |
| RP23-426E16.4 | 0.045882455 | 0.517959912 | 3.724945049 | 0.139051692 |
| Slc2a9 | 46.88057285 | -0.543027725 | 0.1999461 | -2.715870548 |
| Gm15796 | 0.265425963 | -0.889500044 | 2.075801272 | -0.428509249 |
| Wdr1 | 2375.805282 | -0.55089995 | 0.103748956 | -5.309932488 |
| Gm22249 | 0 | NA | NA | NA |
| 4930421P07Rik | 0 | NA | NA | NA |
| Zfp518b | 296.1034572 | -0.093164567 | 0.077748548 | -1.198280474 |
| Gm22779 | 0 | NA | NA | NA |
| Clnk | 4.40624827 | -1.155201635 | 0.507700386 | -2.275360954 |
| Clnkos | 0.326535978 | -0.980071495 | 1.795489104 | -0.545852098 |
| RP24-258B22.1 | 0.156480458 | 0.037059321 | 3.487618632 | 0.010625967 |
| RP24-263A3.3 | 0 | NA | NA | NA |

| | | | | |
|---|---|---|---|---|
| RP24-263A3.2 | 0.133021046 | 0.915476344 | 3.654933187 | 0.250476903 |
| RP24-263A3.1 | 0 | NA | NA | NA |
| Gm25505 | 0.048949532 | 0.037064902 | 3.724945049 | 0.009950456 |
| RP23-477D20.1 | 0.049553047 | -0.443818988 | 3.724945049 | -0.119147795 |
| RP23-477D20.4 | 0.771325962 | 0.680659116 | 1.118030898 | 0.608801704 |
| RP23-477D20.3 | 0.202816535 | 0.262125258 | 3.198091588 | 0.081963024 |
| RP23-477D20.2 | 2.015324298 | 0.08531639 | 0.841434329 | 0.101393997 |
| Gm25377 | 0 | NA | NA | NA |
| RP23-2C16.2 | 158.46767 | 0.355486225 | 0.121443648 | 2.92717017 |
| Hs3st1 | 277.9990456 | 0.269760655 | 0.1051909 | 2.564486612 |
| Gm7816 | 17.85736311 | -0.333896024 | 0.265670364 | -1.256805683 |
| RP23-318O24.1 | 0.038981395 | -0.443816764 | 3.724945049 | -0.119147198 |
| Gm2810 | 23.07760637 | 0.392587245 | 0.263044042 | 1.492477239 |
| Gm23022 | 0 | NA | NA | NA |
| RP23-477O15.2 | 0 | NA | NA | NA |
| RP23-477O15.1 | 3.151818414 | 0.772532412 | 0.878817544 | 0.879058933 |
| RP24-377B20.1 | 0 | NA | NA | NA |
| Rab28 | 662.797046 | -0.078814431 | 0.057396894 | -1.373148027 |
| Nkx3-2 | 0.153659334 | 0.037073358 | 2.839457839 | 0.013056492 |
| Bod1l | 1615.891892 | 0.206134591 | 0.113058551 | 1.823255208 |
| RP24-188B2.2 | 1.139214769 | 0.992130015 | 0.946586808 | 1.048113081 |
| Mir6414 | 0 | NA | NA | NA |
| RP24-188B2.1 | 0.100902008 | -0.876201588 | 3.72093227 | -0.235479048 |
| Gm16223 | 2.960191199 | -0.447561875 | 0.628547301 | -0.712057587 |
| RP23-146B9.1 | 0.099155932 | 0.037073798 | 3.724945049 | 0.009952844 |
| RP24-167K2.2 | 0.184373794 | 0.517963412 | 2.729110023 | 0.18979206 |
| RP24-560A18.4 | 0.659798846 | 0.440718269 | 1.336629408 | 0.329723606 |
| RP24-560A18.1 | 1.041030589 | 0.258820042 | 1.151658019 | 0.22473689 |
| RP24-560A18.3 | 0.172203646 | 0.517964484 | 2.739420462 | 0.189078125 |
| RP24-560A18.2 | 14.93512307 | -0.370790437 | 0.439695171 | -0.843289763 |
| Gm7854 | 28.25369218 | 0.052927042 | 0.280176071 | 0.18890636 |
| Cpeb2 | 1599.741892 | 0.092399269 | 0.065119329 | 1.418922329 |
| RP24-230L5.2 | 66.96472975 | 0.314341504 | 0.221611311 | 1.41843619 |
| RP24-230L5.3 | 79.78188433 | 0.547034571 | 0.216742433 | 2.523892359 |
| RP24-230L5.4 | 0.343955217 | 0.065208027 | 1.752918828 | 0.037199684 |
| RP24-230L5.5 | 0.150451996 | 0.037073243 | 2.858243881 | 0.012970637 |
| RP23-15G1.1 | 0.062426873 | 0.517959912 | 3.724945049 | 0.139051692 |
| C1qtnf7 | 8.52901898 | 0.074403069 | 0.358654301 | 0.207450654 |
| RP23-15G1.2 | 2.619111478 | 0.807989671 | 0.683971789 | 1.181320171 |
| Gm15866 | 0.088534442 | -0.905414734 | 3.723500899 | -0.243162217 |
| RP23-15G1.3 | 0.396976673 | 1.272869628 | 1.555252228 | 0.818432924 |
| Cc2d2a | 374.1500514 | 0.183274154 | 0.122060648 | 1.501500749 |
| Fbxl5 | 1545.557561 | -0.111979247 | 0.07226097 | -1.549650485 |
| RP23-336O18.1 | 33.11233713 | 0.570590695 | 0.314475208 | 1.81442187 |
| RP23-336O18.2 | 3.500141698 | 0.525069196 | 0.551993537 | 0.95122345 |
| RP23-336O18.3 | 11.15943119 | -1.156969471 | 0.55488786 | -2.085050969 |
| RP23-336O18.4 | 0.062426873 | 0.517959912 | 3.724945049 | 0.139051692 |

| | | | | |
|---|---|---|---|---|
| Bst1 | 59.92677726 | -2.196064629 | 0.368246891 | -5.963565969 |
| Cd38 | 164.7434097 | -0.947591725 | 0.206303097 | -4.593201639 |
| Gm16014 | 2.851290958 | -1.356289826 | 0.637365745 | -2.127961593 |
| Gm16015 | 0 | NA | NA | NA |
| Gm7879 | 31.12801235 | 0.137041717 | 0.216910526 | 0.631789152 |
| Gm7882 | 0 | NA | NA | NA |
| Gm5292 | 0 | NA | NA | NA |
| RP23-58C20.3 | 0 | NA | NA | NA |
| RP23-58C20.4 | 0 | NA | NA | NA |
| Fgfbp1 | 20.48618477 | 1.12103784 | 0.294044623 | 3.812475225 |
| Prom1 | 116.7320341 | 0.132417587 | 0.197475267 | 0.67055277 |
| RP24-230J14.2 | 3.662822882 | 1.15832067 | 0.620673061 | 1.866233197 |
| Gm16401 | 1.371463738 | 2.310018433 | 1.004341316 | 2.300033261 |
| RP24-230J14.1 | 0.385783664 | 1.250478907 | 1.735832494 | 0.720391462 |
| RP24-230J14.3 | 0.273584676 | 0.036504207 | 1.806800045 | 0.020203789 |
| RP24-230J14.4 | 0.127971488 | -0.95589744 | 3.6855341 | -0.259364698 |
| RP24-230J14.5 | 0.770607605 | 0.56466751 | 1.035809421 | 0.545146142 |
| Tapt1 | 762.3494051 | 0.171635932 | 0.074599351 | 2.300769773 |
| RP24-230J14.8 | 46.64492421 | 0.765480334 | 0.286820773 | 2.66884552 |
| RP24-230J14.7 | 0 | NA | NA | NA |
| RP24-230J14.9 | 0 | NA | NA | NA |
| RP24-230J14.10 | 12.44662364 | -0.087986637 | 0.320676275 | -0.274378381 |
| RP24-137F10.3 | 0 | NA | NA | NA |
| RP24-137F10.4 | 0.038981395 | -0.443816764 | 3.724945049 | -0.119147198 |
| RP24-137F10.1 | 0.921219545 | -0.146865709 | 1.2101117 | -0.121365416 |
| RP24-137F10.2 | 0.094097695 | 0.037071574 | 3.724945049 | 0.009952247 |
| Ldb2 | 272.9695285 | 0.405544788 | 0.167160371 | 2.42608212 |
| RP24-188E19.2 | 17.17285275 | 0.700441755 | 0.520066388 | 1.346831426 |
| RP24-188E19.3 | 7.245259879 | 0.901359278 | 0.592123446 | 1.522248923 |
| RP24-188E19.4 | 5.313268904 | 0.563372263 | 0.71651364 | 0.78626872 |
| RP24-188E19.5 | 0 | NA | NA | NA |
| RP24-247A21.1 | 40.44753983 | 0.401447312 | 0.389687096 | 1.030178613 |
| RP24-496C22.1 | 0.159422358 | -0.98086361 | 2.814105052 | -0.348552592 |
| RP24-496C22.2 | 19.70483131 | 0.502722438 | 0.386062164 | 1.302180027 |
| RP24-496C22.3 | 8.581378291 | 0.446458394 | 0.500686831 | 0.891691904 |
| RP24-496C22.4 | 7.00608203 | 0.277067378 | 0.506913983 | 0.546576712 |
| RP24-496C22.5 | 14.99026705 | 0.796924083 | 0.460115116 | 1.732010221 |
| RP24-496C22.6 | 21.60948068 | 0.81959226 | 0.357709107 | 2.291225592 |
| RP23-133N3.1 | 0 | NA | NA | NA |
| 4930431F12Rik | 0.1567526 | 0.008815624 | 2.825354425 | 0.003120183 |
| Gm10205 | 0 | NA | NA | NA |
| 4930435H24Rik | 4.37057135 | -0.417676425 | 0.502578863 | -0.831066437 |
| RP23-194K16.1 | 0.718177591 | -0.563346468 | 1.186779171 | -0.474685166 |
| RP23-33C9.1 | 1.272457209 | -0.568725391 | 0.888948367 | -0.639773255 |
| Qdpr | 1058.229875 | 0.055647159 | 0.119585393 | 0.465334082 |
| AC092711.4 | 2.958230677 | -0.736744562 | 0.695439995 | -1.05939343 |
| Clrn2 | 1.123782183 | 0.860521138 | 1.143572152 | 0.752485216 |

| | | | | |
|---|---|---|---|---|
| Lap3 | 1007.429224 | -0.686400792 | 0.147942526 | -4.639644935 |
| Med28 | 563.4186552 | -0.040540002 | 0.086160279 | -0.470518466 |
| Fam184b | 121.4810835 | 0.592583892 | 0.212891635 | 2.783500124 |
| RP23-7D3.3 | 16.87252161 | 0.789650528 | 0.414291243 | 1.906027564 |
| RP23-118K12.1 | 0.502057302 | -0.492056906 | 1.256542373 | -0.391595952 |
| 9630001P10Rik | 17.90177705 | 0.394732053 | 0.297841781 | 1.325307856 |
| RP23-7E4.8 | 0.664660227 | -0.877466998 | 1.361454943 | -0.644506822 |
| RP23-7E4.7 | 0.154585317 | -1.377635992 | 2.821196527 | -0.488316209 |
| RP23-7E4.9 | 3.048203817 | 0.590067135 | 0.7502694 | 0.786473679 |
| Ncapg | 19.54545852 | -0.623133576 | 0.321059775 | -1.940864675 |
| Lcorl | 806.219256 | 0.150175594 | 0.068060921 | 2.206487834 |
| Gm36401 | 0.944820375 | -0.670194742 | 1.063158601 | -0.630380775 |
| RP23-7E4.3 | 66.02500977 | 0.095658844 | 0.177419066 | 0.539168908 |
| Gm29036 | 24.05605521 | 0.349016788 | 0.273130574 | 1.277838593 |
| RP23-7E4.4 | 46.18654444 | 0.296230147 | 0.214797365 | 1.379114435 |
| AC158786.1 | 0.746606991 | 0.922278788 | 1.103570309 | 0.835722727 |
| RP23-228M24.1 | 0 | NA | NA | NA |
| Gm7931 | 9.771161355 | 0.199668925 | 0.33207983 | 0.601267848 |
| RP24-400K16.1 | 0 | NA | NA | NA |
| RP23-277H1.1 | 0 | NA | NA | NA |
| RP23-287A8.1 | 0.858277593 | 1.532263155 | 1.326833072 | 1.154827376 |
| RP23-319E24.2 | 0 | NA | NA | NA |
| RP23-319E24.1 | 0 | NA | NA | NA |
| RP23-195A18.4 | 0 | NA | NA | NA |
| Slit2 | 695.3750179 | 0.232367477 | 0.090598711 | 2.564798925 |
| 4930518C09Rik | 0 | NA | NA | NA |
| Gm20475 | 5.876575912 | 0.088042699 | 0.465810161 | 0.189009829 |
| RP23-195A18.3 | 20.06045718 | -0.150502577 | 0.35235243 | -0.427136481 |
| RP23-219K16.2 | 8.550972595 | -0.141409634 | 0.440743778 | -0.320843177 |
| RP23-219K16.1 | 10.624249 | 0.947624254 | 0.430030634 | 2.203620346 |
| Mir218-1 | 2.27602632 | 1.069912936 | 0.851127171 | 1.257054142 |
| RP23-79E5.4 | 4.747594266 | 0.96085519 | 0.528680877 | 1.817457812 |
| RP23-79E5.5 | 0 | NA | NA | NA |
| Pacrgl | 220.5840451 | 0.08193865 | 0.145303211 | 0.56391493 |
| Kcnip4 | 719.2995326 | 0.643043227 | 0.086906565 | 7.399248013 |
| RP23-14C20.2 | 34.79027562 | 0.838162299 | 0.331693501 | 2.52691806 |
| Gm20700 | 0 | NA | NA | NA |
| RP24-174D16.1 | 87.81495278 | 0.882528542 | 0.238937081 | 3.693560411 |
| Gm19031 | 0 | NA | NA | NA |
| RP24-170C14.2 | 202.1229481 | 0.811717746 | 0.241564088 | 3.360258358 |
| RP23-309G12.4 | 37.2300334 | 1.105989491 | 0.36134366 | 3.060769055 |
| RP23-309G12.3 | 161.4099445 | 0.67047768 | 0.281377231 | 2.382842698 |
| RP23-309G12.2 | 70.38172967 | 0.786338342 | 0.277121094 | 2.83752612 |
| RP23-79E5.7 | 138.4994086 | 0.678303015 | 0.289423772 | 2.343632694 |
| RP23-309G12.1 | 115.2856778 | 0.487755033 | 0.265754208 | 1.835361468 |
| Gm10025 | 0 | NA | NA | NA |
| Gm10048 | 309.3515181 | 0.910272867 | 0.335771893 | 2.710985901 |

| | | | | |
|---|---|---|---|---|
| RP23-187D22.9 | 187.5255152 | 1.069939349 | 0.375275886 | 2.85107407 |
| RP23-187D22.8 | 194.8703213 | 0.772978277 | 0.223097522 | 3.464755098 |
| RP23-187D22.7 | 88.63220688 | 0.774900737 | 0.31120278 | 2.490018684 |
| RP23-187D22.6 | 375.4133243 | 0.894104243 | 0.294031224 | 3.040847948 |
| RP23-187D22.5 | 181.2108272 | 1.016191612 | 0.246525808 | 4.122049613 |
| RP23-187D22.4 | 188.0951161 | 1.050697828 | 0.360501399 | 2.914545773 |
| RP23-187D22.10 | 319.4630086 | 0.755562626 | 0.296246202 | 2.550455066 |
| RP23-187D22.11 | 0.476944705 | 1.319895926 | 1.564556365 | 0.843623123 |
| RP23-187D22.3 | 106.6585838 | 1.07959791 | 0.432312695 | 2.497261639 |
| RP23-187D22.1 | 388.4487423 | 0.925523204 | 0.299061929 | 3.09475434 |
| RP23-140B23.2 | 209.5621235 | 0.732781604 | 0.347090519 | 2.111211814 |
| RP23-140B23.1 | 603.2557397 | 0.788075799 | 0.254482085 | 3.096783019 |
| RP23-394G5.2 | 44.14745288 | 0.982997793 | 0.383483117 | 2.563340466 |
| RP23-394G5.1 | 396.9657275 | 0.754174519 | 0.310569587 | 2.428359219 |
| RP24-158C2.3 | 251.5456547 | 0.556141476 | 0.298609395 | 1.862437969 |
| Gm19932 | 0 | NA | NA | NA |
| RP24-158C2.4 | 0.083526043 | 0.037073798 | 3.724945049 | 0.009952844 |
| RP24-253A20.1 | 476.3543519 | 0.58978475 | 0.308481303 | 1.911897885 |
| Gm5555 | 28.60310479 | -0.283021001 | 0.21219891 | -1.333753323 |
| RP24-79O7.3 | 0.322897824 | 1.354925539 | 1.546878568 | 0.875909439 |
| RP24-79O7.1 | 1.170209419 | 0.129163801 | 1.016989113 | 0.12700608 |
| Adgra3 | 255.2158286 | 0.025824055 | 0.108910363 | 0.237112931 |
| RP24-315J23.1 | 0.596915583 | 1.666107152 | 1.250827065 | 1.332004399 |
| RP24-315J23.2 | 0 | NA | NA | NA |
| RP24-478I4.1 | 0.047948698 | 0.037064902 | 3.724945049 | 0.009950456 |
| Mir8117 | 0 | NA | NA | NA |
| Gm22618 | 0.389682156 | -0.444221812 | 1.586679288 | -0.279969503 |
| RP23-21O1.1 | 0.042716945 | 0.037062677 | 3.724945049 | 0.009949859 |
| RP23-21O1.3 | 0.04393175 | 0.037062677 | 3.724945049 | 0.009949859 |
| RP23-21O1.2 | 0.038981395 | -0.443816764 | 3.724945049 | -0.119147198 |
| RP23-125B15.1 | 0 | NA | NA | NA |
| RP24-363A3.1 | 0.195174816 | -0.918844321 | 2.236723724 | -0.410799202 |
| RP24-312A3.1 | 0 | NA | NA | NA |
| RP24-441G10.1 | 0 | NA | NA | NA |
| AC119939.1 | 0 | NA | NA | NA |
| Ppargc1a | 1065.622016 | 0.346198206 | 0.062221923 | 5.563926483 |
| RP23-385G21.4 | 13.87657411 | 0.732036783 | 0.344763574 | 2.123300831 |
| RP23-385G21.3 | 24.14107729 | 0.624874536 | 0.264329847 | 2.363995373 |
| RP23-385G21.2 | 0 | NA | NA | NA |
| RP23-385G21.1 | 28.94227565 | 0.238013149 | 0.257265417 | 0.925165736 |
| RP24-296L6.1 | 199.7747531 | 0.545923675 | 0.367706878 | 1.484670827 |
| RP24-296L6.2 | 266.7259097 | 0.512936848 | 0.234882045 | 2.183806126 |
| RP24-296L6.3 | 71.01041919 | 0.665255637 | 0.34923566 | 1.904890344 |
| RP24-296L6.4 | 241.104923 | 0.599891288 | 0.307686398 | 1.949684131 |
| RP24-299M10.1 | 94.15749016 | 0.946664218 | 0.464967051 | 2.035981291 |
| RP24-299M10.2 | 0.645477656 | 0.912032457 | 1.470391528 | 0.620265038 |
| Mir6417 | 1.584881738 | -0.877762709 | 0.986087682 | -0.890146712 |

| | | | | |
|---|---|---|---|---|
| RP24-309H3.1 | 30.5102578 | 0.546769282 | 0.293880669 | 1.860514625 |
| Dhx15 | 1662.063983 | -0.014250494 | 0.098528131 | -0.14463376 |
| RP24-309H3.3 | 27.29790927 | 0.139052576 | 0.250364417 | 0.555400716 |
| 9230114K14Rik | 31.71347572 | -0.223883258 | 0.217968413 | -1.027136249 |
| RP24-309H3.5 | 0.319554274 | 1.068637442 | 1.602441036 | 0.666880976 |
| RP24-309H3.6 | 0.802626749 | -1.01554873 | 1.191760547 | -0.852141592 |
| RP24-237I4.4 | 0 | NA | NA | NA |
| Gm25601 | 0.045882455 | 0.517959912 | 3.724945049 | 0.139051692 |
| Sod3 | 146.4300929 | -0.649130377 | 0.148770634 | -4.363296429 |
| Ccdc149 | 659.1518796 | 0.421151442 | 0.076271947 | 5.521708301 |
| RP24-237I4.3 | 0.09689823 | 0.037062677 | 3.724945049 | 0.009949859 |
| Lgi2 | 1350.453356 | 0.227994161 | 0.076409719 | 2.98383717 |
| RP23-181F22.4 | 0 | NA | NA | NA |
| RP23-181F22.3 | 4.780205594 | 1.298675323 | 0.625903701 | 2.074880404 |
| Gm5866 | 0.405467889 | 0.936137988 | 1.585275784 | 0.590520588 |
| 8030423F21Rik | 2.177346141 | -1.256654507 | 0.722168103 | -1.740113557 |
| Sepsecs | 236.6757126 | -0.031891636 | 0.100551367 | -0.317167602 |
| Pi4k2b | 58.07000918 | -0.920864037 | 0.202768104 | -4.54146396 |
| Zcchc4 | 173.3575901 | 0.244496804 | 0.167639049 | 1.458471674 |
| Anapc4 | 711.191152 | 0.072945394 | 0.052275305 | 1.395408303 |
| RP24-373A18.2 | 0 | NA | NA | NA |
| Gm23532 | 0 | NA | NA | NA |
| RP24-373A18.1 | 0 | NA | NA | NA |
| RP24-373A18.3 | 0.457291341 | -2.716321855 | 1.681641413 | -1.615280067 |
| Slc34a2 | 1.437183928 | -2.010565455 | 0.938488025 | -2.142345348 |
| Gm17182 | 0 | NA | NA | NA |
| Sel1l3 | 908.2236204 | 0.140196826 | 0.12758202 | 1.098876048 |
| n-R5s171 | 0 | NA | NA | NA |
| Smim20 | 129.4033044 | -0.203849171 | 0.141295693 | -1.44271326 |
| RP23-425N12.2 | 79.50346624 | 0.122506072 | 0.158817728 | 0.7713627 |
| n-R5s172 | 0.379461129 | -0.343775449 | 1.795716179 | -0.191441974 |
| RP23-425N12.1 | 0.485522898 | -0.517171694 | 1.514245371 | -0.341537576 |
| Gm26486 | 0 | NA | NA | NA |
| Rbpj | 346.6721826 | -0.388742304 | 0.101295695 | -3.837698195 |
| RP24-540G23.3 | 0.236145685 | 0.050066101 | 2.142401746 | 0.023369147 |
| RP23-137N6.2 | 0.29199547 | 0.06225155 | 1.777339783 | 0.035025126 |
| Cckar | 6.464590576 | -0.851939714 | 0.496966878 | -1.714278661 |
| Tbc1d19 | 456.9575057 | 0.170316116 | 0.104998155 | 1.622086749 |
| RP24-196M24.2 | 141.7804511 | 0.579346012 | 0.21617667 | 2.67996547 |
| RP24-196M24.3 | 10.28263209 | 0.470760712 | 0.389700304 | 1.208007043 |
| Stim2 | 905.9735999 | 0.233111368 | 0.062226402 | 3.74618104 |
| Gm15820 | 0 | NA | NA | NA |
| RP24-482F14.1 | 1.962603442 | 0.885152543 | 0.705647422 | 1.254383584 |
| RP23-148F6.1 | 0 | NA | NA | NA |
| RP23-187K3.1 | 6.300906923 | -0.333516138 | 0.415972626 | -0.801774243 |
| RP24-201P24.1 | 0.044544648 | 0.517957688 | 3.724945049 | 0.139051095 |
| Gm24235 | 0 | NA | NA | NA |

| | | | | |
|---|---|---|---|---|
| RP23-267P6.1 | 0 | NA | NA | NA |
| Gm25513 | 0 | NA | NA | NA |
| RP24-133H21.1 | 0 | NA | NA | NA |
| RP23-317I8.3 | 0 | NA | NA | NA |
| RP23-317I8.2 | 0 | NA | NA | NA |
| Gm15975 | 34.36835703 | -0.194522326 | 0.236388267 | -0.822893318 |
| RP23-317I8.1 | 0.181740481 | 0.51796848 | 2.286843757 | 0.226499287 |
| RP23-162F3.1 | 0.072025462 | 0.517959912 | 3.724945049 | 0.139051692 |
| RP23-162F3.2 | 1.035655884 | 0.534853548 | 1.145079512 | 0.467088567 |
| RP24-301M6.1 | 0.124853745 | 1.05357957 | 3.721447096 | 0.283110183 |
| RP24-394H13.1 | 0.707701802 | 1.097486234 | 1.408567406 | 0.779150667 |
| Gm23896 | 0 | NA | NA | NA |
| Gm24902 | 0 | NA | NA | NA |
| Gm23015 | 0 | NA | NA | NA |
| RP23-17A11.1 | 0 | NA | NA | NA |
| RP24-499N22.1 | 0 | NA | NA | NA |
| RP24-499N22.2 | 0 | NA | NA | NA |
| RP23-297D16.1 | 0 | NA | NA | NA |
| RP23-297D16.2 | 0.038981395 | -0.443816764 | 3.724945049 | -0.119147198 |
| 4932441J04Rik | 57.67652023 | 0.163137265 | 0.158659411 | 1.02822306 |
| Gm8121 | 18.9482281 | -0.453354433 | 0.248209214 | -1.826501225 |
| RP24-189M4.2 | 0 | NA | NA | NA |
| RP24-189M4.1 | 0.956833923 | -1.715356766 | 1.151854573 | -1.489212966 |
| Pcdh7 | 3274.629757 | 0.316433513 | 0.092875353 | 3.407077357 |
| RP23-458G12.1 | 132.1876841 | 0.193666086 | 0.136933987 | 1.414302542 |
| RP23-458G12.4 | 95.53520872 | 0.120401311 | 0.211821208 | 0.568410086 |
| RP23-458G12.3 | 115.8575128 | 0.308399719 | 0.182196165 | 1.692679535 |
| RP23-458G12.2 | 78.02550846 | 0.076824391 | 0.228566312 | 0.336114235 |
| RP23-358D12.4 | 15.63086198 | 0.559878651 | 0.406866935 | 1.376073116 |
| RP23-358D12.1 | 30.61318993 | 0.507760696 | 0.27682132 | 1.834254299 |
| RP23-358D12.3 | 67.73431069 | 0.04673966 | 0.183934913 | 0.25410978 |
| RP23-358D12.2 | 74.52970784 | -0.046352697 | 0.23677939 | -0.195763225 |
| RP24-231P10.2 | 87.86336388 | -0.012489186 | 0.208389942 | -0.059931809 |
| RP24-231P10.3 | 68.50219268 | 0.470610503 | 0.267064721 | 1.762159007 |
| RP24-231P10.4 | 91.67305281 | -0.140665783 | 0.240783108 | -0.584201213 |
| Gm37720 | 43.33085034 | 0.069607078 | 0.19998363 | 0.34806388 |
| RP24-231P10.5 | 54.31474321 | 0.221335336 | 0.21289577 | 1.039641772 |
| RP24-340K5.2 | 33.48457548 | 0.194131702 | 0.234532213 | 0.827740037 |
| RP24-340K5.4 | 0 | NA | NA | NA |
| RP24-340K5.3 | 0 | NA | NA | NA |
| RP24-502C10.1 | 0 | NA | NA | NA |
| Gm22536 | 0 | NA | NA | NA |
| RP24-375D24.2 | 0 | NA | NA | NA |
| 1700029E06Rik | 0 | NA | NA | NA |
| RP24-353N4.4 | 0 | NA | NA | NA |
| Gm6632 | 0 | NA | NA | NA |
| RP24-353N4.3 | 0 | NA | NA | NA |

| | | | | |
|---|---|---|---|---|
| Gm22308 | 0 | NA | NA | NA |
| RP23-324H16.1 | 0 | NA | NA | NA |
| RP23-324H16.2 | 3.023846459 | 0.826168239 | 0.606736452 | 1.361659146 |
| RP23-340N4.1 | 0 | NA | NA | NA |
| RP24-267O6.1 | 1.051957033 | 3.280990704 | 1.145047495 | 2.865375208 |
| RP24-543J12.1 | 0 | NA | NA | NA |
| RP24-543J12.2 | 0.062426873 | 0.517959912 | 3.724945049 | 0.139051692 |
| RP24-543J12.3 | 0 | NA | NA | NA |
| RP24-543J12.4 | 0.075043205 | 0.517957688 | 3.724945049 | 0.139051095 |
| RP23-183O3.1 | 0.136194719 | 0.037062677 | 3.724945049 | 0.009949859 |
| Cbfa2t2-ps1 | 30.26364068 | 0.295108518 | 0.204372963 | 1.443970443 |
| Gm23331 | 0 | NA | NA | NA |
| RP23-419P12.1 | 0 | NA | NA | NA |
| RP23-84N24.1 | 0 | NA | NA | NA |
| Gm23708 | 0 | NA | NA | NA |
| RP24-390N7.1 | 2.35460177 | -0.502369718 | 0.678289705 | -0.740641815 |
| RP24-390N7.2 | 0 | NA | NA | NA |
| RP24-390N7.3 | 0 | NA | NA | NA |
| RP24-300K9.1 | 0 | NA | NA | NA |
| RP23-210A16.1 | 0 | NA | NA | NA |
| RP23-86N20.1 | 0 | NA | NA | NA |
| G6pd2 | 0.044191549 | 0.037062677 | 3.724945049 | 0.009949859 |
| RP23-480E8.1 | 0 | NA | NA | NA |
| Gm22273 | 0 | NA | NA | NA |
| RP24-447I18.1 | 0.236971717 | 0.972316888 | 2.106041627 | 0.461679805 |
| Gm24079 | 0 | NA | NA | NA |
| Arap2 | 2567.739427 | -0.028850967 | 0.072522119 | -0.397822997 |
| RP23-387F14.1 | 16.56268708 | -0.173781896 | 0.293953525 | -0.591188337 |
| RP23-387F14.2 | 57.3555258 | 0.040022585 | 0.190103113 | 0.210530931 |
| Dthd1 | 4.062000091 | -1.452565038 | 0.551068886 | -2.635904648 |
| RP23-37I24.1 | 105.4712107 | 0.719743355 | 0.171176551 | 4.204684299 |
| Nwd2 | 487.7321807 | 0.37065329 | 0.161987495 | 2.288159894 |
| Gm3687 | 4.714620213 | 0.288552422 | 0.575117614 | 0.501727673 |
| 0610040J01Rik | 36.24267069 | 0.134324586 | 0.191656426 | 0.700861371 |
| Rell1 | 246.5447793 | -0.323766924 | 0.119309716 | -2.713667715 |
| Gm15819 | 0.356802427 | 0.221915539 | 3.695077298 | 0.060057076 |
| RP23-192C21.2 | 0.04393175 | 0.037062677 | 3.724945049 | 0.009949859 |
| Gm25608 | 0 | NA | NA | NA |
| RP23-192C21.3 | 1.013052095 | -1.901669418 | 1.053151897 | -1.805693389 |
| Pgm1 | 176.2902017 | -0.705290963 | 0.137535244 | -5.128074377 |
| Gm23967 | 0 | NA | NA | NA |
| Gm22395 | 0 | NA | NA | NA |
| Gm22559 | 0 | NA | NA | NA |
| Tbc1d1 | 746.5571268 | -0.202142668 | 0.098674317 | -2.048584413 |
| Gm15824 | 0.060174537 | 0.517957688 | 3.724945049 | 0.139051095 |
| RP23-339I6.1 | 13.2226886 | -0.557110165 | 0.319027821 | -1.746274551 |
| Gm15823 | 0 | NA | NA | NA |

| | | | | |
|---|---|---|---|---|
| Gm6044 | 4.011017284 | 0.965728076 | 0.567288579 | 1.702357692 |
| RP23-339I6.2 | 0.245047857 | 1.020272391 | 2.092747602 | 0.487527684 |
| RP23-339I6.3 | 0.541557618 | 0.629541784 | 1.428780429 | 0.440614787 |
| RP23-195B3.4 | 0.062426873 | 0.517959912 | 3.724945049 | 0.139051692 |
| 1700027F09Rik | 0 | NA | NA | NA |
| RP23-195B3.3 | 0 | NA | NA | NA |
| RP23-195B3.5 | 10.29678862 | 2.364371885 | 0.752606839 | 3.141576403 |
| RP24-448C16.4 | 0.540262476 | 0.86423623 | 1.336742636 | 0.646524025 |
| RP24-448C16.6 | 0.192501058 | -0.443822989 | 2.674470728 | -0.16594797 |
| RP24-448C16.5 | 1.291588001 | 0.859385306 | 0.949868936 | 0.904740932 |
| RP24-448C16.7 | 3.668071862 | -0.214950286 | 0.554936835 | -0.387341896 |
| RP24-448C16.9 | 0.116217011 | 0.517955464 | 3.724945049 | 0.139050498 |
| RP24-448C16.8 | 0 | NA | NA | NA |
| Gm25306 | 0.049553047 | -0.443818988 | 3.724945049 | -0.119147795 |
| Gm26761 | 0.503421469 | 0.086046642 | 1.480570825 | 0.058117207 |
| Gm20033 | 20.30300836 | 0.311121157 | 0.25546469 | 1.217863641 |
| Klf3 | 642.6073925 | -0.48885607 | 0.110570614 | -4.421211506 |
| RP23-326O2.2 | 26.4782624 | 0.051896099 | 0.236330074 | 0.219591599 |
| Tlr1 | 85.37836272 | -0.626927893 | 0.19811536 | -3.164458789 |
| Tlr6 | 42.58363877 | -0.954707653 | 0.238404663 | -4.004567871 |
| Fam114a1 | 72.52929912 | -0.851773227 | 0.175211862 | -4.861390201 |
| Mir574 | 0.038981395 | -0.443816764 | 3.724945049 | -0.119147198 |
| RP24-87P2.3 | 1.276825249 | 0.269193954 | 0.898940883 | 0.299456793 |
| Tmem156 | 6.703461548 | -1.202186977 | 0.445112652 | -2.700860042 |
| Klhl5 | 474.7177188 | -0.078813237 | 0.109225664 | -0.721563359 |
| Gm25154 | 2.287715794 | 0.496189052 | 0.802224428 | 0.618516509 |
| Wdr19 | 272.9715081 | -0.005784423 | 0.119682414 | -0.048331434 |
| Rfc1 | 611.6308179 | -0.151905791 | 0.076583344 | -1.983535615 |
| RP23-344O11.3 | 13.50952572 | 0.470342941 | 0.406432875 | 1.157246302 |
| Klb | 1.57634325 | -0.180974524 | 0.885864736 | -0.204291374 |
| Rpl9 | 1617.17554 | -0.22346166 | 0.087771254 | -2.545954979 |
| Lias | 326.7940019 | 0.123667267 | 0.085926199 | 1.439226558 |
| RP23-477N9.1 | 7.915089792 | 1.117861758 | 0.522597856 | 2.13904773 |
| RP23-477N9.5 | 5.012299285 | 0.114206059 | 0.513982071 | 0.222198527 |
| Ugdh | 393.579355 | -0.412734801 | 0.091797013 | -4.496168084 |
| AC136513.6 | 0.044544648 | 0.517957688 | 3.724945049 | 0.139051095 |
| RP23-477N9.2 | 5.168412192 | 0.691420379 | 0.504172679 | 1.371395967 |
| Smim14 | 1287.523276 | 0.197198008 | 0.055300515 | 3.565934406 |
| RP23-477N9.4 | 25.33295597 | 0.639341584 | 0.354486151 | 1.803572813 |
| RP23-477N9.6 | 5.250217998 | -0.009189019 | 0.508947424 | -0.018054947 |
| RP23-477N9.3 | 33.76352782 | -0.238667177 | 0.439233098 | -0.543372479 |
| Ube2k | 1596.82342 | 0.270347285 | 0.062364383 | 4.334962903 |
| RP23-403P24.1 | 88.56705528 | 0.358658655 | 0.294655614 | 1.217213038 |
| RP23-403P24.3 | 0 | NA | NA | NA |
| Pds5a | 1522.438719 | -0.100748894 | 0.06571203 | -1.533187977 |
| RP23-403P24.5 | 11.79713159 | -0.052827308 | 0.360716547 | -0.146451025 |
| RP23-403P24.6 | 4.819094623 | -0.155674861 | 0.487347016 | -0.319433291 |

| | | | | |
|---|---|---|---|---|
| Gm24399 | 0.0878635 | 0.037058229 | 3.724945049 | 0.009948665 |
| RP23-403P24.7 | 16.35270107 | 0.283615826 | 0.3713951 | 0.763649886 |
| RP23-403P24.8 | 29.36603598 | 0.243210283 | 0.284965332 | 0.853473232 |
| N4bp2os | 32.02296958 | 0.345430746 | 0.20151017 | 1.714209989 |
| N4bp2 | 387.2265267 | 0.20422617 | 0.117071904 | 1.744450742 |
| Gm24346 | 4.621381795 | 0.514116529 | 0.590848781 | 0.870132165 |
| RP23-42O8.7 | 30.0274551 | 0.205743696 | 0.346382435 | 0.593978433 |
| RP23-42O8.6 | 0.112396584 | -0.917692504 | 3.724434845 | -0.246397787 |
| RP23-42O8.5 | 0.165156969 | -1.323927776 | 3.401541256 | -0.389214087 |
| RP23-42O8.4 | 0 | NA | NA | NA |
| Rhoh | 32.07602456 | -0.349864224 | 0.191979213 | -1.822406803 |
| RP23-42O8.9 | 0.263966226 | -0.443366544 | 1.822623182 | -0.243257382 |
| Chrna9 | 0.40746857 | -0.662823141 | 1.451183083 | -0.456746739 |
| RP23-92N2.1 | 0 | NA | NA | NA |
| RP23-92N2.2 | 0 | NA | NA | NA |
| 9130230L23Rik | 1.235688375 | -2.066857472 | 0.954456825 | -2.165480322 |
| Gm25952 | 0 | NA | NA | NA |
| Rbm47 | 123.9754224 | -1.14509949 | 0.195509443 | -5.857003503 |
| RP24-351O18.5 | 7.439759207 | -0.937607611 | 0.480491614 | -1.951350625 |
| RP24-351O18.4 | 9.722078793 | -1.58441465 | 0.419596955 | -3.776039436 |
| Gm26725 | 0.204218301 | -0.868706194 | 2.577641098 | -0.337015962 |
| Gm22929 | 0 | NA | NA | NA |
| 1700126H18Rik | 0 | NA | NA | NA |
| RP24-351O18.2 | 0 | NA | NA | NA |
| Gm15794 | 0 | NA | NA | NA |
| RP24-351O18.7 | 0.049553047 | -0.443818988 | 3.724945049 | -0.119147795 |
| Nsun7 | 120.513026 | -0.632852859 | 0.176167577 | -3.592334463 |
| Apbb2 | 1193.808036 | 0.079170601 | 0.072720471 | 1.088697578 |
| RP23-21D6.2 | 18.43062459 | 1.229023208 | 0.303027472 | 4.055814476 |
| Gm23183 | 8.321784305 | 1.065771637 | 0.453734798 | 2.348886711 |
| RP23-21D6.1 | 24.68721265 | 0.70097216 | 0.343740989 | 2.039245193 |
| Gm16273 | 0.062426873 | 0.517959912 | 3.724945049 | 0.139051692 |
| RP23-157G2.2 | 24.13982844 | 0.694931801 | 0.284159943 | 2.445565669 |
| RP23-157G2.1 | 9.142804473 | 0.786516459 | 0.420004729 | 1.87263715 |
| RP24-360B3.2 | 21.21818474 | 0.765925069 | 0.370455934 | 2.067520041 |
| RP24-360B3.3 | 23.13608884 | 0.367899714 | 0.337300927 | 1.090716582 |
| RP24-360B3.1 | 33.09598643 | 0.401957597 | 0.370121782 | 1.086014431 |
| Uchl1os | 3.562613421 | 1.037033386 | 0.564641738 | 1.836621909 |
| Uchl1 | 2844.203176 | 0.14390274 | 0.077326918 | 1.860965673 |
| RP23-472J13.3 | 20.85386141 | 1.6139002 | 0.519534278 | 3.106436417 |
| RP23-352L15.1 | 0.061920613 | -0.443816764 | 3.724945049 | -0.119147198 |
| Limch1 | 1533.433457 | 0.075298954 | 0.056204652 | 1.33972814 |
| Gm6517 | 9.113272248 | 0.788205907 | 0.412350011 | 1.91149724 |
| RP24-79P8.1 | 29.56015194 | 0.308827266 | 0.292103919 | 1.05725136 |
| RP24-383P24.2 | 11.74058244 | 0.340207832 | 0.377566916 | 0.901053081 |
| Gm15949 | 0 | NA | NA | NA |
| Gm15948 | 0.038981395 | -0.443816764 | 3.724945049 | -0.119147198 |

| | | | | |
|---|---|---|---|---|
| RP24-383P24.3 | 0.808719243 | -0.347557418 | 1.112507838 | -0.312408961 |
| Gm23841 | 0 | NA | NA | NA |
| Phox2b | 13.13116622 | -0.460058672 | 0.554791409 | -0.829246206 |
| RP24-95M11.1 | 1.073023658 | -1.306053514 | 1.22165391 | -1.069086345 |
| RP24-353A8.1 | 1.943439641 | -1.301089051 | 0.716702961 | -1.815381157 |
| Tmem33 | 1537.582746 | -0.121849158 | 0.065346538 | -1.864661269 |
| Slc30a9 | 1760.179162 | 0.23348149 | 0.057185096 | 4.082908066 |
| RP24-353A8.2 | 0.708786783 | 0.465088504 | 1.21377002 | 0.383176793 |
| Bend4 | 204.8953353 | 0.064083451 | 0.115130405 | 0.556616222 |
| Gm26756 | 4.883969858 | 0.93554501 | 0.534783555 | 1.749390013 |
| RP24-103L16.3 | 7.272572362 | 0.704007641 | 0.5752951 | 1.223733074 |
| RP24-103L16.4 | 0 | NA | NA | NA |
| RP24-103L16.5 | 15.55852251 | -0.385736961 | 0.307390244 | -1.254877044 |
| RP23-213N8.1 | 1.075240245 | 0.090552408 | 1.012963305 | 0.089393572 |
| Shisa3 | 72.51937258 | -1.275477348 | 0.540700104 | -2.358936753 |
| Atp8a1 | 4334.315932 | 0.278222988 | 0.060833876 | 4.573487815 |
| Gm23617 | 2.388208885 | -0.717893516 | 0.766875012 | -0.936128449 |
| RP23-213N8.2 | 52.3886502 | 0.514011626 | 0.284905167 | 1.80414989 |
| Gm15478 | 2.676788521 | -0.060796159 | 0.739335023 | -0.082230865 |
| RP24-274I18.1 | 56.96681602 | 0.610472425 | 0.307422852 | 1.985774385 |
| RP23-130C18.1 | 59.36905083 | 0.412612259 | 0.280086773 | 1.473158676 |
| RP23-130C18.2 | 29.63687175 | 0.745337993 | 0.304264217 | 2.449640643 |
| Gm15477 | 9.54473322 | 0.272071198 | 0.34898362 | 0.779610225 |
| RP23-130C18.3 | 0.310726904 | -1.482285493 | 2.07553168 | -0.714171461 |
| Gm21905 | 0.178178593 | 1.413521986 | 3.716108082 | 0.380376984 |
| RP23-130C18.4 | 1.374506326 | 1.895421952 | 1.155482937 | 1.640372083 |
| Gm5108 | 0.225572436 | 0.037617376 | 2.199023191 | 0.017106403 |
| RP23-387L12.2 | 0.395257705 | 0.635734997 | 1.684836379 | 0.37732744 |
| RP23-387L12.3 | 1.482312138 | 0.781935183 | 0.81327318 | 0.961466826 |
| Grxcr1 | 0.175796035 | -0.45233715 | 2.308680796 | -0.19592884 |
| RP23-387L12.4 | 0.038981395 | -0.443816764 | 3.724945049 | -0.119147198 |
| RP24-224K15.1 | 0 | NA | NA | NA |
| Gm23605 | 0 | NA | NA | NA |
| RP24-441E22.1 | 0.086648695 | 0.037058229 | 3.724945049 | 0.009948665 |
| RP23-72G21.1 | 2.775994898 | -0.441481588 | 0.723896859 | -0.60986808 |
| Kctd8 | 186.6518599 | 0.324211061 | 0.099868956 | 3.246364788 |
| RP23-319P12.6 | 23.8423184 | 0.290582749 | 0.262993038 | 1.104906622 |
| RP23-319P12.5 | 17.40396653 | 0.480668169 | 0.306113603 | 1.570228062 |
| RP23-319P12.4 | 57.79128105 | 0.407751613 | 0.263405058 | 1.548002216 |
| RP23-319P12.3 | 56.51122244 | 0.562555441 | 0.266004432 | 2.11483484 |
| RP23-319P12.2 | 47.23969796 | 0.956688182 | 0.275779525 | 3.469032674 |
| RP23-319P12.1 | 45.02692642 | 0.443811596 | 0.314388633 | 1.411665532 |
| RP23-114M24.1 | 151.2873273 | 0.329783802 | 0.229239994 | 1.438596276 |
| Gm24368 | 0.334883495 | 0.480577618 | 1.563151487 | 0.307441488 |
| Gm23932 | 0 | NA | NA | NA |
| Gm5867 | 0.309731683 | 0.507698094 | 1.747949654 | 0.290453499 |
| RP24-351E22.1 | 0.956841293 | 0.404861246 | 1.092464443 | 0.370594438 |

| | | | | |
|---|---|---|---|---|
| Yipf7 | 2.071883479 | -0.444298882 | 0.649529561 | -0.68403181 |
| RP24-351E22.2 | 0.331853251 | -0.461250081 | 1.772207302 | -0.260268695 |
| Guf1 | 544.3541319 | 0.418637451 | 0.109804366 | 3.812575654 |
| Gnpda2 | 249.4753375 | -0.352217438 | 0.130929171 | -2.690137239 |
| Gm26044 | 0 | NA | NA | NA |
| Gm23661 | 0 | NA | NA | NA |
| RP23-236G19.1 | 0 | NA | NA | NA |
| Gm23067 | 0 | NA | NA | NA |
| Gm26072 | 0 | NA | NA | NA |
| RP23-332O18.1 | 0 | NA | NA | NA |
| RP23-332O18.2 | 0.480717083 | 0.020768627 | 1.625268914 | 0.012778579 |
| Gabrg1 | 384.6081965 | 0.276415968 | 0.136650219 | 2.022799306 |
| Gabra2 | 1383.339494 | 0.446794965 | 0.083561376 | 5.346907705 |
| Cox7b2 | 1.615076682 | -1.472168551 | 0.928036154 | -1.586326724 |
| Gabra4 | 1311.463824 | 0.691697755 | 0.110444899 | 6.262831156 |
| Gabrb1 | 2188.449478 | 0.478889988 | 0.081176043 | 5.899400526 |
| RP24-156O12.1 | 9.874941751 | 0.103678916 | 0.36225246 | 0.286206245 |
| Gm15617 | 0.150086409 | 1.027495679 | 3.723011807 | 0.275985071 |
| Gm15616 | 12.70121426 | 0.353255678 | 0.314890101 | 1.12183799 |
| Commd8 | 452.1455523 | 0.018802171 | 0.073125994 | 0.257120208 |
| RP23-27G7.2 | 0 | NA | NA | NA |
| Atp10d | 139.6844161 | -0.268972457 | 0.140352691 | -1.916403992 |
| Corin | 18.43000873 | -0.640735211 | 0.322623656 | -1.986014353 |
| Gm25768 | 0.088383099 | 0.037058229 | 3.724945049 | 0.009948665 |
| Gm20647 | 0 | NA | NA | NA |
| Gm19560 | 0 | NA | NA | NA |
| Gm23712 | 0.116046057 | 0.037062677 | 3.724945049 | 0.009949859 |
| AU023070 | 0.14021869 | -0.443822075 | 2.92152541 | -0.151914501 |
| Nfxl1 | 313.9107044 | -0.100186039 | 0.075383815 | -1.329012597 |
| RP24-316L21.3 | 23.03118454 | -0.137904949 | 0.233629735 | -0.590271393 |
| RP24-506B15.1 | 0.413335106 | 0.046784666 | 1.437919808 | 0.032536353 |
| Zar1 | 3.771757384 | -0.502553239 | 0.576322927 | -0.871999387 |
| Gm5868 | 0.655263372 | 0.968754013 | 1.328328851 | 0.729302848 |
| Cnga1 | 0.310549208 | -1.917494994 | 3.694478014 | -0.519016485 |
| RP24-506B15.2 | 5.852374332 | -0.662843328 | 0.421415335 | -1.572897976 |
| RP24-506B15.3 | 0 | NA | NA | NA |
| Nipal1 | 7.542068972 | -1.170136337 | 0.606211453 | -1.930244522 |
| Txk | 24.20729952 | -0.866494727 | 0.425176891 | -2.037962895 |
| RP23-415O9.1 | 0 | NA | NA | NA |
| Tec | 81.72636874 | -1.487303678 | 0.280808019 | -5.296514272 |
| RP23-204A8.1 | 6.319016411 | -0.558153876 | 0.425681256 | -1.31120144 |
| Gm9870 | 6.265701809 | -0.255485131 | 0.43553427 | -0.586601672 |
| Slain2 | 498.0036851 | -0.30070101 | 0.076527832 | -3.929302616 |
| Gm10135 | 0.045882455 | 0.517959912 | 3.724945049 | 0.139051692 |
| Gm24776 | 0 | NA | NA | NA |
| Slc10a4 | 54.26526042 | 0.155556491 | 0.188361861 | 0.825838578 |
| Gm15847 | 1.549834172 | 2.252190418 | 1.403691178 | 1.604477148 |

| | | | | |
|---|---|---|---|---|
| AC108848.6 | 1.313571363 | 0.561763933 | 0.859194991 | 0.653825893 |
| Fryl | 1628.389783 | -0.094969239 | 0.06177584 | -1.537320076 |
| RP23-300K5.1 | 35.04837016 | 0.281843392 | 0.268414943 | 1.050028692 |
| 1700071G01Rik | 0 | NA | NA | NA |
| RP23-318I19.2 | 17.43330629 | -0.161174207 | 0.345768389 | -0.466133436 |
| Gm17207 | 5.90520374 | 0.007342112 | 0.458256759 | 0.016021829 |
| RP23-153H17.2 | 0.097139547 | -0.443816764 | 3.724945049 | -0.119147198 |
| 1700025M24Rik | 0.068097359 | 0.037064902 | 3.724945049 | 0.009950456 |
| RP23-153H17.3 | 0 | NA | NA | NA |
| Ociad1 | 3089.545592 | 0.167508054 | 0.04694363 | 3.568280805 |
| RP23-153H17.6 | 0.3552573 | -1.017463789 | 2.010188729 | -0.506153365 |
| RP23-153H17.5 | 470.3152913 | 0.518489703 | 0.088428419 | 5.863383147 |
| Ociad2 | 527.2792456 | 0.045860746 | 0.150082629 | 0.305569977 |
| RP23-153H17.7 | 29.63938725 | -0.538817809 | 0.27820204 | -1.936785973 |
| RP23-153H17.9 | 0.315779812 | -1.504557494 | 3.707348516 | -0.405831145 |
| Cwh43 | 14.92583306 | 0.584812739 | 0.320511339 | 1.82462418 |
| RP24-274B10.1 | 0.149988742 | 0.998859423 | 2.860996859 | 0.349129857 |
| Gm15653 | 0 | NA | NA | NA |
| Dcun1d4 | 1162.429234 | 0.305210638 | 0.067991957 | 4.488922686 |
| Lrrc66 | 2.285793377 | 1.32854723 | 0.803572513 | 1.653300988 |
| Sgcb | 773.7364474 | 0.154595955 | 0.092204562 | 1.676662758 |
| RP23-105O12.2 | 2.772617781 | -0.060989706 | 0.629306428 | -0.096915752 |
| Spata18 | 12.87588817 | -0.170740795 | 0.366473323 | -0.465902385 |
| RP23-276J7.1 | 0 | NA | NA | NA |
| RP23-276J7.2 | 0.066050875 | -0.443818988 | 3.724945049 | -0.119147795 |
| Usp46 | 1284.997462 | 0.112890965 | 0.074208774 | 1.521261689 |
| Usp46os1 | 0.374638785 | 1.017784215 | 1.576841806 | 0.645457402 |
| Usp46os2 | 0.045882455 | 0.517959912 | 3.724945049 | 0.139051692 |
| CH25-194I22.2 | 46.11457756 | 0.853796104 | 0.317952401 | 2.68529535 |
| CH25-194I22.3 | 0 | NA | NA | NA |
| CH25-194I22.4 | 0.285862058 | -1.324059374 | 1.790113888 | -0.739650914 |
| Snora26 | 6.265788682 | -0.002640504 | 0.421048543 | -0.006271257 |
| Dancr | 0 | NA | NA | NA |
| RP23-422P4.2 | 0.766635478 | -1.662860339 | 1.142481857 | -1.455480741 |
| Rasl11b | 390.2960293 | 0.338391935 | 0.139195469 | 2.431055679 |
| Scfd2 | 235.5014393 | 0.218916957 | 0.081688505 | 2.679899189 |
| RP23-422P4.3 | 0.472351266 | 0.106107831 | 1.547431229 | 0.068570305 |
| Gm15981 | 4.746611391 | 0.830788172 | 0.517288339 | 1.606044657 |
| Gm15982 | 6.659240228 | 0.500586699 | 0.458026041 | 1.092921919 |
| RP24-465E5.2 | 15.61428043 | 0.321481196 | 0.413410157 | 0.777632553 |
| RP24-465E5.1 | 9.836186905 | 0.807048663 | 0.467321179 | 1.726967874 |
| Fip1l1 | 1364.710559 | 0.120404934 | 0.07025521 | 1.713822126 |
| RP24-245O20.1 | 42.86441423 | 0.141195598 | 0.215249597 | 0.655962193 |
| Lnx1 | 208.7467131 | 0.520722654 | 0.094747798 | 5.495881318 |
| Gm15984 | 0.100902008 | -0.876201588 | 3.72093227 | -0.235479048 |
| Gm15985 | 0 | NA | NA | NA |
| RP23-88L6.1 | 0 | NA | NA | NA |

| | | | | |
|---|---|---|---|---|
| RP24-245O20.2 | 0 | NA | NA | NA |
| Gm6116 | 0.042716945 | 0.037062677 | 3.724945049 | 0.009949859 |
| RP23-382I10.2 | 10.57589096 | -0.299478499 | 0.374973246 | -0.79866631 |
| Chic2 | 373.5242777 | 0.12190817 | 0.10500394 | 1.160986617 |
| RP23-382I10.1 | 56.62750935 | 0.35044675 | 0.279240646 | 1.254999067 |
| RP23-382I10.3 | 0.095897396 | 0.037062677 | 3.724945049 | 0.009949859 |
| Gsx2 | 1.492637219 | 0.659984879 | 0.919702494 | 0.717606925 |
| RP23-55P22.2 | 0 | NA | NA | NA |
| RP23-55P22.3 | 0.097139547 | -0.443816764 | 3.724945049 | -0.119147198 |
| Olfr1401-ps1 | 0 | NA | NA | NA |
| Pdgfra | 784.1705611 | -0.017910697 | 0.109045987 | -0.164249028 |
| RP23-55P22.6 | 0.141219523 | -0.44382205 | 2.915099384 | -0.152249372 |
| RP23-55P22.5 | 0 | NA | NA | NA |
| Mir7025 | 0.116046057 | 0.037062677 | 3.724945049 | 0.009949859 |
| RP23-55P22.7 | 0 | NA | NA | NA |
| RP23-149H20.2 | 0 | NA | NA | NA |
| Gm22084 | 0 | NA | NA | NA |
| RP23-149H20.3 | 0 | NA | NA | NA |
| Gm24502 | 0 | NA | NA | NA |
| Kit | 814.734425 | 0.17552858 | 0.114528626 | 1.53261753 |
| RP24-273B9.2 | 0.092228164 | 0.037073798 | 3.724945049 | 0.009952844 |
| RP24-273B9.1 | 0.222682676 | -1.363802871 | 2.136262691 | -0.638405977 |
| RP24-273B9.3 | 0 | NA | NA | NA |
| RP23-329A21.3 | 0.435363789 | 0.152422409 | 1.463928345 | 0.104118763 |
| Gm38180 | 0.072025462 | 0.517959912 | 3.724945049 | 0.139051692 |
| Gm28865 | 0.066050875 | -0.443818988 | 3.724945049 | -0.119147795 |
| Gm25374 | 0 | NA | NA | NA |
| Kdr | 411.3836948 | -1.822482984 | 0.288416487 | -6.318927881 |
| Gm24738 | 0 | NA | NA | NA |
| RP24-445F15.2 | 0 | NA | NA | NA |
| RP24-445F15.1 | 2.139707088 | 1.035582077 | 0.701502816 | 1.47623367 |
| Srd5a3 | 137.1421468 | -0.116296964 | 0.111788724 | -1.040328212 |
| Tmem165 | 351.682121 | -0.299929017 | 0.096712998 | -3.101227576 |
| Apc-ps1 | 7.578865732 | 0.700918859 | 0.515620684 | 1.359369167 |
| Clock | 2177.681423 | 0.15440011 | 0.060955492 | 2.53299752 |
| RP23-386C10.8 | 47.58777522 | -0.006503939 | 0.205919102 | -0.031584925 |
| RP23-386C10.5 | 89.03088876 | 0.681474829 | 0.287939248 | 2.366731295 |
| Gm7467 | 7.824312757 | 0.40625039 | 0.419776376 | 0.967778116 |
| Pdcl2 | 0.114443069 | -0.443821212 | 3.724945049 | -0.119148392 |
| 4930432L08Rik | 0.99774422 | -0.879962017 | 1.273297503 | -0.691089094 |
| Nmu | 1.605616479 | -0.274453231 | 0.906507487 | -0.302758923 |
| RP24-186A17.1 | 0 | NA | NA | NA |
| RP24-186A17.2 | 0.068097359 | 0.037064902 | 3.724945049 | 0.009950456 |
| Gm7271 | 1.031658297 | 0.097395969 | 1.092599393 | 0.089141518 |
| Exoc1 | 794.8721072 | 0.108323879 | 0.098189215 | 1.103215655 |
| Cep135 | 200.4137152 | 0.061767932 | 0.114848607 | 0.537820468 |
| A730089K16Rik | 11.31091771 | 0.00155572 | 0.344225158 | 0.004519484 |

| | | | | |
|---|---|---|---|---|
| C530008M17Rik | 974.9986135 | 0.070905053 | 0.073353611 | 0.966619807 |
| Gm7494 | 0.044191549 | 0.037062677 | 3.724945049 | 0.009949859 |
| Aasdh | 129.6955173 | -0.032385154 | 0.17462727 | -0.185453017 |
| Ppat | 241.3568791 | 0.265973142 | 0.086723842 | 3.066897574 |
| 2310040G07Rik | 1.39740727 | -0.064234872 | 0.973584164 | -0.065977728 |
| Paics | 931.5296296 | -0.113936793 | 0.092126351 | -1.23674488 |
| Srp72 | 1194.911166 | 0.016476257 | 0.063518046 | 0.259394903 |
| Arl9 | 3.563487532 | -0.765705378 | 0.542449938 | -1.411568744 |
| 1700112J05Rik | 0 | NA | NA | NA |
| 1700023E05Rik | 5.201511052 | -0.064430761 | 0.469931886 | -0.137106596 |
| Hopx | 352.4399641 | 0.018090045 | 0.139180938 | 0.129975018 |
| Hopxos | 6.015739958 | 0.838391933 | 0.392854425 | 2.134103319 |
| Gm15831 | 0.13681464 | 0.037072711 | 2.943790034 | 0.012593531 |
| RP23-3L23.1 | 0 | NA | NA | NA |
| Spink2 | 0.769070381 | -2.726523555 | 1.225761004 | -2.224351685 |
| RP24-439P5.1 | 0 | NA | NA | NA |
| Rest | 302.1062253 | -0.624706863 | 0.133596514 | -4.676071574 |
| Mir5098 | 0.366116714 | -0.291229932 | 1.532280046 | -0.190063124 |
| Noa1 | 216.3475351 | -0.105014473 | 0.085110817 | -1.233855766 |
| Polr2b | 926.3349238 | 0.046686538 | 0.056912343 | 0.820323587 |
| RP24-439P5.3 | 1.692103223 | 1.210652979 | 0.916424433 | 1.321061437 |
| Igfbp7 | 165.7570609 | -0.825240456 | 0.150264343 | -5.49192471 |
| RP24-89O12.2 | 0 | NA | NA | NA |
| RP24-89O12.4 | 0.126225412 | 0.03707172 | 3.698755067 | 0.010022756 |
| RP24-89O12.3 | 0 | NA | NA | NA |
| RP24-89O12.5 | 0 | NA | NA | NA |
| Pea15b | 0.138038828 | 0.998847613 | 3.63333328 | 0.274912191 |
| Gm22011 | 0 | NA | NA | NA |
| RP24-358F13.1 | 0.17692102 | -0.979939382 | 2.723401576 | -0.359821846 |
| RP23-465P13.1 | 0 | NA | NA | NA |
| Mthfr-ps1 | 0.438166815 | 0.26739387 | 1.470183951 | 0.181877832 |
| RP23-135B1.1 | 0 | NA | NA | NA |
| Gm25756 | 0 | NA | NA | NA |
| RP24-318J14.1 | 0 | NA | NA | NA |
| RP24-187F3.3 | 0 | NA | NA | NA |
| Gm22553 | 0 | NA | NA | NA |
| Gm23429 | 0 | NA | NA | NA |
| Gm24304 | 0 | NA | NA | NA |
| Gm22974 | 0 | NA | NA | NA |
| Gm2199 | 0.805896106 | -3.042056716 | 1.213035935 | -2.507804285 |
| Adgrl3 | 3423.719416 | 0.167099123 | 0.083230086 | 2.007676932 |
| RP24-248M2.5 | 137.1470277 | -0.121125409 | 0.146272499 | -0.828080538 |
| RP24-74E24.5 | 20.74210032 | 0.338394729 | 0.32744458 | 1.033441228 |
| AC104196.1 | 2.384345449 | 1.196293977 | 0.654992324 | 1.826424422 |
| Gm16054 | 1.328319734 | 0.503909793 | 1.117762864 | 0.450819945 |
| RP24-82B9.1 | 52.62095602 | 0.447471266 | 0.170962408 | 2.617366424 |
| RP23-393A24.1 | 0.106465261 | 0.037073798 | 3.724945049 | 0.009952844 |

| | | | | |
|---|---|---|---|---|
| 1700031L13Rik | 0.049553047 | -0.443818988 | 3.724945049 | -0.119147795 |
| RP23-151G22.1 | 9.983061651 | -0.523040694 | 0.321563334 | -1.626555763 |
| Mir1187 | 0 | NA | NA | NA |
| Gm24129 | 0 | NA | NA | NA |
| RP24-558F5.1 | 0 | NA | NA | NA |
| Tecrl | 1.615357459 | -0.496399953 | 0.904375769 | -0.548886834 |
| Gm15626 | 0.088383099 | 0.037058229 | 3.724945049 | 0.009948665 |
| AC120867.1 | 0 | NA | NA | NA |
| Gm25765 | 0 | NA | NA | NA |
| Gm25806 | 0 | NA | NA | NA |
| Gm24792 | 0 | NA | NA | NA |
| RP23-189P23.1 | 0 | NA | NA | NA |
| Epha5 | 1720.415688 | 0.27942967 | 0.095193194 | 2.935395453 |
| Hmgn2-ps1 | 6.552657376 | 0.085463073 | 0.3913488 | 0.218380822 |
| RP24-166L23.1 | 2.846346598 | 0.771297673 | 0.801581175 | 0.962220293 |
| RP24-138J15.1 | 0 | NA | NA | NA |
| RP24-273G7.1 | 0 | NA | NA | NA |
| RP23-227I15.1 | 0.342219405 | 0.037077249 | 1.773858817 | 0.020902029 |
| Gm24626 | 0 | NA | NA | NA |
| RP23-258M22.2 | 1.514822285 | -0.764235817 | 0.909368461 | -0.840402818 |
| Cenpc1 | 463.6409797 | 0.186728575 | 0.063768527 | 2.928224683 |
| RP23-19G21.1 | 0.150451996 | 0.037073243 | 2.858243881 | 0.012970637 |
| Stap1 | 5.819746149 | -1.297353491 | 0.428134473 | -3.030247673 |
| Gm24524 | 0.085327104 | -0.905414734 | 3.723500899 | -0.243162217 |
| Uba6 | 668.4127751 | -0.344213874 | 0.066014318 | -5.214230548 |
| Gnrhr | 0.310823584 | 0.357101396 | 1.70131817 | 0.209896892 |
| Tmprss11c | 0.04393175 | 0.037062677 | 3.724945049 | 0.009949859 |
| Gm25473 | 0 | NA | NA | NA |
| Tmprss11d | 1.632697194 | -0.326674445 | 0.909712273 | -0.359096447 |
| Tmprss11a | 3.616981247 | 3.443862757 | 0.965251801 | 3.567838727 |
| Tmprss11g | 0.130746305 | 0.417046913 | 3.663433875 | 0.113840437 |
| Tmprss11f | 0.18352982 | 1.388044403 | 3.713044316 | 0.373829205 |
| Tmprss11b | 2.079765874 | -0.581975814 | 0.696942065 | -0.835041883 |
| RP23-314N3.1 | 0.090074004 | 0.517955464 | 3.724945049 | 0.139050498 |
| Tmprss11e | 0.166952883 | -1.336542448 | 2.746539428 | -0.486627803 |
| Gm16579 | 0.090277459 | -0.443823436 | 3.724945049 | -0.119148989 |
| RP24-567O8.4 | 0 | NA | NA | NA |
| RP24-567O8.3 | 0.115000406 | -0.443821212 | 3.724945049 | -0.119148392 |
| Ythdc1 | 1204.211079 | 0.113945225 | 0.107935602 | 1.055677861 |
| RP24-158N23.1 | 0 | NA | NA | NA |
| Ugt2b34 | 1.309248412 | -0.525095703 | 0.898984041 | -0.584099027 |
| Gm25211 | 0 | NA | NA | NA |
| Ugt2b1 | 0 | NA | NA | NA |
| Gm24121 | 0 | NA | NA | NA |
| RP23-59B8.1 | 0 | NA | NA | NA |
| RP23-227D7.1 | 0.092228164 | 0.037073798 | 3.724945049 | 0.009952844 |
| RP23-59B8.2 | 33.07073811 | -0.273565427 | 0.237306873 | -1.15279184 |

| | | | | |
|---|---|---|---|---|
| Ugt2b35 | 0.394550875 | 0.394384863 | 1.408673765 | 0.279968913 |
| RP23-59B8.3 | 0 | NA | NA | NA |
| RP23-227D7.2 | 0 | NA | NA | NA |
| Ugt2b36 | 0.256447508 | 0.010978678 | 2.469852793 | 0.004445074 |
| RP23-227D7.3 | 0 | NA | NA | NA |
| RP23-67B9.4 | 0 | NA | NA | NA |
| Ugt2b5 | 0.491974008 | -0.705968569 | 1.282196898 | -0.550592947 |
| RP23-67B9.2 | 0.045882455 | 0.517959912 | 3.724945049 | 0.139051692 |
| RP23-227D7.4 | 0 | NA | NA | NA |
| RP23-67B9.1 | 0.072025462 | 0.517959912 | 3.724945049 | 0.139051692 |
| Ugt2b37 | 0.046345709 | -0.443818988 | 3.724945049 | -0.119147795 |
| RP23-67B9.3 | 0.075043205 | 0.517957688 | 3.724945049 | 0.139051095 |
| Ugt2a3 | 0.072025462 | 0.517959912 | 3.724945049 | 0.139051692 |
| RP23-227D7.7 | 0 | NA | NA | NA |
| RP23-67B9.5 | 0 | NA | NA | NA |
| RP23-227D7.8 | 0.093141081 | 0.037060453 | 3.724945049 | 0.009949262 |
| RP23-227D7.6 | 0 | NA | NA | NA |
| RP23-227D7.5 | 0 | NA | NA | NA |
| Ugt2b38 | 0.123841226 | -1.017823543 | 3.719440177 | -0.273649661 |
| Ugt2a2 | 0.062426873 | 0.517959912 | 3.724945049 | 0.139051692 |
| Ugt2a1 | 0 | NA | NA | NA |
| RP23-227D7.10 | 0 | NA | NA | NA |
| Sult1b1 | 0.042716945 | 0.037062677 | 3.724945049 | 0.009949859 |
| Sult1d1 | 1.935656465 | -1.075871674 | 1.192855121 | -0.901929879 |
| Sult1e1 | 0.061920613 | -0.443816764 | 3.724945049 | -0.119147198 |
| mmu-mir-3969 | 0 | NA | NA | NA |
| RP24-571N6.2 | 0 | NA | NA | NA |
| RP24-571N6.3 | 0 | NA | NA | NA |
| Csn1s1 | 0.044544648 | 0.517957688 | 3.724945049 | 0.139051095 |
| Csn2 | 0 | NA | NA | NA |
| Gm25066 | 0 | NA | NA | NA |
| RP23-440C1.3 | 0.266205562 | 0.428176672 | 1.784322685 | 0.239965941 |
| Gm24314 | 0 | NA | NA | NA |
| Csn1s2a | 0.075043205 | 0.517957688 | 3.724945049 | 0.139051095 |
| Csn1s2b | 0 | NA | NA | NA |
| Prr27 | 0 | NA | NA | NA |
| RP23-440C1.5 | 0 | NA | NA | NA |
| Gm7337 | 0 | NA | NA | NA |
| Odam | 0 | NA | NA | NA |
| 1700066N21Rik | 0 | NA | NA | NA |
| RP23-423C8.4 | 0 | NA | NA | NA |
| Csn3 | 0.488468934 | -2.175398575 | 1.284261912 | -1.693890128 |
| BC051076 | 0.204292078 | 0.037060102 | 3.724945049 | 0.009949167 |
| 2310003L06Rik | 0.63240349 | -0.356508002 | 1.164741297 | -0.306083422 |
| Gm28434 | 0 | NA | NA | NA |
| Cabs1 | 0.109869311 | -0.444802825 | 3.724874622 | -0.119414174 |
| Smr3a | 0 | NA | NA | NA |

| | | | | |
|---|---|---|---|---|
| RP24-324D14.5 | 0 | NA | NA | NA |
| RP24-324D14.6 | 0 | NA | NA | NA |
| Smr2 | 0 | NA | NA | NA |
| Gm11116 | 0 | NA | NA | NA |
| RP23-50G20.2 | 0 | NA | NA | NA |
| Gm11115 | 0 | NA | NA | NA |
| RP23-50G20.3 | 0 | NA | NA | NA |
| RP23-50G20.5 | 0 | NA | NA | NA |
| RP23-50G20.4 | 0 | NA | NA | NA |
| Gm7714 | 0 | NA | NA | NA |
| Prol1 | 0 | NA | NA | NA |
| Amtn | 0 | NA | NA | NA |
| RP23-265M4.3 | 1.532002984 | 0.42777718 | 1.104950657 | 0.387145957 |
| RP23-265M4.4 | 0 | NA | NA | NA |
| Ambn | 0 | NA | NA | NA |
| Enam | 0 | NA | NA | NA |
| Jchain | 15.78987055 | -0.854634366 | 0.574797369 | -1.486844603 |
| RP24-132B11.1 | 0.854816446 | -1.044426381 | 1.07723551 | -0.969543216 |
| Utp3 | 218.2673712 | -0.083459642 | 0.183037503 | -0.455970174 |
| Rufy3 | 2816.589347 | 0.118086644 | 0.051052901 | 2.313025146 |
| Grsf1 | 1026.519618 | 0.090326903 | 0.061451623 | 1.469886376 |
| Mob1b | 844.5626815 | 0.073010361 | 0.120189802 | 0.607458872 |
| Gm15682 | 3.374206779 | -0.93105815 | 0.617147186 | -1.508648458 |
| Dck | 405.0431763 | -0.600499213 | 0.110140146 | -5.452137459 |
| RP24-242L17.1 | 22.06035809 | 0.112395828 | 0.393623307 | 0.285541597 |
| RP24-242L17.2 | 0 | NA | NA | NA |
| Mkrn1-ps1 | 1.85792581 | -0.412386356 | 0.709928708 | -0.58088418 |
| Slc4a4 | 4189.981023 | 0.163585098 | 0.13306907 | 1.22932472 |
| Gc | 1.073168736 | -1.2330677 | 1.181887963 | -1.043303375 |
| RP24-365I5.1 | 0.045882455 | 0.517959912 | 3.724945049 | 0.139051692 |
| RP24-365I5.2 | 0 | NA | NA | NA |
| Npffr2 | 7.733873509 | -0.523229858 | 0.515385253 | -1.015220858 |
| Adamts3 | 373.7060012 | 0.552857459 | 0.211848993 | 2.609677075 |
| Gm25320 | 0.186976534 | 0.517972142 | 2.668791346 | 0.194084915 |
| Gm25758 | 0 | NA | NA | NA |
| Eif5al3-ps | 2.402331814 | 0.074618719 | 0.652482508 | 0.114361256 |
| Cox18 | 236.2894485 | -0.056290344 | 0.107614927 | -0.523071896 |
| Ankrd17 | 4578.10987 | -0.004293319 | 0.06263088 | -0.068549559 |
| Gm9958 | 14.24586082 | -1.145083834 | 0.291814825 | -3.924008432 |
| RP23-279P6.5 | 0 | NA | NA | NA |
| Alb | 7.938965767 | 0.0151894 | 0.594806278 | 0.025536717 |
| Afp | 3.204684021 | -0.848491072 | 0.586899779 | -1.445717145 |
| RP23-279P6.3 | 0.060174537 | 0.517957688 | 3.724945049 | 0.139051095 |
| Afm | 0.397459543 | -1.643336892 | 1.684572947 | -0.97552136 |
| RP24-63O24.1 | 0.290146865 | 0.037076371 | 2.036264892 | 0.01820803 |
| RP24-63O24.4 | 0 | NA | NA | NA |
| 5830473C10Rik | 0.606733601 | 0.433835214 | 1.195448455 | 0.36290583 |

| | | | | |
|---|---|---|---|---|
| Rassf6 | 13.93087578 | 0.182319058 | 0.320965949 | 0.568032398 |
| RP24-168A7.1 | 0.237346126 | 0.538906854 | 2.11802704 | 0.254438137 |
| Cxcl5 | 7.088183414 | -1.311207536 | 0.495646188 | -2.645450659 |
| Ppbp | 1.785008614 | -0.723111347 | 0.808254905 | -0.894657543 |
| Pf4 | 3.421604734 | -1.089198771 | 0.687659235 | -1.583922263 |
| Cxcl3 | 3.225786153 | -3.64527493 | 0.913071232 | -3.992322615 |
| Cxcl15 | 0.068097359 | 0.037064902 | 3.724945049 | 0.009950456 |
| Gm24322 | 0 | NA | NA | NA |
| RP24-168A7.7 | 0 | NA | NA | NA |
| RP23-307P18.1 | 0 | NA | NA | NA |
| Cxcl1 | 39.73533034 | 0.45698981 | 0.826570069 | 0.552874858 |
| Gm22816 | 0.235449268 | 1.013096636 | 2.531872841 | 0.40013725 |
| Cxcl2 | 19.17032838 | -0.559038002 | 0.826565649 | -0.676338296 |
| RP23-307P18.3 | 0 | NA | NA | NA |
| Mthfd2l | 111.8726162 | 0.041880809 | 0.123620331 | 0.338785774 |
| Epgn | 0.633611781 | -0.039068896 | 1.418929221 | -0.02753407 |
| RP23-42K10.4 | 0.186036187 | -1.342920348 | 2.703027462 | -0.49682083 |
| Ereg | 4.701452776 | -2.623726149 | 0.758255197 | -3.46021519 |
| Areg | 0.714414789 | -0.790153683 | 1.149446857 | -0.687420805 |
| Gm23092 | 5.083391071 | 0.019863863 | 0.528841453 | 0.037561093 |
| RP23-258C6.2 | 6.099374268 | 0.492225389 | 0.397242165 | 1.239106603 |
| Btc | 22.83547764 | -1.002517232 | 0.270733944 | -3.702960987 |
| Parm1 | 754.069321 | 0.265423205 | 0.059037356 | 4.495851851 |
| RP23-118H9.3 | 3.970608911 | 0.049763046 | 0.531293447 | 0.093663956 |
| RP23-17D18.3 | 4.213956554 | -0.312484609 | 0.487909431 | -0.640456176 |
| Gm28271 | 0.521165882 | -1.047511125 | 1.505400475 | -0.695835522 |
| 1700063O14Rik | 0.323348989 | -0.382211783 | 1.744740668 | -0.219065097 |
| RP24-266G11.2 | 0.541747793 | -0.124984879 | 1.464416483 | -0.085347905 |
| RP24-266G11.4 | 0 | NA | NA | NA |
| Gm5558 | 5.850622695 | -0.355209901 | 0.500039236 | -0.710364059 |
| RP23-442L7.2 | 0.329047954 | 0.87402129 | 1.721363347 | 0.507749448 |
| Rchy1 | 427.1403147 | -0.280035151 | 0.087104615 | -3.214928991 |
| RP23-442L7.3 | 26.15808167 | 0.006813095 | 0.322540941 | 0.021123193 |
| Thap6 | 30.14759305 | -0.535692932 | 0.18559001 | -2.88643193 |
| Gm24113 | 0.102891481 | 0.51795324 | 3.724945049 | 0.139049901 |
| Gm1045 | 3.240137224 | 0.279389093 | 0.587650582 | 0.475434044 |
| Cdkl2 | 400.6574654 | 0.275947409 | 0.094780377 | 2.911440306 |
| Gm26582 | 7.944179279 | -0.140241902 | 0.382380612 | -0.366759971 |
| G3bp2 | 4286.859093 | -0.007354689 | 0.102547513 | -0.071719818 |
| Gm22728 | 6.520177888 | 0.839407397 | 0.455411725 | 1.843183545 |
| Gm24931 | 1.274348854 | 0.998402439 | 0.977204121 | 1.021692825 |
| Uso1 | 1247.998467 | -0.191037809 | 0.055684967 | -3.430689097 |
| Gm15710 | 147.9457653 | -0.413261772 | 0.156365993 | -2.642913355 |
| RP23-321C24.2 | 0 | NA | NA | NA |
| Gm25290 | 1.111053339 | 0.404232062 | 1.167820445 | 0.346142306 |
| U90926 | 0.097139547 | -0.443816764 | 3.724945049 | -0.119147198 |
| Ppef2 | 8.480238973 | -1.241469198 | 0.432776187 | -2.868617167 |

| | | | | |
|---|---|---|---|---|
| Naaa | 229.011174 | -0.088908914 | 0.143122546 | -0.621208305 |
| Sdad1 | 341.6351534 | 0.027551398 | 0.107254961 | 0.256877607 |
| RP23-447A7.2 | 0.049553047 | -0.443818988 | 3.724945049 | -0.119147795 |
| Gm23031 | 0.045882455 | 0.517959912 | 3.724945049 | 0.139051692 |
| Cxcl9 | 597.6617564 | 0.177639664 | 0.428128586 | 0.414921287 |
| Art3 | 40.01616527 | 0.226881137 | 0.181574736 | 1.249519302 |
| Cxcl10 | 3859.437308 | -0.479228424 | 0.523509854 | -0.915414334 |
| Cxcl11 | 885.873838 | -1.159125274 | 0.470130819 | -2.465537733 |
| Gm2673 | 0.689365755 | -2.150130482 | 1.146833383 | -1.874841206 |
| Nup54 | 228.7748306 | -0.302499624 | 0.122584045 | -2.467691654 |
| Gm26039 | 0 | NA | NA | NA |
| Scarb2 | 2989.989026 | -0.14522574 | 0.085405537 | -1.700425346 |
| Mir6415 | 0 | NA | NA | NA |
| Fam47e | 1.965355678 | 0.072489192 | 0.75495707 | 0.096017634 |
| Stbd1 | 13.78635077 | -0.926052368 | 0.374070919 | -2.475606421 |
| Ccdc158 | 6.083259113 | -0.823201564 | 0.447689479 | -1.838777998 |
| Shroom3 | 115.9678624 | -0.72507329 | 0.182963437 | -3.962940906 |
| Gm20500 | 0.399554685 | 0.531425977 | 1.606659144 | 0.330764605 |
| Mir1961 | 0 | NA | NA | NA |
| 4932430I15Rik | 0 | NA | NA | NA |
| RP23-9D24.2 | 1.173754143 | -0.594261235 | 1.103470677 | -0.538538312 |
| Gm25521 | 0.066050875 | -0.443818988 | 3.724945049 | -0.119147795 |
| Sowahb | 261.7368774 | 0.110824676 | 0.132852914 | 0.83419078 |
| Gm22915 | 1.677596521 | 0.191972158 | 0.964198267 | 0.199100293 |
| RP24-470E5.2 | 19.84950728 | 0.804438487 | 0.391280771 | 2.055911118 |
| RP24-470E5.3 | 3.144773293 | 0.722850761 | 0.706885256 | 1.022585709 |
| 11-Sep | 1923.005761 | -0.337602846 | 0.047529138 | -7.103071078 |
| RP24-470E5.4 | 138.2382822 | 0.777765388 | 0.158526584 | 4.906214278 |
| Ccni | 2707.469122 | 0.126411596 | 0.087550803 | 1.443865636 |
| RP24-470E5.9 | 57.10845697 | 0.266532613 | 0.246347857 | 1.081935991 |
| Gm2986 | 3.888599492 | -0.105618744 | 0.582105621 | -0.181442577 |
| 2010109A12Rik | 6.375499996 | -0.267598249 | 0.590007807 | -0.453550353 |
| Ccng2 | 404.8129049 | 0.115437149 | 0.106898848 | 1.079872712 |
| Kat2b-ps | 1.175757053 | -1.747181963 | 0.940673424 | -1.857373578 |
| RP23-388L12.2 | 0 | NA | NA | NA |
| RP23-173B14.3 | 0 | NA | NA | NA |
| Gm16427 | 0 | NA | NA | NA |
| C87414 | 0 | NA | NA | NA |
| RP23-388L12.3 | 0 | NA | NA | NA |
| RP23-388L12.1 | 0 | NA | NA | NA |
| RP23-57N5.1 | 0 | NA | NA | NA |
| RP23-173B14.4 | 0 | NA | NA | NA |
| RP23-57N5.3 | 0 | NA | NA | NA |
| Gm16429 | 0 | NA | NA | NA |
| RP23-173B14.2 | 0 | NA | NA | NA |
| RP23-173B14.1 | 0 | NA | NA | NA |
| BC061212 | 0 | NA | NA | NA |

| | | | | |
|---|---|---|---|---|
| Gm6351 | 0 | NA | NA | NA |
| Gm3106 | 0 | NA | NA | NA |
| Gm16522 | 0 | NA | NA | NA |
| Gm6502 | 0 | NA | NA | NA |
| AA792892 | 0 | NA | NA | NA |
| RP23-432M17.3 | 0 | NA | NA | NA |
| Gm7682 | 0 | NA | NA | NA |
| Gm3139 | 0 | NA | NA | NA |
| Gm3147 | 0 | NA | NA | NA |
| Gm6509 | 0 | NA | NA | NA |
| Gm6205 | 0 | NA | NA | NA |
| RP23-133M13.3 | 0 | NA | NA | NA |
| RP23-432M17.5 | 0 | NA | NA | NA |
| Gm3183 | 0 | NA | NA | NA |
| RP24-85B14.2 | 0 | NA | NA | NA |
| Gm7647 | 0 | NA | NA | NA |
| Gm6367 | 0 | NA | NA | NA |
| RP23-365E3.3 | 0 | NA | NA | NA |
| RP24-85B14.3 | 0 | NA | NA | NA |
| Gm7942 | 0 | NA | NA | NA |
| Gm16513 | 0 | NA | NA | NA |
| Gm10424 | 0 | NA | NA | NA |
| RP23-320H13.5 | 0 | NA | NA | NA |
| Gm3259 | 0 | NA | NA | NA |
| RP23-365E3.2 | 0 | NA | NA | NA |
| RP23-320H13.4 | 0 | NA | NA | NA |
| RP23-320H13.3 | 0 | NA | NA | NA |
| D5Ertd577e | 0 | NA | NA | NA |
| Gm3286 | 0 | NA | NA | NA |
| Gm3176 | 0 | NA | NA | NA |
| RP23-23P3.2 | 0 | NA | NA | NA |
| RP23-23P3.1 | 0 | NA | NA | NA |
| Gm7978 | 0 | NA | NA | NA |
| Gm7982 | 0 | NA | NA | NA |
| E330014E10Rik | 0 | NA | NA | NA |
| RP24-322E6.3 | 0 | NA | NA | NA |
| Gm5559 | 314.382447 | 0.181245547 | 0.130072143 | 1.393423243 |
| Gm25937 | 0 | NA | NA | NA |
| Cxcl13 | 78.29238741 | 0.627836406 | 0.289962873 | 2.165230323 |
| RP23-57A17.2 | 106.8115065 | -0.838482603 | 0.203580002 | -4.118688459 |
| Cnot6l | 1535.89927 | -0.043310844 | 0.072504797 | -0.597351424 |
| Gm2840 | 2.621041726 | -1.82592319 | 0.765317424 | -2.385837736 |
| Mrpl1 | 300.8320522 | 0.003570763 | 0.077712439 | 0.045948406 |
| Fras1 | 497.6484057 | 0.455318978 | 0.174605551 | 2.60770047 |
| RP23-381C14.2 | 0.206671645 | 0.03622389 | 2.636505289 | 0.013739358 |
| RP23-381C14.5 | 0.680615379 | -1.07195374 | 1.252033496 | -0.856170178 |
| RP23-480P21.2 | 0.07796279 | -0.798301147 | 3.721829328 | -0.214491605 |

| | | | | |
|---|---|---|---|---|
| Anxa3 | 185.5288032 | -1.671511726 | 0.219568031 | -7.612728146 |
| RP23-480P21.3 | 7.010785502 | -0.642754472 | 0.550015579 | -1.168611392 |
| RP24-142B15.2 | 0.147290723 | 0.491450248 | 2.859758244 | 0.171850278 |
| RP24-142B15.3 | 0.270258959 | -0.441879886 | 1.796004173 | -0.246035 |
| RP24-142B15.4 | 0.176646398 | -0.526386109 | 3.35399333 | -0.1569431 |
| Gm24623 | 0 | NA | NA | NA |
| RP24-142B15.5 | 2.750814746 | -0.953365959 | 0.597216511 | -1.596348964 |
| RP24-142B15.6 | 0.520715546 | -1.611054368 | 1.455552669 | -1.10683344 |
| RP24-142B15.7 | 1.115544105 | -0.447787346 | 0.871819847 | -0.513623712 |
| RP24-142B15.8 | 0.895791331 | -0.159176023 | 1.031481772 | -0.154317824 |
| RP24-142B15.9 | 0.252544957 | -1.499921942 | 2.086455967 | -0.718885021 |
| RP24-142B15.11 | 5.79381536 | -0.326338621 | 0.446788873 | -0.730409016 |
| Bmp2k | 887.8164416 | -0.357537752 | 0.10558539 | -3.386242657 |
| RP24-142B15.10 | 11.84488042 | 0.548798087 | 0.425631087 | 1.28937501 |
| Paqr3 | 238.9609811 | 0.415439944 | 0.150858394 | 2.753840431 |
| RP23-45P1.1 | 1.824918576 | 1.791948735 | 0.885923675 | 2.022689749 |
| RP23-28F1.3 | 1.733684669 | -0.238769327 | 0.907264304 | -0.263175048 |
| Gm5560 | 26.0495091 | -0.149616839 | 0.262038147 | -0.570973502 |
| RP23-28F1.4 | 0 | NA | NA | NA |
| Naa11 | 0.23191841 | -1.202715736 | 2.569832397 | -0.468013298 |
| 4930467D21Rik | 11.74611447 | 2.234101058 | 0.414349933 | 5.391821939 |
| Gk2 | 0.04393175 | 0.037062677 | 3.724945049 | 0.009949859 |
| Antxr2 | 68.57005387 | -1.355879909 | 0.19575239 | -6.926505003 |
| RP23-178N8.7 | 1.361532721 | -0.277406704 | 0.816437171 | -0.33977716 |
| RP23-178N8.3 | 0.042716945 | 0.037062677 | 3.724945049 | 0.009949859 |
| RP23-178N8.4 | 0 | NA | NA | NA |
| RP23-178N8.5 | 0.202865515 | -0.460713006 | 2.211509892 | -0.208325094 |
| Gm8048 | 0.123841226 | -1.017823543 | 3.719440177 | -0.273649661 |
| Prdm8 | 199.5149206 | 0.706250664 | 0.138738933 | 5.090500887 |
| A730035I17Rik | 9.47161779 | 0.859887624 | 0.376992289 | 2.28091568 |
| Fgf5 | 19.39898948 | 0.428999577 | 0.291502218 | 1.471685464 |
| 1700007G11Rik | 3.95741196 | -0.266709993 | 0.530933753 | -0.502341378 |
| RP23-425F8.1 | 1.376239274 | -1.414280065 | 1.172852136 | -1.20584686 |
| RP24-240P8.1 | 0.208122295 | 0.01451109 | 2.192995807 | 0.006617017 |
| mmu-mir-703 | 13.41816306 | -0.676933781 | 0.303800134 | -2.228220812 |
| Gm24279 | 0.141779851 | 0.517957264 | 3.596372909 | 0.144022124 |
| Gm11111 | 0.419266243 | 0.426563177 | 1.31479293 | 0.324433732 |
| 1700010H22Rik | 0 | NA | NA | NA |
| Bmp3 | 131.1088595 | 0.384474153 | 0.164966639 | 2.330617601 |
| Prkg2 | 141.6564417 | 0.127623648 | 0.139052562 | 0.917808674 |
| Gm3470 | 1.546431532 | 0.346395571 | 0.811630214 | 0.426789891 |
| RP24-548B7.1 | 0.424196832 | -1.202631734 | 1.595195934 | -0.753908475 |
| Rasgef1b | 664.9745574 | -0.155635106 | 0.072374402 | -2.150416461 |
| RP23-253I14.3 | 18.40400765 | 0.487877309 | 0.290335984 | 1.680388705 |
| A930011G23Rik | 101.3913818 | 0.36198837 | 0.294775943 | 1.228011911 |
| RP23-380D12.1 | 0.13861434 | 0.037059817 | 3.618284367 | 0.010242373 |
| Gm16227 | 6.434094977 | 0.002638029 | 0.47883516 | 0.005509264 |

| | | | | |
|---|---|---|---|---|
| Gm16228 | 3.733916757 | 0.620516894 | 0.62430578 | 0.993931041 |
| 4930405H06Rik | 0 | NA | NA | NA |
| Gm25969 | 0 | NA | NA | NA |
| Gm16226 | 0 | NA | NA | NA |
| 4930522N08Rik | 0 | NA | NA | NA |
| RP23-438O22.1 | 33.98607118 | 1.136673648 | 0.370217649 | 3.070284877 |
| RP23-57P23.3 | 0.038981395 | -0.443816764 | 3.724945049 | -0.119147198 |
| Gm26252 | 0 | NA | NA | NA |
| RP23-57P23.4 | 0.879409403 | 0.198138696 | 1.247659686 | 0.158808286 |
| Gm22838 | 0.115603922 | -0.93550548 | 3.724188147 | -0.251197158 |
| RP23-57P23.5 | 0.082913145 | -0.443821212 | 3.724945049 | -0.119148392 |
| Hnrnpd | 2150.330006 | 0.214875082 | 0.079748355 | 2.694413984 |
| Gm17092 | 3.730625592 | 0.00151628 | 0.518632471 | 0.002923611 |
| 4930524J08Rik | 4.763368001 | -0.642361857 | 0.480967489 | -1.335561909 |
| Hnrnpdl | 2079.483625 | 0.265185488 | 0.084226874 | 3.148466473 |
| Enoph1 | 302.3695605 | 0.082144261 | 0.096215559 | 0.853752365 |
| Gm26363 | 0 | NA | NA | NA |
| Tmem150c | 388.2756464 | 0.255766987 | 0.074176781 | 3.448073437 |
| Tmem150cos | 0.555352798 | 0.158385267 | 1.233826311 | 0.128369176 |
| Gm9932 | 0.452014659 | -1.247058973 | 1.488389666 | -0.837857855 |
| 2310034O05Rik | 0.531171913 | -0.345552833 | 1.479362157 | -0.233582312 |
| Gm8091 | 0.417821482 | 0.03663966 | 1.431356788 | 0.025597853 |
| Sec31a | 1485.020543 | -0.138234316 | 0.040507886 | -3.412528558 |
| 5430416N02Rik | 76.01050414 | -0.027742367 | 0.175726853 | -0.157872096 |
| Gm27553 | 0.170081438 | -0.443823536 | 2.780440947 | -0.159623436 |
| Gm27997 | 0.038981395 | -0.443816764 | 3.724945049 | -0.119147198 |
| Gm28030 | 0 | NA | NA | NA |
| Gm27286 | 0.243228626 | -1.073592254 | 3.022466967 | -0.355203966 |
| RP23-325M7.4 | 4.625370955 | -0.206066855 | 0.554723815 | -0.371476489 |
| Lin54 | 315.1002167 | -0.113734689 | 0.09774013 | -1.163643718 |
| Gm23222 | 0.640027454 | -0.115481887 | 1.534154821 | -0.075273946 |
| RP23-313F5.1 | 0 | NA | NA | NA |
| Gm23173 | 0 | NA | NA | NA |
| Cops4 | 680.1477187 | 0.019292537 | 0.063176819 | 0.305373674 |
| Plac8 | 627.7400569 | -1.725948466 | 0.441156014 | -3.912331261 |
| RP23-313F5.2 | 0 | NA | NA | NA |
| Coq2 | 258.0378681 | 0.004704067 | 0.082933229 | 0.056721132 |
| Gm15777 | 0.230076657 | 0.8885161 | 3.108237376 | 0.285858508 |
| Hpse | 70.70159234 | -1.049039582 | 0.200674537 | -5.227566977 |
| Helq | 242.9045317 | 0.217416091 | 0.15292513 | 1.421715917 |
| Mrps18c | 124.8778561 | -0.246686317 | 0.145323808 | -1.697494169 |
| RP23-7M16.1 | 22.40968198 | 0.273901626 | 0.284730061 | 0.96196947 |
| Fam175a | 72.7916956 | 0.081474825 | 0.151279656 | 0.538570927 |
| RP23-7M16.2 | 5.191824483 | 0.446505307 | 0.618059798 | 0.722430595 |
| RP23-7M16.3 | 1.089839937 | -0.605790671 | 1.005401815 | -0.602535884 |
| Gm26286 | 0.158374819 | -0.443821656 | 2.812678268 | -0.157793254 |
| 4930458D05Rik | 1.568767244 | -0.470379785 | 0.767364667 | -0.612980771 |

| | | | | |
|---|---|---|---|---|
| Agpat9 | 94.53062151 | -1.331637988 | 0.212214749 | -6.274954931 |
| RP23-99A9.1 | 2.620592534 | -0.648919121 | 0.651067252 | -0.9967006 |
| RP23-99A9.2 | 0.075043205 | 0.517957688 | 3.724945049 | 0.139051095 |
| RP23-302F24.1 | 0.045882455 | 0.517959912 | 3.724945049 | 0.139051692 |
| RP23-336N16.1 | 0 | NA | NA | NA |
| Gm20484 | 0 | NA | NA | NA |
| Nkx6-1 | 10.48168021 | -1.176905302 | 0.540983427 | -2.175492342 |
| RP24-292F8.2 | 0 | NA | NA | NA |
| RP24-292F8.1 | 1.593328872 | -1.110732743 | 0.870133795 | -1.276507991 |
| Cds1 | 1169.603167 | 0.361693758 | 0.073206565 | 4.940728454 |
| RP24-292F8.3 | 17.79350931 | 0.396125722 | 0.355935118 | 1.112915533 |
| WI1-1010L24.1 | 36.41421054 | 0.930028238 | 0.294839155 | 3.154357963 |
| Wdfy3 | 5687.595008 | 0.104641227 | 0.060639314 | 1.725633418 |
| Gm20548 | 3.946887117 | 0.085955032 | 0.508014651 | 0.169197939 |
| RP24-491C20.4 | 0.127971488 | -0.95589744 | 3.6855341 | -0.259364698 |
| RP23-409I5.1 | 28.85031586 | 0.637358521 | 0.342166598 | 1.862714025 |
| RP23-409I5.2 | 1.952620281 | -0.22556213 | 0.806142588 | -0.27980426 |
| RP23-409I5.3 | 0.494178821 | 1.166829978 | 1.369238532 | 0.852174366 |
| RP23-409I5.4 | 2.040397007 | -0.13814956 | 0.679027101 | -0.203452204 |
| RP23-260N17.2 | 0.650505631 | -0.391399266 | 1.373327052 | -0.285000769 |
| RP23-260N17.1 | 0.199035941 | 0.998854328 | 3.217008259 | 0.310491689 |
| Arhgap24 | 215.5874552 | -0.247338033 | 0.106799089 | -2.315918937 |
| Mapk10 | 3784.726759 | 0.426350103 | 0.076307426 | 5.587268842 |
| RP23-286M22.2 | 10.7259886 | 0.963056303 | 0.387232762 | 2.487021754 |
| RP23-286M22.1 | 125.4642517 | 1.104104387 | 0.262307608 | 4.209196964 |
| RP24-212G3.4 | 0 | NA | NA | NA |
| RP24-212G3.3 | 0 | NA | NA | NA |
| RP24-270K7.2 | 0 | NA | NA | NA |
| RP24-270K7.1 | 0 | NA | NA | NA |
| RP24-270K7.3 | 2.638949413 | -0.991426018 | 0.667636518 | -1.484978714 |
| Ptpn13 | 508.0720606 | -0.224158963 | 0.094310015 | -2.376830942 |
| RP24-102B13.3 | 0 | NA | NA | NA |
| Gm23085 | 0.072025462 | 0.517959912 | 3.724945049 | 0.139051692 |
| Slc10a6 | 36.57358822 | 0.234345839 | 0.824480193 | 0.28423465 |
| Gm15844 | 0 | NA | NA | NA |
| RP24-102B13.5 | 0.358939267 | 1.029623755 | 1.552089588 | 0.663379075 |
| 1700016H13Rik | 1.060754565 | -1.633553572 | 1.032642955 | -1.581915186 |
| Gm25721 | 0 | NA | NA | NA |
| Aff1 | 1183.051492 | -0.484954741 | 0.15316541 | -3.166215797 |
| Klhl8 | 320.9854992 | 0.103372712 | 0.087619672 | 1.179788839 |
| RP23-337H1.2 | 4.609961798 | 0.103000005 | 0.657281312 | 0.156706121 |
| Gm24897 | 0 | NA | NA | NA |
| Hsd17b13 | 1.339240761 | 0.95557824 | 0.9108502 | 1.049105814 |
| Gm8200 | 0 | NA | NA | NA |
| Hsd17b11 | 306.1707364 | 0.203986187 | 0.097357321 | 2.09523214 |
| RP23-102H7.7 | 6.166014486 | 0.248796142 | 0.447314916 | 0.556199074 |
| Nudt9 | 452.8860333 | -0.201228746 | 0.103134891 | -1.951121917 |

| | | | | |
|---|---|---|---|---|
| Mir5619 | 0.706746895 | 0.640850882 | 1.214669443 | 0.527592824 |
| Gm17660 | 2.469786297 | 1.682831539 | 0.784586626 | 2.144863909 |
| RP23-102H7.9 | 2.342408071 | 4.082796198 | 1.024316856 | 3.985872314 |
| Sparcl1 | 15907.62479 | 0.701300126 | 0.0985496 | 7.116214877 |
| Gm26703 | 13.10610984 | 0.616235293 | 0.375920004 | 1.639272414 |
| Dspp | 1.924149456 | -0.404602233 | 0.866206374 | -0.467096809 |
| Dmp1 | 39.86314599 | -0.033819183 | 0.236708477 | -0.142872715 |
| RP24-191K3.1 | 0.046345709 | -0.443818988 | 3.724945049 | -0.119147795 |
| Gm23616 | 0 | NA | NA | NA |
| Ibsp | 1.789549342 | -0.878351556 | 0.800257688 | -1.097585903 |
| Mepe | 0 | NA | NA | NA |
| Spp1 | 1206.760665 | -0.820502729 | 0.318987802 | -2.572207227 |
| RP24-299A7.2 | 2.453753307 | -2.860756354 | 0.89877983 | -3.182933416 |
| Pkd2 | 529.8119712 | -0.031649501 | 0.114056639 | -0.277489338 |
| BC005561 | 622.8318008 | 0.070085556 | 0.117262836 | 0.597679183 |
| D930016D06Rik | 256.987905 | -0.252516299 | 0.156286114 | -1.615730864 |
| Gm8258 | 4.524171958 | 0.079638899 | 0.480169329 | 0.165855864 |
| Zfp951 | 41.98758719 | 0.22458554 | 0.217053107 | 1.03470318 |
| Abcg3 | 1.390608906 | -1.004741007 | 0.848981075 | -1.183466908 |
| Gm24097 | 0 | NA | NA | NA |
| RP23-370D2.2 | 0 | NA | NA | NA |
| RP23-370D2.5 | 0 | NA | NA | NA |
| RP23-370D2.4 | 0.124853745 | 1.05357957 | 3.721447096 | 0.283110183 |
| Gbp8 | 237.2535119 | -0.8054233 | 0.314364849 | -2.562065395 |
| Gbp9 | 1809.026709 | -0.88037402 | 0.217784874 | -4.042402041 |
| Gbp4 | 4374.309881 | -0.959796227 | 0.311598499 | -3.080233795 |
| Gbp6 | 1754.483529 | -0.513695086 | 0.223326677 | -2.300195801 |
| Gbp10 | 886.6277886 | -1.021505239 | 0.321606329 | -3.176259749 |
| Gbp11 | 1003.541595 | -0.448277631 | 0.286007481 | -1.567363308 |
| Gm26519 | 1.456550475 | -0.201216804 | 0.760874182 | -0.264454766 |
| RP24-63G23.2 | 1.81533978 | 0.102168901 | 0.692378138 | 0.147562286 |
| Lrrc8b | 1659.203879 | 0.408885365 | 0.067480692 | 6.059294202 |
| RP23-41J13.2 | 3.634356604 | 0.229803527 | 0.550161459 | 0.417701973 |
| Lrrc8c | 697.388411 | -0.362439592 | 0.107697966 | -3.365333661 |
| C230066G23Rik | 8.141609263 | 0.404262259 | 0.459772124 | 0.879266571 |
| Lrrc8d | 580.2514977 | 0.08985862 | 0.079613507 | 1.128685617 |
| Rps15a-ps5 | 17.05466322 | -0.440944598 | 0.266083385 | -1.657166976 |
| Lrrc8dos | 0.047948698 | 0.037064902 | 3.724945049 | 0.009950456 |
| Gm10359 | 23.82378434 | 0.116276105 | 0.263151564 | 0.441859827 |
| RP24-354P12.2 | 0.324285546 | 1.406759758 | 3.70249052 | 0.379949591 |
| Zfp326 | 429.4665804 | 0.219601941 | 0.098596374 | 2.227282115 |
| RP24-354P12.3 | 23.94047403 | 0.367040457 | 0.318313281 | 1.153079306 |
| RP24-400H4.1 | 0 | NA | NA | NA |
| Gm25672 | 0 | NA | NA | NA |
| RP24-476K21.1 | 0.34546946 | -2.125679809 | 1.660698724 | -1.279991234 |
| RP23-424O22.1 | 0.204914957 | 0.027628924 | 2.19975699 | 0.012559989 |
| RP23-424O22.2 | 0.075043205 | 0.517957688 | 3.724945049 | 0.139051095 |

| | | | | |
|---|---|---|---|---|
| RP23-424O22.3 | 0 | NA | NA | NA |
| RP24-289G19.1 | 0.04393175 | 0.037062677 | 3.724945049 | 0.009949859 |
| RP24-289G19.3 | 0.147068667 | 0.998853574 | 3.51441668 | 0.284216035 |
| RP24-289G19.2 | 0.089089296 | 0.957155977 | 3.723939561 | 0.257027796 |
| RP24-289G19.4 | 0 | NA | NA | NA |
| Barhl2 | 85.11172079 | 0.054120146 | 0.241088152 | 0.224482811 |
| Gm26872 | 0.927992339 | 0.550696682 | 1.030448033 | 0.534424508 |
| Gm28050 | 8.739896192 | 0.656979949 | 0.368192453 | 1.784338446 |
| Gm28039 | 0.075043205 | 0.517957688 | 3.724945049 | 0.139051095 |
| Gm29464 | 0.340386612 | -0.534950876 | 1.653221841 | -0.323580818 |
| Gm17304 | 0 | NA | NA | NA |
| RP23-167H23.3 | 0.042716945 | 0.037062677 | 3.724945049 | 0.009949859 |
| Zfp644 | 2759.468483 | 0.256998714 | 0.075721836 | 3.393984213 |
| RP23-167H23.2 | 69.78784834 | 0.468329537 | 0.323148288 | 1.449271291 |
| Gm24191 | 0 | NA | NA | NA |
| RP24-512F21.1 | 0 | NA | NA | NA |
| Hfm1 | 28.3914873 | 0.45426227 | 0.209156137 | 2.171881142 |
| Gm27527 | 0.338550938 | 1.353687334 | 1.531337516 | 0.883990184 |
| Olfr719-ps | 0 | NA | NA | NA |
| Cdc7 | 158.8450571 | -0.106302455 | 0.196783274 | -0.540200664 |
| RP23-24K16.1 | 0.066050875 | -0.443818988 | 3.724945049 | -0.119147795 |
| Tgfbr3 | 375.510489 | 0.388068018 | 0.119119474 | 3.257805002 |
| RP23-24K16.3 | 17.70335292 | 1.259345255 | 0.474543604 | 2.65380303 |
| RP23-24K16.2 | 5.270225733 | 1.72067681 | 0.627165402 | 2.74357738 |
| Gm8145 | 0 | NA | NA | NA |
| RP24-357G2.2 | 11.97838418 | 1.523892043 | 0.450226942 | 3.384719795 |
| Gm22557 | 0 | NA | NA | NA |
| Brdt | 172.1838994 | 0.371552636 | 0.108270298 | 3.431713424 |
| Gm8152 | 0.483474337 | -0.870601568 | 1.499028921 | -0.580777032 |
| RP24-140D11.1 | 19.5481605 | 0.53210415 | 0.273965855 | 1.942227984 |
| n-R5s173 | 0.044191549 | 0.037062677 | 3.724945049 | 0.009949859 |
| Ephx4 | 172.3772472 | 0.840764296 | 0.129944209 | 6.47019441 |
| Gm17202 | 1.45035625 | 3.894676922 | 1.055484945 | 3.689940761 |
| Lpcat2b | 2.065408989 | 0.354254072 | 0.796437056 | 0.444798581 |
| Btbd8 | 41.15341184 | 0.202058983 | 0.170690315 | 1.183775326 |
| Gm9727 | 10.68833693 | -0.233425159 | 0.34349146 | -0.67956612 |
| A830010M20Rik | 1635.382598 | 0.680617767 | 0.090251135 | 7.541376301 |
| RP23-32E20.1 | 2.258596541 | 0.24028681 | 0.717802884 | 0.334753196 |
| 1700028K03Rik | 21.85631515 | 1.473548566 | 0.326495938 | 4.513221749 |
| Glmn | 205.8721023 | 0.313046895 | 0.093057419 | 3.364018682 |
| Rpap2 | 173.555107 | 0.200380019 | 0.094618249 | 2.117773494 |
| A930041C12Rik | 0.201858732 | 0.517972533 | 2.604406155 | 0.198883163 |
| Gm26692 | 1.12480443 | 0.643012962 | 1.151038532 | 0.558637217 |
| RP23-32E20.3 | 0.156258025 | 0.955385677 | 2.812062187 | 0.339745572 |
| RP24-140D11.4 | 0.388935867 | -0.900862 | 1.607445793 | -0.560430718 |
| Gfi1 | 6.525819315 | -0.179092353 | 0.468352362 | -0.382388065 |
| A430072P03Rik | 0.11147366 | -0.955981391 | 3.722845977 | -0.25678779 |

| | | | | |
|---|---|---|---|---|
| Evi5 | 1027.982555 | 0.100501145 | 0.062674985 | 1.603528828 |
| RP24-140D11.6 | 51.7496199 | 0.537533361 | 0.217281552 | 2.47390244 |
| Ube2d2b | 0 | NA | NA | NA |
| Gm16007 | 0 | NA | NA | NA |
| RP24-140D11.5 | 1.387064387 | -1.187300687 | 0.934825558 | -1.270077264 |
| Rpl5 | 1390.886114 | -0.141143664 | 0.095497851 | -1.477977391 |
| Snord21 | 0 | NA | NA | NA |
| Gm22270 | 2.785641484 | -0.305678853 | 0.640660631 | -0.477130696 |
| Gm26387 | 2.437479595 | 0.274098023 | 0.659561437 | 0.415576181 |
| Fam69a | 302.41774 | 0.26043401 | 0.089543189 | 2.908473695 |
| Gm9850 | 0.572539054 | 0.493635014 | 1.400445546 | 0.352484262 |
| Rps19-ps8 | 0 | NA | NA | NA |
| Mtf2 | 757.8828192 | 0.294517127 | 0.08805557 | 3.344673452 |
| Tmed5 | 501.0379422 | -0.361421639 | 0.113306155 | -3.18977939 |
| Ccdc18 | 25.54476345 | 0.004402325 | 0.24186497 | 0.018201582 |
| RP23-46J11.4 | 0 | NA | NA | NA |
| RP23-46J11.5 | 0 | NA | NA | NA |
| RP23-46J11.6 | 0 | NA | NA | NA |
| Dr1 | 585.9554827 | -0.391635396 | 0.097398388 | -4.020963828 |
| RP24-317M4.1 | 6.093084322 | 0.097176918 | 0.549378208 | 0.17688528 |
| Pigg | 279.9406767 | -0.248831237 | 0.091010612 | -2.734090376 |
| RP24-317M4.2 | 2.428646146 | 0.195161467 | 0.700168266 | 0.278735093 |
| RP24-317M4.3 | 9.503775188 | -0.023788464 | 0.404257483 | -0.058844834 |
| RP24-317M4.6 | 137.5888963 | -0.164079227 | 0.171660595 | -0.955835125 |
| Gm10419 | 867.0367941 | 0.106086635 | 0.12145835 | 0.873440441 |
| Pde6b | 10.02136167 | -3.550670181 | 0.465336113 | -7.630334459 |
| Atp5k | 305.7650685 | 0.162080827 | 0.183142593 | 0.884997994 |
| Mfsd7a | 18.35894452 | -0.594843281 | 0.250679609 | -2.372922481 |
| Pcgf3 | 699.3355955 | -0.08893527 | 0.070329591 | -1.264549794 |
| Gm10086 | 0.202977511 | -1.329524258 | 2.185177584 | -0.608428472 |
| Cplx1 | 5792.979381 | 0.392447137 | 0.079975084 | 4.907117515 |
| RP24-529E15.1 | 10.79973995 | 1.022736967 | 0.551640926 | 1.853990374 |
| Gak | 1581.201267 | -0.012127985 | 0.057375848 | -0.211377886 |
| RP24-529E15.3 | 21.02397872 | 0.336581202 | 0.300977873 | 1.118292181 |
| Tmem175 | 304.8091594 | 0.264344972 | 0.103427381 | 2.555850966 |
| Dgkq | 523.7346149 | 0.012616089 | 0.102431805 | 0.123165742 |
| Idua | 131.045456 | -0.328741 | 0.128013487 | -2.568018475 |
| Slc26a1 | 12.6609391 | 0.053106744 | 0.33897381 | 0.156669164 |
| Fgfrl1 | 247.8383437 | -0.67831665 | 0.148992524 | -4.552689173 |
| Rnf212 | 0.048949532 | 0.037064902 | 3.724945049 | 0.009950456 |
| 1700047L14Rik | 0.220636191 | -1.749523305 | 2.539538519 | -0.688913868 |
| Tmed11 | 0 | NA | NA | NA |
| Vmn2r8 | 0 | NA | NA | NA |
| Gm18594 | 0 | NA | NA | NA |
| Vmn2r9 | 0 | NA | NA | NA |
| Gm20629 | 0.046345709 | -0.443818988 | 3.724945049 | -0.119147795 |
| Gm10417 | 1.121165163 | -0.200481056 | 1.061795473 | -0.188813252 |

| | | | | |
|---|---|---|---|---|
| Vmn2r-ps21 | 0 | NA | NA | NA |
| RP23-37G13.2 | 0.042716945 | 0.037062677 | 3.724945049 | 0.009949859 |
| RP23-37G13.3 | 0 | NA | NA | NA |
| Gm18595 | 0 | NA | NA | NA |
| Vmn2r10 | 0 | NA | NA | NA |
| Gm5871 | 0.397993176 | -0.942205256 | 1.706525184 | -0.552119163 |
| Vmn2r11 | 0 | NA | NA | NA |
| Vmn2r12 | 0 | NA | NA | NA |
| Vmn2r13 | 0 | NA | NA | NA |
| Vmn2r14 | 0 | NA | NA | NA |
| Gm8473 | 0 | NA | NA | NA |
| Vmn2r15 | 0 | NA | NA | NA |
| Vmn2r16 | 0 | NA | NA | NA |
| Gm7171 | 0 | NA | NA | NA |
| Vmn2r17 | 0.044191549 | 0.037062677 | 3.724945049 | 0.009949859 |
| RP23-410M21.3 | 0 | NA | NA | NA |
| Gm8488 | 0.787704992 | -0.829342986 | 1.051150261 | -0.788986139 |
| RP23-203O7.3 | 2.482507492 | -2.191234234 | 0.875196143 | -2.503706458 |
| Crlf2 | 173.2274311 | -0.488095221 | 0.119605178 | -4.080887039 |
| Vmn2r-ps23 | 0.148650935 | 0.966592573 | 2.859657381 | 0.338009924 |
| Gm28000 | 0 | NA | NA | NA |
| 5430403G16Rik | 36.45249734 | 0.043020164 | 0.264363166 | 0.162731308 |
| 4930522L14Rik | 39.79845386 | -0.146091156 | 0.197185278 | -0.740882676 |
| Gm8508 | 4.624103064 | -0.119307126 | 0.488271736 | -0.244345755 |
| RP23-396K22.3 | 1.154557212 | 1.633949264 | 1.048213709 | 1.558794023 |
| RP23-396K22.4 | 4.399392508 | 1.505791341 | 0.617569359 | 2.438254617 |
| Gm26779 | 37.79669302 | 0.228987194 | 0.204035211 | 1.122292536 |
| Gm26808 | 0.439601053 | -1.584337223 | 1.537103412 | -1.030729104 |
| Gm15446 | 190.6866757 | 0.176177741 | 0.134501882 | 1.309853357 |
| 1010001B22Rik | 0.952639941 | -0.208612397 | 1.166596905 | -0.178821319 |
| Zfp932 | 186.9135822 | -0.035135834 | 0.129976567 | -0.270324369 |
| Gm17655 | 4.402479275 | -0.359765036 | 0.541734584 | -0.664098336 |
| Plcxd1 | 122.7290074 | -0.173893287 | 0.148532395 | -1.170743169 |
| Gtpbp6 | 242.7828586 | -0.020325353 | 0.073342318 | -0.277129957 |
| Zfp605 | 230.5230773 | 0.08543343 | 0.082977067 | 1.029602916 |
| Gm15484 | 2.365693163 | -0.625270361 | 0.632580735 | -0.988443572 |
| Chfr | 523.8452094 | 0.05663337 | 0.113903716 | 0.497203883 |
| RP24-374B14.4 | 1.731816556 | -0.088909347 | 0.716125169 | -0.124153361 |
| Gm15787 | 10.57349902 | -0.785407261 | 0.327157944 | -2.400697511 |
| Golga3 | 1104.05704 | -0.051848973 | 0.065942905 | -0.786270681 |
| RP24-374B14.6 | 9.781179111 | 1.036006132 | 0.390823985 | 2.650825364 |
| RP24-374B14.8 | 3.53955661 | 1.620440247 | 0.55605035 | 2.914196972 |
| Ankle2 | 833.7400944 | -0.303580512 | 0.065113849 | -4.662303249 |
| Pgam5 | 509.9939935 | -0.080809457 | 0.068115013 | -1.186367782 |
| Gm26718 | 13.31072805 | -0.041361299 | 0.330659732 | -0.125087198 |
| Gm15788 | 2.377542658 | -0.230163107 | 0.713836642 | -0.322431062 |
| Pxmp2 | 26.9648925 | 0.242647725 | 0.268943056 | 0.902227141 |

| | | | | |
|---|---|---|---|---|
| Pole | 51.3992184 | -0.498097258 | 0.311769315 | -1.597646831 |
| Gm26277 | 0 | NA | NA | NA |
| Gm15791 | 0.092493346 | 0.517955464 | 3.724945049 | 0.139050498 |
| Gm15792 | 0.061920613 | -0.443816764 | 3.724945049 | -0.119147198 |
| P2rx2 | 1.822501977 | 0.321150436 | 0.801377457 | 0.400748028 |
| Lrcol1 | 0.046345709 | -0.443818988 | 3.724945049 | -0.119147795 |
| Fbrsl1 | 654.9798202 | 0.239324346 | 0.082631806 | 2.896273926 |
| Gm26711 | 1.656227335 | 0.687477833 | 0.8634036 | 0.796241564 |
| RP23-231C16.1 | 2.276862309 | 1.015562399 | 0.685619653 | 1.481232917 |
| RP23-231C16.2 | 0 | NA | NA | NA |
| A630023P12Rik | 2.678264972 | 0.916113611 | 0.673622092 | 1.35998154 |
| Galnt9 | 582.7762908 | 0.344622155 | 0.096174065 | 3.583316925 |
| Noc4l | 266.4007122 | -0.266282464 | 0.152321737 | -1.748158008 |
| Ddx51 | 235.8120389 | -0.222246178 | 0.100865512 | -2.203391172 |
| RP23-393E10.2 | 29.91262942 | 0.323726625 | 0.215816705 | 1.500007267 |
| Ep400 | 2474.898703 | 0.041794557 | 0.0400638 | 1.043200017 |
| Mir7026 | 1.550196443 | 0.931669752 | 0.825595398 | 1.12848225 |
| Gm22716 | 3.382263423 | 0.385105575 | 0.592546613 | 0.649916086 |
| Gm26079 | 0.044544648 | 0.517957688 | 3.724945049 | 0.139051095 |
| Gm22583 | 0 | NA | NA | NA |
| Gm15559 | 13.41128988 | 0.284033837 | 0.26573267 | 1.068870596 |
| Pus1 | 142.3904629 | -0.157079369 | 0.101837573 | -1.542450052 |
| Gm26515 | 2.087315227 | 0.04701225 | 0.724002501 | 0.064933822 |
| Ulk1 | 1100.014041 | 0.158625488 | 0.055531573 | 2.856491891 |
| Hscb | 47.36481145 | -0.175846123 | 0.192301447 | -0.914429533 |
| Chek2 | 15.72251656 | -0.629789043 | 0.287617736 | -2.189673877 |
| RP24-207A18.2 | 0.348192234 | 0.101376188 | 1.565732616 | 0.064746808 |
| Ttc28 | 598.1288249 | -0.000409625 | 0.057561427 | -0.00711631 |
| Gm15989 | 3.684955975 | 0.687053188 | 0.587315042 | 1.169820521 |
| RP23-326H18.1 | 27.43115472 | -0.106723413 | 0.226445712 | -0.471298006 |
| Mir701 | 2.293891663 | 1.339576599 | 0.751770044 | 1.781896752 |
| Gm15988 | 0 | NA | NA | NA |
| RP24-230N16.2 | 6.359331415 | 0.243006393 | 0.511514644 | 0.475072211 |
| RP23-72O23.3 | 4.724619101 | 0.238169565 | 0.562655568 | 0.423295491 |
| RP23-72O23.4 | 1.873630482 | 1.000794027 | 0.960674742 | 1.04176157 |
| Gm27602 | 0 | NA | NA | NA |
| RP23-131F7.2 | 0.716273392 | 1.47367994 | 1.343253636 | 1.097097302 |
| Gm27680 | 0 | NA | NA | NA |
| Pitpnb | 878.8123659 | 0.060856214 | 0.065508301 | 0.928984779 |
| RP23-131F7.5 | 0.255202296 | 0.053489473 | 1.822057531 | 0.029356632 |
| Mn1 | 649.1153886 | 0.593242069 | 0.145007432 | 4.091114909 |
| RP23-131F7.4 | 14.92077947 | 0.144641378 | 0.367896208 | 0.393158111 |
| C130026L21Rik | 34.50140241 | 0.033143101 | 0.276796053 | 0.119738344 |
| RP24-572L22.2 | 3.151250669 | -0.498118359 | 0.747769358 | -0.666139035 |
| Gm26897 | 0.18833836 | -0.244781214 | 2.662785935 | -0.091926734 |
| E130006D01Rik | 3.354483374 | -0.140555243 | 0.55870066 | -0.251575222 |
| Gm16019 | 0 | NA | NA | NA |

| | | | | |
|---|---|---|---|---|
| RP24-340B11.1 | 0 | NA | NA | NA |
| RP23-427D2.1 | 0.576279854 | 0.108720917 | 1.317196045 | 0.082539662 |
| RP24-317K11.4 | 0 | NA | NA | NA |
| 1700028D13Rik | 0 | NA | NA | NA |
| Miat | 1661.236509 | 1.364882676 | 0.491203717 | 2.778648914 |
| Gm27998 | 0 | NA | NA | NA |
| Gm27882 | 0 | NA | NA | NA |
| Gm26953 | 0.344925633 | -0.577355578 | 1.75472911 | -0.329028324 |
| Gm27627 | 0 | NA | NA | NA |
| Gm27357 | 0 | NA | NA | NA |
| Cryba4 | 4.194020461 | 0.160160249 | 0.570353985 | 0.280808504 |
| Crybb1 | 14.29264834 | 2.666660059 | 0.585215162 | 4.556717308 |
| Tpst2 | 280.8629906 | -0.149429155 | 0.093434319 | -1.599296241 |
| Tfip11 | 581.9359828 | -0.031996558 | 0.064115637 | -0.49904453 |
| Srrd | 100.6125371 | 0.033085359 | 0.120814433 | 0.273852706 |
| Hps4 | 178.5209335 | -0.208064151 | 0.136694611 | -1.522109388 |
| Gm6583 | 0.632098713 | -0.987611981 | 1.341675181 | -0.736103638 |
| RP23-459I1.3 | 20.2973942 | 0.110440357 | 0.455609126 | 0.242401547 |
| RP23-459I1.5 | 0 | NA | NA | NA |
| Gm20636 | 0.231217418 | 0.037077131 | 2.131487483 | 0.017394956 |
| Asphd2 | 618.5894786 | 0.48747422 | 0.083214885 | 5.858017083 |
| Sez6l | 3346.420789 | 0.177928926 | 0.089597331 | 1.985873063 |
| Gm6588 | 0 | NA | NA | NA |
| Gm27004 | 3.168227339 | -0.753628554 | 0.652997878 | -1.154105671 |
| 1700034G24Rik | 0.106465261 | 0.037073798 | 3.724945049 | 0.009952844 |
| RP23-156K23.3 | 0.158818843 | -0.47083079 | 2.809622806 | -0.167577936 |
| RP23-156K23.4 | 7.736953527 | 1.232573509 | 0.493077811 | 2.499754568 |
| RP24-304N8.2 | 2.801170497 | 2.171111826 | 0.746576119 | 2.908091714 |
| Myo18b | 28.0640786 | -0.521470552 | 0.225592818 | -2.311556534 |
| 1700095B10Rik | 0 | NA | NA | NA |
| Adrbk2 | 1355.458738 | 0.754674627 | 0.082765151 | 9.118265568 |
| Gm22740 | 0 | NA | NA | NA |
| Crybb2 | 0.278057897 | 0.051030661 | 2.409838576 | 0.021175966 |
| RP23-4P1.1 | 0.92508897 | 0.052259887 | 0.901147736 | 0.057992585 |
| Crybb3 | 5.690583048 | -1.090025085 | 0.462267563 | -2.357996047 |
| 2900026A02Rik | 622.1930781 | 0.50573195 | 0.094541408 | 5.349316859 |
| RP23-4P1.6 | 0.061920613 | -0.443816764 | 3.724945049 | -0.119147198 |
| RP23-4P1.5 | 5.731549506 | -0.164878637 | 0.494173652 | -0.33364514 |
| RP23-4P1.3 | 0.48780114 | -1.364290903 | 1.431605335 | -0.952979756 |
| RP23-4P1.7 | 0.066050875 | -0.443818988 | 3.724945049 | -0.119147795 |
| RP23-4P1.2 | 1.950711236 | 1.682081071 | 1.075808579 | 1.563550528 |
| Tmem211 | 0.590173444 | -1.635596566 | 1.331183438 | -1.228678572 |
| Sgsm1 | 1205.098046 | -0.190701525 | 0.082521133 | -2.310941667 |
| RP24-456F12.1 | 1.004172246 | -0.263207831 | 0.950237324 | -0.276991678 |
| RP24-456F12.2 | 0.236337014 | -0.396301249 | 2.149114859 | -0.184402079 |
| mmu-mir-7229 | 0 | NA | NA | NA |
| mmu-mir-7230 | 0 | NA | NA | NA |

| | | | | |
|---|---|---|---|---|
| Aym1 | 0.330879691 | 1.111490832 | 1.992275864 | 0.557900064 |
| Wscd2 | 877.7252593 | 0.295134974 | 0.131066631 | 2.251793392 |
| Cmklr1 | 152.364205 | -0.894995951 | 0.166481436 | -5.375950462 |
| Gm15736 | 0.075043205 | 0.517957688 | 3.724945049 | 0.139051095 |
| 1700069L16Rik | 1.680905595 | 0.094098028 | 0.931870465 | 0.100977584 |
| Ficd | 177.6330377 | -0.218855386 | 0.106213687 | -2.060519633 |
| Sart3 | 480.3375866 | 0.03745147 | 0.096435365 | 0.388358259 |
| Gm3511 | 0.172107867 | 0.517971699 | 2.739902729 | 0.189047477 |
| Iscu | 446.2315743 | -0.08718048 | 0.091649825 | -0.951234546 |
| Tmem119 | 184.4126201 | 1.274104997 | 0.221838249 | 5.743396394 |
| RP24-555M20.7 | 0 | NA | NA | NA |
| Selplg | 268.6005576 | -0.391924272 | 0.158515982 | -2.472459037 |
| Coro1c | 1185.306564 | -0.308219009 | 0.096842264 | -3.182691063 |
| Gm17122 | 0.291024678 | -0.488041998 | 2.054139567 | -0.237589502 |
| RP24-555M20.4 | 0.261813223 | -0.443823324 | 2.533783979 | -0.175162258 |
| RP24-555M20.3 | 33.32994184 | -0.123521411 | 0.362741585 | -0.340521783 |
| RP24-555M20.5 | 0.345648411 | -1.234846767 | 1.551700493 | -0.795802265 |
| Gm22056 | 0 | NA | NA | NA |
| Ssh1 | 648.036513 | -0.263101061 | 0.062482752 | -4.210779006 |
| Dao | 29.66175715 | 0.352092743 | 0.334624708 | 1.052201868 |
| Svop | 1130.54196 | 0.36484394 | 0.079482308 | 4.590253472 |
| Usp30 | 451.0626031 | 0.110682529 | 0.106999714 | 1.034418924 |
| RP24-226L20.2 | 8.375339354 | 0.590477465 | 0.431238005 | 1.369261192 |
| RP24-226L20.1 | 11.87863406 | 1.310250556 | 0.577064167 | 2.270545687 |
| Alkbh2 | 30.67206188 | -0.707061437 | 0.195175867 | -3.62268885 |
| Ung | 32.04545787 | -0.323805012 | 0.236408776 | -1.369682705 |
| Acacb | 113.2622031 | -0.273349884 | 0.131764822 | -2.074528549 |
| Gm13784 | 0 | NA | NA | NA |
| Foxn4 | 0.07796279 | -0.798301147 | 3.721829328 | -0.214491605 |
| Myo1h | 18.55275259 | 1.033407451 | 0.317452919 | 3.255309335 |
| Kctd10 | 678.4952583 | -0.443899655 | 0.11944058 | -3.716489436 |
| Ube3b | 1199.562226 | -0.001978524 | 0.075996314 | -0.02603447 |
| Mir7027 | 0.775147523 | 0.139214248 | 1.203657028 | 0.115659399 |
| Mmab | 164.6481927 | 0.459412658 | 0.094623545 | 4.855162205 |
| Mvk | 173.1109817 | -0.082263467 | 0.131347247 | -0.626305227 |
| Gm13790 | 0.110814304 | 0.037060453 | 3.724945049 | 0.009949262 |
| Fam222a | 196.5946661 | -0.231272678 | 0.128986904 | -1.792993478 |
| Trpv4 | 24.56884945 | 0.239781671 | 0.249068138 | 0.962715154 |
| Gltp | 531.1234222 | -0.442553365 | 0.122817895 | -3.603329667 |
| Tchp | 125.3804559 | 0.016470225 | 0.182713158 | 0.090142522 |
| RP23-3M10.11 | 0.104637558 | -0.443821212 | 3.724945049 | -0.119148392 |
| Mir7028 | 0.496352305 | -0.951061612 | 1.522992197 | -0.624469129 |
| Git2 | 916.5459927 | 0.052813239 | 0.099448287 | 0.531062328 |
| 4930515G01Rik | 7.627541774 | -0.054493018 | 0.43754285 | -0.124543272 |
| Ankrd13a | 914.9811008 | -0.082490021 | 0.102972184 | -0.801090338 |
| 1500011B03Rik | 398.2607319 | 0.327431524 | 0.086216493 | 3.797782902 |
| Gm20499 | 1.052193156 | 0.933338712 | 1.006775245 | 0.927057669 |

| | | | | |
|---|---|---|---|---|
| 2610524H06Rik | 87.22289935 | 0.2031123 | 0.130676912 | 1.554308999 |
| Ywhaq-ps2 | 0.12209515 | 0.037073798 | 3.724945049 | 0.009952844 |
| RP24-297N9.13 | 1.537880876 | 0.04714574 | 0.774866909 | 0.060843661 |
| Mir6240 | 0 | NA | NA | NA |
| 4930519G04Rik | 0.461107361 | -0.019181332 | 1.435771925 | -0.013359596 |
| Gm9936 | 3.314927513 | -0.0114231 | 0.548718722 | -0.020817769 |
| Gm13823 | 0.097139547 | -0.443816764 | 3.724945049 | -0.119147198 |
| Oasl2 | 5567.265836 | -0.287233462 | 0.12013674 | -2.390887765 |
| Gm13822 | 5.72139037 | -2.253279236 | 0.591790859 | -3.807560058 |
| Oasl1 | 994.9398756 | -1.55192852 | 0.375932929 | -4.128205862 |
| 2210016L21Rik | 838.2899106 | 0.308218848 | 0.070917961 | 4.346132432 |
| Hnf1a | 0.04393175 | 0.037062677 | 3.724945049 | 0.009949859 |
| Hnf1aos1 | 0.088476398 | 0.51795324 | 3.724945049 | 0.139049901 |
| Hnf1aos2 | 0.044544648 | 0.517957688 | 3.724945049 | 0.139051095 |
| RP24-556H22.2 | 0.075043205 | 0.517957688 | 3.724945049 | 0.139051095 |
| Sppl3 | 907.0167297 | 0.206933489 | 0.06425809 | 3.220349181 |
| RP24-556H22.3 | 18.34444885 | 0.238124329 | 0.331830725 | 0.717607835 |
| Gm13827 | 0.049553047 | -0.443818988 | 3.724945049 | -0.119147795 |
| Gm10401 | 0.066050875 | -0.443818988 | 3.724945049 | -0.119147795 |
| Rpl37rt | 126.6981566 | -0.028478079 | 0.161207517 | -0.176654787 |
| Acads | 60.73886381 | -0.770900645 | 0.196106637 | -3.931027818 |
| Unc119b | 384.4903987 | -0.49883554 | 0.123156502 | -4.050419846 |
| Mlec | 1723.019123 | -0.563312806 | 0.112621022 | -5.001844212 |
| Cabp1 | 800.6070211 | 0.470676331 | 0.125972305 | 3.736347684 |
| Gm13828 | 0.405693472 | 0.517971627 | 1.728451778 | 0.299673751 |
| Pop5 | 169.6778782 | 0.270959486 | 0.111853603 | 2.422447548 |
| Rnf10 | 2262.437902 | -0.188062455 | 0.066360929 | -2.833933417 |
| Gm13831 | 0.044544648 | 0.517957688 | 3.724945049 | 0.139051095 |
| Coq5 | 405.2808921 | -0.246359536 | 0.078936378 | -3.120988602 |
| Gm13829 | 0 | NA | NA | NA |
| Dynll1 | 1360.661757 | -0.450628333 | 0.115240718 | -3.910322165 |
| Gm13830 | 22.05592487 | -0.139094364 | 0.310386635 | -0.448132579 |
| Srsf9 | 360.4762658 | -0.211863313 | 0.085074299 | -2.490332753 |
| Gatc | 179.8265356 | 0.107964553 | 0.105414294 | 1.024192723 |
| Triap1 | 96.42262351 | -0.133518534 | 0.130117807 | -1.02613575 |
| Cox6a1 | 2038.657447 | 0.053504311 | 0.111896598 | 0.478158511 |
| RP24-132L16.11 | 0.287284012 | 0.076406544 | 1.774613161 | 0.043055324 |
| 4930401G09Rik | 0.097899064 | 0.037062677 | 3.724945049 | 0.009949859 |
| Gm13832 | 4.797855873 | -0.105265181 | 0.452888287 | -0.232430787 |
| RP24-132L16.12 | 3.340097371 | 0.626271862 | 0.702068184 | 0.892038516 |
| RP24-132L16.13 | 6.416277231 | 1.050663715 | 0.556202011 | 1.888996614 |
| Msi1 | 624.614844 | -0.244705093 | 0.097454764 | -2.510960794 |
| RP24-132L16.14 | 0.383028346 | 0.023208446 | 1.504327253 | 0.015427791 |
| Pla2g1b | 0.110595523 | 0.037071574 | 3.724945049 | 0.009952247 |
| Sirt4 | 90.54918992 | -0.074163763 | 0.202511253 | -0.366220453 |
| Gm24265 | 261.4064373 | -0.181747802 | 0.214898542 | -0.845737718 |
| Gm24407 | 234.2932087 | -0.085522775 | 0.180796221 | -0.473034086 |

| | | | | |
|---|---|---|---|---|
| Gm13836 | 0.227885852 | -0.042012864 | 2.499238438 | -0.016810266 |
| Pxn | 603.300534 | -0.564568127 | 0.12427839 | -4.542769865 |
| Gm13840 | 1.630127001 | -0.824126585 | 0.798917015 | -1.031554679 |
| Rplp0 | 924.8935804 | -0.592737035 | 0.130095388 | -4.556172537 |
| Gcn1l1 | 1454.726854 | -0.002551431 | 0.073263038 | -0.034825623 |
| Mir7029 | 0.140122821 | 0.517959509 | 3.572128867 | 0.145000231 |
| RP23-228F6.9 | 3.230897769 | -0.237629699 | 0.659904375 | -0.360097172 |
| Rab35 | 708.4090342 | -0.038053728 | 0.067818921 | -0.561107826 |
| Mir7030 | 0.697123821 | -0.395787604 | 1.298230257 | -0.304867031 |
| Ccdc64 | 496.8343264 | 0.306788104 | 0.103812168 | 2.955222985 |
| Gm13841 | 108.5809223 | -0.170302312 | 0.165803861 | -1.027131161 |
| Cit | 2100.160842 | 0.415863676 | 0.13513636 | 3.077363303 |
| RP23-265I19.3 | 0.244575677 | 0.503526871 | 2.091207802 | 0.2407828 |
| RP23-265I19.2 | 0.484822013 | 1.03017484 | 1.59442402 | 0.646110963 |
| Gm14508 | 2.937257927 | 0.777578623 | 0.6474169 | 1.201047768 |
| Prkab1 | 201.4925811 | -0.004245636 | 0.126763343 | -0.033492619 |
| Gm13842 | 0 | NA | NA | NA |
| Gm17420 | 0 | NA | NA | NA |
| Tmem233 | 0.047948698 | 0.037064902 | 3.724945049 | 0.009950456 |
| C330018A13Rik | 4.719781364 | 1.179378041 | 0.592859101 | 1.989305786 |
| Ccdc60 | 12.74564928 | -0.274042936 | 0.326082431 | -0.840410002 |
| Gm14507 | 0.354637581 | -0.44381978 | 1.725759481 | -0.257173601 |
| Gm13837 | 0.143618953 | -0.891120723 | 2.890309379 | -0.308313265 |
| Gm13839 | 0.393020699 | -0.542174022 | 1.679080581 | -0.322899347 |
| Gm13838 | 0.066050875 | -0.443818988 | 3.724945049 | -0.119147795 |
| B230112J18Rik | 0.528863779 | 0.126264809 | 1.753181869 | 0.072020371 |
| RP24-79A6.9 | 0.038981395 | -0.443816764 | 3.724945049 | -0.119147198 |
| 4930562A09Rik | 0.146089079 | -0.444786384 | 3.522464394 | -0.126271364 |
| Hspb8 | 433.011455 | -1.025879485 | 0.204784161 | -5.009564606 |
| Srrm4os | 0.683645101 | 0.494342056 | 1.288871652 | 0.383546379 |
| Srrm4 | 850.8466253 | 0.183145788 | 0.092605831 | 1.97769176 |
| RP24-79A6.8 | 32.11370528 | 0.40407745 | 0.336079743 | 1.20232611 |
| RP23-312N12.2 | 10.26602459 | -0.035344408 | 0.467458078 | -0.075609791 |
| RP23-312N12.1 | 15.57120856 | 0.752192509 | 0.454092605 | 1.656473814 |
| Gstm2-ps1 | 32.88268672 | 0.405202208 | 0.258780165 | 1.565816326 |
| RP23-418H8.3 | 0.072025462 | 0.517959912 | 3.724945049 | 0.139051692 |
| RP23-280O16.1 | 0.060174537 | 0.517957688 | 3.724945049 | 0.139051095 |
| 4930569F06Rik | 0 | NA | NA | NA |
| RP23-73P16.2 | 0.598047226 | -1.224765504 | 1.174252692 | -1.043016987 |
| Suds3 | 810.9948176 | 0.022254958 | 0.059251253 | 0.375603163 |
| 9530046B11Rik | 1.655761812 | 0.264617312 | 0.86426494 | 0.30617615 |
| Taok3 | 787.4816357 | -0.207771786 | 0.069187085 | -3.003042922 |
| RP23-376H8.2 | 15.63613524 | -0.095133867 | 0.368341899 | -0.258275987 |
| n-R5s174 | 1.199099511 | 0.168683064 | 1.050753055 | 0.160535402 |
| Pebp1 | 1757.043143 | 0.096622314 | 0.072841739 | 1.326469082 |
| RP23-76J15.3 | 0 | NA | NA | NA |
| RP23-76J15.4 | 0.215166026 | 1.019404318 | 2.554583188 | 0.399049177 |

| | | | | |
|---|---|---|---|---|
| BC051077 | 1.218454842 | 0.854291067 | 1.039801551 | 0.821590491 |
| Gm10399 | 0.929291888 | -1.136006333 | 1.06647571 | -1.06519663 |
| Vsig10 | 55.95498491 | -0.385334643 | 0.227673854 | -1.692485266 |
| Gm15727 | 0 | NA | NA | NA |
| Wsb2 | 1799.690181 | 0.201458382 | 0.066453668 | 3.031561505 |
| Rfc5 | 71.308591 | 0.320779695 | 0.16780652 | 1.911604477 |
| Gm15728 | 37.43701224 | -0.475823026 | 0.209721337 | -2.268834603 |
| Ksr2 | 1532.422263 | 0.52810682 | 0.078983415 | 6.68630018 |
| RP23-429P17.1 | 4.731857296 | -0.007981426 | 0.583018149 | -0.013689842 |
| Nos1 | 1164.626249 | 0.359142626 | 0.083379346 | 4.307333204 |
| Gm37028 | 0.69250253 | 1.890655832 | 1.363717502 | 1.38639845 |
| Gm26995 | 0.399888529 | 0.047874869 | 1.504648157 | 0.031817983 |
| Gm25343 | 0.046345709 | -0.443818988 | 3.724945049 | -0.119147795 |
| Fbxo21 | 1283.047223 | 0.524575017 | 0.063516962 | 8.25881783 |
| RP23-42D10.2 | 0.853574741 | -1.084973568 | 1.111222842 | -0.976378029 |
| Tesc | 196.9276744 | 0.184785093 | 0.170962159 | 1.080853762 |
| Fbxw8 | 652.4874552 | -0.433081287 | 0.097532547 | -4.440377086 |
| Gm9754 | 24.46512519 | 0.358761243 | 0.452742711 | 0.792417492 |
| Gm26411 | 1.298260455 | 1.216873818 | 0.941179966 | 1.29292363 |
| Hrk | 446.6006905 | 1.534145726 | 0.193481003 | 7.929180149 |
| Rnft2 | 775.8299915 | 0.486388297 | 0.082964033 | 5.862640486 |
| Gm24050 | 1.923134663 | -1.647495082 | 0.928210154 | -1.77491603 |
| 2410131K14Rik | 480.8992711 | -0.128699484 | 0.099151888 | -1.298003356 |
| Gm25076 | 4.628642042 | -0.6609894 | 0.503966158 | -1.311574972 |
| Gm28563 | 0.587040768 | 0.885364158 | 1.26238256 | 0.701343781 |
| Gm26455 | 0.133946075 | 0.036100024 | 3.626570338 | 0.009954315 |
| Gm15754 | 0.066050875 | -0.443818988 | 3.724945049 | -0.119147795 |
| Med13l | 2276.940129 | 0.103523786 | 0.06664442 | 1.553375159 |
| RP24-122N24.1 | 0 | NA | NA | NA |
| RP24-122N24.2 | 76.64565332 | 0.122103757 | 0.302770632 | 0.40328798 |
| RP24-122N24.3 | 27.41095622 | 0.202885811 | 0.312166018 | 0.649929201 |
| RP24-122N24.4 | 64.02437309 | 0.255754255 | 0.322022104 | 0.794213353 |
| RP23-155F13.2 | 26.31192174 | 0.631576293 | 0.346545337 | 1.822492544 |
| RP23-155F13.3 | 28.57824406 | 0.411835997 | 0.363500077 | 1.132973618 |
| RP24-372H3.1 | 0 | NA | NA | NA |
| RP24-372H3.2 | 0 | NA | NA | NA |
| RP24-372H3.4 | 0 | NA | NA | NA |
| RP24-372H3.3 | 0 | NA | NA | NA |
| 1700081H04Rik | 0.049553047 | -0.443818988 | 3.724945049 | -0.119147795 |
| RP23-316P20.1 | 0.048949532 | 0.037064902 | 3.724945049 | 0.009950456 |
| n-R5s175 | 0 | NA | NA | NA |
| RP23-73O10.1 | 0 | NA | NA | NA |
| Tbx3os1 | 2.722982093 | 0.017954898 | 0.772639802 | 0.02323838 |
| RP23-44J10.1 | 0.387852398 | 1.386882811 | 1.563214796 | 0.887199133 |
| Gm16063 | 0 | NA | NA | NA |
| Gm16064 | 0 | NA | NA | NA |
| Tbx3 | 28.477325 | 0.06217176 | 0.237536585 | 0.261735511 |

| | | | | |
|---|---|---|---|---|
| Tbx3os2 | 0.046345709 | -0.443818988 | 3.724945049 | -0.119147795 |
| RP24-150D8.1 | 0 | NA | NA | NA |
| RP24-150D8.2 | 0 | NA | NA | NA |
| Tbx5 | 0.360567166 | -0.269475575 | 1.957488039 | -0.13766397 |
| RP24-150D8.3 | 0 | NA | NA | NA |
| RP24-69A21.1 | 0 | NA | NA | NA |
| n-R5s176 | 0 | NA | NA | NA |
| Rbm19 | 227.3684793 | -0.552132346 | 0.077967609 | -7.081560617 |
| Gm10390 | 1.46581892 | -1.528198845 | 0.915544768 | -1.669168891 |
| RP23-213H20.1 | 1.459375539 | -0.136308063 | 0.771841931 | -0.176601008 |
| Gm26474 | 0 | NA | NA | NA |
| RP23-213H20.2 | 0.303117673 | 2.239712564 | 3.688052852 | 0.60728863 |
| Gm27199 | 3.757776822 | 0.203539119 | 0.612772637 | 0.332160914 |
| Lhx5 | 49.17951797 | -0.080299831 | 0.212786454 | -0.37737285 |
| Sdsl | 34.31511951 | -0.437470187 | 0.206011636 | -2.123521732 |
| Sds | 0.913087876 | 1.351711058 | 1.340914365 | 1.008051739 |
| Gm15690 | 5.787489139 | 1.190761814 | 0.494241979 | 2.40926887 |
| Plbd2 | 910.6907376 | -0.089383636 | 0.080515886 | -1.110136656 |
| Slc8b1 | 149.2518986 | -1.037142207 | 0.163350658 | -6.349176792 |
| Tpcn1 | 461.0027236 | 0.378493379 | 0.152612478 | 2.48009457 |
| RP23-297M4.7 | 22.51234608 | 0.483838164 | 0.279095239 | 1.73359519 |
| Iqcd | 5.452284118 | 0.468417014 | 0.561971713 | 0.833524184 |
| Rita1 | 142.064277 | -0.041014496 | 0.138773324 | -0.295550286 |
| Ddx54 | 525.9668418 | -0.27768211 | 0.090483294 | -3.068877107 |
| RP23-297M4.4 | 3.794821076 | -0.355043914 | 0.591349551 | -0.600396014 |
| Ccdc42b | 3.366237703 | 0.564336738 | 0.536925826 | 1.05105158 |
| RP23-297M4.1 | 0 | NA | NA | NA |
| Gm15749 | 0.639699895 | -1.976011673 | 1.557682206 | -1.26855893 |
| Rasal1 | 506.8043477 | 1.24616916 | 0.133671847 | 9.322600024 |
| Dtx1 | 1090.783309 | 0.082754377 | 0.074376697 | 1.112638499 |
| Oas2 | 477.0806149 | -1.356826859 | 0.252197605 | -5.380014851 |
| RP24-547J21.3 | 0.066050875 | -0.443818988 | 3.724945049 | -0.119147795 |
| Oas3 | 836.3377262 | -1.718249703 | 0.31895596 | -5.387106436 |
| RP24-547J21.4 | 1.429586369 | -0.53045979 | 0.812526973 | -0.652851914 |
| Oas1e | 0.736605981 | -1.691960331 | 1.324968031 | -1.276982003 |
| Oas1c | 32.60080593 | -0.644132575 | 0.21623257 | -2.978887845 |
| Oas1b | 318.3948597 | -0.399332173 | 0.295266393 | -1.352447088 |
| Oas1f | 0.049553047 | -0.443818988 | 3.724945049 | -0.119147795 |
| Gm22907 | 0 | NA | NA | NA |
| Oas1h | 0.100902008 | -0.876201588 | 3.72093227 | -0.235479048 |
| Oas1g | 272.4663882 | -1.102502189 | 0.277352837 | -3.975088916 |
| Oas1a | 611.2994765 | -1.195343244 | 0.276660978 | -4.32060659 |
| Oas1d | 0.674169638 | -2.684059501 | 1.195690701 | -2.244777431 |
| Rph3a | 3838.839031 | 0.535390754 | 0.089268513 | 5.997531868 |
| RP23-39M18.2 | 36.68051303 | 0.86157715 | 0.355029774 | 2.426774353 |
| Ptpn11 | 2663.867196 | -0.016632036 | 0.074614863 | -0.222905139 |
| RP23-257E17.2 | 45.93026392 | 0.42237669 | 0.34719528 | 1.216539262 |

| | | | | |
|---|---|---|---|---|
| Rpl6 | 1338.143399 | -0.20441529 | 0.084569205 | -2.417136247 |
| Gm26205 | 0 | NA | NA | NA |
| Gm15800 | 6892.751409 | 0.547119504 | 0.090336215 | 6.05648025 |
| Gm24671 | 2.5200291 | 0.647172146 | 0.711687727 | 0.90934847 |
| Trafd1 | 1684.87789 | -0.386047594 | 0.151024064 | -2.556199213 |
| RP23-50I16.1 | 5.493864583 | 0.450605509 | 0.418890694 | 1.075711436 |
| RP23-50I16.2 | 19.99381265 | 0.449408136 | 0.244992032 | 1.834378583 |
| Naa25 | 948.0018726 | -0.470556728 | 0.087365014 | -5.386100328 |
| RP23-50I16.3 | 14.51075867 | -0.200707112 | 0.374978695 | -0.535249375 |
| Erp29 | 574.7376244 | -0.040784292 | 0.100862986 | -0.404353403 |
| Tmem116 | 26.92021731 | -0.205737299 | 0.226208515 | -0.90950289 |
| Gm15546 | 0 | NA | NA | NA |
| Adam1b | 0.227186609 | -1.57076438 | 2.549473144 | -0.616113327 |
| RP24-435H23.1 | 2.953870943 | -0.195593757 | 0.737086212 | -0.26536076 |
| Adam1a | 36.9795004 | -0.349056172 | 0.293458588 | -1.18945632 |
| Mapkapk5 | 369.1571913 | -0.166305127 | 0.094027605 | -1.768684072 |
| Gm15547 | 4.040079771 | 1.292735777 | 0.731112609 | 1.768176011 |
| Aldh2 | 693.3685642 | 0.182307513 | 0.118039349 | 1.544463894 |
| RP23-181P9.3 | 1.724244418 | -0.216833148 | 0.782609157 | -0.277064415 |
| Acad12 | 28.57697775 | -0.490012675 | 0.254211575 | -1.927578139 |
| Acad10 | 41.29551714 | -0.531855505 | 0.204136479 | -2.605391783 |
| Brap | 823.207007 | 0.006459275 | 0.079683652 | 0.081061484 |
| Atxn2 | 1833.502759 | 0.04047568 | 0.070132976 | 0.577127655 |
| RP23-18K20.2 | 63.62259654 | 0.110135913 | 0.365260927 | 0.301526675 |
| Gm16552 | 0 | NA | NA | NA |
| RP23-18K20.3 | 37.1706614 | 0.03271294 | 0.340406443 | 0.096099649 |
| Sh2b3 | 323.628142 | 0.087866939 | 0.105355388 | 0.83400517 |
| Mir7031 | 0.877654538 | -1.402580444 | 1.235281438 | -1.13543392 |
| RP23-431M7.2 | 0.523632775 | -0.134303395 | 1.344165495 | -0.099915818 |
| RP23-431M7.4 | 0.25563639 | 0.037075748 | 2.12528669 | 0.017445057 |
| Fam109a | 123.4761975 | -0.18613837 | 0.113274847 | -1.643245385 |
| Cux2 | 1234.512826 | 0.633366627 | 0.129957524 | 4.87364339 |
| AC113302.1 | 0 | NA | NA | NA |
| RP23-254C8.1 | 0 | NA | NA | NA |
| Gm15637 | 0.291530733 | -0.406735179 | 1.800455957 | -0.225906764 |
| Myl2 | 1.403419482 | 1.925022306 | 1.030567853 | 1.867923883 |
| Ccdc63 | 13.49297597 | -0.253329101 | 0.36172517 | -0.700335841 |
| Ppp1cc | 926.7286209 | -0.005863976 | 0.069741325 | -0.084081797 |
| RP24-347B12.1 | 50.60541787 | 0.436274113 | 0.329617534 | 1.323576777 |
| Hvcn1 | 77.26580927 | 0.922028249 | 0.214276931 | 4.302974863 |
| Tctn1 | 251.1741939 | -0.110155535 | 0.102665927 | -1.072951247 |
| Gm15481 | 0 | NA | NA | NA |
| Pptc7 | 900.5757088 | -0.005353265 | 0.062464881 | -0.085700395 |
| Gm15842 | 8.407355864 | 0.245434179 | 0.449794848 | 0.545658048 |
| Rad9b | 39.0160455 | 0.052670448 | 0.209634026 | 0.251249517 |
| Vps29 | 780.350795 | 0.102777052 | 0.087696614 | 1.171961461 |
| Fam216a | 407.2222889 | -0.11128584 | 0.068406681 | -1.626827061 |

| | | | | |
|---|---|---|---|---|
| Gpn3 | 109.4263101 | 0.039977157 | 0.133005348 | 0.300568045 |
| RP23-407I6.1 | 0.395669761 | -0.400894164 | 1.687772913 | -0.237528497 |
| Arpc3 | 620.2922037 | -0.020925849 | 0.092370318 | -0.226543006 |
| RP23-407I6.2 | 78.71060985 | 0.434769198 | 0.159960793 | 2.717973505 |
| Anapc7 | 355.9121168 | -0.074609056 | 0.083369208 | -0.894923409 |
| Gm15846 | 0 | NA | NA | NA |
| Atp2a2 | 11857.11526 | 0.25827884 | 0.061026515 | 4.232239715 |
| RP23-407I6.6 | 93.54004702 | 0.787452494 | 0.412386323 | 1.909501963 |
| RP23-407I6.5 | 69.66663032 | 0.82922459 | 0.313128726 | 2.64819073 |
| RP23-407I6.4 | 0.863835993 | 0.188917618 | 1.180104811 | 0.160085457 |
| Gm22965 | 1.295159943 | 0.916224562 | 0.821922518 | 1.114733496 |
| Ift81 | 401.1137358 | 0.067502681 | 0.0847853 | 0.7961602 |
| RP23-407I6.7 | 1.063558902 | 3.214136398 | 1.07493303 | 2.990080597 |
| RP24-114E20.2 | 0.51473784 | -0.70149808 | 1.3627605 | -0.514762558 |
| P2rx7 | 217.2536391 | -0.947903475 | 0.160535503 | -5.904634523 |
| Mir8115 | 0 | NA | NA | NA |
| Gm10064 | 0 | NA | NA | NA |
| P2rx4 | 293.0467081 | -0.847674173 | 0.144799868 | -5.854108736 |
| Camkk2 | 1870.397667 | 0.936010826 | 0.142160931 | 6.584163598 |
| RP23-31J24.2 | 7.791943915 | 0.719018773 | 0.450075956 | 1.597549844 |
| Anapc5 | 1309.787107 | 0.141628414 | 0.055396501 | 2.556631043 |
| Rnf34 | 645.0189091 | -0.1147044 | 0.067261255 | -1.705356225 |
| Kdm2b | 295.4828908 | 0.205352129 | 0.095780687 | 2.143982628 |
| RP23-214I6.2 | 6.36596584 | 1.536626151 | 0.625712984 | 2.455800327 |
| RP23-214I6.5 | 0 | NA | NA | NA |
| A930024E05Rik | 27.86517634 | -0.270187401 | 0.211268786 | -1.278879883 |
| RP23-214I6.3 | 0.145501641 | -0.443200222 | 2.888011192 | -0.153462086 |
| Orai1 | 101.2846242 | -0.410829854 | 0.149710319 | -2.744165252 |
| Morn3 | 1.434593181 | -0.665287065 | 0.829341999 | -0.802186632 |
| RP23-214I6.4 | 5.158449836 | 1.704077283 | 0.604221426 | 2.820286088 |
| Gm26745 | 0.042716945 | 0.037062677 | 3.724945049 | 0.009949859 |
| Gm6444 | 0.573335652 | 1.169741126 | 1.224932402 | 0.95494341 |
| Tmem120b | 108.3890549 | 0.139694693 | 0.148619051 | 0.939951454 |
| Rhof | 259.5705259 | 0.071891114 | 0.145153525 | 0.495276392 |
| RP23-371A16.4 | 14.62755803 | 1.274435926 | 0.457295317 | 2.786899137 |
| AI480526 | 273.446315 | 0.519115425 | 0.22287768 | 2.329149446 |
| Gm38102 | 1.104086423 | -0.794452904 | 0.916172652 | -0.86714322 |
| RP23-371A16.3 | 8.618846306 | -0.483756118 | 0.383197861 | -1.262418627 |
| Setd1b | 1298.15226 | 0.027220626 | 0.10339676 | 0.263263823 |
| Psmd9 | 209.0408427 | 0.005332444 | 0.08979496 | 0.059384664 |
| Hpd | 0.5402512 | 0.482404968 | 1.348049491 | 0.357854049 |
| RP23-297I15.1 | 0 | NA | NA | NA |
| Wdr66 | 48.81208703 | 0.740005927 | 0.174993195 | 4.22876973 |
| Rpl35a-ps6 | 0.817924277 | -0.147536682 | 1.134095399 | -0.13009195 |
| Gm15857 | 0.981461204 | 1.261385693 | 1.100461358 | 1.146233517 |
| Gm15860 | 13.45882266 | 0.391344449 | 0.293355082 | 1.334029895 |
| Bcl7a | 835.7958974 | 0.770696584 | 0.10855957 | 7.099296601 |

| | | | | |
|---|---|---|---|---|
| Gm23458 | 4.576786753 | 0.504159774 | 0.587401793 | 0.858287768 |
| RP23-370G17.1 | 16.44557487 | 0.597556053 | 0.379756252 | 1.573525252 |
| Gm15747 | 1.272170519 | -0.057295246 | 0.860015584 | -0.066621172 |
| Mlxip | 530.2039296 | -0.006114334 | 0.102643249 | -0.059568786 |
| RP23-370G17.2 | 23.88538577 | 0.239877255 | 0.306252338 | 0.783266689 |
| RP23-370G17.3 | 10.05076668 | -0.518869584 | 0.422100947 | -1.229254725 |
| RP23-370G17.4 | 55.71562266 | 0.029570911 | 0.257281568 | 0.114935989 |
| Rpl31-ps6 | 0.955200391 | -0.40893274 | 1.292374884 | -0.316419597 |
| Il31 | 0.27099349 | -1.015396582 | 2.046398152 | -0.496187206 |
| Lrrc43 | 5.510592113 | -0.98732953 | 0.587070796 | -1.68178955 |
| Gm15751 | 2.413806088 | 0.067152342 | 0.787435604 | 0.085279789 |
| Diablo | 222.7606092 | -0.384349577 | 0.099273002 | -3.871642518 |
| B3gnt4 | 5.682107971 | -0.856241335 | 0.468515147 | -1.827563828 |
| Gm24682 | 0.386251008 | 1.407039264 | 1.785008109 | 0.78825371 |
| Mir7647 | 1.111319392 | 0.741987615 | 1.076874881 | 0.689019336 |
| Vps33a | 1223.976526 | 0.045805141 | 0.053085551 | 0.862855141 |
| RP23-121M13.2 | 3.773545612 | 0.294807948 | 0.613361022 | 0.480643434 |
| Clip1 | 1856.591368 | 0.123428178 | 0.089046779 | 1.386104927 |
| Gm24840 | 0 | NA | NA | NA |
| RP23-121M13.3 | 21.46809472 | 0.667840504 | 0.331171417 | 2.016600676 |
| RP23-121M13.4 | 2.51877741 | 1.031130836 | 0.61784151 | 1.668924506 |
| Zcchc8 | 358.9731995 | -0.272382183 | 0.102820702 | -2.649098656 |
| Rsrc2 | 977.6377703 | 0.123213213 | 0.079495318 | 1.549943015 |
| Kntc1 | 15.85629698 | -0.862098338 | 0.379371158 | -2.272440379 |
| Hcar2 | 55.90105025 | -0.479330434 | 0.314283142 | -1.525154775 |
| Hcar1 | 26.46370296 | -0.072343631 | 0.259591518 | -0.278682567 |
| Denr | 363.9918925 | 0.039915577 | 0.107813083 | 0.370229431 |
| RP24-492O4.5 | 0.261825437 | 1.0360305 | 2.088576849 | 0.496046147 |
| Ccdc62 | 24.55556878 | -0.228847665 | 0.269227262 | -0.850016684 |
| Hip1r | 946.4062031 | 0.167415587 | 0.068994234 | 2.426515615 |
| Mir7032 | 0 | NA | NA | NA |
| Vps37b | 406.419246 | 0.066943571 | 0.104584534 | 0.640090538 |
| Gm37939 | 0.497466187 | -0.38857915 | 1.558137589 | -0.24938693 |
| RP24-492O4.9 | 0.226827357 | 0.036355176 | 2.143469653 | 0.016960901 |
| RP24-492O4.8 | 0.200931026 | -1.675880452 | 2.148510347 | -0.780019726 |
| Abcb9 | 408.729264 | 0.426189286 | 0.078428155 | 5.434136303 |
| Gm16001 | 32.41362582 | 0.465662445 | 0.211891816 | 2.197642426 |
| Ogfod2 | 183.0496373 | -0.11544988 | 0.086232465 | -1.338821515 |
| Arl6ip4 | 210.5407291 | 0.129422896 | 0.138412682 | 0.935050852 |
| Pitpnm2 | 2668.846213 | 0.841389745 | 0.142161671 | 5.918541473 |
| Pitpnm2os2 | 1.19134753 | -0.25299302 | 0.984637393 | -0.256940292 |
| Pitpnm2os1 | 0.048949532 | 0.037064902 | 3.724945049 | 0.009950456 |
| RP24-257M14.2 | 0 | NA | NA | NA |
| Mphosph9 | 491.0917071 | 0.252555085 | 0.170319712 | 1.482829452 |
| Gm23171 | 0 | NA | NA | NA |
| RP24-257M14.3 | 50.23293639 | 0.493806571 | 0.353717734 | 1.396046968 |
| RP23-180G16.2 | 6.931404989 | 0.542183621 | 0.472913757 | 1.146474623 |

| | | | | |
|---|---|---|---|---|
| 2810006K23Rik | 85.50115535 | -0.581957621 | 0.21721238 | -2.679210183 |
| Cdk2ap1 | 265.3510822 | -0.428786943 | 0.099928061 | -4.290956317 |
| Sbno1 | 3030.840915 | -0.002740837 | 0.053314344 | -0.051409 |
| Gm37415 | 6.225233596 | 0.350158364 | 0.481151762 | 0.727750351 |
| Gm27486 | 0 | NA | NA | NA |
| Setd8 | 525.0620955 | -0.568214159 | 0.101750365 | -5.584394285 |
| Rilpl2 | 121.8049509 | -0.508339788 | 0.119040531 | -4.270308469 |
| Gm15621 | 0.143485256 | -0.955563601 | 3.547405725 | -0.269369696 |
| Snrnp35 | 68.78650209 | -0.232107691 | 0.212952576 | -1.089950147 |
| Rilpl1 | 373.8418609 | 0.192235284 | 0.096011009 | 2.00222127 |
| RP23-311A21.2 | 18.59305175 | 1.189112684 | 0.302145198 | 3.935567048 |
| Tmed2 | 550.4698307 | -0.342654529 | 0.089280724 | -3.83794522 |
| Ddx55 | 213.2561564 | 0.171837905 | 0.139376045 | 1.232908459 |
| Eif2b1 | 210.495533 | -0.06760654 | 0.098770619 | -0.684480272 |
| Gtf2h3 | 153.3355758 | -0.469048087 | 0.13895589 | -3.375517839 |
| Tctn2 | 238.196889 | -0.040473524 | 0.12690742 | -0.318921653 |
| Atp6v0a2 | 1102.977081 | -0.565456171 | 0.101896054 | -5.549343194 |
| Gm24854 | 0 | NA | NA | NA |
| WI1-2065I1.2 | 0.075043205 | 0.517957688 | 3.724945049 | 0.139051095 |
| Dnah10 | 37.63473061 | -0.875456718 | 0.227277018 | -3.851936838 |
| RP23-437D10.1 | 3.569206524 | 1.273435726 | 0.611756528 | 2.081605455 |
| Ccdc92 | 896.6833926 | 0.320467603 | 0.079394148 | 4.0364134 |
| Zfp664 | 1184.705232 | -0.092768506 | 0.063261227 | -1.466435444 |
| Fam101a | 19.01723143 | -3.158459293 | 0.461344164 | -6.846210566 |
| Ncor2 | 4028.896925 | -0.090971796 | 0.074377297 | -1.223112425 |
| RP24-291G12.3 | 0.185374628 | 0.517968595 | 2.272545264 | 0.227924435 |
| RP24-291G12.2 | 0.590920558 | 1.023339716 | 1.191291755 | 0.859016871 |
| RP23-363M4.1 | 0.852297073 | 0.940106853 | 1.152655487 | 0.8156009 |
| Scarb1 | 198.4544971 | -0.121754888 | 0.096623116 | -1.260101033 |
| RP23-363M4.2 | 6.758329482 | -0.252899469 | 0.483640345 | -0.522908131 |
| RP24-105J12.1 | 0 | NA | NA | NA |
| Ubc | 623.813315 | -0.602752354 | 0.146811857 | -4.105610854 |
| Gm10382 | 3.55196556 | -0.322130401 | 0.590683032 | -0.545352386 |
| Dhx37 | 317.3772995 | -0.373104085 | 0.083088801 | -4.490425686 |
| RP24-105J12.3 | 0.046345709 | -0.443818988 | 3.724945049 | -0.119147795 |
| Bri3bp | 1674.728707 | 0.439555047 | 0.067615907 | 6.50076386 |
| RP24-324M10.2 | 0.427937583 | -0.925057667 | 1.468005681 | -0.630145836 |
| Aacs | 461.1003154 | -0.192069217 | 0.09229508 | -2.081034184 |
| Gm27551 | 0.589769752 | -1.149396207 | 1.350380191 | -0.85116489 |
| Tmem132b | 1925.698646 | 0.732163334 | 0.120558748 | 6.073083442 |
| Gm4868 | 0 | NA | NA | NA |
| RP24-332C17.1 | 1.392497718 | -0.822632842 | 0.990036922 | -0.830911276 |
| RP24-332C17.2 | 0 | NA | NA | NA |
| RP24-332C17.3 | 0.274664082 | -1.993775712 | 2.025447357 | -0.984363136 |
| RP24-332C17.4 | 5.259892424 | 0.402185215 | 0.432756173 | 0.929357548 |
| Gm23151 | 0 | NA | NA | NA |
| Gm37700 | 0 | NA | NA | NA |

| | | | | |
|---|---|---|---|---|
| RP23-414E14.1 | 0.0878635 | 0.037058229 | 3.724945049 | 0.009948665 |
| Gm24839 | 0 | NA | NA | NA |
| RP23-74L20.1 | 1.146741161 | 0.233400115 | 1.009477709 | 0.231208786 |
| RP23-74L20.2 | 0.044191549 | 0.037062677 | 3.724945049 | 0.009949859 |
| RP23-145H8.1 | 0 | NA | NA | NA |
| RP23-145H8.2 | 0.512350924 | 1.362178225 | 1.373650676 | 0.991648203 |
| RP23-282A16.1 | 0.230183376 | 0.812871177 | 2.495000623 | 0.32579999 |
| RP23-37N15.1 | 0.075043205 | 0.517957688 | 3.724945049 | 0.139051095 |
| RP23-37N15.3 | 0 | NA | NA | NA |
| RP23-37N15.4 | 0.066050875 | -0.443818988 | 3.724945049 | -0.119147795 |
| RP23-37N15.2 | 0 | NA | NA | NA |
| Tmem132c | 179.9200518 | 0.068496782 | 0.115046547 | 0.595383204 |
| Tmem132cos | 0 | NA | NA | NA |
| RP23-71M14.1 | 1.788302827 | -0.408887478 | 1.094538882 | -0.373570537 |
| WI1-2285D16.1 | 0.302970058 | 1.304836828 | 1.960954596 | 0.665408995 |
| Slc15a4 | 300.9485309 | -0.176514626 | 0.110639381 | -1.595405038 |
| WI1-1024F3.1 | 0.545152564 | 0.727055388 | 1.240998212 | 0.585863365 |
| Glt1d1 | 207.4632902 | 0.41950804 | 0.127093157 | 3.300791716 |
| Tmem132d | 473.83412 | 0.544976219 | 0.146039057 | 3.731715553 |
| Gm17132 | 0.461821276 | 1.79116509 | 1.574434538 | 1.137656122 |
| Gm17131 | 0.522940275 | -0.663576588 | 1.470058809 | -0.451394586 |
| RP23-378C15.2 | 8.098685304 | 0.649179963 | 0.546611922 | 1.187643257 |
| RP23-378C15.1 | 6.802413532 | 0.422061453 | 0.573844757 | 0.73549762 |
| WI1-2255B14.1 | 0.141177696 | 0.037075614 | 2.915366873 | 0.012717307 |
| CH36-12I15.2 | 13.22303069 | -0.6872941 | 0.309237626 | -2.222543578 |
| CH36-12I15.3 | 7.173227082 | 0.322539386 | 0.403617088 | 0.799122227 |
| CH36-12I15.1 | 0 | NA | NA | NA |
| CH36-12I15.4 | 0.075043205 | 0.517957688 | 3.724945049 | 0.139051095 |
| 5930412G12Rik | 23.11388567 | -0.157447656 | 0.250098818 | -0.629541785 |
| RP24-258A20.1 | 1.894888719 | -0.654357175 | 0.829981259 | -0.78839994 |
| Fzd10 | 39.99764841 | -0.100417521 | 0.186547159 | -0.538295634 |
| Piwil1 | 0.809884792 | 0.149349426 | 1.159599426 | 0.12879398 |
| RP23-156F20.2 | 0 | NA | NA | NA |
| Rimbp2 | 1419.490251 | 0.023535148 | 0.223574354 | 0.105267657 |
| RP23-156F20.4 | 0.134481855 | 0.963696503 | 3.654633248 | 0.263691713 |
| RP23-156F20.5 | 1.664368785 | -2.002554788 | 1.076506621 | -1.860234531 |
| Stx2 | 147.9774137 | -0.351653021 | 0.121633303 | -2.891091597 |
| Ran | 1531.557904 | -0.299303655 | 0.089937413 | -3.327910445 |
| Adgrd1 | 48.98301064 | -0.340827017 | 0.171740085 | -1.984551344 |
| Rps16-ps2 | 75.19513727 | -0.25026801 | 0.159987805 | -1.564294287 |
| RP24-158G19.1 | 0 | NA | NA | NA |
| RP24-81J14.1 | 2.60471737 | 0.064506841 | 0.78803383 | 0.081857959 |
| Sfswap | 880.7266452 | 0.166737331 | 0.137492748 | 1.212699091 |
| RP24-161C20.4 | 61.43406924 | -0.214878902 | 0.362040499 | -0.593521727 |
| Mmp17 | 1263.227049 | 0.350707152 | 0.134936026 | 2.599062404 |
| Gm6139 | 8.767336692 | -0.617997374 | 0.385923866 | -1.601345314 |
| Zfp11 | 110.3957661 | 0.113040098 | 0.136791448 | 0.826368163 |

| | | | | |
|---|---|---|---|---|
| 14-Sep | 0.250988417 | 0.054330346 | 2.476902033 | 0.021934798 |
| Mrps17 | 416.7636034 | 0.009793388 | 0.06470967 | 0.151343495 |
| Gbas | 690.6986552 | 0.069741662 | 0.069579649 | 1.002328447 |
| Gm15903 | 27.46139881 | -0.143496844 | 0.334148052 | -0.429440913 |
| Psph | 139.9382442 | -0.626019434 | 0.149296264 | -4.1931353 |
| Cct6a | 1504.12042 | -0.309755754 | 0.087956992 | -3.521672875 |
| Gm23245 | 2.474741284 | -0.289688982 | 0.652172579 | -0.444190681 |
| Snora15 | 0 | NA | NA | NA |
| Sumf2 | 86.25597758 | -0.381691153 | 0.123730601 | -3.084856545 |
| Phkg1 | 43.76477553 | 0.02982558 | 0.286860139 | 0.103972551 |
| WI1-535J9.1 | 0.523518094 | 0.51797437 | 1.447426982 | 0.357858722 |
| Chchd2 | 843.0146962 | -0.166455199 | 0.11760792 | -1.415340041 |
| Zbed5 | 99.19524039 | -0.393711441 | 0.15918739 | -2.473257715 |
| Nupr1l | 8.851478164 | 0.491858781 | 0.375205091 | 1.310906468 |
| Vkorc1l1 | 758.1542901 | -0.063939594 | 0.064006297 | -0.998957869 |
| RP23-189N6.2 | 51.40024564 | 0.487266692 | 0.323082696 | 1.50817948 |
| RP23-189N6.3 | 26.74839225 | 0.245748173 | 0.375219422 | 0.654945237 |
| Gusb | 191.859854 | -0.521307809 | 0.133655712 | -3.900378085 |
| Asl | 102.1205115 | -0.030138859 | 0.133763808 | -0.225314 |
| Crcp | 187.2081582 | -0.068750203 | 0.109070264 | -0.630329485 |
| Tpst1 | 399.6196954 | -0.228994934 | 0.090988301 | -2.516751407 |
| RP23-353L10.2 | 7.510072057 | -0.164397387 | 0.508864039 | -0.323067409 |
| Kctd7 | 165.5730604 | -0.284756593 | 0.107220703 | -2.655798605 |
| Rabgef1 | 458.137125 | -0.162629704 | 0.071972167 | -2.259619374 |
| Gm15920 | 252.9729714 | -0.00294724 | 0.106710596 | -0.027618996 |
| Gm15921 | 0.048949532 | 0.037064902 | 3.724945049 | 0.009950456 |
| Tmem248 | 679.0584884 | -0.159005342 | 0.103894878 | -1.530444463 |
| AC113029.6 | 6.876401235 | 0.404016953 | 0.421287498 | 0.959005322 |
| RP23-206J17.2 | 14.12832924 | 0.660913246 | 0.428608644 | 1.541997009 |
| Sbds | 553.098637 | -0.126379821 | 0.119160167 | -1.060587816 |
| Tyw1 | 213.1214519 | -0.271948906 | 0.1340789 | -2.028275188 |
| Caln1 | 4353.134745 | 0.444615246 | 0.095396462 | 4.660709999 |
| A330070K13Rik | 2.26626686 | 1.076445078 | 0.912648961 | 1.179473295 |
| RP24-571K14.1 | 0 | NA | NA | NA |
| Gm23761 | 4.136581771 | 0.192475065 | 0.572145075 | 0.336409546 |
| RP24-559P3.1 | 12.49495635 | -0.901498228 | 0.374611993 | -2.406485231 |
| Wbscr17 | 856.0949554 | 0.258300931 | 0.08365845 | 3.087565357 |
| RP23-323D23.1 | 6.092562558 | 0.793022384 | 0.5297051 | 1.49710166 |
| RP23-323D23.2 | 13.43160181 | 0.719864474 | 0.428011274 | 1.681882037 |
| RP23-323D23.3 | 19.70298692 | 1.008447844 | 0.433321463 | 2.327251082 |
| RP23-445H19.1 | 18.79099024 | 0.694394826 | 0.306854144 | 2.262947529 |
| RP23-114L5.4 | 5.83927098 | 0.539309025 | 0.474010433 | 1.137757711 |
| Auts2 | 2085.675582 | 0.516147707 | 0.126806535 | 4.07035575 |
| Gm27266 | 0 | NA | NA | NA |
| RP23-120C8.7 | 31.89874726 | 0.816598622 | 0.410321666 | 1.990142586 |
| RP23-120C8.6 | 19.56297173 | 1.03784169 | 0.373769254 | 2.776690912 |
| RP23-120C8.9 | 0.108767819 | -0.443823436 | 3.724945049 | -0.119148989 |

| | | | | |
|---|---|---|---|---|
| RP23-120C8.5 | 11.30893109 | 0.176039534 | 0.511668937 | 0.34404968 |
| RP23-120C8.4 | 44.54237424 | 0.8240811 | 0.360703732 | 2.28464811 |
| RP23-120C8.3 | 22.2103361 | 0.580567401 | 0.375293375 | 1.546969489 |
| RP23-120C8.2 | 22.94359978 | 0.950958654 | 0.328694024 | 2.89314251 |
| RP24-95O4.7 | 0 | NA | NA | NA |
| RP24-95O4.6 | 73.13605096 | 0.631366154 | 0.304601672 | 2.072759973 |
| RP24-95O4.5 | 20.70201435 | 0.842831517 | 0.524646163 | 1.606476091 |
| RP24-95O4.4 | 58.37291526 | 0.875457979 | 0.349669445 | 2.503673089 |
| RP24-95O4.9 | 0 | NA | NA | NA |
| RP24-95O4.3 | 42.64488512 | 0.37672052 | 0.263382799 | 1.430315579 |
| RP24-95O4.2 | 132.5110994 | 0.72142626 | 0.373005336 | 1.93409099 |
| RP24-95O4.1 | 237.0691924 | 0.685669807 | 0.323448511 | 2.119873126 |
| RP23-378O9.2 | 72.35690436 | 1.053170245 | 0.334519921 | 3.148303521 |
| RP23-378O9.1 | 291.0443093 | 0.673652372 | 0.291952975 | 2.307400271 |
| RP24-147H20.3 | 37.48355553 | 0.595128897 | 0.333488835 | 1.784554186 |
| RP24-147H20.2 | 50.07029278 | 0.64186916 | 0.262918814 | 2.441320764 |
| RP23-276C23.3 | 53.66935453 | 0.601379686 | 0.339562476 | 1.771042823 |
| RP23-276C23.2 | 105.0397251 | 0.482137535 | 0.238087141 | 2.025046519 |
| RP23-276C23.1 | 58.80184518 | 0.10467254 | 0.346721494 | 0.301892272 |
| RP24-399A15.2 | 87.32223591 | 0.441851391 | 0.315542401 | 1.400291652 |
| RP24-399A15.3 | 100.1496488 | 0.486497319 | 0.291609105 | 1.668320056 |
| RP24-399A15.5 | 46.61621938 | 0.647745628 | 0.390953641 | 1.65683488 |
| Gm28616 | 0.118974955 | 0.51795324 | 3.724945049 | 0.139049901 |
| RP24-399A15.6 | 0.098502579 | -0.443821212 | 3.724945049 | -0.119148392 |
| Gm38373 | 0 | NA | NA | NA |
| Gm25722 | 0 | NA | NA | NA |
| RP23-455A10.3 | 0 | NA | NA | NA |
| Gm10051 | 4.533521042 | -0.236846001 | 0.587581792 | -0.403086011 |
| RP23-455A10.2 | 0 | NA | NA | NA |
| RP23-455A10.1 | 0.061920613 | -0.443816764 | 3.724945049 | -0.119147198 |
| RP23-409D19.1 | 5.435068931 | 0.397462877 | 0.456133829 | 0.871373382 |
| RP23-409D19.2 | 0 | NA | NA | NA |
| RP24-199E1.1 | 0.733529363 | -1.292878681 | 1.251873963 | -1.03275467 |
| Gm15627 | 4.57824934 | 0.854706325 | 0.533748964 | 1.601326434 |
| Gatsl2 | 1933.057981 | 0.172119324 | 0.077018253 | 2.23478613 |
| Wbscr16 | 129.6578973 | -0.436420692 | 0.118738154 | -3.675488281 |
| B230377A18Rik | 10.36497119 | -0.455753528 | 0.366359704 | -1.244005614 |
| Gtf2ird2 | 207.0338271 | 0.207666132 | 0.101213186 | 2.05176954 |
| Gm24139 | 2.685875012 | 1.115555466 | 0.667800409 | 1.670492338 |
| Ncf1 | 204.7455163 | -0.011652205 | 0.192244602 | -0.060611351 |
| Gtf2i | 2503.932034 | -0.11906092 | 0.103704028 | -1.148083846 |
| RP23-15K13.2 | 110.8580539 | 0.21518415 | 0.374686056 | 0.574305199 |
| RP23-15K13.3 | 5.451269312 | -0.384322698 | 0.63181966 | -0.608279107 |
| Mir3965 | 0 | NA | NA | NA |
| Gtf2ird1 | 384.0610061 | -0.045850046 | 0.139424672 | -0.328851743 |
| n-R5s178 | 0 | NA | NA | NA |
| Gm26340 | 0 | NA | NA | NA |

| | | | | |
|---|---|---|---|---|
| RP24-359H20.2 | 0.585862523 | 0.645113186 | 1.272657201 | 0.506902554 |
| Gm16020 | 0 | NA | NA | NA |
| RP24-359H20.3 | 3.016053333 | -2.811036318 | 0.897787997 | -3.13106917 |
| Clip2 | 1985.813075 | -0.19509217 | 0.051196179 | -3.810678337 |
| RP23-85H24.1 | 29.77219559 | -0.100064682 | 0.329209245 | -0.303954654 |
| RP23-85H24.2 | 8.735753554 | 0.699646702 | 0.381874909 | 1.832135827 |
| Syna | 7.492197562 | 0.413704962 | 0.40705308 | 1.016341558 |
| mmu-mir-7228 | 0.285083685 | -0.291397978 | 1.792546889 | -0.162560868 |
| Rfc2 | 247.4000499 | -0.250108664 | 0.086320282 | -2.897449551 |
| Lat2 | 113.3855065 | -1.130204195 | 0.218265843 | -5.178108411 |
| Eif4h | 3070.987643 | 0.00414723 | 0.049235863 | 0.084231902 |
| Limk1 | 829.834934 | 0.009484951 | 0.12202139 | 0.077731871 |
| Gm10369 | 1.019349322 | 2.156525792 | 1.242811123 | 1.735199944 |
| Eln | 72.96738645 | -0.051308819 | 0.189901607 | -0.270186336 |
| RP23-315E2.1 | 8.223192194 | 1.508654267 | 0.430364467 | 3.50552702 |
| Wbscr28 | 0.045882455 | 0.517959912 | 3.724945049 | 0.139051692 |
| Cldn13 | 0 | NA | NA | NA |
| RP23-201C9.3 | 0 | NA | NA | NA |
| RP23-201C9.2 | 0.038981395 | -0.443816764 | 3.724945049 | -0.119147198 |
| Wbscr27 | 62.20310215 | 0.658122262 | 0.171863861 | 3.829323147 |
| Cldn4 | 0.565252138 | -1.046827866 | 3.059335261 | -0.342174942 |
| Cldn3 | 2.938076242 | 0.076032415 | 0.621587674 | 0.122319695 |
| Wbscr25 | 2.760772158 | 1.423134744 | 0.668556814 | 2.128666875 |
| Rps29-ps | 0 | NA | NA | NA |
| RP23-38B15.1 | 4.804143474 | -0.306702221 | 0.5938244 | -0.516486392 |
| Abhd11 | 157.6817239 | 0.071616125 | 0.117992888 | 0.606952897 |
| Abhd11os | 7.140605062 | -0.512248002 | 0.557716659 | -0.918473554 |
| Stx1a | 708.4200384 | 1.341400301 | 0.162473844 | 8.256100007 |
| Wbscr22 | 194.0156334 | -0.307298452 | 0.083091082 | -3.698332523 |
| Gm25492 | 29.10926416 | 0.065711603 | 0.21048227 | 0.312195432 |
| Dnajc30 | 252.3546012 | 0.134952452 | 0.088521033 | 1.524524142 |
| Vps37d | 164.5301101 | 0.490265664 | 0.122303677 | 4.008593014 |
| Mlxipl | 13.0557667 | 1.002652971 | 0.393850476 | 2.545770622 |
| RP23-240H6.1 | 0 | NA | NA | NA |
| Tbl2 | 380.9054715 | -0.069359844 | 0.066854848 | -1.037469182 |
| Bcl7b | 442.8768574 | -0.095934727 | 0.057354381 | -1.672666071 |
| Baz1b | 2277.625964 | -0.131825231 | 0.087520363 | -1.506223549 |
| Gm25315 | 0 | NA | NA | NA |
| Fzd9 | 41.32503692 | 0.456080518 | 0.197234965 | 2.312371528 |
| RP23-240H6.2 | 0.095295241 | -0.443821212 | 3.724945049 | -0.119148392 |
| Fkbp6 | 0.371024948 | 0.443712294 | 1.636609752 | 0.271116736 |
| Trim50 | 0.160978641 | 0.03706851 | 2.803723868 | 0.01322117 |
| Nsun5 | 116.6234585 | 0.146678296 | 0.175755896 | 0.834556902 |
| Pom121 | 1177.537542 | -0.078046088 | 0.073360758 | -1.063866971 |
| Mir7033 | 0 | NA | NA | NA |
| Hip1 | 1047.747718 | -0.623331002 | 0.114860425 | -5.426856139 |
| RP23-333I24.3 | 8.728400981 | -0.505494371 | 0.40212912 | -1.257044929 |

| | | | | |
|---|---|---|---|---|
| Ccl26 | 0 | NA | NA | NA |
| Ccl24 | 1.573001602 | -3.072466757 | 1.048148592 | -2.931327467 |
| RP23-423A22.3 | 0.045882455 | 0.517959912 | 3.724945049 | 0.139051692 |
| Rhbdd2 | 635.3255966 | 0.202831584 | 0.056805109 | 3.57065745 |
| Gm15607 | 0.101408268 | 0.037730769 | 3.724898049 | 0.010129343 |
| Gm21839 | 0.540306067 | -0.452148968 | 1.364671893 | -0.331324307 |
| RP23-67P7.1 | 0.422515209 | 0.084760085 | 1.374410745 | 0.061670127 |
| Por | 973.1641497 | -0.600376103 | 0.101365671 | -5.922874077 |
| Gm16061 | 0.898502897 | -0.533445588 | 1.110356784 | -0.480427188 |
| Tmem120a | 119.540289 | -0.653738342 | 0.120533662 | -5.423699323 |
| Mir7034 | 0 | NA | NA | NA |
| Styxl1 | 3.717676903 | -0.192457501 | 0.578620763 | -0.332614232 |
| Mdh2 | 1871.37566 | -0.043491598 | 0.059576751 | -0.730009564 |
| Srrm3 | 617.0965085 | 0.172806656 | 0.145945109 | 1.184052397 |
| Gm23268 | 0 | NA | NA | NA |
| Srrm3os | 0.79881978 | 1.998878328 | 3.576497316 | 0.558892724 |
| Gm22450 | 0.295673938 | -0.431701959 | 1.627923377 | -0.265185675 |
| Hspb1 | 217.0812965 | -1.676058321 | 0.432031173 | -3.879484685 |
| RP23-11P12.2 | 1.422147357 | 3.013711726 | 1.199387242 | 2.512709508 |
| Ywhag | 12717.10653 | -0.021595853 | 0.068670956 | -0.314483064 |
| Ssc4d | 19.88563498 | -0.283610639 | 0.258868334 | -1.095578723 |
| Zp3 | 0.564341384 | 0.479123214 | 1.449363505 | 0.330574913 |
| Dtx2 | 165.5707377 | -0.517135713 | 0.132828313 | -3.893264173 |
| Upk3b | 0.665037802 | 0.416373195 | 1.257995135 | 0.330981562 |
| Upk3bl | 6.950616643 | -1.887283776 | 0.469504782 | -4.019732805 |
| Rasa4 | 403.5442416 | -0.440235867 | 0.129364904 | -3.403054872 |
| Gm22214 | 0 | NA | NA | NA |
| Gm15701 | 0.196892959 | 0.037620624 | 2.233059319 | 0.016847123 |
| Mir7035 | 0 | NA | NA | NA |
| Polr2j | 169.450244 | -0.062968602 | 0.104009474 | -0.605412174 |
| RP23-314O1.1 | 1.911784405 | -0.575415057 | 0.72803602 | -0.790366192 |
| Lrwd1 | 120.9503839 | -0.110864808 | 0.115667557 | -0.958477991 |
| Alkbh4 | 76.72992867 | -0.003773486 | 0.121265257 | -0.031117618 |
| Orai2 | 675.4374974 | 0.448647547 | 0.08725949 | 5.141533012 |
| Prkrip1 | 213.2039114 | -0.063737959 | 0.130302743 | -0.489152857 |
| RP23-284P20.2 | 0 | NA | NA | NA |
| Sh2b2 | 93.36965878 | -0.14417594 | 0.159642607 | -0.903116924 |
| Cux1 | 1881.15298 | 0.003990571 | 0.071309137 | 0.055961566 |
| Gm16599 | 1.050839697 | -0.950619197 | 1.202948283 | -0.790241119 |
| A430110C17Rik | 51.6013035 | 0.183276173 | 0.312609002 | 0.586279258 |
| mmu-mir-721 | 2.23838086 | 0.415031404 | 0.845989897 | 0.490586715 |
| Gm38082 | 75.94798503 | -0.283256765 | 0.288165626 | -0.982965138 |
| Myl10 | 0.304278192 | 2.259449884 | 3.686936899 | 0.612825754 |
| Col26a1 | 25.30413262 | 0.514989084 | 0.240487951 | 2.14143404 |
| Gm20485 | 0.218750444 | -1.048061314 | 2.599907078 | -0.403114912 |
| RP24-456B4.2 | 0 | NA | NA | NA |
| Ift22 | 390.4508874 | -0.069252086 | 0.087312527 | -0.793151779 |

| | | | | |
|---|---|---|---|---|
| 4933404O12Rik | 126.181175 | 0.136459764 | 0.113537564 | 1.201890885 |
| Fis1 | 968.7424033 | -0.052423077 | 0.080310696 | -0.652753359 |
| Cldn15 | 7.096013574 | -1.461511986 | 0.525390536 | -2.781763059 |
| Znhit1 | 217.8120474 | -0.289889892 | 0.100416349 | -2.886879419 |
| Plod3 | 660.8862868 | -0.743094771 | 0.109994788 | -6.755727122 |
| Mir702 | 0 | NA | NA | NA |
| Gm7284 | 0.113979814 | 0.517957688 | 3.724945049 | 0.139051095 |
| Gm26989 | 0 | NA | NA | NA |
| Vgf | 2915.612919 | -1.251083508 | 0.163690341 | -7.642989191 |
| Ap1s1 | 1188.779745 | -0.248602076 | 0.070720567 | -3.51527267 |
| Serpine1 | 1051.265589 | -2.972007808 | 0.496888773 | -5.981233571 |
| Trim56 | 1329.516906 | -0.48987082 | 0.16134209 | -3.036224587 |
| Muc3 | 0.044544648 | 0.517957688 | 3.724945049 | 0.139051095 |
| Gm3054 | 0.061920613 | -0.443816764 | 3.724945049 | -0.119147198 |
| CAAA01077340.1 | 0.185747766 | -0.443821885 | 3.2922738 | -0.1348071 |
| CAAA01154573.1 | 3.791680059 | 0.029070949 | 0.488564863 | 0.059502742 |
| Muc3a | 21.091838 | 1.262115138 | 0.477567484 | 2.642799565 |
| WI1-166E14.1 | 15.34798663 | 0.539559367 | 0.341602906 | 1.579492906 |
| Ache | 1111.230768 | -0.033192615 | 0.105337293 | -0.315107918 |
| Mir8116 | 1.629432974 | -0.163186011 | 0.932009687 | -0.175090466 |
| Ufsp1 | 91.40840618 | 0.06807613 | 0.160422395 | 0.424355528 |
| Srrt | 944.0836702 | -0.195557068 | 0.07508589 | -2.604444952 |
| Mir7036 | 0 | NA | NA | NA |
| RP24-210N1.1 | 0.752770407 | -0.591573126 | 1.155097412 | -0.512141331 |
| Trip6 | 103.7948532 | -0.650908882 | 0.208529898 | -3.12141755 |
| Slc12a9 | 407.3671883 | 0.015603097 | 0.084390672 | 0.184891248 |
| RP24-210N1.3 | 0.151841013 | 0.517963177 | 2.85004976 | 0.181738292 |
| Ephb4 | 94.20008279 | -0.454065235 | 0.146156187 | -3.106712387 |
| Zan | 8.749737854 | 0.79776594 | 0.396877432 | 2.010106584 |
| RP24-210N1.4 | 0 | NA | NA | NA |
| Epo | 0.229076001 | 0.037069684 | 2.563865543 | 0.014458513 |
| Pop7 | 101.0875345 | -0.071565351 | 0.135458533 | -0.528319252 |
| RP23-45G4.2 | 6.33503719 | -0.448028326 | 0.427554312 | -1.047886346 |
| RP23-45G4.3 | 32.98247416 | 0.408121372 | 0.227491259 | 1.794009028 |
| Gigyf1 | 841.5272401 | 0.155503498 | 0.12245177 | 1.269916299 |
| Gnb2 | 2014.783637 | -0.253724319 | 0.062540948 | -4.056931149 |
| Mir6418 | 0 | NA | NA | NA |
| Gm17112 | 0.842992085 | -1.637674292 | 1.156410959 | -1.416169813 |
| Actl6b | 409.7728831 | 0.478716533 | 0.111754796 | 4.283633022 |
| Tfr2 | 33.22771086 | 0.816238673 | 0.231352018 | 3.52812428 |
| Mospd3 | 114.8382337 | 0.009706975 | 0.11321376 | 0.08574024 |
| RP23-286M3.4 | 4.875430065 | -0.553812729 | 0.515205569 | -1.074935448 |
| Pcolce | 87.87249262 | -0.434375402 | 0.191909423 | -2.26343967 |
| Fbxo24 | 2.595989607 | 0.9416079 | 0.725491809 | 1.297889084 |
| RP23-286M3.5 | 4.924586763 | -1.082610743 | 0.55285764 | -1.958208886 |
| Gm16089 | 9.122415521 | -0.426297069 | 0.370153237 | -1.151677271 |
| Lrch4 | 4.483355963 | 0.35594815 | 0.643687301 | 0.552983024 |

| | | | | |
|---|---|---|---|---|
| Gm20605 | 6.061787305 | -0.270876204 | 0.502469881 | -0.539089435 |
| Sap25 | 0.075043205 | 0.517957688 | 3.724945049 | 0.139051095 |
| Irs3 | 1.938977309 | -0.828075052 | 0.779351708 | -1.062517787 |
| Agfg2 | 460.5243141 | 0.100366004 | 0.079657816 | 1.259964294 |
| Gm15498 | 0.332718546 | 0.058807751 | 1.96922122 | 0.029863456 |
| Gm15753 | 9.170347486 | -0.711013259 | 0.33712701 | -2.109036766 |
| Nyap1 | 848.2957187 | 0.244457106 | 0.078254359 | 3.123878455 |
| Tsc22d4 | 1384.328407 | -0.296066174 | 0.145419745 | -2.035942049 |
| Ppp1r35 | 80.27309234 | -0.321833864 | 0.153721218 | -2.093620313 |
| Mepce | 500.0316796 | 0.05210198 | 0.07244518 | 0.719191801 |
| Zcwpw1 | 45.4661166 | 0.020790502 | 0.201930571 | 0.102958664 |
| Pilra | 8.121710499 | -0.058811201 | 0.359185977 | -0.163734679 |
| Pilrb1 | 1.522009338 | -2.315670378 | 0.995862169 | -2.325292044 |
| RP23-425N17.3 | 0.609942678 | -1.072580507 | 1.33909239 | -0.800975731 |
| Pilrb2 | 1.815499725 | -0.348167033 | 0.659109311 | -0.528238681 |
| RP23-425N17.5 | 0.143022002 | 0.03610674 | 3.547735707 | 0.010177404 |
| RP23-425N17.6 | 0 | NA | NA | NA |
| RP23-131D6.3 | 0 | NA | NA | NA |
| Cyp3a13 | 18.8351161 | 1.267273042 | 0.343251489 | 3.691966627 |
| Gm15502 | 0.499480636 | 0.041253837 | 1.487597988 | 0.027731845 |
| RP23-131D6.4 | 4.460458762 | 0.453862229 | 0.51956151 | 0.873548599 |
| Gjc3 | 574.2169864 | -0.001212967 | 0.131191802 | -0.009245751 |
| Azgp1 | 0.218824723 | 0.969703469 | 2.580964346 | 0.375713624 |
| Gm4963 | 0.586951709 | 0.467532634 | 1.284007213 | 0.364119944 |
| Gm29127 | 0 | NA | NA | NA |
| Smok3a | 0.044544648 | 0.517957688 | 3.724945049 | 0.139051095 |
| Smok3b | 0 | NA | NA | NA |
| Smok3c | 0 | NA | NA | NA |
| Zkscan1 | 1601.280236 | 0.11044601 | 0.048114908 | 2.295463416 |
| RP23-147H4.1 | 0 | NA | NA | NA |
| Zscan21 | 197.9428209 | -0.06079579 | 0.093815106 | -0.648038385 |
| Zfp113 | 154.778503 | -0.552174195 | 0.170472469 | -3.239081357 |
| Cops6 | 678.0499843 | -0.180046781 | 0.089234522 | -2.017680784 |
| Mcm7 | 172.4442146 | -0.535688561 | 0.142983126 | -3.74651595 |
| Mir25 | 0 | NA | NA | NA |
| Mir93 | 0 | NA | NA | NA |
| Mir106b | 0.072025462 | 0.517959912 | 3.724945049 | 0.139051692 |
| Ap4m1 | 196.8680612 | 0.136505067 | 0.113682055 | 1.200761783 |
| Taf6 | 295.6377165 | -0.346804969 | 0.106545754 | -3.254986293 |
| Cnpy4 | 86.00518766 | -0.109389819 | 0.143491119 | -0.76234557 |
| Mblac1 | 28.62003678 | -0.503311068 | 0.227841821 | -2.209037244 |
| Gm454 | 0 | NA | NA | NA |
| RP24-86H20.4 | 0 | NA | NA | NA |
| Gm22956 | 0 | NA | NA | NA |
| Nxpe5 | 0.64031546 | -2.696515514 | 1.257136648 | -2.144966117 |
| Lamtor4 | 172.5096594 | -0.149021215 | 0.118954507 | -1.252758038 |
| BC037034 | 543.4878643 | 0.00950711 | 0.05895028 | 0.16127337 |

| | | | | |
|---|---|---|---|---|
| 6330418K02Rik | 28.7450452 | 0.038819895 | 0.226354468 | 0.171500457 |
| Gal3st4 | 0 | NA | NA | NA |
| Gpc2 | 31.36245639 | -0.286681696 | 0.23216324 | -1.234828113 |
| Stag3 | 27.52049443 | 0.702187936 | 0.263469537 | 2.665157969 |
| RP24-95J4.1 | 0.060174537 | 0.517957688 | 3.724945049 | 0.139051095 |
| Gm10874 | 0.888510269 | -1.509827563 | 1.192534557 | -1.26606609 |
| Got2-ps1 | 146.5220016 | 0.088400663 | 0.103913254 | 0.850715956 |
| Gm25525 | 0.061920613 | -0.443816764 | 3.724945049 | -0.119147198 |
| Zfp157 | 295.429195 | 0.104845523 | 0.09776558 | 1.072417546 |
| Gm5564 | 8.712984693 | -0.529867838 | 0.383251551 | -1.382558887 |
| RP24-255F19.2 | 1.435448476 | -0.921359209 | 0.908235468 | -1.014449712 |
| 1700123K08Rik | 2.221046269 | 1.590216457 | 0.834829142 | 1.904840615 |
| Zfp68 | 529.1369275 | 0.18895478 | 0.068001017 | 2.778705201 |
| Gm24082 | 0 | NA | NA | NA |
| Gm15497 | 0.148708979 | -0.443201667 | 2.868467393 | -0.154508177 |
| RP24-255F19.5 | 0.796574019 | -1.357411965 | 1.275302743 | -1.064384102 |
| A430033K04Rik | 179.5683617 | 0.24214499 | 0.117791823 | 2.05570288 |
| Gm38232 | 0.132101749 | -0.93550548 | 3.724188147 | -0.251197158 |
| Fam20c | 570.0411155 | -0.284980874 | 0.09710727 | -2.93470174 |
| Gm5294 | 0.042716945 | 0.037062677 | 3.724945049 | 0.009949859 |
| Pdgfa | 337.774443 | -0.046556397 | 0.084677448 | -0.549808691 |
| 6330403L08Rik | 206.3643988 | 0.207682179 | 0.139893751 | 1.484570815 |
| Prkar1b | 4853.393054 | 0.202092778 | 0.083575998 | 2.418071971 |
| Gm15672 | 16.59824677 | 0.061951422 | 0.319251294 | 0.194052219 |
| Dnaaf5 | 128.0720811 | -0.62611479 | 0.1184942 | -5.283927734 |
| Sun1 | 919.0838976 | 0.212049337 | 0.108934493 | 1.946576622 |
| RP23-430G6.3 | 0.066050875 | -0.443818988 | 3.724945049 | -0.119147795 |
| Get4 | 323.1872879 | -0.07216803 | 0.113421713 | -0.636280553 |
| Adap1 | 1303.85996 | 0.325200785 | 0.069581873 | 4.673642315 |
| Gm27316 | 0 | NA | NA | NA |
| Cox19 | 240.9171869 | 0.081884169 | 0.079003482 | 1.036462781 |
| Cyp2w1 | 0 | NA | NA | NA |
| 3110082I17Rik | 68.15138838 | -0.108113908 | 0.126779954 | -0.852768154 |
| Mir339 | 1.20342686 | -0.73464989 | 1.169253681 | -0.628306673 |
| Gpr146 | 293.4623801 | 0.142185845 | 0.096237398 | 1.477448984 |
| C130050O18Rik | 9.082211208 | -0.4896823 | 0.349668893 | -1.400417109 |
| Gper1 | 14.62948997 | 0.195664837 | 0.323669477 | 0.60452051 |
| Gm26285 | 0.142244466 | -1.207781436 | 3.552423657 | -0.339988006 |
| Zfand2a | 664.2777979 | -0.324361911 | 0.099443061 | -3.261785268 |
| RP23-400D9.3 | 0.170157162 | 0.037073895 | 2.749797593 | 0.013482409 |
| Uncx | 53.72924227 | -0.264869816 | 0.204223983 | -1.29695745 |
| RP23-92L4.4 | 0 | NA | NA | NA |
| Micall2 | 117.2123074 | -0.3475282 | 0.279907165 | -1.241583794 |
| Ints1 | 1337.189342 | 0.133595251 | 0.056411792 | 2.368215006 |
| Mir7037 | 0 | NA | NA | NA |
| Mafk | 342.8849901 | -0.377299946 | 0.200417341 | -1.882571355 |
| Tmem184a | 2.905166453 | -1.031725361 | 0.629870421 | -1.637996207 |

| | | | | |
|---|---|---|---|---|
| Gm26938 | 0.061920613 | -0.443816764 | 3.724945049 | -0.119147198 |
| Psmg3 | 54.0547785 | -0.484232286 | 0.163275089 | -2.965745033 |
| Gm24146 | 2.128603593 | 0.088720984 | 0.685562046 | 0.129413501 |
| RP23-168E11.1 | 0.493577707 | 0.032746709 | 1.504542941 | 0.021765221 |
| Elfn1 | 266.0295 | 0.014563309 | 0.135265026 | 0.107664997 |
| RP23-31L10.2 | 0 | NA | NA | NA |
| Mad1l1 | 200.5910728 | 0.319287722 | 0.098770575 | 3.232619851 |
| RP23-31L10.4 | 9.073372328 | 0.062938156 | 0.388772218 | 0.161889541 |
| Gm16122 | 0.099106094 | -0.93550548 | 3.724188147 | -0.251197158 |
| Gm16121 | 0.641775973 | -0.296879863 | 1.508166252 | -0.196848234 |
| RP24-97G6.1 | 10.02733677 | 0.61276681 | 0.399113191 | 1.535320865 |
| Gm16120 | 0.194043089 | -0.881534944 | 2.227684744 | -0.395717997 |
| RP23-307I19.1 | 0.437638936 | 0.816911687 | 1.839191946 | 0.444168804 |
| Nudt1 | 23.04760792 | -0.543756524 | 0.258676573 | -2.102071007 |
| Ftsj2 | 69.98359942 | -0.06006872 | 0.153454305 | -0.391443692 |
| Snx8 | 233.8617382 | -0.668911569 | 0.126594018 | -5.283911346 |
| Eif3b | 1148.304761 | -0.521629401 | 0.083527637 | -6.244991677 |
| Mir7038 | 0.519085648 | 0.517974509 | 1.557490839 | 0.332569859 |
| RP23-307I19.6 | 0 | NA | NA | NA |
| RP23-307I19.5 | 0.947788321 | -3.362384537 | 1.207091884 | -2.785524931 |
| Chst12 | 156.329386 | 0.208884324 | 0.107007144 | 1.952059619 |
| RP23-462C22.2 | 1.646628032 | -1.32371733 | 0.823716193 | -1.607006564 |
| RP23-462C22.1 | 0.174406792 | 0.541348768 | 3.365276146 | 0.160863104 |
| Grifin | 1.541974883 | -1.83412762 | 0.933961528 | -1.963814959 |
| RP23-462C22.5 | 1.34756889 | 1.837084919 | 1.026977313 | 1.788827168 |
| Lfng | 314.5144813 | -0.3294436 | 0.174469431 | -1.888259725 |
| Ttyh3 | 2824.785642 | 0.062114553 | 0.093001528 | 0.667887449 |
| Iqce | 386.6329369 | 0.190953417 | 0.143165802 | 1.333792111 |
| Brat1 | 216.0259619 | -0.181297805 | 0.111926384 | -1.619795069 |
| Amz1 | 137.7511797 | -0.016039013 | 0.162990188 | -0.098404775 |
| Gna12 | 984.1548664 | -0.136587027 | 0.118655751 | -1.151120161 |
| Card11 | 40.90469028 | -0.779131265 | 0.236483359 | -3.294655781 |
| AC161570.1 | 0 | NA | NA | NA |
| Gm26113 | 0 | NA | NA | NA |
| Sdk1 | 231.3078279 | 0.226742436 | 0.100218711 | 2.262476085 |
| Gm6433 | 1.643128883 | 0.778046653 | 0.813089078 | 0.956902108 |
| RP23-49G7.2 | 0 | NA | NA | NA |
| RP23-286C5.1 | 0 | NA | NA | NA |
| RP23-286C5.2 | 0.061920613 | -0.443816764 | 3.724945049 | -0.119147198 |
| Gm16035 | 0 | NA | NA | NA |
| Gm24131 | 0 | NA | NA | NA |
| Gm16036 | 0 | NA | NA | NA |
| Gm26970 | 0 | NA | NA | NA |
| Foxk1 | 1176.476401 | 0.218142533 | 0.076101972 | 2.866450461 |
| Ap5z1 | 229.1563663 | -0.269312462 | 0.115944182 | -2.322776853 |
| Radil | 157.9503125 | 0.026771444 | 0.13471314 | 0.198729269 |
| Papolb | 13.6158541 | 0.329854605 | 0.354033127 | 0.931705481 |

| | | | | |
|---|---|---|---|---|
| RP23-106N8.2 | 0 | NA | NA | NA |
| Mmd2 | 1010.130574 | 0.395006715 | 0.157274787 | 2.511570488 |
| RP23-106N8.4 | 0 | NA | NA | NA |
| Wipi2 | 1453.522254 | -0.099578858 | 0.058699623 | -1.696413939 |
| Slc29a4 | 200.9815486 | -0.274958069 | 0.122378236 | -2.246788952 |
| Tnrc18 | 2698.838952 | -0.043355666 | 0.067023602 | -0.646871629 |
| Gm15770 | 0.972120366 | 0.415346461 | 1.065092702 | 0.389962733 |
| Fbxl18 | 720.5681582 | 0.368389863 | 0.072364819 | 5.09073152 |
| D430018E03Rik | 19.83389332 | 0.873741975 | 0.343666109 | 2.542415301 |
| Actb | 25647.77243 | -0.329532337 | 0.113793817 | -2.895872091 |
| Fscn1 | 1883.804362 | 0.22932825 | 0.087422776 | 2.623209436 |
| Rnf216 | 737.7813341 | -0.143355604 | 0.04778587 | -2.999958027 |
| RP23-393G8.5 | 4.643209367 | -0.500437602 | 0.541458661 | -0.924239721 |
| RP23-393G8.10 | 0.927313021 | 0.306249402 | 1.07050934 | 0.286078216 |
| Olfr718-ps1 | 0 | NA | NA | NA |
| RP23-301L9.8 | 0.114443069 | -0.443821212 | 3.724945049 | -0.119148392 |
| AC123686.1 | 0 | NA | NA | NA |
| Rbakdn | 0.061920613 | -0.443816764 | 3.724945049 | -0.119147198 |
| Rbak | 178.0893661 | 0.396656704 | 0.142387165 | 2.785761659 |
| 4933411G11Rik | 0.556792772 | -0.010026679 | 1.31430255 | -0.007628897 |
| Spdyb | 0.075043205 | 0.517957688 | 3.724945049 | 0.139051095 |
| 4930448H16Rik | 0 | NA | NA | NA |
| Zfp12 | 444.4098317 | 0.195048471 | 0.093836219 | 2.078605396 |
| Zfp316 | 378.1393857 | 0.115561761 | 0.101179032 | 1.142151281 |
| RP23-301L9.3 | 37.38971463 | 0.206286815 | 0.296181834 | 0.696487059 |
| Zfp853 | 32.64981703 | 1.238484427 | 0.229514265 | 5.396110907 |
| E130309D02Rik | 214.9676355 | -0.064572944 | 0.106993084 | -0.603524469 |
| RP23-301L9.6 | 1.497168834 | 1.66482013 | 0.885765091 | 1.87952782 |
| Zdhhc4 | 177.8279622 | 0.267390918 | 0.121188496 | 2.206405111 |
| 0610040B10Rik | 20.77576342 | 0.769768221 | 0.300352016 | 2.56288681 |
| RP23-301L9.4 | 0.904055358 | 2.596717809 | 1.252795664 | 2.072738504 |
| Grid2ip | 247.2010809 | 0.488945814 | 0.223893587 | 2.183831254 |
| Gm20635 | 4.77646696 | 0.668552576 | 0.508923087 | 1.313661323 |
| RP23-372N19.4 | 0.76879839 | 0.933122984 | 1.294940616 | 0.720591332 |
| RP23-372N19.2 | 152.8195759 | 0.383521039 | 0.21995086 | 1.743666928 |
| 9530056K15Rik | 3.981355527 | 1.078887275 | 0.597278296 | 1.806339326 |
| Kdelr2 | 375.7129238 | -0.630657142 | 0.128449117 | -4.909781855 |
| RP23-158A20.2 | 0 | NA | NA | NA |
| RP23-158A20.4 | 0 | NA | NA | NA |
| Daglb | 512.0781547 | -0.056618358 | 0.080113117 | -0.706730181 |
| Rac1 | 2571.473266 | -0.083285908 | 0.076897182 | -1.083081399 |
| RP23-158A20.7 | 0.049553047 | -0.443818988 | 3.724945049 | -0.119147795 |
| Fam220a | 378.2842679 | 0.001764116 | 0.082737497 | 0.021321838 |
| Cyth3 | 478.6545393 | 0.118757575 | 0.074169464 | 1.601165335 |
| Usp42 | 463.5942125 | -0.185391924 | 0.070472604 | -2.630694964 |
| D130017N08Rik | 140.3508044 | 0.160040518 | 0.157751178 | 1.014512342 |
| RP23-151C5.2 | 0 | NA | NA | NA |

| | | | | |
|---|---|---|---|---|
| RP23-151C5.3 | 4.718276104 | 1.457236332 | 0.569592229 | 2.558385205 |
| Eif2ak1 | 746.7429788 | 0.008450247 | 0.061719192 | 0.136914412 |
| Ankrd61 | 4.124592769 | -0.141724744 | 0.517543765 | -0.27384108 |
| Aimp2 | 134.5489307 | -0.449229079 | 0.124898066 | -3.596765711 |
| Pms2 | 217.1478713 | 0.570565133 | 0.105682246 | 5.398874028 |
| Gm17135 | 0.441552168 | 2.232638212 | 1.382632136 | 1.614773846 |
| PMS2 | 0.78876463 | 1.211980727 | 1.095599086 | 1.106226486 |
| Rsph10b | 55.58878597 | 0.421712655 | 0.197192164 | 2.138587283 |
| Ccz1 | 450.6103287 | 0.013277675 | 0.066109969 | 0.200842256 |
| Gm24311 | 0 | NA | NA | NA |
| Ocm | 1.753517396 | 0.424871926 | 0.690832079 | 0.615014761 |
| RP24-194F23.5 | 0.296409788 | 0.517955413 | 3.724945049 | 0.139050484 |
| Lmtk2 | 3277.074586 | 0.342185527 | 0.067869099 | 5.041845716 |
| Gm26925 | 0 | NA | NA | NA |
| Bhlha15 | 4.240558544 | -1.271201201 | 0.651621381 | -1.950827945 |
| Tecpr1 | 1209.265373 | 0.307773095 | 0.067902417 | 4.532579403 |
| RP24-82B23.5 | 8.247112194 | -0.463251695 | 0.420939918 | -1.100517377 |
| 2900089D17Rik | 4.905083075 | 0.014401164 | 0.537938211 | 0.026771038 |
| Bri3 | 199.49671 | -0.476662622 | 0.12605856 | -3.781279278 |
| Baiap2l1 | 221.2967602 | 0.453331366 | 0.127688937 | 3.550279109 |
| Gm15708 | 7.40733758 | -0.477765367 | 0.433689269 | -1.101630594 |
| RP24-182D2.1 | 0.265735694 | 0.023902141 | 3.705814996 | 0.006449901 |
| Gm15709 | 0 | NA | NA | NA |
| Nptx2 | 612.3205461 | -2.032311369 | 0.3173835 | -6.403330261 |
| Gm6272 | 0.315729261 | 0.030230764 | 1.606157485 | 0.018821793 |
| Tmem130 | 1967.351054 | 0.140598579 | 0.08843213 | 1.58990379 |
| Trrap | 3060.283518 | 0.103897204 | 0.076885528 | 1.351323285 |
| Gm23995 | 1.477838792 | 0.067844821 | 0.921221457 | 0.073646592 |
| Smurf1 | 700.542789 | -0.336421034 | 0.068533902 | -4.90882649 |
| Mir7039 | 0.22969359 | -1.373109043 | 3.129889768 | -0.438708435 |
| Kpna7 | 0.7084488 | -0.916308313 | 1.330050325 | -0.688927551 |
| RP23-424M12.7 | 0 | NA | NA | NA |
| Gm4871 | 0 | NA | NA | NA |
| RP23-424M12.4 | 0 | NA | NA | NA |
| RP23-424M12.5 | 0.075043205 | 0.517957688 | 3.724945049 | 0.139051095 |
| 1700018F24Rik | 0.068097359 | 0.037064902 | 3.724945049 | 0.009950456 |
| Arpc1a | 1136.742611 | 0.022085657 | 0.068739598 | 0.321294533 |
| Arpc1b | 585.2155934 | -0.947853251 | 0.159699418 | -5.935232972 |
| Pdap1 | 676.5616892 | 0.036878861 | 0.131448884 | 0.280556667 |
| Bud31 | 300.4083312 | -0.33593383 | 0.088997472 | -3.774644617 |
| Ptcd1 | 337.0823285 | 0.009972041 | 0.078674928 | 0.126749928 |
| Cpsf4 | 256.7055373 | -0.009204238 | 0.092514012 | -0.099490207 |
| Atp5j2 | 504.0996134 | 0.059877062 | 0.243280243 | 0.246123818 |
| Zkscan14 | 94.39392942 | 0.07092708 | 0.12202152 | 0.581266974 |
| Zkscan5 | 178.3410506 | 0.032774578 | 0.124065301 | 0.264171991 |
| Zfp655 | 776.8726219 | -0.261212184 | 0.099090321 | -2.636101898 |
| RP23-60E24.5 | 0 | NA | NA | NA |

| | | | | |
|---|---|---|---|---|
| Zscan25 | 198.2716498 | 0.279136792 | 0.162299069 | 1.719891519 |
| RP23-60E24.3 | 23.00508971 | 0.83450487 | 0.4170696 | 2.000876761 |
| Cyp3a57 | 13.25184546 | -0.305321809 | 0.336539934 | -0.907237979 |
| RP24-443E20.1 | 0.066050875 | -0.443818988 | 3.724945049 | -0.119147795 |
| RP24-443E20.2 | 0.066050875 | -0.443818988 | 3.724945049 | -0.119147795 |
| Cyp3a16 | 0 | NA | NA | NA |
| RP24-127J3.1 | 0.120974994 | 0.517957688 | 3.724945049 | 0.139051095 |
| Cyp3a41b | 0 | NA | NA | NA |
| RP23-379D24.1 | 0 | NA | NA | NA |
| RP23-379D24.2 | 0 | NA | NA | NA |
| RP24-443E20.4 | 0 | NA | NA | NA |
| Cyp3a41a | 0 | NA | NA | NA |
| RP23-102C14.1 | 0 | NA | NA | NA |
| Cyp3a44 | 0 | NA | NA | NA |
| RP23-102C14.2 | 0.060174537 | 0.517957688 | 3.724945049 | 0.139051095 |
| Cyp3a11 | 0.492793603 | 1.876302863 | 1.545223188 | 1.2142601 |
| RP23-161K8.1 | 0 | NA | NA | NA |
| RP23-161K8.2 | 0 | NA | NA | NA |
| RP23-161K8.3 | 0 | NA | NA | NA |
| Cyp3a25 | 0.261272317 | -1.124290489 | 2.011159594 | -0.559025993 |
| RP23-53G23.1 | 0 | NA | NA | NA |
| RP23-53G23.2 | 0 | NA | NA | NA |
| RP23-53G23.3 | 0 | NA | NA | NA |
| Cyp3a59 | 0 | NA | NA | NA |
| RP23-441I24.9 | 0.282778275 | 0.016089598 | 2.005933837 | 0.008021001 |
| Gm6309 | 0.038981395 | -0.443816764 | 3.724945049 | -0.119147198 |
| 1700001J03Rik | 1.161484436 | 0.889475337 | 0.913616073 | 0.973576717 |
| RP23-441I24.5 | 22.34345423 | 1.374389197 | 0.49566558 | 2.772815488 |
| RP23-441I24.4 | 2.948678684 | 0.423133896 | 0.661200639 | 0.639947802 |
| RP23-441I24.3 | 9.915980733 | 0.981007528 | 0.427936795 | 2.292412196 |
| Rnf6 | 700.4219294 | -0.146865309 | 0.069592003 | -2.110376221 |
| Cdk8 | 479.7397194 | 0.149698779 | 0.10618753 | 1.409758551 |
| Gm15739 | 20.56479697 | 0.541775559 | 0.28893243 | 1.875094318 |
| Wasf3 | 1567.401421 | 0.296941724 | 0.074664024 | 3.977038835 |
| Gm6408 | 0.779579221 | 0.388064576 | 1.127265389 | 0.344253074 |
| Gm6370 | 0.038981395 | -0.443816764 | 3.724945049 | -0.119147198 |
| 4930449I24Rik | 0 | NA | NA | NA |
| Gm3402 | 0 | NA | NA | NA |
| Gm3404 | 0 | NA | NA | NA |
| Gm3409 | 0 | NA | NA | NA |
| Gm3415 | 0 | NA | NA | NA |
| Gm8494 | 20.48103304 | -0.614442554 | 0.233382695 | -2.632768269 |
| Gpr12 | 122.0110685 | 0.255417228 | 0.166422406 | 1.534752643 |
| RP23-134K6.2 | 0.089089296 | 0.957155977 | 3.723939561 | 0.257027796 |
| RP23-134K6.3 | 0 | NA | NA | NA |
| Usp12 | 869.2011401 | -0.021029867 | 0.07047263 | -0.298411838 |
| RP24-126A19.2 | 0.044191549 | 0.037062677 | 3.724945049 | 0.009949859 |

| | | | | |
|---|---|---|---|---|
| Rpl21 | 82.06727034 | -0.206670435 | 0.144147468 | -1.433743089 |
| Gm25091 | 0 | NA | NA | NA |
| Gm23202 | 0.066050875 | -0.443818988 | 3.724945049 | -0.119147795 |
| Rasl11a | 102.228073 | -1.056658549 | 0.301588803 | -3.503639855 |
| Gtf3a | 110.5136172 | -0.242954409 | 0.129193965 | -1.88053992 |
| Mtif3 | 139.0778404 | -0.225362606 | 0.096784614 | -2.328496203 |
| Lnx2 | 154.2635579 | -0.235582646 | 0.128583785 | -1.832133382 |
| Polr1d | 179.4983994 | -0.40061672 | 0.124971313 | -3.20566945 |
| Rps10-ps4 | 0.048949532 | 0.037064902 | 3.724945049 | 0.009950456 |
| Gsx1 | 3.152922583 | 0.05512734 | 0.558435314 | 0.098717503 |
| Gm24556 | 0 | NA | NA | NA |
| 2210019I11Rik | 0 | NA | NA | NA |
| Pdx1 | 0.046345709 | -0.443818988 | 3.724945049 | -0.119147795 |
| Gm27033 | 0 | NA | NA | NA |
| Mir7k | 0 | NA | NA | NA |
| Cdx2 | 0 | NA | NA | NA |
| Urad | 0 | NA | NA | NA |
| RP24-510G5.6 | 0 | NA | NA | NA |
| Flt3 | 155.671954 | 0.176541615 | 0.156888175 | 1.125270373 |
| Gm26597 | 1.472170236 | 0.739242661 | 0.915510052 | 0.807465368 |
| Gm6054 | 0.773509025 | 1.870663267 | 1.211979119 | 1.543478133 |
| RP24-544J6.3 | 0.075043205 | 0.517957688 | 3.724945049 | 0.139051095 |
| RP24-544J6.4 | 1.088238703 | -1.368306275 | 1.021804022 | -1.339108328 |
| RP24-544J6.5 | 2.007455813 | 2.062362363 | 0.996855199 | 2.068868544 |
| RP24-544J6.6 | 0.135435006 | 0.434658561 | 3.63325813 | 0.119633273 |
| Pan3 | 1329.249898 | 0.152981532 | 0.100801555 | 1.517650518 |
| RP24-544J6.7 | 8.762375666 | 1.354124499 | 0.497163051 | 2.723703013 |
| Flt1 | 324.459667 | -1.231635033 | 0.198104783 | -6.217088816 |
| RP23-270L23.1 | 0.088534442 | -0.905414734 | 3.723500899 | -0.243162217 |
| RP23-270L23.2 | 0 | NA | NA | NA |
| Pomp | 809.6499027 | -0.282625416 | 0.094166541 | -3.001335844 |
| RP23-270L23.4 | 0.636753058 | 0.505739448 | 1.336096209 | 0.378520233 |
| Slc46a3 | 67.21071847 | 0.050240685 | 0.144953198 | 0.346599353 |
| Gm15721 | 25.84402214 | 0.520788281 | 0.237151317 | 2.196016818 |
| Mtus2 | 777.696009 | 0.562463097 | 0.109401653 | 5.141266889 |
| Gm10167 | 12.31383881 | 0.386272502 | 0.276968719 | 1.394643059 |
| RP24-71D19.1 | 4.780781177 | 1.342548784 | 0.63954473 | 2.099225779 |
| Slc7a1 | 875.5929286 | -0.417572106 | 0.161944244 | -2.578493043 |
| RP24-307I24.1 | 0.318558491 | -1.608188675 | 1.741757248 | -0.923313899 |
| Ubl3 | 1852.018664 | 0.029219264 | 0.059568175 | 0.490518035 |
| RP24-61O18.3 | 0 | NA | NA | NA |
| RP24-61O18.1 | 0.068097359 | 0.037064902 | 3.724945049 | 0.009950456 |
| RP24-61O18.2 | 0 | NA | NA | NA |
| 2210417A02Rik | 0 | NA | NA | NA |
| n-R5s179 | 0 | NA | NA | NA |
| 4930505K14Rik | 0 | NA | NA | NA |
| Gm15407 | 0.046345709 | -0.443818988 | 3.724945049 | -0.119147795 |

| | | | | |
|---|---|---|---|---|
| Katnal1 | 1032.752 | -0.185133112 | 0.06865159 | -2.696705369 |
| RP23-188F20.1 | 0 | NA | NA | NA |
| Gm15410 | 0.390771115 | -1.847754221 | 1.539168694 | -1.200488437 |
| Gm15406 | 5.088172819 | 0.100032658 | 0.450276192 | 0.222158444 |
| 8430423G03Rik | 0.143485256 | -0.955563601 | 3.547405725 | -0.269369696 |
| Gm15409 | 0.354937333 | 1.223144625 | 1.648061695 | 0.742171624 |
| RP23-134H19.3 | 52.59786348 | -0.527650198 | 0.198988666 | -2.651659557 |
| Gm15411 | 7.066285288 | 0.146313471 | 0.477684895 | 0.306297044 |
| Gm15408 | 2.719793829 | -0.375760828 | 0.624028689 | -0.602153129 |
| Hmgb1 | 340.808951 | 0.082720049 | 0.102002441 | 0.81096147 |
| 5730422E09Rik | 12.70416494 | -0.524039722 | 0.398271653 | -1.315784637 |
| RP23-408A1.6 | 1.653178163 | 1.394454598 | 0.845833673 | 1.648615613 |
| RP23-408A1.4 | 3.467320147 | -0.204891671 | 0.548277952 | -0.373700366 |
| Gm23362 | 0 | NA | NA | NA |
| Gm8615 | 17.8882258 | -0.489010939 | 0.26084196 | -1.87474032 |
| RP23-408A1.8 | 0.278466489 | 1.016160899 | 1.792053099 | 0.567037271 |
| Uspl1 | 565.005295 | -0.071039634 | 0.094686764 | -0.750259398 |
| 2310047D07Rik | 2.953893651 | 0.695642962 | 0.636722541 | 1.092537043 |
| Gm29264 | 0.433555451 | 0.545279248 | 1.474191812 | 0.369883514 |
| AC122546.1 | 0 | NA | NA | NA |
| Gm24105 | 0 | NA | NA | NA |
| Alox5ap | 91.57756619 | -1.214769308 | 0.212481879 | -5.717048966 |
| n-R5s180 | 0 | NA | NA | NA |
| Wl1-1993I8.1 | 5.494432944 | -0.464840002 | 0.455781761 | -1.019874075 |
| BC028471 | 1.239154484 | -0.500601948 | 1.041619766 | -0.480599509 |
| Gm20488 | 0.478946765 | -0.41331309 | 1.586904864 | -0.260452343 |
| RP23-375E13.2 | 0.173926504 | -0.427137108 | 2.308837173 | -0.185000966 |
| Medag | 75.60539664 | -0.663165775 | 0.21108654 | -3.141677215 |
| RP23-375E13.4 | 0 | NA | NA | NA |
| RP23-375E13.3 | 0.045882455 | 0.517959912 | 3.724945049 | 0.139051692 |
| Tex26 | 5.173350472 | -0.64943498 | 0.529416278 | -1.226700062 |
| RP23-375E13.6 | 1.443591694 | -0.119507993 | 0.736897763 | -0.162177169 |
| Gm15997 | 2.38686755 | -0.073926322 | 0.811038181 | -0.091150236 |
| Gm2566 | 0.852004391 | -0.886275528 | 1.233811191 | -0.718323463 |
| Wdr95 | 6.751900483 | 0.372792033 | 0.404287147 | 0.922097166 |
| Hsph1 | 3279.88901 | -0.808120698 | 0.228803007 | -3.531949648 |
| RP23-256B18.1 | 1.058337859 | -1.639403232 | 1.102628329 | -1.486813996 |
| Gm5566 | 1.331862033 | -1.534283355 | 0.887515602 | -1.728739586 |
| B3glct | 236.8677375 | 0.432751028 | 0.110546449 | 3.914653363 |
| RP23-346P15.1 | 0 | NA | NA | NA |
| RP23-346P15.3 | 0.483795822 | -0.396981629 | 1.581022548 | -0.251091694 |
| RP23-346P15.4 | 0 | NA | NA | NA |
| RP23-337K7.2 | 0 | NA | NA | NA |
| RP23-337K7.1 | 0.29963536 | -0.806734913 | 1.588520973 | -0.507852856 |
| RP23-337K7.3 | 0 | NA | NA | NA |
| Rxfp2 | 8.210362686 | -0.773017462 | 0.380010675 | -2.034199332 |
| RP23-301H20.1 | 0 | NA | NA | NA |

| | | | | |
|---|---|---|---|---|
| RP23-301H20.2 | 0 | NA | NA | NA |
| Fry | 3743.235337 | 0.597861375 | 0.079406778 | 7.529097533 |
| RP23-327P24.3 | 2.277130459 | 1.99817021 | 0.889613478 | 2.246110541 |
| RP23-327P24.2 | 0 | NA | NA | NA |
| RP24-333I1.2 | 4.567880012 | 0.470034433 | 0.583473482 | 0.805579769 |
| RP24-333I1.3 | 1.611190043 | -0.311908444 | 0.851391021 | -0.366351577 |
| Zar1l | 2.866525085 | 0.500094963 | 0.620334864 | 0.806169365 |
| Gm15933 | 0.686705509 | 0.11131595 | 1.243298063 | 0.089532794 |
| Brca2 | 165.9286918 | 0.570045138 | 0.16965299 | 3.360065371 |
| Gm25284 | 0.075043205 | 0.517957688 | 3.724945049 | 0.139051095 |
| Gm23641 | 0.146089079 | -0.444786384 | 3.522464394 | -0.126271364 |
| RP24-333I1.6 | 1.253964362 | -0.542147213 | 1.064064482 | -0.509505977 |
| RP24-333I1.5 | 4.525610808 | 0.960093394 | 0.521063457 | 1.842565203 |
| RP24-333I1.7 | 13.83704783 | 0.934581243 | 0.278221533 | 3.359126207 |
| N4bp2l1 | 393.7981301 | 0.218036784 | 0.097691039 | 2.231901578 |
| RP23-34P11.1 | 29.55790751 | 0.57758735 | 0.391880624 | 1.473885961 |
| RP23-34P11.2 | 251.9245356 | -0.335457908 | 0.173266352 | -1.936082252 |
| N4bp2l2 | 1453.675956 | 0.118179978 | 0.081326675 | 1.453151474 |
| Pds5b | 2134.751181 | 0.131051941 | 0.07068514 | 1.854023917 |
| Gm24630 | 4.821230019 | 1.285335449 | 0.56726008 | 2.265866212 |
| Gm26013 | 0.049553047 | -0.443818988 | 3.724945049 | -0.119147795 |
| RP23-336F4.1 | 29.09809687 | 0.306154476 | 0.277207007 | 1.104425457 |
| RP23-336F4.4 | 4.80270818 | 0.501138065 | 0.466274648 | 1.074770132 |
| RP23-336F4.5 | 0 | NA | NA | NA |
| Kl | 209.3353533 | 0.55716367 | 0.219994261 | 2.532628206 |
| RP23-412C8.2 | 0.322046281 | -0.80637164 | 1.727692605 | -0.466733282 |
| Stard13 | 272.414666 | -0.48331143 | 0.084591178 | -5.713496835 |
| RP24-155K17.1 | 0.139883403 | -1.137320593 | 3.551352803 | -0.32024996 |
| RP24-343A12.4 | 0 | NA | NA | NA |
| RP24-343A12.3 | 0 | NA | NA | NA |
| D730045B01Rik | 4.778829002 | 0.109681795 | 0.555576769 | 0.197419693 |
| 1700028E10Rik | 23.33760472 | 0.484485526 | 0.224175624 | 2.161187369 |
| Vmn2r-ps25 | 0.141684195 | 0.037074514 | 3.558854665 | 0.010417541 |
| Vmn2r-ps26 | 0.233449105 | 0.831515026 | 2.493543122 | 0.333467273 |
| Vmn2r18 | 0.476067163 | -1.47132355 | 1.58339442 | -0.929221128 |
| RP24-211L21.1 | 0.060174537 | 0.517957688 | 3.724945049 | 0.139051095 |
| Rfc3 | 87.13724546 | -0.390696271 | 0.189934779 | -2.057002266 |
| RP24-211L21.2 | 0 | NA | NA | NA |
| RP24-211L21.3 | 0.154433973 | -0.443826563 | 3.50312282 | -0.126694548 |
| Gm24242 | 0.09689823 | 0.037062677 | 3.724945049 | 0.009949859 |
| Gm26463 | 0 | NA | NA | NA |
| Gm23551 | 0 | NA | NA | NA |
| Vmn1r-ps6 | 0 | NA | NA | NA |
| Vmn1r-ps7 | 0.046345709 | -0.443818988 | 3.724945049 | -0.119147795 |
| RP24-328P2.7 | 0.256207099 | 1.041017243 | 2.130114171 | 0.488714294 |
| RP24-328P2.8 | 0.219080626 | -0.948136098 | 2.168083125 | -0.437315381 |
| RP23-368D12.2 | 0 | NA | NA | NA |

| | | | | |
|---|---|---|---|---|
| RP23-368D12.3 | 0 | NA | NA | NA |
| RP23-258F1.1 | 0.066050875 | -0.443818988 | 3.724945049 | -0.119147795 |
| Gm18222 | 0 | NA | NA | NA |
| RP23-253G12.8 | 136.7192892 | 0.061471965 | 0.216668816 | 0.283713948 |
| RP24-328P2.5 | 839.9270636 | -0.288922454 | 0.1609523 | -1.795081243 |
| RP23-253G12.10 | 0.128477747 | 0.037074008 | 3.677132019 | 0.010082316 |
| RP23-253G12.9 | 311.8525769 | -0.280211257 | 0.184774688 | -1.516502395 |
| Samd9l | 3774.434653 | -0.549032754 | 0.192665903 | -2.84966227 |
| Hepacam2 | 3.625877892 | -0.795032938 | 0.556953859 | -1.427466433 |
| Ccdc132 | 896.7331518 | 0.109357066 | 0.075720304 | 1.444223805 |
| RP23-318M11.2 | 26.8210992 | 0.727403897 | 0.350245613 | 2.076839421 |
| Calcr | 25.19477301 | -0.034778004 | 0.260113386 | -0.133703246 |
| Mir653 | 0 | NA | NA | NA |
| Mir489 | 0.044191549 | 0.037062677 | 3.724945049 | 0.009949859 |
| Tfpi2 | 9.186610503 | -2.233386471 | 0.45218369 | -4.939113291 |
| Gngt1 | 0 | NA | NA | NA |
| Gng11 | 162.5242613 | -0.578733526 | 0.17545907 | -3.298396178 |
| Bet1 | 205.6506794 | -0.495163456 | 0.131960935 | -3.752348795 |
| Gm8602 | 0 | NA | NA | NA |
| Gm24442 | 0 | NA | NA | NA |
| Gm37883 | 0 | NA | NA | NA |
| Col1a2 | 642.3682694 | -0.69035025 | 0.134777341 | -5.122153676 |
| Gm26696 | 3.592646241 | 0.130439209 | 0.591804501 | 0.220409289 |
| Casd1 | 867.3601419 | 0.278497633 | 0.086754367 | 3.210185764 |
| Sgce | 237.5338859 | -0.096031047 | 0.117426733 | -0.817795442 |
| Peg10 | 1587.476363 | -0.5111409 | 0.220507176 | -2.318023882 |
| Gm20714 | 0 | NA | NA | NA |
| Gm26828 | 0 | NA | NA | NA |
| Gm9835 | 0.856801049 | -1.114444062 | 1.0411649 | -1.070381898 |
| Ppp1r9a | 4137.715075 | 0.516903641 | 0.104557336 | 4.943733853 |
| 2210017G18Rik | 30.57888805 | 0.640505495 | 0.354246749 | 1.808077268 |
| Gm26814 | 84.80524745 | 0.250663882 | 0.279820752 | 0.895801616 |
| Gm26736 | 22.89958731 | 0.564336631 | 0.364190991 | 1.549562303 |
| Gm26556 | 14.27493484 | 0.443531639 | 0.32610791 | 1.360076299 |
| Pon1 | 0.759396657 | 0.137406193 | 1.266621949 | 0.108482403 |
| Pon3 | 56.36395702 | -0.403847013 | 0.22987173 | -1.756836358 |
| Pon2 | 309.9460515 | 0.164292039 | 0.110138784 | 1.491681982 |
| Asb4 | 38.68988062 | -0.676239417 | 0.282758462 | -2.391579765 |
| Pdk4 | 314.1644295 | -0.131268993 | 0.189232203 | -0.693692672 |
| Dync1i1 | 1162.518359 | 0.451943355 | 0.062207892 | 7.265048484 |
| 1700019G24Rik | 1.274692649 | -1.117783447 | 0.925545345 | -1.207702521 |
| Slc25a13 | 101.7915475 | 0.390335824 | 0.175630932 | 2.222477662 |
| Gm20619 | 0.313425824 | 0.764647304 | 2.017617321 | 0.378985299 |
| Gm8649 | 0 | NA | NA | NA |
| Gm27018 | 0 | NA | NA | NA |
| Gm8652 | 0.294205253 | -0.875648025 | 1.610483414 | -0.543717506 |
| Gm20617 | 0.379053103 | 0.887239225 | 1.653221653 | 0.53667288 |

| | | | | |
|---|---|---|---|---|
| Gm20685 | 0 | NA | NA | NA |
| Shfm1 | 204.0907547 | -0.214053436 | 0.248740152 | -0.860550397 |
| Gm20618 | 0.47603017 | 0.027470091 | 1.404234985 | 0.019562318 |
| Gm18006 | 0 | NA | NA | NA |
| Gm5873 | 4.742073203 | -0.102433991 | 0.464774192 | -0.22039518 |
| Dlx6os1 | 337.2803174 | 0.042668656 | 0.264151665 | 0.161530899 |
| Gm27699 | 0.095295241 | -0.443821212 | 3.724945049 | -0.119148392 |
| Gm27905 | 0 | NA | NA | NA |
| Dlx6os3 | 6.490635638 | -0.444395529 | 0.567529248 | -0.783035466 |
| Dlx6 | 64.91335236 | -0.070038062 | 0.210678473 | -0.332440522 |
| Dlx6os2 | 8.021264403 | -0.367098322 | 0.578130795 | -0.634974517 |
| Dlx5 | 52.02918498 | -0.318110366 | 0.223511711 | -1.423238024 |
| Gm22155 | 0 | NA | NA | NA |
| Gm26774 | 0.048949532 | 0.037064902 | 3.724945049 | 0.009950456 |
| Sdhaf3 | 139.5460954 | 0.082841587 | 0.117573626 | 0.704593286 |
| Gm27190 | 0 | NA | NA | NA |
| Gm8669 | 0 | NA | NA | NA |
| Gm19272 | 0.25372768 | -1.447354027 | 2.070088829 | -0.699174841 |
| Tac1 | 258.728839 | 0.52977531 | 0.152946563 | 3.463793489 |
| Gm8676 | 0 | NA | NA | NA |
| Asns | 800.622713 | -0.283461415 | 0.089992659 | -3.149828203 |
| C1galt1 | 215.3562405 | -0.486937465 | 0.122509174 | -3.974702053 |
| Gm22276 | 0.301804762 | -0.965901224 | 2.022293767 | -0.477626564 |
| Gm9825 | 372.9565206 | -0.131788871 | 0.085836296 | -1.535351342 |
| Col28a1 | 8.368185802 | 0.575746886 | 0.402984729 | 1.428706462 |
| Mios | 475.6118824 | 0.0936283 | 0.070220833 | 1.333340777 |
| Rpa3 | 69.13831145 | 0.035092429 | 0.145850976 | 0.240604693 |
| Umad1 | 363.7124902 | 0.295644762 | 0.080719374 | 3.662624553 |
| Gm16055 | 0.775287707 | 2.499345386 | 1.356272656 | 1.842804523 |
| Gm8428 | 0 | NA | NA | NA |
| Gm16043 | 0.136120942 | -0.876404342 | 3.595683781 | -0.24373788 |
| Gm24059 | 4.143222737 | 0.91615985 | 0.553910838 | 1.653984338 |
| A430035B10Rik | 32.4009088 | 0.33043299 | 0.255238109 | 1.294606795 |
| Gm26075 | 0.250670796 | -0.443821463 | 2.486942845 | -0.178460661 |
| Glcci1 | 81.25189137 | 0.323875953 | 0.137594662 | 2.353840984 |
| Gm16042 | 0.476187739 | -1.833893783 | 1.666195017 | -1.100647742 |
| Ica1 | 575.9737311 | 0.469648417 | 0.123441992 | 3.804608227 |
| Nxph1 | 529.823522 | 0.267248687 | 0.077427456 | 3.451601036 |
| Gm26058 | 0.921802213 | -1.616642702 | 1.180096341 | -1.369924341 |
| Gm27894 | 0 | NA | NA | NA |
| Ndufa4 | 1081.345019 | 0.256559619 | 0.141193196 | 1.817082028 |
| Phf14 | 900.5267977 | 0.130239781 | 0.064701006 | 2.012948318 |
| Gm6578 | 0.045882455 | 0.517959912 | 3.724945049 | 0.139051692 |
| Thsd7a | 1278.572731 | 0.261286514 | 0.058115127 | 4.496015521 |
| Gm16123 | 0 | NA | NA | NA |
| n-R5s160 | 0 | NA | NA | NA |
| Gm15529 | 0 | NA | NA | NA |

| | | | | |
|---|---|---|---|---|
| Gm5300 | 0.044544648 | 0.517957688 | 3.724945049 | 0.139051095 |
| Tmem106b | 2433.505759 | 0.305240068 | 0.076950191 | 3.966722685 |
| Gm24985 | 0 | NA | NA | NA |
| Vwde | 1.568186543 | 1.501518116 | 0.754887749 | 1.989061445 |
| 1700016P04Rik | 0.132574648 | 0.037054914 | 3.724945049 | 0.009947775 |
| Gm25737 | 0.154696864 | 0.037074681 | 2.833481357 | 0.013084498 |
| Tmem168 | 343.227461 | -0.299531311 | 0.095784066 | -3.127151775 |
| B630005N14Rik | 527.4785247 | -0.184075992 | 0.091542875 | -2.010817243 |
| Gpr85 | 431.2033423 | 0.122524037 | 0.089684918 | 1.366160993 |
| 2610001J05Rik | 197.506035 | -0.558676405 | 0.10794836 | -5.175404265 |
| 1110019D14Rik | 33.23543665 | 0.252781315 | 0.212972226 | 1.186921504 |
| Gm23760 | 0.097899064 | 0.037062677 | 3.724945049 | 0.009949859 |
| Gm23013 | 0 | NA | NA | NA |
| Ppp1r3a | 6.253343815 | 1.233959499 | 0.450686612 | 2.737954639 |
| Foxp2 | 446.2894329 | 0.46316984 | 0.092208918 | 5.023048182 |
| Gm25470 | 1.281282525 | 1.520794107 | 1.09039846 | 1.3947141 |
| Mdfic | 320.1052487 | -1.385486641 | 0.223231704 | -6.20649584 |
| Gm29589 | 0 | NA | NA | NA |
| RP23-244M15.1 | 0.212513282 | 1.689178857 | 3.159662247 | 0.534607412 |
| Tfec | 15.16942446 | -2.674924382 | 0.564280935 | -4.74041247 |
| Gm5721 | 0 | NA | NA | NA |
| Gm15473 | 0.646370819 | -0.907597454 | 1.440899767 | -0.629882435 |
| Gm4876 | 26.61523984 | -0.080391422 | 0.223243583 | -0.360106307 |
| Tes | 60.86189865 | -1.020039243 | 0.229414823 | -4.446265628 |
| D830026I12Rik | 0 | NA | NA | NA |
| Gm15581 | 0.736762124 | 1.491348703 | 1.538513718 | 0.96934378 |
| Cav2 | 214.4350625 | -0.707891604 | 0.141236188 | -5.012112072 |
| Cav1 | 153.6111711 | -1.852440293 | 0.168697182 | -10.9808609 |
| Gm29591 | 0 | NA | NA | NA |
| Gm25021 | 0 | NA | NA | NA |
| Met | 239.4772955 | -1.103959172 | 0.168830666 | -6.538854554 |
| Capza2 | 2658.372187 | -0.008540959 | 0.079569487 | -0.107339622 |
| St7 | 410.6774712 | 0.103743707 | 0.113238965 | 0.91614849 |
| Gm27755 | 0 | NA | NA | NA |
| Gm27762 | 0 | NA | NA | NA |
| Gm27960 | 2.550499828 | -0.369780689 | 0.632467881 | -0.58466319 |
| Gm27365 | 1.826179369 | -0.036650554 | 0.778274422 | -0.047092071 |
| Gm27542 | 1.845381307 | 0.030862935 | 0.808549335 | 0.038170751 |
| Gm28008 | 1.623521978 | -0.624548545 | 0.816998356 | -0.764442842 |
| Gm13898 | 0.097139547 | -0.443816764 | 3.724945049 | -0.119147198 |
| Gm13899 | 0 | NA | NA | NA |
| RP23-479D21.3 | 13.97715435 | -0.154569579 | 0.350618043 | -0.440848902 |
| RP23-479D21.4 | 0.045882455 | 0.517959912 | 3.724945049 | 0.139051692 |
| Gm26738 | 0.238158741 | 0.051475477 | 2.527426437 | 0.020366756 |
| Gm27444 | 0 | NA | NA | NA |
| Gm27402 | 3.526582245 | 0.896990219 | 0.633600777 | 1.415702523 |
| RP23-86H7.3 | 81.37030569 | 1.002419158 | 0.292903414 | 3.422353952 |

| | | | | |
|---|---|---|---|---|
| Wnt2 | 29.40825172 | -0.169383416 | 0.321770888 | -0.526410008 |
| Asz1 | 0.284445263 | 1.818459697 | 2.007700733 | 0.905742408 |
| RP23-275N13.1 | 0 | NA | NA | NA |
| Gm15593 | 0.075043205 | 0.517957688 | 3.724945049 | 0.139051095 |
| Gm15596 | 0.396988867 | 0.15022666 | 1.44999166 | 0.103605189 |
| Cftr | 19.0807837 | 0.357551223 | 0.319868711 | 1.117806182 |
| Gm15592 | 0 | NA | NA | NA |
| Gm15595 | 0.243349506 | -0.450718271 | 2.4640115 | -0.182920523 |
| Cttnbp2 | 1527.7495 | 0.322288087 | 0.140171128 | 2.299247295 |
| Gm15594 | 1.482657567 | -0.65966809 | 0.728987388 | -0.904910155 |
| RP23-16J8.6 | 154.7156367 | 0.774142879 | 0.353429483 | 2.190374364 |
| Gm26233 | 3.766699628 | 0.184524675 | 0.607351673 | 0.303818502 |
| Gm23648 | 0 | NA | NA | NA |
| RP23-148L8.1 | 0 | NA | NA | NA |
| Gm5874 | 1.63591125 | -0.981136958 | 0.911750863 | -1.076102034 |
| Gm26809 | 2.591679042 | 0.068931729 | 0.593451145 | 0.116154008 |
| RP23-148L8.3 | 35.09407007 | -0.029065383 | 0.193565134 | -0.150158152 |
| Lsm8 | 195.2002428 | 0.182111921 | 0.09209842 | 1.97736205 |
| Ankrd7 | 2.04319782 | 0.077389286 | 0.67474343 | 0.11469439 |
| RP24-230L6.3 | 0.39809594 | -1.473355669 | 1.765107619 | -0.834711523 |
| Gm22718 | 0 | NA | NA | NA |
| RP23-18G14.1 | 0.045882455 | 0.517959912 | 3.724945049 | 0.139051692 |
| RP23-179A12.2 | 0 | NA | NA | NA |
| RP23-179A12.1 | 0 | NA | NA | NA |
| Gm24213 | 0 | NA | NA | NA |
| Mir6370 | 0 | NA | NA | NA |
| Gm22828 | 0 | NA | NA | NA |
| RP23-302D18.3 | 0 | NA | NA | NA |
| RP23-302D18.1 | 0 | NA | NA | NA |
| RP23-302D18.2 | 0.044191549 | 0.037062677 | 3.724945049 | 0.009949859 |
| RP23-441A23.1 | 0 | NA | NA | NA |
| Gm23960 | 0 | NA | NA | NA |
| Gm26373 | 0 | NA | NA | NA |
| Kcnd2 | 2260.738386 | 0.303539627 | 0.105315672 | 2.882188564 |
| RP23-307L24.2 | 1439.777736 | 0.366934772 | 0.330857744 | 1.109040906 |
| RP23-94H23.1 | 58.13546571 | 0.634896612 | 0.314499932 | 2.018749601 |
| RP23-301C7.2 | 101.8565513 | 0.317760214 | 0.246822771 | 1.287402344 |
| Tspan12 | 200.3279207 | 0.101444507 | 0.134660219 | 0.753336862 |
| RP24-169J24.2 | 0.152959369 | -0.443823011 | 2.842356906 | -0.156146123 |
| RP24-169J24.1 | 0.164540623 | 1.05688389 | 3.435283929 | 0.307655469 |
| Ing3 | 280.4244847 | -0.037002508 | 0.075803648 | -0.488136236 |
| Gm26719 | 0 | NA | NA | NA |
| Cped1 | 74.68292297 | -0.628408954 | 0.156269858 | -4.021306243 |
| RP24-279C2.3 | 0 | NA | NA | NA |
| Wnt16 | 10.34745373 | -0.906744839 | 0.336993728 | -2.690687578 |
| Fam3c | 626.0288257 | 0.32455688 | 0.094670269 | 3.428287295 |
| RP23-369P2.2 | 0 | NA | NA | NA |

| | | | | |
|---|---|---|---|---|
| Gm8927 | 0.687801143 | -0.021652777 | 1.194272593 | -0.018130515 |
| RP24-241H23.2 | 1.331472478 | 0.47356753 | 0.927056901 | 0.510828979 |
| RP24-241H23.1 | 0.086648695 | 0.037058229 | 3.724945049 | 0.009948665 |
| Gm23961 | 0 | NA | NA | NA |
| Gm25942 | 0 | NA | NA | NA |
| RP23-369P2.3 | 0.047948698 | 0.037064902 | 3.724945049 | 0.009950456 |
| Ptprz1 | 4928.564296 | 0.158948615 | 0.13483634 | 1.178826236 |
| Aass | 39.94585083 | 0.256329742 | 0.243113612 | 1.054361951 |
| RP24-301O17.2 | 0 | NA | NA | NA |
| RP24-301O17.3 | 0 | NA | NA | NA |
| RP24-301O17.1 | 0 | NA | NA | NA |
| Fezf1 | 7.883156094 | -0.056589555 | 0.438178025 | -0.129147406 |
| RP23-124H2.4 | 3.80210492 | -0.480242185 | 0.555430932 | -0.86462989 |
| Cadps2 | 1566.7278 | 0.55382574 | 0.128599223 | 4.306602534 |
| Gm27959 | 0 | NA | NA | NA |
| Gm24217 | 2.108983506 | 1.551707093 | 0.788315367 | 1.968383668 |
| Rnf133 | 16.93650597 | 1.011654329 | 0.434960779 | 2.325851842 |
| Rnf148 | 13.88597797 | -0.052982527 | 0.42280839 | -0.125310965 |
| Tas2r118 | 0 | NA | NA | NA |
| Slc13a1 | 10.97794222 | -0.418864331 | 0.377632229 | -1.109185866 |
| Gm20706 | 0 | NA | NA | NA |
| Gm26386 | 0.072025462 | 0.517959912 | 3.724945049 | 0.139051692 |
| Iqub | 42.66293401 | -0.187449587 | 0.180885393 | -1.036289243 |
| Ndufa5 | 270.339159 | 0.169770391 | 0.219489962 | 0.773476787 |
| Asb15 | 1.031776346 | -0.391068732 | 1.032562439 | -0.378736159 |
| RP23-392I2.4 | 0 | NA | NA | NA |
| Lmod2 | 0.965679557 | 1.907293083 | 1.132890821 | 1.683563012 |
| Wasl | 1854.805742 | 0.215761894 | 0.055432373 | 3.89234451 |
| RP23-392I2.3 | 73.97553726 | 0.280827454 | 0.185449343 | 1.514308162 |
| Hyal6 | 0.086908494 | 0.037058229 | 3.724945049 | 0.009948665 |
| Hyal4 | 0 | NA | NA | NA |
| Spam1 | 0 | NA | NA | NA |
| RP24-255M7.1 | 0 | NA | NA | NA |
| Hyal5 | 0 | NA | NA | NA |
| Gm22267 | 0 | NA | NA | NA |
| Tmem229a | 986.0139867 | 0.206482605 | 0.162600191 | 1.269879225 |
| RP23-339M5.2 | 0.278190828 | -1.677421167 | 2.414660698 | -0.694681935 |
| Gm24443 | 0 | NA | NA | NA |
| Gm22529 | 0 | NA | NA | NA |
| Gpr37 | 655.1826324 | 0.070540463 | 0.152358191 | 0.462990946 |
| Pot1a | 253.1759545 | -0.219607246 | 0.156990208 | -1.398859513 |
| RP23-302O24.1 | 0.342793059 | 0.782052615 | 1.69622989 | 0.461053433 |
| RP23-302O24.2 | 0 | NA | NA | NA |
| Gm24691 | 0 | NA | NA | NA |
| RP23-308F24.1 | 0 | NA | NA | NA |
| Grm8 | 291.4031537 | 0.006789718 | 0.131620694 | 0.051585489 |
| RP23-414I9.1 | 5.120409944 | 0.530328031 | 0.506784851 | 1.046455966 |

| | | | | |
|---|---|---|---|---|
| Gm26310 | 0 | NA | NA | NA |
| Mir592 | 0 | NA | NA | NA |
| RP23-23C4.6 | 185.154323 | -0.70326692 | 0.141077337 | -4.984974463 |
| 6530409C15Rik | 8.058619829 | 0.903010987 | 0.404766297 | 2.230944112 |
| RP23-23C4.5 | 118.802725 | -0.005432219 | 0.255676164 | -0.021246484 |
| RP23-23C4.4 | 75.79663319 | -0.104466814 | 0.23286799 | -0.448609592 |
| Zfp800 | 452.6169934 | -0.277847753 | 0.069355635 | -4.006130881 |
| Gm5303 | 1.49686415 | 0.531900575 | 0.883793132 | 0.601838322 |
| Gcc1 | 343.3632238 | -0.208285983 | 0.090421051 | -2.303512081 |
| Arf5 | 1016.757531 | -0.118499454 | 0.075749729 | -1.564354836 |
| Fscn3 | 1.623741216 | 0.424459267 | 0.846832035 | 0.501231943 |
| Pax4 | 0.249106506 | 0.02017265 | 1.831973674 | 0.01101143 |
| RP23-434O14.5 | 0 | NA | NA | NA |
| Snd1 | 1136.070214 | -0.152260386 | 0.074550896 | -2.042368286 |
| Gm37321 | 29.66800731 | -0.618447927 | 0.31693631 | -1.951331881 |
| Gm37978 | 39.62155469 | -0.615237526 | 0.314036098 | -1.959129956 |
| Lrrc4 | 977.950627 | 0.707909874 | 0.104803283 | 6.754653601 |
| Gm37940 | 70.33009473 | 0.136284143 | 0.277443702 | 0.491213682 |
| Gm3294 | 93.79479471 | 0.06172019 | 0.284454674 | 0.216977241 |
| Mir129-1 | 0 | NA | NA | NA |
| Lep | 0.088534442 | -0.905414734 | 3.723500899 | -0.243162217 |
| Gm25589 | 0 | NA | NA | NA |
| Rbm28 | 422.2309319 | -0.002730215 | 0.115528587 | -0.023632377 |
| Gm24303 | 1.400035301 | 0.793815197 | 1.194249815 | 0.664697777 |
| Prrt4 | 24.29690244 | 1.097736437 | 0.2534305 | 4.331508786 |
| Impdh1 | 348.4065021 | 0.111259458 | 0.080980837 | 1.373898591 |
| Gm26021 | 0 | NA | NA | NA |
| Hilpda | 32.36285512 | -0.474705657 | 0.207094011 | -2.292223011 |
| Fam71f2 | 3.490009053 | 0.364242487 | 0.546466137 | 0.666541734 |
| Fam71f1 | 0.2407585 | -1.317939465 | 2.075179277 | -0.635096678 |
| Gm17172 | 0.045882455 | 0.517959912 | 3.724945049 | 0.139051692 |
| Calu | 1240.173328 | -0.58335443 | 0.116017178 | -5.028172893 |
| Opn1sw | 0.679991162 | -1.942920073 | 1.454761333 | -1.335559331 |
| Ccdc136 | 845.6284117 | 0.57758003 | 0.101326289 | 5.700199189 |
| Gm26220 | 0.160408561 | 0.517955075 | 3.457615493 | 0.149801236 |
| AC044807.1 | 0.581411716 | 1.392958749 | 1.322126779 | 1.053574265 |
| Flnc | 1393.252646 | -2.164908431 | 0.373636386 | -5.794158465 |
| Atp6v1f | 586.7322244 | -0.084501145 | 0.093145521 | -0.907194936 |
| Gm9047 | 2.295347566 | 0.032808005 | 0.661704611 | 0.049581043 |
| Kcp | 10.30552492 | 0.160037246 | 0.338418758 | 0.47289709 |
| Gm26484 | 0 | NA | NA | NA |
| Gm26627 | 0.072025462 | 0.517959912 | 3.724945049 | 0.139051692 |
| Irf5 | 262.9012068 | -1.21851203 | 0.195633062 | -6.228558798 |
| Tnpo3 | 1004.219965 | 0.024022085 | 0.075938456 | 0.316336229 |
| RP23-446B13.7 | 31.58542221 | 0.344429701 | 0.277304032 | 1.242065246 |
| Tspan33 | 165.1105126 | 0.581071117 | 0.130201923 | 4.462845904 |
| Smo | 219.4811244 | -0.258536425 | 0.144108219 | -1.794043582 |

| | | | | |
|---|---|---|---|---|
| Gm13717 | 1.21856855 | 0.908506863 | 1.060011189 | 0.857072899 |
| Ahcyl2 | 2068.520444 | 0.161127914 | 0.045730617 | 3.523414365 |
| RP23-353E8.7 | 0.12092566 | 0.998850474 | 3.724945049 | 0.268151734 |
| Strip2 | 859.1687429 | 0.158643037 | 0.07740611 | 2.049489845 |
| Gm13724 | 2.453329902 | 0.240735111 | 0.677990747 | 0.355071381 |
| Gm24536 | 0.309201626 | -0.912065724 | 1.625167214 | -0.561213466 |
| Gm13725 | 0.309323177 | -0.44449125 | 2.014555284 | -0.220639887 |
| Smkr-ps | 0.763164319 | -0.015958364 | 1.189592188 | -0.013414988 |
| Gm23049 | 0 | NA | NA | NA |
| Nrf1 | 255.4170055 | -0.07779148 | 0.095284307 | -0.816414404 |
| Gm25580 | 0 | NA | NA | NA |
| Mir182 | 0 | NA | NA | NA |
| Mir96 | 0 | NA | NA | NA |
| Mir183 | 0 | NA | NA | NA |
| Gm37046 | 0.075043205 | 0.517957688 | 3.724945049 | 0.139051095 |
| Ube2h | 1345.992283 | 0.174057208 | 0.059101343 | 2.945063486 |
| 1700095J07Rik | 0 | NA | NA | NA |
| Gm25625 | 0.044191549 | 0.037062677 | 3.724945049 | 0.009949859 |
| Zc3hc1 | 183.2490365 | -0.138712606 | 0.095996596 | -1.444974221 |
| n-R5s161 | 0 | NA | NA | NA |
| Klhdc10 | 1743.124813 | 0.088232096 | 0.042709242 | 2.065878304 |
| 1700012C14Rik | 0.900590653 | -0.824434026 | 1.14399307 | -0.720663479 |
| Tmem209 | 379.778664 | -0.175892847 | 0.091531178 | -1.921671408 |
| Ssmem1 | 1.951291546 | 0.082246458 | 0.782374006 | 0.105124221 |
| Cpa2 | 12.99980896 | 0.704217192 | 0.315393714 | 2.232819367 |
| Cpa4 | 11.88556507 | -1.736158556 | 0.477120633 | -3.638825143 |
| Gm13781 | 0.062426873 | 0.517959912 | 3.724945049 | 0.139051692 |
| Cpa5 | 0.592694123 | -1.000795284 | 1.317657599 | -0.759526059 |
| Cpa1 | 0.176646398 | -0.526386109 | 3.35399333 | -0.1569431 |
| Cep41 | 83.12317599 | -0.77098678 | 0.13542623 | -5.693038789 |
| Gm13782 | 0.049553047 | -0.443818988 | 3.724945049 | -0.119147795 |
| Gm14528 | 0 | NA | NA | NA |
| Mest | 610.9026533 | 0.208331945 | 0.121213985 | 1.718712119 |
| Gm27522 | 0 | NA | NA | NA |
| Gm27568 | 0 | NA | NA | NA |
| Mir335 | 5.501266716 | -0.068503506 | 0.581953598 | -0.117713003 |
| Copg2 | 853.6229713 | 0.096840927 | 0.060001242 | 1.613982052 |
| 4930412F09Rik | 0.271012069 | 1.953303653 | 3.700163235 | 0.527896617 |
| Tsga13 | 1.195665945 | -0.604333074 | 0.836634529 | -0.722338193 |
| Klf14 | 3.978034112 | -0.492129535 | 0.550239606 | -0.894391334 |
| Mir29a | 6.497893077 | 0.868735053 | 0.500645764 | 1.735229008 |
| Mir29b-1 | 0.108266322 | -0.955981391 | 3.722845977 | -0.25678779 |
| Gm27019 | 20.56482463 | -0.213756621 | 0.381412264 | -0.560434576 |
| Gm13834 | 35.42125562 | 0.228698592 | 0.340872475 | 0.670921265 |
| Lincpint | 345.1173544 | 0.592214254 | 0.218429782 | 2.711234008 |
| Gm13833 | 0.042716945 | 0.037062677 | 3.724945049 | 0.009949859 |
| 5330406M23Rik | 159.3696281 | 0.491338601 | 0.381658207 | 1.287378583 |

| | | | | |
|---|---|---|---|---|
| Gm13835 | 176.8303318 | -0.096842144 | 0.129625229 | -0.747093328 |
| Gm37728 | 114.4100306 | 1.18500379 | 0.422948553 | 2.801768159 |
| 2210408F21Rik | 141.5258413 | 0.153962882 | 0.165311829 | 0.931348246 |
| RP23-400L4.7 | 44.53988991 | -0.278338158 | 0.428885652 | -0.648979878 |
| Gm14532 | 12.88187382 | 0.098364969 | 0.352013039 | 0.279435584 |
| Mkln1os | 23.92253865 | 0.143509538 | 0.250894891 | 0.571990675 |
| Gm13844 | 13.1985606 | 0.467718154 | 0.493402078 | 0.947945245 |
| Mkln1 | 2442.640936 | 0.205547196 | 0.087090994 | 2.360142965 |
| Podxl | 278.0855086 | -0.667468619 | 0.124303106 | -5.369685771 |
| Gm6117 | 0.646697505 | 0.599801457 | 1.343160926 | 0.446559638 |
| RP24-414G20.1 | 3.468425366 | 0.087158952 | 0.631359021 | 0.138049746 |
| Gm13846 | 0.441802452 | 2.451866641 | 1.628652532 | 1.505457176 |
| Gm13848 | 0.595326517 | -0.644449322 | 1.54648652 | -0.416718357 |
| RP24-414G20.4 | 0.125911488 | -0.761499402 | 3.703518295 | -0.205615132 |
| Gm13849 | 0.554297541 | 0.003489557 | 1.481999285 | 0.002354628 |
| 1700012A03Rik | 0.048949532 | 0.037064902 | 3.724945049 | 0.009950456 |
| Gm13850 | 0 | NA | NA | NA |
| Plxna4 | 2374.903231 | 0.219663283 | 0.101486475 | 2.164458695 |
| Plxna4os3 | 0 | NA | NA | NA |
| Plxna4os2 | 0.093141081 | 0.037060453 | 3.724945049 | 0.009949262 |
| Plxna4os1 | 3.620095065 | -0.618643286 | 0.585611598 | -1.056405455 |
| Gm13852 | 0.049553047 | -0.443818988 | 3.724945049 | -0.119147795 |
| RP23-225G5.2 | 0.573159608 | 0.717725703 | 1.573587049 | 0.456108039 |
| Chchd3 | 512.9631055 | -0.326172005 | 0.096884664 | -3.366600972 |
| Gm14540 | 0 | NA | NA | NA |
| RP24-246G10.3 | 30.7240258 | 0.477452066 | 0.36958396 | 1.291863603 |
| Gm22321 | 0 | NA | NA | NA |
| Gm1401 | 0 | NA | NA | NA |
| Exoc4 | 1197.152527 | 0.015313767 | 0.068239853 | 0.224410896 |
| Gm13854 | 0 | NA | NA | NA |
| RP23-4H14.1 | 15.19819234 | 0.45148062 | 0.43643687 | 1.034469476 |
| RP24-365C10.2 | 0 | NA | NA | NA |
| Gm13853 | 0.045882455 | 0.517959912 | 3.724945049 | 0.139051692 |
| Lrguk | 29.47287266 | 0.440957336 | 0.221673027 | 1.98922414 |
| Gm13856 | 0.085327104 | -0.905414734 | 3.723500899 | -0.243162217 |
| Slc35b4 | 730.9802596 | 0.102160522 | 0.055991451 | 1.824573588 |
| Gm13855 | 0 | NA | NA | NA |
| Gm14546 | 0.956739943 | 0.339878203 | 1.048797607 | 0.324064624 |
| Akr1b3 | 294.7538566 | 0.072211434 | 0.087010021 | 0.829920896 |
| Akr1b8 | 26.07990468 | -1.426515247 | 0.296557582 | -4.810247089 |
| Gm13859 | 0.259720851 | -1.185320894 | 2.026354886 | -0.584952272 |
| Gm13858 | 0.289603658 | -0.011988997 | 2.028663455 | -0.005909801 |
| Akr1b10 | 145.9333712 | -0.009015337 | 0.101496773 | -0.088823873 |
| Akr1b7 | 1.910658871 | -0.008084697 | 0.739305804 | -0.010935525 |
| Gm13857 | 0.038981395 | -0.443816764 | 3.724945049 | -0.119147198 |
| Bpgm | 500.8527338 | -0.117708215 | 0.080742003 | -1.457831247 |
| Gm13860 | 0 | NA | NA | NA |

| | | | | |
|---|---|---|---|---|
| Cald1 | 388.6951145 | -0.68399693 | 0.120584889 | -5.672327095 |
| Gm13861 | 2.144383846 | 0.397785467 | 0.882941343 | 0.450523095 |
| Npn2 | 0 | NA | NA | NA |
| Gm14547 | 0 | NA | NA | NA |
| Agbl3 | 107.7106235 | -0.257331244 | 0.154523382 | -1.665322369 |
| Gm13864 | 2.929070206 | -1.060470219 | 0.605041288 | -1.752723724 |
| Gm13862 | 0.066050875 | -0.443818988 | 3.724945049 | -0.119147795 |
| Gm13863 | 0 | NA | NA | NA |
| RP23-386A17.7 | 1.526274188 | 0.401500637 | 0.854549596 | 0.469838893 |
| Tmem140 | 313.938631 | -0.927732853 | 0.211418161 | -4.388141715 |
| 3110062M04Rik | 36.44138757 | -1.055861835 | 0.249797166 | -4.226876761 |
| Wdr91 | 325.7975239 | 0.016168983 | 0.08038063 | 0.201155212 |
| Stra8 | 0.124347486 | 0.036092774 | 3.717245565 | 0.009709548 |
| 2010107G12Rik | 6.64236274 | 0.39087145 | 0.412113847 | 0.948455027 |
| Cnot4 | 879.3333849 | -0.124892547 | 0.069776518 | -1.789893647 |
| Nup205 | 597.9465125 | -0.112840521 | 0.086818536 | -1.29972846 |
| Gm23435 | 0.60633683 | 0.459141325 | 1.342564386 | 0.34198831 |
| 1810058I24Rik | 158.214639 | -0.532122223 | 0.133285598 | -3.992346001 |
| Slc13a4 | 187.4589476 | 0.608055026 | 0.207434294 | 2.931313892 |
| Fam180a | 13.25868466 | 0.761068778 | 0.366253338 | 2.07798455 |
| RP23-338E5.5 | 0.361225529 | 0.036442078 | 1.931443901 | 0.01886779 |
| Mtpn | 3362.617321 | 0.076033104 | 0.059419389 | 1.279600908 |
| Gm23053 | 1.826002588 | 0.22488075 | 0.751551059 | 0.299222185 |
| RP23-20C5.1 | 0 | NA | NA | NA |
| RP23-13B24.1 | 13.51087949 | 0.153084971 | 0.306367376 | 0.499677783 |
| Gm23273 | 0 | NA | NA | NA |
| RP23-352D5.1 | 6.269730523 | 0.113637733 | 0.470193243 | 0.241683042 |
| 9330158H04Rik | 22.23221858 | -0.023520909 | 0.23054166 | -0.10202455 |
| Chrm2 | 536.4163306 | 0.172774021 | 0.121427705 | 1.422855033 |
| Mir490 | 3.003392332 | 1.399263606 | 0.693540662 | 2.017565346 |
| RP23-81D20.2 | 0 | NA | NA | NA |
| RP23-5K15.1 | 0 | NA | NA | NA |
| Gm25111 | 0.298614356 | 2.027171135 | 1.958621983 | 1.034998663 |
| Ptn | 1821.983259 | 0.480402083 | 0.134930637 | 3.560363272 |
| Dgki | 1466.320099 | 0.228582883 | 0.115138451 | 1.985287108 |
| 1700111E14Rik | 0.088736197 | 0.51795324 | 3.724945049 | 0.139049901 |
| RP23-151F20.5 | 2.105924665 | -2.549927394 | 0.86598896 | -2.944526445 |
| Creb3l2 | 335.1776498 | -0.663731472 | 0.101353113 | -6.548703365 |
| Akr1d1 | 0.049553047 | -0.443818988 | 3.724945049 | -0.119147795 |
| RP24-549J13.2 | 0 | NA | NA | NA |
| RP23-245J24.1 | 9.865094166 | -0.746945529 | 0.351292493 | -2.126278081 |
| Gm15487 | 1555.447171 | -0.027578771 | 0.071084393 | -0.387972236 |
| RP23-459F14.2 | 0.11197992 | 0.037073798 | 3.724945049 | 0.009952844 |
| Trim24 | 495.6674005 | 0.219356533 | 0.110052923 | 1.993191337 |
| Svopl | 1.88997817 | -1.260043071 | 0.726636247 | -1.734076819 |
| Atp6v0a4 | 3.783295768 | -0.581628589 | 0.529274555 | -1.09891659 |
| Tmem213 | 0 | NA | NA | NA |

| | | | | |
|---|---|---|---|---|
| Gm23231 | 0 | NA | NA | NA |
| D630045J12Rik | 1061.614314 | 0.482664039 | 0.134413538 | 3.590888591 |
| RP23-387J8.3 | 0 | NA | NA | NA |
| Zc3hav1l | 80.97671505 | 0.053168901 | 0.194373372 | 0.273540048 |
| Zc3hav1 | 1446.894863 | -0.639891678 | 0.162132715 | -3.946715366 |
| RP24-142K18.4 | 0.189025308 | -0.908251663 | 2.253764128 | -0.40299322 |
| Rpl30-ps1 | 9.381830078 | -0.636442206 | 0.367120183 | -1.733607236 |
| Ttc26 | 133.6908354 | -0.167094409 | 0.122873632 | -1.359888259 |
| Gm16563 | 1.549622433 | -1.485742932 | 1.098620885 | -1.352370916 |
| Ubn2 | 2325.889383 | 0.157009839 | 0.101742945 | 1.543201239 |
| Mir7670 | 0 | NA | NA | NA |
| 1110001J03Rik | 118.1303116 | -0.254937536 | 0.151773548 | -1.679723106 |
| Gm15697 | 2.902282796 | -0.145683169 | 0.715840059 | -0.203513574 |
| Luc7l2 | 4182.925321 | 0.245161077 | 0.112190958 | 2.18521244 |
| Gm26699 | 1.935704857 | 1.505835295 | 0.875455876 | 1.72005847 |
| Klrg2 | 32.43324731 | 0.211649999 | 0.257920056 | 0.820603105 |
| Clec2l | 411.0156647 | 0.422023731 | 0.0802134 | 5.261262236 |
| Hipk2 | 3020.872203 | -0.244765049 | 0.099845618 | -2.451435069 |
| Tbxas1 | 29.76369512 | -0.486523309 | 0.270133643 | -1.801046709 |
| RP23-346B17.4 | 0.320565116 | 1.43835062 | 3.700650276 | 0.388675101 |
| Gm23010 | 0.282660793 | 0.03705195 | 3.724945049 | 0.009946979 |
| 1700025N23Rik | 9.679120195 | 0.867294614 | 0.44408941 | 1.952972971 |
| Parp12 | 1518.309522 | -0.749817931 | 0.145277581 | -5.161277636 |
| 4930599N23Rik | 26.63738886 | -1.16192376 | 0.245931599 | -4.72458101 |
| RP23-345F4.3 | 0.255942976 | -1.985407028 | 3.017299649 | -0.658007907 |
| Kdm7a | 1749.004419 | -0.393622131 | 0.074663177 | -5.271971353 |
| RP24-173B18.1 | 0.413211764 | 0.174600045 | 1.448650725 | 0.120525978 |
| Gm26008 | 0.222111871 | 1.705571485 | 3.112888666 | 0.547906356 |
| Slc37a3 | 628.5897833 | 0.138731825 | 0.071817609 | 1.931724356 |
| Rab19 | 12.00148128 | -0.988220157 | 0.341473493 | -2.893987899 |
| Mkrn1 | 1161.354 | -0.173868431 | 0.050002887 | -3.477167828 |
| Gm10244 | 4.940723394 | 0.388836392 | 0.447372472 | 0.86915583 |
| Gm25402 | 0.04393175 | 0.037062677 | 3.724945049 | 0.009949859 |
| Dennd2a | 408.9845452 | -0.408709554 | 0.093851302 | -4.354862914 |
| 8030453O22Rik | 22.33756815 | 0.213975989 | 0.306532246 | 0.698053766 |
| Adck2 | 176.9877061 | 0.113546338 | 0.098206632 | 1.156198275 |
| Gm16272 | 1.598725798 | -0.572818941 | 0.784292901 | -0.730363541 |
| RP23-366P9.8 | 0.091666476 | 0.037060453 | 3.724945049 | 0.009949262 |
| Ndufb2 | 352.2685807 | -0.124116987 | 0.095107224 | -1.305021648 |
| Braf | 3301.479618 | 0.313592375 | 0.077715326 | 4.035141994 |
| Fth-ps2 | 26.73179484 | 1.009634196 | 0.405120015 | 2.492185425 |
| Mrps33 | 137.537649 | 0.155186492 | 0.135909973 | 1.141832997 |
| RP23-380O16.6 | 0 | NA | NA | NA |
| RP23-380O16.7 | 0.107078754 | -0.443164241 | 3.724898049 | -0.118973522 |
| Gm26833 | 7.290008566 | -0.22838957 | 0.414247378 | -0.551336188 |
| Tmem178b | 2878.275615 | 0.417171776 | 0.084657484 | 4.92776017 |
| Gm37995 | 95.94723699 | 0.646022313 | 0.315082193 | 2.050329495 |

| | | | | |
|---|---|---|---|---|
| Gm37673 | 72.41137234 | 0.745966174 | 0.291422793 | 2.559738611 |
| Gm26638 | 42.96957209 | 0.863420633 | 0.417205865 | 2.069531387 |
| Gm26807 | 1.057496005 | 3.599415197 | 1.247931148 | 2.88430592 |
| Gm37003 | 29.74312774 | 0.468303076 | 0.3867535 | 1.210856724 |
| Agk | 341.3868807 | 0.338604559 | 0.08514474 | 3.976811232 |
| E330009J07Rik | 1236.299548 | 0.045668427 | 0.141344743 | 0.323099579 |
| Wee2 | 3.520209586 | -0.795616587 | 0.546127353 | -1.456833433 |
| Ssbp1 | 232.2371204 | 0.174248814 | 0.136916053 | 1.272668986 |
| Tas2r137 | 4.276107861 | -0.204331212 | 0.543559173 | -0.375913465 |
| Tas2r108 | 4.194868342 | 0.043395613 | 0.530325255 | 0.081828297 |
| Prss37 | 0 | NA | NA | NA |
| RP24-503P15.7 | 0 | NA | NA | NA |
| Olfr461 | 0 | NA | NA | NA |
| Olfr460 | 0.474871048 | 0.379318199 | 1.361529912 | 0.278597037 |
| Clec5a | 38.15951351 | 0.257150956 | 0.205716406 | 1.250026484 |
| Tas2r138 | 0 | NA | NA | NA |
| Mgam | 2.716524655 | -0.895905298 | 0.708131239 | -1.265168446 |
| Mgam2-ps | 0.962702212 | -0.27727074 | 0.891336448 | -0.311073042 |
| Moxd2 | 0 | NA | NA | NA |
| Gm38276 | 0 | NA | NA | NA |
| Trbv1 | 1.116438897 | -0.09625412 | 0.922345914 | -0.104357941 |
| Prss58 | 0 | NA | NA | NA |
| 1700074P13Rik | 0 | NA | NA | NA |
| Gm4744 | 0 | NA | NA | NA |
| Gm29588 | 0 | NA | NA | NA |
| 1810009J06Rik | 0 | NA | NA | NA |
| Gm2663 | 0 | NA | NA | NA |
| Gm38015 | 0 | NA | NA | NA |
| Gm37878 | 0 | NA | NA | NA |
| Gm38294 | 0 | NA | NA | NA |
| 2210010C04Rik | 0.16656219 | -0.323504422 | 2.779637158 | -0.116383687 |
| Trbv2 | 1.408278968 | -0.312369134 | 0.840793453 | -0.371517086 |
| Trbv3 | 0.461584485 | 0.760207169 | 1.603107037 | 0.474208616 |
| 5830405F06Rik | 0.95870128 | 0.711404537 | 0.903274638 | 0.787583872 |
| Trbv5 | 0.150914725 | -0.410358748 | 2.845788253 | -0.144198623 |
| Trbv6 | 0 | NA | NA | NA |
| Trbv7 | 0 | NA | NA | NA |
| Gm37610 | 0 | NA | NA | NA |
| Trbv8 | 0 | NA | NA | NA |
| Trbv9 | 0 | NA | NA | NA |
| Trbv10 | 0 | NA | NA | NA |
| Trbv11 | 0 | NA | NA | NA |
| Trbv12-1 | 0 | NA | NA | NA |
| Trbv13-1 | 2.762763099 | -2.094503907 | 0.721777211 | -2.901870376 |
| Trbv12-2 | 0.921345027 | -0.489086277 | 0.888010655 | -0.550766226 |
| Trbv13-2 | 3.038727124 | -0.189365475 | 0.604982987 | -0.313009588 |
| Trbv12-3 | 0 | NA | NA | NA |

| | | | | |
|---|---|---|---|---|
| Trbv13-3 | 1.194558986 | -1.409416618 | 0.955006768 | -1.475818461 |
| Trbv14 | 1.5023977 | -0.067412002 | 1.651511345 | -0.04081837 |
| Trbv15 | 0.168389893 | 0.129092083 | 2.761872815 | 0.046740778 |
| Trbv16 | 0.74889667 | 1.515406808 | 1.30069162 | 1.165077706 |
| Trbv17 | 1.696056422 | -0.433601829 | 0.801980543 | -0.540663776 |
| Trbv18 | 0 | NA | NA | NA |
| Trbv19 | 0.811168495 | -1.173373491 | 1.076560939 | -1.08992761 |
| Trbv20 | 0.334925437 | -0.769626375 | 1.973786241 | -0.389923873 |
| Trbv21 | 0 | NA | NA | NA |
| Trbv22 | 0.097139547 | -0.443816764 | 3.724945049 | -0.119147198 |
| Trbv23 | 0 | NA | NA | NA |
| Trbv24 | 0.038981395 | -0.443816764 | 3.724945049 | -0.119147198 |
| Trbv25 | 0 | NA | NA | NA |
| Trbv26 | 0.038981395 | -0.443816764 | 3.724945049 | -0.119147198 |
| Gm2719 | 0.3093278 | -0.429786889 | 1.791769011 | -0.239867353 |
| Trbv27 | 0 | NA | NA | NA |
| Trbv28 | 0 | NA | NA | NA |
| Trbv29 | 1.593432551 | 0.238399397 | 0.895107426 | 0.266336073 |
| Trbv30 | 0.049553047 | -0.443818988 | 3.724945049 | -0.119147795 |
| Try4 | 0.044544648 | 0.517957688 | 3.724945049 | 0.139051095 |
| Gm18964 | 0 | NA | NA | NA |
| Try5 | 0 | NA | NA | NA |
| Try10 | 0 | NA | NA | NA |
| Prss3 | 0 | NA | NA | NA |
| Gm5771 | 0 | NA | NA | NA |
| Gm38306 | 0 | NA | NA | NA |
| Gm5409 | 0 | NA | NA | NA |
| Gm10334 | 0.980132196 | 1.521879099 | 1.064795554 | 1.429268833 |
| Prss1 | 0 | NA | NA | NA |
| Gm37301 | 0 | NA | NA | NA |
| Gm29813 | 0.146823232 | 0.517961571 | 2.878417312 | 0.179946656 |
| Gm37841 | 0 | NA | NA | NA |
| Gm38018 | 0 | NA | NA | NA |
| Prss2 | 0.046345709 | -0.443818988 | 3.724945049 | -0.119147795 |
| Trbd1 | 0.960611118 | -0.228283177 | 0.880223398 | -0.259346862 |
| Trbj1-1 | 0.739650571 | -1.541798697 | 1.119736717 | -1.37692966 |
| Trbj1-2 | 0.363286374 | -2.414535097 | 3.679634322 | -0.656188873 |
| Trbj1-3 | 0.347715489 | -1.806188455 | 1.447103179 | -1.248140755 |
| Trbj1-4 | 0.471163529 | -1.543927596 | 1.408658066 | -1.096027229 |
| Trbj1-5 | 0.450331108 | 0.760392771 | 1.333017027 | 0.570429901 |
| Trbj1-6 | 0.583857846 | -1.930982801 | 1.287914728 | -1.499309511 |
| Trbj1-7 | 0.161076545 | -0.410543693 | 2.787893798 | -0.147259445 |
| Trbc1 | 24.37840666 | -0.521714094 | 0.346430966 | -1.50596842 |
| Trbd2 | 0.440524946 | 0.2904864 | 1.635276903 | 0.177637439 |
| Trbj2-1 | 0.112396584 | -0.917692504 | 3.724434845 | -0.246397787 |
| Trbj2-2 | 0.218053095 | -1.59907095 | 2.131916564 | -0.750062633 |
| Trbj2-3 | 0.696513874 | 0.050765556 | 1.306464471 | 0.038857203 |

| | | | | |
|---|---|---|---|---|
| Trbj2-4 | 0.088736197 | 0.51795324 | 3.724945049 | 0.139049901 |
| Trbj2-5 | 0.202417642 | 0.037074454 | 2.653394779 | 0.01397246 |
| Trbj2-6 | 0.354669972 | -1.725391132 | 1.84203353 | -0.936677375 |
| Trbj2-7 | 1.180533128 | 1.0015379 | 1.049437173 | 0.954357179 |
| Trbc2 | 69.31285373 | -0.173997476 | 0.20137582 | -0.86404354 |
| Trbv31 | 1.290124216 | 1.352942409 | 0.907232769 | 1.491284767 |
| Gm27549 | 0 | NA | NA | NA |
| Ephb6 | 693.8526302 | 0.359976076 | 0.134571886 | 2.674972356 |
| Trpv6 | 9.753117248 | 0.928588127 | 0.377233017 | 2.461577023 |
| Trpv5 | 0 | NA | NA | NA |
| 1700034O15Rik | 0.14896402 | -0.924098152 | 2.866934862 | -0.322329665 |
| Kel | 0.227427828 | 0.51797175 | 2.559553304 | 0.202368026 |
| Gm37527 | 0 | NA | NA | NA |
| Olfr459 | 0 | NA | NA | NA |
| Pip | 0.038981395 | -0.443816764 | 3.724945049 | -0.119147198 |
| Sval2 | 0.163190422 | -0.955151543 | 3.396300892 | -0.281232898 |
| Sval1 | 0.173394273 | -1.466453397 | 3.379053676 | -0.433983458 |
| Sval3 | 0.066050875 | -0.443818988 | 3.724945049 | -0.119147795 |
| Sva | 0.066050875 | -0.443818988 | 3.724945049 | -0.119147795 |
| Tas2r139 | 0 | NA | NA | NA |
| Tas2r144 | 0.049553047 | -0.443818988 | 3.724945049 | -0.119147795 |
| Gstk1 | 46.48690246 | 0.09842077 | 0.196014512 | 0.50210961 |
| Tmem139 | 0.247571123 | 0.536094281 | 2.500000251 | 0.214437691 |
| Casp2 | 320.4543346 | -0.017997084 | 0.12690491 | -0.141815509 |
| Gm25108 | 3.821797685 | 0.370756195 | 0.523579208 | 0.708118637 |
| Clcn1 | 102.7118201 | 0.823417216 | 0.285626285 | 2.882848184 |
| Fam131b | 1482.538637 | 0.529814016 | 0.088802184 | 5.9662273 |
| Gm24611 | 0 | NA | NA | NA |
| Zyx | 868.7508899 | -0.464800157 | 0.168503751 | -2.758396506 |
| Epha1 | 4.796176745 | -0.886787973 | 0.50169302 | -1.767590814 |
| 2010310C07Rik | 14.52291164 | 0.898376025 | 0.40501212 | 2.218146029 |
| Tas2r143 | 4.118545241 | 0.773082938 | 0.549059612 | 1.408012757 |
| Tas2r135 | 5.600410292 | 0.744688603 | 0.519373343 | 1.433821379 |
| Tas2r126 | 2.194003788 | 0.217496974 | 0.730621948 | 0.297687435 |
| Olfr458 | 0.062426873 | 0.517959912 | 3.724945049 | 0.139051692 |
| Olfr457 | 0 | NA | NA | NA |
| Olfr456 | 0 | NA | NA | NA |
| Gm4877 | 0.146848595 | 0.037061235 | 3.724945049 | 0.009949472 |
| Olfr455 | 0 | NA | NA | NA |
| Tcaf3 | 0.038981395 | -0.443816764 | 3.724945049 | -0.119147198 |
| Tcaf2 | 17.75907395 | -0.416651141 | 0.256997272 | -1.621227872 |
| Tcaf1 | 3402.596378 | 0.159090561 | 0.081822524 | 1.944337003 |
| Olfr453 | 0.410820671 | 0.560884299 | 1.465928392 | 0.382613709 |
| Olfr38 | 0.225129614 | 1.580880368 | 3.710727143 | 0.426029807 |
| Gm22363 | 0.272364125 | -1.907656335 | 1.995247297 | -0.956100197 |
| Olfr452 | 0.180050447 | -0.925657836 | 3.327036994 | -0.278222886 |
| Olfr450 | 0.211502678 | -1.570513266 | 2.522003765 | -0.622724394 |

| | | | | |
|---|---|---|---|---|
| Gm23982 | 0.194279094 | -1.017823543 | 3.719440177 | -0.273649661 |
| Olfr449 | 0.134148234 | -0.443820953 | 3.624632483 | -0.122445781 |
| Olfr448 | 0.07796279 | -0.798301147 | 3.721829328 | -0.214491605 |
| Olfr447 | 0 | NA | NA | NA |
| Olfr446 | 0.048949532 | 0.037064902 | 3.724945049 | 0.009950456 |
| Olfr444 | 0 | NA | NA | NA |
| Gm17472 | 0 | NA | NA | NA |
| Gm20730 | 0 | NA | NA | NA |
| Olfr443-ps1 | 0.336338362 | -0.727298433 | 1.692875709 | -0.429623055 |
| Olfr441 | 0 | NA | NA | NA |
| Olfr237-ps1 | 0 | NA | NA | NA |
| Olfr437 | 0 | NA | NA | NA |
| Olfr13 | 0 | NA | NA | NA |
| Olfr435 | 0.038981395 | -0.443816764 | 3.724945049 | -0.119147198 |
| Gm21827 | 0 | NA | NA | NA |
| Olfr434 | 0.061920613 | -0.443816764 | 3.724945049 | -0.119147198 |
| Olfr47 | 0 | NA | NA | NA |
| Arhgef5 | 73.22831166 | -0.858318562 | 0.172228257 | -4.983610573 |
| Nobox | 0.405536581 | 2.024916923 | 1.603612602 | 1.262722007 |
| Tpk1 | 61.31573283 | 0.029149044 | 0.176172074 | 0.165457801 |
| Gm15550 | 0.332944463 | -0.834558865 | 1.566951318 | -0.532600378 |
| Gm22939 | 0 | NA | NA | NA |
| Cntnap2 | 2123.978884 | 0.612427744 | 0.063011296 | 9.719332545 |
| Gm22839 | 8.429271064 | 0.747851881 | 0.378218509 | 1.977301116 |
| Gm25161 | 0 | NA | NA | NA |
| Rbpsuh-rs3 | 97.81620039 | -0.510202558 | 0.131722674 | -3.87330855 |
| Cul1 | 1508.243995 | 0.109506493 | 0.059731029 | 1.83332675 |
| A930035D04Rik | 12.16101713 | -0.277094112 | 0.302923705 | -0.914732348 |
| Ezh2 | 210.0637519 | -0.019677378 | 0.107778601 | -0.182572218 |
| Gm26625 | 0.182019239 | -1.313069409 | 2.712073432 | -0.484157027 |
| Gm25283 | 0 | NA | NA | NA |
| Rny3 | 804.362109 | 0.605697456 | 0.320846177 | 1.887812604 |
| Rpl31-ps7 | 0 | NA | NA | NA |
| Rny1 | 909.9952825 | 0.6044683 | 0.418053768 | 1.445910423 |
| Pdia4 | 818.8143374 | -0.615385835 | 0.125025569 | -4.922079855 |
| Mir704 | 0.123992736 | 0.517955521 | 3.720643114 | 0.139211288 |
| Rpl35a-ps7 | 0.314214724 | 2.273291562 | 1.667198477 | 1.36353985 |
| Zfp786 | 50.22326823 | 0.09367003 | 0.174203396 | 0.53770496 |
| Zfp398 | 324.873339 | 0.382409837 | 0.146264192 | 2.614514404 |
| Zfp282 | 307.4764343 | 0.10300775 | 0.111588048 | 0.923107374 |
| Zfp212 | 200.032767 | 0.196467342 | 0.134765323 | 1.457847889 |
| Zfp783 | 130.5839755 | 0.448166372 | 0.160729619 | 2.788324725 |
| Zfp956 | 173.5071749 | 0.446356789 | 0.14445582 | 3.089919043 |
| Zfp777 | 460.5844228 | 0.172038222 | 0.100409186 | 1.713371341 |
| Gm24563 | 1.500420242 | -0.424972358 | 0.817551406 | -0.519811176 |
| Zfp746 | 384.3199641 | 0.056537372 | 0.088328918 | 0.640077712 |
| Gm16630 | 7.347387118 | 0.683202421 | 0.397133288 | 1.720335318 |

| | | | | |
|---|---|---|---|---|
| Gm22122 | 0 | NA | NA | NA |
| Krba1 | 549.5583407 | 0.354991306 | 0.136213335 | 2.606142093 |
| Gm24325 | 1.916854426 | 0.807926839 | 0.853492271 | 0.946612953 |
| Zfp467 | 330.9730189 | 0.219017636 | 0.091380449 | 2.39676691 |
| Sspo | 38.70866699 | 0.114578475 | 0.356339227 | 0.321543255 |
| Gm10243 | 43.3226042 | -0.389720621 | 0.197546957 | -1.97279992 |
| Gm7887 | 75.72156547 | 0.132549433 | 0.158707958 | 0.83517824 |
| 1700026J14Rik | 0.552687697 | -0.315448064 | 1.466043263 | -0.215169683 |
| Atp6v0e2 | 1764.72768 | 0.380242149 | 0.067912389 | 5.599010079 |
| Lrrc61 | 226.76103 | 0.202738521 | 0.111346024 | 1.820797129 |
| Rarres2 | 195.195546 | -0.819139379 | 0.312880204 | -2.618060742 |
| Gm5111 | 0.049553047 | -0.443818988 | 3.724945049 | -0.119147795 |
| Repin1 | 624.3197349 | 0.012252975 | 0.089998856 | 0.136145894 |
| Zfp775 | 56.68412911 | -0.094717489 | 0.166088925 | -0.570281789 |
| AI854703 | 220.6699541 | 0.231672844 | 0.119216333 | 1.943297849 |
| Gimap8 | 51.49836854 | -1.062179768 | 0.249586847 | -4.255752179 |
| Gimap9 | 17.15776395 | -0.791749291 | 0.26196586 | -3.022337688 |
| Gimap4 | 77.7956503 | -0.848868169 | 0.268802286 | -3.157964841 |
| Gimap6 | 85.26288905 | -0.254263834 | 0.228933835 | -1.110643316 |
| Gm28053 | 0.321006207 | -0.754223757 | 2.352813463 | -0.320562496 |
| Gimap7 | 21.5292461 | -1.364058769 | 0.396879672 | -3.436958012 |
| Gimap1os | 2.672780519 | -0.772023873 | 0.61727035 | -1.250706231 |
| Gimap1 | 14.27405217 | -0.595640835 | 0.316351515 | -1.882844898 |
| Gimap5 | 14.86559326 | -0.299846448 | 0.337159995 | -0.889329851 |
| Gimap3 | 25.51805368 | -0.866910003 | 0.34578479 | -2.50707963 |
| Tmem176b | 421.451915 | -0.577703682 | 0.184092144 | -3.138122406 |
| Tmem176a | 202.5557947 | -0.504808901 | 0.184966932 | -2.729184599 |
| Gm7932 | 3.063139515 | -0.93938361 | 0.702724617 | -1.336773448 |
| Aoc1 | 3.736083012 | -0.802356284 | 0.572394095 | -1.401755 |
| 1600015I10Rik | 0 | NA | NA | NA |
| Gm24348 | 0 | NA | NA | NA |
| Gm10242 | 0.061920613 | -0.443816764 | 3.724945049 | -0.119147198 |
| Doxl2 | 0.106112162 | -0.443821212 | 3.724945049 | -0.119148392 |
| Svs1 | 0.044191549 | 0.037062677 | 3.724945049 | 0.009949859 |
| Gpnmb | 840.8388242 | -2.656109016 | 0.422913085 | -6.280508007 |
| Malsu1 | 255.5915214 | 0.071271771 | 0.097210192 | 0.733171791 |
| Igf2bp3 | 6.6227068 | 0.694765927 | 0.50214868 | 1.383586088 |
| Tra2a | 624.172519 | -0.540053485 | 0.089751485 | -6.017209494 |
| Ccdc126 | 154.9854233 | -0.110986014 | 0.114773333 | -0.967001754 |
| Fam221a | 18.46478838 | 0.178561311 | 0.266108807 | 0.671008648 |
| Stk31 | 5.37787014 | 0.292356356 | 0.436625448 | 0.669581578 |
| Gm17060 | 0.066050875 | -0.443818988 | 3.724945049 | -0.119147795 |
| Npy | 399.7067098 | -0.377069059 | 0.185734446 | -2.030151475 |
| Gm21786 | 0.284351963 | 1.61275334 | 3.709438132 | 0.434770249 |
| Mpp6 | 900.1739491 | -0.037695281 | 0.056186405 | -0.670896826 |
| Dfna5 | 75.89232883 | 0.084947098 | 0.148572112 | 0.571756679 |
| Osbpl3 | 392.5077654 | 0.19544582 | 0.113498474 | 1.722012757 |

| | | | | |
|---|---|---|---|---|
| Gm26289 | 0 | NA | NA | NA |
| Cycs | 696.5473769 | -0.262682456 | 0.146853121 | -1.788742755 |
| 5430402O13Rik | 7.75721647 | -0.140443051 | 0.444575129 | -0.315903976 |
| Mir6371 | 0.990815542 | -1.357972835 | 1.330013824 | -1.021021594 |
| 4921507P07Rik | 15.81567947 | 0.720462516 | 0.25271502 | 2.850889175 |
| Gm4782 | 0.19378329 | -0.88152813 | 2.228559639 | -0.395559587 |
| Gm8038 | 0.04393175 | 0.037062677 | 3.724945049 | 0.009949859 |
| Npvf | 3.377670371 | 0.311237761 | 0.54543258 | 0.570625541 |
| G930045G22Rik | 0 | NA | NA | NA |
| Gm29004 | 0.141094079 | 0.037072457 | 3.563850528 | 0.01040236 |
| Mir148a | 0.208883516 | 0.051821621 | 2.631795872 | 0.019690593 |
| Gm26677 | 0.297812842 | -0.443383998 | 1.769393403 | -0.250585312 |
| Nfe2l3 | 68.43159579 | -0.427452015 | 0.162292942 | -2.633829975 |
| Hnrnpa2b1 | 3001.421392 | -0.141995655 | 0.091700139 | -1.548478081 |
| Cbx3 | 448.3183731 | 0.236268116 | 0.106115132 | 2.226526153 |
| Snx10 | 1492.510951 | -0.180975198 | 0.104177654 | -1.737178658 |
| Gm22914 | 0.310734401 | -0.813262086 | 1.746435744 | -0.46566963 |
| Skap2 | 251.8896519 | -0.675422516 | 0.143160097 | -4.717952338 |
| Halr1 | 0.212468866 | 0.027689645 | 2.176631895 | 0.012721326 |
| Rps8-ps3 | 0 | NA | NA | NA |
| Rps19-ps14 | 0 | NA | NA | NA |
| Hoxa1 | 1.953182484 | -0.825669739 | 0.918112149 | -0.899312507 |
| Hotairm1 | 2.721387227 | -1.163611227 | 0.772250395 | -1.50677971 |
| Gm27867 | 0 | NA | NA | NA |
| Gm27861 | 0.467025382 | -1.513921309 | 1.53940331 | -0.983446832 |
| Gm27868 | 0 | NA | NA | NA |
| Gm27477 | 0.460648582 | -1.277776194 | 1.416174728 | -0.902272981 |
| Hoxa2 | 5.976910094 | -0.506715754 | 0.656817448 | -0.771471213 |
| Gm28308 | 0 | NA | NA | NA |
| Hoxaas2 | 3.211323041 | -0.510281706 | 0.740602159 | -0.689009207 |
| Hoxa3 | 12.50440182 | -1.033940513 | 0.790884935 | -1.307321037 |
| Gm15050 | 0.11193333 | 0.037073798 | 3.724945049 | 0.009952844 |
| Hoxaas3 | 0.798143448 | -0.719451172 | 1.376853018 | -0.522533025 |
| 5730596B20Rik | 0.683443395 | 0.026093492 | 1.445887303 | 0.018046699 |
| Gm29430 | 0.397176565 | 2.134718422 | 1.835656498 | 1.162918238 |
| Hoxa4 | 7.87519976 | -0.182332763 | 0.672905923 | -0.270963232 |
| Hoxa5 | 22.4157456 | -0.761793031 | 0.92713811 | -0.821660789 |
| Hoxa6 | 2.164192679 | -1.028838063 | 1.691866927 | -0.608108148 |
| Hoxa7 | 1.260199239 | -1.135883973 | 1.241285013 | -0.915087157 |
| Hoxa9 | 0.137647365 | 1.167618838 | 3.719469251 | 0.31392082 |
| AC015583.1 | 0 | NA | NA | NA |
| Mir196b | 0 | NA | NA | NA |
| Hoxa10 | 0 | NA | NA | NA |
| Hoxa11 | 0 | NA | NA | NA |
| Hoxa11os | 0 | NA | NA | NA |
| Gm28032 | 0 | NA | NA | NA |
| Gm27492 | 0 | NA | NA | NA |

| | | | | |
|---|---|---|---|---|
| Gm27695 | 0 | NA | NA | NA |
| Gm27817 | 0 | NA | NA | NA |
| Gm27454 | 0 | NA | NA | NA |
| Gm27543 | 0 | NA | NA | NA |
| 9530018H14Rik | 0 | NA | NA | NA |
| Hoxa13 | 0 | NA | NA | NA |
| Gm27280 | 0 | NA | NA | NA |
| Gm27985 | 0 | NA | NA | NA |
| Hottip | 0 | NA | NA | NA |
| Gm27383 | 0 | NA | NA | NA |
| Gm27866 | 0 | NA | NA | NA |
| 5730457N03Rik | 3.810690039 | -0.339372826 | 0.557123018 | -0.609152404 |
| Evx1 | 6.587797916 | -0.987438755 | 0.546778284 | -1.805921677 |
| Gm8129 | 27.45075818 | 0.109381211 | 0.351632602 | 0.31106675 |
| Gm15054 | 0 | NA | NA | NA |
| Gm37351 | 0.106358623 | 0.517955464 | 3.724945049 | 0.139050498 |
| Hibadh | 577.34931 | -0.253186413 | 0.102028822 | -2.48151854 |
| Gm24355 | 3.634340466 | 0.433525317 | 0.606945301 | 0.714274114 |
| Tax1bp1 | 2381.399223 | -0.346397906 | 0.070245495 | -4.931247289 |
| Jazf1 | 226.345128 | -0.214882661 | 0.107616568 | -1.996743288 |
| Gm15572 | 3.883676966 | 1.130827834 | 0.622215489 | 1.817421544 |
| Gm26215 | 6.191745285 | 0.212363091 | 0.462755651 | 0.458909774 |
| Gm15573 | 0.191192637 | -0.443820809 | 2.663013325 | -0.16666113 |
| Gm25818 | 0.106618422 | 0.517955464 | 3.724945049 | 0.139050498 |
| Creb5 | 153.7701789 | -0.081746992 | 0.201758955 | -0.405171568 |
| Tril | 1025.612528 | -0.464554023 | 0.167450084 | -2.7742836 |
| Gm16499 | 0.523988269 | -0.283849882 | 1.347114528 | -0.210709539 |
| Gm22910 | 0 | NA | NA | NA |
| Cpvl | 0.864647981 | -1.522447797 | 1.08245112 | -1.406481798 |
| Chn2 | 1528.702195 | 0.145102061 | 0.220276681 | 0.658726384 |
| 9530036M11Rik | 0.430808853 | 0.286408658 | 1.504366254 | 0.190384926 |
| Gm15526 | 0.194037777 | -0.469933603 | 2.636796485 | -0.178221416 |
| Gm15527 | 0.824096613 | -0.262583427 | 1.120524099 | -0.23433983 |
| 9130019P16Rik | 5.722762008 | 0.719549175 | 0.458733161 | 1.56855714 |
| Prr15 | 15.96290937 | 0.271696312 | 0.295361767 | 0.919876379 |
| Wipf3 | 1686.195845 | 0.987144561 | 0.151479296 | 6.516696268 |
| Scrn1 | 2778.202593 | 0.196922283 | 0.053160172 | 3.704319879 |
| Fkbp14 | 126.0321638 | -0.474336733 | 0.16019137 | -2.961062971 |
| Plekha8 | 312.5409318 | -0.043380222 | 0.079916107 | -0.542822012 |
| Gm15863 | 0.061920613 | -0.443816764 | 3.724945049 | -0.119147198 |
| Gm28402 | 0.435961187 | 0.036426512 | 1.456802757 | 0.025004423 |
| Mturn | 1138.29535 | 0.571267932 | 0.102720604 | 5.561376279 |
| Znrf2 | 429.2306728 | 0.009918734 | 0.072499511 | 0.13681105 |
| Nod1 | 393.3170085 | -0.708198089 | 0.172396205 | -4.107967984 |
| Gm24230 | 0 | NA | NA | NA |
| Ggct | 96.47110576 | 0.070568361 | 0.212448801 | 0.332166435 |
| Gars | 1477.498831 | -0.26552906 | 0.093168812 | -2.849977949 |

| | | | | |
|---|---|---|---|---|
| Crhr2 | 14.43709135 | 0.590201324 | 0.308003045 | 1.916219117 |
| Inmt | 2.148441887 | -0.659766368 | 0.733382433 | -0.899621178 |
| Fam188b | 89.47598892 | 0.04702974 | 0.154149318 | 0.305092103 |
| Gm25458 | 0 | NA | NA | NA |
| Aqp1 | 74.91150286 | -1.662079718 | 0.554215726 | -2.998976103 |
| Ghrhr | 0.426601203 | -2.109322949 | 3.210286541 | -0.657051301 |
| Adcyap1r1 | 2280.821534 | 0.295876685 | 0.049938475 | 5.924824188 |
| AC069309.1 | 0.075043205 | 0.517957688 | 3.724945049 | 0.139051095 |
| Neurod6 | 272.3533864 | 1.104981421 | 0.191284116 | 5.776650165 |
| Ccdc129 | 12.62732475 | -0.143819766 | 0.294490628 | -0.488367887 |
| Ppp1r17 | 219.4288715 | -0.019828414 | 0.298942005 | -0.066328633 |
| Pde1c | 512.5279336 | -0.137803633 | 0.11170337 | -1.233656899 |
| AC079956.1 | 1.707215572 | -0.228385641 | 0.855075461 | -0.267094136 |
| Gm24709 | 0 | NA | NA | NA |
| Gm22493 | 0.217296566 | -0.443821436 | 2.200942151 | -0.201650659 |
| Lsm5 | 26.01436472 | 0.171368686 | 0.209997742 | 0.816050136 |
| Avl9 | 1503.371839 | 0.057957185 | 0.083755966 | 0.691976798 |
| Kbtbd2 | 937.4819014 | -0.146969896 | 0.064277382 | -2.28649475 |
| Fkbp9 | 377.8815204 | -0.203736859 | 0.121841155 | -1.672151399 |
| Nt5c3 | 889.5108846 | -0.191627412 | 0.138102143 | -1.387577397 |
| Vmn1r4 | 1.941638245 | 1.204226506 | 0.701514899 | 1.716608595 |
| Vmn1r5 | 0.356992291 | 1.165887933 | 3.30596007 | 0.352662437 |
| Vmn1r6 | 0.118371171 | 0.037073798 | 3.724945049 | 0.009952844 |
| Vmn1r7 | 0 | NA | NA | NA |
| Vmn1r8 | 0 | NA | NA | NA |
| Vmn1r9 | 0 | NA | NA | NA |
| Vmn1r10 | 0 | NA | NA | NA |
| Vmn1r11 | 0 | NA | NA | NA |
| Vmn1r12 | 0 | NA | NA | NA |
| Vmn1r-ps8 | 0 | NA | NA | NA |
| Vmn1r13 | 0 | NA | NA | NA |
| Vmn1r14 | 0 | NA | NA | NA |
| Vmn1r15 | 0 | NA | NA | NA |
| Vmn1r-ps9 | 0 | NA | NA | NA |
| Vmn1r-ps10 | 0 | NA | NA | NA |
| Vmn1r16 | 0 | NA | NA | NA |
| Vmn1r17 | 0 | NA | NA | NA |
| Vmn1r18 | 0 | NA | NA | NA |
| Vmn1r19 | 0 | NA | NA | NA |
| Vmn1r20 | 0 | NA | NA | NA |
| Vmn1r-ps11 | 0 | NA | NA | NA |
| 4930533I22Rik | 0.062426873 | 0.517959912 | 3.724945049 | 0.139051692 |
| Ppm1k | 1833.8061 | 0.375018613 | 0.106646906 | 3.516450928 |
| Gm27896 | 0 | NA | NA | NA |
| Herc6 | 2713.472928 | -0.520820846 | 0.162955648 | -3.196089571 |
| Gm37397 | 18.0739852 | -0.358009756 | 0.273088427 | -1.310966412 |
| Pyurf | 116.2739024 | -0.157083602 | 0.117699788 | -1.334612446 |

| | | | | |
|---|---|---|---|---|
| Gm26712 | 0 | NA | NA | NA |
| Lancl2 | 881.9561078 | 0.303021368 | 0.063628292 | 4.762368421 |
| Vopp1 | 778.6664814 | -0.038218385 | 0.082868364 | -0.461193911 |
| Vmn1r21 | 0.060174537 | 0.517957688 | 3.724945049 | 0.139051095 |
| Gm25726 | 0 | NA | NA | NA |
| Vmn1r-ps12 | 0 | NA | NA | NA |
| Vmn1r22 | 0 | NA | NA | NA |
| Vmn1r23 | 0 | NA | NA | NA |
| Gm20660 | 0 | NA | NA | NA |
| Vmn1r24 | 0 | NA | NA | NA |
| Vmn1r25 | 0 | NA | NA | NA |
| Vmn1r26 | 0 | NA | NA | NA |
| Rps15-ps2 | 2.860642787 | 0.871937513 | 0.774800196 | 1.125370795 |
| Vmn1r-ps14 | 0 | NA | NA | NA |
| Vmn1r-ps15 | 0.09176491 | 0.921395572 | 3.723061103 | 0.247483333 |
| Gm6695 | 0.119587853 | 0.981063299 | 3.723618439 | 0.263470416 |
| Vmn1r-ps17 | 0 | NA | NA | NA |
| Gm20718 | 0.110595523 | 0.037071574 | 3.724945049 | 0.009952247 |
| Vmn1r-ps18 | 0 | NA | NA | NA |
| Vmn1r27 | 0 | NA | NA | NA |
| Vmn1r-ps19 | 0 | NA | NA | NA |
| Vmn1r28 | 0.068097359 | 0.037064902 | 3.724945049 | 0.009950456 |
| Vmn1r29 | 0.139856492 | -0.443820827 | 3.574411099 | -0.124166139 |
| Gm20674 | 0 | NA | NA | NA |
| Vmn1r-ps20 | 0 | NA | NA | NA |
| Vmn1r30 | 0.075043205 | 0.517957688 | 3.724945049 | 0.139051095 |
| Vmn1r-ps21 | 0 | NA | NA | NA |
| Gm20673 | 0 | NA | NA | NA |
| Vmn1r31 | 0 | NA | NA | NA |
| Vmn1r-ps22 | 0.047948698 | 0.037064902 | 3.724945049 | 0.009950456 |
| Vmn1r-ps23 | 0.066050875 | -0.443818988 | 3.724945049 | -0.119147795 |
| Gm20672 | 0 | NA | NA | NA |
| Abcg2 | 150.9866917 | -0.109762999 | 0.114416033 | -0.959332329 |
| Herc3 | 2335.927333 | 0.115246185 | 0.086898646 | 1.326213814 |
| Nap1l5 | 4015.777592 | 0.354606841 | 0.128728043 | 2.754697683 |
| Fam13a | 397.9236056 | 0.190039897 | 0.070722088 | 2.687136415 |
| BB365896 | 2.654833662 | -0.124212709 | 0.685752957 | -0.181133319 |
| Gm22367 | 0 | NA | NA | NA |
| Tigd2 | 330.9779313 | -0.138043541 | 0.109469623 | -1.261021425 |
| Gprin3 | 236.7557592 | 0.524871288 | 0.127106921 | 4.129368289 |
| Gm22622 | 0 | NA | NA | NA |
| Gm5570 | 0.128070691 | 0.517964374 | 3.723419658 | 0.139109856 |
| Gm22723 | 0 | NA | NA | NA |
| n-R5s162 | 0 | NA | NA | NA |
| A530053G22Rik | 0.106618422 | 0.517955464 | 3.724945049 | 0.139050498 |
| Snca | 1013.675009 | 0.52686317 | 0.126763069 | 4.156282865 |
| Mmrn1 | 0.717105192 | -1.340173755 | 1.134664188 | -1.181119286 |

| | | | | |
|---|---|---|---|---|
| A730020E08Rik | 23.88982311 | 0.362107592 | 0.283183075 | 1.278704922 |
| Ccser1 | 527.4917736 | 0.38622623 | 0.106648911 | 3.621473729 |
| Gm22212 | 0 | NA | NA | NA |
| Grid2 | 1329.763725 | 0.265495695 | 0.118524982 | 2.239997766 |
| Gm25205 | 1.741084885 | -0.13233006 | 1.017888146 | -0.13000452 |
| Atoh1 | 0.260468428 | -0.791890979 | 2.018636357 | -0.392290061 |
| Gm15534 | 0.038981395 | -0.443816764 | 3.724945049 | -0.119147198 |
| Smarcad1 | 613.7764423 | -0.18275682 | 0.067724976 | -2.698514336 |
| Hpgds | 46.4750562 | 0.785660547 | 0.270216506 | 2.907522412 |
| Gm17147 | 2.881706403 | -0.307218259 | 0.717371617 | -0.428255386 |
| C130060K24Rik | 9.901210978 | -0.283149583 | 0.370317956 | -0.76461208 |
| Tnip3 | 4.68769279 | -0.930443423 | 0.627892567 | -1.48185131 |
| Ndnf | 237.9647372 | -0.159714212 | 0.165956944 | -0.962383421 |
| Gm15490 | 0.045882455 | 0.517959912 | 3.724945049 | 0.139051692 |
| Gm16838 | 1.376958878 | 0.952665013 | 0.856332546 | 1.112494226 |
| Prdm5 | 99.54165155 | 0.019063106 | 0.114698052 | 0.166202531 |
| Gm23543 | 0 | NA | NA | NA |
| Gm20445 | 3.495574212 | 1.263001391 | 0.622414152 | 2.02919774 |
| 4930544G11Rik | 0 | NA | NA | NA |
| Gm23625 | 0 | NA | NA | NA |
| Mad2l1 | 71.55826495 | 0.107072092 | 0.176696822 | 0.605965012 |
| Vmn1r32 | 0 | NA | NA | NA |
| Vmn1r33 | 0.066050875 | -0.443818988 | 3.724945049 | -0.119147795 |
| Vmn1r34 | 0.225129614 | 1.580880368 | 3.710727143 | 0.426029807 |
| Vmn1r35 | 0.038981395 | -0.443816764 | 3.724945049 | -0.119147198 |
| Vmn1r36 | 0 | NA | NA | NA |
| Vmn1r37 | 0 | NA | NA | NA |
| Vmn1r-ps24 | 0.066050875 | -0.443818988 | 3.724945049 | -0.119147795 |
| Vmn1r38 | 0 | NA | NA | NA |
| Vmn1r39 | 0.047948698 | 0.037064902 | 3.724945049 | 0.009950456 |
| RP23-263B18.4 | 379.7491701 | -0.199655535 | 0.15539966 | -1.284787465 |
| 4930597O21Rik | 2.946845261 | -0.124914147 | 0.563613099 | -0.221631022 |
| Gng12 | 1315.427679 | -0.151579167 | 0.118169438 | -1.282727326 |
| Gm15644 | 4.54732579 | -0.099252152 | 0.555309858 | -0.17873292 |
| Gm25260 | 0.532344628 | -0.211689272 | 1.546673567 | -0.136867453 |
| Gadd45a | 98.08999083 | -1.022782489 | 0.268292079 | -3.812197864 |
| E230016M11Rik | 92.89630384 | 0.504786673 | 0.193365346 | 2.610533296 |
| A430010J10Rik | 11.54166846 | 0.104650804 | 0.377164536 | 0.277467243 |
| Gm27015 | 0.824269312 | 1.54595226 | 1.229414908 | 1.257469915 |
| Gm8566 | 29.8603066 | 0.187054653 | 0.216953449 | 0.862187966 |
| Serbp1 | 2115.70978 | 0.043209388 | 0.111563768 | 0.387306637 |
| Il12rb2 | 27.958596 | -0.966884496 | 0.477003476 | -2.02699675 |
| Gm4761 | 0 | NA | NA | NA |
| Il23r | 2.075698228 | 0.288834504 | 0.793113809 | 0.364177879 |
| RP23-123I10.3 | 0.062426873 | 0.517959912 | 3.724945049 | 0.139051692 |
| Tacstd2 | 4.381073283 | -0.23278664 | 0.509811789 | -0.456612901 |
| Igkv2-137 | 0.061920613 | -0.443816764 | 3.724945049 | -0.119147198 |

| | | | | |
|---|---|---|---|---|
| RP23-123I10.4 | 0 | NA | NA | NA |
| Igkv1-136 | 0 | NA | NA | NA |
| Igkv1-135 | 0.427013396 | -1.886499358 | 1.539464815 | -1.225425447 |
| RP24-322E13.1 | 0 | NA | NA | NA |
| RP24-322E13.2 | 0 | NA | NA | NA |
| Igkv14-134-1 | 0 | NA | NA | NA |
| Igkv17-134 | 0 | NA | NA | NA |
| Igkv1-133 | 0.216076386 | 1.49879282 | 3.709588964 | 0.404032046 |
| RP23-255N3.3 | 0 | NA | NA | NA |
| RP23-255N3.4 | 0 | NA | NA | NA |
| RP23-255N3.6 | 0 | NA | NA | NA |
| Igkv1-132 | 0.132101749 | -0.93550548 | 3.724188147 | -0.251197158 |
| Igkv1-131 | 0 | NA | NA | NA |
| RP23-255N3.8 | 0 | NA | NA | NA |
| Igkv14-130 | 0 | NA | NA | NA |
| Gm22425 | 0 | NA | NA | NA |
| Gm24911 | 0 | NA | NA | NA |
| Igkv9-129 | 0 | NA | NA | NA |
| Igkv9-128 | 0 | NA | NA | NA |
| Igkv17-127 | 0.143022002 | 0.03610674 | 3.547735707 | 0.010177404 |
| Igkv14-126-1 | 0 | NA | NA | NA |
| RP23-234A12.5 | 0 | NA | NA | NA |
| Igkv14-126 | 0 | NA | NA | NA |
| Gm25910 | 0 | NA | NA | NA |
| Igkv11-125 | 0.062426873 | 0.517959912 | 3.724945049 | 0.139051692 |
| Igkv9-124 | 0 | NA | NA | NA |
| Igkv9-123 | 0 | NA | NA | NA |
| Igkv1-122 | 0 | NA | NA | NA |
| Igkv17-121 | 0.097139547 | -0.443816764 | 3.724945049 | -0.119147198 |
| Igkv9-120 | 0.291418641 | -1.653045791 | 3.70657789 | -0.445976272 |
| Igkv9-119 | 0 | NA | NA | NA |
| Igkv14-118-2 | 0 | NA | NA | NA |
| Igkv14-118-1 | 0 | NA | NA | NA |
| RP23-185M15.10 | 0 | NA | NA | NA |
| RP23-185M15.7 | 0 | NA | NA | NA |
| Igkv11-118 | 0 | NA | NA | NA |
| Igkv1-117 | 0.45022207 | -2.689228249 | 3.671271238 | -0.732506011 |
| RP24-76K4.8 | 0 | NA | NA | NA |
| RP24-76K4.7 | 0.141094079 | 0.037072457 | 3.563850528 | 0.01040236 |
| Igkv2-116 | 0.060174537 | 0.517957688 | 3.724945049 | 0.139051095 |
| Igkv1-115 | 0 | NA | NA | NA |
| Igkv11-114 | 0 | NA | NA | NA |
| Igkv2-113 | 0 | NA | NA | NA |
| RP24-76K4.1 | 0 | NA | NA | NA |
| Igkv2-112 | 0.157314084 | 0.037075025 | 3.435173482 | 0.010792766 |
| Igkv14-111 | 0.94930219 | -1.075261167 | 0.990884059 | -1.085153362 |
| Gm25436 | 0 | NA | NA | NA |

| | | | | |
|---|---|---|---|---|
| Igkv1-110 | 0.099106094 | -0.93550548 | 3.724188147 | -0.251197158 |
| RP24-76K4.12 | 0 | NA | NA | NA |
| RP24-76K4.11 | 0 | NA | NA | NA |
| Igkv2-109 | 0.106971521 | 0.998850474 | 3.724945049 | 0.268151734 |
| Igkv1-108 | 0 | NA | NA | NA |
| Igkv2-107 | 0 | NA | NA | NA |
| Igkv19-93 | 0 | NA | NA | NA |
| Igkv2-105 | 0 | NA | NA | NA |
| RP23-29F15.2 | 0 | NA | NA | NA |
| Igkv16-104 | 0.862499085 | 1.607222283 | 3.647176207 | 0.440675797 |
| Igkv15-103 | 0.066050875 | -0.443818988 | 3.724945049 | -0.119147795 |
| Igkv15-102 | 0 | NA | NA | NA |
| Igkv20-101-2 | 0 | NA | NA | NA |
| Igkv15-101-1 | 0 | NA | NA | NA |
| Igkv15-101 | 0 | NA | NA | NA |
| Igkv14-100 | 0 | NA | NA | NA |
| Gm25833 | 0 | NA | NA | NA |
| Igkv1-99 | 0 | NA | NA | NA |
| Igkv12-98 | 0 | NA | NA | NA |
| Igkv15-97 | 0 | NA | NA | NA |
| Igkv10-96 | 0.163190422 | -0.955151543 | 3.396300892 | -0.281232898 |
| Igkv2-95-2 | 0 | NA | NA | NA |
| Igkv2-95-1 | 0 | NA | NA | NA |
| Igkv10-95 | 0 | NA | NA | NA |
| Igkv10-94 | 0.066050875 | -0.443818988 | 3.724945049 | -0.119147795 |
| Igkv2-93-1 | 0 | NA | NA | NA |
| RP24-405O23.8 | 0 | NA | NA | NA |
| AC120404.1 | 0 | NA | NA | NA |
| Igkv4-92 | 0 | NA | NA | NA |
| RP23-211L5.2 | 0 | NA | NA | NA |
| Igkv4-91 | 0 | NA | NA | NA |
| Igkv4-90 | 0 | NA | NA | NA |
| Igkv13-89-1 | 0 | NA | NA | NA |
| Igkv12-89 | 0.175102337 | -1.16604509 | 3.306428121 | -0.352660045 |
| Igkv1-88 | 0.108772582 | 0.037073798 | 3.724945049 | 0.009952844 |
| RP23-211L5.7 | 0 | NA | NA | NA |
| IGKV1D-42 | 0 | NA | NA | NA |
| RP23-211L5.9 | 0.045882455 | 0.517959912 | 3.724945049 | 0.139051692 |
| Igkv13-87 | 0.049553047 | -0.443818988 | 3.724945049 | -0.119147795 |
| RP23-211L5.12 | 0.088534442 | -0.905414734 | 3.723500899 | -0.243162217 |
| Igkv4-86 | 0 | NA | NA | NA |
| Igkv12-47 | 0.097139547 | -0.443816764 | 3.724945049 | -0.119147198 |
| Igkv13-84 | 0.060174537 | 0.517957688 | 3.724945049 | 0.139051095 |
| Igkv4-56 | 0 | NA | NA | NA |
| Igkv13-82 | 0 | NA | NA | NA |
| Igkv4-81 | 0 | NA | NA | NA |
| Igkv13-80-1 | 0 | NA | NA | NA |

| | | | | |
|---|---|---|---|---|
| Igkv4-80 | 0.097139547 | -0.443816764 | 3.724945049 | -0.119147198 |
| Igkv4-79 | 0 | NA | NA | NA |
| Igkv13-78-1 | 0 | NA | NA | NA |
| Igkv4-78 | 0 | NA | NA | NA |
| RP23-344N8.12 | 0 | NA | NA | NA |
| Igkv4-77 | 0 | NA | NA | NA |
| Igkv13-76 | 0 | NA | NA | NA |
| Igkv4-75 | 0 | NA | NA | NA |
| Igkv13-74-1 | 0 | NA | NA | NA |
| Igkv4-74 | 0.062426873 | 0.517959912 | 3.724945049 | 0.139051692 |
| Igkv13-73-1 | 0 | NA | NA | NA |
| Igkv4-73 | 0.046345709 | -0.443818988 | 3.724945049 | -0.119147795 |
| Igkv4-72 | 0.182573437 | -0.443823707 | 3.311478819 | -0.134025833 |
| Igkv13-71-1 | 0 | NA | NA | NA |
| Igkv4-71 | 0 | NA | NA | NA |
| Igkv4-70 | 0.046345709 | -0.443818988 | 3.724945049 | -0.119147795 |
| Igkv4-69 | 0 | NA | NA | NA |
| Igkv4-68 | 0.060174537 | 0.517957688 | 3.724945049 | 0.139051095 |
| Igkv12-67 | 0 | NA | NA | NA |
| Rprl1 | 0 | NA | NA | NA |
| Igkv12-66 | 0 | NA | NA | NA |
| Igkv4-65 | 0 | NA | NA | NA |
| Igkv13-64 | 0 | NA | NA | NA |
| Igkv4-63 | 0 | NA | NA | NA |
| Igkv13-62-1 | 0 | NA | NA | NA |
| Igkv4-62 | 0 | NA | NA | NA |
| Igkv13-61-1 | 0 | NA | NA | NA |
| Igkv4-61 | 0.266304556 | -0.602115226 | 3.715438719 | -0.162057639 |
| Igkv4-59 | 0 | NA | NA | NA |
| Igkv4-60 | 0 | NA | NA | NA |
| Igkv4-58 | 0 | NA | NA | NA |
| Rn7s6 | 476.0174306 | -0.449078057 | 0.334714426 | -1.341675238 |
| AC156953.1 | 0 | NA | NA | NA |
| Igkv13-57-2 | 0 | NA | NA | NA |
| Igkv4-57-1 | 0.097139547 | -0.443816764 | 3.724945049 | -0.119147198 |
| Igkv13-57-1 | 0 | NA | NA | NA |
| Igkv4-57 | 0.102891481 | 0.51795324 | 3.724945049 | 0.139049901 |
| Igkv13-56-1 | 0 | NA | NA | NA |
| Igkv13-55-1 | 0 | NA | NA | NA |
| Igkv4-55 | 0.097139547 | -0.443816764 | 3.724945049 | -0.119147198 |
| Igkv13-54-1 | 0 | NA | NA | NA |
| Igkv4-54 | 0 | NA | NA | NA |
| Igkv4-53 | 0 | NA | NA | NA |
| Gm11143 | 0 | NA | NA | NA |
| RP23-24I14.7 | 0 | NA | NA | NA |
| Igkv4-50 | 0 | NA | NA | NA |
| Igkv12-49 | 0 | NA | NA | NA |

| | | | | |
|---|---|---|---|---|
| Igkv5-48 | 0.183032319 | 0.647812728 | 3.311598859 | 0.195619323 |
| Igkv12-46 | 0.771993125 | -1.940853668 | 3.503801979 | -0.553927899 |
| Igkv5-45 | 0.049553047 | -0.443818988 | 3.724945049 | -0.119147795 |
| Igkv12-44 | 0.240624804 | -1.61948693 | 3.034969775 | -0.533608916 |
| Igkv5-43 | 0.240624804 | -1.61948693 | 3.034969775 | -0.533608916 |
| Igkv12-42 | 0 | NA | NA | NA |
| Igkv12-41 | 0.194279094 | -1.017823543 | 3.719440177 | -0.273649661 |
| Igkv5-40-1 | 0 | NA | NA | NA |
| Igkv12-40 | 0 | NA | NA | NA |
| Igkv5-39 | 0.975696798 | -0.268004585 | 2.595394805 | -0.103261587 |
| RP24-475M8.10 | 0 | NA | NA | NA |
| Igkv12-38 | 0.099106094 | -0.93550548 | 3.724188147 | -0.251197158 |
| Igkv5-37 | 0 | NA | NA | NA |
| Igkv18-36 | 0 | NA | NA | NA |
| Igkv1-35 | 0 | NA | NA | NA |
| Gm23694 | 0 | NA | NA | NA |
| RP24-314D8.6 | 0 | NA | NA | NA |
| Igkv8-34 | 0.066050875 | -0.443818988 | 3.724945049 | -0.119147795 |
| Igkv7-33 | 0.093831153 | 0.517957688 | 3.724945049 | 0.139051095 |
| Igkv6-32 | 0.998588786 | 0.161334084 | 3.649576267 | 0.044206251 |
| Igkv8-31 | 0 | NA | NA | NA |
| Igkv8-30 | 0.203286941 | -0.771106728 | 2.603555075 | -0.296174541 |
| Amd-ps2 | 0.04393175 | 0.037062677 | 3.724945049 | 0.009949859 |
| Igkv6-29 | 0 | NA | NA | NA |
| Igkv8-28 | 0 | NA | NA | NA |
| Gm24096 | 0 | NA | NA | NA |
| Igkv8-27 | 0.112396584 | -0.917692504 | 3.724434845 | -0.246397787 |
| Igkv8-26 | 0 | NA | NA | NA |
| RP23-64J23.3 | 0.049553047 | -0.443818988 | 3.724945049 | -0.119147795 |
| Igkv6-25 | 0.097139547 | -0.443816764 | 3.724945049 | -0.119147198 |
| Igkv8-24 | 0 | NA | NA | NA |
| Igkv8-23-1 | 0 | NA | NA | NA |
| Igkv6-23 | 1.206331356 | -4.00590024 | 3.64722383 | -1.098342308 |
| RP23-64J23.7 | 0 | NA | NA | NA |
| Gm11065 | 0 | NA | NA | NA |
| Igkv8-22 | 0 | NA | NA | NA |
| Igkv8-21 | 0.277388482 | 0.919670127 | 1.767937427 | 0.520193822 |
| Amd-ps5 | 0.048949532 | 0.037064902 | 3.724945049 | 0.009950456 |
| Igkv6-20 | 0.038981395 | -0.443816764 | 3.724945049 | -0.119147198 |
| Igkv8-19 | 0.269481931 | -0.44382286 | 2.435560099 | -0.182226199 |
| Igkv8-18 | 0 | NA | NA | NA |
| RP23-64J23.14 | 0.089814205 | 0.517955464 | 3.724945049 | 0.139050498 |
| Igkv6-17 | 0.659092009 | -2.877801433 | 3.66790651 | -0.784589636 |
| Igkv8-16 | 0.097139547 | -0.443816764 | 3.724945049 | -0.119147198 |
| RP23-175B24.3 | 0 | NA | NA | NA |
| Igkv6-15 | 0.421418085 | -1.569031085 | 1.664974271 | -0.942375574 |
| RP23-175B24.4 | 0.066050875 | -0.443818988 | 3.724945049 | -0.119147795 |

| | | | | |
|---|---|---|---|---|
| Gm10360 | 0 | NA | NA | NA |
| RP23-175B24.5 | 5.844746325 | -0.523519995 | 0.442671335 | -1.182638119 |
| Igkv6-14 | 0 | NA | NA | NA |
| RP23-175B24.8 | 0 | NA | NA | NA |
| Igkv6-13 | 0.044544648 | 0.517957688 | 3.724945049 | 0.139051095 |
| RP23-175B24.10 | 0 | NA | NA | NA |
| RP23-175B24.13 | 0.042716945 | 0.037062677 | 3.724945049 | 0.009949859 |
| RP23-175B24.14 | 0 | NA | NA | NA |
| Igkv3-12-1 | 0 | NA | NA | NA |
| Igkv3-12 | 0.232585366 | 0.127496527 | 2.533852793 | 0.050317259 |
| Igkv3-11 | 0 | NA | NA | NA |
| Igkv3-10 | 0.300744356 | -1.047539541 | 3.711871559 | -0.282213305 |
| Igkv3-9 | 0 | NA | NA | NA |
| Igkv3-8 | 0 | NA | NA | NA |
| Igkv3-7 | 4.685552657 | -3.744607652 | 1.478344757 | -2.532973201 |
| Igkv3-6 | 0 | NA | NA | NA |
| Igkv3-5 | 0.169165009 | 0.036125373 | 3.351346561 | 0.01077936 |
| Igkv3-4 | 0.266304556 | -0.602115226 | 3.715438719 | -0.162057639 |
| Igkv3-3 | 0 | NA | NA | NA |
| Igkv3-2 | 0.309883016 | -2.192905912 | 3.687548901 | -0.594678463 |
| Igkv3-1 | 0 | NA | NA | NA |
| RP23-435I4.11 | 0.318875346 | -1.114473571 | 3.709405606 | -0.300445324 |
| Igkj1 | 0.169165009 | 0.036125373 | 3.351346561 | 0.01077936 |
| Igkj2 | 0 | NA | NA | NA |
| Igkj3 | 0.062426873 | 0.517959912 | 3.724945049 | 0.139051692 |
| Igkj4 | 0.254453631 | -0.50137576 | 2.971545213 | -0.168725604 |
| Igkj5 | 0.189830966 | -1.394586823 | 3.258054901 | -0.428042763 |
| Igkc | 42.12345985 | -2.15565799 | 0.938229679 | -2.297580259 |
| Rpia | 130.920887 | 0.094058073 | 0.106442695 | 0.883649859 |
| Eif2ak3 | 463.4739881 | -0.026569047 | 0.072060079 | -0.368706883 |
| Gm23346 | 0.975898478 | -0.077683071 | 1.014667229 | -0.076560146 |
| RP24-122N23.3 | 0.09349418 | 0.517955464 | 3.724945049 | 0.139050498 |
| Tex37 | 0 | NA | NA | NA |
| Foxi3 | 0.042716945 | 0.037062677 | 3.724945049 | 0.009949859 |
| Gm23485 | 0 | NA | NA | NA |
| Rpl34-ps1 | 54.38793798 | -0.536706529 | 0.251255897 | -2.136095257 |
| Thnsl2 | 33.95158729 | 0.00383421 | 0.215144493 | 0.017821556 |
| Fabp1 | 0.161948271 | -0.443822416 | 3.446645162 | -0.128769396 |
| Smyd1 | 16.55316792 | 0.447156844 | 0.34194927 | 1.307670122 |
| Mir8112 | 5.618825651 | -0.544786655 | 0.487726006 | -1.116993248 |
| Krcc1 | 217.6208105 | -0.870423904 | 0.173463081 | -5.017920237 |
| Gm1070 | 1.970056041 | -0.264916475 | 0.725139182 | -0.3653319 |
| Cd8b1 | 33.23125073 | -0.565427019 | 0.335111736 | -1.687279069 |
| Cd8a | 74.76319009 | -0.267164742 | 0.318764884 | -0.838124761 |
| Rmnd5a | 2145.404065 | 0.136013044 | 0.062542269 | 2.174737913 |
| Rnf103 | 778.7289262 | 0.086027025 | 0.063640385 | 1.351767834 |
| Chmp3 | 861.0926457 | 0.038630122 | 0.074053378 | 0.521652392 |

| | | | | |
|---|---|---|---|---|
| Kdm3a | 752.2610037 | -0.088273193 | 0.076773096 | -1.149793311 |
| Reep1 | 1109.512451 | -0.114102565 | 0.074306504 | -1.535566314 |
| Mrpl35 | 321.4157534 | -0.298033935 | 0.075584189 | -3.943072471 |
| Immt | 1466.512924 | -0.11696597 | 0.059565742 | -1.963644971 |
| Ptcd3 | 534.5440444 | 0.06223596 | 0.07583459 | 0.820680369 |
| Gm22486 | 336.2636012 | -0.14488063 | 0.33011653 | -0.438877236 |
| Polr1a | 650.2299052 | -0.263892773 | 0.055283344 | -4.773458922 |
| Gm26628 | 0.134276636 | -0.890946624 | 2.951368388 | -0.301875777 |
| St3gal5 | 1041.7803 | 0.395504356 | 0.055661045 | 7.10558628 |
| Gm29438 | 74.24082104 | 0.623036307 | 0.314969229 | 1.978086269 |
| Atoh8 | 37.89726758 | -0.496759943 | 0.239886902 | -2.070808949 |
| Sftpb | 0.285479113 | 0.036514558 | 1.787992441 | 0.020422098 |
| Usp39 | 316.7202377 | -0.315204389 | 0.075204031 | -4.191323067 |
| 0610030E20Rik | 238.8730958 | -0.194171413 | 0.089614268 | -2.166746624 |
| Tmem150a | 75.99670554 | -0.78292217 | 0.162914423 | -4.805726551 |
| Rnf181 | 551.102173 | -0.328013436 | 0.093792143 | -3.497237887 |
| Vamp5 | 21.09757441 | -1.237041893 | 0.307333306 | -4.025082433 |
| Vamp8 | 157.0775941 | -0.793304431 | 0.16856575 | -4.706201762 |
| Ggcx | 228.9491815 | -0.236171918 | 0.094962052 | -2.487013634 |
| Mat2a | 2291.964425 | -0.373999005 | 0.123377488 | -3.031339129 |
| Sh2d6 | 8.896846792 | 0.036157408 | 0.357827257 | 0.101047104 |
| Capg | 296.8370465 | -2.125333368 | 0.395192329 | -5.377972228 |
| Elmod3 | 206.9098534 | 0.189294207 | 0.093561028 | 2.023216406 |
| Retsat | 77.16422045 | 0.286334212 | 0.181326617 | 1.579107455 |
| Tgoln1 | 2850.702626 | -0.267549342 | 0.076030477 | -3.518974935 |
| Gm20536 | 0.787340167 | 2.099207955 | 1.175784399 | 1.785368097 |
| Tcf7l1 | 89.55502137 | -0.451731132 | 0.141797213 | -3.185754669 |
| Gm15401 | 1.324332992 | -1.159559961 | 1.014958848 | -1.142469927 |
| Gm37969 | 4.217504979 | -0.094629632 | 0.559629643 | -0.169093315 |
| Gm38114 | 1.798572027 | -0.322127734 | 0.877736907 | -0.366998051 |
| Gm38320 | 0.230331698 | 0.554783528 | 2.557603663 | 0.216915363 |
| Gm15402 | 0.090074004 | 0.517955464 | 3.724945049 | 0.139050498 |
| Gm37736 | 8.749811602 | -0.142160738 | 0.433182429 | -0.328177527 |
| Kcmf1 | 755.6815892 | -0.246557834 | 0.079192848 | -3.1133851 |
| Gm26640 | 0.344365504 | 0.005801062 | 3.722845902 | 0.001558233 |
| Tmsb10 | 1562.811079 | -0.124503861 | 0.218102036 | -0.570851441 |
| Dnah6 | 180.1945128 | 0.44616802 | 0.17550253 | 2.542231271 |
| Gm24533 | 0.239208914 | 0.998867493 | 2.110573294 | 0.473268328 |
| Suclg1 | 851.7330975 | -0.100100441 | 0.057935322 | -1.727796423 |
| Gm24370 | 0 | NA | NA | NA |
| 4931417E11Rik | 0 | NA | NA | NA |
| Gm26179 | 0 | NA | NA | NA |
| Gm23924 | 0 | NA | NA | NA |
| Gm22254 | 0 | NA | NA | NA |
| Gm8993 | 0 | NA | NA | NA |
| Gm9001 | 4.797213872 | 0.139722141 | 0.517684128 | 0.269898445 |
| AC161363.1 | 0 | NA | NA | NA |

| | | | | |
|---|---|---|---|---|
| Gm22283 | 0 | NA | NA | NA |
| Gm9008 | 0.868283866 | -2.01988912 | 1.260119379 | -1.602934733 |
| Ctnna2 | 1519.198385 | 0.515697079 | 0.060149959 | 8.573523338 |
| Lrrtm1 | 446.8085958 | 0.507254731 | 0.111188594 | 4.562111195 |
| Gm21045 | 0 | NA | NA | NA |
| Gm5576 | 0.929768241 | -1.120815149 | 1.160615561 | -0.965707497 |
| Reg3b | 0.091880448 | 0.037060453 | 3.724945049 | 0.009949262 |
| Reg3d | 0 | NA | NA | NA |
| Reg3a | 0 | NA | NA | NA |
| Reg2 | 0 | NA | NA | NA |
| Reg1 | 0 | NA | NA | NA |
| Reg3g | 0.501377686 | -2.156018712 | 1.397110927 | -1.543197945 |
| Gm21392 | 0 | NA | NA | NA |
| Gm23727 | 0 | NA | NA | NA |
| Gm29005 | 0 | NA | NA | NA |
| Gm20594 | 0 | NA | NA | NA |
| Gm4409 | 0.396083486 | 0.530919639 | 1.599554746 | 0.331917142 |
| Lrrtm4 | 669.3441651 | 0.780267863 | 0.122552564 | 6.366801648 |
| Gm26264 | 0 | NA | NA | NA |
| Gm22530 | 0 | NA | NA | NA |
| mmu-mir-7232 | 0.163166504 | 0.517962677 | 2.787726935 | 0.185801081 |
| mmu-mir-468 | 0 | NA | NA | NA |
| 1700009C05Rik | 0.362568279 | 1.408185468 | 1.587692923 | 0.886938178 |
| Gcfc2 | 109.5450915 | 0.133661912 | 0.184411575 | 0.724802182 |
| Mrpl19 | 289.1779516 | -0.126890536 | 0.070314734 | -1.804608072 |
| Eva1a | 89.53357422 | 0.167933457 | 0.173628789 | 0.967198229 |
| Gm15864 | 0 | NA | NA | NA |
| Tacr1 | 94.67186864 | -0.394177308 | 0.128820365 | -3.059899015 |
| Pole4 | 302.9621574 | -0.297847915 | 0.115294178 | -2.583373417 |
| Hk2 | 352.4423473 | -0.660298347 | 0.195848989 | -3.371466704 |
| Gm17034 | 3.741498389 | -2.077620285 | 0.613816048 | -3.384760455 |
| 2310069B03Rik | 0.29717078 | -0.646468591 | 2.009560731 | -0.321696469 |
| Sema4f | 443.9451415 | 0.113518035 | 0.098755934 | 1.149480654 |
| M1ap | 2.42047183 | 0.422304357 | 0.739424014 | 0.571126105 |
| Dok1 | 94.45580167 | -1.146795672 | 0.260851844 | -4.39634873 |
| Loxl3 | 99.54709547 | -0.897088894 | 0.190691453 | -4.704400113 |
| Mir7040 | 0 | NA | NA | NA |
| Htra2 | 198.8763925 | -0.318555042 | 0.084144457 | -3.785811388 |
| Aup1 | 620.1822418 | -0.495139148 | 0.055421016 | -8.934140498 |
| Dqx1 | 10.03751516 | -0.14981368 | 0.418421458 | -0.358044927 |
| Tlx2 | 0.090665643 | 0.037060453 | 3.724945049 | 0.009949262 |
| Gm37092 | 0.240333628 | 0.053682587 | 1.849956675 | 0.029018294 |
| Pcgf1 | 96.34584019 | -0.152753762 | 0.110131466 | -1.387012885 |
| Lbx2 | 0.810336573 | -0.284408707 | 1.055167821 | -0.269538837 |
| Mir3470a | 0 | NA | NA | NA |
| Ccdc142 | 74.38601428 | 0.143223449 | 0.183870117 | 0.778938149 |
| Ccdc142os | 12.6558528 | -0.440241304 | 0.329731101 | -1.335152501 |

| | | | | |
|---|---|---|---|---|
| Mrpl53 | 0 | NA | NA | NA |
| Mogs | 256.0958102 | -0.382181214 | 0.106837438 | -3.577221817 |
| Wbp1 | 156.9620067 | -0.214816059 | 0.129442813 | -1.659544119 |
| Ino80b | 119.6851225 | -0.194518496 | 0.166744617 | -1.166565373 |
| Rtkn | 581.9726703 | -0.012230703 | 0.081117394 | -0.150777805 |
| Wdr54 | 115.1061269 | 0.279643392 | 0.121610525 | 2.29949992 |
| 1700003E16Rik | 151.7522411 | 0.229379927 | 0.110780064 | 2.070588511 |
| Dctn1 | 3696.021488 | 0.325125009 | 0.059097029 | 5.501545715 |
| Gm15624 | 21.19334217 | 1.879591214 | 0.401908177 | 4.676668264 |
| Slc4a5 | 42.25336671 | 1.451363413 | 0.268829421 | 5.398826546 |
| Mthfd2 | 159.228204 | -0.913824615 | 0.167454929 | -5.457137739 |
| Mob1a | 606.7587231 | -0.722018127 | 0.129260599 | -5.585755688 |
| Bola3 | 181.7266574 | -0.072703528 | 0.094557518 | -0.768881517 |
| Tet3 | 1814.49476 | -0.096144147 | 0.106036304 | -0.90670971 |
| Mir6374 | 7.73301019 | 0.783695194 | 0.552816291 | 1.41764128 |
| B230319C09Rik | 9.836627767 | -0.172487124 | 0.334971999 | -0.514929976 |
| Dguok | 113.566417 | -0.248582527 | 0.171581059 | -1.448776045 |
| Actg2 | 1.148058676 | -1.603312431 | 1.015332394 | -1.579101031 |
| Stambp | 312.4465512 | 0.141657978 | 0.083779746 | 1.690838006 |
| Gm24990 | 0 | NA | NA | NA |
| Clec4f | 0.297646291 | -0.387493147 | 1.792885557 | -0.216128211 |
| Cd207 | 0.357781856 | -0.871043738 | 1.526651002 | -0.570558521 |
| Vax2os | 1.069186981 | 0.939694914 | 1.071638561 | 0.876876727 |
| Gm27772 | 0 | NA | NA | NA |
| Gm27631 | 0 | NA | NA | NA |
| Gm27893 | 0 | NA | NA | NA |
| Vax2 | 0.095898756 | -0.917692504 | 3.724434845 | -0.246397787 |
| Gm25054 | 0 | NA | NA | NA |
| Atp6v1b1 | 0.540009155 | 0.626878497 | 1.629301924 | 0.384752812 |
| 1700124L16Rik | 10.03157174 | 0.719041894 | 0.415990076 | 1.728507325 |
| Ankrd53 | 0 | NA | NA | NA |
| Tex261 | 481.6030204 | -0.401063206 | 0.107004764 | -3.748087379 |
| Nagk | 216.7968119 | -0.138081554 | 0.102550542 | -1.346473179 |
| Paip2b | 685.0019845 | 0.062781491 | 0.075414639 | 0.832484132 |
| Gm23014 | 0 | NA | NA | NA |
| Gm5138 | 12.43742249 | 0.086667887 | 0.288541533 | 0.300365378 |
| Zfp638 | 2492.303382 | -0.064803967 | 0.079606337 | -0.814055373 |
| Dysf | 208.5045205 | -1.448275027 | 0.236200452 | -6.131550615 |
| Gm15475 | 0.044544648 | 0.517957688 | 3.724945049 | 0.139051095 |
| Cyp26b1 | 126.4459904 | 0.257789364 | 0.176383366 | 1.461528771 |
| Exoc6b | 1761.121593 | 0.303321221 | 0.047472433 | 6.389418139 |
| Gm7507 | 0.706207703 | 1.263007659 | 1.13789091 | 1.10995496 |
| Gm26353 | 0.038981395 | -0.443816764 | 3.724945049 | -0.119147198 |
| Npm3-ps1 | 4.293674138 | -1.009870182 | 0.639566595 | -1.578991446 |
| Gm5878 | 0.534626289 | 0.253562505 | 1.268278575 | 0.199926506 |
| Spr | 209.6465649 | -0.279684885 | 0.094620986 | -2.955844104 |
| Gm10444 | 0.134148234 | -0.443820953 | 3.624632483 | -0.122445781 |

| | | | | |
|---|---|---|---|---|
| Gm22893 | 0 | NA | NA | NA |
| Spr-ps1 | 0.674932609 | 0.450082472 | 1.496872271 | 0.300681949 |
| Emx1 | 100.6339851 | 0.150312379 | 0.171771093 | 0.875073774 |
| Sfxn5 | 2038.190871 | 0.174652889 | 0.115190199 | 1.516213097 |
| Rab11fip5 | 1169.630522 | -0.08119792 | 0.081242348 | -0.99945314 |
| Mir705 | 0 | NA | NA | NA |
| Noto | 0 | NA | NA | NA |
| Smyd5 | 326.4950938 | -0.099555551 | 0.075573427 | -1.3173354 |
| Pradc1 | 68.96069805 | 0.509724145 | 0.155398594 | 3.280107818 |
| Cct7 | 1480.602166 | -0.406657114 | 0.076208578 | -5.33610685 |
| Fbxo41 | 1325.807268 | 0.600657469 | 0.119871332 | 5.010851726 |
| Egr4 | 333.4082471 | -0.432226003 | 0.53463806 | -0.808446004 |
| Alms1 | 761.6584805 | 0.074652389 | 0.131357455 | 0.568314828 |
| RP23-27D8.2 | 34.95142274 | 0.051051684 | 0.269634397 | 0.189336689 |
| RP24-478I22.8 | 43.9950079 | 0.250860694 | 0.249375043 | 1.005957497 |
| Gm4477 | 0.754021959 | -0.672812965 | 1.124800185 | -0.59816221 |
| Alms1-ps1 | 10.38534064 | -0.382908524 | 0.341564222 | -1.121044007 |
| Cml3 | 8.402099551 | 0.096708961 | 0.420850718 | 0.22979398 |
| Gm27317 | 0.061920613 | -0.443816764 | 3.724945049 | -0.119147198 |
| Alms1-ps2 | 11.88847144 | 0.052395609 | 0.33438295 | 0.156693422 |
| mmu-mir-6375 | 0.167183452 | 0.517964184 | 2.765153692 | 0.187318407 |
| Gm11128 | 2.945191291 | -0.644743589 | 0.80346534 | -0.802453518 |
| Cml5 | 13.56449452 | -0.113122689 | 0.367700605 | -0.307648907 |
| Nat8 | 14.92973809 | -0.768681759 | 0.322725233 | -2.381845855 |
| Gm9769 | 437.2277849 | -0.433490814 | 0.084566196 | -5.126053142 |
| Cml2 | 0.711965957 | 0.930256898 | 1.309065096 | 0.71062692 |
| RP23-448G18.6 | 0 | NA | NA | NA |
| 1700019G17Rik | 48.23925206 | -0.576041182 | 0.200564075 | -2.872105499 |
| Cml1 | 59.08754969 | -0.151958224 | 0.18629149 | -0.815701371 |
| Tprkb | 600.9053318 | -0.061262592 | 0.139026061 | -0.440655452 |
| RP23-345O17.4 | 0 | NA | NA | NA |
| Nat8b | 0.082913145 | -0.443821212 | 3.724945049 | -0.119148392 |
| RP23-345O17.6 | 0.298750603 | 1.015420725 | 1.761006005 | 0.576614005 |
| Dusp11 | 1203.32624 | -0.340570103 | 0.122768321 | -2.774087801 |
| RP23-345O17.5 | 10.75741223 | -0.035798091 | 0.365212401 | -0.098019922 |
| RP23-345O17.7 | 10.85736135 | -0.122780594 | 0.332793008 | -0.368939825 |
| Figla | 0.579661142 | 2.055291281 | 1.384401906 | 1.484605931 |
| Add2 | 1996.526052 | 0.488813366 | 0.115401111 | 4.235776951 |
| Tgfa | 554.9848714 | -0.190533706 | 0.092146422 | -2.067727675 |
| Gm7545 | 0.358406704 | -0.935500506 | 1.733767171 | -0.539576779 |
| Gm10443 | 50.80569197 | 0.187785816 | 0.232716262 | 0.806930356 |
| Fam136a | 140.0252418 | -0.297771291 | 0.157518216 | -1.890392736 |
| Snrpg | 133.4774109 | -0.25161111 | 0.103127199 | -2.439813285 |
| Pcyox1 | 1206.150805 | 0.387415994 | 0.058253733 | 6.650492138 |
| Tia1 | 1452.959773 | 0.274098836 | 0.126838854 | 2.161000569 |
| C87436 | 200.9232409 | 0.060434828 | 0.11802706 | 0.512042131 |
| A430078I02Rik | 3.729996922 | -0.07603161 | 0.596145353 | -0.127538712 |

| | | | | |
|---|---|---|---|---|
| 2310040G24Rik | 5.093180958 | -0.145830894 | 0.449526571 | -0.324409953 |
| Gm28720 | 0.502304703 | 0.503107336 | 1.326888889 | 0.379163124 |
| Gm28719 | 0.796860523 | -1.766695587 | 1.474363903 | -1.19827648 |
| Pcbp1 | 1630.30914 | -0.381438811 | 0.082931388 | -4.599450476 |
| 1600020E01Rik | 80.68127648 | -0.197450709 | 0.157242124 | -1.255711278 |
| Gm22184 | 0 | NA | NA | NA |
| Asprv1 | 17.06775386 | -0.166492902 | 0.27916734 | -0.596391046 |
| Mxd1 | 914.3853629 | -1.26441739 | 0.188922137 | -6.692796364 |
| Snrnp27 | 334.3660018 | -0.137520772 | 0.110720021 | -1.242058759 |
| Gmcl1 | 364.6549906 | 0.195537127 | 0.094520054 | 2.068736936 |
| Anxa4 | 269.2860603 | -1.501027866 | 0.222986497 | -6.731474252 |
| 2610306M01Rik | 36.58485778 | -0.203377179 | 0.187001362 | -1.087570572 |
| Aak1 | 5511.101378 | 0.289166581 | 0.077542994 | 3.729112935 |
| Nfu1 | 197.0210827 | -0.05644045 | 0.092087122 | -0.612902748 |
| Gfpt1 | 1366.619225 | -0.267805323 | 0.054920387 | -4.876246127 |
| D6Ertd527e | 0.429069343 | 1.259090525 | 1.492671403 | 0.84351487 |
| Antxr1 | 334.4251876 | -0.128601796 | 0.122321922 | -1.051338911 |
| n-R5s164 | 0 | NA | NA | NA |
| Gkn1 | 0.539210579 | -2.254398911 | 1.366772384 | -1.649432588 |
| Gkn2 | 0 | NA | NA | NA |
| Gkn3 | 2.877452801 | -1.679664197 | 1.15902072 | -1.44920981 |
| Bmp10 | 1.435473298 | -1.660753486 | 0.817745912 | -2.030891823 |
| Arhgap25 | 146.1837619 | -0.089317235 | 0.147776101 | -0.604409202 |
| Prokr1 | 11.83858572 | -1.741841738 | 0.400967007 | -4.3441024 |
| Aplf | 74.79263325 | -0.591640811 | 0.156723825 | -3.775053407 |
| E230015B07Rik | 2.634345682 | 0.196507955 | 0.650554892 | 0.302062067 |
| 1810020O05Rik | 22.22824979 | 1.111609259 | 0.264197548 | 4.207492718 |
| Efcc1 | 88.67690409 | 0.620820047 | 0.19098122 | 3.250686357 |
| Gm5879 | 12.16014049 | 0.213783786 | 0.303481241 | 0.704438222 |
| Gp9 | 0.777280298 | 0.352979029 | 1.22959372 | 0.287069642 |
| Rab43 | 434.7988563 | -0.237546374 | 0.083126402 | -2.857652536 |
| Isy1 | 190.4889355 | 0.111500765 | 0.094953839 | 1.174262835 |
| Cnbp | 2353.702497 | -0.071203422 | 0.105992319 | -0.671779077 |
| Copg1 | 2888.377927 | 0.168420603 | 0.032358278 | 5.20486919 |
| Gm26636 | 0.2969049 | 2.044157543 | 1.906817054 | 1.072026044 |
| Hmces | 147.1776525 | -0.478056821 | 0.13945211 | -3.428107469 |
| Rps19-ps9 | 0 | NA | NA | NA |
| H1fx | 161.3903661 | 0.453608118 | 0.124610159 | 3.640217781 |
| Gm5577 | 38.22855657 | 0.640353136 | 0.287502292 | 2.227297496 |
| AC153872.1 | 0.355821277 | -0.374937932 | 1.446370001 | -0.259226845 |
| Rab7 | 2929.399181 | -0.186345836 | 0.07699869 | -2.420116955 |
| Rpn1 | 1428.832432 | -0.320911771 | 0.069939409 | -4.588425561 |
| Mir6376 | 0 | NA | NA | NA |
| Gata2 | 41.94893332 | -0.260402137 | 0.222500349 | -1.170344845 |
| Dnajb8 | 0 | NA | NA | NA |
| 1700031F10Rik | 0.520495925 | 1.142721129 | 1.522404196 | 0.750602982 |
| Eefsec | 187.7376768 | -0.114993191 | 0.105290056 | -1.09215623 |

| | | | | |
|---|---|---|---|---|
| Gm25178 | 0 | NA | NA | NA |
| Ruvbl1 | 250.3254379 | -0.256099638 | 0.098363875 | -2.603594456 |
| Sec61a1 | 1211.826051 | -0.65737771 | 0.101355617 | -6.485853773 |
| Kbtbd12 | 4.967738157 | -0.636605421 | 0.548789044 | -1.16001846 |
| Gm26588 | 16.08933923 | 0.555563443 | 0.39039308 | 1.423087323 |
| Mgll | 1645.494617 | -0.010069901 | 0.090405525 | -0.111385904 |
| Abtb1 | 225.610541 | 0.314032662 | 0.091536266 | 3.430691188 |
| Podxl2 | 1494.724341 | 0.089535163 | 0.083307333 | 1.074757284 |
| Gm15612 | 6.463977267 | -0.038183705 | 0.497122906 | -0.076809386 |
| Mcm2 | 112.1005328 | -0.568215988 | 0.22079026 | -2.5735555 |
| Tpra1 | 240.1461408 | -0.144222345 | 0.120335426 | -1.198502806 |
| 4933427D06Rik | 0.766530834 | 0.87249779 | 1.111318218 | 0.785101671 |
| Gm1965 | 0.219739983 | -1.796767288 | 2.555742498 | -0.703031424 |
| Gm6507 | 0 | NA | NA | NA |
| Gm839 | 0.209172349 | -0.950369329 | 2.585388303 | -0.367592492 |
| Gm25961 | 0.088534442 | -0.905414734 | 3.723500899 | -0.243162217 |
| Gm26811 | 0.627488786 | 0.006190165 | 1.388661138 | 0.004457649 |
| Plxna1 | 1720.837352 | 0.021841293 | 0.07097516 | 0.307731509 |
| Chchd6 | 271.9148547 | 0.072002628 | 0.103033326 | 0.698828532 |
| Txnrd3 | 60.18168024 | 0.265023627 | 0.186031209 | 1.42461917 |
| Vmn1r-ps25 | 0 | NA | NA | NA |
| Vmn1r40 | 0.046345709 | -0.443818988 | 3.724945049 | -0.119147795 |
| Vmn1r41 | 0 | NA | NA | NA |
| Vmn1r-ps26 | 0 | NA | NA | NA |
| Vmn1r-ps27 | 0 | NA | NA | NA |
| Vmn1r42 | 0 | NA | NA | NA |
| Vmn1r43 | 0.049553047 | -0.443818988 | 3.724945049 | -0.119147795 |
| Vmn1r44 | 0 | NA | NA | NA |
| Vmn1r-ps28 | 0 | NA | NA | NA |
| Vmn1r45 | 0 | NA | NA | NA |
| Vmn1r46 | 0 | NA | NA | NA |
| Vmn1r-ps29 | 0 | NA | NA | NA |
| Vmn1r47 | 0 | NA | NA | NA |
| Vmn1r48 | 0 | NA | NA | NA |
| Vmn1r-ps31 | 0 | NA | NA | NA |
| Gm20534 | 0 | NA | NA | NA |
| Vmn1r49 | 0 | NA | NA | NA |
| Vmn1r50 | 0 | NA | NA | NA |
| Vmn1r51 | 0.060174537 | 0.517957688 | 3.724945049 | 0.139051095 |
| Vmn1r-ps32 | 0 | NA | NA | NA |
| Vmn1r52 | 0 | NA | NA | NA |
| V1ra8 | 0 | NA | NA | NA |
| Vmn1r53 | 0.265601376 | 0.037076929 | 1.822732534 | 0.020341399 |
| Gm20426 | 1.074962736 | 0.64953133 | 0.908116275 | 0.715251283 |
| Gm4287 | 0.044544648 | 0.517957688 | 3.724945049 | 0.139051095 |
| Vmn1r-ps34 | 0 | NA | NA | NA |
| Vmn1r54 | 0 | NA | NA | NA |

| | | | | |
|---|---|---|---|---|
| BC048671 | 0 | NA | NA | NA |
| Chst13 | 0.497746636 | -0.439293839 | 1.328785741 | -0.330597948 |
| Uroc1 | 0.105143817 | 0.517955464 | 3.724945049 | 0.139050498 |
| Zxdc | 516.6253474 | 0.048603549 | 0.087678802 | 0.554336366 |
| Ccdc37 | 25.13015435 | 0.67849932 | 0.245497018 | 2.763778258 |
| Klf15 | 358.4959565 | 0.504141036 | 0.135458921 | 3.721726343 |
| Aldh1l1 | 1458.014872 | -0.478077944 | 0.110188728 | -4.338719153 |
| Gm15756 | 0.143022002 | 0.03610674 | 3.547735707 | 0.010177404 |
| Slc41a3 | 95.87477886 | 0.004588813 | 0.144216478 | 0.031818922 |
| Iqsec1 | 4998.243591 | 0.2551846 | 0.064398915 | 3.962560532 |
| Gm24816 | 4.887144399 | 0.314142228 | 0.687520197 | 0.456920726 |
| Rpl21-ps12 | 7.418190298 | 0.0389627 | 0.508932242 | 0.076557735 |
| Gm25185 | 0 | NA | NA | NA |
| Ppp1r2-ps2 | 19.04760772 | 0.093047322 | 0.290435825 | 0.320371368 |
| Nup210 | 552.2987281 | 0.037987695 | 0.087660146 | 0.433351941 |
| Hdac11 | 1873.279407 | 0.594431051 | 0.06784343 | 8.761807137 |
| Gm14573 | 0.08169834 | -0.443821212 | 3.724945049 | -0.119148392 |
| Fbln2 | 117.7950445 | -1.195834226 | 0.244306495 | -4.894811444 |
| Wnt7a | 199.3147353 | -0.571439751 | 0.18777906 | -3.043149486 |
| Gm4575 | 4.420079801 | 0.127724879 | 0.511122595 | 0.249890888 |
| 4930471M09Rik | 1.669552124 | -0.428401285 | 0.763129037 | -0.561374636 |
| 1810044D09Rik | 5.670729865 | 0.244793472 | 0.423909015 | 0.577467011 |
| Chchd4 | 184.8478106 | -0.126713918 | 0.098779466 | -1.282796141 |
| Tmem43 | 388.1633489 | -0.518588145 | 0.088192196 | -5.880204486 |
| Xpc | 207.6063262 | 0.027724686 | 0.10929442 | 0.25366973 |
| Lsm3 | 147.4136989 | 0.049799808 | 0.109018622 | 0.456800929 |
| Slc6a6 | 1683.442733 | 0.085762413 | 0.074414446 | 1.152496833 |
| Grip2 | 311.1559795 | -0.161529098 | 0.113574551 | -1.422229687 |
| Ccdc174 | 303.9255113 | 0.113145749 | 0.111999192 | 1.01023719 |
| 4930590J08Rik | 9.858077027 | -0.171235981 | 0.393775111 | -0.434857299 |
| Fgd5 | 79.82046982 | -0.423248596 | 0.173601919 | -2.438040997 |
| 4930466I24Rik | 0.042716945 | 0.037062677 | 3.724945049 | 0.009949859 |
| Nr2c2 | 2086.98739 | 0.322168178 | 0.057590276 | 5.594141922 |
| Mrps25 | 334.8028407 | -0.013653854 | 0.075288407 | -0.181354007 |
| Rbsn | 899.6634951 | 0.002932406 | 0.091128301 | 0.032178876 |
| Trh | 83.76368859 | -1.202715966 | 0.409528781 | -2.936828917 |
| Prickle2 | 2847.184781 | 0.602501731 | 0.107047922 | 5.628336542 |
| Gm17797 | 0.118974955 | 0.51795324 | 3.724945049 | 0.139049901 |
| Adamts9 | 272.0878833 | -0.169198555 | 0.195720024 | -0.864492817 |
| A730049H05Rik | 9.582230866 | -1.035126154 | 0.403255213 | -2.566925657 |
| Gm15737 | 0 | NA | NA | NA |
| 9530026P05Rik | 37.55723895 | 0.649983826 | 0.214699683 | 3.027409336 |
| Gm5313 | 0 | NA | NA | NA |
| Gm25094 | 0 | NA | NA | NA |
| Gm22312 | 0 | NA | NA | NA |
| Magi1 | 1204.640072 | 0.317439562 | 0.071389456 | 4.446588867 |
| Gm23035 | 0 | NA | NA | NA |

| | | | | |
|---|---|---|---|---|
| Gm22840 | 0 | NA | NA | NA |
| Slc25a26 | 101.4816024 | 0.253287063 | 0.12489364 | 2.028022117 |
| Lrig1 | 1206.313659 | -0.304011308 | 0.124707662 | -2.437791736 |
| Mir7041 | 0.248387756 | 0.037077865 | 1.841821919 | 0.02013108 |
| Gm7833 | 0.121388914 | 0.037071574 | 3.724945049 | 0.009952247 |
| Gm23364 | 0 | NA | NA | NA |
| Gm24601 | 0 | NA | NA | NA |
| Gm26501 | 0 | NA | NA | NA |
| Kbtbd8os | 0.511579953 | 1.38495747 | 1.271470668 | 1.089256327 |
| Kbtbd8 | 179.6004661 | -0.21691144 | 0.126639266 | -1.712829253 |
| AY512915 | 0.311830279 | 1.393323908 | 1.959330409 | 0.711122484 |
| Gm10234 | 0 | NA | NA | NA |
| Suclg2 | 141.9514173 | 0.25946043 | 0.151437818 | 1.713313316 |
| Fam19a1 | 272.9938384 | 0.507396183 | 0.143533 | 3.535048969 |
| 1700123L14Rik | 0.257669488 | -0.91619134 | 2.118994881 | -0.432370719 |
| Gm22971 | 0 | NA | NA | NA |
| Gm26011 | 0 | NA | NA | NA |
| Fam19a4 | 11.35413066 | 0.111791827 | 0.317259355 | 0.352367316 |
| Eogt | 164.467491 | 0.128448304 | 0.132003936 | 0.973064197 |
| Tmf1 | 950.0693584 | 0.103266915 | 0.068387641 | 1.510023048 |
| Gm26175 | 2.7824929 | -0.574214222 | 0.669204657 | -0.85805473 |
| Uba3 | 755.4137509 | -0.108232099 | 0.088211216 | -1.226965277 |
| Arl6ip5 | 1177.848838 | -0.147279259 | 0.063476479 | -2.32021784 |
| Lmod3 | 0.289573199 | 0.561202176 | 2.06116879 | 0.27227376 |
| Frmd4b | 362.2741639 | -0.164513366 | 0.088305814 | -1.862995855 |
| Gm25852 | 1.201575708 | -0.342532436 | 0.911517762 | -0.375782514 |
| Gm23510 | 0.586937805 | 1.923052727 | 1.645983806 | 1.168330284 |
| Gm24565 | 0 | NA | NA | NA |
| Gm23703 | 0 | NA | NA | NA |
| Mitf | 247.2340933 | -0.398007665 | 0.113680579 | -3.501105183 |
| Gm15531 | 0.049553047 | -0.443818988 | 3.724945049 | -0.119147795 |
| Gm765 | 8.927308765 | 0.887335008 | 0.403839727 | 2.197245465 |
| Foxp1 | 1512.202388 | 0.590380816 | 0.123903577 | 4.764840788 |
| AC155839.1 | 0.110870145 | -0.444802825 | 3.724874622 | -0.119414174 |
| Gm20696 | 52.11740566 | 0.355433666 | 0.27461996 | 1.294274697 |
| Gm20705 | 26.36886051 | 0.308458644 | 0.394163619 | 0.782564978 |
| Gm22328 | 2.778696684 | -0.211830635 | 0.679901438 | -0.311560799 |
| Gm24936 | 0 | NA | NA | NA |
| Eif4e3 | 223.9200148 | -0.262188187 | 0.090250365 | -2.905120525 |
| Gpr27 | 69.76717276 | 0.603911397 | 0.198073963 | 3.048918628 |
| Prok2 | 9.10031652 | -1.142949395 | 0.420191764 | -2.72006615 |
| Gm26748 | 2.205804803 | 0.977321582 | 0.769842283 | 1.269508838 |
| Tpt1-ps3 | 354.4684848 | -0.626785721 | 0.166294876 | -3.769122272 |
| RP23-191I2.1 | 0 | NA | NA | NA |
| Gm24248 | 0 | NA | NA | NA |
| Rybp | 419.1411725 | -0.172036352 | 0.082499097 | -2.085311952 |
| Gm23234 | 0.066050875 | -0.443818988 | 3.724945049 | -0.119147795 |

| | | | | |
|---|---|---|---|---|
| 1700049E22Rik | 0 | NA | NA | NA |
| Shq1 | 52.12462995 | -0.288305049 | 0.147937061 | -1.948835861 |
| Gm6565 | 46.92634586 | 0.19437419 | 0.184127046 | 1.055652573 |
| Gxylt2 | 11.76095793 | -0.404564051 | 0.369477893 | -1.094961455 |
| Gm15576 | 0.092881282 | 0.037060453 | 3.724945049 | 0.009949262 |
| Ppp4r2 | 799.647895 | -0.27186146 | 0.080122456 | -3.393074485 |
| Pdzrn3 | 253.9254666 | 0.474231412 | 0.131072351 | 3.618088867 |
| Gm26911 | 0.672570739 | 2.15781533 | 1.360845468 | 1.585643175 |
| Gm9871 | 0.521928645 | 1.856831496 | 1.47473546 | 1.259094629 |
| Cntn3 | 401.4213994 | 0.448207542 | 0.14243984 | 3.146644527 |
| Mir6373 | 0 | NA | NA | NA |
| Gm26022 | 0 | NA | NA | NA |
| Chl1 | 3209.185577 | 0.178272764 | 0.075989361 | 2.34602268 |
| Gm24784 | 4.963107029 | 0.106199732 | 0.589806693 | 0.18005854 |
| Gm25656 | 0 | NA | NA | NA |
| RP24-151A23.1 | 0 | NA | NA | NA |
| Cntn6 | 392.3293275 | 0.299999223 | 0.112642624 | 2.663283318 |
| Gm23339 | 0 | NA | NA | NA |
| Gm25570 | 0 | NA | NA | NA |
| Gm15631 | 17.2682059 | -0.185897544 | 0.375932759 | -0.494496794 |
| Cntn4 | 819.1636369 | 0.39084804 | 0.073906195 | 5.28843408 |
| Il5ra | 0.538620109 | -1.202960677 | 1.264009892 | -0.951701949 |
| Trnt1 | 529.4811257 | -0.04826568 | 0.076438705 | -0.631429852 |
| Crbn | 1050.434189 | 0.234471044 | 0.089685859 | 2.614359133 |
| Gm22418 | 0 | NA | NA | NA |
| Gm27690 | 0 | NA | NA | NA |
| Lrrn1 | 1083.285535 | 0.221722967 | 0.076400518 | 2.902113388 |
| Setmar | 60.36555304 | 0.28268242 | 0.137909503 | 2.049767519 |
| Sumf1 | 349.8000208 | -0.326746277 | 0.088132564 | -3.707440962 |
| Gm4604 | 20.23814071 | -0.082386801 | 0.246298796 | -0.334499406 |
| Itpr1 | 13527.53245 | 0.648041139 | 0.165191571 | 3.92296735 |
| Gm27813 | 4.625054459 | 0.768211424 | 0.538372198 | 1.426915109 |
| mmu-mir-7661 | 0.67439107 | -1.264063881 | 1.294454 | -0.976522828 |
| Gm16433 | 10.96864868 | -0.380513426 | 0.332497345 | -1.144410421 |
| 0610040F04Rik | 4.191962974 | 0.808294393 | 0.541972178 | 1.491394625 |
| Gm38167 | 1.527506968 | -0.342929758 | 1.011961932 | -0.338876145 |
| Gm17055 | 0.461373441 | -0.635466603 | 1.760010856 | -0.361058343 |
| Bhlhe40 | 1442.70058 | -0.006658684 | 0.130063628 | -0.051195586 |
| Arl8b | 2174.485167 | -0.038997859 | 0.055423299 | -0.703636549 |
| Edem1 | 637.9057285 | -0.516944976 | 0.134725392 | -3.837027073 |
| Gm20387 | 22.36903969 | 0.762234861 | 0.262084302 | 2.908357558 |
| Grm7 | 1035.879049 | 0.322273357 | 0.144495929 | 2.230328288 |
| Gm20386 | 13.98619218 | 0.148296001 | 0.348708516 | 0.425272096 |
| Gm22093 | 0.0878635 | 0.037058229 | 3.724945049 | 0.009948665 |
| Gm24005 | 0 | NA | NA | NA |
| Gm26362 | 0 | NA | NA | NA |
| Gm25454 | 0 | NA | NA | NA |

| | | | | |
|---|---|---|---|---|
| 1700054K19Rik | 0 | NA | NA | NA |
| n-R5s165 | 0 | NA | NA | NA |
| Lmcd1 | 50.45369243 | -0.282746217 | 0.214774978 | -1.316476524 |
| 5031434C07Rik | 0.668421634 | 0.466384383 | 1.334686352 | 0.349433695 |
| Ssu2 | 1.179082066 | 1.365345759 | 0.886528421 | 1.540103765 |
| Cav3 | 2.338446252 | -1.159114733 | 0.748947459 | -1.547658276 |
| Oxtr | 56.02683163 | 0.371923884 | 0.152925561 | 2.432058321 |
| Gm27944 | 0 | NA | NA | NA |
| Gm15519 | 37.45729361 | -0.017631039 | 0.17234948 | -0.102298184 |
| Gm23002 | 0.075043205 | 0.517957688 | 3.724945049 | 0.139051095 |
| Gm5578 | 12.94208794 | -0.570498973 | 0.339804693 | -1.678902573 |
| Rad18 | 111.1939575 | 0.598790077 | 0.1757078 | 3.407874185 |
| Gm26799 | 2.930040241 | 0.824480639 | 0.602460464 | 1.368522397 |
| Srgap3 | 4773.252643 | 0.257531872 | 0.072133463 | 3.570213624 |
| Gm6134 | 0.068097359 | 0.037064902 | 3.724945049 | 0.009950456 |
| Gm23244 | 0 | NA | NA | NA |
| Thumpd3 | 249.1100717 | -0.403159502 | 0.086073747 | -4.683884633 |
| Gm22591 | 0.068097359 | 0.037064902 | 3.724945049 | 0.009950456 |
| Gt(ROSA)26Sor | 121.8807241 | -0.309163511 | 0.140969721 | -2.19311998 |
| Setd5 | 2417.574525 | 0.117661007 | 0.073036031 | 1.610999467 |
| Lhfpl4 | 1265.24945 | 0.295094993 | 0.080375944 | 3.671434246 |
| Mtmr14 | 178.5916463 | -0.030158612 | 0.085993426 | -0.350708344 |
| Gm16161 | 0.217535639 | -1.056487397 | 3.177269635 | -0.33251424 |
| Cpne9 | 367.3936013 | 0.702187251 | 0.133686183 | 5.252504311 |
| Brpf1 | 657.1446642 | 0.029778424 | 0.116088413 | 0.256515037 |
| Gm15492 | 1.140702544 | 1.36830023 | 1.17533456 | 1.164179355 |
| Ogg1 | 60.44940537 | -0.385471282 | 0.178167336 | -2.163535081 |
| Camk1 | 512.5663802 | -0.238955995 | 0.092978238 | -2.570020689 |
| Tada3 | 319.0164571 | -0.087356696 | 0.074921083 | -1.165982831 |
| Arpc4 | 1219.326062 | -0.114414205 | 0.082665753 | -1.384058109 |
| Ttll3 | 119.1716164 | 0.055811232 | 0.200421076 | 0.278469873 |
| Rpusd3 | 80.45315769 | 0.290192719 | 0.154731371 | 1.875461437 |
| Gm24387 | 0.061920613 | -0.443816764 | 3.724945049 | -0.119147198 |
| Cidec | 0.392922208 | -0.885416733 | 1.677921112 | -0.527686747 |
| Jagn1 | 177.5032672 | -0.404721375 | 0.141996029 | -2.850230236 |
| Il17re | 7.25222539 | -0.382320058 | 0.482928972 | -0.791669333 |
| Il17rc | 69.78172622 | -0.80966701 | 0.213536638 | -3.791700661 |
| Creld1 | 601.1655518 | -0.032170373 | 0.070671432 | -0.455210435 |
| Prrt3 | 474.5330712 | 0.486099996 | 0.118145719 | 4.114410587 |
| Emc3 | 603.6486607 | -0.27351673 | 0.081062961 | -3.374127035 |
| Gm23365 | 0.654623538 | -0.292848661 | 1.255123162 | -0.233322649 |
| Fancd2 | 38.49030515 | 0.148504676 | 0.174867875 | 0.849239321 |
| Fancd2os | 0.484250087 | -1.56369906 | 1.373764387 | -1.138258551 |
| Brk1 | 853.6369896 | -0.078838605 | 0.06389905 | -1.233799341 |
| Vhl | 414.2383988 | -0.138730526 | 0.086996216 | -1.594673107 |
| 1700015O11Rik | 0.140088945 | 0.037059971 | 3.605664301 | 0.010278264 |
| Irak2 | 312.5390426 | -0.030959231 | 0.145932647 | -0.212147397 |

| | | | | |
|---|---|---|---|---|
| Tatdn2 | 686.0036907 | -0.224426936 | 0.101651022 | -2.207817802 |
| mmu-mir-7042 | 1.150390536 | -1.268407689 | 1.012731856 | -1.252461529 |
| Gm26982 | 73.25146826 | 0.767544114 | 0.223758897 | 3.430228351 |
| Ghrl | 1.379464875 | -0.312214723 | 0.843588007 | -0.370103321 |
| Sec13 | 453.3719348 | -0.510022818 | 0.083693056 | -6.093968129 |
| Atp2b2 | 10812.54371 | 0.704545035 | 0.114799844 | 6.13716021 |
| Gm15082 | 0 | NA | NA | NA |
| Gm15083 | 4.119074061 | 0.631563813 | 0.521862987 | 1.210210014 |
| Slc6a11 | 2799.545472 | 0.662761243 | 0.160605695 | 4.126636012 |
| Slc6a1 | 4290.118131 | 0.606565936 | 0.085944473 | 7.057649112 |
| Hrh1 | 148.4312325 | -0.436850081 | 0.128471315 | -3.400370601 |
| Atg7 | 275.6563131 | -0.124425679 | 0.096465716 | -1.289843527 |
| Vgll4 | 362.9008528 | -0.439964582 | 0.095630665 | -4.600664278 |
| Tamm41 | 94.56214301 | 0.352515148 | 0.154343775 | 2.283960902 |
| Gm24767 | 0 | NA | NA | NA |
| Gm17733 | 5.222249821 | -0.74283379 | 0.471700904 | -1.574798317 |
| Syn2 | 4179.711114 | 0.281198918 | 0.087952381 | 3.197172328 |
| Gm17482 | 8.270587696 | 0.690925101 | 0.376045601 | 1.837343927 |
| Timp4 | 415.701942 | -0.092002815 | 0.201364341 | -0.456897258 |
| Pparg | 54.34159874 | -0.446980585 | 0.211707678 | -2.111310223 |
| Tsen2 | 155.5268573 | -0.139816154 | 0.116904352 | -1.195987587 |
| Mkrn2os | 0.394336369 | 1.223008315 | 1.497292882 | 0.816813016 |
| Mkrn2 | 337.3089779 | 0.086942702 | 0.103998102 | 0.836002775 |
| Raf1 | 1017.633135 | -0.105722031 | 0.091053624 | -1.161096354 |
| Gm14335 | 18.11564472 | 0.269889932 | 0.367567202 | 0.734260103 |
| Gm24008 | 0 | NA | NA | NA |
| Tmem40 | 16.46128129 | -2.349307198 | 0.460397472 | -5.102780403 |
| Gm27853 | 0 | NA | NA | NA |
| Cand2 | 86.32620824 | -0.461026534 | 0.167035753 | -2.760047034 |
| Mir7660 | 0.118974955 | 0.51795324 | 3.724945049 | 0.139049901 |
| Rpl32 | 874.900214 | -0.26009442 | 0.142522028 | -1.824941896 |
| Snora7a | 4.44476562 | 0.266265243 | 0.575384067 | 0.46276089 |
| Efcab12 | 50.83190883 | -0.012930934 | 0.244145783 | -0.052963988 |
| Mbd4 | 111.5763811 | 0.323122238 | 0.179155572 | 1.803584638 |
| Ift122 | 349.0430765 | -0.034394109 | 0.11344985 | -0.303165754 |
| Rho | 4.082856454 | -2.122638948 | 0.653662769 | -3.247299755 |
| H1foo | 0.271840384 | 0.420491273 | 2.036217528 | 0.206506067 |
| Plxnd1 | 1058.405309 | 0.521824583 | 0.09401344 | 5.550531728 |
| Tmcc1 | 1022.727567 | 0.113758308 | 0.056287801 | 2.021011779 |
| Gm20404 | 1.065143116 | -0.146608548 | 0.950042708 | -0.15431785 |
| Gm8203 | 110.3300294 | -0.014841361 | 0.136411194 | -0.108798702 |
| Gm8213 | 4.570295035 | 0.045689966 | 0.470969073 | 0.097012666 |
| Fam21 | 1941.589181 | 0.02071379 | 0.083023525 | 0.249493016 |
| Zfand4 | 120.8501776 | 0.467124144 | 0.122594994 | 3.810303585 |
| 8-Mar | 943.7508335 | -0.137915416 | 0.080049826 | -1.722869656 |
| Gm23527 | 0 | NA | NA | NA |
| Alox5 | 9.062762248 | 0.947831853 | 0.39129958 | 2.422266469 |

| | | | | |
|---|---|---|---|---|
| Olfr211 | 0.287786674 | -0.982516562 | 1.999459422 | -0.491391099 |
| Olfr212 | 0.265357793 | -0.869676833 | 1.797908956 | -0.483715724 |
| Olfr213 | 0.046345709 | -0.443818988 | 3.724945049 | -0.119147795 |
| Gm5580 | 0.092493346 | 0.517955464 | 3.724945049 | 0.139050498 |
| Olfr214 | 0 | NA | NA | NA |
| Olfr215 | 0 | NA | NA | NA |
| Zfp422 | 293.8653895 | -0.181436288 | 0.070143631 | -2.586639513 |
| Rassf4 | 281.0708732 | -0.469498486 | 0.153462923 | -3.05936104 |
| Mir7043 | 0 | NA | NA | NA |
| 8430408G22Rik | 5.938959585 | 0.229598775 | 0.550612779 | 0.416987734 |
| Tmem72 | 34.03285026 | 0.390088238 | 0.249350501 | 1.564417297 |
| Gm22882 | 0 | NA | NA | NA |
| Cxcl12 | 287.0988006 | 0.197179921 | 0.122412893 | 1.610777388 |
| Rpl28-ps4 | 0 | NA | NA | NA |
| Gm9946 | 0.160978641 | 0.03706851 | 2.803723868 | 0.01322117 |
| 1700030F04Rik | 1.580617114 | -0.275132169 | 0.996288421 | -0.276157148 |
| Gm7292 | 1492.722712 | 0.491234105 | 0.076040296 | 6.460181388 |
| Gm6253 | 0.669418093 | -0.236256679 | 1.487806722 | -0.158795276 |
| Zfp637 | 220.273934 | -0.080910309 | 0.087415326 | -0.925584938 |
| Zfp239 | 252.8739187 | 0.335577814 | 0.115239436 | 2.912004997 |
| Gm29509 | 0.066050875 | -0.443818988 | 3.724945049 | -0.119147795 |
| Hnrnpf | 625.6970897 | -0.675692619 | 0.116041106 | -5.822872959 |
| 4933440N22Rik | 4.749668983 | 0.130445151 | 0.412997437 | 0.315849783 |
| Fxyd4 | 0.432778942 | -1.225742088 | 1.412434388 | -0.86782232 |
| Gm25535 | 0.060174537 | 0.517957688 | 3.724945049 | 0.139051095 |
| Gm4875 | 25.05432516 | -0.373538953 | 0.21492051 | -1.738033065 |
| Rasgef1a | 1668.685123 | 0.620796518 | 0.108090664 | 5.743294542 |
| Mir7044 | 0.497274054 | 0.59718595 | 1.33700138 | 0.446660683 |
| Csgalnact2 | 271.8412438 | -0.050313023 | 0.07628433 | -0.659545982 |
| Gm23908 | 2.493933935 | 0.3809645 | 0.689650037 | 0.552402638 |
| Ret | 333.8009791 | -0.315406158 | 0.149624234 | -2.107988463 |
| Bms1 | 452.1647011 | -0.416508723 | 0.119625211 | -3.481780482 |
| Zfp248 | 186.5369703 | 0.051534182 | 0.106172568 | 0.485381322 |
| Gm25953 | 0 | NA | NA | NA |
| Zfp9 | 323.1489127 | 0.083236666 | 0.081166539 | 1.025504678 |
| Ankrd26 | 319.7692467 | -0.200911585 | 0.176891652 | -1.135788957 |
| Cacna1c | 2127.985368 | 0.402868531 | 0.124609924 | 3.233037278 |
| Gm18981 | 18.01703932 | 0.472578264 | 0.425464228 | 1.1107356 |
| Gm25905 | 0 | NA | NA | NA |
| Dcp1b | 230.6682507 | 0.1146457 | 0.121125888 | 0.946500387 |
| Cacna2d4 | 45.15394803 | 0.268831009 | 0.168570842 | 1.594765773 |
| Lrtm2 | 701.9863131 | 0.256463131 | 0.072931856 | 3.516476129 |
| Adipor2 | 1326.364367 | -0.33538844 | 0.110438113 | -3.036890342 |
| Wnt5b | 23.55176808 | -0.753155153 | 0.238565766 | -3.157012698 |
| Fbxl14 | 457.0418174 | -0.121242342 | 0.091244839 | -1.328758356 |
| Erc1 | 1392.520042 | -0.026657499 | 0.084378359 | -0.315928149 |
| 3110021A11Rik | 3.049669005 | 0.247475163 | 0.659906061 | 0.375015744 |

| | | | | |
|---|---|---|---|---|
| Gm15532 | 0 | NA | NA | NA |
| 4930540M05Rik | 1.752268337 | -0.199479428 | 0.708328327 | -0.281620006 |
| Rad52 | 228.0315541 | 0.030711485 | 0.095403265 | 0.321912305 |
| Wnk1 | 6814.715123 | -0.262375781 | 0.062925329 | -4.169637003 |
| Gm16199 | 3.97190282 | 0.779175019 | 0.533255979 | 1.461165086 |
| Mir706 | 1.782601153 | -0.047532783 | 0.738553533 | -0.064359293 |
| Ninj2 | 7.372786058 | -0.19450604 | 0.393795134 | -0.493926976 |
| Gm25327 | 0 | NA | NA | NA |
| B4galnt3 | 7.654890081 | 0.360889317 | 0.418339463 | 0.862670986 |
| D330020A13Rik | 0 | NA | NA | NA |
| Ccdc77 | 82.48745712 | -0.254137968 | 0.180561284 | -1.407488703 |
| Kdm5a | 1440.026259 | 0.06794046 | 0.084406862 | 0.80491632 |
| Gm24592 | 0 | NA | NA | NA |
| Il17ra | 446.1211285 | -0.508096998 | 0.184687577 | -2.751116268 |
| Cecr6 | 417.0501185 | 0.561021662 | 0.169609999 | 3.307715728 |
| Cecr5 | 216.0841903 | 0.124907578 | 0.147772995 | 0.845266609 |
| Gm20692 | 2.176719344 | 1.866415402 | 0.89151909 | 2.093522644 |
| Gm24391 | 0.075043205 | 0.517957688 | 3.724945049 | 0.139051095 |
| Cecr2 | 227.0044623 | 0.086575442 | 0.156231773 | 0.554147472 |
| Slc25a18 | 412.4554696 | 0.588251407 | 0.145087317 | 4.0544647 |
| Atp6v1e1 | 2143.722746 | 0.291177262 | 0.066785757 | 4.359870651 |
| Gm25193 | 1.884175004 | 0.244427396 | 0.884560985 | 0.276326223 |
| Bcl2l13 | 678.3631799 | -0.130109418 | 0.072449859 | -1.79585467 |
| Bid | 179.7781668 | -0.137181248 | 0.162909501 | -0.842070277 |
| Mical3 | 1830.192808 | 0.174573439 | 0.123504813 | 1.413495025 |
| Gm4651 | 0.182574094 | 0.401640784 | 2.702311993 | 0.148628576 |
| Pex26 | 225.9169415 | 0.488770822 | 0.110588823 | 4.41971265 |
| Tuba8 | 200.697007 | -0.087840737 | 0.162829872 | -0.53946328 |
| Gm15856 | 6.178163136 | -0.415337902 | 0.542425837 | -0.765704496 |
| Usp18 | 2014.636858 | -0.753793929 | 0.299532549 | -2.516567674 |
| Slc6a13 | 90.14607231 | 0.087583457 | 0.254579326 | 0.344032086 |
| Gm24855 | 0 | NA | NA | NA |
| Slc6a12 | 25.52923883 | -0.290203498 | 0.294344663 | -0.985930898 |
| Iqsec3 | 2651.332656 | -0.151342802 | 0.097482073 | -1.55251933 |
| A2m | 460.9649802 | -1.113562545 | 0.259294924 | -4.294579038 |
| Mug1 | 1.274577395 | 0.952030515 | 1.019034831 | 0.934247276 |
| Mug2 | 0.087930927 | -0.443164241 | 3.724898049 | -0.118973522 |
| Gm10319 | 0 | NA | NA | NA |
| Mug-ps1 | 0.15956642 | 0.036118649 | 3.418731032 | 0.010564929 |
| Gm27703 | 0 | NA | NA | NA |
| Klrg1 | 4.976324902 | 0.001743371 | 0.464948346 | 0.003749602 |
| Gm27826 | 0.044544648 | 0.517957688 | 3.724945049 | 0.139051095 |
| M6pr | 1445.217025 | -0.195493974 | 0.106042948 | -1.843535828 |
| Phc1 | 481.579932 | 0.024288753 | 0.084825946 | 0.286336364 |
| 1700063H04Rik | 0 | NA | NA | NA |
| Rimklb | 372.2860085 | -0.015774555 | 0.063733592 | -0.247507706 |
| Gm8430 | 34.0404713 | -0.094735812 | 0.282056229 | -0.335875623 |

| | | | | |
|---|---|---|---|---|
| Mfap5 | 3.28087796 | -0.45812803 | 0.650906148 | -0.703831162 |
| Aicda | 0 | NA | NA | NA |
| Gm16556 | 1.225054178 | -1.780838541 | 0.937644776 | -1.89926781 |
| Gm16557 | 0.258216092 | -0.837481139 | 2.032403237 | -0.412064458 |
| Gm8459 | 0.839467773 | -2.019622461 | 2.261176567 | -0.893173267 |
| Apobec1 | 484.8295906 | -1.361256991 | 0.204402989 | -6.659672645 |
| Gdf3 | 0.810901572 | 0.285720697 | 1.078081731 | 0.265026935 |
| Gm26168 | 0 | NA | NA | NA |
| Dppa3 | 0.038981395 | -0.443816764 | 3.724945049 | -0.119147198 |
| Gm10420 | 14.33523795 | -0.154349728 | 0.291683997 | -0.529167625 |
| Gm10224 | 1.550258836 | -0.05455669 | 0.730085878 | -0.074726402 |
| Nanog | 0.725797938 | -1.665504125 | 1.551383149 | -1.073560794 |
| Slc2a3 | 1520.555383 | 0.106554143 | 0.08301779 | 1.283509754 |
| AC163108.1 | 10.5610363 | -0.57843268 | 0.349492291 | -1.655065632 |
| Gm26826 | 2.946188958 | 0.92205591 | 0.583842991 | 1.579287453 |
| Foxj2 | 640.279476 | -0.007279188 | 0.065682027 | -0.110824662 |
| mmu-mir-7231 | 1.544805485 | 0.121635561 | 1.131023839 | 0.107544648 |
| C3ar1 | 498.6296836 | -1.791177198 | 0.285545342 | -6.272829327 |
| Necap1 | 1570.429647 | 0.058056452 | 0.064190986 | 0.9044331 |
| Gm5316 | 0 | NA | NA | NA |
| Clec4a1 | 94.94432573 | -1.244576913 | 0.314620232 | -3.955806995 |
| Clec4a3 | 123.1859983 | -1.662350801 | 0.420845343 | -3.950027789 |
| Clec4a4 | 1.253602477 | -2.17931153 | 0.959047534 | -2.272370714 |
| Clec4b1 | 0.237601816 | 0.517974479 | 2.11470486 | 0.244939371 |
| Clec4a2 | 17.55388936 | -0.732990477 | 0.362640376 | -2.021259978 |
| Clec4b2 | 0.180312491 | -0.891783615 | 2.689605142 | -0.331566742 |
| Gm22946 | 0 | NA | NA | NA |
| Gm5883 | 0.359325923 | 0.553536306 | 1.99954138 | 0.276831633 |
| Clec4n | 6.544164894 | -1.002111812 | 0.402369662 | -2.490525271 |
| Clec4d | 34.33899703 | -2.31544635 | 0.826915496 | -2.800100326 |
| Clec4e | 72.26271789 | -0.717394078 | 0.731281966 | -0.981008847 |
| Vmn2r19 | 0 | NA | NA | NA |
| Vmn2r20 | 0 | NA | NA | NA |
| Gm22833 | 0 | NA | NA | NA |
| Vmn2r-ps31 | 0 | NA | NA | NA |
| Vmn2r21 | 0 | NA | NA | NA |
| Gm25891 | 0 | NA | NA | NA |
| Gm22722 | 0 | NA | NA | NA |
| Vmn2r-ps32 | 0 | NA | NA | NA |
| Vmn2r22 | 0 | NA | NA | NA |
| Gm25065 | 0 | NA | NA | NA |
| Vmn2r23 | 0.060174537 | 0.517957688 | 3.724945049 | 0.139051095 |
| Gm22724 | 0 | NA | NA | NA |
| Vmn2r24 | 0 | NA | NA | NA |
| Gm23079 | 0 | NA | NA | NA |
| Vmn2r25 | 0 | NA | NA | NA |
| A330044P14Rik | 0.221456285 | -1.46956717 | 2.15816903 | -0.680932378 |

| | | | | |
|---|---|---|---|---|
| Gm22763 | 0 | NA | NA | NA |
| Vmn2r26 | 0 | NA | NA | NA |
| Gm24843 | 0.072025462 | 0.517959912 | 3.724945049 | 0.139051692 |
| Vmn2r-ps33 | 0 | NA | NA | NA |
| Gm22916 | 0 | NA | NA | NA |
| Vmn2r-ps34 | 0 | NA | NA | NA |
| Gm23554 | 0 | NA | NA | NA |
| Vmn2r27 | 0.851816096 | -0.613269187 | 0.973054444 | -0.630251669 |
| Gm25311 | 0.355991514 | 0.037077826 | 1.697519568 | 0.021842356 |
| Cd163 | 12.17190331 | 1.681921193 | 0.378363014 | 4.445257946 |
| 1700013D24Rik | 0.329404011 | -1.993596761 | 1.952033783 | -1.021292141 |
| Pex5 | 534.6967276 | -0.175430518 | 0.073496906 | -2.386910226 |
| Clstn3 | 4659.131254 | 0.229537491 | 0.061055883 | 3.759465601 |
| C1rl | 16.99094215 | -0.363836999 | 0.248354037 | -1.464993294 |
| C1ra | 434.4635122 | -0.480937831 | 0.270188994 | -1.780005262 |
| C1s1 | 241.4617169 | -0.795113951 | 0.374816561 | -2.121341566 |
| Gm16567 | 0 | NA | NA | NA |
| C1rb | 42.44241018 | -0.472363224 | 0.318358885 | -1.483744432 |
| C1s2 | 22.96225106 | -0.795368607 | 0.328981014 | -2.41767328 |
| Lpcat3 | 223.387876 | -0.181882014 | 0.097808983 | -1.859563489 |
| Emg1 | 187.4943886 | -0.374818802 | 0.097449147 | -3.846301517 |
| Phb2 | 583.7592251 | -0.189136175 | 0.07565219 | -2.50007535 |
| Gm23547 | 0.819923432 | -0.844708966 | 1.243553308 | -0.67927041 |
| Gm15884 | 0.260697633 | -0.444528694 | 2.112155272 | -0.210462128 |
| Gm29008 | 0.068097359 | 0.037064902 | 3.724945049 | 0.009950456 |
| Mir141 | 0 | NA | NA | NA |
| Mir200c | 0 | NA | NA | NA |
| Ptpn6 | 244.4682648 | -1.110112002 | 0.203247749 | -5.461866167 |
| Gm20531 | 0.660431168 | -0.364883688 | 1.148971514 | -0.317574182 |
| Grcc10 | 122.9669071 | 0.330739951 | 0.118595742 | 2.78880123 |
| Atn1 | 2476.146781 | 0.097760969 | 0.07530738 | 1.298159211 |
| Rnu7 | 0 | NA | NA | NA |
| Eno2 | 8072.573921 | 0.168125356 | 0.057569944 | 2.92036687 |
| Lrrc23 | 22.31127345 | -0.645198769 | 0.272358961 | -2.368927998 |
| Spsb2 | 56.84758583 | -1.130170779 | 0.208207616 | -5.428095299 |
| Tpi1 | 3615.771022 | 0.064186742 | 0.059607439 | 1.076824352 |
| Usp5 | 1925.798522 | 0.047010193 | 0.046131269 | 1.019052677 |
| Cdca3 | 13.51901529 | -0.544973682 | 0.418656032 | -1.301721795 |
| Gnb3 | 5.506721047 | -0.074552687 | 0.577162915 | -0.129170958 |
| P3h3 | 139.641216 | 0.050568522 | 0.185140681 | 0.273135661 |
| Gpr162 | 1473.024775 | 0.210564871 | 0.073310227 | 2.872244154 |
| Cd4 | 50.03247416 | 0.537208995 | 0.22616997 | 2.375244581 |
| Lag3 | 118.9770841 | -0.240649214 | 0.20957392 | -1.148278443 |
| Ptms | 3060.359574 | 0.318371201 | 0.079439899 | 4.007698989 |
| A230083G16Rik | 12.40694247 | 0.564614984 | 0.314923304 | 1.792865047 |
| Mlf2 | 3556.102244 | 0.083413124 | 0.061577747 | 1.354598506 |
| Cops7a | 930.3160726 | 0.136737388 | 0.068565398 | 1.994262308 |

| | | | | |
|---|---|---|---|---|
| Gm23851 | 0 | NA | NA | NA |
| Pianp | 1373.810968 | 0.194449471 | 0.064393952 | 3.019685316 |
| Zfp384 | 433.8857635 | -0.144437505 | 0.073834472 | -1.956234013 |
| Ing4 | 294.8789085 | -0.127896569 | 0.082610299 | -1.548191581 |
| Acrbp | 30.74231455 | -0.51145021 | 0.297930703 | -1.716675067 |
| Lpar5 | 6.566982277 | -0.635349737 | 0.369221673 | -1.720781262 |
| Chd4 | 3499.470299 | -0.078730311 | 0.05802233 | -1.35689674 |
| Mir7045 | 0 | NA | NA | NA |
| Gm23751 | 1.854332816 | -0.850219128 | 0.797668873 | -1.065879787 |
| Nop2 | 279.2923541 | -0.56233016 | 0.137500317 | -4.089664458 |
| Iffo1 | 569.9560514 | 0.097380163 | 0.107965726 | 0.901954415 |
| Gapdh | 740.2095212 | 0.150324327 | 0.076922695 | 1.954225956 |
| Ncapd2 | 168.4681777 | 0.119248258 | 0.158666008 | 0.75156777 |
| Scarna10 | 0.038981395 | -0.443816764 | 3.724945049 | -0.119147198 |
| Gm28967 | 0 | NA | NA | NA |
| Mrpl51 | 347.6131604 | -0.295274543 | 0.083065136 | -3.554734947 |
| Mir3098 | 14.8885832 | 0.284445573 | 0.359932106 | 0.790275633 |
| Vamp1 | 2358.40077 | 0.17711346 | 0.164365646 | 1.077557651 |
| Tapbpl | 330.3516113 | -0.587215382 | 0.160244785 | -3.664489808 |
| Cd27 | 20.08776085 | -0.709834722 | 0.341114612 | -2.080927339 |
| Mir8113 | 0 | NA | NA | NA |
| Gm24175 | 0 | NA | NA | NA |
| Tuba3a | 0.099106094 | -0.93550548 | 3.724188147 | -0.251197158 |
| Ltbr | 255.90804 | -0.868232045 | 0.192740506 | -4.504668279 |
| Scnn1a | 38.07952501 | -0.148695771 | 0.192612456 | -0.771994572 |
| Tnfrsf1a | 469.1495194 | -1.133730533 | 0.18118309 | -6.257374981 |
| Plekhg6 | 3.530240572 | 0.624378155 | 0.516757024 | 1.208262541 |
| Cd9 | 682.5462011 | -0.755093516 | 0.13191169 | -5.72423504 |
| Gm28809 | 1.776315563 | -0.706649712 | 0.705374553 | -1.001807776 |
| Gm26728 | 0.409864138 | 0.594568679 | 1.625508122 | 0.365774044 |
| Vwf | 172.1922304 | -0.786368043 | 0.164791902 | -4.771885232 |
| Ano2 | 81.25229014 | 1.10314242 | 0.159996292 | 6.8947999 |
| Gm16303 | 0.512522738 | -0.989503719 | 1.411224198 | -0.701166916 |
| Gm24947 | 0.271265366 | 1.016555112 | 2.096190681 | 0.48495355 |
| Ntf3 | 14.00482386 | 0.600952917 | 0.332679216 | 1.806403551 |
| Gm29009 | 0 | NA | NA | NA |
| Kcna5 | 77.10056723 | 0.711348927 | 0.165638498 | 4.294586922 |
| Gm33320 | 17.12675634 | -0.003225814 | 0.319290443 | -0.010103072 |
| Kcna1 | 5474.951842 | 0.378272474 | 0.094907673 | 3.985689042 |
| Kcna6 | 1135.188091 | -0.165408183 | 0.094998422 | -1.741167686 |
| Gm26673 | 76.86612292 | -0.022230304 | 0.229999973 | -0.096653507 |
| Ndufa9 | 671.0675705 | -0.007319379 | 0.07145745 | -0.102429897 |
| RP23-222G22.5 | 0.062426873 | 0.517959912 | 3.724945049 | 0.139051692 |
| Akap3 | 1.281806 | -0.087213153 | 0.934021692 | -0.093373799 |
| Dyrk4 | 3.739156486 | 0.606590238 | 0.548552134 | 1.105802348 |
| Rad51ap1 | 20.13187148 | 0.151317207 | 0.329430303 | 0.459329956 |
| RP23-222G22.9 | 0 | NA | NA | NA |

| | | | | |
|---|---|---|---|---|
| D6Wsu163e | 358.0575039 | -0.027813481 | 0.079985633 | -0.34773096 |
| RP23-8L20.9 | 0 | NA | NA | NA |
| RP23-8L20.8 | 0.141823039 | -0.910536728 | 3.596944456 | -0.253141726 |
| Fgf6 | 0.246418769 | -0.923583417 | 1.83870084 | -0.502302167 |
| Fgf23 | 4.436253295 | 0.885097725 | 0.507339441 | 1.744586865 |
| Tigar | 254.7556436 | -0.043298403 | 0.094638951 | -0.457511445 |
| Platr31 | 1.918110815 | 0.456044717 | 0.727800668 | 0.626606621 |
| RP23-8L20.6 | 0.061920613 | -0.443816764 | 3.724945049 | -0.119147198 |
| Ccnd2 | 1274.440512 | -0.346242827 | 0.13519246 | -2.561110487 |
| 9330179D12Rik | 23.50546626 | 0.247235044 | 0.22344401 | 1.106474255 |
| RP23-387P23.5 | 2.073082813 | 1.223326153 | 0.82438473 | 1.483926263 |
| RP23-403P6.1 | 26.03534595 | -0.00947737 | 0.217555361 | -0.043563028 |
| Gm4968 | 0.813324618 | -0.357120046 | 1.148763103 | -0.310873535 |
| RP23-387P23.7 | 0 | NA | NA | NA |
| Gm7308 | 60.57587847 | 0.49094587 | 0.18435283 | 2.66307748 |
| RP23-387P23.3 | 0.11147366 | -0.955981391 | 3.722845977 | -0.25678779 |
| RP23-387P23.4 | 2.402022068 | 1.064702362 | 0.679438323 | 1.56703313 |
| RP23-387P23.9 | 0.145088245 | -0.444787099 | 3.530638562 | -0.125979222 |
| RP23-387P23.8 | 0 | NA | NA | NA |
| RP23-223J11.1 | 0.049553047 | -0.443818988 | 3.724945049 | -0.119147795 |
| Parp11 | 870.7509703 | -1.024130876 | 0.180857749 | -5.662631978 |
| RP23-422D12.3 | 30.48963593 | -0.827250556 | 0.322382678 | -2.566051505 |
| Cracr2a | 10.36781213 | 0.637695294 | 0.332728853 | 1.916561454 |
| RP23-42M8.2 | 6.20278278 | 0.011394318 | 0.488924278 | 0.023304873 |
| Prmt8 | 579.4218534 | 0.297100782 | 0.150677434 | 1.97176693 |
| RP23-42M8.9 | 0 | NA | NA | NA |
| RP23-42M8.10 | 14.99285838 | 0.24456604 | 0.341764483 | 0.715598174 |
| RP23-42M8.5 | 0.045882455 | 0.517959912 | 3.724945049 | 0.139051692 |
| RP23-42M8.7 | 0.12092566 | 0.998850474 | 3.724945049 | 0.268151734 |
| mmu-mir-7233 | 0 | NA | NA | NA |
| RP24-473H20.4 | 0.306891002 | -0.481213925 | 1.743439822 | -0.276014072 |
| Gm23098 | 0.762875166 | -1.234868557 | 1.267570263 | -0.974201267 |
| RP24-473H20.3 | 0.711610403 | -3.031539952 | 1.225015695 | -2.474694785 |
| Tspan11 | 99.16132127 | 0.130931953 | 0.164103068 | 0.797864139 |
| Gm15870 | 0.825566729 | -2.144820728 | 1.190485218 | -1.801635749 |
| Tspan9 | 1288.655582 | -0.232255952 | 0.134555117 | -1.726102717 |
| Gm26338 | 0.326202429 | 0.403422994 | 1.805431838 | 0.223449585 |
| Gm10010 | 121.3586711 | 0.739791816 | 0.189810776 | 3.897522736 |
| Gm4691 | 17.793641 | 0.194227696 | 0.299778728 | 0.647903529 |
| Tead4 | 29.15459458 | -0.341546184 | 0.258592579 | -1.320788811 |
| Gm21893 | 0 | NA | NA | NA |
| Gm26951 | 0.061920613 | -0.443816764 | 3.724945049 | -0.119147198 |
| Rpl18-ps2 | 5.774417445 | -1.040703497 | 0.408263825 | -2.549095544 |
| Tulp3 | 551.9556277 | -0.605123649 | 0.108218331 | -5.591692684 |
| Gm26770 | 0.55622196 | -1.541428821 | 1.332888184 | -1.156457713 |
| Rhno1 | 92.43520098 | -0.132802125 | 0.120172646 | -1.105094456 |
| Foxm1 | 45.1673141 | -0.280897408 | 0.364536638 | -0.770560155 |

| | | | | |
|---|---|---|---|---|
| 4933413G19Rik | 17.62518862 | -0.365806623 | 0.372730515 | -0.981423866 |
| Gm15862 | 1.729221609 | 0.054670444 | 0.78838995 | 0.069344421 |
| Nrip2 | 216.9234121 | 0.356299384 | 0.094497622 | 3.770458724 |
| Itfg2 | 280.4983282 | 0.085919851 | 0.103375709 | 0.831141585 |
| Fkbp4 | 1131.070024 | -0.354908864 | 0.082828469 | -4.28486566 |
| Gm10069 | 14.56716468 | -0.058894053 | 0.342340244 | -0.172033684 |
| Pzp | 1.32803898 | 0.770963708 | 0.941435484 | 0.818923571 |
| Gm8724 | 0.141071297 | -0.443820801 | 3.564043909 | -0.12452731 |
| BC048546 | 30.70108197 | 0.447871026 | 0.202177487 | 2.215236879 |
| Klrb1a | 2.241070021 | -0.745527226 | 0.684672773 | -1.088881076 |
| Gm5884 | 150.0674224 | -0.238343574 | 0.121237706 | -1.965919527 |
| Clec2j | 0 | NA | NA | NA |
| Clec2h | 0.169629266 | -0.804564592 | 2.763788843 | -0.291109284 |
| Klrb1 | 0 | NA | NA | NA |
| Klrb1c | 5.966816195 | -1.412546339 | 0.579937348 | -2.435687827 |
| Klrb1b | 4.841237096 | 0.676981 | 0.579419147 | 1.168378719 |
| Gm26656 | 0.322709507 | 1.558286288 | 1.916398 | 0.813132913 |
| BC035044 | 6.979976624 | 0.612221422 | 0.453590248 | 1.349723511 |
| Clec2i | 12.44756223 | -0.908629207 | 0.487522034 | -1.863770543 |
| Gm27336 | 0.319483352 | -1.826497696 | 1.970254238 | -0.927036553 |
| Clec2g | 0.594455495 | 1.27051194 | 1.440818322 | 0.881798851 |
| Gm15987 | 6.297883917 | -1.001204363 | 0.408631582 | -2.45013946 |
| Mir7649 | 0 | NA | NA | NA |
| Gm8778 | 0.092691418 | -0.908687078 | 3.724560878 | -0.2439716 |
| BC064078 | 21.92509358 | -0.230230906 | 0.284574229 | -0.809036388 |
| Gm16187 | 0.11197992 | 0.037073798 | 3.724945049 | 0.009952844 |
| Clec2f | 1.610080848 | 0.80438899 | 0.739052527 | 1.08840571 |
| Gm16186 | 0 | NA | NA | NA |
| Klrb1f | 4.97974666 | -0.867800235 | 0.467019732 | -1.858166103 |
| Clec2e | 0.752785761 | -1.090304515 | 1.247258751 | -0.874160646 |
| Klrb1-ps1 | 0 | NA | NA | NA |
| Gm26160 | 0.088534442 | -0.905414734 | 3.723500899 | -0.243162217 |
| Clec2d | 395.2909725 | -1.263018984 | 0.251183969 | -5.028262709 |
| Gm27514 | 0.12209515 | 0.037073798 | 3.724945049 | 0.009952844 |
| 2310001H17Rik | 41.45082234 | -1.170118854 | 0.25330796 | -4.619352884 |
| Cd69 | 49.45403992 | -0.949068572 | 0.327615398 | -2.896898546 |
| 4922502D21Rik | 0.201858732 | 0.517972533 | 2.604406155 | 0.198883163 |
| Clec12a | 44.34900859 | -1.405589193 | 0.316693695 | -4.438323885 |
| Clec12b | 2.028096924 | -0.96798901 | 0.836981658 | -1.156523563 |
| Clec1b | 1.612175364 | 0.000457505 | 0.816819763 | 0.000560106 |
| Clec9a | 4.424159844 | 0.207414595 | 0.594840208 | 0.348689602 |
| Clec1a | 25.40223335 | -0.292746716 | 0.271813679 | -1.077012448 |
| Clec7a | 77.62764879 | -0.378559037 | 0.252565442 | -1.498855244 |
| Olr1 | 7.563519298 | -2.063105455 | 0.457873084 | -4.505845677 |
| Tmem52b | 0.046345709 | -0.443818988 | 3.724945049 | -0.119147795 |
| 1700101I11Rik | 14.17165844 | -0.153428995 | 0.279946426 | -0.548065561 |
| Gabarapl1 | 4493.649597 | 0.190106181 | 0.059516171 | 3.194193742 |

| | | | | |
|---|---|---|---|---|
| Klre1 | 4.71579917 | -1.912511147 | 0.528750125 | -3.617041502 |
| Klrd1 | 23.39137982 | -1.311839608 | 0.297491431 | -4.409671916 |
| Klrk1 | 186.6532185 | -0.780522659 | 0.250712711 | -3.11321534 |
| Klrc3 | 5.985488163 | -1.171399425 | 0.479984185 | -2.440495877 |
| Klrc2 | 17.16294782 | -1.397625842 | 0.394570037 | -3.542148953 |
| Klrc1 | 33.05721631 | -0.674163153 | 0.29284716 | -2.302098997 |
| Gm7309 | 0 | NA | NA | NA |
| Mir680-1 | 0.066050875 | -0.443818988 | 3.724945049 | -0.119147795 |
| Klri1 | 2.598038951 | -2.388719255 | 0.751227884 | -3.179753185 |
| Klri2 | 8.227621471 | -2.221561036 | 0.610133026 | -3.641109302 |
| Gm156 | 0.097139547 | -0.443816764 | 3.724945049 | -0.119147198 |
| Gm16242 | 0.044544648 | 0.517957688 | 3.724945049 | 0.139051095 |
| Klra17 | 6.058344776 | -0.581132417 | 0.58409238 | -0.994932373 |
| Gm17631 | 0 | NA | NA | NA |
| Gm23977 | 0 | NA | NA | NA |
| Klra5 | 0.677147933 | -1.399523406 | 1.377703404 | -1.015837953 |
| Gm15854 | 0 | NA | NA | NA |
| Gm22555 | 0 | NA | NA | NA |
| Klra6 | 0 | NA | NA | NA |
| Klra4 | 0.378042144 | -1.109251464 | 1.655312826 | -0.670115912 |
| Gm24676 | 0 | NA | NA | NA |
| Klra11-ps | 0 | NA | NA | NA |
| Gm22621 | 0 | NA | NA | NA |
| Klra8 | 0.110595523 | 0.037071574 | 3.724945049 | 0.009952247 |
| Gm24712 | 0 | NA | NA | NA |
| Klra14-ps | 0 | NA | NA | NA |
| Gm23552 | 0 | NA | NA | NA |
| Klra9 | 0.604622052 | -1.384240747 | 1.19448382 | -1.158861028 |
| Klra7 | 3.947549917 | -2.050472385 | 0.58348098 | -3.514206039 |
| Klra10 | 0.285107744 | -1.283288196 | 2.393663047 | -0.536118982 |
| Klra13-ps | 0.52061208 | -2.347787468 | 1.319404763 | -1.77942928 |
| Gm24072 | 0 | NA | NA | NA |
| Klra3 | 12.4313916 | -1.575396605 | 0.410901888 | -3.833996995 |
| Klra1 | 0.328294527 | -0.654960514 | 1.680981452 | -0.389629828 |
| Gm6584 | 0 | NA | NA | NA |
| Gm15861 | 0 | NA | NA | NA |
| Gm20997 | 0 | NA | NA | NA |
| Gm27013 | 0.044544648 | 0.517957688 | 3.724945049 | 0.139051095 |
| Gm8816 | 0 | NA | NA | NA |
| Gm26996 | 0.113979814 | 0.517957688 | 3.724945049 | 0.139051095 |
| Gm26914 | 0 | NA | NA | NA |
| Gm27047 | 0 | NA | NA | NA |
| Gm26919 | 0.161486377 | -0.444693834 | 2.795640213 | -0.159066904 |
| Gm6600 | 0 | NA | NA | NA |
| Gm8837 | 0 | NA | NA | NA |
| Gm5581 | 0.915657815 | -1.944938253 | 1.310611877 | -1.483992543 |
| Klra2 | 81.98899445 | -2.195750982 | 0.429388323 | -5.113671861 |

| | | | | |
|---|---|---|---|---|
| Gm5582 | 1.579793576 | 0.625408367 | 0.802773955 | 0.779059115 |
| Magohb | 84.69062322 | 0.199197918 | 0.160377963 | 1.242052929 |
| Styk1 | 21.45282092 | -0.524517358 | 0.26749748 | -1.960831023 |
| Ybx3 | 331.3395029 | -0.364396073 | 0.103969114 | -3.504849276 |
| Gm22362 | 7.679914887 | 0.430461089 | 0.413314427 | 1.041485758 |
| 5430401F13Rik | 0.172107867 | 0.517971699 | 2.739902729 | 0.189047477 |
| Gm6619 | 0 | NA | NA | NA |
| Gm22190 | 0 | NA | NA | NA |
| Gm16571 | 0 | NA | NA | NA |
| Hmgb1l | 0 | NA | NA | NA |
| Tas2r130 | 0 | NA | NA | NA |
| Tas2r107 | 0 | NA | NA | NA |
| Tas2r106 | 0 | NA | NA | NA |
| Tas2r104 | 0 | NA | NA | NA |
| Tas2r105 | 0 | NA | NA | NA |
| Tas2r114 | 0 | NA | NA | NA |
| Gm4736 | 0 | NA | NA | NA |
| Prpmp5 | 0 | NA | NA | NA |
| Gm8882 | 0.046345709 | -0.443818988 | 3.724945049 | -0.119147795 |
| n-R5s166 | 0 | NA | NA | NA |
| Prh1 | 0 | NA | NA | NA |
| n-R5s167 | 0 | NA | NA | NA |
| Prp2 | 0 | NA | NA | NA |
| A630073D07Rik | 0 | NA | NA | NA |
| Tas2r120 | 0 | NA | NA | NA |
| Tas2r121 | 0 | NA | NA | NA |
| Tas2r122 | 0 | NA | NA | NA |
| Tas2r115 | 0 | NA | NA | NA |
| Tas2r124 | 0 | NA | NA | NA |
| Tas2r102 | 0 | NA | NA | NA |
| Tas2r136 | 0.061920613 | -0.443816764 | 3.724945049 | -0.119147198 |
| Tas2r117 | 0 | NA | NA | NA |
| Tas2r123 | 0 | NA | NA | NA |
| Tas2r116 | 0.061920613 | -0.443816764 | 3.724945049 | -0.119147198 |
| Tas2r110 | 0 | NA | NA | NA |
| Tas2r113 | 0 | NA | NA | NA |
| Tas2r125 | 0 | NA | NA | NA |
| Tas2r129 | 0 | NA | NA | NA |
| Tas2r131 | 0 | NA | NA | NA |
| Tas2r109 | 0 | NA | NA | NA |
| Tas2r103 | 0 | NA | NA | NA |
| Tas2r140 | 0 | NA | NA | NA |
| Smim10l1 | 1843.017742 | 0.162716999 | 0.060462522 | 2.691204297 |
| Gm23403 | 0 | NA | NA | NA |
| Gm25019 | 0.075043205 | 0.517957688 | 3.724945049 | 0.139051095 |
| 5530400C23Rik | 0 | NA | NA | NA |
| Gm5885 | 0 | NA | NA | NA |

| | | | | |
|---|---|---|---|---|
| Gm23375 | 0.502960085 | 0.440289206 | 1.462959228 | 0.300957947 |
| Kap | 0.042716945 | 0.037062677 | 3.724945049 | 0.009949859 |
| Etv6 | 690.3323098 | -0.043035553 | 0.126791554 | -0.339419717 |
| Gm17088 | 0.202171817 | -1.352528503 | 2.5871409 | -0.522788884 |
| Gm17089 | 0.375570109 | -1.785526507 | 1.622215009 | -1.100671919 |
| Bcl2l14 | 0.379111656 | 0.048347328 | 1.43816857 | 0.033617288 |
| Lrp6 | 1132.17589 | 0.176999372 | 0.065029159 | 2.721846257 |
| Mansc1 | 117.9240851 | 0.22386164 | 0.130040844 | 1.721471764 |
| Loh12cr1 | 180.7292652 | 0.38843849 | 0.093819466 | 4.140276065 |
| Dusp16 | 411.5721016 | -0.309385861 | 0.125208166 | -2.470971906 |
| Crebl2 | 520.8784886 | 0.020477104 | 0.112792558 | 0.181546586 |
| Gpr19 | 174.2410689 | -0.223433502 | 0.132010905 | -1.692538216 |
| 2810454H06Rik | 8.20098561 | 0.001853699 | 0.413635998 | 0.004481475 |
| Cdkn1b | 860.406846 | -0.059299561 | 0.100950194 | -0.587414035 |
| 1190002F15Rik | 1.018292001 | 0.137991217 | 0.960823605 | 0.143617638 |
| Apold1 | 80.86165162 | 0.489136627 | 0.256159941 | 1.909496957 |
| Ddx47 | 401.4348954 | -0.103102155 | 0.070003648 | -1.472811167 |
| Gprc5a | 6.993744504 | -0.453553487 | 0.46436447 | -0.976718754 |
| Gprc5d | 0.543589123 | -1.511702073 | 1.445837288 | -1.045554771 |
| Hebp1 | 102.192585 | -0.367126144 | 0.173307041 | -2.118356766 |
| Gm22881 | 0 | NA | NA | NA |
| 8430419L09Rik | 1464.187594 | 0.31693364 | 0.055953988 | 5.664183268 |
| Gsg1 | 0.811303554 | -0.269180214 | 1.18718925 | -0.226737408 |
| Pbp2 | 0 | NA | NA | NA |
| Dynlt1-ps1 | 35.59685521 | -0.479871759 | 0.26576963 | -1.805592909 |
| Emp1 | 465.3937255 | -1.878009556 | 0.40737159 | -4.610065115 |
| Gm25136 | 0.672405421 | -2.06371688 | 1.295524854 | -1.592958154 |
| Grin2b | 9373.268553 | 1.06932752 | 0.189634222 | 5.638895277 |
| Gm14329 | 0.341881936 | 0.523825218 | 1.606429476 | 0.326080432 |
| Gm22892 | 0.112396584 | -0.917692504 | 3.724434845 | -0.246397787 |
| Gm38202 | 40.04288314 | 0.890433971 | 0.353801764 | 2.516759556 |
| Gm37182 | 48.96267409 | 0.986254148 | 0.416922859 | 2.365555467 |
| Gm14330 | 0.038981395 | -0.443816764 | 3.724945049 | -0.119147198 |
| Gm26653 | 7.145952665 | 0.523231304 | 0.474507737 | 1.102682347 |
| Gm26105 | 0.110595523 | 0.037071574 | 3.724945049 | 0.009952247 |
| Gm8994 | 0.107803068 | 0.036092186 | 3.724874622 | 0.009689503 |
| Gm15476 | 0.04393175 | 0.037062677 | 3.724945049 | 0.009949859 |
| E330021D16Rik | 0 | NA | NA | NA |
| Gm25882 | 0 | NA | NA | NA |
| Gm6728 | 11.37417031 | -0.229244182 | 0.321048903 | -0.71404755 |
| Rpl36a-ps3 | 8.907631286 | -0.617649599 | 0.36043515 | -1.71362199 |
| Atf7ip | 2145.155086 | -0.163534624 | 0.075934672 | -2.153622575 |
| 5830415B17Rik | 15.21012998 | -0.034447971 | 0.312857137 | -0.110107671 |
| Plbd1 | 20.58495972 | -1.374916019 | 0.338189117 | -4.065524138 |
| Gucy2c | 13.73199816 | -0.234577172 | 0.319571005 | -0.734037722 |
| Hist4h4 | 361.153536 | 0.096870867 | 0.16008926 | 0.605105342 |
| H2afj | 164.4299334 | -0.311941579 | 0.131062201 | -2.380103325 |

| | | | | |
|---|---|---|---|---|
| AC122804.1 | 2.891209199 | -0.68206007 | 0.599981181 | -1.136802439 |
| Wbp11 | 918.8633835 | 0.166751372 | 0.086231002 | 1.933775197 |
| BC049715 | 12.24740255 | 0.147228094 | 0.317398916 | 0.463858213 |
| Smco3 | 28.46706308 | 0.385873456 | 0.209617389 | 1.840846587 |
| Art4 | 1.936338333 | -1.115467087 | 0.878080438 | -1.270347269 |
| Mgp | 118.7015475 | -0.677425105 | 0.178382422 | -3.797600103 |
| Erp27 | 1.418067754 | 0.761002516 | 0.808088167 | 0.941732038 |
| Arhgdib | 195.7130349 | -0.521630601 | 0.159882 | -3.262597417 |
| Pde6h | 2.208047999 | 0.803076009 | 0.97834072 | 0.82085514 |
| Rerg | 82.27360041 | 0.39500764 | 0.147322129 | 2.681251237 |
| Ptpro | 585.5542697 | -0.394665275 | 0.162771249 | -2.424662079 |
| Eps8 | 778.7034068 | -0.540878223 | 0.122574988 | -4.41263127 |
| Strap | 1329.576707 | -0.157165395 | 0.083514964 | -1.881883051 |
| Dera | 50.46973726 | -0.43675062 | 0.221603977 | -1.970860932 |
| Gm24308 | 0 | NA | NA | NA |
| Slc15a5 | 0.807800175 | 0.97207437 | 1.221328049 | 0.795915864 |
| Mgst1 | 196.4190543 | -0.984338212 | 0.217652339 | -4.522525322 |
| Lmo3 | 872.8499117 | 1.423452148 | 0.124647875 | 11.41978674 |
| A830011K09Rik | 12.61465114 | 0.955550374 | 0.367523877 | 2.599968147 |
| B230110G15Rik | 1.657582323 | 3.262864639 | 1.039251024 | 3.13963091 |
| Igbp1b | 1.236116855 | 3.40520379 | 1.071456596 | 3.178107076 |
| Gm25434 | 0 | NA | NA | NA |
| Rergl | 0.244560259 | 0.51796868 | 2.149863015 | 0.240931016 |
| Pik3c2g | 12.46784403 | -0.266751764 | 0.293966056 | -0.90742369 |
| Gm37269 | 0.211089261 | 0.053234262 | 2.182449338 | 0.024391981 |
| Plcz1 | 6.588952628 | 0.290010733 | 0.377607226 | 0.768022202 |
| Capza3 | 0.060174537 | 0.517957688 | 3.724945049 | 0.139051095 |
| Gm24174 | 0 | NA | NA | NA |
| n-R5s168 | 0 | NA | NA | NA |
| Plekha5 | 1362.818818 | 0.408364598 | 0.108324433 | 3.769829091 |
| Gm25780 | 21.43207649 | 1.406260039 | 0.329531248 | 4.267455814 |
| Aebp2 | 1104.796083 | -0.168890929 | 0.063460658 | -2.661348522 |
| Gm11077 | 0.333293459 | -1.699789151 | 1.700777639 | -0.999418802 |
| Rpl38-ps2 | 307.4974746 | -0.108945243 | 0.090351684 | -1.205790947 |
| Gm28524 | 1.176308512 | 0.374623307 | 1.057452102 | 0.354269764 |
| Gm28523 | 12.22868002 | 0.323124776 | 0.328068127 | 0.984931937 |
| Pde3a | 315.7616379 | -0.345055781 | 0.102506885 | -3.366171737 |
| Slco1c1 | 209.1116899 | 0.143357788 | 0.231087858 | 0.620360538 |
| Slco1b2 | 3.268651982 | -0.165122216 | 0.650384899 | -0.25388384 |
| Gm5724 | 0.34217064 | 0.497281821 | 1.606835421 | 0.309479001 |
| Gm23069 | 0 | NA | NA | NA |
| Slco1a4 | 155.850488 | 0.544847554 | 0.346913151 | 1.57055895 |
| Slco1a1 | 0.113999573 | -0.443821212 | 3.724945049 | -0.119148392 |
| Gm22036 | 0 | NA | NA | NA |
| Gm6614 | 0.602494748 | 0.64052565 | 1.263116052 | 0.507099604 |
| Gm23775 | 0 | NA | NA | NA |
| Slco1a6 | 0.857960599 | -0.043416122 | 1.042267679 | -0.041655443 |

| | | | | |
|---|---|---|---|---|
| Gm18159 | 0.046345709 | -0.443818988 | 3.724945049 | -0.119147795 |
| Gm20400 | 0.439621597 | 0.275333294 | 1.45935284 | 0.188668077 |
| Slco1a5 | 10.05483302 | -1.134397965 | 0.396025037 | -2.864460221 |
| Gm25430 | 0 | NA | NA | NA |
| Iapp | 0.092228164 | 0.037073798 | 3.724945049 | 0.009952844 |
| Pyroxd1 | 173.2075479 | -3.48E-05 | 0.086281636 | -0.000402994 |
| Recql | 183.7198923 | -0.099739301 | 0.148124377 | -0.673348328 |
| Golt1b | 467.436909 | -0.227083154 | 0.06392709 | -3.552221062 |
| B230216G23Rik | 18.91388396 | -0.422237151 | 0.2951394 | -1.430636341 |
| Gys2 | 1.458192719 | 0.636249661 | 0.95385651 | 0.667028694 |
| Ldhb | 2909.672582 | 0.242757515 | 0.07269609 | 3.339347635 |
| Kcnj8 | 29.13393712 | -0.253008545 | 0.201067967 | -1.258323485 |
| Abcc9 | 80.16356092 | -0.592973403 | 0.175642282 | -3.376028802 |
| 5330439B14Rik | 0 | NA | NA | NA |
| Cmas | 639.4281021 | -0.012743948 | 0.06718408 | -0.189687023 |
| Gm766 | 1.893318208 | 1.549716831 | 0.888987009 | 1.743239007 |
| Gm7457 | 1.3052917 | -1.364571793 | 1.032890998 | -1.321118875 |
| St8sia1 | 1060.037098 | 0.476373405 | 0.064680632 | 7.365008558 |
| C2cd5 | 774.2859654 | 0.099784052 | 0.122663743 | 0.81347633 |
| Etnk1 | 6574.447317 | 0.120675843 | 0.090862105 | 1.328120706 |
| D6Ertd474e | 2.829004364 | 0.876644252 | 0.59282495 | 1.47875735 |
| Gm23272 | 0 | NA | NA | NA |
| 4930579D09Rik | 0 | NA | NA | NA |
| 1700060C16Rik | 0.044191549 | 0.037062677 | 3.724945049 | 0.009949859 |
| Sox5 | 636.5935809 | 0.071832825 | 0.097558156 | 0.736307734 |
| Sox5os1 | 0.085433889 | 0.037058229 | 3.724945049 | 0.009948665 |
| Sox5os2 | 0 | NA | NA | NA |
| Sox5it | 4.538093024 | 1.261069988 | 0.50934981 | 2.47584266 |
| Sox5os3 | 0.137647365 | 1.167618838 | 3.719469251 | 0.31392082 |
| Sox5os4 | 0 | NA | NA | NA |
| Sox5os5 | 0 | NA | NA | NA |
| Gm22792 | 0 | NA | NA | NA |
| Bcat1 | 1548.257675 | -0.124067361 | 0.103405336 | -1.199815855 |
| Gm23071 | 1.299875216 | 1.610960794 | 1.031239185 | 1.562160183 |
| Gm26666 | 7.30636117 | 0.751176496 | 0.424219165 | 1.770727391 |
| Gm15687 | 0.219710209 | -0.46022026 | 2.160637949 | -0.213002026 |
| Lrmp | 42.64411814 | -0.232589518 | 0.191517266 | -1.214457177 |
| Casc1 | 19.07366442 | 0.105688351 | 0.268409533 | 0.393757815 |
| Gm15543 | 0.049553047 | -0.443818988 | 3.724945049 | -0.119147795 |
| Lyrm5 | 128.9520887 | -0.001740554 | 0.126083361 | -0.013804791 |
| Kras | 1419.910694 | -0.006437864 | 0.072159762 | -0.08921681 |
| Gm15706 | 24.72915098 | 0.675761782 | 0.240306979 | 2.812077216 |
| Rps25-ps1 | 12.76755632 | 0.07245407 | 0.309668217 | 0.233973221 |
| Gm22793 | 0 | NA | NA | NA |
| 2010013B24Rik | 0.275148976 | -0.917650448 | 2.066424218 | -0.444076507 |
| Gm15707 | 0.08486385 | 0.037730769 | 3.724898049 | 0.010129343 |
| Lmntd1 | 13.42183498 | 0.479365458 | 0.337074274 | 1.422135998 |

| | | | | |
|---|---|---|---|---|
| Gm23498 | 0 | NA | NA | NA |
| 1700073E17Rik | 0.253253743 | 0.236861236 | 2.454665967 | 0.096494284 |
| Gm15499 | 0 | NA | NA | NA |
| Gm25373 | 0 | NA | NA | NA |
| Tuba3b | 0.095898756 | -0.917692504 | 3.724434845 | -0.246397787 |
| Gm15704 | 2.261169167 | -0.194283463 | 0.717603092 | -0.270739444 |
| Rassf8 | 399.0644075 | -0.255876961 | 0.110437826 | -2.316932257 |
| Bhlhe41 | 914.9264729 | -0.008924162 | 0.063242996 | -0.141109094 |
| Sspn | 140.0205592 | -0.130785471 | 0.19300859 | -0.677614772 |
| Sspnos | 0.044191549 | 0.037062677 | 3.724945049 | 0.009949859 |
| Itpr2 | 644.2115075 | 0.060882861 | 0.09844331 | 0.618456055 |
| Gm15720 | 1.860207608 | 0.309280794 | 0.857863826 | 0.360524345 |
| Asun | 359.4234607 | 0.01127065 | 0.106011743 | 0.106315113 |
| Fgfr1op2 | 883.1771317 | 0.052736198 | 0.052893909 | 0.997018351 |
| Tm7sf3 | 427.7210704 | -0.067513884 | 0.080714861 | -0.836449237 |
| Med21 | 211.1955998 | -0.560888164 | 0.156928979 | -3.574152885 |
| Gm6654 | 0.926048393 | -2.79219897 | 1.096833384 | -2.545691087 |
| 4930479D17Rik | 2.935438062 | 0.036213911 | 0.590201214 | 0.061358585 |
| Stk38l | 426.5635259 | -0.343031229 | 0.083585236 | -4.10396916 |
| Arntl2 | 48.93835431 | 0.335230103 | 0.226403165 | 1.48067764 |
| Smco2 | 1.092136027 | -1.350713099 | 1.011685081 | -1.335112205 |
| Ppfibp1 | 368.3784692 | -0.454889393 | 0.08516731 | -5.341126707 |
| 1700034J05Rik | 0.062426873 | 0.517959912 | 3.724945049 | 0.139051692 |
| Rep15 | 3.085232335 | -0.426604201 | 0.610455211 | -0.698829649 |
| Mrps35 | 175.5694965 | -0.003616152 | 0.105553891 | -0.034258822 |
| Mansc4 | 2.80060554 | -0.9436063 | 0.618460543 | -1.525734036 |
| Klhl42 | 434.0929622 | 0.076128779 | 0.074069648 | 1.027799924 |
| Gm7571 | 0.356315666 | 0.519401209 | 1.56269213 | 0.332375904 |
| E330012B07Rik | 0.576204374 | -0.056805463 | 1.418571444 | -0.040044132 |
| Gm15767 | 0.045882455 | 0.517959912 | 3.724945049 | 0.139051692 |
| Pthlh | 19.04363101 | -0.059297271 | 0.303493678 | -0.195382229 |
| Gm15766 | 2.701807514 | 0.00709073 | 0.614987033 | 0.011529885 |
| Gm6288 | 5.185232385 | -0.206281026 | 0.463121443 | -0.445414543 |
| Ccdc91 | 322.5136679 | 0.022269331 | 0.101881874 | 0.218579913 |
| 1700049E15Rik | 0.04393175 | 0.037062677 | 3.724945049 | 0.009949859 |
| Gm15762 | 0.295243939 | 0.037077765 | 1.783564568 | 0.020788575 |
| Far2os2 | 3.668828368 | -0.688810123 | 0.651586961 | -1.057126928 |
| Far2 | 187.1712709 | 0.380267543 | 0.126059439 | 3.016573346 |
| Far2os1 | 2.690587419 | -1.251808675 | 0.6525243 | -1.918409284 |
| Ergic2 | 794.7085187 | 0.081026716 | 0.076531381 | 1.058738456 |
| Gm17216 | 1.982138501 | 0.652496508 | 0.978068969 | 0.667127298 |
| Tmtc1 | 1255.055725 | 0.177150371 | 0.085987464 | 2.060188346 |
| Gm15763 | 0.045882455 | 0.517959912 | 3.724945049 | 0.139051692 |
| Rps4l | 60.78732413 | -0.11275699 | 0.263011243 | -0.428715475 |
| Gm6313 | 0.072025462 | 0.517959912 | 3.724945049 | 0.139051692 |
| Gm23456 | 0.106358623 | 0.517955464 | 3.724945049 | 0.139050498 |
| 4930528G23Rik | 0.514450355 | 1.574704539 | 1.562662751 | 1.007705942 |

| | | | | |
|---|---|---|---|---|
| Gm15778 | 0 | NA | NA | NA |
| Ipo8 | 948.9724006 | -0.083150484 | 0.053850498 | -1.544098696 |
| Caprin2 | 102.1573326 | 0.358271375 | 0.164484833 | 2.178142318 |
| Gm7618 | 0.320079734 | -0.436550909 | 1.699113796 | -0.256928588 |
| Fam60a | 49.2104772 | -0.127765722 | 0.164745366 | -0.775534541 |
| 3010003L21Rik | 2.014334011 | 0.149149826 | 0.65075871 | 0.229193744 |
| Gm25539 | 0 | NA | NA | NA |
| Dennd5b | 2078.167419 | 0.010829558 | 0.05270737 | 0.205465726 |
| Gm26540 | 0.259164831 | -0.488839867 | 1.831509013 | -0.266905521 |
| Gm15779 | 0 | NA | NA | NA |
| Gm15781 | 0 | NA | NA | NA |
| Gm22578 | 4.906737206 | 0.596800216 | 0.514427979 | 1.160123944 |
| Gm15780 | 0 | NA | NA | NA |
| Gm23462 | 2.97460163 | 0.349933802 | 0.716894059 | 0.488124845 |
| Gm10203 | 0.141071297 | -0.443820801 | 3.564043909 | -0.12452731 |
| Mettl20 | 22.19794556 | -0.631700027 | 0.329831302 | -1.915221581 |
| Gm10011 | 0.371635155 | 1.392086274 | 1.549313016 | 0.898518414 |
| Amn1 | 149.0049197 | -0.175027703 | 0.115103832 | -1.520607082 |
| 2810474O19Rik | 949.1809365 | -0.618925948 | 0.145082745 | -4.266020378 |
| Gm15782 | 0.129871752 | -0.443193064 | 2.991078735 | -0.148171647 |
| Gm15784 | 6.345417981 | -0.292933038 | 0.398573704 | -0.734953248 |
| Gm15786 | 0.200503209 | 0.5388051 | 2.612813776 | 0.206216419 |
| Gm15783 | 0.047948698 | 0.037064902 | 3.724945049 | 0.009950456 |
| Bicd1 | 1658.556008 | 0.155737412 | 0.05106328 | 3.049890468 |
| Gm15785 | 0.570614426 | 1.238852448 | 1.335818282 | 0.927410909 |
| Gm24307 | 1.150914894 | -0.107892236 | 0.901103626 | -0.119733439 |
| Gm21814 | 6.725415989 | -0.591104414 | 0.443444002 | -1.332985476 |
| Gm3104 | 0 | NA | NA | NA |
| Gm7353 | 46.36268977 | -0.232090231 | 0.169500569 | -1.369259305 |
| AU018091 | 0.640879238 | -2.060809503 | 1.200842547 | -1.716136314 |
| Mir290a | 0 | NA | NA | NA |
| mmu-mir-290b | 0 | NA | NA | NA |
| Mir291a | 0 | NA | NA | NA |
| Mir292 | 0 | NA | NA | NA |
| mmu-mir-292b | 0 | NA | NA | NA |
| Mir291b | 0 | NA | NA | NA |
| Mir293 | 0 | NA | NA | NA |
| Mir294 | 0 | NA | NA | NA |
| Mir295 | 0 | NA | NA | NA |
| Nlrp12 | 0.605827841 | 0.534074921 | 1.236849957 | 0.431802514 |
| Myadm | 554.9737066 | -0.269544947 | 0.090943168 | -2.963883423 |
| Prkcg | 1983.083602 | 0.905272821 | 0.159465941 | 5.676903905 |
| Cacng7 | 1812.960324 | 0.374563581 | 0.078466106 | 4.773571695 |
| Cacng8 | 593.3089663 | 0.340865935 | 0.10365996 | 3.288308562 |
| Gm23450 | 0 | NA | NA | NA |
| Cacng6 | 9.245691091 | -0.707295087 | 0.377349134 | -1.874378452 |
| Tarm1 | 15.12490142 | -2.111252251 | 0.890551099 | -2.370725557 |

| | | | | |
|---|---|---|---|---|
| Gm25100 | 0 | NA | NA | NA |
| Oscar | 1.834747314 | 0.342706955 | 0.687671524 | 0.498358509 |
| Ndufa3 | 436.48222 | -0.096925685 | 0.119169311 | -0.813344346 |
| Tfpt | 142.0055102 | -0.03092568 | 0.130399282 | -0.237161427 |
| Prpf31 | 410.4576804 | -0.108318959 | 0.072190667 | -1.500456558 |
| Gm15927 | 8.516043887 | 0.917085466 | 0.392979358 | 2.33367338 |
| Cnot3 | 493.1599569 | -0.128276863 | 0.077980483 | -1.644986773 |
| Mir3572 | 0 | NA | NA | NA |
| Leng1 | 69.24448306 | -0.227054775 | 0.179327803 | -1.266143739 |
| Tmc4 | 26.68951419 | -0.152341308 | 0.28644469 | -0.531834986 |
| Gm15929 | 0.377779179 | -0.772555717 | 3.144540354 | -0.245681604 |
| Mboat7 | 1337.109882 | -0.049314612 | 0.052191666 | -0.944875224 |
| Tsen34 | 403.0156256 | -0.183717154 | 0.122599162 | -1.498518841 |
| Rps9 | 1343.617836 | -0.305713488 | 0.140998234 | -2.168207917 |
| Pirb | 100.508228 | -1.546231044 | 0.269718949 | -5.732749037 |
| Gm15922 | 4.272688704 | -1.26340941 | 0.470827373 | -2.683381389 |
| Gm15930 | 0 | NA | NA | NA |
| Gm15925 | 2.457332247 | -1.26053579 | 0.841681653 | -1.497639619 |
| Gm15448 | 3.00180885 | -2.089047501 | 0.780051067 | -2.678090693 |
| Pira2 | 10.48735136 | -1.443829633 | 0.38394087 | -3.760552072 |
| Gm10693 | 0.291730981 | -0.402646272 | 1.75253492 | -0.229750784 |
| Gm14548 | 29.20745553 | -1.743973889 | 0.460652527 | -3.785877177 |
| Lilra6 | 22.5183229 | -1.717334119 | 0.327812418 | -5.23877079 |
| Gm7363 | 4.253190973 | -0.347291944 | 0.459978537 | -0.755017716 |
| Gm15928 | 0.849955171 | 0.920781541 | 1.30227135 | 0.707058126 |
| Lair1 | 165.5340521 | -0.183095026 | 0.209326086 | -0.874688049 |
| 9430041J12Rik | 62.38036442 | 0.050129627 | 0.159548327 | 0.314197136 |
| Gm23741 | 0.042716945 | 0.037062677 | 3.724945049 | 0.009949859 |
| Ttyh1 | 4104.837726 | 0.449521219 | 0.116343525 | 3.863740768 |
| D030047H15Rik | 2.31889725 | 0.937822871 | 0.752601126 | 1.24610878 |
| Leng8 | 1590.8448 | 0.597375988 | 0.28587313 | 2.089654204 |
| Leng9 | 55.71701833 | -0.10539925 | 0.152508274 | -0.691105126 |
| Cdc42ep5 | 10.88113561 | -0.37325551 | 0.352598527 | -1.058584997 |
| Gm5319 | 0 | NA | NA | NA |
| Gm15926 | 0 | NA | NA | NA |
| Lilra5 | 0.951293484 | -2.256252515 | 1.299696597 | -1.735984013 |
| Gm15923 | 0 | NA | NA | NA |
| Gm15931 | 19.55735248 | -1.693858151 | 0.372775982 | -4.543903665 |
| Ncr1 | 7.234741561 | -2.025534791 | 0.46623266 | -4.34447212 |
| Gp6 | 0.386555381 | -1.81614575 | 3.683872171 | -0.492999123 |
| Gm25349 | 0.061920613 | -0.443816764 | 3.724945049 | -0.119147198 |
| Gm24752 | 0.04393175 | 0.037062677 | 3.724945049 | 0.009949859 |
| Rdh13 | 296.6943263 | -0.078620903 | 0.115921992 | -0.678222499 |
| Gm24605 | 0 | NA | NA | NA |
| Gm15494 | 14.28857811 | 1.491006119 | 0.376122757 | 3.964147583 |
| Eps8l1 | 3.37593979 | 1.600930463 | 0.602186925 | 2.658527438 |
| Ppp1r12c | 945.1906909 | 0.006853709 | 0.079462068 | 0.086251324 |

| | | | | |
|---|---|---|---|---|
| Tnnt1 | 15.65154636 | 0.850380374 | 0.284186139 | 2.992335854 |
| Tnni3 | 10.79028321 | 0.389269706 | 0.455628532 | 0.854357615 |
| Dnaaf3 | 15.48723177 | -0.468972809 | 0.3002222 | -1.562085713 |
| Gm15873 | 0.239904818 | -1.115447677 | 2.531083293 | -0.440699711 |
| Syt5 | 799.4033589 | 0.341968892 | 0.117369634 | 2.913606184 |
| Ptprh | 6.302761948 | -0.143860934 | 0.424350486 | -0.33901442 |
| Gm23577 | 0 | NA | NA | NA |
| Tmem86b | 45.98200731 | 0.286659083 | 0.257965379 | 1.111230831 |
| Ppp6r1 | 1889.707859 | 0.053925314 | 0.06253169 | 0.862367765 |
| Hspbp1 | 559.9998375 | 0.049678824 | 0.08184977 | 0.6069513 |
| Brsk1 | 2659.133262 | 0.609744026 | 0.084327828 | 7.230638375 |
| Tmem150b | 4.110811422 | -0.093177573 | 0.529433491 | -0.17599486 |
| Gm24537 | 0 | NA | NA | NA |
| Suv420h2 | 136.5439044 | -0.04474798 | 0.173258199 | -0.258273375 |
| Cox6b2 | 14.20000006 | 0.574449226 | 0.363261667 | 1.581364834 |
| Fam71e2 | 0.747386132 | -1.602660589 | 1.173093 | -1.366183745 |
| Il11 | 12.48613724 | -2.838071597 | 0.550518851 | -5.155266878 |
| Tmem190 | 0.273106308 | -1.620218587 | 1.760687638 | -0.92021921 |
| Tmem238 | 19.96996773 | -0.206005202 | 0.252103697 | -0.817144709 |
| Rpl28 | 949.3495926 | -0.163239537 | 0.099451611 | -1.641396611 |
| Ube2s | 303.8563974 | -0.209764029 | 0.115172303 | -1.821306197 |
| Shisa7 | 1255.260455 | 1.189727728 | 0.237445356 | 5.010532728 |
| Gm15525 | 0 | NA | NA | NA |
| Isoc2b | 8.838166061 | 0.082315259 | 0.406409783 | 0.202542514 |
| Isoc2a | 117.0217193 | 0.053150523 | 0.125615025 | 0.423122337 |
| Zfp628 | 234.1479967 | 0.335929563 | 0.139052128 | 2.415853453 |
| Nat14 | 284.6791148 | 0.153144622 | 0.116183148 | 1.318131114 |
| Ssc5d | 87.37008735 | -0.49140067 | 0.164272735 | -2.991370841 |
| Sbk2 | 2.02355919 | 0.627590514 | 0.715422701 | 0.877230361 |
| Sbk3 | 14.88936924 | 0.903242812 | 0.387081684 | 2.333468227 |
| Zfp579 | 184.6131844 | -0.092368343 | 0.109619391 | -0.842627767 |
| Fiz1 | 274.5806166 | -0.116425003 | 0.081412569 | -1.430061769 |
| Zfp524 | 43.21558028 | -0.191912973 | 0.207825464 | -0.923433391 |
| Zfp865 | 368.9050981 | -0.008892033 | 0.076605903 | -0.11607504 |
| 4632433K11Rik | 17.57441645 | 0.872148071 | 0.279051796 | 3.125398518 |
| Zfp784 | 56.52918824 | -0.475471982 | 0.167598158 | -2.836976186 |
| Gm15510 | 3.40753209 | -0.127581504 | 0.569484701 | -0.22402973 |
| Zfp580 | 125.3343534 | 0.222581157 | 0.136700613 | 1.62823818 |
| Ccdc106 | 206.5384023 | 0.296138593 | 0.113172246 | 2.616706861 |
| U2af2 | 1303.244093 | -0.225690027 | 0.05526929 | -4.083461657 |
| Epn1 | 2656.292776 | 0.407363269 | 0.073104701 | 5.572326614 |
| Rfpl4 | 0 | NA | NA | NA |
| Rasl2-9 | 0.99583754 | 0.058165427 | 1.061891447 | 0.054775304 |
| Gm26217 | 0 | NA | NA | NA |
| Vmn1r55 | 0.048949532 | 0.037064902 | 3.724945049 | 0.009950456 |
| Vmn1r56 | 0 | NA | NA | NA |
| Vmn1r57 | 0.085327104 | -0.905414734 | 3.723500899 | -0.243162217 |

| | | | | |
|---|---|---|---|---|
| Gm23942 | 0 | NA | NA | NA |
| Nlrp2 | 0 | NA | NA | NA |
| Gm5065 | 0.07796279 | -0.798301147 | 3.721829328 | -0.214491605 |
| Vmn1r58 | 0 | NA | NA | NA |
| Vmn1r59 | 0 | NA | NA | NA |
| Vmn2r28 | 0 | NA | NA | NA |
| Vmn1r60 | 0 | NA | NA | NA |
| Vmn1r-ps40 | 0 | NA | NA | NA |
| Vmn1r61 | 0 | NA | NA | NA |
| AC161762.1 | 0 | NA | NA | NA |
| Vmn1r62 | 0 | NA | NA | NA |
| Vmn1r63 | 0 | NA | NA | NA |
| Vmn1r64 | 0 | NA | NA | NA |
| Gm20399 | 0 | NA | NA | NA |
| Vmn1r-ps45 | 0 | NA | NA | NA |
| Vmn1r-ps46 | 0 | NA | NA | NA |
| Vmn1r65 | 0.115603922 | -0.93550548 | 3.724188147 | -0.251197158 |
| Gm5321 | 0.234623521 | -1.363320459 | 2.103129417 | -0.648234221 |
| Nlrp4c | 0 | NA | NA | NA |
| Zfp787 | 156.8006384 | -0.28727814 | 0.101376665 | -2.833769895 |
| Zfp444 | 319.2682102 | -0.001507455 | 0.063180852 | -0.023859362 |
| Galp | 0.278580731 | -0.697897962 | 1.749523481 | -0.398907457 |
| Zscan5b | 0.398992304 | -0.918202307 | 1.60074676 | -0.573608724 |
| Gm6792 | 0.123841226 | -1.017823543 | 3.719440177 | -0.273649661 |
| Zfp667 | 214.0871925 | 0.004096795 | 0.108730883 | 0.037678306 |
| Zfp583 | 81.78065755 | 0.07139765 | 0.14200913 | 0.502768028 |
| Zfp78 | 26.1290204 | 0.022899453 | 0.236195259 | 0.096951366 |
| Zfp28 | 117.9628295 | 0.186048218 | 0.102195295 | 1.82051647 |
| Smim17 | 178.4907971 | 0.336976536 | 0.174360248 | 1.932645434 |
| Gm16306 | 2.355736216 | 0.896706979 | 0.795554973 | 1.127146469 |
| Gm16307 | 0 | NA | NA | NA |
| Gm20715 | 6.848240459 | 0.214987887 | 0.490567247 | 0.438243458 |
| Olfr1344 | 0 | NA | NA | NA |
| Olfr1336 | 0 | NA | NA | NA |
| Olfr1346 | 0.047948698 | 0.037064902 | 3.724945049 | 0.009950456 |
| Olfr5 | 0.046345709 | -0.443818988 | 3.724945049 | -0.119147795 |
| Olfr1347 | 0.072025462 | 0.517959912 | 3.724945049 | 0.139051692 |
| Olfr1348 | 0 | NA | NA | NA |
| n-R5s150 | 0 | NA | NA | NA |
| Olfr1349 | 0.115957212 | 0.517955464 | 3.724945049 | 0.139050498 |
| Olfr1350 | 0 | NA | NA | NA |
| Zim1 | 102.3587915 | 0.03824969 | 0.189354483 | 0.202000447 |
| Peg3 | 8284.965525 | 0.319135853 | 0.077479927 | 4.118948834 |
| Usp29 | 1064.211711 | 0.075885718 | 0.100379956 | 0.755984767 |
| Gm27675 | 0 | NA | NA | NA |
| Mir3099 | 1.920827858 | 0.393752242 | 0.871671494 | 0.451720911 |
| Gm22649 | 0 | NA | NA | NA |

| | | | | |
|---|---|---|---|---|
| Aurkc | 1.107598099 | 0.908923901 | 1.037653978 | 0.875941229 |
| Zfp954 | 110.343462 | -0.49654513 | 0.143057405 | -3.470950212 |
| Zfp773 | 25.68791501 | -0.649116292 | 0.257661678 | -2.519258185 |
| Zfp418 | 91.99907945 | -0.230668777 | 0.160705315 | -1.435352507 |
| Zfp772 | 42.36616452 | -0.28714682 | 0.179570114 | -1.59907912 |
| Vmn2r29 | 3.485300844 | -0.128231969 | 0.624516335 | -0.205330047 |
| Gm18190 | 0.639457996 | -0.104669814 | 1.35021345 | -0.077520938 |
| Clcn4-2 | 1846.754651 | 0.211846374 | 0.057305644 | 3.696780263 |
| Mir5620 | 0.831589853 | 0.933795557 | 1.104452563 | 0.84548272 |
| Vmn2r30 | 0.25221175 | 0.517972928 | 2.131649059 | 0.242991653 |
| Gm3912 | 1.415212796 | 0.27448875 | 0.914444818 | 0.300169835 |
| Vmn2r31 | 0.094294407 | -0.443821212 | 3.724945049 | -0.119148392 |
| Vmn2r-ps36 | 0 | NA | NA | NA |
| Vmn2r32 | 0 | NA | NA | NA |
| Gm3563 | 0 | NA | NA | NA |
| Vmn2r33 | 0 | NA | NA | NA |
| Gm7648 | 0 | NA | NA | NA |
| Gm7651 | 0 | NA | NA | NA |
| Vmn2r-ps37 | 0.072025462 | 0.517959912 | 3.724945049 | 0.139051692 |
| Vmn2r34 | 0 | NA | NA | NA |
| Vmn2r-ps38 | 0 | NA | NA | NA |
| Vmn2r35 | 0.093141081 | 0.037060453 | 3.724945049 | 0.009949262 |
| Vmn2r-ps40 | 0.038981395 | -0.443816764 | 3.724945049 | -0.119147198 |
| Vmn2r36 | 0 | NA | NA | NA |
| Gm7230 | 0 | NA | NA | NA |
| Gm3978 | 0.160408561 | 0.517955075 | 3.457615493 | 0.149801236 |
| Gm20438 | 0.038981395 | -0.443816764 | 3.724945049 | -0.119147198 |
| Gm18200 | 0.398333997 | -0.467155936 | 1.580999136 | -0.295481462 |
| Gm4073 | 0 | NA | NA | NA |
| Gm3675 | 0 | NA | NA | NA |
| 2810047C21Rik1 | 0.042716945 | 0.037062677 | 3.724945049 | 0.009949859 |
| Gm3665 | 0 | NA | NA | NA |
| Vmn2r41 | 0.042716945 | 0.037062677 | 3.724945049 | 0.009949859 |
| Vmn2r42 | 0 | NA | NA | NA |
| Vmn2r43 | 0 | NA | NA | NA |
| Gm3654 | 0 | NA | NA | NA |
| Vmn2r-ps45 | 0.124853745 | 1.05357957 | 3.721447096 | 0.283110183 |
| Gm7696 | 0.676743848 | 0.367531019 | 1.139209864 | 0.322619239 |
| Vmn2r44 | 0.263039019 | 0.5179731 | 2.106825972 | 0.245854715 |
| Vmn2r-ps46 | 0 | NA | NA | NA |
| Vmn2r45 | 0.068097359 | 0.037064902 | 3.724945049 | 0.009950456 |
| Vmn2r-ps47 | 0 | NA | NA | NA |
| Vmn2r-ps49 | 0 | NA | NA | NA |
| Gm3584 | 0.047948698 | 0.037064902 | 3.724945049 | 0.009950456 |
| Gm3956 | 0 | NA | NA | NA |
| Gm3585 | 0 | NA | NA | NA |
| Gm3962 | 0.333123609 | 0.037077385 | 1.720315301 | 0.021552669 |

| | | | | |
|---|---|---|---|---|
| Gm18191 | 0.264294608 | -1.302884165 | 1.785244199 | -0.729807253 |
| Vmn2r-ps44 | 0 | NA | NA | NA |
| Gm3644 | 0 | NA | NA | NA |
| Gm20482 | 0 | NA | NA | NA |
| Vmn2r-ps43 | 0 | NA | NA | NA |
| Vmn2r40 | 0 | NA | NA | NA |
| Vmn2r-ps42 | 0 | NA | NA | NA |
| Vmn2r39 | 0 | NA | NA | NA |
| Zfp419 | 0.038981395 | -0.443816764 | 3.724945049 | -0.119147198 |
| Vmn2r38 | 0.068097359 | 0.037064902 | 3.724945049 | 0.009950456 |
| Gm5583 | 0 | NA | NA | NA |
| Gm18201 | 0 | NA | NA | NA |
| Vmn2r-ps41 | 0 | NA | NA | NA |
| Vmn2r37 | 0 | NA | NA | NA |
| Gm4011 | 0 | NA | NA | NA |
| Gm3623 | 0 | NA | NA | NA |
| Gm3616 | 0 | NA | NA | NA |
| Gm20287 | 0 | NA | NA | NA |
| Gm3611 | 0 | NA | NA | NA |
| Gm3972 | 0.099106094 | -0.93550548 | 3.724188147 | -0.251197158 |
| Gm18199 | 0.40494807 | 0.575394885 | 1.696067947 | 0.339252261 |
| Gm10302 | 0 | NA | NA | NA |
| Gm3683 | 0 | NA | NA | NA |
| Gm20345 | 0 | NA | NA | NA |
| Vmn2r46 | 0.048949532 | 0.037064902 | 3.724945049 | 0.009950456 |
| Vmn2r47 | 0 | NA | NA | NA |
| Vmn2r-ps51 | 0 | NA | NA | NA |
| Vmn2r-ps52 | 0 | NA | NA | NA |
| Vmn2r48 | 0 | NA | NA | NA |
| Vmn2r49 | 0 | NA | NA | NA |
| Vmn2r50 | 0.038981395 | -0.443816764 | 3.724945049 | -0.119147198 |
| Vmn2r51 | 0 | NA | NA | NA |
| Vmn2r52 | 0 | NA | NA | NA |
| Gm20679 | 0 | NA | NA | NA |
| Vmn1r66 | 0 | NA | NA | NA |
| Vmn1r-ps47 | 0 | NA | NA | NA |
| Vmn1r-ps48 | 0 | NA | NA | NA |
| Nlrp4d | 0.062426873 | 0.517959912 | 3.724945049 | 0.139051692 |
| Vmn1r67 | 0 | NA | NA | NA |
| Zik1 | 251.415385 | 0.049139492 | 0.092735109 | 0.529890918 |
| Vmn1r68 | 0 | NA | NA | NA |
| Vmn1r-ps49 | 0 | NA | NA | NA |
| Vmn1r69 | 0 | NA | NA | NA |
| Vmn1r-ps50 | 0 | NA | NA | NA |
| Vmn1r70 | 0 | NA | NA | NA |
| Vmn1r-ps51 | 0 | NA | NA | NA |
| Gm6900 | 2.818881342 | -0.339997736 | 0.609625923 | -0.557715351 |

| | | | | |
|---|---|---|---|---|
| Vmn1r-ps52 | 0 | NA | NA | NA |
| Nlrp4b | 0 | NA | NA | NA |
| Vmn1r71 | 0 | NA | NA | NA |
| Zscan4b | 0 | NA | NA | NA |
| Mir297-1 | 0 | NA | NA | NA |
| Zscan4c | 0 | NA | NA | NA |
| Zscan4d | 0 | NA | NA | NA |
| Gm26248 | 0 | NA | NA | NA |
| Zscan4e | 0 | NA | NA | NA |
| Gm23087 | 0 | NA | NA | NA |
| Zscan4f | 0 | NA | NA | NA |
| Vmn1r-ps53 | 0 | NA | NA | NA |
| Vmn1r72 | 0 | NA | NA | NA |
| Vmn1r73 | 0 | NA | NA | NA |
| Vmn1r-ps54 | 0 | NA | NA | NA |
| Vmn1r74 | 0 | NA | NA | NA |
| Vmn1r-ps55 | 0 | NA | NA | NA |
| Vmn1r75 | 0 | NA | NA | NA |
| Vmn1r-ps56 | 0 | NA | NA | NA |
| Vmn1r76 | 0 | NA | NA | NA |
| Vmn1r-ps57 | 0 | NA | NA | NA |
| Gm24414 | 1.280135606 | -0.872990883 | 0.95134145 | -0.917642012 |
| Vmn1r-ps58 | 0 | NA | NA | NA |
| Vmn1r-ps59 | 0 | NA | NA | NA |
| Vmn1r-ps60 | 0 | NA | NA | NA |
| Vmn1r77 | 0 | NA | NA | NA |
| Gm10179 | 23.18691661 | 0.288379678 | 0.253834191 | 1.136094697 |
| Vmn1r-ps61 | 0 | NA | NA | NA |
| Vmn1r-ps62 | 0 | NA | NA | NA |
| Vmn1r-ps63 | 0 | NA | NA | NA |
| Vmn1r78 | 0 | NA | NA | NA |
| Vmn1r79 | 0 | NA | NA | NA |
| Vmn1r80 | 0 | NA | NA | NA |
| Vmn1r-ps64 | 0 | NA | NA | NA |
| Vmn1r-ps65 | 0 | NA | NA | NA |
| Vmn1r81 | 0 | NA | NA | NA |
| Vmn1r82 | 0 | NA | NA | NA |
| Vmn1r83 | 0 | NA | NA | NA |
| Gm23521 | 0 | NA | NA | NA |
| Vmn1r84 | 0.047948698 | 0.037064902 | 3.724945049 | 0.009950456 |
| Vmn1r-ps66 | 0 | NA | NA | NA |
| Vmn1r-ps67 | 0 | NA | NA | NA |
| Vmn1r-ps68 | 0 | NA | NA | NA |
| Zfp551 | 75.69228729 | 0.083440834 | 0.127972326 | 0.652022482 |
| Vmn2r-ps53 | 0 | NA | NA | NA |
| Zfp606 | 312.4159778 | 0.050196703 | 0.07379081 | 0.680256835 |
| 2900092C05Rik | 0.326351934 | -1.056578655 | 1.67067204 | -0.632427329 |

| | | | | |
|---|---|---|---|---|
| Gm15877 | 0.593305459 | 0.556410941 | 1.191510324 | 0.466979539 |
| Vmn2r53 | 2.096357237 | 0.747364412 | 0.718293242 | 1.040472566 |
| Gm15773 | 0.596424136 | -0.347239449 | 1.337723452 | -0.259574913 |
| Vmn2r54 | 1.895285101 | 0.135024846 | 0.78638424 | 0.171703397 |
| Vmn2r55 | 1.524911348 | 1.492885324 | 0.917383544 | 1.627329522 |
| Vmn2r56 | 2.422070769 | 1.900989674 | 0.944076332 | 2.013597428 |
| Gm6946 | 0 | NA | NA | NA |
| Zscan18 | 180.8411845 | 0.495337626 | 0.148215069 | 3.342019329 |
| Zfp329 | 519.818062 | -0.225028236 | 0.126152298 | -1.783782304 |
| Zfp110 | 270.5868238 | -0.502325392 | 0.113147404 | -4.439566192 |
| Zfp128 | 118.8763104 | -0.104757317 | 0.138369931 | -0.757081513 |
| Zscan22 | 156.5051589 | -0.432863318 | 0.161294379 | -2.683685081 |
| Rps5 | 1620.953915 | -0.214057704 | 0.12902251 | -1.659072549 |
| Rnf225 | 104.6098484 | -0.774332643 | 0.185288884 | -4.17905612 |
| Gm26325 | 0 | NA | NA | NA |
| 1810019N24Rik | 6.340267528 | -0.302306249 | 0.43875506 | -0.689009145 |
| Gm29638 | 0.046345709 | -0.443818988 | 3.724945049 | -0.119147795 |
| Zfp324 | 270.4925259 | -0.32668911 | 0.10675536 | -3.060165886 |
| Zfp446 | 119.2751431 | 0.083947039 | 0.144575928 | 0.580643266 |
| Slc27a5 | 10.7500977 | -0.138525579 | 0.367884196 | -0.376546699 |
| Zbtb45 | 249.6021384 | 0.119567042 | 0.100504216 | 1.189671904 |
| Trim28 | 1232.544845 | -0.037462599 | 0.072825349 | -0.514417026 |
| Chmp2a | 510.643161 | -0.274928785 | 0.108262275 | -2.539469872 |
| Ube2m | 1024.417635 | -0.086213862 | 0.082682791 | -1.042706237 |
| Mzf1 | 76.8160005 | 0.897223911 | 0.199520522 | 4.496900373 |
| Vmn1r85 | 0 | NA | NA | NA |
| Vmn1r86 | 0 | NA | NA | NA |
| Vmn1r87 | 0 | NA | NA | NA |
| Vmn1r88 | 0.289463044 | 1.87930748 | 2.01170423 | 0.934186772 |
| Gm4879 | 6.771121092 | -0.253480381 | 0.453710746 | -0.558682779 |
| Vmn1r89 | 0.099106094 | -0.93550548 | 3.724188147 | -0.251197158 |
| Vmn1r-ps69 | 0 | NA | NA | NA |
| 6330408A02Rik | 197.7921584 | -0.154297194 | 0.184468985 | -0.836439766 |
| Lig1 | 167.3687248 | -0.114289554 | 0.126023173 | -0.90689316 |
| Pla2g4c | 22.31652208 | -1.386714074 | 0.304845667 | -4.548905309 |
| Rps8-ps4 | 2.214157104 | -0.088032169 | 0.640285145 | -0.137489007 |
| Gm24909 | 0.201760502 | -0.46777956 | 2.220505866 | -0.21066351 |
| Gm23499 | 0 | NA | NA | NA |
| Cabp5 | 0.609661125 | -0.501295288 | 1.390809503 | -0.360434184 |
| Bsph1 | 0.222333927 | 0.918104579 | 2.12834778 | 0.431369623 |
| Bsph2-ps | 0 | NA | NA | NA |
| Bsph2 | 0 | NA | NA | NA |
| Sult2a5 | 0 | NA | NA | NA |
| Zbed4-ps1 | 0 | NA | NA | NA |
| Kat6b-ps1 | 0 | NA | NA | NA |
| Tmem167-ps1 | 0 | NA | NA | NA |
| Sult2a-ps1 | 0 | NA | NA | NA |

| | | | | |
|---|---|---|---|---|
| Sult2a2 | 0 | NA | NA | NA |
| Zbed4-ps2 | 0.047948698 | 0.037064902 | 3.724945049 | 0.009950456 |
| Sult2a1 | 0.169165009 | 0.036125373 | 3.351346561 | 0.01077936 |
| Kat6b-ps2 | 0 | NA | NA | NA |
| Sult2a-ps2 | 0 | NA | NA | NA |
| Tmem167-ps2 | 0 | NA | NA | NA |
| Sult2a4 | 0 | NA | NA | NA |
| Zbed4-ps3 | 0.112642008 | 0.517955464 | 3.724945049 | 0.139050498 |
| Tmem167-ps3 | 0 | NA | NA | NA |
| Sult2a3 | 0 | NA | NA | NA |
| Sult2a-ps3 | 0 | NA | NA | NA |
| Sult2a-ps4 | 0 | NA | NA | NA |
| Rpap3-ps1 | 0 | NA | NA | NA |
| Sult2a6 | 0 | NA | NA | NA |
| Rpap3-ps2 | 0 | NA | NA | NA |
| Phf20-ps | 17.46056627 | -0.093092253 | 0.250261024 | -0.371980629 |
| Obox8 | 0 | NA | NA | NA |
| Mrps36-ps3 | 0 | NA | NA | NA |
| Crs-ps | 0 | NA | NA | NA |
| Sult2a8 | 0.047948698 | 0.037064902 | 3.724945049 | 0.009950456 |
| Sult2a7 | 0.119587853 | 0.981063299 | 3.723618439 | 0.263470416 |
| Obox4-ps1 | 0.155414793 | 0.037075986 | 2.829374052 | 0.013103953 |
| Obox4-ps2 | 0.183158741 | -0.443823653 | 2.738290263 | -0.162080572 |
| Nlrp5-ps | 183.9697913 | 0.303675938 | 0.176060837 | 1.724835249 |
| Obox4-ps3 | 0 | NA | NA | NA |
| Vmn1r90 | 3.490537078 | 0.069929583 | 0.590288396 | 0.118466809 |
| Zfp-ps | 0 | NA | NA | NA |
| Ranbp2-ps1 | 0 | NA | NA | NA |
| Obox3-ps7 | 4.106584469 | 0.278541772 | 0.572448572 | 0.486579557 |
| Obox7 | 0.129647038 | -0.443820923 | 2.992845889 | -0.148293945 |
| Obox3-ps1 | 0 | NA | NA | NA |
| RP24-194K24.10 | 0 | NA | NA | NA |
| Gtpbp4-ps1 | 0 | NA | NA | NA |
| Ranbp2-ps2 | 0 | NA | NA | NA |
| Obox4-ps36 | 0 | NA | NA | NA |
| Obox4-ps37 | 0 | NA | NA | NA |
| Gm26056 | 0 | NA | NA | NA |
| Obox3-ps2 | 0 | NA | NA | NA |
| RP23-349I13.7 | 0 | NA | NA | NA |
| Gtpbp4-ps2 | 0 | NA | NA | NA |
| Ranbp2-ps3 | 0 | NA | NA | NA |
| Obox4-ps38 | 0 | NA | NA | NA |
| Obox4-ps39 | 0 | NA | NA | NA |
| Ranbp2-ps4 | 0 | NA | NA | NA |
| Obox4-ps6 | 0 | NA | NA | NA |
| Gm24701 | 0 | NA | NA | NA |
| Obox3-ps3 | 0 | NA | NA | NA |

| | | | | |
|---|---|---|---|---|
| RP23-263E7.8 | 0 | NA | NA | NA |
| Gtpbp4-ps3 | 0 | NA | NA | NA |
| Ranbp2-ps5 | 0 | NA | NA | NA |
| Obox4-ps8 | 0 | NA | NA | NA |
| Obox4-ps9 | 0 | NA | NA | NA |
| Obox3-ps4 | 0 | NA | NA | NA |
| RP23-263E7.7 | 0 | NA | NA | NA |
| Gtpbp4-ps4 | 0 | NA | NA | NA |
| Ranbp2-ps6 | 0 | NA | NA | NA |
| Obox4-ps11 | 0 | NA | NA | NA |
| Obox4-ps12 | 0 | NA | NA | NA |
| Obox4-ps13 | 0 | NA | NA | NA |
| Obox4-ps14 | 0 | NA | NA | NA |
| Obox4-ps15 | 0 | NA | NA | NA |
| Obox4-ps16 | 0 | NA | NA | NA |
| Obox4-ps17 | 0 | NA | NA | NA |
| Obox4-ps18 | 0 | NA | NA | NA |
| Obox4-ps19 | 0 | NA | NA | NA |
| Obox4-ps20 | 0 | NA | NA | NA |
| Obox4-ps21 | 0 | NA | NA | NA |
| Obox4-ps22 | 0 | NA | NA | NA |
| Obox4-ps23 | 0 | NA | NA | NA |
| Mrip-ps | 35.07510409 | 0.376199666 | 0.350642408 | 1.072886955 |
| Obox3-ps5 | 0 | NA | NA | NA |
| RP24-84K14.12 | 0 | NA | NA | NA |
| Gtpbp4-ps5 | 0 | NA | NA | NA |
| RP24-105I10.4 | 0 | NA | NA | NA |
| Obox4-ps26 | 0 | NA | NA | NA |
| Ranbp2-ps7 | 0 | NA | NA | NA |
| Obox2 | 0 | NA | NA | NA |
| Obox1 | 0 | NA | NA | NA |
| Ranbp2-ps8 | 0 | NA | NA | NA |
| Obox4-ps27 | 0 | NA | NA | NA |
| Obox4-ps28 | 0 | NA | NA | NA |
| Obox4-ps29 | 0 | NA | NA | NA |
| Obox3 | 0 | NA | NA | NA |
| Obox3-ps6 | 0 | NA | NA | NA |
| Obox4-ps30 | 0 | NA | NA | NA |
| Ranbp2-ps9 | 0 | NA | NA | NA |
| Ranbp2-ps10 | 0 | NA | NA | NA |
| Obox4-ps31 | 0 | NA | NA | NA |
| Gm25354 | 0 | NA | NA | NA |
| RP23-140B24.1 | 0 | NA | NA | NA |
| RP23-140B24.2 | 0 | NA | NA | NA |
| Obox4-ps33 | 0 | NA | NA | NA |
| Gm23714 | 0 | NA | NA | NA |
| Obox4-ps34 | 0 | NA | NA | NA |

| | | | | |
|---|---|---|---|---|
| RP24-370G23.12 | 0 | NA | NA | NA |
| Obox5 | 0 | NA | NA | NA |
| RP24-370G23.13 | 0 | NA | NA | NA |
| RP24-370G23.14 | 0 | NA | NA | NA |
| Obox4-ps35 | 0.220980774 | -1.406009479 | 3.104094945 | -0.452953116 |
| Obox6 | 0.123845245 | -0.280519638 | 3.720871772 | -0.075390837 |
| Crx | 0.603215896 | 1.81841149 | 2.549944359 | 0.713118105 |
| Crxos | 0.501748666 | 0.037077518 | 1.443598455 | 0.025684094 |
| RP24-466N11.6 | 0 | NA | NA | NA |
| Sepw1 | 5104.264334 | 0.065651564 | 0.09647921 | 0.680473689 |
| Gltscr2 | 737.8364871 | 0.048571638 | 0.092812654 | 0.523329905 |
| Snord23 | 0.926190085 | -0.643594869 | 1.034213301 | -0.622303802 |
| Gm24576 | 0.169505627 | 0.05502394 | 2.747555859 | 0.020026505 |
| Ehd2 | 172.4468587 | -1.201525572 | 0.181620679 | -6.615576937 |
| Gltscr1 | 438.3260183 | -0.053586148 | 0.077705776 | -0.689603143 |
| Zfp541 | 3.78555653 | 0.208720451 | 0.551930615 | 0.378164293 |
| Napa | 1603.381598 | 0.15690278 | 0.0567928 | 2.762723108 |
| Kptn | 155.1087845 | 0.27400729 | 0.11010337 | 2.488636719 |
| Slc8a2 | 2081.402497 | 1.058609473 | 0.140685086 | 7.524674447 |
| Meis3 | 261.2098945 | 0.186914363 | 0.103509185 | 1.805775633 |
| Dhx34 | 160.4271404 | -0.21892151 | 0.088016985 | -2.487264355 |
| C5ar2 | 2.109974762 | -0.848432107 | 0.747314625 | -1.135307778 |
| Gm26473 | 0 | NA | NA | NA |
| C5ar1 | 78.4729806 | -1.343716775 | 0.28449886 | -4.723100729 |
| Inafm1 | 160.8892379 | -0.210805369 | 0.112540111 | -1.873157638 |
| Ccdc9 | 366.0675831 | 0.000772222 | 0.102880107 | 0.007506034 |
| Bbc3 | 74.98142202 | 0.013771902 | 0.146445823 | 0.094040932 |
| Sae1 | 546.3005839 | -0.193005533 | 0.063309851 | -3.048586091 |
| Zc3h4 | 1014.843218 | -0.144440036 | 0.076992757 | -1.876021084 |
| Tmem160 | 153.6281965 | -0.124241398 | 0.149795637 | -0.829405984 |
| Npas1 | 76.47707439 | 0.210673957 | 0.147163727 | 1.431561716 |
| n-R5s151 | 0 | NA | NA | NA |
| Arhgap35 | 4302.533817 | 0.082317688 | 0.051684705 | 1.592689525 |
| Gm17826 | 0.07796279 | -0.798301147 | 3.721829328 | -0.214491605 |
| Gm17981 | 12.95166994 | 0.118923982 | 0.397885212 | 0.298890177 |
| Gm29443 | 4.384705177 | 0.955566131 | 0.550226662 | 1.73667726 |
| Gm29442 | 0.132101749 | -0.93550548 | 3.724188147 | -0.251197158 |
| Gm24708 | 0 | NA | NA | NA |
| Ceacam15 | 0.205464717 | -0.453194645 | 2.642335508 | -0.171512907 |
| Ceacam9 | 1.215943724 | 0.74187646 | 0.951010126 | 0.780093124 |
| Ap2s1 | 780.4435061 | 0.028700731 | 0.097878129 | 0.293229256 |
| Slc1a5 | 49.69459424 | -0.334258157 | 0.224294409 | -1.490265225 |
| Fkrp | 486.4658568 | 0.255190634 | 0.077789472 | 3.280529193 |
| Strn4 | 1924.374574 | 0.46994607 | 0.078946401 | 5.952723129 |
| Prkd2 | 82.10841754 | -0.751628005 | 0.179412601 | -4.189382465 |
| 9330104G04Rik | 30.06220494 | 0.367801946 | 0.208730374 | 1.762091157 |
| Dact3 | 1250.275623 | 0.837197591 | 0.100042034 | 8.368458346 |

| | | | | |
|---|---|---|---|---|
| Gng8 | 8.57217951 | -0.065171237 | 0.390606026 | -0.166846471 |
| Ptgir | 4.875732241 | -1.825276636 | 0.502265065 | -3.634090368 |
| Calm3 | 11988.4177 | 0.484496683 | 0.069461203 | 6.975068947 |
| Pnmal2 | 4095.934161 | 0.35035404 | 0.074672059 | 4.691902776 |
| Gm25815 | 0 | NA | NA | NA |
| Pnmal1 | 642.282333 | 0.303549583 | 0.105796977 | 2.869170661 |
| Ccdc8 | 31.97129462 | -0.27798591 | 0.26236149 | -1.059553022 |
| Ppp5c | 936.7635144 | 0.114904783 | 0.053832465 | 2.134488583 |
| Hif3a | 169.2274098 | 0.889751757 | 0.201528631 | 4.415014143 |
| Psg16 | 99.26004857 | 0.295647366 | 0.189414918 | 1.5608452 |
| Ceacam3 | 2.261397335 | 0.318167451 | 0.796584602 | 0.399414513 |
| Psg29 | 2.218822718 | 0.11225761 | 0.732560455 | 0.153240063 |
| Gm24520 | 0 | NA | NA | NA |
| Gm25068 | 0 | NA | NA | NA |
| Ceacam5 | 0 | NA | NA | NA |
| Gm6065 | 0.197744397 | -0.443050898 | 2.238155105 | -0.197953617 |
| Ceacam14 | 0 | NA | NA | NA |
| Gm5155 | 0 | NA | NA | NA |
| Ceacam11 | 0 | NA | NA | NA |
| Ceacam13 | 0 | NA | NA | NA |
| Gm22414 | 0 | NA | NA | NA |
| Ceacam12 | 0 | NA | NA | NA |
| Igfl3 | 0 | NA | NA | NA |
| Mill1 | 0 | NA | NA | NA |
| Psg18 | 0 | NA | NA | NA |
| Gm24715 | 0 | NA | NA | NA |
| Psg28 | 0.047948698 | 0.037064902 | 3.724945049 | 0.009950456 |
| Psg26 | 0 | NA | NA | NA |
| Psg25 | 0.049553047 | -0.443818988 | 3.724945049 | -0.119147795 |
| Gm26062 | 0 | NA | NA | NA |
| Psg27 | 0 | NA | NA | NA |
| Psg23 | 0.066050875 | -0.443818988 | 3.724945049 | -0.119147795 |
| Psg21 | 0 | NA | NA | NA |
| Psg20 | 0.124347486 | 0.036092774 | 3.717245565 | 0.009709548 |
| Psg22 | 0 | NA | NA | NA |
| Gm24464 | 0 | NA | NA | NA |
| Psg19 | 0 | NA | NA | NA |
| Psg17 | 1.789359587 | -1.19086875 | 0.897128322 | -1.327422979 |
| Mill2 | 14.66817284 | 0.274500052 | 0.341185362 | 0.804548148 |
| Pglyrp1 | 28.41791276 | -0.084702479 | 0.249917457 | -0.338921817 |
| Ccdc61 | 68.71592428 | -0.416876626 | 0.179074289 | -2.327953549 |
| Nova2 | 1854.132093 | 0.496960126 | 0.106631096 | 4.660555357 |
| Gm26821 | 0.975505809 | -2.183424754 | 1.106751005 | -1.972823829 |
| Nanos2 | 10.4316941 | 1.497953096 | 0.439010662 | 3.41211097 |
| Gm10676 | 6.752775585 | 0.161985559 | 0.408689681 | 0.396353437 |
| Mypop | 377.759842 | 0.136495178 | 0.089859128 | 1.518990668 |
| Mypopos | 3.140367151 | -0.734544378 | 0.616875479 | -1.190749839 |

| | | | | |
|---|---|---|---|---|
| Irf2bp1 | 595.038875 | -0.017379466 | 0.077406422 | -0.224522271 |
| Foxa3 | 0.267510113 | -1.560388921 | 2.390276127 | -0.652806972 |
| Sympk | 1388.597211 | 0.063412563 | 0.059549544 | 1.064870676 |
| Rsph6a | 1.359012152 | -0.769879662 | 0.946299407 | -0.813568788 |
| Dmwd | 973.7957556 | 0.304225916 | 0.078839746 | 3.858788628 |
| Dmpk | 178.6755614 | 0.341558233 | 0.206800511 | 1.651631477 |
| Mir3100 | 1.552752546 | -0.411951427 | 0.916471823 | -0.4494971 |
| Six5 | 17.71455458 | -0.188879841 | 0.298649933 | -0.632445616 |
| Gm4969 | 2.587362481 | 0.718164852 | 0.855414207 | 0.839552168 |
| Fbxo46 | 157.8854747 | -0.444790636 | 0.149905548 | -2.967139248 |
| Qpctl | 177.5083971 | -0.036171691 | 0.124813884 | -0.289805025 |
| Snrpd2 | 249.1707862 | -0.265067818 | 0.123617737 | -2.144253932 |
| Gipr | 9.076618891 | -0.659909518 | 0.451036639 | -1.463095146 |
| Eml2 | 1469.733199 | 0.381984018 | 0.072117422 | 5.296695426 |
| Mir330 | 0 | NA | NA | NA |
| Gpr4 | 26.0798917 | 0.080138515 | 0.222444017 | 0.360263748 |
| Opa3 | 585.1491548 | -0.177734604 | 0.064555504 | -2.753206066 |
| Gm25134 | 1.373839161 | 0.565270356 | 0.998728445 | 0.565990043 |
| Vasp | 344.2888316 | -0.668924725 | 0.139496985 | -4.795262958 |
| 1700058P15Rik | 0 | NA | NA | NA |
| Ppm1n | 0.85008157 | -0.359161485 | 1.144311099 | -0.313866994 |
| Rtn2 | 574.5812076 | 0.367527171 | 0.101414621 | 3.624005764 |
| Gm26802 | 0.561122788 | -0.870286143 | 1.540408904 | -0.56497086 |
| Fosb | 461.8672414 | -1.390351104 | 0.612526982 | -2.269860992 |
| Ercc1 | 157.0107263 | -0.65233038 | 0.120985138 | -5.39182242 |
| Cd3eap | 115.4671044 | -0.264409993 | 0.120812837 | -2.188591871 |
| Ppp1r13l | 62.86165383 | -1.060476369 | 0.167594546 | -6.327630547 |
| Ercc2 | 283.3150772 | -0.014341095 | 0.084269376 | -0.170181577 |
| Gm26852 | 0.604815377 | -1.241174731 | 1.214982496 | -1.021557706 |
| Mir343 | 1.738846634 | 1.219731889 | 0.966513288 | 1.261991846 |
| Klc3 | 30.34969473 | -0.054436261 | 0.202927216 | -0.268255102 |
| Ckm | 1.573181334 | -0.787985316 | 0.845607006 | -0.931857602 |
| A930016O22Rik | 2.771915612 | 0.819021993 | 0.708259844 | 1.156386318 |
| Mark4 | 736.3036542 | 0.042182255 | 0.069163745 | 0.60988968 |
| Gm23099 | 0 | NA | NA | NA |
| Exoc3l2 | 2.258282039 | -0.563814957 | 0.687683226 | -0.819875977 |
| Bloc1s3 | 84.9281982 | -0.304194863 | 0.126177222 | -2.410854026 |
| Trappc6a | 46.60324668 | -0.039300311 | 0.183785257 | -0.213838211 |
| Nkpd1 | 9.973239763 | 0.211739238 | 0.33227123 | 0.637248186 |
| Ppp1r37 | 1313.266029 | -0.05293847 | 0.073560305 | -0.719660827 |
| Gemin7 | 188.0515057 | -0.209761751 | 0.102529759 | -2.045862128 |
| Zfp296 | 23.1978082 | -0.171026718 | 0.249978032 | -0.684166993 |
| Clasrp | 352.6239218 | 0.390608768 | 0.176991701 | 2.206932679 |
| Gm26890 | 51.60349904 | 0.25338778 | 0.356512814 | 0.710739616 |
| Relb | 145.3464235 | -0.189618369 | 0.230801795 | -0.821563667 |
| Clptm1 | 1928.605293 | -0.032180371 | 0.064590739 | -0.498219588 |
| Apoc2 | 34.46812154 | -1.625155908 | 0.3275793 | -4.96110685 |

| | | | | |
|---|---|---|---|---|
| Apoc4 | 0.250169628 | 0.517972894 | 2.136490373 | 0.242441015 |
| Apoc1 | 11.42902807 | -0.086426306 | 0.34601645 | -0.249775136 |
| Apoe | 8648.195503 | 0.157133343 | 0.177201504 | 0.886749489 |
| Tomm40 | 595.2610474 | -0.098274993 | 0.066263093 | -1.483103024 |
| Gm20512 | 1.197749286 | 1.754822651 | 1.164941753 | 1.506360851 |
| Pvrl2 | 109.3428072 | -0.434875165 | 0.161118204 | -2.699106329 |
| Bcam | 112.1762543 | -0.255238273 | 0.151481362 | -1.684948366 |
| Gm26600 | 0 | NA | NA | NA |
| Cblc | 0.325981489 | 0.037763393 | 1.519260843 | 0.024856425 |
| Bcl3 | 136.661948 | -1.225155043 | 0.350268037 | -3.497764318 |
| Gm16175 | 0.542127486 | -0.460326899 | 1.28806225 | -0.357379388 |
| Gm16174 | 1.534203801 | 0.625800107 | 0.729322177 | 0.858057147 |
| Gm19345 | 14.35855545 | 0.162518328 | 0.308617369 | 0.526601367 |
| Ceacam16 | 1.967062318 | 0.448812143 | 0.731188537 | 0.613811788 |
| Ceacam19 | 1.492916405 | -2.02442095 | 0.8878985 | -2.280013931 |
| Pvr | 193.8546322 | -1.119958882 | 0.179030986 | -6.255670624 |
| Gm16184 | 0 | NA | NA | NA |
| Gm16251 | 0.990102919 | 0.760173199 | 1.137687387 | 0.668174059 |
| Igsf23 | 0.400433843 | -1.883636344 | 1.587649108 | -1.186431142 |
| Rpl7a-ps8 | 0.1140246 | 0.037073798 | 3.724945049 | 0.009952844 |
| Ceacam20 | 0.152347716 | 0.503763231 | 2.842792142 | 0.177207198 |
| Nlrp9b | 0 | NA | NA | NA |
| n-R5s152 | 0 | NA | NA | NA |
| Vmn1r91 | 0 | NA | NA | NA |
| Gm16442 | 0 | NA | NA | NA |
| Vmn1r93 | 0 | NA | NA | NA |
| Vmn1r94 | 0 | NA | NA | NA |
| Vmn1r95 | 0 | NA | NA | NA |
| Gm6164 | 0 | NA | NA | NA |
| Gm4498 | 0 | NA | NA | NA |
| Gm4133 | 0 | NA | NA | NA |
| Vmn1r-ps79 | 0 | NA | NA | NA |
| Gm5728 | 0 | NA | NA | NA |
| Vmn1r100 | 0 | NA | NA | NA |
| Vmn1r101 | 0 | NA | NA | NA |
| Gm10670 | 0 | NA | NA | NA |
| Vmn1r103 | 0 | NA | NA | NA |
| Vmn1r104 | 0 | NA | NA | NA |
| Gm5726 | 0 | NA | NA | NA |
| Gm4141 | 0 | NA | NA | NA |
| Vmn1r107 | 0 | NA | NA | NA |
| Vmn1r-ps71 | 0 | NA | NA | NA |
| Vmn1r111 | 0 | NA | NA | NA |
| Gm10668 | 0 | NA | NA | NA |
| Vmn1r112 | 0 | NA | NA | NA |
| Vmn1r113 | 0 | NA | NA | NA |
| Vmn1r114 | 0 | NA | NA | NA |

| | | | | |
|---|---|---|---|---|
| Vmn1r115 | 0 | NA | NA | NA |
| Vmn1r116 | 0 | NA | NA | NA |
| Vmn1r117 | 0 | NA | NA | NA |
| Vmn1r118 | 0 | NA | NA | NA |
| Gm4526 | 0 | NA | NA | NA |
| Vmn1r119 | 0 | NA | NA | NA |
| Vmn1r-ps73 | 0 | NA | NA | NA |
| Vmn1r120 | 0 | NA | NA | NA |
| Vmn1r-ps74 | 0 | NA | NA | NA |
| Vmn1r121 | 0 | NA | NA | NA |
| Vmn1r122 | 0 | NA | NA | NA |
| Vmn1r123 | 0 | NA | NA | NA |
| Gm5157 | 0 | NA | NA | NA |
| Vmn1r124 | 0 | NA | NA | NA |
| Vmn1r125 | 0 | NA | NA | NA |
| Vmn1r126 | 0 | NA | NA | NA |
| Vmn1r127 | 0 | NA | NA | NA |
| Vmn1r128 | 0 | NA | NA | NA |
| Vmn1r129 | 0 | NA | NA | NA |
| Vmn1r-ps75 | 0 | NA | NA | NA |
| Gm6882 | 0 | NA | NA | NA |
| Vmn1r130 | 0 | NA | NA | NA |
| Vmn1r-ps76 | 0 | NA | NA | NA |
| Vmn1r131 | 0 | NA | NA | NA |
| Gm4545 | 0 | NA | NA | NA |
| Vmn1r132 | 0 | NA | NA | NA |
| Gm8600 | 0 | NA | NA | NA |
| Gm4175 | 0 | NA | NA | NA |
| Gm4177 | 0 | NA | NA | NA |
| Vmn1r135 | 0 | NA | NA | NA |
| Gm5725 | 0 | NA | NA | NA |
| Vmn1r-ps77 | 0 | NA | NA | NA |
| Gm5891 | 0 | NA | NA | NA |
| Vmn1r137 | 0 | NA | NA | NA |
| Vmn1r-ps78 | 0 | NA | NA | NA |
| Vmn1r138 | 0 | NA | NA | NA |
| Gm6176 | 0 | NA | NA | NA |
| Vmn1r139 | 0 | NA | NA | NA |
| AC127227.1 | 0 | NA | NA | NA |
| Gm16451 | 0 | NA | NA | NA |
| Gm10666 | 0 | NA | NA | NA |
| Vmn1r142 | 0 | NA | NA | NA |
| Vmn1r143 | 0 | NA | NA | NA |
| Gm8653 | 0 | NA | NA | NA |
| Gm8453 | 0 | NA | NA | NA |
| Gm4557 | 0 | NA | NA | NA |
| Gm4558 | 0 | NA | NA | NA |

| | | | | |
|---|---|---|---|---|
| Gm4187 | 0 | NA | NA | NA |
| Vmn1r-ps70 | 0 | NA | NA | NA |
| Gm8660 | 0 | NA | NA | NA |
| Vmn1r148 | 0 | NA | NA | NA |
| Vmn1r149 | 0 | NA | NA | NA |
| Gm10665 | 0 | NA | NA | NA |
| Vmn1r151 | 0 | NA | NA | NA |
| Vmn1r152 | 0 | NA | NA | NA |
| Gm8677 | 0 | NA | NA | NA |
| Gm4201 | 0 | NA | NA | NA |
| Gm4565 | 0 | NA | NA | NA |
| Vmn1r155 | 0 | NA | NA | NA |
| Vmn1r-ps80 | 0 | NA | NA | NA |
| Gm8693 | 0 | NA | NA | NA |
| Gm4567 | 0 | NA | NA | NA |
| Vmn1r157 | 0 | NA | NA | NA |
| Vmn1r158 | 0 | NA | NA | NA |
| Gm4207 | 0 | NA | NA | NA |
| Vmn1r159 | 0 | NA | NA | NA |
| Vmn1r160 | 0 | NA | NA | NA |
| Gm8708 | 0 | NA | NA | NA |
| Gm4214 | 0 | NA | NA | NA |
| Gm4216 | 0 | NA | NA | NA |
| Vmn1r163 | 0 | NA | NA | NA |
| Gm8720 | 0 | NA | NA | NA |
| Gm16456 | 0 | NA | NA | NA |
| Gm10662 | 0 | NA | NA | NA |
| Vmn1r165 | 0 | NA | NA | NA |
| Vmn1r-ps83 | 0 | NA | NA | NA |
| Vmn1r166 | 0 | NA | NA | NA |
| Gm6902 | 0 | NA | NA | NA |
| Nlrp4e | 0 | NA | NA | NA |
| Nlrp5 | 1.484051332 | -2.16248951 | 1.166740548 | -1.853445064 |
| Gm25196 | 0.066050875 | -0.443818988 | 3.724945049 | -0.119147795 |
| Vmn1r-ps84 | 0 | NA | NA | NA |
| Vmn1r167 | 0 | NA | NA | NA |
| Vmn1r168 | 0.047948698 | 0.037064902 | 3.724945049 | 0.009950456 |
| Vmn1r169 | 0 | NA | NA | NA |
| Vmn1r170 | 0 | NA | NA | NA |
| Vmn1r171 | 0 | NA | NA | NA |
| Vmn1r172 | 0 | NA | NA | NA |
| Vmn1r173 | 0 | NA | NA | NA |
| Vmn1r174 | 0 | NA | NA | NA |
| Vmn1r-ps85 | 0 | NA | NA | NA |
| Vmn1r175 | 0 | NA | NA | NA |
| Vmn1r176 | 0 | NA | NA | NA |
| Vmn1r177 | 0 | NA | NA | NA |

| | | | | |
|---|---|---|---|---|
| Vmn1r178 | 0 | NA | NA | NA |
| Vmn1r179 | 0 | NA | NA | NA |
| Gm10175 | 13.75409139 | -0.510439058 | 0.336562742 | -1.516623777 |
| Vmn1r180 | 0 | NA | NA | NA |
| Vmn1r181 | 0 | NA | NA | NA |
| V1rd19 | 0 | NA | NA | NA |
| Vmn1r183 | 0 | NA | NA | NA |
| Ptma-ps1 | 0 | NA | NA | NA |
| Zfp180 | 620.834939 | -0.041182555 | 0.092627719 | -0.444602924 |
| Zfp112 | 130.5214199 | 0.235337903 | 0.166237317 | 1.415674337 |
| Zfp235 | 143.0420723 | -0.109616514 | 0.127428508 | -0.860219709 |
| Zfp114 | 11.47062778 | 0.178293514 | 0.367844628 | 0.484697888 |
| Zfp111 | 197.4067323 | -0.357983806 | 0.161907539 | -2.21103853 |
| Zfp109 | 90.1039288 | -0.07630234 | 0.156360755 | -0.487989075 |
| Zfp108 | 66.9010371 | -0.037262954 | 0.218128008 | -0.170830671 |
| Zfp93 | 102.2549038 | -0.316043224 | 0.142240429 | -2.221894476 |
| Zfp61 | 139.4922126 | 0.280668547 | 0.131514234 | 2.134130571 |
| Zfp94 | 74.342771 | -0.274261361 | 0.153183129 | -1.790414932 |
| Gm15465 | 0 | NA | NA | NA |
| Tescl | 0 | NA | NA | NA |
| Gm27592 | 0.061920613 | -0.443816764 | 3.724945049 | -0.119147198 |
| Lypd5 | 0.324847777 | 1.299109838 | 3.701671382 | 0.350952233 |
| Kcnn4 | 8.685732437 | -1.100279528 | 0.397944123 | -2.764909606 |
| Smg9 | 288.8311134 | 0.070651885 | 0.07839587 | 0.901219488 |
| Irgc1 | 0.04393175 | 0.037062677 | 3.724945049 | 0.009949859 |
| Plaur | 88.94456294 | -1.219970862 | 0.419516581 | -2.908039676 |
| Cadm4 | 2550.068439 | -0.142720206 | 0.088941581 | -1.604651102 |
| Zfp428 | 224.1959431 | -0.175379103 | 0.102626556 | -1.70890567 |
| Irgq | 1232.318714 | 0.090867499 | 0.050964069 | 1.78297181 |
| Pinlyp | 0.104366086 | 0.51795324 | 3.724945049 | 0.139049901 |
| Xrcc1 | 220.1632784 | -0.135373153 | 0.099351298 | -1.362570561 |
| Zfp575 | 72.63402446 | 0.443788258 | 0.137252633 | 3.233367913 |
| Ethe1 | 83.5924299 | -0.351535767 | 0.159966475 | -2.197558997 |
| Phldb3 | 6.837825038 | -0.863988791 | 0.358798181 | -2.408007727 |
| Gm26550 | 0 | NA | NA | NA |
| Lypd3 | 0.412745228 | 1.579242354 | 3.707691451 | 0.425936833 |
| Tex101 | 0 | NA | NA | NA |
| BC049730 | 0 | NA | NA | NA |
| Gm4763 | 0 | NA | NA | NA |
| Cd177 | 15.47993706 | -2.972000402 | 0.54946552 | -5.408893356 |
| Ceacam10 | 1.306976768 | 1.757868791 | 0.944101224 | 1.861949488 |
| Gm5893 | 0 | NA | NA | NA |
| Gm9844 | 66.59610774 | -0.229013699 | 0.188459679 | -1.215186721 |
| Lypd4 | 0 | NA | NA | NA |
| Dmrtc2 | 0.042716945 | 0.037062677 | 3.724945049 | 0.009949859 |
| Rps19 | 1020.693303 | -0.215006073 | 0.087264415 | -2.463845925 |
| Cd79a | 3.200457292 | -0.874975831 | 0.568693404 | -1.538572145 |

| | | | | |
|---|---|---|---|---|
| Arhgef1 | 583.8558966 | -0.153334738 | 0.119919735 | -1.27864474 |
| Rabac1 | 736.5228606 | -0.032624909 | 0.12110832 | -0.269386191 |
| Mir7046 | 0 | NA | NA | NA |
| Atp1a3 | 26298.87918 | 0.307484824 | 0.062739346 | 4.900988652 |
| Mir7047 | 0.284641305 | -0.443820253 | 1.777165317 | -0.249734928 |
| Grik5 | 1931.46398 | 0.493039009 | 0.108713518 | 4.535213467 |
| Zfp574 | 425.7976918 | 0.02248958 | 0.084379463 | 0.266529079 |
| Pou2f2 | 170.0044398 | -0.147894414 | 0.159827903 | -0.925335383 |
| Mir6537 | 0 | NA | NA | NA |
| D930028M14Rik | 1.834100151 | -1.366598793 | 0.878764337 | -1.555136838 |
| Dedd2 | 172.9180783 | -0.245929412 | 0.133582674 | -1.84102777 |
| Mir7048 | 0.228760906 | -0.435568379 | 2.136840713 | -0.203837551 |
| Zfp526 | 147.8424307 | 0.007545911 | 0.118212384 | 0.063833506 |
| Gsk3a | 1893.258904 | 0.205124159 | 0.048009586 | 4.272566744 |
| 9130221H12Rik | 75.48946259 | -0.014302264 | 0.136330727 | -0.104908584 |
| Erf | 724.7532469 | -0.085646 | 0.126159886 | -0.678868716 |
| Cic | 3271.904299 | -0.022833625 | 0.058574456 | -0.389822228 |
| Pafah1b3 | 85.99027182 | -0.547578939 | 0.143425695 | -3.817858028 |
| Prr19 | 0.750844096 | -1.859774889 | 1.192776279 | -1.559198419 |
| Tmem145 | 351.2431402 | 0.635834311 | 0.148752409 | 4.274447163 |
| Megf8 | 2644.487032 | 0.402588391 | 0.083415988 | 4.826273713 |
| Cnfn | 0.044191549 | 0.037062677 | 3.724945049 | 0.009949859 |
| Lipe | 172.463792 | -0.240785751 | 0.130916635 | -1.839229604 |
| 4732471J01Rik | 9.165417789 | 0.607791932 | 0.422222202 | 1.439507274 |
| Cxcl17 | 0.234440768 | -0.276511415 | 2.115872912 | -0.130684321 |
| Gm15495 | 0.148650935 | 0.966592573 | 2.859657381 | 0.338009924 |
| Ceacam1 | 24.75964592 | -0.600595372 | 0.243123354 | -2.470331883 |
| Ceacam2 | 1.575759893 | -0.286002802 | 1.055039954 | -0.271082437 |
| Erich4 | 0.961427984 | 1.607295838 | 1.242424983 | 1.293676367 |
| Atp5sl | 334.5795391 | 0.277530572 | 0.069118694 | 4.015275122 |
| B3gnt8 | 13.86552076 | -1.535360083 | 0.383401053 | -4.004579723 |
| Bckdha | 305.8692003 | 0.305898535 | 0.123322523 | 2.480475804 |
| Exosc5 | 96.23120801 | -0.327528531 | 0.133306541 | -2.456957695 |
| Tmem91 | 111.1486534 | 0.237791447 | 0.155345678 | 1.530724574 |
| B9d2 | 96.55601536 | 0.046389378 | 0.14225186 | 0.326107355 |
| Tgfb1 | 180.5550002 | -0.934997219 | 0.183934185 | -5.083324881 |
| Ccdc97 | 599.3683796 | -0.366869021 | 0.072205036 | -5.080933948 |
| Hnrnpul1 | 1763.970679 | -0.05176646 | 0.063060343 | -0.820903568 |
| Axl | 1705.534697 | -0.529888073 | 0.130054135 | -4.074365436 |
| Cyp2s1 | 45.56318474 | 0.344660377 | 0.218976743 | 1.573958829 |
| Gm26707 | 1.31640046 | 1.255718321 | 1.169963064 | 1.073297406 |
| Gm4613 | 2.523943987 | -0.133689038 | 0.591936735 | -0.225850214 |
| Gm24865 | 0 | NA | NA | NA |
| Gm23921 | 0 | NA | NA | NA |
| Gm6434 | 0 | NA | NA | NA |
| Cyp2b10 | 0.605952604 | -1.065131802 | 1.226028246 | -0.86876612 |
| Cyp2b13 | 0.046345709 | -0.443818988 | 3.724945049 | -0.119147795 |

| | | | | |
|---|---|---|---|---|
| Rnf170-ps | 17.15945834 | 0.32555708 | 0.255104838 | 1.276169759 |
| Cyp2b9 | 0 | NA | NA | NA |
| Gm8902 | 0.060174537 | 0.517957688 | 3.724945049 | 0.139051095 |
| Vmn1r-ps86 | 0 | NA | NA | NA |
| Vmn1r184 | 0 | NA | NA | NA |
| Cyp2a4 | 0.118974955 | 0.51795324 | 3.724945049 | 0.139049901 |
| Nlrp9c | 0.044544648 | 0.517957688 | 3.724945049 | 0.139051095 |
| Nlrp4a | 0 | NA | NA | NA |
| Gm25947 | 0 | NA | NA | NA |
| Nlrp9a | 0 | NA | NA | NA |
| Vmn1r185 | 0 | NA | NA | NA |
| Cyp2b23 | 0 | NA | NA | NA |
| Cyp2b19 | 0.186596897 | 1.134549183 | 3.282617194 | 0.34562336 |
| Cyp2g1 | 8.028728541 | -1.62533183 | 1.658950104 | -0.979735211 |
| Cyp2a5 | 11.27708144 | -0.329456253 | 1.177485162 | -0.279796522 |
| Cyp2a22 | 0.038981395 | -0.443816764 | 3.724945049 | -0.119147198 |
| Cyp2a12 | 0.191971534 | 0.009775915 | 2.650118572 | 0.003688859 |
| Cyp2f2 | 13.0302823 | -1.034850359 | 0.450120982 | -2.299049367 |
| Gm15883 | 0.044544648 | 0.517957688 | 3.724945049 | 0.139051095 |
| Cyp2t4 | 0.092269992 | -0.443823436 | 3.724945049 | -0.119148989 |
| Egln2 | 888.7193016 | 0.14041108 | 0.07604526 | 1.846414631 |
| Rab4b | 72.36944043 | 0.275281106 | 0.167604102 | 1.642448503 |
| Gm21983 | 0 | NA | NA | NA |
| Mir3101 | 0 | NA | NA | NA |
| Mia | 1.133325952 | -1.053935531 | 1.215261553 | -0.867249958 |
| Snrpa | 438.9269378 | -0.182854411 | 0.08367872 | -2.185196074 |
| BC024978 | 169.2292599 | -0.08925255 | 0.099004548 | -0.901499499 |
| Mir1191 | 8.542722226 | 0.186916624 | 0.382720416 | 0.488389476 |
| Itpkc | 197.4348915 | -0.789683083 | 0.214908333 | -3.674511226 |
| Adck4 | 262.1854901 | -0.827658367 | 0.080618404 | -10.26637006 |
| Gm15567 | 1.049380316 | -0.453960939 | 1.098841339 | -0.413126921 |
| Numbl | 1053.281901 | -0.13750227 | 0.096069834 | -1.431274155 |
| Gm15570 | 0 | NA | NA | NA |
| Ltbp4 | 207.6280549 | 0.378236902 | 0.153601299 | 2.462459006 |
| Shkbp1 | 92.29759634 | -0.628455542 | 0.177679922 | -3.537009335 |
| Gm20479 | 0 | NA | NA | NA |
| Sptbn4 | 1935.821563 | 0.323872004 | 0.133172081 | 2.431981259 |
| Blvrb | 91.00347656 | -0.115396688 | 0.123386596 | -0.935244921 |
| Pgam1-ps2 | 38.75767613 | -0.002529738 | 0.178990563 | -0.014133358 |
| Sertad3 | 55.81009826 | -0.419409175 | 0.186707134 | -2.246347883 |
| Sertad1 | 143.2528654 | -0.948566908 | 0.230013748 | -4.12395744 |
| Prx | 13.03989321 | -0.789094412 | 0.315391918 | -2.501948743 |
| Gm15541 | 1.986167605 | 0.345350879 | 0.663504678 | 0.520495018 |
| Hipk4 | 1.250864467 | -1.515183508 | 0.972350533 | -1.55826881 |
| Pld3 | 3221.190747 | 0.498003492 | 0.075048622 | 6.635744695 |
| 2310022A10Rik | 301.4687265 | -0.634380749 | 0.103526656 | -6.127704442 |
| Akt2 | 915.7419638 | -0.247717315 | 0.076202761 | -3.250765592 |

| | | | | |
|---|---|---|---|---|
| Ttc9b | 367.5397197 | 0.877445448 | 0.095714374 | 9.167332038 |
| Map3k10 | 1510.75745 | 0.344606624 | 0.07894855 | 4.364951913 |
| C030039L03Rik | 80.29137703 | -0.564220229 | 0.173754078 | -3.247234449 |
| Zfp60 | 578.4836708 | -0.328259914 | 0.175409389 | -1.871393062 |
| Zfp626 | 57.40632837 | -0.231834036 | 0.155247135 | -1.493322477 |
| Zfp59 | 63.21053847 | -0.312314312 | 0.170353145 | -1.833334585 |
| Zfp607 | 29.83338546 | -0.314047338 | 0.238538723 | -1.31654657 |
| 1700049G17Rik | 131.7655322 | -0.200606026 | 0.103747557 | -1.933597592 |
| Gm26891 | 2.658590328 | 0.484744485 | 0.765822652 | 0.632972247 |
| Gm10651 | 26.89522568 | -0.127913489 | 0.23902244 | -0.535152635 |
| Zfp780b | 147.3540878 | -0.393185679 | 0.142610254 | -2.757064573 |
| Zfp850 | 137.3735265 | -0.140677914 | 0.110660804 | -1.271253322 |
| Psmc4 | 769.5473873 | -0.394198084 | 0.086233744 | -4.571274116 |
| Fcgbp | 5.334611605 | 0.885952048 | 0.448080729 | 1.977215245 |
| Gm10935 | 0.257434735 | -0.924714583 | 2.132651623 | -0.433598518 |
| Gm24356 | 0 | NA | NA | NA |
| 9530053A07Rik | 2.222860927 | -0.23361648 | 0.627751695 | -0.372147909 |
| Fbl | 235.6745308 | -0.343647216 | 0.155747748 | -2.206434577 |
| Dyrk1b | 419.6037486 | 0.460276034 | 0.105451022 | 4.364832374 |
| Eid2 | 336.2126705 | 0.122172504 | 0.095017584 | 1.285788365 |
| Eid2b | 286.8257227 | 0.40726904 | 0.100011143 | 4.072236614 |
| BC089491 | 5.091535971 | 1.229250832 | 0.52113234 | 2.358807425 |
| Dll3 | 12.77550445 | -0.582237329 | 0.348932699 | -1.668623579 |
| Timm50 | 228.1838604 | -0.553357159 | 0.103439203 | -5.349588405 |
| Supt5 | 1576.058628 | -0.203335454 | 0.05788688 | -3.512634541 |
| Rps16 | 917.1611222 | -0.327022063 | 0.096946153 | -3.373234028 |
| AF357399 | 3.515687709 | -0.771442182 | 0.572074273 | -1.348500043 |
| Plekhg2 | 230.709035 | -1.258433918 | 0.193204944 | -6.513466422 |
| Zfp36 | 478.3772212 | -0.817313999 | 0.322228529 | -2.536442077 |
| Med29 | 93.91206282 | -0.31120233 | 0.126079597 | -2.468300477 |
| Paf1 | 477.8677575 | -0.056474089 | 0.079723555 | -0.70837394 |
| Pak4 | 1507.799238 | -0.419198444 | 0.09556643 | -4.386461275 |
| Gmfg | 19.76928172 | -0.453340391 | 0.289766526 | -1.564502283 |
| Lrfn1 | 754.1318081 | 0.448668737 | 0.10117549 | 4.434559572 |
| 1700028B04Rik | 0.752450377 | 1.185804472 | 1.172289189 | 1.011528967 |
| Ifnl2 | 0.046345709 | -0.443818988 | 3.724945049 | -0.119147795 |
| Ifnl3 | 0.046345709 | -0.443818988 | 3.724945049 | -0.119147795 |
| Sycn | 0 | NA | NA | NA |
| Nccrp1 | 4.109619456 | -0.121505897 | 0.526479266 | -0.230789519 |
| Mir7049 | 0.300003454 | -0.926061081 | 2.03223523 | -0.455685969 |
| Papl | 10.97390351 | -0.08286457 | 0.318608584 | -0.260082666 |
| Fbxo27 | 104.6688318 | 0.145402524 | 0.121471487 | 1.197009503 |
| Fbxo17 | 17.81551388 | -0.32665331 | 0.246338243 | -1.326035725 |
| Mrps12 | 129.792039 | -0.363373194 | 0.127963849 | -2.839655074 |
| Sars2 | 103.3753545 | 0.121453659 | 0.11176654 | 1.086672801 |
| Gm6537 | 10.517279 | 0.802818423 | 0.396236094 | 2.026111289 |
| Nfkbib | 264.1176441 | -0.291553412 | 0.158107924 | -1.844015188 |

| | | | | |
|---|---|---|---|---|
| Sirt2 | 1431.100394 | 0.10942292 | 0.06470186 | 1.691186632 |
| Rinl | 62.25372522 | -0.767779381 | 0.233923969 | -3.28217491 |
| Hnrnpl | 2413.616906 | -0.010836866 | 0.059960632 | -0.180733022 |
| Ech1 | 280.1649813 | -0.184620627 | 0.102735382 | -1.797050088 |
| Lgals4 | 13.26484918 | -0.071091568 | 0.321914727 | -0.220839752 |
| Lgals7 | 1.097858384 | 1.573380942 | 1.339285161 | 1.174791588 |
| Capn12 | 3.372768181 | 1.058705118 | 0.670566731 | 1.578821419 |
| Actn4 | 1770.860706 | -0.699125957 | 0.090008171 | -7.767361044 |
| Eif3k | 526.8853122 | -0.193651732 | 0.11324751 | -1.709986672 |
| Map4k1 | 43.16772284 | -0.464254607 | 0.20185538 | -2.299936755 |
| Ryr1 | 194.7438043 | 0.490983448 | 0.228325804 | 2.150363385 |
| AC165142.1 | 0.214156338 | -0.427704962 | 2.173286072 | -0.196801041 |
| Gm26604 | 0.400939804 | 0.037077062 | 1.56730992 | 0.023656497 |
| Mir1963 | 0 | NA | NA | NA |
| Rasgrp4 | 17.50432713 | -1.867712511 | 0.303582765 | -6.152234992 |
| Gm16282 | 0 | NA | NA | NA |
| Fam98c | 181.7263762 | 0.381384799 | 0.107637057 | 3.543248116 |
| 1110035H17Rik | 2.173831266 | 0.555165023 | 0.7103002 | 0.781592097 |
| Spred3 | 426.8502981 | -0.432859826 | 0.113744037 | -3.805560591 |
| Ggn | 9.283367035 | -1.200584104 | 0.344650945 | -3.48347835 |
| Psmd8 | 837.7336857 | -0.151171832 | 0.104361185 | -1.448544606 |
| Catsperg1 | 14.9802426 | 0.169151556 | 0.352888804 | 0.47933387 |
| Gm17113 | 0 | NA | NA | NA |
| Kcnk6 | 62.8823065 | 0.445484628 | 0.184595977 | 2.413295431 |
| Yif1b | 285.5672007 | -0.149146335 | 0.107778876 | -1.383817871 |
| 2200002D01Rik | 2.362804395 | -1.569106748 | 0.665206547 | -2.358826377 |
| Spint2 | 203.5653366 | 0.144187315 | 0.107363611 | 1.342981232 |
| Ppp1r14a | 171.4189243 | -0.448511566 | 0.147536688 | -3.040000239 |
| Dpf1 | 418.7210972 | 0.360468343 | 0.117721189 | 3.06205148 |
| Sipa1l3 | 1249.288852 | 0.041907 | 0.075613876 | 0.554223677 |
| 4932431P20Rik | 0.682519493 | 1.408051125 | 1.31532683 | 1.07049525 |
| 4930432E11Rik | 0.998548739 | 0.037077121 | 1.356191705 | 0.027339144 |
| Gm29326 | 0.495715849 | 0.058434041 | 1.529446112 | 0.038206015 |
| A330087D11Rik | 0.362932359 | 0.037075137 | 1.702605846 | 0.021775525 |
| n-R5s153 | 0 | NA | NA | NA |
| Catsperg2 | 7.493041037 | 0.21638531 | 0.437102776 | 0.495044465 |
| Gm23103 | 0 | NA | NA | NA |
| Zfp84 | 275.1985705 | -0.334246468 | 0.088516938 | -3.776073569 |
| Mir1964 | 1.817730626 | -0.263914021 | 0.827863775 | -0.318789189 |
| Zfp30 | 158.5141845 | 0.297143743 | 0.104434715 | 2.845258343 |
| Zfp790 | 215.9787552 | -0.360319958 | 0.119080355 | -3.025855591 |
| Zfp940 | 138.395624 | 0.173553891 | 0.112020852 | 1.549299871 |
| Gm26554 | 0 | NA | NA | NA |
| Gm17199 | 0 | NA | NA | NA |
| Zfp420 | 147.2350297 | -0.634836278 | 0.121901561 | -5.207778105 |
| Zfp27 | 203.8758977 | -0.060771174 | 0.084431424 | -0.719769625 |
| Mir7668 | 0.042716945 | 0.037062677 | 3.724945049 | 0.009949859 |

| | | | | |
|---|---|---|---|---|
| Zfp383 | 132.1721319 | 0.033597995 | 0.106661677 | 0.314995936 |
| Zfp74 | 213.3163599 | 0.078094244 | 0.09552917 | 0.817491075 |
| C230062I16Rik | 7.143933271 | -0.32555492 | 0.43327995 | -0.751373149 |
| Gm10169 | 7.433721481 | 0.249778307 | 0.38501172 | 0.648755072 |
| Zfp568 | 112.9697705 | -0.099559721 | 0.131713824 | -0.755879062 |
| Zfp14 | 53.58896386 | -0.193995794 | 0.178647619 | -1.085913126 |
| Zfp82 | 6.950672932 | 0.594344535 | 0.421321203 | 1.410668464 |
| Gm26920 | 0.228152142 | 1.00013942 | 2.550451083 | 0.392142169 |
| Zfp566 | 31.53288666 | 0.019355265 | 0.196921193 | 0.098289395 |
| Zfp260 | 833.8589085 | -0.004773146 | 0.068861115 | -0.069315557 |
| Gm23368 | 0 | NA | NA | NA |
| Zfp382 | 241.7030479 | 0.209561853 | 0.09401474 | 2.229031892 |
| Zfp146 | 303.8588219 | -0.259960233 | 0.07531957 | -3.451430136 |
| Gm5113 | 98.06848606 | 0.307499434 | 0.146887664 | 2.093432669 |
| Cox7a1 | 8.620735387 | -0.914072743 | 0.385027884 | -2.374042975 |
| Capns1 | 1143.806175 | -0.484588316 | 0.113453423 | -4.271253389 |
| Gm26810 | 0.29029136 | 0.412690289 | 1.766469122 | 0.2336244 |
| Tbcb | 367.5355379 | -0.15944137 | 0.086852366 | -1.835774625 |
| Polr2i | 115.9016702 | -0.157542182 | 0.134511012 | -1.171221449 |
| Ovol3 | 0.088736197 | 0.51795324 | 3.724945049 | 0.139049901 |
| Wdr62 | 63.98026938 | 0.26172097 | 0.224938962 | 1.163519949 |
| Rps12-ps5 | 0.252860697 | 0.037077175 | 1.851889217 | 0.020021271 |
| Thap8 | 1.268210335 | -0.689461029 | 0.928608558 | -0.742466805 |
| Clip3 | 6878.216135 | 0.313854205 | 0.079688039 | 3.938535962 |
| Alkbh6 | 168.5814911 | 0.111323461 | 0.09938034 | 1.12017589 |
| Gm24757 | 0 | NA | NA | NA |
| Syne4 | 17.72551575 | -0.175912494 | 0.268502893 | -0.655160517 |
| Sdhaf1 | 75.1581409 | 0.113099508 | 0.124853666 | 0.905856521 |
| E130208F15Rik | 4.251243736 | 0.351240263 | 0.506761482 | 0.693107656 |
| Lrfn3 | 312.5931955 | 0.132126973 | 0.089604744 | 1.474553324 |
| Gm25259 | 0.679915734 | -0.576531272 | 1.214068187 | -0.474875528 |
| Tyrobp | 264.1135694 | -0.957663262 | 0.191060044 | -5.012368074 |
| Hcst | 5.467031529 | -1.239458943 | 0.409767058 | -3.024789132 |
| Nfkbid | 33.79332328 | -0.651178008 | 0.256253493 | -2.541147834 |
| Aplp1 | 8728.071125 | 0.413950203 | 0.066054971 | 6.266753194 |
| Kirrel2 | 45.1116197 | 1.86540561 | 0.286403902 | 6.513199015 |
| Nphs1 | 8.109329026 | 1.720127755 | 0.453308795 | 3.794604853 |
| Nphs1os | 0.917433238 | 2.199622674 | 1.264933592 | 1.738923441 |
| Prodh2 | 0.146848595 | 0.037061235 | 3.724945049 | 0.009949472 |
| Arhgap33os | 1.185039339 | 0.967711797 | 1.112124882 | 0.870146701 |
| Arhgap33 | 1004.610349 | 0.899994037 | 0.160431402 | 5.609837126 |
| Proser3 | 27.82645344 | -0.695374653 | 0.275015118 | -2.528496103 |
| Hspb6 | 57.72339347 | -0.215629051 | 0.16705703 | -1.290751134 |
| mmu-mir-9769 | 0 | NA | NA | NA |
| Lin37 | 148.1215379 | -0.229543279 | 0.127531953 | -1.799888375 |
| Psenen | 209.1857129 | -0.333871655 | 0.097255823 | -3.432922003 |
| U2af1l4 | 195.8717764 | 0.079034337 | 0.078343902 | 1.00881287 |

| | | | | |
|---|---|---|---|---|
| Gm17104 | 0.139856492 | -0.443820827 | 3.574411099 | -0.124166139 |
| Igflr1 | 0 | NA | NA | NA |
| Kmt2b | 1103.022709 | 0.124441162 | 0.137344787 | 0.906049401 |
| Mir3569 | 0.55981669 | -1.771793182 | 1.286837285 | -1.376858755 |
| Zbtb32 | 35.12199024 | -1.118565412 | 0.352544526 | -3.172834431 |
| Upk1a | 5.454212958 | -0.143260309 | 0.526875566 | -0.271905395 |
| Cox6b1 | 1385.504029 | 0.085774601 | 0.070783778 | 1.211783325 |
| Etv2 | 0.288531392 | -1.297798892 | 1.732501356 | -0.749089683 |
| Rbm42 | 1.341866388 | 0.433549441 | 0.857236907 | 0.505752188 |
| Gm21982 | 0.195772763 | 0.517968281 | 2.236713726 | 0.231575581 |
| Gm26610 | 2.644816022 | 0.585606677 | 0.667136619 | 0.877791236 |
| Haus5 | 8.540474638 | 0.285514301 | 0.373701486 | 0.764017033 |
| Pmis2 | 0 | NA | NA | NA |
| 2200002J24Rik | 0.066050875 | -0.443818988 | 3.724945049 | -0.119147795 |
| Gm4883 | 0 | NA | NA | NA |
| Atp4a | 2.461184311 | 1.280896123 | 0.696701222 | 1.838515683 |
| Tmem147 | 361.1132233 | -0.28253873 | 0.083468977 | -3.384954976 |
| Tmem147os | 88.86610623 | 0.396518555 | 0.191202128 | 2.07381873 |
| Gapdhs | 0.221390349 | 0.998859591 | 2.615327495 | 0.381925244 |
| Sbsn | 30.54518145 | 0.810532729 | 0.251185509 | 3.226829172 |
| Gm26935 | 15.53492519 | 0.626602294 | 0.368449584 | 1.700645955 |
| Dmkn | 16.49677372 | -0.797646367 | 0.491630367 | -1.622451379 |
| Krtdap | 1.379211929 | 2.116186077 | 3.658216161 | 0.578474859 |
| Ffar2 | 3.090575541 | -1.645858949 | 0.795851997 | -2.068046516 |
| Ffar3 | 0.43344934 | 0.037074874 | 1.676321983 | 0.022116798 |
| Ffar1 | 0.265062405 | 0.006983243 | 2.439004666 | 0.002863153 |
| Cd22 | 18.32792618 | -0.891404929 | 0.385114755 | -2.314647565 |
| Gm29439 | 0.095898756 | -0.917692504 | 3.724434845 | -0.246397787 |
| Gm29440 | 0.110242424 | -0.443823436 | 3.724945049 | -0.119148989 |
| Mag | 2336.670031 | -0.22366793 | 0.128998714 | -1.733877207 |
| Hamp2 | 0.159795471 | -0.939694907 | 2.805348586 | -0.33496547 |
| Hamp | 0.755877001 | 2.426431287 | 1.228307866 | 1.975425994 |
| Usf2 | 555.4235933 | -0.019133976 | 0.061324094 | -0.312014 |
| Gm17077 | 7.799693549 | 1.348715738 | 0.489381891 | 2.755957594 |
| Gm4673 | 75.79538166 | 0.383645519 | 0.23735084 | 1.616364699 |
| Lsr | 56.83401911 | -0.680861339 | 0.212760523 | -3.200130023 |
| Fam187b | 1.876867459 | 0.590602336 | 0.787509049 | 0.749962602 |
| Fxyd5 | 213.0889357 | -0.992334283 | 0.215500643 | -4.604785711 |
| Mir7050 | 0 | NA | NA | NA |
| Fxyd7 | 269.8762191 | 1.228176483 | 0.144112126 | 8.52236738 |
| Fxyd1 | 309.5145642 | -0.308830364 | 0.173971793 | -1.775174916 |
| Lgi4 | 451.055952 | -0.556096736 | 0.125480443 | -4.431740284 |
| Fxyd3 | 1.806277867 | -1.810990979 | 0.80835801 | -2.240332818 |
| Hpn | 35.43406477 | -0.053514379 | 0.289686312 | -0.184732164 |
| Scn1b | 2784.178309 | 0.236260444 | 0.076768107 | 3.077585892 |
| Gramd1a | 871.9470282 | 0.219804697 | 0.096997614 | 2.266083531 |
| G630030J09Rik | 3.656513056 | -0.148141592 | 0.608657133 | -0.243390874 |

| | | | | |
|---|---|---|---|---|
| Gm10640 | 1.251626804 | -1.232802279 | 1.009528117 | -1.221166858 |
| Gm9569 | 0 | NA | NA | NA |
| Gm37292 | 9.829030887 | -0.471155958 | 0.391173298 | -1.204468611 |
| Scgb1b1-ps | 0 | NA | NA | NA |
| Scgb2b1 | 0 | NA | NA | NA |
| Scgb1b2 | 0 | NA | NA | NA |
| Scgb2b2 | 0 | NA | NA | NA |
| Scgb2b3 | 0 | NA | NA | NA |
| Scgb1b3 | 0 | NA | NA | NA |
| Scgb2b4-ps | 0 | NA | NA | NA |
| Scgb1b4-ps | 0 | NA | NA | NA |
| Scgb1b28-ps | 0 | NA | NA | NA |
| Gm29492 | 0 | NA | NA | NA |
| Scgb2b5-ps | 0 | NA | NA | NA |
| Scgb1b5-ps | 0 | NA | NA | NA |
| Scgb2b28-ps | 0 | NA | NA | NA |
| Rpl23a-ps8 | 0 | NA | NA | NA |
| Scgb2b6 | 0 | NA | NA | NA |
| Scgb1b6-ps | 0 | NA | NA | NA |
| Gm4373 | 0 | NA | NA | NA |
| Rpl23a-ps9 | 0 | NA | NA | NA |
| Scgb2b7 | 0 | NA | NA | NA |
| Scgb1b7 | 0 | NA | NA | NA |
| Gm9114 | 0 | NA | NA | NA |
| Gm29627 | 0 | NA | NA | NA |
| Scgb2b8-ps | 0 | NA | NA | NA |
| Scgb1b8-ps | 0 | NA | NA | NA |
| Gm6669 | 0 | NA | NA | NA |
| Gm9120 | 0 | NA | NA | NA |
| Gm21673 | 0 | NA | NA | NA |
| Scgb2b9-ps | 0 | NA | NA | NA |
| Scgb1b9-ps | 0 | NA | NA | NA |
| Gm5329 | 0 | NA | NA | NA |
| Scgb2b29-ps | 0 | NA | NA | NA |
| Gm9127 | 0 | NA | NA | NA |
| Gm29262 | 0 | NA | NA | NA |
| Gm29263 | 0 | NA | NA | NA |
| Scgb2b10 | 0 | NA | NA | NA |
| Scgb1b10 | 0 | NA | NA | NA |
| Rpl23a-ps10 | 0 | NA | NA | NA |
| Scgb2b30 | 0 | NA | NA | NA |
| Gm9135 | 0 | NA | NA | NA |
| Gm28256 | 0 | NA | NA | NA |
| Gm28257 | 0 | NA | NA | NA |
| Scgb2b11 | 0 | NA | NA | NA |
| Scgb1b11 | 0 | NA | NA | NA |
| Scgb2b12 | 0 | NA | NA | NA |

| | | | | |
|---|---|---|---|---|
| Scgb1b12 | 0 | NA | NA | NA |
| Gm9142 | 0 | NA | NA | NA |
| Gm28862 | 0 | NA | NA | NA |
| Rpl23a-ps11 | 0 | NA | NA | NA |
| Scgb1b29 | 0 | NA | NA | NA |
| Gm9146 | 0 | NA | NA | NA |
| Gm5116 | 0 | NA | NA | NA |
| Scgb2b13-ps | 0 | NA | NA | NA |
| Scgb1b13-ps | 0 | NA | NA | NA |
| Gm5327 | 0 | NA | NA | NA |
| Gm9149 | 0 | NA | NA | NA |
| Gm21135 | 0 | NA | NA | NA |
| Scgb2b14-ps | 0 | NA | NA | NA |
| Scgb1b14-ps | 0 | NA | NA | NA |
| Gm24766 | 0 | NA | NA | NA |
| Gm5058 | 0 | NA | NA | NA |
| Gm29429 | 0 | NA | NA | NA |
| Scgb2b31-ps | 0 | NA | NA | NA |
| Gm4398 | 0 | NA | NA | NA |
| Gm29432 | 0 | NA | NA | NA |
| Gm28408 | 0 | NA | NA | NA |
| Scgb2b15 | 0 | NA | NA | NA |
| Scgb1b15 | 0 | NA | NA | NA |
| Scgb2b16-ps | 0 | NA | NA | NA |
| Scgb1b16-ps | 0 | NA | NA | NA |
| Gm25817 | 0 | NA | NA | NA |
| Gm9156 | 0 | NA | NA | NA |
| Gm28409 | 0 | NA | NA | NA |
| Scgb2b32-ps | 0 | NA | NA | NA |
| Gm4405 | 0 | NA | NA | NA |
| Gm28383 | 0 | NA | NA | NA |
| Gm28384 | 0 | NA | NA | NA |
| Scgb2b17 | 0 | NA | NA | NA |
| Scgb1b17 | 0 | NA | NA | NA |
| Gm28992 | 0 | NA | NA | NA |
| Rpl23a-ps7 | 0 | NA | NA | NA |
| Scgb2b33 | 0 | NA | NA | NA |
| Gm9162 | 0 | NA | NA | NA |
| Gm28559 | 0 | NA | NA | NA |
| Gm6003 | 0 | NA | NA | NA |
| Scgb2b18 | 0 | NA | NA | NA |
| Scgb1b18 | 0 | NA | NA | NA |
| Scgb2b19 | 0 | NA | NA | NA |
| Scgb1b19 | 0 | NA | NA | NA |
| Gm9167 | 0 | NA | NA | NA |
| Scgb2b20 | 0 | NA | NA | NA |
| Scgb1b20 | 0 | NA | NA | NA |

| | | | | |
|---|---|---|---|---|
| Gm5732 | 0 | NA | NA | NA |
| Gm28794 | 0 | NA | NA | NA |
| Gm5731 | 0 | NA | NA | NA |
| Scgb1b34-ps | 0 | NA | NA | NA |
| Scgb2b21 | 0 | NA | NA | NA |
| Scgb1b21 | 0 | NA | NA | NA |
| Gm12770 | 0 | NA | NA | NA |
| Gm12771 | 0 | NA | NA | NA |
| Scgb2b22-ps | 0 | NA | NA | NA |
| Scgb1b22-ps | 0 | NA | NA | NA |
| Scgb2b23-ps | 0 | NA | NA | NA |
| Scgb1b23-ps | 0 | NA | NA | NA |
| Gm12763 | 0.048949532 | 0.037064902 | 3.724945049 | 0.009950456 |
| Rpl23a-ps12 | 0 | NA | NA | NA |
| Scgb2b24 | 0 | NA | NA | NA |
| Scgb1b24 | 0 | NA | NA | NA |
| Gm12782 | 0 | NA | NA | NA |
| Gm28614 | 0 | NA | NA | NA |
| Gm26443 | 0 | NA | NA | NA |
| Gm12768 | 0 | NA | NA | NA |
| Scgb2b25-ps | 0 | NA | NA | NA |
| Scgb1b25-ps | 0 | NA | NA | NA |
| Gm12757 | 0 | NA | NA | NA |
| Gm28615 | 0 | NA | NA | NA |
| Gm12765 | 0 | NA | NA | NA |
| Scgb2b26 | 0 | NA | NA | NA |
| Scgb1b26-ps | 0 | NA | NA | NA |
| Gm28184 | 0 | NA | NA | NA |
| Gm12773 | 0 | NA | NA | NA |
| Gm12755 | 0 | NA | NA | NA |
| Scgb2b27 | 0.046345709 | -0.443818988 | 3.724945049 | -0.119147795 |
| Scgb1b27 | 0 | NA | NA | NA |
| Gm12760 | 0 | NA | NA | NA |
| Gm12779 | 0.044544648 | 0.517957688 | 3.724945049 | 0.139051095 |
| Scgb1b30 | 0 | NA | NA | NA |
| Wtip | 46.54900976 | -0.179795998 | 0.162499141 | -1.106442761 |
| Uba2 | 890.9651073 | -0.15033041 | 0.059851988 | -2.511702889 |
| Gm12778 | 0.649479285 | -1.174838888 | 1.319932294 | -0.890075115 |
| Gm12783 | 0 | NA | NA | NA |
| Pdcd2l | 117.7724749 | -0.370049305 | 0.140196981 | -2.639495534 |
| Gpi1 | 4331.119929 | 0.146088401 | 0.0496698 | 2.941191622 |
| Gm26762 | 2.694167648 | -0.459114321 | 0.709653243 | -0.646955856 |
| Gm12762 | 6.006325422 | 0.128747337 | 0.439831054 | 0.29271998 |
| Gm12758 | 0.954952387 | -1.674398923 | 1.098279686 | -1.52456514 |
| 4931406P16Rik | 686.0018774 | -0.216614934 | 0.060672492 | -3.570232991 |
| Gm6096 | 0.066050875 | -0.443818988 | 3.724945049 | -0.119147795 |
| Lsm14a | 923.1856796 | -0.288098222 | 0.057973738 | -4.969460856 |

| | | | | |
|---|---|---|---|---|
| Gm12774 | 0 | NA | NA | NA |
| Gm12764 | 10.54452331 | 0.457537052 | 0.38740872 | 1.181018981 |
| Gm12766 | 0.212513282 | 1.689178857 | 3.159662247 | 0.534607412 |
| Gm12780 | 0 | NA | NA | NA |
| Gm12784 | 0.530195705 | 1.465632575 | 1.413081472 | 1.037189012 |
| Kctd15 | 124.850857 | -0.421557968 | 0.116357248 | -3.622962682 |
| Gm12781 | 0.50824631 | -0.821712589 | 1.310460402 | -0.627041143 |
| Gm12756 | 2.131963076 | 0.029718723 | 0.877047999 | 0.033884944 |
| Chst8 | 122.9302308 | 0.20795045 | 0.129068805 | 1.611159643 |
| Pepd | 269.5084426 | -0.190492828 | 0.095304023 | -1.998791051 |
| Gm25922 | 0.089089296 | 0.957155977 | 3.723939561 | 0.257027796 |
| Cebpg | 578.773901 | -0.690088871 | 0.094384164 | -7.311489982 |
| Gm25033 | 0 | NA | NA | NA |
| Gm22535 | 0 | NA | NA | NA |
| Cebpa | 259.903502 | -0.727682986 | 0.196549757 | -3.702283819 |
| Slc7a10 | 228.5227835 | 0.474597737 | 0.184965664 | 2.565869396 |
| Lrp3 | 882.6663157 | 0.447024921 | 0.077887715 | 5.739350824 |
| Gm24505 | 0.219162254 | 0.503211735 | 2.159165243 | 0.233058464 |
| Gpatch1 | 413.9756821 | -0.068074658 | 0.096216364 | -0.70751642 |
| Rhpn2 | 165.8571245 | -0.294401598 | 0.161904975 | -1.818360421 |
| C230052I12Rik | 49.69667937 | -0.56246164 | 0.188889116 | -2.977734512 |
| Cep89 | 116.2122036 | -0.269092146 | 0.139847012 | -1.924189452 |
| Gm18254 | 0 | NA | NA | NA |
| Slc7a9 | 0.108767819 | -0.443823436 | 3.724945049 | -0.119148989 |
| Tdrd12 | 4.458748399 | -0.518852736 | 0.528979969 | -0.980855167 |
| Gm18558 | 0.451259361 | 1.048020931 | 1.458926085 | 0.718350945 |
| Nudt19 | 464.3454165 | 0.271039426 | 0.105503151 | 2.569017341 |
| B230322F03Rik | 17.22929159 | 0.224506231 | 0.29612449 | 0.758148139 |
| Rgs9bp | 10.39343307 | 0.96429441 | 0.416877956 | 2.313133609 |
| Ankrd27 | 324.283844 | -0.390073117 | 0.149195467 | -2.614510517 |
| Pdcd5 | 124.9127266 | 0.088608996 | 0.218331985 | 0.405845238 |
| Dpy19l3 | 892.6871314 | -0.022915267 | 0.097643248 | -0.234683579 |
| Zfp507 | 543.6392318 | 0.013776406 | 0.072051181 | 0.191203062 |
| Gm26790 | 0.186516865 | -0.444671952 | 2.670993324 | -0.166481866 |
| E130304I02Rik | 2.861485906 | 0.088835752 | 0.625237235 | 0.142083272 |
| Gm28514 | 1.65996821 | -1.105916846 | 0.839496927 | -1.317356634 |
| Rpl17-ps9 | 11.99013794 | -0.034697862 | 0.296521313 | -0.11701642 |
| Gm9211 | 0.12092566 | 0.998850474 | 3.724945049 | 0.268151734 |
| Gm23292 | 2.206361005 | 0.932020376 | 0.764778999 | 1.218679353 |
| Gm25247 | 0 | NA | NA | NA |
| Gm29129 | 0 | NA | NA | NA |
| Gm38285 | 5.376801637 | 0.57928256 | 0.491417949 | 1.178798133 |
| Tshz3 | 524.5764167 | 0.402342082 | 0.135319098 | 2.9732838 |
| Gm37452 | 12.71943701 | 0.040552483 | 0.449502861 | 0.090216297 |
| Gm38340 | 32.55178755 | 0.46416472 | 0.382384624 | 1.213868683 |
| E230020A03Rik | 24.88402293 | 0.660139341 | 0.334921032 | 1.971029816 |
| Gm37827 | 13.69016405 | 0.312504119 | 0.474392385 | 0.658746071 |

| | | | | |
|---|---|---|---|---|
| Gm24550 | 0 | NA | NA | NA |
| Gm28078 | 0 | NA | NA | NA |
| Gm28076 | 0.144050924 | 1.05357957 | 3.721447096 | 0.283110183 |
| Gm28075 | 0 | NA | NA | NA |
| Gm28077 | 0.313601826 | 2.021562109 | 1.732157512 | 1.167077529 |
| Zfp536 | 718.5400894 | 0.086573694 | 0.077725001 | 1.113846149 |
| 4930505M18Rik | 0 | NA | NA | NA |
| Gm22302 | 0 | NA | NA | NA |
| Uri1 | 620.2418666 | 0.376199553 | 0.075285253 | 4.996988602 |
| Ccne1 | 105.1069268 | 0.026249119 | 0.126182829 | 0.208024494 |
| Gm22203 | 0 | NA | NA | NA |
| 1600014C10Rik | 402.5956352 | -0.651824904 | 0.132342085 | -4.925303269 |
| Plekhf1 | 106.4439667 | -0.170690898 | 0.155074615 | -1.10070174 |
| Pop4 | 57.53508151 | 0.366910926 | 0.189151485 | 1.939772905 |
| Gm28581 | 0 | NA | NA | NA |
| Gm9220 | 0 | NA | NA | NA |
| Gm4451 | 0 | NA | NA | NA |
| Gm28453 | 0.075043205 | 0.517957688 | 3.724945049 | 0.139051095 |
| Gm22638 | 0 | NA | NA | NA |
| Gm6818 | 0 | NA | NA | NA |
| Gm28583 | 0 | NA | NA | NA |
| Gm6605 | 0 | NA | NA | NA |
| Gm6819 | 0 | NA | NA | NA |
| Gm29087 | 0.10503227 | -0.905414734 | 3.723500899 | -0.243162217 |
| Gm5591 | 0 | NA | NA | NA |
| Gm4454 | 0 | NA | NA | NA |
| Gm20749 | 0 | NA | NA | NA |
| Gm29086 | 0 | NA | NA | NA |
| Gm25286 | 0 | NA | NA | NA |
| Gm21115 | 0 | NA | NA | NA |
| Gm21276 | 0 | NA | NA | NA |
| Gm21122 | 0 | NA | NA | NA |
| Gm29085 | 0 | NA | NA | NA |
| 1700010N08Rik | 0.146060149 | -0.762452964 | 3.524367124 | -0.216337554 |
| Gm24797 | 0 | NA | NA | NA |
| Gm21129 | 0 | NA | NA | NA |
| Gm21136 | 0 | NA | NA | NA |
| Gm21142 | 0 | NA | NA | NA |
| Gm21152 | 0 | NA | NA | NA |
| Gm25671 | 0 | NA | NA | NA |
| Gm28451 | 0 | NA | NA | NA |
| Gm1982 | 0 | NA | NA | NA |
| Gm1988 | 0 | NA | NA | NA |
| Gm6124 | 0 | NA | NA | NA |
| Gm16387 | 0 | NA | NA | NA |
| Gm9226 | 0 | NA | NA | NA |
| Gm23307 | 0 | NA | NA | NA |

| | | | | |
|---|---|---|---|---|
| Gm29258 | 0 | NA | NA | NA |
| Gm9271 | 0 | NA | NA | NA |
| Gm5114 | 0 | NA | NA | NA |
| Gm27437 | 0 | NA | NA | NA |
| Gm5590 | 0.783781752 | 0.249749782 | 1.315545935 | 0.189844973 |
| Zfp939 | 78.05214499 | 0.245284261 | 0.156555426 | 1.566756683 |
| Gm28455 | 4.111696988 | -0.196254321 | 0.497720983 | -0.394305901 |
| Zfp619 | 83.85373399 | -0.078225035 | 0.144580459 | -0.541048463 |
| Gm33989 | 0.779068608 | -1.065919911 | 1.242902992 | -0.857605073 |
| Gm37494 | 103.4889581 | 0.081630611 | 0.155820776 | 0.523875012 |
| Gm4790 | 216.6323139 | -0.125675143 | 0.15274764 | -0.822763239 |
| 6230415J03Rik | 105.8274719 | 0.240865405 | 0.194621764 | 1.237607758 |
| Gm28582 | 30.69961378 | 0.352265256 | 0.227944816 | 1.545397089 |
| Gm17983 | 0 | NA | NA | NA |
| Gm26526 | 0.0878635 | 0.037058229 | 3.724945049 | 0.009948665 |
| Gm2058 | 91.57356217 | -0.143956432 | 0.244476587 | -0.588835251 |
| Gm26327 | 0.206849446 | 1.292288023 | 2.582154099 | 0.500468978 |
| Gm2021 | 0.044544648 | 0.517957688 | 3.724945049 | 0.139051095 |
| Gm9230 | 0 | NA | NA | NA |
| Gm37302 | 0.167459142 | -0.410658575 | 2.753748719 | -0.149127105 |
| 4930558N11Rik | 0 | NA | NA | NA |
| 4930435C17Rik | 0 | NA | NA | NA |
| Gm38199 | 0 | NA | NA | NA |
| Gm6851 | 4.478289484 | 0.27996135 | 0.49126564 | 0.569877735 |
| 4933404I11Rik | 0 | NA | NA | NA |
| Gm24910 | 0 | NA | NA | NA |
| Gm28807 | 0.144050924 | 1.05357957 | 3.721447096 | 0.283110183 |
| 4921507H08Rik | 0 | NA | NA | NA |
| 4933402C06Rik | 0.04393175 | 0.037062677 | 3.724945049 | 0.009949859 |
| A230077H06Rik | 28.4887614 | 0.336700709 | 0.308315844 | 1.092064243 |
| Gm28307 | 0.072025462 | 0.517959912 | 3.724945049 | 0.139051692 |
| Gm25784 | 0 | NA | NA | NA |
| Vstm2b | 175.071028 | 0.062531854 | 0.138213359 | 0.452429887 |
| Gm29293 | 0.161114758 | 1.389564196 | 3.426332673 | 0.405554372 |
| 4930433I11Rik | 0.134148234 | -0.443820953 | 3.624632483 | -0.122445781 |
| Gm4884 | 0 | NA | NA | NA |
| 1700110I07Rik | 0.044191549 | 0.037062677 | 3.724945049 | 0.009949859 |
| Gm28061 | 0 | NA | NA | NA |
| Gm9239 | 0 | NA | NA | NA |
| Gm6820 | 0 | NA | NA | NA |
| Gm2128 | 0 | NA | NA | NA |
| Gm28060 | 0 | NA | NA | NA |
| Gm28667 | 0 | NA | NA | NA |
| Gm5896 | 0.091880448 | 0.037060453 | 3.724945049 | 0.009949262 |
| Gm6866 | 0 | NA | NA | NA |
| Gm9244 | 0 | NA | NA | NA |
| Gm5592 | 0 | NA | NA | NA |

| | | | | |
|---|---|---|---|---|
| Gm28059 | 0 | NA | NA | NA |
| Gm2108 | 0 | NA | NA | NA |
| Gm17791 | 0.292366188 | 0.037076625 | 2.065110188 | 0.017953824 |
| Gm9246 | 0 | NA | NA | NA |
| AI987944 | 114.8081952 | -0.656245117 | 0.150738174 | -4.353542963 |
| Gm17102 | 8.180380778 | 0.590514852 | 0.407209668 | 1.450149391 |
| Gm20449 | 4.437436714 | -0.231841345 | 0.630363544 | -0.3677899 |
| Vmn2r57 | 0.538693099 | 1.358006265 | 1.451823059 | 0.935380008 |
| AW146154 | 60.96357997 | -0.187983148 | 0.187859322 | -1.000659141 |
| Gm15470 | 0.332094496 | 1.111372254 | 1.990295163 | 0.558395697 |
| Gm6871 | 0 | NA | NA | NA |
| Gm17217 | 0 | NA | NA | NA |
| 2610021A01Rik | 362.0305132 | -0.028416765 | 0.073145863 | -0.388494495 |
| Zfp788 | 649.6253934 | 0.37096389 | 0.073493367 | 5.047583276 |
| Gm9252 | 0 | NA | NA | NA |
| Vmn2r-ps54 | 6.472406225 | -1.756865743 | 0.496868761 | -3.535874827 |
| Gm6872 | 0.060174537 | 0.517957688 | 3.724945049 | 0.139051095 |
| Cd9-ps | 0.178447459 | -1.389338828 | 3.327416506 | -0.417542807 |
| Vmn2r-ps55 | 0.046345709 | -0.443818988 | 3.724945049 | -0.119147795 |
| Vmn2r-ps56 | 0.066050875 | -0.443818988 | 3.724945049 | -0.119147795 |
| Vmn2r58 | 0 | NA | NA | NA |
| Vmn2r-ps57 | 0 | NA | NA | NA |
| Vmn2r59 | 0 | NA | NA | NA |
| Hmgb1-ps7 | 2.510100352 | -0.273015076 | 0.66929139 | -0.407916611 |
| Gm7224 | 23.28790071 | 0.16451903 | 0.224714717 | 0.732123966 |
| Gm5593 | 0.933648904 | -0.677137829 | 1.213587653 | -0.557963677 |
| Vmn2r60 | 0 | NA | NA | NA |
| Gm2308 | 2.058388305 | 0.038553074 | 0.673973063 | 0.057202692 |
| Vmn2r61 | 0 | NA | NA | NA |
| Vmn2r-ps58 | 0 | NA | NA | NA |
| Gm5594 | 0 | NA | NA | NA |
| Vmn2r-ps59 | 0 | NA | NA | NA |
| Vmn2r-ps60 | 0 | NA | NA | NA |
| 4933421I07Rik | 0 | NA | NA | NA |
| Gm6887 | 0.072025462 | 0.517959912 | 3.724945049 | 0.139051692 |
| Zfp141 | 173.0700645 | 0.208952002 | 0.095242079 | 2.193904261 |
| Gm19246 | 0.428879182 | -1.354704548 | 1.457166151 | -0.929684337 |
| Gm21028 | 1.10003083 | 0.212752632 | 0.954065427 | 0.222995851 |
| Gm7221 | 16.48036244 | -0.579197433 | 0.287978446 | -2.011252724 |
| Gm24938 | 1.200098617 | -0.347098758 | 1.097944144 | -0.316135169 |
| 9830147E19Rik | 96.87745312 | -0.047802936 | 0.184090923 | -0.259670249 |
| Gm5595 | 96.44755503 | -0.177552104 | 0.132431946 | -1.340704482 |
| Gm17067 | 0 | NA | NA | NA |
| Gm18128 | 0 | NA | NA | NA |
| Gm6004 | 0 | NA | NA | NA |
| Vmn2r62 | 0.088383099 | 0.037058229 | 3.724945049 | 0.009948665 |
| Gm2381 | 0.372704184 | 1.348070666 | 1.538562441 | 0.876188467 |

| | | | | |
|---|---|---|---|---|
| Gm7211 | 0 | NA | NA | NA |
| Vmn2r63 | 0 | NA | NA | NA |
| Gm9266 | 0 | NA | NA | NA |
| Gm9294 | 4.49822579 | -0.205526789 | 0.547622412 | -0.375307482 |
| Gm23256 | 0 | NA | NA | NA |
| Gm9268 | 0 | NA | NA | NA |
| RP23-367H8.3 | 0 | NA | NA | NA |
| Gm10351 | 0 | NA | NA | NA |
| Zfp936 | 0.092140247 | 0.037060453 | 3.724945049 | 0.009949262 |
| Gm22707 | 0 | NA | NA | NA |
| EU599041 | 0.135598227 | 0.037065847 | 2.950266134 | 0.01256356 |
| Zfp715 | 456.2127529 | -0.244911399 | 0.081557166 | -3.002941525 |
| Siglecf | 5.415502606 | 1.538878629 | 0.548601457 | 2.805093952 |
| Gm10298 | 0 | NA | NA | NA |
| Siglecg | 1.312064872 | -0.284181827 | 0.858630749 | -0.330970941 |
| Lim2 | 0.046345709 | -0.443818988 | 3.724945049 | -0.119147795 |
| Nkg7 | 96.73496261 | -0.887426895 | 0.451204512 | -1.966795259 |
| Cldnd2 | 1.387261314 | -1.322814698 | 1.01957896 | -1.297412706 |
| Etfb | 181.9463623 | -0.325416583 | 0.131920627 | -2.466760434 |
| Mir7051 | 0.307454254 | 0.036532796 | 1.756035211 | 0.020804136 |
| Vsig10l | 33.09847067 | -0.167037765 | 0.179419635 | -0.930989324 |
| Iglon5 | 396.8950678 | -0.18988091 | 0.102820737 | -1.846718037 |
| 4931406B18Rik | 0.154787072 | 0.037073397 | 2.832964014 | 0.013086434 |
| Cd33 | 51.25040222 | 0.906384416 | 0.333241189 | 2.71990512 |
| Zfp658 | 98.3557729 | -0.07251446 | 0.176930771 | -0.409846516 |
| Zfp719 | 304.4942731 | -0.58655937 | 0.158585098 | -3.698704219 |
| Zfp819 | 3.834671984 | 0.358664651 | 0.623770511 | 0.574994561 |
| Ceacam18 | 0.539507586 | -0.704864633 | 1.220964722 | -0.577301392 |
| Siglece | 5.430287587 | 0.49638241 | 0.537516699 | 0.923473467 |
| Ctu1 | 87.06700889 | -0.185113383 | 0.151703937 | -1.220227943 |
| Klk14 | 0 | NA | NA | NA |
| 2310002F09Rik | 9.427952603 | 0.537356972 | 0.372780206 | 1.441484723 |
| Klk13 | 0.134945109 | 0.037074033 | 2.956294124 | 0.012540712 |
| Klk12 | 0 | NA | NA | NA |
| Klk11 | 0.225356219 | 0.939017547 | 2.130397103 | 0.440771134 |
| Klk10 | 1.594085209 | -1.923803555 | 0.876968782 | -2.193696736 |
| Klk9 | 11.12317148 | -2.185128255 | 0.611257164 | -3.574810055 |
| Klk8 | 23.43242316 | -0.895427801 | 0.297759825 | -3.007214964 |
| Klk7 | 6.053054786 | -1.215326191 | 0.732656608 | -1.658793734 |
| Klk6 | 383.1545088 | -1.415666157 | 0.329460654 | -4.296920258 |
| Klk5 | 0.178695003 | 0.401867126 | 2.707437123 | 0.148430825 |
| Klk4 | 0.195922627 | -0.895145272 | 2.218601832 | -0.4034727 |
| Klk15 | 0 | NA | NA | NA |
| Klk1b7-ps | 0.097139547 | -0.443816764 | 3.724945049 | -0.119147198 |
| Klk1b8 | 0.138087489 | -1.320984914 | 3.611429328 | -0.365778974 |
| Klk1b1 | 0.049553047 | -0.443818988 | 3.724945049 | -0.119147795 |
| Klk1b9 | 0.07796279 | -0.798301147 | 3.721829328 | -0.214491605 |

| | | | | |
|---|---|---|---|---|
| Klk1b11 | 0.276660002 | -0.879130875 | 1.827318905 | -0.481104241 |
| Klk1b26 | 0.046345709 | -0.443818988 | 3.724945049 | -0.119147795 |
| Klk1b27 | 1.612931274 | -0.34601727 | 0.79646209 | -0.434442862 |
| Klk1b21 | 0 | NA | NA | NA |
| Klk1b22 | 0 | NA | NA | NA |
| Klk1b16 | 0.166231529 | 1.489754322 | 3.417706705 | 0.435892969 |
| Klk1b24 | 0.038981395 | -0.443816764 | 3.724945049 | -0.119147198 |
| Klk1b3 | 0 | NA | NA | NA |
| Klk1b4 | 0 | NA | NA | NA |
| Klk1b5 | 0 | NA | NA | NA |
| Gm10109 | 2.250427361 | 0.228276539 | 0.662005537 | 0.344825724 |
| Klk1 | 11.42971091 | -1.170250365 | 0.418422289 | -2.796816509 |
| 1700028J19Rik | 6.192777873 | -0.025402398 | 0.43694441 | -0.058136453 |
| Gm25386 | 0 | NA | NA | NA |
| 2410002F23Rik | 502.3878778 | 0.29192732 | 0.065825982 | 4.434834263 |
| Gm28496 | 4.360017817 | 0.638008899 | 0.622223861 | 1.025368744 |
| Gm26247 | 0 | NA | NA | NA |
| Snord88c | 1.895772021 | -0.345399042 | 1.02145726 | -0.338143411 |
| Snord88a | 0 | NA | NA | NA |
| Gm23991 | 0 | NA | NA | NA |
| Acpt | 0 | NA | NA | NA |
| Gm15517 | 0 | NA | NA | NA |
| Clec11a | 34.26572508 | 0.330504515 | 0.202540572 | 1.631794125 |
| Shank1 | 9409.0605 | 0.895386068 | 0.132647788 | 6.750101769 |
| 1700008O03Rik | 12.39141503 | 0.149734559 | 0.310899562 | 0.481617143 |
| Syt3 | 466.7523524 | 0.650827718 | 0.102404786 | 6.35544237 |
| Lrrc4b | 2121.923621 | 0.575305262 | 0.080475552 | 7.14882035 |
| Aspdh | 27.56526144 | 0.272664871 | 0.259531853 | 1.050602722 |
| Josd2 | 428.3256145 | -0.513742144 | 0.108423615 | -4.738286441 |
| Mir7052 | 5.276354033 | 0.153782196 | 0.481611547 | 0.319307536 |
| Emc10 | 1263.340695 | -0.022608957 | 0.076139306 | -0.296941988 |
| Fam71e1 | 39.06557393 | 0.867877354 | 0.196333886 | 4.420415504 |
| Mybpc2 | 3.531971173 | -0.001240541 | 0.544249809 | -0.002279359 |
| Spib | 15.69420495 | -1.221801112 | 0.377245338 | -3.238744095 |
| Pold1 | 95.92363125 | -0.573074349 | 0.191450242 | -2.993333109 |
| Mir7053 | 0.189266655 | 0.445218525 | 2.684314373 | 0.165859308 |
| Nr1h2 | 489.1957701 | -0.307619009 | 0.101414995 | -3.033269483 |
| Napsa | 53.67902564 | -1.591415759 | 0.325631192 | -4.887172361 |
| Gm15396 | 3.09078215 | -0.449010315 | 0.67033566 | -0.669829075 |
| Kcnc3 | 3773.256925 | 0.235846484 | 0.118099821 | 1.997009672 |
| Myh14 | 523.8678887 | 0.12378814 | 0.159014584 | 0.778470354 |
| 2310016G11Rik | 0.744214146 | -0.90191685 | 1.131598384 | -0.797029108 |
| Izumo2 | 0 | NA | NA | NA |
| Zfp473 | 10.07782813 | 0.179535418 | 0.319494052 | 0.561936651 |
| Vrk3 | 217.5211013 | -0.401190537 | 0.127633236 | -3.143307727 |
| Atf5 | 717.0131176 | -1.049267642 | 0.173793818 | -6.037427871 |
| Nup62 | 303.2451805 | -0.29206711 | 0.11527281 | -2.533703393 |

| | | | | |
|---|---|---|---|---|
| Il4i1 | 16.51100415 | -0.249178136 | 0.300561944 | -0.829040871 |
| Tbc1d17 | 424.275241 | 0.08625567 | 0.081501643 | 1.058330449 |
| Akt1s1 | 507.5705319 | -0.452275887 | 0.066080671 | -6.844299272 |
| Mir707 | 0 | NA | NA | NA |
| Pnkp | 197.282501 | -0.24944254 | 0.088971834 | -2.803612445 |
| Ptov1 | 973.5278823 | 0.184465355 | 0.055946515 | 3.297173311 |
| Med25 | 617.720589 | -0.031788405 | 0.073424938 | -0.432937443 |
| Fuz | 99.72731422 | -0.140325274 | 0.146276204 | -0.959317175 |
| Ap2a1 | 2601.373962 | 0.124801461 | 0.051018822 | 2.446184664 |
| Tsks | 3.95301002 | 1.965241978 | 0.603363586 | 3.257143823 |
| Cpt1c | 718.0796799 | 0.348057796 | 0.074324408 | 4.682954173 |
| Prmt1 | 509.9239553 | -0.271195807 | 0.095616278 | -2.836293285 |
| Gm15545 | 9.287909809 | -0.372042384 | 0.385684783 | -0.964628111 |
| Bcl2l12 | 25.31009217 | -0.119886595 | 0.280434012 | -0.427503762 |
| Irf3 | 149.8132699 | -0.760174462 | 0.17189179 | -4.422401226 |
| Scaf1 | 2253.505891 | 0.237800356 | 0.056721262 | 4.192437673 |
| Mir7054 | 0.567171362 | 0.872308733 | 1.370980839 | 0.636266174 |
| Rras | 111.4151899 | -0.566863881 | 0.181066297 | -3.130697934 |
| Prr12 | 1649.371309 | 0.26504983 | 0.065805333 | 4.027786469 |
| Prrg2 | 52.66422563 | -0.286160619 | 0.15134654 | -1.890764188 |
| Nosip | 442.5713574 | -0.03724279 | 0.088515266 | -0.420749908 |
| Rcn3 | 57.62462159 | -1.00297314 | 0.202144978 | -4.961652526 |
| Fcgrt | 111.4072777 | -0.085272196 | 0.193084991 | -0.441630371 |
| Mir150 | 0 | NA | NA | NA |
| Rps11 | 1103.617122 | -0.263928464 | 0.098539185 | -2.678411276 |
| Snord35b | 0.134597393 | 0.037065773 | 2.957007063 | 0.012534895 |
| Rpl13a | 2064.540611 | -0.270078944 | 0.144454329 | -1.869649368 |
| Snord35a | 0.656197145 | 0.041620533 | 1.400457076 | 0.029719249 |
| Snord34 | 0 | NA | NA | NA |
| Snord33 | 0 | NA | NA | NA |
| mmu-mir-5121 | 0 | NA | NA | NA |
| Snord32a | 0 | NA | NA | NA |
| Flt3l | 50.76228596 | 0.146854926 | 0.202730413 | 0.724385276 |
| Aldh16a1 | 113.740617 | -0.36989036 | 0.128065332 | -2.888294241 |
| Pih1d1 | 223.2642417 | -0.263022478 | 0.091657909 | -2.869610268 |
| Slc17a7 | 6082.783442 | 0.68452685 | 0.121784305 | 5.620813379 |
| mmu-mir-7055 | 0.339810401 | 0.983955461 | 1.770309185 | 0.555809951 |
| Gfy | 10.55721659 | -0.123516437 | 0.645408262 | -0.191377216 |
| Pth2 | 1.157525169 | -2.017151547 | 1.047407961 | -1.925850884 |
| Ccdc155 | 18.99181208 | -0.294227243 | 0.284081718 | -1.035713405 |
| Dkkl1 | 52.36991393 | 0.766224318 | 0.221858952 | 3.453655181 |
| Tead2 | 38.54700613 | -0.640538006 | 0.236047725 | -2.713595335 |
| Cd37 | 67.79623972 | -0.989150579 | 0.222494238 | -4.445735713 |
| Slc6a16 | 3.556493767 | -0.177850526 | 0.531328851 | -0.334727778 |
| Gm23585 | 0 | NA | NA | NA |
| 1700039E15Rik | 0.62192417 | 1.3019641 | 1.311410776 | 0.992796555 |
| Trpm4 | 109.1332828 | 0.39122695 | 0.174888113 | 2.237012812 |

| | | | | |
|---|---|---|---|---|
| Rpl14-ps1 | 281.4107973 | -0.078595861 | 0.132557255 | -0.592920099 |
| Hrc | 0.548344614 | -2.049709967 | 1.290942363 | -1.587762573 |
| Ppfia3 | 1660.177301 | 0.437897432 | 0.104454019 | 4.19225067 |
| Mtag2 | 16.72535298 | -0.188580257 | 0.346460192 | -0.544305699 |
| Lin7b | 211.863161 | 0.999224575 | 0.195136994 | 5.120631166 |
| Snrnp70 | 1046.054046 | 0.15775801 | 0.179157636 | 0.880554206 |
| Gm22478 | 0.046345709 | -0.443818988 | 3.724945049 | -0.119147795 |
| Kcna7 | 0.228985044 | -0.524465112 | 2.565006791 | -0.204469288 |
| Ntf5 | 1.628131443 | -0.243148168 | 0.741328407 | -0.327989817 |
| Lhb | 9.980967548 | 0.008012323 | 0.548622892 | 0.014604427 |
| Ruvbl2 | 254.9844317 | -0.544735908 | 0.110020472 | -4.951223147 |
| Gys1 | 207.1643521 | -0.714948042 | 0.126539846 | -5.649983482 |
| Ftl1 | 3794.633164 | -1.286875548 | 0.193221466 | -6.660106519 |
| Gm22133 | 0 | NA | NA | NA |
| Bax | 270.9969915 | -0.494828573 | 0.098109503 | -5.043635495 |
| Dhdh | 156.4792992 | 0.002787375 | 0.15033561 | 0.018541018 |
| Tulp2 | 0.658348016 | -0.201642196 | 1.367360519 | -0.147468201 |
| Nucb1 | 959.7799318 | -0.280670163 | 0.068078619 | -4.122735823 |
| Ppp1r15a | 417.9443 | -0.546957449 | 0.352483104 | -1.551726714 |
| Plekha4 | 287.7791967 | -0.409478154 | 0.304116297 | -1.346452516 |
| Gm16022 | 0.072025462 | 0.517959912 | 3.724945049 | 0.139051692 |
| Hsd17b14 | 2.84382331 | 0.001605126 | 0.661188789 | 0.002427636 |
| 0610005C13Rik | 1.694978974 | -1.625748388 | 0.921020035 | -1.765160721 |
| Bcat2 | 107.8855267 | -0.028687527 | 0.157368065 | -0.182295735 |
| Fgf21 | 1.959085883 | -2.939234293 | 0.888452302 | -3.308263466 |
| Fut1 | 0.956979021 | 0.063361739 | 1.003428992 | 0.063145215 |
| Izumo1 | 0.702218406 | -1.713547394 | 1.411958638 | -1.213596027 |
| Rasip1 | 78.58288542 | -0.207036905 | 0.210278083 | -0.984586229 |
| Gm16047 | 2.391421094 | 0.123351416 | 0.731931081 | 0.16852873 |
| Mamstr | 31.70904901 | 0.472662246 | 0.261819845 | 1.805295719 |
| Fut2 | 5.566764691 | -0.880254306 | 0.405361248 | -2.171530482 |
| Sec1 | 5.465091639 | 0.897692356 | 0.484177805 | 1.854055154 |
| Ntn5 | 29.63111439 | 1.287668556 | 0.333365036 | 3.862638303 |
| Car11 | 810.7850525 | 0.955643813 | 0.13937233 | 6.856768595 |
| Dbp | 869.3346952 | 1.197903617 | 0.154581183 | 7.749349523 |
| Sphk2 | 380.1522572 | -0.528788391 | 0.109692638 | -4.820637023 |
| Rpl18 | 1024.378528 | -0.439678621 | 0.125116604 | -3.514150848 |
| Fam83e | 2.171234967 | -1.081197087 | 0.815653722 | -1.325558946 |
| Spaca4 | 0 | NA | NA | NA |
| Sult2b1 | 46.17871227 | -0.584565986 | 0.257352191 | -2.271463023 |
| Lmtk3 | 2315.447368 | 0.711091779 | 0.110431605 | 6.439205298 |
| Cyth2 | 643.2901348 | -0.020641502 | 0.078523928 | -0.262868947 |
| Kcnj14 | 17.48480922 | -0.632622288 | 0.40255743 | -1.571508162 |
| Grwd1 | 158.3457012 | -0.279825498 | 0.153150317 | -1.827129735 |
| Grin2d | 305.5974088 | 0.195024527 | 0.094845886 | 2.056225486 |
| Kdelr1 | 538.7074619 | -0.262077698 | 0.074203667 | -3.531869912 |
| Gm22098 | 0 | NA | NA | NA |

| | | | | |
|---|---|---|---|---|
| Syngr4 | 0.321511422 | 0.554954838 | 1.755231339 | 0.316171906 |
| Tmem143 | 157.483315 | 0.118042806 | 0.119866795 | 0.984783202 |
| Emp3 | 69.37531286 | -1.578456355 | 0.265686241 | -5.941054196 |
| Ccdc114 | 21.307866 | -0.468202886 | 0.246616163 | -1.898508518 |
| Abcc6 | 6.348614184 | -1.449763554 | 0.465639809 | -3.11348713 |
| Nomo1 | 1371.14784 | 0.19692514 | 0.066434742 | 2.964189124 |
| Kcnj11 | 116.36677 | 0.422537325 | 0.132828706 | 3.181069346 |
| Abcc8 | 310.3174424 | 0.600660561 | 0.125563182 | 4.783731576 |
| Ush1c | 7.85454541 | 0.738920564 | 0.371911594 | 1.986817768 |
| Otog | 2.40295838 | 1.420982282 | 0.766832807 | 1.853053584 |
| Gm38176 | 0 | NA | NA | NA |
| Myod1 | 0.509027769 | 1.472698319 | 1.43241957 | 1.028119379 |
| Kcnc1 | 3456.691312 | 0.219100873 | 0.075857634 | 2.888316714 |
| Sergef | 366.9267911 | -0.016050962 | 0.10606266 | -0.151334713 |
| Gm22969 | 1.792854059 | 2.013607373 | 0.901587106 | 2.23340303 |
| Gm15700 | 0 | NA | NA | NA |
| Tph1 | 2.962511132 | 1.518405283 | 0.834231354 | 1.820124927 |
| Saal1 | 161.7785274 | -0.07203644 | 0.09458859 | -0.761576419 |
| Saa3 | 441.1512112 | -0.303115545 | 0.744250813 | -0.407276069 |
| Saa4 | 0.385868569 | -1.030051114 | 1.619240247 | -0.636132356 |
| Saa1 | 4.201556626 | 0.423567593 | 0.585744473 | 0.723126914 |
| Saa2 | 1.450828853 | 0.968257155 | 0.837715106 | 1.155831079 |
| Hps5 | 293.9049292 | -0.393764825 | 0.116578583 | -3.377677233 |
| Gtf2h1 | 540.0561687 | -0.007374309 | 0.086098932 | -0.08564925 |
| Gm9392 | 24.65709606 | -0.201328807 | 0.225998692 | -0.890840585 |
| Ldha | 1608.193775 | -0.323919693 | 0.081304212 | -3.984045674 |
| Ldhc | 0.061920613 | -0.443816764 | 3.724945049 | -0.119147198 |
| Tsg101 | 606.9962445 | -0.051213013 | 0.074021265 | -0.691868933 |
| Uevld | 206.960226 | 0.242246721 | 0.117991498 | 2.053086245 |
| Gm9999 | 0.062426873 | 0.517959912 | 3.724945049 | 0.139051692 |
| Spty2d1 | 614.2646612 | -0.29146759 | 0.119296592 | -2.443218083 |
| Tmem86a | 239.3545487 | -0.599008042 | 0.136378376 | -4.392250897 |
| Ptpn5 | 1663.361268 | -0.030310462 | 0.133285627 | -0.22740983 |
| Mir7056 | 0.65404267 | 0.047061765 | 1.281838635 | 0.036714266 |
| Gm14377 | 1.857966922 | -1.078975028 | 0.773588091 | -1.394766853 |
| Mrgpra6 | 0 | NA | NA | NA |
| Mrgpra9 | 0.097139547 | -0.443816764 | 3.724945049 | -0.119147198 |
| Mrgprx3-ps | 0 | NA | NA | NA |
| Mrgpra1 | 0 | NA | NA | NA |
| Gm29479 | 0 | NA | NA | NA |
| Gm25326 | 0.128477747 | 0.037074008 | 3.677132019 | 0.010082316 |
| Mrgpra2a | 0.092691418 | -0.908687078 | 3.724560878 | -0.2439716 |
| Mrgpra2b | 0.175945213 | -0.410809826 | 2.710746807 | -0.151548579 |
| Gm7255 | 0 | NA | NA | NA |
| Mrgprc4-ps | 0 | NA | NA | NA |
| Mrgpra14 | 0 | NA | NA | NA |
| Gm9322 | 0 | NA | NA | NA |

| | | | | |
|---|---|---|---|---|
| Mrgpra3 | 0 | NA | NA | NA |
| Mrgprc1-ps | 0 | NA | NA | NA |
| Gm2707 | 0 | NA | NA | NA |
| Gm28288 | 0 | NA | NA | NA |
| Mrgprc3-ps | 0 | NA | NA | NA |
| Gm5733 | 0 | NA | NA | NA |
| Gm22427 | 0 | NA | NA | NA |
| Gm18348 | 0 | NA | NA | NA |
| Gm29563 | 0 | NA | NA | NA |
| Mrgprc7-ps | 0 | NA | NA | NA |
| Gm18349 | 0 | NA | NA | NA |
| Mrgpra18-ps | 0 | NA | NA | NA |
| Gm18350 | 0 | NA | NA | NA |
| Gm5734 | 0 | NA | NA | NA |
| Mrgprc2-ps | 0 | NA | NA | NA |
| Mrgpra4 | 0 | NA | NA | NA |
| Mrgprx1 | 0 | NA | NA | NA |
| Mrgprb7-ps | 0 | NA | NA | NA |
| Gm28666 | 0 | NA | NA | NA |
| Mrgprb12-ps | 0 | NA | NA | NA |
| Gm18351 | 0 | NA | NA | NA |
| Gm28665 | 0 | NA | NA | NA |
| Mrgprb5 | 0 | NA | NA | NA |
| Gm18352 | 0 | NA | NA | NA |
| Mrgprb4 | 0 | NA | NA | NA |
| Gm26950 | 0 | NA | NA | NA |
| Gm18356 | 0 | NA | NA | NA |
| Mrgprb13 | 0 | NA | NA | NA |
| Gm18357 | 0 | NA | NA | NA |
| Mrgprb8 | 0.066050875 | -0.443818988 | 3.724945049 | -0.119147795 |
| Mrgprb1 | 0.19039241 | 0.517965586 | 2.668133052 | 0.194130344 |
| Mrgprx2 | 0.10503227 | -0.905414734 | 3.723500899 | -0.243162217 |
| Mrgprb2 | 0.044544648 | 0.517957688 | 3.724945049 | 0.139051095 |
| Mrgprb9-ps | 0 | NA | NA | NA |
| Gm18353 | 0 | NA | NA | NA |
| Mrgprb3 | 0 | NA | NA | NA |
| Mrgprb11 | 0.045882455 | 0.517959912 | 3.724945049 | 0.139051692 |
| Zdhhc13 | 253.9183394 | -0.014038287 | 0.093525014 | -0.150101946 |
| Csrp3 | 0.531680062 | -2.451798798 | 1.478343946 | -1.658476571 |
| E2f8 | 18.28425174 | -0.772847378 | 0.289341288 | -2.671058056 |
| Gm2788 | 0.049553047 | -0.443818988 | 3.724945049 | -0.119147795 |
| Nav2 | 2595.787216 | -0.479030854 | 0.102637566 | -4.667207827 |
| Gm37613 | 24.19551003 | 0.315835031 | 0.417334254 | 0.756791535 |
| Gm38059 | 24.62141553 | 0.008221588 | 0.448433824 | 0.018334005 |
| Dbx1 | 0.097139547 | -0.443816764 | 3.724945049 | -0.119147198 |
| Htatip2 | 67.03302061 | -0.687586392 | 0.199323302 | -3.449603665 |
| Prmt3 | 302.2229905 | -0.298831757 | 0.090270351 | -3.310408719 |

| | | | | |
|---|---|---|---|---|
| Gm26856 | 0 | NA | NA | NA |
| Slc6a5 | 485.8678254 | -0.425118643 | 0.562356795 | -0.755958933 |
| Nell1 | 633.6668325 | 0.26678875 | 0.102765622 | 2.596089486 |
| Nell1os | 0.048949532 | 0.037064902 | 3.724945049 | 0.009950456 |
| 4933405O20Rik | 9.680996758 | 0.980761467 | 0.43199671 | 2.270298463 |
| Ano5 | 52.81759069 | 0.204275242 | 0.197966874 | 1.031865775 |
| Slc17a6 | 1077.091468 | 0.116198409 | 0.154756821 | 0.750845158 |
| Gm7336 | 39.57013241 | 0.019763918 | 0.185700586 | 0.106428949 |
| Gm29296 | 0.142694128 | -0.443822014 | 2.905728259 | -0.152740371 |
| Fancf | 37.44894291 | -0.184522379 | 0.239720063 | -0.769741075 |
| Gas2 | 177.6414515 | 0.197668222 | 0.131330395 | 1.505121659 |
| Svip | 477.7009006 | 0.177825315 | 0.097001133 | 1.833229252 |
| 1700015G11Rik | 0.161582804 | 0.539616788 | 2.79775094 | 0.192875206 |
| Gm22211 | 0 | NA | NA | NA |
| Gm6181 | 1.4136546 | -1.975336028 | 1.064678504 | -1.85533569 |
| Gm22444 | 0 | NA | NA | NA |
| Mir6238 | 0 | NA | NA | NA |
| Gm27921 | 0 | NA | NA | NA |
| Gm25956 | 0 | NA | NA | NA |
| Gm5776 | 44.67075326 | 0.200294469 | 0.19296313 | 1.037993472 |
| Luzp2 | 603.6111141 | 0.464578005 | 0.109558526 | 4.240455042 |
| Gm24116 | 0 | NA | NA | NA |
| Gm26482 | 0 | NA | NA | NA |
| Siglech | 79.78094538 | 0.686556081 | 0.305806535 | 2.245066743 |
| Tubgcp5 | 553.9336148 | 0.061882373 | 0.0686796 | 0.901029903 |
| Cyfip1 | 1268.126837 | -0.22568822 | 0.083964885 | -2.687888165 |
| Gm17907 | 0.149576918 | -0.443202057 | 2.863264737 | -0.15478906 |
| Nipa2 | 555.4063321 | -0.079004102 | 0.07885076 | -1.001944701 |
| A230056P14Rik | 25.86046502 | -0.755607113 | 0.246270508 | -3.068199761 |
| Nipa1 | 678.816936 | 0.454459845 | 0.108973961 | 4.170352637 |
| Gm15888 | 0 | NA | NA | NA |
| Rps12l1 | 1.168735097 | 1.591207609 | 0.985062282 | 1.615337057 |
| Herc2 | 5466.518983 | 0.222060035 | 0.083947362 | 2.645229462 |
| Oca2 | 2.497417949 | -0.197618199 | 0.658125081 | -0.30027453 |
| Gabrg3 | 564.6589 | 0.394921482 | 0.099107025 | 3.984798087 |
| Gm9962 | 23.2644548 | 0.311769623 | 0.232301278 | 1.342091725 |
| Gabra5 | 375.6717185 | 0.74981011 | 0.110142146 | 6.807658461 |
| Gabrb3 | 2705.952044 | 0.538617554 | 0.077637809 | 6.937567714 |
| Mir6389 | 0 | NA | NA | NA |
| Gm23911 | 0.328919656 | -0.32464846 | 1.72694665 | -0.187989861 |
| Gm26288 | 0 | NA | NA | NA |
| Atp10a | 221.527901 | -1.132368977 | 0.198006094 | -5.718859233 |
| Gm6226 | 2.39103255 | -1.336346139 | 0.785679697 | -1.700879053 |
| Ube3a | 3913.118757 | 0.008682466 | 0.074086887 | 0.117193024 |
| Snhg14 | 6561.535641 | 0.738723196 | 0.125123321 | 5.903960886 |
| Gm25499 | 5.73832025 | 1.751142947 | 0.68280601 | 2.56462732 |
| Gm24966 | 65.59095034 | 0.807689069 | 0.244854043 | 3.298655233 |

| | | | | |
|---|---|---|---|---|
| Gm22496 | 12.77173823 | 1.500284351 | 0.442046574 | 3.39395086 |
| Gm25648 | 15.8722383 | 0.830891017 | 0.483753292 | 1.71759248 |
| Gm24306 | 4.90255571 | 0.653134108 | 0.476281195 | 1.371320377 |
| Gm25934 | 2.147180949 | 1.015511031 | 0.731559601 | 1.388145313 |
| Gm25156 | 8.000524403 | 1.438152191 | 0.464865446 | 3.093695611 |
| Gm26488 | 4.065985139 | 1.227047972 | 0.612887987 | 2.002075418 |
| Gm24220 | 8.515480277 | 2.017793065 | 0.45751554 | 4.410326834 |
| Gm24653 | 7.944433476 | 1.341868908 | 0.420800013 | 3.188851868 |
| Gm24969 | 3.545771104 | 0.303543346 | 0.675336752 | 0.44946961 |
| Gm24417 | 0.567759171 | 0.525845319 | 1.462320906 | 0.35959639 |
| Gm25121 | 85.00767218 | 0.87156973 | 0.276369255 | 3.153642142 |
| Gm24657 | 0.475242527 | 0.991681247 | 1.77640337 | 0.558252289 |
| Gm26230 | 0.075043205 | 0.517957688 | 3.724945049 | 0.139051095 |
| Gm23922 | 0.156071933 | 0.517956562 | 3.489245341 | 0.148443721 |
| Gm25314 | 2.261769448 | -0.596718327 | 0.922340095 | -0.646961279 |
| Gm26096 | 0.262111692 | 1.147505135 | 2.447786287 | 0.468793024 |
| Gm22996 | 6.119234212 | 0.906250091 | 0.607252354 | 1.492378062 |
| Gm26499 | 0.650021907 | 0.408372781 | 1.473854582 | 0.277078069 |
| Gm25646 | 2.157091062 | 0.555385262 | 0.715030871 | 0.776729069 |
| Gm26498 | 21.92470555 | 0.949374985 | 0.34940952 | 2.717083913 |
| Gm23660 | 2.095785588 | 0.752332504 | 0.787058128 | 0.955879214 |
| Gm23933 | 2.60823589 | 1.049008862 | 0.703396102 | 1.491348701 |
| Gm24862 | 0.637223532 | 0.906117155 | 1.253127516 | 0.723084557 |
| Gm24866 | 6.740698188 | 0.91038885 | 0.488904207 | 1.862100666 |
| Gm22348 | 0 | NA | NA | NA |
| Gm26466 | 4.89789051 | 0.345921031 | 0.612417574 | 0.564845043 |
| Gm24799 | 3.843037622 | 0.858906946 | 0.706477127 | 1.215760445 |
| Gm24088 | 1.480005993 | 2.399313489 | 1.037512966 | 2.312562413 |
| Gm25463 | 3.198438298 | 0.912325551 | 0.770844183 | 1.183540813 |
| Gm25229 | 2.607201442 | 1.339884272 | 0.778069667 | 1.722062084 |
| Gm24528 | 0.50461959 | -0.467637163 | 1.314931758 | -0.355636071 |
| Gm23356 | 0.770457139 | 1.068214174 | 1.307157981 | 0.817203574 |
| Gm25449 | 0.090074004 | 0.517955464 | 3.724945049 | 0.139050498 |
| Gm24872 | 0.860548126 | 1.657058828 | 1.159727385 | 1.428834785 |
| Gm26337 | 0.156716517 | -0.443821692 | 2.821984856 | -0.157272882 |
| Gm23145 | 0.371627904 | 0.449375786 | 1.57030842 | 0.286170399 |
| Gm22130 | 0.047948698 | 0.037064902 | 3.724945049 | 0.009950456 |
| Gm25988 | 0 | NA | NA | NA |
| Gm25221 | 0 | NA | NA | NA |
| Gm26389 | 0.044191549 | 0.037062677 | 3.724945049 | 0.009949859 |
| Gm23471 | 2.996745481 | 0.54737148 | 0.621883681 | 0.880183059 |
| Gm25120 | 0.150248541 | 0.998859444 | 2.859451703 | 0.349318522 |
| Gm24418 | 0 | NA | NA | NA |
| Gm23687 | 0 | NA | NA | NA |
| Gm22393 | 0 | NA | NA | NA |
| Gm24639 | 0.168184286 | 0.517964245 | 2.759948177 | 0.187671729 |
| Gm24566 | 0 | NA | NA | NA |

| | | | | |
|---|---|---|---|---|
| Gm22111 | 0 | NA | NA | NA |
| Gm23076 | 0.042716945 | 0.037062677 | 3.724945049 | 0.009949859 |
| Gm25874 | 0 | NA | NA | NA |
| Gm25088 | 0.11657011 | 0.998850474 | 3.724945049 | 0.268151734 |
| Gm22510 | 5.686408051 | 1.182471266 | 0.539782286 | 2.190644816 |
| Gm24983 | 1.441787318 | 1.073299378 | 0.86558839 | 1.239965081 |
| Gm24027 | 3.476611664 | 0.927588594 | 0.616924607 | 1.503568806 |
| Gm22524 | 0.172163037 | -0.92558091 | 2.73973512 | -0.337835911 |
| Gm22630 | 0.258404708 | 1.265030131 | 2.047008666 | 0.617989631 |
| Gm25741 | 0.657611471 | -0.30490907 | 1.228706823 | -0.248154453 |
| Gm26059 | 0.559779376 | 0.915760131 | 1.479281185 | 0.619057513 |
| Gm25175 | 0.444383067 | 1.315369144 | 1.570304231 | 0.837652423 |
| Gm22511 | 1.321027671 | 0.260765969 | 0.885778486 | 0.294391853 |
| Gm25077 | 0.739785724 | -0.389108352 | 1.233075823 | -0.315559144 |
| Gm22834 | 0.106971521 | 0.998850474 | 3.724945049 | 0.268151734 |
| Gm24926 | 1.408313197 | -0.092598813 | 0.961777244 | -0.096278856 |
| Gm26200 | 0.042716945 | 0.037062677 | 3.724945049 | 0.009949859 |
| Gm22632 | 0.060174537 | 0.517957688 | 3.724945049 | 0.139051095 |
| Gm23254 | 4.913973528 | 1.233305038 | 0.496456025 | 2.484218089 |
| Gm26390 | 0.52711509 | 1.954147477 | 1.326906607 | 1.472709131 |
| Gm26366 | 1.250668422 | 2.96181546 | 1.015909234 | 2.915433153 |
| Gm26336 | 0.636553247 | 1.37183931 | 1.298889886 | 1.056162901 |
| Gm26283 | 0.137685729 | 0.5179621 | 2.938031518 | 0.176295624 |
| Gm25098 | 0.174523186 | 0.88002062 | 3.36983679 | 0.261146363 |
| Gm24702 | 0.239175706 | 1.657305723 | 2.993642531 | 0.553608424 |
| Gm26432 | 0.437372945 | 2.310407966 | 1.403955495 | 1.645641884 |
| Gm25462 | 6.852308181 | 0.857047862 | 0.536540703 | 1.597358516 |
| Gm24495 | 12.03897181 | 1.536737122 | 0.382930239 | 4.013099425 |
| Gm25523 | 2.19067646 | 1.178780399 | 0.901219424 | 1.307983791 |
| Gm22255 | 1.065694062 | 1.50942166 | 1.017296996 | 1.483757119 |
| Gm23449 | 8.027250868 | 1.201720651 | 0.422875541 | 2.841783301 |
| Gm23310 | 0.295362062 | 0.037072486 | 2.038352329 | 0.018187477 |
| Gm25461 | 0.060174537 | 0.517957688 | 3.724945049 | 0.139051095 |
| Gm24310 | 0.210314706 | 0.503099069 | 2.185265535 | 0.230223312 |
| Gm22909 | 0.331212083 | 1.069599541 | 1.992109448 | 0.53691806 |
| Gm25823 | 1.425569217 | -0.279532928 | 1.089844051 | -0.256488924 |
| Gm25647 | 0.32719657 | -0.082146803 | 1.765583474 | -0.046526717 |
| Gm22252 | 2.966653922 | 0.969194564 | 0.682822561 | 1.419394466 |
| Gm26392 | 0.252618675 | 0.019915591 | 2.516083652 | 0.007915313 |
| Gm24219 | 2.893504144 | 0.768439039 | 0.656584822 | 1.170357604 |
| Gm25230 | 1.071417476 | 0.728390756 | 0.92553976 | 0.786990239 |
| Gm22627 | 0.53540973 | 1.26629389 | 1.342945969 | 0.94292244 |
| Gm24585 | 0.104637558 | -0.443821212 | 3.724945049 | -0.119148392 |
| Gm23305 | 0.469128522 | 0.336374832 | 1.567702123 | 0.214565527 |
| Gm24206 | 0.186417525 | 0.50278671 | 2.263344327 | 0.22214327 |
| Gm25085 | 2.147377591 | 1.116393803 | 0.785894269 | 1.420539438 |
| Gm25209 | 0.361503534 | 0.524995236 | 1.607906732 | 0.326508513 |

| | | | | |
|---|---|---|---|---|
| Gm23682 | 0.137685729 | 0.5179621 | 2.938031518 | 0.176295624 |
| Gm24654 | 0.322713358 | 1.443508054 | 1.736550592 | 0.831250216 |
| Gm26334 | 0.362177786 | 1.798030302 | 1.542931679 | 1.165333713 |
| Gm25742 | 0.531759696 | 1.81380779 | 1.345303985 | 1.348251258 |
| Gm25585 | 0 | NA | NA | NA |
| Gm25840 | 0 | NA | NA | NA |
| Gm25089 | 0.347269729 | 1.97020313 | 3.262018551 | 0.60398281 |
| Gm24040 | 0.042716945 | 0.037062677 | 3.724945049 | 0.009949859 |
| Gm22050 | 0.315224781 | 0.418564985 | 1.600387936 | 0.261539703 |
| Gm23286 | 0 | NA | NA | NA |
| Gm24021 | 0 | NA | NA | NA |
| Gm22274 | 0 | NA | NA | NA |
| Gm24652 | 0.045882455 | 0.517959912 | 3.724945049 | 0.139051692 |
| Gm23618 | 0 | NA | NA | NA |
| Gm23688 | 0 | NA | NA | NA |
| Gm24570 | 0 | NA | NA | NA |
| Gm26467 | 0 | NA | NA | NA |
| Gm25194 | 0.147068667 | 0.998853574 | 3.51441668 | 0.284216035 |
| Gm22640 | 0 | NA | NA | NA |
| Gm24952 | 0 | NA | NA | NA |
| Gm25087 | 0 | NA | NA | NA |
| Gm23944 | 0 | NA | NA | NA |
| Gm22629 | 0.292758571 | 0.985421902 | 2.033299402 | 0.48464181 |
| Gm26374 | 0.198152624 | -1.482926123 | 3.70739678 | -0.39999121 |
| Gm26284 | 0 | NA | NA | NA |
| Gm25452 | 0 | NA | NA | NA |
| Gm25984 | 0 | NA | NA | NA |
| Gm25146 | 0.176667124 | 0.037076309 | 2.30438001 | 0.016089494 |
| Gm24759 | 1.076799075 | 0.260852285 | 1.02500974 | 0.254487616 |
| Gm24100 | 1.19629966 | 1.179509576 | 1.08583055 | 1.086274075 |
| Gm22253 | 0.244208214 | 0.537959057 | 2.453152573 | 0.219292947 |
| Gm25017 | 0.060174537 | 0.517957688 | 3.724945049 | 0.139051095 |
| Gm23575 | 0.210511418 | 0.037076858 | 2.190351543 | 0.016927355 |
| Gm22391 | 0.323669746 | 0.517974026 | 1.590787234 | 0.325608614 |
| Gm22449 | 1.252922778 | 0.701687469 | 0.914414766 | 0.767362356 |
| Gm24137 | 0.11657011 | 0.998850474 | 3.724945049 | 0.268151734 |
| Gm22912 | 0.442601433 | 1.463208233 | 1.513621332 | 0.966693718 |
| Gm25710 | 0.427556563 | 0.953651398 | 1.562952708 | 0.610160111 |
| Gm26434 | 7.062431358 | 0.295337354 | 0.45791383 | 0.644962728 |
| Gm23560 | 8.541564187 | 0.438779658 | 0.428487394 | 1.024019993 |
| Gm25870 | 11.09216655 | 0.634322783 | 0.358539666 | 1.769184399 |
| Gm23862 | 2.015428526 | 0.656111855 | 0.861219163 | 0.761840753 |
| Gm26504 | 0.522929342 | 1.177349497 | 1.306876567 | 0.900888061 |
| Gm22285 | 0 | NA | NA | NA |
| Gm22131 | 0.271448477 | -0.857599822 | 2.45148479 | -0.349828735 |
| Gm24153 | 0.725964322 | -0.582196539 | 1.223698865 | -0.475767818 |
| Gm26365 | 0 | NA | NA | NA |

| | | | | |
|---|---|---|---|---|
| Gm24609 | 0.334054859 | 1.369470103 | 3.71433345 | 0.368698751 |
| Gm24264 | 0.41844861 | 0.125207582 | 3.305402135 | 0.037879682 |
| Gm25155 | 0.246032045 | 1.019567397 | 2.096073538 | 0.486417761 |
| Gm25816 | 0.401944266 | 1.944522973 | 1.481797825 | 1.312272795 |
| Gm23141 | 0.141693294 | -0.443822038 | 2.912076071 | -0.152407433 |
| Gm23767 | 0.37340313 | 1.54135709 | 1.75779164 | 0.876871328 |
| Gm22417 | 0.221310845 | 0.515336785 | 3.186476227 | 0.161726229 |
| Gm22584 | 1.21151113 | 0.035818823 | 1.023295014 | 0.035003418 |
| Gm26270 | 0.134148234 | -0.443820953 | 3.624632483 | -0.122445781 |
| Gm23089 | 0 | NA | NA | NA |
| Gm24711 | 0.104637558 | -0.443821212 | 3.724945049 | -0.119148392 |
| Gm26188 | 0 | NA | NA | NA |
| Gm23265 | 0.938116572 | 2.334576642 | 1.070902561 | 2.180008459 |
| Gm24518 | 0.272816007 | 1.014377728 | 2.079538099 | 0.487789923 |
| Gm25944 | 0.061920613 | -0.443816764 | 3.724945049 | -0.119147198 |
| Gm23047 | 0.652779004 | 0.245201299 | 1.313238997 | 0.186714908 |
| Gm23953 | 0 | NA | NA | NA |
| Gm26332 | 0 | NA | NA | NA |
| Gm26032 | 0 | NA | NA | NA |
| Gm22863 | 0 | NA | NA | NA |
| Gm24618 | 0 | NA | NA | NA |
| Gm22046 | 0 | NA | NA | NA |
| Gm24760 | 0.092140247 | 0.037060453 | 3.724945049 | 0.009949262 |
| Gm25350 | 0.472357773 | 1.462746783 | 1.436653624 | 1.018162457 |
| Gm26502 | 0 | NA | NA | NA |
| Gm23313 | 0 | NA | NA | NA |
| Gm25471 | 0.129625439 | 0.037055631 | 3.686164085 | 0.010052627 |
| Gm22776 | 0 | NA | NA | NA |
| Gm23446 | 0.198993444 | -0.443820485 | 2.230108596 | -0.199012948 |
| Gm25210 | 0 | NA | NA | NA |
| Gm22128 | 0 | NA | NA | NA |
| Gm22851 | 0.048949532 | 0.037064902 | 3.724945049 | 0.009950456 |
| Gm25074 | 0.045882455 | 0.517959912 | 3.724945049 | 0.139051692 |
| Gm25157 | 0.086648695 | 0.037058229 | 3.724945049 | 0.009948665 |
| Gm22631 | 0.573189294 | 0.040372685 | 1.334938794 | 0.030243098 |
| Gm25474 | 0.37594025 | 0.500223747 | 1.488963107 | 0.335954427 |
| Gm22941 | 0.150983267 | -0.925599732 | 2.85520831 | -0.324179405 |
| Gm26246 | 0.123992736 | 0.517955521 | 3.720643114 | 0.139211288 |
| Gm22047 | 0 | NA | NA | NA |
| Gm25597 | 0.783837108 | 1.393550673 | 1.258350736 | 1.107442172 |
| Gm26097 | 0.21090123 | 0.51796878 | 2.189410504 | 0.236579106 |
| Gm22258 | 0.307376478 | 0.037077148 | 1.794301171 | 0.020663838 |
| Gm26201 | 0 | NA | NA | NA |
| Gm23619 | 0 | NA | NA | NA |
| Gm22812 | 0.249740077 | 0.517964915 | 2.539847055 | 0.203935475 |
| Gm26433 | 0.126225412 | 0.03707172 | 3.698755067 | 0.010022756 |
| Gm23357 | 0.67569888 | 1.529774046 | 1.227966718 | 1.245778101 |

| | | | | |
|---|---|---|---|---|
| Gm25615 | 0.379570916 | -0.369059408 | 1.566217273 | -0.235637427 |
| Gm22110 | 0.872838561 | 0.026660779 | 1.195623247 | 0.022298646 |
| Gm23724 | 0 | NA | NA | NA |
| Gm22188 | 0.093141081 | 0.037060453 | 3.724945049 | 0.009949262 |
| Gm23549 | 0 | NA | NA | NA |
| Snord116l2 | 0.075043205 | 0.517957688 | 3.724945049 | 0.139051095 |
| Gm26223 | 0 | NA | NA | NA |
| Gm23524 | 0.112288909 | 0.037060453 | 3.724945049 | 0.009949262 |
| Gm26136 | 0 | NA | NA | NA |
| Gm24658 | 0.115000406 | -0.443821212 | 3.724945049 | -0.119148392 |
| Gm23359 | 0.04393175 | 0.037062677 | 3.724945049 | 0.009949859 |
| Gm24742 | 0.049553047 | -0.443818988 | 3.724945049 | -0.119147795 |
| Snord116l1 | 0.083526043 | 0.037073798 | 3.724945049 | 0.009952844 |
| Gm22173 | 0.200209858 | -0.452059255 | 2.223473251 | -0.203312208 |
| Gm23696 | 0.675101473 | 1.911854806 | 1.377320656 | 1.388097098 |
| Gm26094 | 1.506060395 | 0.517655402 | 0.832416997 | 0.621870293 |
| Gm22289 | 0.897799824 | 0.279946751 | 1.160052139 | 0.241322559 |
| Gm22373 | 29.41355203 | 1.051325024 | 0.308362256 | 3.409382972 |
| Gm38393 | 1891.788183 | 0.823969737 | 0.150897251 | 5.46046885 |
| Snord64 | 0 | NA | NA | NA |
| Snord107 | 0 | NA | NA | NA |
| Snrpn | 3523.734492 | 0.498888898 | 0.055032835 | 9.065295214 |
| Snurf | 0 | NA | NA | NA |
| Gm7367 | 30.85968858 | 0.391946067 | 0.227831033 | 1.72033661 |
| A230006K03Rik | 161.3268707 | 0.604541602 | 0.175245382 | 3.449686345 |
| Ube2nl | 15.82980231 | 0.596678916 | 0.412190053 | 1.447582035 |
| Gm25533 | 0.514202509 | -0.175282515 | 1.504904238 | -0.116474199 |
| Gm23962 | 0.605864455 | -1.859903183 | 1.316273998 | -1.413006096 |
| Mir344d-1 | 0.868248373 | -0.539319413 | 1.102160027 | -0.489329498 |
| Mir344d-2 | 0.131862677 | -0.443820876 | 2.977354665 | -0.149065505 |
| Gm23490 | 9.098233414 | 0.761127764 | 0.367999023 | 2.068287458 |
| Gm22466 | 0.385656404 | 0.579331909 | 1.456009577 | 0.397890178 |
| Mir344d-3 | 1.914600676 | 2.991313984 | 1.010816611 | 2.959304339 |
| Gm23917 | 0.903269115 | 0.310590346 | 1.208358869 | 0.257034854 |
| Mir344e | 0.296439348 | -0.443819818 | 1.623318933 | -0.273402724 |
| Mir344h-1 | 0.876086486 | 0.31729711 | 1.218679906 | 0.260361321 |
| Mir344h-2 | 2.928954034 | 0.272409434 | 0.69115464 | 0.394136736 |
| Gm23873 | 4.451125399 | 0.448676505 | 0.525405586 | 0.85396219 |
| Mir344b | 5.840177946 | 1.770436419 | 0.571611735 | 3.097270945 |
| Mir344c | 9.878672328 | 0.424168541 | 0.36855522 | 1.15089549 |
| Mir344 | 15.06924853 | 0.403934227 | 0.360119164 | 1.121668235 |
| Mir344-2 | 22.73106971 | 0.45467479 | 0.284587003 | 1.597665337 |
| Mir344g | 49.18631136 | 0.522290719 | 0.178551477 | 2.925154854 |
| Gm23431 | 7.186751093 | 1.082043193 | 0.450660558 | 2.401015964 |
| Mir344f | 6.212148508 | 0.650673246 | 0.429401457 | 1.515302835 |
| Gm24923 | 13.62120263 | 0.317123609 | 0.364540853 | 0.869926119 |
| Mir344i | 13.24250092 | 0.301343857 | 0.450436668 | 0.669003832 |

| | | | | |
|---|---|---|---|---|
| Gm9801 | 35.25565351 | 0.91910632 | 0.303233591 | 3.031017494 |
| Ndn | 2072.502296 | 0.16048374 | 0.104932363 | 1.529401755 |
| Magel2 | 65.47304857 | -0.018558829 | 0.191592476 | -0.096866167 |
| Mkrn3 | 19.82500287 | -0.267806215 | 0.266038369 | -1.00664508 |
| Peg12 | 5.656883354 | -0.150090342 | 0.447940217 | -0.335067797 |
| Atp5l-ps1 | 1.396333172 | -0.091252501 | 0.887122845 | -0.102863432 |
| Gm22494 | 0 | NA | NA | NA |
| Chrna7 | 175.1769349 | 0.250449099 | 0.087875873 | 2.850032564 |
| 4930554H23Rik | 0.967147681 | -3.319217555 | 1.132903376 | -2.929832874 |
| Otud7a | 560.9803126 | 0.235544555 | 0.12796268 | 1.840728518 |
| Gm5898 | 84.2128684 | -0.069751419 | 0.132366756 | -0.526955717 |
| Gm22228 | 3.911342548 | 0.616386517 | 0.530400226 | 1.162115864 |
| Gm3307 | 7.681223622 | 1.26606594 | 0.504405405 | 2.510016603 |
| Klf13 | 2862.082524 | 0.211527305 | 0.074974588 | 2.821319992 |
| Gm27252 | 12.51525409 | 0.262872827 | 0.288499108 | 0.911173795 |
| E030018B13Rik | 0.724732643 | -1.786279065 | 1.250887837 | -1.428008981 |
| Gm20670 | 0.072025462 | 0.517959912 | 3.724945049 | 0.139051692 |
| Trpm1 | 3.26130212 | -1.140242598 | 0.581828986 | -1.959755573 |
| Mir211 | 0 | NA | NA | NA |
| Mir6416 | 0 | NA | NA | NA |
| Mtmr10 | 533.9198899 | -0.176231696 | 0.092409327 | -1.907076925 |
| Gm20457 | 18.61572913 | 0.438508563 | 0.258885669 | 1.693830965 |
| Fan1 | 158.3085833 | 0.285117821 | 0.11162326 | 2.554286803 |
| Mphosph10 | 195.4413562 | -0.264878482 | 0.105848727 | -2.502424824 |
| Mcee | 82.01444615 | 0.169228626 | 0.153637074 | 1.101482999 |
| Apba2 | 1821.774854 | 0.224509895 | 0.084087154 | 2.669966645 |
| Fam189a1 | 495.1526885 | 0.661278195 | 0.113505891 | 5.825937223 |
| Ndnl2 | 270.3594914 | 0.367827745 | 0.123438201 | 2.979853408 |
| Gm25249 | 1.017734822 | 1.707104112 | 1.270260473 | 1.343900836 |
| Tjp1 | 2187.437935 | -0.228910544 | 0.05015365 | -4.564185191 |
| Gm7546 | 0 | NA | NA | NA |
| BC046251 | 8.000213675 | 0.399454081 | 0.413809189 | 0.965309838 |
| Tarsl2 | 377.8388073 | 0.361190271 | 0.076770924 | 4.704779494 |
| Tm2d3 | 199.3078344 | 0.039533409 | 0.092376072 | 0.427961578 |
| Gm7551 | 1.00142042 | -1.539235516 | 1.114654751 | -1.380907869 |
| 1810008I18Rik | 0.117907917 | 0.967119648 | 3.722483401 | 0.259804959 |
| Gm20684 | 1.680189541 | 1.934837269 | 0.944723757 | 2.048045531 |
| Pcsk6 | 300.8982086 | 0.044088448 | 0.137419168 | 0.320831868 |
| Snrpa1 | 232.9850694 | -0.134001984 | 0.133391668 | -1.004575365 |
| Vimp | 381.3181123 | -0.19425781 | 0.101334439 | -1.916996954 |
| Gm26827 | 0.677567072 | -1.81092281 | 1.139333225 | -1.58945844 |
| Chsy1 | 358.5432631 | -0.558491409 | 0.094464591 | -5.912177294 |
| Gm23042 | 0 | NA | NA | NA |
| Lrrk1 | 209.5656891 | -0.559908435 | 0.140067426 | -3.997420756 |
| Gm10974 | 0 | NA | NA | NA |
| Gm23957 | 0.352640022 | 0.039286252 | 1.706065945 | 0.023027393 |
| Mir7057 | 0.172107867 | 0.517971699 | 2.739902729 | 0.189047477 |

| | | | | |
|---|---|---|---|---|
| Aldh1a3 | 20.34501332 | -0.8442824 | 0.331008915 | -2.550633418 |
| Asb7 | 386.1973764 | 0.157195534 | 0.078170677 | 2.010927115 |
| Lins | 133.4462556 | -0.447349763 | 0.099347439 | -4.502881673 |
| Cers3 | 5.123955725 | 0.08093154 | 0.502345976 | 0.161107172 |
| Adamts17 | 97.65468606 | 0.557414362 | 0.206597638 | 2.69806745 |
| 1700112J16Rik | 8.55332328 | 0.456546983 | 0.424203273 | 1.076245781 |
| Lysmd4 | 213.3922501 | 0.11795988 | 0.09374763 | 1.258270522 |
| Mef2a | 2662.223246 | 0.272745094 | 0.061690072 | 4.421215333 |
| Lrrc28 | 238.448027 | 0.212094941 | 0.100343819 | 2.113682177 |
| Gm23233 | 1.460122852 | -1.688871542 | 1.032937471 | -1.635018179 |
| Ttc23 | 41.9836744 | -0.318168493 | 0.199755184 | -1.592792171 |
| Synm | 824.232063 | 0.398765094 | 0.126542853 | 3.15122572 |
| Gm23844 | 0 | NA | NA | NA |
| Gm7583 | 0.042716945 | 0.037062677 | 3.724945049 | 0.009949859 |
| 4833412C05Rik | 2.535954356 | 0.687555945 | 0.731553475 | 0.939857397 |
| Igf1r | 783.90039 | 0.254134264 | 0.078967 | 3.218233746 |
| Pgpep1l | 5.087731363 | -0.019381475 | 0.451508499 | -0.042926048 |
| Gm16157 | 0.108309328 | 0.967119648 | 3.722483401 | 0.259804959 |
| Fam169b | 17.53390412 | -0.024308753 | 0.259920135 | -0.093523934 |
| Gm16158 | 0 | NA | NA | NA |
| Gm5334 | 0 | NA | NA | NA |
| Arrdc4 | 222.7753849 | -0.595771057 | 0.234814071 | -2.537203387 |
| Gm23795 | 0 | NA | NA | NA |
| Gm5342 | 0 | NA | NA | NA |
| Gm23815 | 0.0885994 | 0.517955464 | 3.724945049 | 0.139050498 |
| Gm7627 | 0 | NA | NA | NA |
| Gm24120 | 0 | NA | NA | NA |
| Gm29327 | 0 | NA | NA | NA |
| Nr2f2 | 456.5175412 | -0.075493394 | 0.114008151 | -0.662175407 |
| B130024G19Rik | 28.43045277 | 0.497883132 | 0.298318674 | 1.668964014 |
| Gm28258 | 0.072025462 | 0.517959912 | 3.724945049 | 0.139051692 |
| Gm7656 | 3.484008338 | 1.018536311 | 0.661548203 | 1.53962524 |
| Gm24880 | 0.045882455 | 0.517959912 | 3.724945049 | 0.139051692 |
| 6030442E23Rik | 0.042716945 | 0.037062677 | 3.724945049 | 0.009949859 |
| Gm10172 | 0 | NA | NA | NA |
| Gm10295 | 0.372348714 | 0.655938675 | 3.303542283 | 0.198556161 |
| Gm29295 | 0.243433062 | 0.492154431 | 1.848042707 | 0.266311178 |
| Gm29328 | 0.046345709 | -0.443818988 | 3.724945049 | -0.119147795 |
| Mctp2 | 119.8191935 | -0.378004329 | 0.149658867 | -2.525773026 |
| Gm37620 | 0 | NA | NA | NA |
| Gm28745 | 0 | NA | NA | NA |
| Rpl17-ps10 | 146.566573 | -0.238276427 | 0.112958302 | -2.109419336 |
| Gm28744 | 0 | NA | NA | NA |
| Kansl2-ps | 68.64857961 | -0.039009414 | 0.140060269 | -0.27851877 |
| Gm7693 | 0 | NA | NA | NA |
| Gm5335 | 0.044544648 | 0.517957688 | 3.724945049 | 0.139051095 |
| Gm28746 | 0.114742407 | 0.517955464 | 3.724945049 | 0.139050498 |

| | | | | |
|---|---|---|---|---|
| A730056A06Rik | 25.12540099 | 0.465414702 | 0.285401752 | 1.630735266 |
| Rgma | 807.0540894 | 0.293016617 | 0.112371577 | 2.607568794 |
| Chd2 | 1727.684492 | 0.174093881 | 0.106072459 | 1.641273162 |
| 1810026B05Rik | 562.1011433 | 0.046296763 | 0.110435585 | 0.419219613 |
| Gm26176 | 0 | NA | NA | NA |
| Gm23020 | 0.068097359 | 0.037064902 | 3.724945049 | 0.009950456 |
| A830073O21Rik | 30.67817576 | 1.115627301 | 0.301825487 | 3.696266055 |
| Fam174b | 232.8015944 | 0.005956863 | 0.126126608 | 0.047229237 |
| Gm10619 | 0 | NA | NA | NA |
| mmu-mir-7234 | 0 | NA | NA | NA |
| St8sia2 | 68.7914267 | 0.006840194 | 0.209291996 | 0.03268254 |
| Gm38143 | 0.044191549 | 0.037062677 | 3.724945049 | 0.009949859 |
| Gm27459 | 0 | NA | NA | NA |
| Slco3a1 | 783.7089768 | -0.350976167 | 0.083069808 | -4.225074995 |
| Gm7580 | 0 | NA | NA | NA |
| Gm22326 | 0 | NA | NA | NA |
| Sv2b | 4509.045788 | 0.512445123 | 0.11482723 | 4.462749161 |
| Gm24858 | 0.075043205 | 0.517957688 | 3.724945049 | 0.139051095 |
| 1500012K07Rik | 0.790435591 | 1.356596588 | 1.207848411 | 1.123151363 |
| Gm10161 | 0 | NA | NA | NA |
| Akap13 | 1197.678966 | -0.313673256 | 0.081510267 | -3.848266815 |
| Gm25282 | 0.642535093 | 1.035944219 | 1.154956495 | 0.896955187 |
| Gm23251 | 0.141684195 | 0.037074514 | 3.558854665 | 0.010417541 |
| AU020206 | 84.50291907 | -0.750351771 | 0.202006604 | -3.714491287 |
| Gm26729 | 0 | NA | NA | NA |
| Klhl25 | 139.2928838 | -0.208182674 | 0.149814599 | -1.389602056 |
| Gm5597 | 1.352145875 | 0.02525768 | 0.865383121 | 0.029186703 |
| Agbl1 | 4.71615247 | 0.845673003 | 0.524290279 | 1.612986235 |
| Gm23239 | 0.182574094 | 0.401640784 | 2.702311993 | 0.148628576 |
| AC134536.1 | 0 | NA | NA | NA |
| Ntrk3 | 1530.084009 | 0.320128207 | 0.112827183 | 2.837332261 |
| Gm9885 | 28.00701669 | -0.035251513 | 0.244041505 | -0.144448842 |
| Mrpl46 | 163.3248249 | -0.118425828 | 0.122813215 | -0.964275937 |
| Mrps11 | 173.3279118 | -0.294414856 | 0.097892415 | -3.007534901 |
| Det1 | 52.30670483 | -0.657090997 | 0.185350965 | -3.545117758 |
| Mir7-2 | 0.175313191 | -0.443822753 | 2.775896189 | -0.159884492 |
| Aen | 320.6797765 | -0.500908369 | 0.122126944 | -4.101538555 |
| Isg20 | 337.9177596 | -0.960657851 | 0.378250324 | -2.539740986 |
| Gm26633 | 15.61936475 | -0.151966042 | 0.285044313 | -0.533131288 |
| Acan | 162.5228664 | -1.804829936 | 0.227811085 | -7.922485155 |
| Hapln3 | 8.674975699 | 0.628077106 | 0.357783296 | 1.755467942 |
| Mfge8 | 724.7290691 | 0.5945173 | 0.197251425 | 3.014007627 |
| Abhd2 | 3069.461115 | -0.096586905 | 0.072697091 | -1.328621316 |
| Gm24224 | 0 | NA | NA | NA |
| Rlbp1 | 85.04677146 | -0.09198783 | 0.17239761 | -0.5335795 |
| Fanci | 15.02295619 | -0.800659878 | 0.32505404 | -2.463159293 |
| Gm16017 | 0 | NA | NA | NA |

| | | | | |
|---|---|---|---|---|
| Polg | 722.9823603 | -0.082678752 | 0.097253156 | -0.85013953 |
| AI854517 | 340.1156097 | 0.608613872 | 0.131562334 | 4.626049515 |
| Mir9-3 | 6.978027697 | 0.553760279 | 0.539845123 | 1.0257762 |
| A330074H02Rik | 52.62118314 | 0.096174084 | 0.315038037 | 0.30527769 |
| 2900037B21Rik | 15.38072168 | 0.262924393 | 0.410915002 | 0.639851045 |
| Gm37608 | 67.71388935 | 0.285001284 | 0.332438092 | 0.85730634 |
| Rhcg | 17.15633722 | -2.446425008 | 0.41456989 | -5.901115993 |
| Ticrr | 11.11653513 | 0.151453915 | 0.350844413 | 0.431683987 |
| Kif7 | 33.31114625 | -0.179559869 | 0.18315848 | -0.980352472 |
| 9330171B17Rik | 15.94460044 | 0.628804448 | 0.315432786 | 1.993465731 |
| Plin1 | 0.616538614 | -1.254895181 | 1.309095414 | -0.958597186 |
| Pex11a | 89.1075553 | -0.256328689 | 0.142173006 | -1.802934997 |
| Wdr93 | 5.435520937 | -0.976495307 | 0.516128973 | -1.891959876 |
| Mesp1 | 0.291110193 | -0.431897156 | 1.638514627 | -0.263590663 |
| Mesp2 | 2.119758866 | 1.139410248 | 0.754329012 | 1.510495061 |
| Anpep | 165.5971786 | -1.361714899 | 0.208640073 | -6.526622044 |
| Ap3s2 | 947.0018652 | 0.242033367 | 0.059160851 | 4.091106957 |
| Gm26646 | 0.14985154 | -0.867648036 | 2.846051027 | -0.304860323 |
| 5430400D12Rik | 13.24663712 | 0.677056715 | 0.337065564 | 2.008679582 |
| 2610034B18Rik | 234.4523726 | -0.280916755 | 0.113645861 | -2.471860862 |
| Gm24541 | 0 | NA | NA | NA |
| Zfp710 | 414.4888839 | -0.238996695 | 0.092327144 | -2.588585361 |
| Idh2 | 278.3167953 | 0.531538136 | 0.138419507 | 3.84005222 |
| Sema4b | 443.5342128 | -0.396814817 | 0.112503076 | -3.527146381 |
| Cib1 | 109.0301844 | -0.549345694 | 0.152374855 | -3.605225368 |
| Gdpgp1 | 63.61861299 | -0.045265359 | 0.174584615 | -0.259274617 |
| Gm15504 | 0 | NA | NA | NA |
| Ttll13 | 6.876810528 | 1.303014824 | 0.501975954 | 2.595771399 |
| Ngrn | 276.3196493 | -0.272928531 | 0.09981287 | -2.734402205 |
| Vps33b | 168.9997345 | -0.194590573 | 0.114233864 | -1.703440342 |
| Prc1 | 39.35090745 | -1.092679697 | 0.422184242 | -2.588158413 |
| 6330403N20Rik | 1.079531651 | 0.020656545 | 0.967492704 | 0.021350595 |
| Rccd1 | 59.77890577 | 0.19827045 | 0.162881223 | 1.217270145 |
| Unc45a | 445.147613 | -0.176965471 | 0.112200021 | -1.577232059 |
| Hddc3 | 174.8477751 | 0.090506166 | 0.150834295 | 0.600037056 |
| Man2a2 | 2375.928341 | 0.440389568 | 0.072885751 | 6.042190125 |
| Fes | 119.6055901 | -0.873066422 | 0.170382929 | -5.124142582 |
| Furin | 1024.487266 | -0.722477709 | 0.136740722 | -5.283559274 |
| Gm15544 | 0.049553047 | -0.443818988 | 3.724945049 | -0.119147795 |
| Blm | 98.86775975 | 0.463611135 | 0.132415887 | 3.501174552 |
| Gm17257 | 13.53931913 | -0.728397677 | 0.288952732 | -2.520819489 |
| Crtc3 | 380.3991371 | -0.055568876 | 0.111914072 | -0.496531621 |
| Gm15880 | 1.230449842 | -0.530573671 | 0.881129933 | -0.602151455 |
| Iqgap1 | 1218.575044 | -1.398187688 | 0.179623565 | -7.783988093 |
| Zscan2 | 88.28572028 | 0.296419653 | 0.137012647 | 2.163447385 |
| Wdr73 | 209.5163078 | 0.387932092 | 0.098501381 | 3.938341639 |
| Nmb | 36.76065315 | -0.07637214 | 0.319134345 | -0.239310314 |

| | | | | |
|---|---|---|---|---|
| Sec11a | 259.8445855 | -0.263279791 | 0.096405896 | -2.730951146 |
| Gm16171 | 0 | NA | NA | NA |
| Zfp592 | 595.8256381 | -0.047857024 | 0.073281776 | -0.65305492 |
| Alpk3 | 21.69556852 | 1.347441617 | 0.488523826 | 2.758190176 |
| Platr32 | 1.295334173 | 2.291771566 | 1.020871335 | 2.244917149 |
| Slc28a1 | 1.668031982 | -2.267061795 | 0.993186081 | -2.28261535 |
| Gm6415 | 6.092582732 | -1.193835943 | 0.459489656 | -2.598178061 |
| Pde8a | 237.4167335 | -0.111051495 | 0.125207895 | -0.886936841 |
| Gm7180 | 6.599784284 | -0.12123977 | 0.45746527 | -0.265025081 |
| Rps17 | 912.2411316 | -0.169028073 | 0.11478777 | -1.472526847 |
| Cpeb1 | 643.0184324 | 0.432881867 | 0.118482097 | 3.653563516 |
| Rpl7a-ps9 | 0 | NA | NA | NA |
| Ap3b2 | 1542.537639 | 0.0939135 | 0.056654586 | 1.657650462 |
| BC048679 | 0.097899064 | 0.037062677 | 3.724945049 | 0.009949859 |
| 2900076A07Rik | 53.12144243 | 0.220072876 | 0.188080407 | 1.170099955 |
| Mir1839 | 0 | NA | NA | NA |
| mmu-mir-1839 | 0 | NA | NA | NA |
| Gm37829 | 0.980535863 | 0.631057885 | 1.202281223 | 0.524883756 |
| Fsd2 | 3.637652488 | 0.262214336 | 0.620238762 | 0.422763541 |
| Whamm | 307.5779694 | 0.200760779 | 0.113154539 | 1.774217637 |
| Homer2 | 622.3176929 | 0.596716225 | 0.162846728 | 3.664281339 |
| Gm26149 | 0 | NA | NA | NA |
| Fam103a1 | 318.4598396 | 0.221624149 | 0.084582133 | 2.620224172 |
| 3110040N11Rik | 45.08078921 | -0.035129811 | 0.17801524 | -0.197341592 |
| Btbd1 | 1030.704126 | 0.058696014 | 0.102424611 | 0.573065526 |
| Gm10160 | 0.442186748 | -1.030012218 | 1.581109336 | -0.651449077 |
| Tm6sf1 | 282.4071646 | 0.009235423 | 0.098388723 | 0.093866681 |
| Hdgfrp3 | 1850.998309 | 0.287223048 | 0.065316704 | 4.397390445 |
| Gm25907 | 0 | NA | NA | NA |
| Bnc1 | 0.212279271 | 0.069141073 | 2.578148187 | 0.026818114 |
| 4833418N17Rik | 0 | NA | NA | NA |
| Sh3gl3 | 330.4796501 | 0.382790418 | 0.081438019 | 4.700389594 |
| Adamtsl3 | 118.3270371 | -0.398880951 | 0.130608435 | -3.054021373 |
| Saxo2 | 36.73756128 | -0.336083965 | 0.237173249 | -1.417039936 |
| Eftud1 | 227.2937603 | -0.174015786 | 0.081812713 | -2.127001784 |
| 4933406J10Rik | 0.25679625 | 1.077811015 | 2.073194133 | 0.519879445 |
| n-R5s154 | 0 | NA | NA | NA |
| 1700010L04Rik | 0.351906323 | 0.186721751 | 1.508881506 | 0.123748452 |
| Mex3b | 220.6262723 | 0.315264621 | 0.134128098 | 2.350474103 |
| Gm22934 | 0 | NA | NA | NA |
| A530021J07Rik | 0.162691105 | 0.517965151 | 2.788998835 | 0.185717235 |
| Gm10610 | 1.084114685 | 0.914031751 | 0.994265677 | 0.919303333 |
| Tmc3 | 14.78000229 | -0.205936346 | 0.300336213 | -0.685686029 |
| Gm16638 | 33.35336936 | 0.993453078 | 0.26201803 | 3.791544727 |
| Stard5 | 261.421678 | -0.13285522 | 0.132578473 | -1.002087422 |
| Il16 | 509.4594508 | 0.243816366 | 0.19353907 | 1.259778537 |
| 2310044K18Rik | 1.104418013 | 2.386783462 | 1.169266452 | 2.041265665 |

| | | | | |
|---|---|---|---|---|
| Gm7964 | 331.670112 | -0.235330271 | 0.093842842 | -2.507706132 |
| 1700026D08Rik | 9.589069244 | -0.673080371 | 0.406450065 | -1.655997695 |
| Mesdc1 | 387.3285222 | -0.130927568 | 0.098834045 | -1.324721339 |
| Mesdc2 | 527.7692794 | -0.326323801 | 0.097714789 | -3.339553865 |
| Cemip | 295.0602538 | 0.062371766 | 0.082586104 | 0.75523318 |
| Abhd17c | 249.4441419 | -0.07052332 | 0.084124503 | -0.838320795 |
| Gm26708 | 1.414184224 | 1.13379709 | 1.027422735 | 1.103535138 |
| Gm22177 | 0 | NA | NA | NA |
| Arnt2 | 3090.055291 | 0.291152225 | 0.066664944 | 4.367396203 |
| Gm25790 | 1.94490296 | 1.097522887 | 0.808331477 | 1.357763391 |
| Gm2115 | 123.9262083 | -0.331465866 | 0.156036669 | -2.124281857 |
| Fah | 224.6003258 | -0.10908726 | 0.141745405 | -0.769599972 |
| Zfand6 | 399.9024983 | -0.205804674 | 0.119138356 | -1.727442627 |
| 2610206C17Rik | 0.927099338 | -0.232577416 | 0.950332405 | -0.244732701 |
| Gm18958 | 0.18866986 | 0.517973639 | 2.261034892 | 0.229086973 |
| Vmn2r-ps62 | 0 | NA | NA | NA |
| Gm6155 | 2.358756262 | -0.928045743 | 0.673651792 | -1.377634194 |
| Olfr291 | 0.046345709 | -0.443818988 | 3.724945049 | -0.119147795 |
| Gm8018 | 0.038981395 | -0.443816764 | 3.724945049 | -0.119147198 |
| Olfr290 | 0.092881282 | 0.037060453 | 3.724945049 | 0.009949262 |
| Gm18560 | 0 | NA | NA | NA |
| Vmn2r65 | 0 | NA | NA | NA |
| Vmn2r66 | 0 | NA | NA | NA |
| Vmn2r-ps64 | 0 | NA | NA | NA |
| Vmn2r-ps65 | 0 | NA | NA | NA |
| Vmn2r-ps66 | 0 | NA | NA | NA |
| Vmn2r-ps67 | 0 | NA | NA | NA |
| Gm18358 | 0 | NA | NA | NA |
| Vmn2r67 | 0 | NA | NA | NA |
| Vmn2r-ps68 | 0 | NA | NA | NA |
| Vmn2r68 | 0 | NA | NA | NA |
| Gm20414 | 0 | NA | NA | NA |
| Gm20415 | 0 | NA | NA | NA |
| Vmn2r-ps69 | 0 | NA | NA | NA |
| Vmn2r69 | 0 | NA | NA | NA |
| Gm18359 | 0 | NA | NA | NA |
| Vmn2r-ps70 | 0 | NA | NA | NA |
| Gm18360 | 0 | NA | NA | NA |
| Vmn2r70 | 0 | NA | NA | NA |
| Vmn2r-ps71 | 0 | NA | NA | NA |
| Vmn2r-ps72 | 0 | NA | NA | NA |
| Vmn2r71 | 0 | NA | NA | NA |
| Vmn2r-ps73 | 0 | NA | NA | NA |
| Vmn2r-ps74 | 0 | NA | NA | NA |
| Vmn2r72 | 0 | NA | NA | NA |
| Gm18779 | 0 | NA | NA | NA |
| Gm18354 | 0 | NA | NA | NA |

| | | | | |
|---|---|---|---|---|
| Vmn2r-ps75 | 0 | NA | NA | NA |
| Vmn2r-ps76 | 0 | NA | NA | NA |
| Vmn2r-ps77 | 0 | NA | NA | NA |
| Vmn2r73 | 0.08486385 | 0.037730769 | 3.724898049 | 0.010129343 |
| Vmn2r-ps78 | 0 | NA | NA | NA |
| Vmn2r74 | 0 | NA | NA | NA |
| Vmn2r-ps79 | 0 | NA | NA | NA |
| Vmn2r-ps80 | 0 | NA | NA | NA |
| Vmn2r-ps81 | 0 | NA | NA | NA |
| Vmn2r-ps82 | 0 | NA | NA | NA |
| Vmn2r-ps83 | 0 | NA | NA | NA |
| Vmn2r-ps84 | 0 | NA | NA | NA |
| Vmn2r75 | 0 | NA | NA | NA |
| Vmn2r-ps85 | 0 | NA | NA | NA |
| Vmn2r-ps86 | 0 | NA | NA | NA |
| Vmn2r76 | 0 | NA | NA | NA |
| Olfr310 | 0 | NA | NA | NA |
| Gm20480 | 0 | NA | NA | NA |
| Gm23192 | 0 | NA | NA | NA |
| Olfr309 | 0.297212607 | -1.003000548 | 1.773462994 | -0.56556046 |
| Olfr308 | 0 | NA | NA | NA |
| Olfr307 | 0 | NA | NA | NA |
| Olfr306-ps1 | 0 | NA | NA | NA |
| Olfr305 | 0 | NA | NA | NA |
| Olfr304 | 0 | NA | NA | NA |
| Olfr303 | 0 | NA | NA | NA |
| Olfr301 | 0 | NA | NA | NA |
| Olfr300-ps1 | 0 | NA | NA | NA |
| Olfr299 | 0 | NA | NA | NA |
| Olfr298 | 0.066050875 | -0.443818988 | 3.724945049 | -0.119147795 |
| Olfr297 | 0 | NA | NA | NA |
| Gm17919 | 0 | NA | NA | NA |
| Olfr296-ps1 | 0 | NA | NA | NA |
| Olfr295 | 0 | NA | NA | NA |
| Olfr294 | 0 | NA | NA | NA |
| Olfr293 | 0 | NA | NA | NA |
| Olfr292 | 0 | NA | NA | NA |
| Folh1 | 228.0677052 | 0.274161539 | 0.176244804 | 1.555572317 |
| Vmn2r77 | 0.046345709 | -0.443818988 | 3.724945049 | -0.119147795 |
| Vmn2r-ps87 | 0.08486385 | 0.037730769 | 3.724898049 | 0.010129343 |
| Gm19203 | 0 | NA | NA | NA |
| Vmn2r78 | 0.360847685 | 1.618667148 | 2.011699605 | 0.804626667 |
| Vmn2r79 | 0.061920613 | -0.443816764 | 3.724945049 | -0.119147198 |
| Nox4 | 23.51386303 | -2.286232893 | 0.324167995 | -7.052617564 |
| Tyr | 0.679694067 | 1.563116261 | 1.208016207 | 1.293953054 |
| Grm5 | 3152.369327 | 0.84667751 | 0.105590212 | 8.018522707 |
| Gm24249 | 0.097139547 | -0.443816764 | 3.724945049 | -0.119147198 |

| | | | | |
|---|---|---|---|---|
| Ctsc | 748.4761849 | -1.078701769 | 0.17479077 | -6.17138863 |
| Gm26522 | 0.075043205 | 0.517957688 | 3.724945049 | 0.139051095 |
| Gm25345 | 0.066050875 | -0.443818988 | 3.724945049 | -0.119147795 |
| Rab38 | 4.431782484 | 0.260081843 | 0.617885462 | 0.420922419 |
| Gm15661 | 0 | NA | NA | NA |
| Rps13-ps2 | 13.83618781 | -0.261470175 | 0.361157695 | -0.723977858 |
| Gm8183 | 0.155358952 | 0.517963421 | 2.829692699 | 0.18304582 |
| Tmem135 | 828.7159243 | 0.139187065 | 0.058303532 | 2.387283576 |
| Gm24659 | 0.092269992 | -0.443823436 | 3.724945049 | -0.119148989 |
| Gm22171 | 0.104637558 | -0.443821212 | 3.724945049 | -0.119148392 |
| Gm28748 | 6.9950682 | 0.314856777 | 0.441386973 | 0.713335002 |
| Fzd4 | 116.5445208 | 0.381679766 | 0.185488481 | 2.057700638 |
| Prss23 | 214.4898237 | -0.90974 | 0.189250398 | -4.807070469 |
| Prss23os | 0.31866137 | -1.377166914 | 1.982966072 | -0.694498475 |
| Gm2546 | 6.356979297 | 0.004289877 | 0.449591962 | 0.009541713 |
| A230065N10Rik | 4.255075643 | 0.555893393 | 0.47964338 | 1.158972304 |
| Me3 | 830.2853921 | 0.413533541 | 0.123123331 | 3.358693588 |
| Gm15744 | 0.161532967 | -0.444773734 | 3.449701211 | -0.128931089 |
| Ccdc81 | 14.73968261 | -1.001638998 | 0.29717885 | -3.370492212 |
| l7Rn6 | 302.6286143 | -0.235234196 | 0.117016556 | -2.010264218 |
| Eed | 245.5434659 | -0.264889355 | 0.089316182 | -2.965748755 |
| Gm26529 | 7.028223412 | 0.101090776 | 0.395280576 | 0.255744355 |
| Gm5341 | 0.068097359 | 0.037064902 | 3.724945049 | 0.009950456 |
| 2310010J17Rik | 13.38140942 | -0.04675717 | 0.28354537 | -0.164901899 |
| Picalm | 2942.135727 | -0.26793627 | 0.064205422 | -4.173109724 |
| Ccdc83 | 1.068251668 | 0.499462273 | 0.860120001 | 0.580689058 |
| n-R5s155 | 0 | NA | NA | NA |
| Sytl2 | 471.4003508 | 0.449498347 | 0.100994624 | 4.45071558 |
| Ccdc89 | 10.74032172 | -0.411344178 | 0.334436241 | -1.229962926 |
| Crebzf | 1021.356941 | -0.151804895 | 0.056474981 | -2.68800257 |
| Tmem126a | 179.1913439 | -0.556517521 | 0.144783784 | -3.843783502 |
| Tmem126b | 260.3332693 | -0.173640854 | 0.12762643 | -1.360539923 |
| Dlg2 | 4221.663311 | 0.467139703 | 0.101805474 | 4.588551926 |
| Gm27470 | 0.068097359 | 0.037064902 | 3.724945049 | 0.009950456 |
| Gm27660 | 0 | NA | NA | NA |
| Gm24552 | 6.758948741 | 0.524639559 | 0.419927289 | 1.249358096 |
| Gm22080 | 0 | NA | NA | NA |
| Gm23928 | 0 | NA | NA | NA |
| Ccdc90b | 140.9402912 | -0.398244142 | 0.150496428 | -2.646203289 |
| Ankrd42 | 228.9912909 | 0.055715622 | 0.127068599 | 0.438468843 |
| Gm37008 | 21.96798092 | -0.373554315 | 0.265592699 | -1.406493163 |
| Gm26944 | 56.56169118 | -0.489261064 | 0.276051563 | -1.772353899 |
| 6430511E19Rik | 167.4764364 | -0.624172417 | 0.230919166 | -2.702990957 |
| Pcf11 | 1375.530546 | 0.137994791 | 0.079732812 | 1.730715216 |
| Gm26981 | 0 | NA | NA | NA |
| 4632427E13Rik | 515.8795252 | -0.221614151 | 0.180257895 | -1.229428265 |
| Rab30 | 322.4076359 | 0.176148012 | 0.077815178 | 2.263671637 |

| | | | | |
|---|---|---|---|---|
| Ddias | 20.7592005 | -0.036697978 | 0.319932615 | -0.114705335 |
| Prcp | 124.058315 | -0.371810815 | 0.123031536 | -3.022077333 |
| Gm26862 | 2.576837399 | -0.309878063 | 0.638023338 | -0.485684527 |
| Gm9934 | 11.67004998 | -0.428343672 | 0.325627594 | -1.315440338 |
| Fam181b | 200.5250975 | -0.363922017 | 0.171389005 | -2.123368516 |
| Gm25860 | 0 | NA | NA | NA |
| Gm15501 | 19.75181214 | -0.681948204 | 0.275345431 | -2.476700632 |
| Gm5899 | 1.074124899 | 0.328268682 | 1.002428614 | 0.327473376 |
| Gm23884 | 0.681297893 | 1.35173158 | 1.197985715 | 1.128336977 |
| Gm25684 | 0 | NA | NA | NA |
| Gm9966 | 3.042324184 | 1.4346697 | 0.71099187 | 2.017842623 |
| Tenm4 | 3151.27742 | 0.155323595 | 0.125978145 | 1.232940802 |
| Mir708 | 5.493531071 | 1.304532651 | 0.589845135 | 2.211652813 |
| Mir6394 | 3.129376264 | -0.519742496 | 0.776701797 | -0.669166079 |
| Gm15412 | 0 | NA | NA | NA |
| Rps11-ps5 | 0 | NA | NA | NA |
| Gm15414 | 0.304874617 | -1.286812033 | 3.703501268 | -0.34745824 |
| Gm15413 | 2.589705772 | -0.949683741 | 0.747146773 | -1.271080564 |
| Gm15416 | 0.568378176 | -0.973070936 | 1.204937778 | -0.807569447 |
| Gm25712 | 0 | NA | NA | NA |
| Gm15415 | 0.44530367 | -0.457576747 | 1.481151325 | -0.308933151 |
| Gm22226 | 0 | NA | NA | NA |
| C230038L03Rik | 4.597572509 | 0.571083665 | 0.572401195 | 0.99769824 |
| Nars2 | 182.0977735 | 0.063582555 | 0.092941142 | 0.684116351 |
| Gab2 | 640.0412359 | -0.089943275 | 0.121865425 | -0.738054092 |
| Usp35 | 246.8318203 | 0.224599364 | 0.089992476 | 2.495757144 |
| Kctd21 | 137.2874877 | -0.850356127 | 0.229208234 | -3.709971982 |
| Alg8 | 69.82562007 | -0.688503039 | 0.159042697 | -4.329045296 |
| Ndufc2 | 551.0014102 | -0.05781733 | 0.139449802 | -0.414610338 |
| Gm16053 | 0.824460844 | -0.899588412 | 1.281140921 | -0.702177564 |
| Thrsp | 209.116264 | -0.211958006 | 0.136977675 | -1.54739089 |
| Kctd14 | 10.62226215 | 0.097880632 | 0.365732031 | 0.267629366 |
| Ints4 | 375.7085532 | -0.430317064 | 0.098162463 | -4.383723177 |
| Gm24412 | 1.785420374 | 1.072753283 | 1.041315365 | 1.030190583 |
| Aamdc | 120.3980682 | 0.000148454 | 0.106562697 | 0.001393116 |
| Rsf1 | 1617.979498 | 0.019585794 | 0.058504371 | 0.334774892 |
| Rsf1os1 | 0 | NA | NA | NA |
| Rsf1os2 | 1.842452978 | -0.826498539 | 0.845131102 | -0.977953049 |
| Clns1a | 571.5874786 | -0.001554718 | 0.072749639 | -0.021370796 |
| Gm9990 | 8.809819601 | -0.155116216 | 0.490012554 | -0.316555596 |
| Aqp11 | 58.08426219 | 0.787499985 | 0.183982787 | 4.280291637 |
| Pak1 | 1544.109605 | 0.523698325 | 0.076509909 | 6.844843103 |
| Gdpd4 | 2.28172379 | -1.258690304 | 1.038924711 | -1.211531779 |
| Myo7a | 84.81893064 | 0.032815621 | 0.119880146 | 0.273736909 |
| Capn5 | 668.0892524 | 0.292689739 | 0.098593749 | 2.968643965 |
| Omp | 79.7450718 | -0.807152467 | 0.739960247 | -1.090805176 |
| Gm16938 | 29.29590882 | 0.189410607 | 0.191368194 | 0.989770572 |

| | | | | |
|---|---|---|---|---|
| B3gnt6 | 6.185203872 | 0.185213075 | 0.422268816 | 0.438614142 |
| Acer3 | 413.2037177 | -0.067200291 | 0.094343128 | -0.712296621 |
| Tsku | 55.8336496 | -0.035548954 | 0.221042886 | -0.160823788 |
| Gucy2d | 0.893137065 | 0.818635374 | 0.944451829 | 0.866783619 |
| Lrrc32 | 74.30774312 | -0.55889456 | 0.261354242 | -2.138456045 |
| Gm23040 | 0.075043205 | 0.517957688 | 3.724945049 | 0.139051095 |
| A630091E08Rik | 0 | NA | NA | NA |
| 2210018M11Rik | 690.3005469 | -0.035264933 | 0.10507644 | -0.335612176 |
| Gm15506 | 8.670677481 | 0.493057944 | 0.406945485 | 1.211606868 |
| Prkrir | 815.4013897 | 0.107238706 | 0.073497677 | 1.459076114 |
| Wnt11 | 8.862266067 | 0.776532101 | 0.35970891 | 2.158779168 |
| Gm8398 | 0.117760149 | 0.51795324 | 3.724945049 | 0.139049901 |
| Uvrag | 562.8884481 | -0.10864008 | 0.087819667 | -1.237081436 |
| Gm23479 | 0 | NA | NA | NA |
| Dgat2 | 363.7510314 | 0.549190904 | 0.113440095 | 4.841241565 |
| Mogat2 | 0.144988549 | -0.444784414 | 3.575474176 | -0.124398721 |
| Gm26705 | 2.196663442 | -0.822217766 | 0.697193085 | -1.179325762 |
| Map6 | 1290.520615 | 0.054380604 | 0.075231937 | 0.722839338 |
| Serpinh1 | 377.8077043 | -1.536643356 | 0.205341962 | -7.483338246 |
| Gdpd5 | 476.3039452 | -0.221268146 | 0.121843155 | -1.816008014 |
| Klhl35 | 5.303271472 | 0.789146137 | 0.456714685 | 1.727875548 |
| Rps3 | 2214.045192 | -0.318761647 | 0.11018108 | -2.893070644 |
| Snord15b | 0 | NA | NA | NA |
| Snord15a | 45.47276232 | -0.713947181 | 0.208447321 | -3.425072478 |
| n-R5s156 | 0 | NA | NA | NA |
| Arrb1 | 3455.425929 | -0.004406522 | 0.073632699 | -0.059844636 |
| Mir326 | 2.043157693 | 0.764481697 | 0.746848783 | 1.023609752 |
| Gm4980 | 50.35484282 | 0.124890171 | 0.204324894 | 0.611233259 |
| Gm10605 | 21.76352257 | -0.266073006 | 0.251089408 | -1.059674352 |
| Gm25409 | 0 | NA | NA | NA |
| Slco2b1 | 87.4821162 | 0.141046426 | 0.327449407 | 0.430742651 |
| Gm15635 | 0.464401238 | 0.365334329 | 1.519969167 | 0.240356408 |
| Olfr520 | 0.044544648 | 0.517957688 | 3.724945049 | 0.139051095 |
| Olfr521 | 0 | NA | NA | NA |
| F730035P03Rik | 0.645028293 | -0.396280582 | 1.278762628 | -0.309893778 |
| Neu3 | 60.27959619 | -0.529915162 | 0.140337967 | -3.775992878 |
| Spcs2 | 666.5559383 | -0.395076478 | 0.107821044 | -3.664187085 |
| Xrra1 | 7.264420559 | 0.770844546 | 0.416436467 | 1.851049576 |
| Rnf169 | 651.1755701 | 0.41477451 | 0.080253616 | 5.168296873 |
| Chrdl2 | 5.288126676 | 0.421750673 | 0.521679715 | 0.808447523 |
| Pold3 | 335.8617301 | -0.308114658 | 0.072631629 | -4.242155428 |
| Lipt2 | 20.2504011 | 0.295188042 | 0.237961083 | 1.240488731 |
| Kcne3 | 1.196981447 | -0.16085331 | 0.854723114 | -0.188193472 |
| Pgm2l1 | 3618.346017 | 0.673763212 | 0.110780168 | 6.081984005 |
| Gpx2-ps1 | 5.150345274 | 0.973512314 | 0.601950236 | 1.61726378 |
| P4ha3 | 123.5044264 | -1.021118969 | 0.185408576 | -5.507398785 |
| Ppme1 | 1499.055743 | -0.02847057 | 0.059261537 | -0.480422409 |

| | | | | |
|---|---|---|---|---|
| C2cd3 | 640.0813776 | -0.289192326 | 0.138373567 | -2.089939089 |
| Gm10603 | 1.405100988 | 0.270623429 | 0.93675352 | 0.288895023 |
| Gm37716 | 3.584490054 | 1.069305707 | 0.692891287 | 1.543251774 |
| Ucp3 | 15.93899765 | 1.384657911 | 0.364263773 | 3.801250675 |
| Ucp2 | 1246.309197 | -0.569432054 | 0.157079806 | -3.62511304 |
| Dnajb13 | 7.766840755 | -0.843843287 | 0.396940034 | -2.12587095 |
| Coa4 | 46.56779514 | -0.837440724 | 0.211050778 | -3.967958476 |
| Mrpl48 | 372.182467 | 0.148991727 | 0.07466302 | 1.995522376 |
| Cox20-ps | 0.105143817 | 0.517955464 | 3.724945049 | 0.139050498 |
| Rab6a | 4454.768753 | 0.19689705 | 0.051963429 | 3.789146578 |
| Gm14382 | 10.15270512 | 0.027731369 | 0.461941618 | 0.060032194 |
| Plekhb1 | 2340.080905 | -0.040402941 | 0.243590184 | -0.165864407 |
| Fam168a | 4198.767291 | 0.205030671 | 0.046842229 | 4.377047756 |
| Gm3200 | 5.569043595 | 0.248448085 | 0.452131872 | 0.549503585 |
| Relt | 198.9215136 | 0.045391164 | 0.122028888 | 0.371970646 |
| Arhgef17 | 2322.379047 | 0.344174906 | 0.063604027 | 5.41121248 |
| Mir3102 | 0 | NA | NA | NA |
| P2ry6 | 83.22755977 | -0.54232573 | 0.215337371 | -2.518493322 |
| P2ry2 | 41.59914068 | -1.039197079 | 0.337832641 | -3.076070673 |
| Fchsd2 | 883.8523491 | -0.057503291 | 0.065425128 | -0.878917516 |
| Gm6341 | 2.08700676 | 0.943161171 | 0.770281863 | 1.224436425 |
| Gm15673 | 11.22830619 | 0.697360887 | 0.384691627 | 1.812778957 |
| Atg16l2 | 153.6349008 | 0.05075259 | 0.162479697 | 0.312362659 |
| Stard10 | 401.7502125 | -0.013822389 | 0.092396377 | -0.14959882 |
| Gm20476 | 0.089814205 | 0.517955464 | 3.724945049 | 0.139050498 |
| Arap1 | 603.6362813 | -0.744128116 | 0.096006958 | -7.750772772 |
| Pde2a | 2506.890893 | 0.51605783 | 0.124491952 | 4.145310768 |
| Mir139 | 0.438507067 | 1.927405731 | 1.516536397 | 1.270926128 |
| Art2a-ps | 2.02887955 | -0.307435882 | 0.775123272 | -0.396628373 |
| Gm7027 | 32.67737545 | -0.258029229 | 0.236470154 | -1.091170387 |
| Art2b | 0.341803366 | -1.34010461 | 3.695074291 | -0.362673252 |
| Clpb | 506.8075599 | -0.019959089 | 0.073723102 | -0.270730463 |
| Phox2a | 2.085398638 | -0.318805484 | 0.786192285 | -0.405505739 |
| Inppl1 | 617.7038875 | -0.744974519 | 0.112586818 | -6.616889357 |
| Gm10602 | 1.695248615 | 0.362723227 | 0.803711464 | 0.451310256 |
| Folr2 | 8.630795641 | -1.74179698 | 0.419950308 | -4.147626395 |
| Folr1 | 27.39965009 | 0.970278234 | 0.268873342 | 3.608681415 |
| Gm22655 | 0.092269992 | -0.443823436 | 3.724945049 | -0.119148989 |
| Anapc15 | 157.008238 | 0.145989958 | 0.094237484 | 1.549170793 |
| Tomt | 1.048945322 | -1.633315864 | 1.058709516 | -1.542742215 |
| Lamtor1 | 591.6362557 | -0.289017417 | 0.099030023 | -2.918482795 |
| Lrrc51 | 20.02074397 | -0.982254017 | 0.259431123 | -3.786184191 |
| Numa1 | 1753.65198 | -0.211721227 | 0.083446275 | -2.537216049 |
| Il18bp | 580.6338207 | -0.932091612 | 0.187924643 | -4.959922206 |
| Rnf121 | 426.15743 | -0.362424564 | 0.094362091 | -3.840785634 |
| Xndc1 | 64.93653681 | 0.220746403 | 0.214503165 | 1.029105575 |
| Xntrpc | 2.817151494 | -0.049593764 | 0.656024774 | -0.075597395 |

| | | | | |
|---|---|---|---|---|
| Trpc2 | 0 | NA | NA | NA |
| Art5 | 0.775564919 | -1.802755856 | 1.131872209 | -1.592720311 |
| Art1 | 0.377906432 | -0.909307755 | 1.609146667 | -0.565086933 |
| Chrna10 | 6.942902888 | 0.86849844 | 0.429271006 | 2.023193802 |
| Nup98 | 911.2468387 | -0.218605946 | 0.094661807 | -2.30933629 |
| Pgap2 | 160.2875351 | -0.530970385 | 0.148928128 | -3.565279389 |
| Rhog | 475.1594366 | -0.453441194 | 0.147920578 | -3.06543687 |
| Stim1 | 903.1380357 | 0.441970547 | 0.065099024 | 6.789203944 |
| Rrm1 | 306.720304 | -0.233180414 | 0.123219796 | -1.892394093 |
| Olfr543 | 0 | NA | NA | NA |
| Olfr544 | 0 | NA | NA | NA |
| Olfr545 | 0 | NA | NA | NA |
| Olfr547 | 0 | NA | NA | NA |
| Olfr548-ps1 | 0 | NA | NA | NA |
| Olfr549 | 0 | NA | NA | NA |
| Trim21 | 602.064124 | -0.561568252 | 0.14362057 | -3.910082326 |
| Olfr550 | 0.446685102 | 0.930178401 | 1.603416554 | 0.580122738 |
| Olfr551 | 0 | NA | NA | NA |
| Olfr552 | 0.044544648 | 0.517957688 | 3.724945049 | 0.139051095 |
| Olfr553 | 0 | NA | NA | NA |
| Olfr554 | 0.152980968 | -0.89129297 | 2.833643542 | -0.314539552 |
| Olfr555 | 0 | NA | NA | NA |
| Olfr556 | 0.166709681 | 0.517964155 | 2.76763122 | 0.187150713 |
| Trim68 | 38.7507144 | -0.143425224 | 0.184540097 | -0.77720358 |
| Olfr557 | 0.253342923 | 1.270314211 | 2.063655705 | 0.615564994 |
| Olfr558 | 0.693780966 | 0.413394201 | 1.198014774 | 0.345066029 |
| Olfr33 | 0 | NA | NA | NA |
| Olfr559 | 0 | NA | NA | NA |
| Gm24210 | 0 | NA | NA | NA |
| Olfr78 | 4.264554404 | 2.432140347 | 0.909061231 | 2.675441723 |
| Olfr561 | 0 | NA | NA | NA |
| Olfr564 | 0 | NA | NA | NA |
| Olfr566 | 0.049553047 | -0.443818988 | 3.724945049 | -0.119147795 |
| Olfr568 | 0.044544648 | 0.517957688 | 3.724945049 | 0.139051095 |
| Olfr569 | 0 | NA | NA | NA |
| Olfr570 | 0 | NA | NA | NA |
| Olfr571 | 0 | NA | NA | NA |
| Olfr572 | 0 | NA | NA | NA |
| Olfr573-ps1 | 0 | NA | NA | NA |
| Olfr574 | 0.224720342 | -0.401424306 | 2.133697217 | -0.188135553 |
| Olfr575 | 0 | NA | NA | NA |
| Olfr576 | 0 | NA | NA | NA |
| Olfr577 | 0 | NA | NA | NA |
| Olfr578 | 0 | NA | NA | NA |
| Olfr582 | 0 | NA | NA | NA |
| Olfr583 | 0 | NA | NA | NA |
| Olfr584 | 0.046345709 | -0.443818988 | 3.724945049 | -0.119147795 |

| | | | | |
|---|---|---|---|---|
| Olfr585 | 0.062426873 | 0.517959912 | 3.724945049 | 0.139051692 |
| Olfr586 | 0 | NA | NA | NA |
| Olfr587-ps1 | 0.046345709 | -0.443818988 | 3.724945049 | -0.119147795 |
| Gm15120 | 0.07796279 | -0.798301147 | 3.721829328 | -0.214491605 |
| Olfr588-ps1 | 0 | NA | NA | NA |
| Gm15116 | 0 | NA | NA | NA |
| Olfr589 | 0 | NA | NA | NA |
| Olfr590-ps1 | 0 | NA | NA | NA |
| Olfr591 | 0 | NA | NA | NA |
| Olfr592 | 0 | NA | NA | NA |
| Olfr593 | 0.044191549 | 0.037062677 | 3.724945049 | 0.009949859 |
| Olfr594 | 0 | NA | NA | NA |
| Olfr595-ps1 | 0 | NA | NA | NA |
| Usp17ld | 0 | NA | NA | NA |
| Gm25171 | 0 | NA | NA | NA |
| Gm23550 | 0 | NA | NA | NA |
| Gm23973 | 0 | NA | NA | NA |
| Olfr596 | 0 | NA | NA | NA |
| Olfr597 | 0 | NA | NA | NA |
| Olfr598 | 0 | NA | NA | NA |
| Olfr599 | 0.097139547 | -0.443816764 | 3.724945049 | -0.119147198 |
| Olfr600 | 0 | NA | NA | NA |
| Olfr601 | 0 | NA | NA | NA |
| Olfr602-ps1 | 0 | NA | NA | NA |
| Olfr603 | 0 | NA | NA | NA |
| Usp17lc | 0 | NA | NA | NA |
| Gm25774 | 0.116217011 | 0.517955464 | 3.724945049 | 0.139050498 |
| Olfr605 | 0 | NA | NA | NA |
| Olfr606 | 0 | NA | NA | NA |
| Gm15121 | 0 | NA | NA | NA |
| Olfr607 | 0 | NA | NA | NA |
| Olfr608 | 0 | NA | NA | NA |
| Olfr609 | 0 | NA | NA | NA |
| Olfr610 | 0 | NA | NA | NA |
| Olfr611 | 0 | NA | NA | NA |
| Olfr612 | 0 | NA | NA | NA |
| Olfr613 | 0 | NA | NA | NA |
| Olfr615 | 0 | NA | NA | NA |
| Olfr616 | 0.04393175 | 0.037062677 | 3.724945049 | 0.009949859 |
| Gm26506 | 0.044544648 | 0.517957688 | 3.724945049 | 0.139051095 |
| Olfr617 | 0 | NA | NA | NA |
| Olfr618 | 0 | NA | NA | NA |
| Olfr619 | 0 | NA | NA | NA |
| Olfr620 | 0 | NA | NA | NA |
| Olfr621-ps1 | 0 | NA | NA | NA |
| Olfr622 | 0 | NA | NA | NA |
| Olfr623 | 0 | NA | NA | NA |

| | | | | |
|---|---|---|---|---|
| Olfr624 | 0 | NA | NA | NA |
| Olfr625-ps1 | 0 | NA | NA | NA |
| Olfr626-ps1 | 0 | NA | NA | NA |
| Olfr260-ps1 | 0 | NA | NA | NA |
| Olfr243 | 0 | NA | NA | NA |
| Olfr628 | 0.061920613 | -0.443816764 | 3.724945049 | -0.119147198 |
| Olfr629 | 0 | NA | NA | NA |
| Olfr630 | 0 | NA | NA | NA |
| Olfr69 | 0 | NA | NA | NA |
| Olfr68 | 0 | NA | NA | NA |
| Olfr67 | 0.045882455 | 0.517959912 | 3.724945049 | 0.139051692 |
| Hbb-bt | 10.74344224 | 2.370979122 | 0.616148505 | 3.848064391 |
| Hbb-bs | 223.1020574 | 1.482344422 | 0.690648073 | 2.146309358 |
| Gm15115 | 0 | NA | NA | NA |
| Hbb-bh3 | 0 | NA | NA | NA |
| Hbb-bh2 | 0 | NA | NA | NA |
| Hbb-bh1 | 0 | NA | NA | NA |
| Hbb-bh0 | 0 | NA | NA | NA |
| Hbb-y | 0 | NA | NA | NA |
| Olfr66 | 0 | NA | NA | NA |
| Dnajc19-ps | 11.1957499 | -0.294195322 | 0.333498247 | -0.882149532 |
| Olfr64 | 0 | NA | NA | NA |
| Olfr65 | 0.042716945 | 0.037062677 | 3.724945049 | 0.009949859 |
| Olfr631 | 0 | NA | NA | NA |
| Olfr632 | 0.042716945 | 0.037062677 | 3.724945049 | 0.009949859 |
| Olfr633 | 0 | NA | NA | NA |
| Olfr634-ps1 | 0 | NA | NA | NA |
| Gm4886 | 0 | NA | NA | NA |
| Olfr635 | 0.092691418 | -0.908687078 | 3.724560878 | -0.2439716 |
| Olfr636-ps1 | 0.038981395 | -0.443816764 | 3.724945049 | -0.119147198 |
| Olfr637-ps1 | 0 | NA | NA | NA |
| 4930516K23Rik | 0 | NA | NA | NA |
| Olfr638 | 0 | NA | NA | NA |
| Olfr639 | 0 | NA | NA | NA |
| Olfr640 | 0 | NA | NA | NA |
| Olfr641 | 0 | NA | NA | NA |
| Olfr642 | 0 | NA | NA | NA |
| Olfr643 | 0.062426873 | 0.517959912 | 3.724945049 | 0.139051692 |
| Olfr644 | 0 | NA | NA | NA |
| Olfr645 | 0 | NA | NA | NA |
| Olfr646 | 0 | NA | NA | NA |
| 4931431F19Rik | 0.042716945 | 0.037062677 | 3.724945049 | 0.009949859 |
| Ubqln3 | 0 | NA | NA | NA |
| Ubqlnl | 0 | NA | NA | NA |
| E030002O03Rik | 0.160237607 | 0.037068507 | 2.80611751 | 0.013209891 |
| Olfr647-ps1 | 0 | NA | NA | NA |
| Olfr648 | 0 | NA | NA | NA |

| | | | | |
|---|---|---|---|---|
| Olfr649 | 0.060174537 | 0.517957688 | 3.724945049 | 0.139051095 |
| Olfr650-ps1 | 0.524522254 | -1.66828864 | 1.499420804 | -1.112622044 |
| Trim6 | 12.76726646 | -0.5491868 | 0.282309553 | -1.945335517 |
| Trim34a | 130.1339841 | -0.666690372 | 0.13428465 | -4.964754896 |
| Gm15135 | 0 | NA | NA | NA |
| Trim5 | 3.073902706 | -1.83570092 | 0.634663143 | -2.892401963 |
| Gm24846 | 0 | NA | NA | NA |
| Trim12a | 0.470008961 | 0.085812308 | 1.572237976 | 0.05457972 |
| Trim34b | 43.17410611 | -0.460078004 | 0.182118053 | -2.526262471 |
| Gm15133 | 0.044544648 | 0.517957688 | 3.724945049 | 0.139051095 |
| Trim12c | 411.4065939 | -0.926118238 | 0.261450155 | -3.542236335 |
| Gm27995 | 0 | NA | NA | NA |
| Trim30b | 23.44767613 | -1.837433668 | 0.468559113 | -3.921455409 |
| Trim30c | 106.4359115 | -2.184207562 | 0.649465859 | -3.363082959 |
| Trim30a | 1972.4724 | -1.181316331 | 0.250612808 | -4.713710923 |
| Gm25405 | 0.545822006 | 0.497266589 | 1.304130619 | 0.381301215 |
| Trim30d | 381.5932247 | -1.236465148 | 0.322887804 | -3.829395643 |
| Gm16464 | 7.703381671 | 0.022576682 | 0.417416787 | 0.054086664 |
| Gm6574 | 0 | NA | NA | NA |
| Trim30e-ps1 | 0 | NA | NA | NA |
| Olfr651 | 0.060174537 | 0.517957688 | 3.724945049 | 0.139051095 |
| Olfr652 | 0.038981395 | -0.443816764 | 3.724945049 | -0.119147198 |
| Olfr653 | 0 | NA | NA | NA |
| Olfr654 | 0.212522253 | 0.933526038 | 2.159845669 | 0.432218862 |
| Olfr655 | 0.733075257 | -0.057498301 | 1.180717591 | -0.048697759 |
| Olfr656 | 0 | NA | NA | NA |
| Olfr657 | 0 | NA | NA | NA |
| Olfr658 | 7.112035581 | 0.130015689 | 0.423436983 | 0.307048496 |
| Usp-ps | 0 | NA | NA | NA |
| Olfr659 | 0 | NA | NA | NA |
| Olfr660-ps1 | 0 | NA | NA | NA |
| Olfr661 | 0 | NA | NA | NA |
| Olfr663 | 0 | NA | NA | NA |
| Olfr664 | 0 | NA | NA | NA |
| Usp17le | 0 | NA | NA | NA |
| Gm4887 | 2.328839425 | -0.179993848 | 0.810385655 | -0.222108877 |
| Usp17lb | 0 | NA | NA | NA |
| Usp17la | 0 | NA | NA | NA |
| Olfr665 | 0 | NA | NA | NA |
| Olfr666 | 0.045882455 | 0.517959912 | 3.724945049 | 0.139051692 |
| Olfr667 | 0.038981395 | -0.443816764 | 3.724945049 | -0.119147198 |
| Olfr668 | 0.044544648 | 0.517957688 | 3.724945049 | 0.139051095 |
| Olfr669 | 0 | NA | NA | NA |
| Gm5900 | 87.04897631 | -0.370223031 | 0.137905546 | -2.684613064 |
| Olfr670 | 0 | NA | NA | NA |
| Olfr671 | 0.038981395 | -0.443816764 | 3.724945049 | -0.119147198 |
| Olfr672 | 0.101408268 | 0.037730769 | 3.724898049 | 0.010129343 |

| | | | | |
|---|---|---|---|---|
| Olfr675 | 0 | NA | NA | NA |
| Olfr676 | 0 | NA | NA | NA |
| Olfr677 | 0 | NA | NA | NA |
| Olfr678 | 0 | NA | NA | NA |
| Olfr679 | 0 | NA | NA | NA |
| Olfr680-ps1 | 0.149335367 | 0.517962997 | 2.864896976 | 0.180796378 |
| Olfr681 | 0 | NA | NA | NA |
| Olfr682-ps1 | 0.201268616 | 0.569609419 | 2.613944276 | 0.217911845 |
| Olfr683 | 0.362463993 | -1.712239233 | 1.603191331 | -1.068019269 |
| Olfr684 | 0.256199707 | -1.536922003 | 2.973145954 | -0.516934596 |
| Olfr685 | 0.357595806 | 0.551260586 | 3.297772525 | 0.167161495 |
| Olfr686 | 0.169924525 | 0.517959795 | 3.389539412 | 0.152811262 |
| Olfr687 | 0 | NA | NA | NA |
| Olfr688 | 0 | NA | NA | NA |
| Olfr689 | 0 | NA | NA | NA |
| Olfr690 | 5.292052314 | 1.261373342 | 0.568128741 | 2.220224487 |
| Olfr691 | 5.347759076 | 0.657900925 | 0.654386572 | 1.005370454 |
| Olfr692 | 0.049553047 | -0.443818988 | 3.724945049 | -0.119147795 |
| Fam160a2 | 1185.683241 | 0.278131788 | 0.102022323 | 2.72618561 |
| Gm5901 | 0.044544648 | 0.517957688 | 3.724945049 | 0.139051095 |
| Cnga4 | 8.850792061 | 0.56103401 | 0.404228232 | 1.387913972 |
| Cckbr | 263.3216545 | 0.035427883 | 0.1702194 | 0.208130701 |
| Prkcdbp | 205.5345052 | -1.14187065 | 0.199131183 | -5.734263383 |
| Smpd1 | 1173.492348 | -0.081173599 | 0.069501986 | -1.167932082 |
| Apbb1 | 2731.609977 | 0.334965559 | 0.062885366 | 5.326605892 |
| Hpx | 16.70994554 | -0.097445641 | 0.354131022 | -0.275168326 |
| Trim3 | 1334.924561 | 0.350523351 | 0.084426643 | 4.151809637 |
| Arfip2 | 801.3768636 | 0.127581525 | 0.069958621 | 1.823671224 |
| Timm10b | 130.0419406 | 0.045489535 | 0.105383012 | 0.431659093 |
| Dnhd1 | 163.944155 | -0.113473902 | 0.198529877 | -0.571570906 |
| Gm22504 | 0.072025462 | 0.517959912 | 3.724945049 | 0.139051692 |
| Mrps36-ps2 | 0.9406005 | -0.73347965 | 1.001782252 | -0.73217473 |
| Rrp8 | 163.2337327 | -0.534139982 | 0.126542033 | -4.221047888 |
| Ilk | 832.9278343 | -0.30848138 | 0.104977734 | -2.938541045 |
| Taf10 | 390.2000004 | -0.453089175 | 0.112891168 | -4.013504172 |
| Tpp1 | 810.0873032 | -0.461083067 | 0.117992731 | -3.907724335 |
| Dchs1 | 413.6250274 | -0.073192919 | 0.105440356 | -0.694164191 |
| Gm15645 | 4.914941411 | 0.213707779 | 0.470957047 | 0.453773396 |
| Mrpl17 | 334.4169704 | -0.366374479 | 0.114308412 | -3.205140143 |
| 9130208D14Rik | 37.17479923 | -1.185322873 | 0.208891751 | -5.674340254 |
| Gm4070 | 48.02494048 | -1.308001132 | 0.261631448 | -4.999403318 |
| Gm20663 | 0.150903271 | 0.998859496 | 2.855571818 | 0.349793162 |
| Gm18852 | 4.812241163 | -1.467761089 | 0.478292535 | -3.068751823 |
| Gm27228 | 0 | NA | NA | NA |
| Gm8979 | 1.384150549 | -1.828329847 | 0.869498821 | -2.102739879 |
| Gm27149 | 0.066050875 | -0.443818988 | 3.724945049 | -0.119147795 |
| Gm8982 | 0 | NA | NA | NA |

| | | | | |
|---|---|---|---|---|
| Gvin1 | 6.195143366 | -1.545530952 | 0.553097711 | -2.794318111 |
| Gm18853 | 1.133034309 | -2.326018628 | 1.071881157 | -2.170034068 |
| Gm27150 | 0 | NA | NA | NA |
| Gm8989 | 0.700564593 | -1.1660181 | 1.192926254 | -0.977443573 |
| Gm4759 | 67.3121842 | -1.770655182 | 0.430574472 | -4.11230878 |
| Gm22372 | 0 | NA | NA | NA |
| Gm8995 | 675.8586595 | -1.402292878 | 0.181502991 | -7.726004229 |
| Gm1966 | 19.76971248 | -2.152647552 | 0.367975857 | -5.849969533 |
| Olfr693 | 0.090277459 | -0.443823436 | 3.724945049 | -0.119148989 |
| Olfr694 | 0 | NA | NA | NA |
| Olfr695 | 0 | NA | NA | NA |
| Olfr697 | 0 | NA | NA | NA |
| Olfr698 | 0 | NA | NA | NA |
| Olfr699 | 0.049553047 | -0.443818988 | 3.724945049 | -0.119147795 |
| Olfr700 | 0 | NA | NA | NA |
| Olfr701 | 0.038981395 | -0.443816764 | 3.724945049 | -0.119147198 |
| Olfr702 | 0 | NA | NA | NA |
| Olfr703 | 0.085433889 | 0.037058229 | 3.724945049 | 0.009948665 |
| Olfr704 | 0 | NA | NA | NA |
| Olfr705 | 0.136194719 | 0.037062677 | 3.724945049 | 0.009949859 |
| Olfr706 | 0 | NA | NA | NA |
| Olfr707 | 0.068097359 | 0.037064902 | 3.724945049 | 0.009950456 |
| Olfr1532-ps1 | 0 | NA | NA | NA |
| Olfr709-ps1 | 0 | NA | NA | NA |
| Olfr710 | 0.04393175 | 0.037062677 | 3.724945049 | 0.009949859 |
| Olfr6 | 0 | NA | NA | NA |
| Olfr711 | 0 | NA | NA | NA |
| Olfr2 | 0 | NA | NA | NA |
| Olfr713 | 0 | NA | NA | NA |
| Olfr714 | 0 | NA | NA | NA |
| Olfr17 | 0.048949532 | 0.037064902 | 3.724945049 | 0.009950456 |
| Gm10081 | 0 | NA | NA | NA |
| Olfr715 | 0.097139547 | -0.443816764 | 3.724945049 | -0.119147198 |
| Olfr716 | 0.092691418 | -0.908687078 | 3.724560878 | -0.2439716 |
| Nlrp14 | 0 | NA | NA | NA |
| Gm3765 | 0.678124035 | -1.630082123 | 1.327160064 | -1.228248323 |
| Rbmxl2 | 0 | NA | NA | NA |
| Platr28 | 0.049553047 | -0.443818988 | 3.724945049 | -0.119147795 |
| Gm24932 | 0.060174537 | 0.517957688 | 3.724945049 | 0.139051095 |
| Syt9 | 336.6996381 | 0.22813508 | 0.087638097 | 2.603149627 |
| Olfml1 | 105.9701042 | 0.073442989 | 0.1764686 | 0.416181627 |
| Ppfibp2 | 154.4178595 | -0.468819606 | 0.126136878 | -3.716752899 |
| 5330417H12Rik | 0 | NA | NA | NA |
| Cyb5r2 | 9.463141173 | -0.971508401 | 0.365844638 | -2.65552177 |
| Ovch2 | 0 | NA | NA | NA |
| Olfr467 | 0 | NA | NA | NA |
| Olfr469 | 0 | NA | NA | NA |

| | | | | |
|---|---|---|---|---|
| Olfr470 | 0 | NA | NA | NA |
| Gm10156 | 22.50808326 | 0.131446868 | 0.322565631 | 0.407504259 |
| Olfr472 | 0.092881282 | 0.037060453 | 3.724945049 | 0.009949262 |
| Olfr473 | 0.038981395 | -0.443816764 | 3.724945049 | -0.119147198 |
| Olfr474 | 0 | NA | NA | NA |
| Olfr476 | 0 | NA | NA | NA |
| Olfr477 | 0.165236907 | -0.444772966 | 3.378321243 | -0.131655024 |
| Olfr478 | 0 | NA | NA | NA |
| Olfr479 | 0 | NA | NA | NA |
| Olfr480 | 0 | NA | NA | NA |
| Olfr481 | 0 | NA | NA | NA |
| Olfr482 | 0 | NA | NA | NA |
| Olfr483 | 0.062426873 | 0.517959912 | 3.724945049 | 0.139051692 |
| Gm25424 | 0 | NA | NA | NA |
| Olfr484 | 0 | NA | NA | NA |
| Olfr485 | 0 | NA | NA | NA |
| Olfr486 | 0 | NA | NA | NA |
| Olfr487 | 0.123841226 | -1.017823543 | 3.719440177 | -0.273649661 |
| Olfr488 | 0 | NA | NA | NA |
| Olfr490 | 0 | NA | NA | NA |
| Olfr491 | 0 | NA | NA | NA |
| Olfr492 | 0 | NA | NA | NA |
| Olfr493 | 0 | NA | NA | NA |
| Olfr494 | 0 | NA | NA | NA |
| Olfr495 | 0 | NA | NA | NA |
| Olfr497 | 0.049553047 | -0.443818988 | 3.724945049 | -0.119147795 |
| Olfr498 | 0 | NA | NA | NA |
| Olfr502 | 0 | NA | NA | NA |
| Olfr503 | 0 | NA | NA | NA |
| Olfr504 | 0 | NA | NA | NA |
| Olfr506 | 0 | NA | NA | NA |
| Olfr507 | 0 | NA | NA | NA |
| Olfr508 | 0 | NA | NA | NA |
| Olfr509 | 0 | NA | NA | NA |
| Olfr510 | 0.046345709 | -0.443818988 | 3.724945049 | -0.119147795 |
| Olfr512 | 0 | NA | NA | NA |
| Olfr513 | 0 | NA | NA | NA |
| Olfr514 | 0 | NA | NA | NA |
| Olfr516 | 0 | NA | NA | NA |
| Olfr517 | 0 | NA | NA | NA |
| Olfr518 | 0 | NA | NA | NA |
| Olfr519 | 0 | NA | NA | NA |
| Nlrp10 | 10.06809784 | 0.334397219 | 0.316781401 | 1.055608752 |
| Eif3f | 1059.444892 | -0.045948923 | 0.076089787 | -0.603877665 |
| Gm26599 | 4.712628857 | 1.077243658 | 0.609369884 | 1.767799306 |
| Tub | 1801.451592 | 0.858480693 | 0.111763182 | 7.681247775 |
| Ric3 | 1280.403782 | 0.120305385 | 0.072060234 | 1.669511433 |

| | | | | |
|---|---|---|---|---|
| Gm25450 | 3.817416342 | 0.423227747 | 0.606431653 | 0.697898511 |
| Lmo1 | 71.3151019 | -0.000470071 | 0.135587556 | -0.003466922 |
| Stk33 | 14.06609556 | -0.135270206 | 0.303576462 | -0.445588586 |
| Gm9105 | 5.082316306 | 0.074639534 | 0.440675139 | 0.169375414 |
| 1700095J03Rik | 3.157210742 | 0.397895244 | 0.618440692 | 0.643384644 |
| Trim66 | 632.3889432 | 0.05557537 | 0.105798643 | 0.525293786 |
| Rpl27a | 1290.635761 | -0.274960548 | 0.102157811 | -2.691527401 |
| Snora3 | 0 | NA | NA | NA |
| AC124457.1 | 0.875090066 | 0.489695476 | 1.074897763 | 0.455574002 |
| Gm24888 | 1.185922749 | -1.547973444 | 1.034902177 | -1.495767888 |
| St5 | 362.0690004 | -0.133688026 | 0.085799831 | -1.558138567 |
| Akip1 | 31.41517011 | -0.708780215 | 0.220863934 | -3.209126097 |
| BC051019 | 3.117097637 | -0.980048166 | 0.579838347 | -1.690209298 |
| Ascl3 | 1.052193792 | -0.500926463 | 0.891553056 | -0.561858276 |
| Tmem9b | 679.663783 | -0.007269651 | 0.07440358 | -0.097705663 |
| Nrip3 | 1443.710783 | 0.096619902 | 0.112579438 | 0.85823756 |
| Gm22822 | 1.435595059 | 1.207842319 | 0.92295766 | 1.308664927 |
| Scube2 | 22.45075123 | 0.744842379 | 0.315988098 | 2.357184916 |
| Dennd5a | 3399.692085 | 0.123281811 | 0.068987622 | 1.787013501 |
| Tmem41b | 489.61422 | -0.328798651 | 0.100747527 | -3.263590287 |
| Gm24842 | 0.126912322 | -0.76154727 | 3.693829068 | -0.206167437 |
| 1600010M07Rik | 2.455553585 | -1.160363896 | 0.608530327 | -1.906830021 |
| Gm23100 | 0 | NA | NA | NA |
| Ipo7 | 1735.407786 | -0.357676149 | 0.076051032 | -4.703107119 |
| Gm25636 | 6.97985599 | -0.188837139 | 0.402541596 | -0.469112114 |
| Snora23 | 31.16499087 | -0.326403983 | 0.238729335 | -1.367255443 |
| AA474408 | 22.01653641 | -0.422793753 | 0.285027839 | -1.483341957 |
| Zfp143 | 153.6133789 | -0.310463921 | 0.094917786 | -3.270871923 |
| Wee1 | 194.9507975 | 0.134985777 | 0.12173767 | 1.108825037 |
| Swap70 | 161.8502858 | -0.736728839 | 0.108529538 | -6.788279479 |
| Sbf2 | 1399.071276 | 0.042478863 | 0.073013379 | 0.581795599 |
| Gm9064 | 0 | NA | NA | NA |
| Gm17219 | 0.046345709 | -0.443818988 | 3.724945049 | -0.119147795 |
| Gm10087 | 0 | NA | NA | NA |
| Gm28863 | 0.430307537 | -0.389610815 | 1.43346943 | -0.271795691 |
| Adm | 37.32539147 | -0.538971162 | 0.22096823 | -2.439134179 |
| Ampd3 | 786.3935543 | -0.667356742 | 0.160974896 | -4.14571936 |
| Rnf141 | 819.8137337 | 0.179584184 | 0.109063695 | 1.646599125 |
| AC122844.1 | 0.956902809 | -0.00561708 | 1.214180055 | -0.004626233 |
| Lyve1 | 8.973244537 | 0.522387475 | 0.423066837 | 1.234763468 |
| Mrvi1 | 43.82669296 | 0.227610559 | 0.18809786 | 1.210064585 |
| Gm16336 | 0.075043205 | 0.517957688 | 3.724945049 | 0.139051095 |
| Ctr9 | 691.8584163 | -0.174580107 | 0.060476214 | -2.886756523 |
| Eif4g2 | 8499.079975 | -0.329095065 | 0.08236063 | -3.995781272 |
| Gm23262 | 0 | NA | NA | NA |
| Galnt18 | 280.6267264 | 0.252852089 | 0.132475194 | 1.908674984 |
| Gm25787 | 0 | NA | NA | NA |

| | | | | |
|---|---|---|---|---|
| Gm23345 | 0.066050875 | -0.443818988 | 3.724945049 | -0.119147795 |
| Usp47 | 1697.490691 | -0.119753958 | 0.062838865 | -1.90573076 |
| Dkk3 | 1672.01163 | 0.652291121 | 0.122379299 | 5.330077298 |
| Mical2 | 2673.306519 | 0.455920955 | 0.141648732 | 3.218673037 |
| Micalcl | 33.92286921 | 0.556523781 | 0.285242921 | 1.951052033 |
| Parva | 887.2698577 | -0.14090987 | 0.08699841 | -1.619683273 |
| Parvaos | 2.062454624 | -0.228617015 | 0.649241231 | -0.352129538 |
| 2310014F06Rik | 4.100390628 | -0.377291291 | 0.500820808 | -0.753345876 |
| Gm24154 | 0.181894773 | 0.009507524 | 2.695899385 | 0.003526661 |
| Tead1 | 787.7264761 | -0.718504677 | 0.121379139 | -5.919507119 |
| AC126052.1 | 1.51359592 | 0.023303727 | 0.930677358 | 0.025039534 |
| Rassf10 | 10.84181486 | 1.01702509 | 0.341464965 | 2.978417096 |
| AC104930.1 | 0 | NA | NA | NA |
| Gm23662 | 0 | NA | NA | NA |
| Arntl | 352.8719286 | -0.130083099 | 0.128109368 | -1.015406614 |
| Btbd10 | 732.5892807 | 0.164933161 | 0.078792956 | 2.093247545 |
| Gm15994 | 0.277092755 | -1.457993879 | 2.021993301 | -0.721067611 |
| Pth | 0 | NA | NA | NA |
| Far1 | 1848.391941 | 0.046001687 | 0.06453513 | 0.712816207 |
| Far1os | 0.286281128 | 0.998843802 | 3.724945049 | 0.268149943 |
| Gm23700 | 0 | NA | NA | NA |
| Gm5600 | 0.213133906 | 0.037075446 | 2.1812924 | 0.016997009 |
| Gm29507 | 0.046345709 | -0.443818988 | 3.724945049 | -0.119147795 |
| Spon1 | 1904.191878 | 0.044051102 | 0.107621722 | 0.409314229 |
| Rras2 | 177.862388 | -0.150073003 | 0.11956638 | -1.25514382 |
| Copb1 | 1041.818964 | -0.305071721 | 0.086349586 | -3.532984195 |
| Psma1 | 653.5961011 | -0.319772326 | 0.106613906 | -2.999349127 |
| 4933406I18Rik | 91.43073966 | 0.216186619 | 0.132317181 | 1.633851457 |
| Pde3b | 366.815521 | 0.121808944 | 0.097156923 | 1.253734064 |
| Cyp2r1 | 3.701896043 | -0.496460846 | 0.503882145 | -0.985271756 |
| Calca | 61.65923974 | -1.424057351 | 0.232142131 | -6.134420093 |
| Gm15500 | 180.1198848 | -0.199884299 | 0.163152905 | -1.225134783 |
| Calcb | 5.761881877 | -0.441527819 | 0.511039575 | -0.863979701 |
| Insc | 59.233201 | -0.847712129 | 0.261042962 | -3.247404646 |
| Gm24982 | 0 | NA | NA | NA |
| Rps4l-ps | 0.061920613 | -0.443816764 | 3.724945049 | -0.119147198 |
| Gm22669 | 0 | NA | NA | NA |
| Gm27684 | 0.263733274 | 1.006679293 | 2.101824621 | 0.478954944 |
| Sox6 | 489.52169 | -0.31419982 | 0.087077876 | -3.60826233 |
| Sox6os | 0.611953183 | 0.405812275 | 1.332114548 | 0.304637672 |
| 1700003G18Rik | 0.77082754 | -0.312614945 | 1.205299352 | -0.259367056 |
| Gm4353 | 19.37501129 | -0.431665275 | 0.299908473 | -1.439323375 |
| 1110004F10Rik | 588.1908301 | 0.054119798 | 0.092621873 | 0.584309043 |
| Plekha7 | 339.2168702 | -0.115326839 | 0.102114809 | -1.129384076 |
| Rps13 | 763.6410431 | -0.313921837 | 0.101666993 | -3.087745875 |
| Snord14a | 3.97447326 | 0.54066658 | 0.549756711 | 0.983465175 |
| Pik3c2a | 1083.338475 | -0.097782769 | 0.066937362 | -1.460810022 |

| | | | | |
|---|---|---|---|---|
| Nucb2 | 157.5399997 | 0.076618386 | 0.169725485 | 0.451425346 |
| n-R5s157 | 0 | NA | NA | NA |
| Gm25423 | 0 | NA | NA | NA |
| RP24-246N21.1 | 393.097115 | -0.382846841 | 0.120962916 | -3.164993484 |
| Gm26054 | 0 | NA | NA | NA |
| Xylt1 | 443.4184742 | -0.225231999 | 0.065647215 | -3.430945241 |
| Gm24063 | 0 | NA | NA | NA |
| Rps15a | 1296.320441 | -0.18538437 | 0.074830636 | -2.477386023 |
| Arl6ip1 | 2631.753197 | 0.07938603 | 0.118229964 | 0.671454403 |
| Smg1 | 4666.36924 | -0.012699041 | 0.103838443 | -0.122296146 |
| Gm25683 | 0 | NA | NA | NA |
| 4930583K01Rik | 0.241683003 | 0.945879413 | 2.082187342 | 0.454272002 |
| Gm23229 | 0.044544648 | 0.517957688 | 3.724945049 | 0.139051095 |
| Syt17 | 179.5473353 | 0.684742823 | 0.157369369 | 4.351182365 |
| Itpripl2 | 154.921355 | -0.745285124 | 0.156332537 | -4.767306531 |
| Coq7 | 126.5384451 | 0.004586771 | 0.15117754 | 0.030340294 |
| Tmc7 | 194.611435 | 0.025867403 | 0.125500977 | 0.206113158 |
| Tmc5 | 2.207900702 | 0.720482248 | 0.806164745 | 0.893715897 |
| Gde1 | 1198.036269 | 0.030203731 | 0.059232945 | 0.509914392 |
| Ccp110 | 802.7760504 | 0.074902101 | 0.082653981 | 0.906212863 |
| 9030624J02Rik | 685.4574909 | -0.180635244 | 0.070174825 | -2.574074719 |
| Gm26147 | 1.083234348 | -0.320808871 | 1.191707188 | -0.269201088 |
| Knop1 | 359.1550831 | -0.339092747 | 0.070635989 | -4.800566262 |
| Iqck | 21.26371758 | -0.223966501 | 0.23314431 | -0.960634642 |
| Gprc5b | 1912.548548 | 0.114106079 | 0.094878883 | 1.202649906 |
| Gpr139 | 17.82217913 | -0.301213445 | 0.318059804 | -0.947033988 |
| Gm23321 | 0 | NA | NA | NA |
| Gp2 | 0.244487964 | 0.506965616 | 1.853142184 | 0.273570814 |
| Umod | 0 | NA | NA | NA |
| Pdilt | 0.092881282 | 0.037060453 | 3.724945049 | 0.009949262 |
| Gm37518 | 0.130524232 | 0.517958497 | 3.657871415 | 0.141601067 |
| Gm26967 | 0.180739367 | 0.517958781 | 3.322505153 | 0.155894049 |
| Acsm5 | 5.692917749 | 0.456007122 | 0.473830376 | 0.962384737 |
| Acsm2 | 0.062426873 | 0.517959912 | 3.724945049 | 0.139051692 |
| Acsm1 | 0.038981395 | -0.443816764 | 3.724945049 | -0.119147198 |
| Acsm4 | 1.49381288 | -0.01464104 | 0.88554911 | -0.01653329 |
| Thumpd1 | 343.5482368 | 0.071229683 | 0.105743312 | 0.673609345 |
| Gm5601 | 215.6914571 | -0.058349436 | 0.095628052 | -0.610170705 |
| Acsm3 | 0.623781035 | 0.525244934 | 1.249192954 | 0.420467416 |
| Eri2 | 94.72177517 | -0.268373129 | 0.153102093 | -1.752903076 |
| 2610020H08Rik | 22.09157058 | -0.47027194 | 0.278861977 | -1.686396779 |
| Dcun1d3 | 429.5927485 | -0.41167913 | 0.092891527 | -4.431826466 |
| Lyrm1 | 64.24767704 | 0.138550252 | 0.148521542 | 0.932863004 |
| Dnah3 | 28.90499188 | -0.716002789 | 0.236730313 | -3.024550507 |
| Gm25217 | 0.280224789 | -0.81348717 | 1.80107603 | -0.451667312 |
| Tmem159 | 54.2618588 | -0.302596565 | 0.156538404 | -1.933050022 |
| Zp2 | 1.125736089 | 0.738415033 | 0.961012645 | 0.768371817 |

| | | | | |
|---|---|---|---|---|
| Anks4b | 4.77461678 | 0.573597501 | 0.453398067 | 1.265107956 |
| Crym | 493.688269 | -0.172444495 | 0.12296733 | -1.402360245 |
| Abca14 | 0.666404725 | -1.112467323 | 1.140724097 | -0.97522909 |
| Abca15 | 0.612941558 | -1.751906976 | 1.289508332 | -1.358585232 |
| Abca16 | 0.254958917 | -1.89165757 | 2.048075216 | -0.923627001 |
| E130201H02Rik | 1.649955311 | 0.165219229 | 0.768643466 | 0.214949111 |
| Uqcrc2 | 1539.021604 | 0.084200213 | 0.078736151 | 1.069397114 |
| Pdzd9 | 1.255624142 | -1.366235515 | 0.953526114 | -1.432824435 |
| 9030407P20Rik | 49.63574211 | 0.354467207 | 0.183296674 | 1.933844187 |
| BC030336 | 1048.447558 | 0.051001011 | 0.065561169 | 0.777914917 |
| Vwa3a | 56.37564545 | 0.325784592 | 0.17721078 | 1.838401658 |
| Eef2k | 574.705251 | 0.493340869 | 0.1060663 | 4.651249897 |
| Gm15774 | 0.046345709 | -0.443818988 | 3.724945049 | -0.119147795 |
| Polr3e | 380.9951259 | 0.329072123 | 0.108121494 | 3.043540292 |
| Cdr2 | 189.8166021 | -0.580776435 | 0.177525643 | -3.271507297 |
| 4933427G17Rik | 3.26880693 | 0.996976792 | 0.609116036 | 1.636760049 |
| Gm9234 | 7.527412146 | -0.555079368 | 0.378275395 | -1.467394852 |
| Gm25924 | 0 | NA | NA | NA |
| Gm9905 | 21.38643968 | -0.403445122 | 0.248937458 | -1.620668603 |
| Mettl9 | 576.9325256 | -0.182777784 | 0.11224014 | -1.628452915 |
| Gm27991 | 0 | NA | NA | NA |
| Igsf6 | 72.32097277 | -0.69220695 | 0.320693567 | -2.158468464 |
| Otoa | 4.95839639 | 0.357126602 | 0.48200838 | 0.740913679 |
| Hs3st2 | 198.3223821 | -0.917758518 | 0.314530691 | -2.917866348 |
| Usp31 | 2536.972672 | -0.112127287 | 0.079154082 | -1.416569859 |
| 1700069B07Rik | 1.411393777 | -0.110043182 | 0.783014773 | -0.14053781 |
| Scnn1g | 16.54571043 | 0.239689868 | 0.269309195 | 0.890017395 |
| Scnn1b | 3.3334194 | -2.2248023 | 0.678603197 | -3.278502535 |
| Cog7 | 494.7223238 | 0.169249226 | 0.067145197 | 2.520645303 |
| Gga2 | 393.4606856 | -0.10868656 | 0.097252107 | -1.117575385 |
| Ears2 | 110.4529885 | -0.473460975 | 0.12367493 | -3.828269598 |
| Ubfd1 | 1136.037908 | -0.231935489 | 0.063503697 | -3.652314727 |
| Ndufab1 | 625.6134519 | -0.16579548 | 0.081007231 | -2.046675078 |
| Palb2 | 22.09319308 | -0.231070758 | 0.257952496 | -0.895788029 |
| Gm16326 | 0.136194719 | 0.037062677 | 3.724945049 | 0.009949859 |
| Dctn5 | 636.704053 | -0.045702488 | 0.056066892 | -0.81514217 |
| Gm15489 | 0.496441948 | -0.404692556 | 1.59849342 | -0.253171237 |
| Plk1 | 22.12491667 | -0.392729221 | 0.294048247 | -1.335594501 |
| Ern2 | 3.478004289 | 0.651077717 | 0.602142562 | 1.081268386 |
| Chp2 | 6.555594633 | -0.045008567 | 0.44313413 | -0.10156872 |
| Gm23614 | 0 | NA | NA | NA |
| Prkcb | 4097.896829 | 0.819093035 | 0.120218859 | 6.813348923 |
| Gm14388 | 23.59541605 | 0.177230844 | 0.25734097 | 0.68870046 |
| Gm26414 | 4.07614265 | 0.786328009 | 0.687048493 | 1.144501469 |
| Gm14389 | 0 | NA | NA | NA |
| Cacng3 | 479.8047837 | 1.014167413 | 0.16766816 | 6.048658336 |
| Rbbp6 | 1656.438353 | 0.155702662 | 0.073084655 | 2.130442587 |

| | | | | |
|---|---|---|---|---|
| Tnrc6a | 1721.479868 | 0.000583341 | 0.094631538 | 0.006164337 |
| Slc5a11 | 15.52195762 | -0.18617625 | 0.282045281 | -0.660093476 |
| Arhgap17 | 309.4984358 | -0.643242782 | 0.139552616 | -4.609320858 |
| Lcmt1 | 343.5567309 | 0.124085695 | 0.071115591 | 1.744845158 |
| Gm23559 | 0 | NA | NA | NA |
| Aqp8 | 0.525182132 | 1.759191797 | 1.456755771 | 1.207609286 |
| Zkscan2 | 230.1962317 | 0.715131835 | 0.161505919 | 4.427898608 |
| Gm23788 | 0.167772976 | -0.985237982 | 3.413460593 | -0.288633179 |
| Gm26974 | 9.009306514 | -0.111648526 | 0.329901199 | -0.338430193 |
| Gm27040 | 8.361238006 | 0.166314377 | 0.401814363 | 0.413908493 |
| Hs3st4 | 521.3318301 | 0.456719096 | 0.087803503 | 5.201604505 |
| Gm25759 | 6.10632452 | 0.891147406 | 0.554465527 | 1.607218775 |
| Gm15338 | 14.24273356 | 0.563583056 | 0.456963913 | 1.233320706 |
| Gm15339 | 0 | NA | NA | NA |
| Gm23168 | 0.097139547 | -0.443816764 | 3.724945049 | -0.119147198 |
| 3100003L05Rik | 1.759517327 | -0.284067134 | 0.676396619 | -0.41997125 |
| Gm21957 | 0 | NA | NA | NA |
| 4930533L02Rik | 0 | NA | NA | NA |
| 4933440M02Rik | 0.422952957 | 0.661479879 | 3.716378055 | 0.177990471 |
| 4930571K23Rik | 0 | NA | NA | NA |
| Kdm8 | 38.64691656 | 0.517188611 | 0.245891105 | 2.103323794 |
| Nsmce1 | 119.9047304 | -0.466437741 | 0.166259518 | -2.805479925 |
| Il4ra | 356.9425988 | -1.019581052 | 0.184223445 | -5.534480422 |
| Il21r | 78.43361135 | -0.513432919 | 0.186985791 | -2.74583922 |
| Gtf3c1 | 2212.631377 | 0.190228681 | 0.063090531 | 3.01517007 |
| D430042O09Rik | 320.9984056 | -0.149419167 | 0.077072857 | -1.938674296 |
| Gsg1l | 413.6031104 | 0.5767697 | 0.162043681 | 3.559347073 |
| Xpo6 | 1490.157562 | -0.000662841 | 0.066916051 | -0.00990556 |
| Gm17137 | 0.048949532 | 0.037064902 | 3.724945049 | 0.009950456 |
| Gm17136 | 0.044544648 | 0.517957688 | 3.724945049 | 0.139051095 |
| Sbk1 | 1551.156001 | -0.023573611 | 0.144015156 | -0.163688401 |
| Gm10155 | 8.990861825 | -0.227949228 | 0.356176238 | -0.639989993 |
| Lat | 33.16436472 | -0.556261138 | 0.257953972 | -2.156435638 |
| Mir7058 | 0 | NA | NA | NA |
| Spns1 | 414.4773662 | -0.059885595 | 0.081470503 | -0.73505862 |
| Nfatc2ip | 98.35474023 | -0.342860216 | 0.138724305 | -2.471522323 |
| Cd19 | 0.620398479 | -2.522263255 | 1.469376006 | -1.716553997 |
| Rabep2 | 121.2584971 | -0.13353596 | 0.110134265 | -1.212483327 |
| Atp2a1 | 15.93822915 | 0.318460196 | 0.276435112 | 1.152025131 |
| Sh2b1 | 913.839647 | -0.267918683 | 0.058643755 | -4.568579928 |
| Tufm | 383.5821207 | -0.370776456 | 0.10722884 | -3.457805344 |
| Atxn2l | 1896.105242 | -0.14922368 | 0.084453162 | -1.766940111 |
| Eif3c | 2058.465012 | -0.390305584 | 0.084564969 | -4.615452339 |
| Gm24596 | 0.884909101 | -0.109956626 | 1.15851614 | -0.094911604 |
| G730046D07Rik | 2.910496466 | -2.08295508 | 0.63751166 | -3.267320757 |
| Gm25579 | 0 | NA | NA | NA |
| Cln3 | 171.9378215 | -0.871625395 | 0.104252399 | -8.360722689 |

| | | | | |
|---|---|---|---|---|
| Mir7059 | 0.228481116 | 0.037074694 | 2.191567996 | 0.016916972 |
| Apobr | 49.34344806 | -1.646069621 | 0.328189304 | -5.015610202 |
| Il27 | 13.85132785 | -0.679134244 | 0.371082203 | -1.830145014 |
| Nupr1 | 177.6816824 | -1.907943887 | 0.260647364 | -7.320019878 |
| 2510046G10Rik | 4.55499157 | -0.23882956 | 0.523620375 | -0.456112044 |
| Ccdc101 | 124.2951638 | -0.001776958 | 0.103895886 | -0.017103253 |
| Sult1a1 | 88.10259738 | 1.507554421 | 0.25462659 | 5.920648049 |
| Gm6939 | 1.140676537 | 0.392198437 | 0.948388889 | 0.413541788 |
| Slx1b | 109.1217166 | 0.170615493 | 0.125393781 | 1.3606376 |
| Bola2 | 141.4468845 | -0.377685891 | 0.147240482 | -2.565095446 |
| Coro1a | 641.2179666 | 0.100656075 | 0.087689608 | 1.147867778 |
| Mapk3 | 1099.213407 | 0.279171831 | 0.093757784 | 2.977585625 |
| Gdpd3 | 17.57179256 | 0.036854942 | 0.301102192 | 0.122400113 |
| Gm9967 | 7.556454724 | 0.98586314 | 0.460493546 | 2.140883731 |
| Ypel3 | 1100.394795 | 0.141790391 | 0.072403142 | 1.958345817 |
| Tbx6 | 9.328334444 | -0.169918782 | 0.444342725 | -0.38240478 |
| Ppp4c | 390.6480823 | -0.495265907 | 0.121994768 | -4.059730726 |
| Aldoa | 12436.05844 | -0.048731649 | 0.049845915 | -0.977645808 |
| Gm15676 | 14.57694585 | 0.310733543 | 0.284548403 | 1.092023502 |
| Fam57b | 780.1775602 | 0.250654091 | 0.069513327 | 3.605842251 |
| 4930451I11Rik | 0.843446786 | -0.374088199 | 1.245746864 | -0.300292307 |
| Doc2a | 204.7161822 | 0.509839235 | 0.168773049 | 3.020856932 |
| Ino80e | 331.245082 | -0.248732357 | 0.073379333 | -3.389678655 |
| Hirip3 | 134.4899788 | 0.215191912 | 0.143570264 | 1.498861302 |
| Taok2 | 2099.502443 | 0.280424116 | 0.048498457 | 5.782124449 |
| Tmem219 | 154.4928956 | -0.537083731 | 0.155434679 | -3.455366157 |
| Gm20650 | 0.099106094 | -0.93550548 | 3.724188147 | -0.251197158 |
| Gm23445 | 0.097899064 | 0.037062677 | 3.724945049 | 0.009949859 |
| Kctd13 | 647.9350141 | 0.645671196 | 0.088086538 | 7.329964465 |
| Gm21984 | 106.5590713 | 0.109796928 | 0.19792366 | 0.554743821 |
| Asphd1 | 163.222168 | 0.017287086 | 0.115908681 | 0.149144017 |
| Sez6l2 | 4188.453118 | 0.373098838 | 0.072559111 | 5.141998506 |
| D830044I16Rik | 2.339863245 | -0.864550831 | 0.658814213 | -1.312283211 |
| Cdipt | 964.396228 | 0.061877486 | 0.045082502 | 1.372538884 |
| Mvp | 599.1504578 | -0.929515098 | 0.260433172 | -3.569111764 |
| Pagr1b | 0.194631057 | 1.456047977 | 3.23213671 | 0.450490839 |
| Pagr1a | 269.1231763 | 0.110171288 | 0.147296267 | 0.747957097 |
| Prrt2 | 12.36009762 | 0.683537692 | 0.421228921 | 1.622722606 |
| Maz | 1509.464906 | 0.1362273 | 0.073675482 | 1.849018118 |
| Kif22 | 26.64433543 | -0.15296996 | 0.248937825 | -0.614490625 |
| Gm25333 | 0 | NA | NA | NA |
| AC122863.1 | 0 | NA | NA | NA |
| Zg16 | 0.046345709 | -0.443818988 | 3.724945049 | -0.119147795 |
| Gm22747 | 0 | NA | NA | NA |
| AI467606 | 37.52684094 | -0.735082522 | 0.237849974 | -3.09053017 |
| Qprt | 3.756627285 | 0.33674666 | 0.50262605 | 0.669974547 |
| Spn | 82.89016059 | -0.84214469 | 0.24861452 | -3.387351195 |

| | | | | |
|---|---|---|---|---|
| Gm25735 | 0 | NA | NA | NA |
| Cd2bp2 | 785.0302282 | 0.016151113 | 0.059968299 | 0.26932752 |
| Tbc1d10b | 882.4067378 | -0.29343257 | 0.067764139 | -4.33020434 |
| Mylpf | 4.837556069 | -0.448998716 | 0.488233363 | -0.91963956 |
| 1-Sep | 35.87249771 | -0.387225565 | 0.241165432 | -1.605642908 |
| Zfp553 | 281.7915856 | 0.129588371 | 0.083306651 | 1.555558525 |
| Zfp771 | 217.3863754 | -0.182337384 | 0.134586308 | -1.354798919 |
| Dctpp1 | 50.21803819 | 0.020066768 | 0.16572368 | 0.1210857 |
| Sephs2 | 242.8260877 | 0.094789094 | 0.104308479 | 0.908738145 |
| Gm17511 | 0.08486385 | 0.037730769 | 3.724898049 | 0.010129343 |
| Itgal | 262.0288419 | -0.864251376 | 0.193800006 | -4.459501281 |
| Zfp768 | 163.5362928 | -0.444439222 | 0.099115728 | -4.484043358 |
| Zfp747 | 52.03124678 | -0.208796245 | 0.156547002 | -1.333760737 |
| 9130019O22Rik | 50.28528677 | -0.540968462 | 0.180184714 | -3.002299413 |
| E430018J23Rik | 67.77893546 | 0.033877987 | 0.148845309 | 0.227605341 |
| Smt3h2-ps2 | 0 | NA | NA | NA |
| Zfp764 | 37.42744171 | -0.309141369 | 0.189049411 | -1.635241111 |
| Zfp688 | 66.55993122 | -0.070887569 | 0.140601973 | -0.50417194 |
| Zfp689 | 64.00074658 | -0.184265427 | 0.142983121 | -1.288721538 |
| B130055M24Rik | 80.27483843 | -0.38183773 | 0.194842629 | -1.959723764 |
| Prr14 | 312.8086992 | -0.282781344 | 0.072883366 | -3.879916096 |
| Fbrs | 544.6972203 | -0.044410848 | 0.097875319 | -0.453749202 |
| Mir7060 | 0.754347455 | -0.967204316 | 1.134538265 | -0.852509206 |
| AC136146.1 | 0 | NA | NA | NA |
| 1700008J07Rik | 27.76791627 | -0.290167765 | 0.213555315 | -1.358747566 |
| Srcap | 3720.548078 | 0.096072568 | 0.059445738 | 1.616138863 |
| Snora30 | 11.72211257 | 0.264695072 | 0.317816802 | 0.832854243 |
| Gm28198 | 0.198870357 | 0.037074371 | 2.667920915 | 0.013896353 |
| Tmem265 | 0 | NA | NA | NA |
| Phkg2 | 467.0988482 | 0.207404237 | 0.072793509 | 2.84921333 |
| Gm166 | 30.45493143 | 0.018849269 | 0.208614365 | 0.090354609 |
| Rnf40 | 789.9745989 | -0.235282776 | 0.066175672 | -3.555427076 |
| 1700120K04Rik | 1.43115425 | 2.185075057 | 0.910754289 | 2.399192716 |
| Zfp629 | 457.8583134 | -0.115584081 | 0.066173742 | -1.746675903 |
| Bcl7c | 293.7258915 | -0.129111484 | 0.079905232 | -1.615807635 |
| Mir762 | 1.435335593 | -0.643948215 | 0.990558468 | -0.650086024 |
| Ctf1 | 11.31378371 | -0.216796645 | 0.318121802 | -0.681489427 |
| Ctf2 | 0 | NA | NA | NA |
| Fbxl19 | 610.6258965 | 0.008554911 | 0.099803908 | 0.08571719 |
| Orai3 | 179.2087718 | -0.232923556 | 0.146374701 | -1.591282883 |
| Setd1a | 391.9559766 | 0.162102198 | 0.126873441 | 1.277668491 |
| Hsd3b7 | 112.3828733 | -0.686749263 | 0.138005076 | -4.976260888 |
| Stx1b | 6333.347927 | 0.054371708 | 0.074311818 | 0.731669739 |
| Stx4a | 301.4532526 | -0.374953246 | 0.144001642 | -2.603812294 |
| Zfp668 | 218.1486246 | 0.033477729 | 0.1089135 | 0.307379053 |
| Zfp646 | 586.462955 | -0.252309021 | 0.073324678 | -3.440983708 |
| Prss53 | 5.985399039 | -0.51159688 | 0.463834178 | -1.102973658 |

| | | | | |
|---|---|---|---|---|
| Gm21974 | 0.049553047 | -0.443818988 | 3.724945049 | -0.119147795 |
| Vkorc1 | 34.89639478 | -0.687762923 | 0.26344627 | -2.610638307 |
| Bckdk | 313.1884233 | 0.030737202 | 0.086856578 | 0.353884559 |
| Kat8 | 146.4017955 | 0.163118185 | 0.10114162 | 1.612770147 |
| Prss8 | 2.192463596 | 1.079466479 | 0.778992772 | 1.38572079 |
| Prss36 | 34.47699352 | -0.076162221 | 0.195259553 | -0.390056309 |
| Fus | 2383.412294 | 0.152231455 | 0.118743975 | 1.282014138 |
| Gm26690 | 0.089089296 | 0.957155977 | 3.723939561 | 0.257027796 |
| B230325K18Rik | 116.590507 | 0.680772672 | 0.276255487 | 2.464286517 |
| Pycard | 48.5114067 | -1.132711923 | 0.282973996 | -4.002883439 |
| Gm15533 | 0 | NA | NA | NA |
| Trim72 | 39.26137666 | 0.330922954 | 0.36701573 | 0.901658776 |
| Gm22809 | 0 | NA | NA | NA |
| Itgam | 625.9394924 | -0.724293701 | 0.149016103 | -4.860506261 |
| Itgax | 181.6100497 | -0.201086466 | 0.333941661 | -0.602160464 |
| Itgad | 28.13752969 | 0.455865808 | 0.272060678 | 1.675603442 |
| Cox6a2 | 20.40538094 | 0.761344495 | 0.228868075 | 3.32656485 |
| 9130023H24Rik | 70.01739671 | -0.603908721 | 0.150695673 | -4.007472213 |
| Armc5 | 244.6664953 | -0.001214509 | 0.0950507 | -0.012777488 |
| Tgfb1i1 | 119.9274198 | -0.609236789 | 0.16983534 | -3.587220361 |
| Slc5a2 | 1.211082793 | -2.133973107 | 1.106425973 | -1.928708435 |
| BC017158 | 206.5939799 | 0.003537076 | 0.086261193 | 0.041004261 |
| Mir3103 | 0 | NA | NA | NA |
| Rgs10 | 178.444261 | 0.46671322 | 0.128633621 | 3.628236672 |
| Gm15503 | 2.556904923 | 1.551870299 | 0.741653028 | 2.092447872 |
| Tial1 | 1499.062975 | 0.02929519 | 0.071052254 | 0.412304868 |
| Gm24365 | 0 | NA | NA | NA |
| Bag3 | 352.4742549 | -1.019896427 | 0.156135978 | -6.532103893 |
| Gm7258 | 0 | NA | NA | NA |
| Inpp5f | 1743.240792 | -0.157761817 | 0.073893439 | -2.134990852 |
| RP24-143L4.2 | 0 | NA | NA | NA |
| Gm16044 | 15.72144794 | 1.065429644 | 0.392370953 | 2.715363192 |
| Mcmbp | 481.2577654 | -0.360201445 | 0.068350163 | -5.269942775 |
| Gm23783 | 0 | NA | NA | NA |
| Sec23ip | 696.8616121 | -0.343038926 | 0.074132327 | -4.627386472 |
| n-R5s158 | 0 | NA | NA | NA |
| Gm25778 | 0.04393175 | 0.037062677 | 3.724945049 | 0.009949859 |
| Ppapdc1a | 43.3421754 | -0.251072736 | 0.250720651 | -1.001404291 |
| Wdr11 | 633.8080906 | 0.098723605 | 0.065978521 | 1.496299162 |
| Gm23847 | 0 | NA | NA | NA |
| Fgfr2 | 819.5441903 | -0.3019833 | 0.097213552 | -3.106390964 |
| Gm5903 | 24.16642656 | -0.200870705 | 0.221721029 | -0.905961452 |
| Ate1 | 1165.905196 | 0.068917629 | 0.057524125 | 1.198064785 |
| Gm4275 | 0.372589196 | -0.91620467 | 1.563097316 | -0.586146916 |
| Gm27884 | 0.4630892 | 1.729471756 | 1.490371298 | 1.160430128 |
| Nsmce4a | 299.1185009 | -0.028409665 | 0.11604261 | -0.244820975 |
| Tacc2 | 858.9102761 | 0.442472673 | 0.076264033 | 5.801852525 |

| | | | | |
|---|---|---|---|---|
| Etos1 | 0 | NA | NA | NA |
| Btbd16 | 23.45110209 | -1.22646996 | 0.257553794 | -4.761995316 |
| Gm5602 | 13.63298114 | 0.343661257 | 0.303097941 | 1.133829075 |
| Plekha1 | 1609.865415 | 0.091365633 | 0.071644234 | 1.275268481 |
| mmu-mir-7061 | 2.359287033 | 0.829938586 | 0.774388717 | 1.071733831 |
| Htra1 | 1343.622786 | 0.254927537 | 0.147633652 | 1.72675764 |
| Dmbt1 | 0.462783582 | -0.634467323 | 3.366247751 | -0.188479093 |
| 4933402N03Rik | 0.061920613 | -0.443816764 | 3.724945049 | -0.119147198 |
| 1700029B22Rik | 0 | NA | NA | NA |
| 5430419D17Rik | 1.767964381 | -0.123960687 | 0.854317947 | -0.145099008 |
| Cuzd1 | 2.321753028 | -0.77238482 | 0.663743775 | -1.163679193 |
| 1700007K09Rik | 3.665048522 | 1.699233663 | 0.610103586 | 2.785156 |
| Fam24a | 0 | NA | NA | NA |
| 2310057M21Rik | 207.8650457 | 0.032479296 | 0.130105205 | 0.249638714 |
| Pstk | 158.2756739 | 0.086223715 | 0.116244476 | 0.741744622 |
| Ikzf5 | 441.795929 | -0.107522375 | 0.082372777 | -1.305314434 |
| Acadsb | 717.8726284 | 0.070806992 | 0.086271417 | 0.820746833 |
| Hmx3 | 4.229700681 | -0.594101992 | 0.610881658 | -0.972532052 |
| Hmx2 | 1.376538491 | 0.691044202 | 0.945704983 | 0.73071858 |
| Bub3 | 556.6642844 | -0.194502903 | 0.081751252 | -2.379203965 |
| Gm16477 | 27.01530151 | -0.279281553 | 0.222268118 | -1.256507483 |
| Gpr26 | 2155.295409 | 0.499786779 | 0.17829866 | 2.803087691 |
| Cpxm2 | 34.2720618 | -0.312020951 | 0.200431815 | -1.556743624 |
| n-R5s159 | 0.155551505 | 0.539747996 | 2.83133048 | 0.19063405 |
| Gm15677 | 0.494008362 | -0.795521604 | 1.519749242 | -0.523455832 |
| Chst15 | 526.7601878 | 0.273694371 | 0.09042272 | 3.026831887 |
| Oat | 670.5788141 | -0.097171872 | 0.098574503 | -0.985770854 |
| Nkx1-2 | 0.886311283 | -0.609585092 | 1.245833978 | -0.489298817 |
| Gm16764 | 0 | NA | NA | NA |
| Lhpp | 114.0965916 | 0.078010237 | 0.127094029 | 0.613799387 |
| Gm15582 | 0.853586773 | -0.929549321 | 0.96398717 | -0.964275616 |
| Fam53b | 689.2067909 | -0.011290304 | 0.085174355 | -0.132555202 |
| Mettl10 | 199.6988259 | 0.137611418 | 0.097487804 | 1.411575727 |
| Fam175b | 351.9015338 | -0.252359846 | 0.073606505 | -3.428499232 |
| 1500002F19Rik | 8.219991273 | 0.368238938 | 0.367194716 | 1.002843783 |
| Zranb1 | 1066.949998 | 0.23717995 | 0.076944992 | 3.082461188 |
| Gm15718 | 1.397387052 | 1.669307332 | 0.855616041 | 1.951000509 |
| Ctbp2 | 328.6048763 | -0.301091482 | 0.092234078 | -3.264427739 |
| Tex36 | 0.64913551 | -0.047798439 | 1.186138607 | -0.040297516 |
| Gm28747 | 0.061920613 | -0.443816764 | 3.724945049 | -0.119147198 |
| Edrf1 | 475.154792 | -0.128274888 | 0.124504654 | -1.03028187 |
| Mmp21 | 0.96439918 | 0.37008986 | 1.09076992 | 0.339292323 |
| Uros | 150.8894574 | -0.023840873 | 0.117408659 | -0.203058896 |
| Bccip | 345.0185436 | -0.24695861 | 0.081661247 | -3.024183694 |
| Dhx32 | 391.5252559 | -0.027815793 | 0.075696494 | -0.36746475 |
| Gm15483 | 5.172158321 | -0.085277915 | 0.523020968 | -0.16304875 |
| Fank1 | 22.86474296 | 0.453048989 | 0.254812337 | 1.777971167 |

| | | | | |
|---|---|---|---|---|
| Adam12 | 174.7221536 | -1.568521638 | 0.18741949 | -8.369042299 |
| Gm23631 | 0.152314784 | 0.51796321 | 2.84727539 | 0.181915389 |
| A130023I24Rik | 1.325949706 | -2.46225697 | 0.968391192 | -2.542626358 |
| D7Ertd443e | 78.15056778 | 0.461157968 | 0.194296011 | 2.373481392 |
| Dock1 | 717.8821439 | -0.527495858 | 0.079988155 | -6.594674666 |
| Fam196a | 171.3787744 | -0.334108959 | 0.128205246 | -2.606047492 |
| Gm24581 | 0 | NA | NA | NA |
| 1700120G07Rik | 0.35282422 | -2.312219052 | 3.683638228 | -0.62769982 |
| Nps | 2.222986028 | 0.352581055 | 0.671451983 | 0.525102411 |
| 4930544L04Rik | 0.117650406 | -0.443821212 | 3.724945049 | -0.119148392 |
| Foxi2 | 0.120349074 | 1.027495679 | 3.723011807 | 0.275985071 |
| Clrn3 | 0 | NA | NA | NA |
| Ptpre | 396.1280751 | -0.050875992 | 0.097668735 | -0.52090356 |
| Mir1962 | 1.121758522 | -0.008787344 | 1.056255896 | -0.008319333 |
| 5830432E09Rik | 5.577809048 | -0.927275566 | 0.411911811 | -2.251150711 |
| Mki67 | 177.6667962 | -0.035941665 | 0.244279867 | -0.147133145 |
| Gm9341 | 0 | NA | NA | NA |
| Gm24444 | 0 | NA | NA | NA |
| Mgmt | 8.081835135 | -0.208810643 | 0.343007301 | -0.608764428 |
| Ebf3 | 140.1280894 | -0.030851764 | 0.166915991 | -0.184834085 |
| 9430038I01Rik | 31.06055986 | 0.006919315 | 0.244999536 | 0.028242154 |
| Glrx3 | 666.4075028 | -0.300707124 | 0.105937081 | -2.838544536 |
| Gm25798 | 0 | NA | NA | NA |
| Tcerg1l | 114.589252 | -0.048672163 | 0.137631564 | -0.353640991 |
| Mapk1ip1 | 278.2027142 | 0.26131559 | 0.112092904 | 2.331241148 |
| Ppp2r2d | 650.2346655 | -0.032372848 | 0.116274843 | -0.278416613 |
| Bnip3 | 859.8363811 | 0.359208928 | 0.0868102 | 4.137865443 |
| Gm9358 | 0.765055553 | 0.55164643 | 1.334961536 | 0.413230205 |
| Jakmip3 | 408.0351068 | 0.15552873 | 0.141328315 | 1.100478203 |
| Dpysl4 | 522.9033322 | 0.886008687 | 0.108607217 | 8.157917209 |
| Stk32c | 865.5787329 | 0.261211534 | 0.076100839 | 3.432439612 |
| Gm27975 | 0 | NA | NA | NA |
| Lrrc27 | 79.21060963 | 0.292300387 | 0.146065558 | 2.001158876 |
| Pwwp2b | 187.2201883 | 0.153890637 | 0.101932812 | 1.509726202 |
| Inpp5a | 914.9916766 | -0.05618241 | 0.105280887 | -0.533643012 |
| Nkx6-2 | 248.995156 | -0.509182983 | 0.168873737 | -3.015169751 |
| Cfap46 | 121.7543508 | 0.349072324 | 0.172282196 | 2.02616598 |
| Gm4459 | 68.82999537 | -0.169362706 | 0.161090308 | -1.051352547 |
| Rpl21-ps13 | 0 | NA | NA | NA |
| Mir6401 | 0 | NA | NA | NA |
| Adgra1 | 707.4917962 | 0.442416939 | 0.07252611 | 6.100105769 |
| Kndc1 | 2147.099969 | 0.453958752 | 0.071686047 | 6.332595664 |
| Utf1 | 1.176348267 | -0.48446547 | 0.953476704 | -0.508104149 |
| Gm16201 | 8.156929821 | -0.012351497 | 0.377157685 | -0.032748893 |
| Gm2044 | 0.046345709 | -0.443818988 | 3.724945049 | -0.119147795 |
| mmu-mir-7686 | 0 | NA | NA | NA |
| Mir202 | 0 | NA | NA | NA |

| | | | | |
|---|---|---|---|---|
| 6430531B16Rik | 3.025363873 | 1.465460233 | 0.639708528 | 2.290824911 |
| Adam8 | 325.8439531 | -1.917743981 | 0.48220828 | -3.977003423 |
| Mir7062 | 0 | NA | NA | NA |
| Tubgcp2 | 416.699423 | 0.245408828 | 0.074295999 | 3.303123066 |
| Zfp511 | 95.06049784 | -0.060866436 | 0.121554207 | -0.500734918 |
| Msx3 | 0.762104533 | -0.962470017 | 1.34045209 | -0.718018961 |
| Caly | 767.6017605 | 0.801640416 | 0.075390779 | 10.63313614 |
| Prap1 | 0 | NA | NA | NA |
| Fuom | 141.6570441 | -0.063796087 | 0.115771257 | -0.551052905 |
| Echs1 | 365.0393526 | 0.186050681 | 0.099790427 | 1.864414126 |
| Paox | 52.9831637 | 0.059924199 | 0.152888633 | 0.391946721 |
| Mtg1 | 103.7910862 | 0.146418095 | 0.128362094 | 1.140664591 |
| Sprn | 1168.69905 | 0.727923918 | 0.143854696 | 5.060133163 |
| Olfr522 | 0 | NA | NA | NA |
| Olfr523 | 0.038981395 | -0.443816764 | 3.724945049 | -0.119147198 |
| Olfr524 | 0.042716945 | 0.037062677 | 3.724945049 | 0.009949859 |
| Cd163l1 | 0.514333851 | 2.268962106 | 3.210765874 | 0.706673173 |
| 5830411N06Rik | 0.560014729 | -0.340272536 | 1.297702043 | -0.262211605 |
| Olfr525 | 0.068097359 | 0.037064902 | 3.724945049 | 0.009950456 |
| Olfr527 | 0.1567526 | 0.008815624 | 2.825354425 | 0.003120183 |
| Olfr60 | 0.072025462 | 0.517959912 | 3.724945049 | 0.139051692 |
| Olfr530 | 0.095435502 | 0.037073798 | 3.724945049 | 0.009952844 |
| Olfr531 | 0.095898756 | -0.917692504 | 3.724434845 | -0.246397787 |
| Olfr532 | 0.042716945 | 0.037062677 | 3.724945049 | 0.009949859 |
| Olfr533 | 0.120974994 | 0.517957688 | 3.724945049 | 0.139051095 |
| Olfr535 | 0 | NA | NA | NA |
| Olfr536 | 0.638507312 | -0.056758407 | 1.391912522 | -0.04077728 |
| Olfr538 | 0 | NA | NA | NA |
| Olfr46 | 0.111376404 | 0.517957688 | 3.724945049 | 0.139051095 |
| Olfr61 | 0.224318018 | 0.037067975 | 2.613625264 | 0.01418259 |
| Olfr53 | 0.178643884 | 1.02014301 | 2.710504853 | 0.376366421 |
| Olfr539 | 0.207883488 | 0.517969368 | 2.199582986 | 0.235485259 |
| Olfr45 | 0.066050875 | -0.443818988 | 3.724945049 | -0.119147795 |
| Olfr541 | 0 | NA | NA | NA |
| Cyp2e1 | 1.071126076 | 1.036794874 | 1.159095402 | 0.894486228 |
| Syce1 | 4.365775959 | -1.087960729 | 0.628349572 | -1.731457738 |
| Zfp941 | 465.5985334 | 0.415416594 | 0.102503454 | 4.052708256 |
| Urah | 8.499830305 | 0.039783981 | 0.360050497 | 0.11049556 |
| C330022C24Rik | 1.303324972 | -0.958871345 | 0.943485431 | -1.016307526 |
| Scgb1c1 | 1.058089029 | -0.928103038 | 1.187659431 | -0.781455537 |
| Odf3 | 0.418471408 | -0.849067166 | 1.742807888 | -0.487183454 |
| Bet1l | 215.3921604 | -0.270636614 | 0.09842978 | -2.749539967 |
| Ric8 | 517.4466835 | -0.051037941 | 0.07388084 | -0.6908143 |
| Sirt3 | 296.3129455 | 0.160779186 | 0.113079011 | 1.421830493 |
| Psmd13 | 560.5188265 | -0.284453669 | 0.097189025 | -2.926808538 |
| Gm15542 | 83.64766115 | -0.280660024 | 0.171776047 | -1.633871712 |
| Cox8b | 4.291398722 | 1.45376476 | 0.651978287 | 2.229774807 |

| | | | | |
|---|---|---|---|---|
| Nlrp6 | 8.710158728 | 1.491912675 | 0.408605306 | 3.651231769 |
| Athl1 | 114.0507651 | -0.501696938 | 0.152783092 | -3.283720286 |
| Ifitm5 | 0.496272132 | -0.152289464 | 1.233284292 | -0.123482854 |
| Ifitm2 | 214.7202724 | -0.715063306 | 0.22346384 | -3.1999061 |
| Ifitm1 | 100.7816241 | -0.714236332 | 0.244541783 | -2.920712866 |
| Ifitm3 | 2374.688831 | -0.774646639 | 0.256730908 | -3.017348573 |
| Ifitm6 | 40.88379849 | -1.669581648 | 0.527532425 | -3.164889152 |
| Gm17387 | 0.276680467 | -1.318302158 | 1.752697987 | -0.752155915 |
| B4galnt4 | 713.1671909 | 0.356015952 | 0.108459234 | 3.28248631 |
| Pkp3 | 31.38673572 | -0.335385033 | 0.249942612 | -1.341848157 |
| Sigirr | 8.814058262 | -1.401314911 | 0.408990625 | -3.426276364 |
| Ano9 | 0.759896141 | -1.051005132 | 1.097878868 | -0.957305184 |
| Gm24962 | 0 | NA | NA | NA |
| Ptdss2 | 397.5875048 | -0.000307134 | 0.083024234 | -0.003699326 |
| Rnh1 | 785.7956867 | -1.201208556 | 0.191039213 | -6.287759127 |
| Gm22019 | 0.061920613 | -0.443816764 | 3.724945049 | -0.119147198 |
| Hras | 652.7768577 | 0.061166249 | 0.082246037 | 0.74369843 |
| Lrrc56 | 44.78247708 | 0.146881697 | 0.218090819 | 0.673488675 |
| Lmntd2 | 1.504148252 | -0.10297365 | 0.79348644 | -0.129773673 |
| Rassf7 | 15.32468764 | -0.440015977 | 0.282106456 | -1.559751532 |
| Mir210 | 0.042716945 | 0.037062677 | 3.724945049 | 0.009949859 |
| Phrf1 | 827.2765508 | -0.210028526 | 0.056608973 | -3.710163162 |
| Irf7 | 4351.088714 | -0.88782771 | 0.229890025 | -3.861967089 |
| Cdhr5 | 0.766219994 | -1.566554293 | 1.084201338 | -1.444892418 |
| Sct | 31.79202062 | 0.007928829 | 0.399750428 | 0.019834447 |
| Drd4 | 5.233666748 | -0.954379118 | 0.486296984 | -1.962543772 |
| Deaf1 | 397.411066 | 0.515038077 | 0.083199423 | 6.190404437 |
| Tmem80 | 167.3778283 | -0.001780241 | 0.1110577 | -0.016029871 |
| Eps8l2 | 83.71994531 | 0.221849773 | 0.149851167 | 1.480467441 |
| B230206H07Rik | 17.64932275 | 0.087158286 | 0.398323721 | 0.218812692 |
| Gm25725 | 0 | NA | NA | NA |
| Taldo1 | 760.0746265 | -0.5534965 | 0.115007139 | -4.812714273 |
| Gm4535 | 4.201928798 | -0.806963828 | 0.543411123 | -1.484996889 |
| Pddc1 | 201.5225707 | 0.228843293 | 0.111843507 | 2.046102619 |
| Cend1 | 1948.915175 | 0.325542164 | 0.105811565 | 3.076621755 |
| Slc25a22 | 2538.937697 | 0.317167204 | 0.125443485 | 2.528367291 |
| Pidd1 | 23.70766093 | -0.438831867 | 0.256872949 | -1.708361539 |
| Rplp2 | 1141.878145 | -0.307308117 | 0.093401121 | -3.290197301 |
| Snora52 | 0 | NA | NA | NA |
| Pnpla2 | 482.1622727 | 0.199545137 | 0.19277458 | 1.035121627 |
| Cracr2b | 10.14241748 | -0.073049115 | 0.340158869 | -0.214749993 |
| Gm10575 | 13.15716283 | -1.136213222 | 0.318178535 | -3.570992683 |
| Cd151 | 360.834311 | -1.016939703 | 0.194470077 | -5.229286262 |
| Polr2l | 159.7689273 | -0.329353041 | 0.157896247 | -2.085882635 |
| Tspan4 | 266.6277742 | -0.534754583 | 0.21845957 | -2.447842329 |
| Chid1 | 475.4024328 | -0.092693963 | 0.120528684 | -0.76906144 |
| Ap2a2 | 2741.755715 | 0.018477658 | 0.057575128 | 0.320931263 |

| | | | | |
|---|---|---|---|---|
| Gm16982 | 0.330744509 | 0.506643087 | 1.766158217 | 0.286861665 |
| Mir7063 | 2.38820333 | 0.845958836 | 0.661566458 | 1.27872087 |
| Muc6 | 0.236992836 | -0.793181525 | 2.067263361 | -0.383686733 |
| Muc2 | 8.969982396 | -1.603144021 | 0.513734739 | -3.120567678 |
| Muc5ac | 0.957274486 | -2.681844018 | 1.204415943 | -2.226675954 |
| Muc5b | 2.416483532 | 0.832691536 | 1.060886131 | 0.784901896 |
| Tollip | 1722.402167 | -0.007303626 | 0.063483649 | -0.115047352 |
| Gm20501 | 7.216052579 | 0.369212287 | 0.426921499 | 0.864824771 |
| Brsk2 | 1833.13049 | 0.513108383 | 0.086726722 | 5.916381642 |
| Mir3104 | 0.562632007 | -0.444875629 | 1.372852036 | -0.324052132 |
| Mob2 | 221.1798062 | -0.004641032 | 0.081791822 | -0.056742011 |
| Gm25416 | 0.290864751 | -0.924714556 | 2.059811327 | -0.448931678 |
| Dusp8 | 1691.653108 | 0.328105846 | 0.079910686 | 4.105907001 |
| Gm25370 | 0.286235171 | -1.374375549 | 1.619419991 | -0.848683823 |
| Gm4553 | 0 | NA | NA | NA |
| Krtap5-2 | 0 | NA | NA | NA |
| Gm10153 | 0 | NA | NA | NA |
| Krtap5-3 | 0.115000406 | -0.443821212 | 3.724945049 | -0.119148392 |
| Gm7579 | 0 | NA | NA | NA |
| Krtap5-5 | 0 | NA | NA | NA |
| Gm2431 | 0.097139547 | -0.443816764 | 3.724945049 | -0.119147198 |
| Gm4559 | 0 | NA | NA | NA |
| Gm10013 | 0 | NA | NA | NA |
| Krtap5-1 | 0.044544648 | 0.517957688 | 3.724945049 | 0.139051095 |
| Krtap5-4 | 0 | NA | NA | NA |
| Gm5054 | 0.042716945 | 0.037062677 | 3.724945049 | 0.009949859 |
| Ifitm10 | 182.7874866 | -0.388734759 | 0.117198082 | -3.316903757 |
| Ctsd | 5752.611281 | -0.659647845 | 0.170225301 | -3.8751457 |
| Gm26143 | 0.044544648 | 0.517957688 | 3.724945049 | 0.139051095 |
| Syt8 | 0.543884849 | -1.350783789 | 1.228137019 | -1.099864077 |
| Tnni2 | 2.708583151 | -0.932901377 | 0.585098563 | -1.594434573 |
| Lsp1 | 166.316609 | -0.717928936 | 0.168776005 | -4.253738187 |
| Prr33 | 0.642141737 | 0.066961442 | 1.336158257 | 0.050114904 |
| Tnnt3 | 1.402656864 | -0.047180472 | 0.864683597 | -0.054563857 |
| Mrpl23 | 104.8815584 | -0.08418894 | 0.125009015 | -0.67346295 |
| Nctc1 | 0 | NA | NA | NA |
| H19 | 64.97075824 | -0.941911423 | 0.305269686 | -3.085505919 |
| Gm27483 | 0.184422046 | -0.444673742 | 2.680605813 | -0.16588554 |
| Mir675 | 0 | NA | NA | NA |
| Gm27786 | 0.110595523 | 0.037071574 | 3.724945049 | 0.009952247 |
| Igf2 | 848.6181956 | 0.400471766 | 0.180475006 | 2.218987405 |
| Mir483 | 0 | NA | NA | NA |
| Igf2os | 4.725164502 | 0.993197852 | 0.48477858 | 2.048765959 |
| Ins2 | 0.386429078 | 0.155227166 | 1.421971407 | 0.109163353 |
| Gm7290 | 0.08486385 | 0.037730769 | 3.724898049 | 0.010129343 |
| Gm6471 | 0 | NA | NA | NA |
| Th | 127.7817552 | -0.379343333 | 0.263392086 | -1.440222973 |

| | | | | |
|---|---|---|---|---|
| Ascl2 | 2.369287799 | 0.063470427 | 0.883096925 | 0.071872549 |
| Tspan32os | 6.17140379 | -0.580284803 | 0.465769134 | -1.245863585 |
| Tspan32 | 1.376815522 | -0.821141592 | 0.844736993 | -0.972067754 |
| R74862 | 40.83695903 | 0.827683002 | 0.206410726 | 4.009883688 |
| Cd81 | 3734.509123 | 0.155012685 | 0.106462519 | 1.456030592 |
| Trpm5 | 0.502574849 | -1.805905727 | 3.387646004 | -0.533085725 |
| Tssc4 | 336.1484664 | -0.134796075 | 0.085377031 | -1.578833012 |
| Kcnq1 | 3.259092159 | -0.587124545 | 0.557926567 | -1.052333013 |
| Gm38321 | 0 | NA | NA | NA |
| Gm37235 | 0.046345709 | -0.443818988 | 3.724945049 | -0.119147795 |
| Gm28821 | 0 | NA | NA | NA |
| Kcnq1ot1 | 4980.818381 | 0.14372192 | 0.250703605 | 0.573274246 |
| Gm27395 | 0.113999573 | -0.443821212 | 3.724945049 | -0.119148392 |
| Gm27389 | 0.179789776 | 0.037071311 | 2.267559906 | 0.016348548 |
| Gm27803 | 0 | NA | NA | NA |
| Gm27539 | 0.072025462 | 0.517959912 | 3.724945049 | 0.139051692 |
| Gm37364 | 1.783218599 | 0.5439807 | 0.831412035 | 0.654285333 |
| 4933417O13Rik | 0.382744327 | 0.037076162 | 1.50992271 | 0.024555007 |
| Gm38095 | 0.038981395 | -0.443816764 | 3.724945049 | -0.119147198 |
| Gm38065 | 0 | NA | NA | NA |
| Gm27901 | 0.136963818 | 0.037074342 | 3.599544405 | 0.010299732 |
| Cdkn1c | 70.29508147 | 0.672249086 | 0.150261777 | 4.473852883 |
| Slc22a18 | 7.798117052 | -0.433968777 | 0.458625033 | -0.946238748 |
| Phlda2 | 0.124308499 | -1.207977564 | 3.71482282 | -0.325177706 |
| Nap1l4 | 877.3075028 | 0.04413372 | 0.083876686 | 0.526173866 |
| Gm23297 | 5.403433764 | 0.169405786 | 0.563374949 | 0.300698116 |
| Cars | 570.4979151 | -0.530968634 | 0.113425407 | -4.681214252 |
| Mir7064 | 0.434125078 | 0.037077926 | 1.399370129 | 0.026496153 |
| Gm15580 | 0 | NA | NA | NA |
| Gm15579 | 0.072025462 | 0.517959912 | 3.724945049 | 0.139051692 |
| Tnfrsf26 | 16.36698174 | -1.709243007 | 0.531066645 | -3.218509435 |
| Tnfrsf22 | 17.56781374 | -0.873068301 | 0.261134812 | -3.343362357 |
| Tnfrsf23 | 21.91112602 | -1.685691705 | 0.297304614 | -5.669914369 |
| Osbpl5 | 343.6236498 | -0.239317071 | 0.075116241 | -3.185956418 |
| Mrgprg | 0.045882455 | 0.517959912 | 3.724945049 | 0.139051692 |
| Mrgpre | 90.26427259 | -0.00021576 | 0.140202971 | -0.001538909 |
| Gm22064 | 0.123841226 | -1.017823543 | 3.719440177 | -0.273649661 |
| Nadsyn1 | 56.75566904 | -0.560676941 | 0.195456465 | -2.868551529 |
| Dhcr7 | 321.2309411 | -0.271985348 | 0.106422632 | -2.555709655 |
| Gm498 | 0.049553047 | -0.443818988 | 3.724945049 | -0.119147795 |
| Mir6404 | 0 | NA | NA | NA |
| Shank2 | 2746.162572 | 0.560682095 | 0.117332961 | 4.778555764 |
| Mir3105 | 0.102891481 | 0.51795324 | 3.724945049 | 0.139049901 |
| Gm14372 | 0.097899064 | 0.037062677 | 3.724945049 | 0.009949859 |
| Cttn | 1215.2151 | -0.379796832 | 0.075533372 | -5.028199074 |
| Gm14376 | 6.097514919 | 0.52784732 | 0.618755113 | 0.853079529 |
| Ppfia1 | 744.1652023 | -0.018835453 | 0.082681836 | -0.227806425 |

| | | | | |
|---|---|---|---|---|
| Faddos | 0.049553047 | -0.443818988 | 3.724945049 | -0.119147795 |
| Fadd | 59.54792491 | -0.830628509 | 0.177424135 | -4.681598176 |
| Ano1 | 32.22433977 | 0.191196389 | 0.245616824 | 0.7784336 |
| Gm10152 | 0 | NA | NA | NA |
| Fgf3 | 4.236074083 | -0.44696394 | 0.60365918 | -0.740424323 |
| Fgf4 | 0 | NA | NA | NA |
| Gm26793 | 3.941161089 | 1.193736755 | 0.546507213 | 2.18430192 |
| Fgf15 | 0.844302062 | 0.786551944 | 1.13243645 | 0.694566078 |
| Oraov1 | 196.1845682 | -0.14966463 | 0.081735178 | -1.831091992 |
| Ccnd1 | 896.4298962 | -1.470655938 | 0.242730259 | -6.058807601 |
| Tpcn2 | 87.43800306 | -0.786619663 | 0.199636819 | -3.940253443 |
| Mrgprf | 52.34489822 | -0.384553778 | 0.259520856 | -1.481783711 |
| Mrgprd | 0.203212381 | -0.402877374 | 2.635685714 | -0.152854861 |
| Gm7346 | 0 | NA | NA | NA |
| Gm25775 | 0 | NA | NA | NA |
| Insr | 1190.786105 | 0.077027582 | 0.056937925 | 1.352834379 |
| Gm16180 | 0 | NA | NA | NA |
| A430078G23Rik | 4.533747209 | 0.032681907 | 0.513701478 | 0.063620427 |
| Arhgef18 | 750.8432807 | -0.124008721 | 0.073416173 | -1.689119935 |
| Pex11g | 11.94680139 | -0.536852071 | 0.381377764 | -1.407664845 |
| 1700019B03Rik | 3.816282625 | 0.613188546 | 0.548983176 | 1.116953255 |
| Zfp358 | 245.1757097 | 0.290560755 | 0.110794471 | 2.622520355 |
| Mcoln1 | 510.2888465 | 0.06485854 | 0.059901325 | 1.082756352 |
| Pnpla6 | 618.1933405 | 0.10982498 | 0.092003141 | 1.193709032 |
| C330021F23Rik | 0.687005404 | 0.955886881 | 1.335866491 | 0.715555699 |
| Rps23-ps2 | 34.30525878 | -0.165548868 | 0.185411875 | -0.892870899 |
| Camsap3 | 321.1724317 | 0.184350518 | 0.098652106 | 1.868693178 |
| Xab2 | 460.0137195 | -0.122067324 | 0.070729735 | -1.725827538 |
| Gm22204 | 0.066050875 | -0.443818988 | 3.724945049 | -0.119147795 |
| Pet100 | 155.4280518 | 0.049190094 | 0.115254371 | 0.426795904 |
| Pcp2 | 337.4770539 | 0.0684063 | 0.413548708 | 0.165412922 |
| Stxbp2 | 289.3398245 | -0.039864646 | 0.141857985 | -0.281017994 |
| Retn | 3.230460487 | -2.096860639 | 0.721429925 | -2.906534048 |
| Mcemp1 | 19.01625202 | -0.933924119 | 0.484474717 | -1.927704555 |
| Fcor | 19.48329625 | 0.043576832 | 0.322720539 | 0.13502962 |
| Trappc5 | 363.4451168 | 0.00130153 | 0.087230872 | 0.014920517 |
| Fcer2a | 1.327801032 | 0.93192488 | 0.971078171 | 0.959680598 |
| Clec4g | 0.624219342 | -1.236458103 | 1.308984789 | -0.944593178 |
| Gm16553 | 0.238000031 | 1.464118599 | 2.464171798 | 0.59416255 |
| Gm23886 | 0 | NA | NA | NA |
| Cd209a | 1.799118211 | -1.86217124 | 0.846812261 | -2.199036701 |
| Gm24934 | 0 | NA | NA | NA |
| Rprl3 | 2178.394221 | -0.267107002 | 0.18336546 | -1.456692023 |
| Gm7381 | 0.061920613 | -0.443816764 | 3.724945049 | -0.119147198 |
| Cd209e | 0 | NA | NA | NA |
| Cd209d | 0.062426873 | 0.517959912 | 3.724945049 | 0.139051692 |
| Cd209b | 0.371584567 | 0.03707491 | 1.722170753 | 0.021528011 |

| | | | | |
|---|---|---|---|---|
| Cd209c | 0.190986542 | -0.934215115 | 2.702582435 | -0.345674975 |
| Gm7389 | 8.537660985 | -0.512240011 | 0.404029645 | -1.267827785 |
| Gm16589 | 85.35553407 | -0.09186744 | 0.230716245 | -0.398183663 |
| Gm26750 | 0 | NA | NA | NA |
| Cd209f | 0.631752858 | 0.028428521 | 1.33999871 | 0.021215334 |
| Gm23888 | 0 | NA | NA | NA |
| Cd209g | 0.34920187 | -0.976494745 | 1.727287267 | -0.565334304 |
| 4932443L11Rik | 1.299081245 | 0.73792139 | 0.849166828 | 0.868994603 |
| Evi5l | 1072.773425 | 0.231405392 | 0.07592017 | 3.048009386 |
| Prr36 | 298.1056952 | 0.759718064 | 0.125383613 | 6.059149543 |
| Lrrc8e | 1.436219828 | -2.131381823 | 0.973270905 | -2.189916305 |
| Map2k7 | 1090.223813 | 0.170531481 | 0.059445822 | 2.868687412 |
| Tgfbr3l | 14.69813157 | 0.875170005 | 0.429788663 | 2.036279873 |
| Snapc2 | 202.0178337 | -0.286383632 | 0.11820635 | -2.422743212 |
| Ctxn1 | 1372.549575 | 1.109374853 | 0.124669169 | 8.898550156 |
| Timm44 | 376.9137615 | 0.131960536 | 0.104636991 | 1.261127017 |
| Gm26102 | 0 | NA | NA | NA |
| Elavl1 | 1614.161581 | -0.128124624 | 0.053614578 | -2.38973482 |
| Ccl25 | 51.47610217 | 0.406120238 | 0.202252201 | 2.007989212 |
| Cers4 | 639.9032004 | 0.380245315 | 0.124912389 | 3.044096081 |
| Gm36956 | 0.066050875 | -0.443818988 | 3.724945049 | -0.119147795 |
| Zfp958 | 156.442319 | -0.132065378 | 0.115634212 | -1.142096056 |
| Gm7461 | 2.204253537 | -0.437731255 | 0.742815197 | -0.589286887 |
| Gm6410 | 0.983694817 | 1.546669534 | 1.095172635 | 1.41226094 |
| Shcbp1 | 11.59037295 | -0.128236477 | 0.437933608 | -0.29282173 |
| Gm9457 | 0.283539495 | 0.050971799 | 2.395469754 | 0.021278415 |
| Rpl21-ps14 | 0.417309641 | 0.359149646 | 1.764721593 | 0.203516321 |
| n-R5s93 | 0 | NA | NA | NA |
| Slc10a2 | 0 | NA | NA | NA |
| n-R5s94 | 0 | NA | NA | NA |
| Gm1840 | 36.14079938 | -0.092058518 | 0.177050367 | -0.519956663 |
| Gm22565 | 0 | NA | NA | NA |
| Gm24288 | 0 | NA | NA | NA |
| Gm17215 | 0.061920613 | -0.443816764 | 3.724945049 | -0.119147198 |
| Gm26427 | 0.673557475 | 0.875147265 | 1.201699965 | 0.728257711 |
| Gm27585 | 0 | NA | NA | NA |
| Gm27545 | 0 | NA | NA | NA |
| Gm25169 | 0 | NA | NA | NA |
| Gm28262 | 0 | NA | NA | NA |
| Efnb2 | 602.2040592 | 0.411200099 | 0.074744753 | 5.501390848 |
| 4921522P10Rik | 0.133946075 | 0.036100024 | 3.626570338 | 0.009954315 |
| Arglu1 | 500.6182152 | 0.22368664 | 0.07148336 | 3.129212706 |
| Mir7654 | 0 | NA | NA | NA |
| Fam155a | 942.9312957 | 0.585135503 | 0.068802551 | 8.504561098 |
| n-R5s95 | 14.75670795 | 0.638781815 | 0.547846363 | 1.165987143 |
| AC099760.1 | 0 | NA | NA | NA |
| Gm10217 | 0.045882455 | 0.517959912 | 3.724945049 | 0.139051692 |

| | | | | |
|---|---|---|---|---|
| Gm10067 | 0 | NA | NA | NA |
| Lig4 | 157.1064013 | -0.451082703 | 0.139775376 | -3.227197212 |
| Abhd13 | 722.6949606 | 0.101943757 | 0.08432533 | 1.208933976 |
| Tnfsf13b | 20.67774298 | 0.046313585 | 0.23161578 | 0.199958675 |
| Myo16 | 882.108668 | -0.086225148 | 0.113002603 | -0.763036833 |
| Rps16-ps3 | 0 | NA | NA | NA |
| 3930402G23Rik | 10.57984639 | -1.187555238 | 0.37712138 | -3.149000029 |
| Irs2 | 1258.437026 | 0.098192437 | 0.130505277 | 0.752402046 |
| 9530052E02Rik | 11.90754286 | 0.629714857 | 0.367042806 | 1.71564419 |
| Gm15418 | 2.963241567 | 1.608571561 | 0.786092589 | 2.046287656 |
| Col4a1 | 1092.321152 | -0.541869171 | 0.162415589 | -3.336312558 |
| Col4a2 | 772.5542793 | -0.301280121 | 0.158949626 | -1.895444047 |
| Gm15419 | 0.932741312 | -0.624699131 | 1.061367836 | -0.588579294 |
| Rab20 | 71.74611719 | -0.995372132 | 0.318228436 | -3.127854139 |
| E230013L22Rik | 6.9128239 | -0.828936822 | 0.4100387 | -2.021606306 |
| Carkd | 279.2439548 | -0.057285125 | 0.073600916 | -0.778320814 |
| Cars2 | 87.5486135 | -0.170090861 | 0.132304288 | -1.285603547 |
| Ing1 | 228.1766961 | 0.117911384 | 0.104582532 | 1.127448167 |
| Ankrd10 | 320.9466642 | -0.098173879 | 0.116058972 | -0.845896504 |
| mmu-mir-7238 | 0 | NA | NA | NA |
| 1700016D06Rik | 0.194626871 | 1.57110533 | 2.644262968 | 0.594156235 |
| Gm27854 | 0 | NA | NA | NA |
| 1700128E19Rik | 0.068097359 | 0.037064902 | 3.724945049 | 0.009950456 |
| Arhgef7 | 1845.245703 | -0.102515007 | 0.067832266 | -1.511301526 |
| Gm15875 | 0 | NA | NA | NA |
| Tex29 | 5.082148535 | -1.23527789 | 0.634207081 | -1.947751652 |
| A230072I06Rik | 0.719797104 | -0.956766153 | 1.453375937 | -0.658306037 |
| Gm29175 | 0.569306118 | 0.370817555 | 1.515091316 | 0.24474931 |
| Gm22207 | 0.197462999 | 0.037067291 | 2.656914844 | 0.013951253 |
| Gm5607 | 118.2838747 | 0.646381272 | 0.142449924 | 4.537603482 |
| Sox1 | 235.5645892 | 0.168406795 | 0.159049798 | 1.058830614 |
| Gm25239 | 0 | NA | NA | NA |
| D630011A20Rik | 0 | NA | NA | NA |
| Spaca7 | 0 | NA | NA | NA |
| Tubgcp3 | 535.0497003 | 0.162323318 | 0.083993966 | 1.932559277 |
| Gm26483 | 0 | NA | NA | NA |
| 4931415C17Rik | 18.23396489 | 0.073924617 | 0.293508942 | 0.251864958 |
| Gm15348 | 0.194797293 | 0.037065655 | 3.222608474 | 0.011501756 |
| Atp11a | 1406.61443 | -0.096205689 | 0.071669139 | -1.342358666 |
| Gm15350 | 0.251121349 | -1.362888131 | 2.061297508 | -0.661179731 |
| Gm15347 | 0.159763836 | 0.037068446 | 2.810388006 | 0.013189797 |
| Mcf2l | 2158.382309 | 0.245450502 | 0.0721024 | 3.404193215 |
| Gm15353 | 0.04393175 | 0.037062677 | 3.724945049 | 0.009949859 |
| Gm15351 | 6.732411598 | 0.111223267 | 0.510502019 | 0.217870377 |
| mmu-mir-9768 | 0 | NA | NA | NA |
| Gm15349 | 0 | NA | NA | NA |
| Gm15352 | 1.728865685 | -0.324210519 | 1.030097133 | -0.314737813 |

| | | | | |
|---|---|---|---|---|
| F7 | 20.47438508 | -2.360172429 | 0.383099393 | -6.160731324 |
| F10 | 60.6296221 | -1.657152694 | 0.552439641 | -2.999699097 |
| Proz | 0.405129103 | -1.555351005 | 1.52782059 | -1.018019403 |
| Gm17023 | 0.048949532 | 0.037064902 | 3.724945049 | 0.009950456 |
| Pcid2 | 338.1239794 | 0.028111794 | 0.094124923 | 0.298664726 |
| Gm17022 | 0 | NA | NA | NA |
| Cul4a | 779.7913125 | -0.132441879 | 0.054399362 | -2.434621893 |
| Mir7065 | 0.26402779 | 0.50810766 | 3.724225114 | 0.136433122 |
| Lamp1 | 3957.92154 | -0.531199713 | 0.124400387 | -4.27008088 |
| Grtp1 | 32.55675301 | -0.180929707 | 0.223201474 | -0.810611615 |
| Mir1968 | 0.779555773 | -0.458904163 | 1.293098034 | -0.354887372 |
| 2810030D12Rik | 0.867078918 | 0.523069725 | 1.31171066 | 0.398769135 |
| Adprhl1 | 5.011785424 | 0.446708532 | 0.508798227 | 0.877967941 |
| Dcun1d2 | 379.2848758 | -0.003780329 | 0.098243497 | -0.038479174 |
| Gm25014 | 4.915973584 | -0.267495169 | 0.513503307 | -0.520921998 |
| Tmco3 | 370.903991 | -0.072600775 | 0.093830326 | -0.773745311 |
| Tfdp1 | 412.9010695 | -0.005118411 | 0.0853863 | -0.059944167 |
| Atp4b | 0 | NA | NA | NA |
| Grk1 | 1.091106207 | 1.646571306 | 1.070536161 | 1.538080978 |
| Tmem255b | 12.33555406 | -0.18488921 | 0.351974213 | -0.52529192 |
| Gas6 | 898.867002 | 0.280278366 | 0.079229207 | 3.537563677 |
| 1700029H14Rik | 5.973066547 | 1.062409433 | 0.604396564 | 1.757801907 |
| Rasa3 | 1272.496422 | -0.310220076 | 0.075612086 | -4.10278424 |
| 4932443I19Rik | 8.590972356 | -1.37072848 | 0.41701061 | -3.287035025 |
| Cdc16 | 437.6639685 | -0.347108189 | 0.082910195 | -4.186556182 |
| Upf3a | 175.6020734 | -0.053522457 | 0.167474276 | -0.31958614 |
| AF366264 | 0.110595523 | 0.037071574 | 3.724945049 | 0.009952247 |
| Champ1 | 447.8929496 | -0.015055263 | 0.069274496 | -0.217327651 |
| AC139186.1 | 0 | NA | NA | NA |
| Gm24698 | 0.093141081 | 0.037060453 | 3.724945049 | 0.009949262 |
| Coprs | 191.5374814 | -0.344784253 | 0.12336083 | -2.794924892 |
| Gm7676 | 0.12936564 | 0.037055616 | 3.688232414 | 0.010046985 |
| Fbxo25 | 406.2625769 | 0.145927472 | 0.073261232 | 1.99187848 |
| Tdrp | 156.0986126 | -0.252218847 | 0.114184965 | -2.208862158 |
| Erich1 | 48.89166254 | -0.497002092 | 0.161772163 | -3.072234935 |
| Gm10699 | 61.82645713 | 0.575758971 | 0.180658018 | 3.187010341 |
| Dlgap2 | 896.9164479 | 0.354382242 | 0.183589146 | 1.930300617 |
| Gm3160 | 8.920971897 | 1.089143397 | 0.604945357 | 1.80039963 |
| Gm26184 | 0 | NA | NA | NA |
| AC130213.1 | 3.509323303 | 1.489101032 | 0.753338487 | 1.976669262 |
| C030037F17Rik | 0 | NA | NA | NA |
| Cln8 | 468.4529787 | -0.440546746 | 0.0982936 | -4.481947428 |
| Arhgef10 | 845.0217701 | -0.116512913 | 0.111367134 | -1.046205549 |
| Gm16350 | 0.676711058 | -1.244379464 | 1.326007809 | -0.938440525 |
| Kbtbd11 | 1707.888606 | 0.285489735 | 0.093513605 | 3.052921931 |
| 5830468F06Rik | 0.645565975 | -2.077587624 | 1.32017012 | -1.573727198 |
| Gm37844 | 22.02823964 | 0.074612022 | 0.244978588 | 0.304565484 |

| | | | | |
|---|---|---|---|---|
| BB014433 | 0 | NA | NA | NA |
| Myom2 | 7.622785529 | -0.133591017 | 0.400706251 | -0.333388903 |
| Gm15678 | 0 | NA | NA | NA |
| Gm24231 | 0 | NA | NA | NA |
| Gm24222 | 0 | NA | NA | NA |
| Gm25871 | 1.401691129 | 1.394943074 | 0.985332705 | 1.415707676 |
| Gm25255 | 0 | NA | NA | NA |
| Csmd1 | 2110.968194 | 0.649847063 | 0.239711624 | 2.710953485 |
| Mir3106 | 1.89102523 | 1.401189768 | 0.951151517 | 1.473150958 |
| Gm16346 | 0 | NA | NA | NA |
| Gm16347 | 0 | NA | NA | NA |
| Gm25278 | 0 | NA | NA | NA |
| Gm25665 | 0 | NA | NA | NA |
| Gm24469 | 0 | NA | NA | NA |
| Mcph1 | 236.5966961 | 0.154911551 | 0.103060091 | 1.503118717 |
| Gm25763 | 0 | NA | NA | NA |
| Gm16725 | 0 | NA | NA | NA |
| Gm17149 | 0.323327586 | 1.619732913 | 1.997767806 | 0.810771356 |
| Angpt2 | 48.84955761 | -0.90702579 | 0.29191231 | -3.107185822 |
| Agpat5 | 782.015753 | 0.206344213 | 0.070234771 | 2.937921072 |
| Xkr5 | 3.737801717 | -1.503364359 | 0.811269836 | -1.853100278 |
| Gm22551 | 0.097139547 | -0.443816764 | 3.724945049 | -0.119147198 |
| Defb40 | 0 | NA | NA | NA |
| Defb37 | 0 | NA | NA | NA |
| Defb38 | 0 | NA | NA | NA |
| Gm15366 | 0.11193333 | 0.037073798 | 3.724945049 | 0.009952844 |
| Defb39 | 0 | NA | NA | NA |
| Defb12 | 0 | NA | NA | NA |
| Defb34 | 0 | NA | NA | NA |
| Spag11b | 0.25135723 | 0.037075689 | 2.135305801 | 0.017363175 |
| Spag11a | 0.093744596 | -0.443823436 | 3.724945049 | -0.119148989 |
| Defb14 | 0 | NA | NA | NA |
| Defb4 | 0 | NA | NA | NA |
| Defa-ps12 | 0 | NA | NA | NA |
| Defb6 | 0.199888145 | 1.310060772 | 2.620047342 | 0.500014161 |
| Defb46 | 0 | NA | NA | NA |
| Defb5 | 0 | NA | NA | NA |
| Gm15357 | 0 | NA | NA | NA |
| Defb3 | 0 | NA | NA | NA |
| Defa-ps13 | 0 | NA | NA | NA |
| Defa-ps14 | 0 | NA | NA | NA |
| Defb8 | 0 | NA | NA | NA |
| Defa-ps15 | 0 | NA | NA | NA |
| Rpl19-ps11 | 69.27011738 | 0.215654746 | 0.218225597 | 0.988219298 |
| Gm27288 | 0 | NA | NA | NA |
| Defb7 | 0 | NA | NA | NA |
| Gm15358 | 0 | NA | NA | NA |

| | | | | |
|---|---|---|---|---|
| Gm27852 | 0 | NA | NA | NA |
| Gm16425 | 10.9074589 | -0.210264873 | 0.340749684 | -0.617065495 |
| Gm15359 | 1.904847302 | -0.261789278 | 0.664684853 | -0.393854737 |
| Gm6483 | 149.5723312 | 0.21855713 | 0.126446333 | 1.728457633 |
| 4930467E23Rik | 2.550781979 | 0.157666452 | 0.598255463 | 0.26354369 |
| Gm26853 | 52.27869009 | 0.128662069 | 0.231178003 | 0.55654979 |
| Gm21769 | 152.8336227 | 0.169885436 | 0.169391369 | 1.002916721 |
| Gm21811 | 150.012751 | 0.092113651 | 0.216523511 | 0.425421014 |
| 6820431F20Rik | 191.9164182 | 0.367254323 | 0.119333662 | 3.077541711 |
| Gm26804 | 0 | NA | NA | NA |
| Gm15319 | 17.21563751 | -0.129005135 | 0.350165775 | -0.368411605 |
| 2610005L07Rik | 198.0277139 | 0.076287881 | 0.214324664 | 0.355945414 |
| Gm21092 | 212.4801284 | 0.211761409 | 0.100332703 | 2.110592079 |
| Gm21182 | 0.374378724 | -1.386564292 | 1.745000566 | -0.79459246 |
| Gm21119 | 1.839005393 | -0.078424022 | 0.874779082 | -0.089650088 |
| Gm20946 | 1.715564851 | 0.053379385 | 0.815806019 | 0.065431467 |
| Gm20797 | 0.04393175 | 0.037062677 | 3.724945049 | 0.009949859 |
| Gm20945 | 0.044191549 | 0.037062677 | 3.724945049 | 0.009949859 |
| Gm20796 | 3.51668492 | -0.925757739 | 0.652606692 | -1.418553856 |
| Defb33 | 0.682587184 | -1.772139305 | 1.369318696 | -1.294175937 |
| Gm15056 | 287.1653067 | -1.173727792 | 0.704908523 | -1.665078168 |
| Gm6040 | 1.098746849 | -0.365867134 | 0.918759257 | -0.398218718 |
| AY761185 | 1.275844484 | -0.587350535 | 0.840029907 | -0.699201933 |
| Defa21 | 0 | NA | NA | NA |
| Gm23142 | 0 | NA | NA | NA |
| Defa23 | 0 | NA | NA | NA |
| Gm10104 | 0 | NA | NA | NA |
| Defa-ps3 | 0 | NA | NA | NA |
| Defa25 | 0 | NA | NA | NA |
| Gm6689 | 0 | NA | NA | NA |
| Gm14851 | 0 | NA | NA | NA |
| Gm24568 | 0 | NA | NA | NA |
| Gm2869 | 0.109727584 | 0.037071574 | 3.724945049 | 0.009952247 |
| Defa-ps4 | 0 | NA | NA | NA |
| Gm15284 | 0.075043205 | 0.517957688 | 3.724945049 | 0.139051095 |
| Defa22 | 0 | NA | NA | NA |
| Gm26240 | 0 | NA | NA | NA |
| Defa-rs7 | 0 | NA | NA | NA |
| Gm15293 | 0 | NA | NA | NA |
| Defa-ps5 | 0 | NA | NA | NA |
| Defa-ps6 | 0 | NA | NA | NA |
| Defa-ps7 | 0 | NA | NA | NA |
| Gm15303 | 0 | NA | NA | NA |
| Gm15292 | 0 | NA | NA | NA |
| Defa-ps8 | 0 | NA | NA | NA |
| Defa3 | 0 | NA | NA | NA |
| Defa5 | 0 | NA | NA | NA |

| | | | | |
|---|---|---|---|---|
| Gm15300 | 0 | NA | NA | NA |
| Gm15299 | 0 | NA | NA | NA |
| Defa-rs1 | 0 | NA | NA | NA |
| Gm23561 | 0 | NA | NA | NA |
| Gm15302 | 0 | NA | NA | NA |
| Defa-ps16 | 0 | NA | NA | NA |
| Defa2 | 0 | NA | NA | NA |
| Gm21002 | 0 | NA | NA | NA |
| Gm21498 | 0 | NA | NA | NA |
| Gm15305 | 0 | NA | NA | NA |
| Defa-ps18 | 0 | NA | NA | NA |
| Gm15309 | 0 | NA | NA | NA |
| Gm7849 | 0 | NA | NA | NA |
| Gm23568 | 0 | NA | NA | NA |
| Gm15304 | 0 | NA | NA | NA |
| Defa-ps17 | 0 | NA | NA | NA |
| Defa20 | 0 | NA | NA | NA |
| Gm15308 | 0 | NA | NA | NA |
| Gm24423 | 0 | NA | NA | NA |
| Gm14850 | 0 | NA | NA | NA |
| Gm15311 | 0 | NA | NA | NA |
| Gm6696 | 0 | NA | NA | NA |
| Gm15312 | 0 | NA | NA | NA |
| Gm7861 | 0 | NA | NA | NA |
| Gm24829 | 0 | NA | NA | NA |
| Gm15314 | 0 | NA | NA | NA |
| Defa26 | 0 | NA | NA | NA |
| Gm15313 | 0 | NA | NA | NA |
| Defa17 | 0 | NA | NA | NA |
| Gm15315 | 0 | NA | NA | NA |
| Defa-ps9 | 0 | NA | NA | NA |
| Defa-ps1 | 0 | NA | NA | NA |
| AY761184 | 0.101408268 | 0.037730769 | 3.724898049 | 0.010129343 |
| Gm22447 | 0 | NA | NA | NA |
| Gm7869 | 0.109869311 | -0.444802825 | 3.724874622 | -0.119414174 |
| Defa-ps10 | 0 | NA | NA | NA |
| Defa24 | 0.046345709 | -0.443818988 | 3.724945049 | -0.119147795 |
| Defa-ps11 | 0 | NA | NA | NA |
| Defb1 | 2.378549416 | -0.008821633 | 0.691085836 | -0.012764887 |
| Gm15316 | 0.049553047 | -0.443818988 | 3.724945049 | -0.119147795 |
| Defb50 | 0 | NA | NA | NA |
| Defb2 | 0.062426873 | 0.517959912 | 3.724945049 | 0.139051692 |
| Defb10 | 0.392953962 | -0.401377384 | 1.564359513 | -0.256576177 |
| Defb9 | 0.27226464 | 0.037075733 | 2.061622565 | 0.017983764 |
| Defb11 | 0.092881282 | 0.037060453 | 3.724945049 | 0.009949262 |
| Defb15 | 0 | NA | NA | NA |
| Defb35 | 0 | NA | NA | NA |

| | | | | |
|---|---|---|---|---|
| Defb13 | 0 | NA | NA | NA |
| C86695 | 0 | NA | NA | NA |
| Ccdc70 | 0 | NA | NA | NA |
| Atp7b | 61.90632721 | 0.036402443 | 0.224078879 | 0.162453701 |
| Alg11 | 609.1938378 | -0.075417045 | 0.076941018 | -0.980192967 |
| Nek5 | 10.77738955 | -0.436310216 | 0.330846132 | -1.318770795 |
| Nek3 | 31.0393675 | -0.353121006 | 0.272003699 | -1.298221335 |
| Ckap2 | 17.47088672 | -0.624184581 | 0.503393154 | -1.239954449 |
| Vps36 | 310.0223612 | 0.035536295 | 0.089316399 | 0.397869766 |
| Thsd1 | 30.08606779 | -0.258281559 | 0.263428175 | -0.980462922 |
| Rpl30-ps2 | 0 | NA | NA | NA |
| Tpte | 0.493569458 | 1.003919721 | 1.418578758 | 0.707694032 |
| Slc25a15 | 143.2256164 | -0.26700627 | 0.121038668 | -2.205958423 |
| 1810012K16Rik | 2.758229591 | -0.038665991 | 0.596680693 | -0.064801813 |
| Mrps31 | 170.4153543 | -0.109505854 | 0.089973819 | -1.21708576 |
| Smim19 | 208.6627446 | 0.154130551 | 0.105244252 | 1.464503274 |
| Slc20a2 | 881.8761026 | -0.154313835 | 0.108946221 | -1.416422099 |
| Gm17491 | 71.03296188 | 0.128630771 | 0.259519678 | 0.495649395 |
| Vdac3 | 786.9911644 | -0.073435614 | 0.087291969 | -0.841264261 |
| Dkk4 | 0.121388914 | 0.037071574 | 3.724945049 | 0.009952247 |
| Polb | 409.3894603 | -0.020124343 | 0.073536279 | -0.273665504 |
| Gm26909 | 0.178821284 | -0.427208183 | 2.290359651 | -0.186524497 |
| Ikbkb | 547.644006 | -0.211762713 | 0.097858758 | -2.163962804 |
| Gm15346 | 0 | NA | NA | NA |
| Plat | 1098.82342 | -0.843735176 | 0.183604942 | -4.595383805 |
| Ap3m2 | 1163.088985 | 0.436612485 | 0.077436678 | 5.638316327 |
| 1700041G16Rik | 3.442441086 | -0.693344584 | 0.490765366 | -1.412782223 |
| Gm24735 | 0.372097677 | -1.137037891 | 1.510760496 | -0.752626173 |
| Kat6a | 2202.941112 | -0.323950819 | 0.070843381 | -4.572774671 |
| Ank1 | 2360.60079 | 0.072355505 | 0.091770575 | 0.788439046 |
| AC126445.1 | 0 | NA | NA | NA |
| Gm15815 | 3.535656746 | 1.491810285 | 0.721719656 | 2.067021832 |
| Gm15816 | 4.564354496 | -0.307732741 | 0.537686961 | -0.572326955 |
| Mir486 | 1.840863337 | 0.938155911 | 0.837239044 | 1.120535308 |
| mmu-mir-486b | 0 | NA | NA | NA |
| Nkx6-3 | 1.853806923 | 0.342218907 | 0.763795569 | 0.448050396 |
| Agpat6 | 875.7847179 | -0.128814184 | 0.065080472 | -1.979306242 |
| B230112G18Rik | 0.914342362 | -0.560290666 | 1.220966323 | -0.458891171 |
| Gins4 | 158.5957338 | -0.022194513 | 0.096056108 | -0.231057805 |
| Golga7 | 765.1713379 | -0.234139637 | 0.104568794 | -2.239096657 |
| Gm24335 | 0 | NA | NA | NA |
| Sfrp1 | 332.0038061 | 0.025363762 | 0.186823005 | 0.135763593 |
| Zmat4 | 554.6012764 | 0.362454193 | 0.099784543 | 3.632368142 |
| A730045E13Rik | 1.027651671 | 0.678781245 | 1.246209546 | 0.544676653 |
| Gm26714 | 0.777682211 | 1.638074706 | 1.257571205 | 1.302570144 |
| 1810011O10Rik | 28.96321082 | -0.165247549 | 0.293500314 | -0.563023413 |
| Ido2 | 21.23577249 | 0.063946717 | 0.237439466 | 0.269317979 |

| | | | | |
|---|---|---|---|---|
| Ido1 | 30.3065116 | 0.547173076 | 0.237146694 | 2.307319018 |
| Adam18 | 1.533690469 | -1.190284767 | 0.825018032 | -1.442737881 |
| Adam3 | 2.60343246 | -0.76721326 | 0.62219348 | -1.23307827 |
| Gm26208 | 0 | NA | NA | NA |
| Adam5 | 3.408984752 | 0.008533353 | 0.539501227 | 0.015817115 |
| Adam32 | 15.27834067 | 0.066525163 | 0.281883134 | 0.236002636 |
| Gm23898 | 0 | NA | NA | NA |
| Adam9 | 978.4277452 | -0.481120844 | 0.088237405 | -5.45257246 |
| Tm2d2 | 316.9476182 | -0.046774029 | 0.080867902 | -0.578400426 |
| Htra4 | 16.56292703 | -0.385153764 | 0.443664385 | -0.868119635 |
| Gm16933 | 1.72427518 | -1.233542509 | 0.855761033 | -1.44145674 |
| Plekha2 | 504.2642388 | -0.701505061 | 0.107162011 | -6.546210304 |
| Tacc1 | 2873.630983 | 0.17738226 | 0.089775287 | 1.975847312 |
| Mir8108 | 0 | NA | NA | NA |
| Gm10689 | 1.990043323 | 0.420061453 | 0.745408552 | 0.563531839 |
| Fgfr1 | 1712.090942 | -0.586023581 | 0.111616352 | -5.25033805 |
| Gm16159 | 0.092228164 | 0.037073798 | 3.724945049 | 0.009952844 |
| Letm2 | 106.132572 | 0.101803826 | 0.146704575 | 0.693937632 |
| D830025C05Rik | 3.021300136 | -0.177401513 | 0.599264497 | -0.296032075 |
| Whsc1l1 | 2360.967649 | -0.021998573 | 0.080281184 | -0.274019037 |
| Gm23184 | 0 | NA | NA | NA |
| Gm24992 | 3.122516278 | 0.164158126 | 0.553166662 | 0.296760701 |
| Gm23094 | 7.166830998 | 1.09081337 | 0.583995674 | 1.867844949 |
| Ppapdc1b | 162.0488265 | -0.112228084 | 0.135575626 | -0.827789536 |
| Ddhd2 | 814.1538713 | 0.436571098 | 0.079981446 | 5.458404696 |
| Gm17484 | 32.71518712 | -1.295573938 | 0.309275314 | -4.189063521 |
| Bag4 | 936.8776659 | 0.549664045 | 0.064862422 | 8.474306468 |
| Lsm1 | 143.0297487 | -0.053218702 | 0.118902601 | -0.447582323 |
| Star | 22.62371583 | -0.208205404 | 0.290694382 | -0.716234701 |
| Ash2l | 496.3350927 | -0.139515608 | 0.061826782 | -2.256556204 |
| Kcnu1 | 15.88801623 | 0.356249519 | 0.287455445 | 1.239320825 |
| Hgsnat | 490.0921398 | 0.171851477 | 0.073380053 | 2.34193723 |
| Pomk | 265.4581434 | 0.158131443 | 0.082875847 | 1.908052203 |
| Fnta | 475.5451638 | -0.216636312 | 0.071550081 | -3.027757765 |
| Hook3 | 2198.855902 | 0.037003975 | 0.042696358 | 0.866677543 |
| 1700047A11Rik | 0.066050875 | -0.443818988 | 3.724945049 | -0.119147795 |
| Rnf170 | 300.8605897 | 0.284493791 | 0.104449224 | 2.723752077 |
| Thap1 | 73.51708713 | -0.395678617 | 0.132638719 | -2.983130572 |
| AC154102.1 | 0 | NA | NA | NA |
| Gm22227 | 0 | NA | NA | NA |
| 2310008N11Rik | 0.04393175 | 0.037062677 | 3.724945049 | 0.009949859 |
| Zfp703 | 588.8755344 | -0.328852865 | 0.090302898 | -3.641664582 |
| Erlin2 | 723.3406109 | -0.098266963 | 0.081381971 | -1.207478293 |
| Proscos | 5.115524656 | 0.48141254 | 0.433493937 | 1.110540423 |
| Prosc | 626.0658266 | -0.366708163 | 0.066835731 | -5.486708302 |
| Adgra2 | 197.3133489 | -0.531423515 | 0.1243885 | -4.272288151 |
| Brf2 | 100.3354801 | 0.011608156 | 0.137171577 | 0.08462508 |

| | | | | |
|---|---|---|---|---|
| Rab11fip1 | 53.92462846 | -0.401922977 | 0.20830256 | -1.929515302 |
| 4933416M07Rik | 0.124596252 | 0.037071632 | 3.714681146 | 0.009979761 |
| Got1l1 | 2.891827676 | 0.373328651 | 0.628808248 | 0.59370826 |
| Adrb3 | 6.872911924 | 0.355519844 | 0.465542425 | 0.763667982 |
| Eif4ebp1 | 90.4572564 | -1.558617359 | 0.237114388 | -6.573271951 |
| Tex24 | 0.451016279 | -1.066934664 | 1.481785002 | -0.72003338 |
| Chrnb3 | 25.45046697 | 0.472513197 | 0.251134932 | 1.881511235 |
| Chrna6 | 16.6337043 | 1.238750947 | 0.340942752 | 3.633310699 |
| Gm24410 | 0 | NA | NA | NA |
| Poteg | 0 | NA | NA | NA |
| Gm1698 | 3.756750159 | -0.097913206 | 0.527461238 | -0.185631093 |
| AC111034.1 | 0 | NA | NA | NA |
| Gm24202 | 0 | NA | NA | NA |
| Gm8100 | 1.189584463 | -0.526470849 | 1.064336224 | -0.494647121 |
| Gm26795 | 1.493046057 | 0.15612886 | 0.758565537 | 0.205821188 |
| Unc5d | 798.2940023 | 0.181487171 | 0.078120531 | 2.32316869 |
| 4933433F19Rik | 0 | NA | NA | NA |
| Dusp26 | 500.6484086 | -0.268028079 | 0.108981039 | -2.459401037 |
| Rnf122 | 133.8921804 | -0.16751683 | 0.134950501 | -1.241320545 |
| Snord13 | 7.690137191 | -0.267086298 | 0.481545274 | -0.554644208 |
| Tti2 | 237.0865655 | 0.134902256 | 0.099453955 | 1.356429277 |
| Mak16 | 240.9495093 | -0.228236128 | 0.096192823 | -2.372693945 |
| Fut10 | 112.6731054 | 0.158294448 | 0.126666683 | 1.249692846 |
| Gm8051 | 0 | NA | NA | NA |
| Gm23539 | 0.319751656 | -0.590284186 | 1.799580981 | -0.328012016 |
| Gm24184 | 0.141071297 | -0.443820801 | 3.564043909 | -0.12452731 |
| Gm5117 | 3.857935733 | -0.17947026 | 0.476922272 | -0.376309245 |
| Nrg1 | 264.3991212 | -0.177369286 | 0.143490537 | -1.236104416 |
| Gm26578 | 5.176536363 | -1.513376532 | 0.542765245 | -2.788270886 |
| Gm22140 | 0 | NA | NA | NA |
| Gm5908 | 9.444744683 | -0.149300407 | 0.355652616 | -0.419792798 |
| Gm3985 | 0.861087095 | 1.317537033 | 1.212005459 | 1.087071863 |
| Gm6877 | 13.36504497 | 0.114635449 | 0.303231523 | 0.378045948 |
| Wrn | 282.207455 | -0.057657915 | 0.109205855 | -0.527974578 |
| Purg | 493.6010051 | 0.309144602 | 0.091680546 | 3.371976014 |
| 5930422O12Rik | 8.011746571 | 0.079403089 | 0.43077805 | 0.184324826 |
| Tex15 | 53.01275634 | -0.768920635 | 0.253118839 | -3.037785085 |
| Ppp2cb | 866.8510396 | -0.017785012 | 0.083571581 | -0.212811724 |
| Ubxn8 | 210.3585288 | 0.118787743 | 0.084628211 | 1.403642377 |
| Gsr | 438.4005393 | -0.424948573 | 0.087439372 | -4.859922543 |
| Gm24727 | 28.80332305 | -0.079991799 | 0.329508853 | -0.242760697 |
| 1700104B16Rik | 0.424137699 | 0.081885491 | 1.564760054 | 0.052331021 |
| Gtf2e2 | 144.048718 | -0.336212673 | 0.103876208 | -3.236666808 |
| Smim18 | 19.2016788 | 0.252097036 | 0.287003963 | 0.878374754 |
| Rbpms | 30.315901 | -0.363913085 | 0.247064259 | -1.472949129 |
| Gm26632 | 0 | NA | NA | NA |
| Gm26978 | 0.712265815 | -0.388421437 | 1.16397963 | -0.333701233 |

| | | | | |
|---|---|---|---|---|
| Gm9951 | 0.166952883 | -1.336542448 | 2.746539428 | -0.486627803 |
| Dctn6 | 437.7144974 | 0.33327298 | 0.075153343 | 4.434572918 |
| Gm26768 | 0.185862605 | 0.866959667 | 2.679501836 | 0.323552556 |
| Gm10131 | 44.98302574 | -0.363732798 | 0.199376843 | -1.824348267 |
| Mboat4 | 2.274616701 | -0.534906395 | 0.686184798 | -0.779536937 |
| Leprotl1 | 852.1230496 | -0.306996282 | 0.088054816 | -3.486422393 |
| Saraf | 1822.362344 | -0.116526653 | 0.095819602 | -1.216104546 |
| Mir6395 | 0 | NA | NA | NA |
| Gm29243 | 0.741879394 | -2.184128261 | 1.159842898 | -1.883124227 |
| Gm23177 | 0.047948698 | 0.037064902 | 3.724945049 | 0.009950456 |
| Gm6100 | 0 | NA | NA | NA |
| Gm8254 | 0 | NA | NA | NA |
| Gm4889 | 0 | NA | NA | NA |
| Gm29174 | 0.095435502 | 0.037073798 | 3.724945049 | 0.009952844 |
| Dusp4 | 233.7081911 | -1.599932879 | 0.206548769 | -7.746029615 |
| Tnks | 1939.237959 | 0.038664943 | 0.055678367 | 0.694433859 |
| Gm9939 | 2.193369661 | -1.221273242 | 0.74932922 | -1.629821993 |
| Rpl31-ps23 | 0 | NA | NA | NA |
| Ppp1r3b | 56.58455372 | -1.167591079 | 0.251924363 | -4.634689011 |
| Eri1 | 199.30024 | -0.63603481 | 0.129713118 | -4.903396204 |
| Gm16793 | 8.177702489 | 0.025819045 | 0.36771699 | 0.070214448 |
| Mfhas1 | 794.683006 | 0.096472185 | 0.085377397 | 1.129949951 |
| Gm22216 | 0.304980231 | 1.457189963 | 1.742700559 | 0.836167726 |
| AC129211.1 | 0 | NA | NA | NA |
| Cldn23 | 10.064079 | 0.317732206 | 0.412230355 | 0.77076373 |
| Gm22030 | 0 | NA | NA | NA |
| Rps12l2 | 3.933964785 | 0.165049546 | 0.789930746 | 0.208941792 |
| D8Ertd82e | 662.5180749 | -0.216299039 | 0.095640451 | -2.261585305 |
| Lonrf1 | 282.0614182 | 0.438718235 | 0.12088317 | 3.629274737 |
| Gm5787 | 0.203967006 | -0.467709549 | 2.21351496 | -0.211297216 |
| Gm26150 | 0.099155932 | 0.037073798 | 3.724945049 | 0.009952844 |
| 6430573F11Rik | 301.8566133 | 0.152999903 | 0.184439341 | 0.829540497 |
| Dlc1 | 1005.579271 | 0.00249358 | 0.067679448 | 0.036843978 |
| AI429214 | 46.9821826 | -0.111149106 | 0.162308128 | -0.684803084 |
| Gm25126 | 0 | NA | NA | NA |
| Sgcz | 92.85252959 | 0.697003283 | 0.167012398 | 4.173362523 |
| Gm16205 | 2.502004683 | 2.208732534 | 0.889389479 | 2.483425527 |
| Gm16204 | 0 | NA | NA | NA |
| Mir383 | 2.557294913 | 1.21277647 | 0.696778772 | 1.740547386 |
| Gm23128 | 0 | NA | NA | NA |
| Tusc3 | 992.2313355 | 0.343289847 | 0.07761588 | 4.422933113 |
| Msr1 | 385.467979 | -1.377710575 | 0.353239632 | -3.900215179 |
| Fgf20 | 6.705059015 | 0.268350769 | 0.405793651 | 0.661298591 |
| Micu3 | 1360.461354 | 0.359784413 | 0.066327029 | 5.424401171 |
| Mir7666 | 6.340121748 | 1.192693661 | 0.474552517 | 2.513301728 |
| Zdhhc2 | 665.2779763 | -0.143075966 | 0.100629694 | -1.421806629 |
| Cnot7 | 1005.736534 | 0.049945736 | 0.057938839 | 0.862042392 |

| | | | | |
|---|---|---|---|---|
| Vps37a | 864.8152951 | -0.245809164 | 0.071981479 | -3.414894619 |
| Mtmr7 | 611.4343167 | 0.416719679 | 0.065323589 | 6.379313929 |
| Adam24 | 0.11193333 | 0.037073798 | 3.724945049 | 0.009952844 |
| Adam25 | 0 | NA | NA | NA |
| Adam20 | 0 | NA | NA | NA |
| Adam39 | 0 | NA | NA | NA |
| Gm5345 | 0.200503209 | 0.5388051 | 2.612813776 | 0.206216419 |
| Gm26584 | 0.132101749 | -0.93550548 | 3.724188147 | -0.251197158 |
| Slc7a2 | 630.8632614 | -0.312043616 | 0.135364615 | -2.305208169 |
| Pdgfrl | 37.56964625 | -0.847021736 | 0.245718149 | -3.447127288 |
| Mtus1 | 1041.21851 | -0.255759057 | 0.080739252 | -3.16771644 |
| B430010I23Rik | 0.577003193 | -0.29799096 | 1.266190301 | -0.235344529 |
| Gm16192 | 0.177745132 | 0.037630679 | 2.300135837 | 0.016360198 |
| Gm16193 | 1.147365236 | 0.213635301 | 0.962617579 | 0.221931643 |
| Fgl1 | 0.121388914 | 0.037071574 | 3.724945049 | 0.009952247 |
| Gm16348 | 0.652886586 | -2.576687705 | 3.663061211 | -0.703424692 |
| Pcm1 | 2899.225793 | 0.089366874 | 0.060914569 | 1.467085374 |
| Asah1 | 694.5391705 | -0.421674242 | 0.091943612 | -4.586226615 |
| Frg1 | 216.5532014 | -0.135401148 | 0.107882939 | -1.255074702 |
| Gapdh-ps14 | 0.166952883 | -1.336542448 | 2.746539428 | -0.486627803 |
| Smarce1-ps1 | 16.22242323 | -0.492151642 | 0.276182657 | -1.781978806 |
| 2810404M03Rik | 0.402944817 | -1.158552029 | 1.735325251 | -0.667628174 |
| Zfp353-ps | 0 | NA | NA | NA |
| Gm6180 | 51.37583421 | -0.067337394 | 0.147653751 | -0.456049329 |
| Gm23531 | 0 | NA | NA | NA |
| Gm26181 | 0 | NA | NA | NA |
| Triml1 | 0.169875191 | 0.984813711 | 2.746838256 | 0.358526283 |
| Triml2 | 0.045882455 | 0.517959912 | 3.724945049 | 0.139051692 |
| Zfp42 | 0.483255338 | -2.049950886 | 1.307650721 | -1.567659355 |
| Gm22986 | 0 | NA | NA | NA |
| Adam26b | 0 | NA | NA | NA |
| Adam26a | 0 | NA | NA | NA |
| Gm5346 | 0.068097359 | 0.037064902 | 3.724945049 | 0.009950456 |
| Adam34 | 0.137685729 | 0.5179621 | 2.938031518 | 0.176295624 |
| Gm24591 | 0 | NA | NA | NA |
| Gm37972 | 0 | NA | NA | NA |
| Gm26241 | 0 | NA | NA | NA |
| Gm26089 | 0 | NA | NA | NA |
| Fat1 | 3170.327197 | -0.254796845 | 0.080412571 | -3.168619561 |
| Gm2366 | 0.106971521 | 0.998850474 | 3.724945049 | 0.268151734 |
| AY512931 | 0.183958792 | 0.504050916 | 2.678154971 | 0.188208271 |
| Mtnr1a | 1.009506168 | 0.199848815 | 1.106326088 | 0.180641871 |
| F11 | 1.357931586 | 0.923543167 | 0.787193924 | 1.173209217 |
| Klkb1 | 0.521565614 | -0.402708328 | 1.308105148 | -0.307856237 |
| Cyp4v3 | 176.7598727 | -0.391266082 | 0.145916589 | -2.681436605 |
| Fam149a | 309.9705733 | -0.199254212 | 0.103743844 | -1.920636476 |
| Tlr3 | 1099.900133 | -0.619454588 | 0.188148767 | -3.292365909 |

| | | | | |
|---|---|---|---|---|
| Sorbs2 | 2803.02876 | 0.231040389 | 0.13138125 | 1.758549176 |
| Gm23604 | 0 | NA | NA | NA |
| Gm16351 | 0.27308476 | -0.837846707 | 1.998903769 | -0.419153098 |
| Sorbs2os | 1.437005151 | -0.603875631 | 0.964753448 | -0.62593778 |
| Gm16352 | 7.186401275 | 0.534923153 | 0.422713078 | 1.265452101 |
| Gm25309 | 0 | NA | NA | NA |
| Pdlim3 | 13.56838278 | -0.025608796 | 0.379018613 | -0.067566063 |
| Ccdc110 | 5.316494593 | -0.932022138 | 0.498122515 | -1.871070089 |
| Gm15878 | 0.075043205 | 0.517957688 | 3.724945049 | 0.139051095 |
| 1700029J07Rik | 30.24045965 | 0.169484155 | 0.280335149 | 0.604576897 |
| Ufsp2 | 315.566229 | -0.096845906 | 0.120363207 | -0.804613871 |
| Ankrd37 | 31.55324993 | 0.527349662 | 0.217594614 | 2.423541886 |
| Lrp2bp | 13.15399262 | 0.886407447 | 0.298862939 | 2.965932976 |
| Snx25 | 564.8783107 | 0.185746277 | 0.056688604 | 3.27660701 |
| Gm23030 | 0 | NA | NA | NA |
| Cfap97 | 360.4521644 | 0.059818545 | 0.076000458 | 0.787081376 |
| Gm15634 | 3.396395574 | 0.308100207 | 0.603020768 | 0.51092802 |
| Slc25a4 | 4403.300027 | 0.196414156 | 0.084333065 | 2.329029026 |
| Actg-ps1 | 1.057130107 | 0.389915083 | 0.867258166 | 0.44959517 |
| Gm23812 | 0 | NA | NA | NA |
| Gm10313 | 0.79470004 | -1.976981793 | 1.099495588 | -1.798080697 |
| Gm24669 | 0 | NA | NA | NA |
| Helt | 1.906358267 | -2.506226059 | 0.968812272 | -2.586905773 |
| Acsl1 | 635.4438796 | 0.193439924 | 0.090290945 | 2.142406675 |
| Cenpu | 23.99853257 | 0.266806421 | 0.242791729 | 1.098910666 |
| Primpol | 71.59947947 | -0.282533886 | 0.153106766 | -1.845338996 |
| Casp3 | 141.296778 | -0.586598097 | 0.147114562 | -3.987355763 |
| Gm16675 | 21.12817838 | -0.560378526 | 0.237935954 | -2.355165395 |
| Irf2 | 579.6276244 | -0.473346688 | 0.148687133 | -3.183508075 |
| Enpp6 | 117.2534975 | 0.230884852 | 0.128810107 | 1.792443606 |
| 4930579M01Rik | 0.29175089 | -0.029903536 | 1.768577765 | -0.016908239 |
| Stox2 | 2896.450262 | 0.133007594 | 0.063639746 | 2.090008229 |
| 4930448N21Rik | 21.17633594 | -0.267822141 | 0.353159914 | -0.758359401 |
| Gm10083 | 4.406530871 | 2.302813425 | 0.796431101 | 2.891415744 |
| Gm22857 | 0 | NA | NA | NA |
| Trappc11 | 834.2127417 | 0.037769197 | 0.0579609 | 0.651632333 |
| Rwdd4a | 348.2097546 | -0.435374628 | 0.085521797 | -5.090803097 |
| Gm27409 | 0 | NA | NA | NA |
| Gm8623 | 3.204879861 | -0.124298463 | 0.580506477 | -0.214120718 |
| Ing2 | 227.0032211 | 0.331327724 | 0.108447147 | 3.055200013 |
| AA386476 | 7.480509175 | 1.117851871 | 0.432249927 | 2.586123908 |
| Cdkn2aip | 219.8046481 | -0.162426477 | 0.105939166 | -1.533205167 |
| E030037K01Rik | 3.293407775 | -0.216117248 | 0.613194465 | -0.352444878 |
| n-R5s96 | 0.212081199 | 1.33343622 | 2.576983575 | 0.517440714 |
| Cldn24 | 0.048949532 | 0.037064902 | 3.724945049 | 0.009950456 |
| Cldn22 | 1.967947156 | 1.662350352 | 0.788687969 | 2.107741488 |
| Wwc2 | 675.3607201 | 0.265343424 | 0.065422858 | 4.055821338 |

| | | | | |
|---|---|---|---|---|
| n-R5s97 | 0 | NA | NA | NA |
| Dctd | 53.07648255 | -0.24728895 | 0.197451277 | -1.25240492 |
| Tenm3 | 1960.745608 | 0.567858514 | 0.113469879 | 5.004486833 |
| Gm19744 | 64.58441548 | 0.115403324 | 0.260273609 | 0.443392337 |
| Gm25830 | 0.249706104 | 0.51797477 | 1.848218418 | 0.280256254 |
| 4930555F03Rik | 2.112369234 | 0.008142182 | 0.822945207 | 0.009893954 |
| Gm23986 | 0.048949532 | 0.037064902 | 3.724945049 | 0.009950456 |
| AC140437.1 | 0.038981395 | -0.443816764 | 3.724945049 | -0.119147198 |
| Gm24929 | 0 | NA | NA | NA |
| Gm9892 | 89.20159694 | -0.288697542 | 0.161062184 | -1.792460121 |
| Gm6463 | 0 | NA | NA | NA |
| Gm25813 | 1.893990199 | -0.31820614 | 0.841567304 | -0.378111339 |
| Gm23666 | 0 | NA | NA | NA |
| Gm27964 | 0 | NA | NA | NA |
| Gm22841 | 0 | NA | NA | NA |
| Aga | 79.49866868 | -0.644810833 | 0.148326463 | -4.347240663 |
| Neil3 | 7.05498645 | -0.009558219 | 0.639034185 | -0.014957289 |
| Vegfc | 39.65073296 | 0.088816514 | 0.172196706 | 0.515785207 |
| Spcs3 | 857.5164723 | -0.214994677 | 0.078763216 | -2.729633034 |
| Asb5 | 7.025391348 | -0.093113292 | 0.403152848 | -0.230962753 |
| 5330439A09Rik | 2.898308479 | -0.08434359 | 0.806996215 | -0.104515472 |
| Spata4 | 0.398944595 | 1.390255389 | 1.634224796 | 0.850712456 |
| Wdr17 | 405.2573486 | 0.958957906 | 0.135218343 | 7.091921757 |
| Gm15758 | 0.38984224 | 1.063615912 | 1.454664335 | 0.731176181 |
| n-R5s98 | 6.806331161 | -0.224221767 | 0.459297144 | -0.488184543 |
| Gpm6a | 9377.570103 | 0.71100288 | 0.08442387 | 8.421822899 |
| Adam29 | 0.15906016 | -1.015882374 | 3.433492635 | -0.295874342 |
| Glra3 | 117.655716 | 0.092441847 | 0.131689136 | 0.701970182 |
| Hpgd | 129.4286017 | -1.167369588 | 0.242368827 | -4.816500538 |
| Gm24375 | 0.075043205 | 0.517957688 | 3.724945049 | 0.139051095 |
| Cep44 | 122.059907 | 0.457594962 | 0.151426341 | 3.02189804 |
| Fbxo8 | 283.448663 | 0.235818968 | 0.118465775 | 1.990608417 |
| Gm25992 | 6.166697826 | 1.007164573 | 0.425357173 | 2.367809075 |
| Gm29173 | 0.088736197 | 0.51795324 | 3.724945049 | 0.139049901 |
| AV026068 | 0.061920613 | -0.443816764 | 3.724945049 | -0.119147198 |
| Hand2 | 0.062426873 | 0.517959912 | 3.724945049 | 0.139051692 |
| Gm28063 | 0 | NA | NA | NA |
| 5033428I22Rik | 0 | NA | NA | NA |
| Scrg1 | 178.3231663 | 0.630175568 | 0.213291184 | 2.954531719 |
| Sap30 | 157.6272062 | -0.411693219 | 0.146289672 | -2.814232979 |
| 2500002B13Rik | 14.59964785 | -0.430839574 | 0.326767565 | -1.318489408 |
| Hmgb2 | 86.7028358 | 0.263469213 | 0.275284702 | 0.957079019 |
| Galnt7 | 420.2620518 | -0.478101504 | 0.072771854 | -6.569868392 |
| Gm25379 | 0 | NA | NA | NA |
| Gm10284 | 11.17616707 | 0.577083533 | 0.309030534 | 1.867399717 |
| Galntl6 | 211.6445591 | 0.554197501 | 0.112104763 | 4.943567854 |
| Gm26905 | 1.736582332 | 0.928893641 | 0.915548231 | 1.014576414 |

| | | | | |
|---|---|---|---|---|
| Gm15882 | 0 | NA | NA | NA |
| Gm15881 | 19.66782202 | 0.948303199 | 0.326296152 | 2.906265348 |
| BC030500 | 65.02402097 | 0.276659761 | 0.17419619 | 1.588207882 |
| Gm24847 | 0 | NA | NA | NA |
| Gm26164 | 0 | NA | NA | NA |
| Gm10283 | 0 | NA | NA | NA |
| Aadat | 2.615189709 | -0.305028495 | 0.661893629 | -0.460842167 |
| Mfap3l | 976.8064727 | -0.311496259 | 0.083319499 | -3.738575742 |
| Gm23329 | 0.066050875 | -0.443818988 | 3.724945049 | -0.119147795 |
| 2700029M09Rik | 270.2446424 | 0.066123476 | 0.119153022 | 0.554945862 |
| Clcn3 | 3034.91923 | 0.224888507 | 0.043604516 | 5.15745911 |
| B230317F23Rik | 9.640489829 | -0.011379695 | 0.441287415 | -0.02578749 |
| Nek1 | 677.7359105 | 0.013886065 | 0.152158938 | 0.091260265 |
| Gm16179 | 0 | NA | NA | NA |
| Gm16178 | 0.095898756 | -0.917692504 | 3.724434845 | -0.246397787 |
| Sh3rf1 | 531.7020943 | 0.120774263 | 0.102661456 | 1.176432405 |
| 1700001D01Rik | 0 | NA | NA | NA |
| Gm10015 | 0.445602023 | 0.517788291 | 1.404866202 | 0.36856769 |
| Cbr4 | 83.07961058 | 0.140003425 | 0.148541197 | 0.942522532 |
| Palld | 202.5425458 | -1.099928979 | 0.200096613 | -5.496989496 |
| 4930512H18Rik | 0.155551505 | 0.539747996 | 2.83133048 | 0.19063405 |
| Ddx60 | 1763.240931 | -0.864296365 | 0.175379945 | -4.928136814 |
| Anxa10 | 0.200514362 | -0.862293442 | 2.64038825 | -0.326578276 |
| Gm22324 | 0 | NA | NA | NA |
| Spock3 | 1216.4572 | 0.496471221 | 0.099335656 | 4.997915537 |
| Gm24236 | 0 | NA | NA | NA |
| Gm5350 | 0 | NA | NA | NA |
| Gm24580 | 0 | NA | NA | NA |
| Sgol2b | 2.142608394 | -0.458056849 | 0.804354274 | -0.569471517 |
| Tll1 | 166.2460114 | -0.749524294 | 0.194596601 | -3.851682348 |
| Gm22586 | 0 | NA | NA | NA |
| Mir710 | 0.327710148 | 0.702650715 | 1.935306605 | 0.363069455 |
| Cpe | 12511.93844 | 0.280645358 | 0.093955228 | 2.987011626 |
| Msmo1 | 801.7074196 | -0.683355338 | 0.142311074 | -4.801842308 |
| Klhl2 | 1600.801606 | -0.059928709 | 0.091712907 | -0.653438113 |
| Gm26210 | 0 | NA | NA | NA |
| Tmem192 | 214.6384359 | -0.184092895 | 0.135443316 | -1.35918774 |
| Trim75 | 0 | NA | NA | NA |
| Trim60 | 0 | NA | NA | NA |
| Apela | 0.083526043 | 0.037073798 | 3.724945049 | 0.009952844 |
| Gm10663 | 0 | NA | NA | NA |
| BC030870 | 0 | NA | NA | NA |
| Gm24333 | 0 | NA | NA | NA |
| Gm25403 | 0 | NA | NA | NA |
| 1-Mar | 492.5184607 | 0.228228137 | 0.067494288 | 3.381443728 |
| Gm23515 | 0 | NA | NA | NA |
| Gm15355 | 0.044191549 | 0.037062677 | 3.724945049 | 0.009949859 |

| | | | | |
|---|---|---|---|---|
| Gm26123 | 1.969797064 | 0.609445679 | 0.746900623 | 0.815966221 |
| Gm24205 | 4.278889304 | 0.065821724 | 0.536096911 | 0.122779525 |
| Gm15354 | 0.20829534 | -1.557373471 | 2.531443919 | -0.615211524 |
| Gm15356 | 0.306626524 | -1.242970224 | 1.739684372 | -0.714480307 |
| Tma16 | 219.0930399 | 0.389118127 | 0.181695876 | 2.141590311 |
| Tktl2 | 2.437974107 | 2.455701444 | 0.792366808 | 3.099197771 |
| Gm16330 | 0 | NA | NA | NA |
| Npy5r | 36.55808565 | 0.189412261 | 0.213824798 | 0.885829249 |
| Npy1r | 157.9670103 | 0.540819016 | 0.151321409 | 3.573975552 |
| Naf1 | 250.5788205 | -0.503196124 | 0.116812434 | -4.307727403 |
| Gm25275 | 0 | NA | NA | NA |
| Gm24274 | 0 | NA | NA | NA |
| Gm24677 | 0 | NA | NA | NA |
| Gm10660 | 0 | NA | NA | NA |
| Nat1 | 7.474725557 | -0.201450206 | 0.485674762 | -0.414784177 |
| Nat2 | 24.83237878 | -1.037525846 | 0.232164754 | -4.468920581 |
| Gm9755 | 12.27831489 | -0.434995223 | 0.309739206 | -1.404391872 |
| Nat3 | 0.254414262 | -0.030998796 | 2.456890324 | -0.012617086 |
| AC141868.1 | 0 | NA | NA | NA |
| Psd3 | 7016.150575 | 0.567942324 | 0.104957937 | 5.411142234 |
| Gm15992 | 24.57258576 | 1.189986699 | 0.327365448 | 3.635040618 |
| Gm15991 | 12.26391693 | 0.789787739 | 0.332453061 | 2.375636839 |
| Gm22018 | 0.176406687 | 0.955797087 | 2.705670714 | 0.353256988 |
| Gm21807 | 5.900355672 | -1.030653863 | 0.445392381 | -2.314035681 |
| Gm26659 | 0.106520246 | 0.037071574 | 3.724945049 | 0.009952247 |
| Sh2d4a | 4.03121477 | -1.92183783 | 0.67177789 | -2.860823287 |
| Csgalnact1 | 201.9711671 | -0.014352972 | 0.118028344 | -0.121606148 |
| 1700125H03Rik | 0.107078754 | -0.443164241 | 3.724898049 | -0.118973522 |
| Gm15656 | 0 | NA | NA | NA |
| Gm15654 | 0 | NA | NA | NA |
| Ints10 | 560.1978323 | 0.062751679 | 0.121065161 | 0.518329787 |
| Gm15549 | 5.491253898 | 0.749917653 | 0.618928256 | 1.211639065 |
| Lpl | 272.7124532 | 0.719117289 | 0.132608429 | 5.422862584 |
| Slc18a1 | 6.521337389 | -0.179034056 | 0.37443679 | -0.478142267 |
| Gm15716 | 0 | NA | NA | NA |
| Atp6v1b2 | 5572.862093 | -0.037153424 | 0.048850236 | -0.760557723 |
| Gm15717 | 14.06246178 | 0.257806691 | 0.415387287 | 0.620641746 |
| Lzts1 | 982.0473404 | 0.890041231 | 0.163702176 | 5.436954193 |
| Zfp930 | 119.9920175 | -0.102021687 | 0.13498417 | -0.755804826 |
| D130040H23Rik | 96.03445661 | 0.367582593 | 0.149044587 | 2.466259261 |
| Gm10033 | 37.55209856 | -0.4486696 | 0.243437617 | -1.843057801 |
| Gm10311 | 21.9424117 | -0.638789706 | 0.252907873 | -2.525780233 |
| Gm25661 | 0.130524232 | 0.517958497 | 3.657871415 | 0.141601067 |
| RP23-468H22.1 | 1.045653151 | -0.150996216 | 1.047285912 | -0.1441786 |
| RP23-468H22.3 | 0.584793824 | 0.434856017 | 1.143092128 | 0.380420796 |
| RP23-468H22.2 | 0.794752679 | -0.792785844 | 1.174143596 | -0.675203482 |
| Zfp868 | 254.8017484 | 0.114612691 | 0.086857917 | 1.319542247 |

| | | | | |
|---|---|---|---|---|
| Zfp964 | 24.06935087 | -0.207261259 | 0.243287843 | -0.851917864 |
| Zfp869 | 250.7494571 | -0.071126429 | 0.087864268 | -0.809503458 |
| Zfp963 | 46.44832303 | 0.388792213 | 0.161978434 | 2.400271469 |
| Gm20422 | 0 | NA | NA | NA |
| Zfp866 | 322.7321497 | 0.061141293 | 0.115173877 | 0.530860771 |
| Atp13a1 | 689.8934044 | 0.007084717 | 0.086173133 | 0.08221492 |
| Gmip | 150.8857527 | -0.462825531 | 0.140526429 | -3.293512364 |
| Lpar2 | 13.32151877 | 0.047172779 | 0.291150514 | 0.162021966 |
| Pbx4 | 59.57232151 | 0.7995257 | 0.154080475 | 5.189013719 |
| Cilp2 | 10.39023466 | -0.408949512 | 0.356158338 | -1.148223888 |
| Yjefn3 | 43.79218575 | 0.646614389 | 0.197382756 | 3.275941637 |
| Ndufa13 | 488.3269015 | 0.190234789 | 0.343643987 | 0.55358102 |
| Tssk6 | 2.109268871 | -1.49877008 | 0.775166818 | -1.933480699 |
| Gatad2a | 755.2849878 | -0.560746102 | 0.10143651 | -5.528050001 |
| Mau2 | 1031.702982 | 0.108813213 | 0.065871386 | 1.651904111 |
| Sugp1 | 554.3624155 | 0.076932321 | 0.060598017 | 1.269551792 |
| Tm6sf2 | 17.28368832 | -1.237175044 | 0.507792682 | -2.436378246 |
| Hapln4 | 1233.584627 | 0.204126076 | 0.075593497 | 2.70031263 |
| Ncan | 3935.767652 | -0.928979988 | 0.169820952 | -5.470349656 |
| Mir7066 | 0.192965486 | 1.021055211 | 2.249093311 | 0.45398526 |
| Gm25906 | 0 | NA | NA | NA |
| Rfxank | 136.35992 | -0.934127348 | 0.111584101 | -8.371509403 |
| Nr2c2ap | 176.3814956 | -1.064819607 | 0.15051939 | -7.074301906 |
| 2310045N01Rik | 134.2125858 | -0.201550386 | 0.10275728 | -1.961421956 |
| Mef2b | 0.295756179 | 0.478513462 | 2.097084398 | 0.228180355 |
| Tmem161a | 299.8727589 | -0.249798384 | 0.096131831 | -2.598498134 |
| Slc25a42 | 283.822492 | -0.00664902 | 0.164782516 | -0.040350278 |
| Armc6 | 187.6904792 | -0.286414212 | 0.103757785 | -2.760411778 |
| Sugp2 | 1115.657998 | 0.360801159 | 0.095484239 | 3.778646225 |
| Homer3 | 488.6342012 | -0.077845102 | 0.201835999 | -0.385684922 |
| Ddx49 | 235.0236882 | -0.133885083 | 0.087083332 | -1.537436382 |
| Cope | 659.244897 | -0.195995382 | 0.075444895 | -2.597861418 |
| Cers1 | 345.2784473 | 0.020122183 | 0.093498225 | 0.215214593 |
| Gdf1 | 0 | NA | NA | NA |
| Upf1 | 1210.834756 | -0.370716822 | 0.060362049 | -6.141554623 |
| Comp | 10.12622545 | 0.587294807 | 0.471893918 | 1.244548372 |
| Crtc1 | 1987.206553 | 0.694508098 | 0.100564171 | 6.906118664 |
| Klhl26 | 487.3158746 | 0.106646135 | 0.075016731 | 1.421631325 |
| Tmem59l | 1446.494139 | 0.378923097 | 0.056153756 | 6.747956451 |
| Crlf1 | 37.87736275 | -1.199711742 | 0.216757519 | -5.534810262 |
| 2810428I15Rik | 262.175928 | -0.017606194 | 0.106468521 | -0.165365254 |
| Uba52 | 19.53769797 | -0.384360686 | 0.272269365 | -1.41169274 |
| Kxd1 | 196.5815617 | -0.006552715 | 0.084402149 | -0.07763683 |
| Fkbp8 | 2500.961044 | 0.159184777 | 0.076542678 | 2.079686531 |
| Ell | 288.2178684 | -0.136552077 | 0.121413731 | -1.124683969 |
| Isyna1 | 251.2867603 | -0.179453914 | 0.144710801 | -1.240086524 |
| Ssbp4 | 876.1567567 | 0.299157113 | 0.065138162 | 4.592655113 |

| | | | | |
|---|---|---|---|---|
| Lrrc25 | 65.01020252 | -0.987669962 | 0.3408581 | -2.897598624 |
| Gdf15 | 12.20491556 | -2.16956173 | 0.488035455 | -4.445500238 |
| Pgpep1 | 166.5791501 | -0.776825763 | 0.169733295 | -4.576743558 |
| Lsm4 | 218.808746 | -0.251193021 | 0.096365051 | -2.606681758 |
| Jund | 1476.72973 | -0.105684407 | 0.137854011 | -0.766640062 |
| Gm11175 | 17.16041749 | -0.055135649 | 0.300052954 | -0.183753063 |
| Gm3336 | 0.36285418 | -0.150786639 | 1.711488923 | -0.088102609 |
| Pde4c | 6.152153976 | 0.72080309 | 0.467301699 | 1.542479069 |
| Rab3a | 3789.848654 | 0.596422292 | 0.056072623 | 10.63660408 |
| Mpv17l2 | 124.5250829 | -0.234537706 | 0.145374214 | -1.613337744 |
| Ifi30 | 138.4088315 | -0.378802763 | 0.204082772 | -1.856123175 |
| Pik3r2 | 1054.806414 | 0.398630378 | 0.076623498 | 5.20245599 |
| 2010320M18Rik | 39.80095612 | -0.253382567 | 0.19158463 | -1.322562084 |
| Mast3 | 1971.583413 | 0.999696606 | 0.118396569 | 8.443628206 |
| Mir7240 | 0.07796279 | -0.798301147 | 3.721829328 | -0.214491605 |
| Il12rb1 | 265.2067133 | -0.009542889 | 0.206547145 | -0.046201987 |
| Arrdc2 | 123.0827081 | 0.814727717 | 0.179525811 | 4.538220516 |
| Kcnn1 | 121.7377044 | -0.0513121 | 0.135380962 | -0.379020057 |
| A230052G05Rik | 2.786635864 | -0.448772947 | 0.757968385 | -0.592073437 |
| Ccdc124 | 301.1606489 | -0.072176305 | 0.114823848 | -0.628582875 |
| Slc5a5 | 42.07648303 | 0.657605369 | 0.201464788 | 3.264120617 |
| Rpl18a | 1403.238223 | -0.155338656 | 0.133376968 | -1.164658771 |
| Mir7067 | 0 | NA | NA | NA |
| Snora68 | 19.80479055 | -0.350393073 | 0.289538083 | -1.210179569 |
| Map1s | 619.2559287 | 0.0007386 | 0.06954267 | 0.010620812 |
| Mir1969 | 0.386072962 | -0.360393323 | 3.258545016 | -0.110599461 |
| Gm10654 | 0.749724794 | -1.260733717 | 1.111011008 | -1.134762579 |
| 1700026F02Rik | 0.167183452 | 0.517964184 | 2.765153692 | 0.187318407 |
| Gm17576 | 0.045882455 | 0.517959912 | 3.724945049 | 0.139051692 |
| Gm27628 | 0.044544648 | 0.517957688 | 3.724945049 | 0.139051095 |
| Haus8 | 58.62861715 | -0.682815097 | 0.139912166 | -4.880312538 |
| Myo9b | 1047.53723 | -0.5890571 | 0.093721004 | -6.285219688 |
| Use1 | 300.4227806 | 0.125709157 | 0.097443306 | 1.290074826 |
| Ocel1 | 128.1701314 | -0.320184728 | 0.140698499 | -2.275679772 |
| Nr2f6 | 250.0485222 | -0.407314975 | 0.12091811 | -3.368519204 |
| Ushbp1 | 11.95745292 | -1.175054088 | 0.376186066 | -3.123598115 |
| Babam1 | 547.0408888 | 0.018559407 | 0.075834801 | 0.244734697 |
| Ankle1 | 4.029650605 | 1.083458111 | 0.531546017 | 2.038314797 |
| Abhd8 | 1731.379142 | 0.41491992 | 0.083539479 | 4.966752562 |
| Mrpl34 | 112.3878888 | -0.136709597 | 0.123377051 | -1.108063421 |
| Dda1 | 548.7093606 | -0.115910053 | 0.068830219 | -1.683999485 |
| Ano8 | 471.5065914 | 0.323942947 | 0.100024574 | 3.238633612 |
| Gtpbp3 | 220.5550811 | -0.078257941 | 0.085017339 | -0.920493884 |
| Plvap | 140.3657839 | 0.444247413 | 0.126376284 | 3.515275158 |
| Bst2 | 1856.786934 | -0.640049232 | 0.257996761 | -2.480842123 |
| Mvb12a | 194.6062646 | 0.050518505 | 0.095652208 | 0.528147822 |
| Tmem221 | 4.789566606 | -0.885491491 | 0.59718438 | -1.482777382 |

| | | | | |
|---|---|---|---|---|
| Nxnl1 | 1.098494234 | 0.186780514 | 0.995016013 | 0.187716089 |
| Slc27a1 | 744.5497192 | -0.000234676 | 0.111684768 | -0.002101239 |
| Pgls | 165.2612445 | -0.31853147 | 0.110889018 | -2.872524954 |
| Fam129c | 18.63060221 | -0.796740352 | 0.301291831 | -2.644414052 |
| Colgalt1 | 425.2055677 | -0.393617402 | 0.108795963 | -3.617941239 |
| Unc13a | 2849.774368 | 0.363106458 | 0.134903801 | 2.691595464 |
| Mir6769b | 0.250814404 | 0.51797199 | 1.827301929 | 0.283462728 |
| Jak3 | 179.3964878 | -0.480999452 | 0.107340422 | -4.481065392 |
| Insl3 | 0.723819272 | -0.860840371 | 1.126744171 | -0.764006944 |
| B3gnt3 | 7.857962099 | -1.703404847 | 0.389877531 | -4.369076727 |
| Fcho1 | 729.7161296 | 0.457300098 | 0.102095034 | 4.47916104 |
| Zfp709 | 191.7633267 | -0.357719206 | 0.144122968 | -2.482041627 |
| Zfp882 | 121.9521505 | -0.279125178 | 0.156268846 | -1.786185701 |
| Zfp617 | 199.055627 | -0.316151234 | 0.125689437 | -2.515336545 |
| Ccnb2-ps | 2.364122105 | -1.173290518 | 0.876199751 | -1.339067394 |
| Zfp961 | 151.8196963 | -0.202607382 | 0.145727135 | -1.390320213 |
| Cyp4f18 | 43.06846795 | -2.546629108 | 0.375683596 | -6.778653997 |
| Gm25052 | 0 | NA | NA | NA |
| Olfr372 | 0 | NA | NA | NA |
| Olfr373 | 0 | NA | NA | NA |
| Olfr374 | 0 | NA | NA | NA |
| Gm24747 | 0 | NA | NA | NA |
| Tpm4 | 485.8967539 | -0.992929696 | 0.174173085 | -5.700821672 |
| Gm26586 | 0.363579626 | -0.457448505 | 1.734104313 | -0.263795264 |
| Gm16091 | 1.601615687 | -0.770530131 | 1.009595045 | -0.763207124 |
| Rab8a | 470.4426857 | -0.419350199 | 0.083797567 | -5.004324266 |
| Hsh2d | 15.58486664 | -1.035155614 | 0.307947911 | -3.36146334 |
| Cib3 | 0.321217965 | 0.69343402 | 3.705199858 | 0.187151583 |
| Gm11034 | 0 | NA | NA | NA |
| Fam32a | 524.8981796 | -0.130858974 | 0.119561728 | -1.09448882 |
| Ap1m1 | 552.8521804 | -0.125653398 | 0.059248941 | -2.120770353 |
| Gm10282 | 57.72901061 | -0.292728042 | 0.157414166 | -1.859604188 |
| Klf2 | 122.111689 | -0.467072766 | 0.200614504 | -2.328210354 |
| Eps15l1 | 756.2958603 | 0.142013338 | 0.080378252 | 1.766812975 |
| Calr3 | 22.05166792 | -0.610993589 | 0.238188784 | -2.565165236 |
| Gm3835 | 0.038981395 | -0.443816764 | 3.724945049 | -0.119147198 |
| 1700030K09Rik | 65.49681279 | -0.335020335 | 0.16833206 | -1.990234865 |
| Cherp | 754.5952941 | -0.081540606 | 0.070840145 | -1.151050811 |
| Mir7068 | 0 | NA | NA | NA |
| Slc35e1 | 803.4949084 | -0.172574518 | 0.06559283 | -2.630996682 |
| Gm17435 | 24.35316493 | -0.563174249 | 0.266045083 | -2.116837648 |
| Med26 | 154.280037 | -0.125941759 | 0.124105842 | -1.014793154 |
| Gm28019 | 1.785542434 | 1.479946246 | 0.842386382 | 1.75684968 |
| Smim7 | 545.215953 | -0.067353845 | 0.106946761 | -0.629788546 |
| Tmem38a | 713.1160059 | 0.484659404 | 0.084667395 | 5.724274444 |
| Nwd1 | 810.0404047 | 0.37523821 | 0.1931407 | 1.94282308 |
| Gm26237 | 0.846376972 | 0.001789795 | 1.313788798 | 0.001362316 |

| | | | | |
|---|---|---|---|---|
| Sin3b | 1202.221482 | -0.337939676 | 0.08935597 | -3.781948476 |
| F2rl3 | 3.0074759 | -1.651654107 | 0.616582129 | -2.678725232 |
| Large | 1767.595343 | 0.413014354 | 0.0739185 | 5.587428799 |
| Mir28b | 0 | NA | NA | NA |
| Gm22532 | 8.167798302 | 1.138931519 | 0.432010824 | 2.636349497 |
| Gm23255 | 0 | NA | NA | NA |
| Crry-ps | 1.276549371 | -0.721654405 | 0.919690004 | -0.784671359 |
| Gm27864 | 0 | NA | NA | NA |
| Gm11033 | 0 | NA | NA | NA |
| Isx | 0.612261413 | -0.541128138 | 1.552048477 | -0.348654147 |
| Hmgxb4 | 272.0156998 | -0.266158852 | 0.133801339 | -1.989209175 |
| Tom1 | 187.7576606 | -0.255652072 | 0.097879977 | -2.611893469 |
| Hmox1 | 1627.230529 | -2.63419886 | 0.451255676 | -5.837486368 |
| Mcm5 | 102.0398356 | -0.623063082 | 0.207846571 | -2.997706812 |
| Rasd2 | 1068.740531 | 1.060118472 | 0.146122208 | 7.255012699 |
| Gm7984 | 22.42806742 | -0.319222014 | 0.257939866 | -1.237583083 |
| Gm10358 | 5.720807359 | -0.337806323 | 0.447381044 | -0.755075183 |
| 1700007B14Rik | 4.206508402 | 0.177758648 | 0.590619909 | 0.300969617 |
| Gm38270 | 0 | NA | NA | NA |
| Gm7901 | 51.21551474 | -0.53127906 | 0.259060369 | -2.05079249 |
| Gm22544 | 0 | NA | NA | NA |
| Gm22509 | 0 | NA | NA | NA |
| Gm7069 | 0.097139547 | -0.443816764 | 3.724945049 | -0.119147198 |
| Gm10649 | 41.3073234 | 0.417732468 | 0.222041502 | 1.881326077 |
| Gm27355 | 0 | NA | NA | NA |
| Gm5353 | 7.21028118 | 0.438312354 | 0.504395608 | 0.868985271 |
| Nr3c2 | 828.7814193 | 0.430268683 | 0.0644513 | 6.675872789 |
| Gm24838 | 0 | NA | NA | NA |
| Gm23260 | 3.058609062 | 0.726817848 | 0.61294994 | 1.185770323 |
| Arhgap10 | 375.9267285 | 0.076171361 | 0.076493796 | 0.995784818 |
| 0610038B21Rik | 3.891239497 | -0.697330044 | 0.55303944 | -1.260904728 |
| Prmt10 | 268.6822526 | 0.193466332 | 0.079684212 | 2.427912971 |
| Gm23468 | 0 | NA | NA | NA |
| Tmem184c | 547.9950958 | 0.279233591 | 0.061713841 | 4.52465094 |
| 4933431K23Rik | 0.309777098 | 0.517975643 | 1.619634555 | 0.319810195 |
| 1700092C02Rik | 0 | NA | NA | NA |
| Ednra | 46.53120417 | -0.342805362 | 0.170729652 | -2.007884143 |
| Gm23981 | 0 | NA | NA | NA |
| Ttc29 | 3.962620848 | 0.320009522 | 0.622093812 | 0.514407176 |
| Pou4f2 | 9.952480325 | -0.008566921 | 0.350425792 | -0.024447177 |
| Rbmxl1 | 229.4117921 | -0.208737545 | 0.091481263 | -2.281751896 |
| Slc10a7 | 127.4608844 | -0.730557882 | 0.122855002 | -5.946504984 |
| Lsm6 | 350.1356414 | 0.243097516 | 0.094949157 | 2.560291459 |
| Gm8054 | 0.763271679 | 0.762275055 | 1.09792297 | 0.694288284 |
| Gm22044 | 0 | NA | NA | NA |
| Zfp827 | 1018.675046 | 0.186830713 | 0.068958037 | 2.709339207 |
| 1700011L22Rik | 0.586370721 | 0.741425455 | 1.190652365 | 0.62270523 |

| | | | | |
|---|---|---|---|---|
| Mmaa | 175.5439761 | 0.140844492 | 0.116071834 | 1.213425237 |
| Gm4890 | 7.177265976 | 0.114908354 | 0.439070791 | 0.261708036 |
| Smad1 | 552.5127933 | 0.126202829 | 0.099267657 | 1.271338852 |
| Gm8079 | 0.101408268 | 0.037730769 | 3.724898049 | 0.010129343 |
| Otud4 | 1259.849304 | -0.151834594 | 0.085185034 | -1.782409263 |
| Tpd52-ps | 12.01690052 | -0.074460769 | 0.315595747 | -0.235937174 |
| Abce1 | 1227.066757 | -0.264706785 | 0.078464731 | -3.373576656 |
| Gm27006 | 0.04393175 | 0.037062677 | 3.724945049 | 0.009949859 |
| Anapc10 | 137.3995736 | -0.344922014 | 0.112388856 | -3.06900548 |
| Hhip | 182.8551312 | 0.21495007 | 0.110466252 | 1.945843789 |
| Gm27820 | 0 | NA | NA | NA |
| Gm5909 | 4.376335103 | -0.224226032 | 0.465752497 | -0.48142744 |
| Gypa | 2.705694399 | 0.686267695 | 0.726536556 | 0.944574213 |
| 4930505O20Rik | 0.533083002 | -1.16217708 | 1.311925034 | -0.885856318 |
| Frem3 | 59.61918439 | 0.163462323 | 0.257707667 | 0.634293596 |
| Smarca5 | 1563.50047 | -0.202048501 | 0.07459864 | -2.708474312 |
| Gab1 | 641.4208176 | -0.469214031 | 0.101201085 | -4.636452591 |
| Gm27048 | 0 | NA | NA | NA |
| Gm4891 | 0.083526043 | 0.037073798 | 3.724945049 | 0.009952844 |
| Usp38 | 593.7032144 | -0.115447679 | 0.093912011 | -1.229317499 |
| Gm9725 | 0.99585445 | -1.250166065 | 0.994242833 | -1.257405157 |
| Gm4899 | 0.173022395 | 0.517971728 | 2.735311408 | 0.189364811 |
| Gm5354 | 0 | NA | NA | NA |
| Inpp4b | 528.8835549 | -0.13851887 | 0.084305092 | -1.643066472 |
| 4930579O11Rik | 0 | NA | NA | NA |
| Gm17169 | 0 | NA | NA | NA |
| Gm17072 | 10.76767044 | 0.739812968 | 0.358706275 | 2.062447804 |
| Gm25267 | 0 | NA | NA | NA |
| Il15 | 30.29114704 | -1.776161947 | 0.248754362 | -7.140224311 |
| Gm25138 | 0 | NA | NA | NA |
| Gm8167 | 0 | NA | NA | NA |
| Zfp330 | 179.4385362 | -0.219679162 | 0.118186903 | -1.85874371 |
| Rnf150 | 857.5833068 | 0.552378749 | 0.122969968 | 4.491980898 |
| Gm10645 | 5.503045566 | -0.436488427 | 0.457941212 | -0.953153845 |
| Tbc1d9 | 1881.448684 | -0.076741695 | 0.073240768 | -1.04780026 |
| Ucp1 | 2.960476708 | 1.001062418 | 0.688271806 | 1.454457976 |
| Elmod2 | 319.9756207 | -0.222201242 | 0.076209639 | -2.915657971 |
| Mgat4d | 0.674779921 | -0.576545474 | 1.25505246 | -0.459379582 |
| Gm15660 | 0 | NA | NA | NA |
| Clgn | 175.9442896 | 0.170316755 | 0.123392862 | 1.380280456 |
| Scoc | 643.4231515 | 0.241797663 | 0.101397438 | 2.38465258 |
| Olfr370 | 0.049553047 | -0.443818988 | 3.724945049 | -0.119147795 |
| Ndufb7 | 290.6417338 | 0.142079431 | 0.207729014 | 0.68396527 |
| Tecr | 2387.710836 | 0.246228242 | 0.067490217 | 3.648354553 |
| Dnajb1 | 549.0730292 | -0.171196609 | 0.173304788 | -0.987835428 |
| Gipc1 | 557.0990177 | -0.138087907 | 0.065116937 | -2.120614283 |
| Ptger1 | 17.78461194 | 0.212762259 | 0.32713543 | 0.65037975 |

| | | | | |
|---|---|---|---|---|
| Pkn1 | 501.3936003 | -0.339865964 | 0.097084627 | -3.500718644 |
| Ddx39 | 165.8939195 | -0.710709109 | 0.140714376 | -5.050721383 |
| Adgre5 | 155.9202038 | -1.223813839 | 0.126744591 | -9.655748089 |
| Mir1668 | 0 | NA | NA | NA |
| Gm22024 | 0 | NA | NA | NA |
| Gm26721 | 9.148635249 | 1.93116285 | 0.565735209 | 3.413545453 |
| Adgrl1 | 7570.249026 | 0.641262829 | 0.102717464 | 6.242977646 |
| AC156028.1 | 0.257982025 | 0.532522206 | 1.834375624 | 0.290301615 |
| AC156028.2 | 0.523560496 | -0.84444421 | 1.443140453 | -0.585143469 |
| Gm10644 | 10.70655375 | -0.078551746 | 0.433966041 | -0.181008971 |
| Asf1b | 15.09267688 | -0.440902168 | 0.345986977 | -1.274331685 |
| Prkaca | 1912.644388 | 0.117589587 | 0.052328454 | 2.2471443 |
| Samd1 | 227.0946354 | -0.185935583 | 0.10657885 | -1.744582372 |
| 1700067K01Rik | 1.769403265 | -0.335649178 | 0.897526972 | -0.373971132 |
| Mir8111 | 0 | NA | NA | NA |
| 2210011C24Rik | 4.04060031 | 0.179131115 | 0.543411023 | 0.329642034 |
| Mir1199 | 0 | NA | NA | NA |
| Palm3 | 9.849211078 | -0.247095102 | 0.390839211 | -0.632216767 |
| Il27ra | 17.46056049 | -0.558191741 | 0.267234522 | -2.08877108 |
| Rln3 | 9.200686208 | -0.874398289 | 0.864891747 | -1.010991597 |
| Gm10643 | 25.86145265 | 1.073813768 | 0.2471058 | 4.345562773 |
| C330011M18Rik | 25.69760837 | 0.725498494 | 0.223295545 | 3.249050463 |
| Rfx1 | 271.1011036 | -0.174031018 | 0.121596498 | -1.431217348 |
| Mir709 | 0.78198097 | 0.87812813 | 1.32703597 | 0.661721422 |
| Dcaf15 | 175.8113705 | 0.0459452 | 0.120265463 | 0.382031543 |
| Podnl1 | 7.966600571 | -0.62175141 | 0.373555234 | -1.664416272 |
| Cc2d1a | 577.8296229 | -0.012126413 | 0.049957887 | -0.2427327 |
| 4930432K21Rik | 28.89408785 | 0.498233207 | 0.208102611 | 2.394170858 |
| Gm25110 | 0 | NA | NA | NA |
| Nanos3 | 8.912972197 | 1.07786779 | 0.401224884 | 2.686443025 |
| Mir181d | 2.185676227 | 1.035866596 | 0.794455371 | 1.30387009 |
| Mir181c | 0.986137052 | -0.210476609 | 1.003152979 | -0.209815066 |
| Gm26887 | 0.282313603 | -0.453491188 | 1.791441399 | -0.253143188 |
| Gm37352 | 38.80286291 | -1.048092053 | 0.221503349 | -4.731721013 |
| Mir23a | 1.248589509 | -1.517609249 | 0.99797835 | -1.52068354 |
| Mir27a | 0.444870878 | -2.657912451 | 1.355941138 | -1.960197516 |
| Mir24-2 | 0 | NA | NA | NA |
| mmu-mir-3074-2 | 0.551097043 | -1.497219309 | 1.595975764 | -0.938121582 |
| Gm26532 | 9.112776141 | -1.09527649 | 0.410104729 | -2.670723874 |
| Zswim4 | 345.6336222 | -0.436270153 | 0.153738943 | -2.837733516 |
| D8Ertd738e | 305.0049009 | -0.276453751 | 0.122880174 | -2.249783188 |
| Mri1 | 76.67094331 | -0.119487053 | 0.148377495 | -0.805290949 |
| Ccdc130 | 95.20490353 | -0.585195377 | 0.124039515 | -4.717814142 |
| Gm23626 | 0 | NA | NA | NA |
| Cacna1a | 3274.65666 | 0.371653815 | 0.126547372 | 2.936875026 |
| Gm16183 | 0.17506888 | 1.020218006 | 2.727910183 | 0.373992521 |
| Ier2 | 148.7573661 | -0.620803877 | 0.273810984 | -2.267271634 |

| | | | | |
|---|---|---|---|---|
| Gm26664 | 2.727852729 | -0.118431872 | 0.790531773 | -0.149812918 |
| Nacc1 | 1589.037921 | -0.020885101 | 0.062172522 | -0.335921734 |
| Trmt1 | 439.8140685 | 0.119347614 | 0.094879324 | 1.257888531 |
| Nfix | 4316.174095 | 0.233898839 | 0.090002712 | 2.59879768 |
| Lyl1 | 28.85883109 | -0.46450828 | 0.30261437 | -1.534984208 |
| G430095P16Rik | 2.042153182 | 0.461928431 | 0.793889168 | 0.581855061 |
| Dand5 | 53.8269615 | -0.095151361 | 0.15023251 | -0.633360652 |
| Gadd45gip1 | 231.7276792 | -0.235593497 | 0.108694713 | -2.167478909 |
| Rad23a | 572.6689497 | -0.111827362 | 0.062066063 | -1.80174731 |
| Calr | 4607.859221 | -0.70125501 | 0.107965458 | -6.49517934 |
| 1700122E12Rik | 0.913679305 | -3.153855919 | 1.288394314 | -2.447896491 |
| Farsa | 355.8983591 | -0.151231703 | 0.106737589 | -1.416855147 |
| Mir7069 | 0 | NA | NA | NA |
| Syce2 | 15.66997929 | -0.046377882 | 0.299434061 | -0.154885126 |
| Gcdh | 224.1995156 | 0.199218862 | 0.084425597 | 2.359697408 |
| Klf1 | 0.981329651 | 0.229706686 | 1.039757935 | 0.220923235 |
| Dnase2a | 91.10452649 | -0.737386082 | 0.176730318 | -4.172380215 |
| Mast1 | 1321.949102 | 0.621901828 | 0.082120471 | 7.573042609 |
| Gm24197 | 0 | NA | NA | NA |
| Rtbdn | 11.67166441 | 1.924824155 | 0.435070956 | 4.424161454 |
| Rnaseh2a | 132.5774289 | 0.074769602 | 0.098244999 | 0.761052499 |
| Prdx2 | 1409.970294 | 0.040496432 | 0.073815487 | 0.54861701 |
| Junb | 1137.476461 | -0.409819357 | 0.331351131 | -1.236812913 |
| Hook2 | 101.5563122 | 0.142171184 | 0.147677231 | 0.962715669 |
| Best2 | 0.602809315 | -0.203681119 | 1.190862442 | -0.171036647 |
| Asna1 | 647.8422268 | -0.145141111 | 0.081909725 | -1.771964336 |
| 2310036O22Rik | 524.574075 | -0.215273056 | 0.106690357 | -2.017736764 |
| Tnpo2 | 1895.024094 | 0.107983025 | 0.054330566 | 1.987518888 |
| Gm25506 | 0.455189762 | 1.418639302 | 1.444283825 | 0.982244125 |
| A230103J11Rik | 33.92778836 | -0.030754403 | 0.199896901 | -0.153851327 |
| Fbxw9 | 120.3613756 | -0.089604871 | 0.111416857 | -0.804230829 |
| Mir7070 | 0 | NA | NA | NA |
| Gm5741 | 3.225606799 | 1.152987667 | 0.620357876 | 1.858584718 |
| Dhps | 207.0919154 | -0.423085058 | 0.093946809 | -4.503453203 |
| Wdr83 | 132.9258923 | 0.102201857 | 0.11140977 | 0.917350942 |
| Wdr83os | 100.509984 | -0.225276652 | 0.142711571 | -1.578545104 |
| Man2b1 | 495.4554757 | -0.208015398 | 0.143214805 | -1.452471323 |
| Zfp791 | 32.04972151 | 0.302282761 | 0.206015868 | 1.467279019 |
| Gm23388 | 0 | NA | NA | NA |
| Cks1brt | 0 | NA | NA | NA |
| Olfr371 | 0 | NA | NA | NA |
| Vps35 | 2002.604974 | -0.234313873 | 0.087918906 | -2.665113608 |
| Orc6 | 141.8270405 | 0.258384221 | 0.12436304 | 2.077660867 |
| Gm22336 | 0.044544648 | 0.517957688 | 3.724945049 | 0.139051095 |
| Mylk3 | 3.257485277 | -0.363050876 | 0.647904813 | -0.560346008 |
| AA672651 | 0 | NA | NA | NA |
| 4921524J17Rik | 80.96275031 | 0.053505788 | 0.147592709 | 0.362523249 |

| | | | | |
|---|---|---|---|---|
| 1700051O22Rik | 0.111006857 | 0.037730769 | 3.724898049 | 0.010129343 |
| Gpt2 | 699.6609475 | 0.003010582 | 0.150943839 | 0.019945046 |
| Dnaja2 | 1599.493447 | 0.056558287 | 0.078692242 | 0.718727604 |
| Neto2 | 1594.195169 | -0.149747317 | 0.092322387 | -1.622004394 |
| Mir8109 | 0.42925394 | 0.065791374 | 1.636092943 | 0.040212492 |
| Itfg1 | 2429.683998 | 0.324677704 | 0.081338709 | 3.991675173 |
| Phkb | 740.2327041 | -0.171289769 | 0.103114724 | -1.661157223 |
| Gm16167 | 0 | NA | NA | NA |
| Hm629797 | 14.11744488 | 0.355030939 | 0.374850815 | 0.947125963 |
| Gm23904 | 0 | NA | NA | NA |
| Gm24490 | 0.060174537 | 0.517957688 | 3.724945049 | 0.139051095 |
| Abcc12 | 27.6985328 | -0.48427422 | 0.244326909 | -1.982074848 |
| AC132147.1 | 0 | NA | NA | NA |
| Lonp2 | 859.6725007 | -0.14324095 | 0.080605372 | -1.777064564 |
| Gm16177 | 0.509424666 | 0.885145892 | 1.247964487 | 0.709271699 |
| Gm22305 | 0.045882455 | 0.517959912 | 3.724945049 | 0.139051692 |
| Siah1a | 351.8844398 | 0.288875104 | 0.113293378 | 2.549796905 |
| Gm10638 | 6.090970552 | -0.158052702 | 0.409174498 | -0.386272123 |
| N4bp1 | 1537.795251 | -0.347319544 | 0.093256934 | -3.724329427 |
| 2010110E17Rik | 5.305919521 | -0.274363733 | 0.430693968 | -0.637027107 |
| Gm27167 | 3.969393191 | -0.254174324 | 0.470723587 | -0.539965132 |
| Gm27168 | 0.090665643 | 0.037060453 | 3.724945049 | 0.009949262 |
| Gm24781 | 0 | NA | NA | NA |
| Rps13-ps1 | 58.84124857 | -0.258576957 | 0.181423456 | -1.425267511 |
| Gm5118 | 1.461301642 | 1.104915572 | 0.869760056 | 1.270368263 |
| Gm27246 | 0.76513295 | -0.556307139 | 1.103853834 | -0.503968117 |
| Gm10637 | 0.117650406 | -0.443821212 | 3.724945049 | -0.119148392 |
| Gm27169 | 0.24523014 | -0.989122537 | 2.508599194 | -0.394292775 |
| Gm27207 | 0.273479446 | 0.03707295 | 2.478031715 | 0.014960644 |
| Gm27208 | 0 | NA | NA | NA |
| 4930535O05Rik | 0 | NA | NA | NA |
| Gm19872 | 0 | NA | NA | NA |
| Cbln1 | 1077.308355 | -0.002478495 | 0.238707358 | -0.010382983 |
| Gm2694 | 162.596044 | 0.215801585 | 0.255805316 | 0.843616498 |
| Gm27016 | 0.046345709 | -0.443818988 | 3.724945049 | -0.119147795 |
| 1700120C18Rik | 0 | NA | NA | NA |
| 4933402J07Rik | 1.368974137 | 2.611072877 | 1.095725568 | 2.382962445 |
| Gm24841 | 0.060174537 | 0.517957688 | 3.724945049 | 0.139051095 |
| Zfp423 | 479.0759597 | 0.375058062 | 0.087365531 | 4.292975226 |
| Gm2716 | 0.511053402 | -1.971253888 | 1.322183398 | -1.490908063 |
| Cnep1r1 | 300.1426235 | -0.199101431 | 0.08453307 | -2.355308168 |
| Heatr3 | 504.4058116 | -0.206521831 | 0.062506053 | -3.304029285 |
| Gm9988 | 0.95664278 | 2.003180235 | 1.166951618 | 1.716592362 |
| Mir7071 | 0 | NA | NA | NA |
| 9430002A10Rik | 3.207016025 | -0.01270976 | 0.654926636 | -0.019406388 |
| Papd5 | 693.6320357 | -0.124469468 | 0.072779617 | -1.71022427 |
| Adcy7 | 236.1657784 | -0.209807324 | 0.123715553 | -1.695884785 |

| | | | | |
|---|---|---|---|---|
| Brd7 | 454.0573306 | -0.04469135 | 0.093569051 | -0.477629619 |
| Nkd1 | 108.148034 | 0.344932041 | 0.147106847 | 2.344772169 |
| Snx20 | 40.8375519 | -0.971124156 | 0.223045727 | -4.353924062 |
| Nod2 | 50.86176278 | -0.841438996 | 0.206346856 | -4.077789276 |
| Cyld | 2133.072429 | 0.518160081 | 0.063792157 | 8.122629937 |
| Rps6-ps2 | 0.516489969 | -0.074248603 | 1.44700745 | -0.051311832 |
| Mir8110 | 0 | NA | NA | NA |
| Sall1 | 352.191393 | -0.55686012 | 0.155005417 | -3.592520383 |
| Gm6625 | 0 | NA | NA | NA |
| Gm5356 | 0.530154252 | -1.382838427 | 1.265979285 | -1.092307309 |
| Gm26331 | 1.106943244 | -0.648342025 | 0.962486399 | -0.673611623 |
| Gm24212 | 0 | NA | NA | NA |
| Tox3 | 156.4269144 | 0.706378781 | 0.115505002 | 6.11556876 |
| AC129321.1 | 0 | NA | NA | NA |
| Chd9 | 2318.475297 | -0.127111214 | 0.071947926 | -1.766711304 |
| Gm6658 | 1.730564267 | -0.724158804 | 0.873520508 | -0.829011795 |
| Gm22428 | 0.075043205 | 0.517957688 | 3.724945049 | 0.139051095 |
| Rbl2 | 619.2188564 | 0.221336594 | 0.075123241 | 2.946313183 |
| Aktip | 891.7499773 | 0.247271937 | 0.063596901 | 3.888113022 |
| Gm23408 | 0 | NA | NA | NA |
| Rpgrip1l | 501.0190372 | -0.069818043 | 0.150031684 | -0.465355325 |
| Fto | 1709.590573 | 0.248734169 | 0.048503055 | 5.128216572 |
| Gm3235 | 0.204751453 | -0.427575452 | 2.20193428 | -0.19418175 |
| Irx3 | 54.79817873 | -0.51923698 | 0.227317727 | -2.284190443 |
| Irx3os | 5.187223665 | 0.276239526 | 0.48979823 | 0.563986372 |
| Gm28511 | 0.044544648 | 0.517957688 | 3.724945049 | 0.139051095 |
| Gm28515 | 0.165933562 | 0.486011036 | 2.762106923 | 0.175956634 |
| Gm28516 | 0.048949532 | 0.037064902 | 3.724945049 | 0.009950456 |
| Gm24254 | 0 | NA | NA | NA |
| Crnde | 9.401908757 | -0.156299998 | 0.361298919 | -0.432605773 |
| Gm21817 | 2.549468396 | -1.162432232 | 0.785504374 | -1.479854563 |
| Irx5 | 30.17646328 | -0.346646744 | 0.218255795 | -1.588259062 |
| Gm29177 | 0 | NA | NA | NA |
| Irx6 | 3.738898168 | -0.501271723 | 0.562601804 | -0.890988474 |
| Gm3272 | 14.76791741 | 0.136527029 | 0.302717623 | 0.451004564 |
| Mmp2 | 39.1239749 | 0.074447383 | 0.177242283 | 0.420031734 |
| Lpcat2 | 68.00775929 | 1.198878537 | 0.218412809 | 5.489048657 |
| Capns2 | 4.884769305 | 1.728435829 | 0.648533292 | 2.665145876 |
| Gm22151 | 0 | NA | NA | NA |
| Slc6a2 | 13.59669148 | 0.729957696 | 0.320501058 | 2.277551599 |
| Ces1a | 0.156716517 | -0.443821692 | 2.821984856 | -0.157272882 |
| Ces1b | 0 | NA | NA | NA |
| Ces1c | 0.230645793 | 0.517974543 | 1.885418563 | 0.274726553 |
| Ces1d | 1.445763283 | 0.310599884 | 1.190613006 | 0.260873922 |
| Ces1e | 0 | NA | NA | NA |
| Ces1f | 0.742894727 | 1.936921861 | 1.302061456 | 1.487580983 |
| Ces1g | 0.379165397 | -2.082933692 | 1.488806031 | -1.399063174 |

| | | | | |
|---|---|---|---|---|
| Ces1h | 0 | NA | NA | NA |
| Ces5a | 9.541882804 | 0.786260439 | 0.374713387 | 2.098298235 |
| Gm26843 | 19.49403685 | 0.122737156 | 0.315972474 | 0.388442558 |
| Gnao1 | 5592.419096 | 0.349949738 | 0.063470738 | 5.513560241 |
| Gm24159 | 0.183166439 | 0.998861705 | 2.686322676 | 0.371832362 |
| Amfr | 1992.652925 | -0.121399102 | 0.070214675 | -1.728970492 |
| Nudt21 | 505.4581536 | 0.145765132 | 0.079930996 | 1.823637117 |
| Ogfod1 | 1369.192929 | 0.226620143 | 0.055141617 | 4.109784123 |
| Bbs2 | 229.083479 | 0.280311958 | 0.10701905 | 2.619271587 |
| Mt4 | 0.564343484 | 0.605918808 | 1.413394234 | 0.428697665 |
| Mt3 | 1914.641927 | -0.346132274 | 0.124399983 | -2.782414151 |
| Mt2 | 2941.273043 | 0.212950033 | 0.318920584 | 0.667721194 |
| Mt1 | 3127.581112 | 0.147405196 | 0.231584557 | 0.63650702 |
| Nup93 | 580.310508 | -0.153389829 | 0.129107698 | -1.188076549 |
| Mir138-2 | 5.437333865 | 0.655438086 | 0.558805753 | 1.172926518 |
| Slc12a3 | 11.0278551 | -1.408272769 | 0.416198257 | -3.383658495 |
| Gm15889 | 0.042716945 | 0.037062677 | 3.724945049 | 0.009949859 |
| Gm15890 | 0.82545695 | -0.666546236 | 1.050307202 | -0.634620266 |
| Herpud1 | 583.7203636 | -0.058340413 | 0.095123587 | -0.613311738 |
| Ap3s1-ps2 | 8.423014622 | -0.066253807 | 0.36255095 | -0.182743437 |
| 9330175E14Rik | 54.28032715 | -0.463654394 | 0.22926948 | -2.022311883 |
| Nlrc5 | 1448.823358 | -0.413263559 | 0.123441667 | -3.347844925 |
| Mir7072 | 0.934615587 | 0.115328135 | 1.203367751 | 0.095837814 |
| Cpne2 | 360.9580537 | -0.871512053 | 0.145689432 | -5.981985394 |
| Fam192a | 433.1676574 | -0.2258298 | 0.077726512 | -2.905441047 |
| Rspry1 | 440.5534813 | -0.326807567 | 0.092777321 | -3.522494129 |
| Arl2bp | 939.3975919 | -0.108799687 | 0.074777059 | -1.454987506 |
| Pllp | 348.7112223 | -0.098631305 | 0.175127476 | -0.563197203 |
| Gm22846 | 0 | NA | NA | NA |
| Ccl22 | 3.029817251 | 0.66799236 | 0.649997528 | 1.027684463 |
| Cx3cl1 | 4526.862607 | 0.453195311 | 0.094472563 | 4.797110385 |
| Ccl17 | 8.845676815 | 1.264028254 | 0.456948288 | 2.766239171 |
| Ciapin1 | 565.2127423 | -0.242598024 | 0.084941427 | -2.856062496 |
| Gm7418 | 0.075043205 | 0.517957688 | 3.724945049 | 0.139051095 |
| Coq9 | 367.4866646 | 0.065174265 | 0.076145875 | 0.855913266 |
| Polr2c | 334.1966378 | -0.16848806 | 0.105851378 | -1.59174177 |
| Dok4 | 119.6154433 | -0.281559327 | 0.132142039 | -2.13073242 |
| Ccdc102a | 31.06208967 | -0.39261746 | 0.275072152 | -1.427325366 |
| Adgrg5 | 33.49054987 | -0.063189062 | 0.225800251 | -0.279844961 |
| Gm10286 | 0.154272232 | 0.485804557 | 2.82648974 | 0.171875578 |
| Adgrg1 | 915.8714062 | -0.046839937 | 0.137865398 | -0.339751223 |
| Adgrg3 | 4.584657955 | -0.420957331 | 0.514129217 | -0.818777299 |
| Drc7 | 36.74700655 | 0.065283448 | 0.224893156 | 0.290286504 |
| Katnb1 | 374.5637383 | 0.070021489 | 0.08619089 | 0.812400116 |
| Kifc3 | 519.7752177 | -0.287097566 | 0.09431421 | -3.044054203 |
| Cngb1 | 16.48700249 | 0.013354755 | 0.298426204 | 0.044750611 |
| Tepp | 2.391530947 | 1.420751214 | 0.809146541 | 1.755863915 |

| | | | | |
|---|---|---|---|---|
| Zfp319 | 384.2111316 | -0.180281198 | 0.084063969 | -2.14457158 |
| Usb1 | 184.5297308 | -0.982213454 | 0.143908116 | -6.825281866 |
| Mmp15 | 359.2212613 | -0.416315242 | 0.112234174 | -3.709344742 |
| Cfap20 | 234.0850033 | 0.06795711 | 0.089789937 | 0.756845502 |
| Csnk2a2 | 377.4618607 | -0.150902284 | 0.101472332 | -1.487127391 |
| Ccdc113 | 14.24627225 | 0.030462263 | 0.284050538 | 0.107242407 |
| Prss54 | 0.189486273 | 0.037075562 | 2.65748706 | 0.013951361 |
| Gm5912 | 6.84788527 | -0.281902437 | 0.401178949 | -0.702685017 |
| Gins3 | 30.02162402 | -0.408749493 | 0.232669763 | -1.756779597 |
| Ndrg4 | 14045.15733 | 0.497213377 | 0.062339673 | 7.975873984 |
| Setd6 | 77.58215835 | -0.141438514 | 0.166237122 | -0.850823886 |
| Cnot1 | 2334.865287 | -0.033897552 | 0.045594979 | -0.743449229 |
| Gm26493 | 9.543263168 | 0.537427776 | 0.406357256 | 1.32254997 |
| mmu-mir-7073 | 0.68581662 | -1.072959314 | 1.28291239 | -0.836346521 |
| SNORA76 | 0 | NA | NA | NA |
| Gm26265 | 0 | NA | NA | NA |
| 4930513N10Rik | 11.81172818 | -0.128228998 | 0.314521319 | -0.407695728 |
| Gm10094 | 48.65651831 | 0.027481797 | 0.234208171 | 0.117339188 |
| Slc38a7 | 365.4018507 | -0.352660848 | 0.074434977 | -4.737837774 |
| Got2 | 2359.910276 | 0.116552304 | 0.052979186 | 2.199964028 |
| Gm22780 | 0 | NA | NA | NA |
| Gm24132 | 0 | NA | NA | NA |
| Gm25160 | 0 | NA | NA | NA |
| Gm26399 | 0.072025462 | 0.517959912 | 3.724945049 | 0.139051692 |
| Gm5913 | 8.128038844 | -0.179200958 | 0.354146566 | -0.506007892 |
| Gm7191 | 0 | NA | NA | NA |
| Gm23494 | 0 | NA | NA | NA |
| Gm15679 | 0.097899064 | 0.037062677 | 3.724945049 | 0.009949859 |
| Cdh8 | 694.165117 | 0.56477843 | 0.097517781 | 5.791543056 |
| Gm24693 | 0.415711858 | -0.881917486 | 1.526368473 | -0.577788065 |
| Gm15681 | 35.39573601 | 0.710968694 | 0.338429571 | 2.100787744 |
| Gm15680 | 7.35783165 | -0.063902537 | 0.459455971 | -0.139083048 |
| A330008L17Rik | 88.53438246 | 0.544034681 | 0.224297641 | 2.425503352 |
| Gm8688 | 2.373068043 | 1.84730945 | 0.782775082 | 2.359949227 |
| Gm36942 | 0 | NA | NA | NA |
| Gm15210 | 118.5129343 | -0.497765482 | 0.162201429 | -3.068810701 |
| Gm5742 | 0.038981395 | -0.443816764 | 3.724945049 | -0.119147198 |
| Gm22223 | 0 | NA | NA | NA |
| Gm24602 | 0 | NA | NA | NA |
| Cdh11 | 712.29886 | 0.070080636 | 0.087446467 | 0.801411863 |
| Gm26301 | 4.260983332 | 0.628495715 | 0.639083698 | 0.983432558 |
| Gm8730 | 715.9366467 | -0.594354578 | 0.1279014 | -4.646974765 |
| Cdh5 | 215.0038206 | -0.66205009 | 0.149487302 | -4.428804852 |
| Bean1 | 97.56300219 | -0.158589176 | 0.120243167 | -1.318903857 |
| Gm10631 | 0.136963818 | 0.037074342 | 3.599544405 | 0.010299732 |
| Tk2 | 352.362705 | 0.114132016 | 0.081496517 | 1.400452691 |
| Cklf | 22.85197058 | -0.99457569 | 0.24589782 | -4.044670636 |

| | | | | |
|---|---|---|---|---|
| Cmtm2a | 0.047948698 | 0.037064902 | 3.724945049 | 0.009950456 |
| Cmtm1 | 0.461873605 | -1.589815269 | 1.499189088 | -1.060450134 |
| Gm11020 | 0 | NA | NA | NA |
| Cmtm2b | 0.088534442 | -0.905414734 | 3.723500899 | -0.243162217 |
| Cmtm3 | 162.243504 | -1.15933793 | 0.180446424 | -6.42483182 |
| Cmtm4 | 1562.719887 | 0.209072745 | 0.07768443 | 2.691308219 |
| Dync1li2 | 3022.19108 | 0.222943798 | 0.051230349 | 4.351791474 |
| Ccdc79 | 18.23009573 | -0.547248814 | 0.298604445 | -1.832688105 |
| Gm22148 | 0 | NA | NA | NA |
| Nae1 | 544.9477949 | -0.01484541 | 0.077370015 | -0.191875496 |
| Car7 | 137.94085 | -0.2271368 | 0.129079784 | -1.759662073 |
| Pdp2 | 131.2365206 | 0.113108074 | 0.117152802 | 0.965474763 |
| Cdh16 | 0 | NA | NA | NA |
| Rrad | 32.51512905 | -1.193622093 | 0.289129732 | -4.128327047 |
| Fam96b | 134.4805842 | -0.596079192 | 0.140058218 | -4.255938718 |
| 1700082M22Rik | 0.890885745 | 2.552921749 | 1.108078335 | 2.303918115 |
| AC156564.1 | 0 | NA | NA | NA |
| Ces2a | 0.086648695 | 0.037058229 | 3.724945049 | 0.009948665 |
| 4932416K20Rik | 0 | NA | NA | NA |
| Ces2b | 0.087930927 | -0.443164241 | 3.724898049 | -0.118973522 |
| Matr3-ps2 | 0 | NA | NA | NA |
| Ces2c | 0 | NA | NA | NA |
| Ces2d-ps | 0 | NA | NA | NA |
| Ces2e | 0.208319007 | -0.427624805 | 2.190873284 | -0.195184636 |
| Ces2f | 0 | NA | NA | NA |
| Ces2g | 3.472075296 | 0.246577128 | 0.546006188 | 0.451601344 |
| Ces2h | 0 | NA | NA | NA |
| Ces3a | 0.090890357 | 0.037071574 | 3.724945049 | 0.009952247 |
| Ces3b | 0.093831153 | 0.517957688 | 3.724945049 | 0.139051095 |
| Ces4a | 0.088534442 | -0.905414734 | 3.723500899 | -0.243162217 |
| Gm3830 | 0 | NA | NA | NA |
| Cbfb | 387.4469817 | -0.572659083 | 0.132043671 | -4.336891563 |
| Gm22063 | 3.344407031 | 0.63475123 | 0.559234275 | 1.13503635 |
| Gm24629 | 1.944273076 | 0.087821982 | 0.745263214 | 0.117840221 |
| D230025D16Rik | 379.8537354 | 0.157262616 | 0.086898658 | 1.809724341 |
| B3gnt9 | 40.27416652 | 0.322169162 | 0.191174842 | 1.685206892 |
| Tradd | 49.19104427 | -0.809990737 | 0.215163947 | -3.764528158 |
| Fbxl8 | 12.45441639 | -1.287396289 | 0.358347637 | -3.592590421 |
| Hsf4 | 105.9523221 | 0.27175351 | 0.138820288 | 1.957592169 |
| Nol3 | 174.8745832 | 0.205084548 | 0.122151161 | 1.678940627 |
| 4931428F04Rik | 216.122387 | 0.0563169 | 0.105796752 | 0.532312178 |
| Exoc3l | 19.59593397 | 0.025988917 | 0.260242457 | 0.099864245 |
| E2f4 | 317.7432688 | -0.436288639 | 0.073402578 | -5.94377818 |
| Elmo3 | 30.0446705 | -0.009078757 | 0.197919289 | -0.045871006 |
| Gm27320 | 4.749146687 | 0.516577406 | 0.616046928 | 0.838535803 |
| Mir328 | 3.399306995 | 0.598478306 | 0.672720758 | 0.889638529 |
| Lrrc29 | 48.99782354 | 0.695412355 | 0.239186248 | 2.90740943 |

| | | | | |
|---|---|---|---|---|
| Tmem208 | 199.915556 | -0.592057315 | 0.11020397 | -5.372377377 |
| Fhod1 | 83.53890257 | -0.620203534 | 0.210882215 | -2.940994975 |
| Slc9a5 | 310.0134594 | 0.715510809 | 0.156236782 | 4.579656584 |
| Plekhg4 | 37.98167613 | -0.311561852 | 0.367265618 | -0.848328396 |
| Kctd19 | 6.042255962 | -0.575471406 | 0.437981889 | -1.313915986 |
| Lrrc36 | 11.57992328 | 0.276489867 | 0.304670184 | 0.907505501 |
| Tppp3 | 487.5931413 | -0.413298557 | 0.15725229 | -2.628251441 |
| Zdhhc1 | 228.3370938 | 0.269729291 | 0.098602837 | 2.735512488 |
| Hsd11b2 | 0.385063269 | -0.443378452 | 1.489378307 | -0.297693641 |
| Atp6v0d1 | 1618.375995 | -0.102757125 | 0.049463226 | -2.077444891 |
| Gm38250 | 12.77996492 | -0.279649184 | 0.378581402 | -0.738676496 |
| Agrp | 1.967041363 | -1.269781981 | 0.919330418 | -1.381203055 |
| Fam65a | 1008.594529 | 0.111140035 | 0.095264719 | 1.166644229 |
| Mir1966 | 0 | NA | NA | NA |
| Gm5914 | 15.31483393 | 0.598656206 | 0.325094933 | 1.841481195 |
| Gm38384 | 0 | NA | NA | NA |
| Ctcf | 822.3735919 | 0.065966437 | 0.067543711 | 0.976648094 |
| Gm5915 | 1.873087375 | -0.964424124 | 0.809015153 | -1.19209649 |
| Rltpr | 568.0764324 | 0.465046353 | 0.086146533 | 5.398317679 |
| Acd | 255.1203144 | -0.175228915 | 0.084889765 | -2.064193658 |
| Pard6a | 178.3545304 | -0.004508021 | 0.136086267 | -0.033126198 |
| Enkd1 | 47.5336699 | 0.022556372 | 0.176565939 | 0.127750413 |
| 4933405L10Rik | 1.137565759 | -0.563283962 | 1.081285997 | -0.520938923 |
| Gfod2 | 210.1736066 | -0.16275886 | 0.100457014 | -1.620184124 |
| Ranbp10 | 548.0493242 | -0.304763811 | 0.064190517 | -4.747801201 |
| Tsnaxip1 | 3.0628478 | 0.175428755 | 0.586418658 | 0.299152751 |
| Cenpt | 114.9710339 | -0.469767844 | 0.118192551 | -3.974597709 |
| Thap11 | 254.668566 | -0.106925562 | 0.084150278 | -1.270650126 |
| Nutf2 | 250.6042831 | -0.114739692 | 0.110080367 | -1.042326572 |
| Edc4 | 794.3036961 | 0.100644918 | 0.106677817 | 0.94344748 |
| Nrn1l | 0.575698224 | 0.068826077 | 1.379178446 | 0.049903678 |
| Pskh1 | 139.9944564 | 0.011364496 | 0.107592304 | 0.105625546 |
| Ctrl | 5.684503014 | -0.526108026 | 0.44556122 | -1.180776069 |
| Gm16156 | 0.885244197 | 0.274372583 | 1.178926893 | 0.232730786 |
| Psmb10 | 1108.11009 | -0.157881052 | 0.180298881 | -0.875662965 |
| Lcat | 338.7334527 | -0.295406532 | 0.180164142 | -1.6396522 |
| Slc12a4 | 501.7697974 | -0.539153164 | 0.117725078 | -4.579764751 |
| Mir7074 | 0.042716945 | 0.037062677 | 3.724945049 | 0.009949859 |
| Dpep3 | 0.127971488 | -0.95589744 | 3.6855341 | -0.259364698 |
| Dpep2 | 27.40815167 | -1.923208329 | 0.473809183 | -4.059035575 |
| Gm22972 | 0 | NA | NA | NA |
| Dus2 | 119.8929111 | -0.363999389 | 0.134639327 | -2.703514628 |
| Ddx28 | 135.9796744 | -0.233815549 | 0.120940103 | -1.933316937 |
| Nfatc3 | 441.1062357 | -0.403709681 | 0.074934298 | -5.387515318 |
| Gm27956 | 0 | NA | NA | NA |
| Esrp2 | 1.734636384 | 1.208800518 | 0.865269959 | 1.397021247 |
| 1810019D21Rik | 1.035937776 | -0.126287825 | 1.107521496 | -0.114027425 |

| | | | | |
|---|---|---|---|---|
| Pla2g15 | 252.7616184 | -0.333882038 | 0.114450586 | -2.917259323 |
| Slc7a6 | 332.464804 | 0.136109715 | 0.132099073 | 1.03036086 |
| Slc7a6os | 176.7031225 | -0.060044825 | 0.129236631 | -0.464611501 |
| Prmt7 | 349.6739077 | -0.176646818 | 0.074402536 | -2.374204272 |
| Gm26786 | 0.213709657 | 0.538542576 | 2.560135098 | 0.210357093 |
| Smpd3 | 843.1766415 | 0.993651929 | 0.148076481 | 6.710396706 |
| Gm10629 | 2.127429481 | 0.031667068 | 0.960140279 | 0.03298171 |
| Zfp90 | 174.4767697 | 0.110348523 | 0.092823606 | 1.188798055 |
| 6030452D12Rik | 0.119587853 | 0.981063299 | 3.723618439 | 0.263470416 |
| Cdh3 | 29.60771109 | -0.191703088 | 0.253493578 | -0.756244357 |
| Gm10073 | 77.24332829 | -0.709427994 | 0.204148022 | -3.475066698 |
| 1110028F18Rik | 0.166497232 | -1.455331858 | 3.388676174 | -0.429469145 |
| Cdh1 | 87.82624501 | -0.485637942 | 0.173887044 | -2.792835681 |
| Tango6 | 76.05254567 | -0.105508367 | 0.179798173 | -0.586815567 |
| Has3 | 26.28930354 | 0.415848501 | 0.227748882 | 1.825907978 |
| Chtf8 | 670.8970226 | -0.488395022 | 0.108801053 | -4.488881405 |
| Cirh1a | 288.9954583 | -0.235603841 | 0.083648844 | -2.816582139 |
| Rpl10-ps2 | 1.60060964 | -1.237626216 | 0.802890698 | -1.54146289 |
| Sntb2 | 331.1315353 | 0.448872955 | 0.105093535 | 4.271175718 |
| Gm22085 | 0 | NA | NA | NA |
| Gm16208 | 0.439742618 | 1.063179052 | 1.777195598 | 0.598234124 |
| Vps4a | 870.3055482 | 0.164472268 | 0.05444522 | 3.020876203 |
| Gm16209 | 1.711796903 | 0.517084663 | 0.768815473 | 0.672573175 |
| Pdf | 11.11514947 | 0.632397601 | 0.317378605 | 1.992565318 |
| Cog8 | 288.9214874 | -0.163123989 | 0.096994007 | -1.681794513 |
| Nip7 | 304.7822817 | 0.012495084 | 0.094796038 | 0.131810194 |
| Tmed6 | 0.075043205 | 0.517957688 | 3.724945049 | 0.139051095 |
| Terf2 | 412.9827996 | 0.069244746 | 0.068769507 | 1.006910605 |
| Mir7075 | 0 | NA | NA | NA |
| C630050I24Rik | 0 | NA | NA | NA |
| Cyb5b | 1556.047896 | 0.028449803 | 0.05146984 | 0.552747066 |
| Rps18-ps3 | 31.42787796 | -0.258350212 | 0.203198275 | -1.271419321 |
| Nfat5 | 2785.630128 | -0.017401283 | 0.081942785 | -0.212358936 |
| Nqo1 | 48.27701717 | 0.079402737 | 0.222695459 | 0.356553014 |
| Nob1 | 223.6934552 | -0.145034788 | 0.098885352 | -1.466696379 |
| Wwp2 | 543.3421393 | -0.151333708 | 0.080878734 | -1.871118657 |
| Rps26-ps1 | 30.55227829 | -0.097350326 | 0.202413848 | -0.480946968 |
| Mir140 | 0.882355932 | 1.673935794 | 1.101341981 | 1.519905554 |
| Psmd7 | 734.7468457 | -0.166073085 | 0.094644466 | -1.754704654 |
| Gm23646 | 0.066050875 | -0.443818988 | 3.724945049 | -0.119147795 |
| Gm22825 | 0.0885994 | 0.517955464 | 3.724945049 | 0.139050498 |
| Zfhx3 | 1136.418799 | 0.147153132 | 0.091458354 | 1.608963255 |
| Gm38042 | 32.44911505 | 0.32586964 | 0.359280644 | 0.907005834 |
| Gm38318 | 5.263687666 | -0.225041965 | 0.522710794 | -0.430528635 |
| Gm20686 | 0 | NA | NA | NA |
| Mir3108 | 0.47188914 | -0.012619856 | 1.353300409 | -0.009325244 |
| D030068K23Rik | 6.705867478 | 0.098611716 | 0.390918635 | 0.252256371 |

| | | | | |
|---|---|---|---|---|
| Gm23163 | 0 | NA | NA | NA |
| Pmfbp1 | 9.435034997 | 0.281738498 | 0.475878396 | 0.59203885 |
| Dhx38 | 483.563221 | -0.082468194 | 0.097505816 | -0.845777179 |
| Txnl4b | 126.0385784 | 0.222562994 | 0.111567928 | 1.99486535 |
| Hp | 55.13014879 | -1.491676258 | 0.671820103 | -2.220350733 |
| Dhodh | 150.7597164 | 0.010470906 | 0.117918965 | 0.088797476 |
| Pkd1l3 | 15.56061734 | 0.589037866 | 0.31993522 | 1.841116045 |
| Ist1 | 750.1340784 | -0.370263484 | 0.077871746 | -4.754785924 |
| Zfp821 | 241.0196306 | 0.05058203 | 0.101782229 | 0.496963274 |
| Atxn1l | 914.3276386 | -0.00933559 | 0.088828898 | -0.105096313 |
| Ap1g1 | 1809.349129 | -0.151374642 | 0.047517659 | -3.185650218 |
| Snord71 | 7.711840907 | 0.45114007 | 0.401610307 | 1.123327916 |
| Gm25321 | 0 | NA | NA | NA |
| Gm17344 | 3.933162639 | -1.935253364 | 0.692869919 | -2.793097682 |
| Phlpp2 | 846.0254909 | 0.105121108 | 0.074780091 | 1.405736568 |
| Gm16349 | 3.039183673 | 0.617429423 | 0.814920452 | 0.757656065 |
| Marveld3 | 4.627756683 | -0.583500394 | 0.480115027 | -1.215334578 |
| Tat | 1.816527333 | 0.122089156 | 0.765535511 | 0.159482029 |
| Chst4 | 0.843638514 | -2.730277469 | 1.147092304 | -2.380172423 |
| Zfp612 | 420.3106796 | -0.003964589 | 0.093158854 | -0.042557295 |
| Gm17720 | 0 | NA | NA | NA |
| Gm21964 | 0.038981395 | -0.443816764 | 3.724945049 | -0.119147198 |
| Calb2 | 675.5295577 | 0.082547655 | 0.135692754 | 0.608342395 |
| Cmtr2 | 86.97394878 | -0.167420369 | 0.126372937 | -1.324811889 |
| Gm26832 | 0 | NA | NA | NA |
| Hydin | 49.41383929 | 0.049430108 | 0.216107621 | 0.228729128 |
| Gm23627 | 0 | NA | NA | NA |
| Gm22701 | 0 | NA | NA | NA |
| AC069308.1 | 0 | NA | NA | NA |
| AC069308.2 | 0 | NA | NA | NA |
| Vac14 | 425.0913073 | -0.264988888 | 0.071900134 | -3.685513104 |
| Mtss1l | 3340.251647 | -0.150933516 | 0.106546989 | -1.416591104 |
| Il34 | 286.7182791 | 0.361058994 | 0.11315844 | 3.190738513 |
| Gm15894 | 3.21488869 | -0.334627431 | 0.614956059 | -0.544148523 |
| Gm15895 | 1.066917243 | 1.29220025 | 0.944688504 | 1.367858552 |
| Sf3b3 | 1283.257394 | -0.480982565 | 0.069363417 | -6.934239737 |
| Snord111 | 4.735066032 | 0.632685426 | 0.51859939 | 1.219988759 |
| Gm22193 | 2.867962316 | 0.933164798 | 0.619637952 | 1.505983929 |
| Cog4 | 549.572635 | -0.41971967 | 0.075687002 | -5.545465625 |
| Gm26816 | 0.327007429 | 0.036418815 | 1.984158792 | 0.018354788 |
| Fuk | 171.2700549 | 0.255110737 | 0.134282169 | 1.899810969 |
| St3gal2 | 1026.922896 | 0.193938851 | 0.066273979 | 2.926319736 |
| Gm26132 | 2.688917129 | -0.82165577 | 0.798421917 | -1.029099717 |
| Ddx19a | 222.5239366 | -0.21569843 | 0.097001539 | -2.223659873 |
| Ddx19b | 400.244091 | 0.090298693 | 0.094457329 | 0.955973388 |
| Mir3473d | 0.67306248 | 0.713573438 | 1.205641991 | 0.591861799 |
| Aars | 2524.454962 | -0.48583002 | 0.100717677 | -4.823681758 |

| | | | | |
|---|---|---|---|---|
| Gm24103 | 0 | NA | NA | NA |
| Clec18a | 9.750147804 | 0.221723972 | 0.352581035 | 0.628859611 |
| Pdpr | 569.8637507 | 0.184545982 | 0.063283871 | 2.91616141 |
| 9430091E24Rik | 22.80186376 | -0.100504855 | 0.267682663 | -0.3754627 |
| Glg1 | 3651.900435 | 0.055636562 | 0.060392936 | 0.921242867 |
| Gm23193 | 0 | NA | NA | NA |
| Rfwd3 | 350.9863053 | -0.236512262 | 0.090589825 | -2.610803816 |
| Mlkl | 209.3845513 | -1.256689674 | 0.378789043 | -3.317650538 |
| Fa2h | 504.9529093 | -0.233312378 | 0.168797759 | -1.382200684 |
| Gm24793 | 0 | NA | NA | NA |
| Wdr59 | 307.9043311 | -0.116024154 | 0.073859284 | -1.570881099 |
| Mir1957b | 0.047948698 | 0.037064902 | 3.724945049 | 0.009950456 |
| Znrf1 | 1233.211656 | 0.260387415 | 0.080208236 | 3.246392487 |
| Ldhd | 179.3875016 | 0.190206516 | 0.10002901 | 1.90151353 |
| Zfp1 | 213.1490368 | 0.252881476 | 0.11117486 | 2.274628242 |
| Ctrb1 | 0.649033156 | -1.615026602 | 1.292187576 | -1.249839135 |
| Mir7076 | 0 | NA | NA | NA |
| Bcar1 | 823.9151968 | 0.088427656 | 0.106217188 | 0.832517387 |
| Cfdp1 | 449.2752954 | -0.251951403 | 0.130937184 | -1.924215828 |
| Tmem170 | 50.89811066 | -0.648485395 | 0.168851866 | -3.840558067 |
| Gm23853 | 0 | NA | NA | NA |
| Chst5 | 6.122444543 | -1.00337901 | 0.477649225 | -2.100660814 |
| Tmem231 | 128.7512586 | -0.072770858 | 0.114046234 | -0.638082079 |
| Gabarapl2 | 905.821491 | -0.035841086 | 0.081765534 | -0.438339772 |
| Adat1 | 132.6219706 | -0.372804233 | 0.217566812 | -1.713516094 |
| Kars | 1033.948333 | -0.205085126 | 0.119408255 | -1.71751213 |
| Terf2ip | 508.2144975 | 0.051784576 | 0.081480452 | 0.635546009 |
| Gm6793 | 13.31818528 | -0.110307805 | 0.421642356 | -0.26161462 |
| Gm22827 | 0 | NA | NA | NA |
| Gm26994 | 1.945104103 | 0.084548037 | 0.679912279 | 0.12435139 |
| Cntnap4 | 605.8576847 | 0.344477177 | 0.078777586 | 4.37278157 |
| Gm22664 | 1.761579563 | -0.410995294 | 0.940229969 | -0.437122095 |
| Gm25766 | 0 | NA | NA | NA |
| Gm24291 | 0 | NA | NA | NA |
| Mon1b | 489.7840867 | 0.009911265 | 0.074202202 | 0.133571039 |
| Syce1l | 2.596219419 | 1.048399191 | 0.675326834 | 1.552432301 |
| Adamts18 | 110.6812341 | 0.222185822 | 0.207797256 | 1.069243292 |
| Nudt7 | 34.66451103 | -1.008264869 | 0.238039321 | -4.235707214 |
| Vat1l | 1139.282187 | -0.154452653 | 0.109346886 | -1.41250162 |
| 4933408N05Rik | 0.707695957 | 0.25285561 | 1.382235323 | 0.182932389 |
| Clec3a | 0.137470077 | 0.998852408 | 3.59509949 | 0.277837209 |
| Wwox | 259.2877687 | 0.607888772 | 0.074960016 | 8.109506999 |
| Gm16116 | 0 | NA | NA | NA |
| Gm16117 | 0.066050875 | -0.443818988 | 3.724945049 | -0.119147795 |
| Gm16118 | 0.567921522 | -1.159170259 | 1.202452025 | -0.964005411 |
| Gm22556 | 0 | NA | NA | NA |
| Gm23677 | 1.922491501 | 0.001436931 | 0.801550932 | 0.001792688 |

| | | | | |
|---|---|---|---|---|
| Maf | 623.4348012 | -0.506337144 | 0.10545753 | -4.801337022 |
| Gm15655 | 1.118566861 | -1.806584828 | 1.050270348 | -1.720114094 |
| Dynlrb2 | 11.99619067 | -0.008395256 | 0.288946545 | -0.029054703 |
| Cdyl2 | 447.4685583 | -0.172750109 | 0.090898665 | -1.900469146 |
| Gm10620 | 0 | NA | NA | NA |
| Cmc2 | 85.51804082 | -0.168183469 | 0.132325537 | -1.270982713 |
| Cenpn | 11.50529104 | -0.110565976 | 0.393401349 | -0.28105134 |
| Atmin | 1056.148415 | 0.19412588 | 0.076661042 | 2.532262479 |
| 1700030J22Rik | 85.33078503 | -0.128295141 | 0.206142647 | -0.622360987 |
| Gcsh | 322.5962855 | 0.077726132 | 0.113818608 | 0.68289477 |
| Pkd1l2 | 5.568506049 | -1.018956141 | 0.409468172 | -2.488486799 |
| Gm15395 | 0 | NA | NA | NA |
| Bco1 | 1.165159028 | -0.13863782 | 0.877176728 | -0.158050043 |
| Gm20694 | 0 | NA | NA | NA |
| Gan | 644.4361812 | 0.158148484 | 0.084577812 | 1.869857831 |
| Cmip | 3352.599209 | -0.130620031 | 0.074488311 | -1.753564127 |
| Mir7077 | 0.151026272 | 0.984639628 | 2.850563064 | 0.345419345 |
| Mir7078 | 0.769887329 | -1.773631119 | 1.277155319 | -1.388735648 |
| Plcg2 | 185.0113621 | -0.648950569 | 0.167198366 | -3.881321249 |
| Gm27361 | 0 | NA | NA | NA |
| Sdr42e1 | 58.48922315 | 0.193304875 | 0.181704711 | 1.063840741 |
| Hsd17b2 | 4.082108608 | -0.906114905 | 0.599350478 | -1.511828117 |
| Mphosph6 | 123.4766612 | 0.357438244 | 0.117789571 | 3.034549163 |
| Gm10617 | 0.340027382 | 0.037076093 | 1.556176516 | 0.02382512 |
| Gm25200 | 0 | NA | NA | NA |
| Cdh13 | 744.3496955 | 0.539221309 | 0.066671536 | 8.087728912 |
| Gm24035 | 0 | NA | NA | NA |
| Gm22999 | 0.35553296 | 0.037076692 | 1.796281132 | 0.020640807 |
| Hsbp1 | 1254.7106 | 0.028324657 | 0.053950915 | 0.525007905 |
| Mlycd | 201.1556644 | -0.033288191 | 0.099384598 | -0.334943157 |
| Osgin1 | 24.0618504 | -0.911408395 | 0.329779505 | -2.763690226 |
| Necab2 | 468.9419142 | 0.577775405 | 0.150423812 | 3.840983655 |
| Slc38a8 | 4.05865859 | 0.763354351 | 0.633098961 | 1.205742543 |
| Mbtps1 | 1032.591658 | 0.205961544 | 0.069166445 | 2.977766806 |
| Hsdl1 | 556.8575977 | 0.061018322 | 0.081276547 | 0.75074944 |
| Dnaaf1 | 4.031485904 | -0.218852462 | 0.570994203 | -0.38328316 |
| Taf1c | 112.5897251 | -0.464821274 | 0.112076125 | -4.147371029 |
| Rps13-ps4 | 4.918628032 | 0.489571893 | 0.469137738 | 1.043556835 |
| Adad2 | 0.577193238 | 0.29538182 | 1.278342574 | 0.231066246 |
| Kcng4 | 392.3468948 | 0.119599921 | 0.174784833 | 0.684269449 |
| Wfdc1 | 23.0470312 | -0.164577746 | 0.294485406 | -0.558865544 |
| Atp2c2 | 14.74911433 | -0.028534696 | 0.327821812 | -0.087043311 |
| Tldc1 | 91.32362537 | -0.455838681 | 0.120348142 | -3.787666961 |
| Cotl1 | 860.7212699 | -0.337482931 | 0.111380019 | -3.030013236 |
| Klhl36 | 75.98809112 | 0.218775483 | 0.132086404 | 1.656305843 |
| Usp10 | 645.8447964 | -0.232157081 | 0.058311591 | -3.981319642 |
| Gm20388 | 0 | NA | NA | NA |

| | | | | |
|---|---|---|---|---|
| Crispld2 | 132.7086816 | -0.236168217 | 0.286817984 | -0.823407982 |
| Gm15684 | 0.208379788 | 1.004197562 | 2.623627207 | 0.382751619 |
| Zdhhc7 | 282.094203 | -0.093087169 | 0.102623076 | -0.907078336 |
| Gm15898 | 0 | NA | NA | NA |
| A130014A01Rik | 33.99075599 | -0.058093087 | 0.179785378 | -0.323124649 |
| Gm38379 | 3.088015912 | -0.085846894 | 0.72710392 | -0.118066884 |
| 6430548M08Rik | 3271.759844 | -0.183143752 | 0.095610929 | -1.915510648 |
| Fam92b | 3.336165937 | -0.224418675 | 0.596463063 | -0.376249073 |
| A330074K22Rik | 139.4201663 | 0.316072464 | 0.152583021 | 2.071478609 |
| Gse1 | 1501.615146 | -0.10891647 | 0.09394859 | -1.159319908 |
| Gm22715 | 0 | NA | NA | NA |
| Gm26971 | 9.453064617 | 0.086411386 | 0.532458323 | 0.162287605 |
| 1700016A09Rik | 5.586702209 | -0.284527487 | 0.494262545 | -0.575660628 |
| Mir7687 | 1.495561828 | 2.086383371 | 1.14736438 | 1.81841393 |
| Gins2 | 20.25107559 | -0.777775484 | 0.355870684 | -2.185556492 |
| Gm10614 | 0.913620126 | -0.145447879 | 1.077967794 | -0.134927852 |
| Gm26739 | 0.232719486 | 0.539032328 | 2.13015624 | 0.253048259 |
| 1190005I06Rik | 8.331890292 | 0.406903594 | 0.390446216 | 1.04215018 |
| Gm27021 | 0.176744647 | 1.475421612 | 2.709446342 | 0.544547271 |
| Gm26537 | 0 | NA | NA | NA |
| Emc8 | 634.0511647 | 0.218057091 | 0.072936797 | 2.989671873 |
| Cox4i1 | 2787.69123 | 0.196051517 | 0.092138292 | 2.127796295 |
| Gm33142 | 0.833731302 | 1.238716459 | 1.015177991 | 1.22019633 |
| Gm24239 | 0 | NA | NA | NA |
| Gm18709 | 0.119587853 | 0.981063299 | 3.723618439 | 0.263470416 |
| Irf8 | 490.9940863 | -0.8071888 | 0.174471122 | -4.626489414 |
| Gm26878 | 0.075043205 | 0.517957688 | 3.724945049 | 0.139051095 |
| Gm28324 | 0 | NA | NA | NA |
| Fendrr | 2.814059987 | 0.374777723 | 0.574843572 | 0.651964711 |
| Foxf1 | 17.00115138 | 0.341701796 | 0.282593717 | 1.209162748 |
| Gm27530 | 0.136963818 | 0.037074342 | 3.599544405 | 0.010299732 |
| Mthfsd | 357.8554343 | -0.057979772 | 0.115265894 | -0.503008911 |
| Foxc2 | 30.92999064 | -0.458056444 | 0.261093707 | -1.754375657 |
| AC127554.1 | 0 | NA | NA | NA |
| Foxl1 | 1.257454774 | -1.586635387 | 1.170093458 | -1.355990307 |
| Gm37609 | 0.320352182 | -0.887321112 | 1.997197594 | -0.444283087 |
| 5033426O07Rik | 1.128616957 | 0.789077707 | 1.070751183 | 0.736938441 |
| Gm26815 | 0 | NA | NA | NA |
| Gm26747 | 6.573009254 | -0.395417532 | 0.434766915 | -0.909493152 |
| Gm26784 | 0.109869311 | -0.444802825 | 3.724874622 | -0.119414174 |
| 1700018B08Rik | 0.229212118 | 1.46143958 | 2.536135019 | 0.576246757 |
| Gm27045 | 0.361384634 | 8.12E-05 | 1.820805106 | 4.46E-05 |
| 1700030M09Rik | 0.55198573 | -2.20584773 | 1.272157838 | -1.733941862 |
| Fbxo31 | 1046.295583 | 0.598935154 | 0.075117865 | 7.973271801 |
| Gm17786 | 1.168706595 | -2.097257853 | 1.097675796 | -1.910635053 |
| Map1lc3b | 1461.563508 | 0.078688231 | 0.077324748 | 1.017633214 |
| Zcchc14 | 1352.76978 | 0.254006098 | 0.097403729 | 2.607765648 |

| | | | | |
|---|---|---|---|---|
| Gm27030 | 11.36160826 | 1.145911269 | 0.350969323 | 3.26498983 |
| Gm23299 | 0.157818009 | -0.472760259 | 3.474481451 | -0.136066422 |
| Jph3 | 2592.754649 | 0.668868561 | 0.080972551 | 8.260435807 |
| Klhdc4 | 215.8889794 | -0.291325601 | 0.142039376 | -2.051020001 |
| Gm20406 | 0 | NA | NA | NA |
| Gm24283 | 0.323040168 | 0.068099188 | 1.731577475 | 0.039327832 |
| Gm26812 | 0.227390921 | 0.934164399 | 2.117045321 | 0.441258574 |
| 9530085P06Rik | 0.295432452 | -0.534324477 | 1.770898282 | -0.301725109 |
| Slc7a5 | 545.5188095 | -0.622482162 | 0.135701108 | -4.587156085 |
| BC048644 | 2.632535955 | -0.647710131 | 0.620039175 | -1.044627753 |
| Car5a | 0.305395298 | 0.499654642 | 1.816673536 | 0.275038213 |
| Gm26999 | 0.075043205 | 0.517957688 | 3.724945049 | 0.139051095 |
| Gm37497 | 0 | NA | NA | NA |
| Gm27011 | 0.490536136 | 0.751514889 | 1.489637245 | 0.504495233 |
| Banp | 309.1391948 | -0.671033216 | 0.176122333 | -3.810040463 |
| Gm17709 | 4.900990937 | -0.314551541 | 0.538532421 | -0.584090258 |
| Mir7237 | 0.780673905 | -0.422651027 | 1.281845402 | -0.32972075 |
| Gm22 | 32.71914304 | -0.855240996 | 0.209333213 | -4.085548502 |
| Zfpm1 | 722.8421094 | 0.552042701 | 0.17343269 | 3.183037186 |
| Trhr2 | 33.68978649 | -0.425191629 | 0.272319851 | -1.561368471 |
| Gm20681 | 0.36185533 | -0.594795688 | 1.514841812 | -0.392645412 |
| Zc3h18 | 664.5062256 | -0.262818256 | 0.086793143 | -3.028099314 |
| Il17c | 0.855647594 | 0.772367541 | 1.269264681 | 0.608515743 |
| Cyba | 233.8961265 | -1.112728033 | 0.211506656 | -5.260959896 |
| Mvd | 137.2979585 | -0.34386637 | 0.123519361 | -2.783906658 |
| 9330133O14Rik | 42.62923582 | 0.451406095 | 0.175423705 | 2.573233154 |
| Snai3 | 6.952624755 | -2.152271109 | 0.519764558 | -4.140857773 |
| Rnf166 | 241.1004881 | -0.236265798 | 0.092290019 | -2.560036294 |
| Gm24366 | 2.393340753 | 0.812524253 | 0.771974628 | 1.052527147 |
| Ctu2 | 170.7851 | -0.358934605 | 0.100763612 | -3.562145079 |
| Piezo1 | 341.3561795 | -0.985804482 | 0.156782486 | -6.287720693 |
| Gm26497 | 0.280709254 | 0.037077421 | 1.804674228 | 0.020545216 |
| Cdt1 | 76.18453122 | -1.073186313 | 0.272439755 | -3.939169273 |
| Aprt | 172.1584049 | -0.594149829 | 0.136999536 | -4.336874741 |
| Galns | 112.6437773 | -0.428693447 | 0.149185534 | -2.873559081 |
| Trappc2l | 257.9503243 | -0.02211977 | 0.079840785 | -0.277048499 |
| Pabpn1l | 0.259153158 | 0.45127052 | 2.095748918 | 0.215326615 |
| Mir6396 | 0 | NA | NA | NA |
| Cbfa2t3 | 329.1625399 | -0.19050338 | 0.166410056 | -1.144782862 |
| Gm15899 | 22.41855246 | -0.241415241 | 0.226903437 | -1.063955858 |
| Gm10612 | 0.141684195 | 0.037074514 | 3.558854665 | 0.010417541 |
| Acsf3 | 163.1806245 | 0.217429621 | 0.116708844 | 1.863008945 |
| Gm16378 | 9.858353745 | -0.333326896 | 0.332569343 | -1.002277879 |
| Cdh15 | 97.85341112 | 0.00323292 | 0.238593148 | 0.01354993 |
| Ankrd11 | 2871.409642 | 0.106481311 | 0.079884774 | 1.332936254 |
| Gm24445 | 2.473368846 | 0.330086064 | 0.746756209 | 0.442026541 |
| 2810013P06Rik | 123.5382447 | -0.001931953 | 0.11494636 | -0.01680743 |

| | | | | |
|---|---|---|---|---|
| Spg7 | 394.1502427 | 0.064290863 | 0.087968187 | 0.730842198 |
| Rpl13 | 1506.813565 | -0.318157316 | 0.113254791 | -2.809217275 |
| Snord68 | 0 | NA | NA | NA |
| Mir7079 | 0 | NA | NA | NA |
| Cpne7 | 380.383942 | 1.018732271 | 0.173471961 | 5.872604816 |
| Mir7080 | 0 | NA | NA | NA |
| Sult5a1 | 0.948467964 | 0.134997458 | 1.039169805 | 0.129908949 |
| Dpep1 | 2.460632423 | 0.213359166 | 0.627801501 | 0.339851316 |
| Chmp1a | 416.7948105 | -0.19958369 | 0.089518813 | -2.22951671 |
| Rps12-ps9 | 5.02878481 | 0.198741792 | 0.53419507 | 0.372039734 |
| Spata33 | 6.260054198 | 0.07932651 | 0.439751054 | 0.180389584 |
| Cdk10 | 231.2422657 | 0.375961155 | 0.095620087 | 3.931821947 |
| Spata2l | 264.1685552 | 0.40747135 | 0.161604767 | 2.521406749 |
| 4933417D19Rik | 2.606279457 | -0.048451551 | 0.702426164 | -0.068977429 |
| Vps9d1 | 418.2806154 | 0.369435909 | 0.099460562 | 3.714395933 |
| Zfp276 | 246.8177323 | -0.148953625 | 0.116391732 | -1.279761223 |
| Fanca | 30.7735149 | -1.093434711 | 0.258880646 | -4.223702039 |
| Spire2 | 197.4572324 | -0.096474711 | 0.108028862 | -0.893045703 |
| Tcf25 | 4948.09245 | 0.14356606 | 0.05653856 | 2.539259208 |
| Mc1r | 2.754460718 | -1.227767862 | 0.624998283 | -1.964433977 |
| Gm38092 | 0.099155932 | 0.037073798 | 3.724945049 | 0.009952844 |
| Tubb3 | 2462.821606 | 0.208305177 | 0.098087324 | 2.123670711 |
| Def8 | 517.338147 | 0.03025292 | 0.062360054 | 0.485132997 |
| Gm25428 | 0 | NA | NA | NA |
| Afg3l1 | 423.4960045 | -0.309364947 | 0.063841751 | -4.845809203 |
| Dbndd1 | 119.4149113 | 0.006923136 | 0.116649689 | 0.059349799 |
| Gas8 | 123.9867013 | -0.220141464 | 0.181271234 | -1.214431325 |
| Gm22314 | 2.308872772 | 0.94859477 | 0.686982518 | 1.38081355 |
| n-R5s100 | 0 | NA | NA | NA |
| n-R5s101 | 0 | NA | NA | NA |
| n-R5s102 | 0 | NA | NA | NA |
| n-R5s103 | 0 | NA | NA | NA |
| n-R5s104 | 0 | NA | NA | NA |
| n-R5s105 | 0 | NA | NA | NA |
| n-R5s106 | 0 | NA | NA | NA |
| n-R5s107 | 0 | NA | NA | NA |
| n-R5s108 | 0 | NA | NA | NA |
| n-R5s109 | 0 | NA | NA | NA |
| n-R5s110 | 0 | NA | NA | NA |
| n-R5s111 | 0 | NA | NA | NA |
| n-R5s112 | 0 | NA | NA | NA |
| n-R5s113 | 0 | NA | NA | NA |
| n-R5s114 | 0 | NA | NA | NA |
| n-R5s115 | 0 | NA | NA | NA |
| Gm25018 | 0 | NA | NA | NA |
| n-R5s117 | 0 | NA | NA | NA |
| n-R5s118 | 0 | NA | NA | NA |

| | | | | |
|---|---|---|---|---|
| n-R5s119 | 0 | NA | NA | NA |
| n-R5s120 | 0 | NA | NA | NA |
| n-R5s121 | 0 | NA | NA | NA |
| n-R5s122 | 0 | NA | NA | NA |
| n-R5s123 | 0 | NA | NA | NA |
| n-R5s124 | 0 | NA | NA | NA |
| n-R5s125 | 0 | NA | NA | NA |
| Gm24312 | 0 | NA | NA | NA |
| n-R5s127 | 0 | NA | NA | NA |
| n-R5s128 | 0 | NA | NA | NA |
| n-R5s129 | 0 | NA | NA | NA |
| n-R5s130 | 0 | NA | NA | NA |
| n-R5s131 | 0 | NA | NA | NA |
| Gm26391 | 0 | NA | NA | NA |
| n-R5s133 | 0 | NA | NA | NA |
| n-R5s134 | 0 | NA | NA | NA |
| Rn5s | 0 | NA | NA | NA |
| n-R5s136 | 0 | NA | NA | NA |
| Gm24089 | 0 | NA | NA | NA |
| n-R5s138 | 0 | NA | NA | NA |
| n-R5s139 | 0 | NA | NA | NA |
| Gm22109 | 0 | NA | NA | NA |
| n-R5s141 | 0 | NA | NA | NA |
| n-R5s142 | 0 | NA | NA | NA |
| n-R5s143 | 0 | NA | NA | NA |
| n-R5s144 | 0 | NA | NA | NA |
| Gm25212 | 0 | NA | NA | NA |
| n-R5s146 | 0 | NA | NA | NA |
| Gm22291 | 0 | NA | NA | NA |
| Gm23284 | 0 | NA | NA | NA |
| n-R5s149 | 0 | NA | NA | NA |
| Gm23647 | 0 | NA | NA | NA |
| Rhou | 659.2759545 | 0.034986853 | 0.124734346 | 0.280490932 |
| 4732419C18Rik | 5.763193032 | -1.285155664 | 0.522708651 | -2.458646249 |
| Rab4a | 299.8235972 | 0.466148396 | 0.079858566 | 5.837174652 |
| 2810455O05Rik | 58.83641319 | 0.245719071 | 0.267637856 | 0.918102822 |
| Ccsap | 351.2028742 | 0.3992407 | 0.149318259 | 2.673756724 |
| Gm4342 | 0.738388108 | -0.895946784 | 1.194062058 | -0.750335192 |
| AC147266.1 | 3.758891091 | 0.063440264 | 0.543506603 | 0.116723999 |
| Acta1 | 38.57868714 | 0.013436952 | 0.178822154 | 0.075141427 |
| Nup133 | 281.5212268 | 0.200790425 | 0.072201666 | 2.780966668 |
| Abcb10 | 231.5059475 | -0.031289264 | 0.112066114 | -0.279203618 |
| Taf5l | 237.4112607 | -0.054106446 | 0.111599944 | -0.484825027 |
| Urb2 | 291.7584138 | -0.406146714 | 0.089804813 | -4.522549496 |
| Mir1967 | 0.881693066 | 0.279334625 | 1.023304616 | 0.272973092 |
| Galnt2 | 11.54263694 | -0.403755109 | 0.326869971 | -1.235216278 |
| 1810008B01Rik | 0.299839612 | -0.907736138 | 1.63458901 | -0.555329892 |

| | | | | |
|---|---|---|---|---|
| Gm15775 | 0.11147366 | -0.955981391 | 3.722845977 | -0.25678779 |
| Gm24459 | 0 | NA | NA | NA |
| Pgbd5 | 1529.511435 | 0.528004567 | 0.126402496 | 4.177168841 |
| Cog2 | 399.9630909 | -0.101462443 | 0.082251358 | -1.23356557 |
| Agt | 480.964286 | 0.721769309 | 0.259745224 | 2.778758734 |
| Capn9 | 3.44454021 | 0.183312534 | 0.493444824 | 0.371495504 |
| 2310022B05Rik | 690.1050064 | 0.578133827 | 0.186826637 | 3.094493575 |
| Ttc13 | 501.1809921 | 0.147174132 | 0.133386627 | 1.103364973 |
| Arv1 | 50.9902722 | 0.090256521 | 0.175229552 | 0.515075912 |
| Fam89a | 4.128209337 | -0.065063361 | 0.496468167 | -0.131052432 |
| Trim67 | 432.3924619 | 0.119516246 | 0.133710725 | 0.893841882 |
| Gm16163 | 3.256114358 | 0.586421003 | 0.713110265 | 0.822342675 |
| 2810004N23Rik | 135.240311 | -0.404353178 | 0.115888941 | -3.489143779 |
| Gnpat | 451.9781084 | -0.079074308 | 0.077723004 | -1.017386143 |
| Exoc8 | 423.6774759 | -0.156399808 | 0.071852353 | -2.17668317 |
| Sprtn | 158.0541678 | -0.055931958 | 0.118596282 | -0.471616458 |
| Egln1 | 684.0866431 | 0.056181664 | 0.085898832 | 0.65404456 |
| Tsnax | 775.1924979 | 0.165148976 | 0.080578211 | 2.04954881 |
| Disc1 | 23.96939849 | -0.00553992 | 0.227303685 | -0.02437233 |
| Gm22197 | 0.208494586 | 0.03707459 | 2.629256472 | 0.014100789 |
| Gm16237 | 0.184167273 | 0.998861763 | 2.681673502 | 0.372477023 |
| Sipa1l2 | 1159.737547 | -0.081822114 | 0.084744891 | -0.965510883 |
| Map10 | 80.06219714 | -0.326795149 | 0.126714928 | -2.578979078 |
| Ntpcr | 128.5931328 | 0.108823239 | 0.164452915 | 0.661728853 |
| Pcnxl2 | 1292.263944 | 0.365680964 | 0.122266265 | 2.990857393 |
| A730098A19Rik | 4.29871648 | -0.00112827 | 0.524185561 | -0.002152424 |
| BC021891 | 32.34379101 | 0.843549589 | 0.297572913 | 2.83476604 |
| Kcnk1 | 670.0428367 | -0.009445915 | 0.09349841 | -0.101027549 |
| Slc35f3 | 269.0447631 | 0.21813769 | 0.172426466 | 1.265105611 |
| Coa6 | 108.6067078 | -0.347975008 | 0.154609259 | -2.250673801 |
| Gm17296 | 160.9155127 | -0.049901259 | 0.189220024 | -0.263720814 |
| Gm6091 | 0 | NA | NA | NA |
| Gm26759 | 0.554140925 | 1.478080968 | 1.567603583 | 0.942892057 |
| Irf2bp2 | 1675.480812 | -0.148073356 | 0.116411564 | -1.271981503 |
| Tomm20 | 2682.24209 | 0.481312512 | 0.047207051 | 10.19577584 |
| Gm26397 | 173.2653863 | -0.03935988 | 0.17589338 | -0.223771239 |
| Rbm34 | 225.3370361 | -0.009235887 | 0.1170034 | -0.07893691 |
| Gm22730 | 0 | NA | NA | NA |
| Pard3 | 532.7837827 | -0.0367926 | 0.070331965 | -0.523127711 |
| Gm6921 | 2.797109942 | -0.18650645 | 0.643508893 | -0.289827307 |
| Gm22968 | 0.822547242 | 0.46443197 | 1.212358898 | 0.383081257 |
| Mir21c | 0.121578509 | 0.037073798 | 3.724945049 | 0.009952844 |
| Nrp1 | 639.8097654 | -0.825194935 | 0.104098587 | -7.927052218 |
| Mir1903 | 0 | NA | NA | NA |
| Gm26002 | 0 | NA | NA | NA |
| Itgb1 | 1279.915761 | -0.570357691 | 0.1007218 | -5.662703546 |
| Gm3952 | 0.143618953 | -0.891120723 | 2.890309379 | -0.308313265 |

| | | | | |
|---|---|---|---|---|
| Gm26100 | 0.368403345 | 0.046335683 | 1.696795358 | 0.027307761 |
| Ccdc7 | 0.294450523 | -0.66232641 | 1.735144243 | -0.381712594 |
| Gm22545 | 0 | NA | NA | NA |
| 1700008F21Rik | 0.249972158 | 0.875373185 | 2.089027157 | 0.419033894 |
| Gm22374 | 0.159888962 | 0.51795502 | 3.461347823 | 0.149639691 |
| Gm10999 | 0.583900451 | -1.571784353 | 1.365029172 | -1.15146576 |
| 2610044O15Rik8 | 322.1898291 | 0.025120925 | 0.133110712 | 0.18872204 |
| Gm21399 | 59.65666709 | -0.761270428 | 0.164257071 | -4.63462804 |
| Gm26870 | 1.94439448 | 0.484531656 | 0.800741788 | 0.605103496 |
| Gm10722 | 0.242739364 | -0.401870401 | 2.085877753 | -0.19266249 |
| Gm11168 | 0.318248637 | -1.392959726 | 2.053024126 | -0.67849165 |
| Gm10721 | 0.187213551 | 0.037076654 | 2.667715625 | 0.013898278 |
| Gm10720 | 0 | NA | NA | NA |
| Gm10719 | 0.044544648 | 0.517957688 | 3.724945049 | 0.139051095 |
| Gm10718 | 0.690752925 | 1.105117827 | 1.268381947 | 0.87128158 |
| Gm10717 | 0.158171364 | 0.51796488 | 2.813813721 | 0.184079307 |
| Gm17535 | 0 | NA | NA | NA |
| Gm10715 | 0 | NA | NA | NA |
| mmu-mir-101c | 0 | NA | NA | NA |
| 4930433N12Rik | 0 | NA | NA | NA |
| Gm27679 | 0 | NA | NA | NA |
| Alkbh8 | 460.6917347 | 0.12235304 | 0.060319344 | 2.028421274 |
| Gm24918 | 2.468298956 | -0.908443104 | 0.726399964 | -1.250610062 |
| Gm23275 | 1.098824885 | 2.165643273 | 1.191217368 | 1.818008477 |
| Cwf19l2 | 426.6056693 | 0.209246217 | 0.114279198 | 1.831008799 |
| Gucy1a2 | 1402.939796 | 0.404290424 | 0.096626257 | 4.184063786 |
| Gm22195 | 0.192501058 | -0.443822989 | 2.674470728 | -0.16594797 |
| Aasdhppt | 467.0250061 | -0.031419587 | 0.089622931 | -0.350575312 |
| Kbtbd3 | 165.3891345 | -0.116554632 | 0.136401751 | -0.854495133 |
| Gm23992 | 0.044191549 | 0.037062677 | 3.724945049 | 0.009949859 |
| Msantd4 | 720.9495236 | -0.324704372 | 0.07042281 | -4.610784073 |
| Gria4 | 2416.785679 | 0.348072555 | 0.085796281 | 4.056965545 |
| Gm23811 | 8.724062623 | 1.147238807 | 0.429243305 | 2.672700526 |
| Gm24896 | 0 | NA | NA | NA |
| Casp1 | 190.837744 | -0.910543256 | 0.217167723 | -4.192811179 |
| Casp4 | 185.281841 | -0.994284851 | 0.347718252 | -2.859455448 |
| Casp12 | 173.1181621 | -0.732237025 | 0.205548931 | -3.562348986 |
| Pdgfd | 56.87235133 | 0.222037511 | 0.24057423 | 0.92294803 |
| Ddi1 | 0 | NA | NA | NA |
| Gm23097 | 0 | NA | NA | NA |
| Dync2h1 | 1480.682915 | 0.24461036 | 0.101413512 | 2.412009557 |
| Dcun1d5 | 410.911556 | 0.004409126 | 0.091144971 | 0.048374871 |
| Mmp13 | 18.19977551 | -0.995303723 | 0.410370167 | -2.425380311 |
| Mmp12 | 7.267951943 | -2.003350525 | 0.523506328 | -3.826793331 |
| Mmp1b | 0 | NA | NA | NA |
| Mmp3 | 11.53096597 | 0.066407312 | 0.485363885 | 0.13681964 |
| Mmp1a | 0.181654796 | -1.405197014 | 3.311020957 | -0.424399915 |

| | | | | |
|---|---|---|---|---|
| Mmp10 | 0.099106094 | -0.93550548 | 3.724188147 | -0.251197158 |
| Mmp8 | 40.56306267 | -1.796820549 | 0.859903976 | -2.089559531 |
| Mmp27 | 4.159708068 | -3.503656234 | 0.826468544 | -4.239309843 |
| Mmp20 | 0.336931769 | -1.411904722 | 1.762672688 | -0.801002212 |
| AV064505 | 0 | NA | NA | NA |
| Mmp7 | 0.107803068 | 0.036092186 | 3.724874622 | 0.009689503 |
| Gm10709 | 0.713745664 | -1.679808315 | 1.193144708 | -1.407883138 |
| Tmem123 | 239.3847869 | -0.498128338 | 0.13199872 | -3.773736117 |
| Mir5618 | 0.424774107 | 0.165878997 | 1.432010683 | 0.115836424 |
| Birc2 | 772.896626 | 0.058046048 | 0.106750502 | 0.543754326 |
| Birc3 | 154.0540683 | -0.140233257 | 0.249303813 | -0.562499446 |
| Yap1 | 390.0541373 | -0.601177836 | 0.102212061 | -5.881672163 |
| Gm20416 | 0.109982625 | -0.443823436 | 3.724945049 | -0.119148989 |
| 9230110C19Rik | 76.10268791 | -0.330034863 | 0.144244552 | -2.288023076 |
| Cep126 | 298.1066087 | -0.001499968 | 0.087909468 | -0.017062646 |
| 1700128F08Rik | 0.650208918 | -0.599869314 | 1.266682773 | -0.473575016 |
| Mir1899 | 0.060174537 | 0.517957688 | 3.724945049 | 0.139051095 |
| Trpc6 | 122.6815528 | 0.663013465 | 0.158309784 | 4.188076368 |
| Pgr | 476.6556311 | 0.542063559 | 0.091676652 | 5.912776567 |
| Gm16485 | 192.3635005 | 0.628991639 | 0.141139776 | 4.456515774 |
| Arhgap42 | 263.2227951 | 0.277232403 | 0.083956079 | 3.302112309 |
| Gm16833 | 0 | NA | NA | NA |
| Gm22815 | 0 | NA | NA | NA |
| Cntn5 | 249.9584538 | 0.998628466 | 0.129512631 | 7.710664635 |
| Gm26195 | 0.87738752 | 2.671446857 | 1.115658687 | 2.394501911 |
| Mir6237 | 0 | NA | NA | NA |
| Gm24496 | 0.646963711 | 1.006953388 | 1.256336146 | 0.801499974 |
| Gm25086 | 0.47668666 | 1.601687163 | 1.64976295 | 0.970858973 |
| Gm25365 | 0 | NA | NA | NA |
| Gm21070 | 0 | NA | NA | NA |
| Gm29176 | 0 | NA | NA | NA |
| Jrkl | 160.5671298 | -0.188860522 | 0.139909391 | -1.349877382 |
| Ccdc82 | 240.4966967 | -0.007359519 | 0.097890835 | -0.075180879 |
| Gm26189 | 1.402298525 | -0.329663538 | 0.805476755 | -0.409277532 |
| Phxr4 | 43.51084174 | -0.124520854 | 0.331933883 | -0.375137521 |
| Gm17571 | 0 | NA | NA | NA |
| Maml2 | 397.5331643 | -0.254247982 | 0.086454745 | -2.940821608 |
| Gm22643 | 0 | NA | NA | NA |
| Mtmr2 | 890.4922701 | 0.102987742 | 0.103900242 | 0.991217536 |
| Cep57 | 215.6600884 | 0.155313244 | 0.120060936 | 1.293620138 |
| Fam76b | 356.5380025 | 0.386997745 | 0.143643919 | 2.69414638 |
| Sesn3 | 970.3136834 | 0.29218265 | 0.083301763 | 3.507520586 |
| Endod1 | 2711.618671 | -0.239226352 | 0.179531788 | -1.332501362 |
| Kdm4dl | 4.695898157 | -0.088624099 | 0.5836402 | -0.151847147 |
| Kdm4d | 25.20782128 | 0.319288565 | 0.232008635 | 1.376192591 |
| Cwc15 | 455.3458575 | -0.135516987 | 0.130795103 | -1.03610138 |
| Amotl1 | 1488.185642 | -0.750086835 | 0.112318863 | -6.678191136 |

| | | | | |
|---|---|---|---|---|
| Piwil4 | 14.13799992 | -0.700103473 | 0.270226068 | -2.59080657 |
| Gm10706 | 3.101177844 | -1.308475744 | 0.63718108 | -2.053538289 |
| Gm15586 | 0 | NA | NA | NA |
| Fut4 | 18.43119656 | -0.787567716 | 0.269873463 | -2.918285137 |
| 1700012B09Rik | 5.943573362 | -0.372159727 | 0.44996293 | -0.827089749 |
| Ankrd49 | 112.3845942 | -0.129710498 | 0.124557208 | -1.041372875 |
| Mre11a | 230.8957433 | -0.175581106 | 0.091605006 | -1.916719546 |
| Gm16379 | 0.870448589 | -0.27651001 | 1.010739157 | -0.273572077 |
| Gm16302 | 2.36738784 | -0.821479634 | 0.6770066 | -1.213399743 |
| Gpr83 | 316.7124723 | -0.43378172 | 0.126750765 | -3.422320342 |
| Izumo1r | 3.609677315 | -1.017171498 | 0.510678125 | -1.991805502 |
| Panx1 | 176.4686585 | -0.583099429 | 0.114715492 | -5.083005076 |
| Hephl1 | 1.803729066 | -0.999960393 | 1.001856027 | -0.998107878 |
| Gm22658 | 0.042716945 | 0.037062677 | 3.724945049 | 0.009949859 |
| Vstm5 | 75.69295347 | 0.413624317 | 0.176962842 | 2.337351234 |
| Med17 | 334.3072139 | -0.129894283 | 0.075217037 | -1.726926339 |
| 4931406C07Rik | 230.1835539 | -0.084617914 | 0.118214305 | -0.715800968 |
| Taf1d | 188.0883961 | 0.186295901 | 0.15027348 | 1.239712429 |
| Gm25500 | 1.755293934 | 0.649674388 | 0.83907828 | 0.774271487 |
| mmu-mir-7650 | 0 | NA | NA | NA |
| Gm24455 | 11.03006032 | 0.411648507 | 0.31774593 | 1.295527237 |
| Gm24299 | 7.628192187 | 0.749807084 | 0.396149182 | 1.892739196 |
| AC154296.1 | 0.044191549 | 0.037062677 | 3.724945049 | 0.009949859 |
| Gm25791 | 5.041681076 | 0.940556267 | 0.471332657 | 1.995525354 |
| Gm22620 | 6.972143724 | 0.625793274 | 0.39890056 | 1.568795173 |
| Gm24357 | 17.62472295 | -0.41311432 | 0.261939179 | -1.577138333 |
| Gm23455 | 26.8443768 | -0.678973411 | 0.247805638 | -2.739943361 |
| Gm22579 | 6.383805412 | 0.059291949 | 0.468630188 | 0.12652183 |
| Cep295 | 648.5597578 | 0.379803189 | 0.114130808 | 3.327788483 |
| Scarna9 | 8.758301848 | 0.758521713 | 0.391490791 | 1.937521213 |
| Gm24339 | 46.69042643 | 0.460701333 | 0.281418241 | 1.637069901 |
| Gm16568 | 0.41713571 | -0.32394011 | 1.334587559 | -0.242726757 |
| Smco4 | 17.55463808 | -0.56732939 | 0.299478889 | -1.894388589 |
| Ccdc67 | 5.175431821 | 0.275922442 | 0.434279194 | 0.635357268 |
| Slc36a4 | 909.8448179 | 0.287629778 | 0.074440005 | 3.863913991 |
| Mtnr1b | 0.066050875 | -0.443818988 | 3.724945049 | -0.119147795 |
| Fat3 | 3920.904863 | 0.496064354 | 0.132679138 | 3.738827068 |
| Gm5611 | 1.991326778 | 0.187871541 | 0.789871949 | 0.237850631 |
| CT943659.1 | 0 | NA | NA | NA |
| Chordc1 | 659.9625695 | -0.692442056 | 0.111505027 | -6.209962662 |
| Naalad2 | 9.142264245 | 0.28157997 | 0.489004393 | 0.575822985 |
| Ubtfl1 | 0.110595523 | 0.037071574 | 3.724945049 | 0.009952247 |
| Gm5612 | 0.507208347 | -0.826951908 | 1.421322595 | -0.58181859 |
| Mbd3l2 | 0.241190471 | 0.594215967 | 3.027989488 | 0.196241093 |
| Zfp558 | 31.78569487 | 0.178063056 | 0.218268438 | 0.815798463 |
| Gm22854 | 0 | NA | NA | NA |
| Mbd3l1 | 0.181654796 | -1.405197014 | 3.311020957 | -0.424399915 |

| | | | | |
|---|---|---|---|---|
| Muc16 | 0.130524232 | 0.517958497 | 3.657871415 | 0.141601067 |
| Gm25295 | 0 | NA | NA | NA |
| Gm24546 | 0 | NA | NA | NA |
| Olfr24 | 0 | NA | NA | NA |
| Olfr828 | 0 | NA | NA | NA |
| Olfr829 | 0.068097359 | 0.037064902 | 3.724945049 | 0.009950456 |
| Olfr830 | 0 | NA | NA | NA |
| Olfr831-ps1 | 0 | NA | NA | NA |
| Olfr832 | 0 | NA | NA | NA |
| Olfr833-ps1 | 0 | NA | NA | NA |
| Olfr834 | 0 | NA | NA | NA |
| Olfr835 | 0.046345709 | -0.443818988 | 3.724945049 | -0.119147795 |
| Olfr836 | 0 | NA | NA | NA |
| Olfr837 | 0 | NA | NA | NA |
| Olfr843 | 0 | NA | NA | NA |
| Olfr844 | 0 | NA | NA | NA |
| Olfr845 | 0 | NA | NA | NA |
| Olfr846 | 0.049553047 | -0.443818988 | 3.724945049 | -0.119147795 |
| Olfr847 | 0 | NA | NA | NA |
| Gm23048 | 0 | NA | NA | NA |
| Olfr849 | 0 | NA | NA | NA |
| Olfr850 | 0 | NA | NA | NA |
| Olfr851 | 0 | NA | NA | NA |
| Olfr853 | 0 | NA | NA | NA |
| Olfr854 | 0 | NA | NA | NA |
| Olfr855 | 0 | NA | NA | NA |
| Zfp317 | 272.3857193 | -0.164471739 | 0.085104837 | -1.932578043 |
| Olfr857 | 1.183877471 | 0.644660272 | 0.924877399 | 0.697022408 |
| Olfr58 | 0.474412771 | 1.390508678 | 1.505105482 | 0.923861281 |
| Olfr859 | 0.318756133 | 0.10802338 | 1.776083431 | 0.060821118 |
| Olfr860 | 0 | NA | NA | NA |
| Olfr862 | 0 | NA | NA | NA |
| Olfr77 | 0 | NA | NA | NA |
| Gm7808 | 2.442408819 | -0.7231008 | 0.666379233 | -1.08511905 |
| Olfr866 | 0 | NA | NA | NA |
| AC121953.1 | 0 | NA | NA | NA |
| Gm26231 | 0 | NA | NA | NA |
| Olfr867 | 0.092691418 | -0.908687078 | 3.724560878 | -0.2439716 |
| Olfr868 | 0.089062654 | -0.443823436 | 3.724945049 | -0.119148989 |
| Olfr869 | 0 | NA | NA | NA |
| Olfr870 | 0 | NA | NA | NA |
| Gm25914 | 0 | NA | NA | NA |
| Gm26290 | 0 | NA | NA | NA |
| Olfr871 | 0 | NA | NA | NA |
| Olfr872 | 0 | NA | NA | NA |
| Gm25425 | 0.049553047 | -0.443818988 | 3.724945049 | -0.119147795 |
| Olfr39 | 0 | NA | NA | NA |

| | | | | |
|---|---|---|---|---|
| Olfr873 | 0 | NA | NA | NA |
| Olfr18 | 0.954195627 | -2.153708316 | 1.376405782 | -1.564733558 |
| Zfp560 | 108.4598925 | -0.027163429 | 0.146725615 | -0.185130792 |
| Gm26733 | 0 | NA | NA | NA |
| Zfp26 | 822.179251 | 0.110143082 | 0.089851165 | 1.225839224 |
| Zfp426 | 384.7268472 | -0.280361765 | 0.118757898 | -2.36078416 |
| Gm23316 | 0 | NA | NA | NA |
| Zfp266 | 1050.28284 | -0.144120562 | 0.070240934 | -2.051803042 |
| Zfp846 | 184.0291169 | 0.069219524 | 0.124297089 | 0.556887729 |
| Fbxl12os | 22.46068426 | -0.041503637 | 0.280780739 | -0.147815113 |
| Gm26521 | 12.01662456 | 0.198815914 | 0.387684377 | 0.512829317 |
| Fbxl12 | 213.0329372 | 0.196177143 | 0.16353886 | 1.199575088 |
| Ubl5 | 521.6630355 | 0.085136169 | 0.111746537 | 0.761868521 |
| Pin1 | 657.0847573 | -0.023542096 | 0.068621882 | -0.343069811 |
| Olfm2 | 557.2935379 | 0.057253091 | 0.088246542 | 0.648785657 |
| Col5a3 | 212.1757937 | -1.040402177 | 0.225295602 | -4.617942693 |
| Gm26274 | 0 | NA | NA | NA |
| Rdh8 | 0.705140734 | -3.108144125 | 1.442406378 | -2.154832489 |
| A230050P20Rik | 616.357269 | 0.231568452 | 0.125313342 | 1.847915388 |
| Angptl6 | 38.80084079 | -0.18908305 | 0.194482409 | -0.972237291 |
| Ppan | 173.1887125 | 0.020462945 | 0.146787573 | 0.139405162 |
| Gm24067 | 0 | NA | NA | NA |
| Gm23008 | 1.246341358 | 0.671346544 | 0.901032283 | 0.745086005 |
| Eif3g | 495.9541041 | -0.279443733 | 0.105844288 | -2.640139961 |
| Dnmt1 | 868.320093 | -0.324991414 | 0.097354266 | -3.338234946 |
| Mir7081 | 0.867462339 | 1.446371073 | 1.118389873 | 1.293261955 |
| S1pr2 | 119.737217 | -1.742278748 | 0.223277666 | -7.803193126 |
| Mrpl4 | 356.3497304 | -0.170745537 | 0.067717503 | -2.521438782 |
| Icam1 | 316.3122786 | 0.053992454 | 0.348726965 | 0.1548273 |
| Icam4 | 2.361925876 | 1.359175015 | 0.743344743 | 1.828458501 |
| Icam5 | 1112.549133 | 1.483569576 | 0.231510962 | 6.408204456 |
| Mir1900 | 0.298647883 | -0.206332879 | 2.06488889 | -0.099924446 |
| Gm26592 | 23.65836997 | 0.10819287 | 0.224659551 | 0.481585889 |
| Zglp1 | 9.660883422 | -1.604041626 | 0.431686753 | -3.715753646 |
| Fdx1l | 166.5463135 | -0.061427048 | 0.115310049 | -0.53271201 |
| Raver1 | 630.7586862 | -0.19577721 | 0.073935575 | -2.647943293 |
| Mir7082 | 0 | NA | NA | NA |
| 1700084C06Rik | 2.554977676 | -0.247575478 | 0.654499868 | -0.378266658 |
| Tyk2 | 239.0143666 | -0.650759191 | 0.156231831 | -4.165343165 |
| Cdc37 | 916.0704561 | -0.133016897 | 0.093139137 | -1.428152572 |
| Pde4a | 1129.44041 | 0.262481691 | 0.101637656 | 2.58252406 |
| Gm16754 | 3.260813172 | -0.901352903 | 0.615159007 | -1.465235643 |
| Keap1 | 857.7412683 | 0.040956176 | 0.065886566 | 0.62161649 |
| S1pr5 | 222.4130938 | 0.0426461 | 0.14313713 | 0.297938768 |
| Atg4d | 243.847421 | -0.364721352 | 0.086452205 | -4.218762857 |
| Kri1 | 243.0174812 | -0.189905059 | 0.111816077 | -1.6983699 |
| Gm37379 | 20.17791531 | -0.045368026 | 0.306892389 | -0.147830403 |

| | | | | |
|---|---|---|---|---|
| Cdkn2d | 273.9573276 | 0.020353095 | 0.087308157 | 0.233117913 |
| Ap1m2 | 9.647799368 | 0.196867504 | 0.384922046 | 0.511447722 |
| Slc44a2 | 583.7110967 | -0.538565991 | 0.119692728 | -4.499571533 |
| Ilf3 | 1100.802251 | -0.061391445 | 0.053998442 | -1.13691141 |
| Gm16853 | 1.441198087 | -0.072984917 | 0.894761163 | -0.081569161 |
| Qtrt1 | 87.06778592 | -0.02818499 | 0.152402789 | -0.184937495 |
| Dnm2 | 1057.753246 | -0.167792334 | 0.103624164 | -1.619239435 |
| Mir199a-1 | 0.092691418 | -0.908687078 | 3.724560878 | -0.2439716 |
| Tmed1 | 82.01844689 | -0.274207091 | 0.147525251 | -1.858712922 |
| AB124611 | 105.0329745 | -1.448084269 | 0.237761535 | -6.090490081 |
| Gm20519 | 0.907036939 | -0.525039841 | 1.292538992 | -0.406208126 |
| Carm1 | 707.8332732 | 0.048985486 | 0.062171027 | 0.787915008 |
| Yipf2 | 186.1268908 | 0.120665951 | 0.098564085 | 1.224238541 |
| 1810026J23Rik | 718.2922187 | 0.103907497 | 0.113459674 | 0.915809936 |
| Mir1946b | 0.611133411 | 0.977477475 | 1.473311706 | 0.663455989 |
| Smarca4 | 2424.773633 | -0.119231596 | 0.051833799 | -2.300267346 |
| Gm27373 | 1.398170896 | -0.091266467 | 0.842338375 | -0.108348936 |
| Gm26511 | 0.192512644 | -0.443822369 | 2.697849763 | -0.164509668 |
| Ldlr | 610.6285346 | -0.356982843 | 0.181107947 | -1.971105344 |
| Spc24 | 9.830363938 | 0.005596323 | 0.377057134 | 0.014842108 |
| Kank2 | 390.6615536 | -0.313209224 | 0.144405018 | -2.168963566 |
| Dock6 | 147.3513096 | -0.046866628 | 0.141799778 | -0.330512708 |
| Mir7083 | 0 | NA | NA | NA |
| Gm6484 | 0.140389662 | -0.281291492 | 3.571608989 | -0.078757639 |
| Rab3d | 110.9505493 | -0.309256287 | 0.165429991 | -1.869408841 |
| Tmem205 | 101.5694773 | -0.573676154 | 0.19207175 | -2.986780479 |
| Ccdc159 | 62.58867534 | 0.177548546 | 0.205738428 | 0.862981933 |
| Lppr2 | 1319.490486 | 0.101290629 | 0.096375414 | 1.05100072 |
| Swsap1 | 101.2171663 | 0.32918696 | 0.1279305 | 2.573170277 |
| Epor | 34.63035605 | 0.00414285 | 0.24980232 | 0.016584516 |
| Rgl3 | 100.740861 | 0.0679865 | 0.219463384 | 0.309785164 |
| Ccdc151 | 32.36726646 | 0.321815016 | 0.239372987 | 1.344408238 |
| Prkcsh | 759.8445769 | -0.473269717 | 0.076744296 | -6.166838992 |
| Elavl3 | 2565.303786 | 0.266850294 | 0.071303651 | 3.742449253 |
| Zfp653 | 159.2068104 | 0.266750749 | 0.107380845 | 2.484155798 |
| Gm16845 | 10.87916454 | 0.978103199 | 0.357084351 | 2.739137677 |
| Ecsit | 303.4651564 | -0.026589548 | 0.071940458 | -0.369604934 |
| Cnn1 | 3.417651458 | -1.209582363 | 0.624424564 | -1.937115277 |
| Elof1 | 264.1837638 | -0.359508444 | 0.113710359 | -3.161615593 |
| Mir7084 | 0 | NA | NA | NA |
| Acp5 | 37.28268211 | -0.837696237 | 0.244437048 | -3.427042843 |
| Pigyl | 121.7874717 | -0.253513945 | 0.149089552 | -1.700413891 |
| Rpl15-ps3 | 5.54092659 | -0.328460542 | 0.393461403 | -0.834797364 |
| Zfp872 | 0.324168737 | -2.066539376 | 1.674850757 | -1.233864789 |
| 1810064F22Rik | 0.089062654 | -0.443823436 | 3.724945049 | -0.119148989 |
| Zfp809 | 131.6943064 | -0.311671053 | 0.117691417 | -2.648205469 |
| Zfp599 | 85.70332244 | 0.393366589 | 0.171104174 | 2.298988861 |

| | | | | |
|---|---|---|---|---|
| Zfp810 | 217.69277 | 0.590008579 | 0.114321399 | 5.160963594 |
| Gm10182 | 10.2804556 | 0.146727538 | 0.31079359 | 0.472106062 |
| Anln | 682.8292644 | 0.069515864 | 0.182192464 | 0.381551806 |
| 9530077C05Rik | 75.06799202 | -0.164924984 | 0.175046229 | -0.942179594 |
| Rp9 | 229.1720738 | -0.076328683 | 0.076787356 | -0.994026709 |
| Gm17545 | 0 | NA | NA | NA |
| Bbs9 | 315.6451564 | 0.268600646 | 0.09101442 | 2.951187797 |
| Gm2976 | 0 | NA | NA | NA |
| Gm16165 | 1.13143437 | -2.910944076 | 1.097004828 | -2.653538071 |
| Gm16164 | 0 | NA | NA | NA |
| Gm27639 | 0 | NA | NA | NA |
| Bmper | 91.07485983 | 0.116824852 | 0.119848331 | 0.974772454 |
| Npsr1 | 33.2953994 | 0.858335808 | 0.198109839 | 4.332625843 |
| Gm15600 | 0.84374619 | -0.649451409 | 0.930576671 | -0.697902095 |
| Gm10701 | 1.895990101 | -0.570956197 | 0.702338805 | -0.81293557 |
| Gm22380 | 0.229628521 | -0.911764803 | 2.121064568 | -0.42986188 |
| Dpy19l1 | 977.1122373 | 0.11421322 | 0.060899316 | 1.875443408 |
| Dpy19l2 | 3.565532606 | 1.051636256 | 0.630186002 | 1.668771208 |
| Cypt4 | 0 | NA | NA | NA |
| Gm38179 | 0.167183452 | 0.517964184 | 2.765153692 | 0.187318407 |
| Tbx20 | 7.101997061 | -0.15545108 | 0.502065184 | -0.309623301 |
| Gm23893 | 0 | NA | NA | NA |
| Gm10181 | 0 | NA | NA | NA |
| Gm10180 | 2.742928344 | -0.915932819 | 0.762668065 | -1.200958663 |
| Herpud2 | 587.9118386 | -0.143155001 | 0.085386649 | -1.676550162 |
| Gm29642 | 3.768175155 | -0.498591082 | 0.574851486 | -0.867338946 |
| Gm29375 | 1.122258044 | -0.776995696 | 1.026964685 | -0.756594368 |
| 7-Sep | 3678.656398 | 0.186520612 | 0.076391639 | 2.441636476 |
| Eepd1 | 188.7348402 | -0.285159228 | 0.093792711 | -3.040313308 |
| Gm25861 | 0 | NA | NA | NA |
| Gm25346 | 0 | NA | NA | NA |
| Gm26190 | 0 | NA | NA | NA |
| B3gat1 | 1268.762376 | 0.84705819 | 0.114184952 | 7.418299673 |
| Glb1l2 | 10.70062985 | -0.080298307 | 0.383511625 | -0.209376462 |
| Glb1l3 | 0.449952257 | -1.259362099 | 1.34883245 | -0.933668299 |
| Gm1110 | 2.025517295 | 0.626566349 | 0.868049801 | 0.721809219 |
| Acad8 | 225.253486 | 0.138515506 | 0.103704743 | 1.335671851 |
| Thyn1 | 253.8863181 | 0.032329744 | 0.125099891 | 0.258431431 |
| Vps26b | 1272.753477 | 0.231757405 | 0.060228358 | 3.847978123 |
| Ncapd3 | 405.8601585 | 0.330954094 | 0.168642691 | 1.962457376 |
| Jam3 | 390.9541926 | -0.725283068 | 0.131868227 | -5.500059283 |
| Gm29172 | 0.038981395 | -0.443816764 | 3.724945049 | -0.119147198 |
| Igsf9b | 818.461646 | -0.153496368 | 0.239496523 | -0.640912718 |
| Spata19 | 4.643680173 | -0.05755105 | 0.565043934 | -0.101852345 |
| Opcml | 3000.095646 | 0.521154045 | 0.0832599 | 6.259364273 |
| Gm15606 | 43.86298618 | 1.336914503 | 0.531224872 | 2.516663986 |
| AC157511.1 | 4.057004922 | 0.905659499 | 0.577289511 | 1.568813364 |

| | | | | |
|---|---|---|---|---|
| Ntm | 1996.252357 | 0.490505797 | 0.08042262 | 6.099102426 |
| Gm15520 | 1.48836363 | -0.158419256 | 0.934884475 | -0.169453297 |
| Gm15521 | 42.12449345 | 0.967663387 | 0.325571936 | 2.972195322 |
| Gm37605 | 0 | NA | NA | NA |
| Gm26435 | 0 | NA | NA | NA |
| Snx19 | 1050.451712 | -0.0352175 | 0.049141997 | -0.716647718 |
| Adamts15 | 127.7747721 | -0.072307012 | 0.122912619 | -0.588279811 |
| Adamts8 | 29.93733091 | -0.526550204 | 0.241869549 | -2.177000808 |
| Zbtb44 | 1100.679016 | 0.06690255 | 0.076052884 | 0.879684591 |
| Gm17508 | 0 | NA | NA | NA |
| St14 | 17.50112768 | 0.071518794 | 0.247298749 | 0.289199982 |
| Gm16071 | 0.620902317 | -0.186876025 | 1.177639897 | -0.1586869 |
| Aplp2 | 5929.023008 | 0.312062542 | 0.064066992 | 4.870878644 |
| Gm7244 | 0.710933469 | -0.443818936 | 1.378823029 | -0.321882451 |
| Prdm10 | 216.453307 | 0.2242983 | 0.138467612 | 1.61986111 |
| Nfrkb | 530.6418161 | 0.214410046 | 0.141644229 | 1.513722422 |
| Tmem45b | 0.191334258 | -0.81959484 | 2.66041249 | -0.308070588 |
| Barx2 | 29.27701895 | -0.731168784 | 0.309062889 | -2.365760534 |
| Gm3756 | 26.83663404 | -0.145531852 | 0.260477882 | -0.558710977 |
| AC160635.1 | 2.239755095 | 0.682134576 | 0.704575216 | 0.968150114 |
| Arhgap32 | 5811.857867 | 1.059789947 | 0.098309118 | 10.7801796 |
| Kcnj5 | 21.9094618 | -0.401731238 | 0.280055068 | -1.4344723 |
| Kcnj1 | 0.551016524 | -0.607834109 | 1.412166005 | -0.43042681 |
| Fli1 | 127.9880946 | -0.887790378 | 0.193623471 | -4.585138235 |
| Gm22060 | 0.733464214 | 0.367071442 | 1.045383604 | 0.351135641 |
| Gm38292 | 5.098635657 | -0.853452087 | 0.492167724 | -1.734067566 |
| Gm38346 | 11.05146628 | -0.123827194 | 0.425889384 | -0.290749661 |
| Gm27201 | 0.169165009 | 0.036125373 | 3.351346561 | 0.01077936 |
| Gm37474 | 11.13391284 | -0.7279529 | 0.503673176 | -1.445288204 |
| Gm37082 | 13.07759707 | -0.379722576 | 0.45475127 | -0.835011579 |
| Gm27163 | 0 | NA | NA | NA |
| Ets1 | 209.3380529 | -0.91724735 | 0.201591469 | -4.550030583 |
| Gm27240 | 0 | NA | NA | NA |
| Gm3331 | 0 | NA | NA | NA |
| Gm37167 | 0.620529807 | -1.365126247 | 1.455853195 | -0.937681253 |
| Gm27162 | 0 | NA | NA | NA |
| Gm37192 | 0.061920613 | -0.443816764 | 3.724945049 | -0.119147198 |
| Gm27166 | 0 | NA | NA | NA |
| Gm37758 | 2.309104371 | 0.322791643 | 0.605330499 | 0.533248602 |
| Gm10698 | 69.88234991 | -0.175221897 | 0.163113765 | -1.074231212 |
| Gm25439 | 0 | NA | NA | NA |
| Gm27165 | 0 | NA | NA | NA |
| Gm27164 | 0 | NA | NA | NA |
| 7630403G23Rik | 0.458133289 | 0.020497892 | 1.605292097 | 0.012768949 |
| Gm27161 | 0 | NA | NA | NA |
| Gm38354 | 0 | NA | NA | NA |
| Kirrel3 | 816.5410815 | 0.32638796 | 0.069421403 | 4.701546607 |

| | | | | |
|---|---|---|---|---|
| Gm38217 | 71.70747919 | 0.752476898 | 0.304306187 | 2.472762406 |
| Gm37038 | 54.63741103 | 0.548461782 | 0.285509611 | 1.920992363 |
| Gm26662 | 0.042716945 | 0.037062677 | 3.724945049 | 0.009949859 |
| Kirrel3os | 3.153914063 | -0.060656073 | 0.702925393 | -0.086290912 |
| Gm23702 | 2.111071361 | 1.301877123 | 0.788909248 | 1.650224189 |
| St3gal4 | 323.0715017 | -0.353221749 | 0.095904739 | -3.683047901 |
| 4930581F22Rik | 31.67819825 | -0.10258413 | 0.220003584 | -0.466283905 |
| Dcps | 130.3926271 | -0.372929373 | 0.130525904 | -2.85712922 |
| Gm24262 | 0 | NA | NA | NA |
| Tirap | 88.4573936 | -0.921826282 | 0.18753131 | -4.915586007 |
| Srpr | 878.7068951 | -0.441927879 | 0.071690878 | -6.16435297 |
| Foxred1 | 265.905042 | 0.110486837 | 0.096115488 | 1.149521676 |
| Fam118b | 211.7892874 | -0.189401854 | 0.105587021 | -1.793798627 |
| Rpusd4 | 78.94005876 | -0.602872361 | 0.157730704 | -3.822162374 |
| Gm5614 | 0 | NA | NA | NA |
| 2610105M22Rik | 0.044191549 | 0.037062677 | 3.724945049 | 0.009949859 |
| Cdon | 259.9464149 | 0.409885789 | 0.14800983 | 2.769314644 |
| Gm25630 | 1.242022914 | 0.553686659 | 1.023046999 | 0.541213316 |
| Gm24188 | 0 | NA | NA | NA |
| Gm1113 | 0 | NA | NA | NA |
| Ddx25 | 512.2202113 | 0.182405856 | 0.056242332 | 3.24321286 |
| Pus3 | 127.6005767 | -0.12493904 | 0.107818007 | -1.15879568 |
| Hyls1 | 86.77207718 | 0.133804752 | 0.151579149 | 0.882738514 |
| Pate2 | 15.21999782 | 0.531900515 | 0.332628511 | 1.599082753 |
| Gm6762 | 0 | NA | NA | NA |
| Gm10105 | 0 | NA | NA | NA |
| Pate4 | 0 | NA | NA | NA |
| Gm17365 | 0 | NA | NA | NA |
| Gm17252 | 0 | NA | NA | NA |
| Gm17677 | 0.297465678 | 0.989706925 | 1.757296247 | 0.563198679 |
| Gm27235 | 0 | NA | NA | NA |
| Gm17727 | 0 | NA | NA | NA |
| D730048I06Rik | 0 | NA | NA | NA |
| Gm36974 | 1.090715414 | 0.321391815 | 1.025285672 | 0.313465627 |
| 9230110F15Rik | 0 | NA | NA | NA |
| Gm3428 | 0.042716945 | 0.037062677 | 3.724945049 | 0.009949859 |
| 9230113P08Rik | 1.958122126 | -0.039054015 | 0.756042288 | -0.051655861 |
| Gm27187 | 0 | NA | NA | NA |
| Gm5916 | 0 | NA | NA | NA |
| Gm3434 | 0 | NA | NA | NA |
| Gm3867 | 0.061920613 | -0.443816764 | 3.724945049 | -0.119147198 |
| Gm7257 | 0 | NA | NA | NA |
| Gm9513 | 0 | NA | NA | NA |
| Gm27160 | 0 | NA | NA | NA |
| Gm5615 | 0 | NA | NA | NA |
| Gm27218 | 0 | NA | NA | NA |
| Gm17689 | 0.194279094 | -1.017823543 | 3.719440177 | -0.273649661 |

| | | | | |
|---|---|---|---|---|
| A630095E13Rik | 0.217428854 | -1.735499013 | 2.547916594 | -0.681144358 |
| 1700027I24Rik | 0.357600009 | -1.633827421 | 1.647076287 | -0.991956131 |
| Acrv1 | 0.095898756 | -0.917692504 | 3.724434845 | -0.246397787 |
| Chek1 | 39.39245048 | -0.48363275 | 0.202383819 | -2.389680917 |
| Stt3a | 958.0649057 | -0.316301294 | 0.067206302 | -4.706423126 |
| Gm26787 | 0.495189146 | 1.718242957 | 1.259509963 | 1.364215456 |
| Ei24 | 869.5859806 | -0.052996583 | 0.070494013 | -0.751788421 |
| Gm27219 | 6.744631513 | -0.377287468 | 0.40368022 | -0.934619656 |
| Fez1 | 1689.638317 | 0.035377912 | 0.061511169 | 0.575146156 |
| Pknox2 | 477.999823 | 0.531771721 | 0.108848023 | 4.885451352 |
| Gm3896 | 0.132101749 | -0.93550548 | 3.724188147 | -0.251197158 |
| n-R5s82 | 0 | NA | NA | NA |
| Gm25273 | 0 | NA | NA | NA |
| Tmem218 | 64.43814503 | 0.050183151 | 0.147286954 | 0.340716876 |
| Slc37a2 | 140.0947805 | -1.713934669 | 0.255785764 | -6.700664824 |
| Gm10177 | 68.34373515 | -0.65197722 | 0.201793812 | -3.230907895 |
| Ccdc15 | 56.68857134 | 0.239401911 | 0.194769828 | 1.229152964 |
| Hepacam | 1455.687833 | 0.115329677 | 0.127630923 | 0.903618605 |
| Robo4 | 52.91085951 | -0.474232591 | 0.23072654 | -2.055388131 |
| Robo3 | 261.3456688 | 0.913792091 | 0.423966896 | 2.155338308 |
| Msantd2 | 153.0368494 | -0.227603643 | 0.106359851 | -2.139939473 |
| Esam | 83.18252856 | -0.404778307 | 0.156000597 | -2.594722808 |
| Vsig2 | 19.39362468 | -0.273167752 | 0.313970043 | -0.870043998 |
| Nrgn | 5315.665183 | 1.147916009 | 0.205691447 | 5.580766865 |
| Spa17 | 8.072628227 | -0.571245284 | 0.426203988 | -1.34030957 |
| Siae | 198.8133421 | -0.164568325 | 0.134122841 | -1.226997012 |
| Tbrg1 | 650.3765193 | -0.402087111 | 0.122238678 | -3.289360752 |
| Panx3 | 0.252375374 | 0.505588096 | 2.126445696 | 0.237762054 |
| Gm8343 | 0 | NA | NA | NA |
| Tpt1-ps5 | 7.622765194 | -0.698500551 | 0.418512145 | -1.669009037 |
| Olfr160 | 0 | NA | NA | NA |
| Olfr874 | 0 | NA | NA | NA |
| Olfr875 | 0.072025462 | 0.517959912 | 3.724945049 | 0.139051692 |
| Olfr876 | 0.066050875 | -0.443818988 | 3.724945049 | -0.119147795 |
| Gm24764 | 0 | NA | NA | NA |
| Olfr877 | 0.162452565 | 1.424702199 | 2.773995858 | 0.513592043 |
| Olfr145 | 0.104637558 | -0.443821212 | 3.724945049 | -0.119148392 |
| Olfr878 | 0.170434537 | 0.037075321 | 2.328336905 | 0.015923521 |
| Olfr881 | 0 | NA | NA | NA |
| Olfr883 | 0 | NA | NA | NA |
| Olfr884 | 0.092140247 | 0.037060453 | 3.724945049 | 0.009949262 |
| Olfr885 | 0 | NA | NA | NA |
| Olfr887 | 0.072025462 | 0.517959912 | 3.724945049 | 0.139051692 |
| Olfr888 | 0 | NA | NA | NA |
| Olfr889 | 0 | NA | NA | NA |
| Olfr890 | 0 | NA | NA | NA |
| Olfr891 | 0 | NA | NA | NA |

| | | | | |
|---|---|---|---|---|
| Gm17798 | 0 | NA | NA | NA |
| Olfr892-ps1 | 0 | NA | NA | NA |
| Olfr893 | 0 | NA | NA | NA |
| Olfr894 | 0 | NA | NA | NA |
| Olfr143 | 0 | NA | NA | NA |
| Olfr895 | 0 | NA | NA | NA |
| Olfr25 | 0.111376404 | 0.517957688 | 3.724945049 | 0.139051095 |
| Olfr898 | 0 | NA | NA | NA |
| Olfr250 | 0.072025462 | 0.517959912 | 3.724945049 | 0.139051692 |
| Olfr251 | 0 | NA | NA | NA |
| Olfr147 | 0.068097359 | 0.037064902 | 3.724945049 | 0.009950456 |
| Olfr901 | 0.114443069 | -0.443821212 | 3.724945049 | -0.119148392 |
| Olfr902 | 0 | NA | NA | NA |
| Olfr904 | 0 | NA | NA | NA |
| Olfr905 | 0 | NA | NA | NA |
| Olfr906 | 0 | NA | NA | NA |
| Olfr907 | 0 | NA | NA | NA |
| Gm23271 | 0 | NA | NA | NA |
| Olfr908 | 0 | NA | NA | NA |
| Olfr911-ps1 | 0 | NA | NA | NA |
| Olfr910 | 0 | NA | NA | NA |
| Olfr912 | 0 | NA | NA | NA |
| Olfr913 | 0 | NA | NA | NA |
| Olfr914 | 0 | NA | NA | NA |
| Olfr916 | 0 | NA | NA | NA |
| Olfr917 | 0 | NA | NA | NA |
| Olfr918 | 0 | NA | NA | NA |
| Olfr919 | 0 | NA | NA | NA |
| Vwa5a | 1119.696084 | -0.418443704 | 0.162206023 | -2.579705093 |
| Olfr920 | 22.44487698 | 0.934597213 | 0.415785378 | 2.247787591 |
| Olfr921 | 0.704586541 | -0.065739963 | 1.380264762 | -0.047628517 |
| Olfr922 | 0.673070397 | -1.933100077 | 1.170200505 | -1.651939192 |
| Olfr923 | 0.176917541 | -0.93433906 | 2.766967798 | -0.33767616 |
| Olfr924 | 0.326455024 | -0.443820651 | 1.76049138 | -0.25210044 |
| Olfr26 | 0.232584034 | 0.53903602 | 2.130516083 | 0.253007252 |
| Olfr926 | 0.106056992 | 0.998850474 | 3.724945049 | 0.268151734 |
| Olfr930 | 0 | NA | NA | NA |
| Olfr933 | 0 | NA | NA | NA |
| Olfr934 | 0 | NA | NA | NA |
| Olfr935 | 0 | NA | NA | NA |
| Olfr146 | 0 | NA | NA | NA |
| Olfr936 | 0 | NA | NA | NA |
| Gm25529 | 0 | NA | NA | NA |
| Olfr937 | 0 | NA | NA | NA |
| Gm25032 | 0 | NA | NA | NA |
| Olfr938 | 0 | NA | NA | NA |
| Olfr27 | 0 | NA | NA | NA |

| | | | | |
|---|---|---|---|---|
| Olfr943 | 0 | NA | NA | NA |
| Olfr944 | 0 | NA | NA | NA |
| Olfr1537 | 0 | NA | NA | NA |
| Olfr945 | 0 | NA | NA | NA |
| Olfr948 | 0 | NA | NA | NA |
| Olfr951 | 0 | NA | NA | NA |
| Olfr952 | 0 | NA | NA | NA |
| Olfr954 | 0 | NA | NA | NA |
| Olfr955 | 0 | NA | NA | NA |
| Olfr44 | 0.046345709 | -0.443818988 | 3.724945049 | -0.119147795 |
| Olfr957 | 0 | NA | NA | NA |
| Gm24278 | 0 | NA | NA | NA |
| Olfr958 | 0 | NA | NA | NA |
| Olfr959 | 0.090427103 | 0.998850474 | 3.724945049 | 0.268151734 |
| AW551984 | 246.7130081 | -0.320837058 | 0.155790094 | -2.059418857 |
| Olfr148 | 0 | NA | NA | NA |
| Olfr960 | 0 | NA | NA | NA |
| Olfr961 | 0 | NA | NA | NA |
| Olfr963 | 0 | NA | NA | NA |
| Olfr964-ps1 | 0.048949532 | 0.037064902 | 3.724945049 | 0.009950456 |
| Olfr149 | 0 | NA | NA | NA |
| Olfr965 | 0 | NA | NA | NA |
| Olfr150 | 0 | NA | NA | NA |
| Olfr967 | 0 | NA | NA | NA |
| Olfr968 | 0.046345709 | -0.443818988 | 3.724945049 | -0.119147795 |
| Olfr969 | 0 | NA | NA | NA |
| Olfr970 | 0 | NA | NA | NA |
| Olfr971 | 0 | NA | NA | NA |
| Olfr972 | 0 | NA | NA | NA |
| Olfr229 | 0 | NA | NA | NA |
| Olfr974 | 0.049553047 | -0.443818988 | 3.724945049 | -0.119147795 |
| Olfr975 | 0 | NA | NA | NA |
| Olfr976 | 0.111006857 | 0.037730769 | 3.724898049 | 0.010129343 |
| Olfr978 | 0 | NA | NA | NA |
| Olfr979 | 0.046345709 | -0.443818988 | 3.724945049 | -0.119147795 |
| Olfr980 | 0.09176491 | 0.921395572 | 3.723061103 | 0.247483333 |
| Olfr981 | 0 | NA | NA | NA |
| 1700001J11Rik | 0.049553047 | -0.443818988 | 3.724945049 | -0.119147795 |
| Olfr982 | 0 | NA | NA | NA |
| Olfr983 | 0 | NA | NA | NA |
| Olfr984 | 0.072025462 | 0.517959912 | 3.724945049 | 0.139051692 |
| Olfr985 | 0.09176491 | 0.921395572 | 3.723061103 | 0.247483333 |
| Tmem225 | 0 | NA | NA | NA |
| Olfr986 | 0 | NA | NA | NA |
| Zfp202 | 72.13492 | -0.426644807 | 0.146297749 | -2.916277319 |
| Scn3b | 970.5865091 | 0.893426651 | 0.106644486 | 8.377616956 |
| Gramd1b | 3247.828285 | 0.008920684 | 0.081173117 | 0.109897028 |

| | | | | |
|---|---|---|---|---|
| 1700110K17Rik | 1.343476617 | -0.280822033 | 0.955032077 | -0.294044608 |
| Gm17540 | 0 | NA | NA | NA |
| Gm21915 | 0 | NA | NA | NA |
| Clmp | 308.4342746 | -0.378808843 | 0.128939707 | -2.937875787 |
| Gm16095 | 0.148850706 | -0.86762789 | 2.852067438 | -0.304210159 |
| Gm16096 | 0.692884898 | -1.343567221 | 1.247280947 | -1.077196941 |
| Hspa8 | 7490.665187 | -0.550218634 | 0.080073565 | -6.871414249 |
| Snord14c | 0 | NA | NA | NA |
| Snord14d | 0 | NA | NA | NA |
| Snord14e | 0 | NA | NA | NA |
| Bsx | 2.789926683 | 0.078003422 | 0.578140903 | 0.134921126 |
| 4931429I11Rik | 11.72828212 | -0.013709223 | 0.31840507 | -0.043055921 |
| Crtam | 123.3542397 | -0.063430259 | 0.246993676 | -0.256809242 |
| Ubash3b | 620.5872832 | -0.138726092 | 0.093946174 | -1.476655051 |
| 1700063D05Rik | 6.117005653 | 0.303874958 | 0.433298496 | 0.701306282 |
| Gm25401 | 0 | NA | NA | NA |
| 2610203C20Rik | 984.4296156 | 0.156475074 | 0.145640195 | 1.074394839 |
| Mir100 | 2.750782513 | 0.786500577 | 0.759024694 | 1.036198932 |
| Mirlet7a-2 | 2.112107905 | 0.583698786 | 0.71395363 | 0.817558399 |
| Gm28119 | 36.18489172 | 0.218199081 | 0.240305038 | 0.908008767 |
| Mir125b-1 | 0 | NA | NA | NA |
| Gm8707 | 28.79399415 | 0.638086753 | 0.215634205 | 2.959116587 |
| Sorl1 | 7353.79371 | 0.361651856 | 0.075738854 | 4.774984515 |
| Gm16214 | 0 | NA | NA | NA |
| Sc5d | 491.2699012 | -0.185253476 | 0.105740323 | -1.751966234 |
| Gm26272 | 0 | NA | NA | NA |
| Tecta | 9.746868474 | -0.841115398 | 0.421500496 | -1.995526474 |
| Tbcel | 691.6961961 | 0.126810715 | 0.060283736 | 2.1035643 |
| AC136903.1 | 0 | NA | NA | NA |
| Gm16322 | 0.291180489 | 1.058421007 | 3.708688692 | 0.285389553 |
| Grik4 | 287.3585913 | 0.258283208 | 0.100638139 | 2.566454546 |
| Gm24420 | 0 | NA | NA | NA |
| Gm27509 | 1.432503634 | 1.750843336 | 1.107795646 | 1.580475011 |
| Arhgef12 | 4563.170901 | -0.033787955 | 0.038647687 | -0.874255551 |
| Tmem136 | 181.8275258 | 0.174545542 | 0.109290595 | 1.597077421 |
| Pou2f3 | 2.121136815 | -0.094397868 | 0.660843025 | -0.142844615 |
| Oaf | 220.6241672 | -0.976345845 | 0.162017135 | -6.02618882 |
| D630033O11Rik | 2.615308024 | -0.381090465 | 0.60345096 | -0.631518533 |
| Trim29 | 4.261450001 | -1.02696987 | 0.577075252 | -1.779611701 |
| Gm28215 | 0 | NA | NA | NA |
| Pvrl1 | 1033.013847 | 0.268635484 | 0.079995741 | 3.358122321 |
| Gm3898 | 0.044544648 | 0.517957688 | 3.724945049 | 0.139051095 |
| Gm23326 | 0.048949532 | 0.037064902 | 3.724945049 | 0.009950456 |
| Thy1 | 4661.495848 | 0.275018516 | 0.078041649 | 3.523996717 |
| Rnf26 | 340.15711 | 0.12640284 | 0.078523245 | 1.609750583 |
| Usp2 | 947.8979254 | 0.360779532 | 0.063198559 | 5.70866705 |
| Mfrp | 54.45536832 | 1.05891872 | 0.264185713 | 4.008236125 |

| | | | | |
|---|---|---|---|---|
| C1qtnf5 | 0 | NA | NA | NA |
| Gm20444 | 3.364322678 | -0.57812604 | 0.663673918 | -0.871099532 |
| Gm26737 | 7.342448606 | 0.082664296 | 0.393114108 | 0.210280664 |
| Gm10687 | 8.115134709 | 0.015136915 | 0.395675249 | 0.038255906 |
| Mcam | 257.1599992 | -0.35184847 | 0.154505864 | -2.277249948 |
| Cbl | 967.2924709 | -0.083031733 | 0.07337431 | -1.131618586 |
| Ccdc153 | 11.25998896 | 0.374941314 | 0.415572103 | 0.902229268 |
| Pdzd3 | 3.822296293 | -0.204115532 | 0.651069134 | -0.31350823 |
| Nlrx1 | 97.08268411 | 0.141837672 | 0.141730025 | 1.000759521 |
| Abcg4 | 564.6710316 | 0.38723924 | 0.08687061 | 4.457655357 |
| Hinfp | 148.4591195 | -0.191976665 | 0.1363581 | -1.407886031 |
| C2cd2l | 1532.300989 | 0.321611229 | 0.109863278 | 2.927376963 |
| Dpagt1 | 146.9363667 | -0.371101735 | 0.115613009 | -3.209861399 |
| H2afx | 329.2993814 | -0.080720982 | 0.099538494 | -0.810952408 |
| AC122428.1 | 1.088551472 | 1.960941227 | 1.216016195 | 1.612594664 |
| Hmbs | 132.3982402 | -0.113308685 | 0.128962141 | -0.878619757 |
| Vps11 | 770.3516232 | 0.010102359 | 0.066374904 | 0.152201486 |
| Gm22141 | 0.240440413 | 0.99886561 | 2.156343003 | 0.463222043 |
| Gm10080 | 110.1720267 | -0.213355314 | 0.16722378 | -1.275867065 |
| Hyou1 | 2036.023468 | -0.450068653 | 0.081114187 | -5.548581182 |
| Gm26306 | 0 | NA | NA | NA |
| Slc37a4 | 109.665464 | 0.183630186 | 0.137555829 | 1.334950229 |
| Mir7085 | 0.487060322 | 1.44563216 | 1.423230546 | 1.015739975 |
| Trappc4 | 251.412174 | -0.317478497 | 0.126991058 | -2.500006707 |
| Rps25 | 1102.353914 | -0.004744606 | 0.13166197 | -0.036036267 |
| Ccdc84 | 64.93599903 | -0.126762241 | 0.16984776 | -0.746328598 |
| Foxr1 | 0.310849204 | 0.000948915 | 3.715382321 | 0.000255402 |
| Upk2 | 0.044191549 | 0.037062677 | 3.724945049 | 0.009949859 |
| Gm9830 | 5.961436335 | -0.548600737 | 0.407223439 | -1.347173774 |
| Gm22540 | 5.014127157 | 1.272796026 | 0.576271635 | 2.208673738 |
| C030014I23Rik | 47.83992227 | 0.511116692 | 0.175060821 | 2.919652084 |
| Bcl9l | 785.1067627 | 0.150519275 | 0.082571989 | 1.822885432 |
| Cxcr5 | 2.247268683 | 0.037529863 | 0.653198368 | 0.057455537 |
| Ddx6 | 4111.857025 | -0.027306029 | 0.053612159 | -0.509325305 |
| Treh | 0.042716945 | 0.037062677 | 3.724945049 | 0.009949859 |
| Phldb1 | 1542.489038 | -0.128644255 | 0.071553595 | -1.797872716 |
| Gm24166 | 0 | NA | NA | NA |
| Arcn1 | 1826.393337 | -0.224215568 | 0.081234771 | -2.760093549 |
| Ift46 | 313.3153656 | 0.125759557 | 0.076989196 | 1.633470185 |
| Tmem25 | 356.7404894 | 0.414780918 | 0.099030246 | 4.188426634 |
| Ttc36 | 3.625487916 | -0.389765697 | 0.614760727 | -0.634012031 |
| Kmt2a | 3944.583612 | 0.058640774 | 0.093935644 | 0.624265418 |
| Gm26249 | 0 | NA | NA | NA |
| Atp5l | 692.9294976 | -0.0211126 | 0.138728756 | -0.152186187 |
| Ube4a | 1166.455959 | -0.181687768 | 0.058115062 | -3.126345587 |
| Cd3g | 29.68427331 | -0.546470372 | 0.369523392 | -1.478851905 |
| Cd3d | 21.88467821 | -0.370652418 | 0.291750233 | -1.270444288 |

| | | | | |
|---|---|---|---|---|
| Cd3e | 72.02395476 | -0.537093227 | 0.293857779 | -1.827731869 |
| Mpzl2 | 55.99719323 | -0.181113506 | 0.226529969 | -0.799512341 |
| Mpzl3 | 77.05801845 | 0.137507951 | 0.144864495 | 0.949217759 |
| Amica1 | 164.8869608 | -0.857294467 | 0.313519241 | -2.734423779 |
| Gm10684 | 5.682893444 | 0.955894133 | 0.517626456 | 1.846687165 |
| Scn2b | 3837.721104 | 0.431401479 | 0.098132645 | 4.396105677 |
| Scn4b | 2000.069508 | 0.945545765 | 0.126706736 | 7.462474335 |
| Tmprss4 | 3.058856891 | -0.314043853 | 0.574875805 | -0.546281215 |
| Gm17099 | 0.153914375 | -0.443826546 | 3.506087737 | -0.126587404 |
| BC049352 | 7.258084432 | 0.604993387 | 0.512397608 | 1.180710795 |
| Il10ra | 224.5975288 | -0.538672809 | 0.137988062 | -3.903763871 |
| Gm20612 | 0.047948698 | 0.037064902 | 3.724945049 | 0.009950456 |
| Mir7086 | 0.58926593 | 0.725594049 | 1.278627433 | 0.567478868 |
| 1700003G13Rik | 0 | NA | NA | NA |
| Tmprss13 | 0.459044381 | -0.327175895 | 1.450137847 | -0.2256171 |
| Fxyd6 | 526.6237023 | 0.203816664 | 0.111007595 | 1.836060538 |
| Fxyd2 | 9.473000343 | 0.461367709 | 0.3618513 | 1.275020178 |
| 4833428L15Rik | 0.565850957 | -0.434696385 | 1.327462505 | -0.327464153 |
| Dscaml1 | 663.8123879 | 0.630896939 | 0.115571881 | 5.458913823 |
| Gm22069 | 0 | NA | NA | NA |
| Cep164 | 262.2380998 | 0.068600389 | 0.135672358 | 0.505632762 |
| Gm7286 | 0.119587853 | 0.981063299 | 3.723618439 | 0.263470416 |
| Bace1 | 1452.453558 | -0.005904392 | 0.045026301 | -0.131132068 |
| Gm16536 | 127.1699055 | 0.069295247 | 0.10995847 | 0.630194713 |
| Rnf214 | 859.5718938 | 0.067640142 | 0.049216896 | 1.374327658 |
| Gm24646 | 2.996628699 | 0.66707635 | 0.91250953 | 0.731034941 |
| Pcsk7 | 359.8709397 | -0.058438827 | 0.085309108 | -0.685024474 |
| Tagln | 130.3382176 | -2.779682562 | 0.430098166 | -6.462902623 |
| Sidt2 | 826.3607848 | 0.110889283 | 0.085262622 | 1.30056149 |
| Mir7087 | 1.036669434 | 1.776929332 | 1.070132423 | 1.660476119 |
| Pafah1b2 | 2591.382288 | 0.041275166 | 0.06705551 | 0.615537277 |
| Sik3 | 2827.253407 | -0.138806237 | 0.081567828 | -1.701727753 |
| Apoa1 | 4.148602219 | 0.655908079 | 0.52394961 | 1.251853357 |
| Apoc3 | 5.512392282 | 1.089419944 | 0.439798843 | 2.477086877 |
| Apoa4 | 0.183366429 | -0.400355652 | 2.264892306 | -0.176765867 |
| Gm10680 | 0 | NA | NA | NA |
| Apoa5 | 0.33367842 | -2.111472025 | 1.945604613 | -1.085252374 |
| Zpr1 | 322.0139224 | -0.165135011 | 0.073371754 | -2.250661911 |
| Bud13 | 129.6764734 | -0.199338737 | 0.103002003 | -1.935289907 |
| 4931429L15Rik | 0.137762903 | 0.51796349 | 2.937523392 | 0.176326592 |
| 4930448E22Rik | 1.471695147 | -0.878741644 | 0.876940562 | -1.002053825 |
| Gm22659 | 0.246177344 | -0.443821758 | 2.50940469 | -0.176863365 |
| Gm22805 | 0 | NA | NA | NA |
| 2900052N01Rik | 275.6261956 | 0.759556683 | 0.160757141 | 4.724870564 |
| Gm4791 | 0.045882455 | 0.517959912 | 3.724945049 | 0.139051692 |
| Gm22286 | 0.075043205 | 0.517957688 | 3.724945049 | 0.139051095 |
| Cadm1 | 2092.339405 | 0.237323125 | 0.066816767 | 3.551849864 |

| | | | | |
|---|---|---|---|---|
| Nxpe2 | 2.03192395 | 0.028675009 | 0.68552042 | 0.041829548 |
| Nxpe4 | 71.31690674 | 0.378922563 | 0.137979646 | 2.746220719 |
| Gm5616 | 4.81606905 | -0.726648853 | 0.481727699 | -1.508422403 |
| Rexo2 | 393.6205041 | -0.688109171 | 0.102714809 | -6.699220686 |
| Rbm7 | 414.2773601 | -0.243980268 | 0.110183158 | -2.214315448 |
| Gm5617 | 29.90601096 | -0.070583694 | 0.235326641 | -0.299939242 |
| Gm23653 | 0 | NA | NA | NA |
| Nnmt | 5.720995077 | -0.847704011 | 0.40636858 | -2.086047131 |
| Zbtb16 | 1098.476404 | 0.762992372 | 0.176270175 | 4.32853925 |
| Htr3a | 33.96213922 | 0.297802912 | 0.224383718 | 1.327203747 |
| Htr3b | 0.066050875 | -0.443818988 | 3.724945049 | -0.119147795 |
| Usp28 | 333.1237609 | -0.034795781 | 0.121993533 | -0.285226439 |
| Cldn21 | 0 | NA | NA | NA |
| RP23-136A1.2 | 0 | NA | NA | NA |
| Zw10 | 188.7291143 | -0.332609083 | 0.094535355 | -3.518356526 |
| Tmprss5 | 18.34493739 | -0.914893039 | 0.285180211 | -3.208122456 |
| Gm4894 | 0.044544648 | 0.517957688 | 3.724945049 | 0.139051095 |
| Drd2 | 144.2458334 | 0.621179351 | 0.227647673 | 2.728687463 |
| Ankk1 | 0.421296087 | -1.291313615 | 1.42463838 | -0.906415012 |
| Ttc12 | 60.41204789 | -0.482704979 | 0.179045869 | -2.695985005 |
| Ncam1 | 4558.689867 | 0.148087086 | 0.056881125 | 2.603448634 |
| Gm11149 | 1.147320409 | 0.086108167 | 1.086918414 | 0.079222291 |
| Gm25633 | 0 | NA | NA | NA |
| Gm8907 | 0.242574275 | -1.344190426 | 2.526122974 | -0.532115989 |
| Rpl10-ps3 | 4.878838789 | -0.430940416 | 0.426545323 | -1.010303929 |
| Plet1os | 0.85456564 | -0.480352304 | 1.307537891 | -0.367371613 |
| Plet1 | 4.418087894 | 0.696619621 | 0.496414174 | 1.403303245 |
| Pts | 145.2821715 | -0.02790316 | 0.135072512 | -0.206579112 |
| Bco2 | 15.34273135 | -0.150306783 | 0.280173306 | -0.536477887 |
| Tex12 | 0.048949532 | 0.037064902 | 3.724945049 | 0.009950456 |
| Il18 | 136.7701011 | -0.134398311 | 0.135618115 | -0.991005594 |
| Sdhd | 647.1493877 | -0.076331262 | 0.073261986 | -1.041894526 |
| Timm8b | 499.1712965 | 0.007199171 | 0.149386893 | 0.04819145 |
| AU019823 | 218.1162646 | -0.026686522 | 0.117361937 | -0.227386514 |
| Pih1d2 | 29.01466556 | 0.05902196 | 0.195672592 | 0.301636318 |
| Dlat | 1112.83225 | 0.234944478 | 0.059799419 | 3.928875597 |
| Dixdc1 | 1202.736387 | 0.226795457 | 0.076076422 | 2.981153051 |
| 2310030G06Rik | 16.94180656 | 0.347939249 | 0.24699359 | 1.408697486 |
| Hspb2 | 5.620585044 | 0.083120774 | 0.42795468 | 0.194227982 |
| Cryab | 871.6678754 | -1.137902668 | 0.199870861 | -5.693189406 |
| 1110032A03Rik | 144.5010807 | 0.26633822 | 0.112766366 | 2.361858669 |
| Fdxacb1 | 52.09799359 | 0.097337179 | 0.156882982 | 0.620444473 |
| Alg9 | 194.0505543 | -0.033602386 | 0.087646769 | -0.383384193 |
| Ppp2r1b | 243.7699036 | -0.587599062 | 0.102233097 | -5.747640218 |
| Gm25558 | 0.177937505 | 0.517962461 | 2.743421694 | 0.18880162 |
| Sik2 | 790.3337553 | -0.182011024 | 0.105647565 | -1.722813249 |
| Layn | 62.49321233 | -1.133096122 | 0.240990616 | -4.701826732 |

| | | | | |
|---|---|---|---|---|
| 4833427G06Rik | 5.495872342 | -0.396649804 | 0.441125674 | -0.89917642 |
| Mir34c | 0 | NA | NA | NA |
| Mir34b | 0 | NA | NA | NA |
| Btg4 | 0.260468428 | -0.791890979 | 2.018636357 | -0.392290061 |
| Pou2af1 | 4.531271601 | -1.333465201 | 0.638467461 | -2.088540579 |
| Gm684 | 7.066067338 | -1.03565012 | 0.434681965 | -2.382546792 |
| 2010007H06Rik | 9.722359338 | 0.226132701 | 0.386672727 | 0.584816784 |
| 1810046K07Rik | 0.216202823 | 0.037610776 | 2.173238982 | 0.017306323 |
| Gm7293 | 4.644993497 | 0.127166347 | 0.570870696 | 0.222758582 |
| Arhgap20 | 1285.799934 | 0.556435332 | 0.088989168 | 6.25284338 |
| Arhgap20os | 1.805117362 | 0.490370289 | 1.008891505 | 0.486048585 |
| Fdx1 | 101.2393919 | -0.326219831 | 0.150707335 | -2.164591605 |
| Gm6981 | 16.53353035 | -0.042445407 | 0.286809127 | -0.147991829 |
| Rdx | 1573.201998 | -0.274220725 | 0.091243487 | -3.005373135 |
| Gm27686 | 0 | NA | NA | NA |
| Zc3h12c | 704.8556827 | -0.282660458 | 0.121196932 | -2.332241034 |
| Mir6244 | 0.506103766 | -1.195423058 | 3.275116226 | -0.365001721 |
| Gm25562 | 0 | NA | NA | NA |
| AI593442 | 2277.537907 | 0.770346317 | 0.101367777 | 7.599518681 |
| Gm22242 | 0 | NA | NA | NA |
| Ddx10 | 340.9961152 | 0.008538884 | 0.087745535 | 0.097314177 |
| 4930510E17Rik | 0 | NA | NA | NA |
| Exph5 | 843.2123267 | 0.228447113 | 0.185806924 | 1.229486543 |
| Kdelc2 | 102.2218059 | -0.039841629 | 0.125558597 | -0.317315024 |
| Mir130c | 0.049553047 | -0.443818988 | 3.724945049 | -0.119147795 |
| 4930550C14Rik | 3.372172578 | -0.475935273 | 0.535387028 | -0.888955556 |
| Atm | 883.8572749 | -0.100042577 | 0.096838985 | -1.033081643 |
| Npat | 649.966786 | 0.201061564 | 0.095578897 | 2.103618785 |
| Acat1 | 830.5020792 | 0.372884176 | 0.065300115 | 5.710314202 |
| Cul5 | 1233.233245 | -0.238239194 | 0.08167115 | -2.917054497 |
| Gm22304 | 0 | NA | NA | NA |
| Gm16124 | 9.053711666 | 0.672179467 | 0.392000667 | 1.714740621 |
| Rab39 | 22.16151509 | -0.758627207 | 0.229872857 | -3.300203504 |
| Rps6-ps3 | 1.738215179 | 0.557562843 | 0.703656563 | 0.792379226 |
| Slc35f2 | 23.18056578 | 0.0002721 | 0.229036315 | 0.00118802 |
| Sln | 3.22564905 | -0.123521248 | 0.605289039 | -0.204069857 |
| Gm16380 | 0.31602048 | 1.864827949 | 1.560088262 | 1.195334902 |
| Elmod1 | 1587.467376 | 0.499982708 | 0.06406221 | 7.804643406 |
| Tnfaip8l3 | 95.49101603 | 0.064408533 | 0.140979706 | 0.456863861 |
| Cyp19a1 | 1.568920191 | 0.214276412 | 0.852141998 | 0.251456228 |
| Gldnos | 0.038981395 | -0.443816764 | 3.724945049 | -0.119147198 |
| Gldn | 61.66008331 | -0.861867639 | 0.248667852 | -3.465939138 |
| Dmxl2 | 4593.55088 | 0.233564727 | 0.068082182 | 3.430629268 |
| Gm25980 | 0 | NA | NA | NA |
| Sh2d7 | 0.788119377 | 0.789281867 | 1.329502234 | 0.593667199 |
| Cib2 | 155.6054035 | 0.206593777 | 0.138881614 | 1.487553111 |
| Idh3a | 1642.060698 | 0.066382149 | 0.066620584 | 0.996421002 |

| | | | | |
|---|---|---|---|---|
| Acsbg1 | 1565.688373 | -0.090496777 | 0.147523531 | -0.613439609 |
| Dnaja4 | 456.041311 | -0.398399273 | 0.119366419 | -3.337616033 |
| Mir5710 | 0.417007765 | 0.956447169 | 1.449328986 | 0.659924129 |
| Wdr61 | 268.6245129 | -0.389699994 | 0.093828644 | -4.153315846 |
| Crabp1 | 9.084845767 | 0.938806446 | 0.392143474 | 2.394038173 |
| Ireb2 | 1512.434034 | 0.068643307 | 0.069441773 | 0.988501643 |
| Hykk | 65.99315051 | -0.126563484 | 0.182376962 | -0.693966402 |
| AY074887 | 0.108309328 | 0.967119648 | 3.722483401 | 0.259804959 |
| Psma4 | 828.6826824 | -0.185450558 | 0.095335207 | -1.945247349 |
| Chrna5 | 21.98151757 | 0.742061797 | 0.249181675 | 2.977995063 |
| Chrna3 | 28.52374879 | -0.125850651 | 0.257426434 | -0.488880062 |
| Chrnb4 | 16.87880904 | 0.228603666 | 0.328017428 | 0.696925366 |
| Ube2q2 | 800.5171951 | 0.076393998 | 0.07060929 | 1.08192559 |
| Fbxo22 | 362.6766692 | -0.037493162 | 0.069641364 | -0.53837489 |
| Nrg4 | 36.91795082 | -1.52632937 | 0.279015573 | -5.47040925 |
| Gm15625 | 0 | NA | NA | NA |
| AI118078 | 183.6671056 | 0.062979474 | 0.143879407 | 0.437724029 |
| Etfa | 264.841253 | -0.358727507 | 0.138181719 | -2.596056193 |
| Isl2 | 0.471858061 | 0.485564448 | 1.328087749 | 0.365611722 |
| Gm17226 | 0 | NA | NA | NA |
| Scaper | 701.5265708 | 0.430550257 | 0.08206798 | 5.246263588 |
| Gm23407 | 0.627732484 | 1.220037826 | 1.326622664 | 0.919657005 |
| 4930563M21Rik | 0.038981395 | -0.443816764 | 3.724945049 | -0.119147198 |
| Rcn2 | 1128.177669 | 0.152767844 | 0.08947682 | 1.707345474 |
| Pstpip1 | 94.70433045 | -1.143487334 | 0.219448844 | -5.210723887 |
| Tspan3 | 2389.846793 | 0.047117323 | 0.089498526 | 0.526459204 |
| Peak1 | 1406.181793 | -0.249224874 | 0.072482385 | -3.438419894 |
| Gm24270 | 405.7586766 | 0.028953377 | 0.859714097 | 0.033677914 |
| Gm29322 | 6.151976361 | 0.051323752 | 0.490757649 | 0.104580646 |
| Peak1os | 8.730896443 | -2.700122815 | 0.435059429 | -6.206330981 |
| Gm37842 | 2.222650432 | -3.082892207 | 0.970027111 | -3.17815056 |
| Gm36940 | 30.43733313 | -0.038634727 | 0.305775902 | -0.12634981 |
| Hmg20a | 798.0476359 | 0.194655947 | 0.061636486 | 3.158128537 |
| Gm26868 | 0.201261165 | 0.037075933 | 2.219102811 | 0.016707623 |
| Lingo1 | 2194.009453 | 0.406524342 | 0.187915013 | 2.163341479 |
| Odf3l1 | 0.839566529 | 1.005581233 | 1.305188188 | 0.770449229 |
| Cspg4 | 408.2241713 | -0.172264364 | 0.091765818 | -1.877217114 |
| Snx33 | 323.7784099 | -0.608914271 | 0.143643022 | -4.239080065 |
| Imp3 | 174.2311039 | -0.095363337 | 0.14368102 | -0.66371562 |
| Snupn | 97.8417329 | -0.532792865 | 0.131072017 | -4.06488644 |
| Ptpn9 | 554.0054497 | -0.491203377 | 0.091638734 | -5.360215667 |
| Gm10658 | 5.404131218 | 0.640747672 | 0.51621906 | 1.241232108 |
| Sin3a | 642.6567267 | 0.02571898 | 0.06067785 | 0.423861098 |
| 2700012I20Rik | 1.975685927 | -0.213623626 | 0.787905862 | -0.271128362 |
| Man2c1os | 25.71371313 | -0.123996253 | 0.274482579 | -0.451745438 |
| Man2c1 | 469.3784322 | 0.411368749 | 0.102639134 | 4.007913292 |
| Neil1 | 55.964215 | 0.099001752 | 0.200894613 | 0.492804411 |

| | | | | |
|---|---|---|---|---|
| Commd4 | 212.8717519 | -0.030975331 | 0.088624246 | -0.349513053 |
| 1700041C23Rik | 0.044544648 | 0.517957688 | 3.724945049 | 0.139051095 |
| Trcg1 | 0.119587853 | 0.981063299 | 3.723618439 | 0.263470416 |
| 1700017B05Rik | 450.6317575 | -0.400816908 | 0.152881414 | -2.621750396 |
| Gm5121 | 0.778482992 | -1.568543671 | 1.127558861 | -1.391096931 |
| Ppcdc | 133.777483 | 0.057351053 | 0.137530675 | 0.41700554 |
| Scamp5 | 3375.801194 | 0.287993452 | 0.050605706 | 5.690928405 |
| Gm6166 | 226.0029141 | -0.304921789 | 0.122580289 | -2.487527092 |
| Rpp25 | 85.87965751 | 0.212633738 | 0.167594812 | 1.268736992 |
| Cox5a | 985.6266964 | -0.205152386 | 0.101903702 | -2.013198558 |
| Gm26631 | 4.769631826 | 1.052271025 | 0.680733037 | 1.545791034 |
| Fam219b | 230.0016874 | -0.216665014 | 0.087933856 | -2.463954412 |
| Mpi | 358.5215663 | 0.172488193 | 0.072080519 | 2.392993218 |
| Scamp2 | 323.5294745 | -0.373231669 | 0.1182953 | -3.155084522 |
| Ulk3 | 153.3484004 | 0.256549904 | 0.150208018 | 1.707964114 |
| Cplx3 | 72.27239396 | 0.876479026 | 0.235185266 | 3.726759926 |
| Lman1l | 1.399662294 | -2.251724877 | 0.989108547 | -2.276519481 |
| Csk | 512.8013575 | -0.580317469 | 0.11509458 | -5.042092078 |
| Cyp1a2 | 0.530050215 | -1.172530326 | 1.585358925 | -0.739599284 |
| Cyp1a1 | 0.692555501 | -0.088720866 | 1.18060147 | -0.07514887 |
| Edc3 | 308.5919229 | -0.181319225 | 0.103213419 | -1.756740803 |
| Clk3 | 437.4151506 | -0.134943621 | 0.103401064 | -1.305050603 |
| Gm17231 | 0.756247029 | 2.070354908 | 3.154254159 | 0.656369082 |
| Arid3b | 117.0600692 | -0.136272866 | 0.220532208 | -0.617927273 |
| Ubl7 | 525.5396743 | -0.026848936 | 0.079038963 | -0.339692415 |
| Sema7a | 1597.276098 | -0.107767145 | 0.058849701 | -1.831226716 |
| Gm17322 | 37.01146921 | 1.038281205 | 0.256255137 | 4.051747859 |
| Cyp11a1 | 16.97152848 | 0.606709567 | 0.333218683 | 1.820754952 |
| Ccdc33 | 7.670793784 | -1.157569033 | 0.432398075 | -2.677091087 |
| Gm16130 | 0.257325455 | 0.049306256 | 2.094236517 | 0.023543786 |
| Stra6 | 207.0557773 | -1.09364131 | 0.253577503 | -4.312848332 |
| Gm16131 | 0 | NA | NA | NA |
| Islr | 283.8085694 | -0.130847461 | 0.175076696 | -0.747372234 |
| Islr2 | 662.2018967 | 0.572968921 | 0.179509889 | 3.191851575 |
| 1600029O15Rik | 9.217140603 | -0.072082168 | 0.339337923 | -0.212420019 |
| Pml | 1126.656421 | -0.60362037 | 0.171550096 | -3.518624492 |
| Mir6385 | 0.090427103 | 0.998850474 | 3.724945049 | 0.268151734 |
| Stoml1 | 580.7474093 | 0.299141123 | 0.110179961 | 2.715022962 |
| Loxl1 | 78.19031371 | -0.99512481 | 0.208061013 | -4.782850931 |
| Tbc1d21 | 0 | NA | NA | NA |
| 1700072B07Rik | 0.112288909 | 0.037060453 | 3.724945049 | 0.009949262 |
| 4930461G14Rik | 0.112396584 | -0.917692504 | 3.724434845 | -0.246397787 |
| 6030419C18Rik | 1394.926258 | 0.125845964 | 0.085002638 | 1.480494805 |
| Cd276 | 125.0400392 | -0.474781404 | 0.11032039 | -4.303659579 |
| Gm10657 | 8.005046079 | 1.018444445 | 0.379807393 | 2.681476097 |
| Nptn | 4736.976647 | 0.318405521 | 0.04687635 | 6.792455415 |
| Rec114 | 2.938016276 | -0.183086221 | 0.55855928 | -0.327782972 |

| | | | | |
|---|---|---|---|---|
| Gm20620 | 0 | NA | NA | NA |
| Hcn4 | 140.1825926 | -0.313241819 | 0.123499619 | -2.536378819 |
| Neo1 | 1678.641993 | 0.02378013 | 0.04470705 | 0.531910068 |
| Gm7589 | 1.467463761 | 0.710814815 | 0.856981732 | 0.829439869 |
| Adpgk | 228.1330696 | -0.543349769 | 0.079722832 | -6.81548508 |
| Bbs4 | 465.2733797 | 0.098231078 | 0.063204943 | 1.554167645 |
| Arih1 | 1854.328923 | -0.177883226 | 0.047457663 | -3.748250836 |
| Rps11-ps1 | 39.22874844 | 0.164783382 | 0.238550694 | 0.690768823 |
| Tmem202 | 11.87611272 | -0.733729423 | 0.32917035 | -2.229026467 |
| Hexa | 567.217733 | -0.665595475 | 0.149191451 | -4.461351312 |
| Celf6 | 249.5287256 | -0.103159477 | 0.100930175 | -1.022087567 |
| Parp6 | 1004.151157 | 0.225097655 | 0.078799962 | 2.8565706 |
| Pkm | 10594.62968 | -0.117433583 | 0.062130545 | -1.890110307 |
| Gramd2 | 45.41966732 | 0.636913642 | 0.270214576 | 2.357066193 |
| Senp8 | 100.069545 | -0.73759834 | 0.20062344 | -3.676531208 |
| Myo9a | 3366.723381 | 0.227300523 | 0.072167066 | 3.149643391 |
| Gm7613 | 0 | NA | NA | NA |
| Gm2735 | 0.060174537 | 0.517957688 | 3.724945049 | 0.139051095 |
| Nr2e3 | 0.25324161 | 0.037067095 | 3.011402107 | 0.012308916 |
| Gm7616 | 0.135217741 | 1.027273954 | 3.618332993 | 0.283908075 |
| Gm15507 | 0.044544648 | 0.517957688 | 3.724945049 | 0.139051095 |
| Thsd4 | 209.2641097 | -0.092253622 | 0.15660003 | -0.589103471 |
| Gm23668 | 0.21890733 | 0.517964903 | 2.599046757 | 0.199290337 |
| Gm22099 | 0 | NA | NA | NA |
| Lrrc49 | 843.7713801 | 0.291874505 | 0.067243786 | 4.340542394 |
| B930082K07Rik | 9.514292225 | 0.700941456 | 0.327322675 | 2.141438736 |
| Larp6 | 390.5730594 | 0.390041911 | 0.093878914 | 4.154733929 |
| 1700036A12Rik | 5.346398063 | 0.956182098 | 0.481290957 | 1.986702812 |
| Uaca | 260.6470978 | -0.739552457 | 0.232676815 | -3.178453583 |
| Gm9869 | 0.554399409 | -0.75423718 | 1.527097874 | -0.493902318 |
| Gm10655 | 0.545210175 | 1.135261636 | 1.395076446 | 0.81376303 |
| Tle3 | 1433.600019 | -0.37847483 | 0.130110676 | -2.908868368 |
| Rplp1 | 2533.538003 | -0.208682455 | 0.127404822 | -1.637947849 |
| Kif23 | 18.40282969 | -0.105828996 | 0.361966026 | -0.292372732 |
| Paqr5 | 21.19486437 | 0.134756599 | 0.285851186 | 0.471422212 |
| Gm23221 | 0 | NA | NA | NA |
| Glce | 558.6811353 | -0.097402352 | 0.067271966 | -1.447889176 |
| Mir5133 | 0 | NA | NA | NA |
| Gm25603 | 0.463251208 | -1.87961565 | 1.420366164 | -1.323331756 |
| Spesp1 | 1.257510835 | -3.112690542 | 1.069889859 | -2.909356057 |
| Anp32a | 1220.827071 | -0.123593365 | 0.091930682 | -1.3444191 |
| Coro2b | 2114.073909 | 0.242551984 | 0.084369709 | 2.874870452 |
| n-R5s84 | 0 | NA | NA | NA |
| Itga11 | 34.17670704 | 0.262833923 | 0.235120606 | 1.117868519 |
| Fem1b | 2918.112869 | -0.408295606 | 0.072188503 | -5.655964476 |
| Gm26609 | 0.048949532 | 0.037064902 | 3.724945049 | 0.009950456 |
| Cln6 | 139.097147 | 0.096372676 | 0.096392931 | 0.99978987 |

| | | | | |
|---|---|---|---|---|
| Gm10653 | 0.777406778 | 0.021175396 | 1.120858551 | 0.018892121 |
| Calml4 | 9.674253487 | 0.208670279 | 0.40992115 | 0.509049799 |
| Pias1 | 838.3302677 | 0.228501272 | 0.069712296 | 3.277775707 |
| Gm24526 | 0 | NA | NA | NA |
| Gm25064 | 0.490308497 | -1.912929185 | 1.318917833 | -1.45037783 |
| Skor1 | 43.41229914 | -0.024567395 | 0.227139852 | -0.108159772 |
| Map2k5 | 366.7989628 | 0.151632147 | 0.076002484 | 1.995094612 |
| 2300009A05Rik | 131.9294766 | 0.018268331 | 0.10418751 | 0.175340895 |
| Gm16759 | 25.37684091 | 0.40757082 | 0.386874085 | 1.053497343 |
| Iqch | 1.370360641 | 0.566774571 | 0.862566193 | 0.65707951 |
| Gm25995 | 0.525324978 | -1.070673046 | 1.724129084 | -0.620993553 |
| Gm6257 | 8.599489497 | -0.213441458 | 0.417170141 | -0.511641263 |
| Aagab | 585.5754181 | -0.0342342 | 0.066220828 | -0.516970282 |
| Smad3 | 973.5426315 | -0.105848681 | 0.094031136 | -1.125676936 |
| Smad6 | 31.70011363 | -0.483055415 | 0.23652717 | -2.042282986 |
| 1700055C04Rik | 0 | NA | NA | NA |
| Gm25606 | 0 | NA | NA | NA |
| 1110036E04Rik | 0.062426873 | 0.517959912 | 3.724945049 | 0.139051692 |
| Scarletltr | 0 | NA | NA | NA |
| Lctl | 0.650092409 | 1.887558463 | 1.262268148 | 1.495370429 |
| Zwilch | 32.2726367 | 0.39719306 | 0.253033781 | 1.569723451 |
| Rpl4 | 3612.185091 | -0.220708188 | 0.084939798 | -2.598407266 |
| Gm23344 | 0.928033173 | -0.83280729 | 1.113306126 | -0.74804878 |
| Snord16a | 3.328548066 | 0.555189427 | 0.557481571 | 0.995888396 |
| Gm22571 | 2.155635849 | 0.119432743 | 0.7384836 | 0.161727008 |
| Gm23136 | 2.475450049 | 0.733903896 | 0.754831298 | 0.972275391 |
| Gm22455 | 1.074104802 | -0.003694365 | 0.9302801 | -0.003971239 |
| Snapc5 | 90.39216346 | -0.089616491 | 0.138638304 | -0.646404988 |
| Map2k1 | 2292.231272 | 0.077575276 | 0.089456856 | 0.867180888 |
| Gm24106 | 0 | NA | NA | NA |
| Uchl4 | 10.29624034 | -0.660208156 | 0.362464092 | -1.821444306 |
| Tipin | 120.2139083 | -0.002765148 | 0.131565816 | -0.021017223 |
| Gm24289 | 1.279592076 | 0.548663225 | 0.949331485 | 0.577946938 |
| Dis3l | 359.2069684 | 0.000155038 | 0.089936333 | 0.001723867 |
| Megf11 | 632.3694475 | -0.256530523 | 0.118563192 | -2.163660729 |
| Rab11a | 1359.033138 | -0.076032264 | 0.087535278 | -0.86858996 |
| AC147560.1 | 0 | NA | NA | NA |
| Dennd4a | 1982.107441 | 0.174888154 | 0.072225005 | 2.421434999 |
| Gm24347 | 0 | NA | NA | NA |
| Slc24a1 | 2.426791053 | -0.909332734 | 0.669251357 | -1.358731253 |
| Vwa9 | 254.7088933 | -0.1322795 | 0.082948987 | -1.594709054 |
| Gm25640 | 1.475874304 | -1.318944436 | 1.018828997 | -1.294569 |
| Hacd3 | 1635.485628 | -0.020598423 | 0.067739083 | -0.304084766 |
| Dpp8 | 2380.590348 | 0.211800629 | 0.055429842 | 3.821057768 |
| Igdcc4 | 562.3809832 | 0.081324062 | 0.151478642 | 0.536868177 |
| Igdcc3 | 23.91639695 | 0.772490639 | 0.260115394 | 2.969799775 |
| Gm25313 | 3.277280862 | 0.220981855 | 0.567543975 | 0.389365167 |

| | | | | |
|---|---|---|---|---|
| Rnu5g | 4.132676428 | -0.961575463 | 0.518704612 | -1.853801644 |
| Parp16 | 35.8728912 | -0.063343952 | 0.190265506 | -0.332923992 |
| Cilp | 2.578207035 | -1.556970017 | 0.821095598 | -1.896210405 |
| Clpx | 458.2865452 | -0.034267896 | 0.08405928 | -0.407663444 |
| Gm16218 | 0 | NA | NA | NA |
| Pdcd7 | 276.9287299 | 0.21846798 | 0.08113946 | 2.692499823 |
| Ubap1l | 57.59547211 | 0.171879096 | 0.339010541 | 0.507002217 |
| Kbtbd13 | 1.752461279 | 1.872400214 | 0.856702309 | 2.18559025 |
| Rasl12 | 22.15572106 | 0.002267536 | 0.239902675 | 0.0094519 |
| Slc51b | 0.090890357 | 0.037071574 | 3.724945049 | 0.009952247 |
| Mtfmt | 124.7687732 | 0.235320402 | 0.129801307 | 1.812927829 |
| Spg21 | 367.5189745 | -0.487562923 | 0.136403777 | -3.574409258 |
| Ankdd1a | 14.47040722 | 0.978223908 | 0.383135993 | 2.553202846 |
| Plekho2 | 544.0730231 | -1.013813523 | 0.148622273 | -6.821410405 |
| Pif1 | 5.794504373 | 0.3036432 | 0.493715746 | 0.615016236 |
| Rbpms2 | 39.33216484 | -0.841045054 | 0.208491714 | -4.033949542 |
| Oaz2 | 881.559079 | 0.033321967 | 0.081625581 | 0.408229459 |
| Zfp609 | 1323.683813 | -0.024178892 | 0.058382858 | -0.414143682 |
| Trip4 | 581.8480211 | -0.242867342 | 0.057085231 | -4.254468923 |
| n-R5s85 | 0 | NA | NA | NA |
| Gm16073 | 0.045882455 | 0.517959912 | 3.724945049 | 0.139051692 |
| 2810417H13Rik | 14.58011455 | -0.352241012 | 0.407946995 | -0.863447987 |
| Gm23248 | 0 | NA | NA | NA |
| Csnk1g1 | 818.8766249 | 0.076608252 | 0.094419089 | 0.811364025 |
| Gm22657 | 0 | NA | NA | NA |
| Ppib | 745.1195212 | -0.433771656 | 0.110100072 | -3.939794464 |
| Snx22 | 48.15551827 | -0.050388293 | 0.191911323 | -0.262560292 |
| Snx1 | 927.1821691 | -0.52137768 | 0.082787472 | -6.297784765 |
| Fam96a | 202.4742243 | -0.416024929 | 0.109114151 | -3.81274953 |
| Dapk2 | 23.66058682 | 0.252476217 | 0.232930385 | 1.083912762 |
| Herc1 | 7489.340024 | 0.498382377 | 0.095550839 | 5.215886971 |
| Gm23428 | 0 | NA | NA | NA |
| Gm28379 | 0.527778125 | -0.265341155 | 1.37637584 | -0.192782485 |
| Fbxl22 | 15.26684527 | -0.259076459 | 0.346855297 | -0.746929516 |
| Usp3 | 603.065053 | 0.205877864 | 0.078920371 | 2.608678363 |
| Gm15563 | 0 | NA | NA | NA |
| Car12 | 301.1637606 | -0.120375411 | 0.151649061 | -0.793776173 |
| Aph1b | 309.2262609 | 0.205764764 | 0.087396445 | 2.35438369 |
| Gm10647 | 6.118905949 | -0.06491429 | 0.575137469 | -0.112867433 |
| BC050972 | 1.031670571 | 0.342125419 | 1.015756244 | 0.336818426 |
| Ppp1r2-ps4 | 7.934796858 | 0.172931104 | 0.399824435 | 0.432517597 |
| Aph1c | 99.73584743 | 0.356874607 | 0.126458598 | 2.822066765 |
| Gm17098 | 1.018413619 | 0.609763024 | 1.147376257 | 0.531441208 |
| Rab8b | 840.6987422 | -0.276147242 | 0.113836217 | -2.425829391 |
| Rps27l | 178.3460037 | -0.65275451 | 0.172097417 | -3.792936118 |
| Lactb | 129.5314519 | 0.165954201 | 0.111772624 | 1.484748193 |
| Gm24225 | 0 | NA | NA | NA |

| | | | | |
|---|---|---|---|---|
| Tpm1 | 1521.522177 | -0.251815911 | 0.118277874 | -2.129019594 |
| Gm22145 | 0 | NA | NA | NA |
| Tln2 | 1967.481299 | 0.391195739 | 0.092519862 | 4.228235229 |
| Mir190a | 0.401901456 | 0.423556179 | 1.530318702 | 0.276776451 |
| Mir7241 | 0.17692102 | -0.979939382 | 2.723401576 | -0.359821846 |
| Mir7242 | 0.162691105 | 0.517965151 | 2.788998835 | 0.185717235 |
| M5C1000I18Rik | 30.04157137 | 1.242901381 | 0.23856607 | 5.20988328 |
| C2cd4b | 11.79896043 | -0.937203814 | 0.407334987 | -2.300818351 |
| C2cd4a | 1.024692053 | 1.027277498 | 0.922580632 | 1.113482618 |
| Vps13c | 2530.722944 | 0.066628169 | 0.096942825 | 0.687293457 |
| Gm28195 | 0 | NA | NA | NA |
| Gm22382 | 0 | NA | NA | NA |
| Gm22169 | 0 | NA | NA | NA |
| Gm22962 | 0.044191549 | 0.037062677 | 3.724945049 | 0.009949859 |
| Rora | 3475.748826 | 0.237656929 | 0.071675492 | 3.3157349 |
| AC127262.1 | 19.3378465 | 1.867001029 | 0.415934406 | 4.488691013 |
| Gm15511 | 0.358080405 | -0.309846495 | 1.562467366 | -0.198305899 |
| Ice2 | 215.338252 | -0.178395668 | 0.122629478 | -1.454753541 |
| Gm4978 | 0.090277459 | -0.443823436 | 3.724945049 | -0.119148989 |
| Anxa2 | 640.8248345 | -1.751427277 | 0.340945292 | -5.136974515 |
| Mir3109 | 0.728690676 | -1.743189858 | 1.194996647 | -1.458740376 |
| Gm16144 | 0 | NA | NA | NA |
| Foxb1 | 12.10284569 | -0.177286033 | 0.343588739 | -0.515983248 |
| B230323A14Rik | 7.674232682 | 0.843966622 | 0.517600732 | 1.630535991 |
| Gm28731 | 0.659754175 | -0.086605738 | 1.173285423 | -0.073814723 |
| Bnip2 | 412.8066768 | -0.263505633 | 0.114726344 | -2.296818882 |
| Gtf2a2 | 313.4260604 | 0.104375504 | 0.101303145 | 1.030328371 |
| Gcnt3 | 0.134276636 | -0.890946624 | 2.951368388 | -0.301875777 |
| Fam81a | 750.583868 | 1.100825812 | 0.146579281 | 7.510105163 |
| Gm24257 | 0 | NA | NA | NA |
| Gm26849 | 0.04393175 | 0.037062677 | 3.724945049 | 0.009949859 |
| Myo1e | 393.0843238 | -0.931804253 | 0.136470348 | -6.827888021 |
| Mir5626 | 0.091666476 | 0.037060453 | 3.724945049 | 0.009949262 |
| Ccnb2 | 19.87963883 | -0.530679693 | 0.323068936 | -1.642620613 |
| Rnf111 | 823.3856292 | -0.341177713 | 0.066035138 | -5.166608633 |
| Gm24615 | 0 | NA | NA | NA |
| Gm23588 | 0 | NA | NA | NA |
| Gm23730 | 0.04393175 | 0.037062677 | 3.724945049 | 0.009949859 |
| Gm22229 | 0 | NA | NA | NA |
| Sltm | 725.1236 | -0.090087225 | 0.116307021 | -0.774563938 |
| Fam63b | 1775.64688 | -0.066804374 | 0.092808256 | -0.719810677 |
| Gm10642 | 11.00079997 | 0.277796794 | 0.319685485 | 0.868969055 |
| Adam10 | 1465.376635 | -0.261516711 | 0.092916402 | -2.814537637 |
| Lipc | 1.303781876 | -0.423762521 | 1.099568835 | -0.385389716 |
| Gm25149 | 0 | NA | NA | NA |
| Gm3436 | 6.640611746 | -0.685180529 | 0.378736 | -1.809124374 |
| Aqp9 | 16.64346168 | -0.805298524 | 0.311647874 | -2.584001341 |

| | | | | |
|---|---|---|---|---|
| Aldh1a2 | 316.9562088 | -0.529532666 | 0.196922287 | -2.689043857 |
| Polr2m | 1314.180415 | -0.043780621 | 0.081512338 | -0.537104223 |
| Gcom1 | 0 | NA | NA | NA |
| Myzap | 9.380444241 | 0.188636337 | 0.347843102 | 0.542302941 |
| Cgnl1 | 362.6537632 | -0.065007932 | 0.12591621 | -0.516279292 |
| Gm27220 | 0 | NA | NA | NA |
| Gm37660 | 16.37804138 | 0.561807217 | 0.354016778 | 1.58695082 |
| Tcf12 | 1303.134971 | -0.356629386 | 0.068410975 | -5.213043503 |
| Gm38057 | 46.80930091 | -0.056020257 | 0.314044648 | -0.178383097 |
| Gm38111 | 32.02976615 | 0.242967458 | 0.330531478 | 0.735081147 |
| Gm18821 | 0 | NA | NA | NA |
| Gm37879 | 56.51000279 | 0.20501756 | 0.253417892 | 0.809009809 |
| Gm27152 | 0 | NA | NA | NA |
| Gm37663 | 62.49855959 | 0.019796435 | 0.186734066 | 0.10601405 |
| Gm37023 | 32.44644427 | 0.915257552 | 0.335167793 | 2.730744334 |
| A430027C01Rik | 29.11976123 | 0.974202145 | 0.285433576 | 3.413060791 |
| Gm27450 | 0 | NA | NA | NA |
| Gm37137 | 44.47354403 | 0.509976921 | 0.284889316 | 1.790087912 |
| Rpl15-ps2 | 4.494967837 | -0.300596572 | 0.586680948 | -0.512368047 |
| Gm25045 | 0.207243203 | 1.595699843 | 2.596246059 | 0.614618109 |
| Gm7863 | 0 | NA | NA | NA |
| Gm7866 | 2.795202431 | -0.61443452 | 0.675436802 | -0.909684692 |
| Gm27159 | 0.158742293 | -1.289348677 | 3.440567791 | -0.374748807 |
| Gm27188 | 5.84902935 | -0.167369158 | 0.534587086 | -0.313081184 |
| Gm7870 | 0.124347486 | 0.036092774 | 3.717245565 | 0.009709548 |
| Gm25163 | 0.730530645 | 1.949929187 | 1.264217147 | 1.542400522 |
| Gm27253 | 3.522083342 | 0.988437993 | 0.651262595 | 1.517725722 |
| Zfp280d | 1238.243116 | 0.247850498 | 0.107617244 | 2.303074193 |
| Gm19353 | 21.12517525 | 0.379210873 | 0.331103222 | 1.145295026 |
| Gm17828 | 1.657074577 | -0.215639661 | 0.788027976 | -0.273644677 |
| Gm37983 | 73.73482861 | 0.633607785 | 0.242496655 | 2.612851655 |
| Gm37303 | 69.68360701 | 0.453348356 | 0.24384591 | 1.859159158 |
| Gm37581 | 38.67009967 | 0.345284779 | 0.311619784 | 1.108032277 |
| BC065403 | 0.430924493 | 1.392321557 | 1.430692581 | 0.973180106 |
| Gm37150 | 0.370677815 | -0.461578519 | 1.718783101 | -0.268549603 |
| Gm27230 | 0.066050875 | -0.443818988 | 3.724945049 | -0.119147795 |
| Gm27255 | 0.075043205 | 0.517957688 | 3.724945049 | 0.139051095 |
| Gm27254 | 0.194277958 | 1.05585522 | 3.250214283 | 0.32485711 |
| Mns1 | 77.00500091 | 0.499580337 | 0.151982053 | 3.287100856 |
| Gm36948 | 4.487847835 | 0.522688711 | 0.570148374 | 0.916759101 |
| Gm27231 | 0.460076145 | 1.692420597 | 1.61757049 | 1.046273166 |
| Tex9 | 177.6432418 | 0.344453063 | 0.128635181 | 2.677751615 |
| Gm28010 | 0.172182752 | 0.037075442 | 3.331044665 | 0.011130275 |
| 4930509E16Rik | 0.362648911 | -0.995734401 | 1.727061711 | -0.576548246 |
| Rfx7 | 1180.076257 | 0.165195131 | 0.077755182 | 2.124554624 |
| Gm19427 | 12.05712943 | 0.913663126 | 0.361697002 | 2.526045615 |
| Gm37105 | 45.12253491 | 0.559608546 | 0.345492815 | 1.619740038 |

| | | | | |
|---|---|---|---|---|
| Gm18014 | 0 | NA | NA | NA |
| Gm27241 | 0.220148986 | 0.562109768 | 2.591308868 | 0.216921176 |
| Gm27232 | 0.138573873 | -0.444788617 | 3.62882948 | -0.122570823 |
| Gm23057 | 0.072025462 | 0.517959912 | 3.724945049 | 0.139051692 |
| Nedd4 | 6315.797856 | -0.129635512 | 0.048949365 | -2.648359429 |
| A530064N14Rik | 50.37861278 | 0.318733092 | 0.337274278 | 0.945026385 |
| Gm7265 | 195.1401439 | -0.657035463 | 0.100891645 | -6.512288144 |
| Gm37955 | 4.498464866 | 0.188468629 | 0.604679034 | 0.31168375 |
| Gm36936 | 103.529168 | 0.307461782 | 0.29054033 | 1.058241321 |
| Gm27204 | 0 | NA | NA | NA |
| Prtg | 50.92406102 | 0.664654429 | 0.222998233 | 2.980536757 |
| Gm27203 | 0 | NA | NA | NA |
| A730062M13Rik | 2.892325926 | 1.327054836 | 0.830536179 | 1.597829052 |
| Gm37234 | 1.821448544 | 1.355299498 | 0.90471482 | 1.498040563 |
| Pygo1 | 611.8774595 | 0.511701632 | 0.101595051 | 5.036678749 |
| Dyx1c1 | 5.660739534 | 0.270851706 | 0.43397772 | 0.624114312 |
| Dyx1c1Ccpg1 | 0 | NA | NA | NA |
| Gm23567 | 0.220111735 | -0.948104543 | 2.165130622 | -0.437897157 |
| Gm37710 | 0.046345709 | -0.443818988 | 3.724945049 | -0.119147795 |
| Ccpg1os | 38.04226344 | -0.264565283 | 0.220611655 | -1.199235293 |
| Ccpg1 | 158.4319805 | 0.023899925 | 0.117052621 | 0.204181032 |
| Gm27353 | 16.47684853 | 0.603784615 | 0.370399383 | 1.630090767 |
| Pigb | 210.9882647 | -0.056456988 | 0.092032316 | -0.613447438 |
| Gm38192 | 17.09336393 | 0.354681371 | 0.310108879 | 1.143731751 |
| 2310009A05Rik | 48.02464468 | -0.42144489 | 0.229658647 | -1.835092631 |
| Rab27a | 175.4812393 | -0.208268405 | 0.119001338 | -1.750134987 |
| Gm27186 | 0.89763465 | -0.81977877 | 1.343886856 | -0.610005795 |
| Khdc3 | 3.338837625 | 0.188453145 | 0.669462681 | 0.281499104 |
| Gm20509 | 0 | NA | NA | NA |
| Rsl24d1 | 296.3744691 | -0.113823151 | 0.079981938 | -1.423110689 |
| Gm38215 | 2.049787317 | -1.539566198 | 0.95313798 | -1.61526057 |
| Gm5745 | 0.038981395 | -0.443816764 | 3.724945049 | -0.119147198 |
| Gm27211 | 0.328170597 | -2.006319889 | 3.693612668 | -0.543186324 |
| Unc13c | 1536.822218 | 0.509895928 | 0.098835511 | 5.159035696 |
| Gm37611 | 8.958307541 | 1.127748448 | 0.48025848 | 2.348211423 |
| Gm27148 | 1.654347319 | 2.009165995 | 1.096807385 | 1.831831207 |
| Gm25105 | 4.734499335 | 1.110495992 | 0.538092284 | 2.063764945 |
| Gm37326 | 31.32772203 | 0.248614076 | 0.347332803 | 0.715780583 |
| n-R5s86 | 0 | NA | NA | NA |
| Wdr72 | 4.387779186 | 0.114273374 | 0.522233782 | 0.218816512 |
| Gm24141 | 0 | NA | NA | NA |
| Nr1h2-ps | 0 | NA | NA | NA |
| Gm28622 | 0.044544648 | 0.517957688 | 3.724945049 | 0.139051095 |
| Gm27189 | 0 | NA | NA | NA |
| Gm27233 | 0 | NA | NA | NA |
| Gm22315 | 0 | NA | NA | NA |
| Gm16551 | 2.498792093 | 0.280067457 | 0.696043489 | 0.402370629 |

| | | | | |
|---|---|---|---|---|
| Gm20649 | 0 | NA | NA | NA |
| Gm28182 | 0.458414965 | -1.43140337 | 1.443603549 | -0.991548802 |
| Onecut1 | 8.345816899 | 0.237167357 | 0.396903073 | 0.597544773 |
| Fam214a | 701.9545256 | 0.670969871 | 0.086112349 | 7.79179612 |
| Arpp19 | 2304.068667 | 0.328118135 | 0.123342361 | 2.660222589 |
| Myo5a | 9406.867657 | 0.250210068 | 0.065491104 | 3.820519938 |
| Gm9719 | 0.038981395 | -0.443816764 | 3.724945049 | -0.119147198 |
| Gm16353 | 0 | NA | NA | NA |
| Myo5c | 38.56207624 | 0.784626535 | 0.351685716 | 2.23104465 |
| Gnb5 | 588.9032945 | 0.150426667 | 0.078881774 | 1.906988885 |
| Bcl2l10 | 2.578686406 | 0.40112579 | 0.725171595 | 0.553146031 |
| Mapk6 | 1348.202811 | -0.144807659 | 0.095731302 | -1.512646917 |
| 4933433G15Rik | 7.660142694 | 0.042248855 | 0.43411366 | 0.097322105 |
| A130057D12Rik | 0.39127578 | -0.449291277 | 1.567519167 | -0.286625699 |
| Leo1 | 251.7819994 | 0.039230229 | 0.18006516 | 0.217866849 |
| Tmod3 | 491.2591432 | -0.547723229 | 0.093490054 | -5.858625683 |
| Tmod2 | 9543.757063 | 0.419697025 | 0.06128967 | 6.84776117 |
| Lysmd2 | 376.8025865 | -0.205761797 | 0.107498025 | -1.914098384 |
| Scg3 | 2648.530245 | 0.197772671 | 0.079398108 | 2.490899045 |
| Gm37871 | 0.090665643 | 0.037060453 | 3.724945049 | 0.009949262 |
| Gm3671 | 0.843494912 | 0.301102039 | 1.055407346 | 0.285294622 |
| Bmp5 | 56.01127082 | -0.094916269 | 0.21549882 | -0.440449134 |
| Gm26903 | 0.668743806 | 1.333700405 | 1.47362509 | 0.905047297 |
| AC079245.1 | 0 | NA | NA | NA |
| Gm25300 | 0 | NA | NA | NA |
| Hmgcll1 | 224.7683895 | 0.110929171 | 0.121217222 | 0.915127157 |
| Gm38083 | 27.55755119 | 0.209868653 | 0.281062444 | 0.746697602 |
| Gfral | 1.244634631 | -0.389010177 | 1.047616695 | -0.37132873 |
| Hcrtr2 | 58.13001265 | -0.325288524 | 0.239753891 | -1.356760144 |
| Gm27234 | 0.038981395 | -0.443816764 | 3.724945049 | -0.119147198 |
| Fam83b | 2.476850474 | 0.743675804 | 0.653807673 | 1.137453465 |
| Gm27156 | 0 | NA | NA | NA |
| Gm27245 | 0 | NA | NA | NA |
| Gm22938 | 0 | NA | NA | NA |
| Gm37408 | 0 | NA | NA | NA |
| Gm27155 | 0 | NA | NA | NA |
| Gm23791 | 0 | NA | NA | NA |
| Gm18486 | 0 | NA | NA | NA |
| Tinag | 0.568818845 | -1.795856226 | 1.256567889 | -1.429175647 |
| Gm27157 | 0 | NA | NA | NA |
| Gm27158 | 0 | NA | NA | NA |
| Gm19541 | 0.139037128 | -1.190058937 | 3.717743703 | -0.320102469 |
| Gm5746 | 0.044544648 | 0.517957688 | 3.724945049 | 0.139051095 |
| Mlip | 470.9327951 | 0.756666199 | 0.108938298 | 6.945823588 |
| Gm27210 | 1.42879052 | 0.0040239 | 0.838684794 | 0.004797869 |
| Gm38254 | 2.89654368 | 0.813900767 | 0.651934795 | 1.248438913 |
| 4930590H14Rik | 0 | NA | NA | NA |

| | | | | |
|---|---|---|---|---|
| 5730403I07Rik | 4.072737608 | -0.282262179 | 0.606047499 | -0.465742668 |
| Lrrc1 | 172.5600245 | 0.497442775 | 0.128162289 | 3.881350582 |
| Gm19569 | 3.396866737 | 0.576096644 | 0.546880181 | 1.053423884 |
| Gm21013 | 0.284055481 | 0.517965359 | 2.478960587 | 0.208944572 |
| Gm25250 | 0 | NA | NA | NA |
| Klhl31 | 1.14819629 | 0.401313058 | 1.0327783 | 0.388576191 |
| Gclc | 631.4586054 | 0.04921088 | 0.055143426 | 0.892416072 |
| Elovl5 | 1099.939201 | -0.153391104 | 0.101884596 | -1.505537738 |
| Gm23609 | 0 | NA | NA | NA |
| Gcm1 | 0.062426873 | 0.517959912 | 3.724945049 | 0.139051692 |
| Fbxo9 | 758.0697948 | -0.011962483 | 0.091666111 | -0.130500607 |
| Ick | 452.7014818 | -0.15832081 | 0.088071447 | -1.797640625 |
| Rn7sk | 139206.5501 | 0.079348835 | 0.172335013 | 0.460433627 |
| C920006O11Rik | 1.914692995 | 0.395148912 | 0.71969049 | 0.549053958 |
| Gsta4 | 217.5686189 | 0.239671386 | 0.158443467 | 1.512661831 |
| Gsta1 | 0.047948698 | 0.037064902 | 3.724945049 | 0.009950456 |
| Gm10639 | 0.087930927 | -0.443164241 | 3.724898049 | -0.118973522 |
| Omt2a | 0 | NA | NA | NA |
| Gm8074 | 0 | NA | NA | NA |
| Omt2b | 0 | NA | NA | NA |
| Gsta2 | 0.44449992 | -0.453308258 | 1.647431569 | -0.275160599 |
| Dppa5a | 0.045882455 | 0.517959912 | 3.724945049 | 0.139051692 |
| Ooep | 0.251515219 | 0.940213969 | 1.820460072 | 0.516470525 |
| Gm8087 | 0.090074004 | 0.517955464 | 3.724945049 | 0.139050498 |
| 4930542C12Rik | 0 | NA | NA | NA |
| Ddx43 | 4.317706999 | 1.012413135 | 0.478115188 | 2.117508836 |
| Gm8093 | 1.913696796 | -0.044363864 | 0.61951562 | -0.071610565 |
| Mb21d1 | 40.10846614 | -0.720363412 | 0.223602722 | -3.221621835 |
| Gm17324 | 2.766932339 | -0.019912168 | 0.650543289 | -0.030608521 |
| Mto1 | 264.8336796 | -0.103426448 | 0.078437034 | -1.318592032 |
| Eef1a1 | 14473.51984 | -0.223416535 | 0.094516394 | -2.363786068 |
| Gm26377 | 0 | NA | NA | NA |
| Slc17a5 | 273.2142388 | 0.298268879 | 0.1046716 | 2.849568364 |
| Cd109 | 135.8248537 | -1.208653818 | 0.249706245 | -4.840302723 |
| 4921509A06Rik | 0.07796279 | -0.798301147 | 3.721829328 | -0.214491605 |
| Gm8116 | 61.44867496 | -0.620487628 | 0.145725006 | -4.257935172 |
| Gm28620 | 0 | NA | NA | NA |
| Col12a1 | 350.8884372 | -1.391209323 | 0.158912254 | -8.75457547 |
| 4930562D21Rik | 0 | NA | NA | NA |
| Cox7a2 | 993.8036579 | 0.106146722 | 0.132761015 | 0.799532319 |
| Tmem30a | 5117.156797 | 0.179031971 | 0.090151792 | 1.98589477 |
| 4930429F24Rik | 2.804294664 | -0.779016924 | 0.563933663 | -1.381398159 |
| Filip1 | 118.2003987 | 0.855792018 | 0.133399195 | 6.415271227 |
| Gm20477 | 0.402709178 | 0.510743835 | 1.42783638 | 0.357704736 |
| Gm3211 | 0 | NA | NA | NA |
| Gm25433 | 0 | NA | NA | NA |
| Senp6 | 1335.192463 | -0.060139237 | 0.05607025 | -1.07256945 |

| | | | | |
|---|---|---|---|---|
| Gm20648 | 0 | NA | NA | NA |
| Gm17477 | 0 | NA | NA | NA |
| Myo6 | 1685.660187 | 0.078025414 | 0.080599923 | 0.968058169 |
| Gm27227 | 0 | NA | NA | NA |
| Impg1 | 1.92279121 | 0.627833798 | 0.809177783 | 0.775891048 |
| Gm27224 | 0.046345709 | -0.443818988 | 3.724945049 | -0.119147795 |
| Gm27225 | 0.281850154 | 0.431661762 | 1.782357609 | 0.242185833 |
| Gm8151 | 0.792246753 | 0.352198871 | 1.005274633 | 0.350350898 |
| Gm26713 | 0.150086409 | 1.027495679 | 3.723011807 | 0.275985071 |
| Gm27226 | 0 | NA | NA | NA |
| Htr1b | 292.4427632 | 0.517364181 | 0.133458231 | 3.876600018 |
| D430036J16Rik | 57.68366793 | 1.637762828 | 0.186029944 | 8.80375918 |
| Gm9531 | 81.22717314 | -0.295013714 | 0.131463134 | -2.244079434 |
| Gm26907 | 0.36944391 | 0.859474855 | 3.245846333 | 0.26479222 |
| Mei4 | 18.14348477 | 0.368082617 | 0.261392578 | 1.40816017 |
| Gm37837 | 2.253692409 | 1.477900079 | 0.828978791 | 1.782796008 |
| Gm15896 | 1.074470558 | 2.297317666 | 1.198445951 | 1.916913871 |
| Gm23748 | 0.232637982 | 1.023240166 | 2.544854188 | 0.402082041 |
| 1700063H06Rik | 1.312371618 | 2.055553591 | 1.052870789 | 1.952332245 |
| Gm22836 | 0 | NA | NA | NA |
| Gm24393 | 0 | NA | NA | NA |
| Gm27249 | 0.180969744 | 0.959616598 | 2.733014117 | 0.351120249 |
| Irak1bp1 | 163.4822182 | 0.271217525 | 0.103674377 | 2.616051653 |
| Phip | 3482.802041 | 0.066799465 | 0.086615368 | 0.771219556 |
| Gm27248 | 0.875572801 | 0.854314865 | 1.253524614 | 0.681530187 |
| Hmgn3 | 287.8474579 | 0.236458048 | 0.078429563 | 3.014909665 |
| Gm2065 | 1.354044546 | -1.213904699 | 0.993582361 | -1.22174542 |
| Gm29054 | 6.179723687 | 1.246940255 | 0.487774263 | 2.556387967 |
| Gm38279 | 0.200264842 | -0.388652768 | 2.201307895 | -0.176555387 |
| Gm27216 | 0 | NA | NA | NA |
| Lca5 | 153.3277019 | 0.025770827 | 0.116942609 | 0.220371577 |
| Rps27a-ps2 | 369.3263099 | -0.047053674 | 0.140744456 | -0.33431991 |
| Gm2087 | 1.000715493 | 2.535082525 | 1.134952682 | 2.233646006 |
| Gm37307 | 1.831656741 | 1.941786133 | 0.876279453 | 2.215943928 |
| Gm28552 | 0.11657011 | 0.998850474 | 3.724945049 | 0.268151734 |
| Gm28477 | 0.334445592 | 0.534166855 | 1.973878833 | 0.270617854 |
| Sh3bgrl2 | 558.0448667 | -0.062103246 | 0.086139632 | -0.720960199 |
| Gm2109 | 2.113095361 | 1.612911383 | 0.85564323 | 1.885027926 |
| Elovl4 | 771.8863038 | 0.151411206 | 0.101186761 | 1.496353924 |
| Ttk | 15.5422173 | -0.494957662 | 0.382220418 | -1.294953485 |
| Bckdhb | 244.9808363 | -0.11036371 | 0.077083446 | -1.43174333 |
| Gm29163 | 0 | NA | NA | NA |
| Gm29164 | 0 | NA | NA | NA |
| Gm26146 | 0.204782255 | 0.517969162 | 2.213323883 | 0.234023211 |
| Gm8226 | 4.794882048 | -0.084247822 | 0.465132273 | -0.181126589 |
| Gm26126 | 0 | NA | NA | NA |
| 4930554C24Rik | 0.203235021 | 0.053373928 | 2.187927154 | 0.024394747 |

| | | | | |
|---|---|---|---|---|
| Gm28625 | 0 | NA | NA | NA |
| Gm8228 | 8.163884718 | 0.129462259 | 0.392807566 | 0.329581887 |
| Gm28070 | 0 | NA | NA | NA |
| Gm11114 | 0 | NA | NA | NA |
| Fam46a | 425.1094954 | -1.189256284 | 0.18029309 | -6.596238833 |
| Gm36927 | 0 | NA | NA | NA |
| Gm25125 | 0 | NA | NA | NA |
| Ibtk | 540.332865 | -0.079462419 | 0.055585302 | -1.429558093 |
| 4933431K14Rik | 24.95481328 | -0.610077433 | 0.220068156 | -2.772220402 |
| Gm22830 | 0.143846094 | 0.0370613 | 3.724945049 | 0.009949489 |
| Gm5619 | 19.25933968 | 0.015200486 | 0.371110816 | 0.040959426 |
| Gm18766 | 0.416300838 | 2.032856674 | 1.505816959 | 1.350002509 |
| 9330154J02Rik | 12.67333215 | 0.247699978 | 0.285851909 | 0.866532531 |
| Tpbg | 242.3331958 | -0.73301154 | 0.118265086 | -6.198038362 |
| Ube2cbp | 42.39662063 | -0.107493224 | 0.208827894 | -0.514745524 |
| Gm16202 | 0.329441611 | 0.903901444 | 1.69137295 | 0.534418766 |
| Dopey1 | 1570.3345 | 0.369563882 | 0.110025385 | 3.358896522 |
| Gm17530 | 6.903802255 | -0.92583393 | 0.425055098 | -2.178150395 |
| Pgm3 | 195.6218196 | -0.045140027 | 0.091968282 | -0.490821679 |
| Rwdd2a | 134.250853 | 0.888019042 | 0.126328054 | 7.029468253 |
| Me1 | 717.0322805 | -0.556145801 | 0.10370492 | -5.362771589 |
| Gm38230 | 5.852527318 | 1.200719089 | 0.511313003 | 2.348305407 |
| A330041J22Rik | 15.09013069 | -0.29760343 | 0.271663632 | -1.095484986 |
| Gm25949 | 0 | NA | NA | NA |
| Gm22138 | 0.271189253 | 1.406907094 | 3.71626224 | 0.37858122 |
| Gm37484 | 1.311065665 | 0.023103942 | 1.140861666 | 0.020251309 |
| Prss35 | 55.61444896 | 0.324027761 | 0.178644369 | 1.813814582 |
| Snap91 | 4128.306371 | 0.21120282 | 0.063771473 | 3.311869878 |
| Gm37941 | 94.46575361 | 0.430616719 | 0.274919152 | 1.566339475 |
| Ripply2 | 33.98125597 | 0.26655233 | 0.217528974 | 1.225364718 |
| Cyb5r4 | 351.6163489 | -0.23603522 | 0.081128478 | -2.9094003 |
| Gm8282 | 3.540589861 | 0.012992043 | 0.54690062 | 0.023755766 |
| Mrap2 | 94.81455933 | 0.054588427 | 0.204936983 | 0.266366891 |
| Gm37012 | 4.127795301 | -0.297191688 | 0.500043903 | -0.594331191 |
| Cep162 | 341.6360393 | -0.12820217 | 0.107796864 | -1.189294064 |
| Gm24649 | 0.157258859 | 0.955406371 | 2.806391154 | 0.340439489 |
| Gm9540 | 0 | NA | NA | NA |
| Gm37375 | 0.154920219 | 1.020650266 | 2.83493605 | 0.360025852 |
| Tbx18 | 38.37151821 | -0.182468852 | 0.184377047 | -0.989650586 |
| D030062O11Rik | 0.22856443 | 0.536345688 | 2.143946829 | 0.250167439 |
| Gm37374 | 0.748906342 | -0.627564355 | 1.103799455 | -0.568549253 |
| Gm9541 | 0 | NA | NA | NA |
| Gm25528 | 0 | NA | NA | NA |
| Gm23176 | 0 | NA | NA | NA |
| Gm28877 | 0.115603922 | -0.93550548 | 3.724188147 | -0.251197158 |
| Gm2324 | 0 | NA | NA | NA |
| Gm5368 | 0 | NA | NA | NA |

| | | | | |
|---|---|---|---|---|
| Gm28231 | 0 | NA | NA | NA |
| Gm5066 | 2.101151607 | 0.862897917 | 0.808073574 | 1.067845732 |
| Nt5e | 129.2471619 | 0.028328619 | 0.1529864 | 0.185170831 |
| Snx14 | 532.7999784 | 0.023826681 | 0.071975302 | 0.331039678 |
| Gm10163 | 3.877733872 | 0.66413679 | 0.497199679 | 1.335754665 |
| Gm20537 | 1.59229893 | -0.076955491 | 0.811492844 | -0.094832002 |
| Gm28229 | 64.56740978 | 0.348523092 | 0.183484475 | 1.89946911 |
| Syncrip | 1823.979902 | -0.049361804 | 0.066274556 | -0.744807773 |
| Snhg5 | 101.6600101 | -0.023307986 | 0.154467716 | -0.15089228 |
| Snord50a | 0 | NA | NA | NA |
| Snord50b | 0 | NA | NA | NA |
| Gm22985 | 0 | NA | NA | NA |
| Zfp949 | 311.469506 | 0.057815394 | 0.096119195 | 0.601496854 |
| Gm26461 | 289.3437332 | -0.32007731 | 0.199693813 | -1.602840391 |
| Trim43a | 11.4592027 | -0.511866028 | 0.322941534 | -1.585011447 |
| 9430037G07Rik | 40.90789348 | 0.183170534 | 0.220660349 | 0.830101713 |
| Gm29603 | 0.044544648 | 0.517957688 | 3.724945049 | 0.139051095 |
| Gm23017 | 0 | NA | NA | NA |
| Gm26567 | 0 | NA | NA | NA |
| Gm29415 | 1.336334664 | 0.855680172 | 0.919401995 | 0.930692098 |
| Gm38214 | 23.29150933 | 0.063499807 | 0.336426445 | 0.188747964 |
| Gm37226 | 40.31519145 | -0.052446309 | 0.354452105 | -0.147964446 |
| Mthfsl | 134.439119 | -0.102292092 | 0.160376596 | -0.637824315 |
| Gm10634 | 2.499553143 | -0.342969858 | 0.694395472 | -0.493911426 |
| Bcl2a1d | 28.6456985 | 0.093563601 | 0.314728807 | 0.297283244 |
| Gm37560 | 0.141331096 | -0.443820795 | 3.561840904 | -0.124604329 |
| Gm26082 | 0.238313115 | -0.088739579 | 2.143089712 | -0.04140731 |
| Gm37614 | 0.29572682 | 0.037078117 | 1.775942131 | 0.020877998 |
| Gm29562 | 2.519137971 | 0.755289507 | 0.636994271 | 1.185708476 |
| Trim43c | 0 | NA | NA | NA |
| 9330159M07Rik | 56.07294295 | -0.023606058 | 0.203335246 | -0.116094276 |
| Gm9775 | 0 | NA | NA | NA |
| Gm26135 | 0 | NA | NA | NA |
| Gm26611 | 0 | NA | NA | NA |
| Gm2396 | 3.793477816 | 0.419434324 | 0.528872353 | 0.793072887 |
| Gm23707 | 0 | NA | NA | NA |
| Bcl2a1a | 13.58752781 | 0.157547024 | 0.346907525 | 0.454147036 |
| Gm37833 | 3.739270958 | -0.574304554 | 0.696229764 | -0.824877912 |
| Gm37993 | 0.55378357 | 0.073368158 | 1.333013146 | 0.055039336 |
| Gm37471 | 0.168528002 | 0.037073846 | 2.758169225 | 0.013441469 |
| Gm24463 | 0.710913347 | -0.21998986 | 1.144742891 | -0.192174035 |
| Gm19498 | 0 | NA | NA | NA |
| Trim43b | 0.095295241 | -0.443821212 | 3.724945049 | -0.119148392 |
| 4930579C12Rik | 0.595794717 | -0.723758813 | 1.291750387 | -0.560293088 |
| Gm26423 | 0 | NA | NA | NA |
| Bcl2a1b | 8.616394077 | 0.231918306 | 0.39096838 | 0.593189418 |
| Gm29094 | 0.444881751 | -2.086683106 | 1.564378216 | -1.333873794 |

| | | | | |
|---|---|---|---|---|
| Mthfs | 70.61009405 | -0.153561057 | 0.251651062 | -0.610214222 |
| Gm38091 | 0.04393175 | 0.037062677 | 3.724945049 | 0.009949859 |
| AF529169 | 248.5876439 | 0.0735062 | 0.176207151 | 0.417157871 |
| Tmed3 | 223.4678697 | -0.599537575 | 0.119706508 | -5.008395834 |
| Ankrd34c | 157.1958196 | 0.106021125 | 0.132281352 | 0.80148202 |
| Gm29608 | 0.210489921 | -0.951860691 | 2.624963171 | -0.362618684 |
| Mir184 | 0 | NA | NA | NA |
| 1700017I07Rik | 0 | NA | NA | NA |
| 4930524O08Rik | 0.357611442 | 1.138051066 | 1.943831688 | 0.585467905 |
| Rasgrf1 | 5020.678698 | 0.457184607 | 0.076504551 | 5.975913894 |
| Gm23759 | 0.203875117 | 0.922448301 | 2.61967177 | 0.352123618 |
| Gm17944 | 0.117650406 | -0.443821212 | 3.724945049 | -0.119148392 |
| Ctsh | 415.3048284 | -0.960067762 | 0.198143204 | -4.845322676 |
| Morf4l1 | 2238.38623 | 0.153326721 | 0.065716178 | 2.333165517 |
| Gm26882 | 0.372996712 | 1.394948253 | 1.923135059 | 0.725351164 |
| Gm6223 | 7.567416792 | -0.665478142 | 0.361689819 | -1.839913947 |
| 9430062P05Rik | 1.427283908 | -0.052374055 | 0.897930922 | -0.058327488 |
| Adamts7 | 9.932952153 | 0.073867167 | 0.354144951 | 0.208578908 |
| Tbc1d2b | 421.9794416 | -0.197520259 | 0.088128565 | -2.24127398 |
| Gm16200 | 6.214365972 | 0.361139596 | 0.452932046 | 0.79733726 |
| Gm22717 | 0 | NA | NA | NA |
| Gm22866 | 0.060174537 | 0.517957688 | 3.724945049 | 0.139051095 |
| Gm2497 | 0 | NA | NA | NA |
| Gm29605 | 0 | NA | NA | NA |
| Gm29604 | 0.044544648 | 0.517957688 | 3.724945049 | 0.139051095 |
| Gm28703 | 3.32610108 | 1.783436214 | 0.707410983 | 2.521075097 |
| Gm37665 | 0.317495958 | -0.493297316 | 1.764790896 | -0.27952168 |
| Gm20561 | 0.047948698 | 0.037064902 | 3.724945049 | 0.009950456 |
| Gm29408 | 0.136071997 | 0.037065787 | 2.949782085 | 0.012565602 |
| Gm5620 | 30.93367375 | -0.021949673 | 0.230213622 | -0.095344807 |
| Gm36960 | 3.033497949 | 0.178441549 | 0.65327247 | 0.273150266 |
| Gm29602 | 0 | NA | NA | NA |
| Gm37812 | 6.137587838 | -0.36168508 | 0.428460711 | -0.844149932 |
| Gm37457 | 5.572788009 | 0.476320707 | 0.503247473 | 0.946493987 |
| Gm37040 | 5.798168318 | 0.48366071 | 0.48956929 | 0.987931065 |
| Gm28652 | 0 | NA | NA | NA |
| A730094K22Rik | 3.673889323 | -0.958201451 | 0.767327461 | -1.248751673 |
| Zic1 | 2091.234844 | 0.015015277 | 0.184503575 | 0.081382037 |
| Zic4 | 124.4589485 | 0.260806117 | 0.144157937 | 1.809169323 |
| Gm37477 | 5.296492507 | 0.246044775 | 0.478187132 | 0.514536588 |
| Gm29478 | 1.174654913 | 1.006487136 | 0.9663774 | 1.041505249 |
| Gm37987 | 0.060174537 | 0.517957688 | 3.724945049 | 0.139051095 |
| Gm26101 | 0.062426873 | 0.517959912 | 3.724945049 | 0.139051692 |
| Plscr5 | 1.323501177 | 0.31266546 | 0.977936401 | 0.319719625 |
| Plscr1 | 60.43889054 | -0.409329062 | 0.268575393 | -1.524075075 |
| Mir6386 | 0 | NA | NA | NA |
| Plscr2 | 121.3806642 | -0.133174178 | 0.219501542 | -0.606711809 |

| | | | | |
|---|---|---|---|---|
| Gm29139 | 0 | NA | NA | NA |
| 1700057G04Rik | 0.707934375 | -2.331468803 | 1.243296882 | -1.875230958 |
| Gm15493 | 0 | NA | NA | NA |
| Plscr4 | 44.4848455 | 0.016968487 | 0.186397678 | 0.091033789 |
| Gm25417 | 0.083526043 | 0.037073798 | 3.724945049 | 0.009952844 |
| Gm38391 | 19.9576536 | 0.404565588 | 0.244481808 | 1.654788102 |
| Gm37545 | 0 | NA | NA | NA |
| Plod2 | 158.0764622 | -0.584635466 | 0.109151541 | -5.356181508 |
| Gm2495 | 0 | NA | NA | NA |
| Gm29396 | 0 | NA | NA | NA |
| 4933400C23Rik | 0.454057983 | 0.852863997 | 1.53242511 | 0.556545303 |
| Gm28054 | 0.110242424 | -0.443823436 | 3.724945049 | -0.119148989 |
| Gm29397 | 3.384086637 | -1.022513555 | 0.566957908 | -1.803508762 |
| 1700034K08Rik | 0 | NA | NA | NA |
| Gm9621 | 0 | NA | NA | NA |
| Gm24200 | 0 | NA | NA | NA |
| Gm5370 | 0 | NA | NA | NA |
| Gm28934 | 0 | NA | NA | NA |
| 1190002N15Rik | 756.6875991 | -0.023804677 | 0.13994959 | -0.170094652 |
| Gm24178 | 0.049553047 | -0.443818988 | 3.724945049 | -0.119147795 |
| Gm29395 | 0.832501709 | 1.467414438 | 1.19528623 | 1.227667817 |
| Slc9a9 | 216.5651415 | 0.086945143 | 0.124222513 | 0.69991454 |
| Gm16262 | 0.060174537 | 0.517957688 | 3.724945049 | 0.139051095 |
| Mir7656 | 0.276333946 | 1.013227214 | 1.808099618 | 0.560382406 |
| Chst2 | 1637.467997 | -0.085325787 | 0.107359283 | -0.794768601 |
| Gm37805 | 7.969835414 | 0.295371511 | 0.363002116 | 0.813690879 |
| Gm8449 | 2.212778765 | 1.293488462 | 0.73988083 | 1.748238917 |
| Gm19193 | 0 | NA | NA | NA |
| U2surp | 1949.775422 | 0.315755029 | 0.085139544 | 3.708676541 |
| Gm28424 | 12.26829614 | 0.051544759 | 0.302015544 | 0.170669225 |
| Paqr9 | 964.539708 | 0.209858749 | 0.099415991 | 2.110915424 |
| Gm16126 | 0.766553141 | 1.838351088 | 1.305093657 | 1.40859706 |
| Gm28164 | 0.74026972 | 2.404727367 | 1.238219301 | 1.94208519 |
| Gm25582 | 0.857373094 | 1.654245347 | 1.131194753 | 1.462387749 |
| Gm16127 | 0.660063185 | 2.031522988 | 1.350427124 | 1.504355882 |
| Pcolce2 | 60.65372678 | 0.43291617 | 0.173802395 | 2.490852735 |
| Itpa-ps2 | 0.066050875 | -0.443818988 | 3.724945049 | -0.119147795 |
| Trpc1 | 585.9086551 | 0.148819725 | 0.06637818 | 2.241997676 |
| Pls1 | 183.634994 | 0.191324174 | 0.126322688 | 1.514566994 |
| 1700065D16Rik | 3.065206492 | -0.142975499 | 0.592249966 | -0.241410734 |
| Atr | 564.2015883 | 0.003986456 | 0.098000388 | 0.040677958 |
| Gm24004 | 0 | NA | NA | NA |
| Gm16794 | 29.63475761 | 0.031952068 | 0.205912587 | 0.155172971 |
| Xrn1 | 1619.277144 | 0.110685551 | 0.083220814 | 1.33002245 |
| C78334 | 0.042716945 | 0.037062677 | 3.724945049 | 0.009949859 |
| Gm28085 | 0 | NA | NA | NA |
| Gm38375 | 61.21253001 | 0.001618108 | 0.263195518 | 0.006147933 |

| | | | | |
|---|---|---|---|---|
| Gk5 | 59.46593611 | 0.108168086 | 0.15683582 | 0.689689932 |
| Tfdp2 | 637.9968549 | 0.378857445 | 0.087198905 | 4.344750034 |
| Gm37195 | 34.22903916 | 0.188796154 | 0.324842682 | 0.581192572 |
| Gm28924 | 0.049553047 | -0.443818988 | 3.724945049 | -0.119147795 |
| Atp1b3 | 1332.613662 | -0.160081386 | 0.080216038 | -1.995628186 |
| BC043934 | 5.01349092 | -0.239284613 | 0.530910523 | -0.450706103 |
| Rnf7 | 496.7963806 | -0.263253515 | 0.097233268 | -2.707442826 |
| Gm27289 | 0 | NA | NA | NA |
| Gm28729 | 2.066931119 | -0.588251795 | 0.699474127 | -0.8409915 |
| Gm8495 | 0.930891567 | -0.410476041 | 1.244570383 | -0.329813441 |
| Rasa2 | 837.5261912 | -0.363471129 | 0.084709234 | -4.290808818 |
| n-R5s87 | 2.429268922 | -0.119857233 | 0.695342064 | -0.172371613 |
| Gm19280 | 0 | NA | NA | NA |
| Zbtb38 | 1623.281997 | 0.016546815 | 0.040320626 | 0.410380911 |
| A930006L05Rik | 0.095435502 | 0.037073798 | 3.724945049 | 0.009952844 |
| C430002N11Rik | 6.432897433 | -3.2176247 | 0.653637047 | -4.922647383 |
| Gm28688 | 0.279349467 | -2.1111384 | 1.983068606 | -1.064581626 |
| Gm8520 | 7.409839226 | 0.36394128 | 0.440471601 | 0.826253677 |
| Pxylp1 | 198.9385142 | 0.020118639 | 0.123846853 | 0.162447723 |
| Gm10123 | 609.9706294 | 0.166775728 | 0.095051744 | 1.754578313 |
| Gm8524 | 2.21064614 | -0.329087999 | 0.679134092 | -0.484569988 |
| Spsb4 | 74.46297335 | -0.3779074 | 0.137139242 | -2.755647438 |
| Gm16010 | 1.553703116 | -1.497690652 | 0.920990345 | -1.626174107 |
| Gm18291 | 0.371128506 | -0.914488996 | 1.943412291 | -0.470558409 |
| Gm19325 | 18.24486811 | -0.397459046 | 0.354270738 | -1.121907636 |
| Gm26767 | 0.08169834 | -0.443821212 | 3.724945049 | -0.119148392 |
| Slc25a36 | 1229.888798 | 0.14193843 | 0.06145544 | 2.309615393 |
| Gm16185 | 0.038981395 | -0.443816764 | 3.724945049 | -0.119147198 |
| Rpl7a-ps10 | 2.493931419 | 0.060506467 | 0.730490841 | 0.082829878 |
| Gm5371 | 0 | NA | NA | NA |
| Gm29241 | 0 | NA | NA | NA |
| Gm29240 | 0.044544648 | 0.517957688 | 3.724945049 | 0.139051095 |
| A030012G06Rik | 0.136880458 | -0.425048655 | 3.63688887 | -0.116871499 |
| Trim42 | 0.23239932 | 0.014060737 | 2.131268061 | 0.006597357 |
| Gm15891 | 0.046345709 | -0.443818988 | 3.724945049 | -0.119147795 |
| Gm25607 | 0.495675163 | 0.233058013 | 1.59183977 | 0.14640796 |
| Clstn2 | 1155.089609 | 0.241091566 | 0.085809536 | 2.809612755 |
| Gm37661 | 32.39089962 | -0.000868425 | 0.298087835 | -0.00291332 |
| 4921534H16Rik | 0.120974994 | 0.517957688 | 3.724945049 | 0.139051095 |
| 2610303G11Rik | 0 | NA | NA | NA |
| Nmnat3 | 20.17326344 | -0.299767201 | 0.223877515 | -1.338978594 |
| Gm28530 | 3.500655289 | -0.771381792 | 0.534308277 | -1.443701745 |
| Rbp1 | 140.0093858 | -0.996955078 | 0.181835552 | -5.482729141 |
| Rbp2 | 0.656451805 | -0.507986397 | 1.124245889 | -0.451846346 |
| Gm37113 | 0 | NA | NA | NA |
| 4930579K19Rik | 5.920659574 | 0.406692455 | 0.48992324 | 0.830114642 |
| Copb2 | 1299.911645 | -0.312754614 | 0.077039516 | -4.05966483 |

| | | | | |
|---|---|---|---|---|
| Mrps22 | 205.9148466 | -0.351488812 | 0.089834947 | -3.912606679 |
| Gm37637 | 13.96030438 | -0.432033857 | 0.296261571 | -1.458285172 |
| Gm27919 | 0 | NA | NA | NA |
| Pisrt1 | 0 | NA | NA | NA |
| E330023G01Rik | 0.110595523 | 0.037071574 | 3.724945049 | 0.009952247 |
| Gm37282 | 0 | NA | NA | NA |
| Gm37917 | 0.58604645 | -0.09772665 | 1.462086108 | -0.066840557 |
| Gm38226 | 0.474407165 | 0.501459055 | 1.383857916 | 0.362363108 |
| Prr23a | 0.228844266 | 0.037077102 | 2.137880962 | 0.017342921 |
| Gm6406 | 0 | NA | NA | NA |
| 2410012M07Rik | 0 | NA | NA | NA |
| Gm5920 | 2.082893185 | -0.927169583 | 0.736971988 | -1.258079816 |
| 7420426K07Rik | 0 | NA | NA | NA |
| Gm28266 | 0 | NA | NA | NA |
| Foxl2os | 2.17543179 | -0.734637321 | 0.758799116 | -0.968157852 |
| Foxl2 | 2.235327434 | -0.700356274 | 0.71561017 | -0.978684071 |
| Faim | 196.4395141 | 0.218635999 | 0.107254906 | 2.038470853 |
| Gm1123 | 1.879715592 | 0.310398562 | 0.7636215 | 0.40648222 |
| Pik3cb | 544.1368007 | -0.02692465 | 0.066137782 | -0.407099388 |
| Gm2950 | 0.061920613 | -0.443816764 | 3.724945049 | -0.119147198 |
| Gm6432 | 0.11197992 | 0.037073798 | 3.724945049 | 0.009952844 |
| Gm38327 | 0.266212691 | -0.508106736 | 3.720916417 | -0.136554192 |
| Cep70 | 260.9692196 | 0.429580171 | 0.160338037 | 2.679215603 |
| 4930422M22Rik | 0.158279039 | -0.444696717 | 2.813322133 | -0.158068183 |
| Gm37249 | 36.58656883 | 0.215487509 | 0.32405571 | 0.664970566 |
| Esyt3 | 26.59934933 | 0.175796716 | 0.240029923 | 0.732395004 |
| 1700008A23Rik | 0.963690934 | 0.499170296 | 1.107933235 | 0.450541856 |
| Gm28087 | 0 | NA | NA | NA |
| Mras | 2060.728902 | 0.079760018 | 0.063893894 | 1.248319865 |
| Nme9 | 2.199217757 | -0.649488197 | 0.781245457 | -0.831349726 |
| Armc8 | 1137.205985 | 0.373715793 | 0.079077742 | 4.725928977 |
| Dbr1 | 168.1373829 | -0.058541522 | 0.140017062 | -0.418102774 |
| A4gnt | 0.979608328 | 0.122527468 | 1.129249089 | 0.108503491 |
| Dzip1l | 286.0060264 | 0.304846221 | 0.156661659 | 1.945889147 |
| Cldn18 | 0.262323822 | 1.885183911 | 3.701790202 | 0.509262764 |
| Gm16004 | 0 | NA | NA | NA |
| Gm23949 | 0 | NA | NA | NA |
| Sox14 | 8.749592595 | -0.234382956 | 0.408942565 | -0.573143948 |
| Gm23594 | 0 | NA | NA | NA |
| Gm8641 | 0.504639468 | -0.253179982 | 1.321475944 | -0.191588794 |
| 4930519F24Rik | 0.061920613 | -0.443816764 | 3.724945049 | -0.119147198 |
| 4930540E01Rik | 0 | NA | NA | NA |
| Gm28167 | 0 | NA | NA | NA |
| Gm28267 | 0 | NA | NA | NA |
| Gm28166 | 0.08486385 | 0.037730769 | 3.724898049 | 0.010129343 |
| Il20rb | 65.86335234 | 0.062732671 | 0.263081162 | 0.238453678 |
| Nck1 | 358.9705822 | -0.096741141 | 0.079345668 | -1.219236585 |

| | | | | |
|---|---|---|---|---|
| Gm28586 | 8.399474289 | -0.070168566 | 0.523913045 | -0.133931702 |
| Slc35g2 | 192.5839218 | -0.330251966 | 0.155236629 | -2.12741006 |
| Stag1 | 1284.878169 | 0.01386472 | 0.069566173 | 0.199302606 |
| Gm8661 | 0.119234754 | 0.51795324 | 3.724945049 | 0.139049901 |
| Gm37204 | 117.4552302 | 0.463242397 | 0.272031176 | 1.702901864 |
| Gm38297 | 15.65520555 | 0.65729301 | 0.418963986 | 1.568853249 |
| Gm3081 | 62.39629019 | 0.406322979 | 0.24012356 | 1.692141242 |
| Pccb | 410.7052558 | 0.453941371 | 0.077037897 | 5.892442414 |
| Gm3096 | 0 | NA | NA | NA |
| Msl2 | 1119.25172 | 0.082023461 | 0.06752975 | 1.214627054 |
| Gm37553 | 42.26828888 | 0.013051595 | 0.264748846 | 0.049298025 |
| Ppp2r3a | 1068.136217 | 0.107963825 | 0.070439222 | 1.532723125 |
| Gm37962 | 38.30761735 | 0.067919118 | 0.27699434 | 0.245200382 |
| Gm37621 | 78.87355947 | -0.049112762 | 0.226840243 | -0.216508153 |
| Gm37953 | 20.22364659 | -0.260237671 | 0.31108706 | -0.8365429 |
| Gm38344 | 18.31202231 | 0.87991824 | 0.333258607 | 2.640346627 |
| Gm37314 | 38.77458494 | 0.856481642 | 0.24307169 | 3.523576285 |
| Gm24338 | 0.290020514 | 0.030037802 | 1.633453854 | 0.018389134 |
| Gm3121 | 0 | NA | NA | NA |
| Gm28660 | 0 | NA | NA | NA |
| Gm28979 | 0.404019895 | -1.067083506 | 1.457594874 | -0.732085112 |
| Gm24638 | 0 | NA | NA | NA |
| Gm29385 | 0 | NA | NA | NA |
| Gm5161 | 0.106056992 | 0.998850474 | 3.724945049 | 0.268151734 |
| Gm29521 | 6.846922448 | 0.346054556 | 0.48615348 | 0.711821616 |
| Gm37460 | 0.652148592 | 0.508144816 | 1.29391644 | 0.392718417 |
| Ephb1 | 568.496847 | 0.469865432 | 0.074034621 | 6.346563592 |
| 9630041A04Rik | 1.37032714 | 0.067576592 | 0.881752589 | 0.076638949 |
| Gm37683 | 0.138485956 | -0.44382212 | 2.932778741 | -0.151331607 |
| Gm37562 | 15.3341162 | 0.757918477 | 0.389758396 | 1.944585374 |
| n-R5s88 | 0 | NA | NA | NA |
| Gm29388 | 6.105350049 | 0.95579761 | 0.627072771 | 1.524221198 |
| Gm37945 | 0.049553047 | -0.443818988 | 3.724945049 | -0.119147795 |
| Gm37260 | 0.11197992 | 0.037073798 | 3.724945049 | 0.009952844 |
| Mir7243 | 1.350235793 | -0.279949338 | 1.054308188 | -0.265528942 |
| Gm26846 | 1.767979395 | 0.866077318 | 0.932271803 | 0.928996582 |
| Gm22894 | 0.570356529 | -1.468850257 | 1.337457726 | -1.098240511 |
| Ky | 45.80996683 | 0.629547685 | 0.236046416 | 2.667050384 |
| Gm25669 | 0.261093266 | 1.232229444 | 3.710027525 | 0.332134852 |
| Gm7364 | 0.312630927 | 2.314203508 | 3.68475288 | 0.628048497 |
| Gm29143 | 0 | NA | NA | NA |
| Gm29387 | 0.088534442 | -0.905414734 | 3.723500899 | -0.243162217 |
| Cep63 | 192.8660548 | 0.079631489 | 0.101440434 | 0.785007379 |
| Anapc13 | 138.6441374 | 0.087836679 | 0.156996417 | 0.559482062 |
| 4930533D04Rik | 0.164893722 | 1.344872572 | 3.387385485 | 0.397023775 |
| Gm36957 | 0.137470077 | 0.998852408 | 3.59509949 | 0.277837209 |
| Amotl2 | 701.5681414 | -0.414408825 | 0.181214926 | -2.286836046 |

| | | | | |
|---|---|---|---|---|
| Gm5627 | 0.038981395 | -0.443816764 | 3.724945049 | -0.119147198 |
| Gm36958 | 0.813964436 | 1.431708967 | 1.157887269 | 1.236483901 |
| Gm28621 | 0 | NA | NA | NA |
| Ryk | 428.6544998 | -0.223335615 | 0.075352768 | -2.963867452 |
| Slco2a1 | 62.94329372 | -0.320574063 | 0.172999448 | -1.853035176 |
| Gm25842 | 0.189042777 | 0.037075553 | 2.659470858 | 0.013940951 |
| n-R5s89 | 0 | NA | NA | NA |
| Gm22595 | 0 | NA | NA | NA |
| Rab6b | 7282.57726 | 0.529832661 | 0.087251605 | 6.072468956 |
| Gm37166 | 45.91226253 | 0.935456583 | 0.409256323 | 2.285747415 |
| Srprb | 192.082496 | -0.516662374 | 0.104364289 | -4.950566709 |
| Gm20425 | 0 | NA | NA | NA |
| Trf | 281.060478 | -1.012040304 | 0.207080128 | -4.887191801 |
| 1300017J02Rik | 2.855215039 | -0.75782518 | 0.620203094 | -1.22189842 |
| Topbp1 | 318.7285085 | 0.024193886 | 0.092711203 | 0.260959683 |
| Cdv3 | 1213.725262 | -0.429614068 | 0.094450502 | -4.548563089 |
| Gm29600 | 0.038981395 | -0.443816764 | 3.724945049 | -0.119147198 |
| 5830418P13Rik | 5.81979985 | 0.992616705 | 0.463890725 | 2.139764067 |
| Bfsp2 | 26.48732468 | 0.184630968 | 0.252693721 | 0.730651188 |
| Gm37938 | 2.10811511 | 0.301228352 | 0.958992632 | 0.314109142 |
| Gm37546 | 0.280931382 | 0.674704933 | 3.707356873 | 0.181990824 |
| Gm16252 | 0.115957212 | 0.517955464 | 3.724945049 | 0.139050498 |
| Tmem108 | 366.5574461 | 0.271801929 | 0.070109602 | 3.876814594 |
| Gm26360 | 2.674678528 | 0.341561288 | 0.673883508 | 0.506855093 |
| Gm37649 | 24.6588879 | 0.662312151 | 0.428881996 | 1.544275948 |
| 4932413F04Rik | 0.450128165 | -0.955362508 | 1.455349292 | -0.656448945 |
| Gm38032 | 30.93777638 | 0.343265451 | 0.338358282 | 1.014502877 |
| Gm5372 | 0 | NA | NA | NA |
| Gm29154 | 0.341479653 | 1.533022071 | 3.715883266 | 0.412559266 |
| Nphp3 | 143.1369062 | 0.300878062 | 0.142152726 | 2.116583126 |
| Gm28305 | 0 | NA | NA | NA |
| Gm37200 | 6.440447106 | -0.029243241 | 0.420163429 | -0.069599682 |
| Uba5 | 589.0417371 | -0.226996635 | 0.065898344 | -3.444648551 |
| Gm37750 | 0.049553047 | -0.443818988 | 3.724945049 | -0.119147795 |
| Acad11 | 231.7177883 | -0.125837174 | 0.12390644 | -1.015582193 |
| Ackr4 | 2.634801621 | -0.438180652 | 0.653669449 | -0.670339807 |
| Dnajc13 | 1701.736079 | -0.126149184 | 0.06897807 | -1.828830301 |
| Hmgb1-rs16 | 11.38856345 | 1.121330677 | 0.443348573 | 2.529230373 |
| Gm37563 | 1.931745907 | -0.436044576 | 0.681645823 | -0.639693755 |
| Acpp | 6.437954073 | -0.714811615 | 0.409065137 | -1.747427366 |
| Gm37188 | 0.048949532 | 0.037064902 | 3.724945049 | 0.009950456 |
| Gm28996 | 0 | NA | NA | NA |
| Gm28548 | 0 | NA | NA | NA |
| Mir2136 | 0 | NA | NA | NA |
| Gm25904 | 0 | NA | NA | NA |
| Cpne4 | 536.3195774 | 0.658153086 | 0.157540871 | 4.177665648 |
| Gm23391 | 1.103801871 | 1.976528814 | 1.169466628 | 1.690111344 |

| | | | | |
|---|---|---|---|---|
| Mrpl3 | 481.1686626 | 0.088335376 | 0.060171177 | 1.46806795 |
| Gm38077 | 99.57854452 | 0.806664562 | 0.312966856 | 2.577476008 |
| Nudt16 | 104.3868709 | 0.138466798 | 0.128170288 | 1.080334602 |
| 1700080E11Rik | 0.3422147 | -1.553615211 | 1.895262174 | -0.819736305 |
| Nek11 | 22.44919175 | -0.38973331 | 0.273452093 | -1.425234328 |
| Gm37582 | 0.092140247 | 0.037060453 | 3.724945049 | 0.009949262 |
| Gm20643 | 0 | NA | NA | NA |
| Gm15619 | 0 | NA | NA | NA |
| Aste1 | 73.3563129 | 0.109354669 | 0.137349233 | 0.796179683 |
| Atp2c1 | 2093.602576 | -0.012290037 | 0.059297194 | -0.207261691 |
| Gm19667 | 0.638749378 | 1.857261063 | 1.517420059 | 1.223959741 |
| Gm7436 | 0.368385343 | 0.595993778 | 1.778263426 | 0.335154944 |
| Gm5621 | 1.423724791 | -0.854316488 | 0.988464949 | -0.864286072 |
| Gm28555 | 0 | NA | NA | NA |
| Pik3r4 | 491.4021009 | 0.144925175 | 0.074676382 | 1.940709641 |
| Gm22720 | 0 | NA | NA | NA |
| Col6a6 | 14.261268 | -0.375033112 | 0.311079318 | -1.205586775 |
| Gm38314 | 0 | NA | NA | NA |
| Col6a5 | 28.81713568 | 0.22318529 | 0.275151161 | 0.811137009 |
| Col6a4 | 6.820926848 | -0.222837546 | 0.37612078 | -0.592462735 |
| Gm24631 | 0 | NA | NA | NA |
| Gm36993 | 0 | NA | NA | NA |
| Gm9451 | 0.136331796 | 0.037057167 | 3.649540409 | 0.010153927 |
| Gm29208 | 0 | NA | NA | NA |
| D030055H07Rik | 7.771284628 | 1.108192328 | 0.627319534 | 1.766551602 |
| Glyctk | 39.68295307 | 0.147624719 | 0.224125275 | 0.658670555 |
| Mir135a-1 | 0.193036943 | -0.47213964 | 3.248256614 | -0.145351706 |
| Wdr82 | 1508.867638 | -0.130162391 | 0.058536108 | -2.223625659 |
| Mirlet7g | 2.956734474 | -0.084408904 | 0.770558959 | -0.109542434 |
| Ppm1m | 99.17848393 | -0.06536661 | 0.199502931 | -0.327647369 |
| Twf2 | 271.214745 | 0.106508352 | 0.10115527 | 1.052919454 |
| Tlr9 | 362.3896947 | -1.431347627 | 0.336854612 | -4.249155495 |
| Alas1 | 600.2556519 | -0.112377488 | 0.064790529 | -1.734473982 |
| 4930500F10Rik | 7.519553513 | 0.383964338 | 0.441315084 | 0.870045805 |
| Poc1a | 35.43149234 | 0.318194533 | 0.226775838 | 1.403123614 |
| Gm29123 | 0.049553047 | -0.443818988 | 3.724945049 | -0.119147795 |
| Dusp7 | 640.9294056 | 0.007430547 | 0.085158403 | 0.087255598 |
| Rpl29 | 8.095074919 | 0.08867967 | 0.384109092 | 0.230871051 |
| Acy1 | 102.5946641 | 0.050756692 | 0.143296895 | 0.354206504 |
| Abhd14a | 66.26385201 | -0.310120292 | 0.177057402 | -1.751524022 |
| Abhd14b | 60.06866735 | 0.601217536 | 0.226835821 | 2.650452361 |
| Gm28959 | 0.650039631 | -0.426952079 | 1.339684194 | -0.318696064 |
| Pcbp4 | 1192.577122 | 0.109536593 | 0.064881999 | 1.688243192 |
| Gpr62 | 124.4874142 | -0.24638244 | 0.146187551 | -1.685385913 |
| Parp3 | 396.144577 | -0.986840746 | 0.130920687 | -7.537699109 |
| Rrp9 | 173.3168163 | -0.594131893 | 0.097535979 | -6.091412616 |
| Iqcf1 | 0.403414764 | -1.493367752 | 1.480419834 | -1.008746113 |

| | | | | |
|---|---|---|---|---|
| Iqcf3 | 0.731923588 | -0.441957783 | 1.382759507 | -0.319620137 |
| Iqcf5 | 0.458372996 | -0.443821279 | 1.743100615 | -0.254615985 |
| Iqcf4 | 0 | NA | NA | NA |
| Iqcf6 | 0 | NA | NA | NA |
| Gm26185 | 0 | NA | NA | NA |
| Gm28111 | 0 | NA | NA | NA |
| Grm2 | 419.4914956 | 0.577397004 | 0.134307075 | 4.299081063 |
| Tex264 | 547.2087676 | -0.218792494 | 0.080093845 | -2.731701718 |
| Rad54l2 | 692.1787233 | -0.066047662 | 0.09526773 | -0.693284724 |
| Gm22384 | 0.206359549 | 1.023802084 | 2.640765907 | 0.387691344 |
| Vprbp | 629.97184 | -0.329248576 | 0.086405225 | -3.810516948 |
| Manf | 616.8659586 | -0.723355241 | 0.108926681 | -6.640753533 |
| Gm37612 | 1.882200288 | 0.031082642 | 0.864732445 | 0.035944808 |
| Rbm15b | 823.0495333 | 0.01079947 | 0.068838785 | 0.156880596 |
| 4930524O07Rik | 4.402341252 | -0.065892143 | 0.615604878 | -0.107036422 |
| Dock3 | 4718.325882 | 0.534110455 | 0.112872744 | 4.731970165 |
| Gm17205 | 0.109124069 | 0.517955464 | 3.724945049 | 0.139050498 |
| Gm22454 | 0 | NA | NA | NA |
| Gm17117 | 0 | NA | NA | NA |
| Gm23595 | 0 | NA | NA | NA |
| AA543401 | 0.072025462 | 0.517959912 | 3.724945049 | 0.139051692 |
| Gm22238 | 0 | NA | NA | NA |
| 1700039M15Rik | 0.049553047 | -0.443818988 | 3.724945049 | -0.119147795 |
| Mapkapk3 | 63.6118118 | -1.170386921 | 0.212281459 | -5.513373261 |
| Gm17040 | 0.094097695 | 0.037071574 | 3.724945049 | 0.009952247 |
| Cish | 89.05641637 | -0.343958914 | 0.172018418 | -1.999547018 |
| Gm17041 | 0.049553047 | -0.443818988 | 3.724945049 | -0.119147795 |
| Hemk1 | 115.6953326 | -0.220238351 | 0.117552284 | -1.873535281 |
| 6430571L13Rik | 122.123982 | 0.140163524 | 0.169995677 | 0.824512282 |
| 4930429P21Rik | 0.791934515 | 0.550172934 | 1.38344094 | 0.397684439 |
| Cacna2d2 | 1015.043882 | 0.02931515 | 0.07402572 | 0.396013041 |
| Gm17141 | 0.093831153 | 0.517957688 | 3.724945049 | 0.139051095 |
| Gm17118 | 0 | NA | NA | NA |
| Tmem115 | 377.5379067 | -0.279010481 | 0.094607787 | -2.949128089 |
| Cyb561d2 | 55.18192089 | -0.441140966 | 0.206853612 | -2.132623947 |
| Nprl2 | 184.5566623 | -0.077886827 | 0.090485331 | -0.860767442 |
| Zmynd10 | 22.83363798 | 0.404180962 | 0.27463795 | 1.471686491 |
| Gm34106 | 0.182655267 | 0.866774686 | 2.694309069 | 0.321705738 |
| Rassf1 | 176.3853711 | -0.634459118 | 0.131699458 | -4.817477077 |
| Gm9917 | 9.204737304 | 0.319454992 | 0.500615584 | 0.638124345 |
| Tusc2 | 617.3101563 | 0.079207044 | 0.072182284 | 1.097319716 |
| Hyal2 | 95.99414798 | -0.961855671 | 0.130157723 | -7.389923928 |
| Gm16343 | 0.186399948 | 0.575238522 | 3.295513719 | 0.174552003 |
| Hyal1 | 60.01938957 | -0.757792356 | 0.191785962 | -3.951239962 |
| Nat6 | 87.56719576 | -0.383279372 | 0.15846484 | -2.418702927 |
| Hyal3 | 5.312334351 | -0.43487736 | 0.538859297 | -0.807033232 |
| Ifrd2 | 67.19084161 | -0.184584197 | 0.21982776 | -0.839676469 |

| | | | | |
|---|---|---|---|---|
| Lsmem2 | 2.036564983 | -0.811264303 | 0.784999333 | -1.033458589 |
| Gm38150 | 3.547776317 | 1.407470027 | 0.604274882 | 2.329188367 |
| Sema3b | 58.08865837 | -0.093741723 | 0.198893695 | -0.471315712 |
| Gnai2 | 2404.56764 | -0.393091237 | 0.114558163 | -3.431368192 |
| Gm19721 | 19.767715 | -0.019654691 | 0.415748396 | -0.047275447 |
| Slc38a3 | 598.0789621 | 0.129511724 | 0.181205941 | 0.714721181 |
| Gnat1 | 1.440190137 | 1.21080318 | 0.90817715 | 1.333223567 |
| A930036K24Rik | 0.18066559 | -0.410893234 | 2.687923725 | -0.152866404 |
| Sema3f | 139.5279565 | 0.148670846 | 0.114364441 | 1.299974396 |
| Rbm5 | 1611.009065 | -0.007472524 | 0.158790505 | -0.047059013 |
| Gm37850 | 0 | NA | NA | NA |
| Rbm6 | 1030.666851 | 0.101794099 | 0.124004529 | 0.820890169 |
| Gm37436 | 36.96383849 | 0.623728512 | 0.256994377 | 2.427012287 |
| Mon1a | 192.9155728 | -0.258727671 | 0.10032194 | -2.578973973 |
| Gm23856 | 0.487068913 | -0.931679956 | 1.599732583 | -0.582397312 |
| Mst1r | 16.57438961 | -0.169939067 | 0.260082541 | -0.653404362 |
| Actl11 | 0.123841226 | -1.017823543 | 3.719440177 | -0.273649661 |
| Gm37974 | 0.217247303 | 0.998861747 | 2.596047709 | 0.384762477 |
| Camkv | 2202.703855 | 1.202959024 | 0.15681021 | 7.671433046 |
| Traip | 114.0875043 | 0.583850606 | 0.248455706 | 2.349918288 |
| Uba7 | 688.1155965 | -0.48538528 | 0.125600212 | -3.864525966 |
| Gm20661 | 4.55440648 | -0.674590046 | 0.522182728 | -1.291865874 |
| Fam212a | 16.26337642 | 0.007480261 | 0.247654099 | 0.030204471 |
| Gm24944 | 1.177026929 | 0.819388922 | 0.886039001 | 0.924777489 |
| 4921523L03Rik | 0.198550338 | 1.413806266 | 2.20370556 | 0.641558605 |
| Cdhr4 | 13.4482997 | -0.030267466 | 0.347548063 | -0.087088577 |
| Gm20662 | 0.066050875 | -0.443818988 | 3.724945049 | -0.119147795 |
| 4930535L15Rik | 1.219811481 | -0.527367287 | 0.922659081 | -0.571573291 |
| Ip6k1 | 2129.145135 | 0.145595263 | 0.053189193 | 2.737309122 |
| Gm38134 | 0.127971488 | -0.95589744 | 3.6855341 | -0.259364698 |
| Gmppb | 158.7886363 | -0.821774624 | 0.146394932 | -5.613408963 |
| Rnf123 | 811.9292402 | 0.346235299 | 0.072704938 | 4.762197861 |
| Gm20529 | 0.04393175 | 0.037062677 | 3.724945049 | 0.009949859 |
| Amigo3 | 37.0261778 | -0.433974396 | 0.28611702 | -1.516772391 |
| Mst1 | 0.439905485 | -0.615168162 | 1.361239558 | -0.451917635 |
| Mir7088 | 0 | NA | NA | NA |
| Apeh | 282.2906194 | -0.302596311 | 0.084061709 | -3.5996926 |
| Bsn | 11059.17198 | 0.560865101 | 0.121488447 | 4.616612652 |
| AC137678.1 | 2.386774017 | 1.391364467 | 0.922996757 | 1.507442422 |
| 4930447F24Rik | 14.97539824 | -0.894230227 | 0.339501602 | -2.633949949 |
| Gm37201 | 2.168403122 | 0.09209645 | 0.854024501 | 0.107838182 |
| Gm37247 | 41.41364108 | 0.782393566 | 0.366556174 | 2.134443838 |
| Gm37230 | 0.567387455 | 0.475787337 | 1.365525567 | 0.348427996 |
| Dag1 | 1797.643256 | -0.010260927 | 0.082619712 | -0.12419466 |
| Nicn1 | 612.9850749 | 0.353991355 | 0.073492291 | 4.816714118 |
| Amt | 190.2692267 | -0.146815469 | 0.142305349 | -1.031693257 |
| Tcta | 207.060682 | -0.200691759 | 0.109968901 | -1.824986488 |

| | | | | |
|---|---|---|---|---|
| Rhoa | 1859.627478 | -0.328901497 | 0.106583127 | -3.085868348 |
| Gm37568 | 3.725366168 | -0.198613735 | 0.585938494 | -0.338966866 |
| Gm37401 | 0.049553047 | -0.443818988 | 3.724945049 | -0.119147795 |
| Gpx1 | 610.2857368 | -0.569771564 | 0.144715897 | -3.937173292 |
| Usp4 | 1105.767034 | -0.310373469 | 0.054220441 | -5.724288931 |
| Rps2-ps11 | 0.578661668 | -0.778576777 | 1.20425323 | -0.646522474 |
| Gm37963 | 32.08673899 | 0.144530864 | 0.351158361 | 0.41158315 |
| 1700102P08Rik | 15.20131833 | -0.895879366 | 0.308703362 | -2.902071944 |
| Ccdc36 | 5.280767892 | 0.643847072 | 0.471759364 | 1.364778575 |
| Gm25456 | 0.042716945 | 0.037062677 | 3.724945049 | 0.009949859 |
| BC048562 | 0 | NA | NA | NA |
| Klhdc8b | 159.0275307 | 0.106578771 | 0.131291934 | 0.811769375 |
| Ccdc71 | 275.4617965 | 0.046940676 | 0.075536868 | 0.621427355 |
| Gm37475 | 3.601477103 | -1.259453001 | 0.661656617 | -1.903484328 |
| Lamb2 | 362.809454 | -0.222359298 | 0.149163773 | -1.490705779 |
| Usp19 | 1729.388652 | -0.274427117 | 0.050346115 | -5.450810183 |
| Qars | 489.2628777 | -0.092115107 | 0.06315273 | -1.458608476 |
| Qrich1 | 1514.838001 | 0.129085773 | 0.069267958 | 1.863571223 |
| Gm38366 | 45.47164434 | 0.5829305 | 0.34234423 | 1.70276128 |
| Gm37678 | 6.993644233 | 0.632618452 | 0.497059422 | 1.272721979 |
| Impdh2 | 294.2527551 | -0.423452039 | 0.098642683 | -4.292787112 |
| Ndufaf3 | 182.2571026 | -0.02626256 | 0.08429873 | -0.311541581 |
| 4833445I07Rik | 30.38428203 | 0.675377779 | 0.21278725 | 3.173957924 |
| Gm24083 | 0 | NA | NA | NA |
| mmu-mir-191 | 0 | NA | NA | NA |
| Mir425 | 0 | NA | NA | NA |
| Dalrd3 | 336.6293249 | -0.089964838 | 0.078988493 | -1.138961323 |
| Wdr6 | 1862.952255 | 0.61795206 | 0.090285214 | 6.844443641 |
| P4htm | 622.680632 | 0.264225004 | 0.078978493 | 3.345531082 |
| Gm38163 | 3.058745156 | -0.024797334 | 0.732810618 | -0.033838666 |
| Arih2 | 618.9684878 | -0.123973289 | 0.08251149 | -1.502497271 |
| Gm26298 | 0 | NA | NA | NA |
| Slc25a20 | 138.8267023 | -0.555531687 | 0.153936469 | -3.608837406 |
| Prkar2a | 906.3458615 | -0.243498486 | 0.068867389 | -3.535758952 |
| Gm24259 | 0.258327161 | -0.938250986 | 3.021726918 | -0.310501581 |
| Ip6k2 | 843.8946416 | 0.298729645 | 0.086401106 | 3.457474773 |
| Gm23034 | 0.062426873 | 0.517959912 | 3.724945049 | 0.139051692 |
| Nckipsd | 578.0213764 | 0.117770708 | 0.081423604 | 1.446395172 |
| Gm35025 | 6.065554424 | 0.506774896 | 0.469756395 | 1.078803613 |
| Gm37714 | 0.267221877 | -0.822189084 | 1.778545832 | -0.46228164 |
| Celsr3 | 1375.868327 | 0.354323345 | 0.162306492 | 2.183050976 |
| Gm23156 | 0.616331352 | 2.612314762 | 1.59079523 | 1.64214395 |
| Slc26a6 | 70.00938289 | -0.092496137 | 0.165633733 | -0.558437799 |
| Tmem89 | 0.562660809 | 1.996241194 | 1.272020444 | 1.569346785 |
| Uqcrc1 | 1892.13114 | 0.00239838 | 0.067577217 | 0.035490952 |
| Gm37284 | 0 | NA | NA | NA |
| Col7a1 | 41.89192254 | -0.77219098 | 0.264108173 | -2.923767836 |

| | | | | |
|---|---|---|---|---|
| Mir711 | 0 | NA | NA | NA |
| Ucn2 | 10.62182747 | -2.008684308 | 0.804015966 | -2.498313956 |
| Gm38004 | 0.533662682 | -0.100342729 | 1.272549252 | -0.078851745 |
| Pfkfb4 | 187.9455917 | -0.182829387 | 0.106278072 | -1.720292659 |
| Shisa5 | 1519.355689 | -0.683581065 | 0.144977291 | -4.715090629 |
| Trex1 | 520.4087808 | -0.826594482 | 0.189131052 | -4.370485297 |
| Atrip | 28.91987549 | 0.282499411 | 0.398669558 | 0.708605425 |
| Tma7 | 371.0569712 | 0.030513816 | 0.089682095 | 0.34024424 |
| Ccdc51 | 52.13264183 | -0.407014421 | 0.268144319 | -1.517893134 |
| Gm7628 | 6.615983682 | 0.82664356 | 0.448220102 | 1.844280423 |
| Plxnb1 | 1318.803774 | 0.398876258 | 0.124786527 | 3.196468946 |
| Fbxw21 | 0 | NA | NA | NA |
| Fbxw13 | 0.127364494 | -0.44382566 | 3.724945049 | -0.119149586 |
| Fbxw20 | 0.066050875 | -0.443818988 | 3.724945049 | -0.119147795 |
| Fbxw14 | 0.088383099 | 0.037058229 | 3.724945049 | 0.009948665 |
| Gm22821 | 0 | NA | NA | NA |
| Gm25484 | 0 | NA | NA | NA |
| Gm24641 | 0 | NA | NA | NA |
| Fbxw22 | 0.038981395 | -0.443816764 | 3.724945049 | -0.119147198 |
| Fbxw16 | 0.066050875 | -0.443818988 | 3.724945049 | -0.119147795 |
| RP23-62L19.6 | 0 | NA | NA | NA |
| RP23-62L19.5 | 0 | NA | NA | NA |
| Fbxw19 | 0.038981395 | -0.443816764 | 3.724945049 | -0.119147198 |
| RP24-187P11.4 | 0.450885768 | -0.583270664 | 1.594243267 | -0.365860516 |
| RP24-187P11.2 | 9.209079983 | -0.782556617 | 0.46090265 | -1.697878319 |
| RP24-187P11.5 | 0.238621995 | -0.855974175 | 2.496609277 | -0.34285468 |
| Fbxw15 | 0.204053819 | 0.926603406 | 2.621632572 | 0.353445184 |
| RP24-187P11.3 | 0.046345709 | -0.443818988 | 3.724945049 | -0.119147795 |
| Fbxw24 | 0.124596252 | 0.037071632 | 3.714681146 | 0.009979761 |
| Fbxw25 | 0.109982625 | -0.443823436 | 3.724945049 | -0.119148989 |
| Fbxw18 | 0.15906016 | -1.015882374 | 3.433492635 | -0.295874342 |
| RP24-344B20.5 | 0 | NA | NA | NA |
| Fbxw26 | 0.166952883 | -1.336542448 | 2.746539428 | -0.486627803 |
| RP23-357H5.5 | 0.394548002 | 0.34258472 | 1.76791584 | 0.193778862 |
| Spink8 | 16.32657011 | 1.020467341 | 0.293851043 | 3.472736836 |
| 3000002C10Rik | 56.32280498 | 0.425640191 | 0.188329569 | 2.260081581 |
| Nme6 | 38.63028386 | -0.759029074 | 0.21215872 | -3.577647317 |
| Camp | 2.243591458 | -0.765858848 | 0.707856044 | -1.081941526 |
| Cdc25a | 165.9711309 | -0.04447764 | 0.108185298 | -0.411124621 |
| Mir7089 | 0.220081591 | 0.99757292 | 2.15365768 | 0.463199388 |
| RP24-421E18.9 | 0 | NA | NA | NA |
| Map4 | 7553.036783 | -0.046304646 | 0.080672376 | -0.573983911 |
| RP24-421E18.8 | 10.03429169 | 0.215316461 | 0.341736005 | 0.630066653 |
| RP24-421E18.10 | 0 | NA | NA | NA |
| RP24-421E18.5 | 78.64983153 | 0.285314404 | 0.132311222 | 2.156388549 |
| AC161246.1 | 0 | NA | NA | NA |
| RP24-421E18.7 | 15.20321581 | -0.886482658 | 0.369500238 | -2.399139614 |

| | | | | |
|---|---|---|---|---|
| Dhx30 | 1202.629578 | 0.024687112 | 0.051075583 | 0.4833447 |
| Gm27855 | 0.367828087 | 1.354976519 | 1.572008569 | 0.861939652 |
| Smarcc1 | 1075.459101 | -0.125353285 | 0.051098157 | -2.453186044 |
| Gm23377 | 0 | NA | NA | NA |
| Cspg5 | 2416.656234 | 0.569685291 | 0.078578846 | 7.249855627 |
| RP24-285I4.2 | 0.569412722 | 0.132341865 | 1.291415299 | 0.102478161 |
| Mir6236 | 2.610552123 | -0.958934937 | 1.113840671 | -0.860926488 |
| RP24-330A11.9 | 0.916112882 | 2.214599722 | 1.171457085 | 1.890465942 |
| Elp6 | 89.70397602 | 0.100535308 | 0.116399994 | 0.863705436 |
| RP24-330A11.12 | 0.042716945 | 0.037062677 | 3.724945049 | 0.009949859 |
| Scap | 1121.694434 | 0.151753296 | 0.04875741 | 3.112415067 |
| 5830462I19Rik | 0.695310992 | -3.012745261 | 1.429410433 | -2.107683833 |
| Ptpn23 | 992.3070621 | -0.248317092 | 0.092523075 | -2.683839601 |
| Ngp | 7.348605377 | -2.513638254 | 0.784583922 | -3.203785066 |
| Klhl18 | 377.384243 | -0.030014046 | 0.077629637 | -0.386631282 |
| Kif9 | 65.23232709 | -0.428932717 | 0.159332956 | -2.692052708 |
| Setd2 | 2113.532458 | 0.021787813 | 0.071034211 | 0.306722817 |
| RP24-363K20.7 | 41.47908056 | 0.282947407 | 0.235235824 | 1.202824481 |
| Nradd | 21.00591379 | -1.731946265 | 0.323314336 | -5.356849574 |
| Nbeal2 | 229.3785941 | -0.473490703 | 0.15000782 | -3.156440133 |
| Mir8107 | 0 | NA | NA | NA |
| RP24-179L19.4 | 0.417968807 | 1.305928788 | 1.496387989 | 0.87272071 |
| Ccdc12 | 189.7858103 | -0.025891177 | 0.120365759 | -0.215104175 |
| RP24-179L19.2 | 0.211345377 | -0.843199607 | 2.644317213 | -0.318872336 |
| Pth1r | 56.35433901 | 0.062523681 | 0.157956726 | 0.395827911 |
| Myl3 | 1.171543132 | 0.503557827 | 1.078983319 | 0.466696582 |
| Prss42 | 0.145501641 | -0.443200222 | 2.888011192 | -0.153462086 |
| Prss44 | 0.225957929 | 0.053032434 | 2.139541033 | 0.024786827 |
| Prss43 | 1.895398054 | 0.884301036 | 0.743943676 | 1.188666649 |
| Prss45 | 3.856439033 | 0.574901696 | 0.57345823 | 1.002517124 |
| Prss46 | 14.47717373 | -3.075099855 | 0.522371458 | -5.886806803 |
| Prss50 | 6.552872895 | -0.236927248 | 0.423508156 | -0.559439633 |
| Tmie | 70.89016146 | -0.02232543 | 0.172554674 | -0.129381777 |
| Als2cl | 108.0650459 | 0.26530423 | 0.146155768 | 1.815215599 |
| RP23-265F20.14 | 0.087930927 | -0.443164241 | 3.724898049 | -0.118973522 |
| Gm590 | 0.446134451 | -2.043442631 | 1.512507474 | -1.351029774 |
| Gm24269 | 0 | NA | NA | NA |
| Tdgf1 | 2.307514072 | 0.837490764 | 0.70749326 | 1.183743805 |
| Lrrc2 | 11.32380672 | -0.097168103 | 0.391323403 | -0.248306392 |
| RP24-166N8.8 | 0.109982625 | -0.443823436 | 3.724945049 | -0.119148989 |
| Rtp3 | 1.735450405 | -2.404247859 | 0.846619241 | -2.839821898 |
| 1700061E17Rik | 0.467110691 | 1.242722948 | 1.626949614 | 0.763836161 |
| Gm10030 | 2.141449619 | -0.415424888 | 0.706845168 | -0.587716952 |
| Ltf | 6.616994115 | -1.50094759 | 0.501474796 | -2.993066854 |
| RP24-166N8.9 | 4.949110328 | -0.051190598 | 0.434482001 | -0.117819836 |
| Ccrl2 | 163.4421883 | -0.654077726 | 0.349369628 | -1.87216539 |
| Rp31-ps19 | 0 | NA | NA | NA |

| | | | | |
|---|---|---|---|---|
| Gm26409 | 0.044544648 | 0.517957688 | 3.724945049 | 0.139051095 |
| Lrrfip2 | 487.0826255 | -0.025355247 | 0.092031619 | -0.275505818 |
| Mlh1 | 123.8432082 | -0.105381906 | 0.134020869 | -0.786309674 |
| Gm24860 | 1.095639389 | -0.686244349 | 1.119203953 | -0.613153972 |
| Epm2aip1 | 2946.703591 | -0.030968232 | 0.067262722 | -0.460407063 |
| Trank1 | 3806.887753 | 0.466680425 | 0.277354557 | 1.682613147 |
| RP23-361O11.4 | 10.70607882 | 0.11103426 | 0.44175568 | 0.251347669 |
| Dclk3 | 344.3535978 | 0.779707989 | 0.150832633 | 5.169358738 |
| RP24-362N9.3 | 0.122601409 | 0.998850474 | 3.724945049 | 0.268151734 |
| RP24-362N9.4 | 0 | NA | NA | NA |
| Stac | 39.84967809 | -0.29731978 | 0.204128737 | -1.456530738 |
| n-R5s90 | 0.892725456 | 0.280840443 | 0.964315043 | 0.291233083 |
| Arpp21 | 4354.712088 | 0.580896144 | 0.150350279 | 3.863618675 |
| Mir128-2 | 2.542567827 | 0.89510903 | 0.706494347 | 1.266972671 |
| 2310075C17Rik | 2.414579911 | 0.44308132 | 0.604117441 | 0.733435735 |
| 2900079G21Rik | 54.14502989 | 0.102013094 | 0.169827715 | 0.600685782 |
| Gm27847 | 0 | NA | NA | NA |
| Gm24957 | 0 | NA | NA | NA |
| Gm22492 | 0 | NA | NA | NA |
| Pdcd6ip | 1431.871971 | -0.421785541 | 0.060991118 | -6.915524016 |
| Gm25059 | 0 | NA | NA | NA |
| Clasp2 | 4322.484509 | 0.34940126 | 0.055066635 | 6.345062867 |
| Ubp1 | 888.9733183 | 0.09003529 | 0.116630658 | 0.771969323 |
| Fbxl2 | 321.4966191 | 0.443172946 | 0.100938596 | 4.390520195 |
| Susd5 | 65.24506384 | 0.615334171 | 0.181416004 | 3.391840612 |
| 4921528I07Rik | 0.068097359 | 0.037064902 | 3.724945049 | 0.009950456 |
| Bcl2a1c | 0.061920613 | -0.443816764 | 3.724945049 | -0.119147198 |
| 4930520O04Rik | 10.94558613 | 0.998686704 | 0.330118047 | 3.025241164 |
| Crtap | 52.46044785 | -0.741033101 | 0.232552475 | -3.186519955 |
| Glb1 | 239.9047864 | 0.069890572 | 0.089538224 | 0.780566878 |
| Tmppe | 289.5667612 | -0.123446662 | 0.076010231 | -1.624079556 |
| Ccr4 | 0.577241561 | 1.006589216 | 1.360850267 | 0.739676686 |
| Trim71 | 15.97479315 | 0.275719515 | 0.396471601 | 0.695433203 |
| C130032M10Rik | 0.045882455 | 0.517959912 | 3.724945049 | 0.139051692 |
| Cnot10 | 446.9158187 | 0.081658567 | 0.073640183 | 1.108885995 |
| Gm23232 | 2.256171834 | 1.300951217 | 0.881403681 | 1.475999302 |
| Dync1li1 | 1024.902468 | -0.075210112 | 0.053596501 | -1.403265331 |
| Cmtm6 | 352.6464989 | -0.347521744 | 0.094651325 | -3.671599366 |
| Cmtm7 | 42.27447401 | -0.893628445 | 0.261150043 | -3.421896598 |
| Gm9888 | 1.350285312 | 0.275592866 | 0.811518156 | 0.339601603 |
| Cmtm8 | 7.699545734 | 0.735362771 | 0.378836237 | 1.941109901 |
| Gm23889 | 0 | NA | NA | NA |
| Gpd1l | 938.6073185 | 0.621650147 | 0.080524194 | 7.720041862 |
| Osbpl10 | 236.991532 | 0.567277712 | 0.141168359 | 4.018448013 |
| Rps27rt | 78.90118146 | -0.498074096 | 0.225185232 | -2.211841746 |
| Gm4665 | 0.066050875 | -0.443818988 | 3.724945049 | -0.119147795 |
| Gm26962 | 0.061920613 | -0.443816764 | 3.724945049 | -0.119147198 |

| | | | | |
|---|---|---|---|---|
| Gm27002 | 10.03766149 | -0.445110707 | 0.408788991 | -1.088851992 |
| Stt3b | 1311.019778 | -0.250292266 | 0.065753951 | -3.806497748 |
| Mir467h | 0 | NA | NA | NA |
| Gm26303 | 0 | NA | NA | NA |
| Gm25510 | 0 | NA | NA | NA |
| Gm22243 | 0 | NA | NA | NA |
| Gm5921 | 6.413797967 | -0.177623399 | 0.431336521 | -0.411797726 |
| n-R5s92 | 0 | NA | NA | NA |
| Gadl1 | 3.709958051 | -1.067837061 | 0.576222856 | -1.853166793 |
| Gm16142 | 0.404993149 | -0.368443348 | 1.36769235 | -0.269390516 |
| Gm9385 | 43.14016071 | -0.151564486 | 0.246356067 | -0.615225304 |
| Tgfbr2 | 420.264414 | -0.321919965 | 0.104857135 | -3.070081638 |
| Rbms3 | 651.4525007 | 0.002877869 | 0.096964978 | 0.029679467 |
| Gm18489 | 0.042716945 | 0.037062677 | 3.724945049 | 0.009949859 |
| Gm20397 | 0 | NA | NA | NA |
| Gm17396 | 26.24111455 | -0.307205006 | 0.248598028 | -1.235749971 |
| Gm20396 | 0 | NA | NA | NA |
| Gm26614 | 3.2216295 | 0.32929079 | 0.655080297 | 0.502672408 |
| Zcwpw2 | 38.02155974 | -0.036710483 | 0.20469875 | -0.179339065 |
| Gm22008 | 0 | NA | NA | NA |
| Azi2 | 1280.08819 | -0.070865404 | 0.096324742 | -0.735692642 |
| Cmc1 | 102.3559624 | 0.082971455 | 0.116616676 | 0.711488762 |
| Gm17399 | 4.930845136 | 0.186745333 | 0.530914765 | 0.351742587 |
| Gm22198 | 0 | NA | NA | NA |
| 2610509F24Rik | 0.046345709 | -0.443818988 | 3.724945049 | -0.119147795 |
| Platr11 | 0.154746054 | 0.037067119 | 2.831523697 | 0.013090874 |
| Gm22479 | 0.124853745 | 1.05357957 | 3.721447096 | 0.283110183 |
| Eomes | 133.6882393 | -0.604210047 | 0.419818328 | -1.439217887 |
| Golga4 | 1496.655227 | -0.052865988 | 0.129802835 | -0.407279148 |
| Itga9 | 160.656506 | -0.381585429 | 0.105339806 | -3.622423886 |
| Gm10157 | 51.89187838 | -0.23616889 | 0.155863603 | -1.515227962 |
| Gm2415 | 124.2848123 | 0.656335044 | 0.164206726 | 3.997004631 |
| Gm16295 | 0.648502457 | 0.526559548 | 1.246683838 | 0.422368152 |
| Ctdspl | 341.7777662 | 0.51628241 | 0.135620319 | 3.806821976 |
| Mir26a-1 | 1.512289855 | 1.468676295 | 1.008910311 | 1.455705506 |
| Vill | 6.901825543 | -0.401549785 | 0.359946304 | -1.115582464 |
| Plcd1 | 118.7634359 | -0.180304884 | 0.130515833 | -1.381479006 |
| Dlec1 | 29.71330161 | -0.516580649 | 0.250677924 | -2.060734509 |
| Acaa1b | 17.68007261 | -1.044740546 | 0.28740743 | -3.63505059 |
| Gm10608 | 0.106465261 | 0.037073798 | 3.724945049 | 0.009952844 |
| Slc22a14 | 0.191792366 | -0.86082649 | 2.227787707 | -0.386404184 |
| Slc22a13 | 1.365498172 | -0.815603777 | 1.090672405 | -0.74779904 |
| Slc22a13b-ps | 14.19279457 | -0.32086403 | 0.390182967 | -0.822342482 |
| Oxsr1 | 850.5060215 | -0.144625022 | 0.069443943 | -2.082615343 |
| Gm22729 | 1.308834292 | -0.15210955 | 0.902597765 | -0.168524182 |
| Myd88 | 293.492393 | -0.888063978 | 0.213867725 | -4.152398311 |
| Acaa1a | 291.2994478 | -0.154626485 | 0.103019819 | -1.5009392 |

| | | | | |
|---|---|---|---|---|
| Xylb | 75.68413251 | 0.052703674 | 0.150723186 | 0.349671972 |
| Acvr2b | 67.3176329 | 0.444069091 | 0.146336363 | 3.034577894 |
| 4930516B21Rik | 0 | NA | NA | NA |
| Exog | 250.3105798 | -0.20751043 | 0.088632305 | -2.341250503 |
| Scn5a | 99.30242439 | 0.174313403 | 0.139669143 | 1.248045201 |
| Scn10a | 7.212801384 | 3.769495873 | 0.61123617 | 6.167003951 |
| Scn11a | 3.528093251 | -1.279431368 | 0.562334178 | -2.275215377 |
| Gm2449 | 4.853795414 | 0.525968032 | 0.46556578 | 1.129739458 |
| Wdr48 | 797.9104278 | -0.034554622 | 0.050911947 | -0.67871343 |
| Cx3cr1 | 232.9636481 | 1.81830155 | 0.486303854 | 3.739023521 |
| Gorasp1 | 319.6341586 | -0.353562574 | 0.13259415 | -2.666502061 |
| Ttc21a | 29.13279244 | 0.008509751 | 0.286824181 | 0.029668877 |
| Csrnp1 | 561.5797647 | -0.731040323 | 0.345514356 | -2.115803031 |
| Xirp1 | 5.455029321 | -1.469206985 | 0.615119796 | -2.388489192 |
| Ccr8 | 0.637648044 | 0.369984966 | 1.152405138 | 0.321054596 |
| Slc25a38 | 245.895353 | -0.183270899 | 0.101951506 | -1.797628171 |
| Rpsa | 2894.845799 | -0.320141444 | 0.096999281 | -3.300451722 |
| Gm24044 | 12.71611818 | 0.391740384 | 0.365292604 | 1.072401631 |
| Gm26448 | 99.87717075 | -0.608978477 | 0.18649775 | -3.265339538 |
| Snora62 | 1.810539642 | -0.304270854 | 0.693448116 | -0.438779552 |
| Mobp | 12397.77805 | -0.097848405 | 0.158697917 | -0.616570195 |
| Gm37111 | 46.80149452 | 0.234399419 | 0.20004035 | 1.171760693 |
| Myrip | 677.5660742 | 0.282252959 | 0.147154235 | 1.918075673 |
| Gm15566 | 4.5585197 | 1.943108668 | 0.725562708 | 2.678071303 |
| Gm15565 | 3.283342549 | 0.319626631 | 0.801385812 | 0.398842388 |
| Eif1b | 736.4044258 | -0.166709419 | 0.094642742 | -1.76146016 |
| Entpd3 | 153.1227425 | -0.283157732 | 0.113366969 | -2.497709283 |
| Rpl14 | 1081.536881 | -0.171970718 | 0.112536912 | -1.528127214 |
| 5830454E08Rik | 6.315262053 | -0.174296326 | 0.388166833 | -0.449024263 |
| Gm23624 | 0 | NA | NA | NA |
| 1700020M21Rik | 0.275587006 | -1.939426139 | 2.391100683 | -0.81110183 |
| 4930593C16Rik | 1.144931742 | 0.13961019 | 0.966037307 | 0.144518425 |
| Ctnnb1 | 5088.011416 | -0.302213565 | 0.090563864 | -3.33702153 |
| mmu-mir-7090 | 0.464416953 | 0.034495684 | 1.37107369 | 0.025159613 |
| Ulk4 | 49.64989384 | 0.154295321 | 0.204266865 | 0.755361479 |
| Gm22987 | 0 | NA | NA | NA |
| Gm17157 | 0 | NA | NA | NA |
| Trak1 | 1974.275711 | 0.220716789 | 0.072598342 | 3.0402456 |
| Cck | 1108.958497 | 1.000335026 | 0.122714956 | 8.151696086 |
| Lyzl4 | 13.61068275 | -0.878462311 | 0.381707221 | -2.301403437 |
| Lyzl4os | 0.219234697 | -0.496916407 | 2.544014286 | -0.195327679 |
| Vipr1 | 229.8541341 | -0.491502309 | 0.220533739 | -2.228694401 |
| Sec22c | 369.7565343 | 0.200756112 | 0.074792383 | 2.684178564 |
| Deb1 | 212.2253457 | 0.036675642 | 0.087934214 | 0.417080457 |
| Nktr | 2807.808864 | 0.243328628 | 0.171053752 | 1.422527274 |
| E530011L22Rik | 71.35472267 | 0.468033277 | 0.1597445 | 2.929886646 |
| Zfp651 | 995.5133322 | 0.346031015 | 0.083853289 | 4.126624256 |

| | | | | |
|---|---|---|---|---|
| Klhl40 | 81.7128723 | -0.944925043 | 0.224223161 | -4.21421694 |
| Hhatl | 134.4694746 | 0.424126672 | 0.150050136 | 2.826566389 |
| Ccdc13 | 49.28911322 | 0.686001554 | 0.186015264 | 3.68787776 |
| Gm17163 | 0.831375036 | 1.971061971 | 1.10060292 | 1.790892914 |
| Higd1a | 526.1847965 | 0.010317148 | 0.080342121 | 0.128415187 |
| Ackr2 | 29.87282185 | 0.689186743 | 0.312289127 | 2.20688677 |
| Cyp8b1 | 0.044191549 | 0.037062677 | 3.724945049 | 0.009949859 |
| 1700048O20Rik | 89.81562415 | 0.353215034 | 0.174620385 | 2.022759454 |
| Fam198a | 45.49996011 | 0.562498607 | 0.264411891 | 2.127357453 |
| Pomgnt2 | 324.6867476 | 0.202339054 | 0.090286725 | 2.241072028 |
| Snrk | 1696.951144 | -0.034072794 | 0.107386546 | -0.317291089 |
| Ano10 | 333.863595 | -0.114027471 | 0.076876142 | -1.483262142 |
| Mir8106 | 0.421183792 | 1.570337676 | 1.596833442 | 0.983407308 |
| Gm26797 | 96.16516166 | 0.045252177 | 0.25031216 | 0.180782974 |
| Abhd5 | 237.0914825 | -0.320611764 | 0.12114019 | -2.646617652 |
| A730085K08Rik | 3.727958404 | 0.712973012 | 0.534431455 | 1.334077563 |
| 9530059O14Rik | 357.5582499 | -0.044480716 | 0.343969681 | -0.129315803 |
| Mir138-1 | 4.764516023 | 0.722619468 | 0.535533605 | 1.349344768 |
| Topaz1 | 1.867918734 | -2.522839815 | 0.859655529 | -2.934710161 |
| Tcaim | 210.4247602 | -0.051020211 | 0.104721236 | -0.487200237 |
| Gm25341 | 1.808478971 | 0.448368491 | 0.767966464 | 0.583838633 |
| Zfp445 | 1117.035592 | 0.277227359 | 0.10813781 | 2.563648732 |
| A530083I20Rik | 14.49954602 | 0.377809682 | 0.332517511 | 1.136209882 |
| Zkscan7 | 42.10827637 | -0.429268201 | 0.204687009 | -2.097193188 |
| Zfp105 | 88.67340489 | -0.275821889 | 0.123417344 | -2.234871362 |
| 1110059G10Rik | 126.7733861 | 0.256519018 | 0.111150436 | 2.307854356 |
| Kif15 | 30.3785014 | -0.163539482 | 0.316509143 | -0.516697496 |
| Tmem42 | 63.48058778 | 0.113725012 | 0.138696001 | 0.81995884 |
| Tgm4 | 6.449499428 | -0.090081233 | 0.430773481 | -0.209115085 |
| Zdhhc3 | 963.4259801 | 0.047153021 | 0.052131721 | 0.904497673 |
| Exosc7 | 112.7568782 | -0.160735601 | 0.106292299 | -1.512203641 |
| Clec3b | 48.60893226 | 0.049512316 | 0.221712239 | 0.223317921 |
| Gm23323 | 0.148038037 | 0.517951684 | 3.551555643 | 0.145837975 |
| Cdcp1 | 2.585897959 | -0.401082743 | 0.669178194 | -0.599366128 |
| Tmem158 | 284.2320044 | 0.102523998 | 0.107388368 | 0.954703011 |
| Scp2-ps2 | 91.28446377 | -0.258719255 | 0.162565957 | -1.591472529 |
| Lars2 | 5438.700325 | -0.120418375 | 0.817339111 | -0.14732976 |
| Limd1 | 249.5361358 | -0.81690693 | 0.12176669 | -6.7087882 |
| Sacm1l | 1325.845621 | -0.147427648 | 0.0675611 | -2.182138059 |
| Slc6a20b | 4.677638664 | 0.523318328 | 0.448052323 | 1.167984856 |
| Gm17120 | 4.624289095 | 1.722861126 | 0.820347726 | 2.100159568 |
| Slc6a20a | 203.7172933 | 0.237340379 | 0.200534972 | 1.1835361 |
| Gm17200 | 1.771524358 | -0.066244917 | 0.877235376 | -0.075515555 |
| Ccr9 | 18.10541206 | -1.431491482 | 0.394774908 | -3.62609541 |
| Gm10052 | 130.6631812 | -0.017923398 | 0.128908443 | -0.139039754 |
| Lztfl1 | 833.9950152 | 0.159107216 | 0.064232082 | 2.477067682 |
| Gm17020 | 0.049553047 | -0.443818988 | 3.724945049 | -0.119147795 |

| | | | | |
|---|---|---|---|---|
| Gm17021 | 0.832525973 | -0.924112583 | 1.09916654 | -0.84073937 |
| Fyco1 | 809.1842188 | -0.054943908 | 0.085223818 | -0.644701324 |
| Cxcr6 | 20.53480371 | -1.13665775 | 0.363016233 | -3.131148548 |
| Xcr1 | 0.667406772 | 0.531422549 | 2.555323168 | 0.207966865 |
| Ccr1 | 181.3036799 | -2.059055414 | 0.593940251 | -3.46677197 |
| Ccr1l1 | 0.461281393 | -0.454021306 | 1.456029267 | -0.311821552 |
| Ccr3 | 1.785934789 | 0.93560346 | 0.799098416 | 1.17082382 |
| Ccr2 | 315.4917145 | -1.858084939 | 0.377837788 | -4.917678958 |
| Ccr5 | 468.5098695 | -1.495854816 | 0.295435051 | -5.063227303 |
| 2010315B03Rik | 245.0633303 | 0.061389692 | 0.123143218 | 0.49852272 |
| Nlrp4g | 0.042716945 | 0.037062677 | 3.724945049 | 0.009949859 |
| Ppp2r3d | 158.3671903 | 0.137359123 | 0.117275342 | 1.171253232 |
| Gm21836 | 0 | NA | NA | NA |
| SNORA28 | 0 | NA | NA | NA |
| RP24-571H12.4 | 0 | NA | NA | NA |
| RP24-571H12.6 | 0 | NA | NA | NA |
| AC168054.1 | 0 | NA | NA | NA |
| WI1-2477B13.3 | 0 | NA | NA | NA |
| CAAA01180111.1 | 0 | NA | NA | NA |
| CAAA01202134.1 | 0 | NA | NA | NA |
| CAAA01125382.1 | 0 | NA | NA | NA |
| AC109138.1 | 0 | NA | NA | NA |
| AC109138.3 | 0 | NA | NA | NA |
| AC109138.2 | 0 | NA | NA | NA |
| AC116115.1 | 0 | NA | NA | NA |
| mmu-mir-505 | 0 | NA | NA | NA |
| CAAA01000182.1 | 0 | NA | NA | NA |
| RP23-6C3.5 | 0 | NA | NA | NA |
| RP23-6C3.4 | 0 | NA | NA | NA |
| Hspa9-ps1 | 7.785073125 | -0.999653403 | 0.390599382 | -2.559280554 |
| RP23-299A23.1 | 0 | NA | NA | NA |
| RP23-6C3.2 | 0 | NA | NA | NA |
| RP23-6C3.3 | 0 | NA | NA | NA |
| RP23-445P15.4 | 0 | NA | NA | NA |
| RP23-445P15.5 | 0 | NA | NA | NA |
| RP23-445P15.2 | 0 | NA | NA | NA |
| RP23-163F11.2 | 0 | NA | NA | NA |
| RP23-346N8.3 | 0 | NA | NA | NA |
| CU041226.1 | 0 | NA | NA | NA |
| RP23-402O17.1 | 0 | NA | NA | NA |
| RP23-16C4.1 | 0 | NA | NA | NA |
| AL593853.1 | 0 | NA | NA | NA |
| RP23-199A2.7 | 0 | NA | NA | NA |
| SNORA77 | 0 | NA | NA | NA |
| mmu-mir-1306 | 0 | NA | NA | NA |
| mmu-mir-3618 | 0 | NA | NA | NA |
| mmu-mir-185 | 0 | NA | NA | NA |

| | | | | |
|---|---|---|---|---|
| AC084822.2 | 0 | NA | NA | NA |
| U1 | 0 | NA | NA | NA |
| AC084822.1 | 0 | NA | NA | NA |
| RP23-285E22.4 | 0 | NA | NA | NA |
| RP23-348F1.1 | 0 | NA | NA | NA |
| 7SK | 0 | NA | NA | NA |
| RP23-58H5.2 | 0 | NA | NA | NA |
| RP23-58H5.5 | 0 | NA | NA | NA |
| RP23-58H5.6 | 0 | NA | NA | NA |
| RP23-58H5.8 | 0 | NA | NA | NA |
| OTTMUSG0000001( | 0 | NA | NA | NA |
| RP24-364D22.1 | 0 | NA | NA | NA |
| AC132586.1 | 0 | NA | NA | NA |
| AC245272.1 | 0 | NA | NA | NA |
| AC245272.3 | 0 | NA | NA | NA |
| mmu-mir-290a | 0 | NA | NA | NA |
| mmu-mir-291a | 0 | NA | NA | NA |
| mmu-mir-292a | 0 | NA | NA | NA |
| mmu-mir-291b | 0 | NA | NA | NA |
| mmu-mir-293 | 0 | NA | NA | NA |
| mmu-mir-294 | 0 | NA | NA | NA |
| mmu-mir-295 | 0 | NA | NA | NA |
| AC245272.2 | 0 | NA | NA | NA |
| mmu-mir-935 | 0 | NA | NA | NA |
| Mico1 | 0 | NA | NA | NA |
| MEG3_1 | 0 | NA | NA | NA |
| MEG3_2 | 0 | NA | NA | NA |
| mmu-mir-1906-1 | 0 | NA | NA | NA |
| mmu-mir-770 | 0 | NA | NA | NA |
| AC152063.1 | 0 | NA | NA | NA |
| mmu-mir-673 | 0 | NA | NA | NA |
| mmu-mir-493 | 0 | NA | NA | NA |
| mmu-mir-337 | 0 | NA | NA | NA |
| mmu-mir-540 | 0 | NA | NA | NA |
| mmu-mir-665 | 0 | NA | NA | NA |
| mmu-mir-3070-1 | 0 | NA | NA | NA |
| mmu-mir-3070-2 | 0 | NA | NA | NA |
| mmu-mir-431 | 0 | NA | NA | NA |
| mmu-mir-433 | 0 | NA | NA | NA |
| mmu-mir-127 | 0 | NA | NA | NA |
| mmu-mir-434 | 0 | NA | NA | NA |
| mmu-mir-432 | 0 | NA | NA | NA |
| mmu-mir-3071 | 0 | NA | NA | NA |
| MEG8_1 | 0 | NA | NA | NA |
| mmu-mir-341 | 0 | NA | NA | NA |
| mmu-mir-1188 | 0 | NA | NA | NA |
| AC152063.2 | 0 | NA | NA | NA |

| | | | | |
|---|---|---|---|---|
| mmu-mir-370 | 0 | NA | NA | NA |
| SNORD113 | 0 | NA | NA | NA |
| RP23-204I16.2 | 0 | NA | NA | NA |
| RP23-204I16.3 | 0 | NA | NA | NA |
| mmu-mir-882 | 0 | NA | NA | NA |
| CH36-472P4.1 | 0 | NA | NA | NA |
| AC233742.1 | 0 | NA | NA | NA |
| CH36-472P4.2 | 0 | NA | NA | NA |
| CH36-472P4.3 | 0 | NA | NA | NA |
| CH36-169F23.1 | 0 | NA | NA | NA |
| CH36-169F23.2 | 0 | NA | NA | NA |
| AC240971.1 | 0 | NA | NA | NA |
| CH36-169F23.4 | 0 | NA | NA | NA |
| CH36-169F23.3 | 0 | NA | NA | NA |
| CH36-169F23.5 | 0 | NA | NA | NA |
| RP24-152F23.1 | 0 | NA | NA | NA |
| RP24-152F23.4 | 0 | NA | NA | NA |
| RP24-152F23.5 | 0 | NA | NA | NA |
| AC244694.1 | 0 | NA | NA | NA |
| RP24-505E20.1 | 0 | NA | NA | NA |
| RP24-410B18.1 | 0 | NA | NA | NA |
| RP24-410B18.2 | 0 | NA | NA | NA |
| RP24-410B18.3 | 0 | NA | NA | NA |
| RP24-410B18.5 | 0 | NA | NA | NA |
| AC202135.1 | 0 | NA | NA | NA |
| RP24-410B18.4 | 0 | NA | NA | NA |
| Rhox5 | 0.049553047 | -0.443818988 | 3.724945049 | -0.119147795 |
| Rhox6 | 0 | NA | NA | NA |
| Rhox7a | 0 | NA | NA | NA |
| Rhox8 | 0.068097359 | 0.037064902 | 3.724945049 | 0.009950456 |
| Rhox7b | 0 | NA | NA | NA |
| Rhox9 | 0.095435502 | 0.037073798 | 3.724945049 | 0.009952844 |
| RP23-202F23.7 | 0 | NA | NA | NA |
| Rhox11-ps | 0 | NA | NA | NA |
| AC154679.1 | 0 | NA | NA | NA |
| AC182748.2 | 0 | NA | NA | NA |
| AC182748.1 | 0 | NA | NA | NA |
| U6 | 0 | NA | NA | NA |
| RP24-176A1.1 | 0 | NA | NA | NA |
| RP24-176A1.2 | 0 | NA | NA | NA |
| AC127556.1 | 0 | NA | NA | NA |
| mmu-mir-194-2 | 0 | NA | NA | NA |
| mmu-mir-192 | 0 | NA | NA | NA |
| AC127556.5 | 0 | NA | NA | NA |
| mmu-mir-6989 | 0 | NA | NA | NA |
| AC127556.10 | 0 | NA | NA | NA |
| AC167245.6 | 0 | NA | NA | NA |

| | | | | |
|---|---|---|---|---|
| AC167245.5 | 0 | NA | NA | NA |
| AC124394.1 | 0 | NA | NA | NA |
| Gm14423 | 0 | NA | NA | NA |
| AC124394.3 | 0 | NA | NA | NA |
| AC124394.4 | 0 | NA | NA | NA |
| RP23-188E3.2 | 0 | NA | NA | NA |
| RP23-188E3.1 | 0 | NA | NA | NA |
| BX890553.1 | 0 | NA | NA | NA |
| RP23-324J18.1 | 0 | NA | NA | NA |
| RP23-324J18.2 | 0 | NA | NA | NA |
| AC164883.2 | 0 | NA | NA | NA |
| AC164883.1 | 0 | NA | NA | NA |
| CH25-484K21.2 | 0 | NA | NA | NA |
| CH25-484K21.3 | 0 | NA | NA | NA |
| CH25-484K21.5 | 0 | NA | NA | NA |
| CH25-484K21.6 | 0 | NA | NA | NA |
| Gm9919 | 0 | NA | NA | NA |
| FO907130.1 | 0 | NA | NA | NA |
| SNORD103 | 0 | NA | NA | NA |
| RP23-157N11.1 | 0 | NA | NA | NA |
| RP23-157N11.4 | 0 | NA | NA | NA |
| RP23-157N11.7 | 0 | NA | NA | NA |
| RP23-157N11.6 | 0 | NA | NA | NA |
| RP23-146J17.3 | 0 | NA | NA | NA |
| AC125149.4 | 0 | NA | NA | NA |
| AC125149.1 | 0 | NA | NA | NA |
| AC125149.2 | 0 | NA | NA | NA |
| AC125149.3 | 0 | NA | NA | NA |
| AC133103.4 | 0 | NA | NA | NA |
| AC133103.6 | 0 | NA | NA | NA |
| AC133103.7 | 0 | NA | NA | NA |
| AC133103.2 | 0 | NA | NA | NA |
| AC133103.5 | 0 | NA | NA | NA |
| AC133103.1 | 0 | NA | NA | NA |
| AC133103.3 | 0 | NA | NA | NA |
| AC168977.2 | 0 | NA | NA | NA |
| AC168977.1 | 0 | NA | NA | NA |
| DHRSX | 0 | NA | NA | NA |
| U2 | 0 | NA | NA | NA |
| Vmn1r186 | 0 | NA | NA | NA |
| AC102264.1 | 0 | NA | NA | NA |
| AC132444.1 | 0 | NA | NA | NA |
| AC132444.3 | 0 | NA | NA | NA |
| AC132444.5 | 0 | NA | NA | NA |
| AC132444.4 | 0 | NA | NA | NA |
| AC132444.2 | 0 | NA | NA | NA |
| Csprs | 0 | NA | NA | NA |

| | | | | |
|---|---|---|---|---|
| AC132444.6 | 0 | NA | NA | NA |
| Vamp7 | 0 | NA | NA | NA |
| CAAA01165726.1 | 0 | NA | NA | NA |
| Spry3 | 0 | NA | NA | NA |
| Tmlhe | 0 | NA | NA | NA |
| CR524822.1 | 0 | NA | NA | NA |
| AC087559.4 | 0 | NA | NA | NA |
| AC087559.5 | 0 | NA | NA | NA |
| AC087559.1 | 0 | NA | NA | NA |
| AC087559.3 | 0 | NA | NA | NA |
| AC087559.2 | 0 | NA | NA | NA |
| Ccl21c | 0 | NA | NA | NA |
| AC140325.2 | 0 | NA | NA | NA |
| AC140325.1 | 0 | NA | NA | NA |
| AC140325.3 | 0 | NA | NA | NA |
| AC140325.4 | 0 | NA | NA | NA |
| CAAA01098150.1 | 0 | NA | NA | NA |
| CAAA01064564.1 | 0 | NA | NA | NA |
| CAAA01205117.2 | 0 | NA | NA | NA |
| CAAA01205117.1 | 0 | NA | NA | NA |
| Vmn2r122 | 0 | NA | NA | NA |
| Gm2002 | 0 | NA | NA | NA |
| CR974586.2 | 0 | NA | NA | NA |
| CR974586.4 | 0 | NA | NA | NA |
| CR974586.8 | 0 | NA | NA | NA |
| CR974586.5 | 0 | NA | NA | NA |
| CR974586.7 | 0 | NA | NA | NA |
| CR974586.6 | 0 | NA | NA | NA |
| CR974586.3 | 0 | NA | NA | NA |
| CR974586.1 | 0 | NA | NA | NA |
| Scya19 | 0 | NA | NA | NA |
| CT868723.1 | 0 | NA | NA | NA |
| Gm10931 | 0 | NA | NA | NA |
| CT868723.2 | 0 | NA | NA | NA |
| CAAA01147332.1 | 0 | NA | NA | NA |
| AC164084.2 | 0 | NA | NA | NA |
| AC164084.3 | 0 | NA | NA | NA |
| AC164084.1 | 0 | NA | NA | NA |
| AC165294.1 | 0 | NA | NA | NA |
| AC165294.2 | 0 | NA | NA | NA |
| AC165294.3 | 0 | NA | NA | NA |
| AC125178.1 | 0 | NA | NA | NA |
| AC125178.3 | 0 | NA | NA | NA |
| AC125178.2 | 0 | NA | NA | NA |
| AC123873.3 | 0 | NA | NA | NA |
| AC123873.1 | 0 | NA | NA | NA |
| AC123873.2 | 0 | NA | NA | NA |

| | | | | |
|---|---|---|---|---|
| AC126035.1 | 0 | NA | NA | NA |
| Gm16367 | 0 | NA | NA | NA |
| AC163611.1 | 0 | NA | NA | NA |
| AC140365.1 | 0 | NA | NA | NA |
| AC124606.2 | 0 | NA | NA | NA |
| AC124606.1 | 0 | NA | NA | NA |
| AC133095.2 | 0 | NA | NA | NA |
| AC133095.1 | 0 | NA | NA | NA |
| AC234645.1 | 0 | NA | NA | NA |
| PISD | 0 | NA | NA | NA |
| mt-Tf | 0 | NA | NA | NA |
| mt-Rnr1 | 119593.2604 | 0.44830487 | 0.923923373 | 0.48521867 |
| mt-Tv | 0 | NA | NA | NA |
| mt-Rnr2 | 65951.93075 | 0.399480265 | 0.794776848 | 0.502631986 |
| mt-Tl1 | 208.0973662 | 0.765158204 | 0.217766451 | 3.5136643 |
| mt-Nd1 | 34373.33605 | 0.487806464 | 0.157401546 | 3.099121162 |
| mt-Ti | 143.1190915 | -0.153376762 | 0.339010135 | -0.452425301 |
| mt-Tq | 55.05421077 | 0.469162247 | 0.192951738 | 2.431500501 |
| mt-Tm | 149.670499 | -0.168872962 | 0.247776132 | -0.681554599 |
| mt-Nd2 | 20368.9686 | 0.376223308 | 0.224887857 | 1.672937408 |
| mt-Tw | 186.9799014 | 0.675267568 | 0.185845864 | 3.633481809 |
| mt-Ta | 11.02334367 | 0.095016222 | 0.344521686 | 0.275791701 |
| mt-Tn | 0.72655531 | 1.110252985 | 1.161885524 | 0.955561424 |
| mt-Tc | 1.449755302 | 0.248497405 | 0.851417805 | 0.291863059 |
| mt-Ty | 0.807376506 | 0.470980087 | 1.326422112 | 0.355075569 |
| mt-Co1 | 112876.7948 | 0.200600663 | 0.120042819 | 1.67107591 |
| mt-Ts1 | 1.308396679 | 0.557483759 | 0.835645862 | 0.667129205 |
| mt-Td | 0.616049689 | -0.72406952 | 1.33975206 | -0.540450387 |
| mt-Co2 | 1688.977157 | 0.404992406 | 0.128307447 | 3.15642167 |
| mt-Tk | 0.097139547 | -0.443816764 | 3.724945049 | -0.119147198 |
| mt-Atp8 | 110.3540989 | 0.713716748 | 0.429001015 | 1.663671469 |
| mt-Atp6 | 624.2525603 | 0.387245807 | 0.197293784 | 1.962787674 |
| mt-Co3 | 1101.033194 | 0.427443349 | 0.147099292 | 2.90581513 |
| mt-Tg | 0 | NA | NA | NA |
| mt-Nd3 | 119.756487 | 0.10819803 | 0.17557374 | 0.616254059 |
| mt-Tr | 0 | NA | NA | NA |
| mt-Nd4l | 129.7203189 | 0.418941739 | 0.21743473 | 1.926747112 |
| mt-Nd4 | 21602.70211 | 0.321690916 | 0.143073707 | 2.248427903 |
| mt-Th | 0.134005754 | 0.517961793 | 2.962652882 | 0.174830402 |
| mt-Ts2 | 0.119587853 | 0.981063299 | 3.723618439 | 0.263470416 |
| mt-Tl2 | 0.731462286 | 0.996780359 | 1.265340544 | 0.787756595 |
| mt-Nd5 | 23114.44776 | 0.560699684 | 0.210336871 | 2.66572228 |
| mt-Nd6 | 1598.689532 | 0.032131113 | 0.154854699 | 0.207492011 |
| mt-Te | 5.226567033 | 0.110465351 | 0.490466745 | 0.225224956 |
| mt-Cytb | 37159.01855 | 0.331450488 | 0.140703471 | 2.355666747 |
| mt-Tt | 18.73040197 | 0.982726886 | 0.341299365 | 2.879369217 |
| mt-Tp | 56.15899216 | 0.643844013 | 0.213153161 | 3.02056986 |

| | | | | |
|---|---|---|---|---|
| Gm21950 | 0 | NA | NA | NA |
| Gm21364 | 0 | NA | NA | NA |
| Gm14331 | 0 | NA | NA | NA |
| Gm14333 | 0 | NA | NA | NA |
| Gm14332 | 0 | NA | NA | NA |
| Gm14334 | 0 | NA | NA | NA |
| Gm14352 | 0 | NA | NA | NA |
| Gm14346 | 0 | NA | NA | NA |
| Gm14353 | 0 | NA | NA | NA |
| Gm14360 | 0 | NA | NA | NA |
| Gm14361 | 0 | NA | NA | NA |
| Gm14345 | 0 | NA | NA | NA |
| Gm14351 | 0 | NA | NA | NA |
| Gm14357 | 0 | NA | NA | NA |
| Gm14359 | 0 | NA | NA | NA |
| Gm14355 | 0 | NA | NA | NA |
| Spin2-ps1 | 0 | NA | NA | NA |
| Gm3701 | 0 | NA | NA | NA |
| Gm15251 | 0 | NA | NA | NA |
| Gm15257 | 0 | NA | NA | NA |
| Gm3706 | 0 | NA | NA | NA |
| Gm15258 | 0 | NA | NA | NA |
| Gm14350 | 0 | NA | NA | NA |
| Gm14349 | 0 | NA | NA | NA |
| Gm14358 | 0 | NA | NA | NA |
| Gm14362 | 0 | NA | NA | NA |
| Gm14356 | 0 | NA | NA | NA |
| Gm14347 | 0 | NA | NA | NA |
| Gm10921 | 0 | NA | NA | NA |
| Gm14348 | 0 | NA | NA | NA |
| Gm14364 | 0 | NA | NA | NA |
| Gm14368 | 0 | NA | NA | NA |
| Gm15260 | 0 | NA | NA | NA |
| Gm10922 | 0 | NA | NA | NA |
| Gm6019 | 0 | NA | NA | NA |
| Gm9427 | 0 | NA | NA | NA |
| Gm15256 | 0 | NA | NA | NA |
| Gm5925 | 0 | NA | NA | NA |
| Gm4725 | 0 | NA | NA | NA |
| Gm9498 | 0 | NA | NA | NA |
| Gm15252 | 0 | NA | NA | NA |
| Gm3750 | 0 | NA | NA | NA |
| Gm4726 | 0 | NA | NA | NA |
| Gm3757 | 0 | NA | NA | NA |
| Gm15259 | 0 | NA | NA | NA |
| Gm15254 | 0 | NA | NA | NA |
| Gm3763 | 0 | NA | NA | NA |

| | | | | |
|---|---|---|---|---|
| Gm14370 | 0 | NA | NA | NA |
| Gm14367 | 0 | NA | NA | NA |
| Gm14365 | 0 | NA | NA | NA |
| Gm9428 | 0 | NA | NA | NA |
| Gm5924 | 0 | NA | NA | NA |
| Gm14369 | 0 | NA | NA | NA |
| Gm14363 | 0 | NA | NA | NA |
| Mycs | 0 | NA | NA | NA |
| Gm14366 | 0 | NA | NA | NA |
| Gm14375 | 0 | NA | NA | NA |
| Gm14374 | 0 | NA | NA | NA |
| Gm14373 | 4.562379813 | -0.474301734 | 0.518107281 | -0.915450819 |
| Gm14452 | 0.11997416 | 0.517957688 | 3.724945049 | 0.139051095 |
| Gm14451 | 0.112642008 | 0.517955464 | 3.724945049 | 0.139050498 |
| Gm14448 | 2.617244783 | -0.618626813 | 0.76998793 | -0.80342404 |
| Gm14450 | 0 | NA | NA | NA |
| Gm26618 | 11.11537949 | 0.601810735 | 0.347469202 | 1.731982954 |
| Nudt11 | 271.955026 | 0.009751264 | 0.103458622 | 0.094252795 |
| Rpl19-ps12 | 2.084829622 | 0.873243017 | 0.831776162 | 1.049853383 |
| Gm27360 | 0 | NA | NA | NA |
| AU022751 | 17.60759048 | 1.033616569 | 0.402763895 | 2.566308902 |
| n-R5s2 | 0 | NA | NA | NA |
| Nudt10 | 171.3843856 | -0.036857836 | 0.102222984 | -0.360563103 |
| Bmp15 | 3.216309252 | -0.051209649 | 0.679764508 | -0.075334397 |
| Shroom4 | 90.11915289 | -0.389481461 | 0.156924974 | -2.481959705 |
| Dgkk | 132.1695717 | -0.54106399 | 0.216221035 | -2.502365189 |
| Ccnb3 | 0.629652017 | -3.193139546 | 1.36105023 | -2.346085013 |
| Akap4 | 0 | NA | NA | NA |
| Gm14380 | 0.089089296 | 0.957155977 | 3.723939561 | 0.257027796 |
| Clcn5 | 513.7578379 | -0.182297005 | 0.095867711 | -1.901547499 |
| Gm26401 | 0 | NA | NA | NA |
| Mir500 | 0.181074517 | -0.843807695 | 3.346447532 | -0.252150284 |
| Mir501 | 0.726672012 | -1.114031447 | 1.206090173 | -0.923671771 |
| Mir362 | 0.060174537 | 0.517957688 | 3.724945049 | 0.139051095 |
| Mir188 | 0.206496347 | 0.957549283 | 2.187301055 | 0.437776629 |
| Mir532 | 0.516362146 | 0.996210377 | 1.422527174 | 0.700310261 |
| Usp27x | 183.7584339 | 0.063437568 | 0.129826075 | 0.488635028 |
| Gm14379 | 5.282615923 | -0.302662197 | 0.449574425 | -0.673219338 |
| 2010204K13Rik | 54.17774226 | 0.336878328 | 0.159775508 | 2.108447852 |
| Gm14381 | 0.045882455 | 0.517959912 | 3.724945049 | 0.139051692 |
| Gm14395 | 0 | NA | NA | NA |
| Gm14394 | 0.287498921 | -2.096329295 | 1.959903211 | -1.069608583 |
| Gm23082 | 0.046345709 | -0.443818988 | 3.724945049 | -0.119147795 |
| Ppp1r3f | 110.0662308 | 0.239391651 | 0.115611099 | 2.070663213 |
| Gm36995 | 5.062940487 | 0.751617358 | 0.507016684 | 1.482431214 |
| Ppp1r3fos | 7.493797218 | -0.045125944 | 0.389372729 | -0.115893951 |
| Foxp3 | 2.783566837 | 0.410990235 | 0.595998317 | 0.689582879 |

| | | | | |
|---|---|---|---|---|
| Ccdc22 | 22.02494333 | -0.225046085 | 0.22983655 | -0.979157081 |
| Cacna1f | 2.348133422 | -0.153048869 | 0.6258808 | -0.244533574 |
| Syp | 9951.334057 | 0.468868717 | 0.068829296 | 6.812051595 |
| Gm14703 | 9.786129742 | 0.921568125 | 0.566034171 | 1.628113941 |
| Prickle3 | 15.66947633 | -0.217561648 | 0.262322745 | -0.829366314 |
| Plp2 | 46.01570745 | -1.200763137 | 0.205000775 | -5.857359016 |
| Magix | 1.802146662 | -0.341174283 | 0.813125607 | -0.419583739 |
| Gm6079 | 0 | NA | NA | NA |
| Gpkow | 510.2906317 | -0.026039706 | 0.079337326 | -0.328215073 |
| Gm14720 | 0 | NA | NA | NA |
| Wdr45 | 250.0876241 | 0.200344112 | 0.072502802 | 2.763260277 |
| Praf2 | 530.2469731 | -0.002748063 | 0.065798408 | -0.041764883 |
| Ccdc120 | 144.7907492 | 0.139728616 | 0.143615818 | 0.972933334 |
| Smt3h2-ps | 1.463749607 | 1.55294019 | 1.086030281 | 1.429923472 |
| Tfe3 | 940.6091959 | -0.368546938 | 0.060751841 | -6.066432377 |
| Gripap1 | 833.0662841 | 0.0722016 | 0.116342297 | 0.620596313 |
| Mir1198 | 4.250207662 | 0.511720994 | 0.59491689 | 0.860155431 |
| Kcnd1 | 104.6643679 | 0.397390779 | 0.148778858 | 2.671016464 |
| Otud5 | 1030.482185 | -0.083759322 | 0.047545899 | -1.761651881 |
| Pim2 | 309.3189051 | -0.063263956 | 0.092773509 | -0.681918323 |
| Slc35a2 | 203.1775785 | -0.341075161 | 0.105359568 | -3.237249048 |
| Pqbp1 | 363.0226281 | 0.246840495 | 0.074565173 | 3.310399292 |
| Timm17b | 118.9161447 | -0.17585251 | 0.154523801 | -1.138028631 |
| Gm10491 | 0.046345709 | -0.443818988 | 3.724945049 | -0.119147795 |
| Gm10490 | 0.314722824 | -1.437188589 | 1.730395936 | -0.830554765 |
| Pcsk1n | 5132.141306 | 0.320465036 | 0.101774414 | 3.148777997 |
| Eras | 0.25849572 | -0.511594418 | 2.073274411 | -0.246756732 |
| Hdac6 | 414.7095717 | 0.240665794 | 0.085124168 | 2.827232256 |
| Psmb6-ps | 0 | NA | NA | NA |
| Gata1 | 0.556661178 | -0.229668662 | 1.503959565 | -0.152709333 |
| Glod5 | 2.482897828 | 1.156242767 | 0.629404011 | 1.837043849 |
| Gm9429 | 0.112029109 | 0.037060453 | 3.724945049 | 0.009949262 |
| Rpl22-ps1 | 1.865863811 | 0.137425514 | 0.700646774 | 0.196140936 |
| Gm14799 | 1.762166695 | -0.345260901 | 0.9231179 | -0.37401604 |
| Gm14820 | 0.044544648 | 0.517957688 | 3.724945049 | 0.139051095 |
| Gm7079 | 50.37483144 | -0.555582499 | 0.303041114 | -1.83335684 |
| Suv39h1 | 269.5752616 | 0.357422975 | 0.096337915 | 3.710096648 |
| Was | 67.79222777 | -0.77280988 | 0.210889423 | -3.66452651 |
| Gm22083 | 0 | NA | NA | NA |
| Gm6787 | 1.653410964 | -0.042904893 | 0.915904487 | -0.046844287 |
| Psmb7-ps2 | 0.216874018 | 0.517964955 | 2.599194223 | 0.19927905 |
| Wdr13 | 1616.98844 | 0.633946851 | 0.078816558 | 8.043320721 |
| Rbm3 | 813.7615764 | 0.261308441 | 0.131846074 | 1.98192053 |
| Rbm3os | 4.093244608 | -0.89542808 | 0.587791181 | -1.523377874 |
| Tbc1d25 | 154.8391368 | 0.267495499 | 0.105784832 | 2.528675361 |
| Ebp | 209.4144051 | -0.548911018 | 0.096326221 | -5.698458982 |
| Porcn | 473.2143967 | 0.453745727 | 0.115514235 | 3.928050301 |

| | | | | |
|---|---|---|---|---|
| Ftsj1 | 170.553967 | -0.110112262 | 0.089044693 | -1.236595438 |
| Slc38a5 | 13.85939591 | -0.750866382 | 0.334528889 | -2.244548697 |
| Ssxb10 | 0 | NA | NA | NA |
| Ssxb9 | 0 | NA | NA | NA |
| Ssxb1 | 0 | NA | NA | NA |
| Ssxb2 | 0 | NA | NA | NA |
| Gm14459 | 0 | NA | NA | NA |
| Ssxb6 | 0 | NA | NA | NA |
| Ssxb3 | 0.117907917 | 0.967119648 | 3.722483401 | 0.259804959 |
| Gm14500 | 0 | NA | NA | NA |
| Gm6797 | 0 | NA | NA | NA |
| Gm6798 | 0 | NA | NA | NA |
| Gm9430 | 0.048949532 | 0.037064902 | 3.724945049 | 0.009950456 |
| Ssxb8 | 0.176646398 | -0.526386109 | 3.35399333 | -0.1569431 |
| Ssx9 | 0 | NA | NA | NA |
| Ssxb5 | 0 | NA | NA | NA |
| Gm14502 | 0.848591665 | 0.007633976 | 1.310512506 | 0.005825184 |
| Gm6592 | 0 | NA | NA | NA |
| Gm5751 | 0.339985237 | -0.448804915 | 1.762196452 | -0.25468495 |
| B630019K06Rik | 470.2877301 | 0.211161427 | 0.13392685 | 1.576692256 |
| Gm14504 | 0.188390742 | 0.517957661 | 3.311555221 | 0.156409187 |
| Gm5752 | 0 | NA | NA | NA |
| Gm16479 | 0 | NA | NA | NA |
| Gm14510 | 0 | NA | NA | NA |
| Gm5634 | 0 | NA | NA | NA |
| Gm14509 | 0 | NA | NA | NA |
| Gm14511 | 0 | NA | NA | NA |
| Gm14457 | 0 | NA | NA | NA |
| Gm14458 | 0 | NA | NA | NA |
| Gm9431 | 0.150711795 | 0.037073253 | 2.856704404 | 0.01297763 |
| Gm14865 | 0 | NA | NA | NA |
| Fthl17 | 0 | NA | NA | NA |
| Gm14503 | 0 | NA | NA | NA |
| Gm14499 | 0 | NA | NA | NA |
| Gm14864 | 0.066050875 | -0.443818988 | 3.724945049 | -0.119147795 |
| Gm5379 | 0 | NA | NA | NA |
| Gm5635 | 0 | NA | NA | NA |
| Gm16480 | 0 | NA | NA | NA |
| Gm6826 | 0 | NA | NA | NA |
| 4930402K13Rik | 0 | NA | NA | NA |
| Gm5753 | 0 | NA | NA | NA |
| Lancl3 | 181.9738499 | -0.234268037 | 0.101551095 | -2.306898194 |
| Gm14862 | 0.045882455 | 0.517959912 | 3.724945049 | 0.139051692 |
| Xk | 302.6762856 | 0.104921292 | 0.095239313 | 1.101659481 |
| 1700012L04Rik | 1.468848822 | -0.226461216 | 0.965140859 | -0.234640585 |
| Gm25481 | 0.305717649 | 0.037076492 | 1.781807131 | 0.020808364 |
| Gm6829 | 5.233925965 | -0.928758542 | 0.484057961 | -1.918692834 |

| | | | | |
|---|---|---|---|---|
| Gm14863 | 0.303434138 | -0.896928032 | 1.739739764 | -0.515552987 |
| Gm14501 | 0.106056992 | 0.998850474 | 3.724945049 | 0.268151734 |
| Cybb | 408.2087086 | -0.684421884 | 0.167153841 | -4.094562717 |
| Gm25728 | 0.044191549 | 0.037062677 | 3.724945049 | 0.009949859 |
| Gm5132 | 0.273271296 | 0.517974935 | 2.030826441 | 0.25505623 |
| Dynlt3 | 1172.505651 | 0.165084013 | 0.098893279 | 1.669314786 |
| Hypm | 1.020378237 | 1.016838275 | 1.155053998 | 0.880338302 |
| 4930557A04Rik | 0.934106423 | -0.764549732 | 1.232395332 | -0.62037701 |
| Sytl5 | 63.53451429 | -0.112455431 | 0.175074383 | -0.642329443 |
| Gm5754 | 0.681090533 | 0.145435499 | 1.27238854 | 0.11430117 |
| Srpx | 4.924570278 | -0.910687415 | 0.522510115 | -1.742908681 |
| Rpgr | 331.1611277 | 0.495656468 | 0.145530332 | 3.405863643 |
| Otc | 0.279231709 | -0.919458254 | 2.12426913 | -0.432835106 |
| 4933416E14Rik | 0.989536815 | -0.022522052 | 1.340527135 | -0.016800892 |
| Tspan7 | 5607.471727 | 0.575145362 | 0.086608645 | 6.640738508 |
| Gm25885 | 9.492997069 | 0.969415089 | 0.41199659 | 2.35296872 |
| Gm5931 | 6.023005853 | -0.127844571 | 0.438432429 | -0.291594697 |
| Gm14470 | 0.083526043 | 0.037073798 | 3.724945049 | 0.009952844 |
| Gm14480 | 0.946116905 | 1.173053296 | 1.098727078 | 1.067647571 |
| Gm10489 | 4.50082467 | 0.97626939 | 0.477716086 | 2.043618413 |
| Mid1ip1 | 964.1006633 | -0.215665889 | 0.108281651 | -1.991712234 |
| Gm14493 | 2.463509541 | 0.395488445 | 0.731712336 | 0.540497167 |
| Gm14473 | 0 | NA | NA | NA |
| Gm14485 | 0.048949532 | 0.037064902 | 3.724945049 | 0.009950456 |
| Gm14481 | 0.306362211 | 1.435540486 | 1.985737437 | 0.722925629 |
| Gm14483 | 0.04393175 | 0.037062677 | 3.724945049 | 0.009949859 |
| Gm14474 | 0 | NA | NA | NA |
| Gm14477 | 0 | NA | NA | NA |
| Gm14476 | 0 | NA | NA | NA |
| Gm14484 | 0 | NA | NA | NA |
| Gm14479 | 0 | NA | NA | NA |
| Gm14482 | 0 | NA | NA | NA |
| Gm14478 | 0 | NA | NA | NA |
| Gm14475 | 0 | NA | NA | NA |
| Gm4906 | 0 | NA | NA | NA |
| Gm14515 | 0 | NA | NA | NA |
| Gm14513 | 0 | NA | NA | NA |
| Gm26314 | 0 | NA | NA | NA |
| Rpl23a-ps13 | 0 | NA | NA | NA |
| Gm14491 | 1.107727442 | 1.65482534 | 1.02764437 | 1.610309352 |
| Gm14512 | 0 | NA | NA | NA |
| Bcor | 487.1376583 | -0.122636439 | 0.1241991 | -0.987418099 |
| Gm14521 | 0 | NA | NA | NA |
| Gm14635 | 0.042716945 | 0.037062677 | 3.724945049 | 0.009949859 |
| Rpl30-ps11 | 0.439488226 | -1.723267084 | 1.481578799 | -1.163128877 |
| Atp6ap2 | 2034.077482 | 0.02313606 | 0.064553354 | 0.358402139 |
| Gm9432 | 2.504601602 | 0.141619591 | 0.709675423 | 0.19955544 |

| | | | | |
|---|---|---|---|---|
| Gm4984 | 0.075043205 | 0.517957688 | 3.724945049 | 0.139051095 |
| 1810030O07Rik | 260.8927199 | -0.036297605 | 0.075059127 | -0.483586824 |
| Gm16265 | 0.351984013 | -0.778236132 | 1.762173168 | -0.44163431 |
| Med14 | 1262.541672 | 0.081205162 | 0.043735901 | 1.856716326 |
| Gm14634 | 22.48773761 | 1.076064657 | 0.264110457 | 4.074297813 |
| Gm25063 | 0.044544648 | 0.517957688 | 3.724945049 | 0.139051095 |
| Gm14524 | 0 | NA | NA | NA |
| AA414768 | 46.27663079 | -0.166761839 | 0.191840471 | -0.869273507 |
| Gm14534 | 0.163190422 | -0.955151543 | 3.396300892 | -0.281232898 |
| 5730405O15Rik | 64.71272437 | 0.316090493 | 0.182802782 | 1.729133926 |
| Usp9x | 6600.196581 | 0.083157404 | 0.046085672 | 1.804409038 |
| Gm15027 | 0.158569248 | 0.51797124 | 2.811594837 | 0.184226843 |
| Gm14536 | 0 | NA | NA | NA |
| Rpl3-ps1 | 956.7057751 | 0.058148294 | 0.106539033 | 0.545793331 |
| n-R5s3 | 0 | NA | NA | NA |
| Llph-ps2 | 8.542226199 | -0.172181456 | 0.399408029 | -0.431091623 |
| 2010308F09Rik | 1.66644364 | 0.724432095 | 0.786296049 | 0.921322314 |
| Gm16481 | 0 | NA | NA | NA |
| Ddx3x | 4505.87406 | 0.128585718 | 0.100334043 | 1.281576158 |
| Gm5380 | 0.854198329 | -0.51928139 | 1.104616515 | -0.470101056 |
| Gm7129 | 0.097899064 | 0.037062677 | 3.724945049 | 0.009949859 |
| Gm9434 | 0.22969359 | -1.373109043 | 3.129889768 | -0.438708435 |
| Nyx | 9.174112265 | 0.681484312 | 0.437646752 | 1.557156105 |
| Gm14506 | 0 | NA | NA | NA |
| Cask | 1849.363507 | 0.225206791 | 0.068149281 | 3.304609923 |
| Gpr34 | 42.59505678 | 2.167822229 | 0.703767245 | 3.080311345 |
| Gm25246 | 0 | NA | NA | NA |
| Gpr82 | 1.132182296 | -0.121338897 | 0.998770007 | -0.121488327 |
| Gm5381 | 0 | NA | NA | NA |
| Gm25552 | 0 | NA | NA | NA |
| Gm5382 | 0 | NA | NA | NA |
| Gm14867 | 0 | NA | NA | NA |
| Gm25202 | 0 | NA | NA | NA |
| Gm14505 | 0.222175754 | -0.453056135 | 2.579270609 | -0.175652812 |
| Gm9435 | 0 | NA | NA | NA |
| Ppp1r2-ps9 | 0 | NA | NA | NA |
| Gm14519 | 0.126242988 | 0.037073701 | 3.017245199 | 0.012287268 |
| Gm14518 | 0 | NA | NA | NA |
| Gm14517 | 0 | NA | NA | NA |
| Gm25207 | 0 | NA | NA | NA |
| Gm22464 | 0 | NA | NA | NA |
| Gm23516 | 0 | NA | NA | NA |
| Drr1 | 0.510920245 | -0.822406349 | 1.36562607 | -0.60221928 |
| Gm14520 | 0 | NA | NA | NA |
| Gm14516 | 0.07796279 | -0.798301147 | 3.721829328 | -0.214491605 |
| Cypt1 | 0.038981395 | -0.443816764 | 3.724945049 | -0.119147198 |
| Csnk2a1-ps | 0.061920613 | -0.443816764 | 3.724945049 | -0.119147198 |

| | | | | |
|---|---|---|---|---|
| Maoa | 471.3466681 | -0.243117019 | 0.099483992 | -2.443780302 |
| Maob | 199.4044433 | 0.423927416 | 0.139193031 | 3.045608044 |
| Ndp | 53.73532709 | -0.387006346 | 0.297980697 | -1.298763141 |
| Efhc2 | 28.39597078 | 0.624904426 | 0.241624712 | 2.586260406 |
| Gm14514 | 0 | NA | NA | NA |
| Gm22294 | 0.044191549 | 0.037062677 | 3.724945049 | 0.009949859 |
| Fundc1 | 517.0185679 | -0.120455504 | 0.137234748 | -0.877733265 |
| Gm5073 | 0.100902008 | -0.876201588 | 3.72093227 | -0.235479048 |
| Gm9436 | 0 | NA | NA | NA |
| Gm5755 | 0 | NA | NA | NA |
| Gm5123 | 0 | NA | NA | NA |
| Dusp21 | 0 | NA | NA | NA |
| Kdm6a | 813.7693481 | 0.119247212 | 0.083940278 | 1.420619686 |
| n-R5s5 | 0 | NA | NA | NA |
| Gm14883 | 0 | NA | NA | NA |
| 4930578C19Rik | 7.057036865 | -0.355443919 | 0.487644961 | -0.728898989 |
| Gm6923 | 0 | NA | NA | NA |
| Gm5383 | 0 | NA | NA | NA |
| Mir221 | 8.635094361 | -0.660112461 | 0.424150142 | -1.556317908 |
| Mir222 | 2.111772912 | -0.166509352 | 0.77651351 | -0.214432009 |
| Gm14636 | 161.0641105 | -0.340513167 | 0.186852065 | -1.822367699 |
| Gm26652 | 38.71122557 | 0.185646964 | 0.288501936 | 0.643486024 |
| Gm22783 | 0.140090306 | -0.892132352 | 2.912101179 | -0.306353487 |
| Gm14885 | 0 | NA | NA | NA |
| BC049702 | 0 | NA | NA | NA |
| Gm14531 | 0 | NA | NA | NA |
| Gm23678 | 0 | NA | NA | NA |
| Gm14530 | 0.578381701 | 0.520903495 | 1.435483513 | 0.362876683 |
| Rpl36-ps10 | 0.038981395 | -0.443816764 | 3.724945049 | -0.119147198 |
| Gm5384 | 0.309182497 | 1.508887392 | 1.779925508 | 0.847725023 |
| Chst7 | 15.98671876 | 0.027104545 | 0.302131413 | 0.089711112 |
| Slc9a7 | 557.1947993 | 0.162700015 | 0.070337572 | 2.313130939 |
| Gm9083 | 2.845117016 | -1.098480055 | 0.624253535 | -1.759669738 |
| Gm14529 | 0 | NA | NA | NA |
| Rp2h | 271.8265236 | -0.300149028 | 0.135414633 | -2.216518415 |
| Gm9085 | 0.090427103 | 0.998850474 | 3.724945049 | 0.268151734 |
| Gm14537 | 3.667719452 | 0.812967518 | 0.610501711 | 1.331638394 |
| Jade3 | 162.3751509 | -0.348832431 | 0.124967969 | -2.791374734 |
| Gm14884 | 0.062426873 | 0.517959912 | 3.724945049 | 0.139051692 |
| Gm14538 | 0 | NA | NA | NA |
| Rgn | 3.195312812 | 0.392733049 | 0.549014475 | 0.715341884 |
| Gm14539 | 108.3056856 | 0.059735206 | 0.166832966 | 0.358053972 |
| Gm23628 | 1.296764287 | 0.681944589 | 0.922594443 | 0.739159653 |
| Ndufb11 | 499.4740233 | -0.018923798 | 0.138331014 | -0.136800836 |
| Rbm10 | 598.4934086 | -0.176509973 | 0.074001427 | -2.385223909 |
| Uba1 | 4405.966393 | -0.205961938 | 0.072848423 | -2.827266953 |
| Cdk16 | 2633.067354 | 0.273484319 | 0.058483851 | 4.676236463 |

| | | | | |
|---|---|---|---|---|
| Usp11 | 1238.553824 | 0.373129622 | 0.061400946 | 6.076936039 |
| Araf | 1405.997007 | 0.145386021 | 0.052291821 | 2.78028221 |
| Gm26065 | 0 | NA | NA | NA |
| Gm24824 | 0 | NA | NA | NA |
| Gm15028 | 0.262683741 | -1.12332619 | 2.448530534 | -0.458775651 |
| Syn1 | 5420.391119 | 0.398717781 | 0.093993907 | 4.241953475 |
| Mir5617 | 0 | NA | NA | NA |
| Timp1 | 225.2635421 | -1.472444964 | 0.457450242 | -3.218809021 |
| Gm15029 | 0 | NA | NA | NA |
| Cfp | 118.3901945 | -0.798639495 | 0.211437634 | -3.77718707 |
| Elk1 | 876.8854966 | 0.355328547 | 0.132017037 | 2.691535538 |
| Uxt | 53.85454816 | -0.256328888 | 0.161354602 | -1.588605995 |
| A230072C01Rik | 65.61270515 | -0.081341848 | 0.145629844 | -0.558552049 |
| Zfp182 | 201.9227118 | 0.187627673 | 0.11797127 | 1.590452256 |
| Spaca5 | 0.708400758 | -0.792885429 | 1.1585064 | -0.68440315 |
| Zfp300 | 69.84368967 | -0.417720057 | 0.163872688 | -2.549052327 |
| Ssxa1 | 0 | NA | NA | NA |
| Gm1848 | 2.174616668 | -0.910061614 | 0.821087342 | -1.108361519 |
| Gm21876 | 0 | NA | NA | NA |
| 4930453H23Rik | 0.062426873 | 0.517959912 | 3.724945049 | 0.139051692 |
| Gm6938 | 0.624139293 | 2.725895508 | 1.284546917 | 2.122067689 |
| Gm5124 | 229.7670475 | -0.014757157 | 0.118804366 | -0.124213925 |
| Gm26593 | 1.567651568 | -0.976304654 | 0.898449635 | -1.086654851 |
| Agtr2 | 19.19641764 | 0.057697378 | 0.305274023 | 0.189001925 |
| Gm22592 | 0 | NA | NA | NA |
| Gm9439 | 0.758270043 | 0.887700857 | 1.080280437 | 0.821731864 |
| Gm14558 | 0 | NA | NA | NA |
| Slc6a14 | 1.053279316 | 1.074971526 | 0.960351425 | 1.119352248 |
| Gm25553 | 0.068097359 | 0.037064902 | 3.724945049 | 0.009950456 |
| Gm28269 | 0 | NA | NA | NA |
| Gm28268 | 0.530320922 | -0.723996738 | 1.413385946 | -0.512242774 |
| Gm24535 | 0 | NA | NA | NA |
| Gm14563 | 0 | NA | NA | NA |
| Gm14561 | 0 | NA | NA | NA |
| Gm6071 | 0 | NA | NA | NA |
| Gm25812 | 0 | NA | NA | NA |
| Gm14562 | 0 | NA | NA | NA |
| Gm15030 | 0 | NA | NA | NA |
| Ppp1r2-ps7 | 0 | NA | NA | NA |
| Gm26131 | 0 | NA | NA | NA |
| Klhl13 | 551.8407492 | 0.35808941 | 0.098134633 | 3.6489606 |
| Gm14523 | 0.533370488 | 0.954901487 | 1.327940462 | 0.719084563 |
| Gm7164 | 0.047948698 | 0.037064902 | 3.724945049 | 0.009950456 |
| Wdr44 | 445.6653234 | -0.033126776 | 0.069987193 | -0.473326262 |
| Gm4789 | 0 | NA | NA | NA |
| Gm4907 | 0.266401621 | -1.557452237 | 2.481346097 | -0.62766425 |
| Gm4985 | 0 | NA | NA | NA |

| | | | | |
|---|---|---|---|---|
| Gm25299 | 0 | NA | NA | NA |
| Gm4732 | 0 | NA | NA | NA |
| Gm9440 | 0 | NA | NA | NA |
| Gm6956 | 0 | NA | NA | NA |
| Gm27192 | 0 | NA | NA | NA |
| Gm5934 | 0.046345709 | -0.443818988 | 3.724945049 | -0.119147795 |
| Gm4297 | 0 | NA | NA | NA |
| Gm27910 | 0 | NA | NA | NA |
| Gm27839 | 0 | NA | NA | NA |
| Gm5935 | 0 | NA | NA | NA |
| Gm2759 | 0 | NA | NA | NA |
| CR938728.1 | 0 | NA | NA | NA |
| Gm2768 | 0 | NA | NA | NA |
| Gm5169 | 0 | NA | NA | NA |
| Gm1989 | 0 | NA | NA | NA |
| Gm1993 | 0 | NA | NA | NA |
| Gm2003 | 0 | NA | NA | NA |
| Gm27892 | 0 | NA | NA | NA |
| Gm7350 | 0 | NA | NA | NA |
| E330010L02Rik | 0 | NA | NA | NA |
| Gm14533 | 0 | NA | NA | NA |
| Gm5168 | 0 | NA | NA | NA |
| Gm2012 | 0 | NA | NA | NA |
| Gm26302 | 0 | NA | NA | NA |
| Gm2030 | 0 | NA | NA | NA |
| Gm2005 | 0 | NA | NA | NA |
| Slx | 0 | NA | NA | NA |
| Gm27341 | 0 | NA | NA | NA |
| Gm14525 | 0 | NA | NA | NA |
| Gm6113 | 0 | NA | NA | NA |
| Gm6121 | 0 | NA | NA | NA |
| Gm7365 | 0 | NA | NA | NA |
| Gm7375 | 0 | NA | NA | NA |
| Gm10230 | 0 | NA | NA | NA |
| BX813306.1 | 0 | NA | NA | NA |
| Gm2092 | 0 | NA | NA | NA |
| CR589930.1 | 0 | NA | NA | NA |
| Gm2101 | 0 | NA | NA | NA |
| Gm7386 | 0 | NA | NA | NA |
| Gm10058 | 0 | NA | NA | NA |
| Gm2117 | 0 | NA | NA | NA |
| Gm7391 | 0 | NA | NA | NA |
| Gm4836 | 0 | NA | NA | NA |
| Gm7398 | 0 | NA | NA | NA |
| Gm10147 | 0 | NA | NA | NA |
| BX813332.1 | 0 | NA | NA | NA |
| Gm2165 | 0 | NA | NA | NA |

| | | | | |
|---|---|---|---|---|
| Gm7408 | 0 | NA | NA | NA |
| Gm10096 | 0 | NA | NA | NA |
| Gm2200 | 0 | NA | NA | NA |
| Gm7415 | 0 | NA | NA | NA |
| Gm26818 | 0 | NA | NA | NA |
| Gm2066 | 0 | NA | NA | NA |
| Gm7421 | 0 | NA | NA | NA |
| Gm15432 | 1.975319427 | -0.581405384 | 0.766239985 | -0.758777139 |
| Gm14527 | 0 | NA | NA | NA |
| BX294196.1 | 0 | NA | NA | NA |
| Gm3657 | 0 | NA | NA | NA |
| Gm3669 | 0 | NA | NA | NA |
| Gm10488 | 0 | NA | NA | NA |
| BX294192.1 | 0 | NA | NA | NA |
| E330016L19Rik | 0 | NA | NA | NA |
| Gm3681 | 0 | NA | NA | NA |
| Gm14632 | 0 | NA | NA | NA |
| Gm7437 | 0 | NA | NA | NA |
| Gm14974 | 0 | NA | NA | NA |
| Gm7438 | 0 | NA | NA | NA |
| Gm10487 | 0 | NA | NA | NA |
| Gm21447 | 0 | NA | NA | NA |
| Spin2f | 0 | NA | NA | NA |
| Gm2784 | 0 | NA | NA | NA |
| Gm15280 | 0 | NA | NA | NA |
| Gm15276 | 0 | NA | NA | NA |
| Gm2777 | 0 | NA | NA | NA |
| Gm21883 | 0 | NA | NA | NA |
| Spin2e | 0 | NA | NA | NA |
| Gm21608 | 0 | NA | NA | NA |
| Gm21637 | 0 | NA | NA | NA |
| Gm21645 | 0 | NA | NA | NA |
| Gm2799 | 0 | NA | NA | NA |
| Gmcl1l | 0 | NA | NA | NA |
| Gm5926 | 0 | NA | NA | NA |
| Gm21951 | 0 | NA | NA | NA |
| Gm21657 | 0 | NA | NA | NA |
| Gm21789 | 0 | NA | NA | NA |
| Gm2825 | 0 | NA | NA | NA |
| Spin2-ps6 | 0 | NA | NA | NA |
| Gm15275 | 0 | NA | NA | NA |
| Gm15274 | 0 | NA | NA | NA |
| Gm2863 | 0 | NA | NA | NA |
| Gm2182 | 0 | NA | NA | NA |
| Gm2854 | 0 | NA | NA | NA |
| Gm21979 | 0 | NA | NA | NA |
| Gm21990 | 0 | NA | NA | NA |

| | | | | |
|---|---|---|---|---|
| Gm2892 | 0 | NA | NA | NA |
| Gm2223 | 89.01610747 | -0.089690725 | 0.14994764 | -0.598146957 |
| Gm2231 | 0 | NA | NA | NA |
| Gm2913 | 0 | NA | NA | NA |
| Gm15273 | 0 | NA | NA | NA |
| Gm15281 | 0 | NA | NA | NA |
| Gm15279 | 0 | NA | NA | NA |
| Gm7023 | 0 | NA | NA | NA |
| Gm2927 | 0 | NA | NA | NA |
| Gm2933 | 0 | NA | NA | NA |
| Gm2964 | 0 | NA | NA | NA |
| Gm21870 | 0 | NA | NA | NA |
| Gm21681 | 0 | NA | NA | NA |
| Spin2g | 0 | NA | NA | NA |
| Gm21699 | 0 | NA | NA | NA |
| Gm2288 | 0 | NA | NA | NA |
| Gm14552 | 0 | NA | NA | NA |
| Gm10486 | 0 | NA | NA | NA |
| Gm2309 | 0 | NA | NA | NA |
| Gm2318 | 0 | NA | NA | NA |
| Gm14553 | 0 | NA | NA | NA |
| Gm14819 | 0 | NA | NA | NA |
| Gm4764 | 0 | NA | NA | NA |
| Gm14554 | 0.066050875 | -0.443818988 | 3.724945049 | -0.119147795 |
| Gm25624 | 0 | NA | NA | NA |
| Dock11 | 371.5272742 | -0.353330447 | 0.08381287 | -4.215706342 |
| Il13ra1 | 318.8934833 | -1.113500972 | 0.185282595 | -6.009744053 |
| Zcchc12 | 687.811863 | 0.384015192 | 0.132531119 | 2.897547335 |
| Gm14560 | 0 | NA | NA | NA |
| Gm6023 | 0.589180756 | -0.621510169 | 1.434493616 | -0.433261021 |
| Gm14559 | 0 | NA | NA | NA |
| Lonrf3 | 281.4852912 | -0.321740243 | 0.133771415 | -2.405149432 |
| Gm25047 | 24.14149978 | -0.282766571 | 0.247137857 | -1.144165341 |
| Gm14568 | 0.274682661 | 1.058421007 | 3.708688692 | 0.285389553 |
| AL450395.1 | 0 | NA | NA | NA |
| Gm6268 | 0.075043205 | 0.517957688 | 3.724945049 | 0.139051095 |
| Gm22364 | 0 | NA | NA | NA |
| Gm14569 | 4.657189074 | 0.4299526 | 0.428373886 | 1.003685365 |
| Pgrmc1 | 2005.442762 | 0.106905674 | 0.078977826 | 1.35361632 |
| Akap17b | 352.8596161 | 0.176439993 | 0.073972145 | 2.385222066 |
| Gm6274 | 10.81761153 | -0.355115534 | 0.382064648 | -0.929464517 |
| Slc25a43 | 7.740311858 | -0.863118005 | 0.473306338 | -1.823592748 |
| Gm23613 | 0.350819806 | -1.523424452 | 1.934227274 | -0.787613985 |
| Slc25a5 | 1945.709052 | -0.165754211 | 0.074479418 | -2.22550357 |
| Gm14549 | 0 | NA | NA | NA |
| 2310010G23Rik | 9.327370753 | 0.886657593 | 0.38183399 | 2.32210232 |
| C330007P06Rik | 639.7924073 | -0.030032672 | 0.075664121 | -0.396920912 |

| | | | | | |
|---|---|---|---|---|---|
| Gm14541 | | 2.652685225 | 1.008289155 | 0.716918872 | 1.406420161 |
| Ube2a | | 329.5173645 | -0.330994853 | 0.100346673 | -3.29851346 |
| Nkrf | | 449.5931371 | -0.040716787 | 0.080803574 | -0.503898337 |
| Gm15008 | | 0.250168298 | 0.960022715 | 2.09286295 | 0.458712652 |
| | 6-Sep | 721.875096 | 0.306058254 | 0.065984168 | 4.638358901 |
| Gm14542 | | 0 | NA | NA | NA |
| Sowahd | | 0.087930927 | -0.443164241 | 3.724898049 | -0.118973522 |
| Gm26099 | | 0 | NA | NA | NA |
| Gm24809 | | 0 | NA | NA | NA |
| Rpl39 | | 722.6800404 | -0.233873066 | 0.126966613 | -1.842004448 |
| Snora69 | | 1.244281449 | -0.148966811 | 0.956312496 | -0.155772106 |
| Upf3b | | 273.0236683 | 0.15861066 | 0.120573867 | 1.315464661 |
| Nkap | | 229.4254487 | -0.0259159 | 0.084503929 | -0.30668278 |
| Akap14 | | 1.404017228 | -0.915807732 | 1.289337917 | -0.710293027 |
| Ndufa1 | | 279.6891087 | 0.110622498 | 0.134900033 | 0.820033141 |
| Rnf113a1 | | 34.43701755 | 0.017530886 | 0.197205364 | 0.088896596 |
| Gm9 | | 0 | NA | NA | NA |
| Rhox1 | | 0 | NA | NA | NA |
| Rhox2a | | 0 | NA | NA | NA |
| Rhox3a | | 0 | NA | NA | NA |
| Rhox4a | | 0 | NA | NA | NA |
| Rhox3a2 | | 0 | NA | NA | NA |
| Rhox4a2 | | 0 | NA | NA | NA |
| Rhox2b | | 0 | NA | NA | NA |
| Rhox3b-ps | | 0 | NA | NA | NA |
| Rhox4b | | 0 | NA | NA | NA |
| Rhox2c | | 0 | NA | NA | NA |
| Rhox3c | | 0 | NA | NA | NA |
| Rhox4c | | 0 | NA | NA | NA |
| Rhox2d | | 0 | NA | NA | NA |
| Gm26494 | | 0 | NA | NA | NA |
| Rhox3d-ps | | 0 | NA | NA | NA |
| Rhox4d | | 0 | NA | NA | NA |
| Rhox2e | | 0 | NA | NA | NA |
| Rhox3e | | 0 | NA | NA | NA |
| Rhox4e | | 0 | NA | NA | NA |
| Rhox2f | | 0 | NA | NA | NA |
| Rhox3f | | 0 | NA | NA | NA |
| Rhox4f | | 0 | NA | NA | NA |
| Rhox3g | | 0 | NA | NA | NA |
| Dppa3-ps | | 0 | NA | NA | NA |
| Gm24493 | | 0 | NA | NA | NA |
| Rhox2g | | 0 | NA | NA | NA |
| Rhox4g | | 0 | NA | NA | NA |
| Rhox3h | | 0 | NA | NA | NA |
| Gm24147 | | 0 | NA | NA | NA |
| Rhox2h | | 0 | NA | NA | NA |

| | | | | |
|---|---|---|---|---|
| RP23-457N24.12 | 0 | NA | NA | NA |
| Rhox2-ps | 0 | NA | NA | NA |
| Rhox11-ps1 | 0 | NA | NA | NA |
| Rhox11-ps2 | 0 | NA | NA | NA |
| Btg1-ps1 | 0.062426873 | 0.517959912 | 3.724945049 | 0.139051692 |
| Btg1-ps2 | 0 | NA | NA | NA |
| Rhox10 | 0 | NA | NA | NA |
| Rhox11 | 0 | NA | NA | NA |
| Rhox11-ps3 | 0 | NA | NA | NA |
| Rhox12 | 0.066050875 | -0.443818988 | 3.724945049 | -0.119147795 |
| Rhox13 | 0 | NA | NA | NA |
| Zfp36l1-ps | 0.671662539 | -0.524558927 | 1.088445114 | -0.481934202 |
| Zbtb33 | 539.0021361 | 0.037182884 | 0.101291972 | 0.367086194 |
| Tmem255a | 487.0062224 | -0.007447065 | 0.112310567 | -0.066307789 |
| Mir3110 | 0 | NA | NA | NA |
| Rhox7-ps1 | 0.075043205 | 0.517957688 | 3.724945049 | 0.139051095 |
| Atp1b4 | 0.134880151 | -1.305960502 | 2.927087508 | -0.446163806 |
| Gm40 | 0 | NA | NA | NA |
| Gm4853 | 2.301314035 | -0.776040451 | 0.694540403 | -1.117343855 |
| Lamp2 | 2710.25613 | -0.526651047 | 0.126298236 | -4.169900257 |
| Gm7598 | 7.758147147 | 0.247296598 | 0.410242259 | 0.60280625 |
| Cul4b | 778.5921849 | -0.009592977 | 0.05985949 | -0.160258246 |
| Mcts1 | 310.2787056 | -0.014555643 | 0.096596872 | -0.150684415 |
| C1galt1c1 | 195.6813856 | -0.560075747 | 0.191162838 | -2.929835915 |
| Ap3s1-ps1 | 43.23259583 | -0.403623483 | 0.166451296 | -2.424874372 |
| Gm14565 | 0.060174537 | 0.517957688 | 3.724945049 | 0.139051095 |
| Gm25642 | 0 | NA | NA | NA |
| Gm14564 | 0 | NA | NA | NA |
| 6030498E09Rik | 0.374199036 | 1.015116294 | 1.481481015 | 0.685203714 |
| Gm6024 | 0 | NA | NA | NA |
| Gm14567 | 9.440869076 | -0.128783503 | 0.364025425 | -0.353776122 |
| Gm16423 | 0 | NA | NA | NA |
| Gm14551 | 0 | NA | NA | NA |
| Gm14566 | 0 | NA | NA | NA |
| Gm26437 | 0 | NA | NA | NA |
| Cypt15 | 0.048949532 | 0.037064902 | 3.724945049 | 0.009950456 |
| Gm14550 | 0 | NA | NA | NA |
| Gm14570 | 0 | NA | NA | NA |
| Gm14571 | 0.04393175 | 0.037062677 | 3.724945049 | 0.009949859 |
| Cypt14 | 0 | NA | NA | NA |
| Gm14643 | 0.048949532 | 0.037064902 | 3.724945049 | 0.009950456 |
| Gm26194 | 0 | NA | NA | NA |
| Gm7189 | 0 | NA | NA | NA |
| Gm16383 | 0 | NA | NA | NA |
| Gm7190 | 0 | NA | NA | NA |
| Gm4986 | 0.708087976 | 0.946310616 | 1.11465395 | 0.848972559 |
| Gm14575 | 0 | NA | NA | NA |

| | | | | |
|---|---|---|---|---|
| Gm14574 | 0 | NA | NA | NA |
| Gm14576 | 0 | NA | NA | NA |
| Gm5385 | 0 | NA | NA | NA |
| Gm14577 | 0 | NA | NA | NA |
| Gria3 | 2075.810427 | 0.968389193 | 0.100910265 | 9.596537995 |
| Gm22413 | 0 | NA | NA | NA |
| Gm14614 | 0 | NA | NA | NA |
| Gm14613 | 0 | NA | NA | NA |
| Thoc2 | 1453.981551 | 0.082283281 | 0.062654671 | 1.31328247 |
| Gm7657 | 0 | NA | NA | NA |
| Gm5135 | 0 | NA | NA | NA |
| AL954355.1 | 0 | NA | NA | NA |
| Xiap | 1927.944407 | 0.106906899 | 0.077642847 | 1.376905971 |
| Gm14615 | 0.068097359 | 0.037064902 | 3.724945049 | 0.009950456 |
| Gm37564 | 0.632215448 | 0.406376886 | 1.213766391 | 0.334806508 |
| Stag2 | 1734.40788 | -0.06463864 | 0.081559349 | -0.792535014 |
| Gm14619 | 0.123841226 | -1.017823543 | 3.719440177 | -0.273649661 |
| RP23-280E8.2 | 0.589937099 | -0.335271699 | 1.24896788 | -0.268439008 |
| Sh2d1a | 7.357494345 | -0.262483769 | 0.357949905 | -0.733297497 |
| Tenm1 | 2192.810765 | 0.475657165 | 0.102586175 | 4.636659502 |
| Gm23705 | 2.363489277 | 0.836141905 | 0.738660031 | 1.131971231 |
| Gm14578 | 7.50102364 | 0.623496162 | 0.556775726 | 1.119833595 |
| Gm24763 | 0 | NA | NA | NA |
| Gm14587 | 8.400665238 | -0.099241601 | 0.577975192 | -0.171705641 |
| Gm14588 | 0 | NA | NA | NA |
| Gm14579 | 0 | NA | NA | NA |
| Gm362 | 0 | NA | NA | NA |
| Gm14580 | 0 | NA | NA | NA |
| Gm14628 | 0 | NA | NA | NA |
| Gm14603 | 0 | NA | NA | NA |
| Dcaf12l2 | 11.20309874 | 0.174251435 | 0.321869591 | 0.541372779 |
| Itpa-ps3 | 0 | NA | NA | NA |
| Gm14602 | 0 | NA | NA | NA |
| Dcaf12l1 | 439.0135136 | 0.329347243 | 0.077666304 | 4.240542251 |
| Gm7700 | 0 | NA | NA | NA |
| Prr32 | 4.737155157 | 2.536708665 | 0.865892508 | 2.929588421 |
| Gm14606 | 0 | NA | NA | NA |
| Gm14605 | 0 | NA | NA | NA |
| Gm5386 | 0 | NA | NA | NA |
| Gm14652 | 0 | NA | NA | NA |
| Gm25078 | 0 | NA | NA | NA |
| Gm14653 | 0 | NA | NA | NA |
| 4930515L19Rik | 0 | NA | NA | NA |
| Actrt1 | 0 | NA | NA | NA |
| Gm14654 | 0.049553047 | -0.443818988 | 3.724945049 | -0.119147795 |
| Gm4987 | 0 | NA | NA | NA |
| Gm29242 | 0 | NA | NA | NA |

| | | | | |
|---|---|---|---|---|
| Gm7722 | 0.112396584 | -0.917692504 | 3.724434845 | -0.246397787 |
| Gm14609 | 0.13648314 | -0.890987945 | 2.936510238 | -0.303417279 |
| Gm14610 | 0.187229393 | -0.884247293 | 2.705056179 | -0.32688685 |
| Gm4908 | 0 | NA | NA | NA |
| Gm14608 | 0.401923326 | 0.016847014 | 1.485800304 | 0.01133868 |
| Gm6480 | 0.088476398 | 0.51795324 | 3.724945049 | 0.139049901 |
| Smarca1 | 656.7356124 | 0.433408587 | 0.087814919 | 4.935477845 |
| Gm14648 | 4.891543041 | 0.264029363 | 0.489966353 | 0.538872438 |
| Ocrl | 1059.362513 | 0.220779784 | 0.049961168 | 4.419027691 |
| Apln | 136.796465 | 0.318320073 | 0.189582159 | 1.679061337 |
| Xpnpep2 | 2.216825317 | 2.508693809 | 0.959471843 | 2.614661209 |
| Sash3 | 86.39783744 | -0.748552892 | 0.199270319 | -3.756469577 |
| Zdhhc9 | 642.249467 | -0.080957622 | 0.073207112 | -1.105870997 |
| Gm24038 | 0 | NA | NA | NA |
| Utp14a | 243.5098185 | -0.225991406 | 0.09740259 | -2.320178626 |
| 9530027J09Rik | 0.791221454 | -0.918427225 | 1.308038548 | -0.702140794 |
| Gm22528 | 0 | NA | NA | NA |
| Gm7212 | 0 | NA | NA | NA |
| Gm22612 | 0 | NA | NA | NA |
| Bcorl1 | 414.200989 | -0.093410808 | 0.111068247 | -0.841021717 |
| Elf4 | 210.3602389 | -0.981753337 | 0.166925103 | -5.881400202 |
| Aifm1 | 270.3649458 | -0.175519435 | 0.093510408 | -1.87700427 |
| Rab33a | 124.0995529 | 0.615046416 | 0.110965817 | 5.542665592 |
| Zfp280c | 445.3017861 | 0.290148818 | 0.101395042 | 2.861568098 |
| Slc25a14 | 322.987207 | 0.299553759 | 0.080319289 | 3.72953697 |
| Gm27828 | 1.991846454 | 2.288539909 | 0.934675353 | 2.44848642 |
| Gpr119 | 0.257827269 | -0.481706122 | 1.836990257 | -0.262225736 |
| Rbmx2 | 38.78282912 | 0.040738951 | 0.229405426 | 0.177584952 |
| Gm595 | 0.555008095 | 0.073470785 | 1.470566846 | 0.049960861 |
| Gm7219 | 0 | NA | NA | NA |
| Enox2 | 126.1449739 | -0.023938187 | 0.134051422 | -0.178574656 |
| Gm15034 | 0 | NA | NA | NA |
| Gm15036 | 0 | NA | NA | NA |
| Gm14696 | 0.319846959 | 0.558350678 | 1.772189022 | 0.315062711 |
| Gm14697 | 0.131474741 | 0.037075328 | 2.980045356 | 0.012441196 |
| Arhgap36 | 111.713745 | -0.225574439 | 0.231536749 | -0.974248968 |
| Gm15037 | 0 | NA | NA | NA |
| Gm14719 | 6.72125269 | 0.557696812 | 0.392850224 | 1.419616889 |
| Olfr1320 | 0 | NA | NA | NA |
| Olfr1321 | 0 | NA | NA | NA |
| Igsf1 | 120.6751455 | 0.6458838 | 0.170729977 | 3.783072019 |
| n-R5s6 | 0 | NA | NA | NA |
| Olfr1322 | 0.062426873 | 0.517959912 | 3.724945049 | 0.139051692 |
| Gm15033 | 0 | NA | NA | NA |
| Olfr1323 | 0.233510017 | -0.920396783 | 2.527200164 | -0.364196234 |
| Gm23385 | 0 | NA | NA | NA |
| Gm7803 | 2.841226851 | -0.852551896 | 0.552885436 | -1.542004617 |

| | | | | |
|---|---|---|---|---|
| Gm14612 | 0.038981395 | -0.443816764 | 3.724945049 | -0.119147198 |
| Gm22288 | 0.45637343 | -0.810516484 | 1.361156612 | -0.595461593 |
| Olfr1324 | 0.046345709 | -0.443818988 | 3.724945049 | -0.119147795 |
| Gm14620 | 0 | NA | NA | NA |
| Firre | 150.6157184 | 0.391121814 | 0.217976458 | 1.794330534 |
| Stk26 | 85.23139706 | 0.767677589 | 0.142958179 | 5.369945195 |
| Frmd7 | 56.57679548 | -0.918231965 | 0.65703181 | -1.397545675 |
| Rap2c | 496.5610328 | -0.453021328 | 0.111134621 | -4.07632945 |
| Gm5387 | 0 | NA | NA | NA |
| Mbnl3 | 8.934417775 | -0.628209321 | 0.3670693 | -1.711418857 |
| Gm7834 | 0 | NA | NA | NA |
| Gm5388 | 0.810280881 | 0.283061162 | 1.066319722 | 0.265456182 |
| Gm23417 | 0 | NA | NA | NA |
| Gm14621 | 0 | NA | NA | NA |
| Hs6st2 | 613.6901847 | 0.231825219 | 0.086609954 | 2.676657899 |
| Gm14581 | 4.970320586 | 0.535778778 | 0.532046753 | 1.007014469 |
| Usp26 | 0 | NA | NA | NA |
| Gm7840 | 0 | NA | NA | NA |
| Gm22539 | 0.072025462 | 0.517959912 | 3.724945049 | 0.139051692 |
| 1700080O16Rik | 0 | NA | NA | NA |
| Gpc4 | 223.3113565 | -0.178128677 | 0.091099125 | -1.955328066 |
| Gpc3 | 54.37477559 | 0.341775017 | 0.168283856 | 2.030943586 |
| Gm14582 | 0 | NA | NA | NA |
| mmu-mir-6384 | 0.085433889 | 0.037058229 | 3.724945049 | 0.009948665 |
| Mir717 | 0 | NA | NA | NA |
| A630012P03Rik | 15.99725564 | -1.599344693 | 0.422018264 | -3.789752316 |
| Mir363 | 0 | NA | NA | NA |
| Mir92-2 | 0 | NA | NA | NA |
| Mir19b-2 | 0 | NA | NA | NA |
| Mir20b | 0 | NA | NA | NA |
| Mir18b | 0 | NA | NA | NA |
| Mir106a | 0 | NA | NA | NA |
| Kis2 | 0.066050875 | -0.443818988 | 3.724945049 | -0.119147795 |
| Gm14586 | 42.19938222 | -0.22620592 | 0.220745128 | -1.024737994 |
| Gm7851 | 0 | NA | NA | NA |
| Ccdc160 | 20.72630267 | 0.085760985 | 0.260366398 | 0.329385765 |
| Gm14607 | 0.060174537 | 0.517957688 | 3.724945049 | 0.139051095 |
| Gm6539 | 0 | NA | NA | NA |
| Rps2-ps13 | 156.9088428 | -0.004376368 | 0.152641885 | -0.028670818 |
| Phf6 | 556.50205 | -0.10069293 | 0.082324207 | -1.223126636 |
| Hprt | 957.8882762 | 0.096427254 | 0.077244284 | 1.24834162 |
| Mir450b | 0 | NA | NA | NA |
| Mir450-1 | 0 | NA | NA | NA |
| Mir450-2 | 0 | NA | NA | NA |
| Mir542 | 0 | NA | NA | NA |
| Gm28730 | 0.097899064 | 0.037062677 | 3.724945049 | 0.009949859 |
| Mir351 | 0 | NA | NA | NA |

| | | | | |
|---|---|---|---|---|
| Mir503 | 0.047948698 | 0.037064902 | 3.724945049 | 0.009950456 |
| Mir322 | 0 | NA | NA | NA |
| C430049B03Rik | 3.09373086 | -0.144139728 | 0.645381997 | -0.223340176 |
| Plac1 | 0 | NA | NA | NA |
| Fam122b | 69.8401722 | 0.118773597 | 0.218940043 | 0.542493712 |
| Fam122c | 0 | NA | NA | NA |
| Gm23718 | 0 | NA | NA | NA |
| Gm14585 | 0 | NA | NA | NA |
| Mospd1 | 150.5973759 | 0.008990575 | 0.110415402 | 0.081425012 |
| Gm14584 | 111.7814324 | 0.122058256 | 0.119737201 | 1.019384574 |
| Etd | 0 | NA | NA | NA |
| Gm14597 | 0.48133134 | -1.378283072 | 1.477497941 | -0.932849402 |
| Cxx1c | 578.0914899 | 0.41233254 | 0.108442561 | 3.802312819 |
| Gm14592 | 0.289161111 | -0.932396406 | 1.743865153 | -0.534672308 |
| Gm14600 | 0 | NA | NA | NA |
| Cxx1a | 186.6998316 | 0.199498106 | 0.096517537 | 2.066962245 |
| Cxx1b | 332.64126 | 0.183704906 | 0.080574338 | 2.279943114 |
| 4933416I08Rik | 0.129625439 | 0.037055631 | 3.686164085 | 0.010052627 |
| AW822252 | 37.35059821 | 0.316752482 | 0.255021729 | 1.242060755 |
| 4930502E18Rik | 2.580172256 | 0.390167532 | 0.708692353 | 0.550545706 |
| 1700013H16Rik | 0.066050875 | -0.443818988 | 3.724945049 | -0.119147795 |
| Gm6025 | 0 | NA | NA | NA |
| Gm14593 | 0 | NA | NA | NA |
| Zfp36l3 | 0.434206664 | 0.977060587 | 1.558130532 | 0.627072358 |
| Xlr | 8.912034157 | 0.284155036 | 0.351186064 | 0.809129589 |
| Gm14599 | 0.044544648 | 0.517957688 | 3.724945049 | 0.139051095 |
| Gm14589 | 0 | NA | NA | NA |
| Gm1862 | 2.065160235 | -0.492355838 | 0.718855346 | -0.684916431 |
| Gm14591 | 0.119587853 | 0.981063299 | 3.723618439 | 0.263470416 |
| Gm15482 | 0.887798776 | 3.295513614 | 1.137872037 | 2.896207577 |
| Gm14601 | 0 | NA | NA | NA |
| Gm14644 | 0 | NA | NA | NA |
| Gm22550 | 0 | NA | NA | NA |
| Gm16428 | 0 | NA | NA | NA |
| Gm7916 | 1.902020358 | 0.217553277 | 0.806651108 | 0.269699347 |
| Gm14598 | 0.066050875 | -0.443818988 | 3.724945049 | -0.119147795 |
| Gm6591 | 0 | NA | NA | NA |
| Gm14596 | 0 | NA | NA | NA |
| Gm14594 | 0 | NA | NA | NA |
| Gm16405 | 0 | NA | NA | NA |
| Gm16430 | 0 | NA | NA | NA |
| Gm14595 | 0 | NA | NA | NA |
| Gm14590 | 0 | NA | NA | NA |
| Gm16431 | 0 | NA | NA | NA |
| Gm7950 | 0 | NA | NA | NA |
| Gm7958 | 0 | NA | NA | NA |
| Gm7974 | 0 | NA | NA | NA |

| | | | | |
|---|---|---|---|---|
| Slxl1 | 0 | NA | NA | NA |
| Gm6660 | 0 | NA | NA | NA |
| Gm14625 | 0 | NA | NA | NA |
| Gm16404 | 0 | NA | NA | NA |
| Gm14622 | 0 | NA | NA | NA |
| Gm6664 | 0 | NA | NA | NA |
| Gm14626 | 0 | NA | NA | NA |
| Gm7927 | 0 | NA | NA | NA |
| Gm14623 | 0 | NA | NA | NA |
| Gm14624 | 0 | NA | NA | NA |
| Gm22103 | 0.038981395 | -0.443816764 | 3.724945049 | -0.119147198 |
| Gm14638 | 0 | NA | NA | NA |
| 3830403N18Rik | 1.591056397 | 0.027466546 | 0.758066552 | 0.036232367 |
| Gm14639 | 1.176771261 | -0.472162385 | 1.095369136 | -0.431053213 |
| Gm773 | 0 | NA | NA | NA |
| Gm14627 | 0 | NA | NA | NA |
| 1600025M17Rik | 0 | NA | NA | NA |
| Zfp449 | 65.59676692 | -0.299016178 | 0.141603598 | -2.111642522 |
| Gm2155 | 3.901369078 | 1.326098029 | 0.557407467 | 2.379046042 |
| Smim10l2a | 178.3604988 | 0.549818285 | 0.114473839 | 4.80300383 |
| Gm14633 | 2.099444623 | 0.83760228 | 0.728350318 | 1.149999196 |
| Gm2174 | 0.09349418 | 0.517955464 | 3.724945049 | 0.139050498 |
| Ddx26b | 661.1565071 | 0.48156778 | 0.133107987 | 3.617872897 |
| Gm10477 | 0 | NA | NA | NA |
| Gm648 | 0.240593192 | -0.443822681 | 2.570672612 | -0.172648465 |
| Mmgt1 | 503.4834972 | -0.045608233 | 0.065208248 | -0.699424289 |
| Mir5116 | 0 | NA | NA | NA |
| Slc9a6 | 1377.918565 | 0.411612838 | 0.058506801 | 7.035299006 |
| Gm26312 | 0 | NA | NA | NA |
| Fhl1 | 841.3587556 | 0.075287408 | 0.060267735 | 1.249215827 |
| Gm23258 | 1.567065625 | -0.885007007 | 1.00178798 | -0.883427457 |
| Mtap7d3 | 3.192641839 | -0.467034612 | 0.593171641 | -0.787351552 |
| Adgrg4 | 0.589632284 | -0.383942921 | 1.34236337 | -0.286020111 |
| Brs3 | 2.988176321 | 0.855301971 | 0.734038478 | 1.165200458 |
| Htatsf1 | 484.4912738 | 0.056492238 | 0.129523154 | 0.436155515 |
| Vgll1 | 0 | NA | NA | NA |
| Gm14718 | 0.17796667 | -0.471322379 | 2.715209716 | -0.173585995 |
| Cd40lg | 0.63650561 | 0.579302219 | 1.215470353 | 0.476607445 |
| Arhgef6 | 229.7557389 | 0.043396373 | 0.086400388 | 0.502270577 |
| Rbmx | 637.3674608 | 0.128338879 | 0.096164543 | 1.334575873 |
| Snord61 | 0.989940444 | 0.539375105 | 1.045351921 | 0.515974663 |
| Gm364 | 0.352737941 | -2.240684112 | 3.685510415 | -0.607971179 |
| Gpr101 | 120.5546104 | 0.14533234 | 0.157272424 | 0.924080247 |
| Gm14630 | 0.097139547 | -0.443816764 | 3.724945049 | -0.119147198 |
| Gm5389 | 0 | NA | NA | NA |
| Gm14629 | 0 | NA | NA | NA |
| Gm8107 | 0.038981395 | -0.443816764 | 3.724945049 | -0.119147198 |

| | | | | |
|---|---|---|---|---|
| Gm8111 | 0 | NA | NA | NA |
| Gm14631 | 0 | NA | NA | NA |
| Gm25023 | 0 | NA | NA | NA |
| Zic3 | 63.76716955 | -0.154208845 | 0.165564311 | -0.931413563 |
| Gm14649 | 0 | NA | NA | NA |
| Gm14647 | 0 | NA | NA | NA |
| Gm25851 | 0 | NA | NA | NA |
| Gm14645 | 0 | NA | NA | NA |
| Gm16436 | 0.046345709 | -0.443818988 | 3.724945049 | -0.119147795 |
| Gm14650 | 0.045882455 | 0.517959912 | 3.724945049 | 0.139051692 |
| Gm7071 | 0 | NA | NA | NA |
| AL713977.1 | 0 | NA | NA | NA |
| 4930550L24Rik | 0.32299001 | -1.05268023 | 1.925134339 | -0.546808713 |
| Fgf13 | 698.6388779 | 0.867739412 | 0.105565719 | 8.219897717 |
| Gm26351 | 1.389516516 | 0.647060368 | 1.174222563 | 0.551054279 |
| Mir504 | 1.452681833 | 0.328709514 | 0.868862173 | 0.378321815 |
| Gm26487 | 0 | NA | NA | NA |
| Gm14890 | 0.104106287 | 0.51795324 | 3.724945049 | 0.139049901 |
| AL670091.1 | 0 | NA | NA | NA |
| AL672036.1 | 0 | NA | NA | NA |
| Gm14887 | 0 | NA | NA | NA |
| F9 | 3.199446176 | 0.870135054 | 0.623764231 | 1.394974272 |
| Gm5637 | 10.86361093 | -1.637858842 | 0.370929053 | -4.41555825 |
| Gm14891 | 0.066050875 | -0.443818988 | 3.724945049 | -0.119147795 |
| Mcf2 | 115.7996196 | 0.221261404 | 0.189610415 | 1.166926425 |
| Atp11c | 543.5317974 | -0.024587623 | 0.082931298 | -0.296481825 |
| Gm22287 | 0 | NA | NA | NA |
| Mir505 | 2.930256519 | -0.52181736 | 0.691371977 | -0.754756306 |
| Gm7073 | 0 | NA | NA | NA |
| Gm715 | 17.31431649 | 0.388479519 | 0.254134669 | 1.528636448 |
| Gm14661 | 0.27928303 | 1.466539314 | 2.059348333 | 0.712137568 |
| Gm14651 | 0 | NA | NA | NA |
| Gm4909 | 0 | NA | NA | NA |
| Gm8163 | 0.296485438 | 0.014505471 | 2.092361246 | 0.006932584 |
| Gm14660 | 0 | NA | NA | NA |
| Sox3 | 13.97423569 | -0.428912794 | 0.300498868 | -1.427335809 |
| Gm14662 | 1.313401919 | 1.093121166 | 0.993645766 | 1.100111531 |
| Gm7076 | 0 | NA | NA | NA |
| Rpl7l1-ps1 | 0 | NA | NA | NA |
| Gm14664 | 0.089814205 | 0.517955464 | 3.724945049 | 0.139050498 |
| Gm14663 | 0.061920613 | -0.443816764 | 3.724945049 | -0.119147198 |
| C230004F18Rik | 1115.002913 | 0.454285838 | 0.162070634 | 2.803011418 |
| Gm24396 | 21.15961452 | 0.797543992 | 0.464601067 | 1.716621092 |
| Cdr1 | 62.47634879 | -0.140071843 | 0.484996515 | -0.288809999 |
| Gm14665 | 0.120974994 | 0.517957688 | 3.724945049 | 0.139051095 |
| Gm14666 | 0.43984171 | 0.473004071 | 1.394899412 | 0.33909547 |
| Gm14821 | 0.106112162 | -0.443821212 | 3.724945049 | -0.119148392 |

| | | | | |
|---|---|---|---|---|
| Gm14655 | 0.060174537 | 0.517957688 | 3.724945049 | 0.139051095 |
| Ldoc1 | 14.5236453 | 0.443308761 | 0.304993598 | 1.453501855 |
| Gm7327 | 29.50965119 | 0.001005294 | 0.306223635 | 0.003282877 |
| Gm6743 | 0.441426087 | -0.944136948 | 1.516959041 | -0.622387898 |
| Gm25412 | 0 | NA | NA | NA |
| Gm8215 | 0 | NA | NA | NA |
| Gm5390 | 0 | NA | NA | NA |
| Gm14656 | 0 | NA | NA | NA |
| Gm16411 | 0.044191549 | 0.037062677 | 3.724945049 | 0.009949859 |
| 4933402E13Rik | 0 | NA | NA | NA |
| Gm14679 | 0 | NA | NA | NA |
| 4931400O07Rik | 0 | NA | NA | NA |
| 1700019B21Rik | 0 | NA | NA | NA |
| Gm14678 | 0 | NA | NA | NA |
| Gm14677 | 0 | NA | NA | NA |
| Gm14680 | 169.5303447 | 0.136570076 | 0.097259486 | 1.404182586 |
| Gm8244 | 0 | NA | NA | NA |
| Gm6754 | 0 | NA | NA | NA |
| Gm14659 | 0 | NA | NA | NA |
| Gm14657 | 0 | NA | NA | NA |
| Gm8260 | 0 | NA | NA | NA |
| Gm6760 | 0 | NA | NA | NA |
| 3830417A13Rik | 0 | NA | NA | NA |
| Slitrk4 | 1198.015236 | 0.003901673 | 0.09885952 | 0.03946684 |
| Gm24598 | 1.413287166 | 0.594523444 | 0.956140728 | 0.621794916 |
| Gm14676 | 0 | NA | NA | NA |
| Gm26111 | 0 | NA | NA | NA |
| Gm14667 | 0 | NA | NA | NA |
| Gm14673 | 0 | NA | NA | NA |
| Ctag2 | 0 | NA | NA | NA |
| Gm14669 | 0 | NA | NA | NA |
| Gm14670 | 0 | NA | NA | NA |
| Gm14668 | 0 | NA | NA | NA |
| Gm14671 | 0 | NA | NA | NA |
| Gm14674 | 0 | NA | NA | NA |
| 4930447F04Rik | 0.792081435 | 0.394092271 | 1.184689046 | 0.332654609 |
| Gm23028 | 0 | NA | NA | NA |
| Slitrk2 | 714.9978112 | 0.13455563 | 0.097144455 | 1.385108702 |
| Gm23000 | 0.301026439 | 0.775636169 | 2.013213347 | 0.385272714 |
| Gm8310 | 0.044544648 | 0.517957688 | 3.724945049 | 0.139051095 |
| Mir743 | 0 | NA | NA | NA |
| Mir743b | 0 | NA | NA | NA |
| Gm14681 | 55.50849165 | -0.156536557 | 0.14029181 | -1.115792556 |
| Mir742 | 0 | NA | NA | NA |
| Mir883a | 0 | NA | NA | NA |
| Mir883b | 0 | NA | NA | NA |
| Mir471 | 0 | NA | NA | NA |

| | | | | |
|---|---|---|---|---|
| Mir741 | 0 | NA | NA | NA |
| Mir463 | 0 | NA | NA | NA |
| Mir880 | 0 | NA | NA | NA |
| Mir878 | 0 | NA | NA | NA |
| Mir881 | 0 | NA | NA | NA |
| Mir871 | 0 | NA | NA | NA |
| Mir470 | 0 | NA | NA | NA |
| mmu-mir-465d | 0 | NA | NA | NA |
| Mir465c-1 | 0 | NA | NA | NA |
| Mir465b-1 | 0 | NA | NA | NA |
| Mir465c-2 | 0 | NA | NA | NA |
| Mir465b-2 | 0 | NA | NA | NA |
| Mir465 | 0 | NA | NA | NA |
| Gm6783 | 0 | NA | NA | NA |
| Gm14682 | 0.089089296 | 0.957155977 | 3.723939561 | 0.257027796 |
| Gm8312 | 0 | NA | NA | NA |
| 1700036O09Rik | 0 | NA | NA | NA |
| Gm26276 | 0 | NA | NA | NA |
| Gm14687 | 0 | NA | NA | NA |
| Gm1140 | 0 | NA | NA | NA |
| Gm14692 | 0 | NA | NA | NA |
| Rpl7a-ps13 | 0 | NA | NA | NA |
| 4933436I01Rik | 0 | NA | NA | NA |
| Mir201 | 0 | NA | NA | NA |
| Mir547 | 0 | NA | NA | NA |
| Gm14688 | 0 | NA | NA | NA |
| mmu-mir-509 | 0 | NA | NA | NA |
| Gm4910 | 0 | NA | NA | NA |
| Gm5391 | 0 | NA | NA | NA |
| Gm6806 | 0.624564003 | -0.342520575 | 1.254719444 | -0.272985787 |
| Gm22332 | 0 | NA | NA | NA |
| Gm14701 | 0 | NA | NA | NA |
| Gm5638 | 0.108772582 | 0.037073798 | 3.724945049 | 0.009952844 |
| Fmr1os | 1.106201151 | 1.804100006 | 1.002809669 | 1.799045284 |
| Gm27979 | 0 | NA | NA | NA |
| Gm27838 | 0 | NA | NA | NA |
| Fmr1 | 1356.393382 | -0.004690577 | 0.076173204 | -0.061577789 |
| Gm22435 | 0 | NA | NA | NA |
| Fmr1nb | 0.234997993 | 0.064105491 | 2.121791361 | 0.03021291 |
| Gm14699 | 0 | NA | NA | NA |
| AC055766.1 | 0 | NA | NA | NA |
| Gm14698 | 5.615076243 | -0.237777453 | 0.427798007 | -0.555817113 |
| Gm6812 | 0 | NA | NA | NA |
| Gm8344 | 0 | NA | NA | NA |
| n-R5s7 | 0 | NA | NA | NA |
| Gm14705 | 0.713101696 | 0.510431507 | 1.406007968 | 0.363035999 |
| Aff2 | 361.4846116 | 0.293124583 | 0.089339535 | 3.281017565 |

| | | | | |
|---|---|---|---|---|
| Gm8172 | 2.258517424 | -0.1297021 | 0.721400368 | -0.179792118 |
| 1700111N16Rik | 0.153668717 | 0.998859713 | 2.839403574 | 0.351785045 |
| 1700020N15Rik | 0.116217011 | 0.517955464 | 3.724945049 | 0.139050498 |
| Ids | 5255.302793 | 0.55746386 | 0.070804392 | 7.873294928 |
| 1110012L19Rik | 65.12692351 | -0.297110708 | 0.187279551 | -1.586455686 |
| 4930567H17Rik | 0 | NA | NA | NA |
| Gm16407 | 0 | NA | NA | NA |
| Gm14704 | 0 | NA | NA | NA |
| Gm6823 | 0.187945415 | -1.274220658 | 2.675122942 | -0.47632228 |
| Gm16441 | 0 | NA | NA | NA |
| Gm14722 | 0 | NA | NA | NA |
| Gm14721 | 0 | NA | NA | NA |
| BC023829 | 354.2798382 | -0.114138991 | 0.096398748 | -1.184029809 |
| Gm14727 | 0 | NA | NA | NA |
| Gm14729 | 0 | NA | NA | NA |
| Gm6038 | 0 | NA | NA | NA |
| Gm8401 | 0.88643884 | -0.754484427 | 1.011528383 | -0.745885572 |
| Gm14726 | 0 | NA | NA | NA |
| Gm14724 | 0 | NA | NA | NA |
| Gm14725 | 0 | NA | NA | NA |
| Gm14728 | 0.144050924 | 1.05357957 | 3.721447096 | 0.283110183 |
| Gm14732 | 0 | NA | NA | NA |
| Gm14731 | 0 | NA | NA | NA |
| Rplp1-ps1 | 0.046345709 | -0.443818988 | 3.724945049 | -0.119147795 |
| Gm22785 | 0 | NA | NA | NA |
| Gm8410 | 0.503546025 | 2.095164664 | 1.505157367 | 1.39199044 |
| Gm6062 | 0.075043205 | 0.517957688 | 3.724945049 | 0.139051095 |
| Mamld1 | 657.3894617 | -0.077050851 | 0.095733041 | -0.804851183 |
| n-R5s8 | 8.139267235 | 0.782513586 | 0.40949889 | 1.910905268 |
| Gm14730 | 1.043158539 | 0.206613557 | 1.158370637 | 0.178365672 |
| Mtm1 | 330.8783525 | -0.04707277 | 0.136557339 | -0.344710657 |
| Mtmr1 | 659.908289 | 0.176526731 | 0.07515312 | 2.348894245 |
| Cd99l2 | 1057.193581 | -0.008558147 | 0.056245592 | -0.152156764 |
| Gm16189 | 0 | NA | NA | NA |
| Gm8421 | 0 | NA | NA | NA |
| Gm23945 | 0 | NA | NA | NA |
| Hmgb3 | 158.3977876 | 0.487631599 | 0.173768643 | 2.806211702 |
| Gpr50 | 5.432542524 | -0.721995434 | 0.471015961 | -1.532847068 |
| Gm14723 | 0 | NA | NA | NA |
| Rpl30-ps10 | 27.35008078 | -0.0549308 | 0.214227908 | -0.256412902 |
| Vma21 | 809.8468664 | -0.01901948 | 0.101850775 | -0.186738688 |
| Gm1141 | 6.002430894 | -1.126678842 | 0.485622557 | -2.320071065 |
| Prrg3 | 424.0651492 | 0.234546058 | 0.110046296 | 2.131339874 |
| Fate1 | 0.616736355 | -0.807845478 | 1.299611071 | -0.621605568 |
| Cnga2 | 3.751482543 | -0.777432175 | 0.799832691 | -0.971993497 |
| Magea4 | 0 | NA | NA | NA |
| Gabre | 35.49517603 | -0.42592929 | 0.254704954 | -1.6722458 |

| | | | | |
|---|---|---|---|---|
| Mir224 | 0.097139547 | -0.443816764 | 3.724945049 | -0.119147198 |
| Mir452 | 0.132574648 | 0.037054914 | 3.724945049 | 0.009947775 |
| Magea10 | 0 | NA | NA | NA |
| Gabra3 | 880.1056003 | 0.761897974 | 0.108712326 | 7.00838629 |
| Gm14735 | 0 | NA | NA | NA |
| Mir767 | 0.267172041 | 0.037072061 | 2.044803494 | 0.018129889 |
| Mir105 | 0.580314088 | 1.44700775 | 1.556396901 | 0.929716417 |
| Gm23461 | 0 | NA | NA | NA |
| Gm14736 | 0 | NA | NA | NA |
| Gabrq | 110.4557477 | -0.077084904 | 0.179138402 | -0.430309207 |
| Magea9-ps | 0 | NA | NA | NA |
| Cetn2 | 224.1480122 | 0.051615712 | 0.089402872 | 0.577338418 |
| Nsdhl | 258.6958972 | -0.301839937 | 0.095317301 | -3.166685733 |
| Gm14684 | 0.411174732 | 0.517972505 | 1.736230718 | 0.29833161 |
| Zfp185 | 38.8421771 | -0.185534533 | 0.191937246 | -0.966641635 |
| Mir2137 | 0.131193343 | 0.517960218 | 2.980368697 | 0.173790652 |
| Pnma5 | 2.351169846 | 0.733512226 | 0.753121538 | 0.97396262 |
| Pnma3 | 162.7565536 | 0.8239109 | 0.111978549 | 7.357756502 |
| Xlr4a | 1.613791847 | -1.631676286 | 1.066868642 | -1.529406922 |
| Xlr3a | 10.70218797 | -0.383440773 | 0.347242734 | -1.104244194 |
| Gm20455 | 0 | NA | NA | NA |
| Xlr5a | 0 | NA | NA | NA |
| Gm14685 | 1.153352726 | -0.675394696 | 0.957364087 | -0.705473189 |
| DXBay18 | 1.088618379 | -1.479585727 | 1.034655076 | -1.430027999 |
| Xlr5b | 0.044544648 | 0.517957688 | 3.724945049 | 0.139051095 |
| Spin2d | 0 | NA | NA | NA |
| Xlr3b | 10.24179207 | -1.090006631 | 0.861258897 | -1.265596948 |
| Xlr4b | 4.01574399 | -0.752611436 | 0.570043015 | -1.320271306 |
| F8a | 148.8077908 | -0.051716427 | 0.147980015 | -0.349482513 |
| Xlr4c | 1.800570423 | -0.217897038 | 0.761933976 | -0.285978897 |
| Xlr3c | 1.249216686 | -1.906278141 | 1.035823102 | -1.840351057 |
| Xlr5c | 0 | NA | NA | NA |
| Gm5639 | 0.465319851 | 2.394031314 | 1.334722082 | 1.793655283 |
| Gm8501 | 0 | NA | NA | NA |
| Zfp275 | 599.4099204 | -0.174302875 | 0.073008747 | -2.387424543 |
| Gm18336 | 1.728761539 | 0.263177525 | 0.782738067 | 0.336226812 |
| Gm26726 | 0.597994088 | 1.888840393 | 1.468337417 | 1.286380344 |
| Gm37741 | 0.240513512 | 1.834846136 | 2.473954264 | 0.741665342 |
| Zfp92 | 61.40182776 | -0.473681345 | 0.166014618 | -2.853250816 |
| Trex2 | 0.567095806 | -0.728574863 | 1.27787571 | -0.570145326 |
| Haus7 | 77.87449152 | -0.130151792 | 0.119875048 | -1.085728799 |
| Bgn | 275.5973356 | -0.77385174 | 0.180380712 | -4.290102491 |
| Atp2b3 | 1878.820538 | 0.605699666 | 0.093244911 | 6.495793256 |
| Dusp9 | 7.63352917 | -0.568329744 | 0.405508057 | -1.401525158 |
| Pnck | 377.2175805 | 0.688018684 | 0.078524351 | 8.761851251 |
| Slc6a8 | 1645.774035 | -0.638870582 | 0.064236678 | -9.945573125 |
| Bcap31 | 456.597487 | -0.293176936 | 0.106521232 | -2.752286388 |

| | | | | |
|---|---|---|---|---|
| Abcd1 | 224.9704374 | -0.45660137 | 0.117024238 | -3.901767515 |
| Gm25226 | 0 | NA | NA | NA |
| Plxnb3 | 370.4438468 | -0.356331796 | 0.12080641 | -2.94961002 |
| Srpk3 | 30.04135515 | -0.067974219 | 0.250990755 | -0.270823596 |
| Idh3g | 816.0853577 | 0.279949368 | 0.050626005 | 5.529754269 |
| Ssr4 | 382.7763301 | -0.140537857 | 0.100554708 | -1.397625834 |
| Pdzd4 | 1049.261182 | 0.383678549 | 0.084868811 | 4.520842738 |
| Gm14817 | 2.398130883 | 1.510548873 | 0.818421733 | 1.845685192 |
| Gm23615 | 0.046345709 | -0.443818988 | 3.724945049 | -0.119147795 |
| L1cam | 1414.37122 | 0.546791518 | 0.067264653 | 8.128957732 |
| Arhgap4 | 118.1046268 | -0.134024242 | 0.171280865 | -0.78248228 |
| Avpr2 | 0.170941961 | -0.411789146 | 2.735055455 | -0.150559706 |
| Naa10 | 112.0862865 | 0.207972776 | 0.121048733 | 1.71809131 |
| Renbp | 87.59544553 | -0.929589146 | 0.173733668 | -5.350656319 |
| Hcfc1 | 2331.747261 | 0.125574322 | 0.054452667 | 2.306118863 |
| Gm8545 | 0.428564514 | -1.456512781 | 1.490720249 | -0.97705306 |
| Irak1 | 580.922968 | -0.195761739 | 0.067772526 | -2.888511761 |
| Mir5132 | 0 | NA | NA | NA |
| Mir718 | 0.042716945 | 0.037062677 | 3.724945049 | 0.009949859 |
| Mecp2 | 3093.872843 | 0.29276924 | 0.050903017 | 5.75151054 |
| Opn1mw | 3.912128538 | 0.043196904 | 0.572814381 | 0.075411696 |
| Gm6758 | 1.061458158 | -0.316563374 | 0.938418127 | -0.337337233 |
| Tex28 | 0.045882455 | 0.517959912 | 3.724945049 | 0.139051692 |
| Rpl3-ps2 | 16.62506886 | -0.004276032 | 0.258193703 | -0.016561333 |
| Tktl1 | 1.027298454 | 1.584688141 | 1.023900575 | 1.547697285 |
| Flna | 2140.441872 | -1.085207594 | 0.140085405 | -7.74675703 |
| Emd | 326.2260813 | -0.363830841 | 0.115980781 | -3.136992516 |
| Gm14686 | 0 | NA | NA | NA |
| Rpl10 | 1410.009698 | -0.378988672 | 0.096188919 | -3.940045036 |
| Snora70 | 0 | NA | NA | NA |
| Dnase1l1 | 38.9238987 | -1.379873013 | 0.237674644 | -5.805722442 |
| Mir7091 | 0 | NA | NA | NA |
| Gm14846 | 0 | NA | NA | NA |
| Taz | 323.2193373 | 0.16623053 | 0.12943693 | 1.284258901 |
| Atp6ap1 | 2828.059804 | 0.049799737 | 0.055921661 | 0.890526795 |
| Gdi1 | 4514.466658 | 0.287224315 | 0.045176474 | 6.357829468 |
| Fam50a | 152.1449703 | -0.265458942 | 0.105871675 | -2.50736508 |
| mmu-mir-7092 | 0.044191549 | 0.037062677 | 3.724945049 | 0.009949859 |
| Plxna3 | 360.6800962 | 0.044475928 | 0.094123611 | 0.472526795 |
| Lage3 | 80.66907545 | -0.397539013 | 0.151877767 | -2.617493136 |
| Ubl4 | 481.5895494 | -0.135340715 | 0.079729188 | -1.697505255 |
| Slc10a3 | 39.38722088 | -0.892596686 | 0.222209093 | -4.016922416 |
| Fam3a | 178.954657 | -0.099040962 | 0.132251838 | -0.748881556 |
| Ikbkg | 603.2886534 | -0.090814772 | 0.079377285 | -1.144090166 |
| G6pdx | 686.6621862 | -0.942530004 | 0.149113946 | -6.320870921 |
| Gm6880 | 0.066050875 | -0.443818988 | 3.724945049 | -0.119147795 |
| Gm6881 | 0 | NA | NA | NA |

| | | | | |
|---|---|---|---|---|
| Gm15361 | 20.55718455 | -0.144964902 | 0.268718916 | -0.539466683 |
| Gm15362 | 0 | NA | NA | NA |
| Gm15364 | 0 | NA | NA | NA |
| Gm26125 | 0 | NA | NA | NA |
| Gm4989 | 0 | NA | NA | NA |
| Olfr1326-ps1 | 0 | NA | NA | NA |
| Olfr1325 | 0 | NA | NA | NA |
| Gm26417 | 0 | NA | NA | NA |
| Gm5640 | 1.316030826 | 0.961841678 | 0.907684935 | 1.059664694 |
| Xlr5d-ps | 0.066050875 | -0.443818988 | 3.724945049 | -0.119147795 |
| Spin2-ps2 | 0 | NA | NA | NA |
| Xlr3d-ps | 0.16282664 | -0.51179507 | 3.411119175 | -0.150037288 |
| Gm15365 | 0.152853976 | 1.404519388 | 2.820813914 | 0.497912812 |
| Gm22351 | 0 | NA | NA | NA |
| Xlr4d-ps | 0 | NA | NA | NA |
| Gm6890 | 0 | NA | NA | NA |
| Gm14755 | 0 | NA | NA | NA |
| Olfr1327-ps1 | 0 | NA | NA | NA |
| Gm14758 | 0 | NA | NA | NA |
| Gm14695 | 0 | NA | NA | NA |
| Gm8617 | 0 | NA | NA | NA |
| Spin2-ps3 | 0 | NA | NA | NA |
| Xlr5e-ps | 0 | NA | NA | NA |
| Gm5936 | 1.015210553 | -0.524398592 | 0.942975978 | -0.556110234 |
| Gm14694 | 0 | NA | NA | NA |
| Gm6897 | 0 | NA | NA | NA |
| Gm14756 | 0 | NA | NA | NA |
| Spin2-ps4 | 0 | NA | NA | NA |
| Xlr3e-ps | 0.045882455 | 0.517959912 | 3.724945049 | 0.139051692 |
| Xlr4e-ps | 0.114443069 | -0.443821212 | 3.724945049 | -0.119148392 |
| Gm15363 | 0 | NA | NA | NA |
| Gab3 | 34.85498717 | 0.152245912 | 0.198615689 | 0.766535176 |
| Gm25629 | 0.361882898 | 1.243879106 | 1.850340856 | 0.672243226 |
| Gm15367 | 0 | NA | NA | NA |
| Dkc1 | 249.3852457 | -0.321771893 | 0.090155114 | -3.569091976 |
| Gm25520 | 1.152914642 | 0.159695105 | 1.076378222 | 0.148363374 |
| Mpp1 | 513.8380435 | -0.239571731 | 0.09589009 | -2.498399268 |
| Gm15378 | 0.117046891 | 0.037062677 | 3.724945049 | 0.009949859 |
| Fam220-ps | 0.313388536 | 1.116742708 | 2.016859426 | 0.55370379 |
| Smim9 | 2.883149428 | -0.322996455 | 0.590830517 | -0.546682077 |
| Gm8503 | 0.097139547 | -0.443816764 | 3.724945049 | -0.119147198 |
| F8 | 100.3533705 | 0.203405058 | 0.155052533 | 1.311846083 |
| Gm6039 | 5.682746135 | -0.020790099 | 0.412962185 | -0.050343833 |
| Gm8522 | 1.810659408 | 0.679070682 | 0.915064168 | 0.742101708 |
| Fundc2 | 533.8816821 | 0.286375498 | 0.105707864 | 2.709121974 |
| Mtcp1 | 186.8893314 | 0.088689488 | 0.133025185 | 0.666712008 |
| Brcc3 | 516.7646188 | 0.285452176 | 0.074924841 | 3.809846926 |

| | | | | |
|---|---|---|---|---|
| Gm15379 | 11.17773374 | -0.225594692 | 0.326179855 | -0.691626685 |
| Gm15385 | 2.75264805 | 0.314917486 | 0.703229568 | 0.447816047 |
| Gm8666 | 2.699454663 | -0.696804872 | 0.564807418 | -1.23370347 |
| Vbp1 | 703.5174424 | -0.012452766 | 0.094615991 | -0.131613758 |
| Gm15384 | 1.213939271 | -0.354329961 | 1.085765295 | -0.326341211 |
| Rab39b | 757.2087334 | -0.106210542 | 0.089104179 | -1.1919816 |
| Gm15063 | 0.98952171 | -0.328077592 | 1.031916154 | -0.317930474 |
| Gm25421 | 0 | NA | NA | NA |
| Gm15065 | 1.438276361 | -2.260601438 | 0.977435014 | -2.312789501 |
| 4933407K13Rik | 98.06458889 | 0.587743269 | 0.256516589 | 2.291248579 |
| Gm7153 | 0.141331096 | -0.443820795 | 3.561840904 | -0.124604329 |
| Pls3 | 1303.00074 | 0.099372867 | 0.062391136 | 1.592740151 |
| Gm23320 | 0 | NA | NA | NA |
| Oat-rs1 | 0 | NA | NA | NA |
| Gm14715 | 0 | NA | NA | NA |
| Gm7155 | 0 | NA | NA | NA |
| Gm8692 | 1.34351349 | -0.265013321 | 0.878273622 | -0.30174346 |
| Gm14710 | 0 | NA | NA | NA |
| Gm5397 | 0 | NA | NA | NA |
| Gm14711 | 0.062426873 | 0.517959912 | 3.724945049 | 0.139051692 |
| Gm14707 | 0 | NA | NA | NA |
| Gm14717 | 0 | NA | NA | NA |
| Cldn34b3 | 0.209261529 | 0.034789973 | 2.615696522 | 0.013300462 |
| Gm14712 | 0 | NA | NA | NA |
| Magea7-ps | 0.046345709 | -0.443818988 | 3.724945049 | -0.119147795 |
| Gm14709 | 0 | NA | NA | NA |
| Cldn34b4 | 0 | NA | NA | NA |
| Gm5793 | 0.038981395 | -0.443816764 | 3.724945049 | -0.119147198 |
| Cldn34d | 0.044191549 | 0.037062677 | 3.724945049 | 0.009949859 |
| Gm14716 | 0 | NA | NA | NA |
| Gm14713 | 0 | NA | NA | NA |
| Gm26041 | 0.046345709 | -0.443818988 | 3.724945049 | -0.119147795 |
| Gm23280 | 0 | NA | NA | NA |
| Gm14706 | 0.190396633 | 0.541112798 | 3.26794622 | 0.165581917 |
| Gm14714 | 0 | NA | NA | NA |
| Gm5398 | 2.350384682 | 0.828727139 | 0.814329244 | 1.017680681 |
| Gm5757 | 0 | NA | NA | NA |
| Gm5937 | 0.810948363 | 0.287711332 | 1.117849313 | 0.257379352 |
| Tbl1x | 1255.905752 | 0.032686998 | 0.053105417 | 0.615511556 |
| Gm23121 | 6.016926621 | 0.17869175 | 0.419968203 | 0.425488761 |
| Cldn34-ps | 0 | NA | NA | NA |
| Gm14746 | 0.224947322 | 0.929010107 | 2.177449769 | 0.426650534 |
| Gm14748 | 0 | NA | NA | NA |
| Gm8722 | 2.37024928 | -1.565303848 | 0.77054351 | -2.031428242 |
| Prkx | 384.5278709 | -0.832706032 | 0.138493137 | -6.012615845 |
| Gm14742 | 0.093141081 | 0.037060453 | 3.724945049 | 0.009949262 |
| Pbsn | 0 | NA | NA | NA |

| | | | | |
|---|---|---|---|---|
| Gm14744 | 0.130840244 | 0.037064923 | 2.960883856 | 0.012518195 |
| 5430402E10Rik | 0.090665643 | 0.037060453 | 3.724945049 | 0.009949262 |
| Gm14750 | 0 | NA | NA | NA |
| Gm14753 | 0 | NA | NA | NA |
| Gm16458 | 0 | NA | NA | NA |
| Obp1a | 0.143846094 | 0.0370613 | 3.724945049 | 0.009949489 |
| Gm5938 | 0 | NA | NA | NA |
| Obp1b | 0.091880448 | 0.037060453 | 3.724945049 | 0.009949262 |
| Gm14743 | 0.116046057 | 0.037062677 | 3.724945049 | 0.009949859 |
| Gm8749 | 0 | NA | NA | NA |
| 4930480E11Rik | 0.066050875 | -0.443818988 | 3.724945049 | -0.119147795 |
| Gm14749 | 0 | NA | NA | NA |
| Gm8754 | 0 | NA | NA | NA |
| Gm14752 | 0 | NA | NA | NA |
| Gm14745 | 0 | NA | NA | NA |
| Prrg1 | 108.3946446 | 0.055533602 | 0.164311096 | 0.337978407 |
| Gm22843 | 1.022987396 | 2.424394434 | 1.135467908 | 2.135150114 |
| Gm8757 | 0 | NA | NA | NA |
| Gm8761 | 0 | NA | NA | NA |
| Gm14747 | 0 | NA | NA | NA |
| Fam47c | 0 | NA | NA | NA |
| Gm8770 | 0.097139547 | -0.443816764 | 3.724945049 | -0.119147198 |
| Gm5939 | 0.62385056 | 0.456584847 | 1.18595099 | 0.384994702 |
| Gm23556 | 0 | NA | NA | NA |
| Gm14751 | 0 | NA | NA | NA |
| Rps24-ps3 | 11.66239001 | 0.340271465 | 0.285680643 | 1.191090377 |
| Gm7173 | 57.33673308 | 0.016311288 | 0.163263661 | 0.099907648 |
| Mageb16 | 1.972874096 | 0.164411148 | 0.727961641 | 0.225851389 |
| Gm4772 | 0.201649101 | -0.443821113 | 2.217615465 | -0.200134387 |
| Gm4773 | 0.396881693 | 1.473192524 | 1.543636773 | 0.954364751 |
| Gm14760 | 16.61937256 | 0.15978829 | 0.252737091 | 0.632231263 |
| Gm6952 | 0 | NA | NA | NA |
| Gm14768 | 0 | NA | NA | NA |
| Gm5940 | 3.69570757 | -0.307059273 | 0.54625702 | -0.562115016 |
| Gm8806 | 5.085933841 | -0.616928513 | 0.467553775 | -1.319481408 |
| Gm6954 | 0 | NA | NA | NA |
| Gm14733 | 0 | NA | NA | NA |
| Gm26775 | 0.462969528 | 0.870386507 | 1.532386621 | 0.567994066 |
| Tmem47 | 1344.966627 | -0.237017291 | 0.128622177 | -1.842740469 |
| Gm14754 | 0 | NA | NA | NA |
| 4930595M18Rik | 0.072025462 | 0.517959912 | 3.724945049 | 0.139051692 |
| Gm8822 | 164.0860792 | -0.225568556 | 0.129892878 | -1.736573694 |
| Gm26174 | 0 | NA | NA | NA |
| Gm22686 | 0 | NA | NA | NA |
| Rpl10-ps4 | 0 | NA | NA | NA |
| Gm24706 | 0 | NA | NA | NA |
| Gm14740 | 12.83618259 | 1.928385869 | 0.826457083 | 2.333316403 |

| | | | | |
|---|---|---|---|---|
| Gm14739 | 0 | NA | NA | NA |
| Gm24470 | 0 | NA | NA | NA |
| Gm14741 | 0 | NA | NA | NA |
| Gm23671 | 0 | NA | NA | NA |
| Dmd | 2142.472415 | 0.357693466 | 0.100888055 | 3.545449126 |
| n-R5s9 | 0.908782567 | 0.793150022 | 1.029557294 | 0.770379683 |
| Tsga8 | 0 | NA | NA | NA |
| Gm22580 | 0.283050572 | 0.049243668 | 1.643038255 | 0.029971103 |
| Gm14769 | 8.178611007 | 1.99107086 | 0.542320407 | 3.671392104 |
| Gm14770 | 2.258554499 | 0.600855005 | 0.702386072 | 0.855448348 |
| Gm14771 | 0 | NA | NA | NA |
| Gm16463 | 0.470240119 | -1.203872136 | 1.505520317 | -0.799638585 |
| 1600014K23Rik | 0 | NA | NA | NA |
| Gm14765 | 0 | NA | NA | NA |
| Tab3 | 521.751977 | 0.028635266 | 0.090754635 | 0.315524004 |
| AL645586.1 | 2.461055123 | 1.057870685 | 0.810191715 | 1.30570415 |
| Gyk | 504.0204661 | -0.119161316 | 0.057209862 | -2.082880667 |
| Gm14767 | 0.044544648 | 0.517957688 | 3.724945049 | 0.139051095 |
| Gm14764 | 0 | NA | NA | NA |
| Gm14766 | 0.576946983 | 0.35977538 | 1.193794753 | 0.301371219 |
| Gm6964 | 0 | NA | NA | NA |
| Gm14761 | 0.106520246 | 0.037071574 | 3.724945049 | 0.009952247 |
| Gm14762 | 0.702946434 | 0.282966309 | 1.393196994 | 0.203105742 |
| 5430427O19Rik | 37.85834025 | -0.978456693 | 0.221726276 | -4.412903657 |
| Gm14772 | 0 | NA | NA | NA |
| Samt3 | 0.047948698 | 0.037064902 | 3.724945049 | 0.009950456 |
| Nr0b1 | 0.363180662 | 0.137713385 | 1.490795694 | 0.09237576 |
| Mageb4 | 0.097139547 | -0.443816764 | 3.724945049 | -0.119147198 |
| Gm8839 | 0 | NA | NA | NA |
| Mageb10-ps | 0 | NA | NA | NA |
| Gm8844 | 0.25781401 | 0.492010171 | 1.826039546 | 0.269441137 |
| Mageb8-ps | 0 | NA | NA | NA |
| Gm41 | 0 | NA | NA | NA |
| Mageb6-ps | 0 | NA | NA | NA |
| Mageb7-ps | 0 | NA | NA | NA |
| n-R5s10 | 0 | NA | NA | NA |
| Il1rapl1 | 848.3376803 | 0.701861794 | 0.112103365 | 6.260844997 |
| Gm14775 | 0 | NA | NA | NA |
| Gm8855 | 54.40271974 | 0.31106915 | 0.261767603 | 1.188340902 |
| Gm14776 | 77.87756624 | 0.841854869 | 0.303994845 | 2.769306398 |
| Gm24812 | 2.806226544 | 0.80562171 | 0.785280634 | 1.025902939 |
| Gm14774 | 28.40504812 | 0.621921889 | 0.307852828 | 2.020192225 |
| Gm25006 | 0 | NA | NA | NA |
| Gm8858 | 0 | NA | NA | NA |
| Gm14793 | 0 | NA | NA | NA |
| Gm8864 | 0.154920219 | 1.020650266 | 2.83493605 | 0.360025852 |
| Gm14795 | 0 | NA | NA | NA |

| | | | | |
|---|---|---|---|---|
| Gm14792 | 0 | NA | NA | NA |
| Smek3-ps | 0 | NA | NA | NA |
| Gm14794 | 0.838406277 | -1.022464328 | 1.188566853 | -0.860249742 |
| Gm8876 | 0 | NA | NA | NA |
| Gm27000 | 21.60956527 | 0.819820766 | 0.4131412 | 1.984359744 |
| Mir1906-2 | 0 | NA | NA | NA |
| Gm27335 | 0 | NA | NA | NA |
| Gm5759 | 0 | NA | NA | NA |
| Gm8880 | 0.146451277 | -0.443820591 | 2.882358847 | -0.153978257 |
| Gm16412 | 0.045882455 | 0.517959912 | 3.724945049 | 0.139051692 |
| Gm14791 | 0 | NA | NA | NA |
| Gm14796 | 0 | NA | NA | NA |
| Gm14797 | 0 | NA | NA | NA |
| Gm649 | 0 | NA | NA | NA |
| Gm14805 | 0 | NA | NA | NA |
| Pet2 | 0 | NA | NA | NA |
| Gm14804 | 0 | NA | NA | NA |
| Gm6973 | 0 | NA | NA | NA |
| 4932429P05Rik | 0 | NA | NA | NA |
| Gm7108 | 0.816803644 | 2.778453521 | 1.193258725 | 2.328458584 |
| Gm24954 | 0 | NA | NA | NA |
| Nanog-ps1 | 0 | NA | NA | NA |
| 4930415L06Rik | 0 | NA | NA | NA |
| Gm6027 | 0 | NA | NA | NA |
| Gm4746 | 0 | NA | NA | NA |
| Gm6977 | 10.13847199 | 0.104134022 | 0.386661901 | 0.269315446 |
| Gm44 | 0 | NA | NA | NA |
| Gm14773 | 0 | NA | NA | NA |
| Gm24835 | 0 | NA | NA | NA |
| Gm14786 | 0.042716945 | 0.037062677 | 3.724945049 | 0.009949859 |
| Gm8908 | 0 | NA | NA | NA |
| Gm14788 | 0 | NA | NA | NA |
| Mageb2 | 0.045882455 | 0.517959912 | 3.724945049 | 0.139051692 |
| Gm14783 | 0.291614092 | 0.036519762 | 1.778722282 | 0.020531458 |
| Gm5072 | 0 | NA | NA | NA |
| Gm8914 | 0 | NA | NA | NA |
| Gm14787 | 0 | NA | NA | NA |
| Gm8916 | 0 | NA | NA | NA |
| Gm14785 | 0 | NA | NA | NA |
| 1700084M14Rik | 0 | NA | NA | NA |
| Mageb17-ps | 0 | NA | NA | NA |
| Gm14781 | 0 | NA | NA | NA |
| Mageb5 | 0 | NA | NA | NA |
| Gm14782 | 0 | NA | NA | NA |
| Gm7113 | 0 | NA | NA | NA |
| Gm16465 | 0.297592784 | 0.946830872 | 1.608072887 | 0.58879848 |
| Gm5071 | 0 | NA | NA | NA |

| | | | | |
|---|---|---|---|---|
| Gm14784 | 0 | NA | NA | NA |
| Mageb1 | 0.062426873 | 0.517959912 | 3.724945049 | 0.139051692 |
| Gm4911 | 0 | NA | NA | NA |
| Gm4912 | 0.306841164 | -1.303998395 | 1.969528646 | -0.662086534 |
| Mageb18 | 0.274198742 | 0.870356197 | 1.808329727 | 0.481303926 |
| Gm14777 | 0 | NA | NA | NA |
| Gm5941 | 0 | NA | NA | NA |
| Gm6982 | 0 | NA | NA | NA |
| Gm25332 | 0 | NA | NA | NA |
| Gm24973 | 0.33162218 | 0.037076879 | 1.797740957 | 0.02062415 |
| 1700003E24Rik | 0 | NA | NA | NA |
| Gm24123 | 0 | NA | NA | NA |
| BC061195 | 0 | NA | NA | NA |
| Gm23454 | 0 | NA | NA | NA |
| Arx | 155.2305208 | 0.150141121 | 0.174078262 | 0.862492075 |
| Pola1 | 222.1831865 | -0.070648723 | 0.12621939 | -0.559729551 |
| Gm14780 | 0 | NA | NA | NA |
| Gm23557 | 1.675207292 | -0.871226072 | 0.921399675 | -0.94554632 |
| Gm6985 | 0 | NA | NA | NA |
| Pcyt1b | 616.1736765 | -0.039404269 | 0.068677915 | -0.573754589 |
| Pdk3 | 429.9123672 | 0.081240857 | 0.066323792 | 1.224912719 |
| Supt20-ps | 0 | NA | NA | NA |
| Gm15164 | 0 | NA | NA | NA |
| AU015836 | 0.093831153 | 0.517957688 | 3.724945049 | 0.139051095 |
| Gm14790 | 0 | NA | NA | NA |
| Gm15165 | 0.044191549 | 0.037062677 | 3.724945049 | 0.009949859 |
| Gm6987 | 0 | NA | NA | NA |
| Gm14798 | 0.269031123 | 0.976620945 | 1.798199922 | 0.543110326 |
| Zfx | 737.8114658 | -0.131375793 | 0.093490498 | -1.405231493 |
| Eif2s3x | 977.9631196 | -0.141984505 | 0.073857995 | -1.922398589 |
| Klhl15 | 561.3122414 | 0.110728254 | 0.065185888 | 1.698653758 |
| Mir7673 | 0 | NA | NA | NA |
| Gm5400 | 0.277195307 | -1.874824391 | 2.051686219 | -0.913796844 |
| Gm14826 | 0 | NA | NA | NA |
| Gm14825 | 0.060174537 | 0.517957688 | 3.724945049 | 0.139051095 |
| 4932442L08Rik | 2.026572498 | -0.184344764 | 0.707767283 | -0.260459573 |
| Apoo | 208.2198411 | 0.295805653 | 0.113995651 | 2.594885419 |
| AL627302.1 | 0 | NA | NA | NA |
| Gm9003 | 0 | NA | NA | NA |
| Gm14827 | 54.27178075 | -0.178208291 | 0.403166727 | -0.442021327 |
| Gm24595 | 0 | NA | NA | NA |
| Maged1 | 4444.520797 | 0.176688463 | 0.05535892 | 3.191689123 |
| Gm9005 | 0 | NA | NA | NA |
| Gm14831 | 0.038981395 | -0.443816764 | 3.724945049 | -0.119147198 |
| Gm5401 | 0.15966113 | -1.078283588 | 3.444418822 | -0.313052403 |
| Gm9009 | 0.046345709 | -0.443818988 | 3.724945049 | -0.119147795 |
| Gspt2 | 293.6766047 | 0.116954759 | 0.086273133 | 1.355633611 |

| | | | | |
|---|---|---|---|---|
| Gm5402 | 0.046345709 | -0.443818988 | 3.724945049 | -0.119147795 |
| Zxdb | 233.650166 | 0.053810274 | 0.099476302 | 0.540935605 |
| Gm7061 | 8.775414701 | 0.332787905 | 0.483153098 | 0.688783546 |
| Gm14830 | 3.497870509 | 1.889157164 | 0.693681839 | 2.723376999 |
| Rpl13-ps5 | 0.088383099 | 0.037058229 | 3.724945049 | 0.009948665 |
| Gm26617 | 0.179489757 | 0.502693874 | 2.288344912 | 0.219675745 |
| Zxda | 5.296744549 | 0.108085687 | 0.4302894 | 0.251193003 |
| Gm371 | 0.049553047 | -0.443818988 | 3.724945049 | -0.119147795 |
| Spin4 | 40.24504316 | -0.333459778 | 0.226723429 | -1.470777765 |
| Arhgef9 | 4283.077243 | 0.665220954 | 0.10201062 | 6.521095112 |
| Gm6092 | 0.047948698 | 0.037064902 | 3.724945049 | 0.009950456 |
| Gm14807 | 0.075043205 | 0.517957688 | 3.724945049 | 0.139051095 |
| Gm3712 | 0.119587853 | 0.981063299 | 3.723618439 | 0.263470416 |
| Gm14806 | 0.045882455 | 0.517959912 | 3.724945049 | 0.139051692 |
| Amer1 | 313.2545163 | 0.185011729 | 0.100833255 | 1.834828486 |
| Asb12 | 0.245805814 | 0.446449377 | 2.121039425 | 0.210486129 |
| Gm14818 | 0 | NA | NA | NA |
| Gm9043 | 0 | NA | NA | NA |
| Zc4h2 | 203.86252 | 0.327982652 | 0.102457881 | 3.201146143 |
| Zc3h12b | 477.9337351 | 0.246102452 | 0.118904951 | 2.069741007 |
| 1700010D01Rik | 0 | NA | NA | NA |
| Las1l | 521.4809968 | -0.193362028 | 0.117713935 | -1.642643489 |
| Msn | 2032.338517 | -1.246350846 | 0.186279138 | -6.690769889 |
| Gm24718 | 0 | NA | NA | NA |
| Gm14925 | 0.293430937 | 0.998866886 | 2.044607505 | 0.488537229 |
| Gm9051 | 0 | NA | NA | NA |
| F630028O10Rik | 26.3940196 | -1.881423458 | 0.414429484 | -4.539791525 |
| Mir223 | 0 | NA | NA | NA |
| Vsig4 | 0.106465261 | 0.037073798 | 3.724945049 | 0.009952844 |
| Hsf3 | 1.054375976 | 0.99091484 | 0.982204174 | 1.008868489 |
| Gm14933 | 0.042716945 | 0.037062677 | 3.724945049 | 0.009949859 |
| Heph | 82.76888646 | -0.047168765 | 0.186726091 | -0.252609394 |
| Rpl17-ps8 | 13.95711591 | -0.03997624 | 0.267082435 | -0.149677531 |
| Gm14935 | 0.695213882 | 0.367132004 | 1.238860498 | 0.296346525 |
| Gpr165 | 370.7971054 | 0.199956377 | 0.129765683 | 1.540903363 |
| Gm16394 | 0 | NA | NA | NA |
| Gm14803 | 0 | NA | NA | NA |
| Gm9059 | 0 | NA | NA | NA |
| Gm9060 | 0 | NA | NA | NA |
| Pgr15l | 28.47859877 | -0.375562405 | 0.281600979 | -1.333668678 |
| Gm14800 | 0 | NA | NA | NA |
| Eda2r | 27.60206714 | -1.99328449 | 0.353982399 | -5.631027118 |
| Gm9062 | 0 | NA | NA | NA |
| Gm14802 | 0.124347486 | 0.036092774 | 3.717245565 | 0.009709548 |
| Ar | 593.9010294 | 0.72931026 | 0.122737731 | 5.942021698 |
| Gm14811 | 0 | NA | NA | NA |
| Gm14815 | 0.75340593 | 1.492486787 | 1.205198526 | 1.238374222 |

| | | | | |
|---|---|---|---|---|
| Gm14810 | 2.604223114 | 0.72446316 | 0.722548182 | 1.002650312 |
| Ophn1 | 626.2746632 | 0.397488954 | 0.09118898 | 4.358958202 |
| Gm27554 | 3.06361838 | 0.066452298 | 0.757526994 | 0.087722679 |
| Gm26000 | 0.038981395 | -0.443816764 | 3.724945049 | -0.119147198 |
| Yipf6 | 767.4798748 | -0.06690015 | 0.070481696 | -0.949184743 |
| Gm14813 | 0.038981395 | -0.443816764 | 3.724945049 | -0.119147198 |
| Stard8 | 229.0060904 | -0.228824507 | 0.162321523 | -1.409699114 |
| Gm5760 | 0 | NA | NA | NA |
| Efnb1 | 173.9851955 | -0.371679403 | 0.128072062 | -2.902111486 |
| Gm14812 | 1.182411106 | 3.325077256 | 3.662425806 | 0.907889315 |
| Gm14809 | 0.233612452 | 0.979519378 | 2.527723168 | 0.387510543 |
| Gm14808 | 0.179754939 | 0.959593576 | 2.738727739 | 0.350379325 |
| Pja1 | 1595.105195 | 0.170312095 | 0.083105203 | 2.049355386 |
| Gm14816 | 0.266066471 | -0.41425524 | 2.08887553 | -0.198314947 |
| Pcna-ps1 | 0.075043205 | 0.517957688 | 3.724945049 | 0.139051095 |
| Tmem28 | 262.17291 | 0.092641359 | 0.106779202 | 0.867597413 |
| Eda | 61.73840336 | 0.90745724 | 0.172361124 | 5.264860295 |
| Gm14832 | 1.244840331 | 1.360300317 | 1.133068558 | 1.200545463 |
| Gm14829 | 1.581404248 | 1.703517802 | 0.986087648 | 1.727552114 |
| Gm14828 | 1.203205628 | -0.171677012 | 0.953372422 | -0.180073399 |
| Mir676 | 0 | NA | NA | NA |
| Awat2 | 2.157021785 | -0.594181086 | 0.713617355 | -0.832632618 |
| Otud6a | 0 | NA | NA | NA |
| Igbp1 | 392.0809371 | 0.121290094 | 0.07493917 | 1.618513984 |
| Dgat2l6 | 5.685626011 | -1.089534515 | 0.568492899 | -1.91653144 |
| Awat1 | 0.271018715 | -0.995179771 | 1.808828755 | -0.550179097 |
| Gm6175 | 0 | NA | NA | NA |
| P2ry4 | 1.586926032 | 0.298642138 | 0.787541973 | 0.379207901 |
| Arr3 | 2.257394224 | -0.078513463 | 0.882527797 | -0.088964295 |
| Pdzd11 | 261.6832867 | -0.221194635 | 0.083868938 | -2.63738448 |
| Kif4 | 26.59334305 | -0.256560119 | 0.337326818 | -0.760568403 |
| Gdpd2 | 50.75686142 | 0.443279255 | 0.288360572 | 1.537239478 |
| Gm14866 | 1.936928019 | -0.728528467 | 0.787494591 | -0.925121868 |
| Gm14902 | 0.875624081 | 0.193472077 | 0.986731044 | 0.196073772 |
| Gm22160 | 0.092691418 | -0.908687078 | 3.724560878 | -0.2439716 |
| Dlg3 | 1696.534007 | 0.490989245 | 0.089616795 | 5.478763723 |
| Tex11 | 4.291613721 | 0.924062653 | 0.537314234 | 1.719780706 |
| Gm24577 | 0 | NA | NA | NA |
| Gm27510 | 0 | NA | NA | NA |
| Gm14901 | 0 | NA | NA | NA |
| Gm9078 | 0 | NA | NA | NA |
| Slc7a3 | 112.9406466 | -0.413959964 | 0.165602702 | -2.499717453 |
| Snx12 | 585.2370213 | -0.188047669 | 0.069032116 | -2.72406062 |
| Gm24061 | 0.522088832 | -0.638670568 | 1.306646199 | -0.488786152 |
| Gm14842 | 0 | NA | NA | NA |
| Gm22472 | 0.04393175 | 0.037062677 | 3.724945049 | 0.009949859 |
| Foxo4 | 217.5431392 | -0.180198641 | 0.151273329 | -1.191212237 |

| | | | | |
|---|---|---|---|---|
| Gm614 | 0.87838318 | 0.499183206 | 0.999219246 | 0.49957325 |
| Gm20489 | 0 | NA | NA | NA |
| Il2rg | 109.8692796 | -1.00061738 | 0.210439111 | -4.754902149 |
| Med12 | 591.9911033 | -0.279475338 | 0.120397555 | -2.321270878 |
| Gm21986 | 6.701345333 | 0.16878343 | 0.48081149 | 0.351038678 |
| Nlgn3 | 1054.489164 | 0.579475357 | 0.093591075 | 6.191566438 |
| Gjb1 | 235.1220583 | -0.618206729 | 0.28401481 | -2.17667075 |
| Zmym3 | 1188.784781 | 0.399425883 | 0.061101159 | 6.537124464 |
| Nono | 1296.303821 | -0.093124661 | 0.056094397 | -1.660141926 |
| Itgb1bp2 | 18.97263099 | 0.858733876 | 0.429159507 | 2.000966685 |
| Gm14843 | 1.315890906 | 1.041216752 | 1.122635006 | 0.927475757 |
| Gm6184 | 0.809862658 | 0.389777112 | 1.408657209 | 0.27670118 |
| Taf1 | 1383.236683 | -0.00976353 | 0.046425529 | -0.210305207 |
| Ogt | 3519.482034 | 0.244138834 | 0.125558445 | 1.944423838 |
| Gm14845 | 0.98816996 | 1.894879265 | 1.015224654 | 1.866463011 |
| Gm9089 | 0.045882455 | 0.517959912 | 3.724945049 | 0.139051692 |
| Cxcr3 | 9.848512992 | 0.473554817 | 0.359634452 | 1.316767105 |
| Tpt1-ps6 | 6.439785805 | -0.877627948 | 0.413625079 | -2.121795784 |
| Gm9095 | 0 | NA | NA | NA |
| Gm4779 | 0.781258755 | -0.41586533 | 1.275115748 | -0.326139278 |
| 8030474K03Rik | 0 | NA | NA | NA |
| Gm9097 | 0 | NA | NA | NA |
| H3f3a-ps1 | 4.580768557 | 0.459767287 | 0.62367453 | 0.737191058 |
| Nhsl2 | 1474.969298 | 0.834410269 | 0.08568765 | 9.73781248 |
| Gm25828 | 0 | NA | NA | NA |
| Gm27843 | 0 | NA | NA | NA |
| Rgag4 | 516.2445288 | -0.169021084 | 0.134372115 | -1.25785833 |
| Gm5125 | 0 | NA | NA | NA |
| Pin4 | 43.34765536 | -0.113539081 | 0.296462366 | -0.382979744 |
| Gm16471 | 0.089089296 | 0.957155977 | 3.723939561 | 0.257027796 |
| Ercc6l | 15.19884806 | -1.447308578 | 0.400695261 | -3.611993249 |
| Gm9103 | 0.062426873 | 0.517959912 | 3.724945049 | 0.139051692 |
| Rps4x | 1364.482336 | -0.34455467 | 0.115088109 | -2.993833806 |
| Gm14859 | 0 | NA | NA | NA |
| Cited1 | 20.65218563 | -0.183608283 | 0.272726409 | -0.673232505 |
| Gm14858 | 2.504509112 | 3.120867077 | 0.948456014 | 3.290471071 |
| Gm26318 | 0.128271896 | 0.517956073 | 3.679089552 | 0.140783763 |
| Hdac8 | 196.3053901 | 0.079907473 | 0.11928997 | 0.669859108 |
| Phka1 | 482.7901274 | 0.091723196 | 0.084006624 | 1.091856711 |
| Gm9109 | 0 | NA | NA | NA |
| Gm14840 | 0 | NA | NA | NA |
| Gm3880 | 0 | NA | NA | NA |
| Gm9112 | 0 | NA | NA | NA |
| Dmrtc1b | 0 | NA | NA | NA |
| Gm9115 | 4.289267201 | -1.310095703 | 0.502854034 | -2.605320061 |
| Gm14839 | 0 | NA | NA | NA |
| Gm4234 | 0 | NA | NA | NA |

| | | | | |
|---|---|---|---|---|
| Gm9119 | 0 | NA | NA | NA |
| Gm5166 | 0 | NA | NA | NA |
| Gm25358 | 0 | NA | NA | NA |
| Dmrtc1c1 | 0 | NA | NA | NA |
| Gm9126 | 0 | NA | NA | NA |
| Gm16511 | 0 | NA | NA | NA |
| Gm24509 | 0 | NA | NA | NA |
| Dmrtc1c2 | 0 | NA | NA | NA |
| Gm26152 | 0 | NA | NA | NA |
| 1700031F05Rik | 0 | NA | NA | NA |
| Gm9133 | 0 | NA | NA | NA |
| Dmrtc1a | 21.57087845 | 1.030828516 | 0.245646156 | 4.196395876 |
| 1700011M02Rik | 0.046345709 | -0.443818988 | 3.724945049 | -0.119147795 |
| Gm14841 | 0 | NA | NA | NA |
| Gm14837 | 0.193036943 | -0.47213964 | 3.248256614 | -0.145351706 |
| Gm21998 | 0.296810931 | 0.531670854 | 1.771981397 | 0.300043135 |
| Pabpc1l2b-ps | 6.125206409 | -0.438651409 | 0.426048605 | -1.029580672 |
| Gm9143 | 0.044191549 | 0.037062677 | 3.724945049 | 0.009949859 |
| Pabpc1l2a-ps | 3.182304534 | -0.558731375 | 0.623090948 | -0.896709183 |
| Gm3928 | 0.473680182 | -0.006320131 | 1.346608685 | -0.004693369 |
| Gm14838 | 0.200176296 | 0.517955286 | 3.25927971 | 0.158917102 |
| Gm6206 | 202.4849468 | -0.435348969 | 0.123237322 | -3.532606542 |
| 4930519F16Rik | 7.945419585 | 0.099905416 | 0.416700555 | 0.239753498 |
| Nap1l2 | 1056.694576 | 0.216366169 | 0.067951743 | 3.184115071 |
| Gm9050 | 0 | NA | NA | NA |
| Gm14847 | 0 | NA | NA | NA |
| Gm19200 | 0.512704799 | 0.517971807 | 1.64072462 | 0.31569698 |
| Gm14848 | 4.393179665 | 1.71421087 | 1.339974278 | 1.2792864 |
| Gm3931 | 0.286297534 | 1.452087448 | 2.042866465 | 0.710808794 |
| Cdx4 | 2.430558831 | 2.375583933 | 1.099916478 | 2.159785749 |
| Gm9516 | 0 | NA | NA | NA |
| Chic1 | 1013.690449 | 0.111308299 | 0.065342792 | 1.703451829 |
| Gm26952 | 0 | NA | NA | NA |
| Tsx | 2.523036181 | 0.280028834 | 0.601351719 | 0.465665641 |
| Gm26992 | 0 | NA | NA | NA |
| Tsix | 4.036643334 | 0.772698801 | 0.59857941 | 1.290887706 |
| Xist | 7370.143953 | 0.010230013 | 0.217489056 | 0.04703691 |
| Gm27733 | 0 | NA | NA | NA |
| Gm27927 | 0 | NA | NA | NA |
| Gm27520 | 0.255832186 | 0.517972397 | 1.825382711 | 0.283760986 |
| Gm14861 | 0.384616848 | 1.894095468 | 1.423862812 | 1.330251378 |
| Rps12-ps1 | 0.281032441 | 0.069106975 | 2.074010839 | 0.03332045 |
| Jpx | 90.00391533 | 0.170436682 | 0.221784082 | 0.76848023 |
| Gm27875 | 0 | NA | NA | NA |
| Gm9670 | 11.86610392 | 0.341996017 | 0.407627554 | 0.83899141 |
| Gm14860 | 4.132034165 | -0.559417742 | 0.616701608 | -0.907112508 |
| Gm9785 | 0.044544648 | 0.517957688 | 3.724945049 | 0.139051095 |

| | | | | |
|---|---|---|---|---|
| Gm9157 | 5.137711797 | 0.89174016 | 0.566354093 | 1.574527615 |
| Ftx | 498.1238306 | 0.391146792 | 0.2069347 | 1.890194312 |
| Mir421 | 14.61544255 | 0.461274856 | 0.394307841 | 1.169834348 |
| Mir374b | 9.615598131 | 0.311805355 | 0.4801036 | 0.649454316 |
| mmu-mir-374c | 0 | NA | NA | NA |
| Gm38020 | 450.5061 | 0.431874764 | 0.277707386 | 1.555143239 |
| Gm27798 | 0.19090462 | 0.541105795 | 3.265182312 | 0.165719933 |
| Gm9159 | 0.296935004 | 1.018078826 | 3.723620811 | 0.273410983 |
| Gm27987 | 0.094097695 | 0.037071574 | 3.724945049 | 0.009952247 |
| Gm27285 | 0 | NA | NA | NA |
| Gm27321 | 0.436808164 | 0.575597623 | 1.552423352 | 0.370773618 |
| Zcchc13 | 0.559888142 | -0.015851494 | 1.189753016 | -0.013323349 |
| Gm14874 | 0.823502631 | 1.351692722 | 1.233081915 | 1.096190533 |
| Gm14875 | 0 | NA | NA | NA |
| Gm3943 | 0.062426873 | 0.517959912 | 3.724945049 | 0.139051692 |
| Slc16a2 | 413.5769034 | 0.338434608 | 0.089147371 | 3.79634985 |
| Gm6222 | 51.51232969 | -0.066200865 | 0.199205579 | -0.332324354 |
| Gm37956 | 12.17933112 | 1.133495044 | 0.360818986 | 3.141450667 |
| Gm14893 | 0 | NA | NA | NA |
| Gm9164 | 0.119587853 | 0.981063299 | 3.723618439 | 0.263470416 |
| Gm9166 | 0.719693239 | 2.102039589 | 1.454018729 | 1.445675731 |
| Rlim | 2303.194065 | -0.007016871 | 0.098102885 | -0.071525634 |
| Gm14892 | 0 | NA | NA | NA |
| Gm23378 | 0 | NA | NA | NA |
| Gm22266 | 0.086648695 | 0.037058229 | 3.724945049 | 0.009948665 |
| Gm24993 | 0 | NA | NA | NA |
| C77370 | 997.2398713 | 0.410841165 | 0.084298789 | 4.87363067 |
| Mir672 | 0.913602969 | 0.439640609 | 1.109360878 | 0.396300805 |
| Abcb7 | 785.2393539 | -0.018513278 | 0.08185716 | -0.226165653 |
| Vcp-rs | 650.7603801 | -0.315567297 | 0.07954957 | -3.966926516 |
| Gm6238 | 4.907369357 | 0.402069394 | 0.575530869 | 0.698606132 |
| Uprt | 137.0040339 | -0.351630305 | 0.133565588 | -2.632641453 |
| Zdhhc15 | 117.8220007 | -0.09967586 | 0.13204963 | -0.754836344 |
| Gm22430 | 0 | NA | NA | NA |
| Gm14822 | 2.347436586 | 0.091277712 | 0.724278649 | 0.126025684 |
| 1700121L16Rik | 0.182267886 | 0.037075401 | 2.690523935 | 0.013779993 |
| Magee2 | 43.93040372 | 0.195346124 | 0.168986475 | 1.155986736 |
| Gm6285 | 2.055059636 | -0.067138984 | 0.664207948 | -0.101081271 |
| Gm22090 | 0.110870145 | -0.444802825 | 3.724874622 | -0.119414174 |
| Gm9673 | 0 | NA | NA | NA |
| 5530601H04Rik | 228.1153272 | 0.347754861 | 0.107292904 | 3.241172988 |
| Pbdc1 | 243.3133995 | -0.244170467 | 0.10512622 | -2.322640982 |
| Magee1 | 1870.496208 | 0.385911299 | 0.059741531 | 6.459682075 |
| Gm6292 | 2.268236772 | 0.578538532 | 0.715569223 | 0.80850114 |
| E530001F21Rik | 15.1906836 | -0.211026011 | 0.342883378 | -0.615445438 |
| Rpl30-ps9 | 19.84181199 | 0.03514308 | 0.300076845 | 0.117113602 |
| Gm14824 | 57.64436276 | 0.17819718 | 0.235986439 | 0.755116185 |

| | | | | |
|---|---|---|---|---|
| Mir384 | 25.25469777 | -0.322493526 | 0.243275085 | -1.325633188 |
| 5330434G04Rik | 854.7814095 | 0.224332949 | 0.162767789 | 1.378239211 |
| Mir325 | 9.585432933 | 0.141738018 | 0.492423401 | 0.287837698 |
| Rab9b-ps1 | 0.047948698 | 0.037064902 | 3.724945049 | 0.009950456 |
| Cypt2 | 0 | NA | NA | NA |
| Gm14823 | 0 | NA | NA | NA |
| Gm14855 | 0 | NA | NA | NA |
| Gm14854 | 0 | NA | NA | NA |
| Gm7117 | 0.290452866 | 0.976420952 | 1.763948196 | 0.553542873 |
| Fgf16 | 4.769070217 | 0.599349981 | 0.509934197 | 1.175347691 |
| Atrx | 4307.043563 | 0.367583431 | 0.085511917 | 4.298622253 |
| Gm26020 | 0 | NA | NA | NA |
| Gm14853 | 16.35826218 | 0.71801013 | 0.396133398 | 1.812546311 |
| Gm23656 | 0 | NA | NA | NA |
| Magt1 | 450.0307998 | -0.594558227 | 0.120169057 | -4.947681548 |
| Cox7b | 1087.530571 | 0.159704353 | 0.118508117 | 1.347623741 |
| Atp7a | 328.8971482 | -0.654461208 | 0.102988752 | -6.354686296 |
| Tlr13 | 410.2003387 | -1.395981644 | 0.284945315 | -4.899121245 |
| Gm14856 | 0 | NA | NA | NA |
| Pgk1 | 2368.453568 | -0.116478937 | 0.065409309 | -1.780770033 |
| Taf9b | 306.5549042 | -0.282177403 | 0.118653946 | -2.378154398 |
| Fnd3c2 | 0.045882455 | 0.517959912 | 3.724945049 | 0.139051692 |
| Gm14852 | 0 | NA | NA | NA |
| Gm14857 | 0.631966691 | -1.475447783 | 1.224269866 | -1.205165482 |
| Gm5133 | 0 | NA | NA | NA |
| Mir6382 | 0 | NA | NA | NA |
| Gm14876 | 0 | NA | NA | NA |
| Gm6325 | 0.676409176 | -1.258260974 | 1.207944368 | -1.041654738 |
| Fndc3c1 | 1.066871938 | -1.22776953 | 1.004715309 | -1.222007387 |
| Gm27428 | 0.038981395 | -0.443816764 | 3.724945049 | -0.119147198 |
| Cysltr1 | 35.52481597 | -1.199987408 | 0.31343667 | -3.828484421 |
| Gm5127 | 6.92021651 | -1.314492248 | 0.38927392 | -3.376779638 |
| Gm24622 | 0.231961655 | -0.438143924 | 2.127211574 | -0.205971014 |
| Gm14877 | 0.75733687 | -0.154937404 | 1.430131462 | -0.108337875 |
| Gm14878 | 0.046345709 | -0.443818988 | 3.724945049 | -0.119147795 |
| Gm14879 | 0 | NA | NA | NA |
| Gm9200 | 1.650574175 | 0.505201894 | 0.766424574 | 0.65916714 |
| Gm24491 | 0 | NA | NA | NA |
| Gm22102 | 0 | NA | NA | NA |
| Gm14882 | 0 | NA | NA | NA |
| Gm14881 | 0 | NA | NA | NA |
| Gm14880 | 0 | NA | NA | NA |
| Zcchc5 | 3.290042265 | 0.660436265 | 0.618577687 | 1.067669072 |
| Lpar4 | 34.6581693 | -0.61082524 | 0.244967477 | -2.493495243 |
| P2ry10 | 28.76235556 | -0.557803043 | 0.274357486 | -2.033124921 |
| A630033H20Rik | 13.48921018 | -1.361294808 | 0.388078725 | -3.507780043 |
| Gm14834 | 0 | NA | NA | NA |

| | | | | |
|---|---|---|---|---|
| Gm14836 | 0 | NA | NA | NA |
| Gpr174 | 17.71929983 | -0.832809572 | 0.294037458 | -2.832324761 |
| Gm9208 | 0.060174537 | 0.517957688 | 3.724945049 | 0.139051095 |
| Gm14833 | 0 | NA | NA | NA |
| Gm5392 | 0 | NA | NA | NA |
| Itm2a | 152.8149106 | -0.147368154 | 0.250974117 | -0.587184671 |
| Gm14835 | 0 | NA | NA | NA |
| 4933401B06Rik | 0 | NA | NA | NA |
| Tbx22 | 1.078738829 | 0.929208019 | 1.207521841 | 0.769516531 |
| 2610002M06Rik | 1022.333748 | 0.281843299 | 0.092845332 | 3.035621636 |
| Fam46d | 2.160810062 | 0.898059785 | 0.681277118 | 1.318200422 |
| Gm732 | 0 | NA | NA | NA |
| Gm14849 | 0 | NA | NA | NA |
| Gm378 | 0 | NA | NA | NA |
| Gm9218 | 0 | NA | NA | NA |
| Gm379 | 0.046345709 | -0.443818988 | 3.724945049 | -0.119147795 |
| Brwd3 | 697.4322742 | -0.018291988 | 0.081045294 | -0.225700804 |
| 2810403D21Rik | 30.21590391 | -0.163049259 | 0.193631409 | -0.842059975 |
| Gm14870 | 0.48610956 | -0.583140687 | 1.435595774 | -0.406201172 |
| 5730416F02Rik | 0.253264426 | -0.466228238 | 2.078818565 | -0.224275579 |
| Hmgn5 | 36.85669999 | -0.046202653 | 0.256651189 | -0.180021191 |
| Gm23322 | 0.212513282 | 1.689178857 | 3.159662247 | 0.534607412 |
| Gm25797 | 0 | NA | NA | NA |
| Sh3bgrl | 620.2824578 | -0.230241449 | 0.13311984 | -1.729580272 |
| Gm26415 | 0 | NA | NA | NA |
| Gm14869 | 0 | NA | NA | NA |
| Gm6377 | 31.64666742 | -1.928263712 | 0.356810194 | -5.40417214 |
| Gm14868 | 0 | NA | NA | NA |
| Gm4784 | 0.759591101 | -2.753212664 | 1.159883676 | -2.373697226 |
| Gm4991 | 0 | NA | NA | NA |
| Gm7134 | 0 | NA | NA | NA |
| Gm9225 | 0 | NA | NA | NA |
| Gm14886 | 0 | NA | NA | NA |
| Gm26108 | 0.089814205 | 0.517955464 | 3.724945049 | 0.139050498 |
| Pou3f4 | 86.02711921 | -0.15091375 | 0.148408193 | -1.016882874 |
| Cylc1 | 0 | NA | NA | NA |
| Gm10112 | 0 | NA | NA | NA |
| Rps6ka6 | 115.5535552 | -0.33773204 | 0.140630573 | -2.401554894 |
| Hdx | 271.1100276 | 0.23971931 | 0.109328931 | 2.192642951 |
| Gm7340 | 0 | NA | NA | NA |
| Gm14898 | 0 | NA | NA | NA |
| Gm14899 | 0 | NA | NA | NA |
| Tex16 | 4.590268127 | -0.746840981 | 0.492923439 | -1.515125723 |
| Gm14900 | 0 | NA | NA | NA |
| Gm6335 | 3.683185528 | -0.084531227 | 0.60580465 | -0.139535454 |
| 4933403O08Rik | 4.019211381 | -0.450997235 | 0.512722259 | -0.879613139 |
| Gm5393 | 0 | NA | NA | NA |

| | | | | |
|---|---|---|---|---|
| Apool | 63.23518737 | -0.533580702 | 0.173955085 | -3.06734754 |
| Satl1 | 0 | NA | NA | NA |
| Gm6418 | 0.152980968 | -0.89129297 | 2.833643542 | -0.314539552 |
| Gm14908 | 0.684617987 | -0.119981703 | 1.282604023 | -0.093545398 |
| 2010106E10Rik | 0.828534694 | 0.944698529 | 1.051555378 | 0.898382101 |
| Zfp711 | 267.1925573 | 0.153708548 | 0.098200683 | 1.56524927 |
| Pof1b | 1.108407561 | 0.999026338 | 1.3754611 | 0.726321041 |
| Gm14921 | 0 | NA | NA | NA |
| Gm14922 | 0 | NA | NA | NA |
| Gm14923 | 0 | NA | NA | NA |
| Gm6427 | 0 | NA | NA | NA |
| Gm6429 | 0 | NA | NA | NA |
| Gm14937 | 0 | NA | NA | NA |
| Gm14938 | 0 | NA | NA | NA |
| Gm14936 | 0.124596252 | 0.037071632 | 3.714681146 | 0.009979761 |
| Gm14939 | 1.913926814 | 1.174272769 | 0.858608824 | 1.36764582 |
| Chm | 978.6370897 | 0.119425591 | 0.066939641 | 1.784078735 |
| Mir361 | 1.21006551 | 0.843049422 | 0.913818972 | 0.92255627 |
| Gm7405 | 0 | NA | NA | NA |
| Dach2 | 22.20089386 | -0.476354246 | 0.230476347 | -2.066824864 |
| Gm24831 | 0 | NA | NA | NA |
| Gm23985 | 0 | NA | NA | NA |
| Gm4915 | 0 | NA | NA | NA |
| Gm14896 | 0 | NA | NA | NA |
| Gm7416 | 0 | NA | NA | NA |
| Gm7146 | 0.194279094 | -1.017823543 | 3.719440177 | -0.273649661 |
| Klhl4 | 212.9303465 | -0.032649836 | 0.139393592 | -0.23422767 |
| Gm14895 | 3.529479033 | -0.882473492 | 0.600982419 | -1.468384871 |
| Gm1866 | 7.003298379 | 0.101165888 | 0.39585514 | 0.255562901 |
| Gm14894 | 0.088476398 | 0.51795324 | 3.724945049 | 0.139049901 |
| Gm5942 | 0 | NA | NA | NA |
| Gm14897 | 0.045882455 | 0.517959912 | 3.724945049 | 0.139051692 |
| Ube2dnl1 | 0 | NA | NA | NA |
| Ube2dnl2 | 0 | NA | NA | NA |
| 4930555B12Rik | 0 | NA | NA | NA |
| Gm4914 | 0 | NA | NA | NA |
| Gm7429 | 0.963943105 | -2.768439835 | 1.118767267 | -2.474544899 |
| Gm5943 | 0.089062654 | -0.443823436 | 3.724945049 | -0.119148989 |
| Gm14905 | 0 | NA | NA | NA |
| Gm25366 | 0 | NA | NA | NA |
| Gm27414 | 0 | NA | NA | NA |
| Gm6461 | 0 | NA | NA | NA |
| Gm5944 | 0.066050875 | -0.443818988 | 3.724945049 | -0.119147795 |
| Gm14903 | 0 | NA | NA | NA |
| Gm14906 | 0 | NA | NA | NA |
| Gm14907 | 0 | NA | NA | NA |
| Gm24683 | 0 | NA | NA | NA |

| | | | | |
|---|---|---|---|---|
| Gm14904 | 0 | NA | NA | NA |
| Gm24624 | 0 | NA | NA | NA |
| Cpxcr1 | 0 | NA | NA | NA |
| H2afb2 | 0 | NA | NA | NA |
| Gm24438 | 0 | NA | NA | NA |
| Gm16373 | 0.048949532 | 0.037064902 | 3.724945049 | 0.009950456 |
| Gm14919 | 0 | NA | NA | NA |
| Gm14920 | 0 | NA | NA | NA |
| Gm14918 | 0 | NA | NA | NA |
| Gm14913 | 0 | NA | NA | NA |
| Gm6491 | 0 | NA | NA | NA |
| Gm28579 | 0 | NA | NA | NA |
| Gm14911 | 0 | NA | NA | NA |
| Gm14914 | 0.265235714 | 0.517974841 | 2.048392805 | 0.252868903 |
| Gm14917 | 0 | NA | NA | NA |
| Gm6500 | 0 | NA | NA | NA |
| Gm14916 | 0 | NA | NA | NA |
| Gm14912 | 0 | NA | NA | NA |
| Gm14915 | 0 | NA | NA | NA |
| Gm14909 | 0 | NA | NA | NA |
| Gm14910 | 0 | NA | NA | NA |
| Tgif2lx2 | 0 | NA | NA | NA |
| Tgif2lx1 | 0 | NA | NA | NA |
| Gm7485 | 0 | NA | NA | NA |
| Gm14926 | 0 | NA | NA | NA |
| Gm26182 | 0 | NA | NA | NA |
| Gm14929 | 0 | NA | NA | NA |
| Pabpc5 | 33.05411456 | 0.051523317 | 0.206626119 | 0.249355294 |
| Gm4913 | 0.123841226 | -1.017823543 | 3.719440177 | -0.273649661 |
| Gm16420 | 0 | NA | NA | NA |
| Gm5863 | 0.479236126 | 0.761472987 | 1.322393492 | 0.575829351 |
| Gm14928 | 0.622864522 | 0.906545831 | 1.36348748 | 0.664872868 |
| Pcdh11x | 761.4248623 | 0.537620401 | 0.099056576 | 5.427407488 |
| H2afb3 | 0 | NA | NA | NA |
| Gm14931 | 0 | NA | NA | NA |
| Gm14927 | 1.075015726 | -0.496828785 | 1.093931893 | -0.454167931 |
| Gm14930 | 0 | NA | NA | NA |
| Gm4993 | 0.279479172 | -0.790076176 | 2.485743284 | -0.31784303 |
| Gm22590 | 0.042716945 | 0.037062677 | 3.724945049 | 0.009949859 |
| Gm5394 | 0 | NA | NA | NA |
| Gm25825 | 0 | NA | NA | NA |
| Gm5648 | 1.049721047 | -1.056715588 | 1.006709052 | -1.049673275 |
| Gm14932 | 0.047948698 | 0.037064902 | 3.724945049 | 0.009950456 |
| Gm23277 | 0 | NA | NA | NA |
| Gm14943 | 0 | NA | NA | NA |
| Gm14945 | 0 | NA | NA | NA |
| Gm4992 | 0.108772582 | 0.037073798 | 3.724945049 | 0.009952844 |

| | | | | |
|---|---|---|---|---|
| Gm24783 | 0 | NA | NA | NA |
| Gm16421 | 0.491147404 | 1.226005753 | 1.297827613 | 0.944659939 |
| Gm14946 | 0.141684195 | 0.037074514 | 3.558854665 | 0.010417541 |
| Gm14942 | 0.160761659 | 0.998860243 | 2.799488339 | 0.356801001 |
| Nap1l3 | 532.8242286 | -0.046964842 | 0.123948773 | -0.378905256 |
| Gm14947 | 0 | NA | NA | NA |
| Gm14940 | 0 | NA | NA | NA |
| Gm17521 | 0 | NA | NA | NA |
| Cldn34c1 | 0.062426873 | 0.517959912 | 3.724945049 | 0.139051692 |
| Astx6 | 0 | NA | NA | NA |
| Srsx | 0 | NA | NA | NA |
| Gm17577 | 0 | NA | NA | NA |
| Gm14951 | 0 | NA | NA | NA |
| Rps12-ps21 | 0 | NA | NA | NA |
| Astx2 | 0 | NA | NA | NA |
| Gm17412 | 0 | NA | NA | NA |
| Rps12-ps22 | 0.049553047 | -0.443818988 | 3.724945049 | -0.119147795 |
| Cldn34c2 | 0 | NA | NA | NA |
| Gm14950 | 0 | NA | NA | NA |
| Gm17467 | 0 | NA | NA | NA |
| Rps12-ps23 | 0 | NA | NA | NA |
| Cldn34c3 | 0 | NA | NA | NA |
| Astx5 | 0 | NA | NA | NA |
| Rps12-ps20 | 0 | NA | NA | NA |
| Vmn2r121 | 0 | NA | NA | NA |
| Rps12-ps6 | 0.072025462 | 0.517959912 | 3.724945049 | 0.139051692 |
| Astx1a | 0 | NA | NA | NA |
| Rps12-ps11 | 0 | NA | NA | NA |
| Gm17584 | 0 | NA | NA | NA |
| Astx4a | 0 | NA | NA | NA |
| Rps12-ps12 | 0 | NA | NA | NA |
| Gm17469 | 0 | NA | NA | NA |
| Astx4b | 0 | NA | NA | NA |
| Rps12-ps18 | 0 | NA | NA | NA |
| Astx1b | 0 | NA | NA | NA |
| Rps12-ps15 | 0 | NA | NA | NA |
| Gm17361 | 0 | NA | NA | NA |
| Gm21616 | 0 | NA | NA | NA |
| Rps12-ps17 | 0.094831987 | 0.517957688 | 3.724945049 | 0.139051095 |
| Astx4c | 0 | NA | NA | NA |
| Rps12-ps13 | 0 | NA | NA | NA |
| Gm17693 | 0 | NA | NA | NA |
| Astx1c | 0 | NA | NA | NA |
| Rps12-ps14 | 0 | NA | NA | NA |
| Gm17522 | 0 | NA | NA | NA |
| Astx4d | 0 | NA | NA | NA |
| Rps12-ps16 | 0 | NA | NA | NA |

| | | | | |
|---|---|---|---|---|
| Gm17267 | 0 | NA | NA | NA |
| Astx3 | 0 | NA | NA | NA |
| Gm14957 | 0.04393175 | 0.037062677 | 3.724945049 | 0.009949859 |
| Gm14971 | 0 | NA | NA | NA |
| Gm7746 | 0 | NA | NA | NA |
| 4932411N23Rik | 0 | NA | NA | NA |
| Gm23586 | 0 | NA | NA | NA |
| Gm381 | 0 | NA | NA | NA |
| Gm14973 | 0 | NA | NA | NA |
| Gm14972 | 0 | NA | NA | NA |
| Gm382 | 0 | NA | NA | NA |
| Gm22139 | 0 | NA | NA | NA |
| 4921511C20Rik | 0 | NA | NA | NA |
| Gm22636 | 0 | NA | NA | NA |
| Gm14970 | 0.307002753 | 0.686333419 | 1.712993499 | 0.400663178 |
| Cldn34c4 | 0.191938586 | -1.00180283 | 3.270847114 | -0.306282377 |
| Gm25061 | 0 | NA | NA | NA |
| Gm5403 | 0.038981395 | -0.443816764 | 3.724945049 | -0.119147198 |
| Mdm4-ps | 2.28277985 | -0.96221567 | 0.730345207 | -1.317480639 |
| Gm14975 | 0 | NA | NA | NA |
| Gm4916 | 0 | NA | NA | NA |
| Gm38390 | 0 | NA | NA | NA |
| Gm26475 | 0 | NA | NA | NA |
| Gm24522 | 0 | NA | NA | NA |
| 4930558G05Rik | 0 | NA | NA | NA |
| Gm26029 | 0 | NA | NA | NA |
| Gm22225 | 0 | NA | NA | NA |
| Gm14986 | 0 | NA | NA | NA |
| Gm14985 | 0 | NA | NA | NA |
| Gm14983 | 0 | NA | NA | NA |
| Diap2 | 874.9384135 | -0.038946295 | 0.075067626 | -0.518816128 |
| Gm14984 | 0 | NA | NA | NA |
| Gm14987 | 0 | NA | NA | NA |
| Gm7781 | 0.389466168 | -0.924467286 | 1.440206355 | -0.641899185 |
| Gm14982 | 0.044191549 | 0.037062677 | 3.724945049 | 0.009949859 |
| Gm26209 | 0 | NA | NA | NA |
| Gm25170 | 0 | NA | NA | NA |
| Gm14976 | 0 | NA | NA | NA |
| Gm7790 | 0 | NA | NA | NA |
| Gm14977 | 0 | NA | NA | NA |
| Gm4994 | 0 | NA | NA | NA |
| Gm7058 | 0 | NA | NA | NA |
| Gm14981 | 0 | NA | NA | NA |
| Gm26406 | 0 | NA | NA | NA |
| Gm23768 | 0 | NA | NA | NA |
| Gm22143 | 0 | NA | NA | NA |
| Gm7820 | 0 | NA | NA | NA |

| | | | | |
|---|---|---|---|---|
| Gm7821 | 0 | NA | NA | NA |
| Gm7823 | 0 | NA | NA | NA |
| Gm14979 | 0 | NA | NA | NA |
| Gm22552 | 0 | NA | NA | NA |
| Gm14980 | 0.189785611 | 0.998862084 | 2.656151403 | 0.376056155 |
| Pcdh19 | 966.848733 | 0.540886629 | 0.062760557 | 8.618257373 |
| Gm26851 | 2.30478967 | -0.481588753 | 0.780903268 | -0.616707309 |
| Tnmd | 3.21833566 | -0.055208219 | 0.553639731 | -0.099718673 |
| Tspan6 | 133.5839573 | -0.126141893 | 0.14370195 | -0.877802235 |
| 4930570D08Rik | 0 | NA | NA | NA |
| Srpx2 | 13.98697368 | -0.24113772 | 0.284346225 | -0.848042627 |
| Sytl4 | 25.13300853 | -0.651757622 | 0.278810282 | -2.337638402 |
| Gm7841 | 0.106465261 | 0.037073798 | 3.724945049 | 0.009952844 |
| Cstf2 | 1299.774295 | 0.211910749 | 0.075821147 | 2.794876598 |
| Mir3112 | 0 | NA | NA | NA |
| Nox1 | 3.798596485 | 0.839850508 | 0.525539809 | 1.598072103 |
| Xkrx | 14.27994078 | 0.311648272 | 0.275568487 | 1.130928562 |
| Arl13a | 0.299311364 | -2.016985338 | 3.695466772 | -0.545799885 |
| Trmt2b | 186.6964886 | -0.115096326 | 0.122073985 | -0.942840733 |
| Tmem35 | 346.804519 | -0.018836124 | 0.105528521 | -0.178493207 |
| Cenpi | 15.04443857 | -0.103702424 | 0.326487872 | -0.317630248 |
| Drp2 | 836.9736946 | 0.294543788 | 0.111052282 | 2.652298391 |
| Taf7l | 0.599737706 | 0.386833719 | 1.347703777 | 0.287031709 |
| Gm16410 | 0.92854447 | 0.850351465 | 1.02637724 | 0.828497974 |
| Gm23124 | 0.094831987 | 0.517957688 | 3.724945049 | 0.139051095 |
| Timm8a1 | 33.17250175 | -0.285416887 | 0.234269034 | -1.218329551 |
| Btk | 35.98129176 | -1.028237368 | 0.190414297 | -5.400000852 |
| Gm15154 | 0 | NA | NA | NA |
| Rpl36a | 325.2517739 | -0.025646097 | 0.204290518 | -0.125537384 |
| Gla | 403.723937 | -1.193526956 | 0.163987174 | -7.278172573 |
| Hnrnph2 | 1054.6625 | -0.013458948 | 0.092291246 | -0.145831252 |
| Gm7855 | 0.152402701 | 0.517971011 | 2.846761691 | 0.181950956 |
| Gm7857 | 0.072025462 | 0.517959912 | 3.724945049 | 0.139051692 |
| B230119M05Rik | 0.783798686 | 1.083195292 | 1.085502621 | 0.997874414 |
| Armcx4 | 981.8027628 | 0.241214899 | 0.101546631 | 2.37541017 |
| Armcx1 | 541.5903477 | 0.288629552 | 0.081784578 | 3.529143966 |
| Gm16397 | 0.442616626 | 2.15009367 | 1.480144903 | 1.452623771 |
| Gm10344 | 1.729071564 | -0.05487703 | 0.734374983 | -0.07472617 |
| Armcx6 | 82.02743705 | 0.255083574 | 0.203549778 | 1.253175395 |
| Armcx3 | 414.6905456 | 0.17789444 | 0.078985046 | 2.252254704 |
| Mir7093 | 0.745433557 | 0.566779025 | 1.489820312 | 0.380434486 |
| Armcx2 | 966.547524 | 0.046862937 | 0.076018376 | 0.616468533 |
| Gm7222 | 0 | NA | NA | NA |
| Gm15018 | 0 | NA | NA | NA |
| Nxf2 | 1.760497888 | 0.02403409 | 0.686133513 | 0.035028299 |
| Zmat1 | 230.6846648 | -0.055290533 | 0.157277615 | -0.351547375 |
| 3632454L22Rik | 43.94593524 | 0.367465653 | 0.199324242 | 1.843557261 |

| | | | | |
|---|---|---|---|---|
| Gm6207 | 0 | NA | NA | NA |
| Gm15020 | 0 | NA | NA | NA |
| Gm25868 | 0 | NA | NA | NA |
| Gm15023 | 0.072025462 | 0.517959912 | 3.724945049 | 0.139051692 |
| Tceal6 | 310.5221001 | 0.508255359 | 0.128817531 | 3.945544938 |
| Gm15022 | 0 | NA | NA | NA |
| Pramel | 0 | NA | NA | NA |
| Pramel3 | 0.045882455 | 0.517959912 | 3.724945049 | 0.139051692 |
| Gm25936 | 0 | NA | NA | NA |
| Gm15017 | 0 | NA | NA | NA |
| Gm6215 | 0 | NA | NA | NA |
| Gm5128 | 0 | NA | NA | NA |
| Gm22006 | 0 | NA | NA | NA |
| Gm6221 | 0 | NA | NA | NA |
| Gm7903 | 0 | NA | NA | NA |
| Gm24416 | 0 | NA | NA | NA |
| Gm7905 | 0 | NA | NA | NA |
| AV320801 | 0 | NA | NA | NA |
| Gm22233 | 0 | NA | NA | NA |
| Gm15015 | 0 | NA | NA | NA |
| Gm6228 | 0 | NA | NA | NA |
| Nxf7 | 11.01788405 | -1.220133487 | 0.354393168 | -3.442880951 |
| Prame | 0 | NA | NA | NA |
| Gm15021 | 0.141317957 | 0.482895681 | 3.595358959 | 0.13431084 |
| Tcp11x2 | 0.28975322 | -1.041019518 | 2.356940453 | -0.44168257 |
| Tmsb15a | 0.072025462 | 0.517959912 | 3.724945049 | 0.139051692 |
| Gm15019 | 0 | NA | NA | NA |
| Gprasp1 | 6518.488455 | 0.323296051 | 0.064155176 | 5.039282461 |
| Armcx5 | 396.6884651 | 0.964533945 | 0.171429341 | 5.626422746 |
| mmu-mir-7646 | 0 | NA | NA | NA |
| Mir1970 | 0 | NA | NA | NA |
| Gprasp2 | 1597.674091 | 0.288824026 | 0.111466739 | 2.591122948 |
| Bhlhb9 | 800.6765123 | 0.226592168 | 0.090920017 | 2.492214316 |
| Gm15016 | 2.223121331 | 0.90845026 | 0.736903592 | 1.232793909 |
| Arxes2 | 329.4047032 | -0.035484605 | 0.1009395 | -0.351543295 |
| Arxes1 | 156.3845472 | -0.314655715 | 0.143943868 | -2.185961228 |
| Bex2 | 883.5005139 | 0.250989051 | 0.167290597 | 1.500317746 |
| Nxf3 | 2.723995911 | 0.957909663 | 0.613628028 | 1.561059175 |
| 5730412P04Rik | 0.591118678 | 0.443883693 | 1.183488982 | 0.375063647 |
| Bex4 | 82.98433973 | 0.566682909 | 0.155289774 | 3.649196556 |
| Tceal8 | 272.1310813 | -0.064416367 | 0.142003928 | -0.453623836 |
| Mir6383 | 0 | NA | NA | NA |
| Tceal5 | 256.9790592 | 0.512464598 | 0.233677341 | 2.193043603 |
| Bex1 | 254.7050612 | 0.459216516 | 0.144103757 | 3.186707447 |
| Tceal7 | 6.33875668 | 0.904591503 | 0.46953577 | 1.926565686 |
| Wbp5 | 588.6657451 | -0.164081101 | 0.100732838 | -1.628874007 |
| Ngfrap1 | 809.4390861 | 0.09714891 | 0.155962121 | 0.622900674 |

| | | | | |
|---|---|---|---|---|
| n-R5s12 | 0.061920613 | -0.443816764 | 3.724945049 | -0.119147198 |
| Nsa2-ps2 | 0 | NA | NA | NA |
| Gm14997 | 0 | NA | NA | NA |
| Gm4917 | 0.046345709 | -0.443818988 | 3.724945049 | -0.119147795 |
| Gm7973 | 0.061920613 | -0.443816764 | 3.724945049 | -0.119147198 |
| Gm14994 | 0.090427103 | 0.998850474 | 3.724945049 | 0.268151734 |
| Gm14999 | 0 | NA | NA | NA |
| Gm14998 | 0 | NA | NA | NA |
| Gm14993 | 0 | NA | NA | NA |
| Gm14992 | 0 | NA | NA | NA |
| Gm14996 | 0 | NA | NA | NA |
| Gm14990 | 0 | NA | NA | NA |
| Gm14995 | 0.106112162 | -0.443821212 | 3.724945049 | -0.119148392 |
| Kir3dl2 | 0.620158706 | 1.420562141 | 1.20116347 | 1.182655131 |
| Gm14991 | 0.047948698 | 0.037064902 | 3.724945049 | 0.009950456 |
| Kir3dl1 | 0 | NA | NA | NA |
| Gm15006 | 0 | NA | NA | NA |
| Gm15003 | 0 | NA | NA | NA |
| Gm15005 | 0 | NA | NA | NA |
| Gm15004 | 0 | NA | NA | NA |
| Tceal3 | 620.6626587 | 0.294126412 | 0.120659763 | 2.437651173 |
| Gm15010 | 0.784324088 | 2.389894407 | 1.213851394 | 1.968852546 |
| Gm23068 | 0 | NA | NA | NA |
| Tceal1 | 253.1864694 | 0.319973482 | 0.12402092 | 2.579996033 |
| Morf4l2 | 2139.902612 | -0.106573976 | 0.084216191 | -1.265480841 |
| BC065397 | 34.23957864 | -0.559599129 | 0.269544863 | -2.076089015 |
| Glra4 | 6.335166068 | 0.032093763 | 0.535891978 | 0.059888493 |
| Gm15009 | 0 | NA | NA | NA |
| Plp1 | 16662.56891 | 0.178028278 | 0.142959763 | 1.24530339 |
| Rab9b | 322.6084878 | 0.004330053 | 0.111560603 | 0.038813458 |
| Gm15026 | 0 | NA | NA | NA |
| Gm15025 | 0 | NA | NA | NA |
| H2bfm | 0 | NA | NA | NA |
| Gm6275 | 0.048949532 | 0.037064902 | 3.724945049 | 0.009950456 |
| Tmsb15l | 3.858449061 | 0.268964226 | 0.610333648 | 0.440683922 |
| Tmsb15b2 | 0.857879072 | 1.539351405 | 1.228456368 | 1.253077801 |
| Tmsb15b1 | 3.641358966 | 0.645259588 | 0.664396976 | 0.971195853 |
| Slc25a53 | 178.3064943 | 0.36056295 | 0.132884085 | 2.713364427 |
| Zcchc18 | 1940.436389 | 0.594289026 | 0.069485231 | 8.552738759 |
| Gm15038 | 0 | NA | NA | NA |
| Fam199x | 561.8101836 | 0.066901358 | 0.090267511 | 0.741145474 |
| Gm25770 | 0 | NA | NA | NA |
| Gm8019 | 0 | NA | NA | NA |
| Gm5761 | 0.368947036 | 1.414621492 | 1.554455763 | 0.91004294 |
| Gm15039 | 0 | NA | NA | NA |
| Gm15042 | 0 | NA | NA | NA |
| Gm15045 | 0 | NA | NA | NA |

| | | | | |
|---|---|---|---|---|
| Gm15043 | 0 | NA | NA | NA |
| Gm15044 | 0 | NA | NA | NA |
| Esx1 | 0 | NA | NA | NA |
| Gm15040 | 0.045882455 | 0.517959912 | 3.724945049 | 0.139051692 |
| Gm15041 | 0 | NA | NA | NA |
| Gm23679 | 0 | NA | NA | NA |
| Gm8036 | 0.810529001 | -1.92149885 | 1.204965658 | -1.594650302 |
| Il1rapl2 | 78.53886856 | 0.999879466 | 0.159618028 | 6.264201333 |
| Gm14988 | 0 | NA | NA | NA |
| Gm25393 | 0 | NA | NA | NA |
| Gm14989 | 0 | NA | NA | NA |
| Gm24675 | 0 | NA | NA | NA |
| Tex13a | 0 | NA | NA | NA |
| Gm22057 | 0 | NA | NA | NA |
| Gm23958 | 0 | NA | NA | NA |
| Gm15000 | 0 | NA | NA | NA |
| Gm15001 | 0 | NA | NA | NA |
| Nrk | 62.1717717 | 0.109857499 | 0.26203765 | 0.419243185 |
| Serpina7 | 0.108772582 | 0.037073798 | 3.724945049 | 0.009952844 |
| 4930513O06Rik | 0 | NA | NA | NA |
| 4933428M09Rik | 0 | NA | NA | NA |
| Gm15007 | 0 | NA | NA | NA |
| Mum1l1 | 62.92430637 | -0.088925661 | 0.15181882 | -0.585735424 |
| Platr21 | 0.194631057 | 1.456047977 | 3.23213671 | 0.450490839 |
| Gm15002 | 0 | NA | NA | NA |
| Trap1a | 0 | NA | NA | NA |
| Gm8061 | 0 | NA | NA | NA |
| D330045A20Rik | 6.283406809 | -0.03562516 | 0.416227978 | -0.085590497 |
| Rnf128 | 220.8248196 | -0.178529655 | 0.10926702 | -1.633884181 |
| Gm24380 | 0 | NA | NA | NA |
| Gm6322 | 0 | NA | NA | NA |
| Tbc1d8b | 118.5380032 | -0.666703419 | 0.138541213 | -4.812311117 |
| Gm15013 | 2.788702176 | 0.109410112 | 0.612147674 | 0.178731565 |
| Ripply1 | 0.677889948 | -1.261701199 | 1.327242383 | -0.950618526 |
| Gm15012 | 0.066050875 | -0.443818988 | 3.724945049 | -0.119147795 |
| Cldn2 | 23.20106824 | 0.123891056 | 0.294386797 | 0.42084447 |
| Morc4 | 90.84373692 | -0.538553772 | 0.169340723 | -3.180296878 |
| Rbm41 | 210.9730729 | -0.318305766 | 0.103783023 | -3.067031178 |
| Gm15011 | 0 | NA | NA | NA |
| Nup62cl | 3.861479097 | 0.940746069 | 0.540022444 | 1.742049946 |
| Pih1h3b | 0.733953745 | 0.451783915 | 1.185798941 | 0.380995378 |
| Gm6298 | 0.042716945 | 0.037062677 | 3.724945049 | 0.009949859 |
| Gm15046 | 0.104637558 | -0.443821212 | 3.724945049 | -0.119148392 |
| Mir3475 | 0.844812178 | -0.004804671 | 1.390175003 | -0.003456163 |
| Frmpd3 | 67.41637008 | 0.228164999 | 0.160604133 | 1.420667046 |
| Gm8097 | 0.313170507 | 0.948255994 | 2.070888811 | 0.457898072 |
| Prps1 | 548.0358506 | 0.153144349 | 0.063915509 | 2.396043648 |

| | | | | |
|---|---|---|---|---|
| Tsc22d3 | 1422.675399 | 0.05557938 | 0.154337441 | 0.360115985 |
| Gm15047 | 0 | NA | NA | NA |
| Gm22025 | 0 | NA | NA | NA |
| Mid2 | 552.0542967 | 0.100110453 | 0.076256049 | 1.312819823 |
| Gm23199 | 2.503876714 | -1.434886823 | 0.766292999 | -1.872504153 |
| Eif2c5 | 0 | NA | NA | NA |
| Gm6088 | 0.360750703 | -2.093399773 | 1.505854085 | -1.390174383 |
| Tex13 | 0 | NA | NA | NA |
| Gm7091 | 0 | NA | NA | NA |
| Vsig1 | 0.30513904 | 1.126706188 | 3.711809609 | 0.303546331 |
| Psmd10 | 105.8155227 | -0.451667843 | 0.151306904 | -2.985110611 |
| Atg4a | 98.38075259 | 0.257958918 | 0.170207748 | 1.515553327 |
| Col4a6 | 3.411580873 | -0.563698495 | 0.600313487 | -0.939006881 |
| Gm15077 | 0 | NA | NA | NA |
| Gm15078 | 0 | NA | NA | NA |
| Gm15079 | 0 | NA | NA | NA |
| Col4a5 | 123.2002736 | -1.002499852 | 0.204404282 | -4.904495348 |
| Gm15268 | 0.88858162 | -1.90845248 | 1.272106994 | -1.500229532 |
| Gm15282 | 0 | NA | NA | NA |
| Irs4 | 86.40347737 | 0.198250612 | 0.243038335 | 0.815717456 |
| Gm15295 | 0.642988239 | 0.391873783 | 1.323237547 | 0.296147721 |
| Gm15297 | 0 | NA | NA | NA |
| Hmgn2l6 | 0 | NA | NA | NA |
| Gm15294 | 0.045882455 | 0.517959912 | 3.724945049 | 0.139051692 |
| Gm15298 | 0.198041555 | -1.296529483 | 2.587933318 | -0.500990297 |
| Gm6368 | 8.824265348 | -0.316990161 | 0.370619541 | -0.855298025 |
| Gm5762 | 0 | NA | NA | NA |
| Gm6373 | 0 | NA | NA | NA |
| Gucy2f | 26.45992321 | -0.418250725 | 0.227923264 | -1.835050616 |
| Gm6382 | 25.01211358 | -0.128886909 | 0.222526505 | -0.57919801 |
| Gm15014 | 0 | NA | NA | NA |
| Nxt2 | 433.560056 | 0.154631547 | 0.091270091 | 1.694219278 |
| Gm5643 | 2.51713653 | -0.228558949 | 0.649998682 | -0.351629866 |
| Kcne1l | 125.6411286 | -1.069805239 | 0.241803637 | -4.424272742 |
| Acsl4 | 1433.382608 | 0.127206313 | 0.07229313 | 1.7595906 |
| Gm6067 | 0 | NA | NA | NA |
| Gm7123 | 0 | NA | NA | NA |
| Gm15031 | 0 | NA | NA | NA |
| Gm25915 | 0.060174537 | 0.517957688 | 3.724945049 | 0.139051095 |
| Gm4995 | 0.090537259 | -0.443823436 | 3.724945049 | -0.119148989 |
| Gm15032 | 0 | NA | NA | NA |
| Gm5763 | 8.930037149 | 0.206713694 | 0.338594247 | 0.610505629 |
| Tmem164 | 493.7209216 | 0.021498166 | 0.081416045 | 0.264053183 |
| Mir652 | 2.416261644 | -0.271504973 | 0.824285859 | -0.329382058 |
| Gm15049 | 0.046345709 | -0.443818988 | 3.724945049 | -0.119147795 |
| Ammecr1 | 116.0557192 | 0.11745715 | 0.114868133 | 1.022539031 |
| AL731548.1 | 1.863015902 | -0.090933776 | 0.747545286 | -0.121643167 |

| | | | | |
|---|---|---|---|---|
| Rgag1 | 33.41144712 | 0.763250427 | 0.227494493 | 3.355028146 |
| Gm4760 | 0.046345709 | -0.443818988 | 3.724945049 | -0.119147795 |
| Chrdl1 | 139.3962317 | -0.5503833 | 0.179165953 | -3.071919032 |
| Gm23325 | 0 | NA | NA | NA |
| Pak3 | 2356.012352 | 0.134548719 | 0.102284766 | 1.315432637 |
| Capn6 | 31.3985991 | 0.547981807 | 0.223541478 | 2.451365234 |
| BX530055.1 | 0 | NA | NA | NA |
| Dcx | 707.4354374 | 0.442433407 | 0.161138222 | 2.745676358 |
| Gm15048 | 0.045882455 | 0.517959912 | 3.724945049 | 0.139051692 |
| A730046J19Rik | 10.6199982 | -0.41585861 | 0.362575912 | -1.146955977 |
| Gm8199 | 0.207515118 | 0.010180474 | 2.585168385 | 0.003938031 |
| Gm8202 | 0 | NA | NA | NA |
| Gm15067 | 0 | NA | NA | NA |
| Gm15066 | 0.044544648 | 0.517957688 | 3.724945049 | 0.139051095 |
| Mir680-2 | 0 | NA | NA | NA |
| Alg13 | 168.7701936 | 0.041653119 | 0.113163286 | 0.368079795 |
| Gm2214 | 0.090665643 | 0.037060453 | 3.724945049 | 0.009949262 |
| Trpc5 | 161.8271037 | 0.627927553 | 0.100521554 | 6.246695656 |
| Trpc5os | 0.750003179 | 0.638141516 | 2.46740037 | 0.258629092 |
| Gm15071 | 0 | NA | NA | NA |
| Mageb16-ps1 | 3.715911654 | -0.343662868 | 0.703898587 | -0.488227814 |
| Zcchc16 | 32.00016923 | 0.725264197 | 0.218274719 | 3.322712776 |
| Gm26024 | 2.396750964 | 2.480548187 | 0.948980669 | 2.613908025 |
| Gm8216 | 4.713227208 | 1.716138881 | 0.600174156 | 2.859401499 |
| Mir3472 | 0.066050875 | -0.443818988 | 3.724945049 | -0.119147795 |
| Gm25097 | 0 | NA | NA | NA |
| Gm15073 | 0.066050875 | -0.443818988 | 3.724945049 | -0.119147795 |
| Gm6441 | 0.044191549 | 0.037062677 | 3.724945049 | 0.009949859 |
| Lhfpl1 | 2.080192563 | -0.005990742 | 0.709304955 | -0.008445933 |
| Amot | 561.2978999 | -0.037130034 | 0.106044768 | -0.350135461 |
| Gm5644 | 2.782059835 | 1.236754502 | 0.721320661 | 1.714569635 |
| Gm15060 | 0 | NA | NA | NA |
| Gm27659 | 0.042716945 | 0.037062677 | 3.724945049 | 0.009949859 |
| Gm15058 | 0.106618422 | 0.517955464 | 3.724945049 | 0.139050498 |
| Gm6446 | 0.038981395 | -0.443816764 | 3.724945049 | -0.119147198 |
| Gm15057 | 0.867677903 | -0.513393559 | 1.084720873 | -0.473295547 |
| Gm6447 | 0 | NA | NA | NA |
| Gm15062 | 0 | NA | NA | NA |
| Gm4918 | 0 | NA | NA | NA |
| Hmgb1-ps2 | 15.03640688 | 0.257770166 | 0.263096575 | 0.979754928 |
| Gm15061 | 0 | NA | NA | NA |
| Gm26151 | 0.325583935 | -0.921829719 | 2.048250374 | -0.450057147 |
| Gm15076 | 0 | NA | NA | NA |
| Gm8261 | 2.625663271 | -0.138521377 | 0.737947027 | -0.187711817 |
| Gm15075 | 1.268895825 | -1.137428345 | 1.038545923 | -1.095212373 |
| Gm15074 | 0 | NA | NA | NA |
| Gm4996 | 0 | NA | NA | NA |

| | | | | |
|---|---|---|---|---|
| Htr2c | 718.8775726 | 0.606619357 | 0.113086189 | 5.364221401 |
| Snora35 | 4.50470804 | 0.128694241 | 0.559346276 | 0.230079731 |
| Gm22650 | 4.136795173 | 0.636034729 | 0.63382555 | 1.003485468 |
| Mir764 | 3.164642609 | -0.433103676 | 0.688325335 | -0.629213621 |
| Mir1912 | 2.763607753 | -0.884319534 | 0.70933352 | -1.246690744 |
| Mir1264 | 5.146865135 | 0.504209558 | 0.583302253 | 0.864405297 |
| Mir1298 | 1.82841141 | -0.577259723 | 1.074050496 | -0.537460506 |
| Gm24627 | 1.916860434 | 0.009032276 | 0.857244311 | 0.010536408 |
| Mir448 | 0.601491876 | 0.993155171 | 1.543859997 | 0.643293545 |
| Il13ra2 | 7.235467095 | 0.351232991 | 0.37536072 | 0.935721221 |
| Vmn1r239-ps | 0 | NA | NA | NA |
| Lrch2 | 586.4938171 | 0.26651133 | 0.089291216 | 2.984742977 |
| Gm25107 | 111.0918953 | -0.342432132 | 0.21080517 | -1.624401017 |
| Gm16400 | 0 | NA | NA | NA |
| Gm15167 | 0 | NA | NA | NA |
| Gm8289 | 0 | NA | NA | NA |
| Gm15128 | 0 | NA | NA | NA |
| Gm15081 | 0 | NA | NA | NA |
| Gm15080 | 0 | NA | NA | NA |
| Gm15108 | 0 | NA | NA | NA |
| Gm15110 | 0 | NA | NA | NA |
| Gm8303 | 0 | NA | NA | NA |
| Gm15107 | 0 | NA | NA | NA |
| Gm15109 | 0 | NA | NA | NA |
| Gm15111 | 0 | NA | NA | NA |
| Gm15113 | 0 | NA | NA | NA |
| Gm8320 | 0 | NA | NA | NA |
| Gm15114 | 0 | NA | NA | NA |
| Gm15106 | 0 | NA | NA | NA |
| Gm15112 | 0 | NA | NA | NA |
| Gm8331 | 0 | NA | NA | NA |
| Gm8334 | 0 | NA | NA | NA |
| Gm15127 | 0 | NA | NA | NA |
| Gm15125 | 0 | NA | NA | NA |
| Gm15126 | 0 | NA | NA | NA |
| Kars-ps1 | 0 | NA | NA | NA |
| Luzp4 | 0.048949532 | 0.037064902 | 3.724945049 | 0.009950456 |
| Gm15099 | 0 | NA | NA | NA |
| Gm15084 | 0 | NA | NA | NA |
| Gm15094 | 0 | NA | NA | NA |
| Gm15096 | 0 | NA | NA | NA |
| Ott | 0 | NA | NA | NA |
| Gm15092 | 0 | NA | NA | NA |
| Gm15088 | 0 | NA | NA | NA |
| Gm15090 | 0 | NA | NA | NA |
| Gm8346 | 0 | NA | NA | NA |
| Gm15093 | 0 | NA | NA | NA |

| | | | | |
|---|---|---|---|---|
| Gm15098 | 0 | NA | NA | NA |
| Gm15089 | 0 | NA | NA | NA |
| Gm8366 | 0 | NA | NA | NA |
| Gm15100 | 0 | NA | NA | NA |
| Gm15085 | 0 | NA | NA | NA |
| Gm15086 | 0 | NA | NA | NA |
| Gm10439 | 0 | NA | NA | NA |
| Gm15101 | 0 | NA | NA | NA |
| Gm8383 | 0 | NA | NA | NA |
| Gm15097 | 0 | NA | NA | NA |
| Gm15095 | 0 | NA | NA | NA |
| Gm15087 | 0 | NA | NA | NA |
| Gm15091 | 0 | NA | NA | NA |
| Gm15102 | 0 | NA | NA | NA |
| Gm15103 | 0 | NA | NA | NA |
| Gm8402 | 0 | NA | NA | NA |
| Gm5946 | 2.587997239 | -0.452282588 | 0.653209902 | -0.69240008 |
| Gm15104 | 1.895634264 | 0.799839872 | 0.754288817 | 1.060389407 |
| Gm15105 | 0.604613898 | 0.832574288 | 1.582803096 | 0.526012547 |
| Tmem29 | 269.2719752 | 0.391217144 | 0.111060378 | 3.522562698 |
| Gm15129 | 0 | NA | NA | NA |
| 3110067C02Rik | 4.62715315 | -0.807896725 | 0.567240508 | -1.424257812 |
| Apex2 | 300.6728437 | 0.212743937 | 0.184397377 | 1.153725399 |
| Alas2 | 56.13115503 | 0.702898195 | 0.791725935 | 0.887804938 |
| Mir3620 | 3.543203576 | 1.504477508 | 0.604125257 | 2.490340356 |
| Pfkfb1 | 11.63733137 | 0.221193401 | 0.336338438 | 0.657651271 |
| Tro | 2279.729976 | 0.263441911 | 0.092879003 | 2.836398999 |
| Gm15132 | 0 | NA | NA | NA |
| Gm15131 | 0 | NA | NA | NA |
| Maged2 | 434.343474 | 0.003924132 | 0.116297351 | 0.033742227 |
| Gm24907 | 0.178529143 | 0.037066536 | 2.731815954 | 0.01356846 |
| Itih5l-ps | 0.139037128 | -1.190058937 | 3.717743703 | -0.320102469 |
| Gm27191 | 0 | NA | NA | NA |
| Gm8424 | 0.517778811 | -1.488644659 | 1.360857726 | -1.093901758 |
| Gnl3l | 3189.542655 | 0.222394887 | 0.055478636 | 4.008658149 |
| Gm15152 | 0 | NA | NA | NA |
| Fgd1 | 278.5392714 | -0.098927073 | 0.083608581 | -1.183216743 |
| Tsr2 | 790.8463341 | 0.209741638 | 0.063292621 | 3.313840294 |
| Gm15149 | 0 | NA | NA | NA |
| Rps12-ps7 | 0 | NA | NA | NA |
| Gm15138 | 7.688685255 | 0.213734267 | 0.474344634 | 0.450588563 |
| Wnk3 | 948.2720462 | 0.088242977 | 0.068158706 | 1.294669196 |
| Rpl21-ps15 | 49.38234276 | 0.041925153 | 0.208711382 | 0.200876217 |
| Gm26441 | 0.12209515 | 0.037073798 | 3.724945049 | 0.009952844 |
| A230072E10Rik | 43.55964103 | 0.408701296 | 0.171337961 | 2.385351697 |
| Fam120c | 1332.354665 | -0.026267156 | 0.071502948 | -0.367357662 |
| Gm22248 | 2.415551224 | 1.472884316 | 0.804539358 | 1.830717543 |

| | | | | |
|---|---|---|---|---|
| Gm6451 | 2.057373881 | 0.616609995 | 0.650136781 | 0.948431181 |
| Gm15148 | 0.123992736 | 0.517955521 | 3.720643114 | 0.139211288 |
| Phf8 | 763.8243954 | -0.082448634 | 0.092734172 | -0.889085781 |
| Gm24460 | 0 | NA | NA | NA |
| Gm5395 | 1.549805528 | 0.542998361 | 0.843973773 | 0.64338298 |
| Gm15151 | 0.220031754 | 0.051687712 | 2.58991242 | 0.01995732 |
| Gm4919 | 0 | NA | NA | NA |
| n-R5s13 | 0 | NA | NA | NA |
| Gm15161 | 0 | NA | NA | NA |
| Huwe1 | 8349.182081 | 0.094749392 | 0.052268782 | 1.812733871 |
| Mir3113 | 0 | NA | NA | NA |
| Mirlet7f-2 | 0.623206311 | 0.577381606 | 1.258045622 | 0.458951246 |
| Mir98 | 2.834035009 | -0.138759255 | 0.615720582 | -0.225360755 |
| Gm15191 | 3.302051428 | -0.903075928 | 0.576793367 | -1.565683623 |
| n-R5s14 | 0 | NA | NA | NA |
| Hsd17b10 | 170.8953565 | -0.115684966 | 0.122366115 | -0.945400333 |
| Ribc1 | 4.147795559 | -1.41970026 | 0.539282086 | -2.632574487 |
| Smc1a | 891.0140031 | -0.06944396 | 0.104131202 | -0.666889063 |
| Gm15267 | 3.127372856 | -0.294595205 | 0.550350982 | -0.535286053 |
| Gm15266 | 0 | NA | NA | NA |
| Iqsec2 | 2091.297293 | 0.786421811 | 0.0998806 | 7.873619197 |
| Kdm5c | 1433.211927 | -0.409659509 | 0.057197299 | -7.16221774 |
| Kantr | 325.3470963 | 0.321803099 | 0.089291727 | 3.603952019 |
| n-R5s15 | 0 | NA | NA | NA |
| Tspyl2 | 1239.902915 | 0.204543274 | 0.107936469 | 1.895033955 |
| Gpr173 | 222.5226893 | 0.344177087 | 0.088329314 | 3.896521667 |
| 3010001F23Rik | 25.65934517 | 0.518048962 | 0.375432614 | 1.379872026 |
| Gm4750 | 1.060705812 | -0.330463843 | 0.95967342 | -0.344350313 |
| Gm8464 | 0.320494736 | -0.407637782 | 1.757013116 | -0.232006112 |
| Gm7158 | 0 | NA | NA | NA |
| Cldn34a | 0.10503227 | -0.905414734 | 3.723500899 | -0.243162217 |
| Gm8475 | 0.097139547 | -0.443816764 | 3.724945049 | -0.119147198 |
| Shroom2 | 979.6660779 | 0.43511715 | 0.091824491 | 4.738574048 |
| Gpr143 | 0.64528552 | -0.641471189 | 1.37014302 | -0.46817827 |
| Gm24687 | 0 | NA | NA | NA |
| Rpl7a-ps12 | 0.049553047 | -0.443818988 | 3.724945049 | -0.119147795 |
| Gm6472 | 317.0452057 | -0.349366094 | 0.163119863 | -2.141775302 |
| Gm26035 | 140.1127517 | -0.234412011 | 0.247989238 | -0.945250738 |
| Usp51 | 28.56090422 | -0.019405778 | 0.211637952 | -0.091693278 |
| Mageh1 | 200.6419644 | -0.040567633 | 0.139389554 | -0.291037827 |
| 9530051G07Rik | 22.2217983 | 0.290895197 | 0.277559858 | 1.048044915 |
| Gm5396 | 0.044544648 | 0.517957688 | 3.724945049 | 0.139051095 |
| Gm26960 | 1.043034268 | 1.738519808 | 1.134695573 | 1.532146463 |
| Foxr2 | 13.36216574 | 0.796713207 | 0.313417716 | 2.542017142 |
| Rragb | 646.2289516 | 0.23040336 | 0.073451697 | 3.136801052 |
| Gm15153 | 0.202306569 | -0.443823219 | 2.655697143 | -0.167121172 |
| Klf8 | 93.85799211 | -0.011546784 | 0.130910237 | -0.08820383 |

| | | | | |
|---|---|---|---|---|
| Gm7150 | 0 | NA | NA | NA |
| Ubqln2 | 2353.389334 | 0.153388516 | 0.05391317 | 2.845102883 |
| Gm15166 | 0 | NA | NA | NA |
| Cypt3 | 0 | NA | NA | NA |
| Gm25793 | 0 | NA | NA | NA |
| Gm8534 | 0 | NA | NA | NA |
| Kctd12b | 83.33022565 | -0.074108377 | 0.153125312 | -0.483972091 |
| 2210013O21Rik | 216.6134074 | -0.079131337 | 0.099444349 | -0.795734877 |
| Smt3h1-ps | 0.284401297 | 0.998866659 | 2.063318417 | 0.484106889 |
| Spin2c | 37.12736702 | 0.583411559 | 0.216922809 | 2.689489238 |
| Gm25895 | 0 | NA | NA | NA |
| Samt1 | 0 | NA | NA | NA |
| 4921511M17Rik | 0 | NA | NA | NA |
| Gm10057 | 0 | NA | NA | NA |
| Gm15143 | 0 | NA | NA | NA |
| Gm5947 | 0 | NA | NA | NA |
| Gm15140 | 0 | NA | NA | NA |
| Gm5645 | 0 | NA | NA | NA |
| Gm16445 | 0 | NA | NA | NA |
| Gm6620 | 0 | NA | NA | NA |
| Gm15147 | 0 | NA | NA | NA |
| Gm5060 | 0 | NA | NA | NA |
| Gm15141 | 0 | NA | NA | NA |
| 4930524N10Rik | 0 | NA | NA | NA |
| Gm5646 | 0 | NA | NA | NA |
| Gm8569 | 0 | NA | NA | NA |
| Samt4 | 0 | NA | NA | NA |
| Samt2 | 0 | NA | NA | NA |
| Gm15142 | 0 | NA | NA | NA |
| Gm22359 | 0 | NA | NA | NA |
| Gm8577 | 0.253240508 | 1.037116537 | 2.113001681 | 0.490826177 |
| Cldn34b1 | 0 | NA | NA | NA |
| Magea6 | 0 | NA | NA | NA |
| Magea3 | 0 | NA | NA | NA |
| Gm6645 | 1.418926095 | 0.583322935 | 0.914194904 | 0.638072837 |
| Gm25257 | 0 | NA | NA | NA |
| Magea8 | 0 | NA | NA | NA |
| Magea2 | 0 | NA | NA | NA |
| Gm24268 | 0 | NA | NA | NA |
| Magea5 | 0 | NA | NA | NA |
| Magea1 | 0 | NA | NA | NA |
| Cldn34b2 | 0 | NA | NA | NA |
| Gm25429 | 0.066050875 | -0.443818988 | 3.724945049 | -0.119147795 |
| Gm4997 | 0 | NA | NA | NA |
| Amd-ps1 | 0.844289582 | 1.660048331 | 1.250040482 | 1.327995657 |
| Gm8595 | 0.04393175 | 0.037062677 | 3.724945049 | 0.009949859 |
| Gm15157 | 0.250163086 | 0.561077918 | 2.491267246 | 0.225217876 |

| | | | | |
|---|---|---|---|---|
| Sat1 | 845.2085452 | -0.680165138 | 0.15607573 | -4.357917399 |
| Gm15159 | 0 | NA | NA | NA |
| Acot9 | 212.0243755 | -0.208898297 | 0.120949122 | -1.727158441 |
| Gm650 | 0 | NA | NA | NA |
| Prdx4 | 114.043658 | -0.254341352 | 0.213445369 | -1.191599298 |
| Gm15158 | 0 | NA | NA | NA |
| Gm8606 | 0.172182752 | 0.037075442 | 3.331044665 | 0.011130275 |
| n-R5s16 | 0 | NA | NA | NA |
| Ptchd1 | 278.1680322 | 0.050119307 | 0.094071596 | 0.532778321 |
| Gm15156 | 5.651205916 | 0.019608189 | 0.38871225 | 0.050443969 |
| Gm15155 | 12.34800111 | 0.91428013 | 0.352654125 | 2.592568936 |
| Gm15160 | 1.808882573 | 1.342195129 | 0.793691767 | 1.691078559 |
| Gm23404 | 0 | NA | NA | NA |
| 4930503H13Rik | 1.186325201 | 2.39183519 | 1.069024084 | 2.237400659 |
| Gm7312 | 0 | NA | NA | NA |
| Gm15162 | 0.389425067 | 1.501600372 | 3.266995485 | 0.459627318 |
| Gm15163 | 28.67019377 | 0.55619299 | 0.210379583 | 2.643759357 |
| Gm22666 | 0 | NA | NA | NA |
| Phex | 65.77163163 | -0.177578833 | 0.194687702 | -0.912121471 |
| Gm23798 | 0 | NA | NA | NA |
| Gm25795 | 0.325008436 | 0.053676048 | 2.041842748 | 0.026288042 |
| Gm15172 | 0.412251148 | 0.037075403 | 1.779798312 | 0.020831238 |
| Gm1947 | 0.704217346 | -3.247048989 | 1.306174206 | -2.485923375 |
| Gm24281 | 0 | NA | NA | NA |
| Sms | 958.8198068 | 0.314924789 | 0.058383315 | 5.394088838 |
| Mbtps2 | 467.2768864 | 0.206121717 | 0.064933239 | 3.174363699 |
| Gm6568 | 0.666747433 | -2.19833758 | 1.277592865 | -1.720687115 |
| Yy2 | 0 | NA | NA | NA |
| Gm15176 | 0 | NA | NA | NA |
| Gm15175 | 0.320922609 | 0.461777317 | 1.723584881 | 0.267916783 |
| Gm15173 | 0.066050875 | -0.443818988 | 3.724945049 | -0.119147795 |
| Gm23381 | 0 | NA | NA | NA |
| Gm8644 | 0.329848065 | -2.063853169 | 1.872399294 | -1.102250559 |
| Gm15171 | 0 | NA | NA | NA |
| Gm8648 | 0 | NA | NA | NA |
| Smpx | 10.49823735 | 0.553629903 | 0.3402406 | 1.627171782 |
| Gm15169 | 2.861039839 | 0.029372328 | 0.58580655 | 0.050139978 |
| Klhl34 | 1.755040647 | 0.607758168 | 0.739458002 | 0.821896803 |
| Cnksr2 | 3647.587889 | 0.581411413 | 0.126557246 | 4.594058665 |
| Gm8659 | 0.100902008 | -0.876201588 | 3.72093227 | -0.235479048 |
| Gm15170 | 0.046345709 | -0.443818988 | 3.724945049 | -0.119147795 |
| Gm8662 | 0 | NA | NA | NA |
| Gm15177 | 0 | NA | NA | NA |
| Gm22522 | 0 | NA | NA | NA |
| Gm15168 | 0 | NA | NA | NA |
| Gm5764 | 0.044544648 | 0.517957688 | 3.724945049 | 0.139051095 |
| Gm15185 | 0 | NA | NA | NA |

| | | | | |
|---|---|---|---|---|
| Gm15188 | 0 | NA | NA | NA |
| Gm15189 | 0 | NA | NA | NA |
| Rps6ka3 | 1265.248659 | -0.430112803 | 0.073038242 | -5.888871235 |
| Eif1ax | 1820.91249 | 0.297011284 | 0.070087467 | 4.237723198 |
| Gm15187 | 0 | NA | NA | NA |
| Map7d2 | 1767.738311 | 0.220181028 | 0.090347886 | 2.437035743 |
| Gm23806 | 1.294438978 | 0.607666385 | 0.940499588 | 0.646110208 |
| A830080D01Rik | 225.115431 | -0.275455153 | 0.090592028 | -3.040611407 |
| Gm15190 | 9.37392129 | -0.479440894 | 0.348161773 | -1.37706357 |
| Gm15186 | 6.883933299 | -0.298848022 | 0.471303416 | -0.634088386 |
| Sh3kbp1 | 688.8231907 | -0.032236184 | 0.060011542 | -0.537166401 |
| Map3k15 | 25.20550808 | -0.216869409 | 0.248365413 | -0.873186836 |
| Gm15194 | 0 | NA | NA | NA |
| Gm8682 | 0 | NA | NA | NA |
| Pdha1 | 1985.030852 | 0.247469021 | 0.052983532 | 4.670678093 |
| Gm15195 | 0 | NA | NA | NA |
| Gm15192 | 0 | NA | NA | NA |
| Gm15196 | 0.075043205 | 0.517957688 | 3.724945049 | 0.139051095 |
| Gm15193 | 0 | NA | NA | NA |
| Adgrg2 | 34.96479159 | -0.342120048 | 0.239925388 | -1.425943502 |
| Gm15241 | 0.873806335 | 0.490423795 | 0.993823088 | 0.493471927 |
| Phka2 | 591.6410219 | 0.66915532 | 0.131000249 | 5.108046178 |
| Gm15243 | 0.261996957 | 0.517974802 | 2.055677235 | 0.251972826 |
| Ppef1 | 10.0328463 | -0.08005453 | 0.337711105 | -0.237050333 |
| Gm15242 | 0.08486385 | 0.037730769 | 3.724898049 | 0.010129343 |
| Gm15244 | 0.061920613 | -0.443816764 | 3.724945049 | -0.119147198 |
| Rs1 | 1.55354348 | 0.62983778 | 0.931586109 | 0.676091855 |
| Cdkl5 | 3363.729212 | 0.571740631 | 0.104017483 | 5.496582047 |
| Gja6 | 33.35128538 | 0.510906401 | 0.298555122 | 1.711263224 |
| Scml2 | 17.41690607 | -0.212229324 | 0.274420916 | -0.773371532 |
| Gm5419 | 0 | NA | NA | NA |
| Gm15262 | 0 | NA | NA | NA |
| Prkaca-ps1 | 0.150961315 | -0.410359602 | 2.845512226 | -0.144212911 |
| Rai2 | 127.3185232 | 0.118893123 | 0.100581715 | 1.182055031 |
| Scml1 | 0.926668879 | 0.394528611 | 1.09627147 | 0.35988222 |
| Gm15205 | 1.789402658 | -0.586668689 | 0.869560593 | -0.67467258 |
| Nhs | 212.8785834 | -0.032456243 | 0.095421076 | -0.340137053 |
| Gm15202 | 0.888501202 | -0.703667431 | 1.275735425 | -0.551577872 |
| Gm26317 | 0.531781488 | -0.354381973 | 1.406465642 | -0.251966321 |
| Gm23794 | 0.04393175 | 0.037062677 | 3.724945049 | 0.009949859 |
| Gm27490 | 0 | NA | NA | NA |
| Gm23867 | 0.167772976 | -0.985237982 | 3.413460593 | -0.288633179 |
| Gm15204 | 0.395626157 | -1.237702097 | 1.533945954 | -0.806874645 |
| Reps2 | 2262.978211 | 0.507861346 | 0.078824632 | 6.442926952 |
| Gm7331 | 62.14126764 | -0.204940765 | 0.174429004 | -1.174923662 |
| Gm15206 | 2.290494867 | 1.299990593 | 0.788651252 | 1.648371939 |
| Gm15203 | 0 | NA | NA | NA |

| | | | | |
|---|---|---|---|---|
| Gm22257 | 0 | NA | NA | NA |
| Gm25557 | 0 | NA | NA | NA |
| Gm8737 | 0 | NA | NA | NA |
| Gm15201 | 8.230899355 | 0.991678396 | 0.398738695 | 2.487038271 |
| Gm23786 | 0 | NA | NA | NA |
| Rbbp7 | 1339.028074 | -0.122866353 | 0.106781892 | -1.150629105 |
| Txlng | 376.7131286 | 0.184481267 | 0.099260559 | 1.85855559 |
| Gm16459 | 0 | NA | NA | NA |
| Gm15207 | 0 | NA | NA | NA |
| Gm23901 | 0.060174537 | 0.517957688 | 3.724945049 | 0.139051095 |
| Syap1 | 318.0290325 | -0.184910342 | 0.085744514 | -2.156526783 |
| Mir3473a | 0.568149222 | -0.450680627 | 1.556791845 | -0.289493183 |
| Gm15209 | 0 | NA | NA | NA |
| Ctps2 | 433.4122222 | 0.021435791 | 0.064266312 | 0.333546312 |
| S100g | 0.048949532 | 0.037064902 | 3.724945049 | 0.009950456 |
| Gm23501 | 0 | NA | NA | NA |
| Gm22746 | 0 | NA | NA | NA |
| Gm15215 | 0 | NA | NA | NA |
| Gm15212 | 0 | NA | NA | NA |
| Gm15214 | 0 | NA | NA | NA |
| Gm26007 | 0 | NA | NA | NA |
| Gm7199 | 0 | NA | NA | NA |
| Grpr | 12.17739777 | -0.370387401 | 0.314901793 | -1.176199723 |
| Rnf138rt1 | 7.321455992 | -0.026105037 | 0.378682874 | -0.068936408 |
| Gm15213 | 0 | NA | NA | NA |
| Ap1s2 | 658.6125027 | -0.159874093 | 0.106645372 | -1.4991189 |
| Zrsr2 | 430.8776927 | 0.104249091 | 0.086074854 | 1.211144561 |
| Car5b | 61.00096164 | -0.116958399 | 0.190417709 | -0.614220178 |
| Siah1b | 50.98963704 | 0.186625603 | 0.158200548 | 1.179677347 |
| Tmem27 | 1.459315314 | -0.12489474 | 0.828300477 | -0.150784339 |
| n-R5s18 | 0.046345709 | -0.443818988 | 3.724945049 | -0.119147795 |
| Ace2 | 16.8793861 | 0.344499764 | 0.346913198 | 0.993043117 |
| Bmx | 4.886164753 | -0.167798154 | 0.503315843 | -0.333385401 |
| Gm25367 | 0 | NA | NA | NA |
| Pir | 44.18337336 | 0.07617331 | 0.184198494 | 0.413539263 |
| Gm22023 | 0.437539382 | -0.472039931 | 1.729120837 | -0.272994184 |
| Figf | 22.19425402 | 0.413987876 | 0.296952512 | 1.39412148 |
| Piga | 134.018528 | -0.18471728 | 0.119981173 | -1.539552205 |
| Asb11 | 36.5181848 | 0.777133308 | 0.232768097 | 3.338659027 |
| Gm15211 | 0.30015424 | -1.073815688 | 1.987472642 | -0.54029206 |
| Asb9 | 0.209460102 | 0.053256662 | 2.187386167 | 0.02434717 |
| Gm15216 | 0 | NA | NA | NA |
| Mospd2 | 329.2346432 | -0.21758387 | 0.096793864 | -2.247909745 |
| Fancb | 25.83413697 | 0.138354434 | 0.242090437 | 0.571498965 |
| Gm17604 | 0 | NA | NA | NA |
| Glra2 | 135.3189216 | 0.04236687 | 0.145680703 | 0.290820051 |
| Gm15236 | 0.167734623 | -0.390161801 | 3.392587224 | -0.115004206 |

| | | | | |
|---|---|---|---|---|
| Gm22062 | 0 | NA | NA | NA |
| Gm15224 | 0 | NA | NA | NA |
| Gm15225 | 0 | NA | NA | NA |
| Gemin8 | 60.76505143 | -0.047374968 | 0.178417004 | -0.265529443 |
| Gm15227 | 0 | NA | NA | NA |
| Gpm6b | 8313.7276 | 0.216437779 | 0.111834139 | 1.935346223 |
| Ofd1 | 183.0566898 | 0.029510159 | 0.109420261 | 0.269695566 |
| Trappc2 | 241.3367123 | 0.156474414 | 0.105511337 | 1.483010435 |
| Rab9 | 264.7892433 | -0.252873741 | 0.094877508 | -2.665265431 |
| Gm15229 | 2.091380692 | -3.405915574 | 1.02070518 | -3.33682599 |
| Gm15223 | 0.211816017 | 0.952010571 | 2.167791647 | 0.439161472 |
| Tceanc | 44.56431763 | -0.212581577 | 0.198627509 | -1.070252448 |
| Egfl6 | 12.75317809 | 0.275839352 | 0.358043414 | 0.770407558 |
| Gm15226 | 0.151064894 | 0.48574719 | 2.845233496 | 0.170723138 |
| Gm1720 | 0.332179105 | -0.440483005 | 1.623250558 | -0.271358603 |
| Rpl7a-ps11 | 9.849636779 | 0.143233333 | 0.329367306 | 0.434874166 |
| Gm15230 | 0.199910846 | -0.40079097 | 2.208307074 | -0.181492409 |
| Gm15231 | 0 | NA | NA | NA |
| AL671888.1 | 0.061920613 | -0.443816764 | 3.724945049 | -0.119147198 |
| Gm22368 | 0 | NA | NA | NA |
| Gm8814 | 0.110595523 | 0.037071574 | 3.724945049 | 0.009952247 |
| Gm8817 | 0.926280627 | 0.160436248 | 0.954706013 | 0.168047803 |
| Gm15232 | 1.683504739 | -0.249769314 | 0.925297758 | -0.269933989 |
| Gm15228 | 2.037836299 | 0.029396426 | 0.627666531 | 0.046834465 |
| Tmsb4x | 4681.12954 | 0.00919146 | 0.106074786 | 0.08665075 |
| Tlr8 | 77.00154128 | -2.06259765 | 0.496898684 | -4.150942063 |
| Tlr7 | 273.9150659 | -0.571524684 | 0.192927513 | -2.9623804 |
| Prps2 | 362.3824202 | -0.18055074 | 0.100634494 | -1.794123795 |
| Gm15239 | 0.195968864 | 1.553849027 | 3.234710455 | 0.480367269 |
| Gm26368 | 0.238567011 | -1.095339814 | 2.543006771 | -0.430726267 |
| Gm6744 | 0.523620968 | 2.534981125 | 3.095881875 | 0.818823595 |
| Frmpd4 | 1476.981339 | 0.558334441 | 0.071390036 | 7.820901495 |
| Gm15238 | 0.110870145 | -0.444802825 | 3.724874622 | -0.119414174 |
| Gm24772 | 0 | NA | NA | NA |
| Gm15240 | 0 | NA | NA | NA |
| Gm15237 | 0 | NA | NA | NA |
| Gm8832 | 0.092228164 | 0.037073798 | 3.724945049 | 0.009952844 |
| Gm7209 | 0 | NA | NA | NA |
| Gm23197 | 0 | NA | NA | NA |
| Msl3 | 245.0695052 | -0.105729399 | 0.076146235 | -1.388504609 |
| Gm15233 | 0 | NA | NA | NA |
| Arhgap6 | 136.4378488 | 0.605222366 | 0.133421955 | 4.536152727 |
| Gm15261 | 0.130746305 | 0.417046913 | 3.663433875 | 0.113840437 |
| Amelx | 0.092691418 | -0.908687078 | 3.724560878 | -0.2439716 |
| Hccs | 283.4725314 | -0.292582221 | 0.102645749 | -2.850407573 |
| Gm27980 | 0.850639233 | 0.697760256 | 1.204589926 | 0.57925128 |
| Gm15246 | 24.67077973 | 0.47121931 | 0.313438366 | 1.503387462 |

| | | | | |
|---|---|---|---|---|
| BC022960 | 47.81857404 | 0.535817496 | 0.181494729 | 2.952248252 |
| Gm15245 | 0.062426873 | 0.517959912 | 3.724945049 | 0.139051692 |
| Mid1 | 46.97430459 | -0.973201874 | 0.198970981 | -4.891174937 |
| Gm25651 | 0 | NA | NA | NA |
| 4933400A11Rik | 0 | NA | NA | NA |
| G530011O06Rik | 0.809500859 | -0.165773337 | 1.236577978 | -0.134058134 |
| Gm15726 | 1.372866643 | -0.010680085 | 0.858568815 | -0.012439405 |
| Gm15247 | 0.097139547 | -0.443816764 | 3.724945049 | -0.119147198 |
| Gm21887 | 20.27296318 | 0.214647548 | 0.380502553 | 0.564115921 |
| Asmt | 1.358880654 | 0.235448071 | 0.913864269 | 0.257640088 |
| Gm15069 | 0 | NA | NA | NA |
| Gm15068 | 0 | NA | NA | NA |
| Gm29277 | 0 | NA | NA | NA |
| Gm29089 | 0 | NA | NA | NA |
| Gm2098 | 0.060174537 | 0.517957688 | 3.724945049 | 0.139051095 |
| Zfy1 | 0.044191549 | 0.037062677 | 3.724945049 | 0.009949859 |
| Uba1y | 4.871321793 | -0.614719326 | 0.536658107 | -1.145458009 |
| Gm28588 | 0 | NA | NA | NA |
| Gm28587 | 0.444799355 | 1.781863448 | 3.098969343 | 0.574985826 |
| Rhoay-ps3 | 0.088383099 | 0.037058229 | 3.724945049 | 0.009948665 |
| Kdm5d | 320.9905738 | 0.228170133 | 0.187768929 | 1.215164484 |
| n-R5s1 | 0 | NA | NA | NA |
| AC145392.1 | 0 | NA | NA | NA |
| Eif2s3y | 295.8023547 | -0.311371498 | 0.16182037 | -1.924179861 |
| Gm29650 | 9.742440598 | -0.857914771 | 0.614468524 | -1.396189939 |
| Tspy-ps | 2.062938699 | 1.243453281 | 0.98448891 | 1.263044477 |
| Uty | 395.1440632 | 0.052450788 | 0.126520291 | 0.414564238 |
| Ddx3y | 555.3045698 | 0.023915891 | 0.145945457 | 0.163868691 |
| Gm4017 | 4.097680658 | 1.007254025 | 0.632060511 | 1.593603788 |
| Gm18665 | 3.611403245 | 0.656958093 | 0.591843937 | 1.110019132 |
| Usp9y | 1.14946738 | -1.055350607 | 3.69849518 | -0.285345946 |
| Gm25565 | 0 | NA | NA | NA |
| Gm17790 | 0 | NA | NA | NA |
| Gm29167 | 0 | NA | NA | NA |
| Gm8446 | 0.334381441 | 0.03707683 | 1.80310579 | 0.020562759 |
| Gm28534 | 0 | NA | NA | NA |
| Gm28975 | 0 | NA | NA | NA |
| Gm28974 | 0 | NA | NA | NA |
| Gm18797 | 0 | NA | NA | NA |
| Rhoay-ps1 | 0 | NA | NA | NA |
| Zfy2 | 0 | NA | NA | NA |
| Uba1y-ps2 | 0 | NA | NA | NA |
| Gm18796 | 0 | NA | NA | NA |
| Gm29552 | 0 | NA | NA | NA |
| Gm29550 | 0 | NA | NA | NA |
| Gm29551 | 0 | NA | NA | NA |
| Gm8480 | 0 | NA | NA | NA |

| | | | | |
|---|---|---|---|---|
| Gm29553 | 0 | NA | NA | NA |
| Gm22119 | 0 | NA | NA | NA |
| Gm28630 | 0 | NA | NA | NA |
| Gm28629 | 0 | NA | NA | NA |
| Gm6026 | 0 | NA | NA | NA |
| Uba1y-ps1 | 0 | NA | NA | NA |
| Sry | 0 | NA | NA | NA |
| Gm18798 | 0 | NA | NA | NA |
| Gm16501 | 0 | NA | NA | NA |
| Gm28628 | 0 | NA | NA | NA |
| Gm4064 | 0 | NA | NA | NA |
| Gm6137 | 0 | NA | NA | NA |
| Gm8498 | 0 | NA | NA | NA |
| Rbmy | 0 | NA | NA | NA |
| Gm10256 | 0 | NA | NA | NA |
| Gm8502 | 0 | NA | NA | NA |
| Gm8506 | 0 | NA | NA | NA |
| Gm10352 | 0 | NA | NA | NA |
| Gm8510 | 0 | NA | NA | NA |
| Gm29289 | 0 | NA | NA | NA |
| Gm21677 | 0 | NA | NA | NA |
| Gm29288 | 0 | NA | NA | NA |
| Gm21693 | 0 | NA | NA | NA |
| Gm21139 | 0 | NA | NA | NA |
| Gm21704 | 0 | NA | NA | NA |
| Gm21147 | 0 | NA | NA | NA |
| Gm21708 | 0 | NA | NA | NA |
| Gm21156 | 0 | NA | NA | NA |
| Rhoay-ps2 | 0 | NA | NA | NA |
| Gm2316 | 0 | NA | NA | NA |
| Gm3376 | 0 | NA | NA | NA |
| Gm2325 | 0 | NA | NA | NA |
| Gm28649 | 0 | NA | NA | NA |
| Gm8521 | 0 | NA | NA | NA |
| Gm18177 | 0 | NA | NA | NA |
| Gm28429 | 0 | NA | NA | NA |
| Gm28413 | 0 | NA | NA | NA |
| Gm28412 | 0 | NA | NA | NA |
| Gm8525 | 0 | NA | NA | NA |
| Gm28414 | 0 | NA | NA | NA |
| Gm28648 | 0 | NA | NA | NA |
| Gm28647 | 0 | NA | NA | NA |
| Gm29435 | 0 | NA | NA | NA |
| Vmn2r-ps139 | 0.046345709 | -0.443818988 | 3.724945049 | -0.119147795 |
| Gm29038 | 0 | NA | NA | NA |
| Gm28191 | 0 | NA | NA | NA |
| Gm28186 | 0 | NA | NA | NA |

| | | | | |
|---|---|---|---|---|
| Gm21064 | 0 | NA | NA | NA |
| Gm28241 | 0 | NA | NA | NA |
| Gm28242 | 0 | NA | NA | NA |
| Gm37222 | 0 | NA | NA | NA |
| Gm28356 | 0 | NA | NA | NA |
| Gm28357 | 0 | NA | NA | NA |
| Gm29352 | 0 | NA | NA | NA |
| Gm29351 | 0 | NA | NA | NA |
| Gm29350 | 0 | NA | NA | NA |
| Gm29349 | 0 | NA | NA | NA |
| Gm29355 | 0 | NA | NA | NA |
| Gm20918 | 0 | NA | NA | NA |
| Gm29354 | 0 | NA | NA | NA |
| Gm29353 | 0 | NA | NA | NA |
| Gm21820 | 0 | NA | NA | NA |
| Gm29357 | 0 | NA | NA | NA |
| Gm29356 | 0 | NA | NA | NA |
| Gm29161 | 0 | NA | NA | NA |
| Gm28767 | 0 | NA | NA | NA |
| Gm29160 | 0 | NA | NA | NA |
| Gm20915 | 0 | NA | NA | NA |
| Gm28766 | 0 | NA | NA | NA |
| Gm29159 | 0 | NA | NA | NA |
| Gm27663 | 0 | NA | NA | NA |
| Gm29158 | 0 | NA | NA | NA |
| Gm21854 | 0 | NA | NA | NA |
| Gm28768 | 0 | NA | NA | NA |
| Gm28769 | 0 | NA | NA | NA |
| Gm20914 | 0 | NA | NA | NA |
| Gm28171 | 0 | NA | NA | NA |
| Gm28173 | 0 | NA | NA | NA |
| Gm37236 | 0 | NA | NA | NA |
| Gm29194 | 0.066050875 | -0.443818988 | 3.724945049 | -0.119147795 |
| Gm21778 | 0 | NA | NA | NA |
| Gm29195 | 0 | NA | NA | NA |
| Gm29196 | 0 | NA | NA | NA |
| Gm28435 | 0 | NA | NA | NA |
| Gm28436 | 0 | NA | NA | NA |
| Gm29197 | 0 | NA | NA | NA |
| Gm29198 | 0 | NA | NA | NA |
| Gm29193 | 0 | NA | NA | NA |
| Gm28430 | 0 | NA | NA | NA |
| Gm21746 | 0 | NA | NA | NA |
| Gm28593 | 0 | NA | NA | NA |
| Gm20873 | 0 | NA | NA | NA |
| Gm28916 | 0 | NA | NA | NA |
| Gm28915 | 0 | NA | NA | NA |

| | | | | |
|---|---|---|---|---|
| Gm28918 | 0.060174537 | 0.517957688 | 3.724945049 | 0.139051095 |
| Gm28917 | 0 | NA | NA | NA |
| Gm28919 | 0 | NA | NA | NA |
| Gm21719 | 0 | NA | NA | NA |
| Gm28921 | 0 | NA | NA | NA |
| Gm28920 | 0 | NA | NA | NA |
| Gm28444 | 0 | NA | NA | NA |
| Gm28443 | 0 | NA | NA | NA |
| Gm20830 | 0 | NA | NA | NA |
| Gm28442 | 0 | NA | NA | NA |
| Gm28445 | 0 | NA | NA | NA |
| Gm20788 | 0 | NA | NA | NA |
| Gm28999 | 0 | NA | NA | NA |
| Gm21244 | 0.139376635 | 0.984529594 | 2.922770152 | 0.336848107 |
| Gm28997 | 0 | NA | NA | NA |
| Gm28998 | 0 | NA | NA | NA |
| Gm28398 | 0 | NA | NA | NA |
| Gm28397 | 0 | NA | NA | NA |
| Gm28400 | 0 | NA | NA | NA |
| Gm28399 | 0 | NA | NA | NA |
| Gm28395 | 0 | NA | NA | NA |
| Gm21788 | 0 | NA | NA | NA |
| Gm28394 | 0 | NA | NA | NA |
| Gm28396 | 0 | NA | NA | NA |
| Gm28393 | 0 | NA | NA | NA |
| Gm20826 | 0.204257686 | 0.517965843 | 2.633377662 | 0.196692579 |
| Vmn2r-ps149 | 0 | NA | NA | NA |
| Gm28576 | 0 | NA | NA | NA |
| Gm28575 | 0 | NA | NA | NA |
| Gm28574 | 0 | NA | NA | NA |
| Gm28573 | 0 | NA | NA | NA |
| Gm28572 | 0 | NA | NA | NA |
| Gm28571 | 0 | NA | NA | NA |
| Gm28570 | 0 | NA | NA | NA |
| Gm29049 | 0 | NA | NA | NA |
| Gm20825 | 0 | NA | NA | NA |
| Gm29047 | 0 | NA | NA | NA |
| Gm29048 | 0 | NA | NA | NA |
| Gm29045 | 0 | NA | NA | NA |
| Gm29046 | 0 | NA | NA | NA |
| Gm29043 | 0 | NA | NA | NA |
| Gm29044 | 0 | NA | NA | NA |
| Gm37951 | 0 | NA | NA | NA |
| Gm20824 | 0 | NA | NA | NA |
| Gm28146 | 0 | NA | NA | NA |
| Gm28145 | 0 | NA | NA | NA |
| Gm28148 | 0 | NA | NA | NA |

| | | | | |
|---|---|---|---|---|
| Gm28147 | 0 | NA | NA | NA |
| Gm28142 | 0 | NA | NA | NA |
| Gm28141 | 0 | NA | NA | NA |
| Gm28144 | 0 | NA | NA | NA |
| Gm28143 | 0 | NA | NA | NA |
| Gm28150 | 0 | NA | NA | NA |
| Gm28149 | 0 | NA | NA | NA |
| Gm28098 | 0 | NA | NA | NA |
| Gm28330 | 0 | NA | NA | NA |
| Gm28331 | 0 | NA | NA | NA |
| Gm28099 | 0 | NA | NA | NA |
| Gm20815 | 0 | NA | NA | NA |
| Gm28328 | 0 | NA | NA | NA |
| Gm28329 | 0 | NA | NA | NA |
| Gm28270 | 0 | NA | NA | NA |
| Gm28327 | 0 | NA | NA | NA |
| Gm28097 | 0 | NA | NA | NA |
| Gm29365 | 0 | NA | NA | NA |
| Gm29364 | 0 | NA | NA | NA |
| Gm29363 | 0 | NA | NA | NA |
| Gm21292 | 0 | NA | NA | NA |
| Gm21721 | 0 | NA | NA | NA |
| Gm29366 | 0 | NA | NA | NA |
| Gm29281 | 0 | NA | NA | NA |
| Gm29330 | 0 | NA | NA | NA |
| Gm29329 | 0 | NA | NA | NA |
| Gm21812 | 0 | NA | NA | NA |
| Gm29535 | 0 | NA | NA | NA |
| Gm29533 | 0 | NA | NA | NA |
| Gm29534 | 0 | NA | NA | NA |
| Gm21874 | 0 | NA | NA | NA |
| Gm28849 | 0 | NA | NA | NA |
| Gm28847 | 0 | NA | NA | NA |
| Gm28848 | 0 | NA | NA | NA |
| Gm28846 | 0 | NA | NA | NA |
| Gm29471 | 0 | NA | NA | NA |
| Gm29474 | 0 | NA | NA | NA |
| Gm29475 | 0 | NA | NA | NA |
| Gm20821 | 0 | NA | NA | NA |
| Gm28506 | 0 | NA | NA | NA |
| Gm28507 | 0 | NA | NA | NA |
| Gm21301 | 0 | NA | NA | NA |
| Gm29142 | 0 | NA | NA | NA |
| Gm21310 | 0 | NA | NA | NA |
| Gm28509 | 0 | NA | NA | NA |
| Gm28508 | 0 | NA | NA | NA |
| Gm20834 | 0 | NA | NA | NA |

| | | | | |
|---|---|---|---|---|
| Gm28510 | 0 | NA | NA | NA |
| Gm29525 | 0 | NA | NA | NA |
| Gm29526 | 0 | NA | NA | NA |
| Gm20737 | 0.044544648 | 0.517957688 | 3.724945049 | 0.139051095 |
| Gm21775 | 0 | NA | NA | NA |
| Gm21900 | 0 | NA | NA | NA |
| Gm29527 | 0 | NA | NA | NA |
| Gm20775 | 1.978445198 | -0.520653097 | 0.675638385 | -0.770609113 |
| Gm29522 | 0 | NA | NA | NA |
| Gm29523 | 0 | NA | NA | NA |
| Gm29524 | 0 | NA | NA | NA |
| Gm29528 | 0 | NA | NA | NA |
| Gm20777 | 0 | NA | NA | NA |
| Gm28955 | 0 | NA | NA | NA |
| Gm28954 | 0 | NA | NA | NA |
| Gm28953 | 0 | NA | NA | NA |
| Gm20776 | 0 | NA | NA | NA |
| Gm28958 | 0 | NA | NA | NA |
| Gm20828 | 0 | NA | NA | NA |
| Gm28957 | 0 | NA | NA | NA |
| Gm28956 | 0 | NA | NA | NA |
| Gm28952 | 0 | NA | NA | NA |
| Gm28951 | 0 | NA | NA | NA |
| Gm28390 | 0 | NA | NA | NA |
| Gm28391 | 0 | NA | NA | NA |
| Gm28389 | 0 | NA | NA | NA |
| Gm28655 | 0 | NA | NA | NA |
| Gm28654 | 0 | NA | NA | NA |
| Gm28657 | 0 | NA | NA | NA |
| Gm28656 | 0 | NA | NA | NA |
| Gm29554 | 0 | NA | NA | NA |
| Gm29556 | 0 | NA | NA | NA |
| Gm20812 | 0 | NA | NA | NA |
| Gm35108 | 0 | NA | NA | NA |
| Gm37739 | 0 | NA | NA | NA |
| Gm21736 | 0 | NA | NA | NA |
| Gm21379 | 0 | NA | NA | NA |
| Gm36928 | 0 | NA | NA | NA |
| Gm20807 | 0 | NA | NA | NA |
| Gm38084 | 0 | NA | NA | NA |
| Gm37231 | 0 | NA | NA | NA |
| Gm21891 | 0 | NA | NA | NA |
| Gm21425 | 0 | NA | NA | NA |
| Gm37252 | 0 | NA | NA | NA |
| Gm21828 | 0 | NA | NA | NA |
| Gm21440 | 0 | NA | NA | NA |
| Gm21454 | 0 | NA | NA | NA |

| | | | | |
|---|---|---|---|---|
| Gm37147 | 0 | NA | NA | NA |
| Gm38159 | 0 | NA | NA | NA |
| Gm21725 | 0 | NA | NA | NA |
| Gm37870 | 0 | NA | NA | NA |
| Gm37378 | 0 | NA | NA | NA |
| Gm21866 | 0 | NA | NA | NA |
| Gm20773 | 0 | NA | NA | NA |
| Gm36398 | 0 | NA | NA | NA |
| Gm37441 | 0 | NA | NA | NA |
| Gm37561 | 0 | NA | NA | NA |
| Gm36728 | 0 | NA | NA | NA |
| Gm32295 | 0 | NA | NA | NA |
| Gm20836 | 0 | NA | NA | NA |
| Gm37263 | 0 | NA | NA | NA |
| Gm21767 | 0 | NA | NA | NA |
| Gm37130 | 0 | NA | NA | NA |
| Gm37467 | 0 | NA | NA | NA |
| Gm20928 | 0 | NA | NA | NA |
| Gm38363 | 0 | NA | NA | NA |
| Gm30174 | 0 | NA | NA | NA |
| Gm30353 | 0 | NA | NA | NA |
| Gm30686 | 0 | NA | NA | NA |
| Gm30737 | 0 | NA | NA | NA |
| Gm36987 | 0 | NA | NA | NA |
| Gm31186 | 0 | NA | NA | NA |
| Gm37654 | 0 | NA | NA | NA |
| Gm37044 | 0 | NA | NA | NA |
| Gm31511 | 0 | NA | NA | NA |
| Gm37572 | 0 | NA | NA | NA |
| Gm37344 | 0 | NA | NA | NA |
| Gm37059 | 0 | NA | NA | NA |
| Gm31942 | 0 | NA | NA | NA |
| Gm37440 | 0 | NA | NA | NA |
| Gm38185 | 0 | NA | NA | NA |
| Gm37395 | 0 | NA | NA | NA |
| Gm37577 | 0 | NA | NA | NA |
| Gm20822 | 0 | NA | NA | NA |
| Gm38183 | 0 | NA | NA | NA |
| Gm38149 | 0 | NA | NA | NA |
| Gm38145 | 0 | NA | NA | NA |
| Gm32033 | 0 | NA | NA | NA |
| Gm32447 | 0 | NA | NA | NA |
| Gm20770 | 0 | NA | NA | NA |
| Gm38016 | 0 | NA | NA | NA |
| Gm37158 | 0.04393175 | 0.037062677 | 3.724945049 | 0.009949859 |
| Gm36998 | 0 | NA | NA | NA |
| Gm33112 | 0 | NA | NA | NA |

| | | | | |
|---|---|---|---|---|
| Gm21809 | 0 | NA | NA | NA |
| Gm34015 | 0 | NA | NA | NA |
| Gm20877 | 0 | NA | NA | NA |
| Gm33191 | 0 | NA | NA | NA |
| Gm38003 | 0 | NA | NA | NA |
| Gm20781 | 0 | NA | NA | NA |
| Rbm31y | 0 | NA | NA | NA |
| Gm37325 | 0 | NA | NA | NA |
| Gm20833 | 0 | NA | NA | NA |
| Gm37865 | 0 | NA | NA | NA |
| Gm36941 | 0 | NA | NA | NA |
| Gm21904 | 0 | NA | NA | NA |
| Gm37937 | 0 | NA | NA | NA |
| Gm20772 | 0 | NA | NA | NA |
| Gm20831 | 0 | NA | NA | NA |
| Gm37286 | 0 | NA | NA | NA |
| Srsy | 0 | NA | NA | NA |
| Gm38028 | 0 | NA | NA | NA |
| Gm21764 | 0 | NA | NA | NA |
| Ssty1 | 0 | NA | NA | NA |
| Gm38072 | 0 | NA | NA | NA |
| Gm20851 | 0 | NA | NA | NA |
| Gm30312 | 0 | NA | NA | NA |
| Gm37433 | 0 | NA | NA | NA |
| Gm37473 | 0 | NA | NA | NA |
| Gm37656 | 0 | NA | NA | NA |
| Gm38127 | 0 | NA | NA | NA |
| Gm30638 | 0 | NA | NA | NA |
| Gm38361 | 0 | NA | NA | NA |
| Gm31260 | 0 | NA | NA | NA |
| Gm21660 | 0 | NA | NA | NA |
| Gm38209 | 0 | NA | NA | NA |
| Gm21822 | 0 | NA | NA | NA |
| Gm37734 | 0 | NA | NA | NA |
| Gm38013 | 0 | NA | NA | NA |
| Gm21678 | 0 | NA | NA | NA |
| Gm37948 | 0 | NA | NA | NA |
| Gm37442 | 0 | NA | NA | NA |
| Gm34550 | 0 | NA | NA | NA |
| Gm37544 | 0 | NA | NA | NA |
| Gm34716 | 0 | NA | NA | NA |
| Gm37635 | 0 | NA | NA | NA |
| Gm35070 | 0 | NA | NA | NA |
| Gm35134 | 0 | NA | NA | NA |
| Gm37998 | 0 | NA | NA | NA |
| Gm38094 | 0 | NA | NA | NA |
| Gm21763 | 0 | NA | NA | NA |

| | | | | |
|---|---|---|---|---|
| Gm38369 | 0 | NA | NA | NA |
| Gm35843 | 0 | NA | NA | NA |
| Gm38168 | 0 | NA | NA | NA |
| Gm37740 | 0 | NA | NA | NA |
| Gm27701 | 0 | NA | NA | NA |
| Gm37723 | 0 | NA | NA | NA |
| Gm36261 | 0 | NA | NA | NA |
| Gm37898 | 0 | NA | NA | NA |
| Gm36467 | 0 | NA | NA | NA |
| Gm37927 | 0 | NA | NA | NA |
| Gm36891 | 0 | NA | NA | NA |
| Gm36929 | 0 | NA | NA | NA |
| Gm38174 | 0 | NA | NA | NA |
| Gm29866 | 0 | NA | NA | NA |
| Gm36950 | 0 | NA | NA | NA |
| Gm20910 | 0 | NA | NA | NA |
| Gm30598 | 0 | NA | NA | NA |
| Gm20909 | 0 | NA | NA | NA |
| Gm20865 | 0 | NA | NA | NA |
| Gm37462 | 0 | NA | NA | NA |
| Gm37658 | 0 | NA | NA | NA |
| Gm32347 | 0 | NA | NA | NA |
| Gm32419 | 0 | NA | NA | NA |
| Gm38370 | 0 | NA | NA | NA |
| Gm38054 | 0 | NA | NA | NA |
| Gm37476 | 0 | NA | NA | NA |
| Gm37369 | 0 | NA | NA | NA |
| Gm38181 | 0 | NA | NA | NA |
| Gm31335 | 0 | NA | NA | NA |
| Gm37690 | 0 | NA | NA | NA |
| Gm38107 | 0 | NA | NA | NA |
| Gm31571 | 0 | NA | NA | NA |
| Gm32114 | 0 | NA | NA | NA |
| Gm37075 | 0 | NA | NA | NA |
| Gm37921 | 0 | NA | NA | NA |
| Gm32180 | 0 | NA | NA | NA |
| Gm38371 | 0 | NA | NA | NA |
| Gm37657 | 0 | NA | NA | NA |
| Gm37448 | 0 | NA | NA | NA |
| Gm33163 | 0 | NA | NA | NA |
| Gm21065 | 0.044191549 | 0.037062677 | 3.724945049 | 0.009949859 |
| Gm36978 | 0 | NA | NA | NA |
| Gm37721 | 0 | NA | NA | NA |
| Gm37129 | 0 | NA | NA | NA |
| Gm37990 | 0 | NA | NA | NA |
| Gm21366 | 0 | NA | NA | NA |
| Gm33954 | 0 | NA | NA | NA |

| | | | | |
|---|---|---|---|---|
| Gm37538 | 0 | NA | NA | NA |
| Gm21871 | 0 | NA | NA | NA |
| Gm34274 | 0 | NA | NA | NA |
| Gm34217 | 0 | NA | NA | NA |
| Gm21469 | 0 | NA | NA | NA |
| Gm21661 | 0 | NA | NA | NA |
| Gm37952 | 0 | NA | NA | NA |
| Gm21907 | 0 | NA | NA | NA |
| Gm37627 | 0 | NA | NA | NA |
| Gm20925 | 0 | NA | NA | NA |
| Gm20926 | 0 | NA | NA | NA |
| Gm37454 | 0 | NA | NA | NA |
| Gm38138 | 0 | NA | NA | NA |
| Gm29740 | 0 | NA | NA | NA |
| Gm37840 | 0 | NA | NA | NA |
| Gm37574 | 0 | NA | NA | NA |
| Gm21768 | 0 | NA | NA | NA |
| Gm37172 | 0 | NA | NA | NA |
| Gm35670 | 0 | NA | NA | NA |
| Gm20840 | 0 | NA | NA | NA |
| Gm35955 | 0 | NA | NA | NA |
| Gm37687 | 0 | NA | NA | NA |
| Gm38296 | 0 | NA | NA | NA |
| Gm38290 | 0 | NA | NA | NA |
| Gm38121 | 0 | NA | NA | NA |
| Gm36047 | 0 | NA | NA | NA |
| Gm36345 | 0 | NA | NA | NA |
| Gm37157 | 0 | NA | NA | NA |
| Gm38017 | 0 | NA | NA | NA |
| Gm36468 | 0 | NA | NA | NA |
| Gm21845 | 0 | NA | NA | NA |
| Gm20809 | 0 | NA | NA | NA |
| Gm37434 | 0 | NA | NA | NA |
| Gm37765 | 0 | NA | NA | NA |
| Gm38229 | 0 | NA | NA | NA |
| Gm36866 | 0 | NA | NA | NA |
| Gm30045 | 0 | NA | NA | NA |
| Gm21921 | 0 | NA | NA | NA |
| Gm37112 | 0 | NA | NA | NA |
| Gm37287 | 0 | NA | NA | NA |
| Gm30175 | 0 | NA | NA | NA |
| Gm21350 | 0 | NA | NA | NA |
| Gm21412 | 0 | NA | NA | NA |
| Gm37071 | 0 | NA | NA | NA |
| Gm37547 | 0 | NA | NA | NA |
| Gm36968 | 0 | NA | NA | NA |
| Gm38035 | 0 | NA | NA | NA |

| | | | | |
|---|---|---|---|---|
| Gm20857 | 0 | NA | NA | NA |
| Gm30705 | 0 | NA | NA | NA |
| Gm21773 | 0 | NA | NA | NA |
| Gm37451 | 0 | NA | NA | NA |
| Gm37793 | 0 | NA | NA | NA |
| Gm30858 | 0 | NA | NA | NA |
| Gm21791 | 0 | NA | NA | NA |
| Gm21427 | 0 | NA | NA | NA |
| Gm37875 | 0 | NA | NA | NA |
| Gm37237 | 0 | NA | NA | NA |
| Gm38350 | 0 | NA | NA | NA |
| Gm20858 | 0 | NA | NA | NA |
| Gm31422 | 0 | NA | NA | NA |
| Gm37798 | 0 | NA | NA | NA |
| Gm37318 | 0 | NA | NA | NA |
| Gm31572 | 0 | NA | NA | NA |
| Gm21890 | 0 | NA | NA | NA |
| Gm21443 | 0 | NA | NA | NA |
| Gm37346 | 0 | NA | NA | NA |
| Gm38136 | 0 | NA | NA | NA |
| Gm37991 | 0 | NA | NA | NA |
| Gm37808 | 0 | NA | NA | NA |
| Gm33571 | 0 | NA | NA | NA |
| Gm32181 | 0 | NA | NA | NA |
| Gm28662 | 0 | NA | NA | NA |
| Gm28780 | 0 | NA | NA | NA |
| Gm21755 | 0 | NA | NA | NA |
| Gm20895 | 0 | NA | NA | NA |
| Gm28853 | 0 | NA | NA | NA |
| Gm28852 | 0 | NA | NA | NA |
| Gm28855 | 0 | NA | NA | NA |
| Gm28854 | 0 | NA | NA | NA |
| Gm28858 | 0 | NA | NA | NA |
| Gm20890 | 0 | NA | NA | NA |
| Gm28245 | 0 | NA | NA | NA |
| Gm28246 | 0 | NA | NA | NA |
| Gm28247 | 0 | NA | NA | NA |
| Gm28248 | 0 | NA | NA | NA |
| Gm20892 | 0 | NA | NA | NA |
| Gm28249 | 0 | NA | NA | NA |
| Gm28250 | 0 | NA | NA | NA |
| Gm28251 | 0 | NA | NA | NA |
| Gm28243 | 0 | NA | NA | NA |
| Gm28244 | 0 | NA | NA | NA |
| Gm21771 | 0 | NA | NA | NA |
| Gm21476 | 0 | NA | NA | NA |
| Gm29021 | 0 | NA | NA | NA |

| | | | | |
|---|---|---|---|---|
| Gm28880 | 0.044191549 | 0.037062677 | 3.724945049 | 0.009949859 |
| Gm28452 | 0 | NA | NA | NA |
| Gm21488 | 0 | NA | NA | NA |
| Gm28454 | 0 | NA | NA | NA |
| Gm28878 | 0 | NA | NA | NA |
| Gm28879 | 0 | NA | NA | NA |
| Gm29369 | 0 | NA | NA | NA |
| Gm21506 | 0 | NA | NA | NA |
| Gm29399 | 0 | NA | NA | NA |
| Gm29400 | 0 | NA | NA | NA |
| Gm29401 | 0 | NA | NA | NA |
| Gm20891 | 0 | NA | NA | NA |
| Gm20893 | 0 | NA | NA | NA |
| Gm28799 | 0 | NA | NA | NA |
| Gm28798 | 0 | NA | NA | NA |
| Gm28797 | 0 | NA | NA | NA |
| Gm28796 | 0 | NA | NA | NA |
| Gm20894 | 0 | NA | NA | NA |
| Gm28938 | 0 | NA | NA | NA |
| Gm28937 | 0 | NA | NA | NA |
| Gm28936 | 0 | NA | NA | NA |
| Gm21642 | 0 | NA | NA | NA |
| Gm21804 | 0 | NA | NA | NA |
| Gm29122 | 0 | NA | NA | NA |
| Gm28939 | 0 | NA | NA | NA |
| Gm28983 | 0 | NA | NA | NA |
| Gm28985 | 0 | NA | NA | NA |
| Gm20810 | 0 | NA | NA | NA |
| Gm20811 | 0 | NA | NA | NA |
| Gm28725 | 0 | NA | NA | NA |
| Gm28984 | 0 | NA | NA | NA |
| Gm28987 | 0 | NA | NA | NA |
| Gm28986 | 0 | NA | NA | NA |
| Gm29171 | 0 | NA | NA | NA |
| Gm28129 | 0 | NA | NA | NA |
| Gm28127 | 0 | NA | NA | NA |
| Gm28128 | 0 | NA | NA | NA |
| Gm21853 | 0 | NA | NA | NA |
| Gm28132 | 0 | NA | NA | NA |
| Gm28130 | 0 | NA | NA | NA |
| Gm28131 | 0 | NA | NA | NA |
| Gm21679 | 0 | NA | NA | NA |
| Gm29317 | 0 | NA | NA | NA |
| Gm29316 | 0 | NA | NA | NA |
| Gm29315 | 0 | NA | NA | NA |
| Gm28216 | 0 | NA | NA | NA |
| Gm28217 | 0 | NA | NA | NA |

| | | | | |
|---|---|---|---|---|
| Gm28093 | 0 | NA | NA | NA |
| Gm28092 | 0 | NA | NA | NA |
| Gm28091 | 0 | NA | NA | NA |
| Gm28090 | 0 | NA | NA | NA |
| Gm29580 | 0 | NA | NA | NA |
| Gm28089 | 0 | NA | NA | NA |
| Gm28088 | 0 | NA | NA | NA |
| Gm29579 | 0 | NA | NA | NA |
| Gm29582 | 0 | NA | NA | NA |
| Gm21529 | 0 | NA | NA | NA |
| Gm29581 | 0 | NA | NA | NA |
| Gm21539 | 0 | NA | NA | NA |
| Gm29578 | 0 | NA | NA | NA |
| Gm21908 | 0 | NA | NA | NA |
| Gm21780 | 0 | NA | NA | NA |
| Gm28486 | 0 | NA | NA | NA |
| Gm21737 | 0 | NA | NA | NA |
| Gm28487 | 0 | NA | NA | NA |
| Gm28489 | 0 | NA | NA | NA |
| Gm28488 | 0 | NA | NA | NA |
| Gm28491 | 0 | NA | NA | NA |
| Gm28490 | 0 | NA | NA | NA |
| Gm21562 | 0 | NA | NA | NA |
| Gm28298 | 0 | NA | NA | NA |
| Gm21572 | 0 | NA | NA | NA |
| Gm28297 | 0 | NA | NA | NA |
| Gm21753 | 0 | NA | NA | NA |
| Gm21723 | 0 | NA | NA | NA |
| Gm28296 | 0 | NA | NA | NA |
| Gm21821 | 0 | NA | NA | NA |
| Gm28295 | 0 | NA | NA | NA |
| Gm21842 | 0 | NA | NA | NA |
| Gm28170 | 0 | NA | NA | NA |
| Gm21744 | 0 | NA | NA | NA |
| Gm28764 | 0 | NA | NA | NA |
| Gm28762 | 0 | NA | NA | NA |
| Gm28763 | 0 | NA | NA | NA |
| Gm21094 | 0 | NA | NA | NA |
| Gm21588 | 0 | NA | NA | NA |
| Gm28761 | 0 | NA | NA | NA |
| Gm21599 | 0 | NA | NA | NA |
| Gm28765 | 0 | NA | NA | NA |
| Gm21810 | 0 | NA | NA | NA |
| Gm21916 | 0 | NA | NA | NA |
| Gm29191 | 0 | NA | NA | NA |
| Gm21756 | 0 | NA | NA | NA |
| Gm28210 | 0 | NA | NA | NA |

| | | | | |
|---|---|---|---|---|
| Gm29074 | 0 | NA | NA | NA |
| Gm28211 | 0 | NA | NA | NA |
| Gm28518 | 0 | NA | NA | NA |
| Gm28519 | 0 | NA | NA | NA |
| Gm28520 | 0 | NA | NA | NA |
| Gm20838 | 0 | NA | NA | NA |
| Gm21626 | 0 | NA | NA | NA |
| Gm28521 | 0 | NA | NA | NA |
| Gm21633 | 0 | NA | NA | NA |
| Gm28522 | 0 | NA | NA | NA |
| Gm21745 | 0 | NA | NA | NA |
| Gm28517 | 0 | NA | NA | NA |
| Gm29532 | 0 | NA | NA | NA |
| Gm29531 | 0 | NA | NA | NA |
| Gm28464 | 0 | NA | NA | NA |
| Gm29656 | 0 | NA | NA | NA |
| Gm29655 | 0 | NA | NA | NA |
| Gm21647 | 0 | NA | NA | NA |
| Gm29653 | 0 | NA | NA | NA |
| Gm29654 | 0 | NA | NA | NA |
| Gm29651 | 0 | NA | NA | NA |
| Gm29652 | 0 | NA | NA | NA |
| Gm29080 | 0 | NA | NA | NA |
| Gm21184 | 0 | NA | NA | NA |
| Gm21201 | 0 | NA | NA | NA |
| Gm29081 | 0 | NA | NA | NA |
| Gm29077 | 0 | NA | NA | NA |
| Gm29076 | 0 | NA | NA | NA |
| Gm29079 | 0 | NA | NA | NA |
| Gm29078 | 0 | NA | NA | NA |
| Gm20853 | 0 | NA | NA | NA |
| Gm29075 | 0 | NA | NA | NA |
| Gm29225 | 0 | NA | NA | NA |
| Gm29226 | 0 | NA | NA | NA |
| Gm29227 | 0 | NA | NA | NA |
| Gm28339 | 0 | NA | NA | NA |
| Gm20855 | 0 | NA | NA | NA |
| Gm29386 | 0 | NA | NA | NA |
| Gm29384 | 0 | NA | NA | NA |
| Gm21896 | 0 | NA | NA | NA |
| Gm29280 | 0 | NA | NA | NA |
| Gm20799 | 0 | NA | NA | NA |
| Gm29279 | 0 | NA | NA | NA |
| Gm28260 | 0 | NA | NA | NA |
| Gm21795 | 0 | NA | NA | NA |
| Gm28261 | 0 | NA | NA | NA |
| Gm28264 | 0 | NA | NA | NA |

| | | | | |
|---|---|---|---|---|
| Gm21872 | 0 | NA | NA | NA |
| Gm20798 | 0 | NA | NA | NA |
| Gm28463 | 0 | NA | NA | NA |
| Gm28584 | 0 | NA | NA | NA |
| Gm28585 | 0 | NA | NA | NA |
| Gm20896 | 0 | NA | NA | NA |
| Gm28470 | 0 | NA | NA | NA |
| Gm28469 | 0 | NA | NA | NA |
| Gm21898 | 0 | NA | NA | NA |
| Gm28468 | 0 | NA | NA | NA |
| Gm20898 | 0 | NA | NA | NA |
| Gm29333 | 0 | NA | NA | NA |
| Gm28332 | 0 | NA | NA | NA |
| Gm28333 | 0 | NA | NA | NA |
| Gm28334 | 0 | NA | NA | NA |
| Gm28335 | 0 | NA | NA | NA |
| Gm28336 | 0 | NA | NA | NA |
| Gm28337 | 0 | NA | NA | NA |
| Gm28338 | 0 | NA | NA | NA |
| Gm28812 | 0 | NA | NA | NA |
| Gm28811 | 0 | NA | NA | NA |
| Gm28810 | 0 | NA | NA | NA |
| Gm20835 | 0 | NA | NA | NA |
| Gm29027 | 0 | NA | NA | NA |
| Gm29028 | 0 | NA | NA | NA |
| Gm29029 | 0 | NA | NA | NA |
| Gm29030 | 0 | NA | NA | NA |
| Gm29023 | 0 | NA | NA | NA |
| Gm29024 | 0 | NA | NA | NA |
| Gm29025 | 0 | NA | NA | NA |
| Gm29026 | 0 | NA | NA | NA |
| Gm28213 | 0 | NA | NA | NA |
| Gm28212 | 0 | NA | NA | NA |
| Gm21799 | 0 | NA | NA | NA |
| Gm28227 | 0 | NA | NA | NA |
| Gm20905 | 0 | NA | NA | NA |
| Gm29411 | 0 | NA | NA | NA |
| Gm28222 | 0 | NA | NA | NA |
| Gm21835 | 0 | NA | NA | NA |
| Gm28820 | 0 | NA | NA | NA |
| Gm28223 | 0 | NA | NA | NA |
| Gm29412 | 0 | NA | NA | NA |
| Gm29413 | 0 | NA | NA | NA |
| Gm29671 | 0 | NA | NA | NA |
| Gm21878 | 0 | NA | NA | NA |
| Gm28162 | 0 | NA | NA | NA |
| Gm20922 | 0 | NA | NA | NA |

| | | | | |
|---|---|---|---|---|
| Gm20738 | 0 | NA | NA | NA |
| Gm28545 | 0 | NA | NA | NA |
| Gm28547 | 0 | NA | NA | NA |
| Gm28546 | 0 | NA | NA | NA |
| Gm21924 | 0 | NA | NA | NA |
| Gm21735 | 0 | NA | NA | NA |
| Gm28771 | 0 | NA | NA | NA |
| Gm28772 | 0 | NA | NA | NA |
| Gm28773 | 0 | NA | NA | NA |
| Gm20897 | 0 | NA | NA | NA |
| Gm28774 | 0 | NA | NA | NA |
| Gm21732 | 0 | NA | NA | NA |
| Gm28775 | 0 | NA | NA | NA |
| Gm21765 | 0 | NA | NA | NA |
| Gm20899 | 12.70112357 | 0.103508091 | 0.337796972 | 0.306421015 |
| Gm28776 | 0 | NA | NA | NA |
| Gm21787 | 0 | NA | NA | NA |
| Gm28770 | 0 | NA | NA | NA |
| Gm20901 | 0 | NA | NA | NA |
| Gm28176 | 0 | NA | NA | NA |
| Gm28786 | 0 | NA | NA | NA |
| Gm29360 | 0 | NA | NA | NA |
| Gm29496 | 0 | NA | NA | NA |
| Gm29497 | 0 | NA | NA | NA |
| Gm29499 | 0 | NA | NA | NA |
| Gm29498 | 0 | NA | NA | NA |
| Gm36782 | 0 | NA | NA | NA |
| Gm29839 | 0 | NA | NA | NA |
| Gm29495 | 0 | NA | NA | NA |
| Gm28371 | 0.044191549 | 0.037062677 | 3.724945049 | 0.009949859 |
| Gm28595 | 0 | NA | NA | NA |
| Gm37032 | 0 | NA | NA | NA |
| Gm21865 | 0 | NA | NA | NA |
| Gm29433 | 0 | NA | NA | NA |
| Gm29215 | 0 | NA | NA | NA |
| Gm21882 | 0 | NA | NA | NA |
| Gm21801 | 0 | NA | NA | NA |
| Gm28850 | 0 | NA | NA | NA |
| Gm21914 | 0 | NA | NA | NA |
| Gm28617 | 0 | NA | NA | NA |
| Gm28618 | 0 | NA | NA | NA |
| Gm28619 | 0 | NA | NA | NA |
| Gm21852 | 0 | NA | NA | NA |
| Gm20793 | 0 | NA | NA | NA |
| Gm27466 | 0 | NA | NA | NA |
| Gm28018 | 0 | NA | NA | NA |
| Gm28993 | 0 | NA | NA | NA |

| | | | | |
|---|---|---|---|---|
| Gm28994 | 0 | NA | NA | NA |
| Gm21797 | 0 | NA | NA | NA |
| Gm20795 | 0 | NA | NA | NA |
| Gm28596 | 0 | NA | NA | NA |
| Gm20832 | 0 | NA | NA | NA |
| Gm28597 | 0 | NA | NA | NA |
| Gm28428 | 0 | NA | NA | NA |
| Gm20827 | 0 | NA | NA | NA |
| Gm28427 | 0 | NA | NA | NA |
| Gm21800 | 0 | NA | NA | NA |
| Gm28426 | 0 | NA | NA | NA |
| Gm28425 | 0 | NA | NA | NA |
| Gm21729 | 0 | NA | NA | NA |
| Gm28834 | 0 | NA | NA | NA |
| Gm28833 | 0 | NA | NA | NA |
| Gm21774 | 0 | NA | NA | NA |
| Gm29537 | 0 | NA | NA | NA |
| Gm29274 | 0 | NA | NA | NA |
| Gm29275 | 0.04393175 | 0.037062677 | 3.724945049 | 0.009949859 |
| Gm29276 | 0 | NA | NA | NA |
| Gm21805 | 0 | NA | NA | NA |
| Gm29645 | 0 | NA | NA | NA |
| Gm28312 | 0 | NA | NA | NA |
| Gm29165 | 0 | NA | NA | NA |
| Gm29042 | 0 | NA | NA | NA |
| Gm28311 | 0 | NA | NA | NA |
| Gm28310 | 0 | NA | NA | NA |
| Gm28504 | 0 | NA | NA | NA |
| Gm28567 | 0 | NA | NA | NA |
| Gm28568 | 0 | NA | NA | NA |
| Gm29286 | 0 | NA | NA | NA |
| Gm29285 | 0 | NA | NA | NA |
| Gm28280 | 0 | NA | NA | NA |
| Gm21241 | 0 | NA | NA | NA |
| Gm28278 | 0 | NA | NA | NA |
| Gm28279 | 0 | NA | NA | NA |
| Gm21739 | 0 | NA | NA | NA |
| Gm28886 | 0 | NA | NA | NA |
| Gm29426 | 0 | NA | NA | NA |
| Gm28885 | 0 | NA | NA | NA |
| Gm21856 | 0 | NA | NA | NA |
| Gm28887 | 0 | NA | NA | NA |
| Gm29444 | 0 | NA | NA | NA |
| Gm21783 | 0 | NA | NA | NA |
| Gm29445 | 0 | NA | NA | NA |
| Gm28457 | 0 | NA | NA | NA |
| Gm21779 | 0 | NA | NA | NA |

| | | | | |
|---|---|---|---|---|
| Gm28458 | 0 | NA | NA | NA |
| Gm28459 | 0 | NA | NA | NA |
| Gm28461 | 0 | NA | NA | NA |
| Gm28460 | 0 | NA | NA | NA |
| Gm28851 | 0 | NA | NA | NA |
| Gm28456 | 0 | NA | NA | NA |
| Gm29073 | 0 | NA | NA | NA |
| Gm29648 | 0 | NA | NA | NA |
| Gm29647 | 0 | NA | NA | NA |
| Gm29646 | 0 | NA | NA | NA |
| Gm28549 | 0 | NA | NA | NA |
| Gm29211 | 0 | NA | NA | NA |
| Gm29212 | 0 | NA | NA | NA |
| Gm21247 | 0 | NA | NA | NA |
| Gm29210 | 0 | NA | NA | NA |
| Gm29213 | 0 | NA | NA | NA |
| Gm28550 | 0 | NA | NA | NA |
| Gm28604 | 0 | NA | NA | NA |
| Gm28603 | 0 | NA | NA | NA |
| Gm28565 | 0 | NA | NA | NA |
| Gm28566 | 0 | NA | NA | NA |
| Gm28697 | 0 | NA | NA | NA |
| Gm28554 | 0 | NA | NA | NA |
| Gm21256 | 0 | NA | NA | NA |
| Gm29056 | 0 | NA | NA | NA |
| Gm29098 | 0 | NA | NA | NA |
| Gm21170 | 0 | NA | NA | NA |
| Gm29373 | 0 | NA | NA | NA |
| Gm29370 | 0 | NA | NA | NA |
| Gm21751 | 0 | NA | NA | NA |
| Gm29166 | 0 | NA | NA | NA |
| Gm28194 | 0 | NA | NA | NA |
| Gm20792 | 0 | NA | NA | NA |
| Gm28754 | 0 | NA | NA | NA |
| Gm28753 | 0 | NA | NA | NA |
| Gm21191 | 0 | NA | NA | NA |
| Gm21271 | 0 | NA | NA | NA |
| Gm21281 | 0 | NA | NA | NA |
| Gm28193 | 0 | NA | NA | NA |
| Gm20802 | 0 | NA | NA | NA |
| Gm29070 | 0 | NA | NA | NA |
| Gm28553 | 0 | NA | NA | NA |
| Gm28751 | 0 | NA | NA | NA |
| Gm28752 | 0 | NA | NA | NA |
| Gm28726 | 0 | NA | NA | NA |
| Gm21308 | 0 | NA | NA | NA |
| Gm29267 | 0 | NA | NA | NA |

| | | | | |
|---|---|---|---|---|
| Gm29268 | 0 | NA | NA | NA |
| Gm29269 | 0 | NA | NA | NA |
| Gm29270 | 0 | NA | NA | NA |
| Gm29271 | 0 | NA | NA | NA |
| Gm21209 | 0 | NA | NA | NA |
| Gm28682 | 0 | NA | NA | NA |
| Gm28569 | 0 | NA | NA | NA |
| Gm28681 | 0 | NA | NA | NA |
| Gm28684 | 0 | NA | NA | NA |
| Gm21316 | 0 | NA | NA | NA |
| Gm21923 | 0 | NA | NA | NA |
| Gm28683 | 0 | NA | NA | NA |
| Gm28680 | 0 | NA | NA | NA |
| Gm28679 | 0 | NA | NA | NA |
| Gm28870 | 0 | NA | NA | NA |
| Gm28871 | 0 | NA | NA | NA |
| Gm29512 | 0 | NA | NA | NA |
| Gm29511 | 0 | NA | NA | NA |
| Gm20880 | 0 | NA | NA | NA |
| Gm20881 | 0 | NA | NA | NA |
| Gm29121 | 0 | NA | NA | NA |
| Gm29120 | 0 | NA | NA | NA |
| Gm29117 | 0 | NA | NA | NA |
| Gm20883 | 0 | NA | NA | NA |
| Gm29119 | 0 | NA | NA | NA |
| Gm29118 | 0 | NA | NA | NA |
| Gm29116 | 0 | NA | NA | NA |
| Gm20889 | 0 | NA | NA | NA |
| Gm20887 | 0 | NA | NA | NA |
| Gm29320 | 0 | NA | NA | NA |
| Gm21849 | 0 | NA | NA | NA |
| Gm29321 | 0 | NA | NA | NA |
| Gm21117 | 0 | NA | NA | NA |
| Gm29319 | 0 | NA | NA | NA |
| Gm28317 | 0 | NA | NA | NA |
| Gm28318 | 0 | NA | NA | NA |
| Gm20884 | 0 | NA | NA | NA |
| Gm28313 | 0 | NA | NA | NA |
| Gm28314 | 0 | NA | NA | NA |
| Gm28315 | 0 | NA | NA | NA |
| Gm28316 | 0 | NA | NA | NA |
| Gm21387 | 0 | NA | NA | NA |
| Gm21396 | 0 | NA | NA | NA |
| Gm28944 | 0 | NA | NA | NA |
| Gm28943 | 0 | NA | NA | NA |
| Gm28946 | 0 | NA | NA | NA |
| Gm28945 | 0 | NA | NA | NA |

| | | | | |
|---|---|---|---|---|
| Gm28947 | 0 | NA | NA | NA |
| Gm20920 | 0 | NA | NA | NA |
| Gm28949 | 0 | NA | NA | NA |
| Gm28948 | 0 | NA | NA | NA |
| Gm28950 | 0 | NA | NA | NA |
| Gm21245 | 0 | NA | NA | NA |
| Gm29169 | 0 | NA | NA | NA |
| Gm21899 | 0 | NA | NA | NA |
| Gm29063 | 0 | NA | NA | NA |
| Gm28737 | 0 | NA | NA | NA |
| Gm28738 | 0 | NA | NA | NA |
| Gm20923 | 0 | NA | NA | NA |
| Gm28197 | 0 | NA | NA | NA |
| Gm21249 | 0 | NA | NA | NA |
| Gm28736 | 0 | NA | NA | NA |
| Gm21840 | 0 | NA | NA | NA |
| Gm28735 | 0 | NA | NA | NA |
| Gm21258 | 0 | NA | NA | NA |
| Gm28734 | 0 | NA | NA | NA |
| Gm28232 | 0 | NA | NA | NA |
| Gm28732 | 0 | NA | NA | NA |
| Gm28733 | 0 | NA | NA | NA |
| Gm21268 | 0 | NA | NA | NA |
| Gm21275 | 0 | NA | NA | NA |
| Gm28233 | 0 | NA | NA | NA |
| Gm28235 | 0 | NA | NA | NA |
| Gm28234 | 0 | NA | NA | NA |
| Gm21902 | 0 | NA | NA | NA |
| Gm28236 | 0 | NA | NA | NA |
| Gm21285 | 0 | NA | NA | NA |
| Gm20747 | 0 | NA | NA | NA |
| Gm29250 | 0 | NA | NA | NA |
| Gm29251 | 0 | NA | NA | NA |
| Gm21906 | 0 | NA | NA | NA |
| Gm29252 | 0 | NA | NA | NA |
| Gm29248 | 0 | NA | NA | NA |
| Gm29249 | 0 | NA | NA | NA |
| Gm29484 | 0 | NA | NA | NA |
| Gm21920 | 0 | NA | NA | NA |
| Gm28908 | 0 | NA | NA | NA |
| Gm28907 | 0 | NA | NA | NA |
| Gm21749 | 0 | NA | NA | NA |
| Gm20929 | 0 | NA | NA | NA |
| Gm28910 | 0 | NA | NA | NA |
| Gm28909 | 0 | NA | NA | NA |
| Gm28276 | 0 | NA | NA | NA |
| Gm28274 | 0 | NA | NA | NA |

| | | | | |
|---|---|---|---|---|
| Gm28275 | 0 | NA | NA | NA |
| Gm21302 | 0 | NA | NA | NA |
| Gm28290 | 0 | NA | NA | NA |
| Gm28292 | 0 | NA | NA | NA |
| Gm28291 | 0 | NA | NA | NA |
| Gm28293 | 0 | NA | NA | NA |
| Gm21740 | 0 | NA | NA | NA |
| Gm21912 | 0 | NA | NA | NA |
| Gm28698 | 0 | NA | NA | NA |
| Gm29304 | 0 | NA | NA | NA |
| Gm29305 | 0 | NA | NA | NA |
| Gm29303 | 0 | NA | NA | NA |
| Gm21894 | 0 | NA | NA | NA |
| Gm29307 | 0 | NA | NA | NA |
| Gm29557 | 0 | NA | NA | NA |
| Gm23052 | 0 | NA | NA | NA |
| Gm20927 | 0 | NA | NA | NA |
| Gm20930 | 0 | NA | NA | NA |
| Gm29306 | 0 | NA | NA | NA |
| Gm29300 | 0 | NA | NA | NA |
| Gm29301 | 0 | NA | NA | NA |
| Gm20931 | 0 | NA | NA | NA |
| Gm29559 | 0 | NA | NA | NA |
| Gm29558 | 0 | NA | NA | NA |
| Gm29555 | 0 | NA | NA | NA |
| Gm21733 | 0 | NA | NA | NA |
| Gm21330 | 0 | NA | NA | NA |
| Gm28355 | 0 | NA | NA | NA |
| Gm28354 | 0 | NA | NA | NA |
| Gm28353 | 0 | NA | NA | NA |
| Gm28352 | 0 | NA | NA | NA |
| Gm21858 | 0 | NA | NA | NA |
| Gm29244 | 0 | NA | NA | NA |
| Gm28200 | 0 | NA | NA | NA |
| Gm28201 | 0 | NA | NA | NA |
| Gm28202 | 0 | NA | NA | NA |
| Gm28203 | 0 | NA | NA | NA |
| Gm21340 | 0 | NA | NA | NA |
| Gm28204 | 0 | NA | NA | NA |
| Gm28205 | 0 | NA | NA | NA |
| Gm28206 | 0 | NA | NA | NA |
| Gm28207 | 0 | NA | NA | NA |
| Gm28208 | 0 | NA | NA | NA |
| Gm21851 | 0 | NA | NA | NA |
| Gm21819 | 0 | NA | NA | NA |
| Gm29224 | 0 | NA | NA | NA |
| Gm29223 | 0 | NA | NA | NA |

| | | | | |
|---|---|---|---|---|
| Gm29222 | 0 | NA | NA | NA |
| Gm29221 | 0 | NA | NA | NA |
| Gm29220 | 0 | NA | NA | NA |
| Gm21815 | 0 | NA | NA | NA |
| Gm29407 | 0 | NA | NA | NA |
| Gm29405 | 0 | NA | NA | NA |
| Gm29406 | 0 | NA | NA | NA |
| Gm20913 | 0 | NA | NA | NA |
| Gm29404 | 0 | NA | NA | NA |
| Gm29403 | 0 | NA | NA | NA |
| Gm28350 | 0 | NA | NA | NA |
| Gm28964 | 0 | NA | NA | NA |
| Sly | 0 | NA | NA | NA |
| Gm28165 | 0 | NA | NA | NA |
| Gm28709 | 0 | NA | NA | NA |
| Gm28965 | 0 | NA | NA | NA |
| Gm21868 | 0 | NA | NA | NA |
| Gm21943 | 0 | NA | NA | NA |
| Gm28532 | 0 | NA | NA | NA |
| Gm28533 | 0 | NA | NA | NA |
| Gm28966 | 0 | NA | NA | NA |
| Gm28961 | 0 | NA | NA | NA |
| Gm28962 | 0 | NA | NA | NA |
| Gm28963 | 0 | NA | NA | NA |
| Gm29466 | 0 | NA | NA | NA |
| Gm29467 | 0 | NA | NA | NA |
| Gm21724 | 0 | NA | NA | NA |
| Gm20963 | 0 | NA | NA | NA |
| Gm28326 | 0 | NA | NA | NA |
| Gm28325 | 0 | NA | NA | NA |
| Gm21405 | 0 | NA | NA | NA |
| Gm28284 | 0 | NA | NA | NA |
| Gm29368 | 0 | NA | NA | NA |
| Gm21861 | 0 | NA | NA | NA |
| Gm21413 | 0 | NA | NA | NA |
| Gm28073 | 0 | NA | NA | NA |
| Gm21419 | 0 | NA | NA | NA |
| Gm28074 | 0 | NA | NA | NA |
| Gm21428 | 0 | NA | NA | NA |
| Gm28072 | 0 | NA | NA | NA |
| Gm21758 | 0 | NA | NA | NA |
| Gm29672 | 0 | NA | NA | NA |
| Gm29131 | 0 | NA | NA | NA |
| Gm20871 | 0 | NA | NA | NA |
| Gm29130 | 0 | NA | NA | NA |
| Gm20978 | 0 | NA | NA | NA |
| Gm29132 | 0 | NA | NA | NA |

| | | | | |
|---|---|---|---|---|
| Gm21450 | 0 | NA | NA | NA |
| Gm21843 | 0 | NA | NA | NA |
| Gm28889 | 0 | NA | NA | NA |
| Gm28890 | 0 | NA | NA | NA |
| Gm28891 | 0 | NA | NA | NA |
| Gm28691 | 0 | NA | NA | NA |
| Gm28692 | 0 | NA | NA | NA |
| Gm20794 | 0 | NA | NA | NA |
| Gm28690 | 0 | NA | NA | NA |
| Gm28689 | 0 | NA | NA | NA |
| Gm28687 | 0 | NA | NA | NA |
| Gm20987 | 0 | NA | NA | NA |
| Gm21918 | 0 | NA | NA | NA |
| 1700040F15Rik | 0 | NA | NA | NA |
| Gm29207 | 0 | NA | NA | NA |
| Gm29204 | 0 | NA | NA | NA |
| Gm21489 | 0 | NA | NA | NA |
| Gm21832 | 0 | NA | NA | NA |
| Gm21573 | 0 | NA | NA | NA |
| Gm28179 | 0 | NA | NA | NA |
| Gm21577 | 0 | NA | NA | NA |
| Gm29206 | 0 | NA | NA | NA |
| Gm29203 | 0 | NA | NA | NA |
| Gm21497 | 0 | NA | NA | NA |
| Gm29614 | 0 | NA | NA | NA |
| Gm29613 | 0 | NA | NA | NA |
| Gm29612 | 0 | NA | NA | NA |
| Gm29343 | 0 | NA | NA | NA |
| Gm29342 | 0 | NA | NA | NA |
| Gm20803 | 0 | NA | NA | NA |
| Gm20804 | 0 | NA | NA | NA |
| Gm28598 | 0 | NA | NA | NA |
| Gm28599 | 0 | NA | NA | NA |
| Gm28600 | 0 | NA | NA | NA |
| Gm21518 | 0 | NA | NA | NA |
| Gm28185 | 0 | NA | NA | NA |
| Gm29037 | 0 | NA | NA | NA |
| Gm20823 | 0 | NA | NA | NA |
| Gm21877 | 0 | NA | NA | NA |
| Gm28835 | 0 | NA | NA | NA |
| Gm29548 | 0 | NA | NA | NA |
| Gm29547 | 0 | NA | NA | NA |
| Gm29546 | 0 | NA | NA | NA |
| Gm29545 | 0 | NA | NA | NA |
| Gm29544 | 0 | NA | NA | NA |
| Gm29209 | 0 | NA | NA | NA |
| Gm29549 | 0 | NA | NA | NA |

| | | | | |
|---|---|---|---|---|
| Gm21530 | 0 | NA | NA | NA |
| Gm21617 | 0 | NA | NA | NA |
| Gm28589 | 0 | NA | NA | NA |
| Gm29420 | 0 | NA | NA | NA |
| Gm29421 | 0 | NA | NA | NA |
| Gm21627 | 0 | NA | NA | NA |
| Gm29422 | 0 | NA | NA | NA |
| Gm28808 | 0 | NA | NA | NA |
| Gm21634 | 0 | NA | NA | NA |
| Gm28446 | 0 | NA | NA | NA |
| Gm29060 | 0 | NA | NA | NA |
| Gm29061 | 0 | NA | NA | NA |
| Gm20885 | 0 | NA | NA | NA |
| Gm29062 | 0 | NA | NA | NA |
| Gm29057 | 0 | NA | NA | NA |
| Gm29625 | 0 | NA | NA | NA |
| Gm21654 | 0 | NA | NA | NA |
| Gm29058 | 0 | NA | NA | NA |
| Gm29059 | 0 | NA | NA | NA |
| Gm29628 | 0 | NA | NA | NA |
| Gm20882 | 0 | NA | NA | NA |
| Gm20907 | 0 | NA | NA | NA |
| Gm29624 | 0 | NA | NA | NA |
| Gm29623 | 0 | NA | NA | NA |
| Gm29622 | 0 | NA | NA | NA |
| Gm29265 | 0 | NA | NA | NA |
| Gm20908 | 0 | NA | NA | NA |
| Gm29259 | 0 | NA | NA | NA |
| Gm28860 | 0 | NA | NA | NA |
| Gm28252 | 0 | NA | NA | NA |
| Gm28859 | 0 | NA | NA | NA |
| Gm20921 | 0 | NA | NA | NA |
| Gm28253 | 0 | NA | NA | NA |
| Gm21826 | 0 | NA | NA | NA |
| Gm28254 | 0 | NA | NA | NA |
| Gm28861 | 0 | NA | NA | NA |
| Gm28255 | 0 | NA | NA | NA |
| Gm21171 | 0 | NA | NA | NA |
| Gm20924 | 0 | NA | NA | NA |
| Gm28990 | 0 | NA | NA | NA |
| Gm21848 | 0 | NA | NA | NA |
| Gm29457 | 0 | NA | NA | NA |
| Gm20736 | 0 | NA | NA | NA |
| Gm28558 | 0 | NA | NA | NA |
| Gm28560 | 0 | NA | NA | NA |
| Gm28561 | 0 | NA | NA | NA |
| Gm29584 | 0 | NA | NA | NA |

| | | | | |
|---|---|---|---|---|
| Gm20936 | 0 | NA | NA | NA |
| Gm20934 | 0 | NA | NA | NA |
| Gm29586 | 0 | NA | NA | NA |
| Gm21895 | 0 | NA | NA | NA |
| Gm28134 | 0 | NA | NA | NA |
| Gm28135 | 0 | NA | NA | NA |
| Gm20933 | 0 | NA | NA | NA |
| Gm28133 | 0 | NA | NA | NA |
| Gm28137 | 0 | NA | NA | NA |
| Gm28138 | 0 | NA | NA | NA |
| Gm21837 | 0 | NA | NA | NA |
| Gm28136 | 0 | NA | NA | NA |
| Gm21910 | 0 | NA | NA | NA |
| Gm29377 | 0 | NA | NA | NA |
| Gm21889 | 0 | NA | NA | NA |
| Gm29378 | 0 | NA | NA | NA |
| Gm29379 | 0 | NA | NA | NA |
| Gm29380 | 0 | NA | NA | NA |
| Gm29381 | 0 | NA | NA | NA |
| Gm28795 | 0 | NA | NA | NA |
| Gm20852 | 0 | NA | NA | NA |
| Gm29003 | 0 | NA | NA | NA |
| Gm28749 | 0 | NA | NA | NA |
| Gm29001 | 0 | NA | NA | NA |
| Gm29002 | 0 | NA | NA | NA |
| Ssty2 | 0 | NA | NA | NA |
| Gm29000 | 0 | NA | NA | NA |
| Gm28646 | 0 | NA | NA | NA |
| Gm28612 | 0 | NA | NA | NA |
| Gm28613 | 0 | NA | NA | NA |
| Gm28611 | 0 | NA | NA | NA |
| Gm21114 | 0 | NA | NA | NA |
| Gm29633 | 0 | NA | NA | NA |
| Gm21794 | 0 | NA | NA | NA |
| Gm29219 | 0 | NA | NA | NA |
| Gm29632 | 0 | NA | NA | NA |
| Gm29631 | 0 | NA | NA | NA |
| Gm29637 | 0 | NA | NA | NA |
| Gm20874 | 0 | NA | NA | NA |
| Gm29082 | 0 | NA | NA | NA |
| Gm29217 | 0 | NA | NA | NA |
| Gm29636 | 0 | NA | NA | NA |
| Gm29635 | 0 | NA | NA | NA |
| Gm29634 | 0 | NA | NA | NA |
| Gm29639 | 0 | NA | NA | NA |
| Gm29218 | 0 | NA | NA | NA |
| Gm21806 | 0 | NA | NA | NA |

| | | | | |
|---|---|---|---|---|
| Gm20935 | 0 | NA | NA | NA |
| Gm29662 | 0 | NA | NA | NA |
| Gm29660 | 0 | NA | NA | NA |
| Gm29661 | 0 | NA | NA | NA |
| Gm29659 | 0 | NA | NA | NA |
| Gm20937 | 0 | NA | NA | NA |
| Gm29450 | 0 | NA | NA | NA |
| Gm29449 | 0 | NA | NA | NA |
| Gm21867 | 0 | NA | NA | NA |
| Gm29448 | 0 | NA | NA | NA |
| Gm21127 | 0 | NA | NA | NA |
| Gm20816 | 0 | NA | NA | NA |
| Gm29447 | 0 | NA | NA | NA |
| Gm29446 | 0 | NA | NA | NA |
| Gm29452 | 0 | NA | NA | NA |
| Gm29451 | 0.044191549 | 0.037062677 | 3.724945049 | 0.009949859 |
| Gm20817 | 0 | NA | NA | NA |
| Gm28633 | 0 | NA | NA | NA |
| Gm29410 | 0 | NA | NA | NA |
| Gm28819 | 0 | NA | NA | NA |
| Gm28462 | 0 | NA | NA | NA |
| Gm20818 | 0 | NA | NA | NA |
| Gm21151 | 0 | NA | NA | NA |
| Gm29409 | 0 | NA | NA | NA |
| Gm28632 | 0 | NA | NA | NA |
| Gm28101 | 0 | NA | NA | NA |
| Gm33815 | 0 | NA | NA | NA |
| Gm21160 | 0 | NA | NA | NA |
| Gm28637 | 0 | NA | NA | NA |
| Gm28636 | 0 | NA | NA | NA |
| Gm20932 | 0 | NA | NA | NA |
| Gm29134 | 0 | NA | NA | NA |
| Gm28303 | 0 | NA | NA | NA |
| Gm28304 | 0 | NA | NA | NA |
| Gm28824 | 0 | NA | NA | NA |
| Gm28081 | 0 | NA | NA | NA |
| Gm28080 | 0 | NA | NA | NA |
| Gm21524 | 0 | NA | NA | NA |
| Gm28079 | 0 | NA | NA | NA |
| Gm28302 | 0 | NA | NA | NA |
| Gm28082 | 0 | NA | NA | NA |
| Gm21672 | 0 | NA | NA | NA |
| Gm29644 | 0 | NA | NA | NA |
| Gm21118 | 0 | NA | NA | NA |
| Gm29423 | 0 | NA | NA | NA |
| Gm29424 | 0 | NA | NA | NA |
| 4932431L22Rik | 0 | NA | NA | NA |

| | | | | |
|---|---|---|---|---|
| Gm29425 | 0 | NA | NA | NA |
| Gm20888 | 0 | NA | NA | NA |
| Gm28238 | 0 | NA | NA | NA |
| Gm28433 | 0 | NA | NA | NA |
| Gm28431 | 0 | NA | NA | NA |
| Gm28432 | 0 | NA | NA | NA |
| Gm20886 | 0 | NA | NA | NA |
| Gm21808 | 0 | NA | NA | NA |
| Gm28840 | 0 | NA | NA | NA |
| Gm28841 | 0 | NA | NA | NA |
| Gm28842 | 0 | NA | NA | NA |
| Gm28843 | 0 | NA | NA | NA |
| Gm28837 | 0 | NA | NA | NA |
| Gm28838 | 0 | NA | NA | NA |
| Gm28839 | 0 | NA | NA | NA |
| Gm20866 | 0 | NA | NA | NA |
| Gm20868 | 0 | NA | NA | NA |
| Gm28844 | 0 | NA | NA | NA |
| Gm21841 | 0 | NA | NA | NA |
| Gm28668 | 0 | NA | NA | NA |
| Gm20869 | 0 | NA | NA | NA |
| Gm28481 | 0 | NA | NA | NA |
| Gm28482 | 0 | NA | NA | NA |
| Gm28485 | 0 | NA | NA | NA |
| Gm20870 | 0 | NA | NA | NA |
| Gm28483 | 0 | NA | NA | NA |
| Gm28484 | 0 | NA | NA | NA |
| Gm28480 | 0 | NA | NA | NA |
| Gm20842 | 0 | NA | NA | NA |
| Gm29179 | 0 | NA | NA | NA |
| Gm21880 | 0 | NA | NA | NA |
| Gm29180 | 0 | NA | NA | NA |
| Gm29181 | 0 | NA | NA | NA |
| Gm20843 | 0 | NA | NA | NA |
| Gm29182 | 0 | NA | NA | NA |
| Gm20844 | 0 | NA | NA | NA |
| Gm29178 | 0 | NA | NA | NA |
| Gm21757 | 0 | NA | NA | NA |
| Gm28968 | 0 | NA | NA | NA |
| Gm28495 | 0 | NA | NA | NA |
| Gm20846 | 0 | NA | NA | NA |
| Gm20847 | 0 | NA | NA | NA |
| Gm20848 | 0 | NA | NA | NA |
| Gm28969 | 0 | NA | NA | NA |
| Gm21766 | 0 | NA | NA | NA |
| Gm20800 | 0 | NA | NA | NA |
| Gm21683 | 0 | NA | NA | NA |

| | | | | |
|---|---|---|---|---|
| Gm28677 | 0 | NA | NA | NA |
| Gm21855 | 0 | NA | NA | NA |
| Gm29255 | 0 | NA | NA | NA |
| Gm20902 | 0 | NA | NA | NA |
| Gm28678 | 0 | NA | NA | NA |
| Gm20903 | 0 | NA | NA | NA |
| Gm28272 | 0 | NA | NA | NA |
| Gm29256 | 0 | NA | NA | NA |
| Gm28259 | 0 | NA | NA | NA |
| Gm28273 | 0 | NA | NA | NA |
| Gm21715 | 0 | NA | NA | NA |
| Gm29617 | 0 | NA | NA | NA |
| Gm29615 | 0 | NA | NA | NA |
| Gm29616 | 0 | NA | NA | NA |
| Gm21784 | 0 | NA | NA | NA |
| Gm20805 | 0 | NA | NA | NA |
| Gm29618 | 0 | NA | NA | NA |
| Gm29109 | 0 | NA | NA | NA |
| Gm29108 | 0 | NA | NA | NA |
| Gm29110 | 0 | NA | NA | NA |
| Gm28781 | 0 | NA | NA | NA |
| Gm21913 | 0 | NA | NA | NA |
| Gm28174 | 0 | NA | NA | NA |
| Gm22832 | 0 | NA | NA | NA |
| Gm21076 | 0 | NA | NA | NA |
| Gm29416 | 0 | NA | NA | NA |
| Gm25359 | 0 | NA | NA | NA |
| Gm21869 | 0 | NA | NA | NA |
| Gm20813 | 0 | NA | NA | NA |
| Gm28790 | 0 | NA | NA | NA |
| Gm29272 | 0 | NA | NA | NA |
| Gm28789 | 0 | NA | NA | NA |
| Gm20814 | 0 | NA | NA | NA |
| Gm28787 | 0 | NA | NA | NA |
| Gm28788 | 0 | NA | NA | NA |
| Gm29278 | 0 | NA | NA | NA |
| Gm21922 | 0 | NA | NA | NA |
| Gm21111 | 0 | NA | NA | NA |
| Gm28608 | 0 | NA | NA | NA |
| Gm28607 | 0 | NA | NA | NA |
| Gm28606 | 0 | NA | NA | NA |
| Gm20850 | 0 | NA | NA | NA |
| Gm29574 | 0 | NA | NA | NA |
| Gm29575 | 0 | NA | NA | NA |
| Gm21875 | 0 | NA | NA | NA |
| Gm20904 | 0 | NA | NA | NA |
| Gm29576 | 0 | NA | NA | NA |

| | | | | |
|---|---|---|---|---|
| Gm29577 | 0 | NA | NA | NA |
| Gm29573 | 0 | NA | NA | NA |
| Gm28377 | 0 | NA | NA | NA |
| Gm21776 | 0 | NA | NA | NA |
| Gm21155 | 0 | NA | NA | NA |
| Gm21163 | 0 | NA | NA | NA |
| Gm28122 | 0 | NA | NA | NA |
| Gm28121 | 0 | NA | NA | NA |
| Gm21173 | 0 | NA | NA | NA |
| Gm28123 | 0 | NA | NA | NA |
| Gm28120 | 0 | NA | NA | NA |
| Gm29313 | 0 | NA | NA | NA |
| Gm21180 | 0 | NA | NA | NA |
| Gm20859 | 0 | NA | NA | NA |
| Gm29314 | 0 | NA | NA | NA |
| Gm29310 | 0 | NA | NA | NA |
| Gm29311 | 0 | NA | NA | NA |
| Gm21638 | 0 | NA | NA | NA |
| Gm29312 | 0 | NA | NA | NA |
| Gm29308 | 0 | NA | NA | NA |
| Gm29309 | 0 | NA | NA | NA |
| Gm28702 | 0 | NA | NA | NA |
| Gm28701 | 0 | NA | NA | NA |
| Gm28700 | 0 | NA | NA | NA |
| Gm28699 | 0 | NA | NA | NA |
| Gm28705 | 0 | NA | NA | NA |
| Gm28704 | 0 | NA | NA | NA |
| Gm28540 | 0 | NA | NA | NA |
| Gm21650 | 0 | NA | NA | NA |
| Gm28541 | 0 | NA | NA | NA |
| Gm28542 | 0 | NA | NA | NA |
| Gm28537 | 0 | NA | NA | NA |
| Gm28538 | 0 | NA | NA | NA |
| Gm29569 | 0 | NA | NA | NA |
| Gm20867 | 0 | NA | NA | NA |
| Gm21658 | 0 | NA | NA | NA |
| Gm29568 | 0 | NA | NA | NA |
| Gm29567 | 0 | NA | NA | NA |
| Gm21198 | 0 | NA | NA | NA |
| Gm29566 | 0 | NA | NA | NA |
| Gm29565 | 0 | NA | NA | NA |
| Gm29564 | 0 | NA | NA | NA |
| Gm29147 | 0 | NA | NA | NA |
| Gm29146 | 0 | NA | NA | NA |
| Gm28664 | 0 | NA | NA | NA |
| Gm28663 | 0 | NA | NA | NA |
| Gm29149 | 0 | NA | NA | NA |

| | | | | |
|---|---|---|---|---|
| Gm29148 | 0 | NA | NA | NA |
| Gm29145 | 0 | NA | NA | NA |
| Gm29144 | 0 | NA | NA | NA |
| Gm28817 | 0 | NA | NA | NA |
| Gm28816 | 0 | NA | NA | NA |
| Gm20808 | 0 | NA | NA | NA |
| Gm20806 | 0 | NA | NA | NA |
| Gm28815 | 0 | NA | NA | NA |
| Gm28814 | 0 | NA | NA | NA |
| Gm28813 | 0 | NA | NA | NA |
| Gm20919 | 0 | NA | NA | NA |
| Gm20917 | 0 | NA | NA | NA |
| Gm28421 | 0 | NA | NA | NA |
| Gm28420 | 0 | NA | NA | NA |
| Gm28423 | 0 | NA | NA | NA |
| Gm28422 | 0 | NA | NA | NA |
| Gm20916 | 0 | NA | NA | NA |
| Gm29033 | 0 | NA | NA | NA |
| Gm29034 | 0 | NA | NA | NA |
| Gm21823 | 0 | NA | NA | NA |
| Gm21242 | 0 | NA | NA | NA |
| Gm29031 | 0 | NA | NA | NA |
| Gm29032 | 0 | NA | NA | NA |
| Gm29606 | 0 | NA | NA | NA |
| Gm29607 | 0 | NA | NA | NA |
| Gm21248 | 0 | NA | NA | NA |
| Gm29189 | 0 | NA | NA | NA |
| Gm21873 | 0 | NA | NA | NA |
| Gm29190 | 0 | NA | NA | NA |
| Gm21760 | 0 | NA | NA | NA |
| Gm28159 | 0 | NA | NA | NA |
| Gm28160 | 0 | NA | NA | NA |
| Gm28157 | 0 | NA | NA | NA |
| Gm28158 | 0 | NA | NA | NA |
| Gm21257 | 0 | NA | NA | NA |
| Gm28161 | 0 | NA | NA | NA |
| Gm28695 | 0 | NA | NA | NA |
| Gm28696 | 0 | NA | NA | NA |
| Gm20791 | 0 | NA | NA | NA |
| Gm20841 | 0 | NA | NA | NA |
| Gm28899 | 0 | NA | NA | NA |
| Gm21722 | 0 | NA | NA | NA |
| Gm28898 | 0 | NA | NA | NA |
| Gm28897 | 0 | NA | NA | NA |
| Gm28896 | 0 | NA | NA | NA |
| Gm21282 | 0 | NA | NA | NA |
| Gm28895 | 0 | NA | NA | NA |

| | | | | |
|---|---|---|---|---|
| Gm28894 | 0 | NA | NA | NA |
| Gm21829 | 0 | NA | NA | NA |
| Gm29324 | 0 | NA | NA | NA |
| Gm20911 | 0 | NA | NA | NA |
| Gm28739 | 0 | NA | NA | NA |
| Gm28741 | 0 | NA | NA | NA |
| Gm20912 | 0 | NA | NA | NA |
| Gm20801 | 0 | NA | NA | NA |
| Gm28742 | 0 | NA | NA | NA |
| Gm21919 | 0 | NA | NA | NA |
| Gm28743 | 0 | NA | NA | NA |
| Gm28124 | 0 | NA | NA | NA |
| Gm21790 | 0 | NA | NA | NA |
| Gm21728 | 0 | NA | NA | NA |
| Gm28126 | 0 | NA | NA | NA |
| Gm28125 | 0 | NA | NA | NA |
| Gm21825 | 0 | NA | NA | NA |
| Gm21317 | 0 | NA | NA | NA |
| Gm29468 | 0 | NA | NA | NA |
| Gm29469 | 0 | NA | NA | NA |
| Gm29472 | 0 | NA | NA | NA |
| Gm29473 | 0 | NA | NA | NA |
| Gm29470 | 0 | NA | NA | NA |
| Gm21333 | 0 | NA | NA | NA |
| Gm29382 | 0 | NA | NA | NA |
| Gm28224 | 0 | NA | NA | NA |
| Gm28226 | 0 | NA | NA | NA |
| Gm28225 | 0 | NA | NA | NA |
| Gm21344 | 0 | NA | NA | NA |
| Gm20849 | 0 | NA | NA | NA |
| Gm28829 | 0 | NA | NA | NA |
| Gm21901 | 0 | NA | NA | NA |
| Gm28827 | 0 | NA | NA | NA |
| Gm28828 | 0 | NA | NA | NA |
| Gm28831 | 0 | NA | NA | NA |
| Gm28832 | 0 | NA | NA | NA |
| Gm21864 | 0 | NA | NA | NA |
| Gm21802 | 0 | NA | NA | NA |
| Gm28658 | 0 | NA | NA | NA |
| Gm29238 | 0 | NA | NA | NA |
| Gm21088 | 0 | NA | NA | NA |
| Gm29239 | 0 | NA | NA | NA |
| Gm25729 | 0 | NA | NA | NA |
| Gm21741 | 0 | NA | NA | NA |
| Gm20862 | 0 | NA | NA | NA |
| Gm22624 | 0 | NA | NA | NA |
| Gm20863 | 0 | NA | NA | NA |

| | | | | |
|---|---|---|---|---|
| Gm20864 | 0 | NA | NA | NA |
| Gm29476 | 0 | NA | NA | NA |
| Gm21717 | 0 | NA | NA | NA |
| Gm28472 | 0 | NA | NA | NA |
| Gm21095 | 0 | NA | NA | NA |
| Gm28473 | 0 | NA | NA | NA |
| Gm28474 | 0 | NA | NA | NA |
| Gm28475 | 0 | NA | NA | NA |
| Gm28476 | 0 | NA | NA | NA |
| Gm21380 | 0 | NA | NA | NA |
| Gm21782 | 0 | NA | NA | NA |
| Gm21394 | 0 | NA | NA | NA |
| Gm28471 | 0 | NA | NA | NA |
| Gm21824 | 0 | NA | NA | NA |
| Gm29338 | 0 | NA | NA | NA |
| Gm21409 | 0 | NA | NA | NA |
| Gm29339 | 0 | NA | NA | NA |
| Gm28759 | 0 | NA | NA | NA |
| Gm28758 | 0 | NA | NA | NA |
| Gm29071 | 0 | NA | NA | NA |
| Gm21879 | 0 | NA | NA | NA |
| Gm28757 | 0 | NA | NA | NA |
| Gm20856 | 0 | NA | NA | NA |
| Gm28348 | 0 | NA | NA | NA |
| Gm28973 | 0 | NA | NA | NA |
| Gm21734 | 0 | NA | NA | NA |
| Gm28972 | 0 | NA | NA | NA |
| Gm28970 | 0 | NA | NA | NA |
| Gm28971 | 0 | NA | NA | NA |
| Gm24828 | 0 | NA | NA | NA |
| Gm21785 | 0 | NA | NA | NA |
| Gm20854 | 0 | NA | NA | NA |
| Gm28349 | 0 | NA | NA | NA |
| Gm21892 | 0 | NA | NA | NA |
| Gm28346 | 0 | NA | NA | NA |
| Gm28345 | 0 | NA | NA | NA |
| Gm29090 | 0 | NA | NA | NA |
| Gm28152 | 0 | NA | NA | NA |
| Gm23770 | 0 | NA | NA | NA |
| Gm21881 | 0 | NA | NA | NA |
| Gm21435 | 0 | NA | NA | NA |
| Gm29505 | 0 | NA | NA | NA |
| Gm21862 | 0 | NA | NA | NA |
| Gm28823 | 0 | NA | NA | NA |
| Gm28220 | 0 | NA | NA | NA |
| Gm20820 | 0 | NA | NA | NA |
| Gm28219 | 0 | NA | NA | NA |

| | | | | |
|---|---|---|---|---|
| Gm28218 | 0 | NA | NA | NA |
| Gm23863 | 0 | NA | NA | NA |
| Gm20861 | 0 | NA | NA | NA |
| Gm20860 | 0 | NA | NA | NA |
| Gm28057 | 0 | NA | NA | NA |
| Gm21796 | 0 | NA | NA | NA |
| Gm29302 | 0 | NA | NA | NA |
| Gm28440 | 0 | NA | NA | NA |
| Gm29299 | 0 | NA | NA | NA |
| Gm29298 | 0 | NA | NA | NA |
| Gm29092 | 0 | NA | NA | NA |
| Gm29091 | 0 | NA | NA | NA |
| Gm29297 | 0 | NA | NA | NA |
| Gm21462 | 0 | NA | NA | NA |
| Gm21470 | 0 | NA | NA | NA |
| Gm28366 | 0 | NA | NA | NA |
| Gm21752 | 0 | NA | NA | NA |
| Gm28365 | 0 | NA | NA | NA |
| Gm21477 | 0 | NA | NA | NA |
| Gm28368 | 0 | NA | NA | NA |
| Gm28369 | 0 | NA | NA | NA |
| Gm28367 | 0 | NA | NA | NA |
| Gm29115 | 0 | NA | NA | NA |
| Gm21813 | 0 | NA | NA | NA |
| Gm20906 | 0 | NA | NA | NA |
| Gm29111 | 0 | NA | NA | NA |
| Gm28110 | 0 | NA | NA | NA |
| Gm28109 | 0 | NA | NA | NA |
| Gm28108 | 0 | NA | NA | NA |
| Gm28107 | 0 | NA | NA | NA |
| Gm28106 | 0 | NA | NA | NA |
| Gm28105 | 0 | NA | NA | NA |
| Gm28104 | 0 | NA | NA | NA |
| Gm28103 | 0 | NA | NA | NA |
| Gm28102 | 0 | NA | NA | NA |
| Gm28716 | 0 | NA | NA | NA |
| Gm28717 | 0 | NA | NA | NA |
| Gm28718 | 0 | NA | NA | NA |
| Gm29162 | 0.046345709 | -0.443818988 | 3.724945049 | -0.119147795 |
| Gm21777 | 0 | NA | NA | NA |
| Gm28866 | 0 | NA | NA | NA |
| Gm28405 | 0 | NA | NA | NA |
| Gm21792 | 0 | NA | NA | NA |
| Gm28406 | 0 | NA | NA | NA |
| Gm28978 | 0 | NA | NA | NA |
| Gm29436 | 0 | NA | NA | NA |
| Gm20819 | 0 | NA | NA | NA |

| | | | | |
|---|---|---|---|---|
| Gm29437 | 0 | NA | NA | NA |
| Gm28977 | 0 | NA | NA | NA |
| Gm28407 | 0 | NA | NA | NA |
| Gm21287 | 0 | NA | NA | NA |
| Gm29390 | 0 | NA | NA | NA |
| Gm29391 | 0 | NA | NA | NA |
| Gm21720 | 0 | NA | NA | NA |
| Gm29392 | 0 | NA | NA | NA |
| Gm29393 | 0 | NA | NA | NA |
| Gm21294 | 0 | NA | NA | NA |
| Gm28672 | 0 | NA | NA | NA |
| Gm21838 | 0 | NA | NA | NA |
| Gm28670 | 0 | NA | NA | NA |
| Gm28671 | 0 | NA | NA | NA |
| Gm28673 | 0 | NA | NA | NA |
| Gm28674 | 0 | NA | NA | NA |
| Gm28531 | 0 | NA | NA | NA |
| Gm21996 | 0 | NA | NA | NA |
| Gm21803 | 0 | NA | NA | NA |
| Gm28930 | 0 | NA | NA | NA |
| Gm20879 | 0 | NA | NA | NA |
| Gm29504 | 0 | NA | NA | NA |
| Gm20837 | 0 | NA | NA | NA |
| Gm29503 | 0 | NA | NA | NA |
| Gm28300 | 0 | NA | NA | NA |
| Gm28301 | 0 | NA | NA | NA |
| Gm21857 | 2.299789819 | 0.276410625 | 0.599643089 | 0.460958576 |
| Gm21860 | 0.344895107 | 0.384728534 | 1.501234484 | 0.256274778 |
| Erdr1 | 44.24775491 | -0.293367893 | 0.223766428 | -1.311045163 |
| Gm21742 | 0 | NA | NA | NA |
| Gm21748 | 0 | NA | NA | NA |

| pvalue | padj |
|---|---|
| NA | NA |
| NA | NA |
| 2.71E-17 | 5.50E-15 |
| 0.556010657 | NA |
| 0.018583114 | 0.055353582 |
| 0.000911459 | 0.00460744 |
| 0.003535203 | 0.014370513 |
| NA | NA |
| 0.009536906 | 0.032352991 |
| 0.478316408 | NA |
| 0.009019962 | 0.030933834 |
| 0.005552745 | 0.020907531 |
| 3.56E-05 | 0.000289463 |
| NA | NA |
| NA | NA |
| 0.992061292 | NA |
| 0.905158739 | NA |
| 0.124368483 | NA |
| NA | NA |
| 0.003468767 | 0.014142343 |
| 0.990592969 | NA |
| 0.957425904 | NA |
| 0.621419696 | NA |
| NA | NA |
| 0.980649686 | NA |
| 0.889409301 | NA |
| 0.41062358 | 0.573607677 |
| 0.97251228 | NA |
| 0.992061292 | NA |
| 0.766345984 | NA |
| 0.730867972 | 0.834459624 |
| 0.01461006 | 0.045694675 |
| 0.789575947 | 0.872894237 |
| 0.061290422 | 0.142120456 |
| 0.801661681 | NA |
| 0.513255578 | 0.664813262 |
| 0.958349348 | 0.976945979 |
| 0.242996217 | 0.395871647 |
| 1.13E-05 | 0.000108251 |
| 0.912419498 | 0.950465833 |
| 0.854267045 | 0.91497659 |
| 0.185275654 | NA |
| 0.638959034 | NA |
| 0.144193681 | 0.27058187 |
| NA | NA |
| 0.044885063 | 0.111534583 |

| | | |
|---|---|---|
| | 0.197389114 | NA |
| | 2.54E-06 | 2.96E-05 |
| NA | | NA |
| NA | | NA |
| NA | | NA |
| | 0.366325959 | 0.531177478 |
| | 0.905158266 | NA |
| | 0.017379676 | 0.052536664 |
| | 0.023446726 | 0.06688317 |
| | 0.552132486 | NA |
| | 0.76851712 | NA |
| NA | | NA |
| | 0.906905825 | 0.947360419 |
| | 0.818526268 | 0.892010835 |
| NA | | NA |
| | 0.889410245 | NA |
| | 0.128670975 | 0.2485036 |
| NA | | NA |
| NA | | NA |
| NA | | NA |
| NA | | NA |
| | 0.422290323 | 0.583857369 |
| NA | | NA |
| | 0.369655549 | 0.534382239 |
| | 0.542562063 | 0.689674115 |
| NA | | NA |
| NA | | NA |
| | 0.807879735 | NA |
| | 0.083997073 | 0.180408032 |
| NA | | NA |
| NA | | NA |
| NA | | NA |
| | 0.905158266 | NA |
| NA | | NA |
| NA | | NA |
| | 1.20E-08 | 2.77E-07 |
| | 0.02080998 | 0.060710291 |
| | 0.00379144 | 0.015192743 |
| | 1.69E-05 | 0.000153417 |
| | 0.520586783 | NA |
| | 0.075382963 | 0.165981599 |
| | 0.61803313 | NA |
| | 0.805374319 | NA |
| | 0.640281197 | NA |
| | 0.705908456 | NA |
| | 0.985057863 | NA |
| | 0.662540347 | 0.784001912 |

| | |
|---|---|
| 0.090320784 | 0.190688789 |
| 0.903208911 | 0.945177713 |
| NA | NA |
| 0.878390629 | NA |
| NA | NA |
| NA | NA |
| NA | NA |
| 0.922180181 | 0.955294241 |
| 0.000923564 | 0.004658983 |
| 2.71E-05 | 0.000229664 |
| 0.547592488 | 0.693866899 |
| 0.265196345 | 0.421317214 |
| 1.05E-07 | 1.84E-06 |
| NA | NA |
| 0.368272145 | 0.532943722 |
| 0.403345778 | 0.566852477 |
| 0.513782868 | 0.665221581 |
| 0.997443328 | 0.998898985 |
| 0.705479746 | 0.816681414 |
| NA | NA |
| 0.638474614 | 0.765131197 |
| 0.570688167 | 0.713003042 |
| 0.777036502 | 0.865831661 |
| 0.00923636 | 0.031516293 |
| 0.807115012 | NA |
| 0.419511162 | 0.581385017 |
| NA | NA |
| NA | NA |
| 0.139020955 | 0.263134288 |
| 0.389485097 | 0.55285823 |
| NA | NA |
| 0.313211001 | 0.475117462 |
| NA | NA |
| 0.015025996 | 0.046769078 |
| 0.271406097 | 0.428054765 |
| 0.05667516 | 0.133447334 |
| 0.158459613 | 0.290209729 |
| 0.792413 | NA |
| 0.992061292 | NA |
| NA | NA |
| 0.905158266 | NA |
| NA | NA |
| NA | NA |
| 0.003207346 | 0.013283687 |
| NA | NA |
| NA | NA |
| 0.889409773 | NA |

| | |
|---|---|
| 1.75E-17 | 3.89E-15 |
| 0.018605959 | 0.055406597 |
| NA | NA |
| 0.043739932 | 0.109270212 |
| 0.383933072 | NA |
| NA | NA |
| NA | NA |
| 0.489063535 | NA |
| NA | NA |
| 0.99205891 | NA |
| NA | NA |
| 7.13E-07 | 9.79E-06 |
| 0.000245692 | 0.001508063 |
| NA | NA |
| NA | NA |
| NA | NA |
| 0.740879991 | NA |
| 0.783389773 | 0.8695575 |
| 0.083606339 | 0.179795951 |
| 0.436200727 | 0.59664456 |
| NA | NA |
| 0.734423307 | 0.8369018 |
| 0.24486648 | 0.397973727 |
| 0.746485357 | 0.845213569 |
| 0.505349414 | 0.657325422 |
| 0.956084434 | 0.975600256 |
| 0.878745577 | NA |
| 0.905158739 | NA |
| 8.07E-06 | 8.10E-05 |
| 0.003988507 | 0.015855109 |
| 0.3049809 | 0.465649102 |
| 0.840266151 | NA |
| NA | NA |
| 0.467831779 | NA |
| 0.992061292 | NA |
| NA | NA |
| NA | NA |
| 3.38E-05 | 0.000277239 |
| NA | NA |
| 0.019451873 | 0.057465853 |
| 0.018416658 | 0.054957366 |
| 0.422960204 | 0.584394584 |
| 0.003895758 | 0.01553384 |
| 0.125678408 | 0.24424924 |
| 0.338667524 | 0.501639134 |
| 0.147104176 | 0.274682023 |
| NA | NA |

| | |
|---|---|
| 0.986849022 | 0.993180845 |
| 4.87E-05 | 0.00037866 |
| 0.642240745 | 0.767968009 |
| 0.117816033 | 0.232864291 |
| NA | NA |
| 0.989551585 | NA |
| 0.905158266 | NA |
| 0.99205891 | NA |
| 0.028271376 | 0.077698515 |
| 0.310235315 | 0.47145033 |
| 0.90921216 | 0.948458589 |
| 0.905158739 | NA |
| NA | NA |
| 0.706722674 | 0.817578995 |
| 0.09571985 | 0.1992145 |
| 4.96E-07 | 7.16E-06 |
| 0.183736032 | 0.324303474 |
| NA | NA |
| 6.69E-08 | 1.25E-06 |
| 0.842372003 | NA |
| 0.081141383 | 0.175822343 |
| 0.84291473 | 0.907259774 |
| 0.02763413 | 0.07631969 |
| 0.834866031 | NA |
| 0.873034547 | NA |
| 0.002194558 | 0.009678711 |
| 0.051808572 | 0.124743907 |
| NA | NA |
| 0.110873072 | 0.22236385 |
| 0.748443391 | NA |
| NA | NA |
| 7.89E-30 | 5.77E-26 |
| 0.711683952 | NA |
| 6.69E-05 | 0.000496202 |
| 0.51956162 | 0.670489751 |
| 0.905158266 | NA |
| 0.991570563 | NA |
| 0.749263962 | 0.847157103 |
| 0.905158739 | NA |
| NA | NA |
| 0.145288451 | 0.272031813 |
| 0.005629865 | 0.021158001 |
| 9.99E-05 | 0.000697677 |
| 6.98E-16 | 1.09E-13 |
| 0.284469748 | NA |
| NA | NA |
| 0.516074753 | 0.667166226 |

| | | | |
|---|---|---|---|
| NA | | NA | |
| | 0.932991274 | NA | |
| | 0.026872266 | | 0.074571981 |
| NA | | NA | |
| | 0.992060816 | NA | |
| NA | | NA | |
| NA | | NA | |
| | 0.823459502 | NA | |
| NA | | NA | |
| NA | | NA | |
| NA | | NA | |
| NA | | NA | |
| NA | | NA | |
| | 0.193250352 | NA | |
| | 0.004236442 | | 0.01667169 |
| | 0.707738395 | | 0.817787172 |
| | 0.420250056 | | 0.582078402 |
| | 0.992059387 | NA | |
| NA | | NA | |
| | 0.093215129 | | 0.195092548 |
| NA | | NA | |
| | 0.992062245 | NA | |
| | 0.596464945 | | 0.733346695 |
| NA | | NA | |
| | 0.421987204 | NA | |
| | 0.790039101 | NA | |
| NA | | NA | |
| NA | | NA | |
| | 0.762216526 | | 0.855701058 |
| NA | | NA | |
| NA | | NA | |
| | 0.513219154 | NA | |
| | 0.005147936 | | 0.019646077 |
| NA | | NA | |
| | 0.992060816 | NA | |
| NA | | NA | |
| | 0.317417705 | | 0.479676236 |
| | 0.098671596 | | 0.203831569 |
| NA | | NA | |
| | 0.058361311 | | 0.136568205 |
| | 0.729039491 | | 0.833019991 |
| | 2.88E-06 | | 3.29E-05 |
| | 0.986347789 | NA | |
| | 0.06415232 | | 0.146986727 |
| | 1.49E-08 | | 3.33E-07 |
| | 0.266296699 | | 0.42248459 |
| | 0.011548805 | | 0.037736203 |

| | |
|---|---|
| 0.388294517 | 0.551962569 |
| 0.44036824 | 0.6004748 |
| 0.441195041 | NA |
| 0.905158266 | NA |
| 0.889409773 | NA |
| NA | NA |
| NA | NA |
| 0.905158739 | NA |
| NA | NA |
| NA | NA |
| NA | NA |
| NA | NA |
| 3.91E-14 | 3.78E-12 |
| 4.34E-05 | 0.000343361 |
| 0.593729312 | 0.731140757 |
| 0.034987396 | 0.092032611 |
| 0.162352405 | NA |
| NA | NA |
| NA | NA |
| 1.51E-08 | 3.36E-07 |
| 0.992060816 | NA |
| 0.779470453 | NA |
| 0.631057623 | NA |
| 0.397444805 | NA |
| NA | NA |
| 0.992061292 | NA |
| 0.235095692 | 0.386614716 |
| 0.369748167 | 0.534410583 |
| NA | NA |
| 0.181968984 | 0.322199107 |
| 0.914177989 | NA |
| 5.20E-10 | 1.80E-08 |
| 0.905158266 | NA |
| 0.85498696 | NA |
| 1.19E-06 | 1.52E-05 |
| 0.0710911 | 0.158405832 |
| 0.685084625 | 0.800821797 |
| 0.889410717 | NA |
| 0.745803324 | 0.844702661 |
| 0.922061913 | 0.955294241 |
| NA | NA |
| 0.678053076 | 0.795754769 |
| 5.12E-12 | 3.04E-10 |
| 0.181494723 | NA |
| 0.266266629 | 0.422467407 |
| 0.029234307 | 0.079785751 |
| 0.102025015 | 0.209223693 |

| | |
|---|---|
| 0.555379747 | 0.700815071 |
| 0.144316241 | 0.270742447 |
| 0.032561744 | 0.086975227 |
| 0.005339724 | 0.020253146 |
| 0.821368406 | 0.893671826 |
| 8.58E-07 | 1.15E-05 |
| NA | NA |
| 0.671274182 | NA |
| 0.905158266 | NA |
| 0.854956362 | NA |
| 6.28E-07 | 8.73E-06 |
| 0.690088574 | NA |
| 0.171765088 | 0.308429925 |
| 0.169817242 | 0.305882767 |
| 0.059073709 | 0.137882607 |
| 0.037741364 | 0.097722064 |
| 0.088307892 | 0.187519262 |
| 0.129079679 | 0.249103245 |
| 0.007788864 | 0.027450464 |
| 0.165058315 | 0.299299383 |
| NA | NA |
| 0.030417309 | 0.082394259 |
| 0.027791176 | 0.07664738 |
| 0.03176634 | 0.08530721 |
| 0.000752326 | 0.003927801 |
| 0.030626191 | 0.082824274 |
| 0.098775843 | 0.20399997 |
| 0.153316612 | 0.283462111 |
| 0.078225308 | 0.170971388 |
| 0.012540386 | 0.040377469 |
| 0.041695244 | 0.105397246 |
| 0.012910837 | 0.041291738 |
| 0.115154339 | 0.229150351 |
| 0.0202298 | 0.059333002 |
| 0.781247144 | 0.868271012 |
| NA | NA |
| NA | NA |
| NA | NA |
| NA | NA |
| NA | NA |
| NA | NA |
| 0.992061292 | NA |
| NA | NA |
| 0.777092376 | NA |
| NA | NA |
| 0.541575312 | 0.688938261 |
| 0.992060816 | NA |

| | | |
|---|---|---|
| 0.876460948 | NA | |
| NA | NA | |
| NA | NA | |
| NA | NA | |
| NA | NA | |
| NA | NA | |
| 0.992061292 | NA | |
| 0.250931063 | | 0.405046946 |
| NA | NA | |
| NA | NA | |
| 0.992061292 | NA | |
| 0.37327911 | NA | |
| 0.158412291 | | 0.290147259 |
| 0.624069416 | NA | |
| 0.957876568 | | 0.976600035 |
| 0.11985329 | | 0.235912068 |
| 0.025203929 | | 0.071010146 |
| 7.62E-05 | | 0.000554717 |
| NA | NA | |
| NA | NA | |
| 0.905158739 | NA | |
| NA | NA | |
| NA | NA | |
| NA | NA | |
| 0.992060816 | NA | |
| NA | NA | |
| NA | NA | |
| 0.893532551 | NA | |
| NA | NA | |
| NA | NA | |
| NA | NA | |
| 0.905158266 | NA | |
| NA | NA | |
| NA | NA | |
| 0.447606643 | | 0.606803455 |
| NA | NA | |
| NA | NA | |
| 0.009056947 | | 0.031041283 |
| 0.992060816 | NA | |
| 0.043408235 | | 0.108676483 |
| 0.354366317 | NA | |
| 0.212176024 | | 0.359254631 |
| 0.991782148 | NA | |
| 0.001484799 | | 0.006947516 |
| 0.480008094 | NA | |
| NA | NA | |
| 0.286367981 | | 0.445413975 |

| | | |
|---|---|---|
| | 0.626375226 | NA |
| | 0.095048129 | NA |
| | 0.354677195 | NA |
| | 0.930151745 | NA |
| | 0.370075014 | 0.534636661 |
| | 0.517983246 | NA |
| NA | | NA |
| | 0.10161541 | NA |
| NA | | NA |
| NA | | NA |
| | 0.444374475 | 0.603949935 |
| | 0.992062245 | NA |
| NA | | NA |
| | 0.627197571 | 0.756363966 |
| NA | | NA |
| | 0.972975228 | 0.985912235 |
| NA | | NA |
| | 0.3300545 | 0.492438027 |
| | 0.207186951 | NA |
| | 0.357642177 | 0.522345498 |
| | 0.751310255 | 0.848494006 |
| | 0.982434162 | NA |
| | 0.827180077 | 0.89672859 |
| | 0.018004492 | 0.054018396 |
| | 0.00189575 | 0.008551513 |
| | 0.401282688 | 0.564869943 |
| | 0.000487376 | 0.002703279 |
| | 0.274683612 | 0.431921048 |
| NA | | NA |
| NA | | NA |
| | 0.407082823 | 0.570191295 |
| | 0.349344753 | NA |
| | 0.226555617 | 0.376461598 |
| | 0.042461652 | 0.106928373 |
| | 0.577058501 | NA |
| NA | | NA |
| | 0.835724615 | NA |
| NA | | NA |
| | 0.72199964 | 0.827905488 |
| | 0.55966754 | 0.70400066 |
| NA | | NA |
| | 0.640256703 | 0.766582314 |
| NA | | NA |
| | 3.15E-07 | 4.80E-06 |
| | 0.021807625 | 0.06304287 |
| | 0.344468673 | NA |
| | 0.807290849 | NA |

| | | |
|---|---|---|
| | 0.300819848 | NA |
| NA | | NA |
| | 0.704703613 | NA |
| NA | | NA |
| NA | | NA |
| | 0.849661732 | NA |
| | 5.69E-12 | 3.36E-10 |
| | 0.889409301 | NA |
| | 0.866278329 | NA |
| | 0.087333627 | NA |
| | 0.011083935 | 0.036572823 |
| | 1.49E-23 | 1.56E-20 |
| NA | | NA |
| | 0.378455311 | 0.543017572 |
| | 0.788893855 | 0.872608947 |
| | 0.241705971 | 0.394472049 |
| | 0.269979988 | 0.426392452 |
| | 0.45659562 | NA |
| | 0.933706877 | 0.961917397 |
| | 0.401938249 | NA |
| | 0.483618091 | NA |
| | 0.889409301 | NA |
| NA | | NA |
| NA | | NA |
| | 0.721268272 | NA |
| NA | | NA |
| NA | | NA |
| | 0.117266158 | NA |
| NA | | NA |
| NA | | NA |
| | 0.982889007 | NA |
| NA | | NA |
| | 0.992060816 | NA |
| | 0.320283916 | 0.48257956 |
| | 0.74503104 | NA |
| | 0.787785659 | NA |
| | 0.000491881 | 0.002725438 |
| | 0.7676954 | NA |
| | 6.34E-06 | 6.58E-05 |
| | 0.508570026 | 0.660302087 |
| | 0.308643877 | NA |
| | 0.738475793 | 0.839564584 |
| | 0.992061292 | NA |
| | 0.475898665 | NA |
| | 0.026404588 | 0.073526885 |
| | 0.862859858 | 0.920856319 |
| | 0.327797862 | 0.489912234 |

| | |
|---|---|
| NA | NA |
| 2.00E-05 | 0.00017752 |
| 0.859036022 | NA |
| 0.217856141 | 0.366021652 |
| 0.201277182 | 0.346491505 |
| 0.052964562 | 0.126779865 |
| 0.000136388 | 0.000909485 |
| 0.237254976 | 0.389165794 |
| 0.440955261 | NA |
| NA | NA |
| 0.208921279 | 0.355717999 |
| 0.680446236 | 0.797370131 |
| 0.323701892 | NA |
| 0.005215933 | 0.019858755 |
| 0.035050071 | 0.092084778 |
| 0.066599417 | 0.150993635 |
| NA | NA |
| 0.354875416 | 0.519484363 |
| 0.046238747 | 0.114204828 |
| 0.890850987 | 0.938324867 |
| 0.964001249 | 0.980204827 |
| 0.579826721 | NA |
| NA | NA |
| 0.83596241 | 0.902280588 |
| 0.09439449 | NA |
| 0.550207255 | 0.696121036 |
| NA | NA |
| 2.67E-13 | 2.20E-11 |
| 0.889409773 | NA |
| NA | NA |
| 0.05568594 | 0.131654561 |
| 0.456460102 | 0.614210622 |
| 1.88E-09 | 5.47E-08 |
| 0.020235818 | 0.059334801 |
| 7.90E-09 | 1.92E-07 |
| 0.905158266 | NA |
| 0.633692926 | NA |
| 0.004145366 | 0.016371959 |
| 0.220747821 | 0.369697452 |
| 2.93E-07 | 4.51E-06 |
| 0.499362546 | 0.652400845 |
| 0.00088999 | 0.004510606 |
| 0.160129168 | 0.292486808 |
| 0.184194954 | NA |
| 0.400426702 | NA |
| 0.807433834 | NA |
| NA | NA |

| | | |
|---|---|---|
| | 0.003795029 | 0.015201578 |
| | 0.44458773 | 0.604052837 |
| | 0.002878031 | 0.012139586 |
| | 0.001967201 | 0.008828483 |
| | 0.991700267 | NA |
| | 1.33E-08 | 3.02E-07 |
| | 0.951528444 | NA |
| NA | | NA |
| | 0.003909229 | NA |
| NA | | NA |
| NA | | NA |
| NA | | NA |
| | 0.187704615 | 0.32917028 |
| | 0.001085791 | 0.005337561 |
| NA | | NA |
| | 0.662624541 | NA |
| | 0.080167781 | NA |
| | 0.008899211 | 0.030600966 |
| | 0.668066689 | 0.788322669 |
| | 0.342305031 | NA |
| | 0.694063519 | NA |
| NA | | NA |
| | 3.80E-06 | 4.21E-05 |
| | 0.62072456 | NA |
| | 0.888823417 | 0.93717821 |
| | 0.377773793 | 0.542191812 |
| | 0.000143036 | 0.000944069 |
| | 0.411676861 | 0.574442621 |
| NA | | NA |
| | 0.012238692 | 0.039591843 |
| | 0.881315285 | 0.932485417 |
| | 0.393851454 | 0.557254128 |
| | 0.228468934 | 0.378752025 |
| | 0.001240936 | 0.005965862 |
| | 0.398350365 | 0.561990085 |
| | 0.247677527 | NA |
| | 0.03979916 | 0.101716684 |
| | 0.322490502 | 0.484870953 |
| | 0.199688742 | NA |
| | 0.992061292 | NA |
| | 0.635359926 | NA |
| | 5.63E-07 | 7.96E-06 |
| | 0.10482995 | 0.213460764 |
| | 0.56599259 | 0.709032533 |
| | 0.022092142 | 0.063714387 |
| | 0.764254867 | 0.857062789 |
| | 0.163656224 | 0.297370875 |

| | |
|---|---|
| 0.689086566 | 0.804147946 |
| 0.021793319 | 0.063018107 |
| 0.776535092 | NA |
| 0.008022929 | 0.028156544 |
| 0.082849758 | 0.1787832 |
| 0.00030045 | 0.001787964 |
| 0.889409301 | NA |
| 0.888136516 | 0.936678822 |
| 0.801661681 | NA |
| 4.38E-05 | 0.000346415 |
| NA | NA |
| 0.772122873 | NA |
| 0.039224097 | 0.100621722 |
| 0.416384338 | 0.578659792 |
| NA | NA |
| 0.889409773 | NA |
| 0.991879747 | NA |
| 0.207891487 | 0.354542495 |
| NA | NA |
| NA | NA |
| 0.835064872 | NA |
| NA | NA |
| 0.404598697 | 0.567850108 |
| 0.813836939 | NA |
| 0.53447505 | NA |
| 0.000140464 | 0.000930777 |
| NA | NA |
| 0.000112443 | 0.000771361 |
| 0.441174933 | 0.600941867 |
| 0.808059413 | 0.88498742 |
| 5.24E-05 | 0.00040238 |
| 0.001458366 | 0.006848644 |
| NA | NA |
| 0.430919889 | 0.591854506 |
| 0.98534211 | NA |
| 0.342414277 | 0.505892156 |
| 0.115078396 | NA |
| 0.023489208 | 0.066978251 |
| 0.667690251 | 0.787972818 |
| 0.660474517 | 0.782569467 |
| 0.38077973 | 0.545009913 |
| 0.992060816 | NA |
| NA | NA |
| NA | NA |
| 0.19531888 | 0.33905196 |
| 0.250205097 | 0.404197287 |
| 0.419149928 | 0.581138113 |

| | |
|---|---|
| 0.994131777 | 0.997128749 |
| 0.110592239 | 0.222043977 |
| 0.531781026 | NA |
| 0.15709972 | 0.28840447 |
| 0.451827213 | 0.610038967 |
| NA | NA |
| 0.098846847 | NA |
| NA | NA |
| NA | NA |
| 0.986588132 | NA |
| 0.177202631 | 0.315920482 |
| 0.247295451 | NA |
| NA | NA |
| 0.625386083 | 0.755340311 |
| 0.721993285 | 0.827905488 |
| NA | NA |
| 0.447955793 | 0.607088889 |
| 0.4817845 | 0.637396423 |
| 1.62E-05 | 0.000148128 |
| 0.021914866 | NA |
| 0.545377671 | NA |
| NA | NA |
| NA | NA |
| 0.465764098 | 0.622578149 |
| 0.786307836 | 0.871387018 |
| 0.451715182 | 0.610037742 |
| 0.452072449 | NA |
| 0.671274182 | NA |
| 0.220950216 | 0.3699181 |
| 0.788582525 | NA |
| NA | NA |
| NA | NA |
| NA | NA |
| 2.37E-08 | 5.03E-07 |
| NA | NA |
| 6.05E-10 | 2.03E-08 |
| 0.000499796 | 0.002761704 |
| 0.992060816 | NA |
| 0.808198519 | NA |
| NA | NA |
| NA | NA |
| NA | NA |
| 0.877209064 | NA |
| NA | NA |
| 0.003688007 | 0.014865079 |
| NA | NA |
| 0.200471091 | 0.345564384 |

| | |
|---|---|
| 0.989554908 | NA |
| 0.880258492 | NA |
| 0.70461765 | 0.816199766 |
| NA | NA |
| NA | NA |
| 0.812906299 | NA |
| NA | NA |
| NA | NA |
| 0.242812718 | 0.395690381 |
| 0.992061292 | NA |
| 0.063885236 | NA |
| 0.021845928 | 0.063128666 |
| 0.000184286 | 0.001175355 |
| 0.451035249 | 0.609419335 |
| 0.140166748 | 0.264744678 |
| 0.068408053 | NA |
| 0.197472262 | NA |
| 0.768804414 | NA |
| 0.374037628 | 0.537969442 |
| NA | NA |
| NA | NA |
| 0.884330883 | NA |
| 0.234537956 | NA |
| 0.001496005 | 0.006995478 |
| NA | NA |
| NA | NA |
| NA | NA |
| NA | NA |
| NA | NA |
| NA | NA |
| NA | NA |
| NA | NA |
| NA | NA |
| NA | NA |
| NA | NA |
| NA | NA |
| NA | NA |
| NA | NA |
| NA | NA |
| NA | NA |
| NA | NA |
| 0.992060816 | NA |
| NA | NA |
| NA | NA |
| NA | NA |
| NA | NA |
| NA | NA |

| | | |
|---|---|---|
| | 0.807252815 | NA |
| | 0.220865191 | 0.369803943 |
| | 0.245824561 | 0.399146816 |
| | 0.547939399 | 0.694098718 |
| | 0.859302034 | NA |
| | 0.566312623 | 0.709312051 |
| | 0.095383025 | 0.198607608 |
| | 0.779689731 | 0.86734378 |
| | 0.440737684 | 0.600680164 |
| | 0.208277656 | NA |
| | 0.898195764 | NA |
| | 9.43E-08 | 1.67E-06 |
| | 0.786863035 | 0.871650375 |
| | 0.991799276 | NA |
| | 0.298256486 | NA |
| | 0.011463518 | 0.037526443 |
| | 0.443193836 | 0.602850934 |
| | 0.279546525 | 0.437780783 |
| NA | | NA |
| NA | | NA |
| | 0.598757777 | NA |
| | 0.585247562 | 0.72463779 |
| | 0.375380072 | 0.539532072 |
| | 0.523198884 | NA |
| | 0.088036707 | 0.187053512 |
| | 0.99205891 | NA |
| | 0.193924347 | 0.33716427 |
| | 0.992061292 | NA |
| NA | | NA |
| | 0.000148506 | 0.000975192 |
| | 0.905158739 | NA |
| | 0.772899823 | 0.862973875 |
| | 0.165192979 | 0.299444661 |
| | 0.111184327 | NA |
| | 0.946543497 | 0.969547003 |
| | 0.892713357 | 0.939380965 |
| | 0.362722477 | 0.527608523 |
| | 0.061894206 | 0.143142167 |
| | 0.041395168 | 0.104843885 |
| | 0.948517032 | NA |
| | 0.436552313 | 0.596902382 |
| | 0.001507635 | 0.007037876 |
| | 0.032268713 | NA |
| | 0.430300284 | 0.591410398 |
| | 0.416620161 | 0.578884664 |
| NA | | NA |
| | 0.900119204 | 0.94360613 |

| | | |
|---|---|---|
| | 0.169183795 | 0.304941882 |
| NA | NA | |
| | 0.011316603 | 0.037150736 |
| | 0.992061292 | NA |
| | 0.463558766 | 0.620618341 |
| | 0.391595159 | NA |
| | 0.682192194 | 0.798734119 |
| | 0.004191938 | 0.016529136 |
| | 0.905158739 | NA |
| NA | NA | |
| NA | NA | |
| | 0.008734654 | 0.030162645 |
| | 0.889410245 | NA |
| | 3.97E-06 | 4.37E-05 |
| NA | NA | |
| | 0.000455568 | 0.002549506 |
| | 0.992060816 | NA |
| | 0.075463143 | 0.166108175 |
| | 0.051306169 | 0.123887418 |
| | 0.989535731 | NA |
| | 0.036709583 | 0.095646149 |
| | 0.342139867 | 0.505554764 |
| | 0.19064562 | 0.332805301 |
| | 0.749624458 | 0.847238842 |
| | 0.99205891 | NA |
| | 0.409184874 | 0.572311507 |
| | 0.712500612 | NA |
| | 0.00570024 | 0.021385884 |
| NA | NA | |
| | 0.396079881 | 0.559721869 |
| | 0.067564122 | 0.152455868 |
| | 0.616661457 | 0.749171172 |
| | 0.028861627 | 0.07900007 |
| | 0.162766672 | 0.296097535 |
| | 1.84E-08 | 4.03E-07 |
| | 0.012689824 | 0.040798806 |
| | 0.284692489 | NA |
| | 0.899828785 | 0.943542752 |
| | 0.117235275 | 0.232134302 |
| | 0.518459195 | NA |
| | 0.029987384 | 0.081476486 |
| | 0.84520039 | 0.908856342 |
| | 0.005017567 | 0.019235467 |
| | 0.783889588 | NA |
| | 0.135059321 | 0.257311091 |
| NA | NA | |
| | 0.992060816 | NA |

| | |
|---|---|
| 0.046303962 | 0.114273864 |
| 0.10114553 | 0.207808261 |
| 0.766988453 | 0.858600094 |
| 0.076024181 | 0.167165351 |
| 0.917934109 | 0.953180834 |
| NA | NA |
| NA | NA |
| 0.373245185 | NA |
| NA | NA |
| NA | NA |
| 0.002075375 | 0.009234682 |
| 7.24E-13 | 5.30E-11 |
| 0.000143654 | 0.00094758 |
| NA | NA |
| 0.889409773 | NA |
| 0.189275413 | 0.331073665 |
| NA | NA |
| 0.458547558 | 0.615893683 |
| 0.026412588 | 0.073537977 |
| 0.558023167 | 0.702908387 |
| 0.43443125 | 0.595071276 |
| NA | NA |
| NA | NA |
| NA | NA |
| 0.872834076 | NA |
| 0.889409773 | NA |
| 5.75E-07 | 8.11E-06 |
| 0.991918092 | NA |
| 0.142000324 | 0.267402891 |
| 0.426800119 | 0.587960164 |
| 6.45E-05 | 0.000480808 |
| 0.020698623 | 0.060465752 |
| NA | NA |
| 0.637374562 | 0.764085382 |
| 0.883264219 | NA |
| 0.568251493 | 0.710779543 |
| 0.501267287 | 0.654030205 |
| 0.11341773 | 0.226495082 |
| 0.181886314 | 0.322111323 |
| 0.274746469 | 0.431986274 |
| 0.591259772 | 0.729063515 |
| 0.222037939 | 0.371371095 |
| 0.15043369 | 0.279260579 |
| 0.491052391 | 0.644994584 |
| 0.067056806 | 0.151690546 |
| 0.133128763 | 0.254516325 |
| NA | NA |

| | |
|---|---|
| NA | NA |
| 0.53501355 | 0.683377626 |
| 0.247540007 | NA |
| 0.703153914 | 0.814805109 |
| NA | NA |
| 8.73E-10 | 2.79E-08 |
| NA | NA |
| 9.22E-12 | 5.19E-10 |
| 0.028697044 | 0.078642004 |
| 0.026593543 | NA |
| 0.917189117 | 0.952814839 |
| 0.623282916 | 0.754169581 |
| 0.99205891 | NA |
| 0.678256072 | NA |
| 0.989074458 | NA |
| NA | NA |
| 0.697544541 | 0.810911078 |
| 0.610970668 | 0.744853703 |
| 0.961897107 | 0.979153466 |
| 0.889409773 | NA |
| NA | NA |
| 0.996766868 | 0.998494761 |
| 0.032928151 | 0.087740476 |
| NA | NA |
| 0.989301945 | NA |
| 0.00269427 | 0.011517127 |
| 0.992061292 | NA |
| 0.180747606 | 0.320652503 |
| 0.99205891 | NA |
| 0.807433834 | NA |
| 0.644307627 | 0.769852605 |
| 0.809078395 | NA |
| 0.088751085 | 0.188188979 |
| 0.836159543 | NA |
| 0.929510742 | 0.959575308 |
| 0.150551146 | 0.279431377 |
| NA | NA |
| 0.09424731 | 0.196840088 |
| 0.585184805 | NA |
| 0.036302919 | 0.094777767 |
| 0.995650883 | 0.997695434 |
| 0.991052259 | NA |
| 0.000104764 | 0.000727087 |
| 0.031591803 | 0.08491285 |
| NA | NA |
| 1.35E-11 | 7.30E-10 |
| 0.203201149 | 0.348955503 |

| | | |
|---|---|---|
| | 2.19E-07 | 3.51E-06 |
| | 0.287670328 | 0.446743475 |
| | 0.006002253 | 0.022298 |
| NA | | NA |
| | 0.000494421 | 0.002735449 |
| | 0.788247534 | 0.872260008 |
| | 2.75E-11 | 1.36E-09 |
| | 0.628058787 | 0.756654573 |
| | 0.1946856 | 0.338113024 |
| | 0.763841401 | NA |
| | 0.690826315 | 0.805274956 |
| | 0.801006268 | NA |
| NA | | NA |
| NA | | NA |
| | 2.20E-05 | 0.000193287 |
| | 0.010744535 | 0.035705093 |
| | 0.807879735 | NA |
| | 0.713238604 | NA |
| | 0.193949589 | NA |
| | 0.905158266 | NA |
| | 0.87762688 | NA |
| | 0.624874196 | NA |
| | 0.611230755 | 0.744964174 |
| | 0.976058776 | 0.987682942 |
| | 0.992060816 | NA |
| | 0.013517877 | 0.04283392 |
| | 0.142624183 | 0.268209679 |
| | 0.886956634 | NA |
| NA | | NA |
| NA | | NA |
| NA | | NA |
| | 0.795353865 | NA |
| NA | | NA |
| NA | | NA |
| NA | | NA |
| | 0.797342596 | NA |
| NA | | NA |
| | 0.905158266 | NA |
| NA | | NA |
| | 0.080145997 | 0.174163873 |
| | 0.560458082 | 0.704873941 |
| | 0.608856752 | 0.742936509 |
| | 0.046301088 | 0.114273864 |
| NA | | NA |
| | 0.191172022 | 0.333461469 |
| NA | | NA |
| | 0.992061292 | NA |

| | | |
|---|---|---|
| | 0.356206998 | NA |
| | 0.252073022 | 0.406340577 |
| | 0.94977114 | NA |
| | 0.27454314 | 0.431792924 |
| NA | | NA |
| | 0.423509677 | NA |
| | 5.82E-07 | 8.16E-06 |
| | 0.94974788 | 0.97187724 |
| | 0.313745513 | 0.475632566 |
| | 0.545009535 | NA |
| | 0.669009463 | 0.78895785 |
| NA | | NA |
| NA | | NA |
| NA | | NA |
| NA | | NA |
| | 0.178782246 | 0.318065246 |
| | 0.431809763 | 0.592668954 |
| | 0.001023708 | 0.005076693 |
| | 0.583756204 | NA |
| | 0.079074331 | 0.172415175 |
| NA | | NA |
| | 0.991388785 | NA |
| NA | | NA |
| | 0.348620268 | 0.512543684 |
| | 0.717066785 | 0.82431523 |
| | 0.905158266 | NA |
| | 0.905158266 | NA |
| | 0.992061292 | NA |
| | 0.992061769 | NA |
| | 0.889410245 | NA |
| | 0.906936141 | 0.947360419 |
| | 0.71145679 | NA |
| NA | | NA |
| | 0.944849481 | 0.968866415 |
| | 8.64E-18 | 2.08E-15 |
| NA | | NA |
| | 0.611555094 | NA |
| NA | | NA |
| | 0.629145064 | 0.757547443 |
| | 0.025925221 | 0.072567205 |
| | 0.871597746 | NA |
| | 0.226500692 | NA |
| | 0.065986098 | NA |
| | 0.555541483 | 0.700902288 |
| | 0.528524585 | 0.678126145 |
| | 0.520244185 | NA |
| | 0.775993333 | NA |

| | | |
|---|---|---|
| 0.682253026 | NA | |
| 0.687915585 | NA | |
| 0.328388401 | NA | |
| NA | NA | |
| NA | NA | |
| 0.046786502 | | 0.115113142 |
| 0.802262782 | | 0.880888466 |
| 0.468962692 | NA | |
| NA | NA | |
| 0.013624834 | | 0.043123051 |
| 0.008209881 | | 0.028666644 |
| 0.643388322 | | 0.768937427 |
| 0.337013367 | | 0.500032198 |
| 0.01448204 | | 0.045345945 |
| NA | NA | |
| 0.329844856 | | 0.492265991 |
| NA | NA | |
| 0.564448405 | NA | |
| NA | NA | |
| NA | NA | |
| NA | NA | |
| NA | NA | |
| 0.782559532 | NA | |
| 0.535025646 | | 0.683377626 |
| 0.046730604 | NA | |
| 0.537281389 | NA | |
| 0.000226126 | | 0.001402723 |
| 0.984375676 | | 0.992101408 |
| 0.91829992 | | 0.953425435 |
| 0.1936361 | | 0.336716433 |
| NA | NA | |
| NA | NA | |
| NA | NA | |
| NA | NA | |
| NA | NA | |
| NA | NA | |
| NA | NA | |
| NA | NA | |
| NA | NA | |
| NA | NA | |
| NA | NA | |
| NA | NA | |
| 6.26E-06 | | 6.50E-05 |
| 0.079775662 | | 0.173487874 |
| 0.898916132 | NA | |
| 0.397146515 | | 0.560868244 |
| 1.36E-19 | | 5.34E-17 |

| | | |
|---|---|---|
| | 0.345089298 | NA |
| | 0.000238111 | 0.00146997 |
| | 0.531913077 | 0.681026136 |
| | 0.111451061 | 0.223319084 |
| | 0.001639951 | 0.007536981 |
| | 0.132817242 | 0.25403134 |
| | 0.079005045 | 0.172291646 |
| | 0.023625723 | 0.067253988 |
| | 0.821226893 | 0.893667741 |
| | 0.798130512 | 0.878127024 |
| | 0.137921711 | 0.261561694 |
| | 0.000178559 | 0.001143281 |
| | 0.200976228 | 0.346178874 |
| | 0.889409773 | NA |
| | 0.788582525 | NA |
| | 0.002754947 | 0.011723083 |
| | 0.748556243 | 0.846597988 |
| | 0.15687302 | NA |
| | 6.36E-27 | 1.55E-23 |
| | 0.010109782 | 0.033960639 |
| | 0.807879735 | NA |
| NA | | NA |
| NA | | NA |
| NA | | NA |
| NA | | NA |
| NA | | NA |
| NA | | NA |
| NA | | NA |
| | 5.57E-23 | 4.70E-20 |
| | 0.097502763 | 0.201987092 |
| | 0.015429926 | 0.047769033 |
| | 0.143560409 | NA |
| | 0.843653202 | 0.907770514 |
| | 0.873644739 | 0.927758854 |
| | 0.337081097 | 0.5000989 |
| | 0.241703044 | 0.394472049 |
| | 0.304085163 | 0.464729216 |
| NA | | NA |
| NA | | NA |
| | 0.992060816 | NA |
| | 0.015805249 | 0.048772832 |
| NA | | NA |
| | 0.05703389 | 0.134090705 |
| | 0.015874475 | 0.048924574 |
| | 0.826333149 | 0.89642573 |
| | 0.992061292 | NA |
| | 0.711837103 | 0.820537057 |

| | | |
|---|---|---|
| NA | NA | |
| NA | NA | |
| NA | NA | |
| 0.992061292 | NA | |
| 0.379597637 | NA | |
| NA | NA | |
| 0.046502683 | | 0.114581733 |
| 0.044958755 | | 0.11169242 |
| 0.393436929 | NA | |
| NA | NA | |
| 0.195929482 | | 0.339762734 |
| NA | NA | |
| 2.49E-07 | | 3.91E-06 |
| 5.08E-06 | | 5.40E-05 |
| 0.197768521 | NA | |
| 0.778315897 | NA | |
| 0.719750973 | | 0.826239149 |
| NA | NA | |
| NA | NA | |
| 0.826084685 | NA | |
| 0.866781756 | NA | |
| NA | NA | |
| NA | NA | |
| 0.890296709 | NA | |
| 0.99723853 | NA | |
| 0.325619359 | | 0.487781603 |
| 0.00013245 | | 0.000885918 |
| 0.023219762 | | 0.066330526 |
| 0.361009943 | | 0.525707917 |
| 0.176652124 | | 0.315092518 |
| 0.153049944 | | 0.283088251 |
| 0.012038236 | | 0.039058454 |
| 0.05324988 | | 0.127196139 |
| 0.067007992 | | 0.151634914 |
| 0.912161064 | | 0.950331869 |
| 0.989388079 | NA | |
| 0.065480072 | | 0.149215608 |
| 0.634743277 | NA | |
| 0.588256571 | | 0.726928871 |
| 0.001465293 | | 0.006878233 |
| 0.001896632 | | 0.008553737 |
| 0.030714403 | NA | |
| NA | NA | |
| NA | NA | |
| 0.889409773 | NA | |
| NA | NA | |
| 0.839523182 | NA | |

| | | |
|---|---|---|
| NA | | NA |
| NA | | NA |
| | 0.905158266 | NA |
| | 0.076871038 | NA |
| NA | | NA |
| | 0.398708765 | 0.562302104 |
| | 0.50314126 | NA |
| NA | | NA |
| | 0.334556236 | NA |
| | 0.792928888 | 0.875103053 |
| | 0.007239144 | 0.025927967 |
| | 0.545331921 | 0.691793394 |
| | 0.555506173 | NA |
| NA | | NA |
| NA | | NA |
| | 0.020285283 | 0.059440157 |
| | 0.99205891 | NA |
| | 0.529766079 | 0.679069134 |
| | 0.345943504 | NA |
| | 0.000103204 | 0.000718307 |
| | 0.457827163 | 0.615227139 |
| | 3.18E-08 | 6.59E-07 |
| | 0.063026042 | 0.145041799 |
| | 0.022706541 | 0.065195205 |
| | 0.120459692 | 0.236789398 |
| | 5.76E-07 | 8.11E-06 |
| | 0.002555973 | 0.01100763 |
| | 0.986049583 | NA |
| | 0.099993898 | 0.205989596 |
| | 0.007318673 | 0.026170125 |
| | 0.968462854 | 0.983193519 |
| | 1.36E-07 | 2.32E-06 |
| | 0.000222325 | 0.00138184 |
| | 0.139321138 | 0.26357826 |
| | 0.16124351 | 0.293959836 |
| | 0.128812019 | 0.248710377 |
| | 0.173060846 | 0.310160511 |
| | 2.12E-05 | 0.000187374 |
| | 0.237267255 | 0.389165794 |
| | 0.515866295 | 0.667014542 |
| | 0.029780713 | NA |
| NA | | NA |
| | 0.455845642 | 0.613795063 |
| | 0.905157793 | NA |
| | 0.061456978 | 0.142385922 |
| | 0.38938915 | NA |
| | 1.70E-07 | 2.82E-06 |

| | |
|---|---|
| 0.242230755 | 0.39497625 |
| 0.000321515 | 0.001897959 |
| NA | NA |
| 0.005754698 | 0.021553371 |
| 0.992060816 | NA |
| 0.582580294 | 0.722516699 |
| 0.889409773 | NA |
| 0.993977534 | 0.997117284 |
| 0.05643133 | 0.132987292 |
| 0.000785188 | 0.004060367 |
| 0.188332016 | 0.329848245 |
| 0.68210745 | 0.798677482 |
| 0.035699149 | 0.093479325 |
| 0.803377685 | NA |
| 3.93E-09 | 1.04E-07 |
| 0.542304049 | 0.689465876 |
| 0.740374416 | 0.840635046 |
| NA | NA |
| 2.59E-06 | 2.99E-05 |
| 2.70E-12 | 1.75E-10 |
| 0.426214591 | NA |
| 0.000116385 | 0.000795178 |
| 0.271429271 | 0.428059042 |
| 0.164620713 | NA |
| 0.707712246 | 0.817787172 |
| 0.096448334 | NA |
| 0.040335275 | 0.102668633 |
| 0.000126472 | 0.000851642 |
| 0.835737435 | 0.902244262 |
| 0.001630388 | 0.00750402 |
| NA | NA |
| 0.001627368 | 0.007492389 |
| 2.63E-05 | 0.000223453 |
| 0.768213531 | 0.859627816 |
| 0.27877587 | 0.436916661 |
| 0.095840341 | 0.199425459 |
| 0.0162437 | 0.049838676 |
| 9.88E-05 | 0.000691318 |
| NA | NA |
| 0.945832288 | 0.969270637 |
| 0.905157793 | NA |
| 1.17E-09 | 3.60E-08 |
| 0.936825124 | NA |
| 0.801661681 | NA |
| 0.905157793 | NA |
| NA | NA |
| NA | NA |

| | | |
|---|---|---|
| | 0.98997168 | NA |
| | 0.987989835 | 0.993639148 |
| NA | NA | |
| | 0.748715781 | NA |
| NA | NA | |
| NA | NA | |
| | 7.42E-18 | 1.87E-15 |
| | 0.085157916 | 0.182186436 |
| | 0.037234026 | 0.096691342 |
| | 0.992064361 | NA |
| NA | NA | |
| | 0.889409773 | NA |
| | 0.26583673 | 0.421998753 |
| | 0.284618365 | 0.443415018 |
| | 0.079811221 | NA |
| | 0.655063702 | 0.778121472 |
| | 0.000944771 | 0.004750683 |
| NA | NA | |
| | 0.16675843 | 0.301610112 |
| | 0.636436761 | 0.763403089 |
| | 0.503579265 | NA |
| | 0.248265011 | NA |
| NA | NA | |
| | 0.009705155 | 0.032829161 |
| | 0.366448207 | 0.531307244 |
| | 0.244680999 | 0.397731146 |
| | 0.450464537 | NA |
| | 0.494608745 | 0.647921818 |
| | 0.80372867 | NA |
| | 0.899708905 | 0.943494121 |
| | 0.00038174 | 0.002195552 |
| | 0.905158739 | NA |
| NA | NA | |
| | 0.128163016 | 0.247784088 |
| | 0.468894798 | 0.625240519 |
| | 3.15E-11 | 1.54E-09 |
| | 0.249777416 | NA |
| | 0.150920709 | 0.279998974 |
| | 0.331783289 | 0.494209012 |
| | 0.814133542 | 0.889260721 |
| | 0.806895069 | 0.884206238 |
| NA | NA | |
| | 0.799344342 | NA |
| | 0.635514111 | NA |
| NA | NA | |
| | 0.003249607 | 0.0134258 |
| NA | NA | |

| | |
|---|---|
| 0.00752981 | 0.026750864 |
| 0.892573761 | 0.939380965 |
| 0.521251831 | 0.671999586 |
| 0.156480781 | 0.287617097 |
| 0.595353491 | NA |
| 0.089988566 | NA |
| 0.905157793 | NA |
| 0.000344688 | 0.002017322 |
| 0.384195685 | 0.548184083 |
| 0.871356065 | 0.926316278 |
| 0.17335845 | 0.31052926 |
| NA | NA |
| NA | NA |
| 3.31E-11 | 1.59E-09 |
| NA | NA |
| NA | NA |
| 0.025808055 | 0.072352421 |
| NA | NA |
| NA | NA |
| NA | NA |
| 0.190965652 | 0.333180895 |
| 0.725885217 | NA |
| 6.59E-06 | 6.80E-05 |
| 0.645776025 | 0.771061701 |
| 0.836020775 | NA |
| 0.992269016 | NA |
| 0.605313032 | 0.74025779 |
| 0.039467166 | 0.101044596 |
| NA | NA |
| 0.273279988 | 0.430237687 |
| 0.001688341 | 0.00772864 |
| 0.097329591 | 0.201704463 |
| 0.90515732 | NA |
| NA | NA |
| 0.813836939 | NA |
| 0.830466158 | NA |
| NA | NA |
| 0.757112632 | NA |
| 0.00431514 | 0.016914702 |
| 0.903806311 | NA |
| 0.201491463 | 0.346751649 |
| NA | NA |
| NA | NA |
| NA | NA |
| 0.992061292 | NA |
| 0.905158266 | NA |
| 0.667581844 | NA |

| | |
|---|---|
| NA | NA |
| 0.730944415 | 0.834459624 |
| NA | NA |
| NA | NA |
| 3.63E-05 | 0.000294289 |
| 0.992060816 | NA |
| 0.033520771 | 0.088941833 |
| NA | NA |
| 0.38200725 | NA |
| 0.782559532 | NA |
| NA | NA |
| 0.029466875 | 0.080340589 |
| 0.168950547 | 0.304596475 |
| NA | NA |
| 0.356001583 | 0.520537939 |
| 0.233135165 | 0.384670439 |
| 0.051966108 | 0.124999865 |
| 0.048259184 | 0.117979705 |
| 0.599334623 | NA |
| 0.007766241 | 0.027391404 |
| 0.596104378 | 0.733037803 |
| 0.744075744 | 0.843267922 |
| 0.561364397 | 0.705433071 |
| 0.02643657 | 0.073595416 |
| 2.22E-05 | 0.000194795 |
| 0.165946558 | NA |
| 0.000202242 | 0.001273595 |
| 0.786214937 | 0.871372018 |
| NA | NA |
| 0.008749943 | 0.030201194 |
| NA | NA |
| 1.03E-06 | 1.34E-05 |
| 2.31E-07 | 3.68E-06 |
| 0.801661681 | NA |
| 1.29E-06 | 1.63E-05 |
| 0.905158739 | NA |
| 0.807879735 | NA |
| 0.414066631 | NA |
| 1.19E-06 | 1.52E-05 |
| 0.801661681 | NA |
| 0.001777725 | 0.00808723 |
| 0.084920613 | 0.181826134 |
| 0.14367086 | 0.269785224 |
| 0.675848972 | NA |
| 0.052756988 | NA |
| 0.028805392 | NA |
| 0.197350907 | 0.341607171 |

| | |
|---|---|
| 0.001055041 | 0.005206211 |
| NA | NA |
| 2.36E-09 | 6.72E-08 |
| 0.011662726 | 0.038048106 |
| 3.35E-14 | 3.28E-12 |
| 0.009069826 | NA |
| 0.998849415 | 0.999571253 |
| 3.97E-08 | 7.99E-07 |
| NA | NA |
| 0.008631887 | 0.029868832 |
| NA | NA |
| NA | NA |
| 0.168452613 | NA |
| NA | NA |
| 0.751253092 | 0.848494006 |
| 2.64E-09 | 7.41E-08 |
| 0.078669194 | 0.171773 |
| 0.065275218 | NA |
| 0.204439872 | 0.350325642 |
| 0.987989688 | NA |
| 0.041660385 | 0.105321251 |
| 0.612031787 | 0.745485134 |
| 3.06E-14 | 3.04E-12 |
| 0.012237141 | 0.039591843 |
| 0.152024166 | 0.281665429 |
| 1.11E-09 | 3.45E-08 |
| 0.373641827 | 0.53783542 |
| 0.987444763 | NA |
| 0.761971221 | 0.855600636 |
| 0.268676177 | NA |
| 0.00054053 | 0.002958498 |
| 0.334372571 | 0.497180732 |
| NA | NA |
| NA | NA |
| 0.63503626 | 0.762425436 |
| NA | NA |
| 0.507001184 | NA |
| NA | NA |
| 0.147686183 | NA |
| 0.712967154 | 0.821265333 |
| 0.731848122 | 0.83505601 |
| 0.992061292 | NA |
| NA | NA |
| 0.293088408 | 0.45277861 |
| 0.043912079 | 0.109612976 |
| 0.727524406 | NA |
| 0.942590241 | 0.967619128 |

| | |
|---|---|
| NA | NA |
| 0.224135153 | 0.373796903 |
| 0.773772164 | 0.863681178 |
| 0.000196969 | 0.001244314 |
| 0.992061292 | NA |
| 0.230910954 | 0.381762809 |
| 0.067699706 | 0.152584411 |
| 0.439699013 | 0.599897522 |
| 0.562116514 | 0.706029659 |
| 0.156016226 | 0.286955075 |
| 0.010690289 | 0.035551727 |
| 0.09230093 | 0.193733141 |
| 0.311251233 | 0.472667069 |
| 0.889409301 | NA |
| 0.180985303 | NA |
| 0.04380245 | 0.10935175 |
| 0.916855517 | NA |
| NA | NA |
| NA | NA |
| NA | NA |
| NA | NA |
| 0.401938249 | NA |
| 0.046471668 | 0.114543872 |
| 9.05E-09 | 2.16E-07 |
| 0.027946262 | 0.077017063 |
| NA | NA |
| 0.895432589 | NA |
| 0.002035691 | 0.009089415 |
| 0.044645623 | NA |
| 0.093637699 | 0.19578762 |
| 0.60562739 | NA |
| 0.355765016 | 0.520354583 |
| NA | NA |
| 0.001001859 | 0.004982974 |
| NA | NA |
| 0.7235382 | NA |
| 0.008427194 | 0.029302216 |
| 0.296661994 | 0.456817224 |
| 0.976800545 | NA |
| 0.803431872 | NA |
| 0.000693492 | 0.003661552 |
| 0.046384075 | 0.114392956 |
| 0.984110029 | 0.991970261 |
| 0.98591726 | 0.992834867 |
| 0.435715265 | 0.596240513 |
| 0.944600409 | 0.968822063 |
| 0.793161587 | 0.875259833 |

| | |
|---|---|
| 0.220693465 | 0.369657345 |
| 0.042712387 | 0.107411968 |
| 0.049945052 | 0.121240704 |
| NA | NA |
| 0.905157793 | NA |
| 0.905158266 | NA |
| 0.635596573 | NA |
| 0.905158739 | NA |
| 0.905158739 | NA |
| NA | NA |
| 0.010470836 | 0.03495957 |
| 0.992061292 | NA |
| 0.43328308 | NA |
| 0.110316345 | NA |
| 0.98907643 | NA |
| 0.481178931 | NA |
| 0.340060634 | NA |
| 0.631391481 | NA |
| 0.379850583 | NA |
| 0.139061709 | NA |
| 0.815804877 | NA |
| 0.916465093 | NA |
| 0.302410713 | NA |
| 0.058497226 | 0.136798785 |
| 0.767333857 | NA |
| 0.983087895 | 0.99175941 |
| 0.332139565 | 0.494672592 |
| 0.035253894 | 0.092534099 |
| 0.992059387 | NA |
| 0.567261488 | NA |
| 0.606329466 | NA |
| NA | NA |
| 0.037189745 | 0.096610625 |
| 0.905158739 | NA |
| 0.905158739 | NA |
| 0.992062245 | NA |
| 0.740057435 | 0.840362045 |
| NA | NA |
| 0.992060816 | NA |
| NA | NA |
| NA | NA |
| NA | NA |
| NA | NA |
| 0.233620123 | 0.385226725 |
| 0.627148626 | 0.756363966 |
| 0.889409301 | NA |
| 0.905158266 | NA |

| | | |
|---|---|---|
| | 0.979774332 | NA |
| | 0.100010883 | 0.205989848 |
| | 0.3965892 | 0.560225296 |
| | 0.317734812 | 0.480049999 |
| | 0.217291123 | 0.365491824 |
| | 0.56214276 | NA |
| | 0.021123855 | 0.061421971 |
| | 0.074627944 | 0.164597577 |
| | 0.098528757 | 0.203613115 |
| | 0.657450579 | 0.780096747 |
| | 0.00011377 | 0.000779735 |
| | 0.9709981 | 0.984902179 |
| | 0.99205891 | NA |
| | 0.206185216 | 0.352563554 |
| | 0.992061292 | NA |
| | 0.992060816 | NA |
| | 0.597494152 | NA |
| NA | | NA |
| | 0.40032859 | NA |
| | 0.567634561 | NA |
| | 0.329554857 | 0.492057871 |
| | 0.905158739 | NA |
| | 0.705977814 | 0.817042628 |
| NA | | NA |
| NA | | NA |
| | 0.000400587 | 0.00228956 |
| | 0.193079681 | NA |
| | 0.717981439 | NA |
| NA | | NA |
| | 0.046015069 | 0.113813062 |
| | 0.043441666 | NA |
| | 0.935878949 | NA |
| | 0.018492565 | 0.055143704 |
| | 0.341426727 | 0.504806726 |
| | 0.758117268 | 0.85328022 |
| | 0.889410245 | NA |
| | 0.001504741 | 0.00702735 |
| | 0.905296333 | NA |
| | 0.225277503 | 0.374990047 |
| | 0.024299415 | 0.068877094 |
| | 0.006386684 | 0.023424954 |
| | 0.626144742 | 0.755792126 |
| | 0.106953353 | NA |
| | 0.002234712 | 0.009828166 |
| | 0.001204576 | 0.005826273 |
| NA | | NA |
| NA | | NA |

| | | |
|---|---|---|
| | 0.997717813 | NA |
| | 0.503290757 | 0.65557965 |
| | 0.08300121 | 0.178934181 |
| | 0.989246962 | NA |
| | 4.07E-05 | 0.000324605 |
| | 0.910153848 | 0.949096227 |
| | 0.732438088 | NA |
| NA | | NA |
| | 0.269555124 | 0.425976758 |
| | 0.277628564 | NA |
| NA | | NA |
| | 0.293656515 | 0.453314827 |
| NA | | NA |
| | 0.904518808 | NA |
| | 0.007159274 | 0.025696386 |
| | 0.963503009 | NA |
| NA | | NA |
| NA | | NA |
| NA | | NA |
| NA | | NA |
| NA | | NA |
| NA | | NA |
| NA | | NA |
| NA | | NA |
| NA | | NA |
| NA | | NA |
| NA | | NA |
| | 0.632521441 | 0.760251946 |
| | 0.975180397 | NA |
| | 0.446266969 | 0.605647312 |
| NA | | NA |
| | 0.724999475 | NA |
| | 0.853005301 | 0.914290079 |
| | 0.070528748 | 0.157456363 |
| | 0.26386264 | 0.419592999 |
| | 0.896784355 | 0.941953872 |
| | 0.627469778 | 0.75642702 |
| | 0.513718208 | 0.665180533 |
| | 0.016020955 | 0.04932756 |
| | 0.623380435 | 0.754240365 |
| | 0.068370839 | 0.153638483 |
| | 1.03E-06 | 1.35E-05 |
| | 0.272194775 | NA |
| | 0.724343317 | NA |
| | 1.30E-06 | 1.64E-05 |
| | 0.209423165 | NA |
| | 0.99205891 | NA |

| | |
|---|---|
| 0.822064382 | NA |
| 0.411315352 | 0.574062013 |
| 0.107342554 | 0.217167416 |
| NA | NA |
| 0.165426138 | 0.299740049 |
| 0.23687463 | 0.38869601 |
| 0.975898386 | NA |
| 0.120222911 | 0.236472135 |
| 4.44E-06 | 4.81E-05 |
| 0.98978861 | NA |
| 0.479476152 | NA |
| 0.091338515 | 0.192209147 |
| 0.179850918 | 0.319423028 |
| 1.11E-05 | 0.000105976 |
| 0.023533559 | 0.06708829 |
| 4.88E-05 | 0.000379366 |
| 0.392880114 | 0.556347822 |
| 0.425394456 | 0.586613718 |
| 4.19E-06 | 4.58E-05 |
| 0.401049705 | 0.564673664 |
| 0.024708787 | 0.069821162 |
| 0.501014776 | 0.653778432 |
| 0.986947033 | NA |
| 0.290309077 | 0.44969646 |
| 0.033357018 | 0.08864863 |
| 0.0442526 | 0.110312504 |
| NA | NA |
| 0.435698553 | 0.596240513 |
| 0.59266022 | NA |
| 0.578623335 | 0.719478173 |
| 0.745027466 | 0.844041589 |
| 0.237666602 | 0.389558925 |
| 0.798145709 | NA |
| 0.21676722 | 0.364862132 |
| 0.305861674 | 0.466450204 |
| 0.998162589 | NA |
| 0.917743202 | 0.953115904 |
| NA | NA |
| 0.650582502 | NA |
| NA | NA |
| 0.992062245 | NA |
| 0.634552834 | 0.762095039 |
| 0.425935081 | 0.587180169 |
| 0.151669041 | 0.281102335 |
| 0.277236933 | 0.435063661 |
| 0.601222747 | 0.736773832 |
| NA | NA |

| | |
|---|---|
| NA | NA |
| 0.368707756 | NA |
| NA | NA |
| NA | NA |
| NA | NA |
| NA | NA |
| 0.889409773 | NA |
| NA | NA |
| NA | NA |
| 0.635596573 | NA |
| NA | NA |
| 0.687869472 | 0.803114634 |
| 0.918264028 | 0.953425435 |
| 0.677058425 | NA |
| NA | NA |
| NA | NA |
| 0.131824125 | 0.252726206 |
| 0.18121281 | 0.321296091 |
| 0.77599229 | NA |
| 0.991700267 | NA |
| 0.117911146 | 0.233010335 |
| 0.20968461 | 0.356468475 |
| 4.63E-07 | 6.73E-06 |
| 0.730146653 | 0.834085197 |
| 7.87E-07 | 1.07E-05 |
| 0.001328479 | 0.006327999 |
| NA | NA |
| 0.634693536 | 0.762180649 |
| 0.000374889 | 0.00216068 |
| 0.905158266 | NA |
| 0.049889254 | NA |
| 0.522280273 | NA |
| NA | NA |
| 0.39338966 | NA |
| 0.791147885 | NA |
| 0.028202161 | 0.077547115 |
| NA | NA |
| 0.103396851 | 0.211306789 |
| 0.508316702 | NA |
| 0.889409773 | NA |
| 0.889409773 | NA |
| 1.05E-07 | 1.84E-06 |
| 0.683356401 | 0.799628222 |
| 0.392739811 | 0.556290621 |
| 0.129284453 | 0.249381351 |
| NA | NA |
| 0.416385235 | 0.578659792 |

| | |
|---|---|
| 0.011780261 | 0.038363156 |
| NA | NA |
| NA | NA |
| NA | NA |
| NA | NA |
| NA | NA |
| NA | NA |
| 0.877770242 | NA |
| NA | NA |
| 0.889409301 | NA |
| 0.985674256 | NA |
| NA | NA |
| NA | NA |
| NA | NA |
| NA | NA |
| NA | NA |
| NA | NA |
| NA | NA |
| NA | NA |
| 0.533104325 | 0.682034367 |
| 0.876987616 | NA |
| NA | NA |
| 0.992061292 | NA |
| NA | NA |
| NA | NA |
| NA | NA |
| 0.650582502 | NA |
| 0.116708211 | 0.231361886 |
| 0.027000699 | 0.074862165 |
| NA | NA |
| 0.316495489 | 0.478546061 |
| NA | NA |
| 0.797157316 | NA |
| 0.905158266 | NA |
| 0.467418551 | 0.623732563 |
| NA | NA |
| 0.240425816 | 0.392966914 |
| 0.23851312 | NA |
| 0.000221569 | 0.00137753 |
| 0.079795875 | 0.173506052 |
| 0.624613212 | 0.755196648 |
| 0.048211896 | NA |
| NA | NA |
| NA | NA |
| 0.008585784 | 0.029737418 |
| 0.015502317 | 0.047966095 |
| 0.819268891 | NA |

| | | |
|---|---|---|
| | 0.001772416 | 0.008066421 |
| | 0.842896533 | NA |
| NA | | NA |
| | 0.001926248 | 0.008665947 |
| | 0.000412607 | 0.002345584 |
| | 2.78E-05 | 0.000234758 |
| | 0.805374319 | NA |
| | 0.865500856 | 0.922464903 |
| | 0.03152537 | 0.084777127 |
| | 0.596489153 | 0.733346695 |
| | 0.347629303 | 0.511463442 |
| | 0.707055403 | NA |
| | 0.360375087 | NA |
| | 0.008692523 | 0.030045509 |
| | 0.55736237 | NA |
| | 0.729572771 | NA |
| | 0.728880285 | 0.832938526 |
| | 0.598521616 | 0.734793204 |
| | 0.329819837 | 0.492265991 |
| | 0.009635641 | 0.032642481 |
| NA | | NA |
| | 0.945761934 | NA |
| NA | | NA |
| | 0.992060816 | NA |
| | 0.889410245 | NA |
| | 0.889409773 | NA |
| NA | | NA |
| | 0.527419457 | NA |
| | 0.889409301 | NA |
| NA | | NA |
| | 0.996453356 | NA |
| | 0.807879735 | NA |
| | 0.57413509 | NA |
| NA | | NA |
| | 0.396703839 | NA |
| NA | | NA |
| | 9.02E-05 | 0.000639098 |
| | 0.158780234 | NA |
| | 0.003833263 | 0.015332354 |
| NA | | NA |
| NA | | NA |
| | 0.889409773 | NA |
| NA | | NA |
| | 0.797342596 | NA |
| NA | | NA |
| NA | | NA |
| | 0.992061292 | NA |

| | |
|---|---|
| NA | NA |
| NA | NA |
| NA | NA |
| NA | NA |
| 0.992060816 | NA |
| NA | NA |
| 0.991274049 | NA |
| 0.02582787 | 0.072359556 |
| NA | NA |
| NA | NA |
| NA | NA |
| 0.732029619 | 0.835176308 |
| 0.081862664 | 0.177053308 |
| 0.043494749 | 0.108855845 |
| 0.651637346 | NA |
| 0.992060816 | NA |
| 0.287691377 | 0.446744569 |
| NA | NA |
| 0.559033117 | 0.703518549 |
| 0.989649027 | NA |
| NA | NA |
| NA | NA |
| NA | NA |
| 0.568019869 | 0.710680206 |
| NA | NA |
| NA | NA |
| NA | NA |
| NA | NA |
| NA | NA |
| NA | NA |
| NA | NA |
| 0.964299155 | 0.98039303 |
| NA | NA |
| 0.025047042 | 0.070649712 |
| 9.29E-23 | 7.03E-20 |
| NA | NA |
| 0.223701022 | NA |
| 0.038151929 | NA |
| NA | NA |
| NA | NA |
| NA | NA |
| NA | NA |
| NA | NA |
| NA | NA |
| NA | NA |
| NA | NA |
| NA | NA |

| | | |
|---|---|---|
| NA | | NA |
| NA | | NA |
| | 0.976197687 | NA |
| NA | | NA |
| NA | | NA |
| NA | | NA |
| NA | | NA |
| NA | | NA |
| | 0.992061292 | NA |
| NA | | NA |
| | 0.905158266 | NA |
| NA | | NA |
| | 0.714867828 | 0.822734939 |
| | 0.168583905 | 0.304110241 |
| | 0.780841267 | NA |
| | 0.039703296 | 0.101530764 |
| NA | | NA |
| | 0.482124335 | 0.637653934 |
| NA | | NA |
| | 0.10877138 | 0.219512061 |
| NA | | NA |
| NA | | NA |
| NA | | NA |
| NA | | NA |
| | 0.787647957 | 0.871771467 |
| NA | | NA |
| | 0.697716223 | NA |
| NA | | NA |
| NA | | NA |
| NA | | NA |
| | 0.725280735 | 0.830307868 |
| | 0.890491885 | 0.938091737 |
| | 2.88E-07 | 4.45E-06 |
| NA | | NA |
| | 0.248391885 | 0.402246822 |
| | 0.127391317 | 0.246683063 |
| NA | | NA |
| | 0.599577303 | 0.7355378 |
| | 0.991529086 | NA |
| | 0.430498253 | 0.591460373 |
| | 0.123424812 | 0.241042817 |
| | 0.029536898 | 0.080471556 |
| | 0.991699502 | NA |
| | 0.032411191 | 0.08669964 |
| NA | | NA |
| NA | | NA |
| | 0.889409773 | NA |

| | |
|---|---|
| 0.988858365 | NA |
| NA | NA |
| 0.003883137 | 0.015495151 |
| 0.620232542 | 0.75178479 |
| NA | NA |
| 0.067474924 | 0.152324239 |
| 0.002852599 | 0.012060113 |
| 0.897239053 | 0.942074668 |
| 0.893461926 | 0.939722695 |
| 4.01E-05 | 0.000320068 |
| 0.184590358 | 0.325262372 |
| 0.889409773 | NA |
| 0.559901196 | NA |
| 0.05384476 | 0.128351833 |
| 0.375284867 | NA |
| NA | NA |
| 0.99205891 | NA |
| 0.889409773 | NA |
| NA | NA |
| 0.293135318 | NA |
| 0.889409301 | NA |
| NA | NA |
| 0.04138398 | 0.104827641 |
| NA | NA |
| 0.861251874 | 0.919453056 |
| 0.005766972 | 0.021591976 |
| 0.104509221 | 0.212945902 |
| 0.801661681 | NA |
| 0.03555202 | 0.093205206 |
| NA | NA |
| 0.905158739 | NA |
| 0.111247253 | 0.222971744 |
| NA | NA |
| 9.32E-07 | 1.23E-05 |
| NA | NA |
| NA | NA |
| NA | NA |
| NA | NA |
| NA | NA |
| NA | NA |
| NA | NA |
| NA | NA |
| NA | NA |
| NA | NA |
| 0.735355709 | NA |
| 0.807879735 | NA |
| 0.889410717 | NA |

| | |
|---|---|
| 0.992059387 | NA |
| NA | NA |
| 4.25E-13 | 3.34E-11 |
| 0.000688784 | 0.003644424 |
| 0.006103331 | 0.0225628 |
| 0.371450769 | 0.535811597 |
| 0.071829458 | 0.159678382 |
| 0.038138611 | 0.098458238 |
| 0.626694143 | 0.756248461 |
| 0.016999314 | 0.051673303 |
| NA | NA |
| 0.129809848 | 0.249923029 |
| 0.140271315 | 0.264896612 |
| 0.005037615 | 0.019282026 |
| NA | NA |
| 0.013093394 | 0.041741845 |
| 0.091618562 | 0.192687673 |
| 0.029947529 | 0.08142606 |
| 0.905158739 | NA |
| 0.003444259 | 0.01405286 |
| NA | NA |
| 9.62E-05 | 0.000675735 |
| NA | NA |
| 0.355906511 | 0.520468238 |
| 3.11E-31 | 6.82E-27 |
| 0.389934125 | 0.553315894 |
| 0.424237725 | 0.585527923 |
| 0.135986518 | 0.258718416 |
| 0.957508852 | 0.976497162 |
| NA | NA |
| NA | NA |
| 0.017632233 | 0.053126538 |
| 0.198217212 | 0.342620572 |
| 0.736707729 | 0.838335753 |
| 0.145785891 | 0.272715435 |
| 0.99205891 | NA |
| NA | NA |
| 0.027268107 | 0.075517766 |
| 0.448597021 | 0.607582927 |
| 0.026916002 | 0.074674478 |
| 0.897176769 | NA |
| 0.48395334 | 0.639338968 |
| 0.040057893 | 0.102211396 |
| 0.343455101 | 0.506850172 |
| 0.795515297 | 0.876635741 |
| 0.221141033 | 0.37012469 |
| 0.000169075 | 0.001090696 |

| | |
|---|---|
| 0.509400825 | 0.661068065 |
| 0.002058684 | 0.009171564 |
| 0.453737442 | 0.611777366 |
| 0.062543045 | 0.144217444 |
| 0.031994846 | 0.085794946 |
| NA | NA |
| 0.118638224 | 0.234089025 |
| 0.007059447 | 0.02541702 |
| 0.000366791 | 0.00212292 |
| 9.45E-05 | 0.000665715 |
| 0.000245724 | 0.001508063 |
| 0.043679898 | 0.109157491 |
| 2.12E-05 | 0.00018707 |
| 0.00074991 | 0.003917981 |
| NA | NA |
| NA | NA |
| NA | NA |
| NA | NA |
| 0.24869377 | 0.402525723 |
| 1.51E-06 | 1.87E-05 |
| 0.889409301 | NA |
| 0.996116639 | NA |
| NA | NA |
| 6.06E-05 | 0.000456903 |
| 0.817253826 | 0.891341666 |
| 0.853109161 | 0.914290079 |
| 0.112235734 | 0.224529803 |
| 0.085228715 | 0.182304436 |
| 0.709576856 | 0.819049526 |
| 0.001679655 | 0.007696101 |
| 0.008325691 | 0.028996646 |
| 0.045601212 | 0.112956233 |
| 0.045394651 | NA |
| NA | NA |
| 0.872475511 | NA |
| 0.992060816 | NA |
| 0.03191293 | 0.085627569 |
| 0.129894458 | 0.249987063 |
| 0.081361893 | 0.176213216 |
| 0.157644243 | 0.289198825 |
| 0.053042801 | 0.126880923 |
| 0.38094392 | 0.545138321 |
| 0.889409773 | NA |
| 0.052545235 | 0.126020184 |
| 0.655267694 | 0.778205695 |
| 0.595917667 | 0.732931334 |
| 0.008930744 | 0.030697919 |

| | | |
|---|---|---|
| | 0.272462668 | 0.429289321 |
| | 0.905158266 | NA |
| | 0.448161879 | 0.607293225 |
| | 0.905158266 | NA |
| | 0.989195036 | NA |
| | 0.026434868 | NA |
| | 0.992061292 | NA |
| | 0.47801091 | NA |
| | 0.704556655 | 0.816174015 |
| | 0.889410717 | NA |
| | 0.805374319 | NA |
| | 0.41847424 | NA |
| | 0.091007105 | 0.191677059 |
| | 0.001752922 | 0.007989293 |
| | 0.38519615 | 0.549083338 |
| | 0.566154558 | 0.709154523 |
| | 0.905158739 | NA |
| | 0.847987432 | 0.910649258 |
| | 0.694884617 | 0.808422321 |
| | 0.000837941 | 0.004293127 |
| | 0.608843132 | NA |
| | 8.72E-08 | 1.56E-06 |
| | 0.559369023 | NA |
| | 0.784850956 | 0.870431421 |
| | 0.063153313 | 0.145182538 |
| | 0.493313241 | 0.646928273 |
| | 0.092099963 | 0.193403757 |
| | 0.519652637 | 0.670559555 |
| | 0.822869796 | 0.894551581 |
| | 0.915847652 | 0.95229975 |
| | 0.120724068 | 0.23716747 |
| | 0.907964932 | NA |
| | 0.253257806 | NA |
| | 0.008108723 | 0.028380531 |
| | 7.25E-06 | 7.38E-05 |
| | 0.437034895 | 0.597344215 |
| | 0.414802844 | 0.577264443 |
| | 0.954599578 | 0.97468323 |
| | 0.003320084 | 0.013639985 |
| NA | | NA |
| | 0.000449618 | 0.002525752 |
| | 0.047148333 | 0.115775426 |
| | 0.889409301 | NA |
| NA | | NA |
| | 0.145790177 | 0.272715435 |
| | 0.03330048 | 0.088571372 |
| | 5.66E-14 | 5.26E-12 |

| | | |
|---|---|---|
| | 0.941923274 | 0.967070001 |
| | 0.99205891 | NA |
| | 0.15697444 | 0.288210847 |
| | 0.064536957 | 0.147652328 |
| | 0.990892983 | NA |
| | 0.014258335 | 0.044785979 |
| | 0.444523543 | NA |
| NA | | NA |
| | 0.889409773 | NA |
| NA | | NA |
| | 0.625997939 | NA |
| | 0.007198006 | 0.025814309 |
| | 0.002875328 | 0.012135176 |
| | 0.005490987 | 0.020715172 |
| | 6.83E-06 | 7.01E-05 |
| | 0.992060816 | NA |
| | 0.253471753 | 0.407794434 |
| | 0.641102761 | 0.767212746 |
| | 0.040639237 | 0.103215013 |
| | 0.110506608 | 0.221955678 |
| | 0.567033929 | 0.709891565 |
| | 0.486111331 | 0.64097477 |
| | 0.601476504 | NA |
| | 0.797157316 | NA |
| | 2.33E-17 | 4.88E-15 |
| | 0.022141271 | NA |
| | 0.014963342 | 0.046607097 |
| | 0.711304422 | 0.820181359 |
| | 0.371076999 | 0.535520493 |
| | 0.675933527 | 0.794158605 |
| | 0.179605846 | NA |
| | 0.672258983 | 0.791365659 |
| | 0.000248448 | 0.001520532 |
| | 0.061264957 | 0.142090958 |
| | 0.774984148 | 0.864221186 |
| | 0.943047771 | NA |
| | 0.905158266 | NA |
| | 0.589272446 | NA |
| | 0.79012234 | NA |
| | 0.001143311 | 0.005575385 |
| | 0.662908922 | 0.784311262 |
| NA | | NA |
| NA | | NA |
| | 0.191530484 | 0.333954098 |
| | 0.410831157 | NA |
| | 0.778054002 | 0.866657596 |
| | 0.300378746 | 0.460712222 |

| | |
|---|---|
| NA | NA |
| 0.015557958 | 0.048117914 |
| 9.83E-05 | 0.00068815 |
| 0.738757848 | NA |
| 0.774606007 | 0.864060335 |
| 0.277595868 | NA |
| 0.905158739 | NA |
| 0.705938339 | 0.817040003 |
| 0.67984411 | NA |
| 0.343011702 | 0.506365981 |
| 0.989823193 | NA |
| 0.889410245 | NA |
| 2.97E-05 | 0.000248004 |
| 0.093623404 | 0.19578762 |
| 0.756400462 | NA |
| 0.905157793 | NA |
| 0.167625677 | NA |
| 0.55689198 | 0.701997186 |
| 0.88623727 | 0.935484508 |
| 0.011123933 | 0.036666232 |
| 0.889409773 | NA |
| 0.681555387 | NA |
| 9.54E-07 | 1.26E-05 |
| 0.57502747 | 0.716915245 |
| NA | NA |
| 1.79E-16 | 3.10E-14 |
| 3.06E-06 | 3.45E-05 |
| 0.069571133 | 0.155761879 |
| 0.107557729 | 0.21752258 |
| 0.000440056 | 0.002479012 |
| 0.002137253 | 0.009467812 |
| 0.468554846 | NA |
| 0.366563984 | NA |
| 0.518630639 | NA |
| 0.992061292 | NA |
| 0.478966514 | 0.634662424 |
| 0.032903318 | 0.087695589 |
| 0.290363192 | 0.44974856 |
| 0.004448432 | 0.017353548 |
| 0.416135843 | 0.57848123 |
| 0.992061292 | NA |
| 0.3006151 | 0.46097814 |
| 0.798815277 | NA |
| 0.683121904 | 0.799524246 |
| 0.653044123 | 0.776691601 |
| 3.08E-07 | 4.71E-06 |
| 0.992057737 | NA |

| | |
|---|---|
| 0.004736817 | 0.018322311 |
| 0.038985882 | 0.100081313 |
| 0.992060816 | NA |
| 0.541862075 | 0.689103452 |
| 0.123971439 | 0.241852242 |
| 0.347667171 | 0.511484887 |
| 0.75134968 | NA |
| 0.27970846 | 0.437894746 |
| 2.82E-09 | 7.83E-08 |
| 0.464866928 | 0.621860376 |
| 0.607716446 | 0.741833645 |
| 0.992061292 | NA |
| 0.971174275 | 0.984954128 |
| 0.004065793 | 0.01611275 |
| 0.209062343 | 0.355820802 |
| 0.008530512 | 0.029584817 |
| NA | NA |
| 0.029289099 | 0.079925354 |
| 0.605014157 | 0.740057145 |
| 0.323203366 | 0.485545784 |
| 0.016974454 | 0.051626319 |
| 0.104298981 | 0.212596428 |
| 7.69E-05 | 0.00055955 |
| 0.092351226 | 0.193820183 |
| 0.026630115 | 0.074040352 |
| 3.96E-05 | 0.00031758 |
| 0.690644447 | NA |
| 0.754625048 | 0.850788696 |
| 0.830163713 | NA |
| 0.102148779 | 0.209399276 |
| 0.826844719 | 0.89672859 |
| 0.499320129 | 0.652400845 |
| 0.224945875 | 0.374603464 |
| NA | NA |
| 0.375651067 | NA |
| 0.076734838 | 0.168400989 |
| 0.460268911 | 0.617714522 |
| 0.058122733 | 0.136096938 |
| 4.28E-05 | 0.000339196 |
| 0.905158739 | NA |
| NA | NA |
| 0.335343156 | NA |
| 0.61457121 | 0.747707308 |
| 0.236498627 | 0.388337203 |
| 0.000382883 | 0.002200044 |
| 0.632984771 | 0.760538439 |
| 0.721164089 | 0.827249764 |

| | | |
|---|---|---|
| | 0.239161589 | 0.391469569 |
| | 0.905158739 | NA |
| | 0.986014479 | NA |
| | 0.807252815 | NA |
| | 0.340341222 | NA |
| | 0.599743369 | NA |
| | 0.555870993 | 0.701112644 |
| | 0.001188044 | 0.005762813 |
| | 0.135777059 | 0.258442288 |
| | 0.248410391 | NA |
| | 0.905158266 | NA |
| NA | | NA |
| NA | | NA |
| | 0.558137264 | NA |
| NA | | NA |
| | 0.178876489 | 0.318145307 |
| NA | | NA |
| | 0.905158266 | NA |
| | 0.293890378 | NA |
| NA | | NA |
| NA | | NA |
| | 0.553931337 | NA |
| NA | | NA |
| | 0.348745628 | NA |
| NA | | NA |
| | 0.865928339 | 0.92269606 |
| | 0.631518483 | 0.759406077 |
| NA | | NA |
| | 0.635561011 | 0.762888611 |
| | 0.878673438 | 0.931170794 |
| NA | | NA |
| | 0.789206821 | 0.872787041 |
| | 0.26908722 | 0.425508223 |
| | 0.193545141 | 0.336638238 |
| | 0.36395068 | NA |
| | 0.183673101 | 0.324218458 |
| NA | | NA |
| | 0.01832543 | 0.054792501 |
| NA | | NA |
| | 2.00E-07 | 3.24E-06 |
| | 0.009628535 | 0.032628551 |
| NA | | NA |
| NA | | NA |
| NA | | NA |
| | 0.720763007 | NA |
| | 0.610074459 | NA |
| | 0.626795777 | 0.756272366 |

| | | |
|---|---|---|
| | 0.689856969 | NA |
| | 0.305831501 | 0.466436583 |
| | 0.772496502 | 0.862655141 |
| | 0.992061292 | NA |
| NA | | NA |
| | 0.007553952 | 0.026819279 |
| | 0.265558952 | 0.421710228 |
| NA | | NA |
| | 0.889409773 | NA |
| NA | | NA |
| | 0.398996803 | 0.562576781 |
| | 0.706679458 | 0.817578995 |
| | 0.761420212 | 0.855264578 |
| | 0.665343413 | 0.786370068 |
| | 0.889409301 | NA |
| NA | | NA |
| | 0.584348418 | 0.724096096 |
| | 0.212112684 | 0.35917508 |
| | 0.905296333 | NA |
| NA | | NA |
| | 0.221813519 | 0.371108805 |
| | 0.754267899 | 0.850523216 |
| | 0.992061769 | NA |
| NA | | NA |
| | 0.018326332 | 0.054792501 |
| | 0.062541943 | 0.144217444 |
| | 0.064373154 | NA |
| | 0.905157793 | NA |
| NA | | NA |
| NA | | NA |
| | 0.79959716 | NA |
| NA | | NA |
| | 0.948787134 | NA |
| | 0.358489052 | 0.523373518 |
| | 0.086264339 | 0.184035618 |
| | 0.920101399 | NA |
| NA | | NA |
| NA | | NA |
| | 0.767153727 | NA |
| | 0.001476235 | 0.006914815 |
| | 6.89E-13 | 5.10E-11 |
| | 2.56E-07 | 4.01E-06 |
| NA | | NA |
| | 0.626791926 | NA |
| NA | | NA |
| NA | | NA |
| NA | | NA |

| | |
|---|---|
| 0.636915697 | NA |
| NA | NA |
| NA | NA |
| 0.889409773 | NA |
| NA | NA |
| NA | NA |
| 0.211331223 | 0.358432249 |
| 0.019535121 | 0.057680747 |
| 3.40E-27 | 1.25E-23 |
| 0.498790004 | 0.651804909 |
| 8.67E-05 | 0.00061748 |
| 0.215344398 | 0.363079517 |
| 0.032787529 | 0.087488786 |
| 0.094173949 | 0.196724291 |
| NA | NA |
| 0.003554816 | 0.014439549 |
| 0.956607319 | 0.975940351 |
| 0.982104467 | NA |
| 0.092584283 | 0.194215597 |
| 0.853637712 | NA |
| NA | NA |
| NA | NA |
| 0.319843153 | 0.482180131 |
| NA | NA |
| NA | NA |
| NA | NA |
| NA | NA |
| NA | NA |
| NA | NA |
| NA | NA |
| 0.796361596 | NA |
| 0.826503604 | 0.896566351 |
| 0.992060816 | NA |
| 0.989013136 | 0.993992378 |
| 0.001440377 | 0.006784654 |
| 0.903549779 | 0.945355815 |
| 0.704247604 | NA |
| 0.743044688 | NA |
| 0.99205891 | NA |
| NA | NA |
| 0.991782148 | NA |
| 0.889409773 | NA |
| NA | NA |
| NA | NA |
| NA | NA |
| NA | NA |
| 0.992061292 | NA |

| | | | |
|---|---|---|---|
| NA | | NA | |
| NA | | NA | |
| NA | | NA | |
| | 0.962435159 | | 0.979205986 |
| NA | | NA | |
| NA | | NA | |
| NA | | NA | |
| | 0.889409301 | NA | |
| | 0.696432456 | NA | |
| | 0.824638639 | NA | |
| | 0.992062245 | NA | |
| | 0.889409773 | NA | |
| NA | | NA | |
| NA | | NA | |
| | 8.71E-06 | | 8.67E-05 |
| NA | | NA | |
| | 0.587919859 | NA | |
| | 0.889410717 | NA | |
| | 0.007411427 | | 0.026408578 |
| | 0.001069193 | | 0.005268942 |
| | 0.528996296 | NA | |
| | 0.69372943 | | 0.807442869 |
| | 0.092953682 | | 0.194730957 |
| | 0.811412263 | NA | |
| | 0.889410717 | NA | |
| | 0.183874079 | NA | |
| NA | | NA | |
| | 0.519412825 | | 0.670337127 |
| | 0.886781837 | | 0.935832206 |
| | 0.141367443 | | 0.266439591 |
| | 0.706940795 | NA | |
| | 0.013351647 | | 0.042411227 |
| | 0.905158739 | NA | |
| NA | | NA | |
| NA | | NA | |
| | 0.889410245 | NA | |
| NA | | NA | |
| | 0.889409773 | NA | |
| | 0.275048515 | NA | |
| NA | | NA | |
| | 0.136581654 | | 0.259580798 |
| | 0.043782942 | | 0.109315476 |
| | 0.01725362 | | 0.05224383 |
| | 0.001466321 | | 0.006881588 |
| | 0.250207729 | NA | |
| | 0.323174071 | | 0.485545784 |
| | 0.796451674 | | 0.877227382 |

| | | |
|---|---|---|
| | 0.394083937 | 0.557475305 |
| NA | | NA |
| | 0.85535358 | NA |
| | 0.905158266 | NA |
| | 2.30E-05 | 0.00020026 |
| | 0.895623876 | 0.941075508 |
| | 0.00084886 | 0.004330471 |
| | 0.004915401 | 0.018899892 |
| | 0.773271353 | NA |
| | 0.99205891 | NA |
| NA | | NA |
| | 0.992060816 | NA |
| | 0.334936729 | 0.497758232 |
| | 0.30191727 | 0.462425984 |
| | 0.655478926 | NA |
| | 0.98410783 | 0.991970261 |
| | 0.480128089 | 0.635718634 |
| | 0.019389529 | 0.057335668 |
| | 0.001633274 | 0.007515728 |
| | 0.44201521 | NA |
| | 0.001159474 | 0.005644177 |
| | 8.61E-09 | 2.07E-07 |
| | 0.020235288 | 0.059334801 |
| | 0.3600433 | 0.524665924 |
| | 0.570214517 | NA |
| | 0.52175621 | NA |
| | 0.00031032 | 0.001838946 |
| NA | | NA |
| | 2.77E-07 | 4.30E-06 |
| | 0.029978402 | 0.081462161 |
| | 0.112812728 | 0.225491962 |
| | 0.058192075 | NA |
| | 0.37636935 | 0.540741596 |
| | 0.01313332 | 0.041831192 |
| | 0.973515708 | NA |
| | 0.30641014 | 0.467156861 |
| | 0.93961798 | 0.96589388 |
| | 0.543013715 | 0.689888809 |
| | 0.611588402 | 0.745083347 |
| | 0.24207941 | 0.394788099 |
| | 0.654243265 | NA |
| | 0.023967114 | 0.068084586 |
| | 0.829542163 | NA |
| | 0.827110959 | 0.89672859 |
| | 0.568778371 | NA |
| | 0.404487996 | 0.567803612 |
| | 0.234496205 | 0.385975726 |

| | |
|---|---|
| 0.000145334 | 0.000957228 |
| 0.000425777 | 0.002412346 |
| 0.992060816 | NA |
| 0.060978192 | 0.141532244 |
| 0.905158739 | NA |
| 0.856131524 | NA |
| 0.00702085 | 0.025311252 |
| 0.721956165 | NA |
| 0.219216722 | NA |
| 0.159329683 | NA |
| 0.113524862 | 0.226667798 |
| NA | NA |
| 0.493297091 | 0.646928273 |
| 0.374281392 | NA |
| 0.773354288 | NA |
| 0.983083985 | NA |
| 0.838906477 | NA |
| 1.24E-05 | 0.000117339 |
| 0.690040775 | NA |
| 0.889409773 | NA |
| 0.622574551 | 0.753686233 |
| 2.05E-07 | 3.31E-06 |
| 0.41568489 | 0.578162181 |
| 0.04218201 | 0.106407209 |
| 0.020196191 | 0.059272712 |
| 8.23E-18 | 2.01E-15 |
| 0.689985425 | NA |
| 0.372478827 | 0.536837921 |
| NA | NA |
| 0.008973006 | 0.030801663 |
| 0.166716355 | 0.30155885 |
| 0.846965045 | 0.910197225 |
| NA | NA |
| 0.009067478 | 0.031067679 |
| NA | NA |
| 0.618355686 | NA |
| 0.251183507 | 0.405330587 |
| NA | NA |
| 0.795014225 | NA |
| 0.646242334 | 0.771324894 |
| 0.905158739 | NA |
| 0.033216939 | 0.088391997 |
| 6.72E-05 | 0.000498018 |
| 0.466367703 | 0.623043645 |
| 1.38E-05 | 0.00012852 |
| 0.94983635 | 0.971885067 |
| NA | NA |

| | |
|---|---|
| 0.000459373 | 0.002568068 |
| 0.267615103 | NA |
| 0.775156468 | NA |
| 0.287569847 | 0.446682201 |
| 0.047955818 | 0.117426957 |
| 0.581001083 | 0.721169113 |
| 1.41E-10 | 5.79E-09 |
| 0.901183732 | NA |
| 0.234227875 | 0.385684723 |
| NA | NA |
| 0.917784771 | 0.953115904 |
| 0.285211441 | 0.444149921 |
| 0.000358612 | 0.002082722 |
| 0.547086014 | 0.693377685 |
| NA | NA |
| 0.875766665 | NA |
| 0.195627751 | 0.339400314 |
| 0.781765929 | 0.868484723 |
| 0.228798582 | 0.379241248 |
| 0.058812002 | 0.137403208 |
| 0.086184979 | 0.183955672 |
| 0.699915333 | 0.812553043 |
| NA | NA |
| 0.905158266 | NA |
| 0.00879801 | 0.030333726 |
| 0.00010567 | 0.000731993 |
| 0.953936884 | 0.97434513 |
| 0.865161976 | 0.922201294 |
| 0.116986388 | 0.231829627 |
| 0.325372882 | 0.487570104 |
| 0.033937821 | 0.089840084 |
| 0.185725541 | 0.326712101 |
| 0.902937908 | NA |
| 0.745932212 | 0.844805067 |
| NA | NA |
| NA | NA |
| NA | NA |
| NA | NA |
| 0.889410717 | NA |
| 5.12E-07 | 7.37E-06 |
| 3.51E-26 | 5.93E-23 |
| 0.030753077 | 0.083101066 |
| 0.026286063 | NA |
| 0.126123668 | 0.244854533 |
| 0.026070239 | 0.072843413 |
| 0.465273251 | 0.622187157 |
| 0.608765869 | 0.742908177 |

| | | |
|---|---|---|
| | 0.905158266 | NA |
| NA | | NA |
| NA | | NA |
| | 0.905158739 | NA |
| | 0.145434812 | 0.272259424 |
| | 0.325374071 | 0.487570104 |
| | 0.889409773 | NA |
| | 0.004255832 | 0.016735997 |
| NA | | NA |
| | 0.038037036 | NA |
| | 0.905158266 | NA |
| | 9.23E-08 | 1.64E-06 |
| | 0.086247805 | 0.184018224 |
| | 0.100902537 | 0.207425464 |
| | 0.615539064 | NA |
| NA | | NA |
| NA | | NA |
| | 0.863915285 | 0.921490054 |
| | 0.644797034 | NA |
| | 7.52E-09 | 1.83E-07 |
| | 0.262229889 | 0.417692306 |
| | 0.032252957 | 0.086371058 |
| | 0.680186586 | 0.797241196 |
| | 8.83E-07 | 1.18E-05 |
| | 0.905158266 | NA |
| | 0.700856564 | 0.813129883 |
| | 0.626958452 | 0.756287841 |
| | 0.556647478 | 0.701850627 |
| | 0.363447985 | 0.528400047 |
| | 0.413839667 | 0.576370807 |
| | 0.562838092 | 0.706531851 |
| | 0.863141713 | 0.920982 |
| NA | | NA |
| | 0.804289653 | 0.882408019 |
| | 0.165480384 | 0.299792406 |
| | 0.590083856 | 0.728244332 |
| | 0.99205891 | NA |
| | 0.45181579 | NA |
| | 0.784353866 | NA |
| NA | | NA |
| | 0.992060816 | NA |
| | 0.930334738 | 0.959926975 |
| | 0.730959717 | 0.834459624 |
| | 0.800729051 | 0.879908384 |
| | 0.712997335 | 0.821265333 |
| | 0.097607869 | 0.202128556 |
| | 0.992060816 | NA |

| | | |
|---|---|---|
| | 0.616287802 | NA |
| | 0.331686786 | 0.494098782 |
| | 0.678256072 | NA |
| | 0.939898011 | 0.96589388 |
| NA | | NA |
| NA | | NA |
| NA | | NA |
| NA | | NA |
| NA | | NA |
| NA | | NA |
| NA | | NA |
| NA | | NA |
| NA | | NA |
| NA | | NA |
| | 0.107129222 | 0.21690749 |
| | 0.052517022 | 0.126020184 |
| | 0.324745519 | 0.486946842 |
| | 0.164316468 | 0.298299051 |
| | 0.218275851 | 0.366698753 |
| NA | | NA |
| | 0.187308109 | NA |
| | 2.25E-05 | 0.000196339 |
| NA | | NA |
| | 0.176500317 | NA |
| | 0.103030478 | 0.210813107 |
| NA | | NA |
| NA | | NA |
| | 0.810087483 | 0.886200919 |
| NA | | NA |
| | 0.000230937 | 0.001430101 |
| | 0.020624506 | 0.060281444 |
| | 0.846407861 | 0.909709276 |
| NA | | NA |
| | 0.905158266 | NA |
| | 0.846394063 | 0.909709276 |
| | 0.11778671 | 0.232827289 |
| | 0.880735197 | NA |
| | 3.71E-10 | 1.34E-08 |
| | 0.771321946 | NA |
| | 0.016874283 | 0.051371467 |
| | 0.034884707 | 0.091839501 |
| NA | | NA |
| NA | | NA |
| NA | | NA |
| | 0.265361254 | 0.421426757 |
| | 0.02171165 | 0.062815036 |
| | 0.742917918 | 0.842434007 |

| | | |
|---|---|---|
| | 0.905158739 | NA |
| | 0.244377307 | 0.397472875 |
| | 0.603249487 | NA |
| | 0.568159827 | 0.710774319 |
| | 0.20314086 | 0.348879255 |
| | 0.190790326 | 0.332954364 |
| | 0.867541516 | 0.924011454 |
| | 0.90387548 | 0.945602748 |
| | 0.38018876 | NA |
| | 0.817355691 | NA |
| | 0.992061292 | NA |
| | 0.992061292 | NA |
| NA | | NA |
| | 0.88804077 | NA |
| NA | | NA |
| | 2.76E-09 | 7.71E-08 |
| | 0.809053867 | NA |
| | 0.217556592 | 0.365691912 |
| | 5.31E-05 | 0.000407157 |
| | 0.453088695 | 0.611135218 |
| NA | | NA |
| | 0.905158739 | NA |
| | 0.914787278 | 0.951789385 |
| | 4.79E-05 | 0.000373531 |
| NA | | NA |
| | 7.90E-16 | 1.21E-13 |
| | 0.288088002 | 0.44729721 |
| | 0.246540513 | 0.399954427 |
| | 0.083274202 | 0.179334197 |
| NA | | NA |
| | 0.481964441 | 0.63755766 |
| | 8.23E-05 | 0.000590374 |
| | 0.0021907 | 0.009663635 |
| | 0.488554202 | NA |
| | 0.783414809 | NA |
| | 0.88694015 | 0.935866549 |
| | 0.99205891 | NA |
| | 0.785090986 | 0.87056572 |
| | 0.764714833 | 0.857409906 |
| | 0.447475902 | 0.606775555 |
| | 0.01310263 | 0.041759165 |
| | 0.447607011 | 0.606803455 |
| | 0.000242918 | 0.001493348 |
| | 0.384847785 | 0.548781157 |
| NA | | NA |
| | 0.828133983 | NA |
| | 0.718005881 | NA |

| | |
|---|---|
| 0.412954658 | NA |
| NA | NA |
| 0.042239353 | 0.106490696 |
| NA | NA |
| 0.478222406 | 0.634212212 |
| 0.881822858 | 0.932842669 |
| 0.247036751 | 0.400552316 |
| 0.279011325 | 0.437160877 |
| 0.238915719 | NA |
| 0.889409301 | NA |
| NA | NA |
| 0.833500912 | 0.900819299 |
| 0.224048407 | NA |
| 0.575951747 | 0.717619406 |
| 0.599824381 | 0.735717119 |
| 0.572665257 | 0.715078425 |
| 0.08869975 | 0.188098293 |
| 8.24E-10 | 2.66E-08 |
| 3.02E-05 | 0.000251373 |
| 0.905158739 | NA |
| 0.905296333 | NA |
| 0.515958521 | 0.66709451 |
| 0.957433344 | 0.976465505 |
| 0.004294803 | 0.016850024 |
| 0.740792922 | 0.840979774 |
| 0.848780841 | NA |
| 0.143968208 | NA |
| 0.674222909 | NA |
| 0.996165537 | NA |
| 0.443233299 | NA |
| 0.561755665 | 0.70569752 |
| 0.017786364 | 0.053510175 |
| 0.724089048 | NA |
| 0.28395593 | 0.442728504 |
| NA | NA |
| 0.905158739 | NA |
| 0.604149906 | NA |
| 0.014208385 | 0.044654635 |
| NA | NA |
| 0.14344311 | 0.26949583 |
| NA | NA |
| 0.363529508 | 0.528448594 |
| 0.889410717 | NA |
| 0.905157793 | NA |
| 0.881453338 | NA |
| 0.992061292 | NA |
| 0.012534781 | 0.040371261 |

| | |
|---|---|
| NA | NA |
| NA | NA |
| NA | NA |
| 0.992060816 | NA |
| 0.755799747 | 0.851544722 |
| 0.873030153 | NA |
| 0.616644609 | 0.749171172 |
| 0.115062608 | 0.229009318 |
| 0.146015827 | NA |
| 0.939168579 | 0.965550954 |
| 0.029950284 | 0.08142606 |
| 0.921066744 | 0.95471818 |
| 0.722892019 | 0.828712391 |
| NA | NA |
| NA | NA |
| 0.992061292 | NA |
| 0.424593031 | NA |
| 0.374303313 | NA |
| NA | NA |
| 0.889409301 | NA |
| 0.410785164 | 0.573745796 |
| 0.44000242 | 0.600199598 |
| 0.270462493 | 0.426965196 |
| 0.463996429 | NA |
| 0.889409773 | NA |
| 0.000158984 | 0.001034713 |
| 0.992061292 | NA |
| 0.897257621 | NA |
| NA | NA |
| 0.905158739 | NA |
| 4.98E-05 | 0.000385562 |
| 0.767522376 | 0.859022623 |
| 0.725387258 | NA |
| 0.427957943 | 0.589039017 |
| 2.80E-09 | 7.79E-08 |
| 0.790464404 | NA |
| 0.022873493 | 0.065563116 |
| 0.908803026 | 0.94822796 |
| 0.899699328 | NA |
| 0.006039821 | 0.022388778 |
| 0.260299836 | 0.415556378 |
| 0.274664367 | 0.431921048 |
| 0.889409773 | NA |
| 0.893156782 | NA |
| NA | NA |
| NA | NA |
| NA | NA |

| | | |
|---|---|---|
| | 0.854801436 | 0.915504303 |
| | 0.889409773 | NA |
| NA | | NA |
| | 0.078471452 | 0.171418981 |
| | 0.877158331 | 0.92996909 |
| NA | | NA |
| NA | | NA |
| | 0.923755222 | 0.95601569 |
| | 0.897259678 | NA |
| | 0.123591397 | 0.241325225 |
| | 0.079449877 | 0.17299354 |
| | 6.04E-10 | 2.03E-08 |
| | 0.453755543 | 0.611777366 |
| | 0.350492277 | 0.514654485 |
| | 0.851542467 | NA |
| | 0.026177973 | 0.073070053 |
| | 0.285635184 | NA |
| | 0.000993491 | 0.004951443 |
| | 0.305808912 | 0.466436583 |
| | 0.000476081 | 0.00264473 |
| | 0.258241743 | 0.413137873 |
| | 0.001218821 | 0.005877051 |
| NA | | NA |
| | 0.104748289 | NA |
| | 1.32E-06 | 1.66E-05 |
| | 0.018822456 | 0.055935048 |
| | 0.265351347 | 0.421426757 |
| | 0.129022855 | 0.249051765 |
| | 0.018134205 | 0.054311887 |
| | 0.707105203 | 0.817657881 |
| | 0.100322248 | 0.206541931 |
| | 0.001249064 | 0.005999168 |
| | 0.090592964 | 0.191149725 |
| | 0.174752476 | 0.312389256 |
| | 0.889409301 | NA |
| | 0.080946853 | 0.175556736 |
| | 0.889409301 | NA |
| | 0.99205891 | NA |
| | 0.000274524 | 0.001660587 |
| | 0.018260084 | NA |
| | 0.000294109 | 0.001756416 |
| | 0.133273737 | NA |
| | 0.905158266 | NA |
| NA | | NA |
| | 0.687154228 | 0.802532692 |
| | 0.112612555 | 0.225173838 |
| | 0.83899277 | 0.904395573 |

| | |
|---|---|
| 0.169788464 | 0.305856019 |
| 0.121762754 | 0.238636844 |
| 0.627175734 | 0.756363966 |
| 0.465166568 | NA |
| 3.25E-05 | 0.000268379 |
| 0.952177095 | 0.973288471 |
| 5.22E-05 | 0.000401581 |
| 0.987327988 | 0.993435451 |
| 0.4476459 | 0.606818712 |
| 0.444222007 | 0.603780085 |
| 0.308418307 | 0.469501394 |
| 0.403566073 | 0.567088976 |
| NA | NA |
| 0.547648656 | 0.693866899 |
| 0.275792719 | 0.433168752 |
| 0.286198975 | 0.445245717 |
| 0.174256069 | 0.311577968 |
| 0.025652183 | 0.072032772 |
| 1.76E-07 | 2.90E-06 |
| 0.132185798 | NA |
| 0.055197065 | 0.130766247 |
| 3.09E-07 | 4.71E-06 |
| 0.006704136 | 0.024385643 |
| 0.124911849 | 0.243146821 |
| 0.655523977 | NA |
| 5.12E-05 | 0.000394393 |
| 0.402666281 | NA |
| NA | NA |
| 0.242852177 | 0.3956957 |
| 0.889409773 | NA |
| 0.190263262 | NA |
| NA | NA |
| 0.037296105 | 0.096783877 |
| 4.97E-06 | 5.29E-05 |
| 1.63E-05 | 0.000149366 |
| 0.391361315 | NA |
| 0.454060943 | NA |
| 0.566670824 | NA |
| 0.905158739 | NA |
| 4.11E-05 | 0.00032661 |
| 0.211795575 | 0.358859493 |
| 0.712050807 | 0.820740312 |
| NA | NA |
| 0.502287866 | NA |
| NA | NA |
| NA | NA |
| 0.546721914 | 0.693076233 |

| | | |
|---|---|---|
| | 8.83E-10 | 2.81E-08 |
| | 0.008428107 | 0.029302216 |
| | 0.445807878 | NA |
| | 0.001285521 | 0.006150054 |
| | 3.49E-05 | 0.000284714 |
| | 0.000254269 | 0.001552262 |
| | 0.057764037 | 0.135444793 |
| | 0.002817141 | 0.011928785 |
| | 0.205222084 | 0.351365804 |
| NA | | NA |
| | 0.254900924 | 0.409463744 |
| | 0.518478304 | 0.669367104 |
| | 5.72E-09 | 1.43E-07 |
| | 0.035035019 | 0.092069649 |
| | 0.121920685 | 0.238827505 |
| | 0.040959991 | 0.103849491 |
| | 0.039978517 | 0.102056295 |
| | 0.041868944 | NA |
| | 1.45E-05 | 0.000134697 |
| | 0.075850937 | 0.166894861 |
| | 0.006710815 | 0.024397814 |
| | 0.790914317 | NA |
| | 0.45275755 | 0.610906318 |
| | 0.531014154 | 0.680192487 |
| | 0.996181479 | NA |
| | 0.478581516 | NA |
| | 0.001282306 | 0.006136013 |
| | 0.977051134 | 0.988211775 |
| | 5.14E-08 | 9.95E-07 |
| | 0.825898502 | NA |
| | 0.255789491 | 0.410410743 |
| | 0.794303357 | 0.875871619 |
| | 0.403789332 | 0.567172544 |
| | 0.673830581 | 0.792705761 |
| | 0.096061246 | NA |
| | 0.748865831 | 0.846773676 |
| | 0.324867552 | 0.48701301 |
| NA | | NA |
| | 0.735647038 | NA |
| NA | | NA |
| NA | | NA |
| NA | | NA |
| | 0.905158266 | NA |
| | 0.992061292 | NA |
| NA | | NA |
| | 0.905158266 | NA |
| NA | | NA |

| | | | |
|---|---|---|---|
| NA | | NA | |
| | 0.992061292 | NA | |
| NA | | NA | |
| | 0.875757336 | NA | |
| NA | | NA | |
| NA | | NA | |
| | 0.90244696 | NA | |
| NA | | NA | |
| | 0.782195155 | NA | |
| | 0.184954299 | | 0.325759031 |
| | 9.11E-06 | | 8.99E-05 |
| | 0.211176969 | | 0.358253636 |
| | 0.000337082 | | 0.001978084 |
| | 7.05E-05 | | 0.00051842 |
| NA | | NA | |
| | 0.000195585 | | 0.001236994 |
| | 2.37E-05 | | 0.000205236 |
| NA | | NA | |
| | 0.000143007 | | 0.000944069 |
| | 0.000177927 | | 0.001141216 |
| | 0.798551015 | NA | |
| | 6.01E-07 | | 8.40E-06 |
| | 2.28E-06 | | 2.70E-05 |
| | 1.67E-11 | | 8.80E-10 |
| | 0.001465166 | NA | |
| | 0.04823519 | | 0.117937483 |
| | 6.41E-06 | | 6.64E-05 |
| | 0.001191506 | | 0.005777054 |
| | 4.27E-06 | | 4.65E-05 |
| | 1.03E-05 | | 9.92E-05 |
| | 0.154460527 | | 0.284953266 |
| | 0.001257021 | | 0.00603078 |
| | 0.031555857 | | 0.084835341 |
| | 0.175323776 | NA | |
| | 0.12012604 | NA | |
| NA | | NA | |
| | 0.743044688 | NA | |
| | 0.905158266 | NA | |
| NA | | NA | |
| NA | | NA | |
| NA | | NA | |
| NA | | NA | |
| NA | | NA | |
| | 0.905158739 | NA | |
| NA | | NA | |
| NA | | NA | |
| | 0.25544332 | | 0.410062487 |

| | |
|---|---|
| NA | NA |
| 0.905158266 | NA |
| NA | NA |
| NA | NA |
| NA | NA |
| 0.992061292 | NA |
| 0.689514404 | NA |
| 0.000905715 | 0.004582625 |
| 0.017920429 | 0.053832381 |
| 0.201077757 | 0.34625678 |
| 0.369069278 | 0.533716658 |
| 0.513424234 | NA |
| 0.251336519 | 0.405517901 |
| 0.063040475 | 0.145059812 |
| 0.012669418 | 0.040739163 |
| 0.144364663 | 0.270810151 |
| 0.981431341 | NA |
| 0.992061292 | NA |
| 0.401430125 | 0.564945236 |
| NA | NA |
| 0.992059387 | NA |
| 0.714092122 | 0.82205761 |
| 0.959650124 | 0.977681965 |
| 0.482004869 | 0.637560226 |
| 0.072056056 | 0.160101076 |
| 0.992060816 | NA |
| 0.425089743 | 0.586330861 |
| 0.992061292 | NA |
| 0.784353866 | NA |
| 0.966415254 | NA |
| 0.882927242 | NA |
| NA | NA |
| 0.437309126 | 0.597652468 |
| 0.340836976 | 0.504206357 |
| 0.760890464 | 0.854999166 |
| 0.990571437 | NA |
| 0.847273169 | 0.910371949 |
| NA | NA |
| 0.814054783 | 0.889256242 |
| 0.762439782 | NA |
| 0.992060816 | NA |
| 0.46709402 | 0.623525451 |
| 0.604052927 | 0.739253085 |
| 0.558683647 | 0.703440832 |
| 0.046235796 | 0.114204828 |
| 0.780089629 | 0.867560808 |
| 0.025228828 | 0.071043837 |

| | |
|---|---|
| 0.335810632 | 0.498722939 |
| 0.320990784 | 0.483312989 |
| 0.072624682 | 0.161087413 |
| 0.011245891 | 0.03696834 |
| 0.513336133 | 0.664878386 |
| 0.409722098 | 0.572698125 |
| 0.581725004 | 0.721741306 |
| 0.233560059 | 0.385202381 |
| 2.62E-05 | 0.000223407 |
| 9.30E-06 | 9.15E-05 |
| 0.011922579 | 0.038757649 |
| 0.785420943 | NA |
| 0.75677571 | NA |
| 0.789780056 | NA |
| 0.991879747 | NA |
| 0.120973438 | 0.237607846 |
| NA | NA |
| 0.0440699 | 0.109944437 |
| 4.54E-09 | 1.17E-07 |
| 0.903342961 | NA |
| NA | NA |
| 0.028131839 | 0.077392515 |
| 0.059475186 | 0.138687015 |
| 0.484442772 | NA |
| 0.568165685 | NA |
| 0.627631069 | 0.756471301 |
| 0.041620079 | 0.105231469 |
| 0.230300731 | 0.381046901 |
| 0.009014556 | 0.030920125 |
| 0.0687331 | 0.154168243 |
| 0.175391521 | 0.313251111 |
| 0.953032622 | 0.973603891 |
| 0.992061292 | NA |
| 0.889409773 | NA |
| 9.57E-05 | 0.000672577 |
| 0.00722493 | 0.025889727 |
| 0.554522622 | NA |
| 0.014649764 | 0.045799282 |
| 0.042363402 | 0.106754412 |
| 1.13E-08 | 2.64E-07 |
| 0.759076713 | NA |
| 0.03406386 | 0.090070631 |
| NA | NA |
| 0.006664544 | 0.024265746 |
| 7.83E-05 | 0.000568113 |
| 0.367985962 | NA |
| 0.300736281 | 0.461131764 |

| | | |
|---|---|---|
| | 0.278703635 | NA |
| | 0.777728364 | 0.866338819 |
| | 8.12E-10 | 2.64E-08 |
| | 0.01728911 | 0.052336858 |
| | 0.799588413 | NA |
| | 5.70E-07 | 8.05E-06 |
| | 0.270847857 | NA |
| | 0.00580088 | 0.021689345 |
| | 0.989375694 | 0.994220248 |
| | 0.297939909 | 0.457904814 |
| | 0.000169779 | 0.001093952 |
| | 0.06165489 | 0.142739032 |
| | 0.751314607 | 0.848494006 |
| | 0.030432043 | 0.082409326 |
| | 1.02E-12 | 7.13E-11 |
| | 3.08E-05 | 0.000255823 |
| | 0.611947181 | NA |
| | 0.558920415 | 0.703504149 |
| | 0.453245795 | NA |
| | 0.414995028 | 0.577446489 |
| | 0.357319652 | NA |
| NA | | NA |
| | 3.91E-07 | 5.81E-06 |
| | 0.889410245 | NA |
| | 0.543693515 | NA |
| | 0.617872259 | 0.750177822 |
| | 0.732682314 | NA |
| | 0.73859474 | 0.839639257 |
| NA | | NA |
| | 0.000129297 | 0.000867998 |
| | 0.909755526 | NA |
| NA | | NA |
| | 0.000141836 | 0.000937843 |
| | 0.010356633 | 0.034656044 |
| | 0.101820404 | 0.208862611 |
| | 0.024882422 | 0.070239503 |
| | 0.992060816 | NA |
| NA | | NA |
| | 0.03868356 | 0.099506597 |
| | 0.905158266 | NA |
| NA | | NA |
| | 0.889409773 | NA |
| | 0.524223876 | NA |
| NA | | NA |
| | 0.905158739 | NA |
| | 0.572281563 | 0.714749792 |
| NA | | NA |

| | |
|---|---|
| 0.00011216 | 0.000769725 |
| 0.817125331 | 0.891280307 |
| 0.646136067 | 0.771239979 |
| 0.006985915 | 0.025210141 |
| 0.812766598 | NA |
| 4.45E-05 | 0.000350911 |
| 0.290674732 | 0.450135856 |
| NA | NA |
| NA | NA |
| 0.399137113 | 0.562702354 |
| 0.000745341 | 0.00389782 |
| 0.804534188 | NA |
| 0.841943594 | NA |
| 0.065538706 | 0.149306405 |
| 0.05233605 | NA |
| 0.905158266 | NA |
| 0.001129904 | 0.005517358 |
| 0.06033062 | 0.14035386 |
| 0.07637528 | 0.167779588 |
| 0.212492782 | 0.359624586 |
| 0.297534313 | 0.457496243 |
| 0.528205745 | NA |
| 0.865558722 | 0.922481264 |
| 0.889409773 | NA |
| NA | NA |
| 0.167917113 | 0.303156449 |
| 0.001509801 | 0.007043491 |
| 0.004258153 | 0.016742127 |
| 0.992061292 | NA |
| 0.935416416 | 0.962820337 |
| 0.156182881 | 0.28718974 |
| 1.31E-16 | 2.34E-14 |
| 9.97E-07 | 1.30E-05 |
| 0.001040978 | 0.005150705 |
| NA | NA |
| 0.33379394 | NA |
| 0.889409773 | NA |
| NA | NA |
| 0.524708544 | NA |
| 1.61E-10 | 6.49E-09 |
| 0.738380807 | NA |
| 0.028360418 | 0.077894487 |
| 0.390599025 | NA |
| 0.729117187 | NA |
| 0.022125857 | 0.063776531 |
| 0.739574487 | 0.840161208 |
| 0.000245852 | 0.00150843 |

| | | |
|---|---|---|
| | 0.679919819 | 0.797093552 |
| | 0.367295708 | 0.531951356 |
| | 0.763133079 | NA |
| | 0.02150075 | 0.062282734 |
| | 0.006298763 | 0.023179913 |
| | 0.000139462 | 0.000926329 |
| | 0.123386506 | NA |
| | 0.725538412 | NA |
| | 0.905296333 | NA |
| | 0.78717524 | NA |
| | 0.889409773 | NA |
| | 0.797157316 | NA |
| | 0.053433923 | 0.127497068 |
| | 0.028219858 | 0.07758606 |
| | 0.905158266 | NA |
| NA | | NA |
| | 0.992060816 | NA |
| | 0.99205891 | NA |
| NA | | NA |
| | 0.899179329 | NA |
| | 0.110507622 | NA |
| | 0.779486752 | NA |
| NA | | NA |
| | 0.000618208 | 0.003324817 |
| | 0.678824166 | NA |
| | 0.70270711 | NA |
| | 0.440796652 | 0.600723247 |
| NA | | NA |
| NA | | NA |
| | 0.05110409 | 0.123453763 |
| | 0.181831369 | 0.32205477 |
| | 0.589352881 | NA |
| | 0.911297641 | NA |
| NA | | NA |
| NA | | NA |
| | 0.004459676 | 0.017388145 |
| | 0.780138338 | NA |
| NA | | NA |
| NA | | NA |
| | 0.290131258 | 0.449475613 |
| NA | | NA |
| NA | | NA |
| NA | | NA |
| | 0.208951825 | 0.355717999 |
| | 0.090992582 | 0.191674523 |
| | 0.205521901 | NA |
| | 0.037221316 | 0.096681202 |

| | |
|---|---|
| 0.056953657 | 0.133945096 |
| 1.72E-06 | 2.10E-05 |
| 4.76E-05 | 0.000372346 |
| 0.051628449 | NA |
| NA | NA |
| 0.792436369 | NA |
| 0.390552298 | 0.554049869 |
| 0.734208127 | NA |
| NA | NA |
| NA | NA |
| NA | NA |
| NA | NA |
| 0.992059387 | NA |
| 0.574644283 | 0.71660025 |
| 0.905158739 | NA |
| 0.471212388 | 0.627530647 |
| 0.559092044 | 0.703518549 |
| 0.329179221 | 0.49153043 |
| 0.938945563 | 0.965481746 |
| NA | NA |
| 0.43905457 | NA |
| 0.645963697 | NA |
| 0.753581177 | NA |
| 0.904947233 | NA |
| 0.004072551 | 0.016130799 |
| 0.988250096 | NA |
| 0.978614883 | NA |
| 0.025704557 | 0.072142942 |
| 0.199588749 | NA |
| 0.24585378 | 0.399164744 |
| 0.458822527 | 0.616112265 |
| 0.53809438 | 0.686100017 |
| 0.812592592 | NA |
| 0.000186364 | 0.001184817 |
| 0.06495712 | 0.148386038 |
| 0.496976794 | 0.650209331 |
| NA | NA |
| 0.889409301 | NA |
| 0.012236577 | 0.039591843 |
| 0.889409301 | NA |
| 0.005318067 | 0.020183103 |
| NA | NA |
| NA | NA |
| 0.372899973 | 0.537249061 |
| 0.348415479 | NA |
| 0.808815633 | NA |
| 4.60E-05 | 0.000361142 |

| | | |
|---|---|---|
| | 0.004023732 | 0.01596912 |
| | 0.558094983 | 0.702908387 |
| | 0.097780054 | 0.202370614 |
| | 0.992061292 | NA |
| | 0.889409301 | NA |
| | 0.466064614 | 0.622828193 |
| | 0.992061769 | NA |
| | 0.500191109 | 0.653251435 |
| | 0.282659946 | NA |
| | 0.505141607 | NA |
| | 0.976617485 | NA |
| | 0.18113612 | 0.321186051 |
| | 0.095077661 | 0.198196724 |
| | 0.463659587 | 0.620675986 |
| | 0.396499535 | NA |
| | 0.992662638 | 0.996201218 |
| | 0.997263295 | NA |
| | 0.784353866 | NA |
| | 0.889409301 | NA |
| NA | | NA |
| NA | | NA |
| | 0.236510795 | 0.388337203 |
| | 0.765667233 | 0.857920559 |
| | 0.79898587 | 0.878784408 |
| | 0.905158739 | NA |
| | 0.251419955 | 0.405563122 |
| | 0.005902043 | 0.021977777 |
| | 0.349384295 | 0.513426336 |
| | 0.265046544 | NA |
| | 0.8040279 | NA |
| | 0.09202733 | 0.193269715 |
| | 0.627988712 | 0.75661166 |
| NA | | NA |
| | 0.116056328 | 0.230585711 |
| NA | | NA |
| | 0.878360844 | NA |
| | 2.29E-05 | 0.000199281 |
| NA | | NA |
| | 0.182127988 | 0.322320155 |
| | 0.260267763 | 0.415556378 |
| | 0.199228203 | 0.343987753 |
| | 0.047018457 | 0.115606124 |
| | 0.069454869 | 0.155560001 |
| | 0.000528654 | 0.002904354 |
| NA | | NA |
| | 0.013459928 | 0.042687256 |
| | 0.99205891 | NA |

| | |
|---|---|
| 0.702331823 | NA |
| 0.967948002 | 0.982776689 |
| 0.00126926 | 0.006081535 |
| 0.531452917 | 0.680595708 |
| 0.248575116 | 0.402436865 |
| 0.030801507 | 0.083184145 |
| NA | NA |
| NA | NA |
| NA | NA |
| 0.287057219 | NA |
| 0.86948432 | 0.925335282 |
| 0.002696751 | 0.011523243 |
| 0.995078526 | 0.997551495 |
| 0.323674655 | 0.485985809 |
| 0.198785464 | NA |
| 0.41882449 | NA |
| 0.040132433 | 0.102365906 |
| NA | NA |
| 0.315164433 | 0.477388134 |
| NA | NA |
| 0.442322173 | 0.602017639 |
| 0.889410245 | NA |
| 0.023620041 | 0.067246529 |
| 0.005420223 | 0.020492883 |
| 0.149799785 | 0.278398205 |
| 0.346721771 | NA |
| NA | NA |
| 0.059141312 | NA |
| 0.639789818 | NA |
| 0.000213724 | 0.001335179 |
| 0.653691242 | 0.777272827 |
| 0.886534096 | NA |
| 1.11E-13 | 9.79E-12 |
| 0.012907279 | 0.041291738 |
| 0.843754839 | 0.907835391 |
| NA | NA |
| 0.049847242 | 0.12107019 |
| 0.929032869 | 0.959397704 |
| 0.731007313 | 0.834459624 |
| 0.737670267 | 0.838996291 |
| 0.889410245 | NA |
| 0.889409301 | NA |
| 0.003928257 | 0.015643922 |
| 0.101309695 | 0.208085989 |
| 0.288903732 | 0.448284032 |
| 0.941860755 | 0.967051027 |
| 0.930507109 | 0.959926975 |

| | |
|---|---|
| NA | NA |
| 2.93E-06 | 3.34E-05 |
| NA | NA |
| NA | NA |
| 0.992059387 | NA |
| NA | NA |
| NA | NA |
| NA | NA |
| 0.697960712 | NA |
| 1.12E-06 | 1.44E-05 |
| 0.022923336 | 0.065671695 |
| 0.010854303 | 0.035971874 |
| 0.032851039 | 0.087588146 |
| 0.044108297 | 0.110002737 |
| 0.992060816 | NA |
| 0.980804518 | NA |
| 0.689162991 | NA |
| 3.87E-11 | 1.83E-09 |
| 0.411305048 | 0.574062013 |
| 0.347572822 | 0.511448878 |
| 0.889409301 | NA |
| 0.010561587 | 0.035219726 |
| 0.016482622 | 0.050416758 |
| 0.89190765 | NA |
| 0.001173666 | 0.005701888 |
| NA | NA |
| NA | NA |
| 0.001522978 | 0.007097427 |
| 0.000117994 | 0.000805006 |
| 0.406574852 | 0.569798155 |
| 0.640024095 | 0.766486099 |
| 0.905158266 | NA |
| 0.992061292 | NA |
| NA | NA |
| NA | NA |
| 0.762913759 | NA |
| NA | NA |
| NA | NA |
| 0.293728146 | 0.453361662 |
| 0.59481334 | 0.731893199 |
| 0.889410245 | NA |
| 0.096590715 | 0.200627649 |
| 0.000136855 | 0.000912324 |
| 0.889409301 | NA |
| NA | NA |
| 0.905158266 | NA |
| 0.00879031 | 0.0303167 |

| | |
|---|---|
| 0.327360277 | 0.489542142 |
| 0.366178998 | NA |
| 0.702659489 | NA |
| 0.046531053 | 0.114625914 |
| 0.7667602 | 0.85847587 |
| 0.075136495 | 0.165528585 |
| 0.966839713 | 0.982106512 |
| 0.001826619 | 0.008285629 |
| 0.259956353 | 0.415155023 |
| 0.317752481 | 0.480049999 |
| 0.889410717 | NA |
| 0.259709029 | 0.414820379 |
| 1.25E-07 | 2.16E-06 |
| 0.34859743 | 0.512543684 |
| 0.102836819 | NA |
| NA | NA |
| 0.058075154 | 0.136014538 |
| 5.04E-09 | 1.28E-07 |
| 0.052631026 | 0.126212154 |
| 0.188993589 | 0.33076514 |
| 0.98647646 | 0.9930792 |
| 0.889409773 | NA |
| 0.440730873 | 0.600680164 |
| 0.145274357 | 0.272031813 |
| 0.530300571 | 0.67955592 |
| 0.419155863 | 0.581138113 |
| 0.58500651 | 0.724543595 |
| 3.68E-05 | 0.000297852 |
| 0.921596363 | NA |
| 6.38E-05 | 0.000476748 |
| 0.002668152 | 0.011427725 |
| 0.017138846 | 0.052011044 |
| 7.59E-07 | 1.03E-05 |
| 0.856168084 | NA |
| 0.000240525 | 0.001482373 |
| 0.69880734 | NA |
| 0.095594751 | 0.19899186 |
| 0.725538412 | NA |
| 1.46E-06 | 1.81E-05 |
| 0.721912207 | NA |
| NA | NA |
| 0.983547535 | 0.991858568 |
| 0.347917591 | 0.51176861 |
| NA | NA |
| 0.106830096 | 0.216389824 |
| 0.456684322 | NA |
| 0.534047832 | NA |

| | |
|---|---|
| 0.055276534 | 0.130898026 |
| 0.03227989 | 0.086421007 |
| 0.859604212 | 0.918229747 |
| 0.046951614 | 0.115463865 |
| 0.733234478 | 0.836159945 |
| 0.000197326 | 0.001246213 |
| 0.258319124 | 0.413201461 |
| 0.220411899 | 0.36935477 |
| 0.889409773 | NA |
| 3.90E-12 | 2.40E-10 |
| 0.863536531 | NA |
| NA | NA |
| NA | NA |
| 0.002120354 | 0.00940647 |
| 0.035606942 | 0.09331577 |
| 0.254597197 | NA |
| 0.736602883 | 0.838335753 |
| 4.07E-09 | 1.07E-07 |
| 0.992061292 | NA |
| 1.59E-07 | 2.65E-06 |
| NA | NA |
| 0.988536923 | 0.993877675 |
| 0.905158739 | NA |
| NA | NA |
| NA | NA |
| NA | NA |
| NA | NA |
| 5.48E-08 | 1.05E-06 |
| 0.711680423 | NA |
| 0.12821624 | 0.247843346 |
| 0.00554396 | 0.020881614 |
| 0.106946894 | 0.216586447 |
| 0.000156132 | 0.001018569 |
| 0.889409301 | NA |
| 0.791283264 | NA |
| 0.010793588 | 0.035813901 |
| 0.004432506 | 0.017306793 |
| 0.000288665 | 0.001731441 |
| 0.253071677 | 0.407419425 |
| 0.517637636 | 0.668547152 |
| 0.019246342 | 0.056958277 |
| NA | NA |
| 0.648148617 | NA |
| 0.000863294 | 0.004393298 |
| 0.88033013 | NA |
| 0.992062245 | NA |
| 0.516408108 | NA |

| | | |
|---|---|---|
| | 0.825243546 | NA |
| | 0.014046786 | 0.044241735 |
| | 0.363148691 | 0.52802535 |
| | 0.051583471 | 0.124387039 |
| | 0.973417205 | NA |
| | 0.064637055 | 0.147819734 |
| | 0.992061292 | NA |
| | 0.000160987 | 0.001045274 |
| | 0.591544352 | NA |
| NA | | NA |
| | 0.163941937 | 0.297742205 |
| NA | | NA |
| | 0.034462493 | 0.090946025 |
| | 0.253389029 | 0.407794434 |
| | 0.226912849 | 0.376912197 |
| | 0.788582525 | NA |
| | 0.018958811 | 0.05625139 |
| | 0.668119673 | 0.788322669 |
| | 0.90515732 | NA |
| | 0.073786741 | 0.163121216 |
| | 0.639643326 | 0.76628084 |
| | 0.848776475 | 0.911184679 |
| | 0.010778803 | 0.035781063 |
| | 0.002233335 | 0.009824077 |
| | 3.03E-05 | 0.00025218 |
| | 4.65E-05 | 0.000364457 |
| NA | | NA |
| | 0.539726385 | 0.687103698 |
| NA | | NA |
| | 0.275662579 | 0.433057274 |
| | 0.412770392 | NA |
| | 0.071578535 | 0.159215919 |
| | 0.98933516 | NA |
| NA | | NA |
| | 0.881195111 | 0.932448123 |
| | 0.753471393 | NA |
| | 0.005248712 | 0.01995092 |
| | 0.000214692 | 0.001340084 |
| | 8.04E-05 | 0.000580314 |
| | 1.45E-06 | 1.80E-05 |
| NA | | NA |
| | 0.229823062 | 0.38042248 |
| | 0.000492396 | 0.002725616 |
| | 0.003726857 | 0.014983166 |
| | 0.057221455 | 0.134462242 |
| | 0.889200554 | 0.937385064 |
| | 0.846625998 | 0.909899192 |

| | | |
|---|---|---|
| | 0.000746503 | NA |
| | 2.02E-09 | 5.84E-08 |
| | 0.026625514 | 0.074036934 |
| | 0.75277756 | 0.849447196 |
| NA | | NA |
| | 0.138472739 | 0.262289561 |
| | 0.004050388 | 0.016060395 |
| | 0.685990165 | NA |
| | 0.305801314 | NA |
| | 0.889410245 | NA |
| NA | | NA |
| | 0.220462853 | 0.369383778 |
| | 0.543576387 | NA |
| | 0.095978586 | 0.199658371 |
| | 0.905158266 | NA |
| | 0.358819098 | NA |
| NA | | NA |
| NA | | NA |
| | 0.871493696 | NA |
| | 0.670870289 | NA |
| | 0.484775127 | NA |
| NA | | NA |
| | 0.82210119 | 0.89413175 |
| | 4.30E-05 | 0.000340002 |
| | 0.446584202 | 0.60590315 |
| | 0.927234574 | 0.958351878 |
| | 0.510128183 | NA |
| | 0.520300179 | NA |
| | 0.686359475 | NA |
| | 0.180694274 | NA |
| | 0.004443536 | 0.017337527 |
| | 0.539930585 | 0.687217102 |
| | 0.000264464 | 0.001605133 |
| | 0.023816794 | NA |
| | 0.813836939 | NA |
| | 0.905158266 | NA |
| | 0.350502046 | 0.514654485 |
| | 0.196041659 | 0.33987674 |
| | 0.1721672 | 0.309000372 |
| | 0.905158739 | NA |
| | 0.062627618 | 0.144366981 |
| | 0.580158572 | NA |
| | 0.000283909 | 0.001709451 |
| | 0.238336981 | 0.390366358 |
| | 0.454861405 | 0.612892042 |
| | 0.146708239 | 0.274059228 |
| | 0.81627761 | 0.890670005 |

| | |
|---|---|
| 0.287643654 | 0.446733645 |
| 0.718651198 | 0.825445217 |
| 0.708264862 | NA |
| NA | NA |
| 0.905158739 | NA |
| 0.830163713 | NA |
| NA | NA |
| NA | NA |
| NA | NA |
| 0.905158266 | NA |
| NA | NA |
| NA | NA |
| NA | NA |
| NA | NA |
| NA | NA |
| NA | NA |
| NA | NA |
| NA | NA |
| NA | NA |
| NA | NA |
| NA | NA |
| 0.420409598 | 0.582225931 |
| 0.4888669 | 0.643305506 |
| 0.986263419 | NA |
| 0.983263945 | NA |
| 0.094506404 | 0.197249894 |
| 0.548825641 | NA |
| 0.860468292 | 0.918839877 |
| 2.14E-05 | 0.000188419 |
| 0.568536675 | 0.711002725 |
| 0.352610397 | 0.516852794 |
| NA | NA |
| 0.745769302 | NA |
| 8.46E-14 | 7.64E-12 |
| 0.345289804 | NA |
| 1.54E-09 | 4.60E-08 |
| 0.171711943 | NA |
| 0.237447276 | NA |
| 0.001287742 | 0.006157185 |
| 0.986004707 | NA |
| 0.613985326 | NA |
| 0.189672794 | 0.331607081 |
| 0.308876588 | 0.470003533 |
| 0.291154962 | 0.450581169 |
| 0.233730898 | 0.385329731 |
| 0.761760231 | NA |
| NA | NA |

| | |
|---|---|
| NA | NA |
| 0.309455725 | 0.470673272 |
| 0.163323427 | 0.296839855 |
| 0.983573359 | NA |
| 0.992061292 | NA |
| 0.954476153 | 0.974626452 |
| 0.285644986 | 0.444604356 |
| 4.67E-12 | 2.82E-10 |
| NA | NA |
| 0.41930395 | 0.581274247 |
| 0.816429009 | NA |
| NA | NA |
| 0.905158266 | NA |
| 0.110548059 | 0.221975569 |
| 0.015899202 | 0.048993907 |
| 0.006009844 | 0.022314874 |
| 0.488534177 | 0.642994605 |
| 0.220587719 | 0.369536598 |
| 0.007002814 | 0.025259054 |
| 0.16080335 | 0.293408912 |
| NA | NA |
| 0.026666622 | 0.074104322 |
| 0.248992852 | 0.402752208 |
| 0.054817475 | 0.130133723 |
| 0.570272135 | NA |
| 0.158685587 | 0.290526663 |
| 0.38330643 | 0.547270865 |
| 0.198247056 | 0.342645187 |
| 0.457227416 | 0.614722147 |
| 0.963071979 | 0.979577451 |
| NA | NA |
| NA | NA |
| 0.003715417 | 0.01493991 |
| 0.054945444 | 0.130365104 |
| NA | NA |
| 0.003070084 | 0.012794833 |
| 0.463072002 | 0.620187734 |
| 0.54881978 | 0.695013468 |
| 0.617473521 | NA |
| 5.45E-25 | 7.04E-22 |
| 0.985505086 | 0.99264741 |
| 3.94E-05 | 0.000315838 |
| 0.910334964 | 0.949212477 |
| NA | NA |
| 0.076314937 | 0.167714143 |
| 0.604235759 | NA |
| 0.092781687 | 0.194500531 |

| | |
|---|---|
| 0.729841945 | 0.833850118 |
| 0.003444134 | 0.01405286 |
| 0.715379613 | 0.82306235 |
| 0.192613844 | 0.335470131 |
| 0.056491865 | NA |
| 0.323635117 | 0.485985809 |
| 0.00068369 | 0.003621989 |
| 1.69E-05 | 0.000153351 |
| 0.966855438 | 0.982106512 |
| 0.830144184 | 0.898739641 |
| 0.373051295 | 0.537345582 |
| 0.000354396 | 0.002064786 |
| 0.496677871 | 0.649973145 |
| 0.00296805 | 0.012452315 |
| NA | NA |
| 0.060453437 | 0.140580008 |
| 0.506245097 | 0.658139607 |
| NA | NA |
| 0.057526257 | 0.13504587 |
| NA | NA |
| 0.342054395 | 0.505496498 |
| 0.470610621 | 0.626919354 |
| 0.23676226 | NA |
| NA | NA |
| NA | NA |
| NA | NA |
| NA | NA |
| 0.849488079 | NA |
| 0.016771729 | 0.051143277 |
| 0.158105778 | 0.289706674 |
| 0.751599479 | 0.848725928 |
| 0.592638975 | NA |
| 0.492996057 | 0.646773833 |
| NA | NA |
| 0.417690222 | NA |
| 0.172667878 | 0.30967118 |
| 0.076423699 | 0.167835584 |
| 0.00371054 | 0.014925764 |
| 0.096243796 | 0.200085134 |
| 0.392025144 | 0.555522538 |
| NA | NA |
| 0.992060816 | NA |
| 0.912076839 | 0.950289206 |
| 0.153427422 | 0.283595351 |
| 0.82854455 | NA |
| 0.000107061 | 0.000740458 |
| 0.663491392 | 0.784910205 |

| | |
|---|---|
| 0.669898958 | 0.789501803 |
| 0.354029094 | NA |
| 0.00062387 | 0.003351451 |
| NA | NA |
| 0.52680664 | 0.676697689 |
| 0.010676401 | 0.035510919 |
| 0.348158131 | 0.512001366 |
| 0.99205891 | NA |
| NA | NA |
| 0.014875749 | 0.04635399 |
| 0.959437841 | NA |
| 0.710800381 | 0.819858471 |
| NA | NA |
| 0.103341594 | 0.211252844 |
| 0.01837953 | 0.054888631 |
| 3.63E-07 | 5.42E-06 |
| 0.253845738 | 0.408216665 |
| 3.66E-14 | 3.55E-12 |
| 0.000788781 | 0.004077413 |
| 0.042091414 | 0.106215279 |
| NA | NA |
| 0.987962164 | 0.993639148 |
| 0.984281854 | NA |
| NA | NA |
| 0.029968483 | 0.081445288 |
| 0.007826336 | 0.027550259 |
| 0.777253828 | 0.865942 |
| 0.185795619 | 0.32673068 |
| 3.83E-05 | 0.000308453 |
| 0.00108407 | 0.005331487 |
| 0.762584384 | NA |
| 0.037173877 | NA |
| 0.524448413 | NA |
| 0.326610312 | 0.488858008 |
| 0.800839654 | 0.879945431 |
| NA | NA |
| 0.718131222 | 0.825236459 |
| NA | NA |
| 7.73E-10 | 2.53E-08 |
| 0.087945969 | 0.186896886 |
| 0.999841591 | 0.999887125 |
| 1.62E-06 | 1.99E-05 |
| NA | NA |
| 0.021806284 | 0.06304287 |
| 0.886926061 | 0.935866549 |
| NA | NA |
| NA | NA |

| | |
|---|---|
| 1.66E-07 | 2.75E-06 |
| 0.814603153 | 0.889578811 |
| 0.124237771 | 0.242221186 |
| 0.723671499 | NA |
| 0.905158266 | NA |
| 0.105815444 | 0.214861869 |
| 0.923348404 | NA |
| 0.905158266 | NA |
| 0.036079783 | 0.094341028 |
| 1.40E-07 | 2.38E-06 |
| NA | NA |
| 0.672918545 | 0.792014704 |
| 0.739455843 | 0.840113352 |
| 0.335329334 | 0.498240534 |
| 0.121979011 | 0.238899135 |
| NA | NA |
| 0.017524042 | 0.052902177 |
| 0.197712898 | 0.342125889 |
| 0.623196986 | 0.754169581 |
| 0.792565833 | 0.874922236 |
| 1.83E-08 | 4.01E-07 |
| 0.714058399 | NA |
| 0.889409301 | NA |
| 0.633303353 | 0.760802469 |
| 0.000166579 | 0.001077438 |
| 1.41E-06 | 1.76E-05 |
| 0.277355967 | 0.435219357 |
| 0.905158266 | NA |
| 0.000818711 | 0.004214272 |
| 0.462754263 | 0.620037257 |
| 0.008376247 | 0.029163471 |
| NA | NA |
| NA | NA |
| NA | NA |
| NA | NA |
| 0.441216376 | NA |
| 0.003689398 | 0.014867956 |
| NA | NA |
| 0.123680989 | 0.241435758 |
| NA | NA |
| NA | NA |
| 0.905158266 | NA |
| 0.409814569 | 0.572790918 |
| 0.002120407 | 0.00940647 |
| NA | NA |
| 4.06E-08 | 8.13E-07 |
| NA | NA |

| | |
|---|---|
| NA | NA |
| NA | NA |
| NA | NA |
| NA | NA |
| NA | NA |
| 0.005314819 | 0.020174264 |
| 0.086657824 | 0.18460605 |
| 1.21E-09 | 3.69E-08 |
| NA | NA |
| 0.677693847 | 0.795534131 |
| NA | NA |
| 0.006759493 | 0.024538221 |
| 0.003914828 | 0.015601761 |
| NA | NA |
| 0.44815723 | NA |
| 0.722352826 | NA |
| NA | NA |
| NA | NA |
| 0.75615455 | NA |
| 0.992061292 | NA |
| NA | NA |
| NA | NA |
| NA | NA |
| 0.002304875 | 0.010086241 |
| 0.0158357 | 0.048846206 |
| NA | NA |
| NA | NA |
| 0.767341078 | NA |
| 0.191390094 | 0.333762316 |
| 0.899149809 | 0.943310432 |
| 5.05E-10 | 1.76E-08 |
| 0.140421791 | 0.265135177 |
| 0.008863034 | 0.030510012 |
| 0.238659524 | 0.390807196 |
| 0.906330578 | NA |
| 0.421249447 | 0.58296755 |
| 0.001151381 | 0.005607267 |
| 0.157975486 | 0.289634401 |
| 0.00697321 | 0.025191893 |
| 0.00909554 | 0.03113468 |
| NA | NA |
| NA | NA |
| 0.144623594 | 0.271180044 |
| 0.000801877 | 0.004135674 |
| 0.354501261 | 0.519104641 |
| 0.185097357 | 0.325902399 |
| 0.735450974 | 0.837513247 |

| | | |
|---|---|---|
| | 0.780276289 | NA |
| NA | | NA |
| | 0.5871872 | NA |
| | 0.07102193 | 0.158327312 |
| | 6.39E-06 | 6.62E-05 |
| NA | | NA |
| | 0.432580961 | 0.593430706 |
| | 0.772750597 | NA |
| | 0.00595281 | 0.022129295 |
| | 5.37E-06 | 5.66E-05 |
| | 0.792188023 | NA |
| | 0.797157316 | NA |
| | 0.905158739 | NA |
| | 0.683450055 | NA |
| | 0.815370765 | NA |
| | 0.004001449 | 0.015895046 |
| | 1.52E-06 | 1.88E-05 |
| NA | | NA |
| | 0.032273227 | 0.086414802 |
| | 0.826124187 | 0.896319493 |
| | 0.959003266 | 0.977249651 |
| NA | | NA |
| NA | | NA |
| NA | | NA |
| | 0.075192015 | 0.165627592 |
| | 0.231957173 | 0.383002297 |
| | 0.211438019 | 0.35848504 |
| | 0.000285879 | 0.001717074 |
| | 0.905158739 | NA |
| NA | | NA |
| NA | | NA |
| | 4.04E-07 | 5.97E-06 |
| | 0.701022322 | 0.813193299 |
| | 0.887395124 | NA |
| | 0.269093888 | 0.425508223 |
| | 0.592341465 | 0.729922909 |
| | 0.985580432 | NA |
| | 0.203735736 | 0.349572826 |
| | 0.558890326 | 0.703504149 |
| | 0.889409773 | NA |
| | 0.612423209 | 0.745796431 |
| NA | | NA |
| | 0.01940877 | 0.057369389 |
| | 1.15E-05 | 0.000109616 |
| | 0.476984864 | 0.633143393 |
| | 0.262034292 | 0.417411028 |
| | 1.49E-12 | 1.01E-10 |

| | | |
|---|---|---|
| | 0.574643763 | NA |
| | 3.92E-08 | 7.91E-07 |
| | 0.992061292 | NA |
| | 0.003153694 | 0.013096062 |
| | 0.210583808 | 0.357551214 |
| | 0.001678256 | 0.007692092 |
| | 0.002911762 | 0.012251271 |
| | 1.33E-10 | 5.50E-09 |
| | 0.041756299 | 0.105502999 |
| NA | | NA |
| | 0.245348394 | NA |
| | 0.310533132 | 0.471739678 |
| | 0.992062245 | NA |
| | 0.721614968 | NA |
| | 0.905158266 | NA |
| NA | | NA |
| | 0.673406964 | 0.792334755 |
| | 8.06E-09 | 1.95E-07 |
| | 0.166945807 | NA |
| | 0.029174262 | 0.079641674 |
| | 0.002502098 | 0.010822053 |
| | 0.004911015 | 0.018886337 |
| | 0.991724961 | NA |
| | 0.504157952 | 0.656381713 |
| | 0.535683777 | 0.683939768 |
| | 0.117263354 | 0.232159536 |
| | 0.59225818 | 0.729861236 |
| | 0.002786642 | 0.011824516 |
| | 0.309397587 | NA |
| | 0.237423211 | 0.389324758 |
| | 0.066896329 | 0.151456489 |
| | 0.129159676 | 0.249206338 |
| | 0.042591236 | 0.107168665 |
| | 0.005548824 | 0.020896352 |
| | 0.123877947 | 0.241734278 |
| NA | | NA |
| | 0.590054513 | 0.728244332 |
| | 0.142175996 | 0.267596014 |
| | 0.108598489 | 0.21920344 |
| | 0.889410245 | NA |
| | 0.000662166 | 0.003528407 |
| | 0.875630001 | NA |
| | 0.081401011 | 0.176280553 |
| | 0.248023039 | 0.401736218 |
| | 0.061570115 | 0.142572831 |
| | 3.94E-05 | 0.000315838 |
| | 0.205841128 | 0.35203001 |

| | |
|---|---|
| 0.111033693 | 0.222631229 |
| 0.044764966 | 0.111286526 |
| 0.01181725 | 0.038466497 |
| 0.003639508 | 0.014707391 |
| 0.065453718 | 0.149190181 |
| 0.000181637 | 0.001159807 |
| 0.532055577 | 0.68108945 |
| 0.003343329 | 0.013720081 |
| NA | NA |
| 0.073296836 | 0.162250526 |
| 0.658457397 | 0.780788681 |
| 0.899962636 | 0.943542752 |
| NA | NA |
| 0.889410245 | NA |
| 2.15E-06 | 2.56E-05 |
| NA | NA |
| 0.001608881 | 0.007422146 |
| 0.209306154 | 0.356152952 |
| 0.000566053 | 0.003081301 |
| 0.107749664 | 0.217850554 |
| 0.249660436 | 0.403591072 |
| 0.391727985 | 0.555143906 |
| 0.777092376 | NA |
| 0.508906064 | NA |
| 0.873859502 | 0.927840184 |
| 0.045157105 | 0.112145976 |
| 0.237258286 | 0.389165794 |
| 0.352892762 | 0.517028362 |
| 0.992061769 | NA |
| 0.106823365 | 0.216389824 |
| 0.520371322 | NA |
| 0.056581842 | 0.133270478 |
| 0.889409773 | NA |
| 0.010923925 | 0.036174405 |
| 1.17E-06 | 1.50E-05 |
| 2.91E-13 | 2.38E-11 |
| 0.943331565 | NA |
| NA | NA |
| NA | NA |
| 0.549055263 | NA |
| NA | NA |
| NA | NA |
| NA | NA |
| 0.170932346 | 0.307356228 |
| 0.892876436 | 0.939380965 |
| 6.66E-06 | 6.86E-05 |
| 0.522770494 | 0.673167025 |

| | |
|---|---|
| NA | NA |
| 0.742234807 | 0.842044611 |
| 0.002450597 | 0.010639119 |
| 0.583343686 | 0.723218383 |
| 0.107227346 | 0.216991602 |
| 2.71E-05 | 0.000229322 |
| 3.12E-05 | 0.00025875 |
| 0.889409773 | NA |
| 0.000171365 | 0.001102876 |
| 0.009784662 | 0.033025113 |
| 3.20E-13 | 2.58E-11 |
| NA | NA |
| 0.130494861 | 0.250874788 |
| 0.773967799 | 0.863727138 |
| 0.139884355 | 0.264340942 |
| 0.002020112 | 0.009032711 |
| 0.423069524 | 0.584485652 |
| 0.926264101 | 0.957709454 |
| 1.23E-05 | 0.000116784 |
| 0.905158739 | NA |
| 0.99205891 | NA |
| 0.036263992 | 0.094732453 |
| 0.936967448 | NA |
| 0.077695285 | 0.170032963 |
| 0.761109765 | NA |
| 0.980058715 | NA |
| 0.000313776 | 0.001857196 |
| NA | NA |
| 0.073553488 | 0.16263831 |
| 0.533942344 | 0.682720498 |
| 0.178335011 | NA |
| 0.315720717 | 0.477934042 |
| 0.789658938 | 0.872942037 |
| NA | NA |
| NA | NA |
| 0.920048871 | 0.954241358 |
| NA | NA |
| 0.196526511 | 0.340502262 |
| 0.112734103 | 0.225375834 |
| NA | NA |
| NA | NA |
| 0.86691533 | NA |
| 0.511414819 | 0.663054732 |
| 0.261659207 | 0.417128346 |
| NA | NA |
| 0.500194436 | 0.653251435 |
| 0.722348371 | 0.828218875 |

| | | |
|---|---|---|
| | 0.016396118 | 0.050208109 |
| | 0.000665723 | 0.003543045 |
| | 0.044795043 | 0.11132349 |
| | 0.240667517 | 0.393329075 |
| | 9.87E-14 | 8.81E-12 |
| | 0.338897447 | 0.501878138 |
| | 0.450690826 | 0.60910388 |
| | 0.889409773 | NA |
| | 0.22276738 | 0.372202713 |
| | 0.027965895 | 0.077061499 |
| | 0.53435187 | 0.683033513 |
| | 0.026612658 | 0.074016644 |
| | 0.267942111 | NA |
| NA | | NA |
| | 0.052965913 | 0.126779865 |
| | 0.9197826 | 0.95410771 |
| | 0.448412762 | 0.60748324 |
| NA | | NA |
| | 0.437765964 | 0.598152125 |
| | 0.496914844 | 0.650167018 |
| | 0.58432441 | NA |
| | 0.354366845 | 0.518942418 |
| | 0.063630324 | 0.146078302 |
| | 0.318222701 | 0.480601637 |
| | 0.016238744 | 0.049830432 |
| | 9.20E-07 | 1.22E-05 |
| | 0.24263766 | 0.395548954 |
| | 1.17E-06 | 1.50E-05 |
| | 0.091384559 | NA |
| | 0.905157793 | NA |
| | 0.790051649 | NA |
| NA | | NA |
| NA | | NA |
| NA | | NA |
| | 0.124612245 | 0.242778838 |
| | 0.986869548 | NA |
| | 0.626450728 | NA |
| | 0.382848582 | NA |
| | 0.00685886 | 0.024870164 |
| NA | | NA |
| | 0.259453872 | 0.414593769 |
| | 0.878819756 | 0.93128091 |
| | 0.992061292 | NA |
| | 5.98E-07 | 8.37E-06 |
| | 0.265780317 | NA |
| | 0.475816307 | NA |
| | 0.004127884 | 0.016313606 |

| | |
|---|---|
| 0.93066901 | 0.959958702 |
| 0.185334115 | 0.326154178 |
| NA | NA |
| 0.378093088 | NA |
| NA | NA |
| 0.143843773 | NA |
| 0.00036097 | 0.002094198 |
| 0.080544378 | 0.174804705 |
| 0.062816058 | 0.14464952 |
| 0.880107941 | NA |
| 0.889409773 | NA |
| 0.052469304 | 0.125934358 |
| 0.005281773 | 0.020073113 |
| 0.986195367 | 0.992938931 |
| 4.22E-07 | 6.20E-06 |
| 0.025820657 | 0.072359556 |
| 0.000186182 | 0.001184008 |
| 0.523601028 | NA |
| 4.19E-09 | 1.09E-07 |
| 1.65E-06 | 2.01E-05 |
| 0.967384848 | 0.98241705 |
| 0.007144459 | 0.025655794 |
| NA | NA |
| NA | NA |
| 0.843525319 | NA |
| 0.894946485 | NA |
| NA | NA |
| 0.849264444 | NA |
| 0.680226957 | NA |
| NA | NA |
| NA | NA |
| NA | NA |
| 0.889409301 | NA |
| NA | NA |
| NA | NA |
| 0.889409773 | NA |
| NA | NA |
| NA | NA |
| NA | NA |
| 0.706860257 | 0.817632974 |
| 0.88281598 | NA |
| 0.288578959 | NA |
| 8.84E-06 | 8.77E-05 |
| 5.56E-07 | 7.88E-06 |
| 0.905158266 | NA |
| 0.889409773 | NA |
| 7.33E-05 | 0.000535807 |

| | |
|---|---|
| 0.022613608 | 0.064962353 |
| 8.37E-07 | 1.13E-05 |
| 5.78E-05 | 0.000439307 |
| 0.988682613 | 0.993927811 |
| 0.000638854 | 0.003423281 |
| 0.033736512 | 0.089406236 |
| 0.925587835 | 0.957190509 |
| 0.021218934 | 0.061641297 |
| NA | NA |
| 0.14428331 | NA |
| NA | NA |
| 0.935146574 | 0.962768232 |
| 0.820116371 | 0.892909192 |
| 0.927084333 | 0.958331915 |
| 1.38E-09 | 4.16E-08 |
| NA | NA |
| 4.57E-05 | 0.00035862 |
| 0.815673901 | NA |
| 0.848945189 | NA |
| NA | NA |
| 0.992061292 | NA |
| 0.639558248 | 0.76622072 |
| 0.000153521 | 0.001003321 |
| 0.79081386 | NA |
| NA | NA |
| 0.804534188 | NA |
| 0.78107429 | 0.868128276 |
| 0.105812057 | 0.214861869 |
| NA | NA |
| 0.889410245 | NA |
| 0.797342596 | NA |
| NA | NA |
| 0.905158739 | NA |
| NA | NA |
| NA | NA |
| 0.901183732 | NA |
| 0.665682494 | 0.786617978 |
| NA | NA |
| NA | NA |
| 0.905158266 | NA |
| NA | NA |
| 0.00105498 | 0.005206211 |
| 0.992060816 | NA |
| 0.315582456 | 0.477824638 |
| 0.344817318 | 0.508143311 |
| 0.130741173 | 0.251154354 |
| 0.901393659 | 0.944265974 |

| | | | |
|---|---|---|---|
| | 0.173470187 | | 0.310653387 |
| | 0.509782945 | | 0.661368527 |
| | 0.38247315 | | 0.546496771 |
| | 0.002896789 | | 0.01220464 |
| | 0.202850154 | | 0.348516277 |
| NA | | NA | |
| | 0.565191932 | | 0.708474775 |
| | 0.039842259 | | 0.101791293 |
| | 0.853716831 | | 0.914632881 |
| | 0.020864272 | | 0.060844428 |
| | 0.291167485 | | 0.450581169 |
| | 4.49E-08 | | 8.87E-07 |
| | 0.280620035 | | 0.438992331 |
| | 2.88E-06 | | 3.29E-05 |
| | 0.000257783 | | 0.001571093 |
| | 0.003158814 | | 0.013112361 |
| | 4.98E-07 | | 7.19E-06 |
| | 1.13E-09 | | 3.51E-08 |
| NA | | NA | |
| | 0.989975993 | NA | |
| | 0.383189633 | | 0.547202504 |
| | 0.187227328 | NA | |
| | 0.991918092 | NA | |
| | 1.93E-08 | | 4.21E-07 |
| | 0.017063312 | | 0.051824796 |
| | 3.32E-11 | | 1.59E-09 |
| | 0.722279137 | NA | |
| | 0.023445721 | | 0.06688317 |
| | 0.203512761 | | 0.349354027 |
| | 0.081019598 | | 0.175662456 |
| | 0.583932234 | | 0.723702896 |
| | 0.77474095 | | 0.864072143 |
| | 0.030172115 | | 0.081856867 |
| | 5.91E-05 | | 0.000446955 |
| | 0.033018632 | | 0.08793889 |
| | 9.32E-06 | | 9.17E-05 |
| | 0.614434486 | | 0.747592191 |
| | 0.16921534 | | 0.304973707 |
| | 1.46E-05 | | 0.000135098 |
| | 0.002493764 | | 0.010788133 |
| | 0.09201227 | | 0.19325657 |
| | 0.328720939 | | 0.490946276 |
| | 1.26E-06 | | 1.59E-05 |
| | 2.41E-13 | | 2.01E-11 |
| | 0.173378712 | | 0.310540224 |
| | 0.211388685 | | 0.358481798 |
| | 4.77E-05 | | 0.000372561 |

| | |
|---|---|
| 0.484459779 | 0.639740951 |
| 7.80E-05 | 0.000566101 |
| 0.253667164 | 0.408049026 |
| 0.456973325 | NA |
| NA | NA |
| 0.902716808 | 0.94497585 |
| 1.83E-24 | 2.12E-21 |
| 1.69E-06 | 2.07E-05 |
| 0.000102602 | 0.000714571 |
| 0.000419916 | 0.002382672 |
| 0.040371782 | 0.102749647 |
| 0.043548313 | 0.108965065 |
| NA | NA |
| 0.367296974 | NA |
| 0.10345599 | 0.211373603 |
| NA | NA |
| 0.712603105 | 0.820965252 |
| 0.19046289 | 0.332554388 |
| 0.007262428 | 0.025994401 |
| 0.905157793 | NA |
| 0.139443869 | 0.263674152 |
| 0.983653096 | NA |
| 0.042548529 | 0.107098022 |
| 0.008070603 | 0.028272586 |
| 0.173679146 | 0.310926168 |
| 0.373896021 | 0.53789195 |
| 0.010908234 | 0.036128794 |
| 0.560144466 | 0.704519865 |
| 0.823985045 | 0.895116633 |
| 0.212103506 | 0.35917508 |
| 0.586732962 | NA |
| 0.00302365 | 0.01263729 |
| 0.003034238 | 0.012671898 |
| 0.004207079 | 0.01657396 |
| 0.396522084 | NA |
| 3.73E-06 | 4.14E-05 |
| 9.55E-07 | 1.26E-05 |
| 0.000785299 | 0.004060367 |
| 0.01188898 | 0.038665598 |
| 0.797157316 | NA |
| 5.69E-05 | 0.000433304 |
| 0.27449296 | 0.431744926 |
| 0.67306727 | 0.792104839 |
| 0.001528732 | 0.007119708 |
| 0.549977446 | 0.695996806 |
| NA | NA |
| 2.12E-10 | 8.24E-09 |

| | | |
|---|---|---|
| | 0.958173837 | 0.976812409 |
| | 0.691387744 | 0.805772376 |
| | 0.096280246 | 0.200115278 |
| | 0.051152044 | 0.123556015 |
| | 0.001099975 | 0.005395828 |
| | 0.00593805 | 0.022086956 |
| | 3.81E-05 | 0.000306806 |
| | 0.410518708 | 0.573556042 |
| | 0.023084848 | 0.066013827 |
| | 0.471958322 | 0.628295411 |
| | 0.139892804 | 0.264340942 |
| | 0.215402134 | 0.363148979 |
| | 0.000208522 | 0.00130603 |
| | 0.789940753 | 0.873095346 |
| | 0.813836939 | NA |
| NA | | NA |
| | 2.63E-05 | 0.000223453 |
| | 0.829425697 | 0.898360406 |
| | 0.030386844 | 0.082327541 |
| | 0.036489243 | 0.095183553 |
| | 0.066422494 | 0.150756749 |
| | 0.777092376 | NA |
| | 0.055823564 | 0.131908924 |
| | 0.048343647 | 0.118136896 |
| | 0.029877369 | 0.081278139 |
| | 2.53E-05 | 0.000216943 |
| | 0.000373957 | 0.002156439 |
| | 2.43E-06 | 2.84E-05 |
| | 0.040551257 | 0.103050112 |
| | 4.20E-09 | 1.09E-07 |
| | 0.822874916 | NA |
| | 0.149240555 | 0.27758541 |
| | 0.019036979 | 0.056427456 |
| | 0.443969622 | 0.603585022 |
| | 0.059712955 | 0.139182335 |
| | 0.000285414 | 0.001715219 |
| | 0.026355952 | 0.073436158 |
| | 1.06E-06 | 1.37E-05 |
| | 4.04E-07 | 5.97E-06 |
| | 0.203841086 | 0.349719627 |
| | 0.885496077 | 0.93507114 |
| | 0.822547989 | 0.894351916 |
| | 0.758045688 | NA |
| | 0.103101298 | 0.210859775 |
| | 0.163287519 | 0.296823728 |
| | 0.076068228 | NA |
| | 0.22376784 | 0.37340239 |

| | | |
|---|---|---|
| | 0.000159364 | 0.001036009 |
| | 0.721578566 | NA |
| | 5.44E-06 | 5.72E-05 |
| | 0.039286009 | 0.100721714 |
| | 0.93151477 | 0.960425056 |
| | 0.992062245 | NA |
| NA | | NA |
| | 0.000162533 | 0.001054377 |
| | 0.00476338 | 0.018408847 |
| | 0.837002984 | NA |
| | 0.000102047 | 0.000711154 |
| | 0.293418491 | 0.453042936 |
| | 0.100691697 | 0.207088974 |
| | 0.905157793 | NA |
| | 0.321438016 | 0.48375419 |
| | 0.000840722 | 0.00430236 |
| | 0.189774977 | 0.331735443 |
| | 0.905158739 | NA |
| | 0.321999987 | 0.484334387 |
| | 0.313848851 | NA |
| | 0.912003354 | NA |
| | 0.581117344 | NA |
| | 0.164674235 | 0.298800324 |
| | 0.150156245 | 0.278839829 |
| | 0.130602808 | 0.251004405 |
| | 0.47401382 | 0.630382115 |
| | 0.905158266 | NA |
| | 0.578032233 | 0.719196874 |
| NA | | NA |
| | 0.472500902 | 0.628838262 |
| | 0.687532688 | 0.802889295 |
| | 0.868300685 | 0.92446132 |
| | 0.00519897 | 0.01980504 |
| | 0.060451845 | 0.140580008 |
| | 0.11144055 | 0.223319084 |
| | 0.76380433 | NA |
| | 1.66E-10 | 6.69E-09 |
| | 0.05033574 | 0.121998754 |
| | 2.07E-15 | 2.86E-13 |
| | 0.889409773 | NA |
| NA | | NA |
| | 0.007718463 | 0.027253535 |
| | 0.881689493 | 0.932791461 |
| | 3.32E-07 | 5.03E-06 |
| | 0.007397972 | 0.026380653 |
| | 0.201634542 | 0.346943497 |
| | 0.057513724 | 0.135030884 |

| | |
|---|---|
| NA | NA |
| 0.003675869 | 0.014821596 |
| 0.905158739 | NA |
| 3.79E-05 | 0.000306087 |
| 0.875773908 | NA |
| 0.485560828 | 0.640501606 |
| 0.527655867 | NA |
| 0.025227066 | 0.071043837 |
| 2.42E-07 | 3.82E-06 |
| 0.468511479 | 0.624957088 |
| 0.889409773 | NA |
| 0.017202616 | 0.052146915 |
| 0.190184613 | 0.332232957 |
| 0.035359371 | 0.092766599 |
| 0.095587174 | 0.19899186 |
| 0.396745794 | 0.560374405 |
| NA | NA |
| 0.619066999 | 0.751013327 |
| NA | NA |
| 0.000710132 | 0.003735932 |
| 0.002734212 | 0.011649315 |
| 6.90E-05 | 0.000509549 |
| 0.048176461 | 0.117820127 |
| 0.211443975 | 0.35848504 |
| 0.658460371 | 0.780788681 |
| 0.795793874 | 0.876831882 |
| 0.801957113 | 0.880631879 |
| 0.373680439 | NA |
| 0.011308185 | 0.037140034 |
| 0.905158266 | NA |
| 0.000283199 | 0.0017058 |
| 4.20E-11 | 1.97E-09 |
| NA | NA |
| 0.044633257 | NA |
| 0.266471778 | 0.422640214 |
| 0.000365463 | 0.002117465 |
| 0.005158473 | 0.019679449 |
| NA | NA |
| 0.261071389 | 0.416534386 |
| 0.931486948 | 0.960425056 |
| 0.116622178 | 0.231316629 |
| 0.222300122 | 0.371639769 |
| 0.123416224 | 0.241042817 |
| NA | NA |
| 0.218773532 | NA |
| 0.840482277 | NA |
| NA | NA |

| | | |
|---|---|---|
| | 0.794008527 | NA |
| | 0.889409773 | NA |
| NA | | NA |
| | 0.088438025 | 0.187724562 |
| | 0.228351743 | 0.378643504 |
| | 0.831817855 | NA |
| | 0.920713764 | 0.954551282 |
| | 0.000739645 | 0.003873567 |
| | 0.127351449 | 0.246649362 |
| | 0.000206009 | 0.001293979 |
| NA | | NA |
| | 0.632856295 | 0.760473426 |
| | 0.000453708 | 0.002542525 |
| | 1.92E-10 | 7.58E-09 |
| | 0.01811478 | 0.054285028 |
| | 0.513190807 | NA |
| | 0.257690695 | 0.412496718 |
| | 0.334561645 | 0.497369113 |
| | 0.624204875 | 0.754827625 |
| | 0.0144154 | 0.045169488 |
| NA | | NA |
| | 0.980680523 | 0.990422831 |
| | 0.432963744 | 0.59373327 |
| | 3.66E-05 | 0.000296383 |
| | 0.088536693 | 0.187861362 |
| | 0.730685279 | 0.834459624 |
| | 0.002070508 | 0.009214894 |
| | 0.030038995 | 0.081576342 |
| | 0.945411286 | 0.969042261 |
| | 0.489716501 | 0.643836091 |
| | 0.32310905 | 0.485503738 |
| | 0.052319892 | 0.125630674 |
| | 0.030978815 | 0.083570491 |
| | 0.889409301 | NA |
| | 5.71E-06 | 5.98E-05 |
| | 0.310859489 | 0.472170126 |
| | 3.37E-05 | 0.000276438 |
| NA | | NA |
| | 0.033872662 | 0.089712916 |
| | 0.103462198 | 0.211373603 |
| | 0.247912118 | 0.401586176 |
| | 1.74E-05 | 0.000157697 |
| | 0.61248179 | NA |
| | 0.169107659 | 0.304854699 |
| | 0.202583914 | 0.348167815 |
| | 0.947845631 | 0.970440272 |
| NA | | NA |

| | | | |
|---|---|---|---|
| NA | | NA | |
| | 0.496890637 | | 0.650167018 |
| NA | | NA | |
| | 0.992061292 | NA | |
| | 0.801661681 | NA | |
| NA | | NA | |
| | 0.102235314 | | 0.209553109 |
| | 0.823211199 | NA | |
| | 0.506209114 | NA | |
| NA | | NA | |
| NA | | NA | |
| | 5.29E-08 | | 1.02E-06 |
| | 0.835471475 | NA | |
| | 0.001298329 | | 0.006199177 |
| | 9.81E-06 | | 9.54E-05 |
| | 0.951969541 | | 0.973288471 |
| | 5.40E-10 | | 1.85E-08 |
| | 0.873401525 | NA | |
| | 0.529941823 | | 0.679254756 |
| | 2.68E-06 | | 3.09E-05 |
| | 0.003749944 | | 0.015062196 |
| | 0.530887711 | NA | |
| | 0.163787431 | | 0.297563404 |
| | 0.861596748 | NA | |
| | 0.533617039 | | 0.682412006 |
| | 0.851384534 | NA | |
| | 0.846338124 | NA | |
| | 0.699661429 | NA | |
| | 0.005808722 | | 0.021703884 |
| NA | | NA | |
| | 0.892112101 | NA | |
| | 0.196012032 | | 0.339852207 |
| | 0.992061292 | NA | |
| NA | | NA | |
| | 0.663847908 | | 0.785083811 |
| NA | | NA | |
| | 6.04E-06 | | 6.30E-05 |
| | 0.178975208 | | 0.318277732 |
| NA | | NA | |
| | 0.227622198 | | 0.377604884 |
| | 0.992060816 | NA | |
| | 0.477660287 | | 0.633658082 |
| | 0.92430143 | | 0.956400673 |
| | 0.064935325 | | 0.148368741 |
| NA | | NA | |
| | 1.00E-07 | | 1.77E-06 |
| | 0.372216759 | | 0.536601091 |

| | | |
|---|---|---|
| | 0.048760896 | 0.118957952 |
| | 0.739034185 | NA |
| | 0.607238474 | NA |
| | 0.560844119 | 0.705076774 |
| NA | | NA |
| | 0.671090877 | 0.790456717 |
| | 0.821387042 | NA |
| | 0.605671912 | 0.740504215 |
| | 0.094997 | 0.198091033 |
| | 0.345890815 | 0.509349276 |
| | 0.666769987 | 0.787436923 |
| NA | | NA |
| | 0.43753434 | NA |
| | 0.567550163 | NA |
| NA | | NA |
| | 0.001369401 | 0.006500361 |
| NA | | NA |
| NA | | NA |
| | 0.890296709 | NA |
| NA | | NA |
| | 0.005541098 | 0.020874415 |
| NA | | NA |
| NA | | NA |
| | 9.71E-08 | 1.72E-06 |
| | 1.15E-05 | 0.000109933 |
| | 0.889409773 | NA |
| | 0.809102121 | NA |
| | 0.000825674 | 0.004243149 |
| NA | | NA |
| | 0.215836345 | 0.363655196 |
| | 0.520734809 | NA |
| | 0.905158266 | NA |
| | 7.25E-07 | 9.94E-06 |
| | 0.000890043 | 0.004510606 |
| | 0.054431878 | 0.129414206 |
| NA | | NA |
| | 0.905158266 | NA |
| | 0.365153072 | 0.529896663 |
| | 0.437402782 | NA |
| | 0.070458119 | 0.157330673 |
| | 0.309508075 | NA |
| | 0.132611162 | 0.253791921 |
| NA | | NA |
| NA | | NA |
| NA | | NA |
| NA | | NA |
| NA | | NA |

| | |
|---|---|
| NA | NA |
| NA | NA |
| NA | NA |
| NA | NA |
| NA | NA |
| NA | NA |
| NA | NA |
| NA | NA |
| NA | NA |
| 0.000570628 | 0.003103896 |
| 0.424939438 | 0.58618396 |
| 0.697794461 | 0.810948323 |
| 0.824208621 | NA |
| NA | NA |
| 0.992061292 | NA |
| 0.009383307 | 0.031915744 |
| 0.26000453 | NA |
| 0.90515732 | NA |
| 0.120284475 | 0.236550849 |
| 0.030866866 | 0.083329145 |
| 0.609687398 | NA |
| NA | NA |
| 0.373582666 | 0.53783542 |
| NA | NA |
| 2.41E-09 | 6.86E-08 |
| NA | NA |
| NA | NA |
| NA | NA |
| NA | NA |
| NA | NA |
| NA | NA |
| NA | NA |
| NA | NA |
| NA | NA |
| NA | NA |
| NA | NA |
| NA | NA |
| NA | NA |
| NA | NA |
| 0.386494918 | NA |
| 0.98798397 | NA |
| 0.886225687 | 0.935484508 |
| NA | NA |
| 0.345395093 | 0.50882402 |
| NA | NA |
| 0.927217683 | 0.958351878 |
| 0.029933905 | 0.081401687 |

| | |
|---|---|
| 0.130866521 | NA |
| 0.676676881 | 0.794746653 |
| NA | NA |
| NA | NA |
| NA | NA |
| 0.129255785 | 0.249347956 |
| 0.001211568 | 0.005848499 |
| 0.704558251 | 0.816174015 |
| 0.992060816 | NA |
| 0.026144155 | 0.073022067 |
| NA | NA |
| 0.652858565 | NA |
| 0.625325882 | NA |
| 0.889409773 | NA |
| 0.604027147 | 0.739253085 |
| 0.312738277 | 0.474564288 |
| 0.165176622 | 0.299439729 |
| 0.009604201 | 0.032556137 |
| 0.767096791 | NA |
| NA | NA |
| 2.53E-10 | 9.49E-09 |
| 0.701501067 | NA |
| 0.889410245 | NA |
| 0.769142239 | NA |
| 8.29E-07 | 1.12E-05 |
| 0.561569356 | 0.705503833 |
| 0.286842218 | 0.445898928 |
| NA | NA |
| NA | NA |
| NA | NA |
| NA | NA |
| 0.000539547 | 0.002954594 |
| 0.000157302 | 0.001024984 |
| 0.154458814 | 0.284953266 |
| NA | NA |
| 0.210425796 | 0.357338184 |
| 0.365564138 | 0.53033297 |
| 0.003416913 | 0.013978773 |
| 0.614101066 | 0.747383767 |
| 0.000105329 | 0.000730319 |
| 0.50458232 | 0.656794497 |
| NA | NA |
| 0.992059387 | NA |
| 0.45129271 | 0.609617165 |
| 0.117622951 | 0.232676046 |
| 0.879783349 | NA |
| 0.73258203 | NA |

| | |
|---|---|
| NA | NA |
| NA | NA |
| 0.240683218 | 0.393329075 |
| 1.17E-13 | 1.03E-11 |
| 0.634611217 | NA |
| 0.00218709 | 0.009649651 |
| 0.575789269 | NA |
| 0.991955612 | NA |
| NA | NA |
| NA | NA |
| 3.95E-09 | 1.04E-07 |
| NA | NA |
| 0.420336838 | 0.582161881 |
| NA | NA |
| NA | NA |
| 0.463069177 | 0.620187734 |
| 0.943292459 | 0.967842217 |
| 0.889409773 | NA |
| 0.038321141 | 0.098836497 |
| 0.625159902 | 0.755340311 |
| 0.007389404 | 0.026361862 |
| 0.708468813 | 0.818201018 |
| 0.200136871 | 0.345042439 |
| 0.370250188 | 0.534819358 |
| 0.631397429 | 0.759302088 |
| 0.000120616 | 0.000821275 |
| 0.889409301 | NA |
| 0.324436519 | 0.486699107 |
| 0.992061292 | NA |
| 2.48E-05 | 0.000212805 |
| 0.883340501 | NA |
| 0.024061906 | 0.068309684 |
| 0.702835692 | 0.814565311 |
| 0.134579413 | 0.256508057 |
| 0.29319693 | 0.45285813 |
| 0.992061292 | NA |
| 0.905158266 | NA |
| 0.520765719 | 0.671570027 |
| 0.846190918 | 0.909565146 |
| NA | NA |
| 0.397209972 | 0.560921786 |
| 0.261111786 | 0.416545856 |
| 0.737651134 | NA |
| 0.000760435 | 0.00396081 |
| 0.003739499 | 0.01502849 |
| NA | NA |
| 0.481446097 | 0.637102256 |

| | |
|---|---|
| 0.991759839 | NA |
| NA | NA |
| 1.06E-08 | 2.50E-07 |
| 0.632568118 | 0.760251946 |
| 0.094960342 | 0.198046742 |
| NA | NA |
| NA | NA |
| 0.738923841 | 0.839639257 |
| 0.538911204 | 0.686583379 |
| 0.000139741 | 0.0009279 |
| 0.324050147 | 0.486352073 |
| 0.346304265 | 0.509787179 |
| 6.30E-06 | 6.54E-05 |
| NA | NA |
| 0.038336676 | 0.098841736 |
| 0.32465283 | 0.486897565 |
| 0.566155087 | NA |
| NA | NA |
| NA | NA |
| 0.992060816 | NA |
| NA | NA |
| NA | NA |
| 0.992063809 | NA |
| 0.060048177 | 0.139800458 |
| 0.213380729 | 0.360516075 |
| 0.796625353 | NA |
| NA | NA |
| NA | NA |
| 0.816463243 | 0.890743016 |
| 0.390199004 | NA |
| 0.879279243 | NA |
| 0.727921671 | NA |
| 0.479831332 | 0.635424932 |
| NA | NA |
| 0.889409301 | NA |
| 3.37E-09 | 9.15E-08 |
| 0.021945809 | 0.063355437 |
| 7.33E-06 | 7.44E-05 |
| 0.189912241 | 0.331843948 |
| NA | NA |
| 2.27E-05 | 0.00019785 |
| 0.852086253 | NA |
| NA | NA |
| 0.62417377 | NA |
| NA | NA |
| 0.274998385 | NA |
| 0.905158739 | NA |

| | |
|---|---|
| NA | NA |
| 0.009069418 | 0.031069477 |
| 0.776806993 | NA |
| 0.141466764 | 0.266603903 |
| 0.733822356 | NA |
| 0.380321958 | 0.544567676 |
| 0.88914359 | NA |
| 0.787323853 | NA |
| 0.002913828 | 0.012257615 |
| 5.17E-06 | 5.48E-05 |
| 0.001373176 | 0.006512648 |
| 0.242049471 | 0.394788099 |
| 0.95840602 | 0.976958398 |
| 0.527585088 | 0.677341028 |
| 0.702635693 | NA |
| 0.725264826 | 0.830307868 |
| 0.320643104 | 0.482988196 |
| 0.905158266 | NA |
| 3.83E-09 | 1.02E-07 |
| 0.780523266 | 0.867735439 |
| 0.980870765 | NA |
| 0.348596709 | 0.512543684 |
| NA | NA |
| 0.036529146 | 0.095266451 |
| NA | NA |
| 0.202258773 | 0.347772328 |
| 0.063538587 | 0.145900222 |
| 0.213562403 | 0.360712007 |
| NA | NA |
| 5.50E-09 | 1.39E-07 |
| 0.0523393 | 0.125663535 |
| NA | NA |
| 0.068246569 | 0.153453451 |
| 0.279795188 | 0.437951567 |
| NA | NA |
| 0.815100579 | NA |
| 0.342141342 | NA |
| 0.18788778 | 0.32935293 |
| 0.115339917 | 0.229478051 |
| 0.80483186 | NA |
| NA | NA |
| 0.000139852 | 0.000928077 |
| 0.002416608 | 0.010513745 |
| 0.526454424 | 0.676403528 |
| 2.32E-07 | 3.69E-06 |
| 0.666842132 | 0.787437426 |
| 0.035729457 | 0.09353638 |

| | | |
|---|---|---|
| | 1.07E-06 | 1.39E-05 |
| | 1.37E-05 | 0.000128207 |
| | 6.17E-11 | 2.77E-09 |
| | 0.418841109 | 0.580933042 |
| | 0.889409301 | NA |
| | 0.000184347 | 0.001175403 |
| | 0.08954986 | 0.189443679 |
| | 1.22E-15 | 1.81E-13 |
| | 1.11E-06 | 1.43E-05 |
| | 0.867074297 | 0.923648224 |
| | 0.506557512 | 0.658389843 |
| | 0.000761028 | 0.003960989 |
| | 0.467403028 | 0.623732563 |
| | 0.262365454 | 0.417847622 |
| | 0.811599956 | 0.887325041 |
| | 0.036916083 | 0.096092966 |
| | 8.62E-07 | 1.16E-05 |
| | 1.12E-05 | 0.000107044 |
| | 0.170557724 | NA |
| | 1.21E-05 | 0.000115 |
| | 0.038307698 | 0.098825041 |
| | 0.733700307 | NA |
| | 0.889409301 | NA |
| | 0.003644771 | 0.014720531 |
| NA | | NA |
| | 0.073949267 | 0.163431155 |
| | 0.82046713 | 0.893150789 |
| | 7.73E-14 | 7.01E-12 |
| | 0.589769169 | 0.728143837 |
| | 0.276449566 | 0.43398313 |
| | 0.076853629 | NA |
| | 5.41E-07 | 7.71E-06 |
| | 0.000664273 | 0.003537042 |
| | 0.165888632 | NA |
| | 0.07544975 | 0.166095344 |
| | 0.687804192 | NA |
| | 0.884687831 | 0.934702659 |
| | 0.581580629 | 0.721664547 |
| | 0.932861878 | 0.961327957 |
| | 0.051677209 | 0.124523189 |
| | 0.115849179 | 0.230240939 |
| NA | | NA |
| | 0.018824173 | 0.055935048 |
| | 0.059214302 | 0.138181387 |
| | 0.543022639 | NA |
| | 0.905158739 | NA |
| | 0.992060816 | NA |

| | |
|---|---|
| NA | NA |
| 0.861487601 | NA |
| NA | NA |
| 0.73221552 | NA |
| NA | NA |
| 0.992060816 | NA |
| 0.00669808 | 0.024371687 |
| 0.09016509 | 0.190451636 |
| 4.84E-06 | 5.18E-05 |
| 0.576529488 | 0.718033231 |
| 0.196204982 | NA |
| 0.889409773 | NA |
| NA | NA |
| 0.103834131 | 0.211766852 |
| 0.000201715 | 0.00127247 |
| 0.100839123 | 0.207314512 |
| 0.134280517 | 0.256158924 |
| NA | NA |
| 0.018823522 | 0.055935048 |
| 4.27E-05 | 0.000338892 |
| 0.065751985 | NA |
| 0.238764749 | 0.390892054 |
| 0.000242582 | 0.001491758 |
| 0.072170712 | 0.160290954 |
| NA | NA |
| 0.761822537 | 0.855521176 |
| 0.000310357 | 0.001838946 |
| 0.24772231 | 0.401308314 |
| 0.898549122 | NA |
| 0.982113588 | 0.991173786 |
| 0.000402057 | 0.002297364 |
| NA | NA |
| 0.706462791 | NA |
| 0.882110794 | NA |
| 0.783743882 | NA |
| 0.052131556 | NA |
| 0.054055006 | 0.128755166 |
| NA | NA |
| 0.132679981 | 0.253857255 |
| 0.93323625 | 0.961613008 |
| 0.694737798 | NA |
| 0.025569626 | 0.071846886 |
| 0.002260095 | 0.009916944 |
| 0.365874135 | 0.530627445 |
| 0.12726159 | 0.246534248 |
| 0.000143821 | 0.000948398 |
| NA | NA |

| | |
|---|---|
| 0.848193152 | NA |
| 0.331364487 | 0.493752648 |
| 0.042221018 | 0.106456692 |
| 0.016617831 | 0.050747598 |
| NA | NA |
| 2.00E-09 | 5.80E-08 |
| 0.992061292 | NA |
| 0.023161702 | 0.066199117 |
| 0.919012788 | 0.953894962 |
| 0.856385839 | 0.916173469 |
| 0.382472741 | 0.546496771 |
| 0.766639699 | 0.858439143 |
| 0.014293908 | 0.044878456 |
| 0.012352117 | 0.039888256 |
| 0.000852947 | 0.004346686 |
| 0.434354811 | 0.595009188 |
| 0.778392314 | 0.866682731 |
| 0.000122396 | 0.000831329 |
| 0.000646212 | 0.003456803 |
| 0.00066453 | 0.003537554 |
| 0.000845835 | 0.004321474 |
| 0.492722266 | 0.646649025 |
| 0.005934198 | 0.022078796 |
| 2.00E-07 | 3.24E-06 |
| 1.14E-16 | 2.04E-14 |
| 0.759218025 | 0.853818939 |
| 3.23E-11 | 1.57E-09 |
| 0.721865834 | NA |
| 0.905158739 | NA |
| NA | NA |
| 0.992061292 | NA |
| NA | NA |
| NA | NA |
| NA | NA |
| NA | NA |
| NA | NA |
| NA | NA |
| NA | NA |
| NA | NA |
| NA | NA |
| NA | NA |
| 0.889409301 | NA |
| NA | NA |
| NA | NA |
| NA | NA |
| NA | NA |
| NA | NA |

| | | |
|---|---|---|
| NA | | NA |
| NA | | NA |
| | 0.889409773 | NA |
| NA | | NA |
| NA | | NA |
| NA | | NA |
| NA | | NA |
| | 0.905157793 | NA |
| NA | | NA |
| NA | | NA |
| NA | | NA |
| NA | | NA |
| NA | | NA |
| NA | | NA |
| | 0.905158266 | NA |
| NA | | NA |
| NA | | NA |
| NA | | NA |
| NA | | NA |
| NA | | NA |
| NA | | NA |
| NA | | NA |
| NA | | NA |
| NA | | NA |
| NA | | NA |
| NA | | NA |
| | 0.905158266 | NA |
| | 0.905158266 | NA |
| NA | | NA |
| NA | | NA |
| NA | | NA |
| NA | | NA |
| NA | | NA |
| | 0.889409301 | NA |
| NA | | NA |
| NA | | NA |
| NA | | NA |
| NA | | NA |
| NA | | NA |
| NA | | NA |
| NA | | NA |
| | 0.889409301 | NA |
| NA | | NA |
| NA | | NA |
| | 0.889409301 | NA |
| NA | | NA |
| | 0.889409301 | NA |

| | | |
|---|---|---|
| | 0.183953938 | 0.324490955 |
| | 0.98219698 | 0.991179388 |
| | 0.004010609 | 0.015928554 |
| | 0.152294656 | 0.282000029 |
| | 0.003235473 | 0.013374953 |
| | 0.138236521 | 0.261955106 |
| | 0.011400019 | 0.037368715 |
| NA | | NA |
| | 0.740879991 | NA |
| | 0.910720135 | NA |
| | 0.188952432 | 0.330759432 |
| | 0.622174179 | 0.753367676 |
| | 0.912349466 | NA |
| | 0.50609927 | 0.657988979 |
| | 0.005037012 | 0.019282026 |
| | 0.024243087 | 0.068744052 |
| | 0.358307463 | NA |
| | 0.357134977 | 0.521812958 |
| | 0.072673731 | 0.16116365 |
| | 0.012787128 | 0.041015563 |
| | 0.902160737 | 0.944663947 |
| | 0.000449807 | 0.002526169 |
| | 0.002999915 | 0.012564396 |
| | 0.030050729 | 0.081598116 |
| | 0.139421001 | 0.263674152 |
| | 0.976324606 | 0.987728072 |
| | 0.898380039 | NA |
| | 0.650087881 | NA |
| | 0.033497847 | 0.088923988 |
| | 0.090997172 | 0.191674523 |
| | 0.241242615 | 0.39406729 |
| | 0.137332777 | 0.260715004 |
| | 0.914513702 | NA |
| NA | | NA |
| NA | | NA |
| NA | | NA |
| | 0.950711944 | NA |
| | 0.105227712 | 0.214111872 |
| NA | | NA |
| NA | | NA |
| NA | | NA |
| NA | | NA |
| | 0.000137905 | 0.000917928 |
| | 0.991700267 | NA |
| | 0.037527951 | 0.097293538 |
| | 1.28E-06 | 1.62E-05 |
| | 0.99205891 | NA |

| | |
|---|---|
| NA | NA |
| 0.006284111 | 0.023129868 |
| 6.40E-05 | 0.000477832 |
| 0.14021937 | 0.264821291 |
| 0.000626965 | 0.003364498 |
| 0.246618611 | 0.400051541 |
| 0.086203842 | 0.183960173 |
| 2.42E-06 | 2.83E-05 |
| 0.037108924 | 0.096457726 |
| 0.045061901 | NA |
| 0.128390641 | 0.248093109 |
| 0.04805271 | 0.117582956 |
| 2.90E-05 | 0.000243621 |
| 0.001921111 | 0.00864538 |
| NA | NA |
| 0.693982316 | NA |
| 3.01E-05 | 0.000250555 |
| 0.537931006 | 0.685931535 |
| 0.410017827 | 0.573038535 |
| 0.066487525 | 0.150859417 |
| 9.47E-07 | 1.25E-05 |
| 0.028802739 | 0.078882434 |
| 0.889409301 | NA |
| 4.20E-05 | 0.000334016 |
| 4.10E-06 | 4.49E-05 |
| NA | NA |
| NA | NA |
| 0.404589159 | NA |
| 0.421532971 | 0.583156462 |
| 0.466880637 | 0.623311765 |
| 0.455588977 | 0.613495943 |
| NA | NA |
| 0.969086504 | NA |
| 4.54E-18 | 1.23E-15 |
| 0.246009896 | 0.399359157 |
| 0.008210052 | 0.028666644 |
| 0.896500894 | 0.941790776 |
| 0.293278176 | 0.45285813 |
| 0.000151786 | 0.000993561 |
| 0.586078619 | NA |
| 0.044934088 | 0.111643771 |
| 0.051588087 | 0.124387039 |
| 0.489820573 | 0.643887391 |
| 0.014684877 | 0.045889456 |
| 0.776244828 | NA |
| NA | NA |
| 0.000532467 | 0.002920919 |

| | | |
|---|---|---|
| | 0.330282567 | 0.492708892 |
| | 0.113097462 | 0.225958254 |
| | 0.914439921 | 0.951778255 |
| | 0.45049435 | 0.608909233 |
| | 0.000283225 | 0.0017058 |
| NA | NA | |
| | 0.759183531 | 0.853818939 |
| NA | NA | |
| NA | NA | |
| | 2.56E-15 | 3.41E-13 |
| | 0.032981933 | 0.087851803 |
| NA | NA | |
| | 0.992060816 | NA |
| | 0.454205174 | 0.6122191 |
| | 0.992003131 | NA |
| | 0.000453762 | 0.002542525 |
| | 0.010174472 | 0.034151825 |
| | 0.006697732 | 0.024371687 |
| | 0.003150822 | 0.013086608 |
| | 0.00071901 | 0.00377721 |
| | 0.773847952 | 0.863681178 |
| | 0.821939196 | 0.893999842 |
| | 3.39E-05 | 0.000277312 |
| | 0.035618262 | 0.093334298 |
| | 0.413670773 | NA |
| | 0.84726208 | 0.910371949 |
| | 0.592978322 | NA |
| | 0.000681207 | 0.003611449 |
| | 0.000175634 | 0.001128037 |
| | 2.62E-06 | 3.03E-05 |
| | 0.167763853 | 0.302929566 |
| | 0.564287544 | 0.707889708 |
| | 0.778267824 | 0.866682731 |
| | 0.502875689 | 0.655274222 |
| | 0.882717924 | NA |
| | 0.006487793 | 0.02374424 |
| | 0.463000062 | 0.620167035 |
| NA | NA | |
| | 0.436552053 | 0.596902382 |
| | 2.02E-11 | 1.04E-09 |
| | 0.272191062 | 0.428953892 |
| | 0.059411753 | 0.138597969 |
| | 0.778173562 | 0.866682731 |
| | 0.779790837 | NA |
| | 0.821401037 | NA |
| | 0.718290468 | NA |
| | 0.00095508 | 0.004793064 |

| | |
|---|---|
| NA | NA |
| 0.920196385 | 0.954311534 |
| 5.09E-05 | 0.000392247 |
| 0.003843954 | 0.015361124 |
| NA | NA |
| 0.889409301 | NA |
| 0.003227606 | 0.013357522 |
| NA | NA |
| 0.948544534 | 0.970962588 |
| 0.028655457 | 0.078547645 |
| 0.806786545 | NA |
| 0.273872412 | NA |
| 0.905158266 | NA |
| 0.574643683 | NA |
| 0.025527304 | 0.071755512 |
| 0.905158266 | NA |
| NA | NA |
| NA | NA |
| 0.59021913 | NA |
| NA | NA |
| NA | NA |
| 0.685990165 | NA |
| NA | NA |
| 1.16E-06 | 1.48E-05 |
| NA | NA |
| 0.166919767 | 0.301827335 |
| 4.49E-08 | 8.87E-07 |
| 0.742818495 | NA |
| NA | NA |
| NA | NA |
| NA | NA |
| NA | NA |
| 0.301183085 | NA |
| 0.000528806 | 0.002904463 |
| NA | NA |
| 0.554341776 | 0.699746554 |
| NA | NA |
| 0.002554244 | 0.011006406 |
| 0.428098544 | NA |
| 0.97792622 | 0.98868174 |
| 4.91E-05 | 0.000381355 |
| 0.169972835 | 0.306087706 |
| 0.149452281 | 0.277891406 |
| NA | NA |
| 0.90397203 | 0.945658702 |
| NA | NA |
| NA | NA |

| | | |
|---|---|---|
| | 0.875456202 | 0.928844683 |
| | 0.078575698 | NA |
| | 0.051097523 | 0.123451481 |
| | 0.237742513 | NA |
| | 0.600652637 | NA |
| | 0.556856401 | 0.701992635 |
| | 0.777133351 | 0.865851705 |
| | 0.873616163 | NA |
| | 0.889410245 | NA |
| | 0.836681161 | NA |
| | 5.94E-05 | 0.000448831 |
| | 0.889409773 | NA |
| | 9.69E-06 | 9.45E-05 |
| | 0.049341946 | 0.120108613 |
| | 0.842428725 | NA |
| NA | | NA |
| | 0.083391091 | 0.179492743 |
| NA | | NA |
| | 0.905158739 | NA |
| | 0.115974042 | 0.230468234 |
| | 0.057114288 | NA |
| NA | | NA |
| NA | | NA |
| | 0.435032488 | NA |
| NA | | NA |
| | 0.254794311 | NA |
| NA | | NA |
| NA | | NA |
| NA | | NA |
| NA | | NA |
| NA | | NA |
| NA | | NA |
| | 0.083429904 | 0.179538611 |
| NA | | NA |
| NA | | NA |
| NA | | NA |
| NA | | NA |
| NA | | NA |
| | 3.89E-09 | 1.03E-07 |
| | 0.057699678 | NA |
| | 0.002649775 | 0.011362313 |
| | 0.019564394 | 0.057759414 |
| | 0.439941537 | 0.600177563 |
| NA | | NA |
| NA | | NA |
| NA | | NA |
| | 0.862371391 | 0.920513969 |

| | |
|---|---|
| 0.009709888 | 0.032833299 |
| 0.000140471 | 0.000930777 |
| 0.51242026 | 0.664083834 |
| 1.33E-05 | 0.000124493 |
| NA | NA |
| 0.00024146 | 0.001486048 |
| 0.628619339 | 0.757080681 |
| 1.63E-13 | 1.39E-11 |
| 0.002663477 | 0.011412154 |
| 0.00010934 | 0.000753372 |
| 0.899240667 | 0.943360683 |
| NA | NA |
| NA | NA |
| NA | NA |
| 0.625370341 | 0.755340311 |
| 0.691766277 | NA |
| 0.881935857 | NA |
| 0.905157793 | NA |
| NA | NA |
| 0.519076404 | NA |
| NA | NA |
| 0.638379323 | NA |
| 0.410807114 | NA |
| 0.646326058 | NA |
| 0.061306186 | 0.142126535 |
| NA | NA |
| 0.037035657 | 0.096316824 |
| NA | NA |
| 0.000130998 | 0.000877278 |
| 0.801988994 | 0.880631879 |
| 0.302550007 | 0.463265853 |
| 0.443208957 | 0.602850934 |
| 0.533951538 | 0.682720498 |
| 0.966179509 | NA |
| 0.019549513 | NA |
| 0.985236626 | NA |
| 0.369782939 | NA |
| 0.992061292 | NA |
| 0.807879735 | NA |
| 0.784353866 | NA |
| 0.889409773 | NA |
| 0.007579197 | 0.026868017 |
| NA | NA |
| NA | NA |
| 0.012996477 | 0.041509362 |
| 0.917667314 | 0.953109456 |
| NA | NA |

| | | |
|---|---|---|
| | 0.047420027 | NA |
| | 0.542643148 | NA |
| | 0.067725996 | 0.152626759 |
| | 0.718434192 | 0.8253614 |
| NA | | NA |
| | 0.799743672 | 0.879229303 |
| | 0.45349533 | 0.611539176 |
| | 0.079101883 | 0.172458126 |
| | 0.473715508 | 0.630061711 |
| NA | | NA |
| | 0.000675461 | 0.003585315 |
| NA | | NA |
| | 0.735848873 | 0.837836015 |
| | 0.036143403 | 0.094473634 |
| | 0.992061292 | NA |
| NA | | NA |
| NA | | NA |
| | 0.0360934 | 0.094365396 |
| | 0.837128034 | 0.903172899 |
| NA | | NA |
| | 0.746238761 | NA |
| | 0.961788312 | NA |
| | 0.010546891 | 0.035181404 |
| NA | | NA |
| NA | | NA |
| | 0.789176268 | 0.872787041 |
| | 0.013001328 | 0.041509362 |
| | 0.603668118 | 0.73906937 |
| | 0.756624085 | 0.852080019 |
| NA | | NA |
| NA | | NA |
| NA | | NA |
| NA | | NA |
| | 0.0022422 | 0.009853206 |
| | 0.193438893 | 0.33648009 |
| | 0.253020732 | 0.407367275 |
| | 0.336505329 | NA |
| | 0.747070183 | NA |
| | 0.992060816 | NA |
| | 0.278690628 | 0.436845421 |
| | 8.06E-07 | 1.09E-05 |
| NA | | NA |
| | 0.880258492 | NA |
| NA | | NA |
| | 0.031400272 | 0.084489472 |
| | 0.788582525 | NA |
| | 0.941009306 | NA |

| | | |
|---|---|---|
| | 0.853807602 | NA |
| | 0.395867627 | NA |
| NA | | NA |
| | 0.211164761 | 0.358253636 |
| | 0.613162511 | 0.746448363 |
| | 0.889409773 | NA |
| | 0.679877227 | 0.797093552 |
| | 0.801661681 | NA |
| NA | | NA |
| | 0.076057959 | 0.167216332 |
| NA | | NA |
| | 0.908870836 | 0.948233586 |
| | 0.000286768 | 0.001721471 |
| | 3.18E-09 | 8.74E-08 |
| | 0.068317536 | 0.153573061 |
| | 0.986167737 | NA |
| | 0.000695996 | 0.003671307 |
| | 0.905158266 | NA |
| NA | | NA |
| | 8.12E-07 | 1.10E-05 |
| | 0.003529189 | 0.014354039 |
| | 0.635768082 | 0.763012041 |
| | 0.021069386 | 0.06129606 |
| | 0.732116637 | NA |
| | 0.802608826 | NA |
| | 0.031347008 | 0.084377169 |
| | 0.083622025 | 0.179795951 |
| | 0.016402997 | 0.050222171 |
| | 0.412142522 | 0.574764235 |
| | 0.094755051 | 0.197693698 |
| | 0.24557854 | 0.398918058 |
| | 0.005110298 | 0.019511917 |
| | 0.905158739 | NA |
| | 0.300271378 | 0.460644068 |
| | 0.982950839 | NA |
| NA | | NA |
| NA | | NA |
| | 0.11496471 | 0.228835212 |
| | 0.002765169 | 0.011756116 |
| | 6.45E-07 | 8.94E-06 |
| NA | | NA |
| | 0.283442583 | NA |
| NA | | NA |
| | 0.797157316 | NA |
| | 0.945402596 | NA |
| | 0.905158739 | NA |
| NA | | NA |

| | |
|---|---|
| NA | NA |
| NA | NA |
| NA | NA |
| NA | NA |
| 0.246502147 | NA |
| NA | NA |
| 0.860037372 | NA |
| 7.96E-09 | 1.93E-07 |
| NA | NA |
| NA | NA |
| NA | NA |
| 0.353667978 | 0.517953524 |
| 0.580995077 | 0.721169113 |
| 0.00375146 | 0.015065528 |
| 0.99205891 | NA |
| NA | NA |
| 0.100485997 | 0.206705743 |
| 0.87792082 | NA |
| NA | NA |
| NA | NA |
| NA | NA |
| NA | NA |
| NA | NA |
| NA | NA |
| NA | NA |
| NA | NA |
| 0.494910372 | 0.648162275 |
| NA | NA |
| 0.043610407 | 0.109045882 |
| NA | NA |
| 6.85E-23 | 5.37E-20 |
| 0.503054285 | NA |
| NA | NA |
| NA | NA |
| 0.613645158 | NA |
| 0.905158266 | NA |
| NA | NA |
| 0.004599292 | 0.017850097 |
| 0.602579884 | NA |
| 0.006012259 | 0.022320066 |
| 0.329810237 | 0.492265991 |
| 0.000960003 | 0.004811753 |
| 0.211010372 | 0.358109341 |
| NA | NA |
| 0.99205891 | NA |
| NA | NA |
| 0.992037435 | NA |

| | | |
|---|---|---|
| | 0.388697703 | 0.55209033 |
| | 0.007349926 | 0.026256228 |
| | 0.190295988 | 0.33230295 |
| | 0.165718435 | 0.300099878 |
| | 0.594451694 | 0.73170206 |
| | 0.937897107 | 0.964831003 |
| NA | | NA |
| | 0.953462824 | NA |
| | 0.97595898 | 0.987653037 |
| | 0.294029122 | NA |
| | 0.975891371 | NA |
| | 0.847074558 | 0.910247625 |
| | 2.11E-05 | 0.00018613 |
| | 0.014371577 | 0.045051459 |
| | 0.554138566 | NA |
| NA | | NA |
| NA | | NA |
| | 2.43E-09 | 6.90E-08 |
| | 0.889409773 | NA |
| | 0.033726185 | 0.089396192 |
| | 0.056457133 | 0.133033822 |
| | 0.641530662 | 0.767369638 |
| | 0.485865667 | NA |
| | 0.373854994 | 0.537868166 |
| | 0.205676337 | 0.351942154 |
| NA | | NA |
| | 0.23257949 | NA |
| | 0.869918176 | NA |
| | 0.05064371 | 0.122611382 |
| | 0.889409301 | NA |
| | 0.99405864 | 0.997117284 |
| | 0.173331537 | 0.310511076 |
| | 2.63E-05 | 0.000223765 |
| | 0.40884115 | 0.572063478 |
| | 0.12850775 | 0.248253865 |
| NA | | NA |
| | 0.618159112 | 0.750360752 |
| NA | | NA |
| NA | | NA |
| | 0.000118007 | 0.000805006 |
| | 0.003839415 | 0.015345779 |
| | 0.992060816 | NA |
| | 0.812145995 | 0.887745229 |
| NA | | NA |
| | 0.444665979 | 0.604121774 |
| | 0.965960804 | NA |
| | 4.71E-22 | 3.04E-19 |

| | | |
|---|---|---|
| | 0.766017073 | NA |
| | 0.889409773 | NA |
| | 0.693024074 | NA |
| | 0.627525435 | 0.75642702 |
| | 2.62E-05 | 0.000222972 |
| | 0.023393603 | 0.066766328 |
| | 0.876452546 | 0.929490073 |
| | 0.905158266 | NA |
| | 0.056709569 | 0.133485415 |
| | 0.977610367 | NA |
| NA | | NA |
| | 0.045755638 | NA |
| | 6.89E-13 | 5.10E-11 |
| | 0.012802429 | 0.041052838 |
| | 0.769521988 | 0.860690334 |
| | 0.005571738 | 0.020961077 |
| | 0.630499664 | 0.758596357 |
| NA | | NA |
| NA | | NA |
| | 0.905158266 | NA |
| NA | | NA |
| NA | | NA |
| NA | | NA |
| | 0.819989805 | NA |
| | 0.476419994 | 0.632674587 |
| NA | | NA |
| | 0.769065734 | NA |
| | 0.992061292 | NA |
| | 0.808193729 | 0.88498742 |
| | 0.096920501 | 0.201065403 |
| | 0.67702537 | 0.795058564 |
| | 0.155693024 | 0.286552938 |
| | 0.30300986 | 0.463678991 |
| | 9.74E-05 | 0.000682707 |
| | 0.954174304 | 0.974417042 |
| | 0.012307081 | 0.039777919 |
| | 0.830163713 | NA |
| | 0.988042113 | NA |
| NA | | NA |
| | 2.75E-06 | 3.16E-05 |
| | 0.992061292 | NA |
| | 0.889409301 | NA |
| | 0.143574637 | 0.269650655 |
| | 0.228209975 | 0.378437006 |
| | 0.011046523 | 0.036471996 |
| NA | | NA |
| | 0.96976566 | NA |

| | |
|---|---|
| 5.79E-07 | 8.14E-06 |
| 0.889409301 | NA |
| 0.001596858 | 0.007377456 |
| 1.29E-05 | 0.000121564 |
| NA | NA |
| NA | NA |
| NA | NA |
| 0.170762872 | 0.307233257 |
| 0.028631769 | NA |
| NA | NA |
| 0.92365727 | 0.955959371 |
| 0.243664544 | 0.396607348 |
| 0.761808995 | 0.855521176 |
| 0.483866894 | NA |
| 0.880605919 | NA |
| 0.296535778 | 0.456700039 |
| 0.001254439 | 0.006020199 |
| 0.821014528 | 0.89361378 |
| NA | NA |
| 1.20E-11 | 6.51E-10 |
| NA | NA |
| NA | NA |
| NA | NA |
| 0.992060816 | NA |
| 0.992060816 | NA |
| 0.905158266 | NA |
| 0.191875917 | 0.334359234 |
| 0.905157793 | NA |
| 0.702385324 | 0.814301216 |
| NA | NA |
| NA | NA |
| NA | NA |
| NA | NA |
| 0.70739976 | NA |
| NA | NA |
| NA | NA |
| NA | NA |
| 0.242524885 | 0.395426488 |
| 3.29E-07 | 4.98E-06 |
| 0.295723883 | 0.455769283 |
| 0.156883205 | 0.288091511 |
| 0.024149508 | 0.068531796 |
| NA | NA |
| 0.046269056 | 0.114236811 |
| NA | NA |
| NA | NA |
| 0.054938996 | 0.130365104 |

| | |
|---|---|
| NA | NA |
| 6.00E-05 | 0.0004527 |
| NA | NA |
| 1.55E-07 | 2.60E-06 |
| 1.32E-07 | 2.26E-06 |
| 0.684572113 | 0.800411002 |
| 5.14E-07 | 7.38E-06 |
| NA | NA |
| 0.982094475 | NA |
| NA | NA |
| 0.319486764 | 0.481907532 |
| 0.285267198 | NA |
| 0.531775123 | NA |
| NA | NA |
| 0.905158739 | NA |
| NA | NA |
| 0.025770387 | 0.072290743 |
| 0.815310987 | 0.890052894 |
| NA | NA |
| 0.220201824 | 0.369115408 |
| 0.000690556 | 0.003651318 |
| NA | NA |
| 0.005563331 | 0.02094021 |
| NA | NA |
| NA | NA |
| NA | NA |
| 0.780016604 | 0.867560808 |
| 0.000213914 | 0.00133594 |
| 0.005004071 | 0.019197133 |
| 0.387649329 | NA |
| NA | NA |
| 0.905158739 | NA |
| 0.337191467 | 0.500195056 |
| 0.768855135 | 0.860149545 |
| NA | NA |
| 0.201155569 | NA |
| 0.07003993 | 0.156587948 |
| 0.565850109 | NA |
| 0.133611801 | NA |
| NA | NA |
| 0.276150037 | 0.433671113 |
| NA | NA |
| NA | NA |
| NA | NA |
| NA | NA |
| NA | NA |
| 0.121504285 | 0.238281628 |

| | | |
|---|---|---|
| NA | NA | |
| NA | NA | |
| | 0.889409301 | NA |
| | 0.992061292 | NA |
| | 0.330461815 | NA |
| NA | NA | |
| | 0.000804444 | 0.004147638 |
| | 0.217552206 | 0.365691912 |
| | 0.266218453 | 0.42242149 |
| | 0.048012959 | NA |
| NA | NA | |
| | 0.179239658 | 0.318667759 |
| | 0.000292001 | 0.001747145 |
| | 0.429962772 | NA |
| | 0.391349436 | NA |
| | 0.988888173 | NA |
| | 0.045553306 | 0.11286303 |
| | 0.697003588 | NA |
| | 4.10E-13 | 3.25E-11 |
| | 4.70E-08 | 9.23E-07 |
| | 0.099513313 | 0.205241841 |
| | 0.021209082 | 0.061620831 |
| | 0.992253023 | NA |
| | 0.728146312 | NA |
| | 2.53E-08 | 5.34E-07 |
| | 0.000752246 | 0.003927801 |
| | 0.37292053 | 0.537249061 |
| | 0.022153114 | 0.063823175 |
| | 0.74504658 | NA |
| NA | NA | |
| | 0.852144444 | NA |
| NA | NA | |
| | 0.990963629 | NA |
| | 0.991879747 | NA |
| | 0.31718274 | NA |
| NA | NA | |
| | 0.99205891 | NA |
| | 0.58504919 | 0.72455559 |
| | 0.086856263 | NA |
| | 0.050897313 | 0.123103216 |
| NA | NA | |
| | 7.01E-08 | 1.30E-06 |
| | 0.146396913 | NA |
| | 0.081927017 | 0.177157595 |
| | 0.639449696 | 0.766174271 |
| | 0.905157793 | NA |
| NA | NA | |

| | | |
|---|---|---|
| 0.905157793 | NA | |
| NA | NA | |
| 0.185182689 | NA | |
| 0.672037032 | | 0.791231633 |
| 3.79E-07 | | 5.64E-06 |
| NA | NA | |
| NA | NA | |
| NA | NA | |
| 0.341363659 | | 0.504784484 |
| 1.81E-05 | | 0.000162801 |
| 0.135625647 | | 0.258210818 |
| 0.000768112 | | 0.003991505 |
| 4.42E-05 | | 0.000349056 |
| 2.70E-10 | | 1.01E-08 |
| 3.72E-07 | | 5.53E-06 |
| 1.24E-05 | | 0.000117276 |
| 2.43E-05 | | 0.000209402 |
| 0.117062025 | | 0.231916728 |
| 5.25E-06 | | 5.55E-05 |
| 0.30285277 | NA | |
| 0.000195415 | | 0.001236633 |
| 5.76E-07 | | 8.11E-06 |
| 0.119556313 | NA | |
| 0.000530631 | | 0.002912762 |
| 0.638336523 | NA | |
| NA | NA | |
| 0.193931256 | NA | |
| 0.905158266 | NA | |
| 0.633029199 | NA | |
| 0.804248727 | | 0.882408019 |
| 6.52E-05 | | 0.000485197 |
| 0.380527611 | | 0.544720066 |
| 0.752403888 | NA | |
| 0.371776538 | NA | |
| 0.992037435 | NA | |
| 0.632740913 | NA | |
| NA | NA | |
| 0.905158739 | NA | |
| NA | NA | |
| 0.905158266 | NA | |
| NA | NA | |
| NA | NA | |
| 0.889409301 | NA | |
| NA | NA | |
| NA | NA | |
| NA | NA | |
| NA | NA | |

| | | |
|---|---|---|
| | 0.905158266 | NA |
| NA | | NA |
| | 0.000156663 | 0.001021727 |
| | 0.869989105 | 0.925515189 |
| | 0.160493157 | 0.293005423 |
| | 0.293868648 | 0.453514769 |
| NA | | NA |
| NA | | NA |
| | 0.889409773 | NA |
| | 0.001065576 | 0.005254658 |
| | 8.63E-06 | 8.60E-05 |
| | 0.992061292 | NA |
| | 0.370790165 | 0.535351784 |
| | 0.002946498 | 0.012373715 |
| | 0.519676784 | 0.670559555 |
| | 0.830163713 | NA |
| | 0.000549252 | 0.003000254 |
| NA | | NA |
| | 0.000975164 | 0.004872271 |
| | 3.52E-05 | 0.000286616 |
| | 0.02449865 | 0.069352309 |
| | 0.889410717 | NA |
| | 0.000154378 | 0.001008324 |
| | 0.995402821 | 0.997694248 |
| | 0.102367625 | 0.209696459 |
| | 0.007054864 | 0.025412074 |
| | 0.992060816 | NA |
| | 0.826333325 | NA |
| | 0.759956679 | NA |
| | 0.02095847 | 0.061054263 |
| | 0.758032868 | 0.853272695 |
| NA | | NA |
| | 0.031211802 | 0.084033836 |
| | 1.46E-09 | 4.38E-08 |
| NA | | NA |
| | 0.889409301 | NA |
| NA | | NA |
| | 0.000181037 | 0.001156647 |
| | 0.212801207 | 0.359974826 |
| NA | | NA |
| NA | | NA |
| | 0.231548402 | 0.382499915 |
| | 6.85E-07 | 9.45E-06 |
| | 0.816156606 | 0.890666083 |
| | 0.889409773 | NA |
| | 0.924071833 | NA |
| | 0.992060816 | NA |

| | | |
|---|---|---|
| | 0.889409773 | NA |
| NA | | NA |
| | 0.905158266 | NA |
| NA | | NA |
| NA | | NA |
| NA | | NA |
| NA | | NA |
| | 0.194958108 | 0.338505977 |
| | 0.327906319 | NA |
| NA | | NA |
| | 0.905157793 | NA |
| NA | | NA |
| NA | | NA |
| | 0.905158266 | NA |
| | 0.592921357 | NA |
| | 0.369488426 | NA |
| | 0.92058741 | 0.954536733 |
| | 0.495338068 | 0.648606359 |
| | 0.939013327 | 0.965481746 |
| | 0.889446478 | 0.937385064 |
| | 0.096054988 | 0.199760534 |
| | 0.885536211 | 0.93507114 |
| | 0.00012841 | 0.000863897 |
| | 0.08424398 | 0.180641058 |
| | 0.153951519 | 0.28430085 |
| | 0.38389262 | 0.547822852 |
| | 0.016163177 | 0.049641447 |
| | 0.38611809 | 0.549891746 |
| | 0.591723157 | 0.729447528 |
| | 0.253429878 | 0.407794434 |
| | 0.730879845 | 0.834459624 |
| NA | | NA |
| | 0.987576569 | 0.993587391 |
| | 0.19337155 | 0.336416246 |
| | 0.427705506 | NA |
| | 0.109028657 | 0.219918853 |
| NA | | NA |
| | 0.85072595 | 0.912519105 |
| | 0.824204798 | 0.895142918 |
| | 0.360193182 | NA |
| NA | | NA |
| NA | | NA |
| | 0.765315663 | 0.857689428 |
| | 0.905158266 | NA |
| NA | | NA |
| | 0.077185512 | 0.169170241 |
| | 0.684736303 | 0.80056035 |

| | |
|---|---|
| 0.053168565 | 0.12705719 |
| 0.057623595 | 0.13520064 |
| 0.005827105 | 0.021765165 |
| 0.042849132 | 0.10765722 |
| 0.624794911 | 0.755333156 |
| NA | NA |
| NA | NA |
| 0.120624644 | 0.237028774 |
| 0.656400642 | NA |
| 0.391358779 | 0.554708763 |
| 1.10E-05 | 0.000105609 |
| 0.129785525 | 0.249923029 |
| 0.889409301 | NA |
| 0.235663497 | 0.387403408 |
| 0.049327682 | 0.120087202 |
| 0.875697524 | NA |
| 0.720958085 | NA |
| 0.07964787 | 0.173273438 |
| 0.083533897 | 0.179676839 |
| 0.006096712 | NA |
| 0.031150295 | 0.083909386 |
| 0.018819207 | 0.055935048 |
| 0.824127872 | NA |
| 5.77E-23 | 4.70E-20 |
| 0.224105679 | 0.373776136 |
| 0.836911501 | NA |
| 0.360690638 | 0.52533883 |
| 0.266700701 | 0.422911662 |
| 0.00010495 | 0.00072792 |
| 0.797178279 | NA |
| NA | NA |
| 0.386186919 | NA |
| 0.850588513 | 0.912416256 |
| NA | NA |
| 0.12190801 | NA |
| 0.208003968 | NA |
| NA | NA |
| 0.486833294 | NA |
| 0.017315811 | 0.052385306 |
| 0.210284269 | 0.357224495 |
| 0.550296692 | 0.696121036 |
| 0.875232801 | 0.928844683 |
| NA | NA |
| NA | NA |
| NA | NA |
| 0.635453426 | 0.762801169 |
| 0.105690579 | 0.214754581 |

| | | |
|---|---|---|
| | 0.491991522 | 0.646071824 |
| | 2.96E-12 | 1.90E-10 |
| | 0.727164891 | 0.831570349 |
| | 0.84070747 | NA |
| NA | | NA |
| NA | | NA |
| | 0.057902934 | NA |
| | 0.888274886 | NA |
| | 0.992060816 | NA |
| | 0.462766464 | 0.620037257 |
| | 0.358111416 | NA |
| | 0.64089725 | 0.767071603 |
| | 0.317620171 | 0.479916145 |
| NA | | NA |
| | 0.017686357 | 0.053247175 |
| | 0.735262385 | 0.837385338 |
| | 8.02E-05 | 0.000578735 |
| NA | | NA |
| | 0.094393265 | 0.197088686 |
| | 0.581347067 | NA |
| | 0.246377001 | 0.399807313 |
| | 0.797157316 | NA |
| | 0.095673934 | 0.199137813 |
| NA | | NA |
| | 0.074465673 | 0.164340875 |
| | 0.005156552 | 0.01967554 |
| | 0.60692473 | 0.741298247 |
| | 0.99205891 | NA |
| | 0.01695989 | 0.051603468 |
| | 0.984035801 | 0.991970261 |
| | 0.659394359 | 0.781623359 |
| | 0.868009253 | 0.924373662 |
| NA | | NA |
| | 0.992800006 | NA |
| | 0.88853917 | NA |
| | 0.044496107 | 0.110768623 |
| | 0.889410717 | NA |
| | 0.825990791 | 0.896231435 |
| | 0.788418414 | 0.872361178 |
| | 0.022763427 | 0.065304305 |
| | 0.019800427 | 0.058330772 |
| NA | | NA |
| | 5.54E-10 | 1.88E-08 |
| | 0.789530107 | NA |
| NA | | NA |
| NA | | NA |
| | 0.388728901 | 0.55209033 |

| | | |
|---|---|---|
| | 0.991918092 | NA |
| | 0.905158266 | NA |
| | 0.587552261 | 0.726293159 |
| | 0.61131338 | 0.744981992 |
| | 0.74504658 | NA |
| NA | | NA |
| | 0.398656808 | NA |
| NA | | NA |
| | 3.44E-08 | 7.06E-07 |
| | 0.992060816 | NA |
| | 4.23E-08 | 8.42E-07 |
| | 9.26E-06 | 9.12E-05 |
| | 0.252662921 | NA |
| NA | | NA |
| | 0.006167348 | 0.022772623 |
| NA | | NA |
| | 0.8503631 | NA |
| | 0.88388815 | NA |
| | 0.988120282 | 0.993639148 |
| | 0.314576678 | 0.476662247 |
| | 0.889410717 | NA |
| NA | | NA |
| | 0.991918092 | NA |
| NA | | NA |
| NA | | NA |
| | 0.017686821 | 0.053247175 |
| | 0.861071122 | NA |
| | 0.963732711 | NA |
| | 0.862297989 | 0.920480363 |
| | 2.85E-07 | 4.41E-06 |
| | 0.66692388 | 0.787491611 |
| | 0.975189343 | 0.987192641 |
| | 0.00027218 | 0.001648771 |
| | 7.74E-06 | 7.81E-05 |
| | 6.12E-10 | 2.05E-08 |
| | 0.951544297 | NA |
| | 0.410510127 | 0.573556042 |
| | 0.147634582 | 0.275396674 |
| | 0.208194481 | 0.354921405 |
| | 0.000706323 | 0.00371946 |
| | 0.31758576 | NA |
| | 0.938717182 | 0.965414356 |
| NA | | NA |
| | 0.805374319 | NA |
| | 0.000797545 | 0.004116897 |
| NA | | NA |
| NA | | NA |

| | | |
|---|---|---|
| NA | NA | |
| 0.889409301 | NA | |
| NA | NA | |
| 0.2702696 | | 0.426752725 |
| 0.889409301 | NA | |
| NA | NA | |
| 0.992062245 | NA | |
| 0.992060816 | NA | |
| NA | NA | |
| NA | NA | |
| 0.48386455 | NA | |
| 0.905157793 | NA | |
| 0.889410245 | NA | |
| 0.048552026 | NA | |
| NA | NA | |
| NA | NA | |
| NA | NA | |
| NA | NA | |
| NA | NA | |
| NA | NA | |
| NA | NA | |
| NA | NA | |
| 0.905158266 | NA | |
| NA | NA | |
| 0.71665909 | | 0.824028931 |
| NA | NA | |
| NA | NA | |
| NA | NA | |
| 0.233898846 | | 0.385378497 |
| 0.325528127 | | 0.487736909 |
| 0.959133328 | | 0.977283164 |
| 0.18441077 | | 0.324998082 |
| 0.87762688 | NA | |
| 0.992060816 | NA | |
| 0.92121559 | NA | |
| 0.231419006 | | 0.382430009 |
| 0.001076668 | | 0.005299835 |
| 0.000907747 | | 0.004590957 |
| 0.024578956 | | 0.069543281 |
| 0.552525177 | | 0.698057088 |
| 0.094459375 | | 0.197207969 |
| 0.207410039 | | 0.354106442 |
| 0.104361062 | | 0.212683486 |
| 0.589687462 | | 0.728123424 |
| 5.42E-06 | | 5.71E-05 |
| 0.763195925 | NA | |
| 0.529606063 | | 0.67896394 |

| | | |
|---|---|---|
| | 0.069399005 | 0.155455753 |
| | 0.200015056 | 0.344913665 |
| | 0.827184039 | 0.89672859 |
| | 0.801167114 | 0.880155299 |
| | 0.06561873 | 0.149411209 |
| | 0.278954135 | 0.437102458 |
| | 0.111685038 | 0.223665458 |
| | 0.86156518 | NA |
| | 0.10373998 | 0.211732152 |
| NA | | NA |
| | 0.893186486 | 0.93965803 |
| NA | | NA |
| | 0.000225289 | 0.001398283 |
| | 0.813795952 | 0.889061955 |
| | 0.429769121 | 0.590902269 |
| | 0.213290043 | 0.360403712 |
| NA | | NA |
| | 0.50409647 | 0.656381713 |
| | 0.066589486 | 0.150993635 |
| | 0.000267568 | 0.001623071 |
| | 0.80402271 | NA |
| | 0.586842013 | NA |
| NA | | NA |
| NA | | NA |
| | 0.688443794 | 0.8036541 |
| | 0.981340122 | NA |
| | 0.983718166 | NA |
| | 0.792530476 | 0.874922236 |
| | 0.010550579 | 0.035188362 |
| | 0.039762195 | 0.10165225 |
| | 0.718853331 | NA |
| | 0.685484103 | 0.801050791 |
| | 0.355860483 | 0.520435588 |
| | 0.992061292 | NA |
| | 0.897540122 | NA |
| | 0.343040434 | 0.506374354 |
| | 0.432087278 | 0.592975722 |
| | 0.001627519 | 0.007492389 |
| | 0.768526595 | 0.859883598 |
| NA | | NA |
| | 0.221103633 | 0.370090303 |
| | 0.468244815 | 0.624639323 |
| | 0.641458553 | 0.767366984 |
| NA | | NA |
| NA | | NA |
| | 0.004553417 | 0.017693947 |
| | 0.320954093 | NA |

| | |
|---|---|
| 0.476794768 | 0.632968763 |
| NA | NA |
| 0.1658226 | 0.300206825 |
| 0.092181704 | 0.193520024 |
| 0.00972812 | 0.032873718 |
| 0.694182195 | 0.807800181 |
| 0.840730375 | NA |
| 0.011153305 | 0.036746502 |
| 0.730514185 | 0.834357692 |
| 0.005216326 | 0.019858755 |
| NA | NA |
| NA | NA |
| NA | NA |
| 0.992061292 | NA |
| NA | NA |
| NA | NA |
| NA | NA |
| NA | NA |
| NA | NA |
| NA | NA |
| NA | NA |
| NA | NA |
| NA | NA |
| NA | NA |
| NA | NA |
| NA | NA |
| NA | NA |
| 0.007532737 | 0.026754685 |
| 2.90E-10 | 1.08E-08 |
| 0.377013075 | 0.541383083 |
| 0.781622775 | NA |
| 0.992060816 | NA |
| 0.368733412 | 0.533365192 |
| 0.000163891 | 0.00106162 |
| 0.02334488 | 0.066644593 |
| 0.992060816 | NA |
| 0.905158739 | NA |
| 0.801661681 | NA |
| 0.08509794 | 0.182077916 |
| 0.533546541 | 0.682401333 |
| 0.660948215 | 0.782877278 |
| 0.146657899 | 0.27401181 |
| 2.00E-05 | 0.000177999 |
| 2.21E-05 | 0.000194299 |
| 0.323423931 | 0.485792961 |
| 0.01764914 | 0.053162889 |
| NA | NA |

| | | |
|---|---|---|
| | 7.96E-05 | 0.000575303 |
| | 0.754950656 | NA |
| | 0.380437315 | 0.54466182 |
| | 0.044375798 | 0.110563326 |
| | 0.198419701 | NA |
| NA | | NA |
| | 0.03432991 | 0.09067242 |
| | 0.441466751 | NA |
| | 1.39E-08 | 3.16E-07 |
| | 0.696320689 | 0.809706949 |
| | 5.22E-09 | 1.32E-07 |
| | 0.859935388 | 0.918449474 |
| | 0.26116716 | 0.416603928 |
| | 0.005420124 | 0.020492883 |
| | 0.266094198 | NA |
| | 0.64260267 | 0.768317109 |
| | 0.130504245 | 0.250874788 |
| | 8.64E-07 | 1.16E-05 |
| | 0.101742059 | 0.208740902 |
| | 0.002038857 | 0.009098001 |
| | 0.013597533 | 0.043059083 |
| | 0.833834466 | NA |
| | 0.493408385 | 0.646928273 |
| NA | | NA |
| | 0.531931557 | NA |
| | 0.630956756 | 0.758884411 |
| | 0.905158266 | NA |
| | 0.049541124 | 0.120513297 |
| | 0.026664482 | 0.074104322 |
| | 0.001898132 | 0.008556986 |
| | 0.155385687 | 0.286227187 |
| | 0.000259647 | 0.0015807 |
| | 0.055725259 | 0.131733335 |
| | 0.248817142 | 0.402603242 |
| | 0.2977991 | NA |
| NA | | NA |
| | 0.443087733 | 0.602798062 |
| | 0.338544473 | 0.501524528 |
| | 0.28119913 | 0.439634849 |
| NA | | NA |
| | 8.32E-05 | 0.000596285 |
| | 0.014509038 | 0.045417529 |
| | 0.521328736 | 0.672019825 |
| | 0.630356504 | 0.758530175 |
| | 0.692279857 | NA |
| | 0.076373768 | 0.167779588 |
| | 0.456483326 | 0.614210622 |

| | |
|---|---|
| NA | NA |
| 1.94E-07 | 3.16E-06 |
| 0.766104361 | NA |
| 0.943210954 | 0.96780427 |
| 0.918781737 | NA |
| 0.324487918 | 0.486713739 |
| 0.992061292 | NA |
| 0.116078607 | 0.230592558 |
| 0.283999921 | NA |
| NA | NA |
| NA | NA |
| 0.050002504 | 0.121366749 |
| 0.58595332 | 0.725225395 |
| 0.019725722 | 0.058149701 |
| 0.673222335 | 0.792202414 |
| NA | NA |
| 0.212341632 | 0.359479601 |
| NA | NA |
| 0.749458241 | NA |
| 0.180792086 | 0.320683544 |
| 0.553386407 | NA |
| NA | NA |
| 0.047305993 | 0.116109122 |
| 0.184599453 | NA |
| 0.889409773 | NA |
| NA | NA |
| 0.82010098 | 0.892909192 |
| 0.985204901 | 0.992529528 |
| 0.54300678 | 0.689888809 |
| 0.992061292 | NA |
| NA | NA |
| 0.791880097 | 0.874648913 |
| 0.615416324 | NA |
| 0.001973137 | 0.008846084 |
| 0.256498219 | 0.411097321 |
| NA | NA |
| 0.50435988 | NA |
| 0.9630344 | NA |
| 0.349548721 | 0.513599221 |
| 5.24E-07 | 7.51E-06 |
| 0.038127275 | 0.098452895 |
| 0.812186064 | NA |
| 0.005777342 | 0.0216234 |
| 0.905158266 | NA |
| 0.875378438 | 0.928844683 |
| 0.001327748 | 0.006327041 |
| NA | NA |

| | |
|---|---|
| 9.00E-06 | 8.90E-05 |
| 0.684083637 | 0.800052865 |
| 0.171954201 | 0.308693779 |
| 0.003490087 | 0.014223984 |
| 0.041487627 | 0.104993292 |
| 0.020711695 | 0.060478629 |
| 0.507815605 | 0.65967362 |
| NA | NA |
| 0.285573257 | 0.444604356 |
| NA | NA |
| 0.990164483 | NA |
| 0.07943928 | 0.172987619 |
| 0.000281742 | 0.001698732 |
| 0.225991078 | 0.375779045 |
| 0.001875284 | 0.00847312 |
| NA | NA |
| 0.989118263 | NA |
| 0.163053928 | 0.296497285 |
| 0.889409301 | NA |
| 0.01608253 | 0.049461663 |
| 0.000163707 | 0.001061051 |
| 0.033164587 | 0.088263381 |
| 8.89E-05 | 0.000630379 |
| 0.764356561 | NA |
| NA | NA |
| 4.52E-07 | 6.60E-06 |
| 0.989458783 | NA |
| 0.064398005 | 0.147411216 |
| 0.321039657 | 0.483353434 |
| 0.401743822 | NA |
| 0.037575723 | NA |
| 0.222839673 | 0.372234313 |
| 0.754469038 | 0.850659353 |
| 0.021293991 | 0.061810276 |
| 0.035606803 | 0.09331577 |
| 0.889409773 | NA |
| 0.348733926 | NA |
| NA | NA |
| 0.644198949 | 0.769806525 |
| 0.00615167 | 0.022726199 |
| 0.99205891 | NA |
| 0.313786071 | NA |
| 8.55E-10 | 2.75E-08 |
| 0.201878834 | 0.347282203 |
| 0.069598158 | NA |
| NA | NA |
| 0.173822777 | 0.3111051 |

| | |
|---|---|
| 5.71E-05 | 0.000434619 |
| 0.009508185 | 0.032275503 |
| 0.927343333 | NA |
| 0.17360859 | 0.310875889 |
| 0.900897778 | NA |
| 0.475370754 | NA |
| 0.15497975 | 0.285660112 |
| 0.765927595 | 0.858084513 |
| 2.02E-06 | 2.42E-05 |
| 0.001921283 | 0.00864538 |
| NA | NA |
| NA | NA |
| 0.017870953 | 0.053722742 |
| 0.927941996 | 0.958676937 |
| 8.84E-05 | 0.00062745 |
| 0.318155313 | 0.480559398 |
| 0.008654507 | 0.02993767 |
| 0.167649369 | 0.30279754 |
| 0.779435515 | NA |
| 0.98584549 | NA |
| 6.15E-10 | 2.06E-08 |
| 0.001718298 | 0.007847777 |
| 0.110252817 | 0.221605639 |
| 0.049469799 | 0.120393141 |
| 0.118236479 | 0.233527149 |
| 2.48E-14 | 2.56E-12 |
| 0.185759815 | 0.32673068 |
| 0.520608374 | 0.671445982 |
| NA | NA |
| NA | NA |
| 0.378493362 | 0.543017572 |
| 0.521693619 | 0.672332307 |
| 0.000452152 | 0.002536095 |
| 0.80651194 | 0.8840511 |
| 0.038299716 | 0.098816058 |
| 3.68E-10 | 1.33E-08 |
| 0.200821502 | 0.345978296 |
| 0.02092028 | 0.060964639 |
| 0.501968213 | 0.654672448 |
| 0.230553663 | 0.381344372 |
| 0.051588105 | 0.124387039 |
| 0.001613779 | 0.007436933 |
| 0.986672905 | NA |
| 0.235947762 | 0.387667557 |
| 0.667074965 | 0.787551955 |
| 0.905158739 | NA |
| 0.020713986 | 0.060478629 |

| | |
|---|---|
| 0.170014808 | 0.306138186 |
| 7.79E-07 | 1.06E-05 |
| 0.521743296 | 0.672356868 |
| 0.416718572 | 0.578939841 |
| 0.000246394 | 0.001511332 |
| NA | NA |
| NA | NA |
| 0.656783026 | 0.779499431 |
| NA | NA |
| 0.240873434 | NA |
| 0.503644518 | 0.655923727 |
| 0.559490252 | NA |
| 1.58E-08 | 3.50E-07 |
| 0.000588309 | 0.003188224 |
| NA | NA |
| 0.000219137 | 0.001363957 |
| NA | NA |
| 0.000114968 | 0.000786721 |
| 0.304338152 | 0.464965413 |
| 0.08039434 | 0.174634416 |
| 0.212727452 | 0.359910788 |
| 7.76E-06 | 7.82E-05 |
| 0.889409773 | NA |
| 0.009853825 | 0.033217707 |
| 0.755618593 | NA |
| 2.62E-05 | 0.000223289 |
| 0.085927552 | 0.183513238 |
| 0.999610333 | 0.999864481 |
| 0.391370411 | 0.554708763 |
| NA | NA |
| 0.131836503 | 0.252727872 |
| 0.000199479 | 0.001259087 |
| 4.85E-05 | 0.000377656 |
| 7.65E-06 | 7.73E-05 |
| 0.986683334 | NA |
| 0.001405635 | NA |
| 9.97E-10 | 3.14E-08 |
| 0.002661868 | 0.011407484 |
| NA | NA |
| 0.307734167 | NA |
| 0.10404948 | 0.212146938 |
| 0.859288329 | 0.91798163 |
| 0.051415426 | 0.124096651 |
| 0.038944194 | 0.099997141 |
| 0.01649651 | 0.050452208 |
| NA | NA |
| 0.98358726 | NA |

| | | |
|---|---|---|
| | 0.103572542 | 0.211526329 |
| | 0.601143017 | 0.73671724 |
| | 8.45E-08 | 1.52E-06 |
| | 2.95E-06 | 3.35E-05 |
| | 7.22E-13 | 5.30E-11 |
| | 0.156018171 | 0.286955075 |
| | 3.75E-06 | 4.15E-05 |
| NA | | NA |
| | 1.00E-10 | 4.26E-09 |
| | 0.770230233 | 0.861091697 |
| | 0.165624662 | NA |
| NA | | NA |
| | 0.027556864 | 0.076154188 |
| | 0.119447484 | 0.235326332 |
| | 3.10E-05 | 0.000257515 |
| | 0.641516044 | 0.767369638 |
| | 0.068328421 | 0.153574594 |
| | 0.889409301 | NA |
| | 0.606823896 | NA |
| | 0.262300076 | NA |
| | 0.456534398 | 0.614241704 |
| | 0.921355838 | 0.954934663 |
| | 0.281022367 | 0.439402604 |
| | 0.019151282 | 0.056707526 |
| | 0.115756865 | NA |
| | 0.38004608 | 0.544304363 |
| | 0.175302623 | 0.313117806 |
| | 0.787549252 | 0.871771467 |
| NA | | NA |
| | 0.905158266 | NA |
| | 0.030630536 | 0.082824274 |
| NA | | NA |
| | 0.002433203 | 0.01057615 |
| | 0.011077549 | 0.036557244 |
| | 0.001893325 | 0.008544751 |
| | 0.654076138 | 0.777511713 |
| | 0.301678111 | NA |
| | 0.821979411 | NA |
| | 0.973765012 | NA |
| | 0.000862305 | 0.004389282 |
| | 0.412703362 | NA |
| | 0.000673726 | 0.003577835 |
| | 0.000607583 | 0.003275696 |
| | 0.568196972 | NA |
| | 0.446312892 | 0.605647312 |
| | 0.338731814 | NA |
| | 0.956505922 | NA |

| | | |
|---|---|---|
| | 0.797178279 | NA |
| | 0.003605882 | 0.014606449 |
| | 1.10E-05 | 0.000105338 |
| | 7.19E-08 | 1.32E-06 |
| | 0.817761836 | 0.891575997 |
| | 0.005427504 | 0.02051688 |
| | 0.371687001 | 0.536048526 |
| | 4.44E-15 | 5.60E-13 |
| | 0.043306359 | 0.10848327 |
| | 0.006052788 | 0.022432602 |
| NA | | NA |
| | 0.791283264 | NA |
| NA | | NA |
| NA | | NA |
| | 0.912714454 | 0.950637798 |
| | 2.23E-05 | 0.00019523 |
| | 0.445993413 | 0.605438263 |
| | 0.915781804 | 0.95229975 |
| | 0.355580725 | 0.520165021 |
| | 0.698835136 | NA |
| | 0.051973157 | 0.125003127 |
| | 0.028463034 | 0.078137236 |
| | 0.236804096 | 0.388609308 |
| | 5.01E-05 | 0.000387854 |
| | 0.64894602 | 0.773640908 |
| | 2.31E-07 | 3.67E-06 |
| | 8.91E-06 | 8.84E-05 |
| | 0.654311447 | 0.777658858 |
| NA | | NA |
| NA | | NA |
| | 0.585883372 | NA |
| | 2.13E-07 | 3.42E-06 |
| NA | | NA |
| NA | | NA |
| NA | | NA |
| | 0.215665689 | NA |
| | 0.635596573 | NA |
| | 0.138394004 | NA |
| NA | | NA |
| | 0.818110796 | 0.891724796 |
| | 0.005237504 | 0.019918661 |
| NA | | NA |
| | 0.807879735 | NA |
| NA | | NA |
| | 0.994355368 | NA |
| | 0.077339515 | 0.169355644 |
| | 0.026154825 | 0.073033292 |

| | | |
|---|---|---|
| | 0.825988355 | 0.896231435 |
| | 0.124666835 | 0.242820564 |
| | 0.893344989 | 0.939689721 |
| | 0.029565293 | 0.080528934 |
| | 3.18E-05 | 0.000263544 |
| | 0.268412189 | NA |
| NA | | NA |
| | 5.24E-05 | 0.000402459 |
| | 0.598780059 | NA |
| NA | | NA |
| | 0.239559896 | 0.391784893 |
| NA | | NA |
| | 0.651199154 | 0.775597019 |
| | 0.519657343 | NA |
| | 9.12E-12 | 5.15E-10 |
| | 0.024372719 | 0.069049225 |
| | 0.408903336 | 0.572063478 |
| | 0.192857808 | NA |
| | 0.162584799 | 0.295864723 |
| NA | | NA |
| | 0.249877555 | NA |
| | 1.60E-08 | 3.54E-07 |
| | 0.797343143 | 0.877636997 |
| | 0.775847069 | 0.864857393 |
| NA | | NA |
| | 0.674044197 | 0.792829638 |
| | 0.129837306 | 0.249932253 |
| NA | | NA |
| | 0.98881981 | 0.993934551 |
| | 0.109218331 | 0.220211673 |
| | 0.100486534 | 0.206705743 |
| | 3.04E-07 | 4.65E-06 |
| | 0.003950406 | 0.015717877 |
| | 0.67514305 | 0.793442132 |
| | 0.190295131 | 0.33230295 |
| | 0.00210695 | 0.00935244 |
| | 0.005998077 | 0.022286255 |
| | 0.000125089 | 0.000845235 |
| | 3.69E-06 | 4.10E-05 |
| | 0.456942038 | 0.614601874 |
| NA | | NA |
| | 0.8075516 | 0.884660792 |
| | 0.699833633 | 0.81254411 |
| | 0.906588228 | 0.947328653 |
| | 0.997169979 | NA |
| NA | | NA |
| NA | | NA |

| | | |
|---|---|---|
| NA | | NA |
| | 0.795014225 | NA |
| NA | | NA |
| NA | | NA |
| NA | | NA |
| NA | | NA |
| NA | | NA |
| NA | | NA |
| NA | | NA |
| NA | | NA |
| NA | | NA |
| NA | | NA |
| NA | | NA |
| NA | | NA |
| NA | | NA |
| NA | | NA |
| | 0.813836939 | NA |
| NA | | NA |
| | 0.905158739 | NA |
| NA | | NA |
| | 0.905158266 | NA |
| NA | | NA |
| NA | | NA |
| NA | | NA |
| NA | | NA |
| NA | | NA |
| NA | | NA |
| NA | | NA |
| NA | | NA |
| | 0.992061292 | NA |
| NA | | NA |
| NA | | NA |
| NA | | NA |
| | 0.498907403 | NA |
| NA | | NA |
| NA | | NA |
| | 0.992061292 | NA |
| NA | | NA |
| NA | | NA |
| NA | | NA |
| NA | | NA |
| NA | | NA |
| | 0.905158266 | NA |
| | 0.889409301 | NA |
| NA | | NA |
| NA | | NA |
| NA | | NA |

| | |
|---|---|
| NA | NA |
| 0.905158266 | NA |
| 0.905158266 | NA |
| 0.992060816 | NA |
| 0.042589958 | 0.107168665 |
| 0.764791962 | NA |
| 2.46E-05 | 0.000211807 |
| 0.183045777 | 0.323319034 |
| 0.092902465 | 0.194642231 |
| NA | NA |
| 0.889409773 | NA |
| 1.26E-06 | 1.59E-05 |
| 0.055095107 | 0.130581052 |
| 9.80E-07 | 1.29E-05 |
| NA | NA |
| 0.001021064 | 0.005067014 |
| 0.290146037 | 0.449475613 |
| 0.215884564 | 0.363655196 |
| 0.000983294 | 0.004907307 |
| 0.002719118 | 0.011594001 |
| NA | NA |
| NA | NA |
| NA | NA |
| 0.324915358 | 0.487051426 |
| 0.575465976 | 0.717225632 |
| 1.47E-05 | 0.00013589 |
| 0.153218433 | 0.283328301 |
| 0.461171253 | 0.618585275 |
| 0.216456502 | 0.364422934 |
| 0.029009099 | 0.079261228 |
| 0.3373001 | 0.500300794 |
| 0.034977384 | 0.092017297 |
| 0.029400634 | 0.080179896 |
| 0.992060816 | NA |
| 0.004680924 | 0.018147669 |
| 0.011381866 | 0.03731478 |
| 0.90515732 | NA |
| 0.692886405 | NA |
| 0.308941231 | 0.470069324 |
| 3.80E-05 | 0.000306453 |
| 0.90843699 | 0.948154039 |
| 0.114153151 | 0.227694846 |
| NA | NA |
| 0.605198521 | 0.740200196 |
| 0.692571996 | 0.806617113 |
| 1.10E-07 | 1.91E-06 |
| 2.99E-07 | 4.59E-06 |

| | |
|---|---|
| 0.312949271 | 0.474818838 |
| 0.43028002 | 0.591410398 |
| NA | NA |
| 0.241378699 | 0.394172295 |
| 0.000120951 | 0.000823296 |
| 1.59E-05 | 0.00014598 |
| 7.99E-05 | 0.000577466 |
| 0.972979174 | 0.985912235 |
| 0.992060816 | NA |
| 0.0546677 | 0.129896689 |
| 0.113781532 | 0.227077039 |
| 0.689293523 | 0.804261237 |
| 0.031017927 | 0.083655448 |
| 0.047672996 | 0.116888267 |
| 0.289209999 | 0.448563524 |
| 0.483811749 | 0.639338968 |
| 6.46E-06 | 6.69E-05 |
| NA | NA |
| 0.983459885 | 0.991828033 |
| 0.830922139 | 0.899227207 |
| 0.903553342 | 0.945355815 |
| 0.003112327 | 0.012951222 |
| 0.096936437 | 0.201079465 |
| 0.01237946 | 0.039970676 |
| 1.07E-06 | 1.38E-05 |
| 0.52490711 | NA |
| 0.952543747 | 0.973374154 |
| 0.65374779 | NA |
| 0.105263748 | 0.214143106 |
| 0.396076764 | 0.559721869 |
| 0.200077457 | 0.344994179 |
| 0.228436964 | 0.378727618 |
| 0.855979635 | NA |
| 0.561146985 | 0.705296013 |
| 0.444426038 | 0.603982633 |
| 0.667412856 | 0.787772463 |
| NA | NA |
| 0.904415224 | 0.945942077 |
| NA | NA |
| 0.753716766 | NA |
| 0.02122349 | 0.061646378 |
| 0.230398138 | NA |
| 0.792026402 | 0.874678523 |
| 0.009437171 | 0.032066607 |
| NA | NA |
| 0.001515803 | 0.00706699 |
| 2.36E-07 | 3.74E-06 |

| | | |
|---|---|---|
| | 0.528857012 | 0.678379059 |
| | 0.973245434 | 0.985912235 |
| | 0.398596524 | 0.562192888 |
| | 0.05670321 | 0.133484755 |
| | 0.133873739 | 0.255628995 |
| | 0.947127371 | NA |
| | 0.000108949 | 0.000751147 |
| | 0.508928376 | 0.660650166 |
| NA | | NA |
| NA | | NA |
| | 0.761042477 | NA |
| | 0.000349543 | 0.002040843 |
| | 0.851410992 | NA |
| | 0.471563166 | NA |
| | 7.49E-05 | 0.000546124 |
| | 0.826709774 | NA |
| | 0.067158524 | 0.151789488 |
| | 0.144067547 | 0.270414573 |
| | 0.003414487 | 0.013972926 |
| | 0.066129427 | 0.150309086 |
| | 0.521520702 | 0.67218836 |
| | 0.027060471 | 0.074999479 |
| | 0.057289687 | 0.134562439 |
| NA | | NA |
| NA | | NA |
| | 0.739096709 | NA |
| | 0.002759948 | 0.011738468 |
| | 0.01746381 | 0.052749353 |
| | 0.599014773 | NA |
| | 0.082626698 | 0.178389377 |
| | 0.099748166 | 0.205591326 |
| | 3.32E-05 | 0.000272578 |
| NA | | NA |
| | 0.891911585 | 0.938990232 |
| | 0.781128952 | NA |
| | 0.364835561 | 0.529635003 |
| | 0.135744695 | 0.258415064 |
| | 0.573690083 | 0.715776167 |
| | 0.223602978 | 0.373221175 |
| | 0.199554666 | 0.344351331 |
| | 0.119772258 | 0.235839223 |
| | 0.115699632 | 0.230006176 |
| | 0.891248483 | NA |
| | 0.000297987 | 0.001775727 |
| | 0.992061292 | NA |
| | 0.377254977 | 0.541624207 |
| | 0.000271416 | 0.0016449 |

| | |
|---|---|
| 0.198894546 | 0.343547969 |
| 0.232025056 | 0.38308558 |
| 0.75708901 | NA |
| NA | NA |
| NA | NA |
| 0.889409301 | NA |
| 0.431080344 | 0.591970318 |
| 0.005085609 | 0.019435317 |
| 0.472509282 | 0.628838262 |
| 2.23E-05 | 0.000195022 |
| 0.332972034 | 0.495542724 |
| 0.573786302 | 0.715814873 |
| 8.39E-06 | 8.39E-05 |
| 0.000831556 | 0.004266195 |
| 0.785651581 | 0.871119873 |
| 0.286707892 | 0.445736985 |
| 0.706238893 | NA |
| 0.003812308 | 0.015262438 |
| NA | NA |
| 0.282098278 | 0.440616323 |
| NA | NA |
| 0.366628326 | 0.5313703 |
| NA | NA |
| NA | NA |
| 1.53E-07 | 2.57E-06 |
| 0.000790428 | 0.004084003 |
| 0.00538749 | 0.02040074 |
| 0.000157054 | 0.001023672 |
| 0.179326788 | NA |
| NA | NA |
| 0.000126455 | 0.000851642 |
| NA | NA |
| 0.243806096 | 0.396778928 |
| NA | NA |
| NA | NA |
| 0.001901622 | 0.008569201 |
| 1.02E-14 | 1.16E-12 |
| 0.01433093 | 0.044963704 |
| NA | NA |
| 0.001996675 | 0.008944305 |
| 3.10E-06 | 3.50E-05 |
| 0.118744824 | 0.234257263 |
| 0.889410245 | NA |
| NA | NA |
| NA | NA |
| 0.889409301 | NA |
| 0.655643077 | NA |

| | | |
|---|---|---|
| | 0.992061292 | NA |
| | 0.673446469 | NA |
| | 0.4669875 | NA |
| | 0.746238761 | NA |
| NA | | NA |
| | 0.986570185 | NA |
| | 0.048702228 | 0.118867648 |
| | 0.579032787 | 0.719702325 |
| | 0.216316579 | NA |
| | 0.000769094 | 0.003994449 |
| | 0.026894152 | 0.074623286 |
| | 0.029834448 | 0.081181491 |
| | 1.62E-05 | 0.000148264 |
| | 0.645963697 | NA |
| | 8.87E-13 | 6.36E-11 |
| | 0.871288332 | 0.926316278 |
| | 0.488955624 | 0.643305506 |
| | 9.58E-06 | 9.37E-05 |
| | 0.782612462 | 0.869046121 |
| | 5.39E-05 | 0.000412709 |
| | 0.000447747 | 0.00251653 |
| | 0.004835965 | 0.018653251 |
| | 0.888670512 | 0.937061982 |
| | 0.017911034 | 0.05381152 |
| NA | | NA |
| | 0.33823974 | NA |
| | 0.889409301 | NA |
| | 3.71E-13 | 2.97E-11 |
| NA | | NA |
| | 0.992061292 | NA |
| NA | | NA |
| | 0.026405256 | 0.073526885 |
| NA | | NA |
| | 0.244660472 | NA |
| | 0.537772228 | 0.685825679 |
| | 0.905158739 | NA |
| | 0.163977618 | 0.297782378 |
| | 0.003870789 | 0.015462736 |
| | 0.09180431 | 0.193022869 |
| | 0.798616291 | 0.878509926 |
| | 0.000106475 | 0.000736869 |
| | 0.854565982 | NA |
| | 0.584064485 | 0.723825951 |
| NA | | NA |
| NA | | NA |
| | 0.015192191 | 0.047186044 |
| NA | | NA |

| | |
|---|---|
| NA | NA |
| 0.093770747 | NA |
| 0.283499212 | 0.442330757 |
| 0.530124437 | 0.679409437 |
| 2.38E-05 | 0.000205523 |
| 0.11364842 | 0.226893869 |
| 0.38944172 | 0.55285823 |
| 0.873114763 | NA |
| 0.158997236 | 0.29099151 |
| 0.575530409 | 0.717257222 |
| 0.716492379 | 0.823956855 |
| 0.998192065 | NA |
| 5.42E-08 | 1.04E-06 |
| 0.632594352 | NA |
| 1.16E-05 | 0.00011017 |
| 0.10105503 | NA |
| 3.51E-07 | 5.27E-06 |
| 2.35E-08 | 5.00E-07 |
| 4.63E-06 | 4.99E-05 |
| 0.319321019 | NA |
| 0.000179227 | 0.001147083 |
| 0.000395882 | 0.002266208 |
| 2.27E-06 | 2.68E-05 |
| 0.304339724 | 0.464965413 |
| 0.000279207 | 0.001686221 |
| 0.023783024 | 0.067605362 |
| 0.046600358 | 0.114745153 |
| 0.730277291 | NA |
| 0.0652207 | 0.1488754 |
| 0.877984648 | 0.930694292 |
| 0.022239505 | 0.064013276 |
| 0.11820491 | 0.233485799 |
| NA | NA |
| 1.51E-07 | 2.53E-06 |
| 0.328668536 | 0.490934792 |
| 0.685208971 | 0.80089976 |
| 0.000357652 | 0.002080444 |
| 0.201167485 | 0.346358752 |
| 0.000518171 | 0.002851047 |
| 0.786778373 | NA |
| 0.194924727 | 0.33847478 |
| 0.000437575 | 0.002467569 |
| 0.096812228 | 0.200960871 |
| 3.70E-07 | 5.52E-06 |
| NA | NA |
| 0.843530637 | 0.907714959 |
| 0.084122868 | 0.180554595 |

| | | |
|---|---|---|
| | 0.905158266 | NA |
| NA | | NA |
| | 0.905158739 | NA |
| NA | | NA |
| NA | | NA |
| | 2.41E-05 | 0.000207852 |
| | 0.457647846 | 0.61507145 |
| | 0.588421834 | NA |
| | 0.023673732 | 0.067346998 |
| NA | | NA |
| | 0.992061292 | NA |
| | 0.503898272 | NA |
| NA | | NA |
| | 0.793985797 | 0.875697344 |
| | 0.039436258 | 0.100994154 |
| | 0.496634783 | 0.649955492 |
| | 0.203097321 | 0.348879255 |
| | 0.030179599 | 0.081856942 |
| | 0.717223913 | 0.824405742 |
| | 0.788582525 | NA |
| | 0.31976477 | 0.482134779 |
| | 0.720536159 | 0.826659013 |
| | 0.444961267 | 0.604373382 |
| | 0.535528956 | 0.683861383 |
| NA | | NA |
| NA | | NA |
| | 8.01E-05 | 0.000578278 |
| | 0.477718396 | 0.633696886 |
| | 0.678662856 | NA |
| | 0.1966049 | 0.340611203 |
| | 0.316452978 | 0.478514732 |
| | 0.666622906 | 0.787386413 |
| | 0.929508976 | NA |
| | 0.00394581 | 0.015705282 |
| | 0.028725295 | 0.078699781 |
| | 0.174503178 | 0.311994405 |
| | 0.049836764 | 0.121058131 |
| | 0.416093135 | 0.578478925 |
| | 0.001264059 | 0.006060583 |
| NA | | NA |
| | 0.136932669 | NA |
| | 0.326402552 | 0.488680278 |
| | 0.496502014 | NA |
| | 0.457702223 | 0.615096885 |
| NA | | NA |
| | 0.95734063 | 0.976416297 |
| | 4.47E-13 | 3.51E-11 |

| | | |
|---|---|---|
| | 0.453435822 | NA |
| | 0.992060816 | NA |
| NA | | NA |
| NA | | NA |
| | 0.060164943 | 0.140057456 |
| | 0.170066506 | NA |
| | 0.716172507 | 0.823761567 |
| NA | | NA |
| | 0.877504438 | NA |
| NA | | NA |
| | 0.286288051 | 0.445352742 |
| | 0.09239012 | 0.19388328 |
| | 2.43E-06 | 2.84E-05 |
| | 0.391335907 | 0.554708763 |
| | 0.034753245 | 0.091594515 |
| NA | | NA |
| NA | | NA |
| NA | | NA |
| | 0.030780813 | 0.083152692 |
| | 0.183312896 | 0.323686707 |
| | 0.084167954 | 0.180560702 |
| | 3.45E-09 | 9.34E-08 |
| NA | | NA |
| | 0.226886722 | 0.376897309 |
| | 0.033792158 | 0.089542899 |
| | 0.118483916 | 0.233889635 |
| | 0.016710595 | 0.050982068 |
| | 0.410286923 | 0.573306015 |
| NA | | NA |
| | 0.396690221 | 0.560331954 |
| | 0.725538412 | NA |
| | 0.871895345 | 0.926581323 |
| | 0.992061292 | NA |
| | 0.990967281 | 0.995364126 |
| | 0.119234096 | 0.234969176 |
| | 0.712294559 | NA |
| | 0.90515576 | NA |
| | 4.96E-12 | 2.98E-10 |
| | 0.017568043 | 0.052976746 |
| NA | | NA |
| | 0.081826316 | 0.177026229 |
| | 0.000139939 | 0.000928374 |
| | 4.50E-05 | 0.000354409 |
| | 0.513963188 | 0.665337361 |
| | 0.72768377 | 0.831947097 |
| | 0.807433834 | NA |
| NA | | NA |

| | | |
|---|---|---|
| | 0.889409301 | NA |
| NA | | NA |
| | 0.811608988 | NA |
| NA | | NA |
| | 0.829174742 | NA |
| | 0.065871069 | 0.149845934 |
| | 0.065373021 | 0.149083619 |
| | 0.19897866 | 0.343666226 |
| | 0.977344761 | 0.988372185 |
| NA | | NA |
| | 0.863822773 | 0.921444133 |
| | 0.799375391 | 0.879075598 |
| | 0.000595198 | 0.003223171 |
| | 0.949980327 | NA |
| | 0.788582525 | NA |
| | 6.31E-05 | 0.000472153 |
| | 5.87E-05 | 0.000444346 |
| | 0.797246587 | NA |
| | 0.035061668 | 0.092085538 |
| | 0.024976386 | 0.070486629 |
| | 0.288651379 | NA |
| | 0.868634498 | NA |
| | 0.905158739 | NA |
| | 0.366398744 | NA |
| NA | | NA |
| | 0.905158266 | NA |
| | 0.015862709 | 0.048901823 |
| | 0.175877849 | 0.313992008 |
| | 0.239206643 | 0.391469569 |
| | 1.71E-05 | 0.000154978 |
| | 0.10920635 | 0.220207737 |
| | 0.000121021 | 0.000823518 |
| | 0.832646984 | 0.900383864 |
| | 0.009467598 | 0.032152641 |
| | 0.98522123 | NA |
| NA | | NA |
| | 0.076512758 | 0.167997566 |
| NA | | NA |
| | 0.401004881 | NA |
| | 0.883242602 | 0.933693587 |
| | 0.078670298 | 0.171773 |
| NA | | NA |
| | 0.106885084 | 0.216481236 |
| | 0.992062245 | NA |
| NA | | NA |
| NA | | NA |
| | 8.35E-08 | 1.51E-06 |

| | | |
|---|---|---|
| | 0.992060816 | NA |
| | 0.003017852 | 0.012625076 |
| | 0.992061292 | NA |
| | 0.807783094 | NA |
| | 9.71E-11 | 4.16E-09 |
| | 4.53E-05 | 0.000356045 |
| | 0.726513632 | 0.831171869 |
| NA | | NA |
| | 0.905158739 | NA |
| NA | | NA |
| NA | | NA |
| | 0.110154705 | 0.22149266 |
| | 0.617920733 | NA |
| | 0.000995784 | 0.004959498 |
| | 0.004311768 | 0.016904501 |
| | 0.031028753 | 0.08367437 |
| | 0.775000847 | 0.864221186 |
| NA | | NA |
| NA | | NA |
| | 1.61E-11 | 8.51E-10 |
| | 5.88E-26 | 9.23E-23 |
| | 0.182051711 | 0.322237105 |
| | 0.171090694 | 0.307520713 |
| | 0.743935955 | 0.843153014 |
| | 0.592431647 | 0.729926607 |
| | 0.472705997 | 0.629023815 |
| NA | | NA |
| | 0.84276425 | NA |
| NA | | NA |
| NA | | NA |
| NA | | NA |
| NA | | NA |
| NA | | NA |
| NA | | NA |
| NA | | NA |
| | 0.003060265 | 0.012766026 |
| NA | | NA |
| | 0.889409773 | NA |
| NA | | NA |
| | 0.017152109 | 0.052029722 |
| | 0.010641316 | 0.035415982 |
| | 0.187914489 | 0.32935293 |
| | 0.905158266 | NA |
| | 0.002241753 | 0.009853206 |
| | 9.65E-11 | 4.15E-09 |
| | 0.017228914 | 0.052183412 |
| | 0.025992284 | 0.072671832 |

| | | |
|---|---|---|
| | 0.02838265 | 0.077945798 |
| | 0.224721959 | 0.374377474 |
| | 0.992060816 | NA |
| | 0.012494511 | 0.040271095 |
| | 0.108961648 | 0.219831875 |
| | 3.55E-08 | 7.26E-07 |
| | 0.405802337 | 0.569066577 |
| | 0.844175976 | 0.908199512 |
| | 0.068720714 | 0.154156518 |
| | 0.053324357 | 0.12729096 |
| | 0.11845766 | 0.233871323 |
| | 0.140972684 | 0.265855305 |
| | 0.774544634 | 0.864060335 |
| NA | | NA |
| | 2.93E-05 | 0.000245276 |
| | 0.013643086 | 0.043168375 |
| | 0.385257296 | 0.549127935 |
| | 0.013849559 | 0.043727171 |
| | 0.563741682 | 0.707361916 |
| | 0.986480782 | NA |
| | 0.715446545 | 0.82306235 |
| | 0.72868229 | NA |
| | 0.889409773 | NA |
| NA | | NA |
| NA | | NA |
| NA | | NA |
| NA | | NA |
| NA | | NA |
| NA | | NA |
| | 3.54E-06 | 3.96E-05 |
| NA | | NA |
| | 0.582158768 | 0.722116273 |
| NA | | NA |
| | 8.44E-09 | 2.04E-07 |
| | 0.024023693 | 0.068236485 |
| | 4.57E-07 | 6.66E-06 |
| | 0.220588067 | NA |
| | 0.02422934 | 0.068713946 |
| | 0.275125145 | NA |
| | 0.251842045 | 0.406065017 |
| | 0.093394582 | 0.195430877 |
| | 0.31594394 | NA |
| | 0.125247857 | NA |
| | 0.716512545 | NA |
| | 0.885547585 | 0.93507114 |
| | 0.724976943 | 0.830297927 |
| | 0.653947441 | NA |

| | |
|---|---|
| 0.002586462 | 0.011123407 |
| 0.727145196 | NA |
| 0.075227 | 0.165688034 |
| 0.234722431 | 0.386203346 |
| NA | NA |
| 5.61E-08 | 1.07E-06 |
| 0.039954726 | 0.102019282 |
| NA | NA |
| 0.002111647 | 0.009371394 |
| 0.416332684 | 0.578659792 |
| 0.257767098 | 0.412558871 |
| 0.309207486 | 0.470346617 |
| 0.017379596 | 0.052536664 |
| 0.127894044 | 0.247416555 |
| 0.257560948 | 0.412382332 |
| 0.003384218 | 0.013869736 |
| 0.008992611 | 0.03085449 |
| 5.99E-05 | 0.00045206 |
| 0.056117455 | NA |
| 0.189436955 | 0.331303448 |
| 0.106589841 | NA |
| NA | NA |
| NA | NA |
| 0.561538796 | NA |
| NA | NA |
| NA | NA |
| NA | NA |
| NA | NA |
| NA | NA |
| 0.465838106 | 0.622639173 |
| 0.000878259 | 0.004458089 |
| 0.685918706 | 0.801476876 |
| NA | NA |
| 0.20276473 | NA |
| 0.082886932 | 0.178834475 |
| NA | NA |
| NA | NA |
| NA | NA |
| NA | NA |
| 0.650610771 | 0.775148495 |
| 0.90515732 | NA |
| NA | NA |
| 0.889409301 | NA |
| 0.378114453 | 0.542609807 |
| 0.359886207 | 0.524576251 |
| 0.876401577 | NA |
| 0.359523894 | 0.524361397 |

| | | |
|---|---|---|
| | 0.404967905 | 0.568186712 |
| | 0.869016738 | NA |
| | 0.054383504 | 0.129355217 |
| | 0.199691457 | 0.344517969 |
| | 0.852397399 | NA |
| | 0.371264126 | 0.535614542 |
| | 0.19306433 | 0.336014871 |
| | 0.905158739 | NA |
| | 0.889409301 | NA |
| | 0.745804642 | NA |
| | 0.973951977 | 0.986212831 |
| | 0.494595281 | NA |
| | 0.784667491 | NA |
| | 0.000684741 | 0.003626683 |
| | 0.015659699 | 0.048384878 |
| | 0.96796276 | 0.982776689 |
| | 0.883265034 | 0.933693587 |
| | 0.237603118 | 0.38948394 |
| | 0.992061769 | NA |
| | 0.769737978 | NA |
| NA | | NA |
| | 0.634099925 | 0.761676071 |
| | 0.954164555 | 0.974417042 |
| | 0.236169136 | 0.388002248 |
| | 0.940194971 | NA |
| | 0.008574767 | 0.029708631 |
| | 0.175182268 | 0.312979206 |
| | 0.797475246 | 0.877738406 |
| | 0.317883682 | NA |
| | 0.229250033 | 0.37966525 |
| | 0.010527017 | 0.035120444 |
| | 0.97746857 | NA |
| | 0.017769344 | 0.053473622 |
| | 0.973318333 | 0.985912235 |
| | 0.819774545 | 0.892907912 |
| | 0.897217929 | 0.942074668 |
| NA | | NA |
| | 0.043721505 | 0.109249035 |
| | 0.889409773 | NA |
| | 0.9950149 | 0.997551495 |
| NA | | NA |
| | 5.37E-05 | 0.000410986 |
| | 0.972933391 | 0.985912235 |
| | 0.665564898 | 0.786521343 |
| | 0.648769586 | 0.773500453 |
| | 0.000372099 | 0.00214685 |
| | 0.905158266 | NA |

| | |
|---|---|
| 0.153765444 | 0.284037931 |
| 0.003380012 | 0.013857669 |
| 3.97E-05 | 0.000317855 |
| 0.887686749 | 0.936474337 |
| 1.86E-10 | 7.40E-09 |
| 0.368908303 | 0.533547878 |
| 0.355849201 | NA |
| 0.826681219 | NA |
| 1.23E-14 | 1.36E-12 |
| 0.000439083 | 0.002475404 |
| NA | NA |
| 7.86E-10 | 2.56E-08 |
| NA | NA |
| 0.674133638 | 0.792892371 |
| 0.039438265 | 0.100994154 |
| 0.03674808 | 0.095703684 |
| 1.91E-05 | 0.000170921 |
| NA | NA |
| 0.620208385 | NA |
| 0.602152369 | 0.737666045 |
| 0.568817411 | 0.71126829 |
| 0.013047074 | 0.04163044 |
| 0.195608783 | 0.339394222 |
| 0.151309224 | 0.280624936 |
| 0.879383935 | 0.931609023 |
| 0.00025288 | 0.00154464 |
| 5.09E-05 | 0.000392621 |
| 0.610195717 | 0.744156823 |
| 0.983534203 | NA |
| 0.005669814 | 0.021293219 |
| 0.783383701 | NA |
| 0.99205891 | NA |
| 0.587425812 | 0.726253393 |
| 0.110329553 | 0.221698998 |
| 0.262696619 | 0.418162744 |
| 0.052218929 | 0.125498026 |
| NA | NA |
| 0.483954906 | 0.639338968 |
| 0.889409301 | NA |
| 0.000679386 | 0.003603537 |
| NA | NA |
| 0.001856398 | 0.008401615 |
| 0.293786887 | NA |
| 0.369298963 | 0.533972208 |
| 0.904152147 | NA |
| 0.8148411 | 0.889702743 |
| NA | NA |

| | | |
|---|---|---|
| | 0.053852934 | 0.128357384 |
| | 4.98E-06 | 5.30E-05 |
| | 0.000292159 | 0.001747145 |
| | 0.999747964 | 0.999884566 |
| | 8.44E-07 | 1.14E-05 |
| | 0.00024116 | 0.001485067 |
| | 0.91382713 | NA |
| | 0.178116065 | 0.317183502 |
| | 0.000101396 | 0.000707067 |
| NA | | NA |
| | 0.008572469 | 0.029705356 |
| | 0.88395713 | 0.934155379 |
| | 0.93539296 | 0.962820337 |
| | 0.05955606 | 0.138860855 |
| | 0.889409773 | NA |
| | 0.927169724 | NA |
| | 0.002513303 | 0.010859824 |
| NA | | NA |
| | 0.178516676 | 0.31766999 |
| | 0.636915697 | NA |
| NA | | NA |
| | 0.207173086 | 0.35381193 |
| | 0.009411873 | 0.031998673 |
| | 0.714726398 | NA |
| | 0.001480317 | NA |
| | 3.36E-06 | 3.77E-05 |
| | 0.76109879 | NA |
| | 2.37E-19 | 8.67E-17 |
| | 9.22E-07 | 1.23E-05 |
| | 0.141800807 | 0.267118806 |
| | 0.755891116 | 0.851591389 |
| | 0.326720165 | 0.488955776 |
| | 8.30E-10 | 2.68E-08 |
| | 1.93E-05 | 0.00017239 |
| NA | | NA |
| | 7.32E-05 | 0.000535241 |
| | 1.26E-09 | 3.85E-08 |
| | 0.750908282 | NA |
| | 0.698146738 | 0.811228925 |
| | 0.05783701 | 0.135558001 |
| NA | | NA |
| NA | | NA |
| | 0.00832147 | 0.028986543 |
| | 0.488473083 | 0.642952732 |
| | 0.015399637 | 0.047715625 |
| | 0.243214801 | 0.396168965 |
| | 0.000868842 | 0.004416411 |

| | | | |
|---|---|---|---|
| | 0.097157455 | NA | |
| NA | | NA | |
| | 0.820670654 | | 0.893328057 |
| | 0.205748416 | NA | |
| | 0.482913517 | NA | |
| | 0.24652229 | NA | |
| | 0.040474383 | | 0.102903437 |
| | 0.02215923 | | 0.063832419 |
| | 2.30E-07 | | 3.67E-06 |
| | 0.000866501 | | 0.004406554 |
| | 0.463885027 | NA | |
| | 0.635471743 | NA | |
| NA | | NA | |
| NA | | NA | |
| NA | | NA | |
| NA | | NA | |
| NA | | NA | |
| NA | | NA | |
| NA | | NA | |
| NA | | NA | |
| NA | | NA | |
| NA | | NA | |
| NA | | NA | |
| NA | | NA | |
| NA | | NA | |
| NA | | NA | |
| NA | | NA | |
| NA | | NA | |
| NA | | NA | |
| NA | | NA | |
| | 0.889409301 | NA | |
| NA | | NA | |
| NA | | NA | |
| | 0.601121176 | NA | |
| NA | | NA | |
| NA | | NA | |
| NA | | NA | |
| NA | | NA | |
| NA | | NA | |
| NA | | NA | |
| NA | | NA | |
| | 0.905158266 | NA | |
| NA | | NA | |
| NA | | NA | |
| NA | | NA | |
| NA | | NA | |
| NA | | NA | |

| | | |
|---|---|---|
| NA | | NA |
| NA | | NA |
| NA | | NA |
| | 0.889409773 | NA |
| NA | | NA |
| NA | | NA |
| NA | | NA |
| NA | | NA |
| NA | | NA |
| NA | | NA |
| | 0.905158739 | NA |
| NA | | NA |
| | 0.152968719 | 0.282997906 |
| NA | | NA |
| NA | | NA |
| NA | | NA |
| NA | | NA |
| | 0.889409301 | NA |
| | 0.878605957 | NA |
| NA | | NA |
| | 0.37569316 | NA |
| NA | | NA |
| NA | | NA |
| NA | | NA |
| NA | | NA |
| NA | | NA |
| NA | | NA |
| NA | | NA |
| | 0.991570563 | NA |
| NA | | NA |
| | 0.085745957 | NA |
| | 0.185776328 | 0.32673068 |
| | 0.258600899 | NA |
| | 0.102294944 | 0.209581516 |
| | 0.911874994 | NA |
| NA | | NA |
| | 0.628851435 | NA |
| | 0.005792167 | 0.021664146 |
| | 0.031884177 | 0.085560875 |
| NA | | NA |
| | 0.920451329 | NA |
| | 0.155468844 | 0.286284305 |
| | 0.010709876 | 0.035611472 |
| | 0.243398555 | NA |
| | 0.193180662 | 0.336137413 |
| | 0.775917046 | 0.864891494 |
| | 0.72799235 | 0.832213234 |

| | | |
|---|---|---|
| | 2.84E-08 | 5.94E-07 |
| | 0.532045828 | NA |
| | 0.604927772 | 0.740057145 |
| | 2.35E-05 | 0.00020378 |
| | 0.638451394 | 0.765131197 |
| | 0.492517584 | 0.646627966 |
| | 0.296484749 | NA |
| | 0.169941849 | 0.306057004 |
| | 0.040648797 | 0.103227353 |
| | 0.956554094 | NA |
| | 0.013758762 | 0.04347175 |
| | 0.586701618 | 0.725742498 |
| | 0.684249538 | 0.800119049 |
| | 8.48E-09 | 2.04E-07 |
| | 0.320698717 | 0.483005701 |
| | 0.487130366 | 0.64191645 |
| | 0.000503752 | 0.002779367 |
| | 0.105922102 | 0.215046545 |
| | 0.402427386 | NA |
| | 0.001186992 | 0.005758983 |
| | 0.001149298 | 0.005600853 |
| | 0.322064966 | 0.484365768 |
| | 4.39E-08 | 8.69E-07 |
| | 0.992060816 | NA |
| | 0.001128337 | 0.005510932 |
| | 1.18E-05 | 0.000112695 |
| | 0.006558752 | 0.023952043 |
| | 6.51E-10 | 2.17E-08 |
| | 0.000129242 | 0.000867896 |
| | 0.304553229 | 0.465165498 |
| | 7.21E-05 | 0.000528981 |
| | 0.152203217 | 0.281925807 |
| | 0.002938278 | 0.012348642 |
| | 5.37E-07 | 7.68E-06 |
| | 0.305435959 | 0.466059914 |
| | 1.29E-06 | 1.63E-05 |
| | 0.256388469 | 0.410951415 |
| NA | | NA |
| | 8.65E-16 | 1.32E-13 |
| | 0.02780288 | 0.076670029 |
| | 2.32E-15 | 3.12E-13 |
| | 0.609572532 | 0.743603313 |
| | 0.051477408 | 0.124218944 |
| | 0.844322269 | 0.908312399 |
| | 0.237371821 | 0.389295288 |
| | 0.800001483 | 0.879407568 |
| | 0.700533707 | NA |

| | |
|---|---|
| 0.116838821 | 0.231558094 |
| 0.252403527 | 0.406721235 |
| 0.469375394 | 0.62569139 |
| NA | NA |
| 0.874685676 | 0.928423374 |
| 0.032701924 | 0.087285955 |
| 3.49E-05 | 0.000284328 |
| NA | NA |
| NA | NA |
| NA | NA |
| NA | NA |
| 0.628991101 | NA |
| NA | NA |
| 0.64458986 | 0.770147926 |
| 0.632213549 | 0.760033796 |
| 0.209943033 | 0.356739074 |
| 0.889409301 | NA |
| 0.171053515 | 0.307479055 |
| 0.013271345 | 0.04219277 |
| 0.000388191 | 0.002226826 |
| 0.175938344 | NA |
| 8.38E-08 | 1.51E-06 |
| 0.040718023 | 0.103343396 |
| 0.807879735 | NA |
| 0.568261107 | 0.710779543 |
| 0.085398828 | 0.182561611 |
| NA | NA |
| 1.17E-08 | 2.71E-07 |
| NA | NA |
| 0.427887488 | 0.588978961 |
| 0.905158266 | NA |
| 0.12553367 | 0.244032743 |
| 0.892014958 | NA |
| 0.792742292 | 0.874985071 |
| 0.044511179 | 0.110793581 |
| 0.590307577 | 0.728369873 |
| 0.385712403 | 0.549526935 |
| 0.007789236 | 0.027450464 |
| 0.493731642 | 0.647197536 |
| 0.24939477 | 0.403273914 |
| 0.756585257 | NA |
| 0.650771408 | NA |
| 0.784353866 | NA |
| 0.020190728 | 0.059272712 |
| 0.563930302 | NA |
| 0.277109326 | 0.434925572 |
| 0.986369155 | NA |

| | | |
|---|---|---|
| | 0.001443776 | 0.006796117 |
| | 0.712465483 | 0.820959728 |
| | 0.435064514 | NA |
| | 5.45E-08 | 1.05E-06 |
| | 0.905158739 | NA |
| | 0.94923079 | NA |
| | 0.905158266 | NA |
| | 0.801103067 | NA |
| | 0.216561186 | NA |
| NA | | NA |
| | 0.064649634 | 0.147833105 |
| | 0.000168677 | 0.001089083 |
| | 0.558013764 | 0.702908387 |
| | 0.501317745 | 0.654057178 |
| NA | | NA |
| | 0.417910748 | 0.580118978 |
| | 0.000109337 | 0.000753372 |
| | 0.054724897 | 0.12998421 |
| | 4.92E-06 | 5.25E-05 |
| | 0.164142722 | 0.298032911 |
| | 0.233310323 | 0.38486038 |
| | 0.889410717 | NA |
| | 1.89E-05 | 0.000169731 |
| | 2.71E-07 | 4.23E-06 |
| NA | | NA |
| | 4.09E-08 | 8.19E-07 |
| | 0.012945717 | 0.041379184 |
| | 0.000504569 | 0.002783177 |
| | 0.788582525 | NA |
| | 0.073825584 | 0.163190659 |
| | 0.122336775 | 0.239364986 |
| NA | | NA |
| | 0.078784667 | 0.171954329 |
| | 0.000538356 | 0.002948806 |
| | 0.251925542 | 0.406169822 |
| | 0.777163803 | NA |
| | 0.022930888 | 0.065676192 |
| NA | | NA |
| NA | | NA |
| | 0.402944748 | NA |
| | 0.419540087 | 0.581388411 |
| | 0.11768529 | 0.232731564 |
| | 0.341698291 | NA |
| | 4.66E-06 | 5.00E-05 |
| | 0.707798181 | 0.81781322 |
| NA | | NA |
| | 0.269938039 | 0.426382492 |

| | | |
|---|---|---|
| | 0.488955798 | 0.643305506 |
| | 0.929864903 | NA |
| | 0.623410111 | 0.754240365 |
| | 0.766271684 | NA |
| | 2.35E-06 | 2.76E-05 |
| | 0.065398943 | 0.149111763 |
| | 0.128086188 | 0.247679165 |
| | 0.600814337 | 0.736602201 |
| | 0.000415252 | 0.002359862 |
| | 0.992063385 | NA |
| | 0.39628656 | 0.559869818 |
| | 0.12561832 | 0.244175682 |
| | 0.134910691 | 0.257094841 |
| | 0.295003409 | 0.454818497 |
| | 3.42E-05 | 0.000279771 |
| | 0.004275245 | 0.01679129 |
| | 0.987614736 | 0.993587391 |
| | 0.20907931 | 0.355822101 |
| | 0.000109947 | 0.000756842 |
| | 0.009317202 | 0.031759769 |
| | 0.706251334 | 0.817316102 |
| | 0.014419974 | 0.045177374 |
| | 0.464887287 | 0.621860376 |
| | 0.443996032 | 0.603585022 |
| | 0.758268821 | 0.853322916 |
| | 5.54E-20 | 2.38E-17 |
| | 0.060553877 | NA |
| | 0.669150076 | NA |
| | 0.905158739 | NA |
| | 0.992037435 | NA |
| | 0.54815889 | 0.694296653 |
| | 7.83E-06 | 7.88E-05 |
| | 0.388539012 | 0.551978273 |
| | 0.53002617 | 0.679323217 |
| | 2.18E-05 | 0.000191627 |
| | 0.011094312 | 0.036596064 |
| | 1.75E-10 | 6.99E-09 |
| | 0.19175341 | 0.334243536 |
| | 0.006176052 | 0.022797097 |
| | 0.848398047 | 0.910956565 |
| | 0.905157793 | NA |
| | 2.60E-05 | 0.000222225 |
| | 0.889410717 | NA |
| NA | | NA |
| NA | | NA |
| | 0.671395095 | 0.790730216 |
| | 0.671260627 | NA |

| | |
|---|---|
| NA | NA |
| 0.243890196 | 0.396886381 |
| 0.000459915 | 0.002569553 |
| NA | NA |
| 0.623555904 | NA |
| NA | NA |
| 0.714113058 | NA |
| 0.001573048 | 0.007289 |
| 0.992061292 | NA |
| 0.063270469 | 0.145421418 |
| 0.992059387 | NA |
| 0.992061292 | NA |
| 0.889410245 | NA |
| 0.992060816 | NA |
| 0.202737524 | 0.348404546 |
| NA | NA |
| 0.632158762 | 0.76000954 |
| 0.785036702 | NA |
| 0.607049606 | NA |
| 0.725450777 | 0.83034525 |
| 0.096815204 | 0.200960871 |
| 0.030268024 | 0.082025859 |
| 0.00011605 | 0.000793137 |
| 0.436185398 | 0.59664456 |
| 0.355466071 | 0.52006659 |
| 0.613712406 | NA |
| 0.000543058 | 0.002971416 |
| 0.406133388 | 0.569349021 |
| 0.352097695 | 0.516364453 |
| 0.43962362 | 0.599869203 |
| 0.009349826 | 0.031851197 |
| 0.531350007 | 0.680503605 |
| 0.850420738 | NA |
| 3.55E-05 | 0.000288719 |
| 0.797157316 | NA |
| 0.365611136 | 0.530351099 |
| 5.93E-09 | 1.48E-07 |
| 0.931249867 | NA |
| 0.121148653 | 0.237879731 |
| 0.020228548 | 0.059333002 |
| 0.983928585 | 0.991970261 |
| 0.691349929 | NA |
| 2.88E-07 | 4.44E-06 |
| 0.357479328 | NA |
| 0.397529127 | 0.561129073 |
| NA | NA |
| 0.509571991 | 0.661172971 |

| | | | |
|---|---|---|---|
| | 0.321798482 | | 0.484138825 |
| | 9.12E-05 | | 0.000644838 |
| | 0.014670725 | | 0.045851758 |
| | 0.040411478 | | 0.102810043 |
| | 0.388034224 | | 0.551760897 |
| | 0.959927058 | | 0.977727478 |
| NA | | NA | |
| | 0.793334793 | | 0.875331058 |
| | 0.845501414 | NA | |
| | 0.205839276 | | 0.35203001 |
| | 0.000134993 | | 0.000900733 |
| | 1.20E-11 | | 6.51E-10 |
| | 0.110827134 | | 0.222312326 |
| | 0.986742961 | NA | |
| NA | | NA | |
| | 0.786996571 | | 0.871688903 |
| | 4.66E-11 | | 2.15E-09 |
| | 3.10E-06 | | 3.50E-05 |
| | 0.541212013 | NA | |
| | 0.218360176 | | 0.366784296 |
| | 0.081687339 | | 0.176761162 |
| NA | | NA | |
| NA | | NA | |
| | 0.580243845 | | 0.720553013 |
| | 0.619117489 | | 0.751014237 |
| | 0.008880391 | | 0.03055539 |
| | 0.889409773 | NA | |
| | 0.884008599 | NA | |
| | 0.017061829 | | 0.051824796 |
| NA | | NA | |
| | 0.616631712 | | 0.749171172 |
| | 0.005923351 | | 0.022045908 |
| | 0.396284142 | | 0.559869818 |
| | 0.901183732 | NA | |
| | 0.023672077 | | 0.067346998 |
| | 0.109861439 | | 0.221140134 |
| | 0.905158739 | NA | |
| | 0.007781352 | | 0.027440295 |
| NA | | NA | |
| | 0.269629295 | NA | |
| | 0.889409301 | NA | |
| | 0.992060816 | NA | |
| | 0.667104468 | NA | |
| | 0.006477526 | | 0.023714571 |
| | 0.275555667 | | 0.43292029 |
| | 0.764675526 | NA | |
| | 0.734646595 | NA | |

| | |
|---|---|
| NA | NA |
| 0.473049314 | 0.629328116 |
| 0.987363099 | NA |
| 0.017507934 | 0.052868085 |
| 0.90481849 | 0.946228652 |
| 0.479068025 | 0.634758629 |
| NA | NA |
| NA | NA |
| 0.092446944 | 0.193971877 |
| 2.93E-06 | 3.34E-05 |
| NA | NA |
| NA | NA |
| 0.992060816 | NA |
| 0.857888004 | 0.917175946 |
| 0.889409301 | NA |
| NA | NA |
| 0.025070312 | 0.070697185 |
| NA | NA |
| 0.992060816 | NA |
| 0.808660907 | 0.885169476 |
| 0.257719975 | 0.412513516 |
| 0.001422112 | 0.006714287 |
| 0.431217136 | NA |
| 0.270425474 | NA |
| 0.889409773 | NA |
| 0.986203473 | NA |
| 0.889409773 | NA |
| 0.138360559 | 0.262144911 |
| NA | NA |
| 0.3916037 | 0.555003592 |
| 7.65E-05 | 0.000556529 |
| 0.840141985 | 0.905367711 |
| 0.024105395 | 0.068424297 |
| NA | NA |
| 0.524157219 | 0.674241016 |
| 0.585748688 | 0.725094731 |
| 0.118519299 | 0.233917426 |
| NA | NA |
| 0.000662169 | 0.003528407 |
| 0.889409773 | NA |
| 0.982407966 | 0.991346745 |
| 0.002549002 | 0.010990288 |
| NA | NA |
| 0.807879735 | NA |
| NA | NA |
| NA | NA |
| 0.017843011 | 0.053658542 |

| | |
|---|---|
| NA | NA |
| 0.000353078 | 0.002057652 |
| 0.362190393 | 0.526974984 |
| 0.54284518 | 0.689888809 |
| 1.11E-07 | 1.94E-06 |
| 7.52E-08 | 1.38E-06 |
| 0.744305313 | NA |
| 0.204499029 | 0.350389684 |
| 3.23E-05 | 0.000266945 |
| 0.81073739 | NA |
| NA | NA |
| 0.904947233 | NA |
| 0.889409301 | NA |
| NA | NA |
| NA | NA |
| 0.01115184 | 0.036746502 |
| 3.09E-06 | 3.49E-05 |
| 0.409095066 | 0.572247921 |
| NA | NA |
| 0.076502554 | 0.167991959 |
| 0.176890711 | NA |
| 0.004823908 | 0.018616555 |
| 0.048488988 | NA |
| 7.28E-06 | 7.40E-05 |
| 0.241786341 | 0.394573889 |
| 0.023073932 | 0.0659998 |
| 0.384463852 | NA |
| 0.017170774 | 0.052079149 |
| 0.335250944 | 0.498157767 |
| 0.005015295 | 0.019230115 |
| 0.847652584 | NA |
| 0.037401565 | 0.097023151 |
| 0.889410245 | NA |
| 0.452854143 | NA |
| 0.733526821 | NA |
| 0.992061292 | NA |
| 0.992060816 | NA |
| NA | NA |
| 0.811426724 | 0.887268173 |
| NA | NA |
| 0.81486869 | 0.889702743 |
| 0.166201783 | 0.300801528 |
| 0.000300332 | 0.001787744 |
| 0.041581519 | 0.10518242 |
| 0.002552996 | 0.011003186 |
| 0.989921475 | NA |
| 0.411162912 | 0.574054322 |

| | | |
|---|---|---|
| | 0.048223595 | 0.117922264 |
| | 0.36711024 | NA |
| | 4.50E-10 | 1.58E-08 |
| NA | | NA |
| | 0.77334416 | NA |
| | 3.99E-07 | 5.90E-06 |
| | 4.75E-08 | 9.29E-07 |
| | 0.217316288 | NA |
| | 0.286132079 | NA |
| | 0.029720967 | NA |
| | 0.009718874 | 0.032858621 |
| | 0.000636259 | 0.003411869 |
| | 0.006255747 | 0.023037055 |
| | 0.000281825 | 0.001698765 |
| | 0.026614847 | 0.074016644 |
| | 0.089232489 | 0.18895702 |
| | 9.38E-06 | 9.21E-05 |
| | 0.000648692 | 0.003467534 |
| | 0.797157316 | NA |
| | 0.98097011 | 0.990511759 |
| | 0.482988678 | 0.638514838 |
| | 0.703406217 | 0.815011456 |
| | 0.046498103 | 0.114581733 |
| | 0.037619359 | 0.097495987 |
| | 0.018707408 | 0.055678501 |
| | 0.946276359 | 0.969356826 |
| | 0.578763381 | 0.719530379 |
| | 0.852351155 | NA |
| | 0.857405099 | 0.916906524 |
| | 0.692679768 | 0.806669231 |
| | 0.834184755 | NA |
| NA | | NA |
| | 0.340204977 | NA |
| | 0.730151299 | NA |
| | 0.011733681 | 0.038234145 |
| | 0.25961176 | 0.414725348 |
| | 0.0224812 | NA |
| | 0.001122197 | 0.005487912 |
| | 0.267696051 | 0.423984842 |
| | 0.280825162 | 0.439188072 |
| | 0.077729682 | 0.17009129 |
| NA | | NA |
| | 0.000185202 | 0.001178795 |
| | 0.019450703 | 0.057465853 |
| | 0.3597607 | 0.524471524 |
| | 0.992061292 | NA |
| | 0.0553068 | 0.130927341 |

| | |
|---|---|
| 0.125201967 | 0.243646755 |
| 0.456873172 | 0.61454689 |
| NA | NA |
| NA | NA |
| 0.305515954 | 0.466117198 |
| 0.850504856 | NA |
| 0.007788543 | 0.027450464 |
| 0.003263701 | 0.013476423 |
| 0.663771228 | 0.785035407 |
| 0.300175496 | 0.460551614 |
| 0.219758593 | 0.368625693 |
| 0.645989526 | 0.771235488 |
| 0.98322073 | NA |
| 0.889409301 | NA |
| 0.143563989 | 0.269650655 |
| 0.004733929 | 0.018320743 |
| 0.136040181 | 0.258752189 |
| 0.620682031 | 0.752059116 |
| 0.45031448 | 0.608848242 |
| 0.220322263 | 0.369232739 |
| 0.899348167 | 0.943383319 |
| 2.05E-06 | 2.46E-05 |
| 9.92E-07 | 1.30E-05 |
| 1.08E-13 | 9.62E-12 |
| 0.359621459 | 0.524399204 |
| 2.41E-07 | 3.81E-06 |
| 0.000176752 | 0.001134548 |
| 0.485109367 | 0.640199877 |
| 0.872334167 | 0.926688886 |
| 0.283855237 | 0.442602937 |
| 0.305119556 | 0.465716603 |
| 2.60E-06 | 3.01E-05 |
| 0.279052494 | NA |
| 0.324287065 | 0.486508141 |
| 0.378295275 | NA |
| NA | NA |
| 1.33E-08 | 3.02E-07 |
| 5.53E-10 | 1.88E-08 |
| 0.320387438 | 0.482702418 |
| 0.791539524 | 0.87444868 |
| 0.963734373 | NA |
| NA | NA |
| 0.000547283 | 0.002990243 |
| NA | NA |
| 0.110658708 | 0.222096205 |
| 0.003875832 | 0.015471624 |
| 0.00990184 | 0.03334885 |

| | |
|---|---|
| NA | NA |
| 0.041048392 | 0.104049594 |
| 1.67E-06 | 2.04E-05 |
| 2.93E-05 | 0.000245415 |
| 0.062812683 | 0.14464952 |
| 0.656703511 | 0.779447187 |
| NA | NA |
| NA | NA |
| NA | NA |
| 0.002126023 | 0.009425667 |
| 0.905158739 | NA |
| NA | NA |
| 0.574497442 | NA |
| 0.147496665 | 0.275227674 |
| 0.078858179 | 0.172080567 |
| NA | NA |
| 0.150132809 | 0.278819887 |
| 0.010951972 | 0.036229942 |
| 9.47E-06 | 9.27E-05 |
| 2.40E-05 | 0.000207232 |
| 0.896066414 | NA |
| 0.992060816 | NA |
| 0.001299761 | 0.006204665 |
| NA | NA |
| NA | NA |
| 0.899182161 | NA |
| 0.480588259 | 0.636197106 |
| 0.109468467 | 0.220533768 |
| 0.424180435 | 0.585527921 |
| 0.48631782 | 0.641153519 |
| 3.82E-09 | 1.02E-07 |
| 0.327101187 | NA |
| 0.603330951 | NA |
| 0.988471685 | NA |
| 0.898123435 | 0.942684026 |
| 0.820078869 | 0.892909192 |
| 0.226763894 | 0.376721769 |
| 0.462171938 | 0.619549086 |
| 0.00130124 | 0.006210373 |
| 0.012028087 | 0.039031292 |
| 0.045549608 | 0.11286303 |
| 0.889410245 | NA |
| 0.889409773 | NA |
| 0.889409301 | NA |
| 0.889409773 | NA |
| 1.40E-06 | 1.75E-05 |
| 0.250352115 | 0.404325057 |

| | | |
|---|---|---|
| | 0.117126311 | 0.231981298 |
| | 0.860228632 | 0.918673306 |
| | 0.465796709 | NA |
| | 0.857820087 | 0.917175946 |
| | 0.543578742 | 0.69036701 |
| | 0.905158739 | NA |
| | 0.031548044 | 0.084824721 |
| | 5.49E-13 | 4.17E-11 |
| | 0.861431665 | NA |
| | 0.163301211 | 0.296824046 |
| | 0.000806047 | 0.004153951 |
| | 0.026457393 | 0.073644047 |
| | 0.002895152 | 0.012200084 |
| | 0.7555679 | 0.851416872 |
| | 0.766977975 | 0.858600094 |
| | 0.397355329 | 0.56102487 |
| | 0.907667462 | 0.947610543 |
| | 0.026847358 | 0.074521695 |
| | 0.000678931 | 0.003601993 |
| | 0.12841762 | 0.248123407 |
| | 0.983166751 | NA |
| | 0.349566557 | NA |
| | 0.945119554 | 0.968901974 |
| | 3.93E-07 | 5.83E-06 |
| | 0.302767613 | NA |
| | 0.140756504 | NA |
| | 0.742661097 | 0.842360281 |
| | 0.905158739 | NA |
| | 0.014967724 | 0.046614133 |
| | 0.594184935 | 0.731458306 |
| | 0.497455527 | 0.650525602 |
| | 0.000342659 | 0.002007058 |
| | 0.991918092 | NA |
| | 0.310961328 | 0.472292143 |
| | 4.00E-05 | 0.000320068 |
| | 0.596051433 | NA |
| NA | | NA |
| | 0.478853487 | 0.634613227 |
| | 0.992061292 | NA |
| | 0.000674931 | 0.00358337 |
| | 8.50E-05 | 0.000606196 |
| | 0.004291254 | 0.016839107 |
| | 0.245131847 | 0.398316577 |
| | 0.001948305 | 0.008754418 |
| | 0.0166054 | 0.050721654 |
| | 0.510535065 | 0.662070483 |
| | 4.85E-06 | 5.18E-05 |

| | |
|---|---|
| 0.131389502 | 0.252157146 |
| 0.040883771 | 0.103668213 |
| 0.028320771 | 0.077805056 |
| 0.26674276 | 0.422917276 |
| NA | NA |
| 0.137213671 | NA |
| 0.725049767 | NA |
| 0.734456696 | 0.8369018 |
| 0.914947242 | 0.951879779 |
| 0.991925874 | NA |
| 0.025986548 | 0.072665047 |
| 5.88E-08 | 1.11E-06 |
| 0.184352635 | 0.324947785 |
| 0.810509254 | NA |
| 0.006683515 | 0.024326754 |
| 0.545795478 | NA |
| 0.030968064 | 0.083551754 |
| 0.38256931 | 0.546538252 |
| 9.55E-07 | 1.26E-05 |
| 0.58862196 | NA |
| 0.014707915 | 0.045941835 |
| 0.436201547 | NA |
| 0.015483127 | 0.04791347 |
| 0.001699622 | 0.00777218 |
| 0.00848554 | 0.029450604 |
| NA | NA |
| 0.008390912 | 0.02919747 |
| NA | NA |
| NA | NA |
| 0.565898472 | 0.709032533 |
| 0.027832061 | 0.076731227 |
| 0.123848058 | 0.241697432 |
| 1.52E-06 | 1.88E-05 |
| 0.812784918 | NA |
| 0.077309008 | 0.169309052 |
| 0.462942163 | 0.620127308 |
| 1.48E-05 | 0.000136758 |
| 0.170120329 | 0.306303076 |
| 3.44E-07 | 5.18E-06 |
| 0.113771132 | 0.227076921 |
| 0.473363228 | NA |
| 0.989560379 | NA |
| 0.429810622 | 0.59092233 |
| 0.905158739 | NA |
| 0.456319859 | 0.614178841 |
| 0.048277772 | 0.118002182 |
| 0.782559532 | NA |

| | |
|---|---|
| 3.26E-08 | 6.72E-07 |
| 0.000102873 | 0.000716233 |
| 0.266996267 | 0.423166441 |
| 0.913063904 | 0.9507312 |
| 0.916073972 | 0.952375171 |
| 0.560839122 | 0.705076774 |
| 0.759302087 | 0.853826021 |
| 0.002043314 | 0.009112334 |
| NA | NA |
| 0.000372642 | 0.002149419 |
| NA | NA |
| 0.067135309 | 0.151778467 |
| 0.034609294 | 0.091256663 |
| 0.118058484 | 0.23323853 |
| 0.058558919 | NA |
| 0.140582224 | 0.265369643 |
| 0.517612918 | 0.668547152 |
| 0.390614637 | 0.554102507 |
| 0.654914644 | NA |
| 0.707063707 | 0.817652954 |
| 0.425253074 | 0.586458496 |
| 0.624949777 | 0.755340311 |
| 0.940463998 | 0.966159014 |
| 0.436514449 | 0.596902382 |
| 5.81E-07 | 8.16E-06 |
| NA | NA |
| 0.180384164 | 0.320188817 |
| 0.041813761 | 0.105623879 |
| 0.097190175 | 0.201453563 |
| 0.365469894 | 0.530286336 |
| 0.06792844 | 0.152957404 |
| 0.686189154 | NA |
| 4.59E-06 | 4.95E-05 |
| 0.025614834 | 0.0719555 |
| NA | NA |
| 0.905158266 | NA |
| 0.90515732 | NA |
| 0.992061292 | NA |
| 0.962013211 | 0.979205986 |
| 0.135529665 | NA |
| 5.40E-07 | 7.70E-06 |
| NA | NA |
| NA | NA |
| 0.670581018 | NA |
| NA | NA |
| 0.905158739 | NA |
| 0.661602032 | 0.783313691 |

| | |
|---|---|
| 0.004510998 | NA |
| 0.834139596 | 0.901268968 |
| 0.004636745 | 0.017982744 |
| 2.35E-06 | 2.76E-05 |
| 0.000609529 | 0.003284578 |
| 0.016181214 | 0.049681666 |
| 0.009074138 | 0.031080798 |
| 0.005665772 | 0.021282019 |
| 0.543723756 | 0.690471314 |
| NA | NA |
| 0.954717367 | 0.97468323 |
| 0.001485653 | 0.006950035 |
| 0.80725778 | 0.884393976 |
| 0.78297309 | 0.869226799 |
| 0.844374086 | 0.908323645 |
| 2.18E-10 | 8.43E-09 |
| 0.489245648 | NA |
| NA | NA |
| NA | NA |
| NA | NA |
| NA | NA |
| NA | NA |
| NA | NA |
| NA | NA |
| NA | NA |
| NA | NA |
| 0.054670304 | 0.129896689 |
| 0.006019412 | 0.022342844 |
| 0.058413039 | 0.136660123 |
| 3.18E-05 | 0.00026335 |
| 7.40E-07 | 1.01E-05 |
| 0.332338275 | 0.49490141 |
| 1.28E-07 | 2.20E-06 |
| 0.956714569 | 0.975959083 |
| 0.346204097 | NA |
| 0.956493001 | 0.975869062 |
| 0.060578271 | 0.140825561 |
| 0.051508388 | 0.12426639 |
| 0.408232646 | 0.571467918 |
| 1.61E-09 | 4.76E-08 |
| 0.477118339 | 0.633206915 |
| 0.028930879 | 0.079124819 |
| 0.003278703 | 0.013518032 |
| 4.93E-05 | 0.000382622 |
| NA | NA |
| NA | NA |
| 0.450210664 | 0.608829607 |

| | | |
|---|---|---|
| | 0.033362875 | 0.08864863 |
| | 0.99205891 | NA |
| | 0.077632385 | 0.169929182 |
| | 0.006473199 | 0.023702681 |
| NA | | NA |
| NA | | NA |
| | 1.19E-06 | 1.52E-05 |
| | 0.036931723 | 0.096110891 |
| | 0.135400779 | 0.257916873 |
| | 0.000849555 | 0.004332413 |
| | 0.002474401 | 0.0107255 |
| | 0.217536407 | 0.365691912 |
| | 0.588356987 | 0.727012047 |
| | 0.094085955 | 0.196559175 |
| NA | | NA |
| | 0.487004098 | 0.641827091 |
| | 0.037771723 | 0.09776394 |
| | 0.992060816 | NA |
| | 0.313846245 | NA |
| | 0.804790063 | NA |
| | 0.605883655 | 0.740584424 |
| | 0.00205844 | 0.009171564 |
| | 6.09E-05 | 0.000458201 |
| | 0.052047472 | 0.125140747 |
| | 6.71E-19 | 2.20E-16 |
| | 0.261085627 | 0.416534386 |
| | 0.905158739 | NA |
| | 0.083051881 | 0.179008269 |
| | 0.989121229 | NA |
| | 1.76E-15 | 2.48E-13 |
| | 0.723918262 | NA |
| | 0.002783105 | 0.011811788 |
| | 0.831202584 | NA |
| | 0.343987741 | 0.507363435 |
| | 0.049131786 | 0.119703194 |
| | 0.207629992 | 0.354349206 |
| NA | | NA |
| | 0.669263498 | 0.789091843 |
| NA | | NA |
| NA | | NA |
| | 0.969249665 | 0.983776228 |
| | 0.700845654 | 0.813129883 |
| | 0.905257705 | 0.946540766 |
| | 0.437876808 | 0.598229132 |
| NA | | NA |
| | 0.889410717 | NA |
| | 0.534677938 | 0.683260894 |

| | | |
|---|---|---|
| | 0.022892252 | 0.065593381 |
| | 0.639740422 | 0.766313546 |
| | 1.74E-15 | 2.47E-13 |
| | 0.291407951 | NA |
| | 0.000511896 | 0.002818638 |
| NA | | NA |
| | 0.857487904 | NA |
| | 0.022613182 | 0.064962353 |
| NA | | NA |
| | 5.58E-06 | 5.86E-05 |
| | 0.905158266 | NA |
| NA | | NA |
| NA | | NA |
| NA | | NA |
| | 0.889409773 | NA |
| | 0.981392626 | NA |
| | 0.229010401 | 0.379535049 |
| | 0.992061292 | NA |
| | 0.698709834 | 0.811656465 |
| | 0.005960611 | 0.022154546 |
| | 0.690263216 | 0.805030912 |
| NA | | NA |
| | 0.845054642 | NA |
| | 0.621774877 | NA |
| | 1.30E-06 | 1.64E-05 |
| | 0.291540903 | NA |
| | 0.992061292 | NA |
| NA | | NA |
| | 0.905158266 | NA |
| | 0.372934381 | 0.537249061 |
| | 0.414454756 | 0.577022686 |
| | 0.967285688 | NA |
| | 0.203390327 | 0.34921781 |
| | 0.002934691 | 0.012338289 |
| | 5.35E-10 | 1.84E-08 |
| | 0.01147394 | 0.037543772 |
| NA | | NA |
| | 9.30E-05 | 0.000656117 |
| | 0.150006757 | 0.278703818 |
| | 0.708497618 | NA |
| | 7.23E-05 | 0.0005301 |
| | 0.009867235 | 0.033257806 |
| | 0.001642926 | 0.007544336 |
| | 1.13E-10 | 4.75E-09 |
| | 0.000114304 | 0.000782661 |
| | 0.203417506 | 0.34921781 |
| | 0.379756922 | NA |

| | | |
|---|---|---|
| | 1.06E-06 | 1.37E-05 |
| | 0.161049719 | 0.293731153 |
| NA | | NA |
| | 0.695060601 | 0.80858422 |
| | 0.607707445 | 0.741833645 |
| | 0.923135214 | 0.955599216 |
| | 0.655067519 | 0.778121472 |
| | 0.292839696 | NA |
| | 0.640892093 | NA |
| NA | | NA |
| | 0.889409773 | NA |
| NA | | NA |
| | 0.992060816 | NA |
| | 0.889409773 | NA |
| | 0.660933665 | NA |
| | 0.017149443 | 0.052028822 |
| NA | | NA |
| | 0.081341272 | 0.176203314 |
| | 0.208364117 | 0.355045212 |
| | 0.185960944 | 0.326959122 |
| | 0.382490499 | 0.546496771 |
| | 0.299837605 | NA |
| | 0.867127636 | 0.923660236 |
| | 0.984099697 | 0.991970261 |
| | 0.25248894 | 0.406749661 |
| | 0.096382864 | 0.200290651 |
| | 0.976780844 | 0.988115511 |
| | 0.583818585 | 0.723639366 |
| | 6.69E-06 | 6.88E-05 |
| NA | | NA |
| | 0.07784775 | 0.170315706 |
| | 0.789307087 | 0.872787041 |
| | 0.770738128 | 0.861348596 |
| NA | | NA |
| | 0.685256858 | NA |
| NA | | NA |
| | 0.889409301 | NA |
| | 0.832023774 | NA |
| | 0.900897778 | NA |
| | 0.424690189 | NA |
| NA | | NA |
| | 0.602960022 | NA |
| NA | | NA |
| | 0.466691792 | NA |
| NA | | NA |
| | 0.821110275 | NA |
| | 0.992061292 | NA |

| | |
|---|---|
| NA | NA |
| 0.989517374 | NA |
| 0.905158266 | NA |
| 0.712158425 | NA |
| 0.956994974 | 0.976109092 |
| 0.008925498 | 0.030686553 |
| 0.823234689 | 0.894625048 |
| 0.699215275 | 0.812014475 |
| 0.005488918 | 0.020713379 |
| 0.011019609 | 0.036409809 |
| NA | NA |
| 0.001135991 | 0.005542151 |
| 3.64E-08 | 7.42E-07 |
| 0.447758057 | 0.606886206 |
| NA | NA |
| NA | NA |
| 0.218919242 | 0.367582783 |
| 0.239483303 | 0.391741634 |
| 0.006220663 | 0.022938631 |
| 0.034997704 | 0.092037675 |
| 0.001292106 | 0.006172147 |
| 0.006039944 | 0.022388778 |
| NA | NA |
| NA | NA |
| NA | NA |
| 0.526350221 | NA |
| 0.746262054 | NA |
| 0.05044713 | 0.122216298 |
| 0.98889506 | 0.993964692 |
| 0.149990376 | 0.278703818 |
| 0.56924382 | 0.711562876 |
| 0.983251924 | 0.991763981 |
| NA | NA |
| 0.080736927 | 0.175153346 |
| 0.537162792 | 0.685346777 |
| NA | NA |
| 6.50E-08 | 1.22E-06 |
| 0.525419808 | 0.67542279 |
| 0.493361417 | 0.646928273 |
| 0.003342881 | 0.013720081 |
| 0.019944479 | 0.058660703 |
| 0.822788773 | NA |
| NA | NA |
| NA | NA |
| 0.905158739 | NA |
| 0.744551886 | 0.843589849 |
| 1.35E-05 | 0.000126262 |

| | |
|---|---|
| 0.869196632 | 0.925146083 |
| 0.847872191 | 0.910570032 |
| NA | NA |
| 0.013739084 | 0.043428322 |
| 0.992060816 | NA |
| 0.446945875 | NA |
| 0.143852938 | 0.270104034 |
| 0.613941366 | 0.74723082 |
| 0.09824114 | 0.203152575 |
| NA | NA |
| 0.175945893 | 0.314062418 |
| NA | NA |
| NA | NA |
| 0.083015672 | 0.178947791 |
| 0.132258691 | 0.25324979 |
| 0.000292676 | 0.001748932 |
| 0.271199159 | 0.427789837 |
| 0.020240924 | 0.059341849 |
| 0.48539722 | 0.640362725 |
| 0.082188332 | 0.177635196 |
| 0.003092132 | 0.012872061 |
| 0.572386897 | 0.714840691 |
| NA | NA |
| 4.96E-05 | 0.000384643 |
| 0.150515029 | 0.279387956 |
| 0.895231137 | 0.940997584 |
| 0.000314953 | 0.001863162 |
| 0.054355847 | 0.129331461 |
| NA | NA |
| 0.788866115 | 0.872608947 |
| 0.889409773 | NA |
| 0.604087497 | 0.739253085 |
| NA | NA |
| 0.165535851 | 0.299868154 |
| 0.889410245 | NA |
| NA | NA |
| 0.174212224 | 0.31155988 |
| 0.049196507 | 0.119834286 |
| NA | NA |
| 0.241040833 | NA |
| 0.0123034 | 0.039771878 |
| 0.174953513 | 0.312637664 |
| 0.00364688 | 0.014726342 |
| 0.695898713 | 0.809344835 |
| NA | NA |
| NA | NA |
| NA | NA |

| | | |
|---|---|---|
| | 0.889409773 | NA |
| | 0.002942204 | 0.012360411 |
| NA | | NA |
| NA | | NA |
| NA | | NA |
| | 0.001035272 | 0.005125938 |
| | 0.905157793 | NA |
| | 0.666333631 | NA |
| | 0.025621336 | 0.071955727 |
| | 0.417510236 | 0.579746255 |
| | 0.00419694 | 0.01654292 |
| | 0.131009815 | 0.251604383 |
| | 0.996451369 | NA |
| | 2.07E-08 | 4.49E-07 |
| | 0.905158266 | NA |
| | 0.011046735 | 0.036471996 |
| | 0.262510243 | 0.417956962 |
| NA | | NA |
| NA | | NA |
| NA | | NA |
| | 0.981283813 | NA |
| | 0.001208262 | 0.005837675 |
| | 1.19E-07 | 2.05E-06 |
| | 0.049630248 | 0.120703358 |
| NA | | NA |
| NA | | NA |
| | 6.46E-05 | 0.000481006 |
| | 0.030413312 | NA |
| | 0.541698632 | 0.689039122 |
| | 0.963679454 | 0.979968377 |
| | 0.809936444 | NA |
| | 4.11E-06 | 4.50E-05 |
| | 0.64087215 | 0.767071603 |
| | 0.517320254 | 0.668409049 |
| | 0.669668363 | 0.78941634 |
| | 0.819397492 | 0.892630535 |
| | 0.000108465 | 0.000748282 |
| | 0.658010361 | 0.780449904 |
| | 0.776244828 | NA |
| | 2.59E-06 | 3.00E-05 |
| NA | | NA |
| NA | | NA |
| | 0.00338748 | 0.013880514 |
| | 0.992061292 | NA |
| | 0.002037876 | 0.00909547 |
| | 0.622278298 | 0.753452202 |
| | 0.000531596 | 0.002916869 |

| | | |
|---|---|---|
| | 0.801661681 | NA |
| | 0.106792281 | 0.216357544 |
| | 0.564598392 | NA |
| NA | | NA |
| | 0.992060816 | NA |
| NA | | NA |
| | 0.069770478 | 0.156096784 |
| NA | | NA |
| NA | | NA |
| | 0.303887368 | 0.464621267 |
| NA | | NA |
| NA | | NA |
| | 0.013965352 | 0.044029456 |
| | 2.29E-05 | 0.000199333 |
| | 0.000204995 | 0.001288718 |
| | 0.31215253 | 0.473871926 |
| | 0.844908135 | NA |
| NA | | NA |
| | 0.992061292 | NA |
| | 0.031841914 | 0.085469166 |
| NA | | NA |
| | 0.889409773 | NA |
| | 0.936784399 | 0.964002466 |
| | 0.012734569 | 0.04090673 |
| | 7.73E-06 | 7.79E-05 |
| | 0.914812518 | 0.951789385 |
| | 0.009190564 | 0.031396329 |
| | 0.853394264 | NA |
| | 0.477108552 | 0.633206915 |
| | 5.26E-05 | 0.000403519 |
| | 0.856291041 | NA |
| | 0.946313688 | 0.969356826 |
| | 0.860367507 | NA |
| | 0.171505346 | 0.308039088 |
| | 0.982086287 | 0.991173786 |
| | 0.801661681 | NA |
| | 0.898486504 | 0.942868666 |
| | 0.690050448 | 0.804845052 |
| | 2.65E-08 | 5.56E-07 |
| NA | | NA |
| | 0.756222958 | 0.851753406 |
| | 0.830163713 | NA |
| | 0.002979045 | 0.012491284 |
| NA | | NA |
| NA | | NA |
| | 0.004117062 | 0.016289473 |
| NA | | NA |

| | | |
|---|---|---|
| | 0.833234055 | NA |
| | 0.427144108 | 0.588250704 |
| | 0.580459816 | 0.720660209 |
| | 0.931455826 | NA |
| | 0.875633797 | 0.928855436 |
| | 0.993078066 | NA |
| | 0.580233924 | 0.720553013 |
| | 0.944172081 | 0.968563707 |
| | 0.213494534 | 0.360657861 |
| | 0.679751535 | 0.79706648 |
| | 1.46E-07 | 2.45E-06 |
| | 0.006269668 | 0.023080576 |
| | 1.48E-12 | 1.01E-10 |
| NA | | NA |
| NA | | NA |
| NA | | NA |
| | 0.997810509 | 0.998993342 |
| | 0.540171485 | NA |
| | 0.752746491 | 0.849447196 |
| | 0.974819978 | 0.986909724 |
| | 0.690002736 | 0.804839857 |
| | 0.026485094 | 0.073691791 |
| | 0.976986449 | 0.988191867 |
| | 0.001199076 | 0.005806442 |
| | 0.632050358 | NA |
| | 0.943211402 | 0.96780427 |
| | 0.735544534 | NA |
| | 0.485265828 | 0.640266317 |
| | 0.003281923 | 0.013525808 |
| | 0.296139196 | 0.456281265 |
| NA | | NA |
| | 0.872131943 | 0.926653376 |
| NA | | NA |
| NA | | NA |
| | 0.889409773 | NA |
| | 0.756620589 | NA |
| | 0.002147141 | 0.009503945 |
| | 0.636144101 | 0.763266743 |
| | 0.861210161 | 0.919453056 |
| | 0.142503902 | NA |
| NA | | NA |
| | 0.98257181 | NA |
| | 0.271033458 | NA |
| | 0.129719439 | 0.249869225 |
| NA | | NA |
| | 0.836278944 | NA |
| | 0.393450474 | 0.556866192 |

| | |
|---|---|
| 0.006975122 | 0.025191893 |
| 0.002521002 | 0.010886664 |
| NA | NA |
| 0.889409773 | NA |
| NA | NA |
| NA | NA |
| 0.804906886 | 0.882820554 |
| 0.025149276 | 0.070866249 |
| 0.905158739 | NA |
| NA | NA |
| 0.219675421 | 0.368542484 |
| 0.000294972 | 0.001760133 |
| 0.889409301 | NA |
| 0.004772408 | 0.018437247 |
| 0.000753304 | 0.003930102 |
| 0.644255488 | NA |
| 0.46245238 | 0.61973586 |
| 0.340834315 | 0.504206357 |
| 0.345107858 | 0.508469099 |
| 0.349984662 | NA |
| 0.519922229 | 0.670746698 |
| 0.837007813 | 0.9030982 |
| 0.035008429 | 0.092043834 |
| 0.595060939 | 0.732082646 |
| NA | NA |
| 0.992061292 | NA |
| NA | NA |
| NA | NA |
| NA | NA |
| NA | NA |
| 0.976955209 | 0.988191867 |
| NA | NA |
| 4.75E-13 | 3.67E-11 |
| 0.066352953 | 0.150630052 |
| NA | NA |
| NA | NA |
| NA | NA |
| NA | NA |
| NA | NA |
| NA | NA |
| 0.991879747 | NA |
| NA | NA |
| 0.265132274 | 0.421263841 |
| 0.02695409 | 0.074770699 |
| 0.000368071 | 0.002126969 |
| 0.540825362 | 0.688183586 |
| 0.254598725 | 0.409068077 |

| | | |
|---|---|---|
| | 0.164633641 | 0.298776043 |
| | 0.000126073 | 0.000849767 |
| | 0.130005912 | 0.250135794 |
| | 0.727514261 | 0.831796608 |
| | 0.991782148 | NA |
| NA | | NA |
| NA | | NA |
| NA | | NA |
| NA | | NA |
| | 0.905158739 | NA |
| | 5.48E-06 | 5.76E-05 |
| | 0.226172899 | NA |
| NA | | NA |
| NA | | NA |
| NA | | NA |
| | 0.991782148 | NA |
| | 0.989952082 | NA |
| | 0.546845382 | 0.693192735 |
| | 0.13935635 | 0.263622164 |
| NA | | NA |
| NA | | NA |
| | 0.049885324 | 0.121122494 |
| | 0.017955649 | 0.053908682 |
| | 0.000367878 | 0.002126411 |
| | 0.098856787 | 0.204098927 |
| | 0.001939895 | 0.008723768 |
| | 0.077059399 | 0.169011919 |
| | 0.20116852 | 0.346358752 |
| NA | | NA |
| | 0.13181077 | 0.252722666 |
| | 0.69367651 | 0.807442869 |
| NA | | NA |
| | 0.991635679 | 0.995807742 |
| NA | | NA |
| | 0.972174218 | 0.985471871 |
| | 1.31E-08 | 2.99E-07 |
| | 0.905158739 | NA |
| | 0.087022535 | 0.185296429 |
| | 0.028908639 | 0.079083692 |
| | 0.514265002 | NA |
| | 0.975587446 | 0.987504597 |
| | 0.00298815 | 0.0125199 |
| | 2.37E-06 | 2.78E-05 |
| | 0.578438817 | 0.719452732 |
| | 0.277826073 | 0.435806289 |
| | 0.272590004 | 0.429397698 |
| | 0.830163713 | NA |

| | |
|---|---|
| 1.66E-19 | 6.27E-17 |
| 0.296457318 | 0.456675288 |
| 0.161555784 | 0.294382497 |
| 0.75862276 | 0.853542921 |
| 0.313420283 | 0.475303591 |
| NA | NA |
| 0.745747286 | 0.844682759 |
| 0.707149378 | 0.817665905 |
| 0.00042682 | 0.002416226 |
| 0.93793049 | NA |
| 0.010622232 | 0.035368247 |
| 0.007922236 | 0.027847669 |
| 0.889409773 | NA |
| 0.590970823 | 0.728875998 |
| NA | NA |
| 0.480845053 | 0.636498675 |
| 0.030457401 | 0.082467827 |
| 0.479200826 | 0.634896279 |
| 0.16382794 | 0.297589941 |
| 0.786112585 | NA |
| 0.089510796 | 0.18939753 |
| 0.466181847 | 0.622946947 |
| 0.720758493 | 0.826870892 |
| 0.591939749 | NA |
| NA | NA |
| NA | NA |
| 0.310141684 | 0.471405915 |
| 0.196901792 | 0.341044837 |
| 0.002981068 | 0.01249496 |
| 0.305115884 | 0.465716603 |
| 0.985770346 | 0.992788917 |
| 0.155936484 | 0.286880802 |
| NA | NA |
| 3.56E-15 | 4.59E-13 |
| 0.060449401 | 0.140580008 |
| NA | NA |
| 0.603200339 | 0.738661401 |
| 0.263851727 | 0.419592999 |
| 0.001051766 | 0.005195891 |
| 0.00147655 | 0.006914816 |
| 0.627683506 | 0.756492981 |
| NA | NA |
| NA | NA |
| 0.084245415 | 0.180641058 |
| 0.494222625 | 0.647489411 |
| 0.726862809 | 0.83131148 |
| NA | NA |

| | |
|---|---|
| 0.146677618 | 0.274025336 |
| 0.882330306 | 0.933154595 |
| NA | NA |
| 0.549609809 | 0.695733083 |
| 0.08053657 | 0.174804705 |
| 0.003693668 | 0.014882432 |
| 0.079461785 | 0.173002314 |
| NA | NA |
| 0.61614555 | 0.748833725 |
| 1.74E-10 | 6.96E-09 |
| NA | NA |
| 0.322981478 | 0.485411695 |
| 0.099125089 | 0.204575924 |
| 0.732065642 | NA |
| 1.32E-08 | 3.02E-07 |
| 0.080351327 | 0.17455825 |
| 0.056853205 | 0.133751823 |
| 0.05431039 | 0.129279333 |
| 0.648551948 | NA |
| 0.00120557 | 0.005827229 |
| 0.019083889 | 0.05653084 |
| 2.18E-07 | 3.49E-06 |
| 0.992061292 | NA |
| 0.955449063 | 0.975166441 |
| 0.33096184 | 0.493353544 |
| 0.824151396 | NA |
| 0.001132859 | 0.005529329 |
| 1.34E-17 | 3.07E-15 |
| 0.000996229 | 0.004959559 |
| 0.000105403 | 0.000730602 |
| 0.290127167 | 0.449475613 |
| 0.229135702 | 0.379599463 |
| 0.013952259 | 0.04401193 |
| NA | NA |
| 0.330709269 | 0.493244911 |
| 0.992060816 | NA |
| 0.158915158 | 0.290915817 |
| 0.00091867 | 0.004639622 |
| NA | NA |
| 0.637111008 | 0.763997412 |
| 0.177477549 | 0.316307888 |
| 0.541492181 | NA |
| 0.763369186 | 0.856631755 |
| 4.00E-05 | 0.000320068 |
| NA | NA |
| 0.291404488 | 0.450884382 |
| 0.23384633 | 0.385378497 |

| | |
|---|---|
| 0.611600133 | 0.745083347 |
| NA | NA |
| NA | NA |
| NA | NA |
| 0.001242776 | 0.005970273 |
| 0.751773536 | 0.848794029 |
| 0.008663406 | 0.029959013 |
| 0.905158739 | NA |
| 0.000472059 | 0.002626952 |
| 4.35E-08 | 8.62E-07 |
| 0.010712363 | 0.03561435 |
| 0.018816634 | 0.055935048 |
| 0.114312015 | 0.227845832 |
| 0.000303504 | 0.001804668 |
| NA | NA |
| 0.00010972 | 0.000755754 |
| 0.099432498 | 0.205094705 |
| 0.450848276 | 0.60924168 |
| NA | NA |
| 0.001431709 | 0.006750891 |
| 0.699201804 | 0.812014475 |
| 0.034054843 | 0.090065072 |
| 0.741913059 | 0.841859697 |
| NA | NA |
| 5.67E-05 | 0.000431814 |
| 0.510251795 | NA |
| 0.768219694 | 0.859627816 |
| 3.80E-20 | 1.81E-17 |
| 0.001591667 | 0.007361292 |
| NA | NA |
| 0.501765982 | 0.654486439 |
| 7.24E-05 | 0.00053056 |
| 7.31E-05 | 0.000534645 |
| 2.40E-14 | 2.48E-12 |
| 0.830163713 | NA |
| 0.001511214 | 0.007048588 |
| 0.037041558 | 0.096316824 |
| 0.60133646 | NA |
| 0.078993515 | 0.172290285 |
| 0.002938261 | 0.012348642 |
| 0.372206912 | 0.536601091 |
| 0.129323953 | 0.249406383 |
| NA | NA |
| 0.795353865 | NA |
| NA | NA |
| 4.39E-05 | 0.000346598 |
| 0.905158266 | NA |

| | |
|---|---|
| 0.060978941 | 0.141532244 |
| 0.277270954 | NA |
| 0.155544067 | 0.286350785 |
| 0.299711611 | 0.460069055 |
| 0.187980394 | 0.329403676 |
| 0.88933045 | 0.937385064 |
| 0.363918326 | 0.528873761 |
| 0.289073363 | 0.448383271 |
| NA | NA |
| 0.234063479 | 0.385552466 |
| 0.579027825 | NA |
| 0.067167912 | 0.151789665 |
| 0.905158739 | NA |
| 0.567019208 | 0.709891565 |
| 0.925971649 | 0.957497242 |
| 0.004712866 | 0.018248956 |
| 0.690859096 | 0.805274956 |
| 0.252518923 | NA |
| 0.116391889 | 0.230953967 |
| 0.00052921 | 0.002905958 |
| 6.41E-09 | 1.59E-07 |
| 0.052714125 | 0.126356234 |
| NA | NA |
| 9.15E-08 | 1.63E-06 |
| 0.222638636 | NA |
| NA | NA |
| NA | NA |
| 0.156815008 | 0.288038542 |
| NA | NA |
| NA | NA |
| 0.811214529 | NA |
| 0.095539527 | NA |
| 0.889409773 | NA |
| NA | NA |
| 0.607453915 | 0.741636858 |
| 0.674615118 | 0.79317938 |
| 0.116149364 | 0.230712247 |
| 0.090608748 | 0.191149725 |
| 0.992060816 | NA |
| 0.013325816 | 0.042335299 |
| 0.370768951 | 0.535351784 |
| 0.007116228 | 0.025578967 |
| 0.00037568 | 0.002164671 |
| 0.009721409 | 0.032859929 |
| 0.769420914 | 0.860621121 |
| 0.74770977 | 0.846120012 |
| 0.030698625 | NA |

| | |
|---|---|
| 0.55150626 | 0.697129056 |
| 0.510528688 | 0.662070483 |
| 0.194424678 | 0.337740013 |
| 0.015405301 | 0.047719706 |
| 0.019975055 | 0.058720961 |
| 0.0012201 | 0.005881927 |
| NA | NA |
| 0.325236848 | 0.48746679 |
| 1.20E-06 | 1.53E-05 |
| 0.001196407 | 0.0057957 |
| 0.089431102 | 0.189261672 |
| 0.302800399 | 0.463552284 |
| 0.107553544 | 0.21752258 |
| 0.407467493 | 0.570662223 |
| 0.125965451 | 0.244612252 |
| 0.077092511 | 0.16903817 |
| 0.683114255 | 0.799524246 |
| 0.168505602 | 0.304043921 |
| NA | NA |
| NA | NA |
| 8.64E-07 | 1.16E-05 |
| 0.273974901 | 0.431115361 |
| 0.710365054 | 0.819485486 |
| 0.658130871 | 0.780550678 |
| 0.712535705 | NA |
| 0.069589775 | 0.155787732 |
| 9.69E-13 | 6.84E-11 |
| 0.033810173 | 0.089579825 |
| 0.137514479 | 0.261014819 |
| 0.500458623 | 0.65350931 |
| 0.472080957 | NA |
| 0.571408675 | 0.713781392 |
| 0.058014613 | 0.135901737 |
| 0.000442302 | 0.002489113 |
| 0.019080995 | 0.056529892 |
| 0.885023465 | 0.934832376 |
| 0.171435562 | 0.307938937 |
| 0.85801105 | NA |
| 0.025288967 | 0.071194927 |
| 0.247295661 | 0.400878279 |
| 0.778834447 | 0.86693669 |
| 0.012550347 | 0.040401475 |
| 0.989944333 | 0.994655118 |
| 0.444000975 | NA |
| 0.201479454 | 0.346751649 |
| 0.018373254 | 0.054879246 |
| 5.15E-05 | 0.000396584 |

| | | |
|---|---|---|
| | 0.042125111 | 0.106288096 |
| | 0.027722887 | 0.076526319 |
| | 0.323478514 | 0.485792961 |
| | 0.033953641 | 0.089853209 |
| NA | | NA |
| NA | | NA |
| | 0.76085055 | 0.854998067 |
| | 0.00392204 | 0.015627667 |
| | 0.191899878 | 0.334359234 |
| | 0.776480828 | NA |
| | 8.02E-07 | 1.09E-05 |
| | 0.425118442 | NA |
| | 0.023567486 | 0.067145789 |
| | 0.56411286 | NA |
| | 0.263561325 | 0.419384261 |
| | 0.788582525 | NA |
| | 0.616610819 | 0.749171172 |
| NA | | NA |
| NA | | NA |
| | 0.049879759 | 0.121122494 |
| | 0.231516778 | 0.382499915 |
| NA | | NA |
| | 0.842896533 | NA |
| NA | | NA |
| | 3.10E-07 | 4.73E-06 |
| | 0.511526034 | 0.66315977 |
| | 0.505988695 | 0.657923126 |
| | 0.381829598 | 0.545872145 |
| | 0.398750779 | 0.562302104 |
| | 0.978334058 | 0.988994282 |
| | 0.024248533 | 0.068750617 |
| | 0.125464947 | 0.243920741 |
| | 0.32180383 | 0.484138825 |
| | 0.002308096 | 0.010096312 |
| | 0.062728719 | 0.144554511 |
| | 0.980857979 | 0.990510939 |
| | 0.000764507 | 0.003975351 |
| | 0.000315867 | 0.001867536 |
| | 0.000327968 | 0.001930273 |
| | 0.905158739 | NA |
| | 0.619909678 | 0.751537383 |
| | 0.001428488 | 0.006737151 |
| | 0.281311955 | NA |
| | 0.585640571 | 0.725001764 |
| | 0.905296333 | NA |
| | 0.889409773 | NA |
| NA | | NA |

| | |
|---|---|
| 0.010108946 | 0.033960639 |
| 0.191645647 | 0.334114982 |
| NA | NA |
| NA | NA |
| NA | NA |
| NA | NA |
| NA | NA |
| NA | NA |
| NA | NA |
| NA | NA |
| NA | NA |
| NA | NA |
| NA | NA |
| NA | NA |
| NA | NA |
| NA | NA |
| NA | NA |
| NA | NA |
| NA | NA |
| NA | NA |
| 0.893816483 | NA |
| NA | NA |
| NA | NA |
| NA | NA |
| NA | NA |
| 0.88804077 | NA |
| NA | NA |
| 0.077818213 | 0.170268048 |
| 0.007573004 | 0.026860861 |
| 0.836489909 | NA |
| NA | NA |
| 0.109033041 | 0.219918853 |
| 8.04E-08 | 1.46E-06 |
| 2.63E-05 | 0.000223453 |
| 0.05133132 | NA |
| NA | NA |
| NA | NA |
| NA | NA |
| 0.046300235 | 0.114273864 |
| NA | NA |
| NA | NA |
| 0.421372931 | 0.583045254 |
| 0.797342596 | NA |
| 2.27E-05 | 0.000198032 |
| 0.005496643 | 0.020727502 |
| 0.015705649 | 0.048492738 |
| 0.380743274 | 0.544993257 |

| | | |
|---|---|---|
| | 0.807879735 | NA |
| NA | | NA |
| | 0.847358355 | 0.910374388 |
| | 0.792188023 | NA |
| NA | | NA |
| NA | | NA |
| | 0.110607717 | 0.222052669 |
| | 0.905158266 | NA |
| | 0.889409773 | NA |
| | 0.921800932 | NA |
| | 2.91E-05 | 0.000244459 |
| NA | | NA |
| NA | | NA |
| | 0.286637892 | 0.445707511 |
| NA | | NA |
| | 0.08606812 | 0.183724104 |
| | 0.102139546 | 0.209399276 |
| | 0.000335334 | 0.001968879 |
| NA | | NA |
| | 0.000246865 | 0.001513372 |
| | 4.72E-13 | 3.66E-11 |
| | 0.109582791 | 0.220723584 |
| | 0.003024661 | 0.012639113 |
| | 0.795292913 | 0.87663136 |
| | 0.296756955 | 0.456848428 |
| | 0.385765965 | NA |
| | 0.350328072 | 0.514468945 |
| | 0.866012479 | NA |
| | 0.110159697 | 0.22149266 |
| | 0.742103312 | 0.841988562 |
| | 0.376530961 | 0.540928746 |
| | 0.407355149 | NA |
| | 0.769870849 | 0.860905127 |
| | 0.37372596 | 0.53785872 |
| | 3.32E-11 | 1.59E-09 |
| | 0.889409773 | NA |
| | 0.072459904 | 0.16077064 |
| | 0.016774478 | 0.051143277 |
| | 0.207834149 | 0.354519298 |
| | 3.69E-11 | 1.75E-09 |
| | 0.60594319 | NA |
| | 7.35E-07 | 1.01E-05 |
| | 6.76E-13 | 5.03E-11 |
| | 0.670086127 | NA |
| | 0.045848225 | 0.113503294 |
| | 0.091765642 | 0.192960044 |
| | 0.094537125 | 0.19729526 |

| | |
|---|---|
| 0.080123446 | 0.174132101 |
| 0.587479897 | NA |
| 0.454373694 | NA |
| 4.11E-09 | 1.07E-07 |
| 0.337882721 | 0.500746891 |
| NA | NA |
| NA | NA |
| NA | NA |
| 0.992060816 | NA |
| NA | NA |
| 0.015367329 | 0.047628959 |
| NA | NA |
| 0.667106956 | 0.787551955 |
| 6.52E-06 | 6.74E-05 |
| NA | NA |
| NA | NA |
| 0.420704943 | 0.58248801 |
| 0.265143565 | 0.421263841 |
| 0.603737997 | 0.739113714 |
| 0.000167178 | 0.001080361 |
| 9.06E-05 | 0.000641302 |
| NA | NA |
| 0.000432722 | 0.002444594 |
| 1.08E-09 | 3.37E-08 |
| 0.542702608 | 0.689772911 |
| 0.00126589 | 0.006068038 |
| 0.941784361 | 0.967042615 |
| 0.811532096 | 0.887295025 |
| 0.828052773 | NA |
| 0.998500156 | 0.999410408 |
| 0.837943141 | 0.903707747 |
| 0.889409773 | NA |
| NA | NA |
| 0.455981974 | 0.613799312 |
| 0.194171898 | 0.337487788 |
| 0.001122002 | 0.005487912 |
| 0.013228936 | 0.042094508 |
| 0.43925024 | 0.599545998 |
| 0.001656295 | 0.007599371 |
| NA | NA |
| 0.01099516 | 0.036356381 |
| 0.050495452 | 0.122297849 |
| 0.549096574 | NA |
| 0.361972332 | 0.526847647 |
| 0.015061634 | 0.046853438 |
| 0.618486557 | 0.750475012 |
| 0.443734515 | 0.603341561 |

| | | |
|---|---|---|
| | 0.781517391 | NA |
| | 0.052448432 | 0.125898023 |
| | 0.809775847 | 0.885992417 |
| | 0.215546565 | 0.363280914 |
| | 0.470912023 | 0.627274037 |
| | 0.000469779 | 0.002617577 |
| | 0.992061292 | NA |
| | 0.071768018 | 0.159590269 |
| | 0.102072844 | 0.20930223 |
| NA | | NA |
| | 0.87762688 | NA |
| NA | | NA |
| | 0.905158266 | NA |
| NA | | NA |
| NA | | NA |
| | 0.408243419 | 0.571467918 |
| NA | | NA |
| NA | | NA |
| NA | | NA |
| | 0.767418459 | NA |
| NA | | NA |
| | 0.757535471 | NA |
| | 0.807252815 | NA |
| | 0.366044269 | NA |
| | 0.290244515 | NA |
| NA | | NA |
| | 0.978044343 | NA |
| | 0.095199025 | 0.198280886 |
| NA | | NA |
| | 0.000237501 | 0.001467028 |
| | 0.992062245 | NA |
| | 0.638733623 | NA |
| | 0.807879735 | NA |
| | 0.905158266 | NA |
| NA | | NA |
| | 0.509804191 | NA |
| | 0.801661681 | NA |
| | 0.90766557 | 0.947610543 |
| NA | | NA |
| | 0.23036974 | 0.381097568 |
| | 0.135896209 | 0.258569002 |
| NA | | NA |
| | 0.000259429 | 0.001580249 |
| | 0.030120474 | 0.081757167 |
| | 0.297496558 | 0.457496243 |
| NA | | NA |
| | 0.017850589 | 0.053673981 |

| | |
|---|---|
| 0.5712936 | NA |
| NA | NA |
| 0.081980309 | 0.177255376 |
| NA | NA |
| 0.865775134 | NA |
| 0.788582525 | NA |
| 0.058567088 | 0.136947577 |
| 7.80E-12 | 4.47E-10 |
| 0.326648495 | NA |
| 0.307736563 | 0.468723534 |
| 0.000503276 | 0.002778838 |
| 0.347623159 | 0.511463442 |
| NA | NA |
| 0.919311214 | 0.953979252 |
| 0.75829253 | NA |
| 0.788582525 | NA |
| NA | NA |
| NA | NA |
| NA | NA |
| NA | NA |
| 0.149136462 | 0.277415296 |
| 0.523767537 | 0.673976639 |
| 0.000500184 | 0.002763156 |
| 0.492724235 | 0.646649025 |
| 0.239193945 | 0.391469569 |
| NA | NA |
| 0.001014064 | 0.005035691 |
| 0.607016399 | 0.741298247 |
| 1.59E-05 | 0.00014614 |
| NA | NA |
| 0.283054754 | 0.441856782 |
| NA | NA |
| 0.444113355 | NA |
| NA | NA |
| 0.606631086 | NA |
| 0.813836939 | NA |
| 0.99205891 | NA |
| NA | NA |
| NA | NA |
| NA | NA |
| NA | NA |
| 0.741044223 | NA |
| 0.892067941 | NA |
| NA | NA |
| NA | NA |
| 0.842849934 | NA |
| NA | NA |

| | | |
|---|---|---|
| | 0.668607738 | NA |
| NA | | NA |
| NA | | NA |
| | 0.877082314 | NA |
| | 0.991529086 | NA |
| | 1.22E-10 | 5.07E-09 |
| | 0.003199895 | 0.01325533 |
| | 0.051908746 | 0.124916618 |
| | 0.190049065 | 0.332030187 |
| | 0.104453855 | 0.212852841 |
| | 0.264351542 | NA |
| | 2.04E-10 | 8.02E-09 |
| | 0.000492239 | 0.002725438 |
| NA | | NA |
| | 0.3688047 | 0.533433172 |
| | 1.86E-05 | 0.000167266 |
| | 0.506460192 | 0.658341287 |
| | 0.977579575 | 0.98847308 |
| NA | | NA |
| | 0.000382921 | 0.002200044 |
| | 0.375280622 | 0.539424441 |
| | 0.997739693 | 0.998967988 |
| | 0.852052865 | 0.913585393 |
| | 1.99E-09 | 5.77E-08 |
| | 0.980753582 | 0.990451063 |
| | 0.771728957 | 0.862104908 |
| | 4.28E-08 | 8.50E-07 |
| | 0.351224743 | NA |
| NA | | NA |
| NA | | NA |
| | 0.519139107 | 0.670062638 |
| NA | | NA |
| | 0.031592422 | 0.08491285 |
| | 0.000652246 | 0.003484836 |
| NA | | NA |
| NA | | NA |
| NA | | NA |
| | 0.876792175 | NA |
| | 0.090944004 | 0.191636061 |
| | 0.03879001 | 0.099682836 |
| | 0.042197031 | 0.106432875 |
| NA | | NA |
| | 0.023754808 | 0.067533898 |
| | 0.013301724 | 0.042270994 |
| | 0.102242908 | 0.209553109 |
| | 0.718327207 | 0.8253614 |
| | 0.022053937 | 0.063612558 |

| | |
|---|---|
| 0.263654981 | 0.419384261 |
| 0.030819859 | 0.083223473 |
| 0.069869438 | 0.156246285 |
| 0.687018389 | 0.802416722 |
| 0.569592217 | 0.711957849 |
| NA | NA |
| 0.048543347 | 0.118554978 |
| 0.29567193 | NA |
| 9.49E-05 | 0.000667859 |
| 0.127246189 | 0.246532473 |
| 0.163789283 | 0.297563404 |
| 0.079700773 | 0.173367932 |
| 0.339211805 | 0.502174343 |
| 0.814139876 | 0.889260721 |
| NA | NA |
| NA | NA |
| 0.677416716 | 0.795390529 |
| NA | NA |
| 3.40E-07 | 5.13E-06 |
| NA | NA |
| NA | NA |
| 0.118706941 | 0.234203568 |
| NA | NA |
| NA | NA |
| 0.371065971 | 0.535520493 |
| 0.124556055 | 0.242733973 |
| NA | NA |
| 0.008071444 | 0.028272586 |
| NA | NA |
| 0.265222103 | 0.421327655 |
| 0.319903961 | 0.482206541 |
| 0.239445666 | 0.391737301 |
| 0.938142341 | 0.964947664 |
| 0.026989996 | 0.07484379 |
| 0.008036496 | 0.028182695 |
| 0.066245092 | 0.150478533 |
| 0.067773157 | 0.152705394 |
| 0.110166348 | 0.22149266 |
| 0.103727242 | 0.211725833 |
| 0.02816164 | 0.077445391 |
| 0.189192133 | 0.331007095 |
| 0.183083766 | 0.3233466 |
| 0.700238249 | 0.812670139 |
| NA | NA |
| NA | NA |
| 0.549396526 | 0.695543287 |
| 0.121220298 | 0.237931568 |

| | | |
|---|---|---|
| | 0.274385802 | 0.43164765 |
| | 0.339276093 | 0.502235656 |
| | 0.284589534 | 0.443415018 |
| | 0.301464847 | 0.461861897 |
| | 0.000661934 | 0.003528407 |
| | 0.190816673 | 0.332973881 |
| | 0.210500292 | 0.357437049 |
| | 0.231896519 | 0.382930941 |
| | 0.962919341 | 0.979467566 |
| | 0.057668602 | 0.135264348 |
| | 0.151983163 | 0.281613221 |
| | 0.577334975 | 0.71861917 |
| | 0.589226098 | 0.72775836 |
| | 0.992060816 | NA |
| | 0.573573496 | 0.715672133 |
| | 0.922562887 | 0.955374089 |
| | 0.181555402 | 0.321773614 |
| | 0.005711589 | 0.021417483 |
| NA | | NA |
| | 0.556990696 | 0.702041022 |
| | 0.099639043 | 0.205407913 |
| | 0.229241823 | 0.37966525 |
| | 0.432805072 | 0.593657944 |
| | 0.370558944 | 0.53515974 |
| NA | | NA |
| NA | | NA |
| NA | | NA |
| | 0.121185463 | 0.237917888 |
| | 0.564879642 | 0.70828482 |
| | 0.461254681 | 0.618621606 |
| NA | | NA |
| | 0.286968754 | 0.446000912 |
| | 0.979048407 | 0.989366036 |
| NA | | NA |
| | 0.111036806 | 0.222631229 |
| | 0.47623269 | NA |
| NA | | NA |
| | 0.964180044 | NA |
| | 0.900836801 | NA |
| | 0.000166693 | 0.001077857 |
| | 0.001183792 | 0.005744726 |
| | 0.017567197 | 0.052976746 |
| | 0.154263294 | NA |
| | 0.555287541 | NA |
| | 0.545472245 | NA |
| | 0.055768281 | 0.13180666 |
| | 0.507958096 | 0.659741635 |

| | |
|---|---|
| 0.154282445 | 0.284720414 |
| 0.772366419 | NA |
| 0.672540761 | NA |
| 0.707480788 | 0.817747691 |
| 0.272146836 | 0.428914977 |
| 0.08157417 | 0.176551075 |
| 5.20E-08 | 1.00E-06 |
| 0.433925813 | 0.594651468 |
| 0.281210942 | 0.439634849 |
| 1.32E-06 | 1.66E-05 |
| 0.678893424 | NA |
| 0.412337399 | 0.574889965 |
| 3.03E-06 | 3.43E-05 |
| 0.388452727 | 0.551962569 |
| 0.004198275 | 0.016545211 |
| 0.373778074 | 0.537868166 |
| 0.04798852 | 0.117478252 |
| 0.017185778 | 0.052110259 |
| 0.147639239 | 0.275396674 |
| 0.41214004 | 0.574764235 |
| 0.099997682 | 0.205989596 |
| 0.029801742 | 0.0811126 |
| 0.000552872 | 0.003018525 |
| 0.157356934 | 0.288744122 |
| 0.823098491 | 0.894593615 |
| 0.000141759 | 0.000937618 |
| 0.935779851 | 0.963058994 |
| 0.562340725 | 0.70614971 |
| 0.992061292 | NA |
| 0.992061292 | NA |
| 0.27414487 | 0.431320999 |
| 0.131414822 | 0.252183699 |
| 0.971894545 | 0.985310818 |
| 1.35E-07 | 2.30E-06 |
| 0.919158655 | 0.953956182 |
| 0.583222771 | 0.723150132 |
| 0.390547819 | 0.554049869 |
| 6.10E-05 | 0.0004587 |
| 0.982589167 | NA |
| NA | NA |
| NA | NA |
| 0.03579837 | 0.093675629 |
| 0.277758249 | NA |
| 0.769785117 | 0.860853096 |
| NA | NA |
| 0.00013388 | 0.000894392 |
| 0.516246535 | 0.667349012 |

| | | |
|---|---|---|
| | 0.011159558 | 0.036761588 |
| | 0.331158128 | 0.493512136 |
| | 0.895450219 | 0.941047685 |
| | 0.809516396 | NA |
| | 0.035177802 | 0.092356452 |
| | 0.824325859 | 0.895176873 |
| | 5.62E-05 | 0.000428769 |
| | 0.728016651 | NA |
| | 0.001866366 | 0.008438035 |
| | 0.000107527 | 0.000742981 |
| | 0.00045505 | 0.002548445 |
| | 0.037929175 | 0.098054105 |
| | 0.075136336 | 0.165528585 |
| | 0.003511773 | 0.014299097 |
| | 0.623493075 | NA |
| | 0.641307233 | 0.767353179 |
| | 0.030755607 | 0.083101066 |
| | 0.010671975 | 0.035506954 |
| | 0.000284098 | 0.001710117 |
| | 0.471736943 | NA |
| | 0.021881085 | 0.063215887 |
| | 0.527868161 | NA |
| | 0.386927743 | 0.550758171 |
| | 0.018420065 | 0.054957366 |
| | 0.009884281 | 0.033305036 |
| | 2.34E-05 | 0.000202997 |
| | 0.651080289 | 0.775497509 |
| | 0.889409773 | NA |
| | 0.587082117 | 0.726049572 |
| NA | | NA |
| | 0.089560113 | 0.189447118 |
| | 0.807433834 | NA |
| | 0.054364737 | 0.129338597 |
| | 0.341607628 | 0.504972192 |
| | 0.992060816 | NA |
| | 0.620715587 | NA |
| | 0.952165213 | 0.973288471 |
| | 0.905158739 | NA |
| | 0.674835053 | 0.793207523 |
| NA | | NA |
| | 0.905158739 | NA |
| | 0.150680907 | 0.279648582 |
| | 0.273104414 | 0.430022932 |
| NA | | NA |
| NA | | NA |
| NA | | NA |
| NA | | NA |

| | | |
|---|---|---|
| | 0.858476467 | NA |
| | 0.830163713 | NA |
| | 0.800548326 | NA |
| | 0.398650723 | NA |
| NA | | NA |
| NA | | NA |
| NA | | NA |
| NA | | NA |
| NA | | NA |
| NA | | NA |
| NA | | NA |
| NA | | NA |
| NA | | NA |
| NA | | NA |
| NA | | NA |
| NA | | NA |
| NA | | NA |
| NA | | NA |
| NA | | NA |
| NA | | NA |
| NA | | NA |
| NA | | NA |
| NA | | NA |
| NA | | NA |
| | 0.843922708 | NA |
| NA | | NA |
| NA | | NA |
| NA | | NA |
| NA | | NA |
| NA | | NA |
| NA | | NA |
| NA | | NA |
| NA | | NA |
| NA | | NA |
| | 0.231992229 | NA |
| NA | | NA |
| | 0.889409301 | NA |
| NA | | NA |
| | 0.905158266 | NA |
| | 0.08720265 | 0.185586644 |
| NA | | NA |
| NA | | NA |
| NA | | NA |
| | 0.905158739 | NA |
| NA | | NA |
| NA | | NA |
| NA | | NA |

| | | |
|---|---|---|
| NA | | NA |
| NA | | NA |
| | 0.889409773 | NA |
| NA | | NA |
| NA | | NA |
| NA | | NA |
| NA | | NA |
| NA | | NA |
| NA | | NA |
| NA | | NA |
| NA | | NA |
| NA | | NA |
| | 0.905158266 | NA |
| NA | | NA |
| NA | | NA |
| NA | | NA |
| NA | | NA |
| NA | | NA |
| | 0.582567919 | NA |
| NA | | NA |
| NA | | NA |
| | 0.394551482 | NA |
| NA | | NA |
| NA | | NA |
| NA | | NA |
| | 0.987241596 | NA |
| | 0.889409301 | NA |
| NA | | NA |
| NA | | NA |
| NA | | NA |
| | 0.863203912 | NA |
| NA | | NA |
| NA | | NA |
| NA | | NA |
| NA | | NA |
| NA | | NA |
| NA | | NA |
| NA | | NA |
| NA | | NA |
| NA | | NA |
| NA | | NA |
| NA | | NA |
| NA | | NA |
| NA | | NA |
| NA | | NA |
| NA | | NA |
| NA | | NA |

| | | |
|---|---|---|
| NA | | NA |
| NA | | NA |
| NA | | NA |
| NA | | NA |
| NA | | NA |
| | 0.684061721 | NA |
| NA | | NA |
| NA | | NA |
| NA | | NA |
| | 0.905158739 | NA |
| NA | | NA |
| NA | | NA |
| NA | | NA |
| NA | | NA |
| NA | | NA |
| NA | | NA |
| NA | | NA |
| | 0.703816352 | NA |
| NA | | NA |
| NA | | NA |
| NA | | NA |
| NA | | NA |
| | 0.905158266 | NA |
| NA | | NA |
| NA | | NA |
| NA | | NA |
| NA | | NA |
| NA | | NA |
| NA | | NA |
| NA | | NA |
| NA | | NA |
| NA | | NA |
| NA | | NA |
| NA | | NA |
| | 0.623149859 | NA |
| NA | | NA |
| NA | | NA |
| NA | | NA |
| NA | | NA |
| NA | | NA |
| NA | | NA |
| NA | | NA |
| NA | | NA |
| NA | | NA |
| | 0.784353866 | NA |
| NA | | NA |
| NA | | NA |

| | | |
|---|---|---|
| NA | | NA |
| NA | | NA |
| NA | | NA |
| NA | | NA |
| | 0.905158739 | NA |
| NA | | NA |
| NA | | NA |
| NA | | NA |
| NA | | NA |
| NA | | NA |
| NA | | NA |
| NA | | NA |
| NA | | NA |
| NA | | NA |
| | 0.905158739 | NA |
| NA | | NA |
| NA | | NA |
| NA | | NA |
| NA | | NA |
| NA | | NA |
| NA | | NA |
| NA | | NA |
| NA | | NA |
| NA | | NA |
| NA | | NA |
| NA | | NA |
| NA | | NA |
| NA | | NA |
| NA | | NA |
| NA | | NA |
| NA | | NA |
| NA | | NA |
| NA | | NA |
| NA | | NA |
| NA | | NA |
| NA | | NA |
| | 0.889409301 | NA |
| NA | | NA |
| NA | | NA |
| NA | | NA |
| NA | | NA |
| NA | | NA |
| NA | | NA |
| NA | | NA |
| | 0.598749562 | NA |
| NA | | NA |
| NA | | NA |

| | | |
|---|---|---|
| NA | | NA |
| | 0.777092376 | NA |
| NA | | NA |
| NA | | NA |
| NA | | NA |
| NA | | NA |
| NA | | NA |
| NA | | NA |
| NA | | NA |
| NA | | NA |
| NA | | NA |
| NA | | NA |
| NA | | NA |
| NA | | NA |
| NA | | NA |
| | 0.905158266 | NA |
| NA | | NA |
| NA | | NA |
| NA | | NA |
| NA | | NA |
| NA | | NA |
| | 0.838862418 | NA |
| | 0.905158266 | NA |
| NA | | NA |
| NA | | NA |
| | 0.889409773 | NA |
| NA | | NA |
| NA | | NA |
| NA | | NA |
| NA | | NA |
| NA | | NA |
| NA | | NA |
| NA | | NA |
| NA | | NA |
| NA | | NA |
| NA | | NA |
| NA | | NA |
| NA | | NA |
| NA | | NA |
| | 0.807433834 | NA |
| NA | | NA |
| NA | | NA |
| NA | | NA |
| NA | | NA |
| NA | | NA |
| NA | | NA |
| NA | | NA |

| | | | |
|---|---|---|---|
| | 0.905158266 | NA | |
| NA | | NA | |
| NA | | NA | |
| NA | | NA | |
| NA | | NA | |
| | 0.297552773 | | 0.457496243 |
| NA | | NA | |
| NA | | NA | |
| NA | | NA | |
| NA | | NA | |
| NA | | NA | |
| NA | | NA | |
| NA | | NA | |
| | 0.050197258 | | 0.121718374 |
| | 0.889409301 | NA | |
| NA | | NA | |
| | 0.074292601 | | 0.164123867 |
| | 0.416013782 | | 0.578445997 |
| | 0.656919187 | NA | |
| NA | | NA | |
| NA | | NA | |
| | 0.784963279 | | 0.870512026 |
| | 0.992062245 | NA | |
| NA | | NA | |
| | 0.001793081 | | 0.008150337 |
| | 0.000851811 | | 0.004342909 |
| | 0.555518183 | NA | |
| | 0.992061292 | NA | |
| | 1.91E-17 | | 4.12E-15 |
| | 0.986598218 | NA | |
| | 0.224024755 | | 0.373697934 |
| | 0.00691079 | | 0.025029531 |
| | 0.757780339 | NA | |
| | 0.009643994 | NA | |
| | 0.071206504 | | 0.158569349 |
| | 0.364805992 | NA | |
| | 0.066851207 | | 0.15140116 |
| | 0.905158266 | NA | |
| | 0.208662523 | | 0.355443358 |
| | 0.109880231 | | 0.221140134 |
| | 0.905158266 | NA | |
| NA | | NA | |
| | 0.227255146 | | 0.377109934 |
| NA | | NA | |
| | 0.001193596 | | 0.005784632 |
| | 0.083436929 | | 0.179538611 |
| | 0.426823162 | | 0.587960164 |

| | |
|---|---|
| NA | NA |
| 0.004748209 | 0.018356675 |
| 0.587427483 | 0.726253393 |
| NA | NA |
| NA | NA |
| 0.045631907 | NA |
| 0.012500047 | 0.040277115 |
| 0.055767329 | 0.13180666 |
| 0.792532388 | 0.874922236 |
| 0.823847808 | 0.895011825 |
| 0.108010733 | NA |
| 0.607442373 | NA |
| NA | NA |
| 0.889409301 | NA |
| NA | NA |
| 0.000193049 | 0.001223071 |
| 0.905158739 | NA |
| 0.25571327 | 0.410318429 |
| NA | NA |
| 0.016796966 | 0.051185758 |
| 0.693155741 | 0.807052329 |
| 0.00020074 | 0.001266678 |
| 0.889409773 | NA |
| NA | NA |
| 0.759668215 | NA |
| 0.807879735 | NA |
| 0.337887278 | NA |
| NA | NA |
| 0.054064745 | 0.128764397 |
| 0.953059303 | 0.973603891 |
| 0.902545984 | NA |
| NA | NA |
| NA | NA |
| 0.22600069 | NA |
| 0.000162165 | 0.001052299 |
| NA | NA |
| NA | NA |
| 6.38E-05 | 0.000476806 |
| 0.058678812 | 0.137150402 |
| 0.711785802 | 0.820520994 |
| NA | NA |
| 0.593768816 | NA |
| 3.78E-09 | 1.01E-07 |
| 0.244533134 | 0.397695356 |
| 0.006762758 | 0.024546018 |
| 0.004374787 | 0.017102716 |
| 0.660223574 | NA |

| | | |
|---|---|---|
| | 0.3006332 | NA |
| | 0.905157793 | NA |
| | 0.905158266 | NA |
| | 0.801661681 | NA |
| | 0.553095709 | NA |
| | 0.905158266 | NA |
| | 0.032675616 | 0.087236944 |
| | 0.210387835 | 0.357328987 |
| | 0.931682266 | 0.960437209 |
| NA | | NA |
| | 0.626950218 | 0.756287841 |
| | 0.551268989 | 0.696909368 |
| | 0.992060816 | NA |
| | 0.001163099 | 0.005660571 |
| NA | | NA |
| | 6.18E-16 | 9.77E-14 |
| | 0.939084463 | NA |
| NA | | NA |
| NA | | NA |
| | 3.68E-06 | 4.10E-05 |
| | 0.782195155 | NA |
| | 0.016744038 | 0.051066991 |
| | 0.00098147 | 0.004899318 |
| | 0.54990699 | NA |
| | 2.68E-11 | 1.33E-09 |
| | 0.000255538 | 0.001558709 |
| | 0.034346149 | 0.090693493 |
| | 0.238864579 | 0.391026338 |
| | 0.064020417 | 0.146730439 |
| | 0.164394604 | 0.298391561 |
| NA | | NA |
| | 0.997902434 | NA |
| | 0.48947463 | NA |
| | 0.678707244 | NA |
| | 0.905158266 | NA |
| | 0.531998773 | 0.68108945 |
| NA | | NA |
| NA | | NA |
| | 0.007728387 | 0.027279804 |
| NA | | NA |
| NA | | NA |
| | 9.07E-06 | 8.96E-05 |
| | 0.927338916 | 0.95835985 |
| | 0.151741481 | 0.28121286 |
| | 0.294636161 | 0.454475658 |
| | 0.456982912 | 0.614619205 |
| | 0.319637277 | NA |

| | |
|---|---|
| 0.08202662 | 0.177338047 |
| 0.301487991 | 0.461865132 |
| 0.122594436 | 0.239762309 |
| 0.605793578 | 0.740556766 |
| NA | NA |
| 4.66E-06 | 5.00E-05 |
| 0.00620226 | 0.022878451 |
| 0.004201544 | 0.016555125 |
| 0.905158266 | NA |
| 0.094910683 | 0.19796739 |
| 1.16E-08 | 2.69E-07 |
| 2.69E-09 | 7.51E-08 |
| 0.190282688 | 0.33230295 |
| 1.51E-10 | 6.16E-09 |
| 0.98623254 | NA |
| 0.320881734 | 0.483181924 |
| NA | NA |
| NA | NA |
| NA | NA |
| 0.479413373 | NA |
| 0.00380268 | 0.015229446 |
| 0.884934244 | 0.934832376 |
| 0.326278204 | NA |
| 0.992060816 | NA |
| 0.586630379 | NA |
| 0.000791893 | 0.004090607 |
| 0.001580199 | 0.007315958 |
| 0.797157316 | NA |
| 0.335803685 | 0.498722939 |
| 0.052977678 | 0.126794204 |
| 0.001023119 | 0.005075315 |
| 0.905158266 | NA |
| 4.22E-08 | 8.41E-07 |
| NA | NA |
| 0.992269016 | NA |
| 0.905158739 | NA |
| 0.889409773 | NA |
| 0.871992646 | NA |
| NA | NA |
| 0.327982713 | NA |
| NA | NA |
| NA | NA |
| 0.99205891 | NA |
| 0.337696424 | NA |
| NA | NA |
| 0.905158739 | NA |
| NA | NA |

| | |
|---|---|
| NA | NA |
| 0.973335411 | NA |
| 4.62E-07 | 6.72E-06 |
| 0.584195742 | 0.723947759 |
| 0.756780137 | NA |
| 0.222449543 | 0.371817567 |
| 0.001705566 | 0.007794489 |
| 0.733634795 | NA |
| 0.055730306 | NA |
| 1.14E-07 | 1.97E-06 |
| 0.9135733 | 0.951126308 |
| 0.014307927 | 0.04490928 |
| NA | NA |
| 0.4692325 | 0.62557686 |
| 0.882031044 | NA |
| NA | NA |
| NA | NA |
| 0.612551204 | NA |
| NA | NA |
| 1.22E-06 | 1.55E-05 |
| NA | NA |
| NA | NA |
| 0.219947891 | 0.368774203 |
| 0.905158266 | NA |
| 0.189499481 | 0.331386406 |
| 0.983559068 | NA |
| 0.072898047 | 0.161498004 |
| 0.268663352 | 0.425102936 |
| 0.017573486 | 0.052985881 |
| 0.873018337 | NA |
| 0.72480015 | 0.830249686 |
| NA | NA |
| 0.148482529 | 0.276456492 |
| 0.719869184 | 0.826239149 |
| 0.975696524 | NA |
| NA | NA |
| 0.042515528 | 0.107043883 |
| NA | NA |
| NA | NA |
| NA | NA |
| 0.489848339 | 0.643887391 |
| 0.573383192 | 0.715596562 |
| 0.905158266 | NA |
| NA | NA |
| 0.992061292 | NA |
| NA | NA |
| 0.992061769 | NA |

| | | |
|---|---|---|
| | 0.905158739 | NA |
| NA | | NA |
| | 0.889409301 | NA |
| | 0.073395569 | 0.162403597 |
| | 0.415255592 | 0.577711451 |
| | 0.788582525 | NA |
| | 0.109094153 | 0.220021906 |
| NA | | NA |
| | 0.213287381 | NA |
| | 0.916417926 | 0.952448732 |
| | 0.229097615 | 0.379588547 |
| | 0.785032992 | 0.870545373 |
| | 0.701437796 | 0.813586893 |
| | 0.830163713 | NA |
| | 0.133695888 | 0.255356006 |
| | 0.020766958 | 0.06060088 |
| | 0.916888568 | 0.952667215 |
| | 0.35611313 | 0.52062161 |
| | 0.445905189 | 0.605355923 |
| NA | | NA |
| | 0.604051121 | 0.739253085 |
| | 0.057741798 | 0.135407107 |
| | 0.246736668 | 0.400154393 |
| | 0.889409301 | NA |
| | 0.04375496 | 0.109286711 |
| | 0.001582079 | 0.007323118 |
| | 0.871217504 | 0.926316278 |
| | 0.889409773 | NA |
| | 0.307090834 | 0.467869814 |
| NA | | NA |
| | 0.110109881 | NA |
| | 0.905158739 | NA |
| | 0.155633062 | NA |
| | 0.807879735 | NA |
| | 0.715452167 | 0.82306235 |
| NA | | NA |
| | 0.992060816 | NA |
| | 0.281913572 | NA |
| | 0.129528161 | 0.249641283 |
| NA | | NA |
| NA | | NA |
| | 0.992059387 | NA |
| | 0.904947233 | NA |
| | 0.03435033 | 0.090693627 |
| | 0.070423759 | 0.157303634 |
| | 0.000353014 | 0.002057652 |
| | 0.758980131 | 0.853756605 |

| | | | |
|---|---|---|---|
| | 0.905158739 | NA | |
| | 0.460655925 | | 0.618020487 |
| NA | | NA | |
| NA | | NA | |
| NA | | NA | |
| NA | | NA | |
| NA | | NA | |
| | 0.643954822 | NA | |
| NA | | NA | |
| NA | | NA | |
| NA | | NA | |
| NA | | NA | |
| | 0.593304942 | NA | |
| NA | | NA | |
| NA | | NA | |
| NA | | NA | |
| NA | | NA | |
| | 0.889409773 | NA | |
| NA | | NA | |
| | 0.905158266 | NA | |
| NA | | NA | |
| | 0.919037513 | NA | |
| NA | | NA | |
| NA | | NA | |
| | 0.905158266 | NA | |
| NA | | NA | |
| | 0.905158739 | NA | |
| NA | | NA | |
| | 0.452516938 | NA | |
| NA | | NA | |
| NA | | NA | |
| | 0.889409773 | NA | |
| NA | | NA | |
| NA | | NA | |
| NA | | NA | |
| NA | | NA | |
| NA | | NA | |
| | 0.905158739 | NA | |
| NA | | NA | |
| NA | | NA | |
| | 0.992061292 | NA | |
| | 0.700798622 | NA | |
| | 0.764483348 | NA | |
| | 0.950573947 | NA | |
| NA | | NA | |
| | 0.905158739 | NA | |
| NA | | NA | |

| | | | |
|---|---|---|---|
| | 0.862692732 | NA | |
| NA | | NA | |
| | 0.355898184 | NA | |
| NA | | NA | |
| NA | | NA | |
| NA | | NA | |
| | 0.593949056 | NA | |
| | 0.655478926 | NA | |
| NA | | NA | |
| NA | | NA | |
| NA | | NA | |
| NA | | NA | |
| | 0.905158739 | NA | |
| NA | | NA | |
| NA | | NA | |
| NA | | NA | |
| NA | | NA | |
| | 0.889409773 | NA | |
| NA | | NA | |
| | 0.90515732 | NA | |
| NA | | NA | |
| NA | | NA | |
| NA | | NA | |
| NA | | NA | |
| NA | | NA | |
| NA | | NA | |
| NA | | NA | |
| NA | | NA | |
| NA | | NA | |
| NA | | NA | |
| NA | | NA | |
| | 0.905158739 | NA | |
| NA | | NA | |
| NA | | NA | |
| NA | | NA | |
| NA | | NA | |
| NA | | NA | |
| NA | | NA | |
| | 0.807433834 | NA | |
| | 0.889409773 | NA | |
| NA | | NA | |
| | 0.992037435 | NA | |
| | 0.277996441 | | 0.435881745 |
| NA | | NA | |
| | 0.330735171 | | 0.493250042 |
| | 0.097072607 | | 0.20130488 |
| | 0.032520656 | | 0.086897188 |

| | |
|---|---|
| 0.159005744 | 0.29099151 |
| NA | NA |
| 0.995540448 | NA |
| 0.422771394 | 0.584256638 |
| 0.515314731 | 0.666576119 |
| 0.807879735 | NA |
| 0.74955871 | 0.847208138 |
| 0.888407067 | 0.936919161 |
| 0.580648204 | 0.72084278 |
| 0.916664274 | 0.952583927 |
| 0.062354572 | 0.143890063 |
| 0.002102072 | 0.009332673 |
| 0.6239255 | 0.754656021 |
| 0.395541861 | NA |
| 0.911023513 | 0.949687411 |
| 0.667996923 | 0.78829237 |
| 0.029819199 | 0.081150054 |
| 0.992060816 | NA |
| 0.36726455 | 0.531941313 |
| 0.946157941 | 0.969356826 |
| 0.719430349 | 0.826037701 |
| NA | NA |
| 0.027407034 | 0.075797364 |
| NA | NA |
| 0.99205891 | NA |
| 0.438880969 | 0.599228203 |
| 0.992060816 | NA |
| 4.40E-07 | 6.45E-06 |
| 0.222080609 | NA |
| 0.484414367 | 0.639719454 |
| 0.171210459 | 0.307660431 |
| NA | NA |
| 0.369882186 | 0.534463574 |
| 0.555812734 | 0.701079432 |
| 0.992059387 | NA |
| 0.992060816 | NA |
| 0.962367706 | 0.979205986 |
| 0.026001921 | 0.072687223 |
| 0.905158739 | NA |
| 0.435151394 | 0.595840211 |
| 0.968183247 | NA |
| 0.029187971 | 0.079669192 |
| 0.008650198 | 0.029927476 |
| 0.012894918 | 0.041264864 |
| 0.820458745 | 0.893150789 |
| 0.848242857 | NA |
| NA | NA |

| | | |
|---|---|---|
| | 0.256392237 | NA |
| | 0.273988898 | NA |
| | 0.905158266 | NA |
| NA | | NA |
| NA | | NA |
| | 0.411959479 | NA |
| NA | | NA |
| NA | | NA |
| | 0.028954468 | 0.079169614 |
| NA | | NA |
| | 0.037891798 | 0.097985654 |
| NA | | NA |
| | 0.992061292 | NA |
| NA | | NA |
| NA | | NA |
| | 0.047886438 | 0.117319903 |
| | 0.889409301 | NA |
| | 0.990012186 | NA |
| NA | | NA |
| NA | | NA |
| | 0.105700055 | 0.214754581 |
| | 0.611451125 | 0.745067158 |
| | 0.003264381 | 0.013476694 |
| | 0.839414142 | NA |
| | 0.544016771 | 0.690715906 |
| | 1.80E-11 | 9.46E-10 |
| | 0.809530083 | NA |
| | 0.055666784 | 0.131623443 |
| | 0.020376215 | 0.059674755 |
| | 0.000647321 | 0.003461893 |
| | 0.27505635 | 0.432321408 |
| | 2.59E-09 | 7.30E-08 |
| | 0.003440163 | 0.014046587 |
| | 6.11E-07 | 8.52E-06 |
| | 0.427375637 | 0.588458842 |
| | 0.839933411 | 0.905188055 |
| NA | | NA |
| | 5.39E-06 | 5.69E-05 |
| | 0.905158266 | NA |
| | 0.352498148 | 0.51675725 |
| NA | | NA |
| | 0.642333074 | NA |
| | 0.889409773 | NA |
| NA | | NA |
| | 0.344612653 | 0.508012167 |
| NA | | NA |
| NA | | NA |

| | | | |
|---|---|---|---|
| NA | | NA | |
| | 0.133016574 | | 0.254346129 |
| | 0.905158266 | NA | |
| NA | | NA | |
| NA | | NA | |
| NA | | NA | |
| NA | | NA | |
| NA | | NA | |
| NA | | NA | |
| NA | | NA | |
| | 0.905158266 | NA | |
| NA | | NA | |
| NA | | NA | |
| NA | | NA | |
| NA | | NA | |
| NA | | NA | |
| NA | | NA | |
| NA | | NA | |
| NA | | NA | |
| NA | | NA | |
| NA | | NA | |
| NA | | NA | |
| | 0.905158266 | NA | |
| | 0.889409301 | NA | |
| | 0.889409301 | NA | |
| NA | | NA | |
| NA | | NA | |
| NA | | NA | |
| NA | | NA | |
| NA | | NA | |
| NA | | NA | |
| | 0.000290975 | | 0.00174339 |
| | 0.466615135 | | 0.623222538 |
| | 0.528433686 | NA | |
| | 0.889283189 | NA | |
| NA | | NA | |
| | 0.992003131 | NA | |
| | 0.467235171 | | 0.623633426 |
| | 0.028758239 | | 0.078780211 |
| | 0.299689346 | | 0.460069055 |
| | 0.771599363 | NA | |
| | 0.629321069 | | 0.757608883 |
| NA | | NA | |
| | 0.616367503 | | 0.749021251 |
| | 0.992061292 | NA | |
| | 0.905157793 | NA | |
| NA | | NA | |

| | |
|---|---|
| 1.85E-07 | 3.03E-06 |
| NA | NA |
| 0.677271492 | NA |
| 0.572251868 | 0.714749792 |
| 0.807879735 | NA |
| 0.756440911 | 0.851917426 |
| 0.992061292 | NA |
| 0.913054573 | 0.9507312 |
| NA | NA |
| 0.000247298 | 0.001515608 |
| 0.668899185 | 0.78895785 |
| 0.660696451 | 0.782769748 |
| 0.422930148 | NA |
| 0.905158266 | NA |
| NA | NA |
| 0.857141879 | NA |
| NA | NA |
| 0.772863245 | NA |
| 0.100708033 | NA |
| 0.095783569 | NA |
| NA | NA |
| 0.012880738 | NA |
| NA | NA |
| 0.144829561 | 0.271450352 |
| 0.692853708 | 0.806829007 |
| NA | NA |
| 0.452799747 | NA |
| NA | NA |
| 0.148318419 | 0.27626806 |
| 0.889409773 | NA |
| 0.965614238 | NA |
| 0.722062236 | NA |
| 0.063412294 | 0.145686396 |
| 0.641830725 | 0.767603121 |
| 0.889409773 | NA |
| NA | NA |
| 0.259348113 | 0.414485096 |
| NA | NA |
| 0.018743596 | 0.055771086 |
| 0.74440655 | 0.843468701 |
| 0.817885618 | 0.891604826 |
| 0.836074142 | NA |
| 0.005357191 | 0.020307017 |
| 0.021508861 | NA |
| 0.00021246 | 0.00132842 |
| 0.000142023 | 0.000938553 |
| 0.000246727 | 0.001512954 |

| | | |
|---|---|---|
| | 0.074319247 | 0.164166216 |
| NA | | NA |
| NA | | NA |
| | 0.81281518 | NA |
| NA | | NA |
| NA | | NA |
| | 0.905158739 | NA |
| NA | | NA |
| NA | | NA |
| NA | | NA |
| NA | | NA |
| NA | | NA |
| NA | | NA |
| NA | | NA |
| NA | | NA |
| NA | | NA |
| NA | | NA |
| NA | | NA |
| NA | | NA |
| NA | | NA |
| NA | | NA |
| NA | | NA |
| NA | | NA |
| | 0.934437161 | NA |
| | 0.972232917 | 0.985471871 |
| | 1.05E-07 | 1.85E-06 |
| | 0.725397085 | 0.83034525 |
| | 0.001574412 | 0.007290702 |
| | 6.85E-05 | 0.000506392 |
| | 0.745693563 | NA |
| | 0.713383518 | 0.821500271 |
| | 0.044483027 | 0.110748616 |
| | 0.000459873 | 0.002569553 |
| | 5.12E-11 | 2.35E-09 |
| | 0.007529112 | 0.026750864 |
| | 0.000573237 | 0.003114998 |
| | 0.23496049 | 0.386443514 |
| | 0.001539135 | 0.007159169 |
| | 0.852324861 | NA |
| | 0.905158739 | NA |
| | 0.78378004 | 0.869874117 |
| | 0.419894503 | 0.581696007 |
| | 0.992061769 | NA |
| | 0.905158266 | NA |
| | 0.125626645 | NA |
| | 0.503458743 | NA |
| | 0.380272976 | 0.544533047 |

| | | |
|---|---|---|
| | 0.859523899 | NA |
| NA | | NA |
| | 0.653108539 | 0.776691601 |
| NA | | NA |
| | 0.007484866 | 0.026608415 |
| | 0.020089921 | 0.059027892 |
| | 0.902816228 | 0.945034872 |
| | 0.388406195 | 0.551962569 |
| | 0.889409773 | NA |
| | 0.374047882 | 0.537969442 |
| | 4.92E-06 | 5.25E-05 |
| | 0.564468926 | 0.708012403 |
| | 0.905154837 | NA |
| NA | | NA |
| | 0.792526283 | NA |
| NA | | NA |
| NA | | NA |
| | 0.044523965 | 0.110812847 |
| | 0.2656088 | 0.421758886 |
| | 0.886822368 | 0.935832206 |
| | 0.61066525 | 0.744605376 |
| NA | | NA |
| | 0.878369794 | NA |
| | 3.75E-06 | 4.16E-05 |
| | 0.064959014 | 0.148386038 |
| | 0.832554242 | 0.90036919 |
| | 4.50E-05 | 0.000353806 |
| | 0.029161301 | 0.079616188 |
| | 0.000249765 | 0.001526891 |
| | 0.905158266 | NA |
| | 0.210682777 | 0.357636295 |
| | 0.172294291 | 0.309152666 |
| | 0.225925196 | 0.375754857 |
| NA | | NA |
| NA | | NA |
| | 0.992061292 | NA |
| | 0.414147199 | NA |
| | 0.906603745 | 0.947328653 |
| | 0.495862724 | NA |
| NA | | NA |
| | 0.733849903 | NA |
| | 1.38E-05 | 0.000128677 |
| | 0.519098857 | NA |
| | 0.40273034 | 0.56624123 |
| | 0.001873895 | 0.008468585 |
| | 0.000138298 | 0.000920267 |
| | 0.213166055 | 0.360337793 |

| | | |
|---|---|---|
| | 0.95142254 | NA |
| | 0.056408149 | 0.132946929 |
| | 0.021197284 | 0.061597632 |
| | 0.000376243 | 0.002167347 |
| | 0.718980365 | NA |
| | 0.007533322 | 0.026754685 |
| | 0.114236697 | NA |
| | 0.970364869 | NA |
| | 8.12E-05 | 0.000584711 |
| | 0.907793485 | NA |
| | 0.886549646 | 0.935634339 |
| | 0.905157793 | NA |
| | 0.905158266 | NA |
| NA | | NA |
| | 0.784353866 | NA |
| | 0.22446049 | NA |
| | 0.0623552 | 0.143890063 |
| | 0.703732504 | 0.815346492 |
| NA | | NA |
| | 2.63E-08 | 5.54E-07 |
| | 0.704532748 | NA |
| NA | | NA |
| | 0.045375358 | 0.112523714 |
| NA | | NA |
| | 0.99205891 | NA |
| | 0.811240452 | 0.887108665 |
| | 0.963491421 | 0.979822541 |
| | 0.136253577 | 0.259057127 |
| | 0.327773398 | 0.489912234 |
| | 0.612419464 | 0.745796431 |
| | 0.001266531 | 0.006069784 |
| | 0.986894031 | 0.993180845 |
| | 0.019686543 | 0.058054131 |
| | 0.056358895 | 0.132859364 |
| NA | | NA |
| | 0.099290473 | 0.204840237 |
| NA | | NA |
| | 0.905158266 | NA |
| NA | | NA |
| | 0.071422873 | 0.158951543 |
| | 0.992060816 | NA |
| | 0.70924893 | 0.818800129 |
| NA | | NA |
| | 0.039892578 | 0.10189614 |
| | 0.833229682 | NA |
| | 0.735214941 | NA |
| NA | | NA |

| | |
|---|---|
| 0.001897665 | 0.008556638 |
| NA | NA |
| 2.53E-05 | 0.000216682 |
| 3.83E-05 | 0.000308187 |
| 0.942423556 | 0.967538402 |
| 0.905158739 | NA |
| 0.366293418 | 0.531165368 |
| 0.000326237 | 0.001922662 |
| 0.878143796 | 0.930744261 |
| 0.582450898 | 0.722397021 |
| 0.199292926 | 0.344046648 |
| 2.30E-10 | 8.80E-09 |
| 0.453044437 | 0.611135218 |
| 0.462287829 | 0.619612367 |
| NA | NA |
| NA | NA |
| 0.106190291 | 0.215403547 |
| 0.872205609 | 0.926670213 |
| 0.781452957 | 0.868329394 |
| 0.023155288 | 0.0661894 |
| 0.837323826 | 0.903172899 |
| 0.018471245 | 0.055095094 |
| 0.762784935 | NA |
| 7.15E-08 | 1.32E-06 |
| 0.329578642 | 0.492059927 |
| 0.606164057 | 0.740779963 |
| 0.052857744 | 0.126589944 |
| 0.320878587 | 0.483181924 |
| 0.316168552 | 0.47830712 |
| NA | NA |
| NA | NA |
| 0.422329319 | 0.583857369 |
| 0.955287096 | 0.975137102 |
| 0.288134428 | 0.447337663 |
| NA | NA |
| 0.046261433 | 0.114230833 |
| NA | NA |
| 0.537785526 | 0.685825679 |
| 0.312875414 | 0.47473958 |
| 3.67E-05 | 0.000297381 |
| 0.001696503 | 0.007762765 |
| 2.82E-07 | 4.37E-06 |
| 0.002225699 | 0.009798339 |
| 1.37E-05 | 0.000128225 |
| 0.587580554 | 0.726293159 |
| 3.59E-12 | 2.23E-10 |
| 0.004725424 | 0.018294355 |

| | |
|---|---|
| 0.326318321 | 0.488620798 |
| 0.552627687 | 0.698144711 |
| 0.556003409 | 0.701199108 |
| 0.037670168 | 0.097604628 |
| 0.140948566 | 0.265832652 |
| 0.008656635 | 0.029940312 |
| 0.170834995 | 0.307312662 |
| 0.15270087 | 0.282585404 |
| 3.12E-14 | 3.08E-12 |
| 0.559320363 | 0.703695597 |
| 0.803339107 | 0.881673503 |
| 0.635109997 | NA |
| NA | NA |
| NA | NA |
| NA | NA |
| NA | NA |
| NA | NA |
| NA | NA |
| NA | NA |
| 0.000492055 | 0.002725438 |
| 0.677322692 | NA |
| NA | NA |
| 0.992062245 | NA |
| 0.905158739 | NA |
| NA | NA |
| NA | NA |
| NA | NA |
| 0.00030634 | 0.001818577 |
| 0.583896242 | NA |
| 0.812251103 | 0.887771737 |
| 0.886952954 | NA |
| 3.25E-07 | 4.93E-06 |
| 0.002658485 | 0.011395212 |
| 0.224376153 | 0.373999996 |
| 0.915224843 | NA |
| 0.021017653 | 0.061194198 |
| 0.015662972 | 0.048388182 |
| 7.29E-05 | 0.000533233 |
| NA | NA |
| 0.001590006 | 0.00735516 |
| 0.807879735 | NA |
| NA | NA |
| 0.244667484 | 0.397731146 |
| 2.06E-06 | 2.47E-05 |
| 0.259553995 | 0.414663236 |
| 0.489246673 | NA |
| 0.049811402 | 0.1210233 |

| | |
|---|---|
| 0.720312252 | 0.826488491 |
| 1.04E-06 | 1.36E-05 |
| 0.398244912 | 0.561877411 |
| NA | NA |
| 0.027396759 | 0.075778489 |
| 0.991918092 | NA |
| 0.184177058 | 0.324768651 |
| 0.134442432 | 0.256358228 |
| 0.991501493 | 0.995788655 |
| 0.992269016 | NA |
| 4.63E-13 | 3.62E-11 |
| NA | NA |
| NA | NA |
| 0.651317341 | 0.775695709 |
| 0.023853543 | 0.067770728 |
| 0.959961948 | 0.977727478 |
| 0.37742988 | 0.541804455 |
| 0.067196443 | 0.151822894 |
| 0.024372883 | NA |
| 0.00204205 | 0.009108546 |
| 0.263607412 | 0.419384261 |
| 1.27E-08 | 2.91E-07 |
| 5.25E-05 | 0.00040271 |
| 0.827031753 | 0.89672859 |
| 0.441367716 | 0.601053871 |
| 0.007419487 | 0.026427336 |
| 2.99E-12 | 1.91E-10 |
| NA | NA |
| NA | NA |
| 0.52707487 | 0.676989837 |
| NA | NA |
| 0.206385494 | NA |
| 0.41742575 | NA |
| 0.536766709 | NA |
| 1.56E-09 | 4.64E-08 |
| 2.56E-08 | 5.40E-07 |
| 0.89799259 | 0.942591744 |
| 0.256071741 | NA |
| 0.666284561 | 0.787202339 |
| NA | NA |
| 5.72E-06 | 5.99E-05 |
| 0.383692335 | 0.547615109 |
| 5.90E-06 | 6.16E-05 |
| 0.184789335 | 0.325534618 |
| 0.451760949 | 0.610038967 |
| 6.79E-08 | 1.26E-06 |
| 2.31E-18 | 6.60E-16 |

| | |
|---|---|
| 0.725371566 | NA |
| 0.191209997 | NA |
| 0.012562532 | 0.04043099 |
| 0.359178868 | 0.524065719 |
| NA | NA |
| 0.426881684 | 0.587976906 |
| 1.49E-07 | 2.51E-06 |
| NA | NA |
| 6.40E-11 | 2.84E-09 |
| NA | NA |
| 1.94E-07 | 3.16E-06 |
| 0.001517563 | 0.007073695 |
| 0.448507504 | 0.607517727 |
| 0.004861205 | 0.018717727 |
| 0.748848137 | 0.846773676 |
| 0.889410717 | NA |
| 0.002451684 | 0.010641731 |
| 0.02511911 | 0.070816604 |
| 0.65258335 | 0.776306032 |
| 0.080471164 | 0.174714879 |
| 0.930392602 | 0.959926975 |
| NA | NA |
| 0.336899483 | 0.499897003 |
| 0.793359662 | NA |
| 0.992061292 | NA |
| 0.021146217 | 0.061470713 |
| NA | NA |
| 0.000215238 | 0.001343115 |
| NA | NA |
| 0.985982531 | NA |
| 0.000274584 | 0.001660587 |
| NA | NA |
| NA | NA |
| 0.151428026 | 0.280744338 |
| NA | NA |
| 2.97E-05 | 0.000248094 |
| 0.889409301 | NA |
| 1.45E-05 | 0.00013414 |
| NA | NA |
| 0.000767484 | 0.003989862 |
| 0.905158266 | NA |
| 0.611613188 | NA |
| 0.541789214 | 0.689050697 |
| 0.150180122 | 0.278860587 |
| NA | NA |
| 8.01E-12 | 4.58E-10 |
| 0.043330281 | 0.108518437 |

| | | |
|---|---|---|
| | 0.484656715 | 0.639850374 |
| | 0.945829815 | 0.969270637 |
| | 0.634368872 | NA |
| | 0.385199919 | 0.549083338 |
| | 0.000202175 | 0.001273541 |
| | 0.505788537 | 0.65774076 |
| | 0.225187546 | 0.374925569 |
| | 0.145523378 | 0.27235625 |
| | 0.992061292 | NA |
| NA | | NA |
| | 5.37E-07 | 7.68E-06 |
| | 0.533226819 | 0.682080031 |
| | 0.02634278 | 0.073408769 |
| | 0.498019864 | 0.650953464 |
| | 0.689170459 | 0.80420311 |
| | 4.05E-06 | 4.44E-05 |
| | 1.21E-09 | 3.69E-08 |
| | 0.061389016 | 0.142258486 |
| | 0.04024443 | 0.102544378 |
| | 0.726850194 | NA |
| | 0.719378468 | 0.826037701 |
| | 0.815823219 | NA |
| | 0.062296922 | 0.143816034 |
| | 0.889410245 | NA |
| | 0.014694722 | 0.045907571 |
| | 3.29E-08 | 6.78E-07 |
| | 0.180670612 | 0.320541809 |
| | 0.192912998 | 0.335857966 |
| | 0.001560365 | 0.007239394 |
| | 0.262958831 | NA |
| NA | | NA |
| NA | | NA |
| | 4.19E-06 | 4.58E-05 |
| NA | | NA |
| | 0.992060816 | NA |
| NA | | NA |
| | 0.108567964 | 0.21916731 |
| | 0.011584418 | 0.037831981 |
| NA | | NA |
| NA | | NA |
| | 0.378497516 | 0.543017572 |
| NA | | NA |
| NA | | NA |
| NA | | NA |
| NA | | NA |
| NA | | NA |
| NA | | NA |

| | |
|---|---|
| NA | NA |
| NA | NA |
| NA | NA |
| NA | NA |
| 0.127308553 | NA |
| NA | NA |
| NA | NA |
| 0.213090827 | 0.360331239 |
| 0.889409773 | NA |
| 0.67429234 | 0.793036553 |
| 0.379756416 | 0.544134937 |
| 0.988939007 | NA |
| 0.675197686 | NA |
| 0.584523917 | 0.72424086 |
| 0.905158739 | NA |
| NA | NA |
| 0.197469588 | 0.341785662 |
| 0.90515732 | NA |
| 0.524703907 | 0.674774267 |
| 0.517450642 | 0.668472682 |
| 0.736347087 | NA |
| 0.148771847 | NA |
| 0.602570473 | 0.738054719 |
| 0.992061292 | NA |
| 0.370590139 | 0.535169595 |
| NA | NA |
| 0.812855148 | 0.888210898 |
| NA | NA |
| NA | NA |
| NA | NA |
| 0.689623847 | NA |
| 0.157258599 | 0.28863604 |
| 0.404589212 | 0.567850108 |
| 0.136018209 | 0.258752189 |
| 0.956712083 | 0.975959083 |
| 0.292563657 | 0.452295505 |
| 0.808951365 | NA |
| 0.738666023 | 0.839639257 |
| 0.426497236 | 0.587728447 |
| 0.619233544 | NA |
| 0.103899255 | 0.211860316 |
| NA | NA |
| 0.11621355 | 0.230797987 |
| 0.132108689 | 0.253050829 |
| 0.193615287 | 0.336706905 |
| 0.992061292 | NA |
| 4.48E-06 | 4.85E-05 |

| | |
|---|---|
| 0.426405862 | 0.587639408 |
| 0.000820513 | 0.004220579 |
| 0.00069879 | 0.003685096 |
| 0.13902655 | 0.263134288 |
| 0.915958059 | 0.95229975 |
| 0.618842045 | 0.750823386 |
| 1.44E-07 | 2.43E-06 |
| 0.067111875 | 0.151741109 |
| 0.814966959 | NA |
| 0.003499603 | NA |
| 1.84E-06 | 2.22E-05 |
| NA | NA |
| NA | NA |
| 0.991918092 | NA |
| 0.005186809 | 0.019773807 |
| NA | NA |
| NA | NA |
| 0.161057473 | NA |
| 0.855875864 | 0.916035221 |
| 0.565190667 | 0.708474775 |
| NA | NA |
| NA | NA |
| 0.833780744 | NA |
| NA | NA |
| NA | NA |
| NA | NA |
| 0.135578456 | 0.25816574 |
| NA | NA |
| NA | NA |
| NA | NA |
| NA | NA |
| NA | NA |
| NA | NA |
| NA | NA |
| NA | NA |
| NA | NA |
| NA | NA |
| NA | NA |
| NA | NA |
| NA | NA |
| 0.840647937 | NA |
| 0.018125444 | 0.054307084 |
| 0.394003492 | 0.557433328 |
| 0.535586227 | 0.683894747 |
| 0.937051003 | 0.964178959 |
| 0.255196047 | 0.40975797 |
| 0.955731773 | 0.975318989 |

| | | |
|---|---|---|
| | 0.76492434 | NA |
| | 0.098935195 | 0.204222406 |
| | 0.096247519 | 0.200085134 |
| | 0.69619775 | 0.809606862 |
| | 0.060255983 | 0.140254519 |
| | 0.028990006 | 0.079237185 |
| | 0.052747613 | 0.126408909 |
| | 0.657894338 | 0.780354441 |
| | 0.004800369 | 0.018535487 |
| | 0.985996077 | NA |
| | 0.009293761 | 0.031692915 |
| | 0.989580628 | NA |
| | 0.601000994 | 0.73664624 |
| | 5.12E-05 | 0.000394393 |
| | 0.000330318 | 0.001941504 |
| | 0.294275062 | 0.454014339 |
| | 0.00123138 | 0.005927199 |
| | 0.781072271 | NA |
| NA | | NA |
| | 0.054397062 | 0.129359442 |
| | 0.035634957 | 0.093366903 |
| NA | | NA |
| | 0.018550581 | 0.05527917 |
| | 0.029447523 | 0.080297798 |
| | 0.455410719 | 0.613331124 |
| | 0.204734664 | 0.350683969 |
| | 0.465349823 | 0.622238692 |
| | 0.213297812 | 0.360403712 |
| NA | | NA |
| | 0.084266107 | 0.180667785 |
| NA | | NA |
| | 0.009734068 | 0.032883309 |
| | 0.65748298 | 0.780096747 |
| | 0.532756113 | 0.681759781 |
| NA | | NA |
| | 0.136849893 | 0.260000588 |
| | 0.825758215 | 0.896156204 |
| | 0.006087083 | 0.022517901 |
| | 0.901967787 | 0.94462713 |
| | 0.644249192 | 0.769824673 |
| | 0.4018887 | NA |
| | 2.81E-05 | 0.000236485 |
| | 0.003184711 | 0.013202393 |
| | 0.000101436 | 0.000707121 |
| | 7.17E-09 | 1.76E-07 |
| NA | | NA |
| | 0.148252163 | 0.276176252 |

| | |
|---|---|
| NA | NA |
| NA | NA |
| NA | NA |
| 0.005294109 | 0.020109556 |
| 0.006844288 | 0.024825524 |
| 0.682926021 | NA |
| 0.222978667 | 0.372405579 |
| 0.277835272 | 0.435806289 |
| 0.773819407 | 0.863681178 |
| 0.000171348 | 0.001102876 |
| 0.99205891 | NA |
| 0.807252815 | NA |
| 0.802236522 | NA |
| NA | NA |
| NA | NA |
| 0.11030093 | 0.221661765 |
| 0.002700796 | 0.011531555 |
| 0.865605536 | NA |
| 0.862560572 | NA |
| NA | NA |
| 0.823352925 | 0.894651238 |
| 0.019434805 | 0.057430882 |
| 0.819420116 | 0.892630535 |
| 0.793619578 | 0.875469321 |
| 0.775239228 | 0.864399219 |
| 0.268071328 | 0.424410836 |
| 0.464973302 | 0.621924994 |
| 0.014345454 | 0.044983445 |
| 0.000465052 | 0.002593215 |
| 3.59E-05 | 0.000291489 |
| 1.50E-05 | 0.000138017 |
| 0.122862757 | 0.240201502 |
| 0.842343577 | NA |
| 0.208990377 | 0.355753464 |
| 0.992061292 | NA |
| 0.003222627 | 0.013339427 |
| 0.070376706 | 0.15726082 |
| NA | NA |
| 0.413313032 | 0.575884574 |
| 0.807525235 | NA |
| 1.27E-07 | 2.19E-06 |
| 0.747070183 | NA |
| 0.098598734 | NA |
| 0.066337817 | 0.150630052 |
| 2.11E-05 | 0.000186278 |
| 0.935296281 | NA |
| 0.515295469 | 0.666576119 |

| | |
|---|---|
| 0.204206515 | 0.350094613 |
| 0.4083164 | 0.571533646 |
| 0.755665272 | 0.851480589 |
| 0.004851977 | 0.01869531 |
| 0.042736662 | 0.107460709 |
| NA | NA |
| 0.597888905 | 0.734409715 |
| 0.000865271 | 0.004402337 |
| 0.110734005 | 0.222227016 |
| 6.62E-10 | 2.20E-08 |
| NA | NA |
| 0.058634608 | 0.13709086 |
| 0.000168914 | 0.001089975 |
| 0.000262373 | 0.001595085 |
| 0.572699896 | NA |
| 0.179220929 | 0.31866497 |
| 0.166604151 | 0.30138072 |
| NA | NA |
| NA | NA |
| 3.31E-05 | 0.000272517 |
| 0.610000215 | 0.744042367 |
| 0.001000761 | 0.004979767 |
| 0.922613185 | 0.955374089 |
| NA | NA |
| 0.472108951 | 0.62841974 |
| 0.544514462 | 0.691225485 |
| 0.986412 | NA |
| 0.51394601 | 0.665337361 |
| 0.774609949 | 0.864060335 |
| 0.034811378 | 0.091690991 |
| 0.110443093 | 0.22188654 |
| 0.002669671 | 0.011432003 |
| NA | NA |
| 4.91E-21 | 2.70E-18 |
| NA | NA |
| NA | NA |
| 0.03629026 | 0.094776539 |
| 0.089656303 | 0.189541038 |
| 9.46E-05 | 0.00066622 |
| 1.30E-06 | 1.64E-05 |
| 0.992061292 | NA |
| 0.03627636 | 0.094753489 |
| 0.365071185 | 0.529812845 |
| 0.004707713 | 0.018232216 |
| 8.28E-05 | 0.00059346 |
| NA | NA |
| NA | NA |

| | |
|---|---|
| 0.905158266 | NA |
| NA | NA |
| 0.77005898 | NA |
| 0.90515732 | NA |
| NA | NA |
| 0.088945349 | 0.188528081 |
| 0.150729286 | NA |
| 0.123803812 | 0.241632558 |
| 0.00084783 | 0.004327637 |
| 0.536019805 | 0.684289221 |
| NA | NA |
| 0.042392002 | 0.106801971 |
| 0.187707557 | 0.32917028 |
| 0.012994186 | 0.041509362 |
| 1.91E-05 | 0.000171327 |
| 0.085327233 | 0.182479616 |
| 0.102688592 | 0.210270309 |
| NA | NA |
| NA | NA |
| 1.79E-14 | 1.91E-12 |
| 0.067844912 | 0.152832008 |
| 0.012778025 | 0.040992352 |
| 0.515818719 | 0.666992301 |
| NA | NA |
| 0.014501863 | 0.04540154 |
| 0.723552857 | 0.829210229 |
| NA | NA |
| 0.141926914 | 0.267310499 |
| 0.144151048 | 0.270548107 |
| 0.849350233 | NA |
| 0.992061292 | NA |
| 0.718129384 | 0.825236459 |
| 0.326449061 | 0.488716589 |
| 0.104220112 | 0.212475114 |
| NA | NA |
| 0.654101946 | 0.777511713 |
| 0.340151331 | NA |
| NA | NA |
| NA | NA |
| NA | NA |
| NA | NA |
| NA | NA |
| NA | NA |
| 0.016882458 | 0.05138923 |
| 0.000249639 | 0.001526577 |
| 0.643019122 | 0.768699402 |
| 0.01223773 | 0.039591843 |

| | | |
|---|---|---|
| | 0.992060816 | NA |
| | 0.757944946 | 0.853217464 |
| | 0.118446542 | 0.233871323 |
| | 0.801194238 | 0.880155299 |
| | 0.418017881 | 0.580231014 |
| | 0.607403918 | 0.741617049 |
| | 0.871813082 | NA |
| | 0.439296906 | NA |
| NA | | NA |
| | 0.983464937 | NA |
| NA | | NA |
| NA | | NA |
| | 0.00694805 | 0.025128987 |
| | 0.871314136 | 0.926316278 |
| | 0.167360614 | 0.302400372 |
| NA | | NA |
| | 0.000971077 | 0.004856271 |
| | 0.14398965 | NA |
| | 1.02E-07 | 1.79E-06 |
| | 0.000178581 | 0.001143281 |
| | 0.293750413 | NA |
| | 0.322885404 | 0.485366246 |
| NA | | NA |
| NA | | NA |
| | 0.088147014 | 0.187215425 |
| | 0.672546651 | NA |
| | 0.829815144 | 0.898516309 |
| | 1.63E-09 | 4.82E-08 |
| | 0.876557991 | 0.929557006 |
| NA | | NA |
| NA | | NA |
| | 1.05E-09 | 3.30E-08 |
| | 0.120664145 | NA |
| | 0.008874686 | 0.030545334 |
| | 0.257239566 | 0.411984803 |
| | 0.016286799 | 0.049949976 |
| NA | | NA |
| | 0.537898818 | 0.685930322 |
| | 0.17279475 | 0.309848108 |
| | 0.000172805 | 0.001111164 |
| | 0.000866743 | 0.004406765 |
| | 0.876240141 | 0.929309696 |
| | 0.568962278 | NA |
| | 0.164940788 | 0.299135677 |
| | 8.36E-05 | 0.000598294 |
| | 0.026238674 | 0.073220873 |
| | 0.736466147 | NA |

| | | |
|---|---|---|
| | 0.370456646 | NA |
| | 0.464769222 | 0.621816082 |
| | 0.025383253 | 0.071423737 |
| NA | | NA |
| | 9.97E-07 | 1.30E-05 |
| | 0.769522963 | NA |
| | 0.889409301 | NA |
| | 0.804458936 | NA |
| | 0.029194847 | NA |
| | 1.93E-14 | 2.05E-12 |
| | 0.804591588 | NA |
| | 6.17E-11 | 2.77E-09 |
| | 0.00643277 | 0.023582168 |
| | 0.001951075 | 0.008763274 |
| | 0.621703039 | 0.753043545 |
| | 0.52335651 | NA |
| | 0.001278976 | 0.006124081 |
| | 0.912695952 | 0.950637798 |
| | 0.759589449 | 0.854017956 |
| | 0.091988472 | 0.19325657 |
| | 0.889409773 | NA |
| | 0.543337632 | 0.690180544 |
| | 0.366567562 | 0.5313173 |
| | 0.311597754 | 0.473160576 |
| | 0.122202112 | 0.239208074 |
| | 0.002001629 | 0.008962839 |
| NA | | NA |
| | 0.878896172 | NA |
| | 0.467223942 | 0.623633426 |
| | 0.296839401 | 0.456904294 |
| | 0.017665056 | 0.053196239 |
| | 0.788582525 | NA |
| | 4.72E-10 | 1.64E-08 |
| | 0.889410245 | NA |
| | 0.690104949 | NA |
| | 0.362099078 | 0.526974984 |
| NA | | NA |
| | 0.398550785 | 0.562192888 |
| | 0.720898194 | 0.826987955 |
| | 1.97E-06 | 2.37E-05 |
| NA | | NA |
| | 0.8488843 | 0.911255884 |
| | 0.03865512 | 0.099464234 |
| NA | | NA |
| | 0.037746644 | 0.097722064 |
| NA | | NA |
| | 0.000738024 | 0.003867557 |

| | | |
|---|---|---|
| | 0.001979743 | 0.008872076 |
| | 0.255940614 | 0.410533232 |
| | 0.624958404 | 0.755340311 |
| | 0.093645311 | 0.19578762 |
| | 3.92E-07 | 5.82E-06 |
| | 0.782559532 | NA |
| | 0.95777831 | 0.976545197 |
| | 0.502056312 | NA |
| | 0.000124337 | 0.00084165 |
| | 0.455876705 | 0.613795063 |
| | 0.001104448 | 0.005411107 |
| | 3.89E-09 | 1.03E-07 |
| | 0.650764418 | 0.775247429 |
| | 0.021737714 | 0.062882159 |
| | 0.069547778 | NA |
| | 0.026729037 | 0.074258974 |
| | 0.610601733 | 0.744569272 |
| | 0.011350919 | 0.037241122 |
| | 0.629965648 | 0.758244665 |
| NA | | NA |
| NA | | NA |
| | 0.948582032 | NA |
| NA | | NA |
| NA | | NA |
| | 0.992253023 | NA |
| | 2.29E-05 | 0.000199333 |
| | 0.275761392 | 0.43315053 |
| | 5.80E-06 | 6.06E-05 |
| | 0.360986844 | 0.525707917 |
| NA | | NA |
| | 0.992062245 | NA |
| NA | | NA |
| | 0.050427197 | 0.12219497 |
| | 0.000794501 | 0.00410215 |
| | 0.303135929 | 0.463839583 |
| | 0.466675546 | 0.623253524 |
| | 0.382717918 | NA |
| | 0.003747687 | 0.015055884 |
| | 0.000454249 | 0.002544604 |
| | 0.989111921 | NA |
| NA | | NA |
| | 0.212043559 | 0.359141111 |
| | 2.30E-08 | 4.93E-07 |
| | 0.513124358 | 0.664682502 |
| | 0.995121854 | NA |
| | 0.599782766 | 0.73570717 |
| | 0.086468738 | 0.184364211 |

| | |
|---|---|
| 6.75E-08 | 1.26E-06 |
| 0.00050124 | 0.00276829 |
| 0.002306914 | 0.010093152 |
| 0.321073862 | 0.483371791 |
| 0.944917172 | 0.968866415 |
| 0.242682602 | 0.395566156 |
| 0.089233767 | 0.18895702 |
| 0.18430301 | 0.324886393 |
| 0.808548716 | 0.885134915 |
| NA | NA |
| 0.158999845 | 0.29099151 |
| 4.61E-05 | 0.000361257 |
| 0.904641684 | 0.946133871 |
| 0.063642815 | 0.146078455 |
| 0.689693803 | 0.80464277 |
| 0.01395378 | 0.04401193 |
| 0.631954725 | 0.759805837 |
| 0.590752319 | NA |
| 0.252145359 | 0.40640487 |
| 0.103581339 | 0.211526329 |
| 0.816281882 | 0.890670005 |
| 0.984479014 | NA |
| 0.450476285 | 0.608909233 |
| NA | NA |
| 0.214608563 | NA |
| 0.1646254 | NA |
| NA | NA |
| NA | NA |
| 0.237381662 | 0.389295288 |
| 0.324113733 | 0.48638102 |
| NA | NA |
| NA | NA |
| 2.17E-10 | 8.40E-09 |
| 0.018383256 | 0.054892291 |
| 0.897126576 | NA |
| 0.475047623 | NA |
| 0.000285855 | 0.001717074 |
| 0.203414918 | 0.34921781 |
| NA | NA |
| 0.416804094 | NA |
| NA | NA |
| NA | NA |
| 0.158016637 | 0.289664191 |
| 0.009951797 | 0.033483451 |
| 0.460622633 | 0.618020487 |
| 1.72E-08 | 3.79E-07 |
| 0.331358797 | 0.493752648 |

| | | |
|---|---|---|
| | 0.001202401 | 0.005818315 |
| | 0.474819522 | 0.63103322 |
| | 0.616158192 | 0.748833725 |
| | 0.945917737 | 0.969284153 |
| | 0.056164533 | 0.132529226 |
| | 0.992060816 | NA |
| | 0.009375138 | 0.031902939 |
| | 0.596724248 | 0.733553589 |
| | 0.329753826 | NA |
| | 1.10E-05 | 0.000105237 |
| | 0.274505752 | NA |
| | 0.088648195 | 0.188043442 |
| | 0.960971717 | 0.978483628 |
| | 0.002920891 | 0.012284975 |
| NA | | NA |
| | 0.505666387 | NA |
| | 0.766638171 | NA |
| | 0.010768142 | 0.035761893 |
| | 0.851542467 | NA |
| | 0.031168257 | 0.083946952 |
| | 0.525179759 | 0.675220519 |
| | 0.808812149 | 0.885290898 |
| | 0.398394944 | 0.562016868 |
| | 0.040269677 | 0.102573 |
| | 0.51114793 | NA |
| | 0.096268599 | 0.200110012 |
| | 0.637509907 | 0.764141924 |
| | 0.026361755 | 0.073443007 |
| | 0.803442498 | 0.881742907 |
| | 0.027783135 | 0.076644456 |
| | 0.112843404 | 0.225532748 |
| | 0.779313288 | NA |
| | 0.06077064 | 0.141183081 |
| | 2.44E-05 | 0.000210331 |
| NA | | NA |
| NA | | NA |
| | 0.343141175 | 0.506444774 |
| | 0.369288347 | 0.533972208 |
| | 0.037953475 | 0.098095617 |
| | 0.189780179 | NA |
| | 0.000877311 | 0.004454303 |
| | 0.668047043 | NA |
| | 0.015737979 | 0.048585729 |
| | 0.9993606 | 0.999772568 |
| | 0.315606643 | 0.477824638 |
| | 0.438084846 | NA |
| NA | | NA |

| | | | |
|---|---|---|---|
| NA | | NA | |
| | 0.302296568 | | 0.462910065 |
| NA | | NA | |
| NA | | NA | |
| | 0.889409773 | NA | |
| NA | | NA | |
| | 0.021054974 | | 0.061270365 |
| | 0.250752178 | | 0.404872579 |
| | 0.614339618 | NA | |
| | 0.420344588 | NA | |
| | 1.54E-06 | | 1.90E-05 |
| | 0.855373967 | | 0.915804824 |
| NA | | NA | |
| NA | | NA | |
| | 0.020881299 | | 0.060869832 |
| | 0.408737547 | | 0.572013752 |
| | 0.905154857 | NA | |
| | 0.885057861 | NA | |
| NA | | NA | |
| | 0.119407549 | | 0.235289875 |
| | 0.980161628 | NA | |
| | 8.19E-10 | | 2.65E-08 |
| | 0.219445229 | | 0.368184427 |
| | 0.000415904 | | 0.00236296 |
| | 0.062331759 | | 0.14388133 |
| | 0.786150601 | | 0.871344692 |
| | 0.005160515 | | 0.019681806 |
| | 0.841906282 | NA | |
| | 6.24E-09 | | 1.55E-07 |
| | 0.021567474 | | 0.062455513 |
| | 0.383002356 | | 0.547050132 |
| | 0.369947595 | NA | |
| | 0.992061292 | NA | |
| | 0.905158739 | NA | |
| NA | | NA | |
| | 0.209323937 | NA | |
| | 3.42E-08 | | 7.03E-07 |
| | 0.000835415 | | 0.004282186 |
| NA | | NA | |
| | 0.672367757 | | 0.791451277 |
| | 0.297262962 | | 0.457338848 |
| | 0.138302623 | | 0.262057752 |
| NA | | NA | |
| | 0.006479943 | | 0.023719464 |
| | 0.825060807 | | 0.895709213 |
| | 2.70E-08 | | 5.67E-07 |
| | 0.991906727 | NA | |

| | |
|---|---|
| 0.712619822 | 0.820965252 |
| 0.002474105 | 0.0107255 |
| NA | NA |
| NA | NA |
| 0.522317217 | 0.6726711 |
| 0.002098403 | 0.009318267 |
| 8.87E-06 | 8.80E-05 |
| 0.000292597 | 0.001748932 |
| NA | NA |
| NA | NA |
| 0.114963771 | 0.228835212 |
| 0.619793188 | 0.751479135 |
| NA | NA |
| 0.325961571 | 0.488119895 |
| 0.527766172 | 0.677375497 |
| 0.53276551 | 0.681759781 |
| 0.603285 | 0.738723878 |
| 0.841697906 | 0.906332777 |
| NA | NA |
| 0.889409301 | NA |
| 0.031688875 | 0.085130413 |
| 0.875418535 | 0.928844683 |
| NA | NA |
| 0.00401767 | 0.015947943 |
| 0.177399417 | NA |
| NA | NA |
| 0.11559335 | 0.229834243 |
| 0.210773026 | 0.357734184 |
| 0.012712166 | 0.040846714 |
| 0.678102861 | NA |
| 0.00844574 | 0.029348341 |
| 0.272718508 | 0.429522465 |
| 0.159229754 | 0.291208375 |
| 0.000333995 | 0.001962066 |
| NA | NA |
| 0.056000318 | 0.132226988 |
| 0.002554906 | 0.011007097 |
| 0.162039638 | 0.295166189 |
| NA | NA |
| NA | NA |
| 0.88121865 | NA |
| NA | NA |
| 0.844344776 | NA |
| NA | NA |
| NA | NA |
| NA | NA |
| NA | NA |

| | |
|---|---|
| 0.901384322 | 0.944265974 |
| NA | NA |
| NA | NA |
| 0.355500857 | NA |
| NA | NA |
| 0.770008231 | 0.860971065 |
| NA | NA |
| 0.805374319 | NA |
| 0.944887381 | 0.968866415 |
| NA | NA |
| NA | NA |
| 0.739918048 | 0.840281213 |
| 3.61E-05 | 0.000293172 |
| NA | NA |
| NA | NA |
| 0.076161675 | 0.167427592 |
| 0.07647623 | NA |
| NA | NA |
| 0.662026875 | NA |
| NA | NA |
| NA | NA |
| 0.992062245 | NA |
| 0.19663985 | NA |
| 0.761033396 | 0.855105879 |
| NA | NA |
| 0.297931635 | NA |
| 0.608551259 | NA |
| NA | NA |
| NA | NA |
| 0.885644693 | NA |
| 0.8212117 | 0.893667741 |
| NA | NA |
| 0.992061292 | NA |
| NA | NA |
| NA | NA |
| NA | NA |
| NA | NA |
| 0.889409301 | NA |
| 0.73465785 | NA |
| NA | NA |
| NA | NA |
| NA | NA |
| NA | NA |
| 0.905158739 | NA |
| 0.83327265 | 0.900794276 |
| NA | NA |
| NA | NA |

| | | |
|---|---|---|
| | 0.925933599 | NA |
| NA | | NA |
| | 0.992061292 | NA |
| NA | | NA |
| | 0.960776592 | NA |
| NA | | NA |
| | 0.805374319 | NA |
| | 0.994074939 | 0.997117284 |
| NA | | NA |
| | 0.889409301 | NA |
| | 0.554173014 | NA |
| NA | | NA |
| | 0.992062245 | NA |
| | 0.122640638 | 0.23983131 |
| NA | | NA |
| | 0.991918092 | NA |
| NA | | NA |
| | 0.889409773 | NA |
| | 0.996394248 | 0.998242688 |
| NA | | NA |
| | 0.889409773 | NA |
| | 0.934057239 | 0.962142926 |
| | 0.07713558 | 0.169086918 |
| NA | | NA |
| | 0.992060816 | NA |
| NA | | NA |
| | 0.889409773 | NA |
| NA | | NA |
| | 0.905296333 | NA |
| NA | | NA |
| | 0.992061769 | NA |
| NA | | NA |
| | 0.236192877 | NA |
| NA | | NA |
| | 0.905158266 | NA |
| | 0.219882722 | 0.368721243 |
| NA | | NA |
| | 0.792445003 | NA |
| NA | | NA |
| NA | | NA |
| | 0.472623384 | 0.628951997 |
| NA | | NA |
| | 0.982201828 | NA |
| NA | | NA |
| NA | | NA |
| NA | | NA |
| | 0.729653278 | NA |

| | | |
|---|---|---|
| 0.807879735 | NA | |
| 0.061855682 | | 0.143068139 |
| NA | NA | |
| 0.778531773 | NA | |
| NA | NA | |
| 0.038290093 | NA | |
| NA | NA | |
| 0.992060816 | NA | |
| NA | NA | |
| 0.860037372 | NA | |
| NA | NA | |
| NA | NA | |
| NA | NA | |
| 0.25397175 | NA | |
| NA | NA | |
| 0.599089124 | NA | |
| NA | NA | |
| 0.992061292 | NA | |
| 0.915666606 | NA | |
| NA | NA | |
| NA | NA | |
| NA | NA | |
| NA | NA | |
| NA | NA | |
| 0.543846638 | NA | |
| NA | NA | |
| NA | NA | |
| 0.991918092 | NA | |
| 0.687104623 | NA | |
| NA | NA | |
| NA | NA | |
| 0.289431187 | NA | |
| NA | NA | |
| 0.1242218 | | 0.242221186 |
| 0.748890654 | NA | |
| NA | NA | |
| NA | NA | |
| 0.597700311 | | 0.734342368 |
| NA | NA | |
| 0.905158266 | NA | |
| NA | NA | |
| 0.823136937 | | 0.894593615 |
| 0.84949498 | | 0.911822283 |
| 0.772073142 | NA | |
| NA | NA | |
| NA | NA | |
| 0.809101644 | NA | |

| | |
|---|---|
| NA | NA |
| NA | NA |
| NA | NA |
| NA | NA |
| 0.120409265 | 0.236711464 |
| NA | NA |
| NA | NA |
| NA | NA |
| NA | NA |
| 0.54285867 | NA |
| 0.427454969 | 0.588494274 |
| NA | NA |
| 0.889409301 | NA |
| 0.093547061 | NA |
| NA | NA |
| 0.99205891 | NA |
| NA | NA |
| NA | NA |
| 0.977420838 | NA |
| 0.385595546 | 0.549423474 |
| NA | NA |
| 0.905158266 | NA |
| 0.145713231 | 0.272641177 |
| 0.889409301 | NA |
| 0.443079999 | 0.602798062 |
| 0.905157793 | NA |
| 0.233885773 | 0.385378497 |
| 0.90515732 | NA |
| 0.831680604 | 0.899870628 |
| NA | NA |
| 0.911629943 | 0.950003887 |
| 0.768649441 | NA |
| 0.905157793 | NA |
| 0.825823905 | 0.896183203 |
| 0.905158266 | NA |
| 0.905158739 | NA |
| NA | NA |
| 0.99205891 | NA |
| 0.905158266 | NA |
| 0.533805777 | NA |
| 0.941197464 | 0.966641182 |
| 0.637867216 | NA |
| 0.798045894 | 0.878104385 |
| 0.992061769 | NA |
| NA | NA |
| 0.569319905 | NA |
| 0.887861348 | 0.936517531 |

| | | |
|---|---|---|
| | 0.524122644 | NA |
| | 0.992060816 | NA |
| | 0.90515732 | NA |
| | 0.295867517 | 0.455894661 |
| | 0.087191112 | 0.185580074 |
| | 0.243658055 | 0.396607348 |
| | 0.017931811 | 0.053859205 |
| | 6.68E-09 | 1.65E-07 |
| | 0.204332957 | 0.350187107 |
| | 0.138569742 | 0.262428025 |
| | 0.870075908 | 0.925515189 |
| | 0.003231997 | 0.013365616 |
| NA | | NA |
| | 1.15E-13 | 1.02E-11 |
| NA | | NA |
| | 0.286332079 | 0.445389681 |
| | 0.106614087 | 0.216091815 |
| NA | | NA |
| | 0.889412384 | NA |
| NA | | NA |
| NA | | NA |
| | 0.493504354 | NA |
| | 0.461933934 | NA |
| | 0.186009586 | NA |
| NA | | NA |
| NA | | NA |
| NA | | NA |
| | 1.88E-06 | 2.27E-05 |
| | 0.889410717 | NA |
| | 0.407485935 | 0.570662223 |
| | 0.889409773 | NA |
| | 6.86E-05 | 0.000506509 |
| | 1.13E-14 | 1.27E-12 |
| | 0.000547202 | 0.002990243 |
| | 0.129002738 | 0.249034824 |
| | 0.021830949 | 0.063093685 |
| | 0.000411562 | 0.002341926 |
| NA | | NA |
| | 0.205277687 | 0.351394818 |
| | 0.827678692 | NA |
| NA | | NA |
| | 0.096919959 | NA |
| NA | | NA |
| | 0.034100142 | NA |
| | 0.117636025 | 0.232676046 |
| | 0.49846998 | 0.651464248 |
| | 0.016620826 | 0.050747598 |

| | |
|---|---|
| NA | NA |
| 0.889410245 | NA |
| 0.000134729 | 0.000899519 |
| NA | NA |
| 0.627907743 | 0.756597131 |
| 1.74E-12 | 1.17E-10 |
| 0.00119245 | 0.005780353 |
| 0.037169206 | 0.096580121 |
| 0.022123649 | 0.063776531 |
| 0.093009107 | 0.194814287 |
| 0.036314401 | 0.094785205 |
| 2.12E-05 | 0.00018707 |
| 3.78E-05 | 0.000305437 |
| NA | NA |
| NA | NA |
| 0.991879747 | NA |
| NA | NA |
| 3.90E-05 | 0.000313286 |
| 0.032304133 | 0.086455387 |
| 6.82E-05 | 0.000504551 |
| 0.227106787 | 0.377013462 |
| 0.549067139 | NA |
| 0.095991446 | 0.199666209 |
| 6.92E-07 | 9.53E-06 |
| 0.844991276 | 0.908720478 |
| 0.077139794 | 0.169086918 |
| NA | NA |
| NA | NA |
| NA | NA |
| 0.507335821 | NA |
| 0.536744301 | NA |
| NA | NA |
| 0.239615403 | NA |
| NA | NA |
| 0.996927778 | NA |
| 0.285929386 | 0.444920875 |
| 0.745572737 | NA |
| NA | NA |
| 0.905157793 | NA |
| 0.000153754 | 0.001004551 |
| 0.823096235 | 0.894593615 |
| 1.29E-09 | 3.91E-08 |
| 0.000285 | 0.001714137 |
| 0.063360691 | 0.145598307 |
| NA | NA |
| 0.905158739 | NA |
| 0.251745787 | 0.40596943 |

| | |
|---|---|
| 0.005890547 | 0.021942411 |
| 0.081891477 | 0.177098183 |
| 0.200647075 | 0.345786306 |
| 0.01499615 | 0.046696037 |
| 0.497533415 | 0.650588714 |
| NA | NA |
| 0.005758722 | 0.021564765 |
| 0.157339938 | 0.288737064 |
| 1.20E-10 | 5.02E-09 |
| 0.402470891 | 0.56602141 |
| 0.872328233 | 0.926688886 |
| 0.2497238 | 0.403597919 |
| 0.692065284 | 0.806288029 |
| 0.182753342 | 0.322951918 |
| 1.95E-09 | 5.66E-08 |
| 0.578655859 | 0.719478173 |
| 0.719767946 | NA |
| 0.102608519 | 0.210145538 |
| 0.929749753 | 0.959686698 |
| 0.000827304 | 0.004249538 |
| 0.431361527 | 0.592275856 |
| 0.000358951 | 0.002084135 |
| 8.21E-26 | 1.20E-22 |
| 2.53E-07 | 3.97E-06 |
| 3.53E-11 | 1.68E-09 |
| 0.1623924 | 0.295678699 |
| 0.020243887 | 0.059342612 |
| 0.604360809 | 0.739505127 |
| 0.658882406 | NA |
| 0.005370974 | 0.020348728 |
| 0.540359459 | NA |
| 0.070808431 | 0.15800044 |
| 0.302482378 | NA |
| 0.875642624 | 0.928855436 |
| 0.872763175 | NA |
| 0.668907598 | 0.78895785 |
| 0.60086538 | 0.736607052 |
| 0.584595599 | 0.72424086 |
| NA | NA |
| 0.133252161 | 0.254663551 |
| 0.277859973 | 0.435806289 |
| 0.7125986 | 0.820965252 |
| NA | NA |
| NA | NA |
| 0.524064576 | NA |
| 0.000605548 | 0.003267135 |
| 3.97E-06 | 4.37E-05 |

| | |
|---|---|
| NA | NA |
| 0.848744952 | 0.911184679 |
| NA | NA |
| 0.104795047 | 0.21342447 |
| 0.959648272 | 0.977681965 |
| 0.483007431 | NA |
| 0.600609907 | 0.736433801 |
| 0.931050742 | 0.96017204 |
| 0.992060816 | NA |
| 0.471388871 | NA |
| 0.417925746 | NA |
| NA | NA |
| 0.61844199 | 0.750462404 |
| 0.650212631 | 0.774758216 |
| 0.002865041 | 0.012103391 |
| 0.994323272 | 0.99722972 |
| 0.348170528 | NA |
| 0.713258462 | 0.821458447 |
| 0.795759448 | NA |
| 0.116276343 | 0.230885124 |
| NA | NA |
| NA | NA |
| 0.528358132 | 0.678016376 |
| 0.026752684 | 0.074291148 |
| NA | NA |
| NA | NA |
| NA | NA |
| NA | NA |
| 0.905157793 | NA |
| 0.116532544 | NA |
| 0.905158739 | NA |
| 0.372118729 | NA |
| NA | NA |
| NA | NA |
| NA | NA |
| NA | NA |
| 0.889409773 | NA |
| NA | NA |
| NA | NA |
| NA | NA |
| NA | NA |
| NA | NA |
| 0.853835502 | NA |
| NA | NA |
| 8.50E-05 | 0.000606196 |
| NA | NA |
| NA | NA |

| | |
|---|---|
| 0.156746618 | 0.287961096 |
| 0.709029448 | NA |
| 0.014015198 | 0.044148579 |
| 0.916164776 | 0.952379394 |
| NA | NA |
| 0.062125592 | 0.143564638 |
| 0.24201728 | 0.394788099 |
| 0.591291272 | NA |
| 0.000224829 | 0.001395824 |
| NA | NA |
| 0.430189331 | 0.591331927 |
| 1.29E-08 | 2.95E-07 |
| NA | NA |
| 0.031782039 | 0.085338934 |
| 0.032819156 | 0.087535023 |
| 0.561913441 | 0.705855343 |
| NA | NA |
| 2.87E-06 | 3.28E-05 |
| 0.183950393 | 0.324490955 |
| 0.126103875 | 0.244837753 |
| 1.46E-08 | 3.27E-07 |
| 0.02062455 | 0.060281444 |
| 0.973762037 | NA |
| 0.547378602 | NA |
| 0.220028926 | 0.368881904 |
| 0.036455876 | 0.095109252 |
| 0.143922846 | NA |
| 0.120399754 | 0.236711464 |
| 0.13013915 | 0.250326349 |
| 0.028763048 | 0.078783557 |
| 0.000904781 | 0.004578956 |
| 0.460458764 | 0.617924408 |
| 0.982473465 | NA |
| 0.167214023 | 0.302232513 |
| 0.018833763 | 0.055940835 |
| 0.400801118 | 0.564445198 |
| 0.992060816 | NA |
| 0.071077821 | 0.158405832 |
| 0.028440946 | 0.078086363 |
| 1.61E-05 | 0.000147538 |
| 0.477420352 | 0.633454592 |
| 0.002939801 | 0.012352676 |
| NA | NA |
| 0.000744011 | 0.003892715 |
| 0.039974561 | 0.102056295 |
| 0.154523073 | 0.285044703 |
| 0.647182058 | 0.772110769 |

| | | |
|---|---|---|
| | 0.547447533 | 0.693795832 |
| | 0.21322678 | NA |
| | 0.051592384 | 0.124387039 |
| | 0.3907265 | 0.554118558 |
| | 0.769422281 | NA |
| | 0.798859137 | 0.878689029 |
| | 0.184228091 | 0.324832556 |
| | 4.61E-06 | 4.96E-05 |
| | 6.82E-07 | 9.43E-06 |
| | 0.025595753 | 0.071911097 |
| | 1.98E-05 | 0.000176235 |
| | 0.338417911 | 0.50143637 |
| | 0.478395129 | 0.634326348 |
| | 0.536310495 | 0.684415336 |
| | 0.603586248 | 0.739010338 |
| | 0.412370511 | 0.574899628 |
| | 0.617174963 | 0.749670595 |
| | 3.48E-07 | 5.23E-06 |
| | 0.269004734 | 0.425428517 |
| | 0.85148306 | NA |
| | 0.73637332 | 0.838215838 |
| | 0.00886186 | 0.030510012 |
| | 0.030565014 | 0.082687835 |
| | 0.992060816 | NA |
| | 0.261555939 | NA |
| NA | | NA |
| | 0.056024247 | 0.132255046 |
| | 0.366940296 | 0.531682082 |
| | 8.94E-06 | 8.86E-05 |
| NA | | NA |
| | 0.047929537 | 0.117386204 |
| | 0.157800024 | 0.289392101 |
| | 0.512679999 | NA |
| | 0.695381882 | 0.808872272 |
| | 0.864058096 | NA |
| | 0.992061292 | NA |
| | 0.011950508 | 0.038831192 |
| | 0.46314715 | NA |
| | 0.914353407 | 0.951757985 |
| | 0.194354013 | 0.337643969 |
| | 0.459366865 | 0.616616968 |
| | 0.233895151 | 0.385378497 |
| | 0.518464526 | 0.669367104 |
| | 0.315997598 | 0.478188358 |
| | 0.48379732 | NA |
| | 0.350637438 | 0.514785553 |
| | 0.889409301 | NA |

| | |
|---|---|
| 0.148129719 | 0.276033646 |
| 0.6883314 | NA |
| 0.010641407 | 0.035415982 |
| 0.772002404 | 0.862251933 |
| NA | NA |
| 0.004277501 | 0.016797148 |
| 0.875316426 | 0.928844683 |
| 0.000537941 | 0.002947266 |
| 0.538605633 | 0.686393192 |
| NA | NA |
| 0.886029463 | 0.935379622 |
| 0.741856911 | NA |
| 0.705230054 | 0.816569405 |
| 0.168736828 | 0.3043361 |
| NA | NA |
| 0.807252815 | NA |
| NA | NA |
| 0.390676215 | 0.554118558 |
| NA | NA |
| 0.850184155 | NA |
| 0.254145988 | 0.408543271 |
| 0.270706609 | 0.427197012 |
| 0.131003016 | 0.251604383 |
| 0.000292045 | 0.001747145 |
| 0.125767652 | 0.244379423 |
| 0.999591203 | NA |
| 0.690605109 | 0.805274956 |
| 2.98E-05 | 0.00024907 |
| 0.866442027 | 0.923064258 |
| 0.557010388 | NA |
| 0.889409773 | NA |
| 0.003301965 | 0.013590973 |
| 0.061076313 | NA |
| 0.31170394 | 0.47328909 |
| 0.760387367 | NA |
| 3.79E-06 | 4.20E-05 |
| 0.425240478 | NA |
| 0.763438835 | NA |
| NA | NA |
| NA | NA |
| NA | NA |
| NA | NA |
| NA | NA |
| 0.58239241 | NA |
| 0.013000659 | 0.041509362 |
| 0.985861829 | NA |
| 0.791026341 | NA |

| | |
|---|---|
| 0.553455991 | NA |
| 0.112473933 | 0.224937622 |
| 0.01123958 | 0.036953126 |
| 0.185320399 | 0.326154178 |
| NA | NA |
| NA | NA |
| 0.001536352 | 0.007149416 |
| NA | NA |
| NA | NA |
| 0.636404496 | 0.763403089 |
| 0.901183732 | NA |
| NA | NA |
| 0.579822277 | 0.720275917 |
| NA | NA |
| 0.992061292 | NA |
| 0.889409773 | NA |
| NA | NA |
| 0.889410245 | NA |
| 0.200351799 | 0.345385865 |
| 0.015347315 | 0.047573644 |
| 0.011812252 | 0.038455929 |
| 0.841455762 | 0.906236209 |
| 0.76010128 | NA |
| 0.2316817 | 0.382676555 |
| 0.889409301 | NA |
| NA | NA |
| NA | NA |
| NA | NA |
| 0.997402762 | NA |
| 0.946158566 | NA |
| NA | NA |
| NA | NA |
| NA | NA |
| 0.889409773 | NA |
| 0.992060816 | NA |
| NA | NA |
| NA | NA |
| NA | NA |
| NA | NA |
| NA | NA |
| NA | NA |
| NA | NA |
| NA | NA |
| NA | NA |
| NA | NA |
| NA | NA |
| NA | NA |

| | | |
|---|---|---|
| NA | | NA |
| NA | | NA |
| NA | | NA |
| NA | | NA |
| NA | | NA |
| NA | | NA |
| NA | | NA |
| NA | | NA |
| NA | | NA |
| NA | | NA |
| NA | | NA |
| NA | | NA |
| NA | | NA |
| NA | | NA |
| NA | | NA |
| NA | | NA |
| NA | | NA |
| NA | | NA |
| | 0.889409301 | NA |
| NA | | NA |
| NA | | NA |
| NA | | NA |
| NA | | NA |
| NA | | NA |
| NA | | NA |
| NA | | NA |
| NA | | NA |
| NA | | NA |
| NA | | NA |
| NA | | NA |
| NA | | NA |
| NA | | NA |
| NA | | NA |
| NA | | NA |
| | 0.992060816 | NA |
| NA | | NA |
| NA | | NA |
| NA | | NA |
| | 0.992060816 | NA |
| NA | | NA |
| NA | | NA |
| NA | | NA |
| NA | | NA |
| NA | | NA |
| NA | | NA |
| NA | | NA |
| NA | | NA |

| | |
|---|---|
| NA | NA |
| NA | NA |
| NA | NA |
| NA | NA |
| NA | NA |
| NA | NA |
| NA | NA |
| NA | NA |
| NA | NA |
| 0.905296333 | NA |
| 0.888555194 | 0.936985377 |
| 0.173966634 | 0.311237845 |
| NA | NA |
| 0.869399481 | NA |
| 0.056743733 | 0.133537203 |
| NA | NA |
| NA | NA |
| 0.776404041 | NA |
| NA | NA |
| NA | NA |
| 0.355441124 | 0.52006659 |
| NA | NA |
| NA | NA |
| 0.991918092 | NA |
| 0.672719482 | NA |
| NA | NA |
| 0.905158266 | NA |
| NA | NA |
| NA | NA |
| NA | NA |
| NA | NA |
| 0.822251845 | NA |
| NA | NA |
| NA | NA |
| NA | NA |
| 0.801661681 | NA |
| 0.016152006 | 0.049626683 |
| NA | NA |
| 0.466424453 | NA |
| NA | NA |
| NA | NA |
| 0.905157793 | NA |
| NA | NA |
| NA | NA |
| NA | NA |
| 0.889409301 | NA |
| NA | NA |

| | | | |
|---|---|---|---|
| NA | | NA | |
| | 0.08915742 | | 0.188849984 |
| NA | | NA | |
| NA | | NA | |
| NA | | NA | |
| NA | | NA | |
| NA | | NA | |
| NA | | NA | |
| NA | | NA | |
| NA | | NA | |
| | 0.905158266 | NA | |
| NA | | NA | |
| | 0.209830379 | NA | |
| NA | | NA | |
| NA | | NA | |
| | 0.199957477 | NA | |
| | 0.992060816 | NA | |
| | 0.889409301 | NA | |
| NA | | NA | |
| | 0.905158266 | NA | |
| NA | | NA | |
| NA | | NA | |
| NA | | NA | |
| | 0.889409301 | NA | |
| NA | | NA | |
| | 0.564831657 | NA | |
| | 0.782559532 | NA | |
| NA | | NA | |
| NA | | NA | |
| | 0.664340285 | NA | |
| | 0.992061292 | NA | |
| | 0.091187606 | | 0.191983571 |
| NA | | NA | |
| NA | | NA | |
| NA | | NA | |
| NA | | NA | |
| NA | | NA | |
| | 0.889409773 | NA | |
| NA | | NA | |
| NA | | NA | |
| | 0.004308602 | | 0.016901141 |
| | 0.227111162 | NA | |
| NA | | NA | |
| | 0.926389558 | NA | |
| | 0.125842029 | | 0.244437426 |
| | 1.82E-05 | | 0.000164048 |
| | 7.23E-06 | | 7.36E-05 |

| | |
|---|---|
| NA | NA |
| 0.144175617 | 0.270571094 |
| 0.004439313 | 0.017325671 |
| 0.002196284 | 0.009684379 |
| 0.637307099 | 0.764085382 |
| 0.057554967 | 0.135084386 |
| NA | NA |
| 2.27E-06 | 2.69E-05 |
| 0.204231302 | 0.350094613 |
| 0.097103898 | NA |
| 0.583111039 | 0.72305242 |
| NA | NA |
| 0.492783512 | 0.64668817 |
| 0.889410717 | NA |
| 0.542896206 | 0.689888809 |
| 0.797157316 | NA |
| 0.642869938 | NA |
| NA | NA |
| 0.828724638 | NA |
| NA | NA |
| 0.209750337 | 0.356521995 |
| 0.889409301 | NA |
| 0.251363116 | 0.405531017 |
| 0.982859796 | NA |
| 0.889409301 | NA |
| 0.905158266 | NA |
| 0.905158266 | NA |
| 0.127126843 | 0.246388204 |
| 0.007079481 | 0.02548497 |
| 0.06552803 | 0.149297573 |
| 0.419325349 | 0.581274247 |
| 0.939065968 | 0.965490663 |
| 0.076228756 | 0.167558284 |
| NA | NA |
| 0.321402941 | 0.48375419 |
| 0.792188023 | NA |
| 1.91E-07 | 3.12E-06 |
| 0.808420455 | 0.885124587 |
| 0.825749511 | NA |
| 1.26E-06 | 1.59E-05 |
| 0.000991105 | 0.004941795 |
| 1.79E-07 | 2.95E-06 |
| 0.036622057 | 0.095477908 |
| 5.43E-06 | 5.71E-05 |
| 0.437346783 | 0.597653744 |
| 0.235646665 | 0.387403408 |
| 0.980994114 | 0.990511759 |

| | |
|---|---|
| 0.937428192 | 0.964484171 |
| 0.039954371 | 0.102019282 |
| 0.048145209 | 0.117783048 |
| NA | NA |
| 0.561478005 | 0.705433071 |
| 0.259424887 | 0.414577621 |
| NA | NA |
| 0.994949901 | 0.997551495 |
| 0.409113353 | 0.572247921 |
| NA | NA |
| NA | NA |
| 0.353805414 | NA |
| 0.419468627 | 0.581364515 |
| 0.00768413 | 0.027176004 |
| 0.314608437 | 0.476677477 |
| NA | NA |
| 1.49E-06 | 1.85E-05 |
| 0.014761029 | 0.046074974 |
| NA | NA |
| 0.800234366 | 0.87954084 |
| 0.775463534 | 0.864525581 |
| NA | NA |
| NA | NA |
| NA | NA |
| NA | NA |
| NA | NA |
| 0.798546198 | NA |
| NA | NA |
| NA | NA |
| 0.992060816 | NA |
| NA | NA |
| NA | NA |
| 0.905158266 | NA |
| NA | NA |
| 0.992061769 | NA |
| 0.904947233 | NA |
| 0.905158266 | NA |
| 0.873111673 | NA |
| NA | NA |
| NA | NA |
| NA | NA |
| NA | NA |
| 0.992060816 | NA |
| NA | NA |
| NA | NA |
| NA | NA |
| NA | NA |

| | | | |
|---|---|---|---|
| NA | | NA | |
| NA | | NA | |
| NA | | NA | |
| NA | | NA | |
| | 0.873981309 | NA | |
| | 0.289651925 | | 0.449058643 |
| | 0.780509524 | NA | |
| | 0.238715423 | | 0.390869583 |
| | 0.991923231 | | 0.995930211 |
| | 0.415326069 | | 0.577772895 |
| | 8.70E-18 | | 2.08E-15 |
| | 0.669934589 | | 0.789501803 |
| | 0.090606644 | | 0.191149725 |
| | 0.45702723 | | 0.614641165 |
| | 0.013149862 | | 0.041861094 |
| NA | | NA | |
| | 0.674434771 | | 0.79315145 |
| | 0.847455439 | | 0.910389609 |
| | 0.000224689 | | 0.001395344 |
| | 0.905158739 | NA | |
| | 0.070473557 | | 0.157349144 |
| | 0.018095851 | NA | |
| | 0.905158266 | NA | |
| | 0.905158739 | NA | |
| | 0.889409301 | NA | |
| | 0.889410245 | NA | |
| | 0.591893789 | | 0.729616914 |
| | 0.040206367 | | 0.102459281 |
| NA | | NA | |
| | 0.002777865 | | 0.011798673 |
| | 0.890439715 | | 0.938091737 |
| | 0.889410245 | NA | |
| | 0.905158266 | NA | |
| | 0.905158266 | NA | |
| | 0.627815221 | NA | |
| NA | | NA | |
| NA | | NA | |
| NA | | NA | |
| | 0.905158739 | NA | |
| NA | | NA | |
| NA | | NA | |
| NA | | NA | |
| NA | | NA | |
| NA | | NA | |
| | 0.889409773 | NA | |
| NA | | NA | |
| NA | | NA | |

| | |
|---|---|
| NA | NA |
| NA | NA |
| NA | NA |
| 0.991887208 | NA |
| NA | NA |
| 0.99205891 | NA |
| NA | NA |
| NA | NA |
| 0.614339618 | NA |
| NA | NA |
| NA | NA |
| NA | NA |
| 0.337719813 | 0.500649086 |
| 0.003558476 | 0.014449071 |
| 0.00941206 | 0.031998673 |
| 0.866377143 | NA |
| 0.721376616 | NA |
| 5.01E-09 | 1.27E-07 |
| 0.006278735 | NA |
| 0.000640528 | 0.003431332 |
| 0.303311121 | 0.463978328 |
| 0.905158739 | NA |
| 0.889410717 | NA |
| 6.17E-05 | 0.000463066 |
| 0.581286742 | 0.721482906 |
| 0.454193525 | 0.6122191 |
| 0.008609624 | 0.029805887 |
| 0.297440191 | 0.457496243 |
| 0.677771629 | 0.79555202 |
| 0.25108134 | 0.405219813 |
| 0.261374911 | 0.416844482 |
| NA | NA |
| 2.36E-05 | 0.000204669 |
| 7.79E-05 | 0.000566078 |
| 0.731927603 | 0.835103306 |
| 0.855788113 | NA |
| NA | NA |
| 0.405955271 | 0.569171995 |
| NA | NA |
| NA | NA |
| NA | NA |
| 0.889409301 | NA |
| 0.905158739 | NA |
| 0.373266655 | NA |
| 0.751586013 | NA |
| 0.720918531 | NA |
| NA | NA |

| | | | |
|---|---|---|---|
| | 0.986096642 | NA | |
| | 0.889409773 | NA | |
| | 0.873981309 | NA | |
| | 0.008206271 | | 0.028666644 |
| | 0.576617546 | NA | |
| NA | | NA | |
| | 0.223676264 | NA | |
| | 0.320955415 | NA | |
| NA | | NA | |
| | 0.995059079 | NA | |
| | 0.826889831 | NA | |
| NA | | NA | |
| NA | | NA | |
| NA | | NA | |
| | 0.654987089 | NA | |
| | 0.81295892 | NA | |
| | 0.905158739 | NA | |
| NA | | NA | |
| | 0.318878168 | NA | |
| | 0.889409773 | NA | |
| | 0.889409301 | NA | |
| | 0.991918092 | NA | |
| | 0.905158739 | NA | |
| NA | | NA | |
| NA | | NA | |
| NA | | NA | |
| | 0.905158739 | NA | |
| NA | | NA | |
| | 0.883047992 | NA | |
| NA | | NA | |
| NA | | NA | |
| NA | | NA | |
| NA | | NA | |
| | 0.59570208 | NA | |
| NA | | NA | |
| | 0.889409773 | NA | |
| | 0.572482132 | NA | |
| NA | | NA | |
| NA | | NA | |
| NA | | NA | |
| NA | | NA | |
| NA | | NA | |
| NA | | NA | |
| NA | | NA | |
| | 0.797342596 | NA | |
| NA | | NA | |
| NA | | NA | |

| | | |
|---|---|---|
| NA | | NA |
| | 0.905158266 | NA |
| NA | | NA |
| | 0.399282495 | NA |
| NA | | NA |
| | 0.242813836 | NA |
| | 0.692464693 | NA |
| | 0.905158266 | NA |
| | 0.325405598 | NA |
| NA | | NA |
| NA | | NA |
| | 0.991700267 | NA |
| NA | | NA |
| NA | | NA |
| | 0.581954223 | NA |
| NA | | NA |
| NA | | NA |
| NA | | NA |
| | 0.905158266 | NA |
| NA | | NA |
| | 0.849855223 | NA |
| NA | | NA |
| NA | | NA |
| NA | | NA |
| NA | | NA |
| NA | | NA |
| NA | | NA |
| NA | | NA |
| NA | | NA |
| NA | | NA |
| NA | | NA |
| NA | | NA |
| NA | | NA |
| NA | | NA |
| NA | | NA |
| NA | | NA |
| | 0.991918092 | NA |
| | 0.788582525 | NA |
| NA | | NA |
| NA | | NA |
| NA | | NA |
| NA | | NA |
| NA | | NA |
| | 0.889409301 | NA |
| | 0.845136429 | NA |
| | 0.905158266 | NA |
| | 0.778531773 | NA |

| | | |
|---|---|---|
| NA | NA | |
| NA | NA | |
| 0.889409301 | NA | |
| 0.991570563 | NA | |
| 0.752373727 | NA | |
| 0.873514299 | NA | |
| 0.903365571 | NA | |
| 0.992061292 | NA | |
| NA | NA | |
| 0.782595072 | NA | |
| 0.992253023 | NA | |
| 0.889409773 | NA | |
| 0.878605957 | NA | |
| NA | NA | |
| NA | NA | |
| 0.905158739 | NA | |
| 0.777187101 | NA | |
| NA | NA | |
| 0.801661681 | NA | |
| NA | NA | |
| 0.597813989 | NA | |
| NA | NA | |
| NA | NA | |
| NA | NA | |
| 0.983317411 | NA | |
| 0.777092376 | NA | |
| NA | NA | |
| NA | NA | |
| NA | NA | |
| 0.52409521 | | 0.674216152 |
| 0.868497615 | NA | |
| 0.688318891 | NA | |
| NA | NA | |
| NA | NA | |
| 0.804534188 | NA | |
| NA | NA | |
| NA | NA | |
| NA | NA | |
| NA | NA | |
| 0.787645204 | NA | |
| NA | NA | |
| NA | NA | |
| 0.40982459 | NA | |
| 0.889409301 | NA | |
| 0.797342596 | NA | |
| 0.848263876 | NA | |
| NA | NA | |

| | | |
|---|---|---|
| NA | NA | |
| NA | NA | |
| 0.905158739 | NA | |
| 0.687781615 | NA | |
| 0.989320473 | NA | |
| NA | NA | |
| NA | NA | |
| 0.905158266 | NA | |
| 0.992062245 | NA | |
| NA | NA | |
| 0.740136308 | NA | |
| 0.792103663 | NA | |
| NA | NA | |
| NA | NA | |
| 0.859291006 | NA | |
| 0.893967439 | NA | |
| 0.381206805 | NA | |
| 0.354037494 | NA | |
| 0.010620838 | | 0.035368247 |
| 0.986015719 | NA | |
| 0.517037453 | NA | |
| NA | NA | |
| 0.673858572 | NA | |
| 0.889409773 | NA | |
| 0.905158266 | NA | |
| 0.905158266 | NA | |
| 0.905158739 | NA | |
| NA | NA | |
| 0.992424942 | NA | |
| 0.905158266 | NA | |
| 0.88388815 | NA | |
| 0.905158266 | NA | |
| 0.992059387 | NA | |
| 0.808848586 | NA | |
| NA | NA | |
| 0.85535358 | NA | |
| 0.797157316 | NA | |
| 0.685479708 | NA | |
| 0.475047623 | NA | |
| 0.831676497 | NA | |
| 0.889409301 | NA | |
| 0.919604642 | NA | |
| 0.15203502 | NA | |
| 0.259694413 | NA | |
| 0.180156286 | NA | |
| 0.039011879 | NA | |
| 0.797342596 | NA | |

| | | |
|---|---|---|
| | 0.85323362 | NA |
| | 0.8490673 | NA |
| | 0.572813076 | NA |
| | 0.945608762 | NA |
| | 0.97510837 | NA |
| | 0.057326117 | NA |
| | 0.849445598 | NA |
| | 0.998676816 | NA |
| | 0.905158266 | NA |
| | 0.268818511 | NA |
| | 0.574549467 | NA |
| | 0.668340903 | NA |
| | 0.804838841 | NA |
| | 0.332307945 | NA |
| | 0.816313019 | NA |
| | 0.479744781 | NA |
| | 0.696060699 | NA |
| | 0.485530043 | NA |
| | 0.99205891 | NA |
| | 0.786262515 | NA |
| | 0.963520606 | NA |
| | 0.557784914 | NA |
| | 0.850343426 | NA |
| | 0.964671599 | NA |
| | 0.625325882 | NA |
| | 0.848468971 | NA |
| | 0.873616163 | NA |
| | 0.435711952 | NA |
| | 0.312431048 | NA |
| | 0.543343305 | NA |
| NA | | NA |
| | 0.484213259 | NA |
| | 0.67628142 | NA |
| | 0.318430754 | 0.480810075 |
| | 0.05725859 | 0.134503784 |
| | 0.032495951 | 0.086852312 |
| | 0.992061292 | NA |
| | 0.861756416 | NA |
| NA | | NA |
| NA | | NA |
| | 0.90515732 | NA |
| | 0.337339393 | 0.500313097 |
| | 3.98E-07 | 5.89E-06 |
| | 7.37E-05 | 0.000538258 |
| | 8.27E-06 | 8.28E-05 |
| NA | | NA |
| | 2.04E-07 | 3.30E-06 |

| | |
|---|---|
| 0.002227805 | 0.009805648 |
| 7.86E-09 | 1.91E-07 |
| 0.920701926 | 0.954551282 |
| 1.26E-05 | 0.000118876 |
| NA | NA |
| 0.658866331 | 0.781127619 |
| 0.077310527 | 0.169309052 |
| NA | NA |
| 0.000968098 | 0.00484468 |
| 0.000161884 | 0.001050783 |
| 0.889410717 | NA |
| 0.08399891 | NA |
| 0.115430351 | 0.229595568 |
| 0.997764529 | NA |
| 0.088493101 | NA |
| 0.001002549 | 0.004985277 |
| 0.028966174 | NA |
| 0.664348779 | 0.785591537 |
| NA | NA |
| 0.999530733 | 0.999849461 |
| 0.314108097 | 0.476017923 |
| 0.903513411 | 0.945355815 |
| 9.14E-05 | 0.000646312 |
| 0.573656617 | NA |
| 0.494799934 | 0.64805629 |
| 4.80E-20 | 2.17E-17 |
| 0.126636083 | 0.245653864 |
| 0.187293714 | NA |
| 0.988767841 | 0.993927811 |
| 0.211393087 | 0.358481798 |
| NA | NA |
| 0.003229655 | 0.013358446 |
| 0.332224796 | 0.494765975 |
| 0.727524406 | NA |
| NA | NA |
| 0.905158739 | NA |
| 0.002781923 | 0.011809187 |
| 0.60939036 | 0.743435925 |
| 0.889409773 | NA |
| 0.040696511 | 0.103300739 |
| 0.792208442 | 0.874791571 |
| 0.197939343 | 0.342364559 |
| 0.194499099 | 0.337842567 |
| 0.751130303 | 0.848416764 |
| 0.103718915 | 0.211725833 |
| 8.89E-22 | 5.27E-19 |
| NA | NA |

| | | |
|---|---|---|
| | 0.153406814 | 0.283581128 |
| | 0.17682153 | 0.315343456 |
| | 0.179416794 | 0.318909938 |
| | 0.000168231 | 0.001086774 |
| | 5.83E-11 | 2.65E-09 |
| | 0.824683715 | 0.895476918 |
| | 0.707960164 | 0.817880918 |
| | 0.262540177 | 0.417974316 |
| | 0.383067135 | 0.547107071 |
| | 0.665586168 | NA |
| | 0.670870289 | NA |
| | 0.09123432 | 0.192045095 |
| | 0.882372695 | 0.933154595 |
| | 0.00621673 | 0.022927976 |
| | 0.252813843 | 0.407093876 |
| | 0.001221042 | 0.005885174 |
| | 0.479353138 | 0.63502145 |
| | 0.423496419 | 0.584988228 |
| | 3.47E-06 | 3.89E-05 |
| | 0.680428118 | 0.797370131 |
| | 0.128952499 | 0.248959727 |
| | 0.299918515 | 0.460327859 |
| | 0.441240791 | 0.600955562 |
| | 0.457655286 | 0.61507145 |
| | 0.792188023 | NA |
| | 1.08E-06 | 1.40E-05 |
| | 0.052500691 | 0.125995919 |
| | 9.64E-17 | 1.76E-14 |
| | 3.52E-07 | 5.28E-06 |
| | 0.383435066 | 0.547418933 |
| | 0.088695359 | 0.188098293 |
| | 8.80E-05 | 0.000625163 |
| | 0.008461794 | 0.029391418 |
| | 0.001205508 | 0.005827229 |
| | 0.002965742 | 0.012445008 |
| | 0.611273904 | 0.744975228 |
| | 0.004756597 | 0.018385868 |
| | 0.661094176 | 0.782923472 |
| | 0.062745207 | 0.144555794 |
| | 0.257477314 | 0.412305428 |
| | 0.003871158 | NA |
| | 0.573532968 | NA |
| | 0.191757369 | 0.334243536 |
| | 0.274947775 | NA |
| NA | | NA |
| | 0.687605901 | NA |
| | 0.537602699 | NA |

| | | |
|---|---|---|
| NA | NA | |
| NA | NA | |
| 0.758028621 | NA | |
| 0.889409301 | NA | |
| 0.905158266 | NA | |
| NA | NA | |
| NA | NA | |
| 0.070711688 | NA | |
| 0.252599044 | | 0.406837496 |
| NA | NA | |
| NA | NA | |
| 0.092149755 | | 0.193489814 |
| 0.65298378 | NA | |
| 0.465180871 | | 0.622101508 |
| 0.905158266 | NA | |
| 6.47E-06 | | 6.70E-05 |
| NA | NA | |
| 0.002749625 | | 0.011708166 |
| NA | NA | |
| 0.505720677 | | 0.657691463 |
| 0.046353247 | | 0.114346637 |
| 0.048571083 | | 0.118600289 |
| 0.007137657 | | 0.025639754 |
| 0.905157793 | NA | |
| 0.234952869 | | 0.386443514 |
| 0.380895404 | | 0.545138321 |
| NA | NA | |
| NA | NA | |
| NA | NA | |
| 0.013078094 | | 0.041711237 |
| 0.715971181 | NA | |
| 0.017383054 | NA | |
| 2.62E-07 | | 4.10E-06 |
| 0.133182261 | | 0.254596437 |
| 6.92E-06 | | 7.09E-05 |
| 0.000394631 | | 0.002260225 |
| 0.234385199 | NA | |
| 0.024641909 | | 0.069677012 |
| 0.992060816 | NA | |
| 0.000586044 | | 0.003179871 |
| 0.028555196 | | 0.078341272 |
| 0.033313306 | | 0.088594753 |
| 0.042326884 | NA | |
| 0.187061916 | | 0.328510285 |
| 0.003279351 | | 0.013518167 |
| 0.889410717 | NA | |
| 0.718285282 | NA | |

| | |
|---|---|
| 0.491749036 | 0.645832361 |
| NA | NA |
| 0.423591588 | 0.585082884 |
| 0.655218537 | 0.778189402 |
| 0.388501136 | 0.551962569 |
| 0.120193755 | 0.236435963 |
| 0.0004518 | 0.002535117 |
| 0.653792284 | 0.777294248 |
| 0.753075385 | 0.849610222 |
| NA | NA |
| NA | NA |
| 0.844344776 | NA |
| 0.804607538 | 0.882580395 |
| 0.905158739 | NA |
| NA | NA |
| 0.163906502 | 0.297702471 |
| 2.21E-05 | 0.000194267 |
| 0.000738145 | 0.003867557 |
| 0.002045382 | 0.009119706 |
| 0.003375822 | 0.013843077 |
| 0.004471307 | 0.017427305 |
| 0.838001622 | 0.903726433 |
| 0.818141968 | 0.891724796 |
| 0.205092525 | 0.351214751 |
| 0.011223496 | 0.036911301 |
| 0.011265247 | 0.037015347 |
| 0.181631911 | 0.321857258 |
| 0.290974646 | 0.450441471 |
| 0.837130933 | NA |
| NA | NA |
| NA | NA |
| 0.101634084 | 0.208558345 |
| 0.873394521 | NA |
| 0.241292408 | 0.394091204 |
| 0.06348249 | 0.145786656 |
| 0.037403564 | NA |
| 0.718575578 | 0.825444713 |
| 0.85541371 | NA |
| 0.802884986 | NA |
| 0.385062407 | 0.549005579 |
| 0.000188972 | 0.001199662 |
| 0.387362368 | 0.551023791 |
| 0.478494905 | 0.634420336 |
| 0.232020894 | NA |
| 4.20E-07 | 6.17E-06 |
| 0.341369581 | 0.504784484 |
| 0.577856898 | NA |

| | | |
|---|---|---|
| | 0.051560749 | 0.12437905 |
| | 0.549553752 | 0.695702227 |
| | 0.744862465 | 0.843898197 |
| | 0.03061445 | 0.082811371 |
| | 0.263143246 | 0.418782616 |
| | 0.476762149 | 0.632968763 |
| | 0.889410245 | NA |
| | 0.472080957 | NA |
| | 0.789938605 | NA |
| | 0.162574439 | 0.295864723 |
| | 0.788582525 | NA |
| NA | | NA |
| | 0.193628381 | NA |
| | 0.992060816 | NA |
| | 0.565619769 | 0.708848562 |
| | 0.222474203 | 0.371817567 |
| | 0.445413915 | NA |
| NA | | NA |
| | 0.302635801 | 0.463332604 |
| | 0.076330949 | 0.167726484 |
| | 0.805437493 | 0.883217029 |
| | 0.01526853 | NA |
| NA | | NA |
| | 0.25400829 | 0.408390336 |
| NA | | NA |
| | 0.865459178 | NA |
| NA | | NA |
| | 0.001573704 | 0.007290498 |
| | 0.185787952 | 0.32673068 |
| | 0.907924474 | NA |
| | 0.01033908 | 0.034609124 |
| NA | | NA |
| | 0.854376325 | NA |
| | 0.458281454 | NA |
| NA | | NA |
| | 0.787005569 | NA |
| NA | | NA |
| | 0.991918092 | NA |
| | 0.106670551 | 0.216167571 |
| NA | | NA |
| | 7.48E-08 | 1.37E-06 |
| | 0.362842439 | 0.52772931 |
| | 0.466547643 | 0.62318199 |
| | 0.81062689 | NA |
| | 0.732129785 | NA |
| | 0.992059387 | NA |
| | 7.57E-06 | 7.65E-05 |

| | | |
|---|---|---|
| | 0.760318475 | NA |
| | 0.656436801 | NA |
| | 0.31841794 | 0.480810075 |
| | 0.351930611 | 0.516235424 |
| | 0.657153776 | 0.779855154 |
| | 0.461834589 | 0.61932371 |
| | 0.268804225 | 0.425185078 |
| | 0.616807233 | 0.749306818 |
| | 0.883264219 | NA |
| | 0.516546385 | NA |
| | 0.652065948 | 0.775959887 |
| NA | | NA |
| | 0.187112045 | 0.32854577 |
| | 0.701643061 | NA |
| | 0.538841167 | 0.686573819 |
| | 0.006571589 | 0.023975001 |
| | 0.334429441 | 0.497206235 |
| NA | | NA |
| NA | | NA |
| | 0.523642675 | 0.673880783 |
| | 0.407929395 | NA |
| | 0.047437109 | 0.116387875 |
| | 0.509921708 | 0.661470391 |
| NA | | NA |
| | 7.93E-05 | 0.000573679 |
| | 0.035658415 | 0.093398046 |
| | 1.84E-08 | 4.02E-07 |
| | 0.940426798 | 0.966159014 |
| | 0.843216988 | 0.907523494 |
| | 0.248393229 | 0.402246822 |
| | 0.00082625 | 0.004245113 |
| | 0.170669114 | 0.307129749 |
| | 0.00094428 | 0.004749299 |
| | 0.159103979 | 0.291122763 |
| | 0.016143311 | 0.049606909 |
| | 0.132521769 | 0.253642946 |
| | 0.279992571 | 0.438172243 |
| | 0.02021923 | 0.059325772 |
| | 0.880083127 | 0.932034983 |
| | 0.625479625 | NA |
| | 0.893512561 | NA |
| | 0.832166186 | NA |
| | 0.000503695 | 0.002779367 |
| | 0.777448647 | 0.866115105 |
| | 1.83E-05 | 0.000164873 |
| | 0.023688762 | 0.067381028 |
| | 0.192493692 | 0.335287458 |

| | |
|---|---|
| 0.000841112 | 0.004303348 |
| NA | NA |
| 0.002449536 | 0.010636617 |
| 0.731650061 | NA |
| 0.001568586 | 0.007274462 |
| 0.364443034 | 0.529315448 |
| 0.003441405 | 0.014049044 |
| NA | NA |
| NA | NA |
| 0.11632106 | 0.230907083 |
| 0.032549402 | 0.086963416 |
| NA | NA |
| 0.786112585 | NA |
| 0.761666146 | 0.855476799 |
| NA | NA |
| 0.543044645 | NA |
| NA | NA |
| NA | NA |
| NA | NA |
| NA | NA |
| 0.898466362 | NA |
| 0.682291603 | 0.798785785 |
| 0.657720808 | 0.780275052 |
| 0.502157992 | 0.654861676 |
| 4.99E-07 | 7.19E-06 |
| 0.171195736 | NA |
| 0.922067881 | 0.955294241 |
| 0.050143002 | 0.121627105 |
| NA | NA |
| 0.937338027 | NA |
| 0.188758455 | NA |
| 0.650428396 | NA |
| 0.047150582 | 0.115775426 |
| 0.196727043 | NA |
| NA | NA |
| NA | NA |
| 0.042980627 | 0.107913514 |
| 0.996453356 | NA |
| 0.00307271 | 0.012803349 |
| NA | NA |
| 0.905158739 | NA |
| NA | NA |
| NA | NA |
| NA | NA |
| NA | NA |
| 0.906961884 | NA |
| NA | NA |

| | | |
|---|---|---|
| | 0.992061292 | NA |
| NA | | NA |
| NA | | NA |
| NA | | NA |
| | 0.991918092 | NA |
| | 0.905158739 | NA |
| NA | | NA |
| NA | | NA |
| | 0.544646434 | 0.691323182 |
| NA | | NA |
| | 0.889409773 | NA |
| | 0.000350882 | 0.002047574 |
| NA | | NA |
| NA | | NA |
| | 0.424379267 | 0.58561832 |
| | 0.914475598 | 0.951778255 |
| | 0.046232899 | 0.114204828 |
| | 1.20E-06 | 1.53E-05 |
| | 0.004263659 | 0.016757774 |
| | 0.708597376 | NA |
| | 0.53961449 | 0.687077564 |
| | 1.88E-06 | 2.27E-05 |
| | 8.49E-06 | 8.48E-05 |
| | 0.651903816 | 0.775889209 |
| | 0.624255861 | 0.754847712 |
| | 0.11416368 | 0.227694846 |
| | 0.488328625 | 0.642839655 |
| | 0.70259892 | 0.814462846 |
| | 0.533044559 | 0.681997639 |
| | 0.014587025 | 0.045635628 |
| | 0.167808214 | NA |
| | 0.000294797 | 0.001760047 |
| | 0.99205891 | NA |
| | 0.889410717 | NA |
| | 0.039874833 | 0.101862663 |
| | 0.286592073 | 0.445667824 |
| | 6.05E-16 | 9.67E-14 |
| | 1.62E-05 | 0.000148128 |
| | 0.049520359 | 0.12047613 |
| | 9.09E-07 | 1.21E-05 |
| | 0.019257478 | 0.056983555 |
| | 0.253948223 | 0.408351569 |
| | 0.669903636 | 0.789501803 |
| | 0.89262973 | 0.939380965 |
| | 2.41E-07 | 3.82E-06 |
| | 0.353064691 | 0.517207975 |
| | 0.706739127 | 0.817578995 |

| | |
|---|---|
| 0.226013171 | 0.375787327 |
| 8.50E-10 | 2.74E-08 |
| 0.905158266 | NA |
| 0.02197071 | 0.063405652 |
| NA | NA |
| NA | NA |
| NA | NA |
| 2.48E-11 | 1.24E-09 |
| 0.684169265 | NA |
| 0.008278335 | 0.028868344 |
| 0.899529377 | 0.943475839 |
| 0.455897697 | 0.613795063 |
| 3.18E-05 | 0.00026335 |
| 0.392392581 | 0.555906367 |
| 0.000108136 | 0.000746287 |
| 0.70266127 | 0.814492126 |
| 0.566909132 | 0.709816263 |
| 0.080528078 | 0.17480388 |
| 0.068246529 | 0.153453451 |
| 0.369547857 | NA |
| 0.112792426 | 0.225471906 |
| 0.089381508 | 0.189184862 |
| NA | NA |
| 0.186885506 | 0.328298296 |
| 0.224784315 | 0.374447129 |
| 2.22E-15 | 3.03E-13 |
| 0.159202763 | 0.291206453 |
| 0.017532264 | 0.052912448 |
| 0.853284195 | NA |
| 7.38E-06 | 7.49E-05 |
| NA | NA |
| 0.85323362 | NA |
| NA | NA |
| NA | NA |
| 0.089000661 | 0.188608908 |
| 0.806680947 | 0.884103958 |
| 0.889410245 | NA |
| 0.889410717 | NA |
| NA | NA |
| 0.738753701 | 0.839639257 |
| 0.514579296 | 0.665868348 |
| 0.398732201 | 0.562302104 |
| 0.008568348 | 0.029695764 |
| 5.62E-09 | 1.41E-07 |
| 0.241357256 | 0.394166592 |
| 7.94E-05 | 0.000574015 |
| 7.26E-09 | 1.78E-07 |

| | | |
|---|---|---|
| | 0.075484635 | NA |
| | 0.008951347 | 0.030736925 |
| | 0.180986239 | 0.320972123 |
| | 0.519776055 | 0.670648239 |
| | 0.029317892 | 0.079993986 |
| | 0.495919151 | NA |
| | 0.066630477 | 0.151010284 |
| | 0.712288746 | NA |
| | 0.853071951 | 0.914290079 |
| | 0.327002378 | 0.489178092 |
| | 0.87960624 | 0.931709682 |
| | 0.809325949 | 0.885588425 |
| NA | | NA |
| | 0.2809789 | 0.439365923 |
| NA | | NA |
| | 0.792188023 | NA |
| | 0.014809493 | 0.046186856 |
| | 2.40E-06 | 2.81E-05 |
| NA | | NA |
| | 0.005803146 | 0.02169043 |
| NA | | NA |
| NA | | NA |
| | 0.551244302 | NA |
| NA | | NA |
| | 9.68E-07 | 1.27E-05 |
| | 0.032485762 | 0.086835649 |
| | 0.889409301 | NA |
| | 0.01285082 | 0.041165741 |
| NA | | NA |
| | 0.010379932 | 0.034711031 |
| | 0.139103914 | 0.263258023 |
| | 0.046616308 | 0.114771557 |
| | 0.756199667 | 0.851753406 |
| | 0.62532125 | 0.755340311 |
| NA | | NA |
| | 0.997717909 | 0.998967988 |
| | 8.18E-09 | 1.98E-07 |
| NA | | NA |
| | 0.084178459 | 0.180560702 |
| | 0.644738092 | 0.770218041 |
| NA | | NA |
| | 0.545229103 | 0.691702922 |
| | 0.667118444 | 0.787551955 |
| | 0.050700492 | 0.122708268 |
| | 0.889409301 | NA |
| | 0.320021296 | 0.482347284 |
| NA | | NA |

| | |
|---|---|
| NA | NA |
| NA | NA |
| 0.745202542 | 0.84419638 |
| NA | NA |
| 0.889409773 | NA |
| 0.093921425 | 0.196234117 |
| NA | NA |
| NA | NA |
| NA | NA |
| 0.051891003 | 0.124887608 |
| 0.179251522 | 0.318667759 |
| 0.992060816 | NA |
| 0.992060816 | NA |
| 0.024864882 | 0.070199015 |
| NA | NA |
| NA | NA |
| 0.992060816 | NA |
| 0.007999711 | 0.028093021 |
| NA | NA |
| NA | NA |
| NA | NA |
| NA | NA |
| 3.49E-05 | 0.000284349 |
| 0.457302328 | 0.614785222 |
| NA | NA |
| NA | NA |
| 0.816464882 | NA |
| 0.726844271 | 0.83131148 |
| 0.992061292 | NA |
| NA | NA |
| NA | NA |
| NA | NA |
| NA | NA |
| 0.495780152 | NA |
| 0.001574271 | 0.007290702 |
| 0.95640627 | 0.975825912 |
| NA | NA |
| NA | NA |
| NA | NA |
| NA | NA |
| 0.945666732 | 0.969191439 |
| 0.041616225 | 0.105231469 |
| 0.539794681 | 0.687150806 |
| 0.601899063 | 0.737413184 |
| 0.325780118 | 0.487947999 |
| NA | NA |
| 0.000369022 | 0.002131903 |

| | | |
|---|---|---|
| NA | | NA |
| NA | | NA |
| | 0.859350383 | NA |
| NA | | NA |
| | 0.989376207 | NA |
| | 0.000129826 | 0.000870755 |
| | 0.155955072 | NA |
| NA | | NA |
| NA | | NA |
| NA | | NA |
| NA | | NA |
| NA | | NA |
| | 0.970271853 | NA |
| NA | | NA |
| | 0.015161473 | 0.047142417 |
| | 0.22024594 | 0.369155199 |
| | 0.154184541 | 0.284587571 |
| | 0.322126406 | NA |
| | 2.55E-11 | 1.27E-09 |
| NA | | NA |
| NA | | NA |
| | 0.636184127 | 0.763266743 |
| NA | | NA |
| | 0.037695981 | 0.097659988 |
| NA | | NA |
| | 0.002295477 | 0.010055132 |
| | 1.26E-07 | 2.18E-06 |
| | 0.418945635 | 0.580967932 |
| | 0.0001229 | 0.000833724 |
| | 0.707244104 | 0.817689321 |
| | 0.042378586 | 0.10678042 |
| | 0.794771866 | 0.876199276 |
| | 0.88471069 | NA |
| | 0.889410245 | NA |
| | 7.19E-05 | 0.000527455 |
| | 0.905158266 | NA |
| | 0.636477605 | 0.763404616 |
| | 0.969571565 | 0.9839097 |
| NA | | NA |
| NA | | NA |
| NA | | NA |
| | 0.021496446 | 0.062282683 |
| | 0.068003833 | 0.153093506 |
| NA | | NA |
| | 0.586742151 | 0.725751754 |
| NA | | NA |
| | 0.801661681 | NA |

| | |
|---|---|
| 0.789781345 | NA |
| 0.002032118 | 0.009075304 |
| 0.00202276 | 0.009042711 |
| NA | NA |
| 1.27E-07 | 2.18E-06 |
| NA | NA |
| NA | NA |
| NA | NA |
| NA | NA |
| NA | NA |
| NA | NA |
| NA | NA |
| NA | NA |
| NA | NA |
| 0.043142331 | 0.108163285 |
| 8.00E-08 | 1.45E-06 |
| NA | NA |
| NA | NA |
| NA | NA |
| NA | NA |
| NA | NA |
| NA | NA |
| 0.000835106 | 0.004281599 |
| NA | NA |
| 1.19E-05 | 0.000113318 |
| NA | NA |
| NA | NA |
| 1.11E-05 | 0.000106117 |
| NA | NA |
| NA | NA |
| NA | NA |
| 0.034343953 | 0.090693493 |
| NA | NA |
| NA | NA |
| NA | NA |
| NA | NA |
| NA | NA |
| NA | NA |
| 3.40E-06 | 3.81E-05 |
| NA | NA |
| NA | NA |
| 0.992060816 | NA |
| NA | NA |
| 0.086332953 | 0.184159157 |
| NA | NA |
| 0.702954502 | 0.81461702 |
| 0.889409773 | NA |

| | |
|---|---|
| 0.00068629 | 0.003633289 |
| 0.434674242 | 0.595335351 |
| 0.401448448 | 0.564945236 |
| 0.241581937 | 0.394386868 |
| 0.889409773 | NA |
| NA | NA |
| NA | NA |
| 0.905158739 | NA |
| NA | NA |
| 0.010775598 | 0.035781063 |
| NA | NA |
| 0.54459262 | NA |
| NA | NA |
| 0.992061292 | NA |
| 6.29E-05 | 0.000471215 |
| NA | NA |
| 4.64E-10 | 1.63E-08 |
| 3.00E-06 | 3.41E-05 |
| 0.000315393 | 0.001865264 |
| 0.126476373 | 0.245413244 |
| 7.75E-06 | 7.81E-05 |
| 0.084055629 | 0.180475611 |
| 0.856776697 | 0.916457669 |
| 0.243562596 | 0.396595896 |
| 0.003557351 | 0.014447175 |
| 0.000244574 | 0.001502266 |
| 0.921174896 | NA |
| NA | NA |
| 8.52E-06 | 8.51E-05 |
| 0.026754151 | 0.074291148 |
| 1.81E-11 | 9.46E-10 |
| 0.122312205 | 0.239339688 |
| 0.333512763 | NA |
| 1.41E-06 | 1.76E-05 |
| 0.318781803 | 0.481207783 |
| 0.908581849 | 0.948154039 |
| 0.760600317 | 0.854804358 |
| 0.558637619 | 0.703440832 |
| 0.11821106 | NA |
| 0.549188483 | 0.695360082 |
| 0.029066915 | 0.079391863 |
| 0.89536752 | 0.941005809 |
| 0.088849873 | 0.188362074 |
| 0.235808264 | 0.38746735 |
| 0.992062245 | NA |
| 0.339979232 | NA |
| 0.345632645 | 0.509071517 |

| | | |
|---|---|---|
| | 0.03942104 | 0.100973594 |
| | 0.231070418 | 0.381997689 |
| | 0.590060231 | 0.728244332 |
| | 0.038234051 | 0.098681421 |
| | 0.889409773 | NA |
| | 0.000789952 | 0.004082502 |
| | 0.748832409 | 0.846773676 |
| | 0.023725199 | 0.06746719 |
| | 0.977235187 | 0.988306889 |
| | 0.185465731 | 0.326359643 |
| | 6.11E-27 | 1.55E-23 |
| | 0.018167106 | 0.054394801 |
| NA | | NA |
| | 0.992060816 | NA |
| | 2.19E-12 | 1.44E-10 |
| | 0.026335817 | NA |
| | 0.036152528 | 0.094474993 |
| | 0.889409773 | NA |
| | 0.344089611 | NA |
| | 0.022762085 | 0.065304305 |
| | 0.591692443 | NA |
| | 0.231484118 | 0.382499915 |
| NA | | NA |
| | 0.905158266 | NA |
| NA | | NA |
| | 6.81E-05 | 0.000504235 |
| | 0.158150927 | 0.289765224 |
| | 0.832851174 | NA |
| | 0.007636094 | 0.02704532 |
| | 0.138974516 | 0.263081155 |
| NA | | NA |
| | 0.002268998 | 0.009947083 |
| | 0.415229002 | 0.57771106 |
| | 0.70339438 | NA |
| | 0.061711471 | 0.142839907 |
| NA | | NA |
| | 0.992061292 | NA |
| | 0.286196803 | 0.445245717 |
| | 0.520168258 | 0.670917754 |
| | 0.460095145 | 0.617530219 |
| | 0.570190299 | 0.712502633 |
| | 0.006065472 | 0.022472025 |
| | 0.986153321 | 0.992938931 |
| | 0.02452762 | 0.069425369 |
| | 0.048547127 | 0.118554978 |
| | 0.292824638 | 0.452576596 |
| | 0.991151101 | NA |

| | |
|---|---|
| NA | NA |
| 2.48E-05 | 0.000212718 |
| NA | NA |
| 0.884786536 | NA |
| NA | NA |
| 0.000180418 | 0.001153698 |
| 0.631601028 | NA |
| 0.019455401 | 0.057468542 |
| 0.13449485 | 0.256435913 |
| 0.02018837 | 0.059272712 |
| 0.881183543 | NA |
| 0.044423613 | 0.110625778 |
| 0.061738058 | 0.142871326 |
| NA | NA |
| 0.992060816 | NA |
| 0.905158739 | NA |
| 4.24E-05 | 0.000336819 |
| 0.922418516 | 0.955368118 |
| 0.676631267 | NA |
| 0.117187237 | 0.232060107 |
| 0.026004409 | 0.072687223 |
| 0.021694787 | 0.062782791 |
| 0.137268172 | 0.260637422 |
| NA | NA |
| 0.889410717 | NA |
| 0.609905863 | 0.743968606 |
| 0.001369338 | 0.006500361 |
| 0.951439218 | NA |
| NA | NA |
| 0.530806997 | 0.680006466 |
| 0.709387107 | 0.818873547 |
| 0.927935433 | 0.958676937 |
| 3.97E-05 | 0.000317724 |
| 0.782559532 | NA |
| 0.844344776 | NA |
| 0.806212248 | 0.883854955 |
| 4.96E-07 | 7.16E-06 |
| 0.03564962 | 0.09339418 |
| NA | NA |
| 0.003275079 | 0.013505625 |
| NA | NA |
| 0.992037435 | NA |
| 0.286717743 | 0.445736985 |
| 0.001440414 | 0.006784654 |
| 0.000208915 | 0.001308114 |
| 0.77565447 | NA |
| 0.905158739 | NA |

| | | |
|---|---|---|
| | 0.568466029 | NA |
| | 0.022861031 | 0.065544506 |
| | 0.637384137 | 0.764085382 |
| | 0.978781427 | 0.989230452 |
| | 0.064834012 | 0.148177568 |
| | 0.297365457 | NA |
| | 0.012025796 | 0.039029626 |
| | 0.57357411 | 0.715672133 |
| NA | | NA |
| | 2.57E-06 | 2.99E-05 |
| | 4.73E-23 | 4.33E-20 |
| | 0.786112585 | NA |
| NA | | NA |
| | 6.43E-13 | 4.84E-11 |
| | 0.889409773 | NA |
| | 0.733859954 | 0.836569108 |
| NA | | NA |
| | 0.935224483 | 0.962803301 |
| | 1.86E-07 | 3.05E-06 |
| | 0.178130775 | 0.317183502 |
| | 0.002774994 | 0.011793322 |
| | 0.037914377 | 0.098029178 |
| | 0.172703018 | 0.309708908 |
| | 0.088308471 | 0.187519262 |
| | 0.028723766 | 0.078699781 |
| NA | | NA |
| | 0.421825834 | 0.58337785 |
| | 0.397447044 | 0.561076159 |
| NA | | NA |
| | 0.209526282 | NA |
| | 0.062268592 | 0.143780864 |
| NA | | NA |
| NA | | NA |
| NA | | NA |
| | 2.36E-10 | 8.98E-09 |
| NA | | NA |
| NA | | NA |
| | 0.889409773 | NA |
| NA | | NA |
| | 0.470499513 | 0.626849198 |
| | 0.305952616 | 0.466556492 |
| | 1.21E-05 | 0.000114837 |
| NA | | NA |
| NA | | NA |
| | 0.812064714 | NA |
| | 0.004838323 | 0.018659069 |
| | 0.212449686 | 0.35960708 |

| | |
|---|---|
| 3.37E-07 | 5.09E-06 |
| 0.040665874 | 0.103258779 |
| 0.312569893 | 0.474341554 |
| NA | NA |
| NA | NA |
| 0.02059502 | 0.06021918 |
| 0.14484386 | NA |
| 0.269393324 | 0.425821921 |
| 0.189049169 | 0.330836046 |
| NA | NA |
| NA | NA |
| 2.35E-09 | 6.71E-08 |
| 0.936019481 | NA |
| 0.71487258 | NA |
| 0.022942453 | 0.065692179 |
| 0.630980991 | 0.758884411 |
| 3.91E-08 | 7.89E-07 |
| 0.448778733 | NA |
| 0.930085247 | 0.959774534 |
| 0.842046954 | 0.906575263 |
| 0.889888806 | 0.937671223 |
| 0.279996034 | 0.438172243 |
| 0.424633496 | 0.585895504 |
| 0.871978145 | 0.92661241 |
| NA | NA |
| 1.61E-11 | 8.51E-10 |
| NA | NA |
| 0.852584891 | 0.914066576 |
| 0.192776927 | 0.335698208 |
| 0.338439293 | 0.50143637 |
| 0.789319999 | 0.872787041 |
| NA | NA |
| NA | NA |
| NA | NA |
| NA | NA |
| NA | NA |
| 0.031842217 | 0.085469166 |
| 0.449898557 | 0.608669978 |
| 0.154923515 | 0.285615437 |
| 0.581355726 | NA |
| 2.54E-05 | 0.00021723 |
| 0.992060816 | NA |
| 0.000722109 | 0.003791679 |
| 0.248548622 | 0.402436865 |
| 0.322627261 | 0.485012119 |
| 5.80E-08 | 1.10E-06 |
| 0.000911692 | 0.004607558 |

| | |
|---|---|
| 2.93E-06 | 3.34E-05 |
| 0.392644162 | 0.556191017 |
| NA | NA |
| 0.05937665 | 0.138530797 |
| 0.886681316 | NA |
| 0.067599799 | 0.152460044 |
| 0.344189404 | 0.507535347 |
| 0.246429384 | 0.399862769 |
| 0.828758547 | NA |
| 0.559788932 | NA |
| 0.614399693 | 0.747592191 |
| 0.959719309 | 0.977707089 |
| NA | NA |
| 0.870091332 | 0.925515189 |
| NA | NA |
| 0.062010429 | 0.143350563 |
| 0.99025256 | NA |
| 0.156284344 | 0.287348741 |
| NA | NA |
| 2.68E-05 | 0.000227447 |
| 0.177579798 | 0.316438756 |
| 0.397535634 | 0.561129073 |
| 0.844869309 | 0.908678315 |
| 0.963912592 | 0.980160066 |
| 0.027628593 | 0.076313996 |
| 0.37867495 | 0.543162173 |
| 0.889410245 | NA |
| 0.0001089 | 0.000751051 |
| 0.383722104 | 0.547615109 |
| NA | NA |
| NA | NA |
| 0.016190116 | 0.04970205 |
| 0.032510199 | 0.086879818 |
| NA | NA |
| 0.509243424 | 0.660902858 |
| 0.045267688 | 0.11232013 |
| NA | NA |
| 0.379060872 | 0.543446784 |
| 0.104898964 | 0.213541888 |
| 0.393797726 | NA |
| 0.89505636 | 0.94090395 |
| NA | NA |
| 0.932340945 | NA |
| 0.000217419 | 0.001354798 |
| 0.580141263 | 0.720553013 |
| NA | NA |
| 2.03E-05 | 0.000180461 |

| | |
|---|---|
| 0.004858164 | 0.018709298 |
| NA | NA |
| 0.927333793 | NA |
| 0.483897832 | 0.639338968 |
| 0.568058932 | NA |
| 0.989564328 | NA |
| 0.736612451 | 0.838335753 |
| NA | NA |
| 0.759439139 | NA |
| 1.46E-07 | 2.46E-06 |
| 0.978477519 | NA |
| 0.361647578 | 0.526549473 |
| 0.036162955 | 0.094490995 |
| NA | NA |
| 0.992060816 | NA |
| NA | NA |
| 0.73191801 | NA |
| 0.180347503 | NA |
| 0.042844907 | 0.10765722 |
| 0.151279094 | 0.280592755 |
| 0.44403848 | 0.60360536 |
| 0.558067759 | 0.702908387 |
| 0.723384246 | 0.829146814 |
| NA | NA |
| 0.88804077 | NA |
| NA | NA |
| 0.854779873 | NA |
| 9.33E-06 | 9.17E-05 |
| 0.783359127 | NA |
| 0.106972541 | 0.21661841 |
| 0.216146369 | NA |
| 0.807879735 | NA |
| 0.087595717 | 0.186242675 |
| 0.257272828 | NA |
| 0.053503867 | 0.127622357 |
| 2.33E-06 | 2.75E-05 |
| 0.804534188 | NA |
| 0.007797509 | 0.027475209 |
| 0.000498162 | 0.002753373 |
| 0.00552883 | 0.020838927 |
| 0.830163713 | NA |
| 0.380926791 | 0.545138321 |
| 2.11E-06 | 2.52E-05 |
| NA | NA |
| 0.212475061 | 0.359622311 |
| 0.346424482 | NA |
| 0.640798357 | 0.767071603 |

| | |
|---|---|
| 0.277966487 | 0.435881745 |
| NA | NA |
| NA | NA |
| 0.394633375 | 0.558144679 |
| 0.001210718 | 0.005845681 |
| NA | NA |
| 1.01E-20 | 5.26E-18 |
| 0.999236696 | NA |
| 0.173613098 | NA |
| 0.795353865 | NA |
| 0.234231589 | 0.385684723 |
| 0.173948221 | 0.31123026 |
| 0.325589251 | 0.487781603 |
| 0.608144675 | NA |
| 0.553652647 | NA |
| 0.440602099 | 0.600569926 |
| 0.159297333 | 0.291282384 |
| 0.887529033 | 0.936352938 |
| 0.560606846 | 0.704946235 |
| 0.137309413 | 0.260693187 |
| NA | NA |
| NA | NA |
| 0.080729349 | 0.175153346 |
| 1.35E-05 | 0.000126262 |
| 0.646749179 | 0.771720112 |
| 0.679876989 | 0.797093552 |
| 0.959158703 | 0.977283164 |
| 7.91E-05 | 0.000572806 |
| 0.028074231 | 0.077282441 |
| 0.010783279 | 0.035790513 |
| 0.850230014 | NA |
| 0.765238582 | NA |
| 0.889410245 | NA |
| 7.46E-13 | 5.44E-11 |
| 0.54196557 | NA |
| NA | NA |
| NA | NA |
| NA | NA |
| 0.905158266 | NA |
| NA | NA |
| 1.78E-10 | 7.10E-09 |
| 0.002791946 | 0.011842445 |
| 0.889409773 | NA |
| NA | NA |
| NA | NA |
| NA | NA |
| 0.836699776 | 0.902813162 |

| | |
|---|---|
| 0.585906401 | 0.725225395 |
| 0.464683571 | 0.621777255 |
| 0.000279341 | 0.001686567 |
| 0.411201843 | 0.574062013 |
| 0.973202135 | 0.985912235 |
| 0.229497654 | NA |
| 0.603562779 | 0.739010338 |
| NA | NA |
| NA | NA |
| 0.110299004 | 0.221661765 |
| 0.795807403 | 0.876831882 |
| 0.905158266 | NA |
| 2.07E-05 | 0.000183467 |
| 0.337308313 | 0.500300794 |
| 0.318531898 | 0.480929727 |
| NA | NA |
| NA | NA |
| 0.000305171 | 0.001813106 |
| NA | NA |
| 0.977437708 | NA |
| 0.000123872 | 0.000839281 |
| 5.26E-15 | 6.56E-13 |
| NA | NA |
| 3.61E-05 | 0.000293069 |
| 2.27E-05 | 0.000197859 |
| 0.214350879 | 0.361765503 |
| 0.93983131 | 0.96589388 |
| 0.976152371 | 0.987712189 |
| 0.534514795 | 0.683122476 |
| NA | NA |
| 0.889689235 | 0.937550912 |
| 0.889409773 | NA |
| 0.000383236 | 0.002201278 |
| 0.048306083 | 0.118058239 |
| 0.912905447 | 0.9507312 |
| 0.000660724 | 0.003524127 |
| 0.26581429 | 0.421993637 |
| NA | NA |
| 0.993298863 | NA |
| 0.951702142 | 0.973242087 |
| 0.979260089 | 0.989397824 |
| 0.556804475 | NA |
| 3.95E-05 | 0.000317169 |
| 0.181014587 | 0.320996472 |
| 0.019787562 | NA |
| 0.190618675 | 0.332787048 |
| 0.313006486 | 0.474872836 |

| | | |
|---|---|---|
| | 0.801658106 | NA |
| | 0.057871952 | 0.135610947 |
| | 0.119024489 | 0.234703551 |
| | 0.000250504 | 0.001530902 |
| NA | | NA |
| | 0.375087301 | NA |
| | 0.359677582 | 0.524446216 |
| | 0.597034124 | 0.733852355 |
| | 0.010390551 | 0.034739055 |
| NA | | NA |
| | 0.100805018 | 0.2072638 |
| | 0.204445699 | 0.350325642 |
| NA | | NA |
| | 0.001215867 | 0.005867235 |
| | 0.00317426 | 0.013166523 |
| | 0.441304272 | 0.601004745 |
| NA | | NA |
| | 0.003250982 | 0.013428954 |
| | 0.031171886 | 0.083946952 |
| | 0.075262865 | 0.165750401 |
| | 0.743274479 | 0.842577269 |
| | 0.409324447 | 0.5724338 |
| | 0.687837882 | 0.803114634 |
| | 0.642273858 | NA |
| | 0.000169369 | 0.001091948 |
| | 4.68E-06 | 5.02E-05 |
| | 0.294764796 | 0.454546359 |
| | 0.000349323 | 0.002040102 |
| | 0.905158739 | NA |
| NA | | NA |
| NA | | NA |
| | 0.889409773 | NA |
| | 0.992061292 | NA |
| | 0.539094737 | 0.686737518 |
| | 7.59E-06 | 7.67E-05 |
| | 0.68507006 | NA |
| NA | | NA |
| | 0.905158739 | NA |
| | 0.008163801 | 0.028536915 |
| | 0.292859831 | 0.452594063 |
| NA | | NA |
| | 0.183878742 | NA |
| NA | | NA |
| | 0.762447895 | NA |
| | 0.923659585 | NA |
| NA | | NA |
| | 0.977715921 | NA |

| | | |
|---|---|---|
| | 0.755569952 | 0.851416872 |
| | 0.992061292 | NA |
| NA | | NA |
| | 0.141808482 | NA |
| | 0.905158266 | NA |
| NA | | NA |
| | 0.805374319 | NA |
| NA | | NA |
| | 0.317092988 | NA |
| NA | | NA |
| | 0.064924059 | NA |
| NA | | NA |
| | 0.003273776 | 0.013502788 |
| NA | | NA |
| | 0.706972018 | 0.817643694 |
| | 0.789988362 | NA |
| | 0.383655085 | 0.547615109 |
| NA | | NA |
| NA | | NA |
| | 0.905158739 | NA |
| NA | | NA |
| | 0.199864522 | 0.344735294 |
| | 0.001100836 | 0.005398228 |
| | 0.755095856 | NA |
| | 0.04197733 | 0.105976107 |
| | 0.894383111 | 0.940331262 |
| | 5.28E-22 | 3.31E-19 |
| | 0.889409773 | NA |
| | 0.005224684 | 0.019873875 |
| | 0.004855324 | 0.018701641 |
| | 0.628423634 | 0.756928011 |
| | 0.100160652 | 0.206258503 |
| | 0.016514741 | 0.0504939 |
| NA | | NA |
| | 0.023077695 | 0.066001966 |
| | 1.42E-07 | 2.41E-06 |
| | 0.006309223 | 0.02320285 |
| | 0.632704464 | NA |
| | 0.267991857 | NA |
| | 0.000407061 | 0.002320523 |
| | 0.483543082 | NA |
| | 0.798840642 | 0.878689029 |
| | 0.434179511 | 0.594843268 |
| | 0.056918377 | 0.133890803 |
| | 0.131135301 | 0.251757307 |
| | 0.993508881 | 0.996868244 |
| | 0.074408844 | 0.16425295 |

| | |
|---|---|
| 0.002735944 | 0.011652171 |
| NA | NA |
| NA | NA |
| 0.71070915 | 0.819796303 |
| 0.54018115 | 0.687443516 |
| NA | NA |
| NA | NA |
| 0.022720294 | 0.065226165 |
| NA | NA |
| 0.992061292 | NA |
| 0.011175243 | 0.036791178 |
| 0.795014225 | NA |
| 0.271902674 | 0.428622457 |
| NA | NA |
| 4.96E-05 | 0.000385026 |
| 7.21E-10 | 2.38E-08 |
| 0.093812338 | NA |
| 2.57E-08 | 5.43E-07 |
| 0.478621071 | NA |
| 0.280059273 | 0.438208748 |
| 2.04E-05 | 0.000181246 |
| 0.010944338 | 0.036223918 |
| 0.047840607 | 0.117220698 |
| NA | NA |
| 0.261845938 | 0.417262333 |
| 2.60E-05 | 0.000221887 |
| 0.29248516 | NA |
| 0.104661373 | NA |
| 0.428584061 | 0.589620368 |
| 0.939799216 | 0.96589388 |
| 0.406852928 | 0.570055804 |
| 0.497022604 | 0.650230525 |
| 0.368449403 | 0.533110505 |
| 0.35022158 | 0.514381357 |
| 0.999362807 | 0.999772568 |
| 0.150881498 | 0.279949878 |
| NA | NA |
| 0.842343577 | NA |
| 0.005218689 | 0.019864307 |
| 0.038334881 | 0.098841736 |
| 0.284276551 | 0.443149367 |
| 0.153209421 | NA |
| NA | NA |
| 0.014309861 | 0.04490928 |
| 0.180463581 | NA |
| 0.954798077 | 0.97468323 |
| 0.756185622 | NA |

| | | |
|---|---|---|
| | 0.901996524 | 0.94462713 |
| | 0.276265839 | 0.433756726 |
| | 0.905158266 | NA |
| | 0.905158739 | NA |
| | 0.444086852 | NA |
| NA | | NA |
| NA | | NA |
| | 0.904787575 | NA |
| | 0.288895739 | NA |
| | 0.004148686 | 0.016382123 |
| | 0.038610064 | NA |
| | 0.183459567 | NA |
| | 4.65E-07 | 6.76E-06 |
| | 0.016289527 | 0.049951366 |
| | 0.237670589 | NA |
| | 0.293704046 | NA |
| | 2.59E-05 | 0.000220961 |
| | 0.650847793 | NA |
| | 0.889409301 | NA |
| | 6.54E-06 | 6.75E-05 |
| | 0.00073746 | 0.003865811 |
| | 0.879542993 | NA |
| | 7.24E-05 | 0.000530757 |
| | 2.63E-05 | 0.000223863 |
| | 0.126718339 | NA |
| NA | | NA |
| | 0.889409773 | NA |
| NA | | NA |
| NA | | NA |
| NA | | NA |
| NA | | NA |
| | 0.830163713 | NA |
| | 0.905158739 | NA |
| NA | | NA |
| | 0.935712365 | NA |
| NA | | NA |
| | 0.897412177 | NA |
| NA | | NA |
| | 0.000727868 | 0.003821002 |
| | 0.853748659 | NA |
| | 0.040599493 | 0.10313793 |
| | 0.155667238 | 0.286529495 |
| | 0.031650431 | 0.085054004 |
| | 0.980862383 | NA |
| | 0.183975245 | 0.324490955 |
| | 0.914761072 | 0.951789385 |
| | 0.703991338 | 0.815603345 |

| | |
|---|---|
| NA | NA |
| 0.99205891 | NA |
| 0.004410809 | 0.017225139 |
| 0.471470698 | 0.6277985 |
| 0.920886065 | 0.954597349 |
| 0.092833309 | 0.194553028 |
| 0.531816234 | 0.680941847 |
| NA | NA |
| 0.001320474 | 0.006296699 |
| 0.055587252 | 0.131463701 |
| 0.892483767 | 0.939380965 |
| 0.091487096 | 0.192466483 |
| 0.067872061 | 0.152861804 |
| 0.474515479 | 0.630781852 |
| 0.394124494 | 0.557496764 |
| 0.323761847 | 0.485985809 |
| 0.167483308 | NA |
| 0.017381432 | 0.052536664 |
| NA | NA |
| 0.061411036 | 0.142294495 |
| 2.81E-07 | 4.35E-06 |
| 0.439697002 | 0.599897522 |
| 9.39E-06 | 9.21E-05 |
| 0.220086398 | 0.368950088 |
| 0.730805588 | 0.834459624 |
| 0.001217979 | 0.00587428 |
| 0.728942669 | 0.832952702 |
| NA | NA |
| 0.025279826 | NA |
| 0.991782148 | NA |
| NA | NA |
| 0.987870035 | 0.993639148 |
| 0.000253266 | 0.001546568 |
| 0.810898047 | 0.88686673 |
| 0.492619848 | 0.646627966 |
| 0.853779535 | 0.914632881 |
| 0.076595968 | 0.168146643 |
| 0.115742751 | 0.230050242 |
| 0.168454603 | 0.304001859 |
| 0.013311366 | 0.042295513 |
| 0.187984726 | 0.329403676 |
| 0.419729451 | 0.581540729 |
| NA | NA |
| 8.84E-11 | 3.80E-09 |
| 0.655694497 | 0.778628351 |
| 0.086339019 | 0.184159157 |
| 0.046019681 | 0.113813062 |

| | | |
|---|---|---|
| | 1.26E-14 | 1.38E-12 |
| | 0.347929941 | 0.51176861 |
| | 0.025146154 | 0.070866249 |
| | 0.156500626 | 0.287629498 |
| | 0.270562591 | 0.427092513 |
| | 0.995576205 | 0.997694248 |
| | 0.960356072 | 0.977992812 |
| | 0.96219557 | 0.979205986 |
| | 1.68E-21 | 9.72E-19 |
| | 0.175472859 | NA |
| | 0.016050102 | 0.049391359 |
| | 0.517340507 | 0.668409049 |
| | 0.397268119 | 0.560967823 |
| | 0.414614447 | 0.577113327 |
| | 0.370799592 | 0.535351784 |
| | 0.017227082 | 0.052183412 |
| | 0.000505544 | 0.002787156 |
| | 0.889409773 | NA |
| | 0.031389773 | 0.084471571 |
| | 0.037492941 | 0.097214251 |
| | 0.308386234 | 0.469485116 |
| | 0.002029827 | 0.009068762 |
| | 0.509187481 | 0.660877572 |
| | 0.374655302 | 0.538701923 |
| | 0.392610054 | 0.55617858 |
| | 0.105681844 | 0.214754581 |
| | 0.964932641 | 0.980879357 |
| | 0.202783197 | 0.348455766 |
| | 0.992061292 | NA |
| | 0.70290985 | 0.814608265 |
| | 0.058453331 | 0.136725255 |
| | 0.843271115 | NA |
| | 0.004840874 | 0.018663134 |
| | 0.630742082 | NA |
| | 0.88914359 | NA |
| | 0.981291913 | NA |
| | 0.883264219 | NA |
| NA | | NA |
| | 0.467534142 | NA |
| | 0.680063391 | NA |
| | 7.71E-05 | 0.000560279 |
| | 0.976181115 | NA |
| | 0.721675644 | 0.827706856 |
| | 0.813239196 | NA |
| | 0.001835604 | 0.008321229 |
| | 0.368676822 | 0.533318467 |
| | 1.46E-08 | 3.27E-07 |

| | |
|---|---|
| 0.126010649 | 0.244678384 |
| 0.254634899 | 0.409096265 |
| 0.001762566 | 0.008028249 |
| 8.97E-09 | 2.15E-07 |
| 0.272727889 | 0.429522465 |
| 0.00105465 | 0.005206211 |
| 0.15222908 | 0.28192615 |
| 0.055120948 | 0.130614104 |
| 0.987261983 | 0.993414557 |
| 5.00E-07 | 7.20E-06 |
| 7.56E-06 | 7.65E-05 |
| NA | NA |
| 4.82E-08 | 9.40E-07 |
| NA | NA |
| NA | NA |
| 0.99635246 | NA |
| 4.43E-07 | 6.48E-06 |
| 0.984064632 | 0.991970261 |
| 1.90E-05 | 0.000170603 |
| 0.051959751 | 0.124998267 |
| 0.009762278 | 0.032969835 |
| 0.00234302 | 0.010232772 |
| 0.000925487 | 0.004666594 |
| 0.546063974 | NA |
| 0.00018923 | 0.001200608 |
| 1.80E-10 | 7.16E-09 |
| 0.000108142 | 0.000746287 |
| 0.628377465 | 0.756928011 |
| 0.147775089 | 0.275513091 |
| 9.43E-06 | 9.24E-05 |
| 0.001697745 | NA |
| 0.807879735 | NA |
| 0.008816991 | 0.030384857 |
| 0.833673452 | NA |
| 3.06E-15 | 4.04E-13 |
| 0.429300696 | 0.590387326 |
| 1.57E-08 | 3.50E-07 |
| 3.20E-05 | 0.000264423 |
| 3.87E-08 | 7.83E-07 |
| NA | NA |
| 0.056617248 | 0.133339569 |
| 0.073553278 | 0.16263831 |
| 0.480150757 | 0.635718634 |
| 0.00101638 | 0.005046053 |
| NA | NA |
| 3.60E-07 | 5.39E-06 |
| 0.011632269 | 0.037960023 |

| | |
|---|---|
| 0.035916182 | 0.093957998 |
| 5.17E-08 | 9.99E-07 |
| 4.81E-10 | 1.68E-08 |
| 0.065348447 | 0.149074024 |
| 0.000146597 | 0.000964097 |
| NA | NA |
| 0.949844047 | 0.971885067 |
| 0.025342821 | 0.071328249 |
| NA | NA |
| 0.33368783 | 0.496372515 |
| 0.670420272 | 0.789836298 |
| 0.525530543 | 0.675493163 |
| 0.002589541 | 0.01113447 |
| 0.895271155 | NA |
| NA | NA |
| 0.002094975 | 0.009306807 |
| 0.001395377 | 0.006605105 |
| 0.016417687 | 0.05026014 |
| 0.310169784 | 0.471415995 |
| 0.52580386 | NA |
| 0.795994473 | 0.876987739 |
| 0.022574007 | 0.06487405 |
| 0.013978052 | 0.044063172 |
| 0.62197201 | 0.753205943 |
| 0.491825068 | NA |
| 0.209401238 | 0.356204338 |
| 0.780534908 | NA |
| 0.089199629 | 0.188921167 |
| 0.392222481 | NA |
| 0.048934827 | 0.119315996 |
| 0.654562682 | 0.777852614 |
| 2.63E-07 | 4.11E-06 |
| 0.061217845 | NA |
| 8.53E-05 | 0.000608015 |
| 9.23E-07 | 1.23E-05 |
| 0.000147289 | 0.000967777 |
| 0.359102144 | 0.523989899 |
| NA | NA |
| 0.555644282 | 0.70097846 |
| 0.824889905 | 0.895612234 |
| NA | NA |
| 0.378098476 | 0.542609807 |
| 0.960740221 | NA |
| 7.31E-14 | 6.69E-12 |
| 0.758105898 | 0.85328022 |
| 0.009221868 | 0.031483674 |
| 0.004989082 | 0.01914967 |

| | |
|---|---|
| 4.30E-05 | 0.000340002 |
| 4.25E-05 | 0.000337094 |
| 0.090531217 | 0.191077949 |
| 0.919340575 | NA |
| NA | NA |
| 0.017557456 | 0.052959365 |
| NA | NA |
| 0.012628306 | 0.040618861 |
| 0.342489736 | 0.505922037 |
| NA | NA |
| 1.96E-10 | 7.72E-09 |
| 0.3376678 | NA |
| 0.105301574 | 0.214143106 |
| 4.95E-06 | 5.28E-05 |
| NA | NA |
| 0.991962833 | NA |
| 0.618259594 | 0.750365469 |
| 0.809217305 | 0.885534996 |
| 0.010935666 | 0.036203269 |
| 0.005613286 | 0.021106531 |
| 0.902545984 | NA |
| 0.702107165 | NA |
| 0.101048258 | 0.207705578 |
| 0.905158739 | NA |
| 0.343657363 | 0.507012365 |
| 0.089381204 | 0.189184862 |
| NA | NA |
| 0.401919715 | 0.565463546 |
| 0.116477895 | 0.231075303 |
| 0.85992909 | 0.918449474 |
| 0.321673666 | 0.484023138 |
| 0.007798819 | 0.027475415 |
| 0.007420898 | 0.026428073 |
| 0.090630334 | 0.191158534 |
| 2.98E-19 | 1.04E-16 |
| 0.516330063 | NA |
| 0.940257006 | 0.966081957 |
| 0.012023817 | 0.039028971 |
| 0.000564173 | 0.003073349 |
| 0.376993946 | 0.541383083 |
| 0.386496371 | 0.550358201 |
| 0.001030395 | 0.005104092 |
| 0.581759693 | 0.721743565 |
| 0.889409773 | NA |
| 0.005248079 | 0.01995092 |
| 0.005033445 | 0.019272785 |
| 0.05105023 | 0.12336588 |

| | |
|---|---|
| 0.889409301 | NA |
| 0.02814187 | 0.077410413 |
| 0.000115175 | 0.000787643 |
| 0.267413445 | 0.42364417 |
| 0.718290468 | NA |
| 0.934233954 | 0.962234681 |
| 0.756017727 | 0.851659224 |
| 0.755125713 | 0.851134665 |
| 0.492920569 | 0.646713436 |
| 7.45E-19 | 2.41E-16 |
| 0.791006251 | NA |
| 0.008258004 | 0.028815748 |
| 0.776184247 | 0.865101507 |
| 0.421494912 | 0.583140547 |
| NA | NA |
| 0.684987343 | 0.800768583 |
| 0.043780453 | 0.109315476 |
| NA | NA |
| NA | NA |
| NA | NA |
| NA | NA |
| NA | NA |
| 0.905158266 | NA |
| NA | NA |
| NA | NA |
| 0.919705045 | 0.95407233 |
| NA | NA |
| 0.905158266 | NA |
| NA | NA |
| NA | NA |
| 0.789618189 | NA |
| 0.784353866 | NA |
| 0.905158266 | NA |
| 0.790663689 | NA |
| 0.657363464 | NA |
| NA | NA |
| 0.715748135 | 0.823316568 |
| 4.24E-05 | 0.000336819 |
| 0.893288602 | 0.939675421 |
| 0.00951121 | 0.032280782 |
| 0.149516826 | 0.277981542 |
| 0.978365262 | 0.988994282 |
| 2.09E-06 | 2.49E-05 |
| 0.268804825 | 0.425185078 |
| 0.905158266 | NA |
| 0.00351979 | 0.01432377 |
| 0.001749052 | 0.007974964 |

| | |
|---|---|
| 0.345698243 | 0.509133985 |
| 0.000111902 | 0.000768376 |
| 0.000455698 | 0.002549506 |
| 8.70E-08 | 1.56E-06 |
| 0.070418236 | 0.157303634 |
| 7.40E-06 | 7.50E-05 |
| 0.98463043 | 0.992171589 |
| 0.93002996 | 0.959774534 |
| NA | NA |
| 0.319341978 | NA |
| 0.556647868 | 0.701850627 |
| 0.000667441 | 0.00354955 |
| 4.61E-09 | 1.19E-07 |
| 0.001092808 | 0.005368448 |
| 6.71E-14 | 6.19E-12 |
| 0.136720517 | 0.259799742 |
| NA | NA |
| 0.026859641 | 0.074546367 |
| NA | NA |
| 0.007084307 | 0.02549477 |
| 0.214110953 | 0.361527291 |
| 0.833037063 | 0.900716966 |
| 1.07E-09 | 3.33E-08 |
| 0.905158266 | NA |
| 0.598807685 | 0.734962716 |
| 0.156403649 | 0.287523461 |
| 0.121605237 | 0.238422268 |
| 0.981156804 | 0.990539389 |
| 0.457084159 | 0.61464244 |
| 0.098625959 | 0.203756461 |
| 0.404351443 | 0.567648213 |
| 0.068169287 | 0.153389628 |
| 3.23E-05 | 0.000266945 |
| 4.32E-07 | 6.34E-06 |
| NA | NA |
| 0.961936332 | NA |
| 0.009487788 | 0.032211247 |
| 0.864300101 | 0.921826781 |
| 0.097648321 | 0.202155132 |
| 0.786795408 | 0.871619431 |
| 0.004454921 | 0.017372688 |
| 0.118574344 | 0.233984006 |
| NA | NA |
| NA | NA |
| 0.871704019 | NA |
| 0.099585154 | 0.205332432 |
| 0.679482078 | 0.796870563 |

| | |
|---|---|
| 0.446649568 | 0.605916963 |
| 0.842930639 | 0.907259774 |
| 0.01805566 | 0.054134932 |
| 0.143318046 | 0.269353044 |
| 0.052069108 | 0.12517906 |
| 0.062725269 | 0.144554511 |
| 0.009677154 | 0.032762969 |
| 1.92E-07 | 3.13E-06 |
| 0.096307453 | 0.200152883 |
| 0.248053109 | 0.40175529 |
| 0.231535935 | 0.382499915 |
| 1.00E-05 | 9.71E-05 |
| NA | NA |
| 0.012710691 | 0.040846714 |
| NA | NA |
| 0.768377188 | NA |
| 0.470656446 | NA |
| 0.05477371 | 0.130067073 |
| 0.889409773 | NA |
| NA | NA |
| 0.003065805 | 0.012781853 |
| NA | NA |
| NA | NA |
| 0.293035861 | 0.45277861 |
| 0.033977045 | 0.089902631 |
| 0.001483215 | 0.006941588 |
| 6.24E-12 | 3.67E-10 |
| NA | NA |
| 0.160286831 | 0.292701747 |
| 0.885142852 | 0.934871922 |
| 0.097852574 | 0.202501619 |
| 0.747776804 | 0.846133482 |
| 0.673124855 | NA |
| NA | NA |
| 1.16E-05 | 0.000110628 |
| 0.670373661 | 0.789836298 |
| 0.000635325 | 0.003407692 |
| 0.003999794 | 0.015891348 |
| 0.488200411 | 0.642747935 |
| 0.523892376 | 0.674097778 |
| 0.011356365 | 0.037247861 |
| 0.854072368 | 0.914901953 |
| 4.36E-09 | 1.13E-07 |
| NA | NA |
| 0.748422638 | 0.846490482 |
| NA | NA |
| 0.318877516 | NA |

| | | |
|---|---|---|
| | 3.27E-11 | 1.58E-09 |
| | 0.053795209 | 0.128247638 |
| | 0.060261953 | NA |
| | 0.103601447 | NA |
| | 0.99205891 | NA |
| | 9.08E-07 | 1.21E-05 |
| | 0.513651454 | 0.665169091 |
| | 0.545060864 | 0.691592071 |
| | 0.04515708 | 0.112145976 |
| | 0.468839453 | 0.625204686 |
| | 0.626628018 | 0.756217006 |
| NA | | NA |
| | 0.425905027 | 0.587180169 |
| | 0.910567328 | 0.949347069 |
| | 0.001599607 | 0.007385571 |
| | 5.19E-08 | 1.00E-06 |
| | 0.998304631 | 0.999279094 |
| | 0.003315464 | 0.013625222 |
| | 3.22E-07 | 4.89E-06 |
| | 0.179651782 | 0.319172612 |
| | 1.07E-05 | 0.000103062 |
| | 0.906823911 | 0.947360419 |
| | 0.435822854 | 0.596350576 |
| NA | | NA |
| NA | | NA |
| | 0.202849191 | 0.348516277 |
| NA | | NA |
| | 0.340639008 | 0.503981402 |
| NA | | NA |
| | 0.815192469 | 0.889967755 |
| | 0.011001257 | 0.036371064 |
| | 0.07964764 | 0.173273438 |
| | 2.31E-12 | 1.51E-10 |
| | 0.65373887 | 0.777272827 |
| | 6.27E-05 | 0.000469927 |
| | 0.099345249 | 0.20493399 |
| | 0.000843962 | 0.004313912 |
| | 0.030161333 | 0.081837725 |
| | 0.889409773 | NA |
| NA | | NA |
| | 0.188136091 | 0.329579611 |
| | 0.12981566 | 0.249923029 |
| | 0.072688035 | NA |
| | 0.060117896 | NA |
| | 6.35E-20 | 2.68E-17 |
| | 0.197129221 | 0.341304152 |
| | 0.031056315 | 0.083707575 |

| | |
|---|---|
| 0.008746042 | 0.030197223 |
| 0.765678178 | 0.857920559 |
| 0.794353041 | 0.875882422 |
| 0.065870229 | 0.149845934 |
| 0.000367401 | 0.002125331 |
| 0.992061292 | NA |
| 0.504778118 | 0.656816024 |
| 0.208190302 | 0.354921405 |
| 0.015818188 | 0.048799044 |
| 1.99E-05 | 0.000176908 |
| 0.334711851 | 0.497520004 |
| 9.66E-05 | 0.000677457 |
| 2.82E-07 | 4.37E-06 |
| 0.045562991 | 0.11287429 |
| NA | NA |
| NA | NA |
| 0.561433916 | 0.705433071 |
| 2.91E-13 | 2.38E-11 |
| NA | NA |
| 0.0005649 | 0.003075782 |
| 0.47392714 | NA |
| 0.043034819 | 0.108012526 |
| 2.29E-10 | 8.78E-09 |
| 0.781663705 | NA |
| 0.889409773 | NA |
| 0.171348213 | NA |
| 0.691937748 | 0.806189898 |
| 0.060817502 | 0.141242687 |
| 0.320239435 | 0.482571274 |
| NA | NA |
| 0.654959824 | 0.778098311 |
| 0.935291207 | 0.962820337 |
| NA | NA |
| 0.022149054 | 0.063819851 |
| 0.825063978 | NA |
| NA | NA |
| 0.182379638 | 0.322531567 |
| 0.004218869 | 0.016614447 |
| 0.004610645 | 0.017887836 |
| 0.050068003 | 0.121485446 |
| 0.172947418 | 0.3100206 |
| 0.015902375 | 0.048996807 |
| 0.889409301 | NA |
| 0.034572039 | 0.091202236 |
| 0.194155987 | 0.337486845 |
| 0.899874336 | 0.943542752 |
| 0.005871715 | 0.021894546 |

| | |
|---|---|
| NA | NA |
| 0.2743918 | 0.43164765 |
| 0.245445139 | 0.398766633 |
| 0.006172622 | 0.022788267 |
| 0.889409773 | NA |
| 0.032687272 | 0.087257451 |
| 0.352228907 | 0.516465983 |
| 0.807854396 | 0.884904209 |
| 0.128521113 | 0.24825784 |
| 0.186604383 | NA |
| 0.010292298 | 0.034499857 |
| 0.989446001 | NA |
| NA | NA |
| 0.436724132 | 0.597025787 |
| NA | NA |
| NA | NA |
| 0.905157793 | NA |
| 0.889409301 | NA |
| NA | NA |
| 0.905158739 | NA |
| 0.0026048 | NA |
| 0.80209217 | NA |
| 0.009476431 | 0.032177662 |
| 0.822240121 | NA |
| 0.732010594 | NA |
| 0.970082677 | 0.984200957 |
| 0.035239556 | 0.092507521 |
| 0.182599111 | 0.322763735 |
| 0.711955754 | NA |
| 0.90515732 | NA |
| 0.677949322 | 0.795675529 |
| 1.86E-14 | 1.99E-12 |
| 0.683805734 | 0.799813045 |
| 0.178014458 | 0.317058924 |
| 0.389984874 | 0.553352149 |
| 0.24899877 | 0.402752208 |
| 3.15E-12 | 1.99E-10 |
| 0.055376281 | 0.131035318 |
| 0.629980555 | NA |
| 2.64E-14 | 2.68E-12 |
| 0.02270519 | 0.065195205 |
| 0.099842495 | 0.205766433 |
| 0.352854882 | 0.517028362 |
| 0.199243146 | 0.343987753 |
| 0.027772797 | 0.076633543 |
| 0.648442323 | 0.773152249 |
| 0.083433689 | 0.179538611 |

| | | |
|---|---|---|
| | 0.990856128 | 0.995343537 |
| | 0.260299717 | 0.415556378 |
| | 0.889409773 | NA |
| | 6.96E-06 | 7.11E-05 |
| | 0.108140379 | 0.218580134 |
| | 0.749553417 | 0.847208138 |
| NA | | NA |
| | 0.666530328 | 0.787323264 |
| | 0.023295589 | NA |
| | 0.160965857 | 0.293648688 |
| NA | | NA |
| NA | | NA |
| | 4.06E-18 | 1.12E-15 |
| | 0.162758595 | 0.296097535 |
| | 0.956247259 | 0.975709008 |
| NA | | NA |
| | 0.074602863 | 0.16459054 |
| | 0.55521733 | 0.700650385 |
| | 0.355496025 | 0.520075763 |
| | 0.562591009 | NA |
| | 0.003282437 | 0.013525808 |
| | 0.000196262 | 0.001240206 |
| | 0.992061769 | NA |
| | 0.041204494 | 0.104397078 |
| | 1.23E-13 | 1.07E-11 |
| | 0.905158266 | NA |
| | 0.83611965 | NA |
| | 0.551716995 | 0.697239461 |
| | 0.474654702 | 0.630928729 |
| | 0.300364352 | 0.460712222 |
| | 0.669710705 | 0.789423875 |
| | 0.626309624 | 0.755915853 |
| | 3.16E-07 | 4.81E-06 |
| | 0.726137548 | 0.830958071 |
| | 3.99E-06 | 4.39E-05 |
| | 0.209866817 | 0.35669237 |
| | 0.038882592 | 0.099850643 |
| | 0.953996515 | NA |
| | 1.65E-06 | 2.01E-05 |
| | 0.319561677 | NA |
| | 0.884816681 | NA |
| | 0.801661681 | NA |
| | 0.739372356 | NA |
| | 0.57268994 | 0.715078425 |
| | 0.744197804 | NA |
| NA | | NA |
| | 0.124530875 | 0.242706442 |

| | | |
|---|---|---|
| | 0.016391682 | 0.050201526 |
| | 0.651634153 | 0.775786205 |
| | 0.003305331 | 0.013597183 |
| | 0.653334666 | NA |
| | 0.352605615 | 0.516852794 |
| | 0.068955537 | NA |
| NA | | NA |
| | 0.905158266 | NA |
| NA | | NA |
| | 0.11415759 | 0.227694846 |
| | 0.81880867 | NA |
| NA | | NA |
| | 0.507293665 | 0.659110693 |
| | 0.837285066 | 0.903172899 |
| | 0.028914163 | 0.07908895 |
| | 0.066860881 | 0.151407455 |
| | 0.002802865 | 0.011879583 |
| | 0.864982907 | NA |
| | 0.992062245 | NA |
| | 0.037874359 | 0.097983395 |
| | 0.325924336 | 0.488097421 |
| | 0.006400455 | 0.023471541 |
| | 0.045291337 | 0.112353421 |
| NA | | NA |
| | 9.76E-07 | 1.28E-05 |
| | 9.86E-06 | 9.58E-05 |
| | 0.889410717 | NA |
| | 0.001367085 | 0.006493581 |
| | 2.91E-09 | 8.06E-08 |
| | 0.123567295 | 0.241299619 |
| | 0.96263689 | NA |
| | 6.03E-09 | 1.50E-07 |
| | 0.905158739 | NA |
| | 3.29E-08 | 6.77E-07 |
| | 0.162890838 | NA |
| | 0.250602554 | 0.404690505 |
| | 0.222087935 | 0.371426425 |
| | 0.666320912 | 0.787202932 |
| | 0.091313098 | 0.192192497 |
| | 0.633014301 | 0.760538439 |
| | 0.000391648 | 0.002246066 |
| | 0.905158266 | NA |
| | 0.192940158 | 0.335878621 |
| | 0.298536203 | NA |
| | 0.992061292 | NA |
| NA | | NA |
| | 0.805303911 | NA |

| | | |
|---|---|---|
| | 0.992061292 | NA |
| NA | | NA |
| NA | | NA |
| | 0.742373319 | NA |
| NA | | NA |
| | 1.27E-05 | 0.000120223 |
| NA | | NA |
| NA | | NA |
| NA | | NA |
| NA | | NA |
| | 0.674988072 | NA |
| NA | | NA |
| | 2.36E-06 | 2.78E-05 |
| NA | | NA |
| | 0.375045242 | NA |
| NA | | NA |
| | 0.758186928 | NA |
| NA | | NA |
| | 0.761620888 | NA |
| | 0.058917093 | 0.137604813 |
| NA | | NA |
| | 0.130687649 | 0.251117438 |
| | 8.21E-05 | 0.000589787 |
| | 0.801661681 | NA |
| NA | | NA |
| | 0.548289672 | NA |
| | 0.981033859 | NA |
| | 0.742053152 | NA |
| | 0.00587358 | 0.021897781 |
| | 0.349135052 | 0.513195991 |
| | 0.000346715 | 0.002027568 |
| | 0.004398417 | 0.017179801 |
| | 0.011264059 | 0.037015347 |
| | 0.153393875 | 0.283581083 |
| | 3.48E-05 | 0.000283589 |
| | 1.39E-06 | 1.75E-05 |
| | 0.022121542 | 0.063776531 |
| | 1.09E-07 | 1.90E-06 |
| | 0.002630988 | 0.011294959 |
| | 0.706591362 | NA |
| | 0.001087106 | 0.005341632 |
| | 0.022993569 | 0.065812797 |
| | 0.562457136 | 0.706174743 |
| | 0.00973516 | 0.032883309 |
| | 0.534265525 | 0.68303059 |
| | 0.002880196 | 0.012146384 |
| NA | | NA |

| | |
|---|---|
| 0.003636204 | 0.014702604 |
| 0.006367485 | 0.023362736 |
| 0.542455943 | NA |
| 0.905158739 | NA |
| 5.73E-05 | 0.000435816 |
| 0.293212461 | 0.45285813 |
| 0.321435869 | 0.48375419 |
| 9.07E-07 | 1.21E-05 |
| 0.625367384 | 0.755340311 |
| 0.458234996 | 0.615634004 |
| 0.85620984 | 0.9161369 |
| NA | NA |
| 0.009687421 | 0.03278762 |
| NA | NA |
| 0.014855481 | 0.046303976 |
| 0.602808077 | 0.738263389 |
| NA | NA |
| 0.000875533 | 0.004447336 |
| 0.883802575 | NA |
| NA | NA |
| NA | NA |
| NA | NA |
| NA | NA |
| NA | NA |
| NA | NA |
| NA | NA |
| NA | NA |
| 0.307424149 | 0.468312653 |
| NA | NA |
| 0.99205891 | NA |
| 0.813836939 | NA |
| 0.28398469 | NA |
| NA | NA |
| 0.578646396 | 0.719478173 |
| 0.905155648 | NA |
| 0.320494423 | 0.482830477 |
| 0.565664269 | NA |
| 0.427595897 | NA |
| 0.303432945 | NA |
| 0.019350648 | 0.057243821 |
| 0.50164386 | 0.65440489 |
| 0.562002625 | 0.70592699 |
| 0.000260343 | 0.001584025 |
| 0.001472703 | 0.00690416 |
| 0.109229775 | 0.220214528 |
| 2.61E-05 | 0.000222873 |
| 0.593186918 | 0.730636725 |

| | | |
|---|---|---|
| | 3.00E-06 | 3.41E-05 |
| | 0.99205891 | NA |
| | 0.665442516 | NA |
| | 0.349792379 | NA |
| | 0.763691267 | NA |
| | 1.76E-09 | 5.17E-08 |
| | 3.53E-08 | 7.22E-07 |
| | 0.053321614 | 0.12729096 |
| | 0.204209289 | NA |
| | 0.992061292 | NA |
| NA | | NA |
| | 0.86041537 | NA |
| NA | | NA |
| NA | | NA |
| NA | | NA |
| NA | | NA |
| | 0.002416717 | 0.010513745 |
| | 0.872921339 | NA |
| | 0.236749456 | 0.388577719 |
| | 0.905158739 | NA |
| | 0.429153935 | 0.590278188 |
| | 0.259315224 | 0.414462698 |
| | 0.248784663 | 0.402603242 |
| | 0.051664711 | 0.124523189 |
| | 0.074381957 | 0.164238651 |
| | 0.011006752 | 0.036383752 |
| | 0.402337406 | 0.565942418 |
| NA | | NA |
| | 0.975865863 | 0.987642896 |
| | 0.984063132 | 0.991970261 |
| | 0.953978792 | 0.97434513 |
| | 0.269640899 | NA |
| | 0.006742132 | 0.024491394 |
| | 0.000365914 | 0.002119522 |
| | 0.182015571 | 0.322199107 |
| | 0.981739208 | NA |
| | 0.023454167 | 0.066895708 |
| | 0.485740796 | 0.640700513 |
| | 0.918532861 | 0.953532033 |
| | 0.998980691 | 0.99957245 |
| | 0.983543343 | NA |
| | 4.33E-12 | 2.62E-10 |
| | 0.000842591 | 0.004308908 |
| | 0.815642671 | 0.890356124 |
| NA | | NA |
| | 0.238741734 | 0.390883517 |
| | 0.00261353 | 0.011226625 |

| | | |
|---|---|---|
| | 0.083095065 | 0.179031056 |
| | 4.78E-05 | 0.000373381 |
| | 4.20E-07 | 6.17E-06 |
| | 5.11E-06 | 5.42E-05 |
| | 0.865006126 | 0.922118894 |
| | 0.457591888 | 0.615061534 |
| | 0.421260049 | 0.58296755 |
| | 0.018029626 | 0.054071642 |
| | 0.126864017 | 0.246009092 |
| | 0.925495648 | NA |
| | 0.02049755 | 0.059998093 |
| | 0.182319067 | 0.322450418 |
| | 0.050554403 | 0.122408659 |
| | 0.466771544 | 0.623253524 |
| | 0.521146434 | 0.671903156 |
| | 0.878957938 | 0.931359205 |
| | 0.992061292 | NA |
| | 0.590007995 | NA |
| | 0.689388194 | NA |
| | 1.77E-19 | 6.59E-17 |
| | 0.006567446 | 0.023971001 |
| | 0.066997791 | 0.151634914 |
| NA | | NA |
| | 2.52E-07 | 3.96E-06 |
| | 0.010372619 | 0.034694948 |
| | 0.000213967 | 0.00133594 |
| | 0.012474249 | 0.040221819 |
| | 0.905158266 | NA |
| | 0.989534751 | NA |
| | 0.992061292 | NA |
| | 0.040322127 | 0.102658968 |
| | 1.22E-05 | 0.000116116 |
| | 0.444554501 | 0.604045065 |
| NA | | NA |
| | 0.064443742 | 0.147485163 |
| | 5.60E-23 | 4.70E-20 |
| NA | | NA |
| NA | | NA |
| NA | | NA |
| | 0.062108459 | 0.143546959 |
| NA | | NA |
| | 0.076689699 | 0.16831875 |
| NA | | NA |
| | 2.71E-07 | 4.23E-06 |
| NA | | NA |
| | 0.959420018 | NA |
| | 0.733652063 | 0.836442382 |

| | | |
|---|---|---|
| | 0.415806069 | 0.578279711 |
| | 0.007008612 | 0.025275432 |
| | 0.128203247 | NA |
| | 0.146220273 | 0.273333701 |
| NA | | NA |
| | 2.57E-06 | 2.99E-05 |
| NA | | NA |
| | 0.979901272 | NA |
| | 1.07E-07 | 1.88E-06 |
| NA | | NA |
| NA | | NA |
| | 0.586679642 | NA |
| | 0.14767856 | NA |
| NA | | NA |
| NA | | NA |
| | 2.59E-09 | 7.30E-08 |
| | 0.441184763 | 0.600941867 |
| | 0.000818521 | 0.004214272 |
| | 5.57E-08 | 1.06E-06 |
| | 0.022825867 | 0.065452234 |
| | 0.946252066 | 0.969356826 |
| | 0.010445647 | 0.034896693 |
| | 0.778961711 | NA |
| | 0.002260317 | 0.009916944 |
| | 0.571296583 | NA |
| NA | | NA |
| | 0.889464813 | NA |
| | 0.000767597 | NA |
| | 0.146570978 | 0.273872711 |
| | 0.889409773 | NA |
| | 0.048490902 | NA |
| | 0.047790597 | 0.117137373 |
| | 0.08125476 | NA |
| | 7.81E-05 | 0.00056706 |
| NA | | NA |
| NA | | NA |
| | 0.020223147 | NA |
| | 4.77E-13 | 3.67E-11 |
| | 0.73885741 | 0.839639257 |
| | 0.504128161 | 0.656381713 |
| | 0.295149868 | 0.455012352 |
| | 0.665344142 | 0.786370068 |
| | 0.447778601 | 0.606886206 |
| | 0.420793525 | 0.582573924 |
| | 0.170943195 | 0.307356228 |
| | 0.064388623 | 0.147411216 |
| | 9.78E-05 | 0.000684845 |

| | | |
|---|---|---|
| | 0.001982193 | 0.008881245 |
| | 0.256484934 | NA |
| | 0.249233498 | NA |
| | 0.101205413 | 0.207892392 |
| | 0.797157316 | NA |
| | 0.451272401 | NA |
| | 0.927725827 | 0.958583468 |
| | 0.991782148 | NA |
| | 0.003637654 | 0.014702604 |
| | 0.905296333 | NA |
| | 0.798721772 | NA |
| | 0.299728945 | 0.460069055 |
| | 3.78E-08 | 7.69E-07 |
| | 0.991827912 | NA |
| | 0.753248084 | NA |
| | 0.707366904 | 0.817745188 |
| NA | | NA |
| | 0.36015873 | NA |
| | 0.889409773 | NA |
| | 0.021827943 | 0.0630933 |
| | 0.927764218 | 0.958583468 |
| | 0.779545273 | 0.86734378 |
| NA | | NA |
| | 0.992059387 | NA |
| | 0.006497958 | 0.023773517 |
| | 0.118730445 | NA |
| | 0.46423119 | 0.621285513 |
| | 0.001167357 | 0.005676261 |
| | 5.18E-10 | 1.79E-08 |
| | 0.355450878 | NA |
| | 0.998192065 | NA |
| | 0.315599014 | 0.477824638 |
| | 1.25E-07 | 2.16E-06 |
| | 0.905158739 | NA |
| NA | | NA |
| | 0.940887431 | 0.966427446 |
| | 0.001371364 | 0.00650687 |
| NA | | NA |
| | 0.576039739 | NA |
| | 0.057688348 | 0.135296212 |
| | 0.066017102 | 0.150069311 |
| | 0.472985526 | 0.629328116 |
| NA | | NA |
| | 0.197975553 | 0.342364559 |
| | 0.31969978 | 0.482129488 |
| | 0.889409773 | NA |
| | 0.837728054 | NA |

| | |
|---|---|
| NA | NA |
| 0.671872408 | 0.791165069 |
| 0.340540917 | NA |
| 0.003670589 | 0.014805153 |
| NA | NA |
| 0.735214941 | NA |
| NA | NA |
| 0.012205685 | 0.039525829 |
| 0.015182541 | 0.047182765 |
| 0.012146528 | 0.039363283 |
| 0.035345453 | 0.092752248 |
| 0.002237644 | 0.009837348 |
| 1.58E-09 | 4.70E-08 |
| 0.296370314 | NA |
| 1.19E-06 | 1.52E-05 |
| 0.00056305 | 0.003067996 |
| 6.44E-07 | 8.94E-06 |
| NA | NA |
| 0.075915377 | 0.166969727 |
| NA | NA |
| 0.592728881 | NA |
| 0.041773911 | 0.105535356 |
| 0.081479146 | 0.176432361 |
| 0.805374319 | NA |
| 0.889409773 | NA |
| 0.758527358 | NA |
| 0.656098548 | 0.778855444 |
| 0.298030698 | 0.457974534 |
| 2.66E-07 | 4.16E-06 |
| NA | NA |
| 0.011966237 | 0.038865049 |
| 1.25E-06 | 1.59E-05 |
| 0.103783769 | 0.211741645 |
| 0.001908599 | 0.008597114 |
| 0.196470865 | 0.340461804 |
| 0.206267281 | 0.352676417 |
| 0.889409773 | NA |
| 0.889410717 | NA |
| 3.72E-09 | 9.94E-08 |
| 0.000739579 | 0.003873567 |
| NA | NA |
| 0.051717994 | 0.124594124 |
| 0.159934396 | 0.29220396 |
| 0.984040017 | 0.991970261 |
| 0.90515732 | NA |
| 0.04889175 | 0.119237443 |
| 1.55E-05 | 0.000141958 |

| | |
|---|---|
| 0.61168643 | NA |
| 4.48E-11 | 2.08E-09 |
| 0.55172061 | 0.697239461 |
| 0.616179088 | 0.748833725 |
| 0.938727963 | 0.965414356 |
| 1.05E-06 | 1.37E-05 |
| 0.00032806 | 0.001930299 |
| NA | NA |
| 9.89E-08 | 1.75E-06 |
| 0.384326133 | 0.548213004 |
| 1.44E-05 | 0.000133477 |
| 0.125808901 | 0.244416319 |
| 6.99E-06 | 7.15E-05 |
| 0.205228736 | 0.351365804 |
| 0.074115143 | 0.163764784 |
| 0.000451805 | 0.002535117 |
| 0.00030664 | 0.001819866 |
| NA | NA |
| 4.18E-08 | 8.34E-07 |
| 0.000882883 | 0.004478453 |
| 0.611497432 | 0.745082239 |
| 0.157987197 | 0.289634401 |
| 0.933639727 | 0.961917397 |
| 0.889718181 | NA |
| 0.992061292 | NA |
| 0.741306595 | 0.841432416 |
| 0.837294724 | NA |
| 0.654591318 | 0.777852614 |
| 0.005303425 | 0.020134492 |
| 0.992061292 | NA |
| 0.978685157 | 0.989181044 |
| 1.43E-11 | 7.67E-10 |
| 0.386110819 | 0.549891746 |
| 2.43E-05 | 0.000209402 |
| NA | NA |
| 0.992061769 | NA |
| 0.004371891 | 0.017098236 |
| 0.566853558 | 0.709787152 |
| 0.81426816 | 0.889356604 |
| 0.003709823 | 0.014925613 |
| 0.369384668 | 0.534060964 |
| NA | NA |
| NA | NA |
| 0.041487472 | 0.104993292 |
| 0.207751166 | 0.354440825 |
| 0.814342913 | 0.889394013 |
| 0.492583691 | NA |

| | |
|---|---|
| NA | NA |
| 2.57E-05 | 0.000220027 |
| 0.053304027 | 0.127270101 |
| 0.780415831 | 0.867735439 |
| 0.530763605 | 0.680006466 |
| 0.89960823 | 0.943475839 |
| 0.371198713 | 0.535590548 |
| 0.991933202 | NA |
| 0.294857089 | 0.454656753 |
| 2.74E-08 | 5.73E-07 |
| 0.007044153 | 0.025382763 |
| 0.065209568 | 0.148865464 |
| 0.052281198 | 0.125565223 |
| 0.332633192 | 0.495273411 |
| 0.992060816 | NA |
| 0.338807537 | 0.501785298 |
| 0.378697248 | 0.543162173 |
| 0.422043586 | 0.583642239 |
| NA | NA |
| NA | NA |
| 0.809131053 | NA |
| 0.824082341 | NA |
| 0.769327167 | NA |
| NA | NA |
| 0.067458656 | 0.152305638 |
| NA | NA |
| 0.889409301 | NA |
| 0.708305953 | NA |
| NA | NA |
| NA | NA |
| NA | NA |
| 0.067711028 | NA |
| NA | NA |
| 0.905158266 | NA |
| 0.981760909 | NA |
| 0.905158739 | NA |
| 0.774154168 | NA |
| 0.905158739 | NA |
| 0.863406743 | NA |
| 0.664386455 | NA |
| 0.905158266 | NA |
| NA | NA |
| 0.748954397 | NA |
| 0.653954917 | NA |
| NA | NA |
| NA | NA |
| 0.992062245 | NA |

| | |
|---|---|
| NA | NA |
| NA | NA |
| 0.392577832 | NA |
| 0.889409301 | NA |
| 0.016800139 | 0.051188325 |
| 0.442550242 | 0.602253394 |
| NA | NA |
| 0.835932741 | 0.902280588 |
| 0.01035722 | 0.034656044 |
| 0.750989959 | 0.848382546 |
| 0.564846638 | 0.708283881 |
| 2.52E-05 | 0.000215884 |
| 0.534835119 | 0.683371524 |
| 0.804614115 | NA |
| 0.565863202 | 0.709032243 |
| 0.833234055 | NA |
| NA | NA |
| 0.992061292 | NA |
| 0.00248389 | 0.010758002 |
| 0.793097426 | NA |
| 0.039331966 | 0.10079642 |
| 0.256097062 | 0.410634201 |
| 0.988799963 | NA |
| 0.012235193 | 0.039591843 |
| 5.15E-05 | 0.000396029 |
| NA | NA |
| 0.001604349 | 0.007405122 |
| 0.611354606 | 0.744990888 |
| 0.011946781 | 0.038830576 |
| 1.04E-05 | 0.000100797 |
| 0.552192899 | NA |
| 6.30E-10 | 2.10E-08 |
| 0.969725652 | NA |
| 3.61E-17 | 7.15E-15 |
| 3.44E-07 | 5.17E-06 |
| 0.227144696 | 0.377013462 |
| 0.162455724 | 0.295727865 |
| 0.00057782 | 0.003138348 |
| 0.889409773 | NA |
| 0.003313637 | 0.013625222 |
| NA | NA |
| 0.889409773 | NA |
| NA | NA |
| 0.762134225 | 0.855660376 |
| 0.889410245 | NA |
| NA | NA |
| 0.170858342 | 0.307314234 |

| | |
|---|---|
| 0.93368305 | 0.961917397 |
| 0.79819043 | 0.878129441 |
| NA | NA |
| 0.410913863 | 0.57377957 |
| 0.440207439 | 0.600330093 |
| 0.709181644 | 0.818765496 |
| 0.157171267 | 0.288499863 |
| 0.646212439 | NA |
| 0.559649504 | 0.70400066 |
| 0.006338312 | 0.023274749 |
| 0.278353884 | 0.436411034 |
| 0.833441121 | NA |
| 9.49E-17 | 1.75E-14 |
| NA | NA |
| 0.66759422 | NA |
| NA | NA |
| 0.059033898 | 0.137833637 |
| 0.174998999 | NA |
| 0.649882796 | 0.774584162 |
| 0.024534637 | 0.069436279 |
| 0.905157793 | NA |
| 0.881310752 | 0.932485417 |
| 0.692331986 | 0.806431689 |
| NA | NA |
| NA | NA |
| 0.159569531 | NA |
| 0.06442931 | 0.147467503 |
| 0.000242592 | 0.001491758 |
| 0.911705782 | NA |
| 0.090846899 | 0.19150495 |
| 0.955273895 | NA |
| 0.006511133 | 0.023805855 |
| 0.000344779 | 0.002017322 |
| NA | NA |
| 0.358982215 | NA |
| 8.60E-07 | 1.15E-05 |
| NA | NA |
| NA | NA |
| NA | NA |
| NA | NA |
| NA | NA |
| 0.53429854 | 0.68303059 |
| 0.018533913 | 0.055236996 |
| 0.815824901 | NA |
| 0.974491292 | NA |
| 0.889409301 | NA |
| NA | NA |

| | |
|---|---|
| 0.199891311 | 0.344754422 |
| 0.000945697 | 0.004753162 |
| NA | NA |
| NA | NA |
| 0.508558802 | 0.660302087 |
| 0.917577602 | 0.953081054 |
| 4.82E-05 | 0.0003759 |
| 0.230980097 | NA |
| 0.112452599 | 0.224915441 |
| 0.977700334 | 0.988549666 |
| 0.002059107 | 0.009171587 |
| 0.939121767 | NA |
| 0.562504521 | 0.706193859 |
| 0.340968668 | 0.504334135 |
| NA | NA |
| 0.777614015 | 0.866255385 |
| 0.79751955 | NA |
| 0.268055149 | 0.424410836 |
| 0.809861115 | NA |
| 0.858024477 | 0.917175946 |
| NA | NA |
| 0.986727191 | NA |
| 0.209445973 | 0.356207298 |
| 0.710931066 | 0.819939315 |
| 0.289066564 | 0.448383271 |
| 0.487279046 | 0.642035316 |
| NA | NA |
| 0.293780818 | NA |
| 0.823428588 | 0.894689182 |
| 0.571610609 | NA |
| 0.808437377 | 0.885124587 |
| 0.045716108 | NA |
| NA | NA |
| 0.590357967 | NA |
| 0.453004946 | 0.611135218 |
| 1.18E-11 | 6.43E-10 |
| 0.883666561 | 0.933983154 |
| 0.00135288 | 0.006433065 |
| 2.37E-18 | 6.66E-16 |
| 0.000492203 | 0.002725438 |
| 0.5313918 | NA |
| 2.27E-05 | 0.000198194 |
| 4.46E-05 | 0.000351533 |
| 0.945486164 | 0.969051607 |
| 0.365098122 | NA |
| 0.573179975 | NA |
| 0.492832613 | 0.646713436 |

| | | | |
|---|---|---|---|
| | 0.529152458 | | 0.678592999 |
| NA | | NA | |
| NA | | NA | |
| | 0.889410245 | NA | |
| | 0.720011916 | NA | |
| | 0.388009001 | NA | |
| | 0.573088072 | | 0.715342437 |
| | 0.381722332 | | 0.545872145 |
| | 1.07E-10 | | 4.52E-09 |
| | 0.000818941 | | 0.004214465 |
| | 6.06E-05 | | 0.000456903 |
| | 0.642581419 | NA | |
| NA | | NA | |
| | 0.130665507 | | 0.251096864 |
| | 0.813266252 | | 0.888615894 |
| | 0.000626675 | | 0.003363762 |
| | 0.00050365 | | 0.002779367 |
| NA | | NA | |
| | 0.020870387 | | 0.060854179 |
| | 0.666426541 | | 0.787243016 |
| | 0.002812153 | | 0.011912052 |
| | 0.039250082 | | 0.100669419 |
| | 0.315227493 | | 0.477450719 |
| NA | | NA | |
| | 0.046464755 | | 0.114539689 |
| | 0.507057843 | NA | |
| | 0.00093328 | | 0.00469828 |
| | 0.459199104 | | 0.616429462 |
| NA | | NA | |
| | 0.000318831 | | 0.001883562 |
| | 0.022769536 | | 0.065304305 |
| | 0.02798421 | | 0.077082948 |
| | 0.889409301 | NA | |
| | 0.120281231 | | 0.236550849 |
| | 0.669261904 | | 0.789091843 |
| | 2.06E-10 | | 8.05E-09 |
| NA | | NA | |
| | 0.189247169 | | 0.331050633 |
| | 0.04516175 | | 0.112145976 |
| | 0.541328399 | NA | |
| | 0.00359701 | | 0.014578549 |
| | 0.952919513 | NA | |
| | 0.054375987 | | 0.129351349 |
| | 0.016480772 | | 0.050416758 |
| | 0.765189784 | | 0.857619149 |
| NA | | NA | |
| | 0.281972278 | NA | |

| | | |
|---|---|---|
| | 0.227872911 | 0.377992239 |
| | 3.33E-06 | 3.74E-05 |
| | 5.65E-05 | 0.000430136 |
| | 0.002130251 | 0.009438699 |
| | 0.036908593 | 0.096084859 |
| | 4.41E-06 | 4.78E-05 |
| NA | | NA |
| | 0.137599801 | NA |
| NA | | NA |
| | 0.916258364 | 0.952384067 |
| | 1.47E-11 | 7.87E-10 |
| | 0.561606797 | NA |
| NA | | NA |
| | 0.498895175 | NA |
| | 0.572410998 | NA |
| | 0.905158739 | NA |
| NA | | NA |
| | 2.71E-06 | 3.12E-05 |
| | 0.484348496 | 0.639670935 |
| | 0.000478909 | 0.002659679 |
| | 0.887880951 | 0.936517531 |
| | 0.905158739 | NA |
| | 0.903342961 | NA |
| | 0.026473129 | 0.07367851 |
| | 0.135508629 | 0.258077535 |
| | 0.662182308 | 0.783620462 |
| | 0.813371652 | NA |
| | 0.268850884 | NA |
| NA | | NA |
| | 0.000188531 | 0.001197213 |
| | 0.884799037 | 0.934775177 |
| | 0.935636499 | 0.962956594 |
| | 0.001625255 | 0.007486676 |
| | 0.254405536 | 0.408877345 |
| | 1.43E-06 | 1.78E-05 |
| NA | | NA |
| | 0.784353866 | NA |
| | 0.362735365 | 0.527608523 |
| | 0.1619996 | 0.295117739 |
| | 5.97E-11 | 2.69E-09 |
| | 0.675376813 | NA |
| | 2.10E-06 | 2.50E-05 |
| | 0.503713411 | NA |
| | 0.718372689 | 0.8253614 |
| | 0.140791951 | 0.265603017 |
| | 0.701501067 | NA |
| | 0.063085669 | 0.145067723 |

| | |
|---|---|
| 4.58E-10 | 1.60E-08 |
| NA | NA |
| 0.727229588 | 0.831601027 |
| NA | NA |
| 0.905158739 | NA |
| 0.007615172 | 0.026979922 |
| 0.810546612 | NA |
| 0.905158266 | NA |
| 0.359584223 | 0.524386721 |
| 0.893853583 | 0.939999561 |
| 0.000184039 | 0.001174116 |
| 0.084183272 | 0.180560702 |
| 0.000236006 | 0.001459436 |
| NA | NA |
| 2.62E-07 | 4.09E-06 |
| 0.992061292 | NA |
| 3.26E-06 | 3.67E-05 |
| 1.15E-06 | 1.48E-05 |
| 0.747426375 | 0.845886501 |
| 0.098105365 | 0.202910023 |
| 0.678626863 | 0.796130318 |
| 0.001195439 | 0.005792286 |
| 0.832725425 | 0.900424345 |
| NA | NA |
| 0.889409301 | NA |
| NA | NA |
| 0.667892574 | NA |
| NA | NA |
| 0.952742909 | NA |
| NA | NA |
| 0.854422988 | NA |
| 0.99205891 | NA |
| NA | NA |
| 0.962998974 | NA |
| NA | NA |
| NA | NA |
| NA | NA |
| NA | NA |
| NA | NA |
| NA | NA |
| NA | NA |
| NA | NA |
| 0.889409773 | NA |
| 0.905157793 | NA |
| 0.889409773 | NA |
| NA | NA |
| NA | NA |

| | |
|---|---|
| 0.991879747 | NA |
| NA | NA |
| NA | NA |
| 0.890296709 | NA |
| 0.671565374 | 0.790888343 |
| NA | NA |
| 0.911598 | 0.950003887 |
| 0.229186548 | 0.379655055 |
| 0.007477527 | 0.026598774 |
| 0.988933589 | NA |
| NA | NA |
| 0.686778385 | 0.802221743 |
| 0.02777569 | 0.076633543 |
| 0.883624738 | 0.933983154 |
| 0.756395643 | NA |
| 0.000663737 | 0.003535046 |
| 0.357296912 | 0.522006128 |
| 0.627293101 | 0.75642702 |
| 0.000445868 | 0.002506607 |
| 0.020411607 | 0.059770433 |
| 1.31E-05 | 0.00012334 |
| 0.147861819 | 0.275646934 |
| 0.966949621 | 0.982156748 |
| 0.082425622 | 0.178042912 |
| 0.037293285 | 0.096783877 |
| NA | NA |
| 0.054971911 | 0.130387578 |
| NA | NA |
| 0.095478036 | 0.198786592 |
| 0.788582525 | NA |
| 0.000996248 | 0.004959559 |
| NA | NA |
| NA | NA |
| NA | NA |
| 0.866760016 | NA |
| 0.321980016 | 0.484334387 |
| 0.889409301 | NA |
| 0.611580995 | 0.745083347 |
| 0.986262983 | NA |
| 0.661802816 | 0.783509167 |
| 0.014192342 | 0.044632607 |
| 0.114772655 | 0.228535793 |
| 0.889550534 | 0.937449737 |
| 0.021278919 | 0.061774694 |
| 1.67E-05 | 0.00015242 |
| NA | NA |
| 0.301054642 | 0.461422578 |

| | |
|---|---|
| 0.000363808 | 0.002108432 |
| 0.881183543 | NA |
| 4.78E-05 | 0.000373381 |
| 0.517243339 | 0.668362156 |
| 0.728411158 | 0.832518639 |
| NA | NA |
| 0.290921982 | 0.450391695 |
| NA | NA |
| NA | NA |
| NA | NA |
| 0.001611963 | 0.007433176 |
| 0.764177735 | 0.857062789 |
| NA | NA |
| 0.881453338 | NA |
| 1.79E-21 | 1.01E-18 |
| 0.331486604 | 0.493834079 |
| 0.063250579 | 0.14539092 |
| 0.000920626 | 0.00464534 |
| NA | NA |
| NA | NA |
| 0.07848244 | NA |
| NA | NA |
| NA | NA |
| NA | NA |
| 0.170970355 | 0.307379894 |
| 0.879556438 | 0.931701873 |
| 0.54347541 | 0.690315625 |
| 0.426069696 | 0.587323569 |
| 0.724944119 | 0.830297927 |
| 0.361884125 | 0.526754176 |
| 0.108698208 | 0.219384554 |
| NA | NA |
| 0.194867004 | 0.338401309 |
| 0.754103576 | NA |
| 0.905296333 | NA |
| NA | NA |
| 0.016720449 | 0.051002131 |
| 0.239610027 | 0.391837697 |
| 0.881495756 | NA |
| NA | NA |
| 0.017306801 | 0.052378946 |
| 0.235735342 | 0.387408964 |
| 0.710524387 | 0.819626235 |
| 0.207552464 | 0.35426697 |
| 0.280955875 | 0.439361207 |
| NA | NA |
| 0.626172692 | 0.755792126 |

| | | |
|---|---|---|
| | 7.50E-06 | 7.59E-05 |
| | 0.001469563 | 0.006893856 |
| | 8.79E-11 | 3.80E-09 |
| | 0.303993122 | 0.464688762 |
| | 0.160114733 | 0.292484771 |
| | 0.985829152 | 0.99279167 |
| | 0.132494183 | 0.253612252 |
| | 0.396048039 | 0.559721869 |
| | 0.122199298 | 0.239208074 |
| | 0.357587046 | 0.522345498 |
| | 0.001873328 | 0.008467767 |
| | 1.58E-10 | 6.38E-09 |
| | 0.721406781 | NA |
| | 0.736253459 | NA |
| | 6.11E-18 | 1.61E-15 |
| | 0.048588244 | 0.118629002 |
| | 0.001258762 | 0.006036505 |
| | 0.562787325 | 0.70650851 |
| | 0.005040004 | 0.01928781 |
| | 0.906007645 | 0.947073641 |
| | 0.306869192 | NA |
| | 0.714486906 | NA |
| | 0.54569601 | NA |
| | 0.905158266 | NA |
| | 0.905158739 | NA |
| | 0.961590025 | NA |
| | 0.347040053 | 0.510767596 |
| | 0.220751081 | 0.369697452 |
| | 0.369782986 | 0.534411987 |
| | 0.133549074 | 0.255142171 |
| | 0.889409773 | NA |
| | 0.000256353 | 0.001563246 |
| | 0.850003565 | NA |
| | 0.797342596 | NA |
| | 0.221883876 | 0.371185845 |
| NA | | NA |
| | 0.509890809 | 0.661469385 |
| | 0.578102101 | 0.719196874 |
| NA | | NA |
| | 0.007222893 | 0.025886651 |
| NA | | NA |
| NA | | NA |
| NA | | NA |
| NA | | NA |
| NA | | NA |
| | 0.000148416 | NA |
| NA | | NA |

| | |
|---|---|
| 0.251719472 | NA |
| NA | NA |
| 0.09050261 | NA |
| 0.750656241 | 0.848186703 |
| NA | NA |
| 0.889409773 | NA |
| 0.010842281 | 0.035937457 |
| 0.197953552 | 0.342364559 |
| 0.769409507 | 0.860621121 |
| 0.774177521 | 0.863873377 |
| 0.995409255 | 0.997694248 |
| 0.992060816 | NA |
| 0.889409773 | NA |
| 0.992061292 | NA |
| 0.019030584 | 0.056418602 |
| 0.979817876 | NA |
| 0.62375535 | NA |
| 0.358678503 | 0.523510884 |
| NA | NA |
| 0.992060816 | NA |
| 0.700936495 | 0.813136694 |
| 0.607876785 | NA |
| 0.992061292 | NA |
| 0.793078656 | 0.875224355 |
| 0.000968416 | 0.004845167 |
| NA | NA |
| NA | NA |
| NA | NA |
| 0.934329578 | 0.962288036 |
| 0.31990459 | 0.482206541 |
| NA | NA |
| 1.78E-12 | 1.19E-10 |
| NA | NA |
| NA | NA |
| NA | NA |
| NA | NA |
| NA | NA |
| NA | NA |
| 0.468718522 | 0.625119346 |
| NA | NA |
| NA | NA |
| NA | NA |
| 0.003878274 | 0.015478556 |
| 0.001234411 | 0.005937882 |
| 0.889366074 | 0.937385064 |
| 0.89255079 | 0.939380965 |
| NA | NA |

| | | |
|---|---|---|
| | 0.104550184 | 0.213009602 |
| | 0.597380662 | 0.734092615 |
| | 0.288510231 | 0.447844118 |
| | 0.011311793 | 0.037140501 |
| | 0.946858617 | 0.969824543 |
| | 0.834163237 | 0.901268968 |
| | 0.889410245 | NA |
| | 2.31E-05 | 0.000200912 |
| | 0.995453919 | 0.997694248 |
| | 4.81E-07 | 6.98E-06 |
| | 0.782373854 | NA |
| | 0.992061292 | NA |
| | 0.03202968 | 0.085856901 |
| | 0.87564483 | NA |
| NA | | NA |
| | 0.430653459 | 0.591599581 |
| | 0.01555294 | 0.048109173 |
| | 0.523269446 | 0.673651507 |
| | 0.435474049 | 0.596096163 |
| | 0.017317042 | 0.052385306 |
| | 0.389721776 | 0.553050309 |
| | 0.520478425 | NA |
| | 0.103637626 | 0.211621594 |
| | 0.127114949 | 0.246386898 |
| | 0.012918474 | 0.041304131 |
| NA | | NA |
| | 0.889409301 | NA |
| NA | | NA |
| | 0.305479459 | NA |
| | 0.00075253 | 0.003927933 |
| | 0.146155712 | 0.273259539 |
| | 0.365684106 | NA |
| NA | | NA |
| | 0.080484864 | 0.174727349 |
| | 4.55E-05 | 0.000357205 |
| | 0.550599943 | 0.696424408 |
| NA | | NA |
| NA | | NA |
| NA | | NA |
| NA | | NA |
| | 0.905158266 | NA |
| NA | | NA |
| NA | | NA |
| NA | | NA |
| NA | | NA |
| NA | | NA |
| | 0.873981309 | NA |

| | | |
|---|---|---|
| | 0.992059387 | NA |
| NA | | NA |
| NA | | NA |
| | 0.905158266 | NA |
| NA | | NA |
| NA | | NA |
| | 0.905158739 | NA |
| NA | | NA |
| NA | | NA |
| | 0.892552925 | NA |
| NA | | NA |
| NA | | NA |
| NA | | NA |
| NA | | NA |
| | 0.992060816 | NA |
| | 0.905158266 | NA |
| | 0.889409773 | NA |
| | 0.992060816 | NA |
| | 0.246857686 | NA |
| | 0.905158739 | NA |
| | 0.000508138 | 0.002800048 |
| | 0.42365446 | 0.585132919 |
| | 0.186112929 | 0.327157686 |
| | 0.492572847 | 0.646627966 |
| | 5.42E-10 | 1.85E-08 |
| | 0.981675198 | NA |
| | 0.356129933 | 0.52062161 |
| | 0.875202974 | 0.928844683 |
| | 0.992061292 | NA |
| NA | | NA |
| NA | | NA |
| NA | | NA |
| NA | | NA |
| NA | | NA |
| | 0.7070452 | 0.817652954 |
| NA | | NA |
| NA | | NA |
| | 0.009623387 | 0.032616138 |
| | 0.702811832 | 0.814565311 |
| | 0.79535582 | 0.87663136 |
| NA | | NA |
| | 0.889409301 | NA |
| | 7.12E-06 | 7.26E-05 |
| | 0.23774383 | 0.389569194 |
| | 0.124654852 | 0.242818761 |
| | 0.810473185 | 0.886546735 |
| | 0.231707613 | 0.382676555 |

| | |
|---|---|
| 0.171202203 | 0.307660431 |
| 0.000178081 | 0.001141216 |
| 0.596725847 | NA |
| 0.782055177 | 0.868646922 |
| 0.004188908 | 0.016520155 |
| NA | NA |
| NA | NA |
| 0.410658641 | NA |
| 0.59698375 | 0.733831514 |
| 0.000602972 | 0.003254581 |
| 0.889409301 | NA |
| NA | NA |
| 1.80E-18 | 5.52E-16 |
| 0.445992829 | NA |
| NA | NA |
| 0.373692599 | 0.537845958 |
| NA | NA |
| NA | NA |
| 0.905158266 | NA |
| 0.719447872 | NA |
| 0.497709916 | 0.650740505 |
| 0.783254205 | NA |
| NA | NA |
| 6.99E-13 | 5.15E-11 |
| NA | NA |
| 4.72E-09 | 1.21E-07 |
| NA | NA |
| 0.055360997 | 0.131013271 |
| 0.680082131 | 0.797238393 |
| NA | NA |
| 0.991113901 | NA |
| 0.800835548 | NA |
| 2.78E-07 | 4.31E-06 |
| NA | NA |
| 0.000129948 | 0.000871304 |
| NA | NA |
| 0.889409773 | NA |
| 0.055889823 | 0.132022872 |
| NA | NA |
| NA | NA |
| NA | NA |
| NA | NA |
| 0.000123598 | 0.000837682 |
| 0.592195853 | 0.729845919 |
| 0.889409773 | NA |
| 0.905158739 | NA |
| 2.22E-05 | 0.000194795 |

| | | |
|---|---|---|
| | 0.664815027 | 0.785973574 |
| | 0.155698845 | NA |
| | 0.618654004 | 0.750636715 |
| | 0.438925017 | 0.599251084 |
| | 0.381721657 | 0.545872145 |
| | 0.788886803 | 0.872608947 |
| | 0.001508738 | 0.007041524 |
| | 0.424673498 | 0.585913882 |
| | 0.256030974 | 0.410580543 |
| | 0.507163882 | 0.658982941 |
| | 0.083958592 | 0.180343022 |
| | 0.992061292 | NA |
| | 0.901314663 | 0.944265974 |
| | 0.787290149 | 0.871771467 |
| | 0.908894782 | 0.948233586 |
| | 0.333501917 | NA |
| | 0.950622261 | 0.972436561 |
| | 0.350282536 | 0.514436477 |
| | 0.403820923 | 0.567172544 |
| | 0.98283442 | NA |
| | 0.905158266 | NA |
| NA | | NA |
| | 0.773112738 | 0.86312383 |
| | 0.515697755 | 0.666938095 |
| NA | | NA |
| | 0.199126852 | 0.343868083 |
| NA | | NA |
| | 0.965940371 | 0.981494822 |
| | 0.942581374 | NA |
| NA | | NA |
| NA | | NA |
| NA | | NA |
| NA | | NA |
| NA | | NA |
| NA | | NA |
| | 6.51E-05 | 0.0004846 |
| | 0.838770925 | NA |
| NA | | NA |
| | 0.992061292 | NA |
| NA | | NA |
| NA | | NA |
| | 0.524223876 | NA |
| | 0.003460277 | 0.014110348 |
| NA | | NA |
| | 0.970761443 | 0.984707619 |
| | 0.889409301 | NA |
| | 0.172849717 | 0.30989606 |

| | |
|---|---|
| 0.775408623 | NA |
| 0.225835743 | 0.375670077 |
| 2.27E-06 | 2.69E-05 |
| 1.03E-11 | 5.74E-10 |
| NA | NA |
| 4.49E-06 | 4.86E-05 |
| 0.611668751 | 0.745125602 |
| 0.441173069 | 0.600941867 |
| 0.225988518 | 0.375779045 |
| NA | NA |
| 0.06634517 | 0.150630052 |
| 0.000298747 | 0.001779756 |
| 0.482024449 | 0.637560226 |
| 0.435638684 | 0.596240513 |
| 0.183660904 | 0.324218458 |
| 0.016552606 | 0.050588541 |
| 0.093794863 | 0.196044298 |
| NA | NA |
| 0.99205891 | NA |
| 0.541505019 | NA |
| NA | NA |
| 9.54E-08 | 1.69E-06 |
| NA | NA |
| 0.002601982 | 0.011181393 |
| 0.000165934 | 0.001073904 |
| 0.905296333 | NA |
| 0.887800692 | NA |
| NA | NA |
| NA | NA |
| NA | NA |
| NA | NA |
| NA | NA |
| NA | NA |
| 0.801661681 | NA |
| NA | NA |
| NA | NA |
| NA | NA |
| NA | NA |
| NA | NA |
| 0.905158739 | NA |
| NA | NA |
| NA | NA |
| NA | NA |
| NA | NA |
| NA | NA |
| NA | NA |
| NA | NA |

| | |
|---|---|
| NA | NA |
| NA | NA |
| NA | NA |
| NA | NA |
| NA | NA |
| NA | NA |
| NA | NA |
| NA | NA |
| NA | NA |
| NA | NA |
| NA | NA |
| NA | NA |
| NA | NA |
| NA | NA |
| NA | NA |
| NA | NA |
| NA | NA |
| NA | NA |
| NA | NA |
| 0.905158266 | NA |
| NA | NA |
| 0.889410245 | NA |
| 0.889409301 | NA |
| 0.019811373 | 0.058355189 |
| 0.919864418 | 0.954147509 |
| 0.694345748 | 0.807923818 |
| 0.180849448 | 0.320755434 |
| 0.040573384 | NA |
| 0.889410245 | NA |
| 0.869002849 | 0.925004192 |
| 0.546448725 | 0.692889915 |
| 0.327718445 | 0.489908586 |
| 0.417411636 | 0.579645995 |
| 0.136506945 | 0.259461267 |
| 0.017019474 | NA |
| 0.033516691 | 0.088941833 |
| 5.82E-06 | 6.08E-05 |
| NA | NA |
| 0.889409301 | NA |
| 0.632293725 | NA |
| 0.164340045 | 0.298317191 |
| 0.742777151 | 0.84236171 |
| 0.716235606 | 0.823790994 |
| 0.413883482 | 0.57638682 |
| 0.020583392 | 0.060193197 |
| 0.00141615 | 0.006690843 |
| 0.797157316 | NA |

| | |
|---|---|
| 0.047467553 | 0.116444672 |
| NA | NA |
| NA | NA |
| 5.52E-09 | 1.39E-07 |
| NA | NA |
| 3.80E-15 | 4.85E-13 |
| 0.222287811 | 0.371639769 |
| 0.007702765 | 0.027211231 |
| 0.633568388 | NA |
| NA | NA |
| 0.532080917 | NA |
| 0.00051339 | 0.002826156 |
| 0.49742069 | 0.650518786 |
| 0.071034231 | 0.158327312 |
| 0.289284485 | 0.448628534 |
| 4.59E-05 | 0.000359937 |
| 0.451477815 | 0.609792185 |
| 0.103206525 | 0.211035673 |
| 0.982084718 | 0.991173786 |
| 0.707968509 | 0.817880918 |
| 0.994893908 | 0.997551495 |
| 0.017250569 | 0.052241795 |
| 0.736507287 | NA |
| 0.742452911 | NA |
| 0.930441825 | 0.959926975 |
| NA | NA |
| NA | NA |
| 7.45E-16 | 1.15E-13 |
| 2.65E-14 | 2.68E-12 |
| 0.079295036 | 0.172767672 |
| 2.73E-05 | 0.000230916 |
| 0.282328235 | 0.440942085 |
| NA | NA |
| 0.425491015 | NA |
| 0.597166363 | NA |
| 0.000654985 | 0.003497364 |
| 0.70806608 | NA |
| 9.64E-07 | 1.27E-05 |
| 3.56E-12 | 2.23E-10 |
| 5.57E-05 | 0.000424972 |
| NA | NA |
| NA | NA |
| NA | NA |
| 0.905158266 | NA |
| 0.021409455 | 0.062104389 |
| 0.023062388 | 0.065975374 |
| 0.143631275 | 0.269733958 |

| | |
|---|---|
| NA | NA |
| 0.017442738 | 0.052700204 |
| 0.072260663 | 0.160379605 |
| 0.006022153 | 0.02234924 |
| 1.51E-05 | 0.000139049 |
| 0.033618264 | 0.089189739 |
| 0.327596764 | 0.489766975 |
| 0.920038397 | NA |
| 0.000831272 | 0.004265926 |
| 0.768325719 | NA |
| 0.190969128 | NA |
| 0.142063716 | 0.267476391 |
| 0.992060816 | NA |
| 1.85E-17 | 4.03E-15 |
| 0.490163434 | 0.644250844 |
| 6.66E-07 | 9.22E-06 |
| 0.865170038 | 0.922201294 |
| 0.402410337 | 0.565972497 |
| 0.04145966 | 0.104946706 |
| NA | NA |
| 0.989267257 | NA |
| 3.90E-06 | 4.31E-05 |
| 0.178231171 | 0.317264776 |
| 0.83311102 | 0.90075258 |
| 0.905158266 | NA |
| 0.905157793 | NA |
| 0.118101186 | NA |
| NA | NA |
| 0.00015078 | 0.000988762 |
| 0.006061175 | 0.022459895 |
| NA | NA |
| 0.172403271 | 0.309297665 |
| 0.322045045 | 0.484365768 |
| 0.287547097 | NA |
| 2.17E-06 | 2.58E-05 |
| 0.610913871 | 0.744825812 |
| 0.992061292 | NA |
| 0.486269869 | 0.641128794 |
| 0.624453131 | 0.755044673 |
| 0.104845176 | 0.213471972 |
| 0.402585332 | 0.566073593 |
| 0.989026872 | NA |
| 0.002265565 | 0.009935998 |
| 5.13E-08 | 9.95E-07 |
| 0.317942175 | 0.480270497 |
| 0.154086153 | 0.284459925 |
| 0.131220651 | 0.251899141 |

| | |
|---|---|
| 0.355161887 | 0.519794713 |
| NA | NA |
| 0.060037819 | 0.139791164 |
| 0.991276819 | NA |
| 0.90149378 | 0.944325808 |
| 0.021644686 | 0.062646061 |
| 0.000104022 | 0.000723086 |
| 0.992060816 | NA |
| 0.000302889 | 0.001801498 |
| 0.187524274 | 0.32898023 |
| 0.03813509 | 0.098458238 |
| 0.912946309 | 0.9507312 |
| NA | NA |
| NA | NA |
| 0.677425127 | NA |
| 0.231293922 | 0.382266825 |
| 0.088524447 | 0.187853531 |
| 0.860822856 | 0.919084412 |
| 0.303631809 | 0.464339501 |
| 0.483880761 | NA |
| 0.56526696 | NA |
| NA | NA |
| 8.12E-06 | 8.14E-05 |
| 0.564324026 | NA |
| 0.860886929 | NA |
| 0.511140748 | 0.662777661 |
| 0.02411527 | 0.068443481 |
| 0.905158739 | NA |
| 0.674620814 | 0.79317938 |
| 0.40785458 | NA |
| 0.81166246 | NA |
| 0.992061292 | NA |
| NA | NA |
| 3.88E-07 | 5.77E-06 |
| 0.651711286 | 0.775786205 |
| 0.700753308 | 0.813129883 |
| 0.029662853 | 0.080764611 |
| 0.066695011 | 0.151140944 |
| 0.944430558 | 0.968738351 |
| 0.002476536 | 0.010732635 |
| 9.95E-05 | 0.00069574 |
| 0.257180901 | 0.411984803 |
| 0.509492837 | 0.661148399 |
| 0.008124494 | 0.028426668 |
| 0.910139097 | 0.949096227 |
| 0.849806902 | NA |
| 0.268242487 | 0.424651198 |

| | |
|---|---|
| 0.808498974 | 0.885124587 |
| 0.00993825 | 0.033445828 |
| 0.055500226 | 0.131272024 |
| 0.018029346 | 0.054071642 |
| 0.023533909 | 0.06708829 |
| 0.353250568 | 0.517445749 |
| 1.42E-08 | 3.20E-07 |
| 0.05226706 | 0.125558732 |
| 0.992060816 | NA |
| NA | NA |
| 0.745609582 | 0.844590857 |
| 0.134291249 | 0.256158924 |
| 0.019714011 | 0.058122982 |
| 0.155570543 | 0.286375518 |
| NA | NA |
| 0.000839468 | 0.004297941 |
| 0.992059387 | NA |
| 0.83392426 | NA |
| 0.523247624 | 0.673651507 |
| NA | NA |
| NA | NA |
| 0.139732086 | 0.264128163 |
| 0.038624604 | 0.099409011 |
| 0.003995856 | 0.015878573 |
| 2.46E-05 | 0.000211807 |
| 0.013482444 | 0.042740146 |
| NA | NA |
| 0.504725522 | 0.656816024 |
| 0.51257549 | 0.664206608 |
| 0.034608816 | 0.091256663 |
| 4.11E-08 | 8.20E-07 |
| 0.133565018 | 0.255150433 |
| 0.004844562 | 0.018673291 |
| 0.287609241 | 0.446711792 |
| 0.119644768 | 0.235630445 |
| 0.038500218 | 0.099158608 |
| 0.004380168 | NA |
| 4.58E-06 | 4.94E-05 |
| NA | NA |
| 0.992062245 | NA |
| 0.992060816 | NA |
| 0.004014546 | 0.015938422 |
| 0.578809336 | 0.719546774 |
| 0.36705383 | NA |
| 0.775450426 | 0.864525581 |
| NA | NA |
| 0.807879735 | NA |

| | | |
|---|---|---|
| | 0.233905214 | 0.385378497 |
| | 0.317274096 | 0.479525217 |
| | 0.489074133 | 0.643319689 |
| | 0.005029495 | 0.019263686 |
| | 0.792669312 | 0.874948498 |
| | 0.723875952 | 0.829493923 |
| | 1.50E-05 | 0.00013783 |
| | 0.3211509 | 0.483454624 |
| | 0.805374319 | NA |
| | 0.000475609 | 0.002643357 |
| | 0.687781615 | NA |
| NA | | NA |
| | 0.684173531 | NA |
| | 0.000902491 | 0.004569473 |
| | 0.480168917 | 0.635718634 |
| | 0.006068415 | 0.022473273 |
| NA | | NA |
| | 0.288780675 | 0.448214224 |
| | 0.742353537 | 0.842098425 |
| | 3.43E-07 | 5.16E-06 |
| | 0.208592236 | 0.355351195 |
| | 0.340141429 | 0.503313048 |
| | 3.68E-09 | 9.85E-08 |
| | 0.574275617 | 0.716221859 |
| | 0.34288952 | 0.506314652 |
| | 0.94079084 | 0.966404362 |
| | 0.107696392 | NA |
| | 0.018273884 | 0.054669785 |
| NA | | NA |
| | 0.004348576 | 0.017033603 |
| | 0.979173666 | 0.989397824 |
| | 0.003697975 | 0.014897053 |
| | 0.002649484 | 0.011362313 |
| NA | | NA |
| | 0.160915717 | 0.293581608 |
| | 0.758341495 | 0.853322916 |
| | 0.660444954 | 0.782569467 |
| | 0.341388696 | 0.504784484 |
| | 0.722546121 | 0.828359103 |
| | 1.85E-05 | 0.000166414 |
| | 0.118964026 | 0.234605392 |
| | 0.182798386 | 0.322985981 |
| | 0.000706778 | 0.003720074 |
| NA | | NA |
| | 0.905158266 | NA |
| NA | | NA |
| | 0.82843639 | NA |

| | | |
|---|---|---|
| | 0.833668138 | NA |
| | 0.639822655 | 0.766370244 |
| | 0.889409773 | NA |
| | 0.988120316 | NA |
| | 0.011583399 | 0.037831981 |
| | 0.992060816 | NA |
| NA | | NA |
| NA | | NA |
| | 0.586544763 | 0.725630223 |
| NA | | NA |
| | 0.786756902 | 0.871619431 |
| | 0.598288293 | 0.7347359 |
| | 0.379349674 | 0.543735459 |
| | 0.830972274 | 0.899237146 |
| | 0.791607135 | NA |
| | 0.21273999 | NA |
| NA | | NA |
| NA | | NA |
| | 0.889409301 | NA |
| | 0.002825066 | 0.011957521 |
| | 0.093680804 | 0.195824539 |
| | 0.000156884 | 0.001022867 |
| | 0.191316345 | 0.333660202 |
| NA | | NA |
| | 0.361021694 | 0.525707917 |
| NA | | NA |
| | 0.207781835 | 0.354465609 |
| | 0.053347721 | 0.127332893 |
| | 0.287107639 | NA |
| | 0.003835857 | 0.015337141 |
| NA | | NA |
| | 0.210638884 | 0.357589428 |
| | 0.150298588 | 0.279033369 |
| NA | | NA |
| | 0.559002602 | 0.703518549 |
| | 0.019374949 | 0.057307988 |
| | 0.606178567 | 0.740779963 |
| | 0.501417224 | 0.6541481 |
| | 0.018783278 | 0.055851319 |
| | 0.011117933 | 0.036651957 |
| | 0.345158634 | 0.508509791 |
| NA | | NA |
| | 0.052851865 | 0.126589944 |
| | 0.031445519 | 0.084590487 |
| NA | | NA |
| | 0.142172916 | 0.267596014 |
| | 0.797851955 | 0.877977003 |

| | | |
|---|---|---|
| | 0.052544115 | 0.126020184 |
| | 0.004970364 | 0.019084493 |
| | 0.721773251 | 0.827775568 |
| | 0.000370743 | 0.002140156 |
| | 0.10363672 | NA |
| | 2.74E-07 | 4.26E-06 |
| | 0.087188872 | 0.185580074 |
| | 0.905158739 | NA |
| NA | | NA |
| | 0.421409137 | 0.583058613 |
| | 0.905158266 | NA |
| | 0.257237615 | 0.411984803 |
| | 0.764185711 | 0.857062789 |
| NA | | NA |
| | 0.008418555 | 0.029282616 |
| | 5.80E-05 | 0.000440498 |
| | 0.006334546 | 0.023264808 |
| | 0.072887847 | NA |
| | 0.245281099 | NA |
| | 0.905158739 | NA |
| | 0.786675317 | NA |
| | 0.882207307 | NA |
| | 0.885332346 | NA |
| NA | | NA |
| NA | | NA |
| | 0.905158266 | NA |
| NA | | NA |
| NA | | NA |
| NA | | NA |
| NA | | NA |
| NA | | NA |
| NA | | NA |
| NA | | NA |
| NA | | NA |
| NA | | NA |
| NA | | NA |
| | 0.889409301 | NA |
| NA | | NA |
| NA | | NA |
| NA | | NA |
| NA | | NA |
| NA | | NA |
| NA | | NA |
| NA | | NA |
| NA | | NA |
| NA | | NA |
| NA | | NA |

| | |
|---|---|
| NA | NA |
| NA | NA |
| NA | NA |
| NA | NA |
| NA | NA |
| NA | NA |
| NA | NA |
| NA | NA |
| NA | NA |
| NA | NA |
| NA | NA |
| NA | NA |
| NA | NA |
| 0.843825832 | NA |
| 0.190470264 | 0.332554388 |
| 0.992057737 | NA |
| NA | NA |
| NA | NA |
| NA | NA |
| 0.807879735 | NA |
| NA | NA |
| NA | NA |
| 0.90515732 | NA |
| 0.652247247 | NA |
| NA | NA |
| 0.002092582 | 0.00929994 |
| NA | NA |
| NA | NA |
| 0.223767077 | 0.37340239 |
| NA | NA |
| NA | NA |
| 0.992061769 | NA |
| 0.668856269 | NA |
| 0.905158266 | NA |
| NA | NA |
| NA | NA |
| NA | NA |
| NA | NA |
| NA | NA |
| NA | NA |
| 0.99205891 | NA |
| NA | NA |
| 0.005512772 | 0.020781969 |
| NA | NA |
| 0.905158739 | NA |
| NA | NA |
| 6.58E-05 | 0.000488879 |

| | |
|---|---|
| 5.32E-06 | 5.62E-05 |
| 0.000103361 | 0.000719172 |
| 0.450904353 | 0.609279964 |
| 0.011899267 | 0.038687593 |
| 0.937087634 | 0.964178959 |
| 0.019492448 | 0.057570232 |
| 0.205267832 | 0.351394818 |
| 0.020506302 | 0.060010992 |
| 0.930306789 | 0.959926975 |
| 0.704983448 | 0.81640816 |
| 0.168890872 | 0.304513889 |
| 0.042809834 | 0.107583103 |
| NA | NA |
| 0.905158266 | NA |
| 0.11050783 | 0.221955678 |
| 0.414043418 | 0.576572986 |
| 0.922031634 | 0.955294241 |
| 0.198176908 | 0.342604843 |
| NA | NA |
| 0.000410519 | 0.002338417 |
| NA | NA |
| 0.905158739 | NA |
| 0.114286395 | 0.227836128 |
| 0.103299622 | 0.211206368 |
| 0.452517791 | 0.61070851 |
| 4.39E-05 | 0.000346518 |
| 0.870024014 | 0.925515189 |
| 0.086321587 | NA |
| NA | NA |
| 0.821340268 | NA |
| NA | NA |
| 0.88773506 | NA |
| 0.16122298 | 0.293959836 |
| 0.285325137 | 0.444263964 |
| NA | NA |
| 0.889409773 | NA |
| NA | NA |
| NA | NA |
| NA | NA |
| NA | NA |
| NA | NA |
| NA | NA |
| NA | NA |
| 0.757024967 | NA |
| 0.731301858 | 0.834692942 |
| NA | NA |
| NA | NA |

| | |
|---|---|
| NA | NA |
| NA | NA |
| NA | NA |
| 0.854436427 | NA |
| 0.812795989 | NA |
| 0.301075144 | 0.461422578 |
| 0.008834361 | 0.030439938 |
| 0.000110493 | 0.000760123 |
| 0.538236385 | NA |
| 0.003523329 | 0.014335516 |
| 0.527169593 | NA |
| 0.000376862 | 0.002170344 |
| 0.223625283 | 0.373230036 |
| 2.01E-05 | 0.000178911 |
| 0.76910024 | NA |
| 0.540995326 | NA |
| 0.697979851 | NA |
| 0.781928695 | 0.868550368 |
| 0.413375993 | 0.575935755 |
| 0.942590317 | 0.967619128 |
| 0.003137769 | 0.013039792 |
| 0.433618622 | 0.594371492 |
| 0.632920809 | NA |
| 0.026487969 | 0.073691791 |
| 0.243751963 | 0.39672023 |
| 0.063064911 | 0.14506198 |
| 0.021077245 | 0.0613108 |
| 0.482274819 | 0.637737731 |
| 0.03949904 | NA |
| 0.639912557 | 0.766436121 |
| 0.398619186 | NA |
| 0.807879735 | NA |
| NA | NA |
| NA | NA |
| NA | NA |
| NA | NA |
| NA | NA |
| NA | NA |
| 0.001111695 | 0.005441756 |
| 5.63E-05 | 0.000429308 |
| NA | NA |
| NA | NA |
| NA | NA |
| 0.496571621 | NA |
| 2.80E-05 | 0.000235757 |
| 0.359723617 | 0.524471524 |
| 1.44E-08 | 3.24E-07 |

| | |
|---|---|
| NA | NA |
| 0.09700572 | 0.201186164 |
| 0.019957108 | 0.058682129 |
| 0.65248558 | 0.776245035 |
| 0.322141045 | 0.484447008 |
| 1.63E-07 | 2.71E-06 |
| 0.889410717 | NA |
| 0.905158266 | NA |
| 0.669285285 | 0.789091843 |
| NA | NA |
| 0.432864188 | 0.593657944 |
| NA | NA |
| 0.621093185 | 0.752432156 |
| 0.3578939 | 0.522608868 |
| 0.013999118 | 0.044116909 |
| 0.308501399 | 0.469595329 |
| 0.952028148 | 0.973288471 |
| 0.087230114 | 0.185609117 |
| 0.427182992 | 0.58826736 |
| 0.12637736 | 0.245260313 |
| 4.76E-07 | 6.91E-06 |
| 0.014270667 | 0.044818303 |
| 5.93E-08 | 1.12E-06 |
| 0.734559915 | 0.836928054 |
| NA | NA |
| 0.053053959 | 0.126893791 |
| NA | NA |
| 0.341649254 | 0.504999728 |
| 0.011254737 | 0.036991882 |
| 0.00036737 | 0.002125331 |
| 0.000243737 | 0.001497964 |
| 0.005847721 | 0.021823609 |
| 0.000409643 | 0.002334028 |
| 5.57E-05 | 0.000424972 |
| 0.880445331 | 0.932221934 |
| 0.045340872 | 0.112463596 |
| 0.025971782 | 0.072651511 |
| 0.827977179 | 0.897278333 |
| 0.992061292 | NA |
| 0.02191253 | 0.063287813 |
| NA | NA |
| 0.933704817 | 0.961917397 |
| 0.684237221 | 0.800119049 |
| 7.35E-09 | 1.80E-07 |
| 0.665131674 | NA |
| 0.620956308 | NA |
| 0.019066548 | 0.056502339 |

| | | |
|---|---|---|
| | 0.765213898 | 0.857619149 |
| | 0.510927232 | NA |
| | 0.000220082 | 0.001369058 |
| | 0.190659442 | 0.332805301 |
| | 0.007913616 | 0.02782958 |
| | 0.006544232 | 0.023906969 |
| | 0.288804613 | 0.448219697 |
| | 0.905158266 | NA |
| | 7.14E-08 | 1.32E-06 |
| | 0.144999751 | 0.271699764 |
| | 0.202466965 | 0.348021299 |
| | 0.169183638 | 0.304941882 |
| | 0.971424796 | 0.984954128 |
| | 0.011171003 | 0.036791178 |
| | 7.13E-08 | 1.32E-06 |
| | 0.032296092 | 0.086444404 |
| | 0.821406626 | 0.893671826 |
| | 0.034284786 | 0.090575025 |
| | 0.788920507 | 0.872608947 |
| | 0.606295326 | 0.740881417 |
| | 0.264656852 | 0.420612275 |
| NA | | NA |
| | 0.028269871 | 0.077698515 |
| | 0.905158739 | NA |
| NA | | NA |
| NA | | NA |
| NA | | NA |
| | 0.556759193 | NA |
| | 0.306999224 | 0.467762695 |
| | 0.768040089 | NA |
| | 0.992061292 | NA |
| | 0.040677561 | 0.103276513 |
| | 0.151801472 | 0.281300298 |
| | 0.295448759 | 0.455404015 |
| | 4.08E-06 | 4.47E-05 |
| | 0.000109846 | 0.000756386 |
| | 0.066138305 | 0.150313707 |
| NA | | NA |
| | 0.307966587 | 0.468976302 |
| | 0.082952414 | 0.178899239 |
| | 0.000122229 | 0.000830451 |
| | 0.037893081 | 0.097985654 |
| NA | | NA |
| | 0.02333369 | 0.06662131 |
| | 0.106196038 | 0.215403547 |
| | 2.44E-07 | 3.86E-06 |
| NA | | NA |

| | |
|---|---|
| 0.76362037 | 0.85677877 |
| 0.615432617 | 0.748332145 |
| 0.040289026 | 0.102586587 |
| 0.380060624 | 0.544304363 |
| 2.15E-18 | 6.22E-16 |
| 0.105518195 | 0.21450417 |
| 0.574216894 | 0.716189297 |
| 0.35852514 | 0.523391408 |
| 0.597390214 | NA |
| 0.056110855 | 0.132445262 |
| NA | NA |
| 0.091962487 | 0.193225936 |
| 0.078820763 | 0.172016015 |
| 0.448521216 | 0.607517727 |
| 0.296483459 | 0.456683522 |
| 0.004265318 | 0.016761295 |
| 0.014847797 | 0.046286596 |
| 0.233718505 | 0.385329731 |
| 0.80920819 | 0.885534996 |
| 0.213542648 | 0.360706385 |
| 0.456596464 | 0.614287572 |
| 1.52E-06 | 1.88E-05 |
| 0.57475335 | 0.716654864 |
| 0.690999248 | 0.805395567 |
| 0.042158903 | NA |
| 0.000195529 | 0.001236994 |
| 0.014339598 | 0.044983319 |
| 0.599389325 | 0.73542997 |
| 0.083674642 | 0.179873858 |
| 0.194106411 | 0.337427382 |
| 0.007483636 | 0.02660835 |
| 0.000305679 | 0.001815637 |
| 0.90515732 | NA |
| 0.002672019 | 0.011439826 |
| 0.017273428 | 0.052296597 |
| 0.000947543 | 0.00476026 |
| 0.593155374 | 0.730636725 |
| 0.645096842 | 0.770418315 |
| 0.560611872 | 0.704946235 |
| 0.992059387 | NA |
| 1.05E-11 | 5.81E-10 |
| 0.180588578 | 0.320448048 |
| 0.254850913 | NA |
| 0.129884005 | NA |
| 0.803187987 | 0.881595771 |
| NA | NA |
| NA | NA |

| | |
|---|---|
| 0.000176607 | 0.001133952 |
| 0.052788373 | 0.126465184 |
| 0.362186532 | 0.526974984 |
| NA | NA |
| 0.011360139 | 0.037254674 |
| 0.567138367 | NA |
| 0.716469996 | 0.823956855 |
| NA | NA |
| 0.881172739 | 0.932448123 |
| 0.905158739 | NA |
| 0.057551799 | 0.135084386 |
| 1.91E-15 | 2.66E-13 |
| NA | NA |
| NA | NA |
| 0.002680109 | 0.011465354 |
| 3.92E-08 | 7.91E-07 |
| 0.001126514 | 0.005504478 |
| NA | NA |
| NA | NA |
| 0.41087464 | 0.573771571 |
| 2.80E-06 | 3.22E-05 |
| 5.50E-07 | 7.80E-06 |
| 0.905158266 | NA |
| 0.889409301 | NA |
| NA | NA |
| 0.365982384 | NA |
| NA | NA |
| NA | NA |
| 0.23679126 | 0.388609308 |
| NA | NA |
| 0.158577211 | 0.290400882 |
| 2.41E-06 | 2.82E-05 |
| 0.787226179 | 0.871771467 |
| 0.217132919 | 0.365337657 |
| 0.000667592 | 0.00354955 |
| 0.89795365 | NA |
| 0.001271568 | 0.006091265 |
| 0.006571454 | 0.023975001 |
| 0.862029145 | NA |
| 0.209915776 | 0.356739074 |
| 0.643702915 | NA |
| 0.002511702 | 0.010856119 |
| 0.116601977 | 0.231297453 |
| 0.039406034 | 0.100946931 |
| 3.35E-09 | 9.13E-08 |
| NA | NA |
| 0.545487998 | NA |

| | |
|---|---|
| 0.782879506 | 0.869210793 |
| 0.756687841 | 0.852105516 |
| 0.012112277 | 0.039263874 |
| 0.039620141 | 0.101341721 |
| 0.268751339 | 0.425185078 |
| 0.00110127 | 0.005399151 |
| 0.243926554 | 0.396905118 |
| 0.044095869 | 0.109984234 |
| 0.071194939 | 0.158569349 |
| 0.214637593 | NA |
| 0.0023435 | 0.010232834 |
| NA | NA |
| 0.273021029 | 0.429953298 |
| 0.765044426 | 0.857560519 |
| NA | NA |
| 0.531707374 | NA |
| NA | NA |
| 0.807879735 | NA |
| NA | NA |
| 0.182692714 | 0.322882122 |
| 0.028432403 | 0.078072669 |
| 0.000141025 | 0.000933327 |
| 0.789260309 | 0.872787041 |
| 0.862836158 | 0.920856319 |
| 0.882883177 | 0.933424732 |
| 0.789618189 | NA |
| 0.42264675 | 0.584218809 |
| 0.000369867 | 0.002136219 |
| 0.996026571 | NA |
| 0.905158266 | NA |
| 0.992061292 | NA |
| 0.00020584 | 0.001293288 |
| 0.035333811 | 0.092732778 |
| 0.024673705 | 0.069748956 |
| 0.368095705 | NA |
| 0.905158739 | NA |
| 0.03873812 | 0.099584451 |
| 0.176376639 | 0.314728963 |
| 0.992061292 | NA |
| 0.230111953 | 0.38084326 |
| 0.749476143 | NA |
| 0.039333509 | 0.10079642 |
| 0.62983605 | NA |
| 0.013507472 | 0.042807126 |
| 1.54E-05 | 0.000141232 |
| 0.558813631 | 0.703490713 |
| 0.514646277 | 0.665868348 |

| | | |
|---|---|---|
| | 0.006373057 | 0.023378879 |
| | 0.072565904 | 0.160973298 |
| | 0.0189828 | 0.05630735 |
| | 0.958534361 | 0.976998516 |
| | 0.000960203 | 0.004811753 |
| NA | | NA |
| | 0.417397609 | NA |
| | 0.000692186 | 0.003656414 |
| NA | | NA |
| NA | | NA |
| | 0.58983497 | NA |
| | 0.098443558 | 0.203460836 |
| | 0.269594856 | 0.425993628 |
| | 0.003438 | 0.014040366 |
| | 0.215855641 | 0.363655196 |
| | 0.742848261 | 0.842398521 |
| | 0.688412069 | 0.8036541 |
| NA | | NA |
| | 6.95E-06 | 7.11E-05 |
| | 0.238084024 | 0.390068728 |
| NA | | NA |
| | 0.905158739 | NA |
| | 0.564809876 | 0.708283881 |
| | 0.961678002 | NA |
| | 0.000760452 | 0.00396081 |
| NA | | NA |
| | 0.209340504 | 0.356174292 |
| | 0.816734317 | 0.89098658 |
| NA | | NA |
| | 2.64E-05 | 0.00022409 |
| | 0.800081237 | 0.879416552 |
| | 0.335676962 | NA |
| NA | | NA |
| | 0.640545363 | NA |
| | 0.902247138 | NA |
| | 0.873952925 | 0.927840184 |
| | 0.394906027 | 0.55845228 |
| | 0.395188798 | 0.558748574 |
| | 0.233869353 | 0.385378497 |
| | 0.206748972 | 0.353362444 |
| | 0.000379769 | 0.002184793 |
| | 0.015188062 | 0.04718464 |
| | 0.66955424 | NA |
| | 0.992061292 | NA |
| | 1.44E-07 | 2.43E-06 |
| | 0.889410245 | NA |
| | 8.22E-05 | 0.000589853 |

| | |
|---|---|
| 0.000556216 | 0.003034522 |
| 0.345179541 | NA |
| 0.443306875 | 0.60290943 |
| 0.091885485 | 0.193138066 |
| NA | NA |
| NA | NA |
| 0.241029528 | 0.3937778 |
| 0.840284719 | NA |
| 0.00024543 | 0.001507103 |
| 0.396936386 | 0.560607545 |
| 0.465709245 | 0.622578149 |
| 0.94996158 | 0.971960038 |
| 0.447585867 | 0.606803455 |
| 0.013162391 | 0.041894904 |
| 0.068701247 | 0.154128594 |
| 0.185675089 | 0.326649518 |
| 0.777702224 | NA |
| 0.992061292 | NA |
| 0.841329549 | 0.906158307 |
| 0.018761756 | 0.055802439 |
| NA | NA |
| 0.141490932 | 0.266626565 |
| 7.44E-07 | 1.02E-05 |
| 0.130082555 | NA |
| 0.99205891 | NA |
| 7.86E-18 | 1.96E-15 |
| 0.007436154 | 0.026469528 |
| 0.040505137 | 0.102945869 |
| 0.615462912 | 0.748332145 |
| 0.233142624 | 0.384670439 |
| 0.034273007 | NA |
| 0.520782657 | NA |
| 0.752568704 | 0.849429769 |
| 0.489023385 | 0.643305506 |
| 0.000188418 | 0.001197184 |
| 0.1266819 | 0.245713512 |
| 0.967578118 | NA |
| 0.773794138 | NA |
| 0.982146292 | 0.991173786 |
| 0.589984446 | 0.728244332 |
| 0.002534229 | 0.010930886 |
| 0.848936249 | NA |
| 0.915185953 | 0.952049041 |
| 0.889408834 | 0.937385064 |
| 0.889409301 | NA |
| 0.992061292 | NA |
| 0.004702248 | 0.018217479 |

| | |
|---|---|
| 0.438113221 | 0.598477649 |
| 0.461365406 | NA |
| 0.869362942 | 0.925278249 |
| 0.17389918 | 0.31116787 |
| 0.002964325 | 0.012441441 |
| 0.001181991 | 0.005737255 |
| NA | NA |
| NA | NA |
| 0.428595123 | 0.589620368 |
| 9.93E-12 | 5.52E-10 |
| 0.48186068 | 0.637458804 |
| 0.217244766 | 0.36548984 |
| 0.225449786 | NA |
| 0.559842801 | 0.70418078 |
| 0.05798848 | 0.135855013 |
| NA | NA |
| 0.013852842 | 0.043731249 |
| 0.632798597 | 0.760445706 |
| 0.04262401 | 0.107226558 |
| 0.919052651 | NA |
| 0.025001517 | 0.070548485 |
| 0.779696991 | 0.86734378 |
| 0.137456596 | 0.260927507 |
| 0.74063443 | 0.840843317 |
| 0.905158739 | NA |
| 0.624037847 | NA |
| 0.227456869 | 0.377387637 |
| NA | NA |
| 0.166568339 | 0.301340761 |
| 0.284614111 | 0.443415018 |
| 0.000460556 | 0.002572064 |
| 0.10972184 | 0.220963121 |
| 0.00142703 | 0.006734442 |
| 0.013654181 | 0.043191033 |
| 2.83E-07 | 4.37E-06 |
| 0.8581582 | 0.917175946 |
| 2.20E-06 | 2.60E-05 |
| 0.148429318 | 0.27640429 |
| 0.146430208 | 0.273656249 |
| 0.247310808 | 0.400878279 |
| 0.020900043 | NA |
| 0.000824625 | 0.004238752 |
| NA | NA |
| 4.51E-08 | 8.88E-07 |
| 0.424091889 | 0.585442496 |
| 0.992060816 | NA |
| 0.992060816 | NA |

| | | |
|---|---|---|
| | 0.656078748 | NA |
| | 0.753813258 | 0.85017901 |
| | 0.074103683 | NA |
| | 0.283608546 | 0.442375341 |
| | 0.902978376 | 0.945090858 |
| | 0.251707733 | 0.405937875 |
| | 0.988878521 | NA |
| | 0.005271195 | NA |
| NA | | NA |
| | 0.008031432 | 0.02817383 |
| | 0.006097365 | 0.022548338 |
| NA | | NA |
| | 0.005947731 | 0.022114161 |
| | 0.147871983 | 0.275646934 |
| | 0.007417708 | 0.026425282 |
| | 2.10E-05 | 0.000185773 |
| | 0.019975663 | 0.058720961 |
| | 0.00569337 | 0.021365586 |
| | 0.703550964 | NA |
| | 1.57E-06 | 1.93E-05 |
| | 0.663892377 | NA |
| | 0.289866603 | 0.449264592 |
| | 0.901187599 | NA |
| | 0.058255102 | 0.136363264 |
| | 0.388743598 | 0.55209033 |
| | 0.813834996 | NA |
| | 0.215204259 | 0.362926835 |
| | 0.31401393 | NA |
| | 0.212098481 | 0.35917508 |
| | 0.464279677 | NA |
| | 0.919134237 | NA |
| NA | | NA |
| NA | | NA |
| | 0.013708336 | 0.043343608 |
| | 0.261481701 | NA |
| NA | | NA |
| NA | | NA |
| NA | | NA |
| NA | | NA |
| NA | | NA |
| | 0.512567868 | NA |
| | 1.14E-06 | 1.46E-05 |
| NA | | NA |
| | 0.001234855 | 0.005938715 |
| | 0.840577613 | 0.905703818 |
| | 0.013694571 | 0.043306321 |
| | 0.016464448 | 0.050382219 |

| | | |
|---|---|---|
| | 0.889410717 | NA |
| | 0.144271306 | 0.270704402 |
| | 0.567298611 | 0.710098152 |
| NA | | NA |
| | 0.014197311 | 0.044632607 |
| | 3.60E-06 | 4.01E-05 |
| | 0.992061292 | NA |
| | 0.725202069 | 0.830307868 |
| | 2.29E-05 | 0.000199427 |
| | 0.179811721 | 0.319379243 |
| | 0.992060816 | NA |
| | 0.869905141 | NA |
| | 0.000554162 | 0.003024065 |
| | 0.608990543 | NA |
| | 0.90515732 | NA |
| NA | | NA |
| | 0.905158266 | NA |
| | 0.782195155 | NA |
| | 0.002524395 | 0.01089489 |
| | 0.601051536 | 0.73664624 |
| | 0.01377579 | 0.043513028 |
| | 0.33007857 | 0.492438027 |
| NA | | NA |
| NA | | NA |
| | 0.006450631 | 0.023639753 |
| | 0.000114893 | 0.000786453 |
| NA | | NA |
| NA | | NA |
| | 0.020090827 | 0.059027892 |
| | 0.103085999 | 0.210848124 |
| | 0.028999908 | 0.079254385 |
| | 0.331480332 | 0.493834079 |
| | 0.089038595 | 0.188647216 |
| | 2.49E-06 | 2.91E-05 |
| NA | | NA |
| NA | | NA |
| | 0.281806545 | NA |
| | 2.19E-06 | 2.60E-05 |
| | 5.45E-07 | 7.75E-06 |
| | 0.141866782 | NA |
| | 0.008293375 | 0.028907021 |
| | 0.093662977 | 0.195805912 |
| | 0.004358393 | 0.01706597 |
| | 0.620284932 | 0.75178479 |
| | 0.474690268 | 0.630937812 |
| | 0.009116427 | 0.031181872 |
| | 0.514907557 | 0.666167151 |

| | |
|---|---|
| NA | NA |
| 3.19E-10 | 1.18E-08 |
| 0.003767912 | 0.015112252 |
| 0.596828006 | NA |
| 0.156656414 | NA |
| 0.487616431 | 0.64228067 |
| 0.006943629 | 0.025119463 |
| 0.475470625 | 0.631630941 |
| 0.309747739 | 0.470996139 |
| 0.99205891 | NA |
| 0.692345495 | 0.806431689 |
| 0.000304321 | 0.001808549 |
| 0.993572367 | 0.996886393 |
| 0.091369681 | 0.192237886 |
| 0.001652254 | 0.007582414 |
| 0.97632188 | NA |
| 0.069645011 | 0.155879604 |
| 0.341689042 | 0.505024545 |
| 0.632619948 | 0.760272627 |
| 0.246296674 | 0.399736044 |
| 0.207671346 | 0.354378303 |
| 0.30793144 | NA |
| 8.08E-05 | 0.000582362 |
| 0.020511578 | 0.060015156 |
| 0.434754982 | NA |
| 0.199403625 | 0.344183636 |
| 0.001370802 | 0.006505607 |
| NA | NA |
| 0.506771267 | 0.658589705 |
| 0.769795843 | NA |
| 2.68E-13 | 2.20E-11 |
| 0.351501344 | 0.515812485 |
| NA | NA |
| 0.543557728 | NA |
| 0.83436641 | 0.901399783 |
| 0.882820979 | NA |
| 1.92E-06 | 2.31E-05 |
| 0.048392743 | 0.118217404 |
| 0.470504971 | NA |
| 7.70E-05 | 0.000560115 |
| 0.969925195 | 0.98418406 |
| 0.000202402 | 0.001274242 |
| 0.889409773 | NA |
| 5.66E-08 | 1.08E-06 |
| 0.007196226 | 0.025812142 |
| 3.39E-07 | 5.12E-06 |
| 0.533498738 | NA |

| | |
|---|---|
| 0.00019404 | 0.001228492 |
| 2.16E-07 | 3.46E-06 |
| 0.602985568 | NA |
| 0.045503702 | 0.112791036 |
| NA | NA |
| NA | NA |
| 0.74638038 | 0.845164017 |
| 0.48584552 | 0.64076692 |
| 0.821080976 | NA |
| 4.83E-05 | 0.000376402 |
| 0.889410245 | NA |
| NA | NA |
| 0.889410245 | NA |
| 0.889410245 | NA |
| 0.586107771 | NA |
| 0.843559653 | NA |
| 0.016114757 | 0.049533028 |
| 4.22E-05 | 0.000335272 |
| 1.59E-09 | 4.70E-08 |
| 0.937799842 | 0.964776141 |
| 0.046573801 | NA |
| 0.768855196 | NA |
| 0.842354141 | NA |
| 0.991918092 | NA |
| 0.234833338 | 0.386327934 |
| 0.237544376 | 0.389445793 |
| 0.56888729 | 0.711279167 |
| 2.28E-05 | 0.000198839 |
| 0.105317824 | 0.21415632 |
| 0.577343203 | 0.71861917 |
| 0.000505146 | 0.002785658 |
| 0.905157793 | NA |
| 7.63E-06 | 7.71E-05 |
| 0.361275076 | 0.526041999 |
| 0.599771598 | 0.73570717 |
| 0.013554779 | 0.042944655 |
| 0.004549312 | 0.017681123 |
| 0.000887089 | 0.004498749 |
| NA | NA |
| 0.502356024 | 0.654944839 |
| 0.932773822 | NA |
| 0.010946852 | 0.036223918 |
| NA | NA |
| 0.450205789 | 0.608829607 |
| 0.011065874 | 0.036524204 |
| 0.232985 | 0.384468145 |
| 0.000184829 | 0.001177777 |

| | |
|---|---|
| 4.72E-05 | 0.000369493 |
| 0.001971816 | 0.008841967 |
| 0.000125221 | 0.000845362 |
| 0.167160926 | 0.302188751 |
| 2.57E-06 | 2.99E-05 |
| 0.006918777 | 0.025050194 |
| 0.040254312 | 0.102557655 |
| NA | NA |
| 0.255108182 | NA |
| 0.009091162 | 0.031125877 |
| 0.24510797 | 0.398307254 |
| 0.665359201 | NA |
| 0.000861666 | 0.004387046 |
| 0.614442546 | 0.747592191 |
| 0.801661681 | NA |
| 0.098016217 | 0.202763835 |
| 0.858291706 | NA |
| 0.905158739 | NA |
| 0.992253023 | NA |
| 0.395034208 | 0.558567687 |
| 0.217080913 | NA |
| 0.729038527 | NA |
| NA | NA |
| 0.835307234 | NA |
| 0.992060816 | NA |
| NA | NA |
| 0.006235796 | NA |
| 0.597813989 | NA |
| 0.472559708 | NA |
| 0.890552849 | 0.938100835 |
| 0.610683463 | NA |
| NA | NA |
| NA | NA |
| NA | NA |
| NA | NA |
| NA | NA |
| 0.889410245 | NA |
| 0.991879012 | NA |
| 0.354946961 | 0.519514785 |
| 0.956096248 | 0.975600256 |
| 0.992060816 | NA |
| NA | NA |
| NA | NA |
| NA | NA |
| NA | NA |
| NA | NA |
| NA | NA |

| | | |
|---|---|---|
| | 0.889409301 | NA |
| NA | | NA |
| | 0.748890654 | NA |
| | 0.889409301 | NA |
| NA | | NA |
| NA | | NA |
| | 0.905158266 | NA |
| NA | | NA |
| NA | | NA |
| | 0.051359064 | NA |
| | 0.778531773 | NA |
| | 0.673797815 | NA |
| NA | | NA |
| NA | | NA |
| NA | | NA |
| NA | | NA |
| NA | | NA |
| NA | | NA |
| NA | | NA |
| | 0.851428424 | NA |
| | 0.905158266 | NA |
| NA | | NA |
| | 8.42E-05 | 0.000602127 |
| NA | | NA |
| | 0.069004256 | NA |
| | 0.404935733 | 0.568177875 |
| | 0.350524727 | 0.514654485 |
| | 0.948410236 | NA |
| | 0.905158266 | NA |
| | 0.929978106 | 0.959774534 |
| | 0.0132061 | 0.042027936 |
| | 0.074371212 | 0.16423144 |
| | 0.052650406 | 0.126244842 |
| NA | | NA |
| | 0.00607331 | 0.02248589 |
| | 0.011549935 | 0.037736203 |
| | 0.006895702 | 0.024987246 |
| | 0.199562344 | 0.344351331 |
| | 0.613592306 | 0.746847373 |
| | 0.157459089 | NA |
| | 1.55E-08 | 3.45E-07 |
| | 0.905158266 | NA |
| | 0.992059387 | NA |
| | 0.792188023 | NA |
| | 0.905158266 | NA |
| | 0.046107732 | 0.113966645 |
| | 0.81828543 | NA |

| | | |
|---|---|---|
| | 0.210753033 | 0.357727901 |
| | 0.00508302 | 0.019432109 |
| | 0.097560115 | 0.202048718 |
| | 0.905158739 | NA |
| | 0.00403956 | 0.016025821 |
| | 0.785397031 | NA |
| | 0.905157793 | NA |
| | 0.905158266 | NA |
| NA | | NA |
| | 0.564983171 | 0.708374184 |
| | 0.854100628 | NA |
| NA | | NA |
| | 0.889409301 | NA |
| | 0.000113462 | 0.000777869 |
| | 0.214556784 | 0.361973911 |
| NA | | NA |
| NA | | NA |
| | 0.904947233 | NA |
| | 0.992060816 | NA |
| NA | | NA |
| NA | | NA |
| NA | | NA |
| NA | | NA |
| NA | | NA |
| NA | | NA |
| NA | | NA |
| NA | | NA |
| NA | | NA |
| NA | | NA |
| NA | | NA |
| | 0.99205891 | NA |
| | 0.672317347 | NA |
| | 0.256207933 | NA |
| | 0.719507036 | 0.826082558 |
| | 0.889409301 | NA |
| NA | | NA |
| NA | | NA |
| NA | | NA |
| | 0.905158266 | NA |
| NA | | NA |
| | 0.905158266 | NA |
| NA | | NA |
| | 0.905158739 | NA |
| NA | | NA |
| | 0.18730716 | 0.328756928 |
| | 0.898775728 | NA |
| NA | | NA |

| | | | |
|---|---|---|---|
| | 0.991918092 | NA | |
| NA | | NA | |
| | 0.992061292 | NA | |
| NA | | NA | |
| NA | | NA | |
| | 0.738757848 | NA | |
| NA | | NA | |
| | 0.986799563 | NA | |
| NA | | NA | |
| NA | | NA | |
| NA | | NA | |
| NA | | NA | |
| NA | | NA | |
| NA | | NA | |
| NA | | NA | |
| NA | | NA | |
| | 0.739675887 | NA | |
| NA | | NA | |
| | 0.992060816 | NA | |
| NA | | NA | |
| | 0.992060816 | NA | |
| | 0.992061292 | NA | |
| | 0.905158739 | NA | |
| NA | | NA | |
| NA | | NA | |
| NA | | NA | |
| NA | | NA | |
| NA | | NA | |
| | 0.90515732 | NA | |
| NA | | NA | |
| NA | | NA | |
| NA | | NA | |
| NA | | NA | |
| | 0.992061292 | NA | |
| NA | | NA | |
| NA | | NA | |
| NA | | NA | |
| NA | | NA | |
| NA | | NA | |
| NA | | NA | |
| NA | | NA | |
| | 0.003968356 | | 0.015786436 |
| | 0.479436942 | | 0.635055846 |
| | 0.83248296 | | 0.900339505 |
| NA | | NA | |
| NA | | NA | |
| | 0.990950919 | NA | |

| | |
|---|---|
| NA | NA |
| NA | NA |
| 0.848453806 | NA |
| NA | NA |
| NA | NA |
| NA | NA |
| NA | NA |
| NA | NA |
| NA | NA |
| 0.892758428 | NA |
| 0.812819201 | NA |
| NA | NA |
| 0.001908128 | 0.008596754 |
| 0.867552689 | NA |
| 0.390685998 | NA |
| 0.054225392 | 0.129104997 |
| 0.266150255 | 0.422404846 |
| 0.889409301 | NA |
| 0.557662104 | 0.702597125 |
| 0.951594387 | NA |
| 0.000838721 | 0.004296123 |
| 0.50636418 | 0.658255448 |
| 0.552680984 | 0.698164903 |
| 8.72E-05 | 0.000620198 |
| 0.380212706 | NA |
| 0.001668218 | 0.007650876 |
| 0.905158266 | NA |
| 0.047636158 | NA |
| 3.56E-07 | 5.33E-06 |
| 0.000947224 | 0.004759744 |
| 0.99282068 | NA |
| 0.0109253 | 0.036174405 |
| 0.003382388 | 0.013864822 |
| 0.281299671 | 0.439710954 |
| NA | NA |
| 0.017228469 | 0.052183412 |
| 0.889409301 | NA |
| 1.63E-20 | 8.15E-18 |
| 0.00660754 | 0.02409415 |
| 4.43E-05 | 0.000349274 |
| 0.028872361 | 0.07900007 |
| NA | NA |
| 0.673733436 | NA |
| 0.90515732 | NA |
| 0.024489433 | 0.069335156 |
| 0.000584931 | 0.003175402 |
| NA | NA |

| | | |
|---|---|---|
| | 0.000625395 | 0.003357715 |
| | 0.001242698 | 0.005970273 |
| | 0.901141266 | 0.944236842 |
| | 0.680674233 | NA |
| | 0.693006607 | 0.806921474 |
| | 0.328022186 | 0.490136036 |
| | 1.02E-05 | 9.87E-05 |
| | 0.517844869 | 0.668667264 |
| | 0.029553633 | 0.08050716 |
| | 0.096848359 | 0.200973548 |
| | 0.071478008 | 0.159042008 |
| | 0.557325646 | NA |
| NA | | NA |
| NA | | NA |
| | 0.460563603 | 0.617996711 |
| | 0.00010054 | 0.000701991 |
| | 0.769637375 | NA |
| | 0.002830816 | 0.01197955 |
| | 0.998334379 | NA |
| | 0.001594103 | 0.007367905 |
| | 0.007320161 | 0.026171183 |
| | 0.000178017 | 0.001141216 |
| | 0.040436171 | 0.102842006 |
| | 0.000775627 | 0.00402172 |
| | 0.044778358 | 0.111307217 |
| | 0.000484979 | 0.002690926 |
| | 0.017718836 | 0.053328936 |
| | 0.759902502 | NA |
| NA | | NA |
| | 0.005710604 | 0.021417449 |
| | 0.012815525 | 0.041082643 |
| | 1.72E-10 | 6.90E-09 |
| | 9.51E-07 | 1.26E-05 |
| | 0.600710493 | NA |
| | 0.003360223 | 0.01378426 |
| | 0.990759986 | 0.995333376 |
| | 0.001312245 | 0.006258818 |
| | 0.018360578 | 0.054854413 |
| | 0.213203228 | 0.360337793 |
| | 0.732774096 | 0.835736895 |
| | 0.61931944 | 0.751049447 |
| | 0.166238376 | 0.300842961 |
| | 0.805580165 | NA |
| | 0.258265433 | 0.413145673 |
| | 1.17E-09 | 3.61E-08 |
| | 0.712386997 | 0.820959728 |
| | 0.113824474 | NA |

| | |
|---|---|
| 0.141582884 | 0.266772232 |
| 0.172849633 | 0.30989606 |
| 0.339869139 | 0.503011823 |
| 0.505956971 | 0.657920835 |
| 6.05E-10 | 2.03E-08 |
| 0.000411796 | 0.002342647 |
| 0.140802619 | 0.265603017 |
| 0.008160839 | 0.028535648 |
| 0.01819308 | 0.05445772 |
| 0.017541252 | 0.052925027 |
| 0.600083077 | 0.735952208 |
| 0.98140427 | NA |
| 0.110546129 | 0.221975569 |
| 3.03E-09 | 8.38E-08 |
| 0.146826104 | 0.274256074 |
| 0.841046986 | 0.90593575 |
| 0.036420019 | 0.095026995 |
| 0.81258324 | NA |
| 0.012910809 | 0.041291738 |
| 0.146127859 | 0.273230726 |
| 0.441800361 | 0.601493838 |
| NA | NA |
| 1.84E-07 | 3.03E-06 |
| 0.32468584 | NA |
| 0.202179255 | 0.347690051 |
| 0.293525096 | NA |
| 0.993263396 | 0.99666747 |
| 0.82769551 | 0.897105908 |
| 0.764111779 | NA |
| 0.551217767 | 0.696894045 |
| 0.457476611 | 0.614944228 |
| 0.00014066 | 0.000931748 |
| 8.96E-05 | 0.000635134 |
| 0.196312297 | 0.340292211 |
| 0.000238525 | 0.001471285 |
| 0.486732196 | NA |
| 0.939500431 | 0.965846909 |
| 0.406890559 | 0.570055804 |
| 0.05365372 | NA |
| 0.030224301 | 0.081927593 |
| 0.062300083 | NA |
| 0.002237695 | 0.009837348 |
| 0.983162012 | NA |
| 0.532404051 | NA |
| 0.642112502 | 0.767898291 |
| 0.075621788 | 0.166424016 |
| 0.992060816 | NA |

| | | | |
|---|---|---|---|
| NA | | NA | |
| | 3.90E-05 | | 0.000313286 |
| | 0.656529866 | NA | |
| NA | | NA | |
| NA | | NA | |
| NA | | NA | |
| NA | | NA | |
| NA | | NA | |
| | 2.07E-06 | | 2.48E-05 |
| | 0.002611663 | | 0.0112208 |
| | 0.000211979 | | 0.001325793 |
| | 0.354699515 | | 0.519325687 |
| | 8.45E-08 | | 1.52E-06 |
| | 1.05E-06 | | 1.36E-05 |
| | 0.127555019 | | 0.246891199 |
| | 0.918872785 | NA | |
| | 3.20E-10 | | 1.18E-08 |
| | 0.209396373 | | 0.356204338 |
| | 0.529729242 | NA | |
| NA | | NA | |
| | 0.099522525 | | 0.205241841 |
| NA | | NA | |
| NA | | NA | |
| | 0.028095009 | | 0.077320253 |
| | 0.989975993 | NA | |
| NA | | NA | |
| | 0.905158266 | NA | |
| | 0.808141711 | NA | |
| | 0.627434483 | | 0.75642702 |
| | 0.000177011 | | 0.001135882 |
| | 0.486645306 | | 0.641508241 |
| NA | | NA | |
| | 0.889409773 | NA | |
| | 2.09E-09 | | 6.03E-08 |
| NA | | NA | |
| NA | | NA | |
| | 0.011130713 | | 0.036683074 |
| | 0.000419332 | | 0.002379974 |
| | 0.367123919 | | 0.531807781 |
| | 0.12579122 | | 0.244403592 |
| | 0.884293541 | | 0.934420955 |
| | 0.001226673 | | 0.00590713 |
| | 0.594564166 | | 0.731738715 |
| | 0.410287537 | | 0.573306015 |
| | 0.072165144 | | 0.160290954 |
| | 0.227318818 | | 0.377187088 |
| | 0.14740587 | | 0.275105006 |

| | | |
|---|---|---|
| | 0.336431794 | 0.499371756 |
| | 0.331006776 | 0.493387035 |
| | 0.417588266 | 0.579781945 |
| | 0.675468707 | 0.793739891 |
| | 0.284381248 | 0.443265745 |
| | 0.77164481 | NA |
| | 0.403227897 | 0.566723079 |
| | 0.19279066 | 0.335698208 |
| | 0.376338836 | 0.540733135 |
| | 0.403907342 | NA |
| | 0.032789868 | 0.087488786 |
| | 0.99205891 | NA |
| | 0.847356217 | 0.910374388 |
| | 0.755994417 | NA |
| NA | | NA |
| | 0.790020732 | NA |
| NA | | NA |
| NA | | NA |
| NA | | NA |
| | 0.299540336 | NA |
| | 0.529670737 | 0.678986557 |
| NA | | NA |
| NA | | NA |
| NA | | NA |
| NA | | NA |
| NA | | NA |
| | 0.696477781 | NA |
| NA | | NA |
| NA | | NA |
| | 0.774749143 | 0.864072143 |
| | 0.905158266 | NA |
| NA | | NA |
| NA | | NA |
| | 0.802764718 | 0.881307391 |
| | 7.82E-05 | 0.000567589 |
| | 0.248746777 | NA |
| | 0.240316536 | NA |
| | 0.888500738 | 0.936972948 |
| NA | | NA |
| | 0.034876955 | 0.091830103 |
| | 0.216770722 | NA |
| | 0.810064278 | 0.886200919 |
| | 0.005418988 | 0.020492883 |
| | 0.057583181 | 0.135136163 |
| | 0.539540726 | 0.687077564 |
| | 0.453495145 | 0.611539176 |
| | 0.894033697 | 0.940143949 |

| | | |
|---|---|---|
| | 0.324088933 | 0.486377043 |
| | 1.34E-09 | 4.07E-08 |
| | 0.260557062 | 0.415872404 |
| | 0.927663747 | 0.958569866 |
| | 0.000527729 | 0.002900727 |
| | 0.774739233 | NA |
| | 0.058285116 | 0.13641898 |
| | 0.892773624 | 0.939380965 |
| | 0.008207929 | 0.028666644 |
| | 0.014602968 | 0.045679001 |
| | 0.001541389 | 0.007164978 |
| | 0.212833187 | 0.359978737 |
| | 0.103382023 | 0.211305675 |
| | 1.52E-14 | 1.66E-12 |
| | 0.187329081 | 0.328759444 |
| | 0.792188023 | NA |
| | 0.000425873 | 0.002412346 |
| | 4.33E-06 | 4.71E-05 |
| | 0.101377907 | 0.208175461 |
| | 0.047779409 | 0.11712824 |
| | 1.07E-07 | 1.87E-06 |
| | 0.002981629 | 0.01249496 |
| NA | | NA |
| | 5.02E-05 | 0.000387945 |
| | 0.001453516 | 0.006833173 |
| | 9.72E-07 | 1.28E-05 |
| NA | | NA |
| | 0.807252815 | NA |
| | 0.49860666 | 0.651604098 |
| | 0.247702865 | 0.401306419 |
| | 0.738316799 | 0.83947075 |
| | 0.066870981 | 0.15141471 |
| | 4.05E-05 | 0.000323155 |
| | 0.598936041 | 0.73507917 |
| | 0.037394997 | 0.097017572 |
| | 0.000126158 | 0.000850045 |
| | 0.927160835 | 0.958351878 |
| | 0.130061126 | 0.250220104 |
| | 0.000242101 | 0.001489578 |
| | 0.367779347 | 0.532370909 |
| | 6.65E-06 | 6.85E-05 |
| | 0.339665551 | NA |
| | 0.721658969 | NA |
| | 0.313447285 | 0.475311714 |
| | 0.905158266 | NA |
| | 0.011111838 | 0.036637365 |
| | 0.502172948 | 0.654861676 |

| | |
|---|---|
| 0.789163412 | NA |
| 0.399228436 | 0.562794968 |
| 2.10E-06 | 2.50E-05 |
| 0.068940663 | NA |
| 0.991158452 | NA |
| 1.39E-13 | 1.20E-11 |
| 0.409534812 | 0.572655053 |
| 1.53E-05 | 0.000140457 |
| 5.14E-06 | 5.45E-05 |
| 0.101533055 | 0.208389975 |
| 0.369512467 | 0.534210564 |
| 0.259513529 | 0.414658923 |
| 0.182202338 | 0.322373793 |
| 0.030081149 | 0.081670618 |
| NA | NA |
| 0.001115567 | 0.005458276 |
| 0.003198496 | 0.013252032 |
| 2.31E-08 | 4.94E-07 |
| 3.96E-05 | 0.00031728 |
| NA | NA |
| 0.889409773 | NA |
| NA | NA |
| 0.276152064 | 0.433671113 |
| 0.945301969 | 0.969042261 |
| 0.128038609 | 0.247608967 |
| 6.99E-05 | 0.000515221 |
| 0.801661681 | NA |
| 0.000294907 | 0.001760133 |
| NA | NA |
| NA | NA |
| NA | NA |
| NA | NA |
| NA | NA |
| NA | NA |
| 0.156812599 | 0.288038542 |
| 0.862029991 | NA |
| NA | NA |
| 0.374310404 | 0.538311753 |
| 9.77E-07 | 1.28E-05 |
| 0.484827181 | 0.639918259 |
| NA | NA |
| 0.09326535 | 0.195179055 |
| 0.439444722 | 0.599699624 |
| NA | NA |
| 0.858609768 | NA |
| 0.429882862 | 0.590984647 |
| 5.92E-09 | 1.48E-07 |

| | |
|---|---|
| 0.294676924 | 0.454496354 |
| NA | NA |
| NA | NA |
| 1.37E-08 | 3.11E-07 |
| NA | NA |
| NA | NA |
| NA | NA |
| 0.949053595 | 0.971393115 |
| NA | NA |
| 0.009843649 | 0.033188497 |
| 0.623278772 | 0.754169581 |
| 0.000426958 | 0.002416382 |
| 7.73E-10 | 2.53E-08 |
| 0.178988647 | 0.318277732 |
| 0.002521544 | 0.010886862 |
| 5.82E-05 | 0.000441397 |
| 0.804603306 | 0.882580395 |
| 0.990161651 | NA |
| 0.052124355 | 0.125284447 |
| 0.001911837 | 0.008608168 |
| 0.126268994 | 0.24509333 |
| 0.062278672 | 0.143789019 |
| 0.065850306 | 0.149829746 |
| 0.68315851 | NA |
| 0.000880646 | 0.004468141 |
| 0.587464435 | 0.726253393 |
| NA | NA |
| 0.992060816 | NA |
| 0.805604645 | 0.883232933 |
| 0.159184551 | 0.291197397 |
| 0.7641578 | NA |
| 0.143060682 | 0.268915383 |
| 0.889409301 | NA |
| 0.002528705 | 0.010911346 |
| 1.98E-05 | 0.000176235 |
| 0.864528112 | NA |
| 0.545864031 | 0.692308436 |
| 0.447558054 | 0.606803455 |
| 0.917453945 | 0.952997691 |
| 0.83027301 | 0.898790487 |
| NA | NA |
| 3.64E-07 | 5.43E-06 |
| 0.016793747 | 0.051183051 |
| 0.00085214 | 0.004343582 |
| 2.17E-08 | 4.69E-07 |
| 0.500805396 | 0.653547133 |
| NA | NA |

| | | |
|---|---|---|
| | 0.677678174 | NA |
| | 0.682062926 | 0.798667936 |
| | 0.775345711 | NA |
| NA | | NA |
| | 0.059825602 | 0.139400509 |
| | 5.45E-05 | 0.000416482 |
| | 0.640799784 | 0.767071603 |
| | 0.751816375 | 0.848798755 |
| | 1.28E-05 | 0.000120461 |
| | 1.45E-07 | 2.44E-06 |
| | 0.983472088 | 0.991828033 |
| | 0.128552494 | 0.248296616 |
| | 0.922608602 | 0.955374089 |
| | 0.402556722 | 0.566073593 |
| | 0.889409773 | NA |
| | 0.12920088 | NA |
| | 0.001786267 | 0.008122297 |
| | 0.177245408 | 0.315933141 |
| | 0.02220452 | 0.063948624 |
| | 0.999682348 | 0.999864481 |
| | 0.108176779 | 0.218613463 |
| | 0.570785234 | NA |
| | 0.136306303 | 0.259124761 |
| NA | | NA |
| | 0.700480844 | 0.812908723 |
| | 0.549891726 | 0.695945556 |
| | 0.408968028 | 0.572117533 |
| | 0.619832348 | NA |
| NA | | NA |
| | 0.377254986 | 0.541624207 |
| | 4.40E-10 | 1.55E-08 |
| | 0.889409773 | NA |
| | 0.019398512 | 0.057354508 |
| | 0.797157316 | NA |
| | 0.077031901 | 0.168968486 |
| | 0.852276068 | NA |
| | 0.690671051 | 0.805274956 |
| | 0.281419924 | 0.439805003 |
| | 0.000760773 | 0.003960989 |
| | 3.41E-07 | 5.14E-06 |
| NA | | NA |
| | 0.957356758 | NA |
| | 0.159932703 | 0.29220396 |
| | 0.96177799 | 0.979084024 |
| | 0.949384627 | 0.971596077 |
| | 0.10628835 | 0.215531063 |
| | 0.011607523 | 0.037884898 |

| | |
|---|---|
| 0.004396424 | 0.017175073 |
| 0.33094435 | 0.493353544 |
| 0.98379608 | NA |
| 0.385201003 | 0.549083338 |
| 0.000203973 | 0.001282661 |
| 0.904947233 | NA |
| 0.635428607 | 0.762801169 |
| 0.000115154 | 0.000787643 |
| NA | NA |
| NA | NA |
| NA | NA |
| 0.967518832 | 0.982496569 |
| 0.889409301 | NA |
| 7.93E-10 | 2.58E-08 |
| 0.581560186 | NA |
| 0.051611201 | 0.124418747 |
| 0.037932949 | 0.098054105 |
| 0.530784587 | 0.680006466 |
| 0.889410245 | NA |
| 2.19E-06 | 2.60E-05 |
| 0.481156886 | 0.636800293 |
| 0.688483818 | 0.803658099 |
| 0.000686319 | 0.003633289 |
| 0.338179728 | 0.501119417 |
| 0.991782148 | NA |
| 0.971344042 | 0.984954128 |
| 0.130605962 | 0.251004405 |
| 0.042601445 | 0.107182071 |
| 0.021086139 | 0.061328545 |
| 0.80330859 | 0.881673503 |
| 7.40E-05 | 0.000540231 |
| NA | NA |
| 0.621971365 | 0.753205943 |
| 0.719805059 | 0.826239149 |
| 0.257226896 | 0.411984803 |
| 0.224350764 | 0.373986065 |
| 0.349100787 | 0.513181429 |
| 1.71E-06 | 2.08E-05 |
| 0.991782148 | NA |
| 0.015255418 | 0.047348938 |
| 0.532417891 | 0.681513522 |
| NA | NA |
| NA | NA |
| 0.007648886 | 0.027081892 |
| 0.225319677 | 0.375016283 |
| 0.005433289 | 0.020531679 |
| 0.03336586 | 0.08864863 |

| | |
|---|---|
| 0.198475953 | 0.342986814 |
| 0.57478641 | 0.716655393 |
| 0.841597343 | 0.906268931 |
| 0.829599385 | 0.898460579 |
| 0.523772787 | NA |
| 0.487276809 | 0.642035316 |
| 2.58E-23 | 2.52E-20 |
| 0.864539402 | 0.921887084 |
| 0.605981345 | NA |
| 0.451178705 | NA |
| 2.55E-07 | 4.00E-06 |
| 0.392862003 | 0.556347822 |
| 1.01E-05 | 9.79E-05 |
| 0.212498727 | NA |
| NA | NA |
| NA | NA |
| NA | NA |
| NA | NA |
| 0.107213393 | 0.216991602 |
| NA | NA |
| 0.236265546 | 0.388131602 |
| NA | NA |
| 0.475083303 | 0.631287226 |
| 0.761320525 | 0.85525059 |
| 0.187088783 | 0.328531195 |
| 0.000476097 | 0.00264473 |
| 0.005463379 | 0.020624092 |
| 0.217078231 | 0.365273629 |
| 0.199965425 | NA |
| 0.905158739 | NA |
| 0.471575278 | NA |
| 0.000925498 | 0.004666594 |
| 0.119115176 | 0.234819118 |
| 0.611618623 | NA |
| 0.896086954 | NA |
| 0.78861062 | 0.872441967 |
| 0.991937619 | NA |
| 0.564067216 | 0.707670228 |
| 0.858455358 | NA |
| 0.889410717 | NA |
| 0.1876276 | 0.329135202 |
| 0.879507115 | 0.93169457 |
| 0.006938452 | 0.02510487 |
| 0.833607373 | 0.900845684 |
| 6.54E-13 | 4.89E-11 |
| 0.591278844 | 0.729063515 |
| 0.575387549 | 0.717201294 |

| | |
|---|---|
| NA | NA |
| 0.950622466 | 0.972436561 |
| 0.90515732 | NA |
| 0.776822149 | NA |
| 0.515428602 | 0.66666861 |
| 0.812711122 | NA |
| 0.397485715 | NA |
| 0.992061292 | NA |
| 0.992061769 | NA |
| 0.593585625 | NA |
| 0.58476742 | NA |
| 0.371162767 | 0.535573874 |
| 0.310666641 | NA |
| 0.000171705 | 0.001104736 |
| 7.95E-09 | 1.93E-07 |
| NA | NA |
| 0.191288086 | 0.333637417 |
| NA | NA |
| 0.430839415 | 0.591780992 |
| 0.782559532 | NA |
| NA | NA |
| 0.764844882 | NA |
| 7.25E-10 | 2.38E-08 |
| 0.004509063 | 0.017552653 |
| 0.00850648 | 0.029518615 |
| NA | NA |
| 0.522324101 | 0.6726711 |
| 0.372284601 | 0.536663662 |
| 0.992061292 | NA |
| 1.01E-07 | 1.77E-06 |
| 0.430403814 | 0.591448677 |
| 3.54E-07 | 5.30E-06 |
| 0.449306298 | NA |
| 0.305876667 | NA |
| 0.663522389 | 0.784910205 |
| 6.47E-05 | 0.000482141 |
| 0.992061292 | NA |
| 0.677058425 | NA |
| 6.35E-06 | 6.58E-05 |
| 0.027873315 | 0.076831178 |
| 0.00141178 | 0.006678429 |
| 0.662118484 | NA |
| 0.006068879 | 0.022473273 |
| 0.105105137 | 0.213902105 |
| 0.681708866 | NA |
| 0.207287907 | NA |
| 0.801931434 | NA |

| | |
|---|---|
| NA | NA |
| 0.005093224 | 0.019460956 |
| 0.000848922 | 0.004330471 |
| 0.709302452 | 0.81881887 |
| 0.349157435 | 0.513195991 |
| 0.906111259 | 0.947129528 |
| 0.121511802 | 0.238281628 |
| 0.889409301 | NA |
| 0.450519072 | 0.608909233 |
| 1.06E-06 | 1.37E-05 |
| NA | NA |
| 0.978824139 | 0.989230452 |
| 0.940584839 | 0.966237953 |
| 0.204263044 | 0.350094613 |
| 0.992061292 | NA |
| 0.992063544 | NA |
| 0.159332007 | NA |
| NA | NA |
| 0.61577929 | 0.748596437 |
| 0.797157316 | NA |
| NA | NA |
| NA | NA |
| NA | NA |
| NA | NA |
| 0.025916339 | 0.072561123 |
| NA | NA |
| 0.844408486 | NA |
| 0.114628465 | 0.22831079 |
| 0.440584106 | 0.600569926 |
| 0.000778141 | 0.004031903 |
| 9.86E-10 | 3.11E-08 |
| 0.901183732 | NA |
| 0.550413652 | 0.696228882 |
| NA | NA |
| 0.014101594 | 0.044388891 |
| NA | NA |
| 0.807252815 | NA |
| 0.992059387 | NA |
| 2.95E-05 | 0.000247066 |
| 0.790381637 | 0.87347712 |
| 0.59600142 | 0.732993289 |
| 0.877209064 | NA |
| 0.034921533 | 0.091914412 |
| NA | NA |
| NA | NA |
| NA | NA |
| 0.299409782 | 0.459707692 |

| | | |
|---|---|---|
| | 0.001945509 | 0.008743641 |
| | 0.802548925 | NA |
| | 0.68415513 | 0.800093865 |
| | 0.263978905 | 0.41971261 |
| | 0.447441742 | NA |
| | 2.04E-07 | 3.30E-06 |
| | 0.995576259 | 0.997694248 |
| | 0.757514005 | 0.852907247 |
| | 0.002335665 | 0.010208772 |
| | 0.116397265 | 0.230953967 |
| | 0.067180666 | 0.151802865 |
| | 0.189965625 | 0.331910818 |
| NA | | NA |
| | 0.093582945 | 0.195769066 |
| | 0.136258929 | 0.259057127 |
| | 0.139997387 | 0.264493041 |
| | 0.233411072 | NA |
| | 0.050755524 | 0.122827921 |
| NA | | NA |
| | 0.739755581 | 0.840206898 |
| | 0.725016266 | 0.830297927 |
| NA | | NA |
| | 0.889409301 | NA |
| | 0.180574137 | 0.320448048 |
| | 7.74E-07 | 1.05E-05 |
| | 0.011030822 | 0.036435895 |
| | 0.001861712 | 0.008420458 |
| | 0.812075898 | NA |
| NA | | NA |
| | 0.381793846 | 0.545872145 |
| | 0.164849157 | NA |
| | 0.443656824 | 0.60327328 |
| NA | | NA |
| NA | | NA |
| | 0.004250429 | 0.016717746 |
| | 0.587011253 | NA |
| | 8.66E-07 | 1.16E-05 |
| NA | | NA |
| | 0.905158739 | NA |
| | 0.002251677 | 0.009886937 |
| NA | | NA |
| | 0.540720215 | 0.688089662 |
| NA | | NA |
| NA | | NA |
| NA | | NA |
| | 0.034815013 | 0.091690991 |
| | 0.015918503 | 0.049025863 |

| | |
|---|---|
| 0.000330104 | 0.001940762 |
| 0.59604596 | 0.733007014 |
| 0.868092066 | 0.924373662 |
| 8.89E-13 | 6.36E-11 |
| NA | NA |
| 0.269638069 | NA |
| 0.75755762 | 0.852912622 |
| 0.284758284 | 0.44360153 |
| 0.141062834 | 0.266002472 |
| 0.028577654 | 0.078370821 |
| 0.740629448 | NA |
| 0.153783276 | 0.284037931 |
| 0.256363718 | 0.410941739 |
| 0.905158266 | NA |
| NA | NA |
| 0.06183971 | 0.143060327 |
| NA | NA |
| 0.99205891 | NA |
| 0.869139962 | NA |
| 0.000110024 | 0.000757137 |
| 0.122200747 | 0.239208074 |
| 0.000490988 | 0.002721256 |
| 0.539931515 | 0.687217102 |
| 0.34476038 | 0.508093502 |
| 0.001395287 | 0.006605105 |
| 0.90515732 | NA |
| 0.072073275 | 0.160123132 |
| 0.000172972 | 0.001111914 |
| 0.047058646 | 0.115627257 |
| 0.662829333 | 0.784259352 |
| 0.800834704 | NA |
| 0.996365387 | 0.998242688 |
| 0.808635204 | NA |
| NA | NA |
| 0.360163203 | NA |
| 0.905158739 | NA |
| 0.902889488 | 0.945066507 |
| 0.385974052 | 0.549792697 |
| 0.157926407 | 0.289571307 |
| 0.040788822 | 0.103475248 |
| 1.83E-29 | 1.00E-25 |
| 0.734308205 | NA |
| 0.402363319 | 0.565942617 |
| 0.365429307 | 0.530262483 |
| 0.655386544 | NA |
| NA | NA |
| 0.0758938 | 0.166955716 |

| | |
|---|---|
| 0.995098388 | 0.997551495 |
| 0.667378361 | 0.787772463 |
| 9.99E-05 | 0.000697677 |
| 0.526254264 | 0.67622549 |
| 0.25663459 | 0.411225844 |
| 0.495214381 | 0.648521744 |
| 2.15E-12 | 1.41E-10 |
| 0.081550081 | 0.176516335 |
| 0.29715057 | 0.457230003 |
| 0.001808358 | 0.008211277 |
| 0.015863743 | 0.048901823 |
| 0.889409301 | NA |
| 0.934362802 | NA |
| 0.551432518 | 0.697075969 |
| NA | NA |
| 0.00033937 | 0.001989911 |
| 0.229051779 | 0.379546334 |
| NA | NA |
| 0.562266481 | 0.70614971 |
| NA | NA |
| 0.905158266 | NA |
| NA | NA |
| NA | NA |
| 0.992060816 | NA |
| NA | NA |
| NA | NA |
| 0.889850811 | 0.937671223 |
| 0.733088411 | 0.836036791 |
| NA | NA |
| 0.005339152 | 0.020253146 |
| NA | NA |
| 0.180715703 | NA |
| NA | NA |
| 0.859980034 | NA |
| 0.00135421 | 0.006437996 |
| 0.769104712 | NA |
| 1.30E-10 | 5.40E-09 |
| 0.732725292 | 0.835736895 |
| 0.003417189 | 0.013978773 |
| 1.07E-08 | 2.51E-07 |
| 0.83815308 | 0.903800996 |
| 0.291519858 | 0.451031111 |
| 0.098327728 | 0.203274202 |
| NA | NA |
| 0.98954406 | NA |
| 0.014343972 | 0.044983445 |
| 0.992061292 | NA |

| | |
|---|---|
| NA | NA |
| 0.798047689 | 0.878104385 |
| 0.105719392 | 0.214773997 |
| 0.971464805 | NA |
| 0.001103657 | 0.005409645 |
| 4.29E-09 | 1.11E-07 |
| NA | NA |
| 0.017406963 | 0.052599352 |
| 0.057640839 | 0.135213672 |
| 8.87E-05 | 0.000629315 |
| NA | NA |
| NA | NA |
| 0.000115417 | 0.000789052 |
| 0.22336466 | 0.372936783 |
| 4.10E-05 | 0.00032661 |
| 0.100419971 | 0.206646251 |
| 0.990927264 | 0.995364126 |
| 0.819964581 | 0.892909192 |
| 0.312407306 | 0.474160356 |
| 0.61444923 | NA |
| 0.32390222 | 0.486163285 |
| 0.170241841 | 0.306438842 |
| 0.905158739 | NA |
| 0.025480849 | 0.071643273 |
| 9.46E-05 | 0.000666141 |
| 2.43E-11 | 1.22E-09 |
| 0.889409301 | NA |
| 0.408288516 | NA |
| 0.426688168 | 0.587917768 |
| 0.390951401 | NA |
| 0.00416856 | 0.016451728 |
| 1.49E-05 | 0.000137538 |
| 0.020273932 | 0.059414823 |
| 2.99E-05 | 0.00024907 |
| 0.059309991 | 0.13838998 |
| NA | NA |
| NA | NA |
| NA | NA |
| 0.631155003 | 0.759052123 |
| NA | NA |
| 0.000463873 | 0.002587956 |
| 0.621935397 | NA |
| NA | NA |
| 0.848180693 | NA |
| 0.048376669 | 0.118204436 |
| NA | NA |
| 0.988120101 | 0.993639148 |

| | |
|---|---|
| 0.791051906 | 0.873953955 |
| NA | NA |
| 0.479615068 | 0.635215156 |
| 0.617354016 | 0.749805135 |
| 0.462114808 | 0.619510322 |
| 3.42E-11 | 1.64E-09 |
| 0.958966884 | 0.977249651 |
| 0.028692263 | 0.078638717 |
| 0.864348207 | 0.921826781 |
| 0.528322688 | 0.678010514 |
| 0.889409301 | NA |
| 0.315400296 | 0.477679501 |
| 0.246938341 | 0.400422319 |
| 0.06295486 | 0.144908361 |
| 0.575312434 | NA |
| 0.943502397 | NA |
| 0.889409773 | NA |
| 0.836544856 | 0.90277612 |
| 0.992060816 | NA |
| 0.905158266 | NA |
| 0.50029982 | 0.653350208 |
| 0.009294702 | 0.031692915 |
| NA | NA |
| 0.000344243 | 0.002015798 |
| 5.98E-11 | 2.69E-09 |
| 0.321373049 | 0.48375419 |
| NA | NA |
| 0.565438223 | NA |
| 0.255720219 | NA |
| 0.889409301 | NA |
| 0.004372863 | 0.017098236 |
| NA | NA |
| 0.255149975 | 0.409735675 |
| 1.71E-08 | 3.78E-07 |
| NA | NA |
| 0.017716717 | 0.053328936 |
| 0.79581308 | 0.876831882 |
| 0.577043383 | NA |
| 0.352169478 | 0.516413327 |
| 0.717625932 | 0.824742128 |
| 0.000125097 | 0.000845235 |
| NA | NA |
| NA | NA |
| NA | NA |
| 0.698366912 | NA |
| 0.000819815 | 0.004217973 |
| 0.670417691 | 0.789836298 |

| | | |
|---|---|---|
| | 0.011318661 | 0.037151941 |
| | 0.006163619 | 0.022762682 |
| | 0.003127432 | 0.013004218 |
| | 0.861970235 | NA |
| | 0.534552682 | NA |
| | 0.549904428 | 0.695945556 |
| | 0.083381157 | 0.179492743 |
| | 7.47E-05 | 0.00054447 |
| | 0.093594867 | 0.195775356 |
| | 0.901616885 | 0.944380971 |
| NA | | NA |
| NA | | NA |
| | 0.787273705 | 0.871771467 |
| | 0.621082648 | NA |
| | 0.583321518 | NA |
| | 0.717560555 | 0.824710155 |
| | 0.6938221 | 0.807442869 |
| | 0.782782338 | NA |
| | 0.002283745 | 0.010009731 |
| | 0.170169395 | 0.306366299 |
| | 7.47E-06 | 7.57E-05 |
| | 0.002694963 | 0.011517846 |
| | 0.617738806 | 0.75015919 |
| | 0.062137348 | 0.143568395 |
| | 2.64E-08 | 5.55E-07 |
| | 0.311904157 | 0.473560354 |
| | 0.459158555 | 0.616412711 |
| | 0.852251993 | NA |
| | 0.830353458 | 0.898833264 |
| | 0.539174146 | 0.686798832 |
| NA | | NA |
| NA | | NA |
| NA | | NA |
| | 0.992060816 | NA |
| | 0.001056255 | 0.005211032 |
| | 0.623885596 | 0.754649323 |
| | 0.919993634 | 0.954236465 |
| | 0.69838822 | NA |
| | 0.121004497 | 0.237647595 |
| | 0.416626132 | 0.578884664 |
| | 0.855223421 | 0.915732939 |
| | 0.012429775 | 0.040091868 |
| | 0.655781319 | NA |
| | 0.002700716 | 0.011531555 |
| | 8.95E-06 | 8.87E-05 |
| NA | | NA |
| | 0.014891161 | 0.046395431 |

| | |
|---|---|
| 1.09E-05 | 0.000104766 |
| 0.449572026 | 0.608453136 |
| NA | NA |
| 0.19712454 | 0.341304152 |
| 0.14005201 | 0.264573477 |
| 0.127180976 | 0.246471366 |
| 0.072757192 | 0.161307039 |
| 0.004224253 | 0.016632664 |
| 3.44E-10 | 1.25E-08 |
| 0.116684402 | 0.231361886 |
| 0.21885197 | 0.367554143 |
| 0.960055346 | 0.977731905 |
| 0.739858324 | 0.840266301 |
| 0.787435399 | 0.871771467 |
| 0.000206739 | 0.00129745 |
| 0.270194149 | NA |
| 0.001387509 | 0.00657495 |
| 0.076409369 | 0.167820899 |
| 0.390777357 | 0.554118558 |
| 0.85082022 | 0.912565539 |
| 0.009091551 | 0.031125877 |
| 0.000164371 | 0.001064413 |
| 0.339551848 | 0.502609979 |
| 0.647148707 | 0.772110769 |
| 6.23E-07 | 8.66E-06 |
| 0.005492223 | 0.020715172 |
| 0.905158266 | NA |
| 0.421796537 | NA |
| 0.010802817 | 0.035828283 |
| 0.383527441 | 0.547512277 |
| 0.72648771 | 0.831171869 |
| 0.265679001 | 0.421839855 |
| 0.0675299 | 0.152419474 |
| 0.940955558 | 0.966437938 |
| 0.070847648 | 0.158071887 |
| 0.764083386 | 0.857062789 |
| 0.016620205 | 0.050747598 |
| NA | NA |
| 0.008724203 | 0.030140775 |
| NA | NA |
| 0.157601142 | 0.289143911 |
| 0.494128742 | 0.647447523 |
| 0.449010334 | 0.607917747 |
| 0.057209625 | 0.134460683 |
| 0.074634459 | 0.164597577 |
| 0.779828374 | 0.867445989 |
| 0.135446912 | 0.257982369 |

| | |
|---|---|
| 0.110460684 | 0.221901578 |
| 0.038293174 | 0.098810787 |
| 5.03E-05 | 0.000388577 |
| 0.162136302 | 0.295317773 |
| 0.830163713 | NA |
| 0.151408301 | 0.280737556 |
| 0.023113703 | 0.066087736 |
| 0.001542711 | 0.00716809 |
| 0.285610313 | 0.444604356 |
| 0.890989014 | 0.938380229 |
| 0.001858905 | 0.008411227 |
| 2.79E-05 | 0.000235429 |
| 0.005796343 | 0.021676073 |
| 0.00714576 | 0.025656271 |
| 5.06E-05 | 0.000390419 |
| 0.013084615 | 0.041719915 |
| 1.64E-05 | 0.000149803 |
| 0.671120677 | NA |
| 0.01811561 | 0.054285028 |
| 3.01E-06 | 3.42E-05 |
| 2.16E-05 | 0.000190172 |
| 4.84E-05 | 0.000377038 |
| 0.006006277 | 0.022309172 |
| 0.000969367 | 0.004848825 |
| 9.56E-05 | 0.000672526 |
| 6.27E-05 | 0.000469842 |
| 0.000125926 | 0.000849264 |
| 0.019568094 | 0.057762573 |
| 0.009216431 | 0.031474901 |
| 1.32E-05 | 0.000123794 |
| 0.889409773 | NA |
| 0.093438782 | 0.195504737 |
| 0.000282345 | 0.001701432 |
| 0.558356743 | 0.703157408 |
| 0.126542934 | 0.245494857 |
| 0.454250457 | 0.6122191 |
| 0.905158266 | NA |
| 0.000536251 | 0.002938742 |
| 0.102888895 | 0.210582277 |
| 0.419469892 | 0.581364515 |
| 0.019070462 | 0.056506313 |
| 0.030870376 | 0.083329145 |
| 0.316426507 | 0.478514732 |
| 0.070108816 | 0.156717246 |
| 0.141740881 | 0.267028827 |
| 0.758336722 | NA |
| 0.066216978 | 0.150430233 |

| | |
|---|---|
| 0.598538023 | 0.734793204 |
| 0.001611269 | 0.0074316 |
| 0.876005781 | 0.929106016 |
| 0.105961793 | 0.215087356 |
| 0.492344724 | 0.646460045 |
| 0.755918728 | 0.851591389 |
| 0.475367614 | 0.631532303 |
| 0.165974464 | 0.300439638 |
| 0.949531401 | 0.971700999 |
| 0.136404193 | 0.259288407 |
| 0.027766996 | 0.076628814 |
| 0.173311673 | 0.310511076 |
| 0.858009656 | 0.917175946 |
| 0.025906312 | 0.072542297 |
| NA | NA |
| 0.581889104 | 0.721863332 |
| 6.90E-06 | 7.07E-05 |
| 0.453878306 | NA |
| 0.28699813 | 0.446014999 |
| 0.000306791 | 0.001820273 |
| 7.29E-07 | 1.00E-05 |
| 0.484712606 | 0.639882417 |
| NA | NA |
| NA | NA |
| 0.107354063 | 0.217170692 |
| 0.494892213 | NA |
| 0.418299872 | 0.580512349 |
| 0.80622407 | NA |
| 0.101318612 | 0.208085989 |
| 0.116750477 | 0.231405258 |
| 0.01281007 | 0.041071153 |
| 0.093489585 | 0.195592398 |
| 3.69E-25 | 5.06E-22 |
| NA | NA |
| 0.817593383 | 0.891480863 |
| 0.00502948 | 0.019263686 |
| NA | NA |
| NA | NA |
| 0.001179619 | 0.005728275 |
| 0.855356565 | NA |
| NA | NA |
| 0.889409773 | NA |
| 0.257932359 | 0.412753604 |
| 0.000392233 | 0.002248247 |
| 0.889410245 | NA |
| NA | NA |
| 0.004739388 | 0.018329027 |

| | | |
|---|---|---|
| | 0.484491921 | 0.639744924 |
| | 0.905158266 | NA |
| | 8.48E-05 | 0.00060539 |
| | 5.85E-05 | 0.000443394 |
| | 0.001217461 | 0.005873073 |
| | 0.789108266 | NA |
| NA | | NA |
| | 0.905158266 | NA |
| NA | | NA |
| | 0.032582402 | 0.087019822 |
| | 0.990152369 | NA |
| | 0.678403604 | 0.795978181 |
| | 0.31219219 | NA |
| | 0.133542443 | NA |
| | 9.64E-06 | 9.42E-05 |
| | 0.499746841 | NA |
| | 0.679591283 | 0.796921125 |
| NA | | NA |
| | 0.071419373 | NA |
| | 0.647626621 | NA |
| | 0.111817604 | 0.22381934 |
| | 0.243572707 | 0.396595896 |
| | 2.14E-09 | 6.19E-08 |
| | 0.027421381 | 0.075827491 |
| | 0.889409773 | NA |
| NA | | NA |
| | 0.835770959 | 0.902244262 |
| | 0.072002131 | 0.160013642 |
| | 0.004692247 | 0.01818836 |
| | 0.680793109 | 0.797606354 |
| | 0.000265046 | 0.001608217 |
| NA | | NA |
| | 1.07E-05 | 0.000102913 |
| NA | | NA |
| NA | | NA |
| | 0.992060816 | NA |
| | 0.70853139 | NA |
| | 0.466983843 | NA |
| NA | | NA |
| NA | | NA |
| NA | | NA |
| NA | | NA |
| NA | | NA |
| NA | | NA |
| | 0.992060816 | NA |
| NA | | NA |
| | 0.64426466 | NA |

| | | |
|---|---|---|
| | 0.508882302 | NA |
| | 0.683356112 | 0.799628222 |
| | 0.296636051 | 0.456817224 |
| | 0.043095594 | 0.108096855 |
| NA | | NA |
| | 0.039158124 | 0.100475957 |
| NA | | NA |
| | 0.000661643 | 0.003528175 |
| | 0.391028683 | NA |
| | 0.841525686 | 0.906236209 |
| NA | | NA |
| | 0.395986723 | NA |
| | 0.573899487 | 0.715915403 |
| | 0.048972807 | NA |
| | 0.005601114 | 0.021064372 |
| | 0.013147592 | 0.041861094 |
| | 0.920904195 | NA |
| | 0.583810609 | 0.723639366 |
| | 0.085779648 | NA |
| | 0.018998861 | 0.05633977 |
| | 0.146541499 | 0.273840931 |
| | 0.060823771 | 0.141242687 |
| | 0.182764401 | 0.322951918 |
| | 0.035359214 | 0.092766599 |
| | 5.05E-05 | 0.000390317 |
| | 0.000217017 | 0.001353081 |
| | 0.157735534 | 0.289317958 |
| NA | | NA |
| | 0.746510294 | NA |
| | 0.765447758 | 0.857749915 |
| NA | | NA |
| NA | | NA |
| | 0.645095827 | 0.770418315 |
| | 0.005939409 | 0.022086956 |
| NA | | NA |
| NA | | NA |
| NA | | NA |
| | 0.107655586 | 0.217680387 |
| | 0.73571322 | 0.83776844 |
| NA | | NA |
| | 0.464489611 | 0.621555598 |
| | 2.89E-06 | 3.30E-05 |
| | 0.494728575 | 0.648001478 |
| | 0.184161315 | 0.324766971 |
| | 1.84E-11 | 9.63E-10 |
| NA | | NA |
| NA | | NA |

| | | |
|---|---|---|
| | 0.905157793 | NA |
| | 0.000611747 | 0.003294102 |
| NA | | NA |
| NA | | NA |
| | 0.47858625 | 0.63442653 |
| NA | | NA |
| NA | | NA |
| | 0.224264194 | 0.373926913 |
| NA | | NA |
| | 0.125629644 | 0.24417608 |
| | 0.032444397 | 0.086767327 |
| | 0.797342596 | NA |
| NA | | NA |
| | 0.042411045 | 0.10683769 |
| | 1.62E-05 | 0.000147909 |
| | 0.468542629 | 0.624960675 |
| | 0.000861525 | 0.004387046 |
| | 0.034526147 | 0.09110306 |
| | 0.006905991 | 0.025016273 |
| | 0.457424239 | 0.614911471 |
| | 0.273832037 | 0.430952319 |
| | 0.522014058 | NA |
| | 0.348000549 | 0.511803902 |
| | 0.151287782 | NA |
| | 0.072274146 | 0.160390901 |
| NA | | NA |
| | 0.566229676 | NA |
| NA | | NA |
| NA | | NA |
| | 0.681223676 | NA |
| | 0.981136663 | NA |
| NA | | NA |
| | 0.829913943 | NA |
| NA | | NA |
| NA | | NA |
| | 0.992060816 | NA |
| NA | | NA |
| NA | | NA |
| | 0.006982352 | 0.025205567 |
| | 0.053089605 | 0.126951392 |
| NA | | NA |
| | 0.711738065 | 0.820520994 |
| | 0.242187366 | 0.394934825 |
| | 0.000691372 | 0.003653874 |
| | 0.369797811 | 0.534411987 |
| | 0.266179981 | NA |
| | 0.026751391 | 0.074291148 |

| | | |
|---|---|---|
| | 0.004092831 | 0.016202357 |
| | 0.003564177 | 0.014464195 |
| | 0.465396797 | 0.622238692 |
| | 0.988209178 | 0.993639148 |
| | 0.695805244 | NA |
| | 0.446076633 | 0.605513803 |
| NA | | NA |
| | 0.992061292 | NA |
| | 0.927071646 | 0.958331915 |
| | 0.493207096 | 0.646928273 |
| | 0.103772756 | 0.211741645 |
| NA | | NA |
| | 0.100016565 | 0.205989848 |
| NA | | NA |
| | 0.964506831 | 0.980537292 |
| | 0.2913909 | 0.450884382 |
| | 0.003129816 | 0.013011666 |
| | 0.225260784 | 0.374990047 |
| NA | | NA |
| NA | | NA |
| NA | | NA |
| | 3.67E-05 | 0.00029746 |
| | 0.083162578 | 0.179158938 |
| | 0.003149814 | 0.013084899 |
| | 1.98E-05 | 0.000176197 |
| | 3.24E-05 | 0.000267429 |
| | 0.706592681 | NA |
| | 4.04E-06 | 4.43E-05 |
| | 0.26183732 | 0.417262333 |
| | 0.027647865 | 0.076342552 |
| | 0.563352767 | 0.707056665 |
| | 0.889409773 | NA |
| | 0.649890186 | 0.774584162 |
| | 0.007728092 | 0.027279804 |
| | 0.267351461 | 0.423637662 |
| | 0.904947233 | NA |
| | 0.232768814 | 0.384169139 |
| | 0.889409773 | NA |
| | 1.87E-06 | 2.26E-05 |
| | 0.061722779 | 0.142851023 |
| | 1.06E-16 | 1.93E-14 |
| | 0.182853325 | 0.323031068 |
| | 0.06615231 | 0.150329939 |
| | 0.000387374 | 0.0022233 |
| | 0.983335579 | NA |
| | 7.99E-06 | 8.03E-05 |
| | 0.004848444 | 0.018684974 |

| | |
|---|---|
| NA | NA |
| 0.732582998 | 0.835663333 |
| 0.2785039 | 0.436583891 |
| 0.0081842 | 0.028599117 |
| 0.455965857 | 0.613799312 |
| 0.688841802 | 0.803905035 |
| 0.009403956 | 0.031981023 |
| NA | NA |
| 0.790941328 | 0.873919725 |
| 0.905158739 | NA |
| 0.040801284 | 0.103494905 |
| 0.033814423 | 0.089580276 |
| 0.122309436 | 0.239339688 |
| 0.652356559 | NA |
| 0.776244828 | NA |
| 0.10581684 | 0.214861869 |
| 0.671944007 | 0.79118324 |
| 0.787895951 | NA |
| 0.665531556 | 0.786521343 |
| 0.634948186 | 0.762361377 |
| 0.230356866 | 0.381097568 |
| 0.002978514 | 0.012491284 |
| 0.182839021 | 0.323031068 |
| NA | NA |
| 0.03870234 | 0.099539082 |
| 0.167539865 | 0.302649547 |
| 0.33115889 | NA |
| 0.0180094 | 0.05402574 |
| 0.058668691 | 0.137141345 |
| 0.554828154 | 0.70028 |
| 0.801661681 | NA |
| 0.345964897 | NA |
| 0.262425513 | 0.417912963 |
| 0.465986933 | 0.622800186 |
| 0.030212938 | 0.081906901 |
| 0.000299184 | 0.001781878 |
| 0.031142772 | 0.083899415 |
| NA | NA |
| NA | NA |
| NA | NA |
| NA | NA |
| NA | NA |
| 0.002483831 | 0.010758002 |
| 2.12E-12 | 1.40E-10 |
| NA | NA |
| 0.509059617 | NA |
| 4.04E-06 | 4.43E-05 |

| | |
|---|---|
| 0.253010271 | 0.407367275 |
| 0.305557523 | 0.466148232 |
| 0.037284164 | 0.096781306 |
| 0.001215984 | 0.005867235 |
| 4.77E-05 | 0.000372447 |
| NA | NA |
| NA | NA |
| 0.061952116 | 0.143245921 |
| 0.905158266 | NA |
| 0.788587404 | 0.872441967 |
| 0.905158266 | NA |
| 0.379917577 | 0.544304363 |
| 0.419404244 | 0.581346913 |
| 0.68670587 | 0.802215354 |
| 8.78E-05 | 0.000624465 |
| 0.554041668 | 0.699478241 |
| 0.376944594 | 0.541355549 |
| NA | NA |
| 0.457181338 | NA |
| NA | NA |
| 0.71589052 | NA |
| 2.54E-06 | 2.96E-05 |
| 5.16E-08 | 9.98E-07 |
| 0.988153933 | NA |
| 0.103452031 | NA |
| 3.87E-06 | 4.28E-05 |
| NA | NA |
| NA | NA |
| 0.069006218 | 0.154724324 |
| 0.007739364 | 0.027309771 |
| 0.800259511 | NA |
| 0.450115931 | 0.608829607 |
| 0.197066795 | 0.341276794 |
| 0.082326144 | 0.177880528 |
| 0.000777829 | 0.004031903 |
| 1.99E-07 | 3.23E-06 |
| 0.344196748 | 0.507535347 |
| NA | NA |
| 4.42E-14 | 4.25E-12 |
| 1.05E-05 | 0.000101288 |
| 0.788582525 | NA |
| 0.692365619 | 0.806431689 |
| 0.112085125 | 0.22430304 |
| 0.088315689 | 0.187519262 |
| 0.162781756 | 0.296100445 |
| 0.152975324 | 0.282997906 |
| 0.001296621 | 0.006192365 |

| | |
|---|---|
| 0.700018811 | 0.812630211 |
| 0.000101362 | 0.000707056 |
| NA | NA |
| 6.82E-05 | 0.000504424 |
| 0.161077942 | 0.293731153 |
| 0.883570623 | NA |
| NA | NA |
| 0.883047992 | NA |
| NA | NA |
| 0.753690661 | NA |
| NA | NA |
| 0.046499313 | 0.114581733 |
| 0.037545564 | 0.097327712 |
| NA | NA |
| 0.31335038 | 0.475269566 |
| 0.889409301 | NA |
| 0.954271069 | NA |
| 0.046355197 | 0.114346637 |
| 0.889409301 | NA |
| 0.000656233 | 0.003502729 |
| 0.697634717 | 0.810911078 |
| 0.25628244 | 0.410871438 |
| 0.484541896 | NA |
| 3.05E-05 | 0.000253209 |
| 3.31E-07 | 5.01E-06 |
| NA | NA |
| NA | NA |
| 1.06E-10 | 4.47E-09 |
| NA | NA |
| 0.992061292 | NA |
| 0.015196962 | 0.047194185 |
| 5.21E-05 | 0.000400377 |
| 0.004735577 | 0.018320743 |
| 0.108930687 | 0.219793159 |
| 0.081842082 | 0.177026229 |
| 0.000130022 | 0.000871534 |
| 0.080426948 | 0.17467069 |
| 0.731752444 | 0.834990227 |
| 9.52E-05 | 0.00067007 |
| 0.381805766 | 0.545872145 |
| 0.212612633 | 0.359771966 |
| 0.532823206 | 0.681776847 |
| 0.000134921 | 0.000900526 |
| NA | NA |
| 0.922130131 | 0.955294241 |
| 0.261738783 | 0.4171824 |
| 0.754471586 | 0.850659353 |

| | |
|---|---|
| NA | NA |
| NA | NA |
| 5.48E-06 | 5.76E-05 |
| 0.047128906 | 0.115761034 |
| 0.846927881 | NA |
| 0.430641304 | 0.591599581 |
| 0.142356505 | 0.267843928 |
| 0.298029596 | 0.457974534 |
| 0.002877088 | 0.012139412 |
| 0.800082892 | NA |
| NA | NA |
| NA | NA |
| 0.930923658 | 0.96015743 |
| 1.13E-08 | 2.63E-07 |
| 2.01E-07 | 3.25E-06 |
| 0.553707766 | 0.699187397 |
| NA | NA |
| 4.37E-08 | 8.67E-07 |
| NA | NA |
| 0.248825605 | 0.402603242 |
| 0.708832995 | 0.818449034 |
| 0.121337795 | 0.238009803 |
| 0.742232431 | 0.842044611 |
| 0.416151372 | 0.57848123 |
| 0.096489781 | 0.200474888 |
| 0.514641009 | 0.665868348 |
| 0.054897087 | 0.130294545 |
| NA | NA |
| 0.201996671 | 0.347413317 |
| 0.001839057 | 0.008335162 |
| 0.48611713 | 0.64097477 |
| NA | NA |
| 0.969271239 | 0.983776228 |
| 0.364588847 | 0.529392746 |
| 0.597863507 | 0.734409715 |
| 0.12648969 | 0.245413244 |
| 0.458940968 | 0.616195947 |
| 0.004013247 | 0.015936148 |
| 0.905157793 | NA |
| 0.101381137 | 0.208175461 |
| 0.842246309 | NA |
| 0.090657294 | 0.191178673 |
| 0.312549036 | 0.474341554 |
| 0.090450785 | 0.190944894 |
| NA | NA |
| NA | NA |
| 0.240774585 | 0.393419831 |

| | |
|---|---|
| NA | NA |
| 0.801661681 | NA |
| 0.793253454 | 0.875285292 |
| 0.76987628 | NA |
| 0.001769808 | 0.008056796 |
| 0.206773709 | 0.353376655 |
| 0.826974574 | 0.89672859 |
| 1.66E-07 | 2.75E-06 |
| 0.191509628 | 0.333944248 |
| 0.01908744 | 0.056533733 |
| 2.19E-07 | 3.50E-06 |
| 0.00756814 | 0.026850245 |
| 0.920775321 | 0.954551282 |
| 0.055957106 | 0.132139166 |
| 0.80099795 | 0.88007175 |
| 0.000180618 | 0.001154307 |
| 0.811008456 | 0.886943308 |
| 0.308863665 | NA |
| 0.029157543 | 0.079615828 |
| 0.068918811 | NA |
| 0.273076107 | 0.430009195 |
| NA | NA |
| 0.889409301 | NA |
| NA | NA |
| NA | NA |
| 0.181315624 | 0.321426478 |
| NA | NA |
| 0.013620241 | 0.043114727 |
| 0.038544403 | 0.099260766 |
| 0.073875256 | NA |
| 0.07812605 | 0.170792998 |
| 0.003970434 | 0.015791842 |
| 0.805680227 | NA |
| 0.992061292 | NA |
| 0.910538937 | NA |
| 0.165826111 | 0.300206825 |
| NA | NA |
| NA | NA |
| 0.185826491 | 0.326758802 |
| 0.255122582 | 0.409735675 |
| 0.530665715 | 0.679984154 |
| 0.099556683 | 0.205293004 |
| 0.002755264 | 0.011723083 |
| 0.180163521 | NA |
| 1.77E-05 | 0.000160434 |
| 0.531581159 | 0.68072024 |
| 0.008247666 | 0.028784248 |

| | |
|---|---|
| NA | NA |
| NA | NA |
| NA | NA |
| 0.679511877 | 0.796870563 |
| 0.844421466 | NA |
| 0.381819806 | 0.545872145 |
| 0.490842562 | 0.644873269 |
| NA | NA |
| 0.00014461 | 0.000953026 |
| 0.164770362 | 0.298950044 |
| 0.989649027 | NA |
| NA | NA |
| 0.714949663 | 0.822742881 |
| 0.800408884 | 0.879644597 |
| 0.889409773 | NA |
| 0.281405244 | 0.439805003 |
| 0.00018055 | 0.001154208 |
| 2.61E-05 | 0.000222576 |
| 0.599097756 | 0.735236552 |
| NA | NA |
| NA | NA |
| NA | NA |
| NA | NA |
| 0.362185617 | 0.526974984 |
| NA | NA |
| 0.239173605 | 0.391469569 |
| 0.778531773 | NA |
| NA | NA |
| 0.1435456 | 0.269642243 |
| 0.01000962 | 0.033670535 |
| 0.160153034 | NA |
| 0.000315948 | 0.001867536 |
| 0.731664016 | 0.83493271 |
| 1.97E-05 | 0.00017601 |
| 0.33213253 | 0.494672592 |
| NA | NA |
| 0.042704419 | 0.107404232 |
| 0.003287225 | 0.013540458 |
| 0.640272485 | 0.766582314 |
| 0.14865714 | 0.276734668 |
| 0.043659904 | 0.109119945 |
| 2.20E-11 | 1.12E-09 |
| 0.016697568 | 0.050953571 |
| 0.000473231 | 0.002632805 |
| 1.32E-08 | 3.00E-07 |
| 0.840711454 | NA |
| 0.51909392 | 0.670062638 |

| | |
|---|---|
| 0.502486088 | 0.65498414 |
| 0.607520667 | NA |
| 0.938868647 | 0.965468606 |
| 0.591372018 | NA |
| 0.300190139 | 0.460551614 |
| 0.783183042 | 0.86937628 |
| 0.722249368 | 0.828148602 |
| 0.10213697 | 0.209399276 |
| 0.309097009 | NA |
| NA | NA |
| 0.388488672 | 0.551962569 |
| 0.000693278 | 0.0036613 |
| 3.19E-15 | 4.20E-13 |
| 0.228729228 | 0.379154911 |
| 0.787488966 | 0.871771467 |
| 0.731021439 | 0.834459624 |
| 0.007247294 | 0.025948692 |
| 3.07E-10 | 1.13E-08 |
| 0.127076203 | 0.246333539 |
| 0.144662785 | 0.271226176 |
| 0.080443479 | 0.174672042 |
| 0.678803084 | 0.796251972 |
| 0.292827919 | 0.452576596 |
| 0.016163188 | 0.049641447 |
| 1.71E-09 | 5.04E-08 |
| 4.37E-11 | 2.03E-09 |
| 0.905156847 | NA |
| 2.35E-05 | 0.000204064 |
| 0.337610943 | 0.500580601 |
| NA | NA |
| 1.17E-06 | 1.50E-05 |
| 0.008978235 | 0.030812697 |
| 0.050281385 | 0.121881989 |
| 1.88E-09 | 5.47E-08 |
| 0.032873909 | 0.087627844 |
| 0.190721999 | 0.332861578 |
| 0.500507853 | NA |
| 9.53E-06 | 9.32E-05 |
| 0.390127082 | 0.553518162 |
| 0.00057242 | 0.003111327 |
| 0.992061292 | NA |
| 0.000757809 | 0.003951727 |
| 2.21E-13 | 1.86E-11 |
| 0.0578005 | 0.135514772 |
| 7.11E-05 | 0.000522125 |
| 0.053060204 | 0.126894905 |
| 0.713475213 | 0.821519701 |

| | | |
|---|---|---|
| | 0.254160464 | 0.408543271 |
| | 0.086736928 | NA |
| | 0.363709523 | 0.528675277 |
| | 0.894178359 | 0.940251045 |
| | 2.42E-10 | 9.13E-09 |
| | 0.478136282 | 0.634136293 |
| | 0.009307061 | 0.031730126 |
| | 0.492591161 | 0.646627966 |
| NA | | NA |
| | 0.008596864 | 0.029771099 |
| | 0.025657748 | 0.072039187 |
| | 0.447718774 | 0.606880035 |
| | 0.185480802 | 0.326360011 |
| | 0.039267732 | 0.100698367 |
| | 2.14E-17 | 4.56E-15 |
| | 0.019920251 | 0.058605144 |
| | 0.430157459 | 0.59132513 |
| | 0.000239407 | 0.001476307 |
| | 3.48E-05 | 0.000283589 |
| | 0.004678783 | 0.018142574 |
| | 7.90E-08 | 1.44E-06 |
| | 0.973553289 | 0.986011917 |
| | 1.84E-07 | 3.02E-06 |
| | 1.00E-07 | 1.76E-06 |
| | 0.69365669 | NA |
| | 0.429032015 | 0.590147458 |
| | 6.70E-15 | 8.13E-13 |
| NA | | NA |
| | 0.002471531 | 0.010717289 |
| | 0.008419802 | 0.029282616 |
| | 0.039531419 | 0.10117371 |
| | 0.95044979 | 0.97236894 |
| | 0.397153518 | NA |
| | 0.001851519 | 0.00838299 |
| | 0.889409301 | NA |
| | 0.246695403 | 0.400117022 |
| | 0.082442777 | 0.178047912 |
| NA | | NA |
| | 0.000423776 | 0.00240333 |
| | 0.568352254 | 0.710853059 |
| | 0.270371429 | 0.426852126 |
| | 0.301189582 | 0.461536777 |
| | 2.39E-10 | 9.07E-09 |
| | 0.441203369 | 0.600941867 |
| | 0.49936488 | 0.652400845 |
| | 1.80E-05 | 0.000162302 |
| | 0.435865511 | 0.596371784 |

| | |
|---|---|
| 0.375430631 | NA |
| 6.88E-06 | 7.06E-05 |
| 0.884943576 | 0.934832376 |
| 0.812464777 | 0.887961081 |
| 0.060293463 | 0.140312014 |
| 0.001127543 | 0.005508279 |
| 0.817718597 | 0.891573122 |
| NA | NA |
| 0.007993847 | 0.028084003 |
| 0.257833094 | NA |
| 1.52E-05 | 0.000139766 |
| 0.457614132 | NA |
| 0.565740385 | NA |
| 0.797559319 | 0.87774295 |
| 0.873814768 | 0.927840184 |
| 0.18141454 | 0.32157587 |
| 0.699900891 | 0.812553043 |
| 0.865769343 | 0.922571408 |
| NA | NA |
| 0.466334058 | 0.623036601 |
| NA | NA |
| 0.677550626 | 0.795462692 |
| NA | NA |
| NA | NA |
| NA | NA |
| 0.148963145 | 0.277139854 |
| 1.48E-10 | 6.05E-09 |
| 0.000443875 | 0.002496043 |
| 0.140106809 | 0.26465423 |
| 4.77E-06 | 5.11E-05 |
| 0.037126347 | 0.096480173 |
| 0.087111656 | 0.185446907 |
| 0.900897778 | NA |
| 7.17E-05 | 0.000526708 |
| 0.185101996 | 0.325902399 |
| 0.554912902 | 0.700346711 |
| 0.023571726 | 0.067145789 |
| NA | NA |
| 0.027613537 | 0.076282004 |
| 0.145523743 | 0.27235625 |
| 0.466780248 | 0.623253524 |
| 1.82E-05 | 0.000164112 |
| 0.410660153 | 0.573607677 |
| 0.755789783 | NA |
| 0.905158266 | NA |
| 0.300808887 | 0.461178689 |
| 0.137186434 | 0.260504748 |

| | |
|---|---|
| 0.97753218 | NA |
| 0.092822845 | 0.194550533 |
| 0.134215076 | 0.256111945 |
| 0.878054267 | 0.930694292 |
| 0.00101987 | 0.005062232 |
| NA | NA |
| 0.360337507 | 0.524990135 |
| 0.283130628 | 0.441943806 |
| 0.377818055 | NA |
| 7.98E-06 | 8.02E-05 |
| 0.269211961 | 0.425602984 |
| 0.801661681 | NA |
| 0.834083049 | NA |
| 0.654659911 | 0.777862506 |
| 0.017324259 | 0.052399919 |
| 0.409604481 | 0.572675549 |
| 0.255460396 | 0.410062487 |
| 1.71E-05 | 0.000155108 |
| NA | NA |
| 0.550008314 | 0.695996806 |
| 0.835907109 | 0.902280588 |
| 0.024158178 | 0.068547541 |
| 0.000151164 | 0.000990584 |
| 0.021249 | 0.061707195 |
| 0.000125169 | 0.000845362 |
| 0.33585841 | 0.498722939 |
| 0.822234634 | 0.894188309 |
| NA | NA |
| 0.99162063 | NA |
| 0.229653157 | NA |
| NA | NA |
| 0.889409773 | NA |
| 0.26058549 | 0.415887548 |
| NA | NA |
| 7.92E-06 | 7.96E-05 |
| 0.040150014 | 0.102370457 |
| 0.119815336 | 0.235860598 |
| 0.119047259 | 0.234727374 |
| NA | NA |
| 0.01344092 | 0.042642681 |
| 4.07E-06 | 4.46E-05 |
| 0.740223991 | 0.840507711 |
| 0.124307316 | 0.242313746 |
| 0.00488983 | 0.018811452 |
| 1.40E-08 | 3.17E-07 |
| 0.352465013 | 0.516743172 |
| 0.611066695 | 0.744929415 |

| | |
|---|---|
| 0.677681948 | 0.795534131 |
| 0.038762306 | 0.0996233 |
| 0.625403554 | 0.755340311 |
| 0.000289236 | 0.001733918 |
| 7.64E-05 | 0.000555832 |
| 0.016822001 | 0.051240714 |
| 0.098804964 | 0.204030299 |
| 5.85E-05 | 0.00044297 |
| 0.019685869 | 0.058054131 |
| 0.583848019 | 0.723639366 |
| NA | NA |
| NA | NA |
| 6.99E-08 | 1.30E-06 |
| 0.586980927 | 0.725965316 |
| 0.905158266 | NA |
| 0.092823259 | 0.194550533 |
| 0.914546254 | 0.951778255 |
| 0.100165794 | 0.206258503 |
| 0.857775598 | 0.917175946 |
| 0.381616502 | NA |
| 8.17E-18 | 2.01E-15 |
| 0.003611808 | 0.014622361 |
| 0.226111917 | 0.375923046 |
| 0.435966016 | NA |
| 0.021788125 | 0.063011383 |
| 0.957677648 | 0.976545197 |
| NA | NA |
| NA | NA |
| NA | NA |
| 0.992061292 | NA |
| NA | NA |
| NA | NA |
| NA | NA |
| 0.13378007 | NA |
| 0.99205891 | NA |
| 0.076833981 | NA |
| 0.030555762 | 0.082672988 |
| 0.13798353 | 0.261633835 |
| 3.04E-08 | 6.31E-07 |
| 0.889410717 | NA |
| 5.57E-08 | 1.06E-06 |
| 0.643396555 | 0.768937427 |
| 0.59083045 | 0.728838035 |
| 0.02906829 | 0.079391863 |
| 0.49336834 | 0.646928273 |
| 0.244615013 | 0.397718054 |
| 0.662166252 | 0.783620462 |

| | | |
|---|---|---|
| | 0.439332146 | 0.599583256 |
| | 0.042892582 | 0.107729407 |
| | 0.036051714 | 0.094290088 |
| | 0.442747576 | NA |
| | 0.136051404 | 0.258752189 |
| NA | | NA |
| | 0.844490643 | 0.908369941 |
| | 1.78E-05 | 0.000160875 |
| | 0.497072437 | 0.65023168 |
| | 0.000186093 | 0.00118378 |
| | 0.060328237 | 0.14035386 |
| | 0.922007878 | 0.955294241 |
| | 1.33E-05 | 0.000124493 |
| | 0.231216159 | 0.382181079 |
| | 0.002216914 | 0.009763581 |
| | 0.0504975 | 0.122297849 |
| | 0.320634291 | 0.482988196 |
| | 0.298557658 | 0.458591746 |
| | 0.083093461 | 0.179031056 |
| | 0.340643345 | NA |
| | 0.22468455 | 0.374343554 |
| | 0.001452808 | 0.006831309 |
| | 1.40E-05 | 0.000130507 |
| | 0.345628353 | 0.509071517 |
| | 0.044785568 | 0.11131254 |
| | 0.025955261 | 0.072617539 |
| | 2.28E-05 | 0.000198532 |
| | 0.005881501 | 0.021916152 |
| | 5.97E-06 | 6.23E-05 |
| | 2.61E-12 | 1.70E-10 |
| NA | | NA |
| | 0.083347342 | 0.179468944 |
| | 0.785874018 | 0.871195302 |
| NA | | NA |
| | 0.167938523 | 0.303170176 |
| | 0.889410717 | NA |
| | 0.018730409 | 0.055739403 |
| | 0.869111046 | NA |
| | 0.243964848 | 0.396919619 |
| | 0.855649127 | NA |
| NA | | NA |
| | 0.096762056 | 0.200888531 |
| | 0.905158739 | NA |
| | 0.087614121 | 0.186263771 |
| | 0.628162846 | NA |
| | 0.006307425 | 0.023200127 |
| | 0.006852888 | 0.024852613 |

| | | |
|---|---|---|
| | 0.550238967 | 0.696121036 |
| | 0.084760126 | 0.181532001 |
| | 0.60717047 | 0.741414466 |
| | 0.603842913 | 0.739200944 |
| | 0.858815521 | 0.91762014 |
| | 0.030191573 | 0.081879307 |
| | 0.04292036 | 0.107786848 |
| | 2.94E-06 | 3.34E-05 |
| | 4.13E-09 | 1.08E-07 |
| | 0.90983062 | 0.948984284 |
| | 2.11E-07 | 3.40E-06 |
| | 0.052708451 | 0.126356234 |
| | 1.29E-06 | 1.63E-05 |
| | 0.605260532 | 0.74023481 |
| | 0.000346855 | 0.002027846 |
| | 0.054461878 | 0.129471513 |
| | 0.001028358 | 0.005097453 |
| | 0.301671352 | 0.462081558 |
| | 0.008532294 | 0.029584817 |
| | 0.246067826 | 0.399394145 |
| | 0.905158266 | NA |
| | 0.090233349 | 0.190559157 |
| | 0.041864709 | 0.105740412 |
| | 0.109111172 | 0.220036021 |
| | 0.773645488 | 0.86363079 |
| | 0.00066895 | 0.003555913 |
| | 2.50E-08 | 5.29E-07 |
| | 0.043378733 | 0.108615005 |
| | 0.870892765 | 0.926162874 |
| | 0.749450967 | NA |
| | 0.326966941 | 0.489178092 |
| | 0.011948865 | 0.038831192 |
| | 0.889409773 | NA |
| | 0.127784211 | 0.247247643 |
| | 0.411285847 | NA |
| | 0.01119878 | 0.036846569 |
| | 0.007629616 | 0.027026735 |
| | 0.001331156 | 0.006339118 |
| | 0.521869685 | 0.672409591 |
| | 0.990210336 | NA |
| | 0.171227728 | 0.307666286 |
| | 0.446272613 | NA |
| NA | | NA |
| | 0.977425206 | NA |
| | 0.153810226 | NA |
| | 0.100583023 | NA |
| NA | | NA |

| | | |
|---|---|---|
| | 3.38E-05 | 0.000277312 |
| | 8.48E-13 | 6.12E-11 |
| | 0.979093848 | NA |
| | 0.00075906 | 0.003955432 |
| | 0.447148615 | NA |
| | 1.16E-05 | 0.000110628 |
| | 0.319371497 | 0.481866065 |
| | 0.007661144 | 0.027112179 |
| | 0.013001566 | 0.041509362 |
| | 5.25E-06 | 5.55E-05 |
| | 0.187654953 | 0.32915689 |
| | 0.213184211 | 0.360337793 |
| | 0.333574746 | NA |
| | 8.43E-05 | 0.000602496 |
| | 0.00034927 | 0.002040102 |
| | 0.348563598 | NA |
| | 0.001987684 | NA |
| | 2.69E-05 | 0.00022789 |
| NA | | NA |
| | 0.313812811 | NA |
| | 0.717219282 | NA |
| | 0.971598745 | NA |
| | 0.216964115 | NA |
| NA | | NA |
| NA | | NA |
| NA | | NA |
| | 0.961799825 | NA |
| | 0.905158266 | NA |
| | 0.588120069 | 0.726841996 |
| | 0.35147024 | 0.515812485 |
| | 0.889409301 | NA |
| NA | | NA |
| | 0.413742459 | NA |
| | 0.858280913 | 0.917217762 |
| | 0.99205891 | NA |
| | 2.90E-06 | 3.31E-05 |
| | 0.983556749 | NA |
| NA | | NA |
| | 0.082971695 | 0.17892325 |
| NA | | NA |
| | 0.905158266 | NA |
| | 0.889409301 | NA |
| NA | | NA |
| NA | | NA |
| | 0.90515732 | NA |
| | 0.889409301 | NA |
| NA | | NA |

| | |
|---|---|
| NA | NA |
| NA | NA |
| NA | NA |
| 0.889410717 | NA |
| NA | NA |
| NA | NA |
| 0.503749105 | NA |
| 0.889409773 | NA |
| 0.905157793 | NA |
| NA | NA |
| 4.02E-09 | 1.05E-07 |
| 0.001146471 | 0.005588316 |
| 1.58E-07 | 2.63E-06 |
| 0.197748348 | 0.342160269 |
| 0.87661619 | 0.929573832 |
| 0.000920648 | 0.00464534 |
| 0.033365754 | 0.08864863 |
| 0.905158266 | NA |
| 0.36170757 | NA |
| 0.807252815 | NA |
| 0.494060803 | NA |
| NA | NA |
| 0.655090471 | 0.778121472 |
| 0.038460912 | 0.099103868 |
| 1.08E-08 | 2.53E-07 |
| 0.912389956 | NA |
| 0.092141432 | NA |
| 0.905158266 | NA |
| NA | NA |
| NA | NA |
| 0.99205891 | NA |
| 0.905158266 | NA |
| NA | NA |
| NA | NA |
| NA | NA |
| NA | NA |
| NA | NA |
| NA | NA |
| NA | NA |
| 0.905158266 | NA |
| NA | NA |
| NA | NA |
| NA | NA |
| NA | NA |
| NA | NA |
| NA | NA |
| 0.889409301 | NA |

| | | | |
|---|---|---|---|
| NA | | NA | |
| NA | | NA | |
| NA | | NA | |
| NA | | NA | |
| NA | | NA | |
| NA | | NA | |
| NA | | NA | |
| NA | | NA | |
| | 0.992060816 | NA | |
| NA | | NA | |
| NA | | NA | |
| NA | | NA | |
| NA | | NA | |
| NA | | NA | |
| NA | | NA | |
| NA | | NA | |
| | 0.905158739 | NA | |
| NA | | NA | |
| NA | | NA | |
| | 0.992061292 | NA | |
| NA | | NA | |
| NA | | NA | |
| | 0.992061292 | NA | |
| NA | | NA | |
| NA | | NA | |
| NA | | NA | |
| NA | | NA | |
| NA | | NA | |
| | 0.024636621 | | 0.069671032 |
| NA | | NA | |
| NA | | NA | |
| | 0.810070774 | NA | |
| NA | | NA | |
| | 0.071252294 | | 0.158652314 |
| NA | | NA | |
| | 0.025453029 | | 0.071574218 |
| | 0.698736455 | | 0.811656465 |
| | 1.13E-11 | | 6.21E-10 |
| | 0.400346623 | | 0.563909228 |
| | 0.965428279 | | 0.981292334 |
| | 0.506924388 | | 0.65871073 |
| NA | | NA | |
| | 0.052961661 | | 0.126779865 |
| | 1.05E-06 | | 1.37E-05 |
| | 0.532633748 | NA | |
| | 0.810134348 | NA | |
| | 0.770122589 | | 0.861055088 |

| | |
|---|---|
| NA | NA |
| NA | NA |
| 0.009281527 | 0.03165782 |
| 0.045220733 | 0.112239153 |
| 0.640260403 | 0.766582314 |
| NA | NA |
| NA | NA |
| NA | NA |
| 2.18E-08 | 4.72E-07 |
| 0.889409773 | NA |
| 0.592550967 | 0.729935301 |
| 0.448703276 | 0.60765188 |
| 0.000534318 | 0.002929611 |
| 0.758982706 | 0.853756605 |
| 0.002788116 | 0.011828485 |
| NA | NA |
| 0.015086733 | 0.046924869 |
| NA | NA |
| NA | NA |
| 2.49E-09 | 7.04E-08 |
| 0.824867634 | 0.895612234 |
| NA | NA |
| 0.500527404 | 0.65350931 |
| 0.447134944 | 0.606388108 |
| 0.714614473 | 0.822572691 |
| 0.086203713 | 0.183960173 |
| 0.007650317 | 0.027082591 |
| 0.889409301 | NA |
| 0.26741274 | 0.42364417 |
| 0.448827729 | 0.607745459 |
| 0.000107894 | 0.000745281 |
| 0.22380593 | 0.373418008 |
| 3.00E-07 | 4.60E-06 |
| NA | NA |
| 0.119534947 | 0.235456396 |
| 0.0686927 | 0.154125166 |
| 0.345900327 | NA |
| 0.27115587 | 0.427752281 |
| 0.001058871 | 0.005222766 |
| 0.128130933 | 0.247743872 |
| 0.080987026 | 0.175626517 |
| 0.786112585 | NA |
| 0.72702875 | 0.831457962 |
| 0.032318881 | 0.086484316 |
| 0.284088094 | 0.442903121 |
| NA | NA |
| NA | NA |

| | | |
|---|---|---|
| | 0.919460956 | 0.954020567 |
| | 0.796862171 | 0.877547463 |
| | 0.882062051 | 0.933050753 |
| | 0.006562357 | 0.023957736 |
| | 0.576328464 | 0.717947112 |
| | 0.784353866 | NA |
| | 4.41E-06 | 4.78E-05 |
| | 0.342524031 | 0.505922037 |
| | 0.904947233 | NA |
| | 8.57E-05 | 0.000610171 |
| | 0.842458065 | NA |
| | 0.704913359 | 0.816370046 |
| | 0.060646 | 0.140968006 |
| | 2.06E-12 | 1.37E-10 |
| | 0.786171365 | NA |
| | 0.029521987 | 0.080440913 |
| | 0.42444983 | NA |
| | 0.680918345 | 0.797613891 |
| NA | | NA |
| | 0.992062245 | NA |
| | 1.64E-06 | 2.01E-05 |
| NA | | NA |
| | 0.039596088 | 0.101303799 |
| NA | | NA |
| NA | | NA |
| | 0.834614131 | 0.901538111 |
| | 0.87117164 | NA |
| | 0.036864923 | 0.095982552 |
| NA | | NA |
| | 0.905158739 | NA |
| | 4.45E-14 | 4.27E-12 |
| | 0.905158739 | NA |
| | 0.531136287 | NA |
| | 0.905157793 | NA |
| | 0.02976849 | 0.081042187 |
| | 0.716581268 | 0.824015923 |
| | 0.000431891 | 0.002441156 |
| NA | | NA |
| | 0.074404699 | 0.16425295 |
| | 0.777092376 | NA |
| NA | | NA |
| | 0.992060816 | NA |
| | 1.23E-12 | 8.49E-11 |
| | 0.181997347 | 0.322199107 |
| | 0.802811388 | NA |
| | 0.905296333 | NA |
| | 0.342706481 | 0.506119552 |

| | |
|---|---|
| 0.905158739 | NA |
| 0.000379634 | 0.002184584 |
| 0.809055137 | 0.885468589 |
| 6.91E-08 | 1.28E-06 |
| 0.000104881 | 0.000727671 |
| 0.039518487 | 0.101152401 |
| 0.172184208 | 0.30900564 |
| 0.062747164 | 0.144555794 |
| 0.028510178 | 0.078237315 |
| 0.373279808 | 0.537533694 |
| 0.980958692 | NA |
| 0.929463116 | 0.959575308 |
| 2.49E-06 | 2.90E-05 |
| 0.706840175 | 0.817632974 |
| 0.746175546 | 0.844993493 |
| 0.035021461 | 0.092056051 |
| 0.007579916 | 0.026868017 |
| 0.003729382 | 0.014990572 |
| 0.000250561 | 0.001530902 |
| 0.323753451 | 0.485985809 |
| 0.08007119 | 0.174052986 |
| 0.304008322 | 0.464688762 |
| 0.817874269 | 0.891604826 |
| 0.846154741 | 0.909565146 |
| 1.31E-06 | 1.65E-05 |
| NA | NA |
| NA | NA |
| 5.82E-08 | 1.10E-06 |
| 0.039322526 | 0.10079642 |
| NA | NA |
| 0.998074533 | NA |
| NA | NA |
| NA | NA |
| 0.757095053 | NA |
| 0.642222442 | NA |
| 0.000240325 | 0.001481555 |
| NA | NA |
| 0.818767297 | 0.892155344 |
| 0.748902491 | NA |
| 0.816461755 | 0.890743016 |
| 0.461201964 | 0.618588684 |
| 0.327813833 | 0.489912234 |
| 0.610139388 | 0.744129455 |
| 0.003792133 | 0.015192747 |
| 0.003745312 | 0.015049097 |
| 0.330059917 | 0.492438027 |
| 5.44E-09 | 1.37E-07 |

| | | |
|---|---|---|
| | 0.889409301 | NA |
| | 0.084967268 | 0.181904675 |
| NA | | NA |
| | 0.795353865 | NA |
| NA | | NA |
| | 0.830536717 | NA |
| | 1.33E-05 | 0.000125117 |
| NA | | NA |
| NA | | NA |
| | 0.905296333 | NA |
| | 0.873679372 | 0.927758854 |
| | 0.90515732 | NA |
| | 0.922724538 | NA |
| NA | | NA |
| | 0.788582525 | NA |
| | 0.905158266 | NA |
| | 0.992061769 | NA |
| | 0.626595559 | NA |
| NA | | NA |
| | 0.200306131 | NA |
| | 0.594878053 | NA |
| | 0.00557078 | 0.020961064 |
| | 0.000691608 | 0.003654239 |
| | 0.047540983 | 0.116590623 |
| | 0.573349824 | NA |
| NA | | NA |
| | 0.001756698 | 0.008004842 |
| | 0.468222618 | 0.624639323 |
| | 3.67E-09 | 9.83E-08 |
| | 0.056717916 | 0.133490753 |
| | 3.16E-05 | 0.000262174 |
| | 1.09E-05 | 0.000105 |
| | 1.80E-05 | 0.000162768 |
| | 0.002125437 | 0.009424974 |
| | 0.023596592 | 0.067197195 |
| | 0.019761408 | 0.058239264 |
| NA | | NA |
| NA | | NA |
| | 0.89769384 | 0.942458359 |
| | 0.027628577 | NA |
| | 0.393526705 | 0.556938187 |
| NA | | NA |
| NA | | NA |
| NA | | NA |
| NA | | NA |
| NA | | NA |
| | 0.842985448 | NA |

| | | |
|---|---|---|
| | 0.372864832 | 0.537249061 |
| | 0.271895248 | 0.428622457 |
| | 0.015111632 | 0.046995655 |
| | 0.638338712 | NA |
| | 9.27E-07 | 1.23E-05 |
| NA | | NA |
| | 0.001548273 | 0.00718937 |
| | 0.705719056 | 0.816872319 |
| | 0.412719064 | 0.575269994 |
| | 0.172518653 | 0.309454101 |
| | 0.107235888 | 0.216991602 |
| | 0.048263204 | 0.117979705 |
| | 0.026320844 | 0.073370979 |
| | 0.423223947 | 0.584648631 |
| | 0.306550825 | 0.467274023 |
| | 0.866452144 | NA |
| | 0.091336507 | 0.192209147 |
| NA | | NA |
| | 2.14E-05 | 0.000188832 |
| | 0.431974734 | 0.592858325 |
| | 0.11893722 | 0.234573595 |
| | 0.154398017 | 0.284885813 |
| | 0.059969739 | 0.13967708 |
| NA | | NA |
| | 0.046345916 | 0.114346637 |
| | 0.629821974 | 0.758113288 |
| | 0.532035878 | 0.68108945 |
| | 1.01E-09 | 3.17E-08 |
| | 0.979268995 | NA |
| | 0.02404507 | 0.068279541 |
| NA | | NA |
| | 0.554056714 | NA |
| | 0.718465466 | 0.8253614 |
| | 0.565441817 | NA |
| | 0.970390738 | 0.984468016 |
| | 2.20E-09 | 6.33E-08 |
| | 0.949434805 | NA |
| | 0.56296333 | 0.706646527 |
| NA | | NA |
| | 0.211329646 | 0.358432249 |
| NA | | NA |
| | 5.94E-05 | 0.000448675 |
| | 0.244184706 | 0.397189035 |
| | 0.875786116 | 0.928962773 |
| | 0.011634956 | 0.037963151 |
| | 0.272987075 | NA |
| NA | | NA |

| | | |
|---|---|---|
| | 0.807070704 | NA |
| | 0.905158739 | NA |
| | 0.905158739 | NA |
| | 0.586107771 | NA |
| | 0.889409301 | NA |
| | 0.70389033 | NA |
| NA | | NA |
| | 0.889409301 | NA |
| | 0.859727347 | NA |
| | 0.889409773 | NA |
| | 0.889409301 | NA |
| | 0.889409301 | NA |
| | 0.752891351 | NA |
| | 0.989595644 | NA |
| | 0.943781216 | 0.96829845 |
| | 3.89E-05 | 0.000312828 |
| | 0.21244723 | 0.35960708 |
| | 2.36E-10 | 8.98E-09 |
| | 0.068225445 | 0.153439533 |
| | 0.140518569 | NA |
| | 0.007902178 | 0.027799412 |
| | 0.028586083 | 0.078377051 |
| | 0.145107522 | NA |
| | 0.011355176 | 0.037247861 |
| | 0.785896796 | 0.871195302 |
| | 0.713239625 | NA |
| NA | | NA |
| | 0.445330353 | 0.604725077 |
| | 0.946257666 | 0.969356826 |
| | 0.043583576 | NA |
| | 0.845691689 | 0.909251068 |
| | 0.724493693 | 0.830064068 |
| | 0.222528745 | 0.37188042 |
| | 8.95E-09 | 2.15E-07 |
| | 0.889409301 | NA |
| | 0.018151867 | 0.054356585 |
| | 0.008670505 | 0.02997884 |
| | 0.772381006 | 0.862613901 |
| | 0.054023581 | 0.128708236 |
| | 2.17E-17 | 4.58E-15 |
| | 0.992062245 | NA |
| | 0.375109194 | 0.539253963 |
| | 0.456089203 | 0.61390602 |
| | 0.003901987 | 0.01555341 |
| | 6.19E-06 | 6.43E-05 |
| | 0.012473785 | 0.040221819 |
| | 0.095149965 | 0.1982429 |

| | | |
|---|---|---|
| | 0.195646969 | 0.33940684 |
| | 0.855141364 | 0.915689726 |
| NA | | NA |
| | 3.84E-09 | 1.02E-07 |
| | 0.485370522 | 0.640362725 |
| | 0.266504385 | 0.422661403 |
| | 0.050415735 | NA |
| | 0.010885943 | 0.036071289 |
| | 0.022759215 | 0.065304305 |
| | 0.049115303 | 0.119676315 |
| | 0.86984691 | 0.925515189 |
| | 0.289059626 | 0.448383271 |
| | 0.020819684 | 0.060730532 |
| | 0.416097028 | 0.578478925 |
| | 0.62546551 | NA |
| | 0.98997168 | NA |
| | 2.83E-06 | 3.25E-05 |
| | 0.120902894 | 0.237490531 |
| | 0.681898235 | NA |
| | 0.086963365 | 0.185221002 |
| NA | | NA |
| | 0.02304142 | 0.06593257 |
| | 0.796270232 | 0.877115527 |
| | 0.007997677 | 0.028090369 |
| | 0.034771803 | 0.091630149 |
| NA | | NA |
| | 0.123956581 | 0.24184474 |
| | 0.000256667 | 0.001564729 |
| | 0.003435831 | 0.014034118 |
| | 0.830163713 | NA |
| | 0.992061292 | NA |
| NA | | NA |
| | 0.106332989 | 0.215581765 |
| | 0.060287417 | 0.140312014 |
| | 0.231508419 | 0.382499915 |
| | 0.598038339 | 0.734552181 |
| | 0.525609599 | 0.675555235 |
| | 0.6154795 | 0.748332145 |
| | 0.159714785 | 0.29192417 |
| NA | | NA |
| | 0.087689413 | 0.186387748 |
| | 0.003268236 | NA |
| | 0.002861601 | 0.012091188 |
| | 0.000114111 | 0.000781584 |
| | 0.14872118 | 0.27678349 |
| | 0.682824528 | 0.799346616 |
| | 0.008012241 | 0.028128027 |

| | |
|---|---|
| 3.99E-05 | 0.000319183 |
| 0.758777405 | 0.853662127 |
| 0.000434917 | 0.002454469 |
| NA | NA |
| 0.064395957 | 0.147411216 |
| 0.071033507 | 0.158327312 |
| 0.097639017 | 0.202154929 |
| 0.231866844 | 0.382910734 |
| 0.905158739 | NA |
| 0.940901333 | 0.966427446 |
| 7.34E-06 | 7.45E-05 |
| 0.98173229 | NA |
| 0.598327669 | 0.734743165 |
| NA | NA |
| 0.992061292 | NA |
| 0.992061292 | NA |
| 0.587151328 | 0.726094274 |
| 0.381178834 | 0.545367866 |
| 3.07E-06 | 3.46E-05 |
| 0.091079541 | 0.191811224 |
| 0.006718891 | 0.024423128 |
| 0.006101037 | 0.022558119 |
| 0.476588256 | 0.632809379 |
| 0.002402557 | 0.010461582 |
| 0.600374762 | 0.736227699 |
| 0.755049622 | NA |
| NA | NA |
| 0.483007877 | 0.638514838 |
| 0.909873634 | 0.948984284 |
| 0.014403713 | 0.045139308 |
| 0.758897038 | NA |
| 0.051717252 | 0.124594124 |
| 0.000833601 | 0.004274883 |
| 0.742424883 | 0.842135854 |
| 0.34470508 | 0.508071896 |
| 0.287282879 | 0.446425925 |
| 0.062213026 | 0.143713006 |
| 0.985790394 | NA |
| 5.15E-09 | 1.31E-07 |
| 0.201291994 | NA |
| 0.560595302 | 0.704946235 |
| 5.94E-15 | 7.36E-13 |
| 0.400051159 | 0.563737624 |
| 0.038586866 | NA |
| 0.005906267 | 0.021989778 |
| 0.000276256 | 0.001670241 |
| 1.50E-08 | 3.35E-07 |

| | | |
|---|---|---|
| | 0.298334064 | 0.458344484 |
| | 0.009369361 | NA |
| | 0.805112363 | NA |
| | 3.71E-06 | 4.13E-05 |
| | 0.98694158 | NA |
| | 0.400426702 | NA |
| | 0.001046227 | 0.005173468 |
| | 3.16E-19 | 1.08E-16 |
| | 0.486413572 | 0.641241257 |
| | 0.500673881 | 0.653527774 |
| | 0.190804998 | NA |
| | 0.000234377 | 0.001450586 |
| | 0.248366383 | NA |
| | 0.54787236 | 0.694093876 |
| | 0.409219044 | 0.572322844 |
| | 0.171284639 | 0.307743363 |
| | 0.109386015 | 0.220408104 |
| | 0.679900823 | 0.797093552 |
| | 0.833740162 | 0.900944846 |
| | 0.072187073 | 0.160311079 |
| | 0.000398867 | 0.002280915 |
| | 0.983785879 | NA |
| | 0.889409301 | NA |
| | 6.21E-06 | 6.45E-05 |
| | 3.23E-05 | 0.000266945 |
| | 0.047956872 | 0.117426957 |
| | 0.016235303 | 0.049826836 |
| | 0.851107611 | 0.912750148 |
| | 0.544805684 | 0.691405422 |
| | 0.992061292 | NA |
| | 0.049014088 | 0.119495627 |
| | 0.994724118 | 0.997540618 |
| | 1.90E-09 | 5.52E-08 |
| | 0.457146265 | 0.614688312 |
| NA | | NA |
| | 0.600366502 | NA |
| | 0.131698925 | 0.252584854 |
| | 6.78E-08 | 1.26E-06 |
| | 0.000130433 | 0.000873758 |
| | 0.337601442 | NA |
| | 0.008531234 | 0.029584817 |
| | 0.324756888 | 0.486946842 |
| | 0.032597567 | 0.087049734 |
| | 0.598234085 | NA |
| | 0.061338029 | 0.142155333 |
| NA | | NA |
| NA | | NA |

| | | |
|---|---|---|
| | 0.351559082 | 0.515862739 |
| | 3.22E-05 | 0.000266281 |
| | 0.905158266 | NA |
| | 0.12290269 | 0.240258183 |
| | 0.176934439 | 0.315519193 |
| NA | | NA |
| | 0.145708245 | 0.272641177 |
| | 0.016514577 | 0.0504939 |
| NA | | NA |
| NA | | NA |
| | 0.860455057 | NA |
| | 0.728553967 | 0.832638522 |
| NA | | NA |
| | 0.992061292 | NA |
| NA | | NA |
| NA | | NA |
| NA | | NA |
| NA | | NA |
| | 0.016048704 | 0.049391359 |
| | 0.13794051 | NA |
| | 0.807252815 | NA |
| | 3.49E-06 | 3.90E-05 |
| NA | | NA |
| | 8.37E-05 | 0.000598964 |
| | 0.808185712 | NA |
| | 0.180794311 | 0.320683544 |
| | 0.083474593 | 0.179602056 |
| | 0.681035053 | NA |
| | 0.000648415 | 0.003466895 |
| | 0.001892468 | 0.00854373 |
| | 0.200133879 | 0.345042439 |
| | 0.874017176 | 0.927848352 |
| | 0.889409773 | NA |
| | 0.346033298 | 0.509456601 |
| | 2.06E-11 | 1.06E-09 |
| | 0.129351681 | 0.249420804 |
| | 0.036493025 | 0.095183553 |
| | 6.50E-06 | 6.72E-05 |
| | 0.851875055 | 0.913439345 |
| | 0.980049272 | NA |
| NA | | NA |
| | 0.486732586 | 0.64158478 |
| | 1.61E-08 | 3.56E-07 |
| | 1.73E-06 | 2.11E-05 |
| | 0.266208351 | 0.42242149 |
| | 0.090811063 | 0.191466167 |
| | 0.388502822 | 0.551962569 |

| | |
|---|---|
| 0.655778778 | 0.778686324 |
| NA | NA |
| 0.889409773 | NA |
| 0.955401064 | 0.975162776 |
| 0.83182303 | NA |
| 4.56E-08 | 8.95E-07 |
| 0.958118789 | 0.976801638 |
| 0.102707656 | 0.210277454 |
| 0.026137261 | 0.073012101 |
| 0.007638179 | 0.027048342 |
| 0.748346135 | NA |
| 0.736272805 | 0.838159186 |
| 0.465565004 | 0.622425688 |
| 0.830077042 | NA |
| 0.000292697 | 0.001748932 |
| 0.246636812 | 0.400051541 |
| 0.054238303 | 0.129121736 |
| 0.08432027 | 0.180730971 |
| 0.026066463 | 0.072842131 |
| NA | NA |
| 0.000678115 | 0.003598533 |
| 0.833347244 | 0.900819299 |
| 0.905157793 | NA |
| 0.017871838 | 0.053722742 |
| 0.313498927 | 0.475357198 |
| 0.612603405 | NA |
| 0.507322234 | 0.659110693 |
| NA | NA |
| 0.90515732 | NA |
| 0.98694468 | NA |
| 0.520424232 | NA |
| NA | NA |
| NA | NA |
| 2.17E-10 | 8.40E-09 |
| 0.03516732 | 0.09233997 |
| 0.736285382 | 0.838159186 |
| 1.22E-05 | 0.000115523 |
| 0.850415744 | 0.91232006 |
| 0.992061292 | NA |
| 0.492765019 | NA |
| 0.478808239 | 0.634605875 |
| 0.922303236 | 0.955335317 |
| 0.176598439 | 0.31502235 |
| 0.012326983 | 0.039830521 |
| 0.185297046 | 0.326141218 |
| 0.819704986 | NA |
| 0.270864351 | 0.427415226 |

| | | |
|---|---|---|
| | 5.81E-07 | 8.16E-06 |
| | 0.130289747 | 0.250594076 |
| | 0.040275179 | 0.102575116 |
| | 0.759876132 | 0.854271967 |
| | 0.702621673 | NA |
| NA | | NA |
| | 0.928757791 | 0.959203853 |
| | 0.004541282 | 0.017662418 |
| | 0.235703479 | 0.387408964 |
| | 0.002048308 | 0.009130894 |
| | 0.016563141 | 0.050613695 |
| | 0.992835068 | 0.99632873 |
| | 0.999992169 | 0.999992169 |
| | 0.403681481 | 0.56711195 |
| NA | | NA |
| | 0.320041893 | 0.482347284 |
| | 0.19304212 | 0.336002847 |
| | 0.860058417 | 0.918536199 |
| | 0.86219084 | NA |
| NA | | NA |
| | 0.131715887 | 0.252584854 |
| NA | | NA |
| | 0.731218981 | NA |
| NA | | NA |
| | 1.31E-07 | 2.24E-06 |
| NA | | NA |
| NA | | NA |
| NA | | NA |
| | 0.466556397 | 0.62318199 |
| | 1.41E-06 | 1.76E-05 |
| | 0.889409773 | NA |
| | 0.000689088 | 0.003644432 |
| | 0.747064717 | 0.845575712 |
| | 0.29586719 | NA |
| | 0.905158739 | NA |
| | 0.0733406 | 0.162331038 |
| | 0.281371459 | 0.439791861 |
| | 0.780245744 | 0.867646544 |
| | 0.625476766 | 0.755340311 |
| | 0.931205944 | 0.96025939 |
| | 0.937524243 | 0.964537803 |
| | 0.889409773 | NA |
| | 0.512329241 | NA |
| | 0.000571881 | 0.003109167 |
| NA | | NA |
| | 0.266349314 | NA |
| | 0.992061292 | NA |

| | | |
|---|---|---|
| | 0.889410717 | NA |
| | 0.889409301 | NA |
| | 8.45E-09 | 2.04E-07 |
| | 0.874516807 | NA |
| NA | | NA |
| | 1.43E-06 | 1.78E-05 |
| | 0.000798195 | 0.004119284 |
| NA | | NA |
| | 0.013942266 | 0.04399457 |
| | 9.64E-05 | 0.00067708 |
| | 0.010416016 | 0.034808294 |
| | 0.026148431 | 0.073024724 |
| | 0.041090759 | 0.104144966 |
| | 0.82281678 | 0.894551581 |
| NA | | NA |
| NA | | NA |
| | 0.669505156 | NA |
| | 0.400193946 | 0.563838687 |
| | 0.905158266 | NA |
| | 0.017218147 | 0.052179586 |
| NA | | NA |
| | 0.045141738 | 0.112134323 |
| | 0.121269586 | 0.237934136 |
| | 0.382242116 | 0.546319619 |
| NA | | NA |
| | 0.377205092 | 0.541623398 |
| | 0.654616163 | 0.777852614 |
| | 0.125070248 | 0.243433574 |
| | 0.283486592 | 0.442330757 |
| | 0.637867216 | NA |
| | 0.519557395 | NA |
| | 0.001113535 | 0.005449548 |
| | 0.907016392 | NA |
| NA | | NA |
| | 0.992060816 | NA |
| | 2.54E-05 | 0.000217548 |
| | 0.905157793 | NA |
| | 0.62550192 | 0.755340311 |
| | 0.00603773 | 0.022388778 |
| | 0.207931431 | 0.354583077 |
| | 0.698215798 | 0.811266242 |
| | 0.382184213 | NA |
| | 0.00054422 | 0.002976471 |
| | 0.056338147 | 0.132824712 |
| | 0.688780024 | 0.80387566 |
| NA | | NA |
| | 0.884115531 | NA |

| | |
|---|---|
| 0.038858275 | 0.099799867 |
| 0.690477303 | NA |
| 0.315074539 | NA |
| 0.910562697 | 0.949347069 |
| 0.701439309 | NA |
| 0.946966807 | 0.969890117 |
| 0.003017851 | 0.012625076 |
| 8.06E-05 | 0.000581219 |
| 0.934403312 | NA |
| 0.121726765 | 0.238596628 |
| 0.905158266 | NA |
| 0.025792678 | 0.072326962 |
| 0.830571413 | 0.898980563 |
| 0.46997355 | NA |
| NA | NA |
| 0.99103752 | NA |
| 0.078024621 | NA |
| 0.904016291 | NA |
| 0.889409301 | NA |
| NA | NA |
| NA | NA |
| 0.116707351 | 0.231361886 |
| NA | NA |
| NA | NA |
| NA | NA |
| NA | NA |
| NA | NA |
| NA | NA |
| NA | NA |
| 0.217849506 | 0.366021652 |
| 0.905158266 | NA |
| 0.951920067 | NA |
| 0.71008481 | NA |
| NA | NA |
| 0.770266914 | 0.861091697 |
| 0.76852495 | 0.859883598 |
| 0.33272942 | 0.495349515 |
| 0.11393143 | 0.227314217 |
| 0.099940155 | 0.205948373 |
| 0.00549738 | 0.020727502 |
| 0.017948371 | 0.053894201 |
| 0.905158739 | NA |
| 0.067700256 | 0.152584411 |
| 0.462580204 | NA |
| 0.590805313 | NA |
| 0.317201966 | NA |
| 0.822990651 | NA |

| | | | |
|---|---|---|---|
| | 0.084914797 | NA | |
| | 0.451619593 | NA | |
| | 0.855120302 | NA | |
| | 0.726117338 | NA | |
| | 0.655050211 | NA | |
| | 0.500713728 | NA | |
| | 0.441435861 | NA | |
| | 0.856442839 | NA | |
| | 0.197390955 | NA | |
| | 0.181325644 | NA | |
| | 0.737551201 | | 0.838994288 |
| | 0.78016172 | NA | |
| | 0.536231198 | NA | |
| | 0.992060816 | NA | |
| | 0.404159968 | NA | |
| | 0.842051217 | NA | |
| | 0.294130125 | | 0.453854502 |
| | 0.91976905 | NA | |
| | 0.807166075 | NA | |
| | 0.698817464 | NA | |
| | 0.723095004 | NA | |
| | 0.590219845 | NA | |
| | 0.832112589 | NA | |
| | 0.790014018 | NA | |
| | 0.484915047 | NA | |
| | 0.592045242 | | 0.729721681 |
| | 0.922238427 | NA | |
| | 0.358559603 | NA | |
| | 0.721993847 | NA | |
| | 0.258670983 | NA | |
| | 0.257855486 | NA | |
| | 0.341731917 | NA | |
| NA | | NA | |
| NA | | NA | |
| | 0.810318344 | NA | |
| NA | | NA | |
| | 0.785288496 | NA | |
| | 0.560057525 | NA | |
| NA | | NA | |
| NA | | NA | |
| | 0.889409301 | NA | |
| | 0.889409301 | NA | |
| NA | | NA | |
| NA | | NA | |
| NA | | NA | |
| | 0.905158739 | NA | |
| NA | | NA | |

| | | | |
|---|---|---|---|
| | 0.889409773 | NA | |
| NA | | NA | |
| | 0.889409773 | NA | |
| | 0.470831045 | NA | |
| | 0.889411642 | NA | |
| | 0.836813554 | NA | |
| NA | | NA | |
| NA | | NA | |
| NA | | NA | |
| | 0.797342596 | NA | |
| | 0.905158739 | NA | |
| NA | | NA | |
| | 0.738380807 | NA | |
| NA | | NA | |
| | 0.992062245 | NA | |
| | 0.787579664 | NA | |
| | 0.889409773 | NA | |
| | 0.992060816 | NA | |
| | 0.989651219 | NA | |
| | 0.17869794 | NA | |
| | 0.996890041 | NA | |
| | 0.981824196 | NA | |
| | 0.87441858 | NA | |
| | 0.804320614 | NA | |
| | 0.482367533 | NA | |
| | 0.826533418 | NA | |
| | 0.326634822 | NA | |
| NA | | NA | |
| | 0.462561567 | NA | |
| | 0.122672074 | NA | |
| | 0.597544571 | NA | |
| | 0.481569629 | | 0.63718893 |
| | 0.521691264 | NA | |
| | 0.490708306 | NA | |
| | 0.869750705 | NA | |
| | 0.750259564 | NA | |
| | 0.975035471 | NA | |
| | 0.400397643 | NA | |
| | 0.202671278 | NA | |
| | 0.979079564 | NA | |
| | 0.735924078 | NA | |
| | 0.662901591 | NA | |
| | 0.772624519 | NA | |
| | 0.988656237 | NA | |
| | 0.68711511 | NA | |
| | 0.977851132 | NA | |
| | 0.726439718 | NA | |

| | | |
|---|---|---|
| | 0.076675545 | NA |
| | 0.037550619 | NA |
| | 0.817032209 | NA |
| | 0.342617938 | NA |
| | 0.77688773 | NA |
| | 0.754584106 | NA |
| | 0.82315795 | NA |
| | 0.890331309 | NA |
| | 0.330906148 | NA |
| | 0.410392415 | NA |
| | 0.788045095 | NA |
| | 0.554871967 | NA |
| | 0.992982015 | NA |
| NA | | NA |
| | 0.455242829 | NA |
| | 0.085976233 | NA |
| NA | | NA |
| NA | | NA |
| | 0.823049907 | NA |
| NA | | NA |
| NA | | NA |
| | 0.263551231 | NA |
| | 0.131427236 | 0.252185483 |
| | 0.240768475 | 0.393419831 |
| | 0.577701163 | NA |
| | 0.269805243 | 0.42629539 |
| | 0.032861253 | 0.087604742 |
| NA | | NA |
| | 0.125412527 | 0.24386201 |
| | 0.016782113 | 0.051154694 |
| | 0.714695101 | NA |
| | 0.732772445 | NA |
| | 0.889409301 | NA |
| | 0.17340871 | 0.310568621 |
| NA | | NA |
| | 1.04E-08 | 2.46E-07 |
| | 0.013864706 | NA |
| | 0.478578732 | 0.63442653 |
| | 0.237435091 | 0.389324758 |
| | 0.714530011 | NA |
| NA | | NA |
| | 0.005368314 | 0.020345668 |
| | 0.45640383 | NA |
| | 0.045431748 | 0.112625395 |
| | 0.689255948 | 0.804260129 |
| | 0.006146321 | 0.022710257 |
| | 0.652356559 | NA |

| | |
|---|---|
| 0.889409773 | NA |
| 5.88E-09 | 1.47E-07 |
| 0.07760136 | NA |
| 0.035062033 | 0.092085538 |
| 0.789566051 | 0.872894237 |
| 0.067684607 | 0.152584411 |
| 0.219851127 | 0.368696418 |
| 0.115359886 | 0.229487409 |
| NA | NA |
| 0.338688553 | NA |
| 0.022028024 | 0.063546161 |
| 8.83E-15 | 1.04E-12 |
| 0.576242443 | 0.717913895 |
| 0.031678812 | 0.085113793 |
| 0.026196277 | 0.073111851 |
| 0.134838133 | NA |
| 0.000188507 | 0.001197213 |
| 0.083632055 | 0.179799912 |
| 0.045734388 | 0.113260564 |
| NA | NA |
| 0.280792234 | 0.439167853 |
| 0.175924291 | 0.314049387 |
| 0.012905305 | 0.041291738 |
| 0.193136391 | 0.336090283 |
| 1.61E-13 | 1.38E-11 |
| 0.424811238 | 0.586067096 |
| NA | NA |
| 0.625529706 | 0.755340311 |
| 0.433984508 | 0.594687414 |
| 0.018359568 | 0.054854413 |
| 0.118908864 | 0.234538736 |
| 0.319450974 | 0.481886648 |
| 0.253725529 | 0.408113016 |
| 0.000955164 | 0.004793064 |
| 0.598638746 | NA |
| 0.173804543 | 0.311102146 |
| 0.992059387 | NA |
| 0.989446001 | NA |
| 0.00012124 | 0.000824284 |
| 0.200551177 | 0.345675297 |
| 4.72E-06 | 5.06E-05 |
| 0.189686076 | 0.331607081 |
| 0.000805726 | 0.004153269 |
| 0.916362109 | NA |
| 0.030998076 | 0.083612178 |
| 0.001497466 | 0.007000822 |
| 0.68069521 | NA |

| | | |
|---|---|---|
| | 0.184509047 | NA |
| | 0.198204005 | 0.342620572 |
| | 0.009729327 | 0.032873718 |
| | 0.00572207 | 0.021440883 |
| | 0.795447524 | 0.87663136 |
| | 0.881516819 | NA |
| | 0.85292978 | NA |
| | 0.004481125 | 0.017459375 |
| | 3.81E-05 | 0.000306809 |
| | 0.69598658 | 0.809404158 |
| | 0.020018397 | 0.058838706 |
| | 0.397441965 | 0.561076159 |
| | 0.435291516 | 0.595957753 |
| | 0.802836473 | NA |
| NA | | NA |
| NA | | NA |
| | 0.889409773 | NA |
| NA | | NA |
| | 0.471552308 | NA |
| NA | | NA |
| NA | | NA |
| | 0.504778324 | 0.656816024 |
| | 2.41E-08 | 5.11E-07 |
| | 0.889409301 | NA |
| | 0.436458744 | NA |
| NA | | NA |
| | 0.125952892 | 0.244609494 |
| | 0.075748581 | 0.16668635 |
| | 0.889409301 | NA |
| NA | | NA |
| NA | | NA |
| | 0.697117783 | NA |
| | 0.99205891 | NA |
| | 0.003100441 | 0.012904206 |
| | 0.873365128 | NA |
| | 0.253403329 | 0.407794434 |
| | 0.203343099 | NA |
| | 0.855699189 | 0.915974383 |
| | 0.769110699 | NA |
| NA | | NA |
| | 0.772017635 | 0.862251933 |
| | 0.969530912 | NA |
| | 0.001690872 | 0.007738612 |
| | 0.027649361 | 0.076342552 |
| | 0.550716583 | 0.696531819 |
| NA | | NA |
| | 0.905158739 | NA |

| | | |
|---|---|---|
| | 0.55109848 | NA |
| | 0.616893785 | 0.749370505 |
| | 0.013236092 | 0.042105076 |
| | 0.000855715 | 0.004358765 |
| | 0.134243294 | 0.256111945 |
| | 0.992060816 | NA |
| | 0.622860471 | 0.753949236 |
| | 0.416787223 | 0.578998585 |
| | 0.718821157 | 0.825467722 |
| NA | | NA |
| | 0.786063034 | 0.871291614 |
| | 1.80E-06 | 2.18E-05 |
| | 0.956928503 | 0.976086631 |
| NA | | NA |
| NA | | NA |
| | 0.717746494 | NA |
| | 0.429293175 | 0.590387326 |
| | 0.56985823 | NA |
| NA | | NA |
| | 0.899517123 | NA |
| | 0.98838098 | NA |
| | 9.79E-06 | 9.53E-05 |
| | 0.292698201 | NA |
| | 0.981275271 | NA |
| NA | | NA |
| | 0.889283189 | NA |
| | 6.31E-07 | 8.77E-06 |
| | 0.047902693 | 0.117346636 |
| | 0.801661681 | NA |
| | 0.840454681 | 0.905660238 |
| | 0.905158266 | NA |
| | 0.129749794 | 0.249883855 |
| | 0.580803136 | NA |
| | 0.814561557 | NA |
| | 0.117689536 | NA |
| | 0.731846611 | NA |
| | 0.461738114 | NA |
| | 9.75E-06 | 9.50E-05 |
| | 0.19783468 | 0.342282677 |
| | 0.000470558 | 0.002619926 |
| | 0.585648343 | NA |
| | 0.889409773 | NA |
| NA | | NA |
| | 0.02535069 | 0.071341254 |
| | 0.881776279 | NA |
| | 0.699763763 | 0.812505947 |
| | 0.676127498 | NA |

| | |
|---|---|
| 0.874683018 | 0.928423374 |
| 1.61E-08 | 3.57E-07 |
| NA | NA |
| NA | NA |
| 0.915838123 | 0.95229975 |
| 0.800082288 | NA |
| 0.700849354 | 0.813129883 |
| 0.15263445 | 0.282510105 |
| 0.04420858 | 0.110215283 |
| 0.425892192 | 0.587180169 |
| 0.237034236 | 0.388899791 |
| 0.827902427 | 0.897278333 |
| 0.952629266 | 0.973374154 |
| 0.787852663 | NA |
| 0.889409301 | NA |
| 0.991399481 | NA |
| 0.009111581 | 0.031181872 |
| 0.601786841 | 0.737341661 |
| 0.116062452 | 0.230585711 |
| 0.521097293 | 0.671879246 |
| NA | NA |
| 0.497236392 | 0.650355228 |
| 0.031652686 | 0.085054004 |
| 0.044710771 | 0.111170276 |
| 0.173109777 | 0.310210348 |
| 0.779407566 | 0.867285433 |
| 0.232210926 | 0.383344315 |
| 0.23941342 | 0.391734634 |
| 0.5174459 | 0.668472682 |
| 0.077923722 | 0.170464935 |
| 0.876987616 | NA |
| 0.074975261 | 0.165233015 |
| 0.660303221 | 0.782493169 |
| 0.984499036 | NA |
| NA | NA |
| 0.976965767 | NA |
| 0.000272309 | 0.001649102 |
| NA | NA |
| NA | NA |
| 0.781790497 | 0.868484723 |
| 0.056176226 | 0.132542575 |
| NA | NA |
| NA | NA |
| 0.787645204 | NA |
| 0.789316257 | NA |
| NA | NA |
| 0.905158739 | NA |

| | |
|---|---|
| 0.079532528 | 0.173122005 |
| 4.22E-11 | 1.97E-09 |
| 0.964320281 | 0.98039303 |
| 0.763139826 | NA |
| NA | NA |
| NA | NA |
| 0.836648271 | NA |
| NA | NA |
| 0.311153563 | 0.472551427 |
| NA | NA |
| 2.92E-05 | 0.000245276 |
| NA | NA |
| 0.18396008 | 0.324490955 |
| 0.244107931 | 0.397122985 |
| 0.060379262 | 0.140452142 |
| 0.65496467 | 0.778098311 |
| 0.889409773 | NA |
| 0.190697178 | 0.332844713 |
| 0.550768004 | 0.696556736 |
| 0.203933957 | 0.349776284 |
| 0.467370095 | 0.623732563 |
| 0.327945791 | 0.490055231 |
| 0.996722979 | NA |
| NA | NA |
| NA | NA |
| 6.25E-08 | 1.18E-06 |
| NA | NA |
| 0.128755303 | NA |
| 0.167856582 | NA |
| NA | NA |
| 0.705332551 | NA |
| 0.000703543 | 0.003707489 |
| 0.810100352 | NA |
| 7.76E-08 | 1.42E-06 |
| 0.005838173 | 0.021799089 |
| NA | NA |
| 0.905157793 | NA |
| 0.325679948 | NA |
| 0.030682073 | 0.082932995 |
| 0.975339066 | NA |
| 0.889409773 | NA |
| 0.840260116 | NA |
| 5.09E-13 | 3.89E-11 |
| 0.456462517 | 0.614210622 |
| 0.433604067 | 0.594371492 |
| 0.553125046 | 0.698572087 |
| 0.00747852 | 0.026598774 |

| | |
|---|---|
| 0.31367483 | 0.475558243 |
| 0.773202318 | NA |
| 0.683686891 | NA |
| 5.19E-08 | 1.00E-06 |
| 0.22331238 | 0.372877846 |
| 0.805279752 | 0.883130124 |
| 0.29579547 | 0.455815629 |
| 0.885891408 | 0.935299266 |
| 6.40E-05 | 0.000477832 |
| 5.21E-06 | 5.52E-05 |
| 0.412835275 | 0.575364928 |
| 0.474123572 | 0.630417885 |
| 0.033700174 | 0.089363859 |
| 0.443178568 | 0.602850934 |
| 6.66E-05 | 0.0004942 |
| NA | NA |
| 1.05E-05 | 0.00010127 |
| 0.805509698 | 0.883217029 |
| 1.06E-05 | 0.000102327 |
| 0.830163713 | NA |
| 0.581552553 | 0.721664547 |
| 0.000758762 | 0.00395482 |
| 4.10E-06 | 4.49E-05 |
| NA | NA |
| 0.905296333 | NA |
| 0.421625873 | 0.583248239 |
| 0.391037067 | 0.554415221 |
| 0.000387903 | 0.002225753 |
| 1.34E-15 | 1.98E-13 |
| 0.054113446 | 0.128852435 |
| 0.001337123 | 0.00636365 |
| 0.342080257 | 0.505500697 |
| 0.611591216 | NA |
| 0.144411392 | 0.27087467 |
| 0.409694091 | 0.572695433 |
| 0.992061769 | NA |
| 0.991879012 | NA |
| 0.694304151 | NA |
| 0.805374319 | NA |
| 0.85535434 | NA |
| 0.081108122 | 0.175767615 |
| 0.008777495 | 0.030286772 |
| 0.049663893 | NA |
| 0.992060816 | NA |
| NA | NA |
| 0.905158266 | NA |
| 0.905158266 | NA |

| | |
|---|---|
| NA | NA |
| 0.992061292 | NA |
| 0.878671082 | NA |
| 0.001498442 | 0.007003891 |
| 0.821854665 | 0.893952179 |
| 0.305126134 | 0.465716603 |
| 0.351141659 | 0.515387989 |
| 0.333034458 | 0.495568453 |
| 0.992037435 | NA |
| 0.759924475 | 0.854271967 |
| 0.886052713 | 0.935379622 |
| 0.905158739 | NA |
| 0.979258898 | 0.989397824 |
| 0.928282936 | NA |
| 0.003356959 | 0.01377344 |
| 0.211723421 | 0.358820298 |
| 7.13E-08 | 1.32E-06 |
| 0.02356954 | 0.067145789 |
| 0.002003544 | 0.008967759 |
| 0.283603951 | 0.442375341 |
| 0.71939684 | 0.826037701 |
| 0.258403596 | 0.413306473 |
| 0.605751797 | 0.740546917 |
| 0.542921093 | 0.689888809 |
| 0.500570453 | 0.65350931 |
| 0.646850371 | 0.771798918 |
| 1.66E-07 | 2.75E-06 |
| 0.818780757 | 0.892155344 |
| 0.680948127 | 0.797613891 |
| 9.79E-11 | 4.18E-09 |
| 0.835263455 | NA |
| NA | NA |
| 8.06E-09 | 1.95E-07 |
| NA | NA |
| NA | NA |
| NA | NA |
| 9.97E-05 | 0.00069705 |
| 0.002229682 | 0.009811942 |
| 0.989617443 | NA |
| NA | NA |
| 0.02728443 | 0.075534392 |
| 0.977425817 | NA |
| 0.560565612 | NA |
| NA | NA |
| 0.833331695 | NA |
| 0.013039048 | 0.041610878 |
| 0.012050813 | 0.039087711 |

| | |
|---|---|
| 0.096848767 | 0.200973548 |
| 0.815820835 | 0.890476673 |
| NA | NA |
| NA | NA |
| 0.561461648 | 0.705433071 |
| 0.551964056 | 0.697466838 |
| 4.18E-06 | 4.57E-05 |
| 0.572707512 | 0.715078425 |
| 0.016298654 | 0.049972375 |
| 0.29751473 | 0.457496243 |
| 0.046246075 | 0.114205754 |
| 0.000185542 | 0.001180621 |
| 0.363455823 | NA |
| 0.719535408 | NA |
| 0.400344773 | 0.563909228 |
| 0.951589606 | 0.97321765 |
| 1.96E-07 | 3.19E-06 |
| 0.470500864 | 0.626849198 |
| 0.847612585 | 0.910497043 |
| 0.000173984 | 0.001118093 |
| NA | NA |
| 0.697658652 | 0.810911078 |
| NA | NA |
| NA | NA |
| 0.423989446 | 0.585408359 |
| 0.71501856 | NA |
| 0.016343584 | 0.050082161 |
| 0.10625177 | 0.215481871 |
| 0.836645721 | 0.902813162 |
| 0.629180282 | 0.757547443 |
| 0.373817569 | 0.537868166 |
| 0.096166022 | 0.199972507 |
| NA | NA |
| 0.4166295 | NA |
| 0.423141556 | NA |
| 0.127193716 | 0.246474304 |
| 7.49E-05 | 0.000545698 |
| 0.500492858 | 0.65350931 |
| 2.09E-08 | 4.54E-07 |
| 0.329109782 | 0.491460163 |
| 0.003591 | 0.014557592 |
| 0.809541846 | 0.885780528 |
| 0.356915232 | 0.521608296 |
| 0.692927254 | 0.806871862 |
| 0.725417736 | 0.83034525 |
| 0.794922376 | NA |
| 0.905158266 | NA |

| | |
|---|---|
| 0.000131716 | 0.000881545 |
| NA | NA |
| 0.889409301 | NA |
| 0.505108405 | NA |
| 0.400986393 | 0.564620724 |
| 0.88810401 | 0.936678822 |
| 0.527722183 | 0.677375497 |
| NA | NA |
| NA | NA |
| 0.905158266 | NA |
| 0.001414546 | 0.006687194 |
| 0.014331298 | 0.044963704 |
| 0.983273083 | 0.991763981 |
| 0.117978514 | 0.233101511 |
| 0.002535032 | 0.010932203 |
| 0.000400584 | 0.00228956 |
| 0.991759839 | NA |
| 0.782559532 | NA |
| 0.764288152 | 0.857062789 |
| 0.232482382 | 0.383754087 |
| 0.613342664 | 0.746626285 |
| 0.29325098 | 0.45285813 |
| 0.860142408 | NA |
| 0.114294169 | NA |
| NA | NA |
| 0.829760282 | 0.89850121 |
| 8.62E-10 | 2.76E-08 |
| 0.305374351 | NA |
| 0.449420232 | NA |
| 0.905158266 | NA |
| 0.719840098 | 0.826239149 |
| 0.033442564 | 0.08880944 |
| 0.114405871 | 0.227929461 |
| 0.136648618 | 0.25968559 |
| 0.986159033 | 0.992938931 |
| 0.292344047 | 0.452083304 |
| 0.045187203 | 0.11219057 |
| 0.512962906 | 0.664551767 |
| 0.023544182 | 0.067108727 |
| NA | NA |
| 0.000748507 | 0.003911579 |
| 0.001109988 | 0.005434612 |
| 0.17710455 | NA |
| 0.0946162 | 0.197441523 |
| 0.052992406 | 0.126815632 |
| 0.000493223 | 0.002729505 |
| 0.132476514 | 0.253600538 |

| | |
|---|---|
| 5.96E-05 | 0.000450302 |
| 0.015774905 | 0.048686035 |
| 0.042308261 | 0.106639935 |
| 0.757812843 | 0.853156228 |
| 0.907402141 | 0.947610543 |
| 0.082240255 | 0.177712435 |
| 0.224434259 | 0.374040064 |
| 0.301094044 | 0.461422578 |
| 0.152056732 | 0.281701998 |
| 0.635949449 | 0.763183465 |
| NA | NA |
| 0.502422531 | 0.65498414 |
| 4.25E-05 | 0.000337094 |
| NA | NA |
| 4.60E-11 | 2.13E-09 |
| 0.286964695 | 0.446000912 |
| NA | NA |
| 0.135814638 | 0.258480989 |
| 0.332821138 | 0.495452469 |
| 0.456467696 | 0.614210622 |
| 0.452390765 | 0.610612134 |
| 0.052529344 | 0.126020184 |
| 0.122223474 | NA |
| 0.002700565 | 0.011531555 |
| 0.463192055 | 0.620310687 |
| 0.228840223 | NA |
| 0.015683973 | 0.048446246 |
| 0.98149243 | NA |
| 0.442962302 | 0.602739447 |
| 0.905156847 | NA |
| 0.778120127 | 0.866682731 |
| 0.050172063 | 0.121684155 |
| 0.012519806 | 0.040328944 |
| 0.959926147 | 0.977727478 |
| 0.620927445 | NA |
| 0.022526878 | 0.064755558 |
| 0.709956921 | 0.819272901 |
| 0.258504624 | 0.413377733 |
| 5.84E-05 | 0.000442935 |
| 0.334341585 | 0.4971766 |
| 0.435354978 | 0.595970324 |
| 3.15E-05 | 0.00026115 |
| 0.871777261 | 0.926545514 |
| 0.992061292 | NA |
| 0.984671226 | 0.992171589 |
| 8.82E-10 | 2.81E-08 |
| 0.367739438 | 0.532348232 |

| | |
|---|---|
| NA | NA |
| 0.77708508 | 0.865841855 |
| 0.514988954 | NA |
| 0.064014254 | 0.146730439 |
| 0.818249876 | 0.891756051 |
| NA | NA |
| 0.042950482 | 0.107850157 |
| 0.981278741 | NA |
| 0.171363339 | 0.307859573 |
| 0.445434041 | NA |
| 6.40E-09 | 1.59E-07 |
| 0.788582525 | NA |
| 0.785965787 | 0.8712278 |
| 0.001702199 | 0.007780724 |
| 0.673188868 | NA |
| 0.555544621 | 0.700902288 |
| 0.052783493 | 0.126465184 |
| 0.992228864 | NA |
| 0.023303702 | 0.066552996 |
| 0.066071437 | NA |
| 0.873114763 | NA |
| 0.563757043 | 0.707361916 |
| 0.000263293 | 0.001599414 |
| 0.12215262 | 0.239179901 |
| 0.684240653 | NA |
| 0.56088241 | 0.705084546 |
| 0.304767547 | 0.465428094 |
| 0.049391469 | 0.120215836 |
| 0.186896352 | 0.328298296 |
| 0.946170463 | 0.969356826 |
| 0.050782957 | 0.122880767 |
| 0.28890731 | 0.448284032 |
| 7.66E-13 | 5.57E-11 |
| 0.889409773 | NA |
| 0.992061292 | NA |
| 0.075293063 | 0.165800278 |
| 0.1319013 | 0.252766715 |
| 0.732608717 | 0.835663333 |
| 0.027275893 | 0.07552205 |
| 0.443969004 | NA |
| 0.029798296 | 0.0811126 |
| 0.000920075 | 0.00464534 |
| NA | NA |
| NA | NA |
| 0.100225954 | 0.20636303 |
| 0.00014642 | 0.000963226 |
| 0.530806004 | 0.680006466 |

| | | |
|---|---|---|
| | 0.207095299 | 0.353738678 |
| | 1.41E-12 | 9.68E-11 |
| | 0.889409773 | NA |
| | 2.99E-13 | 2.43E-11 |
| | 0.924889643 | 0.956719373 |
| | 9.00E-05 | 0.00063763 |
| | 0.436801597 | NA |
| NA | | NA |
| | 0.889410245 | NA |
| | 0.020716802 | 0.060478629 |
| NA | | NA |
| | 0.125198869 | 0.243646755 |
| NA | | NA |
| NA | | NA |
| | 0.077553774 | 0.169774032 |
| | 0.284583447 | 0.443415018 |
| | 0.045189926 | 0.11219057 |
| | 0.004503023 | 0.01753225 |
| | 9.28E-08 | 1.65E-06 |
| | 0.033365607 | 0.08864863 |
| | 1.56E-09 | 4.65E-08 |
| NA | | NA |
| | 0.482458923 | 0.637904352 |
| | 0.329693169 | 0.49216399 |
| | 5.67E-12 | 3.36E-10 |
| | 0.114352039 | 0.227884239 |
| | 0.017479772 | 0.052790305 |
| NA | | NA |
| | 0.000939732 | 0.004728591 |
| | 4.52E-08 | 8.90E-07 |
| | 0.002180867 | 0.009624704 |
| | 0.000828255 | 0.004251437 |
| | 0.903682055 | NA |
| | 0.637647217 | 0.764264791 |
| | 6.94E-05 | 0.000511932 |
| | 0.905158739 | NA |
| | 0.037189042 | 0.096610625 |
| | 0.73473161 | 0.836954476 |
| | 0.088292324 | NA |
| | 0.247633174 | 0.40128233 |
| | 0.285334797 | NA |
| | 0.273647162 | 0.430723105 |
| NA | | NA |
| | 0.222012418 | 0.371356694 |
| | 0.99205891 | NA |
| | 0.905158266 | NA |
| | 0.41780544 | 0.580046132 |

| | | |
|---|---|---|
| | 0.874482462 | NA |
| | 0.825530258 | 0.896053297 |
| | 0.749901748 | 0.847421391 |
| | 0.743884988 | 0.843153014 |
| | 0.708098216 | NA |
| | 0.23404935 | 0.385552466 |
| | 0.693391168 | NA |
| | 0.054807252 | 0.130123521 |
| NA | | NA |
| NA | | NA |
| | 0.343071916 | 0.506386784 |
| | 0.904174056 | 0.945779931 |
| | 0.007043001 | 0.025382763 |
| | 5.24E-10 | 1.81E-08 |
| | 0.222838949 | 0.372234313 |
| | 0.068482274 | 0.153810194 |
| | 0.889409773 | NA |
| | 0.629078794 | NA |
| | 0.028866439 | 0.07900007 |
| | 0.094215394 | 0.196792146 |
| | 0.271447796 | 0.428059042 |
| NA | | NA |
| NA | | NA |
| | 0.002804831 | 0.011885621 |
| | 0.889409773 | NA |
| | 0.131739051 | 0.252607215 |
| | 0.47070509 | NA |
| | 0.805276708 | 0.883130124 |
| | 0.973477308 | 0.986004992 |
| | 0.926112948 | 0.957598259 |
| NA | | NA |
| | 0.006538877 | 0.02389138 |
| | 0.221097158 | 0.370090303 |
| | 2.37E-05 | 0.000205236 |
| | 0.031981165 | 0.08578969 |
| NA | | NA |
| | 0.807252815 | NA |
| | 6.36E-12 | 3.72E-10 |
| | 0.00662619 | 0.024149498 |
| | 0.992060816 | NA |
| NA | | NA |
| | 0.601121176 | NA |
| | 0.000366469 | 0.002121909 |
| | 0.006266615 | 0.023073204 |
| | 0.98273701 | 0.991609754 |
| | 0.636645863 | 0.76352302 |
| | 0.992060816 | NA |

| | | |
|---|---|---|
| | 0.088122289 | NA |
| | 0.484098777 | 0.639449867 |
| | 0.047801482 | 0.117150977 |
| NA | NA | |
| | 0.043304164 | 0.10848327 |
| NA | NA | |
| | 0.147344723 | 0.275037638 |
| | 0.002666421 | 0.011422538 |
| NA | NA | |
| | 0.725252809 | NA |
| | 0.689148428 | NA |
| | 0.002680592 | 0.011465354 |
| | 0.992060816 | NA |
| | 0.993651873 | NA |
| | 0.992061769 | NA |
| | 0.788582525 | NA |
| | 0.784353866 | NA |
| | 0.872639999 | NA |
| | 0.624020857 | 0.754701939 |
| NA | NA | |
| NA | NA | |
| NA | NA | |
| | 0.784353866 | NA |
| NA | NA | |
| | 0.905158266 | NA |
| NA | NA | |
| NA | NA | |
| NA | NA | |
| NA | NA | |
| NA | NA | |
| NA | NA | |
| | 0.008700381 | 0.030067936 |
| NA | NA | |
| NA | NA | |
| NA | NA | |
| | 0.905158739 | NA |
| NA | NA | |
| NA | NA | |
| NA | NA | |
| NA | NA | |
| NA | NA | |
| NA | NA | |
| NA | NA | |
| NA | NA | |
| NA | NA | |
| NA | NA | |
| NA | NA | |

| | |
|---|---|
| NA | NA |
| NA | NA |
| NA | NA |
| NA | NA |
| NA | NA |
| NA | NA |
| NA | NA |
| NA | NA |
| 0.992060816 | NA |
| NA | NA |
| NA | NA |
| NA | NA |
| 0.013095695 | 0.041743121 |
| 0.668556125 | 0.788697967 |
| 0.824095256 | NA |
| 0.000145279 | 0.000957151 |
| 0.000238475 | 0.001471285 |
| 0.130717186 | 0.251130244 |
| 1.12E-05 | 0.000106799 |
| 0.169375346 | 0.305202449 |
| NA | NA |
| 0.230177135 | 0.380922428 |
| 0.789840597 | 0.873054952 |
| 0.536076272 | 0.684321524 |
| 0.847721301 | 0.910497043 |
| 0.992061292 | NA |
| 0.002692358 | 0.011511195 |
| 0.1039822 | NA |
| NA | NA |
| 0.657892935 | 0.780354441 |
| 0.991879012 | NA |
| 0.889409301 | NA |
| 4.38E-10 | 1.54E-08 |
| 0.00983604 | 0.033167935 |
| 0.243182762 | NA |
| 0.886750571 | NA |
| NA | NA |
| NA | NA |
| 0.446915078 | 0.606173216 |
| 0.792188023 | NA |
| 0.335113119 | 0.497986667 |
| NA | NA |
| 0.393348956 | 0.556794284 |
| 0.879517119 | NA |
| 0.990599468 | NA |
| 0.743477096 | 0.84276345 |
| NA | NA |

| | | |
|---|---|---|
| | 0.992059387 | NA |
| | 0.241543909 | NA |
| | 0.35930866 | 0.524117102 |
| | 0.031993094 | 0.085794946 |
| | 0.808200146 | 0.88498742 |
| | 0.093909697 | 0.196228284 |
| NA | | NA |
| | 8.66E-12 | 4.92E-10 |
| | 0.349345945 | 0.513404336 |
| | 0.004483807 | 0.017466723 |
| | 0.134978416 | 0.257201582 |
| NA | | NA |
| | 0.905158266 | NA |
| NA | | NA |
| NA | | NA |
| NA | | NA |
| | 0.905158266 | NA |
| | 0.188927143 | 0.330754296 |
| NA | | NA |
| | 0.905158739 | NA |
| NA | | NA |
| NA | | NA |
| | 0.011558752 | 0.037759392 |
| | 2.97E-05 | 0.000248094 |
| | 0.812950636 | NA |
| NA | | NA |
| NA | | NA |
| | 0.167248463 | 0.302247469 |
| NA | | NA |
| | 0.086378088 | 0.184224596 |
| NA | | NA |
| | 0.531278239 | 0.680451377 |
| | 0.012776932 | 0.040992352 |
| | 0.55086304 | 0.696581288 |
| | 0.719949502 | NA |
| | 0.992061292 | NA |
| | 1.05E-09 | 3.30E-08 |
| | 0.362484331 | NA |
| | 0.020542652 | 0.060098069 |
| | 0.262026593 | NA |
| | 0.012261571 | 0.039660013 |
| NA | | NA |
| | 0.867473844 | NA |
| NA | | NA |
| NA | | NA |
| | 0.767302482 | NA |
| | 0.016314475 | 0.0500139 |

| | | |
|---|---|---|
| | 0.992061292 | NA |
| | 0.866999183 | NA |
| NA | | NA |
| NA | | NA |
| | 0.992061292 | NA |
| NA | | NA |
| NA | | NA |
| | 0.889409773 | NA |
| NA | | NA |
| | 0.733126346 | NA |
| NA | | NA |
| | 0.33665873 | NA |
| | 0.849992617 | 0.912133503 |
| NA | | NA |
| NA | | NA |
| | 0.99205891 | NA |
| | 0.889409301 | NA |
| | 0.114796354 | NA |
| | 4.49E-20 | 2.10E-17 |
| | 0.884397418 | 0.934485752 |
| | 0.001416232 | 0.006690843 |
| | 0.414237073 | 0.57680583 |
| | 0.917844338 | 0.953132687 |
| NA | | NA |
| | 0.830163713 | NA |
| | 0.00312155 | 0.012984678 |
| | 0.000397024 | 0.002270967 |
| | 0.304476969 | 0.465113723 |
| | 2.74E-05 | 0.00023166 |
| | 0.543665803 | NA |
| NA | | NA |
| | 0.01117757 | 0.036793324 |
| | 0.000945066 | 0.004751076 |
| | 0.4926172 | 0.646627966 |
| | 0.358895218 | NA |
| | 0.902545984 | NA |
| | 0.665375588 | NA |
| | 0.228017299 | NA |
| NA | | NA |
| | 0.197275602 | 0.34150374 |
| | 0.000286134 | 0.001718132 |
| NA | | NA |
| NA | | NA |
| | 0.779112527 | NA |
| | 0.992060816 | NA |
| | 0.53373617 | NA |
| | 0.905158266 | NA |

| | |
|---|---|
| NA | NA |
| 0.889409301 | NA |
| NA | NA |
| 0.992062245 | NA |
| NA | NA |
| 0.880739341 | 0.932225458 |
| 0.248620543 | 0.402436865 |
| 0.411695396 | 0.574442621 |
| 0.105939197 | 0.215061369 |
| 0.064014342 | 0.146730439 |
| 0.110801961 | 0.222282137 |
| 0.015189591 | 0.04718464 |
| NA | NA |
| NA | NA |
| 0.710550414 | NA |
| 0.455922914 | 0.613795063 |
| 0.673134813 | 0.792141873 |
| 1.76E-06 | 2.14E-05 |
| NA | NA |
| 0.454226616 | 0.6122191 |
| 5.67E-13 | 4.29E-11 |
| 0.802637864 | NA |
| 0.099172406 | 0.204654342 |
| 0.871260465 | NA |
| 0.980060592 | 0.990024406 |
| 0.003818374 | 0.015278366 |
| NA | NA |
| NA | NA |
| NA | NA |
| 0.010837158 | 0.035925898 |
| NA | NA |
| 0.000357858 | 0.002081092 |
| 0.988527135 | NA |
| NA | NA |
| NA | NA |
| 0.769602522 | NA |
| 5.64E-05 | 0.000430061 |
| NA | NA |
| 0.101880506 | 0.208966377 |
| 0.577699923 | NA |
| 0.962829401 | 0.979421449 |
| 2.29E-05 | 0.000199333 |
| 9.60E-06 | 9.38E-05 |
| 0.82206316 | NA |
| NA | NA |
| 0.905158739 | NA |
| NA | NA |

| | |
|---|---|
| NA | NA |
| 0.012108305 | 0.039256793 |
| 0.033347546 | 0.08864863 |
| 0.992061292 | NA |
| NA | NA |
| 0.693787389 | 0.807442869 |
| 0.016099389 | 0.049506578 |
| NA | NA |
| NA | NA |
| 0.12180096 | 0.238645073 |
| NA | NA |
| NA | NA |
| 0.377541624 | NA |
| 0.003994053 | 0.015874283 |
| 0.155380779 | NA |
| 0.013584111 | 0.043025169 |
| 0.608604589 | NA |
| 0.092760118 | 0.19447388 |
| 0.148717611 | 0.27678349 |
| 0.905158739 | NA |
| 1.65E-05 | 0.00015068 |
| NA | NA |
| 1.44E-08 | 3.24E-07 |
| 0.022996958 | 0.065813919 |
| 0.261256673 | NA |
| 1.98E-11 | 1.02E-09 |
| 0.992061292 | NA |
| 0.889409301 | NA |
| 0.507557214 | NA |
| 0.255604323 | 0.41020016 |
| 0.002717127 | 0.011587764 |
| 0.717751148 | NA |
| NA | NA |
| NA | NA |
| 0.196245342 | 0.340203006 |
| 0.335642095 | 0.498637763 |
| 0.715001242 | 0.822759119 |
| 0.899097585 | 0.943310432 |
| 0.127855112 | 0.247363031 |
| 0.083080118 | 0.179031056 |
| 0.715096938 | NA |
| 0.97243262 | 0.985628797 |
| 0.063062448 | 0.14506198 |
| 0.92386121 | NA |
| 0.688023371 | 0.803248722 |
| 0.122987938 | 0.240360639 |
| 0.766530371 | 0.858439143 |

| | | |
|---|---|---|
| | 0.000319967 | 0.001889765 |
| | 0.216981973 | NA |
| | 0.697117783 | NA |
| | 0.000448045 | 0.002517561 |
| NA | NA | |
| NA | NA | |
| NA | NA | |
| | 0.4977384 | 0.650740505 |
| | 1.31E-07 | 2.25E-06 |
| | 0.782595072 | NA |
| NA | NA | |
| NA | NA | |
| NA | NA | |
| NA | NA | |
| | 0.000237955 | 0.00146942 |
| | 0.989346336 | NA |
| | 0.005867535 | 0.021882678 |
| NA | NA | |
| | 0.255161698 | 0.409735675 |
| | 3.13E-12 | 1.98E-10 |
| NA | NA | |
| | 1.31E-06 | 1.64E-05 |
| | 0.373660785 | 0.53783542 |
| | 0.511190684 | 0.662803273 |
| NA | NA | |
| | 0.466799536 | NA |
| | 0.171255166 | NA |
| | 0.455231559 | 0.613222273 |
| | 0.0931047 | 0.194954335 |
| | 0.062774923 | 0.144600287 |
| | 0.824343111 | NA |
| | 6.39E-07 | 8.87E-06 |
| | 0.00194918 | 0.008756554 |
| | 0.992060816 | NA |
| NA | NA | |
| | 0.210343104 | 0.357280649 |
| NA | NA | |
| | 0.788348659 | 0.872327952 |
| NA | NA | |
| | 0.82064933 | NA |
| | 0.003579444 | 0.014518102 |
| | 0.001454667 | 0.006837122 |
| | 0.802550285 | NA |
| NA | NA | |
| NA | NA | |
| | 0.008628115 | 0.029860484 |
| NA | NA | |

| | | |
|---|---|---|
| | 0.782559532 | NA |
| | 0.001011504 | 0.00502639 |
| | 0.889409301 | NA |
| | 3.58E-10 | 1.30E-08 |
| | 2.07E-07 | 3.34E-06 |
| | 0.957577123 | 0.976521435 |
| | 0.294957011 | 0.454778894 |
| | 0.503844754 | 0.656145591 |
| | 0.801661681 | NA |
| NA | | NA |
| | 0.46677764 | 0.623253524 |
| | 0.730606056 | NA |
| | 4.65E-13 | 3.62E-11 |
| | 0.428144591 | 0.589148206 |
| | 0.037880215 | 0.097985654 |
| | 0.876062717 | NA |
| | 0.288315025 | 0.447586401 |
| | 0.148045575 | NA |
| | 0.480425569 | NA |
| | 0.361810173 | 0.526681444 |
| | 0.992060816 | NA |
| | 0.90515732 | NA |
| NA | | NA |
| NA | | NA |
| NA | | NA |
| NA | | NA |
| NA | | NA |
| NA | | NA |
| | 0.68100457 | NA |
| | 0.051018574 | 0.123328584 |
| | 0.215682162 | 0.363453657 |
| | 0.643216749 | 0.768883865 |
| | 0.99205891 | NA |
| | 0.889410245 | NA |
| | 0.626811652 | 0.756272366 |
| | 0.533233202 | 0.682080031 |
| | 0.883842258 | 0.934078936 |
| | 0.754854854 | 0.850958599 |
| | 0.005933965 | 0.022078796 |
| | 0.984351729 | NA |
| | 0.992061292 | NA |
| | 0.001419339 | 0.006702636 |
| | 0.782559532 | NA |
| | 0.643740243 | NA |
| | 0.718303953 | NA |
| | 8.42E-09 | 2.03E-07 |
| | 0.050458914 | 0.122231361 |

| | | |
|---|---|---|
| | 0.992496406 | 0.996170975 |
| | 0.660395808 | 0.782560657 |
| | 0.306682998 | 0.467410603 |
| | 0.893245089 | 0.939674663 |
| | 0.770997365 | 0.8614909 |
| | 0.595001094 | 0.732050035 |
| | 0.040874937 | 0.103662709 |
| | 0.638095481 | NA |
| | 0.06263787 | 0.144375457 |
| NA | | NA |
| NA | | NA |
| | 2.15E-06 | 2.56E-05 |
| | 0.007716965 | 0.027252628 |
| | 0.517826336 | 0.668667264 |
| | 0.477401569 | 0.633454592 |
| | 0.176077809 | 0.314272341 |
| | 0.225886924 | 0.375719662 |
| | 0.428847826 | NA |
| | 0.506514379 | 0.65837275 |
| | 0.87762688 | NA |
| | 0.001805911 | 0.008201863 |
| | 0.575918458 | NA |
| | 0.000100579 | 0.000702041 |
| | 0.52856246 | 0.678126145 |
| | 0.006724811 | 0.024440603 |
| | 0.82290605 | 0.894551581 |
| | 0.089263887 | 0.189002573 |
| | 5.25E-11 | 2.40E-09 |
| | 0.963853447 | NA |
| | 0.001544848 | 0.007174981 |
| | 0.0034216 | 0.013988068 |
| | 0.02776339 | 0.076628491 |
| | 0.04393182 | 0.109649789 |
| | 0.000323524 | 0.001908209 |
| | 2.41E-06 | 2.82E-05 |
| | 0.996229563 | NA |
| | 0.2306341 | 0.381448686 |
| | 0.889410245 | NA |
| | 0.965406552 | NA |
| | 0.07351658 | 0.162605819 |
| | 0.001541086 | 0.007164978 |
| | 0.695041301 | NA |
| | 0.992062245 | NA |
| | 0.664277219 | 0.785549219 |
| | 0.764143939 | 0.857062789 |
| | 0.741642892 | 0.841758403 |
| | 0.260182061 | 0.415485264 |

| | |
|---|---|
| 7.99E-08 | 1.45E-06 |
| 0.777092376 | NA |
| 0.006904364 | 0.025014507 |
| 0.614480225 | NA |
| NA | NA |
| 0.002855087 | 0.012065984 |
| 0.884252762 | NA |
| 0.262509584 | 0.417956962 |
| 0.965779388 | 0.981422073 |
| 0.908689 | 0.948154039 |
| 0.991918092 | NA |
| 0.548147457 | NA |
| 0.003309621 | 0.013612282 |
| 0.905158739 | NA |
| 0.786164713 | NA |
| NA | NA |
| NA | NA |
| 1.91E-06 | 2.30E-05 |
| NA | NA |
| 4.62E-06 | 4.97E-05 |
| NA | NA |
| 0.985930184 | NA |
| 0.754400838 | NA |
| 0.033117411 | NA |
| 0.000271041 | 0.001643232 |
| 0.956643634 | NA |
| 0.769413204 | NA |
| 0.004171446 | 0.016457202 |
| NA | NA |
| NA | NA |
| 0.905158739 | NA |
| 0.895646036 | 0.941075508 |
| 0.955597015 | NA |
| 0.120608197 | 0.237017667 |
| 0.057864946 | 0.135609003 |
| 0.984573214 | 0.992171589 |
| NA | NA |
| 0.321977575 | 0.484334387 |
| 0.002285984 | 0.010017547 |
| 0.03674128 | 0.095703684 |
| NA | NA |
| 0.998601186 | NA |
| NA | NA |
| 0.242911183 | 0.395762478 |
| 0.000768792 | 0.003993828 |
| 0.404153963 | 0.567470944 |
| 1.21E-07 | 2.09E-06 |

| | | |
|---|---|---|
| | 0.924104487 | NA |
| | 0.885481135 | NA |
| | 0.907732836 | 0.947610543 |
| | 0.753886894 | NA |
| | 4.25E-05 | 0.000337064 |
| NA | | NA |
| | 0.20388295 | 0.349743434 |
| | 0.827323718 | 0.896835737 |
| | 0.239790041 | 0.392102875 |
| | 1.81E-15 | 2.52E-13 |
| | 0.047143745 | 0.115775426 |
| NA | | NA |
| | 0.092753443 | 0.19447388 |
| NA | | NA |
| | 0.140537836 | 0.265308661 |
| | 0.413231655 | 0.575807723 |
| NA | | NA |
| | 0.889410717 | NA |
| NA | | NA |
| NA | | NA |
| | 0.992061292 | NA |
| | 0.03001506 | 0.081539432 |
| NA | | NA |
| | 0.680581164 | 0.797443135 |
| NA | | NA |
| NA | | NA |
| | 0.261605062 | 0.417120647 |
| | 0.76617043 | NA |
| NA | | NA |
| NA | | NA |
| NA | | NA |
| NA | | NA |
| NA | | NA |
| NA | | NA |
| NA | | NA |
| NA | | NA |
| | 0.985083122 | NA |
| | 0.476820529 | NA |
| | 0.289343741 | 0.4486508 |
| | 0.941735684 | 0.967042615 |
| | 0.992061292 | NA |
| | 0.992060816 | NA |
| | 0.18519274 | NA |
| | 0.905296333 | NA |
| NA | | NA |
| | 0.176516021 | NA |
| | 0.570364658 | 0.712647575 |

| | |
|---|---|
| 0.13485296 | 0.257007129 |
| NA | NA |
| NA | NA |
| 0.473921969 | NA |
| 0.007378303 | 0.026332265 |
| 0.494126311 | NA |
| 0.047435302 | 0.116387875 |
| 0.760860724 | NA |
| NA | NA |
| 0.889409773 | NA |
| NA | NA |
| 0.992061292 | NA |
| 0.992061292 | NA |
| 0.005352574 | 0.020293022 |
| 5.94E-09 | 1.48E-07 |
| 0.430820308 | NA |
| 0.037240665 | NA |
| NA | NA |
| 0.215245114 | NA |
| 0.547670623 | NA |
| 0.363978673 | NA |
| 0.072261752 | 0.160379605 |
| 0.793725782 | 0.875498515 |
| 0.278811232 | 0.436940897 |
| 0.022118982 | 0.063776531 |
| 0.002793185 | 0.011843128 |
| NA | NA |
| NA | NA |
| 0.000614759 | 0.003307887 |
| 0.148631235 | 0.276709901 |
| 0.466856678 | 0.623311765 |
| 0.044386667 | 0.110571393 |
| 0.151727829 | NA |
| 0.297831038 | 0.457795867 |
| 0.002007263 | 0.008982574 |
| 0.646097183 | 0.771235488 |
| 0.427547702 | NA |
| 0.778347789 | 0.866682731 |
| 0.135245535 | NA |
| 0.000854018 | 0.004351133 |
| 0.308528771 | 0.469604443 |
| 0.828974519 | 0.897960313 |
| NA | NA |
| 0.889409301 | NA |
| 0.000777984 | 0.004031903 |
| 0.092270129 | 0.193687005 |
| 0.038997914 | NA |

| | | |
|---|---|---|
| | 0.733112307 | NA |
| | 0.00970717 | 0.032829161 |
| NA | | NA |
| | 0.710780516 | NA |
| | 0.140372342 | NA |
| NA | | NA |
| NA | | NA |
| | 0.647436604 | NA |
| | 0.992061292 | NA |
| NA | | NA |
| | 0.558916441 | 0.703504149 |
| | 0.033139729 | NA |
| | 0.457614132 | NA |
| | 0.905158266 | NA |
| | 0.889409773 | NA |
| | 0.208220578 | 0.354938339 |
| | 0.034588282 | 0.091234124 |
| NA | | NA |
| | 0.905158739 | NA |
| | 0.901289765 | 0.944265974 |
| | 0.648193601 | 0.773023586 |
| | 0.008802966 | 0.030346048 |
| | 0.399630123 | 0.563252752 |
| | 0.000569898 | 0.00310069 |
| NA | | NA |
| NA | | NA |
| | 0.520027545 | 0.670775662 |
| NA | | NA |
| | 3.33E-06 | 3.74E-05 |
| NA | | NA |
| | 0.000110901 | 0.000762691 |
| | 0.108507567 | 0.219099287 |
| NA | | NA |
| | 0.310401045 | 0.471571645 |
| | 0.065176045 | 0.148815551 |
| | 1.07E-07 | 1.88E-06 |
| | 0.544680689 | NA |
| | 0.460294459 | NA |
| | 0.984860399 | NA |
| | 0.012090647 | 0.039205332 |
| NA | | NA |
| NA | | NA |
| | 0.712474803 | NA |
| NA | | NA |
| NA | | NA |
| NA | | NA |
| | 0.554065509 | 0.699478241 |

| | | |
|---|---|---|
| NA | NA | |
| 0.992060816 | NA | |
| 0.991772468 | NA | |
| NA | NA | |
| NA | NA | |
| 0.063189358 | NA | |
| 0.992059387 | NA | |
| 0.366679523 | | 0.531409427 |
| NA | NA | |
| NA | NA | |
| NA | NA | |
| NA | NA | |
| NA | NA | |
| 0.710240538 | | 0.819476542 |
| NA | NA | |
| 0.054418357 | | 0.12939607 |
| 0.905157793 | NA | |
| 0.905158739 | NA | |
| 1.26E-10 | | 5.21E-09 |
| 0.565874456 | NA | |
| NA | NA | |
| NA | NA | |
| NA | NA | |
| NA | NA | |
| 0.992060816 | NA | |
| 0.905158739 | NA | |
| NA | NA | |
| NA | NA | |
| NA | NA | |
| 0.992061292 | NA | |
| 0.618293911 | NA | |
| 0.000508327 | | 0.002800392 |
| 0.006964136 | | 0.0251718 |
| 0.245637285 | | 0.398918058 |
| 0.591319868 | | 0.72907316 |
| NA | NA | |
| 0.905158739 | NA | |
| 0.771018415 | | 0.8614909 |
| 0.438832012 | | 0.599198617 |
| 0.005395776 | | 0.02042155 |
| 3.53E-10 | | 1.28E-08 |
| 0.404115132 | | 0.56746158 |
| 0.889409301 | NA | |
| 0.001169662 | | 0.005683692 |
| 0.002631495 | | 0.011294959 |
| 0.74673718 | NA | |
| 0.578880068 | | 0.719593966 |

| | | |
|---|---|---|
| | 0.00140005 | 0.006625796 |
| | 0.077964638 | 0.170537452 |
| | 0.99205891 | NA |
| | 0.617454212 | 0.749885351 |
| | 0.28994849 | 0.449296373 |
| | 0.766440153 | 0.858439143 |
| | 0.743481574 | NA |
| | 0.268831476 | 0.425185132 |
| | 0.550102626 | NA |
| | 0.847965549 | NA |
| | 0.027591575 | 0.076230926 |
| | 0.003076841 | 0.01281813 |
| | 0.622905081 | 0.753961673 |
| | 0.644280116 | NA |
| | 6.65E-05 | 0.000493701 |
| | 0.966542867 | NA |
| | 0.090041935 | 0.190209798 |
| | 0.009590347 | 0.032514194 |
| | 0.418507421 | NA |
| | 0.606446973 | NA |
| | 0.992060816 | NA |
| | 0.905158266 | NA |
| | 0.992061292 | NA |
| NA | | NA |
| NA | | NA |
| NA | | NA |
| NA | | NA |
| | 0.992061292 | NA |
| NA | | NA |
| | 0.473537125 | 0.629900759 |
| NA | | NA |
| NA | | NA |
| | 0.905158739 | NA |
| NA | | NA |
| NA | | NA |
| | 0.905158266 | NA |
| NA | | NA |
| NA | | NA |
| NA | | NA |
| NA | | NA |
| | 0.992060816 | NA |
| NA | | NA |
| NA | | NA |
| | 0.905158739 | NA |
| | 0.90515732 | NA |
| NA | | NA |
| NA | | NA |

| | | |
|---|---|---|
| NA | | NA |
| | 0.992061292 | NA |
| NA | | NA |
| NA | | NA |
| NA | | NA |
| | 0.889409301 | NA |
| NA | | NA |
| NA | | NA |
| NA | | NA |
| NA | | NA |
| NA | | NA |
| NA | | NA |
| NA | | NA |
| NA | | NA |
| NA | | NA |
| NA | | NA |
| | 0.99205891 | NA |
| NA | | NA |
| NA | | NA |
| NA | | NA |
| NA | | NA |
| NA | | NA |
| NA | | NA |
| NA | | NA |
| NA | | NA |
| | 0.860142408 | NA |
| NA | | NA |
| NA | | NA |
| NA | | NA |
| NA | | NA |
| NA | | NA |
| NA | | NA |
| | 0.99205891 | NA |
| NA | | NA |
| NA | | NA |
| NA | | NA |
| NA | | NA |
| NA | | NA |
| NA | | NA |
| NA | | NA |
| | 0.889409773 | NA |
| NA | | NA |
| | 0.378994961 | 0.543446784 |
| NA | | NA |
| NA | | NA |
| NA | | NA |
| NA | | NA |

| | | |
|---|---|---|
| NA | | NA |
| NA | | NA |
| NA | | NA |
| NA | | NA |
| NA | | NA |
| NA | | NA |
| NA | | NA |
| NA | | NA |
| NA | | NA |
| NA | | NA |
| NA | | NA |
| NA | | NA |
| NA | | NA |
| NA | | NA |
| NA | | NA |
| NA | | NA |
| NA | | NA |
| NA | | NA |
| NA | | NA |
| NA | | NA |
| NA | | NA |
| NA | | NA |
| NA | | NA |
| NA | | NA |
| NA | | NA |
| NA | | NA |
| NA | | NA |
| NA | | NA |
| NA | | NA |
| NA | | NA |
| NA | | NA |
| NA | | NA |
| NA | | NA |
| | 0.992060816 | NA |
| NA | | NA |
| NA | | NA |
| NA | | NA |
| NA | | NA |
| NA | | NA |
| NA | | NA |
| NA | | NA |
| NA | | NA |
| | 0.905158266 | NA |
| NA | | NA |
| NA | | NA |
| NA | | NA |
| NA | | NA |

| | | |
|---|---|---|
| NA | NA | |
| 0.646024413 | NA | |
| NA | NA | |
| NA | NA | |
| NA | NA | |
| NA | NA | |
| NA | NA | |
| NA | NA | |
| NA | NA | |
| NA | NA | |
| 0.992061292 | NA | |
| NA | NA | |
| NA | NA | |
| NA | NA | |
| NA | NA | |
| NA | NA | |
| NA | NA | |
| NA | NA | |
| NA | NA | |
| 0.02579318 | | 0.072326962 |
| NA | NA | |
| NA | NA | |
| NA | NA | |
| NA | NA | |
| NA | NA | |
| NA | NA | |
| NA | NA | |
| 0.911528918 | | 0.950003887 |
| NA | NA | |
| NA | NA | |
| 0.747813307 | NA | |
| NA | NA | |
| 0.485157163 | | 0.640199877 |
| NA | NA | |
| NA | NA | |
| NA | NA | |
| NA | NA | |
| NA | NA | |
| NA | NA | |
| NA | NA | |
| NA | NA | |
| NA | NA | |
| NA | NA | |
| NA | NA | |
| NA | NA | |
| NA | NA | |
| NA | NA | |

| | | |
|---|---|---|
| NA | | NA |
| NA | | NA |
| NA | | NA |
| NA | | NA |
| NA | | NA |
| NA | | NA |
| NA | | NA |
| NA | | NA |
| NA | | NA |
| NA | | NA |
| NA | | NA |
| NA | | NA |
| NA | | NA |
| NA | | NA |
| NA | | NA |
| NA | | NA |
| NA | | NA |
| NA | | NA |
| NA | | NA |
| NA | | NA |
| NA | | NA |
| NA | | NA |
| NA | | NA |
| NA | | NA |
| NA | | NA |
| NA | | NA |
| NA | | NA |
| | 0.905158266 | NA |
| NA | | NA |
| NA | | NA |
| | 0.905158266 | NA |
| NA | | NA |
| NA | | NA |
| NA | | NA |
| NA | | NA |
| NA | | NA |
| NA | | NA |
| NA | | NA |
| NA | | NA |
| | 0.905158739 | NA |
| NA | | NA |
| NA | | NA |
| NA | | NA |
| NA | | NA |
| NA | | NA |
| NA | | NA |
| NA | | NA |

| | |
|---|---|
| NA | NA |
| NA | NA |
| NA | NA |
| NA | NA |
| NA | NA |
| NA | NA |
| NA | NA |
| NA | NA |
| NA | NA |
| 0.889409301 | NA |
| NA | NA |
| NA | NA |
| 0.992061292 | NA |
| 0.881501945 | NA |
| NA | NA |
| NA | NA |
| NA | NA |
| NA | NA |
| NA | NA |
| NA | NA |
| NA | NA |
| NA | NA |
| NA | NA |
| NA | NA |
| NA | NA |
| NA | NA |
| NA | NA |
| 0.992060816 | NA |
| NA | NA |
| NA | NA |
| NA | NA |
| NA | NA |
| NA | NA |
| NA | NA |
| NA | NA |
| NA | NA |
| NA | NA |
| 0.889409301 | NA |
| 0.99205891 | NA |
| NA | NA |
| 0.992060816 | NA |
| 0.992060816 | NA |
| NA | NA |
| NA | NA |
| NA | NA |
| NA | NA |
| NA | NA |

| | | | |
|---|---|---|---|
| | 0.889409301 | NA | |
| NA | | NA | |
| NA | | NA | |
| | 0.905158266 | NA | |
| NA | | NA | |
| | 0.992060816 | NA | |
| | 0.992060816 | NA | |
| NA | | NA | |
| NA | | NA | |
| NA | | NA | |
| NA | | NA | |
| NA | | NA | |
| NA | | NA | |
| | 0.889410245 | NA | |
| NA | | NA | |
| | 0.889409301 | NA | |
| NA | | NA | |
| NA | | NA | |
| NA | | NA | |
| | 0.992061292 | NA | |
| NA | | NA | |
| NA | | NA | |
| NA | | NA | |
| | 0.992061292 | NA | |
| NA | | NA | |
| NA | | NA | |
| | 0.989534413 | NA | |
| NA | | NA | |
| | 0.625450261 | NA | |
| NA | | NA | |
| | 0.992061292 | NA | |
| | 0.616377951 | NA | |
| | 0.700956025 | NA | |
| NA | | NA | |
| | 0.992060816 | NA | |
| | 0.905158739 | NA | |
| | 0.564925359 | NA | |
| | 0.905158266 | NA | |
| | 0.630936361 | NA | |
| | 0.889409301 | NA | |
| | 0.572211767 | NA | |
| NA | | NA | |
| | 0.905158266 | NA | |
| | 0.624185775 | | 0.754827625 |
| NA | | NA | |
| | 0.049826444 | NA | |
| | 0.784353866 | NA | |

| | |
|---|---|
| NA | NA |
| NA | NA |
| 0.889409773 | NA |
| NA | NA |
| 0.797370036 | NA |
| 0.73936521 | NA |
| 0.762356347 | 0.855801684 |
| 0.837761394 | NA |
| 0.162976002 | 0.296404674 |
| 0.013883514 | 0.043821774 |
| 0.690799586 | 0.805274956 |
| 0.142472384 | 0.268016026 |
| 0.544165912 | 0.69083305 |
| 0.325806149 | 0.487953706 |
| 3.63E-08 | 7.41E-07 |
| 0.42426043 | 0.585527923 |
| 0.564331652 | 0.707889708 |
| 0.79918557 | 0.878916008 |
| 0.806777773 | 0.884165472 |
| 0.164906443 | 0.29909809 |
| 0.905158266 | NA |
| 0.286688533 | 0.445736985 |
| 0.672202033 | NA |
| 0.023729972 | 0.067472026 |
| 0.067566852 | 0.152455868 |
| 3.36E-05 | 0.00027608 |
| 0.12826896 | 0.247923431 |
| 0.065768904 | 0.149675548 |
| 1.00E-10 | 4.26E-09 |
| 0.88587376 | 0.935299266 |
| NA | NA |
| 0.74604618 | 0.844890565 |
| 0.009634942 | 0.032642481 |
| 0.001592364 | 0.007362964 |
| 0.304165162 | 0.464799081 |
| 0.867742132 | 0.924180302 |
| 0.00567545 | 0.02130744 |
| 9.28E-06 | 9.14E-05 |
| 0.708927493 | 0.818515107 |
| 0.554739237 | NA |
| 0.702324993 | 0.814274263 |
| 0.659822653 | NA |
| 0.266197266 | 0.42242149 |
| 0.99120764 | NA |
| 0.079594429 | 0.173222405 |
| 0.649771882 | 0.774527237 |
| 0.385377863 | 0.549228485 |

| | |
|---|---|
| 0.04087688 | 0.103662709 |
| 1.20E-06 | 1.53E-05 |
| 0.103792844 | 0.211741645 |
| NA | NA |
| 0.036688554 | 0.0956027 |
| 0.784514522 | 0.870190149 |
| 3.57E-05 | 0.000290585 |
| 0.000209009 | 0.001308331 |
| 0.000107352 | 0.000742112 |
| 0.779378776 | 0.867285433 |
| 0.563203241 | NA |
| 3.56E-05 | 0.000289463 |
| NA | NA |
| 0.446541616 | 0.605882807 |
| 0.433087651 | 0.593829019 |
| 0.383297494 | 0.547270865 |
| 0.670158159 | 0.789696968 |
| 0.074270266 | 0.164091032 |
| 0.714671422 | 0.822595123 |
| 0.275546422 | 0.43292029 |
| 0.400914685 | 0.564555956 |
| 0.386765984 | 0.550609755 |
| 0.735939652 | NA |
| 4.07E-07 | 6.01E-06 |
| 0.711614161 | NA |
| 0.21974892 | 0.368625693 |
| 0.274609373 | NA |
| NA | NA |
| 0.905158266 | NA |
| NA | NA |
| NA | NA |
| 2.14E-07 | 3.44E-06 |
| 0.218519738 | NA |
| 0.855618785 | 0.915974383 |
| 0.251375329 | NA |
| 0.005001577 | 0.019194274 |
| 0.022959698 | 0.065732986 |
| 0.695066769 | NA |
| NA | NA |
| 0.279446894 | 0.437655969 |
| 0.000299849 | 0.001785356 |
| NA | NA |
| NA | NA |
| 0.008758859 | 0.030227216 |
| 0.459977986 | NA |
| 0.889409773 | NA |
| NA | NA |

| | |
|---|---|
| 0.069698708 | 0.15598389 |
| 0.612007241 | 0.745485134 |
| NA | NA |
| 0.617805984 | 0.750177822 |
| 0.263112715 | 0.418764377 |
| 0.453908367 | NA |
| 0.338811904 | 0.501785298 |
| NA | NA |
| 0.543666862 | NA |
| 0.157495262 | NA |
| 0.22218867 | 0.371510014 |
| 0.001501852 | 0.007016846 |
| 0.326958888 | NA |
| 0.824117289 | NA |
| NA | NA |
| 0.096732316 | 0.200864769 |
| 0.607410884 | NA |
| 0.090293166 | 0.190657768 |
| 0.475119425 | 0.631287226 |
| 0.521781037 | 0.672366046 |
| 0.012213769 | 0.039540344 |
| NA | NA |
| NA | NA |
| NA | NA |
| NA | NA |
| NA | NA |
| NA | NA |
| NA | NA |
| NA | NA |
| NA | NA |
| NA | NA |
| NA | NA |
| 2.89E-14 | 2.90E-12 |
| NA | NA |
| 0.619483509 | 0.751186613 |
| 6.82E-05 | 0.000504424 |
| 0.023405952 | 0.066792892 |
| 0.02243261 | 0.06451836 |
| NA | NA |
| NA | NA |
| 0.022349972 | 0.064305952 |
| NA | NA |
| NA | NA |
| 0.540228512 | NA |
| 0.233752782 | 0.385329731 |
| NA | NA |
| NA | NA |

| | |
|---|---|
| NA | NA |
| 0.992059387 | NA |
| NA | NA |
| 0.992060816 | NA |
| NA | NA |
| 0.680190983 | 0.797241196 |
| NA | NA |
| 0.748778853 | NA |
| 0.89959656 | 0.943475839 |
| 0.490368615 | 0.64447043 |
| 4.62E-05 | 0.000361826 |
| 0.99205891 | NA |
| 0.004497515 | 0.01751391 |
| 0.004938123 | 0.018980615 |
| 0.807198471 | 0.884393976 |
| 0.001241043 | 0.005965862 |
| 0.460635838 | NA |
| 0.677585777 | NA |
| 1.95E-05 | 0.000174147 |
| 0.199524796 | 0.344351331 |
| 0.056258427 | 0.132679497 |
| 0.604071201 | 0.739253085 |
| 2.68E-12 | 1.74E-10 |
| 0.892880345 | 0.939380965 |
| 8.32E-08 | 1.50E-06 |
| NA | NA |
| 0.284286403 | 0.443149367 |
| 0.108355173 | 0.218893399 |
| 0.547632131 | 0.693866899 |
| 0.699409217 | NA |
| 0.986369879 | NA |
| NA | NA |
| 0.124346115 | 0.242367863 |
| NA | NA |
| NA | NA |
| 0.516566256 | NA |
| 0.046086263 | 0.113926405 |
| 0.11175037 | 0.223735082 |
| NA | NA |
| 0.002253095 | 0.009891188 |
| 0.165735437 | 0.300105918 |
| 1.93E-11 | 1.01E-09 |
| 0.872971919 | NA |
| NA | NA |
| 0.362202719 | 0.526974984 |
| NA | NA |
| NA | NA |

| | | |
|---|---|---|
| NA | NA | |
| NA | NA | |
| NA | NA | |
| NA | NA | |
| 0.992061292 | NA | |
| 0.001866347 | | 0.008438035 |
| 0.339526521 | NA | |
| 0.007751385 | | 0.027343399 |
| 0.003925493 | | 0.015638587 |
| 0.015563346 | | 0.0481278 |
| 0.992061292 | NA | |
| 0.272250275 | | 0.429016418 |
| 0.012313565 | | 0.03979302 |
| 0.09029477 | NA | |
| 0.000121246 | | 0.000824284 |
| 0.016986807 | | 0.051656737 |
| 0.973340833 | | 0.985912235 |
| NA | NA | |
| 0.557260472 | | 0.702300431 |
| 0.332928872 | | 0.495542724 |
| NA | NA | |
| NA | NA | |
| 0.364652524 | NA | |
| 0.235761361 | | 0.387419271 |
| NA | NA | |
| 0.823578793 | NA | |
| 0.952638475 | | 0.973374154 |
| 0.678100382 | | 0.795767758 |
| 0.870725732 | NA | |
| 0.25552394 | | 0.410134507 |
| 0.15946235 | | 0.291559846 |
| 0.404876602 | | 0.568177875 |
| 0.026322536 | | 0.073370979 |
| 0.461935536 | | 0.619371794 |
| 0.818160539 | NA | |
| 0.838070435 | | 0.903756258 |
| NA | NA | |
| 0.873981309 | NA | |
| NA | NA | |
| NA | NA | |
| NA | NA | |
| NA | NA | |
| NA | NA | |
| 5.23E-11 | | 2.39E-09 |
| 0.779764553 | NA | |
| 0.038438752 | | 0.099058391 |
| NA | NA | |

| | | |
|---|---|---|
| | 0.09827119 | 0.203176449 |
| | 0.889410245 | NA |
| | 0.035791477 | 0.093675629 |
| | 0.211778741 | 0.35885866 |
| | 0.969976384 | 0.98418406 |
| | 0.636441566 | 0.763403089 |
| | 0.477587932 | NA |
| | 0.001279677 | 0.0061261 |
| | 0.069466796 | 0.155560001 |
| | 0.905157793 | NA |
| NA | | NA |
| | 0.833573894 | 0.900845684 |
| | 0.992060816 | NA |
| NA | | NA |
| | 0.714424162 | NA |
| | 0.043607039 | 0.109045882 |
| | 0.889409301 | NA |
| | 0.002231977 | 0.009820073 |
| | 6.52E-05 | 0.000485197 |
| | 0.905158266 | NA |
| | 0.755964139 | NA |
| | 0.035586696 | 0.093284978 |
| | 0.991918092 | NA |
| | 0.333296133 | NA |
| | 0.515656853 | NA |
| NA | | NA |
| NA | | NA |
| | 0.053495446 | 0.127616133 |
| | 0.525234016 | NA |
| NA | | NA |
| | 0.905157793 | NA |
| | 0.015744523 | 0.048599099 |
| | 0.939903749 | NA |
| | 0.905158739 | NA |
| NA | | NA |
| NA | | NA |
| | 0.391670038 | NA |
| | 0.071154974 | 0.158515987 |
| NA | | NA |
| | 0.213700589 | 0.360917646 |
| | 0.405665402 | NA |
| | 0.401434851 | 0.564945236 |
| NA | | NA |
| NA | | NA |
| | 0.98933516 | NA |
| | 4.51E-06 | 4.87E-05 |
| | 0.847244377 | NA |

| | | | |
|---|---|---|---|
| | 0.001146237 | | 0.005588316 |
| NA | | NA | |
| | 0.834617617 | | 0.901538111 |
| | 0.584552453 | | 0.72424086 |
| NA | | NA | |
| | 0.171057472 | NA | |
| | 0.854025748 | NA | |
| | 0.328122752 | NA | |
| | 0.394041128 | | 0.557450659 |
| | 0.213165898 | | 0.360337793 |
| | 0.992061292 | NA | |
| | 0.905158266 | NA | |
| | 0.821948024 | NA | |
| | 2.37E-10 | | 8.99E-09 |
| | 0.000244343 | | 0.001501267 |
| NA | | NA | |
| NA | | NA | |
| NA | | NA | |
| | 0.784353866 | NA | |
| | 0.084694649 | | 0.181427158 |
| NA | | NA | |
| NA | | NA | |
| | 0.889409773 | NA | |
| NA | | NA | |
| NA | | NA | |
| NA | | NA | |
| NA | | NA | |
| NA | | NA | |
| NA | | NA | |
| | 0.905158739 | NA | |
| NA | | NA | |
| NA | | NA | |
| NA | | NA | |
| NA | | NA | |
| | 0.99205891 | NA | |
| NA | | NA | |
| | 0.889409301 | NA | |
| NA | | NA | |
| NA | | NA | |
| NA | | NA | |
| | 0.903394986 | NA | |
| | 0.991918092 | NA | |
| NA | | NA | |
| NA | | NA | |
| NA | | NA | |
| NA | | NA | |
| NA | | NA | |

| | | |
|---|---|---|
| NA | | NA |
| NA | | NA |
| | 0.905158266 | NA |
| NA | | NA |
| | 0.905158266 | NA |
| NA | | NA |
| | 0.992060816 | NA |
| NA | | NA |
| NA | | NA |
| NA | | NA |
| NA | | NA |
| NA | | NA |
| NA | | NA |
| NA | | NA |
| | 0.992060816 | NA |
| NA | | NA |
| NA | | NA |
| | 0.905158266 | NA |
| NA | | NA |
| NA | | NA |
| | 0.905158266 | NA |
| NA | | NA |
| NA | | NA |
| | 2.94E-05 | 0.00024656 |
| | 0.621129159 | NA |
| | 0.305298975 | 0.465883265 |
| | 4.31E-05 | 0.000340843 |
| | 0.642042252 | NA |
| | 0.145486049 | 0.272332124 |
| | 0.078054136 | 0.170665216 |
| NA | | NA |
| | 0.111637163 | 0.223610368 |
| NA | | NA |
| | 0.002871992 | 0.012124451 |
| | 0.00047505 | 0.002641589 |
| | 0.876100462 | NA |
| | 0.210121134 | NA |
| | 0.016432486 | 0.050294978 |
| | 0.226969818 | NA |
| | 0.838236169 | NA |
| | 2.08E-06 | 2.49E-05 |
| | 0.109339344 | 0.220356438 |
| | 0.036830592 | NA |
| | 0.132643898 | 0.253832449 |
| | 0.992061292 | NA |
| NA | | NA |
| | 0.889409773 | NA |

| | | |
|---|---|---|
| | 0.889409301 | NA |
| | 0.599164734 | 0.735277657 |
| | 1.11E-06 | 1.43E-05 |
| | 0.66120557 | 0.782933739 |
| | 0.575472399 | 0.717225632 |
| | 0.628360162 | 0.756928011 |
| | 0.067333609 | 0.152070216 |
| | 0.073071813 | NA |
| | 0.12524848 | 0.243672488 |
| | 8.99E-13 | 6.41E-11 |
| | 0.808297401 | NA |
| | 0.26127022 | 0.416738051 |
| | 0.889409773 | NA |
| | 0.941945658 | NA |
| | 0.392886348 | 0.556347822 |
| | 0.797854497 | NA |
| | 0.992061292 | NA |
| | 0.375585262 | 0.539791659 |
| NA | | NA |
| | 0.977993482 | NA |
| NA | | NA |
| | 0.566429174 | NA |
| | 0.552058754 | NA |
| | 0.889409773 | NA |
| | 0.344913037 | 0.508216157 |
| | 0.715183383 | NA |
| | 0.04815342 | 0.117790014 |
| | 0.682131888 | NA |
| | 0.057227704 | 0.134462242 |
| | 0.902998829 | 0.945090858 |
| | 0.003700599 | 0.014902155 |
| | 0.719437981 | NA |
| | 0.566617061 | 0.709531481 |
| | 0.445670197 | NA |
| | 0.826656544 | 0.896687959 |
| | 0.01274981 | 0.040937722 |
| NA | | NA |
| | 0.795353865 | NA |
| | 0.109721118 | 0.220963121 |
| | 0.795014225 | NA |
| | 0.0293908 | 0.080175696 |
| | 0.708469154 | 0.818201018 |
| | 0.120338364 | NA |
| | 0.829237509 | NA |
| | 0.192715478 | 0.335620524 |
| | 2.03E-06 | 2.44E-05 |
| | 0.878369794 | NA |

| | | |
|---|---|---|
| | 0.626522117 | NA |
| | 0.992059387 | NA |
| NA | | NA |
| | 0.003895027 | 0.01553384 |
| NA | | NA |
| | 0.992060816 | NA |
| | 1.94E-05 | 0.000173641 |
| | 0.880258492 | NA |
| | 0.547931293 | 0.694098718 |
| NA | | NA |
| | 0.905158266 | NA |
| | 0.87063874 | NA |
| | 0.212750143 | 0.359921448 |
| | 0.277601668 | NA |
| NA | | NA |
| | 0.028122146 | 0.077375543 |
| | 0.021430185 | 0.062148102 |
| NA | | NA |
| | 0.008067849 | 0.028272586 |
| NA | | NA |
| | 0.762384104 | 0.855801684 |
| | 0.863102073 | 0.920982 |
| | 0.0041396 | 0.016352129 |
| | 0.908452678 | 0.948154039 |
| | 0.421316309 | 0.583003644 |
| | 0.038270764 | 0.098764567 |
| | 0.752700049 | NA |
| | 0.025956331 | 0.072617539 |
| | 0.943131176 | 0.96780427 |
| | 0.830069175 | NA |
| | 3.12E-06 | 3.51E-05 |
| | 0.032915814 | 0.087718246 |
| | 0.01405962 | 0.044269458 |
| | 0.954300115 | NA |
| | 0.900836801 | NA |
| | 0.30988904 | 0.471119733 |
| | 0.710261113 | 0.819476542 |
| | 0.54004039 | 0.687304214 |
| | 0.078014671 | 0.170629897 |
| | 0.061488847 | 0.14244473 |
| | 0.82987257 | 0.898534183 |
| | 0.178298903 | 0.317359619 |
| | 0.001900014 | 0.008563711 |
| | 0.946197626 | NA |
| | 0.796710347 | 0.877424269 |
| | 0.889410245 | NA |
| | 0.448449089 | 0.607494974 |

| | |
|---|---|
| 3.07E-09 | 8.45E-08 |
| NA | NA |
| 0.001320987 | 0.006297776 |
| 0.416742165 | NA |
| 0.108290754 | 0.218803521 |
| 6.32E-08 | 1.19E-06 |
| 0.182075305 | 0.322252891 |
| 0.99205891 | NA |
| 0.008466463 | 0.029402983 |
| 0.488449077 | 0.642952732 |
| 1.64E-07 | 2.73E-06 |
| 0.066556683 | 0.150952097 |
| 0.796664224 | 0.877417479 |
| NA | NA |
| 0.017039875 | 0.051783454 |
| 0.91585825 | 0.95229975 |
| 0.769650489 | NA |
| 0.000807854 | 0.004161313 |
| 0.003056457 | NA |
| 2.73E-19 | 9.84E-17 |
| 3.04E-12 | 1.94E-10 |
| 0.81586252 | 0.89047791 |
| NA | NA |
| 8.50E-06 | 8.49E-05 |
| NA | NA |
| 0.0217674 | 0.06295974 |
| 0.030649096 | 0.082864257 |
| 0.992062245 | NA |
| NA | NA |
| 0.613550958 | 0.746838442 |
| 0.807879735 | NA |
| 0.579487872 | 0.719901233 |
| 0.09801452 | 0.202763835 |
| 2.22E-06 | 2.63E-05 |
| 6.08E-07 | 8.50E-06 |
| 0.976681734 | NA |
| 0.635374275 | NA |
| 0.423325519 | NA |
| 0.032028242 | 0.085856901 |
| 0.827094968 | 0.89672859 |
| 0.910351782 | 0.949212477 |
| 1.01E-12 | 7.10E-11 |
| 0.000617811 | 0.003323494 |
| 0.865008748 | 0.922118894 |
| 0.001515507 | 0.00706699 |
| 0.067009664 | 0.151634914 |
| 0.226346262 | 0.376198726 |

| | |
|---|---|
| 0.198883788 | 0.343547969 |
| NA | NA |
| 0.082634762 | 0.178389377 |
| 0.655358301 | 0.77827121 |
| 0.75266361 | 0.849447196 |
| 0.017975325 | 0.053960379 |
| 0.323684372 | 0.485985809 |
| 0.234458562 | 0.385942695 |
| 0.22390095 | 0.373548167 |
| 0.639951338 | 0.766440766 |
| 0.913096076 | NA |
| 0.475956054 | NA |
| NA | NA |
| 0.191959046 | NA |
| 0.148026922 | 0.275888913 |
| 0.988961297 | NA |
| 0.359282249 | 0.524113393 |
| 2.45E-10 | 9.22E-09 |
| 0.000255526 | 0.001558709 |
| 0.854172913 | 0.914957097 |
| 3.69E-12 | 2.29E-10 |
| 0.680240075 | 0.797256181 |
| 0.852583254 | 0.914066576 |
| 4.34E-10 | 1.53E-08 |
| 0.484122867 | 0.639449867 |
| 0.229484367 | 0.379976416 |
| 0.478383216 | 0.634326348 |
| NA | NA |
| 1.71E-07 | 2.83E-06 |
| 0.346021754 | NA |
| 0.577868109 | 0.719068725 |
| 0.040568681 | 0.103071581 |
| 0.515051253 | NA |
| 0.006747801 | 0.024503882 |
| 0.039994869 | 0.102074307 |
| 0.706516698 | 0.817537028 |
| 0.54875502 | NA |
| 0.085471094 | 0.182629802 |
| 0.097213164 | 0.201482196 |
| 0.61929992 | 0.751049447 |
| 0.03220884 | 0.08630232 |
| 0.992061292 | NA |
| 0.150197446 | NA |
| 0.02995651 | 0.081432905 |
| NA | NA |
| 0.514214123 | NA |
| 0.600584751 | 0.736433801 |

| | |
|---|---|
| 0.863106891 | 0.920982 |
| NA | NA |
| 0.000259628 | 0.0015807 |
| 0.418152176 | 0.580380737 |
| 1.00E-05 | 9.71E-05 |
| 7.08E-09 | 1.74E-07 |
| 0.748890654 | NA |
| NA | NA |
| 0.992061292 | NA |
| 0.814979756 | NA |
| 0.000640669 | 0.003431332 |
| 0.105740574 | 0.214797157 |
| 0.665588296 | NA |
| 0.903330316 | 0.945257585 |
| 0.065044967 | 0.148551475 |
| 0.11633358 | 0.230911062 |
| NA | NA |
| 3.38E-05 | 0.000277239 |
| NA | NA |
| 0.474499673 | 0.630781852 |
| 1.21E-08 | 2.80E-07 |
| 0.444907388 | 0.60433758 |
| 9.12E-06 | 9.00E-05 |
| 0.000288025 | 0.001728068 |
| NA | NA |
| 0.008265512 | 0.028832783 |
| 0.073353639 | 0.162343535 |
| 0.480096682 | NA |
| 0.042160919 | 0.106366224 |
| 0.00012491 | 0.000844486 |
| 0.905158266 | NA |
| 0.121949194 | 0.238862041 |
| 0.71458973 | NA |
| 0.348624203 | NA |
| NA | NA |
| 0.006224555 | 0.022949126 |
| 0.018131156 | 0.054311887 |
| NA | NA |
| 0.041400487 | 0.104845265 |
| 0.15009813 | 0.278779061 |
| 0.666698497 | 0.787394842 |
| 0.59409015 | 0.731421036 |
| 0.590993995 | 0.728875998 |
| NA | NA |
| 0.782195155 | NA |
| 0.284458532 | 0.443323271 |
| 0.617973163 | 0.75022516 |

| | |
|---|---|
| 0.44300965 | 0.602766523 |
| 0.002140529 | 0.009476589 |
| 1.04E-10 | 4.42E-09 |
| 8.72E-08 | 1.56E-06 |
| 0.619170352 | 0.751014237 |
| 0.924914412 | 0.956719373 |
| 0.004591822 | 0.017827406 |
| 0.598272529 | 0.7347359 |
| 0.222883289 | NA |
| 0.422636304 | 0.584218809 |
| NA | NA |
| NA | NA |
| 2.69E-06 | 3.11E-05 |
| 0.246629871 | NA |
| 0.119171728 | 0.234909513 |
| 0.00860732 | 0.029802608 |
| 0.677461766 | 0.795400893 |
| 0.070697528 | NA |
| 0.681527924 | 0.798130287 |
| 0.006452734 | 0.023643514 |
| 0.703406073 | 0.815011456 |
| NA | NA |
| 0.190195387 | 0.332232957 |
| NA | NA |
| 0.000655069 | 0.003497364 |
| 0.432381606 | 0.593293979 |
| 0.295750858 | 0.455778868 |
| NA | NA |
| 2.36E-05 | 0.000204669 |
| 0.49627728 | 0.649565047 |
| 0.018933968 | 0.056208059 |
| 0.01861977 | 0.055440208 |
| 0.328432837 | 0.4906161 |
| 3.29E-05 | 0.000271081 |
| 0.03976469 | 0.10165225 |
| 0.72940483 | 0.833394072 |
| 1.94E-05 | 0.000173075 |
| 0.116300704 | 0.230887547 |
| 0.623267375 | NA |
| 0.043772184 | 0.109313475 |
| 0.019771535 | 0.058259359 |
| 6.79E-06 | 6.98E-05 |
| 0.019331539 | 0.057194997 |
| 0.731504269 | 0.834880574 |
| 0.608824207 | 0.742936509 |
| NA | NA |
| 0.053006149 | 0.126834699 |

| | | | |
|---|---|---|---|
| | 0.412009616 | | 0.574682628 |
| | 8.53E-07 | | 1.15E-05 |
| NA | | NA | |
| NA | | NA | |
| | 0.992061292 | NA | |
| | 0.435089195 | | 0.595792194 |
| | 0.082129609 | | 0.177525749 |
| | 0.828474401 | | 0.897728565 |
| | 0.155537414 | | 0.286350785 |
| NA | | NA | |
| | 0.992061292 | NA | |
| NA | | NA | |
| | 0.000806503 | | 0.004155328 |
| | 0.889409773 | NA | |
| | 3.25E-05 | | 0.000267701 |
| | 0.862560572 | NA | |
| | 0.471266349 | NA | |
| NA | | NA | |
| | 0.889409773 | NA | |
| | 0.992061292 | NA | |
| NA | | NA | |
| NA | | NA | |
| | 0.213976024 | | 0.361327246 |
| | 0.001565901 | | 0.007263545 |
| NA | | NA | |
| | 0.268268065 | | 0.424661075 |
| | 0.099217654 | | 0.204728479 |
| | 0.039568165 | | 0.101252113 |
| NA | | NA | |
| | 0.352020967 | | 0.516298985 |
| NA | | NA | |
| | 0.423392862 | NA | |
| | 0.478322369 | | 0.634306474 |
| | 1.37E-05 | | 0.000127811 |
| | 0.255163502 | | 0.409735675 |
| | 0.205029401 | | 0.351134036 |
| | 0.915569704 | | 0.952213132 |
| | 0.588919747 | | 0.727502741 |
| | 0.131935984 | | 0.252808226 |
| | 0.376941522 | | 0.541355549 |
| | 0.007914556 | | 0.02782958 |
| | 0.269564781 | | 0.425976758 |
| | 0.008620462 | | 0.029838702 |
| | 0.441893037 | | 0.601545422 |
| | 0.133091774 | | 0.254467764 |
| | 0.655494219 | NA | |
| | 0.448219685 | | 0.607318514 |

| | |
|---|---|
| 0.731650061 | NA |
| 0.147585406 | 0.275369864 |
| 0.773913116 | 0.863710008 |
| 3.63E-06 | 4.04E-05 |
| NA | NA |
| NA | NA |
| 2.61E-05 | 0.000222873 |
| 0.809273879 | NA |
| 0.1788852 | 0.318145307 |
| 0.007941777 | 0.02791189 |
| 0.660487382 | NA |
| 0.880596384 | 0.932221934 |
| 0.012622101 | 0.0406108 |
| 0.004892223 | 0.018817364 |
| 0.905158266 | NA |
| 0.098445803 | 0.203460836 |
| 0.908737052 | NA |
| 0.000336311 | 0.001974086 |
| NA | NA |
| 0.905158266 | NA |
| 0.905158739 | NA |
| 0.905158266 | NA |
| 0.470415172 | NA |
| 0.419174215 | 0.581138113 |
| 0.160430624 | NA |
| 0.543693406 | 0.690471314 |
| 0.710641144 | NA |
| NA | NA |
| NA | NA |
| NA | NA |
| NA | NA |
| 0.056123773 | 0.132461515 |
| 0.001531383 | 0.007130262 |
| 0.31816171 | NA |
| 0.049222203 | 0.119870285 |
| 0.905158266 | NA |
| NA | NA |
| 0.774986494 | NA |
| 0.905158266 | NA |
| 0.2363587 | 0.388255587 |
| 0.280770165 | 0.439164617 |
| 0.381331439 | 0.545444051 |
| NA | NA |
| 0.819215013 | NA |
| 0.931019351 | 0.96017204 |
| 0.776955996 | 0.865785888 |
| 0.594009926 | 0.731363273 |

| | |
|---|---|
| 0.535161658 | 0.683490636 |
| 0.000471442 | 0.002624181 |
| 1.38E-15 | 1.99E-13 |
| 0.421675251 | 0.583279801 |
| 0.352301479 | 0.516537903 |
| 0.215626853 | 0.36338834 |
| 0.207429346 | 0.354111871 |
| 0.078368691 | 0.171250679 |
| 0.980094639 | NA |
| 0.744867138 | NA |
| 0.005196943 | 0.01980213 |
| 8.72E-07 | 1.17E-05 |
| 0.073076779 | 0.161812342 |
| 0.000485026 | 0.002690926 |
| NA | NA |
| 0.257945157 | 0.412753604 |
| 2.64E-09 | 7.42E-08 |
| 0.037744654 | 0.097722064 |
| 0.795353865 | NA |
| 0.089793984 | 0.189777296 |
| 0.270142377 | NA |
| 0.456847214 | 0.61454689 |
| 0.853087414 | 0.914290079 |
| 0.006090451 | 0.022526566 |
| 0.139692159 | 0.264075424 |
| 4.62E-07 | 6.72E-06 |
| 0.054950562 | 0.130365104 |
| 0.630375571 | 0.758530175 |
| 8.12E-05 | 0.000584711 |
| 0.872123837 | 0.926653376 |
| 0.005715026 | 0.021419964 |
| NA | NA |
| NA | NA |
| 0.992060816 | NA |
| 0.000130361 | 0.000873541 |
| 0.001445767 | 0.006802569 |
| 0.021920932 | 0.063303754 |
| 0.358414355 | NA |
| 0.844963237 | 0.908720478 |
| NA | NA |
| 0.889409301 | NA |
| 1.90E-07 | 3.11E-06 |
| NA | NA |
| 0.154159341 | 0.284564977 |
| 0.642070212 | NA |
| 0.395697931 | NA |
| 0.044446889 | 0.110671191 |

| | |
|---|---|
| 5.84E-05 | 0.000442935 |
| 0.889409301 | NA |
| 0.353462937 | NA |
| 0.000235886 | 0.001459107 |
| 0.000994339 | 0.004953422 |
| 0.349896549 | 0.514007113 |
| 1.55E-05 | 0.000141958 |
| 0.00046644 | 0.002600293 |
| 0.7613479 | 0.85525059 |
| NA | NA |
| 0.766852049 | NA |
| 0.185135058 | 0.325934478 |
| 0.83642152 | NA |
| NA | NA |
| NA | NA |
| 0.889409773 | NA |
| 0.924824475 | 0.956719373 |
| 0.640349188 | NA |
| 0.879879833 | NA |
| 0.109870494 | 0.221140134 |
| 0.73256782 | 0.835663333 |
| 0.336716453 | 0.499721817 |
| 0.7343176 | NA |
| 0.148404683 | 0.276381853 |
| 0.206682206 | 0.353275827 |
| 0.463662684 | 0.620675986 |
| 0.889410717 | NA |
| NA | NA |
| NA | NA |
| NA | NA |
| 0.905158266 | NA |
| NA | NA |
| 0.889409773 | NA |
| NA | NA |
| 0.078697252 | 0.17181477 |
| NA | NA |
| NA | NA |
| 0.03898609 | 0.100081313 |
| 0.016539004 | 0.050558403 |
| 0.992061292 | NA |
| 0.081102849 | 0.175767615 |
| 3.60E-09 | 9.67E-08 |
| 0.15792137 | 0.289571307 |
| 0.895067799 | NA |
| 0.105607169 | 0.2146453 |
| 0.801661681 | NA |
| 0.006082999 | 0.022506585 |

| | | |
|---|---|---|
| | 0.206989662 | 0.353608682 |
| | 0.905158266 | NA |
| NA | | NA |
| | 0.851542467 | NA |
| | 3.77E-12 | 2.33E-10 |
| | 0.132181319 | 0.253132448 |
| NA | | NA |
| | 0.279055221 | NA |
| | 0.547941599 | NA |
| | 0.129843214 | 0.249932253 |
| | 0.889409773 | NA |
| | 9.80E-09 | 2.33E-07 |
| | 0.767326046 | NA |
| | 0.003942346 | 0.015694337 |
| | 0.000139798 | 0.000928002 |
| | 4.99E-18 | 1.34E-15 |
| | 0.992059387 | NA |
| | 0.99384137 | NA |
| NA | | NA |
| | 6.25E-06 | 6.50E-05 |
| | 0.046004156 | 0.113800299 |
| | 0.147772438 | 0.275513091 |
| NA | | NA |
| | 0.787645204 | NA |
| NA | | NA |
| NA | | NA |
| | 0.000151785 | 0.000993561 |
| NA | | NA |
| | 0.922188047 | 0.955294241 |
| NA | | NA |
| | 3.37E-06 | 3.78E-05 |
| | 0.202296198 | 0.34779236 |
| NA | | NA |
| | 0.28465797 | NA |
| | 0.905158266 | NA |
| | 0.873151993 | 0.927333364 |
| NA | | NA |
| | 0.708394443 | 0.818201018 |
| NA | | NA |
| | 7.47E-14 | 6.81E-12 |
| NA | | NA |
| | 2.32E-14 | 2.42E-12 |
| | 0.99205891 | NA |
| | 0.043329237 | 0.108518437 |
| NA | | NA |
| NA | | NA |
| | 0.015843012 | 0.048861896 |

| | | |
|---|---|---|
| | 0.041439406 | 0.10490753 |
| | 0.277299651 | NA |
| NA | | NA |
| | 0.73258203 | NA |
| | 0.669845685 | 0.789501803 |
| | 0.797157316 | NA |
| | 0.97428187 | NA |
| | 0.381073913 | 0.545272827 |
| | 0.209283861 | NA |
| NA | | NA |
| NA | | NA |
| NA | | NA |
| | 0.095313372 | 0.198481397 |
| | 6.35E-05 | 0.000474429 |
| | 6.54E-18 | 1.69E-15 |
| | 0.766053056 | NA |
| | 0.926581366 | 0.957992382 |
| | 0.801661681 | NA |
| NA | | NA |
| | 0.992062245 | NA |
| | 0.0872422 | 0.185616846 |
| | 0.001474188 | 0.0069067 |
| NA | | NA |
| | 0.532793811 | NA |
| | 0.320943056 | NA |
| | 0.585089928 | 0.724565177 |
| | 0.992061292 | NA |
| | 4.66E-10 | 1.63E-08 |
| | 0.899961283 | 0.943542752 |
| | 0.128447111 | 0.248158553 |
| | 0.261551089 | 0.417064873 |
| | 4.52E-08 | 8.90E-07 |
| | 0.266127926 | 0.422399937 |
| | 0.546098295 | NA |
| NA | | NA |
| | 0.685915005 | NA |
| | 0.855119927 | 0.915689726 |
| NA | | NA |
| | 0.001257294 | 0.00603078 |
| | 2.45E-17 | 5.02E-15 |
| | 0.669027921 | 0.78895785 |
| | 0.213190996 | 0.360337793 |
| | 1.45E-05 | 0.000134236 |
| | 0.127661786 | 0.247037191 |
| | 0.005111004 | 0.019511917 |
| NA | | NA |
| | 0.204583078 | 0.350479 |

| | | |
|---|---|---|
| | 0.628996176 | NA |
| NA | | NA |
| | 5.18E-13 | 3.95E-11 |
| | 0.172263296 | 0.309122311 |
| | 0.14255196 | 0.268096822 |
| NA | | NA |
| | 0.98371419 | NA |
| | 0.992060816 | NA |
| | 0.378981305 | 0.543446784 |
| | 0.905158266 | NA |
| | 4.10E-07 | 6.05E-06 |
| | 0.329851167 | 0.492265991 |
| NA | | NA |
| | 0.003067898 | 0.012788151 |
| | 0.064937014 | 0.148368741 |
| | 0.117078381 | 0.231928207 |
| | 0.761438249 | 0.855264578 |
| | 0.064585512 | 0.147732632 |
| | 0.797342596 | NA |
| | 0.038736449 | 0.099584451 |
| NA | | NA |
| | 0.719842444 | NA |
| | 0.791988343 | NA |
| | 0.964833442 | 0.980823923 |
| | 0.433800807 | 0.594584103 |
| | 0.211134416 | NA |
| | 0.792188023 | NA |
| | 0.001054909 | 0.005206211 |
| | 0.825950645 | 0.896231435 |
| | 0.183884857 | 0.324487912 |
| NA | | NA |
| | 0.992060816 | NA |
| | 0.747074748 | 0.845575712 |
| NA | | NA |
| NA | | NA |
| | 0.559713694 | NA |
| NA | | NA |
| NA | | NA |
| | 0.905158266 | NA |
| NA | | NA |
| NA | | NA |
| | 0.981036214 | NA |
| | 0.64609008 | 0.771235488 |
| NA | | NA |
| NA | | NA |
| NA | | NA |
| | 1.56E-06 | 1.92E-05 |

| | | |
|---|---|---|
| | 8.06E-06 | 8.10E-05 |
| | 0.992059387 | NA |
| | 3.05E-09 | 8.42E-08 |
| | 0.003715325 | 0.01493991 |
| NA | | NA |
| | 0.002781332 | 0.011809187 |
| | 0.037706328 | 0.09767527 |
| | 1.12E-10 | 4.72E-09 |
| | 0.497901478 | 0.650880741 |
| | 0.992060816 | NA |
| | 0.782195155 | NA |
| | 0.766220616 | NA |
| | 0.801661681 | NA |
| | 0.991759839 | NA |
| | 0.905158739 | NA |
| | 0.385078737 | NA |
| | 0.05348654 | 0.127608749 |
| | 0.889409773 | NA |
| NA | | NA |
| NA | | NA |
| NA | | NA |
| NA | | NA |
| | 0.083965287 | NA |
| | 0.99205891 | NA |
| NA | | NA |
| | 0.992062245 | NA |
| | 0.965913597 | 0.981494822 |
| | 0.616142072 | 0.748833725 |
| | 0.313880422 | 0.475804238 |
| | 0.170691355 | 0.307129749 |
| | 0.361157656 | NA |
| | 0.249861077 | 0.403760349 |
| NA | | NA |
| | 0.8322255 | NA |
| | 0.018467433 | 0.055091205 |
| | 0.890405224 | NA |
| | 0.079742638 | 0.173441816 |
| NA | | NA |
| | 0.336653621 | 0.499667243 |
| | 0.762141322 | 0.855660376 |
| | 0.128905221 | NA |
| | 0.528979368 | 0.678476053 |
| | 0.014165991 | 0.044566046 |
| | 0.868485979 | NA |
| | 0.986917658 | NA |
| | 0.885390557 | 0.93507114 |
| NA | | NA |

| | | |
|---|---|---|
| | 0.016284938 | 0.049949976 |
| | 0.01143448 | 0.037459309 |
| | 0.778445745 | NA |
| | 0.011593272 | 0.037849636 |
| | 0.111658903 | 0.223633515 |
| | 0.991735419 | 0.995816822 |
| | 0.676647011 | NA |
| NA | | NA |
| | 0.672716393 | NA |
| | 0.007213452 | 0.025861257 |
| | 0.426181252 | 0.587410349 |
| | 0.001528147 | 0.007118492 |
| NA | | NA |
| | 0.626036218 | 0.755772827 |
| | 0.99205891 | NA |
| | 0.03501542 | 0.092051192 |
| | 0.413661748 | NA |
| NA | | NA |
| | 0.324824903 | 0.486985999 |
| NA | | NA |
| NA | | NA |
| NA | | NA |
| NA | | NA |
| NA | | NA |
| NA | | NA |
| | 0.673069945 | NA |
| NA | | NA |
| NA | | NA |
| NA | | NA |
| NA | | NA |
| NA | | NA |
| | 0.762746762 | NA |
| NA | | NA |
| NA | | NA |
| | 0.502940862 | 0.655274222 |
| NA | | NA |
| | 0.757568415 | NA |
| | 0.035513726 | 0.093127048 |
| | 0.122497159 | 0.239636091 |
| | 9.09E-06 | 8.98E-05 |
| | 0.004958968 | 0.01905073 |
| | 2.10E-11 | 1.07E-09 |
| | 0.285625515 | 0.444604356 |
| | 0.743108928 | 0.842520082 |
| | 0.038849966 | NA |
| | 0.340005852 | 0.503180247 |
| | 0.681531081 | 0.798130287 |

| | | |
|---|---|---|
| | 0.57885336 | NA |
| NA | | NA |
| NA | | NA |
| | 0.510187246 | NA |
| | 0.71395495 | NA |
| | 0.0017315 | 0.007906426 |
| | 0.117042894 | NA |
| | 0.99205891 | NA |
| | 0.234062446 | 0.385552466 |
| | 9.75E-05 | 0.000683538 |
| | 0.024051548 | 0.068289105 |
| | 0.018751678 | 0.055780018 |
| | 2.73E-26 | 4.99E-23 |
| | 0.19313827 | 0.336090283 |
| | 0.16564241 | 0.300027731 |
| | 0.625078712 | 0.755340311 |
| | 0.01182138 | 0.038474238 |
| | 0.660719229 | 0.782769748 |
| | 4.80E-06 | 5.14E-05 |
| | 0.578592618 | NA |
| | 0.36649638 | 0.5313173 |
| | 0.00068313 | 0.003619898 |
| | 0.058205982 | 0.136262811 |
| | 0.897958322 | 0.942591744 |
| | 0.902065325 | 0.94465413 |
| | 0.91157329 | 0.950003887 |
| | 0.981348231 | NA |
| | 0.905158739 | NA |
| NA | | NA |
| | 0.942254912 | NA |
| | 0.905158739 | NA |
| NA | | NA |
| NA | | NA |
| NA | | NA |
| | 0.99205891 | NA |
| NA | | NA |
| | 0.992061292 | NA |
| NA | | NA |
| | 0.830163713 | NA |
| NA | | NA |
| NA | | NA |
| | 0.815851452 | NA |
| | 0.736269536 | 0.838159186 |
| | 0.382383308 | NA |
| | 8.69E-06 | 8.66E-05 |
| | 0.662055168 | 0.783554459 |
| | 0.596814718 | NA |

| | |
|---|---|
| 0.001812387 | 0.008225352 |
| 0.992061292 | NA |
| 0.127912346 | 0.247430163 |
| 0.199797138 | NA |
| 0.014196005 | 0.044632607 |
| 0.016627733 | 0.050761629 |
| NA | NA |
| 1.46E-05 | 0.000135258 |
| 0.002621949 | 0.011260585 |
| 0.049718997 | 0.120838899 |
| 0.784366884 | 0.870114291 |
| 0.664897419 | NA |
| 2.04E-10 | 8.02E-09 |
| 0.256851767 | 0.411536994 |
| 0.676941041 | 0.795002049 |
| 0.000185132 | 0.001178694 |
| NA | NA |
| 0.020922386 | 0.060964639 |
| 1.45E-06 | 1.80E-05 |
| 0.003422646 | 0.013988068 |
| 0.619155812 | 0.751014237 |
| 0.092610388 | 0.194215597 |
| 0.693317828 | 0.80719825 |
| 0.009820781 | 0.033131746 |
| 0.02541963 | 0.071507771 |
| 0.541034215 | NA |
| 0.488818218 | NA |
| 0.000325711 | 0.001920077 |
| NA | NA |
| 0.443441817 | 0.60301826 |
| 0.062552571 | 0.144224265 |
| 0.143391933 | 0.269468845 |
| 0.434720863 | 0.595362071 |
| 0.277983049 | 0.435881745 |
| 0.992037435 | NA |
| 0.363926611 | NA |
| 0.339321754 | NA |
| 0.284450356 | 0.443323271 |
| 0.699229551 | 0.812014475 |
| 0.743991246 | NA |
| NA | NA |
| NA | NA |
| 0.991529086 | NA |
| NA | NA |
| 0.01933921 | NA |
| 0.722436794 | NA |
| 0.316960837 | NA |

| | | |
|---|---|---|
| | 0.935233323 | NA |
| | 0.672746301 | NA |
| | 0.659302734 | 0.781583908 |
| | 0.860696643 | 0.919039026 |
| | 0.901154598 | 0.944236842 |
| | 0.251619379 | 0.405825189 |
| | 0.37664024 | 0.541024598 |
| | 4.60E-06 | 4.95E-05 |
| | 0.165330491 | 0.299619729 |
| | 0.788267158 | NA |
| | 0.177632659 | 0.316499477 |
| | 0.00060089 | 0.003247588 |
| | 0.273989172 | NA |
| | 0.95550898 | 0.975182268 |
| | 0.873501929 | NA |
| | 0.048610247 | 0.118669529 |
| | 0.251281028 | 0.40545816 |
| NA | | NA |
| | 0.99205891 | NA |
| NA | | NA |
| | 0.012161014 | 0.039398599 |
| | 0.797708749 | NA |
| | 0.041962531 | 0.10596311 |
| | 0.805452888 | NA |
| | 0.176483973 | 0.314869318 |
| | 0.992061292 | NA |
| | 8.69E-07 | 1.16E-05 |
| | 0.900897778 | NA |
| | 0.050823532 | 0.122938306 |
| | 0.578093383 | 0.719196874 |
| | 0.000705588 | 0.003717122 |
| NA | | NA |
| | 0.373853552 | 0.537868166 |
| | 0.577144289 | 0.71849376 |
| | 0.578625045 | 0.719478173 |
| | 0.067485645 | NA |
| | 4.52E-09 | 1.17E-07 |
| | 1.51E-09 | 4.51E-08 |
| | 0.521037903 | 0.671842121 |
| | 0.136164384 | 0.258922213 |
| | 0.633668488 | NA |
| | 0.043604972 | 0.109045882 |
| | 0.538194162 | 0.686136766 |
| | 0.010163064 | 0.034123964 |
| | 0.905158739 | NA |
| | 0.014238683 | 0.04473065 |
| NA | | NA |

| | | |
|---|---|---|
| | 0.53832663 | 0.686196927 |
| | 0.300617417 | NA |
| | 0.007365665 | 0.026299616 |
| NA | | NA |
| | 9.65E-05 | 0.000677457 |
| | 0.920085262 | 0.954241358 |
| | 0.077491685 | 0.169671943 |
| | 7.40E-08 | 1.36E-06 |
| | 0.38673883 | NA |
| | 0.844651527 | NA |
| | 0.430437636 | 0.591448677 |
| | 0.939327552 | NA |
| | 0.03017748 | 0.081856942 |
| | 0.145276361 | 0.272031813 |
| | 0.088406071 | 0.187693021 |
| | 0.958457604 | 0.976965628 |
| | 0.672057754 | NA |
| NA | | NA |
| | 0.626962242 | 0.756287841 |
| | 0.40964123 | 0.572675549 |
| | 0.570309524 | NA |
| | 0.368069644 | 0.532685778 |
| | 0.965683655 | 0.981422073 |
| | 0.719717629 | 0.826239149 |
| | 0.629368425 | 0.757608883 |
| | 0.145238355 | 0.272021834 |
| | 2.15E-08 | 4.65E-07 |
| | 0.035799712 | 0.093675629 |
| | 0.071089772 | 0.158405832 |
| | 0.568139975 | NA |
| NA | | NA |
| | 0.788582525 | NA |
| | 0.651413798 | 0.775751362 |
| | 0.756022289 | NA |
| | 0.955779237 | NA |
| | 0.754336313 | NA |
| | 2.25E-07 | 3.59E-06 |
| | 0.174511669 | NA |
| NA | | NA |
| | 0.22758699 | 0.377575002 |
| NA | | NA |
| | 0.905158739 | NA |
| | 0.674781019 | 0.793196518 |
| | 0.880670348 | 0.932221934 |
| | 0.379363062 | 0.543735459 |
| | 0.154044471 | 0.284448582 |
| | 0.208454301 | NA |

| | |
|---|---|
| NA | NA |
| 0.778575545 | 0.866798844 |
| 0.558709939 | 0.703440832 |
| 0.797342596 | NA |
| 0.08415497 | 0.180560702 |
| 0.032951787 | 0.087792277 |
| 0.738408059 | 0.839531044 |
| 0.639740303 | 0.766313546 |
| 0.345992935 | 0.509431331 |
| NA | NA |
| 0.000142356 | 0.000940433 |
| 0.010396376 | 0.034753238 |
| 0.607872016 | NA |
| NA | NA |
| 0.553379216 | NA |
| 1.85E-08 | 4.05E-07 |
| 0.04014076 | 0.102370457 |
| 0.848601856 | 0.911130851 |
| 0.751635918 | 0.848725928 |
| 0.105395176 | 0.214275105 |
| 0.422325748 | 0.583857369 |
| 2.17E-09 | 6.26E-08 |
| 2.69E-05 | 0.000228061 |
| 0.227142267 | 0.377013462 |
| 0.578377993 | 0.719417828 |
| 5.50E-05 | 0.000420518 |
| 0.519203505 | 0.670106369 |
| 0.440344104 | 0.6004748 |
| 0.504669721 | 0.656816024 |
| 0.816928849 | 0.891154525 |
| 0.700211268 | 0.812670139 |
| 0.006980257 | 0.025205567 |
| 0.030127546 | 0.081766258 |
| 0.85916788 | 0.91789761 |
| 0.401292004 | 0.564869943 |
| 0.797342596 | NA |
| NA | NA |
| NA | NA |
| 0.287344414 | 0.446465013 |
| 0.736858992 | 0.838420986 |
| 0.00558235 | 0.020997402 |
| 0.000695225 | 0.003668937 |
| 0.178663396 | 0.317879559 |
| 0.889409773 | NA |
| 7.95E-05 | 0.000574595 |
| 0.811203348 | 0.887108665 |
| 0.55675706 | 0.701907697 |

| | | |
|---|---|---|
| | 0.846144595 | NA |
| NA | | NA |
| | 0.889409773 | NA |
| | 0.000753104 | 0.003929991 |
| | 0.469303888 | 0.62563405 |
| NA | | NA |
| NA | | NA |
| NA | | NA |
| | 0.006324008 | 0.023233878 |
| NA | | NA |
| | 0.010604294 | 0.035319231 |
| NA | | NA |
| NA | | NA |
| | 0.017888049 | 0.053757174 |
| | 0.851559792 | NA |
| | 0.774241553 | 0.863900928 |
| | 0.06598901 | 0.150036515 |
| | 0.14789478 | 0.275666028 |
| | 0.117526763 | 0.232522767 |
| | 0.084307381 | 0.180720986 |
| | 0.444618589 | NA |
| | 0.661884742 | 0.783554459 |
| | 0.988648207 | NA |
| | 0.030927039 | 0.083451321 |
| | 0.28473403 | NA |
| | 0.861593485 | 0.919773035 |
| | 0.245715454 | 0.398999162 |
| | 0.415725911 | NA |
| NA | | NA |
| | 0.992061769 | NA |
| NA | | NA |
| NA | | NA |
| NA | | NA |
| NA | | NA |
| NA | | NA |
| NA | | NA |
| | 0.223775282 | NA |
| | 6.31E-10 | 2.10E-08 |
| | 0.637425049 | NA |
| NA | | NA |
| | 0.889409301 | NA |
| | 0.002710582 | 0.011562094 |
| | 0.006367593 | 0.023362736 |
| | 0.203082419 | 0.348879255 |
| | 0.001677516 | 0.007690309 |
| | 0.152740168 | 0.282634309 |
| | 0.889410245 | NA |

| | |
|---|---|
| 4.57E-09 | 1.18E-07 |
| 0.481336149 | 0.636995148 |
| 0.536394746 | NA |
| 0.002733521 | 0.011648628 |
| NA | NA |
| 0.992061292 | NA |
| NA | NA |
| 0.889410245 | NA |
| 0.808923109 | NA |
| NA | NA |
| 0.430925983 | NA |
| 0.406029736 | NA |
| 2.66E-08 | 5.58E-07 |
| 0.569448853 | NA |
| 0.000182116 | 0.001162522 |
| 0.01761173 | 0.053079327 |
| NA | NA |
| NA | NA |
| NA | NA |
| 0.992061292 | NA |
| 0.594581442 | 0.731738715 |
| 0.98964564 | NA |
| 0.315908077 | 0.478118786 |
| 0.114418305 | 0.227933554 |
| 0.905158739 | NA |
| 0.006639519 | 0.024190675 |
| 0.045511948 | 0.112798744 |
| 0.033097855 | 0.088117821 |
| 0.905158266 | NA |
| 0.955322344 | NA |
| NA | NA |
| 0.892870545 | 0.939380965 |
| NA | NA |
| 0.11307179 | 0.22592752 |
| 0.393320364 | 0.556789704 |
| 0.001571831 | 0.007284965 |
| 0.058942242 | 0.137648908 |
| 0.385468931 | 0.54932262 |
| 6.31E-05 | 0.000472153 |
| 0.866748093 | NA |
| 0.966589763 | NA |
| NA | NA |
| 0.529812926 | NA |
| NA | NA |
| 0.198161249 | 0.342604745 |
| 0.003982252 | 0.015835978 |
| 0.88149511 | NA |

| | |
|---|---|
| 0.007818559 | 0.027527296 |
| 0.01840837 | 0.054952336 |
| 0.002426711 | 0.010552109 |
| 9.36E-06 | 9.19E-05 |
| 0.043132542 | 0.108158901 |
| 0.88060637 | 0.932221934 |
| 0.008022474 | 0.028156544 |
| 0.219052803 | 0.367638375 |
| 0.000133674 | 0.000893292 |
| 0.000128747 | 0.000865367 |
| 0.563454348 | 0.70714374 |
| NA | NA |
| 0.102449142 | 0.2098387 |
| 0.992061292 | NA |
| 0.715499324 | NA |
| 0.836278944 | NA |
| 0.905158266 | NA |
| NA | NA |
| NA | NA |
| 0.900836801 | NA |
| NA | NA |
| NA | NA |
| 0.905158739 | NA |
| 0.433319472 | NA |
| 4.99E-06 | 5.31E-05 |
| 0.905158739 | NA |
| 0.089971771 | 0.190098154 |
| 0.945432915 | 0.969042261 |
| 0.001961318 | 0.008803879 |
| 0.016468833 | 0.050388616 |
| 0.23619948 | NA |
| 0.552543925 | NA |
| 0.90515732 | NA |
| 0.402906107 | 0.566345339 |
| 0.851224362 | 0.912830774 |
| 0.347163397 | NA |
| 0.894331014 | 0.94032151 |
| NA | NA |
| 0.889409301 | NA |
| 0.889409773 | NA |
| 0.939903749 | NA |
| NA | NA |
| NA | NA |
| NA | NA |
| 0.905158739 | NA |
| NA | NA |
| 0.905158266 | NA |

| | | | |
|---|---|---|---|
| | 0.895432589 | NA | |
| NA | | NA | |
| NA | | NA | |
| NA | | NA | |
| NA | | NA | |
| | 0.034950951 | | 0.091969793 |
| | 0.001232167 | | 0.005929686 |
| | 0.982843289 | NA | |
| | 0.734112819 | | 0.836640201 |
| | 0.164815577 | | 0.298986738 |
| | 0.739153662 | NA | |
| | 0.017206941 | | 0.052152825 |
| | 3.03E-07 | | 4.64E-06 |
| | 0.011511878 | | 0.037645469 |
| | 0.054964175 | | 0.130383312 |
| | 0.832154132 | | 0.900072534 |
| | 0.246802518 | | 0.400231629 |
| | 1.46E-05 | | 0.000135368 |
| | 0.012118364 | | 0.039277809 |
| | 0.042033175 | | 0.106092699 |
| | 0.66457006 | | 0.785810885 |
| | 0.622805784 | | 0.753924598 |
| | 0.362624043 | | 0.527516487 |
| | 3.62E-15 | | 4.65E-13 |
| | 0.000129677 | | 0.000870283 |
| | 0.000543161 | | 0.002971416 |
| NA | | NA | |
| | 0.471087303 | | 0.627402116 |
| NA | | NA | |
| NA | | NA | |
| | 0.207461899 | | 0.354139912 |
| | 0.000394419 | | 0.002259599 |
| | 0.243567463 | NA | |
| | 0.917190687 | | 0.952814839 |
| | 0.13424111 | | 0.256111945 |
| | 0.623254896 | | 0.754169581 |
| NA | | NA | |
| | 0.150808264 | | 0.279837643 |
| | 0.036647883 | | 0.095521972 |
| | 0.297310889 | | 0.45738054 |
| NA | | NA | |
| | 0.522539997 | NA | |
| | 0.780182136 | | 0.867619748 |
| | 0.915375768 | NA | |
| | 0.438827015 | | 0.599198617 |
| | 0.795014225 | NA | |
| NA | | NA | |

| | |
|---|---|
| NA | NA |
| NA | NA |
| 0.978293206 | NA |
| 1.05E-05 | 0.000101737 |
| 0.000915678 | 0.004625574 |
| NA | NA |
| NA | NA |
| NA | NA |
| 0.699079314 | NA |
| 0.287429243 | NA |
| 0.28084832 | NA |
| 0.138729635 | NA |
| 0.262439422 | NA |
| 4.12E-09 | 1.07E-07 |
| 0.389922951 | NA |
| NA | NA |
| NA | NA |
| 0.661025394 | 0.782909176 |
| 0.001373141 | 0.006512648 |
| 0.27263947 | 0.429444812 |
| 0.141361823 | 0.266439591 |
| 0.14764994 | 0.275396674 |
| 0.678523069 | NA |
| 0.184272092 | 0.32485797 |
| 0.000404869 | 0.002311027 |
| 0.577266975 | 0.718605755 |
| NA | NA |
| NA | NA |
| NA | NA |
| NA | NA |
| 0.121753021 | NA |
| 0.889409301 | NA |
| 0.008446719 | 0.029348341 |
| 0.004163659 | 0.016435339 |
| 0.323026324 | 0.485441103 |
| 0.706966394 | 0.817643694 |
| 0.095744408 | 0.199246727 |
| 0.782559532 | NA |
| 3.67E-16 | 6.11E-14 |
| 0.986671427 | 0.993138883 |
| 0.000205472 | 0.001291342 |
| 0.04060538 | 0.103140952 |
| 0.905158266 | NA |
| 0.007696386 | 0.027197448 |
| 0.979161744 | NA |
| NA | NA |
| 0.538787606 | 0.68654541 |

| | |
|---|---|
| NA | NA |
| 0.002948714 | 0.012380652 |
| 0.622420254 | NA |
| 0.423786221 | NA |
| 3.30E-05 | 0.000271387 |
| 0.2138028 | 0.361062499 |
| 0.652717772 | 0.776353026 |
| 0.2618344 | 0.417262333 |
| 0.398135954 | NA |
| 0.992060816 | NA |
| 0.00412242 | 0.016300049 |
| 0.153149798 | 0.283225232 |
| 0.383532615 | NA |
| NA | NA |
| 0.33772553 | 0.500649086 |
| 0.840865114 | NA |
| 0.669523317 | NA |
| 0.068390766 | 0.153639352 |
| 0.442891905 | 0.602681003 |
| 0.708542265 | 0.818242419 |
| 0.013981542 | 0.044067845 |
| 0.004040209 | 0.016025821 |
| 0.889409773 | NA |
| 0.309764845 | 0.470996139 |
| NA | NA |
| 0.889410717 | NA |
| 0.889409773 | NA |
| 0.267086568 | 0.423279008 |
| 7.92E-15 | 9.40E-13 |
| 0.929974183 | NA |
| 0.731172408 | 0.834588569 |
| 0.992061292 | NA |
| 0.782559532 | NA |
| 0.521401609 | 0.672074309 |
| 0.992061292 | NA |
| NA | NA |
| 0.564338869 | 0.707889708 |
| 0.889409773 | NA |
| 0.976634025 | NA |
| NA | NA |
| NA | NA |
| NA | NA |
| 0.618545497 | NA |
| NA | NA |
| 0.92472642 | 0.956719373 |
| 4.36E-06 | 4.74E-05 |
| 0.576625483 | 0.718092155 |

| | |
|---|---|
| NA | NA |
| NA | NA |
| 0.001122374 | 0.005487912 |
| 0.813836939 | NA |
| 0.105690094 | 0.214754581 |
| 0.508334397 | 0.660113248 |
| 0.816470487 | 0.890743016 |
| 0.36068081 | 0.52533883 |
| 0.635663956 | 0.762970473 |
| 0.774260974 | NA |
| 0.933866302 | 0.962036504 |
| 0.579184094 | 0.719754307 |
| 0.211604336 | 0.358646158 |
| 0.99205891 | NA |
| 0.607249607 | NA |
| 3.32E-11 | 1.59E-09 |
| NA | NA |
| 0.817770736 | NA |
| NA | NA |
| 0.807879735 | NA |
| NA | NA |
| NA | NA |
| NA | NA |
| NA | NA |
| 0.129532892 | 0.249641283 |
| 0.596941972 | NA |
| NA | NA |
| NA | NA |
| 0.688747059 | 0.80387566 |
| NA | NA |
| 0.216540138 | NA |
| 0.117634805 | 0.232676046 |
| 0.115602678 | 0.229834243 |
| 0.137921572 | 0.261561694 |
| 0.06163694 | 0.142712523 |
| 0.890501492 | 0.938091737 |
| 0.001083677 | 0.005331487 |
| 0.136225147 | NA |
| 0.297736743 | 0.457682965 |
| 0.187814758 | 0.329279382 |
| 0.873906655 | 0.927840184 |
| NA | NA |
| 0.069395842 | 0.155455753 |
| 0.372707504 | 0.537097 |
| 0.05919472 | 0.138150373 |
| 0.011406531 | 0.03738448 |
| NA | NA |

| | |
|---|---|
| NA | NA |
| 0.612286859 | NA |
| 0.455452833 | NA |
| 0.417020858 | NA |
| 6.51E-06 | 6.73E-05 |
| 1.16E-09 | 3.59E-08 |
| 0.432845983 | NA |
| NA | NA |
| NA | NA |
| 0.866905076 | NA |
| 0.885737607 | 0.935226816 |
| 0.102730385 | 0.210277454 |
| NA | NA |
| 0.989491875 | NA |
| 0.426332324 | 0.587574939 |
| NA | NA |
| NA | NA |
| 0.566267108 | NA |
| 0.479589631 | 0.635215156 |
| 0.080169321 | 0.17419732 |
| 0.001048966 | 0.005184389 |
| 0.321942857 | NA |
| NA | NA |
| 0.842315032 | NA |
| 0.989064839 | NA |
| 0.589519128 | 0.7280159 |
| 0.512028435 | 0.663654374 |
| 0.78103128 | NA |
| 0.802662566 | NA |
| 0.341506902 | NA |
| 0.285291727 | 0.444243444 |
| 0.683312022 | 0.799628222 |
| 2.00E-07 | 3.24E-06 |
| NA | NA |
| 0.176785253 | NA |
| 0.010788857 | 0.035803614 |
| 0.003611452 | 0.014622361 |
| NA | NA |
| 0.992061292 | NA |
| 0.049501811 | 0.120444352 |
| 0.004232225 | 0.016658079 |
| 0.502812701 | NA |
| 0.748923082 | NA |
| NA | NA |
| 0.588658256 | NA |
| 0.079166195 | 0.17252982 |
| 0.991953915 | NA |

| | | |
|---|---|---|
| | 0.155793649 | 0.286666058 |
| | 0.851410992 | NA |
| | 0.356252325 | 0.520765864 |
| | 0.686332959 | 0.801829402 |
| | 0.533594605 | 0.682412006 |
| NA | | NA |
| | 0.662024217 | 0.783554459 |
| | 0.488180889 | 0.642747935 |
| | 0.990050496 | NA |
| | 0.698311687 | 0.811334727 |
| | 0.202209943 | NA |
| NA | | NA |
| | 0.734670303 | 0.836928054 |
| | 0.00208189 | 0.009261794 |
| NA | | NA |
| | 0.432991967 | 0.593734895 |
| | 0.652271982 | NA |
| | 0.711184804 | 0.820117335 |
| NA | | NA |
| | 0.43288178 | 0.593657944 |
| | 0.68559056 | NA |
| NA | | NA |
| | 0.707659678 | 0.817782279 |
| | 0.678870139 | 0.796288093 |
| | 0.992060816 | NA |
| | 0.16058584 | 0.293077166 |
| NA | | NA |
| | 0.060820652 | 0.141242687 |
| | 0.092010563 | 0.19325657 |
| | 0.656053444 | NA |
| | 0.991827912 | NA |
| | 0.067551736 | 0.15245309 |
| | 0.661152455 | NA |
| | 0.470499575 | 0.626849198 |
| | 0.502498277 | 0.65498414 |
| | 0.989766227 | NA |
| | 0.991879012 | NA |
| | 0.79188669 | NA |
| NA | | NA |
| | 0.723883809 | NA |
| | 0.03880119 | NA |
| NA | | NA |
| | 0.365574488 | 0.53033297 |
| | 0.901632454 | 0.944380971 |
| | 0.145245824 | 0.272021834 |
| | 0.660294721 | NA |
| | 0.905158266 | NA |

| | | |
|---|---|---|
| | 0.945933758 | 0.969284153 |
| | 0.533915992 | NA |
| | 0.225841391 | NA |
| | 0.099084179 | 0.204510713 |
| NA | | NA |
| | 0.610683463 | NA |
| | 0.50317562 | 0.655468555 |
| | 0.224417566 | NA |
| | 0.88077242 | 0.932225458 |
| | 0.690369466 | NA |
| | 0.340034126 | 0.503188178 |
| | 0.909240027 | 0.948458589 |
| | 0.404904623 | 0.568177875 |
| | 0.58736063 | 0.726253393 |
| | 0.315583711 | 0.477824638 |
| | 0.164817818 | 0.298986738 |
| | 0.551228655 | NA |
| | 0.01930535 | NA |
| | 0.003272778 | 0.013502788 |
| | 0.498089519 | 0.651005758 |
| | 0.99205891 | NA |
| | 0.352918535 | 0.517028362 |
| | 0.730161653 | 0.834085197 |
| | 0.978972059 | 0.989334413 |
| | 0.992061292 | NA |
| | 0.239515605 | 0.391741634 |
| | 0.903342961 | NA |
| | 0.523180945 | 0.673616556 |
| | 0.237739195 | 0.389569194 |
| | 0.384048687 | 0.54800995 |
| | 0.210143124 | NA |
| | 0.088108459 | 0.187169745 |
| | 0.319225243 | 0.481777808 |
| | 0.520875218 | 0.67167179 |
| | 0.924294849 | NA |
| | 0.817066975 | 0.891260926 |
| | 0.074849623 | NA |
| | 0.0583363 | 0.136524226 |
| | 9.66E-20 | 3.93E-17 |
| | 2.93E-06 | 3.34E-05 |
| | 0.003764313 | 0.015100576 |
| | 1.52E-10 | 6.16E-09 |
| | 0.006966753 | 0.025174087 |
| NA | | NA |
| | 0.591932081 | 0.729623159 |
| NA | | NA |
| NA | | NA |

| | | |
|---|---|---|
| NA | | NA |
| | 0.004311703 | 0.016904501 |
| | 0.992061292 | NA |
| | 0.179792147 | 0.319370308 |
| | 0.215538842 | 0.363280914 |
| | 0.807879735 | NA |
| | 0.006525306 | 0.023853704 |
| | 0.066502079 | 0.150859417 |
| | 1.38E-06 | 1.73E-05 |
| | 1.22E-06 | 1.56E-05 |
| | 2.48E-18 | 6.90E-16 |
| | 0.800622255 | NA |
| | 0.313376203 | 0.475269566 |
| | 0.007578653 | 0.026868017 |
| | 0.484764995 | 0.639913106 |
| NA | | NA |
| | 0.525355153 | 0.675386324 |
| | 0.439223457 | 0.599545998 |
| | 0.558266672 | 0.7030843 |
| | 0.992061292 | NA |
| | 0.992269016 | NA |
| NA | | NA |
| | 0.233859488 | 0.385378497 |
| | 0.889409773 | NA |
| | 0.988558274 | NA |
| | 0.049883228 | 0.121122494 |
| | 3.88E-06 | 4.28E-05 |
| | 0.007814416 | 0.027517121 |
| | 2.95E-05 | 0.000246824 |
| | 0.101086561 | 0.207725977 |
| | 0.604982737 | 0.740057145 |
| | 0.294584896 | 0.454460428 |
| NA | | NA |
| | 0.762119062 | 0.855660376 |
| | 0.077085284 | 0.16903817 |
| NA | | NA |
| | 0.225071588 | NA |
| | 0.393315692 | 0.556789704 |
| | 0.065686602 | 0.149506188 |
| | 0.732831669 | NA |
| | 0.992061769 | NA |
| | 0.630936361 | NA |
| | 0.932047203 | NA |
| | 0.107142155 | 0.21690749 |
| | 0.117425732 | 0.232385693 |
| | 0.964615806 | NA |
| | 7.25E-06 | 7.38E-05 |

| | | |
|---|---|---|
| | 0.406403238 | NA |
| | 0.022679585 | 0.065134844 |
| | 0.4035447 | 0.567088976 |
| | 0.169383954 | 0.305202449 |
| | 7.20E-12 | 4.20E-10 |
| | 0.020423103 | 0.059796123 |
| | 0.104872787 | 0.213508392 |
| NA | | NA |
| | 0.015285032 | 0.04742069 |
| | 0.000766498 | NA |
| | 0.981306443 | 0.990644915 |
| | 0.558015791 | NA |
| | 0.875319722 | NA |
| | 5.02E-08 | 9.74E-07 |
| | 0.978539429 | 0.989124808 |
| | 0.48926604 | 0.643456487 |
| | 0.057157193 | 0.134351831 |
| | 0.449855762 | 0.60864958 |
| | 0.598424191 | 0.734779513 |
| | 0.315862216 | 0.478115283 |
| | 0.247652434 | 0.401283929 |
| | 0.478032145 | 0.634036475 |
| | 0.1495984 | 0.278109657 |
| | 0.736231406 | NA |
| NA | | NA |
| | 0.039007593 | 0.100113107 |
| | 0.067328849 | 0.152070216 |
| | 0.016234592 | 0.049826836 |
| | 0.016284424 | 0.049949976 |
| | 0.441098362 | 0.600941867 |
| | 0.001506028 | 0.007031865 |
| NA | | NA |
| | 0.000102513 | 0.000714176 |
| | 0.764565753 | 0.857286553 |
| NA | | NA |
| | 0.0010748 | 0.00529301 |
| | 5.55E-08 | 1.06E-06 |
| | 4.00E-08 | 8.02E-07 |
| NA | | NA |
| | 1.09E-07 | 1.91E-06 |
| NA | | NA |
| | 0.119109199 | 0.234819118 |
| | 0.386647289 | 0.550537402 |
| | 0.724997103 | 0.830297927 |
| | 0.835649227 | 0.902218025 |
| | 0.905158266 | NA |
| NA | | NA |

| | | |
|---|---|---|
| NA | NA | |
| NA | NA | |
| NA | NA | |
| 0.610722022 | NA | |
| 0.992060816 | NA | |
| NA | NA | |
| 0.839724535 | NA | |
| NA | NA | |
| 0.787523556 | NA | |
| 0.511797629 | | 0.663433538 |
| 0.697339464 | NA | |
| NA | NA | |
| NA | NA | |
| NA | NA | |
| NA | NA | |
| NA | NA | |
| NA | NA | |
| NA | NA | |
| 0.597850711 | | 0.734409715 |
| NA | NA | |
| 0.953217362 | NA | |
| 0.12125797 | | 0.237932603 |
| 0.793954476 | NA | |
| 0.20405059 | | 0.349921663 |
| NA | NA | |
| 0.149735446 | | 0.278340867 |
| 0.620254458 | | 0.75178479 |
| 0.889409301 | NA | |
| NA | NA | |
| NA | NA | |
| 0.32076059 | NA | |
| 0.721010556 | NA | |
| NA | NA | |
| 0.889409773 | NA | |
| NA | NA | |
| NA | NA | |
| NA | NA | |
| 0.992060816 | NA | |
| 0.989597136 | NA | |
| NA | NA | |
| NA | NA | |
| 0.000432576 | | 0.002444397 |
| NA | NA | |
| 0.780939792 | NA | |
| NA | NA | |
| 0.160434985 | | 0.292923575 |
| 0.017109098 | | 0.051927944 |

| | | |
|---|---|---|
| | 0.941368751 | 0.966726671 |
| | 0.277214619 | NA |
| | 0.807433834 | NA |
| | 0.489020605 | NA |
| | 0.22711649 | 0.377013462 |
| NA | | NA |
| | 0.781242538 | NA |
| | 0.002207846 | 0.009729498 |
| | 0.157106165 | 0.28840447 |
| | 0.00568128 | NA |
| | 0.717258149 | 0.824405742 |
| | 0.762527259 | NA |
| NA | | NA |
| | 0.278816995 | NA |
| | 0.988902895 | NA |
| | 0.182476514 | 0.322624941 |
| NA | | NA |
| | 0.217601846 | 0.365691912 |
| | 0.014150851 | 0.044531176 |
| NA | | NA |
| | 0.76780407 | NA |
| | 0.633513262 | NA |
| | 0.858824588 | 0.91762014 |
| | 0.905158739 | NA |
| | 0.535377628 | 0.6837079 |
| | 0.001242644 | 0.005970273 |
| NA | | NA |
| NA | | NA |
| | 0.319410199 | 0.481886648 |
| NA | | NA |
| | 0.374749522 | 0.538802119 |
| NA | | NA |
| | 0.689404206 | NA |
| | 0.260631421 | 0.415930623 |
| | 0.380928304 | NA |
| | 0.361680023 | 0.5265618 |
| NA | | NA |
| NA | | NA |
| NA | | NA |
| NA | | NA |
| | 0.01894494 | 0.056227649 |
| | 0.041640186 | NA |
| NA | | NA |
| NA | | NA |
| | 0.148411311 | NA |
| | 0.889409773 | NA |
| NA | | NA |

| | |
|---|---|
| 0.864868442 | 0.922103618 |
| NA | NA |
| 0.151264158 | NA |
| 0.003610867 | 0.014622361 |
| 0.966760211 | 0.982100642 |
| 0.000227432 | 0.001410387 |
| 0.905158266 | NA |
| NA | NA |
| 0.992061769 | NA |
| NA | NA |
| NA | NA |
| NA | NA |
| 0.992060816 | NA |
| NA | NA |
| NA | NA |
| NA | NA |
| NA | NA |
| 0.821576899 | NA |
| NA | NA |
| 0.253450138 | 0.407794434 |
| 0.889410245 | NA |
| 0.889409773 | NA |
| 0.12751006 | 0.246825935 |
| 0.127050796 | 0.24632778 |
| 0.774525598 | 0.864060335 |
| 0.021603094 | 0.062542168 |
| 0.001373966 | 0.006513584 |
| 0.889409773 | NA |
| 0.608079798 | 0.742194668 |
| 2.20E-07 | 3.52E-06 |
| 0.801661681 | NA |
| NA | NA |
| 0.620168182 | NA |
| 0.349353654 | NA |
| NA | NA |
| 0.905158739 | NA |
| NA | NA |
| 0.827847378 | NA |
| 9.16E-05 | 0.000646946 |
| NA | NA |
| 0.003388192 | 0.013880842 |
| 0.776027085 | 0.864970243 |
| 0.905158739 | NA |
| 2.05E-05 | 0.000181909 |
| NA | NA |
| 0.992061292 | NA |
| 0.286735163 | NA |

| | | |
|---|---|---|
| | 0.597547577 | 0.734195795 |
| | 0.006553915 | 0.023938361 |
| | 0.157654959 | NA |
| NA | | NA |
| | 0.986220783 | NA |
| | 0.002750518 | NA |
| | 8.45E-08 | 1.52E-06 |
| | 0.002708 | 0.01155557 |
| NA | | NA |
| | 0.001971218 | 0.008841967 |
| | 0.905158266 | NA |
| | 0.089017572 | NA |
| | 1.16E-06 | 1.48E-05 |
| NA | | NA |
| NA | | NA |
| NA | | NA |
| | 0.056187404 | 0.132551255 |
| NA | | NA |
| | 0.805907619 | NA |
| | 0.591092102 | NA |
| | 0.789460955 | NA |
| | 0.746927863 | 0.84549662 |
| NA | | NA |
| NA | | NA |
| | 3.84E-08 | 7.78E-07 |
| | 0.616082448 | 0.748833725 |
| NA | | NA |
| | 0.652561599 | NA |
| | 0.90515732 | NA |
| | 0.168546634 | 0.304067987 |
| | 1.86E-08 | 4.06E-07 |
| NA | | NA |
| | 0.05148434 | 0.124222021 |
| | 0.041606626 | 0.105221685 |
| | 0.804902024 | 0.882820554 |
| NA | | NA |
| NA | | NA |
| NA | | NA |
| NA | | NA |
| | 0.01220884 | 0.039530213 |
| | 0.860103816 | NA |
| NA | | NA |
| | 0.977859697 | 0.988665274 |
| | 0.358256169 | 0.523103079 |
| | 0.000234348 | 0.001450586 |
| | 0.069042975 | 0.154784552 |
| | 0.244745637 | 0.397806768 |

| | | |
|---|---|---|
| | 0.321296827 | NA |
| | 0.004332971 | NA |
| | 0.986493522 | NA |
| | 0.992061292 | NA |
| | 0.001310397 | NA |
| | 0.092651181 | 0.19428259 |
| NA | | NA |
| | 0.910992856 | 0.949687411 |
| | 0.782559532 | NA |
| NA | | NA |
| | 0.264968621 | 0.421077287 |
| | 0.987162995 | NA |
| | 0.506923557 | NA |
| | 0.698823335 | 0.811714447 |
| | 0.307284519 | 0.468132424 |
| | 0.600056741 | 0.735952208 |
| | 0.000166138 | 0.001074902 |
| | 0.1289477 | NA |
| NA | | NA |
| | 0.115080562 | NA |
| | 0.992061292 | NA |
| | 0.290696969 | NA |
| | 0.905158266 | NA |
| NA | | NA |
| NA | | NA |
| NA | | NA |
| NA | | NA |
| NA | | NA |
| | 0.788582525 | NA |
| | 0.283638562 | 0.442390737 |
| | 0.991153054 | NA |
| | 0.847002208 | NA |
| NA | | NA |
| NA | | NA |
| NA | | NA |
| | 0.226977343 | 0.376933787 |
| | 0.115494144 | 0.229686463 |
| NA | | NA |
| | 0.327547349 | 0.489726441 |
| | 7.91E-08 | 1.44E-06 |
| | 0.905289366 | 0.946540766 |
| | 0.535938759 | NA |
| NA | | NA |
| NA | | NA |
| NA | | NA |
| | 0.529207355 | 0.678592999 |
| | 0.905158739 | NA |

| | | |
|---|---|---|
| | 0.837796966 | 0.903638861 |
| | 0.30192942 | NA |
| NA | | NA |
| | 0.607824267 | NA |
| | 0.167226424 | 0.302232513 |
| | 0.175201916 | 0.312988135 |
| NA | | NA |
| NA | | NA |
| | 0.742982306 | 0.842463517 |
| | 0.712532487 | NA |
| NA | | NA |
| | 0.367562996 | 0.532141534 |
| | 7.63E-08 | 1.39E-06 |
| NA | | NA |
| | 0.258601725 | 0.413442685 |
| | 0.266919427 | NA |
| | 0.296749923 | 0.456848428 |
| | 0.845403621 | NA |
| | 0.992061292 | NA |
| | 0.312187733 | NA |
| NA | | NA |
| NA | | NA |
| NA | | NA |
| | 0.03255991 | 0.086975227 |
| NA | | NA |
| | 0.313706682 | NA |
| | 0.397359616 | 0.56102487 |
| | 0.385326557 | NA |
| | 0.889409773 | NA |
| | 0.000230772 | 0.001429483 |
| | 0.695466834 | 0.808888262 |
| | 1.65E-19 | 6.27E-17 |
| NA | | NA |
| | 0.467423063 | 0.623732563 |
| | 0.677137923 | 0.795148217 |
| NA | | NA |
| | 0.685304637 | 0.800926314 |
| NA | | NA |
| | 2.69E-05 | 0.000228153 |
| | 0.013410972 | 0.042575039 |
| | 0.068567803 | 0.153924602 |
| | 0.00073306 | 0.003846418 |
| | 0.376548878 | 0.540928746 |
| | 0.971087957 | 0.984947827 |
| | 0.002025335 | 0.009052377 |
| | 0.580572176 | NA |
| | 0.091155505 | 0.191952793 |

| | | |
|---|---|---|
| | 0.99205891 | NA |
| NA | | NA |
| NA | | NA |
| | 0.452621564 | NA |
| | 2.86E-19 | 1.01E-16 |
| | 1.68E-05 | 0.000153095 |
| | 9.09E-05 | 0.000643276 |
| | 0.008042051 | 0.028187613 |
| | 0.485216751 | NA |
| | 0.606804226 | NA |
| | 0.889409301 | NA |
| | 0.425768527 | NA |
| | 0.743698417 | NA |
| | 0.260932912 | 0.416351244 |
| | 0.739429135 | NA |
| | 0.588226542 | NA |
| | 0.529211297 | NA |
| | 0.801314656 | NA |
| | 0.368350685 | NA |
| NA | | NA |
| | 0.801438949 | 0.88029201 |
| | 0.992061292 | NA |
| | 0.889409773 | NA |
| | 0.357823451 | 0.522540744 |
| | 0.113129098 | 0.226000898 |
| | 0.483864807 | 0.639338968 |
| | 0.451145538 | 0.609530848 |
| | 0.748890654 | NA |
| | 0.719743714 | NA |
| | 0.71179105 | NA |
| | 0.274979248 | NA |
| | 0.186357002 | 0.327560506 |
| | 0.806473335 | 0.8840511 |
| | 0.782559532 | NA |
| | 0.573468914 | 0.715622201 |
| | 0.768774524 | 0.860117174 |
| | 0.348773296 | 0.512734337 |
| NA | | NA |
| | 0.093181428 | 0.195040605 |
| | 0.336138185 | NA |
| | 3.74E-05 | 0.000302461 |
| | 0.036178393 | 0.094520088 |
| | 0.109496599 | 0.220570205 |
| | 0.03417491 | 0.09032822 |
| | 0.468262118 | NA |
| NA | | NA |
| | 0.42653237 | 0.587739979 |

| | | |
|---|---|---|
| | 0.056142696 | 0.132491936 |
| NA | NA | |
| | 0.068642338 | 0.154047636 |
| | 0.889409773 | NA |
| | 0.102649536 | 0.210209938 |
| | 0.181481922 | 0.321669346 |
| | 0.616136908 | NA |
| | 0.187780219 | 0.329245115 |
| NA | NA | |
| | 1.18E-07 | 2.05E-06 |
| | 0.266946323 | 0.423148365 |
| | 0.996232771 | NA |
| | 0.929920476 | 0.959774534 |
| | 0.973816689 | NA |
| | 0.000475241 | 0.002641979 |
| | 0.89195306 | 0.938990232 |
| | 0.008972545 | 0.030801663 |
| NA | NA | |
| | 0.687871945 | 0.803114634 |
| | 0.030782287 | 0.083152692 |
| NA | NA | |
| | 0.51974729 | NA |
| | 0.054712526 | 0.129968883 |
| | 0.10732229 | 0.217146427 |
| | 0.905158266 | NA |
| NA | NA | |
| | 0.904947233 | NA |
| NA | NA | |
| | 0.626547424 | NA |
| | 0.534495936 | NA |
| | 0.984334905 | NA |
| | 0.820360253 | 0.893122994 |
| | 0.90515732 | NA |
| | 0.108512099 | 0.219099287 |
| | 0.102716769 | 0.210277454 |
| | 0.143267828 | 0.269281711 |
| NA | NA | |
| NA | NA | |
| NA | NA | |
| | 0.091615493 | 0.192687673 |
| | 0.008055019 | NA |
| | 0.198448924 | 0.342967095 |
| | 0.025547302 | 0.071802534 |
| | 0.418903682 | 0.580967932 |
| | 0.001612626 | 0.007433176 |
| | 0.003605089 | 0.01460593 |
| | 0.37337366 | NA |

| | |
|---|---|
| NA | NA |
| 0.163491217 | 0.297096768 |
| 0.319294046 | 0.481831003 |
| NA | NA |
| NA | NA |
| 0.565127521 | NA |
| 0.763283214 | NA |
| NA | NA |
| NA | NA |
| 0.188924011 | 0.330754296 |
| 1.74E-05 | 0.00015787 |
| NA | NA |
| 0.799635002 | 0.879190116 |
| 0.519403877 | NA |
| NA | NA |
| 0.992061769 | NA |
| NA | NA |
| 0.282230828 | NA |
| 0.812899309 | NA |
| 0.37751337 | 0.541888881 |
| 0.805309787 | 0.883130124 |
| NA | NA |
| 0.881183543 | NA |
| NA | NA |
| 0.024173959 | 0.068583457 |
| 0.759030825 | 0.853756605 |
| 0.614869515 | 0.748028791 |
| 0.096843803 | 0.200973548 |
| 0.090956085 | 0.191643127 |
| 0.341171234 | 0.504564865 |
| 0.033996446 | 0.089943129 |
| 0.065252138 | 0.148900728 |
| 0.796745564 | NA |
| 0.108402907 | 0.218969684 |
| 0.99205891 | NA |
| 0.14772969 | 0.275483223 |
| NA | NA |
| 0.50569731 | 0.657691463 |
| 0.002929181 | 0.012317479 |
| 0.596365463 | 0.733276734 |
| 0.072048162 | 0.160099737 |
| 0.026529686 | NA |
| 0.417587845 | NA |
| 0.905158739 | NA |
| NA | NA |
| 2.57E-09 | 7.27E-08 |
| NA | NA |

| | | |
|---|---|---|
| | 0.551597104 | 0.697163625 |
| | 0.858360921 | NA |
| | 0.001473823 | 0.0069067 |
| | 0.910071343 | 0.949096227 |
| NA | | NA |
| | 0.458051989 | 0.615453932 |
| | 0.793686831 | 0.875498515 |
| | 0.060695831 | 0.141054053 |
| | 0.570110368 | 0.712443294 |
| NA | | NA |
| NA | | NA |
| | 0.002724962 | 0.01161441 |
| | 0.660491829 | NA |
| | 0.905157793 | NA |
| NA | | NA |
| | 0.905158266 | NA |
| NA | | NA |
| | 0.76147686 | NA |
| | 0.358105114 | NA |
| | 0.060312273 | 0.140340915 |
| | 0.323720178 | 0.485985809 |
| | 0.641425671 | 0.767366984 |
| | 0.889409773 | NA |
| | 0.306819536 | 0.467553795 |
| NA | | NA |
| | 0.005019007 | 0.019237628 |
| | 0.840933431 | 0.90586496 |
| | 0.797178279 | NA |
| | 0.524725414 | 0.674774267 |
| | 0.017312632 | 0.052385306 |
| | 0.727643741 | NA |
| | 0.335433647 | 0.498361803 |
| | 0.261420355 | 0.416886679 |
| | 0.20162977 | 0.346943497 |
| NA | | NA |
| | 0.439561788 | NA |
| | 0.992061292 | NA |
| | 0.021021405 | 0.061197008 |
| | 0.435352266 | 0.595970324 |
| | 0.082677835 | 0.178464818 |
| | 0.303489219 | NA |
| | 0.000125869 | 0.000849139 |
| | 0.655143495 | 0.778142362 |
| | 0.557366744 | NA |
| | 0.992062245 | NA |
| NA | | NA |
| | 0.736800273 | NA |

| | | |
|---|---|---|
| | 0.717746494 | NA |
| | 0.936081704 | NA |
| NA | | NA |
| NA | | NA |
| NA | | NA |
| | 0.988778992 | NA |
| | 0.136941866 | 0.260121605 |
| NA | | NA |
| NA | | NA |
| NA | | NA |
| NA | | NA |
| | 0.198155281 | NA |
| | 0.003222505 | 0.013339427 |
| | 0.76573101 | NA |
| | 0.000192548 | 0.001220249 |
| NA | | NA |
| NA | | NA |
| NA | | NA |
| | 0.00694855 | 0.025128987 |
| | 1.19E-08 | 2.76E-07 |
| NA | | NA |
| | 0.889409773 | NA |
| | 0.992060816 | NA |
| | 0.029901035 | 0.081322372 |
| NA | | NA |
| | 0.852666512 | NA |
| NA | | NA |
| NA | | NA |
| NA | | NA |
| | 0.330662596 | NA |
| | 0.294060611 | NA |
| NA | | NA |
| | 0.142534928 | 0.268087749 |
| NA | | NA |
| | 0.586242131 | 0.725460183 |
| | 0.904347686 | 0.945916492 |
| | 0.499653447 | 0.652739011 |
| NA | | NA |
| | 0.890092006 | 0.937840332 |
| | 0.005789618 | 0.0216583 |
| | 0.218288499 | NA |
| | 0.880414379 | 0.932221934 |
| | 0.58427784 | NA |
| | 0.014805065 | 0.046186165 |
| | 0.207887926 | 0.354542495 |
| | 0.049712933 | 0.120837535 |
| | 0.431534607 | NA |

| | | | |
|---|---|---|---|
| | 0.897828342 | | 0.942509445 |
| | 0.568686551 | NA | |
| NA | | NA | |
| NA | | NA | |
| NA | | NA | |
| | 0.889410245 | NA | |
| | 0.992061769 | NA | |
| | 0.559833991 | NA | |
| | 0.987745158 | NA | |
| | 0.040179843 | | 0.102427347 |
| | 0.151400608 | | 0.280737556 |
| | 0.007372885 | | 0.026321114 |
| | 0.705234244 | | 0.816569405 |
| | 0.002395633 | | 0.01043557 |
| | 0.293061018 | | 0.45277861 |
| | 0.066609712 | | 0.150993635 |
| | 0.712194954 | NA | |
| | 0.756256485 | | 0.851753406 |
| NA | | NA | |
| | 0.905158266 | NA | |
| NA | | NA | |
| NA | | NA | |
| NA | | NA | |
| | 0.69823487 | NA | |
| NA | | NA | |
| NA | | NA | |
| NA | | NA | |
| NA | | NA | |
| NA | | NA | |
| NA | | NA | |
| NA | | NA | |
| NA | | NA | |
| NA | | NA | |
| | 0.48698263 | | 0.641827091 |
| | 0.38050713 | | 0.544720066 |
| | 0.628460751 | | 0.756931199 |
| | 0.992819511 | NA | |
| | 1.12E-06 | | 1.44E-05 |
| | 0.934676117 | NA | |
| | 2.30E-06 | | 2.72E-05 |
| | 4.22E-06 | | 4.60E-05 |
| NA | | NA | |
| | 0.381950754 | NA | |
| | 0.905296333 | NA | |
| NA | | NA | |
| NA | | NA | |
| NA | | NA | |

| | |
|---|---|
| NA | NA |
| 0.134323686 | 0.256198542 |
| NA | NA |
| NA | NA |
| 0.58653004 | 0.725630223 |
| NA | NA |
| NA | NA |
| 0.889409773 | NA |
| NA | NA |
| NA | NA |
| 0.871606873 | NA |
| NA | NA |
| NA | NA |
| 0.992061292 | NA |
| NA | NA |
| NA | NA |
| 0.992003131 | NA |
| NA | NA |
| 0.846445294 | NA |
| 0.227135206 | 0.377013462 |
| NA | NA |
| NA | NA |
| 0.786387587 | NA |
| NA | NA |
| NA | NA |
| NA | NA |
| NA | NA |
| 0.985003519 | 0.992446925 |
| 0.905158266 | NA |
| 7.93E-05 | 0.000573679 |
| NA | NA |
| NA | NA |
| NA | NA |
| NA | NA |
| 0.016077188 | 0.049452159 |
| 0.974053063 | NA |
| 0.876012733 | NA |
| 0.889409301 | NA |
| 0.004862804 | 0.018720604 |
| 0.669626777 | NA |
| 0.807879735 | NA |
| 0.279719004 | 0.437894746 |
| 0.007238771 | 0.025927967 |
| 0.525365364 | NA |
| 0.992061292 | NA |
| NA | NA |
| 9.63E-14 | 8.63E-12 |

| | |
|---|---|
| 3.11E-07 | 4.75E-06 |
| 0.024791518 | 0.070018898 |
| 0.437522037 | 0.59785603 |
| NA | NA |
| 0.253817105 | 0.408203365 |
| 0.992061292 | NA |
| 0.869557568 | NA |
| 0.47701281 | 0.633143393 |
| 0.386768971 | 0.550609755 |
| 0.051767747 | 0.12467295 |
| 0.292546835 | 0.452295505 |
| 0.767781626 | 0.859225193 |
| 0.25062231 | 0.404692646 |
| 0.928684352 | 0.959173119 |
| 0.52729723 | 0.67705063 |
| 0.004120353 | 0.016299556 |
| 0.123022492 | 0.240406773 |
| 0.000189087 | 0.001200048 |
| NA | NA |
| 0.588115337 | NA |
| NA | NA |
| 0.726215019 | 0.830985313 |
| 0.602462841 | NA |
| 0.953358687 | 0.973847672 |
| 0.782559532 | NA |
| 0.227236644 | 0.377109934 |
| 0.889410717 | NA |
| 0.745909069 | NA |
| 0.992087589 | 0.995961112 |
| 0.084737155 | 0.181500506 |
| 0.592415916 | 0.729926607 |
| 0.259074834 | 0.414108624 |
| 0.005069959 | 0.01938893 |
| 0.696252008 | NA |
| 0.155516911 | 0.286348805 |
| 0.432854431 | 0.593657944 |
| 0.434106121 | 0.594816942 |
| 0.934622567 | 0.962454369 |
| 0.821167905 | 0.893667741 |
| NA | NA |
| 0.37248355 | NA |
| NA | NA |
| 0.49100381 | 0.644979831 |
| NA | NA |
| 0.001365098 | 0.006485545 |
| 0.408534826 | NA |
| 0.316716656 | 0.47881453 |

| | | |
|---|---|---|
| | 0.978337971 | NA |
| | 0.424644372 | NA |
| | 0.544786532 | 0.691405422 |
| | 0.449463342 | 0.608418538 |
| | 0.049009244 | NA |
| | 0.496418791 | NA |
| | 0.878509626 | NA |
| NA | | NA |
| | 0.041601477 | 0.105220784 |
| | 0.368460227 | 0.533110505 |
| | 0.000926331 | 0.004668649 |
| | 0.763085966 | NA |
| | 0.465759023 | 0.622578149 |
| | 0.62571683 | 0.755491058 |
| | 0.687609768 | 0.802936607 |
| NA | | NA |
| | 0.992609206 | 0.996201218 |
| NA | | NA |
| | 0.791223477 | NA |
| | 0.022235672 | 0.064010635 |
| | 0.879542993 | NA |
| | 0.130410237 | 0.25075993 |
| | 0.987938331 | 0.993639148 |
| | 0.745760904 | NA |
| | 0.905158739 | NA |
| | 0.00155434 | 0.007212966 |
| NA | | NA |
| | 0.86632646 | 0.922985917 |
| | 0.550699662 | NA |
| | 0.719314367 | NA |
| | 0.757315024 | NA |
| NA | | NA |
| NA | | NA |
| | 0.006735885 | 0.024476801 |
| NA | | NA |
| | 0.801661681 | NA |
| NA | | NA |
| | 0.007084525 | 0.02549477 |
| | 0.161070441 | 0.293731153 |
| | 0.395439346 | 0.558960579 |
| NA | | NA |
| NA | | NA |
| | 0.152098364 | 0.281755355 |
| | 0.665563991 | NA |
| | 0.689785668 | 0.804707192 |
| | 0.265311395 | 0.421426757 |
| | 0.011191858 | 0.036834833 |

| | | |
|---|---|---|
| | 0.137737617 | 0.261302837 |
| | 0.011040832 | 0.036463471 |
| | 0.030017978 | 0.081539432 |
| | 0.307838199 | 0.468813303 |
| | 0.199537472 | NA |
| | 0.106086425 | 0.215260746 |
| | 0.326809763 | 0.489056535 |
| | 0.655950599 | 0.778770817 |
| | 0.792188023 | NA |
| | 0.531099335 | NA |
| | 0.251096407 | 0.405219813 |
| | 0.041424878 | 0.104884576 |
| | 0.346850772 | 0.510523232 |
| | 0.1218179 | NA |
| | 0.784353866 | NA |
| | 0.029009631 | 0.079261228 |
| | 0.063384547 | 0.145637886 |
| | 0.88315578 | NA |
| | 0.500713889 | NA |
| | 0.511611258 | 0.6632311 |
| | 4.83E-07 | 6.99E-06 |
| | 0.845443424 | 0.909073163 |
| | 0.889409773 | NA |
| | 0.992003131 | NA |
| | 0.158292328 | 0.290000103 |
| | 0.132914403 | 0.254195033 |
| | 0.03411182 | 0.090183176 |
| | 0.096659757 | 0.200752067 |
| | 0.01568656 | 0.048447422 |
| NA | | NA |
| | 0.797342596 | NA |
| | 0.905158739 | NA |
| | 0.894259403 | 0.940291238 |
| | 0.905158266 | NA |
| NA | | NA |
| | 3.58E-09 | 9.64E-08 |
| | 0.962962959 | NA |
| | 0.985051756 | NA |
| | 3.31E-11 | 1.59E-09 |
| NA | | NA |
| | 0.7628432 | 0.856185731 |
| NA | | NA |
| | 0.199775388 | 0.344635694 |
| | 0.000443404 | 0.002494671 |
| | 0.018945689 | 0.056227649 |
| | 0.493964343 | 0.647309363 |
| NA | | NA |

| | | |
|---|---|---|
| | 0.604041551 | NA |
| | 0.532840942 | 0.681776847 |
| | 0.989560379 | NA |
| | 0.843400599 | NA |
| | 0.000667393 | 0.00354955 |
| | 0.861596748 | NA |
| | 0.889410245 | NA |
| | 0.009375184 | 0.031902939 |
| | 0.459634039 | NA |
| | 0.786645282 | NA |
| | 0.215788711 | NA |
| | 9.43E-06 | 9.24E-05 |
| | 0.828581995 | 0.897767035 |
| NA | | NA |
| NA | | NA |
| NA | | NA |
| | 0.00120506 | 0.005827229 |
| | 0.000960647 | 0.004812879 |
| | 0.008415161 | 0.029275749 |
| | 0.604619054 | 0.739738678 |
| | 0.091856194 | 0.193113467 |
| | 0.89572766 | 0.941075508 |
| | 0.065418606 | 0.149141109 |
| | 0.128373084 | 0.248081013 |
| | 0.441722672 | 0.601425357 |
| | 0.790069731 | 0.873176366 |
| | 0.342389461 | NA |
| | 0.051775547 | 0.12467806 |
| | 0.49545161 | 0.64871635 |
| NA | | NA |
| | 0.797157316 | NA |
| | 0.551225132 | 0.696894045 |
| | 0.905157793 | NA |
| | 0.050977684 | 0.12327045 |
| | 1.40E-06 | 1.75E-05 |
| | 0.006886393 | 0.024957634 |
| | 4.17E-05 | 0.000331995 |
| | 0.905158266 | NA |
| | 0.887898287 | 0.936517531 |
| | 7.79E-07 | 1.06E-05 |
| | 0.010458757 | 0.034929863 |
| | 0.133830931 | 0.255591704 |
| | 0.461955064 | 0.619371794 |
| | 0.905158739 | NA |
| | 0.293285806 | NA |
| NA | | NA |
| NA | | NA |

| | | |
|---|---|---|
| | 0.889409773 | NA |
| NA | | NA |
| NA | | NA |
| NA | | NA |
| NA | | NA |
| NA | | NA |
| | 0.923299006 | 0.955723714 |
| | 0.905158266 | NA |
| NA | | NA |
| | 0.905158266 | NA |
| | 0.905158266 | NA |
| | 0.905158266 | NA |
| NA | | NA |
| NA | | NA |
| | 0.992060816 | NA |
| | 0.652115544 | NA |
| NA | | NA |
| | 0.090739774 | NA |
| | 0.724524704 | 0.830064068 |
| | 0.452777009 | 0.610906318 |
| | 0.435986299 | 0.596499884 |
| | 0.889409301 | NA |
| | 0.992062081 | NA |
| NA | | NA |
| | 0.74226382 | NA |
| | 0.621970294 | 0.753205943 |
| | 0.290715264 | NA |
| | 0.145050295 | 0.271771281 |
| | 0.905158266 | NA |
| | 0.905158739 | NA |
| | 0.019940507 | 0.058656879 |
| | 0.453112277 | 0.611135218 |
| | 0.841081658 | 0.90593575 |
| | 0.90760829 | 0.947610543 |
| | 0.214153475 | NA |
| | 0.883264219 | NA |
| | 0.055795934 | 0.131857826 |
| | 0.000715602 | 0.003760208 |
| | 0.440113185 | 0.600265306 |
| | 0.001820121 | 0.00825786 |
| | 0.859615715 | NA |
| | 0.951786494 | NA |
| NA | | NA |
| | 0.038469012 | 0.099113109 |
| | 9.67E-13 | 6.84E-11 |
| NA | | NA |
| | 0.788780899 | NA |

| | |
|---|---|
| NA | NA |
| NA | NA |
| 0.431085084 | 0.591970318 |
| NA | NA |
| NA | NA |
| NA | NA |
| 0.807879735 | NA |
| 0.064173709 | 0.147005058 |
| 0.003285723 | 0.01353681 |
| 0.286139386 | 0.445216096 |
| 0.016072942 | 0.049446026 |
| 0.804534188 | NA |
| 0.352695289 | 0.516886225 |
| 0.801661681 | NA |
| 0.828824426 | 0.897886313 |
| 0.955767509 | NA |
| 0.905158266 | NA |
| 0.989978842 | NA |
| 0.905158266 | NA |
| 0.777092376 | NA |
| 0.898195764 | NA |
| 0.878033887 | NA |
| NA | NA |
| NA | NA |
| 0.356116006 | NA |
| NA | NA |
| 0.766789031 | NA |
| 0.863142176 | NA |
| 0.974187146 | NA |
| 0.788885483 | NA |
| NA | NA |
| 0.984480814 | NA |
| NA | NA |
| 0.264466948 | 0.420401745 |
| 0.889409773 | NA |
| NA | NA |
| 0.905158266 | NA |
| NA | NA |
| 0.067385433 | 0.152171608 |
| 2.74E-05 | 0.000231668 |
| 0.015598251 | 0.04822216 |
| 0.911015711 | 0.949687411 |
| 0.057812395 | 0.135514772 |
| 0.054697018 | 0.129946102 |
| 0.868497615 | NA |
| 0.09329131 | NA |
| 0.004745493 | 0.018349407 |

| | |
|---|---|
| NA | NA |
| 0.001697443 | 0.007765448 |
| 0.491127004 | NA |
| NA | NA |
| 0.893967439 | NA |
| 0.502883399 | 0.655274222 |
| 0.478919248 | 0.634638088 |
| NA | NA |
| 0.626522117 | NA |
| 0.882927242 | NA |
| 0.257706023 | NA |
| 0.7954714 | 0.87663136 |
| 0.291914416 | 0.451546116 |
| 0.330913746 | NA |
| 0.258153493 | 0.41302678 |
| NA | NA |
| NA | NA |
| 0.154594624 | 0.285152738 |
| NA | NA |
| NA | NA |
| 0.905158266 | NA |
| NA | NA |
| NA | NA |
| 0.042750444 | 0.107465832 |
| 0.419621102 | NA |
| NA | NA |
| 0.989162833 | NA |
| NA | NA |
| 0.332697648 | 0.495335795 |
| 1.32E-05 | 0.000124196 |
| 0.698041933 | 0.811150066 |
| NA | NA |
| 0.081515729 | 0.176476771 |
| 0.147278488 | 0.274938327 |
| 0.663198051 | 0.784611066 |
| 0.586344518 | 0.725488303 |
| 0.627825655 | 0.756581252 |
| 0.902937908 | NA |
| NA | NA |
| NA | NA |
| 4.38E-07 | 6.42E-06 |
| 0.004130941 | 0.016320859 |
| NA | NA |
| 0.991955612 | NA |
| 0.889409301 | NA |
| 0.010762047 | 0.035747056 |
| 0.000876078 | 0.004449076 |

| | |
|---|---|
| 0.042770544 | 0.107496667 |
| 0.012802488 | 0.041052838 |
| NA | NA |
| 0.38926255 | 0.552742792 |
| 0.98825791 | NA |
| NA | NA |
| 0.992061292 | NA |
| NA | NA |
| 0.00287184 | 0.012124451 |
| 0.036770963 | 0.095749268 |
| 0.579205718 | 0.719754307 |
| 0.397606145 | 0.561162581 |
| 0.992062245 | NA |
| 0.042866109 | 0.107675232 |
| 0.01006749 | 0.033844461 |
| 0.020578637 | 0.060187307 |
| 0.783906671 | NA |
| 0.559414568 | NA |
| 0.330868453 | 0.493350431 |
| 7.85E-10 | 2.56E-08 |
| 0.439958934 | 0.600177563 |
| 0.378646505 | 0.543160348 |
| 0.616780727 | NA |
| 0.786112585 | NA |
| 0.000341533 | 0.002000992 |
| NA | NA |
| NA | NA |
| NA | NA |
| 0.410342717 | NA |
| 0.099990882 | 0.205989596 |
| 0.905158739 | NA |
| NA | NA |
| 0.905158739 | NA |
| 0.951104856 | 0.972767188 |
| 0.576660741 | 0.71809534 |
| NA | NA |
| NA | NA |
| 0.77922419 | 0.867169267 |
| 0.748778853 | NA |
| 0.787645204 | NA |
| NA | NA |
| NA | NA |
| NA | NA |
| NA | NA |
| NA | NA |
| 0.988774692 | NA |
| NA | NA |

| | |
|---|---|
| NA | NA |
| NA | NA |
| NA | NA |
| NA | NA |
| NA | NA |
| NA | NA |
| NA | NA |
| NA | NA |
| NA | NA |
| 0.000405771 | 0.002315575 |
| 4.50E-06 | 4.86E-05 |
| 5.20E-14 | 4.90E-12 |
| 6.87E-09 | 1.69E-07 |
| 0.156293701 | NA |
| 0.002891155 | 0.012187918 |
| NA | NA |
| 0.007667164 | 0.027124737 |
| NA | NA |
| 0.810524717 | 0.886546735 |
| NA | NA |
| NA | NA |
| NA | NA |
| NA | NA |
| NA | NA |
| 0.153742317 | 0.284033951 |
| 0.909415347 | 0.94859641 |
| 0.919341063 | NA |
| 0.003473345 | 0.014158377 |
| NA | NA |
| 0.052532636 | 0.126020184 |
| 6.15E-18 | 1.61E-15 |
| 0.000358021 | 0.00208149 |
| 0.520236877 | NA |
| NA | NA |
| 0.889409301 | NA |
| NA | NA |
| 0.093891309 | 0.196208532 |
| NA | NA |
| 0.992060816 | NA |
| NA | NA |
| 0.006779503 | 0.024601826 |
| 0.139375037 | 0.263634803 |
| 0.889409773 | NA |
| 0.815615096 | NA |
| NA | NA |
| 0.889409773 | NA |
| 0.889409301 | NA |

| | |
|---|---|
| NA | NA |
| NA | NA |
| NA | NA |
| NA | NA |
| 0.889409301 | NA |
| NA | NA |
| 0.69079418 | 0.805274956 |
| 0.868085492 | 0.924373662 |
| NA | NA |
| NA | NA |
| 0.556888053 | NA |
| 0.905158266 | NA |
| 0.904986279 | 0.946313985 |
| 0.924274082 | 0.956400673 |
| NA | NA |
| 1.29E-11 | 6.95E-10 |
| NA | NA |
| 0.929693654 | NA |
| 0.770517631 | 0.861278018 |
| 0.532536757 | NA |
| 4.80E-06 | 5.14E-05 |
| 0.367113123 | 0.531807781 |
| 0.499388977 | NA |
| 0.889409773 | NA |
| 0.070418645 | 0.157303634 |
| 0.007479834 | 0.026599137 |
| 0.42397674 | 0.585408359 |
| 0.341153705 | 0.504564865 |
| 5.60E-07 | 7.93E-06 |
| 0.336315861 | 0.499267172 |
| 0.018398103 | 0.054929155 |
| NA | NA |
| 0.805334324 | NA |
| 0.144470873 | 0.270963094 |
| 0.082755252 | NA |
| 0.889409301 | NA |
| 0.992061769 | NA |
| 0.889410717 | NA |
| 0.124632895 | NA |
| 0.187867716 | 0.329345934 |
| 0.405977251 | NA |
| 3.22E-14 | 3.17E-12 |
| NA | NA |
| 0.007547987 | 0.026802433 |
| NA | NA |
| 0.937972663 | NA |
| 0.738757848 | NA |

| | |
|---|---|
| 0.947049156 | 0.969924394 |
| 0.761042477 | NA |
| NA | NA |
| NA | NA |
| NA | NA |
| 0.942752955 | 0.967650376 |
| NA | NA |
| 0.453266444 | NA |
| 0.601479102 | 0.737046852 |
| 0.78186651 | 0.868525227 |
| 0.204606008 | NA |
| 4.03E-06 | 4.43E-05 |
| 6.89E-16 | 1.08E-13 |
| 0.373554753 | NA |
| 2.01E-11 | 1.04E-09 |
| 0.046559585 | 0.114683335 |
| NA | NA |
| 0.881992598 | NA |
| 0.889409301 | NA |
| NA | NA |
| NA | NA |
| NA | NA |
| 0.454626472 | 0.612650672 |
| 0.798019462 | NA |
| 0.673386867 | 0.792334755 |
| 0.79812283 | NA |
| 0.563231801 | NA |
| 0.118182112 | 0.233461767 |
| 0.439519962 | NA |
| 0.002601212 | 0.011180274 |
| 8.90E-05 | 0.000630924 |
| 4.55E-08 | 8.95E-07 |
| 0.273940993 | NA |
| 0.720622849 | NA |
| 0.905157793 | NA |
| NA | NA |
| NA | NA |
| 0.497082709 | 0.65023168 |
| 0.381325705 | 0.545444051 |
| NA | NA |
| 0.800275352 | 0.879541865 |
| 0.117383794 | 0.232360073 |
| 0.353392789 | 0.517619548 |
| 0.526537117 | 0.676430619 |
| 0.000124748 | 0.000843915 |
| 0.988171651 | NA |
| 0.889409773 | NA |

| | |
|---|---|
| 3.39E-08 | 6.97E-07 |
| 0.35174924 | 0.51603832 |
| 0.188063347 | 0.329478461 |
| 0.71094515 | 0.819939315 |
| 0.176670092 | 0.315098973 |
| 0.603270434 | NA |
| 0.038029164 | 0.098247796 |
| 0.000744252 | 0.003893052 |
| 0.639339939 | 0.766084565 |
| NA | NA |
| NA | NA |
| NA | NA |
| 0.986721446 | NA |
| 0.352112633 | 0.516364453 |
| NA | NA |
| 0.067058913 | 0.151690546 |
| 0.770811593 | 0.861386859 |
| 0.998938482 | 0.99957245 |
| 1.55E-06 | 1.91E-05 |
| 0.053415709 | 0.127467459 |
| 0.732650715 | NA |
| NA | NA |
| NA | NA |
| 0.336429544 | 0.499371756 |
| NA | NA |
| NA | NA |
| NA | NA |
| 0.010954578 | 0.036233106 |
| 0.889410717 | NA |
| 0.499187678 | NA |
| 0.1776427 | 0.316499477 |
| 0.942036307 | NA |
| 0.731601317 | 0.834904549 |
| 0.674750623 | 0.793196518 |
| NA | NA |
| NA | NA |
| 0.000164466 | 0.001064714 |
| NA | NA |
| NA | NA |
| 0.523662356 | 0.673880783 |
| 0.201265253 | 0.346491505 |
| 0.248833335 | 0.402603242 |
| 0.106060679 | 0.215235729 |
| 0.646741715 | 0.771720112 |
| 0.337658321 | 0.500617046 |
| 0.905158266 | NA |
| NA | NA |

| | | |
|---|---|---|
| | 0.889409301 | NA |
| | 0.10662465 | NA |
| | 0.005223785 | 0.019873875 |
| NA | | NA |
| NA | | NA |
| NA | | NA |
| NA | | NA |
| | 1.29E-06 | 1.63E-05 |
| | 0.755603359 | NA |
| | 0.13421575 | 0.256111945 |
| | 0.02422671 | 0.068713946 |
| | 0.006596876 | 0.024059259 |
| | 0.030328821 | NA |
| | 0.098012634 | NA |
| | 0.005403951 | 0.020448968 |
| | 0.972591613 | NA |
| NA | | NA |
| NA | | NA |
| | 0.054905614 | 0.130300701 |
| | 0.808141269 | 0.88498742 |
| | 0.001203001 | 0.00581994 |
| | 0.256021855 | 0.410580543 |
| | 0.02354719 | 0.067108727 |
| | 0.338002828 | 0.500891086 |
| | 0.215474043 | 0.36321444 |
| | 5.30E-05 | 0.000406001 |
| | 2.65E-06 | 3.06E-05 |
| | 0.539670324 | 0.687077564 |
| | 1.47E-05 | 0.000136077 |
| | 1.54E-09 | 4.60E-08 |
| NA | | NA |
| | 2.89E-10 | 1.08E-08 |
| | 1.76E-14 | 1.90E-12 |
| | 0.359927322 | 0.52459998 |
| | 0.097488769 | 0.201977156 |
| | 0.324143207 | 0.48639201 |
| | 0.422780544 | 0.584256638 |
| | 0.992061292 | NA |
| | 0.323729685 | 0.485985809 |
| | 7.51E-05 | 0.000547312 |
| | 0.26998318 | 0.426392452 |
| | 0.078721338 | 0.171850269 |
| | 0.423809927 | 0.58531083 |
| | 0.183274218 | 0.323644436 |
| | 0.000592565 | 0.003210494 |
| | 0.50339802 | NA |
| | 0.162411291 | 0.295678699 |

| | | |
|---|---|---|
| | 0.123258417 | 0.24080352 |
| | 0.569132104 | 0.71146373 |
| | 0.124281703 | 0.242285325 |
| | 0.573038547 | 0.715342437 |
| | 3.32E-10 | 1.21E-08 |
| | 4.03E-05 | 0.000321763 |
| | 0.467097484 | 0.623525451 |
| | 0.517516926 | 0.668518982 |
| | 0.05463385 | 0.129852226 |
| | 0.015165123 | 0.047142417 |
| | 0.905158739 | NA |
| | 0.271682826 | NA |
| | 0.992060816 | NA |
| | 0.293646927 | NA |
| | 0.407917195 | 0.571120485 |
| | 0.209351135 | 0.356174292 |
| | 0.129545516 | 0.249643702 |
| | 1.55E-06 | 1.91E-05 |
| NA | | NA |
| NA | | NA |
| | 3.56E-06 | 3.97E-05 |
| | 0.968159142 | 0.98293063 |
| | 0.006962985 | 0.0251718 |
| | 0.905154761 | NA |
| | 0.649382894 | NA |
| | 0.993798762 | 0.997067986 |
| | 0.211064577 | 0.358139002 |
| | 0.009351841 | 0.031853123 |
| | 0.477244149 | 0.633299266 |
| | 0.043484922 | 0.108843658 |
| | 0.240338017 | NA |
| NA | | NA |
| | 0.320777378 | 0.48309104 |
| | 0.393655336 | 0.557048429 |
| | 0.432620065 | 0.593447277 |
| | 0.000151764 | 0.000993561 |
| | 0.826194401 | 0.896319493 |
| | 0.022671826 | 0.065121075 |
| | 5.37E-14 | 5.02E-12 |
| | 0.016995205 | 0.051667964 |
| | 0.577130516 | 0.71849376 |
| | 0.971436704 | 0.984954128 |
| | 0.848945189 | NA |
| | 0.379771941 | 0.544134937 |
| | 0.319685677 | 0.482129488 |
| | 0.759031831 | NA |
| | 0.272114058 | 0.428894099 |

| | |
|---|---|
| 0.010301064 | 0.034518705 |
| NA | NA |
| NA | NA |
| NA | NA |
| 6.34E-11 | 2.82E-09 |
| 0.300592995 | 0.46097814 |
| 0.091975947 | NA |
| NA | NA |
| 0.452511742 | NA |
| 0.004128362 | 0.016313606 |
| 0.897611556 | 0.942417029 |
| 0.001660636 | 0.007617694 |
| 0.412095653 | NA |
| 0.000206667 | 0.001297368 |
| NA | NA |
| 3.96E-10 | 1.41E-08 |
| 0.405186672 | 0.568312093 |
| 0.001539161 | 0.007159169 |
| 0.284865909 | 0.443737709 |
| 0.648135597 | NA |
| 0.820044889 | 0.892909192 |
| 0.08889525 | 0.18844008 |
| 2.39E-08 | 5.08E-07 |
| 8.35E-06 | 8.35E-05 |
| 0.121805819 | 0.238645073 |
| 0.52403506 | 0.67420236 |
| NA | NA |
| 0.051353222 | 0.123960142 |
| 0.6719599 | 0.79118324 |
| 0.055248447 | 0.130859741 |
| 0.630046829 | NA |
| 0.420567471 | 0.582371111 |
| 0.688245193 | NA |
| 0.000706691 | 0.003720074 |
| 0.002378453 | 0.010368961 |
| 0.402250314 | 0.565856159 |
| 0.978855102 | NA |
| 0.021476875 | 0.062242404 |
| 0.180146074 | 0.319817904 |
| 5.85E-07 | 8.19E-06 |
| 0.00780031 | 0.027476262 |
| 8.58E-06 | 8.55E-05 |
| 0.642578561 | 0.768317109 |
| 0.000341465 | 0.002000992 |
| 0.049670263 | 0.120775353 |
| 1.60E-05 | 0.000147081 |
| 0.367885666 | 0.532489706 |

| | |
|---|---|
| 0.131902809 | 0.252766715 |
| NA | NA |
| 0.003161157 | 0.013119608 |
| 3.16E-13 | 2.56E-11 |
| 0.507476248 | NA |
| 0.046923564 | 0.115424502 |
| 0.85535358 | NA |
| 0.905158266 | NA |
| 6.13E-13 | 4.63E-11 |
| 0.83326192 | 0.900794276 |
| 0.606527858 | 0.741083087 |
| 0.008311442 | 0.028956205 |
| 0.150792136 | 0.279831363 |
| 0.063468395 | 0.145769531 |
| 0.208953343 | 0.355717999 |
| 0.784353866 | NA |
| 0.000598138 | 0.003235313 |
| 0.3569716 | 0.521643665 |
| 0.81800585 | NA |
| 0.421750432 | 0.583310307 |
| NA | NA |
| 0.297003201 | 0.4570673 |
| 0.023309128 | 0.066559835 |
| 0.008037883 | 0.028182695 |
| 0.342930817 | 0.506314652 |
| 1.27E-06 | 1.61E-05 |
| 0.12333986 | 0.240920638 |
| 0.006325621 | 0.023235916 |
| NA | NA |
| 0.584773297 | 0.724336463 |
| 0.005636565 | 0.021179557 |
| 0.378821063 | 0.543268791 |
| 0.000594739 | 0.003221476 |
| 0.766615294 | 0.858439143 |
| 0.206624025 | NA |
| 0.905158266 | NA |
| 0.007831953 | 0.027565612 |
| 0.504175525 | 0.656381713 |
| 0.031806526 | 0.085394242 |
| 0.838348727 | 0.903878809 |
| 0.000452554 | 0.002537051 |
| 0.083575932 | NA |
| 8.96E-06 | 8.88E-05 |
| NA | NA |
| NA | NA |
| 0.60602855 | 0.740679074 |
| 0.391791048 | NA |

| | |
|---|---|
| NA | NA |
| 8.43E-05 | 0.000602496 |
| 0.282004312 | 0.440561518 |
| 2.65E-09 | 7.42E-08 |
| NA | NA |
| 0.001830704 | 0.008300729 |
| 3.14E-05 | 0.000259989 |
| 0.004051916 | 0.016063552 |
| NA | NA |
| NA | NA |
| 0.889409773 | NA |
| 0.507927134 | 0.659740444 |
| NA | NA |
| NA | NA |
| NA | NA |
| NA | NA |
| 0.987031469 | NA |
| NA | NA |
| NA | NA |
| 0.265083061 | 0.421228667 |
| 0.986968087 | NA |
| NA | NA |
| 0.296695079 | 0.456817224 |
| NA | NA |
| NA | NA |
| NA | NA |
| NA | NA |
| NA | NA |
| 0.328061028 | NA |
| NA | NA |
| NA | NA |
| 0.801661681 | NA |
| 0.992060816 | NA |
| 0.185743349 | NA |
| NA | NA |
| NA | NA |
| 0.997017254 | NA |
| NA | NA |
| NA | NA |
| NA | NA |
| 0.874884864 | NA |
| NA | NA |
| 0.905158739 | NA |
| NA | NA |
| NA | NA |
| NA | NA |
| 0.000696009 | 0.003671307 |

| | | |
|---|---|---|
| NA | NA | |
| 0.677738546 | NA | |
| NA | NA | |
| NA | NA | |
| 5.18E-05 | | 0.000398559 |
| NA | NA | |
| 0.644573761 | NA | |
| NA | NA | |
| NA | NA | |
| NA | NA | |
| 0.804534188 | NA | |
| 0.542931594 | NA | |
| 0.70853139 | NA | |
| NA | NA | |
| NA | NA | |
| 0.640597503 | NA | |
| 0.551749102 | NA | |
| 0.103074529 | | 0.210848124 |
| NA | NA | |
| 0.24369659 | NA | |
| NA | NA | |
| 0.520129166 | NA | |
| 0.640597503 | NA | |
| 0.767197858 | NA | |
| 0.804534188 | NA | |
| NA | NA | |
| 0.500005289 | NA | |
| NA | NA | |
| 0.804534188 | NA | |
| 0.627031528 | NA | |
| NA | NA | |
| NA | NA | |
| NA | NA | |
| 0.773869078 | NA | |
| NA | NA | |
| NA | NA | |
| NA | NA | |
| NA | NA | |
| 0.905158739 | NA | |
| 0.42461672 | NA | |
| 0.889409301 | NA | |
| 0.989883573 | NA | |
| NA | NA | |
| NA | NA | |
| NA | NA | |
| NA | NA | |
| NA | NA | |

| | | |
|---|---|---|
| | 0.807252815 | NA |
| NA | | NA |
| | 0.060893587 | 0.14136402 |
| NA | | NA |
| | 0.889409301 | NA |
| | 0.905158739 | NA |
| NA | | NA |
| NA | | NA |
| NA | | NA |
| | 0.992037435 | NA |
| | 0.792188023 | NA |
| | 0.886009757 | NA |
| | 0.607009807 | NA |
| | 0.907077876 | 0.947419271 |
| | 0.786550278 | NA |
| NA | | NA |
| | 6.32E-05 | 0.000472738 |
| NA | | NA |
| | 4.00E-10 | 1.42E-08 |
| NA | | NA |
| NA | | NA |
| NA | | NA |
| NA | | NA |
| NA | | NA |
| NA | | NA |
| NA | | NA |
| NA | | NA |
| NA | | NA |
| NA | | NA |
| NA | | NA |
| NA | | NA |
| | 0.889409773 | NA |
| NA | | NA |
| NA | | NA |
| NA | | NA |
| NA | | NA |
| NA | | NA |
| NA | | NA |
| NA | | NA |
| | 0.889409301 | NA |
| NA | | NA |
| NA | | NA |
| NA | | NA |
| NA | | NA |
| NA | | NA |
| NA | | NA |
| NA | | NA |

| | |
|---|---|
| NA | NA |
| NA | NA |
| NA | NA |
| NA | NA |
| NA | NA |
| NA | NA |
| 0.889409301 | NA |
| NA | NA |
| NA | NA |
| 0.905158739 | NA |
| NA | NA |
| NA | NA |
| NA | NA |
| 0.024356339 | 0.069011724 |
| NA | NA |
| 0.633278338 | NA |
| 0.926779989 | 0.958152626 |
| 0.125389182 | 0.243838209 |
| 0.905158266 | NA |
| 0.008939531 | 0.030705955 |
| 0.992062245 | NA |
| 0.640244569 | 0.766582314 |
| 0.21320957 | 0.360337793 |
| 0.266438767 | 0.422640214 |
| 0.622510952 | 0.753650789 |
| 0.889409301 | NA |
| 0.226555855 | 0.376461598 |
| 0.006078496 | 0.022497504 |
| 0.579730054 | NA |
| 0.108495914 | 0.219099287 |
| 0.000119732 | 0.000815762 |
| 0.110050289 | 0.221421505 |
| 0.365523941 | 0.530329714 |
| 0.445043424 | 0.604416943 |
| 0.020874733 | 0.060858772 |
| 0.889410245 | NA |
| 0.816064948 | 0.890624683 |
| 0.131233792 | 0.251902346 |
| NA | NA |
| 0.529060368 | NA |
| 0.24420391 | NA |
| 0.962253405 | 0.979205986 |
| 0.281865909 | 0.440376628 |
| NA | NA |
| 0.005661894 | 0.021271093 |
| 0.1079615 | NA |
| 0.512812844 | 0.664474966 |

| | |
|---|---|
| 8.35E-05 | 0.000598058 |
| 0.592921357 | NA |
| 0.419126057 | 0.581138113 |
| 0.014377907 | 0.045064868 |
| 0.001920666 | 0.00864538 |
| 0.010741342 | 0.035699883 |
| 0.667270334 | 0.787688918 |
| 0.005450192 | 0.020588469 |
| 0.208284657 | NA |
| 0.007090137 | 0.025510784 |
| 0.905535447 | 0.946752982 |
| 0.004327563 | 0.016957343 |
| 0.062129184 | 0.143564638 |
| 0.292902584 | 0.45262828 |
| NA | NA |
| 0.017806944 | 0.053564751 |
| 0.434143289 | 0.594830754 |
| 0.675872022 | 0.794128831 |
| 0.714926923 | 0.822742881 |
| 0.905158266 | NA |
| 3.96E-06 | 4.36E-05 |
| 0.111789991 | 0.223794003 |
| 0.009159963 | 0.031306403 |
| NA | NA |
| 0.89724257 | 0.942074668 |
| 0.827146035 | 0.89672859 |
| 0.342973597 | 0.506343769 |
| 6.08E-08 | 1.15E-06 |
| 0.585012528 | NA |
| 0.533153833 | 0.682057968 |
| 0.590072332 | 0.728244332 |
| 0.003255791 | 0.013446289 |
| 0.022490254 | 0.064667212 |
| 0.269870711 | 0.426337478 |
| 1.49E-05 | 0.000137538 |
| 0.556412428 | 0.701634347 |
| NA | NA |
| 0.14532708 | NA |
| NA | NA |
| 0.775288276 | NA |
| 0.992060816 | NA |
| 0.891717911 | 0.938832699 |
| 0.785716602 | 0.871119873 |
| 0.009887477 | 0.033308937 |
| 0.000222762 | 0.00138377 |
| 1.31E-07 | 2.25E-06 |
| 0.053014687 | 0.126841307 |

| | |
|---|---|
| 0.797342596 | NA |
| 9.31E-05 | 0.000656518 |
| 0.797206843 | 0.877636997 |
| 0.008275399 | 0.02886269 |
| 0.019526658 | 0.057663514 |
| 0.635753335 | 0.763012041 |
| 2.47E-07 | 3.89E-06 |
| 2.25E-07 | 3.59E-06 |
| 3.03E-10 | 1.12E-08 |
| NA | NA |
| 0.001427301 | 0.006734442 |
| 0.101059792 | 0.207709843 |
| 0.873121745 | NA |
| 0.411280161 | 0.574062013 |
| 0.495878428 | NA |
| 0.020197535 | 0.059272712 |
| 0.697712413 | NA |
| 0.067068788 | 0.151690546 |
| 0.06594838 | 0.149975195 |
| 6.75E-15 | 8.14E-13 |
| 6.12E-05 | 0.000459726 |
| 0.020609117 | 0.060252376 |
| 0.000123532 | 0.000837495 |
| 0.555467495 | 0.700885521 |
| 0.17571917 | 0.313759737 |
| NA | NA |
| 1.26E-05 | 0.000118876 |
| 0.309709583 | 0.470977336 |
| 0.807252815 | NA |
| 0.814633649 | 0.889578811 |
| 2.38E-05 | 0.000205521 |
| 0.00495805 | 0.019050537 |
| 0.001023199 | 0.005075315 |
| 0.681846405 | NA |
| 4.85E-12 | 2.92E-10 |
| 0.803566744 | 0.881835189 |
| 0.571321878 | NA |
| 0.304604145 | 0.465210907 |
| 0.849353009 | 0.911714461 |
| 0.526326173 | NA |
| 0.589554555 | 0.7280159 |
| 0.131713087 | 0.252584854 |
| 0.001470233 | 0.006895524 |
| 0.262023719 | 0.417411028 |
| NA | NA |
| 0.856261312 | 0.9161369 |
| 0.776480828 | NA |

| | | | |
|---|---|---|---|
| | 0.094237587 | NA | |
| NA | | NA | |
| NA | | NA | |
| | 0.002251492 | | 0.009886937 |
| | 0.039994314 | | 0.102074307 |
| | 0.872893094 | | 0.927148082 |
| | 0.830726205 | | 0.899059474 |
| | 0.894840713 | | 0.940722291 |
| | 0.134167077 | | 0.256077779 |
| | 5.24E-10 | | 1.81E-08 |
| NA | | NA | |
| NA | | NA | |
| NA | | NA | |
| | 0.332400174 | | 0.494960021 |
| | 0.538302875 | NA | |
| NA | | NA | |
| NA | | NA | |
| NA | | NA | |
| | 3.10E-07 | | 4.73E-06 |
| | 0.686915061 | NA | |
| | 0.392483946 | NA | |
| | 0.008796254 | | 0.030332433 |
| NA | | NA | |
| | 0.992060816 | NA | |
| NA | | NA | |
| | 0.299132717 | | 0.459378651 |
| NA | | NA | |
| | 0.905158266 | NA | |
| NA | | NA | |
| NA | | NA | |
| NA | | NA | |
| | 0.621090884 | NA | |
| | 0.828662672 | | 0.897799665 |
| | 0.575960232 | NA | |
| | 0.36971337 | | 0.53440858 |
| | 0.719314311 | NA | |
| | 0.494224189 | | 0.647489411 |
| | 0.103386683 | | 0.211305675 |
| | 8.86E-05 | | 0.000628685 |
| | 0.782738268 | | 0.869097923 |
| | 0.063671645 | | 0.146129353 |
| | 0.022768911 | | 0.065304305 |
| | 0.229576317 | | 0.380100003 |
| | 0.889409773 | NA | |
| | 0.019383354 | | 0.057325126 |
| | 0.018496214 | | 0.055147096 |
| | 0.000181123 | | 0.001156855 |

| | |
|---|---|
| 0.13676402 | 0.259859922 |
| 0.183090834 | 0.3233466 |
| 0.050663864 | 0.122646653 |
| 0.388404038 | 0.551962569 |
| 0.253818878 | 0.408203365 |
| 0.062247113 | 0.143761502 |
| 0.019587707 | 0.057812695 |
| 0.782559532 | NA |
| 0.623264663 | 0.754169581 |
| 0.000836418 | 0.004286323 |
| NA | NA |
| NA | NA |
| 0.818730392 | NA |
| 0.889410245 | NA |
| NA | NA |
| 0.900601193 | 0.943931153 |
| 0.151193555 | 0.280457787 |
| 0.889409773 | NA |
| 0.000700366 | 0.003691633 |
| NA | NA |
| NA | NA |
| 0.539674567 | 0.687077564 |
| 0.139436515 | 0.263674152 |
| 0.808629847 | 0.885169476 |
| 0.009378453 | 0.031909117 |
| NA | NA |
| 0.663640045 | 0.784953244 |
| NA | NA |
| 0.757052307 | NA |
| 0.991918092 | NA |
| 0.373623159 | 0.53783542 |
| 0.000520227 | NA |
| 4.68E-09 | 1.20E-07 |
| 0.420223937 | 0.582078402 |
| 0.406438203 | 0.569727606 |
| 0.753842175 | NA |
| 0.905158266 | NA |
| 0.770807091 | NA |
| NA | NA |
| 0.005163765 | 0.019689376 |
| 0.128682489 | 0.248503981 |
| 0.004261206 | 0.016751133 |
| 0.020686449 | 0.06043823 |
| NA | NA |
| 0.991953915 | NA |
| NA | NA |
| NA | NA |

| | | |
|---|---|---|
| NA | | NA |
| NA | | NA |
| | 0.992059387 | NA |
| NA | | NA |
| NA | | NA |
| NA | | NA |
| NA | | NA |
| | 0.566761349 | NA |
| | 0.992060816 | NA |
| NA | | NA |
| NA | | NA |
| | 0.221274609 | 0.370284723 |
| NA | | NA |
| | 0.905158266 | NA |
| | 0.159641046 | NA |
| | 0.473527123 | NA |
| | 0.992059387 | NA |
| NA | | NA |
| | 0.501505187 | NA |
| NA | | NA |
| NA | | NA |
| NA | | NA |
| | 0.245577893 | 0.398918058 |
| | 0.691432619 | 0.805772376 |
| NA | | NA |
| NA | | NA |
| | 0.715047657 | 0.822769414 |
| | 0.97044036 | 0.984472875 |
| NA | | NA |
| NA | | NA |
| | 0.121241715 | 0.237931568 |
| NA | | NA |
| NA | | NA |
| | 0.872671696 | 0.926964083 |
| NA | | NA |
| NA | | NA |
| | 0.143500365 | 0.269580334 |
| | 0.024259511 | 0.068772864 |
| | 0.764565775 | 0.857286553 |
| NA | | NA |
| NA | | NA |
| NA | | NA |
| NA | | NA |
| | 7.43E-07 | 1.02E-05 |
| | 0.460494881 | NA |
| | 0.111488447 | 0.223373613 |
| | 1.28E-06 | 1.62E-05 |

| | |
|---|---|
| 0.029964635 | 0.08144491 |
| 0.001698976 | 0.00777084 |
| 0.008039364 | 0.028182695 |
| 0.071477742 | 0.159042008 |
| NA | NA |
| 0.003561723 | 0.0144583 |
| NA | NA |
| 0.018134467 | 0.054311887 |
| 0.466243927 | 0.622954085 |
| 0.000125655 | 0.000847959 |
| NA | NA |
| 0.426891744 | 0.587976906 |
| 0.011227665 | 0.036919482 |
| 0.010946533 | 0.036223918 |
| NA | NA |
| 0.992060816 | NA |
| 0.889409301 | NA |
| 0.127427829 | 0.246732007 |
| 0.17416395 | 0.311514716 |
| NA | NA |
| 0.028568317 | 0.078367481 |
| 0.369722444 | 0.53440858 |
| 0.982758918 | 0.991609754 |
| NA | NA |
| 0.069718197 | 0.155995709 |
| 0.828920401 | 0.897945986 |
| 0.078281422 | 0.171077005 |
| 0.905158266 | NA |
| 0.992061292 | NA |
| 0.101438717 | 0.20823528 |
| 0.804271358 | 0.882408019 |
| 0.575374512 | 0.717201294 |
| NA | NA |
| NA | NA |
| 0.273932827 | 0.431080046 |
| 0.620164759 | NA |
| 0.550914306 | 0.696581288 |
| 0.000117718 | 0.000803538 |
| 0.049808369 | NA |
| 0.991636109 | NA |
| 0.303862317 | 0.464621267 |
| 0.258533375 | 0.413393606 |
| NA | NA |
| 3.97E-08 | 7.99E-07 |
| 0.006303658 | 0.023190152 |
| 0.991700267 | NA |
| 0.565512757 | NA |

| | | |
|---|---|---|
| NA | | NA |
| | 0.879036601 | NA |
| | 0.231123663 | 0.382056949 |
| | 5.93E-10 | 2.01E-08 |
| | 0.378884225 | NA |
| | 0.992057737 | NA |
| | 0.950199284 | 0.97215795 |
| NA | | NA |
| NA | | NA |
| NA | | NA |
| | 0.429619444 | NA |
| | 0.992061292 | NA |
| | 0.001086714 | 0.005340903 |
| | 0.438631506 | 0.599036585 |
| | 9.90E-08 | 1.75E-06 |
| | 0.338664891 | 0.501639134 |
| | 0.161519868 | 0.294341475 |
| NA | | NA |
| | 0.887609873 | NA |
| | 0.006674399 | 0.024297602 |
| | 0.634385398 | 0.761977297 |
| NA | | NA |
| NA | | NA |
| NA | | NA |
| | 0.935633739 | 0.962956594 |
| | 0.862671691 | 0.920757657 |
| NA | | NA |
| | 0.62921389 | 0.757547443 |
| NA | | NA |
| | 0.00016827 | 0.001086774 |
| | 0.068535095 | 0.153906891 |
| | 0.907697698 | 0.947610543 |
| NA | | NA |
| NA | | NA |
| | 0.000627922 | 0.003368811 |
| NA | | NA |
| | 0.033492288 | 0.08891998 |
| | 0.196521472 | 0.340502262 |
| | 0.804472359 | NA |
| | 0.739947985 | 0.840281213 |
| | 0.13228461 | 0.253270935 |
| | 0.428638681 | NA |
| | 0.733416959 | 0.836281181 |
| | 0.266276066 | NA |
| | 0.493179883 | 0.646928273 |
| | 0.756200829 | 0.851753406 |
| | 0.944857502 | 0.968866415 |

| | | |
|---|---|---|
| | 0.279653683 | 0.43788614 |
| | 0.889409773 | NA |
| | 0.397800624 | 0.561322702 |
| | 0.700681898 | 0.813099076 |
| | 0.601578077 | 0.737127002 |
| | 0.122154473 | 0.239179901 |
| | 0.019887871 | 0.058533409 |
| | 0.081544762 | 0.176516335 |
| NA | | NA |
| | 0.319645301 | 0.482113549 |
| | 0.488631107 | 0.643083636 |
| | 0.991700267 | NA |
| | 0.040450703 | 0.102867052 |
| | 0.440536498 | 0.600569926 |
| | 0.12369581 | 0.241443227 |
| | 0.000977321 | 0.004880828 |
| | 0.206455253 | 0.35294285 |
| | 0.319730651 | 0.482134779 |
| | 0.216987899 | 0.365149612 |
| NA | | NA |
| | 0.293961005 | NA |
| | 0.316126369 | 0.478284342 |
| | 0.992061292 | NA |
| | 0.56544854 | NA |
| | 0.28209959 | 0.440616323 |
| | 0.00134147 | 0.006382955 |
| | 0.120089037 | 0.236293473 |
| | 0.904999809 | NA |
| | 0.792260102 | 0.874804625 |
| | 0.033272918 | 0.088519506 |
| | 0.389330894 | 0.552781398 |
| | 0.14341506 | 0.269489243 |
| NA | | NA |
| | 0.702505648 | NA |
| NA | | NA |
| NA | | NA |
| | 0.750231565 | 0.847750472 |
| | 0.655478926 | NA |
| | 0.076013876 | 0.167165351 |
| | 0.018995285 | 0.056336773 |
| | 5.26E-07 | 7.54E-06 |
| | 1.29E-07 | 2.21E-06 |
| | 2.39E-09 | 6.81E-08 |
| | 0.039251859 | 0.100669419 |
| | 0.014203141 | 0.044644543 |
| | 0.005708949 | 0.021414897 |
| | 0.807879735 | NA |

| | | |
|---|---|---|
| | 0.00068229 | 0.003616317 |
| | 0.122557952 | NA |
| | 0.83186829 | 0.899940673 |
| | 1.21E-15 | 1.80E-13 |
| | 0.889409301 | NA |
| | 0.917691754 | 0.953109456 |
| | 0.000758327 | 0.003953489 |
| | 0.040467392 | 0.102897575 |
| | 0.477808538 | 0.633778175 |
| | 0.094999621 | 0.198091033 |
| | 0.074608782 | 0.16459054 |
| | 0.195300304 | 0.339046515 |
| | 0.103886777 | 0.211854544 |
| | 0.800033015 | 0.879407568 |
| | 0.720256346 | 0.826488491 |
| | 0.52723316 | 0.677023466 |
| | 0.683635203 | 0.799744261 |
| | 0.455311207 | 0.613244864 |
| | 0.824790667 | NA |
| | 0.010503766 | 0.035053527 |
| | 0.950648739 | 0.972436561 |
| | 0.219051805 | 0.367638375 |
| | 0.051933534 | 0.124962576 |
| | 0.055045982 | 0.130520971 |
| | 0.630482699 | 0.758596357 |
| | 0.01059812 | 0.035304023 |
| NA | | NA |
| | 0.013033138 | 0.041598065 |
| | 0.303443085 | 0.464083208 |
| NA | | NA |
| | 0.021570608 | NA |
| | 0.001733704 | 0.007913199 |
| | 0.003008698 | 0.012593977 |
| NA | | NA |
| | 0.807252815 | NA |
| | 0.217266927 | NA |
| NA | | NA |
| | 0.696731144 | NA |
| NA | | NA |
| | 0.941295466 | 0.966696621 |
| | 0.381294077 | 0.545444051 |
| | 1.35E-05 | 0.000126177 |
| | 0.016165389 | 0.049641447 |
| NA | | NA |
| | 0.327207352 | 0.489365566 |
| | 0.604734622 | NA |
| | 0.025395357 | 0.07144864 |

| | | |
|---|---|---|
| | 0.619854065 | 0.751511451 |
| | 3.21E-06 | 3.62E-05 |
| | 0.571539109 | NA |
| | 0.066864876 | NA |
| | 0.427431504 | 0.588494274 |
| | 0.004841077 | 0.018663134 |
| | 0.067598413 | 0.152460044 |
| | 0.905158266 | NA |
| | 8.11E-07 | 1.10E-05 |
| | 0.807879735 | NA |
| | 0.552946973 | NA |
| NA | | NA |
| | 0.797291949 | 0.877636997 |
| | 0.067602232 | 0.152460044 |
| | 0.587710101 | 0.726376209 |
| | 0.006921506 | 0.025055944 |
| | 0.992060816 | NA |
| | 0.797157316 | NA |
| | 0.000775269 | 0.00402101 |
| | 0.252472574 | 0.406749661 |
| | 0.560072168 | NA |
| | 0.021936988 | 0.063341792 |
| | 0.183496302 | 0.323984185 |
| | 0.989651219 | NA |
| | 0.000961836 | 0.004817736 |
| | 0.001812582 | 0.008225352 |
| | 0.991157284 | NA |
| | 0.905158266 | NA |
| | 0.739419838 | NA |
| NA | | NA |
| | 0.981886711 | NA |
| NA | | NA |
| | 7.17E-10 | 2.37E-08 |
| | 8.33E-05 | 0.000596748 |
| | 0.905158739 | NA |
| NA | | NA |
| | 0.124735474 | 0.242889623 |
| | 0.736214175 | 0.838159186 |
| | 0.043681437 | NA |
| | 0.520883381 | NA |
| NA | | NA |
| | 0.823057871 | 0.894593615 |
| | 0.889409773 | NA |
| | 0.003756426 | 0.015079955 |
| | 0.890885574 | NA |
| | 0.577080135 | 0.71849376 |
| NA | | NA |

| | |
|---|---|
| NA | NA |
| 0.412433832 | 0.574914906 |
| 0.713472544 | 0.821519701 |
| 0.519962475 | 0.670746698 |
| 0.125648844 | NA |
| 4.16E-06 | 4.54E-05 |
| 0.007578274 | 0.026868017 |
| 0.594589915 | NA |
| 0.004029047 | 0.015987321 |
| 0.207138715 | 0.353780745 |
| 0.002969664 | 0.012456703 |
| 0.014555399 | 0.045547991 |
| 0.016509399 | NA |
| 0.982635243 | NA |
| 0.06704081 | 0.151689763 |
| 0.000344611 | 0.002017322 |
| 0.831648085 | NA |
| 2.06E-16 | 3.50E-14 |
| 0.048389265 | 0.118217404 |
| 0.364852165 | 0.529635003 |
| 0.755156885 | NA |
| 0.000285227 | 0.001715033 |
| 0.027676638 | 0.07640826 |
| 3.77E-09 | 1.01E-07 |
| 0.000900906 | 0.0045625 |
| NA | NA |
| 0.991979299 | NA |
| 0.03796432 | 0.0981121 |
| NA | NA |
| 0.720367553 | 0.826508756 |
| 0.924819459 | 0.956719373 |
| 0.313558135 | 0.475414147 |
| NA | NA |
| 0.005855579 | 0.021845507 |
| 0.992060816 | NA |
| 0.027995378 | 0.077099582 |
| 0.003291791 | 0.01355418 |
| NA | NA |
| 0.889253628 | 0.937385064 |
| 0.001107284 | 0.005422579 |
| NA | NA |
| 0.709530357 | NA |
| 2.50E-05 | 0.000214777 |
| 0.01284902 | 0.041165741 |
| 0.807879735 | NA |
| 0.247223019 | 0.400795148 |
| 0.817910088 | 0.891604826 |

| | | | |
|---|---|---|---|
| | 0.443104023 | NA | |
| | 0.497904945 | | 0.650880741 |
| NA | | NA | |
| NA | | NA | |
| NA | | NA | |
| NA | | NA | |
| | 0.469070044 | | 0.625410624 |
| | 2.36E-10 | | 8.98E-09 |
| NA | | NA | |
| | 3.89E-11 | | 1.83E-09 |
| | 2.15E-07 | | 3.46E-06 |
| NA | | NA | |
| NA | | NA | |
| | 0.785749412 | | 0.871119873 |
| | 0.289025097 | | 0.448383271 |
| | 0.129656639 | | 0.249813998 |
| NA | | NA | |
| | 0.854687224 | NA | |
| NA | | NA | |
| NA | | NA | |
| | 0.735162184 | NA | |
| | 0.889409773 | NA | |
| NA | | NA | |
| NA | | NA | |
| NA | | NA | |
| | 0.788582525 | NA | |
| NA | | NA | |
| NA | | NA | |
| | 0.687780605 | NA | |
| NA | | NA | |
| NA | | NA | |
| | 0.982114889 | | 0.991173786 |
| NA | | NA | |
| | 0.905158739 | NA | |
| NA | | NA | |
| | 0.905158739 | NA | |
| NA | | NA | |
| | 0.991879747 | NA | |
| | 0.797157316 | NA | |
| | 0.889409773 | NA | |
| NA | | NA | |
| | 0.866220478 | NA | |
| | 0.000404532 | | 0.002310304 |
| | 3.14E-07 | | 4.78E-06 |
| NA | | NA | |
| | 0.992062245 | NA | |
| | 0.889409773 | NA | |

| | | | |
|---|---|---|---|
| | 2.58E-06 | | 2.99E-05 |
| NA | | NA | |
| NA | | NA | |
| NA | | NA | |
| | 0.116394231 | | 0.230953967 |
| | 0.905158266 | NA | |
| NA | | NA | |
| NA | | NA | |
| | 0.788582525 | NA | |
| | 0.000769591 | | 0.003996085 |
| | 0.09648121 | | 0.200474888 |
| | 0.738894262 | | 0.839639257 |
| | 0.071806297 | | 0.159643058 |
| | 0.845803592 | NA | |
| | 0.760320343 | NA | |
| | 0.443287039 | NA | |
| | 0.502543356 | | 0.654994632 |
| | 0.0928575 | | 0.194585155 |
| | 0.54927209 | | 0.695425843 |
| NA | | NA | |
| | 0.003143162 | | 0.013059735 |
| NA | | NA | |
| | 3.20E-05 | | 0.000264656 |
| | 0.444608109 | NA | |
| | 0.742082749 | NA | |
| | 0.823957777 | NA | |
| | 0.503057376 | NA | |
| | 4.18E-16 | | 6.91E-14 |
| | 1.74E-05 | | 0.000157504 |
| NA | | NA | |
| NA | | NA | |
| NA | | NA | |
| | 0.319439145 | | 0.481886648 |
| | 0.606667751 | NA | |
| | 0.895347594 | | 0.941005809 |
| | 0.458312995 | | 0.61569161 |
| | 0.000484736 | | 0.002690678 |
| | 0.209057499 | | 0.355820802 |
| | 0.213158967 | | 0.360337793 |
| | 0.541263953 | | 0.688621967 |
| | 0.845662059 | | 0.909251068 |
| | 2.56E-14 | | 2.62E-12 |
| | 0.285902752 | | 0.444910959 |
| | 0.634610966 | | 0.762123172 |
| | 0.077293455 | | 0.169309052 |
| NA | | NA | |
| | 0.988916036 | NA | |

| | |
|---|---|
| 0.487400001 | 0.642118024 |
| 0.057616445 | 0.135199778 |
| 0.732207762 | NA |
| 0.981858817 | NA |
| NA | NA |
| 0.905158739 | NA |
| 2.77E-07 | 4.30E-06 |
| 0.639533425 | 0.76622072 |
| 0.009808545 | 0.033095551 |
| 0.788582525 | NA |
| NA | NA |
| NA | NA |
| 0.905158739 | NA |
| NA | NA |
| 0.588831087 | NA |
| 0.015563214 | NA |
| 0.350938552 | 0.515124309 |
| 0.789572576 | NA |
| 0.451812281 | 0.610038967 |
| 0.453686352 | 0.611759202 |
| 0.586289582 | NA |
| NA | NA |
| NA | NA |
| 0.006997219 | 0.025246786 |
| NA | NA |
| 0.008111676 | 0.028386342 |
| 0.018950146 | 0.056233279 |
| 0.000255381 | 0.001558616 |
| 0.00020214 | 0.001273541 |
| 0.122312946 | 0.239339688 |
| 1.09E-05 | 0.000104977 |
| 0.003677631 | 0.014825979 |
| NA | NA |
| 0.061300302 | 0.142126535 |
| 4.68E-05 | 0.000366264 |
| 0.000128166 | 0.00086252 |
| 0.000105509 | 0.000731106 |
| NA | NA |
| NA | NA |
| NA | NA |
| NA | NA |
| NA | NA |
| 1.03E-06 | 1.34E-05 |
| 0.889409773 | NA |
| 0.170228038 | NA |
| 0.742591973 | NA |
| 0.905158266 | NA |

| | | |
|---|---|---|
| | 0.164267058 | 0.298234009 |
| | 0.889409773 | NA |
| NA | | NA |
| | 5.97E-07 | 8.36E-06 |
| | 0.46644674 | 0.623111325 |
| | 0.698378281 | NA |
| | 0.194208381 | 0.337502825 |
| | 0.89308898 | NA |
| NA | | NA |
| | 0.03992362 | 0.10196357 |
| | 1.18E-08 | 2.74E-07 |
| | 0.346507503 | 0.51005217 |
| | 0.054392133 | 0.129359442 |
| | 0.153799963 | NA |
| | 0.473494561 | 0.629882296 |
| NA | | NA |
| | 0.807433834 | NA |
| NA | | NA |
| | 0.316817899 | 0.478934617 |
| | 0.053031478 | 0.126867657 |
| | 0.707876893 | 0.817861134 |
| | 0.373106054 | 0.53735396 |
| | 0.657777403 | 0.780300038 |
| | 0.76501065 | 0.857560519 |
| | 0.64770341 | NA |
| NA | | NA |
| | 2.63E-08 | 5.54E-07 |
| | 0.045976744 | 0.113768045 |
| | 0.009722253 | 0.032859929 |
| | 0.508235229 | NA |
| NA | | NA |
| | 0.992061292 | NA |
| | 0.036133742 | 0.094459625 |
| | 0.992062245 | NA |
| | 0.034593959 | 0.091238139 |
| | 0.011469764 | 0.037535925 |
| | 0.005003873 | 0.019197133 |
| | 0.084061412 | 0.180475611 |
| | 0.500513265 | NA |
| NA | | NA |
| | 0.620682516 | 0.752059116 |
| | 0.44557207 | NA |
| | 0.788582525 | NA |
| | 0.450430174 | NA |
| | 0.062033304 | 0.143388349 |
| NA | | NA |
| | 0.992060816 | NA |

| | | |
|---|---|---|
| | 0.08378889 | 0.180048951 |
| | 0.702280011 | NA |
| | 0.889409773 | NA |
| | 0.492019632 | 0.646071824 |
| | 0.8323326 | 0.900221226 |
| | 0.653914579 | NA |
| | 0.793305355 | NA |
| | 0.404407396 | NA |
| NA | | NA |
| | 0.608686591 | NA |
| | 0.000455513 | 0.002549506 |
| | 0.680866984 | 0.797613891 |
| | 0.830533409 | NA |
| | 0.574979138 | NA |
| | 0.889409773 | NA |
| NA | | NA |
| | 0.00289939 | 0.012210915 |
| | 0.889409773 | NA |
| | 0.46623512 | 0.622954085 |
| | 0.037104483 | 0.096457599 |
| | 0.546528306 | 0.692910801 |
| | 4.96E-09 | 1.26E-07 |
| | 0.611230908 | 0.744964174 |
| | 0.905158266 | NA |
| | 0.10871075 | NA |
| | 0.009554995 | 0.032409351 |
| | 0.545238898 | NA |
| | 0.98372463 | 0.991970261 |
| NA | | NA |
| | 0.905158266 | NA |
| | 0.66901675 | 0.78895785 |
| | 0.665158666 | 0.786295174 |
| NA | | NA |
| | 0.792188023 | NA |
| | 0.889409773 | NA |
| NA | | NA |
| | 0.784353866 | NA |
| NA | | NA |
| | 0.142848811 | 0.268600815 |
| | 0.527199441 | 0.677023466 |
| NA | | NA |
| NA | | NA |
| | 0.415725911 | NA |
| | 0.905158266 | NA |
| NA | | NA |
| NA | | NA |
| NA | | NA |

| | |
|---|---|
| NA | NA |
| 0.748375765 | NA |
| 0.602512294 | NA |
| 0.992062245 | NA |
| NA | NA |
| 0.119749742 | 0.235816033 |
| 0.127005472 | 0.246261648 |
| 0.763165081 | NA |
| NA | NA |
| 0.989451431 | NA |
| 0.898446913 | 0.942868666 |
| 0.979708749 | NA |
| 0.847686177 | 0.910497043 |
| 0.110114516 | 0.221483804 |
| 0.03392774 | 0.089826288 |
| 0.482183253 | 0.63769345 |
| 0.16241525 | 0.295678699 |
| 0.192422473 | 0.335189996 |
| 0.372051639 | 0.536468707 |
| 0.153774171 | 0.284037931 |
| NA | NA |
| 0.126830111 | 0.245965064 |
| 0.770568918 | 0.861278018 |
| 0.00358768 | 0.014548821 |
| 0.00770269 | 0.027211231 |
| 0.193327398 | 0.336366082 |
| 0.07942021 | 0.172963243 |
| NA | NA |
| 0.036965293 | 0.096186854 |
| 0.277928385 | 0.435868406 |
| 0.992222885 | 0.996033021 |
| 0.676649716 | 0.794746653 |
| 0.685276179 | 0.800926314 |
| 0.914817012 | 0.951789385 |
| 0.493487096 | 0.646954217 |
| 0.661986283 | 0.783554459 |
| 0.385531323 | 0.549375881 |
| 0.059640354 | 0.139042631 |
| 0.618187366 | 0.750360752 |
| 0.364612996 | 0.529392805 |
| NA | NA |
| NA | NA |
| 0.537478484 | 0.685633207 |
| NA | NA |
| NA | NA |
| NA | NA |
| NA | NA |

| | |
|---|---|
| NA | NA |
| 0.046708965 | 0.11494813 |
| NA | NA |
| 0.489857587 | 0.643887391 |
| 0.775837846 | 0.864857393 |
| 0.677899902 | 0.795660054 |
| 0.999207555 | 0.999708343 |
| 0.89997281 | 0.943542752 |
| NA | NA |
| 0.011485823 | 0.037571458 |
| 3.77E-05 | 0.000304524 |
| 0.024758635 | 0.069944021 |
| 0.794276649 | 0.875871619 |
| 0.414297906 | NA |
| 0.875285295 | NA |
| 7.84E-05 | 0.000568498 |
| 0.184091166 | NA |
| 0.992060816 | NA |
| NA | NA |
| 0.412862092 | NA |
| NA | NA |
| NA | NA |
| NA | NA |
| 0.889409301 | NA |
| NA | NA |
| 0.222772451 | 0.372202713 |
| NA | NA |
| 0.5053401 | 0.657325422 |
| 0.972252336 | NA |
| 0.905158266 | NA |
| 0.692598281 | 0.806617113 |
| NA | NA |
| NA | NA |
| NA | NA |
| 0.992061292 | NA |
| 0.489730599 | 0.643836091 |
| 0.872201222 | NA |
| 0.001175704 | 0.005710526 |
| 0.010577965 | 0.035258278 |
| 0.01346324 | 0.042691593 |
| 0.905157793 | NA |
| 0.836307219 | NA |
| NA | NA |
| 0.552947374 | NA |
| 0.691865425 | 0.806148406 |
| 0.025977122 | 0.072653078 |
| 0.990220335 | 0.994886911 |

| | | |
|---|---|---|
| | 0.736664305 | NA |
| | 0.536336769 | 0.684415336 |
| | 0.565134514 | 0.708474775 |
| | 1.30E-05 | 0.0001225 |
| | 0.031407485 | 0.084498524 |
| | 0.364139188 | 0.529125212 |
| NA | | NA |
| | 6.57E-11 | 2.91E-09 |
| | 0.776244828 | NA |
| | 0.908582635 | 0.948154039 |
| | 0.000374142 | 0.002156941 |
| | 0.000217151 | 0.001353508 |
| | 0.005283046 | 0.020074479 |
| | 1.28E-08 | 2.93E-07 |
| | 5.86E-05 | 0.000443631 |
| | 0.430408703 | 0.591448677 |
| | 0.006503811 | 0.023787007 |
| | 0.309536202 | 0.470746275 |
| | 0.643178562 | NA |
| | 0.701759026 | NA |
| NA | | NA |
| | 0.992060816 | NA |
| | 0.992062245 | NA |
| NA | | NA |
| | 8.08E-05 | 0.000582183 |
| | 0.905158266 | NA |
| NA | | NA |
| | 0.889409773 | NA |
| | 0.626722961 | 0.756248461 |
| | 0.905158266 | NA |
| | 0.002583353 | 0.011112212 |
| | 0.010813605 | 0.035858645 |
| | 0.214284368 | 0.361681049 |
| | 0.786687428 | 0.871587752 |
| | 0.55857981 | NA |
| | 0.023572425 | 0.067145789 |
| | 0.717224331 | 0.824405742 |
| | 0.043568619 | 0.109003453 |
| NA | | NA |
| | 0.527642732 | 0.677375431 |
| | 0.992060816 | NA |
| | 0.99205891 | NA |
| NA | | NA |
| NA | | NA |
| | 0.992060816 | NA |
| | 0.140601267 | 0.265382777 |
| NA | | NA |

| | |
|---|---|
| NA | NA |
| NA | NA |
| 0.380512585 | NA |
| 0.372420956 | 0.536789746 |
| 0.070444267 | 0.157315739 |
| 0.053131152 | 0.127011965 |
| NA | NA |
| 0.336817754 | 0.499809505 |
| 0.488664151 | NA |
| 0.486119245 | 0.64097477 |
| 0.000121803 | 0.000827813 |
| NA | NA |
| 0.002734865 | 0.011649837 |
| NA | NA |
| 0.975710589 | 0.987583721 |
| 0.80708346 | 0.88432442 |
| 6.07E-05 | 0.000457139 |
| 0.038102107 | 0.098410275 |
| 0.418921966 | 0.580967932 |
| 0.091183224 | 0.191983571 |
| 0.889409773 | NA |
| 0.030038008 | 0.081576342 |
| 0.371536535 | 0.535866717 |
| NA | NA |
| NA | NA |
| NA | NA |
| NA | NA |
| 0.889409773 | NA |
| 0.992061292 | NA |
| 0.001812327 | 0.008225352 |
| 0.00321687 | NA |
| NA | NA |
| NA | NA |
| 0.266395456 | 0.422610736 |
| 0.026652629 | 0.074093565 |
| NA | NA |
| 0.975671527 | NA |
| 0.37556139 | NA |
| 0.889410245 | NA |
| NA | NA |
| 0.905158266 | NA |
| 0.994688521 | 0.997540618 |
| 0.860328012 | NA |
| 0.889409773 | NA |
| NA | NA |
| NA | NA |
| NA | NA |

| | |
|---|---|
| NA | NA |
| NA | NA |
| 0.010652681 | 0.035448133 |
| 0.581655001 | 0.721695229 |
| 0.700101474 | 0.812670139 |
| 0.737071561 | NA |
| 0.080899544 | 0.175471462 |
| 0.037784939 | 0.097786622 |
| 0.000925764 | 0.004666862 |
| 0.067603417 | 0.152460044 |
| 0.170863859 | 0.307314234 |
| NA | NA |
| 0.022967916 | 0.065747944 |
| 0.491389628 | 0.645398937 |
| 0.444480739 | 0.604011882 |
| 0.037818399 | 0.09785015 |
| 0.632284784 | 0.760077822 |
| 0.044713142 | 0.111170276 |
| 0.98894168 | NA |
| 0.915479011 | 0.952207446 |
| 0.985456279 | 0.99264741 |
| NA | NA |
| 0.454025851 | NA |
| 0.760702996 | NA |
| 0.656218666 | NA |
| 0.705554301 | 0.816724665 |
| 0.992060816 | NA |
| 0.889409773 | NA |
| 0.363801966 | NA |
| 5.85E-11 | 2.65E-09 |
| 0.403603524 | 0.567088976 |
| 0.992061292 | NA |
| NA | NA |
| 0.992060816 | NA |
| 0.99205891 | NA |
| NA | NA |
| NA | NA |
| 0.801661681 | NA |
| 0.046310083 | 0.114273864 |
| 0.839965616 | NA |
| 0.889409773 | NA |
| 0.003032403 | 0.012666641 |
| 0.992062955 | NA |
| 0.889410717 | NA |
| NA | NA |
| NA | NA |
| NA | NA |

| | |
|---|---|
| NA | NA |
| 0.980466138 | NA |
| 0.536202518 | 0.68440311 |
| 7.05E-05 | 0.00051832 |
| NA | NA |
| 0.288521824 | 0.447844118 |
| 0.784102521 | 0.870084757 |
| NA | NA |
| 0.724814669 | NA |
| 0.027461783 | 0.075910537 |
| 7.05E-05 | 0.00051832 |
| 0.352066825 | NA |
| 0.909651962 | NA |
| 0.11367431 | 0.226924925 |
| 0.257264212 | 0.411994226 |
| 0.495641586 | NA |
| 0.146096646 | 0.273195628 |
| 0.905158739 | NA |
| NA | NA |
| 0.634473106 | NA |
| 0.328315642 | 0.490474398 |
| 0.905158266 | NA |
| 0.839555 | NA |
| 0.486876496 | NA |
| NA | NA |
| NA | NA |
| 0.992061829 | NA |
| 0.215450963 | 0.363203416 |
| 7.07E-06 | 7.22E-05 |
| 0.992060816 | NA |
| 0.964004507 | NA |
| 0.08809749 | 0.187164549 |
| NA | NA |
| NA | NA |
| 0.980424109 | NA |
| NA | NA |
| NA | NA |
| 0.905158739 | NA |
| 0.402582248 | 0.566073593 |
| 0.631786912 | NA |
| 0.002127186 | 0.009427017 |
| 0.9239937 | 0.956217431 |
| NA | NA |
| 0.268359274 | 0.424774837 |
| 0.992061769 | NA |
| 0.326559361 | NA |
| 0.01015421 | 0.034104665 |

| | | |
|---|---|---|
| | 0.805374319 | NA |
| | 0.889410245 | NA |
| | 0.871260465 | NA |
| NA | | NA |
| | 0.99205891 | NA |
| NA | | NA |
| NA | | NA |
| NA | | NA |
| NA | | NA |
| NA | | NA |
| | 0.656053444 | NA |
| | 0.859523899 | NA |
| NA | | NA |
| | 0.043086449 | 0.108096855 |
| | 0.32572241 | 0.487894844 |
| | 0.475125642 | 0.631287226 |
| | 0.103047474 | 0.210828238 |
| | 0.836272544 | 0.902570962 |
| | 0.992059387 | NA |
| | 1.36E-15 | 1.98E-13 |
| | 0.413821066 | 0.576370807 |
| | 0.00058645 | 0.003180501 |
| NA | | NA |
| | 0.141258409 | 0.266302662 |
| | 8.10E-05 | 0.000583804 |
| | 0.000954103 | 0.004789929 |
| | 0.075342296 | NA |
| NA | | NA |
| | 0.992061769 | NA |
| | 7.87E-07 | 1.07E-05 |
| | 0.992060816 | NA |
| | 0.269286117 | 0.425689571 |
| NA | | NA |
| | 0.000451021 | 0.002532339 |
| NA | | NA |
| | 0.633320475 | NA |
| | 0.40883296 | 0.572063478 |
| NA | | NA |
| | 0.747975988 | 0.846215915 |
| | 0.279544383 | NA |
| | 0.642772455 | NA |
| | 0.646077814 | 0.771235488 |
| | 0.231453853 | NA |
| | 0.230460135 | NA |
| | 0.905158266 | NA |
| | 0.165592094 | NA |
| | 0.248064173 | NA |

| | | |
|---|---|---|
| | 0.810984441 | NA |
| NA | | NA |
| | 0.992060816 | NA |
| | 0.036928549 | 0.096110891 |
| NA | | NA |
| | 0.75610447 | NA |
| | 0.756598974 | NA |
| | 0.698835136 | NA |
| | 0.514322595 | 0.665763373 |
| | 0.889409773 | NA |
| | 0.905158739 | NA |
| | 0.359053681 | 0.523953999 |
| | 0.04739943 | 0.116321421 |
| NA | | NA |
| | 0.127241856 | NA |
| | 1.69E-05 | 0.000153931 |
| | 0.172621625 | 0.309613515 |
| | 0.181788617 | 0.322030998 |
| | 0.353629807 | 0.517932163 |
| | 0.016587385 | 0.050673678 |
| NA | | NA |
| | 0.889409301 | NA |
| | 0.008031703 | 0.02817383 |
| | 0.733687394 | NA |
| | 0.333521049 | 0.496190921 |
| | 0.566769876 | NA |
| | 0.143955742 | NA |
| | 0.500558834 | NA |
| | 0.577615954 | NA |
| | 0.992061769 | NA |
| | 0.821870818 | NA |
| | 0.483469642 | NA |
| NA | | NA |
| | 0.984194106 | NA |
| NA | | NA |
| | 0.88471069 | NA |
| | 0.764848394 | NA |
| | 0.621636416 | NA |
| | 0.500014099 | 0.6530936 |
| | 0.127269559 | 0.246534248 |
| NA | | NA |
| | 0.905158266 | NA |
| | 0.655956329 | 0.778770817 |
| | 0.352003451 | 0.516298985 |
| | 0.001137922 | 0.005550338 |
| NA | | NA |
| | 1.11E-07 | 1.93E-06 |

| | |
|---|---|
| 0.780888801 | 0.867966045 |
| 5.56E-05 | 0.000424336 |
| 0.048903816 | 0.119253625 |
| 0.673520554 | 0.792425946 |
| NA | NA |
| 0.30874403 | 0.469899512 |
| NA | NA |
| 0.446499374 | NA |
| 4.90E-05 | 0.000380812 |
| 0.069228007 | 0.155120184 |
| NA | NA |
| 0.538867497 | NA |
| NA | NA |
| NA | NA |
| 7.02E-05 | 0.000517151 |
| 0.359500809 | 0.524361397 |
| 0.240015511 | 0.392370627 |
| 2.59E-09 | 7.30E-08 |
| 0.129331504 | 0.249406383 |
| NA | NA |
| 1.06E-06 | 1.37E-05 |
| 0.905158739 | NA |
| 0.266241953 | NA |
| 0.008299389 | 0.02892339 |
| 0.254653163 | NA |
| 0.905158739 | NA |
| 3.86E-09 | 1.02E-07 |
| 0.889409773 | NA |
| 0.049316101 | 0.120072321 |
| 0.000202021 | 0.001273303 |
| 0.159815827 | 0.292035929 |
| NA | NA |
| NA | NA |
| NA | NA |
| 0.887103057 | NA |
| NA | NA |
| 0.992061292 | NA |
| NA | NA |
| NA | NA |
| NA | NA |
| NA | NA |
| 3.04E-06 | 3.44E-05 |
| 0.370947989 | 0.535436717 |
| 0.604985598 | 0.740057145 |
| 0.351536671 | NA |
| 0.905157793 | NA |
| 0.042444421 | 0.106897241 |

| | | |
|---|---|---|
| | 3.78E-05 | 0.000304991 |
| | 0.994449773 | NA |
| | 0.779652341 | NA |
| NA | | NA |
| | 0.014128214 | 0.044466311 |
| | 0.113489727 | 0.226618252 |
| NA | | NA |
| NA | | NA |
| NA | | NA |
| | 0.82028214 | NA |
| | 0.633269999 | NA |
| | 0.077966687 | NA |
| | 0.905158266 | NA |
| | 0.493320352 | 0.646928273 |
| | 0.25857131 | NA |
| | 0.003022457 | 0.012634711 |
| | 0.000324008 | 0.001910552 |
| | 0.114224183 | 0.22777414 |
| | 0.989401795 | NA |
| NA | | NA |
| NA | | NA |
| | 0.805374319 | NA |
| | 0.881329215 | NA |
| | 0.889409773 | NA |
| | 0.757700567 | NA |
| | 0.719947926 | 0.826286339 |
| | 0.904092239 | 0.945739399 |
| | 0.904947233 | NA |
| | 0.644065695 | NA |
| | 0.744166522 | 0.843327278 |
| | 0.849651462 | 0.911856529 |
| | 0.79397576 | 0.875697344 |
| | 0.070712275 | 0.157801915 |
| | 2.17E-07 | 3.48E-06 |
| | 0.015918265 | 0.049025863 |
| | 0.115105901 | 0.229074722 |
| | 0.707472018 | 0.817747691 |
| | 0.025222393 | 0.071043837 |
| | 0.822203821 | 0.894188309 |
| | 0.272484749 | 0.429289321 |
| | 2.76E-05 | 0.000233006 |
| | 0.436053653 | 0.596517702 |
| | 0.000671913 | 0.003569933 |
| | 0.316099751 | NA |
| | 3.85E-05 | 0.000309834 |
| | 0.043575054 | 0.109007132 |
| | 0.010250793 | 0.034392233 |

| | | |
|---|---|---|
| | 0.757281062 | NA |
| | 0.905158739 | NA |
| | 0.006562493 | 0.023957736 |
| | 0.072828471 | 0.161376428 |
| | 0.381933652 | 0.545985356 |
| | 0.134540363 | 0.256500419 |
| | 0.905158266 | NA |
| | 0.581439707 | 0.721591191 |
| | 0.668211218 | 0.788375786 |
| NA | | NA |
| NA | | NA |
| NA | | NA |
| NA | | NA |
| NA | | NA |
| NA | | NA |
| | 0.266167448 | NA |
| | 0.992061292 | NA |
| | 0.735844279 | 0.837836015 |
| NA | | NA |
| | 0.000219415 | 0.001365297 |
| NA | | NA |
| | 0.5688462 | 0.71126829 |
| NA | | NA |
| | 0.967019327 | NA |
| | 0.019027213 | 0.056416227 |
| | 1.18E-08 | 2.74E-07 |
| | 0.57090186 | 0.713229445 |
| | 0.003240259 | 0.013392217 |
| NA | | NA |
| | 0.838640528 | 0.904149025 |
| | 0.111823033 | 0.22381934 |
| | 0.002645134 | 0.011346845 |
| | 0.012913834 | 0.041295309 |
| | 0.776501806 | 0.865367599 |
| | 0.405099819 | 0.568276942 |
| | 0.992061292 | NA |
| | 0.761069933 | NA |
| NA | | NA |
| | 0.00020769 | 0.001301931 |
| NA | | NA |
| | 0.447165277 | 0.606391794 |
| | 0.314868417 | NA |
| NA | | NA |
| | 0.992060816 | NA |
| | 0.124075125 | NA |
| | 0.989530264 | NA |
| NA | | NA |

| | | |
|---|---|---|
| | 0.992061292 | NA |
| | 0.976162111 | NA |
| | 0.992061292 | NA |
| NA | | NA |
| NA | | NA |
| NA | | NA |
| | 2.08E-06 | 2.49E-05 |
| | 1.18E-14 | 1.32E-12 |
| | 0.06406066 | 0.146807351 |
| | 0.061056953 | 0.141668388 |
| | 0.094913278 | 0.19796739 |
| | 0.048856369 | 0.119177628 |
| | 0.004170936 | 0.016457202 |
| | 0.018602494 | 0.055403793 |
| | 0.03351784 | 0.088941833 |
| | 0.000147185 | 0.000967384 |
| | 0.889409773 | NA |
| | 0.085029193 | 0.18200176 |
| NA | | NA |
| | 0.992062245 | NA |
| NA | | NA |
| | 0.393716654 | NA |
| | 0.975795756 | NA |
| | 0.055842513 | 0.131939502 |
| | 0.121795868 | 0.238645073 |
| | 0.263964249 | 0.41971261 |
| | 0.889409301 | NA |
| | 0.850670341 | NA |
| NA | | NA |
| | 4.72E-08 | 9.24E-07 |
| | 0.84223314 | 0.906642361 |
| | 0.420168804 | NA |
| | 0.000332061 | 0.001951223 |
| | 0.256552009 | 0.411153522 |
| | 0.087383437 | 0.185860337 |
| | 0.035045229 | 0.092084778 |
| | 0.007712746 | 0.02724211 |
| | 0.005011004 | 0.019220374 |
| | 0.016113606 | 0.049533028 |
| | 0.860107653 | NA |
| | 0.988097884 | NA |
| | 0.835581025 | NA |
| NA | | NA |
| NA | | NA |
| | 0.004887439 | 0.018808846 |
| | 0.62447134 | NA |
| | 0.849593512 | 0.911838902 |

| | | |
|---|---|---|
| | 0.045747041 | 0.113279125 |
| | 0.000229555 | 0.00142275 |
| | 0.832057656 | 0.900056851 |
| NA | | NA |
| NA | | NA |
| NA | | NA |
| NA | | NA |
| NA | | NA |
| NA | | NA |
| | 0.865177701 | NA |
| | 0.992061292 | NA |
| NA | | NA |
| NA | | NA |
| | 0.889409773 | NA |
| | 0.418760297 | 0.580933042 |
| NA | | NA |
| | 0.557431919 | 0.702435872 |
| | 0.439569643 | 0.599832823 |
| | 0.799253662 | NA |
| | 0.6444183 | NA |
| | 1.03E-09 | 3.23E-08 |
| | 0.001881736 | 0.008500524 |
| | 0.083036815 | NA |
| | 0.452002307 | 0.610200336 |
| | 0.984369149 | 0.992101408 |
| | 0.244675792 | 0.397731146 |
| NA | | NA |
| NA | | NA |
| | 0.704535257 | NA |
| | 9.84E-05 | 0.000689149 |
| | 0.000140152 | 0.000929507 |
| | 0.835526064 | 0.902208735 |
| NA | | NA |
| | 0.99205891 | NA |
| | 0.804534188 | NA |
| | 0.127663941 | 0.247037191 |
| | 0.487640755 | 0.64228067 |
| | 0.992062245 | NA |
| NA | | NA |
| | 0.040152864 | 0.102370457 |
| NA | | NA |
| NA | | NA |
| NA | | NA |
| NA | | NA |
| | 0.000274238 | 0.001659406 |
| NA | | NA |
| NA | | NA |

| | | |
|---|---|---|
| | 0.67965497 | NA |
| | 0.902594645 | 0.944966123 |
| NA | | NA |
| | 0.83565826 | NA |
| | 7.34E-05 | 0.000536333 |
| | 4.51E-07 | 6.59E-06 |
| | 0.41786632 | 0.580093976 |
| | 0.952181048 | 0.973288471 |
| | 0.458435642 | 0.615781029 |
| NA | | NA |
| NA | | NA |
| NA | | NA |
| | 0.000178102 | 0.001141216 |
| | 0.166622801 | NA |
| NA | | NA |
| | 0.625879805 | NA |
| | 0.851542467 | NA |
| | 0.648480345 | NA |
| NA | | NA |
| NA | | NA |
| | 0.721664761 | 0.827706856 |
| | 0.174945203 | 0.312637664 |
| NA | | NA |
| | 0.905158266 | NA |
| | 0.986575046 | NA |
| | 0.06426699 | 0.147157334 |
| NA | | NA |
| | 0.717000672 | 0.82428238 |
| | 0.461455261 | NA |
| | 0.889409773 | NA |
| | 0.620786298 | NA |
| | 0.008892809 | 0.030588533 |
| | 0.35669969 | 0.521350406 |
| | 0.495473613 | NA |
| | 0.042002587 | 0.106027683 |
| | 0.992061292 | NA |
| NA | | NA |
| | 0.595155243 | 0.732157646 |
| | 0.035167218 | 0.09233997 |
| NA | | NA |
| NA | | NA |
| NA | | NA |
| | 0.468698184 | NA |
| NA | | NA |
| | 0.199274746 | NA |
| NA | | NA |
| | 0.992061292 | NA |

| | | |
|---|---|---|
| NA | | NA |
| | 0.097142534 | 0.201392834 |
| | 0.164032505 | NA |
| | 0.905296333 | NA |
| | 0.051951709 | 0.124992612 |
| | 0.670870289 | NA |
| | 1.06E-08 | 2.50E-07 |
| | 0.723997191 | 0.829589559 |
| | 0.04250063 | NA |
| NA | | NA |
| | 0.224392543 | NA |
| | 0.767326046 | NA |
| | 0.015167411 | 0.047142417 |
| | 0.313368855 | 0.475269566 |
| | 0.007234347 | 0.025919241 |
| | 0.005391784 | 0.020413479 |
| | 0.455318742 | 0.613244864 |
| | 0.988593283 | NA |
| | 0.722824744 | NA |
| | 0.545709827 | NA |
| NA | | NA |
| | 0.000610804 | 0.003289834 |
| NA | | NA |
| | 0.248722162 | 0.402542008 |
| | 0.009529517 | 0.03233292 |
| NA | | NA |
| NA | | NA |
| | 0.00098735 | 0.004925311 |
| | 0.009115902 | 0.031181872 |
| | 0.905158739 | NA |
| NA | | NA |
| | 0.01473162 | 0.046002793 |
| | 0.593519213 | 0.731005014 |
| | 0.992060816 | NA |
| | 0.682882553 | 0.799371927 |
| | 0.992060816 | NA |
| | 0.983328328 | NA |
| | 0.529588762 | 0.67896394 |
| | 0.000496125 | 0.002743491 |
| | 0.146916118 | 0.27440087 |
| | 0.507415168 | 0.659192432 |
| | 0.044537917 | NA |
| | 0.296926476 | 0.45698125 |
| | 0.039272947 | 0.100699981 |
| | 0.024712825 | 0.069823587 |
| | 0.992061292 | NA |
| | 0.226312802 | NA |

| | |
|---|---|
| NA | NA |
| 1.64E-05 | 0.000149803 |
| 0.649303718 | NA |
| NA | NA |
| NA | NA |
| 0.716667631 | 0.824028931 |
| 0.889409773 | NA |
| NA | NA |
| 0.014011324 | 0.04414458 |
| 0.430226138 | NA |
| NA | NA |
| 0.082897698 | 0.178834475 |
| NA | NA |
| NA | NA |
| 0.925777423 | 0.957341485 |
| 0.979967018 | 0.989975422 |
| 0.25180775 | 0.406039535 |
| NA | NA |
| 0.149782721 | 0.278398205 |
| 0.000595472 | 0.003223859 |
| 0.759392525 | 0.853840269 |
| 8.88E-10 | 2.83E-08 |
| NA | NA |
| NA | NA |
| 0.514471793 | 0.665838752 |
| NA | NA |
| NA | NA |
| 0.983974857 | NA |
| NA | NA |
| 0.505939219 | NA |
| 0.170216578 | 0.306426121 |
| NA | NA |
| 0.003753763 | 0.015072022 |
| 0.905158739 | NA |
| 0.001910963 | 0.008605998 |
| 0.007523017 | NA |
| 0.12526848 | 0.243689807 |
| 0.905158266 | NA |
| 0.000306209 | 0.001818292 |
| 0.732787385 | 0.835736895 |
| 0.089629366 | 0.189502335 |
| 0.787818382 | 0.871842829 |
| 0.934967812 | NA |
| 0.007030708 | 0.025342632 |
| 0.799684472 | NA |
| 0.991782148 | NA |
| 0.905158266 | NA |

| | | |
|---|---|---|
| | 0.905157793 | NA |
| | 0.14702859 | NA |
| NA | | NA |
| NA | | NA |
| | 0.992061292 | NA |
| NA | | NA |
| | 0.447774101 | NA |
| | 0.097502948 | NA |
| | 0.805374319 | NA |
| | 0.002777857 | 0.011798673 |
| NA | | NA |
| | 0.889409773 | NA |
| | 0.797342596 | NA |
| NA | | NA |
| NA | | NA |
| NA | | NA |
| NA | | NA |
| | 0.000474463 | 0.00263899 |
| NA | | NA |
| NA | | NA |
| NA | | NA |
| | 0.528299446 | 0.678010514 |
| NA | | NA |
| | 0.798013948 | NA |
| | 4.98E-08 | 9.67E-07 |
| | 0.753755134 | 0.850157121 |
| | 0.870600473 | 0.925967054 |
| | 0.374505506 | 0.538557066 |
| NA | | NA |
| | 0.479170487 | NA |
| | 0.042780721 | NA |
| | 0.889409773 | NA |
| | 0.65228354 | 0.776091825 |
| | 0.365562828 | NA |
| | 1.48E-05 | 0.000136816 |
| | 0.002015797 | 0.009018926 |
| | 0.624032201 | 0.754701939 |
| | 0.594473819 | NA |
| NA | | NA |
| NA | | NA |
| NA | | NA |
| NA | | NA |
| NA | | NA |
| | 0.481904797 | NA |
| | 0.339047244 | 0.502066116 |
| | 0.467672058 | 0.623951073 |
| | 0.000158376 | 0.001031369 |

| | | |
|---|---|---|
| | 0.012626195 | 0.040618021 |
| | 0.840856927 | 0.905826984 |
| | 0.000847712 | 0.004327637 |
| | 0.905158266 | NA |
| NA | | NA |
| NA | | NA |
| | 0.005144321 | 0.019635693 |
| | 0.091869526 | 0.193123007 |
| | 0.039516471 | 0.101152401 |
| | 0.001827525 | 0.008288026 |
| | 0.58548612 | NA |
| NA | | NA |
| NA | | NA |
| NA | | NA |
| NA | | NA |
| | 0.889409773 | NA |
| NA | | NA |
| | 0.816078111 | 0.890624683 |
| | 0.992062245 | NA |
| | 0.296290794 | NA |
| | 0.046047962 | 0.113857359 |
| | 0.65101764 | NA |
| NA | | NA |
| | 0.006317569 | 0.023219905 |
| | 0.564546044 | 0.708068687 |
| | 0.700222582 | 0.812670139 |
| | 0.591663239 | NA |
| | 0.011991929 | 0.038942734 |
| | 0.667529797 | 0.787868144 |
| | 0.435179068 | 0.595840949 |
| | 0.617900008 | 0.750177822 |
| | 0.480348306 | 0.635917794 |
| | 0.450307081 | 0.608848242 |
| | 0.992061292 | NA |
| NA | | NA |
| | 0.889409773 | NA |
| | 0.889410245 | NA |
| NA | | NA |
| | 0.109991898 | 0.221344583 |
| NA | | NA |
| | 6.18E-07 | 8.62E-06 |
| NA | | NA |
| NA | | NA |
| NA | | NA |
| NA | | NA |
| NA | | NA |
| | 0.971434837 | 0.984954128 |

| | |
|---|---|
| NA | NA |
| NA | NA |
| 0.77720069 | NA |
| NA | NA |
| NA | NA |
| NA | NA |
| NA | NA |
| 0.889409301 | NA |
| NA | NA |
| 0.249668073 | 0.403591072 |
| NA | NA |
| NA | NA |
| 0.721880852 | NA |
| NA | NA |
| NA | NA |
| NA | NA |
| NA | NA |
| NA | NA |
| NA | NA |
| 0.074854029 | 0.164998958 |
| 0.319808494 | 0.482160983 |
| 0.992003131 | NA |
| NA | NA |
| NA | NA |
| NA | NA |
| NA | NA |
| NA | NA |
| 0.998010723 | NA |
| 0.905158266 | NA |
| NA | NA |
| 0.99120764 | NA |
| 0.889409773 | NA |
| 0.783906671 | NA |
| 0.996090885 | 0.998090794 |
| NA | NA |
| NA | NA |
| 0.601867093 | NA |
| NA | NA |
| NA | NA |
| 0.322511278 | 0.484870953 |
| 0.425109233 | 0.586330861 |
| 0.56232363 | 0.70614971 |
| 8.11E-11 | 3.53E-09 |
| 0.889409301 | NA |
| 0.698378281 | NA |
| 0.189185579 | 0.331007095 |
| NA | NA |

| | | | |
|---|---|---|---|
| | 0.159713599 | | 0.29192417 |
| | 0.871436548 | | 0.926317899 |
| | 0.817343949 | | 0.891341666 |
| NA | | NA | |
| | 0.085977762 | | 0.183602613 |
| | 0.064137222 | | 0.14696747 |
| | 0.889409301 | NA | |
| | 0.526438891 | | 0.676403528 |
| | 0.944399419 | | 0.968738351 |
| NA | | NA | |
| | 0.147680733 | | 0.275430713 |
| | 0.186830219 | | 0.32823464 |
| | 2.38E-05 | | 0.000205729 |
| NA | | NA | |
| | 4.02E-05 | | 0.000321425 |
| | 4.22E-07 | | 6.20E-06 |
| | 0.728013469 | NA | |
| | 0.015328266 | | 0.047521303 |
| | 0.606905291 | | 0.741298247 |
| NA | | NA | |
| | 0.777092376 | NA | |
| NA | | NA | |
| | 0.012283341 | | 0.039724577 |
| NA | | NA | |
| | 0.968505885 | NA | |
| NA | | NA | |
| | 0.75606103 | NA | |
| | 0.371229057 | | 0.535599137 |
| | 0.807879735 | NA | |
| | 0.905158266 | NA | |
| NA | | NA | |
| NA | | NA | |
| | 0.113252801 | | 0.226186289 |
| | 0.000292117 | | 0.001747145 |
| | 0.449563937 | | 0.608453136 |
| | 0.710328576 | | 0.819485486 |
| | 0.586363969 | | 0.725488303 |
| | 0.529763811 | NA | |
| | 0.617866611 | | 0.750177822 |
| | 0.228371193 | NA | |
| | 0.129997065 | | 0.250135794 |
| | 0.008678966 | | 0.030003373 |
| NA | | NA | |
| | 0.007901953 | | 0.027799412 |
| NA | | NA | |
| | 0.992061292 | NA | |
| | 0.90515732 | NA | |

| | |
|---|---|
| 0.271219009 | 0.427790419 |
| NA | NA |
| 0.905158739 | NA |
| 0.059065175 | 0.137877345 |
| 0.880564166 | 0.932221934 |
| 0.740857779 | 0.841009924 |
| NA | NA |
| NA | NA |
| 8.80E-09 | 2.11E-07 |
| 4.55E-05 | 0.000357205 |
| NA | NA |
| 0.401359335 | 0.564928507 |
| 0.306631788 | 0.467364991 |
| NA | NA |
| NA | NA |
| NA | NA |
| 5.06E-12 | 3.02E-10 |
| 0.481238962 | NA |
| NA | NA |
| 0.640278984 | 0.766582314 |
| 0.992061292 | NA |
| NA | NA |
| NA | NA |
| NA | NA |
| 0.593281011 | NA |
| NA | NA |
| NA | NA |
| NA | NA |
| NA | NA |
| NA | NA |
| 0.98914314 | NA |
| NA | NA |
| 0.889409301 | NA |
| 0.889410717 | NA |
| NA | NA |
| NA | NA |
| 0.905158739 | NA |
| NA | NA |
| 0.889409301 | NA |
| NA | NA |
| 0.089299607 | 0.189059975 |
| 0.056964783 | 0.133956915 |
| 0.100401765 | 0.206628149 |
| 0.226957239 | 0.376928908 |
| 0.221893373 | 0.371185845 |
| NA | NA |
| 0.490986661 | NA |

| | | |
|---|---|---|
| | 9.04E-08 | 1.61E-06 |
| | 0.965877311 | NA |
| NA | | NA |
| | 0.096552903 | 0.200568083 |
| | 0.358789741 | NA |
| | 0.612632319 | 0.745968342 |
| | 0.044686698 | 0.111129695 |
| | 0.992062903 | NA |
| | 0.224973027 | 0.374603464 |
| | 0.058867794 | 0.137518925 |
| | 5.36E-05 | 0.000410684 |
| | 0.995001397 | NA |
| | 0.779829019 | NA |
| | 0.893105785 | NA |
| | 0.072703814 | 0.161214081 |
| | 0.889409773 | NA |
| | 0.672992368 | 0.792059139 |
| | 0.386976525 | 0.550791905 |
| | 0.079009831 | 0.172291646 |
| | 0.994416638 | NA |
| | 0.84780343 | NA |
| NA | | NA |
| | 0.678652186 | NA |
| | 2.97E-05 | 0.000248275 |
| | 0.864106518 | NA |
| | 0.14186091 | 0.267209103 |
| | 0.535879105 | 0.684149376 |
| NA | | NA |
| | 5.04E-06 | 5.36E-05 |
| | 0.347965662 | 0.51178687 |
| | 0.49272314 | 0.646649025 |
| | 0.989734052 | NA |
| | 0.701501067 | NA |
| | 1.79E-06 | 2.18E-05 |
| NA | | NA |
| | 0.674666627 | 0.79317938 |
| | 0.981529786 | NA |
| | 1.09E-07 | 1.90E-06 |
| | 0.344097749 | 0.50745752 |
| | 0.778503097 | 0.866762131 |
| | 0.542047584 | 0.689219624 |
| | 0.108516956 | 0.219099287 |
| | 0.6641117 | NA |
| | 0.00387254 | 0.01546462 |
| | 0.916960377 | NA |
| | 0.615046495 | 0.7481612 |
| | 0.330869885 | 0.493350431 |

| | | |
|---|---|---|
| | 0.97837748 | NA |
| | 0.551922343 | NA |
| | 0.006922731 | 0.025056247 |
| | 0.296506901 | 0.456687595 |
| | 0.001068483 | 0.005266626 |
| | 0.207097975 | 0.353738678 |
| | 0.614206542 | NA |
| | 2.21E-08 | 4.77E-07 |
| | 7.26E-05 | 0.000531738 |
| | 0.002595625 | 0.011158444 |
| | 0.628125261 | NA |
| | 0.788582525 | NA |
| NA | | NA |
| NA | | NA |
| NA | | NA |
| NA | | NA |
| NA | | NA |
| NA | | NA |
| NA | | NA |
| | 0.905158739 | NA |
| | 0.733555284 | NA |
| NA | | NA |
| NA | | NA |
| | 0.905158266 | NA |
| | 0.905157793 | NA |
| | 0.797157316 | NA |
| | 0.983816875 | NA |
| | 0.889409301 | NA |
| NA | | NA |
| | 0.716788373 | NA |
| NA | | NA |
| NA | | NA |
| NA | | NA |
| NA | | NA |
| NA | | NA |
| NA | | NA |
| | 0.259768267 | 0.41488482 |
| NA | | NA |
| | 0.867767434 | NA |
| NA | | NA |
| NA | | NA |
| | 0.889410245 | NA |
| | 0.5043585 | NA |
| NA | | NA |
| NA | | NA |
| | 0.382756506 | 0.546734542 |
| | 0.939068401 | NA |

| | | |
|---|---|---|
| | 0.034460883 | 0.090946025 |
| | 1.33E-07 | 2.27E-06 |
| | 0.006081963 | 0.022506544 |
| | 0.952896391 | 0.973556593 |
| | 0.146339241 | 0.273532805 |
| | 0.520670722 | NA |
| | 0.108074396 | 0.218466874 |
| | 0.259214842 | NA |
| | 1.57E-07 | 2.62E-06 |
| | 0.068780245 | 0.154242791 |
| | 2.08E-06 | 2.49E-05 |
| | 9.36E-05 | 0.000660065 |
| | 0.416701655 | 0.578939841 |
| | 0.629350663 | 0.757608883 |
| | 0.114256034 | 0.227816967 |
| | 0.763544212 | NA |
| | 3.71E-09 | 9.91E-08 |
| | 0.546492344 | 0.692905213 |
| | 0.947496181 | 0.970206045 |
| NA | | NA |
| NA | | NA |
| | 0.001528124 | 0.007118492 |
| | 0.001231071 | 0.005927011 |
| | 0.090922563 | 0.191609267 |
| | 0.112930104 | 0.225685489 |
| | 0.276683516 | 0.434319346 |
| | 0.605683146 | 0.740504215 |
| | 0.020983221 | 0.061110151 |
| | 0.456721085 | 0.614417589 |
| | 0.652607557 | 0.776306032 |
| | 0.730864976 | 0.834459624 |
| | 0.749352347 | 0.847157273 |
| | 0.578505906 | 0.719478173 |
| | 2.30E-09 | 6.57E-08 |
| | 0.015420938 | 0.047754673 |
| | 3.79E-06 | 4.20E-05 |
| | 0.011775249 | 0.03835252 |
| | 0.986730039 | NA |
| | 0.753593746 | NA |
| | 0.615493732 | 0.748332145 |
| | 0.844144589 | 0.908199512 |
| | 0.405993256 | 0.56918891 |
| | 0.828767634 | 0.897869084 |
| | 0.664648358 | 0.785861151 |
| | 0.405220808 | 0.568323671 |
| | 0.055088191 | 0.130581052 |
| | 2.20E-14 | 2.32E-12 |

| | |
|---|---|
| 0.341840252 | 0.505214033 |
| NA | NA |
| NA | NA |
| 0.195923126 | 0.339762734 |
| 0.81201211 | 0.887643067 |
| 0.908016252 | NA |
| NA | NA |
| NA | NA |
| NA | NA |
| NA | NA |
| 0.992060816 | NA |
| NA | NA |
| 0.905157793 | NA |
| 0.077679691 | 0.170015782 |
| 0.543079185 | 0.689932069 |
| 0.001083834 | 0.005331487 |
| 0.724898489 | NA |
| 0.165272215 | 0.299538838 |
| 0.072789418 | 0.161347586 |
| 0.855194509 | NA |
| 0.310699001 | NA |
| 0.512579241 | NA |
| 0.095086411 | 0.198196724 |
| 0.791971404 | 0.874678523 |
| 0.106323987 | 0.215581765 |
| NA | NA |
| NA | NA |
| NA | NA |
| 0.026300958 | 0.073338761 |
| 0.992061769 | NA |
| 0.544704196 | 0.691356537 |
| 0.723298978 | NA |
| 0.61687665 | NA |
| 0.272517921 | NA |
| 0.98063721 | 0.990422831 |
| 0.129210055 | 0.249281638 |
| NA | NA |
| NA | NA |
| 0.126689211 | 0.245713512 |
| 0.023650475 | 0.067306995 |
| 0.212282321 | 0.359406899 |
| 0.613084155 | 0.74643127 |
| 2.61E-06 | 3.01E-05 |
| 3.64E-08 | 7.42E-07 |
| 0.268409254 | 0.424792708 |
| NA | NA |
| 0.038833422 | 0.099776198 |

| | |
|---|---|
| 0.854991458 | NA |
| 0.954726869 | 0.97468323 |
| 0.00392762 | 0.015643922 |
| NA | NA |
| 0.438146274 | 0.59848557 |
| 0.063581055 | 0.145982474 |
| NA | NA |
| 0.009834145 | 0.03316664 |
| 0.989224503 | NA |
| NA | NA |
| 0.948175709 | 0.970720798 |
| 0.370039074 | 0.534619911 |
| 0.689404206 | NA |
| 0.992061292 | NA |
| 0.937637375 | NA |
| 0.889409773 | NA |
| 0.882289743 | 0.933154595 |
| 2.60E-06 | 3.01E-05 |
| 1.38E-10 | 5.68E-09 |
| 0.12122884 | 0.237931568 |
| 3.43E-05 | 0.000280648 |
| 0.768881799 | 0.860149545 |
| 7.78E-08 | 1.42E-06 |
| 0.99934608 | NA |
| 0.09868307 | 0.203836095 |
| 0.011590936 | 0.037847639 |
| 0.487081415 | 0.641890463 |
| NA | NA |
| 0.992061769 | NA |
| 0.103692123 | 0.211710849 |
| 0.571554684 | NA |
| 0.010951748 | 0.036229942 |
| 0.885401792 | NA |
| 0.889410717 | NA |
| 0.000490573 | 0.002719638 |
| 0.000799783 | 0.004126512 |
| 1.14E-07 | 1.97E-06 |
| 0.987944112 | NA |
| 0.657812991 | NA |
| 0.355198901 | 0.51981424 |
| 0.012874489 | 0.04122352 |
| 2.80E-06 | 3.22E-05 |
| 0.961915674 | NA |
| 0.188978406 | 0.33076493 |
| NA | NA |
| 0.036260201 | NA |
| 0.026327666 | 0.073375963 |

| | |
|---|---|
| 0.691441585 | NA |
| NA | NA |
| 0.861367689 | NA |
| 0.996513337 | 0.998286331 |
| 0.261661208 | 0.417128346 |
| 0.886416471 | 0.935538753 |
| 0.074486446 | 0.164370201 |
| 4.83E-07 | 6.99E-06 |
| NA | NA |
| NA | NA |
| 0.905158266 | NA |
| NA | NA |
| NA | NA |
| 0.492883219 | 0.646713436 |
| 0.110626905 | 0.222052669 |
| 1.65E-06 | 2.02E-05 |
| 0.718182814 | NA |
| 0.000284913 | 0.001714085 |
| 0.012828365 | 0.041107197 |
| 0.017146617 | 0.052027437 |
| 0.916677905 | 0.952583927 |
| 0.083592535 | 0.179785355 |
| 0.037014074 | 0.096279561 |
| 0.30399931 | 0.464688762 |
| NA | NA |
| NA | NA |
| NA | NA |
| 0.889409773 | NA |
| NA | NA |
| 7.99E-10 | 2.60E-08 |
| 0.259852791 | NA |
| 0.760105039 | NA |
| 0.889410245 | NA |
| 0.984321992 | NA |
| 0.666946636 | NA |
| 0.889409773 | NA |
| 0.65762435 | 0.780202772 |
| NA | NA |
| 0.162491456 | 0.295768392 |
| 0.325130096 | 0.487340053 |
| 7.09E-07 | 9.74E-06 |
| 1.10E-06 | 1.42E-05 |
| 0.149307779 | 0.277686925 |
| 0.715195955 | 0.822896933 |
| 2.41E-11 | 1.21E-09 |
| 0.161500304 | 0.294330249 |
| 0.034065148 | 0.090070631 |

| | | |
|---|---|---|
| | 0.992060816 | NA |
| NA | | NA |
| | 0.99205891 | NA |
| NA | | NA |
| | 0.36458691 | 0.529392746 |
| NA | | NA |
| NA | | NA |
| NA | | NA |
| | 0.718569988 | NA |
| NA | | NA |
| | 0.875718555 | NA |
| NA | | NA |
| | 0.777092376 | NA |
| NA | | NA |
| | 0.877691128 | NA |
| | 0.536397293 | 0.684452796 |
| | 0.2306915 | 0.381514885 |
| NA | | NA |
| | 0.905158739 | NA |
| NA | | NA |
| NA | | NA |
| | 0.256348196 | NA |
| NA | | NA |
| | 0.889409301 | NA |
| | 0.331794905 | NA |
| | 0.905158266 | NA |
| | 0.889410717 | NA |
| NA | | NA |
| NA | | NA |
| | 0.008264247 | 0.028832783 |
| | 0.006036764 | 0.022388778 |
| | 0.643299553 | NA |
| | 0.905158739 | NA |
| | 0.208262918 | 0.354975639 |
| NA | | NA |
| | 0.551345977 | NA |
| | 0.063862325 | 0.14644832 |
| | 0.822594576 | NA |
| | 0.408886061 | 0.572063478 |
| | 0.084082383 | 0.18049783 |
| | 0.834841151 | 0.901690858 |
| NA | | NA |
| | 0.807267765 | 0.884393976 |
| NA | | NA |
| | 0.025875765 | 0.072466001 |
| NA | | NA |
| | 0.992060816 | NA |

| | | |
|---|---|---|
| | 0.992060816 | NA |
| NA | | NA |
| NA | | NA |
| | 0.889409773 | NA |
| NA | | NA |
| NA | | NA |
| NA | | NA |
| NA | | NA |
| | 0.905296333 | NA |
| NA | | NA |
| NA | | NA |
| NA | | NA |
| | 0.400810327 | 0.564445198 |
| | 0.905158739 | NA |
| | 0.72868229 | NA |
| NA | | NA |
| | 0.989299202 | NA |
| NA | | NA |
| NA | | NA |
| | 0.022747917 | 0.065296928 |
| NA | | NA |
| NA | | NA |
| | 0.992060816 | NA |
| NA | | NA |
| NA | | NA |
| NA | | NA |
| | 0.68984072 | 0.804728664 |
| | 0.981504209 | NA |
| | 0.355698691 | NA |
| | 0.537160651 | NA |
| NA | | NA |
| | 0.080289958 | NA |
| NA | | NA |
| | 0.992059387 | NA |
| | 0.669727145 | NA |
| NA | | NA |
| | 0.992061292 | NA |
| | 0.002342806 | 0.010232772 |
| | 0.013001367 | 0.041509362 |
| | 0.0497714 | 0.12093949 |
| | 0.86624789 | 0.922946988 |
| | 2.09E-07 | 3.36E-06 |
| | 0.727348445 | NA |
| | 0.992060816 | NA |
| | 0.905158739 | NA |
| | 1.81E-07 | 2.98E-06 |
| | 2.31E-09 | 6.61E-08 |

| | |
|---|---|
| 0.006226927 | 0.022954019 |
| 0.000363284 | 0.002106508 |
| 0.630839415 | 0.758838823 |
| NA | NA |
| 0.36798002 | 0.532591172 |
| 0.464628566 | NA |
| 0.011381363 | 0.03731478 |
| NA | NA |
| 0.000104145 | 0.000723708 |
| 0.855881171 | 0.916035221 |
| 0.976528977 | NA |
| 0.864609224 | 0.92191677 |
| NA | NA |
| 0.657499458 | 0.780096747 |
| 2.77E-05 | 0.000233869 |
| NA | NA |
| 0.509735232 | 0.661345699 |
| 0.224413506 | 0.374033866 |
| 0.11172593 | 0.223720501 |
| 0.728310573 | 0.832447006 |
| 0.987468916 | NA |
| 0.064583429 | 0.147732632 |
| NA | NA |
| 0.992061292 | NA |
| 0.085954328 | NA |
| 0.007327523 | 0.026193241 |
| 0.204654665 | 0.350574287 |
| NA | NA |
| 3.43E-17 | 6.85E-15 |
| 0.000131154 | 0.000878054 |
| 0.995508618 | 0.997694248 |
| NA | NA |
| 0.389628249 | 0.553024802 |
| 0.872064438 | NA |
| 0.71487258 | NA |
| NA | NA |
| 0.269509077 | NA |
| 0.000535544 | 0.002935599 |
| 0.602524011 | NA |
| 0.018335995 | 0.054795879 |
| 0.480822888 | NA |
| 2.22E-08 | 4.78E-07 |
| NA | NA |
| 0.379225763 | NA |
| 0.21758402 | 0.365691912 |
| 0.303900691 | 0.464621267 |
| 0.050090582 | 0.121526804 |

| | |
|---|---|
| 0.382736946 | 0.546734542 |
| 0.113208353 | 0.226124191 |
| 0.905158739 | NA |
| NA | NA |
| 0.991074926 | NA |
| 0.99205891 | NA |
| 0.889409301 | NA |
| NA | NA |
| NA | NA |
| NA | NA |
| NA | NA |
| NA | NA |
| NA | NA |
| NA | NA |
| NA | NA |
| NA | NA |
| NA | NA |
| NA | NA |
| NA | NA |
| NA | NA |
| NA | NA |
| NA | NA |
| NA | NA |
| NA | NA |
| NA | NA |
| NA | NA |
| NA | NA |
| NA | NA |
| NA | NA |
| NA | NA |
| NA | NA |
| NA | NA |
| NA | NA |
| NA | NA |
| NA | NA |
| NA | NA |
| NA | NA |
| NA | NA |
| NA | NA |
| NA | NA |
| NA | NA |
| NA | NA |
| NA | NA |
| NA | NA |
| NA | NA |
| 0.98997168 | NA |
| NA | NA |

| | |
|---|---|
| NA | NA |
| NA | NA |
| NA | NA |
| NA | NA |
| NA | NA |
| NA | NA |
| 0.678720252 | NA |
| NA | NA |
| NA | NA |
| 0.889410717 | NA |
| 0.232894317 | 0.384347384 |
| 2.13E-05 | 0.000187723 |
| NA | NA |
| 0.842218492 | 0.906642361 |
| 0.002556533 | 0.01100763 |
| 0.697075828 | 0.81041339 |
| 0.000783224 | 0.004052499 |
| 0.983232732 | NA |
| 0.067160921 | 0.151789488 |
| 0.013605812 | 0.043075263 |
| 0.294016762 | 0.453711459 |
| 0.957705101 | 0.976545197 |
| 0.279154313 | 0.437353708 |
| 0.256012889 | 0.410580543 |
| 0.77836134 | 0.866682731 |
| 0.883979983 | NA |
| 0.754043018 | NA |
| 0.905158266 | NA |
| NA | NA |
| NA | NA |
| NA | NA |
| NA | NA |
| 0.131879822 | 0.252766715 |
| 0.883334782 | NA |
| 0.800942274 | NA |
| 0.478709229 | 0.634518038 |
| 0.980052281 | NA |
| 0.753581177 | NA |
| NA | NA |
| 0.991388785 | NA |
| 0.85219599 | NA |
| 1.81E-06 | 2.19E-05 |
| 0.412716082 | NA |
| 0.000339688 | 0.001991247 |
| 0.079298937 | 0.172767672 |
| 0.047635115 | 0.11680843 |
| NA | NA |

| | |
|---|---|
| NA | NA |
| NA | NA |
| NA | NA |
| NA | NA |
| 0.935949453 | NA |
| NA | NA |
| 0.005842661 | 0.021812137 |
| 0.000126852 | 0.000853938 |
| 0.068226404 | 0.153439533 |
| 1.05E-08 | 2.48E-07 |
| 0.85714868 | NA |
| 0.905158739 | NA |
| NA | NA |
| NA | NA |
| 0.889409773 | NA |
| NA | NA |
| 1.39E-07 | 2.35E-06 |
| 0.000762121 | 0.00396517 |
| 1.56E-07 | 2.61E-06 |
| NA | NA |
| NA | NA |
| NA | NA |
| 2.27E-08 | 4.88E-07 |
| NA | NA |
| NA | NA |
| NA | NA |
| 0.386773326 | 0.550609755 |
| 0.992003131 | NA |
| NA | NA |
| 0.991879747 | NA |
| NA | NA |
| 0.039571275 | 0.101252113 |
| NA | NA |
| 0.067889638 | 0.152885711 |
| 0.76339664 | 0.856631755 |
| 0.557622191 | NA |
| 8.84E-05 | 0.00062745 |
| 0.367562448 | 0.532141534 |
| 0.992061292 | NA |
| 3.97E-07 | 5.88E-06 |
| 0.049030667 | 0.119496606 |
| 0.844103523 | NA |
| 2.79E-12 | 1.79E-10 |
| NA | NA |
| NA | NA |
| NA | NA |
| 0.931565986 | NA |

| | | | |
|---|---|---|---|
| | 0.768168575 | NA | |
| | 0.905157793 | NA | |
| NA | | NA | |
| NA | | NA | |
| | 0.004475244 | | 0.017439555 |
| NA | | NA | |
| | 0.001643646 | | 0.007546064 |
| | 0.984466551 | NA | |
| | 0.524204836 | | 0.674262769 |
| NA | | NA | |
| NA | | NA | |
| NA | | NA | |
| NA | | NA | |
| NA | | NA | |
| NA | | NA | |
| | 0.905158739 | NA | |
| | 0.043224455 | | 0.108339893 |
| NA | | NA | |
| NA | | NA | |
| NA | | NA | |
| NA | | NA | |
| NA | | NA | |
| NA | | NA | |
| | 0.470426713 | | 0.626849198 |
| | 0.992060816 | NA | |
| | 0.654411961 | NA | |
| NA | | NA | |
| NA | | NA | |
| NA | | NA | |
| NA | | NA | |
| | 0.889410245 | NA | |
| | 0.701711656 | | 0.813761835 |
| | 0.837213362 | | 0.903172899 |
| | 0.085012137 | | 0.181982991 |
| | 0.198079326 | | 0.342517042 |
| | 0.071557682 | | 0.159203155 |
| NA | | NA | |
| | 0.905158266 | NA | |
| NA | | NA | |
| NA | | NA | |
| NA | | NA | |
| | 0.723659144 | NA | |
| | 0.395213027 | | 0.558748574 |
| NA | | NA | |
| NA | | NA | |
| NA | | NA | |
| NA | | NA | |

| | | |
|---|---|---|
| | 0.889409773 | NA |
| | 1.19E-09 | 3.66E-08 |
| NA | | NA |
| | 0.807879735 | NA |
| NA | | NA |
| | 0.01418656 | 0.044624363 |
| NA | | NA |
| | 0.874391574 | NA |
| | 0.016943829 | 0.051561743 |
| | 0.003020901 | 0.012630609 |
| | 0.724908537 | NA |
| | 0.992060816 | NA |
| NA | | NA |
| | 0.991782148 | NA |
| NA | | NA |
| | 0.726732747 | 0.831249317 |
| NA | | NA |
| | 0.140947708 | NA |
| | 0.664742791 | NA |
| | 0.74913451 | NA |
| NA | | NA |
| | 0.245628422 | 0.398918058 |
| | 5.22E-07 | 7.49E-06 |
| | 0.110521279 | 0.221962389 |
| | 0.206406613 | 0.35288717 |
| | 0.945040746 | 0.968866415 |
| NA | | NA |
| | 0.905158266 | NA |
| | 0.770879609 | NA |
| NA | | NA |
| NA | | NA |
| | 0.661225037 | NA |
| NA | | NA |
| | 0.234181487 | 0.385684723 |
| | 0.082036067 | 0.177341011 |
| NA | | NA |
| | 0.413010249 | 0.575535732 |
| | 0.293578791 | NA |
| | 0.988864899 | NA |
| | 0.889409773 | NA |
| | 0.479913521 | 0.635495447 |
| | 0.187621451 | NA |
| | 0.000636676 | 0.003413275 |
| NA | | NA |
| NA | | NA |
| | 0.80003058 | 0.879407568 |
| | 0.832101923 | 0.900060397 |

| | |
|---|---|
| 0.673996159 | NA |
| 0.618387877 | 0.750438209 |
| NA | NA |
| 0.619971209 | 0.751570486 |
| 0.085467643 | 0.182629802 |
| 0.889410245 | NA |
| 0.620576982 | 0.752014235 |
| 0.621735027 | 0.753043545 |
| NA | NA |
| 0.395807823 | NA |
| 0.905158266 | NA |
| 0.985944351 | NA |
| NA | NA |
| NA | NA |
| 0.070597762 | 0.157562379 |
| 0.90515732 | NA |
| 0.835251642 | 0.901956764 |
| 0.314926192 | NA |
| NA | NA |
| 0.992060816 | NA |
| 0.000587243 | 0.003184018 |
| 0.273995265 | 0.431116511 |
| 0.195683835 | NA |
| NA | NA |
| 1.00E-08 | 2.38E-07 |
| NA | NA |
| 0.144300043 | 0.270735189 |
| 0.976935263 | NA |
| NA | NA |
| 0.121059131 | 0.237733631 |
| 0.889409773 | NA |
| 0.257453886 | NA |
| NA | NA |
| NA | NA |
| 0.494249169 | 0.647489411 |
| 0.241502776 | 0.394298377 |
| 0.503107629 | NA |
| 0.076674924 | NA |
| 9.48E-10 | 3.01E-08 |
| 6.83E-07 | 9.43E-06 |
| 0.120317976 | 0.236595544 |
| 0.017071151 | 0.051827097 |
| 0.383177685 | 0.547202504 |
| 2.01E-13 | 1.69E-11 |
| 0.724387644 | NA |
| NA | NA |
| 0.889409773 | NA |

| | | |
|---|---|---|
| | 0.942918632 | 0.967684733 |
| | 0.422332094 | 0.583857369 |
| | 0.703479224 | NA |
| | 0.000321527 | 0.001897959 |
| | 0.02532124 | 0.071276644 |
| | 0.580267042 | 0.720553013 |
| | 3.14E-08 | 6.51E-07 |
| | 0.43666298 | 0.596979354 |
| | 0.575184088 | NA |
| | 0.845397003 | NA |
| NA | | NA |
| | 0.241343222 | NA |
| | 0.723274045 | 0.829063776 |
| | 0.71331291 | NA |
| | 0.890573765 | NA |
| | 0.690523596 | NA |
| | 0.889409773 | NA |
| | 0.905158266 | NA |
| | 0.886681316 | NA |
| | 0.554154781 | 0.699550724 |
| | 0.803394287 | NA |
| | 0.871549666 | NA |
| | 0.172053725 | 0.308847196 |
| | 0.829685048 | 0.898465186 |
| | 0.646341912 | 0.771359893 |
| | 0.992061292 | NA |
| | 0.991918092 | NA |
| NA | | NA |
| NA | | NA |
| | 0.901376118 | NA |
| NA | | NA |
| NA | | NA |
| NA | | NA |
| NA | | NA |
| NA | | NA |
| NA | | NA |
| NA | | NA |
| NA | | NA |
| NA | | NA |
| | 0.90515732 | NA |
| NA | | NA |
| NA | | NA |
| NA | | NA |
| NA | | NA |
| | 0.889409301 | NA |
| NA | | NA |
| NA | | NA |

| | | |
|---|---|---|
| NA | NA | |
| NA | NA | |
| 0.731333061 | NA | |
| 0.759335416 | NA | |
| 0.57431887 | NA | |
| 0.751019897 | | 0.848382546 |
| 0.783660966 | | 0.869814554 |
| 0.850494042 | | 0.912359486 |
| 0.489150246 | NA | |
| 0.99205891 | NA | |
| 0.10594459 | NA | |
| 0.992060816 | NA | |
| NA | NA | |
| 0.002754019 | | 0.011723083 |
| NA | NA | |
| NA | NA | |
| 0.905158266 | NA | |
| 0.774095591 | | 0.863825851 |
| 0.83162375 | | 0.899853451 |
| 0.896521609 | NA | |
| 0.067781169 | NA | |
| NA | NA | |
| 0.788582525 | NA | |
| 0.187353508 | | 0.328759444 |
| NA | NA | |
| NA | NA | |
| 0.69262113 | NA | |
| NA | NA | |
| NA | NA | |
| NA | NA | |
| NA | NA | |
| 0.457972223 | NA | |
| NA | NA | |
| 0.889410245 | NA | |
| 0.993956541 | | 0.997117284 |
| NA | NA | |
| 0.732201996 | | 0.835329574 |
| 0.992060816 | NA | |
| 0.692886405 | NA | |
| 0.660223574 | NA | |
| 0.256606564 | | 0.411210942 |
| 0.487429544 | | 0.642118024 |
| NA | NA | |
| NA | NA | |
| 0.305335785 | NA | |
| NA | NA | |
| NA | NA | |

| | | |
|---|---|---|
| NA | | NA |
| NA | | NA |
| NA | | NA |
| NA | | NA |
| | 0.812000296 | 0.887643067 |
| | 0.845897924 | NA |
| NA | | NA |
| NA | | NA |
| NA | | NA |
| NA | | NA |
| NA | | NA |
| NA | | NA |
| | 0.509257996 | NA |
| NA | | NA |
| NA | | NA |
| | 0.905158739 | NA |
| | 0.562329243 | 0.70614971 |
| NA | | NA |
| NA | | NA |
| | 0.005104279 | 0.0194998 |
| | 0.739320043 | NA |
| | 0.005085629 | 0.019435317 |
| | 0.725304682 | 0.830307868 |
| | 0.227145503 | 0.377013462 |
| | 0.591180396 | NA |
| | 0.539628057 | NA |
| | 0.000957228 | 0.00480013 |
| | 0.018885482 | 0.056071702 |
| | 0.774313205 | 0.863936978 |
| | 0.855841474 | 0.916035221 |
| | 0.855806618 | NA |
| | 0.147188939 | NA |
| | 0.836660123 | NA |
| | 8.21E-05 | 0.000589787 |
| | 0.473027088 | 0.629328116 |
| | 0.100639512 | NA |
| | 0.905158266 | NA |
| | 0.399544377 | NA |
| | 0.025767696 | 0.072290743 |
| NA | | NA |
| | 0.771540524 | NA |
| NA | | NA |
| NA | | NA |
| | 0.008546076 | 0.02962325 |
| NA | | NA |
| | 0.043018658 | 0.107984306 |
| | 0.001598878 | 0.007383758 |

| | | |
|---|---|---|
| | 0.009277935 | 0.031650487 |
| | 0.992061292 | NA |
| | 0.37361253 | 0.53783542 |
| NA | | NA |
| NA | | NA |
| NA | | NA |
| | 0.992060816 | NA |
| NA | | NA |
| | 0.504932512 | NA |
| NA | | NA |
| | 0.073453141 | 0.162498239 |
| | 0.388100584 | 0.551819524 |
| | 0.011761352 | 0.038312939 |
| | 0.428245623 | NA |
| | 0.477470556 | NA |
| NA | | NA |
| NA | | NA |
| NA | | NA |
| | 0.096749432 | 0.200881315 |
| | 0.453935246 | 0.611944507 |
| | 0.683522973 | 0.799695294 |
| | 0.915218512 | NA |
| | 0.627470883 | 0.75642702 |
| | 0.902288608 | NA |
| | 0.532998238 | NA |
| NA | | NA |
| NA | | NA |
| | 0.986093117 | NA |
| NA | | NA |
| | 0.187212452 | 0.32866951 |
| NA | | NA |
| | 0.783889569 | 0.869936374 |
| | 0.992061292 | NA |
| | 0.801333715 | 0.880259664 |
| | 0.041974537 | 0.105976107 |
| | 0.740893231 | NA |
| | 0.905158266 | NA |
| | 0.79634602 | 0.877155008 |
| | 0.651613898 | NA |
| | 0.889409301 | NA |
| NA | | NA |
| | 0.222186256 | 0.371510014 |
| | 0.898380742 | 0.942863964 |
| | 0.976347991 | 0.987728072 |
| | 0.182428933 | 0.322566789 |
| | 0.889410245 | NA |
| | 0.413155481 | NA |

| | |
|---|---|
| 0.018261268 | 0.054646931 |
| 0.093072553 | 0.194905606 |
| 0.911453358 | 0.950003887 |
| NA | NA |
| NA | NA |
| NA | NA |
| NA | NA |
| NA | NA |
| NA | NA |
| NA | NA |
| NA | NA |
| 0.611161932 | 0.744964174 |
| 0.109437892 | 0.2204924 |
| NA | NA |
| NA | NA |
| 0.38688661 | 0.550735321 |
| 0.095157381 | 0.1982429 |
| 0.002416931 | 0.010513745 |
| 0.808770739 | NA |
| 0.084455764 | 0.180950738 |
| NA | NA |
| 0.047104772 | 0.115727644 |
| 0.287457284 | 0.446538943 |
| 0.824477825 | NA |
| 0.794384971 | NA |
| NA | NA |
| 0.299576535 | NA |
| NA | NA |
| 0.186642302 | 0.327956975 |
| NA | NA |
| NA | NA |
| 0.052247786 | 0.125536502 |
| 0.905158739 | NA |
| 0.004462584 | 0.017396393 |
| 0.608843132 | NA |
| 0.584698651 | 0.724284857 |
| NA | NA |
| NA | NA |
| 0.976661135 | NA |
| 0.905158266 | NA |
| 0.296780879 | 0.45685323 |
| NA | NA |
| 0.607041535 | 0.741298247 |
| NA | NA |
| 0.378311511 | 0.542857118 |
| 0.791606848 | 0.874479061 |
| 5.88E-11 | 2.66E-09 |

| | | |
|---|---|---|
| | 0.037229783 | 0.096691342 |
| | 6.91E-05 | 0.000510119 |
| | 0.06536314 | 0.149083619 |
| | 0.889410717 | NA |
| | 0.503879276 | 0.656151635 |
| | 0.992060816 | NA |
| NA | | NA |
| NA | | NA |
| | 0.807252815 | NA |
| | 0.860026936 | NA |
| | 0.992060816 | NA |
| | 0.784353866 | NA |
| | 0.202417574 | 0.347963637 |
| | 0.049143761 | 0.119719087 |
| NA | | NA |
| | 1.72E-08 | 3.79E-07 |
| | 0.509298089 | NA |
| | 0.192159226 | 0.334784548 |
| | 0.000430076 | 0.002431522 |
| | 0.267813249 | 0.424118224 |
| | 0.00547897 | 0.020679393 |
| | 0.889409301 | NA |
| | 0.178648211 | 0.317878298 |
| | 0.243621024 | 0.396595896 |
| | 0.860328012 | NA |
| | 0.905158739 | NA |
| | 0.000772837 | 0.004010099 |
| | 0.82836996 | NA |
| NA | | NA |
| | 0.889409773 | NA |
| NA | | NA |
| | 0.013425184 | 0.042607836 |
| NA | | NA |
| NA | | NA |
| | 6.46E-13 | 4.84E-11 |
| NA | | NA |
| | 0.000470191 | 0.002618545 |
| | 0.016072692 | 0.049446026 |
| | 0.430839053 | NA |
| | 0.670086127 | NA |
| | 5.17E-07 | 7.42E-06 |
| | 0.90515732 | NA |
| | 1.43E-17 | 3.24E-15 |
| | 0.043455229 | 0.108781735 |
| | 0.675346975 | 0.793639314 |
| NA | | NA |
| | 0.905157793 | NA |

| | | |
|---|---|---|
| | 0.847834011 | NA |
| NA | | NA |
| | 0.905296333 | NA |
| | 0.978439742 | NA |
| NA | | NA |
| | 0.765118395 | NA |
| | 0.857194446 | NA |
| NA | | NA |
| | 0.889409301 | NA |
| NA | | NA |
| | 0.239837638 | 0.392151503 |
| | 0.049024824 | 0.119495627 |
| | 0.992061292 | NA |
| NA | | NA |
| | 0.439466696 | NA |
| | 0.565771911 | 0.708958308 |
| | 0.618205811 | NA |
| | 0.736674122 | 0.838335753 |
| | 0.782425 | 0.868948491 |
| | 0.04375649 | 0.109286711 |
| | 0.855550651 | 0.915949332 |
| | 0.00705587 | 0.025412074 |
| | 0.022000711 | 0.063475708 |
| | 0.004193874 | 0.016533801 |
| NA | | NA |
| | 0.992060816 | NA |
| | 0.458347168 | 0.61569985 |
| | 0.991918092 | NA |
| | 0.586182922 | 0.725460183 |
| | 0.770675906 | 0.861322894 |
| | 0.620971519 | NA |
| | 0.289229715 | NA |
| | 9.29E-05 | 0.000655701 |
| | 0.06548526 | 0.149215608 |
| | 0.778594615 | NA |
| | 5.52E-07 | 7.82E-06 |
| | 0.003315018 | 0.013625222 |
| | 0.535176374 | 0.683490636 |
| | 0.598757777 | NA |
| | 0.161231409 | 0.293959836 |
| | 0.984275148 | NA |
| NA | | NA |
| | 0.744287956 | 0.843421366 |
| NA | | NA |
| | 0.448796654 | 0.607740857 |
| | 0.818976783 | 0.892324659 |
| NA | | NA |

| | | |
|---|---|---|
| | 0.800694245 | NA |
| NA | | NA |
| | 0.347305487 | NA |
| NA | | NA |
| | 0.394079883 | NA |
| | 0.299315003 | 0.459594305 |
| | 0.03176204 | 0.08530721 |
| | 1.40E-08 | 3.17E-07 |
| | 0.905158266 | NA |
| | 0.003038999 | 0.012682131 |
| | 0.254329716 | 0.408785406 |
| | 0.073534615 | 0.16262933 |
| | 0.81185189 | NA |
| | 0.205427983 | NA |
| | 0.282682626 | 0.441338627 |
| | 0.298969144 | NA |
| | 0.719793148 | 0.826239149 |
| NA | | NA |
| | 0.360464622 | 0.525140492 |
| | 0.432175145 | NA |
| | 1.35E-07 | 2.30E-06 |
| NA | | NA |
| NA | | NA |
| | 0.155334609 | 0.286157104 |
| | 0.298111807 | NA |
| | 0.007443993 | 0.026493136 |
| | 0.905158266 | NA |
| | 0.078484741 | 0.171418981 |
| | 0.873967228 | 0.927840184 |
| | 0.90515732 | NA |
| | 0.096914936 | 0.201065403 |
| | 0.630820125 | 0.758838823 |
| | 0.835917847 | NA |
| | 0.863679941 | NA |
| | 0.309022467 | 0.470127779 |
| | 0.428923657 | 0.590035368 |
| | 0.627373757 | 0.75642702 |
| | 0.063636096 | 0.146078302 |
| | 0.291944454 | 0.45156077 |
| | 0.336272911 | 0.499237161 |
| | 0.057809231 | 0.135514772 |
| | 0.022939381 | 0.065691948 |
| NA | | NA |
| | 5.00E-06 | 5.31E-05 |
| | 0.000326381 | 0.001922995 |
| | 0.985034871 | 0.992446925 |
| | 0.097545521 | 0.202048718 |

| | |
|---|---|
| 0.002492781 | 0.01078717 |
| NA | NA |
| 0.968169251 | NA |
| 0.991918092 | NA |
| 0.020106815 | 0.059062179 |
| 0.772780163 | NA |
| 0.00168137 | 0.007698332 |
| 0.747671577 | NA |
| 0.280053267 | 0.438208748 |
| 0.170682104 | 0.307129749 |
| 0.412740961 | 0.575269994 |
| 0.002085082 | 0.009274119 |
| 0.025938747 | 0.072586842 |
| 0.38797875 | 0.551717743 |
| NA | NA |
| NA | NA |
| 0.889409773 | NA |
| NA | NA |
| 0.306925253 | 0.46768244 |
| 0.24965889 | 0.403591072 |
| 0.03063023 | 0.082824274 |
| NA | NA |
| 0.329112184 | NA |
| 0.044378394 | 0.110563326 |
| 0.049038632 | 0.119502754 |
| 0.000827583 | 0.004249569 |
| 0.031090782 | 0.083779909 |
| 0.67688319 | NA |
| 0.590208542 | 0.728316346 |
| 0.504721891 | 0.656816024 |
| 0.000416156 | 0.002363779 |
| NA | NA |
| 0.000146098 | 0.00096143 |
| 0.905158266 | NA |
| 0.726302422 | 0.831016824 |
| 0.207982755 | NA |
| 0.860328012 | NA |
| 0.855241833 | NA |
| NA | NA |
| 0.42624645 | NA |
| NA | NA |
| NA | NA |
| NA | NA |
| NA | NA |
| NA | NA |
| NA | NA |
| 0.787645204 | NA |

| | | |
|---|---|---|
| | 0.905157793 | NA |
| NA | | NA |
| NA | | NA |
| NA | | NA |
| NA | | NA |
| NA | | NA |
| | 0.875960897 | NA |
| | 0.327827787 | 0.489912234 |
| | 0.887056542 | NA |
| NA | | NA |
| NA | | NA |
| NA | | NA |
| NA | | NA |
| NA | | NA |
| | 0.18426498 | 0.32485797 |
| | 0.889409773 | NA |
| | 0.26388883 | NA |
| | 0.656400642 | NA |
| | 0.889409301 | NA |
| NA | | NA |
| | 0.135012508 | 0.257244223 |
| | 0.275081624 | 0.432330188 |
| | 0.611234786 | NA |
| | 0.570371197 | 0.712647575 |
| | 0.618165901 | NA |
| | 0.298518271 | 0.458563324 |
| | 0.669637648 | 0.78941634 |
| | 0.125697026 | 0.244263804 |
| | 0.000357479 | 0.002079989 |
| | 0.234394501 | 0.38586617 |
| NA | | NA |
| NA | | NA |
| | 0.905157793 | NA |
| | 0.992062903 | NA |
| | 0.905158266 | NA |
| NA | | NA |
| | 0.534389495 | 0.683041846 |
| NA | | NA |
| NA | | NA |
| NA | | NA |
| NA | | NA |
| | 0.702091191 | NA |
| | 0.797342596 | NA |
| NA | | NA |
| NA | | NA |
| NA | | NA |
| NA | | NA |

| | | | |
|---|---|---|---|
| | 0.521304087 | | 0.672019825 |
| NA | | NA | |
| NA | | NA | |
| | 0.002041175 | | 0.009106495 |
| | 0.851672944 | NA | |
| | 3.22E-11 | | 1.57E-09 |
| | 0.167722666 | | 0.302905011 |
| | 0.071960531 | | 0.159937379 |
| | 0.53500735 | | 0.683377626 |
| | 0.609531107 | NA | |
| | 0.898978351 | | 0.943310432 |
| | 0.033046029 | | 0.088001182 |
| | 0.366524689 | | 0.5313173 |
| | 1.81E-18 | | 5.52E-16 |
| | 0.027997271 | | 0.077099582 |
| | 0.680291437 | | 0.797273825 |
| | 0.002806134 | | 0.011888849 |
| | 0.038836215 | | 0.099776198 |
| | 0.877230405 | | 0.930000601 |
| | 0.195777193 | | 0.339605924 |
| | 0.340969287 | | 0.504334135 |
| | 0.079097731 | NA | |
| | 0.905296333 | NA | |
| | 0.196248149 | NA | |
| | 0.983110898 | NA | |
| | 0.05377766 | | 0.12821972 |
| | 0.011479216 | | 0.037555438 |
| NA | | NA | |
| | 0.059425227 | | 0.138601914 |
| | 0.006985711 | | 0.025210141 |
| | 0.362302996 | | 0.52708437 |
| | 0.87314935 | | 0.927333364 |
| | 0.967288007 | | 0.982404998 |
| | 0.036057174 | | 0.094293137 |
| | 0.076806387 | | 0.168541166 |
| | 0.004339652 | | 0.01700168 |
| NA | | NA | |
| | 8.80E-06 | | 8.74E-05 |
| | 0.047038177 | | 0.115625552 |
| | 0.067692147 | | 0.152584411 |
| | 0.08961097 | | 0.189481684 |
| | 0.704219819 | NA | |
| | 0.005802339 | | 0.02169043 |
| | 0.42946635 | | 0.590522955 |
| | 0.369997374 | | 0.534594837 |
| | 0.201876782 | | 0.347282203 |
| NA | | NA | |

| | | |
|---|---|---|
| | 0.992060816 | NA |
| | 0.120709883 | 0.23716747 |
| | 0.920906701 | 0.954597349 |
| | 0.308127921 | 0.469156914 |
| | 0.916918875 | 0.952667215 |
| | 0.992062245 | NA |
| | 0.638218523 | NA |
| | 0.119862969 | 0.235912068 |
| | 0.051318421 | 0.123889755 |
| | 0.905157793 | NA |
| NA | | NA |
| | 0.882781554 | 0.933407172 |
| | 0.782667874 | NA |
| | 0.000249289 | 0.001525254 |
| | 0.924720216 | 0.956719373 |
| | 0.462892274 | 0.620127308 |
| | 8.43E-05 | 0.000602496 |
| | 0.981235318 | NA |
| | 0.811319626 | NA |
| | 0.027787284 | 0.076646272 |
| | 0.557145935 | NA |
| NA | | NA |
| NA | | NA |
| | 0.999804105 | 0.999887125 |
| NA | | NA |
| | 0.565380396 | 0.708589014 |
| | 0.962639912 | 0.979279397 |
| | 0.275586185 | NA |
| | 0.616254514 | NA |
| | 0.88376167 | 0.934038722 |
| NA | | NA |
| | 0.343622374 | 0.506994808 |
| | 0.001571846 | 0.007284965 |
| | 0.941808523 | 0.967042615 |
| | 0.992059387 | NA |
| | 0.360516484 | NA |
| | 0.993459113 | NA |
| | 0.000169299 | 0.001091819 |
| | 0.003412168 | 0.01396604 |
| | 0.797179953 | 0.877636997 |
| | 0.559136761 | NA |
| | 0.875289692 | NA |
| | 0.359201837 | 0.524065719 |
| | 0.674719494 | NA |
| | 0.984401075 | NA |
| | 0.195947464 | 0.339767085 |
| | 0.905158266 | NA |

| | | |
|---|---|---|
| | 0.301000427 | 0.461375707 |
| | 1.86E-07 | 3.05E-06 |
| | 0.825707987 | 0.896145984 |
| | 0.000366519 | 0.002121909 |
| | 5.39E-07 | 7.70E-06 |
| | 0.087257827 | 0.185632109 |
| | 0.270338322 | 0.426830544 |
| | 0.219186561 | 0.36780664 |
| | 0.808281038 | NA |
| | 0.061939702 | 0.143232299 |
| NA | | NA |
| | 9.14E-09 | 2.18E-07 |
| NA | | NA |
| | 0.797176643 | 0.877636997 |
| | 0.307710442 | 0.46871626 |
| | 0.003422383 | 0.013988068 |
| | 2.36E-08 | 5.03E-07 |
| | 0.248606338 | 0.402436865 |
| | 0.905156847 | NA |
| | 0.216426809 | 0.364400882 |
| | 0.002169775 | 0.009582884 |
| NA | | NA |
| NA | | NA |
| | 0.889409301 | NA |
| | 0.799106931 | 0.87887354 |
| | 0.487824215 | 0.642406713 |
| | 0.640862315 | 0.767071603 |
| NA | | NA |
| | 0.594176069 | NA |
| NA | | NA |
| | 0.889409301 | NA |
| | 0.991759839 | NA |
| | 0.759031831 | NA |
| | 0.889409773 | NA |
| NA | | NA |
| NA | | NA |
| NA | | NA |
| NA | | NA |
| NA | | NA |
| NA | | NA |
| NA | | NA |
| NA | | NA |
| NA | | NA |
| | 0.869377566 | NA |
| | 0.440053771 | NA |
| | 0.980094639 | NA |
| | 0.735355709 | NA |

| | | |
|---|---|---|
| NA | | NA |
| NA | | NA |
| | 0.606570688 | NA |
| NA | | NA |
| | 0.894061516 | NA |
| | 0.922528152 | NA |
| | 0.011669269 | 0.038063796 |
| | 0.825955501 | NA |
| | 0.807879735 | NA |
| | 0.887395124 | NA |
| | 0.004068572 | 0.016120854 |
| | 0.037433287 | 0.097082504 |
| | 0.083614376 | 0.179795951 |
| | 0.293594552 | 0.453251039 |
| | 0.766031885 | 0.858141729 |
| | 0.905158739 | NA |
| | 0.000624076 | 0.003351451 |
| | 0.881501945 | NA |
| | 0.62605345 | 0.755772827 |
| | 0.026764231 | 0.074309742 |
| | 0.905158266 | NA |
| | 0.008456471 | 0.029377575 |
| | 0.271964107 | 0.428688524 |
| | 0.387152241 | 0.550934876 |
| | 0.065587847 | 0.149387359 |
| | 0.263648672 | 0.419384261 |
| | 0.631654805 | 0.759528412 |
| | 0.777092376 | NA |
| | 0.54506724 | NA |
| | 0.41476256 | NA |
| | 0.905158266 | NA |
| NA | | NA |
| | 0.004806774 | 0.018556953 |
| | 0.471447024 | 0.6277985 |
| NA | | NA |
| | 0.249342511 | NA |
| | 0.689162991 | NA |
| | 0.015846304 | 0.048865187 |
| | 0.570757182 | NA |
| | 0.064228432 | 0.147099723 |
| | 0.176919613 | NA |
| | 0.072097667 | 0.160161121 |
| | 0.183768139 | 0.324334075 |
| | 0.025827652 | 0.072359556 |
| | 0.776244828 | NA |
| | 0.564905634 | NA |
| | 2.36E-05 | 0.000204348 |

| | |
|---|---|
| 0.131509113 | 0.252276457 |
| 0.686114365 | 0.801616671 |
| 0.549779835 | 0.695868084 |
| NA | NA |
| 0.20570962 | 0.351942154 |
| 0.717446518 | NA |
| 1.04E-06 | 1.35E-05 |
| 0.551384774 | NA |
| 0.619538394 | NA |
| 0.037260442 | 0.096741369 |
| 0.070112037 | 0.156717246 |
| 0.156005377 | 0.286955075 |
| 0.36697088 | NA |
| 0.004987733 | 0.019147838 |
| 0.000171134 | 0.001102034 |
| NA | NA |
| 0.809687584 | NA |
| 0.716507584 | NA |
| NA | NA |
| 0.889409773 | NA |
| NA | NA |
| NA | NA |
| 0.084295835 | 0.180713877 |
| NA | NA |
| NA | NA |
| NA | NA |
| NA | NA |
| NA | NA |
| 0.889409773 | NA |
| NA | NA |
| NA | NA |
| NA | NA |
| NA | NA |
| NA | NA |
| 0.700936353 | 0.813136694 |
| NA | NA |
| NA | NA |
| NA | NA |
| NA | NA |
| 0.905158266 | NA |
| NA | NA |
| NA | NA |
| 0.905158739 | NA |
| NA | NA |
| NA | NA |
| 0.81062689 | NA |
| 0.905731663 | 0.946867964 |

| | |
|---|---|
| 3.52E-12 | 2.21E-10 |
| NA | NA |
| 0.017990562 | 0.053998735 |
| 0.531683354 | 0.680811404 |
| 1.21E-05 | 0.00011471 |
| 0.761919709 | 0.855586545 |
| 0.739096709 | NA |
| 0.879102344 | NA |
| 0.01419482 | 0.044632607 |
| 0.122852726 | NA |
| 3.38E-05 | 0.000277239 |
| 0.0302028 | 0.081889528 |
| 0.396160612 | 0.559799928 |
| 0.209590227 | 0.356360185 |
| 0.99205891 | NA |
| 3.43E-05 | 0.000280214 |
| NA | NA |
| 0.567121113 | 0.709919765 |
| 0.195587885 | 0.339384778 |
| 0.595743653 | 0.732758354 |
| 0.239065977 | NA |
| 0.951868374 | 0.973288471 |
| 0.889409773 | NA |
| NA | NA |
| NA | NA |
| NA | NA |
| 0.105812008 | 0.214861869 |
| 0.455123985 | 0.61313298 |
| 0.912280462 | 0.950411171 |
| 0.00318431 | 0.013202393 |
| 0.737592091 | 0.838994288 |
| 0.01631977 | 0.050023148 |
| 0.98880908 | NA |
| 0.713976697 | NA |
| 0.71642609 | 0.823956855 |
| 0.833478308 | 0.900819299 |
| NA | NA |
| 0.650584012 | 0.775148495 |
| 0.155067271 | 0.285764396 |
| 0.012936764 | 0.041356588 |
| 0.384284604 | NA |
| 0.041370366 | 0.104805247 |
| 0.905696566 | NA |
| 0.792017481 | NA |
| 0.788582525 | NA |
| 0.962719915 | NA |
| NA | NA |

| | |
|---|---|
| 0.460946255 | 0.618359022 |
| 0.087990641 | 0.186973727 |
| NA | NA |
| 0.432119997 | 0.592983566 |
| 0.445665732 | 0.605105658 |
| 2.34E-10 | 8.94E-09 |
| NA | NA |
| 0.004512383 | 0.017562462 |
| 0.044632344 | 0.111044826 |
| 0.857833775 | 0.917175946 |
| 0.550048721 | NA |
| 0.337759695 | 0.500665934 |
| 0.889410717 | NA |
| 0.224247131 | 0.373926856 |
| 0.653418555 | NA |
| 0.02223382 | 0.064010635 |
| 0.270216167 | NA |
| 0.622398668 | 0.753556396 |
| 0.239459438 | 0.391737301 |
| 0.001637978 | 0.007532552 |
| 8.35E-05 | 0.000598058 |
| 0.713611819 | NA |
| 0.028580242 | 0.078370821 |
| 0.409653788 | 0.572675549 |
| 0.792313171 | NA |
| 0.153594945 | 0.283802967 |
| 0.629128649 | 0.757547443 |
| NA | NA |
| 0.002161831 | 0.009552223 |
| 8.60E-05 | 0.000612418 |
| 0.651668767 | 0.775786205 |
| 2.07E-05 | 0.000183467 |
| 0.000235097 | 0.001454636 |
| NA | NA |
| 0.06374598 | 0.146238819 |
| 0.09303928 | 0.194854512 |
| 0.669094809 | NA |
| 0.083266355 | 0.179334197 |
| 1.67E-09 | 4.92E-08 |
| 0.370838198 | 0.535351784 |
| 7.11E-05 | 0.000522347 |
| 0.597114522 | 0.733910096 |
| 5.76E-08 | 1.10E-06 |
| 7.37E-08 | 1.36E-06 |
| 0.40469755 | NA |
| 0.000124891 | 0.000844486 |
| 0.824951436 | 0.895634757 |

| | | |
|---|---|---|
| | 0.743222332 | 0.842561651 |
| | 0.746327291 | NA |
| | 0.002345028 | 0.010237467 |
| | 0.225772291 | 0.375614346 |
| | 0.027276532 | 0.07552205 |
| | 0.424040501 | 0.585408359 |
| | 0.471020094 | 0.627350654 |
| | 0.957800403 | NA |
| | 0.534685332 | 0.683260894 |
| | 0.452770355 | 0.610906318 |
| | 0.181439196 | NA |
| | 0.690283121 | 0.805030912 |
| | 0.131063884 | 0.251664202 |
| | 0.905158739 | NA |
| | 0.38008338 | 0.544304363 |
| | 0.124596952 | 0.242778838 |
| NA | | NA |
| | 0.905158266 | NA |
| | 1.35E-15 | 1.98E-13 |
| | 0.847730278 | NA |
| | 0.047836992 | 0.117220698 |
| | 0.84753701 | NA |
| | 2.36E-10 | 8.98E-09 |
| | 0.358586905 | 0.523411981 |
| NA | | NA |
| NA | | NA |
| | 0.09839707 | 0.203398405 |
| | 0.068347262 | 0.153601221 |
| NA | | NA |
| | 0.010300934 | 0.034518705 |
| | 0.366549808 | 0.5313173 |
| | 0.062824018 | 0.144652679 |
| | 0.179556808 | 0.319098117 |
| | 0.605224671 | NA |
| | 0.479385036 | 0.635025396 |
| NA | | NA |
| | 0.066608052 | 0.150993635 |
| | 0.981525278 | NA |
| | 3.33E-09 | 9.07E-08 |
| | 0.460665756 | 0.618020487 |
| NA | | NA |
| NA | | NA |
| | 0.525056733 | 0.675141361 |
| | 0.054891764 | 0.130294545 |
| | 0.012537883 | 0.04037533 |
| | 0.119110914 | NA |
| NA | | NA |

| | | |
|---|---|---|
| | 0.889409301 | NA |
| | 0.803765645 | NA |
| | 0.491895075 | NA |
| | 0.99205891 | NA |
| | 0.904940095 | 0.946310755 |
| | 0.923116368 | 0.955599216 |
| | 0.788582525 | NA |
| | 0.552058421 | NA |
| | 0.808073433 | 0.88498742 |
| | 0.119231282 | 0.234969176 |
| | 0.077251839 | 0.169281822 |
| NA | | NA |
| | 0.000260409 | 0.001584025 |
| | 0.025130548 | 0.070839756 |
| | 0.229261861 | 0.37966525 |
| | 0.889409301 | NA |
| | 0.875266552 | NA |
| | 0.008788817 | 0.03031631 |
| | 0.1979489 | 0.342364559 |
| | 0.905158266 | NA |
| | 3.50E-09 | 9.45E-08 |
| | 0.57449124 | 0.716450087 |
| | 0.001076124 | 0.005298341 |
| | 0.01442214 | 0.045177715 |
| | 7.83E-19 | 2.49E-16 |
| | 0.625936184 | NA |
| | 0.032476415 | 0.086830142 |
| | 0.030914882 | 0.083428769 |
| | 0.000586225 | 0.00318007 |
| | 0.001415248 | 0.006689072 |
| | 0.594744973 | 0.731893199 |
| NA | | NA |
| | 0.326490499 | 0.488745305 |
| | 0.48920763 | 0.643418204 |
| | 0.196472152 | 0.340461804 |
| | 0.318651525 | 0.481077266 |
| | 0.021130675 | 0.061433667 |
| | 0.889409773 | NA |
| | 0.034958111 | 0.091977614 |
| NA | | NA |
| | 0.387221471 | 0.550991138 |
| | 0.034016218 | 0.089975154 |
| | 0.390875598 | NA |
| NA | | NA |
| | 0.103782492 | 0.211741645 |
| | 7.85E-05 | 0.000568573 |
| | 0.063458422 | 0.145761872 |

| | | |
|---|---|---|
| | 0.854436427 | NA |
| | 0.056333336 | 0.132824712 |
| | 0.034815769 | 0.091690991 |
| | 0.064503679 | 0.147591569 |
| | 0.801844417 | 0.88056119 |
| | 0.48512611 | 0.640199877 |
| | 0.222108648 | 0.371432777 |
| | 0.001254539 | 0.006020199 |
| | 0.976078488 | 0.987682942 |
| | 0.315095279 | 0.477325529 |
| | 0.876062717 | NA |
| | 0.31069861 | 0.47195841 |
| NA | | NA |
| NA | | NA |
| | 0.044563873 | 0.110899602 |
| | 0.505633626 | 0.657656151 |
| | 0.889409773 | NA |
| | 0.014750721 | 0.046055891 |
| | 0.851418405 | NA |
| | 0.197540242 | 0.341854061 |
| | 0.245057994 | 0.398255513 |
| | 0.000425967 | 0.002412346 |
| | 0.069781823 | 0.15610626 |
| | 1.81E-07 | 2.98E-06 |
| | 0.00023658 | 0.001462161 |
| | 0.00361512 | 0.014633073 |
| | 0.002247963 | 0.009876554 |
| | 0.234497864 | NA |
| | 1.87E-10 | 7.40E-09 |
| | 0.29426973 | 0.454014339 |
| | 4.12E-07 | 6.08E-06 |
| | 0.900307157 | 0.943713058 |
| | 0.947461184 | 0.970206045 |
| | 0.000358417 | 0.00208214 |
| | 0.257580665 | 0.412382332 |
| | 0.597833391 | 0.734409715 |
| | 0.944003959 | 0.968455992 |
| | 0.200818137 | 0.345978296 |
| | 0.633656551 | NA |
| | 0.132955804 | NA |
| | 0.98404083 | NA |
| | 0.089725474 | 0.18966901 |
| | 0.881501945 | NA |
| | 0.419618827 | NA |
| NA | | NA |
| | 0.001066786 | 0.005259443 |
| | 0.173334158 | 0.310511076 |

| | |
|---|---|
| 0.992061292 | NA |
| 0.881048892 | NA |
| NA | NA |
| 0.940575801 | NA |
| 0.905158266 | NA |
| 0.905158266 | NA |
| 1.50E-07 | 2.52E-06 |
| 0.207992997 | 0.354660523 |
| 0.298597686 | 0.458621151 |
| 0.124235446 | 0.242221186 |
| 0.620860523 | 0.752233294 |
| NA | NA |
| 0.57681311 | 0.718244348 |
| 0.082412226 | 0.178031487 |
| 0.311214006 | NA |
| 0.416925809 | NA |
| 0.590257235 | NA |
| NA | NA |
| 0.929488443 | 0.959575308 |
| 0.291620706 | 0.451155353 |
| 0.369073612 | 0.533716658 |
| 0.224917856 | 0.374603464 |
| 0.729700761 | NA |
| 0.60750252 | 0.741654963 |
| 0.866775201 | 0.923374407 |
| 0.996865436 | 0.998547948 |
| 0.685958214 | 0.801476876 |
| 0.889409773 | NA |
| 0.536187223 | 0.68440311 |
| 7.02E-11 | 3.09E-09 |
| 0.083383053 | 0.179492743 |
| 0.113737231 | 0.227029894 |
| NA | NA |
| NA | NA |
| NA | NA |
| NA | NA |
| NA | NA |
| 0.065687677 | 0.149506188 |
| 0.001357063 | 0.006448765 |
| 0.114881051 | 0.228710154 |
| 0.526445412 | NA |
| 0.012406135 | 0.040045028 |
| 0.674218089 | NA |
| 0.38200104 | 0.546010599 |
| NA | NA |
| NA | NA |
| 0.35979698 | 0.524481007 |

| | |
|---|---|
| 1.32E-06 | 1.66E-05 |
| 0.091923781 | 0.193181578 |
| 0.003288632 | 0.01354371 |
| 0.214553939 | 0.361973911 |
| 0.222842219 | 0.372234313 |
| NA | NA |
| 5.04E-05 | 0.00038959 |
| 1.81E-09 | 5.31E-08 |
| 0.044045214 | 0.109907825 |
| 0.013956204 | 0.044013254 |
| 0.579397311 | 0.719901233 |
| 1.86E-05 | 0.000166931 |
| NA | NA |
| 0.055092792 | 0.130581052 |
| 0.727346679 | 0.831648306 |
| 0.990693559 | 0.995316826 |
| 0.451691409 | 0.610037742 |
| 0.031075531 | 0.08374909 |
| NA | NA |
| 1.08E-06 | 1.40E-05 |
| 0.017555186 | 0.052959365 |
| 0.24321071 | 0.396168965 |
| 0.005687898 | 0.021350521 |
| NA | NA |
| 0.678102861 | NA |
| 0.008537385 | 0.029597796 |
| 0.046414553 | 0.114428783 |
| 0.043010803 | 0.107976931 |
| 0.835174966 | 0.901955981 |
| 0.652662848 | 0.776329749 |
| 0.719816765 | NA |
| 0.001288135 | 0.006157195 |
| 0.52216364 | 0.67263804 |
| NA | NA |
| 0.736327639 | NA |
| 0.604541543 | 0.739685058 |
| 0.90690505 | 0.947360419 |
| 0.758259185 | 0.853322916 |
| 0.797328626 | 0.877636997 |
| 0.148350316 | NA |
| 0.905158266 | NA |
| 0.716939825 | 0.824279988 |
| 0.005029699 | 0.019263686 |
| 0.013441989 | 0.042642681 |
| 2.63E-14 | 2.68E-12 |
| 0.992061292 | NA |
| NA | NA |

| | |
|---|---|
| 0.066297118 | 0.150565562 |
| 0.654748631 | 0.777925829 |
| 0.895587395 | 0.941075508 |
| NA | NA |
| 0.991529086 | NA |
| 0.296201026 | 0.456312497 |
| NA | NA |
| 0.129467548 | 0.249559155 |
| 0.25345467 | 0.407794434 |
| 0.911787871 | 0.950078288 |
| 0.040097916 | 0.102289747 |
| 0.609489028 | NA |
| NA | NA |
| 0.09588959 | NA |
| 0.022470072 | 0.064617642 |
| 0.026090927 | 0.072891941 |
| 0.118463988 | 0.233871323 |
| 0.00026436 | 0.001604946 |
| 6.22E-07 | 8.66E-06 |
| 0.992060816 | NA |
| NA | NA |
| 0.434306502 | NA |
| 0.531420641 | NA |
| 0.129807906 | 0.249923029 |
| 0.011221376 | 0.036909855 |
| 0.423814695 | NA |
| 5.60E-07 | 7.93E-06 |
| 0.961784302 | 0.979084024 |
| 0.634041665 | 0.761647753 |
| NA | NA |
| 0.518924263 | NA |
| 5.50E-10 | 1.88E-08 |
| 0.003003182 | 0.012575684 |
| 0.62699677 | 0.756287947 |
| 6.67E-17 | 1.24E-14 |
| 0.491205715 | NA |
| NA | NA |
| 0.011963755 | 0.038862737 |
| 0.960280496 | NA |
| 0.021583665 | 0.062494158 |
| 8.95E-08 | 1.60E-06 |
| 0.006008912 | 0.022314874 |
| 0.233528179 | NA |
| 0.003228628 | 0.013357774 |
| 0.000338535 | 0.00198608 |
| 0.639835364 | NA |
| 0.651505587 | 0.775751609 |

| | |
|---|---|
| 0.869542945 | 0.925335282 |
| 0.258433688 | 0.413324498 |
| NA | NA |
| 0.038331351 | 0.098841736 |
| NA | NA |
| 0.000301913 | 0.001796183 |
| 0.889409301 | NA |
| NA | NA |
| NA | NA |
| 0.014082351 | 0.044334672 |
| 0.000195839 | 0.00123789 |
| NA | NA |
| 0.794120391 | 0.875761355 |
| 0.17291149 | 0.309981501 |
| 0.399135354 | NA |
| 0.551754372 | 0.697242001 |
| 2.17E-06 | 2.58E-05 |
| 0.992060816 | NA |
| 0.008731113 | 0.030155159 |
| NA | NA |
| 0.907406003 | 0.947610543 |
| 0.80209217 | NA |
| 0.581158616 | NA |
| 0.199187089 | 0.343945057 |
| 0.235510762 | 0.387210304 |
| 0.625945944 | 0.755726137 |
| 0.760377806 | NA |
| 0.000104591 | 0.000726347 |
| NA | NA |
| 0.746403105 | 0.845164017 |
| 0.1958742 | 0.339720524 |
| NA | NA |
| 0.109351882 | 0.220359545 |
| 0.027447147 | 0.07587963 |
| 1.77E-06 | 2.15E-05 |
| 0.006982278 | 0.025205567 |
| 2.68E-05 | 0.000227704 |
| 0.082585831 | 0.178318807 |
| 0.680327897 | NA |
| 0.003031264 | 0.012664295 |
| 0.339414876 | NA |
| 0.748140303 | 0.846345522 |
| 0.079156085 | 0.17252491 |
| NA | NA |
| NA | NA |
| 0.000994161 | 0.004953422 |
| 0.18643107 | 0.327664468 |

| | |
|---|---|
| 0.005444348 | 0.020569932 |
| 0.694664655 | 0.808252062 |
| 0.001080388 | 0.00531695 |
| NA | NA |
| 0.142091123 | 0.267482124 |
| NA | NA |
| 0.25407317 | 0.408462753 |
| 0.003330338 | 0.013679554 |
| 0.61933619 | 0.751049447 |
| 0.010157917 | 0.034111898 |
| 0.962372858 | 0.979205986 |
| NA | NA |
| NA | NA |
| NA | NA |
| 0.555834195 | NA |
| 0.985963296 | NA |
| 0.729963021 | 0.833945059 |
| 5.75E-08 | 1.09E-06 |
| 6.90E-05 | 0.000509265 |
| 0.000142655 | 0.000942122 |
| 7.91E-05 | 0.000572954 |
| 0.000151801 | 0.000993561 |
| 0.992059387 | NA |
| NA | NA |
| 0.601228656 | NA |
| 0.905158266 | NA |
| NA | NA |
| NA | NA |
| 0.980991335 | NA |
| 0.123173622 | NA |
| 0.086571962 | 0.184512639 |
| 0.002372851 | 0.01034659 |
| NA | NA |
| 0.607402052 | 0.741617049 |
| 0.215029899 | NA |
| NA | NA |
| NA | NA |
| 0.992059387 | NA |
| NA | NA |
| 1.86E-11 | 9.69E-10 |
| NA | NA |
| 0.871386967 | NA |
| 0.016135592 | 0.049590129 |
| 1.19E-09 | 3.66E-08 |
| 0.835766104 | NA |
| 0.03387836 | 0.089717187 |
| 1.11E-08 | 2.58E-07 |

| | |
|---|---|
| 0.771185126 | NA |
| 0.75311676 | 0.849610222 |
| NA | NA |
| 0.363865497 | 0.528843746 |
| 0.27971149 | 0.437894746 |
| 0.45036304 | 0.608848242 |
| 0.42436313 | 0.58561832 |
| 0.711659647 | 0.820520994 |
| 9.77E-07 | 1.28E-05 |
| 0.006504924 | 0.023787113 |
| NA | NA |
| NA | NA |
| 0.734003329 | 0.836640201 |
| 0.538642724 | 0.686400625 |
| 0.004310515 | 0.016904501 |
| 0.927593832 | NA |
| 0.404089897 | 0.56746158 |
| 1.01E-14 | 1.16E-12 |
| 0.013148004 | 0.041861094 |
| 0.992061292 | NA |
| 0.934190245 | 0.962234681 |
| 0.025209876 | 0.071017788 |
| 0.052606394 | NA |
| 0.701994325 | 0.814019823 |
| 0.101781804 | 0.208802937 |
| 0.992003131 | NA |
| NA | NA |
| 0.939903749 | NA |
| 0.003195836 | 0.013243509 |
| 0.117669002 | 0.23272031 |
| 0.734014572 | NA |
| 5.24E-05 | 0.00040238 |
| 0.006499136 | 0.023773869 |
| 0.565946677 | 0.709032533 |
| 0.919692378 | 0.95407233 |
| NA | NA |
| 0.969260324 | 0.983776228 |
| 0.065449009 | 0.149190181 |
| 0.280451402 | 0.438759785 |
| 0.239405557 | 0.391734634 |
| NA | NA |
| 0.258600089 | NA |
| NA | NA |
| 0.126657885 | NA |
| 0.255909143 | 0.410525236 |
| 0.98819809 | 0.993639148 |
| 6.73E-07 | 9.31E-06 |

| | | |
|---|---|---|
| | 0.305206803 | 0.465774981 |
| | 0.645216424 | 0.770444549 |
| NA | | NA |
| | 0.58120914 | NA |
| | 0.766220629 | 0.858298804 |
| | 0.778531773 | NA |
| | 0.859370554 | NA |
| | 0.376059102 | 0.540401926 |
| | 0.9801228 | NA |
| | 0.905157793 | NA |
| | 0.473178621 | NA |
| | 0.483630403 | 0.639222344 |
| NA | | NA |
| | 5.34E-09 | 1.35E-07 |
| | 0.983023836 | NA |
| | 0.603163263 | NA |
| NA | | NA |
| | 0.696449117 | NA |
| | 0.447963431 | NA |
| NA | | NA |
| | 0.905158266 | NA |
| | 0.253279105 | 0.407723471 |
| | 0.950305224 | NA |
| NA | | NA |
| | 1.56E-07 | 2.60E-06 |
| NA | | NA |
| | 0.711672963 | NA |
| | 0.685667372 | NA |
| | 0.280422788 | NA |
| | 0.991918092 | NA |
| | 0.913250302 | 0.950880199 |
| | 0.889409301 | NA |
| | 0.299021539 | 0.459240031 |
| | 0.985459718 | NA |
| | 0.947857505 | 0.970440272 |
| | 0.252315153 | 0.40661885 |
| | 2.17E-05 | 0.000191152 |
| | 0.399560624 | 0.563190946 |
| | 6.08E-07 | 8.49E-06 |
| | 3.43E-05 | 0.000280275 |
| NA | | NA |
| | 0.067256059 | 0.151926324 |
| | 2.74E-05 | 0.000231724 |
| | 0.538873497 | NA |
| | 0.483802155 | 0.639338968 |
| | 0.000929368 | 0.004681804 |
| | 0.367904319 | NA |

| | |
|---|---|
| 0.592212513 | 0.729845919 |
| 0.889409301 | NA |
| 0.805374319 | NA |
| 0.061767005 | 0.142923252 |
| NA | NA |
| NA | NA |
| NA | NA |
| 0.381087576 | 0.545272827 |
| 0.580897096 | NA |
| 0.905158266 | NA |
| 0.992061292 | NA |
| 0.010822121 | 0.035881467 |
| 0.010736062 | 0.035687737 |
| 0.017782197 | 0.053504969 |
| 0.728257436 | 0.8324296 |
| 0.65206921 | 0.775959887 |
| 2.47E-07 | 3.89E-06 |
| 0.56131851 | 0.705430852 |
| 0.907176621 | NA |
| 0.055495983 | 0.131272024 |
| 0.911504912 | 0.950003887 |
| 0.002799628 | 0.011868153 |
| NA | NA |
| 0.179888868 | 0.31946459 |
| 0.002180998 | 0.009624704 |
| 1.37E-07 | 2.33E-06 |
| 0.830144033 | NA |
| NA | NA |
| NA | NA |
| 2.06E-10 | 8.05E-09 |
| 0.680954868 | 0.797613891 |
| 0.95608011 | NA |
| 0.172425949 | 0.30931308 |
| NA | NA |
| NA | NA |
| 0.009783759 | 0.033025113 |
| 0.000279443 | 0.001686719 |
| 0.546267944 | 0.692700686 |
| 0.891643168 | 0.938832699 |
| 0.179610245 | 0.319150458 |
| 0.992269016 | NA |
| 0.163830994 | 0.297589941 |
| NA | NA |
| 0.65188389 | 0.775889209 |
| 0.87040193 | NA |
| NA | NA |
| 0.979188107 | NA |

| | |
|---|---|
| NA | NA |
| 0.159748927 | 0.291962271 |
| 3.19E-08 | 6.59E-07 |
| NA | NA |
| 0.035005588 | 0.092043834 |
| 0.151343974 | 0.280665681 |
| 0.815665474 | NA |
| 0.804534188 | NA |
| 0.003163335 | 0.013126169 |
| NA | NA |
| 0.130885623 | NA |
| NA | NA |
| NA | NA |
| NA | NA |
| NA | NA |
| NA | NA |
| NA | NA |
| NA | NA |
| 0.304518775 | 0.465145226 |
| NA | NA |
| NA | NA |
| NA | NA |
| 0.902146925 | 0.944663947 |
| NA | NA |
| NA | NA |
| NA | NA |
| NA | NA |
| NA | NA |
| NA | NA |
| NA | NA |
| NA | NA |
| NA | NA |
| NA | NA |
| NA | NA |
| NA | NA |
| NA | NA |
| NA | NA |
| NA | NA |
| NA | NA |
| NA | NA |
| NA | NA |
| NA | NA |
| NA | NA |
| 0.573114417 | 0.715342437 |
| NA | NA |
| NA | NA |
| NA | NA |

| | | |
|---|---|---|
| NA | NA | |
| NA | NA | |
| NA | NA | |
| NA | NA | |
| NA | NA | |
| NA | NA | |
| 0.836512434 | NA | |
| 0.992060816 | NA | |
| NA | NA | |
| NA | NA | |
| 0.992060816 | NA | |
| 0.992061292 | NA | |
| 0.905158266 | NA | |
| NA | NA | |
| 0.962130991 | | 0.979205986 |
| NA | NA | |
| NA | NA | |
| 0.889409301 | NA | |
| NA | NA | |
| 0.865472004 | NA | |
| 0.992061292 | NA | |
| 3.77E-20 | | 1.81E-17 |
| 0.749364909 | | 0.847157273 |
| 0.012729179 | | 0.040895397 |
| 1.34E-05 | | 0.000125216 |
| 0.007307524 | | 0.026134515 |
| 0.968030798 | NA | |
| NA | NA | |
| NA | NA | |
| 0.992061292 | NA | |
| 0.992061769 | NA | |
| 0.550822651 | | 0.696581288 |
| NA | NA | |
| 0.821979411 | NA | |
| 0.889409773 | NA | |
| NA | NA | |
| 0.89221045 | | 0.939171145 |
| NA | NA | |
| 0.003448082 | | 0.014065844 |
| NA | NA | |
| 0.559512339 | NA | |
| 0.49767928 | NA | |
| NA | NA | |
| 0.782559532 | NA | |
| NA | NA | |
| NA | NA | |
| 0.604059 | NA | |

| | |
|---|---|
| NA | NA |
| 0.545010998 | 0.691586045 |
| 0.889409301 | NA |
| 0.252688817 | 0.406952235 |
| 0.938397668 | 0.965165077 |
| NA | NA |
| 0.837932936 | 0.903707747 |
| NA | NA |
| 0.535668489 | 0.683939768 |
| NA | NA |
| NA | NA |
| NA | NA |
| 0.175024629 | NA |
| NA | NA |
| NA | NA |
| NA | NA |
| 0.699326867 | NA |
| 0.905158739 | NA |
| 0.992061769 | NA |
| NA | NA |
| 0.205431326 | 0.351630406 |
| 0.140747646 | 0.265590578 |
| 0.78401825 | NA |
| 0.02857853 | 0.078370821 |
| NA | NA |
| NA | NA |
| 0.822346297 | NA |
| 0.905158266 | NA |
| 0.889409773 | NA |
| 0.784353866 | NA |
| 0.083062281 | 0.179013114 |
| 0.905158266 | NA |
| 0.877160953 | NA |
| 0.432400239 | 0.593293979 |
| NA | NA |
| NA | NA |
| NA | NA |
| 0.905158266 | NA |
| NA | NA |
| NA | NA |
| 0.684371666 | NA |
| 0.20969379 | 0.356468475 |
| 0.63011294 | NA |
| 0.773176934 | NA |
| 0.92231471 | 0.955335317 |
| 0.268383304 | 0.424782253 |
| 0.496095813 | 0.64944366 |

| | |
|---|---|
| NA | NA |
| 0.258216991 | NA |
| 0.983871219 | NA |
| 0.690471071 | 0.805192261 |
| NA | NA |
| 0.598384741 | 0.734772159 |
| 0.428194528 | NA |
| NA | NA |
| 0.855659532 | 0.915974383 |
| 0.100370775 | 0.206583733 |
| 0.984630534 | 0.992171589 |
| NA | NA |
| 0.390765093 | 0.554118558 |
| NA | NA |
| 0.661993291 | 0.783554459 |
| 0.669278565 | NA |
| 0.005109379 | 0.019511917 |
| 0.992061769 | NA |
| 0.464610209 | NA |
| 0.926150161 | NA |
| NA | NA |
| 0.03119688 | 0.084003958 |
| 0.291139966 | NA |
| 0.001302698 | 0.006215982 |
| 0.752904354 | 0.849546617 |
| 0.208902811 | 0.355717999 |
| 0.177252894 | 0.315933141 |
| 0.003700347 | 0.014902155 |
| 0.117508875 | 0.232522767 |
| 0.028155526 | 0.077438277 |
| 0.56764598 | NA |
| 1.29E-07 | 2.21E-06 |
| 0.000386515 | 0.002218947 |
| 0.428638014 | NA |
| 0.509181928 | 0.660877572 |
| 0.290105031 | 0.449475613 |
| 0.489034055 | 0.643305506 |
| 0.889409301 | NA |
| 2.79E-12 | 1.79E-10 |
| 0.669956583 | 0.789501803 |
| 0.36896156 | NA |
| 0.038127575 | 0.098452895 |
| 0.848685642 | 0.911176258 |
| 0.003662016 | 0.014779309 |
| 2.30E-08 | 4.93E-07 |
| 0.989838244 | NA |
| 3.48E-05 | 0.000283589 |

| | | |
|---|---|---|
| | 0.087026321 | 0.185296429 |
| | 0.010309654 | 0.034542216 |
| | 0.992060816 | NA |
| | 0.021198293 | 0.061597632 |
| | 0.000194073 | 0.001228492 |
| | 0.241845657 | 0.394609743 |
| | 3.04E-05 | 0.000253132 |
| | 0.242707566 | 0.395577484 |
| | 0.491900404 | NA |
| | 0.484244953 | NA |
| | 0.228960044 | 0.379480233 |
| NA | | NA |
| | 0.31978759 | NA |
| | 4.75E-05 | 0.000371395 |
| | 0.001859662 | 0.008412922 |
| | 0.805485953 | 0.883217029 |
| | 0.682334052 | NA |
| NA | | NA |
| | 0.370123694 | 0.53467181 |
| NA | | NA |
| | 0.028112613 | 0.077359006 |
| | 0.36762903 | NA |
| | 0.306141613 | 0.466779869 |
| NA | | NA |
| | 0.234851198 | 0.386328374 |
| | 0.001209929 | 0.005843155 |
| | 0.418520736 | 0.580672069 |
| | 0.464177347 | 0.621251318 |
| | 6.80E-09 | 1.68E-07 |
| | 1.38E-06 | 1.73E-05 |
| | 0.028073707 | 0.077282441 |
| | 0.788582525 | NA |
| | 0.905158739 | NA |
| | 0.744957774 | NA |
| | 0.999815835 | NA |
| | 0.070962453 | 0.158247641 |
| | 0.098838173 | 0.204079685 |
| | 0.992061292 | NA |
| | 1.82E-05 | 0.000164102 |
| | 0.098780987 | 0.20399997 |
| NA | | NA |
| | 0.170987356 | 0.307385292 |
| | 0.986808628 | NA |
| | 0.847353223 | NA |
| | 4.03E-15 | 5.12E-13 |
| | 2.10E-08 | 4.55E-07 |
| NA | | NA |

| | | |
|---|---|---|
| | 0.040152844 | 0.102370457 |
| | 0.885451361 | 0.93507114 |
| | 0.989109034 | NA |
| | 0.33914481 | NA |
| | 0.984050102 | NA |
| | 0.889409773 | NA |
| | 0.782559532 | NA |
| NA | | NA |
| NA | | NA |
| | 0.16543235 | 0.299740049 |
| | 0.889409301 | NA |
| | 0.000219125 | 0.001363957 |
| | 0.636042449 | 0.763216183 |
| | 0.739096709 | NA |
| | 0.767107778 | 0.858646126 |
| | 0.746774777 | 0.84536691 |
| | 0.173067842 | 0.310160511 |
| | 0.508798811 | 0.660560074 |
| NA | | NA |
| | 0.001573353 | NA |
| | 0.855342185 | NA |
| | 5.15E-06 | 5.46E-05 |
| | 5.76E-05 | 0.000437515 |
| | 0.907955991 | 0.947794524 |
| | 0.428545662 | 0.589620368 |
| | 2.87E-06 | 3.28E-05 |
| | 0.056945721 | 0.133940776 |
| NA | | NA |
| NA | | NA |
| | 0.90515732 | NA |
| | 0.512945661 | 0.664551767 |
| | 0.869725914 | 0.92548514 |
| | 0.942694774 | 0.967644677 |
| | 0.011724313 | 0.038209289 |
| | 9.74E-10 | 3.08E-08 |
| NA | | NA |
| | 0.528432523 | NA |
| | 6.89E-09 | 1.69E-07 |
| | 0.000328891 | 0.001934668 |
| | 0.484510429 | NA |
| | 0.889409301 | NA |
| | 0.905158739 | NA |
| | 0.168811759 | 0.30439624 |
| | 0.797126775 | 0.877636997 |
| | 0.004727367 | 0.018298653 |
| | 5.96E-06 | 6.22E-05 |
| | 0.27940813 | 0.437626471 |

| | | |
|---|---|---|
| | 0.450735865 | 0.60912726 |
| | 9.33E-06 | 9.17E-05 |
| | 0.003089328 | 0.012862827 |
| | 0.050340667 | 0.121998754 |
| | 1.99E-05 | 0.000176642 |
| | 0.824253828 | 0.895142918 |
| | 0.807980726 | NA |
| | 0.794524957 | 0.87598401 |
| NA | | NA |
| | 0.997536236 | 0.998946471 |
| | 0.64475377 | 0.770218041 |
| | 0.781281794 | 0.868271012 |
| | 0.772673843 | 0.862809302 |
| | 2.58E-05 | 0.000220547 |
| | 0.090653379 | 0.191178673 |
| | 2.70E-06 | 3.11E-05 |
| | 0.156991493 | NA |
| | 1.31E-17 | 3.04E-15 |
| | 0.87215209 | NA |
| | 0.079577049 | 0.173201746 |
| | 0.132163755 | 0.253132448 |
| NA | | NA |
| | 0.783033794 | NA |
| NA | | NA |
| | 0.831817855 | NA |
| | 0.705678528 | NA |
| | 0.004818781 | 0.018600036 |
| NA | | NA |
| | 0.992061292 | NA |
| | 0.134426464 | 0.256350042 |
| | 0.000460759 | 0.002572541 |
| | 0.00077063 | 0.004000327 |
| | 0.00104843 | 0.005182909 |
| | 0.804076334 | 0.882306227 |
| | 0.965775297 | 0.981422073 |
| | 0.977717358 | NA |
| NA | | NA |
| NA | | NA |
| | 0.027058094 | 0.074999479 |
| | 0.900897778 | NA |
| NA | | NA |
| | 0.712444696 | 0.820959728 |
| | 0.002624205 | 0.01126807 |
| | 0.357038548 | 0.521706778 |
| NA | | NA |
| | 0.23221668 | 0.383344315 |
| | 0.021418289 | 0.062121807 |

| | |
|---|---|
| 0.153251323 | NA |
| 0.040378427 | NA |
| 0.262191798 | NA |
| 0.208061419 | 0.354749647 |
| NA | NA |
| 0.766602991 | 0.858439143 |
| NA | NA |
| 0.635202685 | 0.762583553 |
| 0.064758414 | 0.148066433 |
| 0.19876162 | 0.343399403 |
| 0.778899578 | 0.866939826 |
| 0.001622646 | 0.007476224 |
| 0.000955547 | 0.004793892 |
| 0.760364982 | 0.854583614 |
| 0.167738431 | 0.302908569 |
| 0.607017266 | 0.741298247 |
| 0.000679877 | 0.003605266 |
| 0.000981295 | 0.004899318 |
| 0.475840873 | 0.632046318 |
| 1.50E-07 | 2.52E-06 |
| 0.026252238 | 0.073230804 |
| 0.840622492 | 0.905707732 |
| 3.02E-06 | 3.42E-05 |
| 0.001739758 | 0.007939184 |
| 0.128072017 | NA |
| 6.63E-05 | 0.000492093 |
| 0.286655486 | NA |
| 0.055017682 | 0.130481087 |
| 0.302619117 | 0.463332604 |
| 0.003245919 | 0.013413086 |
| 0.889409301 | NA |
| 5.01E-05 | 0.000387854 |
| 0.327379496 | 0.489542142 |
| 0.004453255 | 0.017369275 |
| 0.90515732 | NA |
| 0.175015691 | 0.312707043 |
| 0.074674062 | 0.164668381 |
| 0.813983435 | 0.889222539 |
| 0.862683548 | 0.920757657 |
| NA | NA |
| NA | NA |
| 0.002261598 | 0.00992058 |
| 0.992060816 | NA |
| 0.799560257 | 0.87917949 |
| 0.096500558 | 0.200478312 |
| 1.17E-05 | 0.000111068 |
| 0.179924099 | 0.319501317 |

| | |
|---|---|
| 8.08E-08 | 1.46E-06 |
| 0.016837228 | 0.051279983 |
| 0.019841229 | 0.058435292 |
| NA | NA |
| 0.758491725 | NA |
| 0.236534383 | 0.388340823 |
| 0.585925732 | 0.725225395 |
| 0.117838569 | 0.232887871 |
| 0.889410717 | NA |
| 0.711084478 | 0.820056933 |
| 0.079118539 | 0.172460194 |
| 0.863344972 | 0.921150198 |
| 0.139552487 | 0.263839463 |
| 3.58E-12 | 2.23E-10 |
| 0.024186589 | 0.068610426 |
| 6.14E-07 | 8.56E-06 |
| NA | NA |
| 0.059976795 | 0.139678698 |
| 0.001540429 | 0.007163549 |
| 0.643027429 | 0.768699402 |
| 0.011433341 | 0.037459309 |
| 0.496261085 | 0.649565047 |
| 0.522823406 | 0.673195683 |
| 0.789452945 | 0.872846142 |
| 0.336946078 | NA |
| 0.145868978 | 0.272816361 |
| 0.843320323 | 0.907590226 |
| 0.814477595 | NA |
| NA | NA |
| 0.889409773 | NA |
| 0.992061292 | NA |
| 0.160685615 | 0.293234889 |
| 0.113854838 | 0.227182044 |
| 0.009320145 | 0.03176487 |
| 0.026316396 | 0.073370979 |
| 0.523571434 | 0.673880783 |
| 2.83E-05 | 0.000237926 |
| 0.076266915 | 0.16762538 |
| NA | NA |
| 0.098095852 | 0.202909458 |
| 0.845962891 | 0.909490665 |
| 0.870442592 | NA |
| NA | NA |
| NA | NA |
| NA | NA |
| NA | NA |
| NA | NA |

| | |
|---|---|
| NA | NA |
| NA | NA |
| NA | NA |
| NA | NA |
| NA | NA |
| NA | NA |
| 0.162525443 | 0.295805736 |
| NA | NA |
| 0.000623938 | 0.003351451 |
| 0.905157793 | NA |
| 0.803180384 | NA |
| 0.172086392 | 0.308880585 |
| 0.388946793 | 0.552343183 |
| NA | NA |
| 0.007408235 | 0.026404388 |
| 0.132656426 | 0.253834303 |
| 0.088420359 | 0.187705207 |
| 0.743901767 | 0.843153014 |
| 0.574943912 | NA |
| 0.117042237 | 0.231898446 |
| 0.125447519 | 0.243908453 |
| 0.876987616 | NA |
| 0.782559532 | NA |
| 0.565349973 | 0.708589014 |
| NA | NA |
| 0.153044884 | 0.283088251 |
| 0.971453003 | 0.984954128 |
| 0.083731886 | 0.179961679 |
| 0.201856254 | 0.347282203 |
| 0.881745865 | 0.93280616 |
| 0.285718258 | 0.444655367 |
| 0.004954385 | 0.019039787 |
| 0.634824271 | 0.762254275 |
| 0.106374967 | 0.215646963 |
| 0.708161626 | 0.818060979 |
| 0.660036259 | 0.78221902 |
| 0.53720788 | NA |
| 0.011469833 | 0.037535925 |
| NA | NA |
| 2.84E-22 | 1.95E-19 |
| 0.002299302 | 0.010065864 |
| 0.020964767 | 0.061064508 |
| 0.797169572 | NA |
| 0.049761102 | 0.120927849 |
| 0.008857988 | 0.030506988 |
| 0.057088063 | 0.134203701 |
| 0.136949127 | 0.260121605 |

| | |
|---|---|
| 0.064253301 | 0.147141333 |
| 0.303245203 | 0.46390981 |
| 0.931918757 | 0.960525885 |
| 0.002064449 | 0.009193517 |
| 0.419765873 | 0.581554499 |
| NA | NA |
| 0.542352553 | 0.689487623 |
| 0.643401625 | 0.768937427 |
| 0.108436599 | 0.219017593 |
| 0.299817252 | NA |
| 0.747926965 | 0.846215915 |
| 0.585134977 | 0.724580102 |
| 1.75E-07 | 2.89E-06 |
| NA | NA |
| NA | NA |
| 0.982021405 | NA |
| NA | NA |
| NA | NA |
| 0.635596573 | NA |
| 0.803127554 | NA |
| 0.846086633 | NA |
| 0.043851099 | NA |
| 6.34E-05 | 0.000473905 |
| 0.878022179 | 0.930694292 |
| 0.178205934 | 0.31724557 |
| 0.017060711 | 0.051824796 |
| 0.530155322 | 0.679409437 |
| NA | NA |
| NA | NA |
| 0.619838724 | NA |
| 0.903342961 | NA |
| 0.889409773 | NA |
| NA | NA |
| NA | NA |
| 0.793190522 | 0.875259833 |
| 0.875562717 | NA |
| 0.019215951 | 0.056883672 |
| 0.889409301 | NA |
| 4.01E-12 | 2.45E-10 |
| 0.905157793 | NA |
| 0.124604695 | 0.242778838 |
| NA | NA |
| 0.05970265 | 0.139173088 |
| 0.218300777 | 0.366712574 |
| 0.559495044 | 0.703864318 |
| 0.000896462 | 0.004541039 |
| 0.038854459 | 0.099799867 |

| | |
|---|---|
| 0.0731888 | 0.162044046 |
| 0.670388787 | 0.789836298 |
| 0.434431247 | NA |
| 0.614053427 | NA |
| 0.451897177 | 0.610095918 |
| 0.887248776 | NA |
| 6.15E-08 | 1.16E-06 |
| 0.000865547 | 0.004402723 |
| 0.286465083 | 0.445521964 |
| 0.463563096 | 0.620618341 |
| 0.992061292 | NA |
| NA | NA |
| 0.782559532 | NA |
| 0.068568262 | 0.153924602 |
| 0.031385685 | 0.084470922 |
| 0.454595539 | NA |
| 0.103477673 | 0.211373603 |
| 0.769650869 | NA |
| 2.82E-06 | 3.23E-05 |
| 0.980436241 | 0.990358315 |
| 0.39278999 | NA |
| 7.90E-15 | 9.40E-13 |
| 0.006317142 | 0.023219905 |
| NA | NA |
| 0.385970362 | NA |
| 0.813279315 | NA |
| 0.968655498 | NA |
| 0.996735696 | NA |
| 0.866836407 | NA |
| 0.276436447 | NA |
| 0.662005058 | NA |
| 0.992060816 | NA |
| 0.046744986 | NA |
| 0.084274717 | NA |
| NA | NA |
| NA | NA |
| NA | NA |
| NA | NA |
| NA | NA |
| NA | NA |
| NA | NA |
| NA | NA |
| NA | NA |
| NA | NA |
| NA | NA |
| NA | NA |
| NA | NA |

| | |
|---|---|
| NA | NA |
| NA | NA |
| NA | NA |
| NA | NA |
| NA | NA |
| NA | NA |
| NA | NA |
| NA | NA |
| NA | NA |
| NA | NA |
| NA | NA |
| NA | NA |
| 0.962318383 | NA |
| NA | NA |
| NA | NA |
| NA | NA |
| NA | NA |
| NA | NA |
| NA | NA |
| NA | NA |
| NA | NA |
| NA | NA |
| NA | NA |
| NA | NA |
| NA | NA |
| NA | NA |
| NA | NA |
| NA | NA |
| NA | NA |
| NA | NA |
| 0.841476108 | NA |
| NA | NA |
| NA | NA |
| 0.989597136 | NA |
| 0.905158266 | NA |
| 0.21303174 | 0.360259066 |
| NA | NA |
| 0.951897698 | 0.973288471 |
| NA | NA |
| NA | NA |
| 0.97310295 | 0.985912235 |
| NA | NA |
| 0.287846019 | NA |
| 4.10E-05 | 0.000326424 |
| NA | NA |
| 0.164833381 | 0.298990271 |
| 0.858082414 | 0.917175946 |

| | |
|---|---|
| 0.342228853 | 0.505652226 |
| 0.875963076 | 0.929105598 |
| NA | NA |
| NA | NA |
| 0.991918092 | NA |
| 0.995001397 | NA |
| NA | NA |
| NA | NA |
| NA | NA |
| NA | NA |
| NA | NA |
| NA | NA |
| 0.000410685 | 0.002338754 |
| 0.234705927 | 0.386203346 |
| NA | NA |
| 0.770552175 | 0.861278018 |
| 2.07E-05 | 0.000183467 |
| 0.635105497 | NA |
| 0.144610121 | 0.271177938 |
| 0.116288969 | 0.230885124 |
| 0.51654054 | 0.667571875 |
| 0.207845596 | 0.354519298 |
| 0.101517197 | 0.208376904 |
| 0.198116059 | 0.342553586 |
| 0.761896767 | NA |
| 0.016050673 | 0.049391359 |
| 0.219087047 | 0.367667747 |
| 0.905158739 | NA |
| 0.889409301 | NA |
| 0.361019763 | NA |
| 0.905158266 | NA |
| 0.889409301 | NA |
| 0.88886015 | NA |
| 0.376449068 | NA |
| 0.675957888 | NA |
| 0.805374319 | NA |
| 0.549522765 | NA |
| NA | NA |
| NA | NA |
| NA | NA |
| 0.898195764 | NA |
| 0.707060809 | NA |
| NA | NA |
| NA | NA |
| NA | NA |
| 0.889410245 | NA |
| NA | NA |

| | |
|---|---|
| 0.008882661 | 0.030558412 |
| 6.30E-09 | 1.57E-07 |
| 0.992061292 | NA |
| 0.357627817 | 0.522345498 |
| NA | NA |
| NA | NA |
| 0.038565272 | 0.099291219 |
| 0.782537787 | 0.869007144 |
| 0.875583816 | 0.928855436 |
| 0.350113287 | 0.514256701 |
| 0.833260648 | NA |
| 0.752888743 | NA |
| NA | NA |
| NA | NA |
| NA | NA |
| NA | NA |
| NA | NA |
| 0.006318336 | 0.023219905 |
| 0.17189996 | 0.308621636 |
| 0.905158739 | NA |
| 6.97E-07 | 9.60E-06 |
| NA | NA |
| 0.905158739 | NA |
| NA | NA |
| 0.889409773 | NA |
| 0.103812485 | 0.21176204 |
| 0.257581678 | 0.412382332 |
| NA | NA |
| 6.53E-06 | 6.75E-05 |
| 0.992061769 | NA |
| NA | NA |
| 0.097140379 | 0.201392834 |
| NA | NA |
| 0.992060816 | NA |
| 0.571177422 | 0.71353311 |
| 0.001447471 | 0.006809131 |
| 0.070592461 | 0.157562379 |
| 0.807675443 | NA |
| 0.913012263 | 0.9507312 |
| 0.983512969 | NA |
| 0.936146081 | 0.963390749 |
| 0.578237741 | 0.719324868 |
| 0.05079042 | 0.122885284 |
| 3.38E-12 | 2.13E-10 |
| 0.905158739 | NA |
| NA | NA |
| NA | NA |

| | | |
|---|---|---|
| | 0.753640931 | NA |
| | 0.581386097 | NA |
| | 2.90E-06 | 3.31E-05 |
| | 0.422761332 | 0.584256638 |
| NA | | NA |
| | 0.171537539 | 0.308060369 |
| | 1.68E-06 | 2.05E-05 |
| | 0.482359915 | 0.637811849 |
| | 0.337117837 | 0.500119618 |
| | 0.477506164 | 0.633491896 |
| | 0.971295925 | NA |
| | 0.89503558 | NA |
| | 0.470278971 | 0.626743699 |
| | 0.001848122 | 0.008374519 |
| | 0.488715515 | NA |
| | 0.249017282 | 0.402752486 |
| | 0.889409773 | NA |
| | 0.085096415 | 0.182077916 |
| | 0.000904491 | 0.004578541 |
| | 0.055190241 | 0.130764189 |
| | 0.07328289 | 0.162236011 |
| NA | | NA |
| | 0.898638038 | NA |
| | 0.101136088 | 0.207808261 |
| | 0.331133036 | 0.493508236 |
| | 0.508094348 | 0.659879571 |
| NA | | NA |
| NA | | NA |
| NA | | NA |
| | 0.99205891 | NA |
| NA | | NA |
| | 0.499692898 | 0.652751715 |
| | 0.804558367 | 0.882580395 |
| | 0.17833956 | 0.317380533 |
| | 0.711777179 | 0.820520994 |
| | 0.70765437 | 0.817782279 |
| | 0.98794914 | 0.993639148 |
| | 0.773695314 | 0.863642507 |
| NA | | NA |
| | 0.424025546 | 0.585408359 |
| | 0.165651269 | 0.300027731 |
| | 0.029502106 | 0.080404991 |
| | 0.759031831 | NA |
| | 0.929613309 | 0.959636078 |
| | 0.387361619 | 0.551023791 |
| | 0.905158266 | NA |
| NA | | NA |

| | | |
|---|---|---|
| | 0.850172692 | NA |
| NA | | NA |
| | 0.671274182 | NA |
| NA | | NA |
| | 0.013961476 | 0.044023558 |
| NA | | NA |
| | 0.462543179 | 0.619819714 |
| | 0.905158266 | NA |
| | 0.043229672 | 0.108340602 |
| | 0.006121098 | 0.022624673 |
| | 2.00E-12 | 1.33E-10 |
| | 0.824657519 | NA |
| | 0.999445948 | 0.999810195 |
| | 0.009928671 | 0.033418713 |
| | 0.798709671 | NA |
| | 0.915196649 | 0.952049041 |
| | 0.178314596 | 0.317361826 |
| | 0.766750286 | 0.85847587 |
| | 0.507547655 | NA |
| | 0.503718423 | NA |
| | 0.784300484 | 0.870114291 |
| | 0.611216558 | 0.744964174 |
| | 0.400195943 | 0.563838687 |
| | 0.334698636 | 0.497520004 |
| | 0.995377171 | 0.997694248 |
| NA | | NA |
| | 0.142048792 | 0.267471225 |
| | 0.387093004 | 0.550921983 |
| | 0.000321989 | 0.001900175 |
| | 0.133204442 | 0.254616673 |
| | 0.712459212 | 0.820959728 |
| NA | | NA |
| | 0.99205891 | NA |
| | 0.812227822 | 0.887771737 |
| | 0.988721799 | 0.993927811 |
| NA | | NA |
| | 0.006253746 | 0.023037055 |
| | 0.569118182 | 0.71146373 |
| | 0.124824617 | 0.243020105 |
| | 0.005878594 | 0.021909037 |
| | 0.018044767 | 0.05410966 |
| | 2.42E-06 | 2.83E-05 |
| | 0.018639011 | 0.055482451 |
| | 0.290788297 | 0.450248217 |
| NA | | NA |
| | 0.780380112 | 0.867735439 |
| | 0.250000674 | 0.40392648 |

| | | |
|---|---|---|
| | 0.824217541 | 0.895142918 |
| | 1.27E-08 | 2.91E-07 |
| | 0.915938326 | 0.95229975 |
| | 0.981118192 | NA |
| | 0.828981888 | NA |
| | 0.578719418 | 0.719516459 |
| | 0.254498027 | 0.408996061 |
| | 0.853523592 | 0.914492538 |
| | 0.001096127 | 0.005381144 |
| NA | | NA |
| NA | | NA |
| | 0.113995797 | 0.227421978 |
| | 0.290857018 | 0.450322869 |
| | 0.052252089 | 0.125536502 |
| | 0.413498082 | NA |
| | 1.90E-05 | 0.000170746 |
| | 0.694761169 | 0.808321528 |
| | 0.977486538 | 0.98842452 |
| | 0.790842629 | NA |
| NA | | NA |
| | 0.20996534 | 0.356749373 |
| | 0.12728232 | 0.24653722 |
| | 0.610500642 | NA |
| NA | | NA |
| | 0.992059387 | NA |
| NA | | NA |
| | 0.062167406 | 0.143622732 |
| NA | | NA |
| | 0.704788734 | 0.816268766 |
| | 0.992061769 | NA |
| | 0.905158739 | NA |
| | 0.701487036 | 0.813586893 |
| NA | | NA |
| | 0.005844485 | 0.02181524 |
| | 0.236254712 | NA |
| | 0.874215194 | NA |
| | 0.348263869 | 0.512122568 |
| | 0.992061292 | NA |
| | 0.015468742 | 0.047875703 |
| NA | | NA |
| | 0.015049283 | 0.046828286 |
| | 0.583782778 | 0.723639366 |
| | 0.811213603 | NA |
| | 5.26E-20 | 2.31E-17 |
| NA | | NA |
| | 0.694899232 | NA |
| | 0.949878339 | NA |

| | | |
|---|---|---|
| | 0.099741237 | 0.205591326 |
| | 0.757219941 | NA |
| NA | | NA |
| | 0.816827192 | NA |
| | 0.75310421 | 0.849610222 |
| NA | | NA |
| | 0.673461443 | NA |
| | 0.555506173 | NA |
| | 0.864514552 | 0.921887084 |
| | 0.90244696 | NA |
| NA | | NA |
| | 0.797342596 | NA |
| | 0.431591784 | 0.592517911 |
| | 0.905158739 | NA |
| NA | | NA |
| NA | | NA |
| | 1.27E-10 | 5.25E-09 |
| NA | | NA |
| NA | | NA |
| NA | | NA |
| | 0.168700801 | 0.304296114 |
| | 0.006189309 | 0.022838353 |
| | 0.565578181 | NA |
| | 0.99205891 | NA |
| | 6.53E-05 | 0.000485835 |
| | 0.904947233 | NA |
| NA | | NA |
| | 0.17800538 | 0.317058924 |
| | 0.782445398 | 0.868948491 |
| | 0.72444734 | NA |
| | 0.847402695 | 0.910377484 |
| | 0.508864458 | 0.660606245 |
| | 0.623445394 | NA |
| | 0.489441012 | 0.643648052 |
| | 0.370961989 | 0.535436717 |
| | 0.889409773 | NA |
| | 0.234220805 | 0.385684723 |
| | 0.000887689 | 0.004500755 |
| | 0.567433045 | 0.710188785 |
| | 0.011453973 | 0.037510516 |
| | 0.005385597 | 0.02039709 |
| | 0.393081455 | NA |
| | 0.878885557 | NA |
| | 0.042203071 | 0.106435883 |
| | 0.048871007 | 0.119200094 |
| | 1.15E-06 | 1.48E-05 |
| | 0.020107876 | 0.059062179 |

| | | |
|---|---|---|
| | 0.171345351 | NA |
| | 0.283522545 | 0.442330757 |
| | 0.158412086 | 0.290147259 |
| | 0.780841267 | NA |
| NA | | NA |
| NA | | NA |
| | 0.557654429 | 0.702597125 |
| | 0.349572384 | NA |
| | 0.616746062 | NA |
| NA | | NA |
| | 0.781462983 | NA |
| | 0.003783722 | 0.015164583 |
| | 0.605850368 | NA |
| NA | | NA |
| NA | | NA |
| | 0.992059387 | NA |
| | 0.761491057 | NA |
| | 0.992061292 | NA |
| | 0.337825742 | 0.500696239 |
| | 0.373103945 | 0.53735396 |
| | 0.320144615 | NA |
| | 0.446421583 | 0.605757371 |
| | 0.064208859 | 0.147070235 |
| | 3.41E-07 | 5.14E-06 |
| | 0.412415114 | 0.574914906 |
| | 0.451353549 | 0.609661842 |
| | 0.044014157 | 0.10984281 |
| | 0.483864825 | 0.639338968 |
| | 0.125360958 | NA |
| | 0.00013611 | 0.000907911 |
| | 0.502174828 | NA |
| | 9.69E-21 | 5.19E-18 |
| | 0.540901098 | 0.688240074 |
| | 0.905158739 | NA |
| NA | | NA |
| | 0.98996066 | NA |
| | 0.906372871 | 0.947222629 |
| | 0.206890774 | 0.353494748 |
| | 1.16E-09 | 3.58E-08 |
| NA | | NA |
| | 0.529137506 | 0.678592999 |
| | 0.163993977 | 0.297787458 |
| | 0.046193234 | 0.114152288 |
| | 1.07E-05 | 0.000103369 |
| | 0.770272973 | 0.861091697 |
| NA | | NA |
| | 1.15E-05 | 0.000109972 |

| | | |
|---|---|---|
| | 0.139151582 | 0.263325542 |
| | 0.467536758 | 0.623846368 |
| | 0.148240172 | 0.276176252 |
| | 0.009660142 | 0.032710417 |
| | 3.29E-09 | 8.98E-08 |
| | 0.124146668 | 0.242172576 |
| | 0.004735554 | 0.018320743 |
| | 0.054110985 | 0.128852435 |
| | 0.023797148 | 0.067636757 |
| | 0.98620147 | 0.992938931 |
| | 0.514645418 | 0.665868348 |
| | 0.650659159 | 0.775164089 |
| | 2.90E-10 | 1.08E-08 |
| | 5.71E-10 | 1.94E-08 |
| | 0.645036238 | 0.770418315 |
| | 0.372529697 | NA |
| | 0.329417156 | NA |
| NA | | NA |
| | 0.545017883 | NA |
| | 0.444716457 | 0.604152975 |
| | 0.226371641 | 0.376212432 |
| | 0.012400354 | 0.040032251 |
| | 0.000595733 | 0.003224479 |
| | 0.012551521 | 0.040401475 |
| | 0.542184385 | 0.689353651 |
| | 0.010976843 | 0.03630128 |
| | 0.069111747 | 0.154922913 |
| | 0.507028593 | NA |
| | 0.773007641 | 0.863050375 |
| | 0.010035938 | 0.033753892 |
| | 0.415012546 | 0.577446489 |
| | 0.167507514 | 0.302616002 |
| | 0.206805565 | 0.353376655 |
| NA | | NA |
| NA | | NA |
| | 0.804933315 | NA |
| | 0.027142401 | 0.07521706 |
| | 0.787830833 | 0.871842829 |
| | 0.967452994 | NA |
| | 0.868790481 | NA |
| | 0.415628478 | 0.578120336 |
| | 0.829212332 | 0.898173611 |
| | 0.254563461 | 0.409041346 |
| | 0.000168733 | 0.00108913 |
| | 3.02E-07 | 4.63E-06 |
| | 0.002848914 | 0.012046851 |
| | 0.000206385 | 0.001295971 |

| | |
|---|---|
| NA | NA |
| 0.051252843 | 0.123772261 |
| 0.736715552 | NA |
| 0.846986206 | 0.910197225 |
| 0.700223271 | 0.812670139 |
| 0.955675344 | 0.975306728 |
| 0.590952453 | 0.728875998 |
| 0.832308402 | NA |
| 0.342607414 | 0.506007278 |
| 0.004371978 | 0.017098236 |
| NA | NA |
| NA | NA |
| 0.986669997 | 0.993138883 |
| NA | NA |
| 0.889409773 | NA |
| NA | NA |
| 0.9794289 | NA |
| 0.562418283 | 0.706174743 |
| 0.109837735 | 0.221115403 |
| 0.026266372 | 0.073260924 |
| NA | NA |
| 0.016220501 | NA |
| 0.889409773 | NA |
| 0.745638301 | NA |
| 0.07036582 | 0.157252496 |
| 0.025823673 | 0.072359556 |
| 0.80123444 | NA |
| 0.767071281 | 0.858646126 |
| 0.350809407 | 0.515003594 |
| 0.059458466 | 0.138662751 |
| NA | NA |
| NA | NA |
| 0.467583792 | 0.623871217 |
| 0.005199511 | 0.01980504 |
| NA | NA |
| 0.045207054 | 0.112220405 |
| 0.327045699 | 0.489209571 |
| 0.072223559 | 0.160354756 |
| 0.002173761 | 0.009596628 |
| NA | NA |
| 0.261666878 | 0.417128346 |
| 0.549163156 | 0.695360082 |
| 0.905158739 | NA |
| 0.891862982 | NA |
| 0.819792551 | 0.892907912 |
| 0.57770712 | 0.718990629 |
| NA | NA |

| | |
|---|---|
| NA | NA |
| 0.000801935 | 0.004135674 |
| 0.000125232 | 0.000845362 |
| 0.128672764 | NA |
| 0.905158739 | NA |
| 0.850852354 | 0.912565539 |
| 0.434285629 | 0.594951534 |
| 0.112320689 | 0.224672072 |
| 0.905158266 | NA |
| 2.87E-07 | 4.44E-06 |
| 0.757872105 | 0.853179204 |
| NA | NA |
| 0.002161036 | 0.009551969 |
| 0.83993405 | 0.905188055 |
| 0.74200776 | 0.84192365 |
| 0.847685183 | 0.910497043 |
| NA | NA |
| 0.919308106 | 0.953979252 |
| 0.790750881 | 0.873753263 |
| NA | NA |
| 0.966151099 | 0.9816181 |
| 0.426826039 | 0.587960164 |
| 0.004832365 | 0.018642641 |
| 0.000406733 | 0.002319254 |
| 4.03E-11 | 1.90E-09 |
| 0.360696459 | 0.52533883 |
| 0.587791958 | NA |
| 0.992061292 | NA |
| 0.788582525 | NA |
| 0.08979319 | 0.189777296 |
| NA | NA |
| 2.17E-05 | 0.000190675 |
| 0.608843132 | NA |
| 0.002872234 | 0.012124451 |
| 0.044160806 | 0.110108679 |
| NA | NA |
| 0.876979881 | NA |
| 0.621164219 | NA |
| 0.000347929 | 0.002033041 |
| 0.485494394 | 0.640452445 |
| 0.917441565 | 0.952997691 |
| 0.904947233 | NA |
| 0.515015009 | 0.666266913 |
| 0.625269212 | 0.755340311 |
| 1.52E-10 | 6.16E-09 |
| 0.148616145 | NA |
| 0.000670395 | 0.003562732 |

| | |
|---|---|
| NA | NA |
| 0.254529785 | 0.409017164 |
| 0.646599269 | 0.771625093 |
| 0.083514739 | 0.179653227 |
| 0.889283189 | NA |
| 0.323461843 | 0.485792961 |
| 0.992061292 | NA |
| NA | NA |
| 0.43826655 | 0.598612625 |
| 0.071582904 | 0.159215919 |
| 5.06E-05 | 0.000390317 |
| 0.567759935 | NA |
| 0.765094843 | NA |
| 0.858158249 | 0.917175946 |
| 5.69E-05 | 0.000432902 |
| 0.929990993 | 0.959774534 |
| 3.71E-11 | 1.76E-09 |
| 0.905158739 | NA |
| NA | NA |
| 0.905158739 | NA |
| NA | NA |
| NA | NA |
| 0.003064002 | 0.012779187 |
| 0.147090275 | 0.274679424 |
| NA | NA |
| 0.476911075 | 0.633084892 |
| 0.161671593 | 0.294569077 |
| 0.968387088 | NA |
| 0.006792106 | 0.02464032 |
| 0.003491924 | 0.014227783 |
| 0.612500989 | 0.745849787 |
| 0.828427593 | NA |
| 1.33E-07 | 2.27E-06 |
| 0.013110699 | 0.041772758 |
| 0.727500258 | NA |
| 0.079814693 | 0.173529786 |
| 0.441761827 | NA |
| 0.522559856 | NA |
| 0.297943621 | 0.457904814 |
| 0.031766333 | 0.08530721 |
| 0.089572197 | 0.189454428 |
| 0.007562645 | 0.026837125 |
| 0.878564773 | 0.931100572 |
| NA | NA |
| 0.090884169 | 0.191546738 |
| NA | NA |
| 0.283803616 | 0.442593235 |

| | |
|---|---|
| NA | NA |
| 0.00045999 | 0.002569553 |
| 0.556500177 | NA |
| 0.391125666 | 0.554505035 |
| 0.046758578 | 0.115057331 |
| 0.037120084 | 0.096475312 |
| 0.637867216 | NA |
| 0.81929321 | 0.892580849 |
| 0.009825348 | 0.033142062 |
| 0.001641369 | 0.007540339 |
| 0.719240485 | NA |
| NA | NA |
| 0.283808705 | 0.442593235 |
| 0.446997718 | 0.606239447 |
| 4.10E-08 | 8.20E-07 |
| 0.168070659 | 0.303383773 |
| NA | NA |
| 0.523651676 | 0.673880783 |
| NA | NA |
| 0.045224829 | 0.112239153 |
| 0.899072773 | 0.943310432 |
| NA | NA |
| 0.433543224 | 0.594371492 |
| 0.722013582 | NA |
| 0.457975061 | NA |
| 0.905158266 | NA |
| 0.0873905 | 0.185860337 |
| 0.532752129 | 0.681759781 |
| 0.014828129 | 0.046238409 |
| 0.004827935 | 0.018628822 |
| 0.444151044 | 0.603721001 |
| 0.00173276 | 0.007910538 |
| 0.693917595 | NA |
| 0.027443467 | 0.075879009 |
| 0.090041001 | 0.190209798 |
| 0.889409301 | NA |
| 0.006610176 | 0.02409976 |
| 0.668280401 | NA |
| 1.10E-07 | 1.91E-06 |
| NA | NA |
| NA | NA |
| 0.230807846 | 0.381649811 |
| NA | NA |
| 0.022884286 | 0.065585491 |
| 0.585167608 | NA |
| 0.991521864 | NA |
| NA | NA |

| | | |
|---|---|---|
| | 0.802218565 | NA |
| NA | | NA |
| | 0.992060816 | NA |
| | 0.905158266 | NA |
| | 0.542655884 | NA |
| | 0.934676117 | NA |
| | 0.919237702 | 0.953979252 |
| NA | | NA |
| | 0.003420617 | 0.013987583 |
| | 0.010332857 | 0.034593566 |
| | 0.208824008 | 0.355635675 |
| | 0.905158739 | NA |
| | 0.13557408 | 0.25816574 |
| NA | | NA |
| NA | | NA |
| | 0.379369323 | 0.543735459 |
| NA | | NA |
| | 0.169706333 | 0.30573315 |
| | 0.989582723 | NA |
| | 0.068264747 | 0.15347861 |
| | 0.29458651 | NA |
| NA | | NA |
| | 0.813836939 | NA |
| | 0.476429116 | 0.632674587 |
| | 0.99205891 | NA |
| | 0.849472078 | NA |
| | 0.741608815 | NA |
| | 0.822183965 | NA |
| | 0.850031586 | NA |
| | 0.399066407 | 0.562638794 |
| | 0.850166209 | 0.912186054 |
| | 0.155921662 | 0.286877568 |
| | 0.156063456 | 0.286994174 |
| | 0.011606346 | 0.037884898 |
| | 0.97032579 | NA |
| | 0.989651219 | NA |
| | 0.889409301 | NA |
| | 0.835657927 | 0.902218025 |
| | 0.237475557 | 0.389362037 |
| | 0.807879735 | NA |
| | 0.413110029 | NA |
| | 0.133226093 | 0.254635893 |
| | 0.121225429 | 0.237931568 |
| | 0.069612801 | 0.155823395 |
| | 0.341490957 | 0.50483371 |
| | 0.037064674 | 0.09636552 |
| | 0.889409301 | NA |

| | |
|---|---|
| 2.47E-09 | 6.99E-08 |
| 4.36E-06 | 4.74E-05 |
| 0.033340268 | 0.08864863 |
| NA | NA |
| 0.527524663 | 0.677303051 |
| NA | NA |
| NA | NA |
| NA | NA |
| NA | NA |
| 0.000137582 | 0.000916337 |
| 0.502505479 | 0.65498414 |
| 0.062008746 | 0.143350563 |
| 0.021446336 | NA |
| 0.471284006 | NA |
| 0.983880805 | NA |
| 0.795353865 | NA |
| 0.58565301 | NA |
| 0.021404648 | 0.062098647 |
| 0.007611245 | 0.026970361 |
| NA | NA |
| NA | NA |
| 0.783793855 | 0.869874117 |
| NA | NA |
| 0.905158739 | NA |
| 0.903401609 | NA |
| 0.992059387 | NA |
| 0.015262816 | 0.047365204 |
| 0.178034531 | 0.317068959 |
| 0.127946721 | 0.247474856 |
| 0.431710074 | 0.592569165 |
| NA | NA |
| 0.302926168 | 0.463583226 |
| 0.727425224 | NA |
| 0.192854851 | 0.335783355 |
| 0.372558094 | 0.536916925 |
| 0.584669572 | 0.724284857 |
| 0.083271743 | 0.179334197 |
| 0.02195037 | 0.063355437 |
| NA | NA |
| 0.997510458 | NA |
| NA | NA |
| 0.405936099 | 0.569171995 |
| 0.635011392 | NA |
| 0.522320022 | NA |
| 0.641692231 | 0.767479287 |
| 0.289420641 | 0.448731755 |
| 0.451759317 | NA |

| | | |
|---|---|---|
| | 3.49E-06 | 3.90E-05 |
| | 0.637984644 | 0.764627486 |
| | 0.005377584 | 0.020370256 |
| | 0.05664663 | 0.13339446 |
| | 0.69535678 | NA |
| | 0.185069064 | 0.325896679 |
| | 0.519246835 | NA |
| | 0.625325882 | NA |
| | 0.431590034 | 0.592517911 |
| | 0.052274694 | 0.125563334 |
| | 0.027349861 | 0.075683843 |
| | 0.528445486 | NA |
| | 0.589770312 | 0.728143837 |
| | 0.201306343 | 0.346514541 |
| | 0.167859474 | 0.303077305 |
| | 0.403310907 | NA |
| NA | | NA |
| | 0.547661601 | 0.693866899 |
| NA | | NA |
| NA | | NA |
| | 0.248161124 | NA |
| NA | | NA |
| NA | | NA |
| NA | | NA |
| | 0.010323561 | 0.034572988 |
| NA | | NA |
| | 0.850085113 | 0.912150258 |
| | 0.669279923 | 0.789091843 |
| | 0.748329237 | 0.846438134 |
| | 0.027551055 | 0.076147718 |
| | 0.208734031 | 0.355515694 |
| | 0.069147042 | 0.154982829 |
| NA | | NA |
| | 0.572812029 | 0.715127599 |
| | 1.37E-13 | 1.19E-11 |
| | 0.011506835 | 0.037634582 |
| NA | | NA |
| | 0.000221136 | 0.001375226 |
| NA | | NA |
| | 0.000778696 | 0.004033826 |
| | 0.002207693 | 0.009729498 |
| | 0.017179533 | 0.052098519 |
| | 0.004546463 | 0.017676306 |
| | 0.019096968 | 0.056554325 |
| | 0.066452149 | 0.150808468 |
| NA | | NA |
| | 0.006708348 | 0.024395266 |

| | | |
|---|---|---|
| | 0.004357182 | 0.017064269 |
| | 0.000530715 | 0.002912762 |
| | 0.012773638 | 0.040990255 |
| | 0.002359129 | 0.010296983 |
| | 3.76E-05 | 0.000303383 |
| | 0.003562066 | 0.0144583 |
| | 0.010758239 | 0.035739814 |
| | 0.398880039 | NA |
| | 0.012515658 | 0.040321497 |
| | 0.00196976 | 0.00883816 |
| | 0.03475411 | 0.091594515 |
| | 0.00195633 | 0.008783286 |
| | 0.010367033 | 0.034681548 |
| | 0.015167312 | 0.047142417 |
| | 0.062541381 | 0.144217444 |
| NA | | NA |
| | 0.99205891 | NA |
| | 0.055889293 | 0.132022872 |
| | 0.182284713 | 0.322441599 |
| | 0.381079268 | NA |
| | 0.898935586 | NA |
| | 0.812569181 | 0.887986794 |
| | 0.182858743 | NA |
| NA | | NA |
| | 0.992060816 | NA |
| NA | | NA |
| | 0.779500903 | NA |
| | 0.992061292 | NA |
| | 0.992061292 | NA |
| | 0.905158739 | NA |
| NA | | NA |
| | 0.681219778 | NA |
| NA | | NA |
| NA | | NA |
| NA | | NA |
| | 2.64E-08 | 5.55E-07 |
| | 0.033728651 | 0.089396192 |
| | 0.018079038 | 0.054197625 |
| NA | | NA |
| | 0.354879703 | 0.519484363 |
| | 0.137631043 | 0.261145777 |
| | 0.028976497 | 0.079210121 |
| | 0.056794327 | 0.133641944 |
| | 0.051213779 | 0.123691528 |
| | 0.041752234 | 0.105502999 |
| | 0.535083309 | NA |
| | 0.373387113 | NA |

| | | |
|---|---|---|
| | 0.062812751 | 0.14464952 |
| | 0.885000041 | 0.934832376 |
| | 0.578620603 | 0.719478173 |
| | 0.304356312 | 0.464965413 |
| | 0.504848163 | NA |
| | 0.39413551 | NA |
| NA | | NA |
| | 0.889409301 | NA |
| | 1.28E-05 | 0.000120795 |
| | 3.36E-08 | 6.91E-07 |
| | 0.992061292 | NA |
| | 0.002846582 | 0.01203931 |
| NA | | NA |
| | 0.03799762 | 0.098186602 |
| | 0.554841687 | NA |
| | 0.08183908 | 0.177026229 |
| | 0.751116432 | 0.848416764 |
| | 5.59E-06 | 5.86E-05 |
| | 0.144710577 | 0.271273653 |
| | 0.162892749 | 0.296277799 |
| NA | | NA |
| NA | | NA |
| NA | | NA |
| | 0.106250058 | NA |
| | 0.032165702 | NA |
| NA | | NA |
| | 0.271822135 | 0.428557027 |
| NA | | NA |
| | 0.149101262 | 0.277373316 |
| | 0.440491977 | 0.600568939 |
| | 0.848179338 | NA |
| | 0.732698922 | NA |
| NA | | NA |
| | 0.000124193 | 0.000840935 |
| | 0.981355815 | NA |
| | 0.972059705 | NA |
| | 0.08647756 | 0.184365121 |
| | 0.104784775 | 0.21342447 |
| | 0.007362976 | 0.026298565 |
| | 0.227044553 | 0.377013462 |
| | 0.000179547 | 0.001148463 |
| NA | | NA |
| | 0.209702616 | 0.356468475 |
| NA | | NA |
| | 0.42268356 | 0.584232914 |
| | 0.889409773 | NA |
| NA | | NA |

| | | | |
|---|---|---|---|
| NA | | NA | |
| NA | | NA | |
| NA | | NA | |
| NA | | NA | |
| NA | | NA | |
| | 0.410568663 | | 0.573589341 |
| | 0.820813114 | NA | |
| | 0.889409301 | NA | |
| | 0.640436514 | NA | |
| | 0.777092376 | NA | |
| | 0.435890966 | NA | |
| NA | | NA | |
| NA | | NA | |
| NA | | NA | |
| NA | | NA | |
| NA | | NA | |
| NA | | NA | |
| NA | | NA | |
| | 0.905158739 | NA | |
| | 0.303844913 | | 0.464621267 |
| | 0.067774797 | | 0.152705394 |
| NA | | NA | |
| | 0.136431297 | NA | |
| | 0.000656625 | | 0.003503969 |
| | 0.157273091 | | 0.288638512 |
| | 0.569756543 | | 0.712122713 |
| | 0.090516483 | | 0.191065216 |
| | 0.736784724 | | 0.838379923 |
| | 0.168798998 | | 0.30439624 |
| | 0.066616255 | | 0.150993635 |
| | 0.799410748 | | 0.879075598 |
| | 0.844795495 | | 0.908643431 |
| | 0.952209946 | | 0.973288471 |
| | 0.078042436 | | 0.170665216 |
| | 0.559084946 | | 0.703518549 |
| | 0.72779221 | | 0.832027756 |
| | 0.298506362 | | 0.458563324 |
| | 0.40781774 | | 0.571017647 |
| NA | | NA | |
| NA | | NA | |
| NA | | NA | |
| NA | | NA | |
| NA | | NA | |
| NA | | NA | |
| NA | | NA | |
| NA | | NA | |
| NA | | NA | |

| | |
|---|---|
| NA | NA |
| NA | NA |
| 0.17330548 | 0.310511076 |
| NA | NA |
| 0.004165154 | NA |
| NA | NA |
| 0.889409301 | NA |
| NA | NA |
| 0.889409773 | NA |
| 0.992061292 | NA |
| 0.14874729 | 0.276808622 |
| NA | NA |
| NA | NA |
| NA | NA |
| NA | NA |
| 0.458910647 | 0.616192913 |
| NA | NA |
| NA | NA |
| NA | NA |
| NA | NA |
| NA | NA |
| 0.992061292 | NA |
| NA | NA |
| NA | NA |
| 0.644310955 | NA |
| NA | NA |
| 0.690760664 | 0.805274956 |
| 0.554394237 | 0.699772549 |
| 0.833253313 | 0.900794276 |
| 0.008391331 | 0.02919747 |
| 2.61E-05 | 0.000222873 |
| 0.022128211 | 0.063776531 |
| 0.615859096 | 0.748652009 |
| 0.483389534 | 0.638942441 |
| 0.006654288 | 0.024240461 |
| 0.952110177 | NA |
| 0.992061292 | NA |
| NA | NA |
| 0.070966246 | NA |
| 2.93E-07 | 4.51E-06 |
| NA | NA |
| NA | NA |
| NA | NA |
| 0.04050277 | 0.102945869 |
| 0.889409773 | NA |
| 0.080763256 | 0.175193158 |
| NA | NA |

| | | |
|---|---|---|
| | 0.088688336 | 0.188098293 |
| | 0.625884429 | NA |
| | 0.659491896 | NA |
| | 0.889409301 | NA |
| NA | | NA |
| NA | | NA |
| | 0.00168041 | 0.007697145 |
| | 0.517940043 | NA |
| | 0.868197896 | NA |
| | 0.365602651 | NA |
| | 0.698503113 | 0.811501586 |
| | 0.889410245 | NA |
| NA | | NA |
| | 0.905158266 | NA |
| | 0.953655268 | NA |
| | 0.223275795 | 0.372845109 |
| | 9.81E-06 | 9.54E-05 |
| | 0.826189235 | 0.896319493 |
| | 0.001553717 | 0.007211597 |
| | 6.21E-05 | 0.000465802 |
| | 1.17E-06 | 1.49E-05 |
| | 0.905158739 | NA |
| | 0.764591534 | NA |
| | 0.006916044 | 0.025044426 |
| | 0.470562973 | 0.626893911 |
| | 0.536234919 | 0.684404684 |
| | 0.961452103 | 0.978881988 |
| | 0.047307633 | 0.116109122 |
| | 0.247171746 | 0.40074161 |
| | 0.838125811 | NA |
| | 0.010897924 | 0.03610554 |
| | 0.150086343 | 0.278779061 |
| | 0.032431803 | 0.08674421 |
| | 0.824159341 | 0.895142918 |
| | 6.92E-06 | 7.09E-05 |
| | 0.889409773 | NA |
| | 0.170251554 | 0.306438842 |
| | 0.000362562 | 0.002102879 |
| | 0.071298301 | 0.158722567 |
| | 0.985595019 | 0.99269246 |
| | 0.58687337 | 0.725873174 |
| | 1.46E-05 | 0.000135075 |
| | 0.223523171 | 0.373144694 |
| NA | | NA |
| | 0.125229546 | 0.243657238 |
| | 0.883565349 | 0.933966088 |
| | 0.74939797 | 0.847157273 |

| | | |
|---|---|---|
| | 0.992062245 | NA |
| | 0.445075912 | 0.604416943 |
| | 0.393396926 | 0.556826292 |
| | 0.086490167 | 0.184374097 |
| | 0.081080524 | 0.175742495 |
| | 0.384228182 | 0.548192334 |
| | 0.552526538 | 0.698057088 |
| | 0.805374319 | NA |
| | 0.697117783 | NA |
| NA | | NA |
| | 0.068393283 | 0.153639352 |
| | 0.807806017 | NA |
| | 0.647853092 | NA |
| NA | | NA |
| NA | | NA |
| | 0.030350921 | NA |
| NA | | NA |
| | 4.71E-09 | 1.20E-07 |
| | 0.051015351 | 0.123328584 |
| | 0.000159342 | 0.001036009 |
| | 0.736104867 | NA |
| NA | | NA |
| NA | | NA |
| NA | | NA |
| NA | | NA |
| | 0.905158266 | NA |
| | 0.000327729 | 0.001929383 |
| | 0.27628727 | 0.43375936 |
| | 5.00E-05 | 0.000386837 |
| | 0.018829635 | 0.055936141 |
| | 0.041425563 | 0.104884576 |
| | 0.889409301 | NA |
| | 0.014462511 | 0.045291254 |
| | 0.061118508 | 0.141796229 |
| | 0.038685178 | 0.099506597 |
| | 0.275397612 | 0.432733893 |
| | 0.277472608 | 0.435371275 |
| | 0.066265731 | 0.150509845 |
| | 0.062749024 | 0.144555794 |
| | 0.00189357 | 0.008544751 |
| | 0.905158739 | NA |
| | 0.180333745 | 0.320125199 |
| | 0.055940711 | 0.132114656 |
| | 0.290396885 | 0.449769023 |
| | 0.367560098 | 0.532141534 |
| NA | | NA |
| | 0.905158739 | NA |

| | |
|---|---|
| 0.754729741 | NA |
| NA | NA |
| 0.406965104 | 0.570114623 |
| 0.285030764 | NA |
| 0.069465355 | 0.155560001 |
| 0.062228899 | 0.143734553 |
| 4.45E-05 | 0.000350553 |
| 0.701588684 | NA |
| 0.577789668 | 0.719052611 |
| 0.080223626 | 0.174298071 |
| 0.221052939 | 0.370061865 |
| NA | NA |
| 0.209523399 | 0.356274145 |
| 0.928769132 | NA |
| 0.018327379 | 0.054792501 |
| 4.80E-06 | 5.14E-05 |
| 0.349207057 | 0.513234573 |
| 0.071207812 | 0.158569349 |
| 0.934463129 | 0.96234837 |
| 0.047058381 | 0.115627257 |
| 0.140708255 | 0.265542992 |
| 0.014299885 | 0.044890805 |
| 0.435620336 | 0.596240513 |
| 0.475121162 | NA |
| 0.703665597 | NA |
| 0.100927826 | NA |
| 0.986351731 | NA |
| 0.705930283 | NA |
| 0.336317498 | NA |
| 0.844665864 | NA |
| 0.905158739 | NA |
| NA | NA |
| NA | NA |
| 0.992062245 | NA |
| 0.541949198 | 0.689134429 |
| 0.00116889 | 0.005681197 |
| 0.269200053 | 0.425602984 |
| 0.116362063 | 0.230946723 |
| 0.121621763 | 0.23843338 |
| 0.034444026 | 0.090919146 |
| 0.000522336 | 0.002872521 |
| 0.158048467 | 0.289674174 |
| 0.150264942 | 0.278994492 |
| 0.758507354 | NA |
| NA | NA |
| 0.77146932 | NA |
| 0.710939626 | NA |

| | |
|---|---|
| 0.493955079 | 0.647309363 |
| 0.794656519 | NA |
| 0.000137526 | 0.000916243 |
| 0.007142264 | 0.025652106 |
| NA | NA |
| NA | NA |
| NA | NA |
| NA | NA |
| NA | NA |
| NA | NA |
| 0.989804447 | NA |
| 0.043093843 | 0.108096855 |
| 8.95E-08 | 1.60E-06 |
| 0.112665213 | NA |
| 3.78E-10 | 1.36E-08 |
| 3.65E-09 | 9.79E-08 |
| 0.774720161 | 0.864072143 |
| 0.782559532 | NA |
| 0.26193133 | 0.417337838 |
| 0.797085978 | 0.877636997 |
| NA | NA |
| 0.055313693 | 0.130929545 |
| 0.04703173 | 0.115625552 |
| 0.992062245 | NA |
| NA | NA |
| NA | NA |
| 0.992061292 | NA |
| 0.879254371 | NA |
| 0.183843814 | 0.324441558 |
| 0.555008715 | 0.700427378 |
| 0.974044326 | NA |
| 0.383208712 | 0.547202504 |
| 0.465816431 | NA |
| 0.603749245 | NA |
| 0.115742444 | 0.230050242 |
| NA | NA |
| 0.053576547 | 0.127767963 |
| 0.041553644 | 0.105124017 |
| NA | NA |
| 1.18E-07 | 2.05E-06 |
| 0.189789716 | 0.331735443 |
| 0.55747126 | 0.702445136 |
| 8.52E-05 | 0.000607582 |
| 0.889409301 | NA |
| NA | NA |
| 0.408895667 | 0.572063478 |
| 0.108608918 | NA |

| | | |
|---|---|---|
| | 0.513223983 | NA |
| | 0.124214948 | 0.242221186 |
| | 0.293704921 | 0.453357681 |
| NA | | NA |
| | 0.641119992 | 0.767212746 |
| | 0.987216977 | 0.993414557 |
| | 0.905158739 | NA |
| | 0.992060816 | NA |
| NA | | NA |
| | 0.000359331 | 0.002085793 |
| | 0.61274899 | NA |
| | 4.54E-09 | 1.17E-07 |
| | 0.759932082 | 0.854271967 |
| | 0.052771511 | 0.126452381 |
| | 0.684866673 | NA |
| | 0.068057762 | 0.153185772 |
| | 0.726991823 | NA |
| NA | | NA |
| | 7.16E-06 | 7.30E-05 |
| | 0.098269622 | 0.203176449 |
| | 0.093608447 | 0.195785112 |
| | 0.922793299 | 0.955421158 |
| | 0.641285374 | 0.767353179 |
| NA | | NA |
| | 0.905158266 | NA |
| | 0.12819418 | 0.24782252 |
| | 0.518630815 | NA |
| | 0.889409301 | NA |
| | 0.007246574 | 0.025948692 |
| NA | | NA |
| | 0.45951184 | NA |
| | 0.994996294 | 0.997551495 |
| NA | | NA |
| | 0.145536218 | NA |
| | 0.015054901 | 0.046839128 |
| | 0.007364434 | 0.026299495 |
| | 0.945331656 | NA |
| | 0.108264098 | 0.218769793 |
| | 0.274428084 | 0.431673803 |
| | 0.436785664 | 0.597072734 |
| | 0.084173433 | 0.180560702 |
| | 0.086561399 | 0.184508033 |
| | 0.51184845 | 0.663460251 |
| | 3.89E-08 | 7.85E-07 |
| | 0.813836939 | NA |
| NA | | NA |
| NA | | NA |

| | |
|---|---|
| NA | NA |
| 0.992061292 | NA |
| 0.424483927 | 0.585725936 |
| 0.245647342 | 0.398918058 |
| 0.209479099 | 0.356226402 |
| 0.992061292 | NA |
| 0.472999686 | NA |
| NA | NA |
| 0.905158739 | NA |
| NA | NA |
| 0.869535106 | 0.925335282 |
| 0.878990255 | NA |
| NA | NA |
| 0.992061292 | NA |
| NA | NA |
| NA | NA |
| NA | NA |
| NA | NA |
| NA | NA |
| 0.12537012 | 0.243822731 |
| 0.99205891 | NA |
| 0.523209441 | NA |
| NA | NA |
| 0.917075101 | NA |
| 0.889409301 | NA |
| 0.905158266 | NA |
| NA | NA |
| 2.63E-10 | 9.85E-09 |
| NA | NA |
| NA | NA |
| 0.139881174 | 0.264340942 |
| 0.298187441 | 0.458151275 |
| 0.001927201 | 0.008668458 |
| 0.174029636 | 0.311325197 |
| 0.011309175 | 0.037140034 |
| 0.974803066 | 0.986909724 |
| 0.017945957 | 0.053894201 |
| 0.333155196 | 0.495714524 |
| 0.905157793 | NA |
| 0.489509554 | NA |
| 0.76207359 | NA |
| NA | NA |
| 0.992060816 | NA |
| 0.928969442 | NA |
| 0.269933524 | 0.426382492 |
| 0.590701 | 0.728760226 |
| 0.996393986 | 0.998242688 |

| | | |
|---|---|---|
| | 0.333734129 | 0.496374136 |
| | 0.992061292 | NA |
| | 0.852873728 | 0.914286965 |
| | 0.00216293 | 0.009554571 |
| | 0.947395557 | NA |
| | 0.216181811 | 0.364072121 |
| | 0.795330562 | 0.87663136 |
| | 0.158076981 | 0.289678079 |
| NA | | NA |
| | 0.890946539 | 0.938380229 |
| | 0.896586191 | 0.941790776 |
| | 0.032834324 | 0.087554211 |
| | 0.989952082 | NA |
| NA | | NA |
| | 0.026124792 | NA |
| NA | | NA |
| | 2.92E-06 | 3.33E-05 |
| | 0.849259666 | NA |
| | 0.217256655 | 0.36548984 |
| | 0.412031664 | 0.574682628 |
| | 0.18648088 | 0.327725781 |
| | 3.98E-08 | 7.99E-07 |
| NA | | NA |
| | 0.992003131 | NA |
| NA | | NA |
| NA | | NA |
| | 0.783383701 | NA |
| NA | | NA |
| | 0.718980365 | NA |
| NA | | NA |
| | 0.855678602 | NA |
| NA | | NA |
| NA | | NA |
| NA | | NA |
| NA | | NA |
| NA | | NA |
| NA | | NA |
| NA | | NA |
| NA | | NA |
| | 0.012148393 | NA |
| | 0.044677633 | 0.111119736 |
| | 0.407624891 | 0.570820418 |
| | 0.301397461 | 0.461790877 |
| | 0.067786356 | 0.152715768 |
| | 0.652119325 | NA |
| | 0.008861116 | 0.030510012 |
| | 0.99205891 | NA |

| | | |
|---|---|---|
| | 0.905158266 | NA |
| | 0.103831481 | 0.211766852 |
| NA | | NA |
| NA | | NA |
| NA | | NA |
| | 0.583083114 | NA |
| | 0.992062245 | NA |
| NA | | NA |
| NA | | NA |
| NA | | NA |
| NA | | NA |
| NA | | NA |
| | 0.003331228 | 0.013680652 |
| | 0.827132407 | 0.89672859 |
| | 0.335938958 | 0.498808815 |
| NA | | NA |
| NA | | NA |
| | 0.983323808 | NA |
| NA | | NA |
| | 0.40068257 | NA |
| | 0.003409036 | 0.01395582 |
| | 0.989651219 | NA |
| | 0.002443533 | 0.010614745 |
| | 0.807879735 | NA |
| | 1.85E-07 | 3.03E-06 |
| | 0.833748148 | NA |
| | 0.992061292 | NA |
| NA | | NA |
| | 0.719522939 | NA |
| | 0.000359938 | 0.002088762 |
| | 0.909364283 | NA |
| | 0.70853139 | NA |
| | 0.403694127 | 0.56711195 |
| | 0.889410245 | NA |
| | 0.626522117 | NA |
| | 0.90515732 | NA |
| NA | | NA |
| | 0.905157793 | NA |
| | 0.291115415 | 0.450581169 |
| NA | | NA |
| | 0.559153689 | NA |
| NA | | NA |
| NA | | NA |
| NA | | NA |
| NA | | NA |
| | 0.99205891 | NA |
| | 0.248995835 | 0.402752208 |

| | | |
|---|---|---|
| | 0.779501355 | NA |
| NA | | NA |
| NA | | NA |
| | 0.996453356 | NA |
| NA | | NA |
| NA | | NA |
| | 0.581912745 | NA |
| | 0.889409301 | NA |
| NA | | NA |
| | 0.889409301 | NA |
| | 0.905158266 | NA |
| | 0.889409773 | NA |
| | 0.889409301 | NA |
| NA | | NA |
| NA | | NA |
| | 0.992061769 | NA |
| NA | | NA |
| NA | | NA |
| | 0.784353866 | NA |
| | 0.889409301 | NA |
| NA | | NA |
| NA | | NA |
| | 0.992061292 | NA |
| | 0.367094118 | 0.531807781 |
| | 0.905158739 | NA |
| NA | | NA |
| NA | | NA |
| NA | | NA |
| | 0.889409773 | NA |
| NA | | NA |
| NA | | NA |
| | 0.81035666 | NA |
| NA | | NA |
| | 0.889409773 | NA |
| NA | | NA |
| NA | | NA |
| NA | | NA |
| NA | | NA |
| NA | | NA |
| NA | | NA |
| NA | | NA |
| | 0.090286163 | NA |
| | 0.992061844 | NA |
| | 0.75954113 | NA |
| NA | | NA |
| | 0.904947233 | NA |
| NA | | NA |

| | |
|---|---|
| NA | NA |
| NA | NA |
| NA | NA |
| NA | NA |
| NA | NA |
| NA | NA |
| NA | NA |
| NA | NA |
| NA | NA |
| NA | NA |
| NA | NA |
| NA | NA |
| 0.698648156 | NA |
| NA | NA |
| NA | NA |
| NA | NA |
| 0.137055858 | 0.260279304 |
| 0.332274229 | NA |
| 0.648411426 | 0.773152249 |
| 0.020721257 | 0.060483594 |
| 0.14159253 | 0.266772232 |
| 0.543546429 | 0.690365899 |
| 0.131388644 | 0.252157146 |
| 4.98E-08 | 9.67E-07 |
| 0.77522924 | 0.864399219 |
| NA | NA |
| 0.561318515 | 0.705430852 |
| 0.218950083 | 0.367582897 |
| 0.296807808 | NA |
| 0.889409301 | NA |
| NA | NA |
| 0.31000056 | 0.471224027 |
| 0.009062773 | 0.031056403 |
| 0.846109393 | NA |
| NA | NA |
| 0.908951426 | 0.94824763 |
| 0.600924222 | 0.736613542 |
| 0.94534817 | 0.969042261 |
| 8.71E-05 | 0.000619688 |
| NA | NA |
| 0.979626862 | 0.989677322 |
| 0.148256554 | 0.276176252 |
| 0.889409773 | NA |
| 0.329301712 | NA |
| 0.985472897 | NA |
| NA | NA |
| 0.716675231 | NA |

| | | |
|---|---|---|
| | 0.570012969 | 0.712362118 |
| | 0.799157091 | NA |
| | 0.008158218 | 0.028531027 |
| | 0.370970181 | 0.535436717 |
| | 0.11321141 | 0.226124191 |
| | 6.54E-05 | 0.000486378 |
| | 0.992060816 | NA |
| NA | | NA |
| NA | | NA |
| NA | | NA |
| | 0.580349104 | 0.72060424 |
| | 0.689055429 | NA |
| | 0.498825884 | 0.651813007 |
| NA | | NA |
| | 0.734771119 | 0.836956064 |
| | 0.978033767 | NA |
| | 0.619315405 | NA |
| | 0.000539744 | 0.002954933 |
| | 0.491817591 | NA |
| | 0.970037629 | 0.984200725 |
| | 0.215306022 | 0.363042689 |
| | 0.000213098 | 0.001332027 |
| | 6.93E-06 | 7.09E-05 |
| | 0.925376103 | 0.957061693 |
| | 0.521876071 | 0.672409591 |
| | 0.486531841 | NA |
| | 0.826599339 | NA |
| | 0.931984807 | NA |
| NA | | NA |
| | 0.477478405 | 0.633491896 |
| | 0.611629066 | NA |
| | 0.001304768 | 0.006224506 |
| | 0.983147384 | 0.991763981 |
| | 0.00389637 | 0.01553384 |
| | 0.889410717 | NA |
| | 0.634477634 | 0.762046401 |
| | 0.003597666 | 0.014578549 |
| | 0.713798068 | 0.821762178 |
| | 0.942824884 | 0.967678974 |
| | 0.065302217 | 0.148999519 |
| | 0.306926308 | NA |
| | 0.00060205 | 0.003251373 |
| | 0.008219607 | 0.028695444 |
| NA | | NA |
| | 0.729235771 | NA |
| | 0.905158739 | NA |
| | 0.004122705 | 0.016300049 |

| | |
|---|---|
| 0.534462576 | 0.683095495 |
| 0.797273258 | 0.877636997 |
| 0.905158266 | NA |
| 0.889409301 | NA |
| 0.678199526 | 0.795841575 |
| 0.211475198 | 0.358510297 |
| 0.359974156 | 0.52459998 |
| 0.013680776 | 0.043268929 |
| 0.060814573 | NA |
| 0.01359874 | 0.043059083 |
| NA | NA |
| 0.089050948 | 0.188647216 |
| NA | NA |
| 0.923506567 | 0.95584845 |
| 0.013301011 | 0.042270994 |
| 0.065947846 | 0.149975195 |
| 7.40E-05 | 0.00054027 |
| 0.740822298 | NA |
| NA | NA |
| NA | NA |
| 0.590205463 | NA |
| 0.905158266 | NA |
| 0.404173486 | 0.567470944 |
| 0.842184292 | 0.906642361 |
| 0.039791084 | 0.101707883 |
| 0.306503769 | 0.467267167 |
| 1.22E-12 | 8.48E-11 |
| 9.29E-07 | 1.23E-05 |
| 0.148776775 | 0.27684003 |
| 0.279280971 | 0.437465749 |
| 0.856020202 | 0.916094722 |
| 0.650152494 | 0.774728599 |
| 0.280198866 | 0.438395932 |
| 0.063258023 | NA |
| NA | NA |
| NA | NA |
| NA | NA |
| NA | NA |
| NA | NA |
| NA | NA |
| NA | NA |
| NA | NA |
| NA | NA |
| NA | NA |
| NA | NA |
| NA | NA |
| NA | NA |

| | | | |
|---|---|---|---|
| NA | | NA | |
| NA | | NA | |
| NA | | NA | |
| NA | | NA | |
| NA | | NA | |
| NA | | NA | |
| NA | | NA | |
| NA | | NA | |
| NA | | NA | |
| NA | | NA | |
| NA | | NA | |
| NA | | NA | |
| NA | | NA | |
| NA | | NA | |
| NA | | NA | |
| NA | | NA | |
| NA | | NA | |
| NA | | NA | |
| NA | | NA | |
| NA | | NA | |
| NA | | NA | |
| NA | | NA | |
| NA | | NA | |
| NA | | NA | |
| NA | | NA | |
| NA | | NA | |
| NA | | NA | |
| NA | | NA | |
| NA | | NA | |
| NA | | NA | |
| NA | | NA | |
| NA | | NA | |
| NA | | NA | |
| NA | | NA | |
| NA | | NA | |
| NA | | NA | |
| | 0.163491841 | | 0.297096768 |
| NA | | NA | |
| | 0.030370051 | | 0.082292196 |
| | 3.81E-05 | | 0.000306826 |
| | 0.550272778 | | 0.696121036 |
| | 0.017040268 | | 0.051783454 |
| | 0.963351373 | | 0.979770867 |
| | 0.009115268 | | 0.031181872 |
| | 0.989037923 | NA | |
| | 0.391903656 | NA | |
| | 0.830163713 | NA | |

| | | |
|---|---|---|
| | 2.68E-14 | 2.70E-12 |
| | 0.242560236 | 0.395454765 |
| | 0.863555238 | NA |
| | 0.805655139 | NA |
| | 0.875289692 | NA |
| NA | | NA |
| | 0.110410904 | 0.221842167 |
| | 0.268365947 | NA |
| | 0.607515098 | NA |
| | 0.877359144 | NA |
| | 0.472211766 | NA |
| | 0.465140209 | 0.622085014 |
| | 0.000708567 | 0.003728594 |
| | 0.197267745 | 0.34150374 |
| | 0.005890048 | 0.021942411 |
| | 0.043105144 | 0.108102543 |
| | 0.792415662 | 0.874888407 |
| | 0.568017605 | 0.710680206 |
| NA | | NA |
| | 0.63977508 | NA |
| | 6.97E-08 | 1.29E-06 |
| | 0.992061292 | NA |
| | 4.31E-12 | 2.62E-10 |
| | 0.734024349 | NA |
| | 0.992061292 | NA |
| NA | | NA |
| | 0.834975139 | NA |
| | 0.784353866 | NA |
| | 3.57E-07 | 5.34E-06 |
| | 0.022553437 | 0.064823419 |
| | 0.141105835 | 0.266037866 |
| | 0.615427408 | 0.748332145 |
| | 0.227876543 | NA |
| | 0.994720423 | NA |
| | 0.025865794 | 0.072447317 |
| | 0.885483 | NA |
| | 0.745609685 | NA |
| NA | | NA |
| | 0.019773533 | 0.058259359 |
| | 0.35871904 | 0.523535252 |
| | 0.66953237 | NA |
| | 0.450904182 | NA |
| | 0.031522287 | 0.084777127 |
| | 0.092881712 | 0.19461732 |
| | 0.219442497 | 0.368184427 |
| | 0.991827912 | NA |
| | 0.995604265 | 0.997694248 |

| | |
|---|---|
| 0.320256441 | 0.482571274 |
| NA | NA |
| NA | NA |
| NA | NA |
| NA | NA |
| 0.002138547 | 0.009471632 |
| 0.905158739 | NA |
| NA | NA |
| 0.87381992 | NA |
| 0.801661681 | NA |
| 0.905157793 | NA |
| 0.007051252 | 0.025404176 |
| 0.997667299 | 0.998967988 |
| 0.181692507 | 0.32188671 |
| 0.001641295 | 0.007540339 |
| 0.393242213 | 0.556750855 |
| NA | NA |
| 0.000564601 | 0.003074916 |
| 0.897856823 | NA |
| 0.402110541 | NA |
| 0.815309261 | NA |
| 0.979578099 | NA |
| 0.000643632 | 0.003444677 |
| 0.874557586 | 0.928377165 |
| 0.873177717 | NA |
| 0.905158739 | NA |
| NA | NA |
| 0.722436794 | NA |
| 0.710282665 | 0.819476542 |
| 0.244568427 | 0.397695356 |
| 0.939996751 | NA |
| NA | NA |
| NA | NA |
| 0.76008157 | 0.854396273 |
| 9.14E-05 | 0.000646252 |
| NA | NA |
| 0.95476734 | 0.97468323 |
| 0.774986494 | NA |
| 1.72E-07 | 2.84E-06 |
| 0.155108737 | 0.285788955 |
| 0.089603273 | 0.189481684 |
| 0.33606494 | 0.498962138 |
| 0.590182953 | 0.728316346 |
| 0.470029783 | 0.626449623 |
| 0.546817485 | NA |
| 0.874619715 | NA |
| 0.539889054 | NA |

| | |
|---|---|
| 3.50E-10 | 1.27E-08 |
| 0.318909858 | 0.481367994 |
| 0.889409773 | NA |
| 0.889409301 | NA |
| NA | NA |
| NA | NA |
| 0.029593241 | 0.080595062 |
| NA | NA |
| 0.201776004 | NA |
| 7.78E-07 | 1.06E-05 |
| 0.265744708 | 0.421913676 |
| 0.001608517 | 0.007422025 |
| 0.084413383 | 0.180877584 |
| 0.865640951 | 0.922524127 |
| 0.795353865 | NA |
| 0.062502512 | 0.144169396 |
| 0.779627684 | 0.86734378 |
| 0.394117322 | NA |
| 0.8387816 | 0.90421233 |
| 0.775643553 | NA |
| 0.756187079 | NA |
| 0.020562695 | 0.060148691 |
| 2.31E-08 | 4.94E-07 |
| 0.012881752 | 0.041229683 |
| 2.56E-05 | 0.000219316 |
| NA | NA |
| NA | NA |
| NA | NA |
| NA | NA |
| 0.137549463 | 0.261036095 |
| 0.017462089 | 0.052749353 |
| NA | NA |
| 0.889409301 | NA |
| 0.776230563 | 0.865109219 |
| NA | NA |
| 0.507087804 | NA |
| 0.113668934 | NA |
| NA | NA |
| 0.001544362 | 0.007174242 |
| 0.23808421 | 0.390068728 |
| 0.875476463 | 0.928844683 |
| NA | NA |
| 0.294129421 | NA |
| NA | NA |
| 0.036150361 | 0.094474993 |
| 0.578074772 | 0.719196874 |
| 0.051042545 | 0.12336588 |

| | | |
|---|---|---|
| | 0.597781975 | NA |
| | 0.03196373 | 0.085753395 |
| | 6.72E-05 | 0.000498266 |
| | 1.11E-12 | 7.76E-11 |
| | 0.101156538 | 0.207811435 |
| | 0.640430617 | 0.766722054 |
| | 0.886390708 | 0.935538753 |
| | 0.905158266 | NA |
| NA | | NA |
| | 0.272385351 | 0.429198474 |
| NA | | NA |
| | 0.010105239 | 0.033955766 |
| | 0.001457911 | 0.006847973 |
| | 0.781404391 | 0.868329394 |
| | 0.550054018 | 0.696014532 |
| | 0.106152515 | 0.215355051 |
| | 0.868270381 | 0.92446132 |
| | 0.30080755 | 0.461178689 |
| | 0.236624154 | NA |
| NA | | NA |
| NA | | NA |
| NA | | NA |
| | 0.777092376 | NA |
| | 0.010405172 | 0.034777349 |
| | 5.29E-05 | 0.000405647 |
| | 0.002068382 | 0.009207296 |
| | 0.02143713 | 0.062151826 |
| | 0.001491873 | 0.006977644 |
| | 0.117029795 | 0.231894719 |
| | 0.791429516 | NA |
| | 0.882688221 | 0.933353428 |
| | 1.37E-09 | 4.13E-08 |
| | 0.676165029 | 0.794388095 |
| | 0.000764511 | 0.003975351 |
| | 0.379256757 | 0.54368058 |
| | 0.259030479 | 0.414067867 |
| | 0.097485721 | 0.201977156 |
| | 0.992060816 | NA |
| | 0.658590643 | 0.780885093 |
| | 0.703982835 | NA |
| | 0.025928425 | 0.072567205 |
| | 0.248877835 | 0.402645573 |
| NA | | NA |
| NA | | NA |
| | 0.200548219 | NA |
| | 0.989978842 | NA |
| | 0.889409773 | NA |

| | | |
|---|---|---|
| NA | NA | |
| 0.992061292 | NA | |
| 0.776244828 | NA | |
| 0.797157316 | NA | |
| NA | NA | |
| 0.822381641 | | 0.894259604 |
| 0.593047867 | NA | |
| 0.074368687 | | 0.16423144 |
| 0.889409773 | NA | |
| 0.746255412 | NA | |
| NA | NA | |
| 0.992061292 | NA | |
| 0.000688837 | | 0.003644424 |
| 0.147261835 | | 0.274938327 |
| NA | NA | |
| NA | NA | |
| 0.029864629 | | 0.081253549 |
| 0.376701512 | NA | |
| NA | NA | |
| 0.589058655 | | 0.727633403 |
| 0.905158266 | NA | |
| 0.001122775 | | 0.005488652 |
| 0.007959027 | | 0.027968039 |
| 0.006077372 | | 0.022497137 |
| NA | NA | |
| 0.000712509 | | 0.00374575 |
| NA | NA | |
| 0.000599781 | | 0.003243189 |
| 0.561390737 | NA | |
| 0.052109507 | | 0.12526247 |
| 0.992061292 | NA | |
| 9.79E-11 | | 4.18E-09 |
| 0.000224306 | NA | |
| 0.65646532 | | 0.779248714 |
| 0.236502012 | | 0.388337203 |
| 0.496779228 | | 0.650067044 |
| 4.65E-14 | | 4.44E-12 |
| 0.737811269 | | 0.839048097 |
| 6.39E-06 | | 6.62E-05 |
| 0.000768163 | | 0.003991505 |
| 0.034194255 | | 0.090368474 |
| 0.842354141 | NA | |
| 0.576409335 | NA | |
| 0.734048139 | NA | |
| 0.575185684 | NA | |
| 0.702173542 | | 0.814141655 |
| 0.797342596 | NA | |

| | |
|---|---|
| 0.10881795 | 0.219585863 |
| 0.013364619 | 0.042446292 |
| NA | NA |
| NA | NA |
| 0.204057109 | NA |
| 0.13941383 | 0.263674152 |
| NA | NA |
| 0.633269058 | 0.760802469 |
| 0.677720159 | 0.795534131 |
| 0.003631977 | 0.014693182 |
| 0.724475119 | NA |
| NA | NA |
| 0.000823795 | 0.004235475 |
| 0.001423814 | 0.006720881 |
| 0.985478041 | 0.99264741 |
| NA | NA |
| NA | NA |
| NA | NA |
| 5.80E-05 | 0.000440095 |
| 0.859598501 | 0.918229747 |
| 0.006255287 | 0.023037055 |
| 0.780448129 | 0.867735439 |
| 0.953075698 | 0.973603891 |
| 0.339155539 | 0.502124898 |
| 0.382423057 | 0.546496771 |
| 2.34E-14 | 2.44E-12 |
| 0.376157713 | 0.54050826 |
| 0.017647967 | 0.053162889 |
| 0.20603276 | 0.3523303 |
| 0.542903333 | NA |
| 9.24E-07 | 1.23E-05 |
| 0.063740537 | 0.146238819 |
| 0.832592411 | 0.90036919 |
| 0.263442225 | 0.419228048 |
| 0.010592848 | 0.035297169 |
| 0.901975853 | 0.94462713 |
| 0.01022817 | 0.034321576 |
| 0.87550559 | 0.928844683 |
| 5.30E-06 | 5.60E-05 |
| 0.992060816 | NA |
| 0.490877472 | NA |
| NA | NA |
| NA | NA |
| NA | NA |
| NA | NA |
| 0.905158266 | NA |
| 0.850239185 | NA |

| | | |
|---|---|---|
| NA | | NA |
| | 0.992061292 | NA |
| NA | | NA |
| NA | | NA |
| NA | | NA |
| | 0.580866713 | NA |
| NA | | NA |
| NA | | NA |
| NA | | NA |
| NA | | NA |
| NA | | NA |
| NA | | NA |
| NA | | NA |
| NA | | NA |
| | 0.992061292 | NA |
| NA | | NA |
| | 0.430120108 | NA |
| | 0.012289995 | 0.039738142 |
| | 4.49E-05 | 0.000353235 |
| | 0.735355709 | NA |
| NA | | NA |
| | 0.870729995 | 0.926059958 |
| | 0.45876458 | 0.616072127 |
| | 0.80696305 | 0.884236607 |
| | 0.119045139 | NA |
| | 0.014758375 | 0.046073237 |
| | 0.261738079 | 0.4171824 |
| | 0.302667873 | NA |
| | 0.190245449 | 0.332293971 |
| | 0.858078005 | NA |
| | 0.786910719 | 0.871659225 |
| | 0.506627375 | 0.658441674 |
| | 0.241702027 | 0.394472049 |
| | 0.78168032 | 0.868484723 |
| | 0.303196446 | 0.463883519 |
| | 0.322935457 | 0.485375749 |
| | 0.619045283 | 0.751013327 |
| | 0.901193849 | 0.944236842 |
| | 0.016363857 | 0.050130293 |
| | 0.431708925 | 0.592569165 |
| | 0.008029535 | 0.02817383 |
| | 0.003566048 | 0.014469116 |
| | 3.13E-06 | 3.52E-05 |
| | 0.235477089 | 0.387183932 |
| | 0.900454518 | 0.943822471 |
| | 0.747126156 | 0.845590313 |
| | 0.36693622 | 0.531682082 |

| | | | |
|---|---|---|---|
| | 0.110121598 | | 0.221483804 |
| NA | | NA | |
| | 0.889410245 | NA | |
| | 0.905158739 | NA | |
| | 0.688605646 | | 0.803757582 |
| | 0.905158266 | NA | |
| | 0.003776226 | | 0.015140068 |
| | 0.425891643 | NA | |
| | 0.138544517 | | 0.262402886 |
| NA | | NA | |
| | 0.173835765 | | 0.3111051 |
| | 0.000339259 | | 0.001989791 |
| | 0.080436671 | | 0.174672042 |
| | 0.02756719 | | 0.076173137 |
| | 0.13361252 | | 0.255218975 |
| | 0.296855666 | | 0.456904294 |
| | 0.259116309 | NA | |
| | 0.515746427 | | 0.666938095 |
| | 0.889409773 | NA | |
| NA | | NA | |
| | 0.285127983 | | 0.444051445 |
| | 0.122964267 | | 0.240335766 |
| | 0.948226691 | | 0.970727735 |
| | 0.004283509 | | 0.016814727 |
| | 0.360491189 | | 0.525144356 |
| | 0.028547897 | | 0.078331034 |
| | 0.948375594 | NA | |
| | 0.994322054 | | 0.99722972 |
| | 0.242073204 | | 0.394788099 |
| | 0.637427939 | | 0.764085382 |
| | 0.07476607 | | 0.164821618 |
| NA | | NA | |
| | 0.634735504 | | 0.762189366 |
| | 0.672079683 | | 0.791239426 |
| | 0.297522236 | | 0.457496243 |
| NA | | NA | |
| | 0.272598859 | NA | |
| NA | | NA | |
| | 0.352896975 | | 0.517028362 |
| | 0.97658016 | NA | |
| | 4.29E-05 | | 0.000339972 |
| | 0.694202706 | | 0.807800181 |
| | 0.904690425 | | 0.946139784 |
| | 0.505322236 | | 0.657325422 |
| | 0.926756251 | NA | |
| | 0.801369413 | | 0.880259664 |
| NA | | NA | |

| | |
|---|---|
| NA | NA |
| 0.93421758 | NA |
| NA | NA |
| NA | NA |
| 0.005458549 | 0.0206094 |
| NA | NA |
| NA | NA |
| 0.742134279 | NA |
| NA | NA |
| NA | NA |
| 0.778857281 | 0.86693669 |
| 5.20E-06 | 5.50E-05 |
| 0.109754794 | 0.221009219 |
| 0.617748014 | 0.75015919 |
| 0.784197818 | 0.870112479 |
| 0.127981674 | 0.24752066 |
| 0.461667629 | NA |
| 0.808469036 | 0.885124587 |
| NA | NA |
| 0.986121533 | NA |
| 4.68E-09 | 1.20E-07 |
| 0.04704742 | 0.115625552 |
| NA | NA |
| 0.248456849 | 0.402320179 |
| 0.99205891 | NA |
| 0.86691533 | NA |
| 0.012427937 | 0.040091868 |
| 0.003636417 | 0.014702604 |
| 0.020802134 | 0.060695465 |
| NA | NA |
| 7.63E-20 | 3.16E-17 |
| NA | NA |
| 0.983105286 | NA |
| 0.953754535 | NA |
| 0.018373889 | 0.054879246 |
| 8.83E-08 | 1.58E-06 |
| 0.905158739 | NA |
| 0.738647352 | 0.839639257 |
| 0.340600328 | NA |
| 0.905158266 | NA |
| 0.117923161 | 0.23301311 |
| 0.219192343 | NA |
| 0.020836077 | 0.060770277 |
| 0.781786496 | NA |
| 0.853698039 | NA |
| NA | NA |
| NA | NA |

| | | |
|---|---|---|
| | 0.576912628 | NA |
| | 0.024335332 | 0.068961097 |
| | 7.62E-08 | 1.39E-06 |
| | 0.889409773 | NA |
| | 0.919568255 | 0.954020567 |
| | 0.039348891 | 0.100824072 |
| | 0.697750929 | 0.810940651 |
| | 0.850055606 | NA |
| | 0.341485325 | 0.50483371 |
| | 9.28E-09 | 2.21E-07 |
| NA | | NA |
| | 0.013418709 | 0.042593442 |
| | 0.001459132 | 0.006850777 |
| | 0.812199493 | NA |
| | 0.860952139 | NA |
| | 0.733463622 | 0.836290963 |
| | 0.426146975 | NA |
| NA | | NA |
| | 2.54E-05 | 0.000217871 |
| | 0.292706943 | 0.452453312 |
| | 4.43E-06 | 4.80E-05 |
| | 0.300940361 | 0.461315839 |
| | 0.170917644 | 0.307356228 |
| | 0.023174495 | 0.06622706 |
| | 0.000291556 | 0.00174592 |
| | 0.17078597 | 0.30724964 |
| | 0.03803025 | 0.098247796 |
| NA | | NA |
| | 0.830163713 | NA |
| | 0.001132689 | 0.005529329 |
| | 0.00020201 | 0.001273303 |
| | 0.979229845 | 0.989397824 |
| | 0.907922485 | NA |
| | 1.20E-06 | 1.53E-05 |
| | 0.531114758 | 0.680281672 |
| | 0.992061769 | NA |
| | 0.072973967 | 0.161649888 |
| | 0.33569049 | 0.498642188 |
| | 0.000314167 | 0.00185901 |
| | 0.928173959 | 0.958781257 |
| | 0.905157793 | NA |
| | 0.532319538 | NA |
| | 0.595375588 | 0.732387684 |
| | 0.900885141 | 0.944183696 |
| | 0.423079349 | 0.584485652 |
| | 0.000145996 | 0.000961298 |
| | 0.353896593 | NA |

| | | |
|---|---|---|
| | 0.120110728 | 0.236314979 |
| | 0.99205891 | NA |
| | 0.951483718 | NA |
| NA | | NA |
| | 0.989340902 | NA |
| | 0.983391023 | 0.991828033 |
| | 0.905158739 | NA |
| | 0.01680769 | 0.051204227 |
| | 0.000140345 | 0.000930504 |
| | 3.66E-05 | 0.000296383 |
| | 1.39E-05 | 0.000129143 |
| | 0.992061292 | NA |
| | 0.889410717 | NA |
| | 0.889409773 | NA |
| | 0.889409773 | NA |
| | 0.001280345 | 0.006127965 |
| | 0.472999125 | 0.629328116 |
| | 0.905158266 | NA |
| | 0.905158266 | NA |
| | 0.859779557 | 0.918372376 |
| | 8.46E-05 | 0.000604022 |
| | 5.11E-05 | 0.00039392 |
| | 5.68E-07 | 8.02E-06 |
| | 0.000186712 | 0.001186691 |
| | 0.764426023 | NA |
| | 0.015416351 | 0.047747201 |
| | 0.004597891 | 0.017847815 |
| | 0.889409773 | NA |
| | 0.00180245 | 0.008189531 |
| NA | | NA |
| | 9.22E-05 | 0.000651023 |
| | 0.654057517 | 0.777511713 |
| | 0.012762355 | 0.040960033 |
| | 0.305744255 | 0.466400702 |
| | 0.304827601 | 0.465487434 |
| | 0.63253739 | 0.760251946 |
| | 0.965657433 | NA |
| | 0.992061292 | NA |
| | 0.816203439 | 0.890670005 |
| | 0.372372288 | 0.536754829 |
| | 0.058892282 | 0.137561497 |
| | 0.012040307 | 0.039059403 |
| | 0.987690892 | NA |
| | 0.992059387 | NA |
| | 0.714200582 | 0.822139368 |
| | 0.397699077 | 0.561215477 |
| | 0.636188861 | 0.763266743 |

| | | |
|---|---|---|
| | 0.98658798 | NA |
| | 5.55E-06 | 5.83E-05 |
| | 0.302280781 | NA |
| | 5.21E-06 | 5.52E-05 |
| | 0.972218789 | 0.985471871 |
| | 0.88471069 | NA |
| | 0.718774468 | 0.825457274 |
| | 0.574724033 | 0.716654864 |
| | 0.760467427 | NA |
| | 0.003124431 | 0.012994202 |
| | 0.304358707 | 0.464965413 |
| | 0.002088406 | 0.009283261 |
| | 0.809723461 | NA |
| | 0.518207496 | NA |
| | 0.229732672 | 0.380330198 |
| | 0.973281751 | 0.985912235 |
| NA | | NA |
| NA | | NA |
| | 0.992060816 | NA |
| | 0.046667461 | 0.114884617 |
| | 0.400678543 | 0.564331994 |
| | 0.797044762 | NA |
| | 0.757843973 | NA |
| | 0.746771471 | NA |
| | 0.905158266 | NA |
| | 0.942585696 | NA |
| | 0.905158739 | NA |
| | 0.899517123 | NA |
| | 5.46E-07 | 7.75E-06 |
| | 0.701314689 | NA |
| | 0.04796349 | 0.117430069 |
| | 0.229237205 | 0.37966525 |
| | 0.939729547 | 0.96589388 |
| | 0.097625904 | 0.202146839 |
| | 0.117391623 | 0.232360073 |
| | 0.889409301 | NA |
| | 0.889409773 | NA |
| NA | | NA |
| | 0.296940426 | NA |
| | 0.707211939 | 0.817689321 |
| | 0.759470531 | NA |
| | 0.002672947 | 0.011441569 |
| | 0.796193918 | 0.877077681 |
| | 0.872459336 | NA |
| | 0.18468437 | 0.325401916 |
| NA | | NA |
| | 0.689856969 | NA |

| | | |
|---|---|---|
| | 0.411310003 | NA |
| | 0.286786958 | NA |
| | 0.090553487 | 0.19108822 |
| NA | | NA |
| | 0.002432923 | 0.01057615 |
| | 0.055926945 | 0.132096351 |
| | 0.023278386 | 0.066489344 |
| | 2.29E-11 | 1.16E-09 |
| | 0.989077428 | 0.994011498 |
| | 1.65E-05 | 0.000150681 |
| | 0.165625269 | NA |
| | 0.974617206 | NA |
| | 0.905158266 | NA |
| | 1.47E-16 | 2.58E-14 |
| | 0.328877165 | NA |
| | 0.279762169 | 0.437931101 |
| | 8.98E-06 | 8.89E-05 |
| | 0.428117284 | 0.589148206 |
| | 0.196037474 | NA |
| | 2.21E-15 | 3.03E-13 |
| | 4.56E-09 | 1.17E-07 |
| | 0.075911753 | 0.166969727 |
| | 0.194286182 | 0.337579543 |
| | 0.189663582 | 0.331607081 |
| | 0.483088497 | NA |
| | 0.992057737 | NA |
| | 0.905158266 | NA |
| | 0.120333534 | 0.236604949 |
| NA | | NA |
| | 0.686736383 | 0.802215354 |
| | 0.515737955 | 0.666938095 |
| | 0.427071236 | 0.588187235 |
| | 0.068380284 | 0.153639352 |
| | 0.257225333 | 0.411984803 |
| NA | | NA |
| NA | | NA |
| NA | | NA |
| NA | | NA |
| | 0.905158266 | NA |
| | 0.992060816 | NA |
| NA | | NA |
| NA | | NA |
| | 0.981460124 | 0.990708967 |
| | 0.374971699 | NA |
| NA | | NA |
| NA | | NA |
| | 0.793525361 | 0.875469321 |

| | | |
|---|---|---|
| | 0.905158266 | NA |
| NA | | NA |
| NA | | NA |
| | 0.890505996 | NA |
| NA | | NA |
| NA | | NA |
| NA | | NA |
| | 1.43E-12 | 9.75E-11 |
| | 0.095083913 | NA |
| | 0.859821802 | NA |
| NA | | NA |
| | 0.543659383 | NA |
| | 0.739767756 | 0.840206898 |
| | 0.70589654 | 0.817034689 |
| | 0.033710156 | 0.089379537 |
| | 0.313429625 | NA |
| | 0.015984517 | 0.049222271 |
| | 0.266940144 | 0.423148365 |
| | 2.16E-10 | 8.40E-09 |
| | 0.013134754 | 0.041831192 |
| | 0.082989935 | 0.17892744 |
| | 0.404549164 | 0.567850108 |
| | 0.767573545 | 0.859036108 |
| | 0.00214865 | 0.009506789 |
| | 0.548242343 | 0.694362303 |
| | 0.293234901 | 0.45285813 |
| NA | | NA |
| | 0.20459842 | NA |
| | 1.14E-20 | 5.80E-18 |
| | 0.265863719 | 0.422011089 |
| | 7.45E-08 | 1.37E-06 |
| | 0.905158266 | NA |
| | 7.16E-08 | 1.32E-06 |
| | 0.513851751 | NA |
| | 0.201608601 | NA |
| | 0.002892966 | 0.012193215 |
| | 0.176232335 | 0.31449702 |
| | 0.905158266 | NA |
| NA | | NA |
| | 0.813836939 | NA |
| | 7.04E-05 | 0.000517895 |
| | 1.56E-05 | 0.000142785 |
| | 0.024782424 | NA |
| | 2.00E-09 | 5.80E-08 |
| | 0.015233727 | 0.047294984 |
| | 0.823609337 | 0.894841296 |
| | 0.223779564 | 0.37340239 |

| | |
|---|---|
| 0.015643161 | 0.04834058 |
| NA | NA |
| 1.39E-09 | 4.19E-08 |
| 0.363166213 | 0.52802535 |
| 0.010582252 | 0.035267212 |
| 0.282056331 | 0.440611438 |
| 0.066597817 | 0.150993635 |
| 7.20E-08 | 1.32E-06 |
| 0.592477428 | 0.729926607 |
| 0.685952861 | 0.801476876 |
| 0.363084733 | 0.527984271 |
| NA | NA |
| 0.537819727 | NA |
| 0.790731562 | 0.873753263 |
| 0.23426015 | 0.385702829 |
| 0.076946612 | 0.168815131 |
| 0.077031483 | 0.168968486 |
| 0.122475992 | 0.239616028 |
| 0.781730645 | 0.868484723 |
| 0.05390762 | 0.128459841 |
| 0.009176926 | 0.031359495 |
| 0.935393056 | 0.962820337 |
| 0.563853224 | 0.707442175 |
| 0.763012912 | 0.856288676 |
| NA | NA |
| 0.92344143 | 0.955826083 |
| 0.404278071 | 0.567581494 |
| 0.256193554 | NA |
| 0.920411158 | NA |
| 0.986081564 | NA |
| 0.100332173 | 0.206543001 |
| 1.10E-06 | 1.41E-05 |
| NA | NA |
| NA | NA |
| 0.821273937 | NA |
| 0.061772681 | NA |
| 0.483717594 | 0.639299106 |
| 0.932991398 | 0.961405823 |
| 0.185643642 | 0.326620362 |
| 1.69E-05 | 0.000153351 |
| 0.283292991 | 0.44216581 |
| NA | NA |
| 0.931704588 | 0.960437209 |
| 0.585301014 | 0.724663113 |
| 0.801621203 | 0.880404121 |
| 0.241212529 | 0.394047458 |
| 0.103773833 | 0.211741645 |

| | | |
|---|---|---|
| | 0.763743902 | 0.856846285 |
| | 0.812246814 | NA |
| | 0.820779114 | 0.893401833 |
| | 0.00656831 | 0.023971001 |
| | 0.370828032 | 0.535351784 |
| NA | | NA |
| | 2.31E-05 | 0.000200912 |
| | 0.056197369 | 0.132551255 |
| | 0.008092385 | 0.028332381 |
| | 0.872813757 | NA |
| | 0.264964642 | NA |
| | 0.425938928 | 0.587180169 |
| | 0.002789038 | NA |
| | 0.606718953 | NA |
| | 3.53E-09 | 9.55E-08 |
| NA | | NA |
| NA | | NA |
| | 4.80E-09 | 1.22E-07 |
| | 4.57E-11 | 2.12E-09 |
| | 0.110143196 | 0.22149266 |
| | 0.010569125 | 0.035239512 |
| | 0.088128007 | 0.187193163 |
| | 0.032034279 | 0.085858749 |
| | 0.014057124 | 0.044267945 |
| NA | | NA |
| | 0.200939358 | 0.346154182 |
| | 0.878033887 | NA |
| | 0.006066499 | 0.02247204 |
| | 0.422445008 | NA |
| | 0.004798085 | 0.018529925 |
| | 0.992061292 | NA |
| | 0.339606315 | NA |
| | 0.347242463 | 0.511031247 |
| | 0.620405033 | 0.751888853 |
| | 0.005321504 | 0.020192657 |
| | 0.01985115 | 0.058448834 |
| | 0.385863542 | NA |
| | 0.20679819 | 0.353376655 |
| | 0.792347241 | 0.874856852 |
| | 0.952645728 | 0.973374154 |
| | 0.72045254 | NA |
| NA | | NA |
| | 2.35E-05 | 0.000203782 |
| | 0.896493679 | NA |
| | 0.251698543 | NA |
| | 0.182194059 | 0.322373793 |
| | 1.25E-12 | 8.66E-11 |

| | | |
|---|---|---|
| | 0.390733582 | 0.554118558 |
| | 0.115597246 | 0.229834243 |
| | 0.94688329 | NA |
| | 0.95249908 | 0.973374154 |
| | 0.43347052 | 0.59431688 |
| | 0.218976317 | 0.367594292 |
| | 0.908495859 | 0.948154039 |
| | 0.751684044 | NA |
| | 0.619762337 | NA |
| | 0.092609656 | 0.194215597 |
| | 0.932038959 | 0.960604689 |
| | 0.000108104 | 0.000746287 |
| | 0.067615039 | 0.152470593 |
| | 0.430548318 | NA |
| | 0.4908111 | NA |
| | 0.388217117 | 0.551913743 |
| | 0.630769941 | 0.75883839 |
| | 0.165714866 | 0.300099878 |
| NA | | NA |
| | 0.043737202 | 0.109270212 |
| | 0.095132342 | 0.198236013 |
| | 0.008070677 | 0.028272586 |
| | 0.121155194 | 0.237879731 |
| | 0.023059925 | 0.065975374 |
| | 0.127220498 | 0.246504448 |
| | 0.780488442 | 0.867735439 |
| | 0.711211549 | 0.820117335 |
| | 0.619861853 | NA |
| | 0.395315811 | 0.558821932 |
| | 0.015244594 | 0.047322031 |
| NA | | NA |
| | 0.52211374 | 0.67263804 |
| | 0.803061494 | NA |
| | 0.986467808 | NA |
| | 0.435379264 | NA |
| | 5.51E-08 | 1.06E-06 |
| | 0.027974597 | 0.077075807 |
| | 0.180628785 | 0.320493495 |
| | 0.349762093 | 0.513843969 |
| | 3.25E-09 | 8.88E-08 |
| | 0.797224866 | NA |
| | 0.992060816 | NA |
| NA | | NA |
| | 0.138119729 | 0.261801565 |
| NA | | NA |
| | 0.162700352 | 0.296025936 |
| | 0.251598822 | 0.40582184 |

| | |
|---|---|
| 0.007379606 | 0.026332265 |
| 1.78E-05 | 0.000160766 |
| 0.958999614 | 0.977249651 |
| 0.466766422 | 0.623253524 |
| NA | NA |
| 2.35E-08 | 5.00E-07 |
| 1.95E-05 | 0.000174175 |
| 0.787645204 | NA |
| 0.275735105 | 0.433140223 |
| 0.045260939 | 0.112316076 |
| 8.30E-05 | 0.00059504 |
| 0.000124068 | 0.000840349 |
| 0.217609919 | 0.365691912 |
| 0.493671942 | 0.647157912 |
| 0.000736769 | 0.003863112 |
| 0.749785924 | 0.847377721 |
| 2.87E-08 | 5.99E-07 |
| NA | NA |
| 0.889409773 | NA |
| 0.000117187 | 0.000800409 |
| 0.037378523 | 0.096986291 |
| 5.43E-05 | 0.000415266 |
| 0.142529696 | 0.268087749 |
| 7.58E-12 | 4.39E-10 |
| 0.221287236 | 0.370284723 |
| 0.819704986 | NA |
| 0.390331208 | NA |
| 0.414728434 | NA |
| 0.207632918 | 0.354349206 |
| 0.601038186 | 0.73664624 |
| NA | NA |
| 4.03E-05 | 0.000321763 |
| 0.585511181 | 0.724882451 |
| 7.11E-06 | 7.25E-05 |
| 0.905158266 | NA |
| 7.99E-11 | 3.48E-09 |
| 0.528599173 | NA |
| 0.037430777 | 0.097082504 |
| 0.394677763 | NA |
| 1.25E-09 | 3.83E-08 |
| NA | NA |
| 0.406023753 | NA |
| NA | NA |
| 0.324936997 | NA |
| 0.352703821 | 0.516886225 |
| NA | NA |
| NA | NA |

| | | |
|---|---|---|
| | 0.992062245 | NA |
| NA | | NA |
| | 0.817152605 | NA |
| | 0.992061292 | NA |
| NA | | NA |
| | 0.321369166 | NA |
| | 0.744575687 | NA |
| | 0.889409773 | NA |
| NA | | NA |
| | 0.905158266 | NA |
| NA | | NA |
| | 0.55158735 | 0.697163625 |
| NA | | NA |
| | 0.708723857 | 0.818366051 |
| | 0.505788933 | NA |
| | 0.11062169 | 0.222052669 |
| | 0.557967338 | NA |
| | 0.000964124 | 0.004828097 |
| | 0.00019018 | 0.001206287 |
| | 0.255264102 | NA |
| | 0.651705185 | NA |
| | 0.234973988 | 0.386443514 |
| | 0.462036489 | 0.619443144 |
| | 0.989853331 | NA |
| | 0.026246592 | 0.073230804 |
| | 0.424219542 | 0.585527923 |
| NA | | NA |
| | 0.889409773 | NA |
| | 0.528994422 | 0.678476053 |
| | 0.430462806 | 0.591448677 |
| | 0.590372965 | 0.728396445 |
| | 0.897520669 | NA |
| NA | | NA |
| | 0.916163426 | 0.952379394 |
| | 0.792017481 | NA |
| | 0.062852352 | NA |
| | 0.003839062 | 0.015345779 |
| | 0.000875 | 0.004446684 |
| | 0.04719441 | 0.115870087 |
| | 0.117748473 | 0.232772661 |
| NA | | NA |
| | 0.934759666 | 0.962529563 |
| | 0.225244888 | 0.374990047 |
| | 0.552832047 | 0.698282365 |
| | 0.009347878 | 0.031849503 |
| | 0.109300458 | 0.220324998 |
| | 0.408595278 | 0.571887546 |

| | | |
|---|---|---|
| | 0.982499967 | NA |
| | 0.87970476 | 0.931731488 |
| | 0.316184988 | 0.47830712 |
| | 0.667602385 | 0.787911468 |
| | 2.75E-05 | 0.000232543 |
| | 0.000428833 | 0.002425121 |
| | 0.656904729 | 0.779601737 |
| NA | | NA |
| | 0.002036504 | 0.009091194 |
| | 0.917191132 | 0.952814839 |
| | 0.720449043 | NA |
| | 0.15696882 | 0.288210847 |
| | 0.013388755 | 0.042516801 |
| | 0.189889366 | 0.331830382 |
| | 0.317815101 | 0.480111572 |
| | 0.131508598 | 0.252276457 |
| | 0.512503013 | 0.664151884 |
| | 9.60E-05 | 0.00067488 |
| | 0.821735009 | 0.893910585 |
| | 0.528479036 | 0.678126145 |
| | 0.011844237 | 0.038537204 |
| | 0.746644191 | 0.845262659 |
| | 0.007912085 | 0.02782958 |
| | 0.023844884 | 0.067754893 |
| | 0.977966031 | 0.98868174 |
| | 0.992060816 | NA |
| | 0.12590675 | 0.244541512 |
| | 0.337556064 | 0.500533028 |
| | 0.123074321 | 0.240465255 |
| | 0.288877262 | 0.448284032 |
| | 0.04253217 | 0.107069118 |
| | 3.15E-06 | 3.55E-05 |
| | 0.238209764 | 0.390216204 |
| NA | | NA |
| | 0.736562052 | 0.838335753 |
| | 0.016106855 | 0.049522604 |
| | 0.002018034 | 0.009025256 |
| | 0.134366808 | 0.256258532 |
| | 0.092591717 | 0.194215597 |
| | 0.019951906 | 0.058674688 |
| | 0.023638924 | 0.067282843 |
| | 0.255221668 | 0.409769146 |
| | 4.69E-05 | 0.00036722 |
| NA | | NA |
| | 0.046575231 | 0.114696141 |
| | 0.005491537 | 0.020715172 |
| | 0.90515732 | NA |

| | |
|---|---|
| 0.730808925 | 0.834459624 |
| 0.02233347 | 0.064266893 |
| 0.121870603 | 0.238750698 |
| 0.003814082 | 0.015266757 |
| NA | NA |
| 0.038194623 | 0.098591246 |
| 0.108169344 | 0.218613463 |
| 0.012291154 | 0.039738142 |
| NA | NA |
| 0.152626465 | 0.282510105 |
| 0.053101942 | 0.126967066 |
| 0.034016746 | 0.089975154 |
| 0.001642211 | 0.007542629 |
| 0.021032516 | 0.061221238 |
| 0.074333658 | 0.164181533 |
| 0.01463365 | 0.045755422 |
| 0.076553581 | 0.168070394 |
| 0.042862588 | 0.107675232 |
| 0.762734187 | 0.856107136 |
| 0.161426 | 0.29421925 |
| 0.095252209 | 0.198372842 |
| 0.097552863 | 0.202048718 |
| 0.889410717 | NA |
| 0.905157793 | NA |
| NA | NA |
| NA | NA |
| NA | NA |
| 0.686884951 | 0.802303545 |
| NA | NA |
| 0.905158739 | NA |
| 0.383550315 | 0.547512277 |
| NA | NA |
| 0.301718722 | NA |
| 0.109304637 | 0.220324998 |
| 0.025431393 | 0.071522535 |
| 0.000237395 | 0.001466785 |
| 0.213497497 | 0.360657861 |
| 0.040192065 | 0.10243472 |
| 0.094821994 | 0.197814571 |
| 0.951668733 | 0.973242087 |
| 0.250933952 | 0.405046946 |
| 0.565761292 | 0.708958308 |
| 0.543002376 | 0.689888809 |
| NA | NA |
| 0.742267751 | 0.842044611 |
| NA | NA |
| NA | NA |

| | | |
|---|---|---|
| | 0.612223194 | NA |
| NA | | NA |
| | 0.001741711 | 0.007944794 |
| | 0.000138586 | 0.000921905 |
| | 0.761162439 | 0.855173497 |
| | 0.066931187 | 0.151504992 |
| | 0.309466765 | 0.470673272 |
| | 0.8708642 | NA |
| | 0.003762102 | 0.015094465 |
| | 2.24E-07 | 3.58E-06 |
| | 0.932872054 | 0.961327957 |
| | 0.938041342 | 0.964934174 |
| | 0.082705396 | NA |
| | 0.787016904 | 0.871688903 |
| | 0.000455704 | 0.002549506 |
| | 0.889409301 | NA |
| NA | | NA |
| NA | | NA |
| | 0.905158739 | NA |
| | 0.000128496 | 0.000863946 |
| | 0.732219241 | NA |
| | 0.902645834 | 0.944966123 |
| | 0.033281831 | 0.088532493 |
| NA | | NA |
| | 0.605514741 | 0.740421995 |
| | 0.543882172 | 0.690593957 |
| | 0.358370998 | 0.523235954 |
| | 1.51E-16 | 2.62E-14 |
| | 0.00021702 | 0.001353081 |
| | 0.754892003 | 0.850958599 |
| | 0.127377831 | 0.246678701 |
| | 6.11E-05 | 0.000459346 |
| | 0.010903681 | 0.036119163 |
| NA | | NA |
| | 0.299517251 | 0.459840545 |
| | 0.094393052 | 0.197088686 |
| | 0.132009796 | 0.25292759 |
| NA | | NA |
| | 0.020757218 | 0.060580509 |
| | 0.905157793 | NA |
| | 0.78630125 | NA |
| | 0.98945134 | NA |
| | 0.403967237 | 0.567326462 |
| | 0.287388967 | 0.446482866 |
| NA | | NA |
| | 5.74E-08 | 1.09E-06 |
| | 0.208737367 | 0.355515694 |

| | | |
|---|---|---|
| NA | | NA |
| | 0.003375168 | NA |
| | 0.889409301 | NA |
| | 0.000356086 | 0.002073535 |
| | 0.991918092 | NA |
| | 0.740399533 | NA |
| | 0.95082553 | NA |
| | 3.16E-09 | 8.69E-08 |
| | 0.630923666 | NA |
| | 5.84E-08 | 1.11E-06 |
| NA | | NA |
| | 0.739425499 | 0.840113352 |
| | 0.465384339 | 0.622238692 |
| | 0.236392322 | 0.388267415 |
| NA | | NA |
| | 0.576234935 | NA |
| | 0.79086643 | NA |
| | 0.000104678 | 0.000726723 |
| | 0.011980794 | NA |
| | 0.753154177 | 0.849610222 |
| | 0.273263177 | 0.430237687 |
| | 0.740965598 | NA |
| | 9.89E-05 | 0.000691889 |
| | 0.740658413 | NA |
| | 5.83E-05 | 0.000441942 |
| | 0.000666369 | 0.003545625 |
| NA | | NA |
| | 0.986558577 | NA |
| NA | | NA |
| | 0.544905164 | 0.691491707 |
| | 0.42931394 | 0.590387326 |
| | 0.337821784 | 0.500696239 |
| | 0.975175739 | 0.987192641 |
| | 2.73E-07 | 4.24E-06 |
| | 0.624733482 | 0.755300476 |
| NA | | NA |
| | 0.366463843 | 0.531307244 |
| | 0.955372425 | 0.975162776 |
| | 0.429386967 | NA |
| | 0.557687868 | 0.702597125 |
| | 0.623718785 | 0.754530674 |
| | 0.325624597 | 0.487781603 |
| | 0.53999156 | NA |
| | 0.032239052 | 0.086344353 |
| | 0.686863691 | NA |
| NA | | NA |
| | 0.427689398 | 0.588780107 |

| | |
|---|---|
| 0.229405806 | 0.379874979 |
| 0.513915297 | 0.665337361 |
| 0.00540645 | 0.020454898 |
| 0.003890833 | 0.015523039 |
| 1.42E-11 | 7.65E-10 |
| NA | NA |
| 0.889409773 | NA |
| NA | NA |
| 2.12E-14 | 2.24E-12 |
| 0.000439303 | 0.002475404 |
| 2.21E-09 | 6.36E-08 |
| 0.002395609 | 0.01043557 |
| 0.889409773 | NA |
| 0.905158739 | NA |
| 0.892764392 | NA |
| 0.952551682 | 0.973374154 |
| 0.008222369 | 0.028700526 |
| 0.114223042 | 0.22777414 |
| 0.752679717 | 0.849447196 |
| 0.861008549 | NA |
| 0.671306549 | 0.790668338 |
| 0.00920232 | 0.031431599 |
| NA | NA |
| 0.6085521 | NA |
| 0.001799826 | 0.008179302 |
| 0.853314336 | 0.914402182 |
| 0.855788113 | NA |
| 0.001891804 | 0.008542487 |
| 0.044419909 | 0.110625778 |
| NA | NA |
| 0.988464177 | NA |
| 0.597277771 | 0.734069658 |
| 0.294690974 | 0.454496354 |
| 0.072811728 | 0.161355609 |
| 0.204114447 | 0.349999154 |
| 4.97E-05 | 0.0003854 |
| NA | NA |
| 0.156725792 | NA |
| 1.84E-05 | 0.000165336 |
| 0.000418516 | 0.002375953 |
| 0.931672913 | 0.960437209 |
| 0.28240363 | 0.441028469 |
| 0.0236086 | 0.067222671 |
| 0.194325452 | 0.337621061 |
| 0.050205507 | 0.121724934 |
| 0.249453716 | 0.403339529 |
| 0.580275035 | 0.720553013 |

| | | |
|---|---|---|
| | 0.589825145 | 0.728170594 |
| | 0.889409773 | NA |
| | 0.288000688 | 0.447193261 |
| | 0.207682243 | 0.354378303 |
| | 0.976175951 | NA |
| | 0.034941408 | 0.091955703 |
| | 0.001784842 | 0.008117924 |
| | 0.041756175 | 0.105502999 |
| | 0.036293817 | 0.094776539 |
| | 0.47202275 | 0.628343088 |
| | 0.917995778 | 0.953199796 |
| | 0.869940016 | 0.925515189 |
| | 0.020056349 | NA |
| | 0.423145696 | NA |
| | 0.597333685 | 0.734092615 |
| | 0.991879747 | NA |
| NA | | NA |
| NA | | NA |
| | 0.000222527 | 0.0013827 |
| | 0.977876025 | NA |
| | 0.382364129 | 0.546458438 |
| | 0.992623063 | 0.996201218 |
| | 0.707129808 | NA |
| | 0.715768445 | NA |
| NA | | NA |
| | 0.889409773 | NA |
| NA | | NA |
| NA | | NA |
| | 0.021706581 | 0.062808645 |
| NA | | NA |
| | 0.516960124 | 0.668074821 |
| | 0.001199154 | 0.005806442 |
| | 0.043624517 | 0.109056327 |
| | 0.000179308 | 0.001147266 |
| NA | | NA |
| NA | | NA |
| | 0.889409301 | NA |
| | 0.229843621 | 0.380427833 |
| | 0.001133978 | 0.005533563 |
| | 0.445853783 | 0.605323558 |
| | 0.02717205 | 0.075289723 |
| NA | | NA |
| NA | | NA |
| NA | | NA |
| | 0.031955556 | NA |
| | 0.210293777 | 0.357224495 |
| | 0.871878097 | 0.926581323 |

| | | |
|---|---|---|
| | 0.863830271 | 0.921444133 |
| NA | | NA |
| | 0.216894482 | 0.365048359 |
| | 0.007695217 | 0.027197448 |
| | 0.889409773 | NA |
| | 0.205489422 | NA |
| | 0.394927157 | 0.55845228 |
| | 0.905158739 | NA |
| | 0.28353253 | 0.442330757 |
| | 0.166800163 | 0.301660745 |
| | 0.310368214 | NA |
| | 0.056800793 | 0.133642838 |
| | 0.005457603 | 0.020609373 |
| NA | | NA |
| | 0.877209064 | NA |
| | 0.287154733 | NA |
| | 0.039811164 | 0.101735524 |
| | 0.801661681 | NA |
| | 0.003338684 | 0.013706145 |
| | 0.992061292 | NA |
| | 0.582450597 | 0.722397021 |
| | 0.137657551 | 0.261173507 |
| | 0.01560299 | 0.04823002 |
| | 0.846134994 | 0.909565146 |
| | 1.26E-07 | 2.18E-06 |
| | 0.051585514 | 0.124387039 |
| | 0.905158266 | NA |
| | 0.524593578 | 0.674683752 |
| | 2.96E-06 | 3.36E-05 |
| NA | | NA |
| | 0.299986295 | 0.460399709 |
| NA | | NA |
| | 0.393787887 | 0.557200091 |
| | 0.529803061 | NA |
| | 0.139555324 | 0.263839463 |
| | 0.16138845 | 0.294175227 |
| | 0.545497637 | 0.691923672 |
| | 0.73386556 | NA |
| | 0.00110713 | 0.005422579 |
| | 0.98924292 | NA |
| | 0.194645828 | 0.338070689 |
| NA | | NA |
| | 0.214390164 | 0.361803997 |
| | 0.017874145 | 0.053722742 |
| NA | | NA |
| | 0.059758481 | 0.139258887 |
| | 0.101422485 | 0.20823528 |

| | | |
|---|---|---|
| | 0.905158739 | NA |
| | 0.003019507 | 0.012627186 |
| | 0.897030465 | 0.942074668 |
| | 0.982635237 | NA |
| | 0.914261437 | 0.951707366 |
| NA | | NA |
| | 0.001226607 | 0.00590713 |
| | 0.87139284 | 0.926316278 |
| | 0.801661681 | NA |
| | 0.843946304 | NA |
| | 0.124705032 | 0.242851879 |
| | 0.692313078 | NA |
| | 0.656920545 | NA |
| | 0.035547056 | 0.09320332 |
| | 0.695469302 | 0.808888262 |
| | 1.26E-07 | 2.18E-06 |
| | 4.24E-10 | 1.50E-08 |
| | 0.739458998 | NA |
| NA | | NA |
| | 0.005344113 | NA |
| | 0.050931127 | 0.123171433 |
| | 0.108052928 | NA |
| | 0.872201222 | NA |
| | 0.049551555 | NA |
| | 0.073642654 | NA |
| | 0.058991091 | 0.137748338 |
| | 0.50420544 | 0.656381744 |
| | 0.182271997 | 0.322441599 |
| | 0.105276306 | 0.214143106 |
| | 0.921610883 | 0.955146941 |
| | 0.249682805 | 0.403591072 |
| | 0.000985423 | 0.004916816 |
| NA | | NA |
| NA | | NA |
| | 0.023668005 | 0.067346998 |
| | 0.338616667 | NA |
| NA | | NA |
| NA | | NA |
| | 0.905158739 | NA |
| NA | | NA |
| NA | | NA |
| NA | | NA |
| NA | | NA |
| | 0.004151031 | 0.016388438 |
| | 0.020191143 | 0.059272712 |
| | 0.842474527 | 0.906857752 |
| | 0.351488753 | 0.515812485 |

| | |
|---|---|
| NA | NA |
| 0.012019528 | 0.039020818 |
| NA | NA |
| 0.089807516 | 0.189787628 |
| 0.024653518 | 0.069700863 |
| 0.517715029 | 0.668578235 |
| 0.696564105 | NA |
| 3.57E-07 | 5.34E-06 |
| 0.011008928 | 0.036385468 |
| 0.003781065 | 0.0151567 |
| 0.008710571 | 0.030098415 |
| 0.002700168 | 0.011531555 |
| 0.355361519 | 0.520017567 |
| 0.774818216 | NA |
| NA | NA |
| 0.905157793 | NA |
| NA | NA |
| 0.905158739 | NA |
| 0.005340212 | 0.020253146 |
| 0.99391308 | NA |
| 0.889409773 | NA |
| NA | NA |
| 0.037653635 | 0.097573302 |
| 0.253391143 | 0.407794434 |
| 0.486123859 | 0.64097477 |
| 6.81E-08 | 1.27E-06 |
| 0.546159843 | 0.692603603 |
| 0.060172459 | NA |
| 0.027355648 | 0.075683843 |
| 0.010380588 | 0.034711031 |
| 0.038196622 | NA |
| 0.02897465 | 0.079210121 |
| 0.188960202 | 0.330759432 |
| 0.471160988 | NA |
| 0.081217185 | 0.175969233 |
| 0.070865353 | 0.158079264 |
| 9.12E-07 | 1.21E-05 |
| NA | NA |
| NA | NA |
| 0.47973417 | 0.635334578 |
| 0.278772294 | 0.436916661 |
| 0.905158266 | NA |
| 0.982988923 | 0.99175069 |
| 0.109340304 | 0.220356438 |
| 0.008521048 | 0.029559825 |
| 0.310338343 | 0.471515865 |
| NA | NA |

| | |
|---|---|
| 0.010515953 | 0.035088865 |
| 0.891098447 | 0.938410003 |
| 0.784206752 | 0.870112479 |
| 0.000322198 | 0.001900902 |
| 6.71E-08 | 1.25E-06 |
| 0.106359679 | NA |
| 0.268628502 | NA |
| 0.032469108 | 0.086822838 |
| 0.84082192 | 0.905826984 |
| NA | NA |
| 0.538544973 | 0.68635572 |
| 0.889410256 | NA |
| 4.61E-07 | 6.72E-06 |
| NA | NA |
| 0.051077516 | 0.123416723 |
| 5.83E-06 | 6.08E-05 |
| 0.271106763 | 0.427732452 |
| 0.978642353 | 0.989181044 |
| 0.000156025 | 0.00101817 |
| 0.000384823 | 0.002209813 |
| 0.270622302 | 0.42712536 |
| 0.994853759 | NA |
| NA | NA |
| 1.52E-10 | 6.16E-09 |
| 0.984983269 | NA |
| 0.111856493 | 0.223865907 |
| 0.176591896 | 0.31502235 |
| 0.941291596 | NA |
| 9.16E-07 | 1.22E-05 |
| 0.660872814 | NA |
| 0.490868861 | NA |
| NA | NA |
| NA | NA |
| NA | NA |
| 0.889409773 | NA |
| 0.992060816 | NA |
| 0.747987199 | 0.846215915 |
| 2.93E-09 | 8.12E-08 |
| 0.779050457 | 0.86706381 |
| 0.000160236 | 0.001041012 |
| 0.899138328 | 0.943310432 |
| 0.920749063 | 0.954551282 |
| 0.805586386 | 0.883232933 |
| 0.561060535 | 0.705268092 |
| 0.791647392 | 0.874479858 |
| 0.008386454 | 0.029189753 |
| NA | NA |

| | | |
|---|---|---|
| | 0.085452162 | 0.182629802 |
| | 0.045405673 | 0.112573463 |
| | 0.364280964 | 0.529258711 |
| | 0.905158266 | NA |
| | 0.905158266 | NA |
| NA | | NA |
| | 0.889409773 | NA |
| NA | | NA |
| NA | | NA |
| NA | | NA |
| NA | | NA |
| NA | | NA |
| NA | | NA |
| NA | | NA |
| | 0.889409773 | NA |
| | 0.224648417 | NA |
| NA | | NA |
| NA | | NA |
| NA | | NA |
| | 0.576143981 | NA |
| NA | | NA |
| NA | | NA |
| NA | | NA |
| NA | | NA |
| | 0.993600236 | NA |
| | 0.905158739 | NA |
| | 0.330266751 | NA |
| | 0.005557362 | 0.020921329 |
| | 0.522206535 | 0.67263804 |
| | 0.021881869 | 0.063215887 |
| | 0.034825963 | 0.091706838 |
| | 0.158610989 | 0.290438518 |
| | 0.060779749 | 0.141189306 |
| | 6.98E-05 | 0.000514434 |
| | 0.730655971 | NA |
| | 0.905158739 | NA |
| NA | | NA |
| NA | | NA |
| NA | | NA |
| NA | | NA |
| NA | | NA |
| | 0.008469211 | 0.029407875 |
| | 0.124844599 | 0.243037461 |
| | 0.797157316 | NA |
| NA | | NA |
| | 0.765388855 | 0.857727679 |
| | 0.992061292 | NA |

| | | | |
|---|---|---|---|
| | 0.151645591 | | 0.281082597 |
| NA | | NA | |
| | 0.905158266 | NA | |
| | 0.000458946 | | 0.002566332 |
| | 0.060034531 | | 0.139791164 |
| | 0.019885771 | | 0.058533409 |
| | 0.066931551 | | 0.151504992 |
| | 0.001347487 | | 0.006410196 |
| | 0.992060816 | NA | |
| | 0.921362571 | | 0.954934663 |
| NA | | NA | |
| NA | | NA | |
| | 0.905158266 | NA | |
| NA | | NA | |
| NA | | NA | |
| NA | | NA | |
| NA | | NA | |
| NA | | NA | |
| | 0.26047448 | | 0.415770815 |
| | 0.419398419 | NA | |
| | 0.122714809 | NA | |
| | 0.889409773 | NA | |
| | 0.18053541 | NA | |
| | 0.038558424 | | 0.099285229 |
| | 0.904773661 | NA | |
| | 0.129102524 | | 0.249117955 |
| | 0.006455453 | | 0.023645586 |
| | 5.06E-10 | | 1.76E-08 |
| | 0.807879735 | NA | |
| NA | | NA | |
| | 0.002687979 | | 0.01149471 |
| | 0.705044166 | NA | |
| | 0.728892332 | | 0.832938526 |
| | 0.02809074 | | 0.077318194 |
| | 2.73E-07 | | 4.25E-06 |
| | 0.163123504 | | 0.296574683 |
| | 0.035797003 | | 0.093675629 |
| | 0.00992323 | | 0.033405519 |
| | 0.355843647 | NA | |
| | 0.623767371 | | 0.754547882 |
| NA | | NA | |
| | 0.992060816 | NA | |
| NA | | NA | |
| NA | | NA | |
| NA | | NA | |
| NA | | NA | |
| | 0.905158266 | NA | |

| | | |
|---|---|---|
| | 0.00700292 | 0.025259054 |
| NA | | NA |
| | 0.229949701 | NA |
| | 0.824190542 | 0.895142918 |
| | 0.787645204 | NA |
| | 0.457983359 | NA |
| | 0.008009726 | 0.028123692 |
| | 0.759378489 | 0.853840269 |
| | 0.547072212 | 0.693377685 |
| | 0.417387798 | 0.579645995 |
| | 0.188246337 | 0.329746435 |
| | 0.099226408 | NA |
| | 0.708627253 | 0.818297531 |
| NA | | NA |
| | 0.060828457 | 0.141242687 |
| | 0.57068886 | NA |
| | 0.453098491 | 0.611135218 |
| | 0.274597118 | 0.431846889 |
| | 0.711469285 | NA |
| NA | | NA |
| NA | | NA |
| | 1.08E-08 | 2.54E-07 |
| NA | | NA |
| | 0.307788186 | 0.468769648 |
| | 0.630801165 | NA |
| | 0.794514872 | NA |
| | 0.853228278 | NA |
| | 0.001679831 | 0.007696101 |
| NA | | NA |
| | 0.889409301 | NA |
| | 0.219935345 | 0.368774203 |
| | 0.871166339 | NA |
| | 0.927373216 | 0.95835985 |
| | 0.472557866 | NA |
| | 0.356477895 | 0.521060913 |
| | 0.000412508 | 0.002345584 |
| | 0.137063944 | NA |
| | 0.083855713 | NA |
| | 9.05E-05 | 0.000641096 |
| NA | | NA |
| | 0.801743218 | NA |
| NA | | NA |
| NA | | NA |
| | 0.611556539 | NA |
| NA | | NA |
| | 0.041931502 | 0.105896935 |
| NA | | NA |

| | |
|---|---|
| NA | NA |
| 5.11E-14 | 4.84E-12 |
| 0.024696928 | 0.069796634 |
| NA | NA |
| 0.420485194 | 0.582293901 |
| 0.714102748 | NA |
| 0.4201452 | 0.582006589 |
| 0.928658492 | NA |
| 0.00077924 | 0.004035693 |
| 0.889409773 | NA |
| 0.899517123 | NA |
| 0.61039761 | NA |
| 0.065392518 | 0.149111763 |
| 0.000781893 | 0.004048479 |
| 0.025621469 | 0.071955727 |
| 0.140512298 | 0.265283257 |
| 0.052857628 | 0.126589944 |
| 0.146181702 | 0.273284862 |
| 0.063735738 | 0.146238819 |
| 0.023459576 | 0.066902445 |
| 0.905158266 | NA |
| 0.269408623 | 0.425821921 |
| 0.282477656 | 0.441090321 |
| NA | NA |
| 0.011321096 | 0.037154379 |
| 0.640690715 | NA |
| 1.11E-08 | 2.59E-07 |
| 0.748778853 | NA |
| NA | NA |
| NA | NA |
| 0.84349912 | 0.907714959 |
| 0.030680869 | 0.082932995 |
| 0.991688155 | NA |
| 0.738781589 | NA |
| 0.3527745 | NA |
| 0.889409773 | NA |
| 0.039686008 | 0.101498375 |
| NA | NA |
| 0.899182161 | NA |
| 0.992061292 | NA |
| NA | NA |
| NA | NA |
| NA | NA |
| 0.905158266 | NA |
| 0.625043983 | NA |
| 0.661882642 | NA |
| NA | NA |

| | | |
|---|---|---|
| NA | | NA |
| | 0.905158266 | NA |
| NA | | NA |
| | 0.776629605 | 0.865466099 |
| | 0.072640757 | 0.161106795 |
| | 0.991955612 | NA |
| | 0.129392365 | 0.249458029 |
| | 0.004376567 | 0.017106629 |
| | 0.153445494 | 0.283604882 |
| | 0.148676031 | 0.276746373 |
| | 0.037816382 | 0.09785015 |
| | 0.893637238 | 0.939862067 |
| NA | | NA |
| | 0.992061292 | NA |
| | 7.85E-07 | 1.07E-05 |
| NA | | NA |
| | 0.000972388 | 0.004859507 |
| | 0.000175185 | 0.001125482 |
| NA | | NA |
| NA | | NA |
| NA | | NA |
| | 3.02E-07 | 4.63E-06 |
| | 0.825552412 | 0.896053297 |
| | 0.001326492 | 0.006322648 |
| | 0.413474001 | 0.576035756 |
| | 0.020448023 | 0.059861104 |
| NA | | NA |
| NA | | NA |
| | 0.284447444 | NA |
| | 7.66E-07 | 1.04E-05 |
| | 0.070594476 | 0.157562379 |
| | 0.370358726 | 0.53491625 |
| | 0.121246607 | 0.237931568 |
| | 0.17380578 | 0.311102146 |
| | 0.913613038 | NA |
| | 0.078945706 | 0.172237333 |
| | 0.135782535 | 0.258442288 |
| | 0.016776038 | 0.051143277 |
| | 0.487874962 | 0.642435014 |
| | 3.73E-13 | 2.98E-11 |
| | 0.227161708 | NA |
| | 0.026251042 | 0.073230804 |
| | 0.704698781 | NA |
| NA | | NA |
| NA | | NA |
| | 0.586635884 | NA |
| | 0.591493593 | NA |

| | |
|---|---|
| NA | NA |
| 0.389485715 | 0.55285823 |
| 0.984392524 | NA |
| NA | NA |
| 0.825563399 | 0.896053297 |
| 0.871675277 | 0.926481965 |
| 0.905157793 | NA |
| NA | NA |
| 0.433606286 | 0.594371492 |
| 0.739556644 | 0.840161208 |
| 0.525445035 | 0.67542279 |
| 0.154667166 | 0.285262582 |
| NA | NA |
| 0.992060816 | NA |
| 0.481063378 | 0.63674929 |
| NA | NA |
| NA | NA |
| 0.484442772 | NA |
| 0.000532615 | 0.002921003 |
| NA | NA |
| 0.001633665 | 0.007515954 |
| 7.05E-05 | 0.00051832 |
| 0.63291602 | NA |
| 0.124697549 | 0.242851879 |
| 0.15308862 | 0.283135939 |
| 0.182419998 | 0.322566789 |
| 0.809861512 | 0.886041999 |
| 0.000249644 | 0.001526577 |
| 0.065357556 | NA |
| NA | NA |
| 0.807433834 | NA |
| 0.098130695 | 0.202943298 |
| 0.195455907 | 0.339209378 |
| 0.858361213 | NA |
| 0.018580558 | 0.055353475 |
| 0.271049998 | NA |
| 0.000142029 | 0.000938553 |
| 0.000557271 | 0.003039522 |
| 0.170710521 | NA |
| NA | NA |
| 0.069204554 | 0.155083458 |
| 0.044120067 | 0.110019594 |
| 0.889409301 | NA |
| 6.92E-06 | 7.09E-05 |
| NA | NA |
| NA | NA |
| NA | NA |

| | | |
|---|---|---|
| | 0.889409773 | NA |
| | 7.29E-05 | 0.000533233 |
| NA | | NA |
| | 0.046694421 | NA |
| | 0.992062955 | NA |
| | 0.989560379 | NA |
| | 0.001765088 | 0.008038068 |
| | 0.044344764 | 0.110504616 |
| | 0.171888438 | 0.308621636 |
| | 2.27E-07 | 3.62E-06 |
| | 0.235258584 | 0.386853621 |
| | 0.992061292 | NA |
| NA | | NA |
| | 0.00618226 | 0.022816177 |
| | 5.09E-07 | 7.32E-06 |
| | 0.163102071 | NA |
| | 5.42E-10 | 1.85E-08 |
| NA | | NA |
| | 0.592921357 | NA |
| | 2.13E-06 | 2.54E-05 |
| NA | | NA |
| | 0.528771506 | NA |
| | 0.718767636 | 0.825457274 |
| | 8.74E-06 | 8.69E-05 |
| NA | | NA |
| | 0.332373693 | NA |
| | 5.38E-07 | 7.69E-06 |
| | 4.72E-28 | 2.07E-24 |
| NA | | NA |
| NA | | NA |
| | 6.20E-11 | 2.77E-09 |
| | 0.914519553 | 0.951778255 |
| | 0.359589018 | 0.524386721 |
| NA | | NA |
| NA | | NA |
| | 0.55877421 | 0.703481417 |
| | 0.962439847 | 0.979205986 |
| | 0.969551542 | 0.9839097 |
| | 0.444603386 | NA |
| | 0.905158739 | NA |
| NA | | NA |
| | 0.659322388 | 0.781583908 |
| | 0.889409301 | NA |
| | 0.983750803 | NA |
| NA | | NA |
| | 0.156862619 | 0.288091511 |
| | 0.000620814 | 0.003337613 |

| | |
|---|---|
| 0.598603364 | 0.734794078 |
| 0.36507221 | NA |
| NA | NA |
| 0.889409773 | NA |
| 0.917482669 | NA |
| 0.263649781 | 0.419384261 |
| NA | NA |
| 0.85486038 | NA |
| 0.021490901 | 0.062274834 |
| 0.365512987 | NA |
| 0.028497098 | 0.078211195 |
| 0.76126617 | 0.855246282 |
| NA | NA |
| NA | NA |
| 0.281881626 | NA |
| 0.907530487 | 0.947610543 |
| 0.880639841 | 0.932221934 |
| 0.04800072 | 0.117495019 |
| 0.908687361 | 0.948154039 |
| 0.403880152 | NA |
| NA | NA |
| 0.889409301 | NA |
| NA | NA |
| NA | NA |
| NA | NA |
| NA | NA |
| NA | NA |
| NA | NA |
| NA | NA |
| 0.992061292 | NA |
| NA | NA |
| NA | NA |
| NA | NA |
| 0.003949233 | 0.015716059 |
| 0.267412536 | 0.42364417 |
| 0.043513252 | 0.108889743 |
| 0.197954088 | 0.342364559 |
| 0.451247513 | 0.609593611 |
| 0.875917843 | NA |
| 0.758344507 | NA |
| 0.625453346 | 0.755340311 |
| NA | NA |
| 5.79E-05 | 0.000439608 |
| NA | NA |
| 0.007130494 | 0.025618212 |
| 0.000607402 | 0.003275527 |
| NA | NA |

| | | |
|---|---|---|
| | 0.985534734 | NA |
| | 0.609470816 | NA |
| | 0.992062245 | NA |
| NA | | NA |
| NA | | NA |
| | 0.992060816 | NA |
| | 0.238467375 | 0.390540001 |
| | 0.291717235 | 0.451272897 |
| NA | | NA |
| NA | | NA |
| NA | | NA |
| | 0.897241012 | 0.942074668 |
| | 0.387241962 | 0.550991138 |
| | 1.66E-05 | 0.000151116 |
| NA | | NA |
| | 0.04902391 | 0.119495627 |
| | 0.020026456 | 0.058854516 |
| | 0.900277372 | 0.943713058 |
| NA | | NA |
| | 0.267350009 | 0.423637662 |
| NA | | NA |
| | 0.889409301 | NA |
| | 0.300067223 | 0.460491729 |
| | 0.439240261 | 0.599545998 |
| | 0.704883799 | NA |
| NA | | NA |
| | 0.092266152 | NA |
| | 9.93E-05 | 0.000694297 |
| | 0.1299477 | 0.250067613 |
| | 0.992062245 | NA |
| NA | | NA |
| NA | | NA |
| NA | | NA |
| NA | | NA |
| NA | | NA |
| | 0.204127654 | 0.349999154 |
| | 0.487254651 | NA |
| NA | | NA |
| NA | | NA |
| | 0.643370857 | 0.768937427 |
| | 0.161855116 | 0.294878991 |
| | 0.64476027 | NA |
| NA | | NA |
| NA | | NA |
| NA | | NA |
| | 0.958858982 | 0.97723867 |
| | 0.295350565 | 0.45528979 |

| | |
|---|---|
| NA | NA |
| NA | NA |
| 6.20E-07 | 8.63E-06 |
| 0.025684831 | 0.072096793 |
| 0.983049034 | 0.99175941 |
| 0.653713313 | 0.777272827 |
| 6.17E-05 | 0.000463066 |
| 0.547281764 | NA |
| 0.021250048 | 0.061707195 |
| 0.117734261 | 0.23276552 |
| 0.616207896 | NA |
| 0.991214328 | NA |
| NA | NA |
| 0.041115013 | 0.104194411 |
| 0.051017579 | 0.123328584 |
| 0.050097569 | 0.121530328 |
| 1.43E-11 | 7.67E-10 |
| 0.623275323 | 0.754169581 |
| 0.828226081 | 0.897503776 |
| NA | NA |
| 0.807879735 | NA |
| NA | NA |
| 0.981145846 | 0.990539389 |
| 0.506243816 | NA |
| 1.48E-05 | 0.000136758 |
| 0.169473167 | 0.305338142 |
| NA | NA |
| 0.021892778 | 0.063239083 |
| 0.505064897 | 0.657072106 |
| 0.525365364 | NA |
| 0.889409301 | NA |
| 4.95E-07 | 7.15E-06 |
| 0.18169335 | NA |
| 1.20E-08 | 2.76E-07 |
| 0.880921434 | NA |
| 0.292077877 | NA |
| 6.87E-09 | 1.69E-07 |
| 0.364303721 | 0.529258711 |
| 0.960456253 | 0.978049474 |
| 0.636286602 | 0.763342301 |
| NA | NA |
| 0.889409301 | NA |
| 4.71E-10 | 1.64E-08 |
| 0.751747315 | 0.848794029 |
| 0.21421249 | 0.361595192 |
| 8.09E-06 | 8.12E-05 |
| 0.072806213 | 0.161355609 |

| | | |
|---|---|---|
| | 0.391404597 | NA |
| | 0.000426025 | 0.002412346 |
| | 0.788582525 | NA |
| | 0.04041424 | 0.102810043 |
| | 0.722536116 | 0.828359103 |
| | 0.574652024 | NA |
| | 0.825372841 | NA |
| | 0.98929671 | NA |
| NA | | NA |
| | 0.414263161 | 0.57680583 |
| NA | | NA |
| NA | | NA |
| NA | | NA |
| NA | | NA |
| | 0.889409773 | NA |
| | 0.003228884 | 0.013357774 |
| NA | | NA |
| | 0.992061292 | NA |
| | 0.148465127 | 0.27644753 |
| NA | | NA |
| | 0.038839972 | 0.099776198 |
| | 0.471116588 | NA |
| | 0.054647117 | 0.129869702 |
| | 0.91627724 | 0.952384067 |
| | 0.025560859 | 0.071831443 |
| | 0.000273885 | 0.001657727 |
| | 0.889409301 | NA |
| | 0.447537931 | NA |
| | 0.875289692 | NA |
| | 1.25E-08 | 2.87E-07 |
| | 0.905158266 | NA |
| NA | | NA |
| | 0.085666804 | 0.183027569 |
| NA | | NA |
| NA | | NA |
| | 0.906295064 | 0.947186393 |
| | 0.106531319 | 0.21594399 |
| | 0.597571093 | NA |
| | 0.470086577 | NA |
| | 0.371112547 | 0.535536599 |
| | 0.08270025 | 0.178495655 |
| | 0.797342596 | NA |
| | 0.575183053 | 0.717068506 |
| | 0.502270689 | 0.654892766 |
| | 0.00670333 | 0.024385643 |
| | 0.992061292 | NA |
| | 0.197962366 | 0.342364559 |

| | | |
|---|---|---|
| | 0.455007227 | 0.613050908 |
| | 0.005082338 | 0.019432109 |
| | 0.351673453 | 0.515961606 |
| | 0.516351383 | 0.667445256 |
| | 0.779910566 | 0.867493472 |
| | 0.567328292 | 0.710098152 |
| | 0.34315733 | 0.506444774 |
| | 0.018267893 | 0.054659309 |
| | 7.89E-08 | 1.43E-06 |
| | 0.655193043 | NA |
| | 0.890201103 | 0.937910278 |
| | 0.132206579 | NA |
| | 0.676884426 | NA |
| | 0.837091557 | NA |
| | 0.998121281 | NA |
| | 0.992060816 | NA |
| NA | | NA |
| | 0.030429163 | 0.082409326 |
| NA | | NA |
| | 0.992061769 | NA |
| | 0.290783015 | 0.450248217 |
| | 0.905158266 | NA |
| | 0.64831229 | NA |
| | 0.000761007 | 0.003960989 |
| NA | | NA |
| | 0.196404384 | 0.340398096 |
| NA | | NA |
| NA | | NA |
| | 0.822437593 | 0.894276163 |
| NA | | NA |
| | 0.300916739 | 0.461311832 |
| NA | | NA |
| | 0.889409301 | NA |
| | 0.046676469 | 0.114885096 |
| | 0.807879735 | NA |
| | 0.068065402 | 0.153187267 |
| NA | | NA |
| | 0.745889109 | NA |
| | 0.40658351 | 0.569798155 |
| | 1.51E-06 | 1.87E-05 |
| | 0.55857981 | NA |
| | 0.995284688 | NA |
| | 0.929221884 | 0.959434081 |
| | 0.991274888 | 0.995627556 |
| | 0.905158739 | NA |
| | 0.144887063 | 0.271511779 |
| NA | | NA |

| | |
|---|---|
| 1.41E-08 | 3.18E-07 |
| 0.652333305 | 0.776105924 |
| NA | NA |
| NA | NA |
| 0.095848458 | 0.199425459 |
| 0.079649441 | 0.173273438 |
| 0.905158266 | NA |
| NA | NA |
| 0.638470125 | NA |
| 1.14E-05 | 0.000109244 |
| 2.37E-05 | 0.000205018 |
| 0.840577211 | 0.905703818 |
| 0.992253023 | NA |
| 0.342897857 | 0.506314652 |
| 0.07347101 | 0.162521398 |
| 0.193694053 | 0.336790538 |
| 0.73235969 | NA |
| 6.54E-05 | 0.000486378 |
| 0.003375315 | 0.013843077 |
| 0.037710782 | 0.097675285 |
| 0.984946575 | NA |
| 0.20068553 | 0.345825438 |
| 0.764770523 | 0.857428567 |
| NA | NA |
| 0.617301978 | 0.749783403 |
| NA | NA |
| 0.80902577 | 0.885468589 |
| 0.918737188 | 0.95369906 |
| 0.154778184 | 0.285395427 |
| 0.043636549 | 0.109073987 |
| NA | NA |
| NA | NA |
| 0.300669535 | NA |
| 0.000370342 | 0.002138402 |
| 0.047112538 | 0.115733775 |
| 0.889410717 | NA |
| 0.003234493 | 0.01337342 |
| 5.80E-11 | 2.64E-09 |
| 0.905158266 | NA |
| NA | NA |
| 0.033480112 | 0.088898401 |
| 0.69803658 | 0.811150066 |
| 0.99205891 | NA |
| 0.046240499 | 0.114204828 |
| 0.082904454 | 0.178834475 |
| 0.27180445 | 0.428557027 |
| NA | NA |

| | |
|---|---|
| NA | NA |
| 0.000329552 | 0.00193804 |
| NA | NA |
| 0.784438112 | 0.870149349 |
| 7.92E-05 | 0.000573632 |
| 0.686953212 | NA |
| 0.082987796 | 0.17892744 |
| 0.173865287 | 0.311132576 |
| 0.176256689 | NA |
| 0.122781957 | 0.240064909 |
| NA | NA |
| 0.093011202 | 0.194814287 |
| 0.838733637 | 0.904205014 |
| 0.028873274 | 0.07900007 |
| 0.085421814 | 0.182592973 |
| 0.411872378 | 0.574533449 |
| 1.43E-07 | 2.42E-06 |
| 0.014228786 | 0.04471235 |
| 0.071695518 | 0.159445198 |
| 0.697516504 | NA |
| 0.99206359 | NA |
| 0.050822798 | 0.122938306 |
| 2.45E-07 | 3.87E-06 |
| 2.31E-06 | 2.72E-05 |
| 0.51053305 | NA |
| 1.35E-07 | 2.30E-06 |
| 0.904066502 | NA |
| 0.583756204 | NA |
| 0.053393537 | 0.127428397 |
| 0.003803828 | 0.015231266 |
| 0.000506741 | 0.002793051 |
| 0.384761903 | 0.548742393 |
| 0.992061292 | NA |
| 1.33E-05 | 0.000124951 |
| 0.48514357 | 0.640199877 |
| 0.247600065 | 0.40125829 |
| 0.465167998 | NA |
| 0.992061769 | NA |
| 0.191885477 | 0.334359234 |
| 5.46E-05 | 0.000417231 |
| 0.012695974 | 0.040806629 |
| 0.253523444 | 0.407847714 |
| NA | NA |
| 0.905296333 | NA |
| 0.581403235 | 0.721586709 |
| 8.32E-07 | 1.12E-05 |
| 0.040332288 | 0.102668633 |

| | | | |
|---|---|---|---|
| | 0.010475092 | | 0.034968462 |
| | 0.038496249 | | 0.099158608 |
| | 0.003922775 | NA | |
| | 0.225950321 | | 0.375768183 |
| | 6.98E-05 | | 0.000514677 |
| | 0.746619829 | | 0.845262659 |
| | 0.145162365 | | 0.271958055 |
| | 0.203135527 | | 0.348879255 |
| | 0.70698123 | | 0.817643694 |
| | 0.934783253 | | 0.962529563 |
| NA | | NA | |
| NA | | NA | |
| NA | | NA | |
| | 0.780554087 | NA | |
| | 0.211289873 | | 0.358417483 |
| NA | | NA | |
| | 0.205810951 | | 0.35203001 |
| | 0.755745095 | NA | |
| NA | | NA | |
| NA | | NA | |
| | 0.916885299 | NA | |
| NA | | NA | |
| NA | | NA | |
| NA | | NA | |
| NA | | NA | |
| NA | | NA | |
| NA | | NA | |
| NA | | NA | |
| NA | | NA | |
| NA | | NA | |
| | 0.907348463 | NA | |
| | 0.710252432 | NA | |
| | 0.63535115 | NA | |
| | 0.430940152 | NA | |
| | 0.885343628 | NA | |
| NA | | NA | |
| NA | | NA | |
| NA | | NA | |
| NA | | NA | |
| NA | | NA | |
| NA | | NA | |
| NA | | NA | |
| NA | | NA | |
| | 0.00370942 | | 0.014925613 |
| | 0.581793939 | NA | |
| | 0.754273378 | | 0.850523216 |
| NA | | NA | |

| | | |
|---|---|---|
| 0.139992634 | NA | |
| 0.967440694 | NA | |
| 0.96271983 | NA | |
| 0.24398754 | NA | |
| 0.588739351 | | 0.727361731 |
| NA | NA | |
| 0.275745033 | NA | |
| 0.69659284 | NA | |
| NA | NA | |
| 0.905158739 | NA | |
| NA | NA | |
| 0.905158739 | NA | |
| NA | NA | |
| 0.905158739 | NA | |
| 0.810433091 | NA | |
| NA | NA | |
| NA | NA | |
| 0.789980394 | NA | |
| 0.905158266 | NA | |
| 0.889409773 | NA | |
| NA | NA | |
| NA | NA | |
| NA | NA | |
| NA | NA | |
| NA | NA | |
| NA | NA | |
| NA | NA | |
| 0.15292698 | NA | |
| NA | NA | |
| NA | NA | |
| 0.857194446 | NA | |
| NA | NA | |
| NA | NA | |
| 0.905158266 | NA | |
| 0.795367625 | NA | |
| 0.168533996 | NA | |
| 0.511702607 | NA | |
| 0.211979516 | NA | |
| 0.273066859 | NA | |
| 0.568386154 | NA | |
| 0.133793356 | NA | |
| 0.882927242 | NA | |
| 0.132075281 | | 0.253008906 |
| 0.85900772 | NA | |
| 0.805374319 | NA | |
| 0.453216984 | NA | |
| 0.969004238 | NA | |

| | | |
|---|---|---|
| | 0.889410717 | NA |
| | 0.988851952 | NA |
| | 0.348924538 | NA |
| | 0.339902874 | NA |
| | 0.387563973 | 0.551270714 |
| | 0.135886747 | NA |
| NA | | NA |
| | 0.007473541 | 0.026589678 |
| | 0.013832769 | 0.04368044 |
| NA | | NA |
| | 0.747202962 | NA |
| | 0.839629023 | NA |
| NA | | NA |
| NA | | NA |
| | 0.905158739 | NA |
| | 0.778531773 | NA |
| | 0.664300454 | NA |
| | 0.905158266 | NA |
| | 0.905158266 | NA |
| NA | | NA |
| | 0.905158266 | NA |
| | 0.615590421 | 0.748408263 |
| | 0.830205752 | NA |
| | 0.887225733 | 0.936077926 |
| | 0.478871587 | 0.634613227 |
| | 0.003940973 | 0.015691719 |
| | 2.43E-09 | 6.90E-08 |
| NA | | NA |
| | 0.005808569 | 0.021703884 |
| | 0.077129335 | 0.169086918 |
| | 0.026544877 | 0.07384076 |
| | 0.159127293 | 0.291141161 |
| | 0.151623242 | 0.281064898 |
| | 0.765941734 | 0.858084513 |
| | 0.889409301 | NA |
| NA | | NA |
| NA | | NA |
| | 0.992061601 | NA |
| NA | | NA |
| | 0.905158739 | NA |
| | 0.104968777 | 0.213664199 |
| | 0.051854825 | 0.124814218 |
| | 0.702006204 | NA |
| | 0.670086127 | NA |
| | 0.339021615 | NA |
| | 0.780841267 | NA |
| | 0.53346565 | NA |

| | | |
|---|---|---|
| | 0.784353866 | NA |
| | 0.902545984 | NA |
| | 0.830163713 | NA |
| NA | | NA |
| | 0.992060816 | NA |
| NA | | NA |
| NA | | NA |
| NA | | NA |
| | 0.667469864 | NA |
| NA | | NA |
| NA | | NA |
| NA | | NA |
| NA | | NA |
| | 0.905158739 | NA |
| NA | | NA |
| | 0.905158739 | NA |
| NA | | NA |
| | 6.24E-07 | 8.68E-06 |
| | 0.206689102 | NA |
| | 0.868583662 | 0.924705647 |
| | 0.59431024 | NA |
| NA | | NA |
| | 2.49E-22 | 1.77E-19 |
| | 0.048007604 | 0.11749877 |
| NA | | NA |
| | 0.000107368 | 0.000742112 |
| | 0.066753994 | 0.151227788 |
| | 0.360332158 | 0.524990135 |
| | 0.855133685 | 0.915689726 |
| | 0.628274432 | NA |
| NA | | NA |
| | 0.059051112 | 0.137859173 |
| NA | | NA |
| | 0.148202328 | 0.276145517 |
| | 8.56E-07 | 1.15E-05 |
| | 0.889283189 | NA |
| | 0.172712434 | NA |
| | 0.590780754 | 0.728817673 |
| | 0.008935439 | 0.030701853 |
| | 0.355951252 | 0.520499005 |
| | 0.144882475 | 0.271511779 |
| | 0.005298141 | 0.020121254 |
| | 0.002002111 | 0.00896317 |
| | 0.086644245 | 0.18460605 |
| | 0.603195189 | NA |
| | 0.522122078 | 0.67263804 |
| | 0.085371507 | 0.182537543 |

| | |
|---|---|
| NA | NA |
| 0.009156847 | 0.031305495 |
| 0.343836041 | 0.507241896 |
| 0.016540442 | 0.050558403 |
| 0.747798738 | 0.846133482 |
| 0.048518362 | 0.118511092 |
| 0.40361736 | 0.567088976 |
| 0.829635027 | NA |
| 2.16E-08 | 4.66E-07 |
| 0.068637701 | 0.154047636 |
| 0.008843106 | 0.030465292 |
| 0.905158266 | NA |
| 0.891705947 | 0.938832699 |
| 0.568486591 | 0.710980581 |
| 0.051980181 | 0.12500633 |
| 2.08E-05 | 0.000184331 |
| 0.002508306 | 0.010846767 |
| 0.001588747 | 0.007350885 |
| 0.26672191 | 0.422914754 |
| 0.748541963 | NA |
| 0.000588287 | 0.003188224 |
| 0.211041676 | 0.358134789 |
| 0.059721391 | 0.139187223 |
| 0.373825833 | 0.537868166 |
| 0.012173328 | 0.039426861 |
| 0.001700339 | 0.007773836 |
| 0.006349115 | 0.023306623 |
| 0.181296599 | 0.3214187 |
| 0.160988421 | 0.293665453 |
| NA | NA |
| NA | NA |
| 0.905158739 | NA |
| 0.905157793 | NA |
| 0.992061292 | NA |
| 3.37E-10 | 1.23E-08 |
| 0.463453658 | 0.620547492 |
| 0.166485231 | 0.301215224 |
| 1.77E-09 | 5.21E-08 |
| 0.333543157 | 0.496190921 |
| 0.50221502 | 0.65487765 |
| 0.503124561 | 0.655440925 |
| 0.905158266 | NA |
| 0.042341145 | 0.10671057 |
| 0.663729202 | NA |
| 0.502286259 | 0.654892766 |
| 0.56748683 | 0.710215622 |
| 0.085067208 | 0.182047638 |

| | |
|---|---|
| NA | NA |
| 0.073656254 | 0.162849142 |
| 0.752075394 | 0.849003885 |
| 0.307244208 | NA |
| 0.004359716 | 0.017068106 |
| 0.692429955 | NA |
| 0.992061292 | NA |
| 0.568253501 | 0.710779543 |
| NA | NA |
| 0.991700267 | NA |
| 0.984290195 | NA |
| 0.802134739 | NA |
| 0.008442775 | 0.029343923 |
| 0.121507235 | 0.238281628 |
| 0.025978959 | 0.072653078 |
| 0.082355641 | 0.177926753 |
| 0.641451994 | NA |
| 2.38E-06 | 2.80E-05 |
| 0.989850124 | NA |
| NA | NA |
| NA | NA |
| 0.368486227 | 0.533112997 |
| 0.131867132 | 0.252764521 |
| NA | NA |
| 0.325387571 | NA |
| NA | NA |
| 0.366911875 | NA |
| 0.440427679 | 0.600518559 |
| NA | NA |
| 0.490817474 | 0.644873269 |
| 0.191103702 | 0.33336878 |
| 0.99205891 | NA |
| 0.601299259 | NA |
| 0.985601599 | NA |
| 0.244862677 | NA |
| 0.786419312 | 0.871466576 |
| 0.411269981 | 0.574062013 |
| 0.543115749 | 0.689938605 |
| 0.360145877 | NA |
| 0.753581177 | NA |
| NA | NA |
| NA | NA |
| NA | NA |
| NA | NA |
| NA | NA |
| NA | NA |
| NA | NA |

| | |
|---|---|
| NA | NA |
| NA | NA |
| NA | NA |
| NA | NA |
| NA | NA |
| NA | NA |
| NA | NA |
| NA | NA |
| NA | NA |
| NA | NA |
| NA | NA |
| 0.542423425 | 0.689537802 |
| 0.070930575 | 0.158208683 |
| 0.755749878 | 0.851532227 |
| NA | NA |
| 0.889410245 | NA |
| 0.01308239 | 0.041718881 |
| 0.475057696 | 0.631287226 |
| 8.17E-07 | 1.10E-05 |
| 0.045853077 | 0.113503294 |
| 0.069152591 | 0.154982829 |
| 0.64629896 | 0.771350554 |
| 0.867636692 | NA |
| 0.889410245 | NA |
| 0.685351403 | 0.800938343 |
| 0.005532343 | 0.020848586 |
| 0.83311393 | NA |
| NA | NA |
| 0.159581106 | NA |
| 0.510071487 | 0.66158652 |
| 0.849007509 | NA |
| 0.858549092 | NA |
| 0.814721167 | NA |
| 0.116751166 | 0.231405258 |
| 0.357637364 | 0.522345498 |
| 7.19E-11 | 3.16E-09 |
| 0.000211959 | 0.001325793 |
| 0.003065792 | 0.012781853 |
| 0.587252356 | 0.726178314 |
| 0.905158739 | NA |
| 0.980051436 | NA |
| 2.68E-08 | 5.61E-07 |
| 0.891180149 | 0.938410003 |
| 3.99E-05 | 0.00031931 |
| NA | NA |
| 0.739763587 | 0.840206898 |
| 0.004372226 | 0.017098236 |

| | |
|---|---|
| 0.055337211 | 0.130971095 |
| 0.368321883 | 0.532980575 |
| 0.760296005 | 0.854549828 |
| NA | NA |
| 0.002708886 | 0.011557105 |
| 0.51114793 | NA |
| 3.13E-09 | 8.60E-08 |
| 0.889409773 | NA |
| 7.62E-09 | 1.86E-07 |
| 0.625289284 | 0.755340311 |
| 0.947116188 | 0.969924394 |
| 0.217330781 | 0.365502536 |
| 0.789396683 | 0.872827883 |
| NA | NA |
| 0.840189838 | NA |
| 0.414471462 | 0.577022686 |
| 0.4889519 | 0.643305506 |
| 0.022225331 | 0.063997645 |
| 0.094494471 | 0.197243735 |
| 0.165265768 | 0.299538838 |
| 0.086050711 | 0.183704799 |
| 0.724341523 | NA |
| 0.99205891 | NA |
| NA | NA |
| NA | NA |
| NA | NA |
| NA | NA |
| NA | NA |
| NA | NA |
| NA | NA |
| NA | NA |
| NA | NA |
| NA | NA |
| NA | NA |
| NA | NA |
| NA | NA |
| NA | NA |
| NA | NA |
| NA | NA |
| NA | NA |
| NA | NA |
| 0.889409301 | NA |
| 0.000437357 | 0.002466975 |
| NA | NA |
| 0.001393039 | 0.00659688 |
| 0.189869112 | 0.331821395 |
| 0.182003223 | 0.322199107 |

| | |
|---|---|
| NA | NA |
| 1.91E-06 | 2.31E-05 |
| 0.64465949 | 0.770189213 |
| 0.889409773 | NA |
| NA | NA |
| NA | NA |
| NA | NA |
| NA | NA |
| NA | NA |
| NA | NA |
| NA | NA |
| NA | NA |
| 0.260431941 | 0.415733135 |
| NA | NA |
| 0.804534188 | NA |
| 0.792188023 | NA |
| NA | NA |
| 0.992059387 | NA |
| NA | NA |
| NA | NA |
| NA | NA |
| 0.992060816 | NA |
| 0.901183732 | NA |
| NA | NA |
| NA | NA |
| 0.889409773 | NA |
| NA | NA |
| NA | NA |
| NA | NA |
| 0.992060816 | NA |
| 0.905158266 | NA |
| NA | NA |
| 0.337391354 | 0.500333896 |
| 0.184768885 | 0.325524706 |
| 0.005874639 | 0.021898013 |
| 0.007206751 | 0.025841452 |
| 0.856262933 | 0.9161369 |
| NA | NA |
| 0.207301128 | 0.353975542 |
| 3.64E-05 | 0.00029519 |
| NA | NA |
| 0.88936334 | NA |
| NA | NA |
| NA | NA |
| 0.889410245 | NA |
| 3.23E-05 | 0.000267031 |
| 0.237555339 | NA |

| | |
|---|---|
| 0.201000986 | 0.346178874 |
| 0.00029293 | 0.001749848 |
| NA | NA |
| 0.025091068 | 0.070746631 |
| 0.896562851 | 0.941790776 |
| 0.694843907 | NA |
| 0.905158739 | NA |
| 0.006964974 | 0.0251718 |
| 0.003643042 | 0.014718964 |
| 0.668465197 | 0.788633066 |
| 0.444502574 | 0.604011882 |
| 0.138379863 | 0.262158865 |
| 0.335857039 | 0.498722939 |
| 0.889409301 | NA |
| 0.265925711 | NA |
| 0.867997568 | 0.924373662 |
| NA | NA |
| 0.042438154 | 0.106893716 |
| NA | NA |
| NA | NA |
| 0.544537987 | 0.691225485 |
| NA | NA |
| 0.905158266 | NA |
| 0.670086127 | NA |
| 0.905158739 | NA |
| NA | NA |
| NA | NA |
| 0.905158266 | NA |
| NA | NA |
| 0.992060816 | NA |
| 0.198866564 | 0.343547969 |
| 0.824601131 | 0.895431522 |
| 0.199587623 | 0.344365885 |
| 0.858147419 | 0.917175946 |
| 0.891135565 | NA |
| 0.000137737 | 0.000917088 |
| 0.009040117 | 0.030988438 |
| 0.781421355 | 0.868329394 |
| 0.208583526 | NA |
| 0.388584091 | 0.552006602 |
| 0.698529212 | 0.811501586 |
| 0.042662745 | 0.107311709 |
| NA | NA |
| 0.715725185 | 0.823316568 |
| 0.889409301 | NA |
| 0.647949305 | 0.772774212 |
| 0.905158739 | NA |

| | | |
|---|---|---|
| NA | | NA |
| NA | | NA |
| | 0.220414964 | NA |
| NA | | NA |
| NA | | NA |
| NA | | NA |
| NA | | NA |
| | 0.686189154 | NA |
| NA | | NA |
| NA | | NA |
| NA | | NA |
| | 0.801661681 | NA |
| NA | | NA |
| NA | | NA |
| NA | | NA |
| NA | | NA |
| NA | | NA |
| NA | | NA |
| NA | | NA |
| | 0.991879747 | NA |
| NA | | NA |
| NA | | NA |
| NA | | NA |
| NA | | NA |
| | 0.889409301 | NA |
| NA | | NA |
| NA | | NA |
| NA | | NA |
| | 0.905158739 | NA |
| | 0.655614386 | NA |
| NA | | NA |
| NA | | NA |
| NA | | NA |
| NA | | NA |
| NA | | NA |
| NA | | NA |
| | 0.463859775 | NA |
| NA | | NA |
| | 0.991700267 | NA |
| | 0.889409773 | NA |
| NA | | NA |
| NA | | NA |
| NA | | NA |
| NA | | NA |
| | 0.991388785 | NA |
| | 0.277853737 | NA |
| NA | | NA |

| | | |
|---|---|---|
| | 0.801661681 | NA |
| NA | | NA |
| NA | | NA |
| | 0.788582525 | NA |
| NA | | NA |
| NA | | NA |
| NA | | NA |
| NA | | NA |
| NA | | NA |
| | 0.65944772 | NA |
| | 0.905158266 | NA |
| NA | | NA |
| NA | | NA |
| NA | | NA |
| NA | | NA |
| NA | | NA |
| NA | | NA |
| NA | | NA |
| NA | | NA |
| NA | | NA |
| | 0.778531773 | NA |
| NA | | NA |
| NA | | NA |
| NA | | NA |
| | 0.905158266 | NA |
| NA | | NA |
| NA | | NA |
| NA | | NA |
| NA | | NA |
| NA | | NA |
| NA | | NA |
| NA | | NA |
| NA | | NA |
| | 0.724343317 | NA |
| | 0.99205891 | NA |
| NA | | NA |
| NA | | NA |
| | 0.889409301 | NA |
| | 0.905158266 | NA |
| | 0.807879735 | NA |
| NA | | NA |
| | 0.905158739 | NA |
| | 0.889409773 | NA |
| NA | | NA |
| NA | | NA |
| NA | | NA |
| NA | | NA |

| | | | |
|---|---|---|---|
| | 0.905158739 | NA | |
| NA | | NA | |
| NA | | NA | |
| NA | | NA | |
| NA | | NA | |
| NA | | NA | |
| NA | | NA | |
| NA | | NA | |
| NA | | NA | |
| | 0.889409301 | NA | |
| NA | | NA | |
| | 0.905158266 | NA | |
| | 0.893382147 | NA | |
| NA | | NA | |
| NA | | NA | |
| | 0.905158266 | NA | |
| NA | | NA | |
| | 0.889409773 | NA | |
| NA | | NA | |
| NA | | NA | |
| NA | | NA | |
| NA | | NA | |
| NA | | NA | |
| NA | | NA | |
| NA | | NA | |
| NA | | NA | |
| NA | | NA | |
| | 0.871260465 | NA | |
| NA | | NA | |
| NA | | NA | |
| NA | | NA | |
| | 0.179701319 | | 0.319234792 |
| NA | | NA | |
| NA | | NA | |
| | 0.905158739 | NA | |
| NA | | NA | |
| | 0.889410717 | NA | |
| NA | | NA | |
| NA | | NA | |
| | 0.905158739 | NA | |
| NA | | NA | |
| NA | | NA | |
| NA | | NA | |
| NA | | NA | |
| NA | | NA | |
| NA | | NA | |
| NA | | NA | |

| | | |
|---|---|---|
| | 0.844908135 | NA |
| | 0.579628191 | NA |
| | 0.905158266 | NA |
| | 0.593612139 | NA |
| | 0.593612139 | NA |
| NA | | NA |
| | 0.784353866 | NA |
| NA | | NA |
| NA | | NA |
| | 0.917755362 | NA |
| NA | | NA |
| | 0.801661681 | NA |
| NA | | NA |
| NA | | NA |
| NA | | NA |
| NA | | NA |
| NA | | NA |
| | 0.905158266 | NA |
| | 0.889409773 | NA |
| | 0.96474 | NA |
| NA | | NA |
| | 0.767096791 | NA |
| | 0.992061292 | NA |
| NA | | NA |
| NA | | NA |
| NA | | NA |
| | 0.805374319 | NA |
| NA | | NA |
| | 0.905158266 | NA |
| | 0.905158739 | NA |
| NA | | NA |
| NA | | NA |
| | 0.272055044 | NA |
| NA | | NA |
| NA | | NA |
| NA | | NA |
| | 0.602928491 | NA |
| | 0.992060816 | NA |
| | 0.905158739 | NA |
| | 0.855405213 | NA |
| NA | | NA |
| | 0.889410245 | NA |
| | 0.432694209 | NA |
| | 0.905158739 | NA |
| NA | | NA |
| | 0.346000388 | NA |
| | 0.905158266 | NA |

| | |
|---|---|
| NA | NA |
| 0.236952598 | 0.388794897 |
| NA | NA |
| NA | NA |
| 0.889409773 | NA |
| NA | NA |
| 0.992061292 | NA |
| NA | NA |
| NA | NA |
| 0.959869571 | NA |
| NA | NA |
| 0.777779957 | NA |
| NA | NA |
| NA | NA |
| 0.01130996 | 0.037140034 |
| NA | NA |
| 0.991399481 | NA |
| 0.871260465 | NA |
| NA | NA |
| 0.552058421 | NA |
| NA | NA |
| 0.763837496 | NA |
| 0.991399481 | NA |
| NA | NA |
| 0.889409301 | NA |
| 0.866012479 | NA |
| 0.668619987 | NA |
| NA | NA |
| 0.376885255 | 0.541341138 |
| 0.712346218 | 0.820959728 |
| 0.938973465 | NA |
| 0.889410245 | NA |
| NA | NA |
| 0.992061292 | NA |
| NA | NA |
| 0.03267165 | 0.087236944 |
| 0.985781208 | 0.992788917 |
| 0.897540122 | NA |
| 0.190985221 | 0.333188566 |
| 0.263997212 | 0.41971261 |
| 5.22E-07 | 7.49E-06 |
| 0.714863669 | 0.822734939 |
| 0.091549704 | 0.192579744 |
| 0.401960638 | 0.56548489 |
| 0.029649762 | 0.080738979 |
| 0.176449597 | 0.314833566 |
| 0.601912378 | 0.737413184 |

| | | |
|---|---|---|
| | 0.250229011 | 0.404206183 |
| | 0.124644781 | 0.242818761 |
| | 8.04E-05 | 0.000580314 |
| | 0.049571278 | 0.120573294 |
| | 0.411828356 | 0.574508536 |
| | 0.660750492 | 0.782769748 |
| | 1.81E-06 | 2.19E-05 |
| | 0.762746762 | NA |
| | 1.20E-12 | 8.35E-11 |
| | 0.047918975 | 0.117373428 |
| | 0.038376654 | 0.098933193 |
| | 0.983706656 | NA |
| | 2.77E-05 | 0.000233869 |
| | 0.030254179 | 0.08199846 |
| | 1.54E-06 | 1.90E-05 |
| | 0.000470102 | 0.002618545 |
| | 5.70E-05 | 0.000433362 |
| | 2.52E-06 | 2.94E-05 |
| | 0.012882046 | 0.041229683 |
| | 0.002434716 | 0.010580632 |
| | 0.919513068 | 0.954020567 |
| | 7.53E-08 | 1.38E-06 |
| | 0.043050841 | 0.10803389 |
| | 0.114311412 | 0.227845832 |
| | 0.000433218 | 0.002446766 |
| | 0.07420163 | NA |
| | 0.00144377 | 0.006796117 |
| | 0.253258739 | NA |
| | 0.865723243 | 0.922567053 |
| | 0.713620472 | 0.821600794 |
| | 0.828274304 | NA |
| | 0.889410245 | NA |
| | 0.742777439 | 0.84236171 |
| | 0.001849545 | 0.008377508 |
| | 0.99875671 | NA |
| | 0.568100349 | 0.710740404 |
| | 0.011014726 | 0.036399154 |
| | 0.636021755 | NA |
| | 0.084024729 | 0.180432136 |
| NA | | NA |
| NA | | NA |
| NA | | NA |
| NA | | NA |
| NA | | NA |
| NA | | NA |
| | 0.787238384 | 0.871771467 |
| NA | | NA |

| | |
|---|---|
| NA | NA |
| 0.108949064 | NA |
| 1.00E-17 | 2.37E-15 |
| 5.06E-06 | 5.38E-05 |
| NA | NA |
| 0.334190567 | NA |
| 0.992061769 | NA |
| NA | NA |
| NA | NA |
| NA | NA |
| NA | NA |
| 0.122782756 | NA |
| NA | NA |
| NA | NA |
| NA | NA |
| NA | NA |
| 0.739951826 | NA |
| 1.93E-10 | 7.62E-09 |
| NA | NA |
| NA | NA |
| 0.852600757 | NA |
| NA | NA |
| 0.375112186 | NA |
| 0.468573397 | 0.624963753 |
| 0.071136034 | 0.158489871 |
| 0.333444947 | 0.49611204 |
| NA | NA |
| 0.002214116 | 0.009753216 |
| 0.009783936 | 0.033025113 |
| 0.000747691 | 0.003908246 |
| 0.000712404 | 0.00374575 |
| 0.747682653 | NA |
| 0.250357839 | 0.404325057 |
| 0.567914164 | 0.710628932 |
| 1.10E-05 | 0.000105609 |
| 2.55E-06 | 2.96E-05 |
| NA | NA |
| 0.000153208 | 0.001001575 |
| 4.10E-19 | 1.39E-16 |
| 0.720309692 | 0.826488491 |
| 0.992061769 | NA |
| 0.97685 | NA |
| 0.165437834 | 0.299740049 |
| 0.78751506 | NA |
| NA | NA |
| 0.43601615 | 0.59650356 |
| 0.181826439 | 0.32205477 |

| | |
|---|---|
| NA | NA |
| 0.000347265 | 0.002029705 |
| 0.097006198 | 0.201186164 |
| 0.243385934 | 0.396359517 |
| 0.880150995 | 0.932061907 |
| 0.021476568 | 0.062242404 |
| 0.038397267 | 0.098963096 |
| 3.76E-08 | 7.66E-07 |
| 2.92E-06 | 3.33E-05 |
| 6.71E-08 | 1.25E-06 |
| 4.84E-08 | 9.43E-07 |
| 2.33E-08 | 4.97E-07 |
| 0.441963651 | 0.601604253 |
| 0.364560327 | 0.529392746 |
| 0.156295522 | 0.287348741 |
| 0.606601953 | 0.741132381 |
| 0.147400136 | 0.275105006 |
| 0.114312885 | NA |
| 0.090867746 | 0.191530508 |
| NA | NA |
| 0.82888781 | NA |
| 0.568298942 | NA |
| 0.380553598 | NA |
| NA | NA |
| NA | NA |
| NA | NA |
| 0.805374319 | NA |
| NA | NA |
| 0.700420563 | NA |
| 0.083897309 | 0.18024665 |
| NA | NA |
| 0.000178188 | 0.001141434 |
| 0.178149965 | 0.317183502 |
| 0.405135731 | 0.568276942 |
| NA | NA |
| 0.763898469 | 0.856975911 |
| 0.41561323 | 0.578120336 |
| 8.70E-10 | 2.79E-08 |
| 0.889409773 | NA |
| 0.143870386 | 0.270113697 |
| 1.67E-10 | 6.70E-09 |
| 0.267018436 | NA |
| 0.905158739 | NA |
| 0.114338019 | 0.227876979 |
| 0.84153806 | NA |
| 0.003118147 | 0.012972978 |
| 0.902545984 | NA |

| | |
|---|---|
| NA | NA |
| 0.763657035 | NA |
| 0.381533766 | 0.545697907 |
| 0.129465477 | 0.249559155 |
| 0.317575229 | 0.479881258 |
| NA | NA |
| NA | NA |
| 0.187726224 | 0.329176728 |
| 0.001037674 | 0.005135517 |
| 9.50E-08 | 1.68E-06 |
| 5.42E-07 | 7.72E-06 |
| 0.418833878 | 0.580933042 |
| 0.569821212 | 0.712163005 |
| 0.849828941 | 0.91200243 |
| 0.314436016 | 0.476481988 |
| 0.549731704 | NA |
| 0.262269131 | 0.417724512 |
| 0.818251863 | 0.891756051 |
| 0.905158739 | NA |
| 0.875486472 | 0.928844683 |
| 0.851410992 | NA |
| 0.422290666 | 0.583857369 |
| 0.758349502 | 0.853322916 |
| 0.017226106 | 0.052183412 |
| 2.96E-07 | 4.55E-06 |
| 0.477315457 | NA |
| NA | NA |
| 0.004077468 | 0.016144451 |
| 0.414670955 | 0.577117473 |
| 0.659462452 | 0.781623359 |
| NA | NA |
| 0.905157793 | NA |
| 0.564200237 | NA |
| 0.005535674 | 0.020857559 |
| 0.921916474 | 0.955294241 |
| 0.712172578 | 0.820837583 |
| 0.137648243 | NA |
| 2.28E-05 | 0.000198532 |
| 0.038665638 | 0.099479642 |
| 0.589488934 | NA |
| 0.419706608 | 0.581540729 |
| 0.058705454 | 0.13719807 |
| 0.014694855 | 0.045907571 |
| 2.92E-11 | 1.44E-09 |
| 0.030695295 | 0.082958522 |
| 0.608621524 | 0.742773303 |
| 0.898514036 | 0.942868666 |

| | | |
|---|---|---|
| | 0.745627686 | 0.844590857 |
| | 0.704566733 | NA |
| | 0.2308094 | NA |
| | 4.24E-06 | 4.62E-05 |
| | 0.209220669 | 0.356035079 |
| NA | | NA |
| | 0.550914021 | 0.696581288 |
| | 2.19E-11 | 1.11E-09 |
| | 0.214214883 | 0.361595192 |
| | 0.038570779 | 0.099293756 |
| | 1.68E-11 | 8.84E-10 |
| | 0.276784728 | 0.434447165 |
| | 0.000192155 | 0.00121811 |
| | 0.539940646 | 0.687217102 |
| | 1.08E-06 | 1.40E-05 |
| | 0.398940553 | NA |
| | 0.293102962 | 0.45277861 |
| NA | | NA |
| | 0.099059043 | NA |
| NA | | NA |
| | 0.147278999 | 0.274938327 |
| | 0.042265969 | NA |
| | 0.545571619 | 0.691977542 |
| | 1.40E-05 | 0.000130122 |
| | 0.000159973 | 0.001039614 |
| | 0.762604749 | 0.856005607 |
| | 2.58E-05 | 0.000220718 |
| | 0.001151268 | 0.005607267 |
| | 0.48115991 | 0.636800293 |
| | 0.774058997 | NA |
| | 0.004267874 | 0.016768341 |
| | 0.240289765 | 0.392773779 |
| | 0.501724351 | 0.654471012 |
| | 1.94E-07 | 3.16E-06 |
| | 0.283708334 | NA |
| | 0.000607805 | 0.003276089 |
| NA | | NA |
| | 0.000272408 | 0.001649241 |
| | 0.025927398 | 0.072567205 |
| | 0.79546022 | NA |
| | 0.015515516 | 0.048000171 |
| | 4.47E-06 | 4.84E-05 |
| NA | | NA |
| | 0.241862222 | 0.394609743 |
| NA | | NA |
| | 0.452891626 | NA |
| | 0.274764438 | 0.431986274 |

| | |
|---|---|
| NA | NA |
| 0.009225184 | 0.031490102 |
| 8.82E-11 | 3.80E-09 |
| 0.24604129 | 0.399380596 |
| 0.154710843 | 0.285295213 |
| 0.911310337 | 0.949941313 |
| 0.000602046 | 0.003251373 |
| 0.28248341 | 0.441090321 |
| 0.938775184 | 0.965417706 |
| 0.01006595 | 0.033844461 |
| 0.230721331 | 0.381535486 |
| 0.43239396 | NA |
| 0.482036166 | NA |
| NA | NA |
| 0.713177114 | NA |
| 0.807879735 | NA |
| 0.996443322 | NA |
| 0.758286641 | 0.853322916 |
| 0.484659195 | 0.639850374 |
| 0.154267316 | 0.284716422 |
| NA | NA |
| 0.905158266 | NA |
| NA | NA |
| NA | NA |
| NA | NA |
| NA | NA |
| 0.905158266 | NA |
| NA | NA |
| NA | NA |
| NA | NA |
| NA | NA |
| NA | NA |
| NA | NA |
| NA | NA |
| NA | NA |
| NA | NA |
| NA | NA |
| NA | NA |
| 0.889409773 | NA |
| NA | NA |
| NA | NA |
| NA | NA |
| 0.983771031 | NA |
| 0.474453786 | NA |
| 0.889409773 | NA |
| NA | NA |
| NA | NA |

| | |
|---|---|
| NA | NA |
| 0.740948197 | NA |
| 0.889410245 | NA |
| 0.579348667 | 0.719891205 |
| 0.005713635 | 0.021419964 |
| 0.000197865 | 0.00124926 |
| 1.43E-05 | 0.000133181 |
| 0.991879747 | NA |
| 0.974616457 | 0.986794669 |
| 7.42E-05 | 0.000540952 |
| 0.647728026 | 0.772642982 |
| 0.938975383 | 0.965481746 |
| NA | NA |
| 0.748686828 | 0.846702068 |
| 0.664759109 | 0.785949781 |
| 1.92E-18 | 5.70E-16 |
| 0.905157793 | NA |
| 9.84E-07 | 1.29E-05 |
| 0.00234116 | 0.010228717 |
| 0.80267173 | 0.881293426 |
| 0.574542171 | NA |
| 0.563624018 | 0.707316254 |
| 0.199563507 | 0.344351331 |
| 4.10E-09 | 1.07E-07 |
| 0.799750721 | 0.879229303 |
| 0.647814138 | 0.77265497 |
| 0.24911697 | 0.402884044 |
| 0.154959583 | 0.285657949 |
| 0.312381665 | 0.474154212 |
| 0.663666011 | 0.784953244 |
| 0.014767099 | 0.046087369 |
| 0.992061292 | NA |
| 2.22E-08 | 4.78E-07 |
| 0.856089717 | 0.916124469 |
| 0.974329402 | 0.986549515 |
| 0.00331587 | 0.013625222 |
| 1.82E-08 | 4.00E-07 |
| 0.889410717 | NA |
| 0.387317225 | 0.551023791 |
| 0.010260461 | 0.03441941 |
| NA | NA |
| 0.002466597 | 0.010700119 |
| NA | NA |
| NA | NA |
| NA | NA |
| 8.72E-06 | 8.68E-05 |
| NA | NA |

| | |
|---|---|
| NA | NA |
| 0.042557991 | 0.107109561 |
| 0.014777285 | 0.046112605 |
| 0.983938807 | NA |
| 0.992059387 | NA |
| NA | NA |
| NA | NA |
| NA | NA |
| 0.276040866 | NA |
| 0.086743957 | 0.184771612 |
| 0.477008338 | NA |
| NA | NA |
| 0.086654914 | 0.18460605 |
| 0.0004077 | 0.00232356 |
| 0.665471996 | NA |
| NA | NA |
| NA | NA |
| 0.72456281 | 0.830064417 |
| 0.330521397 | 0.49303168 |
| 0.131037543 | 0.251635628 |
| 0.390862243 | 0.554203138 |
| 0.219835642 | 0.368696418 |
| 0.020329096 | 0.059558855 |
| 0.785411531 | NA |
| 0.062462849 | 0.144093046 |
| 0.707078588 | NA |
| 0.242673564 | NA |
| NA | NA |
| NA | NA |
| 0.000463333 | 0.002585598 |
| 0.905158266 | NA |
| 0.02800292 | 0.07710547 |
| 1.89E-06 | 2.28E-05 |
| 0.904947233 | NA |
| 0.195570553 | 0.339381522 |
| 0.433882614 | 0.594651468 |
| 0.75537433 | 0.851300265 |
| NA | NA |
| 0.003671117 | 0.014805153 |
| 0.002296667 | 0.010058338 |
| 0.006526886 | 0.023855507 |
| 0.204259641 | 0.350094613 |
| 0.000163823 | 0.001061487 |
| NA | NA |
| NA | NA |
| 0.037040993 | 0.096316824 |
| 0.905158266 | NA |

| | |
|---|---|
| NA | NA |
| 0.05131503 | 0.123889755 |
| 0.291126972 | 0.450581169 |
| 0.273533523 | 0.4305751 |
| 0.992061769 | NA |
| 0.000691128 | 0.003653464 |
| 0.000296786 | 0.001770479 |
| 0.112820276 | NA |
| 0.207996154 | NA |
| 0.001651556 | 0.007580795 |
| NA | NA |
| NA | NA |
| 0.018974953 | 0.056291676 |
| 0.85710661 | 0.916707965 |
| NA | NA |
| NA | NA |
| 0.007738223 | 0.027309771 |
| NA | NA |
| NA | NA |
| 0.62095537 | 0.752306701 |
| 1.23E-07 | 2.13E-06 |
| 0.341248161 | NA |
| 0.527759502 | 0.677375497 |
| 0.008939502 | 0.030705955 |
| NA | NA |
| NA | NA |
| 0.003706544 | 0.01491986 |
| 0.040387123 | 0.102776779 |
| 0.000209364 | 0.001310182 |
| 0.738002739 | 0.839200567 |
| 8.75E-05 | 0.00062197 |
| 0.153604365 | 0.283802967 |
| 0.328805441 | NA |
| 0.252453469 | 0.406749661 |
| 0.135857918 | 0.258540951 |
| 0.734703036 | NA |
| 0.718055833 | NA |
| 0.959169669 | 0.977283164 |
| 0.481659144 | 0.637268973 |
| 0.000124533 | 0.000842716 |
| 0.003633327 | 0.014695932 |
| 0.025725657 | 0.072192932 |
| 0.670638333 | 0.789966052 |
| 0.992062245 | NA |
| NA | NA |
| NA | NA |
| NA | NA |

| | | | |
|---|---|---|---|
| NA | | NA | |
| NA | | NA | |
| | 0.188014153 | | 0.329418557 |
| | 0.726763741 | NA | |
| | 0.123535062 | NA | |
| | 0.121704593 | | 0.238574464 |
| | 0.01501329 | | 0.04674278 |
| NA | | NA | |
| | 0.918519996 | | 0.953532033 |
| | 0.889409773 | NA | |
| | 0.093171033 | | 0.195037437 |
| | 0.000654711 | | 0.003497152 |
| | 0.171148616 | | 0.307599645 |
| | 0.00035669 | | 0.00207595 |
| | 0.992060816 | NA | |
| NA | | NA | |
| | 2.81E-06 | | 3.23E-05 |
| | 0.992060816 | NA | |
| | 0.028298736 | | 0.077763977 |
| | 0.107179839 | | 0.216958156 |
| | 0.000241193 | | 0.001485067 |
| | 0.725807165 | | 0.830666573 |
| | 0.739500989 | NA | |
| | 1.50E-07 | | 2.52E-06 |
| | 0.797553171 | | 0.87774295 |
| | 0.244351335 | NA | |
| | 0.030500049 | | 0.082552763 |
| | 0.010169244 | | 0.034139494 |
| | 0.243621387 | | 0.396595896 |
| | 0.166340662 | | 0.301003264 |
| | 0.780651689 | | 0.867790343 |
| | 0.06072927 | | 0.141107656 |
| | 0.905158739 | NA | |
| | 0.597716773 | NA | |
| | 0.004368759 | | 0.017097413 |
| | 0.428553508 | | 0.589620368 |
| | 0.000149619 | | 0.000981915 |
| | 0.648957854 | | 0.773640908 |
| | 3.88E-05 | | 0.000312114 |
| | 0.000740502 | | 0.003877129 |
| | 0.815510873 | NA | |
| | 0.395748137 | | 0.559361055 |
| | 0.255012536 | NA | |
| | 0.217277685 | | 0.365491824 |
| | 0.110785355 | | 0.222269128 |
| | 0.991799276 | NA | |
| | 0.831992043 | | 0.900030212 |

| | | |
|---|---|---|
| | 0.027256977 | 0.075496462 |
| | 0.210401738 | NA |
| | 0.000603073 | 0.003254581 |
| | 0.711305507 | NA |
| | 1.10E-09 | 3.43E-08 |
| | 8.40E-10 | 2.71E-08 |
| NA | | NA |
| | 0.226198317 | 0.376029168 |
| | 3.68E-05 | 0.000297852 |
| | 1.69E-12 | 1.14E-10 |
| | 0.000672946 | 0.003574556 |
| | 0.197104992 | 0.341304152 |
| | 4.21E-06 | 4.59E-05 |
| | 0.022373829 | 0.06435773 |
| NA | | NA |
| | 0.115303005 | 0.229425398 |
| | 0.00138782 | 0.006575003 |
| | 0.066159139 | 0.150329939 |
| | 0.647744895 | 0.772642982 |
| | 0.034745656 | 0.091594221 |
| | 0.231701403 | 0.382676555 |
| | 0.414035293 | NA |
| | 0.403153342 | 0.566654563 |
| | 0.245602717 | 0.398918058 |
| | 0.462790229 | 0.620037257 |
| NA | | NA |
| | 3.35E-07 | 5.06E-06 |
| NA | | NA |
| | 0.005779304 | 0.0216234 |
| | 0.889410717 | NA |
| | 0.068009798 | 0.153093506 |
| | 0.643535768 | 0.769055887 |
| | 0.957760601 | 0.976545197 |
| | 0.071296446 | 0.158722567 |
| | 0.761763546 | 0.855521176 |
| | 0.001165056 | 0.005667583 |
| | 0.836395626 | NA |
| | 2.85E-08 | 5.95E-07 |
| | 0.043278546 | 0.108438337 |
| | 0.877359124 | NA |
| | 0.913362153 | 0.950951568 |
| | 0.922716336 | 0.955421158 |
| | 0.802979442 | 0.881454988 |
| | 0.000138796 | 0.000922745 |
| | 0.084912099 | 0.181826134 |
| NA | | NA |
| | 0.015424036 | 0.04775753 |

| | | |
|---|---|---|
| | 0.623149859 | NA |
| | 0.628587632 | NA |
| | 0.905158266 | NA |
| | 0.889410245 | NA |
| NA | | NA |
| NA | | NA |
| | 0.009691693 | 0.032797023 |
| | 0.002218097 | 0.009766831 |
| NA | | NA |
| | 0.67668738 | 0.794746653 |
| | 0.117719601 | 0.232757493 |
| NA | | NA |
| | 0.107228251 | 0.216991602 |
| NA | | NA |
| | 0.98945134 | NA |
| | 0.782427369 | NA |
| | 1.05E-10 | 4.42E-09 |
| | 0.873830171 | NA |
| | 0.354661724 | 0.51930498 |
| | 0.003591169 | 0.014557592 |
| | 0.905158266 | NA |
| | 5.78E-09 | 1.45E-07 |
| | 0.75211653 | 0.849006676 |
| | 0.385491609 | NA |
| | 0.889409773 | NA |
| | 0.082204988 | 0.177653708 |
| | 9.29E-09 | 2.21E-07 |
| | 0.655120074 | NA |
| | 0.509545229 | 0.661172971 |
| | 0.580672523 | 0.72084278 |
| | 0.035031987 | 0.092069649 |
| | 0.000498092 | 0.002753373 |
| | 0.627405879 | 0.75642702 |
| NA | | NA |
| | 0.305125119 | 0.465716603 |
| | 0.2560449 | 0.410580543 |
| | 0.001224816 | 0.005900774 |
| | 0.266682174 | 0.422911662 |
| NA | | NA |
| | 0.343893425 | 0.507263207 |
| | 0.110764624 | 0.22226815 |
| | 0.000437316 | 0.002466975 |
| | 0.002390324 | 0.010416576 |
| | 0.001593944 | 0.007367905 |
| | 0.183927705 | 0.324490955 |
| | 0.752057046 | 0.849003885 |
| | 0.707648758 | 0.817782279 |

| | |
|---|---|
| NA | NA |
| 0.778234898 | 0.866682731 |
| 0.747519131 | 0.845947877 |
| 3.05E-05 | 0.000253476 |
| 0.143970142 | 0.270277879 |
| 0.948684143 | 0.971060229 |
| 0.621357752 | 0.752682924 |
| NA | NA |
| 0.388318389 | 0.551962569 |
| NA | NA |
| 0.159282525 | 0.29127956 |
| 0.42086797 | 0.582640257 |
| NA | NA |
| 0.005939256 | 0.022086956 |
| 0.000940602 | 0.004731887 |
| 0.397962 | 0.561514332 |
| 0.036302525 | 0.094777767 |
| 0.889409773 | NA |
| 0.579477924 | 0.719901233 |
| 5.02E-05 | 0.000388529 |
| 1.30E-05 | 0.000122544 |
| 0.782297516 | 0.868872144 |
| 0.072517633 | 0.160882472 |
| 0.399748617 | 0.563355571 |
| 0.157510223 | 0.289001253 |
| 0.881846723 | NA |
| 9.88E-06 | 9.60E-05 |
| 0.589567227 | 0.7280159 |
| 0.443852151 | 0.603464144 |
| 0.011850414 | 0.038545881 |
| 0.730822156 | 0.834459624 |
| NA | NA |
| 0.324167024 | 0.486394511 |
| 0.120538011 | 0.236922144 |
| 1.75E-05 | 0.000158424 |
| 0.350176354 | NA |
| 0.905296333 | NA |
| NA | NA |
| 0.991570563 | NA |
| NA | NA |
| 0.997008258 | 0.998640084 |
| 0.889409773 | NA |
| 0.065250816 | 0.148900728 |
| 0.774620509 | 0.864060335 |
| NA | NA |
| 0.804515329 | 0.882580395 |
| 0.73696466 | 0.838454327 |

| | |
|---|---|
| 0.481537925 | 0.637185375 |
| NA | NA |
| 0.057529273 | NA |
| 0.680292179 | NA |
| 0.371764392 | NA |
| 2.74E-11 | 1.36E-09 |
| 0.790988712 | NA |
| NA | NA |
| 0.905158739 | NA |
| 0.596689173 | 0.733551537 |
| 0.940432401 | NA |
| 0.283019577 | NA |
| 0.199313545 | 0.344055194 |
| 0.097911185 | 0.202603818 |
| 0.114270138 | 0.227824403 |
| 0.911755388 | 0.950078288 |
| 0.914356907 | NA |
| 3.55E-10 | 1.29E-08 |
| 0.365765794 | 0.530505354 |
| NA | NA |
| 7.63E-05 | 0.000555358 |
| 7.81E-05 | 0.00056706 |
| 0.023064129 | NA |
| 0.806503376 | NA |
| 0.04325286 | 0.108386346 |
| 0.740216436 | NA |
| NA | NA |
| 0.78190939 | NA |
| 0.012755442 | 0.040949819 |
| 0.005108673 | 0.019511917 |
| 0.326588379 | 0.488858008 |
| NA | NA |
| NA | NA |
| NA | NA |
| NA | NA |
| NA | NA |
| NA | NA |
| NA | NA |
| NA | NA |
| NA | NA |
| NA | NA |
| 0.889409773 | NA |
| NA | NA |
| NA | NA |
| NA | NA |
| NA | NA |
| 0.495914279 | NA |

| | |
|---|---|
| NA | NA |
| NA | NA |
| 0.889409301 | NA |
| NA | NA |
| NA | NA |
| NA | NA |
| NA | NA |
| 0.528529939 | NA |
| 0.982573707 | NA |
| 8.78E-06 | 8.72E-05 |
| 0.307116049 | NA |
| 0.01699064 | 0.051661238 |
| 0.000170277 | 0.001096834 |
| 0.142922747 | 0.268702107 |
| 0.0750751 | 0.165436439 |
| 0.033893069 | 0.089745315 |
| NA | NA |
| 0.137876734 | 0.261521568 |
| 0.015620095 | 0.048276096 |
| 0.062947309 | 0.14490617 |
| 0.000119914 | 0.000816748 |
| 0.012416689 | 0.040073204 |
| 0.496966541 | NA |
| 0.833307007 | NA |
| 0.992060816 | NA |
| NA | NA |
| NA | NA |
| 4.71E-08 | 9.24E-07 |
| 0.750807961 | NA |
| 0.005290352 | 0.020098762 |
| 0.194232631 | 0.337513203 |
| NA | NA |
| 0.003496195 | 0.014240949 |
| 0.017839726 | 0.053656012 |
| 5.70E-08 | 1.09E-06 |
| 0.281558744 | 0.439959329 |
| 0.308177959 | 0.469200569 |
| 0.193011506 | 0.335976192 |
| 0.897222376 | 0.942074668 |
| 0.784748935 | 0.870362233 |
| 0.004075679 | 0.016140279 |
| 0.017537325 | 0.052920451 |
| 0.250853635 | 0.405006615 |
| 6.13E-05 | 0.000460457 |
| 0.072994505 | 0.161679075 |
| 0.17554551 | 0.313475144 |
| 0.046123396 | 0.113992533 |

| | |
|---|---|
| NA | NA |
| 0.002530374 | 0.010916403 |
| 0.050437593 | 0.122206677 |
| 0.121576178 | 0.238386578 |
| 0.086038558 | 0.183704799 |
| 0.085290524 | 0.182418878 |
| 0.174814023 | 0.31247384 |
| NA | NA |
| 0.286477987 | 0.445521964 |
| 4.32E-05 | 0.0003416 |
| 0.367081084 | 0.531807781 |
| 0.050674501 | 0.122658881 |
| 0.452311031 | 0.610542042 |
| 0.905158739 | NA |
| NA | NA |
| 0.00037836 | 0.002177828 |
| 0.429366814 | 0.590423062 |
| 0.28123121 | 0.439635234 |
| 0.000247832 | 0.001518032 |
| 0.037440557 | 0.097089891 |
| NA | NA |
| NA | NA |
| 0.801661681 | NA |
| 6.65E-06 | 6.85E-05 |
| 0.440117642 | 0.600265306 |
| 3.92E-10 | 1.40E-08 |
| 0.226946292 | 0.376928908 |
| 1.04E-08 | 2.45E-07 |
| 0.316436441 | 0.478514732 |
| 0.71453369 | NA |
| 1.83E-06 | 2.21E-05 |
| 5.39E-12 | 3.20E-10 |
| 0.483198854 | NA |
| 0.627709295 | NA |
| 0.070855327 | 0.158072959 |
| NA | NA |
| 1.75E-05 | 0.000158424 |
| 0.991939052 | 0.995930211 |
| 6.73E-05 | 0.000498483 |
| 0.08165419 | 0.176706845 |
| 0.923001563 | 0.955550956 |
| 0.918415454 | 0.953500305 |
| 0.889409301 | NA |
| 0.925606605 | NA |
| 0.268812078 | 0.425185078 |
| 0.645997239 | 0.771235488 |
| NA | NA |

| | |
|---|---|
| 0.728042243 | 0.832226945 |
| NA | NA |
| 0.800158696 | NA |
| 0.615454985 | NA |
| 0.081056809 | 0.175710133 |
| 0.647303478 | 0.772213673 |
| 0.530917127 | 0.680107875 |
| 0.905158739 | NA |
| 0.010433817 | 0.034862475 |
| 0.268521301 | 0.424930348 |
| 0.137828484 | 0.261452632 |
| 0.965252723 | 0.981159309 |
| 0.755896765 | NA |
| NA | NA |
| 0.007742958 | 0.027318063 |
| 0.797342596 | NA |
| 0.117106948 | 0.231963871 |
| 0.899748372 | NA |
| NA | NA |
| 0.905158266 | NA |
| 1.49E-08 | 3.33E-07 |
| 0.010286355 | 0.034490468 |
| 0.055293669 | 0.130919405 |
| 0.981407085 | 0.990700969 |
| 0.048636219 | 0.118719735 |
| NA | NA |
| 0.474239501 | 0.630529499 |
| 0.889409301 | NA |
| 0.788582525 | NA |
| NA | NA |
| 0.782537258 | NA |
| 0.329956616 | NA |
| 0.013335008 | NA |
| 0.424949336 | 0.58618396 |
| 0.071602733 | NA |
| 0.084328935 | 0.180731904 |
| 0.823185617 | NA |
| 9.72E-05 | 0.000681576 |
| 0.517047348 | 0.668148221 |
| 0.186571792 | 0.327859313 |
| NA | NA |
| 0.905158739 | NA |
| 0.010800271 | 0.035828283 |
| 2.25E-08 | 4.84E-07 |
| 0.247493987 | NA |
| 0.269118663 | 0.425516757 |
| 0.440967686 | 0.600844467 |

| | |
|---|---|
| 0.326383762 | 0.488680278 |
| 0.944715468 | 0.968866415 |
| 0.000162948 | 0.001056754 |
| 0.40589365 | 0.56915828 |
| 1.83E-05 | 0.000164557 |
| 0.863411048 | 0.92117594 |
| 0.412830021 | NA |
| 0.900897778 | NA |
| 0.02674381 | 0.074290616 |
| 0.276206332 | 0.433714446 |
| 0.049307907 | 0.120065683 |
| NA | NA |
| 0.770967739 | NA |
| NA | NA |
| 0.014863506 | 0.046322413 |
| 0.242654035 | 0.395548954 |
| 0.416141858 | NA |
| 0.177104688 | 0.315771504 |
| 0.062353934 | 0.143890063 |
| 0.353907556 | NA |
| 0.377885593 | NA |
| 0.014280089 | 0.044841482 |
| NA | NA |
| 0.807252815 | NA |
| 0.418494216 | 0.580672069 |
| 0.99205891 | NA |
| 0.276416041 | NA |
| NA | NA |
| 0.06314543 | 0.145179615 |
| 0.382030774 | NA |
| NA | NA |
| 0.807879735 | NA |
| 4.95E-07 | 7.15E-06 |
| 0.99205891 | NA |
| 3.85E-06 | 4.26E-05 |
| 0.003768716 | 0.015112717 |
| 0.842354141 | NA |
| 9.07E-06 | 8.96E-05 |
| 0.247467067 | 0.401101958 |
| 0.9995531 | NA |
| 0.727322352 | 0.831648306 |
| 0.281474705 | 0.439859311 |
| 0.133911189 | 0.255656043 |
| 6.61E-06 | 6.82E-05 |
| 0.905158266 | NA |
| 0.583646887 | 0.723553435 |
| 0.00140222 | 0.006634636 |

| | |
|---|---|
| 0.00029799 | 0.001775727 |
| 1.04E-05 | 0.000100016 |
| 0.001850609 | 0.0083806 |
| 0.014667113 | 0.045846995 |
| 0.000396881 | 0.002270744 |
| 0.02132959 | 0.061905428 |
| NA | NA |
| 0.905158266 | NA |
| 0.001474005 | 0.0069067 |
| 0.000271466 | 0.0016449 |
| 0.905158739 | NA |
| 0.889409773 | NA |
| 0.319769157 | 0.482134779 |
| NA | NA |
| NA | NA |
| 0.30970656 | NA |
| NA | NA |
| NA | NA |
| NA | NA |
| 0.502783902 | NA |
| NA | NA |
| NA | NA |
| NA | NA |
| 0.992059387 | NA |
| NA | NA |
| NA | NA |
| NA | NA |
| 0.246512837 | NA |
| 0.000441071 | 0.002483969 |
| 0.591876322 | NA |
| 0.07516941 | NA |
| NA | NA |
| 0.000126078 | 0.000849767 |
| 0.696810292 | NA |
| NA | NA |
| NA | NA |
| NA | NA |
| 0.889409773 | NA |
| NA | NA |
| 0.889409773 | NA |
| NA | NA |
| NA | NA |
| 0.873616163 | NA |
| NA | NA |
| NA | NA |
| 0.137810899 | NA |
| 3.16E-07 | 4.80E-06 |

| | | | |
|---|---|---|---|
| | 0.435944893 | NA | |
| | 0.214217034 | | 0.361595192 |
| | 0.049898738 | | 0.121141668 |
| | 0.000456866 | | 0.002555353 |
| | 0.29765016 | | 0.457613937 |
| | 0.850055606 | NA | |
| NA | | NA | |
| NA | | NA | |
| NA | | NA | |
| NA | | NA | |
| NA | | NA | |
| NA | | NA | |
| NA | | NA | |
| NA | | NA | |
| NA | | NA | |
| NA | | NA | |
| NA | | NA | |
| NA | | NA | |
| | 0.905158266 | NA | |
| NA | | NA | |
| NA | | NA | |
| NA | | NA | |
| NA | | NA | |
| NA | | NA | |
| NA | | NA | |
| NA | | NA | |
| NA | | NA | |
| NA | | NA | |
| NA | | NA | |
| NA | | NA | |
| | 0.905158739 | NA | |
| NA | | NA | |
| NA | | NA | |
| | 0.905158739 | NA | |
| NA | | NA | |
| NA | | NA | |
| NA | | NA | |
| NA | | NA | |
| NA | | NA | |
| NA | | NA | |
| NA | | NA | |
| NA | | NA | |
| | 0.007119458 | | 0.02558275 |
| NA | | NA | |
| | 0.889409773 | NA | |
| NA | | NA | |
| NA | | NA | |

| | |
|---|---|
| 0.763446562 | NA |
| 0.992061292 | NA |
| 0.734293567 | 0.836802763 |
| 0.601121176 | NA |
| 0.271039472 | NA |
| 0.973182336 | NA |
| 0.006491834 | 0.02375507 |
| 0.085165252 | 0.182186436 |
| 3.47E-05 | 0.000283277 |
| 0.01347464 | 0.042721575 |
| 0.85593857 | 0.916052006 |
| 0.0905434 | 0.191085297 |
| 0.996424312 | 0.998242688 |
| 0.55692567 | 0.701999356 |
| 0.885802412 | NA |
| 0.056198014 | 0.132551255 |
| 0.14080196 | 0.265603017 |
| 0.328708408 | 0.490946276 |
| 0.295766641 | NA |
| 0.034144864 | 0.09025967 |
| NA | NA |
| 1.48E-08 | 3.31E-07 |
| 0.820627937 | NA |
| NA | NA |
| 0.070981951 | 0.158275047 |
| 4.03E-06 | 4.43E-05 |
| 0.111169579 | NA |
| 1.71E-08 | 3.78E-07 |
| 0.744363501 | NA |
| 0.805374319 | NA |
| 0.011843963 | 0.038537204 |
| 0.018003046 | 0.054018396 |
| 0.905158739 | NA |
| 0.270165136 | 0.426618453 |
| 0.992059387 | NA |
| 0.992269016 | NA |
| 0.992061292 | NA |
| NA | NA |
| NA | NA |
| 0.475197777 | 0.631344869 |
| 0.086598172 | 0.184550589 |
| 0.031269782 | 0.084179619 |
| 0.912323985 | 0.950411423 |
| 4.79E-05 | 0.000373713 |
| 0.462925748 | 0.620127308 |
| 0.545109005 | 0.691592071 |
| 0.017307784 | 0.052378946 |

| | | |
|---|---|---|
| | 0.255620884 | 0.41020016 |
| | 0.053140774 | 0.127018422 |
| | 0.642749333 | 0.768450624 |
| | 0.065644044 | 0.149453349 |
| | 0.203960957 | 0.349795272 |
| | 0.000146104 | 0.00096143 |
| | 0.346329848 | NA |
| | 0.001103962 | 0.005409932 |
| | 0.411728786 | 0.574442621 |
| | 0.007334743 | 0.02620625 |
| | 0.015322642 | 0.047510575 |
| | 1.02E-05 | 9.88E-05 |
| | 0.059851896 | 0.13944698 |
| | 0.048739785 | 0.11891966 |
| NA | | NA |
| | 0.42608094 | NA |
| | 6.11E-06 | 6.36E-05 |
| | 3.33E-30 | 3.66E-26 |
| | 0.009323241 | 0.031770493 |
| | 0.001691608 | NA |
| | 0.0014824 | NA |
| NA | | NA |
| | 0.809608583 | NA |
| | 0.364182786 | 0.529153034 |
| | 0.980539945 | NA |
| | 0.442473995 | 0.60218695 |
| | 0.889409773 | NA |
| NA | | NA |
| NA | | NA |
| | 0.000163359 | 0.001059111 |
| | 1.98E-05 | 0.000176197 |
| | 0.007782835 | 0.027441117 |
| | 0.317591855 | NA |
| | 0.227898107 | 0.378005478 |
| | 0.723136725 | NA |
| | 0.324657505 | 0.486897565 |
| | 0.000762192 | 0.00396517 |
| | 0.535020447 | 0.683377626 |
| | 0.799585299 | 0.87917949 |
| | 0.756957183 | NA |
| NA | | NA |
| | 0.116285131 | 0.230885124 |
| | 0.905157793 | NA |
| NA | | NA |
| | 0.612084933 | NA |
| NA | | NA |
| | 0.966773374 | NA |

| | | |
|---|---|---|
| | 0.905158266 | NA |
| | 0.850352973 | NA |
| | 0.004177206 | 0.016476966 |
| NA | | NA |
| | 0.99205891 | NA |
| | 0.999678457 | 0.999864481 |
| | 0.500725714 | 0.653547133 |
| | 0.000381994 | 0.002196439 |
| | 0.152534466 | 0.282372648 |
| | 0.504753805 | NA |
| | 0.000839754 | 0.004298405 |
| | 0.208274791 | 0.354975639 |
| | 0.000735402 | 0.003856866 |
| NA | | NA |
| | 0.84955439 | 0.911838902 |
| | 0.081291874 | 0.176113681 |
| | 0.186461731 | NA |
| | 1.77E-13 | 1.51E-11 |
| | 0.415945012 | 0.578414066 |
| | 0.184138234 | 0.324752348 |
| | 0.139205178 | 0.263381571 |
| NA | | NA |
| NA | | NA |
| | 0.992061292 | NA |
| | 0.461543441 | 0.618971077 |
| | 0.992062245 | NA |
| NA | | NA |
| | 0.013292214 | 0.042253001 |
| | 0.753581177 | NA |
| NA | | NA |
| NA | | NA |
| NA | | NA |
| | 0.230210865 | 0.380949539 |
| | 0.118250258 | NA |
| | 0.076606045 | 0.168151954 |
| | 0.831325383 | NA |
| | 0.224573193 | 0.374202853 |
| | 0.693759852 | 0.807442869 |
| | 0.905158266 | NA |
| | 0.98898572 | 0.993992378 |
| | 0.928909606 | 0.959315527 |
| | 0.004922269 | 0.018922986 |
| | 0.815005768 | 0.889808167 |
| NA | | NA |
| | 0.656987271 | NA |
| | 0.991918092 | NA |
| | 0.154986774 | 0.285660112 |

| | | |
|---|---|---|
| NA | | NA |
| | 0.923128013 | NA |
| NA | | NA |
| NA | | NA |
| | 0.805374319 | NA |
| | 0.786591437 | 0.871533027 |
| | 0.020507422 | 0.060010992 |
| | 0.887783759 | 0.936517531 |
| | 0.498015978 | 0.650953464 |
| | 0.992061292 | NA |
| | 0.536274757 | 0.684415336 |
| | 0.718455054 | 0.8253614 |
| | 0.915332341 | 0.952099989 |
| | 0.318755602 | 0.481201311 |
| | 0.402902223 | 0.566345339 |
| | 0.000351364 | 0.002049294 |
| | 0.010906165 | NA |
| | 0.951073635 | 0.972767188 |
| | 4.06E-05 | 0.000323821 |
| | 0.138692495 | 0.262637849 |
| | 0.181839626 | NA |
| | 9.24E-08 | 1.64E-06 |
| | 0.889409301 | NA |
| | 0.484658497 | 0.639850374 |
| | 0.97267076 | 0.985824667 |
| | 0.127076114 | 0.246333539 |
| | 0.304043952 | 0.464710875 |
| | 0.739605431 | NA |
| | 0.968057942 | NA |
| | 0.889409301 | NA |
| | 0.845093727 | 0.908786149 |
| | 0.990800687 | 0.995333376 |
| | 0.656020204 | 0.778804545 |
| | 0.826977301 | 0.89672859 |
| | 0.992061292 | NA |
| | 0.983414312 | NA |
| | 0.290453663 | 0.449825234 |
| | 0.002556494 | 0.01100763 |
| | 0.055059135 | 0.130538063 |
| | 0.289718909 | 0.449099077 |
| | 0.504690825 | 0.656816024 |
| | 0.039380539 | 0.10089339 |
| | 0.889409301 | NA |
| | 0.668130297 | 0.788322669 |
| | 0.889409301 | NA |
| | 0.889410245 | NA |
| | 0.313595652 | NA |

| | |
|---|---|
| NA | NA |
| 0.122564425 | 0.239724967 |
| 0.029395443 | 0.080175696 |
| 0.797233901 | NA |
| 0.438023863 | 0.598392809 |
| 0.818718334 | 0.892155344 |
| NA | NA |
| 0.837208274 | 0.903172899 |
| 0.789541906 | NA |
| NA | NA |
| NA | NA |
| 0.245998347 | 0.399359157 |
| NA | NA |
| 0.625461414 | 0.755340311 |
| 0.900897778 | NA |
| 0.055464251 | 0.131215201 |
| 0.368909233 | NA |
| 0.128358468 | 0.2480746 |
| 1.99E-05 | 0.000176908 |
| 0.882207307 | NA |
| 0.462367968 | 0.619660555 |
| 0.836621863 | NA |
| 0.992060816 | NA |
| 0.002289248 | 0.010029849 |
| 0.353713229 | NA |
| 0.904694311 | NA |
| 0.18253657 | 0.32270514 |
| NA | NA |
| 0.170918234 | 0.307356228 |
| 0.086137096 | NA |
| NA | NA |
| NA | NA |
| NA | NA |
| NA | NA |
| NA | NA |
| NA | NA |
| NA | NA |
| NA | NA |
| NA | NA |
| 0.665884952 | NA |
| 0.003037833 | 0.012680336 |
| 1.37E-08 | 3.11E-07 |
| 1.81E-06 | 2.19E-05 |
| 0.001007913 | 0.005009679 |
| NA | NA |
| 0.060878281 | 0.141343432 |
| NA | NA |

| | |
|---|---|
| NA | NA |
| 0.618231376 | 0.750365469 |
| 0.416020658 | 0.578445997 |
| 0.812531559 | 0.887986794 |
| 0.133496179 | 0.255085502 |
| 0.019612831 | 0.057879069 |
| 0.099972539 | 0.205989596 |
| NA | NA |
| 0.205461626 | 0.351654859 |
| 0.594840289 | 0.731893199 |
| 0.805928714 | NA |
| 0.344722582 | 0.508071896 |
| 0.133998501 | 0.255800494 |
| 0.030142871 | 0.081797738 |
| 9.88E-09 | 2.34E-07 |
| 0.007288182 | 0.026069587 |
| NA | NA |
| 0.134226907 | 0.256111945 |
| 0.007404317 | 0.026396234 |
| 0.000169539 | 0.001092721 |
| 0.81828543 | NA |
| 0.000153167 | 0.001001575 |
| 1.62E-07 | 2.69E-06 |
| 0.450238353 | 0.608829607 |
| 0.479530356 | NA |
| 0.381743664 | 0.545872145 |
| 0.753371318 | 0.849811516 |
| 0.992061292 | NA |
| 0.000111664 | 0.000767216 |
| 0.212724462 | 0.359910788 |
| 0.036648874 | 0.095521972 |
| 0.489499479 | 0.643686392 |
| 0.289788823 | 0.449175743 |
| NA | NA |
| NA | NA |
| 0.082566661 | NA |
| NA | NA |
| 5.52E-06 | 5.80E-05 |
| 1.40E-05 | 0.000129958 |
| 0.622013207 | NA |
| 0.905158739 | NA |
| 0.992061292 | NA |
| 0.497630629 | 0.650677087 |
| NA | NA |
| 7.37E-05 | 0.000538081 |
| 0.007848296 | 0.027618707 |
| 0.931266632 | 0.96025939 |

| | |
|---|---|
| 0.002768515 | 0.011768065 |
| 0.392906873 | 0.556347822 |
| 0.118267798 | 0.233567999 |
| 0.659430406 | NA |
| 0.003572803 | 0.014493845 |
| 0.734598867 | 0.836928054 |
| NA | NA |
| 0.266469003 | 0.422640214 |
| 0.388485173 | 0.551962569 |
| 0.543883232 | 0.690593957 |
| 4.81E-13 | 3.69E-11 |
| 0.86029798 | 0.918702687 |
| NA | NA |
| 0.796195933 | 0.877077681 |
| 0.113794643 | 0.227082567 |
| 0.171881298 | NA |
| 2.53E-07 | 3.97E-06 |
| 0.357458218 | NA |
| 0.413845736 | 0.576370807 |
| 0.100715113 | 0.207117734 |
| 0.068560325 | 0.153924602 |
| 5.43E-07 | 7.72E-06 |
| NA | NA |
| 0.839492626 | 0.904845559 |
| 0.672206006 | 0.791345718 |
| 0.015698378 | 0.048477104 |
| 0.187459761 | 0.328893327 |
| 0.00277728 | 0.011798673 |
| 0.380361529 | 0.544588825 |
| 0.019623584 | 0.057903021 |
| 0.399436657 | 0.563052352 |
| 0.152699291 | 0.282585404 |
| 0.355781398 | 0.520354583 |
| 0.90759307 | 0.947610543 |
| 0.001775643 | 0.008079434 |
| 0.004554301 | 0.017694249 |
| 0.82273417 | 0.894510058 |
| 0.103474392 | 0.211373603 |
| 0.008878255 | 0.030552829 |
| 4.44E-05 | 0.000349844 |
| 2.51E-08 | 5.32E-07 |
| NA | NA |
| 0.956317475 | NA |
| NA | NA |
| 0.992060816 | NA |
| NA | NA |
| 0.807879735 | NA |

| | | |
|---|---|---|
| NA | | NA |
| NA | | NA |
| | 0.830163713 | NA |
| NA | | NA |
| NA | | NA |
| NA | | NA |
| NA | | NA |
| NA | | NA |
| NA | | NA |
| NA | | NA |
| NA | | NA |
| NA | | NA |
| NA | | NA |
| NA | | NA |
| NA | | NA |
| NA | | NA |
| | 0.801661681 | NA |
| | 0.516833472 | NA |
| NA | | NA |
| | 0.004600244 | 0.017850639 |
| | 0.980964789 | 0.990511759 |
| | 0.689961395 | NA |
| | 0.566232607 | NA |
| | 0.784353866 | NA |
| | 0.969944173 | 0.98418406 |
| | 0.615127375 | 0.748218137 |
| | 0.922765017 | 0.955421158 |
| | 0.06868039 | 0.154113294 |
| | 0.053279891 | 0.127251672 |
| | 0.259680559 | 0.414805077 |
| NA | | NA |
| | 0.661209809 | 0.782933739 |
| NA | | NA |
| NA | | NA |
| | 0.992060816 | NA |
| | 0.905158266 | NA |
| | 0.889409301 | NA |
| NA | | NA |
| NA | | NA |
| | 0.889410245 | NA |
| NA | | NA |
| | 0.839916375 | 0.905188055 |
| | 3.81E-05 | 0.000306806 |
| | 0.449658329 | 0.608515892 |
| NA | | NA |
| | 0.651470052 | 0.775751362 |
| NA | | NA |

| | | | |
|---|---|---|---|
| | 0.381061985 | NA | |
| | 0.00051862 | | 0.002852801 |
| | 0.011760239 | | 0.038312939 |
| | 0.151186672 | | 0.280457787 |
| | 0.109803024 | | 0.221065792 |
| | 0.83731427 | | 0.903172899 |
| | 0.938209135 | NA | |
| | 0.000218351 | | 0.001360219 |
| | 0.397841376 | NA | |
| | 0.808011864 | NA | |
| | 0.764047613 | NA | |
| | 0.905157793 | NA | |
| NA | | NA | |
| NA | | NA | |
| NA | | NA | |
| NA | | NA | |
| NA | | NA | |
| NA | | NA | |
| | 0.889409301 | NA | |
| NA | | NA | |
| NA | | NA | |
| | 0.992061769 | NA | |
| | 0.905158739 | NA | |
| NA | | NA | |
| NA | | NA | |
| | 0.880921434 | NA | |
| | 0.905158739 | NA | |
| | 0.767626112 | NA | |
| NA | | NA | |
| NA | | NA | |
| | 0.992061292 | NA | |
| NA | | NA | |
| | 0.992061292 | NA | |
| NA | | NA | |
| NA | | NA | |
| NA | | NA | |
| | 0.777092376 | NA | |
| | 0.746983622 | NA | |
| | 0.8057947 | NA | |
| NA | | NA | |
| | 0.992060816 | NA | |
| NA | | NA | |
| NA | | NA | |
| | 0.992060816 | NA | |
| NA | | NA | |
| NA | | NA | |
| | 0.98280479 | NA | |

| | | |
|---|---|---|
| | 0.46550801 | NA |
| NA | | NA |
| NA | | NA |
| NA | | NA |
| NA | | NA |
| NA | | NA |
| NA | | NA |
| NA | | NA |
| | 0.905158739 | NA |
| | 0.992060816 | NA |
| NA | | NA |
| NA | | NA |
| NA | | NA |
| NA | | NA |
| NA | | NA |
| NA | | NA |
| NA | | NA |
| NA | | NA |
| NA | | NA |
| | 0.801661681 | NA |
| | 0.734419703 | NA |
| NA | | NA |
| NA | | NA |
| NA | | NA |
| | 0.992060816 | NA |
| NA | | NA |
| NA | | NA |
| NA | | NA |
| NA | | NA |
| NA | | NA |
| | 0.905158739 | NA |
| NA | | NA |
| NA | | NA |
| NA | | NA |
| NA | | NA |
| NA | | NA |
| NA | | NA |
| | 0.889409301 | NA |
| NA | | NA |
| | 0.596187564 | 0.733099043 |
| NA | | NA |
| NA | | NA |
| NA | | NA |
| NA | | NA |
| NA | | NA |
| NA | | NA |
| | 0.577038774 | 0.718484601 |

| | |
|---|---|
| NA | NA |
| NA | NA |
| NA | NA |
| NA | NA |
| NA | NA |
| NA | NA |
| NA | NA |
| NA | NA |
| NA | NA |
| NA | NA |
| NA | NA |
| NA | NA |
| NA | NA |
| NA | NA |
| NA | NA |
| NA | NA |
| NA | NA |
| NA | NA |
| NA | NA |
| NA | NA |
| NA | NA |
| 0.358806315 | NA |
| NA | NA |
| NA | NA |
| NA | NA |
| NA | NA |
| 0.255916934 | 0.410525236 |
| NA | NA |
| NA | NA |
| NA | NA |
| NA | NA |
| NA | NA |
| NA | NA |
| NA | NA |
| NA | NA |
| NA | NA |
| NA | NA |
| NA | NA |
| NA | NA |
| 0.992060816 | NA |
| NA | NA |
| NA | NA |
| NA | NA |
| 0.51438667 | 0.665807066 |
| NA | NA |
| 0.496341851 | 0.649610841 |
| 0.527107681 | NA |

| | | |
|---|---|---|
| | 0.640514517 | NA |
| | 0.298120403 | 0.458080326 |
| | 0.79519169 | NA |
| | 0.863670715 | 0.921363449 |
| | 0.103667131 | NA |
| | 0.044051817 | 0.10991181 |
| NA | | NA |
| | 0.000831713 | 0.004266195 |
| | 0.074459046 | 0.164340875 |
| | 9.01E-06 | 8.92E-05 |
| | 0.449001029 | 0.607917747 |
| | 0.007281566 | 0.026050164 |
| | 0.097101174 | 0.201345107 |
| | 2.93E-05 | 0.000245339 |
| NA | | NA |
| | 0.490817512 | 0.644873269 |
| | 0.905158266 | NA |
| | 0.002212144 | 0.009746484 |
| | 0.561480906 | 0.705433071 |
| | 0.706510505 | 0.817537028 |
| | 0.234175372 | 0.385684723 |
| | 0.606960458 | 0.741298247 |
| | 0.011102061 | 0.036616124 |
| | 0.297084368 | 0.457160171 |
| | 6.90E-06 | 7.07E-05 |
| NA | | NA |
| NA | | NA |
| NA | | NA |
| | 0.350207558 | NA |
| | 0.576378234 | 0.717947112 |
| | 0.801661681 | NA |
| NA | | NA |
| | 0.40290755 | 0.566345339 |
| | 0.364463298 | 0.529315448 |
| | 5.39E-06 | 5.69E-05 |
| | 0.890644281 | 0.938152145 |
| | 0.83314985 | NA |
| NA | | NA |
| | 0.718522467 | NA |
| | 0.666199634 | NA |
| NA | | NA |
| NA | | NA |
| NA | | NA |
| NA | | NA |
| NA | | NA |
| NA | | NA |
| NA | | NA |

| | | |
|---|---|---|
| NA | NA | |
| 0.992060816 | NA | |
| 0.991399481 | NA | |
| NA | NA | |
| NA | NA | |
| NA | NA | |
| NA | NA | |
| 0.889410245 | NA | |
| NA | NA | |
| NA | NA | |
| NA | NA | |
| NA | NA | |
| NA | NA | |
| NA | NA | |
| NA | NA | |
| 0.709907272 | | 0.819267238 |
| NA | NA | |
| NA | NA | |
| NA | NA | |
| 0.992060816 | NA | |
| 0.792188023 | NA | |
| 0.989544857 | NA | |
| 0.871242405 | NA | |
| 0.084557167 | | 0.181150326 |
| NA | NA | |
| 0.905697792 | | 0.946867964 |
| NA | NA | |
| NA | NA | |
| 0.626556314 | | 0.756172031 |
| 0.882110794 | NA | |
| NA | NA | |
| NA | NA | |
| NA | NA | |
| NA | NA | |
| NA | NA | |
| NA | NA | |
| NA | NA | |
| NA | NA | |
| NA | NA | |
| NA | NA | |
| NA | NA | |
| NA | NA | |
| NA | NA | |
| NA | NA | |
| NA | NA | |
| NA | NA | |
| NA | NA | |

| | | | |
|---|---|---|---|
| NA | | NA | |
| NA | | NA | |
| NA | | NA | |
| NA | | NA | |
| NA | | NA | |
| NA | | NA | |
| NA | | NA | |
| NA | | NA | |
| NA | | NA | |
| NA | | NA | |
| NA | | NA | |
| NA | | NA | |
| NA | | NA | |
| NA | | NA | |
| NA | | NA | |
| NA | | NA | |
| NA | | NA | |
| NA | | NA | |
| NA | | NA | |
| NA | | NA | |
| NA | | NA | |
| NA | | NA | |
| | 0.28332184 | | 0.442179408 |
| NA | | NA | |
| NA | | NA | |
| NA | | NA | |
| NA | | NA | |
| NA | | NA | |
| NA | | NA | |
| NA | | NA | |
| NA | | NA | |
| NA | | NA | |
| NA | | NA | |
| NA | | NA | |
| NA | | NA | |
| NA | | NA | |
| NA | | NA | |
| NA | | NA | |
| NA | | NA | |
| NA | | NA | |
| NA | | NA | |
| NA | | NA | |
| NA | | NA | |
| NA | | NA | |
| NA | | NA | |
| NA | | NA | |
| NA | | NA | |

| | | |
|---|---|---|
| NA | | NA |
| NA | | NA |
| NA | | NA |
| NA | | NA |
| | 0.650582502 | NA |
| | 0.939903749 | NA |
| | 0.475772677 | NA |
| | 0.979509311 | NA |
| NA | | NA |
| | 0.496204576 | 0.64954732 |
| | 0.600744693 | 0.736557941 |
| | 0.533742122 | NA |
| | 0.984022229 | NA |
| | 3.70E-11 | 1.76E-09 |
| | 0.49044379 | 0.644481784 |
| | 0.705308549 | 0.816612382 |
| | 0.005732137 | 0.021472535 |
| | 0.012823392 | 0.041101864 |
| | 5.29E-14 | 4.96E-12 |
| | 0.070953393 | 0.158243506 |
| | 0.01287297 | 0.04122352 |
| | 0.256246384 | 0.41084363 |
| NA | | NA |
| | 2.32E-06 | 2.74E-05 |
| | 0.06104662 | 0.141659382 |
| | 0.994011108 | 0.997117284 |
| | 0.925076642 | 0.956842109 |
| | 0.00229921 | 0.010065864 |
| | 0.060652385 | 0.140968006 |
| | 0.406874717 | 0.570055804 |
| | 0.152269293 | 0.281976843 |
| NA | | NA |
| | 0.111229859 | 0.222957231 |
| | 0.830163713 | NA |
| | 0.765023842 | 0.857560519 |
| | 0.082444154 | 0.178047912 |
| | 0.801661681 | NA |
| NA | | NA |
| | 0.863820482 | NA |
| | 0.435336064 | NA |
| | 0.769346924 | 0.860621121 |
| | 0.136154512 | 0.258922213 |
| | 0.001036125 | 0.005129008 |
| | 2.64E-09 | 7.41E-08 |
| | 2.80E-05 | 0.000235606 |
| | 0.078053897 | 0.170665216 |
| | 5.84E-17 | 1.11E-14 |

| | |
|---|---|
| 0.867490853 | 0.92400231 |
| 0.000278963 | 0.001685213 |
| 3.06E-12 | 1.94E-10 |
| 2.71E-06 | 3.12E-05 |
| NA | NA |
| 0.004115497 | 0.016286213 |
| 0.289347995 | 0.4486508 |
| 0.032802807 | 0.087511807 |
| 1.01E-05 | 9.79E-05 |
| 0.11856028 | 0.233977279 |
| 0.689587802 | 0.804561848 |
| 0.878208965 | 0.930768409 |
| NA | NA |
| NA | NA |
| NA | NA |
| 0.843081354 | NA |
| NA | NA |
| NA | NA |
| NA | NA |
| NA | NA |
| NA | NA |
| NA | NA |
| NA | NA |
| NA | NA |
| NA | NA |
| NA | NA |
| 0.992060816 | NA |
| NA | NA |
| 0.905158266 | NA |
| NA | NA |
| NA | NA |
| 0.905158266 | NA |
| NA | NA |
| 0.992253023 | NA |
| NA | NA |
| NA | NA |
| NA | NA |
| 0.184368802 | 0.324950199 |
| 0.421080474 | 0.582824212 |
| 0.734668628 | 0.836928054 |
| 0.01991457 | 0.058596282 |
| 3.15E-06 | 3.55E-05 |
| 0.048515637 | NA |
| 0.000644619 | 0.003449118 |
| 0.691844311 | 0.806148406 |
| 0.128764845 | 0.248641157 |
| 0.233751806 | 0.385329731 |

| | | |
|---|---|---|
| | 0.822350939 | 0.894259604 |
| | 0.513880728 | NA |
| | 0.286934457 | 0.446000912 |
| | 0.415892025 | NA |
| | 0.00011395 | 0.000780729 |
| | 0.098609699 | 0.203742038 |
| | 0.6530731 | NA |
| | 0.527095735 | 0.676989837 |
| | 0.401159527 | 0.56475587 |
| | 0.003005848 | 0.012584445 |
| | 0.771965404 | 0.862251933 |
| | 0.032012546 | 0.085831928 |
| | 0.143441346 | 0.26949583 |
| | 1.18E-07 | 2.05E-06 |
| NA | | NA |
| | 0.718649907 | 0.825445217 |
| | 0.005901474 | 0.021977777 |
| | 0.571400549 | NA |
| | 1.62E-06 | 1.99E-05 |
| NA | | NA |
| | 0.753622059 | NA |
| | 0.000290075 | 0.001738472 |
| | 0.572093585 | NA |
| | 0.023216019 | 0.066328461 |
| | 6.97E-08 | 1.29E-06 |
| | 0.028626517 | 0.078478114 |
| | 2.49E-10 | 9.36E-09 |
| | 0.86486734 | 0.922103618 |
| | 0.306990284 | NA |
| | 0.20695172 | 0.353571371 |
| | 0.788502963 | 0.872410771 |
| | 0.351410121 | NA |
| | 0.247523175 | 0.401163289 |
| | 0.541934889 | 0.689134429 |
| NA | | NA |
| | 0.412286813 | 0.574855935 |
| | 0.015915217 | 0.049025863 |
| | 0.830673226 | 0.899046448 |
| | 0.523963195 | 0.6741494 |
| | 0.471733851 | 0.628034657 |
| | 0.040769937 | 0.10343929 |
| | 0.493869722 | 0.647301255 |
| | 0.027318763 | 0.075619254 |
| | 0.477245606 | 0.633299266 |
| | 0.411325275 | 0.574062013 |
| | 0.618329278 | 0.750408567 |
| | 7.01E-07 | 9.64E-06 |

| | | |
|---|---|---|
| | 0.808438457 | NA |
| | 0.802761249 | 0.881307391 |
| | 0.375213788 | 0.539363679 |
| | 0.138047041 | 0.261708967 |
| | 0.131974565 | NA |
| | 0.006952596 | 0.025139478 |
| | 0.091998534 | 0.19325657 |
| NA | | NA |
| | 0.980169484 | NA |
| | 0.000469176 | 0.00261488 |
| | 0.720807805 | NA |
| | 0.390860908 | NA |
| | 0.598470443 | 0.734793204 |
| | 0.539339707 | 0.686940467 |
| | 0.022606862 | NA |
| | 3.96E-10 | 1.41E-08 |
| NA | | NA |
| | 0.504022492 | NA |
| | 0.235452079 | NA |
| | 0.99205891 | NA |
| | 0.859345642 | NA |
| NA | | NA |
| NA | | NA |
| NA | | NA |
| NA | | NA |
| NA | | NA |
| NA | | NA |
| NA | | NA |
| NA | | NA |
| NA | | NA |
| NA | | NA |
| NA | | NA |
| NA | | NA |
| NA | | NA |
| NA | | NA |
| NA | | NA |
| NA | | NA |
| NA | | NA |
| NA | | NA |
| NA | | NA |
| NA | | NA |
| NA | | NA |
| NA | | NA |
| NA | | NA |
| NA | | NA |
| NA | | NA |

| | |
|---|---|
| NA | NA |
| NA | NA |
| NA | NA |
| NA | NA |
| NA | NA |
| NA | NA |
| NA | NA |
| NA | NA |
| NA | NA |
| NA | NA |
| NA | NA |
| NA | NA |
| NA | NA |
| NA | NA |
| NA | NA |
| NA | NA |
| NA | NA |
| NA | NA |
| NA | NA |
| NA | NA |
| NA | NA |
| NA | NA |
| NA | NA |
| NA | NA |
| NA | NA |
| NA | NA |
| NA | NA |
| NA | NA |
| NA | NA |
| NA | NA |
| NA | NA |
| NA | NA |
| NA | NA |
| NA | NA |
| NA | NA |
| NA | NA |
| NA | NA |
| NA | NA |
| NA | NA |
| NA | NA |
| NA | NA |
| NA | NA |
| NA | NA |
| NA | NA |
| NA | NA |
| NA | NA |
| NA | NA |

| | |
|---|---|
| NA | NA |
| NA | NA |
| NA | NA |
| NA | NA |
| NA | NA |
| NA | NA |
| NA | NA |
| NA | NA |
| NA | NA |
| NA | NA |
| NA | NA |
| NA | NA |
| NA | NA |
| NA | NA |
| NA | NA |
| NA | NA |
| NA | NA |
| NA | NA |
| NA | NA |
| NA | NA |
| NA | NA |
| NA | NA |
| NA | NA |
| NA | NA |
| NA | NA |
| NA | NA |
| NA | NA |
| NA | NA |
| NA | NA |
| NA | NA |
| NA | NA |
| NA | NA |
| 0.063818593 | NA |
| 0.905158266 | NA |
| NA | NA |
| NA | NA |
| 0.992060816 | NA |
| NA | NA |
| NA | NA |
| NA | NA |
| NA | NA |
| NA | NA |
| NA | NA |
| NA | NA |
| NA | NA |
| NA | NA |
| NA | NA |

| | |
|---|---|
| NA | NA |
| NA | NA |
| 0.129361699 | 0.249420804 |
| NA | NA |
| NA | NA |
| NA | NA |
| NA | NA |
| NA | NA |
| 0.656606734 | 0.779374447 |
| 0.156870876 | 0.288091511 |
| 0.389667942 | 0.553045394 |
| 0.627890665 | 0.756597131 |
| 0.027033169 | 0.074942729 |
| 0.625557579 | 0.755340311 |
| 0.864356912 | 0.921826781 |
| 0.026290438 | 0.073318736 |
| 0.032832094 | 0.087554211 |
| 0.073387232 | 0.162401513 |
| NA | NA |
| NA | NA |
| 0.905158739 | NA |
| 0.725624184 | NA |
| 0.005693857 | 0.021365586 |
| 0.367471631 | 0.532135951 |
| 0.992061292 | NA |
| 0.003637022 | 0.014702604 |
| 0.108570609 | 0.21916731 |
| 0.087468424 | 0.18600805 |
| 0.074590892 | 0.164584144 |
| 0.889410717 | NA |
| 0.173017892 | 0.310121614 |
| 0.001223399 | 0.005895242 |
| 0.027980548 | 0.077082531 |
| 0.016039842 | 0.049378786 |
| NA | NA |
| 0.670153875 | NA |
| NA | NA |
| NA | NA |
| NA | NA |
| 6.34E-08 | 1.19E-06 |
| 0.062610211 | NA |
| NA | NA |
| 0.224294884 | 0.373935435 |
| NA | NA |
| 0.992061292 | NA |
| 0.013745518 | 0.043436155 |
| 0.123908783 | 0.241772967 |

| | |
|---|---|
| 0.201022187 | 0.346188237 |
| 0.787632512 | 0.871771467 |
| NA | NA |
| 9.54E-07 | 1.26E-05 |
| 0.802792345 | NA |
| 5.75E-06 | 6.02E-05 |
| 0.789831767 | 0.873054952 |
| 0.354791479 | 0.519425701 |
| NA | NA |
| 0.119913483 | 0.235969189 |
| 0.065617489 | 0.149411209 |
| 0.838480447 | NA |
| 0.949102798 | 0.971398198 |
| 1.93E-05 | 0.000172912 |
| 0.916448347 | 0.952448732 |
| 0.497221049 | 0.650355228 |
| 0.696668006 | 0.810025032 |
| 0.000134615 | 0.000899032 |
| 0.118949425 | NA |
| 1.92E-05 | 0.000171599 |
| 1.39E-06 | 1.74E-05 |
| 0.992061292 | NA |
| 0.065881422 | 0.149853962 |
| 0.15000685 | 0.278703818 |
| 0.896025036 | NA |
| 0.735355709 | NA |
| 0.013498775 | 0.04278574 |
| 0.786327629 | NA |
| 0.195777232 | NA |
| 5.94E-05 | 0.000448675 |
| 6.21E-05 | 0.000465802 |
| 0.013120717 | 0.04179861 |
| 0.014011919 | 0.04414458 |
| 0.125837478 | 0.244437426 |
| 0.744343133 | 0.843440366 |
| 3.71E-07 | 5.53E-06 |
| 3.76E-07 | 5.59E-06 |
| 0.411701199 | 0.574442621 |
| 4.61E-05 | 0.0003617 |
| 0.115496968 | 0.229686463 |
| 0.283137699 | NA |
| 0.821317921 | 0.893671826 |
| NA | NA |
| NA | NA |
| NA | NA |
| 0.384975068 | NA |
| 0.905158266 | NA |

| | | |
|---|---|---|
| | 0.201895516 | 0.347283694 |
| NA | NA | |
| | 0.889409773 | NA |
| NA | NA | |
| NA | NA | |
| | 0.889410717 | NA |
| | 0.889409773 | NA |
| NA | NA | |
| NA | NA | |
| NA | NA | |
| NA | NA | |
| NA | NA | |
| | 0.729625788 | NA |
| | 0.327216841 | 0.489365566 |
| | 0.77963362 | 0.86734378 |
| | 0.905158739 | NA |
| | 0.997056723 | NA |
| | 0.021502136 | 0.062282734 |
| | 0.889409773 | NA |
| | 0.90515732 | NA |
| | 0.064832024 | 0.148177568 |
| | 0.10049709 | 0.206708093 |
| NA | NA | |
| NA | NA | |
| | 0.385805069 | NA |
| | 0.028874473 | 0.07900007 |
| | 0.3673228 | 0.531955508 |
| | 0.625273995 | 0.755340311 |
| | 0.000238305 | 0.001470754 |
| | 9.99E-25 | 1.22E-21 |
| | 0.679513627 | NA |
| | 0.152351658 | 0.282081793 |
| NA | NA | |
| | 0.013798795 | 0.043579423 |
| | 0.000404685 | 0.002310579 |
| NA | NA | |
| | 0.015016484 | 0.046746098 |
| | 0.349662093 | 0.513731427 |
| | 0.988723587 | 0.993927811 |
| | 0.024681734 | 0.069762671 |
| | 3.72E-05 | 0.000301213 |
| | 0.012351181 | 0.039888256 |
| | 0.602718599 | 0.738194976 |
| | 0.119169539 | NA |
| | 3.23E-11 | 1.57E-09 |
| | 8.92E-10 | 2.83E-08 |
| | 0.001150947 | 0.005607267 |

| | | |
|---|---|---|
| | 4.85E-20 | 2.17E-17 |
| | 1.27E-05 | 0.00012009 |
| | 0.001165324 | 0.005667628 |
| | 0.06129062 | 0.142120456 |
| | 0.135352795 | 0.257847837 |
| | 0.066752829 | 0.151227788 |
| | 0.187990659 | 0.329403676 |
| | 0.053162624 | 0.127056819 |
| | 0.526751767 | 0.676666786 |
| | 0.592544315 | 0.729935301 |
| | 0.005832283 | 0.021780801 |
| | 0.203638542 | 0.349487984 |
| | 4.85E-06 | 5.18E-05 |
| | 0.048017305 | 0.117509417 |
| | 0.664580011 | NA |
| NA | | NA |
| | 0.709782728 | 0.819244096 |
| | 0.027353586 | 0.075683843 |
| | 1.27E-05 | 0.000120104 |
| | 0.198516932 | 0.343030635 |
| | 4.66E-05 | 0.000364526 |
| | 0.018333767 | 0.054795879 |
| | 0.095192003 | 0.198280886 |
| | 8.82E-08 | 1.58E-06 |
| | 0.000443687 | 0.002495627 |
| | 0.000742908 | 0.003887872 |
| | 0.177497606 | 0.316317963 |
| | 7.34E-11 | 3.22E-09 |
| | 0.011198527 | 0.036846569 |
| | 0.013575631 | 0.043004513 |
| | 0.478713071 | 0.634518038 |
| | 1.15E-05 | 0.000109947 |
| | 0.117699657 | 0.232739016 |
| | 9.23E-06 | 9.09E-05 |
| | 0.311763324 | NA |
| | 0.905158266 | NA |
| | 0.905158266 | NA |
| NA | | NA |
| | 0.817478324 | 0.891399668 |
| | 0.648615813 | NA |
| | 0.794800009 | 0.876199276 |
| | 0.231302851 | 0.382266825 |
| | 0.184827864 | 0.325576373 |
| | 0.004516234 | 0.017571222 |
| | 0.277181438 | 0.435007662 |
| | 0.042753382 | 0.107465832 |
| | 0.065180927 | 0.148815551 |

| | | |
|---|---|---|
| | 0.090801163 | 0.191463678 |
| | 0.001030097 | 0.005103767 |
| | 0.856577137 | 0.916288843 |
| | 0.072327668 | 0.160493459 |
| | 0.82521721 | 0.89583472 |
| | 0.240078106 | NA |
| | 0.114377024 | 0.227892666 |
| | 8.01E-15 | 9.46E-13 |
| | 0.087268338 | 0.185636484 |
| | 0.021451803 | 0.062186158 |
| | 0.031526482 | 0.084777127 |
| | 0.843983236 | NA |
| | 0.981126607 | NA |
| NA | | NA |
| | 7.64E-10 | 2.51E-08 |
| NA | | NA |
| | 0.000395231 | 0.002263069 |
| | 0.434454333 | 0.595071276 |
| | 0.000141483 | 0.000936075 |
| | 0.000494943 | 0.002737647 |
| | 0.1474648 | 0.2751916 |
| | 0.63170113 | 0.759542524 |
| NA | | NA |
| | 0.015809 | 0.04877755 |
| | 0.16641423 | 0.301111574 |
| | 0.018332831 | 0.054795879 |
| | 0.179278051 | 0.318689122 |
| | 0.00236578 | 0.010317806 |
| | 0.002198256 | 0.009691129 |
| | 0.579425776 | 0.719901233 |
| | 0.284396446 | NA |
| | 0.978189236 | NA |
| | 0.969523425 | NA |
| | 0.982627018 | NA |
| NA | | NA |
| | 0.620568734 | 0.752014235 |
| NA | | NA |
| | 0.00015932 | 0.001036009 |
| | 0.749886377 | 0.847421391 |
| | 0.004437542 | 0.017323374 |
| | 0.002479307 | 0.010742521 |
| | 0.121309651 | 0.237991479 |
| NA | | NA |
| NA | | NA |
| | 1.91E-07 | 3.12E-06 |
| | 0.47166685 | 0.62798353 |
| | 0.992061292 | NA |

| | | |
|---|---|---|
| | 0.75276474 | 0.849447196 |
| | 0.413647848 | 0.576204839 |
| | 0.452428118 | 0.610625018 |
| | 0.516496701 | 0.667554506 |
| | 0.449721709 | 0.608543197 |
| | 0.277517429 | 0.43541049 |
| | 0.158342394 | 0.290067626 |
| | 0.694953171 | NA |
| | 0.921702499 | 0.955196809 |
| | 0.944738443 | 0.968866415 |
| NA | | NA |
| | 0.025811783 | 0.072352421 |
| | 0.000557624 | 0.003040692 |
| | 0.036310549 | 0.094785205 |
| | 0.017594503 | 0.053034685 |
| | 1.94E-05 | 0.000173641 |
| | 0.815276583 | NA |
| | 0.066390996 | 0.150700835 |
| | 0.241509776 | 0.394298377 |
| | 0.889410717 | NA |
| | 0.244618609 | 0.397718054 |
| | 0.984026404 | NA |
| | 0.457804557 | NA |
| | 8.20E-05 | 0.000589265 |
| | 0.262638816 | 0.418101041 |
| NA | | NA |
| | 0.512364412 | 0.664050648 |
| | 0.3650118 | 0.529761673 |
| | 0.488241994 | 0.642764145 |
| | 0.140332672 | 0.264989693 |
| | 0.634875695 | NA |
| | 5.38E-07 | 7.68E-06 |
| | 0.002488065 | 0.010771969 |
| | 0.011048919 | 0.036473724 |
| | 3.69E-10 | 1.33E-08 |
| | 7.36E-11 | 3.22E-09 |
| | 0.000147879 | 0.000971364 |
| | 0.08204823 | NA |
| | 0.992061601 | NA |
| | 0.384220239 | NA |
| | 2.03E-08 | 4.40E-07 |
| | 0.011455236 | 0.037510516 |
| | 0.196789987 | 0.340878073 |
| NA | | NA |
| | 0.071878266 | 0.159770709 |
| | 0.000597114 | 0.003231154 |
| | 0.313064388 | 0.474927868 |

| | | |
|---|---|---|
| | 0.901183732 | NA |
| NA | | NA |
| | 0.364909706 | 0.529683516 |
| | 0.168555921 | NA |
| | 0.001509586 | 0.007043491 |
| | 0.785694763 | 0.871119873 |
| | 0.225595333 | 0.375433718 |
| | 0.453803153 | NA |
| | 0.613030627 | NA |
| | 0.816867675 | NA |
| | 0.380057021 | 0.544304363 |
| | 0.444857092 | 0.604306643 |
| NA | | NA |
| | 0.905158266 | NA |
| NA | | NA |
| | 0.065986453 | 0.150036515 |
| | 0.000711899 | 0.003744335 |
| | 0.038096149 | 0.098406462 |
| | 0.702516815 | NA |
| | 0.001251702 | 0.006009208 |
| | 0.089009489 | 0.188609416 |
| | 0.104706736 | 0.21328898 |
| | 0.562943567 | NA |
| | 0.038635649 | 0.099425785 |
| | 0.982354787 | NA |
| | 0.997715538 | NA |
| | 0.020632228 | 0.060295859 |
| | 0.805374319 | NA |
| | 0.90515732 | NA |
| | 0.082939873 | 0.178889762 |
| | 0.737651134 | NA |
| | 0.048219825 | NA |
| | 0.755029883 | 0.851070336 |
| | 0.005852058 | 0.021836083 |
| | 0.106015486 | 0.215176454 |
| | 0.001373656 | 0.00651352 |
| | 0.453277229 | 0.611320149 |
| | 4.13E-06 | 4.52E-05 |
| NA | | NA |
| | 1.56E-17 | 3.50E-15 |
| | 0.07586901 | 0.166917903 |
| | 9.35E-06 | 9.18E-05 |
| | 0.025069324 | 0.070697185 |
| | 0.853439117 | 0.914446646 |
| | 0.002086847 | 0.009279584 |
| | 0.023446269 | 0.06688317 |
| | 0.807702613 | 0.884782085 |

| | | | |
|---|---|---|---|
| | 0.22202285 | NA | |
| NA | | NA | |
| | 0.228408507 | | 0.378709031 |
| NA | | NA | |
| NA | | NA | |
| NA | | NA | |
| NA | | NA | |
| NA | | NA | |
| NA | | NA | |
| NA | | NA | |
| NA | | NA | |
| NA | | NA | |
| NA | | NA | |
| NA | | NA | |
| NA | | NA | |
| NA | | NA | |
| NA | | NA | |
| NA | | NA | |
| NA | | NA | |
| NA | | NA | |
| NA | | NA | |
| NA | | NA | |
| NA | | NA | |
| NA | | NA | |
| NA | | NA | |
| NA | | NA | |
| NA | | NA | |
| NA | | NA | |
| NA | | NA | |
| NA | | NA | |
| NA | | NA | |
| NA | | NA | |
| NA | | NA | |
| NA | | NA | |
| NA | | NA | |
| NA | | NA | |
| NA | | NA | |
| NA | | NA | |
| NA | | NA | |
| NA | | NA | |
| NA | | NA | |
| NA | | NA | |
| NA | | NA | |
| NA | | NA | |
| NA | | NA | |
| NA | | NA | |
| NA | | NA | |

| | |
|---|---|
| NA | NA |
| NA | NA |
| NA | NA |
| NA | NA |
| NA | NA |
| NA | NA |
| NA | NA |
| NA | NA |
| NA | NA |
| NA | NA |
| NA | NA |
| NA | NA |
| NA | NA |
| NA | NA |
| NA | NA |
| NA | NA |
| NA | NA |
| NA | NA |
| NA | NA |
| NA | NA |
| NA | NA |
| NA | NA |
| NA | NA |
| NA | NA |
| NA | NA |
| NA | NA |
| NA | NA |
| NA | NA |
| NA | NA |
| NA | NA |
| NA | NA |
| NA | NA |
| NA | NA |
| NA | NA |
| NA | NA |
| NA | NA |
| NA | NA |
| NA | NA |
| NA | NA |
| NA | NA |
| NA | NA |
| NA | NA |
| NA | NA |
| NA | NA |
| NA | NA |
| NA | NA |
| NA | NA |

| | | |
|---|---|---|
| NA | | NA |
| NA | | NA |
| NA | | NA |
| NA | | NA |
| NA | | NA |
| NA | | NA |
| NA | | NA |
| NA | | NA |
| NA | | NA |
| NA | | NA |
| NA | | NA |
| NA | | NA |
| | 0.992060816 | NA |
| NA | | NA |
| NA | | NA |
| NA | | NA |
| NA | | NA |
| NA | | NA |
| NA | | NA |
| NA | | NA |
| NA | | NA |
| NA | | NA |
| NA | | NA |
| NA | | NA |
| NA | | NA |
| NA | | NA |
| NA | | NA |
| NA | | NA |
| NA | | NA |
| NA | | NA |
| | 0.905158266 | NA |
| NA | | NA |
| NA | | NA |
| | 0.889409773 | NA |
| NA | | NA |
| | 0.268534926 | 0.424930348 |
| | 0.01201502 | 0.03901195 |
| | 0.373425554 | NA |
| NA | | NA |
| | 0.008302952 | 0.028931215 |
| | 0.003269522 | 0.013495383 |
| | 0.517660514 | 0.668547152 |
| | 0.769736196 | 0.860842227 |
| | 0.127367598 | NA |
| | 0.000356664 | 0.00207595 |
| | 0.905158266 | NA |
| | 6.71E-07 | 9.29E-06 |

| | |
|---|---|
| NA | NA |
| 0.237595183 | 0.38948394 |
| 0.592921357 | NA |
| NA | NA |
| 0.299647778 | NA |
| 0.000291248 | 0.001744548 |
| 0.530632266 | NA |
| 0.972968899 | 0.985912235 |
| 0.107144931 | 0.21690749 |
| 0.045630966 | 0.113017189 |
| 0.797157316 | NA |
| 2.64E-13 | 2.19E-11 |
| NA | NA |
| NA | NA |
| 0.000213667 | 0.001335179 |
| 0.010291757 | 0.034499857 |
| 9.50E-09 | 2.26E-07 |
| 0.815716008 | NA |
| 0.479245612 | 0.634917309 |
| 0.069009064 | 0.154724324 |
| 0.002903874 | 0.012222766 |
| 0.054330841 | 0.12928883 |
| NA | NA |
| 0.90515732 | NA |
| 0.326664169 | 0.488905295 |
| 0.472540926 | NA |
| 0.010198735 | 0.034228035 |
| 0.448362304 | 0.607452359 |
| 0.020715294 | 0.060478629 |
| 0.008935541 | 0.030701853 |
| 0.684856317 | 0.800658035 |
| 0.814454334 | 0.889471464 |
| 0.848366505 | 0.910956565 |
| 0.867777753 | NA |
| 0.887014231 | 0.935899745 |
| 0.18771911 | NA |
| 0.906847042 | 0.947360419 |
| 0.788582525 | NA |
| 0.222965918 | 0.372405579 |
| NA | NA |
| NA | NA |
| 0.238478568 | 0.390540001 |
| 0.002946319 | 0.012373715 |
| 0.928115334 | 0.958765801 |
| 0.224797874 | 0.374447129 |
| 0.048720466 | 0.118885732 |
| 0.510058843 | 0.66158652 |

| | | |
|---|---|---|
| NA | NA | |
| NA | NA | |
| 0.777092376 | NA | |
| NA | NA | |
| 0.243179062 | NA | |
| 0.265345195 | | 0.421426757 |
| NA | NA | |
| NA | NA | |
| 5.82E-07 | | 8.17E-06 |
| 0.835209842 | | 0.901955981 |
| NA | NA | |
| 8.42E-07 | | 1.13E-05 |
| 0.271026489 | | 0.427640344 |
| 0.052407295 | | 0.12581303 |
| NA | NA | |
| NA | NA | |
| NA | NA | |
| 0.889409773 | NA | |
| NA | NA | |
| NA | NA | |
| NA | NA | |
| NA | NA | |
| NA | NA | |
| 0.807879735 | NA | |
| NA | NA | |
| NA | NA | |
| NA | NA | |
| NA | NA | |
| NA | NA | |
| NA | NA | |
| NA | NA | |
| NA | NA | |
| NA | NA | |
| 0.828724638 | NA | |
| NA | NA | |
| NA | NA | |
| NA | NA | |
| NA | NA | |
| NA | NA | |
| NA | NA | |
| NA | NA | |
| NA | NA | |
| NA | NA | |
| NA | NA | |
| NA | NA | |
| NA | NA | |
| NA | NA | |

| | | | |
|---|---|---|---|
| NA | | NA | |
| NA | | NA | |
| NA | | NA | |
| NA | | NA | |
| | 0.849430613 | NA | |
| | 0.117171576 | | 0.232050021 |
| | 0.693355207 | | 0.807198971 |
| | 0.588474179 | | 0.727115941 |
| | 0.391110573 | NA | |
| | 0.600365476 | | 0.736227699 |
| | 0.410642645 | | 0.573607677 |
| | 0.215861536 | | 0.363655196 |
| | 0.122250244 | | 0.239280961 |
| NA | | NA | |
| | 0.992062245 | NA | |
| | 0.555971796 | | 0.701199108 |
| | 0.616744894 | NA | |
| | 0.889409773 | NA | |
| NA | | NA | |
| | 0.881453338 | NA | |
| NA | | NA | |
| NA | | NA | |
| NA | | NA | |
| | 0.568760627 | | 0.71124229 |
| NA | | NA | |
| NA | | NA | |
| | 0.777092376 | NA | |
| NA | | NA | |
| | 0.992061292 | NA | |
| | 0.274804865 | | 0.432016124 |
| | 0.889409301 | NA | |
| NA | | NA | |
| | 0.650959321 | | 0.775399417 |
| | 0.68507006 | NA | |
| | 0.902545984 | NA | |
| NA | | NA | |
| | 0.992061292 | NA | |
| NA | | NA | |
| NA | | NA | |
| NA | | NA | |
| NA | | NA | |
| NA | | NA | |
| NA | | NA | |
| | 0.992061769 | NA | |
| NA | | NA | |
| NA | | NA | |
| NA | | NA | |

| | |
|---|---|
| NA | NA |
| NA | NA |
| 0.98567569 | NA |
| NA | NA |
| 1.34E-05 | 0.000125491 |
| 0.147016864 | 0.274565685 |
| 0.7130299 | 0.821265333 |
| 0.349592496 | NA |
| 0.316991628 | 0.479131274 |
| 0.576574211 | NA |
| NA | NA |
| NA | NA |
| 0.697650127 | 0.810911078 |
| 4.47E-07 | 6.55E-06 |
| NA | NA |
| 0.000406427 | 0.002318715 |
| 0.889409773 | NA |
| 0.67628142 | NA |
| 0.905158266 | NA |
| 0.905158266 | NA |
| NA | NA |
| NA | NA |
| NA | NA |
| 0.683334897 | 0.799628222 |
| 0.464092907 | 0.621176164 |
| 0.576869188 | NA |
| NA | NA |
| 0.954383734 | 0.9745774 |
| NA | NA |
| NA | NA |
| NA | NA |
| NA | NA |
| NA | NA |
| NA | NA |
| 0.889409301 | NA |
| 0.028242293 | 0.077638021 |
| 0.352534546 | NA |
| 0.823538735 | NA |
| 0.044298769 | 0.11041506 |
| 0.751899913 | NA |
| 0.795118144 | 0.87650599 |
| 0.180016419 | 0.319639407 |
| NA | NA |
| NA | NA |
| NA | NA |
| 0.992062245 | NA |
| 0.380927585 | NA |

| | | | |
|---|---|---|---|
| NA | | NA | |
| NA | | NA | |
| NA | | NA | |
| | 0.707431802 | | 0.817747691 |
| NA | | NA | |
| NA | | NA | |
| NA | | NA | |
| NA | | NA | |
| | 0.992061769 | NA | |
| NA | | NA | |
| | 0.989975993 | NA | |
| | 0.002673838 | | 0.011443151 |
| | 0.005030192 | | 0.019263686 |
| NA | | NA | |
| | 0.740666435 | NA | |
| | 0.905158266 | NA | |
| | 0.049206819 | | 0.119846112 |
| | 0.194489223 | NA | |
| | 0.013634154 | | 0.043146331 |
| | 0.983401898 | NA | |
| | 0.351859087 | | 0.516164986 |
| | 0.064788016 | | 0.148113507 |
| | 0.989558834 | NA | |
| | 0.006530065 | | 0.023863155 |
| | 0.681918542 | | 0.798541449 |
| | 0.000216703 | | 0.001351869 |
| | 0.565294975 | | 0.708562838 |
| | 0.563735874 | NA | |
| | 0.355760521 | | 0.520354583 |
| | 0.222378477 | | 0.371742461 |
| NA | | NA | |
| | 0.14944779 | | 0.277891406 |
| | 0.989994222 | NA | |
| NA | | NA | |
| | 0.659378693 | NA | |
| | 0.028257219 | NA | |
| | 0.000350482 | | 0.002045784 |
| | 0.002636532 | | 0.011312157 |
| | 0.097157363 | | 0.201404564 |
| | 1.73E-05 | | 0.000157085 |
| | 0.882002775 | NA | |
| | 0.686600514 | NA | |
| NA | | NA | |
| | 0.905158739 | NA | |
| | 0.714530011 | NA | |
| | 0.905158266 | NA | |
| | 0.830163713 | NA | |

| | | |
|---|---|---|
| | 0.630442414 | NA |
| | 0.905158266 | NA |
| | 0.663966878 | NA |
| NA | | NA |
| NA | | NA |
| | 0.662914387 | NA |
| | 0.905158739 | NA |
| NA | | NA |
| NA | | NA |
| NA | | NA |
| | 0.730225393 | 0.834114617 |
| | 0.005160883 | 0.019681806 |
| | 0.953639938 | 0.974089655 |
| NA | | NA |
| | 9.21E-06 | 9.09E-05 |
| | 0.305189231 | 0.465774981 |
| NA | | NA |
| | 0.735255118 | 0.837385338 |
| NA | | NA |
| NA | | NA |
| NA | | NA |
| NA | | NA |
| | 0.102722863 | 0.210277454 |
| | 1.48E-11 | 7.87E-10 |
| | 0.630077946 | 0.758323426 |
| | 2.08E-10 | 8.11E-09 |
| | 8.75E-13 | 6.30E-11 |
| | 0.293441091 | 0.453045975 |
| | 2.16E-06 | 2.56E-05 |
| | 0.749493319 | 0.847208138 |
| | 0.766510799 | 0.858439143 |
| | 9.85E-06 | 9.58E-05 |
| | 0.998181336 | 0.999273488 |
| | 0.001200572 | 0.00581203 |
| | 0.002759484 | 0.011738468 |
| | 0.868267671 | NA |
| | 0.002419195 | 0.010521508 |
| | 1.02E-06 | 1.34E-05 |
| | 0.502966757 | 0.655274222 |
| | 0.045824132 | 0.113457224 |
| | 0.436291777 | 0.596724387 |
| | 0.425434126 | NA |
| NA | | NA |
| | 0.574159181 | 0.716157993 |
| | 0.001670501 | 0.007659749 |
| | 1.57E-09 | 4.65E-08 |
| | 0.011286422 | 0.037079375 |

| | |
|---|---|
| 0.40708128 | 0.570191295 |
| 0.289904816 | 0.449292106 |
| 7.69E-12 | 4.42E-10 |
| NA | NA |
| 0.005053361 | 0.019335555 |
| 0.000976632 | 0.004878495 |
| 0.665060217 | 0.78622112 |
| 0.337398986 | 0.500333896 |
| 0.014437703 | 0.045220015 |
| 0.001125394 | 0.005500228 |
| 2.83E-06 | 3.24E-05 |
| 0.004564051 | 0.017725858 |
| 0.334731114 | 0.497520004 |
| 0.669012446 | 0.78895785 |
| 9.76E-06 | 9.51E-05 |
| 2.76E-05 | 0.000233006 |
| 0.524602948 | NA |
| 0.001743914 | 0.007953191 |
| 5.63E-05 | 0.000429308 |
| 0.05865583 | 0.137125877 |
| 0.673937713 | 0.792746853 |
| 6.99E-07 | 9.62E-06 |
| 0.658756703 | 0.781039816 |
| NA | NA |
| 0.007397233 | 0.026380653 |
| 0.989998863 | NA |
| 0.061532525 | 0.142515844 |
| 0.97629096 | NA |
| NA | NA |
| NA | NA |
| NA | NA |
| NA | NA |
| 0.468829237 | 0.625204686 |
| 0.003873374 | 0.01546462 |
| 0.00410978 | 0.016266521 |
| 1.90E-08 | 4.14E-07 |
| 0.578340782 | NA |
| 0.848230069 | 0.910865279 |
| 0.054122996 | NA |
| 0.300335867 | 0.460710814 |
| 0.000553044 | 0.003018716 |
| 0.006655741 | 0.024241736 |
| 8.76E-06 | 8.71E-05 |
| 0.737830445 | 0.839048097 |
| NA | NA |
| 0.320809109 | NA |
| 0.025285503 | 0.071194301 |

| | |
|---|---|
| 0.553234626 | 0.698670299 |
| 0.112340038 | NA |
| 2.76E-05 | 0.000233093 |
| 0.586231126 | 0.725460183 |
| 3.05E-07 | 4.66E-06 |
| 0.378559154 | 0.543070521 |
| 0.905158266 | NA |
| 0.83798679 | NA |
| 0.742919362 | NA |
| 0.988347767 | 0.993732996 |
| 7.37E-07 | 1.01E-05 |
| 1.60E-08 | 3.56E-07 |
| 2.74E-11 | 1.36E-09 |
| NA | NA |
| 4.57E-07 | 6.66E-06 |
| 0.985207256 | 0.992529528 |
| 0.882762477 | NA |
| 3.74E-05 | 0.000302592 |
| 0.120727628 | 0.23716747 |
| 0.178156624 | 0.317183502 |
| 0.889409301 | NA |
| 0.998063029 | 0.999200604 |
| 0.077536758 | 0.169753706 |
| 0.855350645 | 0.915804824 |
| 0.000938764 | 0.004724806 |
| 0.94965087 | NA |
| 0.224902025 | NA |
| 0.324827357 | 0.486985999 |
| 0.866167343 | 0.922905948 |
| 0.071028419 | 0.158327312 |
| 0.029891096 | 0.081305409 |
| 0.06373127 | 0.146238819 |
| 0.000112169 | 0.000769725 |
| 7.04E-12 | 4.11E-10 |
| 9.24E-15 | 1.07E-12 |
| 1.43E-06 | 1.78E-05 |
| 0.000441162 | 0.002483969 |
| 0.184985832 | 0.325776237 |
| NA | NA |
| 0.023118963 | 0.066094168 |
| 1.20E-10 | 5.01E-09 |
| 0.792651591 | 0.874948498 |
| 0.11606466 | 0.230585711 |
| 0.067680265 | 0.152584411 |
| 0.039760787 | 0.10165225 |
| 0.000412632 | 0.002345584 |
| NA | NA |

| | | |
|---|---|---|
| | 0.751872031 | NA |
| | 0.324730573 | 0.486946842 |
| | 2.83E-09 | 7.85E-08 |
| | 0.057629127 | 0.13520064 |
| | 0.001848906 | 0.00837634 |
| | 0.003034817 | 0.012671905 |
| | 0.001467325 | 0.006884827 |
| | 1.72E-06 | 2.10E-05 |
| | 0.046942595 | 0.115458383 |
| | 0.063874679 | 0.14644832 |
| NA | | NA |
| | 0.303893676 | NA |
| | 0.003873097 | 0.01546462 |
| | 0.879711688 | 0.931731488 |
| | 0.025522379 | 0.071750853 |
| NA | | NA |
| | 0.068739982 | 0.154168243 |
| | 0.446312854 | 0.605647312 |
| | 0.683805237 | 0.799813045 |
| | 0.524690158 | NA |
| | 0.469601916 | 0.625955352 |
| | 0.247750258 | NA |
| | 0.000731008 | 0.00383657 |
| | 0.931745247 | 0.960437209 |
| | 0.373014699 | 0.537328114 |
| | 6.78E-05 | 0.000501606 |
| | 0.905158739 | NA |
| | 0.489019642 | 0.643305506 |
| | 0.040064214 | 0.102215646 |
| | 0.889409301 | NA |
| | 0.014556938 | 0.045547991 |
| | 1.12E-05 | 0.000107339 |
| | 0.820105077 | 0.892909192 |
| | 0.970712833 | NA |
| | 0.163086158 | 0.296531335 |
| NA | | NA |
| | 0.905158739 | NA |
| NA | | NA |
| NA | | NA |
| NA | | NA |
| | 0.991955612 | NA |
| | 0.807252815 | NA |
| | 0.879542993 | NA |
| NA | | NA |
| NA | | NA |
| NA | | NA |
| NA | | NA |

| | | |
|---|---|---|
| NA | NA | |
| NA | NA | |
| NA | NA | |
| NA | NA | |
| NA | NA | |
| NA | NA | |
| NA | NA | |
| NA | NA | |
| NA | NA | |
| NA | NA | |
| NA | NA | |
| NA | NA | |
| NA | NA | |
| NA | NA | |
| NA | NA | |
| NA | NA | |
| NA | NA | |
| NA | NA | |
| NA | NA | |
| NA | NA | |
| NA | NA | |
| NA | NA | |
| NA | NA | |
| NA | NA | |
| NA | NA | |
| NA | NA | |
| NA | NA | |
| NA | NA | |
| NA | NA | |
| 0.905158266 | NA | |
| 0.846073823 | NA | |
| 0.807879735 | NA | |
| 0.889409773 | NA | |
| NA | NA | |
| NA | NA | |
| NA | NA | |
| 0.889409301 | NA | |
| 0.880684185 | | 0.932221934 |
| 0.097221313 | NA | |
| 0.007561257 | | 0.026837125 |
| 0.905158266 | NA | |
| 3.05E-06 | | 3.45E-05 |
| 0.449174765 | | 0.608065389 |
| 0.9853724 | | 0.99264741 |
| 0.905158739 | NA | |
| 0.00056141 | | 0.003059818 |
| 0.000931598 | | 0.004691047 |

| | |
|---|---|
| NA | NA |
| 0.449673818 | 0.608515892 |
| 0.00942915 | 0.032049958 |
| 0.992060816 | NA |
| 0.02318948 | 0.066261262 |
| 0.302135001 | 0.462727192 |
| 0.452745848 | 0.610906318 |
| 0.915242025 | 0.952051143 |
| 0.878603022 | NA |
| 0.441453499 | 0.601133412 |
| 0.132292802 | 0.253270935 |
| 0.066768486 | 0.151245015 |
| 0.847056703 | NA |
| NA | NA |
| 0.063548305 | NA |
| NA | NA |
| NA | NA |
| NA | NA |
| NA | NA |
| 0.299273094 | 0.459562089 |
| 2.23E-05 | 0.00019523 |
| NA | NA |
| NA | NA |
| 0.024763848 | 0.069949748 |
| 0.36757242 | 0.532141534 |
| 0.007190547 | 0.025795986 |
| 0.876987616 | NA |
| 0.316370302 | 0.478488564 |
| 0.002153526 | 0.00952261 |
| 3.04E-05 | 0.000253064 |
| NA | NA |
| 0.106237723 | NA |
| 0.008163554 | 0.028536915 |
| 0.76396776 | 0.857009861 |
| 6.75E-05 | 0.000500188 |
| 0.179566257 | 0.319098117 |
| 9.92E-12 | 5.52E-10 |
| 3.99E-12 | 2.45E-10 |
| NA | NA |
| 0.850884599 | NA |
| NA | NA |
| 1.07E-08 | 2.52E-07 |
| 0.088965701 | 0.188553019 |
| 0.906707095 | 0.947360419 |
| 3.55E-09 | 9.57E-08 |
| 0.010328668 | 0.034584815 |
| 0.000971491 | 0.004857236 |

| | | |
|---|---|---|
| | 0.000688921 | 0.003644424 |
| | 0.085870973 | 0.183428083 |
| | 0.170275106 | 0.306456115 |
| | 0.165092801 | 0.299337199 |
| | 0.001976802 | 0.008860705 |
| | 0.045276621 | 0.112329603 |
| | 1.03E-05 | 9.98E-05 |
| | 0.001428391 | 0.006737151 |
| | 0.653092926 | 0.776691601 |
| | 0.719148984 | NA |
| | 0.001612467 | 0.007433176 |
| | 0.57667212 | NA |
| | 0.889409773 | NA |
| | 0.881992598 | NA |
| | 0.517657004 | 0.668547152 |
| | 0.639217587 | NA |
| | 0.135600063 | 0.258184495 |
| | 0.781720161 | NA |
| | 0.437318633 | 0.597652468 |
| | 0.006585991 | 0.02402355 |
| | 0.339133262 | 0.502124898 |
| | 0.135869969 | 0.258541477 |
| | 0.469627936 | NA |
| | 0.062588902 | 0.144292882 |
| NA | | NA |
| | 0.572179167 | 0.714703204 |
| | 0.224076191 | 0.373755342 |
| | 0.02074671 | NA |
| | 0.236594881 | 0.388385138 |
| | 0.085058273 | 0.182046258 |
| | 0.722113127 | NA |
| | 0.4138121 | NA |
| | 0.889410245 | NA |
| | 0.153051725 | NA |
| | 0.875029791 | NA |
| | 0.774747615 | NA |
| | 0.992060816 | NA |
| NA | | NA |
| NA | | NA |
| | 0.992061292 | NA |
| | 0.37876015 | 0.543216911 |
| | 0.726850194 | NA |
| NA | | NA |
| NA | | NA |
| NA | | NA |
| | 0.851133994 | NA |
| NA | | NA |

| | | |
|---|---|---|
| NA | | NA |
| | 0.992061292 | NA |
| NA | | NA |
| | 0.788582525 | NA |
| | 0.028477505 | 0.078167192 |
| | 0.21498831 | NA |
| | 0.132692428 | 0.25385895 |
| | 0.735486846 | NA |
| | 0.536582174 | NA |
| | 0.804014905 | NA |
| | 0.535878472 | NA |
| | 0.402225941 | NA |
| | 0.768458492 | NA |
| | 0.752337152 | NA |
| | 0.788582525 | NA |
| | 0.923299104 | NA |
| | 0.992061292 | NA |
| | 0.889409773 | NA |
| | 0.012983626 | 0.041494317 |
| | 0.140829483 | NA |
| | 0.003551952 | NA |
| | 0.290893797 | NA |
| | 0.8600617 | NA |
| | 0.79397964 | NA |
| | 0.57984686 | NA |
| | 0.09983751 | NA |
| | 0.110185814 | 0.221511516 |
| | 5.99E-05 | 0.00045206 |
| | 0.190878807 | 0.333055838 |
| | 0.137873367 | NA |
| | 0.004486197 | 0.017472935 |
| | 0.985489293 | NA |
| | 0.889409773 | NA |
| | 0.817918248 | NA |
| | 0.591324215 | NA |
| | 0.797573332 | NA |
| | 0.96289044 | NA |
| | 0.155784045 | 0.286666058 |
| | 0.99368456 | NA |
| | 0.24185709 | 0.394609743 |
| | 0.43128758 | NA |
| | 0.345720576 | NA |
| | 0.905157793 | NA |
| | 0.830106074 | NA |
| | 0.824202355 | NA |
| | 0.155450695 | 0.286274893 |
| | 0.744039649 | NA |

| | | |
|---|---|---|
| 0.8600617 | NA | |
| 0.405832292 | NA | |
| 0.243883938 | NA | |
| 0.177577583 | NA | |
| NA | NA | |
| NA | NA | |
| 0.545855071 | NA | |
| 0.992061292 | NA | |
| 0.793676336 | NA | |
| NA | NA | |
| NA | NA | |
| NA | NA | |
| 0.889409301 | NA | |
| NA | NA | |
| NA | NA | |
| NA | NA | |
| NA | NA | |
| 0.776244828 | NA | |
| NA | NA | |
| NA | NA | |
| NA | NA | |
| NA | NA | |
| 0.62793045 | NA | |
| 0.689162991 | NA | |
| NA | NA | |
| NA | NA | |
| NA | NA | |
| 0.987162995 | NA | |
| 0.79911887 | NA | |
| 0.277357753 | NA | |
| 0.826421856 | NA | |
| 0.889409773 | NA | |
| 0.98649457 | NA | |
| 0.744720494 | NA | |
| 0.442866104 | NA | |
| 0.788582525 | NA | |
| 0.333697168 | NA | |
| 0.541755751 | NA | |
| 0.518951339 | | 0.669938416 |
| 0.305825832 | | 0.466436583 |
| 0.076863107 | | 0.168648777 |
| 0.446155054 | | 0.605545389 |
| 0.36764784 | NA | |
| NA | NA | |
| 0.726467233 | NA | |
| 0.6342398 | NA | |
| NA | NA | |

| | |
|---|---|
| 0.71235228 | NA |
| 0.969783613 | NA |
| 0.626671 | NA |
| 0.189428108 | NA |
| 0.878865595 | NA |
| 0.380556531 | NA |
| 0.871521449 | NA |
| 0.972077015 | NA |
| 0.902545984 | NA |
| NA | NA |
| 0.905157793 | NA |
| NA | NA |
| 0.029256835 | NA |
| 0.62569865 | NA |
| 0.905158739 | NA |
| 0.851884166 | NA |
| NA | NA |
| NA | NA |
| NA | NA |
| NA | NA |
| NA | NA |
| NA | NA |
| 0.992061769 | NA |
| 0.308600758 | NA |
| NA | NA |
| NA | NA |
| 0.991979299 | NA |
| NA | NA |
| 0.842252617 | NA |
| NA | NA |
| NA | NA |
| 0.992060816 | NA |
| 0.889409301 | NA |
| 0.992062245 | NA |
| 0.975873177 | NA |
| 0.736905233 | NA |
| 0.745802213 | NA |
| 0.889283189 | NA |
| NA | NA |
| 0.268102801 | NA |
| 0.812983331 | NA |
| 0.983513816 | NA |
| NA | NA |
| NA | NA |
| 0.838403923 | NA |
| 0.992003131 | NA |
| 0.212845871 | NA |

| | | |
|---|---|---|
| | 0.813714029 | NA |
| | 0.982209729 | NA |
| NA | | NA |
| | 0.992061769 | NA |
| NA | | NA |
| | 0.889409773 | NA |
| NA | | NA |
| | 0.992061769 | NA |
| NA | | NA |
| | 0.905157793 | NA |
| | 0.992061292 | NA |
| | 0.905158266 | NA |
| | 0.99205891 | NA |
| | 0.838891015 | NA |
| | 0.16510748 | NA |
| | 0.534027161 | NA |
| | 0.809305128 | NA |
| | 0.0006511 | 0.003479559 |
| | 4.75E-08 | 9.29E-07 |
| NA | | NA |
| NA | | NA |
| | 1.24E-19 | 4.96E-17 |
| NA | | NA |
| | 0.085371272 | 0.182537543 |
| | 0.000561238 | 0.00305964 |
| | 0.147733978 | 0.275483223 |
| | 0.907276733 | NA |
| | 0.157653932 | NA |
| | 0.624608441 | NA |
| | 0.881501945 | NA |
| | 0.038613 | 0.099390795 |
| | 0.690711141 | NA |
| | 0.003083344 | 0.012842784 |
| | 0.797151867 | NA |
| | 0.784543659 | NA |
| | 0.794585076 | NA |
| | 0.69348009 | 0.807301559 |
| | 0.393125939 | 0.556622123 |
| | 0.001953113 | 0.008770634 |
| | 0.249775233 | 0.403651336 |
| | 0.262003527 | 0.417411028 |
| | 0.110117477 | 0.221483804 |
| | 0.003442849 | 0.014052325 |
| | 0.016349623 | 0.050093676 |
| | 0.129695731 | 0.249867393 |
| | 0.384340781 | 0.548213004 |
| | 0.503493033 | 0.655765333 |

| | | |
|---|---|---|
| | 0.002437311 | 0.010589814 |
| | 0.126164878 | 0.244912885 |
| | 0.922832678 | 0.955421158 |
| | 0.314105363 | 0.476017923 |
| | 0.737573944 | 0.838994288 |
| | 0.918071361 | NA |
| NA | | NA |
| | 0.004371475 | 0.017098236 |
| | 0.003391444 | NA |
| | 0.065661353 | 0.149477259 |
| | 0.59822434 | 0.7347359 |
| | 0.245188405 | 0.398378999 |
| | 0.012072549 | 0.039152429 |
| | 0.004782647 | 0.018473552 |
| | 0.362203809 | 0.526974984 |
| | 0.153289274 | NA |
| | 0.889409301 | NA |
| | 0.050024366 | 0.121402005 |
| NA | | NA |
| NA | | NA |
| | 0.056510627 | 0.133145585 |
| | 0.090297408 | 0.190657768 |
| | 0.010640557 | 0.035415982 |
| | 0.012334582 | 0.03984921 |
| | 0.2706865 | 0.427195979 |
| | 0.007585878 | 0.026884813 |
| | 5.68E-09 | 1.43E-07 |
| | 0.002883864 | 0.012159516 |
| | 0.178980456 | NA |
| | 5.01E-06 | 5.33E-05 |
| NA | | NA |
| | 0.334389646 | 0.497180732 |
| | 2.54E-06 | 2.96E-05 |
| | 0.668679094 | 0.788800656 |
| | 0.16730729 | NA |
| | 0.795014225 | NA |
| | 0.040555538 | 0.103050112 |
| | 0.748337808 | 0.846438134 |
| | 0.315101365 | 0.477325529 |
| | 0.055238319 | 0.130849864 |
| | 0.11195693 | NA |
| | 3.38E-09 | 9.15E-08 |
| NA | | NA |
| | 6.40E-05 | 0.000477832 |
| NA | | NA |
| | 0.981628422 | NA |
| | 0.850055606 | NA |

| | |
|---|---|
| 0.010752736 | 0.035726937 |
| 0.044333156 | 0.110488227 |
| 6.70E-06 | 6.89E-05 |
| 0.872008992 | 0.92661241 |
| 0.006974331 | 0.025191893 |
| 0.28181735 | 0.44033209 |
| 0.208293938 | 0.354980721 |
| 9.81E-06 | 9.54E-05 |
| 0.034542422 | 0.091135052 |
| 0.102045237 | NA |
| 0.111206822 | 0.222951758 |
| 0.001625868 | 0.007487926 |
| NA | NA |
| 0.992061292 | NA |
| 0.347290713 | 0.511068004 |
| 0.001289827 | 0.006162598 |
| 0.965760485 | 0.981422073 |
| 0.795014225 | NA |
| 0.925487337 | 0.957131655 |
| NA | NA |
| NA | NA |
| 0.011174201 | 0.036791178 |
| NA | NA |
| NA | NA |
| 0.889410245 | NA |
| NA | NA |
| NA | NA |
| NA | NA |
| 0.507858814 | 0.659690724 |
| 0.095124512 | 0.198236013 |
| 0.889409301 | NA |
| 0.12365173 | 0.241421564 |
| 0.889409301 | NA |
| 0.992061292 | NA |
| NA | NA |
| 0.842609949 | NA |
| 0.789999565 | NA |
| 0.905158266 | NA |
| 0.011544404 | 0.037729359 |
| NA | NA |
| NA | NA |
| 0.034908402 | 0.091890865 |
| NA | NA |
| 0.780614158 | 0.867790343 |
| NA | NA |
| 0.889409773 | NA |
| 0.889410245 | NA |

| | | |
|---|---|---|
| | 0.102946192 | 0.210679909 |
| | 0.009118775 | 0.031185044 |
| | 0.100740725 | 0.207151004 |
| | 0.675055639 | 0.793381866 |
| NA | | NA |
| | 0.992060816 | NA |
| | 0.000218794 | 0.001362589 |
| | 0.962330526 | 0.979205986 |
| NA | | NA |
| NA | | NA |
| | 0.973927747 | 0.986212831 |
| | 0.992061292 | NA |
| NA | | NA |
| | 2.39E-05 | 0.000206177 |
| NA | | NA |
| NA | | NA |
| | 8.09E-06 | 8.12E-05 |
| | 0.889409773 | NA |
| | 0.261373215 | NA |
| NA | | NA |
| | 0.000118956 | 0.000811231 |
| | 0.36974283 | NA |
| | 0.991688155 | NA |
| | 0.000203613 | 0.001281128 |
| NA | | NA |
| | 0.164649751 | 0.298780587 |
| | 0.976715686 | NA |
| | 0.106747498 | 0.216302327 |
| | 0.881846723 | NA |
| NA | | NA |
| | 0.004549224 | 0.017681123 |
| | 0.885146051 | 0.934871922 |
| | 0.334907601 | 0.497748631 |
| | 0.002633759 | 0.011302464 |
| | 0.000392438 | 0.002248836 |
| | 0.872972065 | NA |
| | 4.10E-05 | 0.00032661 |
| | 0.011093459 | 0.036596064 |
| | 0.593942702 | 0.731321509 |
| | 2.33E-15 | 3.12E-13 |
| | 0.079179299 | 0.172541256 |
| | 0.002578213 | 0.011092278 |
| | 0.183972936 | 0.324490955 |
| NA | | NA |
| | 0.593632495 | 0.731062529 |
| | 0.013771872 | 0.043506909 |
| NA | | NA |

| | | |
|---|---|---|
| | 0.395247516 | 0.55876136 |
| | 3.73E-06 | 4.14E-05 |
| | 0.304997087 | 0.465649102 |
| | 0.760154666 | 0.854434701 |
| | 0.522269443 | 0.6726711 |
| | 0.391275606 | 0.554681796 |
| | 3.61E-09 | 9.70E-08 |
| | 0.665971107 | 0.786895717 |
| | 0.326912162 | 0.489143102 |
| | 0.046210472 | 0.114182037 |
| | 0.337761711 | NA |
| | 0.071398424 | 0.158913237 |
| | 0.058496327 | 0.136798785 |
| | 0.792095354 | NA |
| | 0.130917149 | 0.251470405 |
| | 6.73E-11 | 2.97E-09 |
| | 4.29E-05 | 0.000339972 |
| | 0.760472536 | NA |
| | 0.044571125 | 0.110905081 |
| | 0.013441182 | 0.042642681 |
| NA | | NA |
| | 0.009637106 | 0.032642481 |
| | 0.000123008 | 0.000834199 |
| | 0.000420064 | 0.002382897 |
| | 0.000311882 | 0.001846985 |
| | 0.795423363 | 0.87663136 |
| NA | | NA |
| | 0.009437883 | 0.032066607 |
| | 0.006249366 | 0.023025138 |
| | 0.088485694 | 0.187789441 |
| | 0.009649059 | 0.03267793 |
| | 0.982965984 | NA |
| | 0.223501449 | 0.3731368 |
| | 0.114742139 | 0.228516473 |
| | 0.54848154 | 0.694625188 |
| | 1.52E-09 | 4.55E-08 |
| | 2.99E-07 | 4.59E-06 |
| | 1.27E-07 | 2.18E-06 |
| | 0.905158266 | NA |
| | 0.000463212 | 0.002585582 |
| | 0.01170819 | 0.038168073 |
| | 0.619519383 | 0.751188632 |
| | 0.547073326 | NA |
| | 7.03E-15 | 8.43E-13 |
| | 0.030506786 | 0.082560824 |
| | 8.20E-05 | 0.00058955 |
| | 0.810864969 | 0.88686673 |

| | |
|---|---|
| 0.006315183 | 0.023219905 |
| NA | NA |
| 0.513720873 | 0.665180533 |
| 0.005812237 | 0.021713324 |
| 0.024773452 | NA |
| 0.022453036 | NA |
| 0.009371988 | 0.031901951 |
| 0.375112905 | 0.539253963 |
| 0.79099014 | 0.873929685 |
| 0.140878663 | 0.265723635 |
| 0.000258626 | 0.001575795 |
| NA | NA |
| 0.097388038 | 0.201806542 |
| 0.992061292 | NA |
| 0.241960746 | 0.394741161 |
| NA | NA |
| NA | NA |
| 0.599664002 | NA |
| 0.67246779 | 0.791526598 |
| 0.076027158 | 0.167165351 |
| 0.000248034 | 0.001518845 |
| NA | NA |
| 0.008787198 | 0.030315489 |
| 0.843560233 | 0.907714959 |
| 0.566600328 | 0.709531481 |
| 0.514756639 | NA |
| 0.925215062 | 0.956940207 |
| 1.10E-05 | 0.000105242 |
| NA | NA |
| 0.978604805 | NA |
| NA | NA |
| 2.60E-06 | 3.00E-05 |
| 0.00225796 | 0.009910562 |
| 0.156471252 | 0.287617097 |
| 0.033419935 | 0.088760081 |
| 0.603147603 | NA |
| NA | NA |
| 0.901514447 | NA |
| 0.018749512 | 0.055780018 |
| NA | NA |
| 0.852666512 | NA |
| 0.357936935 | NA |
| 0.492911118 | 0.646713436 |
| 0.000149713 | 0.000982238 |
| 0.316301372 | 0.478417264 |
| 0.207749265 | 0.354440825 |
| 0.041224428 | NA |

| | |
|---|---|
| 0.012151768 | 0.039374454 |
| 0.097722286 | 0.202270117 |
| 0.185263593 | 0.326108476 |
| 0.000839131 | 0.004297218 |
| 0.450109084 | 0.608829607 |
| 0.40185056 | 0.565402476 |
| 0.269794841 | NA |
| NA | NA |
| 1.26E-05 | 0.000118876 |
| 0.174538775 | 0.312032643 |
| 0.033646582 | 0.089243301 |
| 0.441537222 | 0.601210135 |
| 0.084088201 | 0.18049783 |
| 0.806663406 | NA |
| 0.818801318 | NA |
| NA | NA |
| 0.168316258 | 0.303777161 |
| 0.905158266 | NA |
| 0.905158739 | NA |
| 0.992061769 | NA |
| NA | NA |
| NA | NA |
| NA | NA |
| NA | NA |
| NA | NA |
| NA | NA |
| NA | NA |
| NA | NA |
| NA | NA |
| NA | NA |
| NA | NA |
| NA | NA |
| NA | NA |
| NA | NA |
| NA | NA |
| NA | NA |
| NA | NA |
| NA | NA |
| NA | NA |
| NA | NA |
| NA | NA |
| NA | NA |
| NA | NA |
| NA | NA |
| NA | NA |
| NA | NA |
| NA | NA |

| | | |
|---|---|---|
| NA | NA | |
| NA | NA | |
| NA | NA | |
| 0.991918092 | NA | |
| NA | NA | |
| NA | NA | |
| NA | NA | |
| NA | NA | |
| NA | NA | |
| NA | NA | |
| NA | NA | |
| NA | NA | |
| NA | NA | |
| NA | NA | |
| NA | NA | |
| NA | NA | |
| NA | NA | |
| NA | NA | |
| NA | NA | |
| 0.571692613 | NA | |
| NA | NA | |
| NA | NA | |
| NA | NA | |
| NA | NA | |
| NA | NA | |
| NA | NA | |
| NA | NA | |
| NA | NA | |
| NA | NA | |
| 0.905158266 | NA | |
| NA | NA | |
| NA | NA | |
| NA | NA | |
| NA | NA | |
| NA | NA | |
| NA | NA | |
| NA | NA | |
| 0.119809826 | | 0.235860598 |
| 0.905158266 | NA | |
| 0.991918092 | NA | |
| NA | NA | |
| 0.421035149 | NA | |
| 0.905158739 | NA | |
| 1.76E-12 | | 1.18E-10 |
| 0.195681637 | NA | |
| 1.07E-15 | | 1.61E-13 |
| 0.905158739 | NA | |

| | | |
|---|---|---|
| | 6.77E-10 | 2.25E-08 |
| | 0.889409773 | NA |
| | 0.905158266 | NA |
| | 0.673811734 | 0.792705761 |
| NA | | NA |
| | 0.469079329 | 0.625410624 |
| | 0.854762067 | NA |
| | 0.016973393 | 0.051626319 |
| | 0.90515732 | NA |
| | 0.905157793 | NA |
| | 0.475638483 | 0.631815707 |
| | 0.039618879 | 0.101341721 |
| | 1.53E-06 | 1.89E-05 |
| | 0.487369657 | NA |
| | 0.99238693 | 0.996152158 |
| | 0.246467475 | 0.399895026 |
| | 0.000783118 | 0.004052499 |
| | 0.897412177 | NA |
| | 0.00075034 | 0.003919296 |
| | 0.044403232 | 0.11060011 |
| | 0.00301947 | 0.012627186 |
| | 0.798148243 | 0.878127024 |
| | 0.992060816 | NA |
| | 0.869021198 | 0.925004192 |
| | 3.00E-05 | 0.000250494 |
| | 0.561450037 | NA |
| NA | | NA |
| | 8.56E-06 | 8.54E-05 |
| | 0.218710988 | 0.367345464 |
| | 0.007188084 | 0.025791363 |
| | 0.000121152 | 0.000824156 |
| | 0.173659128 | 0.310915678 |
| | 4.46E-06 | 4.84E-05 |
| | 0.992060816 | NA |
| NA | | NA |
| | 0.211534128 | 0.358582516 |
| NA | | NA |
| NA | | NA |
| | 0.008140088 | 0.028472156 |
| | 0.661046451 | 0.782909176 |
| | 0.159577734 | 0.29174652 |
| | 0.076335829 | 0.167726484 |
| | 0.006871862 | 0.02490908 |
| | 0.08350257 | 0.179644649 |
| NA | | NA |
| | 0.218911278 | 0.367582783 |
| | 0.023594318 | 0.067197195 |

| | |
|---|---|
| 0.908678685 | 0.948154039 |
| 0.002510464 | 0.010853963 |
| 0.627190856 | 0.756363966 |
| 0.188361956 | 0.329848245 |
| 0.033722983 | 0.089396192 |
| NA | NA |
| 0.013260307 | 0.042163783 |
| 0.743309877 | NA |
| 0.259177633 | NA |
| NA | NA |
| 0.043607654 | 0.109045882 |
| 0.217597851 | 0.365691912 |
| 0.026990663 | 0.07484379 |
| 0.503389542 | 0.655669432 |
| NA | NA |
| NA | NA |
| 0.728247086 | NA |
| 0.203699989 | 0.349566119 |
| 0.419338481 | NA |
| NA | NA |
| 0.757372378 | NA |
| NA | NA |
| 0.318425707 | 0.480810075 |
| 0.493901698 | 0.647304529 |
| 0.46048158 | 0.617924408 |
| 0.012568862 | 0.040445434 |
| 0.000207282 | 0.001299746 |
| 1.50E-05 | 0.000137941 |
| 0.678427179 | 0.795978181 |
| 0.482568435 | NA |
| 0.121769017 | 0.238636844 |
| 0.788984625 | 0.872635911 |
| 1.17E-05 | 0.000111068 |
| 0.302920549 | 0.463583226 |
| 0.998888454 | 0.999571253 |
| 0.737794902 | 0.839048097 |
| NA | NA |
| 0.328097546 | 0.490215282 |
| 0.98294987 | 0.99175069 |
| 0.751580834 | 0.848725928 |
| 1.87E-05 | 0.000167564 |
| 7.66E-12 | 4.42E-10 |
| 0.225691663 | 0.375508654 |
| 0.784286811 | 0.870114291 |
| 0.002991169 | 0.012530158 |
| 0.275358618 | 0.432703585 |
| 0.322286272 | 0.484632215 |

| | | |
|---|---|---|
| | 0.660941148 | 0.782877278 |
| | 0.476281116 | 0.632554556 |
| | 0.872232189 | 0.926670213 |
| | 0.386060579 | NA |
| | 0.032479748 | 0.086830142 |
| | 0.889409773 | NA |
| NA | | NA |
| | 0.737163341 | 0.838636919 |
| | 0.225662904 | 0.375489256 |
| | 0.14454416 | 0.271077394 |
| | 0.030867303 | 0.083329145 |
| | 0.889410717 | NA |
| | 0.216056851 | 0.363917495 |
| NA | | NA |
| | 1.29E-06 | 1.63E-05 |
| | 0.900999586 | NA |
| | 0.238268484 | 0.390283279 |
| | 0.469778596 | 0.626152849 |
| | 7.25E-14 | 6.66E-12 |
| | 0.069369143 | 0.15542057 |
| | 0.084010539 | 0.180419308 |
| | 0.003814955 | 0.015267467 |
| NA | | NA |
| | 0.000614635 | 0.003307887 |
| NA | | NA |
| | 0.952279375 | 0.973288471 |
| | 0.306019637 | 0.466626291 |
| | 0.541045168 | 0.688383501 |
| | 0.289292775 | 0.448628534 |
| NA | | NA |
| | 0.666655503 | 0.787386413 |
| | 0.810053969 | NA |
| | 0.889409773 | NA |
| NA | | NA |
| | 0.756641735 | NA |
| | 0.000159372 | 0.001036009 |
| | 0.000248125 | 0.001518981 |
| | 0.064162421 | 0.146994534 |
| | 2.36E-07 | 3.75E-06 |
| | 0.418833004 | 0.580933042 |
| | 2.21E-05 | 0.000194299 |
| | 0.21479468 | 0.362347421 |
| | 0.85072499 | NA |
| | 1.19E-09 | 3.65E-08 |
| | 0.105821354 | 0.214861869 |
| | 3.64E-08 | 7.42E-07 |
| | 0.630927064 | 0.758884411 |

| | | |
|---|---|---|
| | 0.036623272 | 0.095477908 |
| | 0.772661713 | NA |
| | 0.1227697 | 0.240062318 |
| | 0.000143968 | 0.000949079 |
| | 0.000288835 | 0.001731985 |
| | 0.033514006 | 0.088941833 |
| | 7.25E-05 | 0.000531141 |
| | 0.045985936 | 0.113768045 |
| | 0.889410245 | NA |
| | 0.000151166 | 0.000990584 |
| | 0.952129994 | 0.973288471 |
| | 0.868263659 | 0.92446132 |
| | 1.20E-05 | 0.000114356 |
| | 0.582659905 | 0.722574623 |
| | 0.709914705 | 0.819267238 |
| | 6.26E-08 | 1.18E-06 |
| NA | | NA |
| | 0.011785812 | 0.03837554 |
| | 0.002097481 | 0.009316058 |
| | 0.379446001 | 0.543809864 |
| | 0.220787635 | 0.369730472 |
| | 0.069865921 | 0.156246285 |
| | 0.754764926 | 0.850902711 |
| | 0.881081139 | 0.932417385 |
| | 0.889410245 | NA |
| | 9.13E-15 | 1.07E-12 |
| | 3.39E-05 | 0.000277743 |
| | 0.20375493 | NA |
| | 0.691641527 | 0.805973057 |
| | 0.275197916 | 0.432482003 |
| | 0.716848983 | NA |
| | 0.786598345 | 0.871533027 |
| | 0.685105801 | 0.800821797 |
| | 3.67E-11 | 1.75E-09 |
| | 0.651765954 | 0.775809225 |
| | 3.36E-05 | 0.000275979 |
| | 0.000307757 | 0.001825018 |
| | 0.90515732 | NA |
| | 0.121340668 | 0.238009803 |
| | 0.122893335 | NA |
| | 0.003517393 | 0.014319323 |
| | 0.000152978 | 0.00100067 |
| | 0.011173797 | 0.036791178 |
| | 7.05E-07 | 9.69E-06 |
| | 0.000122641 | 0.000832481 |
| | 0.303430067 | 0.464083208 |
| | 0.939739409 | 0.96589388 |

| | |
|---|---|
| NA | NA |
| 0.111222949 | NA |
| 0.572014636 | NA |
| 0.04305317 | 0.10803389 |
| 0.020924927 | 0.060964639 |
| 0.000363469 | 0.002107025 |
| 0.002173523 | 0.009596628 |
| 1.13E-11 | 6.21E-10 |
| 0.058438492 | 0.136705109 |
| NA | NA |
| NA | NA |
| NA | NA |
| NA | NA |
| NA | NA |
| NA | NA |
| 9.23E-05 | 0.00065146 |
| 0.561831851 | NA |
| NA | NA |
| 0.889409773 | NA |
| NA | NA |
| 0.753111281 | NA |
| NA | NA |
| 0.851542467 | NA |
| 0.43703867 | 0.597344215 |
| 0.538181661 | NA |
| 0.730044732 | NA |
| NA | NA |
| NA | NA |
| NA | NA |
| 0.007463085 | 0.026556778 |
| NA | NA |
| NA | NA |
| 0.905158266 | NA |
| 0.889409773 | NA |
| NA | NA |
| NA | NA |
| NA | NA |
| NA | NA |
| NA | NA |
| 0.850770391 | NA |
| NA | NA |
| NA | NA |
| NA | NA |
| NA | NA |
| NA | NA |
| NA | NA |
| 0.905158266 | NA |

| | | |
|---|---|---|
| | 0.889409301 | NA |
| NA | | NA |
| | 0.905158266 | NA |
| | 0.830163713 | NA |
| NA | | NA |
| NA | | NA |
| NA | | NA |
| NA | | NA |
| NA | | NA |
| NA | | NA |
| | 0.992061292 | NA |
| NA | | NA |
| NA | | NA |
| NA | | NA |
| NA | | NA |
| NA | | NA |
| NA | | NA |
| NA | | NA |
| NA | | NA |
| NA | | NA |
| | 0.905158739 | NA |
| NA | | NA |
| NA | | NA |
| NA | | NA |
| NA | | NA |
| NA | | NA |
| | 0.889410245 | NA |
| NA | | NA |
| NA | | NA |
| NA | | NA |
| NA | | NA |
| NA | | NA |
| NA | | NA |
| NA | | NA |
| NA | | NA |
| NA | | NA |
| NA | | NA |
| NA | | NA |
| | 0.992061292 | NA |
| | 0.889409773 | NA |
| NA | | NA |
| NA | | NA |
| NA | | NA |
| NA | | NA |
| NA | | NA |
| NA | | NA |
| NA | | NA |

| | |
|---|---|
| NA | NA |
| NA | NA |
| NA | NA |
| NA | NA |
| NA | NA |
| 0.905158739 | NA |
| NA | NA |
| NA | NA |
| NA | NA |
| NA | NA |
| 0.889409301 | NA |
| 0.000119055 | 0.00081165 |
| 0.031848305 | 0.08547506 |
| NA | NA |
| NA | NA |
| NA | NA |
| NA | NA |
| NA | NA |
| NA | NA |
| NA | NA |
| 0.37769595 | 0.542115521 |
| NA | NA |
| 0.992061292 | NA |
| NA | NA |
| 0.992061292 | NA |
| NA | NA |
| NA | NA |
| NA | NA |
| 0.807252815 | NA |
| 0.905158739 | NA |
| NA | NA |
| NA | NA |
| NA | NA |
| NA | NA |
| NA | NA |
| NA | NA |
| NA | NA |
| 0.889409301 | NA |
| NA | NA |
| NA | NA |
| NA | NA |
| 0.992061292 | NA |
| NA | NA |
| NA | NA |
| 0.989460338 | NA |
| NA | NA |
| NA | NA |

| | | |
|---|---|---|
| | 0.889409773 | NA |
| | 0.265870789 | NA |
| | 0.05173461 | 0.124620481 |
| | 6.88E-07 | 9.48E-06 |
| NA | | NA |
| | 0.003823085 | 0.015294429 |
| NA | | NA |
| | 0.956473296 | NA |
| | 0.011528332 | 0.037693662 |
| | 0.889409773 | NA |
| | 0.00039675 | 0.002270584 |
| NA | | NA |
| | 8.80E-05 | 0.000625279 |
| | 0.000770772 | 0.004000327 |
| | 2.43E-06 | 2.84E-05 |
| | 0.702979752 | NA |
| | 0.000128458 | 0.000863946 |
| | 0.956866117 | 0.976068336 |
| NA | | NA |
| NA | | NA |
| | 0.889409773 | NA |
| | 0.905158739 | NA |
| NA | | NA |
| | 0.665582351 | NA |
| | 0.961160162 | NA |
| NA | | NA |
| NA | | NA |
| | 0.75880646 | 0.853662127 |
| NA | | NA |
| NA | | NA |
| NA | | NA |
| NA | | NA |
| NA | | NA |
| NA | | NA |
| NA | | NA |
| | 0.824229127 | 0.895142918 |
| NA | | NA |
| NA | | NA |
| NA | | NA |
| | 0.889409301 | NA |
| | 0.905158739 | NA |
| | 0.889409773 | NA |
| NA | | NA |
| | 0.007261382 | 0.025994401 |
| NA | | NA |
| | 0.905158739 | NA |
| | 0.991918092 | NA |

| | |
|---|---|
| NA | NA |
| NA | NA |
| NA | NA |
| NA | NA |
| NA | NA |
| 0.856527406 | NA |
| NA | NA |
| 0.827497798 | NA |
| 0.28551182 | NA |
| 0.605201816 | NA |
| 0.867242981 | NA |
| 0.878547116 | NA |
| NA | NA |
| NA | NA |
| NA | NA |
| 0.026403533 | 0.073526885 |
| 0.314718501 | 0.476811341 |
| 0.905158266 | NA |
| 0.006407096 | 0.023491971 |
| 0.889409773 | NA |
| 0.165163243 | 0.299439729 |
| 0.835126916 | 0.901955142 |
| 9.79E-09 | 2.33E-07 |
| 0.242834162 | 0.3956957 |
| 1.00E-07 | 1.76E-06 |
| 0.78318692 | 0.86937628 |
| 3.30E-05 | 0.000271387 |
| 0.068201792 | 0.153439533 |
| 0.665989203 | 0.786895717 |
| 0.56761271 | 0.710332678 |
| 0.889409301 | NA |
| 0.464061926 | NA |
| 2.43E-05 | 0.000209402 |
| 0.00329761 | 0.013575594 |
| 5.98E-05 | 0.00045159 |
| 9.32E-05 | 0.000657004 |
| 0.487579247 | 0.642276705 |
| 0.649991934 | 0.774642098 |
| 0.001349967 | 0.006420605 |
| 1.39E-08 | 3.16E-07 |
| 5.75E-07 | 8.11E-06 |
| 0.726493931 | NA |
| 0.002149551 | 0.009506946 |
| NA | NA |
| 0.035488514 | NA |
| 0.905158266 | NA |
| NA | NA |

| | | |
|---|---|---|
| | 0.005200929 | 0.019807006 |
| | 0.030004265 | NA |
| NA | | NA |
| | 0.3283496 | NA |
| | 3.92E-05 | 0.000314509 |
| NA | | NA |
| | 1.11E-14 | 1.26E-12 |
| | 4.92E-09 | 1.25E-07 |
| | 0.90515732 | NA |
| NA | | NA |
| NA | | NA |
| NA | | NA |
| NA | | NA |
| | 0.905158266 | NA |
| NA | | NA |
| | 0.905158739 | NA |
| NA | | NA |
| | 0.992062245 | NA |
| NA | | NA |
| | 0.992061292 | NA |
| NA | | NA |
| | 0.992060816 | NA |
| NA | | NA |
| NA | | NA |
| | 0.992061292 | NA |
| NA | | NA |
| NA | | NA |
| NA | | NA |
| NA | | NA |
| NA | | NA |
| | 0.992060816 | NA |
| NA | | NA |
| | 0.905158739 | NA |
| | 0.807252815 | NA |
| NA | | NA |
| | 0.219353764 | NA |
| NA | | NA |
| | 0.905158266 | NA |
| | 0.889409773 | NA |
| | 0.009237163 | 0.031516293 |
| | 0.677277103 | 0.795269125 |
| | 0.0002018 | 0.001272636 |
| NA | | NA |
| | 0.007918582 | 0.027839281 |
| NA | | NA |
| NA | | NA |
| NA | | NA |

| | | | |
|---|---|---|---|
| NA | | NA | |
| | 0.683637667 | | 0.799744261 |
| | 0.992061769 | NA | |
| | 0.905158739 | NA | |
| NA | | NA | |
| NA | | NA | |
| | 0.895257161 | NA | |
| NA | | NA | |
| NA | | NA | |
| NA | | NA | |
| NA | | NA | |
| NA | | NA | |
| | 0.889409301 | NA | |
| NA | | NA | |
| NA | | NA | |
| NA | | NA | |
| NA | | NA | |
| | 0.784353866 | NA | |
| NA | | NA | |
| NA | | NA | |
| NA | | NA | |
| NA | | NA | |
| NA | | NA | |
| NA | | NA | |
| NA | | NA | |
| | 0.905158266 | NA | |
| NA | | NA | |
| NA | | NA | |
| NA | | NA | |
| NA | | NA | |
| NA | | NA | |
| NA | | NA | |
| NA | | NA | |
| NA | | NA | |
| | 0.905158266 | NA | |
| NA | | NA | |
| NA | | NA | |
| NA | | NA | |
| NA | | NA | |
| NA | | NA | |
| NA | | NA | |
| NA | | NA | |
| | 0.291147001 | | 0.450581169 |
| | 0.545924979 | | 0.692345747 |
| | 0.077094461 | | 0.16903817 |
| | 1.58E-14 | | 1.71E-12 |
| | 0.095016067 | | 0.198106514 |

| | |
|---|---|
| 0.485240666 | 0.640266317 |
| 0.997233802 | 0.998780254 |
| 0.655894457 | 0.778770817 |
| 0.865501361 | 0.922464903 |
| 0.519974545 | 0.670746698 |
| 0.599378976 | 0.73542997 |
| 0.007112566 | 0.025570535 |
| NA | NA |
| 0.648696342 | NA |
| 0.134714152 | NA |
| 0.119200404 | 0.234944948 |
| 0.001331391 | 0.006339118 |
| 0.09098792 | 0.191674523 |
| 0.574212583 | NA |
| 0.922166019 | 0.955294241 |
| 0.3907613 | 0.554118558 |
| 0.190647877 | NA |
| 0.018414075 | 0.054957366 |
| 0.073935306 | 0.163416748 |
| 0.001100101 | 0.005395828 |
| 0.836660123 | NA |
| 0.056542602 | 0.133192341 |
| NA | NA |
| 2.56E-06 | 2.98E-05 |
| 0.6389895 | 0.765706436 |
| 0.171545252 | 0.308060369 |
| 0.13798358 | 0.261633835 |
| 0.001072165 | 0.005281216 |
| 0.267505668 | 0.423729132 |
| 1.13E-11 | 6.21E-10 |
| 0.560704369 | 0.704981806 |
| NA | NA |
| 0.905158266 | NA |
| NA | NA |
| 0.785779118 | NA |
| 0.014722501 | 0.045980855 |
| 3.39E-05 | 0.000277352 |
| 0.09964047 | 0.205407913 |
| 0.996308813 | NA |
| 0.216918547 | 0.365060881 |
| 0.226254111 | 0.376074031 |
| 0.889409773 | NA |
| 0.003892353 | 0.015526282 |
| 6.45E-05 | 0.000480796 |
| NA | NA |
| 0.056304032 | 0.132772792 |
| NA | NA |

| | |
|---|---|
| 0.905158266 | NA |
| 0.056685146 | 0.133456538 |
| 9.82E-08 | 1.74E-06 |
| 0.001287852 | 0.006157185 |
| 0.051050856 | 0.12336588 |
| 0.105300331 | 0.214143106 |
| 0.724741117 | 0.830225374 |
| 0.451242098 | 0.609593611 |
| 0.997186137 | NA |
| 3.23E-09 | 8.84E-08 |
| 0.98002343 | NA |
| 0.002897414 | 0.012204933 |
| NA | NA |
| NA | NA |
| 0.309912042 | 0.471122085 |
| 0.036327063 | 0.094806984 |
| 0.470867917 | NA |
| NA | NA |
| 0.475959496 | 0.632165642 |
| 0.788583904 | NA |
| NA | NA |
| 0.986439002 | NA |
| 0.905158266 | NA |
| 0.682309075 | 0.798785785 |
| 0.209426555 | 0.356207298 |
| 0.000410897 | 0.002339354 |
| 0.002705571 | 0.01154745 |
| 0.102290045 | 0.209581516 |
| 0.209938682 | 0.356739074 |
| 0.324490602 | 0.486713739 |
| 8.55E-10 | 2.75E-08 |
| 0.220524442 | 0.369458779 |
| 0.387599042 | 0.551284886 |
| 0.001164627 | 0.005666749 |
| NA | NA |
| 0.905158739 | NA |
| NA | NA |
| 0.631970682 | NA |
| 0.000308255 | 0.001827474 |
| 0.760642125 | NA |
| 0.795352046 | NA |
| 0.150058923 | 0.278776979 |
| 0.559012409 | 0.703518549 |
| 0.258735847 | 0.413627 |
| 0.002016809 | 0.009021613 |
| 0.325378547 | 0.487570104 |
| 0.144067585 | 0.270414573 |

| | |
|---|---|
| 0.651683019 | 0.775786205 |
| NA | NA |
| NA | NA |
| 0.001550864 | 0.007199879 |
| NA | NA |
| 0.000601482 | 0.003249986 |
| NA | NA |
| 0.013234867 | 0.042105076 |
| 0.501931098 | 0.654662923 |
| 0.902664484 | 0.944966123 |
| NA | NA |
| 0.649633065 | NA |
| 0.889409773 | NA |
| 1.35E-05 | 0.00012658 |
| 1.87E-06 | 2.26E-05 |
| 0.975795661 | 0.987624305 |
| 0.836702521 | 0.902813162 |
| 0.371473918 | 0.535811597 |
| 0.610111439 | 0.744129455 |
| 0.364823196 | 0.529635003 |
| 0.010050856 | 0.033798891 |
| 0.787774943 | NA |
| 1.58E-06 | 1.94E-05 |
| 0.336735904 | 0.499721817 |
| 0.229111827 | 0.379588547 |
| 0.34362146 | 0.506994808 |
| NA | NA |
| 0.784414466 | NA |
| NA | NA |
| 0.992061769 | NA |
| 0.887395124 | NA |
| 0.876116536 | NA |
| 0.335856378 | 0.498722939 |
| 0.889409301 | NA |
| 0.905158739 | NA |
| 0.986808944 | NA |
| 0.50055971 | 0.65350931 |
| 0.541748733 | 0.689039122 |
| 0.674144028 | NA |
| 0.079618645 | 0.173257936 |
| 0.091719401 | 0.192881281 |
| 9.34E-06 | 9.18E-05 |
| 0.350890706 | 0.51508851 |
| 0.002490029 | 0.010778345 |
| 0.65150867 | NA |
| 0.053230036 | 0.127162572 |
| 0.442266319 | NA |

| | | |
|---|---|---|
| | 0.205832631 | 0.35203001 |
| | 0.160807698 | 0.293408912 |
| | 0.329446636 | NA |
| | 0.174278058 | NA |
| | 0.355680551 | NA |
| | 0.829806995 | NA |
| | 0.284890767 | 0.443744954 |
| | 0.151908016 | NA |
| | 0.053132288 | 0.127011965 |
| | 0.436619174 | 0.59695663 |
| | 0.066003241 | 0.150053336 |
| | 3.30E-06 | 3.71E-05 |
| | 0.905158266 | NA |
| | 0.002338121 | 0.010217474 |
| | 0.001069758 | 0.005270546 |
| | 0.101680616 | 0.208634335 |
| | 0.142268674 | 0.267747499 |
| NA | | NA |
| | 0.105088731 | 0.213888538 |
| | 0.103428885 | 0.211352586 |
| NA | | NA |
| | 0.030891428 | 0.08337572 |
| | 0.458745781 | 0.616072127 |
| | 0.003524354 | 0.014337029 |
| | 0.156608726 | 0.287755921 |
| | 0.888235081 | NA |
| | 0.373456546 | 0.537752937 |
| | 0.001043594 | 0.005162489 |
| | 0.011713986 | 0.038181301 |
| | 0.263748386 | 0.419502449 |
| | 0.000129047 | 0.000866856 |
| | 0.000259887 | 0.001581724 |
| | 0.040689998 | 0.103296146 |
| | 0.370365985 | 0.53491625 |
| | 0.992061292 | NA |
| | 0.414990936 | 0.577446489 |
| | 0.800135892 | NA |
| | 0.181681848 | 0.32188671 |
| | 0.279577746 | 0.437798454 |
| | 0.919099008 | 0.953939364 |
| NA | | NA |
| | 9.54E-12 | 5.34E-10 |
| | 0.491011787 | 0.644979831 |
| | 0.252415729 | 0.406721235 |
| NA | | NA |
| | 1.46E-09 | 4.38E-08 |
| | 0.033135091 | 0.088195571 |

| | |
|---|---|
| 0.995081602 | 0.997551495 |
| 0.509193845 | 0.660877572 |
| 4.04E-06 | 4.43E-05 |
| 0.081011826 | 0.175662456 |
| NA | NA |
| 0.227197586 | NA |
| 9.52E-06 | 9.31E-05 |
| 0.772862102 | NA |
| 0.735039024 | 0.837217798 |
| 0.678941124 | 0.796328819 |
| 1.98E-07 | 3.21E-06 |
| 0.108006385 | 0.218349494 |
| 0.217456136 | 0.365657346 |
| NA | NA |
| 0.905158739 | NA |
| 0.674506456 | 0.793160933 |
| NA | NA |
| NA | NA |
| 0.85873046 | NA |
| NA | NA |
| 0.035437475 | 0.092938196 |
| 0.005024171 | 0.019254062 |
| 3.12E-08 | 6.48E-07 |
| 0.006035633 | 0.022388778 |
| 0.002568352 | 0.011054194 |
| 0.05254101 | 0.126020184 |
| 0.000371778 | 0.002145564 |
| 0.992096636 | 0.995961112 |
| 0.992060816 | NA |
| 0.889409773 | NA |
| 0.869976449 | 0.925515189 |
| 0.522179105 | 0.67263804 |
| 0.031049664 | 0.083707575 |
| NA | NA |
| 0.462303794 | 0.619612367 |
| 0.013453915 | 0.04267435 |
| 0.086060694 | NA |
| 0.225327421 | 0.375016283 |
| 0.249310746 | 0.403167735 |
| 4.91E-06 | 5.24E-05 |
| 0.000544595 | 0.002977778 |
| 0.077238259 | 0.169268954 |
| 3.92E-06 | 4.33E-05 |
| 0.924385039 | NA |
| 0.001085706 | 0.005337561 |
| NA | NA |
| 6.23E-17 | 1.17E-14 |

| | | |
|---|---|---|
| | 0.986502853 | NA |
| | 5.29E-07 | 7.57E-06 |
| | 0.067228257 | 0.151879147 |
| | 2.48E-13 | 2.06E-11 |
| | 0.648309409 | 0.773119709 |
| | 0.986354244 | 0.993034302 |
| | 3.21E-09 | 8.79E-08 |
| | 0.679209712 | NA |
| | 0.173628243 | 0.31088573 |
| | 0.010314743 | 0.034548724 |
| | 0.251023152 | 0.405161146 |
| | 0.002905285 | 0.01222636 |
| | 0.902582149 | 0.944966123 |
| | 0.032283415 | 0.086421007 |
| | 0.050189448 | 0.121712876 |
| | 0.702161145 | 0.814141655 |
| | 4.91E-05 | 0.000381725 |
| | 0.328249533 | 0.490409001 |
| | 0.274822771 | 0.432016124 |
| | 0.000311142 | 0.001843099 |
| | 0.763954201 | NA |
| | 0.002520604 | 0.010886664 |
| | 0.000699746 | 0.003689249 |
| | 0.133909617 | 0.255656043 |
| | 7.38E-09 | 1.80E-07 |
| | 0.000549546 | 0.003001114 |
| | 0.801661681 | NA |
| | 0.992061292 | NA |
| | 2.30E-13 | 1.93E-11 |
| | 0.579069898 | 0.719707714 |
| | 0.881439994 | 0.932572432 |
| | 2.72E-07 | 4.24E-06 |
| | 0.189424595 | 0.331303448 |
| | 0.169895743 | 0.305999066 |
| | 0.000358194 | 0.002081547 |
| | 0.652356559 | NA |
| | 0.454486036 | 0.61249901 |
| | 0.104648717 | 0.213190572 |
| | 0.064455197 | 0.147496005 |
| | 0.538891166 | 0.686583379 |
| NA | | NA |
| NA | | NA |
| | 0.905158266 | NA |
| NA | | NA |
| | 0.001997995 | 0.008948394 |
| | 0.502874016 | 0.655274222 |
| | 0.00070571 | 0.003717122 |

| | |
|---|---|
| NA | NA |
| 0.787677656 | 0.871771467 |
| 1.49E-05 | 0.000137448 |
| 0.357761146 | 0.522484505 |
| 0.108352392 | 0.218893399 |
| 0.119813108 | 0.235860598 |
| 0.175481631 | 0.31338656 |
| 0.903623155 | 0.945383814 |
| 0.363488364 | 0.528423766 |
| 0.991918092 | NA |
| 8.22E-06 | 8.23E-05 |
| 7.32E-06 | 7.44E-05 |
| 0.182282282 | 0.322441599 |
| 0.002679485 | 0.011465084 |
| 0.819953067 | 0.892909192 |
| NA | NA |
| 0.101998513 | 0.209188881 |
| 0.614140557 | 0.747390406 |
| 0.197494912 | 0.341802551 |
| 0.050028086 | 0.121402005 |
| 0.000104493 | 0.000725894 |
| 0.650009349 | 0.774642098 |
| 0.393931531 | NA |
| NA | NA |
| 0.174226594 | 0.31155988 |
| 0.106064294 | 0.215235729 |
| 0.404926938 | 0.568177875 |
| 0.988912671 | NA |
| NA | NA |
| 0.004382748 | 0.017124692 |
| 0.928005426 | 0.958697363 |
| 0.000377366 | 0.002172672 |
| 0.016431264 | NA |
| 0.080693573 | 0.175093889 |
| 0.10613591 | 0.21534126 |
| 0.515636657 | NA |
| 0.495561855 | 0.648783328 |
| NA | NA |
| 0.931691237 | 0.960437209 |
| 0.111545927 | 0.223448004 |
| 0.20136634 | 0.346590645 |
| 6.48E-07 | 8.98E-06 |
| 0.464370171 | 0.621433639 |
| 0.009219323 | 0.031479882 |
| 0.75855487 | 0.853510268 |
| 0.000579603 | 0.003147258 |
| 0.270038605 | 0.426449315 |

| | | |
|---|---|---|
| | 0.905158266 | NA |
| | 0.009037342 | 0.030983762 |
| | 0.723425399 | 0.829150704 |
| | 0.106794454 | 0.216357544 |
| | 0.165832177 | 0.300206825 |
| | 0.696494909 | 0.809866651 |
| | 0.199837687 | 0.344716085 |
| | 0.797157316 | NA |
| | 0.013728632 | 0.04340153 |
| | 6.26E-05 | 0.000468972 |
| NA | | NA |
| | 0.367238158 | 0.531938174 |
| NA | | NA |
| | 1.17E-06 | 1.50E-05 |
| | 0.54706733 | 0.693377685 |
| | 0.093815894 | 0.196069594 |
| | 0.000879236 | 0.004462014 |
| | 6.14E-05 | 0.000460742 |
| | 0.989805317 | 0.994571049 |
| | 0.000334222 | 0.001962872 |
| | 0.053767068 | NA |
| | 0.967292499 | 0.982404998 |
| NA | | NA |
| | 0.000285364 | 0.001715219 |
| | 0.03639847 | 0.094982057 |
| | 0.680115981 | 0.797238393 |
| NA | | NA |
| | 6.49E-11 | 2.88E-09 |
| NA | | NA |
| | 0.032761757 | 0.08743503 |
| NA | | NA |
| | 0.006620312 | 0.024132709 |
| | 1.36E-07 | 2.32E-06 |
| NA | | NA |
| | 3.70E-06 | 4.12E-05 |
| NA | | NA |
| | 0.992061292 | NA |
| | 0.31663139 | 0.478718583 |
| | 0.134575713 | 0.256508057 |
| NA | | NA |
| | 0.001893861 | 0.008544751 |
| | 0.364956256 | 0.52971607 |
| | 0.230891797 | 0.38175988 |
| | 0.557776791 | NA |
| | 0.245873726 | NA |
| | 0.806595052 | 0.884098071 |
| | 6.56E-09 | 1.62E-07 |

| | |
|---|---|
| NA | NA |
| 1.92E-06 | 2.31E-05 |
| 0.256866253 | 0.411536994 |
| 0.20221423 | 0.347722966 |
| 0.283839601 | 0.442602937 |
| 0.084211199 | 0.180602961 |
| 0.850501103 | NA |
| 0.905158739 | NA |
| NA | NA |
| 0.884632702 | 0.934689385 |
| 0.244554044 | 0.397695356 |
| 0.005350198 | 0.020287513 |
| NA | NA |
| 0.802866756 | 0.881375349 |
| 0.458242078 | 0.615634004 |
| 0.191785801 | 0.334266561 |
| 0.411790488 | 0.574492207 |
| 0.330785927 | 0.493292236 |
| 0.464951065 | NA |
| 0.017350073 | 0.052470768 |
| 0.208932035 | 0.355717999 |
| 0.00506159 | 0.01936367 |
| 0.119531367 | 0.235456396 |
| 0.848812311 | NA |
| 0.60065708 | NA |
| 0.002471314 | 0.010717289 |
| 0.324245571 | 0.486479127 |
| 0.624630159 | NA |
| NA | NA |
| 0.539347902 | 0.686940467 |
| 0.334907763 | NA |
| 0.894545163 | 0.940456611 |
| 0.158074924 | 0.289678079 |
| 0.000606928 | 0.003273775 |
| 0.31593624 | 0.478128456 |
| 0.002052965 | 0.009149799 |
| 0.051056985 | NA |
| 0.001096854 | 0.00538351 |
| 0.967855934 | NA |
| 0.905158739 | NA |
| 0.302877707 | 0.463573679 |
| 0.734389525 | NA |
| 0.839088999 | 0.904454905 |
| 0.00249305 | 0.01078717 |
| 0.713272381 | 0.821458447 |
| 0.870480051 | 0.925883819 |
| 0.075408591 | 0.16602138 |

| | | |
|---|---|---|
| | 5.81E-17 | 1.11E-14 |
| | 0.855649127 | NA |
| | 0.011002281 | NA |
| | 0.017621281 | 0.053100824 |
| | 4.26E-11 | 1.99E-09 |
| | 0.009159376 | 0.031306403 |
| NA | | NA |
| | 0.530200603 | NA |
| | 0.599512 | 0.7355378 |
| | 0.905157793 | NA |
| | 0.782559532 | NA |
| NA | | NA |
| | 0.602433956 | 0.737928668 |
| | 0.993362209 | NA |
| | 0.024375993 | 0.069049592 |
| | 0.883026929 | 0.933531767 |
| NA | | NA |
| NA | | NA |
| | 0.542680595 | 0.689772911 |
| | 0.853359172 | 0.914405605 |
| | 0.977469017 | 0.98842452 |
| | 0.004531979 | 0.01762936 |
| NA | | NA |
| | 0.723607951 | 0.82923009 |
| | 0.019740648 | 0.05818589 |
| | 0.780692568 | 0.867791855 |
| | 3.51E-05 | 0.000285526 |
| | 0.67943796 | NA |
| | 0.271123822 | 0.427732452 |
| | 3.41E-16 | 5.76E-14 |
| | 0.000598177 | 0.003235313 |
| NA | | NA |
| | 0.045375271 | 0.112523714 |
| | 0.131113303 | 0.251737082 |
| | 0.593588544 | 0.731049402 |
| | 0.002568355 | 0.011054194 |
| | 0.042747779 | 0.107465832 |
| | 0.293096701 | 0.45277861 |
| NA | | NA |
| NA | | NA |
| | 1.06E-09 | 3.32E-08 |
| | 2.41E-10 | 9.11E-09 |
| | 0.611380306 | NA |
| | 0.973874834 | 0.986212831 |
| | 0.905158266 | NA |
| NA | | NA |
| NA | | NA |

| | |
|---|---|
| 0.021973542 | 0.063405652 |
| 6.98E-05 | 0.000514434 |
| NA | NA |
| 0.000956144 | 0.00479579 |
| 0.616557694 | 0.749171172 |
| 0.472745596 | NA |
| 2.09E-26 | 4.17E-23 |
| NA | NA |
| 0.581597408 | 0.721664547 |
| 0.062263571 | 0.143780864 |
| 0.695097581 | 0.808584403 |
| 0.254009527 | 0.408390336 |
| 4.19E-07 | 6.17E-06 |
| NA | NA |
| 0.905158739 | NA |
| 0.992061292 | NA |
| 0.479769605 | NA |
| 0.793158307 | NA |
| 0.992060816 | NA |
| 0.997510458 | NA |
| 0.889409301 | NA |
| 0.99205891 | NA |
| 0.805374319 | NA |
| 0.992061292 | NA |
| 0.889409773 | NA |
| NA | NA |
| 0.967473452 | NA |
| NA | NA |
| 0.889409773 | NA |
| 0.98868431 | NA |
| 0.706644506 | NA |
| 0.813832119 | NA |
| 0.905158266 | NA |
| NA | NA |
| 0.371061795 | NA |
| 0.083370156 | 0.179492743 |
| 5.06E-05 | 0.000390633 |
| 0.912016371 | 0.950271293 |
| 0.309482966 | NA |
| 0.434534618 | NA |
| 0.62612833 | NA |
| 0.005967898 | 0.022177878 |
| 0.489682248 | 0.643836091 |
| 0.155075465 | 0.285764396 |
| 0.003424596 | 0.013993433 |
| 0.102285791 | 0.209581516 |
| 0.025762397 | 0.072286796 |

| | |
|---|---|
| 0.000260986 | 0.00158709 |
| 0.001024465 | 0.005079304 |
| 0.901724751 | NA |
| 0.001374724 | 0.006515769 |
| 0.003492315 | 0.014227783 |
| 0.002549964 | 0.010992277 |
| NA | NA |
| 0.451957302 | NA |
| 0.00102896 | 0.005099284 |
| 0.179645234 | 0.319172612 |
| 0.000611917 | 0.003294213 |
| 0.338413239 | NA |
| NA | NA |
| 0.997048372 | 0.998640084 |
| 3.22E-10 | 1.18E-08 |
| 0.905158739 | NA |
| 0.45705894 | 0.61464244 |
| 0.500636451 | 0.653527774 |
| 0.896745493 | NA |
| 0.118818609 | 0.234381768 |
| 0.992061292 | NA |
| 0.000207126 | 0.001299477 |
| 0.000112478 | 0.000771361 |
| 0.148488106 | NA |
| 0.984175439 | 0.991990657 |
| 0.04969921 | 0.120830928 |
| 6.00E-10 | 2.02E-08 |
| 0.987210561 | 0.993414557 |
| 0.138748544 | 0.262698678 |
| 0.82679596 | 0.89672859 |
| NA | NA |
| 1.49E-06 | 1.85E-05 |
| 0.137544648 | 0.261036095 |
| 0.040746275 | 0.103391202 |
| 0.002093608 | 0.009302617 |
| 0.01145944 | 0.037518688 |
| 0.087569279 | 0.186204493 |
| 0.001001172 | 0.004980681 |
| NA | NA |
| 0.300612114 | 0.46097814 |
| 0.829962243 | 0.898586968 |
| 0.000355631 | 0.002071432 |
| 1.70E-07 | 2.82E-06 |
| 0.036989256 | 0.096226403 |
| 0.014371453 | 0.045051459 |
| 0.441856838 | 0.601533435 |
| 0.748262482 | 0.846438134 |

| | | |
|---|---|---|
| | 0.774218244 | NA |
| | 0.200995368 | 0.346178874 |
| | 0.701210647 | NA |
| | 0.001805028 | 0.008199548 |
| | 0.025968939 | NA |
| | 0.43251109 | 0.593371925 |
| | 0.90840759 | 0.948154039 |
| | 0.387134973 | 0.550934876 |
| | 3.29E-09 | 8.98E-08 |
| | 0.745898566 | NA |
| | 0.954750708 | 0.97468323 |
| | 0.653480943 | NA |
| | 4.03E-05 | 0.000321585 |
| | 0.39605725 | NA |
| NA | | NA |
| NA | | NA |
| NA | | NA |
| | 0.905157793 | NA |
| NA | | NA |
| NA | | NA |
| | 0.905158739 | NA |
| NA | | NA |
| NA | | NA |
| | 0.889409773 | NA |
| NA | | NA |
| | 0.992061292 | NA |
| | 0.00091021 | 0.004602186 |
| | 0.000106561 | 0.000737231 |
| | 0.889409773 | NA |
| | 0.271391348 | NA |
| | 0.110838734 | 0.222315287 |
| | 2.10E-05 | 0.000185773 |
| | 0.960030823 | NA |
| | 0.956485934 | NA |
| | 0.500652812 | 0.653527774 |
| NA | | NA |
| | 0.002032062 | 0.009075304 |
| | 0.868247027 | NA |
| NA | | NA |
| | 0.992059387 | NA |
| | 0.026487581 | 0.073691791 |
| NA | | NA |
| | 0.040485006 | 0.102918528 |
| | 0.913072927 | NA |
| | 0.991918092 | NA |
| NA | | NA |
| | 0.149804326 | 0.278398205 |

| | | |
|---|---|---|
| | 0.942703337 | 0.967644677 |
| | 0.212814481 | 0.359974826 |
| | 0.331016843 | NA |
| | 6.07E-05 | 0.000457469 |
| | 0.145384163 | 0.272187811 |
| | 0.59397424 | NA |
| | 0.114374364 | 0.227892666 |
| | 0.29264679 | 0.452392177 |
| NA | | NA |
| | 0.905158266 | NA |
| NA | | NA |
| | 0.56645902 | 0.709454949 |
| | 0.905157793 | NA |
| | 0.986956327 | NA |
| NA | | NA |
| | 0.889409301 | NA |
| | 0.512927994 | 0.664551767 |
| | 0.980409908 | NA |
| | 0.905158739 | NA |
| NA | | NA |
| | 0.991782148 | NA |
| | 7.68E-06 | 7.75E-05 |
| | 0.344026827 | 0.507387003 |
| | 0.74504658 | NA |
| | 0.59876741 | 0.734954363 |
| | 0.763644705 | 0.85677877 |
| | 2.85E-06 | 3.27E-05 |
| | 0.978861602 | NA |
| NA | | NA |
| | 0.889409301 | NA |
| | 0.001288587 | 0.006158017 |
| | 0.000827697 | 0.004249569 |
| | 1.43E-08 | 3.22E-07 |
| | 0.001442763 | 0.006794261 |
| | 0.889409301 | NA |
| | 0.998772129 | 0.999545947 |
| | 0.784353866 | NA |
| | 0.00412356 | 0.016300497 |
| | 0.01059715 | 0.035304023 |
| | 0.905158266 | NA |
| NA | | NA |
| | 1.77E-06 | 2.15E-05 |
| | 0.889410717 | NA |
| | 0.992061292 | NA |
| | 4.95E-07 | 7.15E-06 |
| | 0.393615203 | 0.557027534 |
| | 0.819796731 | 0.892907912 |

| | | |
|---|---|---|
| | 0.905158266 | NA |
| | 2.85E-06 | 3.26E-05 |
| | 0.436313437 | 0.596724387 |
| NA | | NA |
| | 0.459042565 | 0.616294674 |
| NA | | NA |
| | 0.02894007 | 0.079140099 |
| | 0.487327277 | NA |
| | 0.067086814 | 0.151715689 |
| | 1.37E-09 | 4.14E-08 |
| | 8.14E-05 | 0.000585829 |
| | 0.138397855 | 0.262170332 |
| | 0.878512733 | NA |
| NA | | NA |
| NA | | NA |
| | 0.176108547 | 0.314301657 |
| NA | | NA |
| | 0.949272466 | 0.97152657 |
| | 0.091196448 | 0.19198378 |
| | 0.159230336 | 0.291208375 |
| | 0.264014312 | 0.41971261 |
| | 0.008728205 | 0.03014986 |
| | 0.278916584 | 0.437074807 |
| | 0.232591806 | 0.383905854 |
| | 0.474265737 | NA |
| | 0.371926313 | 0.536323214 |
| | 0.061665514 | 0.142748578 |
| | 0.084378444 | 0.180820362 |
| | 0.905158266 | NA |
| | 0.66952799 | 0.789335613 |
| | 0.868618983 | 0.924705647 |
| | 0.778696599 | 0.866845717 |
| | 0.003654571 | 0.014754685 |
| | 0.053891885 | 0.128436283 |
| | 0.892588455 | 0.939380965 |
| | 0.988095592 | 0.993639148 |
| | 0.337215991 | NA |
| | 0.344866611 | NA |
| | 0.552403398 | NA |
| NA | | NA |
| | 0.027875313 | 0.076831178 |
| NA | | NA |
| | 0.145201413 | 0.272008004 |
| | 0.905158739 | NA |
| NA | | NA |
| | 0.889409301 | NA |
| | 0.982824459 | NA |

| | | |
|---|---|---|
| | 0.729586993 | NA |
| | 0.204859457 | 0.350870355 |
| | 0.690494807 | 0.805192261 |
| NA | | NA |
| | 0.983073882 | NA |
| NA | | NA |
| | 0.5718464 | NA |
| | 0.384850083 | NA |
| | 0.002303627 | 0.010082789 |
| | 1.37E-09 | 4.13E-08 |
| | 0.028530307 | NA |
| | 0.004121789 | 0.016300049 |
| | 0.04172226 | 0.105453396 |
| | 0.015403811 | 0.047719706 |
| | 5.66E-19 | 1.88E-16 |
| | 0.207263088 | 0.35393811 |
| NA | | NA |
| | 0.016860542 | 0.051343868 |
| | 0.044644438 | 0.111062333 |
| | 0.002333806 | 0.010204097 |
| | 0.905158266 | NA |
| | 0.253414095 | 0.407794434 |
| | 0.55566884 | 0.70097846 |
| | 0.157873143 | NA |
| | 0.769658418 | 0.860799083 |
| | 0.983023562 | NA |
| | 0.83873149 | NA |
| NA | | NA |
| NA | | NA |
| NA | | NA |
| | 0.603093781 | 0.7385721 |
| NA | | NA |
| NA | | NA |
| | 0.905158739 | NA |
| | 0.466455844 | NA |
| NA | | NA |
| NA | | NA |
| NA | | NA |
| NA | | NA |
| | 3.77E-08 | 7.66E-07 |
| | 0.992057737 | NA |
| | 0.001752754 | 0.007989293 |
| NA | | NA |
| | 1.82E-17 | 4.00E-15 |
| | 0.243619644 | 0.396595896 |
| NA | | NA |
| | 0.889409301 | NA |

| | |
|---|---|
| NA | NA |
| 0.001250093 | 0.006002796 |
| 0.226688212 | 0.376624532 |
| 0.841512901 | 0.906236209 |
| 0.44544143 | 0.604838509 |
| NA | NA |
| 0.001638302 | 0.007532552 |
| 0.451809317 | 0.610038967 |
| 0.086227185 | 0.183992106 |
| 0.040728077 | 0.103356969 |
| 0.000848977 | 0.004330471 |
| 0.058033598 | 0.13593171 |
| 0.556143527 | NA |
| 0.001760875 | 0.008022212 |
| 0.04321704 | 0.108333674 |
| 0.436379908 | 0.596778128 |
| 0.198581458 | 0.343115134 |
| 0.259553044 | 0.414663236 |
| 0.397610483 | 0.561162581 |
| NA | NA |
| 0.552407621 | NA |
| NA | NA |
| 0.992060816 | NA |
| 0.130711645 | 0.251130244 |
| NA | NA |
| 0.051444686 | 0.124153629 |
| 0.5103415 | NA |
| 0.806650545 | NA |
| 0.988868872 | NA |
| 5.69E-06 | 5.96E-05 |
| 0.289676929 | 0.449065703 |
| NA | NA |
| NA | NA |
| NA | NA |
| 0.053290513 | 0.127251672 |
| NA | NA |
| 0.801145446 | 0.880155299 |
| 0.990823129 | NA |
| 0.17947973 | 0.318995985 |
| 0.508497058 | NA |
| 0.98947637 | NA |
| 0.000663598 | 0.003535046 |
| 0.992061292 | NA |
| 0.827530109 | 0.896973889 |
| NA | NA |
| NA | NA |
| 0.752960731 | 0.849566576 |

| | | |
|---|---|---|
| | 7.24E-10 | 2.38E-08 |
| | 0.002702464 | 0.011536434 |
| | 0.308668735 | NA |
| | 0.992060816 | NA |
| | 0.765195874 | 0.857619149 |
| NA | | NA |
| | 0.014907358 | 0.046439307 |
| | 0.891478891 | NA |
| | 1.95E-05 | 0.000174282 |
| | 0.417588745 | 0.579781945 |
| | 0.722673969 | NA |
| | 0.690063321 | NA |
| | 0.379961117 | 0.544304363 |
| | 0.969305636 | 0.983776228 |
| | 0.60242111 | 0.737928668 |
| | 0.439081419 | 0.599427347 |
| | 0.952200103 | 0.973288471 |
| NA | | NA |
| | 0.124028816 | NA |
| | 0.599380274 | 0.73542997 |
| | 0.000403837 | 0.002306933 |
| | 0.078781221 | 0.171954329 |
| | 4.08E-05 | 0.000325248 |
| | 0.001012482 | 0.005028976 |
| | 2.83E-05 | 0.000237926 |
| | 0.749282083 | 0.847157103 |
| | 0.992059387 | NA |
| | 0.82795301 | 0.897278333 |
| NA | | NA |
| | 0.992061769 | NA |
| | 0.005191177 | 0.019786549 |
| | 0.991983801 | NA |
| | 0.046384393 | 0.114392956 |
| | 0.02718423 | 0.075313968 |
| | 0.002124624 | 0.009423273 |
| | 0.001437516 | 0.006776819 |
| | 0.053569601 | 0.127765272 |
| | 0.071797559 | 0.159639794 |
| NA | | NA |
| | 0.048079027 | 0.117634247 |
| NA | | NA |
| | 7.40E-06 | 7.50E-05 |
| | 0.295466142 | 0.455404015 |
| | 0.348018068 | NA |
| | 0.002266249 | 0.009937013 |
| | 0.115550531 | NA |
| | 0.760697111 | 0.854869389 |

| | |
|---|---|
| NA | NA |
| 0.738840739 | 0.839639257 |
| NA | NA |
| NA | NA |
| NA | NA |
| 0.15686111 | NA |
| NA | NA |
| 0.006709004 | 0.024395266 |
| 0.140710336 | 0.265542992 |
| NA | NA |
| NA | NA |
| NA | NA |
| NA | NA |
| NA | NA |
| 0.132808433 | 0.25403134 |
| NA | NA |
| NA | NA |
| 0.417496987 | NA |
| 0.001888776 | 0.008530572 |
| 0.003304211 | 0.01359711 |
| 0.063867987 | 0.14644832 |
| 0.905158739 | NA |
| NA | NA |
| NA | NA |
| NA | NA |
| 0.99205891 | NA |
| NA | NA |
| NA | NA |
| NA | NA |
| 0.986146886 | NA |
| 0.90515732 | NA |
| NA | NA |
| NA | NA |
| NA | NA |
| 0.617065107 | NA |
| NA | NA |
| NA | NA |
| NA | NA |
| NA | NA |
| NA | NA |
| NA | NA |
| NA | NA |
| NA | NA |
| 0.323045259 | 0.485441103 |
| NA | NA |
| NA | NA |
| NA | NA |

| | |
|---|---|
| NA | NA |
| 0.537191527 | 0.685346777 |
| 0.693688289 | 0.807442869 |
| 0.08390621 | 0.180248139 |
| 0.792131554 | 0.874750655 |
| 0.577835067 | 0.719068358 |
| 0.315901044 | 0.478118786 |
| 0.67052979 | 0.789880568 |
| 0.002087156 | 0.009279584 |
| NA | NA |
| 0.712566346 | 0.820965252 |
| 0.721881445 | 0.827856415 |
| 0.034807389 | 0.091690991 |
| 0.426850606 | NA |
| 0.92856528 | 0.959095249 |
| 0.94783047 | 0.970440272 |
| 0.992061292 | NA |
| 0.992061292 | NA |
| 0.156029127 | 0.286955075 |
| 0.195604657 | NA |
| 0.095897162 | 0.19950789 |
| 0.690468969 | NA |
| 0.484425843 | NA |
| NA | NA |
| NA | NA |
| NA | NA |
| NA | NA |
| NA | NA |
| NA | NA |
| NA | NA |
| NA | NA |
| NA | NA |
| 0.992059387 | NA |
| NA | NA |
| 0.889409773 | NA |
| NA | NA |
| NA | NA |
| NA | NA |
| NA | NA |
| NA | NA |
| NA | NA |
| NA | NA |
| NA | NA |
| NA | NA |
| NA | NA |
| NA | NA |
| NA | NA |

| | |
|---|---|
| NA | NA |
| NA | NA |
| NA | NA |
| NA | NA |
| NA | NA |
| NA | NA |
| NA | NA |
| NA | NA |
| NA | NA |
| NA | NA |
| NA | NA |
| NA | NA |
| NA | NA |
| NA | NA |
| NA | NA |
| NA | NA |
| NA | NA |
| NA | NA |
| NA | NA |
| NA | NA |
| NA | NA |
| NA | NA |
| NA | NA |
| NA | NA |
| NA | NA |
| NA | NA |
| NA | NA |
| NA | NA |
| NA | NA |
| NA | NA |
| NA | NA |
| NA | NA |
| 0.991918092 | NA |
| NA | NA |
| 0.904947233 | NA |
| NA | NA |
| 0.905158266 | NA |
| NA | NA |
| 0.98981537 | 0.994571049 |
| 0.905158266 | NA |
| NA | NA |
| 0.889409301 | NA |
| 0.797505967 | NA |
| 0.985651806 | NA |
| 0.992061769 | NA |
| NA | NA |
| NA | NA |

| | |
|---|---|
| NA | NA |
| NA | NA |
| NA | NA |
| 0.870948585 | 0.926162874 |
| 0.326990875 | 0.489178092 |
| 0.187245751 | 0.32870169 |
| 0.194211289 | 0.337502825 |
| 0.214992243 | 0.362624983 |
| 0.690726187 | 0.805274956 |
| 0.326857663 | 0.489094884 |
| NA | NA |
| 0.479135284 | NA |
| 0.027386911 | 0.075760793 |
| 0.948331798 | 0.970790078 |
| 0.223571588 | 0.373197149 |
| 0.143056489 | 0.268915383 |
| 0.156651958 | 0.287811273 |
| 0.620141796 | 0.751735783 |
| 0.400199908 | 0.563838687 |
| 0.992059387 | NA |
| 0.78434169 | 0.870114291 |
| 0.85203347 | NA |
| 0.030467204 | 0.082484199 |
| NA | NA |
| 4.32E-06 | 4.70E-05 |
| 1.72E-08 | 3.79E-07 |
| 0.157719767 | 0.289313204 |
| 0.451674585 | NA |
| 4.81E-06 | 5.15E-05 |
| 0.430439939 | 0.591448677 |
| NA | NA |
| 0.038732097 | 0.099584451 |
| 0.567100491 | 0.709919765 |
| 0.262485715 | 0.417956962 |
| NA | NA |
| 0.654116826 | 0.777511713 |
| 0.047781537 | 0.11712824 |
| 0.646312319 | NA |
| 0.817269895 | 0.891341666 |
| 0.025149628 | 0.070866249 |
| NA | NA |
| 0.892008173 | 0.939003235 |
| 0.000280832 | 0.001693708 |
| 0.585975925 | NA |
| 0.192721556 | NA |
| 0.573418941 | 0.715600507 |
| 0.787684998 | 0.871771467 |

| | |
|---|---|
| 0.021037042 | 0.061226297 |
| 0.149094324 | NA |
| 0.217546564 | 0.365691912 |
| NA | NA |
| 0.987380295 | 0.993442561 |
| 0.813430625 | 0.888751273 |
| NA | NA |
| 4.96E-08 | 9.66E-07 |
| 0.562993806 | 0.706646527 |
| 0.385328843 | 0.549194266 |
| 0.14945569 | 0.277891406 |
| 5.90E-11 | 2.67E-09 |
| 0.048172073 | 0.117820127 |
| NA | NA |
| 0.573072786 | 0.715342437 |
| 1.52E-07 | 2.54E-06 |
| 0.99205891 | NA |
| 0.487721322 | 0.642309735 |
| 0.767205586 | 0.858711834 |
| 0.784069992 | 0.870084757 |
| NA | NA |
| 0.766649211 | 0.858439143 |
| 0.061783686 | 0.142946788 |
| 0.407789702 | 0.571014798 |
| 4.80E-08 | 9.38E-07 |
| 2.80E-05 | 0.000235757 |
| 2.36E-17 | 4.90E-15 |
| 0.654454663 | 0.777786976 |
| 0.473846439 | 0.630197684 |
| 0.02403582 | 0.068262101 |
| 0.215226709 | 0.362936823 |
| 0.019183941 | 0.05679657 |
| 0.05638447 | 0.132905385 |
| 0.002463755 | 0.010689902 |
| 0.386118713 | 0.549891746 |
| 0.905158266 | NA |
| 0.006454494 | 0.023645586 |
| 0.002853162 | 0.012060172 |
| NA | NA |
| NA | NA |
| 0.992061292 | NA |
| 0.000270881 | 0.001642716 |
| 0.227248 | 0.377109934 |
| 0.266766219 | 0.422923934 |
| 4.09E-08 | 8.19E-07 |
| 1.93E-05 | 0.000173058 |
| 0.932559459 | 0.961096033 |

| | | |
|---|---|---|
| | 0.053666922 | 0.127969589 |
| | 0.992037435 | NA |
| | 0.552707258 | 0.698164903 |
| | 0.445065125 | 0.604416943 |
| | 4.92E-11 | 2.27E-09 |
| | 0.471504444 | NA |
| | 0.059902408 | 0.139549861 |
| | 0.000279808 | 0.001687994 |
| NA | | NA |
| NA | | NA |
| | 0.852734069 | 0.914181878 |
| NA | | NA |
| NA | | NA |
| | 0.620849234 | NA |
| | 0.83693059 | NA |
| | 0.020170092 | 0.059237 |
| NA | | NA |
| | 0.013916906 | 0.043920859 |
| | 0.214487358 | 0.361940205 |
| | 0.579138044 | 0.719751673 |
| | 0.174962624 | 0.312637664 |
| | 0.01765889 | 0.053184962 |
| | 0.211411772 | 0.35848504 |
| NA | | NA |
| | 0.742902577 | NA |
| | 0.900897778 | NA |
| | 0.706687007 | 0.817578995 |
| | 0.216419754 | 0.364400882 |
| | 0.005299021 | 0.020121254 |
| NA | | NA |
| | 0.674636826 | 0.79317938 |
| | 0.277005051 | NA |
| | 0.705396461 | 0.816628051 |
| | 0.597516981 | 0.734195795 |
| | 0.000746309 | 0.003901955 |
| | 0.85375864 | 0.914632881 |
| | 0.002383239 | 0.01038776 |
| | 0.831473819 | 0.899735554 |
| | 0.160425369 | 0.292923575 |
| | 1.17E-06 | 1.50E-05 |
| | 0.808190784 | 0.88498742 |
| | 0.958264936 | NA |
| | 0.001209345 | 0.005841622 |
| | 0.37974038 | 0.544134937 |
| | 0.140764753 | 0.265600036 |
| NA | | NA |
| | 0.738605019 | NA |

| | |
|---|---|
| 0.626522117 | NA |
| 9.23E-06 | 9.09E-05 |
| 0.746276812 | NA |
| 0.068099438 | 0.153248161 |
| 0.435663488 | 0.596240513 |
| 0.000489527 | 0.002714528 |
| 0.223945099 | 0.373593439 |
| NA | NA |
| 0.059683535 | NA |
| 0.992060816 | NA |
| NA | NA |
| NA | NA |
| NA | NA |
| 0.99205891 | NA |
| 9.48E-15 | 1.09E-12 |
| 0.487410166 | 0.642118024 |
| 0.103139124 | 0.210917492 |
| NA | NA |
| 3.57E-06 | 3.99E-05 |
| 9.42E-07 | 1.25E-05 |
| 0.944022975 | 0.968455992 |
| 0.258497314 | 0.413377733 |
| 0.403060552 | NA |
| NA | NA |
| 0.440846989 | 0.600754564 |
| NA | NA |
| 0.834493679 | 0.901492925 |
| 0.023723037 | 0.06746719 |
| 0.000284219 | 0.001710375 |
| 0.832655355 | NA |
| 0.99205891 | NA |
| 0.406798631 | 0.570055804 |
| 0.970609409 | 0.984598882 |
| 0.493468189 | 0.646954217 |
| NA | NA |
| 3.00E-05 | 0.000250312 |
| 0.013012554 | 0.041538403 |
| NA | NA |
| 0.081762947 | 0.176907336 |
| NA | NA |
| 9.74E-06 | 9.49E-05 |
| 9.61E-05 | 0.000675118 |
| 0.508420845 | 0.660186467 |
| 5.81E-08 | 1.10E-06 |
| 0.011960699 | 0.038858557 |
| 0.155082395 | 0.285764396 |
| 0.38866419 | 0.552084673 |

| | | |
|---|---|---|
| | 0.000638067 | 0.003419895 |
| | 1.78E-10 | 7.10E-09 |
| | 0.99205891 | NA |
| NA | | NA |
| NA | | NA |
| NA | | NA |
| | 0.836621863 | NA |
| | 0.801661681 | NA |
| | 0.021154918 | 0.061487867 |
| | 0.000566582 | 0.003083411 |
| | 0.001536413 | 0.007149416 |
| | 0.813941336 | NA |
| | 0.986947033 | NA |
| | 0.824367097 | NA |
| | 0.992059387 | NA |
| | 0.481791123 | NA |
| | 0.142352832 | 0.267843928 |
| | 4.51E-06 | 4.87E-05 |
| | 0.209451643 | 0.356207298 |
| | 0.626522117 | NA |
| | 0.074752688 | 0.164808662 |
| | 0.504370969 | NA |
| NA | | NA |
| | 0.648354506 | 0.773131502 |
| NA | | NA |
| NA | | NA |
| | 0.719949502 | NA |
| | 0.889409301 | NA |
| | 0.116960652 | NA |
| NA | | NA |
| NA | | NA |
| NA | | NA |
| | 0.992060816 | NA |
| | 0.8600617 | NA |
| NA | | NA |
| NA | | NA |
| NA | | NA |
| NA | | NA |
| | 0.001531647 | 0.007130262 |
| | 0.788582525 | NA |
| | 0.850713389 | NA |
| | 0.856648688 | NA |
| | 0.240711917 | NA |
| | 0.758191728 | NA |
| | 0.00733068 | 0.026195998 |
| | 0.054777553 | 0.130067073 |
| | 0.000993483 | 0.004951443 |

| | |
|---|---|
| 0.078654112 | 0.171771819 |
| NA | NA |
| 0.675104247 | NA |
| 0.531355753 | NA |
| 0.205709308 | 0.351942154 |
| NA | NA |
| 0.946131071 | 0.969356826 |
| 0.061335367 | 0.142155333 |
| 0.889409773 | NA |
| 0.545460161 | 0.691916104 |
| 0.421042535 | 0.582808436 |
| 0.015369981 | 0.047630455 |
| 0.003017662 | 0.012625076 |
| 0.001050625 | 0.00519142 |
| NA | NA |
| 0.431234231 | 0.592138098 |
| 0.60940146 | 0.743435925 |
| 0.019857528 | 0.058459775 |
| 0.653002371 | NA |
| NA | NA |
| 0.072164221 | NA |
| NA | NA |
| 0.009684207 | 0.032781795 |
| 0.032160771 | 0.086187256 |
| 0.271807034 | 0.428557027 |
| 0.064988241 | 0.148437361 |
| 6.68E-05 | 0.000495664 |
| 0.018514463 | 0.05518652 |
| 0.001455021 | 0.00683732 |
| 0.073061935 | 0.161795788 |
| 0.98650982 | NA |
| 0.03661706 | 0.095477908 |
| 0.448235855 | 0.607318514 |
| 0.003835104 | 0.015336924 |
| NA | NA |
| 0.514638384 | 0.665868348 |
| 3.57E-07 | 5.34E-06 |
| NA | NA |
| 0.83045292 | 0.898896617 |
| 0.002249105 | 0.009879594 |
| 0.009706203 | 0.032829161 |
| 0.125225312 | 0.243657238 |
| 0.724504651 | 0.830064068 |
| 0.604848545 | NA |
| 0.992060816 | NA |
| 0.035053357 | 0.092084778 |
| 5.00E-05 | 0.000386837 |

| | |
|---|---|
| NA | NA |
| 0.210422354 | 0.357338184 |
| 5.60E-07 | 7.93E-06 |
| 0.657481978 | 0.780096747 |
| 0.779280911 | NA |
| 0.992105895 | 0.995961112 |
| 0.992060816 | NA |
| 0.905158739 | NA |
| NA | NA |
| 0.073059292 | 0.161795788 |
| NA | NA |
| 0.705347885 | 0.816614868 |
| NA | NA |
| NA | NA |
| NA | NA |
| 1.38E-05 | 0.000128602 |
| 0.988066255 | 0.993639148 |
| 0.606004435 | 0.740679074 |
| 0.006340486 | 0.023278839 |
| 0.817343739 | 0.891341666 |
| 0.916760291 | 0.952592591 |
| 0.394929102 | NA |
| 1.32E-12 | 9.10E-11 |
| 0.464671547 | NA |
| 0.625419132 | 0.755340311 |
| 3.71E-17 | 7.27E-15 |
| 0.767326046 | NA |
| 0.482697759 | 0.638181715 |
| 1.46E-06 | 1.81E-05 |
| 0.889409773 | NA |
| 0.002511951 | 0.010856119 |
| 0.046523954 | 0.114621285 |
| 0.017893767 | 0.053767 |
| 0.889410717 | NA |
| 0.905158739 | NA |
| 0.889409301 | NA |
| NA | NA |
| NA | NA |
| 0.00313144 | 0.013015954 |
| 0.004889378 | 0.018811452 |
| 0.187339868 | 0.328759444 |
| 0.338527372 | 0.501524528 |
| 5.04E-11 | 2.31E-09 |
| NA | NA |
| 0.061845789 | 0.143060327 |
| 7.67E-07 | 1.04E-05 |
| 0.310307787 | 0.471515865 |

| | |
|---|---|
| NA | NA |
| 0.003657711 | 0.014764647 |
| 0.112239339 | 0.224529803 |
| NA | NA |
| NA | NA |
| NA | NA |
| 0.644911846 | 0.770323065 |
| 0.000185066 | 0.001178613 |
| 0.905158266 | NA |
| 0.578931691 | 0.719617401 |
| 2.50E-07 | 3.93E-06 |
| 0.97942684 | 0.989520772 |
| 0.92728579 | 0.958359706 |
| NA | NA |
| 0.805374319 | NA |
| 0.239422129 | 0.391734634 |
| NA | NA |
| 0.712449982 | NA |
| 0.345925181 | 0.509365724 |
| 3.86E-08 | 7.83E-07 |
| 0.848812311 | NA |
| 8.30E-07 | 1.12E-05 |
| 0.743986877 | NA |
| NA | NA |
| 5.80E-07 | 8.15E-06 |
| NA | NA |
| NA | NA |
| NA | NA |
| 0.569036195 | 0.711424835 |
| 0.000117309 | 0.000800992 |
| NA | NA |
| 0.716553001 | NA |
| 0.00281719 | 0.011928785 |
| 1.57E-06 | 1.93E-05 |
| 0.513473875 | 0.665006516 |
| NA | NA |
| 0.174087105 | 0.311402635 |
| NA | NA |
| NA | NA |
| 0.99205891 | NA |
| NA | NA |
| NA | NA |
| NA | NA |
| NA | NA |
| 0.00072106 | 0.003787074 |
| NA | NA |
| 0.992061292 | NA |

| | | |
|---|---|---|
| | 0.414519456 | 0.577052918 |
| | 0.902281688 | 0.944745546 |
| | 0.538415038 | NA |
| | 0.474930229 | NA |
| | 0.032226464 | 0.086331698 |
| | 0.001940454 | 0.008724496 |
| NA | | NA |
| | 0.375709546 | 0.539934943 |
| | 0.000351602 | 0.002050139 |
| | 1.65E-05 | 0.000150537 |
| NA | | NA |
| NA | | NA |
| NA | | NA |
| NA | | NA |
| | 0.678299904 | 0.795916832 |
| | 7.86E-06 | 7.91E-05 |
| | 0.16020219 | 0.292595849 |
| | 0.989933289 | NA |
| NA | | NA |
| | 6.26E-08 | 1.18E-06 |
| | 0.000277937 | 0.001679476 |
| | 0.017518695 | 0.052893308 |
| | 0.723895788 | NA |
| | 0.020665765 | 0.060385833 |
| | 0.992059387 | NA |
| | 0.004225425 | 0.016634297 |
| | 0.903210945 | 0.945177713 |
| | 0.905296333 | NA |
| NA | | NA |
| NA | | NA |
| | 0.604228194 | 0.739384057 |
| | 0.225650574 | 0.375489256 |
| | 5.87E-08 | 1.11E-06 |
| | 0.63254895 | 0.760251946 |
| NA | | NA |
| | 0.446921279 | 0.606173216 |
| | 0.534835366 | 0.683371524 |
| | 5.42E-08 | 1.04E-06 |
| | 0.449766223 | 0.60856593 |
| | 0.013653247 | 0.043191033 |
| | 0.065320572 | 0.149025915 |
| | 0.011544167 | 0.037729359 |
| | 0.887395124 | NA |
| | 0.885359439 | NA |
| | 0.70363308 | NA |
| | 0.499546495 | NA |
| | 0.186987896 | 0.328432827 |

| | | | |
|---|---|---|---|
| | 0.394259681 | | 0.557652067 |
| | 0.418225615 | | 0.580445979 |
| | 0.016382917 | | 0.050181681 |
| NA | | NA | |
| | 0.595515264 | | 0.732518468 |
| | 0.934475809 | | 0.96234837 |
| | 0.00098944 | | 0.004934615 |
| | 0.871288556 | | 0.926316278 |
| | 2.11E-07 | | 3.40E-06 |
| | 0.25087615 | | 0.405013188 |
| | 0.001053103 | | 0.005201326 |
| | 0.579865619 | | 0.72028901 |
| | 0.053177008 | | 0.12706354 |
| | 3.24E-08 | | 6.69E-07 |
| | 0.0985541 | | 0.203646323 |
| | 0.204244331 | | 0.350094613 |
| | 0.014835163 | | 0.046253775 |
| | 0.006927435 | | 0.025069139 |
| | 4.49E-08 | | 8.87E-07 |
| | 0.649839436 | NA | |
| NA | | NA | |
| | 5.69E-17 | | 1.10E-14 |
| | 1.50E-12 | | 1.02E-10 |
| | 0.049829821 | | 0.121054656 |
| | 0.819506033 | NA | |
| | 0.009363255 | | 0.03188211 |
| | 0.96781387 | | 0.982716396 |
| | 0.005772855 | | 0.021610317 |
| | 0.000157683 | | 0.001027163 |
| | 0.699730031 | | 0.812505947 |
| | 0.124186483 | | 0.242221186 |
| | 0.009380634 | | 0.031911596 |
| | 0.829600014 | | 0.898460579 |
| NA | | NA | |
| | 8.17E-10 | | 2.65E-08 |
| | 0.213297808 | | 0.360403712 |
| | 4.98E-12 | | 2.98E-10 |
| | 0.155133306 | | 0.285810242 |
| | 1.50E-11 | | 7.97E-10 |
| | 3.12E-08 | | 6.47E-07 |
| | 0.8686565 | | 0.924705647 |
| | 0.158040452 | | 0.289674174 |
| | 0.938116945 | | 0.964947664 |
| | 0.037554294 | | 0.097338851 |
| | 0.260722977 | | 0.416046498 |
| | 0.214943393 | | 0.362570438 |
| | 4.38E-06 | | 4.75E-05 |

| | | |
|---|---|---|
| | 0.003760315 | 0.015092518 |
| | 8.77E-06 | 8.72E-05 |
| | 4.72E-06 | 5.06E-05 |
| | 0.009142429 | 0.03126107 |
| | 0.443295549 | 0.60290943 |
| | 0.854207179 | 0.914957097 |
| | 0.929795123 | NA |
| | 0.122957221 | 0.240335766 |
| | 2.01E-26 | 4.17E-23 |
| | 0.106671151 | 0.216167571 |
| | 0.063436003 | 0.145725619 |
| | 1.97E-07 | 3.19E-06 |
| | 0.18598105 | 0.326959122 |
| | 3.08E-17 | 6.20E-15 |
| | 0.830163713 | NA |
| | 0.963149259 | 0.97961068 |
| | 5.67E-06 | 5.95E-05 |
| | 0.70467297 | 0.816220791 |
| | 0.553801413 | 0.699254474 |
| | 0.529622174 | 0.67896394 |
| | 0.001098044 | 0.005388145 |
| | 0.244157144 | 0.397173622 |
| NA | | NA |
| | 0.226209996 | 0.376029168 |
| | 0.991525978 | 0.995788655 |
| | 0.911933975 | NA |
| | 0.256474809 | NA |
| | 0.851410992 | NA |
| | 0.889409301 | NA |
| | 0.889409773 | NA |
| | 1.06E-06 | 1.37E-05 |
| | 3.27E-10 | 1.20E-08 |
| | 0.197024679 | 0.34123077 |
| | 0.022865183 | 0.065547853 |
| | 0.000755731 | 0.00394183 |
| | 0.001786543 | 0.008122297 |
| | 0.80666186 | 0.884103958 |
| | 0.041518461 | 0.105047112 |
| | 6.81E-07 | 9.41E-06 |
| | 0.267834427 | 0.424118224 |
| | 0.092181768 | 0.193520024 |
| | 0.001201038 | 0.005813 |
| | 0.357314728 | 0.522006128 |
| | 0.000439299 | 0.002475404 |
| | 0.013107241 | 0.0417678 |
| | 0.597396741 | 0.734092615 |
| | 0.138133567 | 0.261805196 |

| | |
|---|---|
| 0.851099218 | NA |
| 0.998323455 | 0.999279094 |
| 0.004072059 | 0.016130799 |
| 0.008183249 | 0.028599117 |
| 0.000296956 | 0.001770995 |
| 0.007111114 | 0.025570535 |
| 0.776822149 | NA |
| 7.43E-06 | 7.53E-05 |
| 0.444863104 | NA |
| 1.25E-05 | 0.000118048 |
| 7.49E-06 | 7.59E-05 |
| 0.013063201 | 0.041675845 |
| 0.074069196 | 0.163679729 |
| 0.011891881 | 0.038669303 |
| 0.180548734 | 0.320448048 |
| 0.164431663 | 0.298434158 |
| 1.21E-11 | 6.58E-10 |
| NA | NA |
| NA | NA |
| NA | NA |
| NA | NA |
| NA | NA |
| 1.19E-08 | 2.76E-07 |
| 0.791937684 | NA |
| 0.445339881 | NA |
| 5.61E-07 | 7.93E-06 |
| 0.000775306 | 0.00402101 |
| 0.851541785 | NA |
| NA | NA |
| 0.273740648 | 0.43083937 |
| 0.033941133 | 0.089840084 |
| 0.062941546 | 0.14490617 |
| 0.019900937 | 0.058564013 |
| 0.077259555 | 0.169281837 |
| 0.010312668 | 0.034547045 |
| 0.905158739 | NA |
| 0.046565069 | 0.114683979 |
| 0.249711333 | 0.403597919 |
| NA | NA |
| 0.008513487 | 0.029538261 |
| 0.034273629 | 0.090556445 |
| 0.310204438 | 0.471436034 |
| 0.078943435 | 0.172237333 |
| 0.528832941 | 0.678379059 |
| 1.04E-08 | 2.45E-07 |
| 0.052037538 | 0.125130563 |
| 0.99891303 | NA |

| | |
|---|---|
| 0.000155606 | 0.001015738 |
| 0.007390301 | 0.026361862 |
| 2.30E-08 | 4.94E-07 |
| NA | NA |
| 0.008380337 | 0.029173084 |
| NA | NA |
| 0.43264628 | NA |
| NA | NA |
| NA | NA |
| 0.727348972 | NA |
| 0.04667812 | 0.114885096 |
| 0.009004231 | 0.030889533 |
| 5.30E-09 | 1.34E-07 |
| 0.002720192 | 0.011596331 |
| 4.02E-13 | 3.20E-11 |
| 0.21587069 | 0.363655196 |
| 0.450203873 | 0.608829607 |
| 0.763437663 | 0.856634014 |
| NA | NA |
| 0.040287159 | 0.102586587 |
| NA | NA |
| NA | NA |
| 0.905158739 | NA |
| 0.059927576 | 0.139593682 |
| NA | NA |
| 0.384855187 | 0.548781157 |
| 2.46E-11 | 1.23E-09 |
| NA | NA |
| 0.235713014 | 0.387408964 |
| 0.319354709 | 0.481866065 |
| 0.207343175 | 0.35401981 |
| 0.015185987 | 0.04718464 |
| NA | NA |
| 6.05E-06 | 6.30E-05 |
| 0.749112219 | NA |
| NA | NA |
| 0.044655605 | 0.111077529 |
| NA | NA |
| 0.606967343 | 0.741298247 |
| 0.980495918 | 0.990373039 |
| 0.022503992 | 0.064698241 |
| 2.74E-09 | 7.65E-08 |
| 0.010458441 | 0.034929863 |
| 0.487501437 | NA |
| NA | NA |
| 0.006741737 | 0.024491394 |
| 0.533478245 | NA |

| | | |
|---|---|---|
| | 0.224967282 | 0.374603464 |
| | 0.793546545 | 0.875469321 |
| | 0.203608126 | 0.349463095 |
| | 0.991918092 | NA |
| | 0.074682516 | 0.164670486 |
| | 0.813481421 | 0.888762552 |
| | 0.000741984 | 0.003883963 |
| | 0.992061292 | NA |
| | 0.002147727 | 0.00950462 |
| | 0.051673507 | 0.124523189 |
| NA | | NA |
| | 0.630212738 | 0.758417357 |
| | 0.344876297 | 0.508196122 |
| | 0.375694958 | NA |
| | 0.525889235 | 0.675875086 |
| | 0.006759334 | 0.024538221 |
| | 3.54E-06 | 3.96E-05 |
| NA | | NA |
| | 0.99205891 | NA |
| | 0.218952789 | 0.367582897 |
| | 0.208606963 | NA |
| | 0.849806902 | NA |
| NA | | NA |
| | 0.100369181 | 0.206583733 |
| NA | | NA |
| NA | | NA |
| | 0.039165122 | 0.100482173 |
| NA | | NA |
| | 9.32E-13 | 6.62E-11 |
| NA | | NA |
| NA | | NA |
| | 0.063063475 | 0.14506198 |
| | 7.06E-06 | 7.21E-05 |
| | 0.340512128 | 0.503827627 |
| | 0.294730643 | 0.454525611 |
| | 0.145819376 | 0.272746822 |
| | 0.003549394 | 0.014420192 |
| | 0.645961608 | NA |
| NA | | NA |
| | 0.16750031 | 0.302616002 |
| | 0.017095257 | 0.051893109 |
| | 0.905158266 | NA |
| | 0.493997097 | 0.647313656 |
| | 0.000263925 | 0.001602747 |
| | 0.323233254 | 0.485557465 |
| | 0.033954275 | 0.089853209 |
| | 0.515446954 | 0.66666861 |

| | |
|---|---|
| 0.000464005 | 0.002588035 |
| 4.40E-07 | 6.45E-06 |
| 4.65E-22 | 3.04E-19 |
| NA | NA |
| NA | NA |
| 0.000641235 | 0.003433521 |
| 4.29E-10 | 1.52E-08 |
| 0.771585503 | NA |
| 0.558451253 | NA |
| 0.856360535 | 0.916173469 |
| 0.202545892 | 0.348129715 |
| 0.024630807 | 0.06966356 |
| 0.081057591 | 0.175710133 |
| 0.708425794 | 0.818201018 |
| NA | NA |
| 0.741670458 | 0.841758403 |
| NA | NA |
| 0.527245243 | 0.677023466 |
| 0.036728335 | 0.095683652 |
| 0.312020452 | 0.473704169 |
| 1.39E-05 | 0.000129424 |
| 0.001157909 | 0.005637812 |
| 0.152367933 | 0.282088141 |
| 0.508149773 | NA |
| 0.702437968 | 0.814319256 |
| 0.096029271 | 0.199725966 |
| 0.808212474 | 0.88498742 |
| 0.016657987 | 0.050846919 |
| NA | NA |
| 0.007221726 | 0.025886651 |
| 0.19227788 | 0.334964694 |
| 0.833812013 | NA |
| 0.800157566 | NA |
| 2.23E-06 | 2.64E-05 |
| 0.128339271 | NA |
| 0.049972709 | NA |
| NA | NA |
| 0.348181932 | NA |
| 0.007568788 | 0.026850245 |
| 0.00454351 | 0.017667953 |
| 0.024462712 | 0.069268432 |
| 0.420651804 | 0.58245116 |
| 2.38E-06 | 2.80E-05 |
| NA | NA |
| 0.003315377 | 0.013625222 |
| 0.70840988 | NA |
| 0.023373637 | 0.066718016 |

| | | |
|---|---|---|
| | 0.880912217 | 0.932328484 |
| | 0.736929887 | 0.838454327 |
| | 0.208432072 | 0.355124505 |
| | 0.009355089 | 0.031859244 |
| | 0.124787707 | 0.242969788 |
| | 0.560664312 | 0.704971807 |
| | 0.526498154 | 0.676420136 |
| | 0.030198356 | 0.08188759 |
| | 0.071585171 | 0.159215919 |
| | 8.29E-11 | 3.59E-09 |
| | 0.014369293 | NA |
| | 0.156525283 | 0.287650739 |
| NA | | NA |
| | 0.87691188 | 0.929797584 |
| | 0.018289847 | 0.054702635 |
| | 0.825152206 | NA |
| | 3.01E-05 | 0.000251107 |
| | 3.65E-14 | 3.55E-12 |
| NA | | NA |
| | 9.68E-06 | 9.45E-05 |
| | 0.446625711 | 0.605916963 |
| | 0.58326831 | 0.723165762 |
| | 0.216156547 | 0.364057494 |
| | 0.335690231 | 0.498642188 |
| | 0.86419495 | NA |
| | 0.076400478 | 0.167818154 |
| | 0.043618684 | 0.109054159 |
| | 0.046864925 | 0.115293176 |
| | 0.325979594 | NA |
| | 0.877726963 | 0.930482107 |
| | 0.421263677 | 0.58296755 |
| NA | | NA |
| | 0.063086025 | 0.145067723 |
| | 6.69E-06 | 6.88E-05 |
| | 0.358958771 | 0.523850313 |
| | 0.114440435 | 0.227956958 |
| | 0.146370597 | 0.273568129 |
| | 0.142300169 | 0.267783821 |
| NA | | NA |
| NA | | NA |
| NA | | NA |
| | 0.007696232 | 0.027197448 |
| | 0.037740606 | 0.097722064 |
| | 0.889409773 | NA |
| | 0.575243452 | 0.717103091 |
| NA | | NA |
| | 0.716961053 | 0.824279988 |

| | | |
|---|---|---|
| | 0.991918092 | NA |
| | 0.984087211 | 0.991970261 |
| | 0.472308772 | 0.628647613 |
| | 0.104802407 | 0.21342447 |
| | 0.967923719 | NA |
| | 6.56E-05 | 0.000487543 |
| | 0.096681874 | 0.200779012 |
| NA | | NA |
| | 0.343574596 | 0.506992444 |
| NA | | NA |
| | 0.889409773 | NA |
| | 0.047470864 | 0.116444672 |
| NA | | NA |
| | 0.075557618 | 0.166299461 |
| | 0.478155887 | NA |
| | 0.889409301 | NA |
| | 0.010778569 | 0.035781063 |
| | 0.699295144 | 0.812047703 |
| | 0.000195835 | 0.00123789 |
| | 0.524107187 | 0.674216152 |
| | 0.589221079 | 0.72775836 |
| | 0.992061769 | NA |
| NA | | NA |
| | 0.154079888 | 0.284459925 |
| | 0.203953483 | NA |
| | 0.614283782 | NA |
| | 0.905157793 | NA |
| | 0.693364897 | NA |
| | 0.988063578 | NA |
| NA | | NA |
| NA | | NA |
| NA | | NA |
| | 0.991715727 | 0.995816822 |
| | 0.398883742 | 0.562453483 |
| | 0.905158266 | NA |
| NA | | NA |
| | 0.017173946 | NA |
| | 0.889409773 | NA |
| | 1.76E-05 | 0.000159508 |
| | 0.135985634 | NA |
| | 0.01850735 | 0.055172808 |
| | 0.000953059 | 0.00478578 |
| | 0.086053679 | NA |
| NA | | NA |
| | 0.984516914 | 0.992171589 |
| | 0.087224409 | 0.185609117 |
| | 0.089907702 | 0.189981064 |

| | | |
|---|---|---|
| | 0.632913846 | 0.760500965 |
| | 0.019038709 | 0.056427456 |
| | 1.34E-05 | 0.000125327 |
| | 4.55E-05 | 0.000357205 |
| | 4.56E-16 | 7.42E-14 |
| | 0.95907704 | NA |
| NA | | NA |
| | 0.000327495 | 0.001928523 |
| NA | | NA |
| | 0.274698049 | NA |
| | 0.500558262 | NA |
| NA | | NA |
| | 9.62E-10 | 3.05E-08 |
| NA | | NA |
| | 0.077276589 | 0.169302267 |
| | 0.407097733 | 0.570191295 |
| | 0.889409773 | NA |
| | 0.003215866 | 0.013316463 |
| | 0.000101027 | 0.000704939 |
| NA | | NA |
| | 0.641677021 | 0.767479287 |
| | 2.92E-07 | 4.51E-06 |
| | 0.846033572 | NA |
| | 0.022360341 | 0.064327359 |
| | 0.572763407 | 0.715107554 |
| | 0.889409773 | NA |
| | 0.860328012 | NA |
| | 0.992060816 | NA |
| NA | | NA |
| | 0.665301195 | 0.786370068 |
| | 0.138912064 | 0.262985603 |
| | 0.112227771 | 0.224529803 |
| NA | | NA |
| | 0.372935354 | 0.537249061 |
| | 0.65198626 | 0.775945276 |
| | 0.674462272 | 0.79315145 |
| | 4.04E-08 | 8.10E-07 |
| | 0.007695493 | 0.027197448 |
| NA | | NA |
| | 0.022753307 | 0.065303865 |
| | 0.875029791 | NA |
| NA | | NA |
| | 0.783526329 | NA |
| | 0.794189736 | NA |
| NA | | NA |
| | 0.136861432 | NA |
| | 0.16179404 | NA |

| | |
|---|---|
| NA | NA |
| 0.035878808 | 0.093871411 |
| 0.697688555 | 0.810911078 |
| 3.52E-08 | 7.21E-07 |
| 0.710017667 | NA |
| 0.083814376 | 0.180086095 |
| 0.068206951 | 0.153439533 |
| 3.96E-05 | 0.000317387 |
| 0.008811776 | 0.030371652 |
| 0.66814326 | NA |
| 0.005395614 | 0.02042155 |
| 0.504311579 | 0.656480998 |
| 0.524446007 | 0.674533466 |
| 0.234803253 | 0.386307382 |
| 0.240825274 | 0.393473378 |
| 0.000715269 | 0.003759357 |
| 0.992061292 | NA |
| 0.525676107 | NA |
| 0.539670234 | 0.687077564 |
| 0.854999332 | 0.915671592 |
| 0.043144142 | 0.108163285 |
| 0.000814426 | 0.004194178 |
| 0.923649384 | NA |
| 2.20E-09 | 6.34E-08 |
| 0.003667359 | 0.014795433 |
| 0.000427507 | 0.002418242 |
| 0.14567272 | 0.272611834 |
| 0.573300607 | 0.715534161 |
| NA | NA |
| 0.304098279 | 0.464729216 |
| 1.61E-06 | 1.97E-05 |
| 0.005670693 | 0.021293219 |
| 0.004289307 | 0.016834476 |
| 0.889409773 | NA |
| 0.392045757 | 0.555522538 |
| 0.111442739 | 0.223319084 |
| 0.033111193 | 0.088142645 |
| 0.153486132 | 0.283632246 |
| 0.779596455 | 0.86734378 |
| 0.863535347 | NA |
| 0.734043884 | 0.836640201 |
| 0.412913485 | 0.575437407 |
| 0.771597063 | 0.862001419 |
| 0.41656208 | 0.578868922 |
| 0.002334131 | 0.010204097 |
| 0.964306093 | 0.98039303 |
| 0.079111648 | 0.172460194 |

| | |
|---|---|
| 0.031987117 | 0.085794946 |
| 8.78E-12 | 4.98E-10 |
| 0.000207796 | 0.001302226 |
| 0.449142429 | 0.608059099 |
| 0.136981168 | 0.260159962 |
| 0.914596674 | 0.95178562 |
| 0.988868785 | NA |
| 0.482252067 | 0.637737731 |
| 0.078955384 | 0.172238655 |
| 1.51E-15 | 2.17E-13 |
| 0.394867191 | 0.558439405 |
| 0.457209749 | 0.614722147 |
| 0.185985081 | 0.326959122 |
| 0.402959989 | NA |
| NA | NA |
| NA | NA |
| 0.683497074 | 0.799695294 |
| 0.906591272 | 0.947328653 |
| 2.16E-06 | 2.57E-05 |
| 0.027809447 | 0.07667851 |
| NA | NA |
| NA | NA |
| NA | NA |
| 0.889409301 | NA |
| 0.61285111 | 0.746193375 |
| NA | NA |
| NA | NA |
| 0.992061292 | NA |
| 6.97E-09 | 1.71E-07 |
| 0.563407214 | NA |
| 0.035659603 | 0.093398046 |
| 0.889384523 | 0.937385064 |
| 0.015287175 | 0.04742069 |
| 0.018277436 | 0.054672964 |
| NA | NA |
| 0.002149127 | 0.009506946 |
| 0.905158739 | NA |
| NA | NA |
| NA | NA |
| 0.42289325 | 0.584338842 |
| 0.325394593 | 0.487570104 |
| 3.37E-06 | 3.78E-05 |
| 9.48E-06 | 9.28E-05 |
| 0.187201256 | 0.32866951 |
| 0.991782148 | NA |
| 0.161377801 | 0.294175227 |
| 5.24E-05 | 0.00040238 |

| | |
|---|---|
| 0.992060816 | NA |
| 0.288939865 | NA |
| NA | NA |
| 0.807879735 | NA |
| 1.32E-10 | 5.45E-09 |
| 0.007117241 | 0.025578967 |
| 1.35E-05 | 0.000126283 |
| 0.066848967 | 0.15140116 |
| NA | NA |
| 0.847839733 | 0.910570032 |
| 0.07846512 | 0.171418981 |
| 0.334307071 | 0.497158944 |
| NA | NA |
| 3.65E-05 | 0.000296383 |
| 2.08E-05 | 0.000184252 |
| 0.021227241 | NA |
| NA | NA |
| 0.992062245 | NA |
| NA | NA |
| 0.905296333 | NA |
| NA | NA |
| NA | NA |
| NA | NA |
| 0.845248406 | NA |
| NA | NA |
| 0.651556201 | 0.775769811 |
| NA | NA |
| 0.992059387 | NA |
| 0.889409773 | NA |
| 0.807879735 | NA |
| NA | NA |
| 1.45E-05 | 0.000134133 |
| 0.256360088 | 0.410941739 |
| 0.906194259 | 0.947143 |
| 0.070338546 | 0.157207545 |
| 0.091948662 | 0.193215375 |
| 0.000166864 | 0.001078647 |
| 0.000327407 | 0.001928522 |
| 0.050277887 | 0.121881989 |
| 0.093163616 | 0.195037437 |
| 0.594509799 | 0.731732564 |
| 0.920452103 | 0.954486576 |
| 2.79E-09 | 7.76E-08 |
| 0.963413064 | 0.97978823 |
| 0.401729849 | 0.565268855 |
| 0.373660011 | 0.53783542 |
| 0.003644359 | 0.014720531 |

| | |
|---|---|
| 7.77E-08 | 1.42E-06 |
| 0.003271598 | 0.013501415 |
| 4.66E-06 | 5.00E-05 |
| 0.396255107 | 0.559869818 |
| 0.188874478 | 0.330714829 |
| 0.364139542 | 0.529125212 |
| 0.008582505 | 0.029730751 |
| 0.006228322 | 0.022955309 |
| 0.765936997 | NA |
| 0.037760517 | 0.097746456 |
| 0.460103461 | 0.617530219 |
| 0.167216536 | 0.302232513 |
| 0.243354074 | 0.396337026 |
| NA | NA |
| 0.065551074 | 0.14931909 |
| NA | NA |
| 0.328743401 | 0.490946432 |
| 0.233223413 | 0.384774826 |
| 6.73E-08 | 1.26E-06 |
| 0.038999359 | 0.100103672 |
| 0.973573951 | 0.986011917 |
| 0.898346493 | 0.942863964 |
| 0.60240932 | NA |
| 0.105192731 | 0.214060529 |
| 2.06E-06 | 2.46E-05 |
| 0.764823498 | 0.857444183 |
| 7.05E-05 | 0.00051832 |
| 0.203853146 | 0.349719627 |
| 0.297260297 | 0.457338848 |
| 0.345452067 | 0.508873814 |
| 0.960199146 | NA |
| 0.915879456 | 0.95229975 |
| 0.237691693 | 0.389569194 |
| 0.815970462 | NA |
| 0.381213285 | 0.545381622 |
| 0.101077503 | 0.207725977 |
| 4.65E-06 | 5.00E-05 |
| 0.992061292 | NA |
| 0.795353865 | NA |
| 4.93E-05 | 0.000382619 |
| NA | NA |
| 0.006861043 | 0.02487397 |
| 0.053197167 | 0.127097877 |
| 7.14E-08 | 1.32E-06 |
| NA | NA |
| 0.162407182 | 0.295678699 |
| 0.909216053 | NA |

| | |
|---|---|
| 0.003531221 | 0.014359643 |
| 0.30284064 | 0.463572591 |
| 0.642209695 | 0.767968009 |
| 0.017586818 | 0.053018799 |
| 0.833388978 | NA |
| 1.94E-11 | 1.01E-09 |
| 0.973689173 | 0.986083132 |
| 0.234519144 | 0.385984551 |
| 0.792188023 | NA |
| 0.449502703 | 0.608434312 |
| 0.000510726 | 0.002812902 |
| 0.667581844 | NA |
| 0.005224823 | 0.019873875 |
| 0.557327581 | 0.702344697 |
| 0.067864124 | 0.152859606 |
| 7.16E-06 | 7.30E-05 |
| 0.004853763 | 0.018698909 |
| 0.123204168 | NA |
| 1.94E-05 | 0.000173641 |
| NA | NA |
| 0.549683725 | NA |
| 0.002520444 | 0.010886664 |
| 0.501218873 | 0.654005896 |
| 0.046309066 | 0.114273864 |
| 0.092608693 | 0.194215597 |
| 0.895134423 | 0.940940967 |
| 0.889409773 | NA |
| 0.313977735 | 0.47588605 |
| NA | NA |
| NA | NA |
| 0.58043662 | 0.720660209 |
| 0.203579513 | 0.349441297 |
| 0.83182702 | 0.899940362 |
| 0.721426458 | 0.8275075 |
| 0.142458668 | 0.268013185 |
| 0.061328636 | 0.142155333 |
| 0.630554191 | 0.758620396 |
| 0.128534714 | NA |
| 0.07930984 | 0.172774283 |
| 0.905158266 | NA |
| 0.889410245 | NA |
| 0.107624377 | 0.217637323 |
| 0.364403712 | 0.529298923 |
| 0.666811141 | 0.787437426 |
| NA | NA |
| 0.99255964 | NA |
| 0.800842909 | 0.879945431 |

| | |
|---|---|
| NA | NA |
| 0.553824573 | 0.699254474 |
| 0.397677059 | 0.561215477 |
| 0.046057568 | 0.113868287 |
| 0.026394969 | 0.07352621 |
| 0.929242863 | 0.959434081 |
| 0.065604555 | 0.149409917 |
| 1.99E-06 | 2.39E-05 |
| 0.61921495 | 0.751026849 |
| 0.916299385 | 0.952384067 |
| 0.001444291 | 0.006797082 |
| 0.261298252 | 0.416752492 |
| NA | NA |
| 0.005220592 | 0.019868109 |
| 0.159802364 | 0.292035628 |
| 0.448656907 | 0.607626559 |
| 0.224238509 | 0.373926856 |
| 0.873289116 | 0.927434139 |
| 0.017304539 | NA |
| 0.966054438 | 0.981565307 |
| NA | NA |
| 0.905158739 | NA |
| 0.542960408 | 0.689888809 |
| 0.185233551 | 0.326081733 |
| NA | NA |
| 0.819079454 | 0.892392246 |
| NA | NA |
| NA | NA |
| NA | NA |
| NA | NA |
| 0.000228242 | 0.001415012 |
| 0.156602511 | 0.287755921 |
| 0.001419097 | 0.006702636 |
| 0.586339272 | 0.725488303 |
| 0.171356356 | NA |
| 4.08E-12 | 2.49E-10 |
| 0.222469136 | 0.371817567 |
| 0.132071299 | 0.253008906 |
| 2.93E-08 | 6.11E-07 |
| 0.98535582 | NA |
| 0.057457931 | 0.134928746 |
| 0.003429982 | 0.014012833 |
| 0.303432819 | 0.464083208 |
| 0.026171337 | 0.073067248 |
| 0.33908568 | 0.502089173 |
| 0.553943136 | NA |
| 1.41E-06 | 1.76E-05 |

| | |
|---|---|
| NA | NA |
| 0.52944097 | 0.678850535 |
| 0.003543671 | 0.014399607 |
| 0.707316382 | 0.817729832 |
| 0.356923642 | 0.521608296 |
| NA | NA |
| 0.00903297 | 0.030973607 |
| 0.00090778 | 0.004590957 |
| 0.16691009 | 0.301827335 |
| NA | NA |
| 0.11621027 | 0.230797987 |
| 0.992060816 | NA |
| 0.001168776 | 0.005681197 |
| 0.057234782 | 0.134462242 |
| 0.02292824 | 0.065676192 |
| 0.211358317 | NA |
| NA | NA |
| 0.405116968 | 0.568276942 |
| 0.054327536 | 0.12928883 |
| 0.000122755 | 0.000832996 |
| NA | NA |
| 0.035670751 | 0.093416104 |
| 0.523420251 | 0.673766651 |
| 0.661139999 | 0.782933739 |
| 0.086617635 | 0.184574153 |
| 0.08588564 | 0.183441569 |
| 0.52507237 | 0.675141361 |
| 0.793618571 | 0.875469321 |
| NA | NA |
| 0.901037059 | 0.944236842 |
| 1.23E-05 | 0.000116312 |
| 0.662022804 | 0.783554459 |
| NA | NA |
| NA | NA |
| 0.893741787 | 0.939927007 |
| 0.12055882 | 0.236941835 |
| 0.284960056 | 0.443821396 |
| 2.28E-05 | 0.000198532 |
| 0.157802314 | 0.289392101 |
| 0.854851069 | NA |
| 0.781137327 | NA |
| 5.08E-16 | 8.21E-14 |
| NA | NA |
| 0.905158266 | NA |
| 0.335043213 | NA |
| NA | NA |
| 0.998569643 | 0.999434402 |

| | | |
|---|---|---|
| | 1.58E-06 | 1.94E-05 |
| | 0.085411704 | NA |
| | 0.976820963 | 0.988115511 |
| | 0.05737158 | 0.134740378 |
| NA | | NA |
| | 0.203734799 | 0.349572826 |
| | 0.778671023 | 0.866845717 |
| | 0.011332912 | 0.0371876 |
| | 0.533704527 | 0.682484144 |
| | 0.494673339 | 0.647967779 |
| | 0.0128288 | 0.041107197 |
| NA | | NA |
| | 0.874417365 | NA |
| NA | | NA |
| | 0.061503563 | 0.142463791 |
| | 0.079505222 | 0.173079724 |
| | 0.729779136 | NA |
| | 0.16491315 | NA |
| | 0.00010389 | 0.000722398 |
| NA | | NA |
| | 0.287400852 | 0.446482866 |
| | 0.130577595 | 0.25099382 |
| | 0.002408955 | 0.010485281 |
| | 0.980992103 | NA |
| NA | | NA |
| | 6.08E-16 | 9.67E-14 |
| NA | | NA |
| | 0.983532188 | NA |
| | 0.599577696 | 0.7355378 |
| | 0.737667966 | 0.838996291 |
| | 0.005715177 | 0.021419964 |
| | 0.000122542 | 0.000832067 |
| | 0.227916776 | 0.378007891 |
| | 0.002903568 | 0.012222766 |
| | 0.452803463 | 0.610906318 |
| | 0.701509824 | 0.813586893 |
| | 3.36E-05 | 0.00027608 |
| | 0.296690473 | 0.456817224 |
| | 0.817263337 | NA |
| | 0.493805035 | 0.647255104 |
| | 0.576253486 | 0.717913895 |
| | 0.930637085 | 0.959958702 |
| | 0.000152068 | 0.000995016 |
| | 0.00244543 | 0.010620886 |
| | 0.097659898 | 0.202160039 |
| | 6.85E-05 | 0.000506392 |
| NA | | NA |

| | | |
|---|---|---|
| | 0.410276026 | 0.573306015 |
| | 0.701903937 | NA |
| | 0.364365373 | 0.529278245 |
| NA | | NA |
| | 0.746600844 | 0.845262659 |
| | 0.906014663 | 0.947073641 |
| | 0.055427413 | 0.131142178 |
| | 0.706731736 | 0.817578995 |
| | 0.038314091 | 0.098829923 |
| | 0.24632581 | 0.399753785 |
| NA | | NA |
| | 0.871079375 | 0.926252094 |
| | 0.564844593 | 0.708283881 |
| | 0.069000891 | NA |
| | 0.028848069 | 0.078977028 |
| | 0.892668918 | NA |
| | 0.800230921 | NA |
| | 0.297342057 | 0.457396443 |
| | 0.586064928 | NA |
| NA | | NA |
| | 0.002792773 | 0.011843128 |
| | 0.033353977 | 0.08864863 |
| | 0.222390458 | NA |
| NA | | NA |
| | 0.792188023 | NA |
| | 3.72E-06 | 4.13E-05 |
| | 0.889409773 | NA |
| NA | | NA |
| | 0.514423938 | 0.665816059 |
| | 0.226600326 | 0.376507004 |
| | 0.991782148 | NA |
| | 0.614958007 | 0.748095002 |
| | 0.07936616 | 0.172862676 |
| NA | | NA |
| | 0.175102252 | NA |
| | 0.656837928 | NA |
| | 0.461159786 | NA |
| NA | | NA |
| | 0.363089871 | 0.527984271 |
| | 0.904947233 | NA |
| | 0.564448405 | NA |
| | 0.999964426 | NA |
| | 0.082928399 | NA |
| | 1.55E-15 | 2.20E-13 |
| | 0.056051496 | NA |
| | 0.308852297 | 0.46999914 |
| | 0.009113532 | 0.031181872 |

| | |
|---|---|
| 0.00109468 | 0.005375241 |
| 0.891768772 | NA |
| 1.45E-16 | 2.57E-14 |
| 0.040264999 | 0.102572983 |
| NA | NA |
| 0.968629017 | NA |
| 0.659025813 | NA |
| 0.762861626 | NA |
| 4.49E-06 | 4.86E-05 |
| 0.296195048 | 0.456312497 |
| 0.78328688 | NA |
| 0.889409773 | NA |
| NA | NA |
| 0.613913411 | NA |
| 0.000138944 | 0.000923448 |
| 0.559159589 | 0.703563225 |
| 0.741610974 | NA |
| 4.40E-05 | 0.000347088 |
| 0.001457389 | 0.006846984 |
| 0.118436837 | 0.233871323 |
| 0.694581394 | NA |
| 0.002460972 | 0.010679937 |
| 0.542845468 | NA |
| 1.43E-07 | 2.42E-06 |
| 0.005370849 | 0.020348728 |
| 0.010075331 | 0.033865634 |
| 3.46E-05 | 0.000282665 |
| 0.010466123 | 0.034949148 |
| 0.292557762 | 0.452295505 |
| 0.000367837 | 0.002126411 |
| 3.22E-10 | 1.18E-08 |
| 0.983608443 | NA |
| 8.18E-05 | 0.000587905 |
| 1.45E-05 | 0.000134133 |
| 0.004058752 | 0.016087748 |
| 0.781742866 | 0.868484723 |
| 0.829512681 | NA |
| NA | NA |
| 0.252299116 | 0.40661885 |
| 0.287348697 | 0.446465013 |
| 0.991688155 | NA |
| 0.062461008 | 0.144093046 |
| 0.316209403 | 0.478311103 |
| 0.989189051 | 0.994078183 |
| 0.182552724 | 0.322707718 |
| 0.65847 | 0.780788681 |
| 0.986590243 | 0.993138883 |

| | |
|---|---|
| 0.464875545 | 0.621860376 |
| 0.004966212 | 0.019071886 |
| NA | NA |
| NA | NA |
| 4.29E-09 | 1.11E-07 |
| NA | NA |
| 0.896638464 | NA |
| 0.733968501 | 0.836640201 |
| 0.025779545 | 0.072307194 |
| 0.709863266 | 0.819267238 |
| 0.856846731 | 0.916487938 |
| 8.43E-05 | 0.000602496 |
| 0.011688665 | 0.038115741 |
| 0.945007586 | 0.968866415 |
| 0.00020369 | 0.001281244 |
| 0.200629125 | 0.345782511 |
| 2.40E-05 | 0.000207275 |
| 0.371832698 | 0.536223434 |
| 0.011108749 | 0.036632679 |
| 0.049479786 | 0.120404102 |
| 0.99205891 | NA |
| 0.033697688 | 0.089363859 |
| 0.627582007 | 0.756453688 |
| NA | NA |
| 1.26E-06 | 1.60E-05 |
| 0.952673497 | 0.973374154 |
| 0.224583059 | 0.374202853 |
| 0.167336296 | 0.302381314 |
| NA | NA |
| NA | NA |
| NA | NA |
| NA | NA |
| NA | NA |
| NA | NA |
| NA | NA |
| NA | NA |
| NA | NA |
| NA | NA |
| NA | NA |
| NA | NA |
| NA | NA |
| NA | NA |
| NA | NA |
| NA | NA |
| NA | NA |
| NA | NA |
| NA | NA |

| | |
|---|---|
| NA | NA |
| NA | NA |
| NA | NA |
| NA | NA |
| NA | NA |
| NA | NA |
| NA | NA |
| NA | NA |
| NA | NA |
| NA | NA |
| NA | NA |
| NA | NA |
| NA | NA |
| NA | NA |
| NA | NA |
| NA | NA |
| NA | NA |
| NA | NA |
| NA | NA |
| NA | NA |
| NA | NA |
| NA | NA |
| NA | NA |
| NA | NA |
| NA | NA |
| NA | NA |
| NA | NA |
| NA | NA |
| NA | NA |
| NA | NA |
| NA | NA |
| NA | NA |
| 0.779100881 | 0.867075985 |
| 0.013946196 | 0.044000649 |
| 5.31E-09 | 1.34E-07 |
| 0.358565037 | 0.523411981 |
| 0.007500684 | 0.026660331 |
| 0.453052852 | NA |
| 0.907078772 | 0.947419271 |
| 0.940102187 | 0.965968084 |
| 0.00541973 | 0.020492883 |
| 0.780088568 | 0.867560808 |
| 0.627800468 | 0.756581252 |
| 6.11E-06 | 6.36E-05 |
| 0.7848739 | NA |
| 0.216750023 | 0.364861153 |
| 0.578669037 | NA |

| | |
|---|---|
| 0.797342596 | NA |
| NA | NA |
| 2.95E-05 | 0.000246911 |
| 0.217364829 | 0.365531803 |
| 0.005456704 | 0.020609373 |
| 0.710268504 | 0.819476542 |
| 0.001971492 | 0.008841967 |
| 0.269868701 | 0.426337478 |
| 0.606499974 | 0.741083087 |
| 0.89573383 | 0.941075508 |
| 0.371406497 | 0.535784737 |
| 0.410881895 | 0.573771571 |
| 0.00048457 | 0.002690438 |
| 0.308969771 | 0.470080178 |
| 0.029504215 | 0.080404991 |
| 0.637200578 | 0.764063096 |
| 0.513083098 | 0.664668264 |
| 0.04040848 | 0.102810043 |
| 0.980555619 | 0.990387785 |
| 0.988749571 | NA |
| 0.709537716 | NA |
| 0.334288988 | 0.497158944 |
| 0.009909279 | 0.033364163 |
| 0.50814501 | 0.659906339 |
| 0.002781954 | 0.011809187 |
| 0.998282615 | 0.999279094 |
| 0.004585926 | 0.017807664 |
| 0.919528591 | 0.954020567 |
| 0.205833472 | 0.35203001 |
| 0.024406205 | 0.069119475 |
| 0.791995056 | 0.874678523 |
| NA | NA |
| 0.345736118 | NA |
| 0.203379692 | 0.34921781 |
| 2.07E-24 | 2.27E-21 |
| 0.82293531 | 0.894551581 |
| 0.937082805 | 0.964178959 |
| NA | NA |
| 0.600885383 | 0.736607052 |
| 0.771948356 | 0.862251933 |
| 0.701659516 | NA |
| 0.99205891 | NA |
| 2.24E-15 | 3.04E-13 |
| NA | NA |
| NA | NA |
| 1.49E-08 | 3.33E-07 |
| 0.757843973 | NA |

| | |
|---|---|
| 0.978214266 | NA |
| 0.702674557 | NA |
| NA | NA |
| 0.675191361 | NA |
| 0.881048892 | NA |
| 0.249540674 | NA |
| 0.850310676 | 0.912251912 |
| 3.58E-06 | 3.99E-05 |
| 0.545110231 | 0.691592071 |
| 0.847223302 | NA |
| 0.497460016 | NA |
| 0.988911136 | NA |
| NA | NA |
| 0.889409773 | NA |
| 0.383600426 | NA |
| 0.85395124 | NA |
| NA | NA |
| NA | NA |
| NA | NA |
| NA | NA |
| NA | NA |
| 0.042517271 | 0.107043883 |
| 0.211076778 | 0.358139002 |
| 0.069062837 | NA |
| 0.06709923 | 0.151728142 |
| 2.86E-05 | 0.000240452 |
| 0.868197896 | NA |
| 0.72590698 | 0.830737512 |
| 0.392830705 | 0.556347822 |
| 0.992061292 | NA |
| 4.01E-06 | 4.41E-05 |
| 4.97E-05 | 0.0003854 |
| 0.007524339 | 0.026740081 |
| NA | NA |
| 2.76E-05 | 0.000232786 |
| 0.00424369 | 0.01669707 |
| 0.000367551 | 0.002125641 |
| 0.356034289 | 0.520551099 |
| NA | NA |
| NA | NA |
| 0.015864866 | 0.048901823 |
| 0.961417486 | 0.978881988 |
| 0.015292357 | 0.047430067 |
| 0.000129823 | 0.000870755 |
| NA | NA |
| 0.891173359 | 0.938410003 |
| 0.671274182 | NA |

| | |
|---|---|
| 0.801661681 | NA |
| 0.036657385 | 0.095532817 |
| 2.24E-05 | 0.000196151 |
| 0.423130365 | NA |
| NA | NA |
| 0.992269016 | NA |
| 0.159165677 | NA |
| 0.000160821 | 0.001044503 |
| 0.907782182 | NA |
| 0.586610543 | 0.725670717 |
| 0.573775781 | 0.715814873 |
| 4.06E-09 | 1.07E-07 |
| 0.90515732 | NA |
| 0.022136177 | 0.063791118 |
| 0.986386639 | 0.993034302 |
| 0.635802995 | NA |
| 0.889409773 | NA |
| 2.81E-05 | 0.000236512 |
| 3.36E-09 | 9.14E-08 |
| 8.33E-06 | 8.33E-05 |
| 0.000959596 | 0.004810908 |
| NA | NA |
| NA | NA |
| 1.25E-14 | 1.38E-12 |
| 0.016642957 | NA |
| NA | NA |
| 0.422842259 | NA |
| 0.331618511 | NA |
| NA | NA |
| NA | NA |
| NA | NA |
| 0.177055318 | 0.315709113 |
| 0.940070798 | 0.965968084 |
| 0.682336002 | NA |
| 0.707558194 | 0.817782279 |
| NA | NA |
| 0.00327343 | 0.013502788 |
| NA | NA |
| 0.321579369 | 0.483933756 |
| 0.195796663 | 0.339612869 |
| 0.007056916 | 0.025412074 |
| 0.000452303 | 0.002536295 |
| 0.182695494 | 0.322882122 |
| 0.879307496 | 0.931572987 |
| 0.168762015 | 0.304356529 |
| 0.300154849 | 0.460551614 |
| 2.42E-11 | 1.22E-09 |

| | |
|---|---|
| 0.009575129 | 0.032467614 |
| 0.0400204 | 0.102127597 |
| NA | NA |
| 0.003519623 | 0.01432377 |
| 0.408186194 | 0.571460672 |
| 0.297702527 | 0.457662404 |
| 0.055273571 | 0.130898026 |
| 0.784413495 | NA |
| 0.224977025 | 0.374603464 |
| 0.000620891 | 0.003337613 |
| 0.046392402 | 0.114399861 |
| 3.72E-07 | 5.53E-06 |
| 0.31822705 | 0.480601637 |
| 0.992061292 | NA |
| 0.019420929 | 0.0573976 |
| 0.084180893 | 0.180560702 |
| 0.474114255 | 0.630417885 |
| 0.215081778 | 0.362748139 |
| 0.438770261 | 0.599188816 |
| NA | NA |
| 0.195138412 | 0.33879225 |
| 0.058392561 | 0.136626772 |
| 0.992061292 | NA |
| 0.045985612 | 0.113768045 |
| 0.116695674 | 0.231361886 |
| 0.114763699 | 0.228535793 |
| 0.006144977 | 0.022709114 |
| 0.89931887 | 0.943383319 |
| 0.000875383 | 0.004447336 |
| 0.052681663 | 0.126305998 |
| 0.101615865 | 0.208540446 |
| 0.808217078 | NA |
| 0.05817345 | 0.136201171 |
| 0.525195431 | 0.675220519 |
| 0.000111585 | 0.000766912 |
| 0.905158266 | NA |
| 0.000184881 | 0.001177777 |
| 0.811996947 | 0.887643067 |
| NA | NA |
| 5.30E-10 | 1.82E-08 |
| 0.564734836 | 0.708265021 |
| 0.992059387 | NA |
| 0.560688881 | NA |
| 0.844421466 | NA |
| 0.414615413 | 0.577113327 |
| NA | NA |
| 0.671274182 | NA |

| | | | |
|---|---|---|---|
| | 0.887395124 | NA | |
| NA | | NA | |
| NA | | NA | |
| NA | | NA | |
| NA | | NA | |
| | 0.992060816 | NA | |
| NA | | NA | |
| NA | | NA | |
| NA | | NA | |
| NA | | NA | |
| NA | | NA | |
| | 0.905158266 | NA | |
| NA | | NA | |
| NA | | NA | |
| NA | | NA | |
| NA | | NA | |
| NA | | NA | |
| | 0.905158266 | NA | |
| NA | | NA | |
| NA | | NA | |
| NA | | NA | |
| NA | | NA | |
| NA | | NA | |
| NA | | NA | |
| NA | | NA | |
| NA | | NA | |
| | 0.053288199 | | 0.127251672 |
| | 0.48578877 | NA | |
| | 0.355558544 | NA | |
| | 0.951501672 | NA | |
| NA | | NA | |
| NA | | NA | |
| NA | | NA | |
| | 0.277868931 | | 0.435806289 |
| NA | | NA | |
| NA | | NA | |
| NA | | NA | |
| | 0.807252815 | NA | |
| | 0.90515732 | NA | |
| NA | | NA | |
| NA | | NA | |
| NA | | NA | |
| NA | | NA | |
| NA | | NA | |
| NA | | NA | |
| | 0.905158266 | NA | |
| NA | | NA | |

| | |
|---|---|
| NA | NA |
| 0.117645396 | NA |
| 0.85312645 | 0.914290079 |
| NA | NA |
| 0.220259184 | 0.369155199 |
| 0.018236341 | 0.054579776 |
| NA | NA |
| 0.040188808 | 0.10243472 |
| 0.577604139 | 0.718903207 |
| 0.882488682 | 0.933187372 |
| 0.608070721 | 0.742194668 |
| 0.230304407 | 0.381046901 |
| 0.446138479 | 0.605545389 |
| 0.731545942 | 0.834884743 |
| 0.516476929 | 0.667554506 |
| 3.88E-06 | 4.28E-05 |
| NA | NA |
| 0.031174945 | NA |
| 0.064614584 | 0.147783736 |
| 0.330932513 | 0.493353544 |
| 0.889129993 | 0.937385064 |
| NA | NA |
| 0.456219729 | NA |
| 0.008287179 | 0.02889001 |
| 0.000843124 | 0.004310632 |
| 0.195920474 | NA |
| 6.04E-15 | 7.40E-13 |
| 0.011687601 | 0.038115741 |
| 0.876957468 | 0.929801026 |
| 0.067480768 | 0.152324239 |
| 1.47E-10 | 6.02E-09 |
| 0.920404308 | NA |
| 0.630100152 | 0.758323426 |
| 0.000202599 | 0.001275113 |
| 0.594232951 | 0.731473814 |
| 0.008098311 | 0.028348606 |
| NA | NA |
| 0.705232512 | 0.816569405 |
| 3.11E-05 | 0.000258001 |
| 0.153247948 | 0.283359018 |
| 0.009808052 | 0.033095551 |
| 0.142856638 | 0.268600815 |
| 0.534194075 | 0.682990842 |
| 0.765749898 | 0.857957141 |
| 2.46E-05 | 0.00021137 |
| 0.089437969 | 0.189261672 |
| 0.882476615 | 0.933187372 |

| | | |
|---|---|---|
| | 0.815669846 | 0.890356124 |
| | 0.609037586 | 0.743074583 |
| | 6.81E-06 | 6.99E-05 |
| | 0.255575322 | 0.410186997 |
| | 0.934989326 | NA |
| | 0.853278061 | 0.914402182 |
| | 0.105395756 | 0.214275105 |
| | 0.807252815 | NA |
| | 0.063067841 | 0.14506198 |
| | 1.13E-09 | 3.49E-08 |
| | 0.684589684 | NA |
| | 0.430746422 | 0.591690272 |
| | 0.220862254 | 0.369803943 |
| | 0.359766591 | 0.524471524 |
| | 0.507038557 | NA |
| | 0.021433078 | 0.062148286 |
| | 0.913718906 | NA |
| | 0.869329937 | NA |
| | 0.048711828 | 0.118877864 |
| | 0.988158146 | 0.993639148 |
| | 0.03008545 | 0.081672197 |
| | 0.741012592 | 0.841142182 |
| NA | | NA |
| | 0.937225398 | NA |
| | 0.061565956 | 0.142572831 |
| | 0.002819321 | 0.011935505 |
| | 0.3881474 | 0.551850357 |
| | 0.293258259 | 0.45285813 |
| | 0.010077162 | 0.033866604 |
| | 0.986768078 | 0.993163733 |
| | 0.756724335 | 0.852105516 |
| | 0.178816411 | 0.31807449 |
| | 6.97E-10 | 2.31E-08 |
| | 0.000182235 | 0.001162949 |
| | 0.012985898 | 0.041495538 |
| | 0.006160057 | 0.022753355 |
| | 0.711676874 | 0.820520994 |
| | 0.052731254 | 0.126383497 |
| | 0.001568965 | 0.007274685 |
| NA | | NA |
| | 0.000610193 | 0.003287346 |
| | 0.089053101 | 0.188647216 |
| | 0.40383181 | 0.567172544 |
| | 0.217253292 | NA |
| | 0.90515732 | NA |
| | 0.008092033 | 0.028332381 |
| | 0.021505572 | 0.06228447 |

| | |
|---|---|
| 2.46E-07 | 3.88E-06 |
| 0.636851086 | 0.763727432 |
| 0.702793838 | 0.814565311 |
| 0.3461007 | 0.509521672 |
| 0.320209864 | 0.48256732 |
| NA | NA |
| 0.003165544 | 0.01313285 |
| NA | NA |
| 0.007965278 | NA |
| NA | NA |
| NA | NA |
| 0.329673144 | 0.49216399 |
| 1.47E-05 | 0.000136173 |
| 0.485238424 | NA |
| 0.416254999 | 0.578588652 |
| 0.667296116 | NA |
| 0.060731748 | 0.141107656 |
| 0.095162731 | 0.1982429 |
| NA | NA |
| 0.851410992 | NA |
| 0.756847435 | 0.852200432 |
| NA | NA |
| NA | NA |
| 0.229767236 | 0.380358745 |
| 0.093630479 | 0.19578762 |
| 0.385756324 | 0.549553854 |
| 0.449292917 | NA |
| 0.014620861 | 0.045721943 |
| 0.002363322 | 0.010311182 |
| NA | NA |
| NA | NA |
| NA | NA |
| 1.19E-13 | 1.04E-11 |
| 0.834154366 | 0.901268968 |
| 0.350475022 | NA |
| 0.470411781 | 0.626849198 |
| 0.181656554 | 0.321874951 |
| 0.796073962 | 0.877031313 |
| 0.000119097 | 0.000811683 |
| 0.049709258 | 0.120837535 |
| 3.80E-08 | 7.71E-07 |
| 0.905158739 | NA |
| 0.521579393 | 0.67222455 |
| 0.918873876 | 0.953795862 |
| 3.87E-10 | 1.38E-08 |
| 0.011847175 | 0.038541056 |
| 0.116691434 | 0.231361886 |

| | |
|---|---|
| 1.07E-09 | 3.33E-08 |
| 0.865440105 | NA |
| 0.002956785 | 0.012412167 |
| NA | NA |
| NA | NA |
| 0.473591493 | 0.62993492 |
| 0.556344494 | 0.701588971 |
| 0.029480507 | 0.080367779 |
| 0.379030199 | 0.543446784 |
| NA | NA |
| 0.772428346 | 0.862622898 |
| 0.87391556 | NA |
| 1.11E-06 | 1.43E-05 |
| 0.747541748 | NA |
| 0.105262115 | 0.214143106 |
| 0.130096256 | 0.250265762 |
| 0.758028621 | NA |
| 0.017993078 | 0.053998908 |
| 0.576358986 | 0.717947112 |
| 0.332969405 | 0.495542724 |
| 4.27E-27 | 1.34E-23 |
| 0.151437528 | 0.280744338 |
| 0.666885197 | NA |
| 4.54E-06 | 4.90E-05 |
| 0.725486591 | NA |
| 0.082906096 | 0.178834475 |
| 0.771242787 | 0.861649472 |
| 0.991399481 | NA |
| 0.14837695 | 0.276353642 |
| 0.403711189 | 0.56711195 |
| NA | NA |
| 5.36E-06 | 5.66E-05 |
| NA | NA |
| NA | NA |
| 0.348408241 | NA |
| NA | NA |
| 0.905158739 | NA |
| NA | NA |
| 0.593861502 | 0.731262531 |
| 0.282719067 | 0.441364139 |
| NA | NA |
| NA | NA |
| NA | NA |
| 0.98981213 | NA |
| NA | NA |
| NA | NA |
| 2.58E-06 | 2.99E-05 |

| | |
|---|---|
| 0.013407325 | 0.042569612 |
| 0.05473267 | 0.129988612 |
| 0.992061292 | NA |
| 0.931235163 | 0.96025939 |
| 0.098897091 | 0.204162943 |
| 0.000230462 | 0.001427966 |
| 0.641012298 | 0.767167487 |
| 0.004274918 | 0.01679129 |
| NA | NA |
| 8.85E-07 | 1.18E-05 |
| 7.08E-10 | 2.34E-08 |
| 0.250340934 | 0.404325057 |
| 0.072845316 | 0.161397467 |
| 0.000132287 | 0.000885094 |
| NA | NA |
| 0.992061292 | NA |
| 0.005617436 | 0.021115439 |
| 0.58836056 | NA |
| NA | NA |
| NA | NA |
| 0.001181899 | 0.005737255 |
| 0.246539479 | 0.399954427 |
| 0.377377568 | 0.541764776 |
| 0.109802216 | 0.221065792 |
| NA | NA |
| NA | NA |
| NA | NA |
| NA | NA |
| NA | NA |
| 0.573299602 | NA |
| NA | NA |
| NA | NA |
| NA | NA |
| 0.753926932 | NA |
| NA | NA |
| 0.992061292 | NA |
| 0.958802908 | 0.977226877 |
| NA | NA |
| NA | NA |
| NA | NA |
| 0.905158739 | NA |
| NA | NA |
| NA | NA |
| NA | NA |
| NA | NA |
| NA | NA |
| 0.784353866 | NA |

| | | |
|---|---|---|
| | 0.495780152 | NA |
| | 0.321218926 | NA |
| | 0.805374319 | NA |
| | 0.016863016 | 0.051344284 |
| | 2.52E-06 | 2.94E-05 |
| | 0.172499765 | NA |
| | 0.452178307 | 0.610400408 |
| | 0.349984346 | 0.514101696 |
| | 0.565192454 | 0.708474775 |
| | 1.03E-06 | 1.35E-05 |
| | 0.801661681 | NA |
| NA | | NA |
| NA | | NA |
| | 0.733316733 | 0.83621032 |
| | 2.07E-11 | 1.06E-09 |
| | 0.001233977 | 0.005937095 |
| | 0.21901446 | 0.36763022 |
| | 0.366197672 | 0.531061596 |
| | 0.039841531 | 0.101791293 |
| | 0.03113537 | 0.083889766 |
| | 0.032359659 | 0.086582886 |
| | 0.009466723 | 0.032152641 |
| | 0.38427636 | 0.548192334 |
| | 2.39E-08 | 5.09E-07 |
| | 0.180144722 | 0.319817904 |
| | 0.219823714 | 0.368696418 |
| | 0.001004152 | 0.004992117 |
| | 0.812065651 | NA |
| NA | | NA |
| | 0.095115589 | 0.198236013 |
| NA | | NA |
| NA | | NA |
| | 0.889409301 | NA |
| | 0.905158266 | NA |
| NA | | NA |
| | 0.607537244 | NA |
| | 0.905157793 | NA |
| | 0.987295406 | NA |
| NA | | NA |
| NA | | NA |
| | 0.992061769 | NA |
| NA | | NA |
| | 0.889409301 | NA |
| NA | | NA |
| NA | | NA |
| NA | | NA |
| NA | | NA |

| | |
|---|---|
| NA | NA |
| NA | NA |
| NA | NA |
| NA | NA |
| NA | NA |
| NA | NA |
| 0.889409773 | NA |
| NA | NA |
| 0.889409301 | NA |
| NA | NA |
| 0.992060816 | NA |
| 0.905157793 | NA |
| NA | NA |
| NA | NA |
| NA | NA |
| NA | NA |
| NA | NA |
| NA | NA |
| NA | NA |
| NA | NA |
| NA | NA |
| NA | NA |
| NA | NA |
| NA | NA |
| NA | NA |
| NA | NA |
| NA | NA |
| NA | NA |
| 0.009888472 | 0.033308937 |
| 0.024589738 | 0.069556363 |
| 0.962012305 | NA |
| 0.098546947 | NA |
| 0.735607242 | NA |
| 0.80096343 | NA |
| 0.800262609 | NA |
| 0.788582525 | NA |
| NA | NA |
| NA | NA |
| NA | NA |
| NA | NA |
| NA | NA |
| NA | NA |
| NA | NA |
| NA | NA |
| NA | NA |
| NA | NA |
| NA | NA |

| | |
|---|---|
| NA | NA |
| NA | NA |
| NA | NA |
| NA | NA |
| NA | NA |
| NA | NA |
| NA | NA |
| NA | NA |
| NA | NA |
| 0.905158266 | NA |
| NA | NA |
| NA | NA |
| NA | NA |
| 0.788582525 | NA |
| 0.039454131 | 0.101023001 |
| NA | NA |
| NA | NA |
| NA | NA |
| NA | NA |
| 0.992060816 | NA |
| NA | NA |
| NA | NA |
| NA | NA |
| NA | NA |
| 0.905158266 | NA |
| NA | NA |
| NA | NA |
| NA | NA |
| NA | NA |
| NA | NA |
| 0.905158266 | NA |
| NA | NA |
| 0.991918092 | NA |
| NA | NA |
| 0.905158266 | NA |
| 0.804534188 | NA |
| NA | NA |
| 0.905158266 | NA |
| NA | NA |
| NA | NA |
| 0.889409301 | NA |
| 0.804534188 | NA |
| NA | NA |
| NA | NA |
| 0.003542355 | 0.014396922 |
| 5.40E-17 | 1.05E-14 |
| 0.912491039 | 0.950495267 |

| | | |
|---|---|---|
| | 0.768723817 | NA |
| NA | | NA |
| NA | | NA |
| | 0.003304694 | 0.01359711 |
| | 0.760967785 | NA |
| | 0.281392291 | NA |
| | 6.36E-12 | 3.72E-10 |
| NA | | NA |
| NA | | NA |
| NA | | NA |
| | 0.892674236 | 0.939380965 |
| | 0.965656957 | 0.981422073 |
| | 0.797326035 | 0.877636997 |
| | 0.139768127 | 0.264173549 |
| | 0.483111894 | 0.638613898 |
| NA | | NA |
| | 0.282645755 | 0.441312438 |
| | 0.300109345 | 0.460524187 |
| | 0.41360939 | 0.57618782 |
| | 0.36387359 | 0.528843746 |
| NA | | NA |
| | 0.003085224 | 0.012848174 |
| | 1.80E-06 | 2.18E-05 |
| NA | | NA |
| | 0.079779614 | 0.173487874 |
| NA | | NA |
| | 0.04598549 | 0.113768045 |
| | 0.035416471 | 0.092894205 |
| NA | | NA |
| | 0.775345711 | NA |
| | 0.010274409 | 0.034460936 |
| NA | | NA |
| | 0.113998125 | NA |
| | 0.3819791 | 0.546010599 |
| | 0.11024845 | 0.221605639 |
| | 0.8864129 | 0.935538753 |
| | 1.68E-09 | 4.95E-08 |
| | 0.527701535 | 0.677375497 |
| | 0.07513953 | 0.165528585 |
| NA | | NA |
| | 0.000784739 | 0.004059383 |
| | 0.889409773 | NA |
| | 0.992060816 | NA |
| | 0.00042509 | 0.002410158 |
| | 0.107452317 | 0.217349432 |
| | 1.14E-08 | 2.65E-07 |
| | 6.12E-05 | 0.000459726 |

| | |
|---|---|
| NA | NA |
| 0.383699809 | 0.547615109 |
| 0.833448626 | 0.900819299 |
| 0.969483647 | 0.9839097 |
| 0.022771304 | 0.065304305 |
| 0.257794823 | 0.412573174 |
| 0.36693509 | 0.531682082 |
| 0.753894569 | 0.850227048 |
| 0.316943084 | 0.479090877 |
| 8.29E-06 | 8.29E-05 |
| 0.15916482 | 0.29118556 |
| 0.003418343 | 0.013980888 |
| 0.00132799 | 0.006327041 |
| 0.417393002 | 0.579645995 |
| 0.106832599 | NA |
| 0.37960748 | 0.544005786 |
| 0.879028022 | 0.931359205 |
| 0.643205218 | NA |
| 0.202002515 | 0.347413317 |
| 2.88E-08 | 6.01E-07 |
| NA | NA |
| 0.181892637 | 0.322111323 |
| 0.309753226 | NA |
| 0.012419096 | 0.04007508 |
| 0.971253441 | 0.984954128 |
| 0.45546894 | 0.613371915 |
| 0.999796219 | NA |
| 0.992061292 | NA |
| 0.177924272 | 0.316949712 |
| 0.027197342 | 0.07534079 |
| 0.003504224 | 0.014271003 |
| 0.068320745 | 0.153573061 |
| 0.954182324 | 0.974417042 |
| 0.610524224 | 0.744516101 |
| 0.992061292 | NA |
| 0.072197181 | 0.160317311 |
| NA | NA |
| 0.005778481 | 0.0216234 |
| 0.102370146 | 0.209696459 |
| 2.81E-05 | 0.000236329 |
| 0.52607297 | 0.67603209 |
| 0.532453291 | 0.681519108 |
| NA | NA |
| 0.879040088 | 0.931359205 |
| 0.001769934 | 0.008056796 |
| 0.1391799 | 0.263356434 |
| 0.203926416 | 0.349776284 |

| | | |
|---|---|---|
| | 0.067589803 | 0.152460044 |
| | 0.423993394 | 0.585408359 |
| | 0.342509871 | 0.505922037 |
| | 0.006248956 | 0.023025138 |
| | 0.064792493 | 0.148113507 |
| | 1.10E-05 | 0.000105682 |
| | 8.49E-14 | 7.64E-12 |
| | 0.584872647 | 0.724418662 |
| | 0.899266967 | NA |
| | 0.237717632 | 0.389569194 |
| | 9.47E-05 | 0.000666783 |
| | 0.992060816 | NA |
| | 0.57038888 | NA |
| NA | | NA |
| | 0.82149924 | NA |
| | 0.066348704 | 0.150630052 |
| | 0.2023021 | 0.34779236 |
| | 0.743316852 | NA |
| | 4.79E-08 | 9.36E-07 |
| NA | | NA |
| | 0.613114478 | 0.74643127 |
| | 0.792188023 | NA |
| | 0.895670834 | 0.941075508 |
| | 0.528567198 | 0.678126145 |
| | 0.169339983 | 0.305173303 |
| | 0.464757811 | 0.621816082 |
| | 0.493328477 | 0.646928273 |
| | 1.03E-10 | 4.36E-09 |
| | 0.193408596 | 0.336454041 |
| | 0.096818708 | NA |
| | 0.538199959 | 0.686136766 |
| | 0.088806415 | 0.188288122 |
| | 0.210623303 | 0.357589428 |
| | 0.013245966 | 0.042126546 |
| | 0.859692301 | NA |
| NA | | NA |
| | 0.277809893 | NA |
| | 0.024406959 | 0.069119475 |
| | 0.05295473 | 0.126779865 |
| | 0.860037372 | NA |
| | 0.316317598 | NA |
| | 0.859615715 | NA |
| NA | | NA |
| | 2.30E-06 | 2.72E-05 |
| | 0.889409301 | NA |
| | 0.889409773 | NA |
| | 0.000382533 | 0.002198964 |

| | | |
|---|---|---|
| | 0.96663458 | 0.982063882 |
| | 0.006028619 | 0.022369457 |
| | 0.131446453 | 0.25220032 |
| | 2.10E-11 | 1.07E-09 |
| | 0.026807087 | 0.074419319 |
| | 0.764223501 | 0.857062789 |
| NA | | NA |
| | 0.036974354 | 0.096199033 |
| | 1.50E-05 | 0.000138143 |
| | 0.184441293 | 0.325025789 |
| | 0.905158266 | NA |
| | 0.775470666 | 0.864525581 |
| NA | | NA |
| NA | | NA |
| | 0.000434229 | 0.002451215 |
| | 0.001336046 | 0.006359903 |
| | 0.889409773 | NA |
| | 0.006358694 | 0.023337883 |
| | 0.36471623 | NA |
| | 0.007018083 | 0.025305431 |
| | 0.00922911 | 0.031498604 |
| | 0.936855815 | NA |
| NA | | NA |
| | 0.594645664 | NA |
| | 0.312349699 | 0.474138466 |
| | 0.713341843 | NA |
| | 0.160526431 | 0.293038077 |
| | 0.836338575 | 0.902597865 |
| | 0.591628315 | 0.729371557 |
| | 0.992060816 | NA |
| | 0.321682849 | 0.484023138 |
| | 0.297460582 | 0.457496243 |
| | 0.961563664 | 0.978950183 |
| | 0.820123206 | 0.892909192 |
| | 0.762929319 | 0.856238624 |
| | 8.53E-05 | 0.000608266 |
| | 0.002871652 | 0.012124451 |
| | 0.158924639 | 0.290915817 |
| | 0.845997374 | 0.909490665 |
| | 1.25E-08 | 2.87E-07 |
| | 0.018183573 | 0.054436684 |
| | 0.534965201 | 0.683377626 |
| | 0.701434921 | 0.813586893 |
| | 9.05E-09 | 2.16E-07 |
| | 0.850248302 | NA |
| | 0.084922302 | 0.181826134 |
| | 2.58E-06 | 2.99E-05 |

| | |
|---|---|
| 0.368558698 | 0.533182717 |
| NA | NA |
| NA | NA |
| 0.694843907 | NA |
| 0.036749099 | 0.095703684 |
| 0.017193347 | 0.052126012 |
| 0.558670917 | 0.703440832 |
| 0.986192241 | NA |
| 0.823723405 | 0.89492095 |
| 4.03E-10 | 1.43E-08 |
| 0.626932719 | 0.756287841 |
| 0.030418975 | 0.082394259 |
| 0.882349217 | 0.933154595 |
| 0.002652552 | 0.011372002 |
| NA | NA |
| 0.019688015 | 0.058054131 |
| 0.715110122 | NA |
| NA | NA |
| 2.97E-14 | 2.97E-12 |
| NA | NA |
| 0.922476898 | 0.955368118 |
| NA | NA |
| 0.21888944 | 0.367582783 |
| 0.751004579 | 0.848382546 |
| 0.905158266 | NA |
| 0.374026966 | 0.537969442 |
| 0.301565693 | 0.46195194 |
| 0.035411714 | 0.092892824 |
| 1.13E-08 | 2.63E-07 |
| 0.00353354 | 0.014366416 |
| NA | NA |
| 0.086392793 | 0.184238063 |
| 0.000966147 | 0.004836023 |
| 0.428139588 | 0.589148206 |
| 0.999052097 | 0.999598351 |
| 0.838298909 | 0.903869476 |
| 0.231956209 | NA |
| 5.97E-15 | 7.36E-13 |
| 0.647768901 | 0.772642982 |
| 0.8014614 | NA |
| 0.905158739 | NA |
| 0.000528383 | 0.002903593 |
| 0.000602183 | 0.003251373 |
| NA | NA |
| 0.552734726 | NA |
| 0.136868787 | 0.260013988 |
| 0.319045633 | 0.481539835 |

| | | |
|---|---|---|
| | 0.539585703 | 0.687077564 |
| | 0.000845004 | 0.004318234 |
| | 0.509302519 | NA |
| | 3.28E-05 | 0.000270011 |
| | 0.016664014 | 0.050858248 |
| | 0.32290703 | 0.485366246 |
| | 0.487703279 | 0.642309735 |
| | 0.795014225 | NA |
| | 0.051745216 | 0.124632356 |
| | 0.002901407 | 0.012217064 |
| | 0.624926612 | 0.755340311 |
| | 0.485849502 | 0.64076692 |
| | 0.279285593 | 0.437465749 |
| | 0.590318258 | 0.728369873 |
| | 4.49E-08 | 8.87E-07 |
| NA | | NA |
| | 0.661586349 | 0.783313691 |
| | 0.009430064 | 0.032049958 |
| | 0.714654827 | NA |
| NA | | NA |
| | 1.55E-07 | 2.60E-06 |
| | 0.357752027 | NA |
| | 0.905158739 | NA |
| | 0.087757859 | 0.186515178 |
| | 1.88E-07 | 3.08E-06 |
| | 0.59856919 | 0.734793204 |
| | 0.00058512 | 0.003175641 |
| | 0.973133991 | 0.985912235 |
| | 0.916708573 | 0.952583927 |
| | 5.42E-10 | 1.85E-08 |
| | 0.001482178 | 0.006938209 |
| | 0.89945503 | 0.943450346 |
| | 0.001587855 | 0.007348306 |
| | 0.986669866 | NA |
| | 0.030514925 | 0.082572673 |
| | 0.441033461 | NA |
| | 0.060488343 | 0.140646286 |
| | 2.24E-05 | 0.000196273 |
| | 0.50687234 | 0.658682076 |
| | 4.81E-05 | 0.000374604 |
| | 8.31E-08 | 1.50E-06 |
| | 0.214520017 | 0.361967501 |
| | 0.671667117 | 0.790965743 |
| | 0.786292308 | 0.871387018 |
| | 0.65145238 | 0.775751362 |
| | 6.13E-05 | 0.000460197 |
| | 0.622150794 | 0.753367676 |

| | |
|---|---|
| 0.726704173 | 0.831249317 |
| 0.889409773 | NA |
| 0.792188023 | NA |
| 0.008747947 | 0.030199053 |
| 0.164196035 | NA |
| 0.676674355 | 0.794746653 |
| 1.26E-08 | 2.90E-07 |
| 0.012863466 | 0.041200239 |
| 0.204534884 | 0.350423775 |
| 0.044093745 | 0.109984234 |
| 0.122155043 | 0.239179901 |
| 0.013741359 | 0.04342926 |
| 0.01671155 | 0.050982068 |
| 0.001604516 | 0.007405122 |
| 0.087643001 | 0.186307131 |
| 0.000193957 | 0.001228469 |
| 0.022814936 | NA |
| 4.60E-07 | 6.71E-06 |
| 0.459543176 | NA |
| 0.940096265 | NA |
| 0.078961999 | 0.172238655 |
| 0.191875617 | 0.334359234 |
| 0.511586678 | NA |
| 0.53662328 | 0.684701371 |
| 0.734088175 | 0.836640201 |
| 0.067066711 | 0.151690546 |
| 5.08E-05 | 0.000392103 |
| 0.068644115 | 0.154047636 |
| 0.007426444 | 0.026443536 |
| 0.981216512 | NA |
| 1.61E-05 | 0.000147582 |
| NA | NA |
| 0.454838901 | 0.612892042 |
| 0.00141364 | 0.006685787 |
| 0.831779369 | 0.899933148 |
| 0.00043379 | 0.00244937 |
| 0.788582525 | NA |
| 0.006627118 | 0.024149498 |
| 1.73E-06 | 2.10E-05 |
| NA | NA |
| 0.992061769 | NA |
| 0.805374319 | NA |
| 0.138741246 | 0.262698678 |
| 1.68E-05 | 0.000153011 |
| 0.007329815 | 0.026195998 |
| 1.10E-11 | 6.08E-10 |
| 0.743075763 | 0.842520082 |

| | |
|---|---|
| NA | NA |
| 0.01120055 | 0.036846873 |
| 0.594788284 | 0.731893199 |
| 0.406855549 | NA |
| 9.39E-12 | 5.28E-10 |
| 0.120144433 | 0.236360115 |
| 0.000178072 | 0.001141216 |
| 0.489710831 | 0.643836091 |
| 0.025812144 | 0.072352421 |
| 8.14E-06 | 8.16E-05 |
| 0.306739457 | 0.467464206 |
| 0.004282448 | 0.016813564 |
| 0.058743209 | 0.137257089 |
| 0.018419963 | 0.054957366 |
| 0.000236427 | 0.001461627 |
| 0.001634699 | 0.007519134 |
| NA | NA |
| 0.889409773 | NA |
| 0.990179154 | NA |
| 0.776480828 | NA |
| 0.889409773 | NA |
| 0.555791865 | 0.701079432 |
| 0.842035638 | NA |
| NA | NA |
| 1.42E-05 | 0.000132137 |
| 0.032238674 | 0.086344353 |
| 3.26E-05 | 0.000268442 |
| 0.04695534 | 0.115463865 |
| 0.001480629 | 0.006932439 |
| 0.621375167 | NA |
| 0.415780718 | NA |
| 0.003627396 | 0.01467735 |
| 0.101432573 | 0.20823528 |
| 0.770001654 | 0.860971065 |
| 0.637339257 | 0.764085382 |
| NA | NA |
| 0.147648047 | 0.275396674 |
| NA | NA |
| 0.185725075 | NA |
| 0.003621741 | NA |
| 0.178812901 | 0.31807449 |
| 0.004041933 | 0.016029765 |
| NA | NA |
| 0.263623152 | 0.419384261 |
| 1.55E-08 | 3.45E-07 |
| 0.992060816 | NA |
| 0.317412209 | 0.479676236 |

| | | |
|---|---|---|
| | 0.984927165 | NA |
| | 0.610717318 | 0.744627518 |
| | 0.001046285 | 0.005173468 |
| NA | | NA |
| | 0.146953186 | NA |
| | 0.913868955 | 0.95138901 |
| | 0.046032562 | 0.113832098 |
| | 0.86081178 | 0.919084412 |
| | 0.292113112 | 0.451789817 |
| | 0.511129793 | NA |
| | 0.534603865 | NA |
| | 0.608902101 | 0.742950561 |
| | 0.605176904 | 0.740200196 |
| | 0.26030229 | 0.415556378 |
| | 0.041123469 | 0.104203814 |
| NA | | NA |
| NA | | NA |
| | 0.889409301 | NA |
| NA | | NA |
| | 0.134817793 | NA |
| | 0.116479463 | 0.231075303 |
| | 0.009365734 | 0.031885605 |
| | 0.454430735 | NA |
| | 0.319304374 | 0.481831003 |
| | 0.871520836 | 0.926362652 |
| | 0.330913563 | 0.493353544 |
| | 0.996831418 | NA |
| | 0.518017111 | 0.668850341 |
| | 0.385842906 | 0.549641542 |
| NA | | NA |
| | 0.068539345 | 0.153906891 |
| | 0.983231917 | 0.991763981 |
| | 0.563299944 | NA |
| | 0.998624554 | 0.999443807 |
| | 0.030490397 | 0.082536813 |
| | 0.385071448 | 0.549005579 |
| NA | | NA |
| | 0.015459366 | 0.047853427 |
| NA | | NA |
| | 0.174231765 | 0.31155988 |
| | 0.110777313 | 0.222269128 |
| | 0.195468952 | NA |
| | 0.761063313 | 0.855105879 |
| | 0.000132881 | 0.000888527 |
| | 0.591358674 | 0.72908007 |
| | 0.002979939 | 0.012492646 |
| | 0.697006036 | 0.810375155 |

| | |
|---|---|
| 0.063767543 | 0.146273005 |
| 0.739191659 | 0.839900116 |
| 0.057932228 | 0.135737707 |
| 0.683520779 | 0.799695294 |
| NA | NA |
| 0.007091857 | 0.025512794 |
| 0.612153263 | 0.745591741 |
| 0.028845597 | 0.078977028 |
| 0.992458588 | 0.996170975 |
| 0.992059387 | NA |
| 0.069842953 | 0.156227097 |
| 0.00035102 | 0.002047831 |
| 0.010673731 | 0.035507418 |
| 9.02E-12 | 5.10E-10 |
| 0.538543998 | 0.68635572 |
| 5.48E-05 | 0.000419209 |
| 0.683105222 | 0.799524246 |
| 0.678768884 | 0.796251972 |
| 2.10E-05 | 0.000185317 |
| NA | NA |
| 0.889409301 | NA |
| 0.387891205 | 0.551628973 |
| NA | NA |
| 0.417156652 | 0.57943848 |
| NA | NA |
| 8.16E-05 | 0.000586567 |
| 0.792889509 | 0.875103053 |
| 3.02E-10 | 1.12E-08 |
| 0.000137429 | 0.000915876 |
| 0.278403481 | 0.436457631 |
| 1.83E-07 | 3.01E-06 |
| NA | NA |
| 0.847129321 | NA |
| 0.455106108 | 0.61313298 |
| 0.009089263 | 0.031125877 |
| NA | NA |
| 0.427325746 | 0.588427043 |
| 0.018553449 | 0.055280215 |
| 0.910135656 | 0.949096227 |
| 0.736253782 | NA |
| 0.665365265 | 0.786370068 |
| 0.004771524 | 0.018437076 |
| 0.595113072 | NA |
| 0.015273448 | 0.047391501 |
| 0.000148876 | 0.000977333 |
| 0.13761054 | 0.261129438 |
| NA | NA |

| | |
|---|---|
| 0.033252639 | 0.088476276 |
| NA | NA |
| 2.36E-05 | 0.000204106 |
| 0.781951764 | NA |
| 0.718980365 | NA |
| 0.852666512 | NA |
| 1.89E-07 | 3.09E-06 |
| 0.021401901 | 0.062098647 |
| 0.265501209 | NA |
| 0.491897821 | 0.64598913 |
| NA | NA |
| NA | NA |
| NA | NA |
| 0.992061292 | NA |
| 0.000914025 | 0.004618286 |
| 7.17E-06 | 7.30E-05 |
| 0.842805737 | NA |
| 0.145737504 | 0.27266336 |
| 0.90515732 | NA |
| 2.79E-07 | 4.33E-06 |
| 0.144636579 | NA |
| NA | NA |
| 0.605866109 | 0.740584424 |
| 0.102988265 | 0.210746371 |
| 0.941157812 | NA |
| 0.021629104 | 0.062609213 |
| 0.302855885 | 0.463572591 |
| 0.762746762 | NA |
| 5.91E-14 | 5.47E-12 |
| NA | NA |
| 0.992061292 | NA |
| 8.62E-12 | 4.92E-10 |
| 0.992061769 | NA |
| 0.100461453 | 0.206692875 |
| 2.38E-07 | 3.78E-06 |
| NA | NA |
| NA | NA |
| 0.992061292 | NA |
| NA | NA |
| 0.438597374 | 0.599027225 |
| 0.47164157 | 0.62798353 |
| 0.384864057 | 0.548781157 |
| 0.004884746 | 0.018801776 |
| 0.699948701 | NA |
| NA | NA |
| 0.070431683 | 0.157303634 |
| 0.009766141 | 0.03297781 |

| | |
|---|---|
| 0.0071657 | 0.02571525 |
| 0.591195624 | 0.729042775 |
| NA | NA |
| 0.58760983 | 0.726293159 |
| 0.605659357 | 0.740504215 |
| NA | NA |
| 0.112523784 | 0.225016826 |
| 1.86E-07 | 3.05E-06 |
| 0.858422123 | 0.917279387 |
| 0.462290076 | 0.619612367 |
| NA | NA |
| 0.418509504 | 0.580672069 |
| NA | NA |
| 0.915571204 | 0.952213132 |
| 0.006319147 | 0.023219905 |
| 0.000642376 | 0.003438795 |
| NA | NA |
| 0.07343978 | 0.16248505 |
| 0.608393449 | 0.742536224 |
| 0.538806952 | NA |
| NA | NA |
| 0.362988819 | 0.527907244 |
| 0.70784729 | NA |
| 0.754218984 | 0.850523216 |
| 0.992253023 | NA |
| 0.122976295 | NA |
| 0.129083557 | 0.249103245 |
| 0.021274669 | 0.061770522 |
| 0.252086966 | 0.406340577 |
| 0.784357696 | NA |
| 0.008979027 | 0.030812697 |
| 0.063004582 | 0.145007611 |
| 0.267847877 | 0.424118224 |
| 0.330463796 | NA |
| 0.788276299 | NA |
| 0.905158266 | NA |
| 0.889409773 | NA |
| 0.745289219 | NA |
| 0.001012246 | 0.005028938 |
| 0.359268888 | 0.524113393 |
| 0.295434931 | NA |
| 0.007411816 | 0.026408578 |
| 0.991119509 | NA |
| 0.564244669 | NA |
| 0.033623794 | 0.089193632 |
| 0.01153545 | 0.037711322 |
| 0.105288132 | 0.214143106 |

| | |
|---|---|
| NA | NA |
| 0.828269774 | NA |
| 0.90244696 | NA |
| 0.889409301 | NA |
| 0.008088348 | 0.028327278 |
| 0.344645406 | 0.508026345 |
| 7.40E-11 | 3.23E-09 |
| 0.755280879 | 0.851265864 |
| 0.289945438 | 0.449296373 |
| NA | NA |
| 0.002877437 | 0.012139412 |
| NA | NA |
| 0.110081027 | 0.221463058 |
| 0.134122712 | 0.256015354 |
| 4.74E-07 | 6.87E-06 |
| 0.532552515 | 0.681606381 |
| NA | NA |
| 0.661460837 | NA |
| 0.905158266 | NA |
| 0.230436468 | 0.381179238 |
| 0.838212033 | 0.903820183 |
| 0.103082315 | 0.210848124 |
| 0.539580527 | 0.687077564 |
| 0.252734901 | 0.406996604 |
| 0.066491966 | 0.150859417 |
| 0.080095024 | 0.174087561 |
| 0.541857969 | NA |
| 0.778327615 | 0.866682731 |
| NA | NA |
| 0.15470407 | 0.285295213 |
| 0.106254279 | 0.215481871 |
| 0.905158739 | NA |
| 0.587001522 | NA |
| 2.48E-07 | 3.91E-06 |
| 0.01886381 | 0.056014931 |
| 0.066976572 | NA |
| 0.039040008 | 0.100184591 |
| 0.474126844 | 0.630417885 |
| NA | NA |
| 0.826792984 | 0.89672859 |
| NA | NA |
| NA | NA |
| 0.889409773 | NA |
| NA | NA |
| NA | NA |
| NA | NA |
| 0.687411282 | 0.802790211 |

| | |
|---|---|
| NA | NA |
| 0.321417675 | NA |
| 0.550143724 | 0.69608793 |
| 6.61E-15 | 8.06E-13 |
| 0.007808903 | 0.027502116 |
| 0.000133171 | 0.000890196 |
| 0.905158739 | NA |
| NA | NA |
| 0.025678172 | 0.072087314 |
| 0.056522027 | 0.133158158 |
| 0.580163419 | 0.720553013 |
| 0.13036937 | 0.250703301 |
| 0.922470602 | 0.955368118 |
| 0.774398935 | NA |
| 0.827532857 | 0.896973889 |
| 4.67E-09 | 1.20E-07 |
| 7.50E-12 | 4.36E-10 |
| 0.055607586 | 0.131497629 |
| 0.012742032 | 0.040924064 |
| 0.992061769 | NA |
| 0.775418433 | NA |
| 0.659611845 | 0.781758231 |
| 0.365440331 | NA |
| NA | NA |
| NA | NA |
| 0.36012488 | 0.524749983 |
| 0.455246118 | 0.613222273 |
| 0.710392702 | NA |
| 0.174857435 | 0.312525999 |
| 0.905158739 | NA |
| 0.255348768 | 0.409943236 |
| NA | NA |
| NA | NA |
| NA | NA |
| NA | NA |
| NA | NA |
| NA | NA |
| NA | NA |
| 0.152953755 | NA |
| NA | NA |
| NA | NA |
| 0.748890654 | NA |
| 0.889409773 | NA |
| 3.76E-12 | 2.33E-10 |
| 0.996171869 | NA |
| 0.211870366 | 0.358958519 |
| NA | NA |

| | | |
|---|---|---|
| | 0.641399707 | 0.767366984 |
| | 0.000103878 | 0.000722398 |
| | 0.292146763 | 0.45181004 |
| | 0.834491509 | NA |
| NA | | NA |
| | 0.697589682 | NA |
| | 0.37216996 | 0.536601091 |
| | 0.132185882 | 0.253132448 |
| NA | | NA |
| | 0.889409301 | NA |
| | 0.896170374 | 0.941489104 |
| | 0.072233976 | 0.160354756 |
| | 0.645205003 | 0.770444549 |
| | 0.582968421 | 0.722916397 |
| | 0.13036558 | 0.250703301 |
| | 0.992060816 | NA |
| | 0.905296333 | NA |
| NA | | NA |
| NA | | NA |
| NA | | NA |
| | 0.783192856 | NA |
| | 0.889409301 | NA |
| | 0.60552582 | NA |
| | 0.889410245 | NA |
| NA | | NA |
| | 0.034216686 | 0.090416873 |
| | 0.942911832 | 0.967684733 |
| | 0.001274673 | 0.006104806 |
| | 0.975581746 | 0.987504597 |
| | 0.187305539 | 0.328756928 |
| | 0.018089255 | 0.05422085 |
| NA | | NA |
| | 0.00437786 | 0.017108635 |
| | 1.30E-06 | 1.63E-05 |
| | 0.830163713 | NA |
| | 2.06E-05 | 0.000182909 |
| NA | | NA |
| | 2.05E-18 | 6.00E-16 |
| NA | | NA |
| | 0.423981814 | 0.585408359 |
| | 0.04704501 | 0.115625552 |
| | 0.167156572 | 0.302188751 |
| | 1.41E-10 | 5.77E-09 |
| | 0.720564289 | NA |
| NA | | NA |
| NA | | NA |
| | 0.283464338 | 0.442330757 |

| | |
|---|---|
| NA | NA |
| NA | NA |
| 0.33301532 | 0.495568453 |
| NA | NA |
| 0.437813319 | 0.598179609 |
| 0.905158266 | NA |
| 0.808636172 | NA |
| 0.726075371 | NA |
| 0.782559532 | NA |
| NA | NA |
| 0.000105926 | 0.000733302 |
| 1.32E-18 | 4.15E-16 |
| 0.024827291 | 0.070101901 |
| 0.791169531 | NA |
| 0.159083647 | 0.291109818 |
| 0.074619516 | 0.164597577 |
| 0.055248875 | NA |
| 0.687623649 | NA |
| 0.050898772 | NA |
| NA | NA |
| NA | NA |
| 0.725498137 | NA |
| 0.00889531 | 0.030592343 |
| 0.440576805 | 0.600569926 |
| 0.495536073 | NA |
| 0.002570558 | 0.01106151 |
| 0.221803914 | NA |
| 0.010576512 | 0.035258278 |
| 0.859857639 | NA |
| NA | NA |
| 0.825581779 | 0.896053297 |
| 0.738138168 | 0.839311103 |
| 0.025506374 | NA |
| 0.026695343 | 0.074174747 |
| 0.788582525 | NA |
| 0.786684962 | NA |
| 0.470934003 | 0.627274037 |
| 0.059426068 | 0.138601914 |
| 0.13456145 | 0.256508057 |
| 0.195336276 | 0.339055355 |
| 0.152217291 | 0.28192615 |
| NA | NA |
| NA | NA |
| 0.814966959 | NA |
| 0.856268215 | 0.9161369 |
| NA | NA |
| 0.980537739 | NA |

| | |
|---|---|
| NA | NA |
| 0.741715911 | 0.841766484 |
| NA | NA |
| NA | NA |
| 4.22E-11 | 1.97E-09 |
| NA | NA |
| NA | NA |
| 0.152843891 | 0.282802409 |
| 0.005567533 | 0.020952433 |
| 0.992061587 | NA |
| 0.967328242 | 0.982404998 |
| 0.177015178 | NA |
| 0.386198195 | 0.549969271 |
| 5.72E-10 | 1.94E-08 |
| 0.606730857 | 0.741248631 |
| 0.593051839 | NA |
| 0.000782544 | 0.00405089 |
| 0.029394842 | 0.080175696 |
| 0.623552575 | 0.754371164 |
| 2.07E-12 | 1.37E-10 |
| 8.20E-08 | 1.48E-06 |
| 0.018859051 | 0.056008371 |
| 0.273304219 | 0.430244989 |
| NA | NA |
| 0.70499887 | NA |
| 0.983842897 | NA |
| 0.069706277 | 0.155984931 |
| 0.000926746 | 0.004669671 |
| 0.117269164 | 0.232159536 |
| 0.220437834 | 0.369370042 |
| 0.003621229 | 0.014655098 |
| 0.981047423 | 0.990520041 |
| 0.789956662 | 0.873095346 |
| 0.552290621 | 0.697839332 |
| 0.234323971 | 0.385778983 |
| 0.733525585 | NA |
| NA | NA |
| 0.718827801 | NA |
| 0.322344935 | 0.484687239 |
| 0.802457864 | NA |
| 0.569662072 | NA |
| NA | NA |
| NA | NA |
| NA | NA |
| 0.801661681 | NA |
| NA | NA |
| NA | NA |

| | |
|---|---|
| NA | NA |
| 0.285590105 | 0.444604356 |
| 0.853095047 | 0.914290079 |
| 0.740614515 | 0.840843317 |
| 0.181629475 | 0.321857258 |
| 0.924448268 | NA |
| 0.057502824 | 0.135019731 |
| 0.456387935 | 0.614210622 |
| 0.88006069 | 0.932034983 |
| NA | NA |
| NA | NA |
| NA | NA |
| 0.547509107 | 0.693833822 |
| 0.108969897 | 0.219831875 |
| 0.112963733 | 0.225732152 |
| 0.406481279 | 0.569727606 |
| 0.889409773 | NA |
| NA | NA |
| NA | NA |
| 0.352012859 | NA |
| 0.850290357 | 0.912251912 |
| 0.882370827 | 0.933154595 |
| 0.523588049 | 0.673880783 |
| 0.621368735 | 0.752682924 |
| 0.766250273 | 0.858298804 |
| 0.900836801 | NA |
| 0.966971186 | NA |
| 0.983342978 | NA |
| 0.235737446 | 0.387408964 |
| NA | NA |
| 0.907577825 | 0.947610543 |
| NA | NA |
| NA | NA |
| NA | NA |
| 0.427735358 | 0.588806465 |
| NA | NA |
| 0.649723001 | 0.774511014 |
| 0.409440908 | 0.572560205 |
| 0.956107126 | NA |
| 0.989275582 | NA |
| 0.847605884 | NA |
| NA | NA |
| 0.905157793 | NA |
| 0.575279541 | NA |
| NA | NA |
| 0.553054394 | 0.698523032 |
| 0.182245221 | NA |

| | |
|---|---|
| 0.54171991 | 0.689039122 |
| 0.992061292 | NA |
| 0.676562938 | 0.794746653 |
| 5.49E-07 | 7.80E-06 |
| 0.422852649 | 0.584319509 |
| 0.716889751 | NA |
| NA | NA |
| NA | NA |
| 0.558233141 | NA |
| 2.29E-09 | 6.56E-08 |
| 0.724745557 | NA |
| 0.905157793 | NA |
| 1.26E-06 | 1.60E-05 |
| 0.01963946 | 0.057934296 |
| 0.468236625 | NA |
| 0.065780872 | 0.149687271 |
| 0.953487772 | NA |
| 0.834776978 | 0.901665895 |
| 0.025008333 | 0.070558652 |
| 0.425255186 | 0.586458496 |
| NA | NA |
| 0.889409773 | NA |
| NA | NA |
| NA | NA |
| 0.889409773 | NA |
| 0.011699688 | 0.038146021 |
| 0.779844503 | NA |
| 0.992060816 | NA |
| 0.989974364 | NA |
| 0.924040954 | 0.956221268 |
| 0.784737708 | 0.870362233 |
| NA | NA |
| 0.398585629 | 0.562192888 |
| 0.343896688 | 0.507263207 |
| 0.32318641 | 0.485545784 |
| NA | NA |
| 0.211755915 | 0.358847672 |
| 0.935138134 | 0.962768232 |
| 0.070424702 | 0.157303634 |
| 0.606876886 | 0.741298247 |
| 0.297641118 | NA |
| 0.889409773 | NA |
| 0.889409301 | NA |
| 0.749180881 | NA |
| 0.127489958 | 0.24680878 |
| NA | NA |
| 0.544042184 | 0.690715906 |

| | | |
|---|---|---|
| NA | | NA |
| | 0.060760957 | NA |
| NA | | NA |
| | 0.927465743 | 0.958410365 |
| | 0.99205891 | NA |
| | 0.097967488 | 0.202701222 |
| NA | | NA |
| | 8.50E-08 | 1.52E-06 |
| NA | | NA |
| NA | | NA |
| | 0.577838134 | NA |
| | 0.90515732 | NA |
| | 0.071308351 | 0.158728847 |
| NA | | NA |
| NA | | NA |
| NA | | NA |
| NA | | NA |
| NA | | NA |
| | 0.8649357 | 0.922118894 |
| | 0.905158266 | NA |
| | 0.219571692 | NA |
| | 0.483980676 | 0.639338968 |
| | 0.889409773 | NA |
| | 0.57521863 | NA |
| | 0.426748119 | 0.587960164 |
| | 0.415822061 | 0.578279711 |
| | 0.080422665 | 0.17467069 |
| NA | | NA |
| | 0.000208345 | 0.001305294 |
| | 0.864483862 | 0.921887084 |
| | 0.034779582 | 0.09163965 |
| | 0.158954349 | NA |
| | 0.052126788 | NA |
| | 0.143634985 | NA |
| | 0.132489757 | NA |
| | 0.012743693 | 0.040924064 |
| | 0.905158266 | NA |
| | 0.024961522 | 0.070453737 |
| | 0.129882096 | 0.249985182 |
| | 0.809236794 | 0.885534996 |
| | 0.967552634 | 0.982496569 |
| NA | | NA |
| | 0.876684956 | 0.929601862 |
| | 0.183510874 | 0.323984185 |
| | 0.992061292 | NA |
| NA | | NA |
| | 0.99509469 | 0.997551495 |

| | | |
|---|---|---|
| | 0.490389199 | 0.64447043 |
| | 1.39E-05 | 0.000129849 |
| | 0.561110674 | 0.705290743 |
| | 0.905158266 | NA |
| | 0.045974409 | 0.113768045 |
| | 0.652201383 | 0.776075115 |
| | 0.006780375 | 0.024601826 |
| NA | | NA |
| | 0.400352696 | 0.563909228 |
| | 0.741540931 | NA |
| | 1.78E-05 | 0.000160807 |
| | 0.863145388 | 0.920982 |
| NA | | NA |
| | 0.681526547 | 0.798130287 |
| | 0.99205891 | NA |
| | 8.54E-07 | 1.15E-05 |
| | 0.287065298 | NA |
| | 0.408660205 | 0.571941965 |
| | 0.870953292 | 0.926162874 |
| | 0.079331464 | 0.172804247 |
| | 0.627981407 | 0.75661166 |
| | 0.00585761 | 0.021849372 |
| | 0.103912623 | NA |
| | 0.637956114 | NA |
| | 0.261901714 | 0.41732093 |
| | 0.905157793 | NA |
| | 0.020909457 | 0.060943822 |
| | 0.905158739 | NA |
| | 0.933986812 | 0.962115513 |
| NA | | NA |
| NA | | NA |
| | 0.889409773 | NA |
| | 0.906961884 | NA |
| | 0.994736109 | NA |
| | 0.905158266 | NA |
| | 0.883599344 | NA |
| | 0.004960114 | 0.019051801 |
| | 0.99767551 | 0.998967988 |
| | 0.889409773 | NA |
| NA | | NA |
| | 0.180577642 | 0.320448048 |
| | 0.14882289 | 0.276902377 |
| | 4.19E-08 | 8.35E-07 |
| | 0.651379679 | NA |
| NA | | NA |
| | 0.406473969 | 0.569727606 |
| | 4.91E-05 | 0.000381725 |

| | |
|---|---|
| 9.13E-05 | 0.000645723 |
| 0.144761956 | 0.271346802 |
| NA | NA |
| NA | NA |
| 0.992059387 | NA |
| NA | NA |
| 0.946708635 | NA |
| 0.717080705 | NA |
| 0.986163044 | NA |
| NA | NA |
| NA | NA |
| 0.208362892 | 0.355045212 |
| NA | NA |
| NA | NA |
| 0.332965541 | 0.495542724 |
| 0.327736105 | 0.489908586 |
| 0.041502867 | 0.105019757 |
| 0.684388319 | 0.800238716 |
| 0.683934993 | 0.799921625 |
| 0.905158739 | NA |
| 0.99205891 | NA |
| 0.891383187 | NA |
| 0.007379486 | 0.026332265 |
| 0.874403072 | NA |
| 0.50606932 | 0.657988979 |
| 0.463927507 | 0.620992632 |
| 0.652319781 | NA |
| NA | NA |
| 0.211913942 | 0.358976954 |
| 0.405776091 | 0.569066112 |
| 2.29E-06 | 2.70E-05 |
| 0.675871977 | 0.794128831 |
| 0.913596312 | NA |
| 0.051668057 | 0.124523189 |
| 0.610568054 | NA |
| NA | NA |
| NA | NA |
| 0.566547233 | 0.709524962 |
| NA | NA |
| 0.848064321 | NA |
| 0.905158739 | NA |
| NA | NA |
| NA | NA |
| NA | NA |
| 0.991918092 | NA |
| 0.811529239 | 0.887295025 |
| 0.22275441 | 0.372202713 |

| | |
|---|---|
| 0.893456581 | 0.939722695 |
| 0.033386027 | 0.088691479 |
| 0.842026041 | 0.906575263 |
| 0.889410717 | NA |
| 0.088586435 | 0.187948747 |
| 0.116682138 | 0.231361886 |
| 0.090619047 | 0.191153088 |
| 3.81E-09 | 1.01E-07 |
| NA | NA |
| 0.224508366 | 0.374135175 |
| 0.960681794 | 0.978233782 |
| 0.125344089 | 0.243793698 |
| 0.806301281 | 0.883908433 |
| 0.828591671 | 0.897767035 |
| 0.402849553 | 0.566345339 |
| 0.008282127 | 0.028876981 |
| 0.000425765 | 0.002412346 |
| 0.985328421 | NA |
| NA | NA |
| NA | NA |
| 0.464116621 | NA |
| NA | NA |
| NA | NA |
| 0.788582525 | NA |
| 0.476575245 | 0.632809379 |
| 0.694527467 | NA |
| 2.20E-10 | 8.48E-09 |
| 0.938910773 | NA |
| 0.879714138 | NA |
| 0.051824901 | 0.124769543 |
| NA | NA |
| 0.127453464 | 0.246759886 |
| 0.905158266 | NA |
| 0.99205891 | NA |
| 0.790602017 | NA |
| 0.352890858 | 0.517028362 |
| 0.272099481 | NA |
| 0.00765202 | 0.027084253 |
| 0.739787434 | NA |
| 0.529972171 | NA |
| NA | NA |
| 0.807879735 | NA |
| 0.432449242 | 0.593324143 |
| 0.57583277 | 0.717547948 |
| 0.691349929 | NA |
| 0.781137327 | NA |
| 0.022205395 | 0.063948624 |

| | | |
|---|---|---|
| | 0.905158739 | NA |
| | 0.216278746 | NA |
| NA | | NA |
| | 0.003037991 | 0.012680336 |
| | 0.063877317 | 0.14644832 |
| | 0.988877091 | NA |
| NA | | NA |
| NA | | NA |
| | 1.26E-09 | 3.84E-08 |
| | 0.02226904 | 0.064089887 |
| | 7.40E-07 | 1.01E-05 |
| NA | | NA |
| | 1.02E-06 | 1.34E-05 |
| | 0.221746043 | 0.371024182 |
| | 0.794123599 | 0.875761355 |
| | 5.40E-06 | 5.69E-05 |
| | 0.905158739 | NA |
| | 0.032373839 | 0.086610273 |
| | 0.464992238 | 0.621924994 |
| | 0.753438147 | 0.849843243 |
| | 0.855589927 | NA |
| | 0.889410245 | NA |
| | 0.000105833 | 0.000732887 |
| | 0.612256497 | 0.745676119 |
| | 0.122521497 | 0.23966235 |
| | 0.511535307 | NA |
| | 0.310342857 | 0.471515865 |
| NA | | NA |
| | 0.679929551 | NA |
| | 0.034295243 | 0.090591754 |
| NA | | NA |
| | 0.94451229 | 0.968776934 |
| | 0.000571802 | 0.003109167 |
| | 0.905158266 | NA |
| | 0.309828388 | 0.471060138 |
| | 0.502641197 | 0.655083272 |
| | 0.067425034 | 0.15224538 |
| | 0.011431297 | 0.037459309 |
| | 0.522371717 | 0.67269297 |
| | 0.080563234 | 0.174828348 |
| | 0.992060816 | NA |
| NA | | NA |
| NA | | NA |
| NA | | NA |
| NA | | NA |
| | 2.95E-05 | 0.00024656 |
| | 0.091006655 | NA |

| | |
|---|---|
| 0.142085766 | 0.267482124 |
| 0.009952479 | 0.033483451 |
| 0.279993206 | 0.438172243 |
| 0.412366449 | NA |
| 0.154089476 | 0.284459925 |
| 0.992061769 | NA |
| NA | NA |
| NA | NA |
| 0.425927605 | 0.587180169 |
| 0.835805491 | 0.902244262 |
| 0.220967415 | NA |
| 0.737508208 | NA |
| 0.38743076 | NA |
| NA | NA |
| 0.052293506 | 0.125581048 |
| NA | NA |
| 0.227976861 | 0.378078989 |
| NA | NA |
| 0.417286995 | 0.579582866 |
| 0.553540767 | 0.699016718 |
| NA | NA |
| NA | NA |
| 0.991898478 | NA |
| NA | NA |
| 0.077303352 | 0.169309052 |
| 0.510107345 | 0.661593951 |
| 0.884433193 | NA |
| 0.026173641 | 0.073067248 |
| 0.912772267 | 0.950652922 |
| 0.743178302 | 0.842555234 |
| 0.292377909 | 0.452103832 |
| 2.15E-05 | 0.00018893 |
| 0.082834018 | 0.178766802 |
| 0.384275373 | 0.548192334 |
| 0.160579981 | 0.293077166 |
| 0.905158266 | NA |
| 0.930468347 | 0.959926975 |
| 0.817414982 | 0.891374861 |
| 0.72318413 | 0.829003984 |
| 0.079855687 | 0.173601726 |
| 0.008038406 | 0.028182695 |
| 0.749957 | NA |
| 0.091364555 | 0.192237886 |
| 0.09191414 | 0.193179804 |
| 4.78E-14 | 4.55E-12 |
| 1.12E-09 | 3.48E-08 |
| 0.313096411 | NA |

| | | |
|---|---|---|
| | 0.749256307 | NA |
| | 0.799019712 | NA |
| NA | | NA |
| NA | | NA |
| NA | | NA |
| NA | | NA |
| | 1.72E-05 | 0.00015569 |
| | 0.006300816 | 0.023183581 |
| | 0.488130888 | 0.642733475 |
| | 0.698244462 | NA |
| | 0.000138677 | 0.000922228 |
| | 3.12E-11 | 1.53E-09 |
| | 0.971326367 | 0.984954128 |
| | 0.875338953 | 0.928844683 |
| | 0.914760085 | 0.951789385 |
| | 2.22E-06 | 2.63E-05 |
| | 0.889410245 | NA |
| NA | | NA |
| NA | | NA |
| NA | | NA |
| | 0.889409301 | NA |
| NA | | NA |
| | 0.905158266 | NA |
| | 3.52E-08 | 7.21E-07 |
| | 0.992059387 | NA |
| | 0.0455492 | 0.11286303 |
| | 0.905158266 | NA |
| | 0.060994506 | 0.141553408 |
| | 0.40964854 | 0.572675549 |
| | 0.690862809 | NA |
| | 0.692095408 | 0.806288029 |
| | 0.889409773 | NA |
| NA | | NA |
| | 0.003186719 | 0.013208221 |
| | 0.032955587 | 0.087792277 |
| | 0.38936614 | 0.552795698 |
| | 0.141105557 | 0.266037866 |
| | 0.747675631 | NA |
| | 1.45E-06 | 1.81E-05 |
| | 0.52339274 | 0.673766651 |
| | 0.272501655 | 0.429289321 |
| | 1.47E-13 | 1.27E-11 |
| | 0.861431665 | NA |
| | 7.77E-05 | 0.000564942 |
| | 0.015575955 | 0.048160011 |
| | 0.419647338 | 0.58150034 |
| | 0.401089811 | 0.564693926 |

| | | |
|---|---|---|
| | 0.30138934 | 0.461790877 |
| | 0.019849089 | 0.058448834 |
| | 0.637415297 | 0.764085382 |
| | 0.000600545 | 0.00324652 |
| | 0.962293697 | 0.979205986 |
| | 0.474781347 | 0.631020676 |
| | 0.182458448 | NA |
| | 0.87850363 | NA |
| | 0.193609745 | 0.336706905 |
| | 0.962466194 | 0.979205986 |
| NA | | NA |
| | 0.411708832 | 0.574442621 |
| | 0.01522374 | 0.047270661 |
| | 0.009909425 | 0.033364163 |
| | 0.560299092 | NA |
| | 0.513495627 | 0.665006516 |
| | 0.784353866 | NA |
| | 0.700413402 | NA |
| | 1.70E-14 | 1.84E-12 |
| | 0.018777533 | 0.055841799 |
| | 0.000111305 | 0.000765232 |
| | 0.196403598 | 0.340398096 |
| | 0.975903983 | 0.987642896 |
| | 0.355081663 | NA |
| | 0.521159817 | NA |
| | 0.930601105 | 0.959958702 |
| | 0.905158266 | NA |
| | 0.567611095 | NA |
| | 0.006194406 | 0.022853319 |
| | 0.795353865 | NA |
| | 1.98E-08 | 4.32E-07 |
| | 1.91E-06 | 2.31E-05 |
| | 0.992061292 | NA |
| | 0.12932416 | 0.249406383 |
| | 0.651328319 | NA |
| NA | | NA |
| | 0.000318594 | 0.001882669 |
| | 3.90E-06 | 4.31E-05 |
| | 0.131697292 | 0.252584854 |
| | 0.008439793 | 0.0293382 |
| | 0.914124055 | 0.951609468 |
| | 0.032806466 | 0.087511807 |
| | 0.72751878 | NA |
| | 0.90116115 | 0.944236842 |
| | 1.46E-06 | 1.81E-05 |
| | 0.302215837 | 0.462818716 |
| | 0.068003068 | 0.153093506 |

| | | |
|---|---|---|
| | 0.002029587 | 0.009068762 |
| | 0.734634691 | 0.836928054 |
| | 0.905158266 | NA |
| | 8.24E-05 | 0.000591459 |
| | 1.04E-08 | 2.45E-07 |
| | 0.517941047 | NA |
| | 0.680644984 | 0.79747536 |
| | 0.003707034 | 0.01491986 |
| | 0.172322655 | 0.309178297 |
| | 0.992061292 | NA |
| NA | | NA |
| | 0.416923978 | 0.579151925 |
| | 0.534318478 | 0.68303059 |
| | 0.056977377 | 0.133972183 |
| | 0.136038758 | 0.258752189 |
| | 5.01E-08 | 9.74E-07 |
| | 0.144672899 | 0.271226176 |
| | 0.062381942 | 0.143936646 |
| | 0.088612752 | 0.187986418 |
| | 0.203116715 | 0.348879255 |
| | 1.76E-05 | 0.000159577 |
| | 0.755388937 | 0.851300265 |
| | 0.001503755 | 0.00702424 |
| NA | | NA |
| NA | | NA |
| NA | | NA |
| | 0.254719286 | 0.409201903 |
| | 7.68E-12 | 4.42E-10 |
| | 0.000821252 | 0.004223387 |
| | 0.973005802 | 0.985912235 |
| | 0.132968732 | 0.254276791 |
| NA | | NA |
| | 0.000307572 | 0.001824414 |
| | 0.000406606 | 0.00231913 |
| | 0.756179557 | NA |
| | 0.000545263 | 0.002980691 |
| | 0.889409301 | NA |
| | 0.148066395 | 0.275939063 |
| | 0.280675284 | 0.439047483 |
| | 0.643879364 | NA |
| | 0.029032056 | 0.07931263 |
| | 0.100560175 | NA |
| | 0.576545468 | 0.718033231 |
| | 0.116567157 | NA |
| | 0.971688261 | 0.985147169 |
| NA | | NA |
| | 0.003458226 | 0.014104602 |

| | | |
|---|---|---|
| NA | | NA |
| | 0.012478562 | 0.040225595 |
| | 0.937150545 | NA |
| | 0.085379257 | 0.182537543 |
| | 2.42E-06 | 2.83E-05 |
| | 1.24E-05 | 0.000117339 |
| | 0.478569369 | 0.63442653 |
| | 0.733672605 | 0.836442382 |
| | 0.129041341 | 0.249065554 |
| | 0.065142283 | 0.148758257 |
| | 0.001391208 | 0.006589633 |
| NA | | NA |
| | 0.905156847 | NA |
| | 0.905158266 | NA |
| | 0.992062245 | NA |
| NA | | NA |
| NA | | NA |
| NA | | NA |
| | 0.905158739 | NA |
| | 0.905158266 | NA |
| NA | | NA |
| NA | | NA |
| | 0.905158739 | NA |
| | 0.714469161 | NA |
| | 0.08953073 | 0.189421457 |
| | 0.731707788 | NA |
| | 0.723754717 | NA |
| | 0.905158266 | NA |
| | 0.992037435 | NA |
| | 0.90515732 | NA |
| | 0.767326046 | NA |
| NA | | NA |
| | 0.626522117 | NA |
| | 0.846349039 | NA |
| | 0.00051518 | 0.002835299 |
| | 0.023816188 | 0.067682111 |
| | 0.000346701 | 0.002027568 |
| | 0.279278512 | 0.437465749 |
| | 0.680981157 | 0.797613891 |
| | 0.643221455 | NA |
| NA | | NA |
| | 0.565978689 | 0.709032533 |
| | 0.528650976 | 0.678194005 |
| NA | | NA |
| | 0.031053338 | 0.083707575 |
| NA | | NA |
| | 0.016433648 | 0.050294978 |

| | | |
|---|---|---|
| | 0.628851006 | 0.757276625 |
| | 0.388720728 | NA |
| | 0.014159708 | 0.044552663 |
| NA | | NA |
| | 4.17E-13 | 3.30E-11 |
| | 0.918377146 | NA |
| | 0.389278532 | 0.552742792 |
| | 0.058695671 | NA |
| | 0.387749727 | 0.551463488 |
| | 0.992061292 | NA |
| | 0.001855634 | 0.00839989 |
| | 0.035058347 | NA |
| | 0.007278202 | 0.02604237 |
| | 0.001356337 | 0.00644671 |
| | 0.699029196 | 0.811910616 |
| | 0.007101372 | 0.025542839 |
| | 0.759054361 | 0.853756605 |
| | 0.229044249 | 0.379546334 |
| | 8.47E-08 | 1.52E-06 |
| | 0.001597077 | 0.007377456 |
| NA | | NA |
| | 0.382815326 | NA |
| | 0.829686099 | 0.898465186 |
| | 0.749823323 | NA |
| | 0.692231985 | 0.806404358 |
| | 0.640716976 | NA |
| | 0.878033887 | NA |
| | 0.980224998 | NA |
| | 0.23457087 | 0.38604075 |
| | 0.3160939 | 0.478268171 |
| | 3.94E-09 | 1.04E-07 |
| | 0.575861719 | 0.717547948 |
| | 0.897055566 | 0.942074668 |
| | 0.069490783 | 0.15559785 |
| | 0.905296333 | NA |
| | 0.176685895 | NA |
| NA | | NA |
| | 0.236514493 | 0.388337203 |
| | 0.803897353 | 0.882153915 |
| | 0.90515732 | NA |
| | 0.004513873 | 0.017565148 |
| | 0.444964884 | NA |
| | 0.556722295 | 0.701904167 |
| | 0.002761892 | 0.011744461 |
| | 0.906210412 | 0.947143 |
| | 0.061183726 | 0.141917548 |
| NA | | NA |

| | | |
|---|---|---|
| | 0.889409773 | NA |
| | 0.782927656 | 0.869220305 |
| | 0.431686087 | 0.592569165 |
| | 0.539774536 | NA |
| | 0.645224066 | 0.770444549 |
| | 0.092450005 | 0.193971877 |
| | 0.801545323 | 0.880364815 |
| | 2.35E-07 | 3.73E-06 |
| | 0.788582525 | NA |
| NA | | NA |
| | 0.145245959 | 0.272021834 |
| | 0.770873062 | NA |
| | 0.00011172 | 0.00076736 |
| | 0.205165062 | 0.351311571 |
| | 0.463292711 | 0.620407649 |
| | 0.54804929 | 0.694197875 |
| NA | | NA |
| NA | | NA |
| NA | | NA |
| | 4.66E-12 | 2.82E-10 |
| NA | | NA |
| | 2.22E-10 | 8.55E-09 |
| | 0.440132597 | 0.600265306 |
| | 1.13E-05 | 0.00010815 |
| | 0.000694248 | 0.003664662 |
| | 0.992060816 | NA |
| | 0.905158739 | NA |
| | 0.002484349 | 0.010758002 |
| | 0.001439956 | 0.006784654 |
| | 0.435057279 | 0.595785643 |
| | 0.104358831 | 0.212683486 |
| | 0.459496198 | NA |
| | 0.486783859 | 0.641613851 |
| | 0.889409301 | NA |
| | 0.267479366 | 0.423718035 |
| | 0.13994408 | 0.264415079 |
| | 0.160537732 | 0.293038077 |
| | 0.000241037 | 0.001485067 |
| | 0.00062186 | 0.003342001 |
| | 0.734156571 | NA |
| | 0.052244948 | 0.125536502 |
| NA | | NA |
| | 1.16E-14 | 1.30E-12 |
| | 5.86E-05 | 0.000443592 |
| | 0.026977602 | 0.074826469 |
| | 0.905158266 | NA |
| | 0.905158739 | NA |

| | |
|---|---|
| 0.27621916 | 0.433714446 |
| 0.000140949 | 0.000933099 |
| NA | NA |
| NA | NA |
| NA | NA |
| NA | NA |
| 0.680487688 | 0.797376155 |
| NA | NA |
| 0.063858456 | 0.14644832 |
| 0.787629184 | NA |
| 0.538405939 | 0.686258185 |
| 0.002140003 | 0.009476168 |
| 0.976322688 | 0.987728072 |
| 0.992061292 | NA |
| NA | NA |
| 0.216551517 | 0.364554949 |
| NA | NA |
| 0.615194612 | 0.748251601 |
| 0.857671474 | 0.917146721 |
| NA | NA |
| 0.461917769 | 0.619371794 |
| 0.476781413 | 0.632968763 |
| 0.725031319 | 0.830297927 |
| NA | NA |
| 0.905158266 | NA |
| 0.989555292 | NA |
| 0.777092376 | NA |
| 0.150088799 | 0.278779061 |
| 0.683802976 | 0.799813045 |
| 0.000291855 | 0.001747145 |
| 0.129714684 | 0.249869225 |
| 6.41E-05 | 0.000478481 |
| 0.67275632 | NA |
| 0.000140764 | 0.000932158 |
| 0.145474047 | NA |
| 0.264600902 | 0.420553793 |
| 0.167131729 | 0.302185725 |
| 0.039328375 | 0.10079642 |
| 0.000277926 | 0.001679476 |
| 0.99205891 | NA |
| 0.699197352 | NA |
| 0.454581381 | NA |
| 0.410882005 | 0.573771571 |
| 0.0372863 | 0.096781306 |
| 0.866170921 | NA |
| 3.29E-05 | 0.000270994 |
| 0.133371288 | 0.254869038 |

| | |
|---|---|
| 0.72658489 | 0.831210086 |
| 0.002408725 | 0.010485281 |
| NA | NA |
| 0.019219267 | 0.056885818 |
| 0.212014505 | 0.359119601 |
| 6.96E-10 | 2.31E-08 |
| 0.022893013 | 0.065593381 |
| 0.258586025 | 0.413442685 |
| 0.497319455 | 0.650425129 |
| 0.000184736 | 0.001177541 |
| 0.007664514 | 0.027119732 |
| 0.976331133 | 0.987728072 |
| 0.034361563 | 0.090712379 |
| 0.016917806 | 0.051489688 |
| 0.748169018 | NA |
| 0.072235951 | 0.160354756 |
| 0.000965293 | 0.004832848 |
| 0.283539676 | 0.442330757 |
| 0.001093329 | 0.00536981 |
| 0.660821278 | 0.782811374 |
| 0.537518259 | 0.685644115 |
| 0.241293148 | 0.394091204 |
| 0.055101417 | 0.130581913 |
| 0.007404745 | 0.026396234 |
| 0.690009342 | 0.804839857 |
| 0.07816055 | 0.170846857 |
| 0.012499866 | 0.040277115 |
| 0.126480956 | 0.245413244 |
| 0.653414154 | 0.777012965 |
| NA | NA |
| 0.417307195 | NA |
| 0.885091108 | NA |
| 0.000846814 | 0.004325468 |
| 0.979927651 | NA |
| 0.450032119 | 0.608813166 |
| NA | NA |
| NA | NA |
| 0.002363853 | 0.010311452 |
| 3.59E-16 | 6.02E-14 |
| 0.021368838 | 0.062011142 |
| 0.845136429 | NA |
| 0.025834245 | 0.072368183 |
| 0.007270826 | 0.02602022 |
| 0.676619559 | 0.794746653 |
| 0.15487324 | 0.285546723 |
| 0.003390857 | 0.013888844 |
| 3.68E-05 | 0.000297852 |

| | |
|---|---|
| 2.51E-05 | 0.000215082 |
| 0.004704999 | 0.01822492 |
| 0.000226132 | 0.001402723 |
| 0.073310481 | NA |
| 0.897820413 | 0.942509445 |
| 0.027321971 | 0.075619254 |
| 0.992061292 | NA |
| 0.043097954 | 0.108096855 |
| 0.033390394 | 0.088692352 |
| 0.02502141 | 0.070586477 |
| 0.751022738 | 0.848382546 |
| 0.138004777 | 0.261651432 |
| 0.325407015 | NA |
| 0.856537928 | 0.916288843 |
| 0.008130122 | 0.028441827 |
| 0.182178438 | 0.322373793 |
| 0.897107767 | 0.942074668 |
| 0.177226252 | 0.315933141 |
| 0.003338594 | 0.013706145 |
| 0.626116437 | 0.755792126 |
| 0.559328883 | 0.703695597 |
| 0.010357831 | 0.034656044 |
| 0.255868736 | 0.410507896 |
| 0.035976477 | 0.094104522 |
| 0.025425795 | 0.071515952 |
| 0.021007238 | 0.061171986 |
| 0.605367344 | 0.740282982 |
| 0.412239572 | 0.574826566 |
| 0.834358399 | 0.901399783 |
| 0.365731568 | 0.530490752 |
| 0.130482065 | 0.250874788 |
| 0.823288081 | 0.894625048 |
| 0.884049297 | NA |
| 0.548928759 | 0.695111391 |
| 0.339727907 | 0.502836688 |
| 0.111503273 | 0.223382937 |
| 0.882871744 | 0.933424732 |
| 1.96E-11 | 1.02E-09 |
| 0.029099345 | 0.079466797 |
| 0.242812873 | 0.395690381 |
| 0.035714807 | 0.093509174 |
| 0.236596748 | 0.388385138 |
| 0.939804521 | 0.96589388 |
| 0.000287739 | 0.001726828 |
| 0.889418735 | 0.937385064 |
| 0.013246678 | 0.042126546 |
| 0.905158266 | NA |

| | |
|---|---|
| 0.40049396 | NA |
| 0.519120758 | 0.670062638 |
| 0.001741241 | 0.007944297 |
| 0.835254839 | NA |
| 0.000526749 | 0.002896062 |
| 0.755176144 | NA |
| 0.241669601 | 0.394472049 |
| 8.76E-07 | 1.17E-05 |
| 4.12E-07 | 6.08E-06 |
| 0.61811566 | 0.750356668 |
| 0.992061292 | NA |
| 0.241497004 | 0.394298377 |
| NA | NA |
| NA | NA |
| NA | NA |
| NA | NA |
| NA | NA |
| NA | NA |
| NA | NA |
| NA | NA |
| NA | NA |
| NA | NA |
| NA | NA |
| NA | NA |
| NA | NA |
| NA | NA |
| NA | NA |
| NA | NA |
| NA | NA |
| 0.010488905 | 0.035009253 |
| NA | NA |
| NA | NA |
| NA | NA |
| NA | NA |
| NA | NA |
| NA | NA |
| NA | NA |
| NA | NA |
| NA | NA |
| NA | NA |
| NA | NA |
| NA | NA |
| NA | NA |
| NA | NA |
| NA | NA |
| NA | NA |
| NA | NA |

| | |
|---|---|
| NA | NA |
| NA | NA |
| NA | NA |
| NA | NA |
| NA | NA |
| NA | NA |
| NA | NA |
| NA | NA |
| NA | NA |
| NA | NA |
| NA | NA |
| NA | NA |
| NA | NA |
| NA | NA |
| NA | NA |
| NA | NA |
| NA | NA |
| NA | NA |
| NA | NA |
| NA | NA |
| NA | NA |
| NA | NA |
| NA | NA |
| NA | NA |
| NA | NA |
| NA | NA |
| NA | NA |
| NA | NA |
| NA | NA |
| NA | NA |
| NA | NA |
| NA | NA |
| NA | NA |
| NA | NA |
| NA | NA |
| NA | NA |
| NA | NA |
| NA | NA |
| NA | NA |
| NA | NA |
| NA | NA |
| NA | NA |
| NA | NA |
| NA | NA |
| NA | NA |
| NA | NA |
| NA | NA |

| | |
|---|---|
| NA | NA |
| NA | NA |
| NA | NA |
| NA | NA |
| NA | NA |
| NA | NA |
| NA | NA |
| NA | NA |
| NA | NA |
| NA | NA |
| NA | NA |
| NA | NA |
| NA | NA |
| NA | NA |
| NA | NA |
| NA | NA |
| NA | NA |
| NA | NA |
| NA | NA |
| NA | NA |
| NA | NA |
| NA | NA |
| NA | NA |
| NA | NA |
| NA | NA |
| NA | NA |
| 0.905158266 | NA |
| NA | NA |
| NA | NA |
| 0.992060816 | NA |
| NA | NA |
| 0.99205891 | NA |
| NA | NA |
| NA | NA |
| NA | NA |
| NA | NA |
| NA | NA |
| NA | NA |
| NA | NA |
| NA | NA |
| NA | NA |
| NA | NA |
| NA | NA |
| NA | NA |
| NA | NA |
| NA | NA |
| NA | NA |

| | |
|---|---|
| NA | NA |
| NA | NA |
| NA | NA |
| NA | NA |
| NA | NA |
| NA | NA |
| NA | NA |
| NA | NA |
| NA | NA |
| NA | NA |
| NA | NA |
| NA | NA |
| NA | NA |
| NA | NA |
| NA | NA |
| NA | NA |
| NA | NA |
| NA | NA |
| NA | NA |
| NA | NA |
| NA | NA |
| NA | NA |
| NA | NA |
| NA | NA |
| NA | NA |
| NA | NA |
| NA | NA |
| NA | NA |
| NA | NA |
| NA | NA |
| NA | NA |
| NA | NA |
| NA | NA |
| NA | NA |
| NA | NA |
| NA | NA |
| NA | NA |
| NA | NA |
| NA | NA |
| NA | NA |
| NA | NA |
| NA | NA |
| NA | NA |
| NA | NA |
| NA | NA |
| NA | NA |
| NA | NA |

| | |
|---|---|
| NA | NA |
| NA | NA |
| NA | NA |
| NA | NA |
| NA | NA |
| NA | NA |
| NA | NA |
| NA | NA |
| NA | NA |
| NA | NA |
| NA | NA |
| NA | NA |
| NA | NA |
| NA | NA |
| NA | NA |
| NA | NA |
| NA | NA |
| NA | NA |
| NA | NA |
| NA | NA |
| NA | NA |
| NA | NA |
| NA | NA |
| NA | NA |
| NA | NA |
| NA | NA |
| NA | NA |
| NA | NA |
| NA | NA |
| NA | NA |
| NA | NA |
| NA | NA |
| NA | NA |
| NA | NA |
| NA | NA |
| NA | NA |
| NA | NA |
| NA | NA |
| NA | NA |
| NA | NA |
| NA | NA |
| NA | NA |
| NA | NA |
| NA | NA |
| NA | NA |
| NA | NA |
| NA | NA |

| | |
|---|---|
| NA | NA |
| NA | NA |
| NA | NA |
| NA | NA |
| NA | NA |
| NA | NA |
| NA | NA |
| NA | NA |
| NA | NA |
| NA | NA |
| NA | NA |
| 0.62752124 | 0.75642702 |
| NA | NA |
| 0.615223037 | 0.748251601 |
| 0.000441971 | 0.002487885 |
| 0.001940956 | 0.008724966 |
| 0.650962624 | 0.775399417 |
| 0.015036428 | 0.046794914 |
| 0.495520631 | 0.648768039 |
| 0.094339619 | 0.19701414 |
| 0.000279622 | 0.001687338 |
| 0.782708058 | 0.869097923 |
| 0.339293859 | NA |
| 0.770391329 | NA |
| 0.722532979 | NA |
| 0.094706685 | 0.197611564 |
| 0.504689607 | NA |
| 0.588886467 | NA |
| 0.001597178 | 0.007377456 |
| 0.905158739 | NA |
| 0.096178101 | 0.199978688 |
| 0.049670851 | 0.120775353 |
| 0.003662978 | 0.014780473 |
| NA | NA |
| 0.537726855 | 0.685825679 |
| NA | NA |
| 0.054011153 | 0.128692591 |
| 0.024548918 | 0.069467742 |
| 0.861212899 | NA |
| 0.792188023 | NA |
| 0.430839094 | NA |
| 0.007682313 | 0.027173954 |
| 0.835625631 | 0.902218025 |
| 0.821804276 | 0.89394165 |
| 0.018489497 | 0.055142042 |
| 0.003984715 | 0.015842905 |
| 0.002522995 | 0.010890986 |

| | |
|---|---|
| NA | NA |
| NA | NA |
| NA | NA |
| NA | NA |
| NA | NA |
| NA | NA |
| NA | NA |
| NA | NA |
| NA | NA |
| NA | NA |
| NA | NA |
| NA | NA |
| NA | NA |
| NA | NA |
| NA | NA |
| NA | NA |
| NA | NA |
| NA | NA |
| NA | NA |
| NA | NA |
| NA | NA |
| NA | NA |
| NA | NA |
| NA | NA |
| NA | NA |
| NA | NA |
| NA | NA |
| NA | NA |
| NA | NA |
| NA | NA |
| NA | NA |
| NA | NA |
| NA | NA |
| NA | NA |
| NA | NA |
| NA | NA |
| NA | NA |
| NA | NA |
| NA | NA |
| NA | NA |
| NA | NA |
| NA | NA |
| NA | NA |
| NA | NA |
| NA | NA |
| NA | NA |
| NA | NA |

| | |
|---|---|
| NA | NA |
| NA | NA |
| NA | NA |
| NA | NA |
| NA | NA |
| NA | NA |
| NA | NA |
| NA | NA |
| NA | NA |
| NA | NA |
| NA | NA |
| 0.35995501 | 0.52459998 |
| 0.889409773 | NA |
| 0.889410245 | NA |
| 0.421729685 | 0.583310307 |
| NA | NA |
| 0.083276597 | 0.179334197 |
| 0.924908347 | 0.956719373 |
| 0.293785527 | 0.453418358 |
| NA | NA |
| 0.010278725 | 0.034470145 |
| NA | NA |
| 0.718426076 | 0.8253614 |
| 0.939948655 | 0.965900721 |
| 0.013066205 | 0.041679372 |
| 0.01233666 | 0.039850061 |
| 0.01897178 | NA |
| NA | NA |
| 0.797157316 | NA |
| 0.057230337 | 0.134462242 |
| NA | NA |
| 0.800924904 | NA |
| 0.355657234 | NA |
| 0.889409773 | NA |
| 0.661548214 | NA |
| 0.483733565 | NA |
| 0.62510011 | 0.755340311 |
| 0.500807761 | 0.653547133 |
| 0.034992269 | 0.092034404 |
| 0.889409301 | NA |
| NA | NA |
| 0.28479553 | NA |
| 0.905158266 | NA |
| 0.03839028 | 0.098956704 |
| 0.138225594 | 0.261955106 |
| 0.90773659 | 0.947610543 |
| 0.490456537 | 0.644481784 |

| | |
|---|---|
| 0.327502372 | 0.489692535 |
| 0.806817603 | 0.884165472 |
| 9.62E-12 | 5.38E-10 |
| 0.103500728 | 0.211401031 |
| 0.406897157 | 0.570055804 |
| 4.70E-09 | 1.20E-07 |
| 0.674789569 | 0.793196518 |
| NA | NA |
| 0.742749052 | 0.84236171 |
| NA | NA |
| 0.005722711 | 0.021440883 |
| 0.96668613 | 0.982070818 |
| 0.330586436 | 0.493095202 |
| 0.152738985 | NA |
| 1.31E-09 | 3.97E-08 |
| 0.534865266 | 0.683371524 |
| 0.389703369 | 0.553050309 |
| 0.007562194 | 0.026837125 |
| 0.078128133 | 0.170792998 |
| 0.495290599 | 0.648582876 |
| 0.00120688 | 0.005832281 |
| 0.00093163 | 0.004691047 |
| 0.255108527 | 0.409735675 |
| 0.905158266 | NA |
| 0.406225198 | NA |
| 0.001639547 | 0.007536699 |
| 0.805096498 | NA |
| 0.004695226 | 0.018193483 |
| NA | NA |
| 0.878627501 | NA |
| 0.066203422 | 0.150414998 |
| 0.992061769 | NA |
| 0.844499856 | 0.908369941 |
| 0.708392383 | 0.818201018 |
| 0.889409773 | NA |
| 0.066749522 | 0.151227788 |
| 0.00020718 | 0.001299477 |
| 0.000247797 | 0.001518032 |
| NA | NA |
| 0.962637332 | NA |
| 0.842044466 | NA |
| 8.74E-16 | 1.32E-13 |
| 0.047488136 | 0.116474028 |
| 0.127664194 | 0.247037191 |
| 0.011449387 | 0.037502551 |
| 1.21E-08 | 2.79E-07 |
| 8.56E-05 | 0.000610159 |

| | |
|---|---|
| 0.216237314 | 0.364137668 |
| 0.024797119 | 0.070025713 |
| NA | NA |
| NA | NA |
| NA | NA |
| NA | NA |
| NA | NA |
| NA | NA |
| 0.795014225 | NA |
| NA | NA |
| NA | NA |
| NA | NA |
| 0.992060816 | NA |
| 0.875289692 | NA |
| NA | NA |
| NA | NA |
| 0.995352202 | NA |
| NA | NA |
| 0.798966441 | NA |
| 0.114866353 | 0.228701628 |
| 0.875710496 | NA |
| NA | NA |
| NA | NA |
| NA | NA |
| NA | NA |
| NA | NA |
| NA | NA |
| NA | NA |
| NA | NA |
| 0.98964564 | NA |
| NA | NA |
| NA | NA |
| NA | NA |
| NA | NA |
| 0.905158266 | NA |
| NA | NA |
| NA | NA |
| NA | NA |
| NA | NA |
| NA | NA |
| NA | NA |
| 0.021060496 | 0.061278314 |
| 0.889409301 | NA |
| 0.270609739 | 0.42712536 |
| 0.814487706 | NA |
| 0.983398525 | NA |
| 0.055023219 | 0.130481087 |

| | | |
|---|---|---|
| | 0.606166653 | NA |
| | 0.788582525 | NA |
| | 4.23E-05 | 0.000336031 |
| | 0.992061292 | NA |
| | 0.79867967 | NA |
| | 0.095055011 | 0.198168897 |
| | 0.378676069 | NA |
| | 0.535009604 | NA |
| | 0.520659301 | 0.671472226 |
| | 0.908999055 | NA |
| | 0.081349567 | 0.176203901 |
| | 0.000659551 | 0.003518728 |
| | 0.665134567 | NA |
| | 0.986595458 | NA |
| | 3.12E-11 | 1.53E-09 |
| | 0.018624201 | 0.055445882 |
| | 0.77059653 | 0.861278018 |
| | 0.99205891 | NA |
| | 0.285679517 | NA |
| | 0.040991259 | 0.103916769 |
| | 0.046402641 | 0.11441226 |
| | 0.588854214 | 0.727462711 |
| NA | | NA |
| | 0.992060816 | NA |
| | 0.469725577 | NA |
| | 0.992061292 | NA |
| NA | | NA |
| NA | | NA |
| NA | | NA |
| NA | | NA |
| NA | | NA |
| NA | | NA |
| NA | | NA |
| NA | | NA |
| NA | | NA |
| NA | | NA |
| NA | | NA |
| NA | | NA |
| NA | | NA |
| | 0.107330339 | NA |
| NA | | NA |
| | 0.323437715 | 0.485792961 |
| NA | | NA |
| | 0.992061292 | NA |
| | 0.244777218 | NA |
| | 0.720042391 | 0.826351566 |
| | 0.841828281 | 0.906428715 |

| | | |
|---|---|---|
| | 0.889409773 | NA |
| | 0.628679127 | 0.757111164 |
| | 0.658753852 | NA |
| | 0.063351523 | 0.145592476 |
| | 4.62E-05 | 0.0003617 |
| | 0.889409773 | NA |
| NA | | NA |
| | 0.38469755 | 0.548686249 |
| | 0.778531773 | NA |
| | 0.083785128 | 0.180048951 |
| | 0.071167207 | 0.158527156 |
| | 0.853835502 | NA |
| NA | | NA |
| | 0.585208008 | 0.724629675 |
| NA | | NA |
| | 0.666401751 | 0.787243016 |
| | 0.356882175 | NA |
| NA | | NA |
| | 0.199991368 | 0.344899902 |
| | 0.63828282 | NA |
| | 0.992061292 | NA |
| | 0.660872814 | NA |
| | 0.119433427 | 0.235319751 |
| NA | | NA |
| | 0.000951087 | 0.00477697 |
| | 0.002067843 | 0.009206765 |
| NA | | NA |
| | 0.90330426 | NA |
| NA | | NA |
| NA | | NA |
| NA | | NA |
| NA | | NA |
| NA | | NA |
| | 0.860566709 | NA |
| NA | | NA |
| NA | | NA |
| | 0.990196425 | NA |
| NA | | NA |
| NA | | NA |
| NA | | NA |
| NA | | NA |
| NA | | NA |
| | 0.547028184 | NA |
| NA | | NA |
| | 0.830163713 | NA |
| | 0.905158739 | NA |
| | 0.905158739 | NA |

| | |
|---|---|
| 0.014534274 | 0.045490041 |
| 0.002322103 | 0.01015556 |
| 0.194025228 | 0.337312958 |
| 0.00970236 | 0.032828062 |
| NA | NA |
| 0.992061292 | NA |
| 0.380088488 | 0.544304363 |
| 0.813836939 | NA |
| NA | NA |
| NA | NA |
| NA | NA |
| NA | NA |
| 0.155427352 | 0.286255911 |
| NA | NA |
| NA | NA |
| 0.466063453 | 0.622828193 |
| NA | NA |
| NA | NA |
| 0.119632515 | 0.235627446 |
| 0.830210183 | 0.898766782 |
| 0.068399212 | 0.153639352 |
| 0.519908781 | 0.670746698 |
| 0.759335519 | NA |
| NA | NA |
| NA | NA |
| NA | NA |
| NA | NA |
| 0.716697006 | NA |
| 0.905158739 | NA |
| 0.396591127 | NA |
| 0.928516785 | 0.959090272 |
| 0.020715441 | 0.060478629 |
| 0.07846382 | 0.171418981 |
| NA | NA |
| 0.026656018 | 0.074093608 |
| 0.788582525 | NA |
| 0.182979042 | 0.323227157 |
| 0.005248467 | 0.01995092 |
| 0.889409301 | NA |
| NA | NA |
| 0.474397813 | 0.630701797 |
| 0.720302923 | 0.826488491 |
| 0.459810057 | NA |
| 0.891188223 | 0.938410003 |
| 0.017068729 | 0.051827097 |
| 0.004694717 | 0.018193483 |
| 2.92E-06 | 3.33E-05 |

| | |
|---|---|
| 1.23E-09 | 3.74E-08 |
| 0.005431168 | 0.020527196 |
| NA | NA |
| NA | NA |
| 0.646395282 | NA |
| 2.22E-05 | 0.000194396 |
| NA | NA |
| 0.001287242 | 0.006156947 |
| NA | NA |
| 0.00015861 | 0.001032585 |
| 0.007112392 | 0.025570535 |
| 0.112149374 | 0.224411163 |
| 0.576467473 | 0.718017541 |
| 0.111732899 | 0.223720501 |
| 0.493720628 | NA |
| 0.010801609 | 0.035828283 |
| NA | NA |
| 0.267705717 | 0.423984842 |
| NA | NA |
| 0.889409301 | NA |
| 0.033832054 | NA |
| 0.901145897 | 0.944236842 |
| 0.277189374 | NA |
| 0.850091308 | 0.912150258 |
| NA | NA |
| 0.41122952 | NA |
| NA | NA |
| 0.262989894 | NA |
| 0.992060816 | NA |
| NA | NA |
| 0.60848111 | NA |
| NA | NA |
| NA | NA |
| NA | NA |
| NA | NA |
| NA | NA |
| NA | NA |
| NA | NA |
| NA | NA |
| NA | NA |
| 0.000263303 | 0.001599414 |
| 0.472088818 | NA |
| 0.992060816 | NA |
| 0.635980429 | 0.763183465 |
| NA | NA |
| 0.530223909 | NA |
| NA | NA |

| | | |
|---|---|---|
| NA | | NA |
| NA | | NA |
| NA | | NA |
| NA | | NA |
| NA | | NA |
| | 0.905158266 | NA |
| NA | | NA |
| NA | | NA |
| NA | | NA |
| NA | | NA |
| NA | | NA |
| NA | | NA |
| NA | | NA |
| NA | | NA |
| NA | | NA |
| NA | | NA |
| NA | | NA |
| NA | | NA |
| NA | | NA |
| NA | | NA |
| NA | | NA |
| NA | | NA |
| NA | | NA |
| NA | | NA |
| NA | | NA |
| NA | | NA |
| NA | | NA |
| NA | | NA |
| NA | | NA |
| NA | | NA |
| NA | | NA |
| NA | | NA |
| NA | | NA |
| NA | | NA |
| NA | | NA |
| NA | | NA |
| NA | | NA |
| NA | | NA |
| NA | | NA |
| NA | | NA |
| NA | | NA |
| NA | | NA |
| NA | | NA |
| NA | | NA |
| NA | | NA |
| NA | | NA |
| NA | | NA |

| | | | |
|---|---|---|---|
| NA | | NA | |
| NA | | NA | |
| NA | | NA | |
| NA | | NA | |
| NA | | NA | |
| NA | | NA | |
| NA | | NA | |
| | 0.447985883 | | 0.6070922 |
| NA | | NA | |
| NA | | NA | |
| NA | | NA | |
| NA | | NA | |
| NA | | NA | |
| NA | | NA | |
| NA | | NA | |
| NA | | NA | |
| NA | | NA | |
| NA | | NA | |
| NA | | NA | |
| NA | | NA | |
| NA | | NA | |
| NA | | NA | |
| NA | | NA | |
| NA | | NA | |
| NA | | NA | |
| NA | | NA | |
| NA | | NA | |
| NA | | NA | |
| NA | | NA | |
| NA | | NA | |
| NA | | NA | |
| NA | | NA | |
| NA | | NA | |
| NA | | NA | |
| NA | | NA | |
| NA | | NA | |
| NA | | NA | |
| NA | | NA | |
| NA | | NA | |
| NA | | NA | |
| NA | | NA | |
| NA | | NA | |
| NA | | NA | |
| NA | | NA | |
| NA | | NA | |
| NA | | NA | |
| NA | | NA | |

| | | | |
|---|---|---|---|
| NA | | NA | |
| | 0.54974188 | | 0.695860154 |
| NA | | NA | |
| NA | | NA | |
| NA | | NA | |
| NA | | NA | |
| NA | | NA | |
| NA | | NA | |
| NA | | NA | |
| NA | | NA | |
| NA | | NA | |
| NA | | NA | |
| NA | | NA | |
| NA | | NA | |
| NA | | NA | |
| NA | | NA | |
| NA | | NA | |
| NA | | NA | |
| NA | | NA | |
| NA | | NA | |
| NA | | NA | |
| NA | | NA | |
| NA | | NA | |
| | 0.905158266 | NA | |
| NA | | NA | |
| | 2.49E-05 | | 0.000213834 |
| | 1.86E-09 | | 5.43E-08 |
| | 0.00376093 | | 0.015092518 |
| NA | | NA | |
| | 0.664825152 | NA | |
| NA | | NA | |
| | 0.016165854 | | 0.049641447 |
| | 0.252555074 | | 0.406826354 |
| | 0.775345711 | NA | |
| NA | | NA | |
| | 0.889409773 | NA | |
| NA | | NA | |
| | 0.315530293 | | 0.477824638 |
| | 0.175858817 | | 0.313983557 |
| | 0.017068815 | | 0.051827097 |
| | 0.352648406 | | 0.516874005 |
| | 0.068213664 | | 0.153439533 |
| | 0.430922532 | NA | |
| | 0.026047452 | | 0.072798268 |
| NA | | NA | |
| | 0.020227427 | | 0.059333002 |
| | 0.691425783 | | 0.805772376 |

| | |
|---|---|
| 0.159599397 | 0.291761835 |
| 0.000971982 | 0.004858583 |
| 0.614332819 | 0.747582952 |
| 0.646440527 | NA |
| 3.51E-06 | 3.93E-05 |
| NA | NA |
| 0.905296333 | NA |
| NA | NA |
| NA | NA |
| 0.065474499 | 0.149215608 |
| 0.876212658 | NA |
| 0.188353787 | 0.329848245 |
| 0.759084838 | 0.853756605 |
| 0.477522443 | NA |
| 0.412197215 | 0.574804003 |
| 0.929164089 | 0.959434081 |
| NA | NA |
| NA | NA |
| NA | NA |
| NA | NA |
| NA | NA |
| NA | NA |
| NA | NA |
| NA | NA |
| NA | NA |
| NA | NA |
| NA | NA |
| NA | NA |
| NA | NA |
| NA | NA |
| NA | NA |
| NA | NA |
| NA | NA |
| NA | NA |
| NA | NA |
| NA | NA |
| NA | NA |
| NA | NA |
| NA | NA |
| NA | NA |
| NA | NA |
| NA | NA |
| NA | NA |
| NA | NA |
| NA | NA |
| NA | NA |

| | |
|---|---|
| NA | NA |
| NA | NA |
| NA | NA |
| NA | NA |
| 0.889409301 | NA |
| NA | NA |
| NA | NA |
| NA | NA |
| NA | NA |
| 0.905158266 | NA |
| NA | NA |
| 0.629852689 | NA |
| 0.713554725 | 0.821568174 |
| 0.947132783 | 0.969924394 |
| NA | NA |
| 0.889409773 | NA |
| 0.655478926 | NA |
| NA | NA |
| 0.263847331 | 0.419592999 |
| 3.05E-05 | 0.000253279 |
| 0.546637588 | 0.693009341 |
| 0.872677649 | 0.926964083 |
| 0.880224668 | 0.932094974 |
| 0.00339141 | 0.013888844 |
| 0.015313685 | 0.047489508 |
| 0.889409773 | NA |
| NA | NA |
| NA | NA |
| 0.493215381 | NA |
| NA | NA |
| 0.72350667 | 0.829200572 |
| NA | NA |
| NA | NA |
| NA | NA |
| NA | NA |
| 0.992060816 | NA |
| NA | NA |
| NA | NA |
| 0.992061292 | NA |
| NA | NA |
| 0.992060816 | NA |
| NA | NA |
| NA | NA |
| NA | NA |
| NA | NA |
| 0.395896562 | NA |
| NA | NA |

| | |
|---|---|
| NA | NA |
| NA | NA |
| NA | NA |
| NA | NA |
| 8.27E-22 | 5.04E-19 |
| NA | NA |
| NA | NA |
| NA | NA |
| 0.189087783 | 0.330877252 |
| NA | NA |
| NA | NA |
| NA | NA |
| 0.168541321 | 0.304067987 |
| 0.992060816 | NA |
| 0.737771051 | NA |
| 0.428048784 | 0.589127123 |
| 0.784353866 | NA |
| 0.788361417 | NA |
| 0.463377001 | 0.620482686 |
| 3.54E-06 | 3.96E-05 |
| 0.25764653 | 0.412456087 |
| 0.26278468 | 0.4182726 |
| NA | NA |
| 0.863668951 | 0.921363449 |
| NA | NA |
| NA | NA |
| NA | NA |
| NA | NA |
| NA | NA |
| NA | NA |
| 0.588250666 | 0.726928871 |
| NA | NA |
| NA | NA |
| 2.23E-05 | 0.00019523 |
| NA | NA |
| 0.003394112 | 0.013897317 |
| NA | NA |
| NA | NA |
| NA | NA |
| NA | NA |
| NA | NA |
| NA | NA |
| NA | NA |
| NA | NA |
| 0.905158266 | NA |
| NA | NA |
| NA | NA |

| | | |
|---|---|---|
| | 0.805374319 | NA |
| | 0.761571879 | NA |
| | 0.743753467 | NA |
| NA | | NA |
| | 0.990953236 | NA |
| | 0.889410717 | NA |
| | 8.00E-07 | 1.08E-05 |
| | 0.589974876 | 0.728244332 |
| | 9.91E-06 | 9.62E-05 |
| | 0.09314009 | 0.195009844 |
| | 0.0089316 | 0.030697919 |
| | 0.000172327 | 0.001108416 |
| | 0.26878236 | 0.425185078 |
| NA | | NA |
| | 0.020331216 | 0.059558855 |
| | 0.482591364 | NA |
| NA | | NA |
| NA | | NA |
| NA | | NA |
| | 0.400335768 | 0.563909228 |
| | 4.07E-09 | 1.07E-07 |
| | 0.060517509 | 0.140699203 |
| | 2.98E-08 | 6.20E-07 |
| | 0.004215509 | 0.016604192 |
| | 0.000191832 | 0.001216413 |
| | 0.014345784 | 0.044983445 |
| | 0.793147413 | NA |
| | 0.859048944 | 0.917815197 |
| | 0.960153578 | NA |
| NA | | NA |
| | 0.858271697 | 0.917217762 |
| NA | | NA |
| NA | | NA |
| | 0.75271404 | NA |
| | 0.990073618 | NA |
| | 0.329932936 | 0.49235456 |
| NA | | NA |
| | 0.155719246 | 0.286577181 |
| NA | | NA |
| NA | | NA |
| | 0.000154905 | 0.001011463 |
| NA | | NA |
| | 0.889409301 | NA |
| NA | | NA |
| | 0.715711479 | NA |
| NA | | NA |
| | 0.123072472 | 0.240465255 |

| | | |
|---|---|---|
| | 0.905158739 | NA |
| | 0.551534965 | NA |
| | 0.905158266 | NA |
| NA | | NA |
| | 0.072760427 | 0.161307039 |
| | 7.88E-08 | 1.43E-06 |
| | 0.162249542 | 0.295475011 |
| | 4.58E-05 | 0.000359196 |
| NA | | NA |
| | 0.087003819 | 0.185289192 |
| NA | | NA |
| | 0.790658061 | NA |
| NA | | NA |
| NA | | NA |
| | 0.007436052 | 0.026469528 |
| | 0.313927821 | 0.47584324 |
| NA | | NA |
| NA | | NA |
| | 0.889409301 | NA |
| NA | | NA |
| | 0.050544361 | 0.12239784 |
| | 0.042260718 | 0.106532327 |
| NA | | NA |
| | 0.992062245 | NA |
| NA | | NA |
| | 0.000150798 | 0.000988762 |
| NA | | NA |
| NA | | NA |
| NA | | NA |
| NA | | NA |
| NA | | NA |
| NA | | NA |
| | 0.905158266 | NA |
| | 0.305486831 | 0.46610515 |
| NA | | NA |
| | 0.741864126 | 0.841850853 |
| | 0.889409773 | NA |
| NA | | NA |
| | 0.977127131 | 0.988243122 |
| | 0.22128187 | 0.370284723 |
| | 0.211905979 | 0.358976954 |
| NA | | NA |
| NA | | NA |
| NA | | NA |
| NA | | NA |
| | 0.992061292 | NA |
| NA | | NA |

| | | | |
|---|---|---|---|
| | 0.992060816 | NA | |
| NA | | NA | |
| | 0.823270761 | | 0.894625048 |
| NA | | NA | |
| | 0.587478438 | | 0.726253393 |
| NA | | NA | |
| NA | | NA | |
| NA | | NA | |
| | 0.935103958 | | 0.962768232 |
| | 0.308020427 | | 0.469025765 |
| NA | | NA | |
| | 0.35089773 | NA | |
| | 0.000143352 | | 0.00094587 |
| | 0.592876474 | NA | |
| NA | | NA | |
| | 0.038737712 | | 0.099584451 |
| | 0.022611063 | | 0.064962353 |
| | 0.991979299 | NA | |
| | 0.214214147 | | 0.361595192 |
| | 0.581945137 | | 0.72189206 |
| | 0.905158266 | NA | |
| NA | | NA | |
| NA | | NA | |
| | 0.530611805 | NA | |
| | 0.41844061 | | 0.580670966 |
| | 0.889409773 | NA | |
| NA | | NA | |
| | 0.493396657 | | 0.646928273 |
| | 0.792188023 | NA | |
| | 0.003777025 | NA | |
| NA | | NA | |
| NA | | NA | |
| NA | | NA | |
| NA | | NA | |
| | 0.787391563 | | 0.871771467 |
| | 0.905158266 | NA | |
| NA | | NA | |
| NA | | NA | |
| NA | | NA | |
| NA | | NA | |
| NA | | NA | |
| NA | | NA | |
| NA | | NA | |
| NA | | NA | |
| NA | | NA | |
| NA | | NA | |
| NA | | NA | |

| | |
|---|---|
| NA | NA |
| NA | NA |
| NA | NA |
| NA | NA |
| NA | NA |
| NA | NA |
| NA | NA |
| NA | NA |
| NA | NA |
| NA | NA |
| 0.905158739 | NA |
| NA | NA |
| 0.971097077 | NA |
| 0.666429679 | NA |
| NA | NA |
| NA | NA |
| NA | NA |
| 0.034717122 | 0.091529991 |
| 0.017357508 | 0.052486025 |
| 1.56E-06 | 1.92E-05 |
| 0.250144203 | 0.404128645 |
| 0.889410245 | NA |
| 0.000297034 | 0.001770995 |
| NA | NA |
| 0.862927754 | NA |
| 0.484286913 | 0.639628072 |
| NA | NA |
| 1.99E-12 | 1.33E-10 |
| NA | NA |
| 0.211586145 | 0.358643007 |
| 0.377005361 | NA |
| 0.431076099 | 0.591970318 |
| 0.774862718 | NA |
| 0.24393786 | 0.396905118 |
| 0.662723901 | 0.784176859 |
| NA | NA |
| 0.86219084 | NA |
| 0.633641681 | NA |
| 0.615477203 | 0.748332145 |
| 0.182015199 | 0.322199107 |
| 0.605872105 | NA |
| 0.543206589 | NA |
| 0.355444537 | 0.52006659 |
| 0.905158739 | NA |
| NA | NA |
| NA | NA |
| 0.905158739 | NA |

| | |
|---|---|
| NA | NA |
| NA | NA |
| NA | NA |
| 0.351639678 | 0.515946524 |
| NA | NA |
| NA | NA |
| NA | NA |
| NA | NA |
| 0.905158266 | NA |
| 0.889409301 | NA |
| NA | NA |
| NA | NA |
| 0.584510157 | NA |
| 2.04E-16 | 3.49E-14 |
| 0.581596466 | NA |
| 0.705191542 | NA |
| NA | NA |
| 0.889410717 | NA |
| NA | NA |
| NA | NA |
| NA | NA |
| 0.163023598 | 0.296466682 |
| 1.01E-05 | 9.77E-05 |
| 0.905158266 | NA |
| 0.243240084 | 0.396180759 |
| 0.766862145 | 0.858546235 |
| NA | NA |
| 0.450395215 | 0.608854256 |
| NA | NA |
| 0.126354595 | 0.245237807 |
| 0.476379592 | NA |
| NA | NA |
| NA | NA |
| 0.994468642 | NA |
| NA | NA |
| 0.153483124 | 0.283632246 |
| 0.27128353 | NA |
| NA | NA |
| NA | NA |
| 0.889410245 | NA |
| 0.905158739 | NA |
| 0.005062788 | 0.019364876 |
| 0.086048426 | 0.183704799 |
| 0.77272678 | 0.862824537 |
| 0.889409773 | NA |
| 0.734537811 | NA |
| 0.905157793 | NA |

| | | | |
|---|---|---|---|
| | 0.889409773 | NA | |
| | 0.146084465 | | 0.273195628 |
| | 0.997380648 | | 0.998881768 |
| | 0.533686836 | NA | |
| NA | | NA | |
| NA | | NA | |
| NA | | NA | |
| NA | | NA | |
| | 0.992061292 | NA | |
| NA | | NA | |
| NA | | NA | |
| NA | | NA | |
| NA | | NA | |
| NA | | NA | |
| NA | | NA | |
| | 0.160264486 | | 0.292685283 |
| NA | | NA | |
| NA | | NA | |
| NA | | NA | |
| NA | | NA | |
| NA | | NA | |
| NA | | NA | |
| NA | | NA | |
| | 0.96851819 | | 0.983204241 |
| | 0.534076729 | NA | |
| NA | | NA | |
| NA | | NA | |
| NA | | NA | |
| NA | | NA | |
| NA | | NA | |
| NA | | NA | |
| NA | | NA | |
| NA | | NA | |
| NA | | NA | |
| NA | | NA | |
| | 0.739395017 | NA | |
| NA | | NA | |
| | 0.166019231 | | 0.300495903 |
| | 0.700035375 | NA | |
| | 0.889409773 | NA | |
| NA | | NA | |
| NA | | NA | |
| | 0.264510941 | | 0.420441241 |
| NA | | NA | |
| NA | | NA | |
| NA | | NA | |
| NA | | NA | |

| | | |
|---|---|---|
| NA | NA | |
| NA | NA | |
| NA | NA | |
| NA | NA | |
| NA | NA | |
| NA | NA | |
| NA | NA | |
| NA | NA | |
| NA | NA | |
| NA | NA | |
| NA | NA | |
| NA | NA | |
| NA | NA | |
| NA | NA | |
| 0.797157316 | NA | |
| NA | NA | |
| NA | NA | |
| NA | NA | |
| NA | NA | |
| NA | NA | |
| NA | NA | |
| NA | NA | |
| NA | NA | |
| NA | NA | |
| NA | NA | |
| NA | NA | |
| NA | NA | |
| NA | NA | |
| NA | NA | |
| 0.784864141 | NA | |
| NA | NA | |
| NA | NA | |
| 0.99205891 | NA | |
| 0.072011518 | NA | |
| NA | NA | |
| NA | NA | |
| 0.950899065 | | 0.972647316 |
| NA | NA | |
| 0.975897253 | NA | |
| NA | NA | |
| NA | NA | |
| 0.578335886 | | 0.719406204 |
| NA | NA | |
| NA | NA | |
| NA | NA | |
| 0.716577993 | NA | |
| 0.001034333 | | 0.005122444 |

| | |
|---|---|
| 0.857315771 | 0.916855648 |
| 0.724999475 | NA |
| 0.889410245 | NA |
| 3.45E-15 | 4.49E-13 |
| 0.112635977 | 0.225200165 |
| NA | NA |
| NA | NA |
| NA | NA |
| 0.633844795 | NA |
| NA | NA |
| NA | NA |
| NA | NA |
| 0.236401263 | 0.388267415 |
| NA | NA |
| NA | NA |
| NA | NA |
| 0.455736542 | NA |
| NA | NA |
| NA | NA |
| NA | NA |
| 0.777092376 | NA |
| NA | NA |
| NA | NA |
| 0.905158266 | NA |
| NA | NA |
| 0.163925293 | NA |
| 0.889409773 | NA |
| 0.420905561 | 0.582655565 |
| 0.056016756 | 0.13225158 |
| 0.858435807 | NA |
| 0.730311906 | 0.834170046 |
| 0.018829254 | 0.055936141 |
| 0.879063294 | 0.931359205 |
| NA | NA |
| NA | NA |
| NA | NA |
| 0.005012775 | 0.019223807 |
| 0.12531354 | 0.243755872 |
| NA | NA |
| 0.797632026 | 0.877778975 |
| 0.851865521 | 0.913439345 |
| 0.020337033 | 0.059567949 |
| 0.033061152 | 0.08803078 |
| 0.534201259 | NA |
| 0.331053784 | 0.493423609 |
| NA | NA |
| 0.094475862 | 0.197223639 |

| | |
|---|---|
| 0.905158739 | NA |
| 0.992062955 | NA |
| NA | NA |
| 2.41E-12 | 1.57E-10 |
| NA | NA |
| 0.985535234 | NA |
| 0.352517932 | NA |
| NA | NA |
| NA | NA |
| 0.666970731 | 0.787504585 |
| NA | NA |
| 0.563710866 | 0.707361916 |
| 0.001541868 | 0.007165691 |
| 0.765450081 | NA |
| 0.333723213 | 0.496374136 |
| 0.862029991 | NA |
| 0.330075099 | 0.492438027 |
| 1.87E-13 | 1.59E-11 |
| 0.126163598 | NA |
| 0.269487224 | 0.425915499 |
| NA | NA |
| NA | NA |
| 0.480515811 | NA |
| 0.152708983 | NA |
| 0.889409773 | NA |
| NA | NA |
| 0.205657419 | 0.351942154 |
| 0.186744452 | 0.328110211 |
| 0.726727097 | 0.831249317 |
| 0.774894285 | 0.864190127 |
| 0.065716714 | NA |
| NA | NA |
| 0.072868207 | NA |
| NA | NA |
| 0.016966885 | 0.0516176 |
| 0.736699836 | 0.838335753 |
| 0.198310355 | NA |
| 0.458290123 | NA |
| 0.004327445 | 0.016957343 |
| 0.568579135 | NA |
| 0.277598995 | 0.435507347 |
| 1.79E-05 | 0.000161187 |
| 8.26E-11 | 3.58E-09 |
| 0.16105709 | 0.293731153 |
| 1.92E-18 | 5.70E-16 |
| 2.64E-23 | 2.52E-20 |
| 0.005918074 | 0.022030003 |

| | | |
|---|---|---|
| | 9.55E-05 | 0.000671662 |
| NA | | NA |
| | 0.003181753 | 0.01319511 |
| | 0.786526711 | 0.871533027 |
| | 3.21E-08 | 6.63E-07 |
| | 0.162225457 | 0.295455652 |
| | 6.16E-06 | 6.41E-05 |
| | 0.064937929 | 0.148368741 |
| | 0.905158266 | NA |
| | 4.33E-16 | 7.10E-14 |
| | 0.433931195 | 0.594651468 |
| | 0.880323049 | NA |
| | 0.085779962 | 0.183251501 |
| | 8.76E-08 | 1.57E-06 |
| | 0.021103992 | 0.061372342 |
| | 0.328542885 | NA |
| | 0.003870696 | 0.015462736 |
| NA | | NA |
| | 0.992061292 | NA |
| | 8.84E-09 | 2.12E-07 |
| | 0.939887154 | 0.96589388 |
| | 0.735862695 | NA |
| | 0.889409301 | NA |
| | 0.986786572 | 0.993163733 |
| | 0.121695196 | NA |
| | 9.43E-15 | 1.09E-12 |
| | 0.001706905 | 0.007798985 |
| NA | | NA |
| | 8.15E-05 | 0.000586146 |
| NA | | NA |
| | 6.41E-09 | 1.59E-07 |
| NA | | NA |
| NA | | NA |
| | 0.199051382 | 0.343764789 |
| | 0.373183088 | 0.537429658 |
| | 2.05E-10 | 8.02E-09 |
| | 0.012163501 | 0.039400843 |
| | 0.992061292 | NA |
| | 0.636550821 | 0.763450733 |
| | 0.008857827 | 0.030506988 |
| | 0.089601178 | 0.189481684 |
| | 5.90E-05 | 0.000446011 |
| | 0.453928598 | 0.611944507 |
| | 0.252586245 | 0.406837496 |
| | 2.60E-10 | 9.75E-09 |
| | 0.905158266 | NA |
| NA | | NA |

| | | |
|---|---|---|
| | 0.58956488 | 0.7280159 |
| NA | | NA |
| NA | | NA |
| NA | | NA |
| NA | | NA |
| NA | | NA |
| NA | | NA |
| NA | | NA |
| | 0.289297171 | NA |
| | 0.905158266 | NA |
| NA | | NA |
| | 0.880735197 | NA |
| | 0.618545497 | NA |
| NA | | NA |
| NA | | NA |
| NA | | NA |
| NA | | NA |
| NA | | NA |
| NA | | NA |
| NA | | NA |
| NA | | NA |
| NA | | NA |
| NA | | NA |
| | 0.578135499 | NA |
| NA | | NA |
| NA | | NA |
| NA | | NA |
| NA | | NA |
| | 0.889409301 | NA |
| | 0.905157793 | NA |
| NA | | NA |
| | 0.44335793 | 0.602941524 |
| | 0.501428867 | NA |
| NA | | NA |
| | 0.000358221 | 0.002081547 |
| | 0.882056004 | NA |
| | 0.012475559 | 0.040221819 |
| | 0.992061292 | NA |
| | 0.579781581 | NA |
| | 0.58459717 | 0.72424086 |
| | 0.905158739 | NA |
| | 0.189572072 | 0.331486952 |
| | 0.959848394 | 0.977727478 |
| | 0.458025715 | 0.615453932 |
| | 0.006746153 | 0.024501949 |
| | 0.504956107 | 0.656969496 |
| | 0.000139053 | 0.000923892 |

| | | |
|---|---|---|
| | 0.489171799 | 0.643409616 |
| | 0.654285961 | 0.777658858 |
| | 0.21731342 | 0.365501333 |
| | 0.895289803 | 0.941005809 |
| | 0.744166212 | NA |
| | 0.23326846 | 0.384820233 |
| | 0.750537676 | NA |
| NA | | NA |
| | 0.020734215 | NA |
| | 0.021949041 | 0.063355437 |
| | 0.900836801 | NA |
| | 0.111218496 | 0.222954807 |
| NA | | NA |
| NA | | NA |
| NA | | NA |
| NA | | NA |
| | 0.762847635 | NA |
| NA | | NA |
| NA | | NA |
| | 0.889409301 | NA |
| NA | | NA |
| NA | | NA |
| | 0.989388079 | NA |
| NA | | NA |
| | 0.905158266 | NA |
| NA | | NA |
| NA | | NA |
| | 0.905158739 | NA |
| | 0.992061292 | NA |
| NA | | NA |
| NA | | NA |
| | 0.905158266 | NA |
| NA | | NA |
| | 0.868485979 | NA |
| NA | | NA |
| | 0.308829731 | 0.469997371 |
| NA | | NA |
| | 0.796885941 | NA |
| | 0.53821694 | 0.686136766 |
| | 0.670480413 | 0.789864774 |
| NA | | NA |
| | 0.669633885 | NA |
| NA | | NA |
| | 0.042211572 | 0.106445097 |
| | 1.83E-09 | 5.34E-08 |
| | 0.992061769 | NA |
| NA | | NA |

| | |
|---|---|
| 0.990012186 | NA |
| 0.992061769 | NA |
| NA | NA |
| NA | NA |
| NA | NA |
| 0.992061587 | NA |
| NA | NA |
| 0.992061769 | NA |
| 0.992061292 | NA |
| NA | NA |
| 0.905158266 | NA |
| NA | NA |
| NA | NA |
| NA | NA |
| NA | NA |
| 0.73537946 | 0.837475239 |
| 0.03274875 | NA |
| NA | NA |
| NA | NA |
| NA | NA |
| NA | NA |
| 0.905158739 | NA |
| 0.700241341 | NA |
| NA | NA |
| NA | NA |
| 0.233618107 | 0.385226725 |
| 0.920417645 | 0.954486576 |
| 0.821317007 | 0.893671826 |
| 0.84137548 | NA |
| 0.339899042 | NA |
| 0.527235772 | 0.677023466 |
| NA | NA |
| NA | NA |
| 0.574037658 | 0.716047088 |
| 0.187008222 | 0.328442257 |
| NA | NA |
| NA | NA |
| 0.570038997 | NA |
| 0.065366912 | 0.149083619 |
| NA | NA |
| 0.889409301 | NA |
| 0.082462446 | 0.178069904 |
| NA | NA |
| NA | NA |
| NA | NA |
| NA | NA |
| 0.019631545 | 0.057918729 |

| | | |
|---|---|---|
| NA | NA | |
| NA | NA | |
| NA | NA | |
| NA | NA | |
| 0.000391945 | | 0.002247183 |
| 0.441074702 | NA | |
| NA | NA | |
| 0.9760901 | NA | |
| 0.000241233 | | 0.001485067 |
| 0.392302987 | | 0.555851162 |
| NA | NA | |
| 0.423920225 | NA | |
| NA | NA | |
| NA | NA | |
| 0.752363829 | | 0.849242177 |
| 0.191653186 | | 0.334114982 |
| 0.037262102 | | 0.096741369 |
| 0.889409773 | NA | |
| NA | NA | |
| 0.763131439 | NA | |
| NA | NA | |
| 0.992059387 | NA | |
| 0.839052384 | NA | |
| 1.02E-05 | | 9.87E-05 |
| NA | NA | |
| 0.992060816 | NA | |
| 0.926399498 | NA | |
| 0.905158739 | NA | |
| NA | NA | |
| NA | NA | |
| 0.787590234 | NA | |
| NA | NA | |
| NA | NA | |
| NA | NA | |
| NA | NA | |
| NA | NA | |
| 3.83E-10 | | 1.37E-08 |
| NA | NA | |
| 0.23469913 | | 0.386203346 |
| 0.005617578 | | 0.021115439 |
| 0.304937337 | | 0.465622626 |
| 0.043363452 | | 0.108589127 |
| NA | NA | |
| NA | NA | |
| NA | NA | |
| 0.718827801 | NA | |
| NA | NA | |

| | | |
|---|---|---|
| NA | NA | |
| NA | NA | |
| 0.389651391 | NA | |
| NA | NA | |
| 0.04721575 | | 0.115909521 |
| NA | NA | |
| NA | NA | |
| NA | NA | |
| 0.87762688 | NA | |
| 0.889409301 | NA | |
| NA | NA | |
| NA | NA | |
| NA | NA | |
| NA | NA | |
| NA | NA | |
| NA | NA | |
| NA | NA | |
| NA | NA | |
| NA | NA | |
| 0.019887766 | NA | |
| NA | NA | |
| NA | NA | |
| NA | NA | |
| NA | NA | |
| NA | NA | |
| 0.787686947 | | 0.871771467 |
| NA | NA | |
| NA | NA | |
| NA | NA | |
| 0.992061292 | NA | |
| NA | NA | |
| NA | NA | |
| 0.889409301 | NA | |
| 0.983619417 | NA | |
| NA | NA | |
| NA | NA | |
| NA | NA | |
| NA | NA | |
| NA | NA | |
| NA | NA | |
| NA | NA | |
| NA | NA | |
| NA | NA | |
| NA | NA | |
| NA | NA | |
| 0.555996466 | NA | |
| NA | NA | |

| | | | |
|---|---|---|---|
| NA | | NA | |
| | 0.889409301 | NA | |
| NA | | NA | |
| | 0.507915765 | NA | |
| | 0.630300507 | NA | |
| NA | | NA | |
| NA | | NA | |
| NA | | NA | |
| NA | | NA | |
| | 0.983545476 | NA | |
| NA | | NA | |
| NA | | NA | |
| NA | | NA | |
| NA | | NA | |
| | 0.388416792 | | 0.551962569 |
| | 0.575663923 | | 0.7173829 |
| NA | | NA | |
| | 0.344380035 | | 0.507771516 |
| NA | | NA | |
| | 0.566133883 | | 0.709154523 |
| | 0.220608108 | | 0.369542561 |
| NA | | NA | |
| NA | | NA | |
| | 0.889409773 | NA | |
| NA | | NA | |
| | 0.992061292 | NA | |
| NA | | NA | |
| | 0.587053844 | NA | |
| | 0.159952452 | | 0.292212637 |
| | 0.054555623 | | 0.129680334 |
| | 0.089384441 | | 0.189184862 |
| NA | | NA | |
| | 0.360823605 | NA | |
| NA | | NA | |
| | 0.889409773 | NA | |
| | 0.794509295 | | 0.87598401 |
| | 0.009462246 | | 0.032144408 |
| NA | | NA | |
| NA | | NA | |
| | 0.658473773 | | 0.780788681 |
| NA | | NA | |
| | 0.001414435 | | 0.006687194 |
| NA | | NA | |
| | 0.905158739 | NA | |
| | 0.75424085 | NA | |
| | 0.905158266 | NA | |
| | 0.175215772 | | 0.312988135 |

| | | |
|---|---|---|
| | 0.905158266 | NA |
| | 0.588551969 | 0.727171141 |
| | 0.490959491 | 0.644979831 |
| | 0.006461827 | 0.023664988 |
| | 0.992062245 | NA |
| | 0.826123696 | NA |
| | 0.801664894 | 0.880408072 |
| | 0.905158266 | NA |
| | 0.141351227 | 0.266439591 |
| | 6.98E-11 | 3.08E-09 |
| | 0.992060816 | NA |
| | 0.889409773 | NA |
| | 0.792188023 | NA |
| | 0.889409301 | NA |
| | 0.066531106 | 0.150909675 |
| | 0.833288277 | NA |
| NA | | NA |
| NA | | NA |
| | 0.001368821 | 0.006500361 |
| | 0.038476604 | 0.099121041 |
| NA | | NA |
| | 0.100456717 | 0.206692875 |
| | 2.22E-11 | 1.12E-09 |
| NA | | NA |
| | 0.625169362 | NA |
| NA | | NA |
| | 5.63E-06 | 5.91E-05 |
| NA | | NA |
| | 0.99205891 | NA |
| | 0.313037709 | NA |
| | 0.992061292 | NA |
| | 0.80057007 | 0.879777708 |
| | 0.881019036 | 0.932396598 |
| | 0.766965459 | NA |
| | 0.123340307 | 0.240920638 |
| NA | | NA |
| NA | | NA |
| NA | | NA |
| NA | | NA |
| | 0.182312464 | 0.322450418 |
| NA | | NA |
| | 1.79E-08 | 3.94E-07 |
| NA | | NA |
| | 0.992253023 | NA |
| | 2.82E-09 | 7.83E-08 |
| NA | | NA |
| | 0.215577334 | NA |

| | | |
|---|---|---|
| | 0.316029612 | 0.478203848 |
| | 1.31E-05 | 0.000123003 |
| | 0.930097093 | 0.959774534 |
| | 0.905158739 | NA |
| | 0.34252666 | 0.505922037 |
| | 0.905158739 | NA |
| | 0.158628546 | 0.290446448 |
| NA | | NA |
| | 0.003706566 | 0.01491986 |
| | 0.363936703 | NA |
| | 0.698378281 | NA |
| | 0.726054041 | NA |
| | 0.040427376 | 0.102831549 |
| | 0.842798658 | NA |
| | 0.889409773 | NA |
| | 0.385614763 | 0.549423474 |
| | 1.40E-07 | 2.38E-06 |
| | 0.229927567 | NA |
| | 0.084068554 | NA |
| | 0.857094946 | NA |
| NA | | NA |
| | 0.405051957 | 0.568268332 |
| NA | | NA |
| | 0.105551876 | 0.214552776 |
| | 0.055297485 | 0.130919405 |
| | 0.582196539 | NA |
| NA | | NA |
| | 0.704533485 | NA |
| | 0.929110286 | 0.959432531 |
| | 0.008354808 | 0.029093438 |
| | 0.446914898 | 0.606173216 |
| | 0.124234676 | 0.242221186 |
| | 0.354902519 | 0.519484363 |
| | 0.844552425 | NA |
| | 0.807252815 | NA |
| | 4.28E-08 | 8.50E-07 |
| | 0.085472311 | 0.182629802 |
| NA | | NA |
| NA | | NA |
| NA | | NA |
| NA | | NA |
| | 0.012429239 | 0.040091868 |
| | 0.006448467 | 0.023635768 |
| | 0.624993103 | NA |
| NA | | NA |
| | 0.992061292 | NA |
| | 0.233570277 | 0.385202381 |

| | | |
|---|---|---|
| | 0.617375597 | NA |
| NA | | NA |
| | 1.99E-06 | 2.39E-05 |
| | 0.020272229 | 0.059414823 |
| | 0.725559332 | 0.830426215 |
| | 5.96E-10 | 2.01E-08 |
| | 0.029505142 | 0.080404991 |
| | 6.27E-11 | 2.80E-09 |
| | 0.096885904 | 0.201031614 |
| | 0.04539598 | 0.112562142 |
| | 0.353679573 | NA |
| | 0.782009566 | NA |
| | 0.833429472 | 0.900819299 |
| | 0.051844362 | 0.124802712 |
| | 0.061976615 | NA |
| | 0.889409301 | NA |
| | 0.187916702 | 0.32935293 |
| | 0.033854891 | 0.089676664 |
| NA | | NA |
| | 0.74431898 | NA |
| NA | | NA |
| NA | | NA |
| | 0.461006171 | 0.618401619 |
| | 2.08E-22 | 1.52E-19 |
| NA | | NA |
| NA | | NA |
| | 0.208442996 | 0.355124505 |
| NA | | NA |
| | 0.701734783 | 0.813761835 |
| | 0.797157316 | NA |
| | 0.000303852 | 0.001806252 |
| | 0.889409301 | NA |
| | 0.00275496 | 0.011723083 |
| NA | | NA |
| | 0.500799385 | 0.653547133 |
| | 0.001000198 | 0.004978093 |
| | 0.88804077 | NA |
| | 0.502947609 | 0.655274222 |
| | 0.274896087 | 0.432100443 |
| NA | | NA |
| NA | | NA |
| NA | | NA |
| NA | | NA |
| NA | | NA |
| | 0.009178847 | 0.03136118 |
| NA | | NA |
| NA | | NA |

| | |
|---|---|
| NA | NA |
| NA | NA |
| NA | NA |
| NA | NA |
| NA | NA |
| NA | NA |
| NA | NA |
| NA | NA |
| NA | NA |
| NA | NA |
| NA | NA |
| 2.71E-05 | 0.000229664 |
| 0.905158266 | NA |
| NA | NA |
| 0.884433193 | NA |
| 0.764144253 | NA |
| 0.303206892 | 0.463883519 |
| 0.992061292 | NA |
| 0.369874119 | 0.534463574 |
| 0.996255247 | NA |
| 0.873734187 | NA |
| 0.000411484 | 0.002341926 |
| 0.810521359 | 0.886546735 |
| 0.001451973 | 0.006828844 |
| NA | NA |
| NA | NA |
| 0.752232519 | NA |
| 0.20079622 | 0.345978296 |
| 0.477202731 | NA |
| 0.030789259 | 0.083161297 |
| NA | NA |
| 0.088483546 | 0.187789441 |
| NA | NA |
| 0.64145485 | 0.767366984 |
| NA | NA |
| 0.196742619 | 0.340822907 |
| 0.962483811 | 0.979205986 |
| NA | NA |
| NA | NA |
| 0.776593555 | NA |
| 0.183435461 | NA |
| 0.973419046 | NA |
| 0.442201932 | 0.601891292 |
| NA | NA |
| 0.40147413 | 0.564945236 |
| 0.364347304 | 0.529278245 |
| 0.889409773 | NA |

| | |
|---|---|
| 0.115365522 | 0.229487409 |
| 0.058731977 | 0.137245449 |
| 0.242067638 | 0.394788099 |
| 0.516044766 | 0.667166226 |
| NA | NA |
| 0.119911959 | 0.235969189 |
| 0.868377358 | NA |
| 0.784537311 | NA |
| 0.992059387 | NA |
| NA | NA |
| 0.710806153 | NA |
| 0.98936982 | NA |
| 0.272995402 | NA |
| NA | NA |
| 0.889409301 | NA |
| 0.000146842 | 0.000965422 |
| 0.739644352 | 0.84019711 |
| 0.001681131 | 0.007698332 |
| NA | NA |
| 0.792188023 | NA |
| 0.148268175 | NA |
| 0.942979424 | 0.967701896 |
| NA | NA |
| NA | NA |
| 0.992062245 | NA |
| NA | NA |
| 1.10E-06 | 1.41E-05 |
| 0.691883133 | NA |
| 0.821072584 | 0.893632676 |
| 7.28E-05 | 0.000533089 |
| 0.484798211 | 0.639918259 |
| 0.008472373 | 0.029414205 |
| 0.450347184 | 0.608848242 |
| NA | NA |
| 0.899711594 | 0.943494121 |
| 0.989005504 | NA |
| 0.247686582 | 0.401306419 |
| 0.919485945 | 0.954020567 |
| 0.904947233 | NA |
| NA | NA |
| 0.001190389 | 0.005772915 |
| 0.020198447 | 0.059272712 |
| 1.05E-10 | 4.43E-09 |
| 0.41880215 | 0.580933042 |
| 0.538260592 | 0.68615258 |
| 0.906770031 | 0.947360419 |
| 0.450179273 | 0.608829607 |

| | | |
|---|---|---|
| | 0.184961228 | 0.325759031 |
| | 0.168129445 | 0.303464942 |
| | 0.773470979 | 0.863479879 |
| | 0.992060816 | NA |
| NA | | NA |
| NA | | NA |
| NA | | NA |
| NA | | NA |
| | 0.579891732 | NA |
| | 0.239855642 | 0.392151741 |
| | 1.72E-05 | 0.000155948 |
| NA | | NA |
| | 0.069901825 | 0.1562951 |
| NA | | NA |
| | 7.51E-07 | 1.02E-05 |
| | 0.177779431 | 0.316717388 |
| | 2.09E-10 | 8.13E-09 |
| | 9.63E-07 | 1.27E-05 |
| NA | | NA |
| | 0.074950027 | 0.165193982 |
| | 0.017399539 | 0.052584155 |
| | 0.889409301 | NA |
| NA | | NA |
| | 0.228139425 | NA |
| NA | | NA |
| NA | | NA |
| NA | | NA |
| | 0.297571781 | NA |
| | 0.221704833 | NA |
| | 0.905158739 | NA |
| | 0.000128935 | 0.000866364 |
| | 0.000733398 | 0.003847272 |
| | 0.836813554 | NA |
| | 0.913727679 | NA |
| | 0.905158266 | NA |
| NA | | NA |
| | 0.509788446 | NA |
| NA | | NA |
| NA | | NA |
| NA | | NA |
| NA | | NA |
| NA | | NA |
| | 0.285669814 | 0.444611486 |
| | 0.012649228 | 0.040680199 |
| | 0.042039904 | 0.106097488 |
| | 0.000451862 | 0.002535117 |
| NA | | NA |

| | |
|---|---|
| NA | NA |
| 0.004621089 | 0.017925188 |
| 0.889409773 | NA |
| NA | NA |
| NA | NA |
| 0.557079686 | 0.702112887 |
| NA | NA |
| NA | NA |
| 0.441586735 | NA |
| 0.002400404 | 0.010454277 |
| 0.187436564 | 0.328878906 |
| NA | NA |
| NA | NA |
| NA | NA |
| NA | NA |
| 0.905158266 | NA |
| 0.821434133 | 0.893671826 |
| 0.399754383 | 0.563355571 |
| 0.684594793 | NA |
| 0.822542877 | NA |
| 0.857135932 | 0.916707965 |
| 0.592921357 | NA |
| NA | NA |
| 0.083705293 | 0.179922135 |
| NA | NA |
| NA | NA |
| 6.51E-08 | 1.22E-06 |
| NA | NA |
| 0.017610985 | NA |
| NA | NA |
| NA | NA |
| NA | NA |
| NA | NA |
| 0.889410245 | NA |
| 0.309209152 | 0.470346617 |
| NA | NA |
| NA | NA |
| 0.016325559 | 0.050033909 |
| 0.028333114 | 0.077829228 |
| NA | NA |
| NA | NA |
| NA | NA |
| 0.129740569 | 0.249883855 |
| NA | NA |
| 0.889027042 | 0.937347905 |
| 0.379068919 | 0.543446784 |
| NA | NA |

| | | |
|---|---|---|
| | 0.002159676 | 0.009547881 |
| NA | | NA |
| | 0.753111281 | NA |
| | 0.925470286 | NA |
| | 0.368981876 | NA |
| | 0.117524473 | 0.232522767 |
| | 0.467641983 | NA |
| NA | | NA |
| NA | | NA |
| NA | | NA |
| NA | | NA |
| NA | | NA |
| NA | | NA |
| NA | | NA |
| | 0.992037435 | NA |
| | 0.171422967 | 0.307938937 |
| | 0.074410872 | 0.16425295 |
| | 0.356238494 | NA |
| NA | | NA |
| | 0.03875066 | 0.099605026 |
| NA | | NA |
| NA | | NA |
| NA | | NA |
| NA | | NA |
| NA | | NA |
| | 0.784353866 | NA |
| | 0.814808235 | 0.889702743 |
| | 0.141999709 | 0.267402891 |
| | 0.798288368 | 0.87819319 |
| | 0.889410717 | NA |
| NA | | NA |
| | 0.889409301 | NA |
| NA | | NA |
| NA | | NA |
| NA | | NA |
| NA | | NA |
| | 0.013340605 | NA |
| | 0.90515732 | NA |
| NA | | NA |
| NA | | NA |
| NA | | NA |
| NA | | NA |
| | 0.905158266 | NA |
| NA | | NA |
| NA | | NA |
| NA | | NA |
| NA | | NA |

| | | |
|---|---|---|
| NA | NA | |
| NA | NA | |
| NA | NA | |
| NA | NA | |
| NA | NA | |
| 0.992060816 | NA | |
| NA | NA | |
| NA | NA | |
| NA | NA | |
| NA | NA | |
| NA | NA | |
| NA | NA | |
| NA | NA | |
| 0.800369521 | NA | |
| NA | NA | |
| NA | NA | |
| NA | NA | |
| NA | NA | |
| NA | NA | |
| NA | NA | |
| NA | NA | |
| NA | NA | |
| NA | NA | |
| NA | NA | |
| NA | NA | |
| NA | NA | |
| NA | NA | |
| 0.803085963 | | 0.881527852 |
| 0.784353866 | NA | |
| NA | NA | |
| 0.564730532 | NA | |
| 0.506131811 | NA | |
| 5.72E-08 | | 1.09E-06 |
| NA | NA | |
| NA | NA | |
| 0.649707963 | NA | |
| NA | NA | |
| 0.750604009 | NA | |
| 0.992061292 | NA | |
| NA | NA | |
| NA | NA | |
| 0.293868355 | NA | |
| 0.992060816 | NA | |
| NA | NA | |
| NA | NA | |
| NA | NA | |
| 0.99205891 | NA | |

| | |
|---|---|
| NA | NA |
| 0.344832516 | NA |
| 0.991688155 | NA |
| 0.721240786 | NA |
| 0.704758221 | 0.816268766 |
| NA | NA |
| NA | NA |
| NA | NA |
| 0.889409301 | NA |
| NA | NA |
| NA | NA |
| NA | NA |
| NA | NA |
| NA | NA |
| NA | NA |
| NA | NA |
| 0.905158266 | NA |
| NA | NA |
| NA | NA |
| NA | NA |
| NA | NA |
| NA | NA |
| NA | NA |
| NA | NA |
| NA | NA |
| 0.889409301 | NA |
| NA | NA |
| NA | NA |
| NA | NA |
| NA | NA |
| NA | NA |
| NA | NA |
| NA | NA |
| NA | NA |
| NA | NA |
| NA | NA |
| NA | NA |
| NA | NA |
| 0.889409773 | NA |
| NA | NA |
| NA | NA |
| NA | NA |
| NA | NA |
| NA | NA |
| NA | NA |
| NA | NA |
| NA | NA |

| | | |
|---|---|---|
| NA | NA | |
| NA | NA | |
| 0.992061292 | NA | |
| NA | NA | |
| NA | NA | |
| NA | NA | |
| NA | NA | |
| NA | NA | |
| NA | NA | |
| NA | NA | |
| NA | NA | |
| NA | NA | |
| NA | NA | |
| NA | NA | |
| 0.688668124 | NA | |
| 0.759389656 | NA | |
| NA | NA | |
| 0.905158739 | NA | |
| 0.187677567 | | 0.329170263 |
| NA | NA | |
| NA | NA | |
| NA | NA | |
| NA | NA | |
| NA | NA | |
| NA | NA | |
| NA | NA | |
| NA | NA | |
| NA | NA | |
| NA | NA | |
| NA | NA | |
| 0.60388897 | | 0.739216115 |
| NA | NA | |
| NA | NA | |
| 0.520938643 | NA | |
| 0.992061292 | NA | |
| NA | NA | |
| NA | NA | |
| NA | NA | |
| NA | NA | |
| NA | NA | |
| NA | NA | |
| NA | NA | |
| NA | NA | |
| NA | NA | |
| NA | NA | |
| NA | NA | |
| NA | NA | |

| | |
|---|---|
| NA | NA |
| NA | NA |
| NA | NA |
| NA | NA |
| 0.706875149 | NA |
| 6.80E-18 | 1.74E-15 |
| 0.537427799 | 0.68560838 |
| 0.920567676 | 0.954536733 |
| 0.380051051 | 0.544304363 |
| NA | NA |
| 0.396414231 | 0.56001416 |
| 0.019406014 | 0.057368964 |
| 0.99205891 | NA |
| 0.005191953 | 0.019786549 |
| NA | NA |
| 0.110026931 | 0.221394794 |
| 0.258085161 | 0.41294754 |
| 0.585203503 | NA |
| 0.345762373 | 0.509194283 |
| 0.858335656 | 0.917231625 |
| 0.750765427 | 0.848266427 |
| 0.007994586 | 0.028084003 |
| 0.774088041 | NA |
| 0.407388542 | NA |
| 0.889409773 | NA |
| 0.223098765 | 0.372577822 |
| 6.66E-08 | 1.25E-06 |
| NA | NA |
| 0.900098131 | 0.94360613 |
| 3.38E-13 | 2.72E-11 |
| 0.884054604 | 0.934213429 |
| 0.855621215 | NA |
| 0.889409301 | NA |
| 0.31834026 | NA |
| 0.017529459 | 0.052911255 |
| 0.000416906 | 0.002367428 |
| 0.146328242 | NA |
| 0.940432585 | 0.966159014 |
| 0.210141882 | 0.35702171 |
| 0.024306181 | 0.06888738 |
| 0.703622919 | NA |
| 0.537585334 | 0.685689844 |
| NA | NA |
| NA | NA |
| 0.972057175 | 0.985430198 |
| 0.725177738 | 0.830307868 |
| 0.065247689 | 0.148900728 |

| | |
|---|---|
| NA | NA |
| NA | NA |
| NA | NA |
| 0.889409301 | NA |
| 7.96E-05 | 0.000575303 |
| NA | NA |
| NA | NA |
| 0.889409301 | NA |
| NA | NA |
| NA | NA |
| NA | NA |
| NA | NA |
| NA | NA |
| NA | NA |
| NA | NA |
| NA | NA |
| NA | NA |
| NA | NA |
| NA | NA |
| NA | NA |
| NA | NA |
| 0.000575553 | 0.003126808 |
| NA | NA |
| 0.893156782 | NA |
| 0.658718924 | NA |
| 0.889409301 | NA |
| NA | NA |
| 4.67E-07 | 6.79E-06 |
| 1.84E-08 | 4.03E-07 |
| NA | NA |
| NA | NA |
| 0.009566331 | 0.032442789 |
| 0.012694941 | 0.040806629 |
| 0.217652664 | 0.365735756 |
| 0.725180798 | 0.830307868 |
| 0.028818442 | 0.078915597 |
| 0.133532115 | 0.255131968 |
| 0.118509789 | 0.233917426 |
| 0.707613133 | NA |
| 0.000263062 | 0.00159883 |
| 0.650099594 | 0.774707602 |
| NA | NA |
| 0.028304238 | 0.077769364 |
| 0.001439023 | 0.006782463 |
| 0.054033778 | 0.128718566 |
| 0.103339694 | 0.211252844 |
| 0.533349795 | 0.682189431 |

| | | |
|---|---|---|
| | 0.905158739 | NA |
| NA | | NA |
| NA | | NA |
| | 0.905158266 | NA |
| | 0.905158739 | NA |
| | 0.788582525 | NA |
| NA | | NA |
| NA | | NA |
| NA | | NA |
| NA | | NA |
| NA | | NA |
| NA | | NA |
| | 0.905157793 | NA |
| | 0.236945853 | NA |
| | 0.992060816 | NA |
| NA | | NA |
| NA | | NA |
| NA | | NA |
| NA | | NA |
| NA | | NA |
| | 0.014783032 | 0.046123983 |
| | 0.048970026 | NA |
| NA | | NA |
| | 0.009880145 | 0.03329621 |
| | 0.205699012 | 0.351942154 |
| | 0.037885719 | 0.097985654 |
| | 0.952244445 | 0.973288471 |
| NA | | NA |
| | 0.21302025 | 0.360259066 |
| | 0.969039115 | 0.983687589 |
| NA | | NA |
| NA | | NA |
| NA | | NA |
| | 0.992060816 | NA |
| | 0.659441838 | 0.781623359 |
| | 0.210177394 | NA |
| | 0.331450759 | 0.493834079 |
| | 0.006660382 | 0.024254613 |
| | 1.20E-17 | 2.81E-15 |
| NA | | NA |
| | 0.45860524 | 0.615933484 |
| NA | | NA |
| NA | | NA |
| | 0.362799865 | NA |
| NA | | NA |
| NA | | NA |
| NA | | NA |

| | |
|---|---|
| NA | NA |
| NA | NA |
| NA | NA |
| 0.889409301 | NA |
| NA | NA |
| NA | NA |
| 0.110790457 | NA |
| 3.75E-10 | 1.35E-08 |
| NA | NA |
| NA | NA |
| NA | NA |
| NA | NA |
| NA | NA |
| NA | NA |
| NA | NA |
| NA | NA |
| NA | NA |
| 0.675038414 | 0.793381866 |
| 0.99205891 | NA |
| NA | NA |
| NA | NA |
| NA | NA |
| 0.558053325 | 0.702908387 |
| 0.652356559 | NA |
| NA | NA |
| NA | NA |
| NA | NA |
| 0.931791953 | 0.960440269 |
| 0.102283172 | 0.209581516 |
| NA | NA |
| NA | NA |
| 1.49E-06 | 1.85E-05 |
| 0.858148483 | 0.917175946 |
| 0.34179806 | NA |
| 0.905158266 | NA |
| 0.673868656 | 0.792708085 |
| 0.001471242 | 0.006898785 |
| 0.002161963 | 0.009552223 |
| NA | NA |
| 0.081499705 | 0.176459478 |
| 0.703206678 | NA |
| 0.992061292 | NA |
| 0.905157793 | NA |
| 0.997242387 | NA |
| 0.155413577 | 0.286254549 |
| 0.647025672 | NA |
| 0.016573117 | 0.050637132 |

| | |
|---|---|
| 0.718760399 | 0.825457274 |
| NA | NA |
| NA | NA |
| 0.189243669 | 0.331050633 |
| 0.061136888 | 0.141823888 |
| NA | NA |
| 0.16447593 | NA |
| NA | NA |
| NA | NA |
| 0.761473545 | NA |
| 0.002834758 | 0.011993921 |
| 0.129632336 | 0.24978909 |
| 0.347727212 | 0.511538945 |
| NA | NA |
| NA | NA |
| NA | NA |
| 9.37E-07 | 1.24E-05 |
| 0.133554956 | NA |
| NA | NA |
| 0.414661753 | 0.577117473 |
| 0.767117272 | NA |
| NA | NA |
| NA | NA |
| 0.889409301 | NA |
| 0.616377951 | NA |
| 0.392386181 | 0.555906367 |
| NA | NA |
| NA | NA |
| 0.06649819 | 0.150859417 |
| 0.562455573 | 0.706174743 |
| NA | NA |
| 0.090223623 | 0.190556943 |
| 0.725115864 | 0.830307868 |
| 9.68E-06 | 9.45E-05 |
| 0.078477246 | 0.171418981 |
| NA | NA |
| NA | NA |
| NA | NA |
| 0.889409773 | NA |
| 0.90515732 | NA |
| NA | NA |
| 0.541526916 | 0.688916606 |
| 0.791738938 | 0.874536989 |
| 0.741866927 | 0.841850853 |
| 0.905158266 | NA |
| 0.306525849 | 0.467268387 |
| 0.903181625 | 0.945177713 |

| | | |
|---|---|---|
| | 0.000793569 | 0.004098303 |
| | 0.905158266 | NA |
| | 0.002126874 | 0.009427017 |
| NA | | NA |
| | 0.188364543 | 0.329848245 |
| | 0.014231547 | 0.044714628 |
| NA | | NA |
| | 0.00603863 | 0.022388778 |
| | 0.889409301 | NA |
| | 0.251399816 | 0.405560429 |
| | 0.996857914 | NA |
| NA | | NA |
| NA | | NA |
| | 0.889409773 | NA |
| NA | | NA |
| | 0.712813736 | 0.821145569 |
| | 0.992061769 | NA |
| | 4.19E-10 | 1.49E-08 |
| | 0.795921435 | NA |
| NA | | NA |
| | 0.625388486 | 0.755340311 |
| | 0.000891467 | 0.004516778 |
| | 0.008951313 | 0.030736925 |
| | 0.004244412 | 0.01669707 |
| | 0.905158266 | NA |
| NA | | NA |
| | 0.905158266 | NA |
| | 0.992061292 | NA |
| | 0.993261201 | 0.99666747 |
| | 0.726237039 | 0.830985313 |
| | 0.08642416 | 0.184287058 |
| NA | | NA |
| | 0.992061292 | NA |
| | 0.889410245 | NA |
| | 0.905158739 | NA |
| | 0.636002339 | NA |
| NA | | NA |
| NA | | NA |
| NA | | NA |
| | 0.327207105 | 0.489365566 |
| NA | | NA |
| | 0.652669235 | NA |
| NA | | NA |
| | 0.851102566 | 0.912750148 |
| | 0.273423606 | NA |
| NA | | NA |
| NA | | NA |

| | |
|---|---|
| 8.13E-08 | 1.47E-06 |
| 0.818029815 | 0.891691075 |
| 0.315626685 | 0.477824638 |
| 0.529209213 | 0.678592999 |
| 0.212510914 | 0.359627555 |
| 0.387365284 | 0.551023791 |
| 0.590949558 | 0.728875998 |
| 0.991593318 | 0.995807742 |
| 0.520033644 | NA |
| 0.349416742 | 0.513439657 |
| NA | NA |
| 0.002838167 | 0.012006031 |
| 0.10429025 | 0.212596428 |
| NA | NA |
| NA | NA |
| NA | NA |
| NA | NA |
| NA | NA |
| NA | NA |
| NA | NA |
| NA | NA |
| NA | NA |
| NA | NA |
| NA | NA |
| NA | NA |
| NA | NA |
| NA | NA |
| NA | NA |
| NA | NA |
| NA | NA |
| NA | NA |
| NA | NA |
| NA | NA |
| NA | NA |
| NA | NA |
| NA | NA |
| 0.992060816 | NA |
| NA | NA |
| NA | NA |
| NA | NA |
| NA | NA |
| NA | NA |
| NA | NA |
| NA | NA |
| NA | NA |
| NA | NA |
| NA | NA |

| | |
|---|---|
| NA | NA |
| NA | NA |
| NA | NA |
| NA | NA |
| NA | NA |
| NA | NA |
| NA | NA |
| NA | NA |
| NA | NA |
| NA | NA |
| NA | NA |
| NA | NA |
| NA | NA |
| NA | NA |
| NA | NA |
| NA | NA |
| 0.488686115 | 0.643117488 |
| 0.288967479 | 0.448345713 |
| 0.598879489 | NA |
| 0.000427396 | 0.002418239 |
| NA | NA |
| 0.154371855 | 0.284861476 |
| 0.248612767 | 0.402436865 |
| 0.374645686 | 0.538701923 |
| 0.012762082 | 0.040960033 |
| 0.510762241 | 0.662325974 |
| 0.004562541 | 0.017723126 |
| NA | NA |
| NA | NA |
| 0.973082706 | 0.985912235 |
| 0.989174267 | NA |
| 0.748890654 | NA |
| NA | NA |
| 0.273998068 | NA |
| 6.11E-05 | 0.000459346 |
| NA | NA |
| 0.236723259 | 0.388563765 |
| 0.000920241 | 0.00464534 |
| NA | NA |
| NA | NA |
| 0.652286111 | 0.776091825 |
| 0.195434371 | 0.33919881 |
| 0.840795364 | 0.905826984 |
| 0.99205891 | NA |
| 0.0170628 | 0.051824796 |
| 0.713352248 | 0.821500271 |
| 0.067142713 | 0.15177958 |

| | |
|---|---|
| 0.342909992 | 0.506314652 |
| 0.889283189 | NA |
| 0.37395698 | 0.537944404 |
| NA | NA |
| 0.519975625 | NA |
| 0.984077419 | NA |
| NA | NA |
| NA | NA |
| NA | NA |
| 0.069872878 | 0.156246285 |
| NA | NA |
| 0.646269177 | NA |
| 0.821698639 | 0.893910585 |
| 0.117422702 | 0.232385693 |
| NA | NA |
| 0.344454532 | 0.507847259 |
| 0.008474044 | 0.029415354 |
| 0.504842998 | 0.656861254 |
| 0.592452069 | 0.729926607 |
| NA | NA |
| 3.45E-15 | 4.49E-13 |
| 7.94E-13 | 5.75E-11 |
| 0.000313415 | 0.001855562 |
| NA | NA |
| 0.058087903 | 0.136029889 |
| 9.76E-05 | 0.000683742 |
| 0.167626051 | 0.302780329 |
| 0.730582851 | NA |
| 0.816533267 | NA |
| NA | NA |
| 0.807879735 | NA |
| 0.905158739 | NA |
| 2.15E-06 | 2.56E-05 |
| 0.63965711 | NA |
| NA | NA |
| 0.905158266 | NA |
| 0.032211568 | 0.08630232 |
| 0.344530881 | 0.507925722 |
| 0.926941738 | 0.958274734 |
| 0.77102239 | 0.8614909 |
| 0.294617914 | 0.454475658 |
| 0.889409773 | NA |
| 0.125486296 | NA |
| 0.011021477 | 0.036410502 |
| 0.00170802 | 0.007802456 |
| 0.867274707 | NA |
| 0.929714673 | 0.959686698 |

| | | | |
|---|---|---|---|
| NA | | NA | |
| | 0.004439708 | | 0.017325671 |
| NA | | NA | |
| NA | | NA | |
| NA | | NA | |
| NA | | NA | |
| | 0.628405676 | | 0.756928011 |
| | 0.426186151 | | 0.587410349 |
| | 0.628310013 | NA | |
| | 0.007156145 | | 0.025689357 |
| NA | | NA | |
| NA | | NA | |
| NA | | NA | |
| NA | | NA | |
| NA | | NA | |
| NA | | NA | |
| NA | | NA | |
| NA | | NA | |
| NA | | NA | |
| NA | | NA | |
| NA | | NA | |
| NA | | NA | |
| NA | | NA | |
| NA | | NA | |
| NA | | NA | |
| NA | | NA | |
| NA | | NA | |
| NA | | NA | |
| NA | | NA | |
| NA | | NA | |
| | 0.623549394 | NA | |
| NA | | NA | |
| NA | | NA | |
| NA | | NA | |
| | 0.523426268 | NA | |
| NA | | NA | |
| NA | | NA | |
| NA | | NA | |
| NA | | NA | |
| NA | | NA | |
| NA | | NA | |
| NA | | NA | |
| | 0.905158266 | NA | |
| NA | | NA | |
| | 0.184179542 | NA | |
| | 0.992061292 | NA | |
| | 0.821809783 | NA | |

| | |
|---|---|
| 1.31E-05 | 0.000123484 |
| NA | NA |
| 0.084139212 | 0.180560702 |
| NA | NA |
| 0.233418401 | 0.385009739 |
| NA | NA |
| 0.991119509 | NA |
| NA | NA |
| 0.594187042 | 0.731458306 |
| 0.959768598 | 0.977711943 |
| 0.009526209 | 0.032326691 |
| 0.0908218 | 0.19147042 |
| NA | NA |
| 0.025260166 | NA |
| NA | NA |
| 0.645783748 | NA |
| 0.008199092 | 0.028646598 |
| NA | NA |
| 0.361704777 | 0.526562927 |
| NA | NA |
| 0.979027593 | NA |
| 0.983380278 | NA |
| 0.012921578 | NA |
| NA | NA |
| 6.89E-08 | 1.28E-06 |
| 0.001501654 | 0.007016846 |
| 0.085307616 | NA |
| NA | NA |
| NA | NA |
| 0.788763375 | NA |
| 0.905158266 | NA |
| NA | NA |
| 0.270352757 | NA |
| NA | NA |
| NA | NA |
| 0.103700619 | 0.211710849 |
| 0.960010842 | 0.977731905 |
| 0.411135633 | 0.574052735 |
| 4.35E-06 | 4.73E-05 |
| 0.813836939 | NA |
| 0.905158266 | NA |
| NA | NA |
| NA | NA |
| NA | NA |
| NA | NA |
| 0.889409773 | NA |
| NA | NA |

| | |
|---|---|
| NA | NA |
| NA | NA |
| 3.89E-09 | 1.03E-07 |
| 2.26E-05 | 0.000197306 |
| NA | NA |
| 0.014808216 | 0.046186856 |
| 0.518207985 | NA |
| 0.002360983 | 0.010303027 |
| 0.168492595 | 0.304043921 |
| 0.526023142 | 0.676007618 |
| 0.591152677 | 0.729030756 |
| 0.382561249 | 0.546538252 |
| NA | NA |
| NA | NA |
| 3.00E-06 | 3.41E-05 |
| NA | NA |
| NA | NA |
| 0.889409773 | NA |
| NA | NA |
| 0.153884651 | 0.284201266 |
| 0.621679171 | NA |
| 3.26E-07 | 4.94E-06 |
| 0.801062068 | NA |
| 0.812617742 | 0.887995671 |
| 0.991918092 | NA |
| 0.905158739 | NA |
| 0.498982328 | NA |
| 3.87E-08 | 7.83E-07 |
| 0.087032532 | 0.185296429 |
| 0.439302533 | 0.599580105 |
| NA | NA |
| NA | NA |
| 0.885332346 | NA |
| 0.237183866 | 0.389116213 |
| 0.718935214 | NA |
| 0.499883809 | 0.652962261 |
| 0.73375332 | 0.836490974 |
| 0.581237601 | NA |
| 0.801067096 | NA |
| 0.992061292 | NA |
| NA | NA |
| 0.772862102 | NA |
| 0.419738708 | NA |
| 1.17E-10 | 4.90E-09 |
| 0.240025258 | 0.392370627 |
| 0.099276371 | 0.204830388 |
| NA | NA |

| | |
|---|---|
| NA | NA |
| NA | NA |
| NA | NA |
| 0.012881154 | 0.041229683 |
| NA | NA |
| 0.249884854 | 0.403769059 |
| 0.063090157 | 0.145067723 |
| NA | NA |
| NA | NA |
| 0.889409773 | NA |
| 0.031042555 | 0.083701309 |
| 0.772203995 | NA |
| NA | NA |
| 0.738721937 | 0.839639257 |
| 0.992060816 | NA |
| NA | NA |
| NA | NA |
| NA | NA |
| NA | NA |
| NA | NA |
| NA | NA |
| NA | NA |
| 0.239515075 | 0.391741634 |
| 0.945040238 | 0.968866415 |
| NA | NA |
| 0.133842789 | 0.255592122 |
| 0.225840006 | 0.375670077 |
| 0.539069842 | 0.686737518 |
| 0.238128567 | 0.390112295 |
| 0.880145841 | NA |
| 0.905158266 | NA |
| 0.320688943 | 0.483005701 |
| 0.738843381 | 0.839639257 |
| NA | NA |
| 0.679211562 | 0.796603466 |
| 0.784857687 | NA |
| 0.163280924 | 0.296823728 |
| 0.123669543 | 0.241434877 |
| 0.000841838 | 0.004306063 |
| 0.588995633 | NA |
| 0.980575688 | NA |
| NA | NA |
| 0.024581946 | 0.069543281 |
| 0.567661462 | 0.710353205 |
| NA | NA |
| 0.771188949 | 0.86163316 |
| 0.908441788 | NA |

| | |
|---|---|
| NA | NA |
| NA | NA |
| NA | NA |
| 0.790601632 | 0.873676273 |
| NA | NA |
| 0.052947824 | 0.126779865 |
| 0.787394472 | 0.871771467 |
| 0.138071638 | 0.261733003 |
| 0.007692758 | 0.027197448 |
| 0.00084741 | 0.004327505 |
| 0.66054454 | NA |
| 0.284505692 | 0.443365304 |
| 0.441058171 | 0.60093047 |
| 0.864441469 | NA |
| 0.786115238 | NA |
| 0.663653767 | 0.784953244 |
| 0.855981091 | NA |
| NA | NA |
| 0.905158739 | NA |
| NA | NA |
| 0.992059387 | NA |
| 0.866545673 | NA |
| 0.787211038 | 0.871771467 |
| 0.96264516 | 0.979279397 |
| 0.930949125 | 0.96015743 |
| 3.31E-05 | 0.000272316 |
| 0.003052704 | 0.012736903 |
| 0.072793412 | 0.161347586 |
| 0.630966264 | NA |
| 0.666667417 | NA |
| 0.412887067 | NA |
| 5.24E-15 | 6.56E-13 |
| 0.904947233 | NA |
| NA | NA |
| NA | NA |
| NA | NA |
| 0.99205891 | NA |
| NA | NA |
| NA | NA |
| 0.164983442 | 0.299188323 |
| NA | NA |
| 5.73E-06 | 5.99E-05 |
| 0.909364283 | NA |
| 0.807252815 | NA |
| 0.004366324 | 0.017090928 |
| 0.562419634 | NA |
| 0.132739158 | 0.253926228 |

| | |
|---|---|
| 0.003154692 | 0.013097727 |
| 0.889409301 | NA |
| 1.00E-06 | 1.31E-05 |
| NA | NA |
| NA | NA |
| 0.893356605 | NA |
| 0.990075047 | NA |
| 0.905158739 | NA |
| 0.572675243 | 0.715078425 |
| 0.796684688 | NA |
| NA | NA |
| NA | NA |
| NA | NA |
| NA | NA |
| 0.889409773 | NA |
| 0.992061292 | NA |
| 0.25201948 | 0.406291445 |
| NA | NA |
| 0.565300883 | NA |
| 0.992062245 | NA |
| 0.224303363 | 0.373935435 |
| NA | NA |
| NA | NA |
| 0.054332043 | 0.12928883 |
| 0.162657306 | 0.295972139 |
| 0.206573196 | 0.353116987 |
| 0.678460932 | 0.795978181 |
| 0.869834515 | 0.925515189 |
| 0.111024806 | 0.222631229 |
| 0.266990782 | 0.423166441 |
| NA | NA |
| NA | NA |
| NA | NA |
| NA | NA |
| 0.983594448 | NA |
| NA | NA |
| NA | NA |
| NA | NA |
| NA | NA |
| NA | NA |
| NA | NA |
| NA | NA |
| NA | NA |
| NA | NA |
| NA | NA |
| NA | NA |
| NA | NA |

| | |
|---|---|
| NA | NA |
| NA | NA |
| NA | NA |
| NA | NA |
| NA | NA |
| NA | NA |
| NA | NA |
| NA | NA |
| NA | NA |
| NA | NA |
| NA | NA |
| NA | NA |
| NA | NA |
| NA | NA |
| NA | NA |
| NA | NA |
| NA | NA |
| NA | NA |
| NA | NA |
| NA | NA |
| NA | NA |
| NA | NA |
| NA | NA |
| NA | NA |
| NA | NA |
| NA | NA |
| NA | NA |
| NA | NA |
| NA | NA |
| NA | NA |
| NA | NA |
| NA | NA |
| NA | NA |
| NA | NA |
| NA | NA |
| NA | NA |
| NA | NA |
| NA | NA |
| NA | NA |
| NA | NA |
| NA | NA |
| NA | NA |
| NA | NA |
| 0.905158266 | NA |
| NA | NA |
| NA | NA |
| NA | NA |

| | | |
|---|---|---|
| NA | | NA |
| NA | | NA |
| NA | | NA |
| NA | | NA |
| NA | | NA |
| NA | | NA |
| NA | | NA |
| NA | | NA |
| NA | | NA |
| NA | | NA |
| NA | | NA |
| NA | | NA |
| NA | | NA |
| NA | | NA |
| NA | | NA |
| NA | | NA |
| NA | | NA |
| NA | | NA |
| NA | | NA |
| NA | | NA |
| NA | | NA |
| NA | | NA |
| NA | | NA |
| NA | | NA |
| NA | | NA |
| NA | | NA |
| NA | | NA |
| NA | | NA |
| NA | | NA |
| NA | | NA |
| NA | | NA |
| NA | | NA |
| | 0.905158266 | NA |
| NA | | NA |
| NA | | NA |
| NA | | NA |
| NA | | NA |
| NA | | NA |
| NA | | NA |
| NA | | NA |
| NA | | NA |
| NA | | NA |
| NA | | NA |
| NA | | NA |
| NA | | NA |
| NA | | NA |
| NA | | NA |

| | | |
|---|---|---|
| | 0.889409773 | NA |
| NA | | NA |
| NA | | NA |
| NA | | NA |
| NA | | NA |
| NA | | NA |
| NA | | NA |
| NA | | NA |
| NA | | NA |
| NA | | NA |
| NA | | NA |
| NA | | NA |
| NA | | NA |
| | 0.736231406 | NA |
| NA | | NA |
| NA | | NA |
| NA | | NA |
| NA | | NA |
| NA | | NA |
| NA | | NA |
| NA | | NA |
| NA | | NA |
| NA | | NA |
| NA | | NA |
| NA | | NA |
| | 0.844068118 | NA |
| NA | | NA |
| NA | | NA |
| NA | | NA |
| NA | | NA |
| NA | | NA |
| NA | | NA |
| NA | | NA |
| NA | | NA |
| NA | | NA |
| NA | | NA |
| NA | | NA |
| NA | | NA |
| NA | | NA |
| NA | | NA |
| NA | | NA |
| NA | | NA |
| NA | | NA |
| NA | | NA |
| NA | | NA |
| NA | | NA |
| NA | | NA |

| | |
|---|---|
| NA | NA |
| NA | NA |
| NA | NA |
| NA | NA |
| NA | NA |
| NA | NA |
| NA | NA |
| NA | NA |
| NA | NA |
| NA | NA |
| NA | NA |
| NA | NA |
| NA | NA |
| NA | NA |
| NA | NA |
| NA | NA |
| NA | NA |
| NA | NA |
| NA | NA |
| NA | NA |
| NA | NA |
| NA | NA |
| NA | NA |
| NA | NA |
| NA | NA |
| NA | NA |
| NA | NA |
| NA | NA |
| NA | NA |
| NA | NA |
| NA | NA |
| NA | NA |
| NA | NA |
| NA | NA |
| NA | NA |
| NA | NA |
| NA | NA |
| NA | NA |
| NA | NA |
| NA | NA |
| NA | NA |
| NA | NA |
| NA | NA |
| NA | NA |
| NA | NA |
| NA | NA |
| NA | NA |

| | |
|---|---|
| NA | NA |
| NA | NA |
| NA | NA |
| 0.889409773 | NA |
| NA | NA |
| NA | NA |
| NA | NA |
| 0.440938658 | 0.600842196 |
| NA | NA |
| NA | NA |
| NA | NA |
| NA | NA |
| NA | NA |
| NA | NA |
| NA | NA |
| NA | NA |
| NA | NA |
| NA | NA |
| NA | NA |
| NA | NA |
| NA | NA |
| NA | NA |
| NA | NA |
| NA | NA |
| NA | NA |
| NA | NA |
| NA | NA |
| NA | NA |
| NA | NA |
| NA | NA |
| NA | NA |
| NA | NA |
| NA | NA |
| NA | NA |
| NA | NA |
| NA | NA |
| NA | NA |
| NA | NA |
| NA | NA |
| NA | NA |
| NA | NA |
| NA | NA |
| NA | NA |
| NA | NA |
| NA | NA |
| NA | NA |
| NA | NA |

| | | |
|---|---|---|
| NA | | NA |
| NA | | NA |
| NA | | NA |
| NA | | NA |
| NA | | NA |
| NA | | NA |
| NA | | NA |
| NA | | NA |
| NA | | NA |
| NA | | NA |
| NA | | NA |
| NA | | NA |
| NA | | NA |
| NA | | NA |
| NA | | NA |
| NA | | NA |
| NA | | NA |
| NA | | NA |
| NA | | NA |
| NA | | NA |
| NA | | NA |
| NA | | NA |
| NA | | NA |
| NA | | NA |
| NA | | NA |
| NA | | NA |
| NA | | NA |
| NA | | NA |
| NA | | NA |
| NA | | NA |
| NA | | NA |
| NA | | NA |
| NA | | NA |
| NA | | NA |
| NA | | NA |
| NA | | NA |
| NA | | NA |
| NA | | NA |
| NA | | NA |
| NA | | NA |
| NA | | NA |
| NA | | NA |
| NA | | NA |
| NA | | NA |
| | 0.992061292 | NA |
| NA | | NA |
| NA | | NA |

| | |
|---|---|
| NA | NA |
| NA | NA |
| NA | NA |
| NA | NA |
| NA | NA |
| NA | NA |
| NA | NA |
| NA | NA |
| NA | NA |
| NA | NA |
| NA | NA |
| NA | NA |
| NA | NA |
| NA | NA |
| NA | NA |
| NA | NA |
| NA | NA |
| NA | NA |
| NA | NA |
| NA | NA |
| NA | NA |
| NA | NA |
| NA | NA |
| NA | NA |
| NA | NA |
| NA | NA |
| NA | NA |
| NA | NA |
| NA | NA |
| NA | NA |
| NA | NA |
| NA | NA |
| NA | NA |
| NA | NA |
| NA | NA |
| NA | NA |
| NA | NA |
| NA | NA |
| NA | NA |
| NA | NA |
| NA | NA |
| NA | NA |
| NA | NA |
| NA | NA |
| NA | NA |
| NA | NA |
| NA | NA |

| | | |
|---|---|---|
| NA | | NA |
| NA | | NA |
| NA | | NA |
| NA | | NA |
| NA | | NA |
| NA | | NA |
| NA | | NA |
| NA | | NA |
| NA | | NA |
| NA | | NA |
| NA | | NA |
| NA | | NA |
| NA | | NA |
| NA | | NA |
| NA | | NA |
| NA | | NA |
| NA | | NA |
| NA | | NA |
| NA | | NA |
| NA | | NA |
| NA | | NA |
| NA | | NA |
| NA | | NA |
| NA | | NA |
| NA | | NA |
| NA | | NA |
| NA | | NA |
| NA | | NA |
| NA | | NA |
| NA | | NA |
| NA | | NA |
| NA | | NA |
| NA | | NA |
| NA | | NA |
| NA | | NA |
| NA | | NA |
| NA | | NA |
| NA | | NA |
| NA | | NA |
| NA | | NA |
| | 0.992061292 | NA |
| NA | | NA |
| NA | | NA |
| NA | | NA |
| NA | | NA |
| NA | | NA |
| NA | | NA |

| | |
|---|---|
| NA | NA |
| NA | NA |
| NA | NA |
| NA | NA |
| NA | NA |
| NA | NA |
| NA | NA |
| NA | NA |
| NA | NA |
| NA | NA |
| NA | NA |
| NA | NA |
| NA | NA |
| NA | NA |
| NA | NA |
| NA | NA |
| NA | NA |
| NA | NA |
| NA | NA |
| NA | NA |
| NA | NA |
| NA | NA |
| NA | NA |
| NA | NA |
| NA | NA |
| NA | NA |
| NA | NA |
| NA | NA |
| NA | NA |
| NA | NA |
| NA | NA |
| NA | NA |
| NA | NA |
| NA | NA |
| NA | NA |
| NA | NA |
| NA | NA |
| NA | NA |
| NA | NA |
| NA | NA |
| NA | NA |
| NA | NA |
| NA | NA |
| NA | NA |
| NA | NA |
| NA | NA |
| NA | NA |

| | |
|---|---|
| NA | NA |
| NA | NA |
| NA | NA |
| NA | NA |
| NA | NA |
| NA | NA |
| NA | NA |
| NA | NA |
| NA | NA |
| NA | NA |
| NA | NA |
| NA | NA |
| NA | NA |
| NA | NA |
| NA | NA |
| NA | NA |
| NA | NA |
| NA | NA |
| NA | NA |
| NA | NA |
| NA | NA |
| NA | NA |
| NA | NA |
| NA | NA |
| NA | NA |
| NA | NA |
| NA | NA |
| NA | NA |
| NA | NA |
| NA | NA |
| NA | NA |
| NA | NA |
| NA | NA |
| NA | NA |
| NA | NA |
| NA | NA |
| NA | NA |
| NA | NA |
| NA | NA |
| NA | NA |
| NA | NA |
| NA | NA |
| NA | NA |
| NA | NA |
| NA | NA |
| NA | NA |
| NA | NA |

| | | |
|---|---|---|
| | 0.992061292 | NA |
| NA | | NA |
| NA | | NA |
| NA | | NA |
| NA | | NA |
| NA | | NA |
| NA | | NA |
| NA | | NA |
| NA | | NA |
| NA | | NA |
| NA | | NA |
| NA | | NA |
| NA | | NA |
| NA | | NA |
| NA | | NA |
| NA | | NA |
| NA | | NA |
| NA | | NA |
| NA | | NA |
| NA | | NA |
| NA | | NA |
| NA | | NA |
| NA | | NA |
| NA | | NA |
| NA | | NA |
| NA | | NA |
| NA | | NA |
| NA | | NA |
| NA | | NA |
| NA | | NA |
| NA | | NA |
| NA | | NA |
| NA | | NA |
| NA | | NA |
| NA | | NA |
| NA | | NA |
| NA | | NA |
| NA | | NA |
| NA | | NA |
| NA | | NA |
| NA | | NA |
| NA | | NA |
| NA | | NA |
| NA | | NA |
| NA | | NA |
| NA | | NA |
| NA | | NA |

| | |
|---|---|
| NA | NA |
| NA | NA |
| NA | NA |
| NA | NA |
| NA | NA |
| NA | NA |
| NA | NA |
| NA | NA |
| NA | NA |
| NA | NA |
| NA | NA |
| NA | NA |
| NA | NA |
| NA | NA |
| NA | NA |
| NA | NA |
| NA | NA |
| NA | NA |
| NA | NA |
| NA | NA |
| NA | NA |
| NA | NA |
| NA | NA |
| NA | NA |
| NA | NA |
| NA | NA |
| NA | NA |
| NA | NA |
| NA | NA |
| NA | NA |
| NA | NA |
| NA | NA |
| NA | NA |
| NA | NA |
| NA | NA |
| NA | NA |
| NA | NA |
| NA | NA |
| NA | NA |
| NA | NA |
| NA | NA |
| NA | NA |
| NA | NA |
| NA | NA |
| NA | NA |
| NA | NA |
| NA | NA |

| | |
|---|---|
| NA | NA |
| NA | NA |
| NA | NA |
| NA | NA |
| NA | NA |
| NA | NA |
| NA | NA |
| NA | NA |
| NA | NA |
| NA | NA |
| NA | NA |
| NA | NA |
| NA | NA |
| NA | NA |
| NA | NA |
| NA | NA |
| NA | NA |
| NA | NA |
| NA | NA |
| NA | NA |
| NA | NA |
| NA | NA |
| NA | NA |
| NA | NA |
| NA | NA |
| NA | NA |
| NA | NA |
| NA | NA |
| NA | NA |
| NA | NA |
| NA | NA |
| NA | NA |
| NA | NA |
| NA | NA |
| NA | NA |
| NA | NA |
| NA | NA |
| NA | NA |
| NA | NA |
| NA | NA |
| NA | NA |
| NA | NA |
| NA | NA |
| NA | NA |
| NA | NA |
| NA | NA |
| NA | NA |

| | |
|---|---|
| NA | NA |
| NA | NA |
| NA | NA |
| NA | NA |
| NA | NA |
| NA | NA |
| NA | NA |
| NA | NA |
| NA | NA |
| NA | NA |
| NA | NA |
| NA | NA |
| NA | NA |
| NA | NA |
| NA | NA |
| NA | NA |
| NA | NA |
| NA | NA |
| NA | NA |
| NA | NA |
| NA | NA |
| NA | NA |
| NA | NA |
| NA | NA |
| NA | NA |
| NA | NA |
| NA | NA |
| NA | NA |
| NA | NA |
| NA | NA |
| NA | NA |
| NA | NA |
| NA | NA |
| NA | NA |
| NA | NA |
| NA | NA |
| NA | NA |
| NA | NA |
| NA | NA |
| NA | NA |
| NA | NA |
| NA | NA |
| NA | NA |
| NA | NA |
| NA | NA |
| NA | NA |
| NA | NA |

| | |
|---|---|
| NA | NA |
| NA | NA |
| NA | NA |
| NA | NA |
| NA | NA |
| NA | NA |
| NA | NA |
| NA | NA |
| NA | NA |
| NA | NA |
| NA | NA |
| NA | NA |
| NA | NA |
| NA | NA |
| 0.75928411 | 0.853826021 |
| NA | NA |
| NA | NA |
| NA | NA |
| NA | NA |
| NA | NA |
| NA | NA |
| NA | NA |
| NA | NA |
| NA | NA |
| NA | NA |
| NA | NA |
| NA | NA |
| NA | NA |
| NA | NA |
| 0.992061292 | NA |
| NA | NA |
| NA | NA |
| NA | NA |
| NA | NA |
| NA | NA |
| NA | NA |
| NA | NA |
| NA | NA |
| NA | NA |
| NA | NA |
| NA | NA |
| NA | NA |
| NA | NA |
| NA | NA |
| NA | NA |
| NA | NA |
| NA | NA |

| | | |
|---|---|---|
| NA | | NA |
| NA | | NA |
| NA | | NA |
| NA | | NA |
| NA | | NA |
| NA | | NA |
| NA | | NA |
| NA | | NA |
| NA | | NA |
| NA | | NA |
| NA | | NA |
| NA | | NA |
| NA | | NA |
| NA | | NA |
| NA | | NA |
| NA | | NA |
| NA | | NA |
| NA | | NA |
| | 0.992061292 | NA |
| NA | | NA |
| NA | | NA |
| NA | | NA |
| NA | | NA |
| NA | | NA |
| NA | | NA |
| NA | | NA |
| NA | | NA |
| NA | | NA |
| NA | | NA |
| NA | | NA |
| NA | | NA |
| NA | | NA |
| NA | | NA |
| NA | | NA |
| NA | | NA |
| NA | | NA |
| NA | | NA |
| NA | | NA |
| NA | | NA |
| NA | | NA |
| NA | | NA |
| NA | | NA |
| NA | | NA |
| NA | | NA |
| NA | | NA |
| NA | | NA |

| | |
|---|---|
| NA | NA |
| NA | NA |
| NA | NA |
| NA | NA |
| NA | NA |
| NA | NA |
| NA | NA |
| NA | NA |
| NA | NA |
| NA | NA |
| NA | NA |
| NA | NA |
| NA | NA |
| NA | NA |
| NA | NA |
| NA | NA |
| NA | NA |
| NA | NA |
| NA | NA |
| NA | NA |
| NA | NA |
| NA | NA |
| NA | NA |
| NA | NA |
| NA | NA |
| NA | NA |
| NA | NA |
| NA | NA |
| NA | NA |
| NA | NA |
| NA | NA |
| NA | NA |
| NA | NA |
| NA | NA |
| NA | NA |
| NA | NA |
| NA | NA |
| NA | NA |
| NA | NA |
| NA | NA |
| NA | NA |
| NA | NA |
| NA | NA |
| NA | NA |
| NA | NA |
| NA | NA |
| NA | NA |

| | |
|---|---|
| NA | NA |
| NA | NA |
| NA | NA |
| NA | NA |
| NA | NA |
| NA | NA |
| NA | NA |
| NA | NA |
| NA | NA |
| NA | NA |
| NA | NA |
| NA | NA |
| NA | NA |
| NA | NA |
| NA | NA |
| NA | NA |
| NA | NA |
| NA | NA |
| NA | NA |
| NA | NA |
| NA | NA |
| NA | NA |
| NA | NA |
| NA | NA |
| NA | NA |
| NA | NA |
| NA | NA |
| NA | NA |
| NA | NA |
| NA | NA |
| NA | NA |
| NA | NA |
| NA | NA |
| NA | NA |
| NA | NA |
| NA | NA |
| NA | NA |
| NA | NA |
| NA | NA |
| NA | NA |
| NA | NA |
| NA | NA |
| NA | NA |
| NA | NA |
| NA | NA |
| NA | NA |
| NA | NA |

| | |
|---|---|
| NA | NA |
| NA | NA |
| NA | NA |
| NA | NA |
| NA | NA |
| NA | NA |
| NA | NA |
| NA | NA |
| NA | NA |
| NA | NA |
| NA | NA |
| NA | NA |
| NA | NA |
| NA | NA |
| NA | NA |
| NA | NA |
| NA | NA |
| NA | NA |
| NA | NA |
| NA | NA |
| NA | NA |
| NA | NA |
| NA | NA |
| NA | NA |
| NA | NA |
| NA | NA |
| NA | NA |
| NA | NA |
| NA | NA |
| NA | NA |
| NA | NA |
| NA | NA |
| NA | NA |
| NA | NA |
| NA | NA |
| NA | NA |
| NA | NA |
| NA | NA |
| NA | NA |
| NA | NA |
| NA | NA |
| NA | NA |
| NA | NA |
| NA | NA |
| NA | NA |
| NA | NA |
| NA | NA |

| | |
|---|---|
| NA | NA |
| NA | NA |
| NA | NA |
| NA | NA |
| NA | NA |
| NA | NA |
| NA | NA |
| NA | NA |
| NA | NA |
| NA | NA |
| NA | NA |
| NA | NA |
| NA | NA |
| NA | NA |
| NA | NA |
| NA | NA |
| NA | NA |
| NA | NA |
| NA | NA |
| NA | NA |
| NA | NA |
| NA | NA |
| NA | NA |
| NA | NA |
| NA | NA |
| NA | NA |
| NA | NA |
| NA | NA |
| NA | NA |
| NA | NA |
| NA | NA |
| NA | NA |
| NA | NA |
| NA | NA |
| NA | NA |
| NA | NA |
| NA | NA |
| NA | NA |
| NA | NA |
| NA | NA |
| NA | NA |
| NA | NA |
| NA | NA |
| NA | NA |
| NA | NA |
| NA | NA |
| NA | NA |

| | |
|---|---|
| NA | NA |
| NA | NA |
| NA | NA |
| NA | NA |
| NA | NA |
| NA | NA |
| NA | NA |
| NA | NA |
| NA | NA |
| NA | NA |
| NA | NA |
| NA | NA |
| NA | NA |
| NA | NA |
| NA | NA |
| NA | NA |
| NA | NA |
| NA | NA |
| NA | NA |
| NA | NA |
| NA | NA |
| NA | NA |
| NA | NA |
| NA | NA |
| NA | NA |
| NA | NA |
| NA | NA |
| NA | NA |
| NA | NA |
| NA | NA |
| NA | NA |
| NA | NA |
| NA | NA |
| NA | NA |
| NA | NA |
| NA | NA |
| NA | NA |
| NA | NA |
| NA | NA |
| NA | NA |
| NA | NA |
| NA | NA |
| NA | NA |
| NA | NA |
| NA | NA |
| NA | NA |
| NA | NA |

| | |
|---|---|
| NA | NA |
| NA | NA |
| NA | NA |
| NA | NA |
| NA | NA |
| NA | NA |
| NA | NA |
| NA | NA |
| NA | NA |
| NA | NA |
| NA | NA |
| NA | NA |
| NA | NA |
| NA | NA |
| NA | NA |
| NA | NA |
| NA | NA |
| NA | NA |
| NA | NA |
| NA | NA |
| NA | NA |
| NA | NA |
| NA | NA |
| NA | NA |
| NA | NA |
| NA | NA |
| NA | NA |
| NA | NA |
| NA | NA |
| NA | NA |
| NA | NA |
| NA | NA |
| NA | NA |
| NA | NA |
| NA | NA |
| NA | NA |
| NA | NA |
| NA | NA |
| NA | NA |
| NA | NA |
| NA | NA |
| NA | NA |
| NA | NA |
| NA | NA |
| NA | NA |
| NA | NA |
| NA | NA |

| | |
|---|---|
| NA | NA |
| NA | NA |
| NA | NA |
| NA | NA |
| NA | NA |
| NA | NA |
| NA | NA |
| NA | NA |
| NA | NA |
| NA | NA |
| NA | NA |
| NA | NA |
| NA | NA |
| NA | NA |
| NA | NA |
| NA | NA |
| NA | NA |
| NA | NA |
| NA | NA |
| NA | NA |
| NA | NA |
| NA | NA |
| NA | NA |
| NA | NA |
| NA | NA |
| NA | NA |
| NA | NA |
| NA | NA |
| NA | NA |
| NA | NA |
| NA | NA |
| NA | NA |
| NA | NA |
| NA | NA |
| NA | NA |
| NA | NA |
| NA | NA |
| NA | NA |
| NA | NA |
| NA | NA |
| NA | NA |
| NA | NA |
| NA | NA |
| NA | NA |
| NA | NA |
| NA | NA |
| NA | NA |

| | |
|---|---|
| NA | NA |
| NA | NA |
| NA | NA |
| NA | NA |
| NA | NA |
| NA | NA |
| NA | NA |
| NA | NA |
| NA | NA |
| NA | NA |
| NA | NA |
| NA | NA |
| NA | NA |
| NA | NA |
| NA | NA |
| NA | NA |
| NA | NA |
| NA | NA |
| NA | NA |
| NA | NA |
| NA | NA |
| NA | NA |
| NA | NA |
| NA | NA |
| NA | NA |
| NA | NA |
| NA | NA |
| NA | NA |
| NA | NA |
| NA | NA |
| NA | NA |
| NA | NA |
| NA | NA |
| NA | NA |
| NA | NA |
| NA | NA |
| NA | NA |
| NA | NA |
| NA | NA |
| NA | NA |
| NA | NA |
| NA | NA |
| NA | NA |
| NA | NA |
| NA | NA |
| NA | NA |
| NA | NA |

| | | |
|---|---|---|
| NA | | NA |
| NA | | NA |
| NA | | NA |
| NA | | NA |
| NA | | NA |
| NA | | NA |
| NA | | NA |
| NA | | NA |
| NA | | NA |
| NA | | NA |
| NA | | NA |
| NA | | NA |
| NA | | NA |
| NA | | NA |
| NA | | NA |
| | 0.992061292 | NA |
| NA | | NA |
| NA | | NA |
| NA | | NA |
| NA | | NA |
| NA | | NA |
| NA | | NA |
| NA | | NA |
| NA | | NA |
| NA | | NA |
| NA | | NA |
| NA | | NA |
| NA | | NA |
| NA | | NA |
| NA | | NA |
| NA | | NA |
| NA | | NA |
| NA | | NA |
| NA | | NA |
| NA | | NA |
| NA | | NA |
| NA | | NA |
| NA | | NA |
| NA | | NA |
| NA | | NA |
| NA | | NA |
| NA | | NA |
| NA | | NA |
| NA | | NA |
| NA | | NA |
| NA | | NA |
| NA | | NA |

| | |
|---|---|
| NA | NA |
| NA | NA |
| NA | NA |
| NA | NA |
| NA | NA |
| NA | NA |
| NA | NA |
| NA | NA |
| NA | NA |
| NA | NA |
| NA | NA |
| NA | NA |
| NA | NA |
| NA | NA |
| NA | NA |
| NA | NA |
| NA | NA |
| NA | NA |
| NA | NA |
| NA | NA |
| NA | NA |
| NA | NA |
| NA | NA |
| NA | NA |
| NA | NA |
| NA | NA |
| NA | NA |
| NA | NA |
| NA | NA |
| NA | NA |
| NA | NA |
| NA | NA |
| NA | NA |
| NA | NA |
| NA | NA |
| NA | NA |
| NA | NA |
| NA | NA |
| NA | NA |
| NA | NA |
| NA | NA |
| NA | NA |
| NA | NA |
| NA | NA |
| NA | NA |
| NA | NA |
| NA | NA |

| | |
|---|---|
| NA | NA |
| NA | NA |
| NA | NA |
| NA | NA |
| NA | NA |
| NA | NA |
| NA | NA |
| NA | NA |
| NA | NA |
| NA | NA |
| NA | NA |
| NA | NA |
| NA | NA |
| NA | NA |
| NA | NA |
| NA | NA |
| NA | NA |
| NA | NA |
| NA | NA |
| NA | NA |
| NA | NA |
| NA | NA |
| NA | NA |
| NA | NA |
| NA | NA |
| NA | NA |
| NA | NA |
| NA | NA |
| NA | NA |
| NA | NA |
| NA | NA |
| NA | NA |
| NA | NA |
| NA | NA |
| NA | NA |
| NA | NA |
| NA | NA |
| NA | NA |
| NA | NA |
| NA | NA |
| NA | NA |
| NA | NA |
| NA | NA |
| NA | NA |
| NA | NA |
| NA | NA |
| NA | NA |

| | |
|---|---|
| NA | NA |
| NA | NA |
| NA | NA |
| NA | NA |
| NA | NA |
| NA | NA |
| NA | NA |
| NA | NA |
| NA | NA |
| NA | NA |
| NA | NA |
| NA | NA |
| NA | NA |
| NA | NA |
| NA | NA |
| NA | NA |
| NA | NA |
| NA | NA |
| NA | NA |
| NA | NA |
| NA | NA |
| NA | NA |
| NA | NA |
| NA | NA |
| NA | NA |
| NA | NA |
| NA | NA |
| NA | NA |
| NA | NA |
| NA | NA |
| NA | NA |
| NA | NA |
| NA | NA |
| NA | NA |
| NA | NA |
| NA | NA |
| NA | NA |
| NA | NA |
| NA | NA |
| NA | NA |
| NA | NA |
| NA | NA |
| NA | NA |
| NA | NA |
| NA | NA |
| NA | NA |
| NA | NA |

| | |
|---|---|
| NA | NA |
| NA | NA |
| NA | NA |
| NA | NA |
| NA | NA |
| NA | NA |
| NA | NA |
| NA | NA |
| NA | NA |
| NA | NA |
| NA | NA |
| NA | NA |
| NA | NA |
| NA | NA |
| NA | NA |
| NA | NA |
| NA | NA |
| NA | NA |
| NA | NA |
| NA | NA |
| NA | NA |
| NA | NA |
| NA | NA |
| NA | NA |
| NA | NA |
| NA | NA |
| NA | NA |
| NA | NA |
| NA | NA |
| NA | NA |
| NA | NA |
| NA | NA |
| NA | NA |
| NA | NA |
| NA | NA |
| NA | NA |
| NA | NA |
| NA | NA |
| NA | NA |
| NA | NA |
| NA | NA |
| NA | NA |
| NA | NA |
| NA | NA |
| NA | NA |
| NA | NA |
| NA | NA |

| | |
|---|---|
| NA | NA |
| NA | NA |
| NA | NA |
| NA | NA |
| NA | NA |
| NA | NA |
| NA | NA |
| NA | NA |
| NA | NA |
| NA | NA |
| NA | NA |
| NA | NA |
| NA | NA |
| NA | NA |
| NA | NA |
| NA | NA |
| NA | NA |
| NA | NA |
| NA | NA |
| NA | NA |
| NA | NA |
| NA | NA |
| NA | NA |
| NA | NA |
| NA | NA |
| NA | NA |
| NA | NA |
| NA | NA |
| NA | NA |
| NA | NA |
| NA | NA |
| NA | NA |
| NA | NA |
| NA | NA |
| NA | NA |
| NA | NA |
| NA | NA |
| NA | NA |
| NA | NA |
| NA | NA |
| NA | NA |
| NA | NA |
| NA | NA |
| NA | NA |
| NA | NA |
| NA | NA |
| NA | NA |

| | |
|---|---|
| NA | NA |
| NA | NA |
| NA | NA |
| NA | NA |
| NA | NA |
| NA | NA |
| NA | NA |
| NA | NA |
| NA | NA |
| NA | NA |
| NA | NA |
| NA | NA |
| NA | NA |
| NA | NA |
| NA | NA |
| NA | NA |
| NA | NA |
| NA | NA |
| NA | NA |
| NA | NA |
| NA | NA |
| NA | NA |
| NA | NA |
| NA | NA |
| NA | NA |
| NA | NA |
| NA | NA |
| NA | NA |
| NA | NA |
| NA | NA |
| NA | NA |
| NA | NA |
| NA | NA |
| NA | NA |
| NA | NA |
| NA | NA |
| NA | NA |
| NA | NA |
| NA | NA |
| NA | NA |
| NA | NA |
| NA | NA |
| NA | NA |
| NA | NA |
| NA | NA |
| NA | NA |
| NA | NA |

| | | |
|---|---|---|
| NA | | NA |
| NA | | NA |
| NA | | NA |
| NA | | NA |
| NA | | NA |
| NA | | NA |
| NA | | NA |
| NA | | NA |
| NA | | NA |
| NA | | NA |
| NA | | NA |
| NA | | NA |
| NA | | NA |
| NA | | NA |
| NA | | NA |
| NA | | NA |
| NA | | NA |
| NA | | NA |
| NA | | NA |
| NA | | NA |
| NA | | NA |
| NA | | NA |
| NA | | NA |
| NA | | NA |
| NA | | NA |
| NA | | NA |
| NA | | NA |
| NA | | NA |
| NA | | NA |
| NA | | NA |
| NA | | NA |
| NA | | NA |
| NA | | NA |
| NA | | NA |
| NA | | NA |
| NA | | NA |
| NA | | NA |
| NA | | NA |
| | 0.905158266 | NA |
| NA | | NA |
| NA | | NA |
| NA | | NA |
| NA | | NA |
| NA | | NA |
| NA | | NA |
| NA | | NA |
| NA | | NA |

| | | | |
|---|---|---|---|
| NA | | NA | |
| NA | | NA | |
| NA | | NA | |
| NA | | NA | |
| NA | | NA | |
| NA | | NA | |
| NA | | NA | |
| NA | | NA | |
| NA | | NA | |
| NA | | NA | |
| NA | | NA | |
| NA | | NA | |
| NA | | NA | |
| NA | | NA | |
| NA | | NA | |
| NA | | NA | |
| NA | | NA | |
| NA | | NA | |
| NA | | NA | |
| NA | | NA | |
| NA | | NA | |
| NA | | NA | |
| NA | | NA | |
| NA | | NA | |
| NA | | NA | |
| NA | | NA | |
| | 0.644828325 | | 0.770265201 |
| | 0.797738672 | NA | |
| | 0.189842506 | | 0.331801304 |
| NA | | NA | |
| NA | | NA | |
