## Supplementary material for "Comparative pathogenesis of two lineages of Powassan virus reveals distinct clinical outcome, neuropathology, and inflammation": Table S1

| Gene | baseMean | log2FoldChange | lfcSE | stat |
|---|---|---|---|---|
| 4933401J01Rik | 0 | NA | NA | NA |
| Gm26206 | 0 | NA | NA | NA |
| Xkr4 | 1236.723264 | 0.361649927 | 0.098370628 | 3.676401543 |
| Gm18956 | 0.257781492 | 0.751420611 | 2.506666059 | 0.299768933 |
| Gm37180 | 183.2012087 | -0.199743832 | 0.240317933 | -0.831164906 |
| Gm37363 | 156.7123429 | -0.107246198 | 0.238001244 | -0.450611921 |
| Gm37686 | 55.26816088 | 0.04652138 | 0.277949635 | 0.167373416 |
| Gm1992 | 0 | NA | NA | NA |
| Gm37329 | 74.22829483 | -0.015915381 | 0.237459366 | -0.067023596 |
| Gm7341 | 0.472128458 | 0.292213542 | 1.616290581 | 0.180792702 |
| Gm38148 | 143.524156 | -0.168694583 | 0.26731953 | -0.631059701 |
| Gm19938 | 181.8288108 | -0.156331223 | 0.25402851 | -0.615408178 |
| Gm10568 | 17.47471811 | 2.514650463 | 0.331767358 | 7.5795596 |
| Gm38385 | 0 | NA | NA | NA |
| Gm27396 | 0 | NA | NA | NA |
| Gm37381 | 0.042716945 | -0.206490061 | 3.724945049 | -0.055434391 |
| Gm6101 | 0.038981395 | 0.274393828 | 3.724945049 | 0.073663859 |
| Rp1 | 1.19669966 | 1.037778894 | 0.962693886 | 1.077994687 |
| Gm37483 | 0 | NA | NA | NA |
| Sox17 | 14.01852502 | 0.257878972 | 0.285791501 | 0.902332542 |
| Gm37587 | 0.590622753 | 1.167837115 | 1.439861439 | 0.811076041 |
| Gm7357 | 0.431020911 | 0.609073142 | 1.516617494 | 0.401599707 |
| Gm22307 | 0.205539566 | -0.206487548 | 2.613407335 | -0.079010855 |
| Gm38076 | 0 | NA | NA | NA |
| Gm37323 | 0.424160584 | -1.797444836 | 1.403428972 | -1.280752266 |
| Gm7369 | 0.062426873 | 0.755288839 | 3.724945049 | 0.202765096 |
| Gm6085 | 7.026365196 | -0.173403683 | 0.437247322 | -0.39658032 |
| Gm6119 | 1.007284973 | -0.346558273 | 1.03958283 | -0.333362829 |
| Gm25493 | 0.04393175 | -0.206490061 | 3.724945049 | -0.055434391 |
| Gm2053 | 0.509840029 | -1.940301895 | 1.353081268 | -1.4339877 |
| Gm6123 | 6.523572364 | 0.2775134 | 0.387748839 | 0.715704013 |
| Mrpl15 | 339.6237885 | -0.094079666 | 0.09927659 | -0.947652063 |
| Gm37144 | 10.84448109 | -0.900261075 | 0.409406974 | -2.198939279 |
| Lypla1 | 301.4400782 | -0.121431795 | 0.10511088 | -1.15527332 |
| Gm37988 | 0.099106094 | 0.274387056 | 3.724945049 | 0.073662041 |
| Tcea1 | 660.8047814 | 0.468356093 | 0.097916648 | 4.783212094 |
| Gm6104 | 7.397713871 | -0.334139094 | 0.362466238 | -0.921848878 |
| Gm37277 | 41.2031627 | 0.783514309 | 0.180141413 | 4.349440234 |
| Rgs20 | 298.6651195 | -0.025099303 | 0.103879169 | -0.24162018 |
| Gm17100 | 2.600587843 | -0.368544035 | 0.646492293 | -0.570067175 |
| Gm37079 | 17.87741427 | -0.454284644 | 0.345273077 | -1.315725651 |
| Gm16041 | 0.448921586 | -0.997970525 | 1.454094169 | -0.686317672 |
| Gm17101 | 1.407567115 | -0.486942136 | 0.889653864 | -0.547338865 |
| Atp6v1h | 1710.406335 | 0.161826844 | 0.09228553 | 1.753545156 |
| Gm7182 | 0 | NA | NA | NA |
| Gm37567 | 5.040767136 | 0.031763892 | 0.507229155 | 0.062622371 |

| | | | | |
|---|---|---|---|---|
| Gm38264 | 0.54045421 | 0.545733941 | 1.247687386 | 0.437396376 |
| Oprk1 | 98.2787742 | -0.235301229 | 0.146701846 | -1.603941837 |
| Gm36965 | 0 | NA | NA | NA |
| Gm22463 | 0 | NA | NA | NA |
| Gm37429 | 0 | NA | NA | NA |
| Npbwr1 | 11.64878932 | 0.37768382 | 0.31695174 | 1.191613021 |
| Gm19214 | 0.066050875 | 0.274391604 | 3.724945049 | 0.073663262 |
| Rb1cc1 | 2300.690646 | -0.040915632 | 0.053603228 | -0.763305364 |
| 4732440D04Rik | 117.8176336 | -0.723579909 | 0.168230801 | -4.301114343 |
| Fam150a | 0.409266314 | 1.077045044 | 1.4291331 | 0.753635224 |
| Gm2147 | 0.280898025 | 1.183514123 | 1.624766026 | 0.728421265 |
| Gm7417 | 0 | NA | NA | NA |
| Gm19026 | 17.7062556 | -0.54368701 | 0.24417958 | -2.226586716 |
| St18 | 573.9728649 | -0.542516928 | 0.130356739 | -4.161786589 |
| Gm37405 | 0 | NA | NA | NA |
| Gm7449 | 0.090074004 | 0.274402725 | 3.724945049 | 0.073666248 |
| Gm37108 | 10.32889666 | 0.091383536 | 0.325125363 | 0.281071694 |
| Gm37275 | 0 | NA | NA | NA |
| Gm37225 | 0 | NA | NA | NA |
| Gm37489 | 0 | NA | NA | NA |
| Gm5694 | 0 | NA | NA | NA |
| Pcmtd1 | 1844.737977 | 0.074519046 | 0.081281873 | 0.91679785 |
| Gm26983 | 0 | NA | NA | NA |
| Gm38372 | 224.9946318 | -0.819227157 | 0.235900269 | -3.472769065 |
| Gm9826 | 6.912980464 | 0.296504133 | 0.3902158 | 0.759846559 |
| Nras-ps2 | 0 | NA | NA | NA |
| Gm23274 | 0 | NA | NA | NA |
| Gm19002 | 0.085327104 | 0.274389469 | 3.724945049 | 0.073662689 |
| Gm26901 | 31.62112565 | -0.58724381 | 0.295668446 | -1.986156516 |
| Gm18984 | 0 | NA | NA | NA |
| Gm30414 | 0 | NA | NA | NA |
| Gm37791 | 0 | NA | NA | NA |
| Gm7470 | 0.066050875 | 0.274391604 | 3.724945049 | 0.073663262 |
| Rps2-ps2 | 0 | NA | NA | NA |
| Gm38216 | 0 | NA | NA | NA |
| Sntg1 | 622.888287 | -0.022341422 | 0.112042056 | -0.199402111 |
| Gm38024 | 20.53816256 | 0.213486825 | 0.382728504 | 0.557802262 |
| Gm16284 | 4.895741945 | 0.504565501 | 0.499092935 | 1.010965024 |
| Gm38259 | 46.04207795 | -0.021629277 | 0.243954499 | -0.088661112 |
| Gm23358 | 1.395067866 | 0.48456001 | 0.887345854 | 0.546077955 |
| Gm37005 | 8.035261982 | 0.255103599 | 0.365839801 | 0.697309584 |
| Gm15452 | 1.564680506 | -0.401540831 | 0.752591688 | -0.533544069 |
| Gm36964 | 0.112396584 | 0.27438722 | 3.724945049 | 0.073662085 |
| Gm24276 | 0.547110457 | -0.86177769 | 1.585046948 | -0.543692217 |
| Gm6152 | 0.180148697 | 0.776554961 | 2.703307687 | 0.287261034 |
| Gm7445 | 0.326379708 | 0.899393523 | 3.712005061 | 0.242293183 |
| Gm7493 | 10.60824579 | -0.585608488 | 0.483921686 | -1.210130699 |

| | | | | |
|---|---|---|---|---|
| Gm37629 | 21.84260411 | -0.675605845 | 0.281108759 | -2.403361061 |
| Gm38008 | 8.876683074 | -0.702314572 | 0.406862501 | -1.726171793 |
| Gm37143 | 0 | NA | NA | NA |
| Gm6187 | 0.1709406 | -0.20648082 | 3.38513154 | -0.060996395 |
| Gm18299 | 0 | NA | NA | NA |
| Gm7512 | 0 | NA | NA | NA |
| Gm24765 | 0 | NA | NA | NA |
| Gm18300 | 1.926091363 | -0.648335584 | 0.693189853 | -0.935292951 |
| Rrs1 | 218.6880439 | 0.213915799 | 0.124073444 | 1.724106244 |
| Adhfe1 | 178.1879691 | 0.797678037 | 0.144510625 | 5.519857341 |
| 2610203C22Rik | 5.506898143 | -0.453898957 | 0.445152076 | -1.019649198 |
| Gm6161 | 3.14279713 | 0.897872932 | 0.612520472 | 1.465865997 |
| 3110035E14Rik | 1247.772309 | -0.219076052 | 0.181825978 | -1.204866626 |
| Gm29520 | 0 | NA | NA | NA |
| Mybl1 | 100.6319816 | -0.152593819 | 0.163650972 | -0.932434539 |
| Vcpip1 | 2191.495749 | 0.391166264 | 0.077543636 | 5.044466345 |
| 1700034P13Rik | 2.716773814 | -0.39759133 | 0.580514069 | -0.684895252 |
| Sgk3 | 387.7590365 | 0.64208727 | 0.110386719 | 5.816707646 |
| Gm6195 | 2.037763675 | 0.224746892 | 0.862165725 | 0.260677136 |
| Gm22607 | 0 | NA | NA | NA |
| Mcmdc2 | 2.919172438 | 0.407394476 | 0.620904347 | 0.656130817 |
| Snhg6 | 57.04786074 | 0.408697369 | 0.197453889 | 2.069837012 |
| Snord87 | 1.794878382 | -0.073039848 | 0.750275419 | -0.097350714 |
| Tcf24 | 4.573727287 | -0.420126629 | 0.571253857 | -0.735446464 |
| E330040D14Rik | 0.234971822 | 0.75530893 | 2.121545891 | 0.356018191 |
| Ppp1r42 | 4.644333044 | -0.300944225 | 0.457372618 | -0.657984787 |
| Gm15818 | 0 | NA | NA | NA |
| Gm24674 | 0 | NA | NA | NA |
| Cops5 | 641.9122966 | 0.208503678 | 0.099061767 | 2.104784563 |
| Cspp1 | 856.404923 | -0.656935995 | 0.168995667 | -3.887294902 |
| 1700047N06Rik | 0 | NA | NA | NA |
| Arfgef1 | 2686.960645 | 0.04359209 | 0.050249261 | 0.867517035 |
| Gm26348 | 0 | NA | NA | NA |
| Gm37569 | 39.97338864 | 0.225664303 | 0.296835509 | 0.760233518 |
| Cpa6 | 18.52223898 | -1.25981028 | 0.52290272 | -2.409263199 |
| Gm38005 | 2.757436252 | -4.004332872 | 1.085674693 | -3.688335831 |
| Gm15604 | 1.883799777 | -2.278111856 | 1.057467654 | -2.154308785 |
| Gm25253 | 0.347911031 | -1.611046413 | 3.228564737 | -0.498997711 |
| Gm15603 | 0.042716945 | -0.206490061 | 3.724945049 | -0.055434391 |
| Gm37133 | 0 | NA | NA | NA |
| Gm5522 | 0.046345709 | 0.274391604 | 3.724945049 | 0.073663262 |
| Gm28659 | 0 | NA | NA | NA |
| Gm22963 | 0 | NA | NA | NA |
| Prex2 | 1191.400917 | -0.609289305 | 0.156340973 | -3.897182504 |
| Gm24173 | 0 | NA | NA | NA |
| Gm28686 | 0 | NA | NA | NA |
| Gm37410 | 0.060174537 | 0.755286614 | 3.724945049 | 0.202764498 |

| | | | | |
|---|---|---|---|---|
| A830018L16Rik | 657.3086267 | -0.258779434 | 0.104675135 | -2.472214959 |
| Gm38178 | 28.55896715 | -0.376032404 | 0.2989412 | -1.257880828 |
| Gm29273 | 0 | NA | NA | NA |
| Gm38069 | 9.696124784 | -0.003830896 | 0.440924825 | -0.00868832 |
| Gm7560 | 0.344328494 | 1.693954327 | 1.6729816 | 1.012536137 |
| Gm22633 | 0 | NA | NA | NA |
| Mir6341 | 0 | NA | NA | NA |
| Gm2383 | 0.481037375 | -0.247541299 | 1.360763293 | -0.181913563 |
| Gm6216 | 0 | NA | NA | NA |
| Gm17644 | 0.118371171 | 0.75528439 | 3.724945049 | 0.202763901 |
| Gm29663 | 0 | NA | NA | NA |
| Sulf1 | 425.5919252 | 0.226743975 | 0.129888632 | 1.745679901 |
| Slco5a1 | 118.4898984 | -0.279534059 | 0.150158135 | -1.861597834 |
| Gm37162 | 0 | NA | NA | NA |
| Gm5250 | 0 | NA | NA | NA |
| Gm29283 | 0 | NA | NA | NA |
| Prdm14 | 0.178837887 | -0.20648716 | 2.700576645 | -0.076460396 |
| Ncoa2 | 2049.993035 | -0.102241787 | 0.046118431 | -2.21693986 |
| Gm38376 | 24.50757996 | -0.237662407 | 0.371557545 | -0.63963822 |
| Gm37409 | 22.73165489 | -0.588169258 | 0.319361331 | -1.841704684 |
| Gm27881 | 0 | NA | NA | NA |
| Gm38223 | 5.5083264 | -1.215034674 | 0.424707754 | -2.860872355 |
| Gm37702 | 18.1938087 | -0.906566437 | 0.282345841 | -3.210836875 |
| Gm23169 | 4.580170235 | -0.74969263 | 0.564640709 | -1.327733934 |
| Gm38120 | 65.50301172 | -0.658367119 | 0.250063891 | -2.632795627 |
| Gm38380 | 51.36537569 | -0.587580671 | 0.300748621 | -1.9537269 |
| Gm29570 | 1.379700947 | -0.430873245 | 0.84594184 | -0.509341452 |
| Gm7593 | 0.038981395 | 0.274393828 | 3.724945049 | 0.073663859 |
| Tram1 | 601.6593407 | 0.326146925 | 0.102160765 | 3.192487092 |
| Lactb2 | 271.8818834 | -0.268788586 | 0.096675764 | -2.78030992 |
| Gm38319 | 9.470931737 | 0.259846963 | 0.51386227 | 0.505674338 |
| Xkr9 | 0.562487033 | 2.359697676 | 1.367404372 | 1.725676563 |
| Gm26273 | 0 | NA | NA | NA |
| Gm5523 | 0.58821479 | 0.255572398 | 1.280256116 | 0.199625993 |
| Gm36947 | 0.09689823 | -0.687371727 | 3.724945049 | -0.184532045 |
| Gm25491 | 0 | NA | NA | NA |
| Gm37400 | 0 | NA | NA | NA |
| Eya1 | 126.1615464 | -0.033913191 | 0.126541019 | -0.268001566 |
| Gm22616 | 0 | NA | NA | NA |
| Gm37444 | 2.632415511 | -0.342774637 | 0.702828278 | -0.48770752 |
| Gm9947 | 16.04594573 | -0.174127912 | 0.268931148 | -0.647481382 |
| Msc | 6.16192073 | -0.13257933 | 0.443878079 | -0.298684113 |
| Gm37412 | 7.68759231 | 0.096274114 | 0.344600111 | 0.279379231 |
| Smt3h2-ps4 | 2.907781834 | 0.673432778 | 0.540859646 | 1.245115593 |
| Rpl5-ps1 | 8.684837921 | -0.353092873 | 0.37040935 | -0.953250432 |
| Trpa1 | 3.284363746 | 0.072217219 | 0.515784619 | 0.140014294 |
| Gm26345 | 0 | NA | NA | NA |

| | | | | |
|---|---|---|---|---|
| Gm10566 | 6.613104402 | 0.343189978 | 0.404937894 | 0.847512627 |
| Kcnb2 | 762.7240551 | -0.307411079 | 0.104283485 | -2.947840481 |
| Gm38116 | 50.40214469 | -1.302349658 | 0.346191136 | -3.7619382 |
| Gm37138 | 26.46315223 | -0.569349836 | 0.269461673 | -2.112915832 |
| Gm25227 | 0 | NA | NA | NA |
| Gm25168 | 0.155005853 | -1.144125604 | 2.822657929 | -0.405336259 |
| Gm6060 | 0.066050875 | 0.274391604 | 3.724945049 | 0.073663262 |
| Gm28669 | 0.095435502 | 0.75528439 | 3.724945049 | 0.202763901 |
| Terf1 | 191.9488994 | 0.061095896 | 0.128267565 | 0.476316019 |
| Gm37509 | 44.48996355 | -0.017982236 | 0.321707983 | -0.055896146 |
| Sbspon | 3.624262949 | -1.011126014 | 0.560749822 | -1.803167784 |
| Gm23259 | 0.061920613 | 0.274393828 | 3.724945049 | 0.073663859 |
| Gm7634 | 0 | NA | NA | NA |
| 4930444P10Rik | 4.660125666 | -0.246899213 | 0.462779415 | -0.533513818 |
| Rpl7 | 1699.234633 | 0.565993519 | 0.113949248 | 4.967066716 |
| Rdh10 | 199.9939912 | 0.133760017 | 0.160446844 | 0.833671849 |
| Gm38249 | 4.612676316 | -0.52661206 | 0.501806129 | -1.049433297 |
| Gm28095 | 0 | NA | NA | NA |
| Stau2 | 1742.3238 | -0.068652129 | 0.075460403 | -0.909776864 |
| Gm7568 | 0.218065207 | 1.222571812 | 2.536190686 | 0.48205043 |
| Ube2w | 820.6053904 | 0.003054338 | 0.058063094 | 0.052603774 |
| Gm38342 | 3.77249287 | 1.391224084 | 0.560933205 | 2.48019563 |
| Tceb1 | 826.0456009 | 0.050909679 | 0.083301023 | 0.611153109 |
| D030040B21Rik | 0.250367046 | 0.755305742 | 2.484698294 | 0.303982879 |
| Gm7654 | 0.171036444 | -1.081088773 | 2.766397645 | -0.390792978 |
| Tmem70 | 222.9577256 | -0.146960896 | 0.107419946 | -1.368096907 |
| Ly96 | 21.7470159 | 0.475273259 | 0.264482971 | 1.796990021 |
| Gm29649 | 0 | NA | NA | NA |
| Gm5828 | 11.26054785 | 0.409441056 | 0.317205279 | 1.290776301 |
| Gm28376 | 0.229484813 | 0.274412335 | 3.114678745 | 0.088102934 |
| 4921511E07Rik | 0 | NA | NA | NA |
| Jph1 | 403.3093643 | 0.062530723 | 0.091153346 | 0.68599482 |
| Gm28783 | 0.484663642 | 1.140981888 | 1.352032803 | 0.843901039 |
| Gdap1 | 1402.76127 | -0.126047554 | 0.060904479 | -2.06959416 |
| Gm28784 | 4.433742602 | -0.732956018 | 0.488088891 | -1.501685516 |
| Gm28785 | 0.049553047 | 0.274391604 | 3.724945049 | 0.073663262 |
| Gm5251 | 0.15956642 | 0.75529148 | 3.418807262 | 0.22092251 |
| Rbm6-ps1 | 6.916369844 | 0.005318297 | 0.418406533 | 0.012710837 |
| Gm6075 | 0.038981395 | 0.274393828 | 3.724945049 | 0.073663859 |
| Gm25166 | 0 | NA | NA | NA |
| Pi15 | 14.69231755 | -0.021923006 | 0.291828692 | -0.075122859 |
| Gm28154 | 8.974735632 | -0.283243556 | 0.33611655 | -0.842694466 |
| Gm16070 | 24.47860644 | -0.411484652 | 0.22306806 | -1.84465966 |
| Crispld1 | 57.83831356 | -0.131856261 | 0.13935282 | -0.946204469 |
| Gm28153 | 1.598622982 | 0.198929531 | 0.818282152 | 0.243106281 |
| Gm7690 | 0 | NA | NA | NA |
| Gm18775 | 13.00822407 | -0.69778887 | 0.27762916 | -2.513384655 |

| | | | | |
|---|---|---|---|---|
| Gm17969 | 0 | NA | NA | NA |
| Gm28755 | 1.336776882 | -0.231039545 | 0.864729815 | -0.267181194 |
| Gm28756 | 10.18614479 | -0.451085836 | 0.34273538 | -1.316134436 |
| Crisp4 | 0 | NA | NA | NA |
| Defb44-ps | 0.047948698 | -0.206487837 | 3.724945049 | -0.055433794 |
| Defb18 | 0 | NA | NA | NA |
| Gm15386 | 0 | NA | NA | NA |
| Gm17837 | 0.172125443 | 0.755307234 | 2.321733346 | 0.325320406 |
| Gm29006 | 0 | NA | NA | NA |
| Gm5252 | 0 | NA | NA | NA |
| Gm24075 | 0 | NA | NA | NA |
| Gm28343 | 0 | NA | NA | NA |
| Gm28344 | 0 | NA | NA | NA |
| Gm15825 | 1.225430203 | 1.127599417 | 0.861144744 | 1.309419148 |
| Tfap2d | 23.18306498 | 0.308458465 | 0.26299733 | 1.172857781 |
| Gm10075 | 1.913880918 | 0.562781243 | 0.716244625 | 0.785738872 |
| Tfap2b | 140.1785569 | -0.145923183 | 0.153274301 | -0.952039466 |
| Gm28340 | 0.106520246 | 0.755282166 | 3.724945049 | 0.202763304 |
| Gm28341 | 0 | NA | NA | NA |
| Gm4849 | 4.195570756 | 0.548386643 | 0.545744369 | 1.004841596 |
| Gm24633 | 0 | NA | NA | NA |
| Gm38382 | 0.088383099 | -0.687376175 | 3.724945049 | -0.184533239 |
| Pkhd1 | 2.807822083 | -0.484006785 | 0.707853375 | -0.683767008 |
| Gm37315 | 0 | NA | NA | NA |
| 4930486I03Rik | 0.427990203 | -0.745466702 | 1.654135983 | -0.45066833 |
| Gm15795 | 0.244548831 | 0.79518811 | 2.102348598 | 0.378237991 |
| Gm24162 | 0 | NA | NA | NA |
| Mir206 | 0 | NA | NA | NA |
| Gm28653 | 1.730304082 | -1.0447606 | 0.72358462 | -1.443867893 |
| Mir133b | 0 | NA | NA | NA |
| Il17a | 0 | NA | NA | NA |
| Il17f | 0.328867834 | -1.507932147 | 2.052544452 | -0.734664794 |
| Mcm3 | 85.70957654 | 1.645372499 | 0.274352532 | 5.997292931 |
| Gm20587 | 0 | NA | NA | NA |
| Gm28065 | 0.068097359 | -0.206487837 | 3.724945049 | -0.055433794 |
| Gm24723 | 0 | NA | NA | NA |
| 6720483E21Rik | 3.174408629 | -0.458207286 | 0.636777747 | -0.719571762 |
| Paqr8 | 1990.654714 | -0.157079484 | 0.112403676 | -1.397458597 |
| Gm24171 | 0 | NA | NA | NA |
| Gm28064 | 5.60213491 | 0.154607637 | 0.409709806 | 0.377358889 |
| Efhc1 | 14.45078694 | 0.096480848 | 0.281492245 | 0.3427478 |
| Tram2 | 100.5415102 | 1.599533736 | 0.190779357 | 8.38420762 |
| Gm28287 | 0.218494371 | -0.206481361 | 2.166806322 | -0.095292947 |
| Gm2693 | 2.478342139 | 0.365977045 | 0.60030818 | 0.609648605 |
| Tmem14a | 140.3427173 | -0.145805915 | 0.149759741 | -0.973598874 |
| Gm38224 | 3.730747895 | -1.003988767 | 0.55133492 | -1.82101429 |
| Gsta3 | 15.33756154 | -0.22361704 | 0.332721232 | -0.672085272 |

| | | | | |
|---|---|---|---|---|
| Gm28836 | 1.863919813 | -0.443458804 | 0.689361971 | -0.643288755 |
| Gm7094 | 1.910334788 | 0.685566607 | 0.780420326 | 0.878458164 |
| Gm4956 | 0.357620736 | -0.20648172 | 1.569311684 | -0.131574704 |
| Gm21909 | 0.066050875 | 0.274391604 | 3.724945049 | 0.073663262 |
| Khdc1a | 0.075043205 | 0.755286614 | 3.724945049 | 0.202764498 |
| Gm16329 | 0 | NA | NA | NA |
| Gm16327 | 0 | NA | NA | NA |
| Khdc1c | 0.097139547 | 0.274393828 | 3.724945049 | 0.073663859 |
| Gm2709 | 0 | NA | NA | NA |
| Gm16533 | 0 | NA | NA | NA |
| Khdc1b | 0 | NA | NA | NA |
| Gm16328 | 0 | NA | NA | NA |
| Kcnq5 | 996.2770014 | 0.010868846 | 0.143364195 | 0.075812837 |
| Gm36949 | 26.09595903 | 0.003677549 | 0.303778171 | 0.012106034 |
| Gm7658 | 100.9699313 | 0.487010382 | 0.162853911 | 2.990473966 |
| Gm38243 | 62.35611887 | -0.801626039 | 0.339719402 | -2.359671052 |
| Gm26580 | 1.456296832 | 0.939145576 | 0.783855705 | 1.198110277 |
| Gm5696 | 0 | NA | NA | NA |
| Gm28239 | 0 | NA | NA | NA |
| Rims1 | 3306.597451 | -0.36940116 | 0.094830899 | -3.895367051 |
| Gm26866 | 0.048949532 | -0.206487837 | 3.724945049 | -0.055433794 |
| Gm7761 | 0.676990728 | -0.473710213 | 1.182875132 | -0.400473558 |
| Gm29506 | 0.481916195 | -1.437288089 | 1.50080793 | -0.957676235 |
| 4933415F23Rik | 0.213028513 | 2.035187398 | 2.124635671 | 0.957899477 |
| Gm24176 | 0 | NA | NA | NA |
| Gm6420 | 0.044191549 | -0.206490061 | 3.724945049 | -0.055434391 |
| Mir30a | 2.056819932 | -0.48653321 | 0.778574474 | -0.624902596 |
| Gm27028 | 24.51280018 | -0.624923344 | 0.223638706 | -2.794343404 |
| Mir30c-2 | 0 | NA | NA | NA |
| Gm29107 | 3.847956237 | -1.053203026 | 0.572875975 | -1.838448584 |
| Gm20954 | 0.386554245 | 1.153470216 | 3.254673954 | 0.35440423 |
| Ogfrl1 | 2085.851771 | 0.024308306 | 0.074192764 | 0.327637156 |
| Gm7784 | 0.046345709 | 0.274391604 | 3.724945049 | 0.073663262 |
| Gm28822 | 1.28195289 | 0.415080841 | 0.877424998 | 0.473067034 |
| B3gat2 | 201.421169 | -0.299929229 | 0.102722823 | -2.919791523 |
| Smap1 | 968.2258995 | -0.306228626 | 0.061358229 | -4.990832247 |
| Sdhaf4 | 66.50993585 | -0.301110411 | 0.171307475 | -1.757719042 |
| Gm26524 | 0.087261593 | 0.274400501 | 3.724945049 | 0.073665651 |
| Fam135a | 882.3529123 | -0.100315792 | 0.074869614 | -1.339873238 |
| Gm37580 | 2.220335296 | -0.662584337 | 0.665342436 | -0.995854618 |
| Gm26607 | 0 | NA | NA | NA |
| Col9a1 | 27.58382149 | 1.081979662 | 0.314061603 | 3.445119212 |
| Col19a1 | 256.8225277 | -0.260725695 | 0.198130331 | -1.315930243 |
| Gm26418 | 0.607053417 | -0.716305421 | 1.230289514 | -0.582225089 |
| Tubb4b-ps2 | 1.899816927 | -0.499112634 | 0.775299763 | -0.643767298 |
| Gm28439 | 270.7744438 | -0.091629808 | 0.161578923 | -0.567090103 |
| Gm10222 | 81.6198627 | 0.017023546 | 0.18247904 | 0.09329042 |

| | | | | |
|---|---|---|---|---|
| Gm28438 | 233.7213687 | 0.131614565 | 0.185782229 | 0.708434633 |
| Gm28437 | 654.6201061 | 0.142851893 | 0.185693403 | 0.769289003 |
| Gm10925 | 568.5256472 | 0.269044782 | 0.191647847 | 1.403849745 |
| Gm28661 | 361.8513258 | 0.142602722 | 0.139902235 | 1.019302673 |
| Gm29216 | 452.3609317 | 0.14168015 | 0.179019598 | 0.791422567 |
| Lmbrd1 | 1834.054128 | -0.123654385 | 0.057989696 | -2.132350992 |
| Gm28594 | 0 | NA | NA | NA |
| Gm28237 | 0.181654796 | 0.274390172 | 3.327599044 | 0.082458904 |
| Gm9443 | 0.049553047 | 0.274391604 | 3.724945049 | 0.073663262 |
| Gm29414 | 0.167977605 | 1.222129108 | 2.756630383 | 0.443341667 |
| Adgrb3 | 2707.39029 | -0.24625233 | 0.121328924 | -2.029625919 |
| Gm15702 | 0.267089184 | -0.183014427 | 1.805598049 | -0.101359451 |
| Gm20172 | 56.45506021 | -0.739410763 | 0.284064533 | -2.602967559 |
| Gm22260 | 10.97169895 | -0.585900101 | 0.345882482 | -1.693928232 |
| Gm37543 | 46.77977661 | -0.532512182 | 0.330810644 | -1.609719006 |
| Gm37331 | 188.7440919 | -0.600244242 | 0.20212169 | -2.969717122 |
| Gm37211 | 123.4678815 | -0.67269366 | 0.241488444 | -2.785614289 |
| Gm37862 | 145.3838556 | -0.601680467 | 0.236375849 | -2.545439684 |
| Gm38286 | 155.9317435 | -0.583301576 | 0.230125788 | -2.534707565 |
| Gm37880 | 213.8342296 | -0.715528579 | 0.297847502 | -2.402331981 |
| Gm25294 | 0 | NA | NA | NA |
| Gm37011 | 137.8405266 | -0.696583062 | 0.181876235 | -3.829983956 |
| Gm29214 | 43.40613233 | -0.539045439 | 0.220931195 | -2.43987925 |
| Gm37051 | 172.0940709 | -0.656415985 | 0.216508564 | -3.031824567 |
| Gm38260 | 342.1641184 | -0.520381914 | 0.174502339 | -2.982091336 |
| Gm37628 | 460.1944707 | -0.664765209 | 0.246346903 | -2.698492247 |
| 8030445P17Rik | 222.3518729 | -0.742649992 | 0.244702154 | -3.034914003 |
| Gm38198 | 365.9544784 | -0.771038206 | 0.294501089 | -2.618116655 |
| Gm37077 | 292.4722539 | -0.839800201 | 0.18580655 | -4.519755638 |
| Gm37895 | 566.8750103 | -0.65183922 | 0.181700005 | -3.587447447 |
| Gm36952 | 499.6234323 | -0.731119466 | 0.208478624 | -3.50692772 |
| Gm37602 | 362.6410622 | -0.590879288 | 0.149886011 | -3.942191033 |
| 2900022M07Rik | 83.257532 | -0.930422566 | 0.270068123 | -3.445140273 |
| 2900002M20Rik | 64.13250466 | -0.770549857 | 0.245930004 | -3.133208008 |
| Gm9884 | 9.752194563 | -1.017456227 | 0.352507505 | -2.886339191 |
| Gm18443 | 0 | NA | NA | NA |
| Gm19123 | 0 | NA | NA | NA |
| Gm24064 | 0 | NA | NA | NA |
| 4931408C20Rik | 0 | NA | NA | NA |
| n-R5s209 | 0 | NA | NA | NA |
| Gm19641 | 0 | NA | NA | NA |
| Gm5524 | 0.04393175 | -0.206490061 | 3.724945049 | -0.055434391 |
| Gm18776 | 0 | NA | NA | NA |
| Gm19028 | 0.124853745 | 1.290908285 | 3.721447108 | 0.346883416 |
| Gm38089 | 0 | NA | NA | NA |
| Gm7846 | 6.28322175 | 0.265622382 | 0.414161145 | 0.641350319 |
| Gm597 | 0.068097359 | -0.206487837 | 3.724945049 | -0.055433794 |

| | | | | |
|---|---|---|---|---|
| Gm5697 | 0.151010504 | -0.687376735 | 2.854938236 | -0.240767638 |
| Gm6462 | 0 | NA | NA | NA |
| Gm18033 | 0 | NA | NA | NA |
| Gm25586 | 0 | NA | NA | NA |
| Gm22679 | 0 | NA | NA | NA |
| Gm23785 | 0 | NA | NA | NA |
| Mir6342 | 0.042716945 | -0.206490061 | 3.724945049 | -0.055434391 |
| Gm5525 | 5.440920671 | 0.795126735 | 0.462155422 | 1.720474751 |
| Gm7858 | 0 | NA | NA | NA |
| Gm23771 | 0 | NA | NA | NA |
| Gm27430 | 0.117046891 | -0.687371727 | 3.724945049 | -0.184532045 |
| Gm9898 | 0.52260535 | -0.192488626 | 1.275665364 | -0.150892727 |
| Gm6473 | 7.058970522 | 0.059613252 | 0.370109896 | 0.161069058 |
| Gm2914 | 0.311886478 | 2.048824048 | 1.733757825 | 1.18172447 |
| Phf3 | 1888.771435 | -0.124116127 | 0.058770114 | -2.111891872 |
| Gm37522 | 62.34479475 | -0.671109153 | 0.230960735 | -2.905728341 |
| Gm37906 | 105.3489438 | -0.322087344 | 0.222469574 | -1.447781548 |
| Ptp4a1 | 625.4291984 | 0.244059196 | 0.115979172 | 2.104336424 |
| Gm5698 | 0 | NA | NA | NA |
| Gm9458 | 0 | NA | NA | NA |
| Gm5699 | 0.061920613 | 0.274393828 | 3.724945049 | 0.073663859 |
| Gm29669 | 0 | NA | NA | NA |
| Gm28644 | 0 | NA | NA | NA |
| Gm27362 | 0 | NA | NA | NA |
| 4931428L18Rik | 0.048949532 | -0.206487837 | 3.724945049 | -0.055433794 |
| Gm29128 | 0 | NA | NA | NA |
| Gm7893 | 0 | NA | NA | NA |
| Lgsn | 0.247864911 | 1.241966195 | 2.517676371 | 0.493298587 |
| Pih1d3 | 0 | NA | NA | NA |
| Gm26145 | 0 | NA | NA | NA |
| Gm4850 | 0 | NA | NA | NA |
| Gm28645 | 0.066050875 | 0.274391604 | 3.724945049 | 0.073663262 |
| Gm6489 | 0 | NA | NA | NA |
| Gm10307 | 0 | NA | NA | NA |
| Gm19680 | 8.167021386 | 0.633716816 | 0.388711747 | 1.630300143 |
| Gm23032 | 0 | NA | NA | NA |
| Gm7910 | 0 | NA | NA | NA |
| Khdrbs2 | 245.3494503 | -0.141982082 | 0.137127808 | -1.035399634 |
| Gm25814 | 0.068097359 | -0.206487837 | 3.724945049 | -0.055433794 |
| Gm37999 | 19.71845274 | -0.159334917 | 0.345440481 | -0.461251433 |
| Gm37110 | 1.003269596 | -0.654360082 | 0.926416652 | -0.706334542 |
| Gm37911 | 37.09896806 | -0.204400769 | 0.296673314 | -0.688975919 |
| Gm9839 | 0.136963818 | 0.755286583 | 3.59954433 | 0.209828388 |
| Gm37792 | 46.79344322 | -0.282558793 | 0.233907046 | -1.207996075 |
| Gm5415 | 0.486008019 | 0.659817138 | 1.289552062 | 0.511663823 |
| Gm24732 | 0 | NA | NA | NA |
| Gm37426 | 6.59084447 | -0.249334542 | 0.434673938 | -0.573612817 |

| | | | | |
|---|---|---|---|---|
| Gm23024 | 0.184258256 | 1.06537733 | 2.688660402 | 0.396248381 |
| Gm37151 | 1.344706689 | 1.403292365 | 0.994392825 | 1.41120524 |
| Gm28624 | 0.510604427 | 0.297446887 | 1.529772331 | 0.194438664 |
| Gm37724 | 1.002429578 | 0.33405064 | 1.11319759 | 0.300082073 |
| Gm7114 | 3.168220283 | 0.392341091 | 0.531684729 | 0.737920557 |
| Gm4785 | 0.341712212 | 0.785314789 | 1.555097541 | 0.504993911 |
| Gm24901 | 0 | NA | NA | NA |
| Gm37591 | 0.710426184 | 2.891844909 | 1.327744297 | 2.178013429 |
| Gm2987 | 0 | NA | NA | NA |
| Gm25792 | 0 | NA | NA | NA |
| Gm18799 | 2.333919785 | 0.272834517 | 0.716038846 | 0.381033122 |
| Gm7933 | 0.088383099 | -0.687376175 | 3.724945049 | -0.184533239 |
| Gm23453 | 0 | NA | NA | NA |
| Prim2 | 97.92590214 | -0.11719652 | 0.12665491 | -0.925321565 |
| Gm29228 | 0 | NA | NA | NA |
| Gm37618 | 13.20070015 | -1.119199638 | 0.306145677 | -3.655774762 |
| Gm17813 | 0 | NA | NA | NA |
| 1700001G17Rik | 1.688998503 | -1.395951202 | 0.827350251 | -1.687255428 |
| Gm38173 | 0.528487055 | 2.354941994 | 1.404144698 | 1.677136265 |
| Rab23 | 419.6998972 | -0.190228583 | 0.059530059 | -3.195504674 |
| Bag2 | 54.75618287 | -0.272496711 | 0.165240735 | -1.6490892 |
| Gm37905 | 0.392625049 | -1.310578273 | 1.46196489 | -0.896449895 |
| Zfp451 | 1940.361675 | -0.165788633 | 0.068894389 | -2.406417077 |
| Gm15455 | 3.94145278 | 0.815324943 | 0.546567179 | 1.491719544 |
| Bend6 | 1529.651517 | -0.206283231 | 0.129242862 | -1.596089933 |
| Gm37067 | 37.0527924 | -1.123846841 | 0.365742495 | -3.072781689 |
| Dst | 13187.49407 | 0.048046118 | 0.06210017 | 0.773687384 |
| Gm37354 | 31.03348255 | -0.891147494 | 0.320644875 | -2.779235109 |
| Gm28631 | 0 | NA | NA | NA |
| Gm23792 | 0 | NA | NA | NA |
| Gm38157 | 47.50224476 | -0.464749019 | 0.324749143 | -1.431101602 |
| mmu-mir-6896 | 1.465090628 | -0.673497334 | 0.882986077 | -0.762749664 |
| Gm26788 | 4.676338759 | 0.724575064 | 0.455997002 | 1.58899085 |
| D630036G22Rik | 69.39949552 | 0.448056923 | 0.196869736 | 2.275905543 |
| A930005N03Rik | 0.933986728 | -0.249696651 | 1.013890201 | -0.24627583 |
| Gm37958 | 0 | NA | NA | NA |
| Gm37385 | 0.204111068 | 1.256955863 | 2.597989506 | 0.483818684 |
| Gm5266 | 0 | NA | NA | NA |
| Gm3052 | 10.39265568 | -0.966167821 | 0.34625297 | -2.790352448 |
| Gm37233 | 84.28593999 | -0.40138973 | 0.271884106 | -1.476326573 |
| Mir5103 | 0 | NA | NA | NA |
| Ccdc115 | 140.6358465 | 0.254152587 | 0.119272386 | 2.130858584 |
| Gm28306 | 0 | NA | NA | NA |
| Imp4 | 379.5447932 | 0.009769667 | 0.085842503 | 0.113809202 |
| Ptpn18 | 38.27817853 | 1.701805519 | 0.273923735 | 6.212698287 |
| Gm28417 | 1.241452806 | 1.899843318 | 1.086853249 | 1.748021933 |
| 4930568A12Rik | 0.934086492 | 1.472634997 | 1.115617108 | 1.320018299 |

| | | | | |
|---|---|---|---|---|
| Prss39 | 0.301806123 | -0.206484774 | 1.759603476 | -0.117347332 |
| Gm28416 | 0 | NA | NA | NA |
| Cfc1 | 0.191490881 | 0.299687168 | 2.702348657 | 0.11089878 |
| Gm28419 | 0 | NA | NA | NA |
| Prss40 | 0 | NA | NA | NA |
| 1700101I19Rik | 0.559566189 | -0.708550064 | 3.596332518 | -0.197020175 |
| Amer3 | 415.894219 | 0.232383663 | 0.092139061 | 2.522097151 |
| Gm28418 | 0.045882455 | 0.755288839 | 3.724945049 | 0.202765096 |
| Gm36984 | 0.129408498 | 1.221765824 | 2.99039852 | 0.408562877 |
| Gm29730 | 0.628347421 | 3.244593563 | 1.196571547 | 2.71157506 |
| AA619741 | 4.444019085 | 5.523907397 | 1.027797719 | 5.374508322 |
| Arhgef4 | 1837.032545 | 0.003530075 | 0.050109382 | 0.070447387 |
| AC107766.1 | 0 | NA | NA | NA |
| Gm37224 | 4.445654744 | -0.209327239 | 0.455981072 | -0.459070018 |
| Gm37845 | 4.660398994 | -0.449656412 | 0.632433621 | -0.710993845 |
| Gm38336 | 10.69144326 | -1.061581803 | 0.421085202 | -2.521061767 |
| Fam168b | 3287.598993 | -0.04743587 | 0.048061948 | -0.986973508 |
| Gm37127 | 1.263832448 | 0.299604075 | 1.044314416 | 0.286890682 |
| Plekhb2 | 2551.489122 | -0.070354159 | 0.072019523 | -0.976876207 |
| Gm19430 | 0.332159689 | 0.755304491 | 1.991498696 | 0.379264366 |
| Gm37670 | 0.517156391 | 0.411716847 | 1.293883105 | 0.318202506 |
| Gm8009 | 0.062426873 | 0.755288839 | 3.724945049 | 0.202765096 |
| Gm38027 | 0 | NA | NA | NA |
| Gm25096 | 0 | NA | NA | NA |
| Gm37068 | 0.62070299 | 0.898267789 | 1.182995429 | 0.759316365 |
| Gm25634 | 0 | NA | NA | NA |
| Gm28415 | 0 | NA | NA | NA |
| 1110002O04Rik | 0.877137428 | -2.391699921 | 1.115802317 | -2.143479973 |
| Gm8022 | 0 | NA | NA | NA |
| Gm37146 | 0 | NA | NA | NA |
| Gm33222 | 0.331946588 | -0.690773685 | 1.717255376 | -0.40225449 |
| 4930535G08Rik | 0 | NA | NA | NA |
| Gm38049 | 0.068097359 | -0.206487837 | 3.724945049 | -0.055433794 |
| Hs6st1 | 777.2491163 | 0.060671252 | 0.07528733 | 0.805862712 |
| Gm37335 | 0.364365233 | -0.557861579 | 1.542152934 | -0.36174206 |
| Gm33280 | 0.253548292 | 0.255732865 | 1.825859719 | 0.140061617 |
| Uggt1 | 1213.9728 | -0.199536799 | 0.059623306 | -3.346624202 |
| Mir6897 | 0.539626406 | -0.857629997 | 1.419296455 | -0.604264172 |
| Neurl3 | 71.81858134 | 3.70552378 | 0.368271285 | 10.06194056 |
| Arid5a | 533.3445611 | 1.974954119 | 0.248195098 | 7.957264792 |
| 4930403P22Rik | 0.909075398 | 2.938193824 | 2.152865161 | 1.364783023 |
| Kansl3 | 1110.948051 | 0.26455923 | 0.070624147 | 3.746016617 |
| Mir6898 | 0.044191549 | -0.206490061 | 3.724945049 | -0.055434391 |
| Fer1l5 | 1.367163391 | 1.669094389 | 1.082803444 | 1.541456484 |
| Lman2l | 453.4304442 | -0.358418799 | 0.091841461 | -3.902581651 |
| Cnnm4 | 233.328149 | -0.08200973 | 0.107456114 | -0.763192772 |
| Gm37909 | 10.35866414 | -0.707790047 | 0.371094137 | -1.907305928 |

| | | | | |
|---|---|---|---|---|
| AC084391.1 | 0 | NA | NA | NA |
| Cnnm3 | 394.0709263 | 0.034025262 | 0.079305728 | 0.429039142 |
| Gm38033 | 0.17077006 | 1.204318064 | 2.737003855 | 0.440013288 |
| Ankrd23 | 35.9631227 | -0.283036495 | 0.19047126 | -1.485980062 |
| Ankrd39 | 86.73678357 | 0.095119192 | 0.138248374 | 0.688031181 |
| Sema4c | 276.9910831 | 0.420073586 | 0.11833604 | 3.549836419 |
| D430040D24Rik | 19.3189591 | 1.069877997 | 0.241578045 | 4.428705425 |
| Fam178b | 2.063825205 | -0.386832141 | 0.748845108 | -0.516571634 |
| Gm6428 | 1.366437448 | -0.923620879 | 0.901140081 | -1.024947062 |
| Gm24133 | 0 | NA | NA | NA |
| Cox5b | 1330.17609 | 0.04737155 | 0.118364024 | 0.400219156 |
| Actr1b | 2273.379145 | -0.155496909 | 0.039983356 | -3.889040922 |
| 4933424G06Rik | 1.049948633 | -0.099368891 | 1.207597881 | -0.082286407 |
| Gm33533 | 12.17216712 | -0.498194712 | 0.416504994 | -1.196131425 |
| RP23-247A1.6 | 1.790960893 | 0.597976919 | 0.780305109 | 0.766337311 |
| Zap70 | 42.79799715 | 1.935617962 | 0.295295824 | 6.554843674 |
| Gm18828 | 0 | NA | NA | NA |
| Tmem131 | 1521.534467 | -0.097989755 | 0.071243343 | -1.375423319 |
| Gm37506 | 4.051657186 | -0.441701899 | 0.482690907 | -0.915082287 |
| Gm38115 | 35.47402398 | -0.441840334 | 0.312333175 | -1.414644262 |
| F830112A20Rik | 59.60486576 | -0.339388637 | 0.257371872 | -1.318670272 |
| Rpl12-ps1 | 0.348012853 | 1.319432372 | 1.954522607 | 0.675066314 |
| n-R5s210 | 0 | NA | NA | NA |
| Vwa3b | 17.13175727 | 0.28961147 | 0.313748633 | 0.923068467 |
| Gm37020 | 0.834730223 | -0.876100005 | 1.092310075 | -0.802061635 |
| Cnga3 | 3.017996072 | 0.972665797 | 0.751356594 | 1.294546164 |
| 1700074A21Rik | 0 | NA | NA | NA |
| Inpp4a | 3160.431147 | -0.226204888 | 0.060708849 | -3.726061243 |
| 4930439A04Rik | 0.093831153 | 0.274404949 | 3.724945049 | 0.073666845 |
| Gm37135 | 0 | NA | NA | NA |
| Coa5 | 899.7403321 | 0.032352571 | 0.106913273 | 0.30260575 |
| Unc50 | 302.5469516 | 0.137888206 | 0.102336737 | 1.347396937 |
| Mgat4a | 840.1121684 | 0.043368904 | 0.049979752 | 0.867729466 |
| 4930594C11Rik | 47.43194901 | -0.27091275 | 0.314966078 | -0.860133105 |
| 2010300C02Rik | 1765.094823 | 0.018381842 | 0.173337636 | 0.106046456 |
| 4930556I23Rik | 0.066050875 | 0.274391604 | 3.724945049 | 0.073663262 |
| 4930470B04Rik | 0.188205214 | -0.206489193 | 2.682879169 | -0.076965521 |
| Gm19863 | 38.19950049 | -0.809793778 | 0.405977769 | -1.994675178 |
| Tsga10 | 318.6599432 | -0.61864652 | 0.123233279 | -5.020125449 |
| Gm26805 | 21.25672272 | -0.283194426 | 0.283967734 | -0.997276777 |
| Lipt1 | 33.80092537 | -0.424460787 | 0.187306704 | -2.266127041 |
| Mitd1 | 202.445721 | 1.907125517 | 0.182250481 | 10.4643099 |
| Mrpl30 | 491.4665078 | 0.59796255 | 0.104952729 | 5.697446424 |
| Lyg2 | 0.741337718 | -1.223793129 | 1.185708098 | -1.032120073 |
| Gm15457 | 0.21650593 | 0.274406903 | 2.171399783 | 0.126373276 |
| Lyg1 | 0.136120942 | 0.274393632 | 3.599844381 | 0.076223748 |
| Gm23409 | 0 | NA | NA | NA |

| | | | | |
|---|---|---|---|---|
| Txndc9 | 544.4411505 | 0.402274194 | 0.089525285 | 4.493414285 |
| Eif5b | 954.1717507 | 0.248242518 | 0.116760212 | 2.126088282 |
| Rev1 | 696.4174186 | -0.042008253 | 0.111618041 | -0.376357199 |
| Gm37062 | 35.92510916 | -0.032016121 | 0.216236888 | -0.148060405 |
| Gm5099 | 0.141094079 | 0.755285157 | 3.563850454 | 0.211929531 |
| Aff3 | 992.3491355 | -0.127948176 | 0.074190573 | -1.72458806 |
| Gm16150 | 1.482000077 | 0.52479031 | 0.89113256 | 0.588902632 |
| Gm16151 | 0 | NA | NA | NA |
| Gm16152 | 1.435447043 | -1.522808592 | 1.114356697 | -1.366536044 |
| Gm34727 | 0 | NA | NA | NA |
| Gm38135 | 0 | NA | NA | NA |
| 2610300A13Rik | 0 | NA | NA | NA |
| Lonrf2 | 2520.190482 | -0.396390979 | 0.079358349 | -4.994949907 |
| Chst10 | 707.3698007 | -0.383545809 | 0.055359057 | -6.928329868 |
| Gm37707 | 0 | NA | NA | NA |
| Gm6621 | 0.169398393 | 1.717931299 | 3.393596623 | 0.506227313 |
| Nms | 1.222583469 | 0.504194102 | 1.107041236 | 0.455442928 |
| Pdcl3 | 219.8699854 | 0.118891277 | 0.130248747 | 0.912801695 |
| Gm37309 | 20.51143474 | -0.775829477 | 0.34185793 | -2.269449992 |
| Gm37821 | 0.54509763 | 2.579122665 | 1.291986204 | 1.996246288 |
| Gm37091 | 0.205567437 | 0.776036775 | 2.592100219 | 0.299385328 |
| Mir6349 | 0 | NA | NA | NA |
| Npas2 | 366.6556838 | -0.096170886 | 0.157816406 | -0.609384588 |
| Gm20428 | 0.263795272 | -1.166794934 | 1.810819015 | -0.644346522 |
| Gm3617 | 6.543142137 | -1.152948312 | 0.5677172 | -2.030849712 |
| Rpl31 | 1558.50621 | 0.547616722 | 0.125747741 | 4.354883172 |
| Tbc1d8 | 368.5922422 | -0.264271524 | 0.104983345 | -2.517270943 |
| Gm37265 | 6.813130323 | -0.985893953 | 0.458976785 | -2.148025751 |
| Gm24826 | 0 | NA | NA | NA |
| ACEA_U3 | 3.059522033 | -0.951213787 | 0.664874703 | -1.430666233 |
| Cnot11 | 371.6623297 | 0.24399532 | 0.09879053 | 2.469825001 |
| Gm15832 | 37.25351844 | -0.012401063 | 0.304711311 | -0.040697745 |
| Snord89 | 3.528574751 | -0.267697577 | 0.565990752 | -0.472971646 |
| Rnf149 | 381.317406 | 0.741313732 | 0.17676672 | 4.19374039 |
| Gm23722 | 10.82063883 | 0.762531493 | 0.397516599 | 1.918238118 |
| Gm5100 | 0.671689889 | -0.828973761 | 1.34738268 | -0.615247452 |
| Creg2 | 1118.864952 | -0.0324011 | 0.149930879 | -0.216106917 |
| D930019O06Rik | 42.32280546 | -0.190841321 | 0.218402945 | -0.873803786 |
| Rfx8 | 1.59726488 | 0.565473264 | 0.779199749 | 0.725710275 |
| Gm3646 | 0.042716945 | -0.206490061 | 3.724945049 | -0.055434391 |
| 1700066B17Rik | 0.337370548 | 0.672142303 | 1.991998983 | 0.337421007 |
| Map4k4 | 1991.375243 | -0.1055006 | 0.047399928 | -2.225754428 |
| 4930456G14Rik | 5.719135214 | -0.091375929 | 0.480112412 | -0.190321946 |
| Il1r2 | 32.73985817 | 5.042235739 | 0.836149004 | 6.030307653 |
| Gm16894 | 21.14797858 | -0.424326764 | 0.244084079 | -1.738445069 |
| Gm37347 | 2.446674312 | 3.562344216 | 0.96847972 | 3.678284783 |
| Il1r1 | 110.9761096 | 1.136277125 | 0.160929464 | 7.060715271 |

| | | | | |
|---|---|---|---|---|
| Gm38070 | 5.586992304 | 0.375610489 | 0.435395136 | 0.862688757 |
| Il1rl2 | 14.87794066 | 0.201856353 | 0.338637478 | 0.596083913 |
| Gm17970 | 0.509421185 | -0.813541319 | 1.376950584 | -0.590828261 |
| Il1rl1 | 3.917850035 | -0.647672949 | 0.619835464 | -1.044911087 |
| Il18r1 | 13.77975115 | 5.705985972 | 0.884445988 | 6.451480417 |
| Il18rap | 55.22165596 | 7.487449732 | 0.871609347 | 8.590373384 |
| Gm35801 | 0.072025462 | 0.755288839 | 3.724945049 | 0.202765096 |
| Slc9a4 | 29.46342029 | -0.243903166 | 0.22479638 | -1.084995972 |
| Gm37435 | 0.132101749 | 0.274387056 | 3.724945049 | 0.073662041 |
| Slc9a2 | 85.94852822 | 0.397890612 | 0.169460818 | 2.347980001 |
| Gm37623 | 0 | NA | NA | NA |
| Gm37915 | 0.583248015 | 0.216158446 | 1.252319159 | 0.172606515 |
| Mfsd9 | 54.36389778 | -0.11779531 | 0.196900597 | -0.598247602 |
| Tmem182 | 1.853692403 | -1.515940038 | 0.997745827 | -1.519364949 |
| Gm5973 | 0 | NA | NA | NA |
| 4930448I06Rik | 0.044544648 | 0.755286614 | 3.724945049 | 0.202764498 |
| Gm28634 | 0.143022002 | 0.755289479 | 3.547809333 | 0.212888971 |
| Gm29260 | 4.366569511 | -0.156219903 | 0.489144387 | -0.319373802 |
| Gm5267 | 0 | NA | NA | NA |
| Gm8176 | 0 | NA | NA | NA |
| Gm9915 | 0.24066848 | 0.274411084 | 2.105493127 | 0.130331028 |
| Gm29664 | 0 | NA | NA | NA |
| Gm28175 | 3.438849202 | -0.362392068 | 0.519846168 | -0.69711405 |
| Gm28140 | 0.100902008 | 0.274391828 | 3.724945049 | 0.073663322 |
| Gm37047 | 0.333556256 | -0.859286027 | 1.509941237 | -0.569085741 |
| Pantr1 | 157.9407554 | -0.265042105 | 0.157095044 | -1.687144915 |
| 4930420N18Rik | 0 | NA | NA | NA |
| Gm20646 | 38.88834011 | -0.533654021 | 0.180912275 | -2.949794425 |
| Pou3f3 | 1367.814843 | -0.271071282 | 0.089649949 | -3.023663521 |
| Pantr2 | 14.1555736 | -0.923449903 | 0.357031448 | -2.586466569 |
| Dalir | 16.94524304 | -0.232577066 | 0.326700534 | -0.711896804 |
| Mrps9 | 398.9447759 | 0.071310012 | 0.08521488 | 0.836825824 |
| Gm23126 | 0 | NA | NA | NA |
| Gpr45 | 153.2010998 | -0.350635939 | 0.1368721 | -2.561778032 |
| Gm24001 | 0.318202806 | -0.769457448 | 2.004696787 | -0.383827347 |
| Tgfbrap1 | 851.4724595 | 0.042317145 | 0.056550713 | 0.748304354 |
| Gm28782 | 1.198752213 | -2.962888331 | 1.211017544 | -2.446610577 |
| 8430432A02Rik | 7.526368882 | 0.194061267 | 0.436747796 | 0.444332563 |
| AI597479 | 352.8821411 | -0.302863945 | 0.080791831 | -3.748695153 |
| Fhl2 | 255.1157235 | 0.55723614 | 0.120491624 | 4.624687791 |
| Gm8210 | 95.32441064 | 0.424300556 | 0.190005251 | 2.233099103 |
| Gm29040 | 0.048949532 | -0.206487837 | 3.724945049 | -0.055433794 |
| Gm29610 | 0 | NA | NA | NA |
| Gm29041 | 0 | NA | NA | NA |
| Nck2 | 373.3238872 | -0.061648087 | 0.081494145 | -0.756472591 |
| Gm37536 | 17.59062222 | -0.620846814 | 0.341400655 | -1.818528476 |
| Gm38228 | 18.03991603 | -0.847020369 | 0.364364509 | -2.324651134 |

| | | | | |
|---|---|---|---|---|
| 1500015O10Rik | 35.22777033 | -0.530779488 | 0.278610524 | -1.905094898 |
| Gm29155 | 11.99474671 | 0.131635438 | 0.343375813 | 0.383356758 |
| Gm29157 | 0.378942021 | -1.120070302 | 1.499217723 | -0.747103163 |
| Gm29156 | 12.6916125 | 0.097660765 | 0.331561328 | 0.294548118 |
| Uxs1 | 508.6202783 | -0.077098053 | 0.082335012 | -0.936394509 |
| Gm16103 | 0 | NA | NA | NA |
| Gm19587 | 1.101887088 | 1.178782961 | 0.852615955 | 1.38254856 |
| Gm8241 | 0 | NA | NA | NA |
| Gm26352 | 0 | NA | NA | NA |
| Gm24180 | 0.220091977 | -0.206480603 | 2.162429275 | -0.095485483 |
| Tpp2 | 1340.114203 | 0.200199326 | 0.083666117 | 2.392836348 |
| Mettl21c | 0.903763266 | 1.311233399 | 1.043143586 | 1.257001832 |
| Gm15838 | 0 | NA | NA | NA |
| Gm8251 | 2.971797373 | 0.477456011 | 0.623315083 | 0.765994637 |
| Tex30 | 39.96396815 | 0.406490466 | 0.233540411 | 1.740557294 |
| Gm29247 | 0 | NA | NA | NA |
| Kdelc1 | 74.7610229 | -0.32909321 | 0.130654257 | -2.518809707 |
| Bivm | 338.2242815 | -0.175455693 | 0.072697892 | -2.4134908 |
| Ercc5 | 350.2066079 | -0.012647127 | 0.059010141 | -0.214321247 |
| Mettl21e | 0.63532651 | 1.616064647 | 1.180420473 | 1.36905847 |
| Gm15833 | 0.607153663 | -0.083252447 | 1.26131315 | -0.066004582 |
| Gm5700 | 0 | NA | NA | NA |
| Gm28893 | 0 | NA | NA | NA |
| Txn-ps1 | 3.667605082 | 1.12173834 | 0.629945566 | 1.780690906 |
| Gulp1 | 83.35613432 | 0.021999974 | 0.152797289 | 0.14398144 |
| Gm37626 | 20.77771962 | -0.399576926 | 0.298062943 | -1.340579015 |
| Gm16102 | 1.145880322 | -1.693052113 | 0.942557489 | -1.796232201 |
| 4930521E06Rik | 0.181654796 | 0.274390172 | 3.327599044 | 0.082458904 |
| Gm38110 | 3.191642098 | -0.050273505 | 0.546878719 | -0.091928069 |
| Gm3489 | 0.12092566 | 1.236179401 | 3.724945049 | 0.331865138 |
| Gm19480 | 0 | NA | NA | NA |
| Gm3496 | 0 | NA | NA | NA |
| Gm5268 | 0 | NA | NA | NA |
| Col3a1 | 232.7682847 | 0.908932798 | 0.191497135 | 4.746456382 |
| AC125167.1 | 0 | NA | NA | NA |
| Col5a2 | 180.1321098 | 0.477581254 | 0.190492083 | 2.5070924 |
| Gm38211 | 9.579960985 | -0.361584557 | 0.433288922 | -0.834511427 |
| Gm8296 | 0.048949532 | -0.206487837 | 3.724945049 | -0.055433794 |
| Gm7063 | 0.241951302 | 0.274412273 | 2.123051817 | 0.129253686 |
| Gm8304 | 0 | NA | NA | NA |
| Gm23216 | 0 | NA | NA | NA |
| Gm23974 | 0 | NA | NA | NA |
| Gm8307 | 0.148708979 | 0.755295232 | 2.868529146 | 0.263304012 |
| Gm23984 | 0 | NA | NA | NA |
| Wdr75 | 417.7461979 | 0.15551619 | 0.067535427 | 2.302734964 |
| Gm28906 | 0 | NA | NA | NA |
| Gm5526 | 49.19564311 | 0.018674466 | 0.219125828 | 0.085222571 |

| | | | | |
|---|---|---|---|---|
| Gm5269 | 0.154938607 | 0.274408374 | 2.832095734 | 0.096892337 |
| Gm18303 | 0.136201742 | -0.67532486 | 2.944131489 | -0.229379993 |
| Slc40a1 | 53.73490417 | -1.301192026 | 0.261045688 | -4.984537527 |
| Gm9496 | 0 | NA | NA | NA |
| Rpl23a-ps1 | 0 | NA | NA | NA |
| Gm3852 | 0.218641893 | 0.274411858 | 2.185623723 | 0.125553111 |
| Stk-ps2 | 0 | NA | NA | NA |
| Gm18849 | 0 | NA | NA | NA |
| Dnah7b | 166.4217659 | -0.604194283 | 0.14781275 | -4.087565407 |
| n-R5s211 | 0.044191549 | -0.206490061 | 3.724945049 | -0.055434391 |
| Gm8326 | 1.175435645 | 0.845666044 | 0.852153181 | 0.992387358 |
| Dnah7c | 35.29196087 | -0.388612086 | 0.2272773 | -1.709858779 |
| Slc39a10 | 1977.950324 | 0.120648805 | 0.116605292 | 1.034676922 |
| Gm28151 | 62.07798039 | -0.04053154 | 0.230700366 | -0.175689101 |
| Gm37393 | 2.121468972 | -1.628891457 | 1.039313671 | -1.567276081 |
| Gm28527 | 0.844069991 | 0.857607178 | 1.093022225 | 0.784620073 |
| Gm28825 | 0.476041195 | -1.46950433 | 1.450962556 | -1.012778947 |
| Gm28826 | 0.77428951 | 2.966889009 | 1.180077284 | 2.514148057 |
| Gm4852 | 3.57865669 | 0.580033083 | 0.508736757 | 1.140143846 |
| Gm38090 | 0 | NA | NA | NA |
| Mir6350 | 0 | NA | NA | NA |
| Gm18117 | 0.885425998 | 3.151703885 | 1.472495442 | 2.140382779 |
| Gm37196 | 1.287564413 | 0.626051459 | 1.236045366 | 0.506495535 |
| Gm8337 | 21.49979967 | -0.647241559 | 0.298556985 | -2.167899569 |
| Gm22419 | 0 | NA | NA | NA |
| Gm23829 | 0 | NA | NA | NA |
| Gm19206 | 0 | NA | NA | NA |
| Gm5974 | 0 | NA | NA | NA |
| Gm23240 | 0 | NA | NA | NA |
| Gm28830 | 0 | NA | NA | NA |
| Gm8354 | 0 | NA | NA | NA |
| Gm19124 | 0 | NA | NA | NA |
| Gm8357 | 0 | NA | NA | NA |
| Gm22693 | 0 | NA | NA | NA |
| Gm28740 | 0 | NA | NA | NA |
| Gm8367 | 0 | NA | NA | NA |
| C230029F24Rik | 0 | NA | NA | NA |
| Gm6757 | 0 | NA | NA | NA |
| Gm3605 | 0 | NA | NA | NA |
| Gm28388 | 0 | NA | NA | NA |
| Gm8384 | 0 | NA | NA | NA |
| 4933411E06Rik | 0.068097359 | -0.206487837 | 3.724945049 | -0.055433794 |
| Gm8391 | 0 | NA | NA | NA |
| Gm27330 | 0 | NA | NA | NA |
| Gm29665 | 0 | NA | NA | NA |
| Gm20118 | 0 | NA | NA | NA |
| Gm25843 | 0 | NA | NA | NA |

| | | | | |
|---|---|---|---|---|
| Gm28321 | 0.092691418 | 0.274387304 | 3.724945049 | 0.073662108 |
| Tmeff2 | 912.3017426 | -0.250112899 | 0.073791638 | -3.389447723 |
| Gm28322 | 2.545969209 | -1.930761994 | 0.821330623 | -2.350773173 |
| Gm25736 | 12.09809119 | -1.09932092 | 0.354644489 | -3.099782894 |
| Gm28319 | 0.231621402 | 0.683942533 | 2.519475169 | 0.271462303 |
| Gm28320 | 83.01456838 | -0.730279958 | 0.22412077 | -3.258421605 |
| Gm29325 | 37.76672809 | -0.274161174 | 0.261116192 | -1.049958533 |
| 9330175M20Rik | 4.588839173 | 0.197764349 | 0.544822415 | 0.362988642 |
| Sdpr | 92.2652614 | 0.54573629 | 0.148987702 | 3.66296199 |
| Gm5253 | 0.982135049 | -1.91930018 | 1.13789072 | -1.686717491 |
| 1700072G22Rik | 0.174149299 | 0.755285184 | 3.362794904 | 0.224600431 |
| Nabp1 | 240.5774001 | -0.080595741 | 0.195933339 | -0.411342662 |
| Gm17767 | 3.178614829 | 1.494443889 | 0.658866868 | 2.268203126 |
| Gm37535 | 0.140088945 | -1.111948817 | 3.601861891 | -0.308715006 |
| Gm28055 | 0.781209455 | -1.097411741 | 1.299411499 | -0.844545197 |
| Myo1b | 450.2003702 | -0.031569445 | 0.130290629 | -0.242300193 |
| Gm29453 | 3.059357988 | 0.467699508 | 0.634805591 | 0.736760221 |
| Gm28323 | 3.764028883 | -1.819610507 | 0.571626373 | -3.183216507 |
| Gm8419 | 0 | NA | NA | NA |
| Gm27607 | 0 | NA | NA | NA |
| Gm8420 | 0.200957741 | 0.274391329 | 3.253433744 | 0.084338994 |
| Stat4 | 19.2643103 | 2.015546291 | 0.310747962 | 6.486112657 |
| Gm28177 | 2.000698716 | 3.396174647 | 0.992864456 | 3.420582364 |
| Gm3940 | 0.598154365 | -0.681555592 | 1.354185325 | -0.503295656 |
| Stat1 | 2376.29098 | 4.844654695 | 0.114373669 | 42.35812976 |
| Gm18301 | 0.092228164 | 0.75528439 | 3.724945049 | 0.202763901 |
| Gls | 3738.856511 | -0.819509731 | 0.068387717 | -11.98328831 |
| Gm5975 | 0.042716945 | -0.206490061 | 3.724945049 | -0.055434391 |
| Gm23460 | 0 | NA | NA | NA |
| Nab1 | 590.3308437 | 0.031722215 | 0.115274967 | 0.275187374 |
| Gm5976 | 0.038981395 | 0.274393828 | 3.724945049 | 0.073663859 |
| Rps27a-ps1 | 3.694411362 | -0.423864663 | 0.62013918 | -0.683499248 |
| Gm553 | 3.129775993 | -0.019286423 | 0.548660766 | -0.035151816 |
| Gm28178 | 0.452230046 | 0.274413841 | 1.435248485 | 0.19119605 |
| Gm5527 | 9.620821135 | 0.37998176 | 0.373819995 | 1.016483242 |
| Tmem194b | 84.55902287 | -0.136941817 | 0.184074555 | -0.743947564 |
| Gm37242 | 11.17186488 | -0.387267307 | 0.308404754 | -1.255711212 |
| Mfsd6 | 1706.135027 | -0.021728289 | 0.078517995 | -0.27673006 |
| Gm29670 | 25.59481537 | -0.535902285 | 0.29263146 | -1.831321501 |
| Gm28650 | 0.805100971 | -0.324133085 | 1.097088565 | -0.295448422 |
| Inpp1 | 600.5457346 | 0.759561188 | 0.106204934 | 7.151844611 |
| Hibch | 147.5834649 | -0.625761346 | 0.166980585 | -3.747509608 |
| Gm16107 | 1.228146034 | 3.025052058 | 1.02313701 | 2.956644153 |
| 1700019D03Rik | 237.2523422 | -0.025600152 | 0.088465216 | -0.289380995 |
| Mstn | 4.314567637 | 0.823419308 | 0.485236701 | 1.696943586 |
| Gm24349 | 0 | NA | NA | NA |
| 1700019A02Rik | 5.126622909 | -0.69550028 | 0.539135933 | -1.290027684 |

| | | | | |
|---|---|---|---|---|
| Pms1 | 234.6633514 | -0.1528701 | 0.092559129 | -1.651593988 |
| Gm28777 | 0 | NA | NA | NA |
| Ormdl1 | 92.39055438 | 0.174376877 | 0.157388886 | 1.107936405 |
| Gm28778 | 0.044191549 | -0.206490061 | 3.724945049 | -0.055434391 |
| Osgepl1 | 175.8191109 | -0.126362375 | 0.108413788 | -1.165556318 |
| Gm28551 | 1.277925253 | 0.302505329 | 0.860422718 | 0.351577571 |
| Gm18800 | 2.110212028 | -0.732150053 | 0.666892603 | -1.097853013 |
| Asnsd1 | 39.89699009 | -0.332337504 | 0.182914894 | -1.8168969 |
| Gm28779 | 0.097139547 | 0.274393828 | 3.724945049 | 0.073663859 |
| Gm6799 | 0 | NA | NA | NA |
| Gm6801 | 0 | NA | NA | NA |
| Dnah7a | 93.9630857 | 0.125305455 | 0.169167139 | 0.74071983 |
| Gm25240 | 0.108123235 | 0.274402725 | 3.724945049 | 0.073666248 |
| Stk17b | 400.9618622 | 0.872775821 | 0.145056678 | 6.016791725 |
| Gm37863 | 0 | NA | NA | NA |
| Hecw2 | 1282.188723 | -0.309659741 | 0.076283789 | -4.05931251 |
| Mir7681 | 0.068097359 | -0.206487837 | 3.724945049 | -0.055433794 |
| Gm24251 | 2.00647346 | 0.669725866 | 0.738702816 | 0.906624222 |
| Gm37633 | 69.5255469 | -0.835609928 | 0.345835859 | -2.416203832 |
| Gm25829 | 0.15586508 | 0.274407134 | 2.82680955 | 0.097073088 |
| Ccdc150 | 3.346209518 | -0.727734862 | 0.529932592 | -1.373259302 |
| Gtf3c3 | 606.9969703 | 0.066921213 | 0.06497758 | 1.029912359 |
| Pgap1 | 1020.253931 | -0.437214664 | 0.080015921 | -5.464095889 |
| Gm36955 | 30.15051617 | -1.064680741 | 0.24361773 | -4.370292513 |
| Gm19637 | 0.11197992 | 0.75528439 | 3.724945049 | 0.202763901 |
| Ankrd44 | 326.5543126 | 0.344481747 | 0.096088769 | 3.585036509 |
| 4930444A19Rik | 0.477440416 | 1.682558895 | 1.523111478 | 1.104685322 |
| Gm37370 | 15.65747994 | 0.610939506 | 0.284771163 | 2.145369986 |
| Gm24053 | 0 | NA | NA | NA |
| A130048G24Rik | 23.27167967 | 0.144152546 | 0.315729005 | 0.456570489 |
| Sf3b1 | 4398.794146 | 0.288090967 | 0.072655588 | 3.965159105 |
| E330011M16Rik | 2.812432955 | 0.50261149 | 0.660431997 | 0.761034432 |
| Coq10b | 351.6495304 | -0.244149327 | 0.131499754 | -1.856652345 |
| Gm17971 | 11.17290908 | -0.744335082 | 0.49717808 | -1.49711967 |
| Hspd1 | 1608.718732 | -0.085815232 | 0.102515036 | -0.837098979 |
| Hspe1 | 313.2638214 | 0.194332764 | 0.150140898 | 1.294335968 |
| Gm5147 | 0.483827566 | 0.820062608 | 1.390614049 | 0.589712587 |
| Gm16581 | 13.32551551 | 0.020057354 | 0.35687708 | 0.056202415 |
| Mob4 | 696.0879921 | 0.00156997 | 0.090743306 | 0.017301226 |
| Gm6822 | 0.477893029 | -0.191217155 | 1.417188568 | -0.134927108 |
| Rftn2 | 338.1297183 | -0.098605399 | 0.126790934 | -0.777700705 |
| Gm10561 | 20.71569524 | -0.443334825 | 0.253774878 | -1.746961041 |
| Mars2 | 418.0161997 | -0.064326911 | 0.111418188 | -0.577346595 |
| Boll | 0.988924806 | 1.483489183 | 1.193412624 | 1.243064764 |
| Gm8292 | 51.67441755 | 0.352685213 | 0.19465356 | 1.811861101 |
| Gm29138 | 0 | NA | NA | NA |
| Gm8531 | 0.047948698 | -0.206487837 | 3.724945049 | -0.055433794 |

| | | | | |
|---|---|---|---|---|
| Plcl1 | 931.6952418 | 0.224066103 | 0.093774696 | 2.389408993 |
| Gm38056 | 155.7691939 | -0.063592789 | 0.366397115 | -0.173562472 |
| Gm6644 | 4.857481711 | 0.981970999 | 0.494362978 | 1.986336037 |
| Gm37382 | 65.4083701 | -0.08337614 | 0.261442158 | -0.318908554 |
| 9130227L01Rik | 1.675708655 | -0.866756471 | 0.826206026 | -1.049080307 |
| Gm36986 | 0 | NA | NA | NA |
| 1700003I22Rik | 0 | NA | NA | NA |
| Gm28900 | 0.298162442 | 0.755308087 | 1.755280571 | 0.430306185 |
| Gm28240 | 0 | NA | NA | NA |
| Hsfy2 | 0 | NA | NA | NA |
| Satb2 | 608.4974551 | -0.184801617 | 0.197157783 | -0.937328541 |
| 9130024F11Rik | 127.5850177 | 0.14292463 | 0.23146776 | 0.617471004 |
| Gm4081 | 9.66177043 | -0.445321242 | 0.49830262 | -0.893676301 |
| Gm23943 | 0 | NA | NA | NA |
| 1700126A01Rik | 0.044544648 | 0.755286614 | 3.724945049 | 0.202764498 |
| BC055402 | 2.819616475 | 0.212592028 | 0.704883366 | 0.301598872 |
| Gm5254 | 0 | NA | NA | NA |
| 4930558J18Rik | 2.344527321 | -0.424412156 | 0.728915481 | -0.582251532 |
| 1700066M21Rik | 269.4751219 | -0.051194888 | 0.100254198 | -0.510650809 |
| Tyw5 | 138.2526831 | 0.448825444 | 0.093770854 | 4.786406687 |
| 9430016H08Rik | 289.7250437 | 0.430979067 | 0.114782809 | 3.754735336 |
| Gm22371 | 0 | NA | NA | NA |
| Gm24014 | 0 | NA | NA | NA |
| Gm8581 | 0 | NA | NA | NA |
| Gm37373 | 0.354472284 | -1.544478058 | 3.188679565 | -0.484362893 |
| Gm17929 | 0.044544648 | 0.755286614 | 3.724945049 | 0.202764498 |
| Spats2l | 531.7349419 | 0.910192649 | 0.088956232 | 10.2319155 |
| Gm17234 | 0.08486385 | 0.755286617 | 3.724945049 | 0.202764499 |
| Kctd18 | 231.8874669 | 0.251206686 | 0.124713123 | 2.014276288 |
| Sgol2a | 34.52874643 | 0.796849747 | 0.274355445 | 2.904442984 |
| Aox1 | 275.5358162 | 0.469306818 | 0.120284674 | 3.901634366 |
| Aox3 | 36.77314627 | -0.195155422 | 0.230112853 | -0.84808571 |
| Gm24548 | 0 | NA | NA | NA |
| Aox4 | 64.4352454 | -0.137822755 | 0.225107786 | -0.61225228 |
| Gm15759 | 0.15100991 | 1.236179172 | 3.527597384 | 0.350430913 |
| Aox2 | 2.732211411 | -1.090787057 | 0.659598162 | -1.653714518 |
| Gm37607 | 38.38692177 | -0.498043485 | 0.260965954 | -1.908461531 |
| Bzw1 | 2353.558553 | 0.217356722 | 0.094549598 | 2.298864567 |
| Clk1 | 1464.941916 | -0.255574797 | 0.088260216 | -2.895696501 |
| Ppil3 | 142.411199 | 0.020798745 | 0.094960829 | 0.219024467 |
| Nif3l1 | 211.5522702 | 0.195324658 | 0.091490627 | 2.134914413 |
| Orc2 | 434.4142678 | 0.055050307 | 0.093633384 | 0.587934614 |
| Gm23966 | 4.595236341 | 0.369215315 | 0.597495965 | 0.617937754 |
| Gm15834 | 26.01032292 | -0.069561535 | 0.256135105 | -0.271581416 |
| Fam126b | 3027.102502 | -0.185101516 | 0.085895344 | -2.154965666 |
| Ndufb3 | 313.2837583 | -0.020970894 | 0.110226827 | -0.19025218 |
| Gm29389 | 0 | NA | NA | NA |

| | | | | |
|---|---|---|---|---|
| Gm18802 | 0 | NA | NA | NA |
| Gm10068 | 3.8142007 | -0.172155565 | 0.513713402 | -0.335119863 |
| Als2cr12 | 0.590977814 | 1.670362995 | 1.33755992 | 1.248813582 |
| Gm20257 | 14.75086306 | 1.887218372 | 0.396270358 | 4.762451526 |
| Gm28803 | 0 | NA | NA | NA |
| Cflar | 951.4956611 | 1.521431473 | 0.131587063 | 11.56216602 |
| Gm3841 | 0 | NA | NA | NA |
| Casp8 | 157.9166945 | 2.915533068 | 0.239438481 | 12.17654346 |
| Gm37760 | 18.59560002 | 2.04529955 | 0.390577522 | 5.236603331 |
| Gm28802 | 1.046132847 | -0.931843686 | 1.185408323 | -0.78609511 |
| Trak2 | 2825.777811 | 0.011717537 | 0.069684188 | 0.168152014 |
| Stradb | 332.0964291 | -0.426904363 | 0.142670356 | -2.99224293 |
| Gm33589 | 0.118371171 | 0.75528439 | 3.724945049 | 0.202763901 |
| Gm29018 | 0.22549826 | 0.755304379 | 2.143385654 | 0.352388464 |
| Als2cr11 | 0.178692882 | 0.274408127 | 2.707499347 | 0.101351133 |
| Gm25502 | 0 | NA | NA | NA |
| Tmem237 | 159.7002594 | 0.022370829 | 0.10513554 | 0.212780844 |
| G730003C15Rik | 13.69276046 | -0.114036319 | 0.301562044 | -0.378152096 |
| Mpp4 | 26.50791038 | -0.807089005 | 0.258118742 | -3.126812868 |
| Gm29017 | 0.060174537 | 0.755286614 | 3.724945049 | 0.202764498 |
| Gm19125 | 0 | NA | NA | NA |
| Als2 | 1234.116172 | -0.238361181 | 0.107233108 | -2.222831974 |
| Cdk15 | 4.87154909 | -1.649091226 | 0.551568121 | -2.989823314 |
| Gm29016 | 0 | NA | NA | NA |
| Gm37737 | 0.176324247 | -0.171429079 | 2.757307263 | -0.062172642 |
| Fzd7 | 304.579202 | -0.090108303 | 0.155593807 | -0.579125252 |
| Gm26813 | 0.044191549 | -0.206490061 | 3.724945049 | -0.055434391 |
| Gm973 | 103.013939 | 0.267832899 | 0.152761954 | 1.753269665 |
| Gm28411 | 0.118371171 | 0.75528439 | 3.724945049 | 0.202763901 |
| 1700122D07Rik | 0.136120942 | 0.274393632 | 3.599844381 | 0.076223748 |
| Sumo1 | 630.9605461 | 0.496592838 | 0.1283615 | 3.868705488 |
| Gm24545 | 0.226675736 | 0.755308749 | 2.143799133 | 0.352322537 |
| D430013B06Rik | 32.24759346 | -0.156811263 | 0.296108263 | -0.529574088 |
| Gm29596 | 1.33134391 | -1.198248106 | 0.887187844 | -1.350613755 |
| Nop58 | 820.0043818 | 0.323023348 | 0.106344475 | 3.037518848 |
| Snord70 | 4.644160981 | 0.19067529 | 0.449043227 | 0.42462569 |
| Gm26293 | 0 | NA | NA | NA |
| Gm26287 | 2.814965633 | 0.062603188 | 0.546106529 | 0.114635487 |
| Snord11 | 1.304845187 | 0.520368884 | 1.074359578 | 0.484352627 |
| Bmpr2 | 5482.687723 | -0.044483854 | 0.055797851 | -0.797232399 |
| Gm38125 | 29.65508468 | -0.849901097 | 0.25257967 | -3.364883228 |
| Gm37977 | 0.18514425 | -1.164033542 | 3.292450568 | -0.353546247 |
| Fam117b | 1173.743163 | 0.112881563 | 0.069216155 | 1.630855746 |
| Gm37531 | 63.74214748 | -0.214797128 | 0.270498935 | -0.794077539 |
| Gm18829 | 0 | NA | NA | NA |
| Ica1l | 384.5618481 | -0.110319492 | 0.074459291 | -1.481608148 |
| Wdr12 | 444.9492766 | -0.100055878 | 0.077129856 | -1.297239259 |

| | | | | |
|---|---|---|---|---|
| Carf | 373.7108772 | -0.175883874 | 0.10339997 | -1.701005081 |
| Gm15464 | 5.596760458 | 0.251312088 | 0.451451323 | 0.556675936 |
| Nbeal1 | 1755.709698 | -0.398438076 | 0.083324244 | -4.781778486 |
| Gm29332 | 0 | NA | NA | NA |
| Cyp20a1 | 130.1781194 | 0.369608116 | 0.15893701 | 2.325500629 |
| 6030460B20Rik | 3.025787973 | -0.301054684 | 0.592941602 | -0.507730749 |
| Abi2 | 3457.462493 | -0.122063355 | 0.075702702 | -1.612404206 |
| Raph1 | 1729.282084 | 0.0486567 | 0.104583258 | 0.465243684 |
| Gm11575 | 9.518548027 | -0.470314587 | 0.354524702 | -1.326605973 |
| Gm11576 | 0.448283844 | 1.673601795 | 1.538701121 | 1.087671785 |
| Gm11578 | 18.19495742 | -1.257112866 | 0.395276762 | -3.180335873 |
| Gm38388 | 0.262256756 | 0.755309459 | 2.055088433 | 0.367531366 |
| Gm23762 | 0 | NA | NA | NA |
| Gm11577 | 0 | NA | NA | NA |
| Eif4a-ps4 | 428.6153442 | 0.407211467 | 0.116942459 | 3.482152428 |
| Cd28 | 53.32452365 | 4.074791101 | 0.3603677 | 11.30731499 |
| Gm11579 | 0.085327104 | 0.274389469 | 3.724945049 | 0.073662689 |
| 2310016D23Rik | 0.256420178 | 0.755299254 | 2.467917879 | 0.306047158 |
| Gm37198 | 0.546045768 | 1.20984264 | 1.436527639 | 0.842199348 |
| Gm11609 | 0.066050875 | 0.274391604 | 3.724945049 | 0.073663262 |
| Gm11581 | 0.146451277 | -0.68737672 | 2.882358805 | -0.238477152 |
| Gm38137 | 1.155240215 | -1.143478979 | 0.976619215 | -1.170854475 |
| Ctla4 | 27.96364358 | 3.439501113 | 0.474823599 | 7.243745087 |
| Rpl18-ps1 | 12.04120041 | 0.636722664 | 0.365615306 | 1.741509871 |
| Gm11582 | 0.048949532 | -0.206487837 | 3.724945049 | -0.055433794 |
| Icos | 90.9935698 | 6.546303006 | 0.753477296 | 8.688122446 |
| Gm37039 | 2.679819182 | 3.937332998 | 1.104887885 | 3.563558849 |
| Rpl17-ps1 | 1.010478028 | 2.147941499 | 1.202071464 | 1.786866724 |
| Gm11590 | 0 | NA | NA | NA |
| Gm11591 | 0 | NA | NA | NA |
| Gm28083 | 0 | NA | NA | NA |
| Gm11587 | 0.127971488 | 0.274389832 | 3.687653711 | 0.0744077 |
| 9530026F06Rik | 0 | NA | NA | NA |
| Gm11588 | 0 | NA | NA | NA |
| Gm25839 | 0 | NA | NA | NA |
| Gm28449 | 0.11147366 | 0.274389235 | 3.724945049 | 0.073662626 |
| Gm36943 | 0 | NA | NA | NA |
| Gm28094 | 0.046345709 | 0.274391604 | 3.724945049 | 0.073663262 |
| 4930587A21Rik | 0 | NA | NA | NA |
| Pard3b | 166.061435 | 0.402882539 | 0.133499125 | 3.017866507 |
| Gm37205 | 11.98801516 | 0.587390962 | 0.356836875 | 1.646104993 |
| Gm38012 | 7.816280804 | -0.462920147 | 0.391048628 | -1.183791769 |
| Gm38278 | 3.411970267 | -0.993945771 | 0.645491064 | -1.539828862 |
| Pard3bos1 | 0 | NA | NA | NA |
| Gm38075 | 5.595486472 | -0.044931576 | 0.472068613 | -0.09518018 |
| Gm29640 | 0 | NA | NA | NA |
| Gm29641 | 0.044191549 | -0.206490061 | 3.724945049 | -0.055434391 |

| | | | | |
|---|---|---|---|---|
| Pard3bos2 | 0.453587956 | -1.071538798 | 1.355740532 | -0.790371589 |
| Gm38351 | 3.154252535 | 1.163803781 | 0.754921652 | 1.541621939 |
| Pard3bos3 | 0.304629397 | 0.780354813 | 2.020941452 | 0.3861343 |
| Gm29084 | 2.918337729 | -0.754909588 | 0.670239954 | -1.126327345 |
| Gm11600 | 0 | NA | NA | NA |
| Gm11599 | 0.770699074 | 1.288299025 | 1.203577337 | 1.070391562 |
| Nrp2 | 974.5911931 | 0.126372585 | 0.091033321 | 1.388201413 |
| Gm29083 | 5.840962313 | 0.479205014 | 0.445915017 | 1.074655476 |
| Gm37121 | 13.64577828 | -0.875875403 | 0.362902899 | -2.413525506 |
| Gm4208 | 0.515886961 | -0.19889936 | 1.478299294 | -0.13454607 |
| Ino80d | 1352.80268 | -0.065041951 | 0.065815701 | -0.988243695 |
| AL645685.1 | 0 | NA | NA | NA |
| Gm24942 | 0 | NA | NA | NA |
| Rps13-ps7 | 0 | NA | NA | NA |
| Rps27-ps1 | 0 | NA | NA | NA |
| Ino80dos | 70.0990274 | -0.276840867 | 0.138081507 | -2.00490908 |
| Gm20342 | 231.0873436 | -0.67744746 | 0.227320516 | -2.980142184 |
| Ndufs1 | 1507.181369 | -0.106586313 | 0.052849316 | -2.016796452 |
| Gm24557 | 0.222111871 | 1.942901157 | 3.112888699 | 0.624147326 |
| Eef1b2 | 871.8357536 | 0.270152187 | 0.133534039 | 2.023096045 |
| Gm27390 | 0 | NA | NA | NA |
| Gm26457 | 0.157314084 | 0.755289343 | 3.435173412 | 0.219869349 |
| snoZ196 | 0 | NA | NA | NA |
| Snora41 | 4.156744695 | 0.359639415 | 0.546268471 | 0.658356531 |
| Gpr1 | 4.89974049 | -0.724240959 | 0.447907303 | -1.61694385 |
| Gm27512 | 0.049553047 | 0.274391604 | 3.724945049 | 0.073663262 |
| Platr12 | 0.049553047 | 0.274391604 | 3.724945049 | 0.073663262 |
| 1700039I01Rik | 0.097899064 | -0.687371727 | 3.724945049 | -0.184532045 |
| Gm11608 | 0.112029109 | -0.687373951 | 3.724945049 | -0.184532642 |
| Gm24513 | 0.109124069 | 0.274402725 | 3.724945049 | 0.073666248 |
| Zdbf2 | 969.9858277 | -0.605657676 | 0.152044848 | -3.983414666 |
| Gm23448 | 0.180187022 | 0.755304779 | 2.70035346 | 0.279705894 |
| Gm11606 | 0 | NA | NA | NA |
| Gm11605 | 3.581982469 | 0.529967987 | 0.546345894 | 0.970022824 |
| Adam23 | 3251.808784 | -0.245167094 | 0.05879671 | -4.16974169 |
| Gm22281 | 0 | NA | NA | NA |
| Dytn | 1.029483559 | 0.046694525 | 0.928483945 | 0.05029115 |
| Gm12748 | 0 | NA | NA | NA |
| Mdh1b | 8.408068065 | -0.499544873 | 0.360271536 | -1.386578799 |
| Fastkd2 | 291.2035401 | -0.134719573 | 0.126798389 | -1.062470695 |
| 4933402D24Rik | 1.548786521 | 0.311315453 | 0.803127629 | 0.387628867 |
| Gm13751 | 0.449839025 | 0.274413195 | 1.580760945 | 0.173595632 |
| Gm13754 | 0.863190071 | 0.269906728 | 1.041158821 | 0.259236845 |
| Gm13749 | 1.958865323 | -1.995769014 | 0.756684598 | -2.637517692 |
| Klf7 | 1205.650868 | 0.020888161 | 0.103050465 | 0.202698367 |
| Gm26649 | 0.52188035 | 1.714548857 | 1.554453422 | 1.102991465 |
| Mir6899 | 0.842388825 | 0.28993969 | 1.135882725 | 0.255254952 |

| | | | | |
|---|---|---|---|---|
| Gm13748 | 0.622058045 | -0.184820192 | 1.448234678 | -0.127617571 |
| Gm13750 | 1.235703043 | 0.460427132 | 0.957746221 | 0.480740223 |
| Gm28981 | 0.770914299 | -1.368646596 | 1.21794311 | -1.123736063 |
| Gm25748 | 0 NA | NA | NA | |
| Gm28980 | 0 NA | NA | NA | |
| Creb1 | 1118.909764 | -0.335447115 | 0.084921283 | -3.950094741 |
| Mettl21a | 95.46113856 | -0.109741171 | 0.141010132 | -0.778250252 |
| Gm28982 | 1.624436933 | 0.332669621 | 0.869146743 | 0.382754263 |
| Gm29152 | 0 NA | NA | NA | |
| 2810408I11Rik | 4.24223269 | 0.181250264 | 0.556609131 | 0.325632933 |
| Ccnyl1 | 180.7599231 | -0.358822635 | 0.134235561 | -2.673081796 |
| Fzd5 | 105.8313593 | 0.699205696 | 0.188840534 | 3.70262507 |
| Gm38058 | 2.181430594 | 0.244996813 | 0.645604564 | 0.379484326 |
| Plekhm3 | 907.4639202 | -0.077871537 | 0.057563902 | -1.352784212 |
| Rpl31-ps24 | 0 NA | NA | NA | |
| Rpl10a-ps1 | 39.84853431 | 0.457987497 | 0.201016345 | 2.278359495 |
| Gm24788 | 0 NA | NA | NA | |
| Akr1cl | 0.229212118 | 1.217882219 | 2.536134947 | 0.480211915 |
| Gm8550 | 0 NA | NA | NA | |
| Cryge | 0 NA | NA | NA | |
| Crygd | 0 NA | NA | NA | |
| Crygc | 0 NA | NA | NA | |
| Crygb | 0.150086409 | 1.264824188 | 3.723011833 | 0.339731445 |
| Gm7329 | 4.40840556 | -0.049741654 | 0.481446365 | -0.103317124 |
| Cryga | 0.97358459 | 0.834871536 | 1.1038183 | 0.7563487 |
| Gm28845 | 0.23654427 | 1.324606751 | 2.541886324 | 0.521111719 |
| D630023F18Rik | 33.29031821 | 0.795028563 | 0.196985816 | 4.035968579 |
| Idh1 | 461.6797512 | -0.087839625 | 0.126404123 | -0.694911075 |
| Pikfyve | 2046.792619 | 0.32314939 | 0.071130886 | 4.543024938 |
| Pth2r | 15.63521041 | -0.5000181 | 0.336587322 | -1.485552387 |
| Gm28927 | 0 NA | NA | NA | |
| Gm28926 | 0 NA | NA | NA | |
| Gm28925 | 0 NA | NA | NA | |
| Gm28923 | 0 NA | NA | NA | |
| Gm28922 | 0 NA | NA | NA | |
| Gm28728 | 0 NA | NA | NA | |
| Gm5255 | 0 NA | NA | NA | |
| Gm8805 | 0 NA | NA | NA | |
| Crygf | 0 NA | NA | NA | |
| Gm8809 | 0 NA | NA | NA | |
| Gm15659 | 0 NA | NA | NA | |
| Gm8812 | 0 NA | NA | NA | |
| Map2 | 21973.53413 | 0.200174625 | 0.096446517 | 2.075498742 |
| Gm24729 | 8.150369374 | -0.107369605 | 0.43916758 | -0.244484362 |
| Gm10558 | 1.106847758 | 0.742535496 | 1.053706772 | 0.704688928 |
| Gm25411 | 10.74259572 | -0.137259105 | 0.346079972 | -0.396610945 |
| Unc80 | 7728.076732 | -0.226801677 | 0.065775902 | -3.448096783 |

| | | | | |
|---|---|---|---|---|
| Gm37181 | 0.701668669 | -0.825928712 | 1.28610935 | -0.64219167 |
| Rpe | 507.539684 | 0.186816103 | 0.083963494 | 2.224968191 |
| Kansl1l | 470.4693895 | 0.042873753 | 0.08132829 | 0.527168994 |
| Gm37570 | 61.3256198 | -0.294129506 | 0.241017907 | -1.220363706 |
| Gm38387 | 64.00509308 | -0.07046421 | 0.185449932 | -0.379963527 |
| Gm38162 | 96.75562574 | 0.603444754 | 0.190820112 | 3.162375017 |
| Gm15789 | 30.55906383 | 0.78787165 | 0.218468582 | 3.60633847 |
| 6820402A03Rik | 78.68479837 | 0.059428594 | 0.183534217 | 0.323801166 |
| Gm37285 | 31.02313678 | -0.007982743 | 0.224567737 | -0.035547151 |
| Gm37755 | 15.79173624 | 1.331485007 | 0.372067988 | 3.578606737 |
| Gm38218 | 8.540597391 | 1.021717512 | 0.357488117 | 2.858046084 |
| Acadl | 460.9663652 | 0.435236439 | 0.127617225 | 3.410483492 |
| Gm15793 | 6.435362979 | 1.350135756 | 0.45977586 | 2.936508577 |
| Gm27934 | 0.075043205 | 0.755286614 | 3.724945049 | 0.202764498 |
| Gm25832 | 0.12092566 | 1.236179401 | 3.724945049 | 0.331865138 |
| Myl1 | 8.248283288 | 1.30642847 | 0.39413881 | 3.314640522 |
| Rpl31-ps14 | 4.920430513 | 0.222063604 | 0.4813984 | 0.461288621 |
| Gm15826 | 0.731126313 | 2.961744549 | 1.392274918 | 2.127269916 |
| Lancl1 | 1237.017238 | -0.063494777 | 0.046148485 | -1.375879976 |
| Cps1 | 8.332986231 | 0.573629581 | 0.335912033 | 1.707677979 |
| Gm15668 | 0.088534442 | 0.274389469 | 3.724945049 | 0.073662689 |
| Gm26342 | 0 | NA | NA | NA |
| Kif22-ps | 0 | NA | NA | NA |
| Gm22548 | 0 | NA | NA | NA |
| Gm15669 | 0 | NA | NA | NA |
| Gm37957 | 0 | NA | NA | NA |
| Gm23189 | 0 | NA | NA | NA |
| Gm15671 | 0 | NA | NA | NA |
| Erbb4 | 1732.331576 | -0.265044546 | 0.065654497 | -4.036959532 |
| Gm37100 | 8.451353278 | -0.433747821 | 0.372850249 | -1.16332984 |
| Gm15670 | 8.648476067 | 0.290273855 | 0.441930277 | 0.656831792 |
| Gm25542 | 1.041370757 | 0.932866332 | 0.980537146 | 0.951382959 |
| Gm19335 | 10.47773709 | -0.507270402 | 0.370159247 | -1.370411264 |
| Gm37735 | 8.967735426 | -1.056953658 | 0.342864612 | -3.08271435 |
| Gm37769 | 40.95679204 | -0.469960205 | 0.227745335 | -2.06353384 |
| Gm37061 | 10.97852267 | -0.401821529 | 0.437694854 | -0.918040332 |
| Gm16076 | 36.27563093 | -0.816666236 | 0.266754193 | -3.061493535 |
| Gm8840 | 0 | NA | NA | NA |
| Pced1c-ps | 0 | NA | NA | NA |
| Gm28497 | 0.047948698 | -0.206487837 | 3.724945049 | -0.055433794 |
| Gm29113 | 9.644466874 | 0.495949507 | 0.313804891 | 1.58043906 |
| Gm29114 | 0 | NA | NA | NA |
| Ikzf2 | 269.8286907 | 0.330395884 | 0.106922836 | 3.090040403 |
| Gm29112 | 12.57536217 | -0.339578853 | 0.371291056 | -0.914589371 |
| Gm28112 | 19.23386575 | -0.042841495 | 0.306266875 | -0.139882887 |
| Gm37930 | 0.044191549 | -0.206490061 | 3.724945049 | -0.055434391 |
| Spag16 | 19.54478478 | 0.154914055 | 0.237250043 | 0.652956912 |

| | | | | |
|---|---|---|---|---|
| Gm22665 | 0 | NA | NA | NA |
| Gm23879 | 0 | NA | NA | NA |
| Gm28580 | 0 | NA | NA | NA |
| Gm38272 | 0.042716945 | -0.206490061 | 3.724945049 | -0.055434391 |
| Gm37456 | 0.272865106 | 2.35370433 | 2.408694973 | 0.977169943 |
| Gm23422 | 0 | NA | NA | NA |
| Vwc2l | 194.8593581 | -0.095444779 | 0.090453874 | -1.055176251 |
| Bard1 | 62.05484921 | 1.35454942 | 0.229669555 | 5.897818795 |
| Gm16236 | 1.002654625 | -0.485098535 | 1.095939693 | -0.442632508 |
| Gm8870 | 0 | NA | NA | NA |
| Abca12 | 5.342748304 | 0.922302163 | 0.54566239 | 1.690243233 |
| Gm6947 | 0 | NA | NA | NA |
| Atic | 353.5973081 | -0.223810047 | 0.110675553 | -2.022217563 |
| Fn1 | 1063.525137 | 0.02914646 | 0.2247876 | 0.129662225 |
| Apol7d | 1.07675232 | 0.018352921 | 1.194752585 | 0.015361274 |
| Gm5829 | 1.23306525 | 0.948739116 | 0.947550033 | 1.001254903 |
| Gm5256 | 3.462762875 | -0.402328532 | 0.540124903 | -0.744880544 |
| Gm37217 | 0 | NA | NA | NA |
| Gm8883 | 0 | NA | NA | NA |
| Gm28818 | 0.213332449 | 0.713631543 | 2.167688374 | 0.329213162 |
| 4933417E11Rik | 0.198105924 | -1.020160065 | 2.625732037 | -0.388524058 |
| 4930556G22Rik | 0 | NA | NA | NA |
| Gm5528 | 0 | NA | NA | NA |
| Gm8885 | 0.131209559 | 0.755295049 | 2.976009072 | 0.253794606 |
| Gm4319 | 0.520182433 | 0.256709746 | 1.293821216 | 0.19841207 |
| Mreg | 141.6531737 | -0.022216779 | 0.132285631 | -0.167945522 |
| Gm25360 | 67.27497613 | 0.139928068 | 0.219832697 | 0.636520725 |
| D230017M19Rik | 46.21119991 | -0.407651627 | 0.156130631 | -2.610965097 |
| Gm23472 | 572.9451334 | 0.125475508 | 0.804214934 | 0.156022355 |
| Gm24497 | 54.97402124 | -0.066011719 | 0.36930759 | -0.178744549 |
| Gm25939 | 1027.212574 | 0.082079227 | 0.171873041 | 0.477557311 |
| Pecr | 29.32302085 | 0.015460269 | 0.186134359 | 0.083059728 |
| Tmem169 | 181.7981465 | -0.113525966 | 0.111378758 | -1.019278431 |
| Xrcc5 | 253.261693 | 0.137006005 | 0.079194398 | 1.72999618 |
| 4-Mar | 295.8236201 | -0.461894716 | 0.124003262 | -3.724859409 |
| Gm15843 | 0.209261529 | 1.220404626 | 2.616217222 | 0.466476796 |
| Smarcal1 | 689.8282747 | -0.082597818 | 0.054339587 | -1.520030286 |
| Ankar | 0.251200408 | -0.687377909 | 2.100374883 | -0.327264392 |
| Rpl37a | 999.4688314 | 0.274620833 | 0.127360104 | 2.156254782 |
| Igfbp2 | 490.8609578 | 0.377046406 | 0.191599443 | 1.967888842 |
| Igfbp5 | 3859.554287 | 0.68681566 | 0.140166743 | 4.899990152 |
| 1700027A15Rik | 1.147127115 | -1.396266827 | 1.134884656 | -1.230316067 |
| Tnp1 | 0 | NA | NA | NA |
| Gm38106 | 0 | NA | NA | NA |
| Gm29185 | 0.044544648 | 0.755286614 | 3.724945049 | 0.202764498 |
| Gm29184 | 0 | NA | NA | NA |
| Pinc | 0.243249674 | 0.27441246 | 2.557831606 | 0.107283239 |

| | | | | |
|---|---|---|---|---|
| Gm27315 | 0 | NA | NA | NA |
| D530049I02Rik | 0 | NA | NA | NA |
| Gm9553 | 0.046345709 | 0.274391604 | 3.724945049 | 0.073663262 |
| C530043A13Rik | 1.12921268 | 0.64987188 | 1.160844876 | 0.559826634 |
| Gm29186 | 0 | NA | NA | NA |
| 6030407O03Rik | 6.071023221 | -0.914571611 | 0.405636622 | -2.2546574 |
| Gm29183 | 1.411578533 | -1.224242948 | 0.900297991 | -1.359819705 |
| Gm25329 | 0 | NA | NA | NA |
| Mir6351 | 0.439338454 | 1.148794202 | 1.440792567 | 0.797334903 |
| Tns1 | 929.0676011 | 0.387219439 | 0.115905107 | 3.340831548 |
| Rufy4 | 2.528566672 | 0.487627749 | 0.859954831 | 0.567038793 |
| Cxcr2 | 2.628643724 | 2.914101873 | 0.938528315 | 3.104969585 |
| Cxcr1 | 0.182828849 | -0.206486331 | 2.266107739 | -0.09111938 |
| Mir1928 | 0 | NA | NA | NA |
| Gm25524 | 0 | NA | NA | NA |
| Arpc2 | 1724.327588 | 0.477923722 | 0.085975438 | 5.558840228 |
| Gpbar1 | 0.083526043 | 0.75528439 | 3.724945049 | 0.202763901 |
| Aamp | 1126.602311 | 0.110985795 | 0.081484709 | 1.362044442 |
| Gm28364 | 0.299075359 | -0.687378978 | 1.759595324 | -0.390646058 |
| Pnkd | 1381.963466 | -0.309826538 | 0.091431868 | -3.388605567 |
| Gm37503 | 2.04875246 | 0.727979881 | 0.719035964 | 1.012438762 |
| Tmbim1 | 824.4000618 | 0.599875203 | 0.15325033 | 3.914348531 |
| Gm29358 | 41.10390762 | -0.82813349 | 0.332322663 | -2.491956112 |
| Gm37931 | 25.45680512 | -0.516025041 | 0.436438118 | -1.182355574 |
| Catip | 16.47013026 | -0.430088496 | 0.25734262 | -1.671268035 |
| Slc11a1 | 156.1085199 | 3.111764934 | 0.246253793 | 12.63641423 |
| Ctdsp1 | 480.9035942 | 0.52964985 | 0.122052369 | 4.339529453 |
| Mir26b | 0.460912027 | -0.452773024 | 1.395924102 | -0.324353611 |
| Vil1 | 4.671085706 | 0.090001856 | 0.496688574 | 0.181203798 |
| Usp37 | 682.5650065 | -0.143113823 | 0.072512445 | -1.973644991 |
| Gm37733 | 34.22973971 | -0.511296689 | 0.292029245 | -1.750840706 |
| Rqcd1 | 401.5594137 | -0.060777862 | 0.063748559 | -0.953399775 |
| Plcd4 | 276.691242 | 0.302023791 | 0.095837065 | 3.151429884 |
| Zfp142 | 808.2705847 | 0.060137643 | 0.081059113 | 0.741898603 |
| Gm25035 | 2.10081791 | 0.20706218 | 0.654594247 | 0.316321417 |
| Bcs1l | 123.2340065 | -0.587390345 | 0.126166773 | -4.655665913 |
| Rnf25 | 329.113652 | 0.14476348 | 0.092957724 | 1.557304471 |
| Stk36 | 70.93351887 | -0.519299995 | 0.171944252 | -3.020164906 |
| Ttll4 | 457.8466587 | -0.371524088 | 0.110483218 | -3.362719642 |
| Cyp27a1 | 60.63009196 | -0.296502866 | 0.166003608 | -1.786123022 |
| Prkag3 | 1.196546384 | -2.021882538 | 1.018440375 | -1.985273354 |
| Gm27507 | 0 | NA | NA | NA |
| Wnt6 | 20.88210294 | -0.510417054 | 0.29819108 | -1.711711343 |
| Gm29539 | 0.08169834 | -0.206492286 | 3.724945049 | -0.055434988 |
| Wnt10a | 51.88315531 | -0.114736288 | 0.23992102 | -0.478225242 |
| Gm16582 | 1.058309834 | -1.583582073 | 1.15546037 | -1.370520456 |
| Cdk5r2 | 2117.926955 | -0.209829196 | 0.113756111 | -1.84455318 |

| | | | | |
|---|---|---|---|---|
| Fev | 2.243593674 | 0.027962265 | 0.988902127 | 0.02827607 |
| Cryba2 | 3.578913291 | -1.15556868 | 0.762078822 | -1.516337479 |
| Mir375 | 0 | NA | NA | NA |
| Ccdc108 | 54.38524299 | -0.53618332 | 0.179109114 | -2.993612705 |
| Gm15841 | 0.048949532 | -0.206487837 | 3.724945049 | -0.055433794 |
| Ihh | 2.70016216 | -0.567846335 | 0.647020501 | -0.877632678 |
| Gm37744 | 0.044544648 | 0.755286614 | 3.724945049 | 0.202764498 |
| Nhej1 | 18.60744933 | 0.207941365 | 0.270425868 | 0.768940362 |
| Slc23a3 | 6.004310424 | 0.406085917 | 0.526126217 | 0.771841251 |
| Cnppd1 | 349.0250262 | 0.326199754 | 0.096172801 | 3.391808825 |
| Fam134a | 1705.406746 | -0.072575655 | 0.048293068 | -1.502817252 |
| Zfand2b | 100.1118125 | 0.20705583 | 0.140447423 | 1.474258669 |
| Abcb6 | 14.09945832 | -0.018886875 | 0.301041559 | -0.06273843 |
| Gm29253 | 0.323844909 | -0.613806781 | 2.032339249 | -0.302019843 |
| Atg9a | 756.6561104 | -0.068906933 | 0.055345158 | -1.245039949 |
| Ankzf1 | 160.4436551 | -0.398153077 | 0.144495257 | -2.755475067 |
| Glb1l | 98.7643794 | 0.062822352 | 0.128829979 | 0.487637674 |
| Gm27943 | 0 | NA | NA | NA |
| Stk16 | 338.2768 | -0.013259239 | 0.068432864 | -0.193755426 |
| Tuba4a | 2899.871314 | -0.330280536 | 0.070706435 | -4.671152454 |
| A630095N17Rik | 0.826064167 | 0.56597843 | 1.22852712 | 0.460696733 |
| Dnajb2 | 784.3513935 | -0.324527298 | 0.081616623 | -3.976240207 |
| Ptprn | 4437.838423 | -0.063146455 | 0.069430427 | -0.909492544 |
| Gm28294 | 1.007042841 | 0.3073756 | 0.95012283 | 0.323511435 |
| Resp18 | 711.8094129 | 0.451076102 | 0.154492303 | 2.919731885 |
| Gm28902 | 0.705697236 | -2.721869391 | 1.192818601 | -2.281880405 |
| Dnpep | 296.9380932 | 0.255921767 | 0.112071914 | 2.283549543 |
| Des | 14.87844252 | 0.677510215 | 0.320255987 | 2.11552709 |
| Gm15179 | 4.05260616 | -0.627139549 | 0.546190612 | -1.148206386 |
| Speg | 1539.141981 | -0.366693574 | 0.141373897 | -2.593785568 |
| Gm15178 | 0 | NA | NA | NA |
| Gmppa | 265.8934165 | -0.132001007 | 0.083327156 | -1.584129514 |
| Asic4 | 155.5102748 | -0.59871355 | 0.129021433 | -4.640419313 |
| Gm37264 | 4.269864839 | 0.57425496 | 0.51926441 | 1.105900864 |
| Chpf | 816.6335401 | 0.082086987 | 0.087439111 | 0.938790273 |
| Obsl1 | 195.5669334 | -0.264591539 | 0.092741073 | -2.85301355 |
| Tmem198 | 376.0167297 | -0.360541869 | 0.128587392 | -2.80386641 |
| Inha | 132.6233029 | -0.396756195 | 0.180608335 | -2.196776776 |
| Gm15181 | 0 | NA | NA | NA |
| Stk11ip | 336.2431366 | -0.080723857 | 0.080922991 | -0.997539212 |
| Gm15183 | 0.113999573 | -0.206492286 | 3.724945049 | -0.055434988 |
| Slc4a3 | 1347.377089 | -0.454029256 | 0.082056284 | -5.533144228 |
| Gm15180 | 0.245985165 | 1.3331421 | 2.494894433 | 0.5343481 |
| Gm15182 | 0.115603922 | 0.274387056 | 3.724945049 | 0.073662041 |
| Gm5257 | 0.104637558 | -0.206492286 | 3.724945049 | -0.055434988 |
| Gm29065 | 0 | NA | NA | NA |
| Gm29068 | 0 | NA | NA | NA |

| | | | | |
|---|---|---|---|---|
| Gm29069 | 0.135858026 | -1.143903624 | 2.941468349 | -0.388888639 |
| Gm17751 | 4.194728469 | 0.713348534 | 0.502633449 | 1.41922217 |
| Mir6343 | 0 | NA | NA | NA |
| Gm816 | 0.297786538 | -1.404255044 | 1.531524621 | -0.916900078 |
| Gm38265 | 0 | NA | NA | NA |
| Gm25974 | 0 | NA | NA | NA |
| Epha4 | 2103.014444 | 0.174474401 | 0.128356274 | 1.359297807 |
| Mir6352 | 3.045583208 | -0.77817336 | 0.647589701 | -1.20164567 |
| Gm28386 | 2.146862605 | 0.332194752 | 0.694723801 | 0.478168089 |
| Gm28385 | 0.301967822 | -1.7572062 | 3.695942541 | -0.475441969 |
| Gm26263 | 0 | NA | NA | NA |
| Gm28387 | 0.044544648 | 0.755286614 | 3.724945049 | 0.202764498 |
| Pax3 | 26.95793292 | -0.368201982 | 0.226026346 | -1.629022407 |
| Sgpp2 | 367.8063322 | -0.474667672 | 0.109546878 | -4.333009583 |
| Gm28410 | 0.413627519 | 2.263341232 | 1.431057257 | 1.581586775 |
| Gm37884 | 3.006274965 | -0.957177027 | 0.61578152 | -1.554410121 |
| Farsb | 522.3359989 | 0.013531216 | 0.081645504 | 0.165731301 |
| Gm28995 | 0 | NA | NA | NA |
| 4833412K13Rik | 33.12266117 | -0.408027004 | 0.265862399 | -1.534730019 |
| BC035947 | 16.4961278 | -0.797499322 | 0.292845642 | -2.723275364 |
| Mogat1 | 1.455068488 | 0.212151571 | 0.855988801 | 0.247843862 |
| 5730419F03Rik | 0.372487069 | -1.045123078 | 1.742562645 | -0.599762127 |
| Gm29611 | 0 | NA | NA | NA |
| Acsl3 | 1660.961095 | -0.004856681 | 0.07736743 | -0.06277423 |
| Utp14b | 706.5540839 | -0.052999396 | 0.089109464 | -0.594767316 |
| Gm6159 | 2.778261606 | -0.034522178 | 0.631776743 | -0.054643003 |
| Gm29536 | 0.417935612 | 1.236203912 | 1.775007936 | 0.696449794 |
| Kcne4 | 23.89478986 | 0.390033111 | 0.289518979 | 1.347176312 |
| Gm29187 | 0.218130886 | 0.776770355 | 2.170555632 | 0.357867057 |
| Gm5830 | 3.384143917 | -0.115806424 | 0.562986499 | -0.20570018 |
| Scg2 | 2880.675548 | 0.011409431 | 0.14635267 | 0.077958476 |
| Gm19027 | 0.061920613 | 0.274393828 | 3.724945049 | 0.073663859 |
| Gm24160 | 0 | NA | NA | NA |
| Ap1s3 | 62.26140866 | 1.242465543 | 0.205248091 | 6.053481599 |
| Gm37886 | 2.695407072 | 0.378302182 | 0.681947281 | 0.554738162 |
| Wdfy1 | 946.0370216 | 0.196711679 | 0.065349245 | 3.010159917 |
| Gm10555 | 0.378505399 | 0.755307664 | 1.693257481 | 0.446067815 |
| Mrpl44 | 178.4373768 | 0.150105905 | 0.093370508 | 1.607637222 |
| Serpine2 | 1567.911986 | 0.054640377 | 0.124440328 | 0.439088985 |
| Gm5529 | 18.26108814 | -0.168920635 | 0.259828606 | -0.650123318 |
| Gm28071 | 2.944880665 | 1.331201772 | 0.719631304 | 1.849838611 |
| Gm28058 | 0 | NA | NA | NA |
| Gm23934 | 0.310678561 | -0.689509619 | 1.600929012 | -0.430693437 |
| Gm38339 | 0.456870417 | 1.225659773 | 3.440999511 | 0.356192952 |
| 9830004L10Rik | 0 | NA | NA | NA |
| Fam124b | 3.258857179 | 3.389360025 | 0.97544924 | 3.474665709 |
| 2310015K22Rik | 0 | NA | NA | NA |

| | | | | |
|---|---|---|---|---|
| Gm37932 | 4.562962678 | -1.365162053 | 0.556207696 | -2.45441058 |
| Cul3 | 2324.447608 | 0.129754775 | 0.080121753 | 1.619470002 |
| Gm37902 | 63.67772732 | 0.022543693 | 0.163037279 | 0.138273241 |
| Gm37262 | 28.6341495 | -0.315208893 | 0.234310335 | -1.34526244 |
| Gm29125 | 1.604780936 | 0.491695726 | 0.849268856 | 0.578963567 |
| Gm6189 | 0.828249129 | 1.544473667 | 1.44784499 | 1.066739656 |
| 1700016L21Rik | 0.083172944 | -0.206492286 | 3.724945049 | -0.055434988 |
| Dock10 | 1627.450511 | 0.171419769 | 0.093246612 | 1.838348509 |
| Gm37645 | 36.84305094 | 0.178241342 | 0.243463465 | 0.732107143 |
| Gm38062 | 55.87989744 | 0.274451023 | 0.337683329 | 0.812746736 |
| A630081D01Rik | 65.65902623 | -0.04233258 | 0.227711185 | -0.185904703 |
| Gm29124 | 0 | NA | NA | NA |
| Gm28066 | 0 | NA | NA | NA |
| Nyap2 | 954.1496905 | -0.080145108 | 0.093015524 | -0.861631533 |
| Gm37210 | 0 | NA | NA | NA |
| Gm6198 | 0 | NA | NA | NA |
| Gm5530 | 3.003564229 | 2.484494076 | 0.951352919 | 2.611537765 |
| Gm7516 | 0 | NA | NA | NA |
| Gm37114 | 0 | NA | NA | NA |
| Mir6344 | 0 | NA | NA | NA |
| Irs1 | 784.393355 | -0.375204951 | 0.07848623 | -4.780519496 |
| Gm9747 | 0.131672813 | 0.274386859 | 3.668399824 | 0.074797425 |
| Rhbdd1 | 192.9071811 | 0.462129711 | 0.166248046 | 2.779760255 |
| Gm19552 | 8.274522055 | 0.594921441 | 0.395004119 | 1.506114525 |
| Gm28940 | 0.217873223 | -0.473161552 | 2.535311671 | -0.186628554 |
| Gm22112 | 0.107803068 | 0.75528661 | 3.724945049 | 0.202764497 |
| Col4a4 | 19.40116672 | 0.030717245 | 0.262581599 | 0.116981711 |
| Col4a3 | 9.983396163 | -0.173081422 | 0.307038352 | -0.563712712 |
| n-R5s213 | 0 | NA | NA | NA |
| C430014B12Rik | 2.511728143 | 0.091142349 | 0.648356234 | 0.140574493 |
| Mff | 1852.754622 | 0.031135008 | 0.044772907 | 0.695398394 |
| Gm28941 | 40.41534512 | -0.122531303 | 0.25710006 | -0.476589943 |
| Tm4sf20 | 0.089062654 | -0.20649451 | 3.724945049 | -0.055435586 |
| Gm19257 | 0 | NA | NA | NA |
| Gm36999 | 0.100902008 | 0.274391828 | 3.724945049 | 0.073663322 |
| Gm28942 | 0.253304709 | 1.227096947 | 1.826620851 | 0.671785251 |
| Gm17764 | 0 | NA | NA | NA |
| Gm22396 | 0.22174029 | -0.641270223 | 3.158407895 | -0.20303591 |
| Agfg1 | 1873.69111 | 0.012434279 | 0.06051399 | 0.205477751 |
| Gm7553 | 0.943831312 | -0.089147563 | 1.044122176 | -0.085380395 |
| Slc19a3 | 12.62212474 | -0.102613032 | 0.316545139 | -0.324165558 |
| Gm7539 | 0 | NA | NA | NA |
| Krtap28-10 | 0 | NA | NA | NA |
| A030005K14Rik | 0 | NA | NA | NA |
| Krtap28-13 | 0.044191549 | -0.206490061 | 3.724945049 | -0.055434391 |
| Gm29518 | 0.066050875 | 0.274391604 | 3.724945049 | 0.073663262 |
| Ccl20 | 0.166497232 | 0.27439225 | 3.399989659 | 0.080703848 |

| | | | | |
|---|---|---|---|---|
| Gm29519 | 0 | NA | NA | NA |
| Daw1 | 3.764882787 | -0.241756237 | 0.56723834 | -0.42619869 |
| Gm6244 | 0 | NA | NA | NA |
| n-R5s214 | 0 | NA | NA | NA |
| Sphkap | 3563.73903 | -0.099436557 | 0.101515841 | -0.979517638 |
| Gm6374 | 0.068097359 | -0.206487837 | 3.724945049 | -0.055433794 |
| Gm37517 | 48.29894096 | -0.81695506 | 0.354944832 | -2.301639539 |
| Gm37989 | 0 | NA | NA | NA |
| 4933436I20Rik | 1.025359323 | 0.188511753 | 1.301021482 | 0.144895189 |
| Gm25754 | 0.150086409 | 1.264824188 | 3.723011833 | 0.339731445 |
| Gm24555 | 0 | NA | NA | NA |
| Pid1 | 538.7136745 | 0.068146577 | 0.081947113 | 0.831592162 |
| Gm37058 | 25.37561462 | -0.761551561 | 0.325673919 | -2.338386698 |
| Gm18304 | 0 | NA | NA | NA |
| Mir6353 | 3.815514299 | -0.798741524 | 0.705852605 | -1.131598181 |
| Dner | 2450.532823 | -0.153787487 | 0.108551158 | -1.41672821 |
| Gm37959 | 19.69556321 | -0.340899113 | 0.269957826 | -1.262786556 |
| Gm37925 | 43.42924116 | -0.6213466 | 0.265268387 | -2.342331879 |
| Mir5126 | 0.291915594 | 0.007704182 | 1.993755218 | 0.003864156 |
| Trip12 | 5110.806766 | 0.280505328 | 0.052510323 | 5.341908278 |
| Gm38374 | 71.20713703 | -0.244046644 | 0.192108644 | -1.270357433 |
| Gm37738 | 93.58139767 | -0.332895232 | 0.247248654 | -1.346398563 |
| Fbxo36 | 28.2317926 | -0.096689959 | 0.247420864 | -0.390791455 |
| Slc16a14 | 112.1372938 | 0.035895238 | 0.145830606 | 0.246143382 |
| Gm15433 | 3.38607774 | 2.916109014 | 1.043870984 | 2.793553091 |
| Gm29284 | 0.509711911 | 2.221928681 | 1.432560716 | 1.551018855 |
| C130026I21Rik | 27.88048967 | 4.860145572 | 0.669937022 | 7.25463053 |
| Rpl19-ps1 | 2.447118844 | 0.153246345 | 0.656323856 | 0.233491962 |
| Gm27907 | 0 | NA | NA | NA |
| Gm7582 | 5.427132103 | 4.290069241 | 0.967143977 | 4.435812396 |
| Gm2389 | 0 | NA | NA | NA |
| Gm18342 | 8.221653517 | 4.446560418 | 0.916446306 | 4.851959562 |
| A530032D15Rik | 14.11902256 | 5.979197911 | 0.88534824 | 6.753498389 |
| Gm10553 | 0.099106094 | 0.274387056 | 3.724945049 | 0.073662041 |
| A530040E14Rik | 19.7285084 | 6.047451934 | 0.890180837 | 6.793509455 |
| Gm16038 | 0.038981395 | 0.274393828 | 3.724945049 | 0.073663859 |
| Gm2635 | 0.088534442 | 0.274389469 | 3.724945049 | 0.073662689 |
| Gm6264 | 1.445224006 | 1.082687622 | 1.231447711 | 0.879199021 |
| Gm7609 | 9.276052769 | 5.323590717 | 0.894702295 | 5.950125249 |
| Gm16028 | 0.132101749 | 0.274387056 | 3.724945049 | 0.073662041 |
| Gm16026 | 7.688870986 | 4.967156367 | 0.898324801 | 5.529354595 |
| Gm2619 | 4.165531338 | 4.783199311 | 0.925860312 | 5.16622135 |
| Gm16025 | 3.376741903 | -0.065941269 | 0.565084314 | -0.116692797 |
| Gm7281 | 0.301876039 | 1.219313272 | 1.767463114 | 0.689866319 |
| Gm2666 | 0.742190307 | 1.236207578 | 1.383804329 | 0.89333987 |
| Gm2427 | 1.636398745 | 2.464861484 | 1.237960208 | 1.991066811 |
| Gm16092 | 29.37301029 | -0.109509588 | 0.26687262 | -0.410344037 |

| | | | | |
|---|---|---|---|---|
| Gm7592 | 2.43285919 | 2.081717495 | 1.232440902 | 1.689101272 |
| G530012D18Rik | 0 | NA | NA | NA |
| Sp110 | 246.5019786 | 5.636814645 | 0.287788583 | 19.58665138 |
| Gm16094 | 4.539762321 | 4.515637708 | 0.981866205 | 4.599035677 |
| Sp140 | 215.8469477 | 4.893295345 | 0.284476294 | 17.20106544 |
| Gm10552 | 1.59243937 | 2.386173397 | 1.161807068 | 2.053846514 |
| Gm17017 | 6.466014695 | 0.652702274 | 0.448393978 | 1.455644604 |
| Sp100 | 678.1539777 | 5.376708504 | 0.29081573 | 18.48836895 |
| n-R5s215 | 0 | NA | NA | NA |
| A630001G21Rik | 22.19689395 | 2.190925573 | 0.331759922 | 6.603948894 |
| Mir8096 | 0 | NA | NA | NA |
| Gm25472 | 0 | NA | NA | NA |
| 9930111H07Rik | 0.389848623 | -0.206484845 | 1.742185097 | -0.118520612 |
| Gm22906 | 0 | NA | NA | NA |
| Cab39 | 2445.880552 | 0.115137719 | 0.088868378 | 1.295598292 |
| Itm2c | 5468.611581 | 0.2652169 | 0.073578391 | 3.604548774 |
| 4933407L21Rik | 23.38036317 | -0.814273374 | 0.252807786 | -3.220918892 |
| Gm28884 | 0.751763609 | 0.826804109 | 1.1175383 | 0.739844093 |
| Gpr55 | 7.561829528 | 2.762670784 | 0.604950697 | 4.566770152 |
| Gm28100 | 0.240280111 | 0.274411002 | 2.463083575 | 0.111409538 |
| Gm18180 | 25.54930956 | -0.25145502 | 0.244226833 | -1.029596202 |
| Spata3 | 2.778035728 | -0.72739589 | 0.624192478 | -1.165339084 |
| 2810459M11Rik | 146.6634441 | -0.025884125 | 0.162676112 | -0.159114478 |
| Psmd1 | 1536.480455 | 0.174768401 | 0.063801383 | 2.739257243 |
| Gm22786 | 4.654196711 | 0.473506895 | 0.469048157 | 1.009505929 |
| Htr2b | 18.41870533 | 2.181420724 | 0.565987489 | 3.854185411 |
| Gm37914 | 41.69621495 | -0.407507627 | 0.267002557 | -1.526231179 |
| Gm21972 | 0.320475213 | -2.192921172 | 2.280147898 | -0.961745146 |
| Armc9 | 417.345602 | -0.20089444 | 0.177187597 | -1.13379516 |
| Gm16341 | 0.537632031 | 1.25710143 | 1.652563186 | 0.760697951 |
| B3gnt7 | 21.27627369 | 0.901825102 | 0.309528232 | 2.913547163 |
| Gm28626 | 1.81028421 | 2.862479031 | 1.084407352 | 2.639671362 |
| Gm5258 | 0 | NA | NA | NA |
| Gm28627 | 0 | NA | NA | NA |
| Ncl | 3170.074209 | 0.394527117 | 0.101072103 | 3.903422444 |
| Snora75 | 0 | NA | NA | NA |
| mmu-mir-3535 | 0.451658936 | 1.068885701 | 1.288965815 | 0.829258378 |
| Gm24148 | 0 | NA | NA | NA |
| Snord82 | 0.870629472 | 0.961409502 | 1.03136635 | 0.93217071 |
| C130036L24Rik | 32.9831117 | -0.287053209 | 0.236886446 | -1.211775575 |
| Nmur1 | 2.498311769 | 0.686593059 | 0.607633403 | 1.129946206 |
| 1700019O17Rik | 0.04393175 | -0.206490061 | 3.724945049 | -0.055434391 |
| Rpl30-ps6 | 0 | NA | NA | NA |
| Ptma | 1852.709718 | 0.622071381 | 0.137147535 | 4.535782449 |
| Pde6d | 314.3105157 | -0.253633541 | 0.10898974 | -2.327132258 |
| Gm37152 | 0.163519603 | 0.723301248 | 2.774973957 | 0.260651545 |
| Cops7b | 225.9968041 | 0.006742593 | 0.081751984 | 0.082476201 |

| | | | | |
|---|---|---|---|---|
| Gm22955 | 0 | NA | NA | NA |
| Gm37022 | 4.10661082 | 1.12860269 | 0.594690131 | 1.897799596 |
| Nppc | 53.44822697 | 0.506132535 | 0.241950966 | 2.091880615 |
| Dis3l2 | 456.7534953 | 0.048040492 | 0.078921694 | 0.608710855 |
| Gm29097 | 0.04393175 | -0.206490061 | 3.724945049 | -0.055434391 |
| Gm29055 | 12.11508896 | 0.233669995 | 0.415658368 | 0.562168388 |
| Gm37485 | 23.25905464 | -0.402228921 | 0.297608841 | -1.351535525 |
| Gm38142 | 24.33645793 | -0.624683428 | 0.306720416 | -2.036654217 |
| Gm38010 | 20.30950254 | -0.709850322 | 0.278981248 | -2.544437403 |
| Gm37541 | 10.5940186 | -0.047610057 | 0.414081374 | -0.114977538 |
| Gm38021 | 12.72797742 | -0.352794188 | 0.295990796 | -1.191909319 |
| Gm6136 | 179.3671153 | 0.199797622 | 0.112165902 | 1.781268799 |
| Gm37967 | 11.04668228 | -0.728828349 | 0.332830649 | -2.189787361 |
| Gm28375 | 0.045882455 | 0.755288839 | 3.724945049 | 0.202765096 |
| Gm37017 | 0.873848832 | 0.096462871 | 1.08101835 | 0.089233333 |
| Gm37447 | 21.46227591 | -0.542486031 | 0.288785771 | -1.878506786 |
| Akp-ps1 | 0.752438969 | -0.120932202 | 1.13566078 | -0.106486201 |
| Gm29371 | 0 | NA | NA | NA |
| Alppl2 | 0 | NA | NA | NA |
| Alpi | 0 | NA | NA | NA |
| Gm29372 | 0 | NA | NA | NA |
| Akp3 | 0.332159689 | 0.755304491 | 1.991498696 | 0.379264366 |
| Gm29374 | 5.217580992 | 0.777270943 | 0.498961463 | 1.557777506 |
| Ecel1 | 298.3676729 | 0.15983697 | 0.143281151 | 1.115547778 |
| Prss56 | 20.4462505 | 0.869227384 | 0.793607251 | 1.095286596 |
| Chrnd | 0 | NA | NA | NA |
| Chrng | 0.171131556 | 0.755286499 | 3.384058469 | 0.223189554 |
| Eif4e2 | 578.568997 | -0.082070245 | 0.062538019 | -1.312325632 |
| Gm26187 | 0.955307282 | 3.305493832 | 1.134904773 | 2.912573735 |
| Efhd1 | 197.7653228 | -0.70345975 | 0.149600178 | -4.702265461 |
| Efhd1os | 0.337798032 | -0.299923267 | 1.879843707 | -0.159546916 |
| Gigyf2 | 1430.757424 | -0.313395667 | 0.082443041 | -3.801359877 |
| Gm22549 | 0 | NA | NA | NA |
| Kcnj13 | 41.11332588 | -0.323099479 | 0.251976974 | -1.282257955 |
| Gm23504 | 0 | NA | NA | NA |
| 3110079O15Rik | 0.366579243 | -0.544565889 | 1.550378439 | -0.351247073 |
| Ngef | 1590.707428 | -0.571277826 | 0.128841007 | -4.433975152 |
| Neu2 | 18.78655172 | -0.890751017 | 0.272497485 | -3.268841236 |
| Gm17181 | 0.432611478 | -0.589593144 | 1.424641648 | -0.413853648 |
| Gm6153 | 1.379575143 | 1.015744106 | 0.879588079 | 1.154795217 |
| Inpp5d | 416.1094285 | 1.735021012 | 0.132185569 | 13.12564618 |
| Atg16l1 | 673.2924239 | -0.117540853 | 0.122712868 | -0.957852708 |
| Gm25395 | 742.8621918 | -0.135299976 | 0.184946683 | -0.731562057 |
| Scarna6 | 154.5847017 | 0.13066535 | 0.156070829 | 0.837218275 |
| Sag | 17.39870792 | 0.49787063 | 0.292105901 | 1.704418253 |
| Dgkd | 1519.408834 | -0.17316537 | 0.072053275 | -2.403296312 |
| Gm19582 | 6.959264896 | 0.541042163 | 0.484421684 | 1.116882628 |

| | | | | |
|---|---|---|---|---|
| 4833421G17Rik | 11.9590053 | -0.179167376 | 0.319510428 | -0.560755959 |
| Gm38365 | 57.83653722 | -0.180277501 | 0.311280663 | -0.579147765 |
| Usp40 | 453.472211 | -0.154661949 | 0.114991139 | -1.344990148 |
| Gm25198 | 0 | NA | NA | NA |
| Gm5832 | 0.105852363 | -0.206492286 | 3.724945049 | -0.055434988 |
| Gm15375 | 0.049553047 | 0.274391604 | 3.724945049 | 0.073663262 |
| Ugt1a10 | 0.161949631 | 0.274399427 | 2.77761947 | 0.098789424 |
| Gm20528 | 0.038981395 | 0.274393828 | 3.724945049 | 0.073663859 |
| Ugt1a9 | 0.038981395 | 0.274393828 | 3.724945049 | 0.073663859 |
| Ugt1a8 | 0 | NA | NA | NA |
| Ugt1a7c | 12.7903233 | 1.480390123 | 0.385082105 | 3.844349306 |
| Gm15373 | 0.042716945 | -0.206490061 | 3.724945049 | -0.055434391 |
| Ugt1a6b | 1.167440347 | 1.879969037 | 1.167937179 | 1.609649107 |
| Gm15372 | 1.333048035 | 0.921074735 | 1.255216671 | 0.733797404 |
| Gm15370 | 0.204364325 | -0.206481308 | 2.645762338 | -0.078042274 |
| Ugt1a6a | 1.585662245 | 1.41995315 | 0.981168897 | 1.447205629 |
| Gm15369 | 0.593490712 | 1.137529912 | 1.347127182 | 0.844411669 |
| Gm15374 | 1.464328245 | 0.918455814 | 1.135174292 | 0.809087926 |
| Gm15371 | 0.512448221 | 2.695880629 | 1.264625966 | 2.131761248 |
| Ugt1a5 | 1.213945837 | 1.396318405 | 0.938352845 | 1.488052615 |
| Gm15368 | 0.624217637 | 1.133902732 | 1.463391877 | 0.774845583 |
| Gm15376 | 0.474048072 | 1.532961881 | 1.51589145 | 1.011260984 |
| Ugt1a2 | 0.875486306 | -0.406539279 | 1.303899649 | -0.311787245 |
| Dnajb3 | 7.809597006 | 0.128983741 | 0.402849596 | 0.320178405 |
| Ugt1a1 | 0.279519214 | 0.288479884 | 1.798254797 | 0.160422141 |
| Mroh2a | 3.442503238 | -0.056103297 | 0.605349466 | -0.092679188 |
| Hjurp | 549.4911097 | 0.112654356 | 0.083570594 | 1.348014303 |
| Trpm8 | 6.249743756 | 0.822007158 | 0.396384772 | 2.073760691 |
| 4930453O03Rik | 0.121388914 | 0.755282166 | 3.724945049 | 0.202763304 |
| Spp2 | 1.584204494 | 1.597514291 | 0.983676529 | 1.624023999 |
| Gm29538 | 0.330861516 | -1.880082377 | 1.719154108 | -1.093608984 |
| Gm28888 | 0 | NA | NA | NA |
| Glrp1 | 2.778605519 | 3.481122324 | 0.968108812 | 3.595796549 |
| Gm19589 | 0.038981395 | 0.274393828 | 3.724945049 | 0.073663859 |
| Gm38130 | 0.038981395 | 0.274393828 | 3.724945049 | 0.073663859 |
| Gm29336 | 0.085433889 | -0.687376175 | 3.724945049 | -0.184533239 |
| Arl4c | 850.2294767 | 0.249885076 | 0.100988669 | 2.474387255 |
| Gm29335 | 0 | NA | NA | NA |
| Platr5 | 0.048949532 | -0.206487837 | 3.724945049 | -0.055433794 |
| Gm29337 | 0 | NA | NA | NA |
| Gm4753 | 0 | NA | NA | NA |
| 1700067G17Rik | 0 | NA | NA | NA |
| Gm18699 | 0 | NA | NA | NA |
| Sh3bp4 | 304.2424635 | 0.164441418 | 0.139630164 | 1.177692653 |
| Gm38312 | 5.824867419 | -0.767492666 | 0.46402538 | -1.653988553 |
| Gm5259 | 0.045882455 | 0.755288839 | 3.724945049 | 0.202765096 |
| Gm28342 | 0.049553047 | 0.274391604 | 3.724945049 | 0.073663262 |

| | | | | |
|---|---|---|---|---|
| C030007H22Rik | 0.466539735 | 0.575252146 | 3.683462988 | 0.156171556 |
| Agap1 | 3469.23624 | -0.053215907 | 0.065314384 | -0.814765497 |
| Gm37297 | 49.83626593 | -0.539716747 | 0.335893847 | -1.606807481 |
| Gm25180 | 3.378117529 | -0.249047929 | 0.724216958 | -0.343885802 |
| Gm28499 | 9.617358175 | -0.229588331 | 0.455158279 | -0.50441427 |
| 4933400F21Rik | 0.445470297 | 1.660209805 | 1.628906354 | 1.019217465 |
| Gm26037 | 7.202871343 | -0.226297449 | 0.463566763 | -0.48816582 |
| Gm37521 | 23.73742709 | -0.361924548 | 0.339787831 | -1.065148647 |
| Gbx2 | 22.87581777 | -0.151502359 | 0.281946935 | -0.537343523 |
| D130058E05Rik | 7.849306383 | -0.109915246 | 0.444473901 | -0.247292913 |
| Asb18 | 37.68452797 | 0.075267968 | 0.178785131 | 0.420996801 |
| Iqca | 7.094191665 | -0.710878268 | 0.401201501 | -1.7718734 |
| 4930434B07Rik | 0.106465261 | 0.75528439 | 3.724945049 | 0.202763901 |
| Ackr3 | 127.6655458 | 0.778715941 | 0.242064393 | 3.216978467 |
| Gm38277 | 0.04393175 | -0.206490061 | 3.724945049 | -0.055434391 |
| Gm28721 | 0.047948698 | -0.206487837 | 3.724945049 | -0.055433794 |
| Gm28723 | 1.104344843 | 1.854013855 | 1.150748065 | 1.61113793 |
| Gm27648 | 0 | NA | NA | NA |
| Gm28722 | 0.261620403 | -0.206487147 | 2.095026356 | -0.098560644 |
| Gm35048 | 0.834783721 | 1.92127496 | 1.138544554 | 1.687483334 |
| Cops8 | 733.5162779 | 0.151586105 | 0.075533577 | 2.006870454 |
| Gm9991 | 0.061920613 | 0.274393828 | 3.724945049 | 0.073663859 |
| Col6a3 | 94.47715798 | -0.265265818 | 0.152427976 | -1.740269895 |
| Gm36970 | 0 | NA | NA | NA |
| 4930474B08Rik | 0 | NA | NA | NA |
| Mlph | 41.52328994 | -0.018896893 | 0.198088103 | -0.095396405 |
| Prlh | 0.989340548 | -0.571962619 | 1.339136303 | -0.427113071 |
| Rab17 | 0.340708601 | 1.308962897 | 1.755116152 | 0.745798445 |
| Gm28199 | 0 | NA | NA | NA |
| Lrrfip1 | 605.1451411 | 0.055005397 | 0.118889547 | 0.462659652 |
| Rbm44 | 1.28530837 | -0.091414243 | 1.265918006 | -0.07221182 |
| Gm7776 | 0.372084114 | 1.845455805 | 1.963807302 | 0.939733651 |
| Ramp1 | 339.6276143 | -0.207105791 | 0.090808896 | -2.280677326 |
| Ube2f | 531.3694709 | 0.236093045 | 0.122509183 | 1.927145694 |
| Gm7785 | 19.25179799 | 0.75760509 | 0.323190741 | 2.34414231 |
| Gm26683 | 0.0885994 | 0.274402725 | 3.724945049 | 0.073666248 |
| Scly | 285.0654636 | 0.247042787 | 0.094134577 | 2.62435754 |
| Gm17090 | 0.087930927 | -0.206490058 | 3.724945049 | -0.05543439 |
| Espnl | 4.028718013 | 0.232918505 | 0.567379756 | 0.410516066 |
| Klhl30 | 7.180205943 | -0.911805243 | 0.605654009 | -1.505488661 |
| Fam132b | 9.846500243 | 1.094875796 | 0.564967293 | 1.937945451 |
| Ilkap | 381.4142211 | 0.018664065 | 0.105552024 | 0.176823372 |
| 1700020N18Rik | 1.184503545 | 1.861245889 | 0.995689952 | 1.869302673 |
| Hes6 | 86.06443169 | 0.262990703 | 0.135585543 | 1.939666255 |
| Per2 | 585.5330981 | -0.605146581 | 0.195421093 | -3.096628783 |
| Gm28381 | 0 | NA | NA | NA |
| Gm28382 | 0 | NA | NA | NA |

| | | | | |
|---|---|---|---|---|
| Gm19085 | 0.292057199 | 0.777972334 | 2.437401157 | 0.31918108 |
| Traf3ip1 | 223.3191922 | -0.370433149 | 0.107134433 | -3.457647901 |
| Asb1 | 423.9897668 | -0.357967137 | 0.081043394 | -4.416981071 |
| Gm28380 | 0.180091664 | -0.206483904 | 2.750740754 | -0.075064836 |
| Gm29100 | 8.346887551 | -0.581745388 | 0.342844553 | -1.696819692 |
| Twist2 | 5.328893884 | -0.463571443 | 0.418971964 | -1.106449793 |
| Gm29101 | 0.685283607 | -1.010591157 | 1.209528 | -0.835525227 |
| Gm37600 | 0 | NA | NA | NA |
| Hdac4 | 1369.404079 | -0.091926338 | 0.080598115 | -1.140551958 |
| Gm37048 | 0.281888464 | 1.660505073 | 1.753827889 | 0.946789068 |
| Gm26720 | 0 | NA | NA | NA |
| Gm37036 | 19.75259723 | -0.537890527 | 0.319526034 | -1.683401258 |
| Gm29099 | 0 | NA | NA | NA |
| Gm7837 | 0.143650742 | 0.755280773 | 3.584970442 | 0.210679777 |
| Ndufa10 | 1126.271048 | -0.053236247 | 0.054803982 | -0.971393784 |
| Mir6900 | 0.79482971 | -1.113237852 | 1.137828364 | -0.978388206 |
| Olfr1416 | 0 | NA | NA | NA |
| Olfr1415 | 0 | NA | NA | NA |
| Olfr1414 | 0 | NA | NA | NA |
| 1700054K02Rik | 0 | NA | NA | NA |
| Olfr1413 | 0 | NA | NA | NA |
| Gm28478 | 0 | NA | NA | NA |
| Gm28479 | 0 | NA | NA | NA |
| Olfr1412 | 0 | NA | NA | NA |
| Olfr1411 | 0 | NA | NA | NA |
| Olfr1410 | 0 | NA | NA | NA |
| Olfr12 | 0 | NA | NA | NA |
| Myeov2 | 377.7320121 | 0.180632626 | 0.113479343 | 1.591766581 |
| Otos | 0.964645659 | -0.850341207 | 1.040424522 | -0.817302158 |
| Gm29483 | 1.804989916 | -0.024880079 | 0.690167776 | -0.036049321 |
| Gm29482 | 0 | NA | NA | NA |
| Gm29481 | 0.153668717 | 0.755303435 | 2.839403532 | 0.266007782 |
| Gpc1 | 1142.72074 | 0.272237608 | 0.079930812 | 3.405915707 |
| Gm29480 | 6.109138398 | 0.917279567 | 0.426876098 | 2.148819227 |
| Mir149 | 1.829161676 | -0.10072916 | 0.694915105 | -0.14495175 |
| Ankmy1 | 5.039844727 | 1.669163207 | 0.54712085 | 3.050812647 |
| Dusp28 | 267.5008543 | 1.178809204 | 0.1670289 | 7.057516402 |
| 5033417F24Rik | 38.62813476 | -0.239637001 | 0.179375381 | -1.335952564 |
| Rnpepl1 | 468.7456855 | 0.510291554 | 0.093906599 | 5.434032986 |
| 9430060I03Rik | 3.610602274 | 0.527593222 | 0.625676562 | 0.843236352 |
| Capn10 | 256.0341623 | -0.16154939 | 0.079877707 | -2.022459031 |
| Gpr35 | 139.3197398 | 3.789864018 | 0.392772691 | 9.649000828 |
| Gm18697 | 0.51839282 | 1.232437079 | 1.720508678 | 0.716321338 |
| Aqp12 | 0.175102337 | 0.27439644 | 3.315835544 | 0.082753332 |
| Kif1a | 18866.61453 | -0.052817046 | 0.054053818 | -0.977119624 |
| Gm28086 | 0.753737496 | 1.623391549 | 1.136429038 | 1.428502348 |
| Agxt | 0.11197992 | 0.75528439 | 3.724945049 | 0.202763901 |

| | | | | |
|---|---|---|---|---|
| 2310007B03Rik | 0.800599448 | 1.373243307 | 1.179270035 | 1.164485882 |
| Crocc2 | 17.59480686 | -0.253734527 | 0.278891309 | -0.909797185 |
| Sned1 | 257.5400578 | -0.760843938 | 0.123249056 | -6.173223266 |
| Mir6901 | 0 | NA | NA | NA |
| Mterf4 | 142.5738671 | 0.150211622 | 0.10669679 | 1.407836376 |
| Gm28535 | 1.935100933 | 0.148098017 | 0.628633694 | 0.235587145 |
| Gm7867 | 1.11968618 | -0.165607904 | 0.905411277 | -0.182909036 |
| Pask | 68.98058163 | -0.440394223 | 0.212958143 | -2.067984894 |
| Ppp1r7 | 1320.763185 | -0.003963468 | 0.061699941 | -0.06423779 |
| mmu-mir-6902 | 0.22467814 | -0.52320819 | 2.139632498 | -0.244531802 |
| Mir6345 | 0.94748184 | -0.652244986 | 0.967744963 | -0.673984377 |
| Gm26446 | 2.680156834 | -0.4943467 | 0.592566024 | -0.83424746 |
| Ano7 | 2.740637547 | 0.82259131 | 0.639252452 | 1.286801963 |
| Hdlbp | 3229.635038 | 0.049508339 | 0.048485134 | 1.021103484 |
| Gm17415 | 4.04170504 | -0.521198718 | 0.67100864 | -0.776739206 |
| 2-Sep | 1408.822543 | 0.384013024 | 0.084553704 | 4.541646378 |
| Farp2 | 211.4310669 | -0.193998861 | 0.115132626 | -1.68500336 |
| Gm37250 | 18.96035902 | -1.111094017 | 0.33713328 | -3.295711473 |
| Stk25 | 1506.774651 | -0.017462285 | 0.060675862 | -0.287796242 |
| Gm28536 | 1.8285836 | -1.843268231 | 0.816623861 | -2.257181452 |
| Bok | 205.8186137 | -0.183359578 | 0.090897447 | -2.017213738 |
| Thap4 | 221.3880966 | -0.154127497 | 0.079976656 | -1.927156044 |
| Gm10550 | 0.177745132 | -0.206480788 | 2.300214322 | -0.089765891 |
| Atg4b | 616.7307566 | -0.145313632 | 0.048130054 | -3.019186998 |
| Dtymk | 176.2881088 | -0.35745493 | 0.112133903 | -3.187750716 |
| Ing5 | 203.0410209 | 0.164730066 | 0.083434239 | 1.974370086 |
| Gm22089 | 0 | NA | NA | NA |
| D2hgdh | 49.47669206 | -0.549234842 | 0.21781145 | -2.52160684 |
| Gal3st2 | 0.259214904 | 1.206848987 | 1.81222609 | 0.665948357 |
| Gm18666 | 2.840417843 | 0.01016563 | 0.733862502 | 0.013852227 |
| Gm9994 | 1.809180368 | -1.006525121 | 0.711798563 | -1.41405894 |
| Gm7889 | 3.175371776 | -0.764206909 | 0.615293485 | -1.242020154 |
| Gm6086 | 0.846372671 | -0.530229822 | 1.085797193 | -0.488332283 |
| Neu4 | 89.65460407 | -0.349190436 | 0.194266188 | -1.797484362 |
| Pdcd1 | 42.57420289 | 6.159671143 | 0.874863204 | 7.040724896 |
| Gm7891 | 0.255524584 | 1.286900982 | 2.493790686 | 0.5160421 |
| Gm7895 | 67.17479994 | 0.011105836 | 0.195503333 | 0.056806378 |
| Gm23389 | 0 | NA | NA | NA |
| 4930440C22Rik | 0.220980774 | 0.274398268 | 3.11985971 | 0.087952118 |
| Gm23181 | 0 | NA | NA | NA |
| Gm5264 | 0.086908494 | -0.687376175 | 3.724945049 | -0.184533239 |
| Fam174a | 321.8174294 | -0.068676999 | 0.140070193 | -0.490304163 |
| BC055308 | 22.09994013 | -0.228946325 | 0.258947525 | -0.884141777 |
| Gm15427 | 268.0268351 | 0.53949067 | 0.128885376 | 4.185817548 |
| St8sia4 | 341.5027806 | -0.132149881 | 0.157571466 | -0.838666314 |
| Gm37642 | 12.46481921 | 0.246676034 | 0.284885108 | 0.865879005 |
| 4930598F16Rik | 0 | NA | NA | NA |

| | | | | |
|---|---|---|---|---|
| Gm37761 | 0 | NA | NA | NA |
| Gm22360 | 1.558628739 | -0.89924258 | 0.917073015 | -0.980557234 |
| 1700063A18Rik | 0 | NA | NA | NA |
| Gm24586 | 0 | NA | NA | NA |
| Gm22917 | 0 | NA | NA | NA |
| Gm37076 | 0 | NA | NA | NA |
| Gm29601 | 0.044544648 | 0.755286614 | 3.724945049 | 0.202764498 |
| 4930533P14Rik | 0 | NA | NA | NA |
| Gm26245 | 0 | NA | NA | NA |
| Gm1833 | 0.161949631 | 0.274399427 | 2.77761947 | 0.098789424 |
| Gm26078 | 0 | NA | NA | NA |
| Slco4c1 | 49.5126318 | -0.303398996 | 0.190798137 | -1.590157012 |
| Panct2 | 37.25735819 | -0.138666001 | 0.228436022 | -0.607023359 |
| Slco6b1 | 0.161026707 | 0.274389649 | 3.463604896 | 0.079220828 |
| Gm9499 | 0 | NA | NA | NA |
| Gm28901 | 0 | NA | NA | NA |
| Gm6430 | 14.31094521 | 0.852520862 | 0.306020067 | 2.785833196 |
| Slco6c1 | 4.084609871 | -0.117363339 | 0.458642664 | -0.255892764 |
| Gm7133 | 0.216806889 | 0.975799517 | 2.595246896 | 0.375994869 |
| Gm7135 | 0.141094079 | 0.755285157 | 3.563850454 | 0.211929531 |
| Gm37171 | 2.848103185 | 0.806876381 | 0.564047902 | 1.430510384 |
| D1Ertd622e | 880.9560972 | 0.008722401 | 0.062031317 | 0.140612868 |
| Ppip5k2 | 911.1116556 | 0.230216893 | 0.095432368 | 2.412356491 |
| Gin1 | 94.83341611 | -0.086062049 | 0.122662788 | -0.701614977 |
| Pam | 2285.737651 | -0.032827131 | 0.06211438 | -0.528494855 |
| Gm37140 | 76.97524024 | 0.118054905 | 0.235598626 | 0.50108486 |
| Gm25413 | 0 | NA | NA | NA |
| Gm3531 | 31.35057007 | 0.640101943 | 0.207372256 | 3.086728934 |
| B230216N24Rik | 130.1992663 | -0.346218525 | 0.228406346 | -1.515800813 |
| Gm24465 | 0.049553047 | 0.274391604 | 3.724945049 | 0.073663262 |
| 1810006J02Rik | 0.496068496 | -1.109259686 | 1.41404162 | -0.784460422 |
| Gm29461 | 0.196785726 | 1.170169724 | 2.209521717 | 0.529603179 |
| Gm7952 | 0 | NA | NA | NA |
| Gm29462 | 0.477857232 | -2.202167305 | 1.428254104 | -1.541859603 |
| Slco6d1 | 0.368527898 | 0.681394544 | 1.575143789 | 0.432591963 |
| Gm3555 | 28.33472675 | 0.110002982 | 0.240277479 | 0.457816447 |
| Gm26134 | 0 | NA | NA | NA |
| Gm7967 | 10.59292213 | 0.598413102 | 0.342470031 | 1.747344435 |
| Gm29460 | 1.272524668 | 1.296767574 | 1.000720594 | 1.295833804 |
| Gm23434 | 0.075043205 | 0.755286614 | 3.724945049 | 0.202764498 |
| Gm18248 | 0.075043205 | 0.755286614 | 3.724945049 | 0.202764498 |
| Cntnap5b | 377.3845069 | -0.180042674 | 0.092548623 | -1.945384681 |
| Tdpx-ps1 | 13.77002096 | 0.489075183 | 0.276003247 | 1.771990685 |
| Gm37877 | 23.5485973 | -0.575180753 | 0.278453424 | -2.065626434 |
| Gm19029 | 3.564592169 | 0.086031692 | 0.619876974 | 0.138788334 |
| Gm23602 | 0 | NA | NA | NA |
| Gm29668 | 2.625920046 | -0.494436779 | 0.565985426 | -0.873585707 |

| | | | | |
|---|---|---|---|---|
| Gm29667 | 14.43481637 | 0.133715086 | 0.310225436 | 0.43102554 |
| Gm18700 | 0 | NA | NA | NA |
| Gm18185 | 0 | NA | NA | NA |
| Gm29334 | 0 | NA | NA | NA |
| Gm25537 | 0 | NA | NA | NA |
| Gm20268 | 0 | NA | NA | NA |
| Gm28801 | 0 | NA | NA | NA |
| Psmb6-ps2 | 0.576768135 | -0.119891795 | 1.301919974 | -0.092088452 |
| Gm18969 | 0 | NA | NA | NA |
| Gm28188 | 0.072025462 | 0.755288839 | 3.724945049 | 0.202765096 |
| Gm28187 | 1.345496586 | -0.0288598 | 0.859162728 | -0.03359061 |
| Gm24122 | 0 | NA | NA | NA |
| Gm28623 | 0 | NA | NA | NA |
| Gm23036 | 0 | NA | NA | NA |
| Gm3116 | 0 | NA | NA | NA |
| Gm20281 | 0 | NA | NA | NA |
| Gm22034 | 0 | NA | NA | NA |
| Gm18728 | 0 | NA | NA | NA |
| Gm23965 | 0 | NA | NA | NA |
| Gm29141 | 1.908747754 | 3.948723058 | 1.033945568 | 3.819082145 |
| Gm37789 | 0.149576918 | 0.755295313 | 2.863326609 | 0.263782452 |
| Gm18447 | 0 | NA | NA | NA |
| Gm29140 | 0.04393175 | -0.206490061 | 3.724945049 | -0.055434391 |
| Gm23537 | 0 | NA | NA | NA |
| Gm5260 | 0 | NA | NA | NA |
| Gm29592 | 0 | NA | NA | NA |
| Gm29011 | 0.220980774 | 0.274398268 | 3.11985971 | 0.087952118 |
| Cdh20 | 327.0013875 | -0.063919808 | 0.107807291 | -0.592908028 |
| Gm38031 | 85.93314083 | -0.541949779 | 0.197025355 | -2.750660085 |
| n-R5s216 | 0 | NA | NA | NA |
| Gm36969 | 21.4981108 | -0.330232052 | 0.341492483 | -0.967025833 |
| Gm15699 | 0 | NA | NA | NA |
| Gm29012 | 0.089089296 | 1.194485492 | 3.723939576 | 0.320758559 |
| Gm29013 | 0.046345709 | 0.274391604 | 3.724945049 | 0.073663262 |
| Rnf152 | 731.5992459 | -0.238483701 | 0.089233454 | -2.672581764 |
| Gm29014 | 0 | NA | NA | NA |
| Gm17634 | 9.160375891 | 0.328885276 | 0.350278551 | 0.938924965 |
| Gm28403 | 1.125166341 | 0.646887882 | 0.910285723 | 0.710642677 |
| Gm3608 | 115.3994121 | 0.112069453 | 0.23114633 | 0.484842016 |
| Pign | 477.5995311 | 0.008661232 | 0.113154639 | 0.076543323 |
| Gm28404 | 2.874152792 | -0.708641171 | 0.625275148 | -1.133326941 |
| Gm20302 | 1.560084669 | -1.009628922 | 0.962013116 | -1.049496005 |
| Gm22426 | 0 | NA | NA | NA |
| Gm8004 | 0 | NA | NA | NA |
| 2310035C23Rik | 1589.964662 | 0.088320393 | 0.079128884 | 1.116158707 |
| Gm37566 | 68.8298285 | 0.100289259 | 0.184514109 | 0.543531653 |
| Gm10193 | 1.224645236 | 1.163318158 | 1.059379305 | 1.098112973 |

| | | | | |
|---|---|---|---|---|
| Tnfrsf11a | 174.3773544 | 1.033104896 | 0.160651331 | 6.430727273 |
| Gm37779 | 0.176451472 | 1.257521603 | 2.721126865 | 0.462132662 |
| Gm26214 | 0 | NA | NA | NA |
| Gm18801 | 2.226781727 | 2.652386758 | 1.117099002 | 2.374352456 |
| Gm7160 | 28.83074711 | 1.116721303 | 0.29258454 | 3.81674747 |
| Zcchc2 | 822.9552979 | 0.560769696 | 0.110774603 | 5.062258705 |
| Gm37372 | 0.112396584 | 0.27438722 | 3.724945049 | 0.073662085 |
| Gm26509 | 3.937004826 | -0.031559007 | 0.519482546 | -0.060750851 |
| Gm20753 | 4.418263327 | -0.064744563 | 0.463138166 | -0.139795353 |
| Phlpp1 | 1704.681168 | -0.270657481 | 0.075963931 | -3.562973583 |
| Gm38235 | 158.3680684 | -0.643755628 | 0.243361006 | -2.645270246 |
| A530053M12Rik | 0.224329914 | 0.755297126 | 2.590150996 | 0.291603512 |
| Gm6651 | 0.568438163 | 1.707603161 | 1.415346443 | 1.206491294 |
| Bcl2 | 493.9864849 | -0.082963375 | 0.114299723 | -0.725840562 |
| D630008O14Rik | 0.626527047 | 2.729440267 | 1.589023789 | 1.717683704 |
| Mir3473f | 0.393472992 | 0.273911338 | 1.818420811 | 0.150631436 |
| Gm37053 | 17.84686388 | -0.478691757 | 0.37741805 | -1.268332974 |
| Kdsr | 626.2639973 | -0.061977797 | 0.06127298 | -1.011502905 |
| Vps4b | 768.3375885 | 0.318710614 | 0.076606721 | 4.160347922 |
| Serpinb5 | 4.060285521 | 1.050213637 | 0.997834957 | 1.052492328 |
| Pou2f3-rs1 | 0 | NA | NA | NA |
| Serpinb12 | 0.340899075 | 0.930018992 | 1.983191877 | 0.468950586 |
| Serpinb13 | 0 | NA | NA | NA |
| Gm8080 | 0.048949532 | -0.206487837 | 3.724945049 | -0.055433794 |
| Serpinb3a | 0.112642008 | 0.274402725 | 3.724945049 | 0.073666248 |
| Serpinb3d | 0.044544648 | 0.755286614 | 3.724945049 | 0.202764498 |
| Gm15388 | 0 | NA | NA | NA |
| Serpinb3b | 0.36088792 | 2.084836651 | 3.684451194 | 0.56584727 |
| Gm8084 | 0.045882455 | 0.755288839 | 3.724945049 | 0.202765096 |
| Gm15390 | 0 | NA | NA | NA |
| Serpinb3c | 0.256447508 | 1.236196794 | 2.47295682 | 0.499886122 |
| Gm8089 | 0.10503227 | 0.274389469 | 3.724945049 | 0.073662689 |
| Serpinb11 | 0.313959609 | -1.534900332 | 1.551611917 | -0.989229533 |
| Gm15391 | 0 | NA | NA | NA |
| Serpinb7 | 0.394543937 | 0.274412389 | 1.72533481 | 0.159048776 |
| Gm24553 | 0 | NA | NA | NA |
| Serpinb2 | 8.118383424 | 1.989825536 | 0.818712897 | 2.430431404 |
| Serpinb10 | 0.745572437 | 1.075018426 | 1.380070355 | 0.778959147 |
| Serpinb8 | 106.9109276 | 0.155981698 | 0.180893624 | 0.86228411 |
| Gm15389 | 0 | NA | NA | NA |
| Gm24267 | 0 | NA | NA | NA |
| Gm22438 | 0.044544648 | 0.755286614 | 3.724945049 | 0.202764498 |
| D830032E09Rik | 0 | NA | NA | NA |
| Gm18448 | 0.11147366 | 0.274389235 | 3.724945049 | 0.073662626 |
| Gm28282 | 0 | NA | NA | NA |
| Gm5702 | 0 | NA | NA | NA |
| Gm38212 | 0.044191549 | -0.206490061 | 3.724945049 | -0.055434391 |

| | | | | |
|---|---|---|---|---|
| Gm28283 | 0 | NA | NA | NA |
| Gm6679 | 0 | NA | NA | NA |
| Gm17847 | 0 | NA | NA | NA |
| Gm28281 | 0 | NA | NA | NA |
| Gm22159 | 0.068097359 | -0.206487837 | 3.724945049 | -0.055433794 |
| Gm25293 | 0 | NA | NA | NA |
| Gm8141 | 0.177100366 | -1.454105447 | 3.367753361 | -0.431773141 |
| Cdh7 | 599.3121521 | -0.233738648 | 0.1193192 | -1.958935759 |
| Gm18552 | 0 | NA | NA | NA |
| Gm3332 | 0 | NA | NA | NA |
| Gm6693 | 0 | NA | NA | NA |
| Cdh19 | 348.6925766 | 0.221767418 | 0.131306454 | 1.688930069 |
| Gm38255 | 57.38585182 | 0.050649496 | 0.205844032 | 0.246057637 |
| Gm36944 | 56.47739328 | -0.113538098 | 0.241182278 | -0.47075639 |
| Gm29088 | 1.097863377 | 0.58783723 | 1.073230404 | 0.547726963 |
| Gm19927 | 0.068097359 | -0.206487837 | 3.724945049 | -0.055433794 |
| Gm8173 | 15.82173517 | -1.02539348 | 0.377876666 | -2.713566545 |
| Gm37934 | 0 | NA | NA | NA |
| Dsel | 373.0186971 | -0.19780648 | 0.080206499 | -2.466215116 |
| Gm6028 | 0.150520013 | 0.274408177 | 2.857840489 | 0.096019417 |
| Gm17752 | 0 | NA | NA | NA |
| Gm6717 | 0 | NA | NA | NA |
| Gm19086 | 0 | NA | NA | NA |
| Gm18406 | 9.163048501 | -0.446670728 | 0.340200986 | -1.312961297 |
| Gm22331 | 0 | NA | NA | NA |
| Gm8204 | 0 | NA | NA | NA |
| Gm28190 | 0 | NA | NA | NA |
| Gm28189 | 0 | NA | NA | NA |
| Gm17946 | 0 | NA | NA | NA |
| Gm18181 | 0 | NA | NA | NA |
| Gm24937 | 0 | NA | NA | NA |
| 9330185C12Rik | 1.904432073 | 0.976647211 | 0.776174432 | 1.258283153 |
| Gm4854 | 0 | NA | NA | NA |
| Gm17916 | 16.58061403 | -0.311763931 | 0.253671007 | -1.229008925 |
| Cntnap5a | 1043.802266 | 0.071172229 | 0.083457748 | 0.852793555 |
| Gm23393 | 0 | NA | NA | NA |
| Gm28263 | 0.417031551 | 0.648344486 | 1.490254389 | 0.43505625 |
| Gm19965 | 1.062057355 | 2.629303614 | 1.020180314 | 2.577293031 |
| Gm17777 | 0 | NA | NA | NA |
| Gm24030 | 0 | NA | NA | NA |
| Gm27785 | 0 | NA | NA | NA |
| Gm28361 | 0 | NA | NA | NA |
| Zfp813-ps | 0 | NA | NA | NA |
| Gm28362 | 0 | NA | NA | NA |
| Gm28363 | 0 | NA | NA | NA |
| Gm28359 | 0 | NA | NA | NA |
| Gm18182 | 0 | NA | NA | NA |

| | | | | |
|---|---|---|---|---|
| B020011L13Rik | 0 | NA | NA | NA |
| Gm28360 | 0 | NA | NA | NA |
| Gm28169 | 0.218500166 | 1.257331768 | 2.168191684 | 0.579898806 |
| Gm7195 | 0 | NA | NA | NA |
| Gm28168 | 0 | NA | NA | NA |
| Gm7145 | 0 | NA | NA | NA |
| Gm28358 | 0 | NA | NA | NA |
| Gm29489 | 0 | NA | NA | NA |
| Gm29490 | 0.044191549 | -0.206490061 | 3.724945049 | -0.055434391 |
| Gm29104 | 0 | NA | NA | NA |
| Gm29106 | 0.046345709 | 0.274391604 | 3.724945049 | 0.073663262 |
| Gm7200 | 0 | NA | NA | NA |
| Tsn | 868.3708414 | -0.017870392 | 0.069089939 | -0.258654036 |
| Nifk | 240.6864645 | 0.399706747 | 0.120474293 | 3.317776235 |
| Gm28867 | 0.180050447 | -0.206493189 | 3.327115185 | -0.062063733 |
| Clasp1 | 2304.76087 | -0.256924034 | 0.063573698 | -4.041357407 |
| Gm26080 | 0 | NA | NA | NA |
| Gm22710 | 2.569247428 | 0.522636557 | 1.007945198 | 0.518516839 |
| Gm28466 | 0 | NA | NA | NA |
| Tfcp2l1 | 115.5571692 | 0.675414971 | 0.153286462 | 4.406227152 |
| Gm22094 | 0 | NA | NA | NA |
| Gm28882 | 0 | NA | NA | NA |
| Gm23497 | 0 | NA | NA | NA |
| Gm28467 | 0 | NA | NA | NA |
| Gli2 | 130.8498884 | 0.014865037 | 0.170628559 | 0.08711928 |
| Mir6346 | 0 | NA | NA | NA |
| Gm26831 | 0.564941401 | -1.568340625 | 1.196492227 | -1.310782126 |
| Gm8321 | 0 | NA | NA | NA |
| Gm3508 | 0 | NA | NA | NA |
| Gm29456 | 0 | NA | NA | NA |
| Gm29455 | 2.475683411 | -0.6953733 | 0.667980398 | -1.041008542 |
| Inhbb | 109.178341 | 0.049669965 | 0.189521478 | 0.262080928 |
| Ralb | 386.9036816 | 0.57400109 | 0.098166423 | 5.847224287 |
| Gm29454 | 0 | NA | NA | NA |
| Gm27184 | 4.248472554 | 0.509346359 | 0.522451308 | 0.974916421 |
| Tmem185b | 22.02441361 | -0.153903527 | 0.253304064 | -0.607584119 |
| AC132393.1 | 0 | NA | NA | NA |
| 3830432H09Rik | 2.926677728 | 0.278937763 | 0.613745164 | 0.454484662 |
| Gm25322 | 0.15596086 | -1.15072961 | 3.483969583 | -0.330292668 |
| Epb4.1l5 | 218.5589968 | -0.082777665 | 0.090354276 | -0.91614552 |
| Gm37219 | 2.565042467 | -0.262394193 | 0.562716567 | -0.466299037 |
| Ptpn4 | 3441.616975 | -0.189251984 | 0.089374807 | -2.117509305 |
| Gm3851 | 0.583829782 | -0.796599542 | 1.221896047 | -0.651937245 |
| Gm37745 | 125.9235229 | -0.281147684 | 0.239717596 | -1.17282873 |
| Gm29657 | 0 | NA | NA | NA |
| Gm8338 | 0 | NA | NA | NA |
| Gm29658 | 0.060174537 | 0.755286614 | 3.724945049 | 0.202764498 |

| | | | | |
|---|---|---|---|---|
| Gm3551 | 0.190043611 | 0.274408551 | 2.655002295 | 0.103355297 |
| Gm28209 | 0 | NA | NA | NA |
| Tmem177 | 83.02492162 | -0.673435411 | 0.131554028 | -5.119078601 |
| Cfap221 | 5.416510947 | 0.02267408 | 0.455130977 | 0.049818802 |
| Gm17973 | 0 | NA | NA | NA |
| Sctr | 5.377310654 | -0.093482596 | 0.400509352 | -0.233409273 |
| Tmem37 | 24.45334209 | 0.848447608 | 0.324047917 | 2.618278233 |
| Dbi | 1017.882369 | 0.435562879 | 0.144358505 | 3.017230471 |
| 3110009E18Rik | 15.78515723 | -0.404116858 | 0.273905661 | -1.475387023 |
| Steap3 | 67.01145874 | -0.609955425 | 0.253164405 | -2.409325375 |
| C1ql2 | 92.61537039 | -0.142424089 | 0.166332867 | -0.856259448 |
| 1700012E03Rik | 0.060174537 | 0.755286614 | 3.724945049 | 0.202764498 |
| Marco | 0.34718572 | -0.172820816 | 1.769432356 | -0.097670202 |
| En1 | 26.69343009 | -0.252946406 | 0.221261612 | -1.143200593 |
| RP24-358J4.2 | 0.343046485 | 1.591912089 | 1.527802449 | 1.041961996 |
| 2610027F03Rik | 0 | NA | NA | NA |
| Gm29346 | 0.099155932 | 0.75528439 | 3.724945049 | 0.202763901 |
| Gm29345 | 0.060174537 | 0.755286614 | 3.724945049 | 0.202764498 |
| Gm29344 | 0 | NA | NA | NA |
| Gm29348 | 0.333474911 | -0.687379643 | 1.777064702 | -0.386806199 |
| Gm29347 | 0.045882455 | 0.755288839 | 3.724945049 | 0.202765096 |
| Gm37043 | 0 | NA | NA | NA |
| Celrr | 13.48815634 | -0.973541891 | 0.418044594 | -2.328799141 |
| Gm29359 | 0 | NA | NA | NA |
| Gm38283 | 4.643660642 | -0.436469538 | 0.529396341 | -0.824466481 |
| Insig2 | 471.7823211 | -0.192724163 | 0.083784429 | -2.300238424 |
| Gm37174 | 21.08112134 | 0.121907326 | 0.2508919 | 0.485895822 |
| 2210011K15Rik | 0.099106094 | 0.274387056 | 3.724945049 | 0.073662041 |
| Ccdc93 | 456.8414215 | -0.343254118 | 0.104401167 | -3.287837958 |
| Gm15392 | 0 | NA | NA | NA |
| Gm15393 | 0.061920613 | 0.274393828 | 3.724945049 | 0.073663859 |
| Htr5b | 20.85714652 | -0.020924562 | 0.308862535 | -0.067747169 |
| Gm8217 | 0 | NA | NA | NA |
| Ddx18 | 253.9984606 | 0.263074667 | 0.08289296 | 3.173667189 |
| Gm28591 | 0 | NA | NA | NA |
| Gm37920 | 0 | NA | NA | NA |
| Gm15394 | 0 | NA | NA | NA |
| Gm8392 | 0 | NA | NA | NA |
| Gm26472 | 0 | NA | NA | NA |
| Gm28590 | 0 | NA | NA | NA |
| Gm24547 | 0 | NA | NA | NA |
| Gm36982 | 0 | NA | NA | NA |
| Gm25771 | 0 | NA | NA | NA |
| Gm29192 | 0 | NA | NA | NA |
| Gm37970 | 0.148650935 | 0.723036258 | 2.859657322 | 0.252840175 |
| Gm8403 | 0.085327104 | 0.274389469 | 3.724945049 | 0.073662689 |
| Gm28911 | 0.104106287 | 0.274400501 | 3.724945049 | 0.073665651 |

| | | | | |
|---|---|---|---|---|
| Gm20058 | 0.094097695 | 0.755282166 | 3.724945049 | 0.202763304 |
| Gm28912 | 0 | NA | NA | NA |
| Dpp10 | 1832.665455 | -0.14347624 | 0.086689747 | -1.655054306 |
| Gm37478 | 62.41200189 | 0.016104928 | 0.269018755 | 0.059865448 |
| Gm37221 | 43.7843784 | -0.201850621 | 0.273355842 | -0.738417076 |
| Gm24791 | 5.806203063 | -1.260070814 | 0.516751251 | -2.438447534 |
| Gm37066 | 38.05353759 | -0.847337504 | 0.287517495 | -2.947081545 |
| Gm37781 | 35.98403262 | -0.400774432 | 0.316315545 | -1.267008338 |
| Gm37396 | 18.40949471 | -1.035862775 | 0.459778436 | -2.252960763 |
| Gm38263 | 58.17025342 | -0.653527943 | 0.298373874 | -2.190298815 |
| Gm28929 | 0 | NA | NA | NA |
| Gm18875 | 20.78588443 | -1.00748501 | 0.375744587 | -2.681302789 |
| Gm37551 | 88.74313094 | -0.980535552 | 0.293013109 | -3.346388003 |
| Gm38073 | 157.9840511 | -0.104091915 | 0.442746645 | -0.23510492 |
| Gm25578 | 0 | NA | NA | NA |
| Gm28928 | 91.8867847 | -0.025606549 | 0.417205985 | -0.061376275 |
| Gm28299 | 17.41287975 | -0.370920194 | 0.37731976 | -0.98303941 |
| Gm37717 | 149.0700901 | -0.490032115 | 0.458164802 | -1.06955426 |
| 1700121L03Rik | 0.097139547 | 0.274393828 | 3.724945049 | 0.073663859 |
| Actr3 | 2282.827765 | 0.391856144 | 0.087210769 | 4.49320824 |
| Gm28706 | 0 | NA | NA | NA |
| Slc35f5 | 263.8309605 | 0.247057897 | 0.095519934 | 2.586453811 |
| Gm18444 | 0 | NA | NA | NA |
| Gpr39 | 2.657777857 | 1.747660534 | 0.724417501 | 2.412504573 |
| Lypd1 | 535.5413066 | -0.079090104 | 0.082961441 | -0.953335711 |
| Nckap5 | 192.1177816 | 0.360766123 | 0.14778759 | 2.441112424 |
| Gm37161 | 5.830223875 | 0.220848095 | 0.463205521 | 0.476782087 |
| Gm37936 | 6.414345313 | 0.778473026 | 0.484931273 | 1.605326507 |
| Gm38177 | 4.318687175 | -0.066375404 | 0.539700919 | -0.122985531 |
| 4930599A14Rik | 0 | NA | NA | NA |
| Gm5261 | 0 | NA | NA | NA |
| Mgat5 | 2217.417936 | -0.081224658 | 0.060883474 | -1.334100242 |
| Gm37772 | 11.00123342 | -0.015805666 | 0.384758206 | -0.041079477 |
| Gm37510 | 48.12361392 | -0.574154022 | 0.321434663 | -1.786223108 |
| Gm8451 | 18.99290832 | 0.200437857 | 0.328601056 | 0.609973258 |
| SCARNA4 | 0.095435502 | 0.75528439 | 3.724945049 | 0.202763901 |
| SCARNA15 | 0 | NA | NA | NA |
| Gm23734 | 4.197363489 | 0.633409784 | 0.528052424 | 1.199520646 |
| Gm38301 | 6.512391526 | -0.609452369 | 0.448573086 | -1.358646757 |
| Tmem163 | 261.7619097 | 0.075083269 | 0.078659356 | 0.954537041 |
| Acmsd | 0.917997538 | 1.349922105 | 1.182139744 | 1.141931072 |
| 2900009J06Rik | 4.736596173 | -0.316061644 | 0.502475231 | -0.629009401 |
| Ccnt2 | 1690.484171 | 0.001805674 | 0.132182988 | 0.013660409 |
| Map3k19 | 75.93146868 | -0.410847993 | 0.190375969 | -2.158087466 |
| Rab3gap1 | 1284.44217 | -0.0599613 | 0.046980908 | -1.276290767 |
| Zranb3 | 101.8848498 | 0.027942285 | 0.135673349 | 0.205952641 |
| Gm28800 | 35.95127173 | -0.357993964 | 0.178850193 | -2.001641476 |

| | | | | |
|---|---|---|---|---|
| Gm38248 | 15.05177351 | -0.074677415 | 0.403828601 | -0.184923541 |
| Rpl28-ps1 | 66.91960972 | 0.216776246 | 0.168753825 | 1.284570858 |
| Gm37625 | 7.67879382 | -0.41369827 | 0.37520812 | -1.102583467 |
| Gm38026 | 3.696876593 | -1.908228539 | 0.675530371 | -2.824785713 |
| R3hdm1 | 4415.135964 | -0.63098767 | 0.148117334 | -4.260052856 |
| Gm29383 | 0 | NA | NA | NA |
| Gm37588 | 55.83216539 | -0.544106514 | 0.228103204 | -2.385352349 |
| Gm37868 | 81.33316158 | -0.31402143 | 0.275097901 | -1.141489735 |
| Gm37407 | 115.1470443 | -0.396027887 | 0.197210995 | -2.008143039 |
| Mir128-1 | 17.81336244 | -0.299225965 | 0.273344032 | -1.094686293 |
| Ubxn4 | 978.701728 | 0.156848982 | 0.093551869 | 1.676599123 |
| Lct | 31.88336858 | -0.827754019 | 0.304817842 | -2.715569447 |
| Mcm6 | 72.35290303 | 1.675577128 | 0.268014564 | 6.251813719 |
| Dars | 621.5308942 | 0.080233355 | 0.100391591 | 0.799203939 |
| Gm23902 | 0 | NA | NA | NA |
| Gm29598 | 0 | NA | NA | NA |
| Gm29597 | 0 | NA | NA | NA |
| Gm25384 | 0 | NA | NA | NA |
| Gm6170 | 19.62597736 | 0.396399617 | 0.283959614 | 1.39597181 |
| Cxcr4 | 127.9107672 | 2.22034749 | 0.286467132 | 7.750793176 |
| Gm29599 | 0.045882455 | 0.755288839 | 3.724945049 | 0.202765096 |
| Gm26686 | 0.389802664 | 0.612912087 | 1.442196831 | 0.424985047 |
| Gm23056 | 0 | NA | NA | NA |
| Thsd7b | 229.954111 | 0.497600722 | 0.124068998 | 4.010677366 |
| Gm38240 | 12.58215362 | -0.699655862 | 0.313013522 | -2.235225678 |
| Gm38330 | 4.986765232 | -0.210655372 | 0.512874618 | -0.410734641 |
| Gm36951 | 10.68103066 | 0.305685339 | 0.344334796 | 0.887756168 |
| Gm15790 | 8.257872952 | -0.150830413 | 0.398222627 | -0.378759023 |
| Rpl31-ps9 | 4.393169526 | 0.633066048 | 0.516227191 | 1.226332241 |
| Gm37479 | 19.45303257 | 0.483126232 | 0.254904114 | 1.89532536 |
| Gm37064 | 10.99610456 | 0.398530263 | 0.295714068 | 1.347687873 |
| Gm37278 | 4.9996304 | 0.70877503 | 0.445239461 | 1.591896254 |
| Gm29428 | 1.653651842 | -0.258134883 | 0.731269984 | -0.352995321 |
| Gm16081 | 0 | NA | NA | NA |
| Daf2 | 0.977993353 | 0.228384083 | 1.037697176 | 0.220087409 |
| Gm15674 | 0.068097359 | -0.206487837 | 3.724945049 | -0.055433794 |
| Gm15675 | 15.76810382 | -0.045868888 | 0.338969488 | -0.135318633 |
| Cd55 | 67.64943717 | 0.160530454 | 0.143090983 | 1.121876795 |
| Gm16083 | 14.70625333 | -0.307238318 | 0.26975989 | -1.138932544 |
| Gm37109 | 7.99913161 | -0.625839996 | 0.463216052 | -1.351075793 |
| Gm37283 | 5.739234123 | -0.815327529 | 0.435704038 | -1.87128752 |
| Zp3r | 2.754922463 | 0.278937453 | 0.616382595 | 0.45253947 |
| Gm23623 | 0.044544648 | 0.755286614 | 3.724945049 | 0.202764498 |
| C4bp | 2.027159789 | 0.827939267 | 0.840814051 | 0.984687715 |
| Gm29427 | 1.964410769 | -0.767596419 | 0.759664612 | -1.010441196 |
| C4bp-ps1 | 4.021019734 | -0.246769388 | 0.46539884 | -0.530232064 |
| Pfkfb2 | 323.8491531 | -0.207420018 | 0.118971147 | -1.743448087 |

| | | | | |
|---|---|---|---|---|
| Yod1 | 565.3277476 | 0.072898821 | 0.109718626 | 0.664416096 |
| Gm37163 | 0.049553047 | 0.274391604 | 3.724945049 | 0.073663262 |
| AA986860 | 42.90179849 | 0.010420631 | 0.187610864 | 0.055543856 |
| Gm28857 | 0.066050875 | 0.274391604 | 3.724945049 | 0.073663262 |
| Gm28856 | 0.17254495 | 0.27440788 | 2.737704594 | 0.10023283 |
| Fcamr | 0.566848388 | 0.274411186 | 1.562989394 | 0.175568169 |
| Gm15848 | 0.042716945 | -0.206490061 | 3.724945049 | -0.055434391 |
| Pigr | 0.796384302 | 2.328117832 | 1.247876439 | 1.865663746 |
| Fcmr | 1.860507999 | 1.560134059 | 0.813590678 | 1.917590873 |
| Il24 | 0.119234754 | 0.274400501 | 3.724945049 | 0.073665651 |
| Il20 | 0.112396584 | 0.27438722 | 3.724945049 | 0.073662085 |
| Il19 | 1.495078259 | 3.085527171 | 1.184400859 | 2.605137565 |
| Il10 | 32.26081669 | 7.620631193 | 1.035330214 | 7.360580312 |
| Mapkapk2 | 535.196025 | 0.765725453 | 0.13892191 | 5.511912782 |
| Gm25549 | 2.426982663 | 0.356287124 | 0.608301962 | 0.585707668 |
| Gm37084 | 67.84258375 | 0.017794178 | 0.275501943 | 0.064588212 |
| Gm25717 | 0.038981395 | 0.274393828 | 3.724945049 | 0.073663859 |
| Dyrk3 | 126.7275624 | 1.01023284 | 0.183524713 | 5.504614731 |
| Eif2d | 340.1033352 | -0.356340965 | 0.130028236 | -2.740489106 |
| Rassf5 | 359.522077 | 0.173624366 | 0.088603333 | 1.959569243 |
| Gm17796 | 0.165470308 | 1.646276139 | 2.753318586 | 0.597924318 |
| Ikbke | 98.54742827 | 2.293154934 | 0.251916789 | 9.102826933 |
| Gm28914 | 1.174162655 | -0.706056202 | 0.939706837 | -0.751357948 |
| Gm28913 | 1.688430464 | -0.072628292 | 0.747699861 | -0.097135623 |
| Srgap2 | 1739.750525 | -0.120843786 | 0.079728907 | -1.515683471 |
| Gm38262 | 21.98513846 | -0.378410784 | 0.304368503 | -1.243265252 |
| Gm37954 | 37.75257877 | -0.341487149 | 0.290103543 | -1.17712161 |
| Gm29488 | 96.98125143 | -0.271359082 | 0.231225913 | -1.173566915 |
| Gm29487 | 5.722531327 | 0.227422647 | 0.606322455 | 0.375085311 |
| Gm24273 | 9.213624824 | -0.631904787 | 0.393609863 | -1.605408923 |
| Fam72a | 24.35166078 | 0.547700429 | 0.240188006 | 2.280298834 |
| Avpr1b | 0.995114343 | 0.90281531 | 1.336025232 | 0.6757472 |
| Gm8532 | 1.429318806 | 1.125576809 | 0.998184075 | 1.127624491 |
| Ctse | 45.61320862 | 4.660970052 | 0.462895317 | 10.06916658 |
| Rab7b | 58.99444633 | 0.703873022 | 0.31946587 | 2.203280816 |
| Slc26a9 | 1.868919118 | 0.83177258 | 0.675306329 | 1.231696704 |
| Pm20d1 | 45.2287229 | -0.032759553 | 0.261766193 | -0.125148144 |
| Slc41a1 | 1235.334159 | 0.500026091 | 0.110535999 | 4.52364926 |
| Rab29 | 79.93970694 | 1.457180932 | 0.222677629 | 6.543903587 |
| Gm29105 | 0 | NA | NA | NA |
| Nucks1 | 3423.866732 | 0.210147238 | 0.076361953 | 2.751988795 |
| Gm38067 | 40.11111897 | -0.033658368 | 0.26969005 | -0.124803892 |
| Gm37676 | 0.045882455 | 0.755288839 | 3.724945049 | 0.202765096 |
| Gm29103 | 0 | NA | NA | NA |
| Slc45a3 | 23.9080149 | -0.626754823 | 0.243875141 | -2.569982416 |
| Elk4 | 760.1203314 | 0.339799077 | 0.072045502 | 4.716450957 |
| Mfsd4 | 533.3513531 | 0.023206322 | 0.093553403 | 0.248054283 |

| | | | | |
|---|---|---|---|---|
| Gm26892 | 3.660297346 | 1.066196866 | 0.667957944 | 1.596203586 |
| Gm26034 | 0.106465261 | 0.75528439 | 3.724945049 | 0.202763901 |
| Gm37120 | 6.573103098 | 1.524337008 | 0.508330194 | 2.998714273 |
| Cdk18 | 350.945801 | -0.326872396 | 0.127896929 | -2.555748589 |
| Gm29629 | 1.14102893 | 0.896821854 | 0.974972072 | 0.919843634 |
| Lemd1 | 70.07703057 | -1.405786166 | 0.22457812 | -6.259675548 |
| Mir135b | 0.432115038 | -1.845250318 | 1.61152577 | -1.145033082 |
| 1700037F24Rik | 0 | NA | NA | NA |
| Gm10188 | 0.044544648 | 0.755286614 | 3.724945049 | 0.202764498 |
| Gm29630 | 0 | NA | NA | NA |
| Gm7241 | 0.177688696 | 1.081434539 | 2.718670306 | 0.397780686 |
| Klhdc8a | 280.7923049 | 0.022525164 | 0.151392227 | 0.148786791 |
| Nuak2 | 109.3043444 | 2.924435096 | 0.269118333 | 10.86672567 |
| F730311O21Rik | 4.129176112 | 2.889976507 | 0.931852286 | 3.101324696 |
| Tmcc2 | 2513.213478 | 0.225264455 | 0.097135959 | 2.319063472 |
| Gm15849 | 0.048949532 | -0.206487837 | 3.724945049 | -0.055433794 |
| Dstyk | 977.2630414 | -0.030794489 | 0.051243838 | -0.600940341 |
| 6030442K20Rik | 15.5449471 | -0.891837873 | 0.332511456 | -2.682126755 |
| Rbbp5 | 612.646602 | -0.033870344 | 0.058473949 | -0.579238185 |
| Tmem81 | 45.06443145 | 0.121415581 | 0.163838146 | 0.741070282 |
| Cntn2 | 3239.191646 | -0.027372515 | 0.084214572 | -0.325032998 |
| Nfasc | 5772.548432 | -0.032358185 | 0.057370853 | -0.564017847 |
| Gm10538 | 0.168483864 | 1.669410905 | 2.741982635 | 0.608833508 |
| Gm22609 | 0.089089296 | 1.194485492 | 3.723939576 | 0.320758559 |
| Lrrn2 | 1233.035506 | 0.186826274 | 0.094656734 | 1.973724063 |
| AC124095.1 | 0.988133447 | 1.961419558 | 1.266554774 | 1.548625924 |
| Gm38308 | 6.071337244 | 1.030043872 | 0.579578378 | 1.777229639 |
| Mdm4 | 1609.398272 | -0.160735017 | 0.1175915 | -1.366893162 |
| Gm29257 | 3.367574844 | 0.436104298 | 0.566895952 | 0.769284552 |
| Pik3c2b | 418.6707706 | -0.879699145 | 0.130832133 | -6.723876816 |
| Gm28609 | 1.26125615 | 0.913070778 | 1.052344539 | 0.867653838 |
| Gm26616 | 7.818633818 | 0.736170409 | 0.420054546 | 1.752559082 |
| Ppp1r15b | 1586.837848 | 0.574277363 | 0.122372268 | 4.692871763 |
| Plekha6 | 1798.908558 | -0.329345537 | 0.087152072 | -3.778975417 |
| Gm19461 | 22.93070562 | 0.235886521 | 0.239114292 | 0.98650114 |
| Golt1a | 0.589467498 | -0.423116196 | 1.549181796 | -0.273122365 |
| Gm28040 | 0.066050875 | 0.274391604 | 3.724945049 | 0.073663262 |
| Kiss1 | 0.282483437 | 1.193257874 | 1.770585626 | 0.673934012 |
| Ren1 | 0.441022512 | -0.992423539 | 1.462759572 | -0.678459781 |
| Etnk2 | 45.75414376 | -0.447199741 | 0.197820082 | -2.260638736 |
| Sox13 | 89.13343506 | -0.177241959 | 0.141261916 | -1.254704482 |
| Gm8596 | 0 | NA | NA | NA |
| Gm23939 | 0 | NA | NA | NA |
| Snrpe | 146.4574117 | 0.373253962 | 0.146820285 | 2.542250631 |
| Gm26706 | 0.926160546 | 0.200635594 | 0.983080179 | 0.204088738 |
| Zc3h11a | 237.5869313 | -0.123536404 | 0.120540316 | -1.024855481 |
| Gm38394 | 135.9246581 | -0.104744578 | 0.159630081 | -0.656170675 |

| | | | | |
|---|---|---|---|---|
| Zbed6 | 0 | NA | NA | NA |
| Lax1 | 23.20725604 | 3.557226113 | 0.413944177 | 8.593492331 |
| Atp2b4 | 2855.857049 | -0.41055039 | 0.10006847 | -4.102694764 |
| Mir6903 | 0.148659141 | 0.274384287 | 3.724945049 | 0.073661298 |
| Gm37368 | 12.07769997 | -0.584427465 | 0.405834679 | -1.440062901 |
| Gm10537 | 0.578745683 | 0.286523275 | 1.32834724 | 0.215699078 |
| Gm28441 | 0.061920613 | 0.274393828 | 3.724945049 | 0.073663859 |
| Optc | 6.651424553 | 0.283600065 | 0.40247786 | 0.704635194 |
| Gm15851 | 0.236932206 | 0.321096127 | 2.533571261 | 0.126736568 |
| Prelp | 162.3998482 | -0.395518164 | 0.214844342 | -1.8409522 |
| Gm1627 | 0.141089779 | -1.120093354 | 2.901102929 | -0.386092249 |
| Gm18970 | 0.109124069 | 0.274402725 | 3.724945049 | 0.073666248 |
| Fmod | 152.7762662 | -0.636921084 | 0.360793256 | -1.765335335 |
| Btg2 | 283.4370204 | 0.132116537 | 0.292089566 | 0.452315154 |
| Gm38140 | 0.628069627 | 0.559895044 | 1.376344774 | 0.40679854 |
| Gm8618 | 0.104637558 | -0.206492286 | 3.724945049 | -0.055434988 |
| Chit1 | 1.035276703 | 1.046114613 | 0.955545727 | 1.094782367 |
| Chil1 | 168.3470407 | 1.493768183 | 0.252502661 | 5.915851242 |
| Mybph | 3.909958598 | 2.600003318 | 0.722509293 | 3.598574222 |
| Adora1 | 2485.958102 | 0.066483065 | 0.073198944 | 0.908251703 |
| Gm24517 | 0.075043205 | 0.755286614 | 3.724945049 | 0.202764498 |
| Myog | 0.249503106 | 0.274409577 | 2.135059485 | 0.128525495 |
| Ppfia4 | 1225.389552 | -0.375749711 | 0.11269777 | -3.334136158 |
| Tmem183a | 645.1063955 | 0.094854294 | 0.084815665 | 1.118358202 |
| 4933406M09Rik | 0 | NA | NA | NA |
| Cyb5r1 | 155.8595938 | 0.120166006 | 0.1183685 | 1.015185684 |
| Adipor1 | 881.11371 | 0.141472468 | 0.074768597 | 1.89213754 |
| Platr1 | 1.832214876 | 0.353003013 | 1.179690203 | 0.299233657 |
| Gm37935 | 2.337573936 | 0.624725322 | 0.687510839 | 0.908677052 |
| Klhl12 | 493.9899022 | -0.036098095 | 0.079232108 | -0.455599324 |
| Rabif | 301.4519563 | -0.200100195 | 0.083562052 | -2.394629987 |
| Gm25612 | 0.720424393 | 0.552466303 | 1.114176868 | 0.495851529 |
| Gm15454 | 0.589661845 | -0.206000507 | 1.382592188 | -0.148995857 |
| Gm29376 | 0.461892809 | -1.365600366 | 1.424199967 | -0.958854373 |
| Mgat4e | 0.04393175 | -0.206490061 | 3.724945049 | -0.055434391 |
| Kdm5b | 1415.896103 | -0.290725306 | 0.081739953 | -3.556709978 |
| Gm26783 | 4.085067263 | 0.335268417 | 0.478072216 | 0.701292411 |
| Syt2 | 2912.71677 | -0.206589871 | 0.134256716 | -1.538767498 |
| Ppp1r12b | 1746.350506 | -0.43061063 | 0.095084625 | -4.528709353 |
| Gm37949 | 36.84559847 | -0.834472292 | 0.325254621 | -2.565597038 |
| Gm22232 | 0.044191549 | -0.206490061 | 3.724945049 | -0.055434391 |
| Gm37677 | 160.3400663 | -0.836291755 | 0.240477511 | -3.477629788 |
| Gm28892 | 1.029532974 | 0.208131753 | 1.468752285 | 0.141706505 |
| Ube2t | 14.99856726 | 0.479753477 | 0.282549369 | 1.697945665 |
| Lgr6 | 136.8876649 | -0.105823504 | 0.277450244 | -0.381414347 |
| Ptprv | 31.89205934 | 0.174127597 | 0.325077493 | 0.5356495 |
| Gm10535 | 0.133946075 | 0.755288219 | 3.626642455 | 0.20826101 |

| | | | | |
|---|---|---|---|---|
| Ptpn7 | 31.7926034 | 1.613227948 | 0.319273383 | 5.052810642 |
| Arl8a | 2134.195625 | 0.083413458 | 0.061109568 | 1.364981969 |
| Gm26280 | 0.068097359 | -0.206487837 | 3.724945049 | -0.055433794 |
| Gm15445 | 11.41385022 | 0.04962448 | 0.427454264 | 0.116093076 |
| Gpr37l1 | 2282.91187 | 0.199400094 | 0.148785843 | 1.3401819 |
| Gm4204 | 333.7656317 | 0.038438821 | 0.105466606 | 0.364464377 |
| Gm26642 | 0.168999367 | -0.206486166 | 2.746693795 | -0.07517626 |
| Elf3 | 3.18304154 | 0.639497796 | 0.663757175 | 0.963451425 |
| Rnpep | 506.4393182 | 0.25818472 | 0.079211576 | 3.259431692 |
| Timm17a | 833.5513504 | -0.158207583 | 0.065343616 | -2.421163574 |
| Lmod1 | 6.027761192 | -0.046343577 | 0.454452186 | -0.101976794 |
| Gm22796 | 0.04393175 | -0.206490061 | 3.724945049 | -0.055434391 |
| Shisa4 | 891.8069629 | -0.133786842 | 0.086870186 | -1.540077755 |
| Ipo9 | 1823.820265 | -0.125973462 | 0.047083941 | -2.675507996 |
| Gm28277 | 10.85246338 | -0.392100179 | 0.431787587 | -0.908085806 |
| Nav1 | 1957.583854 | -0.472017511 | 0.100811132 | -4.682196301 |
| Mir1231 | 0 | NA | NA | NA |
| 2610012C04Rik | 6.062032631 | -0.451874648 | 0.397700381 | -1.136218796 |
| Gm37333 | 28.22584607 | -1.103670604 | 0.400577213 | -2.755200669 |
| Gm4793 | 13.11457074 | -0.532943828 | 0.310525646 | -1.716263487 |
| Gm38399 | 26.61474392 | 0.954381179 | 0.234699265 | 4.066400378 |
| Csrp1 | 2908.739873 | 0.782170575 | 0.153160385 | 5.106872611 |
| Phlda3 | 300.6871226 | 0.394663614 | 0.157960207 | 2.498500234 |
| Tnni1 | 43.38678691 | 0.495477662 | 0.225184912 | 2.200314652 |
| Lad1 | 13.14689316 | -0.011259509 | 0.366857133 | -0.03069181 |
| Tnnt2 | 21.42389219 | 0.240840842 | 0.279098663 | 0.862923668 |
| Gm28883 | 1.230445023 | 0.43181298 | 0.906822515 | 0.476182464 |
| Pkp1 | 4.016384854 | -0.460418697 | 0.665606654 | -0.691727905 |
| Gm36938 | 0.07796279 | 0.274392189 | 3.724945049 | 0.073663419 |
| Igfn1 | 106.9666211 | -1.561548046 | 0.290305373 | -5.378984305 |
| Tmem9 | 526.3167638 | 0.593143829 | 0.110921689 | 5.347410709 |
| Ascl5 | 1.900208836 | -0.186144965 | 0.73925771 | -0.251799829 |
| Cacna1s | 20.1854872 | 0.768577797 | 0.268344846 | 2.864142195 |
| Gm37014 | 0 | NA | NA | NA |
| Gm37759 | 0.707021654 | -0.904309063 | 1.449050823 | -0.624069942 |
| Gm15850 | 4.717623853 | -1.604546078 | 0.574834715 | -2.791317289 |
| Kif21b | 1190.983809 | -0.439645612 | 0.095254508 | -4.615483516 |
| Mroh3 | 7.987918312 | 0.529096568 | 0.417281888 | 1.267959582 |
| 5730559C18Rik | 56.42049381 | 0.63011324 | 0.189381258 | 3.327220689 |
| Gm26568 | 0.061920613 | 0.274393828 | 3.724945049 | 0.073663859 |
| A430034D21Rik | 0 | NA | NA | NA |
| Gm37552 | 0.476395801 | -1.079250509 | 1.358828785 | -0.79425055 |
| Gm26781 | 4.326125698 | -0.336057158 | 0.465544304 | -0.721858597 |
| Gpr25 | 16.1131376 | -0.377470082 | 0.260931646 | -1.446624385 |
| Camsap2 | 3490.88954 | 0.004365132 | 0.051520103 | 0.084726762 |
| 9230116N13Rik | 6.573994958 | 0.791330836 | 0.516494344 | 1.532119073 |
| Ddx59 | 80.65548895 | 0.322527398 | 0.136605171 | 2.36101896 |

| | | | | |
|---|---|---|---|---|
| Kif14 | 11.46902475 | 1.379879026 | 0.402938171 | 3.424542836 |
| Gm29486 | 0.097139547 | 0.274393828 | 3.724945049 | 0.073663859 |
| Gm29485 | 0.285872948 | -0.987144545 | 2.080135064 | -0.474557909 |
| Gm37799 | 0.092691418 | 0.274387304 | 3.724945049 | 0.073662108 |
| Gm33994 | 0.860407559 | 1.928314294 | 1.199873068 | 1.607098573 |
| 4933409D19Rik | 0.218406194 | 0.755303214 | 2.160775026 | 0.349551992 |
| Platr23 | 2.85951047 | 1.120823231 | 0.676900912 | 1.655815807 |
| Zfp281 | 1236.560802 | 0.396561728 | 0.109446115 | 3.623351358 |
| Gm19705 | 24.95245937 | 0.636468969 | 0.232351743 | 2.739247665 |
| Platr22 | 0.971167388 | 1.9708754 | 1.053877063 | 1.870118886 |
| Gm24086 | 0.066050875 | 0.274391604 | 3.724945049 | 0.073663262 |
| Gm22132 | 0 | NA | NA | NA |
| Gm8762 | 0 | NA | NA | NA |
| Gm29718 | 0.856212859 | -1.101704065 | 1.208320591 | -0.911764703 |
| Gm17781 | 0 | NA | NA | NA |
| Nr5a2 | 2.338360347 | -0.305937711 | 0.744895603 | -0.410712199 |
| Gm23763 | 0 | NA | NA | NA |
| Gm25609 | 0.046345709 | 0.274391604 | 3.724945049 | 0.073663262 |
| Gm37057 | 0.333318405 | 1.056869986 | 1.697104595 | 0.62274888 |
| 1700006P03Rik | 0 | NA | NA | NA |
| Gm23534 | 0 | NA | NA | NA |
| Gm37903 | 0.499362493 | -1.433742827 | 1.56586478 | -0.915623651 |
| Gm22727 | 0 | NA | NA | NA |
| Gm37099 | 0.251787852 | 2.148234503 | 2.03932619 | 1.053404067 |
| Gm37293 | 0 | NA | NA | NA |
| Gm4258 | 79.79539825 | -0.757603335 | 0.248552641 | -3.048059887 |
| Gm37298 | 205.364216 | -0.887621103 | 0.197020514 | -4.505221753 |
| Gm26979 | 0 | NA | NA | NA |
| Gm37124 | 62.91602665 | -0.891513674 | 0.230946853 | -3.860254697 |
| A130050O07Rik | 109.5267668 | -0.939310368 | 0.259698433 | -3.616927363 |
| Gm26936 | 0 | NA | NA | NA |
| Gm26930 | 92.36817408 | -0.877439419 | 0.273722452 | -3.20558074 |
| D130019J16Rik | 127.5229106 | -0.944570881 | 0.213202488 | -4.430393327 |
| Gm37534 | 242.1756396 | -0.930748274 | 0.199647504 | -4.661957969 |
| Mir181a-1 | 0.399769715 | -1.594130933 | 1.544983879 | -1.031810723 |
| A430106G13Rik | 56.85494254 | -0.999574479 | 0.277327507 | -3.604310629 |
| Mir181b-1 | 0 | NA | NA | NA |
| Gm37632 | 17.06609017 | -1.181914399 | 0.278431949 | -4.244895035 |
| Gm23782 | 0 | NA | NA | NA |
| Ptprc | 1013.820159 | 4.001710058 | 0.192769675 | 20.75902267 |
| A130071D04Rik | 74.31188421 | 4.749450753 | 0.423486595 | 11.21511474 |
| Gm18667 | 0 | NA | NA | NA |
| Gm28556 | 0 | NA | NA | NA |
| 1700113B19Rik | 0 | NA | NA | NA |
| Atp6v1g3 | 0.18576184 | 0.274389742 | 3.724945049 | 0.073662762 |
| Gm28500 | 0.232206954 | 0.586111303 | 2.544795705 | 0.230317625 |
| Nek7 | 865.5502947 | 0.528888264 | 0.122140499 | 4.33016294 |

| | | | | |
|---|---|---|---|---|
| Gm28501 | 0.215166026 | 0.775847198 | 2.554583147 | 0.303707945 |
| Gm37080 | 8.379318968 | 0.654750764 | 0.434186033 | 1.507995916 |
| Gm37101 | 16.00512593 | 0.811560631 | 0.366895574 | 2.211966261 |
| Gm5833 | 0.04393175 | -0.206490061 | 3.724945049 | -0.055434391 |
| Gm8790 | 0 | NA | NA | NA |
| Lhx9 | 105.8300051 | 0.096230616 | 0.101244955 | 0.950473195 |
| 2310009B15Rik | 35.43042701 | 0.163698301 | 0.183590311 | 0.89165 |
| 1700019P21Rik | 0 | NA | NA | NA |
| n-R5s217 | 0.075043205 | 0.755286614 | 3.724945049 | 0.202764498 |
| Gm3933 | 0 | NA | NA | NA |
| Gm37045 | 1.708528675 | -0.293349641 | 0.77088639 | -0.380535504 |
| Dennd1b | 1082.456846 | 0.481782741 | 0.07590358 | 6.347299293 |
| Gm37423 | 58.57700844 | -0.178801936 | 0.254546801 | -0.702432464 |
| AC117623.1 | 17.28746858 | 0.010689486 | 0.359738399 | 0.029714609 |
| Gm18553 | 0.072025462 | 0.755288839 | 3.724945049 | 0.202765096 |
| Gm16305 | 0 | NA | NA | NA |
| Gm37470 | 63.54316749 | 0.284829298 | 0.195832838 | 1.454451159 |
| RP23-114L12.4 | 43.15415538 | 0.226454789 | 0.245827046 | 0.921195583 |
| Gm37655 | 0.107078754 | -0.206490058 | 3.724945049 | -0.05543439 |
| Gm16304 | 0 | NA | NA | NA |
| Gm37718 | 31.22466683 | 0.369464932 | 0.250456882 | 1.475163828 |
| Crb1 | 30.89787103 | -0.399804193 | 0.261619734 | -1.528188209 |
| 4933436E23Rik | 0.112288909 | -0.687373951 | 3.724945049 | -0.184532642 |
| Gm34816 | 0 | NA | NA | NA |
| Zbtb41 | 682.7308389 | -0.328612776 | 0.072418677 | -4.537679892 |
| Aspm | 33.81106085 | 0.51895253 | 0.35112338 | 1.477977712 |
| F13b | 0.541494639 | -0.683888701 | 1.538970197 | -0.444380731 |
| Cfhr1 | 0.094294407 | -0.206492286 | 3.724945049 | -0.055434988 |
| Cfhr3 | 0 | NA | NA | NA |
| Gm26271 | 0 | NA | NA | NA |
| Gm4788 | 0.298414574 | -0.676129702 | 1.813350443 | -0.372862126 |
| Gm25619 | 0 | NA | NA | NA |
| Cfhr2 | 0.451902873 | -0.397327185 | 1.309882751 | -0.303330343 |
| Gm16332 | 4.277969054 | 0.657915259 | 0.503264631 | 1.307294847 |
| Gm5837 | 9.46935442 | -0.298344482 | 0.365262968 | -0.816793676 |
| Gm5834 | 1.243360777 | 0.145466028 | 1.054816967 | 0.137906416 |
| Gm38266 | 0 | NA | NA | NA |
| Gm26048 | 0 | NA | NA | NA |
| Gm38019 | 0.981599412 | 0.10388927 | 0.990642766 | 0.104870568 |
| Cfh | 496.518771 | 0.587608254 | 0.132044375 | 4.45008167 |
| Kcnt2 | 523.8644755 | -0.070776784 | 0.116583133 | -0.607092828 |
| Gm37777 | 49.81740225 | -0.385722444 | 0.228504182 | -1.688032317 |
| 4930590L20Rik | 0 | NA | NA | NA |
| Gm37371 | 0.255648062 | -1.236130775 | 1.83531447 | -0.673525325 |
| Gm4845 | 0 | NA | NA | NA |
| Gm25457 | 0 | NA | NA | NA |
| Gm22016 | 0 | NA | NA | NA |

| | | | | |
|---|---|---|---|---|
| Gm37599 | 0.182775946 | 1.806322629 | 3.324025814 | 0.54341414 |
| Gm37142 | 0 | NA | NA | NA |
| Gm37604 | 0 | NA | NA | NA |
| Gm18183 | 0.075043205 | 0.755286614 | 3.724945049 | 0.202764498 |
| Gm28724 | 0 | NA | NA | NA |
| Gm37796 | 0 | NA | NA | NA |
| Gm5835 | 7.977887189 | -0.203537208 | 0.401893621 | -0.50644548 |
| Cdc73 | 1577.910083 | 0.069249753 | 0.05454167 | 1.269666902 |
| B3galt2 | 609.7109376 | 0.165161929 | 0.085020523 | 1.942612479 |
| Gm6397 | 3.497495822 | 0.70281086 | 0.555925971 | 1.264216635 |
| Glrx2 | 603.4570244 | -0.230217527 | 0.062299774 | -3.69531883 |
| Trove2 | 1714.422575 | -0.07590383 | 0.098299008 | -0.772172892 |
| Uchl5 | 265.6573748 | 0.034940425 | 0.139871176 | 0.249804329 |
| Gm29170 | 125.043805 | -0.170045877 | 0.116415779 | -1.460677221 |
| Gm25663 | 0 | NA | NA | NA |
| Rgs2 | 585.4803157 | -0.096674559 | 0.20376871 | -0.474432798 |
| Rgs13 | 2.523201682 | 1.215505618 | 0.700412346 | 1.735414323 |
| Rbm6-ps2 | 0.220340683 | 0.68150599 | 2.19207884 | 0.310894835 |
| Rgs1 | 170.7604979 | 7.404224388 | 0.674181879 | 10.98253248 |
| Gm7266 | 0.253428164 | 1.594735524 | 2.08690747 | 0.764162066 |
| Rgs21 | 0 | NA | NA | NA |
| Gm23672 | 0 | NA | NA | NA |
| Rgs18 | 3.121267796 | 2.199393235 | 0.833273562 | 2.639461199 |
| Gm8856 | 0 | NA | NA | NA |
| Gm22681 | 0 | NA | NA | NA |
| Gm25708 | 0 | NA | NA | NA |
| Gm6550 | 0 | NA | NA | NA |
| Gm5262 | 0 | NA | NA | NA |
| Gm25460 | 0 | NA | NA | NA |
| Gm28392 | 0 | NA | NA | NA |
| Gm38269 | 0.279407529 | 0.291603298 | 2.073798642 | 0.14061312 |
| Gm29514 | 42.66995015 | -0.728729947 | 0.217642584 | -3.348287519 |
| Gm5263 | 0.047948698 | -0.206487837 | 3.724945049 | -0.055433794 |
| Gm38187 | 45.19927449 | -1.062679256 | 0.288213318 | -3.687127521 |
| Brinp3 | 455.9540705 | -0.227357159 | 0.087450192 | -2.59984746 |
| Gm15583 | 12.60536537 | -0.407666228 | 0.389840278 | -1.045726291 |
| A230059L01Rik | 0.205212924 | 1.718778489 | 2.635865588 | 0.652073648 |
| Gm15584 | 0.127515837 | 0.274388643 | 3.69037927 | 0.074352424 |
| Gm37749 | 0.118371171 | 0.75528439 | 3.724945049 | 0.202763901 |
| Gm9931 | 0 | NA | NA | NA |
| Gm17867 | 0.136963818 | 0.755286583 | 3.59954433 | 0.209828388 |
| Gm38284 | 0.044544648 | 0.755286614 | 3.724945049 | 0.202764498 |
| Gm29516 | 0 | NA | NA | NA |
| Gm29515 | 0 | NA | NA | NA |
| Gm22966 | 0 | NA | NA | NA |
| Gm19891 | 0 | NA | NA | NA |
| Gm18505 | 0.044191549 | -0.206490061 | 3.724945049 | -0.055434391 |

| | | | | |
|---|---|---|---|---|
| Gm4034 | 0 | NA | NA | NA |
| Gm29022 | 0 | NA | NA | NA |
| Gm5703 | 0 | NA | NA | NA |
| Gm15443 | 3.976562986 | 0.205843512 | 0.492016757 | 0.418366873 |
| Gm18751 | 0 | NA | NA | NA |
| Gm29020 | 0 | NA | NA | NA |
| Gm18701 | 0 | NA | NA | NA |
| Gm37630 | 0.062426873 | 0.755288839 | 3.724945049 | 0.202765096 |
| Gm38189 | 0.200297358 | 0.776142251 | 2.613669831 | 0.296954972 |
| Gm29398 | 0.199606257 | 0.727999894 | 2.214366496 | 0.328762152 |
| Gm37096 | 0.086648695 | -0.687376175 | 3.724945049 | -0.184533239 |
| Gm37775 | 0.044544648 | 0.755286614 | 3.724945049 | 0.202764498 |
| Gm23535 | 0 | NA | NA | NA |
| n-R5s218 | 0 | NA | NA | NA |
| Pla2g4a | 121.1500737 | 1.038362309 | 0.218395906 | 4.754495302 |
| n-R5s219 | 0 | NA | NA | NA |
| Gm19136 | 0.350167968 | -0.689484978 | 1.604740353 | -0.429655163 |
| AC114655.1 | 0.088736197 | 0.274400501 | 3.724945049 | 0.073665651 |
| Ptgs2os | 10.82272229 | 0.118486622 | 0.395823989 | 0.299341692 |
| Ptgs2 | 389.8698901 | 1.181825911 | 0.353620948 | 3.342069855 |
| Gm9687 | 0.529068976 | -2.518886393 | 1.271744638 | -1.980654227 |
| Ptgs2os2 | 1.911122854 | 1.240284925 | 0.778909224 | 1.59233565 |
| Gm15453 | 44.68224246 | 1.768526034 | 0.235369323 | 7.513834048 |
| Gm4322 | 0.586444394 | -1.19982675 | 1.263787628 | -0.949389536 |
| Pdc | 0.117760149 | 0.274400501 | 3.724945049 | 0.073665651 |
| 2310030A07Rik | 0.378962075 | 2.607491924 | 1.421650037 | 1.834130663 |
| Gm37764 | 0 | NA | NA | NA |
| BC003331 | 544.608875 | 0.204236875 | 0.063724358 | 3.205004826 |
| Tpr | 2374.975998 | 0.136184655 | 0.081541081 | 1.670135515 |
| Prg4 | 160.3573172 | 2.0080674 | 0.389054495 | 5.161403924 |
| Ska2l-ps | 0.636218932 | -0.779273077 | 1.22455852 | -0.63637063 |
| Hmcn1 | 64.89396023 | -0.724912696 | 0.161041297 | -4.501408701 |
| Gm20631 | 0.097139547 | 0.274393828 | 3.724945049 | 0.073663859 |
| Gm29188 | 0 | NA | NA | NA |
| Gm22756 | 0 | NA | NA | NA |
| Gm37671 | 0.106358623 | 0.274402725 | 3.724945049 | 0.073666248 |
| Gm19087 | 0 | NA | NA | NA |
| Gm8941 | 0.044544648 | 0.755286614 | 3.724945049 | 0.202764498 |
| C730036E19Rik | 0.469314954 | 2.764904425 | 1.434505739 | 1.927426534 |
| Gm24402 | 0 | NA | NA | NA |
| Gm10138 | 35.28446059 | -0.96890762 | 0.19363906 | -5.003678584 |
| Ivns1abp | 1648.737111 | -0.143989405 | 0.074710033 | -1.927310161 |
| Swt1 | 331.9812924 | -0.537380189 | 0.114519519 | -4.692476815 |
| Trmt1l | 460.2408708 | -0.474054836 | 0.194843735 | -2.433000154 |
| mmu-mir-7682 | 1.074470705 | -0.591375327 | 1.073884827 | -0.550687851 |
| Gm37893 | 50.46302794 | -1.070715705 | 0.305413663 | -3.50578849 |
| Rnf2 | 494.0673276 | 0.535345108 | 0.097540479 | 5.488440432 |

| | | | | |
|---|---|---|---|---|
| Gm36527 | 43.64518431 | 0.020120877 | 0.346057367 | 0.058143182 |
| Gm29072 | 0 | NA | NA | NA |
| Gm19503 | 0.173853943 | 0.755298194 | 2.74376612 | 0.275277907 |
| Gm28181 | 0.049553047 | 0.274391604 | 3.724945049 | 0.073663262 |
| Fam129a | 103.521438 | 1.07190811 | 0.141712748 | 7.563949762 |
| 2810414N06Rik | 13.51786206 | -0.211102301 | 0.298356053 | -0.707551594 |
| Edem3 | 1232.170216 | 0.036612956 | 0.063643218 | 0.575284487 |
| 1700025G04Rik | 1157.852342 | -0.448298647 | 0.061627241 | -7.274358548 |
| Gm25279 | 0.423681387 | 0.310948283 | 1.754723081 | 0.17720647 |
| Gm28610 | 0.118371171 | 0.75528439 | 3.724945049 | 0.202763901 |
| Gm8964 | 0 | NA | NA | NA |
| Gm38238 | 0.047948698 | -0.206487837 | 3.724945049 | -0.055433794 |
| Tsen15 | 205.9704903 | -0.019422392 | 0.091208688 | -0.21294454 |
| Colgalt2 | 66.2940545 | -0.308028288 | 0.142655736 | -2.159242223 |
| Gm15618 | 0.134880151 | 0.274398319 | 2.937637886 | 0.093407809 |
| Rgl1 | 1379.502362 | 0.226343184 | 0.089972792 | 2.515684791 |
| Gm15479 | 6.574668662 | -0.972170464 | 0.464750856 | -2.091809951 |
| Gm7278 | 19.06484076 | -0.571789222 | 0.297230789 | -1.923721376 |
| Gm28792 | 7.043371214 | -0.433422286 | 0.386275526 | -1.122054743 |
| Apobec4 | 0.396489138 | 1.821102023 | 1.49139203 | 1.221075335 |
| Arpc5 | 1009.34046 | 0.668380211 | 0.120421826 | 5.550324482 |
| Ncf2 | 128.1024071 | 1.873172337 | 0.203762197 | 9.192933553 |
| Smg7 | 2499.293702 | 0.455587634 | 0.086096381 | 5.291600265 |
| Gm28791 | 48.57147184 | 0.019081052 | 0.251992152 | 0.075720818 |
| 1700040K01Rik | 0.217189781 | -0.687379067 | 2.174821597 | -0.316062277 |
| Gm8976 | 0.43436012 | 1.828660786 | 1.682882678 | 1.086624047 |
| Nmnat2 | 3031.207894 | 0.208768148 | 0.076081165 | 2.744018809 |
| Gm28960 | 0 | NA | NA | NA |
| Lamc2 | 154.9877848 | -0.008218726 | 0.141710576 | -0.057996557 |
| Lamc1 | 1030.31094 | 0.615714895 | 0.12206324 | 5.04422869 |
| Gm38009 | 27.75065565 | -0.158433884 | 0.279063351 | -0.567734471 |
| Gm8818 | 0.931703337 | 1.192098965 | 1.356777525 | 0.87862523 |
| E330020D12Rik | 5.955571085 | 0.437642991 | 0.415340708 | 1.053696357 |
| Shcbp1l | 22.81337923 | -0.742799472 | 0.260018142 | -2.856721713 |
| Gm25588 | 0.946605489 | -1.678564125 | 1.305497687 | -1.285765683 |
| Gm37669 | 3.285918002 | -1.145678803 | 0.553114096 | -2.071324546 |
| Dhx9 | 2116.613299 | 0.266630979 | 0.077456715 | 3.442322313 |
| Npl | 38.83715233 | 0.282857019 | 0.211387915 | 1.338094556 |
| Gm6652 | 5.116277746 | 1.273339034 | 0.593151389 | 2.146735315 |
| Rgs8 | 2498.85835 | -0.278912058 | 0.159981771 | -1.743398983 |
| Gm29529 | 0.238359071 | -0.687378567 | 2.119250321 | -0.324349871 |
| 5530400K19Rik | 2.012029913 | 0.298604484 | 0.984173631 | 0.303406304 |
| C230024C17Rik | 0.83423333 | -0.293465879 | 1.1341818 | -0.258746771 |
| Rgs16 | 197.3246427 | 0.120200921 | 0.132733916 | 0.905578052 |
| Gm28513 | 0.329560409 | 2.143332883 | 1.727477455 | 1.240729873 |
| Rnasel | 213.55156 | 1.316285944 | 0.147155202 | 8.944882159 |
| Gm28512 | 4.354440161 | 0.190653084 | 0.51485919 | 0.370301409 |

| | | | | |
|---|---|---|---|---|
| Rgsl1 | 23.12760851 | -0.123625385 | 0.295265257 | -0.418692623 |
| Teddm2 | 33.79161557 | -0.032583791 | 0.283932646 | -0.114758875 |
| Teddm1b | 0.047948698 | -0.206487837 | 3.724945049 | -0.055433794 |
| Gm38352 | 5.642899575 | -1.02474696 | 0.469107075 | -2.184462813 |
| Teddm1a | 0.061920613 | 0.274393828 | 3.724945049 | 0.073663859 |
| Mir8114 | 0.170693195 | 0.755297969 | 2.759623049 | 0.273696065 |
| Glul | 15318.94443 | -0.005153751 | 0.158637417 | -0.032487612 |
| Gm29290 | 0.209920398 | -0.687380217 | 2.625934473 | -0.261765944 |
| Gm29291 | 0.447162105 | -1.833666358 | 1.379414639 | -1.329307596 |
| Gm28286 | 0.825983375 | -1.27747564 | 1.048493086 | -1.21839205 |
| A830008E24Rik | 2.805075798 | 0.288795541 | 0.567338826 | 0.509035392 |
| Gm25336 | 0 | NA | NA | NA |
| Zfp648 | 4.036848141 | 0.337405876 | 0.524049958 | 0.643842959 |
| 4930452N14Rik | 0.785070532 | 0.305626517 | 1.372294762 | 0.222712004 |
| 4930532M18Rik | 0.278668908 | 0.274415848 | 1.798586118 | 0.152573093 |
| n-R5s220 | 0.221251082 | 1.236195239 | 2.577840323 | 0.479546863 |
| Gm18727 | 0.332553061 | 1.236539368 | 1.970583981 | 0.627498944 |
| Cacna1e | 5959.244093 | -0.325975625 | 0.157661049 | -2.067572342 |
| Gm29282 | 1.600249739 | -0.238260148 | 0.794468406 | -0.299898833 |
| Gm29441 | 0.044544648 | 0.755286614 | 3.724945049 | 0.202764498 |
| Gm9530 | 4.308798137 | -0.385790936 | 0.473144521 | -0.815376526 |
| Ier5 | 750.1768822 | -0.061049707 | 0.147203387 | -0.414730313 |
| Mr1 | 88.50137408 | 0.374742141 | 0.154143386 | 2.431126963 |
| Stx6 | 782.513869 | -0.048409926 | 0.05253767 | -0.921432687 |
| Gm8850 | 14.83397756 | -0.004307851 | 0.353343564 | -0.012191678 |
| BC034090 | 156.7399653 | -0.060505085 | 0.110662061 | -0.54675545 |
| Gm37622 | 0.256551671 | 1.697465278 | 2.455008697 | 0.691429436 |
| Xpr1 | 2184.107984 | -0.220182182 | 0.058908855 | -3.737675482 |
| Gm22569 | 0 | NA | NA | NA |
| Gm5532 | 8.533642962 | 1.596741781 | 0.584388348 | 2.732329941 |
| Acbd6 | 499.0889261 | 0.238702224 | 0.082047238 | 2.909326747 |
| Lhx4 | 8.533691663 | -0.539668296 | 0.865475858 | -0.623550953 |
| Gm37089 | 0 | NA | NA | NA |
| Qsox1 | 389.4008561 | 0.254832475 | 0.088222716 | 2.888513145 |
| Gm37539 | 0 | NA | NA | NA |
| Gm37571 | 0.346370079 | 1.236200409 | 2.059371759 | 0.600280354 |
| Cep350 | 1964.651825 | -0.060584267 | 0.075438922 | -0.803090299 |
| Gm38038 | 0 | NA | NA | NA |
| Gm36934 | 0.108309328 | 1.204448679 | 3.72248341 | 0.323560523 |
| Gm37383 | 6.027087886 | -0.843971987 | 0.459984347 | -1.834784146 |
| Gm23744 | 0.061920613 | 0.274393828 | 3.724945049 | 0.073663859 |
| Gm9694 | 3.918626087 | 1.781515852 | 0.632049831 | 2.818631955 |
| Tor1aip1 | 1314.956962 | 1.432242733 | 0.142394253 | 10.05829031 |
| 9430034N14Rik | 86.42678797 | 1.195716581 | 0.279808569 | 4.273337968 |
| Tor1aip2 | 1490.887513 | 1.259127336 | 0.131823671 | 9.551602748 |
| Fam163a | 99.41997446 | -0.018093918 | 0.144637995 | -0.125097956 |
| Gm38113 | 0 | NA | NA | NA |

| | | | | |
|---|---|---|---|---|
| Tdrd5 | 36.39050195 | 0.379992572 | 0.229095166 | 1.658666919 |
| A430050A11Rik | 1.164548605 | 0.538503357 | 0.966438684 | 0.557203852 |
| 4930518J20Rik | 0.382980858 | 0.274415327 | 1.600881824 | 0.171415106 |
| Nphs2 | 3.238706364 | 0.270733909 | 0.630541376 | 0.429367396 |
| Axdnd1 | 8.537287892 | 0.645804319 | 0.368389421 | 1.75304795 |
| Gm2000 | 140.1670121 | 0.665110399 | 0.147491103 | 4.509495061 |
| Soat1 | 740.7127753 | 0.10053544 | 0.149426745 | 0.672807532 |
| Gm37648 | 0.139856492 | -0.206491248 | 3.574411025 | -0.057769307 |
| Gm10031 | 2.652532074 | 0.555603284 | 0.582227718 | 0.954271441 |
| AC114588.1 | 0 | NA | NA | NA |
| Abl2 | 1607.186212 | 0.243570883 | 0.107681002 | 2.261967089 |
| Gm38043 | 100.360574 | -0.313590339 | 0.298691323 | -1.049880982 |
| Tor3a | 2329.182894 | 4.245344713 | 0.248718328 | 17.06888568 |
| Fam20b | 1176.797779 | 0.071372472 | 0.050621344 | 1.409928441 |
| Gm19918 | 0 | NA | NA | NA |
| Gm15428 | 0.152810971 | 0.755295598 | 2.844580432 | 0.265520915 |
| Ralgps2 | 727.6286885 | -0.388557907 | 0.088244071 | -4.40321828 |
| Gm28693 | 10.3212494 | -0.845217242 | 0.405109624 | -2.086391414 |
| Angptl1 | 9.659502383 | -0.120168079 | 0.308990772 | -0.388905072 |
| Gm37802 | 27.9942778 | -0.362200808 | 0.276897559 | -1.308067899 |
| Gm37943 | 22.83945072 | -0.870362317 | 0.38068458 | -2.286308304 |
| 4930439D14Rik | 2.895518095 | -0.631639825 | 0.615221188 | -1.026687373 |
| Gm28694 | 0 | NA | NA | NA |
| Gm10531 | 0.066050875 | 0.274391604 | 3.724945049 | 0.073663262 |
| Tex35 | 6.225377881 | 3.403125162 | 0.90135382 | 3.775570798 |
| Rasal2 | 2052.323456 | -0.06553093 | 0.064066811 | -1.022852999 |
| Gm37788 | 31.13971489 | -0.623234769 | 0.237953427 | -2.619146015 |
| Gm37578 | 17.57312196 | -1.083270531 | 0.337947842 | -3.205437041 |
| Gm6058 | 12.1668676 | -0.138504917 | 0.361208588 | -0.383448572 |
| Gm37060 | 72.32838032 | -0.500292622 | 0.224723215 | -2.226261411 |
| 2810025M15Rik | 74.04682442 | 0.156109023 | 0.154721569 | 1.008967424 |
| BC026585 | 13.64787826 | 0.25157847 | 0.397194862 | 0.633388027 |
| Gm15486 | 0.137416547 | -0.684638423 | 3.63730774 | -0.188226697 |
| Sec16b | 13.26555679 | 0.495183508 | 0.298952343 | 1.656396142 |
| Gm37682 | 0 | NA | NA | NA |
| Gm7412 | 0 | NA | NA | NA |
| Gm37713 | 0 | NA | NA | NA |
| Gm38213 | 0 | NA | NA | NA |
| Gm38256 | 0.119234754 | 0.274400501 | 3.724945049 | 0.073665651 |
| Brinp2 | 1002.677425 | -0.072197188 | 0.124201674 | -0.581289974 |
| Astn1 | 3735.831764 | 0.006292235 | 0.062723553 | 0.100316941 |
| Gm37861 | 45.70823912 | -0.281731317 | 0.338384697 | -0.832577003 |
| Gm37679 | 1.033175386 | 0.087353021 | 0.903090382 | 0.096726776 |
| Mir488 | 2.924473085 | 0.06792053 | 0.689314023 | 0.09853351 |
| Gm37367 | 35.2279183 | -0.671952835 | 0.346039994 | -1.941835761 |
| Gm37848 | 2.466912353 | -2.82116227 | 0.75195965 | -3.751746877 |
| Pappa2 | 44.61674889 | 0.313783648 | 0.257784781 | 1.217231082 |

| | | | | |
|---|---|---|---|---|
| 1600012P17Rik | 0.046345709 | 0.274391604 | 3.724945049 | 0.073663262 |
| 4930562F17Rik | 0 | NA | NA | NA |
| Gm4953 | 0 | NA | NA | NA |
| Gm7430 | 0.038981395 | 0.274393828 | 3.724945049 | 0.073663859 |
| Rfwd2 | 802.379122 | 0.01278901 | 0.103445289 | 0.123630663 |
| Gm37115 | 30.16231669 | -0.532867363 | 0.268299157 | -1.986094063 |
| Gm37088 | 0.075043205 | 0.755286614 | 3.724945049 | 0.202764498 |
| Scarna3a | 3.950984661 | 1.499570687 | 0.515389116 | 2.909589358 |
| mmu-mir-1843b | 0 | NA | NA | NA |
| Gm31256 | 1.235093012 | 1.334475613 | 0.900582295 | 1.48179197 |
| Gm37757 | 0.066050875 | 0.274391604 | 3.724945049 | 0.073663262 |
| Tnr | 2666.999354 | 0.045113884 | 0.079736484 | 0.565787219 |
| Gm37731 | 39.71725266 | -0.90795055 | 0.284553238 | -3.190793245 |
| Gm10530 | 19.32941874 | -0.819854618 | 0.395949468 | -2.070604168 |
| Gm38039 | 0.270352934 | 0.274412105 | 1.818794322 | 0.150875831 |
| Gm24719 | 0 | NA | NA | NA |
| Gm37294 | 0 | NA | NA | NA |
| 4930523C07Rik | 65.68144704 | 0.888174579 | 0.192427586 | 4.615630209 |
| 4930562F07Rik | 0.26443487 | 0.274413825 | 1.808626585 | 0.151724976 |
| Tnn | 13.34929327 | 0.257617223 | 0.424953742 | 0.606224155 |
| Mrps14 | 388.8406474 | 0.001183848 | 0.092453597 | 0.012804785 |
| Cacybp | 851.1120869 | 0.481333353 | 0.1324708 | 3.633505283 |
| 4933417C20Rik | 5.899375495 | -0.059957891 | 0.462002865 | -0.129778179 |
| Rabgap1l | 3165.401473 | -0.126683913 | 0.064014934 | -1.978974365 |
| Gm36975 | 0.049553047 | 0.274391604 | 3.724945049 | 0.073663262 |
| Mir1927 | 6.385949591 | -0.617163413 | 0.408101618 | -1.51227877 |
| 9430087J23Rik | 22.80069575 | -0.945573009 | 0.258440963 | -3.658758258 |
| Gm37529 | 68.10051105 | -0.53635526 | 0.302290788 | -1.774302365 |
| Gm37212 | 65.71378451 | -0.453895496 | 0.248921601 | -1.823447602 |
| Gm37215 | 27.68718647 | -0.480101579 | 0.297563466 | -1.61344262 |
| Gm38302 | 37.68841078 | -0.809926184 | 0.359441773 | -2.253288973 |
| 6720464F23Rik | 73.41064915 | -0.624864421 | 0.252895549 | -2.470839936 |
| Gm23528 | 0 | NA | NA | NA |
| Gm37873 | 11.00141775 | -0.52615633 | 0.362785124 | -1.450324985 |
| Gm37328 | 9.255702839 | -0.682230961 | 0.389630469 | -1.750969225 |
| Gm38329 | 12.82265238 | -0.069132506 | 0.274868151 | -0.251511517 |
| Gm38126 | 0.106465261 | 0.75528439 | 3.724945049 | 0.202763901 |
| Gm37083 | 0.417055923 | -1.068712602 | 1.537219614 | -0.695224412 |
| Gm38060 | 0.123841226 | 0.274391173 | 3.724945049 | 0.073663147 |
| Gm24988 | 0 | NA | NA | NA |
| Gm6177 | 0.068097359 | -0.206487837 | 3.724945049 | -0.055433794 |
| Rc3h1 | 1747.242785 | -0.037741872 | 0.047030487 | -0.802498008 |
| Gm37052 | 30.04310741 | -0.354638823 | 0.25103899 | -1.412684235 |
| Gm37809 | 11.17045412 | -0.742172828 | 0.374219991 | -1.983252752 |
| Gm37653 | 63.56315506 | -0.427406439 | 0.22152414 | -1.929389897 |
| Serpinc1 | 4.738076569 | -0.806149474 | 0.521657735 | -1.545360914 |
| Gm37729 | 0.047948698 | -0.206487837 | 3.724945049 | -0.055433794 |

| | | | | |
|---|---|---|---|---|
| Gm37072 | 1.390520144 | 0.603998662 | 0.910962839 | 0.663033262 |
| Zbtb37 | 638.7082528 | -0.343876704 | 0.096155235 | -3.576266067 |
| Gm37362 | 0.22549826 | 0.755304379 | 2.143385654 | 0.352388464 |
| Gas5 | 565.6631599 | 0.939792264 | 0.117595149 | 7.991760491 |
| Gm26224 | 0 | NA | NA | NA |
| Gm22489 | 0 | NA | NA | NA |
| Gm22357 | 0 | NA | NA | NA |
| Gm23212 | 0 | NA | NA | NA |
| Snord78 | 0 | NA | NA | NA |
| Mir5117 | 0 | NA | NA | NA |
| Snord80 | 0 | NA | NA | NA |
| SNORD77 | 0 | NA | NA | NA |
| Snord47 | 0 | NA | NA | NA |
| Gm25789 | 0 | NA | NA | NA |
| Dars2 | 123.0133515 | -0.426477136 | 0.132088632 | -3.228719469 |
| Gm37634 | 2.14016008 | 1.322063972 | 0.738898763 | 1.789235602 |
| Cenpl | 31.81213049 | 0.61193044 | 0.271482582 | 2.254032042 |
| Klhl20 | 300.3613963 | 0.157011012 | 0.114315318 | 1.373490575 |
| Ankrd45 | 441.8861501 | -0.102923966 | 0.12155181 | -0.846749758 |
| 4930469G21Rik | 0 | NA | NA | NA |
| Gm6185 | 1.432525857 | 0.081783209 | 0.931733795 | 0.087775296 |
| Prdx6 | 2507.892268 | 0.419278106 | 0.222723352 | 1.882506271 |
| Gm19057 | 0 | NA | NA | NA |
| Tnfsf4 | 0.470075119 | 0.75530944 | 1.588921799 | 0.475359731 |
| Tnfsf18 | 2.177622544 | 3.1634383 | 1.0763204 | 2.939123239 |
| Gm25488 | 0 | NA | NA | NA |
| Gm37638 | 0 | NA | NA | NA |
| Gm15429 | 9.641298891 | 0.343903339 | 0.325695493 | 1.055904505 |
| Gm5049 | 0 | NA | NA | NA |
| Fasl | 19.90409191 | 2.6735341 | 0.400067509 | 6.682707387 |
| Gm37357 | 16.51548777 | 0.173309158 | 0.307424921 | 0.563744663 |
| Suco | 1095.050559 | 0.388284572 | 0.093336505 | 4.160050488 |
| Gm7496 | 0 | NA | NA | NA |
| Gm31925 | 0.049553047 | 0.274391604 | 3.724945049 | 0.073663262 |
| 4930558K02Rik | 1.743960802 | 0.818860445 | 0.881153523 | 0.929305079 |
| Pigc | 130.768612 | -0.251646315 | 0.131021028 | -1.920655927 |
| Gm38304 | 0.16282664 | 0.755289343 | 3.41893252 | 0.220913791 |
| Dnm3 | 4093.372687 | -0.005567875 | 0.058988106 | -0.094389795 |
| 2810442N19Rik | 0.321884861 | -0.192291096 | 1.774869743 | -0.108340963 |
| Gm37767 | 5.981304473 | -0.307653387 | 0.442625792 | -0.695064301 |
| Dnm3os | 3.891078412 | -0.621138681 | 0.561200344 | -1.106803814 |
| Mir199a-2 | 0 | NA | NA | NA |
| Mir214 | 0 | NA | NA | NA |
| Gm16101 | 0 | NA | NA | NA |
| Gm10176 | 1.890308503 | 0.805340011 | 0.927885548 | 0.867930331 |
| Mettl13 | 182.4821947 | -0.095278264 | 0.093770776 | -1.016076309 |
| Vamp4 | 850.2108809 | -0.151545679 | 0.055969267 | -2.707658825 |

| | | | | |
|---|---|---|---|---|
| Gm5705 | 0.038981395 | 0.274393828 | 3.724945049 | 0.073663859 |
| Myoc | 87.94067266 | -1.033008652 | 0.44999154 | -2.295617943 |
| 7420461P10Rik | 0.321637403 | 1.207056955 | 2.051235797 | 0.588453535 |
| Prrc2c | 5776.579411 | 0.175353322 | 0.073416229 | 2.38848173 |
| Gm27616 | 4.734205595 | -0.208124345 | 0.476526783 | -0.43675267 |
| Fmo4 | 5.221071454 | 0.642591211 | 0.646372846 | 0.994149453 |
| Fmo1 | 94.46220945 | 0.174408844 | 0.177889999 | 0.980430859 |
| Fmo2 | 41.24708577 | -0.165786166 | 0.269222511 | -0.615796075 |
| Fmo6 | 0.536071372 | -1.406815015 | 1.414984783 | -0.99422625 |
| Fmo3 | 0.266623809 | -0.206481206 | 1.812978894 | -0.113890574 |
| Gm37273 | 0.04393175 | -0.206490061 | 3.724945049 | -0.055434391 |
| Mroh9 | 0.04393175 | -0.206490061 | 3.724945049 | -0.055434391 |
| Gm22671 | 0 | NA | NA | NA |
| Gm37049 | 0.128271896 | 0.274403127 | 3.679089475 | 0.074584521 |
| Gm22434 | 0 | NA | NA | NA |
| Prrx1 | 291.7242276 | 0.205646121 | 0.186680191 | 1.101595836 |
| Gm20471 | 0.226778023 | 0.755308751 | 2.143518328 | 0.352368693 |
| Gm26523 | 7.941957124 | -0.209873741 | 0.417068341 | -0.503211873 |
| Gm37644 | 11.16339375 | -1.163407119 | 0.389015776 | -2.990642517 |
| Gorab | 151.8566572 | 0.551993091 | 0.121822459 | 4.531127461 |
| Gm24940 | 0 | NA | NA | NA |
| Gm29500 | 0.061920613 | 0.274393828 | 3.724945049 | 0.073663859 |
| Rpsa-ps1 | 2.716655785 | 0.605472469 | 0.60613057 | 0.998914258 |
| Mettl11b | 4.382455216 | 0.37615466 | 0.451267845 | 0.833550769 |
| Gm2343 | 0 | NA | NA | NA |
| Kifap3 | 2725.982408 | -0.15524968 | 0.051636037 | -3.006614919 |
| Scyl3 | 425.2181155 | 0.369339579 | 0.10304283 | 3.584330711 |
| BC055324 | 25.44071188 | -0.268917951 | 0.228494123 | -1.176914084 |
| Mettl18 | 30.62632286 | -0.121802519 | 0.20781452 | -0.586111687 |
| Gm37119 | 0 | NA | NA | NA |
| Sele | 15.49838402 | 6.846443134 | 1.159835011 | 5.902945736 |
| Sell | 313.8249993 | 9.744660985 | 0.969961946 | 10.04643639 |
| Selp | 27.22107536 | 7.212155235 | 0.921996499 | 7.822323883 |
| Gm16587 | 0.246813481 | 0.75530715 | 1.830255892 | 0.412678442 |
| Gm16548 | 0.29488824 | -0.264650927 | 1.806883898 | -0.146468142 |
| F5 | 60.57846656 | -0.49956165 | 0.234560397 | -2.129778326 |
| Gm16588 | 0.11197992 | 0.75528439 | 3.724945049 | 0.202763901 |
| Slc19a2 | 224.9421596 | 0.244332024 | 0.111235132 | 2.196536473 |
| Ccdc181 | 152.8998847 | -0.069978667 | 0.151698136 | -0.461302091 |
| Blzf1 | 283.9203218 | 0.141948443 | 0.083487272 | 1.700240526 |
| Nme7 | 157.2285977 | -0.13599188 | 0.104362003 | -1.30307848 |
| Gm37411 | 15.43722212 | -0.510730772 | 0.406126313 | -1.257566317 |
| Atp1b1 | 14543.60903 | -0.109702833 | 0.058652762 | -1.870377934 |
| Gm32391 | 15.06111228 | 0.370564642 | 0.282523045 | 1.311626252 |
| Gm37732 | 0 | NA | NA | NA |
| Gm26685 | 0.956865499 | 2.626073906 | 1.134964207 | 2.31379447 |
| D630023O14Rik | 0.652111363 | 1.364697141 | 1.231678286 | 1.107998052 |

| | | | | |
|---|---|---|---|---|
| Gm37754 | 0.289849351 | 0.721117021 | 1.74806878 | 0.41252211 |
| Gm37853 | 0 | NA | NA | NA |
| Gm37184 | 2.721089389 | 2.45476827 | 0.722142468 | 3.39928529 |
| Gm36945 | 0 | NA | NA | NA |
| Dpt | 2.400487258 | 0.901460821 | 0.791388357 | 1.139087799 |
| Gm32569 | 2.899492925 | -0.060706397 | 0.571797959 | -0.106167566 |
| Gm20743 | 18.05075087 | -0.30085756 | 0.29051525 | -1.035599887 |
| Gm37403 | 2.741653471 | -1.435002465 | 0.640242055 | -2.241343651 |
| Xcl1 | 1.167394941 | 2.854821388 | 1.117095132 | 2.555575891 |
| 1700029M03Rik | 0.062426873 | 0.755288839 | 3.724945049 | 0.202765096 |
| Gm26042 | 0 | NA | NA | NA |
| 4930568G15Rik | 2.644077932 | 0.15541178 | 0.617107701 | 0.25183899 |
| Tbx19 | 0.755699022 | -1.503059279 | 1.386076146 | -1.084398778 |
| Gm16550 | 1.768574416 | -0.05190406 | 0.755573664 | -0.068694904 |
| Sft2d2 | 402.8459024 | 0.117339689 | 0.12640383 | 0.928292197 |
| Tiprl | 413.2944695 | 0.054713217 | 0.09168456 | 0.596754972 |
| Gm38190 | 112.7353609 | -0.498561398 | 0.235281363 | -2.119000808 |
| Gpr161 | 332.1613004 | 0.3375616 | 0.103476678 | 3.262199829 |
| Dcaf6 | 876.2034936 | -0.128352905 | 0.063983236 | -2.006039577 |
| Gm38368 | 0.038981395 | 0.274393828 | 3.724945049 | 0.073663859 |
| Gm24637 | 0.087930927 | -0.206490058 | 3.724945049 | -0.05543439 |
| Gm37864 | 59.27064257 | -0.57784166 | 0.273567005 | -2.1122491 |
| Mpc2 | 532.4916413 | -0.253902064 | 0.056922869 | -4.460457942 |
| Gm37860 | 11.90237977 | -0.230021992 | 0.380208407 | -0.604989232 |
| Adcy10 | 12.42222246 | 0.668414023 | 0.324611976 | 2.059116952 |
| Gm18186 | 0.674049174 | 0.550758447 | 1.477186589 | 0.372842842 |
| Gm38325 | 1.250018007 | -0.823350586 | 0.867091624 | -0.949554307 |
| Gm23599 | 0.39032937 | -0.087173834 | 3.25091475 | -0.026815171 |
| Gm23208 | 0.712615371 | -0.09998909 | 1.13926321 | -0.087766452 |
| Gm38210 | 0.496695103 | 0.77277979 | 1.426443086 | 0.541752978 |
| Mpzl1 | 254.7787603 | -0.204959524 | 0.109906209 | -1.864858459 |
| Gm37559 | 11.16705096 | -0.548188343 | 0.523075896 | -1.048009184 |
| Gm18407 | 0.233595724 | -0.224604699 | 2.527115145 | -0.088877905 |
| Rcsd1 | 77.36566827 | 0.63666767 | 0.134990948 | 4.716373058 |
| AC124587.1 | 0 | NA | NA | NA |
| Gm37073 | 0.097139547 | 0.274393828 | 3.724945049 | 0.073663859 |
| Gm34767 | 0.288888109 | 1.306551378 | 2.062365085 | 0.633520897 |
| Creg1 | 384.0157734 | 0.11908535 | 0.156874762 | 0.759110957 |
| Gm36972 | 0 | NA | NA | NA |
| Gm16565 | 4.973903582 | 1.098951088 | 0.524425706 | 2.095532457 |
| Gm24808 | 0 | NA | NA | NA |
| Cd247 | 27.67774213 | 4.857945482 | 0.543324671 | 8.941146501 |
| Gm23402 | 0.088736197 | 0.274400501 | 3.724945049 | 0.073665651 |
| Gm32999 | 0.098502579 | -0.206492286 | 3.724945049 | -0.055434988 |
| Gm37856 | 0.167459142 | 0.755298223 | 2.757565248 | 0.2739004 |
| Gm37469 | 0.04393175 | -0.206490061 | 3.724945049 | -0.055434391 |
| Pou2f1 | 796.3877359 | -0.160636573 | 0.109141285 | -1.471822255 |

| | | | | |
|---|---|---|---|---|
| Gm17976 | 0 | NA | NA | NA |
| Gm26665 | 0 | NA | NA | NA |
| Gm7626 | 0 | NA | NA | NA |
| Gm37923 | 0.047948698 | -0.206487837 | 3.724945049 | -0.055433794 |
| Dusp27 | 9.458378533 | -0.83399201 | 0.36381248 | -2.292367788 |
| Gpa33 | 0.260737493 | 1.537036514 | 2.048978816 | 0.750147587 |
| Mael | 2.369035115 | 1.158062631 | 0.968569388 | 1.195642403 |
| Ildr2 | 1753.501267 | -0.139633426 | 0.097414053 | -1.433401259 |
| 5330438I03Rik | 1.619090465 | -1.814778519 | 0.908921431 | -1.996628594 |
| Gm15853 | 26.8966923 | 0.056866844 | 0.218161757 | 0.260663671 |
| Tada1 | 290.2844388 | -0.337335712 | 0.083904512 | -4.020471631 |
| Pogk | 1000.722577 | -0.131959566 | 0.069883831 | -1.888270342 |
| Gm16418 | 40.55579905 | -0.439267346 | 0.200841711 | -2.187132061 |
| Gm33354 | 0 | NA | NA | NA |
| Gm4846 | 0 | NA | NA | NA |
| Gm22992 | 0.042716945 | -0.206490061 | 3.724945049 | -0.055434391 |
| Gm4847 | 0.370767229 | 1.56380584 | 1.661248107 | 0.941343941 |
| Fmo9 | 0.364366871 | 0.274413834 | 1.582863764 | 0.173365415 |
| Gm4954 | 0 | NA | NA | NA |
| Gm37904 | 0.072025462 | 0.755288839 | 3.724945049 | 0.202765096 |
| Gm6286 | 20.94810405 | 0.928209552 | 0.271834772 | 3.414609342 |
| Gm16701 | 90.39573473 | -0.314774638 | 0.127325525 | -2.472203734 |
| Fam78b | 369.0147211 | -0.336584534 | 0.086067262 | -3.910715 |
| Gm17868 | 0.454564407 | -0.669278978 | 1.73776936 | -0.385136827 |
| Gm37994 | 0.075043205 | 0.755286614 | 3.724945049 | 0.202764498 |
| Uck2 | 476.1615417 | -0.062538009 | 0.11395367 | -0.548802059 |
| Gm37982 | 0.042716945 | -0.206490061 | 3.724945049 | -0.055434391 |
| Gm2453 | 1.234537134 | -0.6424709 | 0.896840408 | -0.716371491 |
| Mir6347 | 0 | NA | NA | NA |
| Gm37232 | 0.061920613 | 0.274393828 | 3.724945049 | 0.073663859 |
| Tmco1 | 304.4300019 | -0.155885286 | 0.111977535 | -1.392112135 |
| Gm25911 | 23.94632856 | 1.034381088 | 1.064200213 | 0.971979779 |
| Gm24182 | 0.992449309 | -0.509926089 | 0.970476809 | -0.525438717 |
| Aldh9a1 | 305.1290703 | 0.080641568 | 0.115480377 | 0.698314034 |
| Mgst3 | 487.8742445 | -0.12175028 | 0.115090043 | -1.05786979 |
| Lrrc52 | 0.467496845 | 0.652672557 | 1.444178775 | 0.451933354 |
| Rxrg | 90.84925895 | -0.583534977 | 0.138444556 | -4.214936232 |
| Lmx1a | 12.68353485 | 0.185654066 | 0.343382391 | 0.540662745 |
| Gm20711 | 0.175176114 | -0.730240272 | 3.35752178 | -0.21749383 |
| Pbx1 | 3030.795283 | -0.147202391 | 0.082490278 | -1.784481692 |
| Gm37524 | 93.77681393 | -0.661817872 | 0.264277544 | -2.504253149 |
| Gm38381 | 118.8604427 | -0.774486403 | 0.250263907 | -3.094678786 |
| Mir6348 | 0.044544648 | 0.755286614 | 3.724945049 | 0.202764498 |
| 1700063I16Rik | 0.141317957 | 1.201108798 | 3.595358929 | 0.334072014 |
| Mir6354 | 0 | NA | NA | NA |
| Gm37839 | 0 | NA | NA | NA |
| Gm5265 | 0 | NA | NA | NA |

| | | | | |
|---|---|---|---|---|
| Nuf2 | 31.87590655 | 0.416112244 | 0.346623249 | 1.200474133 |
| Gm18900 | 0.075043205 | 0.755286614 | 3.724945049 | 0.202764498 |
| Gm36980 | 0 | NA | NA | NA |
| Rgs5 | 507.967864 | -0.590175911 | 0.170881261 | -3.453719314 |
| Rgs4 | 2513.987794 | -0.327914496 | 0.15110689 | -2.17008302 |
| Gm22114 | 0 | NA | NA | NA |
| Gm37586 | 0 | NA | NA | NA |
| 1700084C01Rik | 205.568575 | 0.792121241 | 0.179522526 | 4.41237797 |
| Gm15852 | 1.264098847 | -0.229679999 | 0.882596494 | -0.260232168 |
| Hsd17b7 | 333.9167156 | 0.333608158 | 0.12933421 | 2.579427044 |
| 3110045C21Rik | 6.064642518 | 0.118833727 | 0.416731 | 0.285156916 |
| Ddr2 | 402.5981112 | 0.681170531 | 0.138587232 | 4.915103087 |
| Uap1 | 505.8478976 | 0.180304179 | 0.069640247 | 2.589080124 |
| Gm37748 | 2.515876782 | -1.188084151 | 0.680125042 | -1.746861352 |
| Gm37502 | 0.166709681 | -0.206481506 | 2.767631179 | -0.074605861 |
| Uhmk1 | 4307.233663 | -0.221472115 | 0.065593777 | -3.376419588 |
| 4930500M09Rik | 0.668413162 | -0.437005523 | 1.279027254 | -0.341670219 |
| Sh2d1b2 | 3.139971286 | 2.735370759 | 1.070088003 | 2.556211033 |
| Gm6345 | 2.814645699 | 0.359356475 | 0.567237113 | 0.633520739 |
| Sh2d1b1 | 8.798894628 | 2.580912144 | 0.631783801 | 4.085119213 |
| Gm7694 | 71.28747091 | -0.025399164 | 0.145471055 | -0.174599434 |
| Nos1ap | 949.0976046 | -0.127240132 | 0.170375782 | -0.746820534 |
| 1700015E13Rik | 0 | NA | NA | NA |
| Gm7299 | 0.686918141 | -1.677967427 | 1.243136595 | -1.349785241 |
| Olfml2b | 107.0665219 | 0.417295871 | 0.116537915 | 3.580773445 |
| Atf6 | 1426.693892 | 0.252130027 | 0.057201184 | 4.407776342 |
| Gm23523 | 2.532308763 | 0.980650538 | 0.764200715 | 1.283236876 |
| Gm25235 | 4.986884554 | 0.535679121 | 0.663280515 | 0.807620771 |
| Gm9929 | 5.148652476 | 0.814578517 | 0.502703687 | 1.620394953 |
| Dusp12 | 141.0307956 | -0.155031483 | 0.124219337 | -1.248046291 |
| Gm26620 | 3.756679203 | 1.40021787 | 0.553350642 | 2.530435071 |
| Fcrlb | 20.36960655 | 1.57217581 | 0.455549955 | 3.45116006 |
| Fcrla | 4.0864204 | 2.762620001 | 1.087793405 | 2.539655039 |
| Gm2962 | 27.33055226 | 0.274983438 | 0.254132682 | 1.082046732 |
| Gm37030 | 0.072025462 | 0.755288839 | 3.724945049 | 0.202765096 |
| Fcgr2b | 292.4194751 | 3.753082481 | 0.213858252 | 17.54939282 |
| Gm38204 | 0.062426873 | 0.755288839 | 3.724945049 | 0.202765096 |
| Gm10522 | 0.106465261 | 0.75528439 | 3.724945049 | 0.202763901 |
| Fcgr4 | 282.2570289 | 10.17219312 | 0.893262989 | 11.38768006 |
| Gm26110 | 0.545665175 | 0.274412676 | 1.437999639 | 0.190829447 |
| Fcgr3 | 124.3411756 | 1.438625352 | 0.154191282 | 9.330134196 |
| Mir6546 | 0.741170898 | 1.164151687 | 1.337036229 | 0.870695693 |
| Gm27552 | 0.046345709 | 0.274391604 | 3.724945049 | 0.073663262 |
| Gm27844 | 0 | NA | NA | NA |
| Cfap126 | 16.16273733 | 0.467221488 | 0.35751979 | 1.306840911 |
| Sdhc | 664.894161 | -0.121778987 | 0.107579679 | -1.131988753 |
| Mpz | 3.16266539 | 0.337780566 | 0.611035567 | 0.552800172 |

| | | | | |
|---|---|---|---|---|
| Pcp4l1 | 1008.326566 | -0.147948861 | 0.109177768 | -1.355118938 |
| Nr1i3 | 2.346028351 | -0.056479416 | 0.578303138 | -0.097664032 |
| Tomm40l | 561.1166494 | 0.332991884 | 0.092902242 | 3.584325579 |
| Apoa2 | 0.57971966 | 0.284255496 | 1.236516516 | 0.229884108 |
| Fcer1g | 255.0199519 | 3.595637641 | 0.252390925 | 14.24630321 |
| Ndufs2 | 1322.178398 | 0.0798279 | 0.073963263 | 1.079291211 |
| Adamts4 | 381.9510903 | -0.075214554 | 0.124281433 | -0.605195419 |
| B4galt3 | 352.3072303 | 0.152458946 | 0.086305579 | 1.76650163 |
| Ppox | 128.0565181 | -0.35980263 | 0.132606927 | -2.713301921 |
| Usp21 | 286.6155179 | -0.326938569 | 0.098631618 | -3.314744056 |
| Ufc1 | 373.6742995 | 0.284212425 | 0.093296798 | 3.046325606 |
| Rpl27-ps1 | 1.899360943 | 0.916064536 | 0.839716942 | 1.090920631 |
| Gm25151 | 0 | NA | NA | NA |
| Gm20045 | 91.15361111 | -0.832378886 | 0.223922891 | -3.717256793 |
| Dedd | 384.4686255 | 0.29930653 | 0.086612575 | 3.455693707 |
| Nit1 | 175.9259044 | -0.349848772 | 0.099255007 | -3.524746854 |
| Pfdn2 | 372.1457708 | 0.170492024 | 0.170211887 | 1.001645814 |
| Gm37154 | 10.37578366 | -0.34820358 | 0.348973984 | -0.997792374 |
| Klhdc9 | 28.71586709 | 0.586843635 | 0.213482167 | 2.748911738 |
| Gm37097 | 0.580816694 | 0.412975677 | 1.249052813 | 0.330631077 |
| Pvrl4 | 89.21914686 | 1.029567505 | 0.136202604 | 7.559088268 |
| Arhgap30 | 261.1304874 | 2.855487152 | 0.186416221 | 15.3178041 |
| Usf1 | 598.6805458 | 0.704907387 | 0.080454904 | 8.761521679 |
| Tstd1 | 8.585507017 | -0.30527004 | 0.370228795 | -0.824544293 |
| Gm26641 | 0.661565226 | -0.302153923 | 1.123296349 | -0.26898861 |
| F11r | 61.91267919 | 0.557314347 | 0.167331778 | 3.33059478 |
| B930036N10Rik | 1.495946937 | 1.143243346 | 0.872333944 | 1.310556987 |
| Gm24871 | 0 | NA | NA | NA |
| Alyref2 | 25.03101443 | 0.356184442 | 0.334845899 | 1.063726458 |
| Gm9765 | 0.094831987 | 0.274404949 | 3.724945049 | 0.073666845 |
| Itln1 | 0.674729533 | 3.05349325 | 1.415761866 | 2.156784502 |
| Gm37450 | 0 | NA | NA | NA |
| Cd244 | 24.4635172 | 4.629986167 | 0.579672941 | 7.987238736 |
| Ly9 | 63.06084566 | 4.782281704 | 0.437532877 | 10.93010823 |
| Slamf7 | 242.896543 | 7.291978226 | 0.698462181 | 10.44004733 |
| Gm37787 | 0.62421388 | 2.340869057 | 1.363756759 | 1.71648576 |
| Mir7683 | 0.465770878 | 1.252713785 | 1.730293408 | 0.723989226 |
| A630035G10Rik | 0.725223345 | 1.524146652 | 1.353786209 | 1.12583999 |
| Gm37981 | 0.097139547 | 0.274393828 | 3.724945049 | 0.073663859 |
| Cd48 | 54.8175684 | 3.996129509 | 0.402832637 | 9.920073852 |
| Gm37010 | 3.46582388 | 1.455233988 | 0.770269548 | 1.889252915 |
| Tma7-ps | 109.4366652 | 0.296695949 | 0.186120776 | 1.594104402 |
| Gm37693 | 0 | NA | NA | NA |
| Gm19030 | 0.512648943 | 0.755281381 | 3.298243301 | 0.228995047 |
| Gm26412 | 0 | NA | NA | NA |
| Gm22950 | 0 | NA | NA | NA |
| Slamf1 | 7.891299907 | 2.574265904 | 0.510928884 | 5.03840355 |

| | | | | |
|---|---|---|---|---|
| Cd84 | 247.5967198 | 3.382109733 | 0.260439405 | 12.98616749 |
| Gm10521 | 6.960180967 | 5.115197521 | 0.984834648 | 5.193965844 |
| Gm37065 | 0.847774459 | 2.432992311 | 1.455209776 | 1.671918613 |
| Slamf6 | 33.56933231 | 3.626821011 | 0.399522402 | 9.077891489 |
| Vangl2 | 264.8998109 | 0.153897685 | 0.098110162 | 1.568621248 |
| Nhlh1 | 18.19327503 | 2.816163544 | 0.484811714 | 5.808777852 |
| Ncstn | 716.1302082 | 0.482958548 | 0.074398187 | 6.491536515 |
| Copa | 2652.049841 | 0.295974278 | 0.066953258 | 4.420610562 |
| Gm38164 | 3.312973942 | 0.439919665 | 0.540502036 | 0.813909359 |
| Gm37756 | 0 | NA | NA | NA |
| Pex19 | 729.4998742 | 0.149829001 | 0.078366233 | 1.911907657 |
| Gm10171 | 8.347475085 | 0.41339763 | 0.378900395 | 1.091045654 |
| Dcaf8 | 1703.591968 | 0.163301577 | 0.051241323 | 3.186911834 |
| Pea15a | 5072.520127 | 0.041535629 | 0.129748437 | 0.320124313 |
| Casq1 | 10.6483708 | 1.313990348 | 0.356678144 | 3.683966535 |
| Gm17224 | 5.574451365 | 0.900737962 | 0.819006457 | 1.09979348 |
| Atp1a4 | 4.799918733 | 1.468418104 | 0.773655381 | 1.898026097 |
| Gm37950 | 1.138245853 | 2.603177467 | 1.336047324 | 1.948417111 |
| Igsf8 | 2030.429245 | 0.613049176 | 0.092885064 | 6.600083462 |
| Atp1a2 | 9906.868086 | -0.297817008 | 0.137361066 | -2.168132624 |
| Kcnj9 | 920.7093185 | -0.42742541 | 0.16703201 | -2.558943098 |
| Gm36937 | 1.071485715 | 0.654667929 | 1.000423933 | 0.654390512 |
| Kcnj10 | 4831.592137 | -0.246292953 | 0.155347309 | -1.585434307 |
| Pigm | 357.594723 | 0.236419975 | 0.126792505 | 1.864621055 |
| Gm37145 | 0.222528853 | 1.219773359 | 2.57372852 | 0.47393241 |
| Gm7804 | 0.35063021 | 0.755304837 | 1.962761046 | 0.384817519 |
| Slamf9 | 59.21569036 | 3.916333375 | 0.332232238 | 11.78793905 |
| Igsf9 | 57.64957815 | -0.004099637 | 0.191260824 | -0.021434799 |
| Tagln2 | 422.7865102 | 2.501859382 | 0.215920783 | 11.5869318 |
| Gm37125 | 0.235978602 | 0.274411603 | 2.125408189 | 0.129110071 |
| Cfap45 | 24.01669692 | -0.243033314 | 0.315789761 | -0.769604793 |
| 4933439K11Rik | 2.636750292 | -0.183467922 | 0.752099669 | -0.243940969 |
| Vsig8 | 4.194719996 | -0.198503847 | 0.482024474 | -0.411812798 |
| Slamf8 | 27.5563302 | 4.20496396 | 0.436620383 | 9.630709247 |
| Fcrl6 | 0.659227985 | 1.205319483 | 1.54256085 | 0.781375647 |
| Dusp23 | 20.89498152 | 0.060783123 | 0.2789361 | 0.217910563 |
| Gm2710 | 3.560626721 | 1.02247288 | 0.552529634 | 1.850530391 |
| Gm22383 | 0 | NA | NA | NA |
| Crp | 0.134618652 | 0.755302229 | 2.958498134 | 0.255299208 |
| Gm38051 | 0 | NA | NA | NA |
| Gm24817 | 0 | NA | NA | NA |
| 9430083B18Rik | 0 | NA | NA | NA |
| Apcs | 0.049553047 | 0.274391604 | 3.724945049 | 0.073663262 |
| Gm26077 | 0.04393175 | -0.206490061 | 3.724945049 | -0.055434391 |
| Olfr16 | 0 | NA | NA | NA |
| RP23-127L7.9 | 0.046345709 | 0.274391604 | 3.724945049 | 0.073663262 |
| RP23-127L7.8 | 0 | NA | NA | NA |

| | | | | |
|---|---|---|---|---|
| Olfr1409-ps1 | 0 | NA | NA | NA |
| Olfr1408 | 0.042716945 | -0.206490061 | 3.724945049 | -0.055434391 |
| Olfr1407-ps1 | 0 | NA | NA | NA |
| Olfr1406 | 0.153801061 | -0.687376744 | 2.838638549 | -0.242150148 |
| Olfr218 | 0 | NA | NA | NA |
| Olfr1404 | 0 | NA | NA | NA |
| Fcer1a | 0.138573873 | 0.75528185 | 3.628901543 | 0.208129606 |
| Olfr418 | 0 | NA | NA | NA |
| Mptx2 | 0.116944185 | 0.274391125 | 3.724945049 | 0.073663134 |
| Ackr1 | 337.5743771 | 0.511814821 | 0.114871046 | 4.455559859 |
| Cadm3 | 5210.19823 | 0.005404541 | 0.102353378 | 0.052802757 |
| Aim2 | 59.97870837 | 2.554201823 | 0.228643982 | 11.17108704 |
| Gm4955 | 118.0231463 | 9.228773384 | 0.92106964 | 10.01962608 |
| BC094916 | 233.0659656 | 8.93433883 | 0.951336491 | 9.39135512 |
| AC084073.1 | 0 | NA | NA | NA |
| Pydc4 | 70.39675328 | 7.631129437 | 0.89756027 | 8.502080237 |
| Pyhin1 | 372.8526333 | 9.409361779 | 0.883439719 | 10.65082493 |
| AC117682.1 | 0 | NA | NA | NA |
| Pydc3 | 163.3167361 | 8.76791587 | 0.907045363 | 9.666457962 |
| Gm18752 | 7.837557856 | 5.128065484 | 0.934163623 | 5.489472464 |
| Gm7871 | 0.290529814 | 1.634278731 | 2.000391364 | 0.816979497 |
| AI607873 | 591.904346 | 7.28535937 | 0.465929829 | 15.63617292 |
| Ifi204 | 992.1935414 | 9.924543751 | 0.741468336 | 13.38498661 |
| Gm16340 | 27.30976778 | 4.577181788 | 0.699628498 | 6.54230324 |
| Gm7897 | 1.063697798 | 0.755310185 | 1.337263208 | 0.564817891 |
| Gm38050 | 43.49727825 | 8.332542954 | 0.882449767 | 9.442512497 |
| Mndal | 289.3519551 | 5.894199555 | 0.385628579 | 15.28465438 |
| Mnda | 254.7490093 | 9.785546543 | 0.954476745 | 10.25226292 |
| Ifi203 | 735.2718426 | 5.497369779 | 0.340665769 | 16.13713581 |
| Ifi202b | 1532.515879 | 8.526963404 | 0.508203346 | 16.77864475 |
| Gm18445 | 3.120173769 | 4.252159355 | 1.101894122 | 3.858954566 |
| Ifi205 | 186.9498831 | 9.606501288 | 1.001811241 | 9.589133059 |
| Olfr433 | 1.889349206 | 2.451953584 | 1.125264961 | 2.179001095 |
| Olfr432 | 0.498029041 | 0.755307172 | 1.709818453 | 0.441747 |
| Olfr431-ps1 | 0.481158361 | 0.755309185 | 1.506747239 | 0.501284598 |
| Olfr430 | 0 | NA | NA | NA |
| Olfr429 | 0.127515837 | 0.274388643 | 3.69037927 | 0.074352424 |
| RP23-343E12.13 | 0.066050875 | 0.274391604 | 3.724945049 | 0.073663262 |
| Olfr427 | 0 | NA | NA | NA |
| Gm37417 | 0 | NA | NA | NA |
| Olfr231 | 0 | NA | NA | NA |
| Olfr424 | 0 | NA | NA | NA |
| Olfr258-ps1 | 0 | NA | NA | NA |
| Olfr421-ps1 | 0.097139547 | 0.274393828 | 3.724945049 | 0.073663859 |
| Olfr420 | 0 | NA | NA | NA |
| RP23-343E12.14 | 0 | NA | NA | NA |
| Spta1 | 4.811948516 | 2.316047933 | 0.625483985 | 3.702809324 |

| | | | | |
|---|---|---|---|---|
| Olfr419 | 0 | NA | NA | NA |
| Mptx1 | 0.066050875 | 0.274391604 | 3.724945049 | 0.073663262 |
| Olfr417 | 0 | NA | NA | NA |
| Olfr416-ps1 | 0 | NA | NA | NA |
| Olfr248 | 0 | NA | NA | NA |
| Olfr414 | 0.117046891 | -0.687371727 | 3.724945049 | -0.184532045 |
| Olfr220 | 0.244461041 | -0.687378099 | 2.522192121 | -0.272532014 |
| Fmn2 | 1375.015257 | -0.314654859 | 0.056604999 | -5.558782121 |
| Grem2 | 314.3592299 | -0.001537261 | 0.170995785 | -0.008990054 |
| Rgs7 | 979.1331399 | 0.064628032 | 0.093963826 | 0.687796944 |
| 5730408A14Rik | 66.38004792 | -0.3896509 | 0.235907738 | -1.651708854 |
| Gm36307 | 0.200008102 | 0.274401468 | 2.638448336 | 0.104001077 |
| Fh1 | 588.3356071 | -0.244242476 | 0.077970817 | -3.132485766 |
| Kmo | 33.53845944 | 3.728508698 | 0.407322569 | 9.153700235 |
| Opn3 | 46.18549766 | -0.568629054 | 0.209004495 | -2.720654659 |
| Chml | 277.8742486 | -0.287584943 | 0.080380275 | -3.577804905 |
| Wdr64 | 0.404309506 | -1.663873624 | 1.482711164 | -1.122183244 |
| Gm25560 | 0.044191549 | -0.206490061 | 3.724945049 | -0.055434391 |
| Exo1 | 11.02950793 | 0.908357596 | 0.351721677 | 2.582603393 |
| Gm23805 | 0 | NA | NA | NA |
| Gm38100 | 0.094097695 | 0.755282166 | 3.724945049 | 0.202763304 |
| Gm37668 | 9.723507462 | -0.312009218 | 0.392399534 | -0.795131469 |
| Pld5 | 273.9533392 | -0.491929116 | 0.098562379 | -4.991043448 |
| B020018G12Rik | 7.849527246 | -0.978216068 | 0.417719006 | -2.341804071 |
| Gm38081 | 26.68633732 | -0.714441929 | 0.314169605 | -2.274064444 |
| Gm37916 | 0.048949532 | -0.206487837 | 3.724945049 | -0.055433794 |
| Gm38101 | 8.572675195 | -0.51792872 | 0.372256249 | -1.391323104 |
| Gm17965 | 0.042716945 | -0.206490061 | 3.724945049 | -0.055434391 |
| Gm38152 | 0.194279094 | 0.274391173 | 3.724945049 | 0.073663147 |
| Gm36536 | 0.394395078 | -0.738359939 | 1.570597864 | -0.470113933 |
| Gm17966 | 0.161076545 | 0.755297744 | 2.79165938 | 0.270555122 |
| Gm26104 | 0 | NA | NA | NA |
| Cep170 | 2565.856711 | -0.149489359 | 0.060639632 | -2.465208877 |
| 3110062G12Rik | 32.76611244 | -0.240430212 | 0.21456098 | -1.120568204 |
| Gm38104 | 128.4296084 | -0.450387882 | 0.185171626 | -2.432272652 |
| Gm36992 | 0.232229192 | -1.817681542 | 3.096265346 | -0.587056127 |
| Mir350 | 3.5612593 | -0.315653517 | 0.535185286 | -0.589802308 |
| Gm25288 | 0 | NA | NA | NA |
| Sdccag8 | 235.1494786 | -0.274937907 | 0.167775858 | -1.638721506 |
| Gm38158 | 1.079846962 | -0.30849942 | 0.915673187 | -0.336909963 |
| Gm38053 | 0.047948698 | -0.206487837 | 3.724945049 | -0.055433794 |
| Gm25993 | 2.439810687 | -0.928642269 | 0.715217237 | -1.298405885 |
| Gm37246 | 8.305594911 | -0.899040981 | 0.459903611 | -1.954846534 |
| Gm37490 | 6.173150445 | -1.31120479 | 0.428603076 | -3.059251938 |
| Gm15423 | 5.069600121 | 0.848204808 | 0.475762898 | 1.782830929 |
| Gm37519 | 17.14460392 | -0.925338429 | 0.447092004 | -2.069682348 |
| Akt3 | 2779.8936 | 0.024815024 | 0.061353771 | 0.404458009 |

| | | | | |
|---|---|---|---|---|
| Gm37960 | 79.23742208 | -0.274969628 | 0.214530026 | -1.281730269 |
| Gm37106 | 32.57544906 | -0.478396682 | 0.253675622 | -1.885859896 |
| Gm37598 | 54.69327838 | 0.330841184 | 0.291110429 | 1.136480014 |
| Gm38146 | 68.83238682 | 0.059916479 | 0.195779281 | 0.306040961 |
| Gm37706 | 171.6006066 | -0.208394995 | 0.241113027 | -0.864304169 |
| Gm37463 | 61.45825268 | -0.350675727 | 0.272447593 | -1.287130943 |
| Gm38328 | 3.391924355 | -0.902051562 | 0.570293212 | -1.581732946 |
| Gm26801 | 7.020008498 | -0.514587366 | 0.382371293 | -1.345779286 |
| Zbtb18 | 3593.895032 | -0.091320714 | 0.103520517 | -0.88215087 |
| Gm29856 | 8.835626795 | 1.638867691 | 0.478092503 | 3.427930119 |
| Gm37306 | 1.917068816 | 1.497210636 | 0.82240034 | 1.820537473 |
| Gm37280 | 0.17486102 | -0.65084711 | 2.71633334 | -0.239605022 |
| Gm37858 | 0.209701617 | 0.755296437 | 2.60475819 | 0.289967967 |
| 2310043L19Rik | 0.252619382 | 0.79499003 | 2.082883734 | 0.381677583 |
| 1700016C15Rik | 0.143022002 | 0.755289479 | 3.547809333 | 0.212888971 |
| Adss | 882.0855769 | -0.020567045 | 0.070489241 | -0.291775661 |
| Gm16585 | 0 | NA | NA | NA |
| Gm7068 | 2.009888605 | -0.159358526 | 0.688273186 | -0.231533828 |
| Gm16432 | 157.8944835 | -0.372649457 | 0.13046598 | -2.856296004 |
| Gm37313 | 0.130017973 | -0.206489706 | 3.662785653 | -0.056375045 |
| Gm25312 | 0 | NA | NA | NA |
| Gm8010 | 62.50190762 | 0.13032462 | 0.137017918 | 0.9511502 |
| Desi2 | 449.7295651 | -0.29223305 | 0.082334657 | -3.549332223 |
| Gm8023 | 0.25372768 | -0.206486202 | 2.095178707 | -0.098553026 |
| 4930527J03Rik | 0.49872596 | -1.575468436 | 1.53830751 | -1.024157021 |
| Gm37486 | 42.26792344 | 1.364277461 | 0.204364651 | 6.675701777 |
| B230369F24Rik | 55.62345512 | -0.663558817 | 0.154271934 | -4.301228361 |
| Cox20 | 107.0345127 | -0.149703189 | 0.191268244 | -0.782687108 |
| Gm16586 | 574.5462637 | -0.104257726 | 0.103863163 | -1.003798872 |
| Gm28062 | 3.424742753 | 0.027786133 | 0.528013311 | 0.052623924 |
| Hnrnpu | 2062.938983 | -0.267094244 | 0.065262402 | -4.092620466 |
| Gm38300 | 25.20762262 | -0.168221572 | 0.241828162 | -0.69562441 |
| Gm27458 | 0.042716945 | -0.206490061 | 3.724945049 | -0.055434391 |
| Gm24919 | 0.075043205 | 0.755286614 | 3.724945049 | 0.202764498 |
| Efcab2 | 203.061846 | 0.053545789 | 0.092306749 | 0.580085304 |
| Kif26b | 843.0367765 | -0.30643026 | 0.109707856 | -2.793147829 |
| Gm24405 | 1.624132796 | -1.25779327 | 0.73963268 | -1.700564757 |
| Gm34176 | 4.619527546 | -0.219179553 | 0.51703822 | -0.423913639 |
| Gm16564 | 2.140983095 | 0.639867958 | 0.730700602 | 0.875691024 |
| Smyd3 | 427.0445891 | -0.094651459 | 0.076780378 | -1.23275584 |
| Gm5561 | 0.317629712 | -0.1757113 | 1.780860294 | -0.098666527 |
| Tfb2m | 166.5859778 | 0.01424538 | 0.092055857 | 0.154747133 |
| Gm15480 | 0 | NA | NA | NA |
| Cnst | 707.9421141 | -0.202881325 | 0.055580125 | -3.650249531 |
| Sccpdh | 630.1321158 | -0.277601737 | 0.096254999 | -2.884024092 |
| Gm1305 | 0.735920534 | -1.346721427 | 1.185360414 | -1.136128228 |
| Ahctf1 | 1340.102284 | -0.172633981 | 0.081378494 | -2.121371055 |

| | | | | |
|---|---|---|---|---|
| Gm10518 | 1.39994924 | -0.013287724 | 0.781252529 | -0.017008232 |
| Gm37070 | 7.2364301 | 0.656756235 | 0.425143682 | 1.544786534 |
| Cdc42bpa | 5990.889057 | -0.200538056 | 0.059431733 | -3.374258927 |
| Gm38169 | 41.0227843 | -0.585568965 | 0.266164946 | -2.20002286 |
| Gm38331 | 0.311669542 | -0.206481588 | 1.75041929 | -0.117961216 |
| Adck3 | 280.2963435 | 0.030515035 | 0.125106874 | 0.243911736 |
| Gm37336 | 1.602826546 | 0.357212357 | 0.861705284 | 0.414541216 |
| Psen2 | 248.7304506 | 0.34547514 | 0.127113304 | 2.717851938 |
| Gm31728 | 3.493469674 | -0.36924338 | 0.61898375 | -0.596531621 |
| Gm5069 | 31.42240588 | -0.168544281 | 0.213319189 | -0.790103704 |
| Itpkb | 1738.483568 | 0.807161843 | 0.13535301 | 5.963383039 |
| Gm37033 | 72.29879847 | 0.567675045 | 0.236567049 | 2.399637005 |
| Gm37390 | 9.932302052 | 0.093148534 | 0.383429779 | 0.242935055 |
| Gm36933 | 22.96547666 | 0.226729937 | 0.329238904 | 0.688648682 |
| 6330403A02Rik | 2006.529945 | 0.019579899 | 0.22963214 | 0.08526637 |
| Gm37984 | 25.25468169 | -1.030104801 | 0.335065604 | -3.074337651 |
| Gm37267 | 0.197212531 | 1.089023091 | 2.626829987 | 0.414576922 |
| Parp1 | 872.0659159 | -0.145124713 | 0.087024245 | -1.66763541 |
| Mir6904 | 1.534744651 | -1.78184783 | 0.840819193 | -2.119180729 |
| Lin9 | 91.26341556 | -0.004129781 | 0.155599897 | -0.026541028 |
| Mixl1 | 0.441773728 | -1.064757232 | 1.359505764 | -0.783194349 |
| Gm8101 | 0 | NA | NA | NA |
| Acbd3 | 598.1054057 | 0.001036392 | 0.081157443 | 0.012770137 |
| Gm37768 | 0.122991902 | 0.274402725 | 3.724945049 | 0.073666248 |
| Gm38293 | 0.815026891 | -0.232547345 | 1.039805402 | -0.223645064 |
| H3f3a | 906.3881747 | 0.110687141 | 0.113581594 | 0.974516537 |
| Gm17275 | 0.368617932 | 0.7553099 | 1.760550776 | 0.429019095 |
| H3f3aos | 16.19441088 | -0.206984414 | 0.292601535 | -0.707393467 |
| Gm24836 | 0 | NA | NA | NA |
| Sde2 | 325.6542778 | 0.03028161 | 0.102270134 | 0.296094358 |
| Gm20305 | 1.024514732 | 0.745231883 | 0.975903188 | 0.763633004 |
| Gm18036 | 0 | NA | NA | NA |
| Lefty2 | 3.909599014 | -0.061329452 | 0.50600352 | -0.121203608 |
| Pycr2 | 201.3612007 | -0.166408081 | 0.109597125 | -1.518361739 |
| Lefty1 | 9.891254432 | -0.556053215 | 0.395642503 | -1.405443579 |
| Tmem63a | 404.5987899 | -0.165001716 | 0.123835133 | -1.332430559 |
| 2210411M09Rik | 0.048949532 | -0.206487837 | 3.724945049 | -0.055433794 |
| Gm36961 | 0 | NA | NA | NA |
| Ephx1 | 210.2340185 | 0.041775549 | 0.130578543 | 0.3199266 |
| Gm37431 | 0.049553047 | 0.274391604 | 3.724945049 | 0.073663262 |
| Gm37086 | 0 | NA | NA | NA |
| 9130409I23Rik | 0.044544648 | 0.755286614 | 3.724945049 | 0.202764498 |
| Gm17967 | 0.193566712 | 0.274401052 | 2.657574471 | 0.103252441 |
| Gm37406 | 0 | NA | NA | NA |
| Gm6606 | 0.061920613 | 0.274393828 | 3.724945049 | 0.073663859 |
| Nvl | 960.5660003 | -0.250421145 | 0.119565516 | -2.094426156 |
| Fgfr3-ps | 0 | NA | NA | NA |

| | | | | |
|---|---|---|---|---|
| Cnih4 | 409.2563578 | 0.065987676 | 0.073076807 | 0.90299068 |
| Wdr26 | 2480.374956 | 0.248802892 | 0.077062422 | 3.228589044 |
| A430110L20Rik | 18.50512536 | -0.790060349 | 0.329978545 | -2.394277932 |
| Rpl35a-ps2 | 28.95887429 | -0.012453692 | 0.226277205 | -0.055037326 |
| Gm8146 | 0.858591824 | 0.11616302 | 1.153103526 | 0.100739455 |
| Cnih3 | 290.3343059 | -0.237107119 | 0.118938545 | -1.993526314 |
| 2900069G24Rik | 4.593070302 | -1.098372754 | 0.525636286 | -2.089606032 |
| Gm16547 | 0 | NA | NA | NA |
| Gm16539 | 0 | NA | NA | NA |
| Ccdc121 | 5.062754143 | -0.269119671 | 0.526867263 | -0.510792168 |
| Dnah14 | 10.40201644 | 1.016109343 | 0.321341062 | 3.162089953 |
| Gm37664 | 0.09176491 | 1.15872478 | 3.72306111 | 0.311229052 |
| Gm23690 | 1.324611736 | 0.27933641 | 1.044066767 | 0.2675465 |
| Lbr | 304.5066716 | 0.403997003 | 0.106963003 | 3.776978882 |
| Gm26004 | 0.608946563 | -0.662701195 | 1.311710593 | -0.505219062 |
| Gm5533 | 0.046345709 | 0.274391604 | 3.724945049 | 0.073663262 |
| Enah | 3997.842318 | 0.076954102 | 0.064363496 | 1.195617183 |
| 2700078F05Rik | 123.0838922 | -0.202310611 | 0.206479508 | -0.979809633 |
| Gm38359 | 39.36611411 | -0.279596409 | 0.244519003 | -1.143454725 |
| Gm37018 | 80.20486681 | -0.290371301 | 0.278262865 | -1.043514377 |
| Gm37258 | 42.3525549 | -0.450418313 | 0.232124535 | -1.940416656 |
| Gm37177 | 0.238685809 | 1.352849929 | 2.530977422 | 0.534516791 |
| Srp9 | 514.3250057 | 0.398272159 | 0.133559741 | 2.981977628 |
| Gm17968 | 0.075043205 | 0.755286614 | 3.724945049 | 0.202764498 |
| Vmn1r1 | 0 | NA | NA | NA |
| Degs1 | 871.076032 | 0.252265259 | 0.092341238 | 2.731880854 |
| 1700047M11Rik | 71.18991338 | -1.153182071 | 0.232810749 | -4.953302525 |
| Fbxo28 | 708.639217 | -0.120181816 | 0.102294534 | -1.174860585 |
| Gm5706 | 0.042716945 | -0.206490061 | 3.724945049 | -0.055434391 |
| Gm10517 | 2.305351242 | -0.735046528 | 0.690080679 | -1.065160279 |
| Trp53bp2 | 449.2976652 | -0.220165186 | 0.080596097 | -2.731710275 |
| Capn2 | 1213.867559 | 0.084113837 | 0.105776235 | 0.79520543 |
| Gm37885 | 64.78973926 | 0.119235261 | 0.271959721 | 0.438429854 |
| Gm37069 | 19.58732916 | -0.626220441 | 0.438303938 | -1.428735605 |
| Gm8197 | 0 | NA | NA | NA |
| Capn8 | 0.352854448 | 0.080802396 | 1.509988585 | 0.053511925 |
| Gm37516 | 0.044544648 | 0.755286614 | 3.724945049 | 0.202764498 |
| Gm37218 | 0 | NA | NA | NA |
| Ccdc185 | 0.590978791 | 0.277273069 | 1.285982497 | 0.215611853 |
| Susd4 | 1048.966416 | -0.116338363 | 0.059734107 | -1.94760363 |
| Gm37542 | 0.158320866 | -0.206490587 | 2.818634005 | -0.073259099 |
| Tlr5 | 15.84910574 | 0.660547616 | 0.495792581 | 1.332306374 |
| Gm38079 | 0.535911413 | -0.180365487 | 1.391180501 | -0.129649234 |
| Gm24975 | 0.165856615 | 0.712026422 | 2.75940713 | 0.258036016 |
| Disp1 | 84.25718148 | -0.447044875 | 0.121155172 | -3.689853812 |
| BC085271 | 7.295759161 | -0.0755459 | 0.42956436 | -0.175866313 |
| Brox | 846.8803349 | 0.080491539 | 0.057875912 | 1.390760601 |

| | | | | |
|---|---|---|---|---|
| Aida | 1155.608412 | 1.750350964 | 0.148930476 | 11.75280578 |
| Mia3 | 1173.748108 | -0.016671541 | 0.062951294 | -0.264832385 |
| Gm37339 | 0.824550338 | 0.2396819 | 1.06499159 | 0.225055205 |
| Taf1a | 110.8777169 | -0.137546926 | 0.149634531 | -0.919219153 |
| Gm37986 | 3.238027032 | -2.247331382 | 0.875470495 | -2.566998425 |
| Hhipl2 | 10.87507933 | -0.687696561 | 0.430266089 | -1.598305277 |
| Gm37214 | 0.761058687 | -1.915737939 | 1.208355895 | -1.585408693 |
| Gm33973 | 0.152667883 | 0.755303377 | 2.845214345 | 0.265464491 |
| Gm8214 | 0.087930927 | -0.206490058 | 3.724945049 | -0.05543439 |
| Gm34068 | 0.293108411 | 1.721877908 | 3.717558119 | 0.463174442 |
| Gm38332 | 0.075043205 | 0.755286614 | 3.724945049 | 0.202764498 |
| Gm38108 | 0.089089296 | 1.194485492 | 3.723939576 | 0.320758559 |
| Dusp10 | 189.900222 | 0.240885414 | 0.16195046 | 1.487401856 |
| 1700056E22Rik | 3.04764041 | 0.484192094 | 0.667584031 | 0.725290109 |
| Gm37223 | 0.046345709 | 0.274391604 | 3.724945049 | 0.073663262 |
| Gm37712 | 0 | NA | NA | NA |
| 2900092O11Rik | 0.068097359 | -0.206487837 | 3.724945049 | -0.055433794 |
| 1700112H15Rik | 0.11193333 | 0.75528439 | 3.724945049 | 0.202763901 |
| Gm23349 | 0 | NA | NA | NA |
| Gm37800 | 0.214735602 | 1.229282501 | 2.178306999 | 0.564329317 |
| Rpl21-ps1 | 1.040400475 | 1.716128717 | 0.993474241 | 1.727401321 |
| Gm38358 | 0.24875112 | 0.274415656 | 1.849989481 | 0.14833363 |
| 4930488B22Rik | 0 | NA | NA | NA |
| Hlx | 28.3948279 | 2.933458384 | 0.329188052 | 8.911193371 |
| Gm34342 | 0.33098618 | 0.755272183 | 3.724945049 | 0.202760624 |
| 1-Mar | 1.626573545 | 0.20552206 | 0.789779827 | 0.260227032 |
| 2-Mar | 436.1785284 | 0.37092502 | 0.109421017 | 3.389888245 |
| Mir1981 | 0 | NA | NA | NA |
| Gm23780 | 0 | NA | NA | NA |
| C130074G19Rik | 294.2439593 | 0.409866668 | 0.174445803 | 2.349535852 |
| Mark1 | 1064.898702 | -0.309965585 | 0.078209514 | -3.963272074 |
| Gm38251 | 0.231207844 | 0.27439312 | 3.067788997 | 0.089443283 |
| Gm34732 | 0.152865955 | 0.755282783 | 3.501818003 | 0.215683049 |
| 4930433J02Rik | 0 | NA | NA | NA |
| Gm37662 | 0 | NA | NA | NA |
| Rab3gap2 | 1548.083402 | -0.238538276 | 0.056803632 | -4.199349017 |
| Snora36b | 0.393059373 | -1.125018133 | 1.537548786 | -0.731695894 |
| mmu-mir-664 | 0 | NA | NA | NA |
| Gm34882 | 0 | NA | NA | NA |
| Iars2 | 594.1626988 | -0.08324155 | 0.054581681 | -1.525082198 |
| 2010103J01Rik | 0 | NA | NA | NA |
| Mir194-1 | 0 | NA | NA | NA |
| Mir215 | 0 | NA | NA | NA |
| Gm37785 | 5.812857154 | 0.464411626 | 0.411683025 | 1.128080581 |
| Bpnt1 | 456.4610138 | -0.316156624 | 0.081470898 | -3.880608077 |
| 4930532G15Rik | 1.624021655 | -0.457119884 | 0.815290574 | -0.560683391 |
| Eprs | 2033.228255 | 0.328874851 | 0.088154033 | 3.730684115 |

| | | | | |
|---|---|---|---|---|
| 9630028B13Rik | 4.198644665 | 0.164653705 | 0.522022223 | 0.315415126 |
| Gm2061 | 20.60515688 | -0.345600111 | 0.212053563 | -1.629777434 |
| Slc30a10 | 131.5005642 | -0.843482435 | 0.176277866 | -4.784959417 |
| 5033404E19Rik | 1.621213957 | -0.349036445 | 0.704930718 | -0.495135815 |
| Gm38093 | 0 | NA | NA | NA |
| Gm9722 | 0.233998759 | 0.795726095 | 2.12938349 | 0.373688487 |
| Lyplal1 | 31.22453688 | -0.28382423 | 0.196789265 | -1.44227496 |
| Gm38268 | 1.323601962 | -3.086848288 | 1.015869583 | -3.038626551 |
| Gm23821 | 0 | NA | NA | NA |
| Gm37134 | 0 | NA | NA | NA |
| Gm37272 | 0 | NA | NA | NA |
| Gm37491 | 0 | NA | NA | NA |
| A730004F24Rik | 0.094097695 | 0.755282166 | 3.724945049 | 0.202763304 |
| Tgfb2 | 267.8778118 | 0.435152102 | 0.119807994 | 3.632079015 |
| Gm26169 | 0.061920613 | 0.274393828 | 3.724945049 | 0.073663859 |
| Gm29491 | 2.076177089 | -0.667817927 | 0.728104781 | -0.917200305 |
| Gm37849 | 4.193292652 | -0.922440876 | 0.575313137 | -1.603371829 |
| Rrp15 | 96.16436869 | 0.168178051 | 0.17559957 | 0.957736125 |
| A430105J06Rik | 9.038545814 | -0.514413708 | 0.400445605 | -1.284603204 |
| 1700023G09Rik | 0 | NA | NA | NA |
| Gm3809 | 0.939714841 | -0.387507417 | 1.090076183 | -0.355486546 |
| Gm37912 | 0.240869569 | -0.206496159 | 3.110743597 | -0.066381607 |
| D1Pas1 | 0.137647365 | 1.404948034 | 3.719469264 | 0.377728093 |
| Gm19058 | 0.109869311 | -0.206490066 | 3.724945049 | -0.055434393 |
| Spata17 | 10.75281961 | -0.888044611 | 0.332278722 | -2.672589459 |
| Gm15508 | 0.503467097 | -0.100454603 | 1.346908512 | -0.074581608 |
| RP23-438D3.4 | 1.299896474 | -1.118670104 | 0.930710806 | -1.201952418 |
| RP23-438D3.3 | 1.714015964 | -2.802969327 | 1.001961344 | -2.797482503 |
| Gm15509 | 0.816619922 | -1.369409782 | 1.153579085 | -1.187096576 |
| 1700007P06Rik | 4.898989025 | -0.321960498 | 0.499973705 | -0.643954862 |
| Gpatch2 | 371.7543322 | -0.017806605 | 0.083267576 | -0.213848004 |
| Gm21710 | 76.9518533 | -0.091309129 | 0.204056688 | -0.447469428 |
| Gm38155 | 0.312806312 | 0.719807827 | 1.780533974 | 0.404265146 |
| Gm37896 | 5.45108928 | 1.537472071 | 0.497346907 | 3.091347408 |
| Esrrg | 699.5845546 | -0.116787343 | 0.099011744 | -1.179530207 |
| Gm36388 | 5.428564864 | -0.953070405 | 0.522212007 | -1.825064137 |
| Gm37929 | 0 | NA | NA | NA |
| 9330162B11Rik | 0.045882455 | 0.755288839 | 3.724945049 | 0.202765096 |
| Gm38315 | 9.475868802 | -0.316418724 | 0.433986847 | -0.729097497 |
| Gm38064 | 0.045882455 | 0.755288839 | 3.724945049 | 0.202765096 |
| Ush2a | 7.7862644 | -0.206617508 | 0.33674472 | -0.613573119 |
| Gm25095 | 0 | NA | NA | NA |
| Gm25269 | 0 | NA | NA | NA |
| Kctd3 | 958.3543934 | -0.166098898 | 0.061993394 | -2.679299949 |
| Gm37178 | 0.897014366 | 0.888142966 | 1.029288558 | 0.862870727 |
| Gm18698 | 0.26837777 | 0.262178897 | 2.090219533 | 0.125431273 |
| Kcnk2 | 433.3427616 | -0.193906923 | 0.101634055 | -1.907893199 |

| | | | | |
|---|---|---|---|---|
| 2900042K21Rik | 2.348091134 | -0.313680773 | 0.700877369 | -0.447554432 |
| Gm3837 | 46.20088099 | 0.464486916 | 0.288102577 | 1.612227564 |
| A430027H14Rik | 10.33216563 | -0.746126629 | 0.297249417 | -2.51010292 |
| Gm24092 | 0.042716945 | -0.206490061 | 3.724945049 | -0.055434391 |
| Gm38122 | 0.072025462 | 0.755288839 | 3.724945049 | 0.202765096 |
| Cenpf | 46.80573964 | 1.071229299 | 0.384360543 | 2.787042838 |
| Gm37239 | 0.092881282 | -0.687373951 | 3.724945049 | -0.184532642 |
| Ptpn14 | 196.8365016 | -0.463876031 | 0.144860843 | -3.202218215 |
| Gm30280 | 0.99636849 | -0.664313936 | 1.20867012 | -0.549623859 |
| Gm26574 | 0.255031674 | 0.274395167 | 3.017653275 | 0.090929985 |
| Gm37422 | 1.509855825 | 0.200159006 | 0.866763256 | 0.230926962 |
| Smyd2 | 468.8475306 | -0.187926017 | 0.074108413 | -2.535825673 |
| Gm38245 | 23.42935459 | -1.007315278 | 0.367463655 | -2.74126506 |
| Prox1 | 753.5982863 | 0.066729467 | 0.072164242 | 0.924688812 |
| Gm28172 | 0.53361457 | 0.566264651 | 1.372310323 | 0.412636006 |
| Prox1os | 9.870830895 | 0.099222976 | 0.316891692 | 0.313113212 |
| Gm29678 | 0.175900428 | 0.27441028 | 2.727499772 | 0.100608727 |
| Gm30725 | 0.194279094 | 0.274391173 | 3.724945049 | 0.073663147 |
| Gm23153 | 0.062426873 | 0.755288839 | 3.724945049 | 0.202765096 |
| AC124613.1 | 0 | NA | NA | NA |
| Gm30932 | 0 | NA | NA | NA |
| Rps6kc1 | 776.7345912 | -0.12515193 | 0.06120013 | -2.044961849 |
| Angel2 | 626.1745066 | 0.024272119 | 0.054379422 | 0.446347506 |
| Vash2 | 94.15710728 | 0.403782551 | 0.173357972 | 2.32918363 |
| Gm31373 | 0.061920613 | 0.274393828 | 3.724945049 | 0.073663859 |
| Gm37095 | 4.062122998 | -0.469266357 | 0.523726498 | -0.896014158 |
| Gm8407 | 6.023452583 | -0.210883984 | 0.409495631 | -0.514984697 |
| Mfsd7b | 308.2483388 | -0.057886394 | 0.101185159 | -0.572083836 |
| A230020J21Rik | 0.378086136 | 0.750253326 | 1.711190538 | 0.438439384 |
| Spata45 | 0.216916081 | 0.776804539 | 2.174076692 | 0.357303191 |
| Tatdn3 | 79.86263131 | -0.220186563 | 0.136609208 | -1.61179884 |
| Nsl1 | 10.37462125 | 0.935461903 | 0.325256914 | 2.876070768 |
| Gm38188 | 0 | NA | NA | NA |
| Batf3 | 9.56190983 | 1.669181981 | 0.357916695 | 4.6636047 |
| Gm38161 | 0 | NA | NA | NA |
| Fam71a | 0.398364559 | 1.887690363 | 1.683200668 | 1.121488601 |
| Atf3 | 253.8533324 | 4.071071905 | 0.462235929 | 8.807346308 |
| Gm24748 | 0 | NA | NA | NA |
| D730003I15Rik | 30.70567579 | -1.347930855 | 0.359481166 | -3.749656406 |
| Gm37074 | 9.818226126 | -1.814351674 | 0.559083341 | -3.245225786 |
| Gm37168 | 4.648304818 | -0.758579302 | 0.479069325 | -1.583443696 |
| Nenf | 240.7507074 | 0.122627822 | 0.105485919 | 1.162504186 |
| Tmem206 | 104.9785531 | -0.371877952 | 0.120543683 | -3.085005714 |
| Ppp2r5a | 813.5882951 | -0.157399429 | 0.08059941 | -1.952860816 |
| Gm2272 | 0 | NA | NA | NA |
| Gm32200 | 9.706590373 | 0.249480015 | 0.354018579 | 0.704708819 |
| Gm37688 | 0.11193333 | 0.75528439 | 3.724945049 | 0.202763901 |

| | | | | |
|---|---|---|---|---|
| Gm37432 | 0.52136618 | 0.523196119 | 1.549991439 | 0.337547748 |
| 9430037O13Rik | 2.32288591 | 1.671415758 | 0.873778665 | 1.912859429 |
| Dtl | 77.14293698 | 0.732139093 | 0.186539179 | 3.924854271 |
| Ints7 | 338.6046142 | 0.15942203 | 0.07297121 | 2.184725033 |
| Gm37349 | 5.323050413 | 0.547019819 | 0.505320885 | 1.082519711 |
| Lpgat1 | 3755.069215 | -0.173702467 | 0.05808112 | -2.990687289 |
| Gm22446 | 0 | NA | NA | NA |
| Gm26203 | 0 | NA | NA | NA |
| Gm20203 | 0 | NA | NA | NA |
| Gm38037 | 1.052686423 | -0.386326636 | 0.927201295 | -0.416658862 |
| Nek2 | 70.81536854 | 0.859411418 | 0.318761972 | 2.696091421 |
| 1700034H15Rik | 43.54077039 | -0.326723992 | 0.177578964 | -1.839880043 |
| Slc30a1 | 668.252428 | 0.452872513 | 0.11543665 | 3.923125914 |
| Rd3 | 12.96523121 | 0.95515154 | 0.291141926 | 3.280707634 |
| Gm26670 | 1.595190335 | -1.209560829 | 0.867838823 | -1.393762064 |
| Gm32460 | 0.281516365 | 0.708145039 | 1.772004464 | 0.399629377 |
| Traf5 | 35.39043388 | 0.665076652 | 0.223278215 | 2.9786903 |
| Mir3473c | 0 | NA | NA | NA |
| Gm3934 | 136.9920633 | -0.033245932 | 0.118224108 | -0.281211109 |
| Gm26879 | 0 | NA | NA | NA |
| Rcor3 | 532.0108606 | -0.122023801 | 0.111059475 | -1.098724808 |
| Gm37055 | 0.109124069 | 0.274402725 | 3.724945049 | 0.073666248 |
| Gm10516 | 32.23956246 | 0.089468408 | 0.183011189 | 0.488868511 |
| Kcnh1 | 1278.299755 | 0.155177835 | 0.14732149 | 1.053327892 |
| Hhat | 36.35158037 | 0.729833009 | 0.236533757 | 3.085534257 |
| Gm22265 | 0.377885927 | 0.274412745 | 1.719526828 | 0.159586196 |
| Gm38360 | 0 | NA | NA | NA |
| A730013G03Rik | 1.204108136 | 1.583212307 | 0.888821849 | 1.781248187 |
| 1700065J18Rik | 0.258344737 | 0.274408982 | 2.151774914 | 0.127526806 |
| Sertad4 | 214.5306265 | 0.058961818 | 0.118396663 | 0.498002361 |
| Gm15867 | 2.355781424 | 0.48423479 | 0.723051486 | 0.669709971 |
| Gm2367 | 0.23837135 | -0.206483062 | 2.548869831 | -0.081009654 |
| Syt14 | 711.6526152 | 0.155993812 | 0.109050545 | 1.430472552 |
| Gm23583 | 5.178783273 | 0.127646919 | 0.446748265 | 0.285724487 |
| Gm31406 | 1.718171857 | 0.475076703 | 0.813483055 | 0.584003194 |
| Gm37103 | 0 | NA | NA | NA |
| Diexf | 261.3112737 | 0.050919641 | 0.094326179 | 0.539825119 |
| Irf6 | 4.847835626 | -0.515091314 | 0.444000519 | -1.160114215 |
| A130010J15Rik | 56.29500924 | 0.045003121 | 0.155012088 | 0.290320078 |
| Traf3ip3 | 31.36915684 | 3.325695131 | 0.314268113 | 10.5823499 |
| Gm15872 | 0.108123235 | 0.274402725 | 3.724945049 | 0.073666248 |
| 4930570N18Rik | 0.062426873 | 0.755288839 | 3.724945049 | 0.202765096 |
| Lamb3 | 38.67870685 | -0.658785798 | 0.223980668 | -2.941261874 |
| Hsd11b1 | 94.82910089 | 0.133390011 | 0.187080266 | 0.713009522 |
| Gm38118 | 1.886096619 | 1.056818112 | 0.742931943 | 1.422496531 |
| G0s2 | 30.35106119 | -0.199442322 | 0.194453326 | -1.025656523 |
| Gm37691 | 3.727288002 | -0.455208955 | 0.526330023 | -0.864873626 |

| | | | | |
|---|---|---|---|---|
| Gm37650 | 0 | NA | NA | NA |
| Camk1g | 290.0946269 | -0.349537689 | 0.147013438 | -2.377590067 |
| Gm37942 | 0 | NA | NA | NA |
| Gm38022 | 0 | NA | NA | NA |
| Gm38322 | 0.094097695 | 0.755282166 | 3.724945049 | 0.202763304 |
| 4631405K08Rik | 0 | NA | NA | NA |
| Mir205 | 0 | NA | NA | NA |
| Gm21362 | 0 | NA | NA | NA |
| 4930503O07Rik | 0.33841178 | 0.451555163 | 3.687300333 | 0.122462268 |
| Plxna2 | 2388.321215 | -0.200592291 | 0.1005552 | -1.994847517 |
| Gm37783 | 41.75688494 | -0.337511492 | 0.285439695 | -1.182426615 |
| Gm37674 | 39.82514733 | -0.490618935 | 0.303429812 | -1.616910782 |
| Gm38036 | 64.68369131 | -0.436675944 | 0.328856866 | -1.327860201 |
| 2900035J10Rik | 6.043773575 | -0.31726419 | 0.422645561 | -0.750662538 |
| Gm19777 | 0.047948698 | -0.206487837 | 3.724945049 | -0.055433794 |
| Mir3962 | 0.624784964 | -0.549503107 | 2.724081254 | -0.201720527 |
| Gm23885 | 0.198152624 | 0.27438216 | 3.724945049 | 0.073660727 |
| Cd34 | 275.1221153 | 0.162483085 | 0.132673368 | 1.224685008 |
| Gm16897 | 7.560911453 | -0.244964767 | 0.389454481 | -0.628994604 |
| A330023F24Rik | 1658.07212 | -0.444236599 | 0.287151801 | -1.547044447 |
| Gm32250 | 0.045882455 | 0.755288839 | 3.724945049 | 0.202765096 |
| Gm37334 | 228.9475402 | -0.391392125 | 0.234329884 | -1.670261249 |
| 9630010A21Rik | 156.1316392 | -0.318765637 | 0.273518656 | -1.165425574 |
| Gm37027 | 0.870290022 | 2.01997061 | 1.281438387 | 1.576330654 |
| Cd46 | 147.5920642 | -0.096048015 | 0.198004344 | -0.485080342 |
| Mir29b-2 | 0 | NA | NA | NA |
| Mir29c | 0 | NA | NA | NA |
| Gm37366 | 35.61469171 | 0.193988731 | 0.238460867 | 0.813503422 |
| Gm37132 | 36.12461116 | -0.321222544 | 0.296763534 | -1.082419191 |
| Cr1l | 392.2221467 | 0.011776145 | 0.083692891 | 0.14070664 |
| Cr2 | 5.163701346 | -0.359028461 | 0.460433192 | -0.779762335 |
| Gm37887 | 0.066050875 | 0.274391604 | 3.724945049 | 0.073663262 |
| Gm24802 | 0.044191549 | -0.206490061 | 3.724945049 | -0.055434391 |
| Gm27940 | 0 | NA | NA | NA |
| Gm38046 | 0 | NA | NA | NA |
| Gm38194 | 0.659436564 | 0.982897083 | 1.308953636 | 0.750902901 |
| 9230019H11Rik | 0 | NA | NA | NA |
| H60c | 0 | NA | NA | NA |
| Gm26752 | 1.987374958 | 0.276406875 | 0.836350518 | 0.330491664 |
| Ppp1r14c | 150.8264208 | -0.247861077 | 0.098566392 | -2.514661156 |
| Gm24482 | 0.106358623 | 0.274402725 | 3.724945049 | 0.073666248 |
| Iyd | 3.589799332 | -0.326322947 | 0.627214679 | -0.520273134 |
| Plekhg1 | 783.4816878 | 0.294593095 | 0.078248249 | 3.764852233 |
| Gm16149 | 0.045882455 | 0.755288839 | 3.724945049 | 0.202765096 |
| Gm23023 | 0 | NA | NA | NA |
| Gm16148 | 0.049553047 | 0.274391604 | 3.724945049 | 0.073663262 |
| Mthfd1l | 264.3319854 | -0.038686079 | 0.096837741 | -0.39949382 |

| | | | | |
|---|---|---|---|---|
| Gm16074 | 3.696842725 | -0.632875025 | 0.713626547 | -0.886843444 |
| Gm25515 | 0.67351111 | 1.437071825 | 1.381631832 | 1.04012646 |
| Gm25369 | 0.211762239 | 0.274414343 | 2.616650133 | 0.104872386 |
| Akap12 | 1387.377891 | 0.786422891 | 0.272560189 | 2.88531826 |
| Zbtb2 | 206.2791963 | 0.431144048 | 0.165896754 | 2.59886971 |
| Gm21781 | 207.0306887 | -0.142868382 | 0.246019742 | -0.580719178 |
| Rmnd1 | 211.8870746 | -0.518794345 | 0.146692901 | -3.536601584 |
| Gm16153 | 6.393807032 | -1.769471478 | 0.532960617 | -3.320079236 |
| Armt1 | 185.8306212 | -0.047194177 | 0.095391975 | -0.494739495 |
| Ccdc170 | 10.84234273 | 0.320208694 | 0.35045984 | 0.913681562 |
| Gm22739 | 0.087261593 | 0.274400501 | 3.724945049 | 0.073665651 |
| Esr1 | 111.3424344 | 0.956205987 | 0.162825288 | 5.872588938 |
| Syne1 | 6019.024116 | -0.254316072 | 0.122782581 | -2.071271582 |
| Gm25694 | 9.729705334 | -1.179703325 | 0.418788352 | -2.816943977 |
| Gm10097 | 0.451409952 | 0.755309338 | 1.502510797 | 0.50269811 |
| Gm23573 | 0 | NA | NA | NA |
| Myct1 | 1.954060925 | 0.142643105 | 0.890248482 | 0.160228417 |
| Vip | 156.4683625 | 0.621526871 | 0.15981796 | 3.888967623 |
| Fbxo5 | 28.97769736 | 0.745205696 | 0.208975196 | 3.56600071 |
| Mtrf1l | 83.2976707 | -0.108329225 | 0.135508566 | -0.799427138 |
| Rgs17 | 1666.499129 | -0.017488092 | 0.083879142 | -0.208491544 |
| Gm10945 | 0.044544648 | 0.755286614 | 3.724945049 | 0.202764498 |
| Oprm1 | 120.7739671 | -0.249507465 | 0.13651841 | -1.827647015 |
| Ipcef1 | 966.4889659 | -0.273070165 | 0.152640689 | -1.788973611 |
| Gm15536 | 9.233390511 | 0.707693707 | 0.346124406 | 2.04462238 |
| Cnksr3 | 268.184003 | -0.3067448 | 0.147836085 | -2.074898022 |
| Ulbp1 | 7.246076538 | 2.994119552 | 0.717673239 | 4.171981608 |
| Lrp11 | 1467.900515 | -0.055772799 | 0.052166448 | -1.069131621 |
| Gm16254 | 0.342362695 | 1.683625948 | 3.707067334 | 0.454166541 |
| Pcmt1 | 1204.251679 | -0.036582401 | 0.05093384 | -0.718233721 |
| Nup43 | 42.06222489 | -0.180146474 | 0.274263693 | -0.656836753 |
| Lats1 | 1258.401443 | 0.210377997 | 0.079513287 | 2.645821905 |
| Katna1 | 226.4998142 | 0.684711339 | 0.116168169 | 5.894139007 |
| Gm20470 | 0.150550172 | -0.206481659 | 2.857661711 | -0.072255459 |
| Ginm1 | 172.1070177 | 0.235876663 | 0.130863995 | 1.802456533 |
| B430219N15Rik | 0.972892922 | 1.253526495 | 1.129304963 | 1.109998217 |
| Ppil4 | 480.6494858 | 0.105356307 | 0.105245829 | 1.001049712 |
| Gm15560 | 0.152667883 | 0.755303377 | 2.845214345 | 0.265464491 |
| Zc3h12d | 64.81458113 | 5.328959076 | 0.616551304 | 8.64317218 |
| Mir5104 | 0.256698001 | 0.755298317 | 2.4675993 | 0.306086291 |
| Gm24726 | 0 | NA | NA | NA |
| Tab2 | 1189.099294 | 0.259189311 | 0.076503005 | 3.38796247 |
| Ust | 495.4936296 | -0.242830815 | 0.09374257 | -2.590400671 |
| Gm24374 | 0 | NA | NA | NA |
| Sash1 | 1877.282141 | 0.069372671 | 0.115960181 | 0.598245625 |
| Gm25410 | 1.528398176 | -0.754492748 | 0.856254617 | -0.881154663 |
| Gm26674 | 0.49731986 | -0.1798632 | 1.403197752 | -0.128180935 |

| | | | | |
|---|---|---|---|---|
| Gm28904 | 9.097189116 | -0.489486213 | 0.346490073 | -1.412699097 |
| Gm9930 | 39.07755842 | -0.675982373 | 0.375076211 | -1.80225339 |
| Samd5 | 168.1922456 | -0.70398556 | 0.117273552 | -6.002935411 |
| Stxbp5 | 2460.054131 | -0.031788353 | 0.081400213 | -0.390519286 |
| Gm25635 | 3.189004039 | -0.915470442 | 0.713938528 | -1.282281885 |
| Gm28905 | 7.03479442 | -0.23750815 | 0.346129378 | -0.686183159 |
| Gm25682 | 1.985393048 | -0.790247041 | 0.80745818 | -0.978684792 |
| Adgb | 11.41173491 | -0.346928317 | 0.351153988 | -0.987966332 |
| Gm15568 | 0.044544648 | 0.755286614 | 3.724945049 | 0.202764498 |
| Rab32 | 75.72339442 | 2.149970249 | 0.251810662 | 8.538042972 |
| Gm5177 | 0.397592075 | 1.442547114 | 1.73346471 | 0.832175645 |
| Gm23601 | 0 | NA | NA | NA |
| Gm10944 | 0 | NA | NA | NA |
| Grm1 | 1592.406977 | -0.1939044 | 0.091948382 | -2.108839717 |
| Shprh | 1486.660042 | -0.176442191 | 0.087918778 | -2.006877213 |
| Gm16577 | 1.038701827 | -0.031806667 | 1.113995162 | -0.028551889 |
| 4930432B10Rik | 21.06911396 | -0.893359938 | 0.379290171 | -2.355346926 |
| Fbxo30 | 323.9844007 | -0.072722313 | 0.096570219 | -0.75305113 |
| Gm25979 | 0.044191549 | -0.206490061 | 3.724945049 | -0.055434391 |
| Epm2a | 429.8223933 | -0.044431957 | 0.102555169 | -0.433249315 |
| Rpl17-ps11 | 0 | NA | NA | NA |
| Gm9797 | 3.625096175 | 0.197455487 | 0.567411347 | 0.347993546 |
| Gm22546 | 0.038981395 | 0.274393828 | 3.724945049 | 0.073663859 |
| Gm22194 | 0 | NA | NA | NA |
| Gm25011 | 0 | NA | NA | NA |
| Gm23680 | 0 | NA | NA | NA |
| Gm25139 | 0 | NA | NA | NA |
| Utrn | 1645.064442 | 0.043574911 | 0.056789322 | 0.767308169 |
| Gm23044 | 1.167901279 | 1.106250493 | 1.003671815 | 1.102203406 |
| B230208H11Rik | 2.934711433 | -1.083474609 | 0.685029972 | -1.581645553 |
| Stx11 | 71.27238607 | 6.467966132 | 0.775854499 | 8.336571019 |
| Sf3b5 | 177.9606754 | 0.194684833 | 0.137122068 | 1.419792129 |
| Plagl1 | 355.2404399 | -0.056480586 | 0.106352936 | -0.531067484 |
| Gm27741 | 0.062426873 | 0.755288839 | 3.724945049 | 0.202765096 |
| Zc2hc1b | 0.135956459 | 0.720160971 | 3.641541792 | 0.197762655 |
| Ltv1 | 143.8273288 | 0.108296938 | 0.135661568 | 0.798287532 |
| Phactr2 | 555.9939026 | -0.157382832 | 0.08045398 | -1.956184526 |
| Fuca2 | 561.995271 | 0.189914022 | 0.098290932 | 1.932162181 |
| Pex3 | 290.4366472 | 0.115163953 | 0.076463393 | 1.50613186 |
| Adat2 | 56.9225531 | -0.037301769 | 0.144331484 | -0.258445132 |
| Aig1 | 695.0458195 | -0.151618364 | 0.06169286 | -2.457632287 |
| Gm22995 | 0 | NA | NA | NA |
| Gm8330 | 0.285805375 | -0.187662268 | 1.810547209 | -0.103649475 |
| Hivep2 | 5288.46622 | -0.244959279 | 0.092785148 | -2.640069946 |
| Gm26835 | 0.195722939 | 0.27440741 | 2.667682423 | 0.102863597 |
| Gm28289 | 0.0878635 | -0.687376175 | 3.724945049 | -0.184533239 |
| Gm24343 | 0.447929572 | -0.089896853 | 1.365891654 | -0.065815508 |

| | | | | |
|---|---|---|---|---|
| Gm37299 | 0.353350569 | 0.721844816 | 1.718542467 | 0.420033156 |
| Adgrg6 | 79.60354276 | 0.611702201 | 0.183076835 | 3.341232118 |
| Gm10335 | 39.8430328 | 0.473281113 | 0.267738851 | 1.767696807 |
| Vta1 | 390.7247282 | 0.290311052 | 0.066571435 | 4.360895209 |
| Gm16234 | 0.304277014 | 0.797671602 | 2.034361095 | 0.39209932 |
| Nmbr | 18.58630088 | -0.252537118 | 0.255045985 | -0.99016308 |
| Gje1 | 0.044191549 | -0.206490061 | 3.724945049 | -0.055434391 |
| Gm8355 | 90.21878376 | -0.105349812 | 0.150886856 | -0.69820404 |
| B230364G03Rik | 0.277885724 | -0.240139972 | 1.77093813 | -0.13560043 |
| Gm23892 | 0 | NA | NA | NA |
| Gm20125 | 6.549262826 | 0.065630658 | 0.495651979 | 0.132412783 |
| Gm25382 | 0 | NA | NA | NA |
| Cited2 | 577.6609852 | 0.378649689 | 0.148643561 | 2.547366913 |
| Gm20655 | 8.589832509 | 0.012186681 | 0.376663455 | 0.032354295 |
| Txlnb | 15.59385469 | -0.878667362 | 0.295289163 | -2.975616694 |
| Gm7584 | 0.106056992 | 1.236179401 | 3.724945049 | 0.331865138 |
| Heca | 505.4761868 | 0.262716296 | 0.091955224 | 2.857002388 |
| Abracl | 74.37190115 | 0.945564176 | 0.187662843 | 5.038632901 |
| Gm17711 | 0.109982625 | -0.20649451 | 3.724945049 | -0.055435586 |
| Reps1 | 606.0807053 | 0.114180415 | 0.092952108 | 1.228378977 |
| Ect2l | 5.746393814 | 0.124371697 | 0.439833696 | 0.282769823 |
| Ccdc28a | 182.412726 | -0.454667449 | 0.142181801 | -3.197789351 |
| Gm10827 | 7.130661091 | -0.392540831 | 0.582431404 | -0.673969207 |
| Nhsl1 | 1061.864252 | 0.319776613 | 0.098390219 | 3.250085384 |
| Hebp2 | 41.95025653 | -0.064048394 | 0.243587783 | -0.262937629 |
| Arfgef3 | 2592.376606 | -0.216110411 | 0.09122199 | -2.369060486 |
| Gm4922 | 0 | NA | NA | NA |
| Perp | 62.70509499 | 0.086915322 | 0.204994577 | 0.423988398 |
| Gm38175 | 0 | NA | NA | NA |
| Tnfaip3 | 387.4893776 | 2.877947393 | 0.375082472 | 7.672838938 |
| Olig3 | 0.674822466 | -1.327374625 | 1.211360687 | -1.095771589 |
| Ifngr1 | 246.1757685 | 0.914979175 | 0.126763319 | 7.218012158 |
| Il22ra2 | 0.165580463 | 0.274406349 | 2.773571271 | 0.098936109 |
| 9230106D20Rik | 0 | NA | NA | NA |
| Il20ra | 10.38114758 | 0.06085192 | 0.345718631 | 0.176015737 |
| Gm25256 | 0.617475372 | 1.513283284 | 1.317505741 | 1.148597108 |
| Slc35d3 | 65.99433708 | -0.630055018 | 0.155100185 | -4.06224544 |
| Pex7 | 193.8965027 | -0.078363175 | 0.080605398 | -0.972182717 |
| Map3k5 | 659.4939879 | -0.126523897 | 0.07434307 | -1.701892278 |
| Map7 | 996.5614196 | -0.137646034 | 0.085371254 | -1.612322978 |
| Mir6413 | 0 | NA | NA | NA |
| 4933406P04Rik | 7.134388926 | -0.579393779 | 0.401882991 | -1.441697688 |
| Bclaf1 | 2858.72397 | 0.31275799 | 0.069416811 | 4.505507869 |
| Mtfr2 | 15.4157451 | 0.165664881 | 0.376282592 | 0.440267195 |
| Pde7b | 825.652375 | -0.001858153 | 0.110968198 | -0.016744919 |
| 4930403O15Rik | 14.60270139 | -0.718822881 | 0.28627052 | -2.510991637 |
| Gm17229 | 23.00707965 | -0.154656705 | 0.34468677 | -0.448687675 |

| | | | | |
|---|---|---|---|---|
| Gm17230 | 1.594890904 | 0.271099553 | 1.260632358 | 0.215050448 |
| Ahi1 | 2789.699633 | -0.261514531 | 0.058605407 | -4.4622936 |
| Myb | 21.21849641 | 0.459006926 | 0.266309961 | 1.723581516 |
| Gm26577 | 5.818997596 | -0.672108727 | 0.421395331 | -1.594960071 |
| Gm2467 | 0 | NA | NA | NA |
| Hbs1l | 606.1812586 | 0.021500662 | 0.08945346 | 0.240355846 |
| Aldh8a1 | 4.539391149 | -0.575116434 | 0.527217755 | -1.090851794 |
| 1700021A07Rik | 0.178864798 | 0.274396414 | 3.277341654 | 0.0837253 |
| 1700020N01Rik | 0.490798134 | 1.232507717 | 1.576270886 | 0.781913647 |
| Gm5420 | 0.09349418 | 0.274402725 | 3.724945049 | 0.073666248 |
| Gm29323 | 0 | NA | NA | NA |
| Platr6 | 2.028201447 | -0.014544233 | 0.65442863 | -0.022224322 |
| Gm23839 | 0.264066224 | 1.236856762 | 2.437241895 | 0.507482152 |
| Sgk1 | 2023.552573 | 1.313492119 | 0.156077728 | 8.415628137 |
| Gm8540 | 0.066050875 | 0.274391604 | 3.724945049 | 0.073663262 |
| Gm7672 | 1.404341276 | 1.052910011 | 0.966849285 | 1.089011522 |
| Gm34267 | 0 | NA | NA | NA |
| 4930444G20Rik | 0 | NA | NA | NA |
| Gm2539 | 0.516867762 | 2.026653512 | 1.669519732 | 1.213914082 |
| Gm19791 | 0.170186936 | 0.27439223 | 3.410018679 | 0.080466489 |
| E030030I06Rik | 0.513032606 | 1.516508858 | 1.376540085 | 1.101681581 |
| Gm7678 | 0 | NA | NA | NA |
| Gm4895 | 29.59563694 | 0.041260144 | 0.341933457 | 0.120667173 |
| Raet1e | 4.132199462 | 0.982119761 | 1.042866155 | 0.941750537 |
| H60b | 4.341397877 | 0.473652409 | 0.510364557 | 0.928066816 |
| Gm26740 | 3.074037044 | 2.243157616 | 0.760056 | 2.951305715 |
| Gm26581 | 0.35312086 | 0.274379465 | 3.724945049 | 0.073660003 |
| Gm1972 | 0.209168331 | 0.274403319 | 2.578781255 | 0.106408141 |
| Hmgb1-ps8 | 0.251652017 | 1.078094959 | 2.082118401 | 0.517787537 |
| Gm26528 | 1.473604354 | 3.094988267 | 1.151130442 | 2.68865122 |
| C920009B18Rik | 12.28007475 | 4.09386849 | 0.717600854 | 5.704938156 |
| Gm5421 | 64.80517792 | -0.089762749 | 0.155383855 | -0.577683881 |
| Raet1d | 2.579584798 | 1.830392834 | 1.15268381 | 1.587940091 |
| Gm10825 | 0.745135914 | 0.755309836 | 1.436467029 | 0.525810771 |
| Gm16531 | 0.100902008 | 0.274391828 | 3.724945049 | 0.073663322 |
| Gm26585 | 0.066050875 | 0.274391604 | 3.724945049 | 0.073663262 |
| Slc2a12 | 91.64299392 | 0.367567753 | 0.168977049 | 2.175252527 |
| Tbpl1 | 649.6904834 | -0.011411009 | 0.049552189 | -0.230282638 |
| Mir7663 | 9.275244636 | 0.326146873 | 0.398854789 | 0.817708305 |
| Tcf21 | 0.038981395 | 0.274393828 | 3.724945049 | 0.073663859 |
| Eya4 | 83.33865471 | 0.580170653 | 0.122001806 | 4.755426748 |
| Gm24655 | 0.526083665 | -0.621572728 | 1.354982501 | -0.458731184 |
| Rps12 | 490.9795335 | 0.51738935 | 0.087945649 | 5.883057936 |
| Snora33 | 5.090834635 | -0.961729921 | 0.512660051 | -1.875960336 |
| Snord100 | 3.200779369 | 1.15033771 | 0.605409811 | 1.900097569 |
| Gm23130 | 6.373645778 | 0.602517044 | 0.519901494 | 1.158906161 |
| Slc18b1 | 157.4498406 | -0.456978612 | 0.117688829 | -3.882939579 |

| | | | | |
|---|---|---|---|---|
| Vnn3 | 3.620663213 | 3.297701365 | 0.8825732 | 3.736462159 |
| Vnn1 | 16.91961705 | 1.411666404 | 0.307785907 | 4.586520603 |
| Taar1 | 0.366161499 | 0.755309942 | 1.748956147 | 0.431863282 |
| Taar2 | 0 | NA | NA | NA |
| Taar3 | 0.061920613 | 0.274393828 | 3.724945049 | 0.073663859 |
| Taar4 | 0.07796279 | 0.274392189 | 3.724945049 | 0.073663419 |
| Taar5 | 0 | NA | NA | NA |
| Taar6 | 0 | NA | NA | NA |
| Taar7a | 0 | NA | NA | NA |
| Taar7b | 0.046345709 | 0.274391604 | 3.724945049 | 0.073663262 |
| Taar7c-ps | 0 | NA | NA | NA |
| Gm15136 | 0 | NA | NA | NA |
| Taar7d | 0 | NA | NA | NA |
| Taar7e | 0 | NA | NA | NA |
| Gm6884 | 0 | NA | NA | NA |
| Taar7f | 0 | NA | NA | NA |
| Taar8a | 0 | NA | NA | NA |
| Taar8b | 0 | NA | NA | NA |
| Taar8c | 0 | NA | NA | NA |
| Taar9 | 0 | NA | NA | NA |
| Gm15137 | 0 | NA | NA | NA |
| Stx7 | 1125.137819 | 0.220786891 | 0.06130734 | 3.60131251 |
| Moxd1 | 76.92434119 | 0.171336505 | 0.135934405 | 1.260435175 |
| Gm15271 | 0.245390278 | -0.693958853 | 2.144183146 | -0.323647191 |
| Gm15272 | 0.315650473 | -0.999578136 | 1.723946956 | -0.579819543 |
| Gm15270 | 6.965063545 | -1.125428279 | 0.444337071 | -2.532825535 |
| Gm6893 | 0.385058506 | -2.033653773 | 1.463958313 | -1.389147324 |
| Ctgf | 300.6738772 | 0.867750226 | 0.206462656 | 4.202940335 |
| Enpp1 | 114.2531846 | -0.426541399 | 0.131082178 | -3.25399994 |
| Enpp3 | 5.598650687 | 0.032270293 | 0.446148471 | 0.072330838 |
| Med23 | 495.4221233 | -0.118562851 | 0.087384078 | -1.356801538 |
| Gm23410 | 0 | NA | NA | NA |
| Mir6905 | 0.793572527 | 0.13652616 | 1.190780534 | 0.114652664 |
| Arg1 | 19.82112316 | 1.445763663 | 0.485489584 | 2.977949913 |
| Gm22566 | 0.337983087 | 0.28091594 | 1.574457448 | 0.178420789 |
| Akap7 | 458.6354203 | 0.044619976 | 0.071538292 | 0.623721569 |
| Gm25016 | 0.821005853 | 0.47839538 | 1.114760972 | 0.429146151 |
| Gm26167 | 1.164232618 | -0.397304423 | 0.846964338 | -0.469092269 |
| Epb4.1l2 | 1228.23948 | 0.1948649 | 0.084336976 | 2.310551192 |
| Smlr1 | 0.237992309 | -0.206481255 | 2.113697579 | -0.097687227 |
| Gm29571 | 1.193153915 | 2.009240644 | 1.119897031 | 1.794129807 |
| Tmem200a | 188.3161123 | -0.226491553 | 0.146845899 | -1.542375742 |
| Gm9767 | 4.750744946 | 0.764669332 | 0.550150636 | 1.389927197 |
| Samd3 | 18.30979764 | -0.022038372 | 0.276388384 | -0.079736969 |
| L3mbtl3 | 344.0952442 | -0.135829751 | 0.08471295 | -1.603411891 |
| Gm27754 | 1.24360625 | -1.434380642 | 0.951719546 | -1.507146351 |
| Gm23757 | 0 | NA | NA | NA |

| | | | | |
|---|---|---|---|---|
| Gm8709 | 0 | NA | NA | NA |
| Arhgap18 | 194.7190015 | 0.382589089 | 0.113297739 | 3.376846638 |
| Lama2 | 360.511942 | -0.237098587 | 0.154530722 | -1.534313588 |
| 4930579H20Rik | 0.409009542 | 1.205720047 | 1.733036458 | 0.695726879 |
| Gm23353 | 0.042716945 | -0.206490061 | 3.724945049 | -0.055434391 |
| Gm36962 | 9.987815949 | -0.775909242 | 0.342968677 | -2.262332669 |
| Gm10145 | 22.9043597 | 0.553080224 | 0.268433144 | 2.060402142 |
| Ptprk | 857.1284641 | 0.189284267 | 0.065648914 | 2.883281006 |
| Gm22370 | 0 | NA | NA | NA |
| Themis | 33.46669582 | 5.998004225 | 0.684119701 | 8.767477702 |
| 4930519F09Rik | 0.493807599 | 0.504291766 | 1.354601018 | 0.372280664 |
| Gm22811 | 0 | NA | NA | NA |
| 2310057J18Rik | 0.049553047 | 0.274391604 | 3.724945049 | 0.073663262 |
| Gm9824 | 52.77535247 | 0.050565326 | 0.188249501 | 0.268608023 |
| Gm9996 | 16.71528001 | 0.382047814 | 0.232219962 | 1.645197988 |
| Soga3 | 1394.424857 | -0.114044577 | 0.071682055 | -1.590978057 |
| 9330159F19Rik | 1284.714598 | -0.196179358 | 0.076004743 | -2.581146262 |
| Echdc1 | 288.0513829 | -0.283828197 | 0.112704177 | -2.518346744 |
| Rnf146 | 973.2977547 | 0.370395175 | 0.112362573 | 3.296428375 |
| Gm25596 | 5.415374794 | 0.16985178 | 0.424007542 | 0.400586695 |
| Gm24286 | 0 | NA | NA | NA |
| Rspo3 | 164.9902427 | 0.357722822 | 0.14938844 | 2.39458168 |
| Gm10275 | 252.6440296 | 0.485170585 | 0.142959802 | 3.393755308 |
| Cenpw | 14.39510704 | 0.424621062 | 0.359888951 | 1.179866903 |
| Gm22623 | 0.372763955 | 0.730578829 | 1.474268088 | 0.49555358 |
| Trmt11 | 140.2023999 | -0.158904483 | 0.097095838 | -1.63657357 |
| Hint3 | 162.1893017 | -0.308127252 | 0.112212389 | -2.745928996 |
| Ncoa7 | 1617.001063 | 0.422963158 | 0.097299029 | 4.347043984 |
| Gm24189 | 3.823607117 | -0.100104378 | 0.513769226 | -0.194843079 |
| Hey2 | 73.53388155 | -0.166137876 | 0.253981342 | -0.654134178 |
| Gm24168 | 0 | NA | NA | NA |
| Gm23790 | 0 | NA | NA | NA |
| Gm5422 | 32.64956114 | -0.22795175 | 0.215736711 | -1.056620119 |
| Hddc2 | 80.54346223 | 0.316426772 | 0.154060743 | 2.053909159 |
| n-R5s76 | 0 | NA | NA | NA |
| Tpd52l1 | 150.8148291 | 0.597336455 | 0.107775725 | 5.542402557 |
| Rnf217 | 787.7636301 | 0.290843135 | 0.087650151 | 3.318227428 |
| Gm7909 | 13.66660191 | -0.297959016 | 0.297925953 | -1.000110977 |
| Gm8793 | 0.447136588 | 0.225469479 | 1.429548651 | 0.157720746 |
| Nkain2 | 470.4928905 | -0.128257919 | 0.074515907 | -1.721215282 |
| Trdn | 7.453187596 | -1.402144321 | 0.720074854 | -1.947220227 |
| Clvs2 | 440.1642126 | -0.078730009 | 0.095363401 | -0.82557887 |
| Gm15939 | 3.414634292 | 0.35092968 | 0.563315253 | 0.622972089 |
| Gm4794 | 0 | NA | NA | NA |
| Gm10327 | 7.81528087 | -0.682088375 | 0.396596269 | -1.719855752 |
| Sult3a1 | 0.245625509 | 1.188101052 | 1.834325955 | 0.647704433 |
| Rsph4a | 69.28623393 | 1.022501366 | 0.178727644 | 5.721002883 |

| | | | | |
|---|---|---|---|---|
| Zufsp | 477.2138068 | 1.65339326 | 0.135318298 | 12.21854899 |
| Gm17567 | 17.61824852 | -0.375876171 | 0.252687465 | -1.487514116 |
| Rwdd1 | 291.9026565 | 0.176863897 | 0.099049451 | 1.785612094 |
| Trappc3l | 1.780324605 | 0.4475721 | 0.697985757 | 0.641233859 |
| Fam26d | 0.532386167 | -0.206485279 | 1.635087532 | -0.12628393 |
| Fam26e | 35.94992537 | 0.552779714 | 0.197810971 | 2.794484606 |
| Fam26f | 40.02314751 | 6.862576052 | 0.902224873 | 7.606281157 |
| Dse | 148.6253122 | 0.481064626 | 0.154750395 | 3.108648774 |
| Tspyl1 | 2089.883738 | -0.098331691 | 0.066431854 | -1.480188859 |
| Nt5dc1 | 71.24363843 | 0.910071973 | 0.162112211 | 5.613839754 |
| Gm26564 | 26.65952218 | -0.328235886 | 0.265922432 | -1.234329435 |
| Tspyl4 | 5615.800342 | -0.078747279 | 0.070454728 | -1.117700415 |
| Col10a1 | 2.713943931 | 0.01188269 | 0.662414493 | 0.017938451 |
| Frk | 10.56880692 | 0.107911028 | 0.375109672 | 0.287678607 |
| Gm26341 | 0 | NA | NA | NA |
| Amd2 | 40.7342373 | -0.449865941 | 0.168657033 | -2.66734172 |
| Hs3st5 | 126.4096297 | -0.6405901 | 0.153152191 | -4.182702812 |
| Gm22554 | 0 | NA | NA | NA |
| Hdac2 | 735.3892852 | -0.256037776 | 0.059744429 | -4.285550612 |
| Gm38231 | 0 | NA | NA | NA |
| Marcks | 2411.516162 | -0.198841256 | 0.115703967 | -1.718534472 |
| 5930403N24Rik | 7.127291511 | -0.691812999 | 0.401647963 | -1.72243622 |
| Gm26535 | 0.348734193 | -0.687380004 | 1.728701328 | -0.397627973 |
| Gm24710 | 0 | NA | NA | NA |
| Gm24889 | 0 | NA | NA | NA |
| Gm24297 | 0 | NA | NA | NA |
| Gm22911 | 0 | NA | NA | NA |
| Rfpl4b | 0.174199136 | 1.236193894 | 2.729447628 | 0.452909915 |
| Lama4 | 161.5765428 | 0.256095491 | 0.109575225 | 2.337166017 |
| Fam229b | 68.30184625 | -0.503566406 | 0.188366089 | -2.673338962 |
| Tube1 | 37.79269275 | -0.102385917 | 0.186329569 | -0.549488295 |
| Wisp3 | 1.507103499 | -0.035855158 | 0.783430802 | -0.045766847 |
| Fyn | 1361.385442 | -0.007695897 | 0.062187459 | -0.123753193 |
| Gm6963 | 0 | NA | NA | NA |
| Gm16364 | 1.605310233 | 2.632099526 | 0.970779972 | 2.711324504 |
| Gm16365 | 6.794706781 | 0.451982852 | 0.480172181 | 0.94129329 |
| Traf3ip2 | 50.94917886 | 1.751358616 | 0.256283871 | 6.833666929 |
| E130307A14Rik | 318.1438882 | -0.402757168 | 0.118796309 | -3.390317202 |
| Rev3l | 1908.960953 | -0.048925513 | 0.096144033 | -0.508877266 |
| Gm22105 | 5.849095356 | -0.242674942 | 0.474715805 | -0.511200468 |
| Gm16210 | 0 | NA | NA | NA |
| Gm26824 | 0.048949532 | -0.206487837 | 3.724945049 | -0.055433794 |
| AA474331 | 80.17615184 | -0.414546908 | 0.171320722 | -2.419712587 |
| 4930547M16Rik | 7.251366017 | -0.095078506 | 0.412653686 | -0.230407504 |
| 2010001E11Rik | 0.234609624 | 1.25636357 | 2.48484488 | 0.505610463 |
| G630090E17Rik | 3.289605183 | 2.022347337 | 1.156188196 | 1.749150652 |
| BC021785 | 2.350275864 | 1.565080437 | 0.785779458 | 1.991755346 |

| | | | | |
|---|---|---|---|---|
| AI317395 | 5.519524843 | -0.217397137 | 0.421958352 | -0.515209938 |
| Gm8055 | 1.673936437 | 0.351108405 | 0.746777198 | 0.470164871 |
| Slc16a10 | 102.3548944 | 0.954661402 | 0.185751054 | 5.139466937 |
| Gm25613 | 0 | NA | NA | NA |
| Gm37786 | 11.97329313 | -0.936599733 | 0.318869733 | -2.937248777 |
| Rpf2 | 140.9629893 | 0.019016679 | 0.125541908 | 0.151476736 |
| Gtf3c6 | 109.1455228 | 0.450057649 | 0.15042094 | 2.991988008 |
| U3 | 0.099155932 | 0.75528439 | 3.724945049 | 0.202763901 |
| Rnu3a | 0 | NA | NA | NA |
| Amd1 | 757.3382757 | -0.384968995 | 0.069185164 | -5.564328719 |
| Cdk19os | 1.037245751 | 1.251867103 | 1.013940948 | 1.234654844 |
| Cdk19 | 1184.871889 | -0.231629794 | 0.095468448 | -2.426244465 |
| Slc22a16 | 0 | NA | NA | NA |
| Ddo | 72.37407717 | -0.34809034 | 0.177931816 | -1.956313082 |
| Mettl24 | 13.24789061 | 0.025688703 | 0.288896376 | 0.08892013 |
| Cdc40 | 528.5442356 | 0.184220102 | 0.06589491 | 2.795665116 |
| Gm22948 | 2.318641632 | -1.220152549 | 0.669330769 | -1.822944058 |
| Wasf1 | 1683.342798 | -0.183822824 | 0.100079345 | -1.836770857 |
| Gpr6 | 59.52113351 | -0.528838468 | 0.254320651 | -2.079416148 |
| Fig4 | 411.9453483 | -0.025858736 | 0.077707695 | -0.332769312 |
| Gm25526 | 0 | NA | NA | NA |
| Ak9 | 11.82815708 | -0.623454635 | 0.430314647 | -1.448834333 |
| Gm25626 | 0.159956389 | 0.274409919 | 2.800363834 | 0.097990809 |
| Gm27783 | 0 | NA | NA | NA |
| Zbtb24 | 171.9864541 | 0.020162305 | 0.089869468 | 0.224350998 |
| Mical1 | 186.6857392 | 0.906443544 | 0.12933547 | 7.008468348 |
| Smpd2 | 91.56976239 | 0.051105115 | 0.146261035 | 0.349410319 |
| Ppil6 | 32.28546686 | -0.362373492 | 0.211143048 | -1.716246382 |
| Cd164 | 1390.831552 | 0.548455886 | 0.129082678 | 4.248872856 |
| Ccdc162 | 21.51099621 | 0.385786544 | 0.267069188 | 1.444519104 |
| Gm17196 | 0.189991899 | -0.687377695 | 2.671422729 | -0.257307721 |
| 5730435O14Rik | 0.634867864 | -1.404226099 | 1.286347423 | -1.091638289 |
| Gm29095 | 0.661465078 | 1.097749068 | 1.291216281 | 0.850166687 |
| Gm29096 | 5.25281502 | 0.125110419 | 0.457954804 | 0.273193813 |
| Gm6976 | 9.172692315 | -0.608111279 | 0.3723526 | -1.633159752 |
| Mir3473b | 0 | NA | NA | NA |
| Cep57l1 | 78.55169042 | -0.164359002 | 0.192901622 | -0.852035355 |
| BC048559 | 0 | NA | NA | NA |
| Sesn1 | 535.6668866 | 0.012224673 | 0.078618825 | 0.155492953 |
| Armc2 | 66.24966393 | -0.674962288 | 0.280837522 | -2.403390699 |
| Gm9855 | 93.44015898 | 0.376637042 | 0.148903881 | 2.529397078 |
| Foxo3 | 2059.889726 | -0.041496964 | 0.066091143 | -0.627874821 |
| Gm23533 | 0 | NA | NA | NA |
| Lace1 | 140.8858273 | -0.328068582 | 0.112808599 | -2.908187725 |
| Gm15197 | 3.036843357 | -1.132501781 | 0.656474921 | -1.725125735 |
| Gm15198 | 0 | NA | NA | NA |
| Gm25277 | 0 | NA | NA | NA |

| | | | | |
|---|---|---|---|---|
| Snx3 | 1059.401976 | -0.060304946 | 0.0722149 | -0.835076234 |
| Nr2e1 | 94.72567818 | -0.519852352 | 0.214334793 | -2.425422143 |
| Ostm1 | 722.7865687 | 0.292223463 | 0.081860029 | 3.569794273 |
| Gm15200 | 0.23382787 | 0.274411136 | 2.130875987 | 0.128778557 |
| Gm15199 | 0.066050875 | 0.274391604 | 3.724945049 | 0.073663262 |
| Sec63 | 1231.43905 | 0.146360781 | 0.064702697 | 2.262050693 |
| Gm26860 | 0.331010926 | 0.247042171 | 1.7529328 | 0.140930771 |
| Gm26464 | 0.049553047 | 0.274391604 | 3.724945049 | 0.073663262 |
| Scml4 | 87.95950352 | 1.210145583 | 0.162640827 | 7.440601515 |
| Sobp | 1030.323232 | -0.299642769 | 0.084030054 | -3.565899997 |
| Gm29245 | 0 | NA | NA | NA |
| Gm29246 | 2.290242257 | -0.959550677 | 0.744252824 | -1.289280533 |
| 9030612E09Rik | 15.14098368 | -0.501942533 | 0.262801586 | -1.909967671 |
| Pdss2 | 173.4444041 | -0.198354148 | 0.093008324 | -2.132649415 |
| Gm16546 | 3.059482485 | -0.192758785 | 0.587492301 | -0.32810436 |
| Rps19-ps11 | 0 | NA | NA | NA |
| Gm3699 | 19.13216094 | 0.323293762 | 0.299931571 | 1.077891739 |
| Bend3 | 116.2620847 | 0.704371293 | 0.177381474 | 3.970940574 |
| Gm9803 | 11.92917661 | 0.482343362 | 0.33698421 | 1.431353005 |
| 1700021F05Rik | 133.4447187 | 0.081913703 | 0.12990911 | 0.630546256 |
| Cd24a | 328.2481463 | 0.763521517 | 0.217669706 | 3.507706839 |
| F930017D23Rik | 0.149263833 | 1.514651884 | 3.486034426 | 0.434491373 |
| Gm9034 | 0.045882455 | 0.755288839 | 3.724945049 | 0.202765096 |
| Qrsl1 | 139.8918007 | -0.25865098 | 0.093610722 | -2.763048666 |
| Rtn4ip1 | 176.0176104 | -0.255118662 | 0.087744139 | -2.907529364 |
| Aim1 | 288.648516 | 2.940993663 | 0.309984175 | 9.487560671 |
| Gm10812 | 1.879239285 | 0.692389285 | 0.852772262 | 0.81192754 |
| Speer5-ps1 | 0.049553047 | 0.274391604 | 3.724945049 | 0.073663262 |
| Atg5 | 276.1062705 | 0.098756411 | 0.099102105 | 0.996511742 |
| Mir1929 | 2.194932386 | -1.10899124 | 0.789210459 | -1.405190754 |
| Prdm1 | 54.05971938 | 1.215794772 | 0.336417619 | 3.613944997 |
| Gm22087 | 0 | NA | NA | NA |
| Gm27977 | 0 | NA | NA | NA |
| Gm22503 | 0 | NA | NA | NA |
| Gm24271 | 0 | NA | NA | NA |
| Gm4795 | 0.55021313 | -0.632649709 | 1.301560295 | -0.486070227 |
| Prep | 326.8201584 | -0.024974965 | 0.068480628 | -0.364701162 |
| Gm23766 | 0 | NA | NA | NA |
| Popdc3 | 11.87580344 | -0.197570245 | 0.307787438 | -0.641904836 |
| Gm27723 | 0 | NA | NA | NA |
| Gm26016 | 0 | NA | NA | NA |
| Gm15934 | 0.066050875 | 0.274391604 | 3.724945049 | 0.073663262 |
| Bves | 10.58168931 | -0.988198415 | 0.351666975 | -2.810040421 |
| Lin28b | 106.2737535 | 0.058446548 | 0.156647236 | 0.373109348 |
| Gm27731 | 0 | NA | NA | NA |
| Hace1 | 690.7874818 | -0.233748747 | 0.102105932 | -2.289276856 |
| Gm27410 | 0 | NA | NA | NA |

| | | | | |
|---|---|---|---|---|
| Gm25740 | 0 | NA | NA | NA |
| Gm23481 | 0 | NA | NA | NA |
| Gm25650 | 0 | NA | NA | NA |
| Gm23584 | 0 | NA | NA | NA |
| Gm24256 | 0 | NA | NA | NA |
| Grik2 | 2090.882152 | -0.051037846 | 0.096328705 | -0.529830087 |
| Gm26543 | 111.823433 | 0.024413534 | 0.191897383 | 0.127221818 |
| Ascc3 | 1896.390272 | 1.132811418 | 0.093992426 | 12.05215635 |
| Gm23883 | 0 | NA | NA | NA |
| Sim1 | 14.16261422 | -0.581398185 | 0.298255296 | -1.949330635 |
| Gm26257 | 0 | NA | NA | NA |
| Gp49a | 332.5075168 | 7.82532227 | 0.666310872 | 11.74425121 |
| Lilrb4 | 459.2260594 | 6.739013517 | 0.485584787 | 13.87813972 |
| Fam162b | 0 | NA | NA | NA |
| Gprc6a | 0.291554621 | 0.189168949 | 1.754890001 | 0.107795331 |
| Rfx6 | 0.232111299 | 1.607789491 | 2.503586656 | 0.642194464 |
| 4933411G06Rik | 0 | NA | NA | NA |
| Gm23906 | 0 | NA | NA | NA |
| Vgll2 | 1.144635215 | 0.767338306 | 0.993473496 | 0.772379242 |
| Gm24084 | 0.042716945 | -0.206490061 | 3.724945049 | -0.055434391 |
| Ros1 | 0 | NA | NA | NA |
| Gm20630 | 0 | NA | NA | NA |
| Gm18513 | 0 | NA | NA | NA |
| Dcbld1 | 120.8113824 | 0.172593747 | 0.12504222 | 1.380283762 |
| Gopc | 773.3070799 | 0.090685386 | 0.06813051 | 1.331053968 |
| Gm26741 | 0 | NA | NA | NA |
| Rps19-ps10 | 0 | NA | NA | NA |
| Nepn | 0.397314471 | -1.311870557 | 1.532927562 | -0.855794227 |
| Nus1 | 1151.136521 | 0.07503154 | 0.08410496 | 0.892117891 |
| Zfa-ps | 7.158863588 | 0.569983403 | 0.489773938 | 1.163768342 |
| Slc35f1 | 918.4030169 | 0.106044853 | 0.063937645 | 1.658566755 |
| Gm25664 | 1.948438448 | -0.385651166 | 0.735748373 | -0.524161766 |
| Cep85l | 359.2181035 | 0.053076441 | 0.122507475 | 0.433250635 |
| Pln | 12.74356708 | 0.929568433 | 0.346763209 | 2.680700857 |
| Gm5423 | 0.736769121 | 0.328056166 | 1.395168263 | 0.235137348 |
| Mcm9 | 170.9008201 | 0.852480578 | 0.108748252 | 7.839027872 |
| Asf1a | 203.4365805 | 0.041816799 | 0.110945843 | 0.376911814 |
| Fam184a | 493.2642014 | -0.313447008 | 0.09121025 | -3.436532699 |
| Man1a | 671.9857947 | -0.159129721 | 0.078490844 | -2.027366676 |
| Gm16998 | 3.462133719 | 0.534231913 | 0.485112194 | 1.10125435 |
| Gm26177 | 0 | NA | NA | NA |
| Gm25602 | 0 | NA | NA | NA |
| Tbc1d32 | 271.3183319 | -0.165029731 | 0.106433827 | -1.550538349 |
| Msl3l2 | 168.4746344 | 0.522591054 | 0.105999832 | 4.930112086 |
| Rps8-ps1 | 3.747707297 | 0.469904223 | 0.48740496 | 0.964094053 |
| Gja1 | 2426.848436 | -0.412059246 | 0.175338295 | -2.350081287 |
| Gm9795 | 5.035917182 | -0.10478995 | 0.455515138 | -0.23004713 |

| | | | | |
|---|---|---|---|---|
| Gm9956 | 0.322113463 | -0.620700844 | 1.772265432 | -0.350230181 |
| Gm24285 | 0 NA | NA | NA | |
| 4930467K11Rik | 0.534824616 | 0.130628404 | 1.272482956 | 0.10265631 |
| Hsf2 | 367.3738673 | -0.056812448 | 0.081353 | -0.698344841 |
| Serinc1 | 9098.13279 | 0.093807138 | 0.069968647 | 1.340702473 |
| Rpl48-ps1 | 0 NA | NA | NA | |
| Pkib | 225.7964083 | -0.090994254 | 0.157400029 | -0.578108243 |
| Gm5777 | 0.168032776 | -0.206492308 | 3.415960319 | -0.06044927 |
| Fabp7 | 502.0611919 | -0.123972927 | 0.33469199 | -0.370409004 |
| Smpdl3a | 299.0070279 | 0.571393031 | 0.176306679 | 3.240904051 |
| AW822073 | 0.119587853 | 1.218392466 | 3.723618458 | 0.327206581 |
| Gm10807 | 0.089089296 | 1.194485492 | 3.723939576 | 0.320758559 |
| Duxf3 | 0.097139547 | 0.274393828 | 3.724945049 | 0.073663859 |
| Gm4981 | 0.424461334 | 1.540633876 | 3.093264643 | 0.498060804 |
| Gm19459 | 0.218736334 | 0.274414121 | 2.218267398 | 0.123706511 |
| Gcc2 | 1409.09042 | -0.064703302 | 0.06311964 | -1.025089844 |
| Lims1 | 608.1636776 | 0.356936486 | 0.106449648 | 3.353101612 |
| Gm23058 | 0 NA | NA | NA | |
| Ranbp2 | 4662.349267 | 0.163526914 | 0.075484704 | 2.166358284 |
| Ccdc138 | 45.03437437 | -0.405818562 | 0.199409784 | -2.03509855 |
| Edar | 4.213958645 | 0.069235389 | 0.498062685 | 0.139009388 |
| BB019430 | 0 NA | NA | NA | |
| Gm25055 | 0 NA | NA | NA | |
| Gm17542 | 0 NA | NA | NA | |
| Gm9987 | 7.178240949 | -0.083144659 | 0.497796141 | -0.167025519 |
| Sh3rf3 | 355.2242412 | -0.372946133 | 0.17535505 | -2.126805774 |
| Gm27672 | 1.813029451 | -1.201740804 | 0.694759525 | -1.729721956 |
| 10-Sep | 84.1994719 | -0.072425831 | 0.185377131 | -0.390694531 |
| Gm23223 | 0.038981395 | 0.274393828 | 3.724945049 | 0.073663859 |
| AC153941.1 | 0 NA | NA | NA | |
| Sowahc | 0 NA | NA | NA | |
| P4ha1 | 725.4105145 | -0.186574649 | 0.101597395 | -1.836411746 |
| Gm10273 | 13.1323639 | 0.293091897 | 0.329844439 | 0.888576136 |
| Pla2g12b | 0.130524232 | 0.274405456 | 3.657871338 | 0.075017799 |
| Oit3 | 3.202472028 | 2.146210197 | 0.815782476 | 2.630860872 |
| Mcu | 514.9080926 | -0.202512954 | 0.063810132 | -3.173680242 |
| Gm10322 | 0.281194119 | -0.206480416 | 2.014383019 | -0.102503056 |
| Micu1 | 809.3638985 | -0.151475103 | 0.059275811 | -2.555428601 |
| Gm17148 | 2.409158253 | 0.691677508 | 0.725105818 | 0.953898716 |
| Gm17059 | 0.044544648 | 0.755286614 | 3.724945049 | 0.202764498 |
| Gm7399 | 5.948605025 | -1.277184844 | 0.479411362 | -2.664068785 |
| Gm23803 | 0 NA | NA | NA | |
| Dnajb12 | 642.9386256 | 0.149370393 | 0.064579523 | 2.312968356 |
| Ddit4 | 632.8851748 | 0.96164234 | 0.17425512 | 5.518588731 |
| Anapc16 | 407.1600245 | 0.222800067 | 0.083780981 | 2.65931557 |
| Ascc1 | 125.3215762 | -0.057282777 | 0.103064237 | -0.555796837 |
| Spock2 | 9371.899727 | -0.10501351 | 0.061795081 | -1.699383006 |

| | | | | |
|---|---|---|---|---|
| mmu-mir-6906 | 1.418442629 | -0.246081988 | 0.849420877 | -0.289705603 |
| Chst3 | 65.59010548 | -0.038382659 | 0.186443551 | -0.205867456 |
| Gm22117 | 0.042716945 | -0.206490061 | 3.724945049 | -0.055434391 |
| Psap | 17356.86525 | 0.366392619 | 0.103862827 | 3.527658841 |
| Cdh23 | 85.39716704 | -0.251785726 | 0.225899282 | -1.114592858 |
| 4632428N05Rik | 287.6489372 | -0.003766441 | 0.179956745 | -0.020929704 |
| Gm17455 | 4.08792871 | -0.279458241 | 0.592683979 | -0.471513067 |
| Slc29a3 | 540.1404158 | 0.719790904 | 0.132588341 | 5.428764715 |
| Unc5b | 525.6811087 | -0.391281176 | 0.099245046 | -3.942576404 |
| Mir6408 | 0 | NA | NA | NA |
| Pldi | 0.297702773 | 0.274408846 | 1.955851176 | 0.140301496 |
| Gm26947 | 9.224316412 | 0.58028641 | 0.407741642 | 1.423171811 |
| Mir466j | 0.085433889 | -0.687376175 | 3.724945049 | -0.184533239 |
| Gm20611 | 0.272151178 | -0.435256579 | 2.454953567 | -0.177297276 |
| Gm20625 | 0.049553047 | 0.274391604 | 3.724945049 | 0.073663262 |
| Gm20610 | 0 | NA | NA | NA |
| Pcbd1 | 47.59642594 | 0.028999072 | 0.182985008 | 0.158477857 |
| Sgpl1 | 896.1568652 | 0.609956126 | 0.133208818 | 4.578947043 |
| AC123530.1 | 1.014434794 | 0.742642527 | 1.003031759 | 0.74039782 |
| Tbata | 34.89983757 | -0.677531292 | 0.365844065 | -1.851967426 |
| Adamts14 | 36.59661118 | 3.025011933 | 0.368867967 | 8.200798666 |
| Prf1 | 187.0444178 | 9.235273112 | 1.022329 | 9.033562691 |
| Rpl27a-ps1 | 0.285326127 | -0.689282622 | 1.779934038 | -0.387251779 |
| Pald1 | 146.0309485 | 0.15471092 | 0.139621092 | 1.108076996 |
| Nodal | 1.922318947 | -1.195763664 | 0.757700536 | -1.578148103 |
| Eif4ebp2 | 324.7426737 | -0.266215842 | 0.095817607 | -2.778360358 |
| D830039M14Rik | 3.979968375 | -0.26342518 | 0.535632902 | -0.491801715 |
| Lrrc20 | 191.831978 | -0.122080193 | 0.096149805 | -1.269687365 |
| Gm28447 | 0.497289099 | 1.580836828 | 1.355219168 | 1.16648057 |
| Npffr1 | 23.64047625 | -0.213107336 | 0.231198934 | -0.921748781 |
| Ppa1 | 944.9657682 | 0.871427284 | 0.138111374 | 6.309598265 |
| Sar1a | 1213.944438 | 0.40076108 | 0.103825096 | 3.859963506 |
| Tysnd1 | 151.8466627 | -0.101685387 | 0.099559845 | -1.021349395 |
| Aifm2 | 86.05177613 | 0.24374004 | 0.159623879 | 1.526964766 |
| H2afy2 | 209.0800445 | 0.10471756 | 0.112651672 | 0.929569512 |
| Mir5108 | 0 | NA | NA | NA |
| Col13a1 | 11.90106664 | -0.80964547 | 0.347667966 | -2.328789389 |
| Gm5424 | 120.3562083 | 0.379708997 | 0.160407523 | 2.367152055 |
| 2010107G23Rik | 182.7260212 | -0.336814779 | 0.105757416 | -3.184786393 |
| Neurog3 | 0.108123235 | 0.274402725 | 3.724945049 | 0.073666248 |
| Tspan15 | 244.3145109 | 0.015844179 | 0.151170318 | 0.104810118 |
| Mir7662 | 1.416932329 | -0.94566605 | 0.836926306 | -1.129927501 |
| Tacr2 | 2.342138215 | 0.231987534 | 0.790171352 | 0.293591426 |
| Hk1 | 4224.69477 | 0.157028786 | 0.055943773 | 2.806903753 |
| Hk1os | 8.888663959 | -0.183184481 | 0.359734991 | -0.509220635 |
| Hkdc1 | 74.93408703 | -0.378901261 | 0.172456155 | -2.197087489 |
| Supv3l1 | 235.6531214 | 0.221972782 | 0.10139301 | 2.1892316 |

| | | | | |
|---|---|---|---|---|
| 4930507D05Rik | 5.319148665 | -0.690368861 | 0.466508831 | -1.479862365 |
| Vps26a | 797.6808432 | 0.515001598 | 0.08177806 | 6.297552136 |
| Srgn | 508.9808035 | 4.382742605 | 0.392300064 | 11.17191406 |
| Kif1bp | 1369.8405 | 0.149704841 | 0.062154504 | 2.408591987 |
| RP24-255E8.1 | 2.889734049 | 1.590691244 | 0.612651242 | 2.596405808 |
| Ddx21 | 587.0568038 | 0.610304249 | 0.117960884 | 5.17378497 |
| Ddx50 | 804.8841463 | 0.025079632 | 0.075173078 | 0.33362519 |
| Stox1 | 37.04183788 | -0.105112769 | 0.230215755 | -0.456583733 |
| mmu-mir-7215 | 0 | NA | NA | NA |
| Ccar1 | 875.9317859 | 0.049473778 | 0.090762589 | 0.545089982 |
| Snord98 | 8.868178155 | 0.540198623 | 0.427909229 | 1.262414051 |
| Tet1 | 553.5602397 | -0.290056532 | 0.120823155 | -2.400670068 |
| Gm26479 | 0 | NA | NA | NA |
| Gm22803 | 0.09349418 | 0.274402725 | 3.724945049 | 0.073666248 |
| Slc25a16 | 401.0961555 | 0.092196755 | 0.090605894 | 1.017558026 |
| Gm25862 | 0 | NA | NA | NA |
| Gm16128 | 5.788109619 | 0.882364966 | 0.506816078 | 1.740996397 |
| Dna2 | 37.65220541 | 0.728834072 | 0.243249095 | 2.996245774 |
| Rufy2 | 990.6841044 | -0.065470254 | 0.109910555 | -0.595668484 |
| Gm16135 | 8.108709463 | 0.119347676 | 0.470447536 | 0.253689661 |
| Hnrnph3 | 390.3370152 | -0.086374626 | 0.135488173 | -0.637506756 |
| Pbld2 | 14.50480443 | -0.236054576 | 0.268498029 | -0.879166887 |
| Gm16143 | 0.144050924 | 1.290908285 | 3.721447108 | 0.346883416 |
| Gm38063 | 0.592709862 | -0.625921917 | 1.283663643 | -0.487605862 |
| Atoh7 | 10.98456152 | 0.086922638 | 0.416824053 | 0.20853556 |
| Mypn | 24.12272005 | -0.852028342 | 0.214558626 | -3.971074743 |
| 1700120B22Rik | 2.541048356 | -0.621739015 | 0.655599967 | -0.9483512 |
| Herc4 | 532.4503219 | -0.079767588 | 0.064951207 | -1.228115565 |
| Gm26789 | 0.091666476 | -0.687373951 | 3.724945049 | -0.184532642 |
| Sirt1 | 314.9020831 | -0.109185542 | 0.083251474 | -1.311514817 |
| Gm16145 | 0.37202309 | 0.311249751 | 1.572703582 | 0.197907447 |
| Dnajc12 | 114.8880949 | -0.014053833 | 0.135195187 | -0.103952166 |
| Gm7075 | 0.075043205 | 0.755286614 | 3.724945049 | 0.202764498 |
| Ctnna3 | 57.98869911 | -0.817952882 | 0.166632479 | -4.90872421 |
| Gm10118 | 118.090575 | -0.401090531 | 0.209982053 | -1.910118151 |
| Lrrtm3 | 650.2187372 | -0.495272344 | 0.079443944 | -6.234236579 |
| AC116138.1 | 1.265617311 | -1.722822788 | 0.905529537 | -1.902558356 |
| Gm23854 | 0 | NA | NA | NA |
| Gm28881 | 0 | NA | NA | NA |
| 1700023F02Rik | 0.746473191 | -0.08912695 | 1.189950702 | -0.0748997 |
| Gm22829 | 0 | NA | NA | NA |
| Gm22594 | 0 | NA | NA | NA |
| Gm28139 | 0 | NA | NA | NA |
| 1110002J07Rik | 9.655946214 | 0.559127145 | 0.434056911 | 1.288142479 |
| Gm26576 | 2.265170197 | 0.60083811 | 0.655048533 | 0.917242127 |
| Reep3 | 781.8290574 | -0.037523166 | 0.086240927 | -0.435096967 |
| Jmjd1c | 3049.812434 | 0.128541387 | 0.093015076 | 1.381941434 |

| | | | | |
|---|---|---|---|---|
| Gm26359 | 0 | NA | NA | NA |
| Nrbf2 | 127.8010227 | 0.245406696 | 0.098141261 | 2.500545578 |
| Egr2 | 185.0575354 | -0.983169329 | 0.529820186 | -1.855666046 |
| Ado | 961.6363525 | -0.162327018 | 0.082247395 | -1.973643261 |
| Gm10797 | 2.84885825 | -0.022145509 | 0.545389108 | -0.040604971 |
| Zfp365 | 3160.688214 | -0.062163989 | 0.126525124 | -0.491317356 |
| Rtkn2 | 179.8255072 | 1.350357203 | 0.156816172 | 8.611083849 |
| Gm16212 | 0.075043205 | 0.755286614 | 3.724945049 | 0.202764498 |
| Arid5b | 664.009088 | 0.154128977 | 0.106579949 | 1.446134839 |
| 1700040L02Rik | 9.415632204 | -0.650795025 | 0.335520876 | -1.939655835 |
| Tmem26 | 24.00413336 | 0.125778434 | 0.315141762 | 0.399116998 |
| Gm24172 | 0 | NA | NA | NA |
| Rhobtb1 | 46.1805193 | -0.069105028 | 0.182999683 | -0.377623756 |
| A930033H14Rik | 22.60880403 | -0.4738991 | 0.236706942 | -2.002049862 |
| Cdk1 | 20.93667416 | 1.976426518 | 0.486879542 | 4.059374745 |
| Ank3 | 10782.92156 | -0.340190625 | 0.09088844 | -3.742947101 |
| Ccdc6 | 1500.046561 | 0.51016905 | 0.088787759 | 5.745939014 |
| Mrln | 8.103814522 | 1.255209694 | 0.454582874 | 2.761234012 |
| Slc16a9 | 87.74711258 | 0.958333019 | 0.194317762 | 4.931782905 |
| Gm7097 | 0.061920613 | 0.274393828 | 3.724945049 | 0.073663859 |
| Gm9877 | 0.095435502 | 0.75528439 | 3.724945049 | 0.202763901 |
| Fam13c | 629.0128233 | -0.590272452 | 0.096749647 | -6.101029507 |
| Gm22320 | 0.300192116 | 1.012530472 | 3.72168013 | 0.272062734 |
| Phyhipl | 1745.320192 | -0.141428003 | 0.057275956 | -2.46923864 |
| 4930533K18Rik | 0.42462922 | -1.356472921 | 1.4976269 | -0.905748235 |
| Gm24486 | 0.340478385 | -0.189920515 | 1.493785414 | -0.127140427 |
| Bicc1 | 210.8407525 | 0.498398726 | 0.133594039 | 3.730695842 |
| Gm15647 | 0 | NA | NA | NA |
| Tfam | 397.4176102 | 0.132290422 | 0.1050535 | 1.259267148 |
| Ube2d1 | 468.1548561 | -0.192564533 | 0.057386207 | -3.355589144 |
| Gm20609 | 0.553453779 | 0.255991098 | 1.113725835 | 0.229851091 |
| Cisd1 | 406.1124699 | -0.150232248 | 0.072711778 | -2.066133586 |
| Ipmk | 440.7198033 | -0.276099197 | 0.071179177 | -3.878932151 |
| Gm22937 | 0 | NA | NA | NA |
| 1700049L16Rik | 0 | NA | NA | NA |
| Gm9923 | 40.21032448 | 0.268407699 | 0.175668782 | 1.527919173 |
| Gm24984 | 0 | NA | NA | NA |
| Zwint | 7079.662798 | 0.137897708 | 0.074623347 | 1.847916413 |
| Pcdh15 | 611.2248178 | -0.242628316 | 0.189634305 | -1.279453715 |
| AC159477.1 | 0 | NA | NA | NA |
| Gm37438 | 0 | NA | NA | NA |
| Gm15398 | 0.158818843 | -0.666974881 | 2.81011023 | -0.237348298 |
| Gm5778 | 18.43137054 | -0.697291041 | 0.517479621 | -1.347475365 |
| Gm15397 | 18.38539889 | -0.515959711 | 0.329650539 | -1.565171748 |
| Gm6407 | 0 | NA | NA | NA |
| Gm6419 | 9.498614017 | 0.0945428 | 0.333056528 | 0.283864126 |
| Rsph14 | 2.362931495 | -1.173630074 | 0.698579366 | -1.68002396 |

| | | | | |
|---|---|---|---|---|
| Gnaz | 847.2906744 | -0.198099485 | 0.094004581 | -2.107338626 |
| Rab36 | 225.5149465 | -0.237780236 | 0.125568767 | -1.893625626 |
| Bcr | 2079.456346 | -0.058938254 | 0.071970434 | -0.818923139 |
| Specc1l | 984.3676726 | -0.026273129 | 0.049667625 | -0.528978979 |
| Adora2a | 245.9031525 | -0.416216567 | 0.171721151 | -2.423793255 |
| Gucd1 | 173.5614678 | 0.106801514 | 0.096770205 | 1.103661139 |
| Upb1 | 7.757773806 | 0.4591301 | 0.396838916 | 1.156968435 |
| Gm25545 | 0.075043205 | 0.755286614 | 3.724945049 | 0.202764498 |
| Snrpd3 | 401.6260286 | 0.228400056 | 0.089632558 | 2.548181841 |
| Lrrc75b | 242.2640389 | -0.431528815 | 0.100967109 | -4.273954368 |
| Ggt1 | 11.84378073 | 0.039423935 | 0.336897767 | 0.117020469 |
| Ggt5 | 50.67091169 | 1.585772289 | 0.272817994 | 5.812564865 |
| Gm9985 | 0.329446336 | -0.687379275 | 1.775395075 | -0.387169754 |
| Gm26402 | 0 | NA | NA | NA |
| Susd2 | 127.9307858 | -0.392436505 | 0.115998033 | -3.383130695 |
| Cabin1 | 1380.689238 | -0.010774552 | 0.071347852 | -0.151014385 |
| Ddt | 134.0537612 | -0.180136937 | 0.110322646 | -1.632819227 |
| Gm20441 | 0 | NA | NA | NA |
| Gstt3 | 21.78233339 | -0.380718505 | 0.260172453 | -1.463331343 |
| Gstt1 | 101.288934 | 0.630909757 | 0.189477581 | 3.329733011 |
| Gstt4 | 0.243832141 | 0.274396967 | 3.044346567 | 0.090133288 |
| Gstt2 | 19.40726938 | 0.23986259 | 0.239082451 | 1.003263058 |
| Mif | 1051.951317 | 0.563486061 | 0.10293212 | 5.474346222 |
| Derl3 | 13.80484017 | -0.595683358 | 0.304124848 | -1.958680335 |
| Smarcb1 | 629.9954016 | -0.003733713 | 0.063078591 | -0.05919144 |
| Mmp11 | 15.83720611 | -0.297232742 | 0.300536189 | -0.989008157 |
| Chchd10 | 1094.33119 | -0.241899988 | 0.091860229 | -2.633348392 |
| Gm867 | 15.27197981 | 1.03922169 | 0.339970659 | 3.056798177 |
| Vpreb3 | 1.181927862 | 0.655164094 | 1.227389817 | 0.533786483 |
| Gm16221 | 0.106112162 | -0.206492286 | 3.724945049 | -0.055434988 |
| Gm5134 | 1.323702316 | 0.384395736 | 0.94888214 | 0.405103774 |
| Gm16219 | 0 | NA | NA | NA |
| Gm16220 | 0 | NA | NA | NA |
| Mir6409 | 0 | NA | NA | NA |
| Zfp280b | 318.8465618 | -0.28900251 | 0.095186269 | -3.036178569 |
| Slc5a4b | 0.19103638 | -0.206481537 | 2.252891137 | -0.091651804 |
| Gm16222 | 0.688368266 | 1.938337174 | 1.48503814 | 1.305244035 |
| Slc5a4a | 0.337090955 | 0.755307964 | 1.766862509 | 0.427485421 |
| Prmt2 | 399.9587037 | -0.366987545 | 0.108877928 | -3.370633065 |
| Mir678 | 0 | NA | NA | NA |
| S100b | 1036.502615 | -0.491853171 | 0.147128063 | -3.343027572 |
| Dip2a | 1191.452903 | -0.63425145 | 0.086174951 | -7.360044204 |
| Gm16240 | 0.04393175 | -0.206490061 | 3.724945049 | -0.055434391 |
| Pcnt | 701.5738582 | 0.270155958 | 0.088677327 | 3.046505412 |
| 1700094J05Rik | 0.456503428 | 0.960660802 | 1.500654409 | 0.64016125 |
| 2610028H24Rik | 0.44299472 | -1.116245979 | 1.408822659 | -0.792325401 |
| Ybey | 77.35572085 | -0.187889282 | 0.131813059 | -1.42542237 |

| | | | | |
|---|---|---|---|---|
| Mcm3ap | 786.2211096 | -0.191691378 | 0.089187978 | -2.149296151 |
| Lss | 497.5698186 | -0.133864591 | 0.071749186 | -1.865729754 |
| 4930483K19Rik | 0 | NA | NA | NA |
| Spatc1l | 0.383664054 | -1.903006744 | 1.526612495 | -1.2465552 |
| Gm15343 | 0 | NA | NA | NA |
| Ftcd | 0.710593519 | -1.660680985 | 1.288300037 | -1.28904831 |
| Col6a2 | 230.4261296 | 0.065801338 | 0.141155789 | 0.466161099 |
| Col6a1 | 323.7855034 | -0.102396758 | 0.186506815 | -0.549024218 |
| Pcbp3 | 627.0954025 | -0.288838919 | 0.115934983 | -2.491387087 |
| Gm37782 | 0.137011352 | -0.687378074 | 2.942485862 | -0.233604546 |
| Gm10787 | 0.044544648 | 0.755286614 | 3.724945049 | 0.202764498 |
| Slc19a1 | 146.9527464 | -0.064833444 | 0.098968869 | -0.65508927 |
| Col18a1 | 325.5916588 | 0.240401794 | 0.166387593 | 1.444830049 |
| Gm10941 | 3.197615884 | -0.752607406 | 0.621793877 | -1.210380858 |
| Pofut2 | 316.8806264 | -0.044913511 | 0.085846918 | -0.523181407 |
| Adarb1 | 3284.804805 | -0.417449287 | 0.073534995 | -5.676879263 |
| Fam207a | 347.3627384 | 0.001627705 | 0.099785037 | 0.016312118 |
| 2810425M01Rik | 0.856869739 | -0.445498052 | 1.019318718 | -0.437054715 |
| Itgb2 | 410.5844244 | 3.444447438 | 0.185101549 | 18.60842039 |
| Pttg1ip | 760.5419193 | 0.501886692 | 0.125279676 | 4.006130201 |
| Sumo3 | 1158.6635 | 0.401872238 | 0.088919567 | 4.519502893 |
| Ube2g2 | 330.4207087 | 0.266245077 | 0.089320456 | 2.980785033 |
| Mir1930 | 0 | NA | NA | NA |
| Gm19402 | 0 | NA | NA | NA |
| Gm10272 | 0.197138355 | 1.236195555 | 2.624156999 | 0.471082925 |
| Gm10024 | 0.182127695 | -1.017794829 | 3.345561342 | -0.304222438 |
| Gm10142 | 0 | NA | NA | NA |
| Krtap12-1 | 0.19999695 | 0.755299832 | 2.625077941 | 0.287724726 |
| Gm10100 | 0.235931372 | 0.321134988 | 2.536876862 | 0.126586747 |
| Gm3233 | 0 | NA | NA | NA |
| Gm3238 | 0 | NA | NA | NA |
| Gm7138 | 0 | NA | NA | NA |
| Gm3250 | 0.062426873 | 0.755288839 | 3.724945049 | 0.202765096 |
| Gm7137 | 0 | NA | NA | NA |
| Gm19668 | 0 | NA | NA | NA |
| Gm9507 | 0.060174537 | 0.755286614 | 3.724945049 | 0.202764498 |
| Gm2696 | 0 | NA | NA | NA |
| Krtap10-4 | 0 | NA | NA | NA |
| AC153874.1 | 0 | NA | NA | NA |
| Tspear | 5.130515031 | -0.524696699 | 0.432690699 | -1.212636878 |
| Gm10318 | 0.127104694 | -0.687378099 | 3.010998535 | -0.228289085 |
| 1700009J07Rik | 0.532715572 | 1.389387355 | 1.443630958 | 0.962425575 |
| Lrrc3 | 123.618813 | 0.104537979 | 0.200495244 | 0.521398795 |
| Trpm2 | 446.088107 | -0.227895086 | 0.137883679 | -1.652806828 |
| Gm22394 | 0.066050875 | 0.274391604 | 3.724945049 | 0.073663262 |
| Gm22871 | 0.044544648 | 0.755286614 | 3.724945049 | 0.202764498 |
| 1810043G02Rik | 263.2113337 | 0.064420689 | 0.082542312 | 0.78045656 |

| | | | | |
|---|---|---|---|---|
| Pfkl | 1143.605665 | -0.12480986 | 0.08210821 | -1.520065524 |
| Aire | 5.516484393 | 3.028482379 | 0.600583094 | 5.042570143 |
| Dnmt3l | 4.167386937 | 1.26845916 | 0.779961531 | 1.626309901 |
| Icosl | 151.2157367 | 2.264333164 | 0.170509928 | 13.27977316 |
| D10Jhu81e | 415.5710078 | -0.162808829 | 0.086311426 | -1.886295206 |
| Pwp2 | 271.1694499 | -0.00111753 | 0.069952645 | -0.015975528 |
| Trappc10 | 1524.4217 | 0.042821553 | 0.042477251 | 1.008105576 |
| Agpat3 | 3193.730264 | -0.025761897 | 0.063663945 | -0.404654426 |
| n-R5s77 | 0 | NA | NA | NA |
| Mir7659 | 1.249419009 | -2.23992087 | 1.055551848 | -2.122037752 |
| Gm22514 | 0 | NA | NA | NA |
| Gm10146 | 20.48723405 | 0.120412565 | 0.251250109 | 0.479253782 |
| Rrp1 | 2027.885291 | 0.023687588 | 0.089042511 | 0.266025609 |
| Mir6907 | 2.924483123 | -0.35697767 | 0.73444092 | -0.486053623 |
| Cstb | 362.581673 | 1.071901656 | 0.202250747 | 5.299865007 |
| Mir6410 | 0 | NA | NA | NA |
| Pdxk | 2399.16992 | 0.053781685 | 0.081714382 | 0.658166705 |
| Mir6908 | 0.341053521 | 1.121014874 | 1.721345712 | 0.651243307 |
| Gm9978 | 0.176220594 | -0.20648024 | 2.719488399 | -0.075926134 |
| Gm25164 | 0 | NA | NA | NA |
| Gm27541 | 0.044191549 | -0.206490061 | 3.724945049 | -0.055434391 |
| Ilvbl | 215.6243573 | 0.018951368 | 0.120366479 | 0.157447225 |
| Syde1 | 248.2945122 | 0.977901421 | 0.162504951 | 6.017671544 |
| Gm37514 | 0.458997541 | 1.155101577 | 1.528617248 | 0.755651278 |
| Olfr1357 | 0 | NA | NA | NA |
| Casp14 | 0.09176491 | 1.15872478 | 3.72306111 | 0.311229052 |
| Ccdc105 | 3.747387982 | 0.400511677 | 0.509749628 | 0.785702734 |
| Slc1a6 | 295.8032966 | 0.37363035 | 0.167162523 | 2.235132276 |
| Olfr1356 | 0 | NA | NA | NA |
| Olfr1355 | 0.089814205 | 0.274402725 | 3.724945049 | 0.073666248 |
| Olfr1354 | 0.11147366 | 0.274389235 | 3.724945049 | 0.073662626 |
| Olfr8 | 0 | NA | NA | NA |
| Olfr1353 | 0.097139547 | 0.274393828 | 3.724945049 | 0.073663859 |
| Olfr1352 | 0 | NA | NA | NA |
| Olfr1351 | 0 | NA | NA | NA |
| Olfr57 | 0.139856492 | -0.206491248 | 3.574411025 | -0.057769307 |
| 2610008E11Rik | 487.1302078 | -0.238439508 | 0.07843058 | -3.040134465 |
| Vmn2r80 | 0 | NA | NA | NA |
| Vmn2r81 | 0 | NA | NA | NA |
| Vmn2r82 | 0.049553047 | 0.274391604 | 3.724945049 | 0.073663262 |
| Vmn2r83 | 0 | NA | NA | NA |
| Ppap2c | 40.71318606 | -0.304564144 | 0.233380508 | -1.30501106 |
| Mir6909 | 0.048949532 | -0.206487837 | 3.724945049 | -0.055433794 |
| Mier2 | 291.2295781 | -0.135739684 | 0.091748361 | -1.479478029 |
| Theg | 1.989322217 | 0.197988123 | 0.864241145 | 0.22908898 |
| C2cd4c | 123.155375 | -0.372777315 | 0.167181191 | -2.229780225 |
| Shc2 | 358.7182919 | -0.69032236 | 0.162533998 | -4.247248999 |

| | | | | |
|---|---|---|---|---|
| Odf3l2 | 1.872511713 | 0.157596309 | 0.838967104 | 0.187845636 |
| Madcam1 | 2.384226188 | 3.518290508 | 1.101889941 | 3.192960003 |
| Tpgs1 | 315.7362571 | 0.216476661 | 0.099165354 | 2.182986814 |
| Cdc34 | 422.4012987 | 0.411400133 | 0.101175073 | 4.066220281 |
| Gzmm | 21.34813532 | 0.251890845 | 0.339918794 | 0.741032415 |
| Gm25794 | 0 | NA | NA | NA |
| Bsg | 2599.444032 | 0.02254964 | 0.082028688 | 0.274899428 |
| Hcn2 | 1794.794068 | -0.147639824 | 0.094623339 | -1.560289728 |
| Polrmt | 300.0243024 | -0.18384554 | 0.106234808 | -1.730558397 |
| Fgf22 | 7.710007984 | -0.48476923 | 0.340536311 | -1.423546371 |
| Rnf126 | 503.0038443 | 0.110160073 | 0.075520724 | 1.458673422 |
| Fstl3 | 34.70897125 | 1.043052863 | 0.345050124 | 3.02290244 |
| Prss57 | 5.311167766 | -0.146125918 | 0.454881526 | -0.32123951 |
| Palm | 1414.659549 | -0.158992549 | 0.081606171 | -1.948290758 |
| Misp | 27.35574496 | 2.010364366 | 0.35295758 | 5.695767649 |
| E130317F20Rik | 72.27215724 | 0.245417058 | 0.144978464 | 1.692782847 |
| Ptbp1 | 656.5387669 | 0.894828202 | 0.120652748 | 7.416558835 |
| Lppr3 | 374.0295548 | -0.55756411 | 0.110385555 | -5.051060444 |
| Mir6910 | 0 | NA | NA | NA |
| Gm17134 | 0.135598227 | -1.141928459 | 2.942835517 | -0.388036794 |
| Prtn3 | 7.905640509 | -0.761084938 | 0.458889475 | -1.658536485 |
| Elane | 0.826815689 | 0.554374486 | 1.274073077 | 0.43511985 |
| Cfd | 0.08486385 | 0.755286617 | 3.724945049 | 0.202764499 |
| Med16 | 415.3832138 | 0.121088264 | 0.068627494 | 1.764427889 |
| Gm24019 | 2.36422992 | -1.058515333 | 0.849084871 | -1.246654333 |
| R3hdm4 | 1784.00259 | -0.016710931 | 0.079551764 | -0.210063619 |
| Gm26602 | 0.222471884 | -0.233884822 | 2.58315595 | -0.090542277 |
| Kiss1r | 45.63901687 | -0.326650709 | 0.161967654 | -2.016765083 |
| Arid3a | 140.4446512 | -0.034732631 | 0.128085207 | -0.271168165 |
| Wdr18 | 517.0323779 | 0.071336312 | 0.068685793 | 1.038589039 |
| Grin3b | 6.451345326 | -1.430866054 | 0.435259807 | -3.287383833 |
| Tmem259 | 829.796043 | -0.001992537 | 0.064371129 | -0.030953889 |
| Gm26243 | 1.926543226 | -1.730687338 | 0.946042746 | -1.829396552 |
| Cnn2 | 130.197291 | 1.623320204 | 0.191045834 | 8.49701966 |
| Abca7 | 370.3957148 | -0.553146612 | 0.098850887 | -5.595767805 |
| Hmha1 | 218.2921494 | 2.235886724 | 0.16397173 | 13.63580617 |
| Polr2e | 305.7771488 | -0.067726056 | 0.076176116 | -0.889072055 |
| Gpx4 | 1189.407151 | 0.143533046 | 0.116897647 | 1.227852309 |
| Sbno2 | 850.3215173 | 2.260158153 | 0.238529564 | 9.475379557 |
| Stk11 | 787.6366792 | 0.266363343 | 0.068404164 | 3.893963881 |
| Dos | 2722.989964 | -0.231337467 | 0.09842428 | -2.350410544 |
| Atp5d | 1392.897457 | -0.135684552 | 0.07966857 | -1.703112681 |
| Midn | 1117.501778 | 0.020186116 | 0.128353432 | 0.15726978 |
| Cirbp | 669.8346141 | 0.862370305 | 0.136349454 | 6.324706719 |
| 1600002K03Rik | 19.50566266 | -0.059832699 | 0.274889045 | -0.217661272 |
| Efna2 | 60.39084732 | -0.809857973 | 0.164878249 | -4.911854522 |
| Mum1 | 291.2372399 | -0.192562382 | 0.078708308 | -2.446531859 |

| | | | | |
|---|---|---|---|---|
| Ndufs7 | 538.4769027 | 0.012448169 | 0.07624457 | 0.1632663 |
| Gamt | 97.19118593 | -0.614514892 | 0.133763266 | -4.594048217 |
| Dazap1 | 600.5946169 | 0.107723666 | 0.092846708 | 1.160231399 |
| Mir6911 | 0.644009716 | -1.129473081 | 1.220503982 | -0.925415318 |
| Gm15122 | 0 | NA | NA | NA |
| Rps15 | 2187.489196 | 0.331846844 | 0.109666693 | 3.025958334 |
| Apc2 | 2618.790862 | -0.157854763 | 0.049570796 | -3.184430674 |
| 2310011J03Rik | 240.725416 | 0.357206989 | 0.098737815 | 3.617732362 |
| Pcsk4 | 31.63242481 | -0.081577449 | 0.240910104 | -0.33862195 |
| Reep6 | 48.59301236 | -0.477922902 | 0.149875506 | -3.188799256 |
| Adamtsl5 | 57.94529312 | -0.379883606 | 0.192891343 | -1.969417597 |
| Plk5 | 149.4888237 | -0.933249633 | 0.21657247 | -4.309179422 |
| Gm15123 | 0 | NA | NA | NA |
| Gm15124 | 0.780980462 | -1.600221004 | 1.139296041 | -1.404569968 |
| Mex3d | 199.4244648 | -0.212285929 | 0.109157042 | -1.944775387 |
| Gm22721 | 0 | NA | NA | NA |
| Mbd3 | 699.1099546 | 0.152549437 | 0.083406003 | 1.828998285 |
| Uqcr11 | 557.7849959 | -0.268836238 | 0.124781174 | -2.154461526 |
| Tcf3 | 302.852696 | -0.006486682 | 0.109124788 | -0.059442792 |
| Gm25044 | 0.104637558 | -0.206492286 | 3.724945049 | -0.055434988 |
| Onecut3 | 5.071639938 | -1.254355614 | 0.497196157 | -2.522858625 |
| 2610034O05Rik | 0.287784626 | -0.966019095 | 1.760443753 | -0.548736131 |
| Atp8b3 | 7.756145753 | -0.461815423 | 0.379309011 | -1.217517669 |
| Rexo1 | 665.8855935 | 0.200017284 | 0.094348982 | 2.119972889 |
| Klf16 | 435.0080941 | -0.404453157 | 0.121998548 | -3.315229284 |
| Gm26710 | 2.145254238 | -0.445967739 | 0.640671966 | -0.696093731 |
| Abhd17a | 822.7097596 | -0.031075382 | 0.079424617 | -0.391256303 |
| Gm29093 | 14.5563475 | -0.573281568 | 0.346826805 | -1.652933279 |
| Adat3 | 37.40970625 | -0.421064313 | 0.279167298 | -1.508286667 |
| Scamp4 | 2.483253496 | -0.52256218 | 0.765498444 | -0.682643034 |
| Mir6912 | 0.975114657 | -1.597555283 | 1.092831594 | -1.461849466 |
| Csnk1g2 | 1099.862112 | 0.001055221 | 0.056794194 | 0.018579727 |
| Btbd2 | 650.9398297 | -0.080033949 | 0.088354662 | -0.905825986 |
| Mknk2 | 631.5413471 | 0.31669566 | 0.100635578 | 3.146955255 |
| Gm26541 | 0.467763849 | -0.727468816 | 1.51666387 | -0.479650653 |
| Mob3a | 177.9043315 | 0.320503008 | 0.170460322 | 1.880220595 |
| Izumo4 | 105.0430274 | -0.294904404 | 0.180177635 | -1.636742567 |
| Ap3d1 | 2190.778951 | -0.006223027 | 0.050800725 | -0.122498778 |
| Dot1l | 1006.566257 | -0.304399209 | 0.154143364 | -1.97477984 |
| Gm17151 | 0.089089296 | 1.194485492 | 3.723939576 | 0.320758559 |
| Plekhj1 | 280.4915415 | 0.058773126 | 0.08740214 | 0.672444931 |
| Sf3a2 | 462.5313837 | 0.331301731 | 0.109215715 | 3.03346208 |
| Amh | 11.52291177 | -0.609126543 | 0.4065711 | -1.498204233 |
| Jsrp1 | 14.91585505 | 0.043057914 | 0.451019284 | 0.095468011 |
| Oaz1 | 1026.280487 | -0.042531199 | 0.106446133 | -0.399556077 |
| Mir1982 | 0 | NA | NA | NA |
| Lingo3 | 479.2193401 | 0.021944845 | 0.12375828 | 0.177320217 |

| | | | | |
|---|---|---|---|---|
| Lsm7 | 91.89504733 | 0.127750186 | 0.142802551 | 0.894593167 |
| Sppl2b | 350.2345351 | -0.031104284 | 0.076489408 | -0.406648252 |
| Tmprss9 | 6.250140971 | -0.730634727 | 0.445170322 | -1.641247609 |
| Timm13 | 280.1181752 | -0.035805837 | 0.112228164 | -0.319045018 |
| Lmnb2 | 221.9066422 | -0.565139859 | 0.097515034 | -5.795412618 |
| Gadd45b | 364.2798403 | 1.989480509 | 0.312604518 | 6.364209073 |
| Gng7 | 1229.082418 | -0.550274011 | 0.240730099 | -2.28585463 |
| Gm3828 | 3.291353755 | -1.308298293 | 0.775783071 | -1.686422845 |
| Diras1 | 1155.444641 | -0.311901968 | 0.067358412 | -4.630482815 |
| Gm16099 | 2.516192357 | -0.383179951 | 0.588683147 | -0.650910346 |
| Slc39a3 | 692.1380104 | -0.121844294 | 0.073668881 | -1.653945221 |
| Sgta | 1016.513989 | 0.103601885 | 0.055442008 | 1.868653203 |
| Thop1 | 406.0563204 | -0.164629442 | 0.066010006 | -2.494007378 |
| Creb3l3 | 2.644382858 | 0.064004226 | 0.663399549 | 0.096479152 |
| mmu-mir-5615-1 | 0.100902008 | 0.274391828 | 3.724945049 | 0.073663322 |
| Mir5615-2 | 0 | NA | NA | NA |
| Map2k2 | 675.7442005 | 0.080786541 | 0.087364722 | 0.924704372 |
| Zbtb7a | 2167.929022 | -0.058979089 | 0.070204858 | -0.840099821 |
| Pias4 | 256.0514789 | -0.024253491 | 0.101850409 | -0.238128561 |
| Eef2 | 12379.31089 | 0.125219276 | 0.074267249 | 1.686063211 |
| Snord37 | 3.244100523 | -1.404540335 | 0.624984797 | -2.247319202 |
| 4930442H23Rik | 0.124853745 | 1.290908285 | 3.721447108 | 0.346883416 |
| Dapk3 | 292.8022491 | 0.033972202 | 0.092477777 | 0.367355309 |
| Atcayos | 2.229235169 | -0.69985456 | 0.873489437 | -0.80121697 |
| Nmrk2 | 2.519283505 | 0.45999921 | 0.806618393 | 0.570281082 |
| Atcay | 1736.526661 | -0.382886089 | 0.073833157 | -5.185828519 |
| Zfr2 | 517.1457258 | -0.407180828 | 0.137871282 | -2.953340418 |
| Matk | 696.4187115 | -0.299431133 | 0.147156982 | -2.034773541 |
| Mrpl54 | 179.3062319 | 0.35083182 | 0.12299132 | 2.852492514 |
| Apba3 | 193.0437716 | -0.325478435 | 0.135289805 | -2.405786848 |
| Mir3057 | 0.237113133 | -0.206481327 | 1.866923053 | -0.110599806 |
| Tjp3 | 12.76057694 | 0.531615561 | 0.306207128 | 1.736130585 |
| Pip5k1c | 3939.793252 | -0.141849337 | 0.090115919 | -1.57407636 |
| Gm16315 | 2.415294132 | 0.212294742 | 0.695275531 | 0.305339009 |
| Cactin | 253.0737681 | 0.193389239 | 0.10534981 | 1.835686654 |
| Tbxa2r | 4.861979082 | -0.832460057 | 0.552410456 | -1.506959269 |
| Gipc3 | 3.688286577 | 0.402465131 | 0.627006085 | 0.641883932 |
| Hmg20b | 157.6554839 | 0.027779685 | 0.10700058 | 0.259621822 |
| Mfsd12 | 595.8008876 | 0.142305118 | 0.111992039 | 1.270671735 |
| 4930404N11Rik | 5.09738268 | 0.846237121 | 0.619693593 | 1.365573456 |
| Fzr1 | 541.0904011 | -0.142271772 | 0.059806327 | -2.37887494 |
| Dohh | 443.5414654 | 0.091535943 | 0.075720484 | 1.20886633 |
| Mir6913 | 0.75335248 | -0.068015762 | 1.152782026 | -0.059001407 |
| Smim24 | 5.77545002 | -0.689840081 | 0.514494969 | -1.340810158 |
| Nfic | 2277.670415 | -0.007079303 | 0.064822254 | -0.109210998 |
| Gm16104 | 0.970068437 | -0.173441196 | 1.187269863 | -0.146084055 |
| mmu-mir-1191b | 3.498764646 | -0.0276792 | 0.639171965 | -0.043304777 |

| | | | | |
|---|---|---|---|---|
| Celf5 | 1725.007547 | -0.396162652 | 0.172347683 | -2.298624763 |
| Gm16105 | 0.342577373 | 1.29974812 | 1.753409315 | 0.741269086 |
| Ncln | 601.4543055 | 0.293795651 | 0.069417324 | 4.232310217 |
| S1pr4 | 11.27433504 | 2.505226137 | 0.457739028 | 5.473044643 |
| Gna15 | 25.30164811 | 1.388390603 | 0.317535341 | 4.372397096 |
| Gm16106 | 0.086908494 | -0.687376175 | 3.724945049 | -0.184533239 |
| Gm25595 | 0 | NA | NA | NA |
| Gna11 | 1114.577472 | 0.02864549 | 0.063913103 | 0.448194328 |
| Aes | 3534.40744 | 0.030731554 | 0.073419106 | 0.418577058 |
| Gm15917 | 0.222899254 | -1.180585499 | 2.149316567 | -0.549284139 |
| Tle2 | 226.3530908 | -0.593209895 | 0.217092959 | -2.732515588 |
| Tle6 | 20.29999438 | -0.284706451 | 0.227014235 | -1.254134794 |
| BC025920 | 34.61278247 | -0.222514218 | 0.206760438 | -1.076193396 |
| Gm26896 | 0.097501745 | -0.206492286 | 3.724945049 | -0.055434988 |
| Sirt6 | 169.5015077 | -0.022287072 | 0.095221253 | -0.234055648 |
| Ankrd24 | 225.4576987 | -0.470955974 | 0.164955866 | -2.855042297 |
| Gm10778 | 8.434511479 | -0.423867762 | 0.387735286 | -1.093188518 |
| Gm8112 | 0.038981395 | 0.274393828 | 3.724945049 | 0.073663859 |
| Zfp433 | 19.00695932 | 0.101082533 | 0.227001586 | 0.445294389 |
| Gm3055 | 1.030732002 | -1.14780218 | 1.088119397 | -1.054849479 |
| Gm4767 | 1.089198015 | -1.567105838 | 1.195572358 | -1.310757838 |
| Zfp781 | 1.228365715 | -0.708385395 | 0.991537544 | -0.714431238 |
| Zfp873 | 74.79504174 | -0.436883784 | 0.162043171 | -2.696095002 |
| AU041133 | 155.8068101 | -0.180856757 | 0.110260736 | -1.640264376 |
| Gm15608 | 10.40952602 | -0.891230199 | 0.388977012 | -2.291215606 |
| Zfp938 | 255.4037447 | -0.024261739 | 0.103165606 | -0.235172745 |
| 4932415D10Rik | 0.046345709 | 0.274391604 | 3.724945049 | 0.073663262 |
| Gm4924 | 47.95002397 | -0.206460578 | 0.172484113 | -1.196983159 |
| Gm1553 | 0 | NA | NA | NA |
| Ap3m1-ps | 10.60277313 | 0.761261246 | 0.344728208 | 2.208294037 |
| 1190007I07Rik | 27.31729701 | -0.361788977 | 0.291983216 | -1.239074568 |
| Tdg | 118.6799162 | 0.075013302 | 0.147613256 | 0.508174563 |
| Glt8d2 | 72.76759065 | -0.03767083 | 0.276249241 | -0.136365371 |
| Hcfc2 | 483.7093496 | -0.39612508 | 0.070151888 | -5.646677388 |
| Nfyb | 378.4988816 | -0.071168171 | 0.074497889 | -0.955304531 |
| 1700028I16Rik | 0.19184735 | 0.755296431 | 2.671497329 | 0.282724008 |
| Txnrd1 | 1189.720619 | 0.350276683 | 0.133728723 | 2.619307774 |
| Gm4799 | 4.084400745 | 0.614217521 | 0.634661145 | 0.967788127 |
| Chst11 | 1110.164625 | 0.099874546 | 0.107225236 | 0.931446273 |
| 1700025N21Rik | 0.075043205 | 0.755286614 | 3.724945049 | 0.202764498 |
| Gm23122 | 0 | NA | NA | NA |
| Slc41a2 | 340.536907 | -0.262160781 | 0.064821094 | -4.044374516 |
| D10Wsu102e | 409.3325013 | -0.193626927 | 0.079122739 | -2.447171698 |
| Aldh1l2 | 203.3414108 | 0.20583312 | 0.134078295 | 1.535171075 |
| A230046K03Rik | 1351.145369 | 0.630313303 | 0.079646088 | 7.913926708 |
| Appl2 | 956.0063235 | -0.837720624 | 0.183176281 | -4.573302935 |
| 1500009L16Rik | 86.40011601 | -0.164053416 | 0.13042535 | -1.257833813 |

| | | | | |
|---|---|---|---|---|
| Gm37908 | 0 | NA | NA | NA |
| Nuak1 | 1233.243768 | -0.011248989 | 0.105933252 | -0.1061894 |
| 4930463O16Rik | 0.038981395 | 0.274393828 | 3.724945049 | 0.073663859 |
| Ckap4 | 392.0257556 | 0.362893631 | 0.117414611 | 3.090702491 |
| Platr7 | 0.182269687 | 1.236183493 | 3.313323614 | 0.373094704 |
| Tcp11l2 | 249.9597353 | -0.028177355 | 0.113899373 | -0.247388146 |
| Gm25657 | 0.602022574 | 0.390398727 | 1.392412294 | 0.280375811 |
| Gm17249 | 4.368992961 | -2.176838816 | 0.604907205 | -3.598632649 |
| Polr3b | 386.4218719 | 0.19116602 | 0.071821838 | 2.661669816 |
| Rfx4 | 218.129252 | -0.225751944 | 0.147825066 | -1.527156051 |
| Gm24226 | 0.075043205 | 0.755286614 | 3.724945049 | 0.202764498 |
| Ric8b | 669.9951763 | -0.227945603 | 0.051699789 | -4.409023865 |
| Fhl4 | 12.95936291 | -0.08460584 | 0.334488692 | -0.252940808 |
| Tmem263 | 569.4293162 | -0.223261007 | 0.081799327 | -2.729374614 |
| Mterf2 | 117.1077398 | -0.075595067 | 0.149917018 | -0.504246071 |
| Cry1 | 350.2670687 | 0.123306924 | 0.152307177 | 0.809593653 |
| Gm27392 | 0 | NA | NA | NA |
| Gm8394 | 52.17199378 | 0.64297936 | 0.206260269 | 3.117320483 |
| Gm25281 | 0 | NA | NA | NA |
| Btbd11 | 507.0576605 | 0.016073142 | 0.097994503 | 0.16402085 |
| Pwp1 | 247.3605164 | 0.203400099 | 0.091354246 | 2.226498567 |
| Prdm4 | 408.5061188 | 0.070751813 | 0.071975212 | 0.983002495 |
| Ascl4 | 2.415104612 | 0.228565868 | 0.59878818 | 0.381714062 |
| Rtcb | 1117.008586 | 0.34264489 | 0.082531299 | 4.15169633 |
| Bpifc | 3.577274519 | 1.313660363 | 0.632225456 | 2.077835288 |
| A230060F14Rik | 5.145365424 | 0.5910135 | 0.47926466 | 1.233167284 |
| Fbxo7 | 420.5312043 | 0.541108908 | 0.086399571 | 6.262865661 |
| Gm24430 | 0.537185458 | 0.285219418 | 1.504512353 | 0.189575989 |
| Syn3 | 338.3405326 | -0.336549258 | 0.168337935 | -1.999247866 |
| 4930486F22Rik | 0.066050875 | 0.274391604 | 3.724945049 | 0.073663262 |
| Gm15990 | 3.87042643 | 3.890791902 | 0.972860407 | 3.999332149 |
| Timp3 | 2834.917203 | 0.717900211 | 0.137587851 | 5.217758744 |
| Gm6729 | 0 | NA | NA | NA |
| Gm5174 | 1.40433107 | 1.489265097 | 0.849983457 | 1.752110684 |
| 1810014B01Rik | 16.79727865 | -0.155461024 | 0.263987196 | -0.588896076 |
| Hsp90b1 | 4072.803122 | 0.259914157 | 0.106247058 | 2.446318617 |
| Gm15344 | 13.86957958 | -0.835421223 | 0.286404132 | -2.916931462 |
| Gm25745 | 0 | NA | NA | NA |
| Ttc41 | 22.08483189 | -0.28624788 | 0.292924766 | -0.977206144 |
| Fabp3-ps1 | 8.356713702 | -0.585853928 | 0.422385172 | -1.387013481 |
| Nt5dc3 | 1819.324601 | 0.156022619 | 0.085288936 | 1.829341827 |
| Gm16268 | 9.292864951 | 0.221149604 | 0.349735427 | 0.632334008 |
| Stab2 | 7.172886002 | 0.644542552 | 0.445310428 | 1.447400537 |
| Gm16280 | 0 | NA | NA | NA |
| Gm16270 | 0.987182834 | 2.321336302 | 1.212769234 | 1.914079149 |
| Gm16271 | 0.570993603 | -0.758273626 | 1.484394989 | -0.51083009 |
| Gm16269 | 0 | NA | NA | NA |

| | | | | |
|---|---|---|---|---|
| 1700113H08Rik | 0.362813327 | -1.232274462 | 1.542316306 | -0.798976485 |
| Gm10764 | 0.093831153 | 0.274404949 | 3.724945049 | 0.073666845 |
| Gm23191 | 0 | NA | NA | NA |
| Ascl1 | 36.60879247 | -0.66952656 | 0.21245635 | -3.151360557 |
| Pah | 2.972707099 | -0.279259796 | 0.563257832 | -0.495793897 |
| Igf1os | 0.184770788 | 1.288632945 | 2.685921747 | 0.479773078 |
| Igf1 | 193.7112454 | -0.203436747 | 0.11552961 | -1.760905688 |
| Pmch | 109.1827471 | -0.136345696 | 0.164881662 | -0.826930624 |
| Parpbp | 16.45101383 | 0.634951917 | 0.326921185 | 1.942217101 |
| Nup37 | 44.56539038 | -0.087753534 | 0.201299033 | -0.435936194 |
| Gm26357 | 0 | NA | NA | NA |
| Ccdc53 | 129.2902162 | 0.207685953 | 0.142819396 | 1.454185904 |
| Dram1 | 46.36865287 | 1.252061172 | 0.264171306 | 4.73958051 |
| Gnptab | 1221.700717 | 0.046123255 | 0.054818999 | 0.841373528 |
| Gm26546 | 4.34695503 | -0.412411115 | 0.528448514 | -0.780418724 |
| Gm16235 | 0.918591783 | 0.162791881 | 1.104349627 | 0.14740973 |
| Chpt1 | 1036.030604 | -0.506690583 | 0.103160968 | -4.911650164 |
| Sycp3 | 7.536531091 | 0.593792713 | 0.436940601 | 1.35897811 |
| Mybpc1 | 139.6857782 | -0.478354304 | 0.227827911 | -2.099629943 |
| Spic | 2.940423805 | 2.916578447 | 0.98269879 | 2.967927178 |
| n-R5s79 | 0 | NA | NA | NA |
| Gm4925 | 6.36087602 | -0.23481062 | 0.387458111 | -0.606028401 |
| Arl1 | 1132.198676 | 0.258357095 | 0.083151393 | 3.107068742 |
| Utp20 | 565.1990458 | 0.031058699 | 0.073733975 | 0.421226431 |
| Slc5a8 | 3.475576094 | 2.200615892 | 0.669554301 | 3.286687707 |
| Ano4 | 411.0563822 | -0.255138822 | 0.101783468 | -2.506682344 |
| Gas2l3 | 106.5758252 | 0.632089554 | 0.254114531 | 2.487419947 |
| Nr1h4 | 10.81026118 | 2.33109105 | 0.58914903 | 3.956708625 |
| Slc17a8 | 21.16662692 | -0.175253623 | 0.320744154 | -0.546396935 |
| Scyl2 | 725.9700778 | 0.062266208 | 0.077527574 | 0.803149193 |
| 1500026H17Rik | 20.55184199 | -1.178987407 | 0.231210196 | -5.099201628 |
| Actr6 | 166.1310719 | -0.059633304 | 0.113262612 | -0.526504753 |
| Uhrf1bp1l | 1814.744137 | 0.106534417 | 0.065879457 | 1.617111351 |
| Gm26180 | 0.045882455 | 0.755288839 | 3.724945049 | 0.202765096 |
| Anks1b | 4400.438249 | -0.121779098 | 0.093328113 | -1.304849028 |
| Gm22602 | 2.708289425 | -0.392104731 | 0.64321566 | -0.609600723 |
| Apaf1 | 344.5692301 | 0.447034691 | 0.116516476 | 3.836665059 |
| Gm25390 | 0 | NA | NA | NA |
| Ikbip | 137.6747652 | -0.355478986 | 0.137625251 | -2.582948876 |
| Slc25a3 | 3200.487187 | -0.056589433 | 0.055853808 | -1.01317054 |
| Gm24119 | 26.8554828 | -0.574578693 | 0.296124623 | -1.940327309 |
| Tmpo | 631.4065621 | 0.251656172 | 0.087464188 | 2.877248129 |
| Gm26765 | 0.307480012 | -1.07248666 | 2.006651716 | -0.534465773 |
| Rmst | 270.1421029 | -0.094774838 | 0.119770861 | -0.7913013 |
| Gm27544 | 10.08854525 | -0.357718552 | 0.319608186 | -1.119240894 |
| Mir135a-2 | 4.853900475 | -0.162455523 | 0.45851194 | -0.354310344 |
| Gm27999 | 0 | NA | NA | NA |

| | | | | |
|---|---|---|---|---|
| Gm27608 | 0 | NA | NA | NA |
| Gm27588 | 11.46356049 | 0.197197529 | 0.321184601 | 0.613969438 |
| Gm27753 | 0 | NA | NA | NA |
| Gm27582 | 0.044191549 | -0.206490061 | 3.724945049 | -0.055434391 |
| Gm27366 | 0.132101749 | 0.274387056 | 3.724945049 | 0.073662041 |
| Gm27419 | 0 | NA | NA | NA |
| Mir1251 | 5.125747786 | 0.312726782 | 0.415096237 | 0.753383804 |
| Gm26700 | 0.516149922 | 0.71617335 | 1.410081913 | 0.507894856 |
| Gm20757 | 0.625729796 | -0.08999801 | 1.16853943 | -0.077017521 |
| Gm25884 | 0 | NA | NA | NA |
| Gm22582 | 0 | NA | NA | NA |
| Nedd1 | 87.64473989 | 0.246982763 | 0.178330213 | 1.38497431 |
| Gm24241 | 0.242925194 | -1.516303534 | 2.100181093 | -0.721987042 |
| Cfap54 | 546.4457364 | -0.491793007 | 0.179845837 | -2.734525382 |
| Cdk17 | 2255.31025 | -0.049908915 | 0.068995412 | -0.723365706 |
| Mir1931 | 9.166638643 | -0.884005656 | 0.374881539 | -2.35809333 |
| Elk3 | 176.0696239 | 1.347444281 | 0.193004372 | 6.981418444 |
| Mir7688 | 0.168606834 | -1.06717182 | 2.775232772 | -0.384534166 |
| Lta4h | 347.4203626 | -0.045543441 | 0.084844503 | -0.536787168 |
| Hal | 122.7623701 | 3.745005466 | 0.590256605 | 6.344707432 |
| Amdhd1 | 3.584129096 | 1.726781361 | 0.838726956 | 2.058812287 |
| Ccdc38 | 1.217260431 | 0.021964646 | 0.911071999 | 0.024108573 |
| Snrpf | 77.19710455 | 0.100008251 | 0.180373849 | 0.554449839 |
| 4933408J17Rik | 0.132790949 | -0.206481859 | 2.970952992 | -0.069500211 |
| Ntn4 | 142.6461492 | -0.53360446 | 0.15792542 | -3.37883832 |
| Gm8580 | 0.167271369 | 1.236193323 | 2.764694848 | 0.44713554 |
| Gm15915 | 0.796817125 | 1.688007909 | 1.166344701 | 1.447263324 |
| Gm15963 | 1.049255278 | -0.313548633 | 0.947361961 | -0.330970258 |
| Gm3571 | 24.10692972 | -0.446038885 | 0.236202466 | -1.888375226 |
| Gm26239 | 0 | NA | NA | NA |
| Usp44 | 0.703834089 | 0.618683381 | 1.12575984 | 0.549569596 |
| Metap2 | 993.8475694 | 0.212434113 | 0.097707599 | 2.174182109 |
| 4930471D02Rik | 0.044191549 | -0.206490061 | 3.724945049 | -0.055434391 |
| Gm25962 | 0 | NA | NA | NA |
| Vezt | 1354.242168 | -0.350729836 | 0.121184464 | -2.894181529 |
| Mir331 | 0 | NA | NA | NA |
| Fgd6 | 264.4343943 | -0.500998346 | 0.11792525 | -4.248439977 |
| Nr2c1 | 165.6055157 | -0.072164441 | 0.11503519 | -0.62732492 |
| Gm26122 | 0 | NA | NA | NA |
| Ndufa12 | 388.6068376 | 0.109855939 | 0.093003613 | 1.181200761 |
| Gm4792 | 0.047948698 | -0.206487837 | 3.724945049 | -0.055433794 |
| Tmcc3 | 859.2376155 | 0.399203799 | 0.115301942 | 3.462246965 |
| Gm16155 | 3.245903696 | -0.176593351 | 0.656498663 | -0.268992705 |
| Tmcc3os | 2.419038749 | 1.875801144 | 0.689400571 | 2.720916147 |
| Mir7211 | 0 | NA | NA | NA |
| Cep83os | 296.969172 | 0.148185652 | 0.082608384 | 1.793833084 |
| Cep83 | 388.069701 | -0.158262782 | 0.085413715 | -1.852896607 |

| | | | | |
|---|---|---|---|---|
| Plxnc1 | 869.6228163 | -0.083209867 | 0.079606403 | -1.045266013 |
| Gm24186 | 1.18741419 | 0.650697415 | 0.87097165 | 0.74709368 |
| Gm25266 | 0.715554978 | 0.148474513 | 1.236967029 | 0.1200311 |
| Cradd | 45.57421372 | 0.419287291 | 0.166801662 | 2.513687727 |
| 2310039L15Rik | 0 | NA | NA | NA |
| Socs2 | 273.8451504 | 0.072402712 | 0.129248199 | 0.560183524 |
| 5730420D15Rik | 1.317555436 | 0.519969552 | 0.88270253 | 0.589065437 |
| Mrpl42 | 211.5063124 | -0.120378339 | 0.113182723 | -1.063575213 |
| Ube2n | 1238.439884 | -0.07496086 | 0.056335134 | -1.330623625 |
| Nudt4 | 1713.099017 | -0.068145912 | 0.119240562 | -0.571499418 |
| Mir3058 | 2.797676517 | -0.133700768 | 0.554244923 | -0.241230479 |
| Gm23776 | 0 | NA | NA | NA |
| Anapc15-ps | 1.031766027 | 1.453537939 | 1.020481113 | 1.424365351 |
| 4732465J04Rik | 0.435690537 | -0.174232469 | 1.407000937 | -0.123832518 |
| Gm24433 | 0 | NA | NA | NA |
| Eea1 | 1239.903993 | 0.016919007 | 0.060810587 | 0.278224702 |
| Gm5426 | 0 | NA | NA | NA |
| Gm22913 | 0 | NA | NA | NA |
| Gm20091 | 0.131209559 | 0.755295049 | 2.976009072 | 0.253794606 |
| Gm8601 | 0 | NA | NA | NA |
| Btg1 | 1120.140172 | 0.513226673 | 0.126988613 | 4.041517267 |
| Gm24587 | 0 | NA | NA | NA |
| mmu-mir-3966 | 0 | NA | NA | NA |
| Dcn | 434.2514662 | 0.932422547 | 0.176948312 | 5.269462808 |
| Lum | 32.29644331 | -0.061477829 | 0.252860577 | -0.243129355 |
| Kera | 0.075043205 | 0.755286614 | 3.724945049 | 0.202764498 |
| Epyc | 1.611010591 | 1.796271752 | 0.922380864 | 1.947429552 |
| Gm10754 | 7.111199995 | -0.631183426 | 0.378961623 | -1.665560278 |
| Ccer1 | 0 | NA | NA | NA |
| C030005K15Rik | 6.350231153 | -0.671240322 | 0.505912868 | -1.32679037 |
| Gm25658 | 1.646274761 | -2.135959148 | 0.831447084 | -2.568965829 |
| Gm37631 | 0.066050875 | 0.274391604 | 3.724945049 | 0.073663262 |
| Atp2b1 | 8738.965953 | -0.010774342 | 0.115361667 | -0.093396205 |
| Poc1b | 99.61505393 | -0.040679127 | 0.122489797 | -0.332102167 |
| Galnt4 | 92.75288412 | 0.227793811 | 0.152726495 | 1.491514689 |
| Phxr2 | 0 | NA | NA | NA |
| Gm16239 | 0.066050875 | 0.274391604 | 3.724945049 | 0.073663262 |
| Dusp6 | 640.3874158 | 0.137646151 | 0.18815794 | 0.731545804 |
| B530045E10Rik | 1.636950402 | 1.173863201 | 0.852024103 | 1.377734735 |
| Gad1-ps | 4.900567814 | -0.381129526 | 0.574902158 | -0.662946765 |
| Csl | 0.893710704 | -1.963423605 | 1.133421793 | -1.732297382 |
| Kitl | 948.0407053 | -0.025978924 | 0.13139315 | -0.197719017 |
| Gm22918 | 0 | NA | NA | NA |
| Gm25287 | 0 | NA | NA | NA |
| Gm9476 | 0 | NA | NA | NA |
| Gm8701 | 0 | NA | NA | NA |
| Gm4301 | 0 | NA | NA | NA |

| | | | | |
|---|---|---|---|---|
| Gm4302 | 0 | NA | NA | NA |
| Gm4303 | 0 | NA | NA | NA |
| Gm4305 | 0 | NA | NA | NA |
| Gm8711 | 0 | NA | NA | NA |
| Gm4307 | 0 | NA | NA | NA |
| Gm4308 | 0 | NA | NA | NA |
| Gm8723 | 0 | NA | NA | NA |
| Gm4312 | 0 | NA | NA | NA |
| Gm4781 | 0 | NA | NA | NA |
| Tmtc3 | 401.2372202 | -0.316658069 | 0.076596331 | -4.134115361 |
| Cep290 | 566.5526866 | -0.151775581 | 0.096798078 | -1.567960686 |
| 4930430F08Rik | 178.6427657 | -0.312613463 | 0.154728652 | -2.020398026 |
| 1700017N19Rik | 0.256989293 | 1.083693938 | 2.09242135 | 0.517913821 |
| Gm38219 | 0 | NA | NA | NA |
| Gm26278 | 0.136194719 | -0.687371727 | 3.724945049 | -0.184532045 |
| Mgat4c | 312.1695497 | -0.207734788 | 0.092488866 | -2.246051835 |
| Mir3059 | 1.49611593 | -0.663824116 | 0.997795198 | -0.665290951 |
| Gm26988 | 0.108767819 | -0.20649451 | 3.724945049 | -0.055435586 |
| Nts | 88.60440578 | 0.689967751 | 0.239581571 | 2.879886576 |
| Rassf9 | 16.58629026 | 0.436318971 | 0.306898362 | 1.421705115 |
| Alx1 | 1.374376221 | -0.367964076 | 0.852351017 | -0.431704859 |
| Gm17028 | 0 | NA | NA | NA |
| Lrriq1 | 58.42473187 | -0.305686362 | 0.180878944 | -1.690005231 |
| Gm25376 | 0 | NA | NA | NA |
| Slc6a15 | 616.6613696 | 0.209594778 | 0.084138507 | 2.491068438 |
| Gm26923 | 0 | NA | NA | NA |
| Gm25143 | 0 | NA | NA | NA |
| Gm6763 | 0 | NA | NA | NA |
| Gm21293 | 0 | NA | NA | NA |
| Gm8764 | 0 | NA | NA | NA |
| Gm21304 | 0 | NA | NA | NA |
| Gm21312 | 0 | NA | NA | NA |
| Gm20765 | 0 | NA | NA | NA |
| Gm4340 | 0 | NA | NA | NA |
| Gm22339 | 0 | NA | NA | NA |
| Mir6411 | 0 | NA | NA | NA |
| Gm22945 | 0 | NA | NA | NA |
| Gm25522 | 0 | NA | NA | NA |
| Gm25587 | 0 | NA | NA | NA |
| Gm25422 | 0.254945911 | 1.540610849 | 2.060262525 | 0.747774048 |
| Gm36997 | 0.361186056 | 0.274416205 | 1.576738903 | 0.174040359 |
| Tmtc2 | 369.308983 | -0.26767715 | 0.102724137 | -2.605786299 |
| Gm15662 | 0 | NA | NA | NA |
| Gm15666 | 12.30954061 | -0.572273986 | 0.391684176 | -1.461059757 |
| Gm15664 | 0 | NA | NA | NA |
| Gm15663 | 93.64214882 | -0.545168181 | 0.132642963 | -4.110042245 |
| Mettl25 | 107.6127449 | 0.151167163 | 0.141291849 | 1.069893019 |

| | | | | |
|---|---|---|---|---|
| n-R5s80 | 1.30631546 | 2.560731698 | 1.079904586 | 2.371257359 |
| Ccdc59 | 282.5902914 | 0.354381459 | 0.113500174 | 3.122298825 |
| Gm24237 | 0 | NA | NA | NA |
| Gm25342 | 0 | NA | NA | NA |
| Rps15a-ps1 | 0 | NA | NA | NA |
| C230021G24Rik | 14.38138754 | -0.730076179 | 0.298814388 | -2.44324306 |
| Ppfia2 | 1751.263262 | -0.101890336 | 0.099197655 | -1.027144601 |
| Acss3 | 21.10431072 | -0.035269743 | 0.260686742 | -0.135295499 |
| Gm24745 | 0.047948698 | -0.206487837 | 3.724945049 | -0.055433794 |
| Lin7a | 1127.089259 | -0.301223325 | 0.105197141 | -2.863417405 |
| Gm38117 | 0 | NA | NA | NA |
| AC155168.1 | 0.358057361 | 0.96476586 | 3.690308502 | 0.261432306 |
| Myf5 | 0.154585317 | 0.274399357 | 2.835007735 | 0.096789632 |
| Myf6 | 0.075043205 | 0.755286614 | 3.724945049 | 0.202764498 |
| Ptprq | 5.567795988 | 0.07155686 | 0.422798582 | 0.169245743 |
| Otogl | 7.088242965 | 0.029584447 | 0.391306985 | 0.075604188 |
| Ppp1r12a | 1620.397248 | 0.131780186 | 0.076329941 | 1.726454706 |
| Pawr | 30.19438579 | 0.960450053 | 0.273620913 | 3.51014855 |
| Gm23105 | 0.203286941 | 0.755296304 | 2.621314208 | 0.288136501 |
| Gm25591 | 0 | NA | NA | NA |
| Syt1 | 12677.56377 | -0.062278114 | 0.113992656 | -0.546334443 |
| Gm22097 | 0.164132501 | 1.537077023 | 3.390982371 | 0.453283696 |
| AC125531.1 | 0.108123235 | 0.274402725 | 3.724945049 | 0.073666248 |
| Gm23041 | 0.2701572 | 0.288686489 | 1.813461493 | 0.159190857 |
| Nav3 | 1988.147118 | -0.05868598 | 0.115730701 | -0.507090852 |
| Gm36935 | 26.04041665 | -0.536670155 | 0.408957676 | -1.312287766 |
| E2f7 | 15.83057167 | 1.185378672 | 0.313683652 | 3.778898473 |
| 1700020G17Rik | 0 | NA | NA | NA |
| Gm23698 | 0 | NA | NA | NA |
| Gm22186 | 0 | NA | NA | NA |
| Gm22236 | 0 | NA | NA | NA |
| Csrp2 | 26.62848996 | 0.070764824 | 0.241676256 | 0.292808341 |
| Zdhhc17 | 1450.600249 | -0.084040098 | 0.05220192 | -1.60990434 |
| Gm5428 | 96.95467252 | 0.298420073 | 0.167816855 | 1.778248506 |
| Gm22101 | 0 | NA | NA | NA |
| Osbpl8 | 2307.979124 | -0.181838368 | 0.051196133 | -3.551798873 |
| Bbs10 | 50.75359481 | -0.24909365 | 0.162980123 | -1.528368275 |
| Nap1l1 | 1669.332245 | 0.05570024 | 0.071327954 | 0.780903367 |
| Gm25117 | 163.1343521 | 0.241456888 | 0.470298022 | 0.513412509 |
| Phlda1 | 275.8361962 | 0.176655196 | 0.182295695 | 0.969058515 |
| Gm28592 | 7.265044534 | 0.078053697 | 0.483591739 | 0.161404115 |
| 4933440J02Rik | 0 | NA | NA | NA |
| Gm37189 | 0.109727584 | 0.755282166 | 3.724945049 | 0.202763304 |
| Krr1 | 496.8011956 | 0.070633597 | 0.072440322 | 0.97505913 |
| Glipr1 | 31.42829398 | 2.520810019 | 0.325505468 | 7.74429392 |
| Glipr1l1 | 0.138226157 | -0.687378071 | 2.934480157 | -0.234241853 |
| Glipr1l2 | 0.144013665 | 0.274391019 | 3.543609481 | 0.077432635 |

| | | | | |
|---|---|---|---|---|
| Gm6808 | 0 | NA | NA | NA |
| Gm15914 | 0 | NA | NA | NA |
| Caps2 | 5.498926737 | -0.152903635 | 0.47476141 | -0.322064161 |
| Kcnc2 | 1574.614437 | 0.181640209 | 0.085103159 | 2.134353292 |
| 1700010J16Rik | 0.193455382 | 1.002139154 | 2.621157884 | 0.382326895 |
| Atxn7l3b | 2117.555783 | -0.045632932 | 0.049433346 | -0.92312044 |
| Gm26596 | 1.459047308 | -0.751896306 | 0.859437808 | -0.874869943 |
| Gm22507 | 0.128477747 | 0.755285235 | 3.677131942 | 0.205400635 |
| Gm38333 | 0 | NA | NA | NA |
| Gm38281 | 0 | NA | NA | NA |
| Trhde | 612.5170607 | -0.325791681 | 0.09168549 | -3.553361414 |
| 4930473D10Rik | 0 | NA | NA | NA |
| Gm15723 | 23.33627371 | -0.323195896 | 0.252110073 | -1.281963437 |
| Gm25319 | 0 | NA | NA | NA |
| Gm22070 | 0 | NA | NA | NA |
| Tph2 | 55.63792861 | -0.263280702 | 0.659529946 | -0.399194462 |
| Tbc1d15 | 653.7365103 | 0.298762553 | 0.097063347 | 3.078016173 |
| Gm22794 | 0.044544648 | 0.755286614 | 3.724945049 | 0.202764498 |
| Rab21 | 1138.336811 | 0.597077176 | 0.094886429 | 6.292545568 |
| Tmem19 | 316.2960843 | -0.091982737 | 0.077904553 | -1.180710668 |
| Thap2 | 234.4484823 | -0.91416671 | 0.239879412 | -3.810942765 |
| Zfc3h1 | 1151.166255 | -0.06434012 | 0.101516774 | -0.633788073 |
| Lgr5 | 129.3954019 | -1.073741587 | 0.277999193 | -3.86239102 |
| A930009A15Rik | 4.998511455 | -1.672455858 | 0.541592137 | -3.088035706 |
| Tspan8 | 5.527738138 | 0.492971126 | 0.473145976 | 1.041900705 |
| Ptprr | 373.9766246 | -0.21900892 | 0.103204897 | -2.122078761 |
| 4933416C03Rik | 0.062426873 | 0.755288839 | 3.724945049 | 0.202765096 |
| Ptprb | 234.5662353 | 0.178737288 | 0.198105441 | 0.902233111 |
| Gm26495 | 0.042716945 | -0.206490061 | 3.724945049 | -0.055434391 |
| Kcnmb4 | 239.9415874 | -0.061688659 | 0.1283091 | -0.480781639 |
| Kcnmb4os1 | 0.534709659 | -0.251230672 | 1.549640255 | -0.162121932 |
| Kcnmb4os2 | 19.7696678 | 0.021884592 | 0.229514774 | 0.09535156 |
| Cnot2 | 821.4440981 | -0.061857794 | 0.056241341 | -1.099863429 |
| Gm25190 | 6.442501395 | -0.137256564 | 0.390071753 | -0.351875168 |
| 5330438D12Rik | 61.66108682 | 0.056598857 | 0.164712561 | 0.343621986 |
| Gm26579 | 0.044191549 | -0.206490061 | 3.724945049 | -0.055434391 |
| 1700030O20Rik | 0.066050875 | 0.274391604 | 3.724945049 | 0.073663262 |
| Myrfl | 6.514367323 | 1.163241033 | 0.515330815 | 2.257270473 |
| Rab3ip | 355.5556215 | 0.210691703 | 0.082514304 | 2.553396117 |
| Gm10271 | 0 | NA | NA | NA |
| D630029K05Rik | 3.06831776 | 2.568557312 | 0.717407018 | 3.580334798 |
| Best3 | 10.8055888 | 1.015392922 | 0.328784608 | 3.088322556 |
| Lrrc10 | 0.159795471 | -0.20649052 | 2.810463622 | -0.073472049 |
| Cct2 | 1508.454392 | 0.154327664 | 0.073067714 | 2.112118404 |
| Frs2 | 1041.72447 | 0.088987989 | 0.063394027 | 1.403728286 |
| Gm26311 | 0 | NA | NA | NA |
| Yeats4 | 230.0577702 | 0.041421627 | 0.115074357 | 0.359955316 |

| | | | | |
|---|---|---|---|---|
| 9530003J23Rik | 0.138076337 | 0.75528675 | 3.589802529 | 0.210397854 |
| Gm10936 | 0 | NA | NA | NA |
| Lyz2 | 917.3809082 | 1.799402183 | 0.451438333 | 3.985931298 |
| Lyz1 | 2.092478253 | 0.274913912 | 0.799336096 | 0.343927809 |
| Cpsf6 | 2549.871472 | -0.100603029 | 0.069513038 | -1.447254106 |
| Gm9002 | 0 | NA | NA | NA |
| Kifc5c-ps | 0 | NA | NA | NA |
| Cpm | 200.0726009 | -0.274539011 | 0.136215051 | -2.015482204 |
| Gm25709 | 2.906247223 | -1.102949386 | 0.645661999 | -1.708245781 |
| Mdm2 | 840.5416714 | 0.6415396 | 0.097141683 | 6.604163919 |
| Slc35e3 | 296.2709981 | -0.178764416 | 0.081282311 | -2.199302814 |
| Nup107 | 192.1877859 | 0.019843661 | 0.101814323 | 0.19490049 |
| Rap1b | 1149.527098 | 0.925627657 | 0.126259508 | 7.331152116 |
| Gm37852 | 0 | NA | NA | NA |
| Mdm1 | 91.75534052 | 0.069181389 | 0.126460911 | 0.547057497 |
| Il22 | 14.70729829 | 1.049298024 | 0.430005849 | 2.440194771 |
| Iltifb | 0.488214936 | 1.795763119 | 1.418668823 | 1.265808545 |
| Gm9040 | 0 | NA | NA | NA |
| Gm9044 | 0 | NA | NA | NA |
| Gm9045 | 0.068097359 | -0.206487837 | 3.724945049 | -0.055433794 |
| Gm9046 | 0 | NA | NA | NA |
| Gm9048 | 0 | NA | NA | NA |
| Gm9049 | 0.219658751 | -1.476801405 | 3.701869585 | -0.398933936 |
| Ifng | 22.72435143 | 7.023739454 | 1.034261076 | 6.791070083 |
| Dyrk2 | 260.899164 | -0.001188355 | 0.110080333 | -0.010795344 |
| 4932442E05Rik | 0.27373663 | 1.659441864 | 1.782130753 | 0.931156068 |
| Gm26976 | 0 | NA | NA | NA |
| 1700064J06Rik | 0 | NA | NA | NA |
| Cand1 | 2796.110183 | -0.05771539 | 0.043176337 | -1.336736601 |
| 1700025F24Rik | 0.942170168 | 0.244704575 | 0.912387554 | 0.268202448 |
| Grip1os3 | 1.395336821 | -1.977447867 | 0.913990708 | -2.163531696 |
| Gm16321 | 1.40953533 | -1.513858923 | 0.871874483 | -1.736326676 |
| Grip1 | 365.1462489 | -0.377101978 | 0.12290644 | -3.068203585 |
| Grip1os2 | 0 | NA | NA | NA |
| Grip1os1 | 0.072025462 | 0.755288839 | 3.724945049 | 0.202765096 |
| Helb | 127.7033369 | 0.681062005 | 0.123611791 | 5.509684775 |
| Irak3 | 30.23155639 | 2.021233297 | 0.286093134 | 7.064948624 |
| Tmbim4 | 275.0985624 | 0.229380248 | 0.125160479 | 1.832689127 |
| Llph | 97.51439187 | 0.201927766 | 0.169497377 | 1.191332688 |
| Gm38141 | 0 | NA | NA | NA |
| Hmga2 | 52.04420527 | 2.897445994 | 0.638022619 | 4.541290396 |
| 1700006J14Rik | 0.548916986 | 0.928014742 | 1.399224064 | 0.663235264 |
| 9230105E05Rik | 0 | NA | NA | NA |
| Mir763 | 0.331210058 | 1.174800763 | 2.02897048 | 0.579013236 |
| Gm24298 | 0 | NA | NA | NA |
| Gm38289 | 0.370157025 | 0.755307989 | 1.705075808 | 0.442976192 |
| Gm37505 | 0.097139547 | 0.274393828 | 3.724945049 | 0.073663859 |

| | | | | |
|---|---|---|---|---|
| 4921513I03Rik | 0 | NA | NA | NA |
| Msrb3 | 98.13132724 | -0.117976155 | 0.124358018 | -0.94868153 |
| Gm15961 | 1.450891998 | -0.261491398 | 0.801540195 | -0.326236164 |
| Gm15910 | 3.363554134 | 0.328339404 | 0.605508735 | 0.542253786 |
| Gm4473 | 0.74372259 | -1.633462283 | 1.22399616 | -1.334532196 |
| Lemd3 | 732.3495551 | -0.115740407 | 0.067218901 | -1.721843205 |
| Gm16166 | 0.157815785 | 1.201305391 | 3.465799399 | 0.346617116 |
| Gm23632 | 0.389039412 | -1.38367347 | 1.51459574 | -0.913559594 |
| Wif1 | 23.54109142 | -0.387158573 | 0.31792938 | -1.217750223 |
| Tbc1d30 | 832.5656543 | -0.625573282 | 0.099065039 | -6.314773493 |
| Gns | 1953.99692 | 0.797092161 | 0.105221907 | 7.575344198 |
| Rassf3 | 242.4931995 | -0.553116782 | 0.122440233 | -4.517443054 |
| Tbk1 | 841.1038649 | 0.509440236 | 0.091707128 | 5.555077879 |
| Xpot | 1103.074494 | -0.107285585 | 0.051591328 | -2.079527493 |
| Gm24330 | 0.750750929 | -0.132953058 | 1.200304432 | -0.110766114 |
| D930020B18Rik | 4.785986039 | -0.044111405 | 0.492372004 | -0.089589589 |
| BC048403 | 125.9765068 | -0.120902139 | 0.102344284 | -1.181327718 |
| C030027H14Rik | 0.046345709 | 0.274391604 | 3.724945049 | 0.073663262 |
| Srgap1 | 808.6398788 | -0.140790025 | 0.079698459 | -1.766533838 |
| Tmem5 | 317.9777111 | 0.455087688 | 0.095438435 | 4.768390109 |
| Gm9081 | 0.90419666 | -0.042430695 | 1.036845161 | -0.040922885 |
| Avpr1a | 7.942965041 | 0.892880188 | 0.407526758 | 2.190973159 |
| Mir6412 | 0 | NA | NA | NA |
| Ppm1h | 1340.411078 | 0.017466636 | 0.079514674 | 0.219665573 |
| Mir8104 | 0 | NA | NA | NA |
| Mirlet7i | 1.909998494 | 2.50685445 | 0.882414759 | 2.840902675 |
| Mon2 | 1822.582098 | -0.198436635 | 0.045472398 | -4.363892036 |
| Usp15 | 1257.977968 | 0.058485537 | 0.056992475 | 1.026197536 |
| Gm27920 | 0 | NA | NA | NA |
| Fam19a2 | 696.2404132 | 0.023394201 | 0.085593604 | 0.273317165 |
| Gm22958 | 7.051847066 | -0.370085264 | 0.470311755 | -0.786893502 |
| Gm23777 | 0 | NA | NA | NA |
| Slc16a7 | 304.2321303 | -0.683922724 | 0.151967656 | -4.500449268 |
| Lrig3 | 67.70650913 | -0.134475633 | 0.142500426 | -0.943685832 |
| Gm22183 | 0 | NA | NA | NA |
| Mir378d | 0.20330765 | -0.685843796 | 2.208071316 | -0.310607629 |
| Gm24179 | 0.30098968 | 1.649518328 | 1.992832736 | 0.827725428 |
| Xrcc6bp1 | 46.49479728 | -0.239409521 | 0.177142348 | -1.351509245 |
| Ctdsp2 | 1189.150957 | 0.291641018 | 0.089485569 | 3.259084347 |
| Mir26a-2 | 0.925632266 | -0.327571187 | 1.118786408 | -0.292791532 |
| Mir546 | 0 | NA | NA | NA |
| Avil | 16.18070083 | -0.523670528 | 0.315955728 | -1.657417423 |
| Tsfm | 123.6160451 | -0.081017445 | 0.113756923 | -0.712197934 |
| Mettl21b | 2.262034746 | 0.590509195 | 0.695983621 | 0.848452719 |
| Mettl1 | 114.9215301 | 0.2147047 | 0.14380479 | 1.493028853 |
| Cyp27b1 | 5.379193204 | 1.104916485 | 0.472369162 | 2.339095298 |
| 9-Mar | 204.9397292 | -0.090115718 | 0.110693708 | -0.814099739 |

| | | | | |
|---|---|---|---|---|
| Cdk4 | 393.057798 | -0.058791891 | 0.130745001 | -0.449668368 |
| Tspan31 | 608.5951114 | 0.437183394 | 0.108788717 | 4.018646484 |
| Agap2 | 8823.335821 | -0.212019516 | 0.139384608 | -1.521111402 |
| Os9 | 1552.713354 | 0.456205728 | 0.079002413 | 5.7745797 |
| Gm23664 | 0.045882455 | 0.755288839 | 3.724945049 | 0.202765096 |
| B4galnt1 | 858.1814942 | -0.061475519 | 0.070773606 | -0.868622117 |
| Slc26a10 | 18.10165712 | -0.334796649 | 0.439443752 | -0.761864626 |
| Arhgef25 | 709.5156223 | -0.189713062 | 0.10391181 | -1.82571223 |
| Dtx3 | 1335.531959 | 0.105409288 | 0.050937023 | 2.069404165 |
| F420014N23Rik | 11.7378327 | -0.033403733 | 0.404226158 | -0.082636247 |
| Pip4k2c | 1116.600625 | -0.129818808 | 0.061584991 | -2.107961803 |
| Kif5a | 22627.74193 | 0.039819771 | 0.105117057 | 0.378813601 |
| Dctn2 | 1693.964189 | -0.00961827 | 0.049080894 | -0.195967695 |
| Mbd6 | 504.0837198 | -0.301301251 | 0.139565665 | -2.158849391 |
| Gm20492 | 1.795058199 | -0.371609198 | 0.716175165 | -0.518880319 |
| Gm4189 | 3.902576099 | -0.075638386 | 0.509635761 | -0.148416559 |
| Ddit3 | 291.5705915 | 1.211432737 | 0.200108767 | 6.053871365 |
| Mars | 495.381841 | 0.082965554 | 0.09987233 | 0.830716114 |
| Gm6283 | 1.594842726 | -0.06180998 | 0.87537519 | -0.070609701 |
| Arhgap9 | 51.31007067 | 3.211058687 | 0.334986296 | 9.585641925 |
| Gli1 | 50.51559619 | -0.336562993 | 0.251367904 | -1.338925881 |
| Inhbe | 0.872322242 | -1.365332731 | 1.229564795 | -1.110419505 |
| Inhbc | 0.072025462 | 0.755288839 | 3.724945049 | 0.202765096 |
| R3hdm2 | 2616.368291 | -0.260364327 | 0.073417191 | -3.546367301 |
| Gm23819 | 0 | NA | NA | NA |
| Stac3 | 24.43958761 | -1.728194821 | 0.421381982 | -4.101254665 |
| Ndufa4l2 | 7.948266496 | 0.059033015 | 0.363345515 | 0.16247074 |
| Shmt2 | 276.9083029 | 0.309575556 | 0.098099188 | 3.155740246 |
| Nxph4 | 143.5244743 | -0.091594905 | 0.227096989 | -0.403329457 |
| Lrp1 | 13094.28162 | -0.020583724 | 0.08422607 | -0.244386607 |
| Gm16217 | 0.962459693 | 1.965374025 | 1.320337677 | 1.488538924 |
| Stat6 | 224.5914028 | 1.156775305 | 0.173836436 | 6.654389207 |
| Nab2 | 546.0189877 | -0.195326433 | 0.14331952 | -1.362873898 |
| Gm16229 | 0.705361133 | -0.545335088 | 1.268962266 | -0.42974886 |
| Tmem194 | 106.2750348 | 0.13726118 | 0.172845189 | 0.79412786 |
| Gm16230 | 1.3230785 | 1.363136111 | 0.992611747 | 1.373282267 |
| Myo1a | 8.599191626 | -0.420334948 | 0.340666671 | -1.233859911 |
| Tac2 | 49.86492223 | 0.459613396 | 0.195396298 | 2.352211388 |
| Zbtb39 | 235.0509989 | 0.180069023 | 0.130055389 | 1.384556409 |
| Gpr182 | 35.51224453 | 0.925222796 | 0.242827465 | 3.810206548 |
| Rdh1 | 7.125358113 | 0.476101516 | 0.494248561 | 0.963283566 |
| Gm28046 | 0 | NA | NA | NA |
| Rdh9 | 1.759257097 | 1.387315489 | 0.831275526 | 1.668899717 |
| Rdh16 | 4.45419811 | 0.664437532 | 0.471489154 | 1.409231849 |
| Rdh18-ps | 0.308514759 | 1.303885431 | 1.753477671 | 0.743599677 |
| Rdh19 | 0.061920613 | 0.274393828 | 3.724945049 | 0.073663859 |
| BC089597 | 0.047948698 | -0.206487837 | 3.724945049 | -0.055433794 |

| | | | | |
|---|---|---|---|---|
| Rdh7 | 0 | NA | NA | NA |
| Sdr9c7 | 0.145038387 | 1.221915589 | 2.886820676 | 0.423273811 |
| Gm15905 | 0 | NA | NA | NA |
| Gm15900 | 1.613670225 | -0.936759442 | 1.088541391 | -0.860563917 |
| Gm23996 | 0 | NA | NA | NA |
| Hsd17b6 | 0.048949532 | -0.206487837 | 3.724945049 | -0.055433794 |
| Prim1 | 70.56909489 | -0.121532913 | 0.142751251 | -0.851361459 |
| Naca | 1262.918694 | 0.190796618 | 0.064378695 | 2.963660873 |
| Ptges3 | 1465.432757 | 0.193612758 | 0.095369126 | 2.030140849 |
| Atp5b | 9103.422248 | -0.102159887 | 0.072627504 | -1.40662809 |
| Snord59a | 1.560301263 | 1.454680828 | 0.9111788 | 1.596482301 |
| mmu-mir-677 | 0.075043205 | 0.755286614 | 3.724945049 | 0.202764498 |
| Mir677 | 0 | NA | NA | NA |
| Baz2a | 1409.129041 | 0.088756268 | 0.085980141 | 1.032288005 |
| Rbms2 | 301.1430177 | 0.479245316 | 0.142975741 | 3.351934468 |
| Gls2 | 183.4405106 | -0.790464608 | 0.125391661 | -6.303964733 |
| Spryd4 | 66.56019931 | 0.170991725 | 0.167733404 | 1.019425591 |
| Mip | 0 | NA | NA | NA |
| Gm26847 | 5.379666823 | 0.722677617 | 0.507476556 | 1.42406109 |
| Timeless | 115.0545271 | 1.185003025 | 0.154430165 | 7.673390936 |
| Apon | 0.725473741 | 0.757863545 | 1.387671555 | 0.546140434 |
| Apof | 3.028410459 | 2.203905046 | 0.817285538 | 2.69661574 |
| Stat2 | 2927.543515 | 3.309826001 | 0.106412596 | 31.10370509 |
| Il23a | 1.801413292 | -2.618675974 | 0.929248021 | -2.818059242 |
| Gm23241 | 0 | NA | NA | NA |
| Pan2 | 211.7421926 | -0.149101453 | 0.12986783 | -1.148101522 |
| Cnpy2 | 199.2616733 | 0.050395491 | 0.129455472 | 0.389288229 |
| Cs | 2382.585503 | -0.081248721 | 0.066317155 | -1.225153905 |
| Gm23182 | 1.59776671 | 0.050724906 | 0.763145231 | 0.066468221 |
| Coq10a | 213.594349 | -0.055084908 | 0.083627743 | -0.658691788 |
| Ankrd52 | 1433.771392 | -0.223255665 | 0.079126975 | -2.82148616 |
| Gm26308 | 0 | NA | NA | NA |
| Mir6914 | 0.577876106 | 1.469936406 | 1.572284205 | 0.934905027 |
| mmu-mir-6915 | 0.129647038 | -0.687376656 | 2.992845844 | -0.229673258 |
| Gm17201 | 0.247536319 | 0.263443005 | 1.847383095 | 0.142603343 |
| Slc39a5 | 1.555265169 | -0.905851151 | 0.71219512 | -1.271914291 |
| Nabp2 | 369.4297074 | 0.083724816 | 0.073749823 | 1.135254464 |
| Gm25361 | 0 | NA | NA | NA |
| Rnf41 | 702.2743574 | -0.024335089 | 0.06365342 | -0.382306073 |
| Gm26347 | 3.130125749 | -0.92914421 | 0.680771114 | -1.364840827 |
| Mir8105 | 0.208009692 | -0.206485804 | 2.605199321 | -0.07925912 |
| Smarcc2 | 3642.728046 | -0.126918875 | 0.071165356 | -1.783436252 |
| Myl6 | 1464.017352 | 0.128961917 | 0.11123489 | 1.159365704 |
| Myl6b | 108.9152618 | -0.358663115 | 0.128398926 | -2.793349809 |
| A430046D13Rik | 90.5350024 | -0.729299291 | 0.162107898 | -4.498851085 |
| Esyt1 | 580.33528 | 1.434316608 | 0.187322732 | 7.656927656 |
| Mir6916 | 0 | NA | NA | NA |

| | | | | |
|---|---|---|---|---|
| Gm29585 | 0.520597108 | 0.909844352 | 1.429064965 | 0.636671092 |
| Zc3h10 | 205.4911794 | 0.185439694 | 0.089040201 | 2.082651342 |
| Rpl41 | 1706.429966 | 0.411537696 | 0.160631801 | 2.561993918 |
| Pa2g4 | 643.314172 | 0.261037856 | 0.091147578 | 2.863903363 |
| AC117232.1 | 0 | NA | NA | NA |
| Erbb3 | 358.7695039 | -0.388603127 | 0.167560314 | -2.319183561 |
| Mir6917 | 0.044191549 | -0.206490061 | 3.724945049 | -0.055434391 |
| Rps26 | 1049.123509 | 0.558858817 | 0.102663212 | 5.443613196 |
| Ikzf4 | 257.6485544 | 0.054951312 | 0.138242662 | 0.397498943 |
| Gm26876 | 8.215402088 | 0.088553827 | 0.366128945 | 0.241865137 |
| Suox | 127.1525145 | 0.176488263 | 0.126631608 | 1.393714148 |
| Rab5b | 1465.571473 | -0.016570911 | 0.062778516 | -0.263958307 |
| Cdk2 | 38.99389103 | 1.278030909 | 0.243244057 | 5.254109502 |
| Pmel | 8.731260839 | 0.150763008 | 0.330769262 | 0.455795097 |
| Dgka | 134.8647648 | 0.264724646 | 0.130564645 | 2.02753698 |
| Wibg | 245.4724351 | 0.541303105 | 0.128670021 | 4.206909289 |
| Rpsa-ps2 | 30.10672328 | 0.574254023 | 0.239139486 | 2.401335024 |
| Mmp19 | 189.6992867 | 3.296244145 | 0.522936577 | 6.303334459 |
| Tmem198b | 157.6632237 | 0.187676204 | 0.152855644 | 1.227800285 |
| Dnajc14 | 675.6783784 | 0.068762075 | 0.055736227 | 1.233705234 |
| Ormdl2 | 72.79417923 | 0.448218562 | 0.218878574 | 2.04779551 |
| Sarnp | 164.5829855 | 0.251041813 | 0.127678872 | 1.966196976 |
| Gdf11 | 478.8249761 | 0.165488708 | 0.133375222 | 1.240775504 |
| Cd63 | 793.3225511 | 1.264762101 | 0.235827362 | 5.363084638 |
| Rdh5 | 84.87448236 | -0.005047594 | 0.186450889 | -0.027071975 |
| Bloc1s1 | 41.20713241 | 0.011932537 | 0.2657537 | 0.044900738 |
| Itga7 | 368.0914646 | 0.423298776 | 0.136805713 | 3.094160086 |
| Gm25494 | 0.763054649 | -0.593523345 | 1.152092564 | -0.515169843 |
| Mettl7b | 0.038981395 | 0.274393828 | 3.724945049 | 0.073663859 |
| 9030616G12Rik | 0 | NA | NA | NA |
| Olfr9 | 0.042716945 | -0.206490061 | 3.724945049 | -0.055434391 |
| Olfr763 | 0 | NA | NA | NA |
| Olfr765 | 0 | NA | NA | NA |
| Olfr767 | 0 | NA | NA | NA |
| Olfr768 | 0 | NA | NA | NA |
| Olfr769 | 0 | NA | NA | NA |
| Olfr770 | 0 | NA | NA | NA |
| Olfr771 | 0 | NA | NA | NA |
| Gm25807 | 0 | NA | NA | NA |
| Olfr772 | 0 | NA | NA | NA |
| Olfr773 | 0 | NA | NA | NA |
| Olfr774 | 0.045882455 | 0.755288839 | 3.724945049 | 0.202765096 |
| Olfr775 | 0 | NA | NA | NA |
| Olfr776 | 0 | NA | NA | NA |
| Olfr777 | 0 | NA | NA | NA |
| Gm25764 | 0 | NA | NA | NA |
| Olfr779 | 0 | NA | NA | NA |

| | | | | |
|---|---|---|---|---|
| Olfr780 | 0 | NA | NA | NA |
| Olfr781 | 0 | NA | NA | NA |
| Olfr782 | 0.044544648 | 0.755286614 | 3.724945049 | 0.202764498 |
| Olfr784 | 0 | NA | NA | NA |
| Olfr786 | 0 | NA | NA | NA |
| Olfr787 | 0 | NA | NA | NA |
| Olfr788 | 0 | NA | NA | NA |
| Olfr790 | 0.08169834 | -0.206492286 | 3.724945049 | -0.055434988 |
| Olfr791 | 0 | NA | NA | NA |
| Olfr792 | 0 | NA | NA | NA |
| Olfr794 | 0 | NA | NA | NA |
| Gm23252 | 0 | NA | NA | NA |
| Olfr796 | 0 | NA | NA | NA |
| Olfr798 | 0 | NA | NA | NA |
| Olfr799 | 0.046345709 | 0.274391604 | 3.724945049 | 0.073663262 |
| Olfr800 | 0 | NA | NA | NA |
| Olfr801 | 0 | NA | NA | NA |
| Olfr802 | 0 | NA | NA | NA |
| Olfr803 | 0 | NA | NA | NA |
| Gm24717 | 0 | NA | NA | NA |
| Olfr804 | 0 | NA | NA | NA |
| Olfr805 | 0 | NA | NA | NA |
| Gm24485 | 0 | NA | NA | NA |
| Olfr806 | 0 | NA | NA | NA |
| Olfr807 | 0 | NA | NA | NA |
| Olfr808 | 0 | NA | NA | NA |
| Olfr809 | 0.046345709 | 0.274391604 | 3.724945049 | 0.073663262 |
| Olfr810 | 0.066050875 | 0.274391604 | 3.724945049 | 0.073663262 |
| Olfr811 | 0 | NA | NA | NA |
| Olfr812 | 0 | NA | NA | NA |
| Olfr813 | 0 | NA | NA | NA |
| Olfr814 | 0 | NA | NA | NA |
| Olfr815 | 0 | NA | NA | NA |
| Olfr816 | 0.045882455 | 0.755288839 | 3.724945049 | 0.202765096 |
| Olfr818 | 0 | NA | NA | NA |
| Olfr819 | 0 | NA | NA | NA |
| Olfr247 | 0 | NA | NA | NA |
| Gm10310 | 0 | NA | NA | NA |
| Olfr820 | 0 | NA | NA | NA |
| Olfr821 | 0 | NA | NA | NA |
| Olfr822 | 0 | NA | NA | NA |
| Olfr823 | 0.045882455 | 0.755288839 | 3.724945049 | 0.202765096 |
| Gm5181 | 0 | NA | NA | NA |
| Olfr824 | 0 | NA | NA | NA |
| Olfr825 | 0.062426873 | 0.755288839 | 3.724945049 | 0.202765096 |
| Olfr826 | 0 | NA | NA | NA |
| Olfr827 | 0.062426873 | 0.755288839 | 3.724945049 | 0.202765096 |

| | | | | |
|---|---|---|---|---|
| Neurod4 | 9.261960518 | 0.872527936 | 0.385226762 | 2.264972277 |
| Tespa1 | 3.163043473 | 4.365845197 | 0.934392916 | 4.672386873 |
| Vmn2r84 | 31.40616345 | -0.117141561 | 0.272952737 | -0.429164266 |
| Gm9770 | 48.13481903 | 0.428541508 | 0.164704545 | 2.601880289 |
| Vmn2r85 | 40.51229935 | -0.25024216 | 0.266984486 | -0.937291015 |
| Vmn2r86 | 19.28575388 | -0.808752194 | 0.33881045 | -2.387034383 |
| Vmn2r87 | 47.38212013 | -0.473376284 | 0.288918914 | -1.638439927 |
| Gm23793 | 0 | NA | NA | NA |
| Gm26445 | 0.178837887 | -0.20648716 | 2.700576645 | -0.076460396 |
| Gm20746 | 0.242679635 | 1.428887174 | 2.49739507 | 0.572151035 |
| Pisd-ps1 | 253.7945557 | -0.190050828 | 0.124050034 | -1.532049786 |
| Sfi1 | 373.2312864 | -0.132841574 | 0.150253393 | -0.884116964 |
| Gm11399 | 1.590475537 | 2.188543496 | 1.066727188 | 2.05164312 |
| Gm11400 | 21.45170634 | -0.003023013 | 0.306322582 | -0.009868724 |
| Drg1 | 586.4306311 | 0.123562233 | 0.074800173 | 1.651897702 |
| Gm12735 | 2.937817048 | -1.618570325 | 0.681606601 | -2.374640037 |
| Fau-ps2 | 1.291849242 | -1.069311054 | 0.843484483 | -1.267730558 |
| Eif4enif1 | 683.4614762 | 0.110992725 | 0.074258525 | 1.494679909 |
| Patz1 | 528.6289566 | 0.016556338 | 0.082397686 | 0.200932081 |
| Gm12592 | 46.92358016 | 0.968194643 | 0.224521811 | 4.312252059 |
| Gm11944 | 38.0840153 | -0.594317844 | 0.229548689 | -2.589070956 |
| Pik3ip1 | 504.3401145 | 0.893481465 | 0.09178237 | 9.734783132 |
| Limk2 | 962.7434137 | -0.167523085 | 0.050216606 | -3.336009701 |
| Gm11948 | 5.24095402 | -0.007338638 | 0.491835991 | -0.014920904 |
| Rnf185 | 379.3246065 | 0.216829818 | 0.083075128 | 2.610044927 |
| 8430429K09Rik | 61.46987694 | -0.365106929 | 0.154710282 | -2.359939649 |
| Gm11945 | 1.421692497 | 0.144356863 | 0.878036659 | 0.164408697 |
| Gm24439 | 0.228704471 | -0.23634757 | 2.18760193 | -0.108039569 |
| Pla2g3 | 97.55501564 | 1.93598091 | 0.239321205 | 8.089466667 |
| Inpp5j | 911.3473185 | -0.372054924 | 0.08131903 | -4.575250367 |
| Selm | 738.0868402 | 0.512926976 | 0.141192758 | 3.632813621 |
| Smtn | 69.98786474 | 0.290976648 | 0.18683683 | 1.557383783 |
| Gm11946 | 0.933952761 | 0.863385489 | 1.167446811 | 0.739550171 |
| Gm11947 | 0 | NA | NA | NA |
| Gm11949 | 0 | NA | NA | NA |
| Gm11950 | 0 | NA | NA | NA |
| Gm11951 | 0.453565911 | 1.370507226 | 1.536245819 | 0.892114536 |
| Tug1 | 1777.476593 | 0.141051297 | 0.071448379 | 1.974170696 |
| Gm27624 | 0 | NA | NA | NA |
| Gm27517 | 0 | NA | NA | NA |
| Gm27937 | 0 | NA | NA | NA |
| Gm27640 | 0 | NA | NA | NA |
| Morc2a | 721.9875827 | -0.068215856 | 0.075007461 | -0.90945427 |
| Gm26393 | 0.141094079 | 0.755285157 | 3.563850454 | 0.211929531 |
| Gm16518 | 10.42668312 | -0.198285478 | 0.354701291 | -0.559021022 |
| Osbp2 | 906.3048646 | -0.347613447 | 0.099919342 | -3.478940531 |
| Gm24013 | 0.11193333 | 0.75528439 | 3.724945049 | 0.202763901 |

| | | | | |
|---|---|---|---|---|
| Gm11952 | 0 | NA | NA | NA |
| 4921536K21Rik | 11.5351327 | 0.551279239 | 0.374293115 | 1.472854338 |
| Dusp18 | 270.8170531 | -0.155406875 | 0.099049528 | -1.568981476 |
| Slc35e4 | 326.5983649 | 0.504081446 | 0.132307521 | 3.809922847 |
| Tcn2 | 256.4564119 | 0.604712076 | 0.145626328 | 4.152491411 |
| 4930556J24Rik | 2.805439532 | 0.212034327 | 0.580940477 | 0.364984598 |
| Gm11953 | 2.130460637 | 0.494175549 | 0.694192748 | 0.711870803 |
| Pes1 | 430.3711271 | 0.27552794 | 0.09400772 | 2.930907598 |
| Gal3st1 | 117.3315211 | -0.298572629 | 0.147470293 | -2.024628987 |
| Gm11954 | 0.875788929 | 1.728408356 | 1.126315068 | 1.53456915 |
| Sec14l4 | 4.148966471 | 0.473833969 | 0.541203458 | 0.87551911 |
| Sec14l3 | 15.8893622 | -0.413582454 | 0.481255003 | -0.859383179 |
| Mtfp1 | 343.8323402 | -0.194780918 | 0.112236512 | -1.735450553 |
| Sec14l2 | 346.4809089 | -0.075441866 | 0.110408474 | -0.683297786 |
| Gm11955 | 0 | NA | NA | NA |
| Gm11957 | 1.061319949 | 0.874840376 | 1.034550491 | 0.845623663 |
| Rnf215 | 199.4153862 | -0.056547427 | 0.095747843 | -0.590586953 |
| Mir3060 | 6.092409896 | -0.005113142 | 0.463904883 | -0.011021963 |
| Ccdc157 | 315.6146368 | -0.069163909 | 0.085205663 | -0.811729011 |
| Sf3a1 | 976.4741568 | -0.094696438 | 0.060091124 | -1.575880613 |
| Tbc1d10a | 286.7296832 | 0.272788908 | 0.111842631 | 2.439042304 |
| Gatsl3 | 28.49626455 | 0.890047117 | 0.301843442 | 2.948704506 |
| Gm11956 | 0.045882455 | 0.755288839 | 3.724945049 | 0.202765096 |
| Osm | 17.76436873 | 3.360939715 | 0.556605214 | 6.038282839 |
| Lif | 163.2509148 | 3.664103905 | 0.517764622 | 7.076775339 |
| Gm11959 | 0 | NA | NA | NA |
| Gm11958 | 0 | NA | NA | NA |
| Gm24803 | 1.135110426 | -0.253369775 | 0.918593608 | -0.275823578 |
| Hormad2 | 2.594629735 | 1.099203686 | 0.722743259 | 1.52087712 |
| Mtmr3 | 1608.347275 | 0.226549133 | 0.069067903 | 3.280092842 |
| Gm11960 | 2.472144487 | 0.420865843 | 0.63710876 | 0.660587122 |
| Gm11032 | 0 | NA | NA | NA |
| Gm11961 | 0.364460012 | 1.109024631 | 1.566734437 | 0.707857442 |
| Ascc2 | 346.5615331 | 0.109030183 | 0.073124972 | 1.491011614 |
| Uqcr10 | 496.5398793 | -0.001387981 | 0.109219981 | -0.012708128 |
| Zmat5 | 89.65500592 | 0.020544277 | 0.13249829 | 0.155053149 |
| Cabp7 | 502.185691 | -0.01130718 | 0.129635823 | -0.087222651 |
| Nf2 | 1204.431154 | 0.109505448 | 0.054210291 | 2.020012184 |
| Nipsnap1 | 497.6595648 | -0.088250049 | 0.08072934 | -1.093159545 |
| Mir6918 | 0.256523886 | -0.184908965 | 1.821889802 | -0.101492947 |
| Thoc5 | 333.4021351 | -0.053700999 | 0.068310653 | -0.786129202 |
| AA413626 | 4.943749683 | -0.177451686 | 0.458284089 | -0.387208918 |
| Nefh | 1154.799543 | -0.420072314 | 0.232532585 | -1.806509457 |
| Ap1b1 | 2045.497515 | 0.065524571 | 0.045599103 | 1.43697059 |
| AL645522.1 | 0.147068667 | 1.236183405 | 3.514416608 | 0.351746404 |
| Gm24958 | 0 | NA | NA | NA |
| Gas2l1 | 586.1682792 | 0.035467718 | 0.067949227 | 0.521973824 |

| | | | | |
|---|---|---|---|---|
| Rasl10a | 112.452118 | -0.838732208 | 0.26470791 | -3.168519628 |
| Ewsr1 | 2343.895997 | 0.173025898 | 0.095787095 | 1.806359178 |
| Rhbdd3 | 147.1104437 | 0.001672218 | 0.123512337 | 0.01353887 |
| Emid1 | 46.41330166 | -0.507026799 | 0.173334907 | -2.925128059 |
| Kremen1 | 346.9264262 | 0.479532042 | 0.117001141 | 4.098524478 |
| Gm25142 | 0 | NA | NA | NA |
| Znrf3 | 372.0960024 | 0.072876414 | 0.093382377 | 0.780408646 |
| Gm11962 | 0 | NA | NA | NA |
| Gm11963 | 0 | NA | NA | NA |
| Xbp1 | 893.6785691 | -0.18924956 | 0.10180586 | -1.858925996 |
| Ccdc117 | 216.8499613 | -0.427303134 | 0.10288063 | -4.153387601 |
| Gm22471 | 0 | NA | NA | NA |
| Ankrd36 | 0.048949532 | -0.206487837 | 3.724945049 | -0.055433794 |
| Gm11964 | 12.76498303 | -0.0900197 | 0.315562314 | -0.285267586 |
| Gm11965 | 0.126225412 | 0.755282661 | 3.698754989 | 0.20419916 |
| Mrps24 | 181.2008852 | -0.023847066 | 0.114884908 | -0.207573534 |
| Urgcp | 807.8473661 | 0.252031301 | 0.068301695 | 3.68997138 |
| 2210015D19Rik | 182.3089558 | 0.029837633 | 0.082468405 | 0.361806841 |
| Dbnl | 1162.267136 | 0.898112082 | 0.125818474 | 7.138157491 |
| Pgam2 | 10.44641268 | -0.588227925 | 0.304473991 | -1.931948019 |
| Gm11966 | 36.99000587 | 0.229510389 | 0.205510788 | 1.11678025 |
| Polm | 78.87651427 | 0.267129594 | 0.136372588 | 1.958821767 |
| Aebp1 | 394.2398871 | 0.102405233 | 0.178848113 | 0.57258213 |
| Pold2 | 110.4180631 | 0.060599758 | 0.146705524 | 0.413070731 |
| Myl7 | 0.376384159 | 1.162507172 | 1.720062231 | 0.67585181 |
| Gck | 15.14292826 | 0.149149229 | 0.324293021 | 0.459921179 |
| Gm11967 | 0.249409806 | -0.691936531 | 2.535662648 | -0.272881935 |
| Ykt6 | 770.8424576 | 0.360207789 | 0.063204949 | 5.699044017 |
| Camk2b | 7644.2209 | -0.336889515 | 0.132788591 | -2.537036606 |
| Nudcd3 | 1857.880843 | 0.052150459 | 0.036167195 | 1.441927107 |
| Rps15a-ps6 | 9.077851534 | 0.212687803 | 0.360306941 | 0.590296157 |
| Gm11969 | 6.242235417 | -0.155267476 | 0.460782639 | -0.336964684 |
| A730071L15Rik | 1.959824955 | 0.829728966 | 0.753581809 | 1.101046968 |
| Npc1l1 | 17.38551341 | 0.383384822 | 0.405597332 | 0.945235069 |
| Gm11401 | 0.198115332 | -0.738034223 | 3.230814308 | -0.228435977 |
| Ddx56 | 347.1312207 | 0.243485847 | 0.083217681 | 2.925890807 |
| Tmed4 | 598.9634342 | 0.223055301 | 0.086262066 | 2.585786695 |
| n-R5s67 | 0 | NA | NA | NA |
| Ogdh | 3550.241053 | -0.263041327 | 0.06348272 | -4.143510643 |
| Zmiz2 | 2387.144131 | 0.166657775 | 0.06767775 | 2.462519447 |
| Mir7651 | 2.015150011 | 0.012619744 | 0.756804488 | 0.016675038 |
| Ppia | 2699.446977 | 0.122650463 | 0.092695092 | 1.323160271 |
| H2afv | 161.7851873 | -0.351550102 | 0.140388704 | -2.504119574 |
| Gm11970 | 0.194008847 | -0.686283198 | 2.639855125 | -0.259970023 |
| Gm11971 | 0.305478853 | 1.113682226 | 1.757003684 | 0.633853097 |
| Gm11972 | 0.155921052 | 0.777072206 | 2.829227972 | 0.274658746 |
| Purb | 8286.574662 | -0.129745592 | 0.084262752 | -1.539773979 |

| | | | | |
|---|---|---|---|---|
| Gm22123 | 0 | NA | NA | NA |
| Gm11973 | 8.076220047 | 0.523473869 | 0.37017568 | 1.4141228 |
| Myo1g | 77.4741113 | 4.740112685 | 0.394241433 | 12.02337523 |
| Gm11974 | 44.13713644 | 1.163754008 | 0.266587334 | 4.365376225 |
| Gm22457 | 0 | NA | NA | NA |
| Gm24000 | 0.045882455 | 0.755288839 | 3.724945049 | 0.202765096 |
| Ccm2 | 367.7041191 | -0.154187399 | 0.115573614 | -1.334105541 |
| Gm11975 | 0 | NA | NA | NA |
| Nacad | 676.8149995 | -0.212392416 | 0.075150528 | -2.826226526 |
| Tbrg4 | 350.0488541 | 0.143171358 | 0.091161246 | 1.570528759 |
| Gm24313 | 0.992467499 | -1.056662303 | 1.019978592 | -1.035965177 |
| Snora5c | 0.582740348 | -0.508823105 | 1.09515181 | -0.464614221 |
| Wap | 0.066050875 | 0.274391604 | 3.724945049 | 0.073663262 |
| Gm11978 | 0.38240426 | -0.128038215 | 1.504880184 | -0.085082 |
| Ramp3 | 89.90826615 | -0.267442606 | 0.143318318 | -1.866074134 |
| Gm11980 | 0.066050875 | 0.274391604 | 3.724945049 | 0.073663262 |
| Gm11981 | 0 | NA | NA | NA |
| Gm11976 | 0 | NA | NA | NA |
| Gm11977 | 0.369792422 | -0.648973593 | 1.563956623 | -0.414956261 |
| Gm11979 | 0 | NA | NA | NA |
| Gm11982 | 0 | NA | NA | NA |
| Gm11983 | 0.178864798 | 0.274396414 | 3.277341654 | 0.0837253 |
| Gm11984 | 0 | NA | NA | NA |
| Adcy1 | 9022.462185 | -0.762509776 | 0.152892748 | -4.98722002 |
| Gm11985 | 0 | NA | NA | NA |
| Igfbp1 | 2.637679718 | 3.07322421 | 0.889329817 | 3.455663073 |
| Igfbp3 | 602.9842508 | 1.705159358 | 0.318954941 | 5.34608228 |
| Gm11986 | 0.276998953 | 1.757664767 | 1.792851877 | 0.980373666 |
| Gm27393 | 0 | NA | NA | NA |
| Gm11987 | 0 | NA | NA | NA |
| Gm24051 | 0 | NA | NA | NA |
| Gm11988 | 0 | NA | NA | NA |
| Gm11989 | 0.796528503 | 1.591267028 | 1.148433934 | 1.38559736 |
| Tns3 | 1722.397234 | 0.161057157 | 0.131766314 | 1.22229386 |
| Gm11990 | 0 | NA | NA | NA |
| Pkd1l1 | 3.97064273 | 0.533057913 | 0.50903043 | 1.047202449 |
| Gm11991 | 0 | NA | NA | NA |
| Hus1 | 255.0186718 | 0.247016969 | 0.119854765 | 2.060969121 |
| Sun3 | 0.968963158 | -0.574023479 | 1.067463917 | -0.537745089 |
| Gm11992 | 17.30127652 | 0.512704219 | 0.305661181 | 1.677361242 |
| Upp1 | 71.12326318 | 2.331229529 | 0.337014939 | 6.917288395 |
| Gm11993 | 27.56449686 | 0.018870525 | 0.22578719 | 0.083576598 |
| Abca13 | 10.03132759 | 0.26376421 | 0.415401521 | 0.63496207 |
| Gm36954 | 0 | NA | NA | NA |
| Eif3s6-ps1 | 4.535504711 | 0.785381831 | 0.525153068 | 1.495529359 |
| Gm11996 | 0 | NA | NA | NA |
| Gm11997 | 0 | NA | NA | NA |

| | | | | |
|---|---|---|---|---|
| Vwc2 | 143.5725646 | -0.349169581 | 0.103521662 | -3.372913205 |
| Gm23065 | 0.966893809 | 0.261950416 | 0.867020226 | 0.302127227 |
| Zpbp | 24.98626369 | -0.239545578 | 0.205949618 | -1.163127077 |
| Gm12598 | 0.344633608 | 1.988332065 | 1.890394607 | 1.051807944 |
| Gm26458 | 0.27289602 | -0.636627682 | 1.799409156 | -0.353798179 |
| Gm12599 | 2.05315699 | 0.646344333 | 0.709835891 | 0.910554596 |
| 4930415F15Rik | 5.574285272 | 0.860882197 | 0.474397279 | 1.814686205 |
| Gm11998 | 0.161486377 | 0.755296344 | 2.795731986 | 0.270160498 |
| 4930512M02Rik | 0.106358623 | 0.274402725 | 3.724945049 | 0.073666248 |
| Gm11999 | 0.204942615 | 0.755300943 | 2.617396202 | 0.288569588 |
| Ikzf1 | 243.3512954 | 2.138222601 | 0.171306329 | 12.48186577 |
| Gm12000 | 0.060174537 | 0.755286614 | 3.724945049 | 0.202764498 |
| Fignl1 | 18.28682388 | 0.301345005 | 0.367030402 | 0.821035543 |
| Ddc | 130.1558318 | 0.338002288 | 0.222469543 | 1.519319384 |
| 1700042O10Rik | 0.263601517 | 1.230388545 | 2.094435763 | 0.587455852 |
| Gm25058 | 0 | NA | NA | NA |
| Grb10 | 839.2894725 | 0.093731987 | 0.070781768 | 1.32423913 |
| Gm24036 | 0 | NA | NA | NA |
| Gm12001 | 0.097139547 | 0.274393828 | 3.724945049 | 0.073663859 |
| Cobl | 1095.669369 | -0.328380622 | 0.127987454 | -2.565725091 |
| Gm12002 | 1.538964163 | -1.17735063 | 0.923453562 | -1.274942974 |
| Gm22156 | 0 | NA | NA | NA |
| Gm22789 | 0 | NA | NA | NA |
| Gm12003 | 0.812725683 | 0.540519441 | 1.209021655 | 0.447071761 |
| Gm12004 | 0 | NA | NA | NA |
| 4930554G24Rik | 0.349921031 | -0.206486673 | 1.999445236 | -0.103271983 |
| Gm12005 | 0 | NA | NA | NA |
| Gm12006 | 0 | NA | NA | NA |
| 1700046C09Rik | 0 | NA | NA | NA |
| Pom121l12 | 0 | NA | NA | NA |
| Gm12007 | 0 | NA | NA | NA |
| Gm12008 | 0 | NA | NA | NA |
| Gm12009 | 7.023367752 | -0.238067195 | 0.409592826 | -0.581228917 |
| Gm24707 | 0 | NA | NA | NA |
| Gm25490 | 0 | NA | NA | NA |
| Gm12010 | 0 | NA | NA | NA |
| Gm37319 | 0 | NA | NA | NA |
| Gm12011 | 0 | NA | NA | NA |
| Vstm2a | 820.6459809 | -0.205613282 | 0.084928203 | -2.42102476 |
| Gm12012 | 0.950406788 | -0.587636129 | 1.115928205 | -0.526589548 |
| Sec61g | 266.2425794 | 0.452855039 | 0.152735123 | 2.964969878 |
| Akt2-ps | 59.61674162 | 0.111320405 | 0.146047133 | 0.762222461 |
| Gm12663 | 89.45054939 | -0.123713593 | 0.111945912 | -1.105119342 |
| Gm12664 | 0 | NA | NA | NA |
| Gm24714 | 0 | NA | NA | NA |
| Gm25698 | 0 | NA | NA | NA |
| Egfr | 584.8862284 | 0.390082947 | 0.124724186 | 3.127564581 |

| | | | | |
|---|---|---|---|---|
| Egfros | 2.163206028 | -0.005530575 | 0.874033532 | -0.006327646 |
| Eldr | 6.358422579 | 0.684933746 | 0.586802607 | 1.167230238 |
| Fbxo48 | 10.03527172 | 0.089338689 | 0.335008792 | 0.266675654 |
| Plek | 727.3535995 | 1.899921349 | 0.122705874 | 15.48354038 |
| Gm12014 | 0 | NA | NA | NA |
| Cnrip1 | 500.2213394 | -0.070858376 | 0.07595018 | -0.932958632 |
| Gm12013 | 1.807586739 | 0.46754258 | 0.797322202 | 0.58639102 |
| Ppp3r1 | 5062.853256 | 0.014111585 | 0.083184756 | 0.169641476 |
| Wdr92 | 87.67747582 | -0.411923313 | 0.157333388 | -2.61815575 |
| Pno1 | 283.54597 | 0.367998009 | 0.123614967 | 2.976969696 |
| C1d | 398.4995549 | 0.256741087 | 0.12564512 | 2.043382883 |
| Gm12015 | 0 | NA | NA | NA |
| Gm23959 | 0 | NA | NA | NA |
| Gm12016 | 0 | NA | NA | NA |
| Etaa1 | 152.4498586 | 0.056351659 | 0.106634009 | 0.528458601 |
| Etaa1os | 0.910758945 | 1.193225792 | 1.166312772 | 1.023075303 |
| Gm12017 | 1.313957777 | -0.058938895 | 0.910740447 | -0.064715359 |
| Gm12018 | 0.106112162 | -0.206492286 | 3.724945049 | -0.055434988 |
| Gm12019 | 0 | NA | NA | NA |
| Gm12020 | 0.546458135 | -0.171635844 | 1.366202921 | -0.125629832 |
| Gm37869 | 0 | NA | NA | NA |
| Gm12022 | 0.803435494 | 1.722325389 | 1.169516106 | 1.472682061 |
| Gm12021 | 0.841174947 | 0.686714295 | 1.10334194 | 0.622394808 |
| Gm12024 | 0.313606012 | 1.456272772 | 3.704357395 | 0.393124263 |
| Gm12023 | 2.23920711 | 2.479136881 | 1.069266431 | 2.318539897 |
| Gm23093 | 0 | NA | NA | NA |
| Gm28401 | 1.961032719 | 1.868827329 | 0.913041424 | 2.046815489 |
| Gm23021 | 0 | NA | NA | NA |
| 8430419K02Rik | 12.33011639 | 1.581558762 | 0.452716267 | 3.49348781 |
| Meis1 | 395.3527126 | -0.238294774 | 0.104810956 | -2.273567415 |
| Gm37818 | 22.45192434 | -0.313340283 | 0.299672158 | -1.045610259 |
| 2900018N21Rik | 18.03296757 | -0.150916345 | 0.35474933 | -0.425416858 |
| 2610001A08Rik | 15.58260202 | -0.944336924 | 0.355000957 | -2.660096842 |
| Gm16140 | 0.421709722 | 1.460934715 | 1.446546501 | 1.009946596 |
| Gm16141 | 1.235942255 | -0.447381561 | 1.123738786 | -0.398118821 |
| 4933406G16Rik | 0.427771506 | -0.615521677 | 1.542049881 | -0.399158085 |
| Gm37439 | 0.296404033 | 0.866813282 | 1.774224298 | 0.488559019 |
| Gm12025 | 0.044191549 | -0.206490061 | 3.724945049 | -0.055434391 |
| Gm12026 | 0.088534442 | 0.274389469 | 3.724945049 | 0.073662689 |
| Gm12027 | 0.123841226 | 0.274391173 | 3.724945049 | 0.073663147 |
| Gm23386 | 0.075043205 | 0.755286614 | 3.724945049 | 0.202764498 |
| Glns-ps1 | 73.63675014 | -0.327020473 | 0.161384817 | -2.026339767 |
| Gm12028 | 0 | NA | NA | NA |
| Gm12029 | 0 | NA | NA | NA |
| Spred2 | 1270.960265 | -0.421407575 | 0.08193175 | -5.143397723 |
| Actr2 | 3727.967054 | 0.254952209 | 0.053972357 | 4.723755364 |
| Gm12030 | 0 | NA | NA | NA |

| | | | | |
|---|---|---|---|---|
| Gm12031 | 1.129362261 | 0.329901715 | 0.863807917 | 0.381915595 |
| Gm26253 | 0.30153761 | 1.313663588 | 1.767889936 | 0.74306865 |
| Rab1 | 1943.110902 | 0.206727212 | 0.080971125 | 2.553097927 |
| Cep68 | 468.5192915 | -0.148551586 | 0.104182128 | -1.425883577 |
| Rpl12-ps2 | 0 | NA | NA | NA |
| Slc1a4 | 846.6499244 | 0.540585862 | 0.083679737 | 6.460176436 |
| Gm12033 | 100.5140019 | 0.053605959 | 0.143507058 | 0.373542318 |
| Gm12034 | 2.661321104 | 0.208292139 | 0.593007375 | 0.351247131 |
| Sertad2 | 498.1303247 | 0.170774517 | 0.092630251 | 1.843614961 |
| Gm12035 | 0 | NA | NA | NA |
| Aftph | 1713.653678 | 0.330415146 | 0.064753185 | 5.102685604 |
| Gm12036 | 0 | NA | NA | NA |
| Lgalsl | 1035.137919 | 0.11174597 | 0.079307074 | 1.409029027 |
| Gm12037 | 2.870345996 | 3.231723904 | 0.843045835 | 3.833390511 |
| Gm12038 | 0.044191549 | -0.206490061 | 3.724945049 | -0.055434391 |
| Gm22807 | 0 | NA | NA | NA |
| Gm23681 | 0 | NA | NA | NA |
| Peli1 | 911.878915 | 0.678614421 | 0.100515308 | 6.751353964 |
| 2610024D14Rik | 21.08894881 | 0.078174676 | 0.310262319 | 0.251963165 |
| Gm22600 | 0 | NA | NA | NA |
| Gm12040 | 0.133633945 | 1.441423131 | 3.720899433 | 0.387385673 |
| Gm12041 | 0.223001177 | 1.203484293 | 2.546337428 | 0.47263347 |
| Vps54 | 967.0089129 | 0.54560764 | 0.087814488 | 6.213184798 |
| Gm23582 | 0 | NA | NA | NA |
| Gm12039 | 0 | NA | NA | NA |
| Gm12042 | 9.247728932 | -0.061827997 | 0.373244414 | -0.165650161 |
| Ugp2 | 931.9428543 | -0.01281168 | 0.113694604 | -0.112685031 |
| Mir1933 | 1.993695893 | -1.708348431 | 0.77519896 | -2.203754802 |
| Gm12043 | 3.417719091 | 0.363251775 | 0.542954095 | 0.66902852 |
| Gm12044 | 0 | NA | NA | NA |
| Gm12045 | 0 | NA | NA | NA |
| Gm12046 | 0 | NA | NA | NA |
| 4932414J04Rik | 0 | NA | NA | NA |
| Mdh1 | 4531.387594 | -0.06209217 | 0.097225358 | -0.63864172 |
| Gm22990 | 2.17550669 | -0.345577296 | 0.680782203 | -0.507617993 |
| Wdpcp | 154.1019773 | 0.066569013 | 0.113161401 | 0.588266071 |
| Pgam1-ps1 | 0.548058809 | -1.547067886 | 1.240574428 | -1.247057695 |
| Gm12047 | 0.149223819 | -0.206489568 | 2.865438769 | -0.072062112 |
| Rpl15-ps1 | 0.048949532 | -0.206487837 | 3.724945049 | -0.055433794 |
| Otx1 | 56.99576563 | 0.361724634 | 0.15456213 | 2.340318638 |
| Ehbp1 | 1086.104915 | -0.032425752 | 0.063797753 | -0.508258528 |
| Rpsa-ps3 | 0 | NA | NA | NA |
| Gm12050 | 0.136201742 | -0.67532486 | 2.944131489 | -0.229379993 |
| Gm12051 | 0 | NA | NA | NA |
| Tmem17 | 31.45379496 | 0.150680915 | 0.221619915 | 0.679906924 |
| Gm26829 | 0.106971521 | 1.236179401 | 3.724945049 | 0.331865138 |
| Gm12052 | 0.878931054 | 0.255773219 | 1.228607985 | 0.208181309 |

| | | | | |
|---|---|---|---|---|
| Gm12053 | 0.158674157 | -0.206481579 | 2.811010873 | -0.073454564 |
| Gm12055 | 0.424314013 | 0.539635785 | 1.496773455 | 0.360532707 |
| Gm12056 | 0 | NA | NA | NA |
| Gm12057 | 7.744039244 | -0.051996772 | 0.335892379 | -0.15480188 |
| Gm12058 | 1.980218008 | -0.128981101 | 0.77554279 | -0.166310748 |
| Gm28048 | 0.044544648 | 0.755286614 | 3.724945049 | 0.202764498 |
| B3gnt2 | 14.36761034 | 0.762764207 | 0.346784738 | 2.199532229 |
| Gm20456 | 0.099106094 | 0.274387056 | 3.724945049 | 0.073662041 |
| Gm23772 | 0 | NA | NA | NA |
| 9130230N09Rik | 6.400237851 | 0.002256824 | 0.432785882 | 0.005214644 |
| Gm24917 | 0 | NA | NA | NA |
| Commd1 | 92.82422243 | 0.218786998 | 0.169656323 | 1.289589412 |
| Zrsr1 | 521.7180302 | 0.185115701 | 0.101024616 | 1.83238212 |
| Cct4 | 1159.74844 | 0.028488289 | 0.097788841 | 0.291324533 |
| Fam161a | 45.44904334 | -0.350367147 | 0.223205716 | -1.569705085 |
| Gm23740 | 0.228592689 | -0.206482037 | 2.138563777 | -0.096551732 |
| Gm12059 | 13.03468439 | -0.892840128 | 0.32538727 | -2.743930723 |
| 1700061J23Rik | 0.046345709 | 0.274391604 | 3.724945049 | 0.073663262 |
| Gm24398 | 0 | NA | NA | NA |
| Xpo1 | 1829.109104 | 0.126137355 | 0.088455462 | 1.425998498 |
| Usp34 | 4434.045314 | -0.190710779 | 0.063378355 | -3.009083741 |
| Gm22753 | 2.309867597 | -1.432898932 | 0.711311832 | -2.014445519 |
| Ahsa2 | 295.3081191 | -0.577630623 | 0.114343768 | -5.051701818 |
| AL672049.1 | 0.184646192 | 0.201506703 | 2.705003829 | 0.074494055 |
| 1700030C12Rik | 0.940659763 | -0.318864177 | 1.033441911 | -0.308545815 |
| 1700093K21Rik | 3.270068486 | 1.709090091 | 0.623086801 | 2.742940611 |
| Gm12060 | 7.122622078 | -0.048047737 | 0.426138023 | -0.112751584 |
| 0610010F05Rik | 731.0037471 | -0.020293535 | 0.066042064 | -0.307281962 |
| Gm23114 | 5.978366796 | -0.630253539 | 0.650518486 | -0.968848007 |
| Pex13 | 566.0191946 | 0.214764286 | 0.087055625 | 2.46697771 |
| Pus10 | 234.206064 | 0.296063368 | 0.10190576 | 2.905266287 |
| Rel | 254.6607661 | 0.897043379 | 0.170498164 | 5.261308159 |
| Gm23827 | 0.061920613 | 0.274393828 | 3.724945049 | 0.073663859 |
| Papolg | 392.4489342 | -0.195865439 | 0.089905359 | -2.17857356 |
| Gm12061 | 0.546436044 | -1.164063167 | 1.502624075 | -0.774686887 |
| Gm12062 | 0 | NA | NA | NA |
| AL662810.1 | 0 | NA | NA | NA |
| A830031A19Rik | 4.718729471 | 0.117045124 | 0.442433282 | 0.264548643 |
| Gm12063 | 6.626652096 | -0.216190551 | 0.373421828 | -0.5789446 |
| Bcl11a | 1084.849886 | -0.210714045 | 0.153478965 | -1.372918072 |
| Gm12064 | 0 | NA | NA | NA |
| 4930538E20Rik | 0.409305803 | 1.390643064 | 1.523564643 | 0.912756194 |
| Gm12065 | 0 | NA | NA | NA |
| Gm12066 | 0.089089296 | 1.194485492 | 3.723939576 | 0.320758559 |
| Gm12068 | 0.5449234 | 1.497466097 | 1.305431301 | 1.147104483 |
| 4933430M04Rik | 0.038981395 | 0.274393828 | 3.724945049 | 0.073663859 |
| Gm12067 | 0 | NA | NA | NA |

| | | | | |
|---|---|---|---|---|
| Gm22787 | 0 | NA | NA | NA |
| Gm25507 | 0 | NA | NA | NA |
| Gm10466 | 0 | NA | NA | NA |
| Gm12665 | 0 | NA | NA | NA |
| 4933427E13Rik | 0.566718995 | 1.696263777 | 1.258903905 | 1.34741323 |
| Gm24255 | 0 | NA | NA | NA |
| Gm12069 | 0.137762903 | -0.206480416 | 2.937523348 | -0.070290647 |
| 5730522E02Rik | 70.17051206 | 0.249276864 | 0.244566967 | 1.019258108 |
| Gm23514 | 0 | NA | NA | NA |
| Mir6372 | 0 | NA | NA | NA |
| Gm22868 | 0 | NA | NA | NA |
| Fancl | 76.3059331 | -0.210806948 | 0.16250136 | -1.297262672 |
| Gm12713 | 5.450982021 | -0.192768076 | 0.495945915 | -0.388687698 |
| Vrk2 | 67.55354626 | 0.783199313 | 0.196256586 | 3.990690609 |
| Gm6899 | 0.12209515 | 0.75528439 | 3.724945049 | 0.202763901 |
| Gm12071 | 0 | NA | NA | NA |
| Gm12070 | 46.42488409 | 0.080852831 | 0.160542277 | 0.503623301 |
| Gm12072 | 0.172942268 | -0.206480807 | 3.372002999 | -0.061233874 |
| Gm12073 | 0 | NA | NA | NA |
| Gm12074 | 0 | NA | NA | NA |
| n-R5s68 | 0 | NA | NA | NA |
| Gm24850 | 0 | NA | NA | NA |
| Gm25387 | 0 | NA | NA | NA |
| Gm12075 | 0 | NA | NA | NA |
| Gm12077 | 0 | NA | NA | NA |
| Gm3810 | 0 | NA | NA | NA |
| Gm12079 | 2.754754232 | -0.348706538 | 0.596905462 | -0.584190563 |
| Gm12080 | 0 | NA | NA | NA |
| Gm6685 | 30.4764154 | -0.074822324 | 0.200961243 | -0.372322159 |
| Gm12081 | 0 | NA | NA | NA |
| Ccdc85a | 925.7833576 | 0.03512425 | 0.07414365 | 0.473732406 |
| 2810471M01Rik | 0.525803784 | -1.687362222 | 1.358382159 | -1.242185206 |
| Mir216b | 0 | NA | NA | NA |
| mmu-mir-216c | 0 | NA | NA | NA |
| Gm12082 | 0.832920528 | -0.351205185 | 1.079884014 | -0.325224914 |
| Mir216a | 0.066050875 | 0.274391604 | 3.724945049 | 0.073663262 |
| Mir217 | 0 | NA | NA | NA |
| Efemp1 | 96.95456267 | 0.127963436 | 0.196870446 | 0.649988043 |
| Rpsa-ps5 | 0.463552209 | 2.257415833 | 1.43177593 | 1.576654409 |
| Pnpt1 | 855.1922142 | 1.475730061 | 0.115273299 | 12.80201121 |
| A630052C17Rik | 26.90946008 | -0.526587544 | 0.259866717 | -2.026375481 |
| Smek2 | 1087.669138 | 0.245237525 | 0.05595324 | 4.382901264 |
| Cfap36 | 532.1389956 | -0.081969784 | 0.083748228 | -0.9787644 |
| Gm12085 | 0 | NA | NA | NA |
| Gm23109 | 0.11197992 | 0.75528439 | 3.724945049 | 0.202763901 |
| Gm12086 | 0 | NA | NA | NA |
| Gm12087 | 0.124596252 | 0.755282361 | 3.714681068 | 0.203323609 |

| | | | | |
|---|---|---|---|---|
| Ccdc88a | 2613.854868 | 0.014239112 | 0.058318207 | 0.24416237 |
| Prorsd1 | 38.98209458 | 0.446983491 | 0.244207842 | 1.830340447 |
| Gm12089 | 10.3364996 | -0.372645546 | 0.413492898 | -0.901213896 |
| Mtif2 | 334.856477 | -0.092361349 | 0.075576211 | -1.222095525 |
| Rps27a | 597.2006918 | 0.451705783 | 0.251876254 | 1.793363909 |
| Clhc1 | 9.730852973 | 0.444846107 | 0.351162238 | 1.266782299 |
| Gm12090 | 0 | NA | NA | NA |
| Gm12088 | 0.547473135 | 0.1776922 | 1.332639384 | 0.133338548 |
| Gm12091 | 2.208204681 | 0.403254962 | 0.699358645 | 0.576606817 |
| Gm12092 | 0.662335159 | -1.17026433 | 1.240119375 | -0.943670709 |
| Gm12093 | 0.303276099 | 0.274416047 | 1.624150591 | 0.168959731 |
| Rtn4 | 7648.216462 | -0.027599756 | 0.071703818 | -0.384913343 |
| Eml6 | 546.8534648 | -0.2154419 | 0.104206112 | -2.067459346 |
| 4931440F15Rik | 25.51154845 | -0.496978997 | 0.284168223 | -1.748890116 |
| Gm12094 | 0.298205325 | -1.134046472 | 2.014259396 | -0.563009151 |
| Gm22588 | 0 | NA | NA | NA |
| Gm12095 | 0 | NA | NA | NA |
| Sptbn1 | 16443.0819 | -0.040250443 | 0.052740818 | -0.763174416 |
| Gm29237 | 0.044544648 | 0.755286614 | 3.724945049 | 0.202764498 |
| Gm8098 | 47.87148983 | 0.003168845 | 0.235667229 | 0.01344627 |
| 4930505A04Rik | 3.929212317 | 0.770689946 | 0.572736081 | 1.345628417 |
| Acyp2 | 181.980473 | 0.175569913 | 0.137287972 | 1.278844098 |
| Gm12096 | 0.263964426 | -0.687380216 | 2.120878983 | -0.324101574 |
| Gm12098 | 9.953545217 | -0.32730087 | 0.470248319 | -0.696017097 |
| Gm12097 | 33.54705527 | -0.139824151 | 0.300285986 | -0.465636617 |
| Gm12099 | 0 | NA | NA | NA |
| Gm12100 | 0.45607993 | 0.762415018 | 1.700136957 | 0.448443294 |
| Gm12101 | 0.170229941 | 1.192332551 | 2.742849982 | 0.434705711 |
| Psme4 | 1642.607637 | 0.270216811 | 0.113793792 | 2.374618207 |
| Gm12103 | 0.045882455 | 0.755288839 | 3.724945049 | 0.202765096 |
| Gpr75 | 147.2383283 | -0.532636019 | 0.112000952 | -4.755638336 |
| Gm12104 | 4.359450978 | 1.131012425 | 0.535249739 | 2.113055537 |
| Erlec1 | 677.4619536 | 0.264475442 | 0.060097565 | 4.400767987 |
| Asb3 | 233.2081696 | 0.044529083 | 0.099217946 | 0.448800692 |
| Chac2 | 85.03464214 | -0.12405853 | 0.169967456 | -0.729895785 |
| Gm12102 | 0 | NA | NA | NA |
| Gm12105 | 4.176201957 | -0.912796518 | 0.526874588 | -1.732473988 |
| Gm12106 | 0 | NA | NA | NA |
| Gm17332 | 0 | NA | NA | NA |
| Stc2 | 26.77438353 | 0.46355936 | 0.343447185 | 1.349725314 |
| Bod1 | 387.4332913 | 0.069201977 | 0.082873512 | 0.835031308 |
| Gm38061 | 0.047948698 | -0.206487837 | 3.724945049 | -0.055433794 |
| D630024D03Rik | 1.773596836 | 0.21243158 | 0.827158973 | 0.256820741 |
| Gm12107 | 0 | NA | NA | NA |
| Cpeb4 | 3230.776281 | 0.163809237 | 0.07017627 | 2.334253971 |
| 4930524B15Rik | 1.33642055 | 2.513015435 | 1.087858519 | 2.310057227 |
| Nsg2 | 3948.965464 | 0.016010113 | 0.068053404 | 0.235258073 |

| | | | | |
|---|---|---|---|---|
| Gm12108 | 0.431234269 | 0.285282978 | 1.575524427 | 0.181071758 |
| Gm12109 | 0.060174537 | 0.755286614 | 3.724945049 | 0.202764498 |
| Il9r | 0.166808115 | 1.571691287 | 3.380229866 | 0.464965801 |
| Snrnp25 | 64.99433264 | -0.089600245 | 0.217974755 | -0.411057901 |
| Rhbdf1 | 175.977258 | 0.774834244 | 0.171591087 | 4.51558561 |
| Nprl3 | 290.7020933 | -0.002547433 | 0.091455025 | -0.027854485 |
| Mpg | 101.0618394 | 0.202869206 | 0.12588677 | 1.611521262 |
| Hba-x | 0.066050875 | 0.274391604 | 3.724945049 | 0.073663262 |
| Hba-a1 | 2.403392355 | 2.5454277 | 1.158253889 | 2.197642265 |
| Hbq1b | 0.787662625 | 0.294476458 | 1.296870153 | 0.227067033 |
| Hba-a2 | 2.268919436 | 3.375910303 | 2.004712925 | 1.6839869 |
| Hbq1a | 1.119917166 | -0.332769023 | 0.973833079 | -0.341710535 |
| Sh3pxd2b | 817.112173 | 0.473848083 | 0.136575448 | 3.469496836 |
| Ubtd2 | 162.9170653 | -0.047775176 | 0.117796473 | -0.405573906 |
| Efcab9 | 6.264832892 | -0.59290107 | 0.389272823 | -1.523099058 |
| Stk10 | 237.0838909 | 0.238901409 | 0.134081452 | 1.781763296 |
| Fbxw11 | 1952.482481 | 0.412860758 | 0.068816596 | 5.999435927 |
| AL669951.2 | 0 | NA | NA | NA |
| AL669951.1 | 0 | NA | NA | NA |
| Gm8192 | 0.049553047 | 0.274391604 | 3.724945049 | 0.073663262 |
| 4930555O08Rik | 0 | NA | NA | NA |
| Smim23 | 0 | NA | NA | NA |
| Gm12110 | 0 | NA | NA | NA |
| Gm12111 | 0.20222093 | 0.74032894 | 2.210400935 | 0.334929708 |
| Gm12112 | 32.64724094 | 0.745073339 | 0.203785565 | 3.656163477 |
| Gm26052 | 0 | NA | NA | NA |
| Rpsa-ps4 | 1.265922642 | 0.766869671 | 1.11159545 | 0.689881981 |
| Gm12114 | 0.044191549 | -0.206490061 | 3.724945049 | -0.055434391 |
| Fgf18 | 14.860062 | -0.772768993 | 0.271871957 | -2.84240052 |
| Npm1 | 1433.536217 | 0.425472037 | 0.082728213 | 5.143010108 |
| Gm12115 | 4.703682786 | 0.346891248 | 0.44117931 | 0.786281768 |
| Tlx3 | 9.956390658 | -0.70554133 | 0.403830854 | -1.747120914 |
| Gm12116 | 10.20532615 | -0.927780488 | 0.529868604 | -1.750963315 |
| Ranbp17 | 205.143616 | 0.118122482 | 0.150516905 | 0.784778839 |
| Gm12117 | 3.560310751 | -0.435971986 | 0.548045857 | -0.795502749 |
| Anp32-ps | 6.77190338 | 0.07625393 | 0.388473292 | 0.196291306 |
| Gm12118 | 0.07796279 | 0.274392189 | 3.724945049 | 0.073663419 |
| Gabrp | 0.357753142 | 0.274416195 | 1.579547554 | 0.173730885 |
| D130052B06Rik | 0 | NA | NA | NA |
| Kcnip1 | 277.5432354 | -0.042510459 | 0.079695789 | -0.5334091 |
| Gm12119 | 0.042716945 | -0.206490061 | 3.724945049 | -0.055434391 |
| Gm12120 | 0.062426873 | 0.755288839 | 3.724945049 | 0.202765096 |
| Kcnmb1 | 5.253596414 | -0.749778183 | 0.478853009 | -1.565779411 |
| 4930469K13Rik | 1.763134642 | -0.819390148 | 0.867983328 | -0.944015999 |
| Lcp2 | 190.2229504 | 3.048233626 | 0.181247815 | 16.81804341 |
| Gm12121 | 0 | NA | NA | NA |
| Foxi1 | 0.275123771 | 0.717838825 | 1.811883765 | 0.396183706 |

| | | | | |
|---|---|---|---|---|
| Dock2 | 455.1128129 | 2.951950116 | 0.164941081 | 17.89699752 |
| 4930403D09Rik | 0.062426873 | 0.755288839 | 3.724945049 | 0.202765096 |
| Fam196b | 116.3261388 | -0.2049849 | 0.162142487 | -1.264226936 |
| Spdl1 | 20.02293662 | 1.184297218 | 0.299382054 | 3.95580564 |
| Gm22022 | 0 | NA | NA | NA |
| Gm12711 | 0 | NA | NA | NA |
| Gm25799 | 0 | NA | NA | NA |
| Slit3 | 590.7003565 | -0.336768123 | 0.145563615 | -2.313546024 |
| Gm26070 | 1.06710716 | 0.812239861 | 1.130224136 | 0.718653792 |
| Gm12123 | 0 | NA | NA | NA |
| Gm12122 | 14.36718598 | -1.320066404 | 0.435699249 | -3.029765159 |
| Mir218-2 | 2.110899703 | -0.314739632 | 0.655582491 | -0.480091578 |
| Pank3 | 1420.981158 | -0.227723081 | 0.066769737 | -3.410573263 |
| Mir103-1 | 0.791513252 | -0.509620792 | 1.132580656 | -0.449964238 |
| Gm12124 | 1.289573466 | -0.080783022 | 0.851097395 | -0.094916307 |
| Fbll1 | 180.3183456 | -0.06804047 | 0.121465958 | -0.560160816 |
| Rars | 540.4176717 | 0.524235554 | 0.101683715 | 5.155550773 |
| Wwc1 | 973.1767011 | -0.224988201 | 0.068845385 | -3.268021535 |
| Gm12125 | 0 | NA | NA | NA |
| Tenm2 | 1701.648525 | -0.081079979 | 0.109199487 | -0.742494136 |
| Gm12126 | 0 | NA | NA | NA |
| Gm12127 | 0 | NA | NA | NA |
| Gm22127 | 0 | NA | NA | NA |
| Gm12128 | 0.048949532 | -0.206487837 | 3.724945049 | -0.055433794 |
| Gm25693 | 0.047948698 | -0.206487837 | 3.724945049 | -0.055433794 |
| Gm12129 | 0.066050875 | 0.274391604 | 3.724945049 | 0.073663262 |
| Gm12130 | 1.911894038 | -0.926041492 | 0.667000474 | -1.388367067 |
| Gm23520 | 0.097501745 | -0.206492286 | 3.724945049 | -0.055434988 |
| Gm12131 | 1.727409906 | -1.845338069 | 0.944526825 | -1.953716951 |
| Gm24192 | 0 | NA | NA | NA |
| Gm12132 | 0 | NA | NA | NA |
| Gm12133 | 0 | NA | NA | NA |
| Gm12134 | 0 | NA | NA | NA |
| 3110004A20Rik | 0.20299191 | -0.206486848 | 2.612742879 | -0.07903068 |
| Gm12135 | 0 | NA | NA | NA |
| Gm12136 | 0 | NA | NA | NA |
| Gm12137 | 0 | NA | NA | NA |
| Gm12138 | 2.171605266 | 0.932198143 | 0.677197211 | 1.376553429 |
| Mat2b | 871.6522152 | -0.110839856 | 0.073645944 | -1.50503679 |
| Hmmr | 18.26964906 | 1.265736655 | 0.386663169 | 3.27348648 |
| Nudcd2 | 124.2665417 | 0.135801175 | 0.125594385 | 1.081267886 |
| Ccng1 | 1174.108608 | 0.261729783 | 0.109958218 | 2.380265767 |
| Gm24446 | 0 | NA | NA | NA |
| Hmgb1-ps1 | 6.658866634 | 0.581389801 | 0.365883485 | 1.58900258 |
| Platr13 | 0 | NA | NA | NA |
| Gm12140 | 0 | NA | NA | NA |
| Hspd1-ps3 | 127.1742578 | 0.377209617 | 0.191504068 | 1.969721172 |

| | | | | |
|---|---|---|---|---|
| Gm12142 | 0 | NA | NA | NA |
| A230004M16Rik | 12.26251621 | 1.500630177 | 0.54421218 | 2.757435854 |
| Gm12143 | 0 | NA | NA | NA |
| Gabrg2 | 1919.016568 | -0.144366485 | 0.074326902 | -1.942318069 |
| Gabra1 | 5062.268846 | -0.049546488 | 0.087611534 | -0.565524717 |
| Gabra6 | 1128.6417 | -0.670513851 | 0.375308749 | -1.786566001 |
| Gabrb2 | 2861.342192 | -0.175930897 | 0.090813818 | -1.937270136 |
| Gm25543 | 0 | NA | NA | NA |
| Gm9972 | 0.444159033 | -2.694740568 | 1.509159056 | -1.785590828 |
| Gm12145 | 0 | NA | NA | NA |
| Atp10b | 25.10547728 | -1.054000233 | 0.20504193 | -5.140413135 |
| Gm12146 | 1.289012822 | 2.330834178 | 1.150578154 | 2.025793875 |
| Gm12144 | 0.970234074 | -0.425845543 | 0.898892423 | -0.473744724 |
| Gm12147 | 0 | NA | NA | NA |
| Mir146 | 0.061920613 | 0.274393828 | 3.724945049 | 0.073663859 |
| Gm12148 | 0 | NA | NA | NA |
| Pttg1 | 36.67629004 | 3.236423314 | 0.333743324 | 9.697342474 |
| Slu7 | 647.400309 | 0.313464793 | 0.070229889 | 4.463410046 |
| Gm12151 | 0 | NA | NA | NA |
| C1qtnf2 | 6.55721845 | 0.182446018 | 0.384606197 | 0.474370979 |
| Ccnjl | 56.86863016 | 0.054125755 | 0.166126967 | 0.325809567 |
| Gm12150 | 0 | NA | NA | NA |
| 4933415A04Rik | 9.074042005 | 0.300385203 | 0.306907669 | 0.978747793 |
| Fabp6 | 0 | NA | NA | NA |
| Gm12149 | 0 | NA | NA | NA |
| Gm12152 | 0 | NA | NA | NA |
| Pwwp2a | 556.3087863 | -0.230077494 | 0.084035846 | -2.737849441 |
| Ttc1 | 454.7934483 | 0.279132715 | 0.106726844 | 2.615393708 |
| Adra1b | 167.6708051 | 0.338395021 | 0.125531907 | 2.695689321 |
| Gm12153 | 0.799763445 | -0.435647241 | 1.175232379 | -0.370690298 |
| Gm12154 | 0 | NA | NA | NA |
| Gm12155 | 0.038981395 | 0.274393828 | 3.724945049 | 0.073663859 |
| Il12b | 80.53366365 | 9.94114545 | 1.268887988 | 7.8345335 |
| Ublcp1 | 282.0163601 | -0.103842019 | 0.10041199 | -1.034159556 |
| Gm22121 | 0 | NA | NA | NA |
| Gm12156 | 0.547961811 | 1.561149322 | 1.406786295 | 1.109727417 |
| Rnf145 | 1154.947274 | -0.066053717 | 0.049546285 | -1.333171961 |
| 4930597A21Rik | 0.434349921 | 2.421788215 | 1.479100499 | 1.637338515 |
| Gm37695 | 1.050344328 | -0.004975597 | 0.982580901 | -0.005063804 |
| AL603913.1 | 0 | NA | NA | NA |
| Ebf1 | 264.2278728 | 0.139276248 | 0.124809678 | 1.115909045 |
| Gm12158 | 0 | NA | NA | NA |
| Gm12159 | 0 | NA | NA | NA |
| Gm12160 | 0 | NA | NA | NA |
| Gm38087 | 0 | NA | NA | NA |
| Gm37312 | 0 | NA | NA | NA |
| Gm12161 | 1.855823375 | -2.15902801 | 0.963622438 | -2.240533143 |

| | | | | |
|---|---|---|---|---|
| Gm12162 | 0 | NA | NA | NA |
| Gm22284 | 0 | NA | NA | NA |
| F630206G17Rik | 0.072025462 | 0.755288839 | 3.724945049 | 0.202765096 |
| Gm12163 | 0.042716945 | -0.206490061 | 3.724945049 | -0.055434391 |
| Gm12164 | 1.520812657 | 1.829272748 | 1.067139347 | 1.714183582 |
| Gm22751 | 0 | NA | NA | NA |
| Clint1 | 800.9757842 | 0.187467646 | 0.064015788 | 2.928459554 |
| Lsm11 | 391.468034 | -0.645563594 | 0.111167579 | -5.80712108 |
| Thg1l | 79.70507417 | 0.173538142 | 0.128475839 | 1.35074535 |
| Sox30 | 1.344747471 | -0.810262759 | 1.005058682 | -0.806184528 |
| Gm12165 | 0 | NA | NA | NA |
| Gm12166 | 16.95436423 | 0.761973738 | 0.273506206 | 2.785946797 |
| Adam19 | 340.8717861 | -0.454598957 | 0.144430069 | -3.147536799 |
| Mir8100 | 0.635485296 | -1.808792314 | 1.512158702 | -1.196165661 |
| Gm16033 | 0.647121192 | -0.294184107 | 1.196984635 | -0.245770997 |
| Gm16034 | 0.200316428 | -0.206483604 | 2.661959317 | -0.077568279 |
| Nipal4 | 54.85333669 | -0.192072219 | 0.203054706 | -0.945913657 |
| Mir7239 | 0.223868884 | -0.687378133 | 2.590860021 | -0.265308866 |
| Cyfip2 | 10488.44967 | -0.070566736 | 0.086389461 | -0.816844266 |
| Fndc9 | 118.2743247 | 0.559330662 | 0.405808946 | 1.378310328 |
| Gm12568 | 12.41784929 | -0.692954168 | 0.372155715 | -1.862000607 |
| Itk | 74.98247743 | 3.193790819 | 0.273597013 | 11.67333952 |
| Gm12167 | 0.124347486 | 0.755286736 | 3.717316019 | 0.203180664 |
| Fam71b | 0.55691522 | 2.353498848 | 1.527182043 | 1.541072892 |
| Med7 | 299.1592794 | 0.356638339 | 0.108145423 | 3.29776636 |
| 2310031A07Rik | 13.18712137 | 5.567002593 | 0.928135113 | 5.998051914 |
| Gm12168 | 5.395875324 | -0.17076647 | 0.473007711 | -0.361022591 |
| Havcr2 | 109.3330086 | 2.320345662 | 0.213632075 | 10.8614105 |
| Gm12169 | 0.124308499 | 0.274388155 | 3.719824689 | 0.073763733 |
| Gm12170 | 0 | NA | NA | NA |
| Gm12171 | 0.288794137 | 0.75530985 | 1.802518342 | 0.419030327 |
| Rpl9-ps2 | 0 | NA | NA | NA |
| BC053393 | 0.265064467 | 0.730016995 | 1.809916268 | 0.403342965 |
| Dppa1 | 0.143022002 | 0.755289479 | 3.547809333 | 0.212888971 |
| Gm4926 | 0.476246927 | 0.364030541 | 1.368048115 | 0.266094837 |
| Gm12173 | 0 | NA | NA | NA |
| Timd2 | 0.095435502 | 0.75528439 | 3.724945049 | 0.202763901 |
| Gm12174 | 30.95181581 | 0.526313337 | 0.263536878 | 1.99711456 |
| Havcr1 | 1.532951352 | 2.58441932 | 1.229298681 | 2.102352634 |
| Timd4 | 7.437687587 | 4.679585084 | 0.899526897 | 5.202273661 |
| Gm12175 | 0 | NA | NA | NA |
| Sgcd | 134.9314582 | -0.277618084 | 0.126006866 | -2.203198066 |
| Gm12176 | 0.97553329 | 0.508628354 | 0.922617152 | 0.551288639 |
| Gm23039 | 4.120196972 | -0.852856061 | 0.510394997 | -1.67097261 |
| Gm12178 | 2.707360892 | 0.155797737 | 0.586765871 | 0.265519425 |
| Gm12179 | 0.104637558 | -0.206492286 | 3.724945049 | -0.055434988 |
| Gm12180 | 0 | NA | NA | NA |

| | | | | |
|---|---|---|---|---|
| Gm12181 | 0.114443069 | -0.206492286 | 3.724945049 | -0.055434988 |
| Gm24422 | 0 | NA | NA | NA |
| Gm12182 | 0.476797581 | -0.207267717 | 1.428133919 | -0.145131849 |
| Gm12183 | 218.1818688 | 0.308788186 | 0.112935637 | 2.734196165 |
| Gnb2l1 | 1496.591963 | 0.450714106 | 0.118286275 | 3.810366882 |
| Gm25296 | 0 | NA | NA | NA |
| Snord96a | 0 | NA | NA | NA |
| Snord95 | 0 | NA | NA | NA |
| Trim41 | 551.7216766 | 0.292052569 | 0.064443069 | 4.531946915 |
| Gm12184 | 52.49590673 | -0.247429047 | 0.170647585 | -1.449941687 |
| Trim7 | 26.45757724 | 0.160720761 | 0.229995205 | 0.698800487 |
| Gm16170 | 11.86762837 | 3.223369156 | 0.533346582 | 6.04366703 |
| Irgm1 | 5470.912599 | 6.531472054 | 0.189073139 | 34.54468509 |
| Gm5431 | 192.2399859 | 5.036111448 | 0.341892676 | 14.73009458 |
| Gm12185 | 296.7910131 | 8.24225362 | 0.822172924 | 10.02496358 |
| Psme2b | 176.4926986 | 2.562196803 | 0.193754817 | 13.22391279 |
| 9930111J21Rik1 | 96.17708593 | 4.063796438 | 0.321533662 | 12.63879003 |
| Olfr56 | 27.72815539 | 7.497739981 | 1.007199079 | 7.444148966 |
| Tgtp1 | 654.3535308 | 8.389218678 | 0.575480023 | 14.57777566 |
| Gm12186 | 0.311786641 | -0.206485711 | 1.982642245 | -0.104146732 |
| 9930111J21Rik2 | 219.1877081 | 4.535642032 | 0.240588261 | 18.85229985 |
| Tgtp2 | 962.1752517 | 9.996337885 | 0.716701057 | 13.94770914 |
| Gm12187 | 0.675604184 | 1.236206545 | 1.489773362 | 0.829795039 |
| Ifi47 | 1143.870919 | 8.292657423 | 0.438577737 | 18.90806744 |
| Gm12188 | 0.333018454 | 1.231950073 | 1.7517077 | 0.703285185 |
| Olfr1396 | 0 | NA | NA | NA |
| Olfr1395 | 0.571466717 | 2.336725878 | 1.616112938 | 1.445892686 |
| Olfr1394 | 0.066050875 | 0.274391604 | 3.724945049 | 0.073663262 |
| Btnl9 | 1.653010361 | 3.644327225 | 1.076158563 | 3.386422179 |
| Zfp62 | 822.4603642 | -0.05764701 | 0.051444196 | -1.120573648 |
| Mgat1 | 662.2687355 | 0.322959377 | 0.077651061 | 4.159110948 |
| Olfr1393 | 2.448580025 | 0.04733384 | 0.730227197 | 0.064820703 |
| Olfr1392 | 1.186358578 | 1.528013081 | 1.079499944 | 1.415482316 |
| Olfr10 | 0.306560087 | 0.226725195 | 1.589446252 | 0.142644141 |
| Olfr1391 | 0.124596252 | 0.755282361 | 3.714681068 | 0.203323609 |
| Olfr1390 | 0.188595572 | 1.807191869 | 3.285103702 | 0.550117145 |
| Gm12189 | 0 | NA | NA | NA |
| Olfr1389 | 0.061920613 | 0.274393828 | 3.724945049 | 0.073663859 |
| Olfr1388 | 0 | NA | NA | NA |
| Olfr1387 | 0.046345709 | 0.274391604 | 3.724945049 | 0.073663262 |
| Olfr1386 | 0 | NA | NA | NA |
| Olfr1385 | 0 | NA | NA | NA |
| Olfr1384 | 0.045882455 | 0.755288839 | 3.724945049 | 0.202765096 |
| Olfr1383 | 0 | NA | NA | NA |
| Olfr1382 | 0 | NA | NA | NA |
| Olfr1381 | 0 | NA | NA | NA |
| Olfr1380 | 0 | NA | NA | NA |

| | | | | |
|---|---|---|---|---|
| 1700024J04Rik | 0.046345709 | 0.274391604 | 3.724945049 | 0.073663262 |
| Gm12190 | 0 | NA | NA | NA |
| Flt4 | 66.9739403 | 1.586566586 | 0.225299131 | 7.042044855 |
| Scgb3a1 | 5.818159514 | 3.894839902 | 0.825227193 | 4.719718319 |
| Cnot6 | 1091.733663 | -0.00452956 | 0.077777627 | -0.058237316 |
| Gm12191 | 162.606518 | 0.413340657 | 0.143646444 | 2.877486181 |
| Gm12192 | 0 | NA | NA | NA |
| Gm23813 | 0 | NA | NA | NA |
| Gm12193 | 0.060174537 | 0.755286614 | 3.724945049 | 0.202764498 |
| Gfpt2 | 101.9890799 | 0.24615632 | 0.135558423 | 1.815868868 |
| Mapk9 | 2585.547055 | -0.073208313 | 0.048779243 | -1.500808708 |
| Gm23492 | 0.042716945 | -0.206490061 | 3.724945049 | -0.055434391 |
| Rasgef1c | 167.346068 | -0.13076172 | 0.135435447 | -0.96549111 |
| Rnf130 | 704.3708933 | -0.170558222 | 0.057002499 | -2.992118323 |
| Mir340 | 5.005408619 | -0.377855732 | 0.489356251 | -0.772148576 |
| Gm12194 | 0.07796279 | 0.274392189 | 3.724945049 | 0.073663419 |
| Tbc1d9b | 2240.773314 | 0.037722627 | 0.079335417 | 0.475482807 |
| Gm12195 | 0 | NA | NA | NA |
| 3010026O09Rik | 75.01584386 | 0.264511207 | 0.14056392 | 1.881785929 |
| Sqstm1 | 5575.809125 | 0.483939427 | 0.098971497 | 4.889684798 |
| Mgat4b | 410.2977067 | -0.081446034 | 0.094909683 | -0.85814251 |
| Mir6919 | 0.119234754 | 0.274400501 | 3.724945049 | 0.073665651 |
| Ltc4s | 8.403515007 | -0.530219349 | 0.389904725 | -1.359869001 |
| Gm26542 | 2.8898106 | -0.875581461 | 0.578641346 | -1.513167814 |
| Maml1 | 333.2785951 | 0.314736541 | 0.127870623 | 2.461367063 |
| Canx | 5951.612133 | 0.100970241 | 0.081946979 | 1.232141107 |
| Gm12196 | 0.048949532 | -0.206487837 | 3.724945049 | -0.055433794 |
| Cby3 | 1.08873821 | 0.481348513 | 1.097931294 | 0.438414057 |
| Mir804 | 0.587766037 | -0.098169305 | 1.242620527 | -0.079001837 |
| Hnrnph1 | 2118.885193 | -0.318951045 | 0.085612235 | -3.725531119 |
| Rufy1 | 315.9868514 | 0.080042507 | 0.084677289 | 0.945265348 |
| Gm23824 | 0 | NA | NA | NA |
| Gm12198 | 7.905317881 | 0.356340662 | 0.354428809 | 1.005394183 |
| Adamts2 | 218.5703262 | 0.133587252 | 0.097136741 | 1.375249472 |
| Gm12199 | 0 | NA | NA | NA |
| Gm12200 | 0.062426873 | 0.755288839 | 3.724945049 | 0.202765096 |
| Gm23392 | 0 | NA | NA | NA |
| Zfp354c | 737.6052082 | -0.045972241 | 0.076449744 | -0.601339373 |
| Zfp879 | 31.60590681 | 0.002503354 | 0.213685139 | 0.011715152 |
| Grm6 | 0 | NA | NA | NA |
| Gm12660 | 0 | NA | NA | NA |
| Zfp454 | 55.6120256 | -0.633226689 | 0.163302867 | -3.877621373 |
| Zfp2 | 299.1331036 | -0.123739929 | 0.079528006 | -1.555928976 |
| Zfp354b | 46.21141538 | -0.879413457 | 0.182423952 | -4.820712658 |
| 4933414I15Rik | 0.060174537 | 0.755286614 | 3.724945049 | 0.202764498 |
| Prop1 | 0.410687608 | 1.908542467 | 1.702160448 | 1.1212471 |
| Olfr1379-ps1 | 0.047948698 | -0.206487837 | 3.724945049 | -0.055433794 |

| | | | | |
|---|---|---|---|---|
| Olfr1378 | 0.11037557 | 0.274404949 | 3.724945049 | 0.073666845 |
| Gm12661 | 0 | NA | NA | NA |
| Olfr1377 | 0.049553047 | 0.274391604 | 3.724945049 | 0.073663262 |
| Olfr1376-ps1 | 0 | NA | NA | NA |
| Olfr51 | 0 | NA | NA | NA |
| Olfr54 | 0 | NA | NA | NA |
| Olfr1375 | 0 | NA | NA | NA |
| Zfp354a | 104.6133705 | -0.338676834 | 0.149689678 | -2.26252631 |
| 9230009I02Rik | 1.062461154 | -0.333881332 | 0.975247472 | -0.342355495 |
| Gm12202 | 0 | NA | NA | NA |
| Gm12201 | 0.113999573 | -0.206492286 | 3.724945049 | -0.055434988 |
| Olfr1374-ps1 | 0 | NA | NA | NA |
| Gm12203 | 0 | NA | NA | NA |
| Platr20 | 0.066050875 | 0.274391604 | 3.724945049 | 0.073663262 |
| Gm12569 | 0.212513282 | 1.926509226 | 3.159662244 | 0.609719988 |
| BC049762 | 1.440851686 | 2.877820106 | 1.036320468 | 2.776959631 |
| Clk4 | 881.9365492 | -0.155553227 | 0.089617041 | -1.735755005 |
| Gm25082 | 22.05723362 | -0.469541113 | 0.282611372 | -1.661437436 |
| Col23a1 | 236.6432467 | 0.441314626 | 0.145948073 | 3.023778365 |
| Phykpl | 90.42005935 | -0.571664115 | 0.183611648 | -3.113441439 |
| Hnrnpab | 1099.124842 | 0.465313916 | 0.124285756 | 3.743903822 |
| Nhp2 | 191.8689779 | 0.259470599 | 0.109145143 | 2.377298629 |
| Rmnd5b | 328.920745 | 0.136391592 | 0.070918199 | 1.923224134 |
| N4bp3 | 118.9340086 | 0.016514004 | 0.104119475 | 0.158606293 |
| D930048N14Rik | 22.3543129 | -0.456970567 | 0.373649513 | -1.222992541 |
| 0610009B22Rik | 109.4926551 | -0.191033128 | 0.154297215 | -1.238085397 |
| Sec24a | 739.2067926 | -0.001860161 | 0.096713418 | -0.019233743 |
| Gm25291 | 176.1642292 | 0.13276585 | 0.306869625 | 0.432645786 |
| Sar1b | 418.9258972 | 0.192598871 | 0.089655628 | 2.148207262 |
| Jade2 | 1658.936098 | 0.726927577 | 0.09583476 | 7.58521828 |
| Gm26551 | 2.891936054 | -1.049416994 | 0.741549691 | -1.415167463 |
| Gm26061 | 0 | NA | NA | NA |
| Cdkn2aipnl | 319.6958028 | 0.035472441 | 0.087868139 | 0.403700829 |
| Ube2b | 1147.001563 | -0.143192646 | 0.061277187 | -2.336801907 |
| Gm12204 | 0.487092162 | -1.261038319 | 1.487365051 | -0.84783377 |
| Cdkl3 | 149.4357486 | 0.191248145 | 0.09415505 | 2.031204322 |
| Gm12205 | 0 | NA | NA | NA |
| Ppp2ca | 3811.689111 | 0.036822215 | 0.080745223 | 0.456029641 |
| Gm24157 | 3.140872897 | 0.200687968 | 0.556713324 | 0.360487093 |
| 9430098F02Rik | 0.363523191 | 2.434569851 | 1.87578847 | 1.297891468 |
| Mir3061 | 0 | NA | NA | NA |
| Olfr1373 | 0 | NA | NA | NA |
| Olfr1372-ps1 | 1.830548353 | 0.169773394 | 0.873871463 | 0.194277307 |
| Gm12206 | 0.049553047 | 0.274391604 | 3.724945049 | 0.073663262 |
| Olfr1371 | 0 | NA | NA | NA |
| Skp1a | 2595.731114 | -0.02674266 | 0.077259159 | -0.346142256 |
| Gm12207 | 10.46687507 | -0.641502132 | 0.402316994 | -1.5945191 |

| | | | | |
|---|---|---|---|---|
| Tcf7 | 44.21850987 | 0.224265808 | 0.204818412 | 1.094949448 |
| A630014C17Rik | 2.221200599 | 0.768636454 | 0.90390033 | 0.85035532 |
| Vdac1 | 3622.033645 | -0.125479223 | 0.054226075 | -2.314001572 |
| 9530068E07Rik | 1167.821488 | 0.595090815 | 0.094391182 | 6.30451704 |
| Gm12208 | 2.679228202 | 0.009192724 | 0.654132964 | 0.014053295 |
| Gm12209 | 0 | NA | NA | NA |
| Gm12210 | 0 | NA | NA | NA |
| Fstl4 | 404.2876281 | -0.075055573 | 0.148466712 | -0.505538056 |
| Gm11186 | 0.223616349 | 0.274411746 | 1.885690919 | 0.145523184 |
| Gm11189 | 2.178677355 | 0.164894493 | 0.723002781 | 0.228068961 |
| Hspa4 | 2862.524315 | -0.20012537 | 0.080079752 | -2.499075788 |
| Zcchc10 | 74.48748572 | 0.107980282 | 0.133782852 | 0.807130959 |
| Gm11188 | 0.045882455 | 0.755288839 | 3.724945049 | 0.202765096 |
| Aff4 | 5179.750789 | 0.035156323 | 0.048765281 | 0.720929366 |
| Gm25930 | 3.800172548 | -0.344185878 | 0.526648611 | -0.653539895 |
| mmu-mir-6920 | 1.519428967 | 0.366307868 | 0.808410883 | 0.453120901 |
| Leap2 | 0.397769051 | -0.539394727 | 1.463568504 | -0.368547646 |
| Uqcrq | 757.3825943 | -0.037973782 | 0.133044705 | -0.285421217 |
| Gdf9 | 9.225389834 | -0.644547697 | 0.378529053 | -1.702769424 |
| Gm10447 | 1.155224236 | 0.90492426 | 0.878955024 | 1.029545579 |
| Shroom1 | 58.79842685 | -0.108520433 | 0.14730039 | -0.736728752 |
| Gm12212 | 3.290890966 | -0.017694447 | 0.494973647 | -0.035748261 |
| Gm9837 | 0.267041986 | -1.477439643 | 1.94646358 | -0.759037908 |
| Sowaha | 702.0530503 | 0.228936252 | 0.138136225 | 1.657322347 |
| Gm9945 | 10.46328265 | -0.757221109 | 0.33699653 | -2.246970051 |
| Rpl9-ps3 | 1.040046165 | -1.071356019 | 1.013546598 | -1.057036767 |
| 8-Sep | 2499.725176 | -0.4304779 | 0.066000934 | -6.522300157 |
| A430108G06Rik | 9.283107748 | -0.472100053 | 0.337809405 | -1.397533777 |
| Kif3a | 1223.698615 | -0.313267273 | 0.07714267 | -4.060881918 |
| Il4 | 0.170190954 | 0.755297539 | 2.753251924 | 0.274329251 |
| Il13 | 0 | NA | NA | NA |
| Rps2-ps4 | 0.4561058 | -0.639736618 | 1.599369667 | -0.399992967 |
| Rad50 | 251.7638459 | 0.261465035 | 0.129456842 | 2.019708126 |
| Gm22275 | 0 | NA | NA | NA |
| Il5 | 1.025767513 | -0.245448789 | 0.881044314 | -0.278588472 |
| Gm12214 | 0.972493339 | 3.327798342 | 1.356932614 | 2.452441858 |
| Gm12213 | 0 | NA | NA | NA |
| Mir7671 | 0.940546779 | 1.231715142 | 1.075062598 | 1.145714811 |
| Irf1 | 1452.897363 | 5.604602154 | 0.280278738 | 19.99652985 |
| Gm17334 | 5.090110731 | 2.844070853 | 0.618620196 | 4.597442617 |
| Gm12216 | 33.70519998 | 3.133441327 | 0.365394194 | 8.575509359 |
| Slc22a5 | 121.7288286 | 0.036645968 | 0.119164136 | 0.307525145 |
| Gm12217 | 0 | NA | NA | NA |
| Gm12218 | 0 | NA | NA | NA |
| Gm12220 | 0 | NA | NA | NA |
| Slc22a21 | 25.5234154 | 0.701522205 | 0.304197503 | 2.30614058 |
| Gm12219 | 2.781362659 | 0.885617903 | 0.709855726 | 1.247602677 |

| | | | | |
|---|---|---|---|---|
| Slc22a4 | 168.0969617 | 0.02271208 | 0.120773546 | 0.188055091 |
| Pdlim4 | 206.5108102 | 1.155659515 | 0.167206203 | 6.911582807 |
| P4ha2 | 113.5551428 | -0.428021112 | 0.155410512 | -2.754132311 |
| Gm12221 | 0.192293863 | 0.274409883 | 2.250822277 | 0.121915393 |
| 4933405E24Rik | 0 | NA | NA | NA |
| Gm12222 | 0 | NA | NA | NA |
| Csf2 | 0.705254363 | 1.239219598 | 1.35525144 | 0.914383532 |
| Gm12223 | 0.986197153 | 0.655452442 | 1.032692876 | 0.634702201 |
| Il3 | 0.048949532 | -0.206487837 | 3.724945049 | -0.055433794 |
| Acsl6 | 2735.937845 | -0.439063158 | 0.0642289 | -6.835912815 |
| Gm12224 | 0.727497348 | -1.476103356 | 1.174997479 | -1.256260871 |
| Meikin | 5.386405023 | 3.378575859 | 0.830290393 | 4.069149647 |
| Gm12226 | 3.723226009 | 0.927849062 | 0.623676769 | 1.48770823 |
| Gm12225 | 0 | NA | NA | NA |
| Fnip1 | 1691.944238 | 0.266770386 | 0.068170565 | 3.913278176 |
| Gm24198 | 6.506667764 | -0.846423264 | 0.509125582 | -1.662503897 |
| Rapgef6 | 1160.993809 | 0.188341404 | 0.067631718 | 2.784808791 |
| Gm26116 | 0 | NA | NA | NA |
| Cdc42se2 | 1261.495251 | -0.074153955 | 0.077595921 | -0.955642431 |
| Gm12227 | 0.36191905 | -0.660450448 | 1.589854568 | -0.415415637 |
| Gm12229 | 9.801174999 | -0.009666869 | 0.362127083 | -0.026694688 |
| Lyrm7os | 0.089089296 | 1.194485492 | 3.723939576 | 0.320758559 |
| Lyrm7 | 91.39166592 | -0.413933711 | 0.127451782 | -3.247767166 |
| Gm25426 | 0 | NA | NA | NA |
| Hint1 | 1051.934254 | 0.317617368 | 0.13552831 | 2.343549979 |
| Gpx3 | 335.6017487 | 0.461333808 | 0.278052647 | 1.659159924 |
| Tnip1 | 367.6412068 | 0.667036256 | 0.095895915 | 6.955835951 |
| Gm12230 | 0.092228164 | 0.75528439 | 3.724945049 | 0.202763901 |
| Anxa6 | 1191.581291 | 0.297253255 | 0.073457924 | 4.046578465 |
| Ccdc69 | 10.64187061 | 1.240603586 | 0.351830361 | 3.526141352 |
| Gm2a | 793.8364917 | 0.241286498 | 0.099645413 | 2.421451126 |
| Gm12231 | 50.21335086 | 0.281873828 | 0.202582541 | 1.391402375 |
| Mup-ps22 | 0 | NA | NA | NA |
| Slc36a3 | 0.361016523 | 1.670171364 | 3.164039747 | 0.527860425 |
| Slc36a3os | 0.686935554 | 2.725048965 | 1.236507189 | 2.203827838 |
| Slc36a2 | 7.128692757 | -1.067676665 | 0.388184102 | -2.750438927 |
| Slc36a1os | 0.119234754 | 0.274400501 | 3.724945049 | 0.073665651 |
| Slc36a1 | 1018.826301 | -0.102326575 | 0.052691743 | -1.94198501 |
| Fat2 | 2711.320312 | -0.510376486 | 0.274448713 | -1.859642478 |
| Sparc | 5655.720685 | 0.432109673 | 0.164974528 | 2.619250848 |
| Atox1 | 160.8036094 | -0.022295286 | 0.136994468 | -0.162745885 |
| Gm12235 | 0 | NA | NA | NA |
| G3bp1 | 943.5798911 | -0.062051991 | 0.075202307 | -0.825134147 |
| Gm22884 | 0.510616014 | 1.114989899 | 1.426483093 | 0.781635551 |
| Gm12238 | 0 | NA | NA | NA |
| Mir7652 | 0 | NA | NA | NA |
| Glra1 | 265.7926779 | -0.822652388 | 0.385980403 | -2.131332011 |

| | | | | |
|---|---|---|---|---|
| Gm12236 | 0.101408268 | 0.755286617 | 3.724945049 | 0.202764499 |
| Gm12237 | 0.066050875 | 0.274391604 | 3.724945049 | 0.073663262 |
| Gm12239 | 4.674577041 | -0.429805095 | 0.473830168 | -0.907086809 |
| Nmur2 | 2.399470382 | -0.06199223 | 0.652122877 | -0.095062192 |
| Gm12240 | 0.124308499 | 0.274388155 | 3.719824689 | 0.073763733 |
| Gm24253 | 0 | NA | NA | NA |
| Gm12241 | 0.724742635 | 0.527555525 | 1.104906206 | 0.477466343 |
| Gm22603 | 0 | NA | NA | NA |
| Gria1 | 3990.825874 | -0.234901612 | 0.110005115 | -2.135369905 |
| Gm12242 | 0.047948698 | -0.206487837 | 3.724945049 | -0.055433794 |
| Fam114a2 | 280.8245509 | -0.305018546 | 0.078161366 | -3.902420845 |
| Mfap3 | 364.5954244 | -0.231344342 | 0.079365796 | -2.914912377 |
| Gm12243 | 0.840033521 | 2.899622461 | 1.438378981 | 2.015896019 |
| 4933424L21Rik | 0 | NA | NA | NA |
| Galnt10 | 170.7549866 | -0.183196533 | 0.129265277 | -1.417213784 |
| 4933426K07Rik | 0 | NA | NA | NA |
| Gm24582 | 0.170988841 | -0.206481469 | 2.745561862 | -0.075205542 |
| n-R5s69 | 0.143140564 | 0.274404261 | 3.546633223 | 0.077370352 |
| 2010001A14Rik | 26.6509321 | -0.213473355 | 0.207320413 | -1.029678419 |
| Sap30l | 239.499411 | -0.0086744 | 0.110897272 | -0.078220142 |
| Hand1 | 0.104366086 | 0.274400501 | 3.724945049 | 0.073665651 |
| Gm12244 | 0 | NA | NA | NA |
| Gm12245 | 0.101408268 | 0.755286617 | 3.724945049 | 0.202764499 |
| Gm26025 | 0 | NA | NA | NA |
| Gm12249 | 0 | NA | NA | NA |
| Larp1 | 2687.565109 | 0.259188461 | 0.066178097 | 3.916529363 |
| Gm12248 | 0.378899548 | 0.616375561 | 1.728045489 | 0.35668943 |
| Gm12247 | 0.394465599 | 0.695364183 | 1.742596176 | 0.399039199 |
| Cnot8 | 374.7702201 | 0.185324633 | 0.061623243 | 3.007382017 |
| Gemin5 | 376.1235968 | 0.099107755 | 0.072609977 | 1.364933025 |
| Gm12246 | 2.686282868 | -0.789191272 | 0.760623468 | -1.037558406 |
| Mrpl22 | 119.8290135 | 0.440831087 | 0.126089236 | 3.496183351 |
| Gm12250 | 1005.570989 | 8.512961339 | 0.456012188 | 18.66827591 |
| Igtp | 4765.845661 | 8.36239413 | 0.277078324 | 30.18061463 |
| Irgm2 | 1836.176375 | 7.425021657 | 0.248438762 | 29.88672784 |
| 4930438A08Rik | 0.296006519 | 0.794927653 | 2.0319577 | 0.391212697 |
| Zfp692 | 130.8218891 | -0.564448594 | 0.201349671 | -2.803325141 |
| Zfp672 | 340.7570754 | -0.2784908 | 0.079700486 | -3.494217079 |
| 1700047K16Rik | 2.233748514 | 1.507337784 | 0.666908294 | 2.260187493 |
| Sh3bp5l | 520.8892961 | -0.228463928 | 0.084387494 | -2.707319733 |
| Lypd8 | 0.598341843 | 1.177701306 | 1.526355837 | 0.771577163 |
| Gm12251 | 32.14483831 | 0.210932001 | 0.217983672 | 0.967650461 |
| Gm12252 | 0 | NA | NA | NA |
| 1810065E05Rik | 0.095898756 | 0.27438722 | 3.724945049 | 0.073662085 |
| Gm12253 | 8.666853529 | 5.479328368 | 0.915254309 | 5.986673118 |
| 4930504O13Rik | 0 | NA | NA | NA |
| Olfr30 | 0 | NA | NA | NA |

| | | | | |
|---|---|---|---|---|
| Olfr333-ps1 | 0 | NA | NA | NA |
| Olfr332 | 0.045882455 | 0.755288839 | 3.724945049 | 0.202765096 |
| Olfr331 | 0 | NA | NA | NA |
| Gm12254 | 27.93282506 | 0.511623683 | 0.221231244 | 2.312619473 |
| Olfr330 | 0.072025462 | 0.755288839 | 3.724945049 | 0.202765096 |
| Gm12255 | 0 | NA | NA | NA |
| Olfr329-ps | 0.0878635 | -0.687376175 | 3.724945049 | -0.184533239 |
| Olfr328 | 0.068097359 | -0.206487837 | 3.724945049 | -0.055433794 |
| Olfr224 | 0 | NA | NA | NA |
| Olfr325 | 0 | NA | NA | NA |
| Olfr324 | 0.389730154 | 2.329117289 | 1.620007911 | 1.4377197 |
| 2210407C18Rik | 0.08169834 | -0.206492286 | 3.724945049 | -0.055434988 |
| Olfr323 | 0.122991902 | 0.274402725 | 3.724945049 | 0.073666248 |
| Trim58 | 1.195637209 | 1.11949657 | 0.903596908 | 1.238933599 |
| Gm22599 | 0 | NA | NA | NA |
| Olfr322 | 0 | NA | NA | NA |
| Olfr320 | 0 | NA | NA | NA |
| Gm12570 | 0 | NA | NA | NA |
| Olfr319 | 0 | NA | NA | NA |
| Olfr318 | 0 | NA | NA | NA |
| Olfr317 | 0 | NA | NA | NA |
| Olfr316 | 0 | NA | NA | NA |
| Olfr315 | 0.049553047 | 0.274391604 | 3.724945049 | 0.073663262 |
| Olfr314 | 0 | NA | NA | NA |
| Fam183b | 11.51123108 | -0.04941927 | 0.455815321 | -0.108419501 |
| Olfr313 | 0 | NA | NA | NA |
| Olfr312 | 0 | NA | NA | NA |
| Olfr311 | 0 | NA | NA | NA |
| Gm12258 | 234.0799246 | -0.254733366 | 0.095718986 | -2.661262681 |
| Gm12259 | 3.878080432 | 0.948097707 | 0.704022876 | 1.346685937 |
| 2810021J22Rik | 159.8131489 | -0.18301021 | 0.089685394 | -2.040579886 |
| Zfp39 | 105.0647483 | 0.635168191 | 0.15411465 | 4.121400461 |
| Gm12256 | 0.146451277 | -0.68737672 | 2.882358805 | -0.238477152 |
| Gm12257 | 0.048949532 | -0.206487837 | 3.724945049 | -0.055433794 |
| Btnl10 | 1.324943677 | -0.469812809 | 1.027099366 | -0.457417096 |
| Rnf187 | 2464.147731 | -0.019631345 | 0.059144538 | -0.33192152 |
| Hist3h2ba | 174.4290419 | -0.052570862 | 0.134328157 | -0.391361451 |
| Hist3h2bb-ps | 13.70298843 | 0.240971651 | 0.263996998 | 0.912781785 |
| Trim17 | 59.87795607 | 0.792603384 | 0.215113407 | 3.684583838 |
| Hist3h2a | 106.0988195 | 0.190023767 | 0.155488319 | 1.222109599 |
| AL662809.1 | 1.743011301 | 1.180126155 | 0.874996299 | 1.348721311 |
| Gm12260 | 5.27939244 | 0.782802218 | 0.457244679 | 1.711998531 |
| 2310058D17Rik | 14.57587431 | -1.111775949 | 0.340782561 | -3.262420312 |
| Trim11 | 219.4000615 | -0.055352586 | 0.118389957 | -0.467544604 |
| Obscn | 127.4374054 | -0.322050567 | 0.280483975 | -1.148195959 |
| Gm10435 | 0.809248464 | -0.463077997 | 1.194034075 | -0.38782645 |
| Iba57 | 77.73534886 | 0.075540942 | 0.143613144 | 0.526002981 |

| | | | | |
|---|---|---|---|---|
| Gjc2 | 99.21955442 | -0.753311413 | 0.178941235 | -4.209825716 |
| Guk1 | 493.4726625 | 0.073278436 | 0.114735211 | 0.638674341 |
| 2610507I01Rik | 3.133265115 | -0.573666057 | 0.584236313 | -0.981907568 |
| Mrpl55 | 89.18665174 | -0.008302327 | 0.176990202 | -0.046908399 |
| Gm10434 | 1.747669216 | -0.402219663 | 0.87413789 | -0.460132969 |
| 2310033P09Rik | 92.90886109 | -0.337411767 | 0.140836891 | -2.39576268 |
| Arf1 | 2962.793734 | 0.187228148 | 0.069516165 | 2.693303754 |
| Wnt3a | 0.23148996 | -1.184170364 | 2.542682818 | -0.465716902 |
| Wnt9a | 76.15545421 | -0.88069366 | 0.161524888 | -5.452371264 |
| Gm24756 | 0 | NA | NA | NA |
| Prss38 | 1.918719673 | 1.097245481 | 0.936059335 | 1.172196505 |
| Gm15755 | 0 | NA | NA | NA |
| Snap47 | 2340.344067 | -0.073896048 | 0.052609188 | -1.404622464 |
| Jmjd4 | 428.2804086 | -0.159578415 | 0.063688321 | -2.505615037 |
| Zfp867 | 99.85436412 | 0.66264399 | 0.17980735 | 3.685299799 |
| Gm12263 | 27.6794227 | 0.32983816 | 0.227049289 | 1.45271611 |
| Gm12262 | 0 | NA | NA | NA |
| Zkscan17 | 266.3947386 | 0.765959907 | 0.124997501 | 6.127801784 |
| 4933439C10Rik | 96.1816591 | -0.739869125 | 0.18158933 | -4.074408579 |
| Nlrp3 | 44.3755037 | 2.300204246 | 0.230479492 | 9.980082074 |
| Olfr222 | 0.26446025 | 0.274412617 | 2.069732024 | 0.132583646 |
| Gm12261 | 0.418705261 | 0.25087784 | 1.425480174 | 0.175995321 |
| Olfr223 | 0 | NA | NA | NA |
| Olfr225 | 0 | NA | NA | NA |
| Mprip | 6005.10193 | 0.08607943 | 0.060106056 | 1.432125745 |
| 1700007J10Rik | 1.636732283 | -1.231974469 | 0.848299065 | -1.452287901 |
| Gm12264 | 0.290485957 | -1.054627115 | 1.772899275 | -0.594860142 |
| Pld6 | 8.31364424 | -0.381607202 | 0.381911805 | -0.999202427 |
| Flcn | 628.2729707 | -0.052218205 | 0.063495941 | -0.822386507 |
| Gm16062 | 8.661166123 | -0.766130437 | 0.371516505 | -2.062170662 |
| Gm24939 | 0.352988078 | -0.494923143 | 1.428969072 | -0.346349794 |
| Cops3 | 618.5280119 | 0.355064951 | 0.082030291 | 4.328461429 |
| Nt5m | 258.4297601 | 0.001771578 | 0.077409602 | 0.022885772 |
| Gm12714 | 0.04393175 | -0.206490061 | 3.724945049 | -0.055434391 |
| 1810063I02Rik | 0.161948271 | -0.68737637 | 3.446645093 | -0.199433464 |
| Med9os | 2.791248013 | -1.121598335 | 0.629296172 | -1.782305989 |
| Med9 | 292.2435254 | -0.031879186 | 0.092937029 | -0.343019211 |
| Rasd1 | 132.7603455 | 1.191827026 | 0.200323986 | 5.949497367 |
| Pemt | 14.2244874 | 0.033341423 | 0.285971446 | 0.116590044 |
| Gm27511 | 0 | NA | NA | NA |
| Rai1 | 1366.993626 | -0.307996612 | 0.079887618 | -3.855373563 |
| 4930412M03Rik | 2.074250591 | 1.168597923 | 0.758644584 | 1.540376017 |
| Srebf1 | 894.7642396 | 0.526789953 | 0.126220049 | 4.173583822 |
| Mir6921 | 0 | NA | NA | NA |
| Mir6922 | 0 | NA | NA | NA |
| Tom1l2 | 3217.981596 | -0.108449195 | 0.068737676 | -1.577725651 |
| Gm12265 | 0.343978253 | 0.27441335 | 1.56586091 | 0.175247589 |

| | | | | |
|---|---|---|---|---|
| Gm27711 | 2.937723515 | -0.978114666 | 0.673350062 | -1.452609454 |
| Gm12266 | 0 | NA | NA | NA |
| Lrrc48 | 90.36745919 | 0.129551948 | 0.119325994 | 1.085697626 |
| Atpaf2 | 218.7768415 | 0.137871951 | 0.075641391 | 1.822705134 |
| Gid4 | 732.8917463 | -0.216465816 | 0.089749386 | -2.411891877 |
| Gm12268 | 1.686071008 | -1.693383958 | 0.900952769 | -1.879547981 |
| Gm12267 | 0.287879534 | 0.983345054 | 1.732894882 | 0.567457994 |
| Drg2 | 463.374575 | 0.107667205 | 0.065478621 | 1.644310819 |
| Myo15 | 13.54234393 | 0.470967276 | 0.454937646 | 1.035234784 |
| Alkbh5 | 1342.8845 | 0.070753834 | 0.06855597 | 1.032059402 |
| Gm12622 | 0 | NA | NA | NA |
| Gm12628 | 0 | NA | NA | NA |
| Gm26837 | 0 | NA | NA | NA |
| Gm12623 | 0.044191549 | -0.206490061 | 3.724945049 | -0.055434391 |
| Gm12621 | 0 | NA | NA | NA |
| Gm12627 | 0 | NA | NA | NA |
| Gm12613 | 0 | NA | NA | NA |
| Gm12614 | 0 | NA | NA | NA |
| Gm12615 | 0 | NA | NA | NA |
| Gm12626 | 0 | NA | NA | NA |
| Gm12625 | 0 | NA | NA | NA |
| Gm12619 | 0 | NA | NA | NA |
| Gm12624 | 0 | NA | NA | NA |
| Gm12612 | 0 | NA | NA | NA |
| Gm12616 | 0 | NA | NA | NA |
| Gm12620 | 0 | NA | NA | NA |
| Gm12617 | 0 | NA | NA | NA |
| Llgl1 | 814.2720307 | 0.073783961 | 0.087641293 | 0.841885814 |
| Flii | 919.6804464 | 0.130882749 | 0.097280345 | 1.345418228 |
| Mief2 | 152.1320646 | 0.228495557 | 0.111272406 | 2.053479073 |
| Mir5100 | 0.418037274 | -2.309060104 | 3.287127819 | -0.702455223 |
| Gm23341 | 0.048949532 | -0.206487837 | 3.724945049 | -0.055433794 |
| Top3a | 117.1519205 | 0.614495756 | 0.117179225 | 5.244067399 |
| Smcr8 | 1137.612935 | 0.357711176 | 0.078088937 | 4.580817589 |
| Shmt1 | 28.85367593 | 1.062273684 | 0.211251951 | 5.028468048 |
| Gm24945 | 0.047948698 | -0.206487837 | 3.724945049 | -0.055433794 |
| Gm12611 | 0.097139547 | 0.274393828 | 3.724945049 | 0.073663859 |
| Gm12618 | 0.132101749 | 0.274387056 | 3.724945049 | 0.073662041 |
| Dhrs7b | 155.5720981 | -0.208614549 | 0.142034065 | -1.468764196 |
| Tmem11 | 223.7050137 | 0.446087095 | 0.106876066 | 4.17387272 |
| Gm26534 | 14.74527245 | -0.539615142 | 0.282968874 | -1.906977028 |
| Natd1 | 186.5605054 | -0.028079496 | 0.126781187 | -0.221479987 |
| Map2k3os | 10.95972877 | 0.077616211 | 0.430980126 | 0.18009232 |
| Map2k3 | 195.4668182 | 0.497238162 | 0.155145255 | 3.204984651 |
| Kcnj12 | 243.7727264 | -0.402893026 | 0.141849272 | -2.840289698 |
| Tnfrsf13b | 25.743611 | 1.348928703 | 0.345544266 | 3.903779735 |
| Gm12269 | 0 | NA | NA | NA |

| | | | | |
|---|---|---|---|---|
| Usp22 | 3131.14818 | 0.121470789 | 0.053143278 | 2.285722569 |
| Rps13-ps5 | 0.882161971 | 0.509650625 | 1.046710238 | 0.48690708 |
| Aldh3a1 | 6.417984165 | 0.045980053 | 0.412987628 | 0.111335182 |
| Aldh3a2 | 629.8388114 | 0.138143559 | 0.098026062 | 1.409253374 |
| Slc47a2 | 0.919894618 | -2.391772391 | 1.481443232 | -1.614488047 |
| Gm25929 | 0 | NA | NA | NA |
| Slc47a1 | 39.2732027 | -0.599384283 | 0.254212193 | -2.357810916 |
| Gm12271 | 0.275857165 | 1.628329138 | 2.43190983 | 0.669568056 |
| Rnf112 | 1605.26594 | -0.292337061 | 0.134656329 | -2.170986424 |
| Mfap4 | 9.173601336 | -0.556092932 | 0.361081582 | -1.540075595 |
| Mapk7 | 210.9337897 | -0.15436508 | 0.107700369 | -1.433282743 |
| B9d1os | 5.30594644 | -0.680583089 | 0.465807546 | -1.461082144 |
| B9d1 | 90.68224864 | -0.090531604 | 0.128442283 | -0.704842687 |
| Epn2 | 1283.27487 | -0.304673354 | 0.061993553 | -4.914597384 |
| Gm12272 | 1.010954397 | 1.492886363 | 1.171521241 | 1.274314379 |
| Grap | 28.95643677 | 1.814285862 | 0.254883336 | 7.11810309 |
| Slc5a10 | 1.756770038 | 1.59847336 | 0.823592512 | 1.940854654 |
| Fam83g | 11.17234595 | 2.736278146 | 0.663184038 | 4.125971057 |
| Prpsap2 | 311.9887896 | -0.000600371 | 0.108578248 | -0.00552939 |
| Ulk2 | 1353.293992 | -0.270732742 | 0.090471777 | -2.992455231 |
| A530017D24Rik | 232.7074356 | -0.249832412 | 0.078532008 | -3.181281359 |
| Gm26964 | 0.259490603 | -0.880776865 | 2.447964515 | -0.359799686 |
| Akap10 | 671.4108366 | -0.162134093 | 0.06034501 | -2.686785428 |
| Gm23721 | 0 | NA | NA | NA |
| Gm12273 | 0.316938377 | 0.274409755 | 2.072901482 | 0.132379545 |
| Specc1 | 1898.855375 | -0.046591437 | 0.065062985 | -0.716097439 |
| Gm12274 | 0.049553047 | 0.274391604 | 3.724945049 | 0.073663262 |
| Adora2b | 44.79440823 | 0.153854961 | 0.205434826 | 0.748923461 |
| Zswim7 | 50.11354397 | 0.410625764 | 0.152180696 | 2.698277608 |
| Ttc19 | 1111.563203 | -0.249621115 | 0.085620505 | -2.915436148 |
| Gm12275 | 2.211543673 | -0.206953378 | 0.626993944 | -0.330072371 |
| Ncor1 | 3691.866875 | -0.071954584 | 0.072380744 | -0.994112249 |
| Gm12276 | 22.41048438 | -0.826166095 | 0.240723924 | -3.432006592 |
| Pigl | 79.7396089 | -0.287382305 | 0.143701579 | -1.99985489 |
| Gm12278 | 0.637771319 | -2.015198466 | 1.1553078 | -1.744295734 |
| Gm12949 | 0 | NA | NA | NA |
| Cenpv | 151.2123072 | -0.025271607 | 0.111174344 | -0.227315097 |
| Gm12279 | 7.228218235 | -0.267200567 | 0.377420422 | -0.707965314 |
| Ubb | 6377.498152 | 0.42772111 | 0.104619285 | 4.088358193 |
| Gm12280 | 0 | NA | NA | NA |
| Trpv2 | 211.9525486 | 0.277950902 | 0.126304074 | 2.200648743 |
| 2410006H16Rik | 257.1498311 | 1.324391106 | 0.184616264 | 7.173750982 |
| Snord49b | 2.860297155 | 0.427050445 | 0.577196825 | 0.739869706 |
| Snord49a | 6.687144773 | 0.245582245 | 0.433637544 | 0.56633068 |
| Snord65 | 1.151518338 | 0.399911229 | 1.041123719 | 0.384114992 |
| Lrrc75a | 93.92612434 | -0.464410406 | 0.168047637 | -2.763564041 |
| Lrrc75aos2 | 8.114142353 | -0.21100776 | 0.418841044 | -0.503789595 |

| | | | | |
|---|---|---|---|---|
| Lrrc75aos1 | 0 | NA | NA | NA |
| Mmgt2 | 82.28640359 | -0.560193203 | 0.125001805 | -4.481480903 |
| Gm12281 | 0.339850366 | -0.589037205 | 1.739294085 | -0.338664525 |
| Zfp287 | 282.6640166 | 0.021990018 | 0.102551561 | 0.214428899 |
| Gm27194 | 0.484088067 | 0.747310105 | 1.481164797 | 0.504542173 |
| Wsb2-ps | 33.81592792 | 0.226345473 | 0.193350288 | 1.170649785 |
| Gm10428 | 0.042716945 | -0.206490061 | 3.724945049 | -0.055434391 |
| Zfp286 | 136.9541373 | -0.093228916 | 0.110918677 | -0.840515935 |
| Zfp286it | 0.804776955 | 0.734943714 | 1.071770126 | 0.685728867 |
| Zfp286os | 0 | NA | NA | NA |
| Gm24151 | 0 | NA | NA | NA |
| Trim16 | 38.08170184 | 0.793031043 | 0.226867685 | 3.495566337 |
| Fbxw10 | 9.908817 | -0.23992716 | 0.338591747 | -0.708603094 |
| Tvp23b | 188.0397332 | -0.347230066 | 0.155901111 | -2.227245617 |
| Tvp23bos | 2.044779979 | 0.24360713 | 0.707556623 | 0.344293477 |
| Cdrt4os2 | 0 | NA | NA | NA |
| Cdrt4 | 2.232638995 | 1.120566888 | 0.832954125 | 1.345292441 |
| Cdrt4os1 | 0 | NA | NA | NA |
| Tekt3 | 0.583929304 | 0.393194918 | 1.291539098 | 0.304439036 |
| Pmp22 | 317.1591218 | 0.13174706 | 0.159420946 | 0.82640998 |
| B430319H21Rik | 0.78030421 | 2.006672561 | 1.216350176 | 1.649749062 |
| Gm12285 | 0 | NA | NA | NA |
| Gm12286 | 16.66800335 | 0.10958449 | 0.235427211 | 0.465470792 |
| B130011K05Rik | 1.258106493 | 1.484095973 | 0.953219009 | 1.556930735 |
| Gm12287 | 0.044544648 | 0.755286614 | 3.724945049 | 0.202764498 |
| Gm12288 | 0 | NA | NA | NA |
| Hmgb1-ps3 | 8.21809902 | 0.351837106 | 0.412205989 | 0.853546806 |
| Hs3st3b1 | 91.21008015 | 0.819681213 | 0.190960467 | 4.292413126 |
| Cox10 | 247.0159409 | 0.060744784 | 0.085373338 | 0.711519373 |
| 9630013K17Rik | 0.09689823 | -0.687371727 | 3.724945049 | -0.184532045 |
| Gm12289 | 0 | NA | NA | NA |
| 2810001G20Rik | 35.83958181 | -0.109476766 | 0.190094708 | -0.575906437 |
| Gm12290 | 1.421699425 | 1.036863177 | 0.928894464 | 1.116233563 |
| F930015N05Rik | 5.181464156 | -0.089226458 | 0.453669447 | -0.196677247 |
| Hs3st3a1 | 37.68402565 | -0.253106114 | 0.24362331 | -1.038924042 |
| Gm24275 | 0 | NA | NA | NA |
| Gm12291 | 0.332139225 | -0.199912414 | 1.775324487 | -0.112606127 |
| Gm12292 | 0.254955332 | 1.159534989 | 2.464216449 | 0.470549164 |
| Elac2 | 339.1600695 | 0.149217678 | 0.087408433 | 1.707131373 |
| Arhgap44 | 1922.5473 | -0.12017871 | 0.073989245 | -1.624272684 |
| Gm12294 | 19.04726638 | -0.437525929 | 0.442415613 | -0.988947759 |
| Gm25991 | 1.580643696 | -0.913245203 | 0.802218779 | -1.138399183 |
| 1700086D15Rik | 40.20348781 | -0.380211429 | 0.382483826 | -0.99405884 |
| Gm12293 | 0.046345709 | 0.274391604 | 3.724945049 | 0.073663262 |
| Myocd | 4.652030534 | 0.866002372 | 0.532356355 | 1.626734353 |
| Gm12295 | 2.004716341 | -0.469249534 | 1.127531644 | -0.416174159 |
| Gm26128 | 0 | NA | NA | NA |

| | | | | |
|---|---|---|---|---|
| Map2k4 | 1997.328949 | -0.108212535 | 0.045536643 | -2.376383667 |
| Mir744 | 16.92311974 | -0.797867177 | 0.289807677 | -2.753091927 |
| Zkscan6 | 133.418627 | 0.055674748 | 0.111024839 | 0.501462097 |
| Dnah9 | 161.9890621 | 0.170619338 | 0.125933354 | 1.354838351 |
| Gm12296 | 7.015874426 | -0.032762037 | 0.456296414 | -0.071799901 |
| Shisa6 | 795.9631443 | -0.027518747 | 0.132117006 | -0.208290722 |
| Gm12297 | 1.996095088 | -0.48919145 | 0.673675408 | -0.726153046 |
| Gm12298 | 0 | NA | NA | NA |
| Pirt | 6.080719795 | 0.265348834 | 0.472032797 | 0.562140673 |
| 9130409J20Rik | 0 | NA | NA | NA |
| Gm12299 | 0.127580795 | 0.274406996 | 3.007567814 | 0.091238839 |
| Tmem220 | 8.211970812 | 0.018866482 | 0.368527457 | 0.051194236 |
| Adprm | 137.339951 | 0.149744537 | 0.126026976 | 1.188194319 |
| Sco1 | 243.6408562 | 1.155087884 | 0.155725556 | 7.417458733 |
| Myh3 | 10.50794026 | 0.671086185 | 0.372161708 | 1.803211267 |
| Mir6923 | 0 | NA | NA | NA |
| 2310065F04Rik | 0.176426445 | 0.274409222 | 2.718482184 | 0.100942071 |
| Myh2 | 5.858603139 | 0.038811607 | 0.511390702 | 0.075894237 |
| 9430073C21Rik | 0.072025462 | 0.755288839 | 3.724945049 | 0.202765096 |
| Myh1 | 7.368640863 | 0.147612734 | 0.402748183 | 0.366513719 |
| Gm12300 | 1.913586201 | 0.793712198 | 0.66578489 | 1.192145106 |
| Myh4 | 5.497083594 | -0.232646122 | 0.432115601 | -0.538388619 |
| Myh8 | 7.157414437 | 0.510979335 | 0.395082859 | 1.293347263 |
| Myh13 | 1.635466931 | 0.108359185 | 0.79811026 | 0.135769693 |
| Gm26476 | 0 | NA | NA | NA |
| Gas7 | 7031.634939 | -0.05701043 | 0.106244575 | -0.536596152 |
| Gm12301 | 0.159545055 | 0.274407309 | 2.806181458 | 0.09778673 |
| Glp2r | 34.73267854 | 1.777181685 | 0.259259539 | 6.854836233 |
| Rcvrn | 8.311551091 | -0.318725198 | 0.458783882 | -0.694717513 |
| Gm12302 | 0.259617586 | 0.274402257 | 2.458491731 | 0.111614065 |
| Dhrs7c | 0.646105857 | -1.077452897 | 1.522728514 | -0.70758043 |
| Usp43 | 27.84987753 | -0.322956596 | 0.200925285 | -1.607346711 |
| 9130017K11Rik | 3.804522088 | -1.012868634 | 0.50186904 | -2.018193099 |
| Cfap52 | 13.68582642 | -0.156408315 | 0.309699197 | -0.505033001 |
| Stx8 | 213.3769252 | 0.133122073 | 0.092368085 | 1.441212872 |
| Gm12303 | 0.044544648 | 0.755286614 | 3.724945049 | 0.202764498 |
| Gm26157 | 1.267602396 | 0.774962027 | 0.961540358 | 0.805958918 |
| Gm12304 | 0 | NA | NA | NA |
| C78197 | 2.497471105 | 0.265551643 | 0.618912654 | 0.429061582 |
| Ntn1 | 541.1663475 | 0.712547496 | 0.154780076 | 4.603612508 |
| Mir6406 | 0.083526043 | 0.75528439 | 3.724945049 | 0.202763901 |
| Gm12305 | 1.238804443 | -2.024998225 | 1.017704604 | -1.989770133 |
| Pik3r5 | 167.1648793 | 2.048611554 | 0.266103384 | 7.69855507 |
| Pik3r6 | 32.88554754 | 0.829297023 | 0.278752352 | 2.975031481 |
| Mfsd6l | 0.872192936 | 0.284268214 | 1.407770726 | 0.20192792 |
| Ccdc42os | 0 | NA | NA | NA |
| Ccdc42 | 2.972495032 | -0.754150569 | 0.551738744 | -1.366861722 |

| | | | | |
|---|---|---|---|---|
| Myh10 | 3525.532232 | -0.341546302 | 0.091237107 | -3.743502086 |
| Ndel1 | 867.2338058 | 0.09887206 | 0.07055793 | 1.401289128 |
| Rnf222 | 1.039519431 | 0.675443213 | 1.141179409 | 0.591881704 |
| Rpl26 | 246.1278693 | 0.428290176 | 0.161126593 | 2.658097379 |
| Gm22608 | 0.141071297 | -0.206491178 | 3.564043836 | -0.057937328 |
| Odf4 | 1.2405255 | -0.662714585 | 1.000176624 | -0.662597554 |
| Arhgef15 | 134.1597504 | -0.109169667 | 0.157743781 | -0.692069547 |
| Slc25a35 | 55.2247238 | -0.641357857 | 0.216642888 | -2.960438093 |
| Rangrf | 38.50784585 | -0.968941618 | 0.223883532 | -4.327882497 |
| Pfas | 417.0734811 | -0.321194463 | 0.07154937 | -4.489130547 |
| Mir3062 | 0 | NA | NA | NA |
| Gm25819 | 0 | NA | NA | NA |
| Ctc1 | 356.1546317 | -0.169196292 | 0.098030866 | -1.725949176 |
| Aurkb | 11.54692778 | 2.606814167 | 0.525760585 | 4.958177244 |
| 9330160F10Rik | 16.80478976 | -0.258109524 | 0.248486974 | -1.038724567 |
| 2310047M10Rik | 219.3535084 | 0.00677496 | 0.103824496 | 0.065253969 |
| Tmem107 | 35.40669845 | -0.887944037 | 0.22897941 | -3.877833538 |
| Snord118 | 16.03095282 | 0.476084057 | 0.377726155 | 1.260394737 |
| Gm12306 | 0.293428113 | -0.684783796 | 1.633506124 | -0.419211037 |
| Gm25371 | 0.515366098 | 0.207211085 | 1.383586831 | 0.149763702 |
| Vamp2 | 8741.82778 | -0.088607545 | 0.07837984 | -1.130488968 |
| Per1 | 1912.220295 | 0.248098218 | 0.164124198 | 1.511649227 |
| 9130213A22Rik | 7.871844516 | -0.111243445 | 0.347166729 | -0.320432334 |
| Hes7 | 8.964305686 | 1.657442674 | 0.421621295 | 3.931117081 |
| Aloxe3 | 69.09659255 | 0.409551151 | 0.181824595 | 2.252451881 |
| Alox12b | 38.03970433 | 1.336182784 | 0.288962927 | 4.624063023 |
| Alox8 | 33.51618146 | -0.148462896 | 0.200871617 | -0.739093449 |
| Gucy2e | 40.20224287 | -0.385247753 | 0.237605767 | -1.621373747 |
| Gm23194 | 0 | NA | NA | NA |
| Cntrobos | 0 | NA | NA | NA |
| Cntrob | 149.0597755 | -0.271756494 | 0.125061732 | -2.172978818 |
| Trappc1 | 165.2164685 | -0.168422738 | 0.124602782 | -1.351677191 |
| Kcnab3 | 622.5251017 | -0.190438112 | 0.104140035 | -1.8286734 |
| Kcnab3os | 3.132524194 | 0.889884814 | 0.649784148 | 1.369508346 |
| Chd3os | 937.6597823 | 0.01941294 | 0.076876605 | 0.252520784 |
| Chd3 | 5496.775121 | -0.541918316 | 0.127984754 | -4.234241181 |
| Gm22442 | 482.6561501 | -0.005609415 | 0.122827577 | -0.045669018 |
| Gm17305 | 8.479929393 | 0.030303932 | 0.376937663 | 0.080395075 |
| Cyb5d1 | 166.3102935 | -0.163213338 | 0.091977044 | -1.774500804 |
| Naa38 | 194.4625342 | 0.058910058 | 0.147038117 | 0.400644806 |
| Tmem88 | 12.86887999 | 0.704359535 | 0.378952538 | 1.858701196 |
| Kdm6b | 1048.172561 | 0.09212495 | 0.16362019 | 0.563041459 |
| Kdm6bos | 0.21690863 | -0.655084719 | 2.210015506 | -0.296416345 |
| Dnah2 | 63.85404554 | -0.59237902 | 0.147333889 | -4.020656928 |
| Dnah2os | 1.999759116 | 1.383662035 | 0.825246004 | 1.676666145 |
| Gm27493 | 0.038981395 | 0.274393828 | 3.724945049 | 0.073663859 |
| Efnb3 | 1089.578534 | -0.23207557 | 0.079967197 | -2.902134601 |

| | | | | |
|---|---|---|---|---|
| Wrap53 | 98.47005554 | 0.016810209 | 0.118054484 | 0.142393649 |
| Trp53 | 281.3127673 | 0.529970534 | 0.129747044 | 4.084644368 |
| Atp1b2 | 7719.462683 | 0.064095945 | 0.113679741 | 0.563829091 |
| Shbg | 1.970655031 | 2.357437102 | 1.00561459 | 2.344274959 |
| Sat2 | 69.39285511 | 0.013510898 | 0.136208458 | 0.099192795 |
| Gm12308 | 0 | NA | NA | NA |
| Gm12309 | 0 | NA | NA | NA |
| Fxr2 | 1268.108725 | -0.106093442 | 0.055923439 | -1.897119423 |
| Mir467f | 0 | NA | NA | NA |
| Sox15 | 1.197362358 | -0.295213623 | 0.873873381 | -0.337821966 |
| Mpdu1 | 282.855519 | 0.283212512 | 0.094691377 | 2.990900756 |
| Mir1934 | 0.521325481 | 0.306584378 | 1.533704038 | 0.199898005 |
| Cd68 | 498.24161 | 2.387707869 | 0.340512465 | 7.012101219 |
| Eif4a1 | 2251.416923 | 0.398782771 | 0.11374106 | 3.506058143 |
| Gm24029 | 0 | NA | NA | NA |
| Gm25835 | 93.03237133 | -0.318042833 | 0.203261056 | -1.564701274 |
| Gm22988 | 0 | NA | NA | NA |
| Senp3 | 708.1528605 | -0.044012535 | 0.058046236 | -0.758232369 |
| Tnfsf13os | 8.601013049 | 0.810155695 | 0.386236665 | 2.097562891 |
| Tnfsfm13 | 5.612199682 | 0.613345345 | 0.548585415 | 1.118048946 |
| Tnfsf13 | 5.550421209 | 0.108152852 | 0.457611431 | 0.236342112 |
| Tnfsf12 | 55.83410333 | 0.107104841 | 0.183661805 | 0.583163392 |
| Gm12307 | 0.075043205 | 0.755286614 | 3.724945049 | 0.202764498 |
| Polr2a | 2028.090813 | 0.371713409 | 0.077653114 | 4.786844842 |
| Slc35g3 | 0.474471242 | -1.468154662 | 1.427873429 | -1.028210647 |
| Zbtb4 | 2349.635844 | -0.160798234 | 0.055420868 | -2.901402328 |
| Chrnb1 | 9.482089685 | 1.329784568 | 0.472487005 | 2.81443628 |
| Fgf11 | 217.0904447 | -0.830216402 | 0.10591739 | -7.838338932 |
| Tmem102 | 2.228356223 | -0.453813374 | 0.801417407 | -0.566263435 |
| 4933402P03Rik | 0 | NA | NA | NA |
| Spem1 | 3.349245703 | -0.261426745 | 0.595945536 | -0.438675566 |
| Nlgn2 | 4433.469855 | 0.32455589 | 0.092798276 | 3.497434475 |
| Tmem256 | 162.3407789 | -0.249403401 | 0.10651093 | -2.341575645 |
| Plscr3 | 159.9402606 | 0.331649653 | 0.117319557 | 2.826891446 |
| Tnk1 | 0.927088552 | -0.305503533 | 1.057593172 | -0.288866779 |
| Tmem95 | 1.006289673 | -2.112854808 | 1.195240543 | -1.76772351 |
| Kctd11 | 45.25365547 | 0.40727341 | 0.299422997 | 1.360194156 |
| Acap1 | 25.63139517 | 6.443467613 | 0.876145267 | 7.35433707 |
| mmu-mir-6924 | 0 | NA | NA | NA |
| 2810408A11Rik | 1.623343953 | -0.564125979 | 0.919653205 | -0.613411638 |
| Neurl4 | 1197.492728 | -0.189798085 | 0.054982629 | -3.451964529 |
| Gps2 | 171.2820456 | 0.436662747 | 0.154883765 | 2.819293206 |
| Eif5a | 3116.497447 | 0.473035251 | 0.09694861 | 4.879237045 |
| Ybx2 | 14.27143314 | -1.083091243 | 0.33420524 | -3.240796716 |
| Slc2a4 | 20.6317304 | 0.121092406 | 0.337184816 | 0.359127695 |
| Gm12310 | 0 | NA | NA | NA |
| Cldn7 | 0.279349812 | -0.691496443 | 1.791276788 | -0.386035507 |

| | | | | |
|---|---|---|---|---|
| Elp5 | 407.5792987 | 0.04356888 | 0.100748692 | 0.432451069 |
| AL596185.1 | 14.58297401 | 0.262536076 | 0.314670522 | 0.834320528 |
| Ctdnep1 | 632.1150543 | 0.075598962 | 0.071876979 | 1.051782687 |
| Gabarap | 1356.650847 | 0.351931222 | 0.127469721 | 2.760900547 |
| Phf23 | 440.621247 | -0.187584868 | 0.092892831 | -2.019368617 |
| Dvl2 | 217.8185947 | -0.347361567 | 0.134071766 | -2.590862918 |
| Acadvl | 388.2749648 | -0.017596147 | 0.093561209 | -0.188070971 |
| Mir324 | 0 | NA | NA | NA |
| Dlg4 | 6142.381317 | -0.165760468 | 0.099494592 | -1.666024906 |
| Asgr1 | 14.73418541 | 0.286259371 | 0.310006965 | 0.923396578 |
| Asgr2 | 0.670311827 | 0.755309886 | 1.551348851 | 0.486873011 |
| Gm12311 | 0 | NA | NA | NA |
| Mgl2 | 3.864736009 | -1.682542912 | 0.57043634 | -2.949571747 |
| Clec10a | 7.893594261 | 0.663608839 | 0.411077864 | 1.614314217 |
| Slc16a11 | 154.4850972 | -0.449144815 | 0.172648004 | -2.601505976 |
| Slc16a13 | 67.2589845 | -0.234641609 | 0.171872563 | -1.365206906 |
| Bcl6b | 16.16416338 | 0.636501186 | 0.295997646 | 2.150358946 |
| Mir497b | 0.072025462 | 0.755288839 | 3.724945049 | 0.202765096 |
| mmu-mir-497a | 0.695279289 | -0.772235401 | 1.200879802 | -0.643058031 |
| Mir195a | 1.190084389 | -1.385231698 | 1.109354226 | -1.248682941 |
| 0610010K14Rik | 9.73724523 | -0.456885707 | 0.359733818 | -1.270066044 |
| Gm21988 | 3.655964835 | 1.375099357 | 0.617480077 | 2.226953399 |
| Rnasek | 9.847312144 | 0.050377467 | 0.322918665 | 0.156006674 |
| Alox12 | 4.024321177 | -0.449542393 | 0.676597514 | -0.664416264 |
| Alox12e | 0.602894331 | -0.203615459 | 1.230279951 | -0.165503354 |
| Alox15 | 1.707423372 | 1.954639524 | 1.03783086 | 1.883389288 |
| Gm12312 | 2.105566461 | -0.71761444 | 0.728924074 | -0.984484483 |
| Pelp1 | 421.1587153 | 0.07684408 | 0.056018022 | 1.371774245 |
| Gm12313 | 0 | NA | NA | NA |
| Gm12314 | 0.049553047 | 0.274391604 | 3.724945049 | 0.073663262 |
| Arrb2 | 535.4570107 | 0.583375708 | 0.077436834 | 7.533568723 |
| Mir7115 | 0 | NA | NA | NA |
| Med11 | 69.16924485 | 0.193102833 | 0.21100087 | 0.91517553 |
| Cxcl16 | 296.0926 | 4.100547805 | 0.210600212 | 19.47076769 |
| Zmynd15 | 30.5256007 | 3.025205672 | 0.347709697 | 8.700377641 |
| Tm4sf5 | 2.723137057 | 0.516419548 | 0.658704148 | 0.783993162 |
| Vmo1 | 1.167525075 | 0.382465308 | 0.971901156 | 0.393522845 |
| Gm12317 | 0.120679735 | -0.206494107 | 3.724945049 | -0.055435477 |
| Gltpd2 | 0.276014283 | 0.799000858 | 2.08966255 | 0.382358797 |
| Psmb6 | 866.9721086 | 0.378001908 | 0.090620408 | 4.171266911 |
| Gm10418 | 0.929389619 | -1.317113196 | 1.084429417 | -1.214567934 |
| Pld2 | 260.2865872 | -0.198000957 | 0.130541617 | -1.516765005 |
| Mink1 | 3061.414012 | -0.144656971 | 0.091450994 | -1.581797683 |
| Gm12316 | 0.706900582 | -0.321901181 | 1.358315029 | -0.236985658 |
| Chrne | 2.081055214 | -0.62089646 | 0.860298944 | -0.721721751 |
| 4930544D05Rik | 1.506657277 | -0.113535249 | 0.875094875 | -0.129740502 |
| Gm12315 | 0.418629604 | 0.756805624 | 1.401871007 | 0.539853967 |

| | | | | |
|---|---|---|---|---|
| Gp1ba | 0.170190954 | 0.755297539 | 2.753251924 | 0.274329251 |
| Slc25a11 | 758.6093433 | -0.089987223 | 0.072509629 | -1.241038246 |
| Rnf167 | 591.5226223 | 0.198604524 | 0.061057028 | 3.252770922 |
| Pfn1 | 1454.63144 | 0.649507306 | 0.130767641 | 4.966880974 |
| Gm12319 | 3.626546672 | 0.423143857 | 0.605990285 | 0.698268384 |
| Eno3 | 42.27077307 | -0.201896594 | 0.193198804 | -1.045019896 |
| Spag7 | 215.7681227 | 0.153557792 | 0.129219415 | 1.18834923 |
| Camta2 | 2839.805525 | -0.063917832 | 0.094522779 | -0.676216181 |
| Inca1 | 17.29699935 | 0.173621067 | 0.248235547 | 0.699420647 |
| Kif1c | 1453.545255 | 0.049141114 | 0.081438714 | 0.603412208 |
| mmu-mir-6925 | 0 | NA | NA | NA |
| Gm12320 | 0.135956459 | 0.720160971 | 3.641541792 | 0.197762655 |
| Gm12318 | 0 | NA | NA | NA |
| Gm22760 | 0 | NA | NA | NA |
| Zfp3 | 139.4854402 | -0.140768031 | 0.105328725 | -1.336463834 |
| Scimp | 44.06491336 | 7.432013485 | 0.882196677 | 8.424440576 |
| Rabep1 | 1978.442608 | -0.085843522 | 0.062501322 | -1.373467301 |
| Gm19967 | 38.13002371 | -0.586075076 | 0.233170575 | -2.513503591 |
| Rpl23a-ps2 | 9.688741338 | -0.430846214 | 0.407067677 | -1.058414213 |
| Gm12322 | 0.177934686 | 0.808082304 | 2.718161593 | 0.297290016 |
| Nup88 | 578.9786411 | 0.254283939 | 0.079711017 | 3.19007271 |
| Rpain | 118.6108747 | -0.210343596 | 0.11833626 | -1.777507556 |
| C1qbp | 471.9490247 | 0.220428849 | 0.124710853 | 1.767519372 |
| Dhx33 | 545.9733439 | 0.256950959 | 0.086602612 | 2.967011661 |
| Gm22297 | 1.938621463 | -0.576325238 | 0.710249026 | -0.811441082 |
| Derl2 | 390.884449 | -0.022244723 | 0.091706284 | -0.242564877 |
| Mis12 | 261.7075808 | -0.431000127 | 0.134046428 | -3.215304835 |
| 6330403K07Rik | 1104.505225 | -0.15784924 | 0.104688037 | -1.507805901 |
| Gm12321 | 0 | NA | NA | NA |
| Gm15377 | 0 | NA | NA | NA |
| Nlrp1a | 0.174317197 | 0.274400285 | 2.750682426 | 0.099757167 |
| Nlrp1b | 15.20331131 | 3.035764574 | 0.518543201 | 5.854410138 |
| Gm23266 | 0 | NA | NA | NA |
| Nlrp1c-ps | 0 | NA | NA | NA |
| Gm23311 | 0 | NA | NA | NA |
| Gm16013 | 0.447195792 | -0.206482858 | 1.615065244 | -0.127847998 |
| 9230020A06Rik | 0.161949631 | 0.274399427 | 2.77761947 | 0.098789424 |
| Gm12324 | 1.538502375 | -3.066377559 | 0.949847451 | -3.228284243 |
| Gm12325 | 0 | NA | NA | NA |
| Gm12326 | 2.668994113 | -1.175241622 | 0.59678173 | -1.96929893 |
| Wscd1 | 774.0552541 | -0.282144889 | 0.083257524 | -3.388821528 |
| Gm22927 | 0 | NA | NA | NA |
| 4930401O10Rik | 0.10503227 | 0.274389469 | 3.724945049 | 0.073662689 |
| Gm23226 | 0 | NA | NA | NA |
| Aipl1 | 1.181230752 | 0.744158197 | 0.919118499 | 0.809643368 |
| Fam64a | 7.10051421 | 1.134502774 | 0.646237653 | 1.755550405 |
| Pitpnm3 | 1123.234306 | -0.455407573 | 0.114073105 | -3.992243162 |

| | | | | |
|---|---|---|---|---|
| 4933427D14Rik | 402.8306143 | -0.322759465 | 0.154000716 | -2.095830939 |
| Txndc17 | 137.0373374 | 0.347455326 | 0.186046562 | 1.867571869 |
| Med31 | 91.72905903 | 0.302869448 | 0.149893822 | 2.020559913 |
| 4930563E22Rik | 22.70421901 | -0.338828867 | 0.245444059 | -1.380472876 |
| Slc13a5 | 115.5056273 | -0.038569924 | 0.178753601 | -0.215771449 |
| 1700051A21Rik | 0.48812072 | 1.578898884 | 1.646710583 | 0.958819904 |
| n-R5s70 | 0 | NA | NA | NA |
| Xaf1 | 848.1965907 | 5.582463565 | 0.169617254 | 32.91212078 |
| Fbxo39 | 1.50133588 | 3.578556454 | 1.041136965 | 3.43716204 |
| Gm23224 | 0 | NA | NA | NA |
| Tekt1 | 14.19528596 | -0.253471446 | 0.290866337 | -0.871436169 |
| Mir6338 | 0 | NA | NA | NA |
| Smtnl2 | 23.5679464 | -0.448114244 | 0.34940589 | -1.282503409 |
| Ggt6 | 0.550474514 | -0.854844094 | 1.234218958 | -0.692619481 |
| Mybbp1a | 930.9934562 | 0.31562217 | 0.086430605 | 3.651740825 |
| Spns2 | 584.9244595 | 0.579609945 | 0.158055433 | 3.66713079 |
| Spns3 | 6.967261194 | 0.870583037 | 0.497899203 | 1.748512615 |
| Gm12712 | 1.554484234 | 0.554999752 | 0.75090826 | 0.739104603 |
| Ube2g1 | 638.5559752 | -0.053226781 | 0.077260335 | -0.688927645 |
| Gm24143 | 0 | NA | NA | NA |
| Gm25678 | 1.584377093 | 0.090055158 | 0.735834965 | 0.122384994 |
| Ankfy1 | 1808.623818 | 1.058173118 | 0.078832367 | 13.42307933 |
| Cyb5d2 | 155.8716509 | -0.19173876 | 0.127474415 | -1.504135236 |
| Zzef1 | 1733.026892 | 0.128976874 | 0.073820666 | 1.747164862 |
| Gm22075 | 0 | NA | NA | NA |
| Gm6733 | 29.33660425 | -0.036006902 | 0.269585648 | -0.13356387 |
| Gm26121 | 4.21018102 | -0.349013184 | 0.520025678 | -0.671146057 |
| AL663082.1 | 0 | NA | NA | NA |
| Atp2a3 | 446.9867994 | -0.109070099 | 0.264113475 | -0.412966807 |
| P2rx1 | 3.999946171 | 0.963628299 | 0.519650397 | 1.854378068 |
| 4732414G09Rik | 8.889423667 | -0.589074282 | 0.346902774 | -1.698096201 |
| Camkk1 | 1051.478754 | 0.155959887 | 0.13137503 | 1.187134922 |
| 1200014J11Rik | 480.4026274 | -0.352046346 | 0.087269446 | -4.034016036 |
| Itgae | 5.448280777 | 1.459145393 | 0.699008433 | 2.087450343 |
| Gsg2 | 7.452195508 | 1.703592512 | 0.50649246 | 3.36351011 |
| P2rx5 | 25.59531151 | 0.160009579 | 0.261243985 | 0.61249096 |
| Emc6 | 323.5210552 | 0.058293515 | 0.108174191 | 0.538885611 |
| Tax1bp3 | 46.14509597 | 0.932735854 | 0.248093555 | 3.759613397 |
| Ctns | 210.1024968 | -0.179090827 | 0.11700245 | -1.530658769 |
| Shpk | 35.71037531 | -0.23262952 | 0.212396175 | -1.095262284 |
| Gm16021 | 0 | NA | NA | NA |
| Trpv1 | 1.771085664 | 1.269224155 | 0.872645223 | 1.454456085 |
| Trpv3 | 13.35021764 | -0.158927827 | 0.325618064 | -0.488080499 |
| Aspa | 279.3669201 | -0.463717033 | 0.189350358 | -2.44898947 |
| Spata22 | 0.720179912 | -0.920600134 | 1.132545786 | -0.812859087 |
| Olfr20 | 0 | NA | NA | NA |
| Olfr376 | 0 | NA | NA | NA |

| | | | | |
|---|---|---|---|---|
| Olfr377-ps1 | 0 | NA | NA | NA |
| Olfr1 | 0.108309328 | 1.204448679 | 3.72248341 | 0.323560523 |
| Olfr75-ps1 | 0 | NA | NA | NA |
| Olfr378 | 0 | NA | NA | NA |
| Olfr379-ps1 | 0 | NA | NA | NA |
| Olfr380 | 0 | NA | NA | NA |
| Olfr381 | 0 | NA | NA | NA |
| Olfr382 | 0 | NA | NA | NA |
| Olfr383-ps1 | 0 | NA | NA | NA |
| Olfr385 | 0 | NA | NA | NA |
| Olfr384 | 0 | NA | NA | NA |
| Gm12327 | 0 | NA | NA | NA |
| Olfr387-ps1 | 0 | NA | NA | NA |
| Zfp735 | 0 | NA | NA | NA |
| Olfr388-ps1 | 0 | NA | NA | NA |
| Gm22381 | 0 | NA | NA | NA |
| Gm12328 | 0.100902008 | 0.274391828 | 3.724945049 | 0.073663322 |
| Olfr389 | 0 | NA | NA | NA |
| Olfr390 | 0.097139547 | 0.274393828 | 3.724945049 | 0.073663859 |
| Olfr391-ps | 0 | NA | NA | NA |
| Olfr392 | 0.046345709 | 0.274391604 | 3.724945049 | 0.073663262 |
| Gm12329 | 0 | NA | NA | NA |
| Olfr393 | 0 | NA | NA | NA |
| Olfr394 | 0 | NA | NA | NA |
| Olfr395 | 0 | NA | NA | NA |
| Olfr396-ps1 | 0 | NA | NA | NA |
| Olfr23 | 0 | NA | NA | NA |
| Olfr22-ps1 | 0 | NA | NA | NA |
| Olfr397 | 0 | NA | NA | NA |
| Olfr398 | 0.042716945 | -0.206490061 | 3.724945049 | -0.055434391 |
| Olfr139 | 0 | NA | NA | NA |
| Olfr399 | 0 | NA | NA | NA |
| Zfp616 | 0 | NA | NA | NA |
| Gm12331 | 0.304339188 | 0.274413212 | 1.767363791 | 0.155266965 |
| Olfr400-ps1 | 0 | NA | NA | NA |
| Olfr401 | 0 | NA | NA | NA |
| Olfr402 | 0.044191549 | -0.206490061 | 3.724945049 | -0.055434391 |
| Olfr403 | 0 | NA | NA | NA |
| Olfr43 | 0 | NA | NA | NA |
| Olfr404-ps1 | 0 | NA | NA | NA |
| Olfr405-ps1 | 0 | NA | NA | NA |
| Olfr406 | 0 | NA | NA | NA |
| Olfr407-ps1 | 0.066050875 | 0.274391604 | 3.724945049 | 0.073663262 |
| Olfr59 | 0.062426873 | 0.755288839 | 3.724945049 | 0.202765096 |
| Olfr408-ps1 | 0 | NA | NA | NA |
| Olfr409-ps1 | 0 | NA | NA | NA |
| n-R5s71 | 0 | NA | NA | NA |

| | | | | |
|---|---|---|---|---|
| Gm12332 | 0 | NA | NA | NA |
| Olfr410 | 0.049553047 | 0.274391604 | 3.724945049 | 0.073663262 |
| Olfr411 | 0.046345709 | 0.274391604 | 3.724945049 | 0.073663262 |
| Olfr412 | 0.048949532 | -0.206487837 | 3.724945049 | -0.055433794 |
| Rap1gap2 | 2227.61315 | -0.153694525 | 0.103702629 | -1.482069711 |
| Gm22993 | 0.552195079 | 1.408422755 | 1.26142654 | 1.116531728 |
| E130309D14Rik | 443.6514258 | 0.028913565 | 0.068582342 | 0.421589054 |
| Cluh | 1336.662119 | -0.081885077 | 0.048607768 | -1.684608877 |
| Pafah1b1 | 8824.289057 | -0.059678123 | 0.057759191 | -1.033222981 |
| Gm24211 | 0 | NA | NA | NA |
| Gm16032 | 0.075043205 | 0.755286614 | 3.724945049 | 0.202764498 |
| Mettl16 | 528.4778054 | -0.433190892 | 0.075224328 | -5.758654202 |
| Mnt | 707.5314908 | 0.116553662 | 0.097310559 | 1.197749387 |
| Sgsm2 | 1130.427896 | 0.335936166 | 0.081456957 | 4.12409423 |
| Mir6926 | 0 | NA | NA | NA |
| Tsr1 | 408.3790304 | 0.335236245 | 0.120580449 | 2.780187397 |
| Snord91a | 2.110678478 | 0.496786227 | 0.687379105 | 0.722725237 |
| Gm22771 | 1.808802678 | 0.116191359 | 0.954313864 | 0.121753821 |
| Srr | 740.5480199 | -0.265397814 | 0.113004142 | -2.348567143 |
| Smg6 | 855.9187194 | 0.121190229 | 0.075680843 | 1.601332971 |
| Gm22733 | 0 | NA | NA | NA |
| Gm28180 | 0 | NA | NA | NA |
| Gm12333 | 0 | NA | NA | NA |
| Gm12335 | 2.169066299 | 0.084197261 | 0.679995203 | 0.123820375 |
| Rps12-ps19 | 2.189863395 | 1.256725737 | 0.753476976 | 1.667901976 |
| Hic1 | 51.99884426 | -0.043419454 | 0.236401798 | -0.183668035 |
| 1700016P03Rik | 48.88953987 | -1.247199513 | 0.496488126 | -2.512042982 |
| Mir212 | 22.07890698 | -0.6079621 | 0.708036802 | -0.858658897 |
| Mir132 | 4.115963635 | 1.57108191 | 0.773270412 | 2.031736745 |
| Ovca2 | 77.25714499 | 0.049919541 | 0.165170321 | 0.302230693 |
| Dph1 | 85.00803555 | -0.142697523 | 0.133922323 | -1.065524552 |
| Rtn4rl1 | 700.6189518 | -0.079510334 | 0.153767359 | -0.517082006 |
| Rpa1 | 558.9030021 | -0.144570466 | 0.083240627 | -1.736777719 |
| AL603834.1 | 0.047948698 | -0.206487837 | 3.724945049 | -0.055433794 |
| Smyd4 | 164.5240954 | -0.270832952 | 0.096890435 | -2.795249618 |
| Serpinf1 | 126.7245348 | -0.25928005 | 0.201510321 | -1.286683719 |
| Gm12336 | 0.089062654 | -0.20649451 | 3.724945049 | -0.055435586 |
| Gm12337 | 0.219222703 | 1.094295263 | 2.169844874 | 0.504319583 |
| Serpinf2 | 5.648606599 | 1.880933744 | 0.577412385 | 3.257522342 |
| Wdr81 | 600.0628896 | 0.283568045 | 0.098989775 | 2.864619552 |
| Mir22hg | 200.1675278 | 0.654892281 | 0.172327629 | 3.800274436 |
| Tlcd2 | 9.434003272 | 0.013246465 | 0.350021449 | 0.037844725 |
| Mir22 | 0 | NA | NA | NA |
| Prpf8 | 4246.36712 | 0.199381085 | 0.058548764 | 3.405385064 |
| Rilp | 13.6021336 | 0.294352047 | 0.287978558 | 1.022131818 |
| Scarf1 | 30.35591607 | 1.275536237 | 0.297812415 | 4.283019023 |
| Slc43a2 | 1420.800711 | -0.122132456 | 0.077305843 | -1.579860609 |

| | | | | |
|---|---|---|---|---|
| Mir3971 | 2.406714335 | -0.248855678 | 0.806603692 | -0.308522859 |
| Pitpna | 3114.811598 | -0.004570246 | 0.062412835 | -0.073226059 |
| 4931413K12Rik | 0 | NA | NA | NA |
| Gm12338 | 94.1563446 | -0.11235574 | 0.134351346 | -0.836282952 |
| Inpp5k | 474.3717951 | 0.34354157 | 0.112778171 | 3.046170786 |
| Myo1c | 416.9620297 | 0.693578235 | 0.147023391 | 4.717468638 |
| Crk | 1895.988957 | -0.035227662 | 0.091187525 | -0.386321072 |
| Ywhae | 6464.299146 | 0.04473889 | 0.06300125 | 0.710127012 |
| Gm26836 | 0.047948698 | -0.206487837 | 3.724945049 | -0.055433794 |
| Doc2b | 509.5744778 | -0.441569498 | 0.13434018 | -3.286950315 |
| Gm12339 | 4.361932539 | -0.661557773 | 0.48845104 | -1.354399354 |
| Rph3al | 23.25949346 | -0.096356075 | 0.253280046 | -0.38043295 |
| Gm16075 | 3.522107247 | 0.048477419 | 0.585489349 | 0.082798123 |
| 1700016K19Rik | 11.13294416 | -0.657412672 | 0.363264784 | -1.809734116 |
| Fam101b | 120.2091784 | -0.261007026 | 0.153239527 | -1.703261756 |
| Vps53 | 790.8122356 | 0.000914333 | 0.066026234 | 0.013848032 |
| Glod4 | 354.7645622 | 0.010380361 | 0.073686182 | 0.140872555 |
| Gm12340 | 0 | NA | NA | NA |
| Fam57a | 179.7700705 | -0.352434459 | 0.099601093 | -3.538459745 |
| Gemin4 | 5.229107669 | -0.321329972 | 0.510776262 | -0.629101223 |
| Dbil5 | 3.336932514 | -0.683724928 | 0.536554944 | -1.274286885 |
| Rnmtl1 | 118.9831452 | -0.075288245 | 0.113897155 | -0.661019535 |
| Nxn | 198.759646 | 0.123250729 | 0.129479273 | 0.951895434 |
| Timm22 | 253.9457515 | 0.271669776 | 0.119335617 | 2.276518798 |
| Abr | 5618.476176 | 0.023029412 | 0.068562823 | 0.335887743 |
| Gm12341 | 0.351219581 | 2.173261185 | 1.618633516 | 1.342651788 |
| Rpl36-ps2 | 4.05888357 | 0.945791598 | 0.622983882 | 1.518163832 |
| Gm26780 | 0.189892101 | 0.274393884 | 3.290457945 | 0.083390789 |
| Bhlha9 | 8.304293226 | 0.068172663 | 0.36576734 | 0.186382587 |
| Tusc5 | 10.29048676 | 0.742014691 | 0.401607397 | 1.847612113 |
| Gosr1 | 767.1062309 | 0.263250341 | 0.065279191 | 4.032683856 |
| Cpd | 2040.908043 | 0.010008286 | 0.04732908 | 0.211461671 |
| Tmigd1 | 0.164253626 | 1.222094934 | 2.776247985 | 0.440196604 |
| Blmh | 568.2821817 | 0.080331925 | 0.068979163 | 1.164582475 |
| Gm22611 | 3.32088423 | -0.211917 | 0.531542039 | -0.398683424 |
| Slc6a4 | 46.2681185 | 0.269858304 | 0.795146007 | 0.339382078 |
| Gm12343 | 0 | NA | NA | NA |
| Ccdc55 | 161.7852798 | 0.373388101 | 0.200046655 | 1.866505091 |
| Gm22772 | 0 | NA | NA | NA |
| Mir423 | 0.516177223 | 1.759039612 | 1.464119928 | 1.201431371 |
| Efcab5 | 66.97098547 | -0.157339761 | 0.172315375 | -0.913091833 |
| Gm12344 | 1.005780181 | -0.453479796 | 1.025101176 | -0.442375647 |
| Gm12345 | 29.19047675 | 0.5108516 | 0.191784386 | 2.663676705 |
| Ssh2 | 1266.750364 | 0.107060131 | 0.057258284 | 1.869775413 |
| Gm24989 | 0 | NA | NA | NA |
| Gm12346 | 262.7722694 | -0.207715899 | 0.105612138 | -1.966780549 |
| Coro6 | 239.6645014 | -0.432022148 | 0.199266379 | -2.168063419 |

| | | | | |
|---|---|---|---|---|
| Ankrd13b | 701.4661961 | -0.087435834 | 0.093809039 | -0.932061926 |
| 2210008F06Rik | 16.34395686 | 0.280158404 | 0.294698905 | 0.950659805 |
| Git1 | 3031.997312 | 0.094832591 | 0.066543075 | 1.42513087 |
| Trp53i13 | 48.23483047 | 0.266060101 | 0.184028093 | 1.445758077 |
| Abhd15 | 33.33330842 | 1.791687885 | 0.239825231 | 7.470806445 |
| Gm10392 | 0.289147236 | 1.674055435 | 2.054353396 | 0.814881918 |
| Taok1 | 6408.541416 | -0.10192292 | 0.047771712 | -2.133541307 |
| Gm12734 | 2.565340877 | 1.196249018 | 0.731885155 | 1.634476406 |
| Gm12732 | 0 NA | NA | NA | |
| Gm12733 | 0 NA | NA | NA | |
| Gm23132 | 0.137483974 | -0.206488593 | 2.933430427 | -0.070391508 |
| Nufip2 | 1887.858992 | 0.027281487 | 0.060985307 | 0.447345249 |
| Cryba1 | 0.167183452 | -0.206481502 | 2.765153652 | -0.074672705 |
| Gm11191 | 0.293994645 | 2.168120992 | 2.01391377 | 1.076570916 |
| Myo18a | 2948.916186 | 0.143123249 | 0.039808198 | 3.595320965 |
| Gm10277 | 0.30796268 | 0.755309885 | 1.79473344 | 0.420847948 |
| Pipox | 16.89892386 | -0.954007245 | 0.278633521 | -3.423878227 |
| Gm11190 | 4.356108356 | -0.870649888 | 0.508787146 | -1.711226188 |
| Sez6 | 1855.373666 | -0.473425101 | 0.135740615 | -3.487718854 |
| Phf12 | 804.2323745 | 0.164773989 | 0.101513221 | 1.623177629 |
| Dhrs13os | 5.524772491 | -0.665367784 | 0.46412481 | -1.433596673 |
| Dhrs13 | 73.22834646 | -0.315204437 | 0.141721906 | -2.224105267 |
| Flot2 | 917.7646301 | 0.096831962 | 0.079791971 | 1.213555207 |
| Mir144 | 0 NA | NA | NA | |
| Mir451a | 0 NA | NA | NA | |
| mmu-mir-451a | 0.164553454 | -0.206488957 | 2.77921914 | -0.074297472 |
| Eral1 | 174.3755178 | 0.043753556 | 0.085695873 | 0.510567834 |
| Fam222b | 474.7107881 | -0.119546327 | 0.071139797 | -1.680442338 |
| Gm12571 | 0.999428373 | 0.14291229 | 1.198826649 | 0.119210137 |
| Traf4 | 100.814411 | -0.160414172 | 0.149692905 | -1.071621745 |
| Nek8 | 29.99850749 | 0.008526272 | 0.236038502 | 0.036122377 |
| Tlcd1 | 129.4641411 | -0.714313742 | 0.157966685 | -4.521926506 |
| Mir7653 | 0 NA | NA | NA | |
| Rpl23a | 798.9907631 | 0.474721825 | 0.125459712 | 3.783858735 |
| Snord42a | 0.955875529 | 0.244834793 | 1.036139412 | 0.236295223 |
| Snord4a | 1.972325891 | -0.456271793 | 0.647169377 | -0.705026858 |
| Snord42b | 3.467268438 | -0.175197274 | 0.647720065 | -0.270483011 |
| Rab34 | 131.9099265 | 0.174134618 | 0.147648461 | 1.179386608 |
| Proca1 | 20.46478121 | -0.61050022 | 0.265056941 | -2.303279506 |
| Supt6 | 2745.970221 | 0.264323533 | 0.090728938 | 2.913332156 |
| Sdf2 | 269.515758 | 0.130081498 | 0.095953379 | 1.355673966 |
| 2610507B11Rik | 4624.687977 | 0.307683009 | 0.056074317 | 5.48705759 |
| Gm11192 | 0.159819677 | -0.671980707 | 3.463442051 | -0.194021063 |
| BC030499 | 82.88575948 | -0.501945575 | 0.455552995 | -1.101837944 |
| Gm11193 | 0.044191549 | -0.206490061 | 3.724945049 | -0.055434391 |
| Spag5 | 131.8137158 | -0.090561341 | 0.173621449 | -0.521602267 |
| Aldoc | 6307.64853 | 0.038943967 | 0.148617011 | 0.262042457 |

| | | | | |
|---|---|---|---|---|
| Pigs | 527.1172141 | -0.195142493 | 0.089513526 | -2.180033579 |
| Unc119 | 149.7412516 | -0.441744371 | 0.126123727 | -3.502468417 |
| Foxn1 | 0.172802443 | 0.802685102 | 3.382542005 | 0.237302331 |
| Slc13a2os | 0 | NA | NA | NA |
| Slc13a2 | 0 | NA | NA | NA |
| Gm11195 | 0.045882455 | 0.755288839 | 3.724945049 | 0.202765096 |
| Slc46a1 | 58.13216195 | -0.306902498 | 0.164640874 | -1.864072329 |
| Sarm1 | 378.1139612 | -0.285941416 | 0.080092498 | -3.570139838 |
| Vtn | 396.8074538 | -0.075719756 | 0.157321623 | -0.48130546 |
| Sebox | 24.6969001 | 0.804979994 | 0.246918544 | 3.260103438 |
| Tmem199 | 242.7545441 | 0.050082246 | 0.089526737 | 0.55941105 |
| Poldip2 | 775.1605727 | -0.056944716 | 0.081762659 | -0.696463609 |
| Tnfaip1 | 559.6315322 | 0.126494106 | 0.078514638 | 1.611089468 |
| Ift20 | 265.7227452 | -0.150344552 | 0.073420318 | -2.047724069 |
| Tmem97 | 90.94084935 | -0.137993939 | 0.147010305 | -0.938668476 |
| D11Bhm181e | 1.727533833 | -0.067331846 | 1.032188296 | -0.065232134 |
| Gm22947 | 0.742175153 | -2.930543661 | 1.159392811 | -2.527653814 |
| Nlk | 1172.726608 | -0.43278204 | 0.097013959 | -4.461028527 |
| Gm17235 | 0 | NA | NA | NA |
| Gm11196 | 6.623448048 | -0.033698392 | 0.48221382 | -0.069882676 |
| Gm11197 | 0 | NA | NA | NA |
| Fam58b | 92.53061276 | 0.280977912 | 0.132421106 | 2.121851421 |
| Gm23840 | 0 | NA | NA | NA |
| Gm23219 | 0 | NA | NA | NA |
| Lyrm9 | 495.4572617 | -0.221759928 | 0.081707811 | -2.714060312 |
| Nos2 | 95.34328637 | 4.630067419 | 0.776393917 | 5.963554479 |
| Lgals9 | 1210.742748 | 5.48658843 | 0.261421295 | 20.98753441 |
| Ksr1 | 731.8760987 | 0.025661592 | 0.087755886 | 0.292420178 |
| Gm11201 | 0.406995362 | -0.687380492 | 1.75904365 | -0.390769434 |
| Gm11200 | 0 | NA | NA | NA |
| Wsb1 | 1011.052735 | -0.00308973 | 0.070353659 | -0.043917125 |
| Gm9964 | 0 | NA | NA | NA |
| Nf1 | 4025.801747 | -0.313212461 | 0.036821497 | -8.506239114 |
| Gm11198 | 0 | NA | NA | NA |
| Gm11199 | 0 | NA | NA | NA |
| AU040972 | 4.210711181 | 0.356882056 | 0.427361816 | 0.835081758 |
| Omg | 735.6960469 | -0.032428873 | 0.080982737 | -0.400441793 |
| Gm21975 | 2.443568159 | 2.51537088 | 1.002185395 | 2.509885789 |
| Evi2b | 0 | NA | NA | NA |
| Evi2a | 264.6184616 | 0.290596154 | 0.123879559 | 2.345795831 |
| Rab11fip4 | 3582.291547 | -0.390653327 | 0.095749154 | -4.079966373 |
| Gm23293 | 1.882325847 | -0.425887548 | 0.779748943 | -0.546185477 |
| Rab11fip4os1 | 0.112642008 | 0.274402725 | 3.724945049 | 0.073666248 |
| Gm25867 | 0 | NA | NA | NA |
| Gm24887 | 0 | NA | NA | NA |
| Rab11fip4os2 | 0.045882455 | 0.755288839 | 3.724945049 | 0.202765096 |
| Gm11205 | 0.304750214 | 1.117631206 | 1.97403922 | 0.566164641 |

| | | | | |
|---|---|---|---|---|
| Mir193a | 0.044191549 | -0.206490061 | 3.724945049 | -0.055434391 |
| Mir365-2 | 0.211497531 | 0.755303694 | 2.189991961 | 0.344888797 |
| 9130204K15Rik | 0.414175607 | 1.238419953 | 1.728334658 | 0.716539443 |
| Gm24454 | 0.133633945 | 1.441423131 | 3.720899433 | 0.387385673 |
| Gm11203 | 0 | NA | NA | NA |
| Gm11204 | 0.206494471 | -0.206482211 | 2.202635613 | -0.093743245 |
| Utp6 | 497.722905 | -0.037972802 | 0.076151695 | -0.498646838 |
| Suz12 | 822.7645743 | 0.261283033 | 0.070691896 | 3.696081864 |
| Gm26563 | 1.315499399 | 1.498786078 | 0.905462829 | 1.655270686 |
| Crlf3 | 219.9408582 | 0.123075574 | 0.099804653 | 1.233164691 |
| Atad5 | 155.6883303 | 0.449927812 | 0.182568215 | 2.464436713 |
| Tefm | 62.86592296 | 0.299119017 | 0.157607017 | 1.897878806 |
| Adap2 | 173.6780421 | 1.830851947 | 0.183657672 | 9.968829128 |
| Adap2os | 0.240869569 | -0.206496159 | 3.110743597 | -0.066381607 |
| Rnf135 | 133.3790234 | 1.101552758 | 0.198197335 | 5.557858563 |
| Rhot1 | 976.0750427 | -0.0972877 | 0.045360908 | -2.144747623 |
| Rhbdl3 | 250.4651554 | -0.199661778 | 0.141825987 | -1.407794037 |
| 5730455P16Rik | 801.0455395 | 0.053597134 | 0.069747116 | 0.768449469 |
| Zfp207 | 1918.849716 | -0.156877671 | 0.077223727 | -2.031469833 |
| Psmd11 | 1260.672455 | 0.263765365 | 0.088107202 | 2.993686766 |
| Cdk5r1 | 2328.335231 | -0.382999622 | 0.098198326 | -3.900266287 |
| Myo1d | 327.8254873 | -0.131704637 | 0.093541252 | -1.407984541 |
| C030013C21Rik | 2.678984608 | -1.015698622 | 0.713665399 | -1.423214049 |
| Tmem98 | 68.95410002 | -0.321492511 | 0.219443468 | -1.465035688 |
| Gm11207 | 0 | NA | NA | NA |
| 4930507D10Rik | 0.549547734 | -1.175918021 | 1.367237974 | -0.860068286 |
| Spaca3 | 0.045882455 | 0.755288839 | 3.724945049 | 0.202765096 |
| Asic2 | 625.5974346 | -0.021514021 | 0.070293323 | -0.306060671 |
| Gm11416 | 0 | NA | NA | NA |
| Gm11417 | 0.04393175 | -0.206490061 | 3.724945049 | -0.055434391 |
| 4930527B05Rik | 0 | NA | NA | NA |
| 1700071K01Rik | 9.922463048 | -0.611191214 | 0.44602152 | -1.370317768 |
| Gm11418 | 0 | NA | NA | NA |
| 5530401A14Rik | 0.732425175 | -0.895538161 | 1.262622587 | -0.709268288 |
| Gm11419 | 2.867374077 | -0.2734212 | 0.654688353 | -0.417635657 |
| Gm17268 | 0.061920613 | 0.274393828 | 3.724945049 | 0.073663859 |
| Ccl2 | 1280.074291 | 10.01558078 | 0.818331246 | 12.23903014 |
| Ccl7 | 524.7321755 | 10.16244958 | 1.031238497 | 9.854606482 |
| Ccl11 | 7.798742698 | 5.280540411 | 0.923652283 | 5.71702199 |
| Ccl12 | 226.9053113 | 8.327799754 | 0.651812546 | 12.77637229 |
| Ccl8 | 52.35493119 | 6.839758122 | 1.026254765 | 6.664775995 |
| Ccl1 | 0.343821097 | 0.791414812 | 2.013041048 | 0.393143901 |
| Tmem132e | 318.4462841 | 0.540143901 | 0.130781997 | 4.130108989 |
| Gm24612 | 0 | NA | NA | NA |
| Gm11426 | 0 | NA | NA | NA |
| Cct6b | 3.620020008 | 1.237752832 | 0.578228304 | 2.140595372 |
| Gm11421 | 0 | NA | NA | NA |

| | | | | |
|---|---|---|---|---|
| Gm11422 | 0 | NA | NA | NA |
| Gm11420 | 0.621419653 | 3.238237916 | 1.223357005 | 2.647009746 |
| Zfp830 | 109.1762647 | -0.110698456 | 0.11514085 | -0.961417747 |
| Gm11423 | 5.437272543 | 0.122947856 | 0.499996794 | 0.245897289 |
| Lig3 | 405.8341476 | -0.094106546 | 0.06782538 | -1.387482769 |
| Rffl | 192.7097897 | -0.244015303 | 0.128528815 | -1.89852605 |
| Rad51d | 202.7225488 | -0.072232322 | 0.081708379 | -0.884025883 |
| Fndc8 | 0.20460011 | -0.600596497 | 3.233930613 | -0.185717187 |
| Nle1 | 99.69246992 | 0.132545751 | 0.118689672 | 1.116742071 |
| Unc45b | 11.03143117 | 1.382333075 | 0.393739912 | 3.510777123 |
| Unc45bos | 2.092676817 | 1.027125343 | 0.811711226 | 1.265382701 |
| Slfn5os | 0.252317246 | 1.28704099 | 2.503377304 | 0.514121857 |
| Slfn5 | 854.2001971 | 4.935503598 | 0.300513353 | 16.42357501 |
| Gm11424 | 0.243199696 | -0.206485386 | 2.459394114 | -0.083957827 |
| Slfn9 | 576.8930042 | 6.543566304 | 0.333963752 | 19.59364233 |
| Mir7679 | 0.98936666 | 2.102199678 | 1.220172633 | 1.7228707 |
| Slfn8 | 1181.313578 | 6.924810432 | 0.288790688 | 23.97864865 |
| Gm11425 | 15.12308696 | 3.028248061 | 0.600582402 | 5.042185803 |
| Slfn10-ps | 120.7759095 | 8.055356723 | 0.914497834 | 8.808502789 |
| Gm27598 | 0.379788221 | 0.755305649 | 1.926284584 | 0.392104913 |
| Slfn2 | 559.7972036 | 6.416850844 | 0.380250773 | 16.87531308 |
| Slfn1 | 299.5776399 | 10.58165978 | 0.921725114 | 11.48027717 |
| Slfn4 | 2011.837015 | 10.9765068 | 0.83916757 | 13.08023234 |
| Slfn3 | 17.21960857 | 3.815730845 | 0.506062575 | 7.54003761 |
| AI662270 | 42.49283015 | 4.515470042 | 0.516508379 | 8.742297757 |
| Gm11427 | 4.559114128 | 3.183103274 | 0.93795766 | 3.393653475 |
| Gm23444 | 10.89494943 | -0.24837143 | 0.412447594 | -0.602189064 |
| Slfn14 | 0.407088326 | 0.80471193 | 1.573746874 | 0.511335046 |
| AA465934 | 34.77868168 | -0.219515809 | 0.182446399 | -1.20317973 |
| Snord7 | 2.473802751 | -1.324160925 | 0.652949255 | -2.02796912 |
| Pex12 | 83.19986102 | 0.295710973 | 0.162314129 | 1.821843693 |
| Ap2b1 | 4452.733103 | -0.04277183 | 0.053596761 | -0.798030126 |
| Gm23951 | 0 | NA | NA | NA |
| Rasl10b | 2095.394555 | -0.362213528 | 0.119860018 | -3.021971238 |
| Gas2l2 | 4.963609251 | -0.648510383 | 0.502351332 | -1.290949863 |
| 1700020L24Rik | 3.62909452 | 0.617335375 | 0.566270875 | 1.090176807 |
| Mmp28 | 45.16190099 | 0.710166994 | 0.222100488 | 3.197503072 |
| Taf15 | 585.2888291 | 0.21576315 | 0.129056061 | 1.67185601 |
| Heatr9 | 16.53926368 | 6.220075031 | 0.952242004 | 6.532031776 |
| Gm23247 | 1.077303893 | 1.675868568 | 1.034017671 | 1.62073494 |
| Ccl5 | 670.2920451 | 11.33196369 | 0.941746033 | 12.03292957 |
| Ccl9 | 101.7341889 | 4.590585151 | 0.563657869 | 8.144275809 |
| E230016K23Rik | 2.428642307 | 0.716209448 | 0.743643198 | 0.963108988 |
| Ccl6 | 71.52556139 | 2.820213154 | 0.391488121 | 7.203828175 |
| Rpl9-ps1 | 0 | NA | NA | NA |
| Ccl3 | 250.7978955 | 8.023870695 | 0.942968431 | 8.509161524 |
| Ccl4 | 181.900904 | 7.97146575 | 0.861304693 | 9.255105436 |

| | | | | |
|---|---|---|---|---|
| Gm11433 | 0.066050875 | 0.274391604 | 3.724945049 | 0.073663262 |
| Gm11438 | 0 | NA | NA | NA |
| Gm11432 | 0.038981395 | 0.274393828 | 3.724945049 | 0.073663859 |
| Gm11430 | 0 | NA | NA | NA |
| Gm11429 | 0 | NA | NA | NA |
| Wfdc17 | 58.81490581 | 4.562762535 | 0.686410186 | 6.647282668 |
| Wfdc18 | 2.91100996 | -0.234765123 | 0.610210615 | -0.384728023 |
| Wfdc21 | 0.36705964 | 2.237258679 | 3.695877945 | 0.605338897 |
| Heatr6 | 295.8490656 | -0.060294084 | 0.087170821 | -0.69167736 |
| Gm22311 | 0 | NA | NA | NA |
| Gm11434 | 0.042716945 | -0.206490061 | 3.724945049 | -0.055434391 |
| Gm12576 | 1.282016488 | -0.940087903 | 0.846991543 | -1.109914154 |
| Gm23211 | 0 | NA | NA | NA |
| Hnf1b | 4.558351918 | -0.171438279 | 0.524562424 | -0.326821501 |
| Ddx52 | 333.6973482 | 0.484900687 | 0.109307857 | 4.436100939 |
| Synrg | 1329.598341 | -0.121830871 | 0.056223442 | -2.166905264 |
| Dusp14 | 294.0225346 | -0.400824138 | 0.179350617 | -2.234863451 |
| Tada2a | 253.1517061 | -0.167821827 | 0.151846367 | -1.105208043 |
| Acaca | 1627.488358 | -0.011599299 | 0.059844407 | -0.193824275 |
| Gm11437 | 0.12092566 | 1.236179401 | 3.724945049 | 0.331865138 |
| Gm37391 | 10.91891655 | -0.19284015 | 0.320454704 | -0.601770383 |
| Aatf | 235.0359788 | 0.458688572 | 0.14166324 | 3.23788001 |
| Lhx1 | 140.9707341 | -0.518612727 | 0.231960729 | -2.235778133 |
| Lhx1os | 31.23601852 | -0.41997067 | 0.223623123 | -1.878028826 |
| 1700109G15Rik | 0 | NA | NA | NA |
| Gm11431 | 0 | NA | NA | NA |
| Mrm1 | 101.0285373 | -0.318869382 | 0.123354928 | -2.584974816 |
| Dhrs11 | 57.07916713 | -0.277069023 | 0.16720193 | -1.65709226 |
| 4930502E09Rik | 0.377064288 | -1.263101192 | 3.710783437 | -0.340386663 |
| Ggnbp2 | 1167.177578 | 0.20524402 | 0.084154293 | 2.438901376 |
| Pigw | 46.32962589 | -0.052886131 | 0.16300223 | -0.32445035 |
| Myo19 | 116.4020599 | -0.467728779 | 0.198277459 | -2.358960935 |
| Gm23564 | 1.319357828 | 0.165244196 | 0.949269029 | 0.1740752 |
| Znhit3 | 108.0352272 | 0.341681127 | 0.117329219 | 2.912157169 |
| Car4 | 225.7880006 | -0.803889253 | 0.201528897 | -3.988952782 |
| Usp32 | 2485.609366 | -0.278106541 | 0.048930082 | -5.683753853 |
| Gm11440 | 7.169804879 | -0.384122091 | 0.353878131 | -1.085464337 |
| Gm11439 | 9.641612845 | -0.300121697 | 0.47867174 | -0.626988544 |
| Rpl13-ps1 | 6.979134821 | 0.46409453 | 0.448876722 | 1.033901975 |
| Gm23506 | 0 | NA | NA | NA |
| 1700125H20Rik | 0.788910745 | 1.23280313 | 1.327071485 | 0.928965126 |
| Appbp2 | 1916.277606 | 0.027485337 | 0.061437807 | 0.447368464 |
| Akirin1-ps | 0.295845421 | 0.208945901 | 2.057543282 | 0.101551157 |
| Appbp2os | 6.2436769 | -0.810864731 | 0.500953218 | -1.618643624 |
| Gm22587 | 0 | NA | NA | NA |
| Ppm1d | 315.9501097 | -0.163657838 | 0.108742626 | -1.505001712 |
| Bcas3 | 671.5506855 | 0.058397255 | 0.068839318 | 0.848312512 |

| | | | | |
|---|---|---|---|---|
| Gm23765 | 0.716442652 | 0.177406067 | 1.128009994 | 0.157273489 |
| Bcas3os1 | 0.047948698 | -0.206487837 | 3.724945049 | -0.055433794 |
| Mir5110 | 0 | NA | NA | NA |
| Bcas3os2 | 0 | NA | NA | NA |
| Gm23473 | 1.868429801 | 0.282387293 | 0.781993889 | 0.361111892 |
| 2610027K06Rik | 0.597123318 | -0.196705668 | 1.48964529 | -0.132048662 |
| Tbx2 | 23.31208816 | 0.605287943 | 0.237622927 | 2.547262387 |
| Gm11444 | 0 | NA | NA | NA |
| Tbx4 | 0.57584922 | -0.626421387 | 1.408273783 | -0.44481506 |
| Gm11443 | 0 | NA | NA | NA |
| Brip1 | 34.57829007 | 3.068097054 | 0.318719606 | 9.626320436 |
| Atp5l-ps2 | 2.273527578 | 0.437503271 | 0.758484654 | 0.57681229 |
| Brip1os | 315.6828442 | 0.80195217 | 0.147856762 | 5.423845052 |
| Ints2 | 398.3495836 | 0.01043934 | 0.128359312 | 0.081329046 |
| Med13 | 3263.787859 | 0.108675729 | 0.050411401 | 2.155776829 |
| Gm25427 | 0 | NA | NA | NA |
| Gm23667 | 0 | NA | NA | NA |
| Gm26439 | 0 | NA | NA | NA |
| Rnft1 | 257.0685552 | -0.150369696 | 0.100955371 | -1.489467022 |
| Rps6kb1 | 899.7579894 | -0.007801425 | 0.057799914 | -0.134972945 |
| Tubd1 | 49.92708075 | -0.44403871 | 0.192550611 | -2.306088294 |
| Vmp1 | 1340.608191 | 0.311056125 | 0.106519341 | 2.920184459 |
| Mir21a | 0 | NA | NA | NA |
| Gm11478 | 49.42339314 | 0.122794322 | 0.171907958 | 0.71430272 |
| Ptrh2 | 123.2383261 | -0.033282278 | 0.119981554 | -0.277394959 |
| Cltc | 12039.16266 | -0.039720555 | 0.056869181 | -0.69845485 |
| Dhx40 | 393.4698973 | 0.135185627 | 0.084947587 | 1.591400445 |
| Ypel2 | 824.3745057 | -0.521966917 | 0.110521373 | -4.722769009 |
| Gm22387 | 0 | NA | NA | NA |
| Gdpd1 | 414.5698482 | -0.280888675 | 0.073814242 | -3.805345261 |
| Gm11479 | 0.152667883 | 0.755303377 | 2.845214345 | 0.265464491 |
| Smg8 | 236.0766196 | 0.34553473 | 0.122185069 | 2.827961995 |
| Gm9378 | 0.044191549 | -0.206490061 | 3.724945049 | -0.055434391 |
| Prr11 | 19.51012886 | 1.163435797 | 0.289521541 | 4.018477496 |
| Ska2 | 149.134786 | -0.038072743 | 0.126986499 | -0.299817251 |
| Mir301 | 1.204075239 | -0.25622756 | 0.818761443 | -0.312945318 |
| Gm11491 | 0.332371241 | -1.816167222 | 3.691775746 | -0.491949497 |
| Trim37 | 3977.710216 | -0.262644827 | 0.077145357 | -3.404544818 |
| Ppm1e | 3157.400212 | -0.181334637 | 0.099241224 | -1.827210812 |
| Gm11490 | 0 | NA | NA | NA |
| Gm22883 | 12.43333056 | -0.302956707 | 0.468153353 | -0.647131342 |
| Rad51c | 45.74186524 | -0.977459196 | 0.20506403 | -4.766604835 |
| Tex14 | 23.09397913 | 0.589533094 | 0.266340194 | 2.213458984 |
| Rnu1a1 | 44.69699138 | -0.610750085 | 0.497986948 | -1.226437935 |
| Gm22068 | 772.7219214 | -0.01314712 | 0.166409619 | -0.079004569 |
| Rnu3b1 | 0.136120942 | 0.274393632 | 3.599844381 | 0.076223748 |
| Gm23137 | 0 | NA | NA | NA |

| | | | | |
|---|---|---|---|---|
| Rnu3b3 | 0.045882455 | 0.755288839 | 3.724945049 | 0.202765096 |
| Rnu3b4 | 0 | NA | NA | NA |
| Gm11489 | 0.231904818 | -1.056019128 | 2.137353648 | -0.494077865 |
| Rnu3b2 | 0 | NA | NA | NA |
| Gm11492 | 0.278403928 | -1.518878349 | 2.018542961 | -0.752462731 |
| 4-Sep | 1581.659409 | -0.383589562 | 0.088544278 | -4.332177854 |
| Mtmr4 | 1614.493203 | -0.161511924 | 0.062118306 | -2.600069704 |
| Hsf5 | 20.47148481 | 0.159264696 | 0.328698102 | 0.48453184 |
| Rnf43 | 59.17937029 | -0.269687445 | 0.170179557 | -1.584722922 |
| Gm25968 | 0 | NA | NA | NA |
| 1110028F11Rik | 2.71851656 | -0.20750233 | 0.676463006 | -0.306746013 |
| Supt4a | 181.1208407 | 0.218154815 | 0.084519326 | 2.581123451 |
| Mir142hg | 12.62452238 | 3.201835132 | 0.557250873 | 5.745769613 |
| Mir142b | 0.614441445 | -1.151842967 | 1.199674109 | -0.960129887 |
| mmu-mir-142a | 0.12092566 | 1.236179401 | 3.724945049 | 0.331865138 |
| Bzrap1 | 2131.147747 | -0.478412367 | 0.154875186 | -3.08901884 |
| Mpo | 4.50923424 | 2.318719397 | 0.978685168 | 2.3692189 |
| Gm11505 | 0.14164095 | -0.20649561 | 3.603178992 | -0.057309284 |
| Lpo | 20.44324603 | 1.041591362 | 0.280877381 | 3.708349028 |
| Mks1 | 36.78772966 | 0.155802591 | 0.200553477 | 0.776863076 |
| Epx | 0.366894703 | 0.755308744 | 1.674168828 | 0.451154466 |
| Olfr462 | 0.183788242 | -0.687377982 | 2.683430441 | -0.256156438 |
| Olfr463 | 0.038981395 | 0.274393828 | 3.724945049 | 0.073663859 |
| Olfr464 | 0.707581183 | -1.403801554 | 1.111223293 | -1.263293852 |
| Gm11504 | 0 | NA | NA | NA |
| Gm11507 | 0.066050875 | 0.274391604 | 3.724945049 | 0.073663262 |
| Dynll2 | 5064.124974 | -0.223296789 | 0.076543856 | -2.917239887 |
| Gm11508 | 7.373820318 | -0.494605412 | 0.42270165 | -1.170105232 |
| 2010015M23Rik | 13.47151829 | 0.114000382 | 0.290481672 | 0.392452925 |
| Srsf1 | 2301.674563 | 0.031921418 | 0.049791092 | 0.641107013 |
| Gm15892 | 24.85441836 | 0.148153652 | 0.28323956 | 0.523068362 |
| Vezf1 | 819.1262531 | 0.098132882 | 0.088392976 | 1.110188679 |
| Cuedc1 | 496.2529045 | 0.239651134 | 0.072243767 | 3.317256907 |
| Mrps23 | 225.4695307 | 0.169750458 | 0.102460138 | 1.656746332 |
| 1700106J16Rik | 0.47539415 | -1.244789406 | 1.358925223 | -0.91601023 |
| Gm11509 | 0 | NA | NA | NA |
| Msi2 | 2918.689029 | 0.187189436 | 0.072710194 | 2.574459291 |
| Mir378b | 0 | NA | NA | NA |
| Gm15893 | 0.758254207 | -0.993782818 | 1.267992993 | -0.783744724 |
| Gm11510 | 2.984155966 | -0.55793368 | 0.591799479 | -0.942774874 |
| C030037D09Rik | 74.7814881 | -0.713412333 | 0.15877028 | -4.493361949 |
| Gm11493 | 0 | NA | NA | NA |
| Akap1 | 422.0526709 | -0.505814665 | 0.097684723 | -5.178032434 |
| 4930556N13Rik | 0.0878635 | -0.687376175 | 3.724945049 | -0.184533239 |
| Gm23507 | 0 | NA | NA | NA |
| Scpep1os | 0 | NA | NA | NA |
| Scpep1 | 257.6478549 | 0.76409211 | 0.189179547 | 4.038978434 |

| | | | | |
|---|---|---|---|---|
| Gm15698 | 0.068097359 | -0.206487837 | 3.724945049 | -0.055433794 |
| Coil | 170.2213894 | 0.257683121 | 0.089376935 | 2.883105357 |
| 2210409E12Rik | 0.042716945 | -0.206490061 | 3.724945049 | -0.055434391 |
| Gm11496 | 0.352842203 | 1.578384772 | 1.997424518 | 0.790209972 |
| Trim25 | 943.4408156 | 4.099586126 | 0.198065021 | 20.69818337 |
| Dgke | 1158.574667 | 0.140136028 | 0.069747854 | 2.0091805 |
| Dgkeos | 4.476524561 | 0.571475607 | 0.484805157 | 1.17877378 |
| Gm525 | 0 | NA | NA | NA |
| Gm22534 | 0.038981395 | 0.274393828 | 3.724945049 | 0.073663859 |
| Gm11497 | 0 | NA | NA | NA |
| Gm24734 | 0 | NA | NA | NA |
| Nog | 67.55626535 | -0.462708284 | 0.173548025 | -2.666168543 |
| Gm2018 | 0.338611762 | 0.361421007 | 2.071994008 | 0.174431492 |
| 4932411E22Rik | 35.54587762 | -0.22846638 | 0.212514035 | -1.0750649 |
| Ankfn1 | 45.04732695 | -0.2824139 | 0.167509743 | -1.685955067 |
| Pctp | 14.03537449 | 0.353113249 | 0.315465066 | 1.119341847 |
| Tmem100 | 84.40838063 | -0.194915095 | 0.145654533 | -1.338201373 |
| Gm11494 | 0 | NA | NA | NA |
| n-R5s72 | 0 | NA | NA | NA |
| Mmd | 1332.487735 | 0.020407015 | 0.099539559 | 0.205014118 |
| Hlf | 2065.341962 | -0.079773931 | 0.131280843 | -0.607658583 |
| Stxbp4 | 295.2899188 | -0.118424934 | 0.098367534 | -1.203902643 |
| Gm11511 | 3.871387691 | 0.731669251 | 0.609673643 | 1.200099857 |
| Gm11512 | 2.280412116 | -0.620303585 | 0.681540396 | -0.910149404 |
| Cox11 | 152.2450534 | 0.018482673 | 0.122827928 | 0.150476145 |
| Tom1l1 | 226.4499419 | -0.380272291 | 0.113754299 | -3.342926752 |
| Gm22762 | 0 | NA | NA | NA |
| 4930405D11Rik | 0.230025872 | -0.657513787 | 2.529213486 | -0.25996769 |
| Gm11516 | 0 | NA | NA | NA |
| Kif2b | 0 | NA | NA | NA |
| Gm11498 | 0 | NA | NA | NA |
| Gm22702 | 0 | NA | NA | NA |
| Gm11500 | 0 | NA | NA | NA |
| Gm11501 | 0 | NA | NA | NA |
| Gm25113 | 0 | NA | NA | NA |
| Car10 | 1148.248468 | -0.450980497 | 0.088449805 | -5.098716674 |
| Gm24856 | 0 | NA | NA | NA |
| Gm11502 | 0.044544648 | 0.755286614 | 3.724945049 | 0.202764498 |
| Gm11506 | 0 | NA | NA | NA |
| Utp18 | 272.3077803 | 0.148083474 | 0.086230913 | 1.717289881 |
| Mbtd1 | 630.0243432 | -0.162840428 | 0.088817097 | -1.833435596 |
| Nme2 | 16.60911457 | 0.141036631 | 0.350835334 | 0.402002358 |
| Gm20390 | 0.049553047 | 0.274391604 | 3.724945049 | 0.073663262 |
| Nme1 | 441.0854408 | -0.012209548 | 0.079152418 | -0.154253628 |
| Spag9 | 5268.156952 | -0.283138418 | 0.049508693 | -5.718963755 |
| Gm11518 | 36.08732431 | 0.588023088 | 0.235392868 | 2.49804972 |
| B230206L02Rik | 10.54869608 | -0.54263895 | 0.316166761 | -1.716306133 |

| | | | | |
|---|---|---|---|---|
| Tob1 | 560.157712 | -0.178561327 | 0.08299864 | -2.151376537 |
| Wfikkn2 | 33.87170838 | 0.156887351 | 0.213490229 | 0.734868998 |
| Gm21885 | 0.047948698 | -0.206487837 | 3.724945049 | -0.055433794 |
| Luc7l3 | 1814.337726 | 0.155150522 | 0.097374276 | 1.593341971 |
| Ankrd40 | 1549.21346 | -0.121877651 | 0.090341683 | -1.349074382 |
| Abcc3 | 101.8740177 | 0.502079383 | 0.312441201 | 1.606956386 |
| Cacna1g | 1923.935302 | -0.609897055 | 0.137071811 | -4.449471062 |
| Spata20 | 1.880027439 | -0.677536206 | 0.740626698 | -0.914814722 |
| Epn3 | 81.88700119 | -0.552558532 | 0.152368247 | -3.626467729 |
| Mycbpap | 22.79270077 | -0.6536035 | 0.254283278 | -2.570375466 |
| Rsad1 | 129.9462632 | -0.610803559 | 0.146459329 | -4.170465354 |
| Acsf2 | 203.4382502 | -0.114356311 | 0.164630887 | -0.694622457 |
| Chad | 2.13663281 | 0.120894934 | 0.688040173 | 0.175709122 |
| Gm11540 | 0 NA | NA | NA | |
| Lrrc59 | 1158.226995 | 0.443823188 | 0.109533369 | 4.051945008 |
| Eme1 | 6.299531046 | 2.164094435 | 0.7619992 | 2.840021926 |
| Mrpl27 | 283.0008542 | 0.74166434 | 0.112533397 | 6.59061541 |
| Xylt2 | 348.7120337 | 0.19436088 | 0.090607688 | 2.145081552 |
| Gm11542 | 2.690072256 | -0.27442865 | 0.578175674 | -0.474645791 |
| Gm11541 | 0.211252453 | -0.206481404 | 2.188118052 | -0.094364837 |
| Cdc34-ps | 2.829894638 | 0.011708241 | 0.582337398 | 0.020105597 |
| Gm11545 | 0.145451803 | 0.274387818 | 3.53800508 | 0.077554388 |
| Tmem92 | 0.060174537 | 0.755286614 | 3.724945049 | 0.202764498 |
| Gm11546 | 0 NA | NA | NA | |
| AA623943 | 0 NA | NA | NA | |
| Gm11543 | 0 NA | NA | NA | |
| Gm11544 | 0 NA | NA | NA | |
| A430060F13Rik | 0 NA | NA | NA | |
| Gm22456 | 0 NA | NA | NA | |
| Col1a1 | 391.7651446 | -0.217434223 | 0.160993744 | -1.350575604 |
| Gm24877 | 0 NA | NA | NA | |
| Sgca | 2.248524686 | 1.428497351 | 0.811059152 | 1.761273944 |
| Hils1 | 0 NA | NA | NA | |
| Ppp1r9b | 8227.147287 | -0.004836014 | 0.100710013 | -0.048019199 |
| Samd14 | 673.600847 | 0.224724576 | 0.116683556 | 1.925931838 |
| Pdk2 | 946.573687 | -0.080072836 | 0.099457686 | -0.805094499 |
| Gm11513 | 1.036123199 | -0.664628313 | 0.966730272 | -0.687501294 |
| Itga3 | 513.8169415 | 0.1080884 | 0.090528607 | 1.193969547 |
| Dlx3 | 0.948391139 | 0.443742738 | 1.003150637 | 0.442349056 |
| Dlx4 | 3.24303925 | -0.034473177 | 0.526546903 | -0.065470287 |
| A730090H04Rik | 3.582880236 | 1.757895801 | 0.627669793 | 2.800669748 |
| Gm11514 | 1.617110251 | 0.238427047 | 0.892321259 | 0.267198663 |
| Gm11515 | 0.488879413 | -1.148542435 | 1.565126008 | -0.733833844 |
| Tac4 | 0.201378205 | 0.235550586 | 2.643723003 | 0.089098058 |
| Kat7 | 1091.977074 | 0.260866821 | 0.077400236 | 3.370362073 |
| Gm11520 | 2.683034273 | 0.420897689 | 0.632110338 | 0.665861106 |
| Gm11521 | 0.240698147 | 1.899486691 | 3.707812888 | 0.512293028 |

| | | | | |
|---|---|---|---|---|
| Fam117a | 84.85769561 | 0.504834597 | 0.15639987 | 3.227845367 |
| Gm29477 | 0.19690755 | -0.687380326 | 2.678289459 | -0.256649005 |
| Slc35b1 | 393.4153695 | 0.40301402 | 0.107889093 | 3.735447278 |
| Spop | 1831.86833 | 0.200760029 | 0.072568253 | 2.766499411 |
| Gm11522 | 0 | NA | NA | NA |
| Nxph3 | 195.7791226 | 0.29932128 | 0.122441826 | 2.444599934 |
| Ngfr | 42.86765523 | 0.143401623 | 0.275694648 | 0.520146562 |
| Gm11528 | 0 | NA | NA | NA |
| Phb | 513.3109458 | -0.073455246 | 0.117504497 | -0.625127103 |
| Gm26830 | 2.519878091 | -0.989975572 | 0.659334359 | -1.501477299 |
| Gm9796 | 6.527712466 | -1.113383399 | 0.435781162 | -2.554914018 |
| Zfp652os | 6.322034818 | -0.211570737 | 0.502798746 | -0.420786127 |
| Zfp652 | 1286.698194 | -0.286323437 | 0.058430511 | -4.900238411 |
| B130006D01Rik | 36.80483133 | -0.384636908 | 0.259192205 | -1.483983314 |
| Polr2k-ps | 4.087166043 | 0.086993198 | 0.468552433 | 0.185663742 |
| Phospho1 | 32.25678491 | -0.549146093 | 0.206330116 | -2.661492671 |
| Abi3 | 109.2635649 | 1.692499383 | 0.188119888 | 8.996918941 |
| Gngt2 | 37.40057289 | 2.019701537 | 0.311543613 | 6.482885392 |
| Gm11527 | 0.940895551 | -0.850857388 | 1.187072807 | -0.716769336 |
| 4833417C18Rik | 16.33142681 | -0.008840157 | 0.296764303 | -0.029788479 |
| B4galnt2 | 0.576309813 | 0.274413038 | 1.553423452 | 0.176650505 |
| Gm11534 | 0 | NA | NA | NA |
| Gm24725 | 0 | NA | NA | NA |
| Igf2bp1 | 1.321944539 | 1.753509141 | 0.981757992 | 1.786091028 |
| Mir3063 | 0 | NA | NA | NA |
| Gm29202 | 0 | NA | NA | NA |
| Gm37179 | 0 | NA | NA | NA |
| Gip | 0 | NA | NA | NA |
| Gm24949 | 0 | NA | NA | NA |
| Snf8 | 318.1547224 | 0.039589869 | 0.101239682 | 0.391050904 |
| Ube2z | 1093.216647 | 0.071772071 | 0.063107751 | 1.137294079 |
| Atp5g1 | 503.7452782 | -0.224682418 | 0.087962741 | -2.554290785 |
| Gm22423 | 0 | NA | NA | NA |
| Calcoco2 | 0.681926656 | 2.093392255 | 1.144692413 | 1.828781452 |
| Ttll6 | 7.585280821 | -1.238378862 | 0.401917046 | -3.08118024 |
| Gm11535 | 0 | NA | NA | NA |
| Hoxb13 | 0 | NA | NA | NA |
| Gm27898 | 0 | NA | NA | NA |
| Gm27605 | 0 | NA | NA | NA |
| Gm11539 | 1.7546716 | -0.65334883 | 0.731988355 | -0.892567245 |
| Gm53 | 0.110242424 | -0.20649451 | 3.724945049 | -0.055435586 |
| Mir196a-1 | 0 | NA | NA | NA |
| Hoxb9 | 0.072025462 | 0.755288839 | 3.724945049 | 0.202765096 |
| Hoxb8 | 30.1408604 | -0.578599152 | 1.733052266 | -0.33386134 |
| Hoxb7 | 3.606077423 | -0.178210335 | 1.461274628 | -0.121955402 |
| Hoxb5os | 0.345588754 | 0.669437753 | 3.23196842 | 0.207130042 |
| Hoxb6 | 10.70947983 | -0.519808096 | 1.246174622 | -0.417122999 |

| | | | | |
|---|---|---|---|---|
| Hoxb5 | 50.63370746 | 0.031279013 | 1.020991383 | 0.030635923 |
| Mir10a | 0.182096932 | -0.687376824 | 2.691326143 | -0.255404506 |
| Hoxb4 | 17.2026822 | -0.468463835 | 0.797750555 | -0.587230973 |
| Hoxb3 | 35.54127179 | -0.318935532 | 0.749379894 | -0.425599265 |
| Gm11536 | 0.751775467 | 1.693293934 | 1.24329889 | 1.361936335 |
| Hoxb3os | 4.383958145 | -0.119067841 | 0.553150624 | -0.215253921 |
| Hoxb2 | 10.86433349 | 0.087929002 | 0.883489744 | 0.099524644 |
| Hoxb1 | 0.038981395 | 0.274393828 | 3.724945049 | 0.073663859 |
| Gm11531 | 0.072025462 | 0.755288839 | 3.724945049 | 0.202765096 |
| Gm11529 | 0.471635551 | -0.183071394 | 1.329187307 | -0.137731825 |
| Skap1 | 41.01891481 | 1.347883138 | 0.251266596 | 5.364354669 |
| 2010300F17Rik | 0.313173757 | -1.179263818 | 2.055799346 | -0.573627879 |
| Gm11537 | 0.487078227 | 2.377527783 | 1.300946005 | 1.827537634 |
| Snx11 | 313.2983275 | 0.265566991 | 0.082027923 | 3.237519365 |
| Cbx1 | 382.9098145 | -0.238994081 | 0.096545393 | -2.475458167 |
| Gm11517 | 5.580449154 | 1.042046013 | 0.476556342 | 2.186616607 |
| Nfe2l1 | 4454.295115 | 0.161765975 | 0.046037257 | 3.513805647 |
| Copz2 | 26.60350595 | 0.234294939 | 0.24024258 | 0.975243184 |
| Mir152 | 0.090665643 | -0.687373951 | 3.724945049 | -0.184532642 |
| Gm11523 | 0.271332793 | 0.380345193 | 2.058157949 | 0.184798836 |
| Gm11524 | 0 | NA | NA | NA |
| Cdk5rap3 | 188.9212374 | 0.14199639 | 0.105113304 | 1.350888847 |
| D030028A08Rik | 76.08496205 | -0.0367797 | 0.142998228 | -0.257203888 |
| Gm11525 | 3.501911378 | 0.710476837 | 0.557938926 | 1.273395355 |
| Prr15l | 1.00014491 | 1.986075752 | 1.193920418 | 1.663490901 |
| Pnpo | 446.5624225 | -0.064869347 | 0.072039793 | -0.900465482 |
| Sp2 | 265.9732879 | 0.696931465 | 0.132822664 | 5.247082428 |
| Sp6 | 5.037085609 | -0.271372794 | 0.440273901 | -0.616372656 |
| Gm11532 | 1.602391015 | 0.626276464 | 0.942580682 | 0.664427435 |
| Scrn2 | 30.51638522 | -0.437908216 | 0.250866359 | -1.745583657 |
| Lrrc46 | 5.763699656 | -0.311719333 | 0.502223836 | -0.620678094 |
| Gm11533 | 0.306021358 | 0.274415327 | 1.802671968 | 0.152226989 |
| Mrpl10 | 403.0972378 | 0.113333015 | 0.084063736 | 1.348179621 |
| Gm11574 | 2.609456233 | 0.029701277 | 0.64094924 | 0.046339516 |
| Osbpl7 | 231.9002821 | 0.142138578 | 0.084928303 | 1.67363026 |
| Mir8103 | 1.700203044 | -1.443072058 | 0.88388791 | -1.632641473 |
| Gm11573 | 0 | NA | NA | NA |
| Tbx21 | 34.3699015 | 0.909599898 | 0.432304339 | 2.104073027 |
| Gm11572 | 0.075043205 | 0.755286614 | 3.724945049 | 0.202764498 |
| Tbkbp1 | 732.199412 | 0.081503577 | 0.07129086 | 1.143254233 |
| Mir7235 | 0 | NA | NA | NA |
| Kpnb1 | 2804.956967 | 0.064653101 | 0.061300883 | 1.054684673 |
| Gm11583 | 2.829785719 | 0.271114417 | 0.659101957 | 0.411339117 |
| Npepps | 2128.553621 | -0.137873569 | 0.057195904 | -2.410549671 |
| Gm11592 | 3.097463869 | -0.303158403 | 0.544476487 | -0.556788788 |
| Mrpl45 | 265.8735872 | -0.13523114 | 0.10696048 | -1.264309401 |
| Gm11594 | 0.046345709 | 0.274391604 | 3.724945049 | 0.073663262 |

| | | | | |
|---|---|---|---|---|
| Gpr179 | 53.52322391 | 1.316282579 | 0.266762107 | 4.934293684 |
| Socs7 | 1787.58141 | -0.024106133 | 0.071454003 | -0.337365748 |
| Arhgap23 | 1777.390419 | -0.178597075 | 0.064181264 | -2.782698013 |
| 4933428G20Rik | 1.863773006 | 0.133963201 | 0.914898951 | 0.146424041 |
| Srcin1 | 3857.360928 | -0.328149512 | 0.090372781 | -3.631065778 |
| Gm11611 | 105.1460556 | -0.641969267 | 0.216487268 | -2.96539041 |
| Gm11612 | 0.519932157 | -0.125941359 | 1.364739906 | -0.092282316 |
| 2410003L11Rik | 0.232084656 | 0.27441153 | 2.13565865 | 0.128490351 |
| E130012A19Rik | 333.7269305 | -0.189977923 | 0.153872373 | -1.234646082 |
| Gm11613 | 75.52422125 | -0.487332392 | 0.164861261 | -2.956015188 |
| Gm24158 | 0 | NA | NA | NA |
| Mllt6 | 2810.779345 | -0.015846656 | 0.061319889 | -0.258426043 |
| Mir6927 | 0 | NA | NA | NA |
| Cisd3 | 84.26726889 | -0.047738536 | 0.166585234 | -0.286571235 |
| Pcgf2 | 208.1768514 | -0.168761596 | 0.145163396 | -1.162563015 |
| Psmb3 | 579.8326605 | 0.559835349 | 0.144067477 | 3.885924583 |
| Pip4k2b | 1705.574018 | -0.143009883 | 0.063756931 | -2.243048429 |
| Gm11614 | 0 | NA | NA | NA |
| Mir8102 | 0.581519725 | 0.926692848 | 1.297457111 | 0.71423775 |
| Cwc25 | 283.5958626 | 0.287578865 | 0.113252335 | 2.539275384 |
| Atp5l2-ps | 17.66472391 | 0.278629969 | 0.326358565 | 0.853754118 |
| 1700001P01Rik | 6.295635024 | -0.925580893 | 0.464273319 | -1.993612071 |
| Rpl23 | 782.0632473 | 0.483897291 | 0.125672128 | 3.850474231 |
| Gm27326 | 2.560534601 | -0.786903833 | 0.749380421 | -1.050072582 |
| Snora21 | 15.9140621 | 0.09376454 | 0.285098904 | 0.328884253 |
| Lasp1 | 1454.162824 | -0.169239281 | 0.081712454 | -2.071156505 |
| B230217C12Rik | 275.8448458 | -0.403264163 | 0.132320113 | -3.047640717 |
| Fbxo47 | 12.48405167 | -0.10480096 | 0.300611386 | -0.348626049 |
| Gm11630 | 0.282072957 | -0.685882003 | 1.785735073 | -0.384089451 |
| Plxdc1 | 370.3130502 | -0.178399229 | 0.127363213 | -1.400712378 |
| Gm22461 | 0.138038828 | 0.755294464 | 3.633333205 | 0.207879218 |
| Gm11633 | 0.1140246 | 0.75528439 | 3.724945049 | 0.202763901 |
| Arl5c | 23.21351478 | 1.388340765 | 0.31309623 | 4.434230224 |
| Cacnb1 | 1086.374005 | -0.450687236 | 0.14214087 | -3.170708309 |
| Gm11629 | 3.893533808 | -0.073336305 | 0.496274341 | -0.147773719 |
| Rpl19 | 1377.971411 | 0.435957756 | 0.144174046 | 3.02382965 |
| Gm27597 | 0 | NA | NA | NA |
| Stac2 | 733.8978338 | -0.124466555 | 0.096886051 | -1.284669505 |
| Gm11631 | 0.045882455 | 0.755288839 | 3.724945049 | 0.202765096 |
| Fbxl20 | 783.1636922 | -0.130161918 | 0.097099989 | -1.340493637 |
| Gm11632 | 0 | NA | NA | NA |
| Med1 | 1253.203097 | -0.020239608 | 0.052043805 | -0.388895629 |
| Gm11604 | 1.649565372 | 1.787619178 | 1.024027856 | 1.745674366 |
| Cdk12 | 1154.320364 | 0.244879974 | 0.094396381 | 2.594166989 |
| Gm27399 | 1.487186809 | 0.118447776 | 0.841996091 | 0.140674971 |
| Mir5119 | 5.066509227 | 0.558412131 | 0.514205666 | 1.085970397 |
| n-R5s73 | 0 | NA | NA | NA |

| | | | | |
|---|---|---|---|---|
| Gm20644 | 4.541882812 | 1.88736749 | 0.749784714 | 2.517212547 |
| Neurod2 | 846.9061386 | 0.195348097 | 0.13504868 | 1.446501342 |
| Ppp1r1b | 2292.267063 | -0.112451594 | 0.257772377 | -0.436243771 |
| 1700003D09Rik | 3.143667968 | -0.078341399 | 0.781475355 | -0.100248073 |
| Stard3 | 334.9821919 | 0.382440395 | 0.10809771 | 3.537913933 |
| Tcap | 15.2625128 | -0.764556885 | 0.259782026 | -2.943070758 |
| Pnmt | 1.554873959 | 1.000507151 | 0.963459559 | 1.03845267 |
| Pgap3 | 82.83766852 | 0.15802549 | 0.161966629 | 0.975666972 |
| Erbb2 | 73.62724615 | 0.862431795 | 0.198210542 | 4.351089436 |
| Gm25048 | 0 | NA | NA | NA |
| Mien1 | 288.359261 | -0.30942558 | 0.134724431 | -2.296729534 |
| Gm12352 | 1.915443829 | 1.420966657 | 0.860642259 | 1.651053784 |
| Grb7 | 32.20120666 | -0.335089259 | 0.203948468 | -1.643009443 |
| Ikzf3 | 75.08787135 | 4.893055715 | 0.419910838 | 11.6526064 |
| Gm25106 | 0.044544648 | 0.755286614 | 3.724945049 | 0.202764498 |
| Zpbp2 | 0.628980615 | 0.331079834 | 1.158565482 | 0.285767045 |
| Ormdl3 | 555.4362379 | 0.253743786 | 0.075330594 | 3.368402839 |
| Gm12 | 0 | NA | NA | NA |
| Gm12355 | 8.225470329 | 0.108413338 | 0.365869171 | 0.296317227 |
| Gsdma3 | 0.182775946 | 1.806322629 | 3.324025814 | 0.54341414 |
| Gsdma2 | 0 | NA | NA | NA |
| Gsdma | 1.768266679 | -0.478568619 | 0.829947379 | -0.576625254 |
| Psmd3 | 1360.534359 | 0.315504241 | 0.09224557 | 3.420264426 |
| Gm12356 | 1.14577023 | 0.455775642 | 0.982983648 | 0.463665538 |
| Csf3 | 0.907385681 | 3.499295554 | 1.177548621 | 2.97167819 |
| Med24 | 900.5465924 | 0.103422271 | 0.071639558 | 1.443647526 |
| Gm22059 | 0.481050399 | 0.745152642 | 1.662884893 | 0.448108372 |
| Thra | 4319.830123 | 0.046376014 | 0.076261663 | 0.608116996 |
| Nr1d1 | 1275.640339 | -0.175875591 | 0.089795919 | -1.958614516 |
| Msl1 | 1734.681355 | -0.267829535 | 0.067466793 | -3.969797898 |
| Gm12359 | 6.105237519 | 0.309824477 | 0.441502914 | 0.701749563 |
| Casc3 | 869.1967834 | 0.008582464 | 0.055317739 | 0.155148504 |
| Rapgefl1 | 1200.68795 | -0.415224778 | 0.143684686 | -2.889833219 |
| Wipf2 | 1511.917858 | 0.008942239 | 0.04839878 | 0.184761659 |
| Gm23640 | 0 | NA | NA | NA |
| Cdc6 | 14.16964528 | 0.302504884 | 0.326386219 | 0.926831055 |
| Rara | 285.8551299 | 0.219051258 | 0.16477201 | 1.32942032 |
| Gm22061 | 0.570559668 | 1.448785963 | 1.316248794 | 1.100693098 |
| Gjd3 | 5.273691302 | 1.082577558 | 0.601798984 | 1.798902268 |
| Top2a | 129.6343859 | 2.190849062 | 0.393874957 | 5.562295912 |
| Gm26394 | 0 | NA | NA | NA |
| Gm23451 | 0 | NA | NA | NA |
| Igfbp4 | 440.1552601 | 0.352300716 | 0.123377612 | 2.85546713 |
| Tns4 | 6.644394917 | 0.763091104 | 0.391813422 | 1.947587963 |
| Ccr7 | 12.21422624 | 3.852409502 | 0.673937559 | 5.716270673 |
| Smarce1 | 477.9356527 | -0.066938869 | 0.071986771 | -0.929877366 |
| Krt222 | 403.0542806 | 0.046959036 | 0.069057553 | 0.67999855 |

| | | | | |
|---|---|---|---|---|
| Krt24 | 0.665888905 | 1.515424454 | 1.44246782 | 1.050577652 |
| Gm12358 | 0 | NA | NA | NA |
| Krt25 | 4.739584563 | -0.376190404 | 0.48330941 | -0.7783635 |
| Krt26 | 0.63227042 | -0.186827013 | 1.515201922 | -0.123301727 |
| Krt27 | 0.305696448 | 2.109902063 | 1.983502693 | 1.063725333 |
| Krt28 | 1.347390254 | 1.270227888 | 0.872477151 | 1.455886708 |
| Krt10 | 43.27398083 | 0.087892965 | 0.399448905 | 0.220035565 |
| Gm11940 | 7.488360699 | -0.670786134 | 0.394064816 | -1.70222285 |
| Krt12 | 43.87784049 | -0.371770996 | 0.195485705 | -1.901780985 |
| A830036E02Rik | 268.3212654 | -0.926014821 | 0.356628572 | -2.596580568 |
| Krt20 | 0.572048112 | 0.151903621 | 1.366669098 | 0.111148793 |
| Krt23 | 0.336553411 | 0.674356583 | 1.992992402 | 0.338363851 |
| Krt39 | 0 | NA | NA | NA |
| Krt40 | 0 | NA | NA | NA |
| Krtap3-3 | 0 | NA | NA | NA |
| Krtap3-2 | 0 | NA | NA | NA |
| Gm11939 | 0 | NA | NA | NA |
| Krtap3-1 | 0 | NA | NA | NA |
| Krtap1-5 | 0 | NA | NA | NA |
| Krtap1-4 | 0 | NA | NA | NA |
| Krtap1-3 | 0 | NA | NA | NA |
| Krtap9-3 | 0 | NA | NA | NA |
| Gm11938 | 0 | NA | NA | NA |
| Gm11937 | 0 | NA | NA | NA |
| Krtap2-4 | 0 | NA | NA | NA |
| Gm11562 | 0 | NA | NA | NA |
| Krtap4-1 | 0 | NA | NA | NA |
| Krtap4-2 | 0 | NA | NA | NA |
| Krtap4-7 | 0 | NA | NA | NA |
| Gm11555 | 0 | NA | NA | NA |
| Gm11563 | 0.072025462 | 0.755288839 | 3.724945049 | 0.202765096 |
| Gm14182 | 0 | NA | NA | NA |
| Krtap4-6 | 0 | NA | NA | NA |
| Gm11557 | 0.202171817 | 0.274403705 | 2.59796949 | 0.105622374 |
| 4930415H17Rik | 0 | NA | NA | NA |
| Gm14190 | 0 | NA | NA | NA |
| Gm14186 | 0 | NA | NA | NA |
| Gm14193 | 0 | NA | NA | NA |
| Gm14180 | 0 | NA | NA | NA |
| Gm14188 | 0 | NA | NA | NA |
| Gm11595 | 0 | NA | NA | NA |
| Krtap4-8 | 0.046345709 | 0.274391604 | 3.724945049 | 0.073663262 |
| Krtap4-9 | 0 | NA | NA | NA |
| Gm11596 | 0 | NA | NA | NA |
| Gm11569 | 0 | NA | NA | NA |
| Gm11554 | 0 | NA | NA | NA |
| Krtap4-13 | 0 | NA | NA | NA |

| | | | | |
|---|---|---|---|---|
| Gm11564 | 0 | NA | NA | NA |
| 2300003K06Rik | 0 | NA | NA | NA |
| Gm11556 | 0 | NA | NA | NA |
| Krtap4-16 | 0.075043205 | 0.755286614 | 3.724945049 | 0.202764498 |
| Gm26324 | 0 | NA | NA | NA |
| Gm11568 | 0 | NA | NA | NA |
| Gm11559 | 0 | NA | NA | NA |
| Krtap9-1 | 0 | NA | NA | NA |
| Gm11567 | 0 | NA | NA | NA |
| Gm11566 | 0 | NA | NA | NA |
| Krtap31-1 | 0.038981395 | 0.274393828 | 3.724945049 | 0.073663859 |
| Gm11565 | 0 | NA | NA | NA |
| Gm11560 | 17.37014342 | 0.399421174 | 0.365629928 | 1.092419255 |
| Krtap31-2 | 0 | NA | NA | NA |
| Krtap9-5 | 0 | NA | NA | NA |
| Gm11561 | 0 | NA | NA | NA |
| Gm11553 | 0 | NA | NA | NA |
| Krtap29-1 | 0.072025462 | 0.755288839 | 3.724945049 | 0.202765096 |
| Krtap16-1 | 0.141781211 | 0.755294552 | 2.911713426 | 0.259398657 |
| Gm11558 | 0 | NA | NA | NA |
| Krtap17-1 | 0.608261712 | 0.524547569 | 1.414086226 | 0.370944543 |
| Krt33a | 0 | NA | NA | NA |
| Krt33b | 0 | NA | NA | NA |
| Krt34 | 0 | NA | NA | NA |
| Krt31 | 0 | NA | NA | NA |
| Gm11571 | 0 | NA | NA | NA |
| Krt32 | 0 | NA | NA | NA |
| Krt35 | 0 | NA | NA | NA |
| Krt36 | 0.15956642 | 0.75529148 | 3.418807262 | 0.22092251 |
| Gm12347 | 0 | NA | NA | NA |
| Krt13 | 0.652975239 | 2.026455391 | 1.598910909 | 1.267397314 |
| Krt15 | 1.82600251 | -2.089671994 | 0.9139943 | -2.286307467 |
| Krt19 | 0.608478225 | -1.432809685 | 1.564307062 | -0.915938897 |
| Krt9 | 26.56616351 | -0.111669335 | 0.299008347 | -0.373465612 |
| Krt14 | 1.447260382 | 2.447818537 | 1.409569365 | 1.73657189 |
| Gm11597 | 0 | NA | NA | NA |
| Gm11598 | 0.706088228 | 0.718474372 | 1.207752946 | 0.594885216 |
| Krt16 | 2.108251394 | 0.787097987 | 1.433023223 | 0.549256965 |
| Krt17 | 3.233276867 | 0.640314694 | 0.922451664 | 0.69414444 |
| Krt42 | 0 | NA | NA | NA |
| Gm14206 | 0.67116138 | 2.004292609 | 1.306889758 | 1.533635562 |
| Gm12349 | 7.416241275 | 4.84491552 | 1.235160787 | 3.92249784 |
| Eif1 | 1981.137352 | 0.176188238 | 0.095959449 | 1.836069708 |
| Gast | 0.589408234 | -1.362454788 | 1.307731152 | -1.041846244 |
| Hap1 | 3690.110612 | 1.305053062 | 0.142378971 | 9.166052091 |
| Gm20394 | 9.564810652 | 1.315126618 | 0.529504737 | 2.483691883 |
| Gm10039 | 471.8830643 | -0.154468177 | 0.089287339 | -1.730012104 |

| | | | | |
|---|---|---|---|---|
| Jup | 876.4788789 | -0.070745741 | 0.099122692 | -0.713718924 |
| Gm12348 | 0.360464465 | -2.091336772 | 1.365601883 | -1.531439578 |
| P3h4 | 107.8090644 | 0.14255455 | 0.116030476 | 1.228595753 |
| Fkbp10 | 103.2641343 | 0.492417808 | 0.165570103 | 2.974074414 |
| Nt5c3b | 220.2515258 | -0.210229314 | 0.086544588 | -2.429144554 |
| Klhl10 | 2.707144304 | -0.513513146 | 0.633189567 | -0.810994325 |
| Gm27374 | 1.132676578 | 0.050428631 | 0.868347631 | 0.058074242 |
| Klhl11 | 985.9766124 | -0.073537307 | 0.062601237 | -1.174694166 |
| Acly | 2346.506125 | -0.166763641 | 0.084249547 | -1.979401031 |
| mmu-mir-7116 | 0.285551684 | -0.582162875 | 2.024829279 | -0.287512079 |
| Ttc25 | 14.3848452 | -0.507956603 | 0.277068561 | -1.833324582 |
| Cnp | 4776.985164 | 0.300488059 | 0.115816097 | 2.59452758 |
| Dnajc7 | 1151.346391 | 0.126155512 | 0.071689473 | 1.75974946 |
| Nkiras2 | 157.3130789 | 0.453006411 | 0.110848814 | 4.086705066 |
| Zfp385c | 87.71940538 | 0.174570037 | 0.226542805 | 0.770583009 |
| Dhx58os | 2.117912561 | 0.544976508 | 0.654764453 | 0.832324518 |
| Dhx58 | 526.9953047 | 5.457528105 | 0.251475143 | 21.70205786 |
| Kat2a | 785.4970274 | 0.381696712 | 0.057474397 | 6.641160776 |
| Hspb9 | 0.468943176 | 0.274387745 | 3.28314321 | 0.083574711 |
| Rab5c | 1897.413452 | 0.781879578 | 0.128363734 | 6.091125219 |
| Kcnh4 | 95.02347477 | -0.659178216 | 0.258590253 | -2.549122439 |
| Hcrt | 38.8486744 | -0.131765807 | 0.201916642 | -0.652575271 |
| Ghdc | 106.0321 | 0.128390776 | 0.182848067 | 0.702171911 |
| Gm24358 | 0.048949532 | -0.206487837 | 3.724945049 | -0.055433794 |
| Stat5b | 602.2188295 | 0.063820398 | 0.106358109 | 0.600052021 |
| Stat5a | 70.6584695 | 1.036437224 | 0.194630092 | 5.32516433 |
| Stat3 | 2401.024981 | 2.012010375 | 0.165925351 | 12.12599742 |
| Ptrf | 400.0738531 | 1.034196185 | 0.181817199 | 5.688109772 |
| Atp6v0a1 | 5069.409655 | -0.069464415 | 0.063070222 | -1.101382114 |
| Mir6928 | 2.18332649 | -0.525843915 | 0.70962728 | -0.741014233 |
| Naglu | 145.2410652 | 0.622894605 | 0.155066406 | 4.016953896 |
| Hsd17b1 | 3.534966268 | -2.232309607 | 0.713173969 | -3.130105283 |
| Coasy | 224.3589791 | 0.125983907 | 0.09435428 | 1.33522196 |
| Mlx | 272.5349141 | -0.035045747 | 0.105716659 | -0.33150638 |
| Psmc3ip | 15.59208128 | -0.165775727 | 0.292016347 | -0.56769331 |
| Fam134c | 741.7829735 | -0.066449177 | 0.066967855 | -0.992254816 |
| Tubg1 | 286.1670221 | -0.202599694 | 0.087204335 | -2.323275476 |
| Gm27626 | 0 | NA | NA | NA |
| Tubg2 | 274.8310085 | 0.180531531 | 0.082090585 | 2.199174625 |
| Plekhh3 | 197.5858469 | -0.218796973 | 0.117780227 | -1.857671518 |
| Cntnap1 | 2524.742535 | -0.284329251 | 0.084620494 | -3.360051888 |
| Ccr10 | 4.101636231 | 1.914870421 | 0.65154731 | 2.93895837 |
| Ezh1 | 709.9648608 | 0.001974641 | 0.079464932 | 0.024849218 |
| Ramp2 | 134.6263258 | 0.654499576 | 0.154668963 | 4.231615472 |
| Vps25 | 18.38762957 | 0.201699496 | 0.268564055 | 0.751029381 |
| Wnk4 | 77.9757232 | -0.070677786 | 0.172468437 | -0.409801278 |
| Gm11615 | 0.280085221 | 0.558328853 | 2.084990456 | 0.267784848 |

| | | | | |
|---|---|---|---|---|
| Coa3 | 343.5032495 | -0.177250444 | 0.142048817 | -1.247813587 |
| Cntd1 | 4.658726077 | -0.828796901 | 0.508121918 | -1.631098505 |
| Becn1 | 989.2074398 | 0.192188917 | 0.068856842 | 2.791137547 |
| Gm11617 | 0 | NA | NA | NA |
| Psme3 | 1184.471621 | 0.230101229 | 0.065878873 | 3.492792419 |
| Aoc2 | 39.91492154 | -0.540904369 | 0.30362401 | -1.781494059 |
| Aoc3 | 5.919817619 | -1.614539222 | 0.52643258 | -3.066943962 |
| Gm28156 | 0 | NA | NA | NA |
| Gm11618 | 0 | NA | NA | NA |
| Gm11619 | 0 | NA | NA | NA |
| Gm25593 | 0 | NA | NA | NA |
| G6pc | 0.591033593 | 0.122422483 | 1.269758699 | 0.096413975 |
| Gm25475 | 0 | NA | NA | NA |
| Gm27029 | 8.009665686 | -1.121254006 | 0.452811263 | -2.476206087 |
| Aarsd1 | 19.81807774 | -0.387629644 | 0.267228755 | -1.450553643 |
| Ptges3l | 9.728067973 | -0.072880983 | 0.350621286 | -0.207862402 |
| Mir6929 | 0.072025462 | 0.755288839 | 3.724945049 | 0.202765096 |
| Rundc1 | 403.5552599 | -0.150619231 | 0.083137453 | -1.811689264 |
| Rpl27 | 526.9415286 | 0.476395048 | 0.128541846 | 3.706147551 |
| Ifi35 | 471.7219268 | 5.308489344 | 0.25225891 | 21.04381302 |
| Gm11626 | 1.005912905 | -0.925594043 | 1.134594468 | -0.81579284 |
| Vat1 | 770.8644348 | -0.176160296 | 0.163542226 | -1.077154811 |
| Rnd2 | 698.8704723 | -0.247796423 | 0.149102536 | -1.661919571 |
| Gm11625 | 0.085327104 | 0.274389469 | 3.724945049 | 0.073662689 |
| Brca1 | 51.12254382 | 0.885804801 | 0.265710995 | 3.333715274 |
| Nbr1 | 2218.562145 | 0.023017795 | 0.038717004 | 0.59451386 |
| Gm27456 | 0 | NA | NA | NA |
| Tmem106a | 300.3975511 | 3.542396288 | 0.32951429 | 10.75035709 |
| Gm11634 | 0 | NA | NA | NA |
| Gm26316 | 17.08146439 | -0.494755442 | 0.342118503 | -1.446152247 |
| Gm11635 | 0.049553047 | 0.274391604 | 3.724945049 | 0.073663262 |
| Rdm1 | 27.87786844 | -0.546175169 | 0.229163063 | -2.38334731 |
| Gm23849 | 1.056059865 | 0.800746565 | 1.119953618 | 0.714981899 |
| Gm24950 | 3.757983618 | 0.360531257 | 0.560282418 | 0.643481297 |
| Gm23971 | 333.7774647 | -0.281428655 | 0.19488708 | -1.444060092 |
| Rnu2-10 | 3.530281436 | 0.319792372 | 0.615979217 | 0.519160976 |
| Arl4d | 211.6319964 | 0.313462308 | 0.336228239 | 0.932290247 |
| Dhx8 | 571.733905 | 0.115463336 | 0.067764296 | 1.703896339 |
| Etv4 | 83.85799785 | -0.472804372 | 0.143887266 | -3.285936177 |
| Meox1 | 5.667363994 | 0.605673667 | 0.469393899 | 1.290331358 |
| Gm11551 | 0.198035107 | 0.755299381 | 3.222442586 | 0.234387227 |
| Rpl27-ps2 | 0.195392278 | 1.699873117 | 3.230774338 | 0.526150371 |
| 4930417O22Rik | 0.123841226 | 0.274391173 | 3.724945049 | 0.073663147 |
| Sost | 2.176193535 | -2.686847859 | 0.95346285 | -2.817989038 |
| Gm20659 | 0.323889686 | 1.887102706 | 3.692557181 | 0.511055784 |
| Dusp3 | 1716.840804 | -0.364301197 | 0.080988783 | -4.498168579 |
| 1700006E09Rik | 0.21733455 | -0.206482122 | 2.170157949 | -0.095146126 |

| | | | | |
|---|---|---|---|---|
| Mpp3 | 656.0571513 | -0.335047539 | 0.11128051 | -3.010837544 |
| E130111B04Rik | 2.471955472 | -0.058853603 | 0.63340845 | -0.092915721 |
| Cd300lg | 0.955408868 | 2.739296032 | 1.157540023 | 2.366480621 |
| Mpp2 | 1306.923634 | -0.483910249 | 0.100652177 | -4.807747468 |
| Ppy | 0.061920613 | 0.274393828 | 3.724945049 | 0.073663859 |
| Pyy | 0.755118542 | 0.53340477 | 1.457691527 | 0.365924313 |
| Gm11584 | 0.066050875 | 0.274391604 | 3.724945049 | 0.073663262 |
| Gm11585 | 0.714893038 | 0.473867914 | 1.181510095 | 0.401069713 |
| Nags | 0.527376817 | -1.110680669 | 1.29536467 | -0.857427021 |
| Gm11586 | 0 | NA | NA | NA |
| Tmem101 | 154.1546923 | -0.168326627 | 0.110129005 | -1.528449549 |
| Lsm12 | 620.6014926 | 0.100804127 | 0.091731628 | 1.09890263 |
| G6pc3 | 331.7313334 | 0.035670535 | 0.100635584 | 0.354452505 |
| Hdac5 | 1648.304379 | -0.438330396 | 0.103998102 | -4.214792247 |
| Mir8101 | 0 | NA | NA | NA |
| BC030867 | 13.51292698 | 0.503365535 | 0.374029064 | 1.34579257 |
| Asb16 | 47.65270219 | -0.01940981 | 0.168354647 | -0.115291206 |
| Tmub2 | 469.4932909 | 0.084889823 | 0.062540162 | 1.357364941 |
| Atxn7l3 | 1951.581005 | -0.113216642 | 0.053454378 | -2.118005019 |
| Ubtf | 1192.509439 | -0.106721378 | 0.069725804 | -1.530586543 |
| Slc4a1 | 10.15205109 | 1.686905368 | 0.639443565 | 2.638083267 |
| Bloodlinc | 0.118974955 | 0.274400501 | 3.724945049 | 0.073665651 |
| Rundc3a | 2248.259499 | 0.055950796 | 0.059263626 | 0.94410011 |
| Slc25a39 | 589.2097923 | -0.020899138 | 0.10194526 | -0.205003527 |
| Mir6930 | 0 | NA | NA | NA |
| Grn | 3148.773445 | 2.642545218 | 0.160723706 | 16.44153983 |
| Fam171a2 | 531.5743851 | -0.386626136 | 0.090030818 | -4.294375475 |
| Itga2b | 26.87028233 | 0.499587572 | 0.243665701 | 2.050299117 |
| Gm23563 | 0.090427103 | 1.236179401 | 3.724945049 | 0.331865138 |
| Gpatch8 | 1536.270915 | -0.075045396 | 0.098196171 | -0.764239537 |
| Gm11628 | 8.708300683 | -0.642813749 | 0.464295329 | -1.384493248 |
| Mdk-ps1 | 0 | NA | NA | NA |
| Gm25337 | 4.232730408 | -0.1669014 | 0.476623449 | -0.350174546 |
| Gm11627 | 10.86367426 | -1.737630451 | 0.406746143 | -4.272026868 |
| Fzd2 | 124.2975267 | -0.890275328 | 0.269941905 | -3.298025658 |
| Gm11636 | 0.35856548 | -0.885835981 | 1.758152834 | -0.503844697 |
| 2810433D01Rik | 2.836445391 | -1.008243156 | 0.667020768 | -1.511561865 |
| Psmb5-ps | 0 | NA | NA | NA |
| Gm22920 | 0 | NA | NA | NA |
| Gm1564 | 0.341943343 | 1.676456225 | 1.720667545 | 0.974305716 |
| Ccdc43 | 214.8599409 | 0.179894947 | 0.087065025 | 2.066213701 |
| Gm11637 | 5.095273339 | 0.710925514 | 0.444830123 | 1.598195529 |
| Gm16342 | 0.648681289 | -0.568822667 | 1.343927268 | -0.423254056 |
| Adam11 | 1411.899054 | -0.216737598 | 0.121113898 | -1.789535317 |
| Gjc1 | 65.81803581 | 0.738273501 | 0.178778428 | 4.129544651 |
| Gm22683 | 0 | NA | NA | NA |
| Higd1b | 5.836578858 | -0.775018096 | 0.452557926 | -1.712527945 |

| | | | | |
|---|---|---|---|---|
| Eftud2 | 907.904142 | 0.12775555 | 0.072015214 | 1.77400777 |
| Gm26668 | 0.400913792 | -0.264518532 | 1.735157565 | -0.152446405 |
| Ccdc103 | 10.14206194 | -0.112263036 | 0.32580191 | -0.344574518 |
| Fam187a | 0.4546328 | -1.680959891 | 1.384035741 | -1.214535031 |
| Gfap | 17257.76253 | 2.426310739 | 0.171395706 | 14.15619324 |
| Kif18b | 11.6304259 | 1.96486279 | 0.576024517 | 3.411074932 |
| C1ql1 | 74.68787927 | -0.475691701 | 0.191888456 | -2.479001133 |
| 2410004I01Rik | 2.828652251 | -0.287227329 | 0.638944648 | -0.449533978 |
| Dcakd | 326.4044415 | 0.089907007 | 0.104324764 | 0.861799282 |
| Mir6931 | 0.170867779 | -1.262680235 | 3.70728864 | -0.340593991 |
| Nmt1 | 1910.096591 | 0.280779313 | 0.046381441 | 6.053699689 |
| Plcd3 | 130.7093725 | -0.080502503 | 0.100182157 | -0.803561282 |
| Acbd4 | 248.1541703 | 0.121254192 | 0.096485815 | 1.256704856 |
| Hexim1 | 470.7862717 | -0.362344194 | 0.104332601 | -3.47297193 |
| Hexim2 | 107.5740266 | 0.610203764 | 0.145429716 | 4.195867134 |
| Fmnl1 | 978.2301766 | -0.166212442 | 0.110416982 | -1.505315923 |
| Gm20511 | 5.853403199 | -0.767633176 | 0.469474667 | -1.635089665 |
| 1700023F06Rik | 21.3606864 | -0.056993489 | 0.25415537 | -0.224246646 |
| Spata32 | 11.37911894 | 0.51122053 | 0.352790588 | 1.449076442 |
| Map3k14 | 131.8227025 | 0.683888406 | 0.190194974 | 3.5957228 |
| 1700028N14Rik | 3.893030208 | -0.09346054 | 0.513184957 | -0.182118627 |
| Arhgap27 | 221.5069098 | 0.632988639 | 0.102295659 | 6.187834785 |
| Arhgap27os3 | 8.082714316 | 2.522704568 | 0.546915799 | 4.612601377 |
| Arhgap27os2 | 8.179740595 | 0.36271966 | 0.367303288 | 0.987520865 |
| Arhgap27os1 | 2.613037336 | -0.21459418 | 0.6219442 | -0.345037673 |
| Plekhm1 | 513.8582558 | -0.045359126 | 0.088457075 | -0.512781208 |
| Lrrc37a | 0.598753221 | 0.755309214 | 1.494272393 | 0.505469564 |
| Gm884 | 1.391014422 | 3.402561298 | 1.115963314 | 3.048990282 |
| Gm11643 | 0.038981395 | 0.274393828 | 3.724945049 | 0.073663859 |
| Lyzl6 | 0.124596252 | 0.755282361 | 3.714681068 | 0.203323609 |
| Rprml | 232.2274179 | 0.180789512 | 0.200005572 | 0.903922374 |
| Gosr2 | 931.0627658 | 0.297986379 | 0.095480202 | 3.120923219 |
| C130046K22Rik | 12.71673352 | -0.301591894 | 0.282822523 | -1.066364483 |
| Gm11642 | 9.924921471 | -0.233000239 | 0.351402789 | -0.663057454 |
| Wnt9b | 31.73347887 | -0.174182064 | 0.306000522 | -0.56922146 |
| Wnt3 | 86.98556824 | 0.489172769 | 0.213959182 | 2.286290137 |
| Nsf | 6788.932109 | -0.145211238 | 0.060918988 | -2.383677767 |
| Gm8738 | 9.128726587 | -0.245355399 | 0.499418435 | -0.491282222 |
| Arf2 | 510.9708085 | 0.270933034 | 0.107423251 | 2.522107928 |
| Crhr1 | 208.0710271 | -0.90206641 | 0.111961159 | -8.056958506 |
| Sppl2c | 0.098502579 | -0.206492286 | 3.724945049 | -0.055434988 |
| Mapt | 4636.496193 | -0.458236852 | 0.067738262 | -6.764815623 |
| Gm11659 | 0.104366086 | 0.274400501 | 3.724945049 | 0.073665651 |
| Gm22743 | 0 | NA | NA | NA |
| AL596383.1 | 0 | NA | NA | NA |
| Kansl1 | 1511.632219 | -0.080880946 | 0.036257841 | -2.230716035 |
| AL593843.1 | 0.534702693 | 1.283737815 | 1.586021667 | 0.809407489 |

| | | | | |
|---|---|---|---|---|
| Gm11660 | 0 | NA | NA | NA |
| Cdc27 | 1303.554971 | 0.059151335 | 0.053704095 | 1.101430621 |
| Myl4 | 65.04662385 | -0.607628125 | 0.21358927 | -2.844843869 |
| Gm11665 | 0 | NA | NA | NA |
| Gm22000 | 0.267335191 | 0.274412688 | 1.813272651 | 0.151335591 |
| Gm11661 | 0 | NA | NA | NA |
| Gm11663 | 0.542039732 | 0.71686547 | 1.552910456 | 0.461627048 |
| Itgb3 | 53.54022327 | -0.107113245 | 0.193712427 | -0.552949782 |
| Gm11662 | 0.044191549 | -0.206490061 | 3.724945049 | -0.055434391 |
| Gm11639 | 8.031358769 | 0.367241963 | 0.358058653 | 1.025647502 |
| Gm11623 | 0.110595523 | 0.755282166 | 3.724945049 | 0.202763304 |
| Gm11621 | 0.116046057 | -0.687371727 | 3.724945049 | -0.184532045 |
| Gm11622 | 0.090074004 | 0.274402725 | 3.724945049 | 0.073666248 |
| Gm11624 | 0.048949532 | -0.206487837 | 3.724945049 | -0.055433794 |
| Efcab3 | 4.279745736 | 0.510209364 | 0.577487545 | 0.883498473 |
| Gm25362 | 0 | NA | NA | NA |
| Mettl2 | 133.1323383 | 0.09053943 | 0.127462443 | 0.710322413 |
| Gm24290 | 0.451210867 | 0.059847184 | 1.433019479 | 0.041762994 |
| Gm10842 | 0.310932149 | -0.206482794 | 1.805403217 | -0.114369351 |
| Gm11620 | 9.032629779 | -0.642792946 | 0.330605489 | -1.944290001 |
| 1700052K11Rik | 167.7122578 | -0.28401418 | 0.096724634 | -2.936316916 |
| Tlk2 | 1033.874952 | 0.519880868 | 0.102012696 | 5.096236924 |
| Mrc2 | 195.2873509 | -0.219566909 | 0.155170309 | -1.415005939 |
| 10-Mar | 11.52093314 | -0.387938572 | 0.331584568 | -1.169953639 |
| Gm11638 | 3.337525259 | -1.820270259 | 0.770086756 | -2.36372103 |
| Gm11640 | 0.456339188 | -0.21443989 | 1.36427717 | -0.157182056 |
| Tanc2 | 3818.814185 | -0.255649566 | 0.105242148 | -2.429155715 |
| Gm23990 | 3.257021581 | -0.86981437 | 0.632696747 | -1.374772945 |
| Gm11652 | 25.69936743 | -0.364147161 | 0.280437578 | -1.298496311 |
| Gm11653 | 35.85399802 | -0.875464887 | 0.290323347 | -3.015482206 |
| Cyb561 | 385.0963259 | -0.074411989 | 0.067767622 | -1.098046326 |
| Gm9910 | 2.885346812 | 0.152312444 | 0.613904008 | 0.248104658 |
| Gm11651 | 0.18135768 | -0.661333131 | 2.732659101 | -0.242010842 |
| Ace | 389.3701917 | 0.339953753 | 0.110226382 | 3.084141451 |
| Ace3 | 0.089089296 | 1.194485492 | 3.723939576 | 0.320758559 |
| Kcnh6 | 35.00512634 | -0.491508878 | 0.182442361 | -2.694050202 |
| Dcaf7 | 2563.358122 | -0.010100054 | 0.061704287 | -0.163684806 |
| Gm11646 | 0.25779132 | 1.200811859 | 2.023395198 | 0.593463828 |
| Taco1os | 4.692172118 | 0.37720028 | 0.502470244 | 0.750691776 |
| Taco1 | 111.1047001 | -0.24008185 | 0.164091693 | -1.463095702 |
| Map3k3 | 378.7212756 | 0.222050851 | 0.119245673 | 1.862129205 |
| Gm11672 | 0.164893722 | 1.582203397 | 3.387385449 | 0.467086908 |
| Limd2 | 471.2960561 | 0.563372161 | 0.107820045 | 5.225115247 |
| Gm10840 | 0.825556236 | 0.106667225 | 0.975045419 | 0.109397185 |
| Strada | 263.1762033 | -0.320948885 | 0.123647114 | -2.595684397 |
| Gm25556 | 0 | NA | NA | NA |
| Ccdc47 | 1290.145382 | 0.043857361 | 0.066799142 | 0.656555755 |

| | | | | |
|---|---|---|---|---|
| Ddx42 | 1058.979334 | 0.041793876 | 0.084950107 | 0.491981437 |
| Ftsj3 | 311.2842433 | 0.198017062 | 0.131702588 | 1.503516867 |
| Psmc5 | 1264.146288 | 0.213465016 | 0.098064928 | 2.176772278 |
| Gm23645 | 2.33541923 | 0.382256044 | 0.611969127 | 0.62463289 |
| Smarcd2 | 283.2968771 | 0.31368928 | 0.115900735 | 2.706534013 |
| Tcam1 | 4.654232259 | 1.10246772 | 0.503039903 | 2.191610873 |
| Gh | 110.6277303 | -2.860354166 | 2.43417786 | -1.175080184 |
| Cd79b | 1.812060818 | 3.450323712 | 1.000560499 | 3.448390893 |
| Scn4a | 1.34423325 | 3.019025971 | 1.108473439 | 2.723588915 |
| 2310007L24Rik | 2.639496584 | 0.117876722 | 0.668192105 | 0.176411425 |
| Icam2 | 17.72490718 | 1.468713357 | 0.341035851 | 4.306624514 |
| Ern1 | 301.964099 | -0.050919447 | 0.106950138 | -0.476104541 |
| Snord104 | 16.93328137 | 0.085981901 | 0.280814871 | 0.306187137 |
| SNORA50 | 0.197850883 | -0.19227119 | 2.220624018 | -0.086584306 |
| Gm22711 | 0 | NA | NA | NA |
| Tex2 | 1392.018177 | -0.212614611 | 0.050634445 | -4.199011368 |
| Pecam1 | 139.8937637 | 1.504186606 | 0.158158084 | 9.510652677 |
| Milr1 | 37.7867777 | 5.17620629 | 0.652433333 | 7.933693808 |
| Gm25889 | 0.53783135 | -0.17273791 | 1.46224392 | -0.118132076 |
| Polg2 | 47.1941341 | -0.543979526 | 0.257874282 | -2.109475681 |
| Ddx5 | 9164.25171 | 0.105965513 | 0.077282673 | 1.371141933 |
| Mir3064 | 0 | NA | NA | NA |
| Gm25994 | 0 | NA | NA | NA |
| Cep95 | 165.3431975 | -0.352500412 | 0.157105732 | -2.243714525 |
| Gm11706 | 1.894273405 | 0.465764785 | 0.860472411 | 0.541289621 |
| Smurf2 | 621.2522263 | 0.105458315 | 0.061394603 | 1.717713139 |
| Gm27322 | 0.075043205 | 0.755286614 | 3.724945049 | 0.202764498 |
| Gm11707 | 0.436105163 | 1.977600229 | 1.683630052 | 1.174604971 |
| Kpna2 | 179.0519059 | 0.049327337 | 0.130510407 | 0.377957113 |
| Gm11708 | 0 | NA | NA | NA |
| Gm11705 | 2.194360983 | 0.204277622 | 0.695666759 | 0.293642925 |
| 1810010H24Rik | 17.13476079 | -0.124795113 | 0.249998011 | -0.499184423 |
| Bptf | 2737.946275 | -0.145969113 | 0.063179239 | -2.310396816 |
| Gm11715 | 0.139856492 | -0.206491248 | 3.574411025 | -0.057769307 |
| Nol11 | 402.0846801 | 0.251685427 | 0.089895862 | 2.799744286 |
| Rps12-ps26 | 3.470784895 | 0.685068598 | 0.58638257 | 1.168296318 |
| Gm11718 | 0.038981395 | 0.274393828 | 3.724945049 | 0.073663859 |
| Pitpnc1 | 980.2405448 | -0.485663952 | 0.065089425 | -7.46148783 |
| Gm11719 | 0 | NA | NA | NA |
| Gm11716 | 0.710007532 | 0.755566991 | 1.315758572 | 0.574244399 |
| Gm11712 | 0.072025462 | 0.755288839 | 3.724945049 | 0.202765096 |
| Gm11713 | 0.068097359 | -0.206487837 | 3.724945049 | -0.055433794 |
| Gm11714 | 1.466365607 | 3.660216296 | 1.064648152 | 3.437958624 |
| Psmd12 | 782.060693 | 0.289585037 | 0.096652341 | 2.996151291 |
| Gm11722 | 4.228535416 | 0.351703458 | 0.494477768 | 0.711262428 |
| C330019F10Rik | 0.3156755 | -0.109950901 | 1.756858421 | -0.062583814 |
| A830035A12Rik | 0.580962004 | 0.644823297 | 1.267882974 | 0.508582661 |

| | | | | |
|---|---|---|---|---|
| Gm11649 | 0 | NA | NA | NA |
| Helz | 1652.750466 | -0.44452786 | 0.089798022 | -4.950307901 |
| Mir6932 | 0.570554915 | 0.651491924 | 1.580743899 | 0.412142615 |
| Cacng1 | 1.900742604 | 0.882780759 | 0.651768918 | 1.354438261 |
| Cacng4 | 438.0588544 | -0.660892464 | 0.108782092 | -6.075379247 |
| Gm11650 | 3.119570997 | -1.146243952 | 0.568771219 | -2.015298795 |
| Gm23120 | 0 | NA | NA | NA |
| Gm27595 | 0 | NA | NA | NA |
| Cacng5 | 366.3860086 | 0.125101958 | 0.155134188 | 0.806411276 |
| Prkca | 1907.502606 | -0.175636407 | 0.094192616 | -1.864651545 |
| Gm11657 | 0 | NA | NA | NA |
| Mir7223 | 0 | NA | NA | NA |
| Gm11655 | 0.048949532 | -0.206487837 | 3.724945049 | -0.055433794 |
| Apoh | 2.789102959 | 1.485605216 | 0.737868635 | 2.013373581 |
| Gm11656 | 0.072025462 | 0.755288839 | 3.724945049 | 0.202765096 |
| Gm11667 | 0 | NA | NA | NA |
| Cep112 | 85.63833006 | -0.284243908 | 0.174461069 | -1.62926841 |
| Cep112os1 | 0 | NA | NA | NA |
| Olfr1397-ps1 | 0.047948698 | -0.206487837 | 3.724945049 | -0.055433794 |
| Rpl23a-ps14 | 5.868082076 | -0.499290265 | 0.414052881 | -1.205861107 |
| Cep112os2 | 13.20886531 | -0.028464054 | 0.451366925 | -0.063061897 |
| Axin2 | 366.2212925 | -0.251235393 | 0.108383852 | -2.31801499 |
| Gm11670 | 1.328031477 | 1.244188721 | 1.074255273 | 1.158187213 |
| Gm11668 | 0.544841187 | -1.340392122 | 1.415271614 | -0.947091787 |
| Gm24149 | 0.044544648 | 0.755286614 | 3.724945049 | 0.202764498 |
| E030025P04Rik | 0.237045706 | -0.568138095 | 2.124314614 | -0.267445364 |
| Gm11658 | 0.044544648 | 0.755286614 | 3.724945049 | 0.202764498 |
| Rgs9 | 522.9399079 | -0.71979952 | 0.245456034 | -2.93249878 |
| 1700096J18Rik | 0 | NA | NA | NA |
| Gm11696 | 30.54970298 | 0.306813524 | 0.208545216 | 1.471208643 |
| Gna13 | 2195.672482 | 1.100170264 | 0.131748643 | 8.350524451 |
| 9930022D16Rik | 1.783663422 | 2.423570524 | 1.031330105 | 2.349946454 |
| Amz2 | 356.9352975 | -0.413882498 | 0.118682853 | -3.487298184 |
| Gm15642 | 0 | NA | NA | NA |
| Slc16a6 | 246.9242258 | 0.540318302 | 0.171896411 | 3.143278545 |
| Gm25540 | 3.251877087 | -0.042379259 | 0.677685356 | -0.062535302 |
| Arsg | 200.3840775 | -0.45248053 | 0.129845444 | -3.484762472 |
| Gm22378 | 0 | NA | NA | NA |
| Wipi1 | 268.8479832 | -0.098026131 | 0.090096988 | -1.088006748 |
| Gm11685 | 0.075043205 | 0.755286614 | 3.724945049 | 0.202764498 |
| Prkar1a | 7255.166353 | -0.01153769 | 0.058053071 | -0.198743837 |
| Fam20a | 97.31167799 | -0.083546288 | 0.131078729 | -0.63737487 |
| Gm11684 | 0 | NA | NA | NA |
| 1700012B07Rik | 0.10503227 | 0.274389469 | 3.724945049 | 0.073662689 |
| Gm11683 | 0 | NA | NA | NA |
| 1700023C21Rik | 0 | NA | NA | NA |
| Abca8b | 635.0819168 | -7.24E-06 | 0.109762601 | -6.60E-05 |

| | | | | |
|---|---|---|---|---|
| Gm11697 | 0 | NA | NA | NA |
| Abca8a | 98.13962934 | -1.183768053 | 0.16806262 | -7.04361298 |
| Abca9 | 172.9692926 | -0.757815145 | 0.218462568 | -3.468855801 |
| Abca6 | 37.25579249 | -0.755500531 | 0.223894101 | -3.374365497 |
| Abca5 | 1374.997336 | -0.147162049 | 0.057476577 | -2.56038296 |
| Map2k6 | 151.8243697 | -0.565774418 | 0.135972419 | -4.16094986 |
| Gm11682 | 0.63216095 | -0.330473407 | 1.151221882 | -0.287063174 |
| Kcnj16 | 546.3597821 | 0.131124758 | 0.156591397 | 0.837368847 |
| Kcnj2 | 326.6825132 | -0.067802816 | 0.099826918 | -0.679203741 |
| Gm11675 | 0.837406104 | 2.49061852 | 1.189276956 | 2.094229193 |
| Gm11676 | 0 | NA | NA | NA |
| Gm11677 | 0.110870145 | -0.206490066 | 3.724945049 | -0.055434393 |
| Gm11674 | 0.062426873 | 0.755288839 | 3.724945049 | 0.202765096 |
| Gm11679 | 0 | NA | NA | NA |
| Gm11680 | 0 | NA | NA | NA |
| BC006965 | 12.81790452 | -0.847270119 | 0.295785604 | -2.864473818 |
| Sox9 | 1126.317198 | 0.706907419 | 0.133951608 | 5.277334323 |
| Gm11681 | 4.401306684 | 0.768295261 | 0.485598358 | 1.582161982 |
| 4933434M16Rik | 0 | NA | NA | NA |
| 2610035D17Rik | 77.49257512 | -0.020170621 | 0.144151192 | -0.139926842 |
| 4732490B19Rik | 0.42613393 | 0.755308324 | 1.653493528 | 0.456795452 |
| Slc39a11 | 151.2282737 | 0.177682229 | 0.104199405 | 1.705213476 |
| Gm16487 | 1.032826873 | -2.684824294 | 1.081560391 | -2.482361887 |
| Sstr2 | 160.0653886 | -1.097678389 | 0.328219892 | -3.344338398 |
| Cog1 | 598.811076 | 0.103367959 | 0.067434263 | 1.532870007 |
| Fam104a | 378.1237319 | 0.149979834 | 0.073036537 | 2.053490487 |
| Gm11693 | 0.456148335 | -0.223856804 | 1.688505072 | -0.132576921 |
| D11Wsu47e | 95.6758204 | -0.226442557 | 0.160822389 | -1.408028809 |
| Cpsf4l | 2.642413763 | 0.088780297 | 0.672433733 | 0.132028321 |
| Cdc42ep4 | 690.6837532 | 0.240348214 | 0.142113319 | 1.691243411 |
| Gm11736 | 0.197525616 | 0.274410924 | 2.227619051 | 0.12318575 |
| Sdk2 | 727.2965037 | -0.231026036 | 0.118511173 | -1.949402995 |
| 1700025D23Rik | 0.122991902 | 0.274402725 | 3.724945049 | 0.073666248 |
| 1700001J04Rik | 0.288935699 | -0.687379378 | 2.086786168 | -0.329396173 |
| 1700092K14Rik | 0.758667428 | 0.011056011 | 1.196130039 | 0.009243151 |
| Gm11690 | 0.042716945 | -0.206490061 | 3.724945049 | -0.055434391 |
| Gm11691 | 0.047948698 | -0.206487837 | 3.724945049 | -0.055433794 |
| Gm11692 | 0 | NA | NA | NA |
| 4932435O22Rik | 2.584340838 | -0.692477235 | 0.684219823 | -1.012068362 |
| Gm11689 | 0 | NA | NA | NA |
| Rpl38 | 617.2880742 | 0.325218192 | 0.198514137 | 1.638262123 |
| Ttyh2 | 758.4082506 | -0.109777051 | 0.096683137 | -1.13543121 |
| Dnaic2 | 33.61650857 | -0.033049017 | 0.238672168 | -0.138470343 |
| Kif19a | 46.14809591 | 0.056398457 | 0.172307077 | 0.327313641 |
| Btbd17 | 82.92747472 | -1.510209113 | 0.260404386 | -5.79947648 |
| Gpr142 | 0.138152447 | 0.274406196 | 2.934963576 | 0.093495605 |
| Gprc5c | 168.3894624 | -0.37641924 | 0.194922709 | -1.931120507 |

| | | | | |
|---|---|---|---|---|
| Cd300a | 55.24117626 | 0.786360528 | 0.269766878 | 2.914963225 |
| Gm11698 | 0.946192865 | 1.352926744 | 1.010165906 | 1.33931143 |
| Cd300lb | 7.999125163 | 0.168615154 | 0.682411533 | 0.247087198 |
| Gm11699 | 0 | NA | NA | NA |
| Cd300c | 0.985558492 | 3.460251747 | 2.187671459 | 1.581705394 |
| Cd300ld | 9.206469351 | 4.413602538 | 0.929855812 | 4.746545087 |
| AF251705 | 73.50869836 | 1.892115435 | 0.221459288 | 8.543852231 |
| Gm11709 | 0.932213635 | 1.712048097 | 1.436376409 | 1.191921621 |
| Gm11710 | 0.854414486 | 1.558607637 | 1.344661666 | 1.15910766 |
| Gm11711 | 0.991446059 | 0.760471892 | 1.464694728 | 0.519201631 |
| Cd300lh | 2.74015677 | 1.997048299 | 1.042570582 | 1.915504171 |
| Cd300e | 5.001744579 | 1.741291837 | 0.703622446 | 2.474753112 |
| Rab37 | 152.6386923 | -0.049632114 | 0.112482622 | -0.441242506 |
| Cd300lf | 291.2418054 | 6.171712528 | 0.424701577 | 14.5318804 |
| Slc9a3r1 | 582.5242563 | 0.602963007 | 0.140800624 | 4.282388746 |
| Nat9 | 95.23938015 | -0.013760432 | 0.133903908 | -0.102763486 |
| Tmem104 | 386.5952224 | 0.455694487 | 0.108291075 | 4.208052109 |
| Grin2c | 638.1417609 | -0.586437846 | 0.162193144 | -3.615675924 |
| Gm25837 | 0.089089296 | 1.194485492 | 3.723939576 | 0.320758559 |
| Fdxr | 101.2810521 | -0.38762433 | 0.116152788 | -3.337193522 |
| Fads6 | 408.9834072 | -0.471706464 | 0.101536665 | -4.645676152 |
| Otop2 | 12.03184821 | -0.700450388 | 0.496083113 | -1.411961766 |
| Ush1g | 18.68407142 | -0.312923248 | 0.342396398 | -0.913920969 |
| Otop3 | 2.248585396 | 0.355037316 | 0.925903249 | 0.383449692 |
| Hid1 | 846.0541876 | -0.175349792 | 0.078443703 | -2.23535842 |
| Cdr2l | 554.9718017 | 0.034641019 | 0.082703001 | 0.418860482 |
| Ict1 | 293.2357144 | 0.078259447 | 0.073105996 | 1.070492867 |
| Ict1os | 0.802129109 | -1.43050158 | 1.143581055 | -1.250896536 |
| Atp5h | 982.123738 | -0.09783837 | 0.088108401 | -1.110431796 |
| Kctd2 | 366.4040735 | -0.088285397 | 0.068963003 | -1.280184921 |
| Trim80 | 1.020577158 | 0.913203683 | 1.26646009 | 0.721067873 |
| Mir3968 | 0 | NA | NA | NA |
| Gm11695 | 0.907486357 | -0.096891956 | 0.951364654 | -0.101845234 |
| Slc16a5 | 0.45393394 | 1.309404029 | 3.371771793 | 0.388343017 |
| Armc7 | 103.9306984 | 0.41242579 | 0.140903156 | 2.927015987 |
| Nt5c | 193.1442848 | -0.137825612 | 0.084878529 | -1.62379831 |
| Gm11694 | 1.069943461 | -0.61902044 | 1.226412237 | -0.504740919 |
| Hn1 | 386.0221868 | -0.103443244 | 0.112484249 | -0.919624253 |
| Sumo2 | 322.8270685 | 0.185316491 | 0.112036359 | 1.654074558 |
| Nup85 | 257.3817318 | -0.30962872 | 0.0780612 | -3.966486806 |
| Gga3 | 719.1270991 | -0.341788745 | 0.077143944 | -4.430532403 |
| Gm25364 | 0 | NA | NA | NA |
| Mrps7 | 278.078163 | 0.18480326 | 0.104463421 | 1.769071491 |
| Mif4gd | 118.7681913 | 0.313435336 | 0.142827507 | 2.1945026 |
| Slc25a19 | 241.7616655 | -0.491514904 | 0.078512456 | -6.260343017 |
| Gm26613 | 0.042716945 | -0.206490061 | 3.724945049 | -0.055434391 |
| Grb2 | 1605.895681 | 0.112616679 | 0.059230291 | 1.901335896 |

| | | | | |
|---|---|---|---|---|
| Gm11702 | 38.96983336 | -0.140534577 | 0.180832672 | -0.777152574 |
| Gm11703 | 12.20134384 | 1.19188224 | 0.366093373 | 3.255678275 |
| Gm23699 | 0 | NA | NA | NA |
| Gm11704 | 0 | NA | NA | NA |
| 2310067B10Rik | 722.2047825 | -0.377266425 | 0.096309039 | -3.917248371 |
| Mir5621 | 0.180265876 | -0.670480839 | 2.743873342 | -0.244355608 |
| Caskin2 | 336.692886 | -0.173104408 | 0.112276757 | -1.541765299 |
| Tsen54 | 70.37095567 | -0.186976882 | 0.131770661 | -1.418956847 |
| Llgl2 | 24.64085341 | 1.13821207 | 0.244196028 | 4.661058909 |
| Myo15b | 2.674434539 | -0.691814854 | 0.886927508 | -0.780012851 |
| Recql5 | 210.8391631 | -0.233591172 | 0.153859104 | -1.51821482 |
| Smim5 | 3.739367707 | 1.176837622 | 0.580599368 | 2.026935761 |
| Smim6 | 0.320213338 | -1.003977014 | 1.71886891 | -0.584091671 |
| Recql5os1 | 0.062426873 | 0.755288839 | 3.724945049 | 0.202765096 |
| Sap30bp | 450.623263 | 0.167430797 | 0.076901157 | 2.177220782 |
| Sap30bpos | 16.7515477 | 1.449740893 | 0.351608752 | 4.123164982 |
| Itgb4 | 300.6627537 | -0.169756574 | 0.139148513 | -1.219966854 |
| B230344G16Rik | 10.95198732 | 0.086985727 | 0.329576629 | 0.26393172 |
| Galk1 | 66.2841132 | -0.147177974 | 0.170160065 | -0.8649384 |
| H3f3b | 3094.200091 | 1.236036943 | 0.17122331 | 7.218859075 |
| Unk | 347.7817187 | 0.251671139 | 0.063895064 | 3.938819758 |
| Unc13d | 61.4735399 | 1.223787134 | 0.259958737 | 4.707620716 |
| Wbp2 | 3196.52038 | 0.159441121 | 0.072414396 | 2.201787633 |
| Trim47 | 116.7135195 | 1.679968335 | 0.233815016 | 7.18503185 |
| Trim65 | 57.48636768 | -0.337295229 | 0.150309253 | -2.244008418 |
| Mrpl38 | 328.9494862 | -0.098504248 | 0.074276859 | -1.3261768 |
| Fbf1 | 655.435408 | 0.046616976 | 0.099839135 | 0.466920873 |
| Acox1 | 2125.616724 | 0.095554814 | 0.05177687 | 1.84551159 |
| Ten1 | 127.8018245 | 0.131578201 | 0.112975666 | 1.164659669 |
| Cdk3-ps | 6.063886428 | 0.05789764 | 0.40990491 | 0.141246516 |
| Evpl | 18.36159365 | -0.46991416 | 0.235812719 | -1.992743069 |
| Srp68 | 664.4255585 | 0.123418272 | 0.071025069 | 1.73767199 |
| Galr2 | 1.093534925 | -0.409175478 | 0.906805356 | -0.45122746 |
| Exoc7 | 686.1507893 | -0.030866631 | 0.049312878 | -0.625934484 |
| Gm26730 | 1.17710504 | -0.655838919 | 1.055555489 | -0.62132112 |
| n-R5s74 | 0 | NA | NA | NA |
| Foxj1 | 234.7792888 | 0.522102645 | 0.165952296 | 3.146100772 |
| Rnf157 | 2678.61245 | -0.144567588 | 0.081303782 | -1.778116392 |
| Rps11-ps2 | 3.484535615 | -0.874219283 | 0.5852103 | -1.493854914 |
| Ubald2 | 148.6109574 | 0.884850076 | 0.134782459 | 6.565023972 |
| Qrich2 | 0.384783943 | 1.214258793 | 1.565787715 | 0.775493882 |
| Gm11739 | 0 | NA | NA | NA |
| Prpsap1 | 399.2746756 | -0.160885733 | 0.073653549 | -2.184358204 |
| Rpl36-ps1 | 0 | NA | NA | NA |
| Gm29292 | 10.23524976 | 2.669009129 | 0.60888636 | 4.383427362 |
| Sphk1 | 151.140251 | 1.544649718 | 0.36571345 | 4.223661222 |
| Ube2o | 1590.155959 | -0.177033819 | 0.073996596 | -2.392458975 |

| | | | | |
|---|---|---|---|---|
| Gm11742 | 0 | NA | NA | NA |
| Aanat | 3.437531617 | -0.698901242 | 0.510897317 | -1.367987693 |
| Rhbdf2 | 162.7293829 | 4.452794451 | 0.330755231 | 13.46250652 |
| Cygb | 534.7165052 | -0.278098759 | 0.085398036 | -3.256500617 |
| Gm11744 | 17.7695872 | -0.388790439 | 0.286730369 | -1.355944406 |
| 1810032O08Rik | 43.33906545 | 0.478578395 | 0.186958519 | 2.559810586 |
| Snord1c | 0 | NA | NA | NA |
| Snord1b | 0 | NA | NA | NA |
| Snord1a | 0 | NA | NA | NA |
| St6galnac2 | 21.17771716 | 0.105161416 | 0.218541133 | 0.48119736 |
| Gm11735 | 0.061920613 | 0.274393828 | 3.724945049 | 0.073663859 |
| BC018473 | 0 | NA | NA | NA |
| St6galnac1 | 0.21290388 | -0.206489223 | 2.636541467 | -0.078318216 |
| Mxra7 | 93.40309728 | -0.475153437 | 0.132668835 | -3.581500041 |
| Jmjd6 | 264.2573602 | -0.026526913 | 0.120303945 | -0.220499114 |
| Gm27401 | 0 | NA | NA | NA |
| Mettl23 | 60.97083361 | -0.275864059 | 0.156770938 | -1.759663255 |
| Srsf2 | 1239.606384 | 0.097160416 | 0.072309797 | 1.343668762 |
| Mfsd11 | 294.976682 | -0.058630635 | 0.080477041 | -0.728538645 |
| Mgat5b | 569.5109985 | -0.456429623 | 0.114371679 | -3.990757396 |
| Gm11728 | 0.813341144 | -0.524943717 | 1.139389863 | -0.460723528 |
| Gm11731 | 0.068097359 | -0.206487837 | 3.724945049 | -0.055433794 |
| Snhg20 | 102.2173952 | 0.028297644 | 0.148544407 | 0.19049956 |
| Gm26083 | 0 | NA | NA | NA |
| mmu-mir-6516 | 0 | NA | NA | NA |
| Gm24323 | 0.154433973 | -0.688825965 | 3.500771551 | -0.196764043 |
| Gm11730 | 1.74719428 | -0.563865861 | 0.750646465 | -0.751173672 |
| Sec14l1 | 2162.074218 | -0.551422817 | 0.100304205 | -5.497504461 |
| Gm22667 | 7.017353583 | -1.181804685 | 0.547218307 | -2.15965853 |
| Gm11745 | 7.216095472 | -0.925311021 | 0.594091033 | -1.557523964 |
| 9-Sep | 1015.394361 | 0.541147451 | 0.115131436 | 4.700257995 |
| Gm16045 | 1.18260799 | -0.811470096 | 0.850840776 | -0.953727323 |
| Gm11729 | 0.295494355 | -1.535152534 | 1.771033766 | -0.866811556 |
| Gm8624 | 0.24265998 | -0.206481094 | 2.509000292 | -0.082296162 |
| Gm11732 | 4.722481653 | -0.758764007 | 0.457772667 | -1.657512696 |
| Gm11733 | 2.291093954 | 1.232772822 | 0.717895667 | 1.717203319 |
| Gm11734 | 3.524723029 | -0.267530635 | 0.572136512 | -0.467599305 |
| 2900041M22Rik | 4.279435866 | -0.047630671 | 0.497219108 | -0.095794127 |
| Tnrc6c | 1645.798208 | -0.23915452 | 0.119440013 | -2.002298177 |
| Tmc6 | 62.04352302 | 0.735806715 | 0.196021699 | 3.753700317 |
| Tmc8 | 13.88221168 | 1.958268045 | 0.375553913 | 5.214346006 |
| Gm11723 | 0.109124069 | 0.274402725 | 3.724945049 | 0.073666248 |
| Gm11724 | 0.044544648 | 0.755286614 | 3.724945049 | 0.202764498 |
| 6030468B19Rik | 0.072025462 | 0.755288839 | 3.724945049 | 0.202765096 |
| Gm20708 | 0.075043205 | 0.755286614 | 3.724945049 | 0.202764498 |
| Syngr2 | 214.0619962 | 1.655155363 | 0.154836711 | 10.68968302 |
| Tk1 | 11.63018472 | 1.737191648 | 0.434247642 | 4.000463056 |

| | | | | |
|---|---|---|---|---|
| Afmid | 13.88472388 | 0.770185991 | 0.306049515 | 2.516540474 |
| Birc5 | 15.65536197 | 1.037854313 | 0.322386591 | 3.21928499 |
| Tmem235 | 0.809771811 | -0.919167165 | 1.173577421 | -0.783218174 |
| Tha1 | 35.97253463 | -0.173275789 | 0.199366175 | -0.869133333 |
| Gm11725 | 4.41486742 | -0.106186523 | 0.509813805 | -0.208284912 |
| Gm11726 | 0.061920613 | 0.274393828 | 3.724945049 | 0.073663859 |
| Socs3 | 775.4504325 | 4.713206368 | 0.414611043 | 11.36777818 |
| Pgs1 | 896.2403674 | 0.284045188 | 0.090173314 | 3.149991668 |
| Mir6933 | 0.186399948 | 1.293453504 | 3.295513693 | 0.392489191 |
| Dnah17 | 7.417164814 | -0.014150614 | 0.379775086 | -0.037260512 |
| Gm11738 | 6.391237767 | 1.981052712 | 0.522483041 | 3.791611507 |
| Cyth1 | 1001.744073 | 0.31250396 | 0.059559243 | 5.246943095 |
| Gm24060 | 17.80235189 | 0.018758523 | 0.265297841 | 0.07070741 |
| Gm11737 | 8.112751975 | 0.221906921 | 0.490443256 | 0.452461968 |
| Usp36 | 634.1888172 | 0.325683058 | 0.151919594 | 2.143785736 |
| Timp2 | 2901.777889 | -0.083133786 | 0.065791768 | -1.26358949 |
| BC100451 | 15.86145379 | -0.496165495 | 0.338130096 | -1.467380456 |
| Lgals3bp | 4154.63823 | 5.507336345 | 0.175224523 | 31.43016883 |
| Cant1 | 623.271791 | 0.236576885 | 0.061116676 | 3.870905641 |
| C1qtnf1 | 107.3234082 | 1.236095472 | 0.200334023 | 6.170172468 |
| Gm11747 | 0.291849543 | 0.755310468 | 1.778233226 | 0.424753321 |
| Gm11748 | 1.545561197 | -0.542345966 | 0.859253572 | -0.631182673 |
| Engase | 86.50548769 | -0.073125209 | 0.144375898 | -0.506491801 |
| Rbfox3 | 2325.205722 | -0.268632225 | 0.104125886 | -2.579879369 |
| Gm11750 | 0.497147205 | 1.222944832 | 1.680550192 | 0.727705033 |
| Gm25040 | 1.127514739 | -1.256708953 | 0.924036821 | -1.360020428 |
| Gm11749 | 17.63452542 | -1.015753271 | 0.346607796 | -2.930555175 |
| Enpp7 | 0.038981395 | 0.274393828 | 3.724945049 | 0.073663859 |
| Cbx2 | 46.53041809 | -0.324612509 | 0.240238584 | -1.351208888 |
| Cbx8 | 139.4173976 | -0.384614363 | 0.115558114 | -3.328319835 |
| Gm26508 | 4.027668366 | 0.330073129 | 0.587362937 | 0.561957707 |
| Cbx4 | 968.7534903 | 0.563929802 | 0.102392937 | 5.507506877 |
| Gm11754 | 0.08486385 | 0.755286617 | 3.724945049 | 0.202764499 |
| Gm11755 | 2.180156761 | -0.908471115 | 0.694113906 | -1.308821372 |
| Tbc1d16 | 764.1409362 | -0.440983435 | 0.120210746 | -3.668419416 |
| Gm26888 | 0.416129427 | -1.404191382 | 1.562437678 | -0.898718331 |
| Mir6934 | 0 | NA | NA | NA |
| Gm11753 | 1.914943955 | -0.975227193 | 0.791357279 | -1.232347535 |
| Gm11752 | 0.044191549 | -0.206490061 | 3.724945049 | -0.055434391 |
| Ccdc40 | 32.62461688 | 0.142344368 | 0.231832273 | 0.613997205 |
| Gaa | 3712.581302 | 0.132723503 | 0.045976916 | 2.886742193 |
| Eif4a3 | 482.0859025 | 0.209060735 | 0.086509878 | 2.416611139 |
| Card14 | 10.20988058 | 0.374997944 | 0.378636976 | 0.990389125 |
| Sgsh | 132.7659391 | -0.264286358 | 0.134598723 | -1.963513119 |
| Slc26a11 | 53.44579997 | -0.498671896 | 0.165222618 | -3.018181788 |
| Mir1932 | 5.939787235 | 4.631397962 | 0.918475842 | 5.042482068 |
| Rnf213 | 13237.76603 | 5.327543707 | 0.164179693 | 32.44946806 |

| | | | | |
|---|---|---|---|---|
| Endov | 287.30136 | -0.637580232 | 0.128255118 | -4.971187432 |
| Nptx1 | 4410.742837 | 0.034175097 | 0.106253136 | 0.321638475 |
| Gm11762 | 16.8669638 | -0.516546254 | 0.292048957 | -1.768697476 |
| Rptor | 1323.078836 | -0.066935966 | 0.040190083 | -1.665484635 |
| Gm11761 | 0 | NA | NA | NA |
| Gm23663 | 1.256631142 | -0.616641556 | 1.066225932 | -0.578340422 |
| Rptoros | 0.455959994 | 1.191823567 | 1.59060326 | 0.749290283 |
| Chmp6 | 176.3066057 | 0.199653923 | 0.109089777 | 1.830179955 |
| Gm11767 | 18.81677128 | -1.26073498 | 0.313322031 | -4.023767423 |
| Gm11766 | 0.898050108 | 1.641639649 | 1.285818623 | 1.27672723 |
| Baiap2 | 2183.791868 | -0.052531258 | 0.134345168 | -0.391017103 |
| Aatk | 3564.404415 | -0.160352199 | 0.045973953 | -3.487892322 |
| Mir338 | 4.047384828 | -0.066116328 | 0.543506973 | -0.121647617 |
| mmu-mir-3065 | 0.136882968 | -0.206497511 | 3.632406564 | -0.056848678 |
| Cep131 | 180.3487011 | -0.311018758 | 0.173202875 | -1.795690499 |
| Gm25948 | 0.585001534 | 0.007324087 | 1.343102643 | 0.005453111 |
| Enthd2 | 135.2998032 | -0.326505793 | 0.123990452 | -2.633313994 |
| 1810043H04Rik | 75.99610051 | 0.011948592 | 0.191172946 | 0.062501478 |
| Slc38a10 | 947.089944 | 0.051983231 | 0.055764921 | 0.932185142 |
| Gm11769 | 0.670691275 | -0.100556916 | 1.223813408 | -0.082166869 |
| 2810410L24Rik | 42.09187015 | -0.488603814 | 0.193753029 | -2.521786717 |
| Gm11770 | 0.567693313 | 1.644244837 | 1.26694754 | 1.297800253 |
| 2900052L18Rik | 38.54445378 | -0.329078206 | 0.271274233 | -1.213083167 |
| Bahcc1 | 551.7191568 | -0.059850127 | 0.09589109 | -0.624146904 |
| Gm11772 | 31.98353754 | 3.28923566 | 0.373445961 | 8.807795507 |
| AL928567.1 | 0 | NA | NA | NA |
| Actg1 | 4258.481376 | 0.30907289 | 0.116615463 | 2.650359418 |
| Gm23419 | 0 | NA | NA | NA |
| Mir6935 | 0 | NA | NA | NA |
| 0610009L18Rik | 11.33403993 | -0.409803681 | 0.30760686 | -1.332231932 |
| Fscn2 | 1.822354792 | 0.605319319 | 0.729235522 | 0.830073825 |
| 2310003H01Rik | 147.9010322 | -0.374196386 | 0.110910855 | -3.373848169 |
| Nploc4 | 1106.700852 | 0.355272999 | 0.077011455 | 4.613248727 |
| Tspan10 | 1.198353109 | 1.30337829 | 1.062512157 | 1.226694943 |
| Pde6g | 3.767627917 | -1.015877398 | 0.576591408 | -1.761867041 |
| Oxld1 | 37.49257177 | 0.015744087 | 0.229785211 | 0.068516537 |
| Ccdc137 | 228.8428678 | 0.213680838 | 0.12783824 | 1.671493898 |
| Arl16 | 155.9283109 | -0.227010755 | 0.106102863 | -2.13953468 |
| Hgs | 966.1986603 | 0.129704185 | 0.062450501 | 2.076911831 |
| Mrpl12 | 411.1473201 | 0.119200437 | 0.122371044 | 0.974090216 |
| Gm16755 | 0.616632713 | 0.859481309 | 1.249115822 | 0.688071749 |
| Gm11788 | 0.154696864 | 0.274407077 | 2.833481316 | 0.096844498 |
| Slc25a10 | 142.1098057 | -0.219527634 | 0.120770889 | -1.817719783 |
| Gm11789 | 0.038981395 | 0.274393828 | 3.724945049 | 0.073663859 |
| Gcgr | 1.708960799 | 0.015587611 | 0.819536524 | 0.019020033 |
| Fam195b | 400.1399544 | 0.09356349 | 0.100388001 | 0.932018666 |
| Ppp1r27 | 0.150086409 | 1.264824188 | 3.723011833 | 0.339731445 |

| | | | | |
|---|---|---|---|---|
| P4hb | 1498.417556 | 0.436218421 | 0.103931348 | 4.197178513 |
| Arhgdia | 2780.717337 | 0.447598821 | 0.095451132 | 4.689298191 |
| Gm3807 | 3.342078322 | -0.656493413 | 0.595863062 | -1.101752156 |
| Alyref | 326.2916262 | 0.354780085 | 0.142335158 | 2.492568178 |
| Anapc11 | 300.5463482 | -0.037908514 | 0.110303985 | -0.343673111 |
| Npb | 4.91649906 | 0.795105458 | 0.476417277 | 1.668926582 |
| Pcyt2 | 527.1846474 | -0.147813674 | 0.075653214 | -1.953832046 |
| Sirt7 | 232.8371362 | -0.14651096 | 0.114004675 | -1.28513116 |
| Mir6936 | 0 | NA | NA | NA |
| Mafg | 947.7400382 | -0.366694459 | 0.08722275 | -4.20411485 |
| AL663030.1 | 0 | NA | NA | NA |
| Gm17586 | 36.02489538 | 0.488619626 | 0.199084355 | 2.454334628 |
| Pycr1 | 25.50189758 | 0.678566337 | 0.245590803 | 2.762995723 |
| Myadml2 | 26.50837914 | -0.084735 | 0.208931671 | -0.405563215 |
| Myadml2os | 1.355090691 | 0.122283511 | 0.959083318 | 0.127500404 |
| Notum | 9.73076196 | -1.088517713 | 0.370706372 | -2.936333973 |
| Notumos | 2.072861717 | -0.18090252 | 0.673004634 | -0.268798327 |
| Aspscr1 | 368.7567786 | 0.230829364 | 0.080520787 | 2.866705265 |
| Gm17178 | 0.221007548 | -0.113778521 | 2.158140354 | -0.052720631 |
| Stra13 | 57.79830129 | -0.091806035 | 0.175497805 | -0.523117856 |
| Lrrc45 | 209.0769499 | -0.507312306 | 0.205818133 | -2.464857192 |
| Rac3 | 88.08912298 | -0.546316141 | 0.136985969 | -3.988117516 |
| Dcxr | 57.27746725 | 0.328438072 | 0.195753003 | 1.677818821 |
| Cbr2 | 7.287723491 | -1.367279238 | 0.656426903 | -2.082911642 |
| Hmga1-rs1 | 524.4408412 | 1.222114668 | 0.262586788 | 4.654136166 |
| Gm11771 | 0 | NA | NA | NA |
| Rfng | 389.6471229 | -0.154176435 | 0.068469032 | -2.251768884 |
| Gps1 | 549.026855 | -0.070386223 | 0.085556637 | -0.822685716 |
| Dus1l | 255.7077501 | 0.080740668 | 0.090045919 | 0.896661051 |
| Fasn | 4709.556278 | -0.122731328 | 0.085514506 | -1.43521064 |
| Ccdc57 | 96.87917321 | -0.140636515 | 0.211333143 | -0.665473069 |
| Gm11773 | 0.216076386 | 1.736121978 | 3.709588981 | 0.468009256 |
| Slc16a3 | 72.70316503 | 1.137968831 | 0.243887204 | 4.665963653 |
| Csnk1d | 1494.74923 | 0.101994111 | 0.057164026 | 1.784235961 |
| Gm28192 | 0 | NA | NA | NA |
| Gm11774 | 0.066050875 | 0.274391604 | 3.724945049 | 0.073663262 |
| Gm11775 | 0.090277459 | -0.20649451 | 3.724945049 | -0.055435586 |
| Gm22152 | 0.044191549 | -0.206490061 | 3.724945049 | -0.055434391 |
| Cd7 | 13.98101131 | 1.830446832 | 0.358527426 | 5.105458325 |
| Sectm1b | 0.607868506 | 1.241623135 | 1.530210492 | 0.811406759 |
| Sectm1a | 17.65934462 | 6.027977667 | 0.920517486 | 6.548466222 |
| Gm11792 | 0 | NA | NA | NA |
| Gm11791 | 0 | NA | NA | NA |
| Gm11793 | 1.517674278 | 0.95990615 | 0.853928707 | 1.124105727 |
| Tex19.2 | 0 | NA | NA | NA |
| Tex19.1 | 0.038981395 | 0.274393828 | 3.724945049 | 0.073663859 |
| Uts2r | 2.195192171 | 1.671738205 | 0.743127727 | 2.249597405 |

| | | | | |
|---|---|---|---|---|
| Gm11790 | 0.802756328 | 3.247767117 | 1.229329682 | 2.641900838 |
| Ogfod3 | 69.70424457 | 0.257433934 | 0.159041687 | 1.61865696 |
| Hexdc | 139.714264 | -0.171893547 | 0.108168604 | -1.589126051 |
| BC017643 | 297.1303026 | 0.667274216 | 0.127573431 | 5.230510856 |
| Narf | 577.2846721 | -0.501437548 | 0.093626399 | -5.355728243 |
| Foxk2 | 1306.171416 | -0.179495167 | 0.048506404 | -3.700442674 |
| Wdr45b | 650.5471699 | 0.201552506 | 0.077366001 | 2.605181916 |
| Rab40b | 176.4499084 | -0.281727369 | 0.157313286 | -1.790868251 |
| Mir7236 | 2.550224353 | -0.208472875 | 0.683894596 | -0.30483188 |
| Gm11866 | 0 | NA | NA | NA |
| Fn3krp | 434.4560457 | -0.356146488 | 0.103597385 | -3.437794175 |
| Fn3k | 227.4287037 | -0.515529479 | 0.128770277 | -4.003481959 |
| Tbcd | 616.9440393 | 0.025947842 | 0.06260554 | 0.414465579 |
| Zfp750 | 17.50032145 | 1.220725883 | 0.313916484 | 3.88869634 |
| B3gntl1 | 72.02965567 | 0.102984084 | 0.179454305 | 0.573873578 |
| Metrnl | 107.2840231 | 0.540201613 | 0.213661803 | 2.52830223 |
| Gm12589 | 0.398050276 | 1.260253297 | 1.673881772 | 0.752892658 |
| Gm29513 | 0 | NA | NA | NA |
| Gm15643 | 0 | NA | NA | NA |
| Ptchd3 | 0 | NA | NA | NA |
| Gm12586 | 0 | NA | NA | NA |
| Gm12590 | 0 | NA | NA | NA |
| Gm12587 | 0 | NA | NA | NA |
| Gm12591 | 0 | NA | NA | NA |
| Gm9071 | 0 | NA | NA | NA |
| Gm9075 | 0 | NA | NA | NA |
| Rab10os | 387.6915989 | -0.173393133 | 0.098529171 | -1.7598152 |
| Rab10 | 2113.636369 | 0.194099031 | 0.078517515 | 2.472047545 |
| Gm26520 | 3.045321195 | 0.513630416 | 0.570447026 | 0.900399849 |
| Kif3c | 2815.319838 | -0.183886412 | 0.05974083 | -3.078069238 |
| 1110002L01Rik | 127.452233 | -0.265768412 | 0.118036006 | -2.251587642 |
| Asxl2 | 1098.710745 | 0.073018576 | 0.076032344 | 0.960362025 |
| Dtnb | 785.8508435 | -0.150816762 | 0.085065087 | -1.772957244 |
| Gm26050 | 2.458093412 | -1.046317498 | 0.766667375 | -1.364760694 |
| Dtnbos | 0.849323088 | -0.174721864 | 0.937176855 | -0.18643425 |
| Dnmt3a | 1911.391238 | -0.419393151 | 0.091926811 | -4.562250607 |
| Dnmt3aos | 1.674671121 | 2.359372117 | 1.114529728 | 2.116921655 |
| Pomc | 33.59521228 | -0.407545753 | 0.91925984 | -0.443341191 |
| Efr3b | 2749.842645 | -0.031627078 | 0.054620012 | -0.579038278 |
| Dnajc27 | 1474.757923 | -0.174504032 | 0.064237164 | -2.716558783 |
| Adcy3 | 597.0799168 | -0.487783871 | 0.080127647 | -6.087585102 |
| Cenpo | 63.78631368 | -0.044099102 | 0.138845334 | -0.317613134 |
| Ptrhd1 | 115.3757405 | 0.104783218 | 0.109868419 | 0.953715533 |
| Ncoa1 | 2595.887549 | 0.154513225 | 0.052950177 | 2.918087044 |
| Gm23756 | 0 | NA | NA | NA |
| Gm22343 | 0 | NA | NA | NA |
| Gm3625 | 2.933921127 | 0.426750602 | 0.568186063 | 0.751075449 |

| | | | | |
|---|---|---|---|---|
| Itsn2 | 887.6567088 | -0.115002628 | 0.064718699 | -1.776961376 |
| Gm17541 | 0.106465261 | 0.75528439 | 3.724945049 | 0.202763901 |
| Fam228a | 28.71218686 | -1.095781632 | 0.330955018 | -3.310968476 |
| Fam228b | 74.82727959 | -0.672445182 | 0.182880448 | -3.676965953 |
| Gm23676 | 0 | NA | NA | NA |
| Gm22003 | 0 | NA | NA | NA |
| Pfn4 | 43.32535029 | -0.247649703 | 0.160152933 | -1.546332613 |
| Gm6682 | 21.88487653 | 0.015523942 | 0.229240661 | 0.067718973 |
| Sf3b6 | 380.7093426 | 0.360041931 | 0.146820428 | 2.452260469 |
| Fkbp1b | 121.655647 | -0.288427572 | 0.1212662 | -2.37846632 |
| BC068281 | 110.397684 | -0.628348956 | 0.135826426 | -4.626117116 |
| Mfsd2b | 23.20891546 | 0.550246214 | 0.407207798 | 1.351266397 |
| Ubxn2a | 760.0083217 | -0.201597859 | 0.0783584 | -2.572766399 |
| Atad2b | 541.029005 | -0.13839189 | 0.092472416 | -1.496574837 |
| Gm23167 | 0 | NA | NA | NA |
| Klhl29 | 743.3499042 | -0.311438689 | 0.120483138 | -2.58491516 |
| 2810032G03Rik | 93.60943587 | -0.496465389 | 0.109078439 | -4.551453003 |
| Gm5652 | 0.048949532 | -0.206487837 | 3.724945049 | -0.055433794 |
| Gm23752 | 0.431854921 | 1.297023935 | 1.529594579 | 0.847952754 |
| 1700101O22Rik | 0.245087052 | 0.755309137 | 2.095767575 | 0.360397377 |
| Gm9144 | 9.249347624 | 0.646337586 | 0.371304672 | 1.740720315 |
| Apob | 11.59973813 | 1.954312174 | 0.464501757 | 4.207329997 |
| Ldah | 329.5450486 | -0.395664985 | 0.111902872 | -3.535789377 |
| Gdf7 | 3.829800073 | 0.401855354 | 0.528473894 | 0.7604072 |
| Hs1bp3 | 177.2730511 | -0.041081227 | 0.118981188 | -0.345274975 |
| Rhob | 3017.567704 | 0.641150735 | 0.115090986 | 5.57081626 |
| Slc7a15 | 0.061920613 | 0.274393828 | 3.724945049 | 0.073663859 |
| Pum2 | 3606.55936 | 0.043408258 | 0.066392237 | 0.653815269 |
| Gm4755 | 0.187019432 | -0.206483795 | 2.718962445 | -0.075942128 |
| Sdc1 | 95.2175769 | 0.639710822 | 0.243062593 | 2.631876894 |
| 9930038B18Rik | 0.162099466 | 0.741185957 | 2.787845873 | 0.265863319 |
| Laptm4a | 2359.716549 | 0.936727642 | 0.119452616 | 7.841834452 |
| Matn3 | 0.908179219 | -0.40370853 | 0.979193841 | -0.412286632 |
| Wdr35 | 500.8882983 | -0.251649525 | 0.070215624 | -3.58395339 |
| Ttc32 | 25.69466073 | -0.269810378 | 0.228424355 | -1.181180431 |
| Gm7099 | 8.593693136 | 0.36864059 | 0.455525561 | 0.809264335 |
| 1700022H16Rik | 0 | NA | NA | NA |
| Osr1 | 21.77471978 | -0.180453199 | 0.280122353 | -0.644194216 |
| Gm23979 | 0 | NA | NA | NA |
| Gm22845 | 0 | NA | NA | NA |
| Nt5c1b | 1.964636517 | 0.38779692 | 0.925167546 | 0.41916399 |
| Rdh14 | 214.0479375 | 0.036293009 | 0.106122876 | 0.341990438 |
| Pgk1-rs7 | 21.9025282 | 0.300204528 | 0.23567203 | 1.273823322 |
| Kcns3 | 56.27639958 | -0.034850751 | 0.171013667 | -0.203789277 |
| Msgn1 | 0 | NA | NA | NA |
| 1700034J04Rik | 0.087261593 | 0.274400501 | 3.724945049 | 0.073665651 |
| Gen1 | 22.65896892 | 0.341580808 | 0.289903178 | 1.178258242 |

| | | | | |
|---|---|---|---|---|
| Smc6 | 783.5087835 | 0.009114366 | 0.065435885 | 0.139286968 |
| Mir7675 | 1.906635116 | -0.172499172 | 0.680000921 | -0.253674909 |
| Vsnl1 | 3374.486255 | -0.156965769 | 0.083826675 | -1.872503812 |
| 4930511A02Rik | 1.096741448 | -0.400905973 | 1.06980249 | -0.374747654 |
| Rad51ap2 | 31.8005175 | -0.01354036 | 0.254248619 | -0.053256378 |
| Gm23781 | 0 | NA | NA | NA |
| Gm25610 | 0.211102819 | 0.274407618 | 2.625543881 | 0.104514581 |
| Gm4294 | 93.43490713 | 0.556806245 | 0.186720741 | 2.982026751 |
| Gm26171 | 0 | NA | NA | NA |
| Fam49a | 2242.155381 | -0.034081145 | 0.073080003 | -0.466353917 |
| Gm37414 | 0.066050875 | 0.274391604 | 3.724945049 | 0.073663262 |
| Gm27952 | 0 | NA | NA | NA |
| Gm25538 | 0 | NA | NA | NA |
| Platr19 | 0 | NA | NA | NA |
| 4930519A11Rik | 0.060174537 | 0.755286614 | 3.724945049 | 0.202764498 |
| Rpl36-ps3 | 0.314164845 | -0.231191423 | 1.632983284 | -0.141576111 |
| Mycn | 47.66468018 | -1.172361044 | 0.189122918 | -6.198936959 |
| Gm24208 | 0.042716945 | -0.206490061 | 3.724945049 | -0.055434391 |
| Ddx1 | 1202.104352 | 0.072790194 | 0.05885234 | 1.236827527 |
| Nbas | 1083.615517 | -0.015569278 | 0.05204558 | -0.299146987 |
| Gm6736 | 3.145663366 | 0.972196858 | 0.563305834 | 1.725877489 |
| Gm24142 | 0 | NA | NA | NA |
| Gm22170 | 0.191089262 | -0.206480188 | 2.650365316 | -0.077906312 |
| Gm4929 | 0.355586071 | -1.704302275 | 1.946909158 | -0.875388699 |
| Fam84a | 632.1102194 | -0.373165821 | 0.116355105 | -3.207128911 |
| Gm9847 | 0.673365293 | -0.579077934 | 1.241320383 | -0.466501591 |
| Gm25414 | 0.044191549 | -0.206490061 | 3.724945049 | -0.055434391 |
| Gm26449 | 0 | NA | NA | NA |
| Gm27682 | 0.049553047 | 0.274391604 | 3.724945049 | 0.073663262 |
| Gm4804 | 2.483041552 | -0.630744696 | 0.627304597 | -1.005483938 |
| Trib2 | 460.7235091 | -1.003434251 | 0.149308243 | -6.720554927 |
| Mir6387 | 0.891236725 | -1.462645994 | 1.141659671 | -1.281157626 |
| Gm5432 | 2.197748528 | -0.773762315 | 0.725568857 | -1.066421618 |
| Lpin1 | 478.9431543 | -0.166017777 | 0.108586525 | -1.528898507 |
| Ntsr2 | 425.5275216 | -0.640728541 | 0.164702625 | -3.890214503 |
| Greb1 | 74.66542959 | -0.181427674 | 0.152364664 | -1.190746395 |
| AC122228.1 | 0 | NA | NA | NA |
| E2f6 | 314.0510662 | -0.359338799 | 0.093041172 | -3.862148258 |
| Rock2 | 3956.78452 | 0.052295692 | 0.076745971 | 0.681412867 |
| Gm22303 | 1.492607829 | -0.605778565 | 0.763901137 | -0.793006497 |
| Pqlc3 | 54.75605864 | 1.202978289 | 0.227554486 | 5.286550526 |
| 2410004P03Rik | 23.25089268 | 0.257444134 | 0.293388231 | 0.877486235 |
| Kcnf1 | 695.0270978 | -0.432884921 | 0.157996982 | -2.739830308 |
| Pdia6 | 1198.63775 | 0.192478979 | 0.119224224 | 1.614428442 |
| Atp6v1c2 | 10.10846665 | -0.403566116 | 0.310940069 | -1.297890353 |
| Nol10 | 367.5541281 | 0.601447951 | 0.12625538 | 4.763741149 |
| Mir3066 | 1.524462984 | 1.314469592 | 1.043033493 | 1.260237183 |

| | | | | |
|---|---|---|---|---|
| Odc1 | 613.484201 | 0.817291326 | 0.170203227 | 4.801855649 |
| Gm22748 | 7.258283802 | 0.351574898 | 0.458548616 | 0.766712374 |
| Gm25925 | 0 | NA | NA | NA |
| Hpcal1 | 649.1744098 | -0.098186862 | 0.142443844 | -0.689302251 |
| B430203G13Rik | 10.4805672 | -0.74512223 | 0.355995639 | -2.09306561 |
| Gm10480 | 62.44942269 | -0.134734022 | 0.143182125 | -0.940997498 |
| 5730507C01Rik | 3.246809683 | 0.431370499 | 0.647403884 | 0.666308173 |
| Gm3944 | 0.500379391 | -0.629885801 | 1.527590079 | -0.412339547 |
| Gm5784 | 1.293338167 | -0.214254394 | 0.987978612 | -0.216861369 |
| Gm6802 | 0 | NA | NA | NA |
| Gm9257 | 0.060174537 | 0.755286614 | 3.724945049 | 0.202764498 |
| Gm22189 | 0 | NA | NA | NA |
| Gm21863 | 0.047948698 | -0.206487837 | 3.724945049 | -0.055433794 |
| Gm3993 | 0.060174537 | 0.755286614 | 3.724945049 | 0.202764498 |
| Gm10479 | 0.270569244 | 0.274410604 | 2.091814456 | 0.131183052 |
| 1700030C10Rik | 18.72189217 | 0.097163285 | 0.251251022 | 0.386717971 |
| Gm28503 | 0 | NA | NA | NA |
| 3110053B16Rik | 90.02746546 | 0.035074511 | 0.158681057 | 0.221037792 |
| Gm10478 | 5.792289935 | -1.072156743 | 0.498878831 | -2.149132569 |
| Zfp125 | 38.28518153 | -0.212013142 | 0.192319215 | -1.102402284 |
| Gm21989 | 3.535470757 | -0.560182842 | 0.546581452 | -1.024884471 |
| Gm5977 | 1.165717276 | -1.97778265 | 0.942928021 | -2.09749059 |
| Asap2 | 769.9183214 | 0.19959046 | 0.086842363 | 2.298307571 |
| Itgb1bp1 | 249.9318875 | -0.099318712 | 0.092676494 | -1.071671007 |
| Gm20535 | 1.863766393 | 0.108024294 | 1.03791548 | 0.104078122 |
| Cpsf3 | 408.1523475 | 0.044205771 | 0.069181527 | 0.638982292 |
| Gm25821 | 2.089596347 | 0.036233831 | 0.674252966 | 0.053739223 |
| Iah1 | 73.05919511 | -0.173083378 | 0.150959127 | -1.146557893 |
| Adam17 | 298.4801547 | 0.343548191 | 0.119354288 | 2.878390018 |
| Gm22766 | 0 | NA | NA | NA |
| Ywhaq | 1407.12046 | -0.127387578 | 0.075415451 | -1.689144286 |
| Gm4419 | 4.781372743 | -0.10595312 | 0.478967419 | -0.22121154 |
| Gm5633 | 5.363150736 | -0.655543063 | 0.454580426 | -1.44208379 |
| Gm22908 | 0 | NA | NA | NA |
| Gm4425 | 1.228836612 | -0.184456235 | 0.875890659 | -0.210592764 |
| Gm20486 | 0 | NA | NA | NA |
| Gm6838 | 0.045882455 | 0.755288839 | 3.724945049 | 0.202765096 |
| Gm10476 | 0.988361678 | -0.382195107 | 1.133491971 | -0.33718378 |
| Gm17391 | 0.141071297 | -0.206491178 | 3.564043836 | -0.057937328 |
| 2410018L13Rik | 0.412847196 | -0.685088896 | 1.6180155 | -0.423413061 |
| Gm26140 | 0 | NA | NA | NA |
| Gm20472 | 0.391014225 | 0.274415707 | 1.486308695 | 0.184629012 |
| Gm20473 | 0 | NA | NA | NA |
| Gm9285 | 0.258012988 | 0.81512789 | 2.115178145 | 0.385370798 |
| Gm9286 | 0 | NA | NA | NA |
| Gm20474 | 0.284703419 | -1.404055673 | 1.983551827 | -0.70784925 |
| Gm10330 | 0.04393175 | -0.206490061 | 3.724945049 | -0.055434391 |

| | | | | |
|---|---|---|---|---|
| Gm17330 | 0 | NA | NA | NA |
| 9030624G23Rik | 0.15672128 | 0.274408451 | 2.821957917 | 0.097240448 |
| Gm9312 | 0.046345709 | 0.274391604 | 3.724945049 | 0.073663262 |
| Gm16372 | 0.480740522 | 0.304813226 | 1.556464 | 0.195836991 |
| Taf1b | 209.6766852 | 0.014143188 | 0.085601704 | 0.165220874 |
| Grhl1 | 110.9007615 | 0.284108314 | 0.169479883 | 1.676354206 |
| Gm6969 | 32.27026886 | -0.032073717 | 0.298796349 | -0.107343068 |
| Klf11 | 102.5281956 | 0.423630475 | 0.156509402 | 2.706741378 |
| Cys1 | 49.24944908 | -0.758643186 | 0.23229155 | -3.265909524 |
| Rrm2 | 38.30661462 | 1.186804154 | 0.420271927 | 2.82389586 |
| Gm26211 | 0 | NA | NA | NA |
| Mboat2 | 425.7860528 | -0.284311618 | 0.103306897 | -2.752106851 |
| Kidins220 | 4110.912429 | -0.108513368 | 0.056938155 | -1.905811098 |
| Id2 | 1035.883774 | 0.918440551 | 0.148857013 | 6.169951516 |
| Gm38052 | 0 | NA | NA | NA |
| Gm22527 | 0 | NA | NA | NA |
| Rnf144a | 371.6551165 | -0.342039826 | 0.089351154 | -3.828040388 |
| Mir6538 | 1.774752459 | -1.023833279 | 0.780638946 | -1.311532411 |
| Rsad2 | 3584.881863 | 9.101178141 | 0.454700946 | 20.01574489 |
| Cmpk2 | 2770.91634 | 3.847196297 | 0.266943202 | 14.41204073 |
| Gm9866 | 90.23738068 | -0.083066457 | 0.137748899 | -0.6030281 |
| Sox11 | 612.1246314 | 0.120236387 | 0.126740056 | 0.948684969 |
| Gm24326 | 0 | NA | NA | NA |
| Gm25923 | 0 | NA | NA | NA |
| Gm25564 | 0 | NA | NA | NA |
| Dcdc2c | 0.829077587 | -0.023307913 | 1.276384795 | -0.018260883 |
| 1700022F17Rik | 0.238210844 | -0.206489586 | 3.042108938 | -0.067877117 |
| Allc | 2.669291208 | -0.827894107 | 0.602736603 | -1.373558705 |
| Colec11 | 3.087887419 | 0.94808105 | 0.583091391 | 1.625956181 |
| Rps7 | 1086.152334 | 0.487346321 | 0.125011787 | 3.898402967 |
| Rnaseh1 | 69.48037499 | -0.075271096 | 0.160521877 | -0.468914874 |
| Adi1 | 240.4881119 | -0.136215762 | 0.110617281 | -1.231414841 |
| Mir6937 | 0 | NA | NA | NA |
| Trappc12 | 622.1510625 | -0.280123547 | 0.080010904 | -3.50106714 |
| Tssc1 | 236.185517 | -0.072332347 | 0.08109057 | -0.891994561 |
| Gm23342 | 0 | NA | NA | NA |
| Myt1l | 2705.158939 | -0.129537878 | 0.126696965 | -1.0224229 |
| Pxdn | 585.0750425 | 0.311017729 | 0.18404735 | 1.689878876 |
| Tpo | 6.12441587 | -0.277927362 | 0.39140024 | -0.710084801 |
| Gm15691 | 3.03177422 | 2.172702051 | 0.717939453 | 3.026302625 |
| Sntg2 | 50.41524398 | 0.043284906 | 0.194221531 | 0.222863581 |
| 2310016D03Rik | 0.251964224 | 0.274411885 | 2.085547615 | 0.131577856 |
| Gm25865 | 0 | NA | NA | NA |
| Gm28806 | 0 | NA | NA | NA |
| Gm25243 | 0.038981395 | 0.274393828 | 3.724945049 | 0.073663859 |
| Tmem18 | 291.1926203 | 0.182061043 | 0.090086474 | 2.020958698 |
| Fam150b | 13.83564331 | 0.343856933 | 0.35048892 | 0.981077899 |

| | | | | |
|---|---|---|---|---|
| Acp1 | 179.9360824 | 0.323780481 | 0.103940507 | 3.115055825 |
| Sh3yl1 | 69.38233294 | -0.285669597 | 0.139031145 | -2.054716562 |
| Gm25408 | 0 NA | NA | NA | |
| Fam110c | 3.004281714 | 0.867520392 | 0.756847411 | 1.146228922 |
| 6030469F06Rik | 0.048949532 | -0.206487837 | 3.724945049 | -0.055433794 |
| Gm24613 | 0.553521388 | -0.462416292 | 1.222196044 | -0.378348706 |
| Lamb1 | 296.7992853 | -0.06603915 | 0.112493498 | -0.587048602 |
| Dld | 1183.816382 | 0.09776909 | 0.084511938 | 1.156867217 |
| Slc26a3 | 2.221929775 | 0.724757641 | 0.674026065 | 1.075266489 |
| Gm9359 | 0 NA | NA | NA | |
| Cbll1 | 375.0362158 | 0.131961974 | 0.073763975 | 1.788975912 |
| Gm29542 | 0 NA | NA | NA | |
| Gm29541 | 0 NA | NA | NA | |
| Slc26a4 | 32.11330933 | -0.213121062 | 0.238971201 | -0.891827387 |
| Bcap29 | 377.0266182 | 0.059314195 | 0.08845693 | 0.670543224 |
| Dus4l | 47.02044193 | -0.060143089 | 0.184274017 | -0.326378564 |
| Cog5 | 397.0078528 | -0.137587285 | 0.076594472 | -1.796308293 |
| Gpr22 | 1006.691692 | 0.38253676 | 0.1012772 | 3.777126139 |
| Gm25913 | 1.857927479 | -0.581678846 | 0.679284422 | -0.856311181 |
| Hbp1 | 434.1401043 | 0.273314088 | 0.103676152 | 2.636229086 |
| Prkar2b | 606.5567347 | -0.586087634 | 0.09781799 | -5.991613925 |
| Pik3cg | 109.1690308 | 1.649268775 | 0.131042718 | 12.58573392 |
| Gm23875 | 0 NA | NA | NA | |
| Ccdc71l | 284.7411437 | 0.219542914 | 0.124193956 | 1.76774234 |
| Nampt | 2161.206624 | 1.94676146 | 0.123793424 | 15.72588756 |
| Gdap10 | 195.362713 | 1.364487198 | 0.195683234 | 6.97293872 |
| 4933406C10Rik | 19.31754882 | 0.296801996 | 0.285849925 | 1.038314057 |
| Sypl | 610.6402324 | 0.11369034 | 0.08739937 | 1.300814189 |
| Gm24922 | 0 NA | NA | NA | |
| Cdhr3 | 9.236620993 | -0.581941791 | 0.325462928 | -1.788043248 |
| Gm22904 | 0.044544648 | 0.755286614 | 3.724945049 | 0.202764498 |
| F730043M19Rik | 202.5005712 | -0.407143026 | 0.13831528 | -2.943586757 |
| Gm16267 | 0 NA | NA | NA | |
| Atxn7l1 | 715.5692364 | 0.316360032 | 0.08770807 | 3.606966061 |
| Atxn7l1os1 | 0.109124069 | 0.274402725 | 3.724945049 | 0.073666248 |
| Atxn7l1os2 | 0 NA | NA | NA | |
| Efcab10 | 8.606127557 | 0.007193843 | 0.345365144 | 0.020829673 |
| Twistnb | 221.7759397 | 0.30248876 | 0.09651816 | 3.134008768 |
| Gm23529 | 0 NA | NA | NA | |
| Ferd3l | 0.671239363 | -1.77557947 | 1.353786918 | -1.311564949 |
| Twist1 | 6.585054892 | -0.268147014 | 0.433181207 | -0.619018115 |
| Gm18025 | 5.060744911 | -0.818952512 | 0.50373059 | -1.625774826 |
| Hdac9 | 424.2109994 | -0.605742889 | 0.128283117 | -4.721922117 |
| 4921508M14Rik | 2.520821878 | -0.641157977 | 0.628111591 | -1.02077081 |
| Gm24429 | 0 NA | NA | NA | |
| Prps1l1 | 0 NA | NA | NA | |
| AC133459.1 | 0 NA | NA | NA | |

| | | | | |
|---|---|---|---|---|
| Gm24359 | 0.075043205 | 0.755286614 | 3.724945049 | 0.202764498 |
| Snx13 | 1558.690179 | -0.131946229 | 0.051293862 | -2.572359023 |
| Mir680-3 | 0 | NA | NA | NA |
| AC132434.1 | 0 | NA | NA | NA |
| Gm26071 | 0 | NA | NA | NA |
| Ahr | 138.7406453 | 0.023312166 | 0.102741785 | 0.226900534 |
| Agr3 | 0.095295241 | -0.206492286 | 3.724945049 | -0.055434988 |
| Agr2 | 0.23039375 | -0.687379257 | 2.132339864 | -0.322359146 |
| Tspan13 | 1131.407557 | -0.107485255 | 0.069443713 | -1.547803975 |
| Gm5434 | 2.449668862 | 0.379658461 | 0.656136905 | 0.578626896 |
| Bzw2 | 293.0080855 | -0.053108032 | 0.093417935 | -0.568499318 |
| Ankmy2 | 458.9611624 | -0.00396825 | 0.064666422 | -0.061364922 |
| Lrrc72 | 0.323815284 | 0.744206468 | 1.740014861 | 0.427701214 |
| Sostdc1 | 41.87300568 | -0.798702323 | 0.238690301 | -3.346186757 |
| Gm25438 | 0.046345709 | 0.274391604 | 3.724945049 | 0.073663262 |
| D630036H23Rik | 7.277711592 | 0.536695657 | 0.352728004 | 1.521556697 |
| Ispd | 147.0734498 | 0.42748876 | 0.09667387 | 4.421968005 |
| Mir5099 | 0 | NA | NA | NA |
| CT010456.1 | 0 | NA | NA | NA |
| Gm29007 | 0 | NA | NA | NA |
| Meox2 | 0.449709139 | -1.891351467 | 1.453760351 | -1.301006363 |
| Agmo | 60.54347211 | -0.092751795 | 0.181496073 | -0.51104023 |
| Dgkb | 1454.497695 | -0.569112475 | 0.149475272 | -3.80740216 |
| Etv1 | 1246.89087 | -0.552018428 | 0.144537211 | -3.819213233 |
| Gm24069 | 0 | NA | NA | NA |
| Gm24434 | 0 | NA | NA | NA |
| Arl4a | 499.8111158 | 0.68757858 | 0.099110063 | 6.937525409 |
| Arl4aos | 1.421581723 | 0.579829802 | 0.78317879 | 0.740354322 |
| Scin | 6.368096109 | 1.040253382 | 0.437810605 | 2.376035139 |
| Lsmem1 | 17.83132388 | 0.809474205 | 0.28861796 | 2.804656387 |
| Gm17056 | 0.249945515 | -0.206481863 | 1.8479485 | -0.111735724 |
| Ifrd1 | 942.5158603 | 0.875784473 | 0.144518865 | 6.060001031 |
| Gm17024 | 29.97646992 | 0.119133644 | 0.315381986 | 0.377743972 |
| Mir1938 | 6.231293264 | 1.233160274 | 0.509045157 | 2.42249682 |
| Gm7008 | 11.68594954 | -0.16554937 | 0.308577389 | -0.536492225 |
| Zfp277 | 160.924537 | 0.035686325 | 0.132026443 | 0.270296795 |
| Dock4 | 2536.949238 | -0.090714929 | 0.10106942 | -0.897550704 |
| Immp2l | 31.50920012 | -0.255225058 | 0.18779074 | -1.359092879 |
| Gm25497 | 0.147068667 | 1.236183405 | 3.514416608 | 0.351746404 |
| Lrrn3 | 669.4920803 | -0.018891084 | 0.073085069 | -0.258480762 |
| Gm25248 | 0 | NA | NA | NA |
| Gm26326 | 0 | NA | NA | NA |
| Dnajb9 | 682.3485262 | 0.343162168 | 0.110072217 | 3.117609294 |
| Mir5627 | 0.044191549 | -0.206490061 | 3.724945049 | -0.055434391 |
| Pnpla8 | 1553.715183 | -0.024473491 | 0.064813538 | -0.377598447 |
| Nrcam | 5819.560715 | -0.059040428 | 0.07910084 | -0.74639445 |
| Stxbp6 | 733.0217604 | -0.384723058 | 0.105586228 | -3.643685991 |

| | | | | |
|---|---|---|---|---|
| Gm15901 | 0.132101749 | 0.274387056 | 3.724945049 | 0.073662041 |
| Gm15902 | 13.4453144 | -0.543032712 | 0.297395677 | -1.825960339 |
| Gm24731 | 0.429113465 | 0.269205552 | 1.483178769 | 0.181505802 |
| Gm22076 | 0 | NA | NA | NA |
| Gm9921 | 0.047948698 | -0.206487837 | 3.724945049 | -0.055433794 |
| Gm24948 | 0 | NA | NA | NA |
| Nova1 | 1267.849016 | -0.251293513 | 0.072110249 | -3.484851535 |
| Gm25051 | 0 | NA | NA | NA |
| Gm36971 | 0 | NA | NA | NA |
| Gm1818 | 26.89079701 | 0.012534667 | 0.22768009 | 0.055053856 |
| Gm26454 | 0 | NA | NA | NA |
| Gm7456 | 0 | NA | NA | NA |
| Foxg1 | 626.865741 | -0.031644704 | 0.135982337 | -0.232711871 |
| 3110039M20Rik | 111.2821837 | -0.080242208 | 0.243149863 | -0.330011324 |
| RP24-490H13.4 | 48.60029554 | 0.124738304 | 0.231883861 | 0.537934395 |
| Gm9804 | 13.1091654 | -0.257843354 | 0.410492827 | -0.628131206 |
| 1810007C17Rik | 0.183032319 | 1.343421106 | 3.313584731 | 0.405428325 |
| Gm25015 | 0 | NA | NA | NA |
| Gm7481 | 0.044191549 | -0.206490061 | 3.724945049 | -0.055434391 |
| Prkd1 | 131.3087976 | -0.380384519 | 0.11113355 | -3.422769449 |
| Gm22088 | 0 | NA | NA | NA |
| Rps11-ps4 | 0.060174537 | 0.755286614 | 3.724945049 | 0.202764498 |
| G2e3 | 315.7949451 | -0.361594983 | 0.103534489 | -3.492507524 |
| Scfd1 | 573.8494792 | 0.477487611 | 0.097541724 | 4.895213965 |
| Coch | 246.9813543 | 0.803066805 | 0.153764145 | 5.222718229 |
| Strn3 | 1743.354882 | 0.032669601 | 0.054811562 | 0.596034851 |
| Ap4s1 | 296.6051223 | -0.017152982 | 0.078947526 | -0.217270672 |
| Gm22663 | 0.270436937 | -1.332849984 | 2.376095253 | -0.56094131 |
| Hectd1 | 2718.891165 | -0.059053201 | 0.040309859 | -1.464981584 |
| Heatr5a | 367.9070692 | -0.013961706 | 0.130273059 | -0.107172625 |
| Gm26517 | 0.276231724 | 1.250988296 | 2.073380336 | 0.603356883 |
| Dtd2 | 62.41787465 | 0.371123942 | 0.164473009 | 2.256442836 |
| Gpr33 | 0.154148777 | 0.755296682 | 2.848943787 | 0.265114631 |
| Nubpl | 61.33971456 | -0.07995722 | 0.146953965 | -0.544097057 |
| Arhgap5 | 3781.700427 | -0.344304405 | 0.104321174 | -3.300426848 |
| 1700031P21Rik | 0.115603922 | 0.274387056 | 3.724945049 | 0.073662041 |
| Gm24859 | 1.957101456 | -0.518445435 | 0.685306566 | -0.756516077 |
| Gm23296 | 1.92086539 | 0.305588011 | 0.812152608 | 0.376269198 |
| Akap6 | 5339.749417 | -0.102743782 | 0.06292521 | -1.632792047 |
| n-R5s58 | 0 | NA | NA | NA |
| Npas3 | 426.0647591 | -0.015951152 | 0.08140002 | -0.195960052 |
| 1700060O08Rik | 0.07796279 | 0.274392189 | 3.724945049 | 0.073663419 |
| Egln3 | 112.0375452 | -0.289893843 | 0.124187579 | -2.334322367 |
| Gm22220 | 0 | NA | NA | NA |
| Gm24692 | 0 | NA | NA | NA |
| Sptssa | 171.7818652 | -0.285366021 | 0.143325259 | -1.991037885 |
| mmu-mir-1892 | 0 | NA | NA | NA |

| | | | | |
|---|---|---|---|---|
| Gm24971 | 0.26923737 | -0.602342381 | 2.112887281 | -0.285080225 |
| Eapp | 505.3675571 | 0.003421459 | 0.074497492 | 0.045927173 |
| Gm22513 | 8.835653036 | -0.334276585 | 0.404680634 | -0.826025655 |
| Rpl31-ps17 | 1.178692824 | 1.328355413 | 0.97892132 | 1.356958303 |
| Gm22634 | 17.48739027 | -1.277136171 | 0.524683313 | -2.434108614 |
| Gm23804 | 1.13792396 | 0.434091631 | 0.97920726 | 0.443309245 |
| Gm25679 | 43.52464816 | -0.019233531 | 0.249662526 | -0.077038119 |
| Gm26444 | 126.6439584 | 0.0602182 | 0.15276865 | 0.39417904 |
| Gm24305 | 91.6640994 | 0.394191564 | 0.206322553 | 1.910559749 |
| Gm22317 | 529.3228771 | -0.319131595 | 0.240019552 | -1.329606658 |
| Snx6 | 482.8085486 | 0.363942344 | 0.088943505 | 4.091837233 |
| Cfl2 | 953.9436151 | 0.17158448 | 0.095747272 | 1.792056078 |
| Baz1a | 191.7919684 | 1.047207715 | 0.186929068 | 5.602166259 |
| Gm27014 | 0 | NA | NA | NA |
| Gm20403 | 0 | NA | NA | NA |
| Gm24296 | 0 | NA | NA | NA |
| 2700097O09Rik | 100.6248846 | -0.641101202 | 0.170470647 | -3.760771789 |
| Srp54a | 0.941593011 | -0.922727262 | 1.105111729 | -0.834962871 |
| Fam177a | 2.46611504 | -0.500832034 | 0.589102916 | -0.850160508 |
| Srp54b | 195.0106852 | 0.176204905 | 0.143265501 | 1.229918604 |
| 1700047I17Rik2 | 3.457287012 | -0.651465031 | 0.613550177 | -1.061795849 |
| Ppp2r3c | 188.8955074 | 0.106201807 | 0.113768054 | 0.933494096 |
| 1110008L16Rik | 73.54196958 | 0.292479658 | 0.123693553 | 2.364550551 |
| Psma6 | 718.3364319 | 0.30332827 | 0.129895713 | 2.335167673 |
| Rpl18a-ps1 | 0.963303439 | -0.524521686 | 0.993905663 | -0.527737899 |
| Nfkbia | 657.3079934 | 2.144828661 | 0.28527025 | 7.518585127 |
| Aldoart2 | 0.61825299 | -0.400946046 | 1.207556372 | -0.332030914 |
| Insm2 | 3.842056669 | 0.173327726 | 0.543628058 | 0.318835137 |
| Ralgapa1 | 2973.912485 | -0.220892886 | 0.065205754 | -3.387628754 |
| Gm9812 | 2.978437241 | 0.578629508 | 0.571742544 | 1.012045568 |
| Gm23394 | 0 | NA | NA | NA |
| Brms1l | 538.8067484 | -0.241715218 | 0.081008212 | -2.983836002 |
| Gm22296 | 0 | NA | NA | NA |
| Gm23799 | 0 | NA | NA | NA |
| Gm25917 | 0.075043205 | 0.755286614 | 3.724945049 | 0.202764498 |
| Gm24685 | 0.291934259 | 0.306467845 | 1.776940518 | 0.172469389 |
| Mbip | 262.0904343 | 0.296817324 | 0.147839522 | 2.007699436 |
| Gm6265 | 9.407196128 | 0.087834008 | 0.561278102 | 0.156489284 |
| Nkx2-1 | 18.9524978 | -0.581484146 | 0.288256596 | -2.017244895 |
| Gm26973 | 0.859979859 | 0.491421648 | 1.225812192 | 0.400894731 |
| Gm25760 | 0 | NA | NA | NA |
| C87198 | 0.421608343 | -1.643746545 | 1.599998564 | -1.027342512 |
| Gm38103 | 1.370820674 | -1.20574282 | 0.993805188 | -1.213258729 |
| Nkx2-9 | 4.229279226 | -2.094712176 | 0.585280522 | -3.578988363 |
| Gm15524 | 0 | NA | NA | NA |
| Pax9 | 0.231631338 | 0.274410131 | 2.134176604 | 0.128578924 |
| Slc25a21 | 5.399330309 | -0.137908276 | 0.411960944 | -0.33476056 |

| | | | | |
|---|---|---|---|---|
| Prps1l3 | 132.0803981 | 0.03616909 | 0.100454159 | 0.360055677 |
| Mipol1 | 126.8360654 | 0.20010276 | 0.109339839 | 1.830099279 |
| Gm11025 | 0 | NA | NA | NA |
| Gm16246 | 0.044544648 | 0.755286614 | 3.724945049 | 0.202764498 |
| Gm2568 | 0 | NA | NA | NA |
| Gm37291 | 0 | NA | NA | NA |
| Foxa1 | 3.942515582 | -0.458054753 | 0.573982786 | -0.798028729 |
| 4921506M07Rik | 10.13732351 | -0.075292256 | 0.354625402 | -0.212314897 |
| 4921518K17Rik | 0.038981395 | 0.274393828 | 3.724945049 | 0.073663859 |
| Rpl26-ps2 | 0 | NA | NA | NA |
| Sstr1 | 99.2471505 | -0.528037105 | 0.168791597 | -3.128337641 |
| Clec14a | 49.14502572 | 1.106336962 | 0.214831916 | 5.149779343 |
| Gm17529 | 0.072025462 | 0.755288839 | 3.724945049 | 0.202765096 |
| Sec23a | 1353.977473 | -0.077085423 | 0.052009999 | -1.482126977 |
| Gemin2 | 59.82869919 | -0.081353039 | 0.146545985 | -0.555136594 |
| Gm22973 | 1.521163403 | 2.896522532 | 1.109635305 | 2.610337396 |
| Trappc6b | 482.938189 | 0.059961504 | 0.070208461 | 0.85404955 |
| Pnn | 591.1787483 | -0.006847109 | 0.154427521 | -0.044338656 |
| Gm5786 | 32.33765274 | 0.482598329 | 0.276286967 | 1.746728537 |
| Rpl21-ps3 | 1.284833043 | 1.910747568 | 0.98652853 | 1.936839644 |
| Mia2 | 5.838823621 | -0.11178386 | 0.519256079 | -0.21527694 |
| Ctage5 | 709.7395958 | 0.14164349 | 0.07629868 | 1.856434346 |
| Gm24233 | 4.255553331 | -0.242915015 | 0.465503933 | -0.521832357 |
| Fbxo33 | 714.5346483 | 0.0881743 | 0.116961918 | 0.753871868 |
| Gm27284 | 0 | NA | NA | NA |
| Gm7719 | 0.04393175 | -0.206490061 | 3.724945049 | -0.055434391 |
| Gm24377 | 0 | NA | NA | NA |
| Gm22673 | 0 | NA | NA | NA |
| Gm23059 | 0 | NA | NA | NA |
| Gm38172 | 0 | NA | NA | NA |
| Gm7745 | 2.081729379 | 0.517916803 | 0.778973078 | 0.664871249 |
| Nanog-ps2 | 0 | NA | NA | NA |
| Lrfn5 | 1144.044751 | -0.096522264 | 0.078831999 | -1.224404619 |
| Gm23213 | 11.15017964 | -0.166687296 | 0.358883808 | -0.464460343 |
| Gm24750 | 0 | NA | NA | NA |
| Gm24590 | 0 | NA | NA | NA |
| Spanxn4 | 0 | NA | NA | NA |
| Gm7763 | 0 | NA | NA | NA |
| Gm5185 | 0 | NA | NA | NA |
| Gm24921 | 0 | NA | NA | NA |
| Fscb | 0.143022002 | 0.755289479 | 3.547809333 | 0.212888971 |
| Gm24066 | 0 | NA | NA | NA |
| Gm6395 | 108.7166347 | 0.301729245 | 0.154491814 | 1.953043571 |
| Gm527 | 45.68336777 | -0.323830984 | 0.147032776 | -2.202440804 |
| Klhl28 | 276.6149016 | -0.360760568 | 0.086366427 | -4.177092667 |
| Fam179b | 1164.381407 | -0.137349322 | 0.073104627 | -1.878804763 |
| Prpf39 | 881.5198811 | 0.075335872 | 0.103156972 | 0.730303246 |

| | | | | |
|---|---|---|---|---|
| Gm25970 | 3.058502859 | -1.073326867 | 0.587542347 | -1.826807671 |
| Fkbp3 | 818.3394673 | 0.045747913 | 0.100778619 | 0.453944632 |
| Fancm | 325.0406225 | -0.268052348 | 0.121901743 | -2.198921373 |
| Mis18bp1 | 20.94898307 | 1.456066735 | 0.34727093 | 4.192884026 |
| Wdr20rt | 0.136963818 | 0.755286583 | 3.59954433 | 0.209828388 |
| Gm26015 | 0 | NA | NA | NA |
| Gm25599 | 0 | NA | NA | NA |
| Gm23948 | 0 | NA | NA | NA |
| Gm22782 | 0 | NA | NA | NA |
| Rpl10l | 0.061920613 | 0.274393828 | 3.724945049 | 0.073663859 |
| Mdga2 | 1092.640628 | -0.449094665 | 0.106183682 | -4.229413192 |
| Gm23826 | 1.550082411 | -0.339541778 | 0.713738339 | -0.475723048 |
| n-R5s60 | 0 | NA | NA | NA |
| Gm26256 | 0 | NA | NA | NA |
| n-R5s61 | 0 | NA | NA | NA |
| Gm15571 | 0.136963818 | 0.755286583 | 3.59954433 | 0.209828388 |
| Gm24499 | 0.13681464 | 0.274404728 | 2.94378999 | 0.093214777 |
| Rps29 | 1758.697591 | 0.258552593 | 0.105115469 | 2.459700709 |
| Rn7s1 | 42650.12024 | 0.224666004 | 0.309752702 | 0.725307649 |
| AC099934.2 | 0 | NA | NA | NA |
| AC099934.1 | 0 | NA | NA | NA |
| Lrr1 | 2.041322758 | -0.165343776 | 1.042688202 | -0.158574515 |
| Rpl36al | 266.3361645 | 0.528152533 | 0.160921256 | 3.282055746 |
| Mgat2 | 450.2752201 | 0.406949938 | 0.112605525 | 3.613942913 |
| Dnaaf2 | 60.5402635 | 0.225356034 | 0.172706463 | 1.304850032 |
| 9330151L19Rik | 197.4609258 | -0.122411053 | 0.096358419 | -1.270372162 |
| Pole2 | 9.423356029 | 0.872288923 | 0.384496421 | 2.268652908 |
| Klhdc1 | 176.7937862 | -0.428964472 | 0.145971029 | -2.93869594 |
| Gm15561 | 0 | NA | NA | NA |
| Klhdc2 | 1197.243616 | -0.066821586 | 0.060108707 | -1.111678971 |
| Nemf | 637.5580402 | 0.035299992 | 0.07912263 | 0.446142807 |
| AC157822.1 | 0 | NA | NA | NA |
| Rn7s2 | 34914.20044 | 0.293119157 | 0.206979543 | 1.416174528 |
| AC157822.2 | 0 | NA | NA | NA |
| Gm9887 | 2.974284523 | 0.043114281 | 0.578638656 | 0.074509853 |
| Arf6 | 1024.329735 | 0.400732268 | 0.094350701 | 4.247263259 |
| Gm23018 | 0.038981395 | 0.274393828 | 3.724945049 | 0.073663859 |
| Vcpkmt | 182.6319952 | 0.593755396 | 0.103294695 | 5.748169309 |
| Sos2 | 821.7158266 | -0.268139694 | 0.058045318 | -4.61948875 |
| Gm24449 | 1.413161897 | -1.076906249 | 0.875883571 | -1.229508447 |
| L2hgdh | 176.5242931 | -0.490020613 | 0.11978471 | -4.090844425 |
| Atp5s | 115.1168851 | -0.245253958 | 0.120111917 | -2.041878647 |
| Cdkl1 | 209.0876337 | 0.295596105 | 0.106156509 | 2.784531143 |
| Mir681 | 2.100525097 | -1.201703016 | 0.745052242 | -1.61291108 |
| 4930512B01Rik | 3.315021913 | 0.166292802 | 0.526962589 | 0.315568516 |
| Map4k5 | 890.2594534 | -0.184617432 | 0.114892473 | -1.606871428 |
| 4931403G20Rik | 0.07796279 | 0.274392189 | 3.724945049 | 0.073663419 |

| | | | | |
|---|---|---|---|---|
| Atl1 | 920.937231 | -0.197248925 | 0.058055847 | -3.397572057 |
| Gm6316 | 3.637700379 | -0.061861025 | 0.529455931 | -0.116838856 |
| Gm22058 | 5.202496254 | -0.134169921 | 0.449296266 | -0.298622381 |
| Gm3086 | 7.899535637 | -0.44824852 | 0.485490554 | -0.923289891 |
| Sav1 | 344.851752 | 0.81131619 | 0.131981727 | 6.147185756 |
| Gm23579 | 0 | NA | NA | NA |
| Nin | 902.9475348 | -0.084118945 | 0.138113124 | -0.609058307 |
| Abhd12b | 8.085658348 | 0.129150482 | 0.407807688 | 0.316694575 |
| Pygl | 47.59505423 | 0.334776414 | 0.183576517 | 1.823634193 |
| F730035M05Rik | 0.381387581 | 0.261098803 | 1.541233689 | 0.169408964 |
| Trim9 | 3338.081355 | -0.166710745 | 0.103888097 | -1.604714592 |
| Tmx1 | 634.4816424 | 0.248471527 | 0.096923242 | 2.563590766 |
| Gm24474 | 0.044544648 | 0.755286614 | 3.724945049 | 0.202764498 |
| Frmd6 | 461.8109092 | -0.268307072 | 0.139073509 | -1.92924643 |
| Gm24857 | 0 | NA | NA | NA |
| Actr10 | 1289.027537 | -0.054022905 | 0.065518048 | -0.82454998 |
| Psma3 | 949.5019891 | 0.416522327 | 0.076346873 | 5.455656636 |
| 3110056K07Rik | 60.20099725 | -0.461490749 | 0.17967066 | -2.568537068 |
| Arid4a | 748.6510582 | 0.003963927 | 0.107423845 | 0.036899882 |
| Tomm20l | 0.146692594 | 0.274392389 | 3.52332538 | 0.07787881 |
| Timm9 | 217.381226 | -0.069542009 | 0.070986745 | -0.979647816 |
| 2700049A03Rik | 196.0398049 | -0.158108982 | 0.124670474 | -1.268215132 |
| Dact1 | 166.182804 | 0.092023576 | 0.158644028 | 0.580063285 |
| 4930474H06Rik | 0.409321926 | -1.073413988 | 1.419380783 | -0.756255122 |
| 1700083H02Rik | 0 | NA | NA | NA |
| Gm22959 | 0 | NA | NA | NA |
| Daam1 | 1425.073008 | -0.025329918 | 0.048068382 | -0.526955913 |
| Gm7985 | 3.193668984 | -0.103738423 | 0.606427727 | -0.171064776 |
| Gpr135 | 77.04711869 | 0.130922425 | 0.133345697 | 0.98182714 |
| L3hypdh | 19.58973956 | -0.450017176 | 0.266756754 | -1.686994497 |
| Jkamp | 337.9089615 | -0.220691885 | 0.100071132 | -2.205350149 |
| Ccdc175 | 2.241016658 | 1.821568487 | 0.946187725 | 1.925166053 |
| Gm24466 | 0 | NA | NA | NA |
| Rtn1 | 7105.395095 | -0.14904112 | 0.073983939 | -2.014506408 |
| Lrrc9 | 75.53725959 | -0.172342714 | 0.149250109 | -1.154724214 |
| AC103353.1 | 0 | NA | NA | NA |
| Pcnxl4 | 483.304744 | -0.195759433 | 0.06826212 | -2.867760832 |
| Dhrs7 | 149.9058334 | 0.108337352 | 0.117426595 | 0.922596383 |
| Ppm1a | 1736.427688 | 0.005404059 | 0.061543246 | 0.087809127 |
| 4930447C04Rik | 136.3889103 | -0.200776119 | 0.189716228 | -1.058297021 |
| Six6 | 2.974111705 | 0.061441654 | 0.883700671 | 0.069527677 |
| Gm26709 | 0 | NA | NA | NA |
| Gm38170 | 0 | NA | NA | NA |
| Six1 | 9.772493715 | 0.47366415 | 0.404009779 | 1.172407636 |
| Six4 | 52.62636027 | -0.466446853 | 0.187214009 | -2.491516819 |
| Mnat1 | 196.833954 | 0.075803461 | 0.085507674 | 0.886510622 |
| Gm23910 | 0 | NA | NA | NA |

| | | | | |
|---|---|---|---|---|
| Trmt5 | 87.0705171 | -0.208326891 | 0.186795407 | -1.115267738 |
| Slc38a6 | 110.6102002 | 0.123666025 | 0.178582994 | 0.692484892 |
| D830013O20Rik | 0 | NA | NA | NA |
| Tmem30b | 3.209381466 | 0.144883328 | 0.585259352 | 0.247554059 |
| Prkch | 135.4372229 | 0.573090108 | 0.145840295 | 3.929573154 |
| Hif1a | 2116.265102 | 0.393318532 | 0.129502986 | 3.037138713 |
| Gm15283 | 3.904436585 | -0.562823761 | 0.473016159 | -1.189861595 |
| Gm5068 | 0 | NA | NA | NA |
| Snapc1 | 313.850414 | -0.055220954 | 0.091123262 | -0.606002825 |
| Syt16 | 456.7941473 | 0.055486824 | 0.144766994 | 0.383283665 |
| Gm29587 | 0 | NA | NA | NA |
| 1700086L19Rik | 251.9530011 | -0.199884805 | 0.174434373 | -1.145902623 |
| Dbpht2 | 497.2679655 | -0.142455006 | 0.156021926 | -0.913044786 |
| Gm11042 | 0.897295135 | -0.961325545 | 1.152023177 | -0.834467191 |
| Kcnh5 | 625.0782806 | -0.437142375 | 0.144675043 | -3.021546535 |
| Rhoj | 239.1716956 | 3.003832602 | 0.331212949 | 9.069188309 |
| Gphb5 | 1.844461247 | 2.518642971 | 1.169347699 | 2.153887141 |
| Ppp2r5e | 1376.244815 | -0.147526789 | 0.046196555 | -3.193458665 |
| Gm29236 | 0.285890061 | 1.207323647 | 1.810472174 | 0.666855677 |
| Wdr89 | 187.0451121 | 0.361091212 | 0.110728647 | 3.261045995 |
| Sgpp1 | 600.9576085 | 0.458162217 | 0.089611878 | 5.112739805 |
| Syne2 | 912.2047069 | -0.205276623 | 0.076039134 | -2.699618098 |
| Mir5101 | 0.042716945 | -0.206490061 | 3.724945049 | -0.055434391 |
| Esr2 | 7.599516194 | 0.221914915 | 0.35960352 | 0.617109963 |
| Gm7862 | 4.308722595 | -0.329152416 | 0.530380719 | -0.620596496 |
| Tex21 | 0.318741927 | 1.247511171 | 1.737368047 | 0.718046573 |
| Mthfd1 | 536.2563463 | -0.200899425 | 0.070798327 | -2.837629548 |
| Akap5 | 895.5134347 | -0.538528575 | 0.184953498 | -2.911697162 |
| Zbtb25 | 191.2371808 | 0.231624778 | 0.088604614 | 2.614139004 |
| Zbtb1 | 407.9327547 | 0.013154626 | 0.087192267 | 0.150869182 |
| 4930426I24Rik | 3.588158775 | -0.418753789 | 0.515664273 | -0.81206671 |
| Hspa2 | 433.2408539 | 0.195392331 | 0.133279226 | 1.466037403 |
| Ppp1r36 | 10.56443581 | -0.106139745 | 0.330013108 | -0.321622816 |
| Gm25563 | 0 | NA | NA | NA |
| Gm10451 | 3.439471239 | 0.196157443 | 0.520124282 | 0.377135715 |
| Mir5135 | 0.047948698 | -0.206487837 | 3.724945049 | -0.055433794 |
| Plekhg3 | 262.9259336 | -0.166952552 | 0.105869086 | -1.576971692 |
| Sptb | 3404.075389 | -0.368688116 | 0.093003112 | -3.964255701 |
| Gm24010 | 0 | NA | NA | NA |
| Churc1 | 76.27497148 | 0.065400719 | 0.147716268 | 0.442745544 |
| Fntb | 334.4021306 | -0.081786497 | 0.082031742 | -0.997010357 |
| Gpx2 | 1.498903582 | -0.103378764 | 0.855815943 | -0.120795558 |
| Rab15 | 1748.719287 | -0.16459604 | 0.124476825 | -1.322302685 |
| Max | 974.4805696 | 0.613386864 | 0.090304053 | 6.792462198 |
| Gm22696 | 0 | NA | NA | NA |
| Fut8 | 862.4398163 | -0.073486973 | 0.072680645 | -1.011094124 |
| Gm24775 | 2.254287688 | -0.859437772 | 0.688764002 | -1.247797169 |

| | | | | |
|---|---|---|---|---|
| Gm25926 | 5.521549126 | -0.332145875 | 0.476104561 | -0.697632206 |
| Gm24070 | 0 | NA | NA | NA |
| Gm24994 | 0 | NA | NA | NA |
| Gm6657 | 0 | NA | NA | NA |
| Gphn | 1083.325677 | -0.020054338 | 0.064313485 | -0.311821664 |
| Fam71d | 7.845798074 | -0.742237609 | 0.447059516 | -1.660265762 |
| Mpp5 | 811.6373095 | 0.136790237 | 0.097675278 | 1.400459151 |
| Gm25225 | 0.097139547 | 0.274393828 | 3.724945049 | 0.073663859 |
| Atp6v1d | 1557.396393 | 0.307514452 | 0.057549659 | 5.34346264 |
| Eif2s1 | 534.2845462 | 0.360858422 | 0.106589473 | 3.385497749 |
| Plek2 | 8.61393615 | 1.924838344 | 0.494406865 | 3.89322738 |
| Tmem229b | 976.9241392 | 0.356974143 | 0.11555898 | 3.089107771 |
| Plekhh1 | 575.4863948 | -0.625005449 | 0.103296393 | -6.050602848 |
| Pigh | 108.6090619 | -0.186575894 | 0.135190273 | -1.380098506 |
| Gm27297 | 0 | NA | NA | NA |
| Arg2 | 76.92116969 | 0.841282126 | 0.293744055 | 2.863997111 |
| Vti1b | 794.0198813 | -0.215711818 | 0.075526704 | -2.856099973 |
| Gm17195 | 12.61275266 | -1.053570882 | 0.34216752 | -3.079108389 |
| Gm17194 | 0 | NA | NA | NA |
| Rdh11 | 314.5614071 | 0.023791351 | 0.072640326 | 0.327522632 |
| Rdh12 | 16.14501922 | 0.071994489 | 0.27116393 | 0.265501715 |
| Zfyve26 | 591.7739187 | 0.18748425 | 0.094978759 | 1.973959792 |
| Rad51b | 1.767988423 | 0.736070221 | 0.839316835 | 0.876987319 |
| Gm28056 | 0 | NA | NA | NA |
| Zfp36l1 | 660.9038404 | 0.670727574 | 0.120486686 | 5.566819006 |
| Gm26669 | 0.480640422 | 1.702010974 | 1.419787414 | 1.198778745 |
| 2310015A10Rik | 112.2249857 | -0.183392709 | 0.10326976 | -1.775860709 |
| Actn1 | 1100.046826 | 0.006300481 | 0.092743255 | 0.067934654 |
| Dcaf5 | 1241.49422 | 0.104278299 | 0.069573649 | 1.498818878 |
| mmu-mir-1843a | 0 | NA | NA | NA |
| Scarna3b | 2.198311352 | -0.916328024 | 0.864564836 | -1.059871955 |
| Exd2 | 554.3707735 | -0.293907162 | 0.100733024 | -2.917684305 |
| Gm26777 | 28.6766377 | -0.179385546 | 0.214252644 | -0.83726176 |
| Galnt16 | 1559.333316 | 0.350851761 | 0.064154583 | 5.468849501 |
| Gm23817 | 0.07796279 | 0.274392189 | 3.724945049 | 0.073663419 |
| Erh | 198.2980328 | 0.150835887 | 0.104576056 | 1.442355852 |
| Slc39a9 | 697.3314638 | 0.120499264 | 0.066110855 | 1.822685011 |
| Plekhd1os | 1.509627474 | 0.22113657 | 0.838915448 | 0.263598161 |
| Plekhd1 | 169.5676397 | -0.265574656 | 0.150669063 | -1.762635614 |
| Ccdc177 | 193.1722448 | -0.539374961 | 0.12117136 | -4.451340328 |
| Gm26796 | 1.873447992 | 0.971620437 | 0.725939077 | 1.338432477 |
| Susd6 | 1200.089806 | 0.673669529 | 0.109395524 | 6.15810869 |
| Gm26545 | 0 | NA | NA | NA |
| Gm23298 | 0.127971488 | 0.274389832 | 3.687653711 | 0.0744077 |
| 1700052I22Rik | 0 | NA | NA | NA |
| Srsf5 | 1958.310891 | 0.177267098 | 0.085654693 | 2.069555005 |
| Slc10a1 | 0.066050875 | 0.274391604 | 3.724945049 | 0.073663262 |

| | | | | |
|---|---|---|---|---|
| Smoc1 | 160.7710477 | 0.330341956 | 0.168564569 | 1.959735415 |
| Mir3067 | 0.784624355 | 1.138423173 | 1.109084803 | 1.026452774 |
| Slc8a3 | 615.5791632 | -0.39866577 | 0.101053851 | -3.945082394 |
| Gm3693 | 13.26815873 | -1.617040571 | 0.449179716 | -3.599985739 |
| Gm20498 | 37.16701504 | -0.117056087 | 0.184525388 | -0.634363045 |
| Cox16 | 98.00354183 | -0.189193913 | 0.12237464 | -1.546022231 |
| Gm3695 | 17.50620631 | -0.33406719 | 0.39701523 | -0.841446788 |
| Gm4787 | 60.98925875 | -0.611506467 | 0.191596341 | -3.191639593 |
| Hspe1-ps2 | 0 NA | NA | NA | |
| Adam4 | 66.87698665 | -0.591754181 | 0.250259653 | -2.364560864 |
| Gm28370 | 1.129831383 | -0.556707861 | 0.948220557 | -0.587107986 |
| Synj2bp | 590.4251372 | -0.491310461 | 0.105687436 | -4.648712099 |
| Gm24853 | 0.038981395 | 0.274393828 | 3.724945049 | 0.073663859 |
| Gm17191 | 9.020757225 | -0.340194385 | 0.364142284 | -0.934234777 |
| 6530401F13Rik | 4.716167235 | -1.399773207 | 0.572006527 | -2.447128032 |
| Adam21 | 15.49915397 | 0.319584605 | 0.245023924 | 1.304299594 |
| Med6 | 165.5733909 | 0.341003522 | 0.12229781 | 2.788304394 |
| Gm16572 | 0.53222664 | -1.855743164 | 1.647754685 | -1.12622539 |
| Gm16570 | 4.946459833 | 0.282831356 | 0.527923175 | 0.535743398 |
| Ttc9 | 455.2096388 | -0.24125213 | 0.087895742 | -2.744753333 |
| Map3k9 | 1178.392681 | -0.156237948 | 0.100126938 | -1.560398744 |
| Pcnx | 2449.47711 | -0.217412571 | 0.069611801 | -3.123214261 |
| n-R5s62 | 0 NA | NA | NA | |
| Sipa1l1 | 4759.581652 | 0.119264592 | 0.127486874 | 0.935504876 |
| Gm5435 | 1.384865981 | 0.698901232 | 0.766167344 | 0.912204412 |
| Gm8358 | 0 NA | NA | NA | |
| n-R5s63 | 0 NA | NA | NA | |
| Rgs6 | 645.1774152 | -0.341186964 | 0.104979472 | -3.250035044 |
| SNORA43 | 0 NA | NA | NA | |
| Gm22149 | 0 NA | NA | NA | |
| 1700085C21Rik | 0.21789665 | 0.755308546 | 2.168530734 | 0.348304284 |
| 4930423C22Rik | 0.847184061 | 1.774670981 | 1.159127028 | 1.53104098 |
| Gm29530 | 0.044544648 | 0.755286614 | 3.724945049 | 0.202764498 |
| 4933436F18Rik | 0 NA | NA | NA | |
| 1700030I03Rik | 2.414365681 | -0.927531287 | 0.635283881 | -1.460026478 |
| Dpf3 | 155.1908164 | -0.482205466 | 0.142699158 | -3.379175276 |
| Dcaf4 | 167.3521746 | -0.003427564 | 0.103509117 | -0.03311364 |
| Zfyve1 | 610.9257158 | -0.188674131 | 0.057352367 | -3.289735749 |
| Gm26623 | 0.048949532 | -0.206487837 | 3.724945049 | -0.055433794 |
| Rbm25 | 2301.160026 | 0.104207267 | 0.110354248 | 0.944297736 |
| Gm29361 | 5.081951061 | -0.661784403 | 0.521799159 | -1.268274185 |
| Psen1 | 837.0310577 | 0.186703464 | 0.065370919 | 2.856063009 |
| Papln | 26.40192091 | -0.563038251 | 0.218875147 | -2.572417463 |
| Numb | 544.7112012 | -0.285841376 | 0.08875796 | -3.22045907 |
| 2410016O06Rik | 126.434428 | 0.07622843 | 0.132465491 | 0.575458781 |
| Gm26571 | 3.09834508 | 0.642329735 | 0.56375282 | 1.139381857 |
| Gm17673 | 1.031598502 | 1.122378165 | 1.062427449 | 1.056428056 |

| | | | | |
|---|---|---|---|---|
| Acot2 | 62.86110075 | 0.648903161 | 0.157212365 | 4.127558039 |
| Acot1 | 68.7388213 | 1.139685042 | 0.253225922 | 4.500664983 |
| Acot4 | 13.42834362 | 0.750764928 | 0.30096752 | 2.494504814 |
| 4732463B04Rik | 11.2321051 | -0.376182681 | 0.31677551 | -1.187537131 |
| Acot3 | 12.40614317 | 0.505555711 | 0.345089309 | 1.464999632 |
| Acot5 | 14.23358867 | 1.026662336 | 0.327504334 | 3.134805343 |
| Gm8385 | 0 | NA | NA | NA |
| Acot6 | 73.66342787 | -0.38427242 | 0.189321017 | -2.02973989 |
| Dnal1 | 1112.279329 | -0.35173133 | 0.094309331 | -3.729549615 |
| Pnma1 | 28.80769789 | -0.35190153 | 0.212556564 | -1.655566514 |
| Elmsan1 | 799.7188921 | 0.87681629 | 0.171179487 | 5.12220422 |
| Gm5436 | 39.89268335 | -0.264348772 | 0.213753501 | -1.236699143 |
| Ptgr2 | 431.9967278 | -0.290644407 | 0.097824369 | -2.971083885 |
| Zfp410 | 288.7236925 | 0.17000304 | 0.08920326 | 1.90579403 |
| Fam161b | 141.580014 | -0.626627034 | 0.183517177 | -3.414541589 |
| Coq6 | 126.6661858 | -0.030054599 | 0.112800323 | -0.266440716 |
| Entpd5 | 391.9413108 | -0.336642704 | 0.071293764 | -4.721909553 |
| Ccdc176 | 64.5472327 | -0.838603128 | 0.184666354 | -4.541179873 |
| AC120402.1 | 0 | NA | NA | NA |
| Rnf113a2 | 0 | NA | NA | NA |
| Aldh6a1 | 576.5073902 | -0.132165546 | 0.127693678 | -1.035020277 |
| Lin52 | 260.6951333 | 0.056462683 | 0.08836898 | 0.638942342 |
| Vsx2 | 9.446677945 | -0.092430123 | 0.481331726 | -0.192029982 |
| Abcd4 | 114.3789344 | -0.236926047 | 0.125757389 | -1.883993045 |
| Vrtn | 1.178380111 | 1.443922041 | 1.078647421 | 1.338641352 |
| Syndig1l | 316.8534421 | -0.62624076 | 0.111417516 | -5.620667061 |
| Npc2 | 1928.697367 | 1.489137241 | 0.159484834 | 9.337171488 |
| Isca2 | 199.1382904 | -0.047236999 | 0.089554383 | -0.527467191 |
| Ltbp2 | 65.48633923 | -0.026545805 | 0.210661509 | -0.126011654 |
| D030025P21Rik | 2.762732937 | -0.173796826 | 0.560341463 | -0.310162353 |
| Gm17139 | 0.226559374 | 2.043498472 | 2.51171147 | 0.813588064 |
| Arel1 | 2266.135036 | 0.474030834 | 0.071180651 | 6.659546175 |
| Gm17193 | 24.019554 | -0.660802977 | 0.331192133 | -1.995225463 |
| Fcf1 | 140.4858461 | 0.566380373 | 0.138843134 | 4.079282539 |
| Ylpm1 | 2692.241911 | -0.340350599 | 0.104947083 | -3.243068704 |
| Prox2os | 42.00672927 | -1.039702569 | 0.298725552 | -3.48046078 |
| Prox2 | 14.36306314 | -0.091708641 | 0.276583967 | -0.331576129 |
| Gm38355 | 1.352228905 | -0.414703271 | 0.87223595 | -0.475448496 |
| Dlst | 1232.377248 | -0.08215182 | 0.056335056 | -1.458271739 |
| Rps6kl1 | 284.8098805 | -0.143714228 | 0.116469644 | -1.233920044 |
| Pgf | 33.44763411 | 1.030058733 | 0.258157157 | 3.990045232 |
| Eif2b2 | 302.5668293 | 0.490652093 | 0.111166085 | 4.413685095 |
| Mlh3 | 324.2556209 | -0.342689812 | 0.106328073 | -3.222947626 |
| Mir6938 | 0.348080155 | -0.68479496 | 1.985741298 | -0.34485608 |
| Acyp1 | 106.7121191 | -0.238486645 | 0.129862804 | -1.836450755 |
| Zc2hc1c | 44.15230839 | -0.00435762 | 0.173460982 | -0.025121612 |
| Nek9 | 985.3989254 | 0.3119566 | 0.070027764 | 4.454756002 |

| | | | | |
|---|---|---|---|---|
| Tmed10 | 1862.85624 | 0.429584911 | 0.097868632 | 4.389403437 |
| Fos | 757.1228931 | 0.467879908 | 0.463273138 | 1.009943962 |
| Jdp2 | 344.9332217 | -0.043520044 | 0.091180896 | -0.477293448 |
| Batf | 16.13625052 | 5.01395028 | 0.777434387 | 6.449354906 |
| n-R5s64 | 0 | NA | NA | NA |
| AC117240.1 | 0 | NA | NA | NA |
| Mfsd7c | 17.4725598 | -0.061865613 | 0.278221562 | -0.222360959 |
| 0610007P14Rik | 325.4729982 | -0.26926429 | 0.064097205 | -4.200874137 |
| Ttll5 | 894.6595886 | -0.333060131 | 0.070905353 | -4.697249487 |
| Gm26531 | 0.135217741 | 1.264602945 | 3.618332936 | 0.349498779 |
| Tgfb3 | 281.890318 | 0.271424323 | 0.129785465 | 2.091330668 |
| Gm37804 | 1.282550361 | 0.926211599 | 0.915695451 | 1.011484329 |
| Ift43 | 63.1959967 | -0.156465422 | 0.150048613 | -1.042764871 |
| Gm805 | 0.516561414 | 0.256963972 | 1.271241896 | 0.202136173 |
| Gpatch2l | 583.809052 | 0.067361971 | 0.067413487 | 0.999235816 |
| 4732487G21Rik | 0.11657011 | 1.236179401 | 3.724945049 | 0.331865138 |
| Esrrb | 65.56798319 | 0.019102758 | 0.154378656 | 0.12373963 |
| Gm27993 | 0 | NA | NA | NA |
| Gm22004 | 0 | NA | NA | NA |
| Vash1 | 371.0212885 | -0.269690886 | 0.070522374 | -3.824188997 |
| Angel1 | 126.5566759 | -0.187976712 | 0.123083227 | -1.527232564 |
| Gm17138 | 0 | NA | NA | NA |
| Lrrc74a | 0.176451472 | 1.257521603 | 2.721126865 | 0.462132662 |
| Gm25888 | 0 | NA | NA | NA |
| Irf2bpl | 1475.011036 | 0.006085865 | 0.111734277 | 0.054467306 |
| Gm26698 | 1.907481327 | 0.496696297 | 0.782008149 | 0.635154886 |
| Cipc | 1257.563404 | -0.216068908 | 0.042471521 | -5.08738334 |
| Zdhhc22 | 154.2953736 | -0.217059143 | 0.115285515 | -1.88279631 |
| Tmem63c | 504.9714176 | -0.176273088 | 0.113363521 | -1.554936596 |
| Gm25847 | 6.00824128 | -0.485957162 | 0.545842139 | -0.890288836 |
| Gm15745 | 1.796415912 | 0.118783336 | 0.752556727 | 0.15783971 |
| Ngb | 44.68760207 | -0.168732453 | 0.221228467 | -0.762706787 |
| Pomt2 | 325.0464884 | -0.173135351 | 0.069352528 | -2.496453358 |
| Gstz1 | 241.2031799 | 0.06545083 | 0.085043587 | 0.769615117 |
| Tmed8 | 929.2418369 | -0.095314802 | 0.074638901 | -1.27701239 |
| Samd15 | 21.85261841 | 0.043914635 | 0.261910977 | 0.167670083 |
| Noxred1 | 12.44760366 | -1.433329368 | 0.391639209 | -3.659820912 |
| Vipas39 | 515.9231004 | -0.021263694 | 0.056620283 | -0.375549055 |
| Ahsa1 | 741.40917 | -0.059461394 | 0.089519224 | -0.664230445 |
| AC151967.1 | 0.038981395 | 0.274393828 | 3.724945049 | 0.073663859 |
| Ism2 | 2.941330987 | -0.535984951 | 0.599720072 | -0.893725216 |
| Sptlc2 | 820.5122325 | 0.578572257 | 0.13159614 | 4.396574683 |
| Gm26764 | 0.618129248 | -0.625482378 | 1.399871298 | -0.446814203 |
| Alkbh1 | 295.3910853 | 0.427822789 | 0.152579811 | 2.803927898 |
| mmu-mir-3068 | 0.086930093 | -0.206490058 | 3.724945049 | -0.05543439 |
| Mir3068 | 0.060174537 | 0.755286614 | 3.724945049 | 0.202764498 |
| Gm29362 | 0 | NA | NA | NA |

| | | | | |
|---|---|---|---|---|
| Slirp | 288.8250465 | 0.408439894 | 0.127372978 | 3.206644768 |
| Snw1 | 789.2603872 | 0.500985503 | 0.081647375 | 6.13596586 |
| Gm8300 | 0 | NA | NA | NA |
| Oog1 | 0 | NA | NA | NA |
| Gm4027 | 0 | NA | NA | NA |
| Gm16381 | 0 | NA | NA | NA |
| Gm21319 | 0 | NA | NA | NA |
| Gm21936 | 0 | NA | NA | NA |
| BB287469 | 0 | NA | NA | NA |
| Gm2001 | 0 | NA | NA | NA |
| Gm2016 | 0 | NA | NA | NA |
| Gm2022 | 0 | NA | NA | NA |
| Gm2035 | 0 | NA | NA | NA |
| Gm2042 | 0 | NA | NA | NA |
| Gm2046 | 0 | NA | NA | NA |
| Gm2075 | 0 | NA | NA | NA |
| Gm6803 | 0 | NA | NA | NA |
| Gm2056 | 0 | NA | NA | NA |
| Gm16368 | 0 | NA | NA | NA |
| RP23-174O6.3 | 0 | NA | NA | NA |
| Gm8587 | 0.789916185 | 0.084510832 | 1.178827381 | 0.071690591 |
| Gm10436 | 0 | NA | NA | NA |
| Gm8332 | 0 | NA | NA | NA |
| Gm5662 | 0 | NA | NA | NA |
| Gm7104 | 0 | NA | NA | NA |
| Gm5039 | 0.128271896 | 0.274403127 | 3.679089475 | 0.074584521 |
| Gm10264 | 0 | NA | NA | NA |
| Adck1 | 270.5190383 | -0.048124727 | 0.082063996 | -0.586429239 |
| Nrxn3 | 5644.729856 | -0.156510899 | 0.105472121 | -1.483907767 |
| Gm37980 | 0.244108046 | -0.843601786 | 2.485561782 | -0.339400852 |
| Gm27313 | 0 | NA | NA | NA |
| Gm23989 | 9.194225744 | 0.160434056 | 0.455471948 | 0.352236964 |
| Dio2 | 725.4291824 | -0.39827257 | 0.186478998 | -2.135750274 |
| Gm26512 | 30.26736896 | 0.257125073 | 0.208680233 | 1.232148678 |
| 4930544I03Rik | 0.534296218 | 1.9731197 | 1.362728961 | 1.447917933 |
| n-R5s65 | 0 | NA | NA | NA |
| Gm23195 | 0 | NA | NA | NA |
| Gm26477 | 0 | NA | NA | NA |
| Cep128 | 104.2602885 | 0.032778231 | 0.122092808 | 0.268469797 |
| 5430427M07Rik | 0 | NA | NA | NA |
| CR974487.1 | 0 | NA | NA | NA |
| Tshr | 7.627433336 | -0.094596328 | 0.372754383 | -0.253776568 |
| Gm18827 | 0.108266322 | 0.274389235 | 3.724945049 | 0.073662626 |
| Gtf2a1 | 1089.805438 | 0.038555804 | 0.062760137 | 0.614335887 |
| Gm25776 | 271.3150006 | -0.565347629 | 0.224530993 | -2.517904642 |
| Ston2 | 295.1567159 | 0.002234792 | 0.122174685 | 0.018291775 |
| Gm8378 | 10.50589885 | -0.485049811 | 0.338515746 | -1.432872228 |

| | | | | |
|---|---|---|---|---|
| Gm16876 | 0.085327104 | 0.274389469 | 3.724945049 | 0.073662689 |
| Gm22015 | 0 | NA | NA | NA |
| Sel1l | 2575.426926 | -0.103842055 | 0.038774024 | -2.678134562 |
| Rpl31-ps1 | 0.104719185 | 1.218392466 | 3.723618458 | 0.327206581 |
| Gm21614 | 0 | NA | NA | NA |
| Gm23249 | 0 | NA | NA | NA |
| Gm9726 | 14.69313774 | -0.112303195 | 0.296367046 | -0.378932802 |
| Mir8099-1 | 0.066050875 | 0.274391604 | 3.724945049 | 0.073663262 |
| Gm26055 | 0.060174537 | 0.755286614 | 3.724945049 | 0.202764498 |
| Gm22246 | 0.682665822 | 0.365025412 | 1.157290634 | 0.315413778 |
| Flrt2 | 1031.326902 | -0.272105545 | 0.073222433 | -3.716150012 |
| 1700019M22Rik | 0 | NA | NA | NA |
| Gm26480 | 0 | NA | NA | NA |
| Gm6863 | 145.6421742 | 0.259388069 | 0.138628921 | 1.871096354 |
| Gm23558 | 0 | NA | NA | NA |
| Galc | 301.800896 | -0.267647803 | 0.096177412 | -2.782855126 |
| Gpr65 | 55.39552026 | 5.246451666 | 0.453169496 | 11.57723923 |
| Kcnk10 | 201.901865 | -0.389047512 | 0.131480025 | -2.958985679 |
| Gm25486 | 0 | NA | NA | NA |
| Spata7 | 282.5135196 | 0.186951364 | 0.141108689 | 1.324874926 |
| Ptpn21 | 244.2151489 | 0.021287667 | 0.092506796 | 0.230120037 |
| Zc3h14 | 1282.947868 | 0.139733528 | 0.058675174 | 2.381476162 |
| Eml5 | 1567.483707 | -0.345618013 | 0.150466565 | -2.296975489 |
| Ttc8 | 210.9206023 | -0.551476556 | 0.153941648 | -3.582373998 |
| Rpl30-ps8 | 10.08383252 | 0.026749979 | 0.334483285 | 0.079974037 |
| Gm28933 | 0.301731016 | -0.687380498 | 2.069751552 | -0.332107734 |
| Foxn3 | 762.8771155 | 0.019274986 | 0.053584317 | 0.359713187 |
| Gm19898 | 0.254453631 | 0.755294773 | 2.976423281 | 0.253759194 |
| Gm20682 | 16.54715445 | -0.301553796 | 0.371970556 | -0.81069265 |
| 3300002A11Rik | 4.616394162 | -0.424212085 | 0.539702189 | -0.786011422 |
| Gm26839 | 2.688780499 | 0.460603558 | 0.633877464 | 0.726644476 |
| 1700064M15Rik | 1.043907264 | 1.224610634 | 1.010806261 | 1.211518648 |
| Efcab11 | 4.011499476 | 0.593519031 | 0.53745737 | 1.104309038 |
| Tdp1 | 173.9666548 | -0.615484456 | 0.160136709 | -3.843493849 |
| Kcnk13 | 123.3766849 | 0.037140246 | 0.153169947 | 0.242477372 |
| Gm23291 | 0.044544648 | 0.755286614 | 3.724945049 | 0.202764498 |
| Psmc1 | 1000.106951 | 0.290259535 | 0.112354158 | 2.583433847 |
| Nrde2 | 211.3513749 | 0.349251851 | 0.102871351 | 3.39503515 |
| Gm10433 | 3.50734834 | 0.453623383 | 0.532283984 | 0.852220613 |
| Calm1 | 27013.61596 | 0.179200085 | 0.082332205 | 2.176549096 |
| Gm26723 | 0.970285468 | 1.652912489 | 1.079234953 | 1.531559448 |
| Ttc7b | 1662.607921 | -0.119386564 | 0.069182237 | -1.725682339 |
| Rps6ka5 | 469.9143002 | 0.011065861 | 0.088900325 | 0.124474924 |
| Gm25801 | 0.225129614 | 1.818209708 | 3.710727189 | 0.489987438 |
| 9030617O03Rik | 99.5661288 | -0.072334855 | 0.168680681 | -0.428827146 |
| Gm8482 | 2.942155206 | 4.653745207 | 1.108337085 | 4.198853642 |
| Gpr68 | 165.6196338 | -0.170794735 | 0.119139507 | -1.433569256 |

| | | | | |
|---|---|---|---|---|
| Ccdc88c | 393.2724972 | 0.103881551 | 0.148374779 | 0.700129443 |
| A630072L19Rik | 0.314467885 | 1.330170726 | 2.014475609 | 0.660306196 |
| Mir1190 | 0.80558184 | -1.503190224 | 1.14608524 | -1.311586757 |
| Smek1 | 938.2289803 | 0.007291205 | 0.053421931 | 0.136483357 |
| D130020L05Rik | 15.63426209 | -0.077959954 | 0.270127135 | -0.288604675 |
| Kif4-ps | 0 | NA | NA | NA |
| Gm18848 | 0 | NA | NA | NA |
| Gm22749 | 0 | NA | NA | NA |
| Catsperb | 0.068097359 | -0.206487837 | 3.724945049 | -0.055433794 |
| Gm24542 | 0 | NA | NA | NA |
| Tc2n | 7.321160109 | 0.187477531 | 0.427714321 | 0.438324185 |
| Gm16285 | 0 | NA | NA | NA |
| Fbln5 | 104.6927819 | 1.364884352 | 0.203645557 | 6.70225451 |
| Trip11 | 1573.263012 | -0.086362681 | 0.048627002 | -1.776023129 |
| Gm20701 | 0 | NA | NA | NA |
| Gm20702 | 0 | NA | NA | NA |
| Gm20036 | 4.340709641 | -0.072351752 | 0.559403478 | -0.12933733 |
| Atxn3 | 451.4130845 | -0.110972443 | 0.072677777 | -1.526910242 |
| Cpsf2 | 797.9273037 | 0.390017902 | 0.077139505 | 5.056007348 |
| Gm16339 | 11.18950982 | 0.964019366 | 0.430297811 | 2.24035387 |
| Slc24a4 | 531.3783664 | 0.331771223 | 0.184414865 | 1.799048156 |
| Gm25203 | 0 | NA | NA | NA |
| Rin3 | 105.1190873 | 2.107888406 | 0.196881418 | 10.70638573 |
| Lgmn | 2861.87464 | 1.359144262 | 0.132827512 | 10.23240024 |
| Golga5 | 336.8030054 | 0.109897924 | 0.088891145 | 1.236320267 |
| Chga | 1338.904208 | -0.252788397 | 0.177403406 | -1.424935415 |
| Itpk1 | 872.6122047 | 0.63674108 | 0.122689924 | 5.189840038 |
| Gm37019 | 35.26459873 | -0.592324648 | 0.310598225 | -1.907044536 |
| Gm20069 | 0.551997393 | 2.367247812 | 1.364747444 | 1.734568416 |
| Gm28373 | 39.99975009 | -0.253063916 | 0.293175089 | -0.863183557 |
| Mir1936 | 2.100202168 | 0.172141615 | 0.773535828 | 0.222538645 |
| Gm28051 | 0.060174537 | 0.755286614 | 3.724945049 | 0.202764498 |
| Gm21971 | 0 | NA | NA | NA |
| Gm20604 | 131.2845531 | 0.127129268 | 0.116483945 | 1.091388754 |
| Moap1 | 4.795281827 | -0.297820592 | 0.54506702 | -0.546392611 |
| Tmem251 | 141.6121926 | 0.623089745 | 0.158373549 | 3.934304368 |
| AK010878 | 58.12541358 | -0.262989266 | 0.153252382 | -1.716053365 |
| Ubr7 | 418.4523783 | 0.129619656 | 0.071798974 | 1.805313502 |
| Btbd7 | 452.2462838 | 0.136695553 | 0.077176604 | 1.771204565 |
| Cox8c | 0 | NA | NA | NA |
| Unc79 | 1662.520299 | -0.38323712 | 0.114880682 | -3.335957894 |
| AC122367.1 | 17.42179249 | -0.681301562 | 0.364811102 | -1.86754613 |
| Gm27976 | 0.999255461 | -1.279781747 | 1.032081869 | -1.240000223 |
| Prima1 | 64.20214066 | -0.042999897 | 0.167044656 | -0.25741558 |
| Fam181a | 11.59340835 | -0.158591141 | 0.316520807 | -0.501044915 |
| Gm29508 | 2.89487079 | -1.526821476 | 0.642964598 | -2.374658699 |
| Asb2 | 49.37814463 | 0.548834159 | 0.220097562 | 2.493594904 |

| | | | | |
|---|---|---|---|---|
| Gm15523 | 0.643207634 | -1.396735377 | 1.263541123 | -1.105413469 |
| Otub2 | 164.6469116 | -0.548743805 | 0.168606176 | -3.254589003 |
| 9330161L09Rik | 14.42659375 | 0.31800907 | 0.28184835 | 1.128298497 |
| Ddx24 | 2634.287795 | 0.823232868 | 0.097772973 | 8.419840834 |
| Ifi27 | 153.7870265 | 0.636450934 | 0.244089204 | 2.607452215 |
| Ifi27l2a | 1103.207706 | 7.987631931 | 0.368177899 | 21.6950337 |
| Ifi27l2b | 0.044191549 | -0.206490061 | 3.724945049 | -0.055434391 |
| Ppp4r4 | 572.2723124 | 0.073957579 | 0.095514267 | 0.774309231 |
| Serpina10 | 1.913476902 | 3.265649848 | 1.142050465 | 2.859461949 |
| Gm18501 | 0 | NA | NA | NA |
| Serpina6 | 0.146451277 | -0.68737672 | 2.882358805 | -0.238477152 |
| Serpina16 | 0 | NA | NA | NA |
| Serpina1f | 0.066050875 | 0.274391604 | 3.724945049 | 0.073663262 |
| Gm5656 | 0 | NA | NA | NA |
| Gm17142 | 0 | NA | NA | NA |
| Serpina1b | 7.892462027 | 0.960040014 | 0.433466146 | 2.214798141 |
| Gm17042 | 0 | NA | NA | NA |
| Serpina1d | 0 | NA | NA | NA |
| Gm17043 | 0 | NA | NA | NA |
| Gm8893 | 0.487995331 | -1.911357535 | 1.320949454 | -1.44695736 |
| Gm6918 | 0 | NA | NA | NA |
| Serpina1a | 0.839309821 | 2.281177265 | 1.164589763 | 1.958781829 |
| Gm8895 | 0.092691418 | 0.274387304 | 3.724945049 | 0.073662108 |
| Serpina1c | 0.721092599 | 0.6575404 | 1.099483785 | 0.598044654 |
| B430119L08Rik | 0.437661561 | -0.655127518 | 1.424835305 | -0.45979175 |
| Gm17198 | 0 | NA | NA | NA |
| Serpina1e | 0.347991053 | -0.092389956 | 1.713772783 | -0.053910272 |
| Serpina11 | 2.314302159 | 3.16979848 | 1.013485117 | 3.12762213 |
| Gm28577 | 0 | NA | NA | NA |
| Serpina9 | 42.26228332 | -0.406415056 | 0.230641819 | -1.762104795 |
| Gm37496 | 0.085433889 | -0.687376175 | 3.724945049 | -0.184533239 |
| Serpina12 | 0.359936507 | 1.493693048 | 3.700439892 | 0.403652834 |
| Serpina4-ps1 | 0.088534442 | 0.274389469 | 3.724945049 | 0.073662689 |
| Serpina5 | 0.066050875 | 0.274391604 | 3.724945049 | 0.073663262 |
| Gm23289 | 0 | NA | NA | NA |
| Serpina3a | 0.228000506 | 0.27440174 | 2.510883511 | 0.109284935 |
| Serpina3b | 0.099106094 | 0.274387056 | 3.724945049 | 0.073662041 |
| Serpina3c | 2.76240905 | 2.244830834 | 0.919948907 | 2.440169031 |
| Serpina3d-ps | 0 | NA | NA | NA |
| Serpina3e-ps | 7.226489367 | 2.559833652 | 0.613976431 | 4.169270225 |
| Serpina3f | 27.21369777 | 5.766399897 | 0.844457731 | 6.828524016 |
| Gm8918 | 0 | NA | NA | NA |
| Serpina3g | 192.2982361 | 5.624644884 | 0.576155925 | 9.762365781 |
| Serpina3h | 81.26495241 | 4.757953399 | 0.523944787 | 9.081020584 |
| Serpina3i | 29.39647476 | 6.426571944 | 0.904605247 | 7.104283294 |
| Serpina3j | 0 | NA | NA | NA |
| Serpina3m | 31.36521844 | 2.551578139 | 0.41603096 | 6.133144848 |

| | | | | |
|---|---|---|---|---|
| Serpina3k | 1.059265352 | 0.367818723 | 1.117654521 | 0.329098765 |
| Serpina3l-ps | 0 | NA | NA | NA |
| Serpina3n | 2233.484408 | 2.924601196 | 0.308536639 | 9.478942947 |
| Gm24334 | 0 | NA | NA | NA |
| Gsc | 0.456948362 | 0.257331922 | 1.459852857 | 0.176272507 |
| Gm10000 | 0.122601409 | 1.236179401 | 3.724945049 | 0.331865138 |
| Dicer1 | 1104.172179 | -0.047110825 | 0.097592241 | -0.482731259 |
| Clmn | 1628.39178 | -0.241456661 | 0.103124343 | -2.341412837 |
| 4930408O17Rik | 1.425419836 | -1.115602239 | 0.964520359 | -1.156639389 |
| Gm28875 | 11.6389744 | 0.228751913 | 0.398369824 | 0.574219981 |
| Syne3 | 100.3522481 | 0.326779509 | 0.152072193 | 2.148844591 |
| Scarna13 | 339.6868988 | 0.324285914 | 0.289904616 | 1.118595209 |
| mmu-mir-3069 | 0 | NA | NA | NA |
| Glrx5 | 236.9685444 | 0.021431445 | 0.10245592 | 0.209177224 |
| Tcl1b2 | 0.273490436 | 0.980875356 | 2.41626262 | 0.405947329 |
| Tcl1b1 | 0.12092566 | 1.236179401 | 3.724945049 | 0.331865138 |
| Tcl1b5 | 0 | NA | NA | NA |
| Tcl1b3 | 0 | NA | NA | NA |
| Tcl1b4 | 0 | NA | NA | NA |
| Tcl1 | 0 | NA | NA | NA |
| Tunar | 293.4716965 | -0.377977995 | 0.102759904 | -3.678263422 |
| D430019H16Rik | 1275.143419 | -0.278245932 | 0.139741797 | -1.991143224 |
| Bdkrb2 | 21.21549069 | 1.478495532 | 0.372722825 | 3.966742668 |
| Bdkrb1 | 6.675953194 | 3.088475389 | 0.76759128 | 4.023593635 |
| Atg2b | 1471.256828 | -0.204356307 | 0.055639012 | -3.672896078 |
| Gm22790 | 0 | NA | NA | NA |
| Gskip | 216.9176705 | -0.187066748 | 0.133501332 | -1.401235073 |
| Ak7 | 31.57741441 | 0.258539712 | 0.206667885 | 1.250991235 |
| Papola | 1915.550768 | -0.208753864 | 0.081688674 | -2.555481123 |
| Gm22120 | 0 | NA | NA | NA |
| Vrk1 | 161.0238951 | 0.028904444 | 0.11260333 | 0.256692617 |
| Gm24986 | 0 | NA | NA | NA |
| Gm17032 | 0.342848402 | -0.687379209 | 1.991277185 | -0.345195141 |
| 1700121N20Rik | 0 | NA | NA | NA |
| Gm3191 | 0.406963084 | 0.755271159 | 3.724945049 | 0.202760349 |
| Gm17031 | 0.100902008 | 0.274391828 | 3.724945049 | 0.073663322 |
| Gm17053 | 0.121578509 | 0.75528439 | 3.724945049 | 0.202763901 |
| Gm17033 | 0 | NA | NA | NA |
| 4933406K04Rik | 0 | NA | NA | NA |
| 1700013N06Rik | 0 | NA | NA | NA |
| Gm9599 | 0 | NA | NA | NA |
| Gm16087 | 0.365883897 | -1.667757981 | 1.455697607 | -1.145676116 |
| Gm16086 | 0.197846125 | -1.013852358 | 3.255083671 | -0.311467372 |
| Gm16085 | 0 | NA | NA | NA |
| Gm16084 | 0 | NA | NA | NA |
| Gm2800 | 0.20562365 | -0.531615553 | 2.592716595 | -0.205041906 |
| Gm5187 | 0 | NA | NA | NA |

| | | | | |
|---|---|---|---|---|
| Gm17052 | 0.203140137 | -0.687378669 | 2.219553625 | -0.309692301 |
| 3110018I06Rik | 0 | NA | NA | NA |
| Gm9006 | 0 | NA | NA | NA |
| 4930465M20Rik | 0.148569308 | -0.687378047 | 2.869495678 | -0.23954664 |
| Gm15208 | 0.15596086 | -1.15072961 | 3.483969583 | -0.330292668 |
| Bcl11b | 1306.155659 | -0.274152498 | 0.151498917 | -1.809600389 |
| Setd3 | 1675.172341 | 0.021916638 | 0.053600133 | 0.408891478 |
| Ccnk | 411.8626785 | 0.213029643 | 0.107593557 | 1.979947957 |
| Ccdc85c | 333.5260322 | -0.467697921 | 0.081903827 | -5.71033049 |
| Hhipl1 | 69.09591632 | -0.289717352 | 0.166165119 | -1.743550954 |
| 4930478K11Rik | 1.119709845 | -1.326290451 | 1.15921056 | -1.144132478 |
| Cyp46a1 | 1040.049922 | -0.173864278 | 0.152481305 | -1.140233411 |
| Eml1 | 662.3640817 | -0.442352923 | 0.098689059 | -4.482289416 |
| Gm15636 | 0 | NA | NA | NA |
| Gm16596 | 7.416129088 | 0.061302372 | 0.361210217 | 0.169713839 |
| Evl | 1049.201664 | -0.02566031 | 0.055344324 | -0.463648443 |
| Mir342 | 1.676187767 | -0.405984548 | 0.976175071 | -0.415893174 |
| Degs2 | 4.353830809 | -0.878687537 | 0.710004582 | -1.237580093 |
| Gm23872 | 0 | NA | NA | NA |
| Yy1 | 536.7063009 | -0.057928541 | 0.081913867 | -0.707188455 |
| Slc25a29 | 103.2788333 | -0.341874235 | 0.129879349 | -2.632244755 |
| Mir345 | 1.700378639 | -0.218312878 | 0.838189995 | -0.260457509 |
| Slc25a47 | 16.0241689 | -0.098248503 | 0.308325468 | -0.318651921 |
| Wars | 1271.197342 | 0.342330795 | 0.116614357 | 2.935580194 |
| Gm22079 | 2.165001819 | -0.49445423 | 0.762411431 | -0.648539895 |
| Wdr25 | 96.15949304 | -0.078159787 | 0.108655843 | -0.71933349 |
| Begain | 792.2631617 | -0.204699027 | 0.107792853 | -1.899003693 |
| Gm28498 | 1.537157928 | -0.05558883 | 0.905462385 | -0.061392755 |
| Gm22234 | 0 | NA | NA | NA |
| Gm25297 | 0 | NA | NA | NA |
| Dlk1 | 383.0032136 | -0.043737882 | 0.175524769 | -0.249183533 |
| Gm27528 | 0 | NA | NA | NA |
| Meg3 | 23754.92684 | -0.537567806 | 0.443912151 | -1.210977902 |
| Gm27596 | 7.046539481 | 0.606714625 | 0.512092672 | 1.184775056 |
| Gm27300 | 0 | NA | NA | NA |
| Mir1906-1 | 0 | NA | NA | NA |
| Mir770 | 0 | NA | NA | NA |
| Gm26906 | 0.900331428 | -0.46159304 | 1.277680598 | -0.361274203 |
| Mir673 | 3.122023216 | 0.148067372 | 0.784766943 | 0.188676872 |
| Mir493 | 16.71408191 | -0.321661086 | 0.467974565 | -0.687347369 |
| Mir337 | 13.9759078 | -0.743827577 | 0.336962276 | -2.207450596 |
| mmu-mir-3544 | 0 | NA | NA | NA |
| Mir540 | 13.29036576 | -0.11501573 | 0.414555192 | -0.27744371 |
| Mir665 | 11.41153131 | -0.214094503 | 0.42317147 | -0.505928489 |
| Mir3070a | 19.33383777 | -0.603158552 | 0.429617268 | -1.403943921 |
| Mir3070b | 15.52429422 | -0.745764747 | 0.397179557 | -1.877651391 |
| Rtl1 | 49.79748458 | 1.118435233 | 0.284813109 | 3.926909257 |

| | | | | |
|---|---|---|---|---|
| Mir431 | 8.591852434 | -0.686142058 | 0.4180517 | -1.641285174 |
| Mir433 | 30.4253769 | -0.64105995 | 0.362222877 | -1.769794209 |
| Mir127 | 8.254821574 | -0.509911297 | 0.406919279 | -1.253101838 |
| Mir434 | 6.186359681 | -0.765555604 | 0.423781606 | -1.806486155 |
| Mir432 | 5.368193358 | -0.776784863 | 0.446041823 | -1.741506788 |
| Mir136 | 0 | NA | NA | NA |
| mmu-mir-136 | 4.439960643 | 0.063978411 | 0.515084853 | 0.124209459 |
| Gm27350 | 1.495241998 | -1.020468851 | 0.973854846 | -1.047865455 |
| Rian | 7362.023447 | -0.559593975 | 0.112592484 | -4.970082874 |
| Mir341 | 46.98560136 | -1.231294225 | 0.304354491 | -4.045592432 |
| Mir1188 | 40.87071727 | -1.125447412 | 0.389622849 | -2.888555986 |
| Gm23912 | 5.84972083 | -1.013402761 | 0.581446843 | -1.74289838 |
| Mir370 | 7.676564023 | -0.583093377 | 0.585575496 | -0.99576123 |
| Gm24564 | 32.85825865 | -1.126564793 | 0.336724629 | -3.34565605 |
| Gm23474 | 0 | NA | NA | NA |
| AF357425 | 0 | NA | NA | NA |
| Gm26945 | 332.9786148 | -1.123060239 | 0.20602837 | -5.450998026 |
| Gm24899 | 0 | NA | NA | NA |
| Gm23600 | 67.41334034 | -0.006104467 | 0.168557149 | -0.036216008 |
| Gm24895 | 0 | NA | NA | NA |
| DQ267100 | 0 | NA | NA | NA |
| DQ267101 | 11.01360181 | -0.364459451 | 0.30602302 | -1.190954362 |
| Gm23508 | 0 | NA | NA | NA |
| Gm26922 | 0 | NA | NA | NA |
| DQ267102 | 24.0816795 | -0.716714414 | 0.262093596 | -2.734574307 |
| Gm25357 | 8.037931983 | -0.991809601 | 0.339395709 | -2.92228091 |
| Gm27604 | 0 | NA | NA | NA |
| Gm37899 | 685.7979278 | -0.369779728 | 0.29247411 | -1.264316108 |
| Gm23347 | 0 | NA | NA | NA |
| AF357428 | 5.570026761 | -0.382067653 | 0.543664315 | -0.702763897 |
| Gm23736 | 2.72795131 | -0.107701697 | 0.552448791 | -0.194953268 |
| AF357341 | 4.666739356 | -0.137348315 | 0.588663486 | -0.233322294 |
| Mir882 | 2.263396373 | -0.944875268 | 0.747094029 | -1.264734065 |
| Gm22981 | 1.907638496 | 0.652021206 | 0.689644836 | 0.945444919 |
| Gm25856 | 14.93485102 | -0.883982171 | 0.310384008 | -2.848027437 |
| Gm23787 | 13.95901734 | -0.374848972 | 0.36095301 | -1.038497981 |
| Gm25854 | 10.14198828 | -0.237982022 | 0.354546192 | -0.671229948 |
| Gm22205 | 21.52538062 | -0.430566023 | 0.272992982 | -1.577205461 |
| Gm25224 | 10.64312297 | 0.088349494 | 0.358282771 | 0.246591522 |
| B830012L14Rik | 195.3433677 | -0.467532227 | 0.354432558 | -1.319100678 |
| Mir379 | 4.424652029 | -0.306060456 | 0.616319995 | -0.496593422 |
| Mir411 | 6.496099058 | -0.434432446 | 0.437245041 | -0.993567464 |
| mmu-mir-299a | 2.459243483 | -1.06545948 | 0.702754398 | -1.516119263 |
| Mir299b | 0 | NA | NA | NA |
| Gm25750 | 0 | NA | NA | NA |
| mmu-mir-380 | 4.04787115 | -0.604380599 | 0.541016115 | -1.11712125 |
| Mir1197 | 3.390250948 | -0.255667681 | 0.562348823 | -0.454642511 |

| | | | | |
|---|---|---|---|---|
| Mir323 | 3.255378368 | -0.623825613 | 0.766392659 | -0.813976498 |
| Mir758 | 6.502832914 | -0.523512181 | 0.451797836 | -1.158731051 |
| Mir329 | 5.909624248 | -0.611711001 | 0.525496468 | -1.164063013 |
| Mir494 | 6.573050005 | -0.410488355 | 0.443116993 | -0.926365637 |
| Mir679 | 11.72323245 | -0.234647187 | 0.342723966 | -0.68465357 |
| Mir1193 | 10.23592663 | -0.463574234 | 0.483203526 | -0.959376762 |
| Mir666 | 14.05147067 | -0.383236504 | 0.386824877 | -0.990723522 |
| Mir543 | 4.380404366 | -0.780084662 | 0.645646904 | -1.208221797 |
| Mir495 | 2.202405632 | -0.486669247 | 0.796813508 | -0.610769323 |
| Mir667 | 9.470118255 | -0.564251396 | 0.349735048 | -1.613368175 |
| Mir376b | 10.35390795 | -0.529827428 | 0.403420126 | -1.313339107 |
| Mir654 | 8.477009777 | -0.452095503 | 0.414988427 | -1.089417134 |
| mmu-mir-376b | 2.209145549 | 0.226046448 | 0.689606498 | 0.327790484 |
| Gm25252 | 0.047948698 | -0.206487837 | 3.724945049 | -0.055433794 |
| mmu-mir-376a | 5.585566604 | -1.13279992 | 0.54786698 | -2.067655033 |
| Mir300 | 6.166566501 | -1.020407218 | 0.51060273 | -1.998436666 |
| Mir381 | 6.601995524 | -0.605335579 | 0.50581613 | -1.196750249 |
| Mir487b | 4.545755139 | 0.006784956 | 0.568454319 | 0.011935797 |
| AC121784.1 | 0 | NA | NA | NA |
| Mir539 | 4.506246945 | -1.441366054 | 0.502226337 | -2.86995314 |
| Gm27379 | 8.227541344 | -0.794019713 | 0.489226913 | -1.623009064 |
| Mir544 | 5.245180705 | -0.945629773 | 0.505514896 | -1.870626918 |
| Mirg | 537.7305624 | -0.709307593 | 0.270220433 | -2.62492212 |
| Mir382 | 2.007426773 | -0.682563881 | 0.630166696 | -1.083148134 |
| Mir134 | 0 | NA | NA | NA |
| Mir668 | 0 | NA | NA | NA |
| Mir485 | 0 | NA | NA | NA |
| Mir453 | 14.69478301 | -0.300649689 | 0.416002963 | -0.722710451 |
| Mir154 | 6.301721876 | -0.868756343 | 0.456806219 | -1.90180498 |
| Mir496a | 7.410274305 | -0.658570256 | 0.411277777 | -1.601278487 |
| Mir377 | 0 | NA | NA | NA |
| Mir541 | 8.228173234 | -0.962381829 | 0.394125427 | -2.441816139 |
| mmu-mir-409 | 3.297680665 | -0.611291615 | 0.552433309 | -1.10654373 |
| Gm24321 | 0 | NA | NA | NA |
| Mir412 | 2.651183465 | -0.729895076 | 0.648265419 | -1.125920117 |
| Mir369 | 1.213203356 | -0.699151884 | 0.885116985 | -0.789897715 |
| Mir410 | 0 | NA | NA | NA |
| Mir3072 | 1.399651712 | -1.51830141 | 1.137545982 | -1.334716516 |
| Mir1247 | 0.141331096 | -0.206491164 | 3.56184083 | -0.057973159 |
| Dio3 | 10.88659069 | -0.145114262 | 0.307252737 | -0.472296078 |
| Ppp2r5c | 1467.571291 | -0.064169154 | 0.054102932 | -1.186056873 |
| Dync1h1 | 16373.17254 | 0.07217142 | 0.045909704 | 1.572029742 |
| Gm17111 | 0.611020101 | 0.880709435 | 1.293883297 | 0.680671461 |
| 1700001K19Rik | 0.241252684 | 0.274396479 | 3.054235743 | 0.08984129 |
| Gm27299 | 0.462939631 | -0.582772237 | 1.362477281 | -0.427729875 |
| Hsp90aa1 | 8408.664402 | -0.010514824 | 0.118845682 | -0.088474603 |
| Wdr20 | 372.1537339 | -0.118314385 | 0.097429859 | -1.214354469 |

| | | | | |
|---|---|---|---|---|
| Mok | 93.83344369 | -0.567572908 | 0.128455616 | -4.418435935 |
| 4921507G05Rik | 0.309276477 | -1.427878544 | 1.712991017 | -0.833558687 |
| Gm26912 | 1.084888343 | 0.879820414 | 1.158120156 | 0.759697005 |
| Zfp839 | 119.0822902 | -0.25145367 | 0.136379301 | -1.843781774 |
| Mpc1-ps | 240.9835416 | 0.237801081 | 0.113315231 | 2.098580034 |
| Cinp | 250.286756 | -0.11421665 | 0.088483413 | -1.290825547 |
| Tecpr2 | 926.178597 | -0.163152044 | 0.078565712 | -2.076631649 |
| Rps19-ps6 | 18.4757316 | 0.004053293 | 0.261174056 | 0.015519509 |
| Ankrd9 | 131.61924 | -0.169380286 | 0.097910049 | -1.729958136 |
| Rcor1 | 394.3624715 | -0.254207812 | 0.090674566 | -2.803518376 |
| 4930595D18Rik | 0.920001838 | 0.88566056 | 1.055272059 | 0.839272254 |
| Traf3 | 916.582985 | 0.010195837 | 0.06205548 | 0.164301967 |
| Amn | 31.33882718 | 0.007754732 | 0.204900051 | 0.037846413 |
| A230087F16Rik | 11.27729028 | -0.996782167 | 0.355417347 | -2.804539998 |
| Cdc42bpb | 2200.324733 | 0.087724561 | 0.063767698 | 1.37568964 |
| A230065H16Rik | 14.31749866 | -0.405752395 | 0.352015445 | -1.152655089 |
| Exoc3l4 | 26.39184743 | 0.580598024 | 0.273001045 | 2.126724545 |
| Gm10425 | 4.124014502 | 3.319615647 | 1.034664186 | 3.208399103 |
| Tnfaip2 | 430.9565934 | 6.435466949 | 0.585738592 | 10.98692665 |
| Gm266 | 27.84625617 | -0.129456813 | 0.237026107 | -0.546171115 |
| Eif5 | 3419.805663 | 0.500847284 | 0.107162247 | 4.673728858 |
| Snora28 | 29.94917886 | -0.109472468 | 0.225662302 | -0.485116331 |
| 2810029C07Rik | 61.53833547 | 0.139816947 | 0.184368047 | 0.758357803 |
| Mark3 | 1177.712359 | 0.11525628 | 0.071421805 | 1.613740786 |
| Ckb | 9080.847841 | -0.238130807 | 0.092989739 | -2.560828864 |
| Trmt61a | 154.0841474 | 0.10309778 | 0.107028181 | 0.963276951 |
| Bag5 | 421.2921291 | 0.179065671 | 0.073700339 | 2.429645157 |
| Apopt1 | 168.4724147 | -0.146181962 | 0.1026165 | -1.424546358 |
| Gm15995 | 0.042716945 | -0.206490061 | 3.724945049 | -0.055434391 |
| Gm15996 | 0.04393175 | -0.206490061 | 3.724945049 | -0.055434391 |
| Klc1 | 4978.643089 | -0.163780615 | 0.075640703 | -2.16524449 |
| Xrcc3 | 62.34707954 | -0.407373585 | 0.166554968 | -2.445880717 |
| Zfyve21 | 80.73079552 | -0.155765493 | 0.156183272 | -0.997325071 |
| Ppp1r13b | 928.8327789 | 0.121456989 | 0.075003787 | 1.619344749 |
| 2010107E04Rik | 287.253024 | -0.213458874 | 0.164198547 | -1.300004645 |
| Tdrd9 | 6.653230718 | 0.276882633 | 0.453825453 | 0.610108207 |
| Rd3l | 8.12228703 | -0.026509212 | 0.458587079 | -0.057806277 |
| Aspg | 140.2133012 | 2.435454545 | 0.212138244 | 11.48050674 |
| Mir3073a | 0.372180926 | 1.212392954 | 1.599605118 | 0.757932655 |
| mmu-mir-3073b | 0 | NA | NA | NA |
| Mir203 | 0 | NA | NA | NA |
| Kif26a | 86.52596024 | 0.055560353 | 0.161730629 | 0.343536368 |
| Gm38123 | 0.492632247 | 0.737114831 | 1.54898083 | 0.475870855 |
| A730018C14Rik | 2.250253578 | -0.111773193 | 0.756057077 | -0.147836978 |
| A530016L24Rik | 0 | NA | NA | NA |
| Tmem179 | 794.5266479 | -0.063261656 | 0.061068173 | -1.035918606 |
| Inf2 | 1281.719595 | -0.329775031 | 0.085429558 | -3.860198258 |

| | | | | |
|---|---|---|---|---|
| Adssl1 | 268.3142132 | 0.17731105 | 0.112252439 | 1.579574149 |
| Siva1 | 44.47611933 | 0.231379587 | 0.203206415 | 1.138643121 |
| Akt1 | 1539.989558 | 0.119807335 | 0.072757838 | 1.646658827 |
| Mir6939 | 0.990373052 | 0.712143145 | 0.985909646 | 0.722320902 |
| Zbtb42 | 29.83589809 | 1.956080967 | 0.29729856 | 6.579517124 |
| Gm30238 | 24.73544283 | -0.570467284 | 0.350409148 | -1.6280034 |
| Cep170b | 3968.143558 | -0.138536591 | 0.076270022 | -1.816396381 |
| Mir6940 | 0.393711421 | -0.367493015 | 1.668424922 | -0.220263441 |
| Pld4 | 304.2279336 | 2.904687683 | 0.160009524 | 18.15321742 |
| Ahnak2 | 140.5418072 | -0.044431234 | 0.25094591 | -0.177055024 |
| BC022687 | 106.0209119 | 0.326309359 | 0.131845648 | 2.474934623 |
| Cdca4 | 92.32272779 | 0.645743073 | 0.159712394 | 4.043161934 |
| Gpr132 | 12.16430624 | 5.492992386 | 0.892836069 | 6.152296684 |
| Jag2 | 271.294481 | -0.13778391 | 0.115387371 | -1.194098698 |
| Mir6941 | 0.653922745 | -1.635306555 | 1.547621897 | -1.056657675 |
| Nudt14 | 37.15232537 | -0.187316931 | 0.192404289 | -0.973559023 |
| Gm26583 | 4.273896503 | 1.682821555 | 0.731091986 | 2.301791823 |
| Brf1 | 375.6428438 | 0.121454185 | 0.076782434 | 1.58179649 |
| Btbd6 | 332.183834 | -0.076894411 | 0.068418759 | -1.123879061 |
| 9230104M06Rik | 1.025204639 | -1.364686952 | 1.199588779 | -1.137628974 |
| Pacs2 | 2506.266713 | 0.180119641 | 0.092674386 | 1.943575233 |
| AC073562.1 | 0.303558719 | 0.274410045 | 2.066409255 | 0.132795594 |
| Tex22 | 3.833641248 | -0.61068855 | 0.518929616 | -1.176823467 |
| Mta1 | 765.704057 | -0.034358806 | 0.059082675 | -0.581537754 |
| Crip2 | 531.6473017 | -0.085476384 | 0.101757488 | -0.840000927 |
| Crip1 | 42.16258074 | 1.024568903 | 0.272338213 | 3.762119508 |
| 4930427A07Rik | 20.00319577 | 0.378778337 | 0.270337023 | 1.401133789 |
| RP24-468O24.5 | 0.060174537 | 0.755286614 | 3.724945049 | 0.202764498 |
| Tmem121 | 107.667371 | -0.00881039 | 0.116493841 | -0.075629665 |
| Gm25622 | 0 | NA | NA | NA |
| Igha | 7.387870444 | 3.458892879 | 1.87284654 | 1.846864014 |
| Ighe | 0.136120942 | 0.274393632 | 3.599844381 | 0.076223748 |
| Ighg2c | 6.374334057 | 1.859459435 | 1.248638503 | 1.48918957 |
| Ighg2b | 4.144711669 | 4.118342931 | 1.524727772 | 2.701034903 |
| Gm37944 | 0.048949532 | -0.206487837 | 3.724945049 | -0.055433794 |
| Ighg1 | 1.048355144 | 1.029510078 | 2.272589011 | 0.453011993 |
| Ighg3 | 2.594242155 | 4.315621349 | 1.814189503 | 2.378815081 |
| Gm17421 | 0.038981395 | 0.274393828 | 3.724945049 | 0.073663859 |
| Gm9134 | 2.929093634 | 1.205283129 | 0.666922084 | 1.807232295 |
| Gm9136 | 0 | NA | NA | NA |
| Gm5190 | 0.038981395 | 0.274393828 | 3.724945049 | 0.073663859 |
| Ighd | 2.086187063 | 0.135756893 | 0.687936556 | 0.197339263 |
| Ighm | 86.67814311 | 0.075806124 | 0.446499598 | 0.169778705 |
| Gm25747 | 0 | NA | NA | NA |
| Gm24022 | 0 | NA | NA | NA |
| AC073553.1 | 0 | NA | NA | NA |
| Gm22175 | 0 | NA | NA | NA |

| | | | | |
|---|---|---|---|---|
| Ighj4 | 0.495312865 | 0.697267679 | 1.693110881 | 0.411826353 |
| Ighj3 | 0.07796279 | 0.274392189 | 3.724945049 | 0.073663419 |
| Ighj2 | 0.566709933 | 0.802843276 | 1.386482455 | 0.579050441 |
| Ighj1 | 0.937655582 | 0.01237045 | 1.026979654 | 0.012045468 |
| Ighd4-1 | 0 | NA | NA | NA |
| Ighd3-2 | 0 | NA | NA | NA |
| Ighd5-6 | 0 | NA | NA | NA |
| Ighd2-8 | 0 | NA | NA | NA |
| Ighd5-5 | 0 | NA | NA | NA |
| Ighd2-7 | 0 | NA | NA | NA |
| Ighd5-8 | 0 | NA | NA | NA |
| Ighd5-4 | 0 | NA | NA | NA |
| Ighd2-6 | 0 | NA | NA | NA |
| Ighd5-7 | 0 | NA | NA | NA |
| Ighd5-3 | 0 | NA | NA | NA |
| Ighd2-5 | 0 | NA | NA | NA |
| Gm37327 | 0 | NA | NA | NA |
| Ighd5-2 | 0 | NA | NA | NA |
| Ighd2-4 | 0 | NA | NA | NA |
| Ighd6-2 | 0 | NA | NA | NA |
| Gm37227 | 0 | NA | NA | NA |
| Ighd2-3 | 0 | NA | NA | NA |
| Ighd6-1 | 0 | NA | NA | NA |
| Ighd1-1 | 0 | NA | NA | NA |
| Adam6b | 0.203125505 | -0.59598799 | 2.619597531 | -0.227511281 |
| Ighd3-1 | 0 | NA | NA | NA |
| Ighd5-1 | 0 | NA | NA | NA |
| Gm16968 | 0 | NA | NA | NA |
| Adam6a | 0 | NA | NA | NA |
| Ighv5-1 | 0 | NA | NA | NA |
| Ighv2-1 | 0 | NA | NA | NA |
| Ighv5-2 | 0 | NA | NA | NA |
| Ighv2-2 | 0 | NA | NA | NA |
| Ighv5-3 | 0 | NA | NA | NA |
| Ighv5-4 | 0.345657073 | 0.274408291 | 1.986933007 | 0.138106463 |
| Ighv6-1 | 0 | NA | NA | NA |
| Ighv2-3 | 0.072025462 | 0.755288839 | 3.724945049 | 0.202765096 |
| Ighv5-5 | 0 | NA | NA | NA |
| Ighv5-6 | 0.066050875 | 0.274391604 | 3.724945049 | 0.073663262 |
| Gm9517 | 109.6899113 | -0.062282052 | 0.132516067 | -0.469996228 |
| Ighv5-7 | 0 | NA | NA | NA |
| Ighv2-4 | 0 | NA | NA | NA |
| Ighv5-8 | 0 | NA | NA | NA |
| Ighv5-9 | 0.097139547 | 0.274393828 | 3.724945049 | 0.073663859 |
| Ighv5-10 | 0 | NA | NA | NA |
| Gm37187 | 0 | NA | NA | NA |
| Ighv2-5 | 0 | NA | NA | NA |

| | | | | |
|---|---|---|---|---|
| Ighv5-11 | 0 | NA | NA | NA |
| Ighv5-12 | 0 | NA | NA | NA |
| Ighv2-6 | 0.060174537 | 0.755286614 | 3.724945049 | 0.202764498 |
| Gm37976 | 0 | NA | NA | NA |
| Gm37418 | 0 | NA | NA | NA |
| Ighv5-9-1 | 0 | NA | NA | NA |
| Gm38203 | 0 | NA | NA | NA |
| Ighv5-12-4 | 0 | NA | NA | NA |
| Gm38205 | 0 | NA | NA | NA |
| Ighv2-9-1 | 0 | NA | NA | NA |
| Gm37722 | 0 | NA | NA | NA |
| Ighv5-13 | 0 | NA | NA | NA |
| Ighv2-6-8 | 0.046345709 | 0.274391604 | 3.724945049 | 0.073663262 |
| Gm7003 | 0 | NA | NA | NA |
| Ighv2-7 | 0 | NA | NA | NA |
| Rpl26-ps3 | 0 | NA | NA | NA |
| Ighv5-15 | 0 | NA | NA | NA |
| Ighv5-16 | 0 | NA | NA | NA |
| Ighv5-17 | 0.427539584 | 0.274386301 | 3.724945049 | 0.073661839 |
| Ighv5-18 | 0 | NA | NA | NA |
| Ighv2-8 | 0 | NA | NA | NA |
| Ighv2-9 | 0.631149539 | 0.274375943 | 3.724945049 | 0.073659058 |
| Ighv5-19 | 0 | NA | NA | NA |
| Ighv7-1 | 0 | NA | NA | NA |
| Ighv7-2 | 0 | NA | NA | NA |
| Ighv14-1 | 0.480014214 | 1.823486035 | 3.702717981 | 0.492472298 |
| Ighv4-1 | 0.072025462 | 0.755288839 | 3.724945049 | 0.202765096 |
| Ighv3-1 | 0 | NA | NA | NA |
| Gm37961 | 0 | NA | NA | NA |
| Ighv11-1 | 0 | NA | NA | NA |
| Ighv14-2 | 0.20773507 | 0.755299473 | 2.580838766 | 0.29265659 |
| Ighv4-2 | 0 | NA | NA | NA |
| Ighv3-2 | 0 | NA | NA | NA |
| Ighv11-2 | 0 | NA | NA | NA |
| Ighv14-3 | 0 | NA | NA | NA |
| Ighv16-1 | 0 | NA | NA | NA |
| Ighv6-2 | 0 | NA | NA | NA |
| Ighv9-1 | 0 | NA | NA | NA |
| Ighv12-1 | 0 | NA | NA | NA |
| Ighv9-2 | 0 | NA | NA | NA |
| Ighv12-2 | 0 | NA | NA | NA |
| Ighv9-3 | 0 | NA | NA | NA |
| Ighv7-3 | 0 | NA | NA | NA |
| Ighv15-1 | 0 | NA | NA | NA |
| Ighv14-4 | 0 | NA | NA | NA |
| Gm37838 | 0 | NA | NA | NA |
| Ighv3-3 | 0 | NA | NA | NA |

| | | | | | |
|---|---|---|---|---|---|
| Ighv7-4 | 0 | NA | NA | NA | |
| Ighv3-4 | 0 | NA | NA | NA | |
| Ighv3-5 | 0 | NA | NA | NA | |
| Gm18551 | 0 | NA | NA | NA | |
| Ighv13-1 | 0 | NA | NA | NA | |
| Ighv3-6 | 0.625300531 | 1.354002705 | | 3.714419835 | 0.364526027 |
| Ighv9-4 | 0 | NA | NA | NA | |
| Ighv3-7 | 0 | NA | NA | NA | |
| Ighv5-21 | 0 | NA | NA | NA | |
| Ighv3-8 | 0.038981395 | 0.274393828 | | 3.724945049 | 0.073663859 |
| Ighv8-1 | 0 | NA | NA | NA | |
| Ighv1-1 | 0 | NA | NA | NA | |
| Ighv13-2 | 0 | NA | NA | NA | |
| Ighv12-3 | 0 | NA | NA | NA | |
| Ighv6-3 | 0 | NA | NA | NA | |
| Ighv6-4 | 0 | NA | NA | NA | |
| Ighv6-5 | 0 | NA | NA | NA | |
| Ighv6-6 | 0.211582616 | -0.206486648 | | 2.578619295 | -0.080076438 |
| Ighv6-7 | 0 | NA | NA | NA | |
| Ighv8-2 | 0 | NA | NA | NA | |
| Ighv1-2 | 0 | NA | NA | NA | |
| mmu-mir-7094-1 | 0 | NA | NA | NA | |
| Ighv10-1 | 0.049553047 | 0.274391604 | | 3.724945049 | 0.073663262 |
| Ighv1-3 | 0 | NA | NA | NA | |
| Ighv1-4 | 0 | NA | NA | NA | |
| Ighv10-2 | 0 | NA | NA | NA | |
| Ighv1-5 | 0 | NA | NA | NA | |
| Mir7094-2 | 0 | NA | NA | NA | |
| Ighv10-3 | 0 | NA | NA | NA | |
| Ighv1-6 | 0 | NA | NA | NA | |
| Ighv1-7 | 0 | NA | NA | NA | |
| Ighv10-4 | 0 | NA | NA | NA | |
| Ighv1-8 | 0 | NA | NA | NA | |
| Ighv15-2 | 0 | NA | NA | NA | |
| Ighv1-9 | 0.287786674 | 0.755304973 | | 2.026686947 | 0.372679646 |
| Ighv1-10 | 0 | NA | NA | NA | |
| Ighv1-11 | 0 | NA | NA | NA | |
| Ighv1-12 | 0 | NA | NA | NA | |
| Ighv1-13 | 0 | NA | NA | NA | |
| Ighv1-14 | 0 | NA | NA | NA | |
| Ighv1-15 | 0 | NA | NA | NA | |
| Ighv1-16 | 0 | NA | NA | NA | |
| Ighv1-17-1 | 0 | NA | NA | NA | |
| Ighv1-17 | 0 | NA | NA | NA | |
| Ighv1-18 | 0.194279094 | 0.274391173 | | 3.724945049 | 0.073663147 |
| Ighv1-19-1 | 0 | NA | NA | NA | |
| Ighv1-19 | 0 | NA | NA | NA | |

| | | | | |
|---|---|---|---|---|
| Ighv1-20 | 0 | NA | NA | NA |
| Ighv1-21-1 | 0 | NA | NA | NA |
| Ighv1-21 | 0 | NA | NA | NA |
| Ighv1-22 | 0 | NA | NA | NA |
| Ighv1-23 | 0.097139547 | 0.274393828 | 3.724945049 | 0.073663859 |
| Ighv1-24 | 0 | NA | NA | NA |
| Ighv1-25 | 0 | NA | NA | NA |
| Ighv1-26 | 0 | NA | NA | NA |
| Ighv1-27 | 0 | NA | NA | NA |
| Ighv1-28 | 0 | NA | NA | NA |
| Ighv1-29 | 0 | NA | NA | NA |
| Ighv1-30 | 0 | NA | NA | NA |
| Ighv1-31 | 0 | NA | NA | NA |
| Ighv1-32 | 0 | NA | NA | NA |
| Ighv1-33 | 0.097139547 | 0.274393828 | 3.724945049 | 0.073663859 |
| Ighv1-34 | 0 | NA | NA | NA |
| Ighv1-35 | 0 | NA | NA | NA |
| Ighv1-36 | 0 | NA | NA | NA |
| Ighv1-37 | 0 | NA | NA | NA |
| Ighv1-38 | 0 | NA | NA | NA |
| Ighv1-39 | 0 | NA | NA | NA |
| Ighv1-40 | 0 | NA | NA | NA |
| Ighv1-41 | 0 | NA | NA | NA |
| Ighv1-42 | 0 | NA | NA | NA |
| Ighv1-43 | 0 | NA | NA | NA |
| Ighv1-44 | 0 | NA | NA | NA |
| Ighv1-45 | 0 | NA | NA | NA |
| Ighv1-46 | 0 | NA | NA | NA |
| Ighv1-47 | 0 | NA | NA | NA |
| Ighv8-3 | 0 | NA | NA | NA |
| Ighv8-4 | 0 | NA | NA | NA |
| Ighv1-48 | 0 | NA | NA | NA |
| Ighv1-49 | 0 | NA | NA | NA |
| Ighv8-5 | 0 | NA | NA | NA |
| Ighv1-50 | 0 | NA | NA | NA |
| Ighv1-51 | 0 | NA | NA | NA |
| Ighv1-52 | 0.072025462 | 0.755288839 | 3.724945049 | 0.202765096 |
| Ighv1-53 | 0 | NA | NA | NA |
| Ighv8-6 | 0 | NA | NA | NA |
| Gm9227 | 0 | NA | NA | NA |
| Ighv1-54 | 0 | NA | NA | NA |
| Ighv1-55 | 0 | NA | NA | NA |
| Gm7142 | 0 | NA | NA | NA |
| Ighv1-56 | 0 | NA | NA | NA |
| Gm7019 | 1.468020652 | 3.434427636 | 1.073920721 | 3.198027162 |
| Ighv8-7 | 0 | NA | NA | NA |
| Gm30948 | 0 | NA | NA | NA |

| | | | | |
|---|---|---|---|---|
| Ighv1-57 | 0 | NA | NA | NA |
| Ighv8-8 | 0.144050924 | 1.290908285 | 3.721447108 | 0.346883416 |
| Ighv1-58 | 0 | NA | NA | NA |
| Ighv1-59 | 0 | NA | NA | NA |
| Ighv1-60 | 0 | NA | NA | NA |
| Ighv1-61 | 0 | NA | NA | NA |
| Ighv1-62 | 0 | NA | NA | NA |
| Ighv1-62-1 | 0 | NA | NA | NA |
| Gm19331 | 0 | NA | NA | NA |
| Gm37511 | 0 | NA | NA | NA |
| Gm38184 | 0 | NA | NA | NA |
| Ighv1-62-2 | 0 | NA | NA | NA |
| Ighv1-62-3 | 0 | NA | NA | NA |
| Ighv8-9 | 0 | NA | NA | NA |
| RP24-350F17.8 | 0 | NA | NA | NA |
| Ighv1-63 | 0.046345709 | 0.274391604 | 3.724945049 | 0.073663262 |
| Gm27722 | 0 | NA | NA | NA |
| Ighv1-64 | 0 | NA | NA | NA |
| Ighv8-10 | 0 | NA | NA | NA |
| RP23-218B2.1 | 0 | NA | NA | NA |
| Ighv1-65 | 0 | NA | NA | NA |
| RP23-218B2.3 | 0.230454321 | -0.206482933 | 2.186279641 | -0.094444887 |
| Ighv8-11 | 0.066050875 | 0.274391604 | 3.724945049 | 0.073663262 |
| RP23-218B2.5 | 0 | NA | NA | NA |
| Ighv1-66 | 0 | NA | NA | NA |
| Ighv1-67 | 0.060174537 | 0.755286614 | 3.724945049 | 0.202764498 |
| Ighv1-68 | 0 | NA | NA | NA |
| RP23-218B2.9 | 0 | NA | NA | NA |
| RP23-218B2.10 | 0 | NA | NA | NA |
| Ighv1-69 | 0 | NA | NA | NA |
| Ighv8-12 | 0 | NA | NA | NA |
| RP23-218B2.13 | 0 | NA | NA | NA |
| Ighv1-70 | 0 | NA | NA | NA |
| Mir6388 | 0 | NA | NA | NA |
| Ighv1-71 | 0 | NA | NA | NA |
| Ighv1-72 | 0 | NA | NA | NA |
| Ighv8-13 | 0 | NA | NA | NA |
| RP23-123N21.1 | 0 | NA | NA | NA |
| Ighv1-73 | 0 | NA | NA | NA |
| Ighv1-74 | 0.136120942 | 0.274393632 | 3.599844381 | 0.076223748 |
| Ighv8-14 | 0 | NA | NA | NA |
| RP23-123N21.5 | 0 | NA | NA | NA |
| RP23-123N21.6 | 0 | NA | NA | NA |
| Ighv1-75 | 0 | NA | NA | NA |
| Ighv8-15 | 0 | NA | NA | NA |
| Ighv1-76 | 0 | NA | NA | NA |
| Ighv8-16 | 0 | NA | NA | NA |

| | | | | |
|---|---|---|---|---|
| Ighv1-77 | 0.046345709 | 0.274391604 | 3.724945049 | 0.073663262 |
| Ighv1-78 | 0 | NA | NA | NA |
| RP23-172K2.2 | 0 | NA | NA | NA |
| RP23-172K2.1 | 0 | NA | NA | NA |
| Ighv1-79 | 0 | NA | NA | NA |
| RP23-172K2.4 | 2.956099666 | -0.199436639 | 0.653957652 | -0.304968737 |
| Ighv1-80 | 0 | NA | NA | NA |
| Ighv1-81 | 0 | NA | NA | NA |
| RP23-172K2.8 | 0 | NA | NA | NA |
| Olfr241-ps1 | 0 | NA | NA | NA |
| Ighv1-82 | 0 | NA | NA | NA |
| Ighv1-83 | 0 | NA | NA | NA |
| Ighv1-84 | 0 | NA | NA | NA |
| RP23-172K2.12 | 5.084652238 | 0.914371303 | 0.583185212 | 1.567891785 |
| Ighv1-85 | 0.072025462 | 0.755288839 | 3.724945049 | 0.202765096 |
| Ighv1-86 | 0 | NA | NA | NA |
| Zfp386 | 523.1915969 | 0.262978593 | 0.082171337 | 3.200368903 |
| Vipr2 | 33.917483 | -0.593430745 | 0.240833583 | -2.46406974 |
| Gm20658 | 0.517478627 | 1.872281989 | 1.345890353 | 1.391110342 |
| AC105950.1 | 0 | NA | NA | NA |
| Gm23732 | 0 | NA | NA | NA |
| Wdr60 | 178.5716657 | -0.26810544 | 0.109935951 | -2.438742167 |
| Gm23190 | 0.086648695 | -0.687376175 | 3.724945049 | -0.184533239 |
| Gm11027 | 0 | NA | NA | NA |
| Esyt2 | 1140.517075 | -0.00584965 | 0.054962134 | -0.106430551 |
| Ncapg2 | 136.9875684 | 0.734551998 | 0.174921141 | 4.1993323 |
| Gm25112 | 0.475792389 | 0.571930434 | 1.419464451 | 0.402919871 |
| Gm27649 | 0.04393175 | -0.206490061 | 3.724945049 | -0.055434391 |
| Ptprn2 | 2936.458257 | -0.026928163 | 0.098835103 | -0.272455458 |
| Gm6750 | 0.877833841 | 0.250594034 | 1.188462535 | 0.210855645 |
| Gm17807 | 4.466807706 | -1.34004921 | 0.514093156 | -2.60662721 |
| Gm10421 | 10.74823256 | 0.143859335 | 0.321183921 | 0.44790329 |
| Gm5441 | 1.251482101 | -0.386296203 | 0.849570888 | -0.454695669 |
| Mir153 | 1.154465371 | -0.340676048 | 0.879798187 | -0.387220676 |
| Rapgef5 | 853.7827578 | -0.204037954 | 0.09667498 | -2.110555958 |
| Gm24741 | 1.59128742 | 0.705276787 | 1.059688239 | 0.665551208 |
| Cdca7l | 18.95984674 | 0.176389958 | 0.315406593 | 0.559246263 |
| Gm20613 | 0.046345709 | 0.274391604 | 3.724945049 | 0.073663262 |
| Dnah11 | 127.3083906 | -0.876830746 | 0.148518773 | -5.903837792 |
| Sp4 | 671.9126227 | 0.073583484 | 0.096412887 | 0.763212123 |
| Gm25577 | 0.046345709 | 0.274391604 | 3.724945049 | 0.073663262 |
| Gm22537 | 0 | NA | NA | NA |
| Sp8 | 103.1593826 | -0.926996165 | 0.574867974 | -1.612537497 |
| Abcb5 | 0 | NA | NA | NA |
| Itgb8 | 2067.75423 | -0.009377087 | 0.147185084 | -0.063709495 |
| Gm6768 | 116.6792349 | 0.16819395 | 0.108854894 | 1.545120704 |
| Macc1 | 2.369395584 | -0.230270554 | 0.698168875 | -0.32982071 |

| | | | | |
|---|---|---|---|---|
| Gm25675 | 0 | NA | NA | NA |
| Tmem196 | 135.2367115 | -0.223102888 | 0.102460518 | -2.177452269 |
| Gm27236 | 2.533080183 | -0.824665076 | 0.654304512 | -1.260368928 |
| Gm17177 | 0 | NA | NA | NA |
| Speer6-ps1 | 0 | NA | NA | NA |
| Gm16505 | 1.0973389 | -0.041224564 | 0.999536268 | -0.04124369 |
| 2810429I04Rik | 28.4214383 | -0.434464965 | 0.199276541 | -2.180211296 |
| Gdi2 | 2568.627456 | 0.15831666 | 0.081946646 | 1.931948018 |
| Fam208b | 1293.848084 | 0.641130157 | 0.092666759 | 6.918663843 |
| Asb13 | 641.9525158 | 0.389354907 | 0.115304018 | 3.37676789 |
| Gm23084 | 0.981575226 | -0.128583036 | 1.20614167 | -0.10660691 |
| Calml3 | 0.554260052 | 2.65137649 | 3.685127339 | 0.719480291 |
| Calm4 | 0.780001735 | 3.036520797 | 3.676122532 | 0.826011856 |
| Calm5 | 0.045882455 | 0.755288839 | 3.724945049 | 0.202765096 |
| AC132239.1 | 0 | NA | NA | NA |
| Net1 | 365.646222 | 0.451681949 | 0.090925914 | 4.967582149 |
| Tubal3 | 0.038981395 | 0.274393828 | 3.724945049 | 0.073663859 |
| Ucn3 | 2.74335279 | 0.334886322 | 0.646708533 | 0.51783192 |
| Gm22460 | 0 | NA | NA | NA |
| Akr1c14 | 23.64608385 | -0.077157428 | 0.381717287 | -0.202132392 |
| Akr1c18 | 8.892648978 | -0.05507097 | 0.351639719 | -0.156611915 |
| Akr1c13 | 6.039600961 | 1.504913974 | 0.599361856 | 2.510860441 |
| Akr1c19 | 0.060174537 | 0.755286614 | 3.724945049 | 0.202764498 |
| Marcksl1-ps4 | 0 | NA | NA | NA |
| Akr1c12 | 0.642786378 | 1.644621764 | 1.414133435 | 1.162989096 |
| Akr1c6 | 0.085327104 | 0.274389469 | 3.724945049 | 0.073662689 |
| Akr1c20 | 0.875592853 | 2.242903763 | 1.562280942 | 1.435659683 |
| Akr1c21 | 0 | NA | NA | NA |
| Akr1e1 | 194.9019369 | -0.293165682 | 0.117784222 | -2.489006404 |
| Rpl29-ps2 | 3.103436511 | -0.441041847 | 0.584564382 | -0.754479508 |
| Gm5444 | 0.134148234 | -0.20649159 | 3.624632408 | -0.056968974 |
| Gm24790 | 0 | NA | NA | NA |
| Gm26043 | 0 | NA | NA | NA |
| 1700016G22Rik | 0.413724333 | 0.755307629 | 1.690033359 | 0.44691877 |
| Klf6 | 1264.230787 | 1.204351237 | 0.170787875 | 7.051737363 |
| Gm26259 | 0 | NA | NA | NA |
| Gm28465 | 0 | NA | NA | NA |
| Pitrm1 | 461.3651909 | -0.050220401 | 0.074718427 | -0.672128726 |
| Pfkp | 2522.968723 | -0.20757122 | 0.074332592 | -2.792465773 |
| Gm10029 | 6.775184329 | 1.119398902 | 0.431260675 | 2.595643346 |
| Gm22689 | 0 | NA | NA | NA |
| Gm9742 | 0.283836568 | 0.274409939 | 2.104902764 | 0.130367038 |
| Adarb2 | 565.2457729 | -0.32194378 | 0.071258537 | -4.517967859 |
| Gm10357 | 4.418419051 | 1.167964178 | 0.492521323 | 2.3713982 |
| Wdr37 | 1368.014662 | -0.029084635 | 0.071554465 | -0.40646849 |
| Idi1 | 182.2969244 | -0.389248442 | 0.17937195 | -2.17006305 |
| Gm22750 | 0.171083144 | 0.27438982 | 3.398950499 | 0.080727807 |

| | | | | |
|---|---|---|---|---|
| Rpl10a-ps2 | 0.785454149 | 1.604633317 | 1.264221508 | 1.269265953 |
| Gm9745 | 0.104637558 | -0.206492286 | 3.724945049 | -0.055434988 |
| Gm25274 | 0.066050875 | 0.274391604 | 3.724945049 | 0.073663262 |
| Gm22115 | 0.115603922 | 0.274387056 | 3.724945049 | 0.073662041 |
| Idi2 | 0.326649557 | -0.670149464 | 1.500963856 | -0.446479415 |
| Gm23923 | 0.066050875 | 0.274391604 | 3.724945049 | 0.073663262 |
| Gtpbp4 | 492.4961443 | 0.313042608 | 0.114860586 | 2.725413643 |
| Larp4b | 1451.804564 | 0.023373436 | 0.05298288 | 0.441150728 |
| Gm23553 | 10.68005917 | -0.119660809 | 0.373293263 | -0.32055443 |
| Gm28155 | 0 | NA | NA | NA |
| Dip2c | 2087.5135 | 0.076277241 | 0.07156153 | 1.065897282 |
| Gm26601 | 155.8395443 | -0.553027449 | 0.387330934 | -1.427790554 |
| Gm23496 | 0.048949532 | -0.206487837 | 3.724945049 | -0.055433794 |
| Zmynd11 | 1907.336184 | -0.105339493 | 0.075347059 | -1.39805713 |
| Gm24187 | 251.7405501 | 0.494827243 | 0.98018866 | 0.504828574 |
| Chrm3 | 417.2410317 | -0.122696971 | 0.106177707 | -1.15558128 |
| Gm26861 | 0.849695755 | -0.516823378 | 1.071306966 | -0.48242324 |
| Gm23415 | 0 | NA | NA | NA |
| Gm25496 | 0 | NA | NA | NA |
| Ryr2 | 4975.297084 | -0.295193014 | 0.152746099 | -1.932573181 |
| Gm26956 | 0.116944185 | 0.274391125 | 3.724945049 | 0.073663134 |
| Mtr | 314.5275728 | -0.195678051 | 0.116461823 | -1.680190514 |
| Actn2 | 215.0662038 | -0.338719105 | 0.202219516 | -1.675007003 |
| Gm5445 | 1.336133173 | -0.416015593 | 0.851905082 | -0.488335615 |
| Heatr1 | 496.059456 | -0.188116878 | 0.09140384 | -2.058085057 |
| Lgals8 | 902.3953984 | 1.305975964 | 0.14231011 | 9.176972517 |
| Edaradd | 6.858033054 | -0.127696228 | 0.380494168 | -0.33560627 |
| Ero1lb | 286.3774935 | -0.238745365 | 0.086550683 | -2.758445761 |
| Gm25483 | 3.19383198 | 0.914133359 | 0.610764808 | 1.496702737 |
| Gpr137b-ps | 100.0559785 | 0.179776067 | 0.155655604 | 1.154960453 |
| Gm24065 | 0 | NA | NA | NA |
| Gm2399 | 0.304316145 | 0.252318154 | 1.601979446 | 0.15750399 |
| Prl2c3 | 1.422091401 | -1.123495038 | 0.824316008 | -1.362942157 |
| Prl2c2 | 0.494725653 | -1.475501929 | 1.503802264 | -0.981180813 |
| Prl2c5 | 0.049553047 | 0.274391604 | 3.724945049 | 0.073663262 |
| Gm2423 | 35.43526429 | 0.031784343 | 0.251937594 | 0.126159587 |
| Gm26133 | 0 | NA | NA | NA |
| Gm25804 | 0 | NA | NA | NA |
| Gpr137b | 160.9115362 | -0.210494707 | 0.115984047 | -1.814859136 |
| Gm23081 | 0 | NA | NA | NA |
| Gm7426 | 0 | NA | NA | NA |
| Nid1 | 422.1371149 | 0.469508192 | 0.146580219 | 3.203080175 |
| Lyst | 2594.837581 | 0.218596089 | 0.100974881 | 2.164856118 |
| Gng4 | 505.6373642 | -0.833439633 | 0.253902394 | -3.282519784 |
| Gm25271 | 3.698276596 | -0.696208965 | 0.627120209 | -1.110168281 |
| B3galnt2 | 227.2140593 | -0.214366382 | 0.108829232 | -1.969750023 |
| Gm18856 | 1.228493641 | 3.302218963 | 1.095579378 | 3.014130266 |

| | | | | |
|---|---|---|---|---|
| Tbce | 525.6659255 | 0.017439399 | 0.103180192 | 0.169018866 |
| Ggps1 | 552.9079473 | 0.149999319 | 0.081120347 | 1.849096115 |
| Arid4b | 1466.024357 | 0.135558895 | 0.08071082 | 1.679562861 |
| Gm26129 | 7.167244977 | -0.348794325 | 0.377708166 | -0.923449257 |
| n-R5s52 | 0 | NA | NA | NA |
| Hecw1 | 1632.790211 | -0.304555475 | 0.096966949 | -3.140817346 |
| Mrpl32 | 170.5037384 | 0.339777759 | 0.115683515 | 2.937132052 |
| Psma2 | 925.2712897 | 0.605294113 | 0.114034215 | 5.308004371 |
| Gm25444 | 0.046345709 | 0.274391604 | 3.724945049 | 0.073663262 |
| AW209491 | 188.160879 | -0.450904718 | 0.146873202 | -3.070027156 |
| Gm6556 | 34.49633554 | 0.692728138 | 0.216693309 | 3.196813702 |
| Gm26645 | 23.14899875 | 0.677542073 | 0.263568763 | 2.570646328 |
| Gli3 | 305.6293553 | -0.265341077 | 0.115015188 | -2.307009027 |
| Inhba | 124.8662631 | -0.370901541 | 0.412500584 | -0.899153977 |
| Gm22628 | 0.250684468 | -0.729055239 | 2.133838007 | -0.341663817 |
| Sugct | 24.65360541 | -0.163041883 | 0.230091069 | -0.708597182 |
| Gm26098 | 0 | NA | NA | NA |
| Gm27043 | 0 | NA | NA | NA |
| Gm25020 | 0 | NA | NA | NA |
| Gm24873 | 1.427842546 | -0.321647576 | 0.938875336 | -0.342588162 |
| Mplkip | 150.6828418 | -0.330515379 | 0.215507309 | -1.533662041 |
| Cdk13 | 1371.430534 | 0.008968437 | 0.059364672 | 0.15107363 |
| Mir466i | 0.78113724 | 1.050649481 | 1.252216816 | 0.839031602 |
| Gm24018 | 0.047948698 | -0.206487837 | 3.724945049 | -0.055433794 |
| Gm18859 | 0.219031842 | -0.206488874 | 2.592141691 | -0.079659563 |
| Rala | 653.2611828 | -0.130029042 | 0.08895368 | -1.461761252 |
| Yae1d1 | 237.983152 | -0.189765816 | 0.093826999 | -2.022507563 |
| Gm37888 | 0.089089296 | 1.194485492 | 3.723939576 | 0.320758559 |
| Vdac3-ps1 | 215.80711 | 0.102512725 | 0.117668155 | 0.871201937 |
| Pou6f2 | 51.8533924 | 0.032737781 | 0.170607163 | 0.191889837 |
| Vps41 | 1555.895699 | -0.008158556 | 0.058640372 | -0.13912865 |
| Gm26172 | 0.049553047 | 0.274391604 | 3.724945049 | 0.073663262 |
| Amph | 1915.694854 | -0.11333762 | 0.07254413 | -1.562326542 |
| RP23-167F18.7 | 0 | NA | NA | NA |
| Tcrg-V7 | 0.107803068 | 0.75528661 | 3.724945049 | 0.202764497 |
| Tcrg-V4 | 0.038981395 | 0.274393828 | 3.724945049 | 0.073663859 |
| Tcrg-V6 | 0.075043205 | 0.755286614 | 3.724945049 | 0.202764498 |
| Tcrg-V5 | 0.192276286 | 0.755286326 | 3.26669843 | 0.231207852 |
| Trgj1 | 0 | NA | NA | NA |
| Tcrg-C1 | 0.641965332 | 1.131951041 | 1.464282204 | 0.773041588 |
| Tcrg-V3 | 0 | NA | NA | NA |
| Trgj3 | 0 | NA | NA | NA |
| Tcrg-C3 | 0.092228164 | 0.75528439 | 3.724945049 | 0.202763901 |
| Tcrg-C2 | 0.588064751 | 2.340526137 | 1.359930646 | 1.721062867 |
| Trgj2 | 0 | NA | NA | NA |
| Trgv2 | 0.097139547 | 0.274393828 | 3.724945049 | 0.073663859 |
| Tcrg-V1 | 0 | NA | NA | NA |

| | | | | |
|---|---|---|---|---|
| Trgj4 | 0 | NA | NA | NA |
| Tcrg-C4 | 0.312934076 | 1.629855866 | 2.043053854 | 0.797754725 |
| Stard3nl | 346.4093214 | -0.10256848 | 0.092695496 | -1.10650986 |
| RP23-306G1.11 | 25.52656834 | -0.611042317 | 0.285321223 | -2.141594344 |
| RP23-306G1.12 | 0.525666135 | -1.755237264 | 1.371009707 | -1.280251522 |
| Epdr1 | 994.4722949 | -0.195200107 | 0.073012029 | -2.673533529 |
| Sfrp4 | 29.48509951 | 0.434831342 | 0.234866223 | 1.85140007 |
| Nme8 | 0.465692922 | 1.591651518 | 1.337905241 | 1.189659379 |
| A530099J19Rik | 0.780625029 | 0.744388422 | 1.378095289 | 0.540157439 |
| Gpr141 | 71.74378744 | 6.527424671 | 0.784705895 | 8.318307171 |
| Gm5446 | 4.31637155 | 0.902807797 | 0.499626293 | 1.806966146 |
| Elmo1 | 1541.907887 | 0.047574564 | 0.056009896 | 0.849395687 |
| Gm7611 | 0 | NA | NA | NA |
| Aoah | 31.82302678 | 4.037583899 | 0.456526123 | 8.844146469 |
| Gm25605 | 0.583760085 | 2.554685931 | 1.323556132 | 1.930168181 |
| Olfr1370 | 0 | NA | NA | NA |
| Olfr1369-ps1 | 0 | NA | NA | NA |
| Olfr263 | 0.213549163 | -0.206489211 | 2.582596912 | -0.0799541 |
| Olfr1368 | 0 | NA | NA | NA |
| Trim27 | 275.0576437 | -0.036536778 | 0.080932142 | -0.451449537 |
| Gpx5 | 0 | NA | NA | NA |
| Gm11270 | 0 | NA | NA | NA |
| Gpx6 | 2.180372029 | -0.916578958 | 0.748531186 | -1.224503367 |
| Olfr1367 | 0.046345709 | 0.274391604 | 3.724945049 | 0.073663262 |
| Zscan12 | 64.42265566 | 0.320528304 | 0.148031727 | 2.165267615 |
| Gm11271 | 0.280341511 | -0.685896002 | 1.78846901 | -0.383510141 |
| Zkscan3 | 573.1062161 | -0.632923525 | 0.13246509 | -4.778040189 |
| Mir6942 | 2.650403497 | -1.64800233 | 0.694111174 | -2.374262787 |
| Pgbd1 | 43.29911835 | -1.213383481 | 0.18613962 | -6.518673879 |
| 4930470G03Rik | 1.381930285 | 0.692527333 | 0.923223287 | 0.750119004 |
| Zscan26 | 980.2046494 | -0.174142509 | 0.057721009 | -3.016969216 |
| Mir1896 | 0 | NA | NA | NA |
| Nkapl | 5.734391589 | -0.271283557 | 0.502948853 | -0.539385975 |
| Zkscan4 | 37.6093419 | -0.03096494 | 0.186339375 | -0.16617497 |
| Gm10065 | 0.298142035 | 0.813465852 | 3.72125273 | 0.218600001 |
| Gm23779 | 0 | NA | NA | NA |
| AK157302 | 180.0808189 | -0.166208179 | 0.088423233 | -1.879689005 |
| AL589742.1 | 0 | NA | NA | NA |
| AL589742.2 | 0 | NA | NA | NA |
| Gm11273 | 0 | NA | NA | NA |
| Zfp389 | 4.935521334 | -0.120823584 | 0.482044001 | -0.250648455 |
| Zkscan8 | 581.7328674 | -0.171235135 | 0.101597102 | -1.685433252 |
| Olfr1366 | 0.049553047 | 0.274391604 | 3.724945049 | 0.073663262 |
| Olfr1535 | 0 | NA | NA | NA |
| Olfr1364 | 0.097899064 | -0.687371727 | 3.724945049 | -0.184532045 |
| Olfr1362 | 0 | NA | NA | NA |
| Olfr11 | 0.091666476 | -0.687373951 | 3.724945049 | -0.184532642 |

| | | | | |
|---|---|---|---|---|
| Olfr1361 | 0.061920613 | 0.274393828 | 3.724945049 | 0.073663859 |
| Olfr1360 | 0 | NA | NA | NA |
| Olfr1359 | 0.062426873 | 0.755288839 | 3.724945049 | 0.202765096 |
| Hist1h2bl | 17.04762427 | 0.928545559 | 0.389450605 | 2.384244749 |
| Hist1h2ai | 8.571363186 | 0.746578559 | 0.407626165 | 1.83152757 |
| AL589651.1 | 0.12092566 | 1.236179401 | 3.724945049 | 0.331865138 |
| Hist1h3h | 40.08083796 | 0.380859005 | 0.235316393 | 1.618497547 |
| AL589651.2 | 0 | NA | NA | NA |
| Hist1h2aj | 0.635935163 | 0.274729986 | 1.366228971 | 0.201086342 |
| Hist1h2bm | 22.0061962 | 1.650143078 | 0.331746959 | 4.974101591 |
| Hist1h4j | 57.33528979 | 0.463820083 | 0.22365561 | 2.073813769 |
| Hist1h4k | 181.1049781 | 0.251298214 | 0.149041869 | 1.686091401 |
| Hist1h2ak | 39.92311535 | 1.093205973 | 0.286459462 | 3.816267628 |
| AL589651.3 | 0.07796279 | 0.274392189 | 3.724945049 | 0.073663419 |
| Hist1h2bn | 10.00898834 | 0.530113812 | 0.397911631 | 1.332240054 |
| Gm11274 | 1.966252851 | -0.198771929 | 0.89684053 | -0.221635756 |
| Hist1h1b | 62.13502671 | 1.010620791 | 0.226019033 | 4.471396847 |
| Hist1h3i | 77.67695009 | 0.607596275 | 0.241021045 | 2.520926239 |
| Hist1h2an | 2.07791248 | 0.778671937 | 0.772040189 | 1.0085899 |
| AL589651.4 | 0 | NA | NA | NA |
| Hist1h2bp | 6.144754156 | 0.608871447 | 0.434156853 | 1.402422748 |
| Hist1h2bq | 12.73633966 | 0.723243978 | 0.337086832 | 2.145571732 |
| Hist1h2ao | 3.983646474 | 0.036555074 | 0.559470819 | 0.065338661 |
| AL589879.1 | 0.072025462 | 0.755288839 | 3.724945049 | 0.202765096 |
| Hist1h4m | 15.4295438 | 0.232930331 | 0.33020661 | 0.70540784 |
| Hist1h4n | 166.7008044 | 0.103094081 | 0.167936242 | 0.613888218 |
| Hist1h2ap | 1.700876845 | 1.044780438 | 0.96749281 | 1.079884447 |
| AL589879.3 | 0.060174537 | 0.755286614 | 3.724945049 | 0.202764498 |
| Hist1h2br | 3.568372849 | 0.556831767 | 0.645712283 | 0.862352755 |
| Gm11278 | 0 | NA | NA | NA |
| Mir1983 | 1.672508104 | -0.88194106 | 0.969936175 | -0.909277418 |
| Gm11279 | 0.061920613 | 0.274393828 | 3.724945049 | 0.073663859 |
| AL589879.2 | 0.514605876 | -0.192395111 | 1.545175731 | -0.124513418 |
| Gm11280 | 0.088534442 | 0.274389469 | 3.724945049 | 0.073662689 |
| Zfp184 | 35.438407 | -0.503741381 | 0.176272012 | -2.857750225 |
| Gm11281 | 0 | NA | NA | NA |
| Gm11282 | 0.068097359 | -0.206487837 | 3.724945049 | -0.055433794 |
| Pom121l2 | 0.476324185 | 1.376533192 | 1.505619648 | 0.914263569 |
| Prss16 | 7.136397789 | 1.488834833 | 0.515082543 | 2.890478147 |
| Gm11290 | 0 | NA | NA | NA |
| Gm11291 | 0 | NA | NA | NA |
| Gm11292 | 0.121388914 | 0.755282166 | 3.724945049 | 0.202763304 |
| AL590614.1 | 0.110870145 | -0.206490066 | 3.724945049 | -0.055434393 |
| Hist1h2ah | 3.414149285 | 0.858957163 | 0.783823122 | 1.095855863 |
| Hist1h2bk | 6.592695331 | 0.545803728 | 0.459172427 | 1.188668342 |
| Hist1h4i | 181.0810821 | 0.341424683 | 0.182465411 | 1.871174817 |
| Hist1h2ag | 7.85075097 | 0.724146002 | 0.401847338 | 1.802042553 |

| | | | | |
|---|---|---|---|---|
| AL590614.2 | 0.038981395 | 0.274393828 | 3.724945049 | 0.073663859 |
| Hist1h2bj | 11.38635759 | 1.19452895 | 0.396339072 | 3.013906612 |
| Vmn1r-ps92 | 0 | NA | NA | NA |
| Vmn1r-ps93 | 0 | NA | NA | NA |
| Gm11285 | 0 | NA | NA | NA |
| Gm26300 | 0 | NA | NA | NA |
| Vmn1r188 | 0 | NA | NA | NA |
| Vmn1r-ps94 | 0.341912768 | 1.195229243 | 2.024046672 | 0.590514665 |
| Vmn1r189 | 0 | NA | NA | NA |
| Vmn1r-ps95 | 0 | NA | NA | NA |
| Vmn1r-ps97 | 0 | NA | NA | NA |
| Vmn1r190-ps | 0 | NA | NA | NA |
| Gm11284 | 0.374772292 | 1.51096677 | 1.516333467 | 0.996460741 |
| Vmn1r-ps99 | 0 | NA | NA | NA |
| Vmn1r191 | 0 | NA | NA | NA |
| Vmn1r192 | 0 | NA | NA | NA |
| Gm25084 | 0 | NA | NA | NA |
| Gm23282 | 0.060174537 | 0.755286614 | 3.724945049 | 0.202764498 |
| Vmn1r193 | 0 | NA | NA | NA |
| Vmn1r194 | 0.049553047 | 0.274391604 | 3.724945049 | 0.073663262 |
| Vmn1r195 | 0 | NA | NA | NA |
| Gm11295 | 0.521690028 | -0.028983489 | 1.278638733 | -0.022667458 |
| Vmn1r196 | 0 | NA | NA | NA |
| Vmn1r-ps100 | 0 | NA | NA | NA |
| Vmn1r197 | 0.066050875 | 0.274391604 | 3.724945049 | 0.073663262 |
| Vmn1r198 | 0 | NA | NA | NA |
| Vmn1r199 | 0.038981395 | 0.274393828 | 3.724945049 | 0.073663859 |
| Vmn1r-ps102 | 0 | NA | NA | NA |
| Vmn1r200 | 0.302366008 | 0.755304438 | 2.043112854 | 0.369683171 |
| Vmn1r-ps103 | 0 | NA | NA | NA |
| Vmn1r-ps104 | 0 | NA | NA | NA |
| Gm11298 | 0.060174537 | 0.755286614 | 3.724945049 | 0.202764498 |
| Vmn1r201 | 0 | NA | NA | NA |
| Vmn1r-ps106 | 0 | NA | NA | NA |
| Vmn1r-ps107 | 0 | NA | NA | NA |
| Vmn1r-ps108 | 0 | NA | NA | NA |
| Vmn1r202 | 0 | NA | NA | NA |
| Gm11300 | 0.038981395 | 0.274393828 | 3.724945049 | 0.073663859 |
| Vmn1r203 | 0 | NA | NA | NA |
| Vmn1r204 | 0 | NA | NA | NA |
| Vmn1r205 | 0.044191549 | -0.206490061 | 3.724945049 | -0.055434391 |
| Vmn1r206 | 1.041252144 | -1.345628619 | 1.041557971 | -1.291938285 |
| Vmn1r-ps109 | 0.733882059 | -1.923332888 | 1.108517832 | -1.735049119 |
| Vmn1r-ps110 | 1.137431431 | -0.705963582 | 0.997054461 | -0.708049169 |
| Gm11307 | 0 | NA | NA | NA |
| Vmn1r-ps111 | 0.097139547 | 0.274393828 | 3.724945049 | 0.073663859 |
| Gm11309 | 0 | NA | NA | NA |

| | | | | |
|---|---|---|---|---|
| Vmn1r207-ps | 0.447844902 | 1.277784483 | 1.565303948 | 0.816317166 |
| Vmn1r-ps112 | 0 | NA | NA | NA |
| Vmn1r-ps113 | 1.019683961 | -0.354100775 | 1.220012844 | -0.290243481 |
| Gm11313 | 0 | NA | NA | NA |
| Vmn1r208 | 0 | NA | NA | NA |
| Gm11314 | 0 | NA | NA | NA |
| Vmn1r209 | 0.344645118 | -0.653526122 | 1.585942002 | -0.412074414 |
| Vmn1r210 | 0.134880151 | 0.274398319 | 2.937637886 | 0.093407809 |
| Vmn1r-ps114 | 0 | NA | NA | NA |
| Vmn1r211 | 0 | NA | NA | NA |
| Vmn1r-ps116 | 0 | NA | NA | NA |
| Gm23774 | 0 | NA | NA | NA |
| Vmn1r212 | 0.038981395 | 0.274393828 | 3.724945049 | 0.073663859 |
| Gm11320 | 0 | NA | NA | NA |
| Gm11321 | 0 | NA | NA | NA |
| Vmn1r-ps120 | 0 | NA | NA | NA |
| Gm11322 | 0 | NA | NA | NA |
| Gm11318 | 0.044544648 | 0.755286614 | 3.724945049 | 0.202764498 |
| Vmn1r-ps123 | 0 | NA | NA | NA |
| Vmn1r213 | 0.109982625 | -0.20649451 | 3.724945049 | -0.055435586 |
| Vmn1r-ps125 | 0 | NA | NA | NA |
| Vmn1r214 | 0 | NA | NA | NA |
| Vmn1r-ps126 | 0 | NA | NA | NA |
| Vmn1r215 | 0 | NA | NA | NA |
| Vmn1r216 | 0 | NA | NA | NA |
| Gm11325 | 0 | NA | NA | NA |
| Vmn1r-ps127 | 0 | NA | NA | NA |
| Vmn1r217 | 0 | NA | NA | NA |
| Vmn1r-ps128 | 0 | NA | NA | NA |
| Vmn1r218 | 0 | NA | NA | NA |
| Vmn1r219 | 0 | NA | NA | NA |
| Vmn1r220 | 0.038981395 | 0.274393828 | 3.724945049 | 0.073663859 |
| Vmn1r-ps132 | 0 | NA | NA | NA |
| Vmn1r221 | 0 | NA | NA | NA |
| Vmn1r222 | 0 | NA | NA | NA |
| Vmn1r223 | 0 | NA | NA | NA |
| Gm11331 | 0 | NA | NA | NA |
| Vmn1r-ps135 | 0 | NA | NA | NA |
| 4930557F10Rik | 0.136120942 | 0.274393632 | 3.599844381 | 0.076223748 |
| 4933404K08Rik | 0.044544648 | 0.755286614 | 3.724945049 | 0.202764498 |
| Gm11333 | 0 | NA | NA | NA |
| 4930586N03Rik | 0.124596252 | 0.755282361 | 3.714681068 | 0.203323609 |
| Zfp322a | 495.523674 | -0.47715963 | 0.112820578 | -4.229367 |
| Gm11334 | 0 | NA | NA | NA |
| Abt1 | 128.8824264 | 0.277291927 | 0.130471498 | 2.125306539 |
| C230035I16Rik | 12.59447301 | 2.343893058 | 0.448107816 | 5.230645337 |
| Btn1a1 | 18.16397336 | 6.395586855 | 0.934317466 | 6.845196724 |

| | | | | |
|---|---|---|---|---|
| Btn2a2 | 7.935836238 | 0.046180669 | 0.421361855 | 0.109598599 |
| AC034285.1 | 0 | NA | NA | NA |
| Gm11335 | 0.44640837 | 0.206268564 | 1.356057929 | 0.152108962 |
| Hist1h4h | 362.5692348 | -0.339471227 | 0.179585853 | -1.890300494 |
| Hist1h2af | 13.35131425 | 2.224737807 | 0.521484821 | 4.266160235 |
| AC034285.3 | 0.10503227 | 0.274389469 | 3.724945049 | 0.073662689 |
| Hist1h3g | 12.55213016 | 1.348818201 | 0.410329216 | 3.287161015 |
| Hist1h2bh | 13.53059552 | 1.917910118 | 0.412250917 | 4.652288297 |
| Hist1h3f | 117.9595612 | 1.005735143 | 0.180428733 | 5.574140704 |
| Hist1h4f | 133.4200694 | 0.08621739 | 0.185080418 | 0.465837451 |
| Hist1h1d | 190.8763881 | 1.204423517 | 0.194221407 | 6.201291294 |
| Hist1h3e | 180.3700906 | 0.867762855 | 0.136182635 | 6.372052155 |
| Hist1h2ae | 13.93887753 | 0.346856307 | 0.39035055 | 0.888576451 |
| AC034285.2 | 0.480269614 | 0.882259631 | 1.307718971 | 0.674655373 |
| Hist1h2bg | 107.7133796 | 0.715096348 | 0.113700163 | 6.289316828 |
| Hist1h2bf | 17.61300926 | 1.091304627 | 0.366747404 | 2.975630138 |
| Hist1h2ad | 7.472743362 | 0.208201305 | 0.447981147 | 0.464754614 |
| AC034285.4 | 0.068097359 | -0.206487837 | 3.724945049 | -0.055433794 |
| Hist1h3d | 38.7205137 | 0.193935822 | 0.24013136 | 0.807623885 |
| Hist1h4d | 957.8385019 | 0.594108565 | 0.126468176 | 4.69769221 |
| Hist1h2be | 153.5786973 | 0.88509567 | 0.148902568 | 5.944126287 |
| Gm11398 | 0 | NA | NA | NA |
| Hist1h1e | 485.7404665 | 0.821167855 | 0.180130872 | 4.558729148 |
| Gm22452 | 0 | NA | NA | NA |
| Gm24991 | 0.1140246 | 0.75528439 | 3.724945049 | 0.202763901 |
| Hist1h2ac | 9.474268641 | 0.155801159 | 0.394800756 | 0.394632373 |
| AL592149.1 | 0.048949532 | -0.206487837 | 3.724945049 | -0.055433794 |
| Hist1h2bc | 147.070612 | 0.525007611 | 0.147264006 | 3.565077602 |
| Hist1h1t | 0.61465116 | 0.248102357 | 1.340623364 | 0.185064921 |
| Hist1h4c | 207.5103062 | 0.198618763 | 0.167836415 | 1.183406846 |
| Hfe | 68.88156831 | 1.630252928 | 0.214710657 | 7.592790011 |
| Gm11338 | 0 | NA | NA | NA |
| Hist1h1c | 665.5210639 | 1.699075791 | 0.166106595 | 10.22882802 |
| Hist1h3c | 20.18455272 | 2.377298642 | 0.375235054 | 6.335491897 |
| Gm11336 | 0.090890357 | 0.755282166 | 3.724945049 | 0.202763304 |
| AL590388.2 | 0.068097359 | -0.206487837 | 3.724945049 | -0.055433794 |
| Hist1h2bb | 38.92068366 | 0.903037499 | 0.293658882 | 3.075124083 |
| Hist1h2ab | 19.95108352 | 1.202109334 | 0.278627456 | 4.314396544 |
| AL590388.1 | 0.038981395 | 0.274393828 | 3.724945049 | 0.073663859 |
| Hist1h3b | 12.81895394 | 1.865891108 | 0.403180051 | 4.627935098 |
| 4930558J22Rik | 0.487944759 | -0.854909297 | 1.376595117 | -0.621031766 |
| Hist1h4b | 52.56315827 | -0.444324508 | 0.241543448 | -1.839522087 |
| Hist1h4a | 71.08590372 | -0.333524696 | 0.150894616 | -2.210315422 |
| Hist1h3a | 21.90840992 | 0.428457563 | 0.279114732 | 1.535058935 |
| Hist1h1a | 77.50845308 | 0.762967288 | 0.20034581 | 3.808251771 |
| Trim38 | 1.17402693 | 0.476622992 | 0.876707464 | 0.543651117 |
| Gm11337 | 0 | NA | NA | NA |

| | | | | |
|---|---|---|---|---|
| Slc17a2 | 0.366957926 | 0.274413833 | 1.47719331 | 0.185767043 |
| Slc17a3 | 0.879126543 | 1.570506545 | 1.228707835 | 1.278177367 |
| Slc17a1 | 0.066050875 | 0.274391604 | 3.724945049 | 0.073663262 |
| Slc17a4 | 0 | NA | NA | NA |
| Snrp1c-ps2 | 0 | NA | NA | NA |
| Hist1h2ba | 0.304971633 | 0.755307569 | 1.778244728 | 0.424748943 |
| Hist1h2aa | 0 | NA | NA | NA |
| AL606464.1 | 0 | NA | NA | NA |
| Scgn | 17.90384641 | -0.353797027 | 0.570515684 | -0.620135496 |
| Gm11339 | 0 | NA | NA | NA |
| Lrrc16a | 976.9628538 | 0.942460462 | 0.091694934 | 10.27821736 |
| Gm11343 | 0 | NA | NA | NA |
| Gm11340 | 0.044191549 | -0.206490061 | 3.724945049 | -0.055434391 |
| Gm23340 | 0 | NA | NA | NA |
| Gm11341 | 0 | NA | NA | NA |
| Cmah | 7.485215113 | 1.127932199 | 0.533220222 | 2.1153215 |
| Gm11342 | 0.062426873 | 0.755288839 | 3.724945049 | 0.202765096 |
| Gm11345 | 0.130840244 | -1.039642477 | 2.947200097 | -0.352755986 |
| Gm11344 | 0 | NA | NA | NA |
| Gm22013 | 0 | NA | NA | NA |
| Fam65b | 791.2498213 | -0.39536491 | 0.115221441 | -3.431348431 |
| Gm11346 | 3.822611367 | 0.785606888 | 0.562488036 | 1.396664173 |
| C530050E15Rik | 25.2429927 | 0.016414592 | 0.32556819 | 0.050418291 |
| 1700016G14Rik | 1.09380521 | 0.063016213 | 0.931889709 | 0.067621964 |
| Gmnn | 28.95427006 | 0.107556261 | 0.254077788 | 0.423320204 |
| BC005537 | 3337.74108 | 0.303394211 | 0.051602531 | 5.879444425 |
| Gm22358 | 0.227031315 | -0.70707812 | 2.14026266 | -0.330369787 |
| Acot13 | 187.4166705 | -0.088351312 | 0.156121033 | -0.565915497 |
| Tdp2 | 199.2904976 | 0.102334477 | 0.090946562 | 1.125215451 |
| D130043K22Rik | 395.2440717 | -0.459213862 | 0.121315482 | -3.785286544 |
| 4932702P03Rik | 1.792617364 | -0.222300211 | 0.669301248 | -0.33213775 |
| Aldh5a1 | 1047.224437 | -0.693161946 | 0.065052184 | -10.65547532 |
| 9330162012Rik | 49.19052179 | 0.201940173 | 0.241038167 | 0.837793349 |
| Gpld1 | 489.1648644 | 0.031584571 | 0.133193713 | 0.237132599 |
| Mrs2 | 387.1535434 | -0.193359675 | 0.085497391 | -2.261585693 |
| Dcdc2a | 119.9340571 | 0.688362351 | 0.164218541 | 4.191745615 |
| Gm11348 | 0 | NA | NA | NA |
| Nrsn1 | 2860.65622 | 0.055315238 | 0.088486913 | 0.625123376 |
| Tpi-rs8 | 0.046345709 | 0.274391604 | 3.724945049 | 0.073663262 |
| Gm11349 | 2.028713949 | -0.614011981 | 0.736804834 | -0.833344127 |
| Gm11350 | 0 | NA | NA | NA |
| Gm11351 | 0.19500485 | 0.274391147 | 3.236821722 | 0.084771782 |
| Gm11352 | 0.044544648 | 0.755286614 | 3.724945049 | 0.202764498 |
| 1700092E19Rik | 0 | NA | NA | NA |
| Gm11353 | 25.4493118 | 0.442670246 | 0.249607878 | 1.773462641 |
| Hdgfl1 | 0 | NA | NA | NA |
| Gm11354 | 0 | NA | NA | NA |

| | | | | |
|---|---|---|---|---|
| Gm11355 | 0 | NA | NA | NA |
| Prl | 159.9440114 | -2.136193715 | 2.888538176 | -0.739541451 |
| Prl3d1 | 0.046345709 | 0.274391604 | 3.724945049 | 0.073663262 |
| Prl3d2 | 0 | NA | NA | NA |
| Prl3d3 | 0 | NA | NA | NA |
| Prl3c1 | 0 | NA | NA | NA |
| Prl3b1 | 0 | NA | NA | NA |
| Prl3a1 | 0 | NA | NA | NA |
| 4930511O05Rik | 0 | NA | NA | NA |
| Gm11356 | 0 | NA | NA | NA |
| Prl6a1 | 0.046345709 | 0.274391604 | 3.724945049 | 0.073663262 |
| Gm11357 | 0 | NA | NA | NA |
| Gm11358 | 0 | NA | NA | NA |
| Prl8a2 | 0 | NA | NA | NA |
| Prl2b1 | 0 | NA | NA | NA |
| Prl8a6 | 0 | NA | NA | NA |
| Prl8a8 | 0 | NA | NA | NA |
| Prl8a9 | 0 | NA | NA | NA |
| Prl8a1 | 0 | NA | NA | NA |
| Prl7b1 | 0 | NA | NA | NA |
| Prl7a1 | 0 | NA | NA | NA |
| Prl7a2 | 0 | NA | NA | NA |
| Gm11359 | 0.049553047 | 0.274391604 | 3.724945049 | 0.073663262 |
| Prl7d1 | 0.045882455 | 0.755288839 | 3.724945049 | 0.202765096 |
| Prl7c1 | 0.062426873 | 0.755288839 | 3.724945049 | 0.202765096 |
| Prl2a1 | 0 | NA | NA | NA |
| Prl2c1 | 0 | NA | NA | NA |
| Gm22476 | 0 | NA | NA | NA |
| Gm11360 | 0 | NA | NA | NA |
| Prl4a1 | 0 | NA | NA | NA |
| Prl5a1 | 0 | NA | NA | NA |
| Gm11361 | 310.4479338 | 0.35594618 | 0.153018207 | 2.326168814 |
| 2610307P16Rik | 11.882246 | -0.361770226 | 0.306682339 | -1.179625234 |
| Gm6081 | 0.66818121 | -1.035613134 | 1.238859542 | -0.835940718 |
| Mir6368 | 0.123992736 | 0.274402763 | 3.720643035 | 0.073751435 |
| Gm17528 | 0 | NA | NA | NA |
| Gm11362 | 0.126225412 | 0.755282661 | 3.698754989 | 0.20419916 |
| Gm26735 | 7.2496826 | -0.205316935 | 0.37833994 | -0.542678458 |
| Sox4 | 422.4467281 | -0.221823475 | 0.150952024 | -1.469496528 |
| A330102I10Rik | 83.11002475 | -0.629892267 | 0.239435508 | -2.630738738 |
| Gm11363 | 0.134452335 | 1.290546799 | 3.628238117 | 0.355695177 |
| Cdkal1 | 296.7585439 | -0.035764108 | 0.110090694 | -0.324860409 |
| Gm11364 | 0 | NA | NA | NA |
| E2f3 | 366.9818336 | -0.047570976 | 0.071398834 | -0.666271041 |
| Gm11365 | 0.042716945 | -0.206490061 | 3.724945049 | -0.055434391 |
| Gm11368 | 0.105852363 | -0.206492286 | 3.724945049 | -0.055434988 |
| Gm11367 | 0 | NA | NA | NA |

| | | | | |
|---|---|---|---|---|
| Mboat1 | 33.58227926 | -0.729010968 | 0.243841552 | -2.9896913 |
| Gm23206 | 0 | NA | NA | NA |
| Gm11366 | 0.322434302 | -0.534071343 | 1.730602793 | -0.308604231 |
| Agtr1a | 5.923613152 | -0.94241342 | 0.448828589 | -2.099717896 |
| Gm11369 | 0.085327104 | 0.274389469 | 3.724945049 | 0.073662689 |
| Uqcrfs1 | 840.2137481 | -0.033012592 | 0.112924289 | -0.292342707 |
| 4930519D14Rik | 0.04393175 | -0.206490061 | 3.724945049 | -0.055434391 |
| Dusp22 | 130.6997309 | -0.163068534 | 0.116521699 | -1.399469249 |
| Gm11370 | 0 | NA | NA | NA |
| Irf4 | 27.40671112 | 2.604223304 | 0.367932989 | 7.077982627 |
| Exoc2 | 757.5578618 | 0.192340284 | 0.045119106 | 4.262945333 |
| Hus1b | 7.907650439 | -0.545509777 | 0.389598001 | -1.400186283 |
| Gm5447 | 0.689547818 | 0.265654711 | 1.390191562 | 0.191092162 |
| Gm11371 | 0.046345709 | 0.274391604 | 3.724945049 | 0.073663262 |
| Gm11372 | 0 | NA | NA | NA |
| 4930401O12Rik | 0.147083031 | -0.196511238 | 2.875092884 | -0.068349527 |
| Gm11376 | 0 | NA | NA | NA |
| Gm11373 | 0.313638404 | -0.196976918 | 1.786883446 | -0.1102349 |
| Gm11374 | 0.506145433 | 0.690467485 | 1.322926792 | 0.521924183 |
| G630018N14Rik | 1.020023321 | 3.146601318 | 1.277656677 | 2.462790962 |
| Gm22330 | 0 | NA | NA | NA |
| Gm11375 | 1.368265809 | 0.134305094 | 0.803068318 | 0.167239936 |
| Gm23351 | 0 | NA | NA | NA |
| A530084C06Rik | 4.406782827 | -0.310316629 | 0.480930663 | -0.64524193 |
| Foxq1 | 10.28519308 | -0.382236529 | 0.341166729 | -1.120380437 |
| Gm11377 | 0 | NA | NA | NA |
| 1700018A04Rik | 0.705754361 | -0.03870943 | 1.236699822 | -0.031300587 |
| Gm11378 | 0 | NA | NA | NA |
| Foxf2 | 22.59941583 | 0.920592492 | 0.336253524 | 2.737792848 |
| Gm27516 | 0.075043205 | 0.755286614 | 3.724945049 | 0.202764498 |
| Gm25186 | 0.382576472 | 1.195308721 | 1.766333606 | 0.67671742 |
| Gm11379 | 0.403183719 | 0.294938388 | 1.412754309 | 0.208768351 |
| Foxc1 | 125.7748194 | 0.863614907 | 0.198367351 | 4.353614161 |
| Gmds | 102.4660629 | -0.107918362 | 0.121970055 | -0.884793912 |
| Gm11380 | 0.044544648 | 0.755286614 | 3.724945049 | 0.202764498 |
| Gm11381 | 0 | NA | NA | NA |
| Mylk4 | 2.177335128 | 0.018278927 | 0.619517712 | 0.029505092 |
| Gm27342 | 0 | NA | NA | NA |
| Wrnip1 | 563.9693568 | 0.025551335 | 0.070723336 | 0.361285774 |
| D930007J09Rik | 3.931524232 | -0.522233063 | 0.495483993 | -1.053985741 |
| Serpinb1a | 58.83320495 | -1.790276467 | 0.218007127 | -8.212008909 |
| Serpinb1-ps1 | 0.259499533 | 1.715206316 | 2.494191293 | 0.68768034 |
| Serpinb1c | 2.097314789 | 1.713055966 | 1.109099056 | 1.544547313 |
| Gm11382 | 0.61025173 | -0.206484367 | 1.490554925 | -0.138528519 |
| Serpinb6b | 64.34938334 | 2.634750829 | 0.352384664 | 7.476916832 |
| Serpinb9 | 320.5775695 | 2.266717368 | 0.228605517 | 9.915409751 |
| Serpinb9b | 16.14145208 | 5.069700163 | 0.966581011 | 5.244982167 |

| | | | | |
|---|---|---|---|---|
| Serpinb1b | 57.17042438 | 0.847052693 | 0.342218971 | 2.475177492 |
| Gm11384 | 0 | NA | NA | NA |
| Gm11383 | 0 | NA | NA | NA |
| Serpinb9c | 0.260452197 | -0.548454096 | 1.825306483 | -0.300472332 |
| Serpinb9d | 0 | NA | NA | NA |
| Gm11385 | 0 | NA | NA | NA |
| Serpinb9e | 0.061920613 | 0.274393828 | 3.724945049 | 0.073663859 |
| Serpinb9f | 0 | NA | NA | NA |
| Gm11397 | 0 | NA | NA | NA |
| Serpinb9g | 0 | NA | NA | NA |
| Gm11387 | 0 | NA | NA | NA |
| Gm11386 | 0 | NA | NA | NA |
| Gm11393 | 0 | NA | NA | NA |
| Gm25043 | 0 | NA | NA | NA |
| Gm11388 | 0 | NA | NA | NA |
| Serpinb6d | 0 | NA | NA | NA |
| Gm11389 | 0 | NA | NA | NA |
| Gm26701 | 0 | NA | NA | NA |
| Gm11392 | 0 | NA | NA | NA |
| Gm11394 | 0 | NA | NA | NA |
| Gm11395 | 0 | NA | NA | NA |
| Vmn2r-ps94 | 0 | NA | NA | NA |
| Gm16566 | 0 | NA | NA | NA |
| Serpinb6e | 0.733278584 | -0.568461948 | 1.305235011 | -0.435524594 |
| Serpinb6c | 3.256068972 | 0.663588454 | 0.579949901 | 1.144216859 |
| Serpinb6a | 261.2870453 | 0.280304161 | 0.111215776 | 2.520363313 |
| 1110046J04Rik | 30.45720878 | -0.298153263 | 0.222559181 | -1.339658341 |
| Nqo2 | 104.6663154 | 0.252261895 | 0.15926887 | 1.583874451 |
| Ripk1 | 212.9648014 | 1.357515629 | 0.167330895 | 8.112761407 |
| Gm17083 | 0.214446769 | 1.546731252 | 2.554573644 | 0.605475304 |
| Gm17084 | 5.185998495 | 0.809658953 | 0.484594212 | 1.67079782 |
| Bphl | 141.617793 | -0.071679516 | 0.098967869 | -0.724270581 |
| Tubb2a | 3268.123541 | 0.129832816 | 0.114447697 | 1.134429254 |
| Tubb2b | 1731.528415 | 0.83414959 | 0.1737043 | 4.802124027 |
| Psmg4 | 78.61758626 | 1.044865783 | 0.190514176 | 5.484451639 |
| Slc22a23 | 1092.32252 | -0.09046459 | 0.04993593 | -1.811613218 |
| Gm24530 | 6.119618873 | -0.562656523 | 0.432490479 | -1.300968577 |
| Pxdc1 | 58.73051673 | 0.103601426 | 0.212009451 | 0.488664186 |
| Gm15908 | 0.16390591 | -0.20648029 | 2.782471951 | -0.074207501 |
| Fam50b | 0.097139547 | 0.274393828 | 3.724945049 | 0.073663859 |
| Prpf4b | 962.083236 | -0.181535246 | 0.098031996 | -1.851795878 |
| Fam217a | 2.489500889 | -0.686072617 | 0.70789317 | -0.969175358 |
| 4933417A18Rik | 0.093141081 | -0.687373951 | 3.724945049 | -0.184532642 |
| Gm8245 | 0.066050875 | 0.274391604 | 3.724945049 | 0.073663262 |
| Eci3 | 1.298423698 | -0.366244889 | 0.976415722 | -0.375091142 |
| Gm16984 | 0.868294842 | -0.960272363 | 1.144888527 | -0.838747476 |
| Eci2 | 255.3897712 | 0.192067099 | 0.131123537 | 1.464779733 |

| | | | | |
|---|---|---|---|---|
| 1700011B04Rik | 0.157356115 | 0.755296948 | 2.830742007 | 0.266819423 |
| Gm22674 | 0 | NA | NA | NA |
| Cdyl | 458.3848708 | 0.110679477 | 0.113037741 | 0.979137376 |
| Gm22126 | 0 | NA | NA | NA |
| Rpp40 | 47.71410153 | -0.575461429 | 0.208011078 | -2.766494141 |
| Ppp1r3g | 58.16805527 | -0.878829749 | 0.287685986 | -3.054822944 |
| Lyrm4 | 177.4291997 | -0.143084203 | 0.110821168 | -1.291127012 |
| Fars2 | 206.9435174 | 0.051467048 | 0.114960337 | 0.447693953 |
| Gm26395 | 0.075043205 | 0.755286614 | 3.724945049 | 0.202764498 |
| Nrn1 | 1029.09811 | -0.343108106 | 0.159515956 | -2.150932828 |
| F13a1 | 144.5358196 | 1.508106404 | 0.427312563 | 3.529281691 |
| Ly86 | 116.6552724 | 1.583392089 | 0.278383239 | 5.687814021 |
| Gm4035 | 0.308888673 | -1.985473465 | 3.685836331 | -0.538676514 |
| Gm29459 | 0 | NA | NA | NA |
| Gm29458 | 0.240304698 | 0.795617473 | 2.11323539 | 0.376492593 |
| Gm29590 | 0 | NA | NA | NA |
| Gm37149 | 0 | NA | NA | NA |
| Rreb1 | 710.6869941 | 0.171589747 | 0.174145365 | 0.985324798 |
| Ssr1 | 2094.214592 | 0.055197626 | 0.045773758 | 1.205879289 |
| Cage1 | 29.07551637 | -0.47253524 | 0.226105932 | -2.089884309 |
| Riok1 | 219.0593976 | 0.347163899 | 0.123121023 | 2.819696348 |
| Gm27387 | 0 | NA | NA | NA |
| Gm10129 | 0.169723681 | 0.755289365 | 3.415329259 | 0.221146867 |
| Dsp | 96.30191792 | 0.303970229 | 0.163731453 | 1.85651702 |
| Snrnp48 | 159.0698413 | -0.402440867 | 0.168515742 | -2.388149985 |
| Bmp6 | 104.867172 | -0.006106229 | 0.19647948 | -0.031078201 |
| Txndc5 | 655.6323935 | 0.174464111 | 0.083300022 | 2.094406542 |
| Bloc1s5 | 130.6301522 | -0.24341998 | 0.106793294 | -2.279356421 |
| Eef1e1 | 223.262343 | 0.171819034 | 0.126445318 | 1.358840626 |
| Gm22035 | 0.046345709 | 0.274391604 | 3.724945049 | 0.073663262 |
| Slc35b3 | 116.4993211 | 0.327114222 | 0.120535183 | 2.713848469 |
| Gm26514 | 39.29548882 | -0.19936149 | 0.222768857 | -0.894925317 |
| 5033403F01Rik | 3.803373425 | 0.760575127 | 0.522285764 | 1.456243267 |
| Gm38206 | 0 | NA | NA | NA |
| Gm23972 | 0 | NA | NA | NA |
| Ofcc1 | 0.044191549 | -0.206490061 | 3.724945049 | -0.055434391 |
| Gm9979 | 0.471418172 | 0.287792624 | 1.563732415 | 0.184042117 |
| Tfap2a | 38.58151316 | -0.022905655 | 0.237386451 | -0.096490994 |
| Gm26688 | 1.413787916 | 1.230292241 | 0.883318248 | 1.392807456 |
| Gm37823 | 0 | NA | NA | NA |
| Gm6245 | 0.59099902 | 1.059519387 | 1.299270694 | 0.815472397 |
| Gcnt2 | 477.1222298 | 0.67525569 | 0.142784091 | 4.729208166 |
| Mir5124a | 0.74668639 | 1.670983097 | 1.355621479 | 1.232632503 |
| A730081D07Rik | 27.84660377 | -0.208941921 | 0.284820241 | -0.733592248 |
| Pak1ip1 | 411.5182475 | 0.201812419 | 0.117159958 | 1.72253748 |
| Tmem14c | 229.1318417 | 0.034762699 | 0.103401184 | 0.336192461 |
| Mak | 31.22020358 | -0.018001909 | 0.209379628 | -0.085977367 |

| | | | | |
|---|---|---|---|---|
| Gcm2 | 0.462408164 | 1.504512291 | 1.426400378 | 1.054761563 |
| Sycp2l | 3.696698795 | 0.283550451 | 0.503573095 | 0.563077047 |
| Elovl2 | 197.6053173 | 0.040423059 | 0.121910623 | 0.331579461 |
| Smim13 | 2327.736401 | -0.332051781 | 0.085561761 | -3.880843242 |
| Gm17364 | 0.17692102 | -0.206487546 | 2.728717236 | -0.07567202 |
| Nedd9 | 334.1506425 | 0.987932105 | 0.185379366 | 5.329245241 |
| Gm26877 | 0.927545067 | -1.595361681 | 1.213302642 | -1.314891789 |
| Gm24819 | 0.43165004 | -0.805925497 | 1.315363026 | -0.612701954 |
| Tmem170b | 1737.683987 | -0.09321875 | 0.048574843 | -1.919074664 |
| Gm5082 | 0.47229425 | -0.906004628 | 3.685326863 | -0.245841051 |
| Adtrp | 1.749374567 | 1.74858634 | 1.052946471 | 1.660660242 |
| 1700061E18Rik | 0.094294407 | -0.206492286 | 3.724945049 | -0.055434988 |
| Gm28564 | 0.066050875 | 0.274391604 | 3.724945049 | 0.073663262 |
| Gm37698 | 0 | NA | NA | NA |
| Gm22157 | 0.123841226 | 0.274391173 | 3.724945049 | 0.073663147 |
| Gm28707 | 0.850789295 | -0.071497072 | 1.377563355 | -0.051901114 |
| Hivep1 | 1475.754828 | -0.099811948 | 0.061393721 | -1.625768014 |
| Edn1 | 31.52376167 | 2.038538346 | 0.335935477 | 6.068243721 |
| Gm23086 | 0 | NA | NA | NA |
| Phactr1 | 3991.264433 | -0.108818815 | 0.117383724 | -0.927034951 |
| Gm15810 | 0.418886444 | 0.282644672 | 1.39140604 | 0.203136011 |
| Gm15812 | 0 | NA | NA | NA |
| Gm15809 | 21.27740935 | -0.16761012 | 0.316459528 | -0.529641566 |
| Gm26086 | 0 | NA | NA | NA |
| Gm15813 | 0.11193333 | 0.75528439 | 3.724945049 | 0.202763901 |
| Tbc1d7 | 105.3200195 | -0.081582237 | 0.115785615 | -0.704597347 |
| Gfod1 | 1583.085298 | -0.37237287 | 0.089698277 | -4.151393811 |
| Sirt5 | 74.72223409 | -0.469580172 | 0.138870361 | -3.381428331 |
| Nol7 | 275.008259 | 0.040254287 | 0.138420118 | 0.290812403 |
| Ranbp9 | 1368.855223 | 0.325675095 | 0.075977615 | 4.286461136 |
| Gm26064 | 7.226595754 | 0.397501195 | 0.427765702 | 0.929249806 |
| Mcur1 | 233.0888101 | 0.963183381 | 0.109641906 | 8.784810592 |
| Rnf182 | 236.6714893 | -0.482400614 | 0.108326041 | -4.453228489 |
| Cd83 | 231.474017 | 0.491504005 | 0.128965186 | 3.811137116 |
| Gm23117 | 0 | NA | NA | NA |
| Gm5083 | 5.023184258 | -0.956751789 | 0.49067675 | -1.949861674 |
| Gm27007 | 0 | NA | NA | NA |
| Gm27036 | 0.066050875 | 0.274391604 | 3.724945049 | 0.073663262 |
| 1700029N11Rik | 0 | NA | NA | NA |
| Jarid2 | 797.878151 | -0.106240776 | 0.088159689 | -1.205094731 |
| Gm22213 | 0.047948698 | -0.206487837 | 3.724945049 | -0.055433794 |
| Dtnbp1 | 383.1011531 | 0.299027915 | 0.111176913 | 2.689658383 |
| Gm23104 | 0 | NA | NA | NA |
| Hsp25-ps1 | 4.708514708 | 1.164613083 | 0.794545754 | 1.465759621 |
| Gm9817 | 0.309768565 | 0.767670325 | 1.794602956 | 0.42776611 |
| Gm8513 | 0.148910734 | 0.723040935 | 2.858091615 | 0.252980321 |
| Gm23755 | 0 | NA | NA | NA |

| | | | | |
|---|---|---|---|---|
| Mylip | 196.9662467 | 0.077406508 | 0.103964127 | 0.744550172 |
| Gm23387 | 0 | NA | NA | NA |
| Gmpr | 269.5668325 | -0.308810007 | 0.085288802 | -3.620756778 |
| Atxn1 | 2899.31005 | 0.097967266 | 0.081839646 | 1.197063652 |
| 5033430I15Rik | 31.65815391 | 0.518307952 | 0.185715271 | 2.790874166 |
| Gm10113 | 0.097139547 | 0.274393828 | 3.724945049 | 0.073663859 |
| Stmnd1 | 2.201457459 | 2.30013515 | 0.85688493 | 2.684298756 |
| Rbm24 | 161.1267455 | -0.195626869 | 0.136759241 | -1.430447161 |
| Cap2 | 1272.593817 | -0.523874719 | 0.134820218 | -3.885728185 |
| Gm24915 | 2.015269402 | -1.715970136 | 0.738917928 | -2.32227433 |
| C78339 | 1105.225121 | -0.105097561 | 0.081217016 | -1.294033765 |
| Nup153 | 1192.694187 | 0.113220481 | 0.060190658 | 1.881030783 |
| Kif13a | 663.6183412 | -0.408330058 | 0.085452959 | -4.778419158 |
| Gm24620 | 4.394585356 | -0.36403172 | 0.473848117 | -0.768245577 |
| Gm22960 | 0 | NA | NA | NA |
| Gm24769 | 0 | NA | NA | NA |
| 2010001K21Rik | 1.984554007 | -0.550337855 | 0.773510076 | -0.711481172 |
| Nhlrc1 | 147.1966146 | -0.271851134 | 0.098201606 | -2.768296216 |
| Tpmt | 47.2986511 | -0.23381345 | 0.173571026 | -1.347076496 |
| Kdm1b | 181.3782971 | -0.271377853 | 0.120500684 | -2.252085587 |
| Dek | 884.7800832 | 0.860747471 | 0.121372668 | 7.091773503 |
| Rnf144b | 226.1613996 | -0.166412083 | 0.116381064 | -1.429889689 |
| 4931429P17Rik | 0.349470464 | -0.478145487 | 1.765781349 | -0.270784085 |
| A330033J07Rik | 16.48077188 | -0.304284562 | 0.292541574 | -1.040141265 |
| Id4 | 466.0183473 | -0.647192265 | 0.161632527 | -4.004096686 |
| A330048O09Rik | 5.209108024 | -1.215070275 | 0.447537451 | -2.715013621 |
| Zfp169 | 346.1617771 | -0.237080247 | 0.095961428 | -2.470578568 |
| A830005F24Rik | 4.046338601 | -1.421475365 | 0.583409305 | -2.436497587 |
| 6720427I07Rik | 280.6546708 | 0.090854883 | 0.101948177 | 0.891186938 |
| Mirlet7d | 0 | NA | NA | NA |
| Gm24111 | 0 | NA | NA | NA |
| mmu-let-7f-1 | 12.7179762 | -0.115236511 | 0.306875937 | -0.37551498 |
| Mirlet7a-1 | 9.148397703 | 0.421170798 | 0.442895366 | 0.950948758 |
| Gm37238 | 83.05504175 | -0.311321438 | 0.232831072 | -1.337112932 |
| Gm25232 | 0 | NA | NA | NA |
| Ptpdc1 | 475.0012653 | 0.184430896 | 0.07789755 | 2.36760844 |
| Barx1 | 0 | NA | NA | NA |
| Phf2 | 871.8668907 | 0.015533197 | 0.06408183 | 0.242396282 |
| Phf2os1 | 14.57948276 | -0.723737445 | 0.329824626 | -2.19430991 |
| Fam120a | 2396.718561 | 0.055311351 | 0.05901292 | 0.937275282 |
| Fam120aos | 129.3350566 | -0.014487614 | 0.145325727 | -0.099690638 |
| Wnk2 | 2363.287208 | -0.107621463 | 0.11185776 | -0.962127821 |
| Ninj1 | 228.5158888 | 0.871869269 | 0.173613766 | 5.021890194 |
| 1110007C09Rik | 111.7269501 | 0.861034301 | 0.188698887 | 4.563006773 |
| Susd3 | 9.113448631 | 0.520961865 | 0.358980982 | 1.451224135 |
| Fgd3 | 292.1672564 | 0.191184043 | 0.162240274 | 1.17840064 |
| Bicd2 | 1037.207963 | -0.183862446 | 0.058444979 | -3.145906628 |

| | | | | |
|---|---|---|---|---|
| Ippk | 334.3509724 | 0.035445496 | 0.091889213 | 0.385741644 |
| Cenpp | 10.13615864 | 0.366086699 | 0.405425253 | 0.90296965 |
| Tes3-ps | 44.94843152 | 0.387095897 | 0.295400174 | 1.310411875 |
| Ecm2 | 93.28506817 | 0.87721566 | 0.183073679 | 4.791599006 |
| Aspn | 20.67132123 | -1.177046833 | 0.295618576 | -3.981640286 |
| Omd | 21.95627783 | 0.997801331 | 0.31510758 | 3.166541818 |
| Ogn | 102.8893265 | -0.117814182 | 0.217677128 | -0.541233627 |
| Nol8 | 394.8682349 | 0.089474052 | 0.064442885 | 1.38842406 |
| Iars | 963.3246765 | -0.052124657 | 0.074683976 | -0.69793629 |
| Gm25394 | 8.321309178 | -0.286525546 | 0.444020663 | -0.645297776 |
| Gm23482 | 2.859218534 | -1.462607146 | 0.592246058 | -2.469593721 |
| Gm7251 | 0.539950272 | 1.358656295 | 1.41270551 | 0.961740635 |
| Gm8674 | 0 | NA | NA | NA |
| Gm10784 | 0 | NA | NA | NA |
| Gm17611 | 0 | NA | NA | NA |
| Gm24300 | 0 | NA | NA | NA |
| Gm906 | 0 | NA | NA | NA |
| Gm24017 | 0 | NA | NA | NA |
| Gm8739 | 0 | NA | NA | NA |
| Fbxw17 | 179.6250207 | 1.092127083 | 0.164123653 | 6.654294267 |
| Nutm2 | 0.276311821 | 1.719880949 | 2.407135304 | 0.714492844 |
| Mir683-1 | 0 | NA | NA | NA |
| Mir683-2 | 0.088123299 | -0.687376175 | 3.724945049 | -0.184533239 |
| Gm904 | 0.038981395 | 0.274393828 | 3.724945049 | 0.073663859 |
| Gm23952 | 0 | NA | NA | NA |
| Gm8765 | 0 | NA | NA | NA |
| Gm22199 | 0 | NA | NA | NA |
| Spin1 | 2716.345326 | 0.236742993 | 0.065248019 | 3.628355267 |
| Nxnl2 | 1.249332652 | 1.561723861 | 0.999522384 | 1.562470122 |
| Hist1h2al | 110.2101923 | 0.573459827 | 0.164864367 | 3.478373388 |
| AC160122.1 | 0.260264958 | 0.704285785 | 2.093597966 | 0.336399727 |
| S1pr3 | 554.2109028 | 2.507651106 | 0.297468223 | 8.429979773 |
| Shc3 | 425.7683225 | -1.084324141 | 0.148653976 | -7.294282811 |
| Cks2 | 12.99324003 | 0.129778748 | 0.383684521 | 0.338243377 |
| Gm22806 | 0.426919779 | 0.71318198 | 1.738149862 | 0.410310984 |
| Secisbp2 | 291.1100637 | 0.054788874 | 0.088653423 | 0.618011944 |
| Gm15440 | 205.1826132 | -0.193304144 | 0.116274846 | -1.662476017 |
| Sema4d | 1108.340496 | 0.284182489 | 0.091498804 | 3.105860156 |
| Gm23566 | 0 | NA | NA | NA |
| Gadd45g | 275.2348046 | 1.693737744 | 0.32597864 | 5.19585499 |
| Gm26651 | 0.10503227 | 0.274389469 | 3.724945049 | 0.073662689 |
| Gm37872 | 0 | NA | NA | NA |
| Diras2 | 2681.114 | -0.584718911 | 0.107204346 | -5.454246353 |
| Syk | 204.4789825 | 1.99528016 | 0.232371612 | 8.586591706 |
| Auh | 575.8122667 | -0.200218458 | 0.05581075 | -3.58745329 |
| n-R5s54 | 0.832946694 | 0.736903946 | 1.387098219 | 0.531255779 |
| Nfil3 | 206.9757982 | 1.290873687 | 0.346033692 | 3.730485543 |

| | | | | |
|---|---|---|---|---|
| Ror2 | 25.4305805 | 0.211213403 | 0.227779663 | 0.92727068 |
| Sptlc1 | 374.6291411 | 0.191113905 | 0.094405611 | 2.02439138 |
| RP23-340E6.2 | 12.21145091 | -0.460964146 | 0.307653929 | -1.49832036 |
| RP23-340E6.3 | 0 | NA | NA | NA |
| Msx2 | 19.08535284 | -0.21154808 | 0.345619109 | -0.612084444 |
| Gm5449 | 0.111006857 | 0.755286617 | 3.724945049 | 0.202764499 |
| Drd1 | 333.3471759 | -0.526348408 | 0.220936128 | -2.382355539 |
| Sfxn1 | 561.8273373 | -0.067412572 | 0.066616871 | -1.01194444 |
| Hrh2 | 63.56667738 | -0.199879367 | 0.174286646 | -1.146842693 |
| Gm16578 | 0.107803068 | 0.75528661 | 3.724945049 | 0.202764497 |
| Cplx2 | 9264.934547 | -0.302313076 | 0.121087039 | -2.49665924 |
| Gm16248 | 0 | NA | NA | NA |
| Gm22668 | 0 | NA | NA | NA |
| Thoc3 | 293.4655856 | 0.165075569 | 0.095152432 | 1.734853918 |
| Gm2830 | 71.59267597 | 0.637229953 | 0.158609017 | 4.017614909 |
| Simc1 | 267.6907852 | -0.273752695 | 0.088666059 | -3.087457564 |
| 4833439L19Rik | 1366.025163 | -0.025929057 | 0.061449035 | -0.421960366 |
| Arl10 | 135.7640415 | -0.338322196 | 0.130855464 | -2.585464797 |
| Gm24195 | 1.207241537 | -0.154354427 | 0.892704347 | -0.172906548 |
| Nop16 | 117.0143181 | -0.055357276 | 0.141900719 | -0.390112722 |
| Higd2a | 413.1629045 | -0.369282197 | 0.093805507 | -3.936679293 |
| Cltb | 2228.333823 | 0.230200421 | 0.093895747 | 2.451659719 |
| Faf2 | 759.1273305 | 0.247765599 | 0.072115808 | 3.435662783 |
| Rnf44 | 1513.683017 | -0.168511953 | 0.063979103 | -2.633859279 |
| Cdhr2 | 4.783323946 | -0.371385967 | 0.431045186 | -0.861594047 |
| Gprin1 | 1231.562756 | -0.249372763 | 0.096821432 | -2.575594642 |
| Sncb | 3503.722137 | -0.082368681 | 0.101428791 | -0.812083829 |
| Gm16249 | 0 | NA | NA | NA |
| Eif4e1b | 0 | NA | NA | NA |
| Tspan17 | 212.3703016 | -0.05073172 | 0.130481525 | -0.388803856 |
| Gm16250 | 0 | NA | NA | NA |
| Gm29431 | 0.504101427 | -2.126512216 | 1.390065848 | -1.529792433 |
| 5330429C05Rik | 1.674398058 | 0.876524279 | 0.88851908 | 0.986500233 |
| 4930526F13Rik | 0.237545104 | 1.324557094 | 2.53859436 | 0.521767918 |
| Unc5a | 875.420532 | 0.146815408 | 0.125328204 | 1.171447474 |
| Hk3 | 302.6549373 | 4.248239515 | 0.376825012 | 11.27377264 |
| Uimc1 | 267.0504781 | 0.144560529 | 0.117432357 | 1.231011048 |
| Gm6344 | 1.345796852 | 0.778528297 | 0.97158674 | 0.80129572 |
| Zfp346 | 338.7158424 | 0.05088843 | 0.100720634 | 0.50524335 |
| Gm17617 | 0 | NA | NA | NA |
| Fgfr4 | 13.86132062 | 1.542698791 | 0.503055167 | 3.06665927 |
| Nsd1 | 4468.214251 | -0.259836545 | 0.069122265 | -3.759086082 |
| Rab24 | 365.9432809 | 0.207933509 | 0.069015661 | 3.012845289 |
| Prelid1 | 846.0882492 | 0.270809403 | 0.091502822 | 2.959574322 |
| Mxd3 | 8.517488434 | -0.005260315 | 0.514851915 | -0.010217143 |
| Lman2 | 1016.200266 | 0.044218172 | 0.08639892 | 0.511790789 |
| Rgs14 | 301.6653793 | 0.130512266 | 0.113827182 | 1.146582595 |

| | | | | |
|---|---|---|---|---|
| Slc34a1 | 0.764795395 | 0.30230694 | 1.093392948 | 0.276485174 |
| Pfn3 | 0.525579064 | 0.062764237 | 1.348082372 | 0.046558162 |
| F12 | 5.68547348 | -0.422912666 | 0.677151479 | -0.624546618 |
| Grk6 | 508.6179697 | 0.007845542 | 0.056264656 | 0.139439975 |
| Mir6943 | 0 | NA | NA | NA |
| Prr7 | 196.075479 | -0.225417052 | 0.124544794 | -1.809927537 |
| Dbn1 | 1526.862046 | -0.099655664 | 0.101672746 | -0.980161034 |
| Mir6944 | 0 | NA | NA | NA |
| Pdlim7 | 426.9912292 | 0.095560549 | 0.089281987 | 1.07032283 |
| Mir6945 | 0 | NA | NA | NA |
| Dok3 | 92.02924505 | 0.640953213 | 0.234791672 | 2.729880527 |
| Ddx41 | 338.5120362 | 0.187169944 | 0.109975839 | 1.701918754 |
| Fam193b | 537.9735023 | -0.574201271 | 0.180436729 | -3.182285971 |
| Tmed9 | 877.8577826 | 0.554089237 | 0.078806121 | 7.031043187 |
| B4galt7 | 189.7572043 | -0.044994181 | 0.085377723 | -0.527001413 |
| Gm15911 | 0.088476398 | 0.274400501 | 3.724945049 | 0.073665651 |
| Caml | 264.5844916 | 0.338324607 | 0.08192046 | 4.12991585 |
| Ddx46 | 1095.580132 | 0.135696625 | 0.079823587 | 1.699956488 |
| B230219D22Rik | 1459.612451 | -0.109298244 | 0.074009219 | -1.476819316 |
| Txndc15 | 462.6097524 | 0.026083408 | 0.100106733 | 0.260555984 |
| Pcbd2 | 43.31228221 | -0.115717876 | 0.191991155 | -0.602725036 |
| 4930451E10Rik | 0 | NA | NA | NA |
| Catsper3 | 9.093479106 | -0.697207288 | 0.348801607 | -1.998864896 |
| Pitx1 | 1.253999855 | 0.025237193 | 0.888755505 | 0.028396104 |
| Gm28760 | 0.04393175 | -0.206490061 | 3.724945049 | -0.055434391 |
| H2afy | 903.7078744 | 0.371077209 | 0.072711221 | 5.103437975 |
| Gm25148 | 0 | NA | NA | NA |
| Tifab | 116.9424394 | 1.792789208 | 0.300263105 | 5.970727604 |
| 4930550C17Rik | 0 | NA | NA | NA |
| Neurog1 | 0.267034963 | -2.003226468 | 2.037186825 | -0.983329778 |
| Cxcl14 | 505.7540529 | -0.129611318 | 0.122309844 | -1.059696534 |
| Gm24716 | 0 | NA | NA | NA |
| Gm10782 | 0 | NA | NA | NA |
| Slc25a48 | 3.125560887 | -0.697836441 | 0.806161314 | -0.865628788 |
| Il9 | 0 | NA | NA | NA |
| Fbxl21 | 83.03568542 | 0.241779072 | 0.128436322 | 1.882482065 |
| Lect2 | 0.045882455 | 0.755288839 | 3.724945049 | 0.202765096 |
| Tgfbi | 1350.337662 | 4.883892146 | 0.322011382 | 15.16683078 |
| Gm27698 | 0 | NA | NA | NA |
| Smad5 | 771.3450184 | -0.10799266 | 0.079470038 | -1.358910388 |
| Gm27301 | 0.049553047 | 0.274391604 | 3.724945049 | 0.073663262 |
| Gm27834 | 0.26282868 | -1.148284131 | 1.811378309 | -0.633928388 |
| Gm27271 | 2.902142703 | -1.139830901 | 0.566136658 | -2.0133494 |
| Trpc7 | 38.7170757 | -0.726690152 | 0.171591325 | -4.235005193 |
| Gm22777 | 0 | NA | NA | NA |
| Spock1 | 3106.932593 | -0.087781187 | 0.072246811 | -1.215018163 |
| Klhl3 | 316.2281292 | -0.593862045 | 0.159574055 | -3.721545116 |

| | | | | |
|---|---|---|---|---|
| Mir874 | 4.532224978 | -1.582478766 | 0.614627916 | -2.574693933 |
| Hnrnpa0 | 1502.786418 | 0.132618493 | 0.100201181 | 1.323522252 |
| 1700066J03Rik | 0.225300421 | 0.497061484 | 2.565566006 | 0.193743401 |
| Idnk | 305.8515994 | 1.617617817 | 0.160318129 | 10.09004927 |
| Ubqln1 | 2320.123517 | -0.025044679 | 0.062840383 | -0.398544349 |
| Gkap1 | 124.6432152 | 0.051908727 | 0.139572799 | 0.371911488 |
| Gm26555 | 4.731241973 | -0.393198983 | 0.539030764 | -0.729455551 |
| Kif27 | 62.59338873 | -0.053894173 | 0.171259417 | -0.314693197 |
| Mir6369 | 0.042716945 | -0.206490061 | 3.724945049 | -0.055434391 |
| Gm24284 | 0 | NA | NA | NA |
| 2210016F16Rik | 128.8048172 | 0.388518394 | 0.167743031 | 2.316152222 |
| Hnrnpk | 4038.157108 | 0.296858201 | 0.076804079 | 3.865135892 |
| Mir7-1 | 15.77224037 | 0.677142936 | 0.266224114 | 2.543507147 |
| Rmi1 | 123.5749391 | 0.294707635 | 0.112263488 | 2.625142337 |
| Slc28a3 | 2.52226614 | -1.280841779 | 0.718968183 | -1.781499945 |
| Ntrk2 | 6695.738885 | -0.432724718 | 0.105611334 | -4.097332176 |
| Agtpbp1 | 2886.668731 | -0.193524471 | 0.068614012 | -2.820480324 |
| Naa35 | 495.813822 | -0.190155503 | 0.055842373 | -3.40521887 |
| Golm1 | 357.7061881 | 0.491473015 | 0.117312101 | 4.189448591 |
| 4921517D22Rik | 0.150248541 | 0.755303235 | 2.859451661 | 0.264142684 |
| Spata31d1a | 2.564266786 | 0.096522162 | 0.672378205 | 0.143553377 |
| Spata31d1b | 1.567374066 | 0.914596649 | 0.897451092 | 1.019104726 |
| Spata31d1d | 3.578253317 | 0.290219747 | 0.563257883 | 0.515251993 |
| 4930528D03Rik | 0.109124069 | 0.274402725 | 3.724945049 | 0.073666248 |
| 1700014D04Rik | 13.37826059 | 0.206100549 | 0.285634028 | 0.721554608 |
| Isca1 | 1058.772678 | 0.134195382 | 0.055046971 | 2.437834072 |
| Etohd2 | 12.41969927 | -1.67065499 | 0.32194718 | -5.189220758 |
| Zcchc6 | 1650.158349 | 1.035039429 | 0.098479746 | 10.51017568 |
| Gas1 | 102.0490903 | -0.524406741 | 0.187980219 | -2.78969109 |
| Gm5084 | 0.588182734 | -2.491410303 | 1.487519075 | -1.674876205 |
| Gm24999 | 0 | NA | NA | NA |
| Gm25000 | 0 | NA | NA | NA |
| Dapk1 | 1289.496349 | 0.268046843 | 0.120656747 | 2.221565309 |
| Gm10312 | 0 | NA | NA | NA |
| BC051665 | 0.048949532 | -0.206487837 | 3.724945049 | -0.055433794 |
| Ctsll3 | 0 | NA | NA | NA |
| 4930486L24Rik | 7.005733943 | 3.26290045 | 0.70808191 | 4.608083336 |
| Ctla2b | 52.79599323 | 3.593107095 | 0.365858725 | 9.8210234 |
| Tpbpb | 0 | NA | NA | NA |
| Gm24594 | 0 | NA | NA | NA |
| Ctla2a | 216.8941751 | 4.317614046 | 0.276791886 | 15.5987739 |
| Tpbpa | 0 | NA | NA | NA |
| Ctsj | 0 | NA | NA | NA |
| Ctsq | 0 | NA | NA | NA |
| Ctsr | 0 | NA | NA | NA |
| Cts6 | 0 | NA | NA | NA |
| Cts8 | 0 | NA | NA | NA |

| | | | | |
|---|---|---|---|---|
| Gm27704 | 0 | NA | NA | NA |
| Cts7 | 0 | NA | NA | NA |
| Ctsm | 0 | NA | NA | NA |
| Cts3 | 0 | NA | NA | NA |
| Gm19961 | 1.330930655 | -0.017554631 | 0.981915856 | -0.017877938 |
| Gm23739 | 0 | NA | NA | NA |
| Gm24193 | 0 | NA | NA | NA |
| Gm20563 | 3.002659297 | 0.593988815 | 0.689701016 | 0.861226533 |
| Gm25649 | 0.044544648 | 0.755286614 | 3.724945049 | 0.202764498 |
| Gm6478 | 2.109479346 | 0.269814972 | 0.644655968 | 0.418541028 |
| Zfp808 | 54.99002886 | -0.170622467 | 0.194235295 | -0.878431835 |
| Gm23713 | 0.185738947 | 0.274409542 | 2.674436203 | 0.102604632 |
| Gm24539 | 0.901339752 | -1.243771748 | 1.088951436 | -1.142173752 |
| Gm3604 | 26.94184347 | 0.534404554 | 0.234846489 | 2.275548402 |
| Gm25933 | 0.061920613 | 0.274393828 | 3.724945049 | 0.073663859 |
| Gm23855 | 0 | NA | NA | NA |
| Zfp935 | 99.08513468 | -0.094624377 | 0.13629517 | -0.694260677 |
| Gm22256 | 0.093484797 | -0.20649451 | 3.724945049 | -0.055435586 |
| Zfp934 | 37.404617 | -0.049801081 | 0.217752554 | -0.228704922 |
| Gm7664 | 14.88251027 | -0.158908671 | 0.27214454 | -0.583912767 |
| Gm24095 | 0.244748102 | -0.929947208 | 2.502495982 | -0.371607873 |
| Gm22615 | 1.125305835 | -2.262388965 | 1.02113161 | -2.215570395 |
| 6720489N17Rik | 53.55286888 | -0.442617966 | 0.185785028 | -2.382419996 |
| Gm24020 | 0.042716945 | -0.206490061 | 3.724945049 | -0.055434391 |
| Platr25 | 30.64885467 | -0.842266369 | 0.302597692 | -2.783452723 |
| Mir713 | 0 | NA | NA | NA |
| Gm5141 | 99.08605519 | -0.305495746 | 0.163696267 | -1.866235269 |
| Gm21963 | 0 | NA | NA | NA |
| Gm24588 | 0 | NA | NA | NA |
| Gm25448 | 0 | NA | NA | NA |
| Fbp2 | 0.981774026 | -1.267143737 | 1.062994434 | -1.19205115 |
| Fbp1 | 2.587088777 | 0.25001685 | 0.73304591 | 0.341065746 |
| 2010111I01Rik | 498.5900927 | 0.008350216 | 0.090669597 | 0.092094999 |
| Gm16907 | 5.155791298 | 0.680657186 | 0.426705829 | 1.595143865 |
| Mir23b | 8.607625547 | 0.285929199 | 0.42961653 | 0.665545153 |
| Mir27b | 11.22797161 | 0.407928824 | 0.360090092 | 1.132852121 |
| Mir3074-1 | 0.352569346 | 0.274415344 | 1.608257285 | 0.170629007 |
| mmu-mir-24-1 | 9.164853348 | 0.096999009 | 0.348802267 | 0.278091681 |
| Fancc | 74.63926839 | -0.223865735 | 0.134341397 | -1.666394277 |
| Gm16133 | 0.546112947 | 2.652953944 | 1.330718843 | 1.993624693 |
| Gm16132 | 1.982187692 | 1.509413758 | 0.708721388 | 2.129770294 |
| Gm16134 | 0 | NA | NA | NA |
| Ptch1 | 1765.336376 | -0.407810109 | 0.102689837 | -3.97128011 |
| A930032L01Rik | 2.69933516 | -1.547213873 | 0.678542715 | -2.280201141 |
| Ercc6l2 | 388.7591555 | 0.123794298 | 0.084002614 | 1.473695776 |
| Hsd17b3 | 0.044191549 | -0.206490061 | 3.724945049 | -0.055434391 |
| Slc35d2 | 16.4056275 | 1.842527682 | 0.411197915 | 4.48087798 |

| | | | | |
|---|---|---|---|---|
| Zfp367 | 307.36151 | 0.068769396 | 0.125360822 | 0.548571669 |
| Habp4 | 1131.608262 | -0.175059421 | 0.079285349 | -2.207966824 |
| Cdc14b | 233.8312342 | 0.011056234 | 0.098140446 | 0.112657261 |
| 1810034E14Rik | 1.766063332 | 1.42487074 | 0.951123348 | 1.498092485 |
| Aaed1 | 90.98287385 | -0.42664546 | 0.141368561 | -3.017965643 |
| Gm25654 | 2.492378667 | 0.120669904 | 0.649392264 | 0.185819744 |
| Ctsl | 1539.723944 | 0.857040865 | 0.180275671 | 4.754057263 |
| Cdk20 | 89.79521824 | -0.061098098 | 0.122326027 | -0.499469322 |
| 1190003K10Rik | 0.204782255 | -0.699723755 | 2.205163063 | -0.317311571 |
| Gm4810 | 1.123243924 | 0.151135766 | 0.807742592 | 0.187108824 |
| Cntnap3 | 108.0176306 | -0.337413167 | 0.126919046 | -2.6584912 |
| Spata31 | 0 | NA | NA | NA |
| Spata31d1c | 0 | NA | NA | NA |
| Prss47 | 0.08486385 | 0.755286617 | 3.724945049 | 0.202764499 |
| Hiatl1 | 800.205226 | -0.009327753 | 0.082197753 | -0.113479411 |
| Olfr466 | 0 | NA | NA | NA |
| Gm24130 | 0 | NA | NA | NA |
| Nlrp4f | 0.716239967 | 0.67384588 | 1.367151432 | 0.492883132 |
| Gm10775 | 2.001003253 | 0.830751837 | 0.834458168 | 0.995558398 |
| Zfp369 | 347.6968427 | -0.388676524 | 0.106905058 | -3.635716865 |
| Gm5792 | 0 | NA | NA | NA |
| Gm10139 | 0 | NA | NA | NA |
| Gm6901 | 0.223727156 | 0.274410664 | 2.154193795 | 0.127384391 |
| Gm26639 | 0 | NA | NA | NA |
| Platr2 | 0 | NA | NA | NA |
| Gm6917 | 0 | NA | NA | NA |
| Gm6910 | 5.181120878 | 0.577965398 | 0.432738196 | 1.33560061 |
| Gm10324 | 0 | NA | NA | NA |
| Gm17514 | 0 | NA | NA | NA |
| Gm10772 | 0 | NA | NA | NA |
| Gm26754 | 0 | NA | NA | NA |
| Gm26715 | 0 | NA | NA | NA |
| Vmn2r-ps104 | 0 | NA | NA | NA |
| 2410141K09Rik | 0 | NA | NA | NA |
| Gm26806 | 0 | NA | NA | NA |
| Gm27501 | 0 | NA | NA | NA |
| Zfp640 | 0 | NA | NA | NA |
| Gm17404 | 0 | NA | NA | NA |
| Gm17449 | 0 | NA | NA | NA |
| Gm10323 | 0 | NA | NA | NA |
| Gm5196 | 0.205773706 | -1.144255305 | 2.189704783 | -0.522561449 |
| Gm5451 | 26.41010186 | 0.270422045 | 0.254371893 | 1.063097189 |
| Uqcrb | 583.1177258 | 0.214679259 | 0.10010284 | 2.144587105 |
| Gm10767 | 12.81661586 | -0.21871915 | 0.27438904 | -0.797113289 |
| Mterf3 | 229.6188801 | -0.014258533 | 0.080439714 | -0.177257381 |
| Ptdss1 | 668.8494159 | -0.032687667 | 0.069611494 | -0.46957284 |
| Zfp712 | 93.19383338 | -0.3688672 | 0.172053582 | -2.143908869 |

| | | | | |
|---|---|---|---|---|
| Zfp708 | 0.458268618 | -1.728105339 | 1.359973036 | -1.270690884 |
| Gm28557 | 4.418616242 | -1.460939293 | 0.568294407 | -2.570743747 |
| Rslcan18 | 4.299918267 | -1.646608137 | 0.548671193 | -3.00108363 |
| Zfp759 | 131.3437174 | -0.097883241 | 0.157661409 | -0.620844641 |
| Rsl1 | 30.30590387 | 0.374958181 | 0.286589232 | 1.308347068 |
| Zfp455 | 45.95318099 | 0.461300744 | 0.280069628 | 1.647093073 |
| Zfp458 | 240.4665658 | -0.366412582 | 0.13443658 | -2.725542274 |
| Zfp457 | 21.39111743 | -0.380876404 | 0.276229058 | -1.378842642 |
| Zfp595 | 65.32548619 | 0.085286917 | 0.20463789 | 0.416769919 |
| Gm28044 | 0.23851432 | -0.206481968 | 2.112354461 | -0.097749678 |
| Zfp953 | 36.77585063 | 0.223881071 | 0.215885413 | 1.037036582 |
| Gm28041 | 0.153723802 | 0.274408321 | 2.839085032 | 0.096653787 |
| Gm17039 | 2.363953117 | -0.591252214 | 0.721654562 | -0.819300874 |
| Zfp456 | 19.86136891 | 0.33425752 | 0.28950215 | 1.15459426 |
| Zfp429 | 33.84957497 | 0.64257821 | 0.229305473 | 2.802280302 |
| Zfp459 | 66.34388881 | 0.024431898 | 0.154472725 | 0.158163183 |
| Zfp874a | 90.85288293 | -0.190660464 | 0.158068873 | -1.206186018 |
| Gm38307 | 0.282611191 | -0.680160452 | 1.64517603 | -0.413427159 |
| Gm7928 | 0 | NA | NA | NA |
| Zfp874b | 171.5607145 | -0.340233986 | 0.158858446 | -2.14174313 |
| Zfp58 | 50.28513376 | -0.395734232 | 0.174167822 | -2.272143204 |
| Gm26965 | 0 | NA | NA | NA |
| Zfp87 | 184.3921584 | -0.229483995 | 0.105013937 | -2.185271798 |
| Zfp748 | 247.3472852 | -0.048899557 | 0.135529344 | -0.360804206 |
| 9430065F17Rik | 2.796353279 | -0.046641741 | 0.60095018 | -0.077613324 |
| Zfp729b | 275.2153557 | 0.323809506 | 0.118830331 | 2.724973523 |
| Gm37276 | 23.42523615 | 0.002413867 | 0.236469857 | 0.010207928 |
| Zfp729a | 293.0100012 | 0.123267913 | 0.087366078 | 1.410935644 |
| Gm26587 | 0 | NA | NA | NA |
| Zfp738 | 303.073871 | -0.252766483 | 0.132660381 | -1.905365267 |
| Gm26875 | 0 | NA | NA | NA |
| Zfp65 | 240.5209492 | 0.055140903 | 0.114907351 | 0.479872717 |
| Zfp85os | 44.02786501 | 0.765509919 | 0.198372405 | 3.85895366 |
| Zfp85 | 74.6551353 | 0.300573813 | 0.163240955 | 1.841289235 |
| Gm9894 | 0 | NA | NA | NA |
| Zfp493 | 42.58231206 | -0.466356157 | 0.282154709 | -1.652838468 |
| 4930525G20Rik | 7.15404337 | -0.446274949 | 0.428741767 | -1.040894504 |
| Gm9625 | 125.9289179 | 0.945078381 | 0.161931648 | 5.836279638 |
| Zfp273 | 60.68720866 | -0.035273179 | 0.16135295 | -0.218608828 |
| Gm10037 | 3.305635379 | -0.361272393 | 0.542637558 | -0.665771079 |
| BC048507 | 0.806474979 | 0.570528067 | 1.085754541 | 0.525466895 |
| Mtrr | 182.8372363 | -0.226825964 | 0.101838132 | -2.227318597 |
| Fastkd3 | 102.6350786 | -0.36380983 | 0.124822615 | -2.914614708 |
| 1700001L19Rik | 42.40692636 | -0.154876105 | 0.161652575 | -0.958080037 |
| Adcy2 | 1089.553893 | -0.453628393 | 0.090036547 | -5.038269544 |
| Gm26929 | 0 | NA | NA | NA |
| Gm37122 | 2.65229942 | -0.642458385 | 0.653332793 | -0.983355484 |

| | | | | |
|---|---|---|---|---|
| Gm26844 | 0 | NA | NA | NA |
| Gm4812 | 0 | NA | NA | NA |
| AC161592.1 | 0 | NA | NA | NA |
| Papd7 | 947.3406657 | 0.644023025 | 0.081895822 | 7.863930191 |
| Nsun2 | 820.5404197 | 0.130808786 | 0.063815686 | 2.049790507 |
| A530095I07Rik | 0.197913552 | -0.68737861 | 2.226349723 | -0.308746916 |
| Srd5a1 | 47.64261977 | -0.061169077 | 0.171359955 | -0.356962496 |
| Gm17108 | 3.053155272 | 0.734450533 | 0.616740829 | 1.190857648 |
| Ube2ql1 | 1424.867787 | -0.254923059 | 0.085293045 | -2.988790687 |
| Med10 | 183.4117316 | 0.082911908 | 0.11212085 | 0.739486969 |
| Gm26819 | 0.108772582 | 0.75528439 | 3.724945049 | 0.202763901 |
| 1700084F23Rik | 0.092691418 | 0.274387304 | 3.724945049 | 0.073662108 |
| AC123920.1 | 0.239188194 | 0.238481171 | 1.824742444 | 0.130693058 |
| Gm26018 | 0 | NA | NA | NA |
| 1700100L14Rik | 0 | NA | NA | NA |
| Ice1 | 1670.349411 | -0.102387683 | 0.110759195 | -0.924417003 |
| Adamts16 | 57.86550676 | -0.67162628 | 0.229303321 | -2.928986276 |
| 8030423J24Rik | 1.260089781 | -0.634866909 | 0.924849567 | -0.686454242 |
| Gm38191 | 0.247225956 | 1.236189465 | 2.992047273 | 0.4131584 |
| Mir466f-4 | 0 | NA | NA | NA |
| Irx1 | 50.49064827 | 0.054159448 | 0.210572691 | 0.257200721 |
| Rpl9-ps4 | 43.5655805 | 0.084512071 | 0.193722061 | 0.43625424 |
| Gm20554 | 3.453678518 | 0.296087929 | 0.523056451 | 0.566072608 |
| Irx2 | 81.72096566 | 0.31977809 | 0.212435697 | 1.50529358 |
| Irx4 | 3.045905106 | 0.481437944 | 0.597682253 | 0.80550818 |
| Gm10263 | 1.909878088 | 0.45787812 | 0.704912849 | 0.649552807 |
| Ndufs6 | 294.0100797 | -0.039483743 | 0.077256561 | -0.511072999 |
| Mrpl36 | 129.8851055 | -0.39309188 | 0.178266842 | -2.205075692 |
| Lpcat1 | 574.2185807 | 0.247089455 | 0.088739246 | 2.78444394 |
| Slc6a3 | 74.17620595 | -0.05505143 | 0.286434486 | -0.192195537 |
| Clptm1l | 878.6178926 | 0.131297117 | 0.064596623 | 2.032569363 |
| Tert | 17.88891691 | -0.729921484 | 0.280943951 | -2.598103576 |
| Slc6a18 | 0.178490723 | 1.257478905 | 2.711241583 | 0.463801866 |
| Slc6a19 | 6.934680317 | 0.72770421 | 0.418377932 | 1.739346546 |
| Slc6a19os | 0.042716945 | -0.206490061 | 3.724945049 | -0.055434391 |
| Slc12a7 | 159.1782715 | 1.348039675 | 0.174494018 | 7.725420545 |
| Nkd2 | 44.99299481 | -0.196810639 | 0.189341888 | -1.039445849 |
| Gm8062 | 0 | NA | NA | NA |
| Trip13 | 13.34211523 | 0.989847628 | 0.304104581 | 3.25495797 |
| Gm25197 | 0.235280811 | -0.699147686 | 2.117696254 | -0.330145404 |
| Brd9 | 453.7906305 | 0.026047469 | 0.090771343 | 0.286956959 |
| Zdhhc11 | 0.149223819 | -0.206489568 | 2.865438769 | -0.072062112 |
| Gm10126 | 0.987133562 | 0.275140695 | 0.938307131 | 0.293230954 |
| Gm15912 | 22.38223563 | -0.286415232 | 0.238879524 | -1.198994483 |
| Tppp | 4530.77903 | -0.316916798 | 0.080633997 | -3.930312391 |
| Gm6263 | 0.459143553 | 1.461224349 | 1.374201252 | 1.063326312 |
| Cep72 | 28.65246828 | -0.271393184 | 0.225603614 | -1.2029647 |

| | | | | |
|---|---|---|---|---|
| Slc9a3 | 15.35249687 | -0.564302791 | 0.321780271 | -1.753689836 |
| Exoc3 | 1219.916005 | -0.00882711 | 0.047341606 | -0.186455654 |
| Ahrr | 8.460366272 | -0.249919196 | 0.35518759 | -0.703625921 |
| Pdcd6 | 440.5530706 | 0.235359464 | 0.084464143 | 2.786501545 |
| Sdha | 3363.70453 | -0.063231585 | 0.061131891 | -1.034346939 |
| Gm16125 | 0 | NA | NA | NA |
| Ccdc127 | 908.4665307 | -0.231846673 | 0.06236461 | -3.717599996 |
| Lrrc14b | 12.94872141 | 1.167160311 | 0.483284577 | 2.415058059 |
| Zfp72 | 58.03635434 | -0.805707479 | 0.212152272 | -3.797779166 |
| Gm10116 | 125.931969 | 0.953570488 | 0.219841011 | 4.337545958 |
| Mir692-3 | 0 | NA | NA | NA |
| Zfp825 | 69.54643972 | -0.142416829 | 0.187992568 | -0.757566271 |
| Erap1 | 747.9723258 | 2.709610574 | 0.123176842 | 21.99772728 |
| Cast | 225.7625238 | 1.108770928 | 0.150008528 | 7.39138598 |
| Hnrnpa1l2-ps | 0.737427016 | -0.456524981 | 1.19476976 | -0.382102892 |
| Pcsk1 | 401.8836238 | -0.055906299 | 0.165364682 | -0.338078833 |
| Gm4149 | 23.46034018 | -0.044000543 | 0.661431675 | -0.066523186 |
| Mir682 | 0 | NA | NA | NA |
| Gm23012 | 0 | NA | NA | NA |
| Ell2 | 509.0942641 | 0.177073457 | 0.100176494 | 1.767614839 |
| Gm23127 | 1.778293916 | 0.324933856 | 0.748568989 | 0.43407336 |
| Glrx | 434.8245854 | 0.797058467 | 0.174732484 | 4.561592944 |
| Rhobtb3 | 441.2117491 | -0.705996525 | 0.128646309 | -5.487887922 |
| Gm24163 | 0 | NA | NA | NA |
| Gm15528 | 4.932600309 | -0.242457054 | 0.512702615 | -0.472899976 |
| Gm6311 | 20.93266756 | 0.256988406 | 0.291695149 | 0.881017073 |
| Gm23319 | 0.214976432 | -0.206481381 | 2.177043894 | -0.094844841 |
| Spata9 | 2.596037066 | 2.056533228 | 0.863476235 | 2.381690595 |
| Rfesd | 170.1520903 | 0.096594676 | 0.100925603 | 0.957087925 |
| Gpr150 | 6.580668916 | -0.287462925 | 0.363766988 | -0.790239177 |
| Arsk | 95.44909683 | -0.500678581 | 0.140617037 | -3.560582644 |
| Ttc37 | 526.0109267 | -0.046914854 | 0.067918208 | -0.690755191 |
| Gm24025 | 0 | NA | NA | NA |
| Mctp1 | 1090.396185 | 0.232631638 | 0.095825619 | 2.427655998 |
| Gm24519 | 0 | NA | NA | NA |
| Gm21959 | 0 | NA | NA | NA |
| Gm15740 | 8.875000737 | 0.450173309 | 0.369071753 | 1.219744685 |
| Gm8185 | 13.50337661 | -0.755773474 | 0.359435565 | -2.102667483 |
| Ankrd32 | 492.724753 | -0.123568599 | 0.081709271 | -1.512295937 |
| 2210408I21Rik | 307.5143814 | 0.129544153 | 0.079915287 | 1.621018435 |
| Gm26088 | 0.042716945 | -0.206490061 | 3.724945049 | -0.055434391 |
| Fam172a | 517.357941 | -0.021084366 | 0.066591599 | -0.316622016 |
| Pou5f2 | 26.05560058 | -0.055371937 | 0.362796059 | -0.15262552 |
| Nr2f1 | 507.987299 | -0.415742916 | 0.105124839 | -3.954754384 |
| A830082K12Rik | 207.7537088 | -0.664764775 | 0.156021652 | -4.260721298 |
| Gm6317 | 0 | NA | NA | NA |
| Gm25383 | 0 | NA | NA | NA |

| | | | | |
|---|---|---|---|---|
| Gm29318 | 0.046345709 | 0.274391604 | 3.724945049 | 0.073663262 |
| Gm38349 | 0 | NA | NA | NA |
| Gm24863 | 0.296104695 | 0.274415945 | 1.772080761 | 0.154855214 |
| Gm28526 | 0.110242424 | -0.20649451 | 3.724945049 | -0.055435586 |
| Gm24635 | 0 | NA | NA | NA |
| Arrdc3 | 1031.103706 | 0.225270273 | 0.098032334 | 2.29791808 |
| 1700023H06Rik | 0.473445911 | -0.188069928 | 1.422750749 | -0.132187545 |
| Gm4211 | 15.12852727 | 0.002894099 | 0.287652817 | 0.010061085 |
| 9330111N05Rik | 6.889709052 | 0.015523833 | 0.402484847 | 0.038569981 |
| Gm8399 | 71.44903874 | 0.24591972 | 0.15217677 | 1.616013531 |
| Gm25241 | 0 | NA | NA | NA |
| Adgrv1 | 98.32611678 | -0.544619733 | 0.158391893 | -3.438431873 |
| Lysmd3 | 266.3986882 | 0.316881188 | 0.106492351 | 2.975623917 |
| Polr3g | 69.8321072 | -0.170599403 | 0.126613027 | -1.347407982 |
| Mblac2 | 357.5721259 | -0.295062946 | 0.100392542 | -2.939092297 |
| Gm17259 | 17.35136828 | -0.72897844 | 0.289131757 | -2.52126728 |
| Cetn3 | 274.7140815 | 0.085892249 | 0.13816186 | 0.621678439 |
| Gm24295 | 0 | NA | NA | NA |
| mmu-mir-3961 | 0 | NA | NA | NA |
| Mef2c | 4656.118685 | -0.048638287 | 0.158402141 | -0.307055742 |
| C130071C03Rik | 839.3578359 | -0.31579781 | 0.126900364 | -2.488549289 |
| Gm26803 | 22.51607708 | -0.133012539 | 0.225177295 | -0.590701378 |
| Gm33366 | 4.40666377 | 0.286824062 | 0.450482941 | 0.636703492 |
| Mir9-2 | 0 | NA | NA | NA |
| 2810049E08Rik | 34.32347081 | 0.676996755 | 0.255524731 | 2.64943731 |
| Gm17750 | 9.902264217 | -0.530874214 | 0.340247166 | -1.560260502 |
| Gm24331 | 0 | NA | NA | NA |
| A230107N01Rik | 12.62433136 | -1.39189315 | 0.3958866 | -3.515888513 |
| Tmem161b | 374.8315931 | 0.249324114 | 0.09819653 | 2.539031807 |
| Gm26927 | 0.174715781 | -0.206480246 | 2.726888653 | -0.075720087 |
| Gm26913 | 0.044191549 | -0.206490061 | 3.724945049 | -0.055434391 |
| Gm4076 | 2.353414094 | -0.214212009 | 0.708612608 | -0.302297767 |
| Ccnh | 371.5577082 | -0.308676144 | 0.098095244 | -3.146698374 |
| Rasa1 | 1059.457239 | -0.259817603 | 0.077438887 | -3.355130927 |
| Gm37641 | 0 | NA | NA | NA |
| Cox7c | 960.0717164 | 0.035590319 | 0.111253047 | 0.319904219 |
| Gm22574 | 1.539931973 | -0.622042752 | 0.885879535 | -0.702175327 |
| Gm27025 | 0 | NA | NA | NA |
| Gm25700 | 0 | NA | NA | NA |
| Gm24935 | 0 | NA | NA | NA |
| Gm21232 | 0 | NA | NA | NA |
| Gm27044 | 0 | NA | NA | NA |
| Gm8526 | 0 | NA | NA | NA |
| Edil3 | 2220.587599 | -0.207704984 | 0.091056011 | -2.281068347 |
| Hapln1 | 201.6502422 | -0.567602269 | 0.213209019 | -2.662186957 |
| Gm8546 | 11.32043437 | -0.0288124 | 0.337401898 | -0.085394897 |
| Vcan | 1385.185601 | 0.891990303 | 0.123655713 | 7.213498546 |

| | | | | |
|---|---|---|---|---|
| Gm16318 | 0.048949532 | -0.206487837 | 3.724945049 | -0.055433794 |
| Gm4117 | 32.71244237 | 2.698991941 | 0.294936956 | 9.151080895 |
| Xrcc4 | 102.7407697 | -0.02814174 | 0.135383596 | -0.207866692 |
| Gm24498 | 3.65796296 | -0.939825207 | 0.554639511 | -1.694479366 |
| Tmem167 | 992.695371 | 0.049443656 | 0.056671923 | 0.872454166 |
| Gm24507 | 5.663741534 | 0.159971571 | 0.410528614 | 0.389672159 |
| Gm37727 | 1.381073991 | -0.130048124 | 0.895144833 | -0.145281657 |
| Gm37054 | 0.576775271 | 2.217838226 | 1.339201504 | 1.656090005 |
| Gm37708 | 0 | NA | NA | NA |
| Gm21726 | 0.255118679 | 1.260676538 | 2.475692927 | 0.509221691 |
| Gm36966 | 0 | NA | NA | NA |
| AC147633.1 | 0 | NA | NA | NA |
| Atp6ap1l | 69.06133185 | -0.19974515 | 0.196262122 | -1.017746814 |
| Rps23 | 1420.870961 | 0.555205745 | 0.091505689 | 6.0674451 |
| Atg10 | 83.32890499 | 0.10471005 | 0.118693127 | 0.882191349 |
| Gm17450 | 0 | NA | NA | NA |
| A830009L08Rik | 17.31106247 | -1.226390903 | 0.271908508 | -4.510307212 |
| Gm29540 | 0.823808628 | 0.043270556 | 1.288100572 | 0.033592529 |
| Ssbp2 | 1111.715029 | 0.058702128 | 0.083763703 | 0.700806275 |
| 4833422C13Rik | 42.21647888 | -0.469869743 | 0.188846973 | -2.488097828 |
| Gm22778 | 1.426922496 | 0.060184856 | 0.86919427 | 0.069242123 |
| Acot12 | 6.478410357 | 0.369788576 | 0.461964769 | 0.800469216 |
| Gm24597 | 0 | NA | NA | NA |
| Gm27656 | 0 | NA | NA | NA |
| Zcchc9 | 113.9382792 | 0.079541086 | 0.116488105 | 0.682825827 |
| Ckmt2 | 1.022167862 | -0.255934908 | 1.004658713 | -0.25474811 |
| 1700119I11Rik | 3.377993944 | -0.490277909 | 0.586166892 | -0.836413512 |
| Rasgrf2 | 1981.271647 | 0.07727893 | 0.094684633 | 0.816171827 |
| Msh3 | 384.3197434 | 0.331807851 | 0.103192994 | 3.215410634 |
| Gm20379 | 0 | NA | NA | NA |
| AC161588.1 | 0 | NA | NA | NA |
| Dhfr | 139.8089927 | 0.805498957 | 0.199085906 | 4.045986837 |
| C030017D09Rik | 1.139049109 | -1.13868152 | 1.122179021 | -1.014705763 |
| Ankrd34b | 374.0828976 | -0.271820502 | 0.100985547 | -2.691677287 |
| Fam151b | 60.4619812 | -0.230662593 | 0.145722904 | -1.582884961 |
| Zfyve16 | 462.5719769 | -0.399127296 | 0.097180664 | -4.107064909 |
| Spz1 | 0 | NA | NA | NA |
| Serinc5 | 1004.030055 | -0.309376983 | 0.08231319 | -3.758534705 |
| Thbs4 | 44.88367789 | -0.184576493 | 0.345074278 | -0.534889166 |
| Mtx3 | 682.3862295 | -0.27022527 | 0.074967793 | -3.604551499 |
| Cmya5 | 96.50354882 | -0.21880693 | 0.14498793 | -1.509138926 |
| Papd4 | 358.1413963 | 0.390571729 | 0.119769417 | 3.261030565 |
| Gm2076 | 0.653050501 | 1.713694209 | 1.266027692 | 1.353599309 |
| Homer1 | 3096.738791 | -0.506057699 | 0.242877234 | -2.083594623 |
| Jmy | 1417.927992 | -0.086186517 | 0.052848877 | -1.630810757 |
| Bhmt | 2.668832678 | 0.467701894 | 0.606399407 | 0.771276964 |
| Gm15622 | 0.30042186 | 1.24727193 | 1.616248366 | 0.771708084 |

| | | | | |
|---|---|---|---|---|
| Bhmt2 | 0.855497761 | 0.346874556 | 1.049499355 | 0.330514311 |
| Dmgdh | 2.316670105 | -1.000708976 | 0.683568012 | -1.463949394 |
| AW495222 | 7.0482006 | -0.232089391 | 0.392799315 | -0.590859969 |
| Gm15620 | 0 | NA | NA | NA |
| Arsb | 966.5384739 | -0.085783255 | 0.127694077 | -0.671787268 |
| Mir5624 | 0.311503589 | -0.64207563 | 1.785305867 | -0.359644609 |
| Gm25534 | 0.045882455 | 0.755288839 | 3.724945049 | 0.202765096 |
| Gm26527 | 0.345934107 | 0.745435541 | 1.751102565 | 0.425694963 |
| Gm24737 | 0.579409632 | 2.080977154 | 1.28639712 | 1.617678648 |
| Lhfpl2 | 534.543113 | 0.420604361 | 0.158440229 | 2.654656363 |
| Gm15907 | 0.284494328 | 1.710715801 | 2.051916821 | 0.833715959 |
| Scamp1 | 2084.156384 | -0.184421834 | 0.05290423 | -3.485956329 |
| Gm9776 | 34.06058694 | 0.478535477 | 0.175346184 | 2.729089769 |
| Ap3b1 | 770.6625569 | 0.074979518 | 0.064952202 | 1.154379936 |
| Gm25109 | 1.803894262 | -1.206585619 | 0.872530611 | -1.382857636 |
| Gm36991 | 0.478351411 | 2.59053752 | 3.687373727 | 0.702542707 |
| Tbca | 287.401201 | 0.310902 | 0.125394494 | 2.479391159 |
| Otp | 27.26391552 | 0.249776464 | 0.333209904 | 0.749606961 |
| Wdr41 | 404.2473427 | 0.041721255 | 0.068915758 | 0.605394991 |
| Pde8b | 739.5660274 | -0.412914926 | 0.138935497 | -2.971990126 |
| Gm16243 | 0.075043205 | 0.755286614 | 3.724945049 | 0.202764498 |
| Zbed3 | 136.959666 | 0.557112756 | 0.156161537 | 3.567541451 |
| Snora47 | 6.619789565 | -0.121844361 | 0.362881183 | -0.335769301 |
| Gm22661 | 5.694343269 | 0.179651798 | 0.40547099 | 0.443069423 |
| Aggf1 | 620.7099307 | -0.12198383 | 0.075604042 | -1.613456454 |
| Crhbp | 115.8235207 | 0.036205039 | 0.155594811 | 0.23268796 |
| S100z | 0 | NA | NA | NA |
| F2rl1 | 1.572027216 | -0.97165058 | 0.757777851 | -1.282236713 |
| F2r | 208.5978153 | 0.29679372 | 0.140610009 | 2.110758133 |
| Iqgap2 | 782.9040417 | 1.261642916 | 0.106206847 | 11.87911095 |
| F2rl2 | 16.42119601 | 1.01089903 | 0.341720265 | 2.958264796 |
| Gm24032 | 0.090427103 | 1.236179401 | 3.724945049 | 0.331865138 |
| Sv2c | 1728.872108 | -0.286199785 | 0.136737808 | -2.093055238 |
| Gm17190 | 0.114742407 | 0.274402725 | 3.724945049 | 0.073666248 |
| Gm23955 | 0.962110563 | -0.016869336 | 1.164406423 | -0.014487498 |
| Gm29543 | 3.164183009 | 0.187675544 | 0.626588881 | 0.29951943 |
| Gm25213 | 0 | NA | NA | NA |
| Poc5 | 406.0730432 | -0.033169492 | 0.078319535 | -0.423514928 |
| Ankdd1b | 3.542632829 | 0.595362641 | 0.593995563 | 1.002301494 |
| Polk | 176.7680094 | -0.196800461 | 0.124181713 | -1.584778109 |
| Gm17622 | 0 | NA | NA | NA |
| Col4a3bp | 1491.199334 | -0.028485982 | 0.051107838 | -0.55737013 |
| Hmgcr | 1361.035643 | -0.306724725 | 0.093871726 | -3.267487846 |
| Gm27792 | 0 | NA | NA | NA |
| Gcnt4 | 101.3429506 | -0.851044315 | 0.148486422 | -5.731462195 |
| 1700029F12Rik | 0 | NA | NA | NA |
| Fam169a | 842.6174546 | -0.158726924 | 0.074520797 | -2.12996814 |

| | | | | |
|---|---|---|---|---|
| Gm6169 | 6.279356964 | 1.185135075 | 0.542593992 | 2.184202354 |
| Nsa2 | 317.8801227 | 0.450969697 | 0.127403373 | 3.539699837 |
| Gfm2 | 387.8031722 | -0.255400895 | 0.122205831 | -2.089923966 |
| Hexb | 1048.089142 | 0.363070729 | 0.170927243 | 2.12412441 |
| Enc1 | 4686.802237 | -0.046290333 | 0.131560112 | -0.351856902 |
| C430039J16Rik | 0.150086409 | 1.264824188 | 3.723011833 | 0.339731445 |
| Lincenc1 | 22.69480111 | 0.57269449 | 0.257114994 | 2.227386592 |
| Gm8756 | 1.313089905 | 0.451690678 | 0.852795028 | 0.529659137 |
| 5330416C01Rik | 28.18081706 | -0.911266987 | 0.328029951 | -2.777999343 |
| Gm26619 | 4.459207448 | 0.176956766 | 0.529299921 | 0.334322299 |
| Gm10260 | 388.9520094 | 0.663781203 | 0.141148644 | 4.702710454 |
| Arhgef28 | 438.5791712 | -0.037169586 | 0.089732769 | -0.41422533 |
| Utp15 | 405.8469762 | 0.1311226 | 0.105228601 | 1.246073778 |
| Ankra2 | 370.3092848 | 0.073287545 | 0.091173889 | 0.803821643 |
| Gm21976 | 0.640962652 | -0.65567245 | 1.28887187 | -0.508718101 |
| Btf3 | 823.2235028 | 0.478393967 | 0.089100853 | 5.369128944 |
| Gm9828 | 25.61986482 | -0.286847399 | 0.238508743 | -1.202670376 |
| Foxd1 | 28.76092711 | -0.201043976 | 0.296566113 | -0.677906096 |
| Gm2445 | 3.175554283 | -0.163868316 | 0.569275431 | -0.287854185 |
| 2310005E17Rik | 0 | NA | NA | NA |
| Gm29501 | 0 | NA | NA | NA |
| Gm10320 | 0.789918879 | 1.781021068 | 1.196985818 | 1.487921613 |
| Gm37425 | 0 | NA | NA | NA |
| Gm22698 | 0 | NA | NA | NA |
| Tmem174 | 0.124347486 | 0.755286736 | 3.717316019 | 0.203180664 |
| Tmem171 | 14.34229216 | 3.503421266 | 0.625835249 | 5.597992878 |
| Fcho2 | 753.4025212 | 0.006005121 | 0.066949255 | 0.089696604 |
| Tnpo1 | 2400.224482 | 0.256299557 | 0.087572415 | 2.926715658 |
| Gm5453 | 2.269219555 | 0.876249941 | 0.730978051 | 1.198736323 |
| 1700024P04Rik | 0 | NA | NA | NA |
| Gm24733 | 0.085433889 | -0.687376175 | 3.724945049 | -0.184533239 |
| 2310020H05Rik | 0 | NA | NA | NA |
| Gm807 | 7.497632429 | 1.424028669 | 0.551478448 | 2.582201852 |
| Zfp366 | 15.07922646 | -0.619885324 | 0.323795645 | -1.91443379 |
| Ptcd2 | 245.1837604 | -0.032252942 | 0.075050602 | -0.429749275 |
| Mrps27 | 194.2719257 | -0.021331839 | 0.084194162 | -0.253364824 |
| 6430562O15Rik | 0.528537426 | 0.282190517 | 1.480017632 | 0.190666997 |
| Map1b | 25160.03304 | -0.044036934 | 0.096011907 | -0.458661174 |
| Gm26559 | 14.25506631 | 0.306684946 | 0.329158218 | 0.931725016 |
| 1700012J22Rik | 0.18753903 | -0.206483787 | 2.716639112 | -0.076007073 |
| Gm24471 | 0 | NA | NA | NA |
| Cartpt | 99.17765322 | -0.357170827 | 0.233772937 | -1.527853615 |
| Mccc2 | 207.1112753 | -0.035084854 | 0.086317137 | -0.406464521 |
| Bdp1 | 1242.860492 | 0.013452094 | 0.080904324 | 0.166271628 |
| BC001981 | 0.782345574 | 1.655814522 | 1.209187902 | 1.369360808 |
| Serf1 | 47.59636433 | -0.311611405 | 0.177654457 | -1.754030893 |
| Smn1 | 181.1517505 | 0.231606255 | 0.093267755 | 2.483240364 |

| | | | | |
|---|---|---|---|---|
| Naip2 | 165.0036826 | 3.509588601 | 0.233774435 | 15.01271342 |
| Naip5 | 40.97982787 | 2.641780863 | 0.316776403 | 8.33957591 |
| Naip6 | 36.81044465 | 1.971016723 | 0.291960425 | 6.750972242 |
| Naip3 | 1.709236637 | 2.119407906 | 0.994368177 | 2.131411639 |
| Naip1 | 6.360266967 | 0.913172856 | 0.480836693 | 1.899133052 |
| Rpl31-ps13 | 5.091748836 | 0.295034065 | 0.562642604 | 0.524372066 |
| Gtf2h2 | 239.663237 | -0.032305489 | 0.125234503 | -0.257959976 |
| Ocln | 88.83136681 | -0.196201789 | 0.157640544 | -1.244615018 |
| Marveld2 | 42.42039179 | -0.364910235 | 0.192476522 | -1.895868806 |
| Gm24261 | 0 NA | NA | NA | |
| Rad17 | 254.2430578 | 0.035040788 | 0.099882331 | 0.350820689 |
| Ak6 | 71.85688038 | -0.330825088 | 0.14668846 | -2.255290487 |
| Taf9 | 6.955395866 | -0.783095635 | 0.384908803 | -2.034496558 |
| Ccdc125 | 6.403646332 | 1.445078896 | 0.471289014 | 3.066226567 |
| Cdk7 | 820.0282203 | -0.245649786 | 0.055775054 | -4.404294911 |
| Mrps36 | 129.039131 | -0.212397584 | 0.130741614 | -1.624559912 |
| Cenph | 6.451376144 | 0.452035596 | 0.599219614 | 0.75437383 |
| Gm16335 | 0.346789506 | 0.755311633 | 1.494913149 | 0.505254525 |
| Ccnb1 | 20.0589336 | 0.718115805 | 0.398740068 | 1.800962239 |
| Slc30a5 | 515.9001532 | -0.187383704 | 0.075873986 | -2.46966996 |
| Gm29502 | 6.014679714 | 0.48911361 | 0.439856872 | 1.111983559 |
| Gm37830 | 0.152961209 | 0.274409656 | 2.839987012 | 0.09662356 |
| Gm10257 | 10.11133075 | 0.533198467 | 0.319596446 | 1.668349176 |
| Gm6114 | 0 NA | NA | NA | |
| Hmgb1-ps9 | 1.368962209 | 1.680532955 | 1.00633697 | 1.669950529 |
| 4932411K12Rik | 0.386457151 | -0.206484855 | 1.753477855 | -0.117757321 |
| Gm36994 | 0 NA | NA | NA | |
| Gm29341 | 0 NA | NA | NA | |
| Pik3r1 | 3272.596283 | 0.330424148 | 0.069053819 | 4.785023549 |
| Gm36638 | 65.83543421 | -0.177190446 | 0.239877373 | -0.738670946 |
| Gm38133 | 0 NA | NA | NA | |
| Gm25808 | 0 NA | NA | NA | |
| Cd180 | 333.6593441 | 5.049782512 | 0.281023611 | 17.9692464 |
| Mast4 | 1713.710742 | -0.128068974 | 0.081310205 | -1.575066415 |
| 1700099I09Rik | 0.905219458 | 1.265835803 | 1.201821712 | 1.053264216 |
| Gm17160 | 0.372057435 | 1.238040298 | 1.559422649 | 0.793909399 |
| Gm6211 | 10.83872605 | -0.188018956 | 0.395627024 | -0.475242955 |
| Gm5454 | 142.1539982 | -0.601299645 | 0.183274264 | -3.280873326 |
| n-R5s56 | 0 NA | NA | NA | |
| Gm16416 | 0.30254151 | 0.300195191 | 1.762573777 | 0.170316383 |
| Gm24441 | 0 NA | NA | NA | |
| Gm24870 | 0 NA | NA | NA | |
| Srek1 | 793.2803317 | -0.077624257 | 0.15199829 | -0.510691643 |
| Erbb2ip | 3344.991394 | 0.585511639 | 0.13241452 | 4.421808417 |
| Nln | 399.6247544 | 0.054139314 | 0.080163557 | 0.67536067 |
| Sgtb | 1145.826265 | -0.220712874 | 0.088261563 | -2.500668098 |
| Trappc13 | 722.8966951 | -0.0085354 | 0.071595206 | -0.119217482 |

| | | | | |
|---|---|---|---|---|
| Trim23 | 1039.038422 | 0.146014737 | 0.090821792 | 1.607705966 |
| Ppwd1 | 182.8657831 | 0.011977453 | 0.098847535 | 0.121170981 |
| Cenpk | 9.293126097 | 0.709027235 | 0.394337503 | 1.798021315 |
| Adamts6 | 345.3432011 | 1.520619655 | 0.150760451 | 10.08632994 |
| Cwc27 | 204.665214 | 0.247883148 | 0.162578243 | 1.524700623 |
| 2610204G07Rik | 0.047948698 | -0.206487837 | 3.724945049 | -0.055433794 |
| Srek1ip1 | 192.1915529 | -0.242650209 | 0.127488783 | -1.903306336 |
| Fam159b | 3.808355539 | 0.911407349 | 0.625637453 | 1.456765967 |
| Gm4938 | 0.726615386 | 0.832467555 | 1.076540547 | 0.773280261 |
| Rgs7bp | 3653.625861 | -0.302767772 | 0.098864082 | -3.0624648 |
| 4933425L06Rik | 0.717517214 | 0.268299387 | 1.218079107 | 0.220264337 |
| Rnf180 | 161.367705 | 0.147350494 | 0.096188373 | 1.531895071 |
| Gm25631 | 2.548041448 | -0.351633563 | 0.633461068 | -0.555098933 |
| Htr1a | 192.2048649 | 0.089255533 | 0.109013429 | 0.81875723 |
| Dph3b-ps | 0 | NA | NA | NA |
| Gm24527 | 0.388660859 | 0.046085866 | 1.550453686 | 0.029724116 |
| Gm28989 | 0.486723951 | 1.428267925 | 1.581581507 | 0.903063116 |
| Ipo11 | 842.5268281 | -0.128602008 | 0.060265408 | -2.133927456 |
| Gm28988 | 0.166709681 | -0.206481506 | 2.767631179 | -0.074605861 |
| Dimt1 | 147.6538309 | -0.287734251 | 0.096260998 | -2.989105208 |
| Kif2a | 1440.730209 | -0.360401144 | 0.065234696 | -5.524684956 |
| Gm24431 | 2.158765044 | -0.569499977 | 0.826814661 | -0.688787952 |
| 3830408C21Rik | 12.03259843 | -0.695533999 | 0.330619836 | -2.103727374 |
| Apoo-ps | 41.16296874 | -0.168136523 | 0.283350536 | -0.593386996 |
| 1700006H21Rik | 0.426601203 | 0.755280747 | 3.26857073 | 0.231073705 |
| Zswim6 | 635.8544846 | -0.142039711 | 0.095698607 | -1.484240113 |
| Gm22118 | 4.848883265 | -0.320531761 | 0.532396964 | -0.602054073 |
| Smim15 | 306.3096422 | 0.016194844 | 0.106634211 | 0.151872877 |
| Ndufaf2 | 61.73980177 | -0.261782021 | 0.255545914 | -1.024403078 |
| Ercc8 | 184.4501039 | -0.01580955 | 0.110714444 | -0.142795727 |
| Elovl7 | 210.6233394 | -0.447412659 | 0.144821719 | -3.089403047 |
| Gm2822 | 0.316450815 | -0.173923092 | 1.756454279 | -0.099019424 |
| Depdc1b | 9.399548456 | -0.398033622 | 0.381208122 | -1.044137306 |
| Pde4d | 1367.797614 | -0.104005588 | 0.086171806 | -1.206956108 |
| Gm27572 | 0 | NA | NA | NA |
| Gm28016 | 0 | NA | NA | NA |
| Rps3a3 | 38.33507003 | 0.366040026 | 0.193672145 | 1.889998308 |
| Mir582 | 2.019407445 | 0.509484267 | 0.684437766 | 0.744383628 |
| AC161585.1 | 9.306437351 | -0.329549256 | 0.376126519 | -0.876165968 |
| Mir1904 | 1.272510498 | 0.647427931 | 0.824231875 | 0.78549247 |
| Rab3c | 4667.9155 | -0.335044666 | 0.092538291 | -3.620605737 |
| Gm5455 | 0.386702458 | -0.684241688 | 1.701273527 | -0.402193814 |
| Gapt | 3.214970131 | 2.64112131 | 1.040961718 | 2.537193506 |
| Plk2 | 1844.679949 | 0.268855364 | 0.140235138 | 1.917175458 |
| Actbl2 | 1.958193336 | -2.494923401 | 0.86540556 | -2.88295282 |
| Gm15285 | 0.507331407 | 0.287934141 | 1.434537364 | 0.200715679 |
| Gm15289 | 0 | NA | NA | NA |

| | | | | |
|---|---|---|---|---|
| Gm6270 | 0 | NA | NA | NA |
| Gpbp1 | 1669.061245 | 0.23170364 | 0.071387698 | 3.245708247 |
| Gm15290 | 0 | NA | NA | NA |
| Gm15288 | 0 | NA | NA | NA |
| Gm15286 | 0.075043205 | 0.755286614 | 3.724945049 | 0.202764498 |
| Gm10198 | 0 | NA | NA | NA |
| Gm10736 | 67.77195831 | 0.40973766 | 0.181771039 | 2.25414159 |
| Mier3 | 595.0131209 | 0.17969327 | 0.091474379 | 1.964410938 |
| Gm15488 | 0.182466651 | 0.274402709 | 2.68257363 | 0.10229084 |
| Gm15287 | 0.237498462 | -0.206486869 | 2.136352402 | -0.096653936 |
| Map3k1 | 549.7994848 | 0.219648169 | 0.157539634 | 1.394240698 |
| Gm15327 | 2.461737376 | -0.12171173 | 0.693460351 | -0.175513611 |
| Gm15325 | 0 | NA | NA | NA |
| Gm15324 | 0 | NA | NA | NA |
| Gm15326 | 3.336705128 | 1.647551108 | 0.7492674 | 2.198882678 |
| Gm9025 | 2.20249356 | 2.402272754 | 0.962419592 | 2.496076321 |
| Gm15322 | 0.293377407 | -1.726578811 | 3.695996407 | -0.467148401 |
| Gm15323 | 0.143781519 | -0.192986053 | 3.581286957 | -0.053887347 |
| Gm37427 | 0 | NA | NA | NA |
| Ankrd55 | 97.25075632 | -0.179824685 | 0.174821738 | -1.028617417 |
| Gm22768 | 0.696415487 | -1.90642413 | 1.18690449 | -1.606215283 |
| Il6st | 2388.948021 | 0.087529908 | 0.071090562 | 1.23124512 |
| Il31ra | 12.44555901 | 0.469066742 | 0.369231251 | 1.270387436 |
| Ddx4 | 13.81354613 | 0.737200715 | 0.370081745 | 1.991994268 |
| Slc38a9 | 402.4228987 | -0.662091484 | 0.126415884 | -5.237407384 |
| Ppap2a | 177.5947723 | 0.118267538 | 0.094851881 | 1.246865499 |
| Skiv2l2 | 776.3773442 | -0.071308152 | 0.058789593 | -1.212938348 |
| Gm26943 | 0.235215884 | 0.755301565 | 2.493330199 | 0.302928816 |
| Dhx29 | 365.0039571 | -0.240559565 | 0.06926747 | -3.472908199 |
| Ccno | 13.15772638 | 0.803339555 | 0.391661625 | 2.051106117 |
| Mcidas | 3.060161834 | 0.575452051 | 0.624013694 | 0.922178563 |
| Cdc20b | 1.498584608 | -1.084690833 | 0.899075356 | -1.206451524 |
| Mir449c | 0.04393175 | -0.206490061 | 3.724945049 | -0.055434391 |
| Mir449b | 0.038981395 | 0.274393828 | 3.724945049 | 0.073663859 |
| Mir449a | 0 | NA | NA | NA |
| Gm10735 | 1.512916718 | 1.406774834 | 0.816842888 | 1.722209809 |
| Gpx8 | 87.13244342 | 0.650004386 | 0.2107729 | 3.083908734 |
| Gzma | 199.9537804 | 7.77327601 | 0.794405254 | 9.785025934 |
| Gm9848 | 0 | NA | NA | NA |
| Gzmk | 39.86666651 | 2.795296468 | 0.358564334 | 7.79580177 |
| Esm1 | 8.383027344 | 2.528343069 | 0.505265292 | 5.003991186 |
| BC067074 | 57.61241419 | -0.314849085 | 0.168514223 | -1.868382856 |
| Gm22385 | 0 | NA | NA | NA |
| Snx18 | 579.4915124 | 0.318334501 | 0.104682972 | 3.04093871 |
| Hspb3 | 12.0017422 | 0.89841609 | 0.316724483 | 2.836585548 |
| Arl15 | 416.4383343 | 0.122506287 | 0.092114285 | 1.329937992 |
| Gm37907 | 0.135812198 | -1.150633786 | 3.646939241 | -0.315506706 |

| | | | | |
|---|---|---|---|---|
| Ndufs4 | 556.7189796 | 0.202789593 | 0.101667428 | 1.994636808 |
| Fst | 27.88029129 | -0.610856585 | 0.291924384 | -2.09251648 |
| Gm25950 | 0 | NA | NA | NA |
| 4930544M13Rik | 0 | NA | NA | NA |
| Mocs2 | 459.8435434 | 0.346448696 | 0.07256121 | 4.774571637 |
| Itga2 | 10.13867756 | 1.791454818 | 0.512063145 | 3.498503721 |
| Itga1 | 103.8467935 | 0.034086 | 0.183745129 | 0.185506958 |
| Pelo | 139.7216595 | -0.067516574 | 0.114436874 | -0.58998968 |
| Gm25985 | 0 | NA | NA | NA |
| Gm6404 | 0 | NA | NA | NA |
| Isl1 | 36.11947267 | 0.16641684 | 0.183353205 | 0.907629837 |
| Gm38193 | 6.0191764 | -0.899161675 | 0.492617901 | -1.825272029 |
| Gm23198 | 0 | NA | NA | NA |
| Parp8 | 540.5637389 | 0.593554967 | 0.113669051 | 5.221781677 |
| Gm17509 | 11.37141038 | -0.636669328 | 0.36071311 | -1.765029632 |
| Emb | 394.0332575 | 0.170694308 | 0.123042042 | 1.387284422 |
| Gm6421 | 12.74783509 | 0.261830742 | 0.346757071 | 0.755084074 |
| Hcn1 | 981.9976838 | 0.028382789 | 0.068343758 | 0.415294531 |
| 4933413L06Rik | 0 | NA | NA | NA |
| Gm26349 | 0.072025462 | 0.755288839 | 3.724945049 | 0.202765096 |
| Mrps30 | 237.5140506 | 0.145470749 | 0.095516196 | 1.522995635 |
| B430218F22Rik | 1.974484148 | 0.394540766 | 0.672592222 | 0.586597276 |
| 1700003P14Rik | 0 | NA | NA | NA |
| Fgf10 | 52.0629943 | -0.322622623 | 0.152043927 | -2.121904044 |
| Gm16263 | 1.381163524 | -0.74347706 | 0.813392667 | -0.914044459 |
| AC154241.1 | 0 | NA | NA | NA |
| Nnt | 359.3284767 | -0.447200019 | 0.112791828 | -3.964826417 |
| Paip1 | 990.7888142 | 0.113994008 | 0.073228849 | 1.556681679 |
| 4833420G17Rik | 380.1582038 | -0.18443883 | 0.102634411 | -1.79704671 |
| Gm9752 | 0.269061364 | 1.295750407 | 3.708688725 | 0.349382357 |
| 3110070M22Rik | 13.70697502 | 0.474111808 | 0.275539886 | 1.720664891 |
| Gm7120 | 31.73718637 | -0.27039184 | 0.206035706 | -1.312354282 |
| Gm21967 | 80.16892809 | -0.342668592 | 0.151286223 | -2.265035012 |
| Ccl28 | 18.99182376 | 0.504874951 | 0.2741834 | 1.841376798 |
| 1700074H08Rik | 0 | NA | NA | NA |
| Hmgcs1 | 2950.403097 | -0.255976194 | 0.130186929 | -1.966220396 |
| Nim1k | 238.1724293 | -0.363347734 | 0.131904901 | -2.754618919 |
| Zfp131 | 574.5633546 | 0.296533536 | 0.08738074 | 3.393580036 |
| Gm23308 | 0 | NA | NA | NA |
| AF067063 | 0 | NA | NA | NA |
| D13Ertd608e | 1.077945179 | 3.219197767 | 1.386641576 | 2.321578858 |
| Gm21358 | 0 | NA | NA | NA |
| Gm37619 | 0.171181394 | 1.257632701 | 2.747344421 | 0.457763028 |
| Tcstv1 | 0 | NA | NA | NA |
| Gm21761 | 0 | NA | NA | NA |
| Gm21181 | 0 | NA | NA | NA |
| B020031M17Rik | 0 | NA | NA | NA |

| | | | | |
|---|---|---|---|---|
| Gm20784 | 1.793435801 | -0.365466268 | 0.728921625 | -0.501379374 |
| Gm22108 | 0 | NA | NA | NA |
| Gm21370 | 0 | NA | NA | NA |
| Gm21188 | 0.521673782 | 1.659008194 | 1.46949802 | 1.128962524 |
| Gm24798 | 0 | NA | NA | NA |
| Gm21378 | 13.07532112 | 0.293638382 | 0.310029852 | 0.947129382 |
| Gm20767 | 0 | NA | NA | NA |
| Gm21818 | 0.095898756 | 0.27438722 | 3.724945049 | 0.073662085 |
| Gm21762 | 1.958730402 | 4.187297759 | 1.091620436 | 3.835855048 |
| Gm21731 | 0 | NA | NA | NA |
| AF067061 | 0 | NA | NA | NA |
| Gm20900 | 10.70701964 | 0.401280878 | 0.361987153 | 1.108550055 |
| BC147527 | 33.1074984 | 6.653546558 | 0.931758591 | 7.140848094 |
| Tcstv3 | 0 | NA | NA | NA |
| Gm2888 | 0 | NA | NA | NA |
| Gm2897 | 10.21470906 | -0.537074298 | 0.344164206 | -1.560517591 |
| Gm2904 | 0.731111494 | -1.117409993 | 1.405219654 | -0.795185286 |
| Gm17051 | 0 | NA | NA | NA |
| Gm10340 | 0.412722675 | 0.291930722 | 1.584250556 | 0.184270551 |
| Gm17221 | 0 | NA | NA | NA |
| Gm17188 | 0 | NA | NA | NA |
| Gm5795 | 0.0878635 | -0.687376175 | 3.724945049 | -0.184533239 |
| Gm17222 | 1.483555279 | 1.645072319 | 1.039288073 | 1.582883862 |
| D830030K20Rik | 12.49083266 | -0.358737212 | 0.30508443 | -1.175862076 |
| Gm4904 | 0 | NA | NA | NA |
| 1700110I01Rik | 0.924198543 | -2.034869147 | 1.127870657 | -1.804168885 |
| Gm2956 | 0.84966665 | -1.282044285 | 1.201549164 | -1.066992782 |
| Gm17223 | 0 | NA | NA | NA |
| Gm10413 | 0 | NA | NA | NA |
| Gm10409 | 0.801287066 | -0.241697674 | 1.445893721 | -0.167161438 |
| Gm3005 | 2.629406015 | 0.49032468 | 0.839306869 | 0.584201916 |
| Gm26817 | 0 | NA | NA | NA |
| Gm10408 | 0.044191549 | -0.206490061 | 3.724945049 | -0.055434391 |
| Gm8323 | 0 | NA | NA | NA |
| Gm8324 | 0 | NA | NA | NA |
| Gm11109 | 0 | NA | NA | NA |
| Gm17107 | 0 | NA | NA | NA |
| Gm3012 | 0.072025462 | 0.755288839 | 3.724945049 | 0.202765096 |
| Gm3020 | 0.88261066 | 0.067782727 | 1.284374055 | 0.052774911 |
| Gm2866 | 0 | NA | NA | NA |
| Gm3043 | 0 | NA | NA | NA |
| Gm2871 | 0 | NA | NA | NA |
| Gm2877 | 0 | NA | NA | NA |
| Gm21965 | 0.038981395 | 0.274393828 | 3.724945049 | 0.073663859 |
| Gm3002 | 2.349117343 | -0.135555617 | 0.622554369 | -0.21774101 |
| Gm26886 | 0 | NA | NA | NA |
| Gm26570 | 0 | NA | NA | NA |

| | | | | |
|---|---|---|---|---|
| Gm3033 | 0.403254479 | 1.698169006 | 1.653272262 | 1.027156292 |
| Gm2907 | 0 | NA | NA | NA |
| Gm2916 | 0.044191549 | -0.206490061 | 3.724945049 | -0.055434391 |
| Gm3029 | 0 | NA | NA | NA |
| Gm3095 | 0.864161856 | -0.118418191 | 1.121945741 | -0.105547164 |
| Gm17129 | 0 | NA | NA | NA |
| Gm3099 | 0.095898756 | 0.27438722 | 3.724945049 | 0.073662085 |
| Gm5796 | 5.645051647 | 2.314459133 | 0.555596352 | 4.165720534 |
| Gm11107 | 0 | NA | NA | NA |
| Gm3115 | 0.045882455 | 0.755288839 | 3.724945049 | 0.202765096 |
| Gm8108 | 0.608376789 | -0.404202632 | 1.238114674 | -0.326466232 |
| Gm17152 | 0 | NA | NA | NA |
| Gm3127 | 0.0878635 | -0.687376175 | 3.724945049 | -0.184533239 |
| Gm2974 | 9.121692315 | 1.321800697 | 0.356178016 | 3.711067604 |
| Gm11105 | 0 | NA | NA | NA |
| Gm3138 | 0.101408268 | 0.755286617 | 3.724945049 | 0.202764499 |
| Gm17085 | 0 | NA | NA | NA |
| Gm3149 | 0.060174537 | 0.755286614 | 3.724945049 | 0.202764498 |
| 2610042L04Rik | 9.926510475 | 1.732167836 | 0.382431187 | 4.529358207 |
| Gm17086 | 0 | NA | NA | NA |
| Gm3159 | 0.075043205 | 0.755286614 | 3.724945049 | 0.202764498 |
| Gm3164 | 5.22863904 | 0.760179364 | 0.542827937 | 1.400405751 |
| Gm3173 | 3.269157551 | 2.478938222 | 0.685889371 | 3.614195418 |
| Gm17170 | 0 | NA | NA | NA |
| Gm3182 | 0.048949532 | -0.206487837 | 3.724945049 | -0.055433794 |
| Gm17074 | 0 | NA | NA | NA |
| Gm3030 | 0.11997416 | 0.274404949 | 3.724945049 | 0.073666845 |
| Gm26655 | 0 | NA | NA | NA |
| Gm3047 | 0.107078754 | -0.206490058 | 3.724945049 | -0.05543439 |
| Gm8164 | 0 | NA | NA | NA |
| Gm26898 | 0.093141081 | -0.687373951 | 3.724945049 | -0.184532642 |
| Gm8159 | 0 | NA | NA | NA |
| Gm3239 | 1.60056093 | 0.914040653 | 0.831244386 | 1.099605205 |
| Gm17073 | 0 | NA | NA | NA |
| Gm7876 | 0.046345709 | 0.274391604 | 3.724945049 | 0.073663262 |
| Gm3252 | 10.87374526 | 1.062251812 | 0.402125688 | 2.641591534 |
| Gm3084 | 0 | NA | NA | NA |
| Gm9602 | 0.3514278 | -0.714905126 | 1.749197382 | -0.408704663 |
| Gm3091 | 0 | NA | NA | NA |
| Gm3269 | 0 | NA | NA | NA |
| Gm3264 | 2.10462685 | 2.579690923 | 0.89310758 | 2.888443656 |
| Gm26650 | 0 | NA | NA | NA |
| Gm3278 | 0.274437282 | -1.081619431 | 1.607404148 | -0.672898245 |
| Gm8305 | 0 | NA | NA | NA |
| Gm10410 | 0 | NA | NA | NA |
| Gm8297 | 0 | NA | NA | NA |
| Gm3298 | 0.938975218 | 0.804232289 | 1.064030931 | 0.755835442 |

| | | | | |
|---|---|---|---|---|
| 4930555G01Rik | 0.088534442 | 0.274389469 | 3.724945049 | 0.073662689 |
| Gm8281 | 11.41715526 | 0.440148913 | 0.326656068 | 1.34743835 |
| Gm17069 | 0 | NA | NA | NA |
| Gm8279 | 0.163190422 | 0.274394252 | 3.398594894 | 0.080737558 |
| Gm26552 | 0 | NA | NA | NA |
| Gm3317 | 1.057189525 | 2.227303692 | 1.069731183 | 2.082115326 |
| Gm3140 | 0 | NA | NA | NA |
| Gm8271 | 0.068097359 | -0.206487837 | 3.724945049 | -0.055433794 |
| Gm3326 | 0 | NA | NA | NA |
| Gm3526 | 0.137762903 | -0.206480416 | 2.937523348 | -0.070290647 |
| Gm3339 | 0 | NA | NA | NA |
| Gm3542 | 0 | NA | NA | NA |
| Gm17126 | 0 | NA | NA | NA |
| Gm3500 | 0.712486263 | -1.20342086 | 1.175959449 | -1.023352345 |
| Gm3348 | 0 | NA | NA | NA |
| Gm3194 | 0.375475346 | 1.551527642 | 1.680009848 | 0.923522945 |
| Gm8246 | 0 | NA | NA | NA |
| Gm3187 | 0.044191549 | -0.206490061 | 3.724945049 | -0.055434391 |
| Gm3488 | 1.283947918 | 0.090624612 | 0.852411767 | 0.106315534 |
| Gm26893 | 0 | NA | NA | NA |
| Gm26746 | 0 | NA | NA | NA |
| Gm3532 | 0 | NA | NA | NA |
| Gm8266 | 0 | NA | NA | NA |
| Gm26731 | 0 | NA | NA | NA |
| Gm3383 | 0.828062179 | 0.622949265 | 1.132683767 | 0.549976334 |
| Gm3344 | 0 | NA | NA | NA |
| Gm8265 | 0 | NA | NA | NA |
| Gm9603 | 0.111006857 | 0.755286617 | 3.724945049 | 0.202764499 |
| Gm3373 | 0.817666292 | 0.315302726 | 1.170609639 | 0.269349162 |
| Gm3170 | 0 | NA | NA | NA |
| Gm26557 | 0 | NA | NA | NA |
| Gm3424 | 0.342449771 | -1.204867977 | 1.5770729 | -0.763990033 |
| Gm17158 | 0 | NA | NA | NA |
| Gm8237 | 12.34732519 | 0.394046101 | 0.398439865 | 0.98897258 |
| Gm17046 | 0.139037128 | 0.274384718 | 3.724945049 | 0.073661414 |
| Gm3242 | 0 | NA | NA | NA |
| Gm17047 | 0 | NA | NA | NA |
| Gm3248 | 20.30228851 | 0.607474533 | 0.235341619 | 2.58124566 |
| Gm17048 | 0 | NA | NA | NA |
| Gm3453 | 0.066050875 | 0.274391604 | 3.724945049 | 0.073663262 |
| Gm3261 | 0 | NA | NA | NA |
| Gm8206 | 2.632186321 | 0.346113738 | 0.606555286 | 0.570621913 |
| Gm3468 | 14.54750696 | 0.674598517 | 0.364175534 | 1.852399334 |
| Gm17045 | 0.385383671 | 0.177164693 | 1.583513041 | 0.111880792 |
| Gm6337 | 0 | NA | NA | NA |
| Gm3275 | 0 | NA | NA | NA |
| Gm3476 | 0.611787247 | 0.336948982 | 1.386304066 | 0.243055611 |

| | | | | |
|---|---|---|---|---|
| Gm26991 | 0 | NA | NA | NA |
| Gm3208 | 0 | NA | NA | NA |
| Gm21560 | 0 | NA | NA | NA |
| Gm21772 | 0 | NA | NA | NA |
| Gm3411 | 2.644118259 | 1.020737148 | 0.621029156 | 1.643621944 |
| Gm21103 | 0 | NA | NA | NA |
| Gm3618 | 0 | NA | NA | NA |
| Gm26630 | 0 | NA | NA | NA |
| Gm8340 | 0 | NA | NA | NA |
| Gm16440 | 0.365173462 | 0.224328948 | 1.392842239 | 0.161058404 |
| Gm3591 | 2.764354947 | 1.188101799 | 0.703455875 | 1.688949999 |
| Gm3594 | 0 | NA | NA | NA |
| Gm8356 | 0.045882455 | 0.755288839 | 3.724945049 | 0.202765096 |
| Gm3629 | 1.032577368 | -0.080079062 | 0.879771676 | -0.091022551 |
| Gm3460 | 0 | NA | NA | NA |
| Gm10251 | 0.095435502 | 0.75528439 | 3.724945049 | 0.202763901 |
| Gm26680 | 0 | NA | NA | NA |
| Gm3466 | 0 | NA | NA | NA |
| Gm8050 | 0.427185557 | -0.592132877 | 1.532078782 | -0.386489836 |
| Gm3636 | 17.74053186 | 0.377943513 | 0.259761038 | 1.454966134 |
| Gm8362 | 0 | NA | NA | NA |
| Gm3642 | 0.049553047 | 0.274391604 | 3.724945049 | 0.073663262 |
| Gm3667 | 13.76281871 | 0.781454761 | 0.306723482 | 2.547750027 |
| Gm6356 | 0.072025462 | 0.755288839 | 3.724945049 | 0.202765096 |
| Gm10406 | 10.70611474 | 1.347337201 | 0.360252063 | 3.739984687 |
| Gm3685 | 0.098502579 | -0.206492286 | 3.724945049 | -0.055434988 |
| Gm3696 | 16.41020923 | 0.93465869 | 0.270053018 | 3.461019236 |
| Gm6676 | 0.092269992 | -0.20649451 | 3.724945049 | -0.055435586 |
| Gm3512 | 4.130618934 | 1.674145534 | 0.590894872 | 2.833237543 |
| Gm16434 | 0 | NA | NA | NA |
| Gm3715 | 1.793529573 | 0.933452912 | 0.757852758 | 1.231707482 |
| Gm5456 | 0.437842326 | -0.201481538 | 1.401874453 | -0.143722954 |
| Gm3727 | 0.094294407 | -0.206492286 | 3.724945049 | -0.055434988 |
| Gm3739 | 19.15242329 | 0.874255154 | 0.302954145 | 2.885767263 |
| Gm5797 | 0.049553047 | 0.274391604 | 3.724945049 | 0.073663262 |
| Gm8374 | 0.038981395 | 0.274393828 | 3.724945049 | 0.073663859 |
| Gm10339 | 0 | NA | NA | NA |
| 4933406F09Rik | 0.083526043 | 0.75528439 | 3.724945049 | 0.202763901 |
| Gm8379 | 0.046345709 | 0.274391604 | 3.724945049 | 0.073663262 |
| Gm3755 | 0.852403115 | -0.039225504 | 0.960049543 | -0.040857792 |
| Gm3752 | 3.599181491 | 1.175797244 | 0.73231636 | 1.605586477 |
| Gm3760 | 0.150455055 | 0.274400633 | 2.848105408 | 0.096344971 |
| Gm3558 | 7.230339464 | -0.3524583 | 0.354677505 | -0.993743033 |
| Gm17121 | 0.092140247 | -0.687373951 | 3.724945049 | -0.184532642 |
| Gm10338 | 0 | NA | NA | NA |
| Gm10128 | 0.356838182 | 2.168027783 | 1.708919403 | 1.268654203 |
| Gm10044 | 5.427300065 | -0.565935595 | 0.449051755 | -1.260290353 |

| | | | | |
|---|---|---|---|---|
| Gm16741 | 0.543837322 | -1.567075365 | 1.339462543 | -1.169928471 |
| Gm26661 | 0.047948698 | -0.206487837 | 3.724945049 | -0.055433794 |
| Gm8396 | 0.090277459 | -0.20649451 | 3.724945049 | -0.055435586 |
| Flnb | 1267.997399 | 0.01645215 | 0.150817289 | 0.109086632 |
| Dnase1l3 | 17.4478056 | 5.701609961 | 0.766535493 | 7.4381552 |
| Abhd6 | 413.5173793 | -0.232414418 | 0.077684227 | -2.991783855 |
| Rpp14 | 225.7839547 | -0.206237114 | 0.081529861 | -2.529589905 |
| Pxk | 710.6699321 | 0.354731537 | 0.08784205 | 4.038288456 |
| Pdhb | 939.7685917 | -0.13519452 | 0.086210223 | -1.568195927 |
| Kctd6 | 337.2401318 | 0.077742241 | 0.143779343 | 0.540705219 |
| Acox2 | 3.533179683 | 0.632769654 | 0.618181217 | 1.023598966 |
| Fam107a | 4664.005083 | 1.041866704 | 0.182668878 | 5.703580783 |
| Oit1 | 0 | NA | NA | NA |
| 4930452B06Rik | 90.32014768 | 0.031507391 | 0.135369762 | 0.232750579 |
| 4930455B14Rik | 0 | NA | NA | NA |
| Gm9800 | 154.1573682 | 0.794185486 | 0.175274395 | 4.531098149 |
| Fhit | 18.766523 | -0.259492913 | 0.238181419 | -1.089475891 |
| Hmgb3-ps | 0 | NA | NA | NA |
| Mnd1-ps | 0.293028276 | -1.308578471 | 3.692614927 | -0.354377182 |
| Gm22053 | 0 | NA | NA | NA |
| Fhitos | 0 | NA | NA | NA |
| Slc25a5-ps | 0.541865536 | 0.745191716 | 1.413341193 | 0.527255358 |
| Rpl21-ps4 | 0.428504882 | -0.644443694 | 1.789526364 | -0.360119698 |
| Gm3839 | 0.806310386 | 0.071639248 | 1.008112759 | 0.071062734 |
| AC158396.1 | 0 | NA | NA | NA |
| Mir5124b | 0 | NA | NA | NA |
| Gm25466 | 0 | NA | NA | NA |
| Ptprg | 1480.140492 | -0.066133113 | 0.060238986 | -1.097845708 |
| Gm3848 | 0.118974955 | 0.274400501 | 3.724945049 | 0.073665651 |
| 3830406C13Rik | 205.9760077 | -0.054700287 | 0.108183684 | -0.505624182 |
| Gm24578 | 0.075043205 | 0.755286614 | 3.724945049 | 0.202764498 |
| Fezf2 | 140.4500694 | -0.117798143 | 0.125840557 | -0.93609044 |
| Cadps | 3883.453482 | -0.131602502 | 0.052676593 | -2.498310842 |
| n-R5s45 | 5.980340237 | -0.021955851 | 0.430803928 | -0.050964835 |
| Gm5457 | 6.294359381 | 0.169500478 | 0.426200926 | 0.397700869 |
| Gm5087 | 8.752821209 | 0.730825648 | 0.358016259 | 2.041319717 |
| Synpr | 975.6483675 | 0.114444908 | 0.167189799 | 0.684520876 |
| Gm15913 | 0 | NA | NA | NA |
| Sntn | 3.353049333 | -0.378647679 | 0.644240807 | -0.587742463 |
| Rpl19-ps4 | 1.071806319 | -0.060962561 | 1.120855614 | -0.054389308 |
| Gm27506 | 0 | NA | NA | NA |
| Gm281 | 0.82274574 | -0.621047799 | 1.224033125 | -0.507378261 |
| Gm25964 | 0 | NA | NA | NA |
| Gm11100 | 1.662603624 | 1.648533391 | 1.13808445 | 1.448515873 |
| Thoc7 | 350.5995921 | 0.077776009 | 0.097622076 | 0.796705136 |
| Atxn7 | 575.3815957 | 0.030038617 | 0.092596008 | 0.324405092 |
| Psmd6 | 699.5893116 | 0.278770024 | 0.088089389 | 3.164626589 |

| | | | | |
|---|---|---|---|---|
| Olfr720 | 0 | NA | NA | NA |
| Olfr31 | 0.115957212 | 0.274402725 | 3.724945049 | 0.073666248 |
| Il3ra | 52.25554781 | 1.99564358 | 0.327866284 | 6.086760599 |
| Olfr721-ps1 | 0 | NA | NA | NA |
| Slc4a7 | 745.5514613 | 0.149287296 | 0.088238279 | 1.691865455 |
| Nek10 | 32.25918065 | -0.075968765 | 0.185288921 | -0.410001661 |
| Lrrc3b | 129.9205039 | -0.165885693 | 0.096895708 | -1.712002481 |
| B230110C06Rik | 12.80781388 | -0.486331014 | 0.323949161 | -1.501257209 |
| Oxsm | 84.03786033 | 0.054927319 | 0.158492249 | 0.346561551 |
| Ngly1 | 371.6785756 | 0.232361153 | 0.069782991 | 3.329767765 |
| Top2b | 2165.3358 | 0.030996783 | 0.067264106 | 0.460822049 |
| Rarb | 169.0811627 | -0.219323341 | 0.111098854 | -1.974127852 |
| Thrb | 548.3553106 | -0.410966685 | 0.097569252 | -4.212051217 |
| Gm24140 | 0 | NA | NA | NA |
| Gm27851 | 0 | NA | NA | NA |
| Gm6773 | 0.143022002 | 0.755289479 | 3.547809333 | 0.212888971 |
| CU024892.1 | 0 | NA | NA | NA |
| Nr1d2 | 2201.828554 | -0.16115294 | 0.06252186 | -2.57754554 |
| Rpl15 | 1078.184859 | 0.293678795 | 0.12099846 | 2.427128363 |
| Nkiras1 | 524.1011176 | -0.312843955 | 0.081159513 | -3.854680043 |
| Ube2e1 | 611.3267082 | 0.1847758 | 0.10010392 | 1.845839801 |
| Gm22350 | 0.766686841 | 1.446672762 | 1.328402905 | 1.089031616 |
| Gm6781 | 3.825407769 | 1.543700798 | 0.573502919 | 2.691705215 |
| Ube2e2 | 649.6422344 | 0.059630949 | 0.082883458 | 0.719455364 |
| Gm20678 | 11.63441953 | -0.300888972 | 0.447371909 | -0.672570106 |
| Gm20699 | 63.74099188 | -0.473740826 | 0.312207109 | -1.517392821 |
| Gm22499 | 0 | NA | NA | NA |
| Gm15916 | 0 | NA | NA | NA |
| Gm20677 | 0 | NA | NA | NA |
| Gm8582 | 0.356401535 | 0.901906232 | 1.720761407 | 0.524132066 |
| Gm8584 | 0.374155823 | 0.823209805 | 1.723270112 | 0.477702131 |
| Gm9637 | 0 | NA | NA | NA |
| Gm21738 | 1.310113234 | 0.563252816 | 0.992480197 | 0.567520458 |
| Gm7352 | 0 | NA | NA | NA |
| Gm2244 | 0.202043434 | 0.274411029 | 2.61025331 | 0.105128122 |
| Gm2237 | 15.97977484 | 1.032973985 | 0.290536624 | 3.555400246 |
| Gm5458 | 0.372874528 | -0.588887666 | 1.542095856 | -0.381874878 |
| 4930469B13Rik | 0 | NA | NA | NA |
| Gm17030 | 0.08169834 | -0.206492286 | 3.724945049 | -0.055434988 |
| Nid2 | 107.7830705 | 0.760896285 | 0.17482667 | 4.352289522 |
| 2700060E02Rik | 556.5044864 | 0.674626527 | 0.100158461 | 6.735591954 |
| Gng2 | 1773.216353 | 0.202076688 | 0.064736098 | 3.121545707 |
| Saysd1 | 59.98590933 | -0.424830675 | 0.156322199 | -2.717660557 |
| Kcnk5 | 24.84306935 | 0.668156458 | 0.268609834 | 2.487460889 |
| Gm24912 | 0 | NA | NA | NA |
| Kcnk16 | 0.038981395 | 0.274393828 | 3.724945049 | 0.073663859 |
| Nudt13 | 124.3202546 | 0.580294406 | 0.129751842 | 4.472340423 |

| | | | | |
|---|---|---|---|---|
| Ecd | 421.6024906 | 0.080052653 | 0.07281726 | 1.099363716 |
| Fam149b | 307.5778511 | -0.051943698 | 0.085551447 | -0.60716329 |
| Dnajc9 | 241.2584147 | 0.114284343 | 0.120091599 | 0.951643109 |
| Mrps16 | 150.6596876 | 0.269356812 | 0.108999642 | 2.471171518 |
| Cfap70 | 17.61224759 | 0.264250054 | 0.258419387 | 1.02256281 |
| 7330404K18Rik | 0 | NA | NA | NA |
| Anxa7 | 995.7442115 | 0.527915763 | 0.084589801 | 6.24089143 |
| Mss51 | 28.46984622 | -0.70927389 | 0.272223153 | -2.605487008 |
| Ppp3cb | 3014.123295 | -0.095683006 | 0.059993799 | -1.59488159 |
| 1810062O18Rik | 11.1715319 | 0.111792281 | 0.322820714 | 0.346298352 |
| Usp54 | 1368.558277 | -0.057954338 | 0.072144469 | -0.803309519 |
| 2810402E24Rik | 26.39841974 | -0.073532054 | 0.204073307 | -0.360321767 |
| Myoz1 | 2.494795275 | 0.032420356 | 0.806369752 | 0.040205323 |
| Synpo2l | 4.120754503 | 1.068789634 | 0.559526919 | 1.910166602 |
| Sec24c | 1144.691689 | 0.047217952 | 0.073476671 | 0.642625085 |
| 6230400D17Rik | 14.90222924 | 0.19977195 | 0.275715726 | 0.724557691 |
| Mir6946 | 0.27962944 | -1.444734298 | 2.469604592 | -0.585006322 |
| Fut11 | 188.4790295 | -0.475756771 | 0.095004119 | -5.007748911 |
| Gm25328 | 3.501233001 | -0.405853286 | 0.588937221 | -0.689128266 |
| Chchd1 | 206.0508832 | 0.3925701 | 0.114916168 | 3.416143312 |
| Zswim8 | 1959.999352 | 0.135038039 | 0.072838852 | 1.853928719 |
| Ndst2 | 264.5758868 | 0.429211828 | 0.112814739 | 3.804572277 |
| Camk2g | 2341.047828 | -0.160906489 | 0.055706144 | -2.88848728 |
| Plau | 140.9153563 | 1.039030339 | 0.294616877 | 3.52671697 |
| Vcl | 823.9348543 | -0.012023959 | 0.114116389 | -0.105365754 |
| Ap3m1 | 731.0242943 | 0.006743611 | 0.068449866 | 0.098518987 |
| Adk | 509.0668211 | -0.138437479 | 0.129439334 | -1.069516316 |
| Gm26372 | 4.248258162 | -0.553142954 | 0.517344713 | -1.069196109 |
| Gm25864 | 0.26743452 | -1.152719122 | 2.086856071 | -0.552371166 |
| Kat6b | 1504.25356 | -0.272285713 | 0.079048774 | -3.444527953 |
| Dupd1 | 0.592353126 | 1.512246641 | 1.266273941 | 1.1942492 |
| Dusp13 | 5.809763333 | 0.129547036 | 0.497098105 | 0.260606578 |
| Gm15935 | 0.68377221 | 1.783359478 | 1.498622377 | 1.189999233 |
| Samd8 | 1631.759024 | 0.130655536 | 0.075245906 | 1.736380658 |
| Gm23502 | 0.371001759 | -1.518173993 | 3.187383283 | -0.476307321 |
| Vdac2 | 1632.626953 | 0.124573577 | 0.068113244 | 1.828918556 |
| Comtd1 | 88.38321911 | 0.052011464 | 0.120646397 | 0.431106649 |
| A430057M04Rik | 12.38941375 | 0.19337696 | 0.347697673 | 0.55616409 |
| Gm37590 | 0 | NA | NA | NA |
| Zfp503 | 121.7214339 | 0.245426136 | 0.156292635 | 1.570298797 |
| C130012C08Rik | 2.040813782 | -0.034121016 | 0.667579935 | -0.051111506 |
| 1700112E06Rik | 23.04170766 | 0.888645397 | 0.313868611 | 2.831265587 |
| Gm27532 | 0 | NA | NA | NA |
| Gm5670 | 0 | NA | NA | NA |
| Gm10248 | 1.061926442 | 1.818430065 | 1.079443511 | 1.684599561 |
| Kcnma1 | 2520.351437 | -0.294885997 | 0.120423055 | -2.448750345 |
| A830039N20Rik | 615.447544 | -0.215618463 | 0.11316752 | -1.905303425 |

| | | | | |
|---|---|---|---|---|
| Gm29626 | 0 | NA | NA | NA |
| Gm6158 | 5.934383314 | -0.163853011 | 0.41272588 | -0.397002028 |
| mmu-mir-7210 | 0 | NA | NA | NA |
| Dlg5 | 927.9578943 | 0.291319047 | 0.083511024 | 3.488390291 |
| Gm17105 | 11.22543593 | -0.337033073 | 0.399432986 | -0.84377877 |
| E330034G19Rik | 0.520501943 | 0.693886091 | 1.44809381 | 0.479172058 |
| Polr3a | 517.7839581 | -0.048600191 | 0.066584329 | -0.729904343 |
| Rps24 | 2358.715317 | 0.813335588 | 0.106663824 | 7.62522438 |
| 4930542C16Rik | 0.048949532 | -0.206487837 | 3.724945049 | -0.055433794 |
| 4931403M11Rik | 0.541236733 | -0.596481041 | 1.627453358 | -0.366511912 |
| 4930428N03Rik | 0.347568383 | 0.274412348 | 1.782599482 | 0.15393943 |
| Gm10398 | 0 | NA | NA | NA |
| Gm26660 | 26.92887363 | -0.577649217 | 0.287165828 | -2.011552773 |
| Zmiz1os1 | 24.20908764 | -0.478001565 | 0.212726721 | -2.247021734 |
| Zmiz1 | 3086.44194 | 0.107441846 | 0.081615448 | 1.316440069 |
| Mir3075 | 1.765392608 | -0.923685808 | 0.804557139 | -1.148067382 |
| Gm26772 | 0.433939115 | -0.684777593 | 1.484356723 | -0.461329533 |
| Ppif | 256.0875963 | 0.007083844 | 0.079866414 | 0.088696161 |
| 1700054O19Rik | 0.226439421 | 0.79634234 | 2.149951938 | 0.370400066 |
| Zcchc24 | 1699.51765 | 0.062538763 | 0.127309228 | 0.491235112 |
| Gm27185 | 0 | NA | NA | NA |
| Gm23701 | 0 | NA | NA | NA |
| Anxa11 | 152.4094582 | 0.349609334 | 0.098825676 | 3.537636636 |
| Anxa11os | 1.923336751 | 2.539352508 | 1.071641111 | 2.369592285 |
| Plac9a | 0 | NA | NA | NA |
| Tmem254a | 0 | NA | NA | NA |
| Cphx1 | 0 | NA | NA | NA |
| Gm19013 | 0 | NA | NA | NA |
| Duxbl1 | 0.094294407 | -0.206492286 | 3.724945049 | -0.055434988 |
| Gm2260 | 1.263942296 | -2.183061561 | 1.348354825 | -1.619055698 |
| Gm9780 | 0.038981395 | 0.274393828 | 3.724945049 | 0.073663859 |
| Tmem254c | 0.63148455 | -0.253169321 | 1.284418485 | -0.197108126 |
| Gm19014 | 0 | NA | NA | NA |
| Cphx2 | 0 | NA | NA | NA |
| Cphx3 | 0 | NA | NA | NA |
| Duxbl2 | 0 | NA | NA | NA |
| Gm2274 | 0.075043205 | 0.755286614 | 3.724945049 | 0.202764498 |
| Plac9b | 0 | NA | NA | NA |
| Tmem254b | 0 | NA | NA | NA |
| Gm19015 | 0 | NA | NA | NA |
| Duxbl3 | 0 | NA | NA | NA |
| 4933413J09Rik | 0 | NA | NA | NA |
| Gm16245 | 0.158569248 | 1.236192552 | 2.811594796 | 0.43967664 |
| Slmapos2 | 0 | NA | NA | NA |
| Slmap | 1697.427193 | -0.170027897 | 0.045099549 | -3.770057619 |
| Gm24031 | 0 | NA | NA | NA |
| Gm2178 | 0 | NA | NA | NA |

| | | | | |
|---|---|---|---|---|
| Dennd6a | 497.1097104 | -0.29211478 | 0.082137891 | -3.556394948 |
| Arf4os | 0.854956099 | 0.062487885 | 1.109202853 | 0.05633585 |
| Arf4 | 1525.733304 | 0.881539431 | 0.129833248 | 6.789781845 |
| Pde12 | 370.8916435 | 0.338356108 | 0.139197087 | 2.430770029 |
| Mir7672 | 0 | NA | NA | NA |
| 9930004E17Rik | 4.759985783 | 0.556803844 | 0.442547716 | 1.258178099 |
| Dnah12 | 39.20255926 | -0.708258202 | 0.223338347 | -3.171234183 |
| Asb14 | 0.828364824 | -1.91506959 | 1.153071285 | -1.660842322 |
| Appl1 | 1598.16168 | -0.083548747 | 0.049681303 | -1.681693953 |
| Hesx1 | 0 | NA | NA | NA |
| Il17rd | 75.9257241 | -0.265230805 | 0.176909008 | -1.499249858 |
| Arhgef3 | 850.6410935 | 0.116520107 | 0.059933506 | 1.944156365 |
| Gm23633 | 0 | NA | NA | NA |
| Fam208a | 1486.670546 | -0.346081829 | 0.120157966 | -2.880223764 |
| Ccdc66 | 229.1061877 | -0.404177493 | 0.12640475 | -3.197486583 |
| Gm22485 | 2.784173388 | 0.242988054 | 0.704530304 | 0.344893687 |
| Gm25697 | 0 | NA | NA | NA |
| Erc2 | 1776.688752 | -0.1986456 | 0.111616884 | -1.779709239 |
| Wnt5a | 214.1936815 | -0.099457332 | 0.090745927 | -1.09599775 |
| Gm2670 | 2.740285017 | 2.632617357 | 0.961921257 | 2.736832497 |
| Cacna2d3 | 929.5729355 | -0.185674121 | 0.115607543 | -1.606072717 |
| Gm23999 | 4.885903765 | -0.320110481 | 0.566046146 | -0.565520114 |
| Lrtm1 | 0.20890592 | -0.149301274 | 2.610374561 | -0.057195345 |
| Gm6043 | 0.823931708 | -1.058673795 | 1.169892401 | -0.90493262 |
| Selk | 760.4563827 | 0.227566074 | 0.108980758 | 2.088130772 |
| Actr8 | 526.6961103 | 0.025557459 | 0.067990734 | 0.375896206 |
| mmu-mir-6947 | 1.07422636 | 0.657713518 | 0.954699754 | 0.688921847 |
| Il17rb | 16.19919013 | -0.320058937 | 0.25755141 | -1.242699224 |
| Chdh | 62.42317444 | -0.65365478 | 0.241769529 | -2.703627633 |
| Cacna1d | 1000.382172 | -0.251562253 | 0.148138278 | -1.698158344 |
| Dcp1a | 375.2765008 | -0.10407879 | 0.068668223 | -1.515676183 |
| Tkt | 1261.009206 | -0.039671108 | 0.075298618 | -0.526850421 |
| Mir3076 | 0.281790215 | 0.725616371 | 2.06832146 | 0.350823789 |
| Prkcd | 783.4000836 | 0.457182281 | 0.099260443 | 4.605885975 |
| Rft1 | 140.5032926 | 0.517542645 | 0.114622348 | 4.515198438 |
| Sfmbt1 | 877.7713785 | 0.177188862 | 0.061481365 | 2.881992973 |
| 1700087M22Rik | 0.048949532 | -0.206487837 | 3.724945049 | -0.055433794 |
| Gm15667 | 10.02380282 | 0.512382681 | 0.357029748 | 1.435126019 |
| Tmem110 | 229.3938144 | -0.15938675 | 0.091457089 | -1.742748991 |
| Mustn1 | 28.82764176 | 0.814764441 | 0.340827419 | 2.390548396 |
| Itih4 | 5.953922204 | 2.875908015 | 0.707137826 | 4.066969561 |
| Itih3 | 692.5237228 | 0.338786256 | 0.165004551 | 2.053193406 |
| Itih1 | 0 | NA | NA | NA |
| Nek4 | 254.004222 | -0.371673766 | 0.091801439 | -4.048670357 |
| Spcs1 | 557.2353825 | 0.290562316 | 0.147231 | 1.973513162 |
| Glt8d1 | 210.2471507 | -0.28539416 | 0.102800277 | -2.776200288 |
| Gnl3 | 342.5965878 | 0.622325919 | 0.168903401 | 3.684507922 |

| | | | | |
|---|---|---|---|---|
| Snord69 | 6.499084164 | 0.47385782 | 0.43991459 | 1.077158681 |
| Gm24916 | 0.901079044 | 0.268877191 | 1.296818858 | 0.207335966 |
| Snord19 | 6.156095358 | 1.01205053 | 0.490521454 | 2.063213591 |
| Pbrm1 | 1674.501212 | 0.060616542 | 0.045072412 | 1.344870169 |
| Gm5459 | 0.740733539 | -1.000631938 | 1.373088293 | -0.728745517 |
| Smim4 | 39.16029245 | -0.13141697 | 0.175266873 | -0.749810662 |
| Nt5dc2 | 56.55209549 | 0.230252635 | 0.179406303 | 1.28341441 |
| Stab1 | 251.2128195 | 0.349819395 | 0.142790468 | 2.449879191 |
| Nisch | 5669.263495 | 0.058951321 | 0.043124069 | 1.367016645 |
| Tnnc1 | 26.93316509 | -0.632490847 | 0.237361393 | -2.66467448 |
| Sema3g | 57.06689575 | 1.281729251 | 0.251682055 | 5.092652521 |
| Phf7 | 44.60374799 | 0.342663932 | 0.187999864 | 1.822681808 |
| Bap1 | 1295.484164 | 0.0245987 | 0.066091636 | 0.372190818 |
| Dnah1 | 99.51919862 | -0.1697693 | 0.124244315 | -1.366415033 |
| Capn7 | 711.6756629 | -0.210022377 | 0.067982953 | -3.089338855 |
| Sh3bp5 | 520.8229502 | 0.020528781 | 0.090743357 | 0.226229019 |
| Mettl6 | 226.22855 | 0.234352011 | 0.105031413 | 2.23125638 |
| Eaf1 | 382.8875853 | 0.218769053 | 0.123390336 | 1.772983683 |
| Colq | 11.36104049 | -1.067042343 | 0.379101812 | -2.814659042 |
| Gm24057 | 0.166923653 | -0.206481504 | 2.766511172 | -0.074636064 |
| Hacl1 | 56.72216686 | -0.573971399 | 0.228013703 | -2.517267121 |
| Btd | 112.7781148 | 0.179201952 | 0.14307131 | 1.252535904 |
| Ankrd28 | 1173.696225 | 0.144884368 | 0.072142915 | 2.008296547 |
| Gm22876 | 0.047948698 | -0.206487837 | 3.724945049 | -0.055433794 |
| D830044D21Rik | 0.364803694 | 1.17339196 | 1.690398086 | 0.694151259 |
| Gm26045 | 0 | NA | NA | NA |
| Galnt15 | 194.1661234 | 1.565719242 | 0.244019229 | 6.416376471 |
| Dph3 | 331.9002246 | -0.010165623 | 0.096905913 | -0.104901992 |
| Oxnad1 | 151.3315786 | -0.561531854 | 0.128746164 | -4.361542393 |
| Msmb | 0 | NA | NA | NA |
| Ncoa4 | 556.7801885 | 0.148666418 | 0.076096567 | 1.953654733 |
| Timm23 | 256.1091139 | 0.304114401 | 0.119651557 | 2.541666893 |
| Gm23946 | 1.072725949 | -0.573838274 | 0.860784174 | -0.666645939 |
| Parg | 736.7172274 | -0.034836115 | 0.053828499 | -0.647168606 |
| Gm7707 | 0.203659793 | 0.755299243 | 2.597234627 | 0.290809015 |
| Gm17210 | 0.136194719 | -0.687371727 | 3.724945049 | -0.184532045 |
| Ogdhl | 1425.208207 | -0.029265833 | 0.068926978 | -0.42459185 |
| 1700024G13Rik | 0.700875826 | -0.564100208 | 1.089077377 | -0.517961552 |
| Chat | 45.18468771 | 0.11618427 | 0.203312227 | 0.57145737 |
| Slc18a3 | 49.59830663 | -0.115910825 | 0.340316024 | -0.340597612 |
| Gm2990 | 1.893473903 | -0.756273728 | 0.749171739 | -1.009479788 |
| Ercc6 | 588.9859623 | -0.404376193 | 0.113619876 | -3.559026876 |
| Prrxl1 | 7.584775115 | -0.322012899 | 0.610386664 | -0.527555595 |
| 3425401B19Rik | 4.622495198 | 0.148773057 | 0.481577618 | 0.308928511 |
| Gm28651 | 0.300197857 | -1.122974879 | 1.743475689 | -0.644101255 |
| 1810011H11Rik | 3.566868347 | -0.084955601 | 0.609473778 | -0.139391724 |
| Fam170b | 0.045882455 | 0.755288839 | 3.724945049 | 0.202765096 |

| | | | | |
|---|---|---|---|---|
| Vstm4 | 53.0402276 | 0.259243274 | 0.261874731 | 0.989951464 |
| Gm7734 | 0.640869663 | 0.054780827 | 1.217104069 | 0.045009156 |
| Wdfy4 | 177.0804989 | 1.931649373 | 0.185443609 | 10.41637069 |
| Lrrc18 | 16.544867 | 1.46561687 | 0.333184746 | 4.39881144 |
| Gm25498 | 0 | NA | NA | NA |
| Arhgap22 | 86.59583876 | 0.272793488 | 0.143970554 | 1.894786674 |
| Gm15512 | 6.43024678 | -1.208520558 | 0.493022016 | -2.451250693 |
| Mapk8 | 1298.322725 | -0.235649404 | 0.06325482 | -3.725398398 |
| Ptpn20 | 4.379735502 | -0.491311591 | 0.485826692 | -1.011289826 |
| Gm26228 | 0 | NA | NA | NA |
| Gdf10 | 323.3389736 | 0.267001923 | 0.354610293 | 0.75294465 |
| Gdf2 | 0.41755988 | 0.274415845 | 1.478488252 | 0.185605699 |
| Rbp3 | 3.927501353 | 1.28733448 | 0.727708711 | 1.76902442 |
| Zfp488 | 48.54553077 | 0.04827111 | 0.18932143 | 0.254969076 |
| Gm5460 | 0 | NA | NA | NA |
| A630023A22Rik | 0.092691418 | 0.274387304 | 3.724945049 | 0.073662108 |
| Antxrl | 0 | NA | NA | NA |
| Anxa8 | 7.103537313 | 1.256274398 | 0.53888415 | 2.331251343 |
| Gm24998 | 0 | NA | NA | NA |
| Npy4r | 1.230904645 | -0.169055302 | 0.979014678 | -0.172679027 |
| Rpl23a-ps3 | 207.5274121 | 0.45992523 | 0.13997268 | 3.285821407 |
| Gprin2 | 18.00709799 | 0.266292831 | 0.291108465 | 0.914754681 |
| Syt15 | 3.163297818 | -0.834876018 | 0.59215691 | -1.409889851 |
| Fam35a | 121.8552786 | 0.150168124 | 0.127849341 | 1.174570965 |
| Glud1 | 2362.144562 | -0.323689301 | 0.10471252 | -3.091218706 |
| Fam25c | 0.591489534 | 0.822441994 | 1.34707968 | 0.610537006 |
| Sncg | 85.14162449 | -0.010096644 | 0.327777164 | -0.030803379 |
| Mmrn2 | 26.86649225 | 1.724061949 | 0.34083323 | 5.058374004 |
| Bmpr1a | 1071.470205 | 0.02658131 | 0.09374246 | 0.283556783 |
| Gm17110 | 1.151183522 | 0.166764044 | 0.943315941 | 0.176784931 |
| 9230112D13Rik | 0.075043205 | 0.755286614 | 3.724945049 | 0.202764498 |
| Ldb3 | 91.38560832 | -0.302526918 | 0.148763197 | -2.033613985 |
| Opn4 | 1.339915753 | 0.218551738 | 1.181174312 | 0.1850292 |
| Wapal | 1670.7941 | 0.222181497 | 0.070571488 | 3.148318167 |
| A930038B10Rik | 0.504415982 | -1.731457198 | 1.355216152 | -1.277624381 |
| Grid1 | 726.7856293 | -0.173974715 | 0.065389047 | -2.660609419 |
| Mir346 | 1.214291556 | -0.212484726 | 0.970473025 | -0.218949646 |
| Gm27617 | 0.732444829 | 0.275298412 | 1.053358458 | 0.261353018 |
| 4930596D02Rik | 0 | NA | NA | NA |
| Gm7853 | 0 | NA | NA | NA |
| 4930474N05Rik | 0 | NA | NA | NA |
| Gm24394 | 0 | NA | NA | NA |
| Gm25204 | 0 | NA | NA | NA |
| n-R5s46 | 0.182014725 | 1.713926057 | 2.685235113 | 0.638277836 |
| Ccser2 | 2137.802388 | -0.006033233 | 0.037788518 | -0.159657824 |
| Rgr | 0.246474914 | -0.206481214 | 2.09234022 | -0.09868434 |
| Lrit1 | 0.184428051 | 0.755298719 | 2.670659372 | 0.282813573 |

| | | | | |
|---|---|---|---|---|
| Lrit2 | 0.338204618 | -0.206504546 | 3.724945049 | -0.05543828 |
| Cdhr1 | 304.1165299 | -0.939565689 | 0.588760466 | -1.59583692 |
| 2610528A11Rik | 2.326728918 | -0.737323517 | 0.800987806 | -0.920517779 |
| Ghitm | 4045.339356 | 0.16285601 | 0.090227976 | 1.804939188 |
| Gm22469 | 0 | NA | NA | NA |
| Gm23909 | 0 | NA | NA | NA |
| Nrg3 | 1077.56296 | -0.125438302 | 0.135451907 | -0.926072619 |
| Gm20641 | 0 | NA | NA | NA |
| Nrg3os | 0 | NA | NA | NA |
| Gm27332 | 2.333541239 | -0.867677802 | 0.64808871 | -1.33882567 |
| Gm20642 | 0.139856492 | -0.206491248 | 3.574411025 | -0.057769307 |
| 1700109I08Rik | 0 | NA | NA | NA |
| Eif1-ps1 | 3.471245013 | -0.965956375 | 0.537622857 | -1.79671746 |
| Gm25012 | 0 | NA | NA | NA |
| Gm22191 | 0.04393175 | -0.206490061 | 3.724945049 | -0.055434391 |
| Gm22726 | 0.094831987 | 0.274404949 | 3.724945049 | 0.073666845 |
| Gm24388 | 0 | NA | NA | NA |
| Gm5205 | 0.0885994 | 0.274402725 | 3.724945049 | 0.073666248 |
| Sh2d4b | 40.5303677 | 0.079422781 | 0.248349138 | 0.319802926 |
| Tspan14 | 417.092454 | 0.231756281 | 0.090206781 | 2.56916697 |
| Fam213a | 601.8361932 | 0.023990587 | 0.113733582 | 0.210936704 |
| Dydc2 | 4.886726711 | 0.425780193 | 0.492495491 | 0.864536227 |
| Dydc1 | 0.704807757 | 0.240872844 | 1.12172898 | 0.214733549 |
| Mat1a | 4.404352952 | 1.448054226 | 0.706566867 | 2.04942277 |
| Sftpa1 | 0.045882455 | 0.755288839 | 3.724945049 | 0.202765096 |
| Mbl1 | 0 | NA | NA | NA |
| Sftpd | 0 | NA | NA | NA |
| Gm7925 | 0 | NA | NA | NA |
| Gm22823 | 0.476740939 | 2.020667866 | 1.647461387 | 1.22653428 |
| Gm2832 | 0.567915064 | 1.084088359 | 1.312238677 | 0.826136569 |
| Gm9671 | 0 | NA | NA | NA |
| Gm5798 | 0 | NA | NA | NA |
| Gm2855 | 0 | NA | NA | NA |
| Gm7945 | 0.044544648 | 0.755286614 | 3.724945049 | 0.202764498 |
| Gm6482 | 0.047948698 | -0.206487837 | 3.724945049 | -0.055433794 |
| Gm7953 | 0 | NA | NA | NA |
| Gm7954 | 0 | NA | NA | NA |
| Gm2878 | 0 | NA | NA | NA |
| Gm3486 | 0 | NA | NA | NA |
| Gm6490 | 0 | NA | NA | NA |
| Gm17061 | 0 | NA | NA | NA |
| Gm7970 | 0 | NA | NA | NA |
| 1700049E17Rik2 | 0 | NA | NA | NA |
| Gm17159 | 0 | NA | NA | NA |
| Gm3072 | 0 | NA | NA | NA |
| Gm3077 | 0 | NA | NA | NA |
| Gm3676 | 0 | NA | NA | NA |

| | | | | |
|---|---|---|---|---|
| Gm8065 | 0 | NA | NA | NA |
| Gm8068 | 0 | NA | NA | NA |
| Gm7929 | 0 | NA | NA | NA |
| Gm17062 | 0 | NA | NA | NA |
| Gm7980 | 0 | NA | NA | NA |
| 1700049E17Rik1 | 0 | NA | NA | NA |
| 1700024B05Rik | 0 | NA | NA | NA |
| Gm9890 | 0 | NA | NA | NA |
| Gm6401 | 0 | NA | NA | NA |
| Gm3543 | 0 | NA | NA | NA |
| Gm17063 | 0 | NA | NA | NA |
| Gm2930 | 0 | NA | NA | NA |
| Gm7991 | 0 | NA | NA | NA |
| 1700091H14Rik | 0 | NA | NA | NA |
| Gm7951 | 0 | NA | NA | NA |
| Gm17064 | 0 | NA | NA | NA |
| Gm17027 | 0 | NA | NA | NA |
| Gm17166 | 0 | NA | NA | NA |
| Gm3573 | 0.045882455 | 0.755288839 | 3.724945049 | 0.202765096 |
| Gm17165 | 0 | NA | NA | NA |
| Gm17038 | 0 | NA | NA | NA |
| Gm17026 | 0 | NA | NA | NA |
| Gm25208 | 0 | NA | NA | NA |
| Gm25892 | 0 | NA | NA | NA |
| Gm9611 | 0 | NA | NA | NA |
| Gm7995 | 0 | NA | NA | NA |
| Gm17164 | 0 | NA | NA | NA |
| Gm2951 | 0 | NA | NA | NA |
| Gm2959 | 0 | NA | NA | NA |
| Gm8005 | 0 | NA | NA | NA |
| Gm8011 | 0 | NA | NA | NA |
| Gm2966 | 0 | NA | NA | NA |
| Gm8020 | 0 | NA | NA | NA |
| Gm8024 | 0 | NA | NA | NA |
| Gm8032 | 0.048949532 | -0.206487837 | 3.724945049 | -0.055433794 |
| Gm17204 | 0 | NA | NA | NA |
| Gm17124 | 0 | NA | NA | NA |
| Gm25228 | 0 | NA | NA | NA |
| Gm23620 | 0.048949532 | -0.206487837 | 3.724945049 | -0.055433794 |
| Gm3633 | 0 | NA | NA | NA |
| Gm10378 | 0 | NA | NA | NA |
| Gm17162 | 0 | NA | NA | NA |
| Gm5929 | 0 | NA | NA | NA |
| Gm3008 | 0 | NA | NA | NA |
| Gm17161 | 0 | NA | NA | NA |
| Gm3015 | 0 | NA | NA | NA |
| Gm3025 | 0 | NA | NA | NA |

| | | | | |
|---|---|---|---|---|
| Gm10377 | 0 | NA | NA | NA |
| Gm7152 | 0 | NA | NA | NA |
| Gm21977 | 0 | NA | NA | NA |
| Gm3102 | 0 | NA | NA | NA |
| Gm8082 | 0 | NA | NA | NA |
| Gm3111 | 0 | NA | NA | NA |
| Gm10376 | 0 | NA | NA | NA |
| Gm6536 | 0 | NA | NA | NA |
| Gm3123 | 0 | NA | NA | NA |
| Gm8094 | 0.086930093 | -0.206490058 | 3.724945049 | -0.05543439 |
| Gm8104 | 2.686914903 | -0.566493947 | 0.694533966 | -0.815646138 |
| Gm7236 | 14.77965937 | 0.192676757 | 0.272156235 | 0.707963778 |
| Gm8114 | 0 | NA | NA | NA |
| Gm3141 | 0.276783799 | -0.541300263 | 2.489059134 | -0.217471837 |
| Gm7233 | 2.888901366 | 0.311091185 | 0.609621371 | 0.510302295 |
| Gm7316 | 0 | NA | NA | NA |
| Gm8122 | 0 | NA | NA | NA |
| Gm8126 | 0.513218277 | 1.098598653 | 1.304179085 | 0.842367942 |
| Gm17153 | 0 | NA | NA | NA |
| Gm8127 | 0 | NA | NA | NA |
| Gm9732 | 2.036409272 | 1.181781353 | 0.864741787 | 1.366629172 |
| Gm17155 | 0 | NA | NA | NA |
| Gm17154 | 0 | NA | NA | NA |
| 1700001F09Rik | 0.08486385 | 0.755286617 | 3.724945049 | 0.202764499 |
| Gm9597 | 0.358995975 | 0.755275353 | 3.724945049 | 0.202761475 |
| Gm8138 | 0 | NA | NA | NA |
| Gm5799 | 0.049553047 | 0.274391604 | 3.724945049 | 0.073663262 |
| Gm3216 | 0 | NA | NA | NA |
| Gm17654 | 0 | NA | NA | NA |
| Gm3221 | 0 | NA | NA | NA |
| Gm10375 | 0 | NA | NA | NA |
| Gm8046 | 0.277407222 | 1.213561779 | 1.783033258 | 0.680616457 |
| Gm8161 | 0 | NA | NA | NA |
| Gm8165 | 0 | NA | NA | NA |
| Gm9672 | 0 | NA | NA | NA |
| Gm17208 | 0.132101749 | 0.274387056 | 3.724945049 | 0.073662041 |
| Gm7324 | 15.17033637 | -0.009287021 | 0.291045604 | -0.031909159 |
| Gm16506 | 0 | NA | NA | NA |
| Gm6526 | 0.404621698 | 1.236204676 | 1.727794934 | 0.715481132 |
| Gm9587 | 0 | NA | NA | NA |
| Gm8180 | 0 | NA | NA | NA |
| Ear1 | 0.104637558 | -0.206492286 | 3.724945049 | -0.055434988 |
| Ear-ps12 | 0 | NA | NA | NA |
| Ang-ps2 | 0 | NA | NA | NA |
| Ear-ps9 | 0 | NA | NA | NA |
| Ear-ps8 | 0.045882455 | 0.755288839 | 3.724945049 | 0.202765096 |
| Ang-ps3 | 0 | NA | NA | NA |

| | | | | |
|---|---|---|---|---|
| Ear10 | 0 | NA | NA | NA |
| Gm8113 | 3.368233017 | 0.856463298 | 0.58496883 | 1.464117837 |
| Gm9550 | 0 | NA | NA | NA |
| Ang5 | 0 | NA | NA | NA |
| Ear-ps7 | 0 | NA | NA | NA |
| Ear-ps10 | 0 | NA | NA | NA |
| Ang6 | 0 | NA | NA | NA |
| Gm8194 | 0 | NA | NA | NA |
| Gm3287 | 0 | NA | NA | NA |
| Ear-ps2 | 0 | NA | NA | NA |
| Ang-ps1 | 0.046345709 | 0.274391604 | 3.724945049 | 0.073663262 |
| Gm16545 | 0 | NA | NA | NA |
| Ear2 | 0.663448651 | 1.616782142 | 1.385331109 | 1.167072717 |
| Gm3327 | 0 | NA | NA | NA |
| Gm8207 | 0 | NA | NA | NA |
| 4930503E14Rik | 0.57934571 | 0.220253299 | 1.427510819 | 0.15429186 |
| Gm8212 | 0.068097359 | -0.206487837 | 3.724945049 | -0.055433794 |
| Gm8220 | 0.072025462 | 0.755288839 | 3.724945049 | 0.202765096 |
| Gm8224 | 0 | NA | NA | NA |
| Gm5930 | 0.049553047 | 0.274391604 | 3.724945049 | 0.073663262 |
| Gm8229 | 0 | NA | NA | NA |
| Gm3370 | 0 | NA | NA | NA |
| Gm3371 | 0 | NA | NA | NA |
| Gm8232 | 0.045882455 | 0.755288839 | 3.724945049 | 0.202765096 |
| Gm21991 | 0 | NA | NA | NA |
| Gm17184 | 0.465126201 | 2.554943879 | 1.685545978 | 1.515796016 |
| BC061237 | 0.120349074 | 1.264824188 | 3.723011833 | 0.339731445 |
| Gm17093 | 0 | NA | NA | NA |
| Gm17183 | 0 | NA | NA | NA |
| Gm5624 | 0.172257197 | -0.206488151 | 2.742791845 | -0.075283931 |
| Gm8247 | 0.04393175 | -0.206490061 | 3.724945049 | -0.055434391 |
| Gm16439 | 6.205147171 | 0.202261818 | 0.380378348 | 0.531738515 |
| Gm8256 | 0 | NA | NA | NA |
| Gm9593 | 0 | NA | NA | NA |
| Gm8257 | 0 | NA | NA | NA |
| Gm21754 | 0 | NA | NA | NA |
| Gm6541 | 0 | NA | NA | NA |
| Gm8267 | 0.060174537 | 0.755286614 | 3.724945049 | 0.202764498 |
| Gm17187 | 0 | NA | NA | NA |
| Gm17070 | 0 | NA | NA | NA |
| Ptgdr | 10.70749769 | -1.095656003 | 0.478322411 | -2.290622344 |
| Gm16976 | 1.330389483 | -0.713249925 | 1.030966576 | -0.69182643 |
| mmu-mir-327 | 0 | NA | NA | NA |
| 4930431P03Rik | 1.669097723 | 1.012033029 | 0.830792728 | 1.218153452 |
| Gm17173 | 1.687842655 | 1.616208938 | 1.070215927 | 1.510170889 |
| Ptger2 | 14.92504073 | 0.80500044 | 0.384692054 | 2.092584009 |
| Txndc16 | 1225.114205 | 0.033728688 | 0.068182689 | 0.494681109 |

| | | | | |
|---|---|---|---|---|
| Gm15765 | 0 | NA | NA | NA |
| Gpr137c | 408.4638035 | -0.454663083 | 0.081898124 | -5.551568972 |
| Ero1l | 547.1939839 | -0.134057085 | 0.055142969 | -2.431082094 |
| Psmc6 | 1042.749598 | 0.337183895 | 0.100740202 | 3.347063923 |
| Styx | 68.64544997 | -0.367844894 | 0.210719248 | -1.74566347 |
| Gnpnat1 | 150.1275212 | 0.241618585 | 0.153469376 | 1.57437654 |
| Gm25353 | 0 | NA | NA | NA |
| Fermt2 | 877.773119 | -0.160104528 | 0.074212998 | -2.157365051 |
| Gm22637 | 3.849995044 | 0.014555658 | 0.532799494 | 0.027319205 |
| Gm15601 | 0.688336999 | 0.737588299 | 1.380275354 | 0.534377649 |
| Ddhd1 | 1001.935963 | -0.316475315 | 0.08091632 | -3.911143189 |
| Mir5131 | 0 | NA | NA | NA |
| A530076I17Rik | 4.579416614 | 2.231955703 | 0.59058602 | 3.779222041 |
| Gm15516 | 0.119234754 | 0.274400501 | 3.724945049 | 0.073665651 |
| Gm7206 | 18.53655595 | 0.278975435 | 0.256313477 | 1.088415009 |
| Gm15218 | 0.089089296 | 1.194485492 | 3.723939576 | 0.320758559 |
| Gm8317 | 0.802853338 | 0.498165963 | 1.177925321 | 0.422918121 |
| Gm15220 | 0 | NA | NA | NA |
| Gm6580 | 0.146060149 | -0.206490268 | 3.528689472 | -0.058517552 |
| Gm37257 | 0 | NA | NA | NA |
| Gm1821 | 160.7341621 | 0.606305352 | 0.166018638 | 3.652031838 |
| Gm15217 | 0.045882455 | 0.755288839 | 3.724945049 | 0.202765096 |
| Bmp4 | 27.37682956 | -0.513338479 | 0.245662651 | -2.089607339 |
| Gm15222 | 0.402965827 | -0.687254588 | 1.538064226 | -0.446830878 |
| Gm15221 | 0.062426873 | 0.755288839 | 3.724945049 | 0.202765096 |
| E130120K24Rik | 0.066050875 | 0.274391604 | 3.724945049 | 0.073663262 |
| Rubie | 0.049553047 | 0.274391604 | 3.724945049 | 0.073663262 |
| Cdkn3 | 4.411638001 | 1.089564115 | 0.565837993 | 1.925576098 |
| Cnih1 | 494.7336958 | -0.063228634 | 0.07690367 | -0.822179664 |
| Gmfb | 1861.18559 | -0.078691016 | 0.091264851 | -0.862226967 |
| Gm10101 | 2.636333195 | 0.079568099 | 0.582879484 | 0.136508663 |
| Cgrrf1 | 236.9675109 | 0.452892063 | 0.095996385 | 4.717803309 |
| Samd4 | 517.3838205 | -0.094482387 | 0.076994154 | -1.227137151 |
| Mir378c | 0 | NA | NA | NA |
| Gm15562 | 5.089503423 | -0.615441595 | 0.456881127 | -1.347049722 |
| Gm10371 | 0.119587853 | 1.218392466 | 3.723618458 | 0.327206581 |
| Gch1 | 83.87125827 | 2.196236984 | 0.321625608 | 6.828551371 |
| Wdhd1 | 97.88412802 | 0.307686435 | 0.169162096 | 1.81888521 |
| Gm24378 | 1.312748754 | -0.761214287 | 0.907057961 | -0.839212399 |
| Socs4 | 478.1944072 | -0.060721414 | 0.07375723 | -0.823260494 |
| Mapk1ip1l | 616.9594476 | 0.42663152 | 0.073669684 | 5.791140942 |
| Lgals3 | 777.9121779 | 5.053444635 | 0.459703274 | 10.9928402 |
| Dlgap5 | 14.65831982 | 2.198787828 | 0.4965637 | 4.428007578 |
| Fbxo34 | 544.4943191 | 0.105474571 | 0.070866175 | 1.488362692 |
| Atg14 | 247.0243887 | 0.038896999 | 0.074786541 | 0.52010694 |
| Ktn1 | 1793.170985 | 0.226441355 | 0.079650026 | 2.842953945 |
| Gm37874 | 52.01025927 | -0.296419136 | 0.223485063 | -1.326348761 |

| | | | | |
|---|---|---|---|---|
| Gm6055 | 178.1871439 | 0.42522599 | 0.109619255 | 3.879117694 |
| Peli2 | 376.3843311 | -0.108008573 | 0.092294883 | -1.170255274 |
| Gm6498 | 16.31733287 | -0.105730766 | 0.330999567 | -0.319428713 |
| Gm23881 | 0 | NA | NA | NA |
| Gm3534 | 2.621659813 | 1.195006127 | 0.681976543 | 1.752268665 |
| Tmem260 | 200.6788583 | -0.537211529 | 0.16856732 | -3.186925723 |
| Gm23469 | 0 | NA | NA | NA |
| Otx2 | 54.10925475 | 0.139200205 | 0.143475756 | 0.970200188 |
| Otx2os1 | 2.535154825 | -1.549234946 | 0.690094707 | -2.244959903 |
| Gm25380 | 0 | NA | NA | NA |
| Gm22699 | 0 | NA | NA | NA |
| Gm23470 | 0.573140788 | 0.981791402 | 1.275504059 | 0.769728168 |
| Exoc5 | 1321.293313 | 0.114170718 | 0.081575886 | 1.399564549 |
| Ap5m1 | 326.8043512 | -0.294513741 | 0.077859683 | -3.782621904 |
| Naa30 | 880.0118064 | 0.238177552 | 0.103981887 | 2.2905677 |
| 1700011H14Rik | 0 | NA | NA | NA |
| Slc35f4 | 116.8448588 | -0.088152357 | 0.120845111 | -0.729465645 |
| Gm22116 | 5.008872367 | -0.794444429 | 0.456248678 | -1.741253109 |
| Gm22949 | 0 | NA | NA | NA |
| 3632451O06Rik | 470.0630739 | -0.723103722 | 0.105135628 | -6.877818086 |
| Gm16082 | 1.838486464 | -1.304098467 | 0.814576854 | -1.600952028 |
| Gm22989 | 0 | NA | NA | NA |
| Olfr722 | 0 | NA | NA | NA |
| Olfr723 | 0 | NA | NA | NA |
| Olfr724 | 0 | NA | NA | NA |
| Olfr725 | 0 | NA | NA | NA |
| Gm37194 | 0.41257556 | -1.129766608 | 1.524729781 | -0.740961856 |
| Olfr726 | 0 | NA | NA | NA |
| Olfr727 | 0 | NA | NA | NA |
| Olfr728 | 0.068097359 | -0.206487837 | 3.724945049 | -0.055433794 |
| Olfr729 | 0 | NA | NA | NA |
| Olfr730 | 0 | NA | NA | NA |
| Olfr731 | 0.046345709 | 0.274391604 | 3.724945049 | 0.073663262 |
| Olfr732 | 0 | NA | NA | NA |
| Olfr733 | 0.092881282 | -0.687373951 | 3.724945049 | -0.184532642 |
| Olfr734 | 0.109869311 | -0.206490066 | 3.724945049 | -0.055434393 |
| Olfr735 | 0.066050875 | 0.274391604 | 3.724945049 | 0.073663262 |
| Tlr11 | 0.527131164 | 1.391740273 | 1.636786493 | 0.85028822 |
| Olfr736 | 0 | NA | NA | NA |
| Olfr738 | 0 | NA | NA | NA |
| Olfr739 | 0 | NA | NA | NA |
| Olfr740 | 0 | NA | NA | NA |
| Olfr741 | 0.048949532 | -0.206487837 | 3.724945049 | -0.055433794 |
| Olfr742 | 0 | NA | NA | NA |
| Olfr743 | 0 | NA | NA | NA |
| Olfr744 | 0 | NA | NA | NA |
| Olfr745 | 0 | NA | NA | NA |

| | | | | |
|---|---|---|---|---|
| Olfr746 | 0 | NA | NA | NA |
| Olfr747 | 0 | NA | NA | NA |
| Olfr748 | 0 | NA | NA | NA |
| Gm24316 | 0 | NA | NA | NA |
| Olfr749 | 0.172571121 | 0.755296067 | 2.73763 | 0.275894137 |
| Ttc5 | 245.1817073 | -0.091798284 | 0.092706085 | -0.990207752 |
| Gm10916 | 0.925454891 | 1.631383419 | 1.086444812 | 1.501579649 |
| Ccnb1ip1 | 5.389104856 | -1.274194904 | 0.494888387 | -2.574711667 |
| Rpph1 | 38912.15563 | 0.371345769 | 0.218597602 | 1.698764147 |
| Parp2 | 165.0646803 | 0.097124183 | 0.118679855 | 0.818371265 |
| Tep1 | 233.8406273 | 0.599898675 | 0.121525124 | 4.936416908 |
| Gm26782 | 38.7123841 | -0.596169936 | 0.228716722 | -2.606586575 |
| Klhl33 | 73.15949532 | 0.438116031 | 0.182180551 | 2.404845237 |
| A930018M24Rik | 44.75620921 | 0.124371465 | 0.181333149 | 0.685872745 |
| Osgep | 153.0925057 | -0.127161299 | 0.097279714 | -1.3071718 |
| Gm24689 | 0 | NA | NA | NA |
| Apex1 | 280.3220763 | 0.263347371 | 0.100031793 | 2.632636719 |
| Tmem55b | 636.4352874 | 0.12438304 | 0.058045457 | 2.142855739 |
| Pnp | 408.8476884 | 1.679958665 | 0.274516888 | 6.119691494 |
| Gm38316 | 0.038981395 | 0.274393828 | 3.724945049 | 0.073663859 |
| Pnp2 | 4.854385997 | 1.33917036 | 0.542550401 | 2.468287476 |
| Rnase10 | 0.957491927 | -1.903767211 | 1.152892627 | -1.651296197 |
| Rnase9 | 0.066050875 | 0.274391604 | 3.724945049 | 0.073663262 |
| Rnase11 | 0.038981395 | 0.274393828 | 3.724945049 | 0.073663859 |
| Rnase12 | 0.062426873 | 0.755288839 | 3.724945049 | 0.202765096 |
| Olfr750 | 0.092493346 | 0.274402725 | 3.724945049 | 0.073666248 |
| Rnase4 | 86.11690773 | 0.329534376 | 0.223593724 | 1.473808699 |
| Ang | 3.984262514 | 0.626258786 | 0.593079509 | 1.055944063 |
| Eddm3b | 0 | NA | NA | NA |
| Rnase6 | 15.68530113 | 6.43148211 | 0.909536582 | 7.071163749 |
| Rnase1 | 3.783653538 | -0.309062902 | 0.510302401 | -0.605646576 |
| Rnase2b | 0.112642008 | 0.274402725 | 3.724945049 | 0.073666248 |
| Ang2 | 0.049553047 | 0.274391604 | 3.724945049 | 0.073663262 |
| Ear14 | 0.049553047 | 0.274391604 | 3.724945049 | 0.073663262 |
| Rnase2a | 0.216167348 | -1.057762314 | 1.870438752 | -0.565515611 |
| Gm21718 | 0 | NA | NA | NA |
| Gm3719 | 0 | NA | NA | NA |
| Gm16289 | 0 | NA | NA | NA |
| Gm7247 | 0.038981395 | 0.274393828 | 3.724945049 | 0.073663859 |
| Vmn2r-ps111 | 0 | NA | NA | NA |
| Vmn2r88 | 0 | NA | NA | NA |
| Vmn2r89 | 0 | NA | NA | NA |
| Gm5622 | 0 | NA | NA | NA |
| Gm16290 | 0 | NA | NA | NA |
| Gm17175 | 0.075043205 | 0.755286614 | 3.724945049 | 0.202764498 |
| Gm17174 | 0 | NA | NA | NA |
| Gm17078 | 0 | NA | NA | NA |

| | | | | |
|---|---|---|---|---|
| Gm4181 | 0 | NA | NA | NA |
| Gm17186 | 0 | NA | NA | NA |
| Gm17079 | 0 | NA | NA | NA |
| Gm5800 | 0.136813032 | -1.150638635 | 3.638145365 | -0.31627066 |
| Ear-ps3 | 0 | NA | NA | NA |
| Ang4 | 0.095435502 | 0.75528439 | 3.724945049 | 0.202763901 |
| Ear-ps4 | 0 | NA | NA | NA |
| AY358078 | 0 | NA | NA | NA |
| Gm6047 | 0.182466651 | 0.274402709 | 2.68257363 | 0.10229084 |
| Gm27375 | 0 | NA | NA | NA |
| Ear-ps5 | 0 | NA | NA | NA |
| Ear6 | 0 | NA | NA | NA |
| Mettl17 | 160.3336998 | -0.28225648 | 0.15675437 | -1.800629097 |
| Slc39a2 | 25.14959081 | -1.171136692 | 0.285959914 | -4.095457557 |
| Ndrg2 | 5788.517411 | 0.323627668 | 0.150587728 | 2.149097221 |
| Tppp2 | 0.205880021 | -1.154755909 | 2.63262206 | -0.438633379 |
| Rnase13 | 0.16050186 | 0.755303814 | 2.800912406 | 0.26966349 |
| Arhgef40 | 469.4959225 | 0.026018903 | 0.087145639 | 0.298568047 |
| Gm16617 | 1.237774572 | 2.153466379 | 1.112567345 | 1.935582945 |
| Zfp219 | 473.6521586 | -0.190770445 | 0.090524908 | -2.10738072 |
| Tmem253 | 3.25900046 | -0.022411675 | 0.604529069 | -0.037072948 |
| Olfr221 | 0.038981395 | 0.274393828 | 3.724945049 | 0.073663859 |
| Gm22354 | 0.087261593 | 0.274400501 | 3.724945049 | 0.073665651 |
| Hnrnpc | 1799.363866 | 0.041711968 | 0.073812618 | 0.565106197 |
| Rpgrip1 | 13.58329738 | -0.102764627 | 0.299894158 | -0.342669653 |
| Supt16 | 1326.937056 | 0.233712382 | 0.080152101 | 2.915860963 |
| Gm26590 | 3.018699778 | -0.581864329 | 0.673815274 | -0.863536864 |
| Chd8 | 2275.929444 | -0.005770235 | 0.076323303 | -0.075602534 |
| Snord8 | 7.001247873 | -0.254366912 | 0.431993734 | -0.588820837 |
| RP24-223I19.6 | 3.473627573 | -0.367576406 | 0.498532587 | -0.737316708 |
| Rab2b | 331.1686235 | 0.011704611 | 0.072569508 | 0.161288274 |
| Gm23758 | 0 | NA | NA | NA |
| Tox4 | 931.9067073 | 0.351170307 | 0.068140642 | 5.153610193 |
| Mettl3 | 190.3858702 | -0.522752235 | 0.139953283 | -3.735190931 |
| Sall2 | 992.2939578 | -0.218810246 | 0.064933174 | -3.369775928 |
| Gm26328 | 0.151841013 | -0.206481646 | 2.850049718 | -0.072448437 |
| Olfr1513 | 0 | NA | NA | NA |
| RP24-461L18.3 | 26.78966518 | 0.66056208 | 0.220363819 | 2.997597718 |
| Olfr1512 | 0 | NA | NA | NA |
| Gm37504 | 0 | NA | NA | NA |
| Olfr1511 | 0 | NA | NA | NA |
| Olfr1510 | 0.062426873 | 0.755288839 | 3.724945049 | 0.202765096 |
| RP24-461L18.7 | 0.061920613 | 0.274393828 | 3.724945049 | 0.073663859 |
| Trav1 | 0.280850934 | 1.541557173 | 2.004912157 | 0.768890132 |
| Olfr1509 | 0.473637868 | -1.317240375 | 1.518815718 | -0.867281237 |
| Olfr1508 | 0.186324499 | 0.244775121 | 2.713146848 | 0.090218162 |
| RP23-117I19.7 | 0 | NA | NA | NA |

| | | | | |
|---|---|---|---|---|
| Olfr1507 | 1.123152174 | -0.306250439 | 0.941348333 | -0.325331684 |
| RP23-117I19.2 | 0.060174537 | 0.755286614 | 3.724945049 | 0.202764498 |
| RP23-117I19.3 | 0.172571121 | 0.755296067 | 2.73763 | 0.275894137 |
| Trav2 | 2.527889093 | 4.175534809 | 0.963532927 | 4.333567326 |
| Trav3-1 | 0.371273269 | 2.140976915 | 1.685231706 | 1.270434746 |
| Trav4-1 | 0 | NA | NA | NA |
| Trav5-1 | 0.4844674 | -0.206481701 | 1.468643431 | -0.140593487 |
| Trav6-1 | 0.59120296 | 1.444611005 | 1.537670213 | 0.939480386 |
| RP24-447O5.2 | 0 | NA | NA | NA |
| Trav7-1 | 0.214323314 | 1.236200973 | 2.185419544 | 0.565658423 |
| Trav6-2 | 0.503770797 | 2.081929109 | 1.521560708 | 1.368285273 |
| Trav7d-2 | 0 | NA | NA | NA |
| Trav4d-2 | 0 | NA | NA | NA |
| Trav6d-3 | 0 | NA | NA | NA |
| Trav7d-3 | 0.231728546 | 0.274380666 | 3.724945049 | 0.073660326 |
| Trav6d-4 | 0.360686062 | 2.023627626 | 1.544913628 | 1.309864571 |
| Trav7d-4 | 0.038981395 | 0.274393828 | 3.724945049 | 0.073663859 |
| RP23-452K12.17 | 0 | NA | NA | NA |
| Trav8d-1 | 0.480868668 | 1.247904498 | 1.673151649 | 0.745840641 |
| Trav9d-1 | 0.075043205 | 0.755286614 | 3.724945049 | 0.202764498 |
| Trav6d-5 | 0.045882455 | 0.755288839 | 3.724945049 | 0.202765096 |
| Trav10d | 0.08486385 | 0.755286617 | 3.724945049 | 0.202764499 |
| Trav6d-6 | 0.038981395 | 0.274393828 | 3.724945049 | 0.073663859 |
| Trav11d | 0 | NA | NA | NA |
| Trav7d-5 | 0 | NA | NA | NA |
| Trav12d-1 | 0 | NA | NA | NA |
| Trav13d-1 | 0.061920613 | 0.274393828 | 3.724945049 | 0.073663859 |
| Trav14d-1 | 0 | NA | NA | NA |
| RP23-452K12.18 | 0.160559904 | 0.755297704 | 2.794491024 | 0.270280955 |
| Trav15d-1-dv6d-1 | 0 | NA | NA | NA |
| Trav3d-2 | 0 | NA | NA | NA |
| Trav9d-2 | 0 | NA | NA | NA |
| Trav4d-3 | 0 | NA | NA | NA |
| Rps19-ps2 | 0.546944188 | 1.090408238 | 1.53308398 | 0.711251472 |
| Trav5d-2 | 0 | NA | NA | NA |
| Trav12d-2 | 0.075043205 | 0.755286614 | 3.724945049 | 0.202764498 |
| Trav9d-3 | 0.216532645 | 0.274393532 | 3.170680374 | 0.0865409 |
| Trav5d-3 | 0 | NA | NA | NA |
| Trav12d-3 | 0 | NA | NA | NA |
| Trav13d-2 | 0 | NA | NA | NA |
| Trav14d-2 | 0 | NA | NA | NA |
| Trav15d-2-dv6d-2 | 0 | NA | NA | NA |
| RP24-548I6.12 | 0 | NA | NA | NA |
| Trav3d-3 | 0 | NA | NA | NA |
| Trav9d-4 | 0.11147366 | 0.274389235 | 3.724945049 | 0.073662626 |
| Trav4d-4 | 0 | NA | NA | NA |
| RP24-334B8.4 | 0 | NA | NA | NA |

| | | | | |
|---|---|---|---|---|
| Trav5d-4 | 0 | NA | NA | NA |
| Trav6d-7 | 0.046345709 | 0.274391604 | 3.724945049 | 0.073663262 |
| Trav7d-6 | 0 | NA | NA | NA |
| RP24-334B8.8 | 0.414202207 | 1.014088922 | 1.489231086 | 0.680947995 |
| Trav13d-3 | 0 | NA | NA | NA |
| Trav8d-2 | 0.305990011 | 0.274407512 | 1.999909996 | 0.137209931 |
| Trav16d-dv11 | 0.583486461 | 1.636092181 | 3.416274413 | 0.47891123 |
| Trav13d-4 | 0.066050875 | 0.274391604 | 3.724945049 | 0.073663262 |
| Trav14d-3-dv8 | 0.35026241 | 0.755307866 | 1.722706575 | 0.438442552 |
| Trav15d-3 | 0 | NA | NA | NA |
| Trav7n-4 | 0 | NA | NA | NA |
| Trav9n-1 | 0.141094079 | 0.755285157 | 3.563850454 | 0.211929531 |
| Trav6n-5 | 0 | NA | NA | NA |
| Trav10n | 0 | NA | NA | NA |
| Trav6n-6 | 0.230315653 | 0.755303602 | 2.138118016 | 0.353256273 |
| Trav11n | 0 | NA | NA | NA |
| Trav7n-5 | 0 | NA | NA | NA |
| Trav12n-1 | 0 | NA | NA | NA |
| Trav13n-1 | 0.049553047 | 0.274391604 | 3.724945049 | 0.073663262 |
| Trav14n-1 | 0 | NA | NA | NA |
| RP23-371K2.5 | 0.606297752 | 0.641157928 | 1.291548949 | 0.496425574 |
| Trav15n-1 | 0 | NA | NA | NA |
| Trav3n-2 | 0 | NA | NA | NA |
| Trav9n-2 | 0 | NA | NA | NA |
| Trav4n-3 | 0 | NA | NA | NA |
| Rps19-ps1 | 0 | NA | NA | NA |
| Trav5n-2 | 0 | NA | NA | NA |
| Trav12n-2 | 0 | NA | NA | NA |
| Trav9n-3 | 0 | NA | NA | NA |
| Trav5n-3 | 0 | NA | NA | NA |
| Trav12n-3 | 0 | NA | NA | NA |
| Trav13n-2 | 0 | NA | NA | NA |
| Trav14n-2 | 0 | NA | NA | NA |
| Trav15n-2 | 0 | NA | NA | NA |
| RP23-125A3.6 | 0 | NA | NA | NA |
| Trav3n-3 | 0 | NA | NA | NA |
| Trav9n-4 | 0.08486385 | 0.755286617 | 3.724945049 | 0.202764499 |
| Trav4n-4 | 0.12092566 | 1.236179401 | 3.724945049 | 0.331865138 |
| Trav5n-4 | 0 | NA | NA | NA |
| Trav6n-7 | 0 | NA | NA | NA |
| Trav7n-6 | 0 | NA | NA | NA |
| RP24-334B8.20 | 0 | NA | NA | NA |
| Trav13n-3 | 0 | NA | NA | NA |
| Trav8n-2 | 0.072025462 | 0.755288839 | 3.724945049 | 0.202765096 |
| Trav16n | 0.219234697 | 0.755300834 | 2.549697479 | 0.296231549 |
| Trav13n-4 | 0.049553047 | 0.274391604 | 3.724945049 | 0.073663262 |
| Trav14n-3 | 0.163190422 | 0.274394252 | 3.398594894 | 0.080737558 |

| | | | | |
|---|---|---|---|---|
| Trav15n-3 | 0 | NA | NA | NA |
| Trav7-2 | 0 | NA | NA | NA |
| Trav4-2 | 0.045882455 | 0.755288839 | 3.724945049 | 0.202765096 |
| Trav6-3 | 0.15956642 | 0.75529148 | 3.418807262 | 0.22092251 |
| Trav7-3 | 0.350945957 | 1.479742684 | 1.736320146 | 0.852229174 |
| Trav6-4 | 0.163577647 | 0.755296524 | 2.784101768 | 0.271289122 |
| Trav7-4 | 0.394584743 | 1.384360599 | 3.712147512 | 0.372927152 |
| RP23-230L15.11 | 0.04393175 | -0.206490061 | 3.724945049 | -0.055434391 |
| Trav8-1 | 0 | NA | NA | NA |
| Trav9-1 | 0.21728767 | 0.75529784 | 2.544699458 | 0.296812198 |
| Trav6-5 | 0.124347486 | 0.755286736 | 3.717316019 | 0.203180664 |
| Trav10 | 0.044544648 | 0.755286614 | 3.724945049 | 0.202764498 |
| Trav6-6 | 0.141781211 | 0.755294552 | 2.911713426 | 0.259398657 |
| Trav11 | 0 | NA | NA | NA |
| Trav7-5 | 0 | NA | NA | NA |
| Trav12-1 | 0.061920613 | 0.274393828 | 3.724945049 | 0.073663859 |
| Trav13-1 | 0.353543486 | 1.960691973 | 1.662396355 | 1.179437123 |
| Trav14-1 | 0 | NA | NA | NA |
| Trav15-1-dv6-1 | 0.099106094 | 0.274387056 | 3.724945049 | 0.073662041 |
| Trav3-2 | 0 | NA | NA | NA |
| Trav9-2 | 0.219744001 | 0.755303984 | 2.154043771 | 0.350644678 |
| Trav4-3 | 0 | NA | NA | NA |
| Trav5-2 | 0 | NA | NA | NA |
| Trav12-2 | 0 | NA | NA | NA |
| Trav12-3 | 0.264301748 | 1.260486074 | 2.448937563 | 0.514707314 |
| Trav13-2 | 0.124853745 | 1.290908285 | 3.721447108 | 0.346883416 |
| Trav14-2 | 0 | NA | NA | NA |
| Trav15-2-dv6-2 | 0 | NA | NA | NA |
| RP24-100P9.4 | 0 | NA | NA | NA |
| Trav3-3 | 2.07315737 | 0.836847831 | 0.696608482 | 1.201317315 |
| Trav9-4 | 0.183309527 | 0.755297975 | 2.685861896 | 0.281212514 |
| Trav4-4-dv10 | 0.315232318 | 2.186312145 | 3.698483093 | 0.591137526 |
| RP24-100P9.5 | 0 | NA | NA | NA |
| Trav5-4 | 0 | NA | NA | NA |
| Trav6-7-dv9 | 0.09176491 | 1.15872478 | 3.72306111 | 0.311229052 |
| Trav7-6 | 0 | NA | NA | NA |
| Trav13-3 | 0 | NA | NA | NA |
| Trav8-2 | 0 | NA | NA | NA |
| Trav16 | 0 | NA | NA | NA |
| Trav13-4-dv7 | 0.143485256 | 0.274391988 | 3.549605256 | 0.077302113 |
| Trav14-3 | 0 | NA | NA | NA |
| Trav15-3 | 0 | NA | NA | NA |
| Trav3-4 | 0.305831651 | 0.938072266 | 2.017130918 | 0.465052743 |
| Trav12-4 | 0.062426873 | 0.755288839 | 3.724945049 | 0.202765096 |
| B230359F08Rik | 0.108266322 | 0.274389235 | 3.724945049 | 0.073662626 |
| Trav17 | 0.511617548 | 2.351474232 | 1.36928864 | 1.71729624 |
| Trav18 | 0 | NA | NA | NA |

| | | | | |
|---|---|---|---|---|
| Trav19 | 0 | NA | NA | NA |
| Trav20 | 0 | NA | NA | NA |
| Trav21-dv12 | 0.038981395 | 0.274393828 | 3.724945049 | 0.073663859 |
| Trdv1 | 0.294928389 | 0.755274557 | 3.724945049 | 0.202761262 |
| RP23-339J4.3 | 0.166286699 | 0.274407613 | 2.769850422 | 0.09906947 |
| Trav22 | 0 | NA | NA | NA |
| Trdv2-1 | 0 | NA | NA | NA |
| Trdv2-2 | 0.066050875 | 0.274391604 | 3.724945049 | 0.073663262 |
| Trav23 | 0.0878635 | -0.687376175 | 3.724945049 | -0.184533239 |
| Trdv3 | 0 | NA | NA | NA |
| RP23-6A14.5 | 0.391491935 | 0.251113572 | 1.377258784 | 0.182328532 |
| Trdv4 | 0.800063227 | 1.648211483 | 1.186001252 | 1.389721536 |
| Trdd1 | 0 | NA | NA | NA |
| Gm23704 | 0 | NA | NA | NA |
| Trdd2 | 0.215047464 | 1.193240132 | 2.545125707 | 0.468833476 |
| Trdj1 | 0.138110619 | 1.201070057 | 3.622723897 | 0.33153784 |
| RP23-6A14.10 | 1.151756246 | -0.244557232 | 1.012980703 | -0.241423386 |
| Trdj2 | 0.506381433 | 1.24242931 | 1.780128007 | 0.697943802 |
| Trdc | 6.909450248 | 4.405172938 | 0.919908767 | 4.788706335 |
| Trdv5 | 0.261680116 | -0.693751059 | 2.10587865 | -0.32943544 |
| Traj61 | 0.355005915 | 0.770696139 | 1.565405626 | 0.492329992 |
| Traj60 | 0 | NA | NA | NA |
| Traj59 | 0.384053089 | -0.585316234 | 1.984477947 | -0.29494721 |
| Traj58 | 0.060174537 | 0.755286614 | 3.724945049 | 0.202764498 |
| Traj57 | 0.066050875 | 0.274391604 | 3.724945049 | 0.073663262 |
| Traj56 | 0.049553047 | 0.274391604 | 3.724945049 | 0.073663262 |
| Traj55 | 0.038981395 | 0.274393828 | 3.724945049 | 0.073663859 |
| Traj54 | 0 | NA | NA | NA |
| Traj53 | 0.235191775 | 1.194607255 | 2.115589888 | 0.564668635 |
| Traj52 | 0.049553047 | 0.274391604 | 3.724945049 | 0.073663262 |
| Traj51 | 0.143140564 | 0.274404261 | 3.546633223 | 0.077370352 |
| Traj50 | 0.066050875 | 0.274391604 | 3.724945049 | 0.073663262 |
| Traj49 | 0.094097695 | 0.755282166 | 3.724945049 | 0.202763304 |
| Traj48 | 0.268211629 | 0.262177368 | 2.090580652 | 0.125408875 |
| Traj47 | 0 | NA | NA | NA |
| Traj46 | 0.173853943 | 0.755298194 | 2.74376612 | 0.275277907 |
| Traj45 | 0.089089296 | 1.194485492 | 3.723939576 | 0.320758559 |
| Traj44 | 0.378197461 | 1.229767908 | 1.844010208 | 0.666898644 |
| Traj43 | 0.372057435 | 1.238040298 | 1.559422649 | 0.793909399 |
| Traj42 | 0.270744093 | 1.093805541 | 2.434245076 | 0.449340763 |
| Traj41 | 0.062426873 | 0.755288839 | 3.724945049 | 0.202765096 |
| Traj40 | 0.373373093 | 1.639822933 | 1.5609011 | 1.050561713 |
| Traj39 | 0.520484338 | 0.755307927 | 1.67274114 | 0.451539039 |
| Traj38 | 0.304511964 | 0.27440773 | 2.020152696 | 0.135835143 |
| Traj37 | 0.321009791 | 0.274410415 | 1.755991433 | 0.15627093 |
| Traj36 | 1.007510561 | 0.249033028 | 1.43035099 | 0.174106237 |
| Traj35 | 0.11147366 | 0.274389235 | 3.724945049 | 0.073662626 |

| | | | | |
|---|---|---|---|---|
| Traj34 | 0.174360202 | 1.222187254 | 2.724220671 | 0.448637391 |
| Traj33 | 0.478372902 | 1.773386211 | 3.500355772 | 0.506630276 |
| Traj32 | 0.292144397 | 0.274379759 | 3.724945049 | 0.073660082 |
| Traj31 | 0.534577455 | 1.041889152 | 1.360510826 | 0.765807322 |
| Traj30 | 0.493327417 | 1.94679307 | 1.369215899 | 1.421830605 |
| Traj29 | 0.56458922 | 0.755309493 | 1.469569008 | 0.513966672 |
| Traj28 | 0.229902237 | 1.536497829 | 2.516990621 | 0.610450359 |
| Traj27 | 0.668611641 | 1.739008611 | 1.272910557 | 1.366167168 |
| Traj26 | 0.066050875 | 0.274391604 | 3.724945049 | 0.073663262 |
| Traj25 | 0.632730142 | 1.035325957 | 1.304188962 | 0.793846587 |
| Traj24 | 0.739637387 | 2.291559167 | 1.350945285 | 1.696263492 |
| Traj23 | 0.722755445 | 1.750480867 | 1.355040021 | 1.291829644 |
| Traj22 | 0.569236379 | 1.61222838 | 1.299639771 | 1.240519424 |
| Traj21 | 0.962101734 | 2.303770357 | 1.09854615 | 2.097108399 |
| Traj20 | 0.661355215 | 2.681370965 | 1.221222791 | 2.195644387 |
| Traj19 | 0.223817141 | 1.277658211 | 2.152523403 | 0.593562983 |
| Traj18 | 0.206905613 | 0.755303967 | 2.185615118 | 0.345579586 |
| Traj17 | 0.414345469 | 1.197877739 | 1.728064805 | 0.693190288 |
| Traj16 | 0.11193333 | 0.75528439 | 3.724945049 | 0.202763901 |
| Traj15 | 0.43445422 | 1.719950146 | 1.735582558 | 0.990992988 |
| Traj14 | 0.377056126 | 1.043216355 | 3.303679098 | 0.315774118 |
| Traj13 | 0.208613207 | 0.274403186 | 2.608118164 | 0.105211179 |
| Traj12 | 0.486308198 | 1.961076459 | 1.527382534 | 1.283945845 |
| Traj11 | 0.728352697 | 2.666182382 | 1.221503139 | 2.182706124 |
| Traj9 | 0.154585317 | 0.274399357 | 2.835007735 | 0.096789632 |
| Traj8 | 0.203604809 | 0.755300031 | 2.610247297 | 0.289359568 |
| Traj7 | 0.161486377 | 0.755296344 | 2.795731986 | 0.270160498 |
| Traj6 | 0.490533913 | 1.577015846 | 1.524654451 | 1.034343123 |
| Traj5 | 0.517714159 | 1.204256231 | 1.489251576 | 0.808631833 |
| Traj4 | 0.260182639 | 0.755303864 | 2.094294743 | 0.360648312 |
| Traj3 | 0 | NA | NA | NA |
| Traj2 | 0.364308502 | 1.225858047 | 1.762806502 | 0.695401365 |
| Traj1 | 0.178447459 | 0.27439001 | 3.343252531 | 0.082072774 |
| Trac | 42.85427748 | 3.405089494 | 0.36342624 | 9.369410117 |
| Dad1 | 331.339338 | -0.090396226 | 0.096221608 | -0.939458701 |
| Abhd4 | 923.6765976 | 0.3542819 | 0.137052319 | 2.585012078 |
| Olfr49 | 0.04393175 | -0.206490061 | 3.724945049 | -0.055434391 |
| RP24-378G4.3 | 0.161488833 | 0.75528682 | 3.43662875 | 0.219775505 |
| RP24-378G4.4 | 0 | NA | NA | NA |
| RP24-378G4.5 | 0 | NA | NA | NA |
| RP24-378G4.6 | 0.093484797 | -0.20649451 | 3.724945049 | -0.055435586 |
| Oxa1l | 446.5440451 | 0.339169818 | 0.062442299 | 5.43173179 |
| Slc7a7 | 63.95572232 | 1.499354133 | 0.199138276 | 7.529211185 |
| Mrpl52 | 179.7423914 | 0.340632239 | 0.118832047 | 2.866501494 |
| Mmp14 | 516.3861542 | 1.612251276 | 0.191506914 | 8.418762742 |
| Gm24424 | 0 | NA | NA | NA |
| Lrp10 | 683.6561126 | 0.776302005 | 0.138676434 | 5.597937456 |

| | | | | |
|---|---|---|---|---|
| Rem2 | 153.9580932 | 0.076482733 | 0.337828291 | 0.226395286 |
| Prmt5 | 476.4345395 | 0.039343211 | 0.062008706 | 0.634478824 |
| Haus4 | 37.03329486 | 0.552583579 | 0.201453064 | 2.742989195 |
| Ajuba | 48.90169317 | 0.920072347 | 0.278344596 | 3.30551539 |
| Gm24705 | 0 | NA | NA | NA |
| 4931414P19Rik | 61.32594599 | 0.292166886 | 0.142150754 | 2.055331248 |
| Psmb5 | 352.4206726 | 0.333650116 | 0.100897703 | 3.306815761 |
| Mir686 | 0 | NA | NA | NA |
| Psmb11 | 6.261784242 | 1.115526883 | 0.451985403 | 2.468059534 |
| Cdh24 | 130.6564516 | 0.571311861 | 0.128198714 | 4.456455474 |
| Gm20726 | 0.104106287 | 0.274400501 | 3.724945049 | 0.073665651 |
| Gm17606 | 1.502658278 | 0.385964435 | 1.253522724 | 0.30790382 |
| Acin1 | 1506.43145 | -0.171055914 | 0.097972892 | -1.745951457 |
| mmu-mir-6948 | 0.536873645 | -1.271502089 | 1.248451656 | -1.018463216 |
| 4930579G18Rik | 0.898085247 | -0.856065483 | 1.231487246 | -0.695147665 |
| 1700123O20Rik | 137.487668 | 0.249994339 | 0.136211694 | 1.835336827 |
| Cebpe | 1.473260831 | 2.223923951 | 1.213264125 | 1.833008909 |
| Slc7a8 | 825.5824292 | 0.144173451 | 0.084880656 | 1.698543087 |
| Gm8865 | 0 | NA | NA | NA |
| Homez | 148.0548322 | -0.399848126 | 0.153763618 | -2.600407891 |
| Ppp1r3e | 55.21987148 | -0.199645135 | 0.176469848 | -1.131327177 |
| Gm37271 | 5.557962796 | -0.930502423 | 0.455898696 | -2.041028918 |
| Bcl2l2 | 1139.670049 | -0.081192972 | 0.064513744 | -1.258537577 |
| Gm20521 | 0.375216023 | 2.316621342 | 3.695373188 | 0.626897805 |
| Pabpn1 | 264.758843 | -0.442292986 | 0.215090336 | -2.056312681 |
| Slc22a17 | 4291.308412 | -0.071549912 | 0.054754828 | -1.306732489 |
| Efs | 132.9613014 | -0.194284376 | 0.181750419 | -1.068962463 |
| Il25 | 0.855745269 | -0.465037893 | 1.149805881 | -0.404449047 |
| Cmtm5 | 177.6437715 | -0.78235478 | 0.189001007 | -4.139421236 |
| Myh6 | 15.5941189 | 1.279983044 | 0.370416897 | 3.455520129 |
| Mir208a | 0 | NA | NA | NA |
| Mhrt | 3.992004508 | 0.678205251 | 0.581981045 | 1.165339072 |
| Myh7 | 58.29767794 | 1.46182802 | 0.28296595 | 5.166091619 |
| Gm29015 | 0.163519603 | 0.723301248 | 2.774973957 | 0.260651545 |
| Mir208b | 0 | NA | NA | NA |
| Gm17428 | 0.038981395 | 0.274393828 | 3.724945049 | 0.073663859 |
| Ngdn | 176.4773903 | 0.345177714 | 0.136790313 | 2.523407586 |
| Gm10364 | 2.051660529 | -0.429958475 | 0.720753953 | -0.596539878 |
| Gm20687 | 0.094831987 | 0.274404949 | 3.724945049 | 0.073666845 |
| Thtpa | 328.5777116 | 0.045442282 | 0.082812553 | 0.548736638 |
| Zfhx2 | 1084.53136 | -0.732685607 | 0.172690576 | -4.242765435 |
| Zfhx2os | 47.4886808 | -0.78872548 | 0.380272879 | -2.074103947 |
| Gm37034 | 35.49877453 | -1.100190758 | 0.346249205 | -3.177453525 |
| Gm38129 | 19.96999586 | -1.40180181 | 0.361138997 | -3.88161296 |
| Ap1g2 | 74.95724327 | 0.684108134 | 0.211645551 | 3.232329387 |
| Jph4 | 3324.850563 | -0.193054978 | 0.12864326 | -1.500700295 |
| Gm23596 | 0 | NA | NA | NA |

| | | | | |
|---|---|---|---|---|
| Dhrs2 | 1.791652276 | -2.695960923 | 0.946877961 | -2.847210554 |
| Gm8894 | 12.48756017 | 0.113510232 | 0.333825207 | 0.340028941 |
| Dhrs4 | 75.51960051 | -0.060040154 | 0.158065957 | -0.379842412 |
| Lrrc16b | 392.3729988 | -0.5597737 | 0.136054666 | -4.114329306 |
| Cpne6 | 868.2829338 | -0.256848606 | 0.126509879 | -2.030265213 |
| Nrl | 2.663415179 | -1.112724079 | 0.632624516 | -1.758901292 |
| Pck2 | 235.7861946 | 0.168872074 | 0.102259845 | 1.65140163 |
| A730061H03Rik | 1.918506749 | 0.895116462 | 0.853135307 | 1.049208086 |
| Dcaf11 | 703.1883374 | 0.119044384 | 0.052519701 | 2.26666151 |
| Gm15932 | 0.28189265 | 1.358627762 | 3.706919369 | 0.36651128 |
| Fitm1 | 0.170186936 | 0.27439223 | 3.410018679 | 0.080466489 |
| Psme1 | 1237.910375 | 2.95960787 | 0.157116964 | 18.8369721 |
| Emc9 | 130.4834388 | -0.592178886 | 0.137821492 | -4.296709309 |
| Psme2 | 385.2943989 | 2.466127229 | 0.165145854 | 14.93302534 |
| Rnf31 | 653.5334504 | 1.324945906 | 0.11045465 | 11.99538372 |
| Irf9 | 958.7618358 | 3.574720616 | 0.141901295 | 25.19159967 |
| Rec8 | 6.735313399 | 0.780316214 | 0.420949606 | 1.853704582 |
| Ipo4 | 859.5313815 | 0.10056806 | 0.047429761 | 2.120357705 |
| Tm9sf1 | 264.973998 | 0.048646576 | 0.087000273 | 0.559154293 |
| Tssk4 | 3.653146442 | -0.305233944 | 0.523879707 | -0.582641281 |
| Mdp1 | 234.6429327 | 0.078910052 | 0.095994587 | 0.822026062 |
| Nedd8 | 570.9432889 | 0.037913487 | 0.11023382 | 0.343936979 |
| Gmpr2 | 130.1104825 | -0.192617091 | 0.133271972 | -1.445293318 |
| Tinf2 | 129.962745 | 0.542095674 | 0.184360276 | 2.940414741 |
| Gm25887 | 0.119587853 | 1.218392466 | 3.723618458 | 0.327206581 |
| Tgm1 | 450.7877214 | 6.335004441 | 0.49991735 | 12.67210358 |
| Rabggta | 331.6413799 | -0.004861845 | 0.066154037 | -0.07349279 |
| Dhrs1 | 1107.416416 | 0.805008678 | 0.131954064 | 6.100673614 |
| Nop9 | 182.2591248 | 0.366943694 | 0.1463425 | 2.50743081 |
| Cideb | 12.60069031 | -1.078742903 | 0.33218704 | -3.247396115 |
| Ltb4r2 | 1.834185785 | 0.361692797 | 0.919028256 | 0.39356004 |
| Ltb4r1 | 13.33054419 | 3.797458527 | 0.871124025 | 4.359262764 |
| Adcy4 | 30.55724621 | 1.781977908 | 0.343416233 | 5.188974014 |
| Ripk3 | 29.79384862 | 4.033220762 | 0.520019338 | 7.755905347 |
| Nfatc4 | 25.04774085 | 0.09746772 | 0.242031779 | 0.40270629 |
| Nynrin | 300.1840239 | -0.146836847 | 0.106096171 | -1.383997614 |
| Gm26751 | 4.016269843 | 0.937102635 | 0.582926784 | 1.60758205 |
| Cbln3 | 2048.884425 | 0.011250013 | 0.317355283 | 0.035449271 |
| Khnyn | 336.7112361 | 0.300435008 | 0.091966201 | 3.266798059 |
| Sdr39u1 | 401.4242335 | -0.196148072 | 0.093922703 | -2.088398928 |
| Cma1 | 1.421829526 | -0.778269081 | 1.239580895 | -0.627848561 |
| Cma2 | 0.212113984 | 2.012468375 | 3.153803206 | 0.638108418 |
| Gm5801 | 19.00002645 | 0.136648271 | 0.255972444 | 0.533839769 |
| Mcpt1 | 0.342615221 | 0.75530919 | 1.575882798 | 0.479292744 |
| Mcpt9 | 0 | NA | NA | NA |
| Mcpt2 | 0.74687904 | 1.697380386 | 3.610585167 | 0.470112269 |
| Mcpt4 | 0.384964691 | -0.669426541 | 1.465961003 | -0.456646895 |

| | | | | |
|---|---|---|---|---|
| Mcpt8 | 0 | NA | NA | NA |
| Mir6949 | 0 | NA | NA | NA |
| Ctsg | 0.191334258 | 0.755296462 | 2.673199576 | 0.282543985 |
| Gzme | 0.045882455 | 0.755288839 | 3.724945049 | 0.202765096 |
| Gzmd | 0.066050875 | 0.274391604 | 3.724945049 | 0.073663262 |
| Gzmg | 0 | NA | NA | NA |
| Gzmn | 0 | NA | NA | NA |
| Gzmf | 1.629358998 | 2.680047186 | 1.33587521 | 2.006210735 |
| Gzmc | 1.857191415 | 3.02192708 | 1.140852956 | 2.648831353 |
| Gm24118 | 0 | NA | NA | NA |
| Gm4819 | 0 | NA | NA | NA |
| Gzmb | 1148.613576 | 12.34853763 | 1.011123466 | 12.21269018 |
| Atp12a | 1.447939701 | -0.34353154 | 0.891707621 | -0.385251322 |
| Rnf17 | 3.730754265 | 0.081157778 | 0.646687041 | 0.125497765 |
| Gm24392 | 0.066050875 | 0.274391604 | 3.724945049 | 0.073663262 |
| Cenpj | 130.4607887 | 0.933603723 | 0.208453532 | 4.47871387 |
| Gm22431 | 0 | NA | NA | NA |
| Parp4 | 406.4978698 | 1.412949429 | 0.111414301 | 12.68193957 |
| Gm16573 | 0 | NA | NA | NA |
| Mphosph8 | 456.601835 | -0.180379746 | 0.122555243 | -1.471823982 |
| Gm16973 | 101.7464204 | -0.019367821 | 0.148295686 | -0.130602728 |
| Pspc1 | 679.2798492 | -0.186635803 | 0.073783496 | -2.529506095 |
| C030013D06Rik | 71.86212202 | -0.418717657 | 0.296081565 | -1.414196988 |
| Gm4479 | 0.082913145 | -0.206492286 | 3.724945049 | -0.055434988 |
| Zmym5 | 534.3723691 | -0.042477667 | 0.079783372 | -0.532412532 |
| 2410022M11Rik | 46.59279127 | 0.350236058 | 0.164203314 | 2.13294147 |
| Gm22498 | 0 | NA | NA | NA |
| Gm27635 | 0 | NA | NA | NA |
| Gm22218 | 0 | NA | NA | NA |
| Zmym2 | 1925.381415 | -0.036258464 | 0.072765439 | -0.498292388 |
| B020004C17Rik | 0.385592413 | 0.775059807 | 3.723584315 | 0.208148854 |
| Gja3 | 0.839270164 | -1.874660644 | 1.446975307 | -1.295571967 |
| Gjb2 | 207.1465136 | 0.817826817 | 0.191955645 | 4.260498918 |
| Gm4491 | 3.32060927 | -1.121139342 | 0.541264867 | -2.071332189 |
| Gjb6 | 476.1042231 | 0.015675384 | 0.16429569 | 0.095409588 |
| Cryl1 | 92.72897768 | -0.172199659 | 0.148369053 | -1.160617089 |
| Gm23593 | 0.852011583 | -1.428098143 | 1.218711484 | -1.171809868 |
| Ift88 | 288.14491 | 0.009656175 | 0.09964555 | 0.096905234 |
| Gm15836 | 0.068097359 | -0.206487837 | 3.724945049 | -0.055433794 |
| Ift88os | 3.791653849 | 0.613159548 | 0.505055531 | 1.214043824 |
| Il17d | 56.83289221 | -0.471903659 | 0.16841752 | -2.801986753 |
| N6amt2 | 87.43391111 | -0.452327714 | 0.152758614 | -2.961061922 |
| 1700039M10Rik | 1.307453931 | 1.240439622 | 0.805050314 | 1.540822481 |
| Xpo4 | 372.4125732 | 0.225471208 | 0.109507479 | 2.05895716 |
| Lats2 | 654.4618862 | 0.291832188 | 0.114106123 | 2.557550643 |
| Gm26440 | 0.104366086 | 0.274400501 | 3.724945049 | 0.073665651 |
| Gm20430 | 0.779952531 | -0.780234292 | 1.030983874 | -0.756786125 |

| | | | | |
|---|---|---|---|---|
| Sap18 | 492.5086436 | 0.165319747 | 0.094121069 | 1.756458464 |
| Mir3077 | 0 | NA | NA | NA |
| Ska3 | 10.9912364 | 1.010338278 | 0.497247679 | 2.031861223 |
| Mrpl57 | 188.5806294 | -0.093960367 | 0.121564081 | -0.772928699 |
| Gm22738 | 2.721618137 | -0.302937009 | 0.595534897 | -0.508680533 |
| Zdhhc20 | 1037.770143 | 0.172596284 | 0.098952766 | 1.744229006 |
| Micu2 | 298.7097166 | 0.494498175 | 0.114545713 | 4.317037825 |
| 3110083C13Rik | 6.409078304 | -1.240237089 | 0.525804686 | -2.358741034 |
| Fgf9 | 261.7121968 | -0.482490705 | 0.082396784 | -5.855698289 |
| Gm17109 | 0 | NA | NA | NA |
| Gm25614 | 0 | NA | NA | NA |
| Gm9022 | 0.171181394 | 1.257632701 | 2.747344421 | 0.457763028 |
| Rpl13-ps3 | 2.686924039 | 1.256644526 | 0.701952132 | 1.790213989 |
| 1700129C05Rik | 0.097139547 | 0.274393828 | 3.724945049 | 0.073663859 |
| Gm5142 | 0 | NA | NA | NA |
| Rcbtb1 | 723.0799953 | -0.069340755 | 0.083226917 | -0.833152995 |
| Gm6904 | 9.706043522 | 5.293616557 | 0.933668565 | 5.669695602 |
| Phf11a | 116.1058508 | 8.822916373 | 0.914327048 | 9.649628536 |
| Phf11b | 380.8323055 | 7.643359374 | 0.524759397 | 14.56545498 |
| Phf11d | 768.3922217 | 8.648674504 | 0.54710659 | 15.80802472 |
| Phf11c | 63.97794148 | 2.386578875 | 0.259522394 | 9.196042159 |
| Setdb2 | 303.1628553 | 2.149253632 | 0.205666821 | 10.45017188 |
| Gm16301 | 4.198859835 | 2.434978173 | 0.793033543 | 3.070460507 |
| Cab39l | 377.7726523 | -0.045855834 | 0.094781581 | -0.483805333 |
| Cdadc1 | 398.4046235 | 0.006160695 | 0.075367089 | 0.081742513 |
| Shisa2 | 57.22810011 | -0.250405996 | 0.270832571 | -0.924578587 |
| Atp8a2 | 620.7074487 | -0.420012905 | 0.093988764 | -4.468756554 |
| Gm9013 | 6.16443421 | -0.683453983 | 0.464464792 | -1.47148717 |
| 4930563I02Rik | 0.208097268 | 0.274410299 | 2.199917172 | 0.12473665 |
| Gm6913 | 0 | NA | NA | NA |
| Gm16261 | 0 | NA | NA | NA |
| Nupl1 | 647.3223968 | -0.030374144 | 0.070649264 | -0.42992867 |
| Mir719 | 0.1633926 | -0.687376774 | 2.785222606 | -0.246794196 |
| Mtmr6 | 1302.815769 | -0.150639913 | 0.052242974 | -2.883448287 |
| Amer2 | 525.7164864 | -0.406737984 | 0.106253707 | -3.827988655 |
| Gm29266 | 1.126110934 | -0.271118564 | 0.910682963 | -0.297709055 |
| Spata13 | 889.6645646 | 1.3646172 | 0.131270442 | 10.39546437 |
| C1qtnf9 | 1.516716675 | 1.34452687 | 0.960815398 | 1.399360245 |
| Mipepos | 0.63216027 | 0.140146963 | 1.296421591 | 0.108102923 |
| Mipep | 261.2781114 | -0.030268405 | 0.083990334 | -0.360379622 |
| Tnfrsf19 | 737.4688333 | 0.293447318 | 0.127483184 | 2.301851189 |
| Sacs | 3375.54935 | -0.249630882 | 0.069503582 | -3.591626148 |
| Sgcg | 2.524458954 | -0.413197872 | 0.717595943 | -0.575808539 |
| 1700109G14Rik | 1.219573449 | 1.093056365 | 0.950414961 | 1.150083289 |
| Arl11 | 45.06214385 | 2.783896682 | 0.333629084 | 8.344286566 |
| Ebpl | 74.2502746 | -0.224055443 | 0.153924123 | -1.455622671 |
| Gm25716 | 0.646183563 | -0.112644551 | 1.301097706 | -0.08657655 |

| | | | | |
|---|---|---|---|---|
| Kpna3 | 1362.917462 | 0.338328896 | 0.071941768 | 4.702815955 |
| Spryd7 | 646.9547196 | 0.339677525 | 0.10972739 | 3.095649374 |
| Trim13 | 93.97118903 | 0.082140267 | 0.14427661 | 0.569324904 |
| Dleu2 | 509.193176 | -0.214649895 | 0.13828991 | -1.552173217 |
| Kcnrg | 4.493413694 | -0.124995226 | 0.442153784 | -0.282696272 |
| Gm27010 | 39.15760298 | -0.433525585 | 0.262498508 | -1.651535425 |
| Gm27845 | 0.122991902 | 0.274402725 | 3.724945049 | 0.073666248 |
| Gm26916 | 0.226559374 | 2.043498472 | 2.51171147 | 0.813588064 |
| Mir16-1 | 0.351804329 | 0.267863388 | 1.493499804 | 0.179352811 |
| Mir15a | 2.378988063 | -1.229202868 | 0.775792791 | -1.584447397 |
| Gm27747 | 0.12092566 | 1.236179401 | 3.724945049 | 0.331865138 |
| Gm27972 | 0 | NA | NA | NA |
| Gm27017 | 1.269792884 | -0.105654375 | 0.94825392 | -0.111419919 |
| Gm27928 | 0.047948698 | -0.206487837 | 3.724945049 | -0.055433794 |
| Gm37420 | 92.02312766 | -0.504643163 | 0.227353934 | -2.219636816 |
| Gm37472 | 129.5225664 | -0.341161391 | 0.238857775 | -1.428303478 |
| Gm37820 | 1.452044762 | 1.371018366 | 0.969234651 | 1.414537093 |
| Gm26969 | 0 | NA | NA | NA |
| Dleu7 | 46.2767513 | 0.004785375 | 0.178090782 | 0.026870428 |
| Rnaseh2b | 84.37813906 | 0.292782409 | 0.1308291 | 2.237899741 |
| Gucy1b2 | 4.464211597 | 1.55622104 | 0.559764625 | 2.780134667 |
| Gm4131 | 0.940148893 | 0.274414895 | 1.294108721 | 0.212049336 |
| Gm24625 | 0 | NA | NA | NA |
| Fam124a | 214.0573691 | -0.341285744 | 0.096252409 | -3.545737183 |
| Gm23816 | 0 | NA | NA | NA |
| Serpine3 | 0.736171263 | 0.072543957 | 1.145900042 | 0.063307404 |
| Gm25162 | 0 | NA | NA | NA |
| Ints6 | 590.4517194 | -0.362022894 | 0.119335092 | -3.033666681 |
| 4931440J10Rik | 4.963742587 | -0.748568612 | 0.488525591 | -1.532301737 |
| Gm26536 | 1.065495214 | 0.275316651 | 0.924989097 | 0.297643131 |
| Wdfy2 | 169.1506073 | -0.020679038 | 0.101459835 | -0.203815018 |
| Mir8098 | 0 | NA | NA | NA |
| Mir6541 | 1.551969969 | 0.04589019 | 0.779936856 | 0.05883834 |
| n-R5s47 | 0.9747039 | 0.032328522 | 1.070667161 | 0.030194745 |
| Defb48 | 0 | NA | NA | NA |
| Defb47 | 0.450831908 | -0.031730801 | 1.35691187 | -0.02338457 |
| Defb43 | 0 | NA | NA | NA |
| Defb30 | 0.101408268 | 0.755286617 | 3.724945049 | 0.202764499 |
| Defb42 | 4.17522043 | -0.285312309 | 0.672823342 | -0.424052335 |
| Gm23629 | 0 | NA | NA | NA |
| Ctsb | 9370.229512 | 0.976249278 | 0.145596102 | 6.705188294 |
| Fdft1 | 819.3075922 | 0.204068449 | 0.117445704 | 1.737555667 |
| Neil2 | 16.96188054 | -0.112434684 | 0.310426714 | -0.362193968 |
| Gata4 | 0.136623169 | -0.206497534 | 3.634681883 | -0.056813097 |
| Blk | 0.439667587 | 2.102396686 | 1.39350534 | 1.508710894 |
| Gm21430 | 0.110595523 | 0.755282166 | 3.724945049 | 0.202763304 |
| Fam167a | 87.58736384 | 1.040537545 | 0.208443674 | 4.991936308 |

| | | | | |
|---|---|---|---|---|
| Tdh | 0.432707378 | -1.351598969 | 1.490785736 | -0.906635297 |
| Gm17232 | 0.288435958 | 1.155695392 | 2.026353017 | 0.570332702 |
| Mtmr9 | 857.9831177 | -0.006847287 | 0.052391087 | -0.130695653 |
| Gm15918 | 7.496844332 | -0.740823151 | 0.380246831 | -1.9482691 |
| Xkr6 | 411.7970283 | -0.700574042 | 0.158620361 | -4.416671588 |
| Mir598 | 1.860426698 | -1.068331348 | 0.715032529 | -1.494101743 |
| Pinx1 | 73.25514319 | 0.092126356 | 0.139351202 | 0.661109161 |
| Sox7 | 14.05727425 | 0.982011309 | 0.334745363 | 2.933606909 |
| 4930578I06Rik | 0.15100991 | 1.236179172 | 3.527597384 | 0.350430913 |
| Rp1l1 | 0.46004984 | 1.691256468 | 1.496368187 | 1.130240861 |
| Prss55 | 1.808854001 | 2.5660714 | 0.990116341 | 2.591686748 |
| Prss51 | 0 | NA | NA | NA |
| 4930471C04Rik | 6.910637616 | -0.028633482 | 0.534719873 | -0.053548566 |
| Prss52 | 0.240888117 | -0.696105019 | 2.53545742 | -0.274548101 |
| Gm17116 | 16.40399705 | -0.594741078 | 0.383444606 | -1.551048233 |
| Msra | 295.1015591 | 0.239045664 | 0.089575138 | 2.668660847 |
| Mir124a-1hg | 1670.155338 | -0.814445088 | 0.360108857 | -2.261663582 |
| Mir124a-1 | 0 | NA | NA | NA |
| Mir3078 | 0 | NA | NA | NA |
| Kif13b | 733.5243985 | -0.306925639 | 0.106798894 | -2.873865338 |
| Gm26225 | 1.137833813 | -0.3353699 | 0.913289507 | -0.367210942 |
| Hmbox1 | 1181.12912 | -0.172527863 | 0.05365737 | -3.215361928 |
| Gm37183 | 36.08914756 | -0.378432402 | 0.322600933 | -1.17306667 |
| Gm20111 | 0 | NA | NA | NA |
| Ints9 | 171.8987803 | 0.11316166 | 0.106682287 | 1.060735235 |
| Extl3 | 2015.132037 | -0.033142097 | 0.063008526 | -0.525993855 |
| Fzd3 | 1027.303391 | -0.116154229 | 0.075538424 | -1.537684045 |
| Gm27263 | 1.700332359 | 0.123623137 | 0.827035235 | 0.149477473 |
| Fbxo16 | 53.77017717 | -0.366103724 | 0.14397942 | -2.542750374 |
| Zfp395 | 203.6050118 | -0.027613432 | 0.122842539 | -0.224787216 |
| Pnoc | 87.06028916 | 0.069443537 | 0.14840347 | 0.467937419 |
| Elp3 | 772.4740043 | 0.029938259 | 0.061393652 | 0.487644209 |
| Nuggc | 2.07464996 | 0.737532349 | 1.287018904 | 0.573054791 |
| Scara5 | 27.21228639 | 1.729858401 | 0.378543666 | 4.56977241 |
| Pbk | 12.44226385 | 0.473958543 | 0.609028272 | 0.77822092 |
| Esco2 | 8.165872688 | 1.235383576 | 0.550811086 | 2.242844431 |
| Ccdc25 | 465.8537132 | 0.769228274 | 0.128477543 | 5.987258611 |
| Scara3 | 275.1443948 | 0.586715416 | 0.099549955 | 5.893678365 |
| Clu | 7658.013756 | 0.538989207 | 0.108757833 | 4.955865624 |
| Gulo | 1.585533927 | -1.194448392 | 1.056917555 | -1.130124471 |
| Adam2 | 0.443182709 | 1.276307657 | 1.764187312 | 0.723453597 |
| Gm10233 | 0.231561134 | 0.274410832 | 2.132800968 | 0.128662185 |
| Ephx2 | 103.3385784 | -0.205719359 | 0.136949725 | -1.502152412 |
| Chrna2 | 9.44470098 | -0.236346065 | 0.343919416 | -0.687213499 |
| Ptk2b | 3860.082922 | 0.389254992 | 0.150898328 | 2.579584527 |
| 1700001G11Rik | 4.137181521 | -0.88578653 | 0.590257335 | -1.500678563 |
| Trim35 | 1878.620081 | 0.283028175 | 0.069217582 | 4.088963638 |

| | | | | |
|---|---|---|---|---|
| Stmn4 | 1213.698151 | -0.023450506 | 0.098271482 | -0.238629815 |
| Gm23899 | 0 | NA | NA | NA |
| Adra1a | 340.4045469 | -0.512354016 | 0.11493078 | -4.45793562 |
| Gm10032 | 0.180493943 | -0.206488401 | 2.699100097 | -0.076502684 |
| Dpysl2 | 11326.80766 | -0.007358628 | 0.054431364 | -0.135190954 |
| Gm5464 | 7.318891932 | -0.355738412 | 0.405048184 | -0.87826196 |
| Pnma2 | 428.5622982 | -0.505603316 | 0.119683955 | -4.224487032 |
| 4930578I07Rik | 0 | NA | NA | NA |
| Gm24258 | 0 | NA | NA | NA |
| Bnip3l | 1147.218713 | -0.179789714 | 0.098044718 | -1.833752169 |
| Ppp2r2a | 1306.363493 | 0.239061625 | 0.072261256 | 3.308295992 |
| Gm27647 | 0 | NA | NA | NA |
| Gm20675 | 0.146692594 | 0.274392389 | 3.52332538 | 0.07787881 |
| Ebf2 | 68.02274029 | 0.040535758 | 0.178365821 | 0.227261913 |
| Gm23178 | 0 | NA | NA | NA |
| Gm6878 | 0.454723033 | -0.136256814 | 1.716718704 | -0.079370496 |
| 1700120O09Rik | 0 | NA | NA | NA |
| 4930438E09Rik | 0 | NA | NA | NA |
| Gm10860 | 0 | NA | NA | NA |
| Gm24981 | 0 | NA | NA | NA |
| Gm20667 | 1.359754244 | 1.035466245 | 1.052829685 | 0.983507836 |
| Cdca2 | 13.24071265 | 1.06985967 | 0.402081949 | 2.66080005 |
| Kctd9 | 293.1358824 | -0.226399108 | 0.138039592 | -1.6401027 |
| Gnrh1 | 6.140945712 | -0.59095585 | 0.462822473 | -1.276852109 |
| Dock5 | 320.7362463 | -0.282923047 | 0.119556321 | -2.366441566 |
| Mir6539 | 0.673941775 | -0.920308361 | 1.166086895 | -0.789227943 |
| Nefm | 2120.125399 | -0.147984942 | 0.220762134 | -0.670336615 |
| Nefl | 4244.358884 | -0.24283673 | 0.134213616 | -1.809330065 |
| Gm22319 | 0 | NA | NA | NA |
| Adam7 | 0.28902587 | 0.806467247 | 2.091842962 | 0.385529537 |
| Adamdec1 | 0.646331757 | -0.95752068 | 1.374904638 | -0.696426977 |
| Adam28 | 1.311804504 | -0.808599583 | 1.149532289 | -0.703416156 |
| Stc1 | 48.95481958 | 0.035190571 | 0.204663098 | 0.1719439 |
| Rps2-ps5 | 0.934890173 | -1.796709785 | 1.144634818 | -1.56967948 |
| 1700092C10Rik | 0 | NA | NA | NA |
| Nkx2-6 | 0 | NA | NA | NA |
| Nkx3-1 | 12.50312542 | 0.961603747 | 0.31672014 | 3.036130717 |
| Gm37762 | 0.256447508 | 1.236196794 | 2.47295682 | 0.499886122 |
| Slc25a37 | 364.4713757 | -0.300852999 | 0.11982707 | -2.51072649 |
| Gm27222 | 0 | NA | NA | NA |
| Gm20236 | 0.061920613 | 0.274393828 | 3.724945049 | 0.073663859 |
| Gm31748 | 0 | NA | NA | NA |
| Gm27223 | 0 | NA | NA | NA |
| Synb | 0 | NA | NA | NA |
| Gm16677 | 0 | NA | NA | NA |
| Entpd4 | 0.136880458 | -0.675115491 | 3.636764192 | -0.185636312 |
| Gm27221 | 0 | NA | NA | NA |

| | | | | |
|---|---|---|---|---|
| Gm28049 | 0.042716945 | -0.206490061 | 3.724945049 | -0.055434391 |
| Gm21451 | 0 | NA | NA | NA |
| Gm38378 | 0 | NA | NA | NA |
| Gm37094 | 0 | NA | NA | NA |
| Gm37847 | 0.111006857 | 0.755286617 | 3.724945049 | 0.202764499 |
| Gm27175 | 0.061920613 | 0.274393828 | 3.724945049 | 0.073663859 |
| Gm27179 | 0 | NA | NA | NA |
| Gm27174 | 0 | NA | NA | NA |
| Gm21685 | 17.74312811 | 0.455373735 | 0.335043607 | 1.35914766 |
| Gm27176 | 0 | NA | NA | NA |
| Gm27177 | 0.075043205 | 0.755286614 | 3.724945049 | 0.202764498 |
| Loxl2 | 21.89650722 | 1.315189078 | 0.356484294 | 3.689332464 |
| Gm37513 | 0 | NA | NA | NA |
| Mir6950 | 0 | NA | NA | NA |
| R3hcc1 | 176.6082625 | 0.072080999 | 0.09708871 | 0.742424104 |
| Chmp7 | 739.9914465 | -0.020353574 | 0.058812936 | -0.346073078 |
| 4930480K23Rik | 35.48654686 | -0.464742359 | 0.174961031 | -2.656262115 |
| Tnfrsf10b | 17.05474737 | 1.631740488 | 0.395982757 | 4.120736215 |
| Rhobtb2 | 1292.582811 | -0.288671599 | 0.099200754 | -2.909973842 |
| Pebp4 | 0.585213848 | 0.016127032 | 1.207694418 | 0.01335357 |
| Egr3 | 501.7132685 | -0.724990879 | 0.321150545 | -2.257479837 |
| Bin3 | 135.9347836 | 0.564636789 | 0.112638165 | 5.012837256 |
| Ccar2 | 987.7091471 | 0.048879108 | 0.051307974 | 0.952661051 |
| 9930012K11Rik | 11.13584856 | 0.17908866 | 0.335117319 | 0.534405862 |
| Pdlim2 | 142.1719157 | -0.263123835 | 0.136096074 | -1.933368294 |
| Sorbs3 | 390.0743824 | 0.315280554 | 0.117989279 | 2.672111892 |
| Gm26908 | 10.16694102 | 0.098674133 | 0.343463388 | 0.287291562 |
| Ppp3cc | 243.8924496 | -0.059209332 | 0.087817201 | -0.674233876 |
| AC139294.1 | 1.827339133 | -0.343247523 | 0.79798174 | -0.430144583 |
| Slc39a14 | 953.2277302 | 0.951610575 | 0.178527379 | 5.330334079 |
| Gm9174 | 1.024625068 | 0.220110979 | 1.154753292 | 0.190612991 |
| Piwil2 | 21.10693623 | 0.790209777 | 0.259588697 | 3.044083916 |
| Gm22725 | 0.106465261 | 0.75528439 | 3.724945049 | 0.202763901 |
| Gm24890 | 0.088476398 | 0.274400501 | 3.724945049 | 0.073665651 |
| Polr3d | 217.1158271 | 0.028999969 | 0.115687742 | 0.25067452 |
| Mir320 | 1.741823475 | -0.746101613 | 0.747031162 | -0.998755676 |
| Phyhip | 4018.25641 | 0.058583046 | 0.112872201 | 0.519021025 |
| Bmp1 | 483.1780716 | -0.289815746 | 0.099934943 | -2.900044124 |
| Sftpc | 3.468929289 | 0.342871125 | 0.501470691 | 0.683731136 |
| Lgi3 | 1208.980655 | 0.172603841 | 0.085233151 | 2.025078727 |
| Reep4 | 37.19800003 | 0.499655015 | 0.208514882 | 2.396255889 |
| Hr | 708.6183582 | -0.223799914 | 0.115627255 | -1.935529072 |
| Nudt18 | 347.76934 | 0.138860751 | 0.076851434 | 1.806872592 |
| Fam160b2 | 415.787422 | -0.218662407 | 0.093621552 | -2.335599049 |
| Dmtn | 2327.269897 | -0.239430886 | 0.084873682 | -2.821026271 |
| Fgf17 | 6.855534712 | -0.33327148 | 0.481452489 | -0.692220911 |
| Npm2 | 6.446150388 | -0.14013299 | 0.419420311 | -0.33411112 |

| | | | | |
|---|---|---|---|---|
| Xpo7 | 1222.808954 | -0.173416228 | 0.053997423 | -3.211564887 |
| Dok2 | 40.29440054 | 1.640511451 | 0.265630896 | 6.175906028 |
| 2410012E07Rik | 0.066050875 | 0.274391604 | 3.724945049 | 0.073663262 |
| Gfra2 | 372.4700337 | -0.510689687 | 0.097093289 | -5.259783568 |
| Gm37455 | 0 | NA | NA | NA |
| Gm9192 | 0 | NA | NA | NA |
| Gm23735 | 0 | NA | NA | NA |
| Fndc3a | 2663.159039 | 0.911545099 | 0.100381906 | 9.08077095 |
| Gm16549 | 0.514076732 | 1.278139183 | 1.592170981 | 0.802765028 |
| Cysltr2 | 20.03900042 | 2.694305194 | 0.444345654 | 6.063534484 |
| Rcbtb2 | 290.6187941 | -0.058345154 | 0.089275323 | -0.653541784 |
| Gm17233 | 4.303707252 | -0.65664919 | 0.54030832 | -1.215323113 |
| Rb1 | 791.2567668 | -0.145433183 | 0.070710804 | -2.056732149 |
| Mir687 | 2.668100091 | -1.044936618 | 0.695208621 | -1.503054747 |
| Lpar6 | 59.93468125 | 0.441293514 | 0.192736931 | 2.28961576 |
| Itm2b | 7614.81607 | 0.774068034 | 0.10271905 | 7.535778725 |
| Med4 | 170.1772693 | 0.085333717 | 0.132761316 | 0.642760405 |
| Nudt15 | 22.3670934 | 0.183939896 | 0.232143861 | 0.792353051 |
| Sucla2 | 1247.988688 | -0.015732267 | 0.086720008 | -0.181414497 |
| Gm21750 | 0.774729477 | 1.138723957 | 1.237635639 | 0.920080128 |
| Gm24540 | 2.261342538 | -0.385696892 | 0.72051413 | -0.535307881 |
| Gm16409 | 3.92403765 | 0.359264428 | 0.550834649 | 0.652218281 |
| Gm22164 | 0 | NA | NA | NA |
| Gm6984 | 34.159564 | 0.023468759 | 0.178700266 | 0.131330296 |
| Htr2a | 177.8320817 | -0.250797171 | 0.178056639 | -1.408524687 |
| Esd | 684.0533462 | 0.374665899 | 0.139559303 | 2.684635783 |
| Lrch1 | 437.4414308 | 0.224363119 | 0.078744724 | 2.849246353 |
| 5031414D18Rik | 62.98482591 | 2.06983978 | 0.232525575 | 8.901557529 |
| Lrrc63 | 0.699855268 | 0.329705786 | 1.395282958 | 0.236300303 |
| Lcp1 | 670.0106953 | 2.697650051 | 0.165332939 | 16.31647064 |
| Gm15628 | 0 | NA | NA | NA |
| Gm15629 | 0.174360202 | 1.222187254 | 2.724220671 | 0.448637391 |
| Cpb2 | 0 | NA | NA | NA |
| AC161877.1 | 0 | NA | NA | NA |
| Zc3h13 | 935.6372034 | -0.029549898 | 0.127617423 | -0.231550658 |
| Siah3 | 24.32705589 | -0.338546095 | 0.22424756 | -1.509698011 |
| Spert | 0.0885994 | 0.274402725 | 3.724945049 | 0.073666248 |
| 4930564B18Rik | 0.087261593 | 0.274400501 | 3.724945049 | 0.073665651 |
| Gm23622 | 0 | NA | NA | NA |
| Cog3 | 540.4936263 | -0.034454367 | 0.073217736 | -0.470574057 |
| Gm24876 | 5.352175477 | -0.269151809 | 0.493461128 | -0.5454367 |
| Slc25a30 | 95.05997151 | 0.259808504 | 0.151514928 | 1.714738649 |
| Gm4285 | 34.11371507 | 0.440049561 | 0.209688002 | 2.098591993 |
| Tpt1 | 4005.741273 | 0.445083359 | 0.098394427 | 4.523461052 |
| Snora31 | 5.414963088 | -0.623764137 | 0.460955205 | -1.353199032 |
| Gtf2f2 | 148.747947 | 0.459946063 | 0.110972147 | 4.144698266 |
| Kctd4 | 189.0479674 | -0.341816723 | 0.138936656 | -2.460234286 |

| | | | | |
|---|---|---|---|---|
| Gm25517 | 1.093745531 | -2.856537514 | 1.055848127 | -2.705443558 |
| Gpalpp1 | 612.613148 | 0.062665893 | 0.056820847 | 1.102867977 |
| Nufip1 | 200.4613363 | 0.28493787 | 0.10988725 | 2.593002106 |
| Rps2-ps6 | 4.539665628 | 0.712158265 | 0.453128386 | 1.571647875 |
| Tsc22d1 | 3622.389764 | -0.089738677 | 0.064891199 | -1.382909814 |
| 4930444M15Rik | 1.224655951 | 0.576094426 | 0.917378978 | 0.627978665 |
| Serp2 | 345.0104461 | -0.093585456 | 0.105437285 | -0.887593576 |
| 1700108F19Rik | 0.956737924 | -0.576855674 | 1.03498316 | -0.557357546 |
| Lacc1 | 58.05462338 | 1.72096311 | 0.196691702 | 8.749546091 |
| Ccdc122 | 14.09176844 | 1.089464104 | 0.312954517 | 3.481221856 |
| Enox1 | 345.9653688 | -0.331809926 | 0.124294328 | -2.669550032 |
| Gm10132 | 18.18438502 | 0.583651605 | 0.290623375 | 2.008274814 |
| Gm6994 | 0 | NA | NA | NA |
| n-R5s48 | 4.967990864 | -0.947874287 | 0.570395255 | -1.661785016 |
| Gm23823 | 0 | NA | NA | NA |
| Gm1587 | 0.104719185 | 1.218392466 | 3.723618458 | 0.327206581 |
| Dnajc15 | 183.7589994 | 0.154788177 | 0.098392934 | 1.573163548 |
| Epsti1 | 187.7081649 | 6.062328122 | 0.39746865 | 15.25234284 |
| Gm19301 | 0 | NA | NA | NA |
| Fam216b | 12.69086337 | -0.482522102 | 0.329091047 | -1.466226769 |
| Mir1971 | 0 | NA | NA | NA |
| Gm23652 | 0 | NA | NA | NA |
| Tnfsf11 | 1.022937004 | 0.755898007 | 1.058628066 | 0.714035487 |
| Gm26197 | 0 | NA | NA | NA |
| Akap11 | 5712.428926 | -0.001396201 | 0.059840883 | -0.023331898 |
| Dgkh | 929.9045385 | -0.044472419 | 0.117944446 | -0.37706243 |
| Gm26251 | 0.072025462 | 0.755288839 | 3.724945049 | 0.202765096 |
| Vwa8 | 1020.041174 | -0.314899536 | 0.064762104 | -4.862404355 |
| Zfp957 | 0 | NA | NA | NA |
| Rgcc | 194.1349241 | 0.012979189 | 0.146929059 | 0.088336436 |
| Naa16 | 214.9779865 | -0.499934434 | 0.133125668 | -3.755357176 |
| Mtrf1 | 71.56468441 | 0.479568143 | 0.133457797 | 3.593406712 |
| Kbtbd7 | 573.16381 | -0.28112937 | 0.077319369 | -3.635950151 |
| Kbtbd6 | 5.414111675 | 0.13038488 | 0.436498293 | 0.298706505 |
| AC124745.1 | 0 | NA | NA | NA |
| Wbp4 | 689.3854996 | 0.195887919 | 0.088699757 | 2.208438053 |
| Elf1 | 298.3952992 | 1.545908968 | 0.163059842 | 9.480623462 |
| Gm27857 | 0 | NA | NA | NA |
| Sugt1 | 598.6834534 | -0.127734618 | 0.083987603 | -1.520874683 |
| Lect1 | 10.35988659 | 0.350118453 | 0.354813751 | 0.986766864 |
| Mir759 | 0 | NA | NA | NA |
| Pcdh8 | 429.8832926 | -0.151219375 | 0.158593715 | -0.953501691 |
| Gm9748 | 3.167342963 | -0.083475845 | 0.528441247 | -0.15796618 |
| Gm6999 | 2.919730593 | 0.513496484 | 0.736871373 | 0.696860406 |
| Gm10845 | 0.072025462 | 0.755288839 | 3.724945049 | 0.202765096 |
| Olfm4 | 11.07339153 | -0.380000548 | 0.339456654 | -1.119437618 |
| Gm9578 | 0 | NA | NA | NA |

| | | | | |
|---|---|---|---|---|
| Gm22891 | 0 | NA | NA | NA |
| Gm24774 | 0 | NA | NA | NA |
| Pcdh17 | 1873.471252 | -0.032971482 | 0.064657722 | -0.509938814 |
| Gm23926 | 0 | NA | NA | NA |
| Gm25115 | 0.075043205 | 0.755286614 | 3.724945049 | 0.202764498 |
| Diap3 | 34.4707947 | 0.417053456 | 0.243342153 | 1.713856195 |
| Gm23438 | 0 | NA | NA | NA |
| Gm37002 | 0 | NA | NA | NA |
| Gm23278 | 0 | NA | NA | NA |
| Tdrd3 | 499.7196116 | 0.027885611 | 0.05639136 | 0.494501482 |
| Rps3a2 | 226.2208805 | 0.395141462 | 0.146845601 | 2.690863455 |
| Gm26405 | 0.047948698 | -0.206487837 | 3.724945049 | -0.055433794 |
| Gm9274 | 0.048949532 | -0.206487837 | 3.724945049 | -0.055433794 |
| Pcdh20 | 350.8125221 | -0.47660547 | 0.157423544 | -3.027536148 |
| Gm36946 | 0 | NA | NA | NA |
| Gm25886 | 0 | NA | NA | NA |
| Gm25415 | 0.048949532 | -0.206487837 | 3.724945049 | -0.055433794 |
| Gm10110 | 26.30868621 | 0.198616166 | 0.233161079 | 0.851840995 |
| Gm25495 | 0 | NA | NA | NA |
| Gm23225 | 0 | NA | NA | NA |
| Gm24127 | 0 | NA | NA | NA |
| Gm37341 | 0 | NA | NA | NA |
| Pcdh9 | 3659.38219 | -0.331516934 | 0.073505068 | -4.51012352 |
| Gm23509 | 13.87461055 | 0.177586067 | 0.40917551 | 0.43400952 |
| Gm23324 | 0 | NA | NA | NA |
| Gm24680 | 0 | NA | NA | NA |
| 4921530L21Rik | 0.197024265 | -0.673766873 | 2.225999129 | -0.302680654 |
| Klhl1 | 133.1425135 | 0.459320948 | 0.194179644 | 2.365443353 |
| Gm15515 | 0.04393175 | -0.206490061 | 3.724945049 | -0.055434391 |
| Gm24043 | 0 | NA | NA | NA |
| Gm25133 | 0 | NA | NA | NA |
| Gm24770 | 0 | NA | NA | NA |
| Gm27441 | 0 | NA | NA | NA |
| Gm23276 | 0.217428854 | 0.274401121 | 2.581304093 | 0.106303291 |
| Dach1 | 251.06346 | -0.400875792 | 0.105586099 | -3.796672058 |
| Gm25831 | 2.466058065 | 0.086207351 | 0.693103695 | 0.124378721 |
| Gm16331 | 0 | NA | NA | NA |
| Gm26006 | 0 | NA | NA | NA |
| Gm26139 | 0 | NA | NA | NA |
| Gm27034 | 0 | NA | NA | NA |
| Rpl36a-ps1 | 11.21814357 | 0.556640236 | 0.319762946 | 1.740790303 |
| Mzt1 | 677.8415215 | -0.054868758 | 0.072313636 | -0.75876088 |
| Bora | 35.47661271 | -0.125448503 | 0.22633059 | -0.554271091 |
| Dis3 | 309.1793791 | 0.261165363 | 0.132109042 | 1.976892415 |
| Pibf1 | 250.2980066 | 0.136431685 | 0.068371664 | 1.995441932 |
| Gm4412 | 0 | NA | NA | NA |
| Klf5 | 148.4473388 | 0.074204545 | 0.171730834 | 0.432097975 |

| | | | | |
|---|---|---|---|---|
| Gm22984 | 0 | NA | NA | NA |
| Gm25130 | 0 | NA | NA | NA |
| Gm37093 | 0.13998015 | 1.236190668 | 2.923064887 | 0.422909075 |
| Gm22970 | 0 | NA | NA | NA |
| Gm5958 | 0.163532861 | 0.274408734 | 2.784469947 | 0.09854972 |
| Klf12 | 318.1322375 | -0.046096957 | 0.08896016 | -0.518175293 |
| 4930517O19Rik | 0.663801701 | 1.029854454 | 1.453551894 | 0.7085089 |
| Gm16260 | 0 | NA | NA | NA |
| Gm26367 | 0 | NA | NA | NA |
| Gm24345 | 0 | NA | NA | NA |
| Gm22401 | 0 | NA | NA | NA |
| Gm23538 | 0 | NA | NA | NA |
| Prr30 | 0.448613728 | 0.687378764 | 1.394970511 | 0.49275505 |
| Gm26173 | 0 | NA | NA | NA |
| Tbc1d4 | 374.3985729 | -0.400939282 | 0.087922184 | -4.560160655 |
| Commd6 | 158.7771116 | -0.150688391 | 0.11497448 | -1.310624683 |
| Uchl3 | 201.0806668 | -0.136797908 | 0.099508407 | -1.374737199 |
| Gm9922 | 0.794139519 | -0.550133036 | 1.116929228 | -0.49254064 |
| Lmo7 | 977.0007035 | 0.168630415 | 0.183200707 | 0.920468144 |
| Gm22347 | 0 | NA | NA | NA |
| Gm23764 | 0 | NA | NA | NA |
| Kctd12 | 489.5908408 | -0.362706935 | 0.14002448 | -2.590310889 |
| AC102815.1 | 0 | NA | NA | NA |
| Gm26778 | 8.453551642 | -0.034399386 | 0.34471581 | -0.099790568 |
| Mir5130 | 0 | NA | NA | NA |
| Irg1 | 221.3397588 | 9.111487172 | 0.966040693 | 9.431784023 |
| Cln5 | 245.9919061 | 0.209735417 | 0.136678732 | 1.534513915 |
| Fbxl3 | 1187.753289 | -0.038116638 | 0.064828443 | -0.587961636 |
| Mycbp2 | 9169.971231 | -0.011011442 | 0.08375245 | -0.131476061 |
| Gm4654 | 6.877366695 | -0.30661957 | 0.37595412 | -0.815577095 |
| Scel | 26.78325797 | -0.308674102 | 0.227711138 | -1.355551181 |
| Slain1 | 561.6860723 | -0.44956966 | 0.080472476 | -5.586626426 |
| Gm22877 | 0.140122821 | 0.274406091 | 3.572128793 | 0.076818644 |
| Slain1os | 0.09349418 | 0.274402725 | 3.724945049 | 0.073666248 |
| Ednrb | 961.6585733 | -1.005596184 | 0.176976395 | -5.68209216 |
| Gm37910 | 0.066050875 | 0.274391604 | 3.724945049 | 0.073663262 |
| Pou4f1 | 42.92036986 | 0.29965847 | 0.338172769 | 0.886110584 |
| Rnf219 | 139.6039557 | -0.078364422 | 0.09575618 | -0.818374567 |
| Gm22021 | 0 | NA | NA | NA |
| Gm25724 | 0 | NA | NA | NA |
| Gm24280 | 0 | NA | NA | NA |
| Rbm26 | 1630.5591 | 0.207007429 | 0.0954071 | 2.169727722 |
| Gm17066 | 187.5789837 | 0.417956774 | 0.124907248 | 3.346137068 |
| Gm22290 | 0 | NA | NA | NA |
| Ndfip2 | 1106.337769 | -0.22959609 | 0.063266053 | -3.629056639 |
| Gm24207 | 0 | NA | NA | NA |
| Gm16259 | 0.115603922 | 0.274387056 | 3.724945049 | 0.073662041 |

| | | | | |
|---|---|---|---|---|
| 4930449E01Rik | 0.297771212 | -0.725501578 | 1.617836294 | -0.448439425 |
| Mir6390 | 7.744732698 | -0.6752459 | 0.357961577 | -1.886364187 |
| Gm10076 | 151.4983943 | 0.358700661 | 0.171251502 | 2.094584024 |
| Gm22406 | 0 | NA | NA | NA |
| Spry2 | 675.0338336 | -0.242085411 | 0.131636914 | -1.839038937 |
| Gm4775 | 0 | NA | NA | NA |
| Trim52 | 0 | NA | NA | NA |
| Gm23459 | 0 | NA | NA | NA |
| 1700128A07Rik | 0 | NA | NA | NA |
| Gm24634 | 0 | NA | NA | NA |
| Gm17366 | 0 | NA | NA | NA |
| Gm23437 | 0 | NA | NA | NA |
| Gm24818 | 0 | NA | NA | NA |
| Gm24229 | 0 | NA | NA | NA |
| Rpl27a-ps2 | 2.255766892 | 1.291898446 | 0.745341764 | 1.733296735 |
| Slitrk1 | 1192.574154 | -0.342164768 | 0.093593957 | -3.655842502 |
| Gm25901 | 0 | NA | NA | NA |
| Gm24369 | 0 | NA | NA | NA |
| n-R5s50 | 0 | NA | NA | NA |
| Gm23242 | 0 | NA | NA | NA |
| Gm25670 | 0 | NA | NA | NA |
| Gm26255 | 0 | NA | NA | NA |
| Slitrk6 | 94.50009073 | 0.202018405 | 0.130520913 | 1.547785717 |
| Gm22802 | 0 | NA | NA | NA |
| Slitrk5 | 1071.418396 | 0.077182974 | 0.099031855 | 0.779375224 |
| Gm22764 | 0 | NA | NA | NA |
| Gm4822 | 0 | NA | NA | NA |
| Tpm3-rs7 | 867.6432377 | 0.876934444 | 0.112729156 | 7.779127188 |
| Gm4487 | 0 | NA | NA | NA |
| Gm37495 | 0 | NA | NA | NA |
| Gm24073 | 0 | NA | NA | NA |
| Mir17hg | 18.52657897 | 0.62547166 | 0.298159293 | 2.097776842 |
| Mir17 | 0 | NA | NA | NA |
| Mir18 | 0 | NA | NA | NA |
| Mir19a | 0 | NA | NA | NA |
| Mir20a | 0 | NA | NA | NA |
| Mir19b-1 | 0 | NA | NA | NA |
| Mir92-1 | 0 | NA | NA | NA |
| Gpc5 | 280.2541914 | -0.283917224 | 0.133425504 | -2.127908191 |
| 4930505G20Rik | 0 | NA | NA | NA |
| Gm18367 | 0 | NA | NA | NA |
| Gm20713 | 0.048949532 | -0.206487837 | 3.724945049 | -0.055433794 |
| Gm38045 | 0 | NA | NA | NA |
| Gpc6 | 374.2491254 | 0.082258149 | 0.082680684 | 0.994889556 |
| Mir6239 | 0 | NA | NA | NA |
| Dct | 6.057780134 | -0.01052253 | 0.445726627 | -0.023607586 |
| Gm26773 | 0.075043205 | 0.755286614 | 3.724945049 | 0.202764498 |

| | | | | |
|---|---|---|---|---|
| Tgds | 62.65483873 | -0.149264553 | 0.203504708 | -0.733469781 |
| Gpr180 | 179.3319749 | -0.178416786 | 0.086609585 | -2.060012012 |
| Sox21 | 98.70143058 | -1.170825632 | 0.233773355 | -5.008379299 |
| Gm4675 | 4.582536418 | -0.186158618 | 0.428783796 | -0.434154975 |
| Mir6241 | 0.075043205 | 0.755286614 | 3.724945049 | 0.202764498 |
| Gm9376 | 0 | NA | NA | NA |
| Gm23669 | 0 | NA | NA | NA |
| Gm26791 | 0.061920613 | 0.274393828 | 3.724945049 | 0.073663859 |
| 1700044C05Rik | 0 | NA | NA | NA |
| Abcc4 | 177.0567019 | 0.075875066 | 0.142769843 | 0.531450227 |
| Gm23302 | 0 | NA | NA | NA |
| Mir6391 | 0.804474276 | 0.933450147 | 1.083467794 | 0.861539358 |
| Gm27198 | 0 | NA | NA | NA |
| Gm22379 | 0.042716945 | -0.206490061 | 3.724945049 | -0.055434391 |
| Cldn10 | 133.9096642 | -0.70186387 | 0.184769432 | -3.798593006 |
| n-R5s51 | 0 | NA | NA | NA |
| Dzip1 | 1031.570717 | -0.036220288 | 0.074447601 | -0.486520553 |
| Dnajc3 | 1082.666445 | 0.164221658 | 0.075833839 | 2.165545871 |
| Uggt2 | 618.685887 | -0.321152422 | 0.088637937 | -3.623193766 |
| Gm16835 | 39.44922461 | -0.21518525 | 0.269118048 | -0.799594275 |
| Hs6st3 | 282.2429297 | -0.26290581 | 0.157966736 | -1.664311209 |
| 1700006F04Rik | 17.63169857 | -0.300423638 | 0.335254348 | -0.896106613 |
| Oxgr1 | 3.717291957 | 0.212180345 | 0.584267296 | 0.363156292 |
| Mbnl2 | 3584.706982 | 0.151373731 | 0.068199799 | 2.219562695 |
| Gm26679 | 5.318848262 | 0.563643134 | 0.518629448 | 1.086793541 |
| Rap2a | 997.0783507 | -0.090910107 | 0.069721359 | -1.303906124 |
| Gm17613 | 0.138137327 | 0.755284675 | 3.610850559 | 0.209170848 |
| Gm25781 | 0 | NA | NA | NA |
| Ipo5 | 1921.943571 | 0.154070346 | 0.062944827 | 2.447704657 |
| Farp1 | 948.5700461 | -0.083159411 | 0.073152172 | -1.136800295 |
| B930095G15Rik | 416.2277983 | -0.267151937 | 0.112405967 | -2.376670422 |
| Stk24 | 646.4051468 | 0.191318444 | 0.083821305 | 2.282456042 |
| Slc15a1 | 1.066426709 | 0.596754543 | 1.125278144 | 0.530317367 |
| Dock9 | 2161.334577 | -0.15394656 | 0.091199436 | -1.688020954 |
| 1810041H14Rik | 1.973916092 | 0.301927023 | 0.701839224 | 0.430194 |
| Ubac2 | 353.8238038 | 0.097069939 | 0.092996745 | 1.043799321 |
| Gpr18 | 9.840232673 | 5.973035682 | 0.935238774 | 6.386642479 |
| Gpr183 | 14.82188085 | 0.51591617 | 0.315984717 | 1.632725073 |
| Timm8a2 | 1.438846906 | 2.590020748 | 0.956357812 | 2.708213094 |
| A330035P11Rik | 10.89891607 | 0.216809859 | 0.310626441 | 0.697976188 |
| Tm9sf2 | 1465.863616 | 0.049962209 | 0.069681458 | 0.71700866 |
| Gm6254 | 34.64912996 | 0.387782472 | 0.324383825 | 1.195443304 |
| Gm17082 | 26.87597815 | 0.152499312 | 0.249109299 | 0.612178319 |
| Clybl | 80.6719109 | -0.255201247 | 0.144431792 | -1.76693264 |
| 1700108J01Rik | 0.0878635 | -0.687376175 | 3.724945049 | -0.184533239 |
| Gm25464 | 0.76305848 | -2.454935243 | 1.157386799 | -2.121101817 |
| Gm5089 | 8.639524202 | -1.097403535 | 0.462044866 | -2.375101675 |

| | | | | |
|---|---|---|---|---|
| Zic5 | 171.9911989 | -0.279316143 | 0.145744497 | -1.91647814 |
| 2610035F20Rik | 11.57667783 | -0.333385149 | 0.306385538 | -1.088122995 |
| Zic2 | 736.6056247 | 0.072151824 | 0.192316141 | 0.375173001 |
| Gm10837 | 14.11353788 | -0.146837509 | 0.4243206 | -0.346053219 |
| Gm9399 | 0.044544648 | 0.755286614 | 3.724945049 | 0.202764498 |
| Pcca | 484.4393937 | -0.056711479 | 0.110346181 | -0.51394148 |
| Ggact | 197.5556307 | 0.23601208 | 0.125144032 | 1.885923568 |
| 4930594M22Rik | 104.9172262 | -0.054333495 | 0.198275209 | -0.274030705 |
| Tmtc4 | 235.3928922 | -0.144153582 | 0.106468206 | -1.353958971 |
| Gm15735 | 9.699810692 | -0.620109803 | 0.401293412 | -1.545277806 |
| Nalcn | 1752.798546 | -0.178519926 | 0.058191983 | -3.067775244 |
| Itgbl1 | 26.65978906 | -0.076310675 | 0.309083722 | -0.246893219 |
| Gm9308 | 0 | NA | NA | NA |
| Gm25272 | 0.047948698 | -0.206487837 | 3.724945049 | -0.055433794 |
| Fgf14 | 940.1745018 | -0.103191566 | 0.090160403 | -1.144533104 |
| Gm28932 | 1.468722268 | 0.585149855 | 0.749782099 | 0.780426548 |
| Gm15422 | 16.80940044 | 0.05365802 | 0.34887475 | 0.153803106 |
| 1700024B18Rik | 0.075043205 | 0.755286614 | 3.724945049 | 0.202764498 |
| Gm17615 | 0.320086762 | 0.3526156 | 1.570515221 | 0.224522243 |
| Sepp1 | 2912.542392 | 0.271637293 | 0.136155581 | 1.995050748 |
| Ccdc152 | 0.495781838 | -0.597541265 | 1.431083436 | -0.417544673 |
| Ghr | 164.1599138 | -0.361552129 | 0.101460067 | -3.56349191 |
| Gm22031 | 0 | NA | NA | NA |
| Gm4823 | 0.066050875 | 0.274391604 | 3.724945049 | 0.073663262 |
| Gm23228 | 0 | NA | NA | NA |
| Fbxo4 | 73.81450778 | 0.053857728 | 0.142459549 | 0.378056285 |
| AW549877 | 1232.520806 | -0.084115093 | 0.072611141 | -1.158432333 |
| Gm2093 | 0.919878397 | 1.573473967 | 1.091774084 | 1.441208387 |
| BC037032 | 34.71017491 | 0.57104976 | 0.259314312 | 2.20215288 |
| Oxct1 | 2597.175127 | -0.272081415 | 0.087904278 | -3.095201058 |
| Gm15632 | 0 | NA | NA | NA |
| Plcxd3 | 320.4411957 | -0.294441939 | 0.108038785 | -2.725335527 |
| C6 | 2.125003216 | 4.587466264 | 1.062480699 | 4.317693739 |
| Mroh2b | 0.19444009 | 0.299620658 | 2.689656738 | 0.111397359 |
| C7 | 8.962534464 | 1.051786134 | 0.389340151 | 2.701458177 |
| Gm25652 | 0 | NA | NA | NA |
| Gm16077 | 0.04393175 | -0.206490061 | 3.724945049 | -0.055434391 |
| Card6 | 43.77270322 | 1.002974054 | 0.177703734 | 5.644079789 |
| Rpl37 | 814.5795284 | 0.31711628 | 0.109856758 | 2.886634253 |
| Snord72 | 5.885017498 | 0.131186933 | 0.464809667 | 0.282237962 |
| Gm10250 | 114.0624678 | -0.17011262 | 0.110722438 | -1.536387961 |
| Prkaa1 | 690.6189185 | 0.279995725 | 0.104954966 | 2.667770145 |
| Gm15938 | 2.619102942 | -0.518633291 | 0.714230106 | -0.72614314 |
| Ttc33 | 464.303568 | 0.170943964 | 0.069853676 | 2.447172062 |
| Ptger4 | 41.58960014 | 1.827312654 | 0.261029896 | 7.000396053 |
| Gm15937 | 14.4175352 | 0.068489484 | 0.323746841 | 0.211552594 |
| Gm27760 | 0.173816988 | -1.279622137 | 3.321126619 | -0.385297606 |

| | | | | |
|---|---|---|---|---|
| Gm23139 | 0 | NA | NA | NA |
| Dab2 | 418.7976734 | 1.093664024 | 0.185648502 | 5.891046847 |
| Gm16311 | 0 | NA | NA | NA |
| C9 | 0.135879625 | -0.66959118 | 2.940650251 | -0.227701741 |
| Gm23218 | 0 | NA | NA | NA |
| Fyb | 220.4543545 | 2.521578033 | 0.189025862 | 13.33985733 |
| Gm2245 | 0.436872751 | 0.853081584 | 1.709325331 | 0.499075026 |
| Gm7666 | 27.5160756 | 0.280828461 | 0.20689208 | 1.357366897 |
| Rictor | 1871.956267 | 0.205836149 | 0.065930314 | 3.122025928 |
| Osmr | 628.1045177 | 3.958124916 | 0.294812674 | 13.42589809 |
| Gm38282 | 0.159330469 | 1.236180454 | 3.465376669 | 0.356723258 |
| Lifr | 1758.792625 | 0.201309007 | 0.071711371 | 2.807211802 |
| Egflam | 43.8423358 | -0.372229423 | 0.170515929 | -2.182959832 |
| Gm16030 | 0 | NA | NA | NA |
| Gm16029 | 0.048949532 | -0.206487837 | 3.724945049 | -0.055433794 |
| Gm37743 | 0.061920613 | 0.274393828 | 3.724945049 | 0.073663859 |
| Gdnf | 23.58641988 | 0.868697021 | 0.276997367 | 3.136120143 |
| Wdr70 | 191.8981038 | 0.082956563 | 0.106152208 | 0.781486932 |
| Gm27529 | 0.871987541 | 3.050621475 | 1.332163978 | 2.289974452 |
| Nup155 | 488.0096314 | 0.128291446 | 0.114563108 | 1.119832105 |
| 2410089E03Rik | 1512.066955 | -0.140998614 | 0.161708771 | -0.871929295 |
| Gm24144 | 14.79135611 | -0.269108326 | 0.433175828 | -0.621245021 |
| Nipbl | 1929.6716 | 0.05560191 | 0.064100381 | 0.867419345 |
| Gm2310 | 0 | NA | NA | NA |
| Gm2348 | 0.104106287 | 0.274400501 | 3.724945049 | 0.073665651 |
| Slc1a3 | 7685.648105 | 0.399594231 | 0.11677836 | 3.421817463 |
| Gm37310 | 119.6237137 | 1.133275728 | 0.298055179 | 3.802234645 |
| RP24-341L14.2 | 0.280881662 | 1.516895756 | 2.085559961 | 0.727332604 |
| Gm5210 | 0 | NA | NA | NA |
| Ranbp3l | 56.33245746 | -0.825727797 | 0.249108884 | -3.31472641 |
| Nadk2 | 494.6776195 | -0.300371744 | 0.110434962 | -2.719897201 |
| Skp2 | 125.2851504 | -0.344820514 | 0.133661696 | -2.579800522 |
| Lmbrd2 | 1052.783199 | -0.239485843 | 0.053461652 | -4.479581805 |
| Ugt3a1 | 0.150086409 | 1.264824188 | 3.723011833 | 0.339731445 |
| Ugt3a2 | 0.171181394 | 1.257632701 | 2.747344421 | 0.457763028 |
| Capsl | 15.44470158 | 1.017189724 | 0.310586032 | 3.275065904 |
| Il7r | 194.8832465 | 2.845420695 | 0.448743649 | 6.340860094 |
| Spef2 | 48.63543957 | -0.361365199 | 0.186354915 | -1.939123522 |
| Gm26350 | 0 | NA | NA | NA |
| Prlr | 148.8562054 | -0.481001333 | 0.176762481 | -2.721173236 |
| Gm21973 | 0 | NA | NA | NA |
| Agxt2 | 0.124596252 | 0.755282361 | 3.714681068 | 0.203323609 |
| Dnajc21 | 254.7197003 | -0.288493957 | 0.107297548 | -2.688728328 |
| Brix1 | 263.1432847 | 0.222087475 | 0.110955593 | 2.001588814 |
| Rad1 | 85.02316572 | -0.560602655 | 0.141231881 | -3.969377533 |
| Ttc23l | 0.267080418 | 0.274415778 | 1.817541942 | 0.150981813 |
| Gm17044 | 0.038981395 | 0.274393828 | 3.724945049 | 0.073663859 |

| | | | | |
|---|---|---|---|---|
| Gm8174 | 0.386177252 | -0.736898538 | 1.581149024 | -0.466052552 |
| Rai14 | 244.3821753 | 0.917781788 | 0.134502633 | 6.823522822 |
| 4930556M19Rik | 7.969462537 | 0.502938765 | 0.387475573 | 1.297988311 |
| C1qtnf3 | 3.38967256 | 1.256108372 | 0.614452012 | 2.044274163 |
| Amacr | 74.0218532 | -0.389134014 | 0.154428811 | -2.519827815 |
| Slc45a2 | 0.883742299 | -1.505373139 | 1.185365137 | -1.269965762 |
| Rxfp3 | 34.39250656 | -0.247991751 | 0.208672566 | -1.188425274 |
| Adamts12 | 17.29016175 | 0.05979175 | 0.279569619 | 0.2138707 |
| Gm24844 | 0 | NA | NA | NA |
| Tars | 425.9557127 | 0.451987833 | 0.121456392 | 3.721400131 |
| Npr3 | 146.1473597 | 0.148091488 | 0.12878877 | 1.149878884 |
| 1700047G03Rik | 0.146692594 | 0.274392389 | 3.52332538 | 0.07787881 |
| Gm24074 | 0 | NA | NA | NA |
| Sub1 | 1754.400611 | 0.26642693 | 0.078730261 | 3.384047329 |
| RP24-105K16.2 | 0.060174537 | 0.755286614 | 3.724945049 | 0.202764498 |
| Zfr | 4001.152899 | 0.100081576 | 0.057339631 | 1.745417157 |
| Gm25713 | 0 | NA | NA | NA |
| Mir1898 | 18.01255643 | -0.03182236 | 0.256356145 | -0.124133401 |
| Mtmr12 | 542.6306912 | 0.038703252 | 0.092908585 | 0.416573474 |
| 1810049J17Rik | 2.812275527 | 0.81859413 | 0.620750402 | 1.318717036 |
| Golph3 | 1317.912574 | 0.171969814 | 0.091870185 | 1.871878383 |
| Pdzd2 | 1799.644411 | 0.29017818 | 0.166606228 | 1.741700683 |
| Gm24135 | 1.998901016 | -0.756018611 | 0.676264727 | -1.117932936 |
| Gm2606 | 25.26788695 | 0.256972674 | 0.241843699 | 1.062556829 |
| F830212C03Rik | 14.03902647 | -0.548088586 | 0.3969275 | -1.38082795 |
| Gm24302 | 0 | NA | NA | NA |
| 6030458C11Rik | 453.2360569 | -0.25761253 | 0.10148696 | -2.538380597 |
| Drosha | 1331.381267 | -0.185407179 | 0.050068782 | -3.70304953 |
| AC133189.1 | 0 | NA | NA | NA |
| Gm24664 | 0.315718849 | 1.856681716 | 1.673673441 | 1.10934527 |
| Cdh6 | 210.1792556 | -0.014474521 | 0.116770578 | -0.123956922 |
| Gm9630 | 0 | NA | NA | NA |
| Gm24080 | 0 | NA | NA | NA |
| Gm25550 | 0 | NA | NA | NA |
| Cdh9 | 185.9667162 | -0.247782845 | 0.129792829 | -1.909064204 |
| Gm22032 | 0 | NA | NA | NA |
| Gm22895 | 0 | NA | NA | NA |
| Gm24713 | 0.044544648 | 0.755286614 | 3.724945049 | 0.202764498 |
| 4921515E04Rik | 0 | NA | NA | NA |
| Gm8318 | 1.912323087 | 0.987126467 | 0.722433326 | 1.366391099 |
| C030047K22Rik | 5.198307323 | -0.704486627 | 0.496754227 | -1.418179433 |
| Cdh10 | 687.055273 | -0.34584882 | 0.118864248 | -2.909611804 |
| Gm25976 | 0 | NA | NA | NA |
| Gm37815 | 0 | NA | NA | NA |
| Acot10 | 1.155605027 | 0.065288104 | 0.997241586 | 0.065468693 |
| Gm25711 | 4.168298335 | -0.455090264 | 0.480655518 | -0.94681169 |
| Gm27397 | 3.423615971 | -1.027714964 | 0.548794807 | -1.872676184 |

| | | | | |
|---|---|---|---|---|
| Cdh12 | 473.520525 | -0.059519188 | 0.108575114 | -0.548184441 |
| Gm25101 | 4.033556537 | -1.100573264 | 0.546228422 | -2.01485902 |
| Gm38234 | 6.221313409 | -0.890030948 | 0.431530416 | -2.062498759 |
| AC166159.1 | 0 | NA | NA | NA |
| Gm27525 | 0 | NA | NA | NA |
| Cdh18 | 540.5763889 | -0.275639121 | 0.102774184 | -2.681987922 |
| Gm5803 | 0.68701006 | -1.884460168 | 1.206841064 | -1.561481643 |
| Gm19840 | 13.48907376 | -0.760860861 | 0.290140434 | -2.622388237 |
| Gm24656 | 2.60150024 | -0.441861733 | 0.583854044 | -0.7568017 |
| Gm23980 | 0 | NA | NA | NA |
| Gm22810 | 0 | NA | NA | NA |
| Gm2824 | 69.97913716 | -0.444377738 | 0.199083358 | -2.232118964 |
| 9230109A22Rik | 0.817550754 | 1.736940987 | 1.165146109 | 1.490749507 |
| Gm38341 | 0.308655657 | 2.071435376 | 3.704213912 | 0.559210517 |
| Basp1 | 2855.029311 | -0.142745123 | 0.107327871 | -1.329991188 |
| Gm5468 | 5.212139954 | -0.479661719 | 0.55666936 | -0.86166359 |
| Myo10 | 988.6576999 | -0.38434787 | 0.120234967 | -3.196639699 |
| Gm2862 | 0.999928428 | 1.425264339 | 1.098752957 | 1.297165418 |
| Fam134b | 559.6848899 | -0.011962327 | 0.082468204 | -0.145053807 |
| Mir7212 | 1.890715168 | -1.235938911 | 0.897352813 | -1.377316584 |
| Zfp622 | 385.825252 | 0.2940811 | 0.097625415 | 3.012341612 |
| 11-Mar | 93.58712996 | -0.489403572 | 0.127067413 | -3.851527009 |
| Fbxl7 | 13.24742036 | -0.061296432 | 0.331345124 | -0.184992709 |
| Gm6576 | 4.198207553 | 0.703091499 | 0.498156489 | 1.411386814 |
| Gm23903 | 0 | NA | NA | NA |
| Ank | 1809.490276 | -0.208560754 | 0.050340063 | -4.143037196 |
| Mir7117 | 0 | NA | NA | NA |
| Otulin | 178.6870836 | 0.421155799 | 0.10317664 | 4.081890991 |
| Fam105a | 144.2450621 | -0.03254497 | 0.121472252 | -0.267921024 |
| Trio | 4183.182376 | -0.089546176 | 0.085577236 | -1.046378451 |
| Dnah5 | 99.5374478 | -0.860262249 | 0.127144915 | -6.765998065 |
| Mir3964 | 0 | NA | NA | NA |
| Gm23033 | 0 | NA | NA | NA |
| Gm24554 | 0 | NA | NA | NA |
| Gm23327 | 0 | NA | NA | NA |
| Gm26163 | 0 | NA | NA | NA |
| Ctnnd2 | 3344.046455 | 0.037338918 | 0.061704459 | 0.605125114 |
| Gm23956 | 8.765840549 | -0.639271375 | 0.42842643 | -1.492138043 |
| Gm24487 | 2.253680491 | 0.025024139 | 0.617780964 | 0.040506492 |
| Gm26416 | 0.503286714 | 2.74708429 | 3.679457924 | 0.746600273 |
| Dap | 102.7869284 | 0.581662447 | 0.200225126 | 2.905042234 |
| Fam136b-ps | 0.047948698 | -0.206487837 | 3.724945049 | -0.055433794 |
| Ankrd33b | 1536.741971 | 0.506903822 | 0.161444183 | 3.139808525 |
| Ropn1l | 5.055244084 | 0.210266224 | 0.452535007 | 0.464640792 |
| Gm6361 | 5.200498561 | 0.374041619 | 0.465233005 | 0.803987711 |
| 6-Mar | 3828.438788 | -0.19795107 | 0.047839437 | -4.137821926 |
| Cmbl | 49.64000691 | -0.352071979 | 0.182225486 | -1.932067718 |

| | | | | |
|---|---|---|---|---|
| Cct5 | 1440.817807 | 0.111355384 | 0.077629312 | 1.434450229 |
| Fam173b | 140.20352 | -0.740796504 | 0.233611842 | -3.171057155 |
| Tas2r119 | 0 | NA | NA | NA |
| Snhg18 | 13.92677096 | -0.49917519 | 0.297522245 | -1.677774342 |
| Snord123 | 0.145446656 | 0.755296437 | 2.882544899 | 0.262024171 |
| Sema5a | 1219.347837 | -0.212722403 | 0.080276051 | -2.649886235 |
| Sdc2 | 591.0431138 | 0.06380052 | 0.104086673 | 0.612955703 |
| Cpq | 169.9525733 | 0.589106227 | 0.172378882 | 3.417508102 |
| 1700084J12Rik | 0.496918936 | -0.513731602 | 3.597050567 | -0.142820233 |
| Gm24476 | 0.306418947 | 0.274378581 | 3.724945049 | 0.073659766 |
| Tspyl5 | 513.6623244 | -0.163086599 | 0.094931891 | -1.717932686 |
| Mtdh | 1529.028053 | 0.161580853 | 0.061727195 | 2.617660727 |
| Gm24500 | 0 | NA | NA | NA |
| Laptm4b | 633.1682788 | -0.077082853 | 0.103517375 | -0.744636856 |
| Gm25809 | 0 | NA | NA | NA |
| Matn2 | 218.3605345 | 0.723217132 | 0.168140429 | 4.301268501 |
| 9430069I07Rik | 0.203825279 | 1.19261434 | 2.627357347 | 0.453921634 |
| Rpl30 | 585.149464 | 0.368543219 | 0.081534603 | 4.52008356 |
| Gm24523 | 0 | NA | NA | NA |
| Erich5 | 14.29563529 | -0.339856814 | 0.332320021 | -1.022679324 |
| Hrsp12 | 208.6478451 | -0.289146692 | 0.138749475 | -2.083948013 |
| Pop1 | 119.2294828 | -0.335790951 | 0.159685453 | -2.102827432 |
| Gm22208 | 2.694090619 | -0.250365856 | 0.583605274 | -0.428998618 |
| Nipal2 | 159.5133862 | -0.153177831 | 0.096199773 | -1.592288895 |
| Kcns2 | 254.6488734 | -0.215284602 | 0.09787883 | -2.199501173 |
| Stk3 | 354.1359327 | 0.290358148 | 0.08973482 | 3.23573555 |
| Osr2 | 0.114742407 | 0.274402725 | 3.724945049 | 0.073666248 |
| Vps13b | 2358.901886 | -0.080329308 | 0.070142083 | -1.145236988 |
| Gm24771 | 4.664523441 | -0.856861587 | 0.502184811 | -1.706267429 |
| Mir599 | 0 | NA | NA | NA |
| Mir875 | 0 | NA | NA | NA |
| Cox6c | 1187.10824 | 0.116455665 | 0.149182759 | 0.780624153 |
| Rgs22 | 32.16090155 | 0.219777768 | 0.187053365 | 1.174946881 |
| Gm25372 | 0 | NA | NA | NA |
| Gm18889 | 2.742579127 | -0.140342753 | 0.598267857 | -0.234581803 |
| Fbxo43 | 1.809171643 | -0.734900108 | 0.911078283 | -0.806626742 |
| Gm22979 | 0 | NA | NA | NA |
| Polr2k | 87.53921573 | 0.260185075 | 0.140015676 | 1.858256743 |
| Spag1 | 177.5961547 | -0.588567813 | 0.133951834 | -4.393876469 |
| Gm23754 | 0.706791778 | -1.418455032 | 1.196784884 | -1.185221381 |
| Rnf19a | 663.4722742 | 0.144211799 | 0.100344393 | 1.437168477 |
| Mir8097 | 0 | NA | NA | NA |
| Rpl7a-ps3 | 1.338063731 | 0.431882646 | 0.894347997 | 0.482902234 |
| Ankrd46 | 1500.096336 | 0.140982619 | 0.058668672 | 2.403030684 |
| Snx31 | 4.226100588 | -0.160946215 | 0.463680999 | -0.347105478 |
| AC137871.1 | 0 | NA | NA | NA |
| Pabpc1 | 2806.56132 | 0.651141237 | 0.076231835 | 8.541592035 |

| | | | | |
|---|---|---|---|---|
| Ywhaz | 10312.43491 | 0.050429596 | 0.066029743 | 0.763740614 |
| Gm25303 | 0 | NA | NA | NA |
| Gm10384 | 1.188161708 | -1.681569258 | 1.002010511 | -1.678195227 |
| Zfp706 | 1437.63576 | -0.198464311 | 0.05654816 | -3.509651109 |
| Gm26766 | 0.392978793 | -0.673024736 | 1.718397406 | -0.391658375 |
| Gm16136 | 0.154527273 | 0.274405781 | 2.834454908 | 0.096810776 |
| Grhl2 | 2.049607807 | 0.913203725 | 0.990833067 | 0.921652451 |
| Gm16137 | 0 | NA | NA | NA |
| Gm16138 | 0.695937234 | 2.097968399 | 2.395895722 | 0.875650964 |
| Ncald | 3437.612725 | -0.111876949 | 0.106745522 | -1.048071592 |
| Gm28221 | 0.522018477 | -0.159155374 | 1.246516988 | -0.127680068 |
| Gm23654 | 2.696581576 | -0.30482709 | 0.63073272 | -0.483290434 |
| Gm15941 | 47.58132438 | -0.551600372 | 0.36994543 | -1.491031723 |
| Gm8668 | 0 | NA | NA | NA |
| 4930447A16Rik | 0.047948698 | -0.206487837 | 3.724945049 | -0.055433794 |
| Gm3267 | 0 | NA | NA | NA |
| Gm24833 | 0.580994137 | -1.71016892 | 1.275356137 | -1.340934402 |
| Gm15942 | 0.537603422 | 0.697202887 | 1.247748592 | 0.558768722 |
| Gm15940 | 9.81108231 | -0.579487135 | 0.378832252 | -1.529666841 |
| Rps19-ps5 | 1.865250533 | 0.410272435 | 0.814490656 | 0.503716565 |
| Rrm2b | 427.3554432 | -0.154863208 | 0.068396945 | -2.264183119 |
| Gm3362 | 13.70112511 | 0.513142935 | 0.307879947 | 1.66669814 |
| Ubr5 | 4115.687366 | 0.039194409 | 0.060034964 | 0.6528597 |
| Gm22353 | 0 | NA | NA | NA |
| Odf1 | 0.128271896 | 0.274403127 | 3.679089475 | 0.074584521 |
| Gm24098 | 0 | NA | NA | NA |
| Gm8705 | 0.498759394 | 1.573649056 | 1.665840162 | 0.944657892 |
| Klf10 | 430.8187853 | -0.026146172 | 0.252778924 | -0.103434935 |
| Gm26863 | 0.301278804 | -0.663485424 | 1.60712777 | -0.41283925 |
| Azin1 | 2466.052833 | 0.168694996 | 0.08219355 | 2.052411601 |
| Mir6951 | 0.490407978 | 0.906683316 | 1.2777157 | 0.709612722 |
| 2310043O21Rik | 0.088534442 | 0.274389469 | 3.724945049 | 0.073662689 |
| Atp6v1c1 | 1712.938684 | 0.019835293 | 0.043892111 | 0.451910209 |
| Gm26704 | 1.33281409 | 0.455596607 | 1.01507476 | 0.448830594 |
| Baalc | 276.5131432 | -0.408503444 | 0.091536816 | -4.46272291 |
| Fzd6 | 96.39112974 | 0.205179675 | 0.139485325 | 1.470976787 |
| Gm24789 | 0.09176491 | 1.15872478 | 3.72306111 | 0.311229052 |
| Cthrc1 | 75.42071906 | -0.553535154 | 0.157079232 | -3.523923227 |
| Slc25a32 | 170.269925 | -0.163933607 | 0.091895809 | -1.783907332 |
| Dcaf13 | 241.4209333 | 0.340735603 | 0.091709777 | 3.71536835 |
| n-R5s39 | 0.07796279 | 0.274392189 | 3.724945049 | 0.073663419 |
| Gm9522 | 11.31287606 | 0.07124228 | 0.323627375 | 0.220136754 |
| Rims2 | 2518.228734 | -0.245346741 | 0.088740159 | -2.764776908 |
| Gm25381 | 0 | NA | NA | NA |
| Dcstamp | 2.546063767 | 0.922088957 | 0.728482115 | 1.265767461 |
| Dpys | 1.614744503 | 0.292218092 | 0.734031421 | 0.39810025 |
| Gm16291 | 9.213264714 | 3.121948328 | 0.614003897 | 5.084574125 |

| | | | | |
|---|---|---|---|---|
| Lrp12 | 640.2551409 | -0.132372311 | 0.079309146 | -1.669067407 |
| 9330182O14Rik | 0 | NA | NA | NA |
| Gm25958 | 0 | NA | NA | NA |
| Gm16294 | 6.25597695 | 0.744543305 | 0.456142857 | 1.632259047 |
| Zfpm2 | 484.4239322 | 0.124406892 | 0.11410637 | 1.090271231 |
| 4930555K19Rik | 0 | NA | NA | NA |
| Oxr1 | 3719.638992 | 0.179966335 | 0.050787977 | 3.543483034 |
| Gm26854 | 0.181780715 | 1.120942148 | 3.312348268 | 0.338413131 |
| Abra | 0.899802893 | 1.54933242 | 1.001153242 | 1.547547724 |
| Gm24751 | 0.127971488 | 0.274389832 | 3.687653711 | 0.0744077 |
| Angpt1 | 263.5197957 | 1.478764406 | 0.164879904 | 8.968736472 |
| Gm17473 | 35.3169963 | 5.35341584 | 0.522969208 | 10.23657943 |
| Rspo2 | 100.8944359 | -0.037348777 | 0.155192567 | -0.240660864 |
| Gm9658 | 1.11165528 | 1.45281077 | 0.995905665 | 1.458783519 |
| Gm22037 | 0.385685737 | -0.206482135 | 1.737808134 | -0.118817567 |
| Eif3e | 970.451907 | 0.560337927 | 0.10407744 | 5.383855763 |
| Gm10373 | 1.964982618 | 0.501504266 | 0.690157772 | 0.726651624 |
| Emc2 | 503.3907252 | -0.044737112 | 0.098331361 | -0.454962812 |
| Tmem74 | 82.76716195 | 0.084863758 | 0.144303687 | 0.588091402 |
| Trhr | 65.62486328 | -0.046395405 | 0.133982692 | -0.34627909 |
| Gm24903 | 0 | NA | NA | NA |
| Gm25093 | 0 | NA | NA | NA |
| Nudcd1 | 289.802079 | -0.183134122 | 0.127909987 | -1.431742161 |
| Eny2 | 530.2164271 | 0.237805867 | 0.075286329 | 3.158685918 |
| Gm22392 | 8.528292636 | -0.080604273 | 0.335627748 | -0.24015974 |
| Pkhd1l1 | 13.16766861 | -0.726207079 | 0.273837005 | -2.651968382 |
| Ebag9 | 211.2230902 | -0.036216917 | 0.08663019 | -0.418063461 |
| Sybu | 684.3366891 | -0.393851326 | 0.064759345 | -6.081768183 |
| 2310069G16Rik | 44.1689834 | 0.110307522 | 0.168676395 | 0.653959448 |
| A930017M01Rik | 111.592861 | 0.145581216 | 0.144082859 | 1.010399275 |
| Gm24768 | 0.351351704 | 0.28109647 | 1.583870974 | 0.177474349 |
| Gm23343 | 0.299746944 | 1.18434574 | 1.603458335 | 0.738619591 |
| Gm5471 | 0.105143817 | 0.274402725 | 3.724945049 | 0.073666248 |
| Kcnv1 | 528.0842402 | -0.623685181 | 0.149677424 | -4.166862062 |
| Gm5472 | 0.382197862 | 0.237087919 | 1.560542782 | 0.151926574 |
| Gm26063 | 0 | NA | NA | NA |
| 1700022A22Rik | 0 | NA | NA | NA |
| Gm23562 | 0 | NA | NA | NA |
| 4930548G14Rik | 0.066050875 | 0.274391604 | 3.724945049 | 0.073663262 |
| Gm23665 | 0 | NA | NA | NA |
| Csmd3 | 1405.512621 | -0.425677351 | 0.114287567 | -3.724616433 |
| Gm16300 | 8.844286263 | -0.466095897 | 0.319329891 | -1.459606223 |
| Gm23530 | 0.060174537 | 0.755286614 | 3.724945049 | 0.202764498 |
| Gm25007 | 0 | NA | NA | NA |
| Gm23185 | 0 | NA | NA | NA |
| Gm23848 | 0 | NA | NA | NA |
| Trps1 | 1156.791181 | -0.127156067 | 0.063147466 | -2.013636895 |

| | | | | |
|---|---|---|---|---|
| Mir1907 | 3.164287563 | -0.660298589 | 0.602448488 | -1.096024974 |
| Eif3h | 1020.195418 | 0.33865251 | 0.083873544 | 4.037655905 |
| Utp23 | 94.48504293 | -0.201747509 | 0.144747251 | -1.393791648 |
| Rad21 | 1630.277696 | 0.160654447 | 0.088987526 | 1.805359179 |
| Aard | 24.65755326 | -0.314902321 | 0.240968366 | -1.306820169 |
| Slc30a8 | 0.5812948 | -1.423524953 | 1.306639394 | -1.089455101 |
| Gm10020 | 9.983419005 | 0.598203358 | 0.446477928 | 1.339827392 |
| Gm23545 | 0 | NA | NA | NA |
| Gm23267 | 0 | NA | NA | NA |
| Med30 | 115.6898845 | 0.216084195 | 0.142274092 | 1.518788072 |
| Ext1 | 627.9625793 | -0.059226353 | 0.064230989 | -0.922083782 |
| Gm27926 | 0.066050875 | 0.274391604 | 3.724945049 | 0.073663262 |
| Samd12 | 154.297153 | -0.387541029 | 0.111750893 | -3.467900946 |
| Gm23200 | 7.643354964 | -1.210602826 | 0.430067138 | -2.814915906 |
| mmu-mir-28c | 7.291715215 | -0.657238897 | 0.607801832 | -1.081337471 |
| Gm7489 | 0 | NA | NA | NA |
| Gm26933 | 0 | NA | NA | NA |
| Tnfrsf11b | 55.23446549 | 0.867216394 | 0.251485577 | 3.448374272 |
| Gm5215 | 0 | NA | NA | NA |
| Colec10 | 0.252184553 | 0.720535611 | 1.821054519 | 0.395669434 |
| Mal2 | 1117.686592 | -0.163148092 | 0.101507791 | -1.607246999 |
| Nov | 659.4404828 | 0.071189007 | 0.138949816 | 0.512336098 |
| Gm7577 | 0 | NA | NA | NA |
| Enpp2 | 3338.021252 | -0.476628922 | 0.137531108 | -3.465608106 |
| Gm26684 | 4.138921123 | 0.466750263 | 0.496810408 | 0.939493731 |
| Taf2 | 560.7641421 | -0.345756251 | 0.072464202 | -4.771407696 |
| Dscc1 | 3.002576494 | 0.66504694 | 0.603191453 | 1.102547022 |
| Gm9920 | 2.459937312 | -0.763475643 | 0.650767606 | -1.173192452 |
| Deptor | 493.9836769 | -0.227515912 | 0.084320567 | -2.698225588 |
| Gm26296 | 0 | NA | NA | NA |
| Col14a1 | 17.16314811 | -0.509535647 | 0.256739014 | -1.984644402 |
| 1700040F17Rik | 0.075043205 | 0.755286614 | 3.724945049 | 0.202764498 |
| Mrpl13 | 304.6669654 | 0.131289319 | 0.083878575 | 1.565230678 |
| Mtbp | 67.98096681 | 0.853265445 | 0.183038508 | 4.66167176 |
| Sntb1 | 66.74833637 | 0.217139641 | 0.163590598 | 1.327335701 |
| Has2 | 80.3213104 | 3.06531865 | 0.521792234 | 5.874596152 |
| Has2os | 3.183229149 | -0.758168065 | 0.650614376 | -1.165310962 |
| Gm26178 | 0 | NA | NA | NA |
| Slc22a22 | 0.334826625 | 1.158547884 | 1.600799583 | 0.723730751 |
| Zhx2 | 102.0661113 | -0.568324043 | 0.148452558 | -3.828320975 |
| Gm16006 | 1.944759997 | 1.400128259 | 0.813948974 | 1.720167115 |
| 9330154K18Rik | 0.788164698 | 0.785083364 | 1.318290382 | 0.595531436 |
| Derl1 | 846.7041548 | 0.337253754 | 0.090585663 | 3.72303677 |
| Tbc1d31 | 114.9980324 | -0.671813768 | 0.197638628 | -3.399202754 |
| Fam83a | 0.697414605 | -1.593143473 | 1.40217046 | -1.136198143 |
| 9130401M01Rik | 105.150685 | -0.147193988 | 0.116577144 | -1.262631623 |
| Gm29394 | 4.265622792 | -1.100403608 | 0.576386042 | -1.909143399 |

| | | | | |
|---|---|---|---|---|
| Gm22299 | 0.817774247 | -0.853716495 | 1.179214557 | -0.723970451 |
| Zhx1 | 1069.677375 | 0.093277312 | 0.06773759 | 1.377039126 |
| Atad2 | 218.8401466 | 0.17485254 | 0.16426699 | 1.064441127 |
| Wdyhv1 | 104.6557226 | -0.220039965 | 0.11534786 | -1.907620685 |
| Gm23475 | 0 | NA | NA | NA |
| Gm22488 | 1.584432859 | 0.95243088 | 0.94689402 | 1.005847391 |
| Fbxo32 | 391.5402532 | -0.361239089 | 0.103263058 | -3.49824125 |
| Gm15943 | 6.572403966 | -0.705156051 | 0.400263549 | -1.76172937 |
| Klhl38 | 13.89074863 | 0.011682539 | 0.352310378 | 0.033159793 |
| Anxa13 | 0 | NA | NA | NA |
| D15Ertd621e | 1138.565379 | -0.038154967 | 0.055110939 | -0.692330185 |
| Fer1l6 | 7.081223484 | 0.70915804 | 0.467066779 | 1.518322586 |
| Gm23789 | 0 | NA | NA | NA |
| Gm20712 | 48.91645952 | -0.139943936 | 0.174094957 | -0.803836816 |
| Tmem65 | 1185.359275 | -0.150016274 | 0.062044382 | -2.417886483 |
| Gm24041 | 11.06577238 | -0.618055341 | 0.321350854 | -1.923303871 |
| Ube2d4 | 0 | NA | NA | NA |
| Trmt12 | 90.91210585 | -0.326690278 | 0.120855926 | -2.703138267 |
| Rnf139 | 607.7861192 | 0.777171766 | 0.108795549 | 7.143415071 |
| Tatdn1 | 116.8025155 | -0.215574191 | 0.13207662 | -1.632190392 |
| Ndufb9 | 863.2536908 | -0.05658955 | 0.123087922 | -0.459749006 |
| Mtss1 | 982.1708213 | -0.284703857 | 0.099339604 | -2.865965277 |
| Gm5045 | 4.498534705 | 0.934101494 | 0.609903827 | 1.53155539 |
| Zfp572 | 0.238889773 | -0.15270574 | 2.141900865 | -0.071294495 |
| Sqle | 640.1137635 | -0.33998073 | 0.114222625 | -2.976474488 |
| E430025E21Rik | 722.4473385 | 0.074211137 | 0.087594452 | 0.847212756 |
| Nsmce2 | 214.2833592 | 0.022141914 | 0.078840231 | 0.280845367 |
| Trib1 | 585.2550097 | 0.703985069 | 0.258376043 | 2.724653033 |
| Gm7083 | 0.061920613 | 0.274393828 | 3.724945049 | 0.073663859 |
| Gm7713 | 0 | NA | NA | NA |
| 4933412E24Rik | 0 | NA | NA | NA |
| Gm25875 | 0.044544648 | 0.755286614 | 3.724945049 | 0.202764498 |
| Gm23835 | 0.042716945 | -0.206490061 | 3.724945049 | -0.055434391 |
| Fam84b | 212.0412458 | 0.203541843 | 0.153300353 | 1.327732383 |
| 9930014A18Rik | 33.67317411 | 0.543233597 | 0.184449396 | 2.945163326 |
| A1bg | 0.161114758 | 1.626894704 | 3.426332624 | 0.474821006 |
| Gm24696 | 0 | NA | NA | NA |
| Gm37947 | 0.038981395 | 0.274393828 | 3.724945049 | 0.073663859 |
| Myc | 144.727581 | 1.89347508 | 0.278470391 | 6.799556216 |
| Pvt1 | 146.0044478 | -0.130955131 | 0.14334165 | -0.913587435 |
| Gm27913 | 0 | NA | NA | NA |
| Gm27782 | 1.314786873 | -1.994913877 | 1.025094074 | -1.94607883 |
| Gm27718 | 0 | NA | NA | NA |
| Gm27957 | 0.583672784 | -1.235825338 | 1.236746278 | -0.999255352 |
| H2afy3 | 0.122507438 | 0.755282569 | 3.724945049 | 0.202763412 |
| Gm24810 | 0 | NA | NA | NA |
| Gm22521 | 0.240278643 | 0.720214332 | 1.843540437 | 0.390669126 |

| | | | | |
|---|---|---|---|---|
| RP23-55P19.1 | 7.30667397 | 0.547846521 | 0.406106737 | 1.349021014 |
| Gm25702 | 0.042716945 | -0.206490061 | 3.724945049 | -0.055434391 |
| Gm37615 | 0 | NA | NA | NA |
| Gm5473 | 0.91428204 | 0.089711331 | 0.960514049 | 0.093399291 |
| Gsdmc | 0.046345709 | 0.274391604 | 3.724945049 | 0.073663262 |
| 1700065I16Rik | 0 | NA | NA | NA |
| Gsdmc2 | 0.112396584 | 0.27438722 | 3.724945049 | 0.073662085 |
| Gsdmcl1 | 0 | NA | NA | NA |
| Gsdmc3 | 0.258721378 | 0.274408032 | 2.055350618 | 0.13350911 |
| Gsdmcl2 | 0 | NA | NA | NA |
| Gsdmc4 | 1.216796211 | -0.31606415 | 1.182390393 | -0.26730947 |
| Gsdmcl-ps | 0 | NA | NA | NA |
| Fam49b | 1750.706656 | 0.110298089 | 0.048916967 | 2.254802257 |
| Gm25628 | 0 | NA | NA | NA |
| Asap1 | 1466.62248 | -0.308000841 | 0.060900736 | -5.057423985 |
| Gm20717 | 0.527708665 | 2.495681685 | 1.462485676 | 1.706465729 |
| Adcy8 | 473.5293802 | -0.785678468 | 0.071617425 | -10.97049318 |
| Gm21798 | 0 | NA | NA | NA |
| Gm21961 | 0 | NA | NA | NA |
| Gm5046 | 0.097139547 | 0.274393828 | 3.724945049 | 0.073663859 |
| Gm23907 | 0 | NA | NA | NA |
| Efr3a | 2113.727247 | -0.067783371 | 0.065059088 | -1.041873988 |
| Gm20405 | 22.08023906 | 0.957055934 | 0.382967188 | 2.499054656 |
| Oc90 | 0.439163765 | 2.35351327 | 1.459322953 | 1.612743269 |
| Hhla1 | 1.7811158 | 0.927795283 | 0.894476348 | 1.037249655 |
| Gm2991 | 7.931298069 | 0.603567827 | 0.478063344 | 1.262526888 |
| Kcnq3 | 2910.472199 | -0.156843615 | 0.091898007 | -1.70671399 |
| Gm27153 | 0.044544648 | 0.755286614 | 3.724945049 | 0.202764498 |
| Gm27242 | 2.088118819 | 1.619218171 | 0.705265922 | 2.295897363 |
| Lrrc6 | 31.55937458 | -0.178223685 | 0.204243972 | -0.872601935 |
| Tmem71 | 8.010071188 | 3.93584179 | 0.705687732 | 5.577313608 |
| Phf20l1 | 1418.308876 | 0.042348526 | 0.070740365 | 0.598647275 |
| Tg | 5.923287473 | 1.619347985 | 0.569913313 | 2.841393505 |
| Gm17140 | 0 | NA | NA | NA |
| Sla | 260.5870413 | 1.420559664 | 0.112361773 | 12.64273092 |
| Wisp1 | 38.31189845 | -1.005885926 | 0.336319339 | -2.990865551 |
| Ndrg1 | 857.5715411 | -0.288431945 | 0.132979828 | -2.168990214 |
| Gm17035 | 0.30380643 | -1.129486131 | 2.020584286 | -0.558989862 |
| Gm2895 | 1.340196514 | 0.870250919 | 1.191730612 | 0.730241306 |
| St3gal1 | 321.0877622 | 0.367967756 | 0.111505235 | 3.300004311 |
| 1700012I11Rik | 0.522753486 | -0.455040644 | 1.308664298 | -0.347713806 |
| Zfat | 173.6083377 | -0.308190744 | 0.130378835 | -2.363809618 |
| Gm10240 | 8.242000484 | 0.383951772 | 0.468885711 | 0.81886004 |
| Gm37320 | 57.92546311 | -1.318486474 | 0.491665034 | -2.681676312 |
| Gm26621 | 67.44066586 | -0.75185966 | 0.312825227 | -2.403449582 |
| Mir30b | 2.936662866 | -0.721617792 | 0.636987039 | -1.132861029 |
| Gm37296 | 72.21542741 | -0.520482361 | 0.208459229 | -2.496806509 |

| | | | | |
|---|---|---|---|---|
| Gm20732 | 11.82845525 | -0.49633491 | 0.384247013 | -1.29170792 |
| Mir30d | 0 | NA | NA | NA |
| Gm25987 | 0 | NA | NA | NA |
| Gm7125 | 2.234104145 | -0.556540664 | 0.660247071 | -0.842927879 |
| Khdrbs3 | 814.1907786 | -0.137321238 | 0.116468804 | -1.179038798 |
| Gm24232 | 0 | NA | NA | NA |
| Gm23987 | 0 | NA | NA | NA |
| Fam135b | 709.6683967 | -0.386004696 | 0.118267602 | -3.263824484 |
| AC157602.1 | 0 | NA | NA | NA |
| Gm22519 | 0.044191549 | -0.206490061 | 3.724945049 | -0.055434391 |
| Col22a1 | 42.46963243 | 0.466229452 | 0.213757831 | 2.181110512 |
| Gm16308 | 0.108309328 | 1.204448679 | 3.72248341 | 0.323560523 |
| Gm23217 | 4.535027026 | -0.429556302 | 0.486198066 | -0.88350064 |
| Gm24390 | 0 | NA | NA | NA |
| Kcnk9 | 245.3309406 | -0.177329092 | 0.093113422 | -1.904441793 |
| Trappc9 | 814.4156458 | -0.075726786 | 0.043089034 | -1.757449143 |
| Gm28020 | 1.075827988 | -0.04587771 | 0.903118347 | -0.050799222 |
| Peg13 | 3399.486185 | -0.310350036 | 0.051908941 | -5.978739444 |
| Gm3150 | 0.456099668 | -0.199777123 | 1.492557364 | -0.133848874 |
| Chrac1 | 76.46489437 | 0.769396918 | 0.164706857 | 4.671310792 |
| Ago2 | 2822.203106 | -0.255325065 | 0.072104585 | -3.541037873 |
| Ptk2 | 1504.870551 | -0.03983162 | 0.06839563 | -0.582370832 |
| Mir151 | 5.981125562 | -0.570909904 | 0.518175148 | -1.101770137 |
| Gm24787 | 0 | NA | NA | NA |
| Dennd3 | 115.7213391 | 1.199768598 | 0.223399706 | 5.370502125 |
| Slc45a4 | 787.9133075 | 0.243137603 | 0.072228393 | 3.366233042 |
| 1700010B13Rik | 0.609775176 | 0.668249572 | 1.158251689 | 0.576946771 |
| Gpr20 | 1.015418215 | 0.842474171 | 0.948732552 | 0.887999647 |
| Ptp4a3 | 340.214372 | -0.255992509 | 0.118186745 | -2.166000169 |
| Gm22106 | 2.287044039 | -1.154118831 | 0.675573996 | -1.708352954 |
| Mroh5 | 17.84213286 | -0.163145867 | 0.244150573 | -0.668218245 |
| Gm3244 | 3.864046566 | -0.120315833 | 0.518852703 | -0.231888227 |
| Gm6569 | 1.915143953 | -0.100316035 | 0.731360144 | -0.137163662 |
| Gm7935 | 7.955384209 | 0.229020067 | 0.371157296 | 0.61704315 |
| Gm15387 | 13.22477098 | 0.530260621 | 0.4936933 | 1.074068901 |
| Adgrb1 | 3078.93178 | -0.358476964 | 0.14578497 | -2.458943218 |
| Gm26117 | 0 | NA | NA | NA |
| Mroh4 | 0.178490723 | 1.257478905 | 2.711241583 | 0.463801866 |
| Arc | 2012.660561 | -1.552983244 | 0.498443232 | -3.115667228 |
| Jrk | 151.5292579 | -0.038570326 | 0.107343892 | -0.359315518 |
| 4933427E11Rik | 4.797225747 | -0.208859902 | 0.571966292 | -0.365161207 |
| Psca | 1.592565788 | 2.66071636 | 1.090980546 | 2.438830251 |
| Gm28976 | 0 | NA | NA | NA |
| Them6 | 413.7288862 | 0.207811493 | 0.078626409 | 2.643024098 |
| Slurp1 | 1.036256916 | 2.075605766 | 1.086693616 | 1.910019288 |
| Lypd2 | 2.552977391 | 0.6175806 | 0.650312968 | 0.949666746 |
| 2300005B03Rik | 0.58378306 | 2.684437206 | 1.604471517 | 1.673097452 |

| | | | | |
|---|---|---|---|---|
| Lynx1 | 4130.871836 | 0.017509712 | 0.070137223 | 0.24964934 |
| Ly6d | 10.57966156 | 3.642985585 | 0.718339623 | 5.071397248 |
| D730001G18Rik | 0.049553047 | 0.274391604 | 3.724945049 | 0.073663262 |
| Gm17189 | 0.097139547 | 0.274393828 | 3.724945049 | 0.073663859 |
| Ly6k | 0.732543458 | 0.911440698 | 1.268310279 | 0.718625965 |
| Gml | 0 | NA | NA | NA |
| Hemt1 | 0 | NA | NA | NA |
| Cyp11b1 | 0.755541319 | 2.281550561 | 1.22808402 | 1.857813084 |
| Cyp11b2 | 1.063164669 | -0.023844534 | 1.167833194 | -0.020417756 |
| 2010109I03Rik | 3.337759678 | 0.678993567 | 0.803346774 | 0.845206067 |
| Gm6610 | 1.171300456 | 2.099623799 | 1.275180161 | 1.646531105 |
| Gm29126 | 0.728772687 | 1.606189101 | 1.288991104 | 1.246082379 |
| Ly6e | 7038.794082 | 2.967976165 | 0.168101953 | 17.65581017 |
| Ly6i | 26.75927167 | 6.62742866 | 1.011099767 | 6.554673313 |
| Gm8000 | 1.59659966 | 3.490622561 | 1.05020722 | 3.323746488 |
| Gm37042 | 1.174124634 | 0.432006395 | 1.063706804 | 0.406132962 |
| Ly6a | 4998.068703 | 8.150018016 | 0.341028506 | 23.89834829 |
| Gm28068 | 1.632511506 | 2.224916397 | 1.000689248 | 2.223383935 |
| Gm28069 | 0.045882455 | 0.755288839 | 3.724945049 | 0.202765096 |
| Ly6c1 | 1373.160618 | 3.973047585 | 0.366551739 | 10.83898167 |
| Gm5960 | 192.4448412 | 9.282135136 | 1.067173217 | 8.697871154 |
| Gm28502 | 0.175945213 | 0.755298819 | 2.714629575 | 0.278232738 |
| Ly6c2 | 840.6845555 | 10.3408706 | 0.922400208 | 11.21082856 |
| I830127L07Rik | 12.15111785 | 5.448058418 | 0.960673242 | 5.671083755 |
| Ly6g | 0.512981325 | 0.274414171 | 1.472370823 | 0.186375719 |
| Gm28117 | 0 | NA | NA | NA |
| BC025446 | 0.044544648 | 0.755286614 | 3.724945049 | 0.202764498 |
| Gm10238 | 0 | NA | NA | NA |
| Gm28115 | 0 | NA | NA | NA |
| Ly6f | 0 | NA | NA | NA |
| Gm28116 | 0 | NA | NA | NA |
| Gm3454 | 0.07796279 | 0.274392189 | 3.724945049 | 0.073663419 |
| Gm28113 | 0.038981395 | 0.274393828 | 3.724945049 | 0.073663859 |
| Gm29517 | 0 | NA | NA | NA |
| Gm28114 | 0.149988252 | 0.755288975 | 3.49731137 | 0.215962748 |
| Gm3142 | 0 | NA | NA | NA |
| 9030619P08Rik | 0.161532967 | 0.755285283 | 3.44977679 | 0.218937435 |
| Gm20654 | 0 | NA | NA | NA |
| Ly6h | 944.0802929 | 0.293483918 | 0.09597472 | 3.057929398 |
| Gpihbp1 | 0.278632317 | 0.755309716 | 1.818253597 | 0.415403945 |
| Zfp41 | 98.87672504 | 0.030814842 | 0.130976067 | 0.235270781 |
| 2810039B14Rik | 19.1597757 | -0.430207377 | 0.262379917 | -1.639635308 |
| Top1mt | 124.7567236 | -0.132370993 | 0.117931038 | -1.12244406 |
| Gm15945 | 0.585634519 | -0.493396463 | 1.331157614 | -0.370652174 |
| Rhpn1 | 84.34622614 | 0.057971722 | 0.14421415 | 0.40198359 |
| Mafa | 7.826514485 | -0.225027663 | 0.40735476 | -0.552412014 |
| Zc3h3 | 159.3577311 | -0.085786294 | 0.152825442 | -0.561335158 |

| | | | | |
|---|---|---|---|---|
| Gm23747 | 0 | NA | NA | NA |
| Gm9568 | 0.083526043 | 0.75528439 | 3.724945049 | 0.202763901 |
| Gsdmd | 138.4714216 | 3.742609699 | 0.231501621 | 16.16666734 |
| Mroh6 | 1.934735662 | -0.605461976 | 0.767299576 | -0.789081598 |
| Naprt | 43.80295543 | 0.179144568 | 0.237960279 | 0.752833914 |
| Eef1d | 688.7842089 | 0.110615207 | 0.066080915 | 1.673935759 |
| Tigd5 | 16.71972297 | -0.162470659 | 0.266229058 | -0.610266438 |
| AC116520.1 | 0 | NA | NA | NA |
| Pycrl | 118.6008582 | -0.195295257 | 0.173254055 | -1.127218972 |
| Tsta3 | 168.0293222 | 0.315568214 | 0.128338826 | 2.458867848 |
| Zfp623 | 159.2088236 | 0.080928247 | 0.133467279 | 0.606352715 |
| Zfp707 | 75.27886443 | 0.176153912 | 0.14088281 | 1.250357738 |
| Ccdc166 | 61.81052859 | 0.376390296 | 0.146350747 | 2.571837208 |
| Mapk15 | 15.79109506 | 0.158143009 | 0.333184108 | 0.474641513 |
| Fam83h | 77.85627554 | 0.309084363 | 0.137306381 | 2.251056067 |
| K230010J24Rik | 12.00267003 | 0.493138556 | 0.326705149 | 1.509430011 |
| Gm23027 | 0.654384519 | -1.121655118 | 1.507247211 | -0.744174618 |
| Scrib | 592.0251915 | -0.200108322 | 0.076224934 | -2.62523444 |
| Mir6952 | 0 | NA | NA | NA |
| Puf60 | 1138.944899 | 0.104446718 | 0.086768195 | 1.20374427 |
| Nrbp2 | 715.0247032 | -0.128055157 | 0.115684095 | -1.106938312 |
| BC024139 | 9.779656096 | 0.21826247 | 0.319832758 | 0.682426877 |
| Plec | 5010.938922 | -0.050104292 | 0.078523677 | -0.638078784 |
| Gm25720 | 3.45116213 | -1.896126078 | 0.684965521 | -2.768206601 |
| Mir1942 | 3.221922816 | -1.101544832 | 0.624324152 | -1.764379654 |
| Parp10 | 866.3362482 | 5.917063062 | 0.213632486 | 27.69739365 |
| Grina | 4908.811971 | 0.613632225 | 0.085495865 | 7.177332168 |
| Mir6953 | 0 | NA | NA | NA |
| Gm10872 | 0.972832988 | 0.755311014 | 1.371317123 | 0.550792374 |
| Spatc1 | 0.136963818 | 0.755286583 | 3.59954433 | 0.209828388 |
| Mir3079 | 0 | NA | NA | NA |
| Smpd5 | 4.917982605 | 0.422147491 | 0.467403525 | 0.903175669 |
| Oplah | 309.3643565 | 0.120308027 | 0.115866124 | 1.038336516 |
| Exosc4 | 178.5313927 | 0.485704564 | 0.113974535 | 4.261518275 |
| Gpaa1 | 246.0551654 | -0.086572526 | 0.087973132 | -0.984079153 |
| Rps6-ps1 | 8.203160127 | 0.419536462 | 0.360977724 | 1.16222258 |
| Cyc1 | 1071.038313 | -0.101798415 | 0.092858783 | -1.09627125 |
| Sharpin | 316.269203 | 0.221095489 | 0.084334382 | 2.621653024 |
| Maf1 | 531.081438 | 0.305128111 | 0.080761899 | 3.778119578 |
| Hgh1 | 82.07062805 | 0.297487794 | 0.191586236 | 1.552761827 |
| Tssk5 | 6.475749014 | 0.928260138 | 0.457966843 | 2.026915598 |
| Mroh1 | 1129.595412 | -0.072166967 | 0.086808165 | -0.831338466 |
| Mir6954 | 0 | NA | NA | NA |
| Bop1 | 410.1160048 | -0.258556008 | 0.064235596 | -4.025120412 |
| Scx | 29.69111678 | 0.061078587 | 0.223085226 | 0.273790371 |
| Hsf1 | 313.4887952 | -0.025005998 | 0.099271413 | -0.25189526 |
| Dgat1 | 157.725041 | -0.388563128 | 0.121211475 | -3.205662902 |

| | | | | |
|---|---|---|---|---|
| Scrt1 | 1878.856055 | -0.422561243 | 0.078158368 | -5.406474777 |
| Fbxl6 | 146.2430035 | -0.012568247 | 0.095714265 | -0.131310069 |
| Slc52a2 | 132.5881942 | -0.014779297 | 0.107411647 | -0.137594917 |
| Adck5 | 90.29773825 | -0.302165317 | 0.122132858 | -2.474070622 |
| Cpsf1 | 643.6551908 | 0.094036227 | 0.078298388 | 1.20099825 |
| Slc39a4 | 6.380351065 | 1.563750836 | 0.49032303 | 3.189225756 |
| Tonsl | 636.9176432 | 0.068720982 | 0.084761869 | 0.810753496 |
| Cyhr1 | 972.3595972 | -0.214015748 | 0.056798845 | -3.767959503 |
| Kifc2 | 2493.40518 | -0.164272078 | 0.139052275 | -1.181369218 |
| Foxh1 | 0.909626782 | 0.398554355 | 0.920232748 | 0.433101687 |
| Ppp1r16a | 515.0001235 | -0.097163623 | 0.069165395 | -1.404801097 |
| Gpt | 84.05780739 | -1.277917873 | 0.190404391 | -6.711598735 |
| Mfsd3 | 34.06178562 | -0.027733298 | 0.222426018 | -0.124685493 |
| Recql4 | 21.17447754 | 0.187103567 | 0.304968605 | 0.613517471 |
| Lrrc14 | 184.1576737 | 0.034718886 | 0.085279259 | 0.407119923 |
| Lrrc24 | 405.4748953 | -0.124032736 | 0.078458391 | -1.580872798 |
| Arhgap39 | 936.4403093 | -0.096266908 | 0.093250779 | -1.032344278 |
| Gm17271 | 0.060174537 | 0.755286614 | 3.724945049 | 0.202764498 |
| Zfp251 | 456.6486897 | -0.256382701 | 0.066033336 | -3.882625288 |
| Zfp7 | 92.43301678 | -0.255583683 | 0.121900018 | -2.096666485 |
| Commd5 | 140.0820505 | 0.24304866 | 0.130493194 | 1.862538974 |
| Rpl8 | 2132.795094 | 0.314596592 | 0.101946158 | 3.085909233 |
| Zfp647 | 28.94240596 | 0.042428607 | 0.200431884 | 0.211685915 |
| 1110038F14Rik | 133.089957 | 0.701288196 | 0.110299643 | 6.358027775 |
| Mb | 1.797034208 | 0.597858692 | 0.793257324 | 0.753675603 |
| Apol6 | 78.23562678 | 5.840396747 | 0.497748713 | 11.733625 |
| Gm24753 | 2.783881763 | 0.750752348 | 0.7284724 | 1.030584477 |
| Rbfox2 | 3954.719732 | 0.075805259 | 0.080486569 | 0.94183738 |
| 1700109K24Rik | 0.04393175 | -0.206490061 | 3.724945049 | -0.055434391 |
| Gm24056 | 9.581094532 | -0.84116995 | 0.421640464 | -1.994993415 |
| Apol7a | 18.60379838 | 3.941679407 | 0.548127809 | 7.191168454 |
| Gm20688 | 0.311586316 | 1.597817682 | 1.967195023 | 0.812231458 |
| Apol9a | 239.6974814 | 8.024779352 | 0.705290974 | 11.37796973 |
| Apol7b | 0.961738709 | 0.884652075 | 1.193861731 | 0.741000445 |
| 4930571N24Rik | 0.157815785 | 1.201305391 | 3.465799399 | 0.346617116 |
| Gm6506 | 0.409269261 | 1.195729019 | 1.461092471 | 0.818380111 |
| Apol10a | 0.15100991 | 1.236179172 | 3.527597384 | 0.350430913 |
| Apol11a | 0 | NA | NA | NA |
| Apol7c | 0.292232717 | 0.755305041 | 2.000916511 | 0.377479538 |
| 1700025B11Rik | 0.199129965 | 0.755302396 | 2.233169354 | 0.338219936 |
| Apol10b | 22.02299047 | 6.282251607 | 0.909509508 | 6.907296242 |
| Gm8221 | 0.475668934 | 0.260822356 | 1.310371564 | 0.199044579 |
| Apol11b | 3.059439512 | 4.614260822 | 0.968715936 | 4.763275435 |
| Apol7e | 1.633903828 | 3.276331882 | 1.06177598 | 3.085709173 |
| Apol9b | 163.6719493 | 8.467166019 | 0.887639979 | 9.53896424 |
| Apol8 | 54.12186927 | 0.085115044 | 0.156497646 | 0.543874277 |
| Myh9 | 2978.335968 | 0.914597555 | 0.130684792 | 6.998500304 |

| | | | | |
|---|---|---|---|---|
| Gm22107 | 5.021758635 | 0.610693244 | 0.492759348 | 1.239333656 |
| Txn2 | 707.7197206 | 0.041045999 | 0.072214192 | 0.568392423 |
| Foxred2 | 319.3024116 | 0.116834291 | 0.093533668 | 1.249114816 |
| Eif3d | 737.7459319 | 0.331377201 | 0.096738301 | 3.425501563 |
| Gm17638 | 11.65945967 | 0.749761627 | 0.375853546 | 1.994823876 |
| Cacng2 | 970.2458113 | -0.281587738 | 0.088702807 | -3.174507642 |
| A730060N03Rik | 73.25342862 | -0.572355962 | 0.168905783 | -3.388610808 |
| Ift27 | 107.8713168 | 0.040984714 | 0.129939486 | 0.315413851 |
| Pvalb | 1103.933019 | -0.047261979 | 0.152062858 | -0.310805544 |
| Ncf4 | 100.2792551 | 3.812482679 | 0.308801751 | 12.34605267 |
| Csf2rb2 | 172.9194105 | 4.414105618 | 0.289807419 | 15.23116845 |
| Mir7676-1 | 0 | NA | NA | NA |
| Csf2rb | 284.283327 | 4.210374874 | 0.256187154 | 16.43476186 |
| Mir7676-2 | 0 | NA | NA | NA |
| Gm26329 | 0 | NA | NA | NA |
| Tex33 | 0.613719218 | 1.911486891 | 1.25822419 | 1.519194199 |
| Tst | 291.9109073 | 0.201699919 | 0.157863869 | 1.277682606 |
| Mpst | 172.564482 | 0.063619532 | 0.13695706 | 0.464521743 |
| Kctd17 | 1094.573162 | 0.04310621 | 0.05909384 | 0.729453534 |
| Tmprss6 | 5.777471174 | 2.239395454 | 0.491268926 | 4.558390193 |
| Il2rb | 153.2540098 | 5.765344733 | 0.487149887 | 11.83484772 |
| C1qtnf6 | 29.05525177 | 0.224225023 | 0.214728858 | 1.044223978 |
| Sstr3 | 170.7394951 | -0.657688075 | 0.166401635 | -3.952413545 |
| Gm6723 | 0.648304949 | -0.150639673 | 1.257168309 | -0.119824587 |
| Rac2 | 192.5000097 | 4.046200982 | 0.246460501 | 16.41723916 |
| Cyth4 | 257.1621641 | 1.810216144 | 0.148741764 | 12.17019415 |
| Elfn2 | 1215.602818 | -0.357232143 | 0.12123928 | -2.946504989 |
| Mfng | 23.1312152 | 0.880019723 | 0.285614745 | 3.081142481 |
| Card10 | 133.9315844 | -0.121643747 | 0.178349824 | -0.682051396 |
| Cdc42ep1 | 428.9155534 | 0.432570856 | 0.167067875 | 2.589192299 |
| Lgals2 | 1.456018128 | -0.899340078 | 1.13323152 | -0.793606657 |
| Gm26634 | 0.085327104 | 0.274389469 | 3.724945049 | 0.073662689 |
| Gga1 | 712.034101 | -0.003989325 | 0.069240215 | -0.057615725 |
| Mir6955 | 1.342239839 | -1.436800174 | 0.942576828 | -1.524332162 |
| Sh3bp1 | 1228.874401 | -0.008225705 | 0.056624866 | -0.145266661 |
| 1700027A07Rik | 1.879021917 | -0.592684636 | 0.681875246 | -0.869198053 |
| Lgals1 | 413.7512805 | 1.637527973 | 0.333590277 | 4.908800058 |
| Nol12 | 155.5073409 | 0.372310804 | 0.115715208 | 3.217475138 |
| Triobp | 275.7571062 | 0.924025546 | 0.17243409 | 5.35871731 |
| Mir6956 | 0 | NA | NA | NA |
| H1f0 | 1010.011615 | 0.378724033 | 0.102779529 | 3.684819688 |
| Gcat | 70.63897891 | -0.020047268 | 0.152603552 | -0.131368292 |
| Ankrd54 | 161.076979 | 0.13946778 | 0.144720326 | 0.963705536 |
| Eif3l | 629.3207194 | 0.259123972 | 0.077294967 | 3.35240419 |
| Gm25072 | 0 | NA | NA | NA |
| Micall1 | 863.8231058 | -0.224131552 | 0.081507059 | -2.749842224 |
| 1700088E04Rik | 67.10927695 | 0.267700026 | 0.149537999 | 1.790180607 |

| | | | | |
|---|---|---|---|---|
| Polr2f | 144.8684693 | 0.123287387 | 0.193491334 | 0.637172654 |
| Sox10 | 508.9917057 | -0.014765479 | 0.108713165 | -0.135820525 |
| Gm10863 | 17.85705234 | -0.954135824 | 0.320863511 | -2.973650134 |
| Pick1 | 359.5554522 | -0.215614952 | 0.089543474 | -2.407935966 |
| Slc16a8 | 3.295224942 | -0.915218959 | 0.515547011 | -1.775238611 |
| Lgals1-ps2 | 0 | NA | NA | NA |
| Baiap2l2 | 8.450121399 | -0.635194151 | 0.436574488 | -1.454950229 |
| Pla2g6 | 246.7582411 | -0.372075291 | 0.106303806 | -3.500112607 |
| Gm20420 | 0 | NA | NA | NA |
| Maff | 106.5092356 | 3.546085994 | 0.482381253 | 7.351210219 |
| Tmem184b | 1712.559106 | 1.197913137 | 0.127933462 | 9.363563854 |
| Mir1943 | 2.514149661 | 0.867171972 | 0.709261002 | 1.222641551 |
| Csnk1e | 892.3901685 | 0.038889585 | 0.078647545 | 0.494479332 |
| Gm16059 | 0.387099777 | -0.877979378 | 1.859292104 | -0.472211642 |
| AC165278.1 | 0 | NA | NA | NA |
| Kcnj4 | 701.1998511 | -0.476203956 | 0.152014754 | -3.132616698 |
| Kdelr3 | 10.19668411 | 0.649457103 | 0.459403669 | 1.413695943 |
| Ddx17 | 3680.454381 | -0.276936062 | 0.063222894 | -4.38031292 |
| Dmc1 | 9.014701878 | -1.076485996 | 0.386152681 | -2.787721152 |
| Gm27903 | 0.358948382 | 0.247046166 | 1.431444914 | 0.172585172 |
| Fam227a | 83.10424768 | -1.068408014 | 0.247135511 | -4.323166708 |
| Cby1 | 177.468213 | -0.273274803 | 0.104967898 | -2.603413136 |
| Tomm22 | 344.3752573 | 0.087099365 | 0.098424016 | 0.884940163 |
| Josd1 | 713.1554615 | 0.00988055 | 0.085994918 | 0.114896908 |
| Gm26884 | 1.046429311 | 0.661377173 | 1.167073265 | 0.566697218 |
| Gtpbp1 | 638.7878775 | 0.125225148 | 0.062828965 | 1.99311174 |
| n-R5s40 | 1.378629347 | -0.511454356 | 0.920421422 | -0.555674111 |
| Sun2 | 786.962264 | 0.369134555 | 0.071340406 | 5.174270475 |
| Gm16575 | 0.364940181 | -1.102041118 | 1.735229899 | -0.635098046 |
| Gm16576 | 14.37465387 | -0.423585645 | 0.332988117 | -1.272074357 |
| Dnal4 | 230.0228909 | -0.223605358 | 0.083143746 | -2.689382787 |
| Nptxr | 4627.776829 | -0.345091921 | 0.152628477 | -2.260993023 |
| Npcd | 0.498288516 | 1.773874898 | 1.327470789 | 1.336281682 |
| Cbx6 | 4151.173169 | -0.299282588 | 0.088463303 | -3.383126992 |
| Apobec3 | 148.4452752 | 2.791047812 | 0.196301845 | 14.21814355 |
| Cbx7 | 316.6582599 | -0.103470572 | 0.085068865 | -1.216315414 |
| Pdgfb | 148.5994786 | 0.296403629 | 0.160914265 | 1.841997223 |
| Mir7213 | 0 | NA | NA | NA |
| Rpl3 | 1750.858766 | 0.305665872 | 0.095484437 | 3.201211449 |
| Snord83b | 1.863983961 | 0.107665236 | 0.723104018 | 0.148893151 |
| Gm22077 | 0 | NA | NA | NA |
| AC161199.1 | 2.341657818 | 0.249298813 | 0.75578594 | 0.329853732 |
| Gm24204 | 0.553939527 | 0.201978117 | 1.406113805 | 0.143642795 |
| Syngr1 | 4833.272579 | -0.267897077 | 0.060019195 | -4.46352335 |
| Tab1 | 241.9134514 | 0.137935904 | 0.085725961 | 1.609033038 |
| Mgat3 | 1733.281968 | -0.220671771 | 0.070811448 | -3.116329042 |
| Mief1 | 547.7891883 | 0.13530813 | 0.080620729 | 1.678329275 |

| | | | | |
|---|---|---|---|---|
| Atf4 | 1610.207181 | 0.792601388 | 0.158435734 | 5.002668082 |
| Rps19bp1 | 105.055002 | 0.046168669 | 0.154686624 | 0.298465816 |
| Cacna1i | 1385.507454 | -0.572769352 | 0.120589845 | -4.749731232 |
| Enthd1 | 1.187168766 | 3.342704841 | 1.08189982 | 3.089662073 |
| Gm23220 | 0 | NA | NA | NA |
| Grap2 | 30.59568962 | 3.679297265 | 0.419673263 | 8.767051864 |
| Mir6957 | 0 | NA | NA | NA |
| Fam83f | 18.37852399 | 1.402570627 | 0.403700912 | 3.474281539 |
| Tnrc6b | 3449.044664 | -0.316652128 | 0.077660107 | -4.077410401 |
| Mir5113 | 0 | NA | NA | NA |
| Adsl | 284.5366327 | 0.196647099 | 0.07997175 | 2.458957059 |
| Gm6612 | 1.14679352 | -1.641340644 | 1.161185279 | -1.413504524 |
| Sgsm3 | 461.0312359 | -0.250894157 | 0.06778535 | -3.701303583 |
| Mkl1 | 795.5391381 | 0.24764723 | 0.084000527 | 2.948162829 |
| Gm25131 | 0 | NA | NA | NA |
| 4930483J18Rik | 0.130580445 | -1.034201839 | 3.667847045 | -0.281964277 |
| Mchr1 | 166.8774582 | 0.289582001 | 0.187027795 | 1.54833671 |
| Slc25a17 | 303.5824259 | 0.185648158 | 0.097304948 | 1.907900489 |
| St13 | 1980.343047 | -0.114006649 | 0.064624838 | -1.764130518 |
| Xpnpep3 | 220.8414576 | -0.100353247 | 0.110282657 | -0.909963993 |
| Dnajb7 | 0.134148234 | -0.20649159 | 3.624632408 | -0.056968974 |
| Gm17025 | 0.441788749 | -1.199706509 | 1.233644925 | -0.972489316 |
| Rbx1 | 792.4993545 | 0.056308859 | 0.081874366 | 0.687747116 |
| Gm5218 | 0.038981395 | 0.274393828 | 3.724945049 | 0.073663859 |
| 1110025M09Rik | 12.66128553 | -0.170399154 | 0.309832544 | -0.549971775 |
| Ep300 | 2415.831866 | -0.069927557 | 0.063711415 | -1.097567161 |
| Gm23880 | 0 | NA | NA | NA |
| Gm22067 | 12.43328101 | -1.035027522 | 0.289756909 | -3.572054674 |
| L3mbtl2 | 382.5292509 | 0.001355963 | 0.063870368 | 0.021229919 |
| Gm18284 | 12.73109718 | 0.009208253 | 0.287708551 | 0.032005489 |
| Chadl | 117.5180505 | -0.06376524 | 0.131761366 | -0.483944894 |
| Rangap1 | 1965.883829 | -0.213243729 | 0.065988721 | -3.231517853 |
| Zc3h7b | 3000.898726 | 0.159051365 | 0.079578083 | 1.998683039 |
| Gm17597 | 7.602086333 | -0.571104577 | 0.343617082 | -1.662037795 |
| Tef | 3374.752106 | 0.01920196 | 0.106993916 | 0.179467776 |
| Gm8444 | 1.234579709 | 0.157910624 | 0.858529564 | 0.183931493 |
| Tob2 | 667.3737093 | 0.190019815 | 0.111046208 | 1.711177889 |
| Phf5a | 193.4131225 | 0.436308864 | 0.136695339 | 3.191834239 |
| Aco2 | 4367.876781 | -0.114808033 | 0.067949658 | -1.689604288 |
| Polr3h | 303.152543 | -0.170206661 | 0.094609038 | -1.799052868 |
| Csdc2 | 982.7828558 | -0.555532899 | 0.112265272 | -4.948394908 |
| Pmm1 | 652.8758253 | -0.032931358 | 0.057409936 | -0.573617752 |
| Gm5805 | 68.93469726 | 0.788917304 | 0.215661336 | 3.658130473 |
| 1700029P11Rik | 0.075043205 | 0.755286614 | 3.724945049 | 0.202764498 |
| Xrcc6 | 153.2673552 | -0.281137682 | 0.119482689 | -2.352957446 |
| Desi1 | 1377.267412 | -0.285481535 | 0.078499338 | -3.63673813 |
| Nhp2l1 | 547.7572345 | 0.139045483 | 0.105873629 | 1.31331554 |

| | | | | |
|---|---|---|---|---|
| 4930407I10Rik | 0.062426873 | 0.755288839 | 3.724945049 | 0.202765096 |
| Mei1 | 19.77925728 | -0.5089842 | 0.291398042 | -1.74669739 |
| Ccdc134 | 79.33616336 | -0.34188554 | 0.148202289 | -2.306884338 |
| Srebf2 | 2183.707136 | -0.434676218 | 0.079150298 | -5.491782452 |
| Mir33 | 0.155921052 | 0.777072206 | 2.829227972 | 0.274658746 |
| Shisa8 | 125.8398763 | -0.039918549 | 0.202699726 | -0.1969344 |
| Tnfrsf13c | 6.674742414 | 1.232695622 | 0.469418155 | 2.626007556 |
| Cenpm | 13.06399243 | 0.55387951 | 0.327739012 | 1.690001766 |
| 1500009C09Rik | 1247.839228 | 0.955357996 | 0.137026666 | 6.972058997 |
| 3-Sep | 4869.520885 | 0.035483214 | 0.069285577 | 0.512129874 |
| Wbp2nl | 0.428891288 | 1.594727963 | 1.540523798 | 1.035185543 |
| Naga | 263.3133539 | 0.535961812 | 0.121886944 | 4.397204449 |
| Fam109b | 4.886605523 | 0.278510961 | 0.480195067 | 0.579995464 |
| Smdt1 | 561.4413452 | 0.084530288 | 0.109248377 | 0.773744102 |
| n-R5s41 | 0 | NA | NA | NA |
| Ndufa6 | 517.8924798 | 0.129013613 | 0.107935151 | 1.195288207 |
| Cyp2d22 | 207.7231852 | -0.017208164 | 0.134104785 | -0.128318789 |
| Cyp2d11 | 0 | NA | NA | NA |
| Mir7684 | 0 | NA | NA | NA |
| Cyp2d10 | 0 | NA | NA | NA |
| Gm27331 | 0 | NA | NA | NA |
| Gm25990 | 0 | NA | NA | NA |
| Cyp2d9 | 0.049553047 | 0.274391604 | 3.724945049 | 0.073663262 |
| Gm27622 | 0 | NA | NA | NA |
| Gm27825 | 0 | NA | NA | NA |
| Gm5806 | 3.903672454 | 1.140556031 | 0.600224148 | 1.900216835 |
| Gm28023 | 0 | NA | NA | NA |
| Cyp2d12 | 0.066050875 | 0.274391604 | 3.724945049 | 0.073663262 |
| Gm27924 | 0 | NA | NA | NA |
| Gm27742 | 0 | NA | NA | NA |
| Cyp2d34 | 0.18989612 | 0.755298873 | 2.65941639 | 0.284009257 |
| Gm27463 | 0.194279094 | 0.274391173 | 3.724945049 | 0.073663147 |
| Cyp2d37-ps | 0.066050875 | 0.274391604 | 3.724945049 | 0.073663262 |
| Cyp2d40 | 0.493912024 | 2.056966279 | 1.529934988 | 1.344479534 |
| Cyp2d41-ps | 0.567195248 | 0.594832084 | 1.385159617 | 0.429432158 |
| Gm27618 | 0 | NA | NA | NA |
| Cyp2d26 | 2.149324495 | 0.752487768 | 0.701353074 | 1.072908633 |
| Tcf20 | 4578.253049 | -0.302944716 | 0.097007297 | -3.122906481 |
| Gm20324 | 2.615112804 | 0.542508759 | 0.678562525 | 0.799497083 |
| Mir3080 | 2.487744732 | -0.302477176 | 0.562304839 | -0.537923836 |
| Gm29019 | 5.11266864 | -0.361389648 | 0.557586812 | -0.648131627 |
| AW121686 | 10.56313198 | -0.305967894 | 0.343096002 | -0.891785077 |
| Nfam1 | 67.01581601 | 1.171501906 | 0.237219147 | 4.938479547 |
| Serhl | 61.58247766 | 0.55998619 | 0.182358714 | 3.07079479 |
| Gm17206 | 0.066050875 | 0.274391604 | 3.724945049 | 0.073663262 |
| Rrp7a | 918.7979819 | 0.174805505 | 0.075571229 | 2.313122443 |
| Poldip3 | 1119.919797 | 0.201465441 | 0.077474882 | 2.600396876 |

| | | | | |
|---|---|---|---|---|
| Rnu12 | 255.543187 | 0.128984744 | 0.165627994 | 0.778761737 |
| Cyb5r3 | 666.3127489 | 0.358831365 | 0.117937554 | 3.042553889 |
| A4galt | 36.02363458 | 0.844368904 | 0.266677579 | 3.166253821 |
| Arfgap3 | 242.8767015 | 0.324369525 | 0.079969331 | 4.056174032 |
| 1700001L05Rik | 36.6738785 | -0.247651517 | 0.197822695 | -1.251886278 |
| Pacsin2 | 643.9682478 | -0.02364809 | 0.089478947 | -0.264286638 |
| Gm5417 | 28.18046227 | 0.05233368 | 0.249583532 | 0.209684027 |
| Ttll1 | 251.422686 | -0.5755591 | 0.111127579 | -5.179264267 |
| Gm24575 | 0 | NA | NA | NA |
| Bik | 0.708533877 | 2.084146954 | 1.330399357 | 1.566557397 |
| Mcat | 123.3968876 | 0.6358109 | 0.120185277 | 5.290256147 |
| Tspo | 459.8898812 | 5.03163464 | 0.344167491 | 14.61972664 |
| Ttll12 | 215.0437835 | -0.20709965 | 0.114187991 | -1.813672764 |
| Scube1 | 480.4921202 | -0.853379758 | 0.172104114 | -4.958508776 |
| Mpped1 | 824.048876 | -0.402066993 | 0.1425541 | -2.820451972 |
| Efcab6 | 85.58806344 | -0.408638719 | 0.294086714 | -1.389517785 |
| Rpl31-ps22 | 0 | NA | NA | NA |
| Sult4a1 | 2771.319555 | -0.281297842 | 0.086070644 | -3.268220502 |
| Pnpla5 | 0 | NA | NA | NA |
| Pnpla3 | 102.6316791 | -0.676747261 | 0.186369388 | -3.631214697 |
| Mir6392 | 3.244976357 | -0.596013909 | 0.510243168 | -1.168097774 |
| Samm50 | 528.8198655 | -0.008355445 | 0.070349096 | -0.118771171 |
| Parvb | 380.2549727 | 0.189685552 | 0.102928161 | 1.842892652 |
| Gm22890 | 0.046345709 | 0.274391604 | 3.724945049 | 0.073663262 |
| Parvg | 82.64255846 | 1.383957871 | 0.182845884 | 7.568985646 |
| 1810041L15Rik | 649.3280647 | -0.532807402 | 0.126675084 | -4.206094723 |
| Ldoc1l | 614.5341085 | -0.127617466 | 0.082771743 | -1.541799917 |
| Prr5 | 124.3176796 | 0.30711573 | 0.134037725 | 2.291263375 |
| Arhgap8 | 19.60055845 | 4.745473354 | 0.575705191 | 8.242887901 |
| Phf21b | 132.294507 | -0.508426493 | 0.174568791 | -2.912470729 |
| Gm4217 | 13.23963404 | 0.24249124 | 0.367336731 | 0.660133388 |
| Gm29666 | 48.12944454 | -0.438954285 | 0.211336175 | -2.077042823 |
| Nup50 | 602.2802685 | 0.017669159 | 0.082585838 | 0.213949013 |
| 5031439G07Rik | 1832.693521 | -0.410895373 | 0.056794972 | -7.234713869 |
| Mir1249 | 0 | NA | NA | NA |
| Upk3a | 0.636985128 | -0.868322264 | 1.328438448 | -0.653641322 |
| Fam118a | 232.4075255 | 0.055849324 | 0.085221059 | 0.655346517 |
| Smc1b | 5.078404989 | 1.313842118 | 0.533398642 | 2.463152346 |
| Gm10923 | 2.764924986 | 0.386148228 | 0.687344409 | 0.561797293 |
| Ribc2 | 3.994063061 | 0.527018301 | 0.502418313 | 1.04896316 |
| Fbln1 | 204.5288485 | 0.817662597 | 0.180170252 | 4.538277489 |
| Atxn10 | 2123.828845 | 0.009697533 | 0.062535375 | 0.155072764 |
| Gm23517 | 0 | NA | NA | NA |
| 7530416G11Rik | 0 | NA | NA | NA |
| Gm4825 | 0.947696482 | 2.611796943 | 1.138357082 | 2.294356476 |
| Wnt7b | 268.9427193 | -0.440749816 | 0.133058582 | -3.312449381 |
| AU022754 | 9.118481155 | -0.313517895 | 0.356108941 | -0.880398829 |

| | | | | |
|---|---|---|---|---|
| D130051D11Rik | 5.759629403 | 0.062416477 | 0.440967097 | 0.141544523 |
| Gm37699 | 45.08239792 | -0.584171197 | 0.252625587 | -2.312399172 |
| Mirlet7c-2 | 5.369208344 | 0.859953311 | 0.442763699 | 1.942239872 |
| Mirlet7b | 13.61717214 | 0.483225909 | 0.357114539 | 1.353139836 |
| Ppara | 39.64814777 | -0.223526251 | 0.207101561 | -1.079307421 |
| Cdpf1 | 48.27012333 | -0.180816249 | 0.191090883 | -0.946231687 |
| Pkdrej | 11.56756557 | 0.831186742 | 0.312967278 | 2.655826346 |
| Ttc38 | 59.42181862 | 0.410648847 | 0.147206636 | 2.789608253 |
| Gtse1 | 17.82322006 | 1.527144533 | 0.351770991 | 4.341303207 |
| Trmu | 131.3669997 | -0.024682405 | 0.123887557 | -0.199232321 |
| Celsr1 | 124.8124239 | -0.416336188 | 0.154322001 | -2.697840788 |
| Gramd4 | 450.6965781 | -0.292355909 | 0.071215982 | -4.105200812 |
| Gm15722 | 0 | NA | NA | NA |
| Cerk | 1311.25289 | -0.273142581 | 0.083350355 | -3.277041586 |
| Gm24011 | 0 | NA | NA | NA |
| Gm15569 | 1.568615446 | 0.340592839 | 0.926573781 | 0.367583074 |
| Gm22818 | 0.772793883 | -1.578257143 | 1.080683282 | -1.460425242 |
| Tbc1d22a | 375.15338 | 0.271944671 | 0.07020513 | 3.873572658 |
| Gm23416 | 0.044544648 | 0.755286614 | 3.724945049 | 0.202764498 |
| n-R5s42 | 0 | NA | NA | NA |
| Fam19a5 | 1108.298912 | -0.085571202 | 0.075875758 | -1.127780522 |
| Mir6393 | 0 | NA | NA | NA |
| Zdhhc25 | 0 | NA | NA | NA |
| Brd1 | 988.6755636 | 0.02057084 | 0.114953635 | 0.178949017 |
| Zbed4 | 103.903103 | -0.085460329 | 0.114085362 | -0.749091099 |
| Alg12 | 119.1075428 | 0.013655061 | 0.114987087 | 0.118752991 |
| Creld2 | 178.5176635 | 0.246950109 | 0.165795513 | 1.489486076 |
| Gm23144 | 0 | NA | NA | NA |
| Pim3 | 592.2908598 | 0.594380259 | 0.171991026 | 3.455879485 |
| Gm20621 | 2.301900616 | -0.727191373 | 0.659142545 | -1.103238409 |
| Ttll8 | 19.07562425 | -0.172122689 | 0.364254404 | -0.47253427 |
| Mlc1 | 1260.237927 | -0.23128473 | 0.129654402 | -1.783855597 |
| Mov10l1 | 0.232306593 | -1.197690402 | 2.17831276 | -0.549824811 |
| Gm23737 | 0 | NA | NA | NA |
| Panx2 | 872.8564831 | -0.334242989 | 0.099196271 | -3.369511637 |
| 1810021B22Rik | 9.773541415 | 0.501065118 | 0.345964211 | 1.448314889 |
| Trabd | 275.7218993 | 0.076719075 | 0.092119678 | 0.832819627 |
| Selo | 248.0954824 | 0.086045909 | 0.089877216 | 0.957371761 |
| Tubgcp6 | 650.1368911 | -0.116115804 | 0.083574816 | -1.389363569 |
| Hdac10 | 82.94525066 | -0.211223988 | 0.156473096 | -1.349906104 |
| Mapk12 | 96.05029267 | -0.161345698 | 0.167018145 | -0.966036941 |
| Mapk11 | 187.7470408 | -0.582682584 | 0.153528296 | -3.795278133 |
| Plxnb2 | 1637.125997 | 0.259945691 | 0.126206487 | 2.059685659 |
| mmu-mir-7118 | 0 | NA | NA | NA |
| Dennd6b | 661.1196248 | 0.356394086 | 0.182526552 | 1.952560228 |
| Mir6958 | 0 | NA | NA | NA |
| Gm26798 | 0.47411084 | 0.241787113 | 1.287202753 | 0.187839183 |

| | | | | |
|---|---|---|---|---|
| Ppp6r2 | 1298.582558 | -0.03290117 | 0.053575916 | -0.614103738 |
| Sbf1 | 2446.17373 | -0.147750555 | 0.052616291 | -2.808076214 |
| Mir6959 | 1.302949929 | 0.29260989 | 0.767291966 | 0.381354038 |
| Adm2 | 0.810290415 | 1.703791387 | 1.375255008 | 1.238891243 |
| Miox | 0.11193333 | 0.75528439 | 3.724945049 | 0.202763901 |
| Lmf2 | 236.2729926 | 0.206553008 | 0.117734566 | 1.754395637 |
| Ncaph2 | 806.3681184 | -0.038963285 | 0.063327133 | -0.615269993 |
| Sco2 | 38.3484847 | 0.063358868 | 0.176520781 | 0.358931495 |
| Tymp | 10.59392881 | -0.564111532 | 0.33141105 | -1.702150646 |
| Odf3b | 13.0856276 | -0.359562734 | 0.287242637 | -1.251773543 |
| Klhdc7b | 7.376514687 | -1.436847055 | 0.384284487 | -3.739019146 |
| Syce3 | 0 | NA | NA | NA |
| Chkb | 231.7816191 | 0.099713869 | 0.095673794 | 1.042227606 |
| Cpt1b | 2.011645178 | -0.134923677 | 0.71692743 | -0.188197118 |
| C730034F03Rik | 13.21654071 | -0.565071971 | 0.298512056 | -1.892961974 |
| Mapk8ip2 | 3763.788627 | -0.062582713 | 0.054707848 | -1.143943978 |
| Arsa | 324.9924385 | -0.080456954 | 0.097863327 | -0.822135894 |
| C230037L18Rik | 26.54613617 | -0.741147585 | 0.268412991 | -2.761220991 |
| Shank3 | 2814.903766 | -0.571755946 | 0.132924153 | -4.301369873 |
| Gm15609 | 24.97572873 | -0.661170864 | 0.346156769 | -1.910033037 |
| Acr | 23.80281401 | 1.101199193 | 0.416603161 | 2.643280936 |
| Rabl2 | 143.4438333 | -0.212884061 | 0.134952476 | -1.577474283 |
| Syt10 | 129.1730868 | -0.591471701 | 0.176167972 | -3.357430379 |
| Gm24643 | 0 | NA | NA | NA |
| Gm24415 | 0 | NA | NA | NA |
| Alg10b | 588.5402971 | 0.20781687 | 0.08087521 | 2.569599151 |
| Mir7648 | 0 | NA | NA | NA |
| Cpne8 | 261.9200795 | 0.219837381 | 0.106668935 | 2.060931618 |
| Gm6746 | 0 | NA | NA | NA |
| Rpl31-ps8 | 113.0537037 | 0.481340531 | 0.242517528 | 1.984765948 |
| Tcea1-ps1 | 127.5401109 | 0.525945002 | 0.110220615 | 4.77174805 |
| Kif21a | 3097.142762 | -0.226811369 | 0.060182333 | -3.768736719 |
| Abcd2 | 509.0775856 | -0.289275018 | 0.096564551 | -2.99566473 |
| CN725425 | 2.738760097 | -0.802658158 | 0.647332735 | -1.239946807 |
| Slc2a13 | 1660.411825 | -0.148477839 | 0.100618264 | -1.475654953 |
| Gm15383 | 16.38285574 | -1.299346537 | 0.370098726 | -3.510810622 |
| Gm15381 | 25.44622861 | -0.915222649 | 0.284620804 | -3.215585911 |
| Gm24647 | 4.156244294 | -0.588709288 | 0.546742981 | -1.076756918 |
| 4933438A12Rik | 0.060174537 | 0.755286614 | 3.724945049 | 0.202764498 |
| Gm25389 | 0 | NA | NA | NA |
| Lrrk2 | 1067.387671 | -0.519483708 | 0.139446713 | -3.725320566 |
| Gm15382 | 4.663180715 | -1.2461494 | 0.514056707 | -2.424147733 |
| Smgc | 0.890372413 | -0.149378657 | 1.362651583 | -0.109623516 |
| Muc19 | 0.988759962 | -0.961769494 | 1.310663996 | -0.733803245 |
| Cntn1 | 4874.212289 | -0.340902336 | 0.066575045 | -5.120572334 |
| Gm26760 | 2.85183316 | -0.131150349 | 0.577777109 | -0.226991251 |
| Pdzrn4 | 94.36282265 | -0.330255055 | 0.112670196 | -2.931166067 |

| | | | | |
|---|---|---|---|---|
| Gm37304 | 37.54844209 | -0.635403203 | 0.306874755 | -2.070561988 |
| Gm4335 | 6.712093293 | 0.198123107 | 0.438949712 | 0.451357186 |
| Gm24447 | 11246.06972 | 0.179634925 | 0.147219046 | 1.220188083 |
| Gxylt1 | 308.8367025 | -0.058445713 | 0.097491491 | -0.599495532 |
| Yaf2 | 648.3196561 | -0.012708559 | 0.075324841 | -0.168716709 |
| Gm26371 | 0.04393175 | -0.206490061 | 3.724945049 | -0.055434391 |
| Zcrb1 | 380.1198483 | 0.025926361 | 0.123936373 | 0.209190897 |
| Pphln1 | 620.7309094 | 0.06297504 | 0.055797797 | 1.128629497 |
| Prickle1 | 831.5948449 | -0.229925286 | 0.107042016 | -2.147991001 |
| Adamts20 | 204.9752347 | -0.081274093 | 0.199011638 | -0.408388644 |
| Pus7l | 33.70876001 | 0.026031414 | 0.190760692 | 0.136461103 |
| Irak4 | 54.36204711 | 1.229456715 | 0.254955699 | 4.822236645 |
| Twf1 | 752.74737 | 0.231392948 | 0.083567159 | 2.768945963 |
| Tmem117 | 131.7499303 | -0.202221139 | 0.105161763 | -1.922953112 |
| Gm25546 | 0 | NA | NA | NA |
| Gm23129 | 0 | NA | NA | NA |
| 1700129L04Rik | 4.754384054 | 0.672882876 | 0.653448433 | 1.029741356 |
| Nell2 | 3266.316947 | -0.009856223 | 0.122989983 | -0.080138422 |
| Gm24668 | 0 | NA | NA | NA |
| Dbx2 | 163.3891668 | 0.024664457 | 0.151341906 | 0.162971764 |
| Gm8849 | 1.166878852 | 2.563827487 | 1.109621386 | 2.310542604 |
| A130051J06Rik | 4.097955438 | 1.018410405 | 0.618050804 | 1.647777818 |
| Ano6 | 622.2228295 | 0.430451226 | 0.101819372 | 4.227596562 |
| Gm17546 | 0.627185525 | -0.390575438 | 1.184275075 | -0.329801282 |
| Gm25070 | 0.044544648 | 0.755286614 | 3.724945049 | 0.202764498 |
| D030018L15Rik | 0.718519848 | -0.135392577 | 1.367537531 | -0.099004652 |
| 2610037D02Rik | 115.5236633 | -0.662657803 | 0.22622616 | -2.929182913 |
| 4833422M21Rik | 10.72830272 | -0.400845028 | 0.370455186 | -1.082033789 |
| Arid2 | 997.3079313 | -0.313693241 | 0.069668619 | -4.502647636 |
| Gm25397 | 0 | NA | NA | NA |
| Scaf11 | 1520.248971 | 0.279686769 | 0.067478424 | 4.144832556 |
| Slc38a1 | 3775.966176 | -0.340455055 | 0.048028797 | -7.088560915 |
| Gm22045 | 0 | NA | NA | NA |
| Slc38a2 | 2617.581037 | -0.058418074 | 0.076729911 | -0.761346821 |
| Gm38144 | 0.240186774 | 0.27441168 | 2.114576897 | 0.129771436 |
| Gm8888 | 0 | NA | NA | NA |
| Slc38a4 | 9.499651179 | -0.43259678 | 0.326176681 | -1.326265198 |
| Amigo2 | 212.2260951 | 0.822643276 | 0.16767198 | 4.906265657 |
| Pced1b | 100.5813245 | 0.809041476 | 0.131743158 | 6.141051161 |
| Rpap3 | 291.5755603 | 0.220173584 | 0.077363278 | 2.845970188 |
| Endou | 46.06882177 | 1.061489085 | 0.207705763 | 5.110542296 |
| Rapgef3 | 390.02648 | -0.040037163 | 0.113023585 | -0.354237241 |
| Rapgef3os1 | 0.072025462 | 0.755288839 | 3.724945049 | 0.202765096 |
| Rapgef3os2 | 3.594151187 | 0.45825953 | 0.522110442 | 0.877706118 |
| Slc48a1 | 1148.386551 | -0.115224694 | 0.077210542 | -1.492344065 |
| Hdac7 | 379.135267 | 0.276630852 | 0.164704878 | 1.679554702 |
| Vdr | 23.71495927 | 1.592492553 | 0.338074352 | 4.710480241 |

| | | | | |
|---|---|---|---|---|
| Gm22202 | 0 | NA | NA | NA |
| Tmem106c | 169.9404391 | -0.264958139 | 0.143664375 | -1.8442856 |
| Col2a1 | 22.89251041 | -0.492280094 | 0.29257978 | -1.682549948 |
| Senp1 | 454.6842185 | 0.351179785 | 0.079603451 | 4.411615083 |
| Gm25668 | 0.044544648 | 0.755286614 | 3.724945049 | 0.202764498 |
| Pfkm | 3358.190513 | -0.108855712 | 0.064786134 | -1.680231636 |
| Gm26513 | 7.485808625 | 0.114118873 | 0.388606257 | 0.293661955 |
| Asb8 | 417.0210594 | 0.118562872 | 0.095254579 | 1.244694731 |
| Ccdc184 | 256.2935464 | -0.621347662 | 0.125570685 | -4.94819045 |
| Olfr288 | 1.13985991 | 0.667230307 | 0.893324108 | 0.74690731 |
| Olfr287 | 28.52921329 | -0.368674228 | 0.219390698 | -1.680446034 |
| Olfr286 | 0.16123844 | -1.17716678 | 2.795194053 | -0.421139555 |
| H1fnt | 0 | NA | NA | NA |
| Zfp641 | 111.9084074 | -0.443879986 | 0.124164685 | -3.57492942 |
| Olfr285 | 0 | NA | NA | NA |
| Olfr284 | 0 | NA | NA | NA |
| Olfr283 | 0.082913145 | -0.206492286 | 3.724945049 | -0.055434988 |
| 1700031M16Rik | 0.045882455 | 0.755288839 | 3.724945049 | 0.202765096 |
| Olfr282 | 0 | NA | NA | NA |
| Olfr281 | 0.061920613 | 0.274393828 | 3.724945049 | 0.073663859 |
| Lalba | 0.845223607 | -1.083738568 | 1.33483603 | -0.811888909 |
| Olfr279 | 0.133280846 | 0.671929825 | 3.652577664 | 0.183960448 |
| Kansl2 | 403.2058431 | 0.040865841 | 0.075273113 | 0.542900898 |
| Snora34 | 1.248930875 | 1.303510274 | 1.003135753 | 1.299435565 |
| mmu-mir-1291 | 1.156326783 | -1.394790263 | 1.037809987 | -1.343974601 |
| Snora2b | 4.632530903 | -1.078233401 | 0.487175617 | -2.213233509 |
| Ccnt1 | 886.6569998 | -0.122531071 | 0.046091352 | -2.658439526 |
| 9330020H09Rik | 18.2002576 | 0.05960565 | 0.280333117 | 0.212624362 |
| 4930415O20Rik | 0.192434788 | -0.206487112 | 2.657301408 | -0.077705567 |
| Gm29331 | 0.089062654 | -0.20649451 | 3.724945049 | -0.055435586 |
| Adcy6 | 518.3618143 | -0.350551167 | 0.097014762 | -3.613379673 |
| Cacnb3 | 1158.604023 | -0.264399495 | 0.096302303 | -2.74551581 |
| Ddx23 | 691.4598963 | 0.094949191 | 0.093216238 | 1.018590678 |
| Rnd1 | 252.5003933 | 0.277338628 | 0.219154888 | 1.265491407 |
| Ccdc65 | 49.36712147 | -0.209915429 | 0.151024349 | -1.389944273 |
| Fkbp11 | 8.263769739 | 0.89719107 | 0.442676122 | 2.026743766 |
| Arf3 | 8081.452134 | 0.119511095 | 0.068045579 | 1.75633888 |
| Wnt10b | 8.994095922 | 1.514602367 | 0.50411842 | 3.004457499 |
| Wnt1 | 0.424046041 | -0.194587129 | 1.614258639 | -0.120542721 |
| Ddn | 6153.970663 | -0.102769957 | 0.183323978 | -0.560592008 |
| B130046B21Rik | 36.69080057 | -0.257805372 | 0.238086548 | -1.082822083 |
| Prkag1 | 322.4183075 | -0.04320466 | 0.084475493 | -0.511446077 |
| Kmt2d | 5538.759192 | -0.071119355 | 0.082164889 | -0.865568694 |
| Rhebl1 | 33.80626636 | -0.133717397 | 0.178871699 | -0.747560388 |
| Dhh | 5.749732932 | 1.89567269 | 0.589473884 | 3.215872223 |
| Lmbr1l | 116.8431874 | 0.116838613 | 0.155894131 | 0.749474093 |
| Tuba1b | 2389.31285 | -0.04342074 | 0.133621909 | -0.324952247 |

| | | | | |
|---|---|---|---|---|
| Tuba1a | 4753.134624 | -0.065906528 | 0.115933807 | -0.568484123 |
| 4930578M01Rik | 10.95583967 | -0.526295806 | 0.358405972 | -1.468434814 |
| Gm8973 | 0.060174537 | 0.755286614 | 3.724945049 | 0.202764498 |
| Tuba1c | 280.0639814 | 0.492459171 | 0.191090065 | 2.577105041 |
| Prph | 45.07156546 | 0.091646506 | 0.334317464 | 0.274130176 |
| Troap | 4.659496152 | 1.994317123 | 0.714072411 | 2.792877994 |
| Mir6960 | 0 | NA | NA | NA |
| C1ql4 | 6.463082896 | -0.201679156 | 0.407558146 | -0.494847564 |
| Dnajc22 | 1.159556269 | 0.85637822 | 0.85821157 | 0.997863755 |
| Spats2 | 344.7749557 | -0.013134891 | 0.063177534 | -0.207904464 |
| Gm25183 | 0 | NA | NA | NA |
| Gm25057 | 0 | NA | NA | NA |
| Kcnh3 | 619.2177828 | -0.42156334 | 0.163769916 | -2.57411953 |
| Mcrs1 | 447.4221575 | 0.119899787 | 0.083069627 | 1.443364943 |
| 1700120C14Rik | 12.4281098 | -0.032856086 | 0.320997156 | -0.102356315 |
| Gm25999 | 0 | NA | NA | NA |
| Fam186b | 17.25303227 | 0.296693454 | 0.277168246 | 1.070445329 |
| Prpf40b | 472.5579373 | -0.424704078 | 0.13172648 | -3.224135938 |
| Fmnl3 | 135.4055067 | 0.667574687 | 0.149407726 | 4.46814035 |
| Gm22951 | 0.744072464 | -0.209814749 | 1.421044122 | -0.1476483 |
| Tmbim6 | 4439.906879 | 0.685983202 | 0.122890081 | 5.582087625 |
| Nckap5l | 263.9144233 | -0.048030751 | 0.105403852 | -0.455683069 |
| Nckap5los | 0.092140247 | -0.687373951 | 3.724945049 | -0.184532642 |
| Bcdin3d | 22.52486403 | 0.022272722 | 0.227922804 | 0.097720462 |
| Faim2 | 4926.385511 | 0.009821074 | 0.068207389 | 0.143988419 |
| Gm5808 | 0.066050875 | 0.274391604 | 3.724945049 | 0.073663262 |
| Aqp2 | 0.235388486 | 0.274410979 | 2.119097237 | 0.129494284 |
| Aqp5 | 2.102526643 | 0.086031489 | 0.804251384 | 0.106970893 |
| Aqp6 | 109.2123799 | -0.189043697 | 0.225837299 | -0.837079159 |
| Racgap1 | 122.2464084 | -0.20793588 | 0.125845237 | -1.652314263 |
| Asic1 | 756.9424049 | -0.388927999 | 0.070786703 | -5.494365225 |
| Gm17349 | 9.498973506 | -0.529365048 | 0.334723978 | -1.581497242 |
| Smarcd1 | 1391.542852 | -0.053419382 | 0.081050714 | -0.659085896 |
| Gm16537 | 2.123900911 | -0.933356391 | 0.699512324 | -1.334295851 |
| Gpd1 | 1414.234084 | -0.04368571 | 0.104202291 | -0.419239436 |
| Cox14 | 343.5443005 | -0.168761236 | 0.092567041 | -1.823124457 |
| Cers5 | 400.0721546 | -0.241479402 | 0.073043265 | -3.305977669 |
| Gm17058 | 0.41865154 | 1.261100797 | 1.554425868 | 0.811296842 |
| Lima1 | 217.8937591 | 0.457124666 | 0.114281548 | 3.999986632 |
| Gm25897 | 0.60093359 | 1.490727129 | 1.253508298 | 1.189243925 |
| Gm4468 | 0.964311207 | 0.66638694 | 1.08129071 | 0.616288417 |
| Gm17057 | 0.099106094 | 0.274387056 | 3.724945049 | 0.073662041 |
| Gm9763 | 0.448689577 | 0.975707882 | 1.378827833 | 0.70763576 |
| Fam186a | 4.175781228 | -0.700348196 | 0.522756405 | -1.339721885 |
| Gm21917 | 0.34674037 | -0.209351869 | 1.577287399 | -0.132729057 |
| Gm23133 | 0 | NA | NA | NA |
| Larp4 | 1360.21083 | 0.265534765 | 0.085103241 | 3.120148682 |

| | | | | |
|---|---|---|---|---|
| 2310068J16Rik | 5.082496618 | 0.600102591 | 0.4422056 | 1.357066917 |
| Dip2b | 2171.940491 | -0.274586467 | 0.044993445 | -6.102810457 |
| Atf1 | 263.8766334 | 0.26116869 | 0.128288689 | 2.035788912 |
| Tmprss12 | 0.281502737 | -1.137283297 | 2.053473602 | -0.553833902 |
| Mettl7a1 | 199.3407439 | -0.50279277 | 0.216817466 | -2.318968014 |
| Mettl7a3 | 1.559777635 | 0.920263527 | 0.792479033 | 1.161246529 |
| Methig1 | 0 | NA | NA | NA |
| Mettl7a2 | 0.046345709 | 0.274391604 | 3.724945049 | 0.073663262 |
| Higd1c | 0 | NA | NA | NA |
| Slc11a2 | 350.6928657 | 0.119870729 | 0.097223696 | 1.232937387 |
| Gm5475 | 0.572323712 | 0.545088516 | 1.422854342 | 0.383095093 |
| n-R5s43 | 0 | NA | NA | NA |
| 5330439K02Rik | 13.54861615 | -0.574795189 | 0.276823264 | -2.0763977 |
| Letmd1 | 357.2468629 | 0.109008913 | 0.080812671 | 1.348908667 |
| Csrnp2 | 483.3671844 | 0.204912378 | 0.128437738 | 1.595421884 |
| Tfcp2 | 253.237872 | -0.397464732 | 0.11012379 | -3.609254035 |
| Pou6f1 | 1003.020766 | -0.07263228 | 0.080814423 | -0.898753933 |
| Gm20680 | 0.523790777 | -0.150493667 | 1.300793181 | -0.11569377 |
| Gm27209 | 0.085433889 | -0.687376175 | 3.724945049 | -0.184533239 |
| C330013E15Rik | 16.45784469 | 0.363374172 | 0.259445128 | 1.400581981 |
| Dazap2 | 1246.964278 | 0.419083041 | 0.107784735 | 3.888148379 |
| Smagp | 10.67246666 | 0.955744204 | 0.375936677 | 2.542301038 |
| Bin2 | 77.96195219 | 1.8689464 | 0.224610691 | 8.320825643 |
| Mir6961 | 0 | NA | NA | NA |
| Cela1 | 27.94372578 | 1.39570909 | 0.298675458 | 4.672995562 |
| Galnt6 | 328.7283006 | 0.46786986 | 0.190991911 | 2.449684164 |
| Galnt6os | 0.119234754 | 0.274400501 | 3.724945049 | 0.073665651 |
| Slc4a8 | 2144.399781 | -0.229178371 | 0.08258221 | -2.775154235 |
| Scn8a | 4962.265033 | -0.354699025 | 0.113378363 | -3.128454285 |
| Fignl2 | 63.27470895 | 0.360736545 | 0.169763119 | 2.124940605 |
| Ankrd33 | 0.864742418 | -0.695071364 | 1.033271266 | -0.672690112 |
| Acvrl1 | 107.1769413 | 1.474908689 | 0.158580464 | 9.300696014 |
| Gm16031 | 0.097139547 | 0.274393828 | 3.724945049 | 0.073663859 |
| Acvr1b | 1299.302991 | -0.069683771 | 0.065359261 | -1.066165225 |
| mmu-mir-6962 | 0.93249843 | -1.371781013 | 1.174692835 | -1.16777848 |
| A330009N23Rik | 60.10913543 | -0.404430996 | 0.180028635 | -2.24648149 |
| Grasp | 286.1441832 | -0.180450028 | 0.216622334 | -0.833016728 |
| Nr4a1 | 763.9124175 | -0.946459124 | 0.422596108 | -2.239630479 |
| Atg101 | 182.2922892 | -0.069242971 | 0.107314364 | -0.645234875 |
| 6030408B16Rik | 2.433334847 | -0.047235292 | 0.687240838 | -0.068731788 |
| Krt80 | 28.28862397 | 0.439978014 | 0.23712557 | 1.855464236 |
| Mir1941 | 0.066050875 | 0.274391604 | 3.724945049 | 0.073663262 |
| Krt7 | 2.609690864 | -0.051302794 | 0.701190196 | -0.073165305 |
| Krt83 | 0.375360954 | 1.360093242 | 1.542600486 | 0.881688587 |
| 1700011A15Rik | 0.04393175 | -0.206490061 | 3.724945049 | -0.055434391 |
| Krt81 | 0 | NA | NA | NA |
| Krt86 | 0.614518174 | 2.420972668 | 1.387598954 | 1.744720736 |

| | | | | |
|---|---|---|---|---|
| 5430421N21Rik | 0.095897396 | -0.687371727 | 3.724945049 | -0.184532045 |
| Krt84 | 0 | NA | NA | NA |
| Krt82 | 0 | NA | NA | NA |
| 4732456N10Rik | 0.738905749 | -1.050919941 | 1.503955753 | -0.698770518 |
| AC103674.1 | 0 | NA | NA | NA |
| Krt75 | 1.991379766 | 0.274412976 | 1.294538739 | 0.211977415 |
| AC103674.2 | 0 | NA | NA | NA |
| Gm5414 | 0 | NA | NA | NA |
| Gm5478 | 0 | NA | NA | NA |
| AC120787.3 | 0 | NA | NA | NA |
| Krt6b | 0.519670895 | 1.307732299 | 1.785672983 | 0.732347026 |
| AC120787.2 | 0 | NA | NA | NA |
| Krt6a | 4.257533425 | 0.748924073 | 1.337483292 | 0.559950227 |
| AC120787.4 | 0 | NA | NA | NA |
| Krt5 | 1.350984256 | 2.818374975 | 1.127629361 | 2.499380622 |
| AC120787.1 | 0 | NA | NA | NA |
| Krt71 | 0.51940401 | -0.2621534 | 1.409364855 | -0.186008186 |
| Krt74 | 0 | NA | NA | NA |
| Krt72 | 0.233626327 | 1.660244406 | 3.105378673 | 0.53463509 |
| Krt73 | 9.672351053 | -0.136792081 | 0.38349982 | -0.35669399 |
| Krt2 | 16.20338829 | 1.922175169 | 0.854350571 | 2.249867016 |
| Krt1 | 24.4159562 | 0.168390347 | 0.302796986 | 0.556116324 |
| Krt77 | 35.13814104 | -0.046474225 | 0.280341977 | -0.165776904 |
| Krt76 | 0.099106094 | 0.274387056 | 3.724945049 | 0.073662041 |
| AC139844.1 | 0 | NA | NA | NA |
| Krt4 | 0.254192696 | 1.239251716 | 2.47001095 | 0.501719119 |
| Krt79 | 0.300336086 | 1.201607644 | 3.722238664 | 0.322818538 |
| Krt78 | 0.207810967 | 0.129934436 | 2.587603752 | 0.050214194 |
| Krt8 | 17.9984126 | 1.396285531 | 0.40159092 | 3.476885211 |
| Krt18 | 21.72902479 | 0.810128396 | 0.349908316 | 2.315259056 |
| Eif4b | 2876.650182 | 0.265900827 | 0.058317404 | 4.559544992 |
| Tns2 | 268.4246283 | 0.635436054 | 0.122025693 | 5.207395559 |
| Spryd3 | 1083.158277 | -0.087994115 | 0.068087658 | -1.292365133 |
| Igfbp6 | 171.4122214 | -0.015692222 | 0.114843834 | -0.136639658 |
| Soat2 | 21.31164465 | 4.043372418 | 0.582959543 | 6.935940003 |
| Csad | 172.5508944 | 0.006781631 | 0.102568077 | 0.066118343 |
| Zfp740 | 877.8255617 | 0.18302235 | 0.066667361 | 2.745306653 |
| Itgb7 | 116.7476434 | 3.096648424 | 0.422185894 | 7.334798409 |
| Rarg | 131.1736488 | -0.179477932 | 0.116379387 | -1.542179737 |
| Gm9918 | 0.827590884 | -0.639130712 | 1.124364925 | -0.568437077 |
| Mfsd5 | 351.832301 | 0.222341374 | 0.103286455 | 2.152667307 |
| Espl1 | 27.92326781 | 0.644914493 | 0.316228116 | 2.039396436 |
| Pfdn5 | 660.9445207 | 0.191014691 | 0.094233541 | 2.027035049 |
| Myg1 | 214.9838279 | -0.128991286 | 0.090157294 | -1.430736003 |
| Aaas | 128.6515988 | -0.001436059 | 0.169669663 | -0.00846385 |
| Sp7 | 27.79878053 | -1.437869835 | 0.352150731 | -4.083109044 |
| Gm23590 | 0 | NA | NA | NA |

| | | | | |
|---|---|---|---|---|
| Sp1 | 861.7402076 | 0.266646217 | 0.098241615 | 2.71418805 |
| Amhr2 | 7.396618114 | -0.787858634 | 0.432854624 | -1.82014605 |
| Gm26518 | 0.703641397 | -1.597612609 | 1.184433252 | -1.348841403 |
| Gm24128 | 0.061920613 | 0.274393828 | 3.724945049 | 0.073663859 |
| Prr13 | 456.4408238 | 0.30716524 | 0.114965785 | 2.671796995 |
| Pcbp2 | 2462.12792 | 0.270769641 | 0.071073382 | 3.809719369 |
| Gm27406 | 2.84356551 | 1.249148252 | 0.600541777 | 2.080035562 |
| Map3k12 | 1206.534751 | -0.336856893 | 0.066880015 | -5.036734686 |
| Gm10337 | 2.690945547 | 0.687184731 | 0.75761026 | 0.907042535 |
| Tarbp2 | 121.0001225 | -0.060564396 | 0.115211992 | -0.525677888 |
| Npff | 8.501785748 | -0.609936273 | 0.579966 | -1.051675913 |
| Gm28047 | 0 | NA | NA | NA |
| Atf7 | 596.7832338 | -0.092791167 | 0.069498984 | -1.335144228 |
| Gm25141 | 0 | NA | NA | NA |
| Rpl39-ps | 40.47241404 | 0.240816779 | 0.248035885 | 0.970894915 |
| Atp5g2 | 389.3075053 | 0.098711219 | 0.113359787 | 0.87077809 |
| Mir688 | 0 | NA | NA | NA |
| Calcoco1 | 660.9432622 | 0.055402317 | 0.095747645 | 0.578628514 |
| Hoxc13 | 0.157352566 | 0.75529745 | 2.812320981 | 0.268567299 |
| Hoxc12 | 0 | NA | NA | NA |
| Hotair | 0 | NA | NA | NA |
| Gm27337 | 0 | NA | NA | NA |
| Gm27443 | 0 | NA | NA | NA |
| Gm28011 | 0 | NA | NA | NA |
| Gm27535 | 0 | NA | NA | NA |
| Hoxc11 | 0 | NA | NA | NA |
| Hoxc10 | 0 | NA | NA | NA |
| Hoxc5 | 4.717838401 | -1.153125458 | 1.24708146 | -0.92465929 |
| Mir196a-2 | 0 | NA | NA | NA |
| Hoxc9 | 0.095435502 | 0.75528439 | 3.724945049 | 0.202763901 |
| Hoxc8 | 0.100902008 | 0.274391828 | 3.724945049 | 0.073663322 |
| Hoxc6 | 1.521766428 | -1.40791378 | 2.099207086 | -0.67068837 |
| Mir615 | 0 | NA | NA | NA |
| Hoxc4 | 16.48418288 | -0.351775444 | 1.027357196 | -0.342408119 |
| 5730585A16Rik | 0.314534502 | -1.789473481 | 3.692614927 | -0.484608744 |
| Gm28876 | 4.308624721 | -0.49538432 | 0.531913777 | -0.931324477 |
| Gm10830 | 0.40206694 | -0.736563636 | 1.56914819 | -0.46940349 |
| Gm28265 | 0.496950623 | 0.705671836 | 1.709011608 | 0.412912255 |
| D930007P13Rik | 1.440959972 | -0.58345009 | 0.943693808 | -0.618262073 |
| Smug1 | 140.5464891 | -0.225078968 | 0.119598908 | -1.881948346 |
| Cbx5 | 3979.323211 | -0.212515653 | 0.065222036 | -3.258341275 |
| Hnrnpa1 | 1468.269564 | 0.096924537 | 0.093096546 | 1.041118506 |
| Nfe2 | 3.16484666 | 2.092737866 | 1.064812762 | 1.965357611 |
| Copz1 | 570.9806916 | 0.27313207 | 0.093113211 | 2.933333184 |
| Mir148b | 1.892187285 | 0.384976736 | 0.695406963 | 0.553599197 |
| Gpr84 | 34.27634392 | 4.768747989 | 0.444499555 | 10.72835267 |
| Zfp385a | 1024.49315 | -0.016043817 | 0.082148396 | -0.195302862 |

| | | | | |
|---|---|---|---|---|
| Itga5 | 497.2109644 | 2.80998919 | 0.34402634 | 8.167947821 |
| Mir6963 | 0.095435502 | 0.75528439 | 3.724945049 | 0.202763901 |
| Gm37322 | 0.248806291 | -1.155962446 | 2.130457273 | -0.54258889 |
| Gtsf1 | 0.666252615 | -0.889142591 | 1.272486888 | -0.69874401 |
| BC048502 | 0.169045294 | 0.778759426 | 3.400490153 | 0.229013875 |
| Nckap1l | 484.3215996 | 1.960018227 | 0.14907943 | 13.14747599 |
| Pde1b | 1849.40123 | -0.316550807 | 0.124129492 | -2.550165987 |
| Ppp1r1a | 376.4709638 | -0.426312561 | 0.135205174 | -3.153078754 |
| Glycam1 | 0.54398204 | 0.249835459 | 1.207993342 | 0.206818573 |
| Mucl1 | 0.04393175 | -0.206490061 | 3.724945049 | -0.055434391 |
| Gm22852 | 0 | NA | NA | NA |
| Spt1 | 0.136548098 | -0.206480423 | 2.945560552 | -0.070098855 |
| Gm23215 | 0 | NA | NA | NA |
| Gm22862 | 0 | NA | NA | NA |
| Gm22598 | 0 | NA | NA | NA |
| Olfr161 | 0 | NA | NA | NA |
| Mefv | 48.54285063 | 7.827383332 | 0.979966308 | 7.987400453 |
| Tdgf1-ps1 | 0.267098211 | 1.236195215 | 2.473599901 | 0.499755524 |
| Zfp263 | 304.9461567 | 0.050827707 | 0.105519989 | 0.481687948 |
| Olfr15 | 0.061920613 | 0.274393828 | 3.724945049 | 0.073663859 |
| Zfp174 | 93.47116396 | -0.265275582 | 0.172492708 | -1.537894471 |
| Zfp597 | 424.0273003 | -0.088061131 | 0.105926649 | -0.831340663 |
| Gm26675 | 8.550105792 | -0.873961843 | 0.432894247 | -2.018880707 |
| Naa60 | 734.2670764 | -0.146761517 | 0.064192118 | -2.286285643 |
| Gm20695 | 1.863767 | 0.29102871 | 0.741267536 | 0.392609545 |
| 1700037C18Rik | 31.49051414 | -0.287887291 | 0.230118594 | -1.251038805 |
| Cluap1 | 161.5850996 | -0.111454958 | 0.101577181 | -1.097244058 |
| 1700016D08Rik | 0 | NA | NA | NA |
| Gm15538 | 10.93859038 | -0.803395918 | 0.438005688 | -1.834213435 |
| Gm23493 | 2.266218546 | -0.041206616 | 0.674000436 | -0.061137373 |
| Gm15539 | 10.65583297 | -0.697311736 | 0.416901722 | -1.672604596 |
| Gm15537 | 0.604736192 | -0.647892844 | 1.289989303 | -0.502246679 |
| Nlrc3 | 40.04733098 | 0.42360545 | 0.214900832 | 1.971167102 |
| Slx4 | 453.7919637 | -0.458986081 | 0.092927261 | -4.939197337 |
| Gm15879 | 36.75047168 | -0.705918125 | 0.203990496 | -3.460544189 |
| Dnase1 | 0.317440118 | -1.12836775 | 1.562758706 | -0.722035811 |
| Trap1 | 710.530572 | -0.010850337 | 0.055468717 | -0.195611834 |
| Gm23413 | 3.363014587 | -0.701787802 | 0.539811389 | -1.300061126 |
| Crebbp | 3534.609602 | -0.052215758 | 0.053034588 | -0.984560456 |
| Gm37186 | 92.75499785 | -0.774002217 | 0.314263176 | -2.462910947 |
| Gm5766 | 0.297064053 | 0.755310525 | 1.770854511 | 0.426523196 |
| Adcy9 | 953.9955112 | 0.053398875 | 0.128519566 | 0.415492186 |
| Gm6142 | 13.94176219 | -0.095005242 | 0.291010886 | -0.326466284 |
| Srl | 63.68776743 | -0.718505025 | 0.156051882 | -4.604270165 |
| Gm15885 | 0 | NA | NA | NA |
| Tfap4 | 61.90644834 | 0.531201291 | 0.17463996 | 3.041693845 |
| Glis2 | 152.6195355 | -0.070940489 | 0.104968491 | -0.675826506 |

| | | | | |
|---|---|---|---|---|
| Pam16 | 156.721799 | -0.09063455 | 0.151544856 | -0.598070776 |
| Coro7 | 608.4531973 | -0.111136632 | 0.107465669 | -1.034159397 |
| Vasn | 199.7417839 | 0.554845506 | 0.217888817 | 2.546461598 |
| Dnaja3 | 623.6357018 | -0.209899433 | 0.086259496 | -2.433348698 |
| Nmral1 | 147.3588111 | 0.206137049 | 0.1370827 | 1.503742262 |
| Hmox2 | 693.4042527 | 0.417704435 | 0.081596493 | 5.119146945 |
| Gm15859 | 12.58934561 | -0.450698166 | 0.388025425 | -1.161517099 |
| Cdip1 | 1931.022938 | -0.185604242 | 0.054000451 | -3.43708691 |
| Gm15835 | 24.15715004 | -0.385653388 | 0.304259128 | -1.267516245 |
| 4930562C15Rik | 0.450624642 | 2.132192396 | 1.519822319 | 1.40292215 |
| Ubald1 | 776.8048839 | -0.054868808 | 0.086358716 | -0.635359242 |
| Mgrn1 | 1884.371733 | -0.04674628 | 0.050910674 | -0.918201944 |
| Gm16861 | 17.52762606 | 0.947388612 | 0.262286726 | 3.61203416 |
| Nudt16l1 | 313.4024066 | 0.125252028 | 0.085950429 | 1.4572589 |
| Anks3 | 314.2732224 | -0.15880668 | 0.097715918 | -1.625187414 |
| 4930451G09Rik | 27.70571581 | -0.583709618 | 0.201987991 | -2.889823382 |
| 12-Sep | 0.042716945 | -0.206490061 | 3.724945049 | -0.055434391 |
| Smim22 | 0.651416456 | 1.287192956 | 1.262685034 | 1.01940937 |
| RP23-450O11.3 | 0.373087082 | 0.762339574 | 1.722682665 | 0.442530472 |
| Rogdi | 663.6085146 | 0.049701711 | 0.076894504 | 0.646362332 |
| Glyr1 | 1981.897547 | 0.155283278 | 0.054703955 | 2.838611569 |
| Ubn1 | 976.2780589 | -0.030947323 | 0.080429874 | -0.384773984 |
| Gm23483 | 0 | NA | NA | NA |
| Ppl | 31.86021987 | 0.74752598 | 0.289071504 | 2.585955276 |
| Sec14l5 | 164.8073848 | -0.315876894 | 0.133110339 | -2.373045523 |
| Nagpa | 106.0130434 | 0.292827154 | 0.125312301 | 2.336778997 |
| AU021092 | 9.404176926 | 0.327890996 | 0.439239939 | 0.746496314 |
| Mir6364 | 0.049553047 | 0.274391604 | 3.724945049 | 0.073663262 |
| Gm23689 | 0.15591167 | 0.274407136 | 2.826544723 | 0.097082184 |
| Gm26095 | 0.09689823 | -0.687371727 | 3.724945049 | -0.184532045 |
| Alg1 | 199.3302287 | 0.064698068 | 0.10945146 | 0.591111965 |
| Eef2kmt | 50.93323876 | -0.051085482 | 0.222063736 | -0.230048736 |
| Gm15983 | 5.210713431 | 0.879972944 | 0.462801145 | 1.901406152 |
| n-R5s30 | 0 | NA | NA | NA |
| 1700123O21Rik | 37.36951086 | -0.406689294 | 0.377849124 | -1.07632721 |
| Rbfox1 | 3132.04572 | -0.054893525 | 0.080432239 | -0.682481626 |
| Gm23476 | 0 | NA | NA | NA |
| Gm26159 | 0 | NA | NA | NA |
| Gm24579 | 0 | NA | NA | NA |
| Tmem114 | 8.994908763 | -0.370224212 | 0.365795256 | -1.012107745 |
| Gm25782 | 0 | NA | NA | NA |
| Mettl22 | 149.3545907 | 0.198425274 | 0.089554427 | 2.215694755 |
| Gm23706 | 0 | NA | NA | NA |
| Abat | 3761.014197 | -0.166222884 | 0.08529601 | -1.948776774 |
| Gm22137 | 0 | NA | NA | NA |
| Gm7541 | 1.437230806 | 0.025796044 | 0.91977121 | 0.028046153 |
| Tmem186 | 191.5203021 | -0.158046081 | 0.098922938 | -1.597668689 |

| | | | | |
|---|---|---|---|---|
| Pmm2 | 257.7422973 | 0.205339818 | 0.101076893 | 2.031520867 |
| Carhsp1 | 736.470531 | 0.822408727 | 0.155907561 | 5.274976534 |
| RP24-183G19.6 | 0.790581521 | 3.351531327 | 1.42893599 | 2.34547338 |
| Usp7 | 2331.28947 | 0.065351736 | 0.068566623 | 0.953112938 |
| Gm25805 | 0 | NA | NA | NA |
| 1810013L24Rik | 1107.486804 | -0.088793511 | 0.067187632 | -1.321575232 |
| Gm22224 | 0 | NA | NA | NA |
| Gm10247 | 1.54124956 | 1.003487564 | 0.922814195 | 1.087421031 |
| Grin2a | 4615.108378 | -0.216756351 | 0.161468693 | -1.342404812 |
| Gm7572 | 0 | NA | NA | NA |
| Gm3890 | 0 | NA | NA | NA |
| Rpl39l | 0.823456907 | 2.659737244 | 1.149539051 | 2.313742401 |
| Atf7ip2 | 0.994634002 | 1.389308937 | 0.987531788 | 1.406849839 |
| Gm24667 | 0 | NA | NA | NA |
| Gm22932 | 0 | NA | NA | NA |
| Emp2 | 291.2677059 | 0.167334821 | 0.135382976 | 1.236010804 |
| Tekt5 | 64.82616325 | -0.564762733 | 0.177608636 | -3.179815711 |
| Nubp1 | 155.2730509 | 0.051092181 | 0.137434381 | 0.371756911 |
| Tvp23a | 456.5008628 | 0.269241948 | 0.108643376 | 2.478217784 |
| Ciita | 53.45720072 | 5.291604279 | 0.520236581 | 10.1715344 |
| Dexi | 379.7312674 | 0.051179966 | 0.07993636 | 0.640258897 |
| Clec16a | 1806.749408 | -0.031335699 | 0.06734037 | -0.465333042 |
| Gm15558 | 0.339464816 | 1.212575972 | 1.746503674 | 0.694287673 |
| Socs1 | 257.5468929 | 4.23044102 | 0.382821895 | 11.05067676 |
| Tnp2 | 0.356459053 | 1.187764872 | 1.650601265 | 0.719595275 |
| Prm3 | 0 | NA | NA | NA |
| Prm2 | 0.134416897 | 0.75529538 | 2.953956728 | 0.255689385 |
| Prm1 | 0.978261292 | 0.274412269 | 1.387128588 | 0.197827563 |
| Gm10343 | 5.069691734 | 1.65809287 | 0.507993455 | 3.264004393 |
| Gm11172 | 0.044544648 | 0.755286614 | 3.724945049 | 0.202764498 |
| Gm26822 | 0.840740177 | 1.899503563 | 1.164467878 | 1.631220232 |
| Rmi2 | 18.46361415 | 1.118532587 | 0.295766578 | 3.781808593 |
| Gm21859 | 0.468571393 | -0.674062617 | 1.523069349 | -0.442568566 |
| Litaf | 694.90215 | 1.626051501 | 0.257766145 | 6.308243088 |
| Gm26268 | 0 | NA | NA | NA |
| Gm24961 | 0 | NA | NA | NA |
| Gm4262 | 0.72304365 | 0.274414141 | 1.370596561 | 0.20021511 |
| Snn | 1735.197761 | 0.115227198 | 0.079614416 | 1.447315737 |
| Txndc11 | 466.704329 | -0.354301195 | 0.092284929 | -3.839209717 |
| Zc3h7a | 853.6995849 | 0.537075694 | 0.126793324 | 4.235835733 |
| Gm23935 | 15251.03865 | 1.526267675 | 0.922168559 | 1.655085354 |
| Gm24145 | 5.507590669 | 0.285144839 | 0.540328336 | 0.527725125 |
| 4930509G22Rik | 5.036420442 | 0.187177548 | 0.424255798 | 0.441190311 |
| Gm7638 | 2.798124148 | 0.106167042 | 0.613385695 | 0.173083661 |
| Gm6305 | 2.348678771 | -0.530452615 | 0.723721943 | -0.732950852 |
| Rsl1d1 | 606.0058677 | 0.522918928 | 0.101739645 | 5.139775431 |
| 2610020C07Rik | 17.10628321 | 0.235407878 | 0.244418429 | 0.963134735 |

| | | | | |
|---|---|---|---|---|
| Gspt1 | 1482.382663 | 0.01415349 | 0.052431719 | 0.269941361 |
| Mir1945 | 0.407802109 | -0.206482002 | 1.469268452 | -0.140533884 |
| Tnfrsf17 | 1.11445967 | 0.07703366 | 1.200972628 | 0.064142727 |
| Snx29 | 234.388902 | -0.439094501 | 0.11988629 | -3.662591457 |
| Gm15897 | 0.089089296 | 1.194485492 | 3.723939576 | 0.320758559 |
| Gm4279 | 0.739397817 | -0.436592423 | 1.245824987 | -0.350444427 |
| Cpped1 | 428.5321132 | 0.220325541 | 0.081541832 | 2.701993996 |
| Gm9961 | 0.136963818 | 0.755286583 | 3.59954433 | 0.209828388 |
| Shisa9 | 635.7404322 | -0.552090556 | 0.100313908 | -5.5036292 |
| Gm6327 | 0.086930093 | -0.206490058 | 3.724945049 | -0.05543439 |
| 1700003L19Rik | 0.364102347 | -2.18564865 | 1.637457783 | -1.334781679 |
| Ercc4 | 327.5978949 | 0.040981422 | 0.115996294 | 0.35329941 |
| Mkl2 | 3224.420774 | -0.156163609 | 0.122215249 | -1.277775159 |
| Gm15738 | 0.13861434 | -1.111919598 | 3.614496194 | -0.307627824 |
| Mir193b | 0.600656357 | -1.189286765 | 1.443646226 | -0.823807622 |
| Mir365-1 | 1.735414367 | -2.084017753 | 0.894444824 | -2.329956749 |
| Mir6365 | 0 | NA | NA | NA |
| 2310015D24Rik | 1.269356065 | 0.023456438 | 0.837195524 | 0.028017873 |
| Gm25276 | 0.11037557 | 0.274404949 | 3.724945049 | 0.073666845 |
| Parn | 231.9657214 | 0.093601402 | 0.08569168 | 1.092304432 |
| Bfar | 419.0822541 | 0.364953514 | 0.096544893 | 3.78014313 |
| 3110001I22Rik | 41.39457374 | 0.179607132 | 0.176271496 | 1.01892329 |
| Pla2g10 | 0 | NA | NA | NA |
| Pla2g10os | 0.090890357 | 0.755282166 | 3.724945049 | 0.202763304 |
| Rrn3 | 908.1970389 | 0.078719866 | 0.065005847 | 1.210965919 |
| Gm27734 | 0.941805692 | -0.068286732 | 0.900663996 | -0.07581821 |
| Ntan1 | 347.9210283 | 0.028563218 | 0.088921895 | 0.321216928 |
| Pdxdc1 | 751.5294376 | 0.133824306 | 0.061206532 | 2.186438317 |
| Mpv17l | 545.4884099 | -0.107458931 | 0.080642294 | -1.332538124 |
| Gm15950 | 0.843513109 | -0.833996425 | 1.383006503 | -0.603031456 |
| Gm26213 | 0.252317246 | 1.28704099 | 2.503377304 | 0.514121857 |
| Ifitm7 | 16.50735806 | 0.555878029 | 0.312720878 | 1.777553302 |
| 2900011O08Rik | 1845.656536 | -0.076043008 | 0.060758999 | -1.25155138 |
| Gm15806 | 0.061920613 | 0.274393828 | 3.724945049 | 0.073663859 |
| Gm15807 | 0 | NA | NA | NA |
| Gm15808 | 7.400018289 | -1.265348055 | 0.600602935 | -2.106796321 |
| Marf1 | 2957.81909 | -0.218219644 | 0.075012525 | -2.909109433 |
| Gm27671 | 0 | NA | NA | NA |
| Mir484 | 0.262798536 | 1.254086688 | 1.818294299 | 0.68970501 |
| Nde1 | 330.4944427 | 0.950326093 | 0.146846673 | 6.471553443 |
| Myh11 | 25.31456723 | 0.583317974 | 0.285244454 | 2.044975692 |
| Fopnl | 265.3373334 | -0.07468386 | 0.093001905 | -0.803035805 |
| Gm15868 | 0 | NA | NA | NA |
| Gm15869 | 19.70131526 | -0.375356587 | 0.268726291 | -1.396798899 |
| Abcc1 | 333.839765 | -0.300412013 | 0.112840383 | -2.662273956 |
| Gm23116 | 0.060174537 | 0.755286614 | 3.724945049 | 0.202764498 |
| Gm1758 | 0.227501076 | 0.274410816 | 2.140181937 | 0.128218452 |

| | | | | |
|---|---|---|---|---|
| A630010A05Rik | 0 | NA | NA | NA |
| Snai2 | 6.714012416 | 0.260839931 | 0.376890458 | 0.6920842 |
| Gm25547 | 1.090696383 | 0.007551551 | 0.994039008 | 0.007596836 |
| Efcab1 | 65.3173956 | -0.167444513 | 0.154305147 | -1.085151833 |
| Gm7731 | 0 | NA | NA | NA |
| Gm21897 | 0.148910734 | 0.723040935 | 2.858091615 | 0.252980321 |
| Gm6031 | 0 | NA | NA | NA |
| Ube2v2 | 393.0830429 | -0.133447274 | 0.09721373 | -1.372720436 |
| Mcm4 | 207.5248428 | 0.447204133 | 0.163555764 | 2.73426092 |
| Prkdc | 719.6145477 | -0.417148663 | 0.122778808 | -3.397562403 |
| Mzt2 | 126.2984278 | 0.329059648 | 0.130634589 | 2.518932019 |
| Cebpd | 477.928814 | 3.375219496 | 0.34822469 | 9.692648437 |
| Spidr | 147.239883 | 1.392815649 | 0.270246532 | 5.153870579 |
| n-R5s31 | 0.50758071 | 0.269574428 | 1.387865186 | 0.194236754 |
| Pkp2 | 124.1564146 | 0.404560652 | 0.133689663 | 3.026117667 |
| Yars2 | 199.1181058 | -0.107607081 | 0.110499801 | -0.973821492 |
| Dnm1l | 3082.769836 | 0.089076052 | 0.043633252 | 2.041471782 |
| Gm7765 | 0 | NA | NA | NA |
| Fgd4 | 1158.129079 | -0.272463063 | 0.085088201 | -3.2021251 |
| Gm16139 | 0 | NA | NA | NA |
| Olfr19 | 0.447268164 | -0.09307593 | 1.452145475 | -0.064095459 |
| Spag6 | 66.08212979 | 0.016833715 | 0.156082097 | 0.107851674 |
| Rpl31-ps12 | 2.913194718 | 1.128279502 | 0.592422721 | 1.904517604 |
| Gm15714 | 0.095898756 | 0.27438722 | 3.724945049 | 0.073662085 |
| Igll1 | 0.075043205 | 0.755286614 | 3.724945049 | 0.202764498 |
| Vpreb1 | 0.913331526 | -1.217213492 | 1.167837831 | -1.042279553 |
| Top3b | 435.3366088 | 0.059748728 | 0.119764476 | 0.498885232 |
| Ppm1f | 789.0102064 | -0.326390104 | 0.089720877 | -3.637838983 |
| Mapk1 | 4968.897064 | -0.133220453 | 0.070167041 | -1.898618662 |
| Gm22558 | 0 | NA | NA | NA |
| 1700056N10Rik | 40.39149775 | 0.084553771 | 0.30999519 | 0.272758331 |
| Ypel1 | 118.1725456 | -0.389720754 | 0.204438416 | -1.906299029 |
| Gm9974 | 6.459321561 | 0.00771288 | 0.397962067 | 0.019380943 |
| Ppil2 | 680.5363805 | 0.016037169 | 0.054485891 | 0.294336183 |
| Gm15585 | 2.078701808 | -0.273942077 | 0.701518471 | -0.390498737 |
| 2610318N02Rik | 4.21806971 | 0.097437766 | 0.448074951 | 0.217458633 |
| Mir130b | 0.075043205 | 0.755286614 | 3.724945049 | 0.202764498 |
| Mir301b | 0.118974955 | 0.274400501 | 3.724945049 | 0.073665651 |
| Sdf2l1 | 144.324589 | 0.192532647 | 0.161624604 | 1.191233531 |
| Ccdc116 | 27.44048453 | 0.189891793 | 0.222524654 | 0.853351704 |
| Gm15646 | 0 | NA | NA | NA |
| Ydjc | 88.53278356 | -0.326380252 | 0.128619202 | -2.537570184 |
| Ube2l3 | 1433.809399 | 0.338702876 | 0.081311262 | 4.165509997 |
| Gm15648 | 2.296535557 | -0.558784139 | 0.677931027 | -0.824249248 |
| Gm22129 | 0.093744596 | -0.20649451 | 3.724945049 | -0.055435586 |
| Rimbp3 | 28.96339831 | -0.308288441 | 0.217955161 | -1.414458095 |
| Hic2 | 126.3818273 | -0.135566186 | 0.175032056 | -0.774522043 |

| | | | | |
|---|---|---|---|---|
| Gm26635 | 0.934835317 | 0.341985061 | 1.081342224 | 0.316259787 |
| Tmem191c | 275.0346229 | -0.272405109 | 0.156121898 | -1.744823201 |
| Pi4ka | 4405.809757 | -0.08390959 | 0.080689078 | -1.039912618 |
| Serpind1 | 61.15174061 | 0.978397928 | 0.2025944 | 4.829343389 |
| Gm24927 | 3.890661782 | 0.031808399 | 0.538467847 | 0.059072048 |
| Snap29 | 384.8317826 | 0.086893445 | 0.145853164 | 0.595759753 |
| Crkl | 918.3467102 | 0.267542576 | 0.083324249 | 3.210860924 |
| Gm22770 | 0 | NA | NA | NA |
| Aifm3 | 673.3284071 | -0.461442522 | 0.17382446 | -2.654646653 |
| Lztr1 | 1143.994835 | -0.183823065 | 0.052026315 | -3.533270928 |
| Thap7 | 186.8558046 | 0.191252751 | 0.107332305 | 1.781875007 |
| Lrrc74b | 11.80310187 | -0.696832581 | 0.316978592 | -2.198358493 |
| P2rx6 | 28.77111211 | -0.374549385 | 0.255662895 | -1.465012689 |
| Slc7a4 | 422.6595786 | 0.032247807 | 0.106365005 | 0.303180605 |
| Smpd4 | 474.152001 | 0.047083163 | 0.142516096 | 0.330370844 |
| Ccdc74a | 148.3892086 | -0.090020663 | 0.191192522 | -0.470837782 |
| Med15 | 779.2021573 | 0.240181908 | 0.067399202 | 3.563571959 |
| Rpl26-ps4 | 0 | NA | NA | NA |
| Klhl22 | 742.0166947 | 0.037893916 | 0.045628869 | 0.830481151 |
| Scarf2 | 44.76671024 | 0.378567042 | 0.177184854 | 2.136565478 |
| Car15 | 16.28727946 | -0.348344613 | 0.313897123 | -1.10974134 |
| Gm20518 | 10.60745708 | -0.136518023 | 0.41083416 | -0.332294721 |
| Dgcr2 | 954.3027326 | 0.000842551 | 0.070877439 | 0.011887432 |
| Tssk1 | 0.674642588 | 0.340638172 | 1.172234197 | 0.290588837 |
| Tssk2 | 0.04393175 | -0.206490061 | 3.724945049 | -0.055434391 |
| Dgcr14 | 286.5719584 | 0.211659764 | 0.069679843 | 3.037603909 |
| Gsc2 | 0.267511063 | -1.228247163 | 2.479702835 | -0.495320304 |
| AA914427 | 7.57215109 | -0.8729833 | 0.383518378 | -2.276248936 |
| Slc25a1 | 386.8663658 | 0.140651501 | 0.117828485 | 1.193696935 |
| Vpreb2 | 0.042716945 | -0.206490061 | 3.724945049 | -0.055434391 |
| Dgcr6 | 244.4688473 | -0.124415184 | 0.109119716 | -1.140171445 |
| Prodh | 187.7866812 | -0.055592446 | 0.16023038 | -0.346953221 |
| Gm15954 | 5.737854463 | -1.043628114 | 0.443498524 | -2.353171559 |
| Rtn4r | 444.6635611 | -0.343217367 | 0.123093237 | -2.788271517 |
| Mir6366 | 0 | NA | NA | NA |
| Gm7873 | 0.093141081 | -0.687373951 | 3.724945049 | -0.184532642 |
| Zdhhc8 | 854.0005545 | -0.449381199 | 0.086171994 | -5.21493326 |
| Ranbp1 | 621.7089953 | 0.811925788 | 0.150022343 | 5.412032447 |
| Gm25777 | 2.388976987 | -0.629645683 | 0.734163836 | -0.857636473 |
| Trmt2a | 346.4940552 | 0.160161471 | 0.076944164 | 2.08152851 |
| Dgcr8 | 527.8985729 | -0.200278056 | 0.104963059 | -1.908081365 |
| Mir1306 | 0 | NA | NA | NA |
| Gm24572 | 0 | NA | NA | NA |
| Tango2 | 629.321088 | -0.095056897 | 0.094358444 | -1.007402124 |
| Mir185 | 2.706681505 | -1.04092295 | 0.706572156 | -1.473201204 |
| Arvcf | 417.1584827 | -0.205816488 | 0.065999051 | -3.118476474 |
| Gm26170 | 1.640007924 | -1.211590809 | 0.827263099 | -1.464577365 |

| | | | | |
|---|---|---|---|---|
| Gm26219 | 0 | NA | NA | NA |
| Comt | 659.6866933 | -0.03971014 | 0.058515499 | -0.678626019 |
| Gm15764 | 1.82265688 | -0.081988976 | 0.86164529 | -0.095153977 |
| Txnrd2 | 113.8682495 | 0.001849152 | 0.136246072 | 0.013572152 |
| Gnb1l | 33.08347225 | 0.118410142 | 0.186344729 | 0.635435961 |
| Gm16314 | 20.96206684 | -0.342014634 | 0.23059594 | -1.483177171 |
| Tbx1 | 2.659687333 | 0.256348205 | 0.629989463 | 0.406908718 |
| 4930588K23Rik | 0.132314849 | -1.044454757 | 3.65395166 | -0.285842522 |
| Gp1bb | 61.84750879 | -0.20820252 | 0.188455329 | -1.104784469 |
| 5-Sep | 2743.444644 | -0.319482377 | 0.110308053 | -2.896274289 |
| Cldn5 | 120.886244 | 1.914928384 | 0.187479402 | 10.21407345 |
| Cdc45 | 22.31723845 | 0.295049959 | 0.286491282 | 1.029874128 |
| Ufd1l | 707.9940102 | 0.364590996 | 0.055221048 | 6.602391866 |
| 2510002D24Rik | 76.81866928 | -0.500200429 | 0.121722277 | -4.109358119 |
| Gm28539 | 0.048949532 | -0.206487837 | 3.724945049 | -0.055433794 |
| Mrpl40 | 175.5484898 | 0.216488181 | 0.103794024 | 2.08574803 |
| Hira | 202.5271098 | -0.028499166 | 0.120157584 | -0.237181583 |
| Gm15799 | 5.251177882 | -0.079244168 | 0.594767253 | -0.133235593 |
| Gm15798 | 0 | NA | NA | NA |
| RP23-374P12.1 | 0 | NA | NA | NA |
| Gm10088 | 0.415109666 | -0.180360617 | 1.542016427 | -0.116964135 |
| RP23-247I11.8 | 0.329535318 | 1.247430042 | 1.721980721 | 0.7244158 |
| Iglc1 | 0.667277359 | 2.083565829 | 2.481966711 | 0.839481779 |
| Iglj1 | 0 | NA | NA | NA |
| Iglc3 | 1.77022427 | 3.157740098 | 1.0745869 | 2.938561876 |
| Iglj3p | 0 | NA | NA | NA |
| Iglj3 | 0.072025462 | 0.755288839 | 3.724945049 | 0.202765096 |
| Iglv1 | 0.611248024 | 1.814566791 | 2.561884616 | 0.708293723 |
| RP23-247I11.7 | 0 | NA | NA | NA |
| Iglc4 | 0 | NA | NA | NA |
| Iglj4 | 0 | NA | NA | NA |
| Iglc2 | 0.938589659 | 1.238273971 | 1.578969602 | 0.784229139 |
| Iglj2 | 0 | NA | NA | NA |
| RP23-272P19.8 | 0.046345709 | 0.274391604 | 3.724945049 | 0.073663262 |
| RP23-272P19.7 | 0.343041433 | 1.696053492 | 1.743117694 | 0.972999986 |
| Iglv3 | 0.097139547 | 0.274393828 | 3.724945049 | 0.073663859 |
| RP23-272P19.10 | 1.305037701 | 3.321036088 | 1.314496067 | 2.52647092 |
| Iglv2 | 0.097139547 | 0.274393828 | 3.724945049 | 0.073663859 |
| Olfr164 | 0.207953851 | -0.687376882 | 2.579759988 | -0.266449935 |
| Gm22158 | 0.391389582 | -0.19325443 | 3.393771449 | -0.056943855 |
| Gm23612 | 0.049553047 | 0.274391604 | 3.724945049 | 0.073663262 |
| Olfr165 | 0 | NA | NA | NA |
| Gm25762 | 0.144848288 | -0.206489769 | 2.892439516 | -0.071389486 |
| Olfr166 | 0.296045541 | 1.155808 | 2.011454234 | 0.574613123 |
| Olfr167 | 0 | NA | NA | NA |
| Olfr168 | 0 | NA | NA | NA |
| Olfr169 | 0.0878635 | -0.687376175 | 3.724945049 | -0.184533239 |

| | | | | |
|---|---|---|---|---|
| Olfr170 | 0 | NA | NA | NA |
| Olfr171 | 0 | NA | NA | NA |
| Lamp3 | 0.505840948 | 1.271526335 | 1.272604085 | 0.999153114 |
| Gm17799 | 0.062426873 | 0.755288839 | 3.724945049 | 0.202765096 |
| B3gnt5 | 145.4656175 | 1.198670744 | 0.153543902 | 7.806697178 |
| A930003A15Rik | 2.53052731 | 1.665309786 | 0.758568255 | 2.195332819 |
| Gm27883 | 0 | NA | NA | NA |
| Klhl6 | 70.61673169 | 3.521395879 | 0.313332524 | 11.23852652 |
| Klhl24 | 1378.234747 | -0.139961339 | 0.07877827 | -1.77664906 |
| Yeats2 | 481.9769426 | -0.353880849 | 0.130324918 | -2.715373649 |
| Gm10241 | 24.31474395 | 0.443541206 | 0.266814581 | 1.662357446 |
| Map6d1 | 369.8382378 | -0.701916789 | 0.102613878 | -6.840369019 |
| Parl | 312.4995396 | 0.248102715 | 0.072464514 | 3.42378223 |
| Cyp2ab1 | 1.187519348 | 3.239726587 | 1.106555217 | 2.927758631 |
| Abcc5 | 1948.385061 | -0.446756017 | 0.137746853 | -3.243311961 |
| Gm20614 | 0.26923737 | -0.602342381 | 2.112887281 | -0.285080225 |
| Gm23332 | 0 | NA | NA | NA |
| Gm20615 | 0.04393175 | -0.206490061 | 3.724945049 | -0.055434391 |
| Eif2b5 | 489.2421831 | 0.080636467 | 0.064008771 | 1.259772139 |
| Gm16618 | 0.359481004 | -1.025775347 | 1.766352822 | -0.58073072 |
| Dvl3 | 617.5796496 | -0.30928165 | 0.073646645 | -4.199534815 |
| Ap2m1 | 2464.108253 | -0.045336585 | 0.040855309 | -1.109686509 |
| Gm15760 | 0.195053328 | -0.206481366 | 2.683801234 | -0.076936162 |
| Abcf3 | 658.1946899 | 0.113622338 | 0.065418938 | 1.736841666 |
| Vwa5b2 | 365.3797502 | -0.361471651 | 0.201668924 | -1.792401342 |
| Mir1224 | 0 | NA | NA | NA |
| Alg3 | 92.27766346 | 0.104332985 | 0.13040634 | 0.800060676 |
| Ece2 | 434.908798 | -0.27684096 | 0.100851688 | -2.745030508 |
| Camk2n2 | 831.1974061 | -0.124119603 | 0.098329298 | -1.262285046 |
| Psmd2 | 2330.871484 | 0.136649945 | 0.082369405 | 1.658989102 |
| Gm4462 | 1.62367863 | 0.534657313 | 0.828922419 | 0.645002838 |
| Eif4g1 | 4413.780657 | 0.043714898 | 0.045436961 | 0.962099946 |
| Snord66 | 1.086592418 | 1.727294319 | 1.176240463 | 1.468487417 |
| Fam131a | 1339.029144 | -0.269090494 | 0.111046293 | -2.423228073 |
| Clcn2 | 427.9328786 | 0.084970605 | 0.108903916 | 0.780234611 |
| Polr2h | 104.4427863 | 0.126366082 | 0.115146927 | 1.097433387 |
| Thpo | 27.14520041 | -0.480357761 | 0.387403738 | -1.239940953 |
| Chrd | 234.488815 | 0.037968848 | 0.208500394 | 0.182104441 |
| Gm22741 | 0.075043205 | 0.755286614 | 3.724945049 | 0.202764498 |
| Rps10-ps2 | 203.4377443 | 0.420366532 | 0.136296809 | 3.084199368 |
| Gm9697 | 0 | NA | NA | NA |
| Teddm3 | 0.044544648 | 0.755286614 | 3.724945049 | 0.202764498 |
| AC168061.1 | 0 | NA | NA | NA |
| Ephb3 | 170.6997336 | -0.801119835 | 0.132032643 | -6.06758917 |
| Gm16863 | 0.106358623 | 0.274402725 | 3.724945049 | 0.073666248 |
| Mir7680 | 0 | NA | NA | NA |
| Vps8 | 810.4907143 | -0.042456696 | 0.056960468 | -0.745371266 |

| | | | | |
|---|---|---|---|---|
| 2510009E07Rik | 1017.889882 | 0.084688493 | 0.08869148 | 0.954866165 |
| Ehhadh | 44.10308811 | 0.104607574 | 0.21717656 | 0.481670648 |
| Gm24898 | 0 | NA | NA | NA |
| 1300002E11Rik | 152.2564682 | -0.606885591 | 0.180624159 | -3.359935879 |
| Gm26744 | 2.147345651 | 2.101078981 | 0.815642353 | 2.575980727 |
| Map3k13 | 489.4826492 | -0.632551847 | 0.081266104 | -7.783710779 |
| Gm6467 | 24.4487381 | 0.033418859 | 0.222371918 | 0.150283631 |
| 2610020F03Rik | 0.233260489 | 0.27439915 | 3.05580847 | 0.089795926 |
| Tmem41a | 97.70389257 | -0.062304529 | 0.131152565 | -0.475053837 |
| Liph | 10.28461881 | 0.507972453 | 0.369245869 | 1.375702467 |
| Senp2 | 1234.002812 | 0.029030115 | 0.072667684 | 0.399491403 |
| Gm4943 | 1.131366322 | -0.124971205 | 0.967930264 | -0.129111786 |
| Igf2bp2 | 94.65434489 | 1.238504652 | 0.303440038 | 4.081546592 |
| Gm25810 | 0 | NA | NA | NA |
| Gm5809 | 0.299370719 | -0.209942342 | 1.769513522 | -0.118644102 |
| Gm26980 | 0 | NA | NA | NA |
| Tra2b | 1042.494832 | 0.233710681 | 0.08324579 | 2.807477495 |
| Gm15776 | 0.868382071 | 1.564041091 | 1.269005739 | 1.232493315 |
| Etv5 | 804.3619116 | -0.624548395 | 0.118818697 | -5.256314107 |
| Dgkg | 917.1223497 | 0.049602436 | 0.101430013 | 0.489031147 |
| 9230117E06Rik | 0.105852363 | -0.206492286 | 3.724945049 | -0.055434988 |
| Mir8095 | 6.066797584 | -1.261673566 | 0.482982588 | -2.612254765 |
| Gm15651 | 2.236386351 | -0.736629816 | 0.674810857 | -1.091609313 |
| Gm22481 | 0 | NA | NA | NA |
| Mir6362 | 0 | NA | NA | NA |
| Crygs | 0.564059227 | 0.742438121 | 1.508105043 | 0.492298679 |
| Tbccd1 | 144.3603587 | -0.031607398 | 0.119678363 | -0.264102864 |
| Dnajb11 | 661.7688257 | 0.346015869 | 0.135185777 | 2.559558245 |
| Ahsg | 1.513507497 | -0.727975667 | 1.016095922 | -0.716443843 |
| Fetub | 1.838490909 | -0.612750427 | 0.817733796 | -0.749327507 |
| Hrg | 0.529344572 | 1.753302419 | 1.415769143 | 1.238409827 |
| Kng2 | 1.792037434 | 1.765119411 | 1.168238337 | 1.510924059 |
| Gm6493 | 33.39922536 | 0.334457078 | 0.210362309 | 1.589909707 |
| Gm23715 | 0 | NA | NA | NA |
| Kng1 | 0.385993897 | 0.274412695 | 1.769042953 | 0.155119295 |
| Eif4a2 | 5039.606027 | 0.110909245 | 0.063202647 | 1.754819634 |
| Snord2 | 0 | NA | NA | NA |
| Gm24201 | 0 | NA | NA | NA |
| AC154540.1 | 0 | NA | NA | NA |
| Snora81 | 0 | NA | NA | NA |
| Gm26447 | 0 | NA | NA | NA |
| Gm24616 | 7.869682849 | 0.373255306 | 0.384372959 | 0.971075869 |
| Rfc4 | 32.64420203 | -0.032464925 | 0.204988769 | -0.158374166 |
| Adipoq | 0.218718758 | 0.740540577 | 2.160441111 | 0.342772859 |
| B630019A10Rik | 0.40993559 | -0.592618102 | 1.549304898 | -0.382505795 |
| Gm6630 | 0.062426873 | 0.755288839 | 3.724945049 | 0.202765096 |
| Gm15746 | 0 | NA | NA | NA |

| | | | | |
|---|---|---|---|---|
| BC106179 | 11.069689 | 0.01528448 | 0.33898004 | 0.045089617 |
| St6gal1 | 316.7496297 | -0.044598993 | 0.10162624 | -0.438853123 |
| Gm23430 | 0 | NA | NA | NA |
| Gm23066 | 0 | NA | NA | NA |
| Rtp1 | 5.214837505 | -0.825325898 | 0.619764521 | -1.33167658 |
| Masp1 | 117.7213065 | 0.454821896 | 0.111744404 | 4.07019841 |
| Rtp4 | 910.9235272 | 6.828517861 | 0.234696816 | 29.0950596 |
| Gm6640 | 0.597217552 | 1.749092207 | 1.262070619 | 1.385890916 |
| Sst | 654.4591436 | -0.118188936 | 0.150591191 | -0.784832999 |
| Rtp2 | 3.004889707 | -1.03217681 | 0.708524126 | -1.456798396 |
| Bcl6 | 818.2508927 | 0.470625849 | 0.134624766 | 3.495834106 |
| Gm37419 | 0.567687411 | 1.63492994 | 1.409802472 | 1.159687241 |
| Lppos | 20.64171431 | -0.948036164 | 0.246461622 | -3.846587376 |
| Lpp | 689.3135003 | 0.551812335 | 0.131982075 | 4.180964243 |
| Gm24440 | 0 | NA | NA | NA |
| Mir28a | 2.122337162 | -0.432998124 | 0.865787267 | -0.500120689 |
| Gm22672 | 0.704522364 | -0.221436908 | 1.073298154 | -0.20631444 |
| A230028O05Rik | 10.09423595 | 2.984432486 | 0.605548234 | 4.928480206 |
| Tprg | 0.366915913 | 0.721691033 | 1.541227604 | 0.468257272 |
| Trp63 | 10.83564004 | -0.457101416 | 0.351728459 | -1.299586098 |
| Gm24928 | 0 | NA | NA | NA |
| Gm27005 | 0.212482393 | -0.206480633 | 2.184635012 | -0.094514933 |
| Gm4524 | 1.767199681 | -1.929478128 | 0.908455017 | -2.123911576 |
| P3h2 | 56.59938758 | -0.115140298 | 0.157943701 | -0.728995821 |
| AU015336 | 6.800499267 | -0.508556061 | 0.395258264 | -1.286642447 |
| Cldn1 | 56.29871389 | 0.940153617 | 0.180142721 | 5.218937573 |
| Cldn16 | 0 | NA | NA | NA |
| Tmem207 | 0.473315337 | 0.323155645 | 1.296989964 | 0.249158169 |
| Il1rap | 1004.716365 | 0.447037174 | 0.069690184 | 6.41463617 |
| Gm20319 | 0.326503072 | -0.540120219 | 1.588169525 | -0.340089777 |
| Gmnc | 8.309823615 | -0.919958079 | 0.4286047 | -2.146402219 |
| Ostn | 1.447793589 | 1.35135105 | 1.108060415 | 1.219564413 |
| Gm15970 | 0 | NA | NA | NA |
| Uts2b | 1.222794617 | -0.420466553 | 1.21471634 | -0.346143819 |
| Ccdc50 | 1385.300587 | 0.260512693 | 0.067014762 | 3.88739267 |
| Fgf12 | 2033.20002 | -0.09396334 | 0.081746186 | -1.149452293 |
| Mir690 | 11.63860447 | -1.168798088 | 0.399668824 | -2.924416464 |
| Mb21d2 | 459.0744955 | 0.14891052 | 0.083196169 | 1.789872309 |
| Hrasls | 202.7297218 | 0.117294645 | 0.082039779 | 1.429728927 |
| Atp13a5 | 116.1853686 | -0.524896554 | 0.152018282 | -3.452851501 |
| Gm15752 | 0.667341687 | -2.149831613 | 1.229716455 | -1.74823359 |
| Atp13a4 | 513.9834346 | 0.7895712 | 0.15338748 | 5.147559625 |
| Opa1 | 2535.740764 | -0.074230389 | 0.055222519 | -1.344205044 |
| Gm26834 | 2.849524632 | -0.047669887 | 0.632282932 | -0.075393285 |
| Gm26569 | 0.312910946 | 0.755307749 | 1.728418962 | 0.436993441 |
| Gm1968 | 0.303184894 | 0.274378537 | 3.724945049 | 0.073659754 |
| 4632428C04Rik | 13.7946314 | -0.015416062 | 0.309292943 | -0.049842916 |

| | | | | |
|---|---|---|---|---|
| Hes1 | 94.25849431 | 0.418415023 | 0.128970013 | 3.244281463 |
| n-R5s32 | 0 | NA | NA | NA |
| 1700025H01Rik | 0 | NA | NA | NA |
| Gm26562 | 0.090074004 | 0.274402725 | 3.724945049 | 0.073666248 |
| Cpn2 | 0.207381971 | -0.206487621 | 2.582242976 | -0.079964443 |
| Lrrc15 | 1.556998443 | 1.475598047 | 0.994396651 | 1.483912928 |
| Gp5 | 14.94000079 | 0.702363099 | 0.306499774 | 2.291561555 |
| Atp13a3 | 1577.269787 | -0.025036426 | 0.054636587 | -0.458235544 |
| Tmem44 | 273.2629563 | -0.711551039 | 0.124041668 | -5.736387211 |
| Lsg1 | 245.3569069 | 0.264533787 | 0.10398245 | 2.544023402 |
| Fam43a | 229.047573 | -0.194206992 | 0.095721237 | -2.028880925 |
| Gm26295 | 0.327620686 | 1.219034923 | 2.061932165 | 0.591210004 |
| Gm25375 | 0 | NA | NA | NA |
| Xxylt1 | 202.0539679 | -0.26936919 | 0.090861109 | -2.964625813 |
| Gm15742 | 2.112839908 | -1.060315537 | 0.827061754 | -1.282027047 |
| Acap2 | 1866.515927 | -0.103059157 | 0.080477962 | -1.28058855 |
| Ppp1r2 | 1509.036798 | 0.336748532 | 0.078904131 | 4.267818769 |
| Gm27415 | 0 | NA | NA | NA |
| Apod | 8482.021723 | 3.199242537 | 0.259694557 | 12.31925142 |
| 1700007E05Rik | 2.68870073 | -0.042382623 | 0.707162225 | -0.059933381 |
| Bdh1 | 477.7842783 | -0.000651073 | 0.064968527 | -0.010021354 |
| Gm15743 | 22.43508857 | 0.656052289 | 0.314160173 | 2.08827326 |
| Dlg1 | 2632.440439 | -0.049722876 | 0.04628358 | -1.074309213 |
| Gm24879 | 0 | NA | NA | NA |
| Mfi2 | 8.341406001 | -1.241190215 | 0.397236764 | -3.124560281 |
| Gm15703 | 0.299449808 | 0.274410379 | 2.052015864 | 0.133727221 |
| Pigz | 89.07597162 | -0.263049904 | 0.150347139 | -1.749616961 |
| NCBP2-AS2 | 84.09966569 | -0.320134463 | 0.129805499 | -2.466262707 |
| 0610012G03Rik | 0 | NA | NA | NA |
| Ncbp2 | 349.8016824 | -0.457541444 | 0.126587711 | -3.61442228 |
| Senp5 | 762.9688528 | -0.008987616 | 0.069321035 | -0.129652078 |
| Gm15696 | 12.05015047 | -0.044162519 | 0.33336733 | -0.132474046 |
| Gm26269 | 0.072025462 | 0.755288839 | 3.724945049 | 0.202765096 |
| Gm15694 | 6.232026559 | -0.156805079 | 0.445595286 | -0.35190022 |
| Hmgb1-ps6 | 11.46318819 | 0.060270168 | 0.356456537 | 0.169081395 |
| Pak2 | 1045.01706 | 0.14510615 | 0.083259597 | 1.742815912 |
| Gm26419 | 0 | NA | NA | NA |
| Pigx | 116.9372743 | 0.068204369 | 0.137453752 | 0.496198671 |
| Cep19 | 336.7429659 | -0.164609823 | 0.090797647 | -1.812930487 |
| Gm15729 | 0.971776759 | -0.711389214 | 1.126120075 | -0.631717017 |
| Nrros | 147.122329 | 1.270583758 | 0.138078805 | 9.201873968 |
| Bex6 | 0.377709955 | 2.006996205 | 1.519258462 | 1.321036714 |
| Fbxo45 | 678.3823491 | -0.079107466 | 0.073272165 | -1.079638718 |
| Gm25848 | 45.86024405 | -0.523883328 | 0.325910128 | -1.607447216 |
| Wdr53 | 71.48528253 | 0.224814983 | 0.148083569 | 1.518162915 |
| Mir1946a | 0 | NA | NA | NA |
| Smco1 | 1.487580774 | 1.518937192 | 0.815201409 | 1.863266151 |

| | | | | |
|---|---|---|---|---|
| Rnf168 | 482.2284669 | -0.003019572 | 0.059327986 | -0.050896257 |
| Ubxn7 | 1865.438353 | 0.20442041 | 0.074759844 | 2.734361098 |
| Tm4sf19 | 8.203405253 | 3.208106917 | 0.980828856 | 3.270812128 |
| Gm20056 | 14.90719541 | 0.385417012 | 0.357929988 | 1.076794416 |
| Tctex1d2 | 113.226245 | -0.105558732 | 0.107595582 | -0.981069387 |
| Pcyt1a | 752.3083677 | 0.259516594 | 0.093474259 | 2.776342899 |
| Slc51a | 0 | NA | NA | NA |
| Zdhhc19 | 0.768268477 | -0.970160146 | 1.277507174 | -0.759416594 |
| Gm5405 | 0 | NA | NA | NA |
| Tfrc | 1390.85657 | 0.04859797 | 0.113792679 | 0.42707466 |
| Tnk2 | 2566.202347 | -0.043940375 | 0.047876112 | -0.917793311 |
| Tnk2os | 0.364228253 | -0.664652072 | 1.597498839 | -0.416057937 |
| Gm9556 | 0 | NA | NA | NA |
| Muc4 | 0.409426173 | 1.239731403 | 1.699187069 | 0.729602659 |
| Muc20 | 0.330893424 | 2.082206921 | 1.543751684 | 1.348796534 |
| Gm933 | 0.097139547 | 0.274393828 | 3.724945049 | 0.073663859 |
| Gm26769 | 0 | NA | NA | NA |
| 1700021K19Rik | 991.8281154 | -0.39485509 | 0.104336111 | -3.784452837 |
| Fyttd1 | 1180.673463 | 0.140754234 | 0.094140194 | 1.495155551 |
| Lrch3 | 977.5154375 | -0.112234296 | 0.065238458 | -1.720370151 |
| Gm6611 | 33.95866455 | -0.535892611 | 0.345096907 | -1.552875729 |
| Gm17106 | 0.097139547 | 0.274393828 | 3.724945049 | 0.073663859 |
| n-R5s33 | 0.130017973 | -0.206489706 | 3.662785653 | -0.056375045 |
| Iqcg | 37.65510366 | -0.004385072 | 0.200305602 | -0.021891909 |
| Rpl35a | 394.3677081 | 0.385268645 | 0.093570231 | 4.117427524 |
| Lmln | 221.22393 | -0.352212809 | 0.089037476 | -3.955781609 |
| Gm22231 | 0.549303492 | -1.05880701 | 1.316131783 | -0.804484037 |
| Mir1947 | 0.769571883 | 1.071096802 | 1.075846258 | 0.995585376 |
| Gm23916 | 0 | NA | NA | NA |
| Osbpl11 | 369.2866785 | -0.04630954 | 0.094341418 | -0.490871783 |
| Snx4 | 448.4376452 | 0.421043781 | 0.08300299 | 5.072633932 |
| 1700007L15Rik | 6.048360371 | -0.341329551 | 0.389474256 | -0.876385398 |
| Zfp148 | 2196.281864 | -0.066555362 | 0.047612428 | -1.397856941 |
| Slc12a8 | 28.22738804 | -0.234439037 | 0.216745002 | -1.081635261 |
| Heg1 | 831.6435367 | 0.291638512 | 0.078590393 | 3.71086719 |
| Gm15658 | 0 | NA | NA | NA |
| Gm15657 | 0.194279094 | 0.274391173 | 3.724945049 | 0.073663147 |
| Muc13 | 4.915050087 | 3.840170154 | 0.908807931 | 4.225502466 |
| Itgb5 | 478.5994184 | 0.124788404 | 0.198723546 | 0.627949766 |
| Umps | 206.1452455 | -0.128056441 | 0.091567406 | -1.39849371 |
| Gm15829 | 0.247806867 | 1.114639832 | 2.094906211 | 0.532071472 |
| Kalrn | 4921.446429 | -0.335149156 | 0.182968595 | -1.8317305 |
| Ropn1 | 0.318826904 | -1.43003122 | 1.74441836 | -0.819775378 |
| Ccdc14 | 47.35523537 | 0.185510694 | 0.174566501 | 1.062693542 |
| Mylk | 164.66998 | 0.6551249 | 0.132402229 | 4.947989959 |
| 1700119H24Rik | 0.164132501 | 1.537077023 | 3.390982371 | 0.453283696 |
| Hacd2 | 451.6459504 | 0.518058841 | 0.09467822 | 5.471784732 |

| | | | | |
|---|---|---|---|---|
| Adcy5 | 2609.480313 | -0.444472552 | 0.131395606 | -3.382704833 |
| Gm25179 | 0 | NA | NA | NA |
| Sec22a | 183.1160017 | 0.022346412 | 0.113441619 | 0.19698601 |
| Gm25079 | 0 | NA | NA | NA |
| Gm23518 | 0.038981395 | 0.274393828 | 3.724945049 | 0.073663859 |
| Pdia5 | 30.59877438 | -0.39330338 | 0.232318849 | -1.692946488 |
| Gm25548 | 0.246683951 | -0.206481881 | 1.854028192 | -0.111369332 |
| 1600019K03Rik | 0.049553047 | 0.274391604 | 3.724945049 | 0.073663262 |
| Sema5b | 130.570813 | -0.540582625 | 0.116868543 | -4.62556142 |
| Gm5963 | 36.49740693 | 0.244219721 | 0.293432286 | 0.832286467 |
| Dirc2 | 652.6475954 | -0.026352837 | 0.077863977 | -0.33844709 |
| Gm25967 | 2.125672366 | 0.670315665 | 0.682780213 | 0.981744422 |
| Hspbap1 | 70.57271391 | -0.844486011 | 0.203802915 | -4.143640493 |
| Gm26838 | 0 | NA | NA | NA |
| Parp14 | 2858.919116 | 6.343073444 | 0.187644633 | 33.80364962 |
| Gm10237 | 0.042716945 | -0.206490061 | 3.724945049 | -0.055434391 |
| Dtx3l | 1910.456235 | 4.981879809 | 0.167350753 | 29.76909102 |
| Parp9 | 1102.792659 | 5.057047617 | 0.168612039 | 29.99220956 |
| Gm15564 | 359.9778991 | 0.517657904 | 0.972551041 | 0.53226811 |
| Kpna1 | 1948.470927 | 0.022755423 | 0.057720603 | 0.394233986 |
| Wdr5b | 49.21408818 | 0.122476118 | 0.167286174 | 0.732135325 |
| Fam162a | 149.3833853 | -0.374301538 | 0.121469862 | -3.081435442 |
| Ccdc58 | 55.1879007 | -0.124823301 | 0.162060414 | -0.770226969 |
| Csta1 | 0 | NA | NA | NA |
| Gm10092 | 0.045882455 | 0.755288839 | 3.724945049 | 0.202765096 |
| Gm23080 | 0 | NA | NA | NA |
| Stfa2l1 | 0.423264953 | 1.709131408 | 1.436647373 | 1.189666608 |
| Gm22885 | 0 | NA | NA | NA |
| Gm5483 | 0.535697641 | 2.343465072 | 1.692053694 | 1.384982687 |
| Gm25024 | 0 | NA | NA | NA |
| Gm10913 | 0.156524568 | 1.236192361 | 2.823069743 | 0.437889416 |
| Gm5416 | 0.092228164 | 0.75528439 | 3.724945049 | 0.202763901 |
| Gm24801 | 0 | NA | NA | NA |
| 2010005H15Rik | 0.699696385 | 2.649592528 | 1.218684162 | 2.174142087 |
| Gm10813 | 0 | NA | NA | NA |
| Gm24569 | 0 | NA | NA | NA |
| Stfa1 | 0 | NA | NA | NA |
| Gm23306 | 0 | NA | NA | NA |
| Gm4758 | 0 | NA | NA | NA |
| Gm25254 | 0 | NA | NA | NA |
| Gm24437 | 0 | NA | NA | NA |
| BC117090 | 0 | NA | NA | NA |
| Gm25465 | 0.060174537 | 0.755286614 | 3.724945049 | 0.202764498 |
| BC100530 | 0.115000406 | -0.206492286 | 3.724945049 | -0.055434988 |
| Gm15845 | 0.075043205 | 0.755286614 | 3.724945049 | 0.202764498 |
| Gm24218 | 0 | NA | NA | NA |
| Gm22349 | 0 | NA | NA | NA |

| | | | | |
|---|---|---|---|---|
| Stfa2 | 0.143022002 | 0.755289479 | 3.547809333 | 0.212888971 |
| Gm22263 | 0 | NA | NA | NA |
| Gm25903 | 0 | NA | NA | NA |
| Stfa3 | 0.131209559 | 0.755295049 | 2.976009072 | 0.253794606 |
| Casr | 17.16410259 | 0.47146026 | 0.287381089 | 1.640540307 |
| Gm25902 | 0 | NA | NA | NA |
| Cd86 | 81.75402155 | 2.966738874 | 0.208301102 | 14.24255009 |
| Ildr1 | 14.35142951 | 1.689390659 | 0.359379567 | 4.700853402 |
| Slc15a2 | 296.0330333 | -0.727429889 | 0.218360796 | -3.331320922 |
| Gm8425 | 0.186025035 | 0.274409551 | 2.673127101 | 0.102654884 |
| Gm22617 | 0 | NA | NA | NA |
| Eaf2 | 14.00843118 | 0.801431025 | 0.295448703 | 2.712589414 |
| Iqcb1 | 366.8902262 | 0.008765688 | 0.083484293 | 0.104998053 |
| Golgb1 | 1930.593704 | -0.194146099 | 0.107899949 | -1.799315952 |
| 4930565N06Rik | 0.233785498 | 0.755298085 | 2.508473841 | 0.301098649 |
| Hcls1 | 216.1493986 | 3.378370055 | 0.199092214 | 16.96887079 |
| Fbxo40 | 25.85214164 | 0.371917316 | 0.31728303 | 1.172194165 |
| Polq | 42.09947402 | -0.106614341 | 0.28825188 | -0.369865205 |
| Stxbp5l | 2108.032249 | -0.156746372 | 0.094960763 | -1.650643551 |
| Gm15665 | 42.97331767 | -0.306612024 | 0.29600775 | -1.035824314 |
| Gtf2e1 | 231.0323693 | -0.078646306 | 0.134360621 | -0.585337466 |
| Rabl3 | 128.7194147 | 0.405603363 | 0.122206176 | 3.319008727 |
| Hgd | 1.882553937 | 0.234818871 | 0.773170434 | 0.303709067 |
| Ndufb4 | 405.4328191 | 0.093329778 | 0.135666011 | 0.687937804 |
| Fstl1 | 650.539503 | 0.710836217 | 0.175369658 | 4.053359201 |
| n-R5s34 | 0 | NA | NA | NA |
| Lrrc58 | 3637.314835 | 0.23358009 | 0.100203684 | 2.331052913 |
| Gm15725 | 0 | NA | NA | NA |
| Gpr156 | 12.90207318 | 0.627309911 | 0.456065926 | 1.375480771 |
| Gm25140 | 0.106056992 | 1.236179401 | 3.724945049 | 0.331865138 |
| Gsk3b | 4829.972571 | -0.129453301 | 0.045467263 | -2.847176026 |
| SNORA17 | 0 | NA | NA | NA |
| Gm25617 | 0 | NA | NA | NA |
| Gm27931 | 0 | NA | NA | NA |
| Nr1i2 | 0.200534954 | -1.090116161 | 2.636631645 | -0.413450306 |
| Maats1 | 18.76607067 | -0.005780676 | 0.248014389 | -0.023307825 |
| Maats1os | 0.072025462 | 0.755288839 | 3.724945049 | 0.202765096 |
| Cox17 | 102.4271693 | -0.36072362 | 0.207585969 | -1.737707136 |
| Gm21987 | 0.2386254 | -0.206480538 | 2.112272161 | -0.09775281 |
| Popdc2 | 3.459091227 | 0.387076163 | 0.526212901 | 0.735588508 |
| Pla1a | 56.89416345 | 3.400364482 | 0.362505424 | 9.380175451 |
| Adprh | 679.8642621 | 0.020658998 | 0.08401107 | 0.24590805 |
| D930030I03Rik | 2.891043216 | -0.538938299 | 0.602105925 | -0.895088848 |
| Cd80 | 14.42150522 | 2.288619599 | 0.478739051 | 4.780515806 |
| Timmdc1 | 207.9193946 | -0.353493453 | 0.134552806 | -2.627172663 |
| Gm15953 | 0 | NA | NA | NA |
| Poglut1 | 522.8527119 | 0.086294454 | 0.062296846 | 1.38521387 |

| | | | | |
|---|---|---|---|---|
| Tmem39a | 165.5822775 | -0.061843821 | 0.13272105 | -0.465968438 |
| Gm17103 | 0.159330469 | 1.236180454 | 3.465376669 | 0.356723258 |
| Arhgap31 | 855.9405782 | 0.388733113 | 0.094642481 | 4.107385087 |
| Gm15530 | 5.747139663 | -0.499502942 | 0.445952039 | -1.120082202 |
| B4galt4 | 168.6399155 | -0.047676243 | 0.098554728 | -0.48375399 |
| Gm15575 | 0 | NA | NA | NA |
| Upk1b | 9.705317616 | 1.573393699 | 0.431981875 | 3.642267857 |
| 4930435E12Rik | 0 | NA | NA | NA |
| Gm15803 | 0 | NA | NA | NA |
| Gm15802 | 0 | NA | NA | NA |
| Igsf11 | 559.2207678 | -0.188223221 | 0.138325975 | -1.360722175 |
| Gm22500 | 7.621091703 | -0.828937238 | 0.514688653 | -1.610560545 |
| Gm24795 | 0 | NA | NA | NA |
| Gm8609 | 0.167459142 | 0.755298223 | 2.757565248 | 0.2739004 |
| Lsamp | 5421.197146 | -0.085933708 | 0.063054397 | -1.362850368 |
| Gm37508 | 3.928946461 | -0.236834757 | 0.56431529 | -0.419685169 |
| Gm22543 | 16.99085758 | -0.780338756 | 0.344381998 | -2.265910416 |
| Gm36939 | 48.56639706 | -0.148063018 | 0.231253382 | -0.640263147 |
| Gm27887 | 0 | NA | NA | NA |
| Gm27717 | 0 | NA | NA | NA |
| Gm26381 | 1.397116197 | -0.16793368 | 0.753805895 | -0.222781064 |
| AC124193.1 | 0 | NA | NA | NA |
| Gm27587 | 0 | NA | NA | NA |
| Gm28750 | 0 | NA | NA | NA |
| Gap43 | 1479.213936 | -0.270688889 | 0.095912171 | -2.822257964 |
| Mir6540 | 2.543989632 | -1.061238716 | 0.694470005 | -1.528127507 |
| 4932412D23Rik | 1.816140692 | -0.814519159 | 0.803210119 | -1.014079802 |
| Zbtb20 | 6323.085067 | -0.00086837 | 0.083917609 | -0.010347884 |
| BC002163 | 27.11636185 | -0.269452147 | 0.215289646 | -1.251579686 |
| Gm25873 | 5.983328485 | 0.432180057 | 0.432653955 | 0.998904671 |
| Gm15712 | 29.69002061 | -0.121898819 | 0.352677727 | -0.345637985 |
| Gm37946 | 0 | NA | NA | NA |
| Gm15711 | 21.32018645 | -0.208539471 | 0.309898674 | -0.672927922 |
| Gm9968 | 1.290099097 | 1.361342875 | 0.994741551 | 1.36853927 |
| Gm15713 | 0.087930927 | -0.206490058 | 3.724945049 | -0.05543439 |
| Mir568 | 0 | NA | NA | NA |
| Tigit | 22.00729436 | 7.128174521 | 0.922702818 | 7.725319987 |
| Drd3 | 8.262393275 | -0.564839074 | 0.421275236 | -1.340783949 |
| Gm25996 | 0.1580736 | -0.206480907 | 3.474509404 | -0.059427356 |
| Gm26074 | 0 | NA | NA | NA |
| Qtrtd1 | 200.210793 | -0.314660983 | 0.110817767 | -2.839445265 |
| Gm25989 | 9.070232471 | -0.588651475 | 0.427709012 | -1.376289625 |
| 2610015P09Rik | 57.77977805 | -0.676795862 | 0.219993986 | -3.076428924 |
| Zdhhc23 | 82.33084375 | -0.322440701 | 0.153526342 | -2.100230471 |
| Gramd1c | 24.07085944 | -0.019274403 | 0.208526706 | -0.09243134 |
| Mir3470b | 0 | NA | NA | NA |
| Atp6v1a | 5242.393468 | -0.115821054 | 0.059926893 | -1.932705794 |

| | | | | |
|---|---|---|---|---|
| Naa50 | 1580.032754 | 0.129811916 | 0.090393054 | 1.436082871 |
| Gm608 | 1399.505967 | -0.145349895 | 0.072787597 | -1.996904705 |
| Sidt1 | 676.9943462 | 0.329793758 | 0.140563956 | 2.346218527 |
| Gm26732 | 6.046421774 | -1.166978041 | 0.405330167 | -2.879080155 |
| Spice1 | 128.0135962 | -0.7811178 | 0.168576696 | -4.633604869 |
| Cfap44 | 93.24835868 | -0.697598165 | 0.173526352 | -4.020128117 |
| Boc | 445.6221661 | -0.474127284 | 0.167160082 | -2.836366666 |
| Mir3081 | 1.772274333 | -1.745796071 | 0.851432079 | -2.050423181 |
| BC027231 | 116.0115484 | 0.094453228 | 0.166262163 | 0.568098153 |
| Gtpbp8 | 357.6076025 | 0.141290954 | 0.084119225 | 1.679651161 |
| Cd200r1 | 51.42872498 | 1.357077216 | 0.306379264 | 4.429402955 |
| Cd200r4 | 69.15376312 | 2.54677654 | 0.413445013 | 6.159891784 |
| Cd200r2 | 1.386349714 | 0.94857069 | 0.829366686 | 1.143728951 |
| Cd200r3 | 0.534659868 | 0.716179269 | 1.393413158 | 0.513974815 |
| Ccdc80 | 217.6073964 | 0.818746 | 0.159772455 | 5.124450274 |
| Slc35a5 | 550.0251731 | -0.163649073 | 0.101668661 | -1.609631435 |
| Atg3 | 539.1722443 | 0.147180754 | 0.070489104 | 2.087992979 |
| Gm6030 | 17.33716352 | 0.021003481 | 0.296467489 | 0.070845815 |
| Btla | 10.07865719 | 1.345826348 | 0.402072484 | 3.347223206 |
| Cd200 | 1252.555305 | 0.810170693 | 0.104419266 | 7.758823838 |
| Gm5406 | 0.966602524 | 0.288383719 | 1.205448436 | 0.239233558 |
| Gm609 | 0.707603298 | 0.022846396 | 1.301196037 | 0.017557997 |
| Gm16011 | 0.581873452 | 0.332857156 | 1.326732312 | 0.250884939 |
| Gm6903 | 0.049553047 | 0.274391604 | 3.724945049 | 0.073663262 |
| Slc9c1 | 0.038981395 | 0.274393828 | 3.724945049 | 0.073663859 |
| Gcsam | 0.711975465 | 0.009701155 | 1.16792252 | 0.008306334 |
| BC016579 | 3.578730896 | -1.04958437 | 0.687845295 | -1.525901793 |
| Tmprss7 | 24.10678365 | -0.837540649 | 0.310522596 | -2.697197111 |
| Tagln3 | 1506.536885 | -0.262468168 | 0.095635374 | -2.744467414 |
| Abhd10 | 203.4530973 | -0.228213471 | 0.09382732 | -2.43227101 |
| Gm15591 | 0.060174537 | 0.755286614 | 3.724945049 | 0.202764498 |
| Phldb2 | 129.9886621 | -0.312698247 | 0.127033998 | -2.461531976 |
| Gm15640 | 0.229952095 | 0.263768843 | 2.124178835 | 0.124174499 |
| Gm15638 | 0.108266322 | 0.274389235 | 3.724945049 | 0.073662626 |
| Plcxd2 | 1552.118851 | -0.615831823 | 0.078676555 | -7.827386751 |
| Gm24754 | 0 | NA | NA | NA |
| Cd96 | 3.576396818 | 3.724145395 | 0.878917233 | 4.237196921 |
| Gm4737 | 112.4227191 | 0.272704613 | 0.165712345 | 1.645650561 |
| Gm26297 | 0 | NA | NA | NA |
| Pvrl3 | 517.9500622 | -0.556245867 | 0.087324381 | -6.369880468 |
| Gm6912 | 0 | NA | NA | NA |
| Gm24651 | 0 | NA | NA | NA |
| Gm25318 | 0 | NA | NA | NA |
| Gm25339 | 0 | NA | NA | NA |
| Gm7275 | 0 | NA | NA | NA |
| Gm7204 | 1.57829385 | -0.68062751 | 1.252955932 | -0.543217437 |
| Dppa4 | 0 | NA | NA | NA |

| | | | | |
|---|---|---|---|---|
| Dppa2 | 0.379773675 | 0.755308329 | 1.683019321 | 0.448781734 |
| n-R5s36 | 0 | NA | NA | NA |
| Morc1 | 1.278798984 | -0.519103942 | 1.012391547 | -0.512750175 |
| Trat1 | 6.136963414 | 1.427798683 | 0.460102365 | 3.103219614 |
| Retnlb | 0 | NA | NA | NA |
| Retnla | 0.060174537 | 0.755286614 | 3.724945049 | 0.202764498 |
| Retnlg | 1.932622373 | 1.497106213 | 1.203737876 | 1.243714469 |
| Dzip3 | 1926.33653 | -0.225631132 | 0.095028756 | -2.374345853 |
| Gm22576 | 7.017922756 | -0.333474062 | 0.428248457 | -0.778692967 |
| C330027C09Rik | 62.11505808 | 0.587774398 | 0.167031971 | 3.518933499 |
| Myh15 | 9.416055846 | -0.286202772 | 0.332642769 | -0.860390782 |
| 1700026J12Rik | 0.047948698 | -0.206487837 | 3.724945049 | -0.055433794 |
| Gm5407 | 0.075043205 | 0.755286614 | 3.724945049 | 0.202764498 |
| Gm6931 | 0.04393175 | -0.206490061 | 3.724945049 | -0.055434391 |
| Ift57 | 340.5122929 | -0.122734958 | 0.085463583 | -1.436108267 |
| Gm15518 | 0.295614739 | 0.274414019 | 2.088175921 | 0.131413266 |
| Gm16619 | 0.436453079 | 1.816507081 | 1.380146455 | 1.316169798 |
| Cd47 | 3446.906187 | 0.758951542 | 0.098711507 | 7.688582258 |
| AC107830.1 | 0 | NA | NA | NA |
| Gm8824 | 0.047948698 | -0.206487837 | 3.724945049 | -0.055433794 |
| Bbx | 1297.513501 | 0.80154005 | 0.112168876 | 7.145832943 |
| G730013B05Rik | 0.417980132 | 0.145228645 | 1.513557514 | 0.095951851 |
| Ccdc54 | 0.042716945 | -0.206490061 | 3.724945049 | -0.055434391 |
| Gm9575 | 16.04186629 | 0.051697306 | 0.274696839 | 0.188197673 |
| 5330426P16Rik | 100.6912749 | -0.862325586 | 0.178198001 | -4.839142883 |
| Mir6363 | 0 | NA | NA | NA |
| 1700116B05Rik | 0 | NA | NA | NA |
| Gm24925 | 0 | NA | NA | NA |
| Gm6767 | 3.585344364 | 0.484989093 | 0.549658898 | 0.882345569 |
| Cblb | 789.0152622 | 0.551239303 | 0.092576458 | 5.954422055 |
| Gm23513 | 0 | NA | NA | NA |
| Alcam | 2206.12951 | 0.02722339 | 0.071329547 | 0.381656565 |
| Gm25723 | 0 | NA | NA | NA |
| Gm22459 | 0 | NA | NA | NA |
| Gm25041 | 0 | NA | NA | NA |
| Gm25853 | 0 | NA | NA | NA |
| Gm23180 | 0 | NA | NA | NA |
| Gm22977 | 0 | NA | NA | NA |
| Zpld1 | 1.833497833 | -0.657668102 | 0.797109326 | -0.825066375 |
| Mir5118 | 0 | NA | NA | NA |
| Gm23003 | 0 | NA | NA | NA |
| Gm26369 | 0 | NA | NA | NA |
| Nfkbiz | 257.9280466 | 1.158800848 | 0.165416076 | 7.005370179 |
| Nxpe3 | 312.8495634 | -0.081970342 | 0.093964311 | -0.872356119 |
| Cep97 | 386.1470673 | -0.656452951 | 0.147333046 | -4.455571701 |
| Rpl24 | 16.95551249 | 0.457820386 | 0.308959353 | 1.481814295 |
| Gm28037 | 0 | NA | NA | NA |

| | | | | |
|---|---|---|---|---|
| Zbtb11os1 | 37.78932254 | 0.509309487 | 0.199521922 | 2.552649268 |
| Zbtb11 | 1088.837139 | -0.053251968 | 0.068976366 | -0.772032088 |
| Pcnp | 1051.54738 | 0.288184799 | 0.078111561 | 3.689400092 |
| Trmt10c | 191.2919896 | 0.301626057 | 0.101813832 | 2.962525348 |
| Gm22422 | 89.64062021 | -0.108518932 | 0.388590841 | -0.279262713 |
| Senp7 | 795.4829463 | -0.287588526 | 0.119359686 | -2.409427631 |
| Gm24047 | 0.105143817 | 0.274402725 | 3.724945049 | 0.073666248 |
| Impg2 | 63.00063838 | -0.023348219 | 0.151797928 | -0.153811183 |
| Abi3bp | 266.089939 | -0.165900152 | 0.205832017 | -0.805997795 |
| Tfg | 756.0033368 | 0.081674072 | 0.057843242 | 1.411989874 |
| Adgrg7 | 0.562361829 | 1.649566784 | 1.402195012 | 1.176417524 |
| Gm15693 | 0.044191549 | -0.206490061 | 3.724945049 | -0.055434391 |
| Tmem45a | 10.44425536 | 1.520729987 | 0.397159414 | 3.829016589 |
| Gm15839 | 0.181290818 | 0.274405526 | 2.740334485 | 0.100135778 |
| Gm15840 | 0 | NA | NA | NA |
| Tmem45a2 | 3.654246218 | 1.761970525 | 0.608644479 | 2.894909237 |
| Tomm70a | 2858.851143 | 0.226695323 | 0.081159752 | 2.7931988 |
| Nit2 | 67.55208235 | -0.016771102 | 0.143169286 | -0.117141757 |
| Gm16892 | 2.540529182 | -0.048920914 | 0.574995875 | -0.085080462 |
| Tbc1d23 | 546.2969052 | 0.066810471 | 0.057904588 | 1.153802723 |
| Tmem30c | 8.630201669 | -0.063550967 | 0.396776173 | -0.160168306 |
| Gm26800 | 0.820767868 | 0.702811874 | 1.343353423 | 0.523177194 |
| Cmss1 | 65.09916204 | 0.189847756 | 0.129611196 | 1.464748121 |
| Filip1l | 99.78162097 | 0.46184908 | 0.286898482 | 1.609799668 |
| Col8a1 | 91.62796141 | 0.671190637 | 0.169590689 | 3.957709241 |
| Gm24039 | 0 | NA | NA | NA |
| Gm37458 | 0.072025462 | 0.755288839 | 3.724945049 | 0.202765096 |
| Gm37825 | 0 | NA | NA | NA |
| 4930461C15Rik | 1.662681963 | -0.565307018 | 0.760403339 | -0.743430478 |
| Dcbld2 | 658.0902064 | 0.147997674 | 0.093534684 | 1.582275882 |
| St3gal6 | 198.8642845 | 0.315013256 | 0.10177281 | 3.095259491 |
| Gm813 | 1.602843074 | 3.531771676 | 1.086939797 | 3.249279937 |
| E330017A01Rik | 0 | NA | NA | NA |
| Cpox | 461.6201025 | -0.101092333 | 0.061242942 | -1.65067729 |
| Gpr15 | 6.100871677 | 3.688636503 | 1.009873869 | 3.652571492 |
| Cldnd1 | 811.4291424 | 0.24173537 | 0.088235967 | 2.739646622 |
| Gm27564 | 0 | NA | NA | NA |
| Gm26019 | 0 | NA | NA | NA |
| Olfr172 | 0 | NA | NA | NA |
| Olfr173 | 0 | NA | NA | NA |
| Olfr175-ps1 | 0.049553047 | 0.274391604 | 3.724945049 | 0.073663262 |
| Olfr177 | 0 | NA | NA | NA |
| Olfr178 | 0 | NA | NA | NA |
| Olfr180 | 0 | NA | NA | NA |
| Olfr181 | 0 | NA | NA | NA |
| Olfr183 | 0 | NA | NA | NA |
| Olfr186 | 0.172571121 | 0.755296067 | 2.73763 | 0.275894137 |

| | | | | |
|---|---|---|---|---|
| Olfr187 | 0.106520246 | 0.755282166 | 3.724945049 | 0.202763304 |
| Olfr190 | 0 | NA | NA | NA |
| Olfr191 | 0 | NA | NA | NA |
| Olfr192 | 0.046345709 | 0.274391604 | 3.724945049 | 0.073663262 |
| Olfr193 | 0 | NA | NA | NA |
| Olfr194 | 0 | NA | NA | NA |
| Olfr195 | 0.097139547 | 0.274393828 | 3.724945049 | 0.073663859 |
| Olfr196 | 0 | NA | NA | NA |
| Olfr197 | 0 | NA | NA | NA |
| Olfr198 | 0.194201709 | -0.992840384 | 3.270763994 | -0.303549992 |
| Olfr199 | 0 | NA | NA | NA |
| Olfr201 | 0 | NA | NA | NA |
| Olfr202 | 0 | NA | NA | NA |
| Olfr203 | 0 | NA | NA | NA |
| Olfr204 | 0.068097359 | -0.206487837 | 3.724945049 | -0.055433794 |
| Olfr205 | 0.046345709 | 0.274391604 | 3.724945049 | 0.073663262 |
| Olfr206 | 0.075043205 | 0.755286614 | 3.724945049 | 0.202764498 |
| Olfr209 | 0.048949532 | -0.206487837 | 3.724945049 | -0.055433794 |
| Gabrr3 | 1.001170369 | 1.990954962 | 1.027970208 | 1.936782744 |
| 1700022E09Rik | 0.097139547 | 0.274393828 | 3.724945049 | 0.073663859 |
| Mina | 105.9311312 | 0.010606296 | 0.138509069 | 0.076574741 |
| Crybg3 | 125.2170434 | 0.101021774 | 0.111240092 | 0.908141773 |
| Gm9581 | 73.34347339 | 0.033762644 | 0.127664425 | 0.264463992 |
| Arl6 | 230.3950377 | 0.017321102 | 0.103413298 | 0.16749395 |
| Epha6 | 866.3468396 | -0.389130404 | 0.12374835 | -3.144530032 |
| Gm24755 | 0.366471122 | -0.22931651 | 1.596060386 | -0.143676588 |
| Gm4660 | 10.78697284 | -0.831843647 | 0.417768027 | -1.991161586 |
| Gm9017 | 5.516248819 | -0.406541116 | 0.407749847 | -0.997035606 |
| Gm27001 | 0.095897396 | -0.687371727 | 3.724945049 | -0.184532045 |
| Gm27784 | 0 | NA | NA | NA |
| Gm17065 | 0 | NA | NA | NA |
| Gm23852 | 0 | NA | NA | NA |
| Gm17054 | 0 | NA | NA | NA |
| n-R5s37 | 0 | NA | NA | NA |
| Gm9027 | 1.878461702 | 0.385058231 | 0.754397458 | 0.510418251 |
| Gm17192 | 0 | NA | NA | NA |
| Gm24164 | 0 | NA | NA | NA |
| Gm9816 | 144.6420699 | -0.089167934 | 0.113364762 | -0.786557769 |
| Nsun3 | 164.1841752 | 0.592402739 | 0.09833199 | 6.024516939 |
| Arl13b | 185.8996067 | 0.151519496 | 0.093066506 | 1.628077625 |
| Gm25986 | 0 | NA | NA | NA |
| Stx19 | 0.062426873 | 0.755288839 | 3.724945049 | 0.202765096 |
| Pros1 | 274.6408184 | 1.34264409 | 0.167114654 | 8.034269038 |
| Epha3 | 257.2072295 | -0.352975538 | 0.156286959 | -2.258509216 |
| Gm22769 | 4.369109362 | -0.324886554 | 0.454523786 | -0.714784493 |
| Csnka2ip | 2.872518017 | 0.583284056 | 0.643209928 | 0.906833104 |
| 4930453N24Rik | 641.6073447 | 0.76163457 | 0.114936542 | 6.626565905 |

| | | | | |
|---|---|---|---|---|
| Zfp654 | 590.4534688 | 0.292113277 | 0.120367893 | 2.426837177 |
| Cggbp1 | 1135.996861 | 0.207959228 | 0.095236989 | 2.183597255 |
| Gm9061 | 0.350048956 | -0.537835696 | 1.555220398 | -0.34582603 |
| Gm23374 | 2.380614889 | -0.269351281 | 0.697972951 | -0.385905042 |
| Htr1f | 54.47759132 | -0.204580106 | 0.15755383 | -1.29847752 |
| Gm22429 | 0 | NA | NA | NA |
| Gm23659 | 0 | NA | NA | NA |
| Pou1f1 | 0.726361874 | -0.206506192 | 3.724945049 | -0.055438722 |
| Chmp2b | 560.9048391 | 0.120141374 | 0.085438774 | 1.406169215 |
| Vgll3 | 46.32731703 | 0.210776345 | 0.153559956 | 1.372599669 |
| Gm22217 | 0.045882455 | 0.755288839 | 3.724945049 | 0.202765096 |
| Gm22448 | 0 | NA | NA | NA |
| Cadm2 | 4980.463628 | 0.051417362 | 0.091643329 | 0.561059513 |
| 1700010K23Rik | 0.914990437 | 1.627653631 | 1.151521408 | 1.413481 |
| 4933411O13Rik | 0 | NA | NA | NA |
| Gm15828 | 81.5457743 | -0.47019036 | 0.296605071 | -1.585240463 |
| Gm24681 | 0 | NA | NA | NA |
| Gm23591 | 0 | NA | NA | NA |
| RP24-377L12.1 | 0.046345709 | 0.274391604 | 3.724945049 | 0.073663262 |
| Speer2 | 0.175920186 | -0.206487576 | 2.733592047 | -0.075537086 |
| Gbe1 | 229.1812539 | 0.586267729 | 0.094274073 | 6.218758869 |
| Gm24968 | 0.168853147 | 0.755296059 | 2.763777273 | 0.273283982 |
| Gm22797 | 0 | NA | NA | NA |
| Robo1 | 968.3837098 | -0.152238519 | 0.091264062 | -1.668110264 |
| Gm25699 | 0 | NA | NA | NA |
| Robo2 | 1762.408431 | -0.216128686 | 0.108161103 | -1.998210819 |
| Gm24272 | 0 | NA | NA | NA |
| Gm22163 | 1.874863541 | -0.710392385 | 0.674441009 | -1.053305443 |
| mmu-mir-691 | 8.12823547 | -0.421335177 | 0.442632839 | -0.951884136 |
| Gm26307 | 0 | NA | NA | NA |
| Gm26937 | 0.185762564 | -1.52982335 | 3.304313696 | -0.462977638 |
| Lipi | 0.34836924 | 0.765398954 | 1.581290856 | 0.484034263 |
| Rbm11 | 76.42304198 | 1.200851864 | 0.182787426 | 6.569663421 |
| Gm15553 | 0 | NA | NA | NA |
| Hspa13 | 738.3975908 | 0.091002298 | 0.076801919 | 1.184896153 |
| Gm22923 | 0 | NA | NA | NA |
| Gm15554 | 0.120974994 | 0.274404949 | 3.724945049 | 0.073666845 |
| Samsn1 | 95.65148686 | 2.857304471 | 0.278075465 | 10.27528436 |
| Gm15555 | 0 | NA | NA | NA |
| Gm37705 | 0 | NA | NA | NA |
| Gm22268 | 0 | NA | NA | NA |
| Gm26915 | 0 | NA | NA | NA |
| Nrip1 | 1272.517009 | 0.333870578 | 0.089953568 | 3.711587925 |
| Gm9843 | 41.81605505 | 0.366962866 | 0.200700402 | 1.828411214 |
| 1700063K16Rik | 0.044544648 | 0.755286614 | 3.724945049 | 0.202764498 |
| 1700041M19Rik | 0.061920613 | 0.274393828 | 3.724945049 | 0.073663859 |
| Usp25 | 1277.058717 | 1.078429347 | 0.111323721 | 9.68732749 |

| | | | | |
|---|---|---|---|---|
| Mir99ahg | 442.0252824 | -0.107860538 | 0.130023446 | -0.829546831 |
| Gm31258 | 0.728787387 | 0.40648799 | 1.148400827 | 0.353960029 |
| Gm37466 | 10.46537345 | -0.434022446 | 0.376835171 | -1.151756734 |
| Gm21816 | 50.06920957 | -0.510091362 | 0.234499007 | -2.175238899 |
| 9430092D12Rik | 19.231994 | -0.130765916 | 0.271140943 | -0.482280229 |
| Gm38071 | 38.74659047 | -0.515195996 | 0.239862252 | -2.147882758 |
| 9430053O09Rik | 25.41324745 | 0.727427156 | 0.209470598 | 3.472693373 |
| Gm37063 | 47.20854318 | 0.029569918 | 0.204922057 | 0.144298366 |
| Gm37241 | 6.880611024 | 0.101972915 | 0.489494371 | 0.208322958 |
| Gm36963 | 34.35472376 | -0.119091538 | 0.228613435 | -0.520929744 |
| Gm37606 | 95.09549254 | 0.064962442 | 0.216194019 | 0.300482141 |
| Gm11146 | 0.04393175 | -0.206490061 | 3.724945049 | -0.055434391 |
| Mir99a | 7.589586068 | 0.185060393 | 0.39741611 | 0.465659011 |
| Mirlet7c-1 | 8.265456032 | 0.336244551 | 0.377468445 | 0.890788503 |
| Gm37311 | 115.4089426 | 0.017885515 | 0.205227993 | 0.087149488 |
| Mir125b-2 | 0.487613095 | -1.055158867 | 1.549435632 | -0.680995612 |
| C130023A14Rik | 179.7393386 | -0.069308209 | 0.147882136 | -0.468671947 |
| 9530003O04Rik | 61.28675002 | 0.254448277 | 0.166843584 | 1.525070788 |
| Gm37694 | 93.40348466 | -0.111721341 | 0.212392734 | -0.526013011 |
| Gm17333 | 0.423282238 | 1.718769033 | 3.704057142 | 0.464023358 |
| Gm25025 | 0.066050875 | 0.274391604 | 3.724945049 | 0.073663262 |
| Gm25916 | 0 | NA | NA | NA |
| Cxadr | 476.9468567 | -0.076004834 | 0.144538954 | -0.525843258 |
| Btg3 | 111.9560118 | 0.890771572 | 0.239193441 | 3.724063541 |
| Gm25038 | 0 | NA | NA | NA |
| D16Ertd472e | 110.9694825 | 0.142530809 | 0.127662105 | 1.116469201 |
| Gm24861 | 0 | NA | NA | NA |
| Chodl | 16.16523681 | 0.040780535 | 0.307069699 | 0.132805467 |
| Tmprss15 | 0.287833757 | 0.650385045 | 2.042040449 | 0.318497631 |
| Gm15692 | 0 | NA | NA | NA |
| Gm37709 | 0 | NA | NA | NA |
| Rps19-ps12 | 0 | NA | NA | NA |
| 1700066C05Rik | 0 | NA | NA | NA |
| Gm23083 | 0 | NA | NA | NA |
| Gm24773 | 0 | NA | NA | NA |
| Ncam2 | 1411.136076 | -0.112071798 | 0.077289885 | -1.450018957 |
| Gm26023 | 0.228677828 | -1.537873234 | 2.143310477 | -0.71752238 |
| Gm23355 | 0 | NA | NA | NA |
| Gm21833 | 0.117046891 | -0.687371727 | 3.724945049 | -0.184532045 |
| Rpl21-ps5 | 0 | NA | NA | NA |
| Gm24508 | 0 | NA | NA | NA |
| Gm25292 | 0.193566712 | 0.274401052 | 2.657574471 | 0.103252441 |
| Gm26667 | 11.04978208 | 5.562728877 | 0.885833766 | 6.279653237 |
| Mir155 | 0 | NA | NA | NA |
| Mrpl39 | 254.7822395 | 0.333499462 | 0.103328803 | 3.227555638 |
| Gm26153 | 0.072025462 | 0.755288839 | 3.724945049 | 0.202765096 |
| Jam2 | 457.3571526 | -0.645919259 | 0.15615337 | -4.136441365 |

| | | | | |
|---|---|---|---|---|
| Gm25908 | 0.500634304 | -1.846718422 | 1.302710034 | -1.41759745 |
| Atp5j | 806.9086276 | -0.021103086 | 0.130551475 | -0.161645712 |
| Gabpa | 471.5751915 | 0.011159067 | 0.060974941 | 0.183010709 |
| App | 15888.22377 | 0.097359405 | 0.058594493 | 1.661579439 |
| Gm27295 | 0 | NA | NA | NA |
| Cyyr1 | 66.70143812 | 1.494200414 | 0.188370288 | 7.932251061 |
| Gm2541 | 17.77359248 | 0.074715568 | 0.270872893 | 0.275832574 |
| Adamts1 | 613.1894008 | 1.84410178 | 0.303358668 | 6.078948696 |
| Adamts5 | 105.3116989 | 0.428520178 | 0.200862386 | 2.133401808 |
| Gm25715 | 0 | NA | NA | NA |
| N6amt1 | 193.0747588 | 0.003689504 | 0.112200313 | 0.032883188 |
| Ltn1 | 1170.966149 | -0.196533323 | 0.041548501 | -4.730214566 |
| Gm25896 | 1.990209358 | -1.171814699 | 0.873683907 | -1.341234159 |
| Rwdd2b | 73.7279705 | 0.067162237 | 0.128577301 | 0.522349096 |
| Usp16 | 561.8276137 | 0.221330576 | 0.122675157 | 1.804200477 |
| Cct8 | 1716.316278 | 0.118903108 | 0.087280084 | 1.36231661 |
| B130034C11Rik | 16.77205567 | -0.247362926 | 0.323246871 | -0.76524461 |
| Gm24891 | 0 | NA | NA | NA |
| Rpl31-ps4 | 0.092228164 | 0.75528439 | 3.724945049 | 0.202763901 |
| Map3k7cl | 1.664220238 | 0.922138161 | 0.822537887 | 1.12108898 |
| Gm22808 | 0 | NA | NA | NA |
| Bach1 | 550.4479081 | 0.667637301 | 0.125044542 | 5.339195855 |
| Gm23994 | 0 | NA | NA | NA |
| 2810407A14Rik | 5.016221076 | 1.434629406 | 0.723106063 | 1.983981991 |
| Grik1 | 248.488386 | -0.297875644 | 0.094722791 | -3.144709313 |
| Cldn17 | 0.086930093 | -0.206490058 | 3.724945049 | -0.05543439 |
| Cldn8 | 0.133745955 | -0.206483268 | 3.671134981 | -0.056245077 |
| Krtap24-1 | 0 | NA | NA | NA |
| Gm23587 | 0 | NA | NA | NA |
| 2310079G19Rik | 0 | NA | NA | NA |
| Krtap26-1 | 0 | NA | NA | NA |
| Krtap27-1 | 0 | NA | NA | NA |
| 2310061N02Rik | 0 | NA | NA | NA |
| Krtap13-1 | 0.115603922 | 0.274387056 | 3.724945049 | 0.073662041 |
| Krtap13 | 0 | NA | NA | NA |
| 2310034C09Rik | 0 | NA | NA | NA |
| AC125199.1 | 0 | NA | NA | NA |
| 2310057N15Rik | 0 | NA | NA | NA |
| Gm5965 | 0 | NA | NA | NA |
| Krtap14 | 0.061920613 | 0.274393828 | 3.724945049 | 0.073663859 |
| Krtap15 | 0 | NA | NA | NA |
| Krtap19-1 | 0 | NA | NA | NA |
| Krtap19-2 | 0 | NA | NA | NA |
| Krtap19-3 | 0 | NA | NA | NA |
| Krtap19-4 | 0 | NA | NA | NA |
| Krtap19-5 | 0 | NA | NA | NA |
| Krtap19-9a | 0 | NA | NA | NA |

| | | | | |
|---|---|---|---|---|
| Krtap19-9b | 0 | NA | NA | NA |
| Krtap16-3 | 0 | NA | NA | NA |
| AC140194.1 | 0 | NA | NA | NA |
| Krtap22-2 | 0 | NA | NA | NA |
| Gm10229 | 0 | NA | NA | NA |
| Krtap6-1 | 0 | NA | NA | NA |
| Gm10228 | 0 | NA | NA | NA |
| Krtap6-5 | 0 | NA | NA | NA |
| 1110025L11Rik | 0 | NA | NA | NA |
| Krtap6-3 | 0 | NA | NA | NA |
| Gm6358 | 0 | NA | NA | NA |
| Gm10061 | 0 | NA | NA | NA |
| Gm9789 | 0 | NA | NA | NA |
| Gm7735 | 0 | NA | NA | NA |
| 1110057P08Rik | 0 | NA | NA | NA |
| Krtap20-2 | 0 | NA | NA | NA |
| Krtap21-1 | 0 | NA | NA | NA |
| Krtap6-2 | 0 | NA | NA | NA |
| AC121818.1 | 0 | NA | NA | NA |
| Gm23655 | 0.049553047 | 0.274391604 | 3.724945049 | 0.073663262 |
| Krtap8-1 | 0 | NA | NA | NA |
| Krtap7-1 | 0.106618422 | 0.274402725 | 3.724945049 | 0.073666248 |
| Krtap11-1 | 0.045882455 | 0.755288839 | 3.724945049 | 0.202765096 |
| Tiam1 | 2404.733508 | -0.09403181 | 0.13029334 | -0.721693142 |
| Sod1 | 1216.13554 | 0.110946992 | 0.094947426 | 1.168509738 |
| Scaf4 | 656.9093113 | -0.04278458 | 0.107575259 | -0.397717659 |
| Hunk | 320.5353761 | -0.017447397 | 0.069397535 | -0.251412343 |
| 1110008E08Rik | 0.670543963 | 0.274413005 | 1.483944559 | 0.184921332 |
| Gm23406 | 0.090074004 | 0.274402725 | 3.724945049 | 0.073666248 |
| Mis18a | 41.5770124 | 0.229436836 | 0.209506965 | 1.095127487 |
| Gm17518 | 6.352540116 | -0.415663557 | 0.394529641 | -1.05356737 |
| Mrap | 10.74916131 | 2.491819429 | 0.625683263 | 3.982557269 |
| Urb1 | 305.5173951 | -0.066854629 | 0.087160583 | -0.767028242 |
| Eva1c | 41.11539447 | 0.702523689 | 0.204907502 | 3.428491794 |
| Gm7831 | 1.072190385 | 1.309767182 | 0.87767751 | 1.492310293 |
| 1110004E09Rik | 190.7191731 | 0.124549878 | 0.117059579 | 1.063987067 |
| Synj1 | 5333.078176 | 0.008112473 | 0.061726775 | 0.131425517 |
| Gm22933 | 0 | NA | NA | NA |
| Gm15965 | 0.045882455 | 0.755288839 | 3.724945049 | 0.202765096 |
| 4930404I05Rik | 0.517542306 | 0.747455263 | 1.390934196 | 0.537376437 |
| Paxbp1 | 967.3107701 | 0.07825968 | 0.135418433 | 0.577910096 |
| 4931406G06Rik | 3.965964068 | 1.460953866 | 0.609575495 | 2.396674206 |
| 4932438H23Rik | 22.83793654 | 0.001057646 | 0.210633681 | 0.005021259 |
| H3f3a-ps2 | 2.549826182 | 0.42876005 | 0.648171224 | 0.661491956 |
| Gm9881 | 5.470790701 | -0.531700546 | 0.443692342 | -1.198354119 |
| Olig2 | 347.143824 | 0.212875978 | 0.162117368 | 1.313097913 |
| Gm37016 | 0.089089296 | 1.194485492 | 3.723939576 | 0.320758559 |

| | | | | |
|---|---|---|---|---|
| Olig1 | 1119.055513 | 0.023360857 | 0.148045706 | 0.1577949 |
| Gm15966 | 0 | NA | NA | NA |
| Gm24695 | 0 | NA | NA | NA |
| Ifnar2 | 243.5630765 | 1.014737391 | 0.122646441 | 8.273679879 |
| Gm21970 | 0 | NA | NA | NA |
| Il10rb | 126.5063742 | 1.386101566 | 0.204102802 | 6.791193207 |
| A930006K02Rik | 5.618572222 | 0.044676743 | 0.429985047 | 0.103903016 |
| Ifnar1 | 638.5446908 | 0.000118014 | 0.068302386 | 0.001727824 |
| Gm7856 | 0.270435483 | 2.00322837 | 3.70434369 | 0.540778215 |
| Mir6367 | 0 | NA | NA | NA |
| Gm7860 | 1.38062972 | 0.278785379 | 1.049632833 | 0.265602762 |
| Ifngr2 | 476.6547008 | 0.05765442 | 0.066974691 | 0.86083891 |
| Tmem50b | 796.4973955 | -0.16685546 | 0.059172335 | -2.819822145 |
| Gm15964 | 4.772112467 | 2.715413327 | 0.762873417 | 3.559454644 |
| Dnajc28 | 95.3526294 | -0.475812234 | 0.177568763 | -2.679594242 |
| Gart | 409.6463015 | 0.044499393 | 0.099604221 | 0.446762115 |
| Son | 6238.367876 | 0.027058744 | 0.060453294 | 0.447597515 |
| Donson | 181.5744247 | -0.243055574 | 0.098514617 | -2.46720315 |
| Atp5o | 1115.945799 | -0.049386432 | 0.076226368 | -0.647891715 |
| Gm10785 | 5.027936496 | 0.056958143 | 0.478962498 | 0.118919839 |
| Cryzl1 | 541.6613726 | -0.260821121 | 0.079453582 | -3.282685514 |
| Itsn1 | 2861.680614 | -0.387385982 | 0.057837284 | -6.697859095 |
| Gm25245 | 0.133144048 | 0.755302125 | 2.968531442 | 0.25443629 |
| Gm15976 | 0.272598396 | 0.331446257 | 1.805525767 | 0.183573263 |
| D430001F17Rik | 1.721321887 | 0.112901759 | 0.750147165 | 0.150506146 |
| Gm27773 | 0 | NA | NA | NA |
| Gm23210 | 0 | NA | NA | NA |
| Mrps6 | 345.9199876 | 0.672896669 | 0.172896246 | 3.891910231 |
| Slc5a3 | 2337.043536 | 0.050793946 | 0.131946088 | 0.384959851 |
| Gm16310 | 32.27711759 | -1.030712491 | 0.326435565 | -3.157476086 |
| Kcne2 | 13.78272213 | -0.899028802 | 0.472355186 | -1.903289787 |
| Smim11 | 43.31674533 | -0.46457906 | 0.222904337 | -2.084208257 |
| 1700048M11Rik | 0 | NA | NA | NA |
| 4930563D23Rik | 0.422874972 | -0.209193911 | 1.734107608 | -0.120634907 |
| Kcne1 | 0.561001133 | 0.890120891 | 1.327084743 | 0.670734025 |
| Rcan1 | 683.9052406 | 0.414371331 | 0.15763988 | 2.628594566 |
| 2410124H12Rik | 1.475924581 | -0.719503616 | 0.881513023 | -0.816214391 |
| Clic6 | 121.7859387 | -0.274509317 | 0.164137743 | -1.672432624 |
| Runx1 | 156.9186044 | 1.66116135 | 0.185974786 | 8.932185834 |
| Gm26626 | 3.202063597 | 2.626130701 | 1.023897951 | 2.564836366 |
| Gm28003 | 0 | NA | NA | NA |
| 1810053B23Rik | 0 | NA | NA | NA |
| Mir802 | 0 | NA | NA | NA |
| 1700029J03Rik | 0.049553047 | 0.274391604 | 3.724945049 | 0.073663262 |
| Setd4 | 64.56323875 | -0.107600604 | 0.161884711 | -0.664674285 |
| Cbr1 | 336.8011914 | -0.133677553 | 0.082713416 | -1.616153209 |
| Gm5678 | 9.467024167 | 0.269591563 | 0.361444621 | 0.74587239 |

| | | | | |
|---|---|---|---|---|
| Gm22344 | 0 | NA | NA | NA |
| Cbr3 | 89.79952323 | 0.638000515 | 0.160020849 | 3.98698369 |
| Dopey2 | 1643.428677 | 0.003964584 | 0.06461254 | 0.061359352 |
| 2310043M15Rik | 4.325146265 | 3.03291044 | 0.883302148 | 3.433604737 |
| Morc3 | 612.5180941 | 0.604025273 | 0.090849556 | 6.648632047 |
| Chaf1b | 23.6865518 | 1.95825127 | 0.341177478 | 5.739685051 |
| Cldn14 | 11.98292911 | 1.006326914 | 0.393522055 | 2.557231294 |
| Sim2 | 1.471461872 | 0.236135179 | 0.80995746 | 0.291540223 |
| Hlcs | 243.6874178 | -0.169678457 | 0.080405022 | -2.11029675 |
| Gm23062 | 1.571274675 | -1.04152183 | 0.946625827 | -1.100246582 |
| Ripply3 | 0.878604591 | -0.48351605 | 1.322054583 | -0.365730777 |
| Pigp | 188.369765 | 0.006661052 | 0.160892773 | 0.041400567 |
| Gm15310 | 0.048949532 | -0.206487837 | 3.724945049 | -0.055433794 |
| Ttc3 | 12670.90535 | -0.2023617 | 0.057638704 | -3.510864859 |
| Gm23692 | 8.647424846 | -1.161356862 | 0.361308479 | -3.214308355 |
| Dscr3 | 356.437628 | -0.104851508 | 0.167349907 | -0.62654058 |
| Dyrk1a | 1886.330775 | 0.23548995 | 0.085731302 | 2.746837429 |
| Gm15971 | 19.22130654 | -0.427522167 | 0.34227605 | -1.249056622 |
| Kcnj6 | 616.7000442 | -0.45098332 | 0.117990359 | -3.822204838 |
| Gm22745 | 0 | NA | NA | NA |
| Gm7976 | 0.168942356 | 0.274406511 | 2.756030556 | 0.099565845 |
| Kcnj15 | 5.024696343 | 1.931768425 | 0.888676183 | 2.173759646 |
| Erg | 22.49114717 | 0.972013477 | 0.261938723 | 3.710842999 |
| Ets2 | 1062.543237 | 0.290075781 | 0.106490089 | 2.723969739 |
| Gm37259 | 0.066050875 | 0.274391604 | 3.724945049 | 0.073663262 |
| 2810404F17Rik | 0.08169834 | -0.206492286 | 3.724945049 | -0.055434988 |
| 1600002D24Rik | 0.782188385 | 1.622715106 | 1.361213306 | 1.192109347 |
| Gm15340 | 0 | NA | NA | NA |
| Psmg1 | 92.11413496 | -0.255433703 | 0.128242005 | -1.991809959 |
| Brwd1 | 2327.039944 | -0.14724607 | 0.115049126 | -1.279853885 |
| Gm15342 | 0 | NA | NA | NA |
| Gm27833 | 5.512382878 | -0.170694453 | 0.476864843 | -0.357951432 |
| 1700093J21Rik | 4.649104205 | -0.512327989 | 0.507059955 | -1.01038937 |
| Gm15341 | 0.325655381 | 1.777339705 | 1.900201532 | 0.935342739 |
| Hmgn1 | 496.6434735 | 0.104598662 | 0.111380531 | 0.939110818 |
| Wrb | 502.1431527 | -0.380551202 | 0.089094695 | -4.271311551 |
| Gm15317 | 0.04393175 | -0.206490061 | 3.724945049 | -0.055434391 |
| Lca5l | 22.78202919 | -0.426717862 | 0.2776926 | -1.536655504 |
| Sh3bgr | 25.544519 | -0.412780992 | 0.233060599 | -1.771131603 |
| B3galt5 | 665.4843436 | 0.157311133 | 0.135680003 | 1.159427546 |
| Igsf5 | 4.211675288 | -0.313518602 | 0.500508292 | -0.626400414 |
| Itgb2l | 0.181155367 | 0.274408223 | 2.695761639 | 0.101792465 |
| Pcp4 | 324.4405224 | -0.546855695 | 0.170047033 | -3.215908486 |
| Dscam | 1116.965166 | -0.163902184 | 0.09031503 | -1.814783028 |
| Bace2 | 46.32598539 | -0.277135463 | 0.170089037 | -1.629355238 |
| Mx1 | 1975.925507 | 9.651263044 | 0.469533423 | 20.55500751 |
| Fam3b | 2.301576675 | 3.914035732 | 0.971330347 | 4.029561872 |

| | | | | |
|---|---|---|---|---|
| Gm9242 | 1.910909441 | 2.194157382 | 1.050602076 | 2.088476153 |
| Gm24777 | 0 | NA | NA | NA |
| Mx2 | 1286.777433 | 6.58166681 | 0.263388722 | 24.98841545 |
| Tmprss2 | 0.798754509 | -1.029170008 | 1.249183844 | -0.823873934 |
| Ripk4 | 12.75005239 | 0.457415643 | 0.302413766 | 1.512549013 |
| Prdm15 | 265.211607 | -0.978284796 | 0.150021091 | -6.52098177 |
| C2cd2 | 212.6215641 | -0.425452329 | 0.09806024 | -4.338683324 |
| mmu-mir-6964 | 0.047948698 | -0.206487837 | 3.724945049 | -0.055433794 |
| Zbtb21 | 472.0135521 | 0.010710077 | 0.093049584 | 0.115100748 |
| C030010L15Rik | 1.895179359 | 0.847489627 | 0.733299061 | 1.155721687 |
| B230307C23Rik | 102.4124531 | 0.573818643 | 0.137879376 | 4.161743836 |
| Gm10232 | 17.35894481 | 0.521593049 | 0.304865634 | 1.710894867 |
| Gm16569 | 0 | NA | NA | NA |
| Gm5479 | 0 | NA | NA | NA |
| Gm25909 | 0.276643538 | -1.443424282 | 2.085476793 | -0.692131549 |
| Pisd-ps2 | 107.0691569 | -0.071467072 | 0.126091373 | -0.56678796 |
| Scaf8 | 1244.518299 | 0.178892946 | 0.095210228 | 1.878925728 |
| Tiam2 | 517.9732827 | -0.377313376 | 0.120144326 | -3.140500999 |
| Tfb1m | 51.82495722 | -0.333898941 | 0.177276424 | -1.883493205 |
| Cldn20 | 1.178313167 | 2.115781035 | 1.106105514 | 1.912820258 |
| Nox3 | 0.272415186 | 1.273327956 | 1.801614001 | 0.706770682 |
| Gm23186 | 0 | NA | NA | NA |
| Arid1b | 2127.270207 | 0.145546175 | 0.063464416 | 2.293350879 |
| Gm15599 | 0.896855871 | -1.317875146 | 1.195911611 | -1.101983737 |
| Gm29050 | 0.223558305 | 0.156567632 | 2.146558956 | 0.072938892 |
| Tmem242 | 341.4840666 | 0.11216104 | 0.097657644 | 1.148512663 |
| Ldhal6b | 8.737325657 | -0.218526237 | 0.428590407 | -0.509871974 |
| Gm22475 | 0.038981395 | 0.274393828 | 3.724945049 | 0.073663859 |
| Zdhhc14 | 490.2302194 | 0.013467122 | 0.113686393 | 0.118458519 |
| 3300005D01Rik | 0.549721405 | 2.784797786 | 1.439664343 | 1.934338237 |
| Gm10231 | 1.299158781 | 0.052553203 | 1.089418923 | 0.048239664 |
| Gm26595 | 0.095897396 | -0.687371727 | 3.724945049 | -0.184532045 |
| Gm26622 | 0 | NA | NA | NA |
| Snx9 | 182.1926714 | -0.012122583 | 0.128383591 | -0.094424703 |
| Synj2 | 857.4048739 | -0.408029055 | 0.091150939 | -4.476410878 |
| Serac1 | 260.8621082 | -0.509020939 | 0.121298922 | -4.196417661 |
| Gtf2h5 | 444.1448567 | -0.020668673 | 0.082393394 | -0.250853523 |
| Tulp4 | 5049.784843 | -0.243538102 | 0.066772725 | -3.6472692 |
| Gm5812 | 12.25279831 | 0.506934069 | 0.319060065 | 1.588835846 |
| Gm22774 | 1.94531079 | -0.383978913 | 1.025251406 | -0.374521713 |
| Gm15590 | 16.93295172 | 1.18477601 | 0.365155256 | 3.244581562 |
| n-R5s26 | 7.164646703 | -0.79699195 | 0.428766077 | -1.858803652 |
| Ppp1r2-ps6 | 4.838948672 | 0.517172092 | 0.492384256 | 1.050342464 |
| Tmem181a | 366.4664287 | -0.362576473 | 0.125913741 | -2.87956238 |
| Dynlt1a | 11.06668227 | 0.064610915 | 0.339132333 | 0.190518299 |
| Gm26848 | 0.173022395 | 1.236193799 | 2.735311368 | 0.451938969 |
| Ppp1r2-ps1 | 3.995967374 | -0.310709292 | 0.510148803 | -0.609056201 |

| | | | | |
|---|---|---|---|---|
| Dynlt1b | 39.38951723 | 0.232617667 | 0.205941938 | 1.129530341 |
| Tmem181b-ps | 797.3566581 | -0.606955445 | 0.258086204 | -2.351754705 |
| Dynlt1c | 50.2770515 | -0.208830896 | 0.215779877 | -0.967795973 |
| Tmem181c-ps | 423.9445639 | -0.436307319 | 0.268235236 | -1.626584658 |
| Dynlt1f | 15.90420978 | 0.596549562 | 0.306975287 | 1.943314617 |
| Sytl3 | 19.32529137 | 4.297868773 | 0.553174957 | 7.769456507 |
| Ezr | 1074.367285 | 0.908856419 | 0.130619284 | 6.958056979 |
| Gm2885 | 11.26381867 | 0.02955251 | 0.296879283 | 0.099543862 |
| Mir692-1 | 0.048949532 | -0.206487837 | 3.724945049 | -0.055433794 |
| Gm29205 | 0 | NA | NA | NA |
| Rsph3b | 25.73551809 | 0.306409401 | 0.243803218 | 1.256789811 |
| Tagap1 | 187.261273 | 0.133478839 | 0.08501699 | 1.570025466 |
| Rnaset2b | 43.79098034 | 0.335106762 | 0.500790209 | 0.669155978 |
| Gm1604b | 6.225126559 | -0.348910764 | 0.574877445 | -0.606930689 |
| Gm25119 | 0 | NA | NA | NA |
| Rps6ka2 | 480.6686781 | -0.248120452 | 0.091652236 | -2.707194752 |
| Tcp10a | 0.109727584 | 0.755282166 | 3.724945049 | 0.202763304 |
| Ttll2 | 0.198340327 | 0.274410802 | 2.234943494 | 0.122781987 |
| Gm9992 | 2.241760946 | 3.180128777 | 1.00505256 | 3.164141761 |
| Gm26057 | 0 | NA | NA | NA |
| Gm25872 | 0 | NA | NA | NA |
| Fndc1 | 124.9694906 | -0.203351119 | 0.170674791 | -1.191453744 |
| E430024P14Rik | 37.33337678 | -0.391922551 | 0.21855242 | -1.793265667 |
| Tagap | 96.63924738 | 1.686456823 | 0.166905306 | 10.10427329 |
| Rsph3a | 115.5642771 | -0.106711842 | 0.105041485 | -1.015901873 |
| Gm1604a | 4.428521402 | 0.038185513 | 0.503878212 | 0.07578322 |
| Gm22416 | 0 | NA | NA | NA |
| Rnaset2a | 127.2262442 | 0.958545121 | 0.441123854 | 2.172961431 |
| Fgfr1op | 286.1447221 | -0.080206812 | 0.101757299 | -0.78821679 |
| Ccr6 | 7.182727363 | -2.221005506 | 0.533778606 | -4.160911434 |
| Mpc1 | 409.8670687 | 0.152604995 | 0.065688149 | 2.323173933 |
| 4930506C21Rik | 16.13295338 | -0.381619137 | 0.260249173 | -1.466360612 |
| Sft2d1 | 202.5780256 | -0.120948507 | 0.13689531 | -0.883510962 |
| Prr18 | 521.3498503 | -0.538855838 | 0.161567151 | -3.335181894 |
| T2 | 5.357486873 | -0.08678793 | 0.40751351 | -0.212969456 |
| Gm16702 | 11.86087499 | 0.124296023 | 0.274466114 | 0.452864733 |
| T | 0.764042974 | -1.159272826 | 1.318229094 | -0.879416811 |
| Pde10a | 2524.291417 | -0.534455141 | 0.183246198 | -2.916596061 |
| Gm15425 | 0 | NA | NA | NA |
| Gm17087 | 0.0878635 | -0.687376175 | 3.724945049 | -0.184533239 |
| Gm15426 | 0.048949532 | -0.206487837 | 3.724945049 | -0.055433794 |
| 1700010I14Rik | 16.46611213 | 0.163148934 | 0.285729636 | 0.57099059 |
| 6530411M01Rik | 2.907305987 | -1.227746911 | 0.586857862 | -2.092068611 |
| Gm8492 | 0.83813809 | 0.882099985 | 1.027144838 | 0.858788316 |
| Gm17748 | 3.088466909 | -0.045249617 | 0.598837528 | -0.075562427 |
| Gm17728 | 0 | NA | NA | NA |
| Pabpc6 | 0.088534442 | 0.274389469 | 3.724945049 | 0.073662689 |

| | | | | |
|---|---|---|---|---|
| Qk | 8468.123674 | -0.322739352 | 0.120853802 | -2.670493991 |
| 1700110C19Rik | 10.49805674 | -1.507131669 | 0.465284963 | -3.23915834 |
| Pacrg | 86.69560762 | -0.254610058 | 0.133439233 | -1.908059959 |
| A230009B12Rik | 11.05179156 | -1.281252178 | 0.398141988 | -3.218078513 |
| Gm16168 | 29.62534574 | -0.646536168 | 0.319695742 | -2.022348384 |
| Gm16169 | 3.038906809 | 1.258603651 | 0.692916356 | 1.81638612 |
| Park2 | 207.7195687 | -0.173608738 | 0.228676369 | -0.759189673 |
| Gm3145 | 3.839486989 | -0.159129544 | 0.528444293 | -0.301128324 |
| Gm28505 | 0.112396584 | 0.27438722 | 3.724945049 | 0.073662085 |
| Gm37402 | 18.41567043 | 0.027188234 | 0.285562867 | 0.095209278 |
| Gm37332 | 0.321071396 | 0.755274557 | 3.724945049 | 0.202761262 |
| Gm28118 | 0 | NA | NA | NA |
| Gm10513 | 0.405560335 | 0.78066265 | 1.522033112 | 0.512907797 |
| Agpat4 | 470.2599501 | -0.105272787 | 0.110707219 | -0.950911676 |
| Tdgf1-ps2 | 2.738925537 | -0.251674945 | 0.603443654 | -0.417064532 |
| Map3k4 | 824.0897494 | -0.09355184 | 0.059417863 | -1.574473315 |
| Gm24794 | 0 | NA | NA | NA |
| 4732491K20Rik | 48.6473752 | -0.159943793 | 0.174394879 | -0.917135834 |
| Plg | 1.917939655 | 1.104522232 | 0.741702954 | 1.489170598 |
| Slc22a3 | 26.84679186 | -0.421407554 | 0.190949783 | -2.206902507 |
| Slc22a2 | 15.39931173 | 0.876297267 | 0.319228606 | 2.74504618 |
| Slc22a1 | 2.772496349 | -0.09638142 | 0.691088801 | -0.139463148 |
| Igf2r | 1261.515538 | 0.00919512 | 0.066122713 | 0.139061443 |
| Airn | 54.16339736 | -0.879390228 | 0.235626358 | -3.732138603 |
| Gm23833 | 6.456263911 | -0.919708519 | 0.480758477 | -1.913036508 |
| Gm16574 | 6.618925518 | -1.426156803 | 0.494503357 | -2.884018445 |
| Gm7162 | 0.709135383 | -0.207189591 | 1.220699124 | -0.169730269 |
| Mas1 | 77.7093898 | -0.422830142 | 0.205557013 | -2.056996911 |
| Gm24085 | 0 | NA | NA | NA |
| Mrgprh | 1.689186202 | -3.32551472 | 0.984899048 | -3.376503131 |
| Pnldc1 | 15.6955681 | -0.77877859 | 0.365221333 | -2.132346933 |
| Mrpl18 | 276.4221612 | -0.129244415 | 0.093317925 | -1.384990246 |
| Tcp1 | 1522.478764 | 0.127077846 | 0.065793961 | 1.931451516 |
| Gm26130 | 0 | NA | NA | NA |
| Snora20 | 1.9528199 | -0.628244158 | 0.649921957 | -0.966645534 |
| Acat3 | 48.71270492 | -0.137544859 | 0.193108161 | -0.712268491 |
| Acat2 | 268.88389 | -0.270879285 | 0.085920849 | -3.152660709 |
| Gm15946 | 1.917568322 | -1.029914471 | 0.754382899 | -1.365241011 |
| Wtap | 592.5866412 | 0.173918467 | 0.074132764 | 2.346040494 |
| Sod2 | 2289.929684 | 0.053714649 | 0.066727666 | 0.804983173 |
| Gpr31b | 8.443869178 | 5.955137892 | 0.908477927 | 6.55507164 |
| Tcp10b | 7.010379163 | 0.949763468 | 0.598520735 | 1.586851404 |
| Unc93a | 6.630768804 | 2.940747377 | 0.731255652 | 4.021503791 |
| Gm23050 | 0 | NA | NA | NA |
| Smok2a | 0.046345709 | 0.274391604 | 3.724945049 | 0.073663262 |
| Smok2b | 0 | NA | NA | NA |
| Gm26500 | 0.250452015 | 0.274411203 | 2.084898498 | 0.131618495 |

| | | | | |
|---|---|---|---|---|
| Tcp10c | 0.224267024 | 0.274410676 | 2.152688503 | 0.127473471 |
| Gm8597 | 4.592470227 | 0.638683459 | 0.563202446 | 1.134021102 |
| Gm25730 | 0.075043205 | 0.755286614 | 3.724945049 | 0.202764498 |
| Rnu6 | 0.503427435 | -0.645034068 | 1.508137562 | -0.427702408 |
| Tcte2 | 120.1207726 | -0.250707911 | 0.206320522 | -1.215138019 |
| Gm16050 | 0.048949532 | -0.206487837 | 3.724945049 | -0.055433794 |
| Gm8603 | 0 | NA | NA | NA |
| Smok4a | 0 | NA | NA | NA |
| Gm26375 | 1.918309794 | -0.495053455 | 0.706871403 | -0.700344437 |
| Gm24867 | 0 | NA | NA | NA |
| Gm16046 | 10.12097898 | -0.656814811 | 0.489899018 | -1.340714692 |
| Gm16052 | 12.43848109 | -0.71842012 | 0.293451797 | -2.448170798 |
| Gm16049 | 10.2923578 | -1.053471937 | 0.357677937 | -2.94530869 |
| Mllt4 | 2109.491735 | -0.209421753 | 0.082140817 | -2.549545529 |
| Gm3355 | 0.441163179 | -0.562162645 | 1.435105221 | -0.391722249 |
| Gm7168 | 0.355971994 | 0.274407688 | 1.941233925 | 0.141357352 |
| Gm7356 | 0 | NA | NA | NA |
| Gm7358 | 0 | NA | NA | NA |
| 4930488N24Rik | 0.072025462 | 0.755288839 | 3.724945049 | 0.202765096 |
| Dact2 | 350.5464997 | -0.161137473 | 0.13841712 | -1.164144099 |
| Smoc2 | 139.6069329 | 0.562759068 | 0.213365394 | 2.637536751 |
| Thbs2 | 224.1080342 | 0.482898166 | 0.183339212 | 2.63390554 |
| Gm3222 | 8.619157282 | 0.118651181 | 0.330109368 | 0.359429912 |
| Gm23352 | 0 | NA | NA | NA |
| Wdr27 | 15.49212616 | -0.407752806 | 0.317272528 | -1.28518157 |
| Gm7393 | 0 | NA | NA | NA |
| 1600012H06Rik | 204.8958293 | 0.243565402 | 0.095751207 | 2.543731917 |
| Phf10 | 695.9833845 | -0.155019096 | 0.077380188 | -2.003343493 |
| Gm3448 | 0.554247143 | 0.277545222 | 1.472562632 | 0.188477703 |
| Ermard | 167.2744434 | 0.950124907 | 0.153047175 | 6.208052553 |
| Tcte3 | 0 | NA | NA | NA |
| Gm28869 | 9.40787293 | 1.988782643 | 0.440502512 | 4.514804319 |
| Gm28873 | 3.819986632 | -0.219667956 | 0.577475681 | -0.380393432 |
| Gm5091 | 0 | NA | NA | NA |
| Dll1 | 136.4076956 | 0.711556824 | 0.146512357 | 4.85663352 |
| A930024N18Rik | 109.5960546 | 0.987687895 | 0.156734576 | 6.301659279 |
| Fam120b | 782.860338 | -0.186332424 | 0.09087838 | -2.050349323 |
| Psmb1 | 1128.534319 | 0.520832218 | 0.119214831 | 4.368854216 |
| Tbp | 458.9414412 | 0.062472046 | 0.130399795 | 0.479080858 |
| Pdcd2 | 107.0321996 | 0.183829181 | 0.14025295 | 1.31069743 |
| Prdm9 | 37.41141876 | -0.16312333 | 0.177478875 | -0.919114063 |
| Gm6686 | 0 | NA | NA | NA |
| Chd1 | 779.4457597 | 0.042155833 | 0.091459172 | 0.460925151 |
| Rgmb | 695.6204429 | 0.036257573 | 0.076047177 | 0.476777361 |
| Gm25598 | 0 | NA | NA | NA |
| BC002059 | 100.3680048 | -0.104543445 | 0.150045638 | -0.696744315 |
| Zfp960 | 12.66666622 | -0.392732815 | 0.441366198 | -0.88981172 |

| | | | | |
|---|---|---|---|---|
| Zfp97 | 21.19205506 | -0.157388318 | 0.272012091 | -0.578607804 |
| Gm6712 | 39.55212394 | -0.472971504 | 0.201809963 | -2.34364794 |
| Oaz1-ps | 23.5739542 | 0.147236693 | 0.242367376 | 0.607493863 |
| Riok2 | 310.94823 | 0.160855994 | 0.077675366 | 2.070875261 |
| Gm26873 | 0.801515891 | 1.610021228 | 1.111283076 | 1.448794878 |
| Lix1 | 332.7540407 | -0.505236306 | 0.160840819 | -3.141219427 |
| Lnpep | 2780.673188 | 0.106262549 | 0.056242984 | 1.889347641 |
| Vmn2r90 | 0.038981395 | 0.274393828 | 3.724945049 | 0.073663859 |
| Gm23837 | 0 | NA | NA | NA |
| Mir99b | 6.303641128 | -1.194421704 | 0.430173598 | -2.776603932 |
| Gm25927 | 0.066050875 | 0.274391604 | 3.724945049 | 0.073663262 |
| mmu-let-7e | 6.339667104 | -0.541538019 | 0.399607842 | -1.355173656 |
| Spaca6 | 577.2041235 | -0.498071033 | 0.291341152 | -1.709580093 |
| Mir125a | 0 | NA | NA | NA |
| Has1 | 11.28386909 | -0.396341248 | 0.363706676 | -1.089727723 |
| Fpr1 | 8.843961322 | 5.167963667 | 0.940363171 | 5.495710406 |
| Fpr2 | 37.5235979 | 2.703706065 | 0.463938346 | 5.827727082 |
| Fpr3 | 0.406865304 | 0.274411005 | 1.629984143 | 0.168351948 |
| Gm7535 | 0.668988238 | -0.912303082 | 1.296321483 | -0.70376299 |
| Fpr-rs4 | 0.097139547 | 0.274393828 | 3.724945049 | 0.073663859 |
| Vmn2r124 | 0.527106987 | 2.076108619 | 1.437276223 | 1.444474336 |
| Vmn2r91 | 0.129871752 | 0.755293287 | 2.991137803 | 0.252510361 |
| Vmn2r92 | 0.150961315 | 0.755296921 | 2.849194963 | 0.265091344 |
| Vmn2r94 | 0 | NA | NA | NA |
| Vmn2r93 | 0 | NA | NA | NA |
| Vmn2r95 | 0.066050875 | 0.274391604 | 3.724945049 | 0.073663262 |
| Vmn2r-ps114 | 0 | NA | NA | NA |
| Vmn2r96 | 0 | NA | NA | NA |
| Vmn2r-ps115 | 0 | NA | NA | NA |
| Vmn2r-ps116 | 0 | NA | NA | NA |
| Vmn2r-ps117 | 0 | NA | NA | NA |
| Vmn2r97 | 0 | NA | NA | NA |
| Vmn2r-ps118 | 0 | NA | NA | NA |
| Vmn2r98 | 0 | NA | NA | NA |
| Vmn2r-ps119 | 0 | NA | NA | NA |
| Vmn2r-ps120 | 0 | NA | NA | NA |
| Vmn2r99 | 0.062426873 | 0.755288839 | 3.724945049 | 0.202765096 |
| Vmn2r-ps121 | 0 | NA | NA | NA |
| Vmn2r100 | 0 | NA | NA | NA |
| Vmn2r101 | 0 | NA | NA | NA |
| Vmn2r102 | 0 | NA | NA | NA |
| Vmn2r-ps122 | 0 | NA | NA | NA |
| Vmn2r103 | 0 | NA | NA | NA |
| Vmn2r-ps123 | 0 | NA | NA | NA |
| Vmn2r-ps124 | 0 | NA | NA | NA |
| Vmn2r-ps125 | 0 | NA | NA | NA |
| Vmn2r104 | 0 | NA | NA | NA |

| | | | | |
|---|---|---|---|---|
| Vmn2r-ps126 | 0 | NA | NA | NA |
| Fpr-rs7 | 0 | NA | NA | NA |
| Fpr-rs6 | 0 | NA | NA | NA |
| Vmn2r105 | 0 | NA | NA | NA |
| Vmn2r106 | 0 | NA | NA | NA |
| Vmn2r-ps127 | 0 | NA | NA | NA |
| Vmn2r107 | 0 | NA | NA | NA |
| Vmn1r-ps141 | 0 | NA | NA | NA |
| Vmn1r224 | 0 | NA | NA | NA |
| Vmn1r-ps142 | 0 | NA | NA | NA |
| Vmn1r-ps143 | 0 | NA | NA | NA |
| Vmn2r108 | 0 | NA | NA | NA |
| Vmn1r225 | 0 | NA | NA | NA |
| Vmn2r109 | 0.225617294 | 0.755301136 | 2.52655979 | 0.298944493 |
| Gm5145 | 2.222558862 | -0.413969544 | 0.741788176 | -0.558069753 |
| Vmn2r110 | 0.133946075 | 0.755288219 | 3.626642455 | 0.20826101 |
| Fpr-rs3 | 0 | NA | NA | NA |
| Vmn1r226 | 0 | NA | NA | NA |
| Vmn1r227 | 0 | NA | NA | NA |
| Vmn1r228 | 0.088534442 | 0.274389469 | 3.724945049 | 0.073662689 |
| Vmn1r229 | 0 | NA | NA | NA |
| Vmn1r230 | 0 | NA | NA | NA |
| Vmn1r231 | 0.108767819 | -0.20649451 | 3.724945049 | -0.055435586 |
| Vmn1r232 | 0.192963196 | -0.206487533 | 2.645912331 | -0.078040202 |
| Vmn1r-ps144 | 0 | NA | NA | NA |
| Ppp2r1a | 4099.317594 | 0.11616625 | 0.060504973 | 1.919945508 |
| Vmn1r-ps145 | 0 | NA | NA | NA |
| Vmn1r233 | 0 | NA | NA | NA |
| Zfp160 | 354.561668 | 0.193965227 | 0.084231143 | 2.302773288 |
| Gm5681 | 0 | NA | NA | NA |
| Vmn1r-ps146 | 0 | NA | NA | NA |
| Gm5092 | 0.092140247 | -0.687373951 | 3.724945049 | -0.184532642 |
| Gm6811 | 0.490429956 | 0.744012719 | 1.379482168 | 0.539342035 |
| Gm7732 | 0.049553047 | 0.274391604 | 3.724945049 | 0.073663262 |
| Gm7736 | 0 | NA | NA | NA |
| Gm7740 | 0 | NA | NA | NA |
| Gm7742 | 0 | NA | NA | NA |
| Vmn1r-ps147 | 0 | NA | NA | NA |
| Vmn1r234 | 0 | NA | NA | NA |
| Vmn1r-ps148 | 0 | NA | NA | NA |
| Vmn1r235 | 0.11193333 | 0.75528439 | 3.724945049 | 0.202763901 |
| Vmn1r-ps149 | 0 | NA | NA | NA |
| Gm26384 | 1.87298797 | -1.065940838 | 0.929598849 | -1.146667554 |
| Vmn1r236 | 0 | NA | NA | NA |
| Vmn1r237 | 0.097139547 | 0.274393828 | 3.724945049 | 0.073663859 |
| Vmn1r-ps150 | 0 | NA | NA | NA |
| Zfp677 | 55.30162814 | 0.15793778 | 0.174526357 | 0.904950879 |

| | | | | |
|---|---|---|---|---|
| Zfp54 | 27.41881594 | -0.448215069 | 0.28052762 | -1.597757359 |
| Zfp51 | 105.8080167 | 0.347778457 | 0.135682663 | 2.563175352 |
| Gm26753 | 1.899580325 | -0.061666884 | 0.796883462 | -0.077385072 |
| Zfp53 | 100.822271 | 0.013414925 | 0.142315152 | 0.0942621 |
| 9330136K24Rik | 11.4954026 | -1.056353027 | 0.435512956 | -2.425537549 |
| Zfp52 | 67.11583658 | -0.200958611 | 0.176603437 | -1.137908836 |
| Zfp948 | 235.9906384 | -0.033210401 | 0.122169573 | -0.271838564 |
| 3110052M02Rik | 172.7372577 | -0.263947312 | 0.106746964 | -2.472644673 |
| Gm10509 | 18.52972673 | 0.358999785 | 0.249332691 | 1.439842419 |
| Gm10226 | 26.0094674 | 0.295608092 | 0.221971584 | 1.331738444 |
| Zfp760 | 279.5658373 | -0.332967439 | 0.149105375 | -2.233101524 |
| Zfp229 | 105.1619278 | -0.124112559 | 0.181205381 | -0.684927559 |
| 4930515G13Rik | 0 | NA | NA | NA |
| Gm9818 | 0.049553047 | 0.274391604 | 3.724945049 | 0.073663262 |
| Zfp820 | 16.99717544 | 0.214366632 | 0.286228889 | 0.748934297 |
| 2210404O09Rik | 62.10937004 | 0.167629804 | 0.147187563 | 1.138885657 |
| Zfp942 | 313.3301051 | -0.072076169 | 0.097136903 | -0.742006043 |
| Zfp943 | 306.7466701 | 0.112523136 | 0.100293702 | 1.121936212 |
| Gm7805 | 0 | NA | NA | NA |
| Gm7809 | 5.578788543 | -0.296167307 | 0.544896319 | -0.543529653 |
| Gm9772 | 0 | NA | NA | NA |
| Gm7052 | 0.038981395 | 0.274393828 | 3.724945049 | 0.073663859 |
| Zfp947 | 19.48152888 | 0.123924183 | 0.259525131 | 0.477503595 |
| Gm4944 | 183.4875709 | 0.290556127 | 0.117206623 | 2.479007739 |
| Zfp944 | 179.0922888 | 0.189721819 | 0.101901283 | 1.861819734 |
| Zfp758 | 147.1029375 | 0.051558338 | 0.107794388 | 0.478302619 |
| Zfp946 | 267.7372296 | 0.131771992 | 0.09404594 | 1.401144931 |
| Vmn2r111 | 0.571153248 | 0.755307651 | 1.639113165 | 0.460802626 |
| Vmn2r112 | 0 | NA | NA | NA |
| Gm9805 | 0.259702268 | 0.274412011 | 2.067438046 | 0.132730464 |
| Gm5224 | 0 | NA | NA | NA |
| Gm5493 | 1.253500652 | -0.111582833 | 0.84741307 | -0.131674666 |
| Gm16386 | 36.63223039 | -0.640989583 | 0.216147902 | -2.96551378 |
| Zfp945 | 660.5015324 | 1.146419975 | 0.121131025 | 9.464296799 |
| Vmn2r-ps128 | 0 | NA | NA | NA |
| Vmn2r113 | 0.044544648 | 0.755286614 | 3.724945049 | 0.202764498 |
| Vmn2r-ps129 | 0 | NA | NA | NA |
| Gm18415 | 0 | NA | NA | NA |
| Gm18396 | 0 | NA | NA | NA |
| Vmn2r-ps130 | 0 | NA | NA | NA |
| Gm18397 | 0 | NA | NA | NA |
| Gm18416 | 0 | NA | NA | NA |
| Vmn2r-ps131 | 0 | NA | NA | NA |
| Vmn2r-ps132 | 0.415262948 | -0.23107806 | 1.394183618 | -0.165744352 |
| Zfp40 | 220.4414794 | -0.107214496 | 0.129981211 | -0.824846109 |
| Vmn2r-ps133 | 0 | NA | NA | NA |
| Vmn2r114 | 0 | NA | NA | NA |

| | | | | |
|---|---|---|---|---|
| Vmn2r115 | 0 | NA | NA | NA |
| Vmn2r116 | 0 | NA | NA | NA |
| Vmn2r-ps134 | 0 | NA | NA | NA |
| Vmn2r117 | 0.156477978 | 1.236192357 | 2.823333319 | 0.437848535 |
| Casp16 | 0.468340293 | -0.503862605 | 1.413037221 | -0.356581269 |
| Gm26695 | 3.060190583 | 0.782055679 | 0.670053796 | 1.167153568 |
| Zfp213 | 78.04727618 | 0.118292422 | 0.142397543 | 0.830719546 |
| Zfp13 | 172.7206419 | 0.264950695 | 0.102515758 | 2.584487498 |
| Zscan10 | 0.304370145 | 2.506421941 | 3.686692052 | 0.679856605 |
| Mmp25 | 15.55909309 | 3.883588251 | 0.632406141 | 6.140971762 |
| Ccdc64b | 0.263919366 | 2.00510328 | 2.035124318 | 0.985248549 |
| Thoc6 | 57.82215382 | 0.337082499 | 0.183624122 | 1.835720137 |
| Hcfc1r1 | 508.7659468 | -0.011304267 | 0.096805063 | -0.116773513 |
| Tnfrsf12a | 409.2070675 | 1.956267272 | 0.394860673 | 4.954322896 |
| Cldn6 | 0.270371171 | 0.288681709 | 1.813105715 | 0.159219458 |
| Cldn9 | 1.372057613 | 0.079547435 | 0.868710198 | 0.091569588 |
| 1520401A03Rik | 0.641047896 | -0.661316939 | 1.330743201 | -0.496953085 |
| Pkmyt1 | 17.10398538 | 0.209311839 | 0.406355334 | 0.515095587 |
| Paqr4 | 569.8115097 | -0.154120769 | 0.099686303 | -1.546057635 |
| Kremen2 | 6.737691786 | 0.016925794 | 0.421488851 | 0.040157157 |
| 9530082P21Rik | 164.7586404 | 3.613821482 | 0.249879867 | 14.46223547 |
| Flywch1 | 2434.68631 | -0.007986266 | 0.05916607 | -0.134980496 |
| Gm16275 | 0.151377979 | 0.274399297 | 2.848448388 | 0.096332901 |
| Flywch2 | 102.393296 | -0.28907341 | 0.137946566 | -2.095546252 |
| D930048G16Rik | 12.89624846 | -1.092838079 | 0.328187329 | -3.329921615 |
| Srrm2 | 8917.247412 | 0.059369444 | 0.097936833 | 0.606201386 |
| Mir5125 | 32.86136734 | -0.523358502 | 0.342347576 | -1.528734357 |
| Tceb2 | 859.0037201 | 0.026690038 | 0.098460177 | 0.271074449 |
| Prss33 | 1.317762752 | 0.3444081 | 0.948097126 | 0.363262466 |
| Prss41 | 25.3916789 | -0.185346641 | 0.2090476 | -0.886624103 |
| Prss32 | 1.380959146 | -1.582053763 | 1.00350122 | -1.57653397 |
| Gm15947 | 0.088534442 | 0.274389469 | 3.724945049 | 0.073662689 |
| Prss21 | 0 | NA | NA | NA |
| Dcpp1 | 0 | NA | NA | NA |
| Dcpp2 | 0 | NA | NA | NA |
| Dcpp3 | 0 | NA | NA | NA |
| Sbp | 0 | NA | NA | NA |
| Sbpl | 0 | NA | NA | NA |
| Prss30 | 3.148853658 | -1.368761815 | 0.687390685 | -1.99124289 |
| Prss22 | 4.912676513 | -1.094492633 | 0.579674282 | -1.888116597 |
| Gm27700 | 0 | NA | NA | NA |
| Gm25092 | 0 | NA | NA | NA |
| Gm5225 | 0 | NA | NA | NA |
| Prss27 | 0.22024148 | -0.206486895 | 2.187052008 | -0.094413345 |
| Kctd5 | 261.9043685 | 0.205288443 | 0.112558997 | 1.823829721 |
| Pdpk1 | 2152.703641 | -0.303177222 | 0.072950743 | -4.155916869 |
| Amdhd2 | 151.6535398 | -0.103686719 | 0.162684883 | -0.637346977 |

| | | | | |
|---|---|---|---|---|
| RP23-27H19.7 | 0 | NA | NA | NA |
| Atp6v0c | 323.8959897 | 0.08182708 | 0.127805559 | 0.640246647 |
| Tbc1d24 | 880.7733529 | -0.204480575 | 0.110307035 | -1.853740116 |
| BC028777 | 3.029716819 | -0.12739804 | 0.544025974 | -0.234176392 |
| 1600002H07Rik | 75.97695628 | 0.146691145 | 0.165240186 | 0.887744979 |
| Ccnf | 89.28741565 | 0.063305017 | 0.185127796 | 0.341953065 |
| Mir5134 | 0.102891481 | 0.274400501 | 3.724945049 | 0.073665651 |
| Mir6965 | 0.049553047 | 0.274391604 | 3.724945049 | 0.073663262 |
| Abca17 | 4.624081096 | 0.221724941 | 0.470564629 | 0.471189137 |
| Gm24427 | 0 | NA | NA | NA |
| Abca3 | 1345.901901 | -0.257440804 | 0.06618032 | -3.889990294 |
| Gm25618 | 0 | NA | NA | NA |
| D330041H03Rik | 54.50038671 | -1.082575096 | 0.211180432 | -5.126304017 |
| Rnps1 | 386.2518119 | 0.050254184 | 0.065223718 | 0.770489413 |
| Eci1 | 99.86578218 | -0.367773367 | 0.174910521 | -2.102637196 |
| Dnase1l2 | 15.71919709 | -1.789923845 | 0.569535304 | -3.142779445 |
| E4f1 | 179.8701777 | -0.077306229 | 0.102648916 | -0.753112964 |
| Pgp | 366.4948428 | -0.267192465 | 0.120860828 | -2.210744955 |
| Mlst8 | 286.6817983 | -0.01316636 | 0.06968779 | -0.18893353 |
| Bricd5 | 7.409630445 | -0.485097549 | 0.354985378 | -1.366528254 |
| Caskin1 | 1583.967537 | -0.545749443 | 0.070260655 | -7.767497243 |
| Traf7 | 402.3587103 | 0.033468911 | 0.067682135 | 0.494501415 |
| Rab26 | 113.4819396 | -0.334504796 | 0.160483514 | -2.084356127 |
| Rab26os | 18.9116278 | -0.149997884 | 0.276343588 | -0.542794877 |
| Snord60 | 0 | NA | NA | NA |
| Pkd1 | 2223.361591 | -0.238530955 | 0.139522898 | -1.709618695 |
| Gm24589 | 0 | NA | NA | NA |
| Tsc2 | 1269.544961 | 0.009978073 | 0.048553248 | 0.205507836 |
| Nthl1 | 19.09256242 | -0.146696131 | 0.302658495 | -0.484691931 |
| Slc9a3r2 | 379.4006015 | 0.018839469 | 0.080526641 | 0.233953244 |
| Npw | 2.00579695 | -0.892615526 | 1.170472811 | -0.762611073 |
| Zfp598 | 484.0735371 | -0.053512781 | 0.113364673 | -0.472041068 |
| Syngr3 | 1091.758285 | -0.247899524 | 0.062836982 | -3.945121398 |
| Gfer | 118.1883123 | 0.176831552 | 0.136225809 | 1.298076724 |
| Noxo1 | 16.22553485 | -0.658664129 | 0.271754565 | -2.423746332 |
| Tbl3 | 217.5084557 | 0.204779674 | 0.105757376 | 1.936315764 |
| Rnf151 | 7.038343901 | -0.968802547 | 0.415715141 | -2.330448068 |
| Rps2 | 374.06437 | 0.46405411 | 0.135072654 | 3.435588894 |
| Snhg9 | 0.044191549 | -0.206490061 | 3.724945049 | -0.055434391 |
| Snora78 | 10.19340988 | -0.677547658 | 0.443782136 | -1.526757395 |
| Snora64 | 0 | NA | NA | NA |
| Ndufb10 | 578.8159123 | -0.172337776 | 0.100878953 | -1.708362059 |
| Rpl3l | 1.739503778 | -0.009939651 | 0.782347276 | -0.012704909 |
| Msrb1 | 303.5260477 | 1.096896103 | 0.199523129 | 5.497588716 |
| Hs3st6 | 1.483788266 | -0.747382708 | 0.944833584 | -0.791020472 |
| Meiob | 0.700302977 | 1.843997083 | 1.256868721 | 1.467135789 |
| Hagh | 454.8144511 | -0.244367218 | 0.081581167 | -2.995387603 |

| | | | | |
|---|---|---|---|---|
| Fahd1 | 88.20798843 | -0.237174099 | 0.140484876 | -1.688253605 |
| Igfals | 0.769314682 | 1.921558157 | 3.606436968 | 0.532813459 |
| Nubp2 | 229.2468458 | 0.415385505 | 0.133464283 | 3.112334587 |
| Spsb3 | 316.2208862 | 0.298415236 | 0.112699798 | 2.647877298 |
| Eme2 | 125.0526026 | -0.135662165 | 0.135354231 | -1.002275023 |
| Mapk8ip3 | 5120.798928 | -0.012188824 | 0.049876987 | -0.244377715 |
| Mrps34 | 292.3664307 | 0.532887905 | 0.102883756 | 5.179514502 |
| Nme3 | 0.049553047 | 0.274391604 | 3.724945049 | 0.073663262 |
| Hn1l | 155.596658 | 0.946129381 | 0.201299755 | 4.700102009 |
| Cramp1l | 709.8548994 | -0.090721591 | 0.099784668 | -0.909173652 |
| Ift140 | 319.2473791 | -0.438226072 | 0.127465268 | -3.438003778 |
| Tmem204 | 25.86076114 | -0.123475605 | 0.215581691 | -0.572755528 |
| Telo2 | 173.173594 | -0.278592691 | 0.120686135 | -2.308406764 |
| Ptx4 | 2.281795086 | 1.232223823 | 0.638254106 | 1.930616368 |
| Clcn7 | 760.3513654 | 0.253269229 | 0.101221353 | 2.502132415 |
| Ccdc154 | 1.912657768 | -0.479473669 | 0.864097482 | -0.554883771 |
| BC003965 | 211.743186 | -0.126656683 | 0.107866742 | -1.174195862 |
| Unkl | 277.8387526 | -0.020690854 | 0.12570135 | -0.164603277 |
| Gnptg | 438.2214292 | -0.066903417 | 0.072643629 | -0.9209812 |
| Tsr3 | 147.8211637 | 0.446078974 | 0.132438143 | 3.368206194 |
| Baiap3 | 567.9275415 | 0.313073609 | 0.159868839 | 1.9583154 |
| Mir3547 | 0 | NA | NA | NA |
| Ube2i | 715.8321189 | 0.354072984 | 0.106467579 | 3.325641373 |
| Prss34 | 0.061920613 | 0.274393828 | 3.724945049 | 0.073663859 |
| Gm25921 | 0 | NA | NA | NA |
| Prss28 | 0 | NA | NA | NA |
| Prss29 | 0 | NA | NA | NA |
| Gm5430 | 0.892004414 | 0.293881143 | 1.018545312 | 0.28853026 |
| Tpsab1 | 3.207720559 | 0.412041522 | 0.590800738 | 0.697428922 |
| Tpsb2 | 1.275377428 | -0.364716857 | 0.830293445 | -0.4392626 |
| Tpsg1 | 0.042716945 | -0.206490061 | 3.724945049 | -0.055434391 |
| Cacna1h | 727.8784213 | -0.206748867 | 0.128971613 | -1.60305715 |
| Tekt4 | 20.49047743 | 0.882007017 | 0.375303036 | 2.350119587 |
| Sstr5 | 11.97499174 | -0.233059629 | 0.30829134 | -0.755972028 |
| Sox8 | 497.4054169 | -0.366939127 | 0.115361629 | -3.180772762 |
| 2810468N07Rik | 100.8375544 | -0.423526666 | 0.186839891 | -2.266789307 |
| Lmf1 | 167.6280357 | -0.065612549 | 0.135609031 | -0.483836134 |
| Gm22645 | 0 | NA | NA | NA |
| Gng13 | 254.443737 | -0.202093342 | 0.202187102 | -0.999536272 |
| Chtf18 | 22.59824114 | 0.061173668 | 0.341045513 | 0.17937098 |
| Rpusd1 | 275.4402358 | -0.12979656 | 0.085967544 | -1.509832121 |
| Mslnl | 3.752562866 | -1.017782473 | 0.598423155 | -1.700773885 |
| Gm25083 | 0 | NA | NA | NA |
| Msln | 26.85201184 | 1.723148819 | 0.35601499 | 4.840101869 |
| Narfl | 208.0663513 | 0.087072769 | 0.078080089 | 1.115172504 |
| Haghl | 395.0534574 | -0.373684488 | 0.097401787 | -3.836526012 |
| Mir6966 | 0 | NA | NA | NA |

| | | | | |
|---|---|---|---|---|
| Ccdc78 | 1.842483344 | -0.895747218 | 0.731803063 | -1.224027698 |
| Fam173a | 248.2616466 | -0.335036972 | 0.12168631 | -2.753284011 |
| Metrn | 475.4136771 | -0.512250584 | 0.260384369 | -1.967286233 |
| Gm26694 | 2.65610448 | 1.492555917 | 0.755627073 | 1.975254686 |
| Fbxl16 | 5262.479825 | -0.208136574 | 0.127632737 | -1.630745991 |
| Wdr24 | 242.1513502 | 0.076291772 | 0.080253397 | 0.950636051 |
| Jmjd8 | 285.0629585 | -0.105348032 | 0.088844117 | -1.185762602 |
| Stub1 | 735.6538842 | -0.108939702 | 0.089991668 | -1.210553209 |
| Mir3082 | 1.578690045 | -0.366193688 | 0.771306911 | -0.474770396 |
| Rhbdl1 | 30.55610411 | -0.669742267 | 0.198286371 | -3.377651534 |
| Gm20683 | 0 | NA | NA | NA |
| Rhot2 | 307.1541338 | -0.130070751 | 0.087186541 | -1.491867307 |
| Wdr90 | 125.7964112 | -0.410792728 | 0.193539392 | -2.122527743 |
| Fam195a | 55.95550801 | -0.079707092 | 0.157941429 | -0.504662345 |
| 0610011F06Rik | 119.2220752 | -0.427305331 | 0.11980215 | -3.56675846 |
| Wfikkn1 | 12.97584509 | 1.121467882 | 0.59025794 | 1.899962383 |
| Rab40c | 356.9302055 | 0.013534625 | 0.106933172 | 0.126570869 |
| Gm10012 | 55.0896093 | -0.07194927 | 0.199837713 | -0.360038497 |
| Gm16278 | 14.17840066 | 0.149784596 | 0.320862359 | 0.466818845 |
| Pigq | 882.6485576 | -0.162934946 | 0.101072572 | -1.612058965 |
| Nhlrc4 | 4.440385187 | -0.423478034 | 0.457853965 | -0.924919441 |
| A930017K11Rik | 39.98571024 | 0.681043373 | 0.208052285 | 3.273424147 |
| D630044L22Rik | 2.092098959 | -0.740395181 | 0.764990621 | -0.967848704 |
| Capn15 | 519.3883034 | -0.361914106 | 0.100770224 | -3.591478613 |
| 1700022N22Rik | 1.65831888 | -0.223377038 | 0.959765084 | -0.232741367 |
| Rab11fip3 | 1324.716031 | -0.247530582 | 0.082200117 | -3.011316675 |
| Decr2 | 128.9572076 | -0.251423261 | 0.116277521 | -2.162268842 |
| Nme4 | 11.32066076 | -0.137779777 | 0.314930129 | -0.43749316 |
| Gm8186 | 9.968334961 | -0.274092109 | 0.382898143 | -0.715835566 |
| Tmem8 | 155.1022499 | -0.094194108 | 0.106956844 | -0.880673968 |
| Mrpl28 | 258.4561109 | -0.09437406 | 0.10128417 | -0.931775026 |
| Axin1 | 402.0340246 | 0.098980169 | 0.090410512 | 1.094786073 |
| Pdia2 | 0.090277459 | -0.20649451 | 3.724945049 | -0.055435586 |
| Arhgdig | 330.2245481 | -0.244085385 | 0.09712511 | -2.513102782 |
| Rgs11 | 126.1618791 | -0.620821522 | 0.287121544 | -2.162225496 |
| Itfg3 | 218.9037408 | -0.156140095 | 0.140325361 | -1.112700465 |
| Gm23123 | 4.679423458 | 0.835147967 | 0.618085571 | 1.351185022 |
| Luc7l | 1216.152592 | -0.100931812 | 0.070988752 | -1.421800068 |
| Gm16580 | 14.51804168 | 0.252863804 | 0.259380892 | 0.974874445 |
| Hba-ps4 | 0.121388914 | 0.755282166 | 3.724945049 | 0.202763304 |
| Neurl1b | 522.8334351 | -0.678168633 | 0.173715228 | -3.903910093 |
| Dusp1 | 584.3483468 | 0.072626496 | 0.31415853 | 0.231177857 |
| Gm8225 | 0.51166646 | -1.103974945 | 1.358469462 | -0.812660848 |
| 1700049J03Rik | 1.147468153 | -0.624190057 | 0.845001933 | -0.738684768 |
| Ergic1 | 1341.076534 | 0.041946703 | 0.054617287 | 0.768011478 |
| Gm22420 | 0 | NA | NA | NA |
| Gm17218 | 0 | NA | NA | NA |

| | | | | |
|---|---|---|---|---|
| Atp6v0e | 321.5954019 | 0.330344085 | 0.130209068 | 2.53702827 |
| Crebrf | 2489.558306 | 0.334529246 | 0.091307757 | 3.663754951 |
| Bnip1 | 161.7726201 | 0.137724461 | 0.157058567 | 0.876898749 |
| Nkx2-5 | 0 | NA | NA | NA |
| Gm17382 | 7.241949452 | 1.109703986 | 0.379017961 | 2.927840103 |
| Gm20468 | 0.607877193 | 0.556130384 | 1.201609801 | 0.462821112 |
| Kifc5b | 20.39272261 | 0.175207733 | 0.265976486 | 0.658733918 |
| Gm27775 | 0 | NA | NA | NA |
| Phf1 | 420.9294313 | -0.053487316 | 0.084307099 | -0.634434312 |
| Mir6968 | 0.061920613 | 0.274393828 | 3.724945049 | 0.073663859 |
| Cuta | 316.9454618 | 0.060598309 | 0.107852456 | 0.561863041 |
| Syngap1 | 4379.193043 | 0.062380407 | 0.135433075 | 0.460599501 |
| Mir3083 | 0 | NA | NA | NA |
| Zbtb9 | 0 | NA | NA | NA |
| Ggnbp1 | 24.84235746 | -0.395576801 | 0.241348214 | -1.639029325 |
| Bak1 | 376.0418335 | 1.511108754 | 0.165434224 | 9.134196765 |
| Itpr3 | 158.6747885 | 0.863550692 | 0.168221124 | 5.133425994 |
| Mir7677 | 0 | NA | NA | NA |
| Itpr3os | 0 | NA | NA | NA |
| Uqcc2 | 404.6628392 | 0.00175248 | 0.106292808 | 0.016487287 |
| Ip6k3 | 19.80013244 | 0.831899179 | 0.266309948 | 3.123800608 |
| Lemd2 | 337.854379 | 0.050201682 | 0.087094359 | 0.57640567 |
| Gm26724 | 0.046345709 | 0.274391604 | 3.724945049 | 0.073663262 |
| mmu-mir-7214 | 0.072025462 | 0.755288839 | 3.724945049 | 0.202765096 |
| Mir7216 | 0 | NA | NA | NA |
| Gm10505 | 1.106753324 | 1.271904412 | 1.111348421 | 1.144469536 |
| mmu-mir-7217 | 0 | NA | NA | NA |
| mmu-mir-7218 | 0 | NA | NA | NA |
| Grm4 | 1362.166969 | -0.660573753 | 0.169374361 | -3.900081148 |
| Gm16005 | 0 | NA | NA | NA |
| Gm9703 | 3.905880329 | 0.217261961 | 0.570516108 | 0.380816524 |
| Hmga1 | 483.005091 | 1.072396295 | 0.264439762 | 4.055351919 |
| AI413582 | 165.7897589 | -0.184607141 | 0.126277286 | -1.461918826 |
| Nudt3 | 2683.122826 | -0.0107864 | 0.064869704 | -0.166277929 |
| Rps10 | 731.2072688 | 0.279541872 | 0.115678078 | 2.416550104 |
| Gm15420 | 1.006594715 | 0.480978135 | 1.058245504 | 0.454505248 |
| Pacsin1 | 4703.755317 | -0.249132232 | 0.0749406 | -3.324396005 |
| Spdef | 9.032787089 | 1.761716618 | 0.562133492 | 3.133982665 |
| Gm15458 | 0.243671637 | -0.206481895 | 2.127579275 | -0.097050154 |
| D17Wsu92e | 3051.297663 | 0.384508744 | 0.078033492 | 4.927483466 |
| Rps2-ps9 | 0.733389001 | 1.163900076 | 1.441699192 | 0.807311319 |
| Snrpc | 219.5654359 | 0.498210967 | 0.090576437 | 5.500447856 |
| Uhrf1bp1 | 565.7822905 | -0.119113612 | 0.099576941 | -1.196196739 |
| Taf11 | 229.7296066 | 0.10867837 | 0.083437602 | 1.302510709 |
| Anks1 | 569.0372512 | -0.22713934 | 0.066195728 | -3.43132929 |
| Gm15597 | 0 | NA | NA | NA |
| Gm15598 | 0.962678808 | 0.15326925 | 1.059983473 | 0.144595886 |

| | | | | |
|---|---|---|---|---|
| Tcp11 | 4.048470876 | 0.058375596 | 0.540885185 | 0.10792604 |
| 4930526A20Rik | 2.494837672 | 0.176190094 | 0.701201705 | 0.251268776 |
| Scube3 | 58.8908513 | -0.397539496 | 0.233822753 | -1.700174563 |
| Zfp523 | 600.1388987 | -0.315860113 | 0.105536799 | -2.992890775 |
| Def6 | 48.95250913 | 2.004833138 | 0.224895597 | 8.914505953 |
| Ppard | 357.9241621 | 0.238435006 | 0.071933231 | 3.314671174 |
| Fance | 139.6977314 | 0.269941004 | 0.110256554 | 2.448298924 |
| Rpl10a | 628.5301939 | 0.429194902 | 0.092451764 | 4.642365731 |
| Tead3 | 55.35341921 | 0.963895209 | 0.259995871 | 3.707348142 |
| Tulp1 | 1.121467505 | 3.825108932 | 1.313539 | 2.912063465 |
| Fkbp5 | 903.5710338 | 1.366948224 | 0.171826178 | 7.955413088 |
| Gm22146 | 0 | NA | NA | NA |
| Gm20109 | 2.702534539 | 0.767488248 | 0.61165582 | 1.254771429 |
| E230001N04Rik | 14.53897155 | 0.545317725 | 0.295617597 | 1.844672747 |
| Gm25195 | 0 | NA | NA | NA |
| Armc12 | 0.464894546 | -1.057076315 | 1.530731217 | -0.690569516 |
| Clpsl2 | 0 | NA | NA | NA |
| Clps | 0.10503227 | 0.274389469 | 3.724945049 | 0.073662689 |
| Mir6969 | 0 | NA | NA | NA |
| Lhfpl5 | 32.4049498 | -0.480744733 | 0.249676782 | -1.925468315 |
| Srpk1 | 762.4953308 | -0.428260261 | 0.083264098 | -5.14339636 |
| Slc26a8 | 26.07395248 | -0.406733129 | 0.215952069 | -1.883441687 |
| Mapk14 | 951.4501698 | -0.026615287 | 0.062660106 | -0.424756488 |
| Mapk13 | 6.079458248 | 0.030691759 | 0.4451441 | 0.068947919 |
| Brpf3 | 576.6212084 | 0.030227686 | 0.080745941 | 0.374355481 |
| Pnpla1os | 0.18989612 | 0.755298873 | 2.65941639 | 0.284009257 |
| Pnpla1 | 2.646101737 | 0.809873302 | 0.609815452 | 1.328062941 |
| 4930539E08Rik | 83.46186464 | 0.841955298 | 0.162111822 | 5.193669933 |
| 1700030A11Rik | 0.307834097 | 0.744009458 | 1.761311287 | 0.422417924 |
| Gm16191 | 0.066050875 | 0.274391604 | 3.724945049 | 0.073663262 |
| Pxt1 | 0.042716945 | -0.206490061 | 3.724945049 | -0.055434391 |
| Kctd20 | 733.6283113 | -0.211377162 | 0.056404676 | -3.74751135 |
| Stk38 | 479.5171983 | -0.301672202 | 0.116288213 | -2.594176961 |
| Gm23887 | 7.551778246 | 0.093656303 | 0.472568999 | 0.198185457 |
| Gm16196 | 0.628440879 | -0.650318237 | 1.445736286 | -0.449818022 |
| Gm16195 | 0.038981395 | 0.274393828 | 3.724945049 | 0.073663859 |
| Srsf3 | 1460.401065 | 0.187305808 | 0.086756899 | 2.15897306 |
| Trp53cor1 | 8.461310733 | 1.154293551 | 0.452656465 | 2.550043226 |
| Gm27599 | 0.04393175 | -0.206490061 | 3.724945049 | -0.055434391 |
| Cdkn1a | 962.23884 | 2.710999995 | 0.358349462 | 7.56524087 |
| Gm16194 | 0.143847454 | -0.206489815 | 2.898675551 | -0.071235918 |
| Rab44 | 2.105186647 | 1.439019514 | 0.936302242 | 1.536917727 |
| Rpl35a-ps3 | 1.427311429 | 1.541041808 | 0.962137915 | 1.60168494 |
| Cpne5 | 630.1572259 | -0.37283448 | 0.246481032 | -1.512629496 |
| Ppil1 | 83.28828306 | -0.077475965 | 0.158462522 | -0.48892296 |
| BC004004 | 500.5361452 | 0.25254933 | 0.062171348 | 4.062149794 |
| Gm10045 | 2.961333848 | -0.10624057 | 0.620815404 | -0.171130693 |

| | | | | |
|---|---|---|---|---|
| Pi16 | 15.76972868 | 0.253169413 | 0.291416658 | 0.868754086 |
| Mtch1 | 2170.561962 | -0.197316425 | 0.065581075 | -3.008740328 |
| Fgd2 | 41.20869392 | 0.65358193 | 0.305812547 | 2.137197887 |
| Gm26885 | 6.791476013 | 1.330888231 | 0.466299663 | 2.85414796 |
| Pim1 | 294.2735208 | 2.302178098 | 0.308277475 | 7.46787645 |
| Gm17657 | 0 | NA | NA | NA |
| Tmem217 | 0.817974253 | -0.251141929 | 1.365986532 | -0.183853884 |
| Tbc1d22b | 597.8773199 | 0.035457416 | 0.063131698 | 0.561642035 |
| Gm28052 | 0 | NA | NA | NA |
| Tbc1d22bos | 0 | NA | NA | NA |
| Mir7667 | 0.28873988 | -0.620058017 | 1.779346547 | -0.348475129 |
| Rnf8 | 73.9322441 | 0.146550657 | 0.163306444 | 0.897396657 |
| Gm28043 | 40.56409608 | 0.746175177 | 0.339408302 | 2.198458828 |
| Cmtr1 | 326.8605501 | 0.58031922 | 0.118133367 | 4.912407343 |
| Ccdc167 | 100.0491434 | -0.930291707 | 0.133831628 | -6.951209671 |
| Mdga1 | 427.208838 | -0.706107834 | 0.142831415 | -4.943645164 |
| Gm16758 | 7.249476259 | -1.127180749 | 0.385635298 | -2.922919022 |
| Gm22327 | 0 | NA | NA | NA |
| Zfand3 | 884.6551979 | 0.281584796 | 0.083713325 | 3.363679528 |
| Btbd9 | 1023.020761 | 0.230541543 | 0.059449493 | 3.877939619 |
| Gm25049 | 0 | NA | NA | NA |
| Gm9874 | 0.097139547 | 0.274393828 | 3.724945049 | 0.073663859 |
| Glo1 | 537.9219683 | -0.19717862 | 0.092479 | -2.132144809 |
| 1700097N02Rik | 0.406736653 | 1.174111142 | 1.562734955 | 0.751318155 |
| Dnah8 | 41.46003794 | 0.555078492 | 0.26503358 | 2.094370427 |
| Gm24661 | 0 | NA | NA | NA |
| Glp1r | 10.69878445 | -0.162512818 | 0.308948546 | -0.52601904 |
| Umodl1 | 10.985597 | -0.235872511 | 0.591865864 | -0.398523593 |
| Gm24026 | 0 | NA | NA | NA |
| Abcg1 | 1340.501293 | 0.322139879 | 0.076435795 | 4.214515966 |
| Gm25447 | 2.050495351 | 0.5680413 | 0.771384013 | 0.736392369 |
| Tff3 | 1.029781442 | -0.443702091 | 0.909668875 | -0.487762199 |
| Tff2 | 0 | NA | NA | NA |
| Gm15318 | 0.237699072 | 0.755303524 | 2.139904856 | 0.352961265 |
| Tff1 | 0.320114016 | 1.415032895 | 1.999100828 | 0.70783468 |
| Tmprss3 | 1.885701583 | -1.837196532 | 0.78426518 | -2.342570574 |
| Ubash3a | 6.422161135 | 2.799679745 | 0.615139584 | 4.551291799 |
| Rsph1 | 24.15190014 | -0.17372645 | 0.257664691 | -0.674234599 |
| Slc37a1 | 77.30229299 | 0.344898948 | 0.137686507 | 2.50495822 |
| Pde9a | 243.102937 | 0.137570635 | 0.139072153 | 0.989203319 |
| Wdr4 | 235.3192353 | -0.031020719 | 0.104530566 | -0.296762183 |
| Ndufv3 | 462.2896515 | 0.108831781 | 0.08811566 | 1.235101466 |
| Gm38099 | 1.2404959 | -0.645530263 | 0.872436991 | -0.7399162 |
| 4833413E03Rik | 0.117046891 | -0.687371727 | 3.724945049 | -0.184532045 |
| Pknox1 | 214.0257755 | -0.282369191 | 0.09264931 | -3.047720389 |
| Gm24970 | 0.09349418 | 0.274402725 | 3.724945049 | 0.073666248 |
| Cbs | 185.6562434 | 0.099010322 | 0.152217532 | 0.650452816 |

| | | | | |
|---|---|---|---|---|
| U2af1 | 321.058565 | 0.305727426 | 0.102854818 | 2.972417177 |
| Cryaa | 0.372981075 | -1.173147092 | 1.577072716 | -0.743876347 |
| Sik1 | 432.2003957 | 0.037863654 | 0.188256328 | 0.201128188 |
| Hsf2bp | 3.787275169 | -0.231861349 | 0.638341368 | -0.363224695 |
| Gm17572 | 0 | NA | NA | NA |
| Gm25883 | 0 | NA | NA | NA |
| Rrp1b | 198.546731 | -0.304010452 | 0.145866827 | -2.084164432 |
| Pdxk-ps | 1.082653786 | -1.540589215 | 0.95452051 | -1.613992784 |
| Gm17276 | 31.48193691 | 0.442053201 | 0.24187355 | 1.827621091 |
| Notch3 | 246.8071195 | -0.727430861 | 0.198957319 | -3.656215629 |
| Ephx3 | 2.938467936 | 0.198049911 | 0.630465916 | 0.314132622 |
| Brd4 | 1978.347052 | 0.078564606 | 0.074545245 | 1.053918399 |
| Gm26549 | 15.20514642 | -1.087748636 | 0.396774569 | -2.741477709 |
| Akap8 | 845.3496397 | 0.127617285 | 0.132292603 | 0.964659259 |
| Akap8l | 594.3929537 | -0.251950002 | 0.138323642 | -1.821452926 |
| Wiz | 755.6427441 | -0.053003132 | 0.062614842 | -0.846494699 |
| Gm26858 | 0.274350536 | 0.340268248 | 2.080539175 | 0.163548109 |
| Gm22201 | 0.044191549 | -0.206490061 | 3.724945049 | -0.055434391 |
| Rasal3 | 63.09606561 | 2.112554927 | 0.262672584 | 8.042540636 |
| Pglyrp2 | 9.032459669 | 5.829519231 | 0.916539041 | 6.360361063 |
| Cyp4f39 | 0 | NA | NA | NA |
| Cyp4f17 | 39.89199084 | 0.426453337 | 0.189944948 | 2.245141777 |
| Cyp4f16 | 69.22286846 | 0.25132191 | 0.191926828 | 1.309467321 |
| Cyp4f37 | 0.44264929 | 0.755309715 | 1.499906351 | 0.503571249 |
| Cyp4f40 | 0.108772582 | 0.75528439 | 3.724945049 | 0.202763901 |
| Cyp4f15 | 57.06117773 | -0.230178842 | 0.201987698 | -1.139568617 |
| Zfp871 | 1423.833684 | -0.084239891 | 0.060214694 | -1.398992261 |
| Gm17115 | 5.126403706 | 0.116737814 | 0.414538484 | 0.281609111 |
| Zfp811 | 109.0107342 | -0.319182799 | 0.166230741 | -1.920118966 |
| Gm23294 | 1.960852514 | 0.040532616 | 0.633716333 | 0.063960189 |
| Gm25555 | 1.60505521 | -0.465068744 | 0.782117304 | -0.594627866 |
| Gm26693 | 0.210884633 | -0.206486093 | 2.191835561 | -0.094206927 |
| Zfp799 | 476.0807173 | -0.493058676 | 0.094951673 | -5.192732916 |
| Zfp870 | 141.8172216 | -0.256706758 | 0.094037137 | -2.729844466 |
| Cyp4f14 | 50.57166942 | -0.013895184 | 0.189051869 | -0.073499323 |
| Cyp4f13 | 93.77020389 | -0.12641 | 0.152951389 | -0.826471736 |
| Cyp4f41-ps | 1.046059986 | 0.496044376 | 1.059152268 | 0.468340947 |
| Gm25667 | 0.310541583 | 1.305237013 | 3.71555195 | 0.351290207 |
| Zfp472 | 71.14659937 | 0.269267795 | 0.147999641 | 1.819381404 |
| Zfp952 | 219.1287848 | -0.04832801 | 0.089100034 | -0.542401699 |
| Zfp763 | 281.1564641 | 0.001196771 | 0.097114185 | 0.012323338 |
| 4921501E09Rik | 0.687686063 | -0.074339035 | 1.213672302 | -0.061251323 |
| Zfp563 | 131.0379676 | -0.400277002 | 0.123819661 | -3.232741878 |
| Morc2b | 49.27469275 | -0.897069813 | 0.361301395 | -2.482884994 |
| Olfr55 | 0.072025462 | 0.755288839 | 3.724945049 | 0.202765096 |
| Olfr239 | 0.04393175 | -0.206490061 | 3.724945049 | -0.055434391 |
| Gm4461 | 9.889894451 | -1.157092151 | 0.339133125 | -3.411911326 |

| | | | | |
|---|---|---|---|---|
| Zfp955a | 185.3926737 | -0.192576979 | 0.130285458 | -1.478115678 |
| Olfr63 | 0.330364319 | 0.274412412 | 1.756994906 | 0.156182816 |
| Zfp955b | 228.1232927 | -0.246935045 | 0.086948905 | -2.84000179 |
| Zfp422-rs1 | 1.757111958 | 0.596670073 | 0.697764019 | 0.855117284 |
| Zfp81 | 498.1008011 | 0.04900341 | 0.084231419 | 0.581771155 |
| Zfp101 | 191.0430263 | -0.242037362 | 0.098911354 | -2.447012939 |
| CT030734.1 | 0 | NA | NA | NA |
| Actl9 | 0 | NA | NA | NA |
| Adamts10 | 190.1840441 | -0.423056089 | 0.146123363 | -2.895198138 |
| Myo1f | 224.7996339 | 2.314437698 | 0.227560947 | 10.17062784 |
| Zfp414 | 163.0997787 | 0.207347442 | 0.103896385 | 1.99571373 |
| Gm20507 | 3.129648303 | -0.114052986 | 0.625059801 | -0.182467318 |
| Pram1 | 0.821047175 | -1.164092427 | 1.210482356 | -0.961676493 |
| Hnrnpm | 845.7194937 | 0.180562219 | 0.145305636 | 1.242637411 |
| 2-Mar | 559.2133353 | -0.090815334 | 0.06249813 | -1.453088814 |
| Gm22943 | 0 | NA | NA | NA |
| Rab11b | 1217.267332 | 0.03906589 | 0.076195211 | 0.512707939 |
| Gm17251 | 14.95472732 | 0.479553592 | 0.268944697 | 1.783093686 |
| Angptl4 | 147.9950521 | 2.880588325 | 0.441926559 | 6.518251201 |
| 4931413I07Rik | 0.260232477 | 1.246526948 | 2.458796504 | 0.506966293 |
| Kank3 | 84.23068527 | -0.745403192 | 0.199098532 | -3.743890952 |
| Rps28 | 433.7597313 | 0.516398812 | 0.169127155 | 3.053316974 |
| Ndufa7 | 437.6705464 | -0.085787436 | 0.096611728 | -0.887960894 |
| Cd320 | 102.4821458 | -0.171977558 | 0.128699612 | -1.336270989 |
| Gm4134 | 0.061920613 | 0.274393828 | 3.724945049 | 0.073663859 |
| Gm7029 | 1.064012846 | 0.557079947 | 1.108754014 | 0.502437817 |
| Kifc1 | 18.16257989 | -0.210493845 | 0.266476763 | -0.789914447 |
| Gm27740 | 0 | NA | NA | NA |
| Platr17 | 2.945475634 | 2.367269679 | 0.962871453 | 2.458552148 |
| BC051226 | 10.35737011 | -0.599035836 | 0.382666439 | -1.56542559 |
| Daxx | 959.7967857 | 2.139735768 | 0.175732191 | 12.17611727 |
| Tapbp | 4153.144377 | 4.102232676 | 0.161597279 | 25.38553064 |
| Zbtb22 | 294.4051857 | 0.156034367 | 0.077552476 | 2.011984346 |
| Gm19412 | 30.80291434 | 1.707250663 | 0.232272678 | 7.350200099 |
| Rgl2 | 188.6076046 | -0.903820893 | 0.135716591 | -6.659619759 |
| Pfdn6 | 222.2854681 | 0.023665446 | 0.140469312 | 0.168474139 |
| Wdr46 | 114.1398418 | 0.383667038 | 0.151080371 | 2.53948965 |
| B3galt4 | 35.09101575 | 0.426622678 | 0.172702564 | 2.470274146 |
| Rps18 | 253.3875219 | 0.513454255 | 0.202239304 | 2.538845046 |
| Vps52 | 738.2973603 | -0.099081644 | 0.064822025 | -1.528518166 |
| Gm23111 | 0.99397625 | 1.881746574 | 0.972553284 | 1.934851904 |
| H2-K2 | 47.37552283 | 6.142795431 | 0.691313604 | 8.885685734 |
| Gm4152 | 0 | NA | NA | NA |
| H2-K1 | 2006.445401 | 6.186838146 | 0.185147698 | 33.41569034 |
| Ring1 | 223.6236612 | 0.268851446 | 0.082687424 | 3.2514188 |
| Gm26940 | 0.048949532 | -0.206487837 | 3.724945049 | -0.055433794 |
| Mir219a-1 | 0.047948698 | -0.206487837 | 3.724945049 | -0.055433794 |

| | | | | |
|---|---|---|---|---|
| mmu-mir-219c | 1.228891946 | -0.994608605 | 0.872370591 | -1.140121658 |
| H2-Ke6 | 1.732078677 | -0.744324206 | 0.743916434 | -1.000548143 |
| Gm20427 | 1.185090727 | 0.251685855 | 1.152560391 | 0.218371078 |
| Slc39a7 | 92.16732074 | 0.503246059 | 0.155947707 | 3.227018009 |
| Rxrb | 612.8230062 | 0.002637509 | 0.05665626 | 0.046552825 |
| Col11a2 | 1625.713743 | 2.810361927 | 0.245397626 | 11.45227836 |
| H2-Pb | 0.204910391 | -1.634333233 | 2.623494331 | -0.622960459 |
| BC051537 | 0.820567722 | 1.69024713 | 1.090457322 | 1.550035105 |
| H2-Pa | 0 | NA | NA | NA |
| H2-Oa | 2.48498024 | 3.952289487 | 0.934544045 | 4.229109915 |
| Brd2 | 3845.362694 | 0.497056792 | 0.080194493 | 6.198141234 |
| Gm20493 | 0 | NA | NA | NA |
| H2-DMa | 136.095749 | 0.720426919 | 0.163544432 | 4.405083745 |
| H2-DMb2 | 12.8997608 | 2.588727975 | 0.516295641 | 5.014041893 |
| H2-DMb1 | 15.23037961 | 4.205150826 | 0.665918363 | 6.314814336 |
| Psmb9 | 510.8582445 | 6.941755942 | 0.323684921 | 21.44602817 |
| Tap1 | 2068.383747 | 6.888831116 | 0.196444487 | 35.06757165 |
| Psmb8 | 1292.910119 | 7.416163261 | 0.29036428 | 25.54089386 |
| Gm20496 | 0.676590931 | 2.226160203 | 1.479149945 | 1.505026729 |
| Tap2 | 1370.634535 | 4.629900261 | 0.174585988 | 26.51931182 |
| Gm15821 | 0.89598866 | 2.663983506 | 1.400415584 | 1.902280678 |
| H2-Ob | 4.820269621 | 0.9593745 | 0.54498854 | 1.7603572 |
| Gm20506 | 0.159280631 | 0.755283203 | 3.464371592 | 0.218014489 |
| H2-Ab1 | 139.2632816 | 3.788009081 | 0.290021093 | 13.06115028 |
| H2-Aa | 90.90688896 | 4.297200321 | 0.348979163 | 12.31363008 |
| Gm20513 | 0.289806465 | 1.089223608 | 1.986122181 | 0.548417221 |
| H2-Eb1 | 122.6882474 | 5.051121593 | 0.52414155 | 9.636941758 |
| H2-Eb2 | 1.375804068 | 0.743898679 | 1.0376095 | 0.716935108 |
| H2-Ea-ps | 95.63606511 | 4.541568273 | 0.38698429 | 11.73579495 |
| Btnl2 | 0.44006443 | 0.76141473 | 1.55709143 | 0.488998086 |
| Btnl1 | 1.318555613 | 3.616022427 | 1.129496339 | 3.201446787 |
| Gm15320 | 0 | NA | NA | NA |
| Gm15321 | 0 | NA | NA | NA |
| BC051142 | 7.899042157 | 0.409428444 | 0.33627895 | 1.217526234 |
| Btnl4 | 0.635806793 | 1.653233459 | 1.458048078 | 1.133867589 |
| Gm15332 | 0 | NA | NA | NA |
| Btnl5-ps | 0 | NA | NA | NA |
| Gm15331 | 0 | NA | NA | NA |
| Btnl6 | 0 | NA | NA | NA |
| Gm15329 | 0 | NA | NA | NA |
| Btnl7-ps | 0.643414179 | 2.122119267 | 1.27255501 | 1.667605134 |
| Notch4 | 57.84156358 | 0.524371264 | 0.193453428 | 2.710581404 |
| Gm20462 | 0 | NA | NA | NA |
| Gpsm3 | 60.20938841 | 2.520240832 | 0.286054588 | 8.810349286 |
| Pbx2 | 551.6966075 | -0.254510642 | 0.069008299 | -3.688116434 |
| Ager | 5.903964042 | 1.372577346 | 0.521058662 | 2.634208865 |
| Rnf5 | 318.1261628 | -0.216490101 | 0.073831785 | -2.932207326 |

| | | | | |
|---|---|---|---|---|
| Gm20463 | 0.104106287 | 0.274400501 | 3.724945049 | 0.073665651 |
| Agpat1 | 1285.566767 | -0.167952504 | 0.061514294 | -2.730300447 |
| Egfl8 | 5.399883896 | -1.16629189 | 0.471346175 | -2.474384966 |
| Gm20460 | 0 | NA | NA | NA |
| Ppt2 | 144.3949312 | -0.446674301 | 0.097761244 | -4.569032485 |
| Prrt1 | 1088.589987 | 0.011571611 | 0.118516605 | 0.097637046 |
| Gm20461 | 0.042716945 | -0.206490061 | 3.724945049 | -0.055434391 |
| Fkbpl | 63.97067545 | 0.411790272 | 0.142486359 | 2.890032941 |
| Atf6b | 618.9161125 | -0.034044511 | 0.080656105 | -0.422094654 |
| Tnxb | 123.7697874 | -0.515653585 | 0.197887328 | -2.605793857 |
| Gm24389 | 0.068097359 | -0.206487837 | 3.724945049 | -0.055433794 |
| Cyp21a2-ps | 0.84670602 | 1.27098295 | 1.308342867 | 0.971444858 |
| C4b | 4516.244341 | 3.760190905 | 0.14300107 | 26.29484457 |
| Stk19-ps1 | 0.301345325 | 0.274411508 | 2.025462448 | 0.135480916 |
| Skiv2l-ps1 | 0.092269992 | -0.20649451 | 3.724945049 | -0.055435586 |
| Gm27471 | 0 | NA | NA | NA |
| Tnxa | 0.049553047 | 0.274391604 | 3.724945049 | 0.073663262 |
| Cyp21a1 | 0.116944185 | 0.274391125 | 3.724945049 | 0.073663134 |
| C4a | 141.065081 | 2.337055996 | 0.216550896 | 10.79217884 |
| Stk19 | 195.7826537 | 0.282310681 | 0.166999071 | 1.690492521 |
| Dxo | 129.6180202 | 0.077310624 | 0.129561525 | 0.596709743 |
| Skiv2l | 714.6748977 | 0.132066762 | 0.055622371 | 2.374346142 |
| Mir6971 | 0 | NA | NA | NA |
| Mir6970 | 0 | NA | NA | NA |
| Nelfe | 167.1258369 | 0.185002303 | 0.090528853 | 2.043572811 |
| Cfb | 85.69436929 | 8.93803566 | 0.884886898 | 10.10076618 |
| Gm20547 | 0 | NA | NA | NA |
| Mir6972 | 0 | NA | NA | NA |
| C2 | 78.60544563 | 3.082896516 | 0.257254191 | 11.98385342 |
| Zbtb12 | 59.55277683 | -0.554170667 | 0.241146847 | -2.298063081 |
| Ehmt2 | 1573.434019 | -0.20313932 | 0.072523779 | -2.801002985 |
| Slc44a4 | 2.383600578 | -0.022024304 | 0.634618601 | -0.034704788 |
| Neu1 | 530.836773 | 0.11850802 | 0.079677148 | 1.487352692 |
| 1110038B12Rik | 128.359516 | 0.698023263 | 0.174650066 | 3.996696256 |
| Snord52 | 0 | NA | NA | NA |
| Gm25744 | 0 | NA | NA | NA |
| Gm8696 | 0.535596869 | 0.755310167 | 1.571798478 | 0.480538808 |
| Hspa1b | 205.5992703 | -0.671306897 | 0.623265353 | -1.077080401 |
| Hspa1a | 168.1854328 | -0.659087304 | 0.577732343 | -1.140817737 |
| Gm20481 | 0.568897959 | -0.199810834 | 1.607106598 | -0.124329546 |
| Hspa1l | 8.258336928 | 1.107414192 | 0.450196451 | 2.459846562 |
| Gm10501 | 57.48590907 | -0.460761648 | 0.170063593 | -2.709349124 |
| Lsm2 | 72.82678909 | -0.124565582 | 0.132723851 | -0.938532007 |
| Vars | 655.1030319 | 0.359050736 | 0.067229586 | 5.34066554 |
| Vwa7 | 91.21471042 | -0.492555279 | 0.193846013 | -2.540961623 |
| Sapcd1 | 2.507914302 | 1.362354019 | 0.606778716 | 2.245223805 |
| Msh5 | 15.17733262 | 0.32336669 | 0.377966987 | 0.855542152 |

| | | | | |
|---|---|---|---|---|
| Gm23421 | 0 | NA | NA | NA |
| Clic1 | 397.1204603 | 2.234195989 | 0.211963851 | 10.54045764 |
| Ddah2 | 89.30682263 | -0.494146462 | 0.162386141 | -3.043033468 |
| G6b | 1.225313711 | -0.893884901 | 0.901562276 | -0.991484366 |
| Ly6g6c | 0.500444145 | 0.274412726 | 1.57174866 | 0.174590717 |
| Ly6g6d | 2.936279196 | -0.903656926 | 0.57568019 | -1.56972038 |
| Ly6g6e | 18.02936065 | 0.013345093 | 0.466754711 | 0.028591233 |
| Ly6g6f | 3.98262746 | -0.881925706 | 0.552991566 | -1.594826684 |
| Abhd16a | 1009.474395 | 0.440810299 | 0.103621215 | 4.254054529 |
| Ly6g5c | 0.083526043 | 0.75528439 | 3.724945049 | 0.202763901 |
| Ly6g5b | 3.278244484 | -0.146764867 | 0.554933114 | -0.264473075 |
| Csnk2b | 658.9798502 | 0.18858053 | 0.068686423 | 2.745528459 |
| Gpank1 | 122.9329401 | 0.21512868 | 0.127554117 | 1.686567903 |
| Gm20522 | 3.138335889 | -0.870131299 | 0.632443785 | -1.375823939 |
| D17H6S53E | 194.6956857 | -0.06595249 | 0.09492814 | -0.694762272 |
| Apom | 0.669558094 | 2.030017307 | 1.242971782 | 1.633196615 |
| Bag6 | 2358.238324 | 0.013113821 | 0.046376699 | 0.282767451 |
| Prrc2a | 6282.854268 | -0.154993033 | 0.076352266 | -2.02997293 |
| Gm23442 | 7.065932701 | 0.038954439 | 0.437464513 | 0.08904594 |
| Gm17705 | 6.475434284 | -0.306426474 | 0.405908279 | -0.754915556 |
| Aif1 | 114.5567538 | 2.321361596 | 0.197871555 | 11.73165896 |
| Ncr3-ps | 0.383232264 | 1.931584503 | 1.661725613 | 1.162396781 |
| Lst1 | 34.76546749 | 1.90239664 | 0.293614918 | 6.479223363 |
| Ltb | 23.43129296 | 3.51468645 | 0.449614099 | 7.817117956 |
| Mir6973a | 0.196800764 | 0.274406863 | 2.203382765 | 0.124538899 |
| Tnf | 84.48636142 | 6.831757108 | 0.731203065 | 9.343173514 |
| Lta | 7.270911819 | 1.95046139 | 0.515429203 | 3.784149947 |
| Mir6974 | 0 | NA | NA | NA |
| Nfkbil1 | 124.1921778 | 0.658642077 | 0.156859911 | 4.198919101 |
| Gm16181 | 0.333516142 | -0.23475915 | 1.743267149 | -0.134666193 |
| Atp6v1g2 | 2556.244658 | -0.184094228 | 0.094074377 | -1.956900858 |
| Ddx39b | 1299.932362 | 0.075898493 | 0.071069233 | 1.067951488 |
| Gm22589 | 0 | NA | NA | NA |
| Mir6975 | 0.574701389 | -0.566521478 | 1.350160199 | -0.419595747 |
| Gm25128 | 2.512166706 | -0.642752775 | 0.674756595 | -0.952569831 |
| Mir8094 | 0.904930538 | -1.37250788 | 1.174508546 | -1.168580581 |
| H2-D1 | 7783.618613 | 4.87805101 | 0.188143676 | 25.92726532 |
| Gm18733 | 1.69076615 | -0.722633659 | 0.888685372 | -0.813149042 |
| Gm18734 | 0.938757343 | 1.126762928 | 1.02343603 | 1.100960778 |
| H2-Q1 | 113.5987717 | 3.863730256 | 0.299122416 | 12.91688636 |
| Gm8741 | 9.465940354 | -0.293676447 | 0.358006148 | -0.820311183 |
| H2-Q2 | 32.27579733 | 4.554785243 | 0.504895664 | 9.021240553 |
| H2-Q3 | 0.044544648 | 0.755286614 | 3.724945049 | 0.202764498 |
| Gm11131 | 30.17597747 | 6.831490334 | 0.865720666 | 7.891102297 |
| H2-Q4 | 2741.587666 | 8.177563921 | 0.277354201 | 29.48418989 |
| H2-Q5 | 203.0857095 | 8.512661074 | 0.737651247 | 11.5402246 |
| Gm8750 | 0.56888582 | 0.236787338 | 1.330949049 | 0.177908642 |

| | | | | |
|---|---|---|---|---|
| H2-Q6 | 1065.781777 | 9.393224931 | 0.506290679 | 18.55302758 |
| H2-Q7 | 469.1606159 | 9.007012058 | 0.625843188 | 14.3918033 |
| Gm8752 | 0.72059279 | 1.125243799 | 1.318706307 | 0.853293711 |
| H2-Q10 | 15.54474821 | 3.013182941 | 0.460971578 | 6.536591585 |
| Pou5f1 | 0 | NA | NA | NA |
| Gm19553 | 0 | NA | NA | NA |
| Tcf19 | 44.18076177 | 0.79265274 | 0.321285809 | 2.467126523 |
| Cchcr1 | 54.80302612 | 0.84528905 | 0.194410478 | 4.347960349 |
| Psors1c2 | 0.771204239 | -0.856508897 | 1.282578292 | -0.667802428 |
| Cdsn | 7.227748083 | 0.633278737 | 0.500029112 | 1.266483735 |
| Gm9613 | 0.0885994 | 0.274402725 | 3.724945049 | 0.073666248 |
| 2300002M23Rik | 0 | NA | NA | NA |
| Sfta2 | 0.116217011 | 0.274402725 | 3.724945049 | 0.073666248 |
| Gm9573 | 0.122991902 | 0.274402725 | 3.724945049 | 0.073666248 |
| Dpcr1 | 0.220032676 | -0.206488847 | 2.588496343 | -0.079771736 |
| Gm20483 | 0.193987248 | -1.189164022 | 2.240590368 | -0.530736916 |
| Vars2 | 218.6187569 | -0.361794579 | 0.09189792 | -3.936918052 |
| Gtf2h4 | 76.61225677 | -0.417932759 | 0.148291654 | -2.818316118 |
| Ddr1 | 1349.901542 | 0.242090709 | 0.11381168 | 2.127116558 |
| Gm23864 | 3.332665857 | -0.187413815 | 0.543295804 | -0.344957228 |
| Gm4577 | 0.622983114 | 2.12824951 | 1.191770981 | 1.785787323 |
| Gm20443 | 0.385495486 | 0.261869444 | 1.563551986 | 0.167483682 |
| Gm20442 | 0.201858732 | 1.23619586 | 2.604406118 | 0.474655566 |
| 4833427F10Rik | 0.08486385 | 0.755286617 | 3.724945049 | 0.202764499 |
| Ier3 | 124.9554162 | 1.065308784 | 0.2526106 | 4.217197476 |
| Flot1 | 628.8586158 | 0.129742543 | 0.089299156 | 1.452897753 |
| Tubb5 | 3619.132628 | -0.036283126 | 0.093182131 | -0.38937858 |
| Mdc1 | 303.644182 | -0.293813409 | 0.115383402 | -2.546409653 |
| Ppp1r18os | 5.57648041 | 0.548314235 | 0.499773903 | 1.097124582 |
| Nrm | 18.32685686 | 0.279321875 | 0.242879883 | 1.150041209 |
| Ppp1r18 | 331.6778556 | 2.060426184 | 0.208574985 | 9.878587253 |
| Gm28033 | 0.098502579 | -0.206492286 | 3.724945049 | -0.055434988 |
| Dhx16 | 338.5688034 | 0.067792388 | 0.100439845 | 0.674955119 |
| 2310061I04Rik | 406.7426439 | -0.289380634 | 0.082926362 | -3.48960967 |
| Gm16279 | 3.489032565 | -0.187932757 | 0.635220774 | -0.295854236 |
| Atat1 | 349.1959485 | -0.325623609 | 0.078033256 | -4.172882536 |
| Mrps18b | 109.9134655 | 0.264363795 | 0.1243249 | 2.126394585 |
| Ppp1r10 | 838.888173 | 0.145384386 | 0.082001457 | 1.772948824 |
| Mir1894 | 0 | NA | NA | NA |
| Gm9840 | 3.211017017 | 0.094948811 | 0.588718472 | 0.161280502 |
| Gm8801 | 0.654072159 | -0.862567083 | 1.179162498 | -0.73150824 |
| Abcf1 | 700.9733728 | 0.036591712 | 0.085676119 | 0.427093474 |
| Mir877 | 0 | NA | NA | NA |
| Prr3 | 210.8648339 | 0.143250147 | 0.127264263 | 1.125611727 |
| Gnl1 | 1616.148979 | 0.157728577 | 0.068154602 | 2.314276254 |
| Gm20508 | 2.081650308 | -1.209328439 | 0.771292491 | -1.56792456 |
| A930015D03Rik | 113.1905163 | 1.120719362 | 0.170866614 | 6.559030683 |

| | | | | |
|---|---|---|---|---|
| H2-T24 | 905.0134114 | 5.877139468 | 0.264394562 | 22.22867004 |
| H2-T23 | 3145.49009 | 7.220466957 | 0.211398613 | 34.15569698 |
| H2-T22 | 1287.107806 | 4.520831361 | 0.167613407 | 26.97177663 |
| Gm6034 | 25.06496248 | 3.920972414 | 0.43637901 | 8.985245227 |
| Gm11127 | 246.9603189 | 5.494485572 | 0.291804812 | 18.82931791 |
| Gm8810 | 46.0581403 | 5.737044992 | 0.59348423 | 9.666718507 |
| H2-Bl | 5.276086801 | 4.888431549 | 0.947139663 | 5.161257352 |
| Gm20530 | 0 | NA | NA | NA |
| Gm8815 | 1.211501154 | 2.954012395 | 1.133491547 | 2.60611771 |
| C920025E04Rik | 19.33663716 | 5.548554348 | 0.853740577 | 6.499110496 |
| Gm7030 | 5.681141433 | 5.405662232 | 0.958295081 | 5.640916187 |
| BC023719 | 0.374687972 | 0.7850974 | 1.56443739 | 0.50184009 |
| H2-T10 | 170.5753034 | 3.71417319 | 0.224922332 | 16.51313659 |
| Gm19684 | 10.16906943 | 5.971395253 | 0.897243593 | 6.655266532 |
| Gm9574 | 0.115603922 | 0.274387056 | 3.724945049 | 0.073662041 |
| Gm10499 | 19.27648584 | 3.360722664 | 0.510018707 | 6.589410585 |
| 2410017I17Rik | 0.142780685 | -0.187625351 | 2.899062601 | -0.064719317 |
| Gm17782 | 0.038981395 | 0.274393828 | 3.724945049 | 0.073663859 |
| Gm6659 | 0.124347486 | 0.755286736 | 3.717316019 | 0.203180664 |
| Gm8909 | 5.891852665 | 4.14832759 | 0.848588392 | 4.888503815 |
| Gm20478 | 1.20453066 | 2.039725145 | 1.465276163 | 1.392041444 |
| Gm6623 | 0.15621502 | -0.206488227 | 2.837166009 | -0.072779748 |
| Gm20495 | 0 | NA | NA | NA |
| H2-T3 | 0.239012627 | 1.142325694 | 2.112402267 | 0.540770909 |
| Gm4246 | 0.068097359 | -0.206487837 | 3.724945049 | -0.055433794 |
| Gm20447 | 0 | NA | NA | NA |
| Gm8835 | 0.696575882 | 0.274412544 | 1.413966574 | 0.194072865 |
| Gm5682 | 0.219744001 | 0.755303984 | 2.154043771 | 0.350644678 |
| Gm20392 | 0.238833577 | 2.005445876 | 2.461453524 | 0.814740501 |
| Gm10074 | 1.752125518 | 1.404921464 | 0.834414608 | 1.683721078 |
| Gm20545 | 0.204225461 | 1.868934307 | 3.204890169 | 0.583150813 |
| Gm29402 | 0 | NA | NA | NA |
| Gm18604 | 0 | NA | NA | NA |
| Gm20391 | 0 | NA | NA | NA |
| Gm22453 | 0 | NA | NA | NA |
| Gm4252 | 0 | NA | NA | NA |
| Gm20546 | 0.114742407 | 0.274402725 | 3.724945049 | 0.073666248 |
| Gm6633 | 0.13733263 | -1.154748075 | 3.63430361 | -0.31773572 |
| Rpp21 | 97.94756067 | -0.058442204 | 0.124334591 | -0.470039786 |
| Trim39 | 333.3226075 | -0.261702093 | 0.130915387 | -1.999017071 |
| H2-M10.2 | 0.048949532 | -0.206487837 | 3.724945049 | -0.055433794 |
| H2-M10.1 | 0 | NA | NA | NA |
| H2-M10.3 | 0 | NA | NA | NA |
| Stk-ps1 | 0 | NA | NA | NA |
| H2-M10.5-ps1 | 0 | NA | NA | NA |
| Lgals1-ps1 | 0 | NA | NA | NA |
| Rpl9-ps9 | 0 | NA | NA | NA |

| | | | | |
|---|---|---|---|---|
| Obox3-ps8 | 0.062426873 | 0.755288839 | 3.724945049 | 0.202765096 |
| H2-M10.4 | 0 | NA | NA | NA |
| Gm20452 | 0.139037128 | 0.274384718 | 3.724945049 | 0.073661414 |
| Gm20451 | 0.072025462 | 0.755288839 | 3.724945049 | 0.202765096 |
| Gm20454 | 0 | NA | NA | NA |
| Gm20453 | 0 | NA | NA | NA |
| H2-M11 | 0.066050875 | 0.274391604 | 3.724945049 | 0.073663262 |
| Gm8868 | 0 | NA | NA | NA |
| Gm20450 | 0 | NA | NA | NA |
| H2-M9 | 0.541282201 | 0.755309719 | 1.46216738 | 0.516568574 |
| Rpl9-ps5 | 0.163190422 | 0.274394252 | 3.398594894 | 0.080737558 |
| H2-M1 | 0.163289894 | 0.274392153 | 3.417209707 | 0.080297136 |
| Gm8877 | 0 | NA | NA | NA |
| Gm20549 | 0 | NA | NA | NA |
| Gm20523 | 0 | NA | NA | NA |
| Gm20525 | 0 | NA | NA | NA |
| Gm20526 | 0 | NA | NA | NA |
| Gm4271 | 0 | NA | NA | NA |
| Gm20401 | 0 | NA | NA | NA |
| Gm8878 | 0.22323753 | 0.716838832 | 2.560144723 | 0.27999934 |
| H2-M10.5 | 0.066050875 | 0.274391604 | 3.724945049 | 0.073663262 |
| H2-M10.6 | 0 | NA | NA | NA |
| Trim26 | 648.1818355 | 1.185083563 | 0.102013637 | 11.61691312 |
| Gm25973 | 0 | NA | NA | NA |
| Trim15 | 1.517773654 | 2.862120732 | 1.163333362 | 2.460275641 |
| Trim10 | 2.069903214 | 3.955748651 | 1.044320994 | 3.787866636 |
| Trim40 | 1.721525315 | -0.345736661 | 0.767355586 | -0.450555997 |
| Trim31 | 0.590394638 | 1.934910294 | 1.25440088 | 1.542497558 |
| 1700031A10Rik | 0.066050875 | 0.274391604 | 3.724945049 | 0.073663262 |
| Rnf39 | 43.61237388 | 0.107300053 | 0.247404997 | 0.433702043 |
| Ppp1r11 | 632.5865216 | 0.233139668 | 0.097365035 | 2.394490672 |
| Znrd1 | 116.024339 | 0.230696978 | 0.138054979 | 1.671051488 |
| Znrd1as | 20.09275304 | -0.03201627 | 0.26230597 | -0.122056962 |
| Gm27735 | 0 | NA | NA | NA |
| H2-M6-ps | 4.48790112 | 0.385558294 | 0.494044313 | 0.780412372 |
| 2410137M14Rik | 2.456754703 | 0.151106118 | 0.603753907 | 0.250277666 |
| H2-M5 | 10.19594007 | -0.507335516 | 0.378777962 | -1.339400828 |
| Zfp57 | 63.14501023 | -0.567697645 | 0.220729049 | -2.57192086 |
| Mog | 603.8852982 | -0.185982426 | 0.156042749 | -1.191868426 |
| Gm22388 | 0.655055538 | -0.005231119 | 1.07814289 | -0.004851972 |
| Gabbr1 | 8020.403664 | -0.071983113 | 0.070679481 | -1.018444269 |
| Olfr90 | 0.046345709 | 0.274391604 | 3.724945049 | 0.073663262 |
| Olfr91 | 0.106520246 | 0.755282166 | 3.724945049 | 0.202763304 |
| Olfr92 | 0.119587853 | 1.218392466 | 3.723618458 | 0.327206581 |
| Gm4831 | 0.046345709 | 0.274391604 | 3.724945049 | 0.073663262 |
| Gm19807 | 2.281184893 | 0.365035234 | 0.834987274 | 0.437174607 |
| Olfr93 | 0.291730981 | 1.227259542 | 1.769789387 | 0.693449487 |

| | | | | |
|---|---|---|---|---|
| Vhl-ps1 | 3.944458201 | 4.783429399 | 0.961555631 | 4.974677746 |
| Olfr753-ps1 | 11.84344878 | 6.019811343 | 0.942983683 | 6.383791628 |
| Gm20497 | 17.12714754 | 6.903117708 | 0.941624614 | 7.331071861 |
| Gm7031 | 0.038981395 | 0.274393828 | 3.724945049 | 0.073663859 |
| Ubd | 60.38255723 | 7.971218485 | 1.043069498 | 7.642078018 |
| Olfr94 | 0.247779669 | 1.668159097 | 1.824981731 | 0.914068929 |
| Olfr95 | 0.082913145 | -0.206492286 | 3.724945049 | -0.055434988 |
| Olfr96 | 0.046345709 | 0.274391604 | 3.724945049 | 0.073663262 |
| Olfr97 | 0 | NA | NA | NA |
| Gm20429 | 1.971464702 | 3.949547442 | 1.05263874 | 3.752044546 |
| Olfr754-ps1 | 0.99187381 | 3.061775217 | 1.174083653 | 2.607799886 |
| Olfr98 | 0 | NA | NA | NA |
| Gm25104 | 0.045882455 | 0.755288839 | 3.724945049 | 0.202765096 |
| H2-M3 | 192.7482628 | 4.021592278 | 0.234520303 | 17.14816254 |
| Olfr755-ps1 | 6.424303794 | 2.735752685 | 0.703861631 | 3.886776271 |
| Olfr99 | 0 | NA | NA | NA |
| Olfr756-ps1 | 0 | NA | NA | NA |
| Olfr101 | 0.110870145 | -0.206490066 | 3.724945049 | -0.055434393 |
| Olfr102 | 0.068097359 | -0.206487837 | 3.724945049 | -0.055433794 |
| Olfr103 | 0 | NA | NA | NA |
| Olfr104-ps | 0 | NA | NA | NA |
| Olfr757-ps1 | 0 | NA | NA | NA |
| Olfr105-ps | 0 | NA | NA | NA |
| Olfr758-ps1 | 0 | NA | NA | NA |
| Olfr106-ps | 0 | NA | NA | NA |
| Olfr107 | 0 | NA | NA | NA |
| Olfr759-ps1 | 0 | NA | NA | NA |
| Gm18808 | 0 | NA | NA | NA |
| Olfr108 | 0 | NA | NA | NA |
| Olfr760-ps1 | 0 | NA | NA | NA |
| Olfr109 | 0.108772582 | 0.75528439 | 3.724945049 | 0.202763901 |
| H2-M2 | 1.126195538 | 3.609844815 | 1.260923868 | 2.862857073 |
| Olfr110 | 0.784115673 | 0.832190028 | 1.112651431 | 0.747934173 |
| Olfr111 | 1.850398219 | 1.336477238 | 0.81970167 | 1.630443473 |
| Olfr112 | 0.062426873 | 0.755288839 | 3.724945049 | 0.202765096 |
| Olfr113 | 0 | NA | NA | NA |
| CR974439.2 | 0 | NA | NA | NA |
| Olfr114 | 0 | NA | NA | NA |
| Olfr115 | 0.046345709 | 0.274391604 | 3.724945049 | 0.073663262 |
| CR974439.1 | 0 | NA | NA | NA |
| Olfr116 | 0.046345709 | 0.274391604 | 3.724945049 | 0.073663262 |
| Olfr117 | 0 | NA | NA | NA |
| Olfr118 | 0.061920613 | 0.274393828 | 3.724945049 | 0.073663859 |
| Olfr119 | 0 | NA | NA | NA |
| Gm7335 | 43.44352658 | -0.10033732 | 0.195685501 | -0.512747851 |
| Olfr120 | 0.155118291 | 0.755284404 | 3.496534233 | 0.216009441 |
| Olfr121 | 0 | NA | NA | NA |

| | | | | |
|---|---|---|---|---|
| Olfr122 | 0.08486385 | 0.755286617 | 3.724945049 | 0.202764499 |
| Olfr123 | 0 | NA | NA | NA |
| Gm20524 | 0.042716945 | -0.206490061 | 3.724945049 | -0.055434391 |
| Olfr124 | 0 | NA | NA | NA |
| Olfr125 | 0 | NA | NA | NA |
| Gm9577 | 0.148659141 | 0.274384287 | 3.724945049 | 0.073661298 |
| Olfr126 | 0 | NA | NA | NA |
| Gm18737 | 0.194588561 | -0.206482317 | 2.240900713 | -0.092142555 |
| Gm18520 | 0 | NA | NA | NA |
| Olfr127 | 0 | NA | NA | NA |
| Olfr128 | 0 | NA | NA | NA |
| Gm20543 | 0 | NA | NA | NA |
| Gm20542 | 0 | NA | NA | NA |
| Gm5967 | 0 | NA | NA | NA |
| Olfr761 | 0 | NA | NA | NA |
| Gm18522 | 0 | NA | NA | NA |
| Gm20541 | 1.213329596 | 0.55300754 | 1.010357517 | 0.547338473 |
| Olfr129 | 0 | NA | NA | NA |
| Olfr130 | 0.068097359 | -0.206487837 | 3.724945049 | -0.055433794 |
| Gm18833 | 0 | NA | NA | NA |
| Olfr131 | 0.048949532 | -0.206487837 | 3.724945049 | -0.055433794 |
| Olfr132 | 0.042716945 | -0.206490061 | 3.724945049 | -0.055434391 |
| Olfr133 | 0.061920613 | 0.274393828 | 3.724945049 | 0.073663859 |
| Olfr134 | 0 | NA | NA | NA |
| Olfr135 | 0 | NA | NA | NA |
| Olfr138 | 0 | NA | NA | NA |
| Olfr137 | 0 | NA | NA | NA |
| Olfr762-ps1 | 0 | NA | NA | NA |
| Olfr136 | 0.089062654 | -0.20649451 | 3.724945049 | -0.055435586 |
| Esp36 | 0 | NA | NA | NA |
| Gm20409 | 0 | NA | NA | NA |
| Gm6726 | 0 | NA | NA | NA |
| Esp34 | 0 | NA | NA | NA |
| Esp31 | 0.04393175 | -0.206490061 | 3.724945049 | -0.055434391 |
| Esp24 | 0 | NA | NA | NA |
| Gm21903 | 0 | NA | NA | NA |
| Esp23 | 0 | NA | NA | NA |
| Esp18 | 0 | NA | NA | NA |
| Gm26429 | 0 | NA | NA | NA |
| Esp16 | 0 | NA | NA | NA |
| Esp15 | 0 | NA | NA | NA |
| Gm26917 | 335746.0181 | 0.452776741 | 0.104829135 | 4.319188005 |
| Rn18s-rs5 | 102.7885986 | 0.931357879 | 0.819016696 | 1.137165925 |
| Yam1 | 128743.8113 | 0.850245942 | 0.635052405 | 1.338859495 |
| Esp38 | 0 | NA | NA | NA |
| Gm26376 | 0 | NA | NA | NA |
| Tgif2-ps2 | 0.524319023 | 0.26289914 | 1.370459655 | 0.19183282 |

| | | | | |
|---|---|---|---|---|
| Pgk2 | 0 | NA | NA | NA |
| Crisp3 | 0 | NA | NA | NA |
| Crisp1 | 0.331267484 | 0.765790488 | 1.746708477 | 0.438419174 |
| Esp8 | 0 | NA | NA | NA |
| Esp6Esp5 | 0 | NA | NA | NA |
| Esp6 | 0 | NA | NA | NA |
| Esp4 | 0 | NA | NA | NA |
| Esp3 | 0 | NA | NA | NA |
| Esp1 | 0 | NA | NA | NA |
| Crisp2 | 0.293737151 | 1.63295758 | 3.709858878 | 0.440167034 |
| Rhag | 0.351655166 | 2.04418537 | 3.703161682 | 0.552010834 |
| Gm20574 | 0 | NA | NA | NA |
| 9130008F23Rik | 2.649375936 | -0.729836224 | 0.808099096 | -0.903151888 |
| Glyatl3 | 0.199876958 | -1.124725491 | 2.647758279 | -0.424784052 |
| Gm23078 | 0.852119224 | -0.033128156 | 1.004340386 | -0.032984989 |
| Cenpq | 63.31617002 | 0.845117996 | 0.171057485 | 4.940549655 |
| Mut | 485.5759982 | -0.182263025 | 0.100193886 | -1.819103257 |
| Gm6771 | 0.062426873 | 0.755288839 | 3.724945049 | 0.202765096 |
| Ptchd4 | 384.0058745 | -0.168108712 | 0.079277615 | -2.120506695 |
| Opn5 | 0.714665336 | 0.58974311 | 1.168888639 | 0.504533187 |
| Adgrf4 | 19.83835518 | -0.221216166 | 0.273775716 | -0.808019678 |
| Adgrf2 | 18.64465502 | -0.190819562 | 0.296024083 | -0.644608237 |
| Cd2ap | 541.0115779 | 0.400065834 | 0.141502497 | 2.827270486 |
| Tnfrsf21 | 965.9785103 | -0.054501078 | 0.06672648 | -0.816783364 |
| Adgrf1 | 1.021366362 | 1.387929656 | 1.038860037 | 1.336012173 |
| Adgrf5 | 194.5698432 | 0.686141699 | 0.183999622 | 3.729038641 |
| Gm25135 | 0.066050875 | 0.274391604 | 3.724945049 | 0.073663262 |
| Mep1a | 1.39389272 | 0.836531298 | 1.259174986 | 0.664348726 |
| Ankrd66 | 10.82149774 | 0.683055633 | 0.521624472 | 1.309477739 |
| Pla2g7 | 1523.964145 | -0.00510361 | 0.129938657 | -0.03927707 |
| 1700071M16Rik | 0.275776889 | -0.603493152 | 2.054707378 | -0.293712457 |
| Tdrd6 | 25.59334993 | -0.186041933 | 0.273804204 | -0.679470695 |
| Slc25a27 | 391.4686422 | -0.33669744 | 0.13353463 | -2.521424144 |
| Cyp39a1 | 17.12697954 | -0.683404438 | 0.295306006 | -2.314224647 |
| Gm25783 | 0 | NA | NA | NA |
| Rcan2 | 1857.55946 | 0.006402173 | 0.10134021 | 0.063175051 |
| Gm24532 | 0.062426873 | 0.755288839 | 3.724945049 | 0.202765096 |
| Enpp5 | 2499.132028 | 0.013583554 | 0.0503842 | 0.269599475 |
| Enpp4 | 696.8122402 | 1.145265813 | 0.167046073 | 6.855987651 |
| Clic5 | 79.94705142 | 1.153157237 | 0.210356776 | 5.481911532 |
| Runx2 | 205.0751855 | 0.515595944 | 0.169907375 | 3.034570713 |
| Runx2os3 | 0 | NA | NA | NA |
| n-R5s27 | 0.766229633 | 0.775216947 | 1.252103833 | 0.619131518 |
| Runx2os2 | 0.092269992 | -0.20649451 | 3.724945049 | -0.055435586 |
| Runx2os1 | 7.863194607 | 0.102918766 | 0.486843878 | 0.211399939 |
| Supt3 | 134.9145071 | -0.325835165 | 0.125636729 | -2.593470608 |
| Gm24979 | 0 | NA | NA | NA |

| | | | | |
|---|---|---|---|---|
| Cdc5l | 544.4221431 | -0.077670484 | 0.06613447 | -1.174432704 |
| B230354K17Rik | 184.8456956 | -0.68877293 | 0.130623508 | -5.272963045 |
| Spats1 | 16.79904519 | -0.082188704 | 0.277352735 | -0.296332771 |
| Gm9104 | 1.30365031 | -0.219627219 | 0.883072557 | -0.248708011 |
| Aars2 | 207.7781551 | -0.287448734 | 0.090598229 | -3.17278536 |
| Tcte1 | 13.23124629 | 0.053711316 | 0.309512145 | 0.173535408 |
| Tmem151b | 1924.460039 | -0.173407227 | 0.086346696 | -2.008267072 |
| Nfkbie | 174.6593782 | 2.487874865 | 0.271620527 | 9.159377224 |
| Slc35b2 | 231.0133132 | 0.225885891 | 0.148570865 | 1.520391574 |
| Hsp90ab1 | 12795.80252 | 0.120748517 | 0.083476641 | 1.446494679 |
| Slc29a1 | 163.1308914 | 0.028445435 | 0.100151638 | 0.284023664 |
| Gm17080 | 1.296318896 | -0.874667638 | 0.934868606 | -0.935604888 |
| Gm7325 | 0 | NA | NA | NA |
| Gm25008 | 0 | NA | NA | NA |
| Capn11 | 4.929061397 | -0.615189329 | 0.550899535 | -1.116699671 |
| Tmem63b | 1144.513863 | -0.368917264 | 0.073033885 | -5.051316435 |
| Gm16172 | 0.25133133 | 1.463094664 | 2.418873113 | 0.604866231 |
| Mrpl14 | 161.5517627 | -0.059664608 | 0.136406022 | -0.4374045 |
| 1600014C23Rik | 0.431841237 | -0.714711469 | 1.403184555 | -0.509349584 |
| F630040K05Rik | 2.147486072 | -0.772221312 | 0.59554655 | -1.296659871 |
| Vegfa | 569.830768 | -0.787217117 | 0.101271542 | -7.773330018 |
| Mrps18a | 383.939429 | 0.030134412 | 0.080568732 | 0.374021182 |
| Rsph9 | 56.95556391 | 0.049849755 | 0.137225789 | 0.363268123 |
| Mad2l1bp | 60.28485607 | -0.019895236 | 0.164742338 | -0.120765773 |
| Gtpbp2 | 597.9990834 | 0.155169483 | 0.083276544 | 1.863303574 |
| Polh | 84.99680479 | -0.154046624 | 0.156014371 | -0.987387399 |
| Xpo5 | 708.0881202 | 0.100716474 | 0.059323732 | 1.697743383 |
| Mir693 | 0.626433637 | -1.427751107 | 1.267865766 | -1.12610589 |
| Gm22474 | 0 | NA | NA | NA |
| Polr1c | 186.1516209 | 0.145506593 | 0.112596991 | 1.292277811 |
| Yipf3 | 497.8450709 | 0.151514453 | 0.072665915 | 2.085082842 |
| Lrrc73 | 196.1398283 | -0.130181605 | 0.113529772 | -1.146673715 |
| Tjap1 | 262.6446001 | -0.129388169 | 0.109453801 | -1.182125872 |
| Gm26785 | 0.440626437 | 2.443453806 | 1.642052545 | 1.488048487 |
| Dlk2 | 119.3259875 | -0.627880453 | 0.177010157 | -3.547143654 |
| Abcc10 | 137.5955981 | -0.47379021 | 0.168992881 | -2.803610457 |
| Zfp318 | 978.2373152 | -0.023376259 | 0.093895503 | -0.248960365 |
| Gm23037 | 4.835503307 | 0.920228076 | 0.5255552 | 1.750963697 |
| Crip3 | 13.66803183 | -2.250304416 | 0.398193443 | -5.651284455 |
| Slc22a7 | 3.396474255 | 0.761699658 | 0.564088404 | 1.350319652 |
| Gm5093 | 4.877282298 | -0.207362169 | 0.565379351 | -0.366766434 |
| Ttbk1 | 1698.649847 | -0.264233597 | 0.089342409 | -2.957538318 |
| Gm26904 | 0.300318634 | -0.604574979 | 1.776140781 | -0.340386858 |
| Dnph1 | 20.31270367 | 0.038155736 | 0.258154763 | 0.147801791 |
| Cul9 | 1048.399803 | -0.157198348 | 0.075504174 | -2.081982211 |
| Srf | 574.4040805 | -0.071016826 | 0.135610773 | -0.523681302 |
| mmu-mir-6976 | 0.656458977 | 1.606970627 | 1.153893737 | 1.392650445 |

| | | | | |
|---|---|---|---|---|
| Ptk7 | 194.7380974 | 0.048552189 | 0.089881625 | 0.540179253 |
| Klc4 | 177.2125805 | -0.339992846 | 0.096587316 | -3.520056883 |
| Mrpl2 | 228.4860238 | 0.368037138 | 0.115472377 | 3.187231 |
| Cul7 | 465.2672236 | -0.205739577 | 0.094487545 | -2.17742537 |
| Rrp36 | 135.9417813 | 0.150677004 | 0.10522022 | 1.432015668 |
| Klhdc3 | 1295.864183 | -0.075074257 | 0.049742917 | -1.509245159 |
| Mea1 | 401.4088604 | 0.028850827 | 0.123722628 | 0.233189574 |
| Ppp2r5d | 1040.331458 | 0.060027543 | 0.066135851 | 0.907639984 |
| Pex6 | 410.5596741 | -0.437170666 | 0.132161033 | -3.307863556 |
| Gnmt | 15.57555665 | 1.011525888 | 0.308936744 | 3.274216839 |
| Cnpy3 | 552.2456984 | 0.408351458 | 0.07940327 | 5.142753674 |
| Ptcra | 0.771865818 | 0.254143613 | 1.10541366 | 0.229908153 |
| 2310039H08Rik | 29.97597163 | 0.33682064 | 0.227272946 | 1.482009389 |
| Rpl7l1 | 408.9885597 | 0.092569504 | 0.105504382 | 0.877399616 |
| Gltscr1l | 1089.761893 | -0.18565023 | 0.082184816 | -2.25893589 |
| A330017A19Rik | 35.0263174 | -1.195589143 | 0.267349403 | -4.472009773 |
| Tbcc | 165.35841 | 0.184500115 | 0.131313803 | 1.405032151 |
| Prph2 | 0.359258722 | 0.297945553 | 1.483187177 | 0.200881964 |
| Ubr2 | 1461.655788 | -0.212151669 | 0.05542076 | -3.828018066 |
| Gm23657 | 4.719513391 | -0.022400839 | 0.454106766 | -0.049329455 |
| Gm16494 | 2.052894931 | 0.375815104 | 0.948205077 | 0.396343695 |
| Gm4945 | 0 | NA | NA | NA |
| Trerf1 | 592.2320734 | -0.460719355 | 0.153761286 | -2.996328706 |
| Gm26216 | 1.145909063 | -0.962091315 | 0.823981304 | -1.167613039 |
| Mrps10 | 197.4832179 | -0.059182096 | 0.100728293 | -0.587541926 |
| Guca1b | 0.605175792 | -0.238263622 | 1.373518782 | -0.173469504 |
| Guca1a | 2.149392939 | 3.245942955 | 1.119329218 | 2.899900138 |
| Gm5814 | 2.49590342 | 1.670147701 | 0.733120474 | 2.278135395 |
| 1700001C19Rik | 0.29699235 | 0.288107838 | 1.771717407 | 0.162615007 |
| AI661453 | 6.189948269 | 1.24138437 | 0.428537586 | 2.896792274 |
| Taf8 | 186.7542863 | 0.256121866 | 0.092532739 | 2.767905372 |
| Ccnd3 | 416.7992574 | 0.691346713 | 0.129497722 | 5.338678589 |
| Bysl | 253.0965139 | 0.020923207 | 0.091732204 | 0.228090095 |
| Med20 | 240.4788844 | -0.049989129 | 0.120676586 | -0.414240499 |
| Gm20517 | 21.19396069 | 0.456821607 | 0.317287157 | 1.439773395 |
| Kpna2-ps | 1.554960773 | 0.091992654 | 0.886919581 | 0.103721528 |
| Usp49 | 129.8291219 | -0.27691902 | 0.126670834 | -2.186130874 |
| Tomm6 | 2.522522481 | -0.574553447 | 0.777968535 | -0.738530443 |
| Gm21981 | 0.57942775 | 2.092549892 | 1.210656956 | 1.728441638 |
| Tomm6os | 39.84955966 | -0.224998917 | 0.167409376 | -1.344004271 |
| Prickle4 | 1.072312625 | 0.439645667 | 0.915215589 | 0.480373883 |
| Frs3 | 230.2853897 | 0.151169104 | 0.109023089 | 1.386578804 |
| Frs3os | 0.363326102 | -0.018816084 | 1.694519498 | -0.011104082 |
| Pgc | 0.329750641 | 2.230692761 | 1.734075649 | 1.286387225 |
| Tfeb | 165.6362251 | -0.042503748 | 0.129412311 | -0.328436667 |
| Mdfi | 11.42660599 | -0.442375858 | 0.333018396 | -1.328382645 |
| Gm25201 | 0.048949532 | -0.206487837 | 3.724945049 | -0.055433794 |

| | | | | |
|---|---|---|---|---|
| n-R5s28 | 0 | NA | NA | NA |
| Foxp4 | 579.0805347 | -0.029384313 | 0.104344001 | -0.281609986 |
| Gm15556 | 0.196245641 | 0.274393285 | 3.243303546 | 0.084603023 |
| Gm5228 | 0 | NA | NA | NA |
| Gm14871 | 0 | NA | NA | NA |
| 1700122O11Rik | 0 | NA | NA | NA |
| 1700067P10Rik | 0 | NA | NA | NA |
| 9830107B12Rik | 0 | NA | NA | NA |
| A530064D06Rik | 30.08457509 | 5.274025453 | 0.923517116 | 5.71080423 |
| Trem1 | 13.84162536 | 5.970969536 | 0.990209682 | 6.030005203 |
| Trem3 | 3.961576269 | 3.790023675 | 0.99551936 | 3.807081838 |
| Treml4 | 2.160387598 | 2.808158797 | 1.170056243 | 2.400020352 |
| Treml2 | 120.9667195 | 5.326316063 | 0.39609561 | 13.44704642 |
| B430306N03Rik | 110.3507723 | 7.281609676 | 0.901143924 | 8.080407006 |
| Trem2 | 83.40596837 | -0.374330084 | 0.490333082 | -0.763420004 |
| Treml1 | 1.368647145 | -0.390118954 | 0.839658617 | -0.464616151 |
| Nfya | 266.8544357 | -0.335746406 | 0.130082045 | -2.58103572 |
| Oard1 | 77.30484208 | 0.060877451 | 0.147323363 | 0.413223333 |
| Apobec2 | 1.336996185 | -0.377967839 | 0.948040008 | -0.398683427 |
| Tspo2 | 0 | NA | NA | NA |
| Unc5cl | 6.179264061 | -0.642461099 | 0.452448693 | -1.419964537 |
| Gm24071 | 0 | NA | NA | NA |
| Gm23642 | 0 | NA | NA | NA |
| Lrfn2 | 299.7160112 | -0.014481325 | 0.136157749 | -0.106356968 |
| AC154247.1 | 0.135598227 | -1.141928459 | 2.942835517 | -0.388036794 |
| 1700008K24Rik | 0 | NA | NA | NA |
| Gm22978 | 0.046345709 | 0.274391604 | 3.724945049 | 0.073663262 |
| Gm20540 | 0.410699371 | -0.19944126 | 1.470536074 | -0.135624868 |
| Mocs1 | 175.5053655 | 0.575387657 | 0.116968883 | 4.919151514 |
| Daam2 | 1383.444506 | 0.003674934 | 0.099418617 | 0.03696424 |
| Kif6 | 28.85323937 | -0.051469235 | 0.238164242 | -0.216108157 |
| Gm16555 | 0 | NA | NA | NA |
| Gm16554 | 0.075043205 | 0.755286614 | 3.724945049 | 0.202764498 |
| Rftn1 | 212.415448 | 0.136279372 | 0.141836592 | 0.960819561 |
| Gm37173 | 0 | NA | NA | NA |
| AY702103 | 0 | NA | NA | NA |
| Dazl | 26.34726813 | -0.144764607 | 0.262794916 | -0.550865326 |
| Plcl2 | 1538.397032 | -0.131907702 | 0.082523655 | -1.598422932 |
| Gm7334 | 7.373441669 | 1.097788112 | 0.455682965 | 2.409105004 |
| Tbc1d5 | 596.1325065 | -0.043557792 | 0.06172844 | -0.70563572 |
| Gm25177 | 2.526566463 | -0.275922611 | 0.68290358 | -0.404043292 |
| Satb1 | 1280.976831 | -0.240266801 | 0.109020122 | -2.203875721 |
| E430014B02Rik | 30.56272651 | -0.389031047 | 0.329609261 | -1.180279478 |
| C230085N15Rik | 40.73930876 | -0.39827765 | 0.334506251 | -1.190643369 |
| Gm36931 | 44.82250833 | -0.16597841 | 0.230027358 | -0.721559435 |
| 5830444F18Rik | 19.30851287 | -0.25923499 | 0.381010092 | -0.680388777 |
| Gm37248 | 39.69177475 | -0.211096395 | 0.331966643 | -0.635896406 |

| | | | | |
|---|---|---|---|---|
| Gm6934 | 12.03475686 | -0.753859313 | 0.439435225 | -1.715518624 |
| Gm37593 | 18.0879042 | -0.898750931 | 0.419081874 | -2.144571233 |
| Gm37266 | 10.80255973 | -1.885623176 | 0.439449699 | -4.290873748 |
| Gm37176 | 9.616469211 | -1.241568482 | 0.521526988 | -2.380640908 |
| Gm19585 | 5.697516218 | 3.576738414 | 0.833705888 | 4.29016811 |
| Gm27217 | 0.396216368 | 0.778355944 | 1.751930835 | 0.444284631 |
| Kcnh8 | 23.78866619 | -0.064080075 | 0.250566789 | -0.255740497 |
| Efhb | 28.27811259 | 2.036039229 | 0.285865675 | 7.122363429 |
| Rab5a | 1148.748334 | 0.267691979 | 0.085744027 | 3.121989828 |
| Pp2d1 | 0.305447682 | 2.152978526 | 1.71895518 | 1.252492532 |
| Kat2b | 803.7724207 | 0.428755557 | 0.100320389 | 4.273862589 |
| Gm6919 | 0.108772582 | 0.75528439 | 3.724945049 | 0.202763901 |
| Sgol1 | 6.962193728 | 2.060945716 | 0.67462952 | 3.05492964 |
| Gm19518 | 0.186767869 | 0.778496236 | 3.289032344 | 0.236694613 |
| 4932415M13Rik | 0 | NA | NA | NA |
| Ck-ps1 | 0.277663158 | 1.707812409 | 3.716337667 | 0.459541775 |
| RP24-173D2.6 | 0 | NA | NA | NA |
| RP24-173D2.8 | 0 | NA | NA | NA |
| RP24-173D2.9 | 0 | NA | NA | NA |
| Sult1c2 | 0.362371572 | 0.605244393 | 1.68816104 | 0.358522901 |
| Mrps36-ps1 | 2.335139537 | 0.756708809 | 0.830684345 | 0.910946274 |
| RP24-173D2.10 | 0 | NA | NA | NA |
| RP24-173D2.11 | 0 | NA | NA | NA |
| RP23-416K11.2 | 0 | NA | NA | NA |
| RP23-416K11.3 | 0 | NA | NA | NA |
| RP23-416K11.4 | 0 | NA | NA | NA |
| Sult1c1 | 1.161929334 | 0.142701247 | 1.13207653 | 0.126052651 |
| RP23-416K11.5 | 0 | NA | NA | NA |
| Gm37966 | 0 | NA | NA | NA |
| Slc5a7 | 146.1741277 | -0.09825811 | 0.24728831 | -0.397342318 |
| 1700025K24Rik | 0.046345709 | 0.274391604 | 3.724945049 | 0.073663262 |
| Trp53-ps | 0 | NA | NA | NA |
| Gm26291 | 0 | NA | NA | NA |
| Gm17377 | 2.328145098 | -0.401102376 | 0.731578672 | -0.548269642 |
| St6gal2 | 374.6218881 | -0.614116536 | 0.159995557 | -3.838334947 |
| Gm26547 | 0.778497894 | -1.138948768 | 1.467292056 | -0.776224994 |
| Vmn2r118 | 1.673678531 | 0.236479189 | 0.856498197 | 0.276100043 |
| Pot1b | 63.94887526 | 0.076677632 | 0.155036317 | 0.49457852 |
| Gm22654 | 0 | NA | NA | NA |
| Adgre4 | 3.084562812 | 2.385605754 | 0.997905764 | 2.390612259 |
| Gm18417 | 1.232924928 | 2.406938537 | 1.062535946 | 2.26527728 |
| Zfp119a | 72.0705982 | 0.337524131 | 0.137763325 | 2.450028935 |
| Zfp959 | 70.36865212 | -0.128036969 | 0.141887263 | -0.902385219 |
| Rpl21-ps6 | 2.92891158 | 1.902149925 | 0.629086286 | 3.023670947 |
| Zfp119b | 52.43127221 | 0.846628033 | 0.195890317 | 4.321949375 |
| Ebi3 | 12.63508742 | 1.292827827 | 0.347562896 | 3.719694602 |
| Gm16712 | 2.434106523 | -0.122565636 | 0.611319892 | -0.200493453 |

| | | | | |
|---|---|---|---|---|
| Ccdc94 | 63.31143036 | 0.33256667 | 0.164101007 | 2.026597367 |
| Shd | 180.526558 | -0.175654635 | 0.083882703 | -2.094050735 |
| Fsd1 | 267.0610619 | -0.331940165 | 0.08452386 | -3.927177087 |
| Stap2 | 23.62948936 | -0.592276465 | 0.215570765 | -2.747480466 |
| Mpnd | 574.8071085 | 0.060707186 | 0.058781917 | 1.032752745 |
| Gm16276 | 0.401552702 | -0.959584859 | 1.493568484 | -0.642477977 |
| Sh3gl1 | 328.504642 | 0.074702847 | 0.085648887 | 0.872198688 |
| Chaf1a | 59.75683498 | 0.528188814 | 0.216419025 | 2.440584012 |
| Ubxn6 | 593.1748788 | 0.012341951 | 0.051681144 | 0.238809555 |
| Hdgfrp2 | 630.2550697 | 0.086094675 | 0.076547359 | 1.124724299 |
| Plin4 | 580.4887041 | 2.457818123 | 0.280883152 | 8.750322337 |
| Plin5 | 6.445794394 | 0.269317298 | 0.385477938 | 0.698658136 |
| Lrg1 | 47.22509705 | 3.778984869 | 0.407329451 | 9.277465364 |
| Sema6b | 1185.955188 | -0.113934081 | 0.093905733 | -1.213281419 |
| Tnfaip8l1 | 28.12050717 | -0.501966164 | 0.217911213 | -2.303535271 |
| AC026385.1 | 0.119587853 | 1.218392466 | 3.723618458 | 0.327206581 |
| Mydgf | 321.3040017 | 0.388371561 | 0.122911543 | 3.159764755 |
| Dpp9 | 687.0448644 | -0.287520873 | 0.056579082 | -5.08175219 |
| Mir7b | 2.668141924 | -1.985024025 | 0.781063736 | -2.541436677 |
| Fem1a | 839.6826125 | 0.232836352 | 0.077473665 | 3.005361282 |
| Ticam1 | 133.3080346 | 0.547536667 | 0.148965152 | 3.675602382 |
| Plin3 | 209.6287355 | 0.293843014 | 0.197821146 | 1.48539739 |
| Arrdc5 | 0 | NA | NA | NA |
| Uhrf1 | 83.37304998 | 2.231207085 | 0.338831976 | 6.584995635 |
| Kdm4b | 809.6209241 | -0.101940064 | 0.08054581 | -1.265615973 |
| Ptprs | 5796.573917 | -0.310856165 | 0.087463452 | -3.554126413 |
| Mir6977 | 0 | NA | NA | NA |
| Znrf4 | 0.092691418 | 0.274387304 | 3.724945049 | 0.073662108 |
| Safb2 | 645.9551208 | -0.101203162 | 0.178707101 | -0.566307443 |
| Safb | 970.0585309 | 0.02823885 | 0.159772014 | 0.176744658 |
| 2410015M20Rik | 237.0366201 | -0.063850808 | 0.141502613 | -0.451234128 |
| Rpl36 | 557.6166036 | 0.324063485 | 0.151521986 | 2.138722524 |
| Lonp1 | 634.6361425 | -0.027373268 | 0.075589899 | -0.362128643 |
| Catsperd | 6.596655121 | -0.447595472 | 0.408575418 | -1.095502695 |
| Ranbp3 | 724.4876957 | 0.064043788 | 0.058860501 | 1.088060541 |
| Vmac | 91.17240515 | 0.032502082 | 0.117817517 | 0.275867994 |
| Ndufa11 | 437.8651622 | -0.096366056 | 0.107011483 | -0.90052071 |
| Nrtn | 25.07553935 | 0.294162179 | 0.225484034 | 1.30458097 |
| Dus3l | 477.2858719 | 0.007271633 | 0.090230907 | 0.080589162 |
| Prr22 | 6.373319375 | 0.081657544 | 0.388962613 | 0.209936743 |
| Rfx2 | 53.12569894 | 0.359407761 | 0.182299882 | 1.97151944 |
| Acsbg2 | 0.382360723 | 0.274413013 | 1.723359121 | 0.159231474 |
| 1700061G19Rik | 1.170313971 | 0.455370047 | 1.167961464 | 0.389884479 |
| Mllt1 | 783.4489317 | -0.064725745 | 0.07002139 | -0.924371034 |
| Acer1 | 0.046345709 | 0.274391604 | 3.724945049 | 0.073663262 |
| Clpp | 295.2646874 | 0.073680621 | 0.088546806 | 0.8321093 |
| Alkbh7 | 82.81251037 | -0.369949045 | 0.18299883 | -2.021592406 |

| | | | | |
|---|---|---|---|---|
| Pspn | 0.342416647 | -0.113073311 | 1.981371761 | -0.057068196 |
| Gtf2f1 | 546.3864564 | 0.681646164 | 0.131674299 | 5.17675941 |
| Khsrp | 1672.094708 | 0.009236742 | 0.041239504 | 0.223978005 |
| Slc25a41 | 1.408439352 | -2.517104043 | 1.026545805 | -2.452013374 |
| Gm17168 | 0.17329739 | -1.27723404 | 3.323993654 | -0.384246835 |
| Slc25a23 | 2717.008405 | -0.263951651 | 0.089394718 | -2.952653781 |
| Crb3 | 6.90059268 | 0.822552763 | 0.4324524 | 1.902065439 |
| Dennd1c | 54.0186804 | 2.538701321 | 0.287821672 | 8.820396682 |
| Tubb4a | 6699.351768 | -0.302594215 | 0.095192645 | -3.178756262 |
| Gm11110 | 11.24476628 | 0.290205582 | 0.319424209 | 0.9085272 |
| Tnfsf9 | 20.88192311 | 1.186012122 | 0.344618703 | 3.441519895 |
| Cd70 | 3.132506103 | -0.022910388 | 0.585715237 | -0.039115234 |
| Tnfsf14 | 6.752466611 | 5.013785027 | 1.016982 | 4.930062702 |
| C3 | 1371.649863 | 5.651689779 | 0.473366409 | 11.93935537 |
| Mir6978 | 0 | NA | NA | NA |
| Gpr108 | 284.67282 | 0.269197064 | 0.098880449 | 2.722449857 |
| Trip10 | 80.25548548 | 0.700002628 | 0.177879924 | 3.935253697 |
| Vav1 | 298.6647182 | 3.159478789 | 0.192844418 | 16.38356357 |
| Adgre1 | 147.8605273 | 1.443060297 | 0.247576317 | 5.82874935 |
| Vmn2r120 | 0 | NA | NA | NA |
| Gm21834 | 0.060174537 | 0.755286614 | 3.724945049 | 0.202764498 |
| Gm25102 | 0 | NA | NA | NA |
| Cntnap5c | 204.371656 | 0.046493284 | 0.148768114 | 0.312521835 |
| Rpl7a-ps5 | 12.52156343 | 0.841614013 | 0.458991812 | 1.833614438 |
| Gm5815 | 49.22368419 | -0.0750073 | 0.185900931 | -0.403479961 |
| Pdzph1 | 37.42674745 | 1.028005148 | 0.228145716 | 4.505914761 |
| Nudt12os | 0.099106094 | 0.274387056 | 3.724945049 | 0.073662041 |
| Nudt12 | 83.70631604 | -0.397446522 | 0.177603308 | -2.237832882 |
| Gm23769 | 0 | NA | NA | NA |
| Gm23203 | 0 | NA | NA | NA |
| Gm29051 | 0 | NA | NA | NA |
| Gm23447 | 0 | NA | NA | NA |
| Gm23773 | 0 | NA | NA | NA |
| Obox6-ps1 | 0 | NA | NA | NA |
| Gm4518 | 4.468190482 | 0.459709421 | 0.478256245 | 0.961219902 |
| Gm27023 | 0.131193343 | -0.194387882 | 2.978303909 | -0.065267981 |
| Gm25800 | 0 | NA | NA | NA |
| Efna5 | 380.5177091 | -0.218761083 | 0.087025259 | -2.513765386 |
| Fbxl17 | 855.8478773 | -0.08467266 | 0.06033969 | -1.403266407 |
| Gm25348 | 2.594574602 | -0.420474127 | 0.59286357 | -0.709225779 |
| Gm24730 | 0 | NA | NA | NA |
| 4930405O22Rik | 44.37612019 | -0.445282903 | 0.274015144 | -1.625030269 |
| Gm17228 | 21.45301415 | -0.625839377 | 0.29755829 | -2.103249677 |
| Gm24813 | 0 | NA | NA | NA |
| A930002H24Rik | 1.337378336 | 0.457424275 | 0.99686584 | 0.458862423 |
| Fer | 419.2667044 | 0.01950359 | 0.062753918 | 0.310794775 |
| Pja2 | 6703.040193 | -0.085679 | 0.059202311 | -1.447223903 |

| | | | | |
|---|---|---|---|---|
| Gm24194 | 3.223858552 | -0.389660403 | 0.653246163 | -0.596498571 |
| Mir6420 | 0 | NA | NA | NA |
| AU016765 | 0 | NA | NA | NA |
| Man2a1 | 771.0017978 | 0.182075994 | 0.110697128 | 1.644812274 |
| Gm17133 | 0 | NA | NA | NA |
| 4930583I09Rik | 0 | NA | NA | NA |
| Gm25344 | 0 | NA | NA | NA |
| Tmem232 | 14.97100193 | -0.02901328 | 0.265424243 | -0.109309079 |
| Gm25731 | 0 | NA | NA | NA |
| Vapa | 1462.977574 | 0.217380894 | 0.090787973 | 2.394379866 |
| Txndc2 | 4.780619236 | 0.820786814 | 0.493221591 | 1.664133989 |
| Rab31 | 1383.126595 | 0.409436231 | 0.129102637 | 3.171401005 |
| Ppp4r1 | 548.0723031 | -0.038041526 | 0.087623231 | -0.434148859 |
| Ralbp1 | 849.2774884 | -0.120179482 | 0.083998205 | -1.430738689 |
| Twsg1 | 633.6219113 | 0.580633686 | 0.106555019 | 5.449144422 |
| Ankrd12 | 2379.847845 | -0.037562993 | 0.072176214 | -0.520434518 |
| Ndufv2 | 1210.518907 | 0.10121521 | 0.095962082 | 1.0547417 |
| Gm23264 | 0.127717592 | 0.274405555 | 3.006584758 | 0.091268192 |
| Wash1 | 277.8590539 | 0.225067572 | 0.078013739 | 2.884973545 |
| Ddx11 | 26.93519898 | -0.708412018 | 0.235938032 | -3.002534237 |
| Mtcl1 | 1748.393845 | -0.770610061 | 0.141072367 | -5.462515977 |
| Gm4705 | 22.01963504 | -0.304105096 | 0.260155425 | -1.168936209 |
| Rab12 | 884.2363581 | 0.204425327 | 0.0855102 | 2.390654288 |
| Themis3 | 0.652038697 | 0.60220072 | 1.369662808 | 0.439670784 |
| Ptprm | 817.855665 | -0.070073304 | 0.066552651 | -1.052900264 |
| Mir5709 | 2.098507799 | 0.267183496 | 0.716638572 | 0.372828796 |
| Gm22277 | 0 | NA | NA | NA |
| Lrrc30 | 0.047948698 | -0.206487837 | 3.724945049 | -0.055433794 |
| Lama1 | 126.6409752 | 0.044135441 | 0.179096318 | 0.246434107 |
| Arhgap28 | 21.82346039 | 1.398617817 | 0.266008478 | 5.257794139 |
| Gm17097 | 0.284087786 | 1.734188951 | 1.783398378 | 0.972406935 |
| L3mbtl4 | 19.07932733 | -0.234424201 | 0.238500784 | -0.982907465 |
| Gm15974 | 0.045882455 | 0.755288839 | 3.724945049 | 0.202765096 |
| Tmem200c | 195.2513106 | -0.125051513 | 0.09773168 | -1.279539175 |
| Epb4.1l3 | 2609.206492 | -0.306714998 | 0.070479662 | -4.351822783 |
| 2410021H03Rik | 0.229803803 | -0.206481986 | 2.163245747 | -0.095450083 |
| Zbtb14 | 266.5595265 | 0.040386679 | 0.085419412 | 0.472804465 |
| A930029G22Rik | 17.70809633 | -0.73830995 | 0.273183576 | -2.702614709 |
| C030034I22Rik | 43.35133415 | 0.274346905 | 0.190488094 | 1.440231248 |
| A330050F15Rik | 100.5892871 | 0.157345344 | 0.129110203 | 1.218690238 |
| Gm22967 | 0 | NA | NA | NA |
| Dlgap1 | 5541.017591 | -0.066795189 | 0.104125133 | -0.64148959 |
| 5031415H12Rik | 2.165235617 | 0.420904204 | 0.848966656 | 0.49578414 |
| Gm20703 | 2.008244415 | 3.451800972 | 0.920556332 | 3.749690109 |
| Tgif1 | 173.2492349 | 3.061195517 | 0.289968731 | 10.55698492 |
| Gm28727 | 2.554427972 | 1.475718408 | 0.744085952 | 1.983263365 |
| Mir1195 | 0 | NA | NA | NA |

| | | | | |
|---|---|---|---|---|
| Gm26510 | 1.053857694 | 2.882720638 | 1.097682332 | 2.626188427 |
| Gm9320 | 7.10377163 | 0.576845774 | 0.435751755 | 1.323794495 |
| Gm26561 | 0.274682661 | 1.776633033 | 3.708688794 | 0.479046135 |
| Gm16519 | 0 | NA | NA | NA |
| Myl12b | 1231.826391 | 0.053803466 | 0.100796658 | 0.53378224 |
| Myl12a | 318.06527 | 1.247784913 | 0.188482502 | 6.620163156 |
| Myom1 | 11.02360487 | -0.315967327 | 0.330772739 | -0.955239926 |
| Lpin2 | 1181.411907 | 0.148132521 | 0.060158762 | 2.462359879 |
| Emilin2 | 338.4431336 | 2.10473824 | 0.294063799 | 7.157420414 |
| Smchd1 | 1270.998503 | 0.242281011 | 0.072264214 | 3.352710794 |
| Gm37639 | 37.94753101 | 0.001285438 | 0.259365918 | 0.00495608 |
| Gm38220 | 18.1195503 | 0.822980082 | 0.407192108 | 2.0211101 |
| Gm4707 | 30.94223545 | 0.035075375 | 0.330063843 | 0.106268456 |
| Ndc80 | 7.673135682 | 2.712224243 | 0.550766181 | 4.924456763 |
| Gm23115 | 0.060174537 | 0.755286614 | 3.724945049 | 0.202764498 |
| Mettl4-ps1 | 0.604171475 | -1.423928378 | 1.223464959 | -1.163848925 |
| Spdya | 9.137166137 | 0.120489261 | 0.316114887 | 0.381156554 |
| Trmt61b | 64.18131499 | 0.170982918 | 0.155510977 | 1.099490993 |
| Wdr43 | 540.8743764 | 0.620415542 | 0.106536323 | 5.823511872 |
| Snord92 | 1.968787839 | -0.322525916 | 0.909106598 | -0.354772385 |
| Snord53 | 3.604104618 | 0.873107487 | 0.661048115 | 1.320792642 |
| Gm22858 | 0.576946659 | -0.035413207 | 3.535657565 | -0.010016017 |
| Fam179a | 77.43913187 | 0.713175856 | 0.181797504 | 3.922913352 |
| Gm15641 | 0 | NA | NA | NA |
| BC027072 | 1.833725576 | 0.266626633 | 0.758136393 | 0.351686893 |
| Clip4 | 625.3662385 | 0.039499313 | 0.055320742 | 0.714005491 |
| Alk | 126.1837875 | 0.211119248 | 0.11113749 | 1.899622241 |
| Gm24736 | 0.908322817 | -1.106149875 | 1.041584134 | -1.061988023 |
| Gm26963 | 0 | NA | NA | NA |
| Ypel5 | 905.935313 | 0.445396705 | 0.079877345 | 5.576007903 |
| Lbh | 1015.797448 | 0.408454893 | 0.136208518 | 2.998747059 |
| Gm23649 | 0.075043205 | 0.755286614 | 3.724945049 | 0.202764498 |
| Lclat1 | 432.1555874 | -0.1343997 | 0.081540763 | -1.648251678 |
| Gm9311 | 0.089089296 | 1.194485492 | 3.723939576 | 0.320758559 |
| Gm25406 | 3.791534426 | -2.18757657 | 0.638073384 | -3.428409057 |
| Capn13 | 0.17782045 | 0.274410407 | 2.296516967 | 0.119489823 |
| Galnt14 | 193.3014098 | -0.217596623 | 0.151479196 | -1.43647859 |
| D630014O11Rik | 13.27744998 | -0.394393791 | 0.421697983 | -0.935251784 |
| Ehd3 | 1261.372702 | 0.254124246 | 0.085744738 | 2.963729935 |
| Xdh | 746.6583068 | 4.590616558 | 0.268530205 | 17.09534525 |
| Gm24475 | 0 | NA | NA | NA |
| Srd5a2 | 1.0712781 | -0.419171494 | 0.974763223 | -0.430023912 |
| Memo1 | 287.5100735 | 0.004968288 | 0.085111606 | 0.058373798 |
| Dpy30 | 145.3608656 | 0.06635679 | 0.120768943 | 0.549452437 |
| Spast | 787.6234006 | -0.095611782 | 0.054701521 | -1.747881595 |
| Slc30a6 | 112.2934621 | -0.000914142 | 0.133187965 | -0.00686355 |
| Nlrc4 | 11.8016311 | 1.593502087 | 0.388897966 | 4.097481155 |

| | | | | |
|---|---|---|---|---|
| Yipf4 | 489.5110183 | -0.151181542 | 0.098077423 | -1.541451006 |
| Birc6 | 5732.824248 | 0.051865404 | 0.064959718 | 0.798424097 |
| Gm26749 | 0.249552943 | 0.755309902 | 1.848392422 | 0.408630707 |
| Ttc27 | 207.7543264 | 0.224184468 | 0.08049017 | 2.785240327 |
| Ltbp1 | 508.8403771 | 1.766542861 | 0.18302155 | 9.652103039 |
| Gm6276 | 0.110870145 | -0.206490066 | 3.724945049 | -0.055434393 |
| Rasgrp3 | 150.64243 | -0.242109029 | 0.188023135 | -1.287655531 |
| Fam98a | 451.495364 | -0.054935493 | 0.089140082 | -0.616282725 |
| Gm24126 | 0 | NA | NA | NA |
| Gm26124 | 0 | NA | NA | NA |
| Gm16391 | 0 | NA | NA | NA |
| Gm17538 | 12.6024622 | -0.502996357 | 0.305290752 | -1.647597754 |
| Gm5229 | 0.072025462 | 0.755288839 | 3.724945049 | 0.202765096 |
| Crim1 | 1289.41108 | 0.388646139 | 0.077069747 | 5.04278465 |
| Gm25069 | 0.582565216 | -1.45273683 | 1.432134883 | -1.014385479 |
| Fez2 | 418.1889862 | 0.323566145 | 0.093657392 | 3.4547849 |
| Gm10093 | 94.11779407 | 0.819940173 | 0.164167875 | 4.994522662 |
| Vit | 19.4303546 | -0.253167852 | 0.235739226 | -1.073931804 |
| Gm22562 | 0.092493346 | 0.274402725 | 3.724945049 | 0.073666248 |
| Strn | 1780.921145 | -0.283351368 | 0.059933706 | -4.727746473 |
| Heatr5b | 1002.913214 | -0.359234321 | 0.059982354 | -5.989000099 |
| Gpatch11 | 308.9091273 | -0.492109552 | 0.124416408 | -3.955342878 |
| Gm6548 | 198.422454 | 3.626521185 | 0.204157139 | 17.76338169 |
| Eif2ak2 | 2234.053102 | 4.837922433 | 0.166109484 | 29.12490192 |
| n-R5s29 | 0.136963818 | 0.755286583 | 3.59954433 | 0.209828388 |
| Gm26637 | 5.907155527 | 4.298261748 | 0.884320502 | 4.860524819 |
| Sult6b1 | 1.803778753 | -0.433315651 | 1.003057899 | -0.431994656 |
| Cebpzos | 71.15640056 | 0.158232246 | 0.150839941 | 1.049007609 |
| Cebpz | 422.2918863 | -0.02831574 | 0.089293428 | -0.317108894 |
| Ndufaf7 | 198.2430765 | -0.381310895 | 0.081624338 | -4.671534325 |
| Prkd3 | 324.5446214 | -0.040682863 | 0.118077398 | -0.344544034 |
| Gm26957 | 0 | NA | NA | NA |
| Qpct | 101.3108535 | -0.278199465 | 0.159922361 | -1.739590786 |
| Cdc42ep3 | 151.9232374 | -0.25131598 | 0.119870664 | -2.096559511 |
| Gm17315 | 8.227418531 | 0.017974844 | 0.353281549 | 0.050879657 |
| Rmdn2 | 100.9181395 | 0.048895242 | 0.111524195 | 0.438427214 |
| 4921513D11Rik | 17.43713135 | -0.310080484 | 0.280677975 | -1.104755315 |
| Cyp1b1 | 282.3238744 | 2.122978409 | 0.231558316 | 9.168223579 |
| Gm37555 | 0.136963818 | 0.755286583 | 3.59954433 | 0.209828388 |
| Atl2 | 973.0030504 | -0.200842485 | 0.068818253 | -2.918447865 |
| Hnrnpll | 974.8453895 | 0.043109089 | 0.085415745 | 0.504697219 |
| Gm22215 | 0 | NA | NA | NA |
| Gm23125 | 0 | NA | NA | NA |
| Galm | 27.71090595 | -0.467394543 | 0.314491343 | -1.486192079 |
| Gm25706 | 3.582960898 | -1.203430219 | 0.611184225 | -1.969013875 |
| Srsf7 | 905.8057965 | -0.005492763 | 0.07251532 | -0.075746237 |
| Ttc39d | 1.912265302 | -1.228962077 | 0.886483731 | -1.386333482 |

| | | | | |
|---|---|---|---|---|
| Gemin6 | 32.80368467 | -0.032603354 | 0.208463947 | -0.156398048 |
| Dhx57 | 544.389297 | -0.362740778 | 0.104362596 | -3.475773799 |
| Morn2 | 84.79043694 | 0.023888133 | 0.165656202 | 0.144203073 |
| Arhgef33 | 272.4084982 | -0.105166349 | 0.302859578 | -0.347244586 |
| Gm10190 | 0.322675615 | 0.263651205 | 1.479355313 | 0.178220339 |
| Sos1 | 1511.796088 | -0.180494879 | 0.052633548 | -3.429274428 |
| Cdkl4 | 159.4655411 | 0.795846798 | 0.155937783 | 5.103617496 |
| Map4k3 | 1163.46542 | 0.006266763 | 0.050858156 | 0.123220412 |
| Gm9959 | 0.271824581 | -1.135676865 | 1.789228915 | -0.634729774 |
| Gm9386 | 0.459282249 | 1.790265827 | 1.442140986 | 1.241394457 |
| Tmem178 | 593.6338004 | -0.0083873 | 0.089684719 | -0.093519838 |
| Thumpd2 | 64.74403033 | 0.361933031 | 0.153577843 | 2.356674784 |
| Slc8a1 | 2319.537813 | 0.097498414 | 0.087381605 | 1.115777333 |
| Gm11096 | 0.461313532 | 0.941443942 | 1.274705679 | 0.738557895 |
| Rpl31-ps25 | 0 | NA | NA | NA |
| Gm27359 | 0 | NA | NA | NA |
| Gm22386 | 0 | NA | NA | NA |
| Gm23400 | 0 | NA | NA | NA |
| Gm6594 | 14.12482822 | 0.455886812 | 0.331895172 | 1.373586754 |
| Gm29052 | 0 | NA | NA | NA |
| Pkdcc | 80.91185254 | -0.208879209 | 0.147545894 | -1.415689748 |
| Gm24240 | 0 | NA | NA | NA |
| Rpl21-ps7 | 2.037653348 | 1.017527402 | 0.722347841 | 1.408639086 |
| Eml4 | 575.2161808 | -0.193920312 | 0.079738229 | -2.431961609 |
| Cox7a2l | 853.8877004 | 0.049917552 | 0.078711009 | 0.634187679 |
| Kcng3 | 81.41433254 | -0.4626809 | 0.181917316 | -2.543358214 |
| Mta3 | 891.3237956 | -0.051829214 | 0.057845116 | -0.895999834 |
| Haao | 4.556506885 | 4.180299591 | 0.94103006 | 4.442259357 |
| C430042M11Rik | 1.362936493 | -0.389468756 | 0.959578054 | -0.405875014 |
| Gm23905 | 0.061920613 | 0.274393828 | 3.724945049 | 0.073663859 |
| Gm37549 | 0.452867449 | 2.479078131 | 1.772692 | 1.398482157 |
| Zfp36l2 | 928.29596 | 1.292403722 | 0.226431705 | 5.707697698 |
| Thada | 445.9157641 | -0.31713031 | 0.091921525 | -3.450011402 |
| Gm24492 | 0.916014489 | -0.584826201 | 1.034245932 | -0.565461447 |
| Gm24722 | 0.550570057 | 0.701064623 | 1.333919739 | 0.525567321 |
| Plekhh2 | 185.6708911 | -0.362254852 | 0.110220925 | -3.286625056 |
| Gm26066 | 0.13205505 | -1.042928876 | 3.656016608 | -0.285263714 |
| Dync2li1 | 99.93135035 | -0.155333871 | 0.142208632 | -1.092295653 |
| Abcg5 | 0.732414602 | 1.403019582 | 1.448950842 | 0.968300333 |
| Abcg8 | 0.118974955 | 0.274400501 | 3.724945049 | 0.073665651 |
| Lrpprc | 1213.010164 | -0.113030358 | 0.072715775 | -1.554413168 |
| 1110020A21Rik | 4.802739388 | -0.497015336 | 0.440162824 | -1.129162458 |
| Ppm1b | 1207.845512 | -0.196020653 | 0.062509517 | -3.135852954 |
| Slc3a1 | 11.21614055 | 0.152444204 | 0.391513413 | 0.3893716 |
| Prepl | 4734.893635 | -0.128795712 | 0.053768816 | -2.395360745 |
| Camkmt | 72.64996259 | -0.096817079 | 0.14051694 | -0.689006458 |
| Rpl31-ps16 | 2.862600439 | 0.44053164 | 0.641305539 | 0.686929418 |

| | | | | |
|---|---|---|---|---|
| Gm27178 | 0 | NA | NA | NA |
| Six3os1 | 52.19109397 | -0.495595706 | 0.225525176 | -2.19751832 |
| Gm27435 | 0.089062654 | -0.20649451 | 3.724945049 | -0.055435586 |
| Gm27801 | 0.250814404 | -1.030920997 | 1.784277208 | -0.577780735 |
| Six3 | 109.2161158 | -0.443931749 | 0.149942954 | -2.960670952 |
| Gm27610 | 0.226876767 | -1.056945463 | 2.148259335 | -0.492000871 |
| Gm27283 | 0.388809915 | -0.114578331 | 1.685363391 | -0.067984348 |
| Gm27275 | 0.04393175 | -0.206490061 | 3.724945049 | -0.055434391 |
| Gm27502 | 0.090665643 | -0.687373951 | 3.724945049 | -0.184532642 |
| Gm27427 | 0.359940881 | -1.83459091 | 1.718336012 | -1.067655509 |
| Gm28528 | 1.48446612 | 0.175825153 | 0.878169414 | 0.200217805 |
| Six2 | 7.803849726 | 0.384200625 | 0.368883413 | 1.041523179 |
| Gm29418 | 0.434220697 | 0.645176124 | 1.341715704 | 0.480859039 |
| Srbd1 | 196.7237653 | -0.51278443 | 0.118353667 | -4.33264504 |
| Prkce | 4692.067451 | 0.018421744 | 0.093206843 | 0.197643682 |
| 2010106C02Rik | 0 | NA | NA | NA |
| Gm10309 | 10.12416963 | 4.622334927 | 0.777120765 | 5.948026529 |
| Gm22235 | 0 | NA | NA | NA |
| Epas1 | 1364.101885 | 0.07190652 | 0.114113503 | 0.630131563 |
| Gm29168 | 0.150086409 | 1.264824188 | 3.723011833 | 0.339731445 |
| Tmem247 | 0 | NA | NA | NA |
| Atp6v1e2 | 0.434133818 | -0.123618703 | 1.574747085 | -0.078500671 |
| Rhoq | 718.01171 | -0.161757441 | 0.106485179 | -1.51906061 |
| Pigf | 100.958299 | -0.329961222 | 0.154578967 | -2.134580327 |
| Cript | 263.4740334 | -0.095743235 | 0.142215781 | -0.673225113 |
| Gm24648 | 0 | NA | NA | NA |
| Gm5499 | 18.0195093 | 0.060407144 | 0.266485649 | 0.226680666 |
| Socs5 | 1211.318638 | -0.372798657 | 0.092705964 | -4.021301752 |
| Gm22346 | 0.044191549 | -0.206490061 | 3.724945049 | -0.055434391 |
| Mcfd2 | 543.0470171 | 0.219812675 | 0.086873319 | 2.530266802 |
| 4833418N02Rik | 13.1105544 | 0.099067052 | 0.28584191 | 0.346579871 |
| Ttc7 | 196.4810743 | 0.16855821 | 0.149622188 | 1.126558914 |
| Gm15978 | 0.6240965 | -0.228420251 | 1.158860323 | -0.197107664 |
| 0610012D04Rik | 0.332525094 | -0.687379803 | 1.751416426 | -0.392470799 |
| 1700011E24Rik | 3.15672782 | 0.640200354 | 0.594113353 | 1.077572741 |
| Calm2 | 8582.912101 | -0.039481873 | 0.104328509 | -0.378438006 |
| Gm26005 | 0.04393175 | -0.206490061 | 3.724945049 | -0.055434391 |
| Gm28676 | 0.161026707 | 0.274389649 | 3.463604896 | 0.079220828 |
| Epcam | 12.12751263 | 1.103494958 | 0.522496236 | 2.111967288 |
| Msh2 | 333.7442204 | -0.143906889 | 0.096961342 | -1.484167681 |
| Kcnk12 | 130.455902 | -0.582982776 | 0.139844431 | -4.168795076 |
| Rpl36-ps4 | 0.224455361 | 1.12570297 | 3.133384974 | 0.359260984 |
| Msh6 | 331.2527847 | -0.164432303 | 0.077836022 | -2.112547627 |
| Fbxo11 | 1810.721455 | -0.011534434 | 0.066316718 | -0.173929509 |
| Gm4832 | 0.217012189 | 0.755304083 | 2.167447184 | 0.34847635 |
| Gm26612 | 0.40569442 | 0.290487826 | 1.345270335 | 0.215932678 |
| Gm22542 | 0.253603277 | 0.316213348 | 2.106677492 | 0.150100501 |

| | | | | |
|---|---|---|---|---|
| Gm23947 | 0 | NA | NA | NA |
| Foxn2 | 537.5500919 | -0.11958206 | 0.093696205 | -1.27627432 |
| Ppp1r21 | 678.815562 | -0.162872989 | 0.056880974 | -2.863400159 |
| Gm38109 | 8.730948186 | -1.14812181 | 0.417684336 | -2.748778711 |
| Ston1 | 93.35916316 | 0.639267026 | 0.175065541 | 3.651586834 |
| Gtf2a1l | 2.669826932 | -0.157393913 | 0.584618071 | -0.269225192 |
| Lhcgr | 3.114871368 | -0.519420155 | 0.580186591 | -0.895263977 |
| Fshr | 0.045882455 | 0.755288839 | 3.724945049 | 0.202765096 |
| Gm10184 | 3.494817459 | 0.254206292 | 0.573822479 | 0.443005113 |
| Nrxn1 | 7760.329357 | -0.27835376 | 0.098129925 | -2.836583846 |
| Gm24510 | 0 | NA | NA | NA |
| Gm10493 | 3.600385196 | -0.519430924 | 0.54034121 | -0.9613017 |
| Gm19815 | 7.90218533 | -0.695719662 | 0.543130009 | -1.280944987 |
| Gm20491 | 88.08114114 | -0.553108098 | 0.31213265 | -1.77202897 |
| 4930480K15Rik | 0.097899064 | -0.687371727 | 3.724945049 | -0.184532045 |
| AC151286.1 | 0.277228681 | -2.028466505 | 3.683251455 | -0.55072713 |
| Gm10308 | 1.537590905 | 0.130766237 | 0.703961895 | 0.18575755 |
| Gm27475 | 0 | NA | NA | NA |
| Gm6741 | 17.12533191 | -0.254369475 | 0.352483222 | -0.72164988 |
| Gm15404 | 0.089089296 | 1.194485492 | 3.723939576 | 0.320758559 |
| Gm15403 | 0 | NA | NA | NA |
| Gm24244 | 0 | NA | NA | NA |
| AC131325.1 | 0 | NA | NA | NA |
| Gm27039 | 0 | NA | NA | NA |
| Adcyap1 | 181.5271029 | -0.188389188 | 0.183731158 | -1.025352422 |
| Gm25935 | 0 | NA | NA | NA |
| Gm22593 | 0.196742721 | 0.274410082 | 2.232417207 | 0.122920609 |
| Mettl4 | 150.9550503 | -0.378950779 | 0.163207004 | -2.321902677 |
| 2700099C18Rik | 6.495084589 | 0.814506735 | 0.4651093 | 1.751215758 |
| Gm27038 | 3.867668655 | 0.365673965 | 0.455032832 | 0.803621055 |
| Gm1976 | 94.15986379 | -0.081348 | 0.11246641 | -0.723309296 |
| Gm23964 | 0.139856492 | -0.206491248 | 3.574411025 | -0.057769307 |
| Gm26734 | 6.887014156 | -0.161052926 | 0.409475481 | -0.393315189 |
| Gm23102 | 0 | NA | NA | NA |
| Gm20939 | 23.37788906 | -0.185123203 | 0.367433792 | -0.503827376 |
| Vmn1r-ps151 | 0 | NA | NA | NA |
| Vmn1r-ps152 | 0.092691418 | 0.274387304 | 3.724945049 | 0.073662108 |
| Vmn1r238 | 0.090890357 | 0.755282166 | 3.724945049 | 0.202763304 |
| Crem | 361.0165932 | 0.864958032 | 0.164648354 | 5.25336579 |
| Gm6225 | 15.32231194 | 0.726518482 | 0.370085985 | 1.963107253 |
| Cul2 | 847.8838813 | 0.249848469 | 0.080817129 | 3.091528639 |
| G430049J08Rik | 0.148708979 | 0.755295232 | 2.868529146 | 0.263304012 |
| Bambi | 60.79279935 | 0.3043015 | 0.181496385 | 1.67662568 |
| Gm23643 | 0 | NA | NA | NA |
| Gm24632 | 0 | NA | NA | NA |
| Lyzl1 | 0 | NA | NA | NA |
| Gm10557 | 201.5185117 | 0.260784109 | 0.14741779 | 1.769013823 |

| | | | | |
|---|---|---|---|---|
| Map3k8 | 99.88469274 | 1.71352436 | 0.18673058 | 9.176452821 |
| Gm26865 | 0.332096027 | -0.809100656 | 1.703305367 | -0.475017969 |
| Mtpap | 376.5909295 | 0.163865078 | 0.095886996 | 1.708939523 |
| 9430020K01Rik | 1287.14579 | -0.536832426 | 0.111611412 | -4.809834559 |
| Gm10556 | 0.381271471 | 0.274412283 | 1.709622201 | 0.160510482 |
| Svil | 295.6710269 | -0.114684916 | 0.106183012 | -1.080068399 |
| Gm26682 | 7.326093239 | 0.734712148 | 0.468223278 | 1.569149129 |
| Zfp438 | 70.78775138 | -0.113190072 | 0.149802254 | -0.755596587 |
| Gm10125 | 71.18097263 | 0.07193055 | 0.182777519 | 0.393541562 |
| Gm26575 | 19.11664666 | -0.117903768 | 0.31997941 | -0.368472983 |
| Gm2238 | 59.71255148 | -0.551960778 | 0.190389947 | -2.899106735 |
| Zeb1 | 1461.053986 | -0.13027047 | 0.051928556 | -2.508648017 |
| Gm16147 | 0 | NA | NA | NA |
| Gm22709 | 4.178956302 | -1.33392068 | 0.593920923 | -2.245956705 |
| Arhgap12 | 782.7285597 | -0.150480378 | 0.075786385 | -1.985585899 |
| Kif5b | 4777.179878 | 0.095074021 | 0.115661887 | 0.821999565 |
| Gm17036 | 1.215361072 | 0.238891583 | 1.148452962 | 0.208011639 |
| Rpl27-ps3 | 9.972102879 | 0.209394645 | 0.391745336 | 0.534517264 |
| Gm6291 | 0 | NA | NA | NA |
| Epc1 | 763.0696826 | 0.060410529 | 0.074504695 | 0.810828483 |
| Gm28529 | 13.22339979 | 0.134285207 | 0.348957931 | 0.384817754 |
| Mir1893 | 0 | NA | NA | NA |
| 4921524L21Rik | 0.072025462 | 0.755288839 | 3.724945049 | 0.202765096 |
| Rab18 | 1241.658341 | 0.034510452 | 0.080618984 | 0.428068563 |
| Mkx | 204.0449348 | 0.200709755 | 0.12038117 | 1.667285297 |
| Armc4 | 20.81306997 | 0.689583508 | 0.233077396 | 2.958603105 |
| Mpp7 | 246.8020213 | -0.146011393 | 0.101102222 | -1.444195683 |
| Fabp5l2 | 0.240849226 | -0.206481911 | 1.864898668 | -0.110720177 |
| Gm24292 | 0 | NA | NA | NA |
| Wac | 1913.852717 | 0.294447027 | 0.063182442 | 4.660266609 |
| Gm7502 | 0.768676318 | -0.503479839 | 1.058954823 | -0.475449781 |
| Gm5819 | 9.13781552 | -0.813839201 | 0.346835575 | -2.346469795 |
| Gm26119 | 0 | NA | NA | NA |
| Gm37148 | 0 | NA | NA | NA |
| Fzd8 | 66.68569282 | -0.264834877 | 0.15816692 | -1.67440117 |
| Gjd4 | 0.815409053 | 0.329981918 | 1.112762557 | 0.296542974 |
| Ccny | 1305.979739 | 0.159725062 | 0.051097949 | 3.125860508 |
| Gm22765 | 0 | NA | NA | NA |
| Cetn1 | 0.038981395 | 0.274393828 | 3.724945049 | 0.073663859 |
| Colec12 | 279.2173052 | 0.64518017 | 0.145269604 | 4.441260621 |
| Gm17430 | 0 | NA | NA | NA |
| Gm7497 | 4.858631947 | 0.146936166 | 0.608147728 | 0.24161262 |
| Gm23350 | 0 | NA | NA | NA |
| Gm23637 | 0 | NA | NA | NA |
| Gm5501 | 0 | NA | NA | NA |
| Thoc1 | 432.2063309 | 0.13187177 | 0.079208804 | 1.664862525 |
| Usp14 | 1395.726703 | -0.008501602 | 0.050220897 | -0.169284148 |

| | | | | |
|---|---|---|---|---|
| Rock1 | 1035.477942 | 0.155204218 | 0.057181172 | 2.714253872 |
| Gm2531 | 25.92851566 | -0.331981172 | 0.313202901 | -1.059955611 |
| Gm26244 | 55.4930898 | 0.390579768 | 0.191381106 | 2.040848112 |
| Greb1l | 215.658039 | 0.39384928 | 0.141301541 | 2.787296418 |
| AC124740.1 | 0 | NA | NA | NA |
| Gm10349 | 1.210176943 | 0.617712216 | 0.840455188 | 0.734973411 |
| Gm24894 | 2.156355716 | -0.847977028 | 0.688565001 | -1.231513405 |
| Esco1 | 664.6422751 | 0.063671703 | 0.081323618 | 0.782942331 |
| Snrpd1 | 236.8850658 | 0.214658127 | 0.132139905 | 1.624476169 |
| Abhd3 | 409.5174636 | -0.421278908 | 0.144398324 | -2.917477818 |
| 4930563E18Rik | 1.299310135 | -0.973762515 | 0.813735791 | -1.196656858 |
| Mib1 | 3835.229447 | -0.243585941 | 0.060322021 | -4.038093181 |
| Gm25116 | 4.118368001 | -0.397823664 | 0.663295583 | -0.599768299 |
| Gm21747 | 0.049553047 | 0.274391604 | 3.724945049 | 0.073663262 |
| Mir133a-1 | 0 | NA | NA | NA |
| Mir1b | 7.73100226 | -0.142689718 | 0.346119583 | -0.412255547 |
| mmu-mir-1a-2 | 0 | NA | NA | NA |
| 1010001N08Rik | 0.099155932 | 0.75528439 | 3.724945049 | 0.202763901 |
| Gata6 | 1.608823659 | 0.595945494 | 0.862635985 | 0.690842376 |
| Rbbp8 | 116.5073145 | 0.460093407 | 0.122219278 | 3.764491293 |
| Gm6277 | 35.32926637 | 0.105611606 | 0.238731012 | 0.442387459 |
| Cables1 | 273.0530706 | -0.310458334 | 0.118479805 | -2.620348127 |
| Mir1901 | 6.131714793 | 0.745128041 | 0.421359893 | 1.768388622 |
| Tmem241 | 88.72054137 | -0.377353621 | 0.158107679 | -2.386687494 |
| Riok3 | 978.9154087 | 0.322159363 | 0.084570776 | 3.809346208 |
| Gm25289 | 0.089062654 | -0.20649451 | 3.724945049 | -0.055435586 |
| 3110002H16Rik | 281.027959 | -0.080219824 | 0.092102605 | -0.870983223 |
| Npc1 | 1148.204684 | -0.1575716 | 0.08726607 | -1.80564565 |
| Gm15956 | 8.077703934 | -0.700982364 | 0.417329555 | -1.679685409 |
| Ankrd29 | 171.8859524 | -0.517635405 | 0.08901824 | -5.814936415 |
| Gm16072 | 0.87006192 | -1.441394796 | 1.148705374 | -1.254799385 |
| Lama3 | 110.321609 | -0.408120167 | 0.166933754 | -2.444803145 |
| Gm29201 | 0.436251183 | -1.705618065 | 1.72723773 | -0.987483098 |
| Gm29199 | 0 | NA | NA | NA |
| Gm29200 | 0.582012116 | 0.683061354 | 1.374020252 | 0.497126118 |
| Gm22340 | 0.038981395 | 0.274393828 | 3.724945049 | 0.073663859 |
| Ttc39c | 360.5212832 | 0.254326421 | 0.10489788 | 2.424514398 |
| Mir1948 | 2.898584899 | -0.512776714 | 0.607770392 | -0.843701373 |
| Cabyr | 18.12948923 | 1.429505525 | 0.436086703 | 3.278030529 |
| Gm7599 | 1.737466599 | 0.517425132 | 0.722285775 | 0.71637176 |
| Osbpl1a | 2161.262522 | -0.015064642 | 0.069182673 | -0.217751661 |
| Gm5687 | 0.269295688 | -0.175237236 | 2.080409373 | -0.084232093 |
| Gm26407 | 27.03744251 | -0.994854013 | 0.282881753 | -3.51685466 |
| Impact | 3627.068778 | -0.100185668 | 0.083351673 | -1.201963497 |
| Hrh4 | 0.228099782 | -0.206496245 | 3.135223884 | -0.065863317 |
| Gm22251 | 0 | NA | NA | NA |
| Zfp521 | 803.5141072 | 0.105187459 | 0.103826899 | 1.013104125 |

| | | | | |
|---|---|---|---|---|
| Gm5160 | 3.237902636 | 0.284465392 | 0.600626361 | 0.473614564 |
| Gm6457 | 44.70599757 | 0.440472847 | 0.159237586 | 2.766136171 |
| Ss18 | 416.8945723 | -0.050014731 | 0.097922246 | -0.510759644 |
| Gm24924 | 13.08384193 | 0.179969173 | 0.287065777 | 0.626926604 |
| Psma8 | 5.559946366 | 0.471506167 | 0.498571266 | 0.945714683 |
| Taf4b | 120.7625759 | 0.652236067 | 0.137961617 | 4.727663243 |
| Kctd1 | 603.8212261 | -0.149671853 | 0.098041326 | -1.526620043 |
| Gm6304 | 6.355730337 | -0.023192052 | 0.405011885 | -0.057262646 |
| Aqp4 | 4262.513783 | 0.891639736 | 0.140923623 | 6.327113347 |
| Chst9 | 8.958957809 | 0.092789043 | 0.3217912 | 0.288351712 |
| Gm10036 | 42.03144848 | 0.236132982 | 0.267176017 | 0.883810548 |
| Gm24228 | 0.072025462 | 0.755288839 | 3.724945049 | 0.202765096 |
| Gm4835 | 1.43143147 | 0.118547386 | 0.834676516 | 0.14202794 |
| Gm22389 | 1.98025533 | 0.32337616 | 0.78389961 | 0.412522415 |
| Gm24385 | 0 | NA | NA | NA |
| Gm7665 | 11.35479647 | 2.934574577 | 0.736755851 | 3.983103189 |
| Cdh2 | 2013.677982 | 0.215352954 | 0.062150686 | 3.465013316 |
| Gm15485 | 0 | NA | NA | NA |
| Gm15328 | 30.8296316 | -1.231913576 | 0.284397371 | -4.331663023 |
| Gm7670 | 0 | NA | NA | NA |
| 1700001G01Rik | 0.046345709 | 0.274391604 | 3.724945049 | 0.073663262 |
| Gm22470 | 0 | NA | NA | NA |
| Gm24015 | 0 | NA | NA | NA |
| Gm26235 | 0.048949532 | -0.206487837 | 3.724945049 | -0.055433794 |
| Gm25392 | 0 | NA | NA | NA |
| Gm24238 | 0 | NA | NA | NA |
| Dsc3 | 13.88762389 | 0.448302216 | 0.303159229 | 1.478768162 |
| Dsc2 | 11.39730211 | -0.019913151 | 0.365820871 | -0.054434158 |
| Dsc1 | 2.506558332 | 4.07456224 | 1.24910315 | 3.261990206 |
| Dsg1c | 4.094471938 | -0.100009705 | 0.613047794 | -0.163135251 |
| Dsg1a | 1.903639553 | 2.323787865 | 2.922897958 | 0.795028734 |
| Dsg1b | 0.835693444 | 1.582708454 | 1.240531622 | 1.275830802 |
| Dsg4 | 0 | NA | NA | NA |
| Dsg3 | 1.159328825 | 1.706624647 | 1.114339854 | 1.531511811 |
| Dsg2 | 36.65644163 | -0.522211823 | 0.206603587 | -2.527602886 |
| Gm16090 | 0.646850989 | 0.351824743 | 1.326002054 | 0.265327449 |
| Ttr | 2265.392118 | -0.659342989 | 0.334646385 | -1.970267778 |
| Gm10269 | 7.797321979 | 0.115782235 | 0.404067072 | 0.286542119 |
| B4galt6 | 2045.964961 | -0.244581309 | 0.062515447 | -3.912334021 |
| Trappc8 | 1142.654653 | -0.063745998 | 0.05396272 | -1.181296981 |
| Gm24959 | 4.935114692 | -0.50116832 | 0.461339428 | -1.086333164 |
| Rnf125 | 36.46586033 | 3.145243682 | 0.404338557 | 7.778737966 |
| Rnf138 | 267.4053105 | 0.096482099 | 0.088325757 | 1.092343867 |
| Mep1b | 0 | NA | NA | NA |
| Gm6378 | 0.155469778 | 0.274407114 | 2.829060383 | 0.096995849 |
| Garem | 494.2667245 | -0.150414037 | 0.099117687 | -1.517529734 |
| Gm24372 | 0.044191549 | -0.206490061 | 3.724945049 | -0.055434391 |

| | | | | |
|---|---|---|---|---|
| Gm22886 | 0 | NA | NA | NA |
| Klhl14 | 23.8882674 | -0.647001455 | 0.243997305 | -2.651674591 |
| 4930426D05Rik | 6.412194933 | -0.242463673 | 0.392157743 | -0.618280979 |
| Ccdc178 | 1.26832265 | 1.47369026 | 1.065552254 | 1.383029556 |
| Asxl3 | 371.9529472 | 0.044381287 | 0.116244777 | 0.381791667 |
| Nol4 | 882.8164278 | -0.418095357 | 0.094282865 | -4.434478701 |
| Gm22495 | 0 | NA | NA | NA |
| Dtna | 1930.500953 | -0.353366511 | 0.08471489 | -4.171244431 |
| Gm15972 | 1.928713869 | 0.460160908 | 0.672084253 | 0.684677414 |
| Mapre2 | 3079.479868 | -0.248914965 | 0.060265597 | -4.13029953 |
| Mir6360 | 0 | NA | NA | NA |
| Gm23207 | 0 | NA | NA | NA |
| Zfp397 | 704.3356492 | -0.365241776 | 0.083831519 | -4.356855018 |
| Zscan30 | 8.824857606 | -0.654793449 | 0.350683123 | -1.867194076 |
| Zfp35 | 243.5789265 | -0.060604621 | 0.096108099 | -0.63058808 |
| Zfp191 | 825.6271123 | 0.274643857 | 0.097388955 | 2.820071919 |
| Ino80c | 294.4170576 | 0.560484347 | 0.109283902 | 5.128699988 |
| Galnt1 | 801.5304355 | 0.318753645 | 0.089072587 | 3.578582992 |
| Gm7701 | 14.47583842 | -0.516319796 | 0.274710354 | -1.879506138 |
| Mir187 | 0.214710016 | 0.27439204 | 3.154113791 | 0.086994972 |
| 2700062C07Rik | 106.228182 | 0.303596688 | 0.113874956 | 2.666053181 |
| Rprd1a | 820.2179591 | -0.062493559 | 0.068868796 | -0.907429246 |
| Gm23606 | 0.061920613 | 0.274393828 | 3.724945049 | 0.073663859 |
| Slc39a6 | 544.4852359 | 0.003277135 | 0.093965118 | 0.034876084 |
| Elp2 | 877.5310991 | -0.011166946 | 0.0492625 | -0.226682496 |
| Mocos | 101.3743352 | 2.225607565 | 0.216304002 | 10.28925747 |
| n-R5s23 | 0.045882455 | 0.755288839 | 3.724945049 | 0.202765096 |
| Gm9955 | 0.957795399 | 0.093254071 | 0.907066991 | 0.102808361 |
| Fhod3 | 695.6586359 | -0.17978943 | 0.130600166 | -1.37664013 |
| Gm24294 | 1.176033076 | -1.800360754 | 1.074911016 | -1.674892831 |
| Tpgs2 | 457.1009357 | -0.379526713 | 0.074482078 | -5.095544077 |
| AW554918 | 270.5192402 | 0.023291985 | 0.076935003 | 0.302748868 |
| Gm16558 | 5.405763959 | -0.218410359 | 0.428406364 | -0.509820528 |
| Celf4 | 4385.110905 | -0.029202561 | 0.090764367 | -0.32174037 |
| Gm25845 | 9.017131998 | -0.9901361 | 0.341500904 | -2.899365973 |
| Gm7847 | 0 | NA | NA | NA |
| Gm23674 | 0 | NA | NA | NA |
| Gm26106 | 0 | NA | NA | NA |
| Gm7729 | 7.595035811 | -0.242978074 | 0.391081732 | -0.621297428 |
| Gm24246 | 0 | NA | NA | NA |
| Pik3c3 | 522.6692759 | -0.052055106 | 0.057174033 | -0.910467622 |
| Rpl29-ps5 | 0.839167439 | -0.936295049 | 1.079016356 | -0.867730173 |
| Gm22870 | 0 | NA | NA | NA |
| Gm7936 | 0.203976984 | -1.052174267 | 2.161973372 | -0.486673093 |
| Rit2 | 621.7940845 | -0.279653056 | 0.091935621 | -3.041835724 |
| Gm23283 | 0 | NA | NA | NA |
| Syt4 | 3383.452968 | 0.079746364 | 0.077639875 | 1.027131537 |

| | | | | |
|---|---|---|---|---|
| Gm26658 | 4.064855943 | 0.105349759 | 0.512779584 | 0.205448427 |
| Gm25396 | 0 | NA | NA | NA |
| Slc25a46 | 1147.053562 | 0.020201978 | 0.080965199 | 0.24951434 |
| B930094E09Rik | 4.156831401 | 0.409955473 | 0.532261549 | 0.77021433 |
| Sap130 | 827.4019423 | -0.11620882 | 0.055485892 | -2.094385 |
| Ammecr1l | 554.7126231 | -0.238122056 | 0.079301206 | -3.002754545 |
| Gm26533 | 2.98239301 | -0.03473922 | 0.574252757 | -0.06049465 |
| Polr2d | 122.288999 | 0.237512998 | 0.120266619 | 1.974887127 |
| Wdr33 | 970.9585418 | 0.041658669 | 0.05365675 | 0.776391956 |
| Gm6665 | 1.827239475 | 0.919591467 | 0.810034111 | 1.135250298 |
| Gm26823 | 0.062426873 | 0.755288839 | 3.724945049 | 0.202765096 |
| Sft2d3 | 54.7958328 | -0.192001695 | 0.153081595 | -1.254244153 |
| Lims2 | 61.79027259 | -0.072831932 | 0.163882447 | -0.4444157 |
| Gpr17 | 643.1742382 | -1.002220156 | 0.285097431 | -3.515360179 |
| Myo7b | 0.405683672 | -1.723741171 | 1.61424135 | -1.067833612 |
| Gm16344 | 0.401548357 | -0.067053563 | 1.53562718 | -0.043665262 |
| Gm16345 | 0.075043205 | 0.755286614 | 3.724945049 | 0.202764498 |
| Iws1 | 844.5236566 | -0.028053098 | 0.077462761 | -0.36214947 |
| Proc | 0.048949532 | -0.206487837 | 3.724945049 | -0.055433794 |
| Gm26717 | 0.046345709 | 0.274391604 | 3.724945049 | 0.073663262 |
| Map3k2 | 1196.837555 | 0.036322602 | 0.088583971 | 0.410035826 |
| Ercc3 | 374.8626789 | 0.157877875 | 0.06734926 | 2.344166439 |
| Gm22325 | 0 | NA | NA | NA |
| A830052D11Rik | 8.708922263 | -0.28376148 | 0.323901682 | -0.876072884 |
| Bin1 | 1973.364051 | -0.101613598 | 0.040721114 | -2.495354066 |
| Gypc | 57.79268309 | 2.604535415 | 0.30283436 | 8.600528085 |
| Gm25826 | 0 | NA | NA | NA |
| 4930455D15Rik | 0.382692223 | 0.752884284 | 1.611123129 | 0.467304001 |
| Tslp | 1.429800265 | 1.07436043 | 1.026225611 | 1.046904714 |
| Gm26378 | 0.072025462 | 0.755288839 | 3.724945049 | 0.202765096 |
| Wdr36 | 397.0988872 | 0.075726952 | 0.063871368 | 1.185616559 |
| Mir6361 | 0 | NA | NA | NA |
| 2310026I22Rik | 3.939202308 | 0.096678798 | 0.486866172 | 0.19857366 |
| Camk4 | 6160.809613 | -0.132554519 | 0.157722212 | -0.840430253 |
| Gm22814 | 0 | NA | NA | NA |
| Stard4 | 406.2855849 | -0.589556656 | 0.100420528 | -5.870877861 |
| Nrep | 1304.95571 | -0.776872385 | 0.156679396 | -4.95835702 |
| Gm10549 | 1.094754251 | -1.746753045 | 1.064787267 | -1.640471387 |
| Epb4.1l4aos | 32.18986031 | 0.442323907 | 0.245584037 | 1.801110168 |
| Gm23639 | 3.703464063 | 0.920788718 | 0.540961995 | 1.70213199 |
| Epb4.1l4a | 142.1446161 | -0.242067899 | 0.132927587 | -1.821050876 |
| Gm22788 | 0 | NA | NA | NA |
| Gm24432 | 0 | NA | NA | NA |
| Gm23411 | 0 | NA | NA | NA |
| Gm10548 | 0.669147434 | 2.249081776 | 1.217202243 | 1.847746987 |
| Apc | 7569.545602 | -0.275206978 | 0.074928952 | -3.672905729 |
| Srp19 | 313.7947171 | 0.203953246 | 0.09110234 | 2.238726755 |

| | | | | |
|---|---|---|---|---|
| Reep5 | 3174.602423 | -0.007681265 | 0.064375357 | -0.119319959 |
| Pkd2l2 | 2.995780043 | -1.820999265 | 0.737278935 | -2.469891894 |
| Fam13b | 1185.027783 | -0.078757338 | 0.064919915 | -1.213146037 |
| n-R5s24 | 0.131453142 | -0.196408214 | 2.976894461 | -0.065977554 |
| Wnt8a | 0.202010919 | -0.206486746 | 2.213058433 | -0.093303793 |
| Nme5 | 50.66745939 | -0.386916849 | 0.161650585 | -2.393538188 |
| 4933408B17Rik | 16.70085405 | -0.075775283 | 0.30752842 | -0.246400912 |
| Brd8 | 925.5748569 | 0.182867238 | 0.074072866 | 2.468747992 |
| Kif20a | 22.72681451 | 1.661103166 | 0.423043727 | 3.92655194 |
| Cdc23 | 555.3324902 | -0.173697313 | 0.072840807 | -2.384615448 |
| n-R5s25 | 0 | NA | NA | NA |
| Gfra3 | 0 | NA | NA | NA |
| Cdc25c | 5.025978301 | 0.968755431 | 0.484166249 | 2.000873527 |
| Gm3550 | 0.207873913 | -0.20649053 | 2.611896668 | -0.079057695 |
| 2010110K18Rik | 0.294784161 | 1.814187379 | 1.997959103 | 0.908020277 |
| Fam53c | 1037.28919 | 0.021930349 | 0.055033754 | 0.398489054 |
| Kdm3b | 1100.794087 | 0.018929563 | 0.051071652 | 0.370647161 |
| Gm26538 | 0.390497984 | 0.249468225 | 1.571852264 | 0.158709715 |
| Reep2 | 1197.302419 | -0.063528665 | 0.067883123 | -0.935853592 |
| Egr1 | 3179.097212 | -0.445566759 | 0.353958014 | -1.258812463 |
| Etf1 | 1390.357339 | 0.430381067 | 0.099977161 | 4.304793817 |
| Hspa9 | 2869.344892 | 0.36933403 | 0.106922194 | 3.454231688 |
| Gm22200 | 4.566833944 | -0.401726232 | 0.524236038 | -0.766307928 |
| Gm26109 | 3.869151513 | 0.430800845 | 0.525720894 | 0.819447828 |
| Ctnna1 | 941.2579512 | 0.379484771 | 0.10288292 | 3.688510889 |
| Lrrtm2 | 1096.201602 | -0.196495334 | 0.078755997 | -2.494988834 |
| Sil1 | 202.6507413 | 0.26814605 | 0.096258905 | 2.785675246 |
| Gm5239 | 0.164165709 | -0.206480289 | 2.781083781 | -0.074244541 |
| Snhg4 | 73.54980766 | -0.494469962 | 0.145801936 | -3.391381332 |
| Gm28285 | 6.985251871 | 0.140777449 | 0.376278707 | 0.374130787 |
| Mir1949 | 0 | NA | NA | NA |
| mmu-mir-1949 | 0 | NA | NA | NA |
| Snora74a | 0 | NA | NA | NA |
| AC121821.1 | 0.048949532 | -0.206487837 | 3.724945049 | -0.055433794 |
| Matr3 | 4954.108798 | 0.06984095 | 0.068599416 | 1.018098323 |
| Gm36989 | 219.4274089 | -0.590074269 | 0.135449699 | -4.356408858 |
| Paip2 | 1058.0764 | -0.072490163 | 0.095688054 | -0.757567527 |
| Slc23a1 | 0.196018198 | 0.776228636 | 2.631724082 | 0.294950615 |
| Mzb1 | 4.18320868 | 0.49173217 | 0.664279339 | 0.740249081 |
| Prob1 | 99.4461201 | -0.491053121 | 0.149070801 | -3.294093261 |
| Spata24 | 30.05154039 | -0.143997328 | 0.206467457 | -0.697433536 |
| Dnajc18 | 1508.902544 | 0.006999349 | 0.044970088 | 0.155644543 |
| 1700066B19Rik | 1.640932998 | 0.160447812 | 0.820316038 | 0.19559268 |
| Ecscr | 35.18133015 | 1.858054738 | 0.273919188 | 6.783222267 |
| Tmem173 | 105.9553805 | 2.06192233 | 0.211814915 | 9.734547406 |
| Gm16490 | 0 | NA | NA | NA |
| Ube2d2a | 1153.052153 | 0.045910426 | 0.076917554 | 0.596878395 |

| | | | | |
|---|---|---|---|---|
| Cxxc5 | 830.8617964 | -0.394162973 | 0.063214452 | -6.235330025 |
| Gm29417 | 3.083648494 | -1.117994134 | 0.580805811 | -1.924901771 |
| Gm23882 | 0 | NA | NA | NA |
| Psd2 | 681.0294117 | -0.759141495 | 0.142775046 | -5.317046049 |
| Nrg2 | 144.6574193 | -0.442539687 | 0.123440304 | -3.585050203 |
| Pura | 8433.573605 | -0.436673362 | 0.077322298 | -5.647444154 |
| Cystm1 | 401.2118129 | -0.110992088 | 0.100047042 | -1.109398995 |
| Pfdn1 | 311.4709643 | 0.258515677 | 0.135357886 | 1.909867873 |
| Hbegf | 582.7227112 | 1.24128351 | 0.317284602 | 3.91220848 |
| Slc4a9 | 2.54483004 | -0.23020032 | 0.638673466 | -0.360435077 |
| Ankhd1 | 1581.43401 | -0.092741217 | 0.084671182 | -1.09531029 |
| Gm6542 | 84.43238 | 0.472807673 | 0.140549276 | 3.36399935 |
| Eif4ebp3 | 7.191909191 | -0.539626609 | 0.39190335 | -1.376937984 |
| Sra1 | 541.6420646 | -0.032871792 | 0.068254102 | -0.481609031 |
| Slc35a4 | 487.0228943 | 0.163784341 | 0.076314286 | 2.146181914 |
| E230025N22Rik | 1.365445969 | -0.744780066 | 0.956683659 | -0.77850192 |
| Cd14 | 75.49450281 | 1.982301123 | 0.33286106 | 5.955341009 |
| Tmco6 | 58.12891111 | -0.377225395 | 0.175924093 | -2.144250905 |
| Ndufa2 | 320.0471109 | 0.128879932 | 0.117199836 | 1.099659655 |
| Ik | 860.0388227 | 0.220128903 | 0.117737654 | 1.86965594 |
| Wdr55 | 90.11495044 | 0.064318478 | 0.142527459 | 0.451270786 |
| Dnd1 | 14.61022796 | -0.721999905 | 0.338153293 | -2.135126051 |
| Hars | 700.2311977 | 0.176420309 | 0.125178416 | 1.409350873 |
| Hars2 | 250.6053811 | -0.101943317 | 0.107178212 | -0.95115709 |
| Zmat2 | 1268.32983 | 0.260871812 | 0.062096626 | 4.201062553 |
| Vaultrc5 | 18.77737098 | 0.438459672 | 0.426042889 | 1.029144443 |
| Gm23424 | 0.049553047 | 0.274391604 | 3.724945049 | 0.073663262 |
| Gm6756 | 1.105122771 | -0.328892169 | 1.081543214 | -0.304095264 |
| Pcdha1 | 15.4569917 | -0.702179147 | 0.246517371 | -2.848396214 |
| Gm37751 | 3.355297506 | -0.333617705 | 0.524304699 | -0.636305006 |
| Pcdha2 | 32.39827103 | -0.749325159 | 0.198948924 | -3.766419769 |
| Pcdha3 | 47.11146249 | -0.912875207 | 0.180647851 | -5.053341082 |
| Pcdha4 | 76.22283242 | -0.128137213 | 0.158572289 | -0.808068127 |
| Gm38225 | 2.632135046 | -0.435315982 | 0.761454237 | -0.571690274 |
| Pcdha5 | 55.26370248 | -0.685632707 | 0.185256498 | -3.700991398 |
| Gm37753 | 3.33123091 | -1.425743802 | 0.587487517 | -2.426849525 |
| Pcdha6 | 52.26043541 | -0.678964847 | 0.161692767 | -4.199104626 |
| Gm37013 | 0 | NA | NA | NA |
| Pcdha7 | 45.39741467 | -0.666175653 | 0.19607352 | -3.397580925 |
| Gm37388 | 0 | NA | NA | NA |
| Gm36858 | 26.49116316 | -0.957309245 | 0.278947345 | -3.43186362 |
| Pcdha8 | 34.08858227 | -0.942229478 | 0.311959739 | -3.020356026 |
| Pcdha9 | 61.46070133 | -0.883154713 | 0.170581344 | -5.177322982 |
| Pcdha11 | 154.8993467 | -0.844924118 | 0.172258633 | -4.904974027 |
| Gm10545 | 8.703390065 | -0.538353851 | 0.319041794 | -1.687408553 |
| Gm18150 | 13.89624275 | -1.048013163 | 0.426498808 | -2.457247576 |
| Pcdha12 | 117.7268123 | -0.690700883 | 0.137668392 | -5.017134806 |

| | | | | |
|---|---|---|---|---|
| Gm19035 | 45.21062936 | -0.791988554 | 0.281566949 | -2.812789483 |
| Pcdhac1 | 92.62921019 | -0.421307303 | 0.123840762 | -3.402008322 |
| Pcdhac2 | 545.5711865 | -0.060346113 | 0.096662665 | -0.624295978 |
| Gm10544 | 2.182918281 | -0.91369495 | 0.778087175 | -1.174283525 |
| Gm38097 | 0.07796279 | 0.274392189 | 3.724945049 | 0.073663419 |
| Pcdhb1 | 5.985971091 | -0.255427513 | 0.400542757 | -0.637703486 |
| Pcdhb2 | 60.01309998 | -0.512532344 | 0.174105467 | -2.94380385 |
| Pcdhb3 | 83.43922558 | -0.361863887 | 0.126695243 | -2.856175809 |
| Gm18529 | 8.138222198 | -1.051441222 | 0.360509682 | -2.916540872 |
| Pcdhb4 | 60.73868439 | -0.685087908 | 0.166954542 | -4.103439773 |
| Pcdhb5 | 51.69657475 | -0.315238328 | 0.169917354 | -1.855245037 |
| Pcdhb6 | 68.6643995 | -0.247318191 | 0.15513485 | -1.594214263 |
| Pcdhb7 | 114.8261978 | -0.279432916 | 0.144428477 | -1.934749445 |
| Pcdhb8 | 89.35877465 | -0.145472163 | 0.158471911 | -0.917968123 |
| Pcdhb9 | 170.4389826 | -0.043930486 | 0.103910895 | -0.422770735 |
| Pcdhb10 | 105.3601154 | -0.126372338 | 0.126073542 | -1.002370013 |
| Pcdhb11 | 148.8037279 | -0.214219716 | 0.118543922 | -1.807091521 |
| Gm37446 | 0.821410149 | 0.325000206 | 1.448322088 | 0.224397742 |
| Gm20162 | 21.72530242 | -0.128565226 | 0.307185586 | -0.418526233 |
| Pcdhb12 | 117.9366013 | 0.015841193 | 0.154976513 | 0.102216737 |
| Pcdhb13 | 90.40139946 | -0.110984602 | 0.166474398 | -0.666676698 |
| Pcdhb14 | 293.0642983 | -0.176059795 | 0.116813007 | -1.507193414 |
| Pcdhb15 | 108.4402084 | -0.355704191 | 0.153571446 | -2.316213079 |
| Pcdhb16 | 347.3551605 | -0.252053308 | 0.07916996 | -3.183698815 |
| Pcdhb17 | 319.6024478 | -0.2212283 | 0.10926237 | -2.024743744 |
| Pcdhb18 | 229.3400325 | -0.172123301 | 0.089413249 | -1.925031287 |
| Pcdhb19 | 338.4815422 | -0.219433632 | 0.138810744 | -1.580811586 |
| Pcdhb20 | 206.2975924 | -0.313870688 | 0.1238812 | -2.533642622 |
| Pcdhb21 | 135.2304553 | -0.145881924 | 0.128235917 | -1.137605813 |
| Pcdhb22 | 238.2686593 | -0.169738908 | 0.07940486 | -2.137638786 |
| 4930517L18Rik | 0.044544648 | 0.755286614 | 3.724945049 | 0.202764498 |
| Slc25a2 | 10.03606488 | -0.675671902 | 0.382267834 | -1.767535331 |
| Taf7 | 183.7126313 | 0.283629919 | 0.118466201 | 2.394184316 |
| Gm8242 | 6.947208237 | 0.567006823 | 0.435635613 | 1.301562145 |
| 3222401L13Rik | 32.8963234 | -0.044886219 | 0.202400201 | -0.221769638 |
| Pcdhga1 | 89.5175488 | 0.009759732 | 0.12828534 | 0.076078309 |
| Gm37118 | 0.066050875 | 0.274391604 | 3.724945049 | 0.073663262 |
| Pcdhga2 | 78.94781358 | -0.138858637 | 0.141660119 | -0.980223917 |
| Pcdhga3 | 73.82529401 | -0.176819418 | 0.148957262 | -1.187047988 |
| Pcdhgb1 | 82.71764577 | -0.274204017 | 0.152772197 | -1.79485549 |
| Pcdhga4 | 165.6903328 | -0.145171702 | 0.094906873 | -1.529622641 |
| Pcdhgb2 | 147.8807373 | 0.20808462 | 0.136127581 | 1.528599998 |
| Pcdhga5 | 172.6956286 | -0.211196488 | 0.113411188 | -1.862219172 |
| Gm24401 | 7.516116141 | -0.162993514 | 0.520778304 | -0.312980615 |
| Pcdhga6 | 188.1446477 | -0.218134523 | 0.111434025 | -1.957521715 |
| Gm37165 | 0.949659951 | 0.561290323 | 1.123991142 | 0.49937255 |
| Pcdhga7 | 242.4294219 | -0.308511391 | 0.103649954 | -2.976473962 |

| | | | | |
|---|---|---|---|---|
| Pcdhgb4 | 169.0788982 | -0.586399913 | 0.156712424 | -3.74188528 |
| Pcdhga8 | 424.9442205 | -0.218936367 | 0.092725661 | -2.361119502 |
| Pcdhgb5 | 111.8894427 | -0.32062292 | 0.133791892 | -2.396430128 |
| Pcdhga9 | 152.843188 | -0.342451483 | 0.102507488 | -3.340746013 |
| Pcdhgb6 | 97.63560197 | -0.506170946 | 0.192248844 | -2.632894618 |
| Gm26672 | 13.40694889 | -0.163115268 | 0.295706446 | -0.551612148 |
| Pcdhga10 | 152.3565402 | -0.064423804 | 0.13858213 | -0.464878149 |
| Pcdhgb7 | 116.7201272 | -0.311701726 | 0.13900719 | -2.242342465 |
| Pcdhga11 | 97.79522649 | -0.541535176 | 0.18403336 | -2.942592447 |
| Pcdhgb8 | 105.1432484 | -0.300665144 | 0.166933181 | -1.80111073 |
| Pcdhga12 | 164.3031905 | -0.506981382 | 0.162637844 | -3.117241169 |
| BC037039 | 16.91683977 | 0.315843335 | 0.282251499 | 1.119013846 |
| Pcdhgc3 | 1154.35759 | -0.100774562 | 0.10720054 | -0.940056473 |
| Pcdhgc5 | 972.9292026 | -0.369079362 | 0.104068511 | -3.546503715 |
| Gm38182 | 0 | NA | NA | NA |
| Gm29994 | 1.935845755 | 1.30941738 | 0.929623041 | 1.408546608 |
| Diap1 | 993.6614968 | 0.2927718 | 0.095573585 | 3.063312938 |
| Mir6979 | 0.451714327 | 0.755310918 | 1.55190433 | 0.486699407 |
| Hdac3 | 567.0461211 | -0.150644607 | 0.070563659 | -2.134875215 |
| Rell2 | 1076.046952 | -0.01182628 | 0.086351991 | -0.136954344 |
| Fchsd1 | 97.53749871 | 0.179354708 | 0.125138043 | 1.433254855 |
| Arap3 | 55.62706146 | 0.628828381 | 0.308666343 | 2.037243108 |
| Mir6981 | 0 | NA | NA | NA |
| Mir6980 | 0 | NA | NA | NA |
| Pcdh1 | 2873.63663 | -0.32140719 | 0.110866919 | -2.899036017 |
| 2010320O07Rik | 1.189163173 | 0.222906876 | 0.944110635 | 0.236102494 |
| 1700086O06Rik | 44.53713014 | 0.035191148 | 0.181816528 | 0.193553074 |
| Gm10268 | 1.309431784 | 0.601954252 | 0.861130424 | 0.69902797 |
| 0610009O20Rik | 346.6488783 | 0.057161177 | 0.077384162 | 0.738667651 |
| Pcdh12 | 4.608338074 | 0.735622641 | 0.545914974 | 1.347504055 |
| Rnf14 | 3142.218856 | 0.329052704 | 0.0718237 | 4.581394472 |
| Gm23205 | 10.03026786 | -0.513360782 | 0.4787381 | -1.072320717 |
| Gnpda1 | 269.096148 | -0.095230381 | 0.114424981 | -0.832251669 |
| Gm24690 | 0 | NA | NA | NA |
| Ndfip1 | 2572.872965 | 0.055883481 | 0.070681068 | 0.79064285 |
| Spry4 | 260.3722342 | -0.42485926 | 0.153964569 | -2.759461231 |
| Gm27369 | 0 | NA | NA | NA |
| Gm27804 | 0 | NA | NA | NA |
| Arhgap26 | 2536.520982 | 0.257002804 | 0.09571894 | 2.684973356 |
| Gm19774 | 0.900419082 | -1.065644597 | 1.0135989 | -1.051347429 |
| Gm15335 | 0 | NA | NA | NA |
| Gm8302 | 0.044544648 | 0.755286614 | 3.724945049 | 0.202764498 |
| Fgf1 | 1013.624655 | -0.271286519 | 0.122718189 | -2.210646361 |
| Gm15334 | 1.714908477 | 2.266880442 | 0.80572259 | 2.813475096 |
| Gm15336 | 0.09349418 | 0.274402725 | 3.724945049 | 0.073666248 |
| Gm15333 | 0 | NA | NA | NA |
| Gm15337 | 14.68375219 | 1.142403909 | 0.325937329 | 3.504980275 |

| | | | | |
|---|---|---|---|---|
| Nr3c1 | 1778.771107 | -0.094389438 | 0.053176632 | -1.775017232 |
| Pabpc2 | 0.066050875 | 0.274391604 | 3.724945049 | 0.073663262 |
| Yipf5 | 514.5136962 | 0.177308971 | 0.085616308 | 2.070971938 |
| 2900055J20Rik | 110.5064999 | -0.632862708 | 0.196221406 | -3.225248053 |
| Kctd16 | 294.0495108 | -0.569315034 | 0.189679224 | -3.001462272 |
| Rps19-ps13 | 0.270136144 | 0.317971004 | 2.09378328 | 0.151864334 |
| Gm22515 | 0 | NA | NA | NA |
| Gm22483 | 0.046345709 | 0.274391604 | 3.724945049 | 0.073663262 |
| AC149064.1 | 0 | NA | NA | NA |
| Prelid2 | 7.690273632 | 0.770487545 | 0.436423222 | 1.765459551 |
| Grxcr2 | 0.311588305 | -0.990050133 | 1.986639063 | -0.498354307 |
| Sh3rf2 | 92.9302299 | -0.53770916 | 0.27805314 | -1.933835958 |
| Gm5689 | 0.669351884 | 1.145907518 | 1.365394912 | 0.839249881 |
| Plac8l1 | 0.74423012 | 0.196470106 | 1.595939048 | 0.123106272 |
| Lars | 747.0783727 | 0.162316549 | 0.114168153 | 1.421732288 |
| Rps19-ps4 | 0.931580594 | 0.330061844 | 1.223462169 | 0.269776911 |
| Rbm27 | 1071.37103 | 0.059941099 | 0.063145113 | 0.949259512 |
| Gm22697 | 0 | NA | NA | NA |
| Pou4f3 | 1.201894167 | -0.234141761 | 1.184755703 | -0.197628726 |
| Gm16415 | 0.834345451 | 0.984103649 | 1.314716666 | 0.748529074 |
| Tcerg1 | 1111.337889 | 0.182370019 | 0.087057263 | 2.094828311 |
| Gm27457 | 2.628954575 | -0.084222524 | 0.596875481 | -0.141105686 |
| Gpr151 | 33.37780463 | 0.866080685 | 0.309910134 | 2.794618794 |
| Ppp2r2b | 978.0231163 | -0.370760679 | 0.066628716 | -5.564577857 |
| 4933407I08Rik | 0.075043205 | 0.755286614 | 3.724945049 | 0.202764498 |
| Gm22264 | 0 | NA | NA | NA |
| Gm8181 | 9.04075607 | -0.089704445 | 0.466632342 | -0.19223795 |
| Stk32a | 162.2987704 | 0.08262472 | 0.142854434 | 0.578384006 |
| Dpysl3 | 999.9963044 | -0.104722566 | 0.068935328 | -1.519142209 |
| Gm3650 | 17.99992091 | 0.165376321 | 0.358669327 | 0.461082978 |
| Eif3j2 | 165.1918275 | 0.147744317 | 0.121271941 | 1.218289368 |
| Gm27325 | 0 | NA | NA | NA |
| Jakmip2 | 846.4346633 | 0.053842745 | 0.077044385 | 0.698853581 |
| Spink1 | 0 | NA | NA | NA |
| Gm23894 | 0 | NA | NA | NA |
| Scgb3a2 | 0.047948698 | -0.206487837 | 3.724945049 | -0.055433794 |
| Gm94 | 0.18352982 | 1.625373558 | 3.713044335 | 0.437746876 |
| Spink5 | 0.975055063 | 2.370123977 | 3.519700185 | 0.673388031 |
| Spink14 | 0 | NA | NA | NA |
| Spink6 | 0 | NA | NA | NA |
| Spink12 | 0 | NA | NA | NA |
| Gm37797 | 0 | NA | NA | NA |
| Gm10267 | 0 | NA | NA | NA |
| Spinkl | 0 | NA | NA | NA |
| Spink11 | 0.047948698 | -0.206487837 | 3.724945049 | -0.055433794 |
| Gm10542 | 0 | NA | NA | NA |
| Npy6r | 0.261451683 | -1.095683512 | 2.093011973 | -0.523496055 |

| | | | | |
|---|---|---|---|---|
| Myot | 0.836501452 | -0.545701943 | 1.200299818 | -0.454638029 |
| Dcp2 | 1781.048542 | 0.740896909 | 0.094911138 | 7.806216641 |
| Mcc | 632.7057759 | -0.11149851 | 0.092443422 | -1.206127031 |
| A930012L18Rik | 13.61962428 | -0.474778472 | 0.272268325 | -1.743788858 |
| RP23-342A15.2 | 0 | NA | NA | NA |
| Ythdc2 | 676.2351253 | -0.141186956 | 0.083660535 | -1.687617183 |
| Gm26928 | 0 | NA | NA | NA |
| Kcnn2 | 557.7293241 | -0.661174132 | 0.108870211 | -6.073049032 |
| Gm10540 | 0.30732292 | 1.573930812 | 1.558251173 | 1.010062331 |
| A330093E20Rik | 2.019082719 | -0.797968299 | 0.905193271 | -0.881544666 |
| Gm26262 | 0 | NA | NA | NA |
| Gm38337 | 0.335887269 | 1.529433438 | 1.522245107 | 1.00472219 |
| Trim36 | 95.57263784 | -0.669253538 | 0.173110855 | -3.86604028 |
| 1700018A14Rik | 0.75323169 | -0.206483523 | 1.508227525 | -0.136904757 |
| Pggt1b | 266.2078042 | 0.162755016 | 0.090484787 | 1.798700333 |
| Ccdc112 | 108.5123134 | 0.147775412 | 0.153536594 | 0.962476816 |
| Gm24617 | 0.282138413 | 1.256526863 | 3.723547076 | 0.337454271 |
| Mospd4 | 3.756273683 | 0.145462445 | 0.520489541 | 0.27947237 |
| Gm24076 | 0.776630353 | 2.770339045 | 1.218179211 | 2.274163785 |
| Fem1c | 864.4434414 | 0.149922048 | 0.084055869 | 1.783599987 |
| Ticam2 | 12.34676471 | 1.1529728 | 0.414513215 | 2.781510358 |
| Tmed7 | 1507.489644 | 0.243667242 | 0.107475202 | 2.267195011 |
| Eif1a | 755.0916563 | 0.995394586 | 0.149636073 | 6.652103099 |
| Cdo1 | 339.5541898 | 0.281289548 | 0.149342866 | 1.88351513 |
| Atg12 | 650.7230075 | 0.050437467 | 0.08736735 | 0.577303389 |
| Ap3s1 | 461.3279815 | -0.092718135 | 0.087705207 | -1.057156558 |
| Lvrn | 10.74167226 | -0.952521761 | 0.385519162 | -2.470750757 |
| Gm23610 | 0 | NA | NA | NA |
| Arl14epl | 1.575196263 | 1.242960371 | 0.914486508 | 1.359189402 |
| Commd10 | 179.0565603 | 0.0247503 | 0.089561003 | 0.276351307 |
| Gm22791 | 0 | NA | NA | NA |
| Gm25036 | 0 | NA | NA | NA |
| Sema6a | 831.0817601 | -0.182516336 | 0.059288341 | -3.078452404 |
| Eno1b | 521.4844324 | -0.074070155 | 0.123567049 | -0.599432909 |
| Gm5839 | 0 | NA | NA | NA |
| 1700044K03Rik | 0 | NA | NA | NA |
| Gm25440 | 0 | NA | NA | NA |
| Dtwd2 | 52.5847649 | 0.055419062 | 0.148709157 | 0.37266745 |
| Gm16283 | 2.090780166 | -0.422045584 | 0.698147355 | -0.604522213 |
| 1700065O20Rik | 0 | NA | NA | NA |
| Dmxl1 | 3104.946172 | -0.075035853 | 0.049820512 | -1.506123688 |
| Tnfaip8 | 137.5382232 | 0.69868413 | 0.149965469 | 4.658966716 |
| C030005K06Rik | 2.787674368 | -3.022412484 | 0.778311415 | -3.88329456 |
| Hsd17b4 | 542.6081984 | -0.372717864 | 0.086213569 | -4.323192628 |
| Cd63-ps | 108.0511914 | 1.052929636 | 0.215354339 | 4.889289176 |
| Gm8529 | 0 | NA | NA | NA |
| Fam170a | 0 | NA | NA | NA |

| | | | | |
|---|---|---|---|---|
| Hdhd1a | 0.113999573 | -0.206492286 | 3.724945049 | -0.055434988 |
| Prr16 | 138.718701 | -0.393560826 | 0.135475638 | -2.905030238 |
| Gm25739 | 0 | NA | NA | NA |
| Gm25785 | 0 | NA | NA | NA |
| Gm4950 | 20.25727402 | 0.406032005 | 0.255858915 | 1.586937101 |
| Gm22259 | 0 | NA | NA | NA |
| Ftmt | 0 | NA | NA | NA |
| Srfbp1 | 140.6912451 | 0.15837724 | 0.11450176 | 1.383186073 |
| Gm27026 | 0 | NA | NA | NA |
| Lox | 47.85031176 | 2.051634429 | 0.34559095 | 5.936597672 |
| Zfp474 | 3.585501729 | -0.654642138 | 0.598098234 | -1.094539494 |
| 1700034E13Rik | 0.108266322 | 0.274389235 | 3.724945049 | 0.073662626 |
| Gykl1 | 0 | NA | NA | NA |
| Sncaip | 113.2002606 | 0.138165035 | 0.108332813 | 1.27537568 |
| Snx2 | 781.8559442 | 0.561560092 | 0.08526235 | 6.586261017 |
| Snx24 | 137.1094418 | 0.162383716 | 0.100194174 | 1.620690203 |
| Ppic | 20.57672658 | 0.115256448 | 0.250626122 | 0.459874044 |
| Prdm6 | 12.21164467 | -0.473716578 | 0.339601327 | -1.394919691 |
| Cep120 | 755.1587019 | -0.318424535 | 0.060622696 | -5.252563111 |
| Csnk1g3 | 932.5335226 | -0.09662908 | 0.060405821 | -1.599665047 |
| Gm5507 | 5.787175505 | -0.23195921 | 0.394084776 | -0.58860231 |
| Redrum | 0.747472414 | 1.9096399 | 1.217502937 | 1.568488948 |
| Zfp608 | 685.0116038 | -0.062514482 | 0.072501083 | -0.862255829 |
| Gm26742 | 0.583001974 | -1.410510138 | 1.266536387 | -1.113675179 |
| Gm4221 | 12.78354649 | -0.74342741 | 0.34660181 | -2.144903426 |
| Gm24183 | 0 | NA | NA | NA |
| Gm34073 | 0.428599116 | 0.47376613 | 3.302571138 | 0.143453724 |
| Gm22597 | 0 | NA | NA | NA |
| Gm37828 | 0 | NA | NA | NA |
| Gm37337 | 0.587544172 | 0.321144926 | 1.302595268 | 0.246542371 |
| Gm24514 | 0.493679557 | -1.938694779 | 1.435516245 | -1.350520961 |
| Gm26959 | 0 | NA | NA | NA |
| Gm8614 | 0.29391207 | -1.655086777 | 2.017889165 | -0.820206979 |
| Gm23335 | 0 | NA | NA | NA |
| Gm6944 | 0 | NA | NA | NA |
| Gm22484 | 0.068097359 | -0.206487837 | 3.724945049 | -0.055433794 |
| Gm23555 | 0 | NA | NA | NA |
| Gm25476 | 0 | NA | NA | NA |
| Gramd3 | 170.2177708 | 0.254801148 | 0.112485861 | 2.265183784 |
| Aldh7a1 | 399.3017201 | -0.284637372 | 0.077231943 | -3.68548766 |
| Mir1258 | 0 | NA | NA | NA |
| Phax | 381.2748711 | 0.317849126 | 0.102019984 | 3.115557481 |
| Tex43 | 2.674966445 | -1.126464076 | 0.638778644 | -1.763465462 |
| Lmnb1 | 214.1019998 | 0.453434868 | 0.14745818 | 3.075006534 |
| 3-Mar | 67.75458388 | 0.424324994 | 0.179407154 | 2.365150907 |
| Gm15345 | 0.999978248 | 0.677018775 | 1.237550583 | 0.547063517 |
| C330018D20Rik | 87.14994236 | -0.564219108 | 0.142043268 | -3.972163666 |

| | | | | |
|---|---|---|---|---|
| Megf10 | 704.5314188 | -0.134376654 | 0.126511159 | -1.062172334 |
| Prrc1 | 265.116747 | 0.073669711 | 0.08317771 | 0.885690536 |
| 4930511M06Rik | 2.334218344 | 1.135465804 | 0.71844081 | 1.580458388 |
| Ctxn3 | 52.35061762 | 0.230791982 | 0.191240899 | 1.206812891 |
| 1700066O22Rik | 0.436302845 | 1.223415768 | 1.44146001 | 0.848733755 |
| 1700011I03Rik | 2.706585767 | 0.150833926 | 0.698889391 | 0.215819453 |
| Gm26038 | 0 | NA | NA | NA |
| AC102232.1 | 0.04393175 | -0.206490061 | 3.724945049 | -0.055434391 |
| Slc12a2 | 1824.003985 | -0.510744913 | 0.088398893 | -5.777729757 |
| Fbn2 | 140.4520598 | -0.625339245 | 0.168586198 | -3.709314605 |
| Gm26507 | 13.7398474 | -0.715793595 | 0.290615698 | -2.463024537 |
| Gm25660 | 0 | NA | NA | NA |
| Slc27a6 | 3.148605881 | 0.159039794 | 0.662593989 | 0.240026014 |
| Gm23270 | 0 | NA | NA | NA |
| Isoc1 | 459.7670982 | 0.712267957 | 0.124942604 | 5.700761262 |
| Adamts19 | 19.03947336 | -0.377736096 | 0.336093815 | -1.123900763 |
| A730017C20Rik | 501.0808718 | 0.738846413 | 0.108986544 | 6.779244365 |
| Chsy3 | 230.6243843 | -0.475366164 | 0.155674357 | -3.053593231 |
| Mir6355 | 0 | NA | NA | NA |
| Gm26028 | 0 | NA | NA | NA |
| Gm23576 | 0 | NA | NA | NA |
| Gm4951 | 925.2780923 | 8.862331992 | 0.448488933 | 19.76042514 |
| Gm5970 | 122.1217523 | 8.490412755 | 0.812047502 | 10.45556169 |
| Gm4841 | 260.7014249 | 10.54665629 | 0.920981953 | 11.45153415 |
| F830016B08Rik | 944.1534938 | 9.555105529 | 0.530476707 | 18.01229987 |
| Iigp1 | 5189.009943 | 9.461832339 | 0.274885794 | 34.42095788 |
| Gm6960 | 0.926812825 | 3.288254982 | 1.147364177 | 2.865920906 |
| BC023105 | 164.7922582 | 9.082842421 | 0.902040521 | 10.06921774 |
| Smim3 | 173.6181185 | 2.029738852 | 0.231871808 | 8.753711246 |
| Dctn4 | 1434.033031 | 0.32242361 | 0.059396111 | 5.428362333 |
| Rbm22 | 390.7352864 | 0.471067496 | 0.142711226 | 3.300844004 |
| Myoz3 | 0.060174537 | 0.755286614 | 3.724945049 | 0.202764498 |
| Synpo | 1700.826908 | -0.221435248 | 0.129266318 | -1.713015819 |
| Ndst1 | 2081.012663 | 0.513355246 | 0.091005944 | 5.640897958 |
| Rps14 | 1385.960924 | 0.60818621 | 0.137554226 | 4.421428744 |
| Gm8731 | 0.109869311 | -0.206490066 | 3.724945049 | -0.055434393 |
| Cd74 | 447.6522266 | 3.899057383 | 0.329717456 | 11.82545027 |
| Mir5107 | 0.060174537 | 0.755286614 | 3.724945049 | 0.202764498 |
| Tcof1 | 457.500707 | 0.238650775 | 0.095736264 | 2.492793913 |
| Arsi | 22.1297461 | 1.141138486 | 0.299349752 | 3.812057569 |
| Camk2a | 16056.95831 | 0.0384056 | 0.175240399 | 0.219159512 |
| Mir7220 | 4.383661021 | -1.369277182 | 0.541163828 | -2.530245207 |
| Mir6982 | 3.753094389 | -0.944684752 | 0.586431844 | -1.61090289 |
| Slc6a7 | 440.8262577 | -0.436314588 | 0.142209125 | -3.068119483 |
| Cdx1 | 0.329049156 | -1.352102056 | 1.701879364 | -0.794475851 |
| Pdgfrb | 307.8251967 | 0.029530737 | 0.092431671 | 0.319487216 |
| Csf1r | 863.5867635 | 0.139540529 | 0.300828774 | 0.463853662 |

| | | | | |
|---|---|---|---|---|
| Mir6983 | 0 | NA | NA | NA |
| Hmgxb3 | 688.8088613 | -0.120050252 | 0.059207357 | -2.027623877 |
| Gm26403 | 2.513420025 | 0.022865407 | 0.675850913 | 0.033832029 |
| Slc26a2 | 293.7967227 | 0.255073873 | 0.102792525 | 2.48144378 |
| Pde6a | 3.821803709 | -0.453202091 | 0.520363658 | -0.870933402 |
| Rps2-ps10 | 1236.213371 | 0.346802554 | 0.129671686 | 2.674466301 |
| Ppargc1b | 407.0618076 | -0.54649904 | 0.126860145 | -4.307885963 |
| Gm25301 | 0 | NA | NA | NA |
| mmu-mir-378a | 3.818950962 | -1.32052512 | 0.516258161 | -2.557877474 |
| Gm25834 | 0.038981395 | 0.274393828 | 3.724945049 | 0.073663859 |
| Arhgef37 | 140.4268379 | 0.262704742 | 0.142199881 | 1.847432925 |
| Csnk1a1 | 4557.884024 | 0.114701562 | 0.066961596 | 1.712945462 |
| Bvht | 5.444855971 | 1.226056426 | 0.609594976 | 2.011263994 |
| Mir145a | 0.194631057 | 1.693379269 | 3.232136674 | 0.52391945 |
| Mir143 | 0.147068667 | 1.236183405 | 3.514416608 | 0.351746404 |
| Mir143hg | 8.471314973 | -0.163413577 | 0.437642696 | -0.373394959 |
| Il17b | 2.212656516 | 0.852751319 | 0.872591753 | 0.977262639 |
| Gm38165 | 0.662591016 | 0.953019154 | 1.302352312 | 0.731767545 |
| Pcyox1l | 207.9956365 | -0.45915414 | 0.136924021 | -3.3533498 |
| Grpel2 | 247.9051696 | -0.104420127 | 0.096615538 | -1.080779852 |
| 1500015A07Rik | 35.15775905 | -0.208924841 | 0.185967061 | -1.123450787 |
| Afap1l1 | 138.1820362 | 0.047440309 | 0.093484353 | 0.507467908 |
| Ablim3 | 445.2700515 | -0.82471833 | 0.115748983 | -7.125058955 |
| Gm37774 | 0 | NA | NA | NA |
| Sh3tc2 | 44.0296305 | -0.309413562 | 0.22485876 | -1.376035171 |
| Adrb2 | 16.87312987 | -0.240213204 | 0.253053178 | -0.949259779 |
| Gm9949 | 1.519386227 | 1.677746754 | 0.878629035 | 1.909505249 |
| Htr4 | 92.71409538 | -0.013796607 | 0.141219332 | -0.097696303 |
| Fbxo38 | 754.8506995 | 0.124207513 | 0.05745026 | 2.16200089 |
| Spink10 | 11.52016113 | -0.06104506 | 0.326825063 | -0.186782065 |
| Spink7 | 0.099106094 | 0.274387056 | 3.724945049 | 0.073662041 |
| Spink13 | 0.546984718 | -1.286696425 | 1.301381396 | -0.988715859 |
| Gm17732 | 5.889985807 | -0.063957041 | 0.419812075 | -0.152346836 |
| 2700046A07Rik | 9.548929167 | 0.105003656 | 0.449246128 | 0.233733024 |
| Apcdd1 | 206.8351242 | -0.08208339 | 0.163677247 | -0.501495424 |
| Napg | 1597.555114 | -0.07896978 | 0.055898547 | -1.412734055 |
| Piezo2 | 47.03296206 | 0.113760961 | 0.204059716 | 0.557488575 |
| Gm26972 | 0 | NA | NA | NA |
| Gm23581 | 0 | NA | NA | NA |
| Gm22560 | 0 | NA | NA | NA |
| Gm22210 | 0 | NA | NA | NA |
| Gm24848 | 0 | NA | NA | NA |
| Txnl1 | 825.0032602 | 0.177166502 | 0.096637508 | 1.833309922 |
| Wdr7 | 3498.348225 | 0.066333335 | 0.044875532 | 1.478162644 |
| Gm24384 | 0 | NA | NA | NA |
| St8sia3os | 1.428069919 | 0.910928186 | 1.017552773 | 0.895214686 |
| St8sia3 | 1740.780446 | -0.001735328 | 0.080871413 | -0.021457865 |

| | | | | |
|---|---|---|---|---|
| Onecut2 | 194.3572374 | -0.046642903 | 0.154787818 | -0.301334459 |
| Amd-ps3 | 30.2089109 | -0.04833069 | 0.193434632 | -0.249855413 |
| Fech | 621.1316025 | 0.061686163 | 0.067245335 | 0.917329992 |
| Nars | 1964.183135 | 0.226196319 | 0.088436751 | 2.557718558 |
| Atp8b1 | 51.70323717 | 0.196689852 | 0.212976273 | 0.923529409 |
| Gm23016 | 0 | NA | NA | NA |
| Gm24504 | 0 | NA | NA | NA |
| Nedd4l | 2403.215903 | 0.025756868 | 0.10503666 | 0.245217892 |
| Mir122 | 0.049553047 | 0.274391604 | 3.724945049 | 0.073663262 |
| Alpk2 | 11.61777028 | -0.204936792 | 0.351283888 | -0.583393657 |
| Gm22567 | 0.046345709 | 0.274391604 | 3.724945049 | 0.073663262 |
| Malt1 | 178.3264262 | 0.864979567 | 0.180692469 | 4.787026095 |
| Zfp532 | 1029.175328 | -0.06661619 | 0.060499565 | -1.101101969 |
| Gm37643 | 15.88324691 | -1.111843211 | 0.360056569 | -3.087968134 |
| Oacyl | 19.6943626 | 0.821273843 | 0.358521366 | 2.290724961 |
| Sec11c | 480.2888366 | 0.031879984 | 0.087025196 | 0.366330509 |
| Grp | 19.80146899 | -0.593280401 | 0.24897322 | -2.382908495 |
| Gm25150 | 0 | NA | NA | NA |
| Rax | 0.423125362 | -1.098735172 | 1.534721057 | -0.715918484 |
| Raxos1 | 0 | NA | NA | NA |
| Cplx4 | 0.598788582 | 0.512032733 | 1.33937673 | 0.382291794 |
| Lman1 | 670.4253144 | 0.166987401 | 0.064301609 | 2.596939708 |
| Ccbe1 | 53.63741432 | 0.130082535 | 0.205218496 | 0.633873344 |
| Mir694 | 0.217748738 | -1.493307638 | 2.460727054 | -0.606856269 |
| Gm15958 | 0.904555577 | -0.342607703 | 0.933526504 | -0.367003724 |
| Pmaip1 | 46.65352456 | 1.864648392 | 0.216756049 | 8.602520691 |
| Gm24679 | 0 | NA | NA | NA |
| Gm22219 | 0 | NA | NA | NA |
| Mc4r | 36.60476633 | 0.542138021 | 0.221643955 | 2.445986045 |
| Gnal | 2244.602178 | -0.328174659 | 0.095062618 | -3.452194639 |
| Gm23512 | 0.291502873 | -1.078693865 | 2.009766309 | -0.536726017 |
| Mppe1 | 130.6858995 | 0.33934319 | 0.109847832 | 3.089211529 |
| Impa2 | 34.6996934 | -0.208186919 | 0.219418266 | -0.948813072 |
| B430212C06Rik | 2.198202818 | 0.90264134 | 0.832637104 | 1.084075326 |
| Cidea | 13.25459769 | 0.436999025 | 0.288264008 | 1.515968044 |
| Tubb6 | 218.1172078 | 2.946667138 | 0.334491894 | 8.809382814 |
| Afg3l2 | 820.858718 | -0.00215794 | 0.077595755 | -0.027810029 |
| Gm24900 | 0 | NA | NA | NA |
| Slmo1 | 256.90097 | 0.221410069 | 0.098627776 | 2.244905816 |
| Spire1 | 2405.877136 | 0.095183541 | 0.071705777 | 1.327418022 |
| Gm17669 | 0.12092566 | 1.236179401 | 3.724945049 | 0.331865138 |
| Cep76 | 301.4880058 | 0.134037785 | 0.160647043 | 0.834361981 |
| Psmg2 | 96.05648597 | -0.152449555 | 0.160262369 | -0.951249851 |
| Ptpn2 | 330.6143394 | 0.204459029 | 0.096349951 | 2.122046009 |
| Gm26910 | 20.45356711 | -0.14591584 | 0.30316833 | -0.481303042 |
| Seh1l | 391.9333236 | 0.420754255 | 0.06546272 | 6.427387298 |
| Cep192 | 279.5064016 | -0.087255321 | 0.12864663 | -0.67825578 |

| | | | | |
|---|---|---|---|---|
| Ldlrad4 | 528.0018019 | -0.138232672 | 0.097785543 | -1.413630975 |
| Fam210a | 679.3179832 | -0.467216401 | 0.07079446 | -6.599618134 |
| Mir7219 | 0 | NA | NA | NA |
| Rnmt | 878.6655392 | 0.031400939 | 0.050162937 | 0.625978878 |
| Mc5r | 18.63850153 | 0.855899052 | 0.32028821 | 2.672277731 |
| Mc2r | 1.528578462 | 2.667004529 | 1.107928219 | 2.407199746 |
| Mir6356 | 0 | NA | NA | NA |
| 4930546C10Rik | 0.160559904 | 0.755297704 | 2.794491024 | 0.270280955 |
| Mir145b | 0 | NA | NA | NA |
| Gm23536 | 0.335787014 | 0.318036574 | 1.722461501 | 0.184640744 |
| Gm4842 | 0 | NA | NA | NA |
| Tcf4 | 6112.811251 | -0.052387908 | 0.087952861 | -0.595636206 |
| Gm24845 | 5.630675305 | -0.266523452 | 0.43294348 | -0.615607958 |
| Gm22508 | 0 | NA | NA | NA |
| Ccdc68 | 4.327574142 | -0.257753172 | 0.531950615 | -0.484543424 |
| 1700061H18Rik | 0.072025462 | 0.755288839 | 3.724945049 | 0.202765096 |
| 4930448D08Rik | 0.605479419 | 0.306065062 | 1.208617367 | 0.253235698 |
| Rab27b | 340.9708684 | -0.144818193 | 0.101047905 | -1.433163736 |
| Dynap | 0.072025462 | 0.755288839 | 3.724945049 | 0.202765096 |
| 4930503L19Rik | 92.03394658 | 0.05019871 | 0.135291691 | 0.371040599 |
| Stard6 | 8.232924272 | -0.198580759 | 0.348903648 | -0.569156443 |
| Poli | 92.01233665 | 0.14705815 | 0.12431265 | 1.182970119 |
| Mir6357 | 0.407891471 | 0.334920279 | 1.413953075 | 0.236868029 |
| Mbd2 | 568.7990903 | 0.709242685 | 0.125994455 | 5.629157925 |
| Dcc | 395.337002 | -0.364236888 | 0.121431561 | -2.999524048 |
| Gm25509 | 0 | NA | NA | NA |
| n-R5-8s1 | 0 | NA | NA | NA |
| Mex3c | 685.3858209 | -0.051307881 | 0.095468579 | -0.537432115 |
| Gm26438 | 0 | NA | NA | NA |
| Gm22065 | 0.042716945 | -0.206490061 | 3.724945049 | -0.055434391 |
| Smad4 | 842.5159027 | 0.062403842 | 0.063421505 | 0.983953966 |
| Elac1 | 466.4413671 | 0.072333816 | 0.084775221 | 0.853242431 |
| Me2 | 344.6597954 | -0.057239752 | 0.072511754 | -0.789385834 |
| Mro | 101.9368696 | -0.480782899 | 0.119108371 | -4.036516438 |
| Mapk4 | 1333.921859 | -0.082657695 | 0.098043077 | -0.843075287 |
| D730045A05Rik | 2.135082683 | -0.613370861 | 1.184212021 | -0.517956962 |
| Gm9925 | 7.46780898 | -0.410204478 | 0.415151288 | -0.98808432 |
| Ska1 | 2.02584455 | 3.178222877 | 0.980933524 | 3.239998225 |
| Cxxc1 | 352.5600067 | -0.234544796 | 0.090685636 | -2.586350002 |
| Mbd1 | 382.0312851 | 0.378877467 | 0.138580753 | 2.733983318 |
| Cfap53 | 11.98097365 | 1.368010203 | 0.480236521 | 2.848617591 |
| Gm14328 | 4.493268881 | 1.106826574 | 0.506204688 | 2.186519799 |
| Myo5b | 428.355463 | 0.171909107 | 0.10841968 | 1.585589498 |
| 1700120E14Rik | 0 | NA | NA | NA |
| Scarna17 | 10.66151498 | -0.24213583 | 0.387839784 | -0.624319216 |
| Gm23119 | 43.96335493 | -0.043450074 | 0.176293671 | -0.246464176 |
| Acaa2 | 104.6941341 | -0.591281094 | 0.174948905 | -3.379735891 |

| | | | | |
|---|---|---|---|---|
| Gm24559 | 0 | NA | NA | NA |
| Gm25594 | 0 | NA | NA | NA |
| Lipg | 453.85713 | 2.656812698 | 0.429110598 | 6.191440408 |
| Rpl17 | 766.1645459 | 0.390744642 | 0.077235936 | 5.059104107 |
| Snord58b | 1.376506768 | -0.029520073 | 0.913252938 | -0.032324094 |
| Gm23301 | 2.714581473 | 0.364844662 | 0.614917827 | 0.59332263 |
| Gm26202 | 2.364995654 | 0.831335028 | 0.649728874 | 1.279510673 |
| BC031181 | 600.9122168 | -0.038406416 | 0.09442981 | -0.406719192 |
| Dym | 677.9260573 | -0.196678673 | 0.058314795 | -3.372706228 |
| Gm27781 | 0.317560373 | -1.261353796 | 3.707204303 | -0.340243939 |
| Gm8807 | 0 | NA | NA | NA |
| 2010010A06Rik | 0 | NA | NA | NA |
| Gm20544 | 9.279688315 | 0.193461619 | 0.336584385 | 0.57477895 |
| Smad7 | 616.9288546 | -0.12548152 | 0.081705222 | -1.535783352 |
| Ctif | 1554.441727 | 0.068895423 | 0.066182876 | 1.040985635 |
| Gm10532 | 5.803090283 | 0.749242264 | 0.528194315 | 1.418497401 |
| 1700034B16Rik | 0 | NA | NA | NA |
| Zbtb7c | 113.4256275 | 0.588395839 | 0.13142556 | 4.477027453 |
| Mir6358 | 0 | NA | NA | NA |
| Smad2 | 378.1534108 | 0.049390208 | 0.080343042 | 0.614741578 |
| Skor2 | 31.77545133 | -0.437647358 | 0.303165455 | -1.443592434 |
| Ier3ip1 | 232.6003113 | 0.147904663 | 0.093877223 | 1.575511704 |
| Hdhd2 | 753.876256 | -0.168597419 | 0.05978452 | -2.820084843 |
| Katnal2 | 40.6261297 | -0.86306452 | 0.208630447 | -4.136810006 |
| Pias2 | 1238.28436 | -0.055007506 | 0.052901751 | -1.039805003 |
| Gm7276 | 15.00176566 | -0.788142004 | 0.355097772 | -2.219507039 |
| St8sia5 | 623.4113925 | -0.331713469 | 0.107159696 | -3.095505868 |
| Gm15957 | 0 | NA | NA | NA |
| Loxhd1 | 23.89937881 | 1.266979461 | 0.642831655 | 1.970935082 |
| Rnf165 | 750.5636098 | -0.552240021 | 0.160133527 | -3.4486221 |
| 8030462N17Rik | 630.218415 | -0.114324275 | 0.067514341 | -1.693333204 |
| Gm26014 | 0 | NA | NA | NA |
| 4930465K10Rik | 2.048990233 | -0.98118527 | 0.669956165 | -1.464551445 |
| A330094K24Rik | 6.372749315 | 1.201198578 | 0.445219763 | 2.697990248 |
| Haus1 | 42.89733926 | -0.063797529 | 0.163736619 | -0.389635072 |
| Atp5a1 | 6848.041245 | 0.004506204 | 0.069070021 | 0.065241093 |
| F830208F22Rik | 1.770308639 | 0.213575825 | 0.812588438 | 0.262833945 |
| Pstpip2 | 100.7389169 | 0.039652123 | 0.130719562 | 0.303337327 |
| Gm23954 | 0.113999573 | -0.206492286 | 3.724945049 | -0.055434988 |
| Epg5 | 1099.858229 | -0.286473033 | 0.088835892 | -3.224744251 |
| Siglec15 | 0.39439978 | 1.239268355 | 1.594060808 | 0.777428533 |
| Slc14a1 | 350.1029007 | 0.618823059 | 0.124607944 | 4.966160561 |
| Slc14a2 | 4.084512964 | -0.670078282 | 0.575891748 | -1.163549026 |
| Gm6133 | 2.464583914 | -0.293042672 | 0.561307223 | -0.5220718 |
| Gm26430 | 0 | NA | NA | NA |
| Gm23895 | 0 | NA | NA | NA |
| Setbp1 | 1103.372977 | -0.252820401 | 0.087895458 | -2.876376164 |

| | | | | |
|---|---|---|---|---|
| Gm25824 | 0 | NA | NA | NA |
| Gm26977 | 0.099106094 | 0.274387056 | 3.724945049 | 0.073662041 |
| Pard6g | 127.4897943 | 0.107240243 | 0.144508424 | 0.742103749 |
| Gm21886 | 0.269331125 | 0.39059162 | 2.059802211 | 0.189625789 |
| Adnp2 | 487.4784465 | 0.039495309 | 0.097585759 | 0.404724109 |
| Rbfa | 151.797274 | -0.006662449 | 0.090777109 | -0.073393497 |
| Gm16286 | 543.6190761 | 0.456431091 | 0.098251526 | 4.64553693 |
| Txnl4a | 164.8076413 | -0.375769867 | 0.115455518 | -3.254672226 |
| Hsbp1l1 | 8.924943958 | -0.134717672 | 0.334069067 | -0.403262933 |
| Pqlc1 | 221.7266294 | 0.106481041 | 0.102625318 | 1.037570883 |
| Kcng2 | 77.54725198 | 0.711013544 | 0.187540631 | 3.791250685 |
| Gm37015 | 4.004740963 | 0.296765929 | 0.531214717 | 0.558655322 |
| Gm26676 | 5.505563926 | 1.566149509 | 0.522926834 | 2.994968714 |
| D330025C20Rik | 2.748297996 | 1.692047706 | 0.62387998 | 2.712136563 |
| Ctdp1 | 381.7265928 | 0.170294307 | 0.09889268 | 1.722011248 |
| Nfatc1 | 215.3281497 | 1.309729275 | 0.137167049 | 9.548424953 |
| Atp9b | 951.6403139 | 0.161826731 | 0.073353752 | 2.206113886 |
| Gm25071 | 1.130492263 | -0.99509794 | 1.019265741 | -0.976289009 |
| Sall3 | 188.0210551 | -0.708947907 | 0.17720049 | -4.00082363 |
| Gm27238 | 1.264061948 | -3.809807716 | 1.068735272 | -3.56478149 |
| Gm27239 | 2.200041218 | -0.857938242 | 0.759639772 | -1.129401427 |
| Gm25718 | 0 | NA | NA | NA |
| 4930594M17Rik | 0.045882455 | 0.755288839 | 3.724945049 | 0.202765096 |
| Gm26012 | 0 | NA | NA | NA |
| Mir5127 | 0 | NA | NA | NA |
| Gm17383 | 86.5616375 | -0.075696235 | 0.174368959 | -0.434115311 |
| Gm25060 | 0 | NA | NA | NA |
| Galr1 | 70.21229872 | -0.338109502 | 0.262042761 | -1.290283698 |
| Mbp | 30062.83706 | -0.057261078 | 0.176841873 | -0.323798185 |
| Rpl21-ps8 | 1.385867572 | -0.266213423 | 0.985684717 | -0.27007969 |
| Zfp236 | 860.9371065 | -0.651212536 | 0.109633401 | -5.939910003 |
| Gm10524 | 101.1240389 | -0.139298748 | 0.121996974 | -1.14182134 |
| Mir5112 | 0.320492979 | -0.94692372 | 1.556450558 | -0.608386636 |
| 2210420H20Rik | 0.04393175 | -0.206490061 | 3.724945049 | -0.055434391 |
| 4930445N18Rik | 0.552122516 | -0.157524466 | 1.423278397 | -0.110677199 |
| Zfp516 | 555.826877 | 0.279018642 | 0.091989081 | 3.033171318 |
| 1700095A13Rik | 0 | NA | NA | NA |
| Gm24720 | 0 | NA | NA | NA |
| Tshz1 | 687.2218015 | -0.335734216 | 0.140786171 | -2.384710188 |
| Zadh2 | 324.0628898 | -0.397002109 | 0.109663641 | -3.620179897 |
| Zfp407 | 728.9401691 | -0.085086406 | 0.098792345 | -0.861265176 |
| Gm37216 | 25.25091243 | 0.241870716 | 0.291693282 | 0.829195361 |
| Cndp1 | 9.704862909 | 2.245107725 | 0.398265761 | 5.63720998 |
| Gm7289 | 1.584876195 | 0.2304351 | 0.965153112 | 0.238754968 |
| Cndp2 | 787.0114979 | 1.015805737 | 0.162290279 | 6.259190261 |
| D030046N08Rik | 4.544185544 | -0.12464881 | 0.472486634 | -0.263814469 |
| Fam69c | 47.9173264 | -0.229265159 | 0.178893272 | -1.281575076 |

| | | | | |
|---|---|---|---|---|
| Gm22014 | 0 | NA | NA | NA |
| Gm25005 | 0 | NA | NA | NA |
| Gm22819 | 0 | NA | NA | NA |
| Cyb5a | 343.7727805 | -0.059723165 | 0.076809469 | -0.777549511 |
| Gm16146 | 0.241190471 | 1.348328565 | 3.031116258 | 0.44482905 |
| Fbxo15 | 2.684576295 | 0.206868785 | 0.647582977 | 0.319447533 |
| Timm21 | 78.28501918 | -0.375833038 | 0.203357835 | -1.848136505 |
| Gm25453 | 0 | NA | NA | NA |
| Neto1 | 920.988223 | 0.078275955 | 0.112617403 | 0.695060909 |
| Cbln2 | 282.7539475 | -0.502578933 | 0.110236873 | -4.559081908 |
| Gm5096 | 0.150520013 | 0.274408177 | 2.857840489 | 0.096019417 |
| Gm24987 | 0 | NA | NA | NA |
| Socs6 | 448.8133279 | 0.277130737 | 0.09687534 | 2.860694343 |
| Rttn | 192.02834 | 0.015084658 | 0.097661568 | 0.154458484 |
| Mir6359 | 0.075043205 | 0.755286614 | 3.724945049 | 0.202764498 |
| Cd226 | 7.086717468 | 3.862790954 | 0.850133672 | 4.543745393 |
| 9330132A10Rik | 749.6793612 | -0.071464248 | 0.097516004 | -0.732846355 |
| Dok6 | 388.457898 | -0.120357527 | 0.104073452 | -1.15646714 |
| Gm16296 | 5.46580188 | -0.147841017 | 0.429182794 | -0.344470978 |
| Gm16297 | 0 | NA | NA | NA |
| Gm22694 | 0 | NA | NA | NA |
| Gm27037 | 0 | NA | NA | NA |
| Gm23859 | 0 | NA | NA | NA |
| Eif3s6-ps2 | 6.153886045 | 0.104193978 | 0.516018906 | 0.201918916 |
| Gm25418 | 0 | NA | NA | NA |
| Tmx3 | 1329.967682 | 0.106040065 | 0.089708427 | 1.182052434 |
| 1700030N03Rik | 0.163190422 | 0.274394252 | 3.398594894 | 0.080737558 |
| Gm25431 | 0 | NA | NA | NA |
| Ighmbp2 | 158.9101558 | 0.35225925 | 0.113736725 | 3.09714606 |
| Mrpl21 | 165.3835846 | 0.087330753 | 0.116559395 | 0.749238216 |
| Mir6984 | 0.950361064 | -0.825858198 | 0.969535752 | -0.851807885 |
| Cpt1a | 573.796439 | 0.618084985 | 0.143825751 | 4.297457044 |
| Mtl5 | 3.167783942 | -0.772011555 | 0.649072159 | -1.189407902 |
| Gal | 59.25410188 | 0.146002724 | 0.37230284 | 0.392161189 |
| Ppp6r3 | 1233.355573 | 0.067969369 | 0.061399248 | 1.10700654 |
| Gm23940 | 0 | NA | NA | NA |
| Lrp5 | 186.6160727 | 0.456146729 | 0.152090831 | 2.999173091 |
| 1810055G02Rik | 195.6912383 | -0.14115292 | 0.099164139 | -1.423427064 |
| Suv420h1 | 939.8295677 | -0.233396567 | 0.073475699 | -3.176513712 |
| Gm16066 | 1.927026077 | 0.290465708 | 0.667008951 | 0.435474977 |
| Chka | 1064.49316 | -0.02821104 | 0.168589337 | -0.16733585 |
| Tcirg1 | 611.8661545 | 2.353718052 | 0.172878004 | 13.61490763 |
| Ndufs8 | 433.8570073 | 0.079585286 | 0.110701413 | 0.718918422 |
| Aldh3b1 | 78.94818307 | 0.490910335 | 0.233085813 | 2.106135626 |
| Unc93b1 | 424.5399235 | 2.026789794 | 0.159481247 | 12.70864021 |
| Aldh3b3 | 1.060607173 | 0.662074551 | 0.989212583 | 0.66929451 |
| Aldh3b2 | 8.359127856 | -0.557610603 | 0.359631963 | -1.550503459 |

| | | | | |
|---|---|---|---|---|
| Acy3 | 20.90646341 | 0.463346022 | 0.242327704 | 1.912063765 |
| Tbx10 | 0.479449896 | -0.227621741 | 1.465617158 | -0.155307776 |
| Nudt8 | 43.46713457 | -0.474899121 | 0.175884848 | -2.700057028 |
| Gm16312 | 0.283672814 | -0.206481676 | 1.641924311 | -0.125755904 |
| Doc2g | 55.67602762 | -1.027651619 | 0.61780712 | -1.663385847 |
| Ndufv1 | 1083.51582 | -0.14545362 | 0.065082743 | -2.23490305 |
| Gstp1 | 541.1525819 | -0.118743407 | 0.138963151 | -0.85449564 |
| Gstp2 | 1.939495377 | -1.98870811 | 0.952100267 | -2.088759114 |
| BC021614 | 1.429393014 | 4.089445455 | 1.073455198 | 3.809609813 |
| Gm23927 | 0 | NA | NA | NA |
| Cabp2 | 3.994848174 | -0.842391021 | 0.721583089 | -1.16742068 |
| Cdk2ap2 | 150.3217526 | -0.009393005 | 0.134851227 | -0.069654577 |
| Pitpnm1 | 2202.207616 | 0.058907552 | 0.069562778 | 0.84682575 |
| Aip | 570.1830399 | 0.094543505 | 0.082068972 | 1.152000601 |
| Tmem134 | 248.6189701 | 0.245489908 | 0.097090719 | 2.528459064 |
| Cabp4 | 3.477720489 | 3.428384451 | 0.92844725 | 3.692600147 |
| Gpr152 | 2.789673889 | -0.838257831 | 0.843039467 | -0.994328099 |
| Coro1b | 812.1320772 | 0.466918036 | 0.093228774 | 5.008303939 |
| Ptprcap | 24.66290501 | 4.149550234 | 0.527528648 | 7.866018739 |
| Rps6kb2 | 211.501265 | -0.1929196 | 0.14790692 | -1.304331128 |
| Carns1 | 117.9629228 | -0.566070744 | 0.121182478 | -4.671226034 |
| Gm17552 | 4.682108124 | 0.370154482 | 0.471769795 | 0.784608269 |
| Tbc1d10c | 30.14122667 | 2.328465811 | 0.259672891 | 8.966919113 |
| Ppp1ca | 1432.944828 | -0.045594754 | 0.084101273 | -0.542141073 |
| Rad9a | 127.978386 | 0.234071787 | 0.125672907 | 1.862547726 |
| Clcf1 | 180.1899159 | 3.000351299 | 0.388207636 | 7.72872818 |
| Gm26115 | 0.127364494 | -0.687378399 | 3.724945049 | -0.184533836 |
| Pold4 | 108.3674615 | 0.308137948 | 0.142082328 | 2.168728173 |
| Ssh3 | 269.6560211 | 0.042395808 | 0.150824997 | 0.281092713 |
| Mir6985 | 0 | NA | NA | NA |
| Ankrd13d | 332.9753029 | -0.422596972 | 0.109531889 | -3.858209481 |
| Adrbk1 | 2117.90304 | -0.098439835 | 0.059357774 | -1.658415222 |
| Kdm2a | 1799.114924 | -0.070638109 | 0.062158229 | -1.136424096 |
| Rhod | 32.92487569 | 0.808782583 | 0.337297213 | 2.397833577 |
| A930001C03Rik | 2.701377055 | -0.256445842 | 0.759573033 | -0.337618412 |
| Syt12 | 590.78774 | 0.0853227 | 0.116790096 | 0.730564517 |
| 2010003K11Rik | 0.302393035 | 0.747822753 | 1.922854408 | 0.388912832 |
| Pcx | 931.2379665 | -0.047959447 | 0.104172433 | -0.460385207 |
| Lrfn4 | 397.610308 | 0.212279868 | 0.093586665 | 2.268270458 |
| Rce1 | 129.1447744 | 0.00861711 | 0.14945259 | 0.05765782 |
| Mir6986 | 0.044191549 | -0.206490061 | 3.724945049 | -0.055434391 |
| Gm960 | 25.50013179 | 0.011850166 | 0.208400695 | 0.056862409 |
| Sptbn2 | 9930.18535 | 0.177368601 | 0.103286741 | 1.71724463 |
| Gm21992 | 41.25034136 | -0.250457983 | 0.178232099 | -1.405234998 |
| Rbm4b | 188.1038022 | -0.5307979 | 0.089824609 | -5.909270345 |
| Gm37206 | 190.853055 | -0.114141572 | 0.19971944 | -0.571509576 |
| Rbm4 | 178.0962253 | -0.432397038 | 0.173702335 | -2.489298933 |

| | | | | |
|---|---|---|---|---|
| Rbm14 | 411.7655184 | -0.327678084 | 0.085390761 | -3.837395061 |
| Gm21844 | 0.416305032 | -0.206483894 | 1.551347348 | -0.133099718 |
| Ccs | 117.7739688 | 0.101022621 | 0.137917656 | 0.732485051 |
| Ccdc87 | 18.82604018 | -0.207323596 | 0.271238273 | -0.764359667 |
| Ctsf | 447.0332469 | -0.121635605 | 0.083577054 | -1.455370814 |
| Actn3 | 10.89038947 | -0.121679321 | 0.316049355 | -0.38500101 |
| Zdhhc24 | 191.9093747 | -0.159173335 | 0.093182734 | -1.708184865 |
| Bbs1 | 425.8384116 | -0.388840577 | 0.077503128 | -5.017095258 |
| Gm25334 | 0 | NA | NA | NA |
| Dpp3 | 394.3142871 | 0.035745823 | 0.08155067 | 0.438326536 |
| Peli3 | 250.3912601 | -0.276705679 | 0.090038622 | -3.07318875 |
| Mrpl11 | 282.4075369 | 0.121199948 | 0.087368368 | 1.387229154 |
| Npas4 | 211.216604 | -1.105632286 | 0.600513732 | -1.841144051 |
| Slc29a2 | 100.667391 | -0.614502226 | 0.121827006 | -5.044055883 |
| B4gat1 | 647.79448 | -0.123928987 | 0.061159698 | -2.026317828 |
| Brms1 | 180.8653628 | 0.568312496 | 0.107315381 | 5.295722676 |
| Rin1 | 659.0041585 | 0.173387447 | 0.180667718 | 0.959703533 |
| Cd248 | 53.27825101 | 0.086796774 | 0.206862717 | 0.419586355 |
| Tmem151a | 1868.678076 | 0.052593532 | 0.06905536 | 0.76161404 |
| Yif1a | 156.5502458 | 0.365607328 | 0.14980442 | 2.440564353 |
| Cnih2 | 897.9476727 | 0.11274781 | 0.098336586 | 1.146549973 |
| Rab1b | 1115.051491 | 0.217986567 | 0.09755551 | 2.234487487 |
| Klc2 | 2105.216524 | -0.044572011 | 0.069684265 | -0.639628067 |
| Pacs1 | 821.0770873 | -0.134932535 | 0.070313943 | -1.919001127 |
| Gm27468 | 0.909851095 | -1.69494434 | 1.037912806 | -1.633031532 |
| Sf3b2 | 2394.762425 | 0.039232997 | 0.061515829 | 0.637770759 |
| Gal3st3 | 390.9376513 | -0.045232101 | 0.108443112 | -0.417104413 |
| Catsper1 | 0 | NA | NA | NA |
| Cst6 | 23.01852267 | -0.03888415 | 0.234305676 | -0.165954793 |
| Banf1 | 258.7929532 | 0.025039252 | 0.095542798 | 0.26207367 |
| Eif1ad | 254.8369414 | 0.335631101 | 0.108855827 | 3.083262611 |
| Sart1 | 490.457391 | 0.236700743 | 0.1479362 | 1.600019083 |
| Tsga10ip | 0.362047852 | 1.059797072 | 1.537031735 | 0.689508907 |
| 4930481A15Rik | 2.199134214 | 1.887402148 | 0.698101623 | 2.703620913 |
| Gm6293 | 0.045882455 | 0.755288839 | 3.724945049 | 0.202765096 |
| Drap1 | 882.2497186 | 0.32077366 | 0.090799862 | 3.532754914 |
| AI837181 | 578.5093112 | 0.811945776 | 0.115095666 | 7.054529546 |
| Gm25432 | 0 | NA | NA | NA |
| Fosl1 | 112.2972541 | 6.204976293 | 0.831649352 | 7.461048674 |
| Ccdc85b | 470.1620099 | -0.059248656 | 0.105523273 | -0.56147477 |
| Fibp | 216.1359764 | -0.095907145 | 0.076886715 | -1.247382519 |
| Ctsw | 32.65133826 | 4.155837872 | 0.421998805 | 9.847984919 |
| Efemp2 | 186.5996474 | 0.575172469 | 0.15907361 | 3.61576298 |
| Mus81 | 195.7583366 | -0.271810986 | 0.115871761 | -2.345791445 |
| Cfl1 | 6985.640524 | 0.252474684 | 0.069338002 | 3.641216621 |
| Snx32 | 463.4435586 | -0.377325683 | 0.082262602 | -4.586843511 |
| 1700020D05Rik | 15.41874029 | -0.77350941 | 0.408652739 | -1.892828156 |

| | | | | |
|---|---|---|---|---|
| Ovol1 | 1.251933375 | 2.292928722 | 1.39904631 | 1.638922676 |
| Ap5b1 | 86.41281867 | 0.226022936 | 0.142808859 | 1.582695488 |
| Rnaseh2c | 82.70313133 | -0.604915691 | 0.174689642 | -3.462802278 |
| Kat5 | 511.7440207 | 0.122002014 | 0.064458155 | 1.892732009 |
| Rela | 372.9586236 | 0.851496097 | 0.163464523 | 5.209057492 |
| Sipa1 | 217.0465711 | 0.945346618 | 0.152964926 | 6.180152809 |
| Pcnxl3 | 996.3058461 | -0.098963866 | 0.072108033 | -1.37243884 |
| Mir6987 | 0 | NA | NA | NA |
| Map3k11 | 329.8028705 | 0.344024616 | 0.110092499 | 3.12486881 |
| Kcnk7 | 0.811426433 | 1.032844955 | 1.355190844 | 0.762139856 |
| Ehbp1l1 | 284.648246 | 0.003807321 | 0.110216722 | 0.034543954 |
| Gm16538 | 0.233003758 | 0.274401717 | 2.562080697 | 0.107101122 |
| Fam89b | 0.325597809 | -0.512278214 | 1.712862321 | -0.299077286 |
| Sssca1 | 172.9191299 | -0.04990878 | 0.101602233 | -0.491217353 |
| Ltbp3 | 514.8847917 | -0.310861799 | 0.074414367 | -4.177443318 |
| Scyl1 | 711.1667025 | 0.026342347 | 0.072750863 | 0.362089823 |
| Malat1 | 158003.9038 | -0.173687295 | 0.226881966 | -0.76554033 |
| Gm20417 | 3.344837808 | 0.372574708 | 0.923177748 | 0.403578519 |
| Gm27694 | 0 | NA | NA | NA |
| Gm37376 | 17.03796044 | 0.554114934 | 0.963945025 | 0.574840805 |
| Gm27702 | 0 | NA | NA | NA |
| Neat1 | 2624.857955 | 1.259980843 | 0.289220651 | 4.356469151 |
| Gm27533 | 0 | NA | NA | NA |
| Gm27505 | 0 | NA | NA | NA |
| Gm28006 | 0 | NA | NA | NA |
| Frmd8os | 2.120407615 | -0.988692499 | 0.824876504 | -1.198594571 |
| Frmd8 | 356.4512706 | 0.425237159 | 0.188678646 | 2.253764104 |
| Slc25a45 | 48.19350137 | 0.444524665 | 0.236602524 | 1.878782429 |
| Tigd3 | 29.86989943 | -0.058922727 | 0.202884064 | -0.290425604 |
| Dpf2 | 712.9918355 | 0.063057568 | 0.067672777 | 0.931801103 |
| Cdc42ep2 | 113.290494 | -0.643329577 | 0.16217003 | -3.967006581 |
| Pola2 | 84.75758802 | 0.666842914 | 0.150873371 | 4.419884765 |
| Slc22a20 | 0.141071297 | -0.206491178 | 3.564043836 | -0.057937328 |
| Capn1 | 318.3485814 | 0.413990295 | 0.102049615 | 4.056755072 |
| Gm10814 | 14.06531991 | 0.205387602 | 0.295029154 | 0.69616036 |
| Gm8034 | 7.330356508 | -0.533169947 | 0.405446952 | -1.315017773 |
| Syvn1 | 680.7779014 | 0.3296585 | 0.083094506 | 3.967271927 |
| Mir6988 | 0 | NA | NA | NA |
| Mrpl49 | 285.2862106 | -0.148814073 | 0.091985887 | -1.617792441 |
| Fau | 1011.116782 | 0.463514363 | 0.100356545 | 4.61867596 |
| Znhit2 | 165.2069863 | 0.310312279 | 0.098158479 | 3.161339512 |
| Tm7sf2 | 103.8839401 | -0.579200255 | 0.140157422 | -4.132497931 |
| Vps51 | 374.0248763 | 0.014918411 | 0.070669801 | 0.21110022 |
| Gm28374 | 0.132725991 | -0.206490349 | 2.971275708 | -0.06949552 |
| BC048609 | 0.046345709 | 0.274391604 | 3.724945049 | 0.073663262 |
| Zfpl1 | 181.431277 | 0.366116486 | 0.111308237 | 3.289212888 |
| Cdca5 | 6.165523022 | 1.815484574 | 0.605940533 | 2.996143144 |

| | | | | |
|---|---|---|---|---|
| Naaladl1 | 0.666154845 | -1.657719944 | 1.201740396 | -1.379432654 |
| Sac3d1 | 83.16707754 | -0.108994793 | 0.143127825 | -0.761520637 |
| Snx15 | 211.7518732 | -0.255318933 | 0.080005075 | -3.191284228 |
| Arl2 | 383.7572647 | -0.029956815 | 0.098025078 | -0.305603581 |
| Batf2 | 180.1949883 | 6.345763372 | 0.536596831 | 11.82594269 |
| n-R5s19 | 0 | NA | NA | NA |
| Gm5510 | 2.351247315 | 0.226470158 | 0.690015797 | 0.328210106 |
| 1700123I01Rik | 7.996023092 | 2.480450782 | 0.499326755 | 4.967590375 |
| Gpha2 | 0.377319533 | 1.107322032 | 1.531744303 | 0.722915718 |
| Ppp2r5b | 1407.304023 | -0.026632001 | 0.052864443 | -0.503779083 |
| Atg2a | 650.2802182 | -0.091927721 | 0.060502498 | -1.519403728 |
| Mir194-2 | 0.099106094 | 0.274387056 | 3.724945049 | 0.073662041 |
| Mir192 | 0.254293348 | 0.755305854 | 2.472801316 | 0.305445427 |
| Gm14963 | 2.026491629 | -0.655303245 | 0.753721439 | -0.869423651 |
| Ehd1 | 703.3125589 | 0.396193796 | 0.118716333 | 3.337314981 |
| Cdc42bpg | 176.7180393 | -0.16001133 | 0.152561273 | -1.048833215 |
| Men1 | 585.1800558 | -0.171731343 | 0.053311637 | -3.221273129 |
| Map4k2 | 638.3859077 | -0.273308037 | 0.097444872 | -2.804745207 |
| Mir6989 | 0 | NA | NA | NA |
| Gm14966 | 12.5179983 | -0.423597595 | 0.306614569 | -1.381531205 |
| Sf1 | 2104.915733 | 0.230485394 | 0.082496157 | 2.793892497 |
| Pygm | 267.9090404 | -0.569372047 | 0.169411914 | -3.360873709 |
| Rasgrp2 | 263.4585251 | -0.739468417 | 0.195645687 | -3.779630554 |
| Gm14965 | 2.017609801 | -1.394961102 | 0.815760688 | -1.710012659 |
| Nrxn2 | 6243.867174 | -0.206520637 | 0.076291937 | -2.706978537 |
| Gm14964 | 23.8586541 | -0.184858463 | 0.264913691 | -0.697806377 |
| Slc22a12 | 5.133294896 | 0.661773039 | 0.568766867 | 1.163522486 |
| Gm14967 | 0 | NA | NA | NA |
| Gm26460 | 0.404356784 | 1.692740899 | 1.536213687 | 1.101891562 |
| Hmgb1-ps4 | 0.663061751 | -1.082192818 | 1.284508298 | -0.842495778 |
| 2410152P15Rik | 0 | NA | NA | NA |
| Gm14968 | 0.11037557 | 0.274404949 | 3.724945049 | 0.073666845 |
| Rps6ka4 | 374.0853682 | -0.127403322 | 0.080048168 | -1.591583243 |
| mmu-mir-5046 | 0 | NA | NA | NA |
| Ccdc88b | 121.8708714 | 2.54977066 | 0.193324499 | 13.1890716 |
| Prdx5 | 1538.036727 | 0.151990364 | 0.086154102 | 1.76416862 |
| Trmt112 | 108.60505 | 0.360450791 | 0.164671518 | 2.188907927 |
| Esrra | 457.3522009 | -0.452462008 | 0.084359909 | -5.363472 |
| Mir6990 | 0 | NA | NA | NA |
| Tex40 | 21.48008408 | -0.414104933 | 0.296988016 | -1.394348965 |
| Kcnk4 | 51.05303909 | -0.526716535 | 0.202285703 | -2.603824821 |
| Gpr137 | 725.8468719 | -0.171689895 | 0.059873323 | -2.867552466 |
| Bad | 198.4137653 | -0.123441279 | 0.120954118 | -1.020562842 |
| Plcb3 | 323.134626 | 0.228417092 | 0.121562754 | 1.879005564 |
| Ppp1r14b | 149.5878577 | 0.12130321 | 0.171009632 | 0.709335539 |
| Fkbp2 | 300.3276835 | 0.114769975 | 0.283079663 | 0.405433488 |
| Vegfb | 398.1291403 | -0.303999296 | 0.106203352 | -2.862426553 |

| | | | | |
|---|---|---|---|---|
| Dnajc4 | 95.01009773 | -0.186503357 | 0.133931219 | -1.392530868 |
| Nudt22 | 67.37727446 | 0.214152606 | 0.149478645 | 1.43266355 |
| Trpt1 | 56.60693119 | -0.371442299 | 0.147110459 | -2.524921077 |
| Fermt3 | 175.2517023 | 2.717575937 | 0.16933632 | 16.04839374 |
| Stip1 | 1604.279865 | -0.142088117 | 0.096587965 | -1.471074751 |
| Macrod1 | 69.5949187 | -0.060469555 | 0.140017298 | -0.431872033 |
| Flrt1 | 608.6636325 | -0.694486936 | 0.125994091 | -5.5120596 |
| Otub1 | 1632.950069 | 0.088880864 | 0.065470996 | 1.357560895 |
| Cox8a | 2092.571017 | -0.141293261 | 0.074521088 | -1.896017163 |
| Naa40 | 276.9429562 | -0.160894585 | 0.122048432 | -1.318284736 |
| Gm22113 | 0 | NA | NA | NA |
| Gm23997 | 0 | NA | NA | NA |
| Rcor2 | 69.31615873 | -0.365567282 | 0.175973083 | -2.077404541 |
| Mark2 | 1468.643762 | -0.190049198 | 0.078471865 | -2.421876905 |
| Gm17227 | 0.066050875 | 0.274391604 | 3.724945049 | 0.073663262 |
| AI846148 | 273.2784663 | -0.185368788 | 0.101462231 | -1.826973313 |
| 1700105P06Rik | 6.754696947 | -0.277548482 | 0.413142264 | -0.671798812 |
| Rab11b-ps2 | 191.3767199 | 0.176656152 | 0.124438482 | 1.419626384 |
| 2700081O15Rik | 415.5788108 | -0.457705715 | 0.103911014 | -4.404785372 |
| Mir6991 | 0.662460401 | 0.319751923 | 1.361225429 | 0.234900051 |
| Rtn3 | 9090.002618 | -0.124105656 | 0.051290132 | -2.419678985 |
| Atl3 | 817.3787042 | 0.0557824 | 0.062697709 | 0.889703957 |
| Pla2g16 | 615.7304892 | 0.780798833 | 0.131544831 | 5.935610161 |
| Lgals12 | 1.476747611 | -1.322682217 | 0.831612528 | -1.590502995 |
| Hrasls5 | 1.687958028 | 3.316894183 | 1.00065497 | 3.314723139 |
| Gm16437 | 26.12384685 | 0.291891048 | 0.261987296 | 1.114141992 |
| Slc22a19 | 0 | NA | NA | NA |
| Gm15967 | 0 | NA | NA | NA |
| Gm15968 | 0 | NA | NA | NA |
| Slc22a26 | 0.04393175 | -0.206490061 | 3.724945049 | -0.055434391 |
| Slc22a27 | 0 | NA | NA | NA |
| Gm24951 | 0 | NA | NA | NA |
| Slc22a28 | 0 | NA | NA | NA |
| Slc22a29 | 1.523828694 | -0.914066611 | 0.908151918 | -1.006512889 |
| Slc22a30 | 0.083526043 | 0.75528439 | 3.724945049 | 0.202763901 |
| Gm6425 | 1.341472408 | -1.394684405 | 1.033003757 | -1.350125201 |
| Slc22a8 | 69.67438683 | -1.16655452 | 0.282698664 | -4.126494638 |
| Slc22a6 | 62.43237234 | -0.747833493 | 0.265796203 | -2.813559727 |
| Chrm1 | 1319.762724 | -0.392078295 | 0.17638618 | -2.222840221 |
| Gm22820 | 0.118974955 | 0.274400501 | 3.724945049 | 0.073665651 |
| Slc3a2 | 1681.539151 | 0.804379325 | 0.142758185 | 5.634558375 |
| Gm25855 | 7.213846394 | 0.519060155 | 0.397510998 | 1.305775581 |
| Gm25894 | 7.96667369 | 0.893380813 | 0.435791195 | 2.05002034 |
| Gm24452 | 7.467891793 | 1.116262183 | 0.398549872 | 2.800809288 |
| Gm24453 | 6.941141185 | 0.979037123 | 0.410567722 | 2.384593505 |
| Gm22744 | 7.168190812 | 0.226623477 | 0.42916299 | 0.528059229 |
| Gm22680 | 5.248670073 | 1.216995935 | 0.489420796 | 2.486604463 |

| | | | | |
|---|---|---|---|---|
| Gm23246 | 6.637239202 | 0.274565729 | 0.397777464 | 0.690249584 |
| Snord22 | 166.9869455 | 0.096523037 | 0.24372418 | 0.396033896 |
| Wdr74 | 196.0682104 | 0.188794817 | 0.115167923 | 1.639300354 |
| 1700092M07Rik | 1.245725704 | -0.633065505 | 0.932186122 | -0.679119212 |
| Stx5a | 534.5973207 | -0.163721394 | 0.081185484 | -2.016633829 |
| Mir6992 | 0 | NA | NA | NA |
| Nxf1 | 945.8212657 | -0.079981294 | 0.068049327 | -1.175342912 |
| Taf6l | 170.5846256 | 0.175288922 | 0.113876541 | 1.539289137 |
| Polr2g | 319.7014694 | 0.25312685 | 0.07864755 | 3.218496302 |
| Zbtb3 | 40.55560979 | -0.454372671 | 0.19965349 | -2.2758063 |
| Ttc9c | 517.0751682 | 0.420047217 | 0.078935653 | 5.321387742 |
| Hnrnpul2 | 2234.773551 | -0.193368113 | 0.111079023 | -1.740815747 |
| Gng3 | 1184.138002 | -0.207273414 | 0.070747777 | -2.929751622 |
| Bscl2 | 767.6870937 | -0.01924508 | 0.068913019 | -0.279266238 |
| Lrrn4cl | 17.94000658 | 0.027083838 | 0.266571465 | 0.101600664 |
| Ubxn1 | 553.7697328 | 0.294668922 | 0.092352906 | 3.190683803 |
| Uqcc3 | 121.6763236 | 0.141054623 | 0.141704223 | 0.995415802 |
| Lbhd1 | 49.53080462 | 0.088296477 | 0.175968701 | 0.501773763 |
| Snora57 | 16.08879982 | 0.735500226 | 0.344694169 | 2.133776236 |
| Mir5136 | 0.568319789 | -0.445706735 | 1.238088246 | -0.359995934 |
| 1810009A15Rik | 2.938865308 | 0.380075003 | 0.579250637 | 0.656149478 |
| Ints5 | 306.8553324 | 0.213263992 | 0.10324807 | 2.065549433 |
| Ganab | 1308.618353 | -0.025897874 | 0.062256472 | -0.415986864 |
| B3gat3 | 910.6532437 | -0.033253634 | 0.091277371 | -0.364314111 |
| Rom1 | 80.3855847 | 0.149464121 | 0.16547205 | 0.903259017 |
| Eml3 | 209.4625949 | 0.136829105 | 0.109606518 | 1.248366497 |
| Mta2 | 687.4573254 | 0.238131844 | 0.080865274 | 2.944797332 |
| Tut1 | 264.2884532 | 0.058353053 | 0.095586065 | 0.610476572 |
| Eef1g | 1677.123862 | 0.158794359 | 0.083888802 | 1.892914837 |
| Ahnak | 4057.702084 | 1.488199509 | 0.17349297 | 8.577866361 |
| Scgb1a1 | 0 | NA | NA | NA |
| Asrgl1 | 1213.125555 | -0.467242366 | 0.119196571 | -3.919931276 |
| Gm9750 | 3.69099669 | 0.197386815 | 0.544271954 | 0.362662109 |
| Gm24543 | 0 | NA | NA | NA |
| Pcna-ps2 | 9.410251089 | 0.570596343 | 0.365292504 | 1.562025875 |
| Gm2617 | 0.087261593 | 0.274400501 | 3.724945049 | 0.073665651 |
| Stxbp3-ps | 23.11005125 | 0.535347298 | 0.249346667 | 2.147000019 |
| Gm29231 | 0.2218918 | -0.566750704 | 3.137147691 | -0.180657961 |
| Gm6445 | 1.943633249 | 0.400043666 | 0.656338199 | 0.609508432 |
| Scgb2a2 | 0.152314784 | -0.206481641 | 2.847275348 | -0.072519028 |
| Gm22369 | 0 | NA | NA | NA |
| Incenp | 144.1045022 | 0.091762735 | 0.125598695 | 0.730602611 |
| Gm24484 | 0.061920613 | 0.274393828 | 3.724945049 | 0.073663859 |
| Fth1 | 13460.26008 | 0.803839643 | 0.167240104 | 4.806500486 |
| Best1 | 1.289165993 | 2.624776569 | 1.037245594 | 2.530525639 |
| Rab3il1 | 131.0561169 | 0.404543381 | 0.173787654 | 2.327802755 |
| Fads3 | 258.367544 | -0.079397423 | 0.089063746 | -0.891467362 |

| | | | | |
|---|---|---|---|---|
| Fads2 | 1025.070797 | -0.086245609 | 0.119155463 | -0.723807421 |
| Fads1 | 1910.999182 | -0.143432153 | 0.09817288 | -1.461016043 |
| Mir6993 | 1.831497397 | -1.128769328 | 0.73501747 | -1.535704081 |
| Gm10143 | 2.039768525 | -0.305525415 | 0.758624948 | -0.402735787 |
| Fen1 | 177.2425644 | 0.863609465 | 0.146735635 | 5.88547878 |
| Tmem258 | 152.1698681 | 0.087635588 | 0.166827822 | 0.525305596 |
| Myrf | 1402.777588 | 0.663044176 | 0.136965237 | 4.840966883 |
| Dagla | 1427.846811 | -0.108933716 | 0.1129455 | -0.964480354 |
| Syt7 | 3169.726564 | -0.377113773 | 0.103630353 | -3.639028158 |
| Gm26859 | 2.155457834 | -0.918837014 | 0.690268806 | -1.331129273 |
| Lrrc10b | 214.7262662 | -0.137633009 | 0.212462132 | -0.647800185 |
| Ppp1r32 | 18.15485456 | 0.230146654 | 0.241743265 | 0.952029229 |
| Sdhaf2 | 369.9013947 | 0.110248194 | 0.073778047 | 1.494322464 |
| Cpsf7 | 982.7429653 | 0.081225462 | 0.079703144 | 1.01909985 |
| Tmem216 | 43.25074293 | -0.139783548 | 0.191378602 | -0.730403226 |
| Tmem138 | 84.89562639 | 0.022596046 | 0.122820262 | 0.183976532 |
| Cyb561a3 | 70.26552882 | 0.278440758 | 0.198210692 | 1.404771637 |
| Tkfc | 98.20098442 | -0.076178863 | 0.130444055 | -0.583996432 |
| Ddb1 | 2897.624334 | 0.035895016 | 0.055276186 | 0.649375774 |
| Vwce | 6.549817523 | 0.270587136 | 0.482580079 | 0.560709295 |
| Pga5 | 0.066050875 | 0.274391604 | 3.724945049 | 0.073663262 |
| Vps37c | 286.9571528 | 0.255517442 | 0.100186674 | 2.55041345 |
| Cd5 | 21.9583943 | 3.389312063 | 0.431109156 | 7.86184199 |
| A430093F15Rik | 5.018136537 | 3.154887831 | 0.843201748 | 3.741557509 |
| Cd6 | 55.03509925 | 1.887971707 | 0.224860773 | 8.396180797 |
| Slc15a3 | 354.5772757 | 4.232665208 | 0.304122323 | 13.91764066 |
| Tmem132a | 2470.423893 | 0.134465197 | 0.09310184 | 1.444280772 |
| Tmem109 | 292.8556633 | -0.384865294 | 0.121564821 | -3.165926552 |
| Prpf19 | 1647.79328 | 0.170280543 | 0.065669533 | 2.592991542 |
| Zp1 | 0.369197529 | 0.71894924 | 1.430393995 | 0.502623223 |
| Ptgdr2 | 2.273986841 | -0.064446821 | 0.718233976 | -0.089729563 |
| Ccdc86 | 181.1409562 | 1.058488527 | 0.139519246 | 7.58668471 |
| Ms4a10 | 0.044544648 | 0.755286614 | 3.724945049 | 0.202764498 |
| Ms4a15 | 28.11515156 | -1.758487318 | 1.139837309 | -1.542752903 |
| Ms4a18 | 0.263774563 | 0.274402418 | 2.424655244 | 0.113171726 |
| Gm28347 | 2.460283742 | 3.428876484 | 1.027473863 | 3.337190956 |
| AW112010 | 268.3498837 | 6.11778596 | 0.45651287 | 13.40112483 |
| Ms4a8a | 62.44774592 | 7.294291613 | 1.064682184 | 6.851144616 |
| Gm336 | 28.43566127 | 0.608713609 | 0.210264663 | 2.894987685 |
| 1700017D01Rik | 0.314055832 | 1.581491647 | 1.736223241 | 0.910880358 |
| Gm6501 | 23.72387051 | 0.43660801 | 0.276516333 | 1.578959208 |
| 1700025F22Rik | 0.715113835 | 2.060397062 | 1.477828511 | 1.394205786 |
| Ms4a13 | 0 | NA | NA | NA |
| 4930526L06Rik | 0.358247177 | 0.286922598 | 1.575819605 | 0.182078327 |
| Ms4a12 | 0.470940457 | 0.755308069 | 1.657003296 | 0.455827741 |
| Ms4a1 | 1.60259925 | 2.577006291 | 1.061429647 | 2.427863494 |
| Ms4a5 | 0 | NA | NA | NA |

| | | | | |
|---|---|---|---|---|
| Ms4a14 | 14.20065461 | 4.083413365 | 0.822370169 | 4.965420099 |
| Ms4a7 | 116.0378029 | 3.867583049 | 0.3942402 | 9.810219882 |
| Gm28935 | 0.610760527 | 2.541714856 | 1.457527538 | 1.743853746 |
| Ms4a4a | 77.63966184 | 3.832192419 | 0.420016964 | 9.123899151 |
| Gm5692 | 0.269752699 | 0.755307134 | 1.822342568 | 0.414470445 |
| Ms4a4c | 622.4266349 | 8.772061636 | 0.672000371 | 13.05365595 |
| Gm5511 | 2.864211081 | 1.824211698 | 0.653582994 | 2.791094191 |
| Ms4a4b | 229.0039815 | 8.661093468 | 0.797017659 | 10.86687776 |
| Gm37387 | 7.980888526 | 4.940347344 | 0.962233196 | 5.134251621 |
| Ms4a6c | 638.4433901 | 5.888618362 | 0.333573797 | 17.6531203 |
| BE692007 | 14.17246462 | 4.910974136 | 0.793126948 | 6.191914358 |
| Gm8369 | 26.70344983 | 4.919097954 | 0.577250485 | 8.52160038 |
| Gm19261 | 0.679959982 | 2.087293355 | 1.293415471 | 1.613784125 |
| Ms4a6b | 404.189003 | 4.854161034 | 0.306547323 | 15.83494838 |
| Ms4a4d | 12.73072494 | 6.044318444 | 0.912785918 | 6.621835771 |
| Gm10212 | 1.028441291 | 1.314939141 | 2.17715269 | 0.603971943 |
| Gm38275 | 1.398451413 | 0.277425148 | 1.285042659 | 0.215887889 |
| Ms4a6d | 780.7598453 | 5.745074487 | 0.381798572 | 15.04739646 |
| Gm24599 | 0 | NA | NA | NA |
| Ms4a2 | 1.450910466 | 1.575600292 | 1.044022671 | 1.50916291 |
| Ms4a3 | 0.922047842 | 1.794503851 | 1.179048386 | 1.521993391 |
| Oosp2 | 0.454235466 | 1.956039177 | 1.38614993 | 1.41113103 |
| Oosp1 | 0.420049603 | 0.755307554 | 1.702880423 | 0.44354703 |
| Gm26183 | 0 | NA | NA | NA |
| Gm25443 | 0 | NA | NA | NA |
| Oosp3 | 0 | NA | NA | NA |
| Gm19224 | 0.308730509 | 1.454182358 | 3.721091989 | 0.39079452 |
| Gif | 0.046345709 | 0.274391604 | 3.724945049 | 0.073663262 |
| Mrpl16 | 136.9670042 | 0.156879901 | 0.117183985 | 1.338748645 |
| Stx3 | 268.7199862 | 0.211926167 | 0.114660186 | 1.848297768 |
| Olfr1417 | 0.045882455 | 0.755288839 | 3.724945049 | 0.202765096 |
| Olfr1418 | 0 | NA | NA | NA |
| Olfr1419 | 0.385885129 | -1.666408644 | 1.510471091 | -1.103237695 |
| Gm6365 | 6.926587766 | 0.998181068 | 0.397823421 | 2.509105839 |
| Olfr1420 | 0.092228164 | 0.75528439 | 3.724945049 | 0.202763901 |
| Patl1 | 512.1726233 | 0.215019496 | 0.08425572 | 2.551986935 |
| mmu-mir-6994 | 0.26492599 | 0.755310125 | 1.821085228 | 0.414758251 |
| Gm22272 | 0 | NA | NA | NA |
| Osbp | 1002.556198 | -0.095504393 | 0.047347272 | -2.017104479 |
| Olfr1423 | 0 | NA | NA | NA |
| Olfr1424 | 0.049553047 | 0.274391604 | 3.724945049 | 0.073663262 |
| Olfr1425 | 0.045882455 | 0.755288839 | 3.724945049 | 0.202765096 |
| Olfr1426 | 0 | NA | NA | NA |
| Olfr1427 | 0 | NA | NA | NA |
| Olfr1428 | 0.090277459 | -0.20649451 | 3.724945049 | -0.055435586 |
| Olfr76 | 0.045882455 | 0.755288839 | 3.724945049 | 0.202765096 |
| Olfr1431 | 0 | NA | NA | NA |

| | | | | |
|---|---|---|---|---|
| Olfr1432 | 0 | NA | NA | NA |
| Olfr262 | 0 | NA | NA | NA |
| Gm26864 | 0 | NA | NA | NA |
| Olfr235 | 0.119234754 | 0.274400501 | 3.724945049 | 0.073665651 |
| Olfr1434 | 0 | NA | NA | NA |
| Olfr1436 | 0 | NA | NA | NA |
| Olfr1437 | 0.435432306 | 0.274381276 | 3.724945049 | 0.07366049 |
| Olfr1440 | 0.060174537 | 0.755286614 | 3.724945049 | 0.202764498 |
| Olfr1441 | 0.105852363 | -0.206492286 | 3.724945049 | -0.055434988 |
| Pfpl | 0 | NA | NA | NA |
| Mpeg1 | 2850.264192 | 3.67183925 | 0.186673327 | 19.66986561 |
| Dtx4 | 733.6416179 | 0.21740186 | 0.095488291 | 2.276738407 |
| Fam111a | 54.9166158 | 1.826653339 | 0.371168478 | 4.921359026 |
| Gm16274 | 2.331142904 | 0.516929806 | 0.632315063 | 0.817519361 |
| A330040F15Rik | 6.392297317 | 0.402175555 | 0.425640173 | 0.944872174 |
| Gm4952 | 2.655904937 | 2.490010493 | 1.045708639 | 2.381170433 |
| Glyat | 2.115680473 | -1.289820771 | 0.959873017 | -1.343741046 |
| Gm24521 | 0.066050875 | 0.274391604 | 3.724945049 | 0.073663262 |
| Olfr1442 | 0.259089178 | 0.274407562 | 2.066703684 | 0.132775474 |
| Olfr1443 | 0.092691418 | 0.274387304 | 3.724945049 | 0.073662108 |
| Keg1 | 0.403815225 | 0.274379694 | 3.724945049 | 0.073660065 |
| Gm15962 | 0 | NA | NA | NA |
| Cntf | 4.709263031 | -0.077230072 | 0.466069822 | -0.165704941 |
| Zfp91 | 2396.758906 | 0.263759806 | 0.061336707 | 4.3001951 |
| Lpxn | 113.3029764 | 3.843201384 | 0.332334967 | 11.5642402 |
| Gm5244 | 0.635603962 | 1.846089651 | 1.329040663 | 1.389039254 |
| Olfr1444 | 0.544199285 | 0.278833682 | 1.381969265 | 0.201765472 |
| Olfr1445 | 0.066050875 | 0.274391604 | 3.724945049 | 0.073663262 |
| Olfr1446 | 0 | NA | NA | NA |
| Olfr1447 | 0 | NA | NA | NA |
| Olfr1448 | 0.092228164 | 0.75528439 | 3.724945049 | 0.202763901 |
| Olfr1449 | 0.066050875 | 0.274391604 | 3.724945049 | 0.073663262 |
| Olfr1450 | 0 | NA | NA | NA |
| Olfr1451 | 0 | NA | NA | NA |
| Olfr1452-ps1 | 0 | NA | NA | NA |
| Olfr1453 | 0 | NA | NA | NA |
| Olfr1454 | 0 | NA | NA | NA |
| Olfr1456-ps1 | 0 | NA | NA | NA |
| Olfr1457 | 0 | NA | NA | NA |
| Olfr1458 | 0.049553047 | 0.274391604 | 3.724945049 | 0.073663262 |
| Olfr1459 | 0 | NA | NA | NA |
| Olfr1461 | 0 | NA | NA | NA |
| Olfr1462 | 0 | NA | NA | NA |
| Olfr1463 | 0 | NA | NA | NA |
| Gm22443 | 0 | NA | NA | NA |
| Olfr1465 | 0 | NA | NA | NA |
| Olfr1466 | 0.062426873 | 0.755288839 | 3.724945049 | 0.202765096 |

| | | | | |
|---|---|---|---|---|
| Olfr1467 | 0 | NA | NA | NA |
| Olfr1469 | 0 | NA | NA | NA |
| Olfr1471 | 0 | NA | NA | NA |
| Olfr1472 | 0 | NA | NA | NA |
| Olfr1474 | 0 | NA | NA | NA |
| Olfr1475 | 0 | NA | NA | NA |
| Olfr1477 | 0 | NA | NA | NA |
| Olfr1480 | 0 | NA | NA | NA |
| Olfr1484 | 0.068097359 | -0.206487837 | 3.724945049 | -0.055433794 |
| Gm22323 | 0 | NA | NA | NA |
| Gm23931 | 0 | NA | NA | NA |
| Olfr1487 | 0 | NA | NA | NA |
| Olfr1489 | 0 | NA | NA | NA |
| Gm24124 | 0 | NA | NA | NA |
| Olfr1490 | 0 | NA | NA | NA |
| Olfr1491 | 0 | NA | NA | NA |
| Olfr1494 | 0.061920613 | 0.274393828 | 3.724945049 | 0.073663859 |
| Olfr1495 | 0 | NA | NA | NA |
| Olfr1496 | 0 | NA | NA | NA |
| Olfr1497 | 0.04393175 | -0.206490061 | 3.724945049 | -0.055434391 |
| Olfr1499 | 0 | NA | NA | NA |
| Olfr1500 | 0 | NA | NA | NA |
| Olfr1501 | 0.044191549 | -0.206490061 | 3.724945049 | -0.055434391 |
| Olfr1502 | 0 | NA | NA | NA |
| Olfr1503-ps1 | 0 | NA | NA | NA |
| Olfr1504 | 0 | NA | NA | NA |
| Olfr1505 | 0 | NA | NA | NA |
| Gm26993 | 0 | NA | NA | NA |
| Tle4 | 1024.20378 | 0.300782864 | 0.069083676 | 4.353892006 |
| Gm37997 | 0 | NA | NA | NA |
| Gm26026 | 0 | NA | NA | NA |
| Gm5513 | 1.595119001 | 0.906946804 | 0.838127367 | 1.082110954 |
| Gm24319 | 0 | NA | NA | NA |
| Psat1 | 926.2091933 | -0.066018387 | 0.126250643 | -0.522915258 |
| Gm3329 | 0 | NA | NA | NA |
| Cep78 | 106.2535858 | -0.247513387 | 0.122246965 | -2.02469964 |
| C130060C02Rik | 7.941562867 | -0.907200338 | 0.470695053 | -1.927363229 |
| Gnaq | 3032.47275 | 0.00919271 | 0.037749768 | 0.243516988 |
| E030024N20Rik | 20.01620722 | -0.163993188 | 0.2203527 | -0.744230445 |
| Gm10819 | 108.4716668 | -0.073409949 | 0.128748979 | -0.570178882 |
| Gm17068 | 25.77906067 | -0.072319152 | 0.372667631 | -0.194058044 |
| Mir496b | 0 | NA | NA | NA |
| Gna14 | 16.24861454 | 1.057442605 | 0.475041257 | 2.226001615 |
| Vps13a | 1858.188761 | -0.087192565 | 0.073091581 | -1.192922124 |
| Gm9806 | 0.46212231 | 0.252139248 | 1.599210812 | 0.157664797 |
| Foxb2 | 1.423250569 | 0.044688634 | 0.953789304 | 0.04685378 |
| Prune2 | 1550.090265 | 0.317145371 | 0.11406252 | 2.780452083 |

| | | | | |
|---|---|---|---|---|
| Gm27970 | 0 | NA | NA | NA |
| Gm27796 | 0 | NA | NA | NA |
| Gcnt1 | 110.511412 | -0.326831238 | 0.192522256 | -1.697628339 |
| Rfk | 726.378061 | 0.026136203 | 0.109071267 | 0.239625008 |
| Pcsk5 | 144.4692259 | -0.104061843 | 0.189445513 | -0.549296953 |
| Gm22880 | 0 | NA | NA | NA |
| Gm23063 | 0 | NA | NA | NA |
| Gm23061 | 0 | NA | NA | NA |
| Ostf1 | 237.4482545 | 0.959797092 | 0.147576353 | 6.503732303 |
| Gm8250 | 0.075043205 | 0.755286614 | 3.724945049 | 0.202764498 |
| Nmrk1 | 21.71250768 | -0.044987101 | 0.23485215 | -0.191554991 |
| 2410127L17Rik | 57.66373482 | 0.103994239 | 0.153841364 | 0.675983601 |
| D030056L22Rik | 161.6691014 | -0.34259048 | 0.106067126 | -3.229940248 |
| Gm23238 | 1902.741476 | 0.307754761 | 0.398685618 | 0.771923408 |
| Trpm6 | 41.27246436 | 1.333433291 | 0.233361902 | 5.714014502 |
| Gm24190 | 0 | NA | NA | NA |
| Rorb | 1457.656088 | 0.000751858 | 0.083893473 | 0.008962055 |
| Gm25437 | 0 | NA | NA | NA |
| Gm22684 | 0 | NA | NA | NA |
| Anxa1 | 128.5436956 | 1.719641956 | 0.309846873 | 5.549973571 |
| 1500015L24Rik | 2.063333161 | -0.701795881 | 0.735776844 | -0.953816211 |
| Gm6684 | 0 | NA | NA | NA |
| Aldh1a1 | 547.0244569 | 0.65486497 | 0.18765713 | 3.489688718 |
| Aldh1a7 | 11.46744072 | -0.478467554 | 0.484383793 | -0.98778605 |
| Tmc1 | 1.743158711 | 0.267020748 | 0.842923478 | 0.316779346 |
| Zfand5 | 1899.320038 | -0.201785316 | 0.07421961 | -2.718760124 |
| Gda | 1770.327303 | -0.232890884 | 0.10457113 | -2.227104984 |
| Gm3443 | 0.062426873 | 0.755288839 | 3.724945049 | 0.202765096 |
| 1110059E24Rik | 203.1103973 | 0.129460974 | 0.13414202 | 0.965103806 |
| Abhd17b | 456.3664533 | 0.15208405 | 0.07651593 | 1.987612912 |
| Tmem2 | 509.6691455 | 0.258633157 | 0.106602946 | 2.426135174 |
| Gm5514 | 10.26669978 | -0.053658378 | 0.349308726 | -0.153613049 |
| Trpm3 | 1663.018852 | -0.264825544 | 0.081201068 | -3.261355451 |
| Gm22506 | 0 | NA | NA | NA |
| Gm27151 | 4.77538895 | -0.043487707 | 0.447141311 | -0.097257189 |
| Mir204 | 3.450472164 | 0.202462988 | 0.629936112 | 0.321402416 |
| C330002G04Rik | 1.481948713 | 1.545299037 | 0.979876115 | 1.577035111 |
| 2410080I02Rik | 2.836846521 | 0.39092858 | 0.622838553 | 0.627656363 |
| Klf9 | 2856.577684 | 0.477268338 | 0.121764931 | 3.91958781 |
| Mir1192 | 10.09841093 | 0.02955036 | 0.385227622 | 0.076708829 |
| Smc5 | 682.7510276 | 0.031917277 | 0.088521297 | 0.360560431 |
| Gm24263 | 0.146692594 | 0.274392389 | 3.52332538 | 0.07787881 |
| Mamdc2 | 41.82610674 | -0.403163531 | 0.224313366 | -1.797322816 |
| 1700028P14Rik | 7.326566073 | 0.029478646 | 0.428129552 | 0.068854499 |
| Gm9493 | 43.04589625 | 0.433652666 | 0.200843841 | 2.159153415 |
| Gm22144 | 0 | NA | NA | NA |
| Gm21542 | 0 | NA | NA | NA |

| | | | | |
|---|---|---|---|---|
| Gm6563 | 117.5227222 | 0.47024311 | 0.145473233 | 3.232506084 |
| Ptar1 | 312.4373579 | -0.189133198 | 0.084089208 | -2.24919704 |
| Gm9938 | 199.0723875 | -0.20364635 | 0.094157529 | -2.162825991 |
| Apba1 | 2979.940385 | -0.186964078 | 0.068108564 | -2.745089107 |
| Fam189a2 | 83.27374125 | -0.004439116 | 0.147304036 | -0.030135742 |
| 1700021P04Rik | 0.123841226 | 0.274391173 | 3.724945049 | 0.073663147 |
| Tjp2 | 802.6063976 | 0.621754344 | 0.109918818 | 5.656486797 |
| Fxn | 57.39510889 | 0.051962382 | 0.158134538 | 0.328596033 |
| Gm16216 | 0.110870145 | -0.206490066 | 3.724945049 | -0.055434393 |
| Pip5k1b | 239.2249923 | -0.043752079 | 0.108650354 | -0.402686948 |
| Pip5k1bos | 0.196777805 | -1.156436132 | 2.227175217 | -0.519238955 |
| Fam122a | 94.19792559 | 0.169483186 | 0.159645096 | 1.061624753 |
| Tmem252 | 56.70084035 | 4.625183044 | 0.434557301 | 10.64343652 |
| Pgm5 | 91.2960709 | 0.544417263 | 0.226060185 | 2.40828461 |
| Gm17800 | 0.452161616 | 0.633574641 | 1.419021311 | 0.446487052 |
| Gm10053 | 118.3751616 | -0.034621082 | 0.161158006 | -0.214826947 |
| Foxd4 | 0.431881386 | 2.493054063 | 1.635564743 | 1.52427721 |
| Cbwd1 | 142.8925874 | 0.335536772 | 0.104744255 | 3.203390693 |
| mmu-mir-3084-1 | 0 | NA | NA | NA |
| Gm23658 | 0.086648695 | -0.687376175 | 3.724945049 | -0.184533239 |
| Dock8 | 246.4602774 | 1.743299066 | 0.175075435 | 9.957416719 |
| Gm24252 | 0 | NA | NA | NA |
| Kank1 | 435.2904436 | 0.357509935 | 0.133237953 | 2.683243978 |
| Gm26207 | 0 | NA | NA | NA |
| Dmrt1 | 0 | NA | NA | NA |
| Dmrt3 | 3.281054849 | 1.198634223 | 0.597727146 | 2.005320037 |
| 2610016A17Rik | 0.172734917 | -0.6873768 | 2.736751389 | -0.251165233 |
| Dmrt2 | 7.322241276 | -0.20114583 | 0.364869834 | -0.551281063 |
| Gm8825 | 0 | NA | NA | NA |
| Gm25176 | 0.061920613 | 0.274393828 | 3.724945049 | 0.073663859 |
| Smarca2 | 2885.991631 | -0.152008036 | 0.069411126 | -2.189966452 |
| Gm20616 | 0.038981395 | 0.274393828 | 3.724945049 | 0.073663859 |
| 4931403E22Rik | 1.142959485 | -0.228678686 | 1.002998482 | -0.227995047 |
| Gm815 | 0.098502579 | -0.206492286 | 3.724945049 | -0.055434988 |
| Vldlr | 1529.815537 | -0.415095581 | 0.062705351 | -6.619779229 |
| Kcnv2 | 1.799299276 | 0.056832 | 0.891093655 | 0.063777808 |
| D19Bwg1357e | 318.8741132 | -0.021078499 | 0.098665671 | -0.213635597 |
| Gm24782 | 0 | NA | NA | NA |
| Rfx3 | 1251.313148 | -0.461683213 | 0.117057548 | -3.944070433 |
| Gm8412 | 0.124853745 | 1.290908285 | 3.721447108 | 0.346883416 |
| Gm20407 | 0 | NA | NA | NA |
| Gm28228 | 0.047948698 | -0.206487837 | 3.724945049 | -0.055433794 |
| Glis3 | 334.0083546 | 1.038580416 | 0.121720989 | 8.532467742 |
| 4933413C19Rik | 10.33626457 | 0.002643393 | 0.389786684 | 0.006781639 |
| D930032P07Rik | 0.131806241 | 0.671889633 | 3.666020728 | 0.183274914 |
| Rps15a-ps2 | 0.107078754 | -0.206490058 | 3.724945049 | -0.05543439 |
| Gm6788 | 2.966347917 | -0.187926831 | 0.600085596 | -0.313166709 |

| | | | | |
|---|---|---|---|---|
| Gm26942 | 0.038981395 | 0.274393828 | 3.724945049 | 0.073663859 |
| Slc1a1 | 1257.835077 | -0.147729169 | 0.080070957 | -1.844978187 |
| Gm5822 | 3.84996234 | 0.359928391 | 0.531865038 | 0.676728804 |
| 4430402I18Rik | 33.49194021 | 0.084865769 | 0.184291452 | 0.460497586 |
| Ppapdc2 | 422.5883542 | -0.01677653 | 0.082675435 | -0.202920371 |
| Cdc37l1 | 944.3468794 | -0.093401032 | 0.059370459 | -1.573190339 |
| Gm10136 | 20.53681307 | 0.242462233 | 0.244631511 | 0.991132466 |
| Ak3 | 872.3625232 | -0.012069139 | 0.118036544 | -0.102249173 |
| 1700018L02Rik | 10.4047421 | 0.836245791 | 0.370692536 | 2.255901348 |
| Rcl1 | 158.5573071 | 0.118973345 | 0.090309122 | 1.317401198 |
| Mir101b | 0.39937024 | -1.564242072 | 1.452789668 | -1.076716131 |
| Gm5518 | 30.36473408 | 0.220314889 | 0.276081697 | 0.798006138 |
| Jak2 | 1349.483123 | 0.692424557 | 0.097761967 | 7.082760101 |
| Insl6 | 6.983240532 | 1.950879477 | 0.624404047 | 3.124386343 |
| Rln1 | 4.858786034 | 1.163446682 | 0.577753844 | 2.013741134 |
| Plgrkt | 208.1764301 | 0.516815983 | 0.106391838 | 4.85766571 |
| Cd274 | 1796.888883 | 6.635891837 | 0.274322992 | 24.190068 |
| Pdcd1lg2 | 5.725591314 | 3.955334525 | 0.92855662 | 4.259658957 |
| A930007I19Rik | 28.01513506 | 0.43949668 | 0.232447867 | 1.890732257 |
| Ric1 | 1103.079595 | -0.051632648 | 0.084862851 | -0.608424629 |
| Ermp1 | 862.3818722 | -0.234899315 | 0.081033316 | -2.898799229 |
| Mlana | 1.722373549 | 4.181580912 | 1.003936837 | 4.165183265 |
| 9930021J03Rik | 1199.788204 | -0.199422098 | 0.04936426 | -4.039807307 |
| Ranbp6 | 784.8181587 | -0.068206506 | 0.091869325 | -0.742429589 |
| Il33 | 391.409633 | 0.664069428 | 0.290508862 | 2.285883549 |
| Gm26046 | 0 | NA | NA | NA |
| Trpd52l3 | 0 | NA | NA | NA |
| Uhrf2 | 805.07859 | -0.191144895 | 0.068080512 | -2.807630116 |
| Gldc | 269.8156168 | -0.554402725 | 0.171281869 | -3.236785831 |
| Rpl31-ps20 | 0 | NA | NA | NA |
| Mbl2 | 0.229843471 | -0.130205387 | 2.548386578 | -0.051093264 |
| Gm24108 | 0 | NA | NA | NA |
| Gm24883 | 0 | NA | NA | NA |
| Ppp1r2-ps3 | 1.44902218 | 0.926748644 | 0.925825154 | 1.000997477 |
| Dkk1 | 0.239409371 | -0.155399504 | 2.141382825 | -0.072569698 |
| Prkg1 | 254.0169383 | -0.268523572 | 0.124756775 | -2.152376677 |
| Gm23426 | 0 | NA | NA | NA |
| Cstf2t | 1157.615467 | -0.016948817 | 0.054557125 | -0.310661843 |
| A1cf | 0.581447236 | 2.152937302 | 1.371598754 | 1.569655336 |
| Asah2 | 367.16758 | 0.34163108 | 0.145147752 | 2.353678073 |
| Sgms1 | 487.606945 | -0.031098122 | 0.093645915 | -0.332081995 |
| 2700046G09Rik | 25.94331831 | -0.159084115 | 0.229368367 | -0.693574782 |
| Rpl9-ps6 | 47.62432018 | 0.389513497 | 0.179129973 | 2.174474149 |
| Minpp1 | 627.6333895 | 0.401101038 | 0.080693705 | 4.970660843 |
| Papss2 | 133.0147862 | -0.301133663 | 0.181091785 | -1.662878647 |
| Atad1 | 1547.98136 | 0.452012203 | 0.101510594 | 4.452857439 |
| Pten | 3804.391786 | 0.075927865 | 0.069761013 | 1.088399685 |

| | | | | |
|---|---|---|---|---|
| Gm22482 | 0.072025462 | 0.755288839 | 3.724945049 | 0.202765096 |
| Rnls | 22.89813653 | 0.158881571 | 0.242303703 | 0.655712518 |
| mmu-mir-3970 | 0 | NA | NA | NA |
| Gm7237 | 0.127971488 | 0.274389832 | 3.687653711 | 0.0744077 |
| Gm8975 | 0 | NA | NA | NA |
| Lipo4 | 0.195256443 | 0.713177197 | 2.223427333 | 0.320755793 |
| Lipo1 | 94.81852167 | 0.397594584 | 0.125967059 | 3.156337749 |
| Gm27440 | 0 | NA | NA | NA |
| Gm8978 | 0 | NA | NA | NA |
| Gm8980 | 0.107078754 | -0.206490058 | 3.724945049 | -0.05543439 |
| Lipo2 | 1.691184024 | 0.74758894 | 0.918231978 | 0.814161299 |
| Gm5097 | 0.090537259 | -0.20649451 | 3.724945049 | -0.055435586 |
| Gm5519 | 0.296824366 | 1.460662515 | 2.36875491 | 0.616637251 |
| Gm8988 | 0 | NA | NA | NA |
| Gm23095 | 0.106971521 | 1.236179401 | 3.724945049 | 0.331865138 |
| Lipf | 0.066050875 | 0.274391604 | 3.724945049 | 0.073663262 |
| Lipk | 0.091880448 | -0.687373951 | 3.724945049 | -0.184532642 |
| Lipn | 0.251288325 | 0.755305259 | 2.115013813 | 0.357115993 |
| Gm23334 | 0 | NA | NA | NA |
| Lipm | 1.074639982 | 1.241334438 | 1.007198904 | 1.232462062 |
| Ankrd22 | 0.527224786 | -0.923535934 | 1.497746911 | -0.616616817 |
| Stambpl1 | 217.0575449 | 0.278848078 | 0.086141994 | 3.237074819 |
| Acta2 | 30.64504911 | 0.261415431 | 0.30177198 | 0.866268071 |
| Fas | 57.72959901 | 3.13139097 | 0.341601736 | 9.166788805 |
| Gm27818 | 0.914190395 | 2.701367992 | 1.310506303 | 2.061316291 |
| Gm23060 | 0 | NA | NA | NA |
| Ch25h | 163.957365 | 5.987828131 | 0.499274634 | 11.99305497 |
| Gm26902 | 14.57275529 | 0.648419825 | 0.376575709 | 1.721884362 |
| Lipa | 315.4234948 | 0.117825852 | 0.154815323 | 0.761073579 |
| Ifit2 | 10029.24711 | 6.511410024 | 0.281724566 | 23.112681 |
| Ifit3 | 3311.123546 | 7.635983054 | 0.263709151 | 28.95607915 |
| Gm14446 | 387.1233819 | 3.382222738 | 0.465269128 | 7.269389989 |
| Ifit3b | 2018.601927 | 6.006139174 | 0.25839178 | 23.24431211 |
| 2010002M12Rik | 153.7706214 | 3.349027093 | 0.252234261 | 13.27744725 |
| Ifit1 | 4083.025541 | 8.656107765 | 0.28626423 | 30.23817462 |
| Slc16a12 | 39.40020055 | -0.508081339 | 0.210162597 | -2.417563103 |
| Gm27476 | 0 | NA | NA | NA |
| Gm22082 | 0 | NA | NA | NA |
| Pank1 | 222.2087571 | -0.563153603 | 0.105693517 | -5.328175472 |
| Mir107 | 0.848910726 | 0.5881629 | 1.186483057 | 0.495719594 |
| Kif20b | 26.85031262 | 0.972661985 | 0.263427568 | 3.692331799 |
| Mir1950 | 0 | NA | NA | NA |
| Gm5248 | 0 | NA | NA | NA |
| Gm24006 | 0 | NA | NA | NA |
| Gm23075 | 0 | NA | NA | NA |
| Gm22853 | 0 | NA | NA | NA |
| Gm25611 | 0.204668643 | -0.526358365 | 2.595258787 | -0.202815368 |

| | | | | |
|---|---|---|---|---|
| A830019P07Rik | 16.97608067 | -0.18233529 | 0.310273562 | -0.587659768 |
| Htr7 | 177.6287066 | 0.264109768 | 0.111263639 | 2.373729383 |
| Rpp30 | 118.3865456 | 0.145147487 | 0.121062034 | 1.198951337 |
| Ankrd1 | 0.384174469 | 2.33093856 | 1.651755651 | 1.411188489 |
| Pcgf5 | 290.0103938 | 0.999897498 | 0.112458555 | 8.891253295 |
| Gm23170 | 0 | NA | NA | NA |
| Hectd2 | 578.0733075 | -0.368172516 | 0.105558693 | -3.487846479 |
| 1500017E21Rik | 5.192905494 | -0.773605923 | 0.454899608 | -1.700608022 |
| Ppp1r3c | 367.6091572 | -0.577743601 | 0.178961044 | -3.228320475 |
| Tnks2 | 2533.779749 | -0.021569068 | 0.059089213 | -0.36502547 |
| Fgfbp3 | 45.89989236 | -0.544510499 | 0.201889821 | -2.697067613 |
| Btaf1 | 981.4939569 | -0.043599566 | 0.090137089 | -0.483702844 |
| Cpeb3 | 1630.611145 | -0.194851459 | 0.050566603 | -3.853362662 |
| Gm22714 | 7.127945666 | -0.642774226 | 0.437210834 | -1.470169939 |
| A330032B11Rik | 4.137844924 | -0.159002884 | 0.608597897 | -0.261260982 |
| Gm23918 | 1.204149766 | 0.456076222 | 1.009235623 | 0.451902619 |
| 5-Mar | 1036.41543 | 1.056260526 | 0.121146076 | 8.71890005 |
| Gm25268 | 0 | NA | NA | NA |
| Ide | 826.1150129 | -0.118749247 | 0.067926403 | -1.748204561 |
| Rpl10-ps6 | 9.176167568 | -0.434225844 | 0.337124252 | -1.288029092 |
| Kif11 | 42.94126884 | 0.81589104 | 0.277255861 | 2.942736857 |
| Gm38345 | 1.850005102 | 1.135759005 | 0.920527896 | 1.233812695 |
| Hhex | 20.51235511 | 0.954336963 | 0.342201103 | 2.788819073 |
| Gm23026 | 0 | NA | NA | NA |
| Exoc6 | 443.4655257 | 0.021259595 | 0.076856062 | 0.276615715 |
| Cyp26c1 | 3.181018617 | -0.451057781 | 0.697988852 | -0.646224907 |
| Cyp26a1 | 1.890572446 | 0.786685017 | 0.710060152 | 1.107913202 |
| Myof | 324.4012162 | 0.918475067 | 0.277143682 | 3.314075422 |
| I830134H01Rik | 0.540537234 | -2.057353608 | 1.269207492 | -1.620974996 |
| Cep55 | 11.79181242 | 1.720968672 | 0.432951607 | 3.974967744 |
| Gm23300 | 0 | NA | NA | NA |
| Ffar4 | 0.430805093 | 0.761628303 | 1.566287069 | 0.486263545 |
| Rbp4 | 74.47465664 | 0.035733481 | 0.184135433 | 0.194060862 |
| Pde6c | 1.452976049 | 0.0519199 | 0.871260789 | 0.059591687 |
| Fra10ac1 | 83.34982365 | -0.239777877 | 0.173570527 | -1.381443504 |
| Lgi1 | 1139.244008 | -0.028435963 | 0.092354369 | -0.307900569 |
| RP24-417P12.1 | 0.567844938 | -0.051059655 | 1.291545717 | -0.039533757 |
| Slc35g1 | 45.27465032 | -0.051630271 | 0.18509598 | -0.278937829 |
| Gm24930 | 0 | NA | NA | NA |
| Plce1 | 890.1324761 | 0.801275552 | 0.135500878 | 5.913434389 |
| Gm28991 | 0 | NA | NA | NA |
| Noc3l | 300.5445623 | -0.002078125 | 0.080135912 | -0.025932509 |
| Tbc1d12 | 321.3981898 | -0.16574755 | 0.09019258 | -1.837707167 |
| Gm25838 | 2.422369998 | -0.840435693 | 0.663193034 | -1.267256514 |
| Hells | 136.3095865 | 0.023376712 | 0.145313415 | 0.160870983 |
| Cyp2c55 | 1.660109277 | -0.060316762 | 0.813747404 | -0.074122217 |
| Gm23800 | 0 | NA | NA | NA |

| | | | | |
|---|---|---|---|---|
| Cyp2c65 | 0.792025016 | 1.639336349 | 1.224230761 | 1.339074626 |
| Cyp2c66 | 0.061920613 | 0.274393828 | 3.724945049 | 0.073663859 |
| Cyp2c53-ps | 0.038981395 | 0.274393828 | 3.724945049 | 0.073663859 |
| Cyp2c29 | 0.220032676 | -0.206488847 | 2.588496343 | -0.079771736 |
| Cyp2c38 | 0.045882455 | 0.755288839 | 3.724945049 | 0.202765096 |
| Cyp2c39 | 0.206165386 | -0.687378167 | 2.656893911 | -0.258714947 |
| Cyp2c67 | 0 | NA | NA | NA |
| Cyp2c68 | 0.045882455 | 0.755288839 | 3.724945049 | 0.202765096 |
| Cyp2c40 | 0.138763737 | -0.206480411 | 2.930963593 | -0.070447962 |
| Cyp2c69 | 0.075043205 | 0.755286614 | 3.724945049 | 0.202764498 |
| Cyp2c37 | 0.045882455 | 0.755288839 | 3.724945049 | 0.202765096 |
| Cyp2c54 | 0.045882455 | 0.755288839 | 3.724945049 | 0.202765096 |
| Cyp2c50 | 0.153424464 | -0.206487971 | 2.834765573 | -0.072841286 |
| Cyp2c70 | 0.309994989 | -0.60702744 | 1.723884649 | -0.352127644 |
| Gm16470 | 3.026502306 | 0.220979977 | 0.620312552 | 0.356239732 |
| Pdlim1 | 63.17543641 | -0.004366424 | 0.188463531 | -0.023168533 |
| Sorbs1 | 1628.853159 | 0.356602625 | 0.09317115 | 3.827393192 |
| Aldh18a1 | 344.8097266 | 0.036791711 | 0.117902217 | 0.312052746 |
| Gm15801 | 18.90733818 | 0.479282223 | 0.263127205 | 1.821484871 |
| Gm27042 | 1.18341529 | -0.504423338 | 1.01159097 | -0.498643576 |
| Tctn3 | 216.4391839 | -0.205563724 | 0.100804447 | -2.039232686 |
| Entpd1 | 233.1810674 | 0.870122974 | 0.184203063 | 4.723716091 |
| Gm16027 | 0.848235736 | 0.369313584 | 1.066125353 | 0.34640728 |
| Gm8783 | 30.04226012 | 0.397078537 | 0.200257444 | 1.982840329 |
| Ccnj | 128.7530246 | -0.031244103 | 0.128475387 | -0.243191351 |
| E030044B06Rik | 1.397451337 | 1.000103349 | 0.9152755 | 1.092680126 |
| Mir8092 | 0 | NA | NA | NA |
| Zfp518a | 366.4925213 | 0.421323025 | 0.114401318 | 3.682851152 |
| Blnk | 124.657933 | 1.430164684 | 0.193612722 | 7.386728864 |
| Dntt | 1.408550033 | 2.063204953 | 1.319901178 | 1.563151081 |
| Opalin | 167.6099803 | -0.009257408 | 0.136918346 | -0.067612617 |
| Tll2 | 11.98457804 | 0.047181162 | 0.305629268 | 0.154373836 |
| Tm9sf3 | 2559.714795 | 0.035228403 | 0.058366375 | 0.603573598 |
| Pik3ap1 | 444.6212228 | 3.985884712 | 0.195629003 | 20.37471263 |
| Gm9788 | 0.072025462 | 0.755288839 | 3.724945049 | 0.202765096 |
| Lcor | 579.5255258 | -0.123595029 | 0.113361367 | -1.090274691 |
| Gm340 | 323.3061025 | 0.29460311 | 0.112842096 | 2.61075538 |
| Mir8091 | 15.23441784 | 0.467084472 | 0.296712215 | 1.574200349 |
| AI606181 | 120.0613128 | 0.168116609 | 0.124532667 | 1.349979997 |
| Slit1 | 868.1752054 | 0.075718524 | 0.098420255 | 0.769338832 |
| Gm22601 | 0.086648695 | -0.687376175 | 3.724945049 | -0.184533239 |
| Arhgap19 | 33.0013039 | -0.132813652 | 0.180418729 | -0.736141155 |
| Frat1 | 29.17957322 | 0.286072074 | 0.212867957 | 1.343894491 |
| Frat2 | 52.5338819 | -0.114694948 | 0.149785419 | -0.765728392 |
| Rrp12 | 552.962244 | 0.143029452 | 0.086116709 | 1.660879212 |
| Pgam1 | 1666.384237 | -0.108389368 | 0.061364489 | -1.766320717 |
| Exosc1 | 140.0755093 | 0.103129704 | 0.098805912 | 1.043760452 |

| | | | | |
|---|---|---|---|---|
| Zdhhc16 | 217.4843809 | 0.177546658 | 0.087872913 | 2.020493582 |
| Mms19 | 424.2345625 | 0.148258814 | 0.116173168 | 1.276188095 |
| Gm23113 | 0 | NA | NA | NA |
| Ubtd1 | 125.2316124 | 0.445874481 | 0.157852338 | 2.824630194 |
| Ankrd2 | 4.166437747 | 0.834980161 | 0.525332087 | 1.589433012 |
| Hoga1 | 9.277047816 | 0.35703549 | 0.416314788 | 0.857609434 |
| 4933411K16Rik | 1.522479514 | -0.55259942 | 0.773499698 | -0.714414526 |
| Morn4 | 316.6990762 | -0.168453744 | 0.100411564 | -1.677632915 |
| Pi4k2a | 1103.365836 | 0.692047312 | 0.106861815 | 6.476095406 |
| Avpi1 | 112.560822 | -0.232663073 | 0.131105164 | -1.774629355 |
| Marveld1 | 84.73861597 | 0.027829117 | 0.125118838 | 0.222421482 |
| Zfyve27 | 786.8275757 | -0.008018325 | 0.074237442 | -0.108009173 |
| Sfrp5 | 34.79157235 | -0.375597092 | 0.3448494 | -1.089162667 |
| Golga7b | 962.3374335 | -0.316361107 | 0.06988765 | -4.52670976 |
| Gm26339 | 1.445692662 | 1.685183379 | 0.953816346 | 1.766779722 |
| Mir3085 | 0.135858026 | -1.143903624 | 2.941468349 | -0.388888639 |
| Crtac1 | 458.2694328 | -0.316143645 | 0.104094892 | -3.037071636 |
| R3hcc1l | 325.8532248 | -0.063380752 | 0.09324914 | -0.679692621 |
| Gm16541 | 0.261817563 | 1.720971234 | 2.44413035 | 0.704124162 |
| Loxl4 | 7.914160501 | 2.127410497 | 0.486801756 | 4.37017835 |
| Gm25216 | 0 | NA | NA | NA |
| Pyroxd2 | 33.56925608 | -0.445079785 | 0.235443525 | -1.890388727 |
| Hps1 | 90.94188656 | 0.279320956 | 0.149108171 | 1.87327733 |
| Gm16244 | 3.388720562 | 0.354085066 | 0.558128426 | 0.634415037 |
| Hpse2 | 1.819823472 | 1.403579318 | 0.702445686 | 1.998132162 |
| Gm22135 | 0 | NA | NA | NA |
| Gm6776 | 3.379576263 | -1.25254759 | 0.581478641 | -2.154073256 |
| Cnnm1 | 1191.472548 | -0.264458976 | 0.08064663 | -3.279231594 |
| Got1 | 2463.067521 | -0.007419856 | 0.062940985 | -0.117885926 |
| Gm20467 | 0.07796279 | 0.274392189 | 3.724945049 | 0.073663419 |
| Nkx2-3 | 0.04393175 | -0.206490061 | 3.724945049 | -0.055434391 |
| Gm22646 | 0 | NA | NA | NA |
| Slc25a28 | 398.2739226 | 1.114311648 | 0.134914328 | 8.259401819 |
| Entpd7 | 424.4745962 | -0.072269151 | 0.095552487 | -0.756329354 |
| Cox15 | 306.6258307 | 0.56054148 | 0.117642929 | 4.764769841 |
| Cutc | 59.7964683 | 1.000251564 | 0.188744042 | 5.299513315 |
| Abcc2 | 1.945998912 | 0.805169837 | 0.695431853 | 1.157798328 |
| Dnmbp | 248.0196757 | -0.092928782 | 0.10881755 | -0.853987076 |
| Gm24400 | 2.333191932 | -1.077385847 | 0.625547802 | -1.72230778 |
| n-R5s21 | 0 | NA | NA | NA |
| Cpn1 | 2.984315308 | -0.638761892 | 0.624602615 | -1.022669256 |
| Cyp2c44 | 1.49985572 | -2.967723751 | 1.011166803 | -2.934949744 |
| Erlin1 | 312.6811813 | -0.019851065 | 0.078730788 | -0.252138529 |
| Chuk | 597.7287007 | -0.124174698 | 0.086240787 | -1.439860438 |
| Cwf19l1 | 148.8001304 | 0.121723231 | 0.099219756 | 1.226804376 |
| Gm24336 | 70.39117647 | 0.200494948 | 0.144519246 | 1.387323512 |
| Bloc1s2 | 164.8635678 | 0.161152655 | 0.121579481 | 1.325492214 |

| | | | | |
|---|---|---|---|---|
| Pkd2l1 | 8.957326903 | 0.943645779 | 0.490332872 | 1.92450034 |
| Scd3 | 92.32162226 | -0.480833188 | 0.151986555 | -3.16365608 |
| Scd2 | 13180.9979 | -0.652608933 | 0.125846411 | -5.185757199 |
| Mir5114 | 0 | NA | NA | NA |
| Scd4 | 17.24753904 | -0.194840319 | 0.276676337 | -0.704217505 |
| Scd1 | 2469.814438 | -0.293123163 | 0.14191889 | -2.065427395 |
| Wnt8b | 3.609855578 | -0.180223942 | 0.492767301 | -0.365738435 |
| Sec31b | 4.76996336 | -0.223607545 | 0.51831555 | -0.431411994 |
| Gm20538 | 0.061920613 | 0.274393828 | 3.724945049 | 0.073663859 |
| Ndufb8 | 791.3026735 | -0.059833932 | 0.119519776 | -0.50061951 |
| Hif1an | 511.8430662 | -0.376660858 | 0.072558541 | -5.191130567 |
| Gm26644 | 1.498020843 | -0.512256631 | 0.838538865 | -0.610891937 |
| Pax2 | 56.35874315 | 0.073377489 | 0.303744199 | 0.241576594 |
| Gm20395 | 0 | NA | NA | NA |
| Mir6405 | 0.042716945 | -0.206490061 | 3.724945049 | -0.055434391 |
| 1700039E22Rik | 0.097899064 | -0.687371727 | 3.724945049 | -0.184532045 |
| Fam178a | 1155.011791 | -0.074350525 | 0.091494725 | -0.812620887 |
| Gm25482 | 3.723338729 | 0.572471536 | 0.643903184 | 0.889064615 |
| Sema4g | 448.4563521 | -0.278905744 | 0.10573994 | -2.637657486 |
| Mrpl43 | 278.1537674 | 0.180172699 | 0.096094032 | 1.874962423 |
| Peo1 | 143.5811533 | -0.459934358 | 0.113674255 | -4.046073195 |
| Lzts2 | 639.0048677 | 0.261944791 | 0.091081783 | 2.875929557 |
| Pdzd7 | 38.25076259 | -0.572915118 | 0.320521576 | -1.78744634 |
| Sfxn3 | 897.777185 | -0.0401231 | 0.042892095 | -0.935442767 |
| Kazald1 | 26.8175025 | 0.037485407 | 0.21419495 | 0.175006026 |
| Tlx1 | 0.675592865 | 0.219745128 | 1.134298118 | 0.193727844 |
| Tlx1os | 0 | NA | NA | NA |
| Lbx1 | 26.55161603 | -0.027837352 | 0.332950574 | -0.083608062 |
| Gm29595 | 9.837264003 | 0.071642838 | 0.321698861 | 0.222701559 |
| Gm29593 | 4.239070863 | -0.966209201 | 0.598193528 | -1.615211725 |
| Gm29594 | 0.57177649 | -0.593562688 | 1.249263829 | -0.475129972 |
| Gm28578 | 19.28251988 | -0.520885586 | 0.378774031 | -1.375188222 |
| Btrc | 1150.587465 | -0.060776043 | 0.064839495 | -0.937330599 |
| Gm6807 | 12.07198287 | 0.52943494 | 0.31473669 | 1.682151961 |
| Poll | 98.22017834 | -0.016017689 | 0.138431981 | -0.115708012 |
| Dpcd | 221.260172 | 0.440846939 | 0.141355684 | 3.11870685 |
| Fbxw4 | 248.8152368 | -0.077550761 | 0.101811334 | -0.761710497 |
| Fgf8 | 0.275408967 | -0.675136508 | 2.070152497 | -0.326128877 |
| Npm3 | 55.30170471 | -0.29319738 | 0.206240437 | -1.421628973 |
| Gm15491 | 0.865681968 | -0.459920386 | 1.157461288 | -0.397352715 |
| Mgea5 | 3029.551855 | 0.203306638 | 0.106226506 | 1.913897435 |
| Kcnip2 | 682.9003887 | -0.332659842 | 0.158695922 | -2.096209141 |
| 9130011E15Rik | 279.4855959 | 0.139590291 | 0.077534948 | 1.800353193 |
| 4930505N22Rik | 0.892093866 | -1.930132014 | 1.111378462 | -1.736700935 |
| Hps6 | 95.93381243 | 0.141095858 | 0.158216998 | 0.891786975 |
| Ldb1 | 957.9423255 | -0.020209927 | 0.076879699 | -0.262877289 |
| Pprc1 | 454.9862607 | -0.000527397 | 0.090430557 | -0.005832063 |

| | | | | |
|---|---|---|---|---|
| Nolc1 | 518.4413524 | 0.069010421 | 0.110817885 | 0.62273721 |
| Elovl3 | 0.899697409 | 0.274413282 | 1.385490196 | 0.198062233 |
| Pitx3 | 0.694683682 | 0.527409734 | 1.232125953 | 0.428048555 |
| Gbf1 | 1560.460353 | -0.167448569 | 0.051120409 | -3.275571766 |
| Gm8974 | 0.178004931 | -0.206480786 | 2.299245362 | -0.08980372 |
| Gm16726 | 0.21650593 | 0.274406903 | 2.171399783 | 0.126373276 |
| Nfkb2 | 234.2002576 | 2.47667215 | 0.242323367 | 10.22052549 |
| Psd | 2382.095847 | -0.235041024 | 0.119698238 | -1.963613073 |
| Fbxl15 | 156.8858204 | 0.031240523 | 0.113256505 | 0.275838658 |
| Cuedc2 | 723.9696748 | 0.001820619 | 0.082282209 | 0.022126525 |
| Gm26792 | 1.450243399 | -1.480218249 | 0.970956764 | -1.524494503 |
| Tmem180 | 197.5786801 | 0.013113732 | 0.105074397 | 0.124804261 |
| Mir146b | 0.495786544 | -0.566657931 | 1.421708874 | -0.398575223 |
| 2310034G01Rik | 6.1931368 | -1.152028808 | 0.507658645 | -2.269298117 |
| Gm24610 | 1.876595299 | -1.322019676 | 0.692794555 | -1.908242013 |
| Actr1a | 1762.40465 | 0.055995092 | 0.051122685 | 1.095308124 |
| Sufu | 283.8186178 | -0.052250052 | 0.094540145 | -0.55267582 |
| Trim8 | 1307.724394 | 0.127628476 | 0.07130399 | 1.789920527 |
| Arl3 | 347.6175142 | -0.306708249 | 0.118168854 | -2.595508364 |
| Sfxn2 | 104.3476033 | 0.611144534 | 0.151787876 | 4.026306633 |
| Wbp1l | 711.1586369 | 0.529779484 | 0.098193704 | 5.395249036 |
| Cyp17a1 | 0.280011045 | 0.755310321 | 1.796631694 | 0.420403538 |
| Gm6967 | 0.062426873 | 0.755288839 | 3.724945049 | 0.202765096 |
| 2010012O05Rik | 189.8383209 | 0.138378822 | 0.090331968 | 1.531892038 |
| As3mt | 170.4653533 | -0.127914142 | 0.13695932 | -0.933957189 |
| Cnnm2 | 423.4727026 | -0.245873956 | 0.068662574 | -3.58090212 |
| Nt5c2 | 724.1452932 | 0.054491355 | 0.078403586 | 0.695011005 |
| Ina | 2102.910469 | -0.196708848 | 0.113233167 | -1.73720168 |
| Pcgf6 | 132.4800137 | -0.600413627 | 0.149915594 | -4.005011171 |
| Gm23140 | 0.042716945 | -0.206490061 | 3.724945049 | -0.055434391 |
| Taf5 | 105.5145098 | -0.088048127 | 0.167296346 | -0.52630036 |
| Usmg5 | 854.0068898 | 0.105910653 | 0.140867168 | 0.751847677 |
| Pdcd11 | 426.8069572 | -0.121552966 | 0.089599573 | -1.356624384 |
| Calhm2 | 55.28273003 | 0.244224583 | 0.24824572 | 0.98380179 |
| Calhm1 | 0 | NA | NA | NA |
| Calhm3 | 0.222639008 | -0.206490445 | 2.573738039 | -0.080229783 |
| Gm6970 | 2.783210482 | 0.284544641 | 0.596851996 | 0.47674238 |
| Neurl1a | 1174.052599 | -0.332905765 | 0.103312586 | -3.222315654 |
| Sh3pxd2a | 901.8376151 | -0.168678875 | 0.079471289 | -2.1225134 |
| Mir6995 | 1.143944469 | -0.54242268 | 0.950469015 | -0.570689493 |
| Obfc1 | 85.46752745 | -0.32939121 | 0.134865788 | -2.442363001 |
| Gm19557 | 1.777092953 | 0.683121963 | 0.890355264 | 0.767246503 |
| Slk | 1543.68933 | 0.195135792 | 0.073869123 | 2.64164218 |
| Gm16068 | 0.225284295 | -0.18591703 | 1.883450619 | -0.09871086 |
| Col17a1 | 13.5201549 | 0.62458467 | 0.297352668 | 2.1004845 |
| Gm26085 | 0 | NA | NA | NA |
| Gm24805 | 0 | NA | NA | NA |

| | | | | |
|---|---|---|---|---|
| Sfr1 | 586.6289326 | 0.153963693 | 0.099683399 | 1.544526912 |
| Cfap43 | 93.17411037 | 1.093232693 | 0.193050751 | 5.662928978 |
| Mir8090 | 0.046345709 | 0.274391604 | 3.724945049 | 0.073663262 |
| Gsto1 | 187.9833352 | 0.168141606 | 0.109843634 | 1.530736015 |
| Gsto2 | 5.405473811 | -0.865244063 | 0.451882344 | -1.914755188 |
| Gm22240 | 0 | NA | NA | NA |
| Cfap58 | 6.680071795 | 1.610256712 | 0.421372334 | 3.821458084 |
| Sorcs3 | 648.2072618 | -0.045280339 | 0.142573176 | -0.317593676 |
| Gm23857 | 0 | NA | NA | NA |
| Gm27935 | 0 | NA | NA | NA |
| Rpl13a-ps1 | 0.376112881 | 1.294864633 | 3.296299608 | 0.392823707 |
| Sorcs1 | 756.9127055 | -0.056901736 | 0.155105244 | -0.366858879 |
| Gm27749 | 0 | NA | NA | NA |
| Gm26629 | 0.136194719 | -0.687371727 | 3.724945049 | -0.184532045 |
| Gm25659 | 0 | NA | NA | NA |
| Gm25644 | 0 | NA | NA | NA |
| Rpl31-ps18 | 0 | NA | NA | NA |
| Ins1 | 0 | NA | NA | NA |
| Xpnpep1 | 575.104864 | 0.345419542 | 0.084706635 | 4.077833375 |
| Mir6407 | 1.567823505 | 0.439928891 | 0.793542054 | 0.55438636 |
| 1700054A03Rik | 0.092691418 | 0.274387304 | 3.724945049 | 0.073662108 |
| Add3 | 1815.758105 | -0.030788558 | 0.094892722 | -0.324456477 |
| Gm24724 | 0 | NA | NA | NA |
| Mxi1 | 685.6681876 | -0.261688579 | 0.069970584 | -3.739979911 |
| Gm10197 | 0.330485341 | 1.196861364 | 1.766489151 | 0.677536776 |
| Smndc1 | 390.3324799 | 0.407587257 | 0.122800443 | 3.3191025 |
| 5830416P10Rik | 35.48233613 | -0.166428454 | 0.249397532 | -0.667321978 |
| 4833407H14Rik | 1.343421229 | 2.27189601 | 1.069490368 | 2.124279075 |
| Dusp5 | 436.4752899 | 0.104499741 | 0.26534666 | 0.393823465 |
| Nutf2-ps1 | 25.95248433 | -0.230205299 | 0.279407578 | -0.823904996 |
| Smc3 | 928.9007826 | -0.023728086 | 0.087819787 | -0.27019066 |
| Rbm20 | 75.31453989 | -0.427282923 | 0.132711799 | -3.219630267 |
| Gm16298 | 0.11037557 | 0.274404949 | 3.724945049 | 0.073666845 |
| Gm16299 | 1.831283359 | 0.503070011 | 0.788952548 | 0.637642925 |
| Pdcd4 | 830.0440107 | -0.054900928 | 0.076174677 | -0.720724133 |
| Bbip1 | 292.1551775 | -0.191718976 | 0.104499206 | -1.834645291 |
| Shoc2 | 875.8097201 | -0.024803886 | 0.056697422 | -0.437478199 |
| Adra2a | 223.1903651 | -0.032489931 | 0.154925437 | -0.209713342 |
| Gpam | 844.7644142 | 0.134909767 | 0.087390484 | 1.543758087 |
| Tectb | 0.98697613 | 0.159412343 | 1.056017983 | 0.150956088 |
| Mir6715 | 0 | NA | NA | NA |
| Gucy2g | 23.72729764 | -1.332606923 | 0.264605666 | -5.036199497 |
| Acsl5 | 698.4865369 | 0.558078133 | 0.07039217 | 7.928127949 |
| Zdhhc6 | 305.077262 | -0.023420886 | 0.085421736 | -0.27417946 |
| Vti1a | 524.7365872 | -0.052734856 | 0.063199482 | -0.83441912 |
| 4930552P12Rik | 3.0018596 | 2.038663626 | 0.80459698 | 2.53376992 |
| Tcf7l2 | 729.2267912 | 0.088674826 | 0.084041044 | 1.055137133 |

| | | | | |
|---|---|---|---|---|
| Ppnr | 24.84417134 | -0.474693227 | 0.397116638 | -1.195349631 |
| Gm22271 | 0 | NA | NA | NA |
| Habp2 | 0.044544648 | 0.755286614 | 3.724945049 | 0.202764498 |
| Nrap | 37.22294834 | 1.93478496 | 0.280729754 | 6.891983952 |
| Gm17197 | 0.54074031 | -0.187325006 | 1.280302004 | -0.146313139 |
| Casp7 | 137.0862569 | 2.238989533 | 0.182529236 | 12.26647074 |
| Plekhs1 | 3.487526041 | 0.854629473 | 0.566226746 | 1.509341405 |
| Dclre1a | 154.6814328 | 0.443342016 | 0.106061256 | 4.180056261 |
| Nhlrc2 | 312.5790796 | 0.1904343 | 0.130551605 | 1.458689847 |
| Adrb1 | 210.0560459 | -0.284579536 | 0.102458485 | -2.777510674 |
| Gm6990 | 0.88041345 | -1.483348247 | 1.07683704 | -1.377504852 |
| Ccdc186 | 1033.921743 | -0.06907104 | 0.060602086 | -1.139746913 |
| Tdrd1 | 1.956606073 | 0.635195583 | 0.725753108 | 0.875222683 |
| Vwa2 | 0.389464624 | 1.176497709 | 1.573647012 | 0.747624913 |
| Afap1l2 | 288.9123761 | -0.052159444 | 0.139837381 | -0.373000719 |
| Ablim1 | 1533.129369 | -0.223258723 | 0.06399787 | -3.48853362 |
| B230217O12Rik | 56.18862831 | -0.363364496 | 0.150320341 | -2.417267647 |
| Fam160b1 | 577.2204722 | -0.00632499 | 0.073382162 | -0.086192475 |
| Trub1 | 145.5468936 | -0.020069099 | 0.147060421 | -0.136468391 |
| Gm26874 | 0 | NA | NA | NA |
| Atrnl1 | 1851.542762 | -0.18763296 | 0.085375855 | -2.197728616 |
| Mir5623 | 3.511196682 | 0.083944536 | 0.787923686 | 0.106538917 |
| Gm16277 | 0.097501745 | -0.206492286 | 3.724945049 | -0.055434988 |
| Gfra1 | 699.0851246 | 0.650804674 | 0.093648119 | 6.949468697 |
| Ccdc172 | 1.84083352 | -0.049864796 | 0.800645773 | -0.062280722 |
| 1810007D17Rik | 0.526835603 | -0.224802525 | 1.593914092 | -0.141038044 |
| Pnlip | 3.471747287 | 0.376144609 | 0.521367387 | 0.721457879 |
| 1810018F18Rik | 0 | NA | NA | NA |
| Pnliprp1 | 0.093484797 | -0.20649451 | 3.724945049 | -0.055435586 |
| Pnliprp2 | 0.200944149 | -0.20648068 | 2.220319025 | -0.092995951 |
| 1700019N19Rik | 1.010445843 | 1.283654229 | 1.116491937 | 1.149721002 |
| Gm22561 | 0 | NA | NA | NA |
| Gm17150 | 0 | NA | NA | NA |
| Hspa12a | 2209.460681 | -0.24943992 | 0.080020454 | -3.117202017 |
| Mir3086 | 1.922482983 | -0.476940099 | 0.670557297 | -0.711259279 |
| Eno4 | 32.39598207 | 0.242229017 | 0.184557098 | 1.31248822 |
| Shtn1 | 427.0917043 | -0.052801105 | 0.105343212 | -0.501229302 |
| Vax1 | 9.031124418 | 0.342215086 | 0.40733947 | 0.840122578 |
| Kcnk18 | 0.044191549 | -0.206490061 | 3.724945049 | -0.055434391 |
| Gm29261 | 1.407425821 | 0.369429697 | 0.925402033 | 0.399209948 |
| Slc18a2 | 109.7658993 | 0.037910859 | 0.300582506 | 0.126124634 |
| Pdzd8 | 1685.906693 | -0.043683704 | 0.063482142 | -0.688125859 |
| Rps12-ps3 | 13.1319493 | 0.474735371 | 0.331927425 | 1.430238464 |
| Emx2os | 17.8875887 | -0.524003903 | 0.308201816 | -1.700197327 |
| Emx2 | 54.25553675 | -0.178205603 | 0.16929659 | -1.052623695 |
| 2700089I24Rik | 0.185374628 | -0.680385936 | 2.266154646 | -0.300238087 |
| Gm17203 | 36.89967017 | -0.231545476 | 0.236392644 | -0.979495268 |

| | | | | |
|---|---|---|---|---|
| Rab11fip2 | 973.2053457 | 0.064163235 | 0.068962853 | 0.930402854 |
| Gm4219 | 3.097868212 | -0.311258932 | 0.620486725 | -0.50163673 |
| E330013P04Rik | 2.49875752 | -1.673638684 | 0.748280415 | -2.236646384 |
| Fam204a | 247.5678764 | 0.116758839 | 0.106124982 | 1.100201263 |
| 4933412A08Rik | 0.53212218 | 1.185158149 | 1.300345847 | 0.911417645 |
| Gm28351 | 0 | NA | NA | NA |
| Prlhr | 4.925174813 | -0.389264638 | 0.439695335 | -0.885305362 |
| Cacul1 | 885.5461787 | 0.218965316 | 0.07926888 | 2.762311203 |
| Gm25238 | 3.512009492 | 0.340981703 | 0.552134244 | 0.617570286 |
| Nanos1 | 296.0425935 | 0.046327425 | 0.104869442 | 0.441762861 |
| Eif3a | 2000.929003 | 0.186295576 | 0.118365209 | 1.573904836 |
| Snora19 | 2.074201168 | 0.210064059 | 0.696320838 | 0.301677112 |
| Gm26158 | 3.094791288 | 1.565446329 | 0.65117338 | 2.404039198 |
| Fam45a | 537.0189958 | 0.357629115 | 0.106601746 | 3.354814785 |
| Gm15959 | 0 | NA | NA | NA |
| Sfxn4 | 143.5137547 | -0.382134163 | 0.143757944 | -2.658177718 |
| Prdx3 | 403.3254737 | 0.071080947 | 0.123900796 | 0.573692415 |
| Grk5 | 237.0149773 | -0.26632705 | 0.091421853 | -2.913166188 |
| Gm9529 | 0.17796667 | -0.684257857 | 2.715150213 | -0.252014733 |
| Gm22520 | 0 | NA | NA | NA |
| 5830428H23Rik | 346.0755543 | -0.83768133 | 0.223811685 | -3.742795327 |
| Gm22365 | 0 | NA | NA | NA |
| Gm7102 | 0.828685883 | -0.951165344 | 1.153340278 | -0.824704871 |
| Gm6020 | 0 | NA | NA | NA |
| Csf2ra | 189.2145181 | 0.521054885 | 0.154492407 | 3.372689288 |
| Gm21060 | 0 | NA | NA | NA |
| Gm37392 | 0 | NA | NA | NA |
| Gm27306 | 0 | NA | NA | NA |
| Fam171a1 | 789.1479404 | -0.179369621 | 0.076863519 | -2.333611863 |
| Nmt2 | 1499.866317 | 0.299051833 | 0.07026782 | 4.255886014 |
| Gm22005 | 0.075043205 | 0.755286614 | 3.724945049 | 0.202764498 |
| Rpp38 | 43.57699851 | -0.402529132 | 0.215064611 | -1.871666059 |
| Acbd7 | 3.006698544 | 0.388014451 | 0.725845608 | 0.534568848 |
| Olah | 0.742270129 | 2.639387976 | 1.116529303 | 2.363921815 |
| Gm37525 | 0.038981395 | 0.274393828 | 3.724945049 | 0.073663859 |
| Meig1 | 9.115918498 | -0.235968016 | 0.347483638 | -0.679076624 |
| Dclre1c | 259.4107841 | 0.47146934 | 0.15323053 | 3.076862949 |
| Suv39h2 | 54.79723866 | -0.430884682 | 0.159760534 | -2.697065866 |
| Gm13184 | 2.921456852 | 1.630373516 | 0.752664736 | 2.166135118 |
| Hspa14 | 484.0326706 | 0.117723737 | 0.114251018 | 1.030395513 |
| Cdnf | 15.49217047 | 0.048148891 | 0.268799056 | 0.179125969 |
| Gm13186 | 0.075043205 | 0.755286614 | 3.724945049 | 0.202764498 |
| Gm13182 | 0.275763028 | -0.628040219 | 1.987791608 | -0.315948722 |
| Fam107b | 448.5992859 | 0.428576509 | 0.176528489 | 2.427803648 |
| Gm13183 | 0 | NA | NA | NA |
| Gm13181 | 0.648853987 | -1.153291849 | 1.347865663 | -0.855643021 |
| Gm13185 | 0.04393175 | -0.206490061 | 3.724945049 | -0.055434391 |

| | | | | |
|---|---|---|---|---|
| Gm13180 | 0.088736197 | 0.274400501 | 3.724945049 | 0.073665651 |
| Gm13191 | 0.045882455 | 0.755288839 | 3.724945049 | 0.202765096 |
| Frmd4a | 1510.085395 | 0.17698243 | 0.063028139 | 2.807990738 |
| Gm2639 | 0.385855529 | -0.433640638 | 3.262688888 | -0.132908976 |
| Gm13188 | 0 | NA | NA | NA |
| Gm10862 | 12.42465393 | 0.76284716 | 0.304248418 | 2.507316767 |
| Gm38085 | 10.53103825 | 0.749748263 | 0.422531283 | 1.774420718 |
| 1700080N15Rik | 0 | NA | NA | NA |
| Gm13189 | 2.288528955 | 0.254318436 | 0.692282817 | 0.367362052 |
| Gm37814 | 13.6296294 | 0.685305031 | 0.426506534 | 1.606786712 |
| Gm37107 | 25.13242011 | -0.645419295 | 0.353057015 | -1.828088007 |
| Gm38348 | 18.53952862 | -0.569513437 | 0.411672522 | -1.383413773 |
| Gm13187 | 1.991338835 | -0.743511899 | 0.822621423 | -0.903832406 |
| Gm36988 | 22.65632828 | -1.29847508 | 0.338842714 | -3.832087948 |
| Gm23691 | 0 | NA | NA | NA |
| Gm13175 | 0 | NA | NA | NA |
| Gm13179 | 0.804355203 | 1.904210715 | 1.213121717 | 1.569678202 |
| Prpf18 | 625.4248675 | 0.102664063 | 0.06503867 | 1.578508031 |
| Gm13196 | 0.049553047 | 0.274391604 | 3.724945049 | 0.073663262 |
| Bend7 | 58.21736025 | -0.287987137 | 0.146352261 | -1.967766915 |
| Gm13176 | 0 | NA | NA | NA |
| Sephs1 | 350.7542985 | 0.097147402 | 0.079887921 | 1.2160462 |
| Phyh | 939.6459375 | 0.73241136 | 0.083699147 | 8.750523543 |
| Gm13194 | 2.056906148 | 1.260111926 | 0.896029887 | 1.406328008 |
| Gm13193 | 0 | NA | NA | NA |
| Gm13192 | 14.54002108 | 0.193253694 | 0.262897268 | 0.73509206 |
| Ucma | 8.601202928 | 0.693875566 | 0.39043873 | 1.777168895 |
| Mcm10 | 26.1128149 | 1.311477927 | 0.408201783 | 3.212817731 |
| Optn | 314.9999699 | 0.329989606 | 0.090886487 | 3.630788421 |
| 4930551O13Rik | 0.150455055 | 0.274400633 | 2.848105408 | 0.096344971 |
| Gm26776 | 1.257166043 | 1.202598214 | 1.105151843 | 1.088174645 |
| Ccdc3 | 115.3722469 | 0.258403798 | 0.115684718 | 2.233690008 |
| Gm13198 | 1.380091768 | -0.693662376 | 0.873329646 | -0.794273249 |
| Gm23118 | 0 | NA | NA | NA |
| Gm13197 | 0.068097359 | -0.206487837 | 3.724945049 | -0.055433794 |
| Camk1d | 1938.800003 | 0.186183117 | 0.091523564 | 2.034264274 |
| Gm13216 | 0.097501745 | -0.206492286 | 3.724945049 | -0.055434988 |
| Cdc123 | 504.6193172 | 0.071065685 | 0.070553288 | 1.00726255 |
| Nudt5 | 187.3923638 | -0.467926175 | 0.129862904 | -3.603232028 |
| Gm13199 | 8.870008164 | -0.488297281 | 0.397486113 | -1.228463749 |
| Sec61a2 | 789.7387573 | -0.269392559 | 0.112755303 | -2.389178623 |
| Gm13267 | 0.3753498 | 0.274413637 | 1.666784745 | 0.164636518 |
| Dhtkd1 | 85.77264844 | -0.000576426 | 0.14776135 | -0.003901062 |
| Upf2 | 873.5434251 | 0.105323051 | 0.079751926 | 1.32063332 |
| Proser2 | 48.48109559 | -0.389389206 | 0.164309306 | -2.369854856 |
| Gm13384 | 0 | NA | NA | NA |
| Gm10857 | 0.297167372 | 1.278704114 | 1.748534056 | 0.731300663 |

| | | | | |
|---|---|---|---|---|
| Echdc3 | 8.401856911 | 0.276850968 | 0.400641881 | 0.691018539 |
| A230108P19Rik | 0.266247585 | -0.155954515 | 2.501021303 | -0.062356332 |
| Gm13383 | 0.729279448 | 0.181295155 | 1.350497001 | 0.134243286 |
| Usp6nl | 655.1216898 | 0.045857159 | 0.07135131 | 0.642695399 |
| Gm13388 | 0.223035371 | 1.698867651 | 2.557900941 | 0.664164755 |
| Gm13391 | 5.144468292 | 1.225749639 | 0.510331035 | 2.401871635 |
| Gm38386 | 2.604362067 | -0.715975399 | 0.858522136 | -0.833962654 |
| Celf2 | 6175.024994 | -0.168866005 | 0.120444623 | -1.402021942 |
| Gm13389 | 0.654952557 | 0.57013022 | 1.400154315 | 0.407190989 |
| Gm10855 | 169.6995702 | -0.773341911 | 0.496927182 | -1.556247956 |
| Gm37340 | 0.066050875 | 0.274391604 | 3.724945049 | 0.073663262 |
| Gm10115 | 101.9444587 | -0.733928721 | 0.467613342 | -1.569520487 |
| Gm26315 | 191.8007175 | -0.22953736 | 0.252074594 | -0.910592997 |
| Gm26565 | 0.531771505 | -0.406673749 | 1.355369006 | -0.300046516 |
| Gm24340 | 0 | NA | NA | NA |
| Gm28641 | 0.190780345 | -1.424321918 | 3.283197437 | -0.433821586 |
| Gm13210 | 0.681349771 | -1.844650486 | 1.557907125 | -1.184056775 |
| Gm13211 | 0.138642343 | 1.152959107 | 3.605171821 | 0.319806978 |
| Gm13254 | 0.062426873 | 0.755288839 | 3.724945049 | 0.202765096 |
| Gm24534 | 0 | NA | NA | NA |
| Gm13255 | 0 | NA | NA | NA |
| Gm13217 | 27.36624143 | 0.023191711 | 0.186551966 | 0.124317696 |
| 1700061F12Rik | 0 | NA | NA | NA |
| Gm23963 | 0 | NA | NA | NA |
| Gm13221 | 0 | NA | NA | NA |
| Gm13219 | 0 | NA | NA | NA |
| Gm13220 | 0 | NA | NA | NA |
| Gm13218 | 0 | NA | NA | NA |
| Gm37866 | 0 | NA | NA | NA |
| Gata3 | 51.8974266 | 0.337886153 | 0.164934926 | 2.048602807 |
| Gm13256 | 0.066050875 | 0.274391604 | 3.724945049 | 0.073663262 |
| 4930412O13Rik | 0.725537781 | 1.488080695 | 1.568503487 | 0.948726418 |
| 9230102O04Rik | 0.685158741 | -1.33973753 | 1.097518236 | -1.220697284 |
| Gm13262 | 0 | NA | NA | NA |
| Taf3 | 355.5580828 | 0.023903006 | 0.065048441 | 0.367464693 |
| C630004M23Rik | 3.262821873 | -0.229327147 | 0.693901122 | -0.330489662 |
| Atp5c1 | 1526.152544 | 0.023579114 | 0.079260578 | 0.297488538 |
| Gm23608 | 6.323166952 | -0.583910126 | 0.384537457 | -1.518473989 |
| Kin | 91.12058367 | -0.092941023 | 0.130920997 | -0.709901586 |
| Itih2 | 90.97624014 | 1.227569972 | 0.234190902 | 5.241749187 |
| Itih5 | 570.7931782 | 0.284969659 | 0.154215823 | 1.847862651 |
| Gm13261 | 0.038981395 | 0.274393828 | 3.724945049 | 0.073663859 |
| Sfmbt2 | 174.706509 | 0.694767667 | 0.126412979 | 5.496015287 |
| Gm22072 | 0.909783392 | 0.440553092 | 1.061600839 | 0.414989397 |
| Gm22677 | 0.580675992 | 1.816320518 | 1.740282076 | 1.043693171 |
| Mir466m | 0.940874839 | -0.120320309 | 0.950351671 | -0.12660609 |
| Mir466f-1 | 0.953625643 | 0.356394274 | 1.140792578 | 0.312409355 |

| | | | | |
|---|---|---|---|---|
| Mir669f | 1.177175292 | -0.135773493 | 1.078550416 | -0.125885161 |
| Mir669e | 0.868322942 | 1.691291718 | 1.084464111 | 1.55956449 |
| Mir669b | 1.131281954 | 0.540866812 | 1.109444001 | 0.487511593 |
| Gm23578 | 1.362203202 | 1.893208379 | 1.068117647 | 1.772471773 |
| Mir669d | 0.802169344 | 0.423158776 | 1.094895718 | 0.386483177 |
| Mir466f-2 | 0.667839854 | 0.722592566 | 1.432652919 | 0.504373778 |
| Mir669l | 0.610120006 | -1.08286495 | 1.159770705 | -0.933688828 |
| Mir669d-2 | 0.709755772 | 0.049756449 | 1.187868037 | 0.041887186 |
| Mir466f-3 | 0.856346278 | 1.335987303 | 1.076093046 | 1.241516529 |
| Mir297a-2 | 0.78950392 | -0.912985851 | 1.252873663 | -0.72871342 |
| Mir466o | 1.891729189 | 0.580837143 | 0.732322856 | 0.793143541 |
| Gm26073 | 1.013961535 | 0.771000204 | 0.987879995 | 0.780459376 |
| Mir467c | 0.194094924 | -0.206487125 | 2.244581821 | -0.091993583 |
| mmu-mir-466b-1 | 0.068097359 | -0.206487837 | 3.724945049 | -0.055433794 |
| Mir669a-3 | 0.890972926 | 1.364994603 | 1.323691871 | 1.031202679 |
| Mir669k | 1.409213138 | -0.795113817 | 0.833673076 | -0.953747746 |
| Gm26491 | 1.682866984 | 1.143876514 | 0.850050468 | 1.345657178 |
| Mir467a-1 | 0.921542032 | 0.409798636 | 1.118024766 | 0.366538066 |
| Mir466b-8 | 0.330565531 | 0.676487137 | 1.742413514 | 0.388247182 |
| Mir669a-1 | 1.495866474 | 0.847307594 | 0.836904398 | 1.012430567 |
| Mir669g | 0.495109938 | 0.745543106 | 1.336634804 | 0.557776218 |
| Gm25701 | 0.561285994 | 0.247403985 | 1.298319929 | 0.190557026 |
| Mir669j | 1.067120961 | 2.138121295 | 1.274821087 | 1.677193229 |
| Mir467a-2 | 0.612405649 | 0.580860597 | 1.314432573 | 0.441909771 |
| mmu-mir-466e | 0.75911489 | 1.682427234 | 1.467803914 | 1.146220702 |
| mmu-mir-669a-4 | 2.460216864 | 0.412834603 | 0.62557085 | 0.659932608 |
| Gm24315 | 0.616248996 | 0.138087988 | 1.265822905 | 0.1090895 |
| Gm25451 | 0.626970746 | -0.799575093 | 1.282039816 | -0.623674151 |
| Mir467b | 1.221212496 | -0.32696181 | 0.899651591 | -0.363431593 |
| mmu-mir-466c-1 | 1.101702777 | -0.376704792 | 1.110940551 | -0.339086364 |
| Gm24761 | 0.825397108 | 1.073349769 | 1.205643159 | 0.890271521 |
| Gm23651 | 1.10380611 | 0.249585923 | 0.930091152 | 0.268345659 |
| Mir467a-3 | 0 | NA | NA | NA |
| Mir466c-2 | 0 | NA | NA | NA |
| Gm23108 | 0.643541246 | 2.049540065 | 1.551577099 | 1.320939879 |
| Gm25680 | 0 | NA | NA | NA |
| Gm22775 | 0.32589515 | 0.731608926 | 1.788364052 | 0.409093957 |
| Mir467a-4 | 0.173861546 | 0.274400692 | 2.720914747 | 0.100848692 |
| Mir466b-4 | 0 | NA | NA | NA |
| Gm26060 | 0 | NA | NA | NA |
| Gm26431 | 0.062426873 | 0.755288839 | 3.724945049 | 0.202765096 |
| Gm26229 | 0.045882455 | 0.755288839 | 3.724945049 | 0.202765096 |
| Mir467a-5 | 0 | NA | NA | NA |
| Mir466b-5 | 0 | NA | NA | NA |
| Mir669p-1 | 0 | NA | NA | NA |
| Gm24167 | 0.038981395 | 0.274393828 | 3.724945049 | 0.073663859 |
| Gm24640 | 0 | NA | NA | NA |

| | | | | |
|---|---|---|---|---|
| Gm23288 | 0.060174537 | 0.755286614 | 3.724945049 | 0.202764498 |
| Mir467a-6 | 0 | NA | NA | NA |
| Gm25285 | 0.094831987 | 0.274404949 | 3.724945049 | 0.073666845 |
| Gm24673 | 0.458955183 | 1.254056413 | 1.427576155 | 0.878451499 |
| Gm22635 | 0.309435895 | -1.149739964 | 3.702082823 | -0.310565705 |
| Gm23569 | 0.231961655 | 0.274410909 | 2.128142741 | 0.128943846 |
| Mir467a-7 | 0 | NA | NA | NA |
| Mir466b-6 | 0 | NA | NA | NA |
| Gm26426 | 0 | NA | NA | NA |
| Gm24488 | 0.108266322 | 0.274389235 | 3.724945049 | 0.073662626 |
| Gm25158 | 0.038981395 | 0.274393828 | 3.724945049 | 0.073663859 |
| Mir467a-8 | 0 | NA | NA | NA |
| Mir466b-7 | 0.158320866 | -0.206490587 | 2.818634005 | -0.073259099 |
| Mir669p-2 | 0 | NA | NA | NA |
| Gm24028 | 0.085433889 | -0.687376175 | 3.724945049 | -0.184533239 |
| Gm22626 | 0.305016863 | -0.179707781 | 2.038473242 | -0.088158028 |
| Gm22166 | 0.060174537 | 0.755286614 | 3.724945049 | 0.202764498 |
| Mir467a-9 | 0.048949532 | -0.206487837 | 3.724945049 | -0.055433794 |
| Mir466b-2 | 0.150451996 | 0.274405555 | 2.858243839 | 0.096004949 |
| Gm26156 | 1.082398122 | -2.175337688 | 1.036686038 | -2.098357274 |
| Gm23571 | 0.95850512 | -0.462965411 | 1.063703053 | -0.435239336 |
| Gm25316 | 0.639171633 | -1.07068008 | 1.166819747 | -0.917605382 |
| Gm25022 | 0.159191956 | -0.687378025 | 2.808135645 | -0.24478092 |
| Gm25943 | 0.342780899 | 0.252270308 | 1.606282233 | 0.157052293 |
| Gm23988 | 0.869956765 | 0.909758574 | 1.202059392 | 0.756833298 |
| Gm25673 | 0.68128338 | 1.975514228 | 1.324022683 | 1.49205467 |
| Gm25753 | 0.533146218 | -1.169364677 | 1.395087588 | -0.838201621 |
| Mir467a-10 | 0 | NA | NA | NA |
| Mir466b-3 | 0.655224956 | -0.074742706 | 1.226681795 | -0.060930803 |
| Gm26398 | 0.832395869 | -0.89945455 | 1.263748921 | -0.711735167 |
| Mir467e | 0.572335319 | 1.305288149 | 1.291991962 | 1.010291231 |
| Mir466p | 1.677881506 | -0.480171893 | 0.828546025 | -0.579535571 |
| Gm22051 | 0.979291814 | 0.487382511 | 1.268275781 | 0.384287485 |
| Gm22678 | 0.741352231 | -0.017113677 | 1.081844488 | -0.015818981 |
| Mir467d | 0.189382193 | -1.154673968 | 2.701715785 | -0.427385432 |
| Mir466 | 0.816872591 | -0.318617218 | 1.111154288 | -0.286744353 |
| Gm22091 | 0.829130969 | -1.487807611 | 1.080145891 | -1.377413573 |
| Mir297c | 0.615674597 | -0.689094481 | 1.366197281 | -0.504388708 |
| Mir669c | 0.542133415 | 1.848490697 | 1.64214011 | 1.12565955 |
| Gm24342 | 0.492872509 | -1.125378582 | 1.272387966 | -0.884461825 |
| Gm25263 | 0.34438493 | 0.714209137 | 1.48136371 | 0.482129495 |
| Mir669a-2 | 0.768366922 | 0.061109929 | 1.191502577 | 0.051288122 |
| Gm22239 | 0.500671887 | 1.661061239 | 1.546857303 | 1.073829652 |
| Gm22925 | 0.21460851 | -0.193563921 | 2.598321842 | -0.074495745 |
| Gm23542 | 0.931192987 | 0.933890769 | 0.999787946 | 0.934088846 |
| Mir297b | 0.635701853 | -0.699180891 | 1.174619982 | -0.595240079 |
| Mir466d | 0.422902843 | 0.81761386 | 1.556047151 | 0.525442857 |

| | | | | |
|---|---|---|---|---|
| Gm26092 | 0.669827454 | 1.668237344 | 1.299738351 | 1.283517827 |
| Mir669m-1 | 0.636524932 | -1.310651731 | 1.318245983 | -0.994239124 |
| Mir669m-2 | 0.70834526 | 0.229509456 | 1.176176681 | 0.195131786 |
| Mir466n | 0.60136891 | -0.430069566 | 1.291472335 | -0.333007184 |
| Mir669o | 1.139979212 | 0.626589168 | 1.048551438 | 0.597575994 |
| Mir466g | 0.512204714 | -0.891479936 | 1.267260997 | -0.703469876 |
| Mir466h | 0.440637317 | 0.993827306 | 3.385008509 | 0.293596693 |
| Mir297a-3 | 0.589191525 | -0.535133295 | 1.39829755 | -0.382703449 |
| Mir466l | 0.747968401 | 2.113287168 | 1.214004587 | 1.740757153 |
| Mir297a-4 | 0.343551435 | 0.274411678 | 1.799476252 | 0.152495304 |
| Gm24806 | 0.654401101 | 0.944322792 | 1.211549323 | 0.779434047 |
| Mir669i | 0.597540007 | -1.140897841 | 1.22212314 | -0.933537549 |
| Mir669h | 0.264963494 | -0.603091194 | 2.078180522 | -0.290201543 |
| Gm18547 | 0 | NA | NA | NA |
| Gm13265 | 0.236919184 | 2.089753366 | 2.480798699 | 0.842371195 |
| Gm13263 | 0.565796537 | 2.650616027 | 1.405508211 | 1.885877298 |
| Gm37343 | 0 | NA | NA | NA |
| Gm23877 | 0 | NA | NA | NA |
| Gm13264 | 0.423046101 | 0.781636429 | 1.677724587 | 0.465890788 |
| Gm26478 | 0 | NA | NA | NA |
| Gm13297 | 0 | NA | NA | NA |
| Gm13294 | 1.544067245 | 0.853378798 | 0.796610443 | 1.071262378 |
| Prkcq | 193.2675698 | 0.502899315 | 0.123185378 | 4.082459496 |
| Gm36932 | 3.868039749 | 0.183454945 | 0.567102123 | 0.32349543 |
| Gm37520 | 0.177524535 | 0.274400303 | 2.74030457 | 0.100134965 |
| Gm37766 | 3.664845353 | 0.499274302 | 0.742926368 | 0.672037397 |
| Gm38171 | 1.860369092 | 2.306634015 | 1.149850691 | 2.006029158 |
| Gm37851 | 0 | NA | NA | NA |
| 8030442B05Rik | 2.076510791 | -0.451117225 | 0.844779768 | -0.534005717 |
| Gm13293 | 46.58195302 | 0.705062477 | 0.18981325 | 3.714506104 |
| Gm37730 | 0.669761723 | -1.721491212 | 1.258552495 | -1.367834253 |
| 4933403L11Rik | 0.631690809 | -2.262438689 | 1.13861945 | -1.987001618 |
| Gm13291 | 0.045882455 | 0.755288839 | 3.724945049 | 0.202765096 |
| Gm13292 | 11.7837924 | 0.387543677 | 0.287675717 | 1.347154638 |
| Gm13296 | 0 | NA | NA | NA |
| Pfkfb3 | 910.8026235 | 0.307187155 | 0.131143465 | 2.342374849 |
| Gm10851 | 1.168248907 | 1.625690081 | 1.268674094 | 1.281408747 |
| Gm37975 | 7.09582318 | -1.45256843 | 0.49613243 | -2.927783677 |
| Rbm17 | 378.7991564 | 0.266870352 | 0.130696813 | 2.041904055 |
| Gm13260 | 0.138087489 | 0.274387362 | 3.620422247 | 0.075788774 |
| Gm17490 | 0 | NA | NA | NA |
| Il2ra | 71.84653869 | 4.90863463 | 0.447768125 | 10.96244766 |
| Gm37881 | 0.909342915 | 1.843117589 | 2.600149921 | 0.708850507 |
| Il15ra | 192.4286151 | 1.79902008 | 0.220971115 | 8.14142644 |
| Fbxo18 | 817.0147377 | -0.313479679 | 0.135771376 | -2.308878998 |
| Ankrd16 | 279.6460949 | -0.004267914 | 0.175440814 | -0.024326803 |
| Gm34768 | 0.194631057 | 1.693379269 | 3.232136674 | 0.52391945 |

| | | | | |
|---|---|---|---|---|
| Gm13310 | 0.075043205 | 0.755286614 | 3.724945049 | 0.202764498 |
| Itga8 | 97.06132719 | -0.314857334 | 0.213558018 | -1.474340965 |
| Gm37701 | 1.141257353 | -1.034830551 | 0.839864335 | -1.232140131 |
| E030013I19Rik | 6.292898879 | 0.080497414 | 0.456535223 | 0.176322461 |
| Fam188a | 576.4656474 | 0.42708865 | 0.088656793 | 4.817325711 |
| Gm37289 | 14.48869372 | 0.878282679 | 0.342643927 | 2.56325185 |
| Gm37255 | 40.61645279 | 0.512872366 | 0.310823049 | 1.650046121 |
| Gm38014 | 40.65078467 | 0.194155376 | 0.322771304 | 0.601526139 |
| Gm13321 | 0 | NA | NA | NA |
| Gm37565 | 0.527338585 | 0.274385874 | 3.217039136 | 0.085291432 |
| Pter | 24.4696158 | -0.400507948 | 0.230833484 | -1.735051351 |
| C1ql3 | 535.3698455 | -0.273865005 | 0.080341684 | -3.408753607 |
| Gm37824 | 3.481665934 | -0.282048307 | 0.555699307 | -0.50755562 |
| Gm37811 | 9.289570475 | -0.30792304 | 0.316500127 | -0.972900209 |
| Gm37356 | 2.692269002 | 0.031905767 | 0.559037907 | 0.057072636 |
| Gm37742 | 0.85289683 | 0.9094986 | 1.111029976 | 0.818608516 |
| Rsu1 | 237.1157615 | -0.280436584 | 0.117951956 | -2.377549246 |
| Gm38156 | 11.10230631 | -1.201131328 | 0.425966527 | -2.819778671 |
| Gm37780 | 17.42481316 | -1.013089963 | 0.292057877 | -3.468798629 |
| Gm27540 | 0 | NA | NA | NA |
| Gm37160 | 10.15377686 | -1.091390146 | 0.332519851 | -3.282180423 |
| Gm13270 | 8.36952636 | -0.386176257 | 0.401413395 | -0.962041281 |
| Cubn | 24.44027687 | -0.578125361 | 0.263364744 | -2.195150922 |
| Trdmt1 | 114.6086781 | 0.537224642 | 0.127931587 | 4.199311947 |
| Vim | 4040.904579 | 2.247716675 | 0.283345188 | 7.932785776 |
| St8sia6 | 68.51491012 | -0.025783173 | 0.159713377 | -0.161434024 |
| Gm37126 | 0 | NA | NA | NA |
| Hacd1 | 81.82130967 | -0.332668324 | 0.138840948 | -2.396038988 |
| D930036K23Rik | 55.51995555 | -0.394301454 | 0.208141987 | -1.894386906 |
| Gm37894 | 18.6674481 | -0.415116187 | 0.289351896 | -1.434641322 |
| Stamos | 3.400319483 | -0.502354409 | 0.597623845 | -0.840586286 |
| Stam | 1143.237138 | 0.095060844 | 0.058679268 | 1.620007324 |
| Gm37499 | 111.4820659 | -0.216118787 | 0.128205213 | -1.685725427 |
| Tmem236 | 7.691974964 | 0.873137819 | 0.491595554 | 1.776130424 |
| Gm13320 | 0.278513079 | 1.333749957 | 2.083018119 | 0.640296858 |
| Mrc1 | 62.46851479 | -0.450061478 | 0.275647465 | -1.632743031 |
| Mir511 | 0.109727584 | 0.755282166 | 3.724945049 | 0.202763304 |
| Gm37697 | 0.16123844 | -1.17716678 | 2.795194053 | -0.421139555 |
| Slc39a12 | 432.812937 | -0.59365248 | 0.174461407 | -3.402772512 |
| Gm13266 | 1.979745462 | 0.620646591 | 0.699503072 | 0.887267856 |
| Cacnb2 | 1007.083002 | 0.370860059 | 0.150970679 | 2.456503881 |
| Gm10849 | 2.270217088 | 0.225693995 | 0.707683362 | 0.318919459 |
| Gm38105 | 15.2328292 | 0.288702073 | 0.29143948 | 0.99060729 |
| Gm13268 | 1.66036403 | 1.924029396 | 0.92894407 | 2.071200472 |
| Gm13312 | 24.04166377 | 0.341469018 | 0.436271259 | 0.782698861 |
| 2900092N22Rik | 49.08839729 | 0.287266316 | 0.312015438 | 0.920679815 |
| Gm13314 | 0.346172528 | -0.128829361 | 1.662683993 | -0.07748277 |

| | | | | |
|---|---|---|---|---|
| Gm13311 | 0.601079571 | 1.45458189 | 1.262875695 | 1.151801318 |
| Gm10848 | 3.555860714 | 1.117779673 | 0.62458115 | 1.789646827 |
| Gm37006 | 66.77607684 | 0.235431098 | 0.346278634 | 0.679889183 |
| Gm13315 | 24.30272673 | 0.341976312 | 0.421973662 | 0.810420988 |
| Gm38273 | 9.78169311 | 0.120652127 | 0.381339291 | 0.316390495 |
| Gm13269 | 0.198547815 | 0.75530139 | 2.608871479 | 0.289512686 |
| Gm13316 | 0.175141309 | 0.27441033 | 2.306610632 | 0.118966906 |
| Gm37437 | 53.34605896 | 0.261570079 | 0.371803731 | 0.703516553 |
| Gm37891 | 58.0044897 | 0.819238164 | 0.255694314 | 3.203974904 |
| Nsun6 | 110.3532385 | -0.157144722 | 0.156517659 | -1.004006339 |
| Gm36930 | 6.501557468 | 0.295539334 | 0.368593584 | 0.801802709 |
| Arl5b | 609.464288 | -0.184908406 | 0.101821229 | -1.81601036 |
| Gm38257 | 38.93688588 | -0.469921501 | 0.291903506 | -1.6098522 |
| Gm13313 | 0.521845371 | 0.289243333 | 1.363713044 | 0.212099851 |
| Gm13367 | 0 | NA | NA | NA |
| Gm13366 | 0 | NA | NA | NA |
| Gm13365 | 0.11997416 | 0.274404949 | 3.724945049 | 0.073666845 |
| Gm13368 | 0 | NA | NA | NA |
| Malrd1 | 1.059665568 | 3.082736779 | 1.108658809 | 2.780600086 |
| Gm37595 | 0 | NA | NA | NA |
| Gm6587 | 0 | NA | NA | NA |
| Gm37416 | 2.356264561 | 0.513194709 | 0.66010084 | 0.777448955 |
| Plxdc2 | 861.0341044 | -0.068278143 | 0.073519087 | -0.928713161 |
| Gm24933 | 0.072025462 | 0.755288839 | 3.724945049 | 0.202765096 |
| Gm13326 | 1.471932881 | -0.862085828 | 0.892378055 | -0.966054492 |
| Gm13325 | 0 | NA | NA | NA |
| Gm13322 | 3.121057985 | -1.06655662 | 0.564008616 | -1.891028948 |
| Nebl | 1622.729757 | -0.109400266 | 0.063571141 | -1.720910848 |
| Gm13324 | 0.045882455 | 0.755288839 | 3.724945049 | 0.202765096 |
| Mir6419 | 0 | NA | NA | NA |
| Gm13323 | 0 | NA | NA | NA |
| H2afb1 | 0.165156969 | 0.274389239 | 3.414282971 | 0.080365114 |
| Gm13329 | 0.121578509 | 0.75528439 | 3.724945049 | 0.202763901 |
| A930004D18Rik | 38.49094849 | 0.027157583 | 0.186811843 | 0.145373992 |
| Gm13330 | 0.172363982 | -1.143861708 | 2.780806472 | -0.411341717 |
| Skida1 | 111.6125948 | -0.035438263 | 0.118206515 | -0.299799574 |
| 1810059C17Rik | 0.960935127 | -0.408900489 | 1.091878165 | -0.37449278 |
| Mllt10 | 761.3719482 | -0.261800266 | 0.095018068 | -2.755268246 |
| Mir7655 | 0.321829426 | -1.632020342 | 1.66780851 | -0.9785418 |
| Gm23638 | 0 | NA | NA | NA |
| Gm25147 | 6.022279029 | -0.996420838 | 0.48252852 | -2.06499885 |
| Gm13351 | 0.9210764 | 2.358740476 | 2.153813453 | 1.095146134 |
| Dnajc1 | 546.8891029 | -0.171389512 | 0.069556677 | -2.464026743 |
| Gm13350 | 4.932017945 | -1.013709241 | 0.46573443 | -2.176582139 |
| Gm13349 | 8.865425504 | -1.065709778 | 0.410956467 | -2.593242503 |
| Gm13353 | 0 | NA | NA | NA |
| Gm13355 | 0.097139547 | 0.274393828 | 3.724945049 | 0.073663859 |

| | | | | |
|---|---|---|---|---|
| Gm13352 | 0.195196415 | -0.206485298 | 2.224677853 | -0.09281582 |
| Commd3 | 408.9733762 | 0.192170126 | 0.105924853 | 1.814211877 |
| Bmi1 | 684.4399484 | 0.136610197 | 0.096314743 | 1.418372659 |
| Spag6l | 18.02665997 | -0.09903958 | 0.271608299 | -0.364641213 |
| Gm20539 | 2.239606267 | -1.006712212 | 0.753951785 | -1.335247468 |
| Gm13333 | 0.068097359 | -0.206487837 | 3.724945049 | -0.055433794 |
| Carlr | 12.90806688 | 0.828018829 | 0.29272934 | 2.82861577 |
| Pip4k2a | 1173.833449 | 0.023026744 | 0.112065298 | 0.205476135 |
| 4930426L09Rik | 9.715138516 | 0.116049273 | 0.323850835 | 0.358341743 |
| Gm13332 | 0 | NA | NA | NA |
| Armc3 | 8.367447745 | -0.608385153 | 0.365106076 | -1.666324372 |
| 4930447M23Rik | 25.2336097 | 0.126787994 | 0.204740767 | 0.619261108 |
| Msrb2 | 108.7981829 | 0.186826232 | 0.128272471 | 1.456479562 |
| Gm13344 | 0 | NA | NA | NA |
| Ptf1aos | 0 | NA | NA | NA |
| Ptf1a | 0.167536551 | 0.755304179 | 2.763312762 | 0.273332859 |
| 1810010K12Rik | 2.426680169 | -1.309164651 | 0.630987339 | -2.074787513 |
| 4921504E06Rik | 0.234904022 | 0.755303265 | 2.138287965 | 0.353228039 |
| Gm13345 | 0 | NA | NA | NA |
| Gm13348 | 0.146089079 | -0.206488916 | 3.522538524 | -0.058619349 |
| Gm13343 | 0.231405101 | -0.206482188 | 2.545874405 | -0.081104625 |
| Otud1 | 292.6909858 | 0.312824066 | 0.164831423 | 1.897842414 |
| Gm13346 | 0.525489152 | 1.142792953 | 1.661510444 | 0.687803653 |
| Gm13347 | 0.240250011 | 1.257488971 | 2.112720328 | 0.595198974 |
| Gm24670 | 0 | NA | NA | NA |
| BX571892.1 | 0.123992736 | 0.274402763 | 3.720643035 | 0.073751435 |
| Etl4 | 1406.655128 | -0.341644393 | 0.095614651 | -3.573138539 |
| Gm25859 | 5.580134033 | -0.491925435 | 0.461581191 | -1.065739778 |
| Gm13361 | 0.132101749 | 0.274387056 | 3.724945049 | 0.073662041 |
| Gm13363 | 9.874902262 | -0.783854012 | 0.381816591 | -2.052959539 |
| Gm13360 | 0.049553047 | 0.274391604 | 3.724945049 | 0.073663262 |
| Gm13328 | 2.887214733 | 0.806080056 | 0.558157914 | 1.444179213 |
| Gm23970 | 0.864953071 | -0.792359168 | 1.020284687 | -0.776605959 |
| Gm17171 | 1.518325143 | 0.628433624 | 0.916573323 | 0.685633772 |
| Gm13335 | 0.728721824 | 0.335992528 | 1.161208753 | 0.289347223 |
| Gm27446 | 0.851828701 | -2.049760625 | 1.11731277 | -1.834545061 |
| Gm13362 | 0.31548088 | -1.115658084 | 1.74004331 | -0.641166848 |
| Arhgap21 | 2959.631058 | -0.080558619 | 0.083306713 | -0.967012331 |
| Gm13380 | 18.50560142 | -0.492296397 | 0.369082833 | -1.333837156 |
| Gm13375 | 61.63895035 | -0.269057239 | 0.157651392 | -1.706659459 |
| Gm13377 | 0.2958732 | -0.477954454 | 1.7132165 | -0.278980767 |
| Gm13376 | 0.075043205 | 0.755286614 | 3.724945049 | 0.202764498 |
| Gm25368 | 0 | NA | NA | NA |
| Enkur | 19.33843805 | -0.262631596 | 0.275221987 | -0.954253687 |
| Gm23007 | 0.130861843 | -0.687376661 | 2.984314958 | -0.230329798 |
| Thnsl1 | 325.2700769 | -0.354738775 | 0.115676404 | -3.066647667 |
| Gm13378 | 1.494280535 | -0.291699072 | 0.86132411 | -0.33866354 |

| | | | | |
|---|---|---|---|---|
| Gm13327 | 3.973417911 | -0.068531885 | 0.52202219 | -0.131281555 |
| Gpr158 | 2965.115641 | 0.128870457 | 0.096643231 | 1.333465943 |
| Gm13379 | 0 | NA | NA | NA |
| Gm24886 | 0 | NA | NA | NA |
| Gm13337 | 5.556964523 | 0.27393739 | 0.481636867 | 0.568763333 |
| Myo3a | 0.925384949 | -0.745700744 | 0.99146619 | -0.752119186 |
| Gm13342 | 2.124844546 | 0.983467873 | 0.739638985 | 1.329659324 |
| Gm13341 | 22.54937325 | 0.161435142 | 0.329586222 | 0.489811562 |
| Gm13340 | 244.019484 | 0.112780311 | 0.142057712 | 0.793904884 |
| Gm13339 | 4.846303826 | 0.489605053 | 0.85771718 | 0.570823419 |
| Gad2 | 3413.524295 | -0.006222615 | 0.088214254 | -0.070539789 |
| mmu-mir-3967 | 2.839739068 | 0.202938586 | 0.633270956 | 0.320460909 |
| Gm23636 | 1.903429752 | 0.458995545 | 0.75920973 | 0.604570157 |
| Gm13338 | 0.254763098 | 0.248560473 | 1.825179947 | 0.136184092 |
| 1700092C17Rik | 0.045882455 | 0.755288839 | 3.724945049 | 0.202765096 |
| Gm13356 | 0.169165009 | 0.755292496 | 3.351424224 | 0.225364635 |
| Apbb1ip | 159.2402503 | 1.791435382 | 0.126160174 | 14.19969016 |
| A130006I12Rik | 3.281667885 | -0.925270996 | 0.563533329 | -1.641909978 |
| Pdss1 | 223.7617087 | 0.120567347 | 0.130476727 | 0.924052513 |
| Gm25544 | 3.664064784 | -0.221931559 | 0.626646218 | -0.354157661 |
| Gm14441 | 0 | NA | NA | NA |
| Abi1 | 1539.731744 | 0.177280238 | 0.049758208 | 3.562834025 |
| Gm13357 | 12.25840608 | 0.379649765 | 0.36885208 | 1.029273755 |
| Acbd5 | 986.3592934 | -0.134460833 | 0.075152425 | -1.789174913 |
| Mastl | 28.61317683 | 0.546850473 | 0.313731502 | 1.743052479 |
| Yme1l1 | 1605.641626 | 0.244958316 | 0.064655234 | 3.788684983 |
| Gm13423 | 14.64943441 | 0.040877789 | 0.282761697 | 0.144566219 |
| 4931423N10Rik | 14.857103 | 0.830531125 | 0.313409153 | 2.649990009 |
| Gm13422 | 10.18410332 | 0.467104001 | 0.321999926 | 1.450633878 |
| Nxph2 | 18.63269461 | -0.174235722 | 0.231523904 | -0.752560402 |
| Gm38309 | 0.149576918 | 0.755295313 | 2.863326609 | 0.263782452 |
| Spopl | 699.9338885 | 0.005242051 | 0.093158541 | 0.056270219 |
| Gm13331 | 22.66616647 | 0.194319205 | 0.323504659 | 0.600668955 |
| Sh3d2c-ps1 | 0.219384069 | -0.666725122 | 1.896297501 | -0.351593103 |
| Gm26323 | 0 | NA | NA | NA |
| Gm13392 | 0.616639959 | -0.789258465 | 1.289808618 | -0.61191905 |
| Hnmt | 147.4068612 | -0.136878969 | 0.121524929 | -1.126344771 |
| Tbpl2 | 0 | NA | NA | NA |
| Il1f8 | 0 | NA | NA | NA |
| Gm13410 | 44.62557519 | 0.1834062 | 0.164246568 | 1.116651638 |
| Il1f9 | 2.482099328 | 2.623738725 | 1.062367274 | 2.469709664 |
| Il1f6 | 0 | NA | NA | NA |
| Gm37703 | 0 | NA | NA | NA |
| Gm13409 | 0.143485256 | 0.274391988 | 3.549605256 | 0.077302113 |
| Il1f5 | 0.291332888 | 1.772594257 | 3.704813872 | 0.478457034 |
| Il1f10 | 0 | NA | NA | NA |
| Gm13411 | 0.061920613 | 0.274393828 | 3.724945049 | 0.073663859 |

| | | | | |
|---|---|---|---|---|
| Il1rn | 704.4699326 | 11.50888128 | 1.063376637 | 10.82295857 |
| Psd4 | 122.4570999 | 2.659162402 | 0.270273231 | 9.838793114 |
| Pax8 | 14.32991156 | 0.879234608 | 0.498526729 | 1.76366593 |
| Gm13415 | 0.371620636 | 1.606184227 | 1.744277252 | 0.920830806 |
| Gm27193 | 0 | NA | NA | NA |
| Gm13418 | 0 | NA | NA | NA |
| Gm13417 | 0.292450921 | -0.665239599 | 2.077829062 | -0.320160888 |
| Gm38383 | 0 | NA | NA | NA |
| Gm13416 | 20.11579464 | -0.536302843 | 0.241173954 | -2.223717923 |
| Gm25455 | 0 | NA | NA | NA |
| Cacna1b | 2031.063943 | -0.2165471 | 0.13032876 | -1.661545005 |
| Gm13459 | 0 | NA | NA | NA |
| Ehmt1 | 682.9022921 | -0.172883278 | 0.104746729 | -1.650488567 |
| Gm38147 | 36.95672247 | -0.501324464 | 0.303373547 | -1.652498934 |
| Gm37139 | 53.01011326 | -0.364024359 | 0.296385777 | -1.228211299 |
| Arrdc1 | 89.73200459 | 0.37166269 | 0.124216669 | 2.992051653 |
| Gm22875 | 0.044544648 | 0.755286614 | 3.724945049 | 0.202764498 |
| Zmynd19 | 269.41162 | 0.139668371 | 0.120516591 | 1.158914052 |
| Dph7 | 184.0627944 | 0.239867186 | 0.088552685 | 2.708751123 |
| Mrpl41 | 329.4748938 | 0.24197615 | 0.119288455 | 2.02849597 |
| Pnpla7 | 332.8704237 | 0.89658059 | 0.111479485 | 8.042561275 |
| Gm13370 | 5.147326582 | 0.633537528 | 0.442189249 | 1.43272938 |
| Gm13369 | 0.218114541 | 0.274411417 | 2.547375155 | 0.107723205 |
| Gm25518 | 0 | NA | NA | NA |
| Gm24875 | 0 | NA | NA | NA |
| Nsmf | 2802.591237 | -0.079673175 | 0.091839016 | -0.867530797 |
| Gm24328 | 0 | NA | NA | NA |
| Mir7664 | 1.286742338 | -0.805081686 | 0.884245024 | -0.910473527 |
| Entpd8 | 0.593920364 | 1.20441992 | 1.4180032 | 0.849377434 |
| Noxa1 | 0 | NA | NA | NA |
| Gm13387 | 1.538924929 | -1.406557836 | 0.779684941 | -1.804007955 |
| Nrarp | 119.8106717 | -0.972603551 | 0.257002112 | -3.784418518 |
| Gm38287 | 0.509714849 | 1.141099403 | 1.582261448 | 0.721182586 |
| Tor4a | 84.46660317 | 1.448297842 | 0.22587961 | 6.411813092 |
| Nelfb | 451.1973938 | 0.032250293 | 0.066374722 | 0.485882158 |
| Gm22572 | 0 | NA | NA | NA |
| 4933433C11Rik | 0.117650406 | -0.206492286 | 3.724945049 | -0.055434988 |
| Fam166a | 0.395870588 | 1.153006434 | 1.420931463 | 0.811444087 |
| Tubb4b | 1827.495376 | -0.044874754 | 0.098919923 | -0.453647275 |
| Slc34a3 | 1.160340464 | 0.460574689 | 0.919330577 | 0.500989199 |
| Rnf224 | 0.060174537 | 0.755286614 | 3.724945049 | 0.202764498 |
| Cysrt1 | 0.419553954 | -0.466378888 | 1.560420081 | -0.298880342 |
| Rnf208 | 937.0051333 | -0.231524596 | 0.065392846 | -3.540518718 |
| Ndor1 | 276.8690761 | 0.086814567 | 0.147053522 | 0.590360341 |
| Tmem203 | 61.6019059 | -0.263241927 | 0.16644176 | -1.581585819 |
| Tprn | 478.8306303 | 0.413029664 | 0.144867243 | 2.851090795 |
| Ssna1 | 171.2042094 | -0.148344529 | 0.122763198 | -1.208379474 |

| | | | | |
|---|---|---|---|---|
| Anapc2 | 780.9388462 | -0.105312084 | 0.073564197 | -1.431567095 |
| Tmem210 | 0.270178086 | 0.755274493 | 3.724945049 | 0.202761244 |
| Lrrc26 | 0.469296467 | 0.293875643 | 3.28873187 | 0.089358347 |
| Grin1 | 5090.601218 | -0.038644904 | 0.120049226 | -0.321908815 |
| Grin1os | 5.022424966 | -0.168526565 | 0.464810489 | -0.362570487 |
| AA543186 | 4.244450911 | -0.885173345 | 0.543117518 | -1.629800763 |
| Man1b1 | 806.4183462 | 0.088151992 | 0.061197837 | 1.44044295 |
| Gm25220 | 2.58962508 | -0.892510342 | 0.673067976 | -1.32603299 |
| Dpp7 | 187.4441202 | 0.031368689 | 0.109180148 | 0.287311288 |
| Uap1l1 | 215.8935137 | 0.497037689 | 0.144117181 | 3.448844089 |
| Sapcd2 | 9.831372002 | -0.426142079 | 0.384154693 | -1.10929812 |
| Entpd2 | 57.81792083 | -0.471453249 | 0.20154091 | -2.33924343 |
| Npdc1 | 1413.945948 | 0.03487808 | 0.04979789 | 0.700392731 |
| Fut7 | 5.453509397 | 1.460915961 | 0.627044246 | 2.329845098 |
| Abca2 | 5084.226177 | -0.024040232 | 0.069987586 | -0.343492804 |
| mmu-mir-3087 | 0 | NA | NA | NA |
| BC029214 | 267.2170635 | 0.451399746 | 0.084696329 | 5.329625839 |
| Clic3 | 1.915678769 | 0.771508096 | 0.797937157 | 0.966878267 |
| Ptgds | 4437.782382 | -0.029143488 | 0.159837718 | -0.18233173 |
| Lcn12 | 0.07796279 | 0.274392189 | 3.724945049 | 0.073663419 |
| C8g | 3.739276094 | -0.289183447 | 0.51389118 | -0.562732849 |
| Fbxw5 | 594.4106511 | 0.07900998 | 0.058514508 | 1.350263085 |
| Traf2 | 170.0261398 | 0.515412584 | 0.134914472 | 3.820291297 |
| Gm13433 | 0 | NA | NA | NA |
| Gm13432 | 0.086930093 | -0.206490058 | 3.724945049 | -0.05543439 |
| Edf1 | 525.3660091 | 0.062517548 | 0.200911328 | 0.31116985 |
| Mamdc4 | 22.73292264 | -1.123045772 | 0.307715657 | -3.64962181 |
| Phpt1 | 63.01003723 | -0.428696431 | 0.152205669 | -2.81656021 |
| Gm996 | 1271.384592 | -0.409505228 | 0.128316754 | -3.191362121 |
| Gm26702 | 5.898309265 | -0.00142969 | 0.546097917 | -0.002618011 |
| Rabl6 | 1439.377497 | -0.108298008 | 0.05620534 | -1.926827742 |
| Fcnaos | 8.104967773 | -0.182518995 | 0.346965846 | -0.526043115 |
| Ccdc183 | 0.262568881 | -1.146829727 | 1.811501988 | -0.633082235 |
| Tmem141 | 79.20590439 | -0.158676113 | 0.17306685 | -0.91684868 |
| Fcna | 2.11389961 | -1.345955265 | 0.647120105 | -2.0799157 |
| Lcn8 | 0.091666476 | -0.687373951 | 3.724945049 | -0.184532642 |
| Lcn5 | 0.13733263 | -1.154748075 | 3.63430361 | -0.31773572 |
| Lcn6 | 0.208472373 | 0.328356647 | 2.629152744 | 0.12489067 |
| Lcn10 | 0.112396584 | 0.27438722 | 3.724945049 | 0.073662085 |
| Obp2a | 0.229116515 | -0.217090614 | 2.177627078 | -0.099691364 |
| A230005M16Rik | 4.640067633 | 0.019584129 | 0.418876014 | 0.046753999 |
| Bmyc | 286.5842141 | -0.039871779 | 0.096751397 | -0.412105454 |
| Obp2b | 1.10611924 | -1.047554395 | 1.331948051 | -0.786482922 |
| Lcn3 | 0.068097359 | -0.206487837 | 3.724945049 | -0.055433794 |
| Lcn11 | 0 | NA | NA | NA |
| Gm13538 | 0.049553047 | 0.274391604 | 3.724945049 | 0.073663262 |
| Gm13539 | 0.066050875 | 0.274391604 | 3.724945049 | 0.073663262 |

| | | | | |
|---|---|---|---|---|
| Glt6d1 | 0 | NA | NA | NA |
| Lcn9 | 0.042716945 | -0.206490061 | 3.724945049 | -0.055434391 |
| Sohlh1 | 0.142434329 | -0.687378061 | 2.907370952 | -0.236425992 |
| Kcnt1 | 1559.190496 | -0.130104902 | 0.182016798 | -0.714796125 |
| Camsap1 | 1832.116154 | -0.038800934 | 0.069167051 | -0.560974249 |
| Gm13542 | 4.306872891 | -0.399321156 | 0.574985134 | -0.694489531 |
| Ubac1 | 350.8632011 | -0.114513267 | 0.066060876 | -1.733450623 |
| Nacc2 | 2614.976181 | -0.193139071 | 0.101697503 | -1.899152533 |
| Gm13554 | 0.124596252 | 0.755282361 | 3.714681068 | 0.203323609 |
| C330006A16Rik | 755.4666773 | 0.010328459 | 0.075278177 | 0.137203892 |
| 1810012K08Rik | 1.703298555 | -1.335077298 | 0.924375204 | -1.44430237 |
| Gm13553 | 0.097139547 | 0.274393828 | 3.724945049 | 0.073663859 |
| Lhx3 | 3.930402932 | 0.629929375 | 0.528865517 | 1.191095571 |
| Gm27196 | 0.679199126 | 0.266809255 | 1.186278457 | 0.224912839 |
| Qsox2 | 316.2840522 | 0.016383542 | 0.08181065 | 0.20026173 |
| C030048H21Rik | 9.452175138 | -0.836532278 | 0.354455986 | -2.360045569 |
| 4932418E24Rik | 35.63244048 | -0.409788604 | 0.220865974 | -1.855372272 |
| Gpsm1 | 740.3008245 | -0.221277296 | 0.090510808 | -2.444761029 |
| Dnlz | 309.3938586 | -0.174379812 | 0.11005072 | -1.584540408 |
| Card9 | 28.78353682 | 1.52116392 | 0.268674185 | 5.661742006 |
| Gm13562 | 6.216571396 | -1.03176178 | 0.430471629 | -2.396817144 |
| Snapc4 | 357.5941975 | 0.070942591 | 0.172720553 | 0.410736243 |
| Gm13563 | 26.0969972 | -0.665565834 | 0.269143669 | -2.472901692 |
| Sdccag3 | 585.8077495 | -0.128205645 | 0.073105298 | -1.753712098 |
| Pmpca | 864.0585262 | 0.047041817 | 0.065977367 | 0.712999308 |
| Inpp5e | 354.6391807 | -0.070618421 | 0.090509409 | -0.780232929 |
| Sec16a | 1481.474974 | 0.058158614 | 0.045680272 | 1.273166967 |
| 0610009E02Rik | 13.44346893 | -0.621896973 | 0.329211872 | -1.889047836 |
| Gm13568 | 2.19974275 | -0.011216743 | 0.693051888 | -0.016184565 |
| Notch1 | 1250.567747 | 0.605985619 | 0.118879605 | 5.097473342 |
| Mir6996 | 0.216707685 | 0.274410505 | 2.17420368 | 0.126211959 |
| Gm13358 | 0 | NA | NA | NA |
| Egfl7 | 168.1436807 | 0.530122134 | 0.127802599 | 4.147976147 |
| Gm20532 | 0 | NA | NA | NA |
| Mir126a | 0 | NA | NA | NA |
| mmu-mir-126b | 0 | NA | NA | NA |
| Agpat2 | 19.91019668 | 1.324615907 | 0.339066453 | 3.90665575 |
| Fam69b | 271.5122921 | -0.111415457 | 0.071443562 | -1.55948911 |
| Snhg7 | 0.151994617 | 0.274408244 | 2.849149087 | 0.09631235 |
| Snora43 | 0 | NA | NA | NA |
| Snora17 | 8.121164419 | -0.613727538 | 0.405330996 | -1.51413917 |
| Snhg7os | 0.566020539 | -0.540657391 | 1.312594624 | -0.411899745 |
| Lcn4 | 0.485200684 | 1.577459198 | 1.697312518 | 0.929386416 |
| Gm13359 | 0 | NA | NA | NA |
| Abo | 0.44885175 | 0.639431553 | 1.380042638 | 0.463341882 |
| Surf6 | 237.2764312 | 0.163319317 | 0.092389221 | 1.76773129 |
| Med22 | 318.481323 | 0.483803916 | 0.106610873 | 4.538035403 |

| | | | | |
|---|---|---|---|---|
| Rpl7a | 773.724984 | 0.497744894 | 0.132407633 | 3.759185793 |
| Gm23969 | 2.942191425 | 0.734235735 | 0.665573081 | 1.103163208 |
| Gm22879 | 0 | NA | NA | NA |
| Gm24134 | 0 | NA | NA | NA |
| Surf1 | 388.1526091 | 0.311587904 | 0.083756904 | 3.720145913 |
| Surf2 | 198.1272666 | 0.033709962 | 0.084923991 | 0.396942747 |
| Surf4 | 907.7278515 | 0.479109472 | 0.105493993 | 4.541580592 |
| Stkld1 | 2.796069476 | -0.026171459 | 0.714580003 | -0.036624952 |
| Rexo4 | 357.7792058 | -0.240978356 | 0.107273397 | -2.246394391 |
| Adamts13 | 7.574428338 | -0.364711025 | 0.362303613 | -1.006644738 |
| Cacfd1 | 416.4720264 | 0.050621691 | 0.072691533 | 0.696390474 |
| Slc2a6 | 213.3264522 | 0.075143201 | 0.100170734 | 0.750151254 |
| Gm13397 | 0.26409243 | 0.274408736 | 2.052976414 | 0.133663852 |
| Gm13398 | 0.795847279 | 1.469006388 | 1.279076784 | 1.148489603 |
| Tmem8c | 3.290439168 | 0.868841552 | 0.628954883 | 1.381405209 |
| Adamtsl2 | 40.03111391 | -0.352336634 | 0.210996996 | -1.669865643 |
| Fam163b | 723.0454923 | -0.564867033 | 0.147397819 | -3.832261817 |
| Dbhos | 1.810426623 | -0.756640734 | 0.79050108 | -0.957165972 |
| Dbh | 21.92637974 | -0.064458155 | 0.347593077 | -0.185441422 |
| Sardh | 177.0436753 | -0.037332831 | 0.167175283 | -0.223315496 |
| Sardhos | 0 | NA | NA | NA |
| Vav2 | 234.323641 | 0.064179233 | 0.094712373 | 0.677622471 |
| AA645442 | 0 | NA | NA | NA |
| Gm24049 | 0 | NA | NA | NA |
| Brd3 | 790.2310845 | -0.215966077 | 0.06777452 | -3.186537915 |
| Mir7578 | 0.414979285 | 0.280875689 | 1.468527103 | 0.191263537 |
| Wdr5 | 327.3142271 | 0.066202204 | 0.10138508 | 0.652977779 |
| Gm25541 | 8.286915501 | 0.380602066 | 0.364609402 | 1.043862458 |
| Gm13421 | 4.157031413 | -0.659979948 | 0.523660953 | -1.26031919 |
| Rxra | 493.0791904 | 0.338153353 | 0.080407389 | 4.205500972 |
| 2810430I11Rik | 3.721301919 | -0.417506662 | 0.613870159 | -0.680122101 |
| Col5a1 | 208.2168104 | -0.123294427 | 0.145831876 | -0.845455952 |
| Gm13372 | 0 | NA | NA | NA |
| Fcnb | 0.737107465 | 0.619361375 | 1.200851412 | 0.515768536 |
| Gm13371 | 0.299853729 | 0.267427398 | 1.770061259 | 0.151083696 |
| F730016J06Rik | 3.372121501 | 0.196700411 | 0.519677477 | 0.378504784 |
| Olfm1 | 4803.122293 | -0.147574271 | 0.095036764 | -1.552812449 |
| Gm13373 | 2.464667239 | 0.571066607 | 0.649540431 | 0.879185621 |
| Ppp1r26 | 345.0378359 | -0.234383909 | 0.065449194 | -3.581158073 |
| 1700007K13Rik | 15.71640074 | -0.438326171 | 0.309279047 | -1.417251429 |
| Mrps2 | 321.9683543 | 0.155646706 | 0.093319428 | 1.667891767 |
| Rpsa-ps9 | 16.7483217 | 0.471502917 | 0.296218726 | 1.591739064 |
| Gbgt1 | 30.60507794 | 0.533729693 | 0.231550181 | 2.305028181 |
| Gm10134 | 5.669552113 | 0.695526139 | 0.469360318 | 1.481859699 |
| Ralgds | 1434.688426 | 0.430996181 | 0.103779247 | 4.153009325 |
| Cel | 0.371662472 | -2.054590509 | 1.612385808 | -1.274254895 |
| Gm13381 | 0.066050875 | 0.274391604 | 3.724945049 | 0.073663262 |

| | | | | |
|---|---|---|---|---|
| Gtf3c5 | 232.8597514 | 0.103192862 | 0.087917307 | 1.173749124 |
| Gm22675 | 0 | NA | NA | NA |
| Gm22824 | 0.072025462 | 0.755288839 | 3.724945049 | 0.202765096 |
| Gfi1b | 0.585889909 | 2.300181247 | 1.470126436 | 1.564614574 |
| Tsc1 | 1195.112856 | -0.223609102 | 0.085775302 | -2.606917114 |
| 1700026L06Rik | 2.561385301 | 0.194549458 | 0.675497612 | 0.288009098 |
| Ak8 | 10.0543632 | -0.046717241 | 0.336341238 | -0.138898345 |
| Mir3088 | 0 | NA | NA | NA |
| Gm13385 | 0 | NA | NA | NA |
| Gm13386 | 0.066050875 | 0.274391604 | 3.724945049 | 0.073663262 |
| Gtf3c4 | 803.8265441 | 0.056421343 | 0.056948961 | 0.990735246 |
| Ddx31 | 110.419689 | -0.126611541 | 0.113759321 | -1.112977292 |
| Barhl1 | 48.18788692 | -0.360658911 | 0.250036754 | -1.442423585 |
| 1700101E01Rik | 8.146087367 | 0.144717832 | 0.362155129 | 0.399601773 |
| Gm13394 | 2432.869465 | 0.134678212 | 0.074850957 | 1.799285097 |
| Ttf1 | 162.3829832 | -0.433187908 | 0.140981166 | -3.0726651 |
| Gm13393 | 0.138076337 | 0.75528675 | 3.589802529 | 0.210397854 |
| Gm13396 | 0.120349074 | 1.264824188 | 3.723011833 | 0.339731445 |
| Setx | 1922.416912 | -0.269377662 | 0.06745147 | -3.993651475 |
| Ntng2 | 269.5695528 | -0.406939353 | 0.125315025 | -3.247330898 |
| 6530402F18Rik | 198.2560696 | -0.388740848 | 0.101720186 | -3.821668679 |
| Med27 | 188.5821441 | -0.060201376 | 0.102936233 | -0.584841454 |
| Gm13402 | 0.13742593 | -0.206481804 | 2.939744481 | -0.07023801 |
| Gm24976 | 1.183229494 | 0.302657294 | 0.929463326 | 0.32562586 |
| Mir133c | 0 | NA | NA | NA |
| Gm13401 | 1.918047756 | -1.642353484 | 0.806461342 | -2.036493751 |
| Rapgef1 | 1745.697021 | 0.253184018 | 0.058783626 | 4.307050053 |
| Gm9823 | 0.449135467 | 1.217729418 | 1.427268342 | 0.853188837 |
| Gm13420 | 1.634298146 | -1.511711424 | 0.931251938 | -1.623310902 |
| Gm13547 | 0.049553047 | 0.274391604 | 3.724945049 | 0.073663262 |
| Trub2 | 328.3662552 | 0.298747387 | 0.083070109 | 3.596328332 |
| Coq4 | 113.9900504 | -0.146601231 | 0.106744948 | -1.373378629 |
| Slc27a4 | 871.144694 | -0.188951372 | 0.052743704 | -3.58244413 |
| Urm1 | 399.0492176 | 0.34554134 | 0.087888908 | 3.931569381 |
| Mir219a-2 | 4.701385445 | -0.140091778 | 0.63980906 | -0.218958728 |
| mmu-mir-219b | 2.33080191 | 0.913961418 | 0.717257334 | 1.274244786 |
| 2600006K01Rik | 2.48771953 | -1.594263156 | 0.793288259 | -2.009689592 |
| Cercam | 101.0078076 | 0.156312236 | 0.146661972 | 1.065799365 |
| Odf2 | 917.3723799 | -0.192782029 | 0.147929554 | -1.303201584 |
| Gm23865 | 0 | NA | NA | NA |
| Gle1 | 489.083346 | 0.005324571 | 0.06830199 | 0.077956314 |
| Sptan1 | 16083.58909 | -0.117660044 | 0.070462798 | -1.669817943 |
| Mir6997 | 1.080983412 | -0.645655141 | 0.993083822 | -0.650151706 |
| Wdr34 | 186.8631446 | 0.089765581 | 0.120531517 | 0.744747797 |
| Set | 1857.674102 | 0.131264613 | 0.079685181 | 1.647290137 |
| Pkn3 | 10.32177187 | 1.940589113 | 0.466679234 | 4.158293261 |
| Zdhhc12 | 49.78225683 | 0.540597541 | 0.155502888 | 3.476446952 |

| | | | | |
|---|---|---|---|---|
| Zer1 | 1722.618374 | -0.144517506 | 0.054716309 | -2.641214443 |
| Tbc1d13 | 421.3439125 | 0.13344742 | 0.088419551 | 1.509252411 |
| Slc39a1-ps | 5.938480901 | 0.528689849 | 0.509626854 | 1.037405789 |
| Endog | 111.3222115 | 0.088899871 | 0.136773399 | 0.649979245 |
| D2Wsu81e | 251.2755507 | 0.015550249 | 0.091608439 | 0.169746901 |
| Ccbl1 | 92.26571604 | -0.578014044 | 0.144702351 | -3.994503466 |
| 1700084E18Rik | 9.307150498 | 0.477712922 | 0.331713567 | 1.440136821 |
| Lrrc8a | 523.7153376 | 0.659659354 | 0.102631564 | 6.427451043 |
| Gm28035 | 4.247830997 | -0.080442038 | 0.465532362 | -0.172795803 |
| Phyhd1 | 42.27143342 | 0.684836211 | 0.254374006 | 2.69224133 |
| Gm28038 | 0 | NA | NA | NA |
| Dolk | 191.5944489 | 0.051338129 | 0.107175599 | 0.479009485 |
| Nup188 | 388.5180733 | 0.052275554 | 0.091579848 | 0.570819405 |
| Gm14471 | 0 | NA | NA | NA |
| Sh3glb2 | 1245.303918 | -0.059424558 | 0.066623175 | -0.891950253 |
| Fam73b | 803.1893246 | -0.090138284 | 0.063600774 | -1.417251369 |
| AL954388.1 | 0 | NA | NA | NA |
| Dolpp1 | 175.3732175 | -0.115903522 | 0.090047059 | -1.287143889 |
| Crat | 430.9212061 | -0.088519679 | 0.080338492 | -1.101833964 |
| Gm16323 | 10.6181011 | -0.153854417 | 0.39832748 | -0.386251075 |
| Ppp2r4 | 1830.296797 | 0.029045011 | 0.073776297 | 0.393690273 |
| Ier5l | 217.6479524 | 0.116964543 | 0.168792708 | 0.692947843 |
| Gm14487 | 1.256366316 | 3.065094653 | 1.356253881 | 2.259971158 |
| Cstad | 12.63152331 | 0.427866325 | 0.298524932 | 1.433268311 |
| 9330198N18Rik | 2.48591285 | 0.726360658 | 0.759223538 | 0.956715146 |
| Gm14486 | 1.55717633 | 2.720984619 | 0.96370088 | 2.82347425 |
| Gm14488 | 4.29828934 | 1.260520978 | 0.619502587 | 2.034730775 |
| Mir3089 | 0.367473112 | -1.433061541 | 1.508726601 | -0.949848396 |
| 1700001O22Rik | 34.98746724 | 0.036446283 | 0.205426517 | 0.177417617 |
| 4930527E20Rik | 0.124415284 | -0.687378399 | 3.724945049 | -0.184533836 |
| Ntmt1 | 195.850967 | 0.308998177 | 0.121154327 | 2.550451028 |
| Asb6 | 292.993291 | 0.257167327 | 0.083896763 | 3.065283045 |
| Prrx2 | 13.36258994 | 0.558779629 | 0.33201355 | 1.683002484 |
| Ptges | 99.72996419 | 1.939989524 | 0.285320258 | 6.799340275 |
| Gm13400 | 1.094212761 | 0.931406325 | 1.111754195 | 0.837780805 |
| Tor1b | 368.7015473 | -0.031576541 | 0.076574173 | -0.412365419 |
| Tor1a | 257.103606 | 0.20524637 | 0.085188683 | 2.409314978 |
| BC005624 | 537.4115427 | 0.041033507 | 0.09934093 | 0.413057411 |
| Usp20 | 792.5786421 | -0.168912254 | 0.069487777 | -2.430819648 |
| Fnbp1 | 2552.528974 | -0.391639014 | 0.071602622 | -5.469618353 |
| D330023K18Rik | 31.26496909 | -0.46898648 | 0.21872695 | -2.144164131 |
| Gpr107 | 862.5963975 | 0.102492032 | 0.048926105 | 2.094833238 |
| Gm22516 | 4.36673596 | -0.227491752 | 0.533725628 | -0.426233518 |
| Gm13405 | 0.116046057 | -0.687371727 | 3.724945049 | -0.184532045 |
| Ncs1 | 3353.155418 | -0.172959533 | 0.10565158 | -1.637074744 |
| Hmcn2 | 32.42021316 | -0.408521424 | 0.303476332 | -1.346139321 |
| Ass1 | 313.9716636 | 0.676679241 | 0.133040785 | 5.086254118 |

| | | | | |
|---|---|---|---|---|
| Gm13426 | 0 | NA | NA | NA |
| Fubp3 | 562.5375217 | 0.202689702 | 0.080618307 | 2.514189521 |
| Mir6998 | 1.273153199 | -0.218568775 | 0.983988615 | -0.222125309 |
| Gm13425 | 3.449187758 | 0.716612911 | 0.519792941 | 1.378650716 |
| Prdm12 | 5.973420824 | -0.332125478 | 0.454242052 | -0.731164093 |
| Gm13424 | 0 | NA | NA | NA |
| Exosc2 | 146.3618478 | 0.278738152 | 0.103653757 | 2.689127345 |
| Gm13427 | 1.245568681 | -0.076031011 | 0.881404601 | -0.086261191 |
| Abl1 | 1015.182759 | 0.049628507 | 0.052704964 | 0.9416287 |
| Qrfp | 1.096023364 | 0.85800456 | 1.055907119 | 0.812575789 |
| Fibcd1 | 171.8501216 | -0.404769718 | 0.138477573 | -2.922998364 |
| Gm13441 | 0.122601409 | 1.236179401 | 3.724945049 | 0.331865138 |
| Lamc3 | 85.86094398 | -0.287664542 | 0.183673229 | -1.566175666 |
| Aif1l | 75.90635778 | -0.480182834 | 0.221488952 | -2.167976455 |
| Gm13442 | 0.192796986 | -0.687378705 | 2.253311075 | -0.305052734 |
| Nup214 | 817.8687593 | 0.191791149 | 0.073500033 | 2.609402229 |
| Gm13608 | 10.74285294 | 0.45710424 | 0.461762587 | 0.989911814 |
| Gm13609 | 0.288698636 | 0.741869676 | 2.060578828 | 0.360029748 |
| Fam78a | 86.39529847 | -0.016868603 | 0.139613438 | -0.120823631 |
| Gm16534 | 0.731853201 | 0.770096604 | 1.084218153 | 0.710278279 |
| Mir7674 | 0.20460011 | -0.600596497 | 3.233930613 | -0.185717187 |
| Ppapdc3 | 209.3998458 | -0.048687567 | 0.093307818 | -0.521795158 |
| Prrc2b | 8454.239442 | -0.126464867 | 0.069748164 | -1.8131641 |
| Gm27805 | 3.918894907 | -0.387945148 | 0.518290229 | -0.748509476 |
| Gm22192 | 0.800855801 | -0.714019278 | 1.112000236 | -0.642103531 |
| Gm25632 | 3.196964654 | -0.342416879 | 0.719910052 | -0.47563842 |
| Gm13610 | 1.740926027 | -1.19545803 | 0.802477062 | -1.489709907 |
| Pomt1 | 154.4461912 | -0.468387908 | 0.169494665 | -2.763437478 |
| Uck1 | 264.0364267 | 0.03625653 | 0.09618547 | 0.376943937 |
| Gm13611 | 4.282612097 | 1.420336489 | 0.587070678 | 2.419361997 |
| Swi5 | 679.3934042 | 0.077202325 | 0.075758751 | 1.019054886 |
| Golga2 | 1066.72792 | 0.061488899 | 0.082631792 | 0.744131251 |
| Dnm1 | 10321.66057 | -0.088873475 | 0.084365179 | -1.05343787 |
| Gm23363 | 0.044544648 | 0.755286614 | 3.724945049 | 0.202764498 |
| Mir199b | 0.042716945 | -0.206490061 | 3.724945049 | -0.055434391 |
| Ciz1 | 515.2909118 | 0.212028974 | 0.077336448 | 2.741643571 |
| 1110008P14Rik | 542.4132862 | -0.565548866 | 0.097115176 | -5.823485968 |
| Lcn2 | 343.4844549 | 7.162416926 | 0.805749927 | 8.889131341 |
| Ptges2 | 252.589667 | -0.1020945 | 0.090200629 | -1.131860172 |
| Gm13413 | 0.473410537 | -0.179746079 | 1.47886471 | -0.121543288 |
| Slc25a25 | 955.9872403 | 0.014185873 | 0.131815762 | 0.107618944 |
| Naif1 | 76.9182076 | -0.103744549 | 0.153627446 | -0.675299575 |
| Gm37169 | 0 | NA | NA | NA |
| Gm13412 | 0 | NA | NA | NA |
| Fam102a | 1133.257023 | -0.04120438 | 0.088736334 | -0.464346211 |
| Dpm2 | 258.7392704 | -0.022916037 | 0.084850071 | -0.270076816 |
| 9430097D07Rik | 4.085120842 | -0.496258418 | 0.544021465 | -0.912203745 |

| | | | | |
|---|---|---|---|---|
| Pip5kl1 | 31.38216028 | -1.334561574 | 0.296523208 | -4.500698549 |
| St6galnac4 | 520.5645504 | 0.195423902 | 0.090741955 | 2.153622347 |
| Gm23369 | 0.044544648 | 0.755286614 | 3.724945049 | 0.202764498 |
| St6galnac6 | 748.2103617 | -0.211115282 | 0.087811899 | -2.404176253 |
| Ak1 | 1088.578713 | -0.165597873 | 0.071399683 | -2.319308238 |
| Eng | 245.6138286 | 1.719204275 | 0.173805474 | 9.891542736 |
| Mir1954 | 0.415365334 | -0.206499749 | 3.28012919 | -0.062954761 |
| AL772271.1 | 0 | NA | NA | NA |
| Gm26236 | 0.112642008 | 0.274402725 | 3.724945049 | 0.073666248 |
| Fpgs | 67.89073458 | 0.151094699 | 0.180288161 | 0.838073331 |
| Mir8093 | 0 | NA | NA | NA |
| Cdk9 | 551.2940997 | 0.056995562 | 0.071359927 | 0.798705444 |
| Mir2861 | 0 | NA | NA | NA |
| mmu-mir-3960 | 0 | NA | NA | NA |
| Sh2d3c | 241.4715273 | -0.070842379 | 0.117198018 | -0.604467382 |
| 6330409D20Rik | 20.79074414 | -0.489831061 | 0.22422064 | -2.184593985 |
| Ttc16 | 12.6100887 | -0.466592972 | 0.35655132 | -1.308627807 |
| Tor2a | 191.8598868 | 0.112131762 | 0.099741989 | 1.124218227 |
| Ptrh1 | 37.42700904 | 0.066372024 | 0.228767641 | 0.290128549 |
| 1700019L03Rik | 4.625372364 | -1.202020014 | 0.534346584 | -2.249513799 |
| Stxbp1 | 9943.087999 | 0.037486215 | 0.062005031 | 0.60456731 |
| Gm24165 | 0 | NA | NA | NA |
| Gm13524 | 5.575737892 | -0.091717991 | 0.478695385 | -0.191599906 |
| Gm13523 | 5.508606278 | 0.759475151 | 0.457538881 | 1.659913908 |
| Fam129b | 816.2277192 | 0.79707474 | 0.173534566 | 4.593175637 |
| Lrsam1 | 401.4005097 | -0.19207235 | 0.074331234 | -2.584005939 |
| Rpl12 | 1113.211754 | 0.461192954 | 0.099152797 | 4.651335818 |
| Snora65 | 0 | NA | NA | NA |
| Slc2a8 | 138.0519622 | -0.390236694 | 0.124080313 | -3.145033106 |
| Garnl3 | 1194.21391 | 0.186155735 | 0.128070204 | 1.453544458 |
| Gm38299 | 48.15575876 | 0.497250864 | 0.407533622 | 1.220146847 |
| Ralgps1 | 1094.760979 | -0.076215457 | 0.097813855 | -0.779188773 |
| Gm13528 | 0.134209209 | -0.684635135 | 3.665582479 | -0.1867739 |
| Gm23546 | 4.82094354 | -1.047456308 | 0.527132029 | -1.987085303 |
| Angptl2 | 62.62522016 | -0.423201212 | 0.170162097 | -2.487047463 |
| Gm13537 | 0.097139547 | 0.274393828 | 3.724945049 | 0.073663859 |
| Gm25998 | 3.056188022 | 0.479504042 | 0.555464522 | 0.863248729 |
| Zbtb34 | 281.4807306 | -0.274108949 | 0.098680187 | -2.777750605 |
| Gm13536 | 1.068960359 | -2.33046957 | 1.159488252 | -2.009912188 |
| Zbtb43 | 400.5586369 | -0.133827366 | 0.104724961 | -1.277893682 |
| Gm13530 | 1.493521525 | -0.70466601 | 0.782107947 | -0.900983059 |
| Lmx1b | 29.823212 | 0.046461954 | 0.381338707 | 0.121839072 |
| Gm38011 | 0 | NA | NA | NA |
| C130021I20Rik | 27.33416534 | -0.245060706 | 0.384715016 | -0.636992828 |
| Gm13529 | 0.108772582 | 0.75528439 | 3.724945049 | 0.202763901 |
| 9430024E24Rik | 0.066050875 | 0.274391604 | 3.724945049 | 0.073663262 |
| Mvb12b | 1603.937987 | 0.020944772 | 0.058571485 | 0.357593324 |

| | | | | |
|---|---|---|---|---|
| Nron | 0.158932157 | -0.687378026 | 2.809576835 | -0.244655358 |
| Gm13403 | 3.213919713 | 0.477676697 | 0.631793113 | 0.756065058 |
| Gm13404 | 2.447136968 | 0.352104055 | 0.608298702 | 0.578834139 |
| C230014O12Rik | 11.75502483 | -0.152805674 | 0.283779034 | -0.538467099 |
| Gm38389 | 0.847399432 | -2.415024777 | 1.150370928 | -2.09934441 |
| Pbx3 | 410.290469 | -0.474686038 | 0.158762957 | -2.989904235 |
| C79798 | 0.252257838 | 0.755299723 | 2.491960718 | 0.303094554 |
| Mapkap1 | 523.1598314 | -0.206520972 | 0.094112551 | -2.194404148 |
| Gm13407 | 0 | NA | NA | NA |
| Gm13406 | 0 | NA | NA | NA |
| Gm13408 | 58.61577562 | 0.337966716 | 0.197359197 | 1.712444723 |
| Gm13414 | 23.60150067 | 0.330539521 | 0.274538953 | 1.20398041 |
| Gapvd1 | 1564.103693 | -0.105132898 | 0.043008615 | -2.444461385 |
| Hspa5 | 4684.153117 | 0.28615875 | 0.123447034 | 2.318069071 |
| Rabepk | 186.7520506 | -0.062814888 | 0.099174673 | -0.633376307 |
| Fbxw2 | 731.5136731 | 0.169702872 | 0.076972317 | 2.20472605 |
| Gm13419 | 0.060174537 | 0.755286614 | 3.724945049 | 0.202764498 |
| RP23-446N16.6 | 0.320849018 | 0.581619982 | 1.721917182 | 0.337774655 |
| Psmd5 | 541.5249382 | 0.092145247 | 0.067565384 | 1.363793727 |
| Cutal | 4.992381895 | -0.212436151 | 0.488069987 | -0.435257559 |
| Phf19 | 22.86602922 | 0.484203274 | 0.388585289 | 1.246066919 |
| Gm13448 | 0 | NA | NA | NA |
| Gm24905 | 0 | NA | NA | NA |
| Traf1 | 68.5058879 | 1.500048017 | 0.234349791 | 6.400893347 |
| Gm13450 | 0.044544648 | 0.755286614 | 3.724945049 | 0.202764498 |
| Hc | 5.496526028 | 0.537294104 | 0.47371508 | 1.134213638 |
| Gm13449 | 0.579463306 | -0.328728363 | 1.576179586 | -0.208560221 |
| AI182371 | 3.278154351 | 0.862762287 | 0.611846471 | 1.410096042 |
| Cntrl | 338.5002484 | 0.630411993 | 0.153576357 | 4.104876588 |
| Rab14 | 2869.966926 | -0.015622577 | 0.038339379 | -0.407481224 |
| Gm13605 | 0 | NA | NA | NA |
| Gsn | 990.1649303 | -0.032526366 | 0.135198581 | -0.240582153 |
| RP23-186B18.8 | 6.833187758 | -0.293739617 | 0.467525409 | -0.628285889 |
| Stom | 265.4412871 | 0.75641664 | 0.145981008 | 5.181609908 |
| Gm13443 | 73.68671835 | -0.242604421 | 0.143062668 | -1.695791247 |
| 4930568D16Rik | 0 | NA | NA | NA |
| 4930402F06Rik | 0.092691418 | 0.274387304 | 3.724945049 | 0.073662108 |
| Ggta1 | 165.7903113 | 1.810506004 | 0.198269522 | 9.13153965 |
| RP23-458A10.4 | 2.488423977 | -0.283739835 | 1.011972465 | -0.280382959 |
| Gm13444 | 0.047948698 | -0.206487837 | 3.724945049 | -0.055433794 |
| Gm13445 | 0 | NA | NA | NA |
| Gm13447 | 0.202171817 | 0.274403705 | 2.59796949 | 0.105622374 |
| Gm13446 | 33.55988347 | -0.75030594 | 0.200048575 | -3.750618774 |
| Dab2ip | 1883.686994 | -0.245389541 | 0.076922863 | -3.190072885 |
| Ttll11 | 316.1783459 | -0.178578298 | 0.114184547 | -1.563944527 |
| RP23-366N13.3 | 20.18499025 | -1.035528763 | 0.409965496 | -2.525892478 |
| 9030204H09Rik | 0.047948698 | -0.206487837 | 3.724945049 | -0.055433794 |

| | | | | |
|---|---|---|---|---|
| RP23-366N13.2 | 0.655197851 | -0.680666332 | 1.439990488 | -0.472688075 |
| Ndufa8 | 520.9048915 | -0.015070916 | 0.104427309 | -0.144319685 |
| Morn5 | 2.567728017 | -1.855984022 | 0.723053187 | -2.566870673 |
| RP23-169L4.6 | 0 | NA | NA | NA |
| Lhx6 | 117.6126763 | -0.21769907 | 0.128249184 | -1.697469436 |
| Gm27953 | 0 | NA | NA | NA |
| Rbm18 | 518.3893294 | 0.08960657 | 0.076850236 | 1.165989529 |
| Mrrf | 178.907885 | -0.061086357 | 0.10191307 | -0.599396689 |
| Gm13428 | 0 | NA | NA | NA |
| Gm13429 | 0.948980887 | 0.012379531 | 1.136167406 | 0.010895868 |
| Gm13431 | 0.366933406 | 1.389237196 | 1.701867807 | 0.816301472 |
| Ptgs1 | 131.6150916 | -0.443053836 | 0.225615921 | -1.963752535 |
| RP23-454A14.10 | 0.048949532 | -0.206487837 | 3.724945049 | -0.055433794 |
| Olfr334-ps1 | 0.427301431 | 1.851649441 | 3.366868605 | 0.549961896 |
| Olfr335-ps | 0.092140247 | -0.687373951 | 3.724945049 | -0.184532642 |
| Olfr336-ps1 | 0.11657011 | 1.236179401 | 3.724945049 | 0.331865138 |
| Olfr337-ps1 | 0.194279094 | 0.274391173 | 3.724945049 | 0.073663147 |
| Olfr338 | 0.195866688 | 0.755291911 | 3.257342435 | 0.23187366 |
| Gm13430 | 9.735155595 | -0.160789553 | 0.379777667 | -0.423378117 |
| Olfr339 | 0 | NA | NA | NA |
| Olfr340 | 0 | NA | NA | NA |
| Olfr341 | 0 | NA | NA | NA |
| Olfr342 | 0.194279094 | 0.274391173 | 3.724945049 | 0.073663147 |
| Olfr343-ps1 | 0 | NA | NA | NA |
| Olfr344 | 0.046345709 | 0.274391604 | 3.724945049 | 0.073663262 |
| Olfr345 | 0 | NA | NA | NA |
| Olfr346 | 0 | NA | NA | NA |
| Olfr347 | 0 | NA | NA | NA |
| Gm13438 | 0 | NA | NA | NA |
| Olfr348 | 0 | NA | NA | NA |
| Olfr50 | 0 | NA | NA | NA |
| Gm13436 | 42.25227907 | 0.578462667 | 0.223026543 | 2.593694273 |
| Olfr3 | 0 | NA | NA | NA |
| Olfr349-ps1 | 0 | NA | NA | NA |
| Olfr350 | 0 | NA | NA | NA |
| Olfr351 | 0.097139547 | 0.274393828 | 3.724945049 | 0.073663859 |
| Olfr352 | 0 | NA | NA | NA |
| Olfr353 | 0 | NA | NA | NA |
| Olfr354 | 0 | NA | NA | NA |
| Gm13440 | 0 | NA | NA | NA |
| Olfr355 | 0 | NA | NA | NA |
| Olfr356 | 0 | NA | NA | NA |
| Olfr357 | 0 | NA | NA | NA |
| Olfr358 | 0 | NA | NA | NA |
| Olfr359-ps1 | 0 | NA | NA | NA |
| Gm13439 | 0 | NA | NA | NA |
| Olfr360 | 0 | NA | NA | NA |

| | | | | |
|---|---|---|---|---|
| Gm13437 | 0 | NA | NA | NA |
| Olfr361 | 0 | NA | NA | NA |
| Gm13434 | 0 | NA | NA | NA |
| Olfr362 | 0 | NA | NA | NA |
| Olfr363-ps | 0 | NA | NA | NA |
| Olfr364-ps1 | 0 | NA | NA | NA |
| Olfr365 | 0 | NA | NA | NA |
| Gm13435 | 0 | NA | NA | NA |
| Olfr366 | 0.068097359 | -0.206487837 | 3.724945049 | -0.055433794 |
| Olfr367-ps | 0 | NA | NA | NA |
| Gm22306 | 0 | NA | NA | NA |
| Olfr368 | 0 | NA | NA | NA |
| Pdcl | 328.2885349 | -0.094788594 | 0.123248957 | -0.76908232 |
| Rc3h2 | 3083.559831 | 0.041734029 | 0.037451397 | 1.114351729 |
| Snord90 | 1.013447778 | -0.18010426 | 1.041295648 | -0.172961695 |
| Zbtb6 | 434.5484348 | 0.13321456 | 0.076623227 | 1.738566296 |
| Zbtb26 | 221.5652181 | -0.480443067 | 0.130578588 | -3.679340344 |
| Rabgap1 | 1677.895686 | -0.099025038 | 0.043003802 | -2.302704249 |
| Strbp | 4912.750144 | -0.812011924 | 0.131728379 | -6.164289957 |
| Gpr21 | 129.8264781 | -0.149259293 | 0.152084808 | -0.981421451 |
| Gm13543 | 0 | NA | NA | NA |
| Mir5128 | 8.038112018 | -1.25931386 | 0.410675967 | -3.066441576 |
| Crb2 | 102.1842241 | 0.190236818 | 0.160560082 | 1.184832595 |
| Dennd1a | 974.3781944 | 0.304304416 | 0.060839418 | 5.001764102 |
| AL929186.1 | 4.630423038 | -0.571378728 | 0.52839994 | -1.081337609 |
| Gm13556 | 0.04393175 | -0.206490061 | 3.724945049 | -0.055434391 |
| Lhx2 | 320.8917483 | -0.132245157 | 0.124061345 | -1.065965852 |
| Gm27197 | 1.480046296 | 0.942465145 | 0.894935515 | 1.053109559 |
| Gm13584 | 0 | NA | NA | NA |
| Gm13589 | 3.470889967 | 0.017663599 | 0.550443082 | 0.032089783 |
| Gm25081 | 0.13733263 | -1.154748075 | 3.63430361 | -0.31773572 |
| Gm13588 | 0.045882455 | 0.755288839 | 3.724945049 | 0.202765096 |
| Nek6 | 512.9966335 | 0.927168164 | 0.139520641 | 6.645383477 |
| Psmb7 | 966.8015143 | 0.265698147 | 0.085850009 | 3.094911113 |
| Gm13586 | 0.536876103 | -0.72996244 | 1.531907332 | -0.476505611 |
| Gm27634 | 1.100154067 | 0.121633612 | 1.138910966 | 0.106798174 |
| Adgrd2-ps | 0 | NA | NA | NA |
| Gm13587 | 0 | NA | NA | NA |
| Nr5a1 | 5.753592802 | 0.404126857 | 0.495527004 | 0.815549615 |
| Nr5a1os | 0.104719185 | 1.218392466 | 3.723618458 | 0.327206581 |
| Nr6a1 | 153.9253641 | -0.371517263 | 0.141835562 | -2.619352007 |
| Rpl15-ps4 | 0 | NA | NA | NA |
| Nr6a1os | 2.499784555 | -1.072809444 | 0.705159092 | -1.521372205 |
| Mir181a-2 | 0.139226991 | -0.687378068 | 2.927945967 | -0.234764602 |
| Mir181b-2 | 0.377368388 | 0.274415619 | 1.725159792 | 0.159066783 |
| Olfml2a | 59.91228772 | -0.321360061 | 0.209674354 | -1.532662697 |
| Gm13473 | 0 | NA | NA | NA |

| | | | | |
|---|---|---|---|---|
| Gm13474 | 0.110595523 | 0.755282166 | 3.724945049 | 0.202763304 |
| Gm13496 | 0.873180464 | -1.612763943 | 1.099680273 | -1.466575315 |
| Wdr38 | 12.8811223 | -0.22162484 | 0.304340877 | -0.72821253 |
| Rpl35 | 950.799867 | 0.538296995 | 0.103389331 | 5.20650426 |
| Arpc5l | 615.8797353 | 0.382945684 | 0.098918209 | 3.8713366 |
| Golga1 | 507.3639855 | -0.330880462 | 0.086151424 | -3.84068476 |
| Gm26127 | 0 | NA | NA | NA |
| Scai | 1810.878198 | 0.079341365 | 0.073596525 | 1.078058575 |
| Gm16523 | 10.92727314 | -0.924680214 | 0.435342115 | -2.124031151 |
| Ppp6c | 892.2623954 | 0.161891991 | 0.083195092 | 1.945931994 |
| Gm13511 | 6.921421994 | -0.516612174 | 0.433827922 | -1.190822785 |
| Gm13504 | 0 | NA | NA | NA |
| Vmn2r-ps2 | 0.066050875 | 0.274391604 | 3.724945049 | 0.073663262 |
| Gm13506 | 0 | NA | NA | NA |
| Gm13454 | 0 | NA | NA | NA |
| Gm13451 | 0 | NA | NA | NA |
| Gm13452 | 0.066050875 | 0.274391604 | 3.724945049 | 0.073663262 |
| Gm13453 | 7.860772947 | -0.134055057 | 0.373966611 | -0.358467984 |
| Gm13457 | 0 | NA | NA | NA |
| Gm13458 | 0.061920613 | 0.274393828 | 3.724945049 | 0.073663859 |
| Gm13455 | 0 | NA | NA | NA |
| Gm24467 | 0 | NA | NA | NA |
| Gm13456 | 339.2160491 | 0.434602396 | 0.115420746 | 3.765375027 |
| Lrp1b | 2840.147083 | -0.378584091 | 0.12267889 | -3.085975836 |
| Gm27348 | 0.290738398 | -0.253190086 | 1.761780102 | -0.143712649 |
| Gm13462 | 0 | NA | NA | NA |
| Gm13460 | 0 | NA | NA | NA |
| Gm13461 | 25.32005373 | 0.041215301 | 0.222586073 | 0.185165677 |
| Gm13463 | 0 | NA | NA | NA |
| Mir6336 | 9.1407384 | -0.182390373 | 0.358709939 | -0.508461999 |
| Gm24350 | 0 | NA | NA | NA |
| Gm13464 | 14.15448784 | 0.208802461 | 0.301642515 | 0.692218271 |
| Kynu | 6.170416941 | 1.801694862 | 0.557888733 | 3.229487812 |
| Arhgap15 | 136.0233683 | 0.940312409 | 0.16333536 | 5.756943305 |
| Arhgap15os | 0.169875191 | 0.741257059 | 2.746838206 | 0.269858289 |
| Gm22867 | 0.042716945 | -0.206490061 | 3.724945049 | -0.055434391 |
| Gtdc1 | 447.6665794 | 0.039700371 | 0.095927573 | 0.413857764 |
| Gm13477 | 0.271375346 | 1.619808885 | 2.046214177 | 0.791612581 |
| Zeb2 | 3739.770028 | -0.111953315 | 0.097115739 | -1.152782399 |
| Mir5129 | 1.526907099 | -1.775262202 | 0.907995599 | -1.95514406 |
| Zeb2os | 56.92489478 | 1.288854647 | 0.210113396 | 6.13409079 |
| Gm27761 | 0 | NA | NA | NA |
| Gm27676 | 0.743251506 | 1.49372147 | 1.197847187 | 1.247005032 |
| Gm27404 | 0 | NA | NA | NA |
| Gm27645 | 0 | NA | NA | NA |
| Gm28643 | 0.172790758 | -0.206490367 | 3.395198804 | -0.060818343 |
| Vis1 | 2.225936112 | 0.662973589 | 0.853842632 | 0.776458758 |

| | | | | |
|---|---|---|---|---|
| Gm13475 | 0.044191549 | -0.206490061 | 3.724945049 | -0.055434391 |
| Gm13479 | 1.673299741 | -0.077774673 | 0.750011567 | -0.103697965 |
| Gm13478 | 0 | NA | NA | NA |
| 1700019E08Rik | 0 | NA | NA | NA |
| Gm13465 | 0.044191549 | -0.206490061 | 3.724945049 | -0.055434391 |
| Gm13467 | 0 | NA | NA | NA |
| Gm13466 | 0 | NA | NA | NA |
| Gm23072 | 0.060174537 | 0.755286614 | 3.724945049 | 0.202764498 |
| Gm13469 | 0 | NA | NA | NA |
| Gm13470 | 0.459694823 | 2.11474383 | 1.484694465 | 1.424362978 |
| Gm25264 | 0 | NA | NA | NA |
| Gm25959 | 0.869297867 | 1.925265562 | 1.17397037 | 1.639960948 |
| Gm13481 | 2.425955829 | 0.815574952 | 0.728005298 | 1.120287111 |
| Gm13471 | 0 | NA | NA | NA |
| Gm13482 | 0 | NA | NA | NA |
| Gm25338 | 0.121578509 | 0.75528439 | 3.724945049 | 0.202763901 |
| Gm13472 | 0.062426873 | 0.755288839 | 3.724945049 | 0.202765096 |
| Gm13489 | 0.524554913 | 0.275382318 | 1.272203105 | 0.21646097 |
| Acvr2a | 849.6809722 | 0.210738273 | 0.062479418 | 3.372923087 |
| Orc4 | 1017.373273 | 0.220473399 | 0.089712171 | 2.457563984 |
| Mbd5 | 1674.173319 | -0.068045782 | 0.096469488 | -0.705360667 |
| Gm13510 | 17.67851426 | -0.32824072 | 0.295828241 | -1.109565195 |
| Gm13509 | 582.147492 | 0.056656908 | 0.097690238 | 0.579964884 |
| Gm13508 | 0 | NA | NA | NA |
| Gm25215 | 0.07796279 | 0.274392189 | 3.724945049 | 0.073663419 |
| Epc2 | 915.7051038 | 0.462929585 | 0.098374784 | 4.705774848 |
| Kif5c | 11473.79799 | -0.07456501 | 0.08649287 | -0.86209429 |
| Gm13525 | 21.45150064 | -0.560594104 | 0.438188383 | -1.279344971 |
| Lypd6b | 138.4529101 | 0.880449627 | 0.158992294 | 5.537687427 |
| Gm13480 | 0.55522199 | 1.94344951 | 1.297839456 | 1.497449858 |
| Gm25223 | 0 | NA | NA | NA |
| Lypd6 | 201.0019211 | -0.074884753 | 0.112931134 | -0.66310105 |
| Mmadhc | 325.9114616 | 0.095883152 | 0.095472554 | 1.004300692 |
| Gm13483 | 20.39339472 | -0.412886598 | 0.261355509 | -1.579789151 |
| Gm13486 | 0.761951221 | -0.237010449 | 1.037565183 | -0.228429455 |
| Gm13488 | 0.134148234 | -0.20649159 | 3.624632408 | -0.056968974 |
| Gm13487 | 0.658756006 | -0.08415912 | 1.322522357 | -0.06363531 |
| Gm13485 | 1.138498193 | 0.662118781 | 0.907767271 | 0.729392656 |
| Gm13484 | 0 | NA | NA | NA |
| Gm13498 | 2.899184114 | 1.38897759 | 0.590195648 | 2.353418894 |
| Rnd3 | 459.0834406 | 0.502200421 | 0.166458749 | 3.016966217 |
| Gm28639 | 0 | NA | NA | NA |
| Gm13500 | 0 | NA | NA | NA |
| Gm13497 | 0.338153501 | 0.274200907 | 1.51243417 | 0.181297747 |
| Gm13499 | 0.047948698 | -0.206487837 | 3.724945049 | -0.055433794 |
| 1700057H21Rik | 0.253674139 | -1.068891966 | 2.126659042 | -0.502615579 |
| Gm23505 | 0.049553047 | 0.274391604 | 3.724945049 | 0.073663262 |

| | | | | |
|---|---|---|---|---|
| Gm13494 | 0 | NA | NA | NA |
| Gm13491 | 0.045882455 | 0.755288839 | 3.724945049 | 0.202765096 |
| Tas2r134 | 0 | NA | NA | NA |
| Gm13490 | 0.088383099 | -0.687376175 | 3.724945049 | -0.184533239 |
| Gm13492 | 0 | NA | NA | NA |
| Gm13493 | 88.56531709 | -0.08055425 | 0.131426182 | -0.612923912 |
| Rbm43 | 483.6855949 | 2.117740258 | 0.156204734 | 13.55746525 |
| Gm13495 | 1.701798027 | 4.290644768 | 1.166846602 | 3.67712839 |
| Nmi | 215.7031363 | 3.940466181 | 0.264519672 | 14.89668481 |
| Tnfaip6 | 134.9876113 | -0.173764589 | 0.178752373 | -0.972096682 |
| Rif1 | 1558.481454 | -0.24000898 | 0.070249208 | -3.416536459 |
| Neb | 246.6161145 | 0.183092939 | 0.160222482 | 1.142741872 |
| Gm22095 | 0 | NA | NA | NA |
| Gm13521 | 0 | NA | NA | NA |
| Gm13522 | 0.21580698 | 0.271959002 | 2.575005099 | 0.105614937 |
| Arl5a | 1365.664992 | -0.007174134 | 0.064720578 | -0.110847811 |
| A430018G15Rik | 10.01027238 | -0.506609893 | 0.356149449 | -1.422464346 |
| Cacnb4 | 3963.510753 | -0.311601429 | 0.067741863 | -4.599835559 |
| Gm13541 | 0 | NA | NA | NA |
| Bloc1s2-ps | 5.732035973 | 0.62151497 | 0.433685249 | 1.433101474 |
| Stam2 | 454.7228158 | -0.035578222 | 0.078234905 | -0.454761492 |
| Gm13548 | 1.65657422 | -0.061168431 | 0.840666384 | -0.072761837 |
| Fmnl2 | 1786.717828 | 0.738201576 | 0.116198192 | 6.352952364 |
| Gm26603 | 0.266304556 | 0.759334413 | 3.724657033 | 0.203866935 |
| Prpf40a | 1253.218375 | 0.414604298 | 0.094979017 | 4.365219928 |
| Arl6ip6 | 155.8469731 | 0.002801879 | 0.104756341 | 0.026746633 |
| Gm25898 | 0 | NA | NA | NA |
| Gm13501 | 0 | NA | NA | NA |
| Gm13503 | 0 | NA | NA | NA |
| Gm13502 | 9.659148329 | 0.27597958 | 0.380926318 | 0.724495963 |
| Gm23315 | 0 | NA | NA | NA |
| Rprm | 258.4970102 | -0.23671847 | 0.093231207 | -2.539047577 |
| Gm22298 | 0.222122807 | -0.206482265 | 2.578296719 | -0.080084757 |
| Gm13505 | 0 | NA | NA | NA |
| Gm14035 | 0 | NA | NA | NA |
| Gm14034 | 0.195609542 | 1.483204386 | 2.611791994 | 0.567887638 |
| Galnt13 | 988.7175182 | -0.165207676 | 0.103685939 | -1.59334696 |
| Gm24549 | 0 | NA | NA | NA |
| Gm25489 | 4.670143882 | -0.591573498 | 0.493134443 | -1.199619102 |
| Gm14032 | 0.804512061 | 0.563668706 | 1.096123675 | 0.514238237 |
| Gm14033 | 39.47582969 | -0.29258119 | 0.239261342 | -1.222851917 |
| Gm26794 | 31.41234798 | -0.094654607 | 0.186431018 | -0.507719196 |
| Kcnj3 | 1639.12147 | -0.471997615 | 0.094013828 | -5.020512702 |
| Gm13517 | 0.255816663 | 0.274411948 | 2.076440269 | 0.132154993 |
| Gm13615 | 0 | NA | NA | NA |
| Gm13518 | 0.061920613 | 0.274393828 | 3.724945049 | 0.073663859 |
| Mir195b | 0 | NA | NA | NA |

| | | | | |
|---|---|---|---|---|
| Gm13527 | 0 | NA | NA | NA |
| Nr4a2 | 294.6002765 | -0.525487158 | 0.192871613 | -2.724543797 |
| BB557941 | 2.213086149 | 0.134553831 | 0.642257079 | 0.209501515 |
| 4930555B11Rik | 0.042716945 | -0.206490061 | 3.724945049 | -0.055434391 |
| Gm13526 | 0 | NA | NA | NA |
| A930012O16Rik | 7.994888912 | 0.503869892 | 0.375851877 | 1.340607626 |
| Gpd2 | 1202.471657 | -0.258365219 | 0.087608389 | -2.949092228 |
| Gm13535 | 0 | NA | NA | NA |
| Gm13531 | 0 | NA | NA | NA |
| Gm13532 | 2.612576842 | 0.926632351 | 0.719018427 | 1.28874632 |
| Gm13533 | 0 | NA | NA | NA |
| Gm25388 | 0 | NA | NA | NA |
| Galnt5 | 0.79124893 | 1.4422845 | 1.214921123 | 1.187142501 |
| Ermn | 1375.435044 | 0.3902068 | 0.176796869 | 2.207091127 |
| Gm13534 | 0.908099955 | -1.781597561 | 1.106363786 | -1.610318037 |
| Cytip | 92.50638567 | 5.001638759 | 0.401163121 | 12.46784289 |
| Gm13546 | 1.506085986 | 2.001937418 | 1.018774076 | 1.965045505 |
| Acvr1c | 451.2463121 | -0.808513679 | 0.170698918 | -4.736489771 |
| Gm13544 | 2.162167769 | 1.392352721 | 0.679762462 | 2.048293041 |
| Gm13545 | 0.038981395 | 0.274393828 | 3.724945049 | 0.073663859 |
| Acvr1 | 465.7691716 | -0.134988524 | 0.095132163 | -1.418957803 |
| Gm17409 | 0 | NA | NA | NA |
| Upp2 | 286.6269545 | -0.290954978 | 0.099616789 | -2.920742374 |
| Gm13558 | 10.32835312 | -0.228609079 | 0.31626852 | -0.722832229 |
| Gm13559 | 1.310073478 | 0.157681718 | 0.895339058 | 0.176113972 |
| Ccdc148 | 215.227496 | -0.063884286 | 0.106629821 | -0.599122137 |
| Gm11099 | 0.095897396 | -0.687371727 | 3.724945049 | -0.184532045 |
| Gm13560 | 0.045882455 | 0.755288839 | 3.724945049 | 0.202765096 |
| Gm13557 | 1.02920101 | -0.189043446 | 1.415569178 | -0.13354589 |
| Gm13550 | 2.191863138 | 0.15901437 | 0.698997519 | 0.227489177 |
| Pkp4 | 2887.099339 | -0.256658327 | 0.058415327 | -4.393681268 |
| 5330411J11Rik | 0.17957102 | -0.206487468 | 2.71597396 | -0.076027043 |
| Gm13551 | 0 | NA | NA | NA |
| Gm13549 | 0 | NA | NA | NA |
| Dapl1 | 3.189629895 | -0.636999625 | 0.584375398 | -1.090052091 |
| Gm13552 | 6.37558536 | -0.059556645 | 0.411812598 | -0.144620746 |
| Tanc1 | 1294.003133 | 0.208418791 | 0.087020716 | 2.39504798 |
| Wdsub1 | 122.8229018 | 0.213196141 | 0.126192286 | 1.689454625 |
| Baz2b | 1280.473641 | -0.123112685 | 0.099695253 | -1.234890146 |
| Gm13574 | 8.429661362 | -0.302753221 | 0.34408346 | -0.879883098 |
| Gm13572 | 1.149049355 | 1.533017985 | 1.140030282 | 1.344716899 |
| Gm13570 | 9.56573459 | -0.261703888 | 0.358785296 | -0.729416426 |
| Gm13620 | 2.37993816 | -1.526804885 | 0.753665463 | -2.025838996 |
| Ldha-ps | 0.208487022 | 0.284960502 | 2.585355588 | 0.110221009 |
| 7-Mar | 1192.427979 | 0.191983994 | 0.095767417 | 2.004690109 |
| Cd302 | 130.1829803 | 0.844999051 | 0.161557611 | 5.230326489 |
| Gm13573 | 2.148767599 | 0.206418023 | 0.659440119 | 0.313020117 |

| | | | | |
|---|---|---|---|---|
| Ly75 | 45.62619954 | 1.020432053 | 0.234967701 | 4.342860953 |
| Gm13571 | 1.148968944 | 0.640478326 | 1.163765369 | 0.550350048 |
| Gm13580 | 0.165156969 | 0.274389239 | 3.414282971 | 0.080365114 |
| Pla2r1 | 18.48570985 | 0.686548636 | 0.322659648 | 2.127779658 |
| Gm13583 | 0 | NA | NA | NA |
| Gm13566 | 0 | NA | NA | NA |
| Gm13567 | 0 | NA | NA | NA |
| Itgb6 | 5.350769152 | 2.045923619 | 0.569896252 | 3.589993112 |
| Rbms1 | 693.4012186 | 0.599321308 | 0.124759107 | 4.803828149 |
| Gm13582 | 0.21728767 | 0.75529784 | 2.544699458 | 0.296812198 |
| Gm13579 | 0 | NA | NA | NA |
| Gm13581 | 0 | NA | NA | NA |
| Gm26422 | 0 | NA | NA | NA |
| Gm22338 | 0 | NA | NA | NA |
| Tank | 472.8096782 | 0.367338959 | 0.085643359 | 4.28917038 |
| Gm13626 | 0.359719716 | 0.299568488 | 1.585130268 | 0.188986668 |
| Psmd14 | 942.4841296 | 0.267134381 | 0.074736422 | 3.574353371 |
| Gm22403 | 0 | NA | NA | NA |
| Gm25966 | 3.561922538 | 0.455148747 | 0.515406943 | 0.883086178 |
| Tbr1 | 776.6407633 | -0.148596162 | 0.180144959 | -0.824869945 |
| Gm13555 | 0 | NA | NA | NA |
| Slc4a10 | 3386.482564 | 0.021090015 | 0.082906246 | 0.254383913 |
| Dpp4 | 80.82233867 | 1.293098338 | 0.170156604 | 7.599460204 |
| Gm13561 | 3.915321939 | 0.430945129 | 0.631753252 | 0.682141529 |
| Gm13565 | 0 | NA | NA | NA |
| Gm13564 | 0.340218975 | 1.671046367 | 1.97997212 | 0.843974695 |
| Gcg | 1.314123019 | -0.081410493 | 1.13187799 | -0.071925149 |
| Fap | 70.46597968 | 1.367289639 | 0.318007681 | 4.299549097 |
| Ifih1 | 2560.45954 | 5.846711433 | 0.240581778 | 24.30238679 |
| Gca | 214.646939 | 0.450693998 | 0.122794684 | 3.670305443 |
| Gm13590 | 0.316156321 | -0.20648233 | 2.002651866 | -0.103104456 |
| Kcnh7 | 1930.150615 | -0.388229795 | 0.15610843 | -2.486923955 |
| Gm22020 | 4.453022388 | 0.670792223 | 0.553534985 | 1.211833473 |
| Gm13569 | 0.048949532 | -0.206487837 | 3.724945049 | -0.055433794 |
| Gm23503 | 0 | NA | NA | NA |
| Fign | 177.2882869 | -0.385156499 | 0.091388734 | -4.214485565 |
| Gm13575 | 0 | NA | NA | NA |
| Gm13577 | 10.38401885 | 0.147348466 | 0.328711121 | 0.448261275 |
| Gm13576 | 0 | NA | NA | NA |
| Gm13578 | 0.248954575 | -0.215825541 | 1.842482071 | -0.117138476 |
| Grb14 | 287.0919978 | -0.13415137 | 0.09427396 | -1.422994961 |
| Cobll1 | 254.6797965 | 0.209150971 | 0.132702313 | 1.576091382 |
| Gm13594 | 1.366457952 | -0.82421612 | 0.879330423 | -0.93732242 |
| Gm13592 | 0 | NA | NA | NA |
| Gm13593 | 0 | NA | NA | NA |
| Slc38a11 | 11.09653691 | -0.206419555 | 0.34453866 | -0.599118705 |
| Mir6337 | 0 | NA | NA | NA |

| | | | | |
|---|---|---|---|---|
| Gm24138 | 0.150086409 | 1.264824188 | 3.723011833 | 0.339731445 |
| Scn3a | 1030.005389 | -0.187514046 | 0.074372927 | -2.521267529 |
| Gm13595 | 0.072025462 | 0.755288839 | 3.724945049 | 0.202765096 |
| Scn2a1 | 6263.540254 | -0.193334612 | 0.11055511 | -1.74876233 |
| Csrnp3 | 2276.632116 | 0.148582852 | 0.080134468 | 1.854169068 |
| Gm13619 | 2.502007469 | -0.630817378 | 0.646326219 | -0.976004622 |
| Gm13617 | 8.016718732 | 3.539525595 | 0.718799381 | 4.924219034 |
| Galnt3 | 26.28681886 | 0.311567787 | 0.241515321 | 1.290053921 |
| Gm28638 | 0.099106094 | 0.274387056 | 3.724945049 | 0.073662041 |
| Gm13616 | 0 | NA | NA | NA |
| Ttc21b | 309.885117 | -0.70448739 | 0.169565194 | -4.154669783 |
| Gm13618 | 0.367269401 | -0.440585242 | 1.530026955 | -0.287959138 |
| Scn1a | 3205.479907 | -0.346040533 | 0.065795618 | -5.259324934 |
| Gm13630 | 13.73615189 | 0.122033554 | 0.311832078 | 0.391343814 |
| Gm13629 | 78.07883946 | -0.289391532 | 0.158801962 | -1.822342293 |
| Rpl35-ps1 | 0.152586256 | -0.68737674 | 2.845690286 | -0.241550089 |
| Scn9a | 113.7734046 | -0.235479502 | 0.17445436 | -1.349805769 |
| Gm13627 | 0.809136868 | -0.35516742 | 1.037940007 | -0.342184922 |
| Gm23046 | 0.7960769 | 1.96745165 | 1.219388171 | 1.613474442 |
| Scn7a | 68.38610524 | 0.046934862 | 0.179948293 | 0.260824158 |
| Gm13600 | 0.068097359 | -0.206487837 | 3.724945049 | -0.055433794 |
| Gm13602 | 0.090277459 | -0.20649451 | 3.724945049 | -0.055435586 |
| Gm13598 | 0 | NA | NA | NA |
| Gm1322 | 0 | NA | NA | NA |
| Gm13599 | 0 | NA | NA | NA |
| Gm21830 | 0 | NA | NA | NA |
| Gm23328 | 0 | NA | NA | NA |
| Gm26727 | 0 | NA | NA | NA |
| Xirp2 | 1.65068616 | -0.317691459 | 0.784253913 | -0.405087503 |
| Gm13601 | 10.77684248 | -0.144286083 | 0.379332214 | -0.380368651 |
| B3galt1 | 1121.981432 | 0.472858582 | 0.085686448 | 5.518475695 |
| Mir7224 | 5.441005428 | -0.129566892 | 0.400096792 | -0.323838868 |
| Gm26743 | 0.083526043 | 0.75528439 | 3.724945049 | 0.202763901 |
| Gm13604 | 0.106618422 | 0.274402725 | 3.724945049 | 0.073666248 |
| Gm13603 | 9.345943703 | 0.008011334 | 0.441041589 | 0.018164578 |
| Mir6335 | 8.512614594 | -0.693483035 | 0.455554397 | -1.522283702 |
| Gm37964 | 115.6859084 | -0.284086195 | 0.233531492 | -1.216479167 |
| Gm13607 | 9.156164614 | -0.517678294 | 0.464040186 | -1.115589359 |
| Stk39 | 1177.70026 | 0.109936551 | 0.058697858 | 1.872922718 |
| Gm13596 | 0.577259679 | -1.377092371 | 1.287190644 | -1.069843365 |
| Gm26321 | 0.044191549 | -0.206490061 | 3.724945049 | -0.055434391 |
| Gm13597 | 10.00332192 | 0.258915331 | 0.393191542 | 0.658496695 |
| 4933409G03Rik | 0.150086409 | 1.264824188 | 3.723011833 | 0.339731445 |
| 4932414N04Rik | 0.339353417 | 0.755310922 | 1.596146134 | 0.473209129 |
| Gm13612 | 0.421611163 | 0.348941459 | 1.441744121 | 0.242027315 |
| Cers6 | 1364.374441 | -0.135161054 | 0.049085134 | -2.753604687 |
| Gm37159 | 76.06150798 | -0.666985656 | 0.238837231 | -2.792636861 |

| | | | | |
|---|---|---|---|---|
| Gm38236 | 46.00811634 | -0.541522138 | 0.308316519 | -1.756383796 |
| Gm37651 | 29.9890416 | -0.580492888 | 0.339144541 | -1.711638601 |
| Gm37747 | 50.62951173 | -0.369804806 | 0.253860359 | -1.456725292 |
| Gm37666 | 22.07432209 | -0.297298543 | 0.316621099 | -0.93897262 |
| Gm38377 | 75.00233876 | -0.2396504 | 0.19734063 | -1.214399692 |
| Nostrin | 27.43524618 | 1.208616136 | 0.253077022 | 4.775684993 |
| Spc25 | 14.006815 | 1.612225258 | 0.43772661 | 3.683178545 |
| G6pc2 | 0.378093966 | 14.95568928 | 3.723504053 | 4.016563178 |
| Abcb11 | 1.943768998 | 0.886236308 | 1.002125753 | 0.884356385 |
| Gm13613 | 0 | NA | NA | NA |
| Gm13614 | 0 | NA | NA | NA |
| Dhrs9 | 8.333878881 | 1.755536574 | 0.755437673 | 2.32386686 |
| Lrp2 | 28.81441089 | -0.394644882 | 0.302137164 | -1.306177886 |
| Bbs5 | 149.8333186 | -0.154645379 | 0.11799047 | -1.310659907 |
| Klhl41 | 8.40861694 | 0.101652322 | 0.377164971 | 0.269516867 |
| Fastkd1 | 167.3175697 | -0.322207738 | 0.090324916 | -3.567207721 |
| Ppig | 1069.962896 | -0.029854232 | 0.08037818 | -0.371422101 |
| Gm13622 | 1.277078827 | 1.269442798 | 0.996574559 | 1.273806146 |
| Ccdc173 | 33.00873947 | -0.444954019 | 0.20500652 | -2.170438378 |
| Gm13623 | 0.146451277 | -0.68737672 | 2.882358805 | -0.238477152 |
| Phospho2 | 316.3809453 | -0.114468015 | 0.124145626 | -0.9220463 |
| Gm13624 | 0.441815039 | 1.636604795 | 3.361288925 | 0.486897982 |
| Klhl23 | 607.9612626 | -0.183836365 | 0.092534141 | -1.986686892 |
| Ssb | 1044.348789 | 0.315173177 | 0.092188779 | 3.418780261 |
| Mettl5 | 79.04134513 | -0.123312688 | 0.133584309 | -0.923107576 |
| Gm13625 | 2.609485436 | -0.423645219 | 0.608986077 | -0.695656658 |
| Mettl5os | 3.090821495 | 0.040033367 | 0.5226019 | 0.076603944 |
| Ubr3 | 4154.724246 | 0.102036435 | 0.068752034 | 1.484122421 |
| Gm13631 | 0.295881152 | 0.721026219 | 2.058284145 | 0.35030451 |
| Myo3b | 31.90249792 | -0.596983609 | 0.275933393 | -2.163506211 |
| Rpl9-ps7 | 51.73231683 | -0.078473649 | 0.209449934 | -0.374665425 |
| Sp5 | 11.17023444 | 0.3489942 | 0.34214582 | 1.020015966 |
| Gad1os | 2.192682401 | 0.035248342 | 0.75572242 | 0.046641916 |
| Erich2os | 3.966806659 | 0.929799567 | 0.714456472 | 1.301408278 |
| Erich2 | 9.894614024 | -0.116930277 | 0.423568414 | -0.276059954 |
| Gm37026 | 0.230844305 | -0.142560319 | 2.12948879 | -0.066945795 |
| Gad1 | 4252.286034 | -0.311056893 | 0.092250134 | -3.371885553 |
| Gm16510 | 0.089814205 | 0.274402725 | 3.724945049 | 0.073666248 |
| Gm26558 | 1.60906251 | 0.768297399 | 0.874571823 | 0.878484052 |
| Gorasp2 | 1047.261539 | 0.231362257 | 0.068835449 | 3.361091734 |
| Tlk1 | 1317.27295 | 0.041809062 | 0.056706289 | 0.737291457 |
| Gm13633 | 0.219273179 | 0.755305126 | 2.150245573 | 0.351264588 |
| Gm13632 | 0.086908494 | -0.687376175 | 3.724945049 | -0.184533239 |
| Mettl8 | 114.1898919 | -0.595192924 | 0.159818423 | -3.724182197 |
| Dcaf17 | 286.7511595 | -0.295980069 | 0.130712385 | -2.264361327 |
| Cybrd1 | 58.06687379 | 0.302612866 | 0.282172048 | 1.072440973 |
| Dync1i2 | 1658.410193 | -0.063485566 | 0.043157568 | -1.471018164 |

| | | | | |
|---|---|---|---|---|
| Slc25a12 | 1679.553867 | 0.140959592 | 0.045824513 | 3.076074019 |
| Gm13641 | 0.124853745 | 1.290908285 | 3.721447108 | 0.346883416 |
| Hat1 | 224.6790855 | 0.492813445 | 0.132233926 | 3.726830613 |
| Gm13642 | 0.843921636 | 2.130013729 | 1.171594106 | 1.818047495 |
| Gm23166 | 0 | NA | NA | NA |
| Metap1d | 226.2128202 | -0.218876358 | 0.092315786 | -2.370952649 |
| Gm23253 | 0.412462835 | -0.033085548 | 1.516170747 | -0.021821782 |
| Dlx1as | 32.54297881 | -1.402763087 | 0.288692958 | -4.859013865 |
| Dlx1 | 196.3712922 | -0.510109928 | 0.299222826 | -1.704782804 |
| Dlx2 | 66.09861157 | -0.430671811 | 0.265989461 | -1.61913111 |
| Platr26 | 0.08486385 | 0.755286617 | 3.724945049 | 0.202764499 |
| Gm13647 | 0.342086031 | 1.971973305 | 3.689232862 | 0.534521235 |
| Itga6 | 388.4782796 | 0.112956672 | 0.140580734 | 0.80350037 |
| Gm13663 | 0.038981395 | 0.274393828 | 3.724945049 | 0.073663859 |
| Gm17250 | 1.158557209 | 0.729238918 | 1.082560524 | 0.673624154 |
| Gm13662 | 0 | NA | NA | NA |
| Gm13739 | 0 | NA | NA | NA |
| Pdk1 | 468.9053974 | 0.379254784 | 0.151328838 | 2.506163324 |
| Gm13746 | 0 | NA | NA | NA |
| Rapgef4 | 3912.521623 | -0.062992488 | 0.112335909 | -0.560751125 |
| Gm26408 | 0 | NA | NA | NA |
| Rapgef4os1 | 0.258731086 | -0.206480476 | 2.063349631 | -0.100070523 |
| Rapgef4os2 | 0.479549184 | -0.169217644 | 1.402828411 | -0.120626046 |
| Rapgef4os3 | 1.315840983 | 1.963066275 | 0.913484104 | 2.148987888 |
| Zak | 235.4261635 | -0.071214782 | 0.089387184 | -0.796700146 |
| Zakit | 0.427144302 | -0.405550287 | 1.386196511 | -0.292563344 |
| Rpl36a-ps4 | 0.431106946 | -0.209555597 | 1.404677678 | -0.149184115 |
| Cdca7 | 32.93658422 | -0.5702804 | 0.205474868 | -2.775426537 |
| Rps13-ps8 | 0 | NA | NA | NA |
| Eif1-ps3 | 0 | NA | NA | NA |
| Pex13-ps | 0.061920613 | 0.274393828 | 3.724945049 | 0.073663859 |
| Ak3l2-ps | 18.80802623 | -0.286013839 | 0.247454644 | -1.155823282 |
| 8430437L04Rik | 0.222986776 | 1.275022439 | 2.151495606 | 0.592621447 |
| 6430710C18Rik | 1.790214168 | -1.376714417 | 0.855059613 | -1.610080041 |
| Sp3 | 1150.979175 | 0.217563822 | 0.087226727 | 2.494233461 |
| Gm25960 | 2.212404147 | -0.631187007 | 0.716165679 | -0.881342161 |
| Hspe1-ps5 | 0 | NA | NA | NA |
| Gm11084 | 3.542390089 | -0.442013058 | 0.542288215 | -0.815088813 |
| Sp3os | 52.06250527 | -0.052692348 | 0.168671326 | -0.312396597 |
| Ola1 | 736.2602298 | -0.06545782 | 0.069211215 | -0.945768976 |
| Gm13665 | 0 | NA | NA | NA |
| Gm13666 | 0.293512495 | 1.285753678 | 2.023886053 | 0.635289559 |
| Gm13664 | 0 | NA | NA | NA |
| Sp9 | 90.07522838 | -0.576380478 | 0.243634451 | -2.365759343 |
| Cir1 | 188.1924414 | 0.176043029 | 0.107672938 | 1.634979336 |
| Scrn3 | 297.9077961 | -0.289802799 | 0.110379509 | -2.625512674 |
| Gpr155 | 760.0828086 | -0.302637433 | 0.069883915 | -4.330573524 |

| | | | | |
|---|---|---|---|---|
| Gm13703 | 0.773921687 | 1.743702875 | 1.268672132 | 1.374431448 |
| Gm13709 | 0.142443712 | -0.20648103 | 3.596298704 | -0.057414872 |
| D230022J07Rik | 3.614518035 | -0.488299443 | 0.523242775 | -0.933217746 |
| Gm13707 | 1.060348673 | -0.373749394 | 0.957361945 | -0.390395081 |
| Wipf1 | 548.8928039 | 0.979662103 | 0.143967935 | 6.804724282 |
| Gm13708 | 0 | NA | NA | NA |
| Chrna1 | 3.856870735 | 0.881731652 | 0.705830835 | 1.249211013 |
| Chrna1os | 4.004686001 | 0.167979585 | 0.509174453 | 0.32990576 |
| Chn1os3 | 3.866965461 | -0.138636826 | 0.547198573 | -0.253357433 |
| Chn1 | 4895.40377 | 0.049826951 | 0.117395958 | 0.424434976 |
| Chn1os1 | 6.863703603 | 0.060810191 | 0.371305063 | 0.163774204 |
| Gm13640 | 0 | NA | NA | NA |
| Atf2 | 1720.92591 | -0.098199956 | 0.056939322 | -1.724642159 |
| Gm27421 | 0 | NA | NA | NA |
| Gm10822 | 3.167618981 | -1.451652312 | 0.623090281 | -2.329762405 |
| Atp5g3 | 1417.808151 | -0.160231285 | 0.094609726 | -1.693602685 |
| Gm13668 | 0 | NA | NA | NA |
| A630050E04Rik | 0.102891481 | 0.274400501 | 3.724945049 | 0.073665651 |
| Gm28640 | 0 | NA | NA | NA |
| Gm13667 | 0 | NA | NA | NA |
| 4930441J16Rik | 0.04393175 | -0.206490061 | 3.724945049 | -0.055434391 |
| Lnp | 1061.691288 | -0.111554376 | 0.072144565 | -1.54626167 |
| 4930445N08Rik | 0 | NA | NA | NA |
| Mrpl23-ps1 | 10.74443234 | 0.553155531 | 0.44155331 | 1.252749144 |
| Gm22975 | 0 | NA | NA | NA |
| Gm13672 | 0 | NA | NA | NA |
| Evx2 | 6.495853999 | -0.439633396 | 0.404961063 | -1.08561893 |
| Hoxd13 | 0.292684482 | 1.194545305 | 1.614636671 | 0.739822974 |
| Hoxd12 | 0 | NA | NA | NA |
| Hoxd11 | 0 | NA | NA | NA |
| Gm28309 | 0 | NA | NA | NA |
| Hoxd10 | 0 | NA | NA | NA |
| Gm14396 | 0 | NA | NA | NA |
| Gm28793 | 0 | NA | NA | NA |
| Hoxd9 | 0 | NA | NA | NA |
| AL928644.1 | 0 | NA | NA | NA |
| Hoxd3os1 | 0.132812548 | 0.274408785 | 2.967372194 | 0.092475351 |
| Hoxd8 | 0.583508756 | -1.215953436 | 3.440502546 | -0.353423205 |
| Hoxd3 | 4.727112919 | -0.810089271 | 0.687998636 | -1.17745767 |
| Hoxd4 | 1.678255395 | -0.447300901 | 0.9321293 | -0.479870015 |
| Mir10b | 0.042716945 | -0.206490061 | 3.724945049 | -0.055434391 |
| Gm28230 | 0.047948698 | -0.206487837 | 3.724945049 | -0.055433794 |
| Haglr | 0.983322447 | -2.693557827 | 1.277162342 | -2.109017577 |
| Gm14424 | 0.086930093 | -0.206490058 | 3.724945049 | -0.05543439 |
| Gm14426 | 0 | NA | NA | NA |
| Hoxd1 | 1.509597968 | 0.187552021 | 1.103925132 | 0.169895598 |
| Mtx2 | 366.8248721 | -0.071690888 | 0.080705266 | -0.888304955 |

| | | | | |
|---|---|---|---|---|
| Gm14425 | 13.47600813 | -0.147493807 | 0.316340724 | -0.466249826 |
| Gm14427 | 0 | NA | NA | NA |
| n-R5s198 | 0 | NA | NA | NA |
| Gm13652 | 1.412897596 | -0.648154591 | 0.853866515 | -0.75908187 |
| 2600014E21Rik | 28.30992819 | -0.279351676 | 0.212818402 | -1.31262933 |
| A330043C09Rik | 1.006727404 | 1.792193322 | 1.032568891 | 1.73566465 |
| Rps6-ps4 | 71.2506606 | 0.449636264 | 0.147167864 | 3.05526119 |
| Gm13653 | 0.886214715 | 0.201064992 | 1.111235525 | 0.180938232 |
| Gm37243 | 0 | NA | NA | NA |
| Gm13658 | 0.044544648 | 0.755286614 | 3.724945049 | 0.202764498 |
| Gm13660 | 0 | NA | NA | NA |
| Gm13655 | 0.042716945 | -0.206490061 | 3.724945049 | -0.055434391 |
| Gm24574 | 0.04393175 | -0.206490061 | 3.724945049 | -0.055434391 |
| Hnrnpa3 | 910.2237922 | 0.182100575 | 0.076679864 | 2.374816088 |
| Nfe2l2 | 396.3091985 | 0.841107964 | 0.157710837 | 5.333228711 |
| Gm24996 | 0.882521852 | -1.087297852 | 1.097527047 | -0.990679779 |
| E030042O20Rik | 0.923975021 | -0.617542152 | 1.223663547 | -0.504666625 |
| Gm13656 | 0 | NA | NA | NA |
| Gm13657 | 0.629646617 | 0.755309849 | 1.577530686 | 0.478792492 |
| Gm13669 | 1.634157898 | 2.532999298 | 1.063258296 | 2.382299115 |
| Gm13670 | 1.659798175 | -2.111160193 | 0.934760166 | -2.258504661 |
| Agps | 976.4418691 | -0.037376063 | 0.058667124 | -0.637087013 |
| Ttc30b | 125.5045713 | 0.14972912 | 0.120450062 | 1.243080481 |
| Ttc30a2 | 2.514664855 | 0.776060583 | 0.608403009 | 1.275569929 |
| Ttc30a1 | 46.83376903 | -0.223059912 | 0.193712608 | -1.151499193 |
| Pde11a | 85.82156519 | -0.04847126 | 0.119901812 | -0.404257948 |
| AL845373.1 | 0 | NA | NA | NA |
| Cyct | 0 | NA | NA | NA |
| Rbm45 | 248.2191173 | -0.05637621 | 0.081250578 | -0.693856111 |
| Osbpl6 | 1518.253489 | -0.357715803 | 0.089173217 | -4.011471328 |
| Gm23164 | 1.915350769 | -0.572493034 | 0.778424811 | -0.73545065 |
| Prkra | 162.1113476 | -0.037667183 | 0.095666119 | -0.393735875 |
| Dfnb59 | 0.622664233 | 0.427202198 | 1.23270216 | 0.346557516 |
| Fkbp7 | 23.5112496 | 0.468584168 | 0.212789546 | 2.202101446 |
| Plekha3 | 466.6346452 | 0.007932825 | 0.077944301 | 0.101775564 |
| Ttn | 182.6205874 | -0.200456182 | 0.222257219 | -0.901910784 |
| Gm13938 | 1.375616079 | 0.405128108 | 0.997081539 | 0.406313919 |
| Ccdc141 | 278.830304 | -0.09343145 | 0.116290607 | -0.803430756 |
| AL928681.1 | 1.008651653 | -0.134331713 | 0.931638166 | -0.144188718 |
| Sestd1 | 1270.998036 | -0.182762928 | 0.0752898 | -2.427459358 |
| Gm13650 | 19.02386222 | -1.216897781 | 0.400336088 | -3.039690444 |
| Gm13651 | 3.760012592 | 0.45550993 | 0.528280916 | 0.862249452 |
| Gm13943 | 0 | NA | NA | NA |
| Gm13649 | 0.072025462 | 0.755288839 | 3.724945049 | 0.202765096 |
| Zfp385b | 633.2440097 | -0.314883012 | 0.09968749 | -3.158701384 |
| Gm13944 | 5.255555274 | -0.809120525 | 0.651884759 | -1.241201783 |
| Gm13661 | 1.194926524 | 1.091569455 | 0.817158486 | 1.335811196 |

| | | | | |
|---|---|---|---|---|
| Gm13727 | 0.386897144 | -0.79453399 | 1.375377069 | -0.577684482 |
| Cwc22 | 371.0030043 | 0.354226072 | 0.105359006 | 3.362086318 |
| Gm13726 | 0 | NA | NA | NA |
| 4930440I19Rik | 0.335959249 | 0.675521024 | 2.042566958 | 0.330721606 |
| Gm14461 | 0.674729951 | 0.278755562 | 1.378103688 | 0.202274738 |
| Gm14460 | 0 | NA | NA | NA |
| Gm23149 | 0 | NA | NA | NA |
| Gm14464 | 0.757266105 | 1.40614882 | 1.273987211 | 1.103738568 |
| Gm14463 | 0.136194719 | -0.687371727 | 3.724945049 | -0.184532045 |
| Gm17014 | 0 | NA | NA | NA |
| Ube2e3 | 827.2373707 | -0.130560875 | 0.083985829 | -1.554558378 |
| Gm37004 | 1.348672834 | 0.390792125 | 1.084692633 | 0.360279136 |
| Gm14466 | 0.233003758 | 0.274401717 | 2.562080697 | 0.107101122 |
| Gm14469 | 0.066050875 | 0.274391604 | 3.724945049 | 0.073663262 |
| Gm14465 | 0.075043205 | 0.755286614 | 3.724945049 | 0.202764498 |
| Itga4 | 452.6472453 | 0.828485783 | 0.139746123 | 5.928506388 |
| Cerkl | 55.52887644 | -0.077006524 | 0.265426212 | -0.290124038 |
| Gm22606 | 0 | NA | NA | NA |
| Gm13673 | 0.097139547 | 0.274393828 | 3.724945049 | 0.073663859 |
| Neurod1 | 741.6134161 | -0.607387221 | 0.284672583 | -2.133634417 |
| Gm26755 | 3.778983902 | 1.498575212 | 0.585467355 | 2.559622152 |
| Ssfa2 | 1354.956421 | 0.894128379 | 0.138357048 | 6.462470789 |
| Ppp1r1c | 7.321389695 | 1.624705023 | 0.425782233 | 3.81581216 |
| Pde1a | 1335.258322 | -0.003192026 | 0.139231789 | -0.022925989 |
| Gm13674 | 0 | NA | NA | NA |
| Prdx6b | 0 | NA | NA | NA |
| Dnajc10 | 1132.405827 | 0.045623259 | 0.050190003 | 0.90901088 |
| Gm13752 | 0 | NA | NA | NA |
| Frzb | 132.7433967 | -0.190798576 | 0.171854018 | -1.110236341 |
| Gm13687 | 0 | NA | NA | NA |
| Nckap1 | 5774.673254 | -0.140854731 | 0.046631035 | -3.02062201 |
| Gm13689 | 37.9418997 | -0.315805546 | 0.298775444 | -1.05699967 |
| Gm24461 | 0 | NA | NA | NA |
| Gm13688 | 12.81351646 | -0.269714116 | 0.31489351 | -0.856524847 |
| Dusp19 | 119.9719043 | -0.566903299 | 0.116814564 | -4.853019021 |
| Nup35 | 181.7556722 | -0.333020713 | 0.131204415 | -2.538182213 |
| mmu-mir-684-1 | 0.866431109 | 0.3728892 | 1.19894921 | 0.311013341 |
| Gm13690 | 0.627304132 | 0.772144398 | 1.24780927 | 0.618800017 |
| Gm25420 | 0.262453343 | 0.722164893 | 2.108606376 | 0.342484449 |
| Gm4735 | 399.2680756 | -0.106475174 | 0.115135143 | -0.924784313 |
| Gm22652 | 0 | NA | NA | NA |
| Gm13700 | 0 | NA | NA | NA |
| Gm13755 | 0.332110845 | 1.261400303 | 1.580142736 | 0.798282506 |
| Gm13756 | 0 | NA | NA | NA |
| Gm23452 | 0 | NA | NA | NA |
| Gm23900 | 0 | NA | NA | NA |
| Zfp804a | 500.2371726 | 0.318585415 | 0.126648044 | 2.515517844 |

| | | | | |
|---|---|---|---|---|
| Gm13677 | 0 | NA | NA | NA |
| A130030D18Rik | 0.048949532 | -0.206487837 | 3.724945049 | -0.055433794 |
| Gm13675 | 0.141830814 | -1.124141163 | 3.589786304 | -0.313149884 |
| Gm13676 | 0 | NA | NA | NA |
| Gm13678 | 0 | NA | NA | NA |
| Fsip2 | 0.617250809 | 1.514294862 | 1.177311908 | 1.286230822 |
| Gm28675 | 0.121388914 | 0.755282166 | 3.724945049 | 0.202763304 |
| Gm13680 | 10.62432231 | 0.472268877 | 0.333150442 | 1.417584422 |
| Gm13679 | 0 | NA | NA | NA |
| Gm13682 | 0 | NA | NA | NA |
| Gm13681 | 0 | NA | NA | NA |
| Gm13683 | 0 | NA | NA | NA |
| Gm13685 | 0 | NA | NA | NA |
| Zc3h15 | 953.3117506 | 0.112685841 | 0.104602778 | 1.077273883 |
| Gm13686 | 0 | NA | NA | NA |
| Itgav | 2001.084212 | 0.113141505 | 0.077589356 | 1.458209098 |
| Gm13684 | 0.113999573 | -0.206492286 | 3.724945049 | -0.055434988 |
| Dnmt3l-ps1 | 0.097139547 | 0.274393828 | 3.724945049 | 0.073663859 |
| Fam171b | 2724.632699 | 0.049562069 | 0.085692732 | 0.578369571 |
| Zswim2 | 0.405375818 | 1.130435139 | 1.555548724 | 0.726711495 |
| Gm13698 | 0 | NA | NA | NA |
| Gm13693 | 0 | NA | NA | NA |
| Gm13695 | 0 | NA | NA | NA |
| Gm13696 | 0 | NA | NA | NA |
| Gm13694 | 0.047948698 | -0.206487837 | 3.724945049 | -0.055433794 |
| Gm13697 | 0.061920613 | 0.274393828 | 3.724945049 | 0.073663859 |
| Gm13691 | 0 | NA | NA | NA |
| Gm13692 | 0 | NA | NA | NA |
| Gm13702 | 0 | NA | NA | NA |
| Gm25972 | 0.044191549 | -0.206490061 | 3.724945049 | -0.055434391 |
| Gm13711 | 0.180501107 | 1.103983082 | 2.70443857 | 0.408211558 |
| Calcrl | 175.101438 | 0.236004486 | 0.125660852 | 1.878106674 |
| Tfpi | 63.47998308 | 0.534128673 | 0.150589271 | 3.546923833 |
| Gm13710 | 7.944981639 | -0.106258809 | 0.486292311 | -0.218508101 |
| Ctnnd1 | 284.3298198 | -0.026383163 | 0.092686027 | -0.284650916 |
| Gm28635 | 0 | NA | NA | NA |
| Btbd18 | 0.061920613 | 0.274393828 | 3.724945049 | 0.073663859 |
| 2700094K13Rik | 93.3688872 | -0.04634537 | 0.111222243 | -0.416691562 |
| Tmx2 | 1256.227034 | 0.078583626 | 0.057735323 | 1.361101352 |
| Med19 | 347.2590899 | 0.382357403 | 0.090845887 | 4.208857602 |
| Zdhhc5 | 1049.31998 | 0.019932872 | 0.064230152 | 0.310335117 |
| n-R5s200 | 2.639998601 | -1.25799829 | 0.749388606 | -1.678699516 |
| Gm27942 | 0.072025462 | 0.755288839 | 3.724945049 | 0.202765096 |
| Clp1 | 87.12695073 | -0.101238821 | 0.164903548 | -0.613927487 |
| Ypel4 | 577.9281882 | 0.058040751 | 0.115185303 | 0.503890245 |
| Mir130a | 0.283042562 | -1.124648394 | 3.709272273 | -0.303199202 |
| Gm19426 | 6.293410423 | -0.182777085 | 0.464434669 | -0.393547461 |

| | | | | |
|---|---|---|---|---|
| Serping1 | 1376.102164 | 4.575117578 | 0.281271005 | 16.26586991 |
| Ube2l6 | 958.9461895 | 5.576493755 | 0.286410498 | 19.47028405 |
| Smtnl1 | 0.1140246 | 0.75528439 | 3.724945049 | 0.202763901 |
| Timm10 | 121.5002785 | 0.278988085 | 0.122095035 | 2.285007627 |
| Slc43a1 | 4.475148185 | 1.316867931 | 0.593897823 | 2.217330791 |
| Rtn4rl2 | 239.9515241 | -0.062113417 | 0.189530866 | -0.327721908 |
| Gm24332 | 0.173129629 | -0.206487326 | 2.728586275 | -0.075675572 |
| Slc43a3 | 68.69465978 | 2.755672538 | 0.33343379 | 8.264526945 |
| Prg2 | 0.484678492 | -0.20650112 | 3.724945049 | -0.05543736 |
| Prg3 | 0.205912129 | 0.755300156 | 2.600938185 | 0.290395274 |
| P2rx3 | 14.94560384 | 0.630808641 | 0.278706689 | 2.26334231 |
| Ssrp1 | 920.1403945 | 0.254978406 | 0.068535754 | 3.72037061 |
| Tnks1bp1 | 1348.991537 | 0.591313026 | 0.08363302 | 7.07032972 |
| 4930443O20Rik | 0.247670153 | -0.117531396 | 2.088306145 | -0.056280731 |
| Aplnr | 18.49192457 | 0.064150558 | 0.287542861 | 0.223099115 |
| Gm13712 | 0.447666499 | 1.545006031 | 1.727861895 | 0.894172177 |
| Gm13713 | 6.805315018 | 1.938995878 | 0.516377188 | 3.754999106 |
| Lrrc55 | 258.0702094 | -0.303202769 | 0.128044045 | -2.367956825 |
| Gm13716 | 0.932468966 | 1.392933881 | 1.24719558 | 1.116852804 |
| Gm13715 | 0.832268342 | 2.161885399 | 1.176378498 | 1.837746441 |
| Gm13714 | 0.047948698 | -0.206487837 | 3.724945049 | -0.055433794 |
| Olfr987 | 0.046345709 | 0.274391604 | 3.724945049 | 0.073663262 |
| Olfr988 | 0.042716945 | -0.206490061 | 3.724945049 | -0.055434391 |
| Olfr990-ps1 | 0 | NA | NA | NA |
| Olfr992 | 0 | NA | NA | NA |
| Olfr993 | 0 | NA | NA | NA |
| Gm23716 | 0 | NA | NA | NA |
| Olfr994 | 0.042716945 | -0.206490061 | 3.724945049 | -0.055434391 |
| Olfr995 | 0 | NA | NA | NA |
| 4833423E24Rik | 3.225753529 | -0.686171059 | 0.56506521 | -1.2143219 |
| Olfr996 | 0 | NA | NA | NA |
| Olfr997-ps1 | 0 | NA | NA | NA |
| Olfr998 | 0.038981395 | 0.274393828 | 3.724945049 | 0.073663859 |
| Olfr999-ps1 | 0 | NA | NA | NA |
| Olfr1000 | 0 | NA | NA | NA |
| Olfr1001-ps1 | 0.046345709 | 0.274391604 | 3.724945049 | 0.073663262 |
| Olfr1002 | 0 | NA | NA | NA |
| Olfr1003-ps1 | 0 | NA | NA | NA |
| Olfr1004-ps1 | 0 | NA | NA | NA |
| Olfr154 | 0 | NA | NA | NA |
| Olfr1006 | 0.048949532 | -0.206487837 | 3.724945049 | -0.055433794 |
| Olfr1008 | 0 | NA | NA | NA |
| Olfr1009 | 0 | NA | NA | NA |
| Olfr1010 | 0.038981395 | 0.274393828 | 3.724945049 | 0.073663859 |
| Olfr1012 | 0.089062654 | -0.20649451 | 3.724945049 | -0.055435586 |
| Olfr1013 | 0 | NA | NA | NA |
| Olfr1014 | 0 | NA | NA | NA |

| | | | | |
|---|---|---|---|---|
| Olfr1015 | 0 | NA | NA | NA |
| Olfr1016 | 0.04393175 | -0.206490061 | 3.724945049 | -0.055434391 |
| Olfr1017-ps1 | 0 | NA | NA | NA |
| Gm13720 | 0 | NA | NA | NA |
| Olfr1018 | 0 | NA | NA | NA |
| Olfr1019 | 0.062426873 | 0.755288839 | 3.724945049 | 0.202765096 |
| Olfr1020 | 0 | NA | NA | NA |
| Olfr1021-ps1 | 0 | NA | NA | NA |
| Olfr1022 | 0 | NA | NA | NA |
| Gm24037 | 0 | NA | NA | NA |
| Olfr1023 | 0 | NA | NA | NA |
| Olfr1024 | 0 | NA | NA | NA |
| Olfr1025-ps1 | 0 | NA | NA | NA |
| Olfr1026 | 0 | NA | NA | NA |
| Olfr1027-ps1 | 0 | NA | NA | NA |
| Olfr1028 | 0 | NA | NA | NA |
| Olfr1029 | 0.108772582 | 0.75528439 | 3.724945049 | 0.202763901 |
| Olfr1030 | 0 | NA | NA | NA |
| Olfr1031 | 0 | NA | NA | NA |
| Olfr1032 | 0 | NA | NA | NA |
| Olfr1033 | 0 | NA | NA | NA |
| Olfr1034 | 0 | NA | NA | NA |
| Olfr1035-ps1 | 0 | NA | NA | NA |
| Olfr1036 | 0 | NA | NA | NA |
| Olfr1037 | 0 | NA | NA | NA |
| Gm13722 | 0.13742593 | -0.206481804 | 2.939744481 | -0.07023801 |
| Olfr1038-ps | 0 | NA | NA | NA |
| Olfr1039 | 0 | NA | NA | NA |
| Olfr1040 | 0 | NA | NA | NA |
| Olfr1042 | 0 | NA | NA | NA |
| Olfr1043 | 0 | NA | NA | NA |
| Olfr1044 | 0 | NA | NA | NA |
| Olfr52 | 0.108772582 | 0.75528439 | 3.724945049 | 0.202763901 |
| Olfr1045 | 0 | NA | NA | NA |
| Olfr1046 | 0 | NA | NA | NA |
| Olfr1047 | 0 | NA | NA | NA |
| Olfr1048 | 0 | NA | NA | NA |
| Olfr1049 | 0 | NA | NA | NA |
| Olfr1050-ps1 | 0 | NA | NA | NA |
| Olfr1051 | 0 | NA | NA | NA |
| Olfr1052 | 0.044544648 | 0.755286614 | 3.724945049 | 0.202764498 |
| Olfr1053 | 0 | NA | NA | NA |
| Gm13719 | 2.60552406 | 0.98856298 | 0.64663826 | 1.528772796 |
| Olfr1054 | 0 | NA | NA | NA |
| Olfr1055 | 0 | NA | NA | NA |
| Olfr1056 | 0 | NA | NA | NA |
| Olfr1057 | 0 | NA | NA | NA |

| | | | | |
|---|---|---|---|---|
| Olfr1058 | 0 | NA | NA | NA |
| Olfr1060 | 0 | NA | NA | NA |
| Olfr1061 | 0 | NA | NA | NA |
| Olfr1062 | 0 | NA | NA | NA |
| Olfr1063-ps1 | 0 | NA | NA | NA |
| Olfr1065 | 0 | NA | NA | NA |
| Olfr1066 | 0 | NA | NA | NA |
| Olfr1069-ps1 | 0 | NA | NA | NA |
| Olfr228 | 0 | NA | NA | NA |
| Olfr1073-ps1 | 0 | NA | NA | NA |
| Olfr1076 | 0 | NA | NA | NA |
| Olfr1077-ps1 | 0 | NA | NA | NA |
| Olfr1078-ps1 | 0 | NA | NA | NA |
| Olfr1079 | 0 | NA | NA | NA |
| Olfr1080 | 0 | NA | NA | NA |
| Olfr1081-ps1 | 0 | NA | NA | NA |
| Olfr1082 | 0 | NA | NA | NA |
| Olfr1083-ps | 0 | NA | NA | NA |
| Olfr1084 | 0 | NA | NA | NA |
| Olfr1085 | 0 | NA | NA | NA |
| Olfr1086 | 0 | NA | NA | NA |
| Olfr1087 | 0 | NA | NA | NA |
| Olfr1089 | 0 | NA | NA | NA |
| Olfr1090 | 0 | NA | NA | NA |
| Olfr1093 | 0 | NA | NA | NA |
| BX284685.1 | 0 | NA | NA | NA |
| Olfr141 | 0 | NA | NA | NA |
| Olfr1094 | 0 | NA | NA | NA |
| Olfr1095 | 0 | NA | NA | NA |
| Olfr1096-ps1 | 0 | NA | NA | NA |
| Gm13723 | 0 | NA | NA | NA |
| Olfr1097 | 0 | NA | NA | NA |
| Olfr1098 | 0 | NA | NA | NA |
| Olfr1099 | 0.048949532 | -0.206487837 | 3.724945049 | -0.055433794 |
| Olfr1100 | 0 | NA | NA | NA |
| Olfr1101 | 0 | NA | NA | NA |
| Olfr1102 | 0 | NA | NA | NA |
| Olfr1103-ps1 | 0 | NA | NA | NA |
| Olfr1104 | 0 | NA | NA | NA |
| Olfr1105 | 0 | NA | NA | NA |
| Olfr1106 | 0 | NA | NA | NA |
| Olfr1107 | 0 | NA | NA | NA |
| Olfr1109 | 0.046345709 | 0.274391604 | 3.724945049 | 0.073663262 |
| Olfr259 | 0 | NA | NA | NA |
| Gm13733 | 0 | NA | NA | NA |
| Olfr1110 | 0 | NA | NA | NA |
| Gm13734 | 0 | NA | NA | NA |

| | | | | |
|---|---|---|---|---|
| Olfr1111 | 0 | NA | NA | NA |
| Nlk-ps1 | 0.21574249 | 0.274415833 | 2.174796322 | 0.126180015 |
| Olfr1112 | 0 | NA | NA | NA |
| Gm13730 | 0 | NA | NA | NA |
| Olfr1113 | 0 | NA | NA | NA |
| Gm13735 | 0 | NA | NA | NA |
| Olfr1115 | 0 | NA | NA | NA |
| Gm27438 | 0 | NA | NA | NA |
| Gm13732 | 0 | NA | NA | NA |
| Olfr1116 | 0 | NA | NA | NA |
| Gm13728 | 0.042716945 | -0.206490061 | 3.724945049 | -0.055434391 |
| Olfr1117-ps1 | 0 | NA | NA | NA |
| Gm13729 | 0 | NA | NA | NA |
| Olfr1118 | 0 | NA | NA | NA |
| Gm13737 | 0 | NA | NA | NA |
| Gm13731 | 0 | NA | NA | NA |
| Olfr1120 | 0 | NA | NA | NA |
| Olfr1121 | 0 | NA | NA | NA |
| Olfr1122 | 0 | NA | NA | NA |
| Gm1826 | 8.12848589 | -0.608632118 | 0.387669411 | -1.569977151 |
| Olfr1123 | 0 | NA | NA | NA |
| Olfr1124 | 0 | NA | NA | NA |
| Olfr1126 | 0 | NA | NA | NA |
| Pramel7 | 0 | NA | NA | NA |
| Pramel6 | 0 | NA | NA | NA |
| Gm13738 | 0 | NA | NA | NA |
| Olfr1127-ps1 | 0 | NA | NA | NA |
| Gm13736 | 22.02889899 | 0.473309103 | 0.274280711 | 1.725637582 |
| Olfr153 | 0 | NA | NA | NA |
| Olfr1128 | 0 | NA | NA | NA |
| Gm13740 | 0.18866314 | 0.169827872 | 2.656144464 | 0.063937739 |
| Gm13744 | 0 | NA | NA | NA |
| Gm13743 | 3.080495725 | 0.684522569 | 0.571166086 | 1.19846501 |
| Olfr1129 | 0 | NA | NA | NA |
| Olfr1130 | 0 | NA | NA | NA |
| Olfr1131 | 0 | NA | NA | NA |
| Olfr1132 | 0 | NA | NA | NA |
| Gm13742 | 0 | NA | NA | NA |
| Olfr1133 | 0 | NA | NA | NA |
| Gm13741 | 0 | NA | NA | NA |
| Olfr1134 | 0 | NA | NA | NA |
| Olfr1135 | 0 | NA | NA | NA |
| Olfr1136 | 0 | NA | NA | NA |
| Olfr1137 | 0 | NA | NA | NA |
| Olfr1138 | 0 | NA | NA | NA |
| Olfr1140 | 0 | NA | NA | NA |
| Olfr1141 | 0 | NA | NA | NA |

| | | | | |
|---|---|---|---|---|
| Olfr152 | 0 | NA | NA | NA |
| Olfr1143 | 0 | NA | NA | NA |
| Olfr1144-ps1 | 0 | NA | NA | NA |
| Olfr1145 | 0 | NA | NA | NA |
| Olfr1146-ps1 | 0 | NA | NA | NA |
| Olfr1147-ps1 | 0 | NA | NA | NA |
| Olfr1148 | 0 | NA | NA | NA |
| Olfr1149-ps1 | 0 | NA | NA | NA |
| Olfr1150-ps1 | 0 | NA | NA | NA |
| Olfr1151 | 0 | NA | NA | NA |
| Olfr1152 | 0 | NA | NA | NA |
| Gm13745 | 0 | NA | NA | NA |
| Olfr1153 | 0 | NA | NA | NA |
| Olfr1154 | 0 | NA | NA | NA |
| Olfr1155 | 0 | NA | NA | NA |
| Olfr1156 | 0 | NA | NA | NA |
| Olfr1157 | 0 | NA | NA | NA |
| Olfr74 | 0 | NA | NA | NA |
| Olfr1158 | 0 | NA | NA | NA |
| Olfr1159-ps1 | 0 | NA | NA | NA |
| Olfr1160 | 0 | NA | NA | NA |
| Olfr1161 | 0 | NA | NA | NA |
| Olfr73 | 0 | NA | NA | NA |
| Gm13747 | 0 | NA | NA | NA |
| Olfr1162 | 0 | NA | NA | NA |
| Olfr1163 | 0 | NA | NA | NA |
| Olfr1164 | 0 | NA | NA | NA |
| Olfr1165-ps | 0.046345709 | 0.274391604 | 3.724945049 | 0.073663262 |
| Olfr1166 | 0 | NA | NA | NA |
| Olfr1167 | 0 | NA | NA | NA |
| Olfr1168 | 0.046345709 | 0.274391604 | 3.724945049 | 0.073663262 |
| Olfr1169-ps1 | 0 | NA | NA | NA |
| Olfr1170 | 0 | NA | NA | NA |
| Olfr1171-ps1 | 0 | NA | NA | NA |
| Gm13753 | 0 | NA | NA | NA |
| Olfr1172-ps1 | 0 | NA | NA | NA |
| Olfr1173 | 0 | NA | NA | NA |
| Olfr1174-ps | 0 | NA | NA | NA |
| Olfr1175-ps | 0 | NA | NA | NA |
| Olfr1176 | 0.061920613 | 0.274393828 | 3.724945049 | 0.073663859 |
| Olfr1177-ps | 0 | NA | NA | NA |
| Olfr1178 | 0 | NA | NA | NA |
| Olfr1179 | 0 | NA | NA | NA |
| Olfr1180 | 0 | NA | NA | NA |
| Olfr1181 | 0 | NA | NA | NA |
| Olfr1182 | 0 | NA | NA | NA |
| Gm13757 | 0 | NA | NA | NA |

| | | | | |
|---|---|---|---|---|
| Olfr1183 | 0 | NA | NA | NA |
| Olfr1184 | 0 | NA | NA | NA |
| Olfr1185-ps1 | 0 | NA | NA | NA |
| Olfr1186 | 0 | NA | NA | NA |
| Olfr1187-ps1 | 0 | NA | NA | NA |
| Olfr1188 | 0 | NA | NA | NA |
| Gm13758 | 0 | NA | NA | NA |
| Olfr1189 | 0 | NA | NA | NA |
| Olfr475-ps1 | 0 | NA | NA | NA |
| Olfr1191-ps1 | 0.062426873 | 0.755288839 | 3.724945049 | 0.202765096 |
| Gm13759 | 0 | NA | NA | NA |
| Olfr1192-ps1 | 0 | NA | NA | NA |
| Olfr1193 | 0.042716945 | -0.206490061 | 3.724945049 | -0.055434391 |
| Olfr1195 | 0.131862677 | -0.687376665 | 2.97735462 | -0.230868255 |
| Olfr1196 | 0 | NA | NA | NA |
| Olfr1197 | 0 | NA | NA | NA |
| Olfr1198 | 0 | NA | NA | NA |
| Olfr1199 | 0 | NA | NA | NA |
| Gm13760 | 0 | NA | NA | NA |
| Olfr1200 | 0 | NA | NA | NA |
| Olfr1201 | 0 | NA | NA | NA |
| Olfr1202 | 0 | NA | NA | NA |
| Olfr1205 | 0 | NA | NA | NA |
| Olfr1204 | 0 | NA | NA | NA |
| Olfr1206 | 0 | NA | NA | NA |
| Olfr1208 | 0 | NA | NA | NA |
| Olfr1209 | 0 | NA | NA | NA |
| Olfr1211 | 0.068097359 | -0.206487837 | 3.724945049 | -0.055433794 |
| Gm13763 | 0 | NA | NA | NA |
| Olfr1212 | 0 | NA | NA | NA |
| Olfr1213 | 0 | NA | NA | NA |
| Gm13762 | 0 | NA | NA | NA |
| Olfr1214 | 0 | NA | NA | NA |
| Olfr1215 | 0 | NA | NA | NA |
| Olfr1216 | 0 | NA | NA | NA |
| Olfr1217 | 0 | NA | NA | NA |
| Olfr1218 | 0 | NA | NA | NA |
| Olfr1219 | 0.045882455 | 0.755288839 | 3.724945049 | 0.202765096 |
| Olfr1220 | 0.095435502 | 0.75528439 | 3.724945049 | 0.202763901 |
| Olfr1221 | 0 | NA | NA | NA |
| Gm13764 | 0.048949532 | -0.206487837 | 3.724945049 | -0.055433794 |
| Olfr1222 | 0.048949532 | -0.206487837 | 3.724945049 | -0.055433794 |
| Olfr1223 | 0 | NA | NA | NA |
| Olfr1224-ps1 | 0 | NA | NA | NA |
| Olfr1225 | 0 | NA | NA | NA |
| Gm13761 | 0 | NA | NA | NA |
| Olfr1226 | 0 | NA | NA | NA |

| | | | | |
|---|---|---|---|---|
| Olfr1228 | 0.062426873 | 0.755288839 | 3.724945049 | 0.202765096 |
| Olfr1229 | 0 | NA | NA | NA |
| Olfr1230 | 0 | NA | NA | NA |
| Olfr1231 | 0.046345709 | 0.274391604 | 3.724945049 | 0.073663262 |
| Olfr1232 | 0 | NA | NA | NA |
| Olfr1233 | 0.048949532 | -0.206487837 | 3.724945049 | -0.055433794 |
| Olfr1234 | 0.048949532 | -0.206487837 | 3.724945049 | -0.055433794 |
| Olfr1235-ps1 | 0 | NA | NA | NA |
| Olfr1238 | 0 | NA | NA | NA |
| Olfr1239 | 0 | NA | NA | NA |
| Olfr1240 | 0 | NA | NA | NA |
| Olfr1241 | 0 | NA | NA | NA |
| Olfr1242 | 0 | NA | NA | NA |
| Olfr1243 | 0.106618422 | 0.274402725 | 3.724945049 | 0.073666248 |
| Olfr1245 | 0 | NA | NA | NA |
| Olfr1246 | 0.045882455 | 0.755288839 | 3.724945049 | 0.202765096 |
| Olfr1247 | 0 | NA | NA | NA |
| Olfr1248 | 0 | NA | NA | NA |
| Olfr1249 | 0 | NA | NA | NA |
| Olfr1250 | 0.042716945 | -0.206490061 | 3.724945049 | -0.055434391 |
| Olfr1251 | 0 | NA | NA | NA |
| Olfr1252 | 0 | NA | NA | NA |
| Olfr1253 | 0 | NA | NA | NA |
| Olfr1254 | 0.04393175 | -0.206490061 | 3.724945049 | -0.055434391 |
| Olfr1255 | 0 | NA | NA | NA |
| Olfr1256 | 0 | NA | NA | NA |
| Olfr48 | 0.156220659 | -1.153238739 | 2.817542717 | -0.409306568 |
| Olfr1257 | 0 | NA | NA | NA |
| Olfr1258 | 0.254495662 | 0.755305347 | 2.107454834 | 0.358396932 |
| Olfr1259 | 0 | NA | NA | NA |
| Olfr1260 | 0.042716945 | -0.206490061 | 3.724945049 | -0.055434391 |
| Olfr1261 | 0.198041555 | 0.274404852 | 2.604148426 | 0.105372201 |
| Olfr1262 | 0.31782415 | 0.755274284 | 3.724945049 | 0.202761188 |
| Olfr1263 | 0 | NA | NA | NA |
| Olfr1264 | 0.068097359 | -0.206487837 | 3.724945049 | -0.055433794 |
| Olfr1265 | 0.097139547 | 0.274393828 | 3.724945049 | 0.073663859 |
| Olfr140 | 0.229124067 | -0.206488753 | 2.577799727 | -0.080102714 |
| Olfr1267-ps1 | 0.049553047 | 0.274391604 | 3.724945049 | 0.073663262 |
| Olfr1268-ps1 | 0.202674043 | -0.206490498 | 2.623803806 | -0.078698909 |
| Olfr1269 | 0.045882455 | 0.755288839 | 3.724945049 | 0.202765096 |
| Olfr32 | 0.631605561 | -1.202009627 | 1.402740921 | -0.856900664 |
| Gm4222 | 0 | NA | NA | NA |
| Olfr1270 | 0.066050875 | 0.274391604 | 3.724945049 | 0.073663262 |
| Gm13768 | 4.397907185 | -0.019004416 | 0.560363328 | -0.033914454 |
| Olfr1506 | 0 | NA | NA | NA |
| Gm18953 | 1.627769934 | -0.419547033 | 0.98332977 | -0.426659546 |
| Olfr142 | 0.123841226 | 0.274391173 | 3.724945049 | 0.073663147 |

| | | | | |
|---|---|---|---|---|
| Olfr1271 | 0 | NA | NA | NA |
| Olfr1272 | 0 | NA | NA | NA |
| Olfr1273-ps | 0.044544648 | 0.755286614 | 3.724945049 | 0.202764498 |
| Gm13769 | 0 | NA | NA | NA |
| Gm13767 | 0.326846603 | 0.75531142 | 1.600550911 | 0.471907151 |
| Olfr1274-ps | 0.163996116 | -0.206488978 | 2.782191723 | -0.074218098 |
| Ptprj | 2106.140895 | 0.140934147 | 0.077299957 | 1.823211212 |
| 4933423P22Rik | 0.57303442 | 2.070639605 | 1.37299065 | 1.508123602 |
| Nup160 | 358.4498607 | 0.021690361 | 0.1080747 | 0.200697861 |
| Fnbp4 | 883.9475744 | 0.71633684 | 0.154881098 | 4.625075931 |
| Rpl30-ps3 | 4.003505758 | 1.097716048 | 0.577419747 | 1.901071196 |
| Agbl2 | 38.0546618 | -0.281578295 | 0.204066742 | -1.379834323 |
| Gm13772 | 2.379780159 | -0.43358388 | 0.733366496 | -0.591224009 |
| Mtch2 | 1065.574097 | -0.143006768 | 0.054282987 | -2.634467558 |
| C1qtnf4 | 729.131802 | -0.348422754 | 0.094926393 | -3.670451836 |
| Ndufs3 | 556.9779417 | -0.020851759 | 0.053797827 | -0.387594808 |
| Kbtbd4 | 295.8165819 | 0.085631086 | 0.078892133 | 1.085419847 |
| Ptpmt1 | 185.7476871 | 0.160477179 | 0.09095727 | 1.764313936 |
| Gm17661 | 14.75789179 | -0.835699259 | 0.346187771 | -2.41400572 |
| Celf1 | 3282.919782 | -0.132078373 | 0.069292674 | -1.906094335 |
| Gm13777 | 0.066050875 | 0.274391604 | 3.724945049 | 0.073663262 |
| Rapsn | 7.482500097 | 0.373703496 | 0.399864035 | 0.934576414 |
| Gm13778 | 1.349943917 | -1.009705351 | 0.946072883 | -1.067259584 |
| Psmc3 | 1077.312818 | 0.174261293 | 0.092085526 | 1.892385277 |
| Slc39a13 | 259.1457734 | -0.138037173 | 0.081724066 | -1.689063947 |
| Spi1 | 202.8166777 | 3.108692891 | 0.228366129 | 13.6127582 |
| Mybpc3 | 140.415035 | -0.129497002 | 0.244027908 | -0.530664724 |
| Madd | 2828.091406 | -0.259322306 | 0.077404675 | -3.350215032 |
| Nr1h3 | 80.38358335 | 0.760573293 | 0.229445812 | 3.314827517 |
| Gm13783 | 1.952547309 | -1.413229475 | 0.750084331 | -1.884094116 |
| Gm23390 | 0 | NA | NA | NA |
| Acp2 | 985.571155 | 0.304690566 | 0.044959968 | 6.776930188 |
| Ddb2 | 90.31751625 | -0.062485849 | 0.208033373 | -0.300364541 |
| A330069E16Rik | 10.08996547 | -0.679863138 | 0.365667025 | -1.859241037 |
| Gm17281 | 10.85400332 | -0.857928083 | 0.355949609 | -2.410251513 |
| Pacsin3 | 255.622615 | -0.48835354 | 0.122578557 | -3.98400465 |
| Arfgap2 | 730.4068622 | 0.244829386 | 0.075003067 | 3.264258339 |
| 1110051M20Rik | 549.9156413 | -0.074224687 | 0.056670694 | -1.309754344 |
| Gm13787 | 4.652905194 | -1.001410561 | 0.544306867 | -1.839790422 |
| Gm22071 | 0.442430462 | -1.489073054 | 3.362438263 | -0.442855136 |
| Lrp4 | 768.9458671 | 0.073039469 | 0.142275894 | 0.513365033 |
| Gm13770 | 1.121024803 | 0.074404829 | 0.975331642 | 0.076286696 |
| Ckap5 | 3392.946938 | 0.225250474 | 0.056544282 | 3.983611882 |
| Snord67 | 0.174512195 | -1.282572172 | 3.342629336 | -0.383701584 |
| F2 | 4.096713608 | 0.525022572 | 0.621796177 | 0.844364425 |
| Gm13771 | 3.006370356 | -0.293055948 | 0.65308039 | -0.448728752 |
| Zfp408 | 162.7132082 | 0.461987866 | 0.172594746 | 2.676720333 |

| | | | | |
|---|---|---|---|---|
| Arhgap1 | 635.3928673 | 0.171303424 | 0.063216595 | 2.709785702 |
| Atg13 | 1068.978477 | 0.210755993 | 0.050739256 | 4.153706773 |
| Harbi1 | 67.29079701 | -0.374148135 | 0.160311375 | -2.333883885 |
| Gm13776 | 0 | NA | NA | NA |
| Ambra1 | 1182.397324 | 0.065033594 | 0.068925228 | 0.943538329 |
| Gm10805 | 2.718864041 | -0.204596582 | 0.720495866 | -0.28396635 |
| Chrm4 | 179.7457586 | -0.191479893 | 0.161602988 | -1.184878419 |
| Mdk | 159.9674338 | 0.115989054 | 0.186083719 | 0.623316506 |
| Dgkz | 3070.276697 | -0.229034197 | 0.091668744 | -2.498498251 |
| Mir6999 | 2.840504449 | -1.075342722 | 0.712316597 | -1.509641536 |
| Gm9821 | 0.506210706 | 0.305044246 | 1.713128459 | 0.178062681 |
| Creb3l1 | 136.2000294 | 1.689107031 | 0.23368831 | 7.228033904 |
| Gm13780 | 0 | NA | NA | NA |
| 2900072N19Rik | 2.504782524 | -0.993442487 | 0.72378456 | -1.372566565 |
| Phf21a | 888.8724875 | -0.206355507 | 0.08261244 | -2.497874495 |
| Mir1955 | 9.919958007 | -0.438220092 | 0.359428688 | -1.219212896 |
| Gm13786 | 17.47248729 | -0.540427465 | 0.306848198 | -1.761220916 |
| Gm13785 | 11.39462224 | -0.312864593 | 0.321086089 | -0.974394732 |
| Gyltl1b | 4.391866736 | -0.792811678 | 0.505643216 | -1.567927052 |
| Pex16 | 172.030486 | 0.204662646 | 0.101410555 | 2.018159212 |
| 1700029I15Rik | 16.98714461 | -0.291044809 | 0.274473516 | -1.060374834 |
| Mapk8ip1 | 3383.018323 | -0.149192769 | 0.083744848 | -1.781515776 |
| Mir7000 | 0.999544483 | -0.553341854 | 1.076375487 | -0.514078833 |
| Cry2 | 1477.051092 | 0.071883234 | 0.123672862 | 0.581236923 |
| D930015M05Rik | 1.112373992 | 0.208042727 | 0.976199563 | 0.213114956 |
| Slc35c1 | 209.4076668 | 0.140477224 | 0.098615799 | 1.424490038 |
| Gm13814 | 0.592656722 | -1.442056816 | 1.300391865 | -1.108940201 |
| Gm13817 | 0 | NA | NA | NA |
| Gm13815 | 0.046345709 | 0.274391604 | 3.724945049 | 0.073663262 |
| Mir7221 | 0 | NA | NA | NA |
| Mir7222 | 0 | NA | NA | NA |
| Chst1 | 1513.4659 | -0.072634779 | 0.103996534 | -0.698434618 |
| Gm13816 | 0.371244445 | -0.687380144 | 1.700509805 | -0.404220041 |
| Gm13791 | 2.292177245 | 0.799780662 | 0.771291869 | 1.036936462 |
| Gm13792 | 0.47588132 | 0.395841677 | 1.674854635 | 0.236343901 |
| Syt13 | 2064.734027 | -0.043913494 | 0.077526591 | -0.566431387 |
| Prdm11 | 417.3168277 | -0.384948337 | 0.112494654 | -3.421925609 |
| 4631405J19Rik | 0.851482706 | -2.41971295 | 1.103094351 | -2.193568436 |
| Gm13802 | 0 | NA | NA | NA |
| Trp53i11 | 736.2819127 | 0.481965046 | 0.113882302 | 4.232132963 |
| Tspan18 | 97.99565433 | 0.318259215 | 0.141807334 | 2.244307161 |
| Gm23380 | 0 | NA | NA | NA |
| Gm13807 | 0 | NA | NA | NA |
| Cd82 | 303.0056944 | 0.527208555 | 0.142629312 | 3.696354888 |
| Mir7001 | 0.756442295 | 0.745579728 | 1.2618217 | 0.590875658 |
| Gm10804 | 0.060174537 | 0.755286614 | 3.724945049 | 0.202764498 |
| Gm10803 | 0 | NA | NA | NA |

| | | | | |
|---|---|---|---|---|
| Alx4 | 48.87403417 | 0.024153913 | 0.361707501 | 0.066777474 |
| Ext2 | 641.1248268 | 0.170305166 | 0.076186021 | 2.235386029 |
| Gm13818 | 0 | NA | NA | NA |
| Accs | 46.96754752 | 0.114818771 | 0.19466096 | 0.589839746 |
| Accsl | 3.620414628 | 1.072073831 | 0.603978717 | 1.77501922 |
| Gm1335 | 0.929738683 | 1.709760279 | 1.126094433 | 1.518309858 |
| Gm13889 | 181.9860749 | 0.671400032 | 0.343220351 | 1.956177806 |
| AL772372.1 | 0 | NA | NA | NA |
| Gm27027 | 1.373972637 | -0.578668382 | 0.822836334 | -0.70326061 |
| 4921507L20Rik | 0.891453265 | 2.147598248 | 1.283677312 | 1.673004756 |
| Alkbh3 | 168.1665116 | 0.308874943 | 0.138639576 | 2.227898784 |
| Hsd17b12 | 914.0604724 | 0.316800465 | 0.127127449 | 2.491991057 |
| Gm23630 | 0.694735342 | -0.188662953 | 1.356994513 | -0.139030004 |
| Mir129-2 | 0.549517402 | 0.394051167 | 1.252419226 | 0.314632001 |
| mmu-mir-129b | 0 | NA | NA | NA |
| E530001K10Rik | 26.89108908 | -1.152628272 | 0.296467084 | -3.887879414 |
| Mir670 | 0.304542382 | -0.290700539 | 2.001182684 | -0.145264369 |
| Ttc17 | 942.8773192 | -0.027933684 | 0.095283399 | -0.293164228 |
| Itpa-ps1 | 137.8875215 | 0.329232119 | 0.121990651 | 2.698830748 |
| 2810002D19Rik | 26.44289622 | -0.283385034 | 0.225086782 | -1.259003445 |
| Api5 | 1200.933468 | 0.119379688 | 0.06477517 | 1.842985314 |
| AL672251.1 | 0 | NA | NA | NA |
| Gm13788 | 0 | NA | NA | NA |
| Gm24318 | 0 | NA | NA | NA |
| Gm13793 | 0 | NA | NA | NA |
| Gm26396 | 0 | NA | NA | NA |
| Gm13794 | 0 | NA | NA | NA |
| Gm13798 | 0 | NA | NA | NA |
| Gm13797 | 0 | NA | NA | NA |
| Gm13799 | 0 | NA | NA | NA |
| Gm37175 | 0 | NA | NA | NA |
| Gm13801 | 0 | NA | NA | NA |
| Lrrc4c | 1313.773726 | 0.087161335 | 0.083585778 | 1.0427771 |
| 4930445B16Rik | 0 | NA | NA | NA |
| Gm17075 | 11.93369344 | -0.255154464 | 0.310008895 | -0.823055299 |
| Gm17076 | 16.91552809 | 0.275701902 | 0.326238039 | 0.845094286 |
| Gm22957 | 5.852693499 | -0.764769354 | 0.462905566 | -1.652106628 |
| Gm22313 | 8.145373654 | -0.393905201 | 0.56094629 | -0.702215539 |
| Gm23317 | 0 | NA | NA | NA |
| BX813336.1 | 0 | NA | NA | NA |
| Gm13803 | 11.5628337 | 0.392112373 | 0.326033242 | 1.202676053 |
| Gm13804 | 0 | NA | NA | NA |
| Gm13806 | 0 | NA | NA | NA |
| Gm10801 | 0.412640929 | -0.144918153 | 1.533171204 | -0.094521833 |
| Gm10800 | 8.185264119 | -0.553584332 | 0.448074172 | -1.235474764 |
| Gm29235 | 0 | NA | NA | NA |
| Gm13809 | 0 | NA | NA | NA |

| | | | | |
|---|---|---|---|---|
| Gm13808 | 0 | NA | NA | NA |
| Gm13811 | 0.109727584 | 0.755282166 | 3.724945049 | 0.202763304 |
| Gm13813 | 0 | NA | NA | NA |
| Gm13812 | 0.047948698 | -0.206487837 | 3.724945049 | -0.055433794 |
| Gm13810 | 0 | NA | NA | NA |
| B230118H07Rik | 178.272976 | -0.055023712 | 0.149017011 | -0.369244501 |
| Gm20693 | 0 | NA | NA | NA |
| Rag2 | 0.139883403 | 0.274393101 | 3.559413475 | 0.077089415 |
| Rag1 | 5.95628265 | 0.467363708 | 0.425055991 | 1.099534455 |
| Traf6 | 437.7135397 | 0.315017999 | 0.107542224 | 2.929249442 |
| Prr5l | 253.5858244 | 0.117482334 | 0.139485099 | 0.842257238 |
| Gm13905 | 0.083526043 | 0.75528439 | 3.724945049 | 0.202763901 |
| Commd9 | 191.447391 | 0.091378688 | 0.118189322 | 0.773155193 |
| Ldlrad3 | 167.5338321 | -0.230938117 | 0.127855443 | -1.806243926 |
| Gm13919 | 2.169032929 | 1.384578419 | 0.882311825 | 1.569262 |
| Gm13920 | 4.969440295 | 1.068787075 | 0.741509937 | 1.441365816 |
| Gm13921 | 0.311841693 | 1.505303941 | 1.718579712 | 0.875899983 |
| Gm13866 | 1.228132247 | -0.235389367 | 0.962437528 | -0.244576256 |
| Trim44 | 3706.544855 | 0.005793887 | 0.046663655 | 0.124162737 |
| Gm13868 | 13.87276866 | 0.065790012 | 0.395755898 | 0.166238866 |
| Fjx1 | 429.7281807 | -0.409587639 | 0.092312518 | -4.436967465 |
| Pamr1 | 172.126028 | -0.554094452 | 0.151175645 | -3.665236241 |
| Slc1a2 | 36923.0076 | -0.282951745 | 0.160589758 | -1.761953865 |
| Gm13872 | 1.821369377 | 0.015079791 | 0.895247505 | 0.01684427 |
| Cd44 | 1546.027959 | 3.406107783 | 0.333812987 | 10.2036407 |
| Gm23731 | 0 | NA | NA | NA |
| Pdhx | 517.8134996 | 0.099209257 | 0.068548922 | 1.447276687 |
| Apip | 121.8280582 | 0.582221316 | 0.113363945 | 5.13585969 |
| Gm10038 | 4.822154624 | 0.787696588 | 0.503053256 | 1.565831408 |
| Gm13874 | 0.503556157 | 1.818388892 | 1.745548227 | 1.041729391 |
| Gm13869 | 0.217279565 | -0.206481368 | 2.170317414 | -0.095138788 |
| Gm13873 | 0 | NA | NA | NA |
| Ehf | 2.553683234 | 0.154920639 | 0.68767654 | 0.225281262 |
| A930006I01Rik | 0 | NA | NA | NA |
| Gm37220 | 0 | NA | NA | NA |
| BC016548 | 0.652016589 | -1.184043212 | 1.238058637 | -0.956370867 |
| Elf5 | 2.42823633 | -1.085877758 | 0.586393179 | -1.851791252 |
| Cat | 444.9583296 | -0.372854318 | 0.115450318 | -3.229565107 |
| Gm13996 | 0 | NA | NA | NA |
| Abtb2 | 322.9374325 | 0.555382741 | 0.142230808 | 3.904799166 |
| Nat10 | 403.567961 | 0.207671626 | 0.170028029 | 1.221396417 |
| Caprin1 | 3734.751293 | 0.15056727 | 0.052557417 | 2.864814892 |
| 4930547E08Rik | 0.171199454 | 1.25961124 | 3.386170909 | 0.371986906 |
| Gm13878 | 0 | NA | NA | NA |
| Gm13880 | 2.206276467 | -0.211225823 | 0.679952178 | -0.310648056 |
| Gm13877 | 0 | NA | NA | NA |
| Gm13875 | 0 | NA | NA | NA |

| | | | | |
|---|---|---|---|---|
| Gm22757 | 0 | NA | NA | NA |
| Gm13879 | 0 | NA | NA | NA |
| Gm22761 | 0 | NA | NA | NA |
| Gm13876 | 0 | NA | NA | NA |
| Gm13881 | 0.097899064 | -0.687371727 | 3.724945049 | -0.184532045 |
| Lmo2 | 263.3668701 | 1.244138342 | 0.110268938 | 11.28276345 |
| 4931422A03Rik | 0.485755142 | 1.492026481 | 1.313672778 | 1.135767221 |
| Fbxo3 | 1097.464659 | -0.077676745 | 0.054184374 | -1.433563584 |
| Gm24644 | 12.13107012 | 0.079086864 | 0.342881587 | 0.230653577 |
| Gm10912 | 3.939700132 | -1.682431629 | 0.570432435 | -2.949396857 |
| Gm10799 | 0.09689823 | -0.687371727 | 3.724945049 | -0.184532045 |
| Cd59b | 12.1589774 | -0.425065083 | 0.312070898 | -1.362078571 |
| Cd59a | 104.2603775 | 0.12896677 | 0.217986315 | 0.591627827 |
| A930018P22Rik | 1.374894827 | -0.559298177 | 1.017122805 | -0.549882644 |
| D430041D05Rik | 2766.772851 | -0.385988859 | 0.150434253 | -2.565830928 |
| Gm22501 | 3.530580087 | 0.018721277 | 0.550093288 | 0.034032913 |
| Gm13883 | 6.270791699 | 0.687493486 | 0.547110977 | 1.256588726 |
| Gm25554 | 0 | NA | NA | NA |
| Gm13882 | 5.232096263 | 0.460799412 | 0.432139682 | 1.066320525 |
| Hipk3 | 2248.948924 | -0.194110469 | 0.062160944 | -3.12270789 |
| Mir1902 | 0 | NA | NA | NA |
| Gm13870 | 0 | NA | NA | NA |
| Gm13884 | 0.326587101 | -0.687380503 | 2.031357508 | -0.3383848 |
| Platr14 | 2.284861264 | 0.121810117 | 0.65426359 | 0.186178963 |
| Gm22439 | 0 | NA | NA | NA |
| Gm13885 | 0.266822755 | -0.708619068 | 2.396167317 | -0.295730212 |
| Cstf3 | 425.9326561 | -0.0407229 | 0.074515522 | -0.546502245 |
| Tcp11l1 | 978.2370623 | -0.358713843 | 0.071528154 | -5.015002094 |
| Pin1rt1 | 0.178918687 | -1.004501303 | 2.708509187 | -0.370868708 |
| Depdc7 | 17.9411153 | -0.456045053 | 0.290710855 | -1.568723851 |
| Qser1 | 622.2252455 | -0.089138255 | 0.065717804 | -1.356379085 |
| Gm13886 | 11.02501732 | -0.398824827 | 0.338834277 | -1.177049828 |
| Prrg4 | 12.12341216 | 0.71248966 | 0.352620682 | 2.020555502 |
| Ccdc73 | 1.808963643 | 0.431098518 | 0.739661369 | 0.582832274 |
| Gm13887 | 0.209356743 | 0.740421159 | 2.188175762 | 0.338373714 |
| Eif3m | 484.4513724 | 0.339756666 | 0.081231597 | 4.18256784 |
| Gm13888 | 0 | NA | NA | NA |
| Wt1os | 0.172571121 | 0.755296067 | 2.73763 | 0.275894137 |
| Gm11060 | 0 | NA | NA | NA |
| Gm27715 | 0 | NA | NA | NA |
| Gm27538 | 0 | NA | NA | NA |
| Gm27636 | 0 | NA | NA | NA |
| Gm27308 | 0 | NA | NA | NA |
| Wt1 | 11.92317521 | 0.174700332 | 0.631181069 | 0.276783226 |
| Them7 | 0.493638219 | -0.167605786 | 1.429343373 | -0.117260687 |
| Rcn1 | 399.3853322 | 0.584852565 | 0.128338274 | 4.557117255 |
| 4930527A07Rik | 0 | NA | NA | NA |

| | | | | |
|---|---|---|---|---|
| Pax6os1 | 4.443791399 | -0.221272913 | 0.496823035 | -0.445375711 |
| Gm13890 | 0.122991902 | 0.274402725 | 3.724945049 | 0.073666248 |
| Rpl10-ps1 | 5.581713243 | -0.074972455 | 0.452720752 | -0.165604193 |
| Paupar | 4.745398909 | 0.117393594 | 0.423806698 | 0.276997966 |
| Pax6 | 311.7227136 | -0.347549009 | 0.161508518 | -2.151892755 |
| Elp4 | 237.3869113 | -0.07786038 | 0.084449727 | -0.921973142 |
| Gm13954 | 0.303403639 | 2.13028887 | 1.988292982 | 1.071415978 |
| Immp1l | 117.1741609 | 0.079447805 | 0.197393935 | 0.402483515 |
| Dnajc24 | 115.5254544 | -0.071901493 | 0.102341277 | -0.702565921 |
| Gm22002 | 0.104637558 | -0.206492286 | 3.724945049 | -0.055434988 |
| n-R5s201 | 0 | NA | NA | NA |
| Dcdc5 | 20.38028449 | -0.811145309 | 0.268774174 | -3.017943637 |
| Gm13992 | 0.047948698 | -0.206487837 | 3.724945049 | -0.055433794 |
| Gm29053 | 0 | NA | NA | NA |
| Gm14013 | 0.459558249 | -1.06105345 | 1.322849286 | -0.802097005 |
| Gm14015 | 1.812101644 | -0.072952994 | 0.648449004 | -0.112503826 |
| Gm14014 | 0.072025462 | 0.755288839 | 3.724945049 | 0.202765096 |
| BB218582 | 7.281320362 | -0.118137335 | 0.377408563 | -0.313022402 |
| Mpped2 | 264.5495323 | -0.340429146 | 0.098695966 | -3.44927113 |
| Gm13901 | 0.066050875 | 0.274391604 | 3.724945049 | 0.073663262 |
| Gm13902 | 0.292336004 | -0.187397027 | 1.609637724 | -0.116421866 |
| Arl14ep | 637.6380912 | 0.182020103 | 0.079561615 | 2.287787938 |
| Gm13900 | 0.111006857 | 0.755286617 | 3.724945049 | 0.202764499 |
| Gm13904 | 0.527697161 | -0.016963203 | 1.323746248 | -0.012814543 |
| Fshb | 0.248616998 | 0.274395128 | 3.034959991 | 0.090411448 |
| Gm24885 | 0 | NA | NA | NA |
| Gm13903 | 0 | NA | NA | NA |
| Kcna4 | 692.1728847 | -0.305731154 | 0.115268492 | -2.652339334 |
| Gm9864 | 1.048421568 | -0.37859294 | 0.996238913 | -0.380022237 |
| Gm23439 | 0 | NA | NA | NA |
| Gm23749 | 0.105852363 | -0.206492286 | 3.724945049 | -0.055434988 |
| Gm13912 | 1.945135077 | 1.95160132 | 1.076626503 | 1.812700425 |
| Gm13916 | 0.123845245 | 0.755285145 | 3.724945049 | 0.202764104 |
| Gm13915 | 0.038981395 | 0.274393828 | 3.724945049 | 0.073663859 |
| Gm13913 | 0 | NA | NA | NA |
| Gm13914 | 0 | NA | NA | NA |
| Rpl35a-ps4 | 1.664460293 | 0.44325441 | 0.892503378 | 0.496641717 |
| Gm13910 | 140.7366961 | -0.295783922 | 0.155806577 | -1.898404606 |
| Gm13923 | 0 | NA | NA | NA |
| Mettl15 | 54.80204305 | -0.338371437 | 0.170725234 | -1.981964992 |
| Gm24115 | 0.713863726 | -0.51621184 | 1.180891859 | -0.437137267 |
| Kif18a | 32.64308567 | 0.146842457 | 0.21531925 | 0.681975517 |
| Gm13924 | 0 | NA | NA | NA |
| Gm13925 | 0 | NA | NA | NA |
| Gm20638 | 0.165580463 | 0.274406349 | 2.773571271 | 0.098936109 |
| Bdnf | 331.5846942 | -0.576198141 | 0.218567494 | -2.636248096 |
| Gm13932 | 0.182620985 | 1.236194542 | 2.688869866 | 0.459745024 |

| | | | | |
|---|---|---|---|---|
| Lin7c | 1365.827837 | 0.138048304 | 0.062920759 | 2.194002537 |
| Platr8 | 0 | NA | NA | NA |
| Lgr4 | 521.05221 | -0.249463465 | 0.099037764 | -2.518872152 |
| Ccdc34 | 111.7137007 | -0.316466513 | 0.147346364 | -2.147772798 |
| Ccdc34os | 0 | NA | NA | NA |
| Gm13936 | 1.575920251 | 1.114820689 | 0.870019927 | 1.281373741 |
| Gm13937 | 0.157911564 | -0.206480316 | 2.815266154 | -0.073343089 |
| Gm22997 | 0.40088837 | 1.140227142 | 1.415056804 | 0.805781887 |
| Bbox1 | 28.0529557 | -0.992537805 | 0.266300868 | -3.727129437 |
| Fibin | 60.99905906 | 0.616630745 | 0.213620199 | 2.886575093 |
| Gm13935 | 0.04393175 | -0.206490061 | 3.724945049 | -0.055434391 |
| Gm10794 | 0.046345709 | 0.274391604 | 3.724945049 | 0.073663262 |
| Slc5a12 | 0.290078826 | -0.688641848 | 1.811650007 | -0.380118591 |
| Ano3 | 1378.79017 | -0.356564685 | 0.16572141 | -2.151590941 |
| Muc15 | 15.98576495 | 1.066519268 | 0.327050272 | 3.261025475 |
| Gm13942 | 0 | NA | NA | NA |
| Gm13941 | 0 | NA | NA | NA |
| Gm15130 | 0 | NA | NA | NA |
| 4930430A15Rik | 0.123841226 | 0.274391173 | 3.724945049 | 0.073663147 |
| Gm13961 | 21.20792997 | -0.32708248 | 0.23207109 | -1.409406401 |
| Olfr1275 | 0 | NA | NA | NA |
| Gm13962 | 0 | NA | NA | NA |
| Olfr1276 | 0.075043205 | 0.755286614 | 3.724945049 | 0.202764498 |
| Olfr1277 | 0 | NA | NA | NA |
| Olfr1278 | 0 | NA | NA | NA |
| Olfr1279 | 0 | NA | NA | NA |
| Olfr1280 | 0 | NA | NA | NA |
| Olfr1281 | 0 | NA | NA | NA |
| Olfr1282 | 0 | NA | NA | NA |
| Olfr1283 | 0.038981395 | 0.274393828 | 3.724945049 | 0.073663859 |
| Olfr1284 | 0 | NA | NA | NA |
| Olfr1285 | 0 | NA | NA | NA |
| Olfr1286 | 0 | NA | NA | NA |
| Olfr1287 | 0 | NA | NA | NA |
| Olfr1288 | 0.11197992 | 0.75528439 | 3.724945049 | 0.202763901 |
| Olfr1289 | 0 | NA | NA | NA |
| Olfr1290 | 0.062426873 | 0.755288839 | 3.724945049 | 0.202765096 |
| Olfr1291-ps1 | 0 | NA | NA | NA |
| Olfr1292-ps1 | 0 | NA | NA | NA |
| Olfr1293-ps | 0 | NA | NA | NA |
| Olfr1294 | 0.134986936 | -0.664675289 | 3.655468937 | -0.181830375 |
| Olfr1295 | 0.08486385 | 0.755286617 | 3.724945049 | 0.202764499 |
| Olfr1296-ps1 | 0 | NA | NA | NA |
| Olfr1297 | 0 | NA | NA | NA |
| Olfr1298 | 0 | NA | NA | NA |
| Olfr1299 | 0 | NA | NA | NA |
| Olfr1300-ps1 | 0 | NA | NA | NA |

| | | | | |
|---|---|---|---|---|
| Gm13922 | 0 | NA | NA | NA |
| Gm13930 | 0 | NA | NA | NA |
| Olfr1301 | 0.046345709 | 0.274391604 | 3.724945049 | 0.073663262 |
| Olfr1302 | 0 | NA | NA | NA |
| Olfr1303 | 0.046345709 | 0.274391604 | 3.724945049 | 0.073663262 |
| Gm26421 | 0 | NA | NA | NA |
| Olfr268-ps1 | 0.068097359 | -0.206487837 | 3.724945049 | -0.055433794 |
| Olfr1305 | 0 | NA | NA | NA |
| Olfr1306 | 0 | NA | NA | NA |
| Gm13928 | 0 | NA | NA | NA |
| Olfr1307 | 0 | NA | NA | NA |
| Olfr1308 | 0 | NA | NA | NA |
| Olfr1309 | 0 | NA | NA | NA |
| Gm13929 | 0 | NA | NA | NA |
| Olfr1310 | 0.048949532 | -0.206487837 | 3.724945049 | -0.055433794 |
| Olfr1311 | 0 | NA | NA | NA |
| Olfr1312 | 0 | NA | NA | NA |
| Olfr1313 | 0.066050875 | 0.274391604 | 3.724945049 | 0.073663262 |
| Olfr1314 | 0 | NA | NA | NA |
| Olfr1315-ps1 | 0 | NA | NA | NA |
| Olfr1316 | 0.049553047 | 0.274391604 | 3.724945049 | 0.073663262 |
| Olfr1317 | 0 | NA | NA | NA |
| Olfr1318 | 0 | NA | NA | NA |
| Lpcat4 | 780.3235412 | -0.351408165 | 0.094252957 | -3.728351633 |
| Nutm1 | 0.463200932 | 0.274415306 | 1.551173734 | 0.176908169 |
| Nop10 | 173.0751968 | -0.089224865 | 0.117064278 | -0.762186951 |
| Slc12a6 | 925.7716682 | -0.366040193 | 0.083234875 | -4.397678184 |
| Gm21985 | 0.288125237 | 1.480087932 | 3.707988866 | 0.399161914 |
| Emc4 | 460.0520838 | 0.164981266 | 0.073899373 | 2.232512381 |
| Katnbl1 | 227.6757466 | 0.102302055 | 0.106086409 | 0.964327633 |
| Gm13940 | 0 | NA | NA | NA |
| Emc7 | 1191.037724 | 0.033322702 | 0.066038562 | 0.504594606 |
| n-R5s202 | 0 | NA | NA | NA |
| Chrm5 | 22.04456264 | 0.04126069 | 0.221539516 | 0.186245284 |
| Aven | 92.20401975 | -0.221142743 | 0.115617719 | -1.91270634 |
| Gm22676 | 0.183828322 | -0.959894242 | 2.667155968 | -0.359894304 |
| AL691423.1 | 0.566015516 | -0.462416804 | 1.39471777 | -0.331548657 |
| Ryr3 | 1324.13524 | -0.22161617 | 0.270289517 | -0.81992144 |
| Tmco5b | 1.585639959 | 4.210504368 | 1.015881589 | 4.144680258 |
| Gm26505 | 0.194120612 | 1.257156926 | 2.639891475 | 0.476215382 |
| Fmn1 | 837.5690741 | -0.113781626 | 0.074935429 | -1.518395602 |
| Gm13967 | 4.347705094 | 0.702208488 | 0.545551917 | 1.287152452 |
| Gm13963 | 0.697976008 | 0.755309058 | 1.478697749 | 0.510793405 |
| Gm13965 | 7.79987822 | -0.267876389 | 0.381192003 | -0.702733497 |
| Gm13966 | 0.044191549 | -0.206490061 | 3.724945049 | -0.055434391 |
| AL691437.1 | 0 | NA | NA | NA |
| Gm13964 | 0.060174537 | 0.755286614 | 3.724945049 | 0.202764498 |

| | | | | |
|---|---|---|---|---|
| 4930533B01Rik | 0.045882455 | 0.755288839 | 3.724945049 | 0.202765096 |
| Grem1 | 36.57688901 | 0.20250291 | 0.223852538 | 0.904626375 |
| Scg5 | 2876.760524 | 0.092432021 | 0.095413297 | 0.968754077 |
| Arhgap11a | 55.95224419 | 0.748412022 | 0.24188682 | 3.094058707 |
| Gm21596 | 8.155034204 | 0.281951752 | 0.384816643 | 0.73269116 |
| Gjd2 | 128.1457705 | -0.33011226 | 0.13190385 | -2.50267342 |
| A530058N18Rik | 102.4745166 | -0.612934972 | 0.235777341 | -2.599634758 |
| Actc1 | 0.709411339 | 1.244996614 | 1.387748022 | 0.89713449 |
| C130080G10Rik | 3.984159091 | 0.822844228 | 0.502432312 | 1.637721556 |
| Aqr | 833.2625079 | -0.174046279 | 0.047085381 | -3.69639738 |
| Mir7002 | 0.230049058 | -0.655550379 | 2.117076221 | -0.309648926 |
| Zfp770 | 474.7123103 | -0.186972019 | 0.088176823 | -2.120421366 |
| Gm25103 | 0.044544648 | 0.755286614 | 3.724945049 | 0.202764498 |
| Gm29234 | 0.986742042 | -0.992971571 | 1.053121598 | -0.942884063 |
| Dph6 | 238.7373716 | -0.161993091 | 0.115665994 | -1.400524786 |
| Gm13972 | 0.263371022 | 0.234975714 | 1.812076923 | 0.129672042 |
| Gm13975 | 0.044191549 | -0.206490061 | 3.724945049 | -0.055434391 |
| Gm13974 | 2.444458607 | -0.281899447 | 0.632242066 | -0.445872652 |
| Gm28494 | 0 | NA | NA | NA |
| Gm28492 | 0.508786284 | -1.165972958 | 1.272945871 | -0.915964288 |
| Gm13973 | 0 | NA | NA | NA |
| Gm28493 | 0.257385133 | 0.755303993 | 2.06610423 | 0.365569162 |
| 4930528P14Rik | 0.479723347 | -0.18305382 | 1.403495505 | -0.130427079 |
| Gm13976 | 0.060174537 | 0.755286614 | 3.724945049 | 0.202764498 |
| 3110099E03Rik | 2.086914652 | 0.105611853 | 0.712175852 | 0.148294628 |
| Gm13977 | 0.219276728 | 0.755299055 | 2.607357526 | 0.289679895 |
| BC052040 | 104.7146314 | -0.453936496 | 0.143270469 | -3.168388433 |
| Mir1951 | 0.245748936 | -0.94905779 | 2.060458615 | -0.460605121 |
| Meis2 | 1352.514286 | -0.329780612 | 0.141353789 | -2.333015714 |
| 2700033N17Rik | 6.560169035 | 0.164495671 | 0.394924113 | 0.416524758 |
| G630016G05Rik | 19.18698179 | -0.480182606 | 0.300583296 | -1.597502634 |
| 2810405F15Rik | 1.233431567 | -0.704601719 | 0.987326357 | -0.713646216 |
| Gm13991 | 44.848571 | 0.010482617 | 0.165868361 | 0.06319841 |
| Gm13990 | 0.730890257 | -1.137388168 | 1.341015559 | -0.848154342 |
| Tmco5 | 9.517001854 | 0.471384061 | 0.370026155 | 1.273920923 |
| D330050G23Rik | 15.28741942 | -1.050232139 | 0.284997492 | -3.685057477 |
| Gm24914 | 0 | NA | NA | NA |
| Gm25514 | 0.127971488 | 0.274389832 | 3.687653711 | 0.0744077 |
| Gm29340 | 5.225310162 | 1.073374738 | 0.544729508 | 1.970472909 |
| Gm13981 | 0.117907917 | 1.204448679 | 3.72248341 | 0.323560523 |
| Spred1 | 3571.650489 | 0.085019044 | 0.08834504 | 0.962352206 |
| Mir674 | 5.596050554 | -0.37779063 | 0.411571266 | -0.917922754 |
| Rap1a-ps2 | 0.847725382 | -0.944527078 | 1.212742337 | -0.778835742 |
| Gm25189 | 0.284781514 | -0.206482516 | 2.061091955 | -0.100181128 |
| Fam98b | 385.3078931 | -0.048590231 | 0.086851312 | -0.559464561 |
| Rasgrp1 | 4351.652765 | -0.053237447 | 0.139082431 | -0.382776215 |
| Gm13982 | 0.169723681 | 0.755289365 | 3.415329259 | 0.221146867 |

| | | | | |
|---|---|---|---|---|
| Gm28183 | 0.166952883 | 0.274401811 | 2.756036668 | 0.099563919 |
| Gm13985 | 0.090890357 | 0.755282166 | 3.724945049 | 0.202763304 |
| 4930412B13Rik | 0 | NA | NA | NA |
| Gm13986 | 9.239123008 | 0.466888884 | 0.449852972 | 1.037869954 |
| Gm36979 | 0 | NA | NA | NA |
| Gm25308 | 0.048949532 | -0.206487837 | 3.724945049 | -0.055433794 |
| Thbs1 | 1597.215543 | 3.623566069 | 0.45802395 | 7.9113026 |
| Gm29233 | 0.136201742 | -0.67532486 | 2.944131489 | -0.229379993 |
| Fsip1 | 6.681599266 | 0.419345754 | 0.382920253 | 1.095125556 |
| Gm22800 | 0 | NA | NA | NA |
| Gm29232 | 0.049553047 | 0.274391604 | 3.724945049 | 0.073663262 |
| A930104D05Rik | 0.722394163 | -0.653614002 | 1.272286313 | -0.51373185 |
| Gpr176 | 129.1472356 | -0.465092356 | 0.120756241 | -3.851497456 |
| 1700054M17Rik | 0.459216712 | -0.192242066 | 1.416210775 | -0.135743965 |
| Gm22651 | 0 | NA | NA | NA |
| Eif2ak4 | 429.1232767 | -0.062959573 | 0.137705985 | -0.457202878 |
| Srp14 | 522.9120673 | 0.037426835 | 0.105352783 | 0.355252453 |
| Gm13983 | 0 | NA | NA | NA |
| Bmf | 64.79741328 | 0.527691754 | 0.148512244 | 3.553186867 |
| Gm14016 | 0 | NA | NA | NA |
| Bub1b | 14.01725363 | 2.784352802 | 0.441332752 | 6.308964816 |
| Pak6 | 596.5307493 | -0.440981456 | 0.150352124 | -2.932991193 |
| Ankrd63 | 286.9989274 | -0.703110887 | 0.331814472 | -2.118988009 |
| Plcb2 | 54.89159527 | 1.296880111 | 0.252508368 | 5.135988636 |
| Inafm2 | 508.6639642 | -0.01998883 | 0.088200443 | -0.22662959 |
| A430105I19Rik | 127.3481097 | 0.362069527 | 0.14235051 | 2.543507058 |
| Phgr1 | 0.264111009 | 1.88737933 | 3.710686682 | 0.50863344 |
| Disp2 | 4917.700072 | -0.137645149 | 0.0590268 | -2.331909374 |
| Knstrn | 21.59877154 | 1.123522178 | 0.389544548 | 2.884194337 |
| Gm14091 | 1.226992653 | -1.66786517 | 1.088736661 | -1.531927077 |
| Ivd | 475.053727 | -0.043426677 | 0.081334506 | -0.533926861 |
| Bahd1 | 560.8711464 | -0.200874073 | 0.0900357 | -2.23104917 |
| Chst14 | 36.12922769 | 0.597058097 | 0.213112564 | 2.80160909 |
| Gm14089 | 0.273591019 | 1.272631092 | 2.02731918 | 0.627740863 |
| Gm14088 | 0.141331096 | -0.206491164 | 3.56184083 | -0.057973159 |
| Ccdc32 | 448.3274695 | 0.599463719 | 0.118824684 | 5.044942667 |
| Rpusd2 | 91.45199017 | 0.153944523 | 0.144256019 | 1.067161871 |
| Casc5 | 31.52527001 | 2.180160928 | 0.36477748 | 5.976687295 |
| Rad51 | 19.29236894 | 0.325915786 | 0.348010788 | 0.936510583 |
| Rmdn3 | 789.494809 | 0.412301178 | 0.102696327 | 4.014760697 |
| Gchfr | 7.98091292 | -0.805009596 | 0.40550068 | -1.985223789 |
| Dnajc17 | 78.13003113 | 0.343909327 | 0.139885254 | 2.458510218 |
| Gm14137 | 2.178995086 | 2.193640802 | 1.110937801 | 1.974584715 |
| Gm14138 | 1.456796327 | -0.336751424 | 0.824558168 | -0.408402265 |
| Zfyve19 | 182.5094088 | 0.19041403 | 0.087808871 | 2.168505608 |
| Ppp1r14d | 0.089814205 | 0.274402725 | 3.724945049 | 0.073666248 |
| Spint1 | 12.20675764 | 0.805858606 | 0.312036292 | 2.582579742 |

| | | | | |
|---|---|---|---|---|
| Rhov | 153.8616767 | 0.862818008 | 0.173230362 | 4.980755104 |
| Gm14175 | 0 | NA | NA | NA |
| Vps18 | 675.7131133 | 0.088091388 | 0.055172617 | 1.596650533 |
| Gm14207 | 0.327809541 | 2.258329075 | 1.896473878 | 1.190804208 |
| Dll4 | 18.4801647 | 0.845929275 | 0.391298684 | 2.161850548 |
| Chac1 | 175.096179 | 0.879715895 | 0.26439732 | 3.327249665 |
| Ino80 | 813.323168 | 0.210799404 | 0.063880458 | 3.299904366 |
| Gm14208 | 0.083526043 | 0.75528439 | 3.724945049 | 0.202763901 |
| Exd1 | 16.27157243 | -0.865145253 | 0.287009417 | -3.014344479 |
| Chp1 | 1163.742042 | -0.146498116 | 0.055177904 | -2.655014139 |
| 1700020I14Rik | 6679.70898 | -0.498338224 | 0.088233184 | -5.647968301 |
| Oip5 | 7.956782189 | 0.378113085 | 0.379147429 | 0.99727192 |
| Nusap1 | 22.39643515 | 1.923131534 | 0.413279303 | 4.653345863 |
| n-R5s204 | 0 | NA | NA | NA |
| Ndufaf1 | 192.2611336 | 0.062053348 | 0.078611237 | 0.789369942 |
| H3f3c | 22.18271704 | 0.209812191 | 0.250429907 | 0.837808045 |
| Gm14383 | 0.267580615 | 0.28874427 | 1.817776807 | 0.158844732 |
| Rtf1 | 860.2757225 | 0.060709849 | 0.079826284 | 0.76052455 |
| Gm25857 | 1.36269325 | 0.896651797 | 0.791455401 | 1.132915128 |
| Itpka | 639.0705542 | 0.121019855 | 0.132438408 | 0.913782161 |
| Ltk | 62.0682925 | -0.115466471 | 0.139717817 | -0.826426245 |
| Rpap1 | 285.1701272 | -0.175096842 | 0.094527485 | -1.852337885 |
| Gm23814 | 0 | NA | NA | NA |
| Tyro3 | 1444.38765 | -0.587730171 | 0.086177448 | -6.819999712 |
| Gm13998 | 0 | NA | NA | NA |
| Mga | 2323.037544 | -0.193493329 | 0.065998024 | -2.931804862 |
| Mapkbp1 | 844.8660816 | -0.217738012 | 0.112043842 | -1.943328683 |
| Mir7665 | 0.088383099 | -0.687376175 | 3.724945049 | -0.184533239 |
| Gm13999 | 0 | NA | NA | NA |
| Jmjd7 | 2.162814427 | 0.027046218 | 0.739192013 | 0.036588894 |
| Gm28042 | 0.10503227 | 0.274389469 | 3.724945049 | 0.073662689 |
| Pla2g4b | 5.643927168 | -0.663317669 | 0.420063505 | -1.579089022 |
| Sptbn5 | 13.1047588 | 0.107775179 | 0.337951825 | 0.318906931 |
| Ehd4 | 458.2061878 | 1.460871775 | 0.158103255 | 9.239985456 |
| Pla2g4e | 246.4736302 | 0.09465322 | 0.138382718 | 0.68399596 |
| Gm13997 | 1.380678606 | -0.265810484 | 0.861608613 | -0.308504906 |
| Pla2g4d | 1.305860104 | 2.228446268 | 1.081395284 | 2.060713877 |
| Pla2g4f | 1.649860538 | 0.247982051 | 0.862866557 | 0.287393281 |
| Vps39 | 1015.694296 | 0.102931026 | 0.055379698 | 1.858641901 |
| Tmem87a | 799.4634507 | 0.321395142 | 0.065325311 | 4.919917494 |
| Ganc | 158.6786428 | 0.256304746 | 0.109927578 | 2.331578219 |
| Capn3 | 66.72612438 | 0.912503273 | 0.230679885 | 3.955712356 |
| Zfp106 | 4291.41446 | -0.073801379 | 0.048122425 | -1.53361722 |
| Gm26899 | 5.004844821 | 0.010459039 | 0.501033823 | 0.020874916 |
| Snap23 | 220.6488251 | -0.048720455 | 0.142264634 | -0.34246357 |
| Lrrc57 | 243.5709634 | 0.23297083 | 0.087741396 | 2.655198588 |
| Haus2 | 232.8508983 | 0.417319231 | 0.11946161 | 3.493333381 |

| | | | | |
|---|---|---|---|---|
| Stard9 | 314.5610356 | -0.296160707 | 0.173081017 | -1.711110278 |
| n-R5s205 | 0 | NA | NA | NA |
| Cdan1 | 293.1784353 | -0.125444094 | 0.156164552 | -0.803281488 |
| Ttbk2 | 1910.365418 | -0.339927182 | 0.065216794 | -5.212264489 |
| AV039307 | 6.926568459 | -0.899461403 | 0.410963696 | -2.188663895 |
| Ubr1 | 1554.190815 | -0.108956862 | 0.046490014 | -2.343661602 |
| Tmem62 | 383.5196607 | -0.059689818 | 0.072736765 | -0.820627893 |
| AL844548.1 | 1.862757 | 0.352162243 | 0.681413451 | 0.516811405 |
| Gm22865 | 3.966952568 | 0.878936984 | 0.644218143 | 1.364346834 |
| Ccndbp1 | 483.2632691 | -0.11709815 | 0.073814685 | -1.586380129 |
| Epb4.2 | 1.132377036 | 1.401574492 | 1.113304935 | 1.258931357 |
| Tgm5 | 0.516597926 | 0.284582554 | 1.529097782 | 0.186111416 |
| Gm14213 | 0 | NA | NA | NA |
| Tgm7 | 0.64993548 | -1.867203868 | 1.163195557 | -1.605236417 |
| Lcmt2 | 75.2947259 | 0.396623018 | 0.162582921 | 2.439512195 |
| Gm23858 | 0.195161473 | -0.171747621 | 2.671768033 | -0.064282385 |
| Adal | 331.7277646 | -0.129061844 | 0.064053823 | -2.014896827 |
| Zscan29 | 154.5557525 | 0.217963463 | 0.134302869 | 1.622924837 |
| Tubgcp4 | 399.137152 | -0.519651884 | 0.089453445 | -5.809188035 |
| Trp53bp1 | 1866.10298 | -0.32511 | 0.07557907 | -4.301587742 |
| Map1a | 23447.45946 | -0.151507373 | 0.068336432 | -2.217080528 |
| Gm14978 | 2.354129177 | -0.565206872 | 0.712727171 | -0.793019959 |
| Ppip5k1 | 1660.739469 | -0.21033242 | 0.084584102 | -2.486666107 |
| Ckmt1 | 1627.233016 | -0.239427219 | 0.066390474 | -3.606348995 |
| Strc | 2.337199581 | 0.854273403 | 0.675276248 | 1.265072488 |
| Catsper2 | 52.60120326 | -0.229651456 | 0.192885978 | -1.190607315 |
| Gm14001 | 3.014229995 | -2.009381524 | 0.63144996 | -3.182170642 |
| Gm14000 | 0 | NA | NA | NA |
| Pdia3 | 2693.142216 | 0.411255464 | 0.103205344 | 3.984827208 |
| Ell3 | 0.594313001 | 0.519763485 | 1.195683296 | 0.434699964 |
| Serinc4 | 3.508801613 | 0.843371499 | 0.539227802 | 1.564035636 |
| Serf2 | 753.224674 | 0.065370977 | 0.070236532 | 0.930726149 |
| Gm22953 | 0.938485331 | 0.629391354 | 1.180834699 | 0.53300547 |
| Hypk | 31.77497691 | 0.077357155 | 0.290251912 | 0.266517298 |
| Mfap1b | 106.0742756 | 0.170291815 | 0.134344831 | 1.267572511 |
| Gm14017 | 2.307785566 | -0.043756002 | 0.697069389 | -0.062771373 |
| Gm14018 | 0.899881821 | -1.341261687 | 1.045010347 | -1.283491298 |
| Mfap1a | 281.266986 | 0.140208168 | 0.063472781 | 2.208949496 |
| Wdr76 | 70.42691052 | -0.263371353 | 0.136823049 | -1.92490486 |
| Frmd5 | 605.5814555 | 0.047561371 | 0.073740409 | 0.644983819 |
| Gm14021 | 44.18792118 | -0.463073527 | 0.256089479 | -1.808248933 |
| Gm14020 | 0.04393175 | -0.206490061 | 3.724945049 | -0.055434391 |
| Gm14019 | 0.838491377 | -0.151765264 | 1.27424693 | -0.119101926 |
| Casc4 | 1001.124057 | -0.043609288 | 0.064551692 | -0.675571583 |
| Gm23983 | 0 | NA | NA | NA |
| Mageb3 | 1.086684801 | -0.850747838 | 0.964455377 | -0.882101814 |
| Ctdspl2 | 624.7685435 | 0.157995543 | 0.086209414 | 1.832694784 |

| | | | | |
|---|---|---|---|---|
| Eif3j1 | 235.1963187 | 0.148080624 | 0.130298618 | 1.136471174 |
| AL845457.1 | 0 | NA | NA | NA |
| Spg11 | 412.0809888 | -0.222315661 | 0.094842801 | -2.344043596 |
| Patl2 | 2.452851275 | 0.318182927 | 0.638128239 | 0.498619099 |
| B2m | 9604.418411 | 5.075159876 | 0.161465836 | 31.43178774 |
| Trim69 | 8.404189488 | 2.973981646 | 0.924410043 | 3.217167175 |
| 4933406J08Rik | 0.139037128 | 0.274384718 | 3.724945049 | 0.073661414 |
| Gm14050 | 0 | NA | NA | NA |
| Gm14058 | 0.04393175 | -0.206490061 | 3.724945049 | -0.055434391 |
| Rps12-ps10 | 0.709725355 | 1.53862068 | 1.361258388 | 1.130292892 |
| Sord | 185.688388 | 0.036107221 | 0.084824449 | 0.425669973 |
| Duox2 | 2.808703517 | -0.196149034 | 0.742609878 | -0.264134695 |
| Duoxa2 | 0.276302848 | 1.114582553 | 2.029676133 | 0.549143055 |
| Duoxa1 | 3.074242624 | 1.580093473 | 0.980424629 | 1.611641963 |
| Duox1 | 2.917555992 | 0.383544119 | 0.579208081 | 0.662187099 |
| Shf | 467.0710296 | 0.05981136 | 0.112445809 | 0.531912752 |
| Gm24409 | 0 | NA | NA | NA |
| Slc28a2 | 188.3325738 | 8.386396452 | 1.05930875 | 7.91685753 |
| Gm14086 | 0 | NA | NA | NA |
| Bambi-ps1 | 12.14750959 | 0.867709751 | 0.36425108 | 2.382174823 |
| Gm14085 | 41.28799092 | 7.053962856 | 0.921782234 | 7.652526374 |
| Gatm | 1118.00039 | 0.247270138 | 0.15471878 | 1.598190849 |
| Spata5l1 | 53.6862279 | 0.04212302 | 0.204849081 | 0.20562953 |
| AA467197 | 88.13508126 | 5.694845996 | 0.647573782 | 8.794126874 |
| Mir147 | 0 | NA | NA | NA |
| Slc30a4 | 1022.722383 | 0.217401736 | 0.08295077 | 2.620852547 |
| 4930417H01Rik | 4.957477421 | 0.349397581 | 0.500495485 | 0.698103364 |
| Bloc1s6os | 0.578202636 | -1.453225663 | 1.287543264 | -1.128681034 |
| Bloc1s6 | 473.7227365 | 0.672436723 | 0.103794368 | 6.478547312 |
| Sqrdl | 62.88662421 | 0.53611065 | 0.22125622 | 2.423030863 |
| Gm13988 | 0.046345709 | 0.274391604 | 3.724945049 | 0.073663262 |
| Sema6d | 1571.33555 | -0.370447571 | 0.077489225 | -4.78063329 |
| 4930517E11Rik | 0.136309558 | 1.557673883 | 3.610467106 | 0.431432786 |
| 4930583P06Rik | 0.286786342 | -0.206487158 | 2.043472028 | -0.101047215 |
| Gm13994 | 0 | NA | NA | NA |
| Slc24a5 | 5.913197406 | -0.011190594 | 0.391431264 | -0.028588912 |
| Myef2 | 1104.874051 | -0.148666339 | 0.087908831 | -1.691142267 |
| Gm22352 | 0 | NA | NA | NA |
| Gm14004 | 11.63604071 | -0.065647364 | 0.299927464 | -0.21887747 |
| Ctxn2 | 55.00778503 | -0.15507074 | 0.17711544 | -0.875534852 |
| Slc12a1 | 4.418703675 | -0.251241472 | 0.454513135 | -0.552770542 |
| Gm14002 | 3.728554256 | -0.087653733 | 0.54795649 | -0.159964769 |
| Dut | 174.499205 | -0.108764157 | 0.103923141 | -1.046582654 |
| A530010F05Rik | 0 | NA | NA | NA |
| Gm14003 | 0.116944185 | 0.274391125 | 3.724945049 | 0.073663134 |
| Fbn1 | 344.1427708 | 1.112839151 | 0.131786864 | 8.444234225 |
| Gm9913 | 2.575840985 | 0.948208924 | 0.676906563 | 1.400797357 |

| | | | | |
|---|---|---|---|---|
| Cep152 | 85.54108715 | -0.168368973 | 0.153889769 | -1.094088153 |
| Shc4 | 125.3363377 | -0.048188388 | 0.145125312 | -0.33204675 |
| Eid1 | 1534.982027 | 0.021214618 | 0.071355477 | 0.297308894 |
| Secisbp2l | 3169.337731 | 0.140893565 | 0.042998521 | 3.276707279 |
| Cops2 | 1452.800378 | 0.405853599 | 0.108315767 | 3.746948467 |
| Galk2 | 221.865611 | -0.013185302 | 0.089437781 | -0.147424298 |
| Fam227b | 26.91957889 | -1.018063611 | 0.295505579 | -3.445158682 |
| Fgf7 | 42.37332393 | -0.551651789 | 0.238858368 | -2.309535117 |
| Gm26496 | 0.514662899 | -1.111958762 | 1.359287093 | -0.818045553 |
| Dtwd1 | 64.96816098 | 0.212743032 | 0.139435696 | 1.52574297 |
| Gm17555 | 0 | NA | NA | NA |
| Gm24739 | 0 | NA | NA | NA |
| Atp8b4 | 80.28755012 | 6.391122106 | 0.639793747 | 9.989347565 |
| Gm26697 | 1.020000067 | -1.152217786 | 1.104046695 | -1.04363139 |
| Slc27a2 | 66.76116308 | -0.578916867 | 0.143884605 | -4.023480264 |
| Hdc | 76.84591766 | 1.698553574 | 0.272904176 | 6.223992616 |
| Gabpb1 | 280.3852881 | 0.22498598 | 0.116894722 | 1.924688958 |
| Gm23753 | 1.578568981 | -0.363203384 | 0.766217201 | -0.474021443 |
| Gm27003 | 13.76199491 | -0.650567494 | 0.351359555 | -1.851571946 |
| Usp8 | 1065.714832 | 0.057617424 | 0.054353669 | 1.060046638 |
| Usp50 | 5.50178608 | 0.10246718 | 0.476927741 | 0.214848438 |
| Gm10774 | 0 | NA | NA | NA |
| Trpm7 | 1310.402208 | -0.008772592 | 0.100334751 | -0.087433231 |
| Gm14212 | 0 | NA | NA | NA |
| Sppl2a | 1346.162496 | 0.846190902 | 0.101404127 | 8.344738337 |
| Mir3473g | 5.710082525 | 0.883834907 | 0.602778985 | 1.466266956 |
| Ap4e1 | 265.198369 | 0.12090097 | 0.126931556 | 0.952489466 |
| Gm24081 | 0 | NA | NA | NA |
| Blvra | 139.0631896 | 0.098721674 | 0.13761903 | 0.717354813 |
| Ncaph | 24.30005576 | 1.221100036 | 0.285237119 | 4.280999754 |
| Itpripl1 | 51.57497973 | 1.188788559 | 0.201361679 | 5.90374774 |
| 1810024B03Rik | 6.848270579 | 0.236309703 | 0.376391647 | 0.627829296 |
| Snrnp200 | 2169.627635 | 0.193184527 | 0.05493092 | 3.516863116 |
| Ciao1 | 447.7927703 | -0.165720395 | 0.099075507 | -1.672667655 |
| Tmem127 | 1631.67063 | 0.105223926 | 0.062161473 | 1.69275149 |
| Gm10766 | 3.235620746 | -1.520536937 | 0.595967312 | -2.551376401 |
| Stard7 | 1005.190084 | 0.027854642 | 0.052591098 | 0.529645573 |
| Dusp2 | 53.83102264 | 3.77967046 | 0.414766267 | 9.112772083 |
| Astl | 1.38321721 | 2.218043532 | 1.106495316 | 2.004566581 |
| Adra2b | 12.20916545 | -1.277064038 | 0.361951202 | -3.528276824 |
| Gpat2 | 10.37835042 | -0.275353931 | 0.321739066 | -0.855829959 |
| Fahd2a | 142.0280488 | -0.197348413 | 0.10481158 | -1.882887496 |
| Kcnip3 | 821.4776332 | -0.064184653 | 0.078878339 | -0.813717095 |
| Gm22613 | 5.411512881 | -0.943218137 | 0.53014605 | -1.779166585 |
| Prom2 | 9.157968272 | -0.291121385 | 0.36994034 | -0.786941443 |
| Gm14229 | 0 | NA | NA | NA |
| Zfp661 | 72.87159097 | -0.416702273 | 0.137018751 | -3.041206193 |

| | | | | |
|---|---|---|---|---|
| Mrps5 | 469.1048793 | -0.008009676 | 0.066856807 | -0.119803442 |
| Mal | 1616.595918 | 0.0481201 | 0.095089745 | 0.506049309 |
| Gm22889 | 0 | NA | NA | NA |
| Gm14245 | 0 | NA | NA | NA |
| Gm14244 | 0.04393175 | -0.206490061 | 3.724945049 | -0.055434391 |
| Mall | 5.752169691 | 0.535172962 | 0.478656047 | 1.118074168 |
| Nphp1 | 102.8958306 | 0.210081649 | 0.108771447 | 1.931404385 |
| 1500011K16Rik | 147.7564642 | 0.145049682 | 0.179839914 | 0.806548886 |
| Bub1 | 15.64735813 | 1.834213794 | 0.452285482 | 4.055433721 |
| Acoxl | 0 | NA | NA | NA |
| Gm22411 | 0.044191549 | -0.206490061 | 3.724945049 | -0.055434391 |
| Gm23101 | 0 | NA | NA | NA |
| Bcl2l11 | 158.1317849 | 0.995433864 | 0.18226596 | 5.46143594 |
| Gm14009 | 0.120974994 | 0.274404949 | 3.724945049 | 0.073666845 |
| Gm14005 | 19.18807827 | 2.918906969 | 0.57592433 | 5.068212643 |
| Gm14012 | 0.247225956 | 1.236189465 | 2.992047273 | 0.4131584 |
| Gm14010 | 0.594047536 | 1.232913393 | 1.435103619 | 0.859111061 |
| Gm29010 | 0 | NA | NA | NA |
| Gm14006 | 0.075043205 | 0.755286614 | 3.724945049 | 0.202764498 |
| Gm14008 | 0.04393175 | -0.206490061 | 3.724945049 | -0.055434391 |
| Gm355 | 0 | NA | NA | NA |
| Gm14007 | 0 | NA | NA | NA |
| Anapc1 | 1911.795479 | 0.0208817 | 0.058805891 | 0.355095375 |
| Mertk | 498.8880327 | 0.321723482 | 0.1350657 | 2.381977672 |
| Gm14011 | 0 | NA | NA | NA |
| Gm23172 | 3.964087094 | -0.079657777 | 0.71240885 | -0.111814693 |
| Tmem87b | 666.3521244 | 0.174952928 | 0.074743752 | 2.340703042 |
| Fbln7 | 48.08116317 | -0.486035223 | 0.206050428 | -2.358816859 |
| Gm22859 | 0 | NA | NA | NA |
| Zc3h8 | 36.68443589 | -0.584092227 | 0.249864623 | -2.337634759 |
| Gm14028 | 0 | NA | NA | NA |
| 4933427J07Rik | 0.358075244 | -0.414335564 | 1.677820589 | -0.246948671 |
| Gm10762 | 30.64394088 | 0.285815331 | 0.206471788 | 1.384282729 |
| Zc3h6 | 404.6004255 | 0.359247623 | 0.109168078 | 3.290775362 |
| Gm14027 | 11.52035073 | -0.278106588 | 0.308245392 | -0.902224642 |
| Gm14025 | 1.764317597 | 0.285965414 | 0.894854651 | 0.319566327 |
| Gm14022 | 5.545450415 | -1.147225301 | 0.445308256 | -2.57624979 |
| Ttl | 570.9256818 | -0.560992904 | 0.096054499 | -5.840360519 |
| Polr1b | 151.5190305 | -0.050967928 | 0.151403293 | -0.336636855 |
| Chchd5 | 87.19880141 | 0.380469249 | 0.157072361 | 2.422254597 |
| AI847159 | 3.26828567 | -0.272340437 | 0.657545494 | -0.414177331 |
| Gm14026 | 27.00074528 | 0.059884561 | 0.23003982 | 0.260322588 |
| Gm14029 | 5.503856369 | 0.23659109 | 0.490930222 | 0.481924069 |
| Gm4430 | 83.58514794 | -0.235909459 | 0.129670611 | -1.819297816 |
| Slc20a1 | 1626.484812 | 0.045033385 | 0.079846159 | 0.564001891 |
| 9830144P21Rik | 0.631002787 | 1.070050066 | 1.327040585 | 0.806343135 |
| A730036I17Rik | 9.097055571 | -0.343701498 | 0.388121604 | -0.885551058 |

| | | | | |
|---|---|---|---|---|
| Gm14024 | 0.180493943 | -0.206488401 | 2.699100097 | -0.076502684 |
| Ckap2l | 17.79315673 | 0.816106433 | 0.483448614 | 1.688093438 |
| Gm14023 | 2.610220779 | -0.327590733 | 0.660759406 | -0.495779145 |
| Il1a | 25.73372477 | 2.221657891 | 0.316710794 | 7.014784257 |
| Gm25703 | 0 | NA | NA | NA |
| Gm14039 | 0 | NA | NA | NA |
| Il1b | 17.42109151 | 3.601921819 | 0.54560998 | 6.601642114 |
| Il1bos | 0.58991373 | -0.120182183 | 1.327044508 | -0.090563792 |
| Spcs2-ps | 12.01699771 | 0.48100976 | 0.304075265 | 1.581877301 |
| F830045P16Rik | 7.997923877 | -0.349883675 | 0.47713871 | -0.733295513 |
| Gm14040 | 0.25468147 | -1.116550492 | 2.104063842 | -0.53066379 |
| Gm14041 | 2.468877162 | -1.140205333 | 0.703682739 | -1.620340062 |
| Sirpa | 3103.836334 | -0.210629549 | 0.083939089 | -2.509314228 |
| Pdyn | 260.8278238 | 0.101075095 | 0.150515472 | 0.671526279 |
| Gm14046 | 0.049553047 | 0.274391604 | 3.724945049 | 0.073663262 |
| Stk35 | 654.0656122 | 0.026802552 | 0.064058558 | 0.418407047 |
| Gm28196 | 0.664320874 | -0.259747225 | 1.556991942 | -0.166826313 |
| Tgm3 | 28.71390692 | 0.468974759 | 0.257138618 | 1.823820793 |
| Mir6339 | 0 | NA | NA | NA |
| Gm22424 | 0.046345709 | 0.274391604 | 3.724945049 | 0.073663262 |
| Gm24665 | 0.038981395 | 0.274393828 | 3.724945049 | 0.073663859 |
| AU015228 | 0.066050875 | 0.274391604 | 3.724945049 | 0.073663262 |
| Tgm6 | 0.648101363 | -0.612874955 | 1.076120978 | -0.569522356 |
| Snrpb | 546.91693 | 0.433047232 | 0.106566121 | 4.063648242 |
| Tmc2 | 0.617199072 | -0.318761259 | 1.223049732 | -0.260628207 |
| Nop56 | 546.9667433 | 0.153202147 | 0.102175117 | 1.499407597 |
| Snord110 | 0.459163063 | -1.188203828 | 1.274644767 | -0.932184291 |
| Gm24451 | 0 | NA | NA | NA |
| Gm23187 | 0 | NA | NA | NA |
| Gm23650 | 0 | NA | NA | NA |
| Snord57 | 0 | NA | NA | NA |
| Idh3b | 1254.967299 | -0.11287982 | 0.057095173 | -1.977046639 |
| Ebf4 | 86.18724845 | -0.509041503 | 0.123999321 | -4.105195885 |
| Gm14044 | 0.90691343 | -0.158251668 | 1.132907599 | -0.139686298 |
| Cpxm1 | 57.79122351 | -0.117126279 | 0.270034613 | -0.433745427 |
| Gm14042 | 0.049553047 | 0.274391604 | 3.724945049 | 0.073663262 |
| Gm28372 | 0 | NA | NA | NA |
| 1700020A23Rik | 0.230076657 | 1.606735026 | 3.108237438 | 0.516928021 |
| Tmem239 | 0.066050875 | 0.274391604 | 3.724945049 | 0.073663262 |
| Pced1a | 228.252696 | -0.126984024 | 0.098434788 | -1.290031972 |
| Vps16 | 502.9920582 | -0.036599592 | 0.06313217 | -0.579729664 |
| Ptpra | 1842.435841 | 0.118261532 | 0.061788793 | 1.913964091 |
| Gm23876 | 0 | NA | NA | NA |
| 4930473A02Rik | 0.931693893 | 0.768279135 | 1.089581471 | 0.705113987 |
| Mrps26 | 372.6054324 | 0.11460437 | 0.126272002 | 0.907599216 |
| Oxt | 55.53220185 | -0.048998546 | 0.528851858 | -0.092650797 |
| Avp | 105.9017596 | -0.058466133 | 0.310325273 | -0.188402744 |

| | | | | |
|---|---|---|---|---|
| Ubox5 | 144.7706665 | -0.048150455 | 0.109700988 | -0.438924539 |
| Fastkd5 | 53.00231664 | -0.571878595 | 0.174586093 | -3.275625148 |
| Lzts3 | 1886.840697 | -0.11387353 | 0.135817729 | -0.838429055 |
| Ddrgk1 | 422.8448348 | 0.327622001 | 0.088282086 | 3.711081335 |
| Itpa | 134.0024633 | 0.244009397 | 0.116690126 | 2.091088637 |
| Slc4a11 | 82.95192658 | 1.983803265 | 0.236100723 | 8.402359994 |
| Gm24893 | 1.856422655 | 0.160941159 | 0.661810509 | 0.243183142 |
| 4930402H24Rik | 1406.23195 | -0.785248182 | 0.111895791 | -7.017673991 |
| Gm14057 | 0.232395771 | 1.236201847 | 2.124734505 | 0.581814737 |
| A730017L22Rik | 0.834338752 | 0.903953759 | 1.10702942 | 0.816558027 |
| Gm14048 | 32.5064405 | 0.488557983 | 0.219921148 | 2.221514339 |
| Atrn | 2962.760329 | -0.287985503 | 0.071654207 | -4.019101127 |
| Gm14049 | 10.85113572 | -0.282579216 | 0.334953132 | -0.843638076 |
| Gfra4 | 152.1835605 | -0.113313063 | 0.133036217 | -0.851745979 |
| Mir6973b | 0 | NA | NA | NA |
| Adam33 | 45.18950367 | -0.281857707 | 0.210885968 | -1.336540832 |
| Siglec1 | 680.3250049 | 5.625297663 | 0.217870232 | 25.81948719 |
| Hspa12b | 36.83173324 | 0.665922133 | 0.211683527 | 3.145838238 |
| Gm11037 | 0.127971488 | 0.274389832 | 3.687653711 | 0.0744077 |
| 1700037H04Rik | 296.061265 | -0.373263479 | 0.087285271 | -4.276362694 |
| Gm14232 | 0.292476601 | 1.67318075 | 1.738169557 | 0.962610778 |
| Spef1 | 54.9579452 | -0.482268661 | 0.154920988 | -3.112997585 |
| Cenpb | 1691.510661 | 0.281571185 | 0.087266575 | 3.226563963 |
| Cdc25b | 96.01317364 | -0.356507723 | 0.152950053 | -2.330876753 |
| Ap5s1 | 93.61776372 | -0.220981771 | 0.117704418 | -1.877429698 |
| Mavs | 109.308156 | -0.333775726 | 0.196085403 | -1.702195683 |
| Pank2 | 601.5473895 | -0.262051153 | 0.067007153 | -3.910793742 |
| Mir103-2 | 2.477700388 | -0.950763653 | 0.647884394 | -1.467489666 |
| Rnf24 | 607.3774196 | 0.047049134 | 0.063300375 | 0.743267866 |
| Gm14233 | 1.848697502 | -1.461433835 | 0.754258006 | -1.937578154 |
| Gm14284 | 25.84299431 | 0.312465008 | 0.22051256 | 1.416994148 |
| Gm22902 | 0 | NA | NA | NA |
| Gm14283 | 0 | NA | NA | NA |
| Gm14304 | 0.048949532 | -0.206487837 | 3.724945049 | -0.055433794 |
| Smox | 612.36022 | 0.448102359 | 0.116148506 | 3.858012251 |
| Adra1d | 160.5552724 | -0.396387168 | 0.122024305 | -3.248427986 |
| Gm14285 | 9.998759481 | 0.224117397 | 0.384875772 | 0.582311005 |
| Gm14280 | 0.350616066 | 1.744903481 | 1.641532238 | 1.062972411 |
| 5330413P13Rik | 17.0164514 | -0.46941423 | 0.30294886 | -1.549483401 |
| Gm14282 | 0 | NA | NA | NA |
| Gm14281 | 0.072025462 | 0.755288839 | 3.724945049 | 0.202765096 |
| Erv3 | 13.96015691 | 1.084813486 | 0.515181353 | 2.105692452 |
| Lamr1-ps1 | 0 | NA | NA | NA |
| Prnp | 7268.086947 | 0.56671285 | 0.091792802 | 6.17382669 |
| Gm28045 | 0.311063295 | 1.224254379 | 1.787616453 | 0.684852937 |
| Prnd | 0.302678705 | 2.189698824 | 1.732506501 | 1.263890682 |
| Rassf2 | 1100.292024 | 0.127584133 | 0.113747662 | 1.121641808 |

| | | | | |
|---|---|---|---|---|
| Slc23a2 | 2945.124527 | 0.377165049 | 0.071179567 | 5.298782611 |
| Gm14052 | 0.072025462 | 0.755288839 | 3.724945049 | 0.202765096 |
| Gm14051 | 0.790290772 | 1.068741695 | 1.213091375 | 0.881006754 |
| Tmem230 | 195.6462545 | -0.166906149 | 0.102503527 | -1.628296645 |
| Pcna | 440.6784249 | 0.48497207 | 0.157342967 | 3.082260858 |
| AV099323 | 3.007737845 | -1.525829509 | 0.690038526 | -2.21122365 |
| Cds2 | 6744.842328 | -0.254400389 | 0.060725977 | -4.189317327 |
| Prokr2 | 44.48347877 | -0.880541854 | 0.21533574 | -4.089157953 |
| 4921508D12Rik | 3.197768018 | 3.33384588 | 0.931448402 | 3.579206184 |
| Gm14095 | 0 | NA | NA | NA |
| 1700026D11Rik | 0.269634638 | 1.734717565 | 1.806416841 | 0.960308565 |
| Gpcpd1 | 1339.978017 | -0.128539166 | 0.093191837 | -1.379296398 |
| AU019990 | 0.221462189 | -0.207202482 | 2.214040853 | -0.093585663 |
| Rpl23a-ps4 | 0 | NA | NA | NA |
| Gm14099 | 0 | NA | NA | NA |
| Gm14098 | 0.075043205 | 0.755286614 | 3.724945049 | 0.202764498 |
| 1110034G24Rik | 59.26919426 | -0.241929981 | 0.185748184 | -1.302462159 |
| Gm14097 | 0.654170826 | 1.141642039 | 1.392093958 | 0.820089788 |
| Gm22245 | 1.105592398 | -1.504161905 | 0.978097249 | -1.537844939 |
| Chgb | 2627.906697 | 0.239604204 | 0.25286968 | 0.947540267 |
| Trmt6 | 322.2692483 | 0.151246784 | 0.081975056 | 1.845034229 |
| Mcm8 | 58.14385559 | 0.040735679 | 0.175890338 | 0.231597027 |
| Gm14104 | 1.507066517 | -0.640289623 | 0.828609293 | -0.772728026 |
| Crls1 | 253.7650353 | 0.166909497 | 0.107542518 | 1.552032632 |
| Lrrn4 | 5.522564292 | 0.124487644 | 0.545579862 | 0.228174924 |
| Fermt1 | 3.449481863 | -0.364735224 | 0.65801318 | -0.554297748 |
| Gm14102 | 0.088736197 | 0.274400501 | 3.724945049 | 0.073665651 |
| Gm14101 | 0 | NA | NA | NA |
| Gm25704 | 0 | NA | NA | NA |
| Gm14103 | 0 | NA | NA | NA |
| Gm14094 | 0.049553047 | 0.274391604 | 3.724945049 | 0.073663262 |
| Gm27449 | 0 | NA | NA | NA |
| A430048G15Rik | 0.11037557 | 0.274404949 | 3.724945049 | 0.073666845 |
| Gm14100 | 0 | NA | NA | NA |
| Bmp2 | 72.0830543 | 0.234576411 | 0.178607093 | 1.313365596 |
| Mir3090 | 0 | NA | NA | NA |
| Gm25258 | 0 | NA | NA | NA |
| Hao1 | 8.182416264 | 4.374734039 | 0.851111295 | 5.140025827 |
| Tmx4 | 4150.07463 | 0.034426354 | 0.066548848 | 0.51730954 |
| Gm14037 | 0.044191549 | -0.206490061 | 3.724945049 | -0.055434391 |
| Gm14038 | 8.079079625 | 0.1728837 | 0.34901789 | 0.495343376 |
| Plcb1 | 2558.946918 | -0.341730842 | 0.097191371 | -3.516061543 |
| Gm14036 | 21.28777894 | 0.195271775 | 0.234526313 | 0.83262203 |
| 4930545L23Rik | 0.686786262 | 1.023164151 | 1.230659845 | 0.831394764 |
| 9630028H03Rik | 3.015362371 | 0.446616713 | 0.591379425 | 0.755211788 |
| Platr3 | 0.132101749 | 0.274387056 | 3.724945049 | 0.073662041 |
| Plcb4 | 1658.213635 | -0.215517707 | 0.059729503 | -3.608228723 |

| | | | | |
|---|---|---|---|---|
| Gm14209 | 16.51388486 | -0.636997531 | 0.335462873 | -1.89886149 |
| Gm14211 | 0.046345709 | 0.274391604 | 3.724945049 | 0.073663262 |
| Gm14210 | 0 | NA | NA | NA |
| Gm26227 | 0.166709681 | -0.206481506 | 2.767631179 | -0.074605861 |
| Lamp5 | 675.5566326 | -0.001371564 | 0.152684157 | -0.008983015 |
| Pak7 | 618.9095195 | -0.53706886 | 0.119699207 | -4.486820542 |
| Gm14218 | 0 | NA | NA | NA |
| Gm14056 | 0.369423471 | 0.274413248 | 1.591049024 | 0.172473157 |
| Gm14054 | 0.515606452 | 2.647449943 | 1.693841514 | 1.562985628 |
| Ankef1 | 28.51623535 | -0.182014895 | 0.196568392 | -0.92596217 |
| Gm36967 | 0.044544648 | 0.755286614 | 3.724945049 | 0.202764498 |
| Snap25 | 16400.01604 | -0.314292422 | 0.076271652 | -4.1206977 |
| Gm14053 | 0.15906016 | 0.274395307 | 3.439462686 | 0.079778539 |
| Mkks | 278.0206276 | -0.622330839 | 0.149696392 | -4.157286842 |
| Slx4ip | 166.0135647 | -0.307324991 | 0.12139811 | -2.531546758 |
| Jag1 | 129.0559854 | -0.708185839 | 0.14074239 | -5.031787774 |
| Gm28214 | 0.110595523 | 0.755282166 | 3.724945049 | 0.202763304 |
| Gm14055 | 0.284116082 | 1.236203209 | 2.014748405 | 0.613576964 |
| Gm14064 | 0 | NA | NA | NA |
| Gm14062 | 10.87638177 | 5.891781841 | 0.874251893 | 6.739226858 |
| Btbd3 | 2584.985912 | -0.398299315 | 0.107916729 | -3.690802335 |
| Gm14061 | 4.211247599 | 0.084291696 | 0.484966312 | 0.173809383 |
| Mir1952 | 0 | NA | NA | NA |
| Gm14065 | 0.143485256 | 0.274391988 | 3.549605256 | 0.077302113 |
| Gm14067 | 0 | NA | NA | NA |
| Gm14069 | 0 | NA | NA | NA |
| Gm14066 | 25.91994233 | -0.148650278 | 0.210337606 | -0.706722307 |
| Gm14071 | 0 | NA | NA | NA |
| Sptlc3 | 1.944739822 | 0.448182417 | 0.860920214 | 0.52058531 |
| Gm27385 | 0 | NA | NA | NA |
| Ism1 | 46.24466388 | 0.152550077 | 0.164156264 | 0.929297935 |
| Tasp1 | 470.1320558 | -0.051163811 | 0.089589952 | -0.571088722 |
| Gm14070 | 0 | NA | NA | NA |
| Rps19-ps7 | 1.19142324 | 0.09521495 | 1.081113213 | 0.088071211 |
| Gm14073 | 0.066050875 | 0.274391604 | 3.724945049 | 0.073663262 |
| Esf1 | 372.9243209 | 0.255320804 | 0.089501721 | 2.85269157 |
| Gm17374 | 0 | NA | NA | NA |
| Ndufaf5 | 256.6486657 | -0.038819862 | 0.080586353 | -0.481717566 |
| Sel1l2 | 0 | NA | NA | NA |
| Macrod2 | 727.1380872 | -0.044525313 | 0.089520186 | -0.497377359 |
| Gm23846 | 0 | NA | NA | NA |
| Flrt3 | 365.9243577 | -0.017688034 | 0.076061052 | -0.232550479 |
| Calr-ps | 0.108772582 | 0.75528439 | 3.724945049 | 0.202763901 |
| Zcchc9-ps | 30.12202849 | -0.557125871 | 0.372890131 | -1.49407513 |
| Macrod2os2 | 0 | NA | NA | NA |
| Macrod2os1 | 0 | NA | NA | NA |
| Gm25583 | 2.242176843 | -0.226630833 | 0.57516839 | -0.394025188 |

| | | | | |
|---|---|---|---|---|
| Fau-ps1 | 8.271865471 | -0.556891451 | 0.483659094 | -1.15141317 |
| Gm23179 | 0.743559953 | -0.747577928 | 1.204427751 | -0.620691384 |
| Rpl26-ps5 | 0 | NA | NA | NA |
| Rbmx2-ps | 0.144013665 | 0.274391019 | 3.543609481 | 0.077432635 |
| Kif16b | 346.7941092 | -0.331385926 | 0.102914163 | -3.220022556 |
| Gm22310 | 0.089089296 | 1.194485492 | 3.723939576 | 0.320758559 |
| Kif16bos | 0.612922179 | 0.699649291 | 1.277631344 | 0.547614376 |
| Snrpb2 | 243.6495243 | 0.609603574 | 0.133870351 | 4.553686215 |
| Otor | 0.348607679 | 0.274407441 | 1.976929533 | 0.138804867 |
| Gm14083 | 0 | NA | NA | NA |
| Gm14082 | 0 | NA | NA | NA |
| Gm11644 | 0 | NA | NA | NA |
| Gm11645 | 3.276776717 | -0.220781921 | 0.655129738 | -0.337004884 |
| Pcsk2os1 | 65.67909268 | -0.085838382 | 0.167295877 | -0.51309323 |
| Pcsk2 | 3501.594654 | -0.074751056 | 0.104880248 | -0.712727679 |
| Pcsk2os2 | 0.594248913 | 0.730791372 | 1.433669193 | 0.509735004 |
| Bfsp1 | 5.877618629 | 0.108727564 | 0.414984497 | 0.262003918 |
| Rps13-ps6 | 0.132101749 | 0.274387056 | 3.724945049 | 0.073662041 |
| Gm11673 | 0 | NA | NA | NA |
| Gm11686 | 0.0878635 | -0.687376175 | 3.724945049 | -0.184533239 |
| Dstn | 1131.896076 | 0.024870697 | 0.091142325 | 0.272877583 |
| Rrbp1 | 1060.324141 | 1.121772797 | 0.171563496 | 6.538528433 |
| Banf2 | 0 | NA | NA | NA |
| Banf2os | 0.710948324 | -0.005788712 | 1.084504685 | -0.005337655 |
| Gm11687 | 1.624027788 | 0.448559406 | 0.878544422 | 0.510571116 |
| Gm5535 | 4.621059661 | -0.008679712 | 0.449308717 | -0.019317924 |
| 4930444E06Rik | 0.044191549 | -0.206490061 | 3.724945049 | -0.055434391 |
| Snx5 | 1040.74114 | 0.438499354 | 0.097813579 | 4.483011038 |
| Gm11688 | 3.796974204 | 0.546478601 | 0.528138657 | 1.034725624 |
| Gm26165 | 17.87795363 | 0.168461384 | 0.300767607 | 0.56010481 |
| Snord17 | 426.2071553 | 0.136030775 | 0.144896844 | 0.938811164 |
| Mgme1 | 58.73843915 | -0.540702052 | 0.165067753 | -3.275637075 |
| Ovol2 | 23.1269052 | 0.424844516 | 0.297442168 | 1.428326452 |
| 1700108N11Rik | 1.600434186 | 1.370781234 | 0.811016624 | 1.690201154 |
| Gm11700 | 0 | NA | NA | NA |
| Gm11701 | 1.140052013 | 0.196745636 | 0.947910949 | 0.207557088 |
| Csrp2bp | 346.2940874 | -0.03706232 | 0.065701464 | -0.564101886 |
| Pet117 | 0 | NA | NA | NA |
| Zfp133-ps | 17.17923113 | -1.488108171 | 0.291905156 | -5.097916713 |
| Dzank1 | 3640.794771 | -0.25028615 | 0.06189686 | -4.043600131 |
| Polr3f | 438.0816624 | 0.283611392 | 0.116378264 | 2.436979057 |
| Rbbp9 | 270.2515903 | -0.369715944 | 0.135715295 | -2.724202478 |
| Sec23b | 663.5913093 | 0.385526859 | 0.091172446 | 4.22854573 |
| Gm561 | 84.95980064 | -0.131774429 | 0.153339698 | -0.859362777 |
| Dtd1 | 350.4218442 | -0.058911958 | 0.072423052 | -0.813442072 |
| Gm24823 | 0 | NA | NA | NA |
| Gm11743 | 12.81133691 | -0.20155684 | 0.401304597 | -0.502254002 |

| | | | | |
|---|---|---|---|---|
| Scp2d1 | 0.431157666 | 2.36711553 | 1.506530558 | 1.57123632 |
| Gm11746 | 0 | NA | NA | NA |
| Slc24a3 | 1355.879502 | -0.030149517 | 0.077056169 | -0.391266759 |
| Gm14093 | 14.45402812 | -0.429604721 | 0.318552174 | -1.348616507 |
| Gm22731 | 3.697861772 | -0.728492796 | 0.555293139 | -1.311906712 |
| Gm22390 | 0 | NA | NA | NA |
| Gm11759 | 0.254720735 | 0.755309979 | 1.838959555 | 0.410726803 |
| Gm14092 | 0.048949532 | -0.206487837 | 3.724945049 | -0.055433794 |
| BC039771 | 5.098733496 | -0.015704083 | 0.522500372 | -0.030055641 |
| Gm11763 | 0.066050875 | 0.274391604 | 3.724945049 | 0.073663262 |
| Rin2 | 761.9091662 | 0.807118448 | 0.124502536 | 6.482747045 |
| Gm25026 | 1.82714634 | -1.32731777 | 0.811073396 | -1.636495262 |
| Gm11764 | 0 | NA | NA | NA |
| Naa20 | 524.4015188 | 0.537058771 | 0.100030836 | 5.368932154 |
| Crnkl1 | 313.9592317 | 0.084659779 | 0.074206125 | 1.140873195 |
| Cfap61 | 40.89814708 | -0.633043848 | 0.182385189 | -3.470916968 |
| Insm1 | 43.73815878 | -0.553967245 | 0.191740686 | -2.889148134 |
| Ralgapa2 | 804.7755793 | -0.170484301 | 0.151300253 | -1.126794558 |
| A930019D19Rik | 0.11147366 | 0.274389235 | 3.724945049 | 0.073662626 |
| Gm14117 | 89.00856018 | 0.103788315 | 0.130729256 | 0.793918039 |
| 4933406D12Rik | 0 | NA | NA | NA |
| Gm14111 | 0.398068774 | 2.480963831 | 3.688640457 | 0.672595733 |
| Gm14114 | 1.234305601 | -0.20936412 | 0.868515094 | -0.241059853 |
| Kiz | 194.5668815 | -0.019832203 | 0.100628732 | -0.197082906 |
| Xrn2 | 1167.834356 | 0.595960797 | 0.087482913 | 6.812310853 |
| Nkx2-4 | 1.324777524 | 0.090336011 | 1.336078838 | 0.067612785 |
| Gm14116 | 0.119587853 | 1.218392466 | 3.723618458 | 0.327206581 |
| Nkx2-2 | 88.16107763 | 0.124093915 | 0.145796108 | 0.851146969 |
| Nkx2-2os | 6.403733378 | -0.133337715 | 0.408757111 | -0.326202803 |
| 6430503K07Rik | 6.531065735 | 0.196655533 | 0.42805088 | 0.45942093 |
| Gm22261 | 0 | NA | NA | NA |
| Gm14115 | 0.048949532 | -0.206487837 | 3.724945049 | -0.055433794 |
| Gm14113 | 3.843754305 | -0.099119523 | 0.482497948 | -0.205429937 |
| Gm14110 | 0 | NA | NA | NA |
| AI646519 | 0 | NA | NA | NA |
| Pax1 | 0.59047672 | -0.215015715 | 1.357923013 | -0.158341609 |
| Gm14109 | 0 | NA | NA | NA |
| Gm14107 | 0 | NA | NA | NA |
| Gm14106 | 0.066050875 | 0.274391604 | 3.724945049 | 0.073663262 |
| A530006G24Rik | 0 | NA | NA | NA |
| Gm25516 | 0 | NA | NA | NA |
| 9030622O22Rik | 0.80144685 | -1.380375643 | 1.253297867 | -1.101394712 |
| Foxa2 | 2.785346695 | 0.694821316 | 0.817233345 | 0.850211657 |
| Gm27420 | 0 | NA | NA | NA |
| Gm14108 | 0 | NA | NA | NA |
| Gm24221 | 0 | NA | NA | NA |
| Sstr4 | 86.59783816 | -0.396526772 | 0.146106421 | -2.713958565 |

| | | | | |
|---|---|---|---|---|
| Thbd | 190.3799444 | 0.463345679 | 0.176198347 | 2.629682334 |
| Gm14119 | 0.109727584 | 0.755282166 | 3.724945049 | 0.202763304 |
| Cd93 | 290.705625 | 2.97171527 | 0.212374417 | 13.99281189 |
| Gm14120 | 2.718509683 | 0.917559674 | 0.730153018 | 1.256667645 |
| Gm25866 | 0 | NA | NA | NA |
| Nxt1 | 62.52770177 | 0.060886869 | 0.195919296 | 0.310775257 |
| Gzf1 | 464.5606654 | -0.115362421 | 0.062906703 | -1.833865299 |
| Napb | 4683.537772 | -0.032205995 | 0.049474106 | -0.650966702 |
| Cstl1 | 7.743612027 | 0.51095846 | 0.398415491 | 1.28247639 |
| Cst11 | 0.068097359 | -0.206487837 | 3.724945049 | -0.055433794 |
| 8030411F24Rik | 0.116944185 | 0.274391125 | 3.724945049 | 0.073663134 |
| Cst12 | 0.585343961 | 2.027049625 | 1.542520782 | 1.314114952 |
| Cst8 | 0.132101749 | 0.274387056 | 3.724945049 | 0.073662041 |
| Cst13 | 0.138076337 | 0.75528675 | 3.589802529 | 0.210397854 |
| Cst9 | 0.038981395 | 0.274393828 | 3.724945049 | 0.073663859 |
| 9230104L09Rik | 0.693909001 | 0.272788455 | 1.298162204 | 0.210134337 |
| Cst3 | 6207.776889 | -0.136222842 | 0.125061703 | -1.089245059 |
| Gm38221 | 0.060174537 | 0.755286614 | 3.724945049 | 0.202764498 |
| Gm14129 | 0 | NA | NA | NA |
| Gm1330 | 0 | NA | NA | NA |
| Gm10750 | 0 | NA | NA | NA |
| Gm14135 | 0 | NA | NA | NA |
| Gm14132 | 0.875345121 | 0.689690438 | 0.931611302 | 0.740319956 |
| Gm14131 | 0.1140246 | 0.75528439 | 3.724945049 | 0.202763901 |
| Cst10 | 0 | NA | NA | NA |
| Gm24551 | 0.086648695 | -0.687376175 | 3.724945049 | -0.184533239 |
| Gm14130 | 2.711335961 | 0.343107248 | 0.748644392 | 0.458304707 |
| Gm14133 | 2.358317775 | 0.151629088 | 0.711836015 | 0.213011262 |
| Syndig1 | 375.012134 | -0.232638396 | 0.086615131 | -2.685886322 |
| C530025M09Rik | 3.362260024 | -1.563952425 | 0.6366101 | -2.45668805 |
| Gm14140 | 1.613620997 | -0.26153341 | 0.793098756 | -0.329761468 |
| Zfp120 | 248.9582029 | 0.011417117 | 0.132781302 | 0.085984373 |
| Gm10770 | 0 | NA | NA | NA |
| Gm14139 | 0.236020217 | -0.546590255 | 2.491798399 | -0.219355729 |
| Zfp937 | 71.56539432 | -0.501177465 | 0.163957967 | -3.056743596 |
| Gm21994 | 0.580893185 | 1.185061074 | 1.539832452 | 0.769603909 |
| Gm14124 | 2.427470934 | 0.601883622 | 0.701035705 | 0.858563433 |
| Gm14125 | 0 | NA | NA | NA |
| 3300002I08Rik | 53.22250888 | -0.262750428 | 0.201868882 | -1.30158955 |
| Gm10130 | 6.196226755 | 0.798952294 | 0.413387083 | 1.932697771 |
| Gm14126 | 0.606327527 | 0.77907949 | 1.205844085 | 0.646086422 |
| Gm14122 | 12.30982907 | -0.470582937 | 0.296806452 | -1.58548756 |
| Zfp442 | 70.96873342 | -0.450472194 | 0.178608602 | -2.522119256 |
| Gm14127 | 0.190396633 | 1.25932864 | 3.267946179 | 0.385357827 |
| Zfp345 | 0.580755766 | 1.109489684 | 1.306990802 | 0.84888867 |
| Gm14121 | 87.55734381 | -0.232832767 | 0.16940531 | -1.374412449 |
| Gm14128 | 0 | NA | NA | NA |

| | | | | |
|---|---|---|---|---|
| Cst7 | 47.58322733 | 6.580172404 | 0.619648515 | 10.61920145 |
| Apmap | 559.647641 | 0.427635015 | 0.090758567 | 4.711786785 |
| Gm14123 | 1.105496479 | 2.109137302 | 1.097606069 | 1.921579482 |
| Acss1 | 208.0155599 | -0.480886478 | 0.11820671 | -4.068182583 |
| E130215H24Rik | 2.738071508 | -0.035473622 | 0.581954475 | -0.060956008 |
| Vsx1 | 2.657225507 | -1.138006124 | 0.661974735 | -1.719108091 |
| Platr15 | 0.232351398 | 0.38005454 | 2.535740505 | 0.149879114 |
| Gm28450 | 0 | NA | NA | NA |
| Gm26681 | 0.300525961 | 0.755271801 | 3.724945049 | 0.202760522 |
| Entpd6 | 438.5268923 | -0.114753858 | 0.072492763 | -1.582969848 |
| Pygb | 1906.36878 | -0.167369785 | 0.105657533 | -1.584078118 |
| Abhd12 | 2275.162201 | 0.143096643 | 0.048469928 | 2.952276757 |
| Gm23598 | 0 | NA | NA | NA |
| Gins1 | 16.63953435 | 1.4680808 | 0.333273628 | 4.405031413 |
| Gm14143 | 0.083526043 | 0.75528439 | 3.724945049 | 0.202763901 |
| Ninl | 67.5515232 | 0.180812191 | 0.162360724 | 1.11364489 |
| Gm14142 | 0.423021739 | 1.12181008 | 1.666025266 | 0.673345178 |
| Gm14141 | 0 | NA | NA | NA |
| Nanp | 8.071585247 | 0.244599998 | 0.467499472 | 0.523209143 |
| Gm22187 | 0 | NA | NA | NA |
| Gm24199 | 0 | NA | NA | NA |
| Gm14151 | 0 | NA | NA | NA |
| Gm14147 | 0 | NA | NA | NA |
| Gm14145 | 0 | NA | NA | NA |
| Gm22517 | 0 | NA | NA | NA |
| Gm14149 | 0.579212071 | -0.518278426 | 1.297720751 | -0.399375926 |
| Gm14144 | 0 | NA | NA | NA |
| Gm23005 | 0 | NA | NA | NA |
| Gm24077 | 0 | NA | NA | NA |
| 4930442J19Rik | 0 | NA | NA | NA |
| Gm23261 | 0 | NA | NA | NA |
| Gm14148 | 0.134276636 | -0.206488731 | 2.954908363 | -0.06987991 |
| Gm14152 | 0 | NA | NA | NA |
| Gm24868 | 0 | NA | NA | NA |
| Gm14150 | 75.86947207 | 0.098027808 | 0.217740745 | 0.450204245 |
| Gm14146 | 0 | NA | NA | NA |
| 4921509C19Rik | 0.322748876 | 0.764644002 | 3.724286181 | 0.205312902 |
| Nsfl1c | 781.3049052 | 0.205048603 | 0.05989416 | 3.423515804 |
| Gm14167 | 3.663554111 | 0.524423216 | 0.492741301 | 1.064297257 |
| Fkbp1a | 3786.330142 | 0.257124483 | 0.067337654 | 3.818435444 |
| Sdcbp2 | 21.61712664 | 2.503196151 | 0.444264973 | 5.634466594 |
| Snph | 3369.871354 | 0.17212671 | 0.092054457 | 1.869835698 |
| Rad21l | 5.277545605 | 0.101833198 | 0.434272138 | 0.234491667 |
| Tmem74bos | 12.83654197 | 0.199314486 | 0.336146648 | 0.592939083 |
| Gm14159 | 1.598224206 | 0.490795621 | 0.707496726 | 0.693707269 |
| Tmem74b | 29.74572494 | -0.674039497 | 0.220298944 | -3.059658323 |
| Psmf1 | 502.2961145 | 0.555388489 | 0.099198164 | 5.598777922 |

| | | | | |
|---|---|---|---|---|
| 4930556L07Rik | 0.192951122 | 1.698434623 | 2.632329555 | 0.64522112 |
| Gm14153 | 0 | NA | NA | NA |
| Gm14155 | 0 | NA | NA | NA |
| Rspo4 | 1.536911596 | -1.210639941 | 0.88986927 | -1.360469433 |
| Rspo4os | 0.417636614 | 0.768697481 | 1.449630623 | 0.530271276 |
| Angpt4 | 4.455728532 | 1.482768956 | 0.764246878 | 1.940170119 |
| Gm14154 | 0.934869149 | 1.995612922 | 1.101341324 | 1.811984058 |
| Mir1953 | 0.11193333 | 0.75528439 | 3.724945049 | 0.202763901 |
| Fam110a | 57.87769406 | 0.045420644 | 0.153716975 | 0.295482294 |
| Slc52a3 | 22.60185597 | 1.698543373 | 0.296904938 | 5.720832355 |
| AA387200 | 2.676545693 | -1.452366708 | 0.748688551 | -1.939881019 |
| Scrt2 | 297.4913887 | -0.054420573 | 0.143954917 | -0.378039002 |
| Srxn1 | 1496.550893 | 0.289136355 | 0.112188494 | 2.577237162 |
| Tcf15 | 3.917975245 | -0.458305976 | 0.55682007 | -0.823077329 |
| Rps15a-ps7 | 16.65743553 | 0.56140649 | 0.274970205 | 2.041699358 |
| Csnk2a1 | 1499.289359 | -0.071955099 | 0.038364778 | -1.875551021 |
| Tbc1d20 | 711.5116405 | 0.159120517 | 0.055483494 | 2.867889261 |
| Rbck1 | 775.7363636 | 0.0661827 | 0.065128158 | 1.0161918 |
| Trib3 | 48.90459884 | 2.688591922 | 0.406742865 | 6.610053073 |
| Gm14164 | 0.458177261 | -1.317290779 | 1.526537663 | -0.862927139 |
| Gm14165 | 3.039451885 | 0.575725783 | 0.640565734 | 0.898777054 |
| Nrsn2 | 894.8007339 | 0.090686748 | 0.086471931 | 1.048742024 |
| Sox12 | 175.956736 | -0.212804444 | 0.14155384 | -1.503346315 |
| Zcchc3 | 147.4086379 | 0.10514412 | 0.119190307 | 0.882153274 |
| 6820408C15Rik | 27.31515061 | -0.09828292 | 0.193375355 | -0.508249459 |
| Gm27343 | 3.046712631 | -0.956403826 | 0.561237712 | -1.704097577 |
| Gm14158 | 0.634981691 | -0.772403679 | 1.215841739 | -0.635283075 |
| Defb23 | 0.097139547 | 0.274393828 | 3.724945049 | 0.073663859 |
| Gm14156 | 0 | NA | NA | NA |
| Defb20 | 0.545971167 | -1.128420404 | 1.320022014 | -0.854849686 |
| Defb22 | 0.061920613 | 0.274393828 | 3.724945049 | 0.073663859 |
| Defb26 | 0 | NA | NA | NA |
| Defb28 | 0 | NA | NA | NA |
| Defb29 | 0 | NA | NA | NA |
| Gm14161 | 0.118371171 | 0.75528439 | 3.724945049 | 0.202763901 |
| Gm17416 | 0 | NA | NA | NA |
| Defb21 | 0.04393175 | -0.206490061 | 3.724945049 | -0.055434391 |
| Defb19 | 0 | NA | NA | NA |
| Defb45 | 0.07796279 | 0.274392189 | 3.724945049 | 0.073663419 |
| Defb36 | 0 | NA | NA | NA |
| Gm14163 | 0 | NA | NA | NA |
| Defb25 | 0.234671876 | 0.261859813 | 2.17025674 | 0.120658449 |
| Rem1 | 3.041196478 | -1.095315764 | 0.634298138 | -1.726815355 |
| 1700030C14Rik | 0.290525896 | 0.834076697 | 1.784208654 | 0.467477105 |
| H13 | 832.2285858 | 0.081109635 | 0.073111414 | 1.109397714 |
| Mcts2 | 80.0283442 | -0.001538592 | 0.123217901 | -0.01248676 |
| Gm26841 | 0.794703889 | -1.134986753 | 1.200050691 | -0.945782341 |

| | | | | |
|---|---|---|---|---|
| Id1 | 158.7587211 | 1.335940091 | 0.207886973 | 6.426280926 |
| Gm14162 | 0.044191549 | -0.206490061 | 3.724945049 | -0.055434391 |
| Cox4i2 | 12.07102929 | 0.533361992 | 0.315022477 | 1.693091858 |
| Gm23802 | 0.93732221 | -0.442152713 | 0.928345356 | -0.476280416 |
| Bcl2l1 | 1656.202641 | 0.539538775 | 0.112343708 | 4.80257225 |
| Tpx2 | 97.61010764 | 1.841764879 | 0.238875812 | 7.710135501 |
| Mir5622 | 0 | NA | NA | NA |
| Mylk2 | 13.81209412 | 5.40034746 | 0.986784397 | 5.472672123 |
| Foxs1 | 7.663944057 | 1.201637862 | 0.517586005 | 2.32161969 |
| Dusp15 | 161.0227663 | -0.670458245 | 0.119332399 | -5.618409173 |
| Ttll9 | 15.9748206 | 0.027270999 | 0.325445896 | 0.083795801 |
| Gm14199 | 0.260717194 | 0.755305441 | 2.054894108 | 0.367564167 |
| Pdrg1 | 314.6393574 | 0.402523198 | 0.092681385 | 4.343085706 |
| Xkr7 | 77.04392738 | -0.928198654 | 0.162691848 | -5.705256064 |
| Ccm2l | 14.10545208 | 0.336227336 | 0.279740692 | 1.201925015 |
| Hck | 217.2183995 | 4.081185861 | 0.220182178 | 18.53549594 |
| Gm25643 | 0 | NA | NA | NA |
| Tm9sf4 | 988.450612 | -0.021075947 | 0.042808599 | -0.49232975 |
| Tspyl3 | 233.7002 | -0.055331058 | 0.092027308 | -0.601246081 |
| Plagl2 | 263.3156821 | 0.740856315 | 0.114596778 | 6.464896558 |
| Pofut1 | 393.8680983 | -0.00782316 | 0.08401568 | -0.093115473 |
| Kif3b | 984.8683119 | -0.038452295 | 0.045734395 | -0.840774115 |
| 2500004C02Rik | 39.85926624 | -0.73144924 | 0.22241908 | -3.288608325 |
| Asxl1 | 918.9589658 | 0.129205752 | 0.08269648 | 1.562409332 |
| Nol4l | 699.5239319 | -0.281365002 | 0.12520767 | -2.247186635 |
| Gm14472 | 0.227745227 | 1.329211109 | 2.57303662 | 0.516592379 |
| 4930404H24Rik | 0.825833058 | -0.759116221 | 1.137961857 | -0.667084065 |
| Commd7 | 204.4885226 | 0.083615087 | 0.094881052 | 0.881262218 |
| Dnmt3bos | 0 | NA | NA | NA |
| Dnmt3b | 18.90429361 | 0.174197791 | 0.281611402 | 0.618575063 |
| Gm14490 | 1.779788623 | 0.196255943 | 0.877887232 | 0.223554844 |
| Mapre1 | 1211.391518 | 0.130441956 | 0.071119053 | 1.834135155 |
| Efcab8 | 3.424510338 | 2.571897001 | 0.877518633 | 2.93087452 |
| Gm14494 | 80.10444224 | 0.272936937 | 0.130498968 | 2.09148733 |
| Sun5 | 0.124596252 | 0.755282361 | 3.714681068 | 0.203323609 |
| Bpifb2 | 0.955311604 | -0.657616337 | 1.138445686 | -0.577644016 |
| Bpifb6 | 1.501918467 | 0.02019397 | 0.894839334 | 0.022567146 |
| Bpifb3 | 1.988936712 | -1.849508095 | 1.565562224 | -1.181369904 |
| Bpifb4 | 3.358904097 | -0.026053523 | 1.220358424 | -0.021349074 |
| Bpifa6 | 0.496470236 | -1.489195296 | 3.697278825 | -0.402781442 |
| Bpifa2 | 0 | NA | NA | NA |
| Gm21850 | 0 | NA | NA | NA |
| Gm14497 | 0.15596086 | -1.15072961 | 3.483969583 | -0.330292668 |
| Bpifa3 | 0 | NA | NA | NA |
| Bpifa1 | 1.270782233 | -3.422064185 | 1.278368282 | -2.676900102 |
| Bpifa5 | 0.243228626 | -0.206488331 | 3.033562892 | -0.068067925 |
| Bpifb1 | 0.700162942 | -0.153499901 | 1.238254319 | -0.123964761 |

| | | | | |
|---|---|---|---|---|
| Bpifb5 | 0.765485319 | -0.580236069 | 1.154824825 | -0.502445095 |
| Bpifb9a | 1.362283058 | -0.27948302 | 0.883040505 | -0.316500793 |
| Gm14197 | 1.834732021 | -0.355586787 | 0.733611939 | -0.484706925 |
| Bpifb9b | 3.934499224 | -1.411595784 | 1.501756693 | -0.939963038 |
| Cdk5rap1 | 105.428526 | -0.002187062 | 0.136275888 | -0.016048784 |
| Snta1 | 269.5638173 | -0.293680319 | 0.113623224 | -2.584685687 |
| Cbfa2t2 | 875.8606125 | -0.074875786 | 0.050543364 | -1.481416746 |
| Necab3 | 257.0570188 | -0.853200391 | 0.170390624 | -5.007320073 |
| 1700003F12Rik | 3.150202034 | 0.315002994 | 0.642732812 | 0.490099444 |
| Actl10 | 0.266983968 | -0.19768134 | 1.823280154 | -0.108420716 |
| E2f1 | 73.04566475 | -0.323297379 | 0.225540133 | -1.433436143 |
| Pxmp4 | 135.3858191 | -0.126768158 | 0.112012829 | -1.131728925 |
| 4930519P11Rik | 1.196740622 | 0.442366034 | 0.825756096 | 0.535710286 |
| Zfp341 | 104.6788915 | -0.523869162 | 0.130935641 | -4.000966874 |
| Gm14198 | 0.162949104 | -0.687376772 | 2.787607662 | -0.24658304 |
| Chmp4b | 2004.387491 | 0.742261579 | 0.136649762 | 5.431854146 |
| Rpl5-ps2 | 5.302412506 | 0.474922158 | 0.450367286 | 1.054521882 |
| Gm14215 | 0 | NA | NA | NA |
| Gm14216 | 0.099155932 | 0.75528439 | 3.724945049 | 0.202763901 |
| Gm14214 | 0 | NA | NA | NA |
| Raly | 850.5641207 | 0.412827404 | 0.087042759 | 4.74281156 |
| a | 0.140640116 | -0.206495687 | 3.611596447 | -0.057175736 |
| Eif2s2 | 723.9443759 | 0.532610597 | 0.115386939 | 4.615865553 |
| Gm14226 | 0.71216146 | 1.058200276 | 1.334325148 | 0.793060281 |
| Ahcy | 112.8096952 | 0.31305872 | 0.149015827 | 2.100842081 |
| 2310005A03Rik | 0.04393175 | -0.206490061 | 3.724945049 | -0.055434391 |
| Itch | 1259.365133 | -0.171805679 | 0.062736312 | -2.738536493 |
| Gm14227 | 0.141071297 | -0.206491178 | 3.564043836 | -0.057937328 |
| Dynlrb1 | 713.8695448 | 0.125301846 | 0.088058893 | 1.422932319 |
| Map1lc3a | 781.5540749 | -0.25350031 | 0.121777782 | -2.081663054 |
| Pigu | 261.7083534 | -0.227957245 | 0.111173212 | -2.050469188 |
| Mir695 | 0 | NA | NA | NA |
| Trp53inp2 | 2971.079894 | -0.052487487 | 0.106908563 | -0.490956812 |
| Ncoa6 | 1310.679018 | -0.111074637 | 0.068866004 | -1.612909572 |
| Gm14239 | 28.68396613 | -0.703674182 | 0.301733367 | -2.332105951 |
| Gm14238 | 0.886238366 | 1.074996566 | 0.938456365 | 1.145494459 |
| Ggt7 | 845.8338809 | -4.54E-05 | 0.0722928 | -0.000628038 |
| Acss2 | 635.0895078 | 0.060386586 | 0.065236806 | 0.925652091 |
| Mipep-ps | 1.837594399 | -0.252435209 | 0.861611742 | -0.292980233 |
| Gm20637 | 2.163212633 | 0.184063209 | 0.619932923 | 0.296908266 |
| Acss2os | 0.277114854 | -1.294816432 | 2.066548853 | -0.626559798 |
| Gss | 143.0865772 | -0.157523675 | 0.12558177 | -1.254351448 |
| Gssos2 | 1.802868249 | 0.294580848 | 0.874687816 | 0.336783985 |
| Gssos1 | 1.992642508 | 0.618241022 | 0.746353744 | 0.828348524 |
| Gm14257 | 0.038981395 | 0.274393828 | 3.724945049 | 0.073663859 |
| Myh7b | 88.10465837 | 0.341957944 | 0.26898386 | 1.271295403 |
| Mir499 | 0 | NA | NA | NA |

| | | | | |
|---|---|---|---|---|
| Trpc4ap | 1703.710218 | 0.087938761 | 0.054804589 | 1.604587528 |
| Gm22334 | 1.228515858 | -1.20166206 | 1.162339632 | -1.033830412 |
| Edem2 | 274.0937995 | 0.128868726 | 0.090611977 | 1.422204119 |
| Gm17581 | 0 | NA | NA | NA |
| Procr | 21.11771009 | 2.922256973 | 0.45942652 | 6.360662355 |
| Mmp24 | 793.763571 | -0.297223331 | 0.133025945 | -2.234326032 |
| BC029722 | 543.0851477 | -0.019105982 | 0.091963636 | -0.207755835 |
| Eif6 | 475.8961878 | 0.432581269 | 0.119432292 | 3.621979145 |
| Gm16098 | 1.335672075 | 1.211574028 | 1.016909497 | 1.191427587 |
| Fam83c | 0.240161549 | 0.753270913 | 3.046061241 | 0.247293424 |
| Uqcc1 | 374.2692931 | 0.183668458 | 0.083968867 | 2.187339949 |
| Gm15557 | 0.53525542 | 0.147381771 | 1.446093747 | 0.101917162 |
| Gdf5 | 4.286069218 | 1.878857365 | 0.56058188 | 3.351619863 |
| Cep250 | 337.5416032 | -0.106634918 | 0.158033025 | -0.674763505 |
| 6430550D23Rik | 7.007522312 | -1.534674686 | 0.45446055 | -3.376915086 |
| Ergic3 | 697.3852742 | 0.179337408 | 0.089307634 | 2.008085986 |
| Fer1l4 | 0.699011511 | 2.028315124 | 2.610454809 | 0.776996835 |
| Gm14240 | 0 | NA | NA | NA |
| Spag4 | 20.11396154 | -0.81218164 | 0.305597867 | -2.657680984 |
| Cpne1 | 7.154347536 | -0.715749943 | 0.436733662 | -1.638870565 |
| Gm28036 | 0.689591474 | -0.679067328 | 1.147004842 | -0.592035275 |
| Gm37274 | 40.70937969 | -1.126621552 | 0.330115131 | -3.412814039 |
| Rbm12 | 333.4415097 | -0.116634902 | 0.083378968 | -1.398852802 |
| Nfs1 | 286.4122944 | -0.059350827 | 0.089474535 | -0.663326462 |
| Romo1 | 128.8625568 | -0.314578495 | 0.179614544 | -1.751408808 |
| Rbm39 | 3275.570639 | 0.263291367 | 0.088124211 | 2.987730209 |
| Phf20 | 1427.32006 | 0.004529229 | 0.051175726 | 0.088503472 |
| Scand1 | 367.1199098 | 0.338741955 | 0.1766532 | 1.917553454 |
| Cnbd2 | 68.49456786 | 0.078859128 | 0.138388457 | 0.569838917 |
| Gm14251 | 0.489024281 | 1.566014413 | 1.678363182 | 0.933060514 |
| Gm22759 | 0 | NA | NA | NA |
| Gm14252 | 0.106056992 | 1.236179401 | 3.724945049 | 0.331865138 |
| Gm14253 | 61.94935748 | 0.662831746 | 0.199369827 | 3.324634207 |
| Gm14170 | 0.179599912 | -0.665371588 | 2.727661601 | -0.2439348 |
| Gm22316 | 0.234323835 | 1.295694608 | 2.538217475 | 0.510474229 |
| Rpl21-ps9 | 0.666172099 | -1.084624294 | 1.17958834 | -0.919493909 |
| 2900097C17Rik | 2086.303339 | -0.037416005 | 0.092032212 | -0.406553357 |
| Gm14173 | 0.760931553 | -1.200676325 | 1.154993653 | -1.039552314 |
| Gm22758 | 0 | NA | NA | NA |
| Epb4.1l1 | 5881.508901 | -0.375506349 | 0.078115571 | -4.807061423 |
| Gm14225 | 0 | NA | NA | NA |
| Gm14224 | 0.038981395 | 0.274393828 | 3.724945049 | 0.073663859 |
| Aar2 | 389.6193864 | 0.273834803 | 0.071532303 | 3.828127868 |
| Gm14168 | 20.52960382 | 0.386634153 | 0.303423628 | 1.274238779 |
| Gm14169 | 37.69901258 | -0.599019314 | 0.203967976 | -2.936830213 |
| Dlgap4 | 2838.730653 | -0.18186898 | 0.100990252 | -1.800856783 |
| Gm14174 | 0.748568137 | -0.633730593 | 1.032235907 | -0.61393969 |

| | | | | |
|---|---|---|---|---|
| Gm14172 | 0 | NA | NA | NA |
| 4930405A21Rik | 7.408513515 | -0.055721772 | 0.368566077 | -0.151185299 |
| Myl9 | 37.34594635 | -0.374328537 | 0.266515874 | -1.404526235 |
| Gm14231 | 0.108266322 | 0.274389235 | 3.724945049 | 0.073662626 |
| Gm14230 | 5.976257995 | 1.213990724 | 0.515238749 | 2.356171242 |
| Tgif2 | 38.14849271 | 1.654700994 | 0.323797565 | 5.110294735 |
| 5430405H02Rik | 85.38075778 | 0.067592056 | 0.173112422 | 0.390451794 |
| Gm37790 | 6.47799121 | 0.20708497 | 0.432129951 | 0.479219203 |
| 4930518I15Rik | 26.40534196 | -0.465506781 | 0.257850687 | -1.805334653 |
| Gm27651 | 0 | NA | NA | NA |
| 1110008F13Rik | 336.8881828 | 0.104726838 | 0.112046008 | 0.934677104 |
| Sla2 | 15.40209609 | 1.434780471 | 0.405379131 | 3.539354544 |
| Gm14278 | 0.888132063 | -1.378276029 | 1.191138768 | -1.157107859 |
| Gm14248 | 0 | NA | NA | NA |
| Gm14247 | 0.534422126 | -1.408533355 | 1.640944256 | -0.858367583 |
| Ndrg3 | 3682.906368 | -0.361071976 | 0.077904707 | -4.63479022 |
| Gm27653 | 9.457828482 | -0.177880068 | 0.424334503 | -0.419197749 |
| Gm14276 | 8.337030673 | -0.455625216 | 0.359224751 | -1.26835697 |
| Dsn1 | 45.4075692 | 0.248057874 | 0.204896353 | 1.210650507 |
| Soga1 | 1303.081876 | -0.099473561 | 0.119955561 | -0.829253435 |
| Tldc2 | 4.0710617 | -0.551475828 | 0.468740188 | -1.176506393 |
| Samhd1 | 1799.230623 | 3.041209999 | 0.185582311 | 16.38739155 |
| Rbl1 | 152.3466098 | 1.85333551 | 0.167138185 | 11.08864206 |
| Gm14277 | 3.35648137 | -0.098425048 | 0.549822432 | -0.179012426 |
| Mroh8 | 16.34303778 | -0.46018526 | 0.280840265 | -1.638601429 |
| Rpn2 | 1339.45037 | 0.270485616 | 0.077177042 | 3.50474195 |
| Ghrh | 3.638464397 | 0.56574908 | 0.629817872 | 0.898274096 |
| Manbal | 220.2483533 | 0.442096477 | 0.128444875 | 3.441916041 |
| Src | 1335.75724 | 0.029838006 | 0.061733694 | 0.483334208 |
| Gm14279 | 2.044822302 | 1.725291185 | 0.766388663 | 2.251196121 |
| Gm14287 | 40.93809312 | -0.055230234 | 0.254516156 | -0.217000897 |
| Gm14286 | 22.25133751 | -0.279745934 | 0.334257965 | -0.836916286 |
| Blcap | 862.3755201 | -0.081771327 | 0.063046221 | -1.297005993 |
| Nnat | 1274.354139 | -0.015816677 | 0.167628025 | -0.094355806 |
| Gm23134 | 19.82322296 | -0.116881329 | 0.444501491 | -0.26294924 |
| Platr27 | 0.11147366 | 0.274389235 | 3.724945049 | 0.073662626 |
| Gm25407 | 0 | NA | NA | NA |
| Gm23463 | 0 | NA | NA | NA |
| Ctnnbl1 | 279.0876754 | -0.12218406 | 0.080509453 | -1.51763619 |
| Gm14176 | 7.679034499 | 0.163306178 | 0.360889079 | 0.452510722 |
| Vstm2l | 466.6791405 | -0.322281979 | 0.097486547 | -3.305912364 |
| Tti1 | 247.1897018 | -0.256248871 | 0.121111379 | -2.115811698 |
| Rprd1b | 501.6807887 | -0.03259838 | 0.065806592 | -0.495366482 |
| 2010009K17Rik | 0.075043205 | 0.755286614 | 3.724945049 | 0.202764498 |
| Tgm2 | 2012.035572 | 4.656823987 | 0.332946033 | 13.98672315 |
| Gm20412 | 1.717954688 | 3.322280346 | 1.378391266 | 2.410259283 |
| D630003M21Rik | 59.49929145 | -0.811340787 | 0.179076232 | -4.530700564 |

| | | | | |
|---|---|---|---|---|
| 1700060C20Rik | 0.29895891 | 1.772698507 | 2.029398503 | 0.87350932 |
| Mir7685 | 0 | NA | NA | NA |
| Bpi | 0.044544648 | 0.755286614 | 3.724945049 | 0.202764498 |
| Lbp | 183.630573 | 1.336988933 | 0.251602487 | 5.313893943 |
| Snhg17 | 148.0843631 | 0.297763504 | 0.187267824 | 1.590040919 |
| Gm24411 | 0 | NA | NA | NA |
| Gm26003 | 0 | NA | NA | NA |
| Gm23925 | 0 | NA | NA | NA |
| Snhg11 | 9158.007078 | -0.350251999 | 0.334549478 | -1.046936321 |
| Gm25187 | 0 | NA | NA | NA |
| Gm25129 | 29.82494218 | -0.135727797 | 0.286478936 | -0.473779326 |
| Gm14201 | 0 | NA | NA | NA |
| Gm14202 | 0 | NA | NA | NA |
| Ralgapb | 2672.82956 | -0.118869501 | 0.052481242 | -2.264990234 |
| Gm14200 | 0.166639798 | 1.236193269 | 2.767997355 | 0.446602041 |
| Gm24379 | 4.859614805 | -0.380701725 | 0.4493815 | -0.847168218 |
| Adig | 1.688935393 | -2.258404108 | 0.929666038 | -2.429263859 |
| Arhgap40 | 1.707670524 | -1.118397706 | 0.933064915 | -1.198627971 |
| Gm14204 | 129.1982071 | -0.840081726 | 0.146445629 | -5.736475268 |
| Slc32a1 | 1229.442361 | -0.223883469 | 0.062055712 | -3.607781791 |
| Gm14205 | 1.579080267 | 0.363588131 | 0.864266103 | 0.420690027 |
| Actr5 | 113.0741844 | 0.125893066 | 0.170644181 | 0.737751883 |
| Mir3474 | 0.201505633 | 0.27440894 | 2.605863055 | 0.105304436 |
| Ppp1r16b | 2425.103286 | -0.129395697 | 0.077279215 | -1.67439198 |
| Gm26141 | 1.537427175 | -1.28460533 | 0.906436677 | -1.417203609 |
| Fam83d | 51.99511303 | 0.219085032 | 0.218638382 | 1.00204287 |
| Dhx35 | 216.8272346 | 0.02410299 | 0.111087895 | 0.216972244 |
| AL669913.1 | 0.255320825 | 2.173795215 | 2.379358213 | 0.913605696 |
| Gm25090 | 0 | NA | NA | NA |
| Gm14219 | 0.112288909 | -0.687373951 | 3.724945049 | -0.184532642 |
| Gm11445 | 0 | NA | NA | NA |
| Gm11446 | 0 | NA | NA | NA |
| Gm14223 | 0 | NA | NA | NA |
| 4933416E03Rik | 0 | NA | NA | NA |
| Gm14220 | 0 | NA | NA | NA |
| 9130015L21Rik | 0 | NA | NA | NA |
| Gm826 | 0.8433133 | 2.352608502 | 1.135910276 | 2.07112177 |
| Mafb | 493.6009209 | 1.400801944 | 0.173289533 | 8.083592357 |
| Gm14221 | 0.304607851 | 1.168147094 | 2.019904409 | 0.578318008 |
| Gm14222 | 0 | NA | NA | NA |
| Gm23968 | 0.062426873 | 0.755288839 | 3.724945049 | 0.202765096 |
| Top1 | 1394.007593 | -0.350191374 | 0.063168051 | -5.54380523 |
| Plcg1 | 938.5103693 | -0.06573098 | 0.067933298 | -0.967581181 |
| Gm27206 | 3.426664286 | 0.983131591 | 0.540324238 | 1.819521543 |
| Zhx3 | 1591.941502 | -0.062791884 | 0.05265406 | -1.19253643 |
| Gm16751 | 10.07322116 | -0.490249418 | 0.331672734 | -1.47811191 |
| Gm11447 | 0.089814205 | 0.274402725 | 3.724945049 | 0.073666248 |

| | | | | |
|---|---|---|---|---|
| Lpin3 | 58.0580359 | 2.59736561 | 0.264693165 | 9.81274153 |
| Emilin3 | 11.35137183 | 0.131138396 | 0.326370456 | 0.401808417 |
| Gm14228 | 0.449436652 | 2.518411052 | 3.689435082 | 0.682600722 |
| Chd6 | 2195.755601 | -0.208496602 | 0.068798662 | -3.030532803 |
| Gm25561 | 0 | NA | NA | NA |
| Gm14243 | 0.04393175 | -0.206490061 | 3.724945049 | -0.055434391 |
| Gm14242 | 0 | NA | NA | NA |
| Gm5270 | 0.106618422 | 0.274402725 | 3.724945049 | 0.073666248 |
| Gm11448 | 0.467761363 | -0.435050408 | 1.406381506 | -0.309340251 |
| Gm14241 | 0 | NA | NA | NA |
| Gm11449 | 0.644287204 | 1.227551691 | 1.205900056 | 1.017954751 |
| Gm29229 | 0.291544834 | -1.117282897 | 1.605693497 | -0.69582576 |
| Ptprt | 2587.731476 | -0.336938243 | 0.076418572 | -4.40911459 |
| Ptprtos | 0.352863859 | 1.127274643 | 1.927690512 | 0.584779889 |
| 9430021M05Rik | 23.91964892 | -0.365863661 | 0.235837264 | -1.551339489 |
| Gm14246 | 8.592019774 | 0.131953116 | 0.391764669 | 0.336817294 |
| Gm22936 | 0 | NA | NA | NA |
| Gm11450 | 0 | NA | NA | NA |
| Gm24478 | 0 | NA | NA | NA |
| Gm14255 | 0.044191549 | -0.206490061 | 3.724945049 | -0.055434391 |
| Gm11451 | 16.98984007 | 0.231072441 | 0.313796025 | 0.736377844 |
| Gm11452 | 0.150711795 | 0.27440557 | 2.856704362 | 0.096056692 |
| Srsf6 | 1346.747418 | 0.186350346 | 0.082123581 | 2.269145399 |
| L3mbtl1 | 55.09161921 | -1.390533506 | 0.363618924 | -3.824150543 |
| Gm11453 | 0.038981395 | 0.274393828 | 3.724945049 | 0.073663859 |
| Sgk2 | 21.73993536 | -0.168084029 | 0.278295 | -0.603977897 |
| Ift52 | 322.3681409 | -0.008402568 | 0.076479131 | -0.109867467 |
| Mybl2 | 17.83251159 | 0.297507573 | 0.306907847 | 0.969371021 |
| Mir7003 | 0.049553047 | 0.274391604 | 3.724945049 | 0.073663262 |
| Gtsf1l | 0.489956056 | 0.721521501 | 1.365638639 | 0.528339987 |
| Gm22174 | 0 | NA | NA | NA |
| Gm11454 | 2.386488514 | -0.112712212 | 0.660223989 | -0.170718141 |
| Gm14254 | 0 | NA | NA | NA |
| Tox2 | 401.2220022 | -0.042736937 | 0.103795638 | -0.411741169 |
| Jph2 | 9.352170419 | -0.38999295 | 0.354409932 | -1.100400736 |
| Oser1 | 111.6600522 | 0.163645901 | 0.13630381 | 1.200596676 |
| 2900093K20Rik | 14.33683216 | -0.685618669 | 0.285646222 | -2.400237129 |
| Gdap1l1 | 668.5251384 | 0.065486538 | 0.090078347 | 0.726995336 |
| Fitm2 | 424.7360197 | -0.301248963 | 0.066514796 | -4.529051879 |
| 2310001K24Rik | 0.198365723 | -1.021572254 | 2.624989092 | -0.389172 |
| R3hdml | 0.349069061 | -0.073252342 | 1.754847664 | -0.041742849 |
| Hnf4aos | 0 | NA | NA | NA |
| Hnf4a | 0.497304619 | -0.06347441 | 1.537657228 | -0.041279948 |
| Gm25881 | 0 | NA | NA | NA |
| Ttpal | 655.7681734 | -0.278426372 | 0.08914504 | -3.12329628 |
| Serinc3 | 3221.802251 | 0.429859234 | 0.072113096 | 5.96090385 |
| 0610039K10Rik | 0.721809532 | 0.956371586 | 1.180940962 | 0.809838609 |

| | | | | |
|---|---|---|---|---|
| Pkig | 413.2230987 | -0.095259852 | 0.077840655 | -1.223780192 |
| Gm16316 | 3.804176175 | 3.400699857 | 0.951535699 | 3.57390675 |
| Ada | 48.39031174 | 2.560747715 | 0.2889122 | 8.863411487 |
| Wisp2 | 23.40200403 | 1.952319911 | 0.509915959 | 3.828709172 |
| Kcnk15 | 8.160985404 | -1.234000541 | 0.52493818 | -2.350754029 |
| Rims4 | 1059.645651 | -0.388903266 | 0.082697251 | -4.702735103 |
| Ywhab | 4440.302886 | -0.024843168 | 0.058921146 | -0.421634162 |
| Pabpc1l | 10.92999753 | 1.413947745 | 0.500683691 | 2.824033957 |
| Tomm34 | 543.6160348 | 0.023876623 | 0.06907388 | 0.345667901 |
| Stk4 | 685.9737304 | 0.044193883 | 0.077031912 | 0.573708767 |
| Gm11455 | 4.303662858 | -0.741957162 | 0.572102013 | -1.296896612 |
| n-R5s207 | 0 | NA | NA | NA |
| Kcns1 | 75.78496726 | -0.389995683 | 0.198132547 | -1.968357493 |
| Wfdc5 | 0.042716945 | -0.206490061 | 3.724945049 | -0.055434391 |
| A730032A03Rik | 0.162822621 | 0.274394889 | 3.407465873 | 0.080527553 |
| Wfdc12 | 0.231631338 | 0.274410131 | 2.134176604 | 0.128578924 |
| Wfdc15a | 0.066050875 | 0.274391604 | 3.724945049 | 0.073663262 |
| Wfdc15b | 0 | NA | NA | NA |
| Svs2 | 0 | NA | NA | NA |
| Svs3b | 0.141331096 | -0.206491164 | 3.56184083 | -0.057973159 |
| Svs4 | 0 | NA | NA | NA |
| Svs3a | 0 | NA | NA | NA |
| Svs6 | 0.038981395 | 0.274393828 | 3.724945049 | 0.073663859 |
| Svs5 | 0.282943117 | -0.687379931 | 2.049488077 | -0.335391037 |
| Slpi | 106.5214453 | 1.696653111 | 0.525753191 | 3.227090469 |
| Mir7678 | 0.061920613 | 0.274393828 | 3.724945049 | 0.073663859 |
| Matn4 | 90.53204317 | -0.865525114 | 0.176028547 | -4.916958806 |
| Rbpjl | 13.41464962 | -0.895674317 | 0.3020956 | -2.964870444 |
| Sdc4 | 2785.760239 | 1.57739514 | 0.252042748 | 6.258442875 |
| Sys1 | 225.5397579 | 0.12875601 | 0.082421039 | 1.562174066 |
| Gm14302 | 1.474490841 | 0.345675587 | 0.779143438 | 0.443661038 |
| Gm20458 | 0.478777135 | 0.558790688 | 1.275804482 | 0.437990849 |
| Trp53tg5 | 0.109982625 | -0.20649451 | 3.724945049 | -0.055435586 |
| Dbndd2 | 527.4245514 | 0.127846176 | 0.092972211 | 1.37510095 |
| Pigt | 917.1375041 | 0.24406406 | 0.083100194 | 2.936985439 |
| Gm14317 | 0.567797178 | 0.615211196 | 1.194351088 | 0.515100796 |
| Wfdc2 | 6.468348977 | 0.685146911 | 0.44238315 | 1.548763581 |
| Spint3 | 0 | NA | NA | NA |
| Wfdc6a | 0.062426873 | 0.755288839 | 3.724945049 | 0.202765096 |
| Eppin | 0.044544648 | 0.755286614 | 3.724945049 | 0.202764498 |
| Wfdc8 | 0.123845245 | 0.755285145 | 3.724945049 | 0.202764104 |
| Gm25512 | 0 | NA | NA | NA |
| Wfdc6b | 0 | NA | NA | NA |
| Gm24473 | 0 | NA | NA | NA |
| Wfdc16 | 0.038981395 | 0.274393828 | 3.724945049 | 0.073663859 |
| Gm11456 | 0 | NA | NA | NA |
| Wfdc9 | 0.046345709 | 0.274391604 | 3.724945049 | 0.073663262 |

| | | | | |
|---|---|---|---|---|
| Wfdc10 | 0.171131556 | 0.755286499 | 3.384058469 | 0.223189554 |
| Wfdc11 | 0 | NA | NA | NA |
| Wfdc13 | 0 | NA | NA | NA |
| Spint4 | 0 | NA | NA | NA |
| Spint5 | 0 | NA | NA | NA |
| Wfdc3 | 9.036228213 | -0.511494515 | 0.364988073 | -1.40140063 |
| Dnttip1 | 245.2841238 | 0.16023303 | 0.108873412 | 1.471737016 |
| Gm11457 | 0.366597908 | -0.0758297 | 1.564654931 | -0.048464169 |
| Gm22596 | 5.545541213 | -0.013108042 | 0.481249485 | -0.027237519 |
| Ube2c | 12.17535567 | 1.604146184 | 0.485498928 | 3.304118899 |
| Tnnc2 | 0.984947528 | 0.885765497 | 1.235935499 | 0.716676151 |
| Snx21 | 316.0422822 | -0.276392125 | 0.069537441 | -3.974723855 |
| Acot8 | 121.1509639 | -0.093256177 | 0.114135091 | -0.817068413 |
| Zswim3 | 75.20523749 | -0.700302492 | 0.179702496 | -3.897010388 |
| Zswim1 | 246.2299447 | 0.112534508 | 0.073338358 | 1.534456347 |
| Spata25 | 2.304449883 | 0.870180922 | 0.865164633 | 1.005798075 |
| Neurl2 | 13.0448324 | 0.147333135 | 0.289116907 | 0.509597091 |
| Ctsa | 1093.099412 | 0.403742668 | 0.092809931 | 4.350209789 |
| Pltp | 377.8729552 | -0.011026811 | 0.180360219 | -0.061137712 |
| Pcif1 | 806.4495592 | 0.324928482 | 0.092265331 | 3.521674686 |
| Zfp335 | 457.5970439 | 0.11532468 | 0.101626206 | 1.134792735 |
| Zfp335os | 35.33703191 | 0.22896469 | 0.198175675 | 1.155362228 |
| Mmp9 | 18.18569822 | 0.339341608 | 0.251506496 | 1.349235958 |
| Slc12a5 | 6336.594774 | -0.326593313 | 0.07333316 | -4.453555732 |
| Ncoa5 | 554.329722 | -0.240237092 | 0.101952472 | -2.356363582 |
| Cd40 | 64.99215734 | 5.528275254 | 0.527928112 | 10.47164401 |
| 1700025C18Rik | 0 | NA | NA | NA |
| Cdh22 | 506.5456601 | -0.260411453 | 0.105381123 | -2.471139469 |
| Gm14397 | 0 | NA | NA | NA |
| Gm25569 | 0 | NA | NA | NA |
| Slc35c2 | 388.3991378 | 0.181748436 | 0.062599889 | 2.903334777 |
| Elmo2 | 1659.397843 | 0.011749419 | 0.071262434 | 0.164875356 |
| Zfp663 | 0.749070597 | -2.102399777 | 1.358949382 | -1.547077326 |
| Zfp334 | 573.5096397 | -0.200250195 | 0.06062301 | -3.30320446 |
| Ocstamp | 4.715999993 | 2.650141937 | 0.949689868 | 2.790534075 |
| Slc13a3 | 348.6971527 | 0.138198505 | 0.129722859 | 1.065336561 |
| Gm14437 | 0 | NA | NA | NA |
| Trp53rka | 121.9068042 | 0.007957024 | 0.141292694 | 0.056315888 |
| Slc2a10 | 29.04143457 | 0.501856435 | 0.279694608 | 1.794301428 |
| Eya2 | 64.95457155 | 0.103966527 | 0.171431747 | 0.606460174 |
| Gm28163 | 0 | NA | NA | NA |
| Gm11460 | 0.325497449 | -0.548564783 | 1.54523698 | -0.35500366 |
| Zmynd8 | 1768.666945 | -0.064762749 | 0.094538871 | -0.685038312 |
| BC046401 | 0.509709817 | -0.062301188 | 1.272115627 | -0.048974469 |
| Gm11465 | 2.089642419 | 0.253939472 | 0.661009264 | 0.384169308 |
| Gm22692 | 0.117907917 | 1.204448679 | 3.72248341 | 0.323560523 |
| Gm11461 | 0 | NA | NA | NA |

| | | | | |
|---|---|---|---|---|
| Gm11462 | 0 | NA | NA | NA |
| Platr29 | 0 | NA | NA | NA |
| Gm11464 | 0.663210203 | 0.57629381 | 1.17141364 | 0.491964401 |
| Ncoa3 | 801.9318103 | -0.099121093 | 0.093095948 | -1.064719735 |
| Sulf2 | 1316.311321 | 0.410362241 | 0.063753592 | 6.436692074 |
| Gm22687 | 0 | NA | NA | NA |
| Gm11466 | 0 | NA | NA | NA |
| Gm11467 | 0 | NA | NA | NA |
| Gm11468 | 0.423089777 | 0.798187492 | 1.451476186 | 0.549914287 |
| Gm14268 | 0.409889595 | 0.274412043 | 1.692240347 | 0.162159024 |
| 5031425F14Rik | 0.951684498 | 0.25471262 | 1.154819974 | 0.220564786 |
| Gm11469 | 0.922271222 | -0.392468459 | 1.197571647 | -0.327720232 |
| Gm14267 | 0.809026523 | 0.445217914 | 1.047099144 | 0.425191746 |
| Prex1 | 2399.621753 | 0.214528056 | 0.071516317 | 2.999707852 |
| Gm23152 | 1.245355593 | -0.554544719 | 1.115415182 | -0.497164399 |
| Gm25265 | 0 | NA | NA | NA |
| Gm24477 | 0 | NA | NA | NA |
| Trp53rkb | 54.71816161 | -0.402405919 | 0.144063192 | -2.793259777 |
| Arfgef2 | 1429.215814 | -0.087950829 | 0.043930862 | -2.002028313 |
| Cse1l | 859.0610123 | -0.034577008 | 0.069452713 | -0.497849646 |
| Gm17096 | 7.764297659 | 0.30405909 | 0.511858259 | 0.59402986 |
| Stau1 | 911.3828237 | 0.121378723 | 0.069738437 | 1.740485279 |
| Gm11470 | 0.342301797 | 2.250724531 | 3.697442713 | 0.608724653 |
| Ddx27 | 291.9835414 | 0.082525488 | 0.121765347 | 0.677741988 |
| Znfx1 | 5280.077714 | 2.526734085 | 0.16706239 | 15.12449378 |
| 1500012F01Rik | 269.2484598 | 1.420116413 | 0.147173173 | 9.64928852 |
| Gm27494 | 0 | NA | NA | NA |
| Gm25878 | 0 | NA | NA | NA |
| Gm23201 | 0 | NA | NA | NA |
| Snord12 | 0 | NA | NA | NA |
| Gm11472 | 0.906216104 | 1.909805512 | 1.115900934 | 1.711447185 |
| Gm11471 | 0.062426873 | 0.755288839 | 3.724945049 | 0.202765096 |
| Gm14290 | 607.9128014 | -0.521910289 | 0.175476717 | -2.974242381 |
| Kcnb1 | 1138.507184 | -0.428704897 | 0.148372693 | -2.889378687 |
| 1110018N20Rik | 50.53414942 | -0.413616349 | 0.210012038 | -1.969488763 |
| Ptgis | 12.61769256 | 0.011703248 | 0.279097236 | 0.041932511 |
| Gm14291 | 10.76029111 | 0.23108342 | 0.366950537 | 0.629739968 |
| Gm11473 | 0.150086409 | 1.264824188 | 3.723011833 | 0.339731445 |
| B4galt5 | 1227.37508 | 0.312449426 | 0.107597995 | 2.903859176 |
| Slc9a8 | 501.9560524 | 0.221111578 | 0.075475642 | 2.929575334 |
| Spata2 | 691.9053702 | 0.133659661 | 0.087962443 | 1.519508281 |
| Rnf114 | 1225.538309 | 1.595577069 | 0.122772199 | 12.99624082 |
| Gm11474 | 0.049553047 | 0.274391604 | 3.724945049 | 0.073663262 |
| Snai1 | 13.66662012 | 0.966244308 | 0.364410742 | 2.651525318 |
| Gm11476 | 0.445249545 | -0.760367061 | 1.377843826 | -0.551852864 |
| Gm11475 | 0 | NA | NA | NA |
| Ube2v1 | 21.15999022 | -0.019331451 | 0.241732954 | -0.079970276 |

| | | | | |
|---|---|---|---|---|
| Gm20431 | 0 | NA | NA | NA |
| Tmem189 | 432.2688695 | 0.264907458 | 0.094023447 | 2.817461674 |
| Gm22355 | 0.959597608 | 0.404851258 | 1.021826152 | 0.396203656 |
| Gm14320 | 1.589793752 | -1.574570027 | 0.84513807 | -1.86309206 |
| Cebpb | 250.3011784 | 2.267788769 | 0.288782121 | 7.852940341 |
| A530013C23Rik | 6.464581192 | 0.190895453 | 0.515572749 | 0.370259005 |
| Gm14321 | 4.54434231 | -0.887911427 | 0.542909142 | -1.635469653 |
| 9230111E07Rik | 2.522272708 | 0.080724992 | 0.597524056 | 0.13509915 |
| 1200007C13Rik | 1.134624916 | 3.536657764 | 1.285543598 | 2.751099044 |
| Gm14319 | 0.048949532 | -0.206487837 | 3.724945049 | -0.055433794 |
| Ptpn1 | 822.4208813 | 0.529938766 | 0.106332474 | 4.983790423 |
| Fam65c | 5.824423608 | 0.465414646 | 0.507181592 | 0.917648933 |
| Gm14236 | 0.314827273 | 1.592408362 | 1.555042093 | 1.024029104 |
| Gm24327 | 0 | NA | NA | NA |
| Pard6b | 76.59336974 | -0.072005896 | 0.205488336 | -0.350413542 |
| Gm14235 | 0.229025006 | 0.274411602 | 2.133783704 | 0.128603289 |
| E130018N17Rik | 1.228602664 | 0.825515073 | 1.131202221 | 0.729767903 |
| Gm24561 | 2.66823802 | -0.143060251 | 0.532728884 | -0.268542322 |
| Adnp | 84.6346395 | -0.146139663 | 0.181141366 | -0.806771339 |
| Gm20716 | 2.018522609 | 0.151625354 | 0.674733709 | 0.224718806 |
| Dpm1 | 240.6952631 | -0.070473155 | 0.190188241 | -0.37054423 |
| Mocs3 | 110.2099467 | -0.105160231 | 0.153346024 | -0.685770835 |
| Gm14237 | 0.075043205 | 0.755286614 | 3.724945049 | 0.202764498 |
| Kcng1 | 163.4779742 | -0.676044342 | 0.199264737 | -3.392694332 |
| Gm14262 | 0 | NA | NA | NA |
| Gm22704 | 0.118974955 | 0.274400501 | 3.724945049 | 0.073665651 |
| Gm14234 | 0.075043205 | 0.755286614 | 3.724945049 | 0.202764498 |
| Nfatc2 | 337.8978929 | 0.126017317 | 0.086626331 | 1.454723018 |
| Atp9a | 3839.633382 | 0.018304195 | 0.038589526 | 0.474330659 |
| mmu-mir-7004 | 0.834414618 | -0.251271065 | 1.217875605 | -0.206319154 |
| Sall4 | 2.654068103 | -0.693203799 | 0.671244816 | -1.032713821 |
| Gm14261 | 0.095897396 | -0.687371727 | 3.724945049 | -0.184532045 |
| Gm25214 | 0.150086409 | 1.264824188 | 3.723011833 | 0.339731445 |
| Zfp64 | 184.749999 | 0.23623241 | 0.110002501 | 2.147518539 |
| 1700017J07Rik | 0.04393175 | -0.206490061 | 3.724945049 | -0.055434391 |
| Gm14260 | 0 | NA | NA | NA |
| Gm9873 | 2.521204246 | 0.074589583 | 0.616075187 | 0.121072208 |
| Gm26469 | 0.075043205 | 0.755286614 | 3.724945049 | 0.202764498 |
| Gm14258 | 0.182681975 | 1.236192873 | 2.725430215 | 0.453577151 |
| 9430093N23Rik | 0 | NA | NA | NA |
| Gm14250 | 0 | NA | NA | NA |
| 4930529I22Rik | 0 | NA | NA | NA |
| Gm14249 | 0.152853976 | 1.641852012 | 2.820813903 | 0.582049036 |
| Gm11011 | 0 | NA | NA | NA |
| Gm34294 | 5.209546091 | -0.504835054 | 0.508800913 | -0.99220548 |
| Tshz2 | 720.7148521 | 0.007120784 | 0.10615656 | 0.067078131 |
| Gm26883 | 2.358426798 | 0.122969305 | 0.671947884 | 0.18300423 |

| | | | | |
|---|---|---|---|---|
| AY702102 | 0 | NA | NA | NA |
| A630075F10Rik | 0 | NA | NA | NA |
| Zfp217 | 142.5419381 | 1.335828778 | 0.160379282 | 8.329185413 |
| Gm17619 | 0.100902008 | 0.274391828 | 3.724945049 | 0.073663322 |
| Gm14270 | 11.75654133 | 0.320035734 | 0.311220644 | 1.028324246 |
| Bcas1 | 1958.212372 | 0.039832306 | 0.141923443 | 0.280660511 |
| Bcas1os2 | 1.937199938 | -0.85191144 | 0.774776656 | -1.099557446 |
| Bcas1os1 | 3.671955489 | 0.030471097 | 0.575889849 | 0.052911328 |
| Gm14269 | 52.69193393 | 0.378997189 | 0.167931302 | 2.256858515 |
| Cyp24a1 | 1.170018213 | 0.326127181 | 1.087520385 | 0.299881441 |
| Pfdn4 | 118.4952192 | 0.493724939 | 0.194000852 | 2.544962739 |
| Gm16796 | 10.39617679 | 0.32517869 | 0.366632425 | 0.8869338 |
| Gm14637 | 2.883279791 | 1.889454734 | 0.667377346 | 2.831164027 |
| 4930470P17Rik | 0.106465261 | 0.75528439 | 3.724945049 | 0.202763901 |
| Gm14264 | 0.597452826 | 0.906623591 | 1.453145949 | 0.623904014 |
| Dok5 | 201.9773433 | -0.486954737 | 0.151007762 | -3.224700036 |
| Gm14263 | 0 | NA | NA | NA |
| Gm14266 | 0.20369824 | 1.453677228 | 3.191915274 | 0.455424754 |
| Gm14640 | 0 | NA | NA | NA |
| 1700007M16Rik | 0.088534442 | 0.274389469 | 3.724945049 | 0.073662689 |
| Gm26489 | 0 | NA | NA | NA |
| 1700028P15Rik | 0 | NA | NA | NA |
| Gm14272 | 0 | NA | NA | NA |
| Gm14641 | 0 | NA | NA | NA |
| Cbln4 | 171.4002324 | -0.328824577 | 0.135964898 | -2.418451976 |
| Gm14271 | 0.25568908 | 0.274412207 | 1.851889213 | 0.148179602 |
| Gm14274 | 0 | NA | NA | NA |
| AL844846.1 | 0 | NA | NA | NA |
| Mc3r | 7.192941036 | 0.499414064 | 0.448617207 | 1.113229847 |
| Gm14275 | 0 | NA | NA | NA |
| Gm14273 | 0.454387946 | -0.206482618 | 1.583396151 | -0.130404901 |
| Fam210b | 297.4704023 | 0.008735214 | 0.097244381 | 0.08982744 |
| Aurka | 15.8043844 | 0.891945127 | 0.332981776 | 2.678660485 |
| Cstf1 | 247.1233043 | 0.130729064 | 0.080842656 | 1.61708027 |
| Cass4 | 5.704398591 | 0.912960171 | 0.564867564 | 1.616237556 |
| Gm14455 | 6.260709098 | -0.348314809 | 0.434811548 | -0.80107074 |
| Rtfdc1 | 1259.401244 | 0.307425576 | 0.064525669 | 4.76439196 |
| Gcnt7 | 0.721712508 | 0.280128967 | 1.410110608 | 0.198657443 |
| Fam209 | 5.869341775 | 0.348688263 | 0.523634445 | 0.665900164 |
| Gm14303 | 72.23225202 | 0.218215624 | 0.194524359 | 1.121790737 |
| Tfap2c | 3.754962934 | -1.185971752 | 0.664185346 | -1.785603611 |
| Gm22773 | 0 | NA | NA | NA |
| Bmp7 | 187.5782872 | 0.068045366 | 0.196262606 | 0.34670571 |
| Spo11 | 3.986507545 | 0.333881254 | 0.506402666 | 0.659319701 |
| Rae1 | 377.0248253 | 0.322949175 | 0.081265511 | 3.974000415 |
| Rbm38 | 50.55568089 | 0.28114001 | 0.195014341 | 1.441637617 |
| Gm20490 | 0 | NA | NA | NA |

| | | | | |
|---|---|---|---|---|
| Gm14453 | 0 | NA | NA | NA |
| Ctcfl | 0.389334344 | 0.261141016 | 1.476228357 | 0.176897439 |
| Ctcflos | 0.803203798 | 2.532459862 | 1.229675771 | 2.05945333 |
| Pck1 | 0.852421781 | 0.527367397 | 1.458495146 | 0.361583238 |
| Zbp1 | 1311.11427 | 9.066213536 | 0.533552663 | 16.99216248 |
| Pmepa1 | 867.4379167 | 0.107232547 | 0.159232334 | 0.673434497 |
| Pmepa1os | 1.650675276 | 1.644958878 | 0.9393589 | 1.751150576 |
| Gm14642 | 0 | NA | NA | NA |
| 1700021F07Rik | 0 | NA | NA | NA |
| Ankrd60 | 0.203084296 | -1.124763467 | 2.634616364 | -0.426917362 |
| Ppp4r1l-ps | 195.0349838 | -0.329603438 | 0.15545445 | -2.12025734 |
| Rab22a | 871.4199496 | 0.109477554 | 0.074086324 | 1.477702606 |
| Mir6340 | 0 | NA | NA | NA |
| 1700010B08Rik | 0.15795635 | 0.274407234 | 2.815015565 | 0.097479828 |
| Vapb | 1573.7862 | -0.204547775 | 0.061624271 | -3.319272926 |
| Gm14371 | 0 | NA | NA | NA |
| Gm14336 | 0 | NA | NA | NA |
| Atp5k-ps2 | 1.191345417 | -0.014349386 | 0.995680186 | -0.014411642 |
| Vamp7-ps | 9.408148121 | 0.379751547 | 0.401666606 | 0.94543968 |
| Stx16 | 905.3860616 | -0.132614809 | 0.070868041 | -1.871292168 |
| Npepl1 | 158.3376538 | 0.066499775 | 0.13718997 | 0.484727677 |
| Gm10714 | 1.585474913 | -1.666707706 | 0.774292407 | -2.152555923 |
| Gm27591 | 0 | NA | NA | NA |
| Mir296 | 0.229181457 | 0.274412217 | 2.603735228 | 0.105391752 |
| Mir298 | 0.18819539 | 1.118023756 | 3.283761112 | 0.34047049 |
| Nespas | 11.25411321 | 0.048093606 | 0.365099767 | 0.1317273 |
| Gnas | 10595.54034 | 0.031535587 | 0.07965094 | 0.395922343 |
| Gm27899 | 0.248714005 | -1.132483845 | 1.8126423 | -0.624769622 |
| Gm27869 | 0.247908139 | 0.29222644 | 2.143973162 | 0.136301352 |
| Gm27849 | 0.638233968 | 0.1692665 | 2.382317657 | 0.071051188 |
| Gm20721 | 37.27970981 | -0.209137081 | 0.269687406 | -0.775479597 |
| Nelfcd | 290.7180538 | 0.005632219 | 0.065598811 | 0.085858559 |
| Ctsz | 1102.778525 | 2.162889984 | 0.149868553 | 14.43191348 |
| Rpl30-ps5 | 0 | NA | NA | NA |
| Tubb1 | 0.687228199 | 0.588171877 | 1.340133637 | 0.438890467 |
| Atp5e | 358.6752708 | -0.38272372 | 0.137497621 | -2.783493404 |
| Slmo2 | 386.0940015 | 0.047323495 | 0.100061352 | 0.472944787 |
| Gm14385 | 0 | NA | NA | NA |
| Vmn1r-ps1 | 0.042716945 | -0.206490061 | 3.724945049 | -0.055434391 |
| Zfp831 | 392.3466882 | -0.422615057 | 0.189607286 | -2.228896714 |
| Edn3 | 25.09549224 | -1.429212437 | 0.395752738 | -3.61137726 |
| Gm14617 | 0.43206172 | -0.687385376 | 3.332226331 | -0.20628412 |
| Gm14618 | 0.490955376 | 0.33213655 | 3.715550159 | 0.089390948 |
| Gm14616 | 0 | NA | NA | NA |
| Gm14444 | 1.336470638 | -0.801382066 | 1.009483008 | -0.793853943 |
| Gm16354 | 2.08236629 | 0.231742394 | 0.808993884 | 0.286457535 |
| Gm14445 | 0.131100043 | -1.037279138 | 3.663654906 | -0.283126868 |

| | | | | |
|---|---|---|---|---|
| Gm14393 | 21.09049984 | -0.136033942 | 0.314216626 | -0.432930438 |
| Gm14439 | 0.167183452 | -0.206481502 | 2.765153652 | -0.074672705 |
| Gm14387 | 3.01954408 | -1.188733231 | 0.620896895 | -1.914542078 |
| Gm14392 | 2.964599863 | -0.388690382 | 0.574965065 | -0.676024346 |
| Gm14399 | 171.4815682 | -0.530439745 | 0.141546292 | -3.747464789 |
| Gm16355 | 0 | NA | NA | NA |
| Gm14443 | 181.6545351 | -0.661780855 | 0.199273082 | -3.320974652 |
| Gm14391 | 24.59774181 | -0.279826121 | 0.256294918 | -1.091812992 |
| Gm14442 | 0.132475575 | 0.274407273 | 2.973122108 | 0.092295998 |
| Gm14438 | 2.078804666 | 0.25475756 | 0.644199251 | 0.395463918 |
| Gm14440 | 1.419637689 | -0.918110865 | 0.8922271 | -1.029010287 |
| Gm14298 | 0 | NA | NA | NA |
| Gm4631 | 1.996170162 | -0.594725516 | 0.710299515 | -0.83728836 |
| Gm4723 | 1.755952801 | 0.623025802 | 0.6519612 | 0.955617913 |
| Gm14297 | 0 | NA | NA | NA |
| Gm4724 | 1.988766949 | -0.86926086 | 0.789549186 | -1.100958466 |
| Gm8923 | 0.572800032 | 1.387969532 | 1.381074981 | 1.004992162 |
| Gm11007 | 1.812554613 | -0.243680159 | 0.721891775 | -0.337557744 |
| Gm16359 | 0 | NA | NA | NA |
| Gm4245 | 2.26497711 | -0.251884388 | 0.772633567 | -0.326007565 |
| Gm2007 | 0.319719969 | 1.507446878 | 2.037703262 | 0.739777428 |
| Gm16358 | 0 | NA | NA | NA |
| Gm14288 | 19.90464905 | -0.69672626 | 0.317294617 | -2.195833851 |
| Rps8-ps2 | 2.50324314 | 0.840494258 | 0.727049477 | 1.156034472 |
| Gm2020 | 0.047948698 | -0.206487837 | 3.724945049 | -0.055433794 |
| Gm6710 | 28.49017795 | 0.102060982 | 0.252820819 | 0.403688995 |
| Gm16356 | 0 | NA | NA | NA |
| Gm8898 | 2.984855315 | -2.012206758 | 0.730277616 | -2.755399745 |
| Gm2026 | 2.773167693 | -1.849898878 | 0.849625163 | -2.177311781 |
| Gm11008 | 0.226297694 | 1.663785031 | 2.123090118 | 0.783661992 |
| Gm16357 | 0.13861434 | -1.111919598 | 3.614496194 | -0.307627824 |
| Gm2004 | 6.050452602 | 0.190808812 | 0.436014781 | 0.437620055 |
| Gm6182 | 0.19897497 | 0.247093634 | 2.216505264 | 0.11147893 |
| 2210418O10Rik | 24.2680911 | -0.915246641 | 0.325768554 | -2.809499657 |
| Gm17258 | 4.979667379 | -0.66179431 | 0.660207117 | -1.002404083 |
| Gm1980 | 0.288549458 | 0.263200517 | 1.63853571 | 0.160631541 |
| Gm11009 | 0.13733263 | -1.154748075 | 3.63430361 | -0.31773572 |
| Gm14435 | 0.646335998 | -0.734704976 | 1.2335135 | -0.595619729 |
| Gm16362 | 0 | NA | NA | NA |
| Gm14434 | 1.597696658 | 0.369036993 | 1.000198652 | 0.368963697 |
| Gm14431 | 1.363863261 | 0.451757544 | 0.769142931 | 0.587351877 |
| Gm14307 | 0 | NA | NA | NA |
| Gm14308 | 2.466484677 | -0.380463811 | 0.861352554 | -0.441705095 |
| Gm14306 | 4.680219553 | -0.333167212 | 0.496507601 | -0.671021373 |
| Gm14305 | 173.9135921 | -0.205080369 | 0.125911634 | -1.628764257 |
| Gm14301 | 0.675521499 | -0.241895749 | 1.493391656 | -0.161977435 |
| Gm14390 | 0.066050875 | 0.274391604 | 3.724945049 | 0.073663262 |

| | | | | |
|---|---|---|---|---|
| Gm14295 | 67.51412393 | -0.208938354 | 0.257182297 | -0.812413436 |
| Gm14408 | 0.284066144 | -1.216217132 | 2.087184509 | -0.582707052 |
| Gm14405 | 1.448471955 | -0.214709213 | 0.754097614 | -0.284723369 |
| Gm14296 | 30.11651545 | 0.035727362 | 0.298083362 | 0.119856949 |
| Mup-ps25 | 0 | NA | NA | NA |
| Gm14417 | 0.204225461 | 1.868934307 | 3.204890169 | 0.583150813 |
| Gm14419 | 4.87970569 | -0.009143117 | 0.450707782 | -0.020286132 |
| Gm14402 | 1.138899903 | -1.697018594 | 1.063218171 | -1.596115116 |
| Gm14421 | 5.323853793 | -0.698908794 | 0.451098248 | -1.549349387 |
| Gm14401 | 0.702992681 | 0.264262324 | 1.124444127 | 0.235015967 |
| Gm14407 | 3.984100659 | -0.448634573 | 0.503965001 | -0.890209781 |
| Gm14415 | 18.81040368 | -0.951175552 | 0.322286536 | -2.951335054 |
| Gm14411 | 16.94595019 | -0.211658709 | 0.316056444 | -0.669686421 |
| Gm14416 | 3.389752464 | -0.334395831 | 0.533625912 | -0.626648414 |
| Gm14413 | 14.03696361 | -0.810248506 | 0.290343648 | -2.790653458 |
| Gm14410 | 143.7066193 | -0.353441291 | 0.149395974 | -2.365801982 |
| Gm14400 | 0.439370512 | -0.191860611 | 1.358832657 | -0.141195172 |
| Gm14409 | 0.902861002 | 0.623824908 | 0.889902912 | 0.701003334 |
| Gm14412 | 64.87765343 | 0.137464439 | 0.234643443 | 0.585843936 |
| Gm14418 | 19.13716893 | -0.186488779 | 0.266949114 | -0.698592985 |
| Gm14404 | 0.1140246 | 0.75528439 | 3.724945049 | 0.202763901 |
| Gm14414 | 1.844525372 | 0.21652087 | 0.757022166 | 0.286016552 |
| Gm14420 | 72.73691108 | -0.338276423 | 0.189413504 | -1.785915024 |
| Gm14403 | 33.59942204 | -0.652357272 | 0.239896394 | -2.719329208 |
| Gm16363 | 0.136194719 | -0.687371727 | 3.724945049 | -0.184532045 |
| Gm14406 | 1.816762255 | -0.491584271 | 0.729657198 | -0.673719484 |
| Gm14324 | 0.130017973 | -0.206489706 | 3.662785653 | -0.056375045 |
| Gm14322 | 38.40290494 | -0.187446743 | 0.200967022 | -0.932723893 |
| Gm14325 | 156.7524774 | -0.122018662 | 0.172009164 | -0.709373029 |
| Gm14429 | 5.50850624 | 0.153664933 | 0.449585986 | 0.341792089 |
| Gm14323 | 1.346967727 | 0.025924685 | 0.903968288 | 0.028678755 |
| Gm14327 | 9.478232069 | 0.342922586 | 0.332454402 | 1.031487576 |
| C330013J21Rik | 6.088359734 | 0.665989251 | 0.551952539 | 1.206606011 |
| Gm14326 | 207.3808432 | -0.137584413 | 0.11401029 | -1.206771892 |
| Gm14428 | 0.751182044 | 1.02893659 | 1.219936102 | 0.843434823 |
| Etohi1 | 168.5832989 | -0.375087036 | 0.123167657 | -3.045337114 |
| Gm26869 | 1.499946283 | -2.164065945 | 0.965289617 | -2.241882546 |
| Zfp931 | 152.8081414 | -0.401511319 | 0.163541119 | -2.455109279 |
| Phactr3 | 1341.248435 | 0.113105316 | 0.099646188 | 1.135069176 |
| Sycp2 | 51.90291932 | 0.013323863 | 0.173276767 | 0.076893532 |
| Ppp1r3d | 90.63451093 | 0.177083555 | 0.165146468 | 1.072281816 |
| Fam217b | 760.5922785 | -0.075554271 | 0.085405975 | -0.884648537 |
| Cdh26 | 2.606367607 | -2.403043521 | 0.84116597 | -2.856800686 |
| Gm14314 | 0 | NA | NA | NA |
| Gm14292 | 70.00844022 | 0.276444334 | 0.129597762 | 2.133094968 |
| Gm14293 | 0 | NA | NA | NA |
| Gm14294 | 0 | NA | NA | NA |

| | | | | |
|---|---|---|---|---|
| 4930591A17Rik | 0 | NA | NA | NA |
| Cdh4 | 374.0015759 | -0.220834547 | 0.089036234 | -2.480277277 |
| Gm14300 | 0.042716945 | -0.206490061 | 3.724945049 | -0.055434391 |
| Taf4a | 305.5847678 | -0.33042541 | 0.098819661 | -3.343721338 |
| 4921531C22Rik | 29.89290028 | -0.620782084 | 0.280774611 | -2.210962315 |
| Gm14299 | 0.10503227 | 0.274389469 | 3.724945049 | 0.073662689 |
| Lsm14b | 804.4044537 | -0.390863765 | 0.072284875 | -5.407269042 |
| Psma7 | 1015.88698 | 0.724703667 | 0.116655226 | 6.212354932 |
| Ss18l1 | 608.8536376 | -0.172812218 | 0.065611729 | -2.633861662 |
| Mtg2 | 112.4777664 | -0.489520304 | 0.118173903 | -4.142372305 |
| Hrh3 | 541.8584486 | -0.202631521 | 0.144356452 | -1.403688705 |
| Gm17180 | 0.40273248 | -0.043050747 | 3.367818266 | -0.012782978 |
| Osbpl2 | 658.7052841 | -0.075124786 | 0.052712938 | -1.425167886 |
| Adrm1 | 690.902229 | 0.378380328 | 0.084047028 | 4.502007227 |
| Lama5 | 192.3097236 | 0.894147015 | 0.194762212 | 4.590967645 |
| Mir7005 | 0 | NA | NA | NA |
| Rps21 | 796.5226521 | 0.199220412 | 0.147360365 | 1.351926704 |
| Mir3091 | 3.756198497 | -0.053153317 | 0.571206913 | -0.0930544 |
| Cables2 | 296.6979083 | -0.154257718 | 0.085024273 | -1.814278593 |
| Rbbp8nl | 0.042716945 | -0.206490061 | 3.724945049 | -0.055434391 |
| Gata5 | 0.107803068 | 0.75528661 | 3.724945049 | 0.202764497 |
| Gata5os | 0 | NA | NA | NA |
| B230312C02Rik | 9.425892428 | -0.97426904 | 0.356912221 | -2.729716111 |
| Mir1a-1 | 0.044544648 | 0.755286614 | 3.724945049 | 0.202764498 |
| Mir133a-2 | 0.230754588 | -1.812433634 | 2.536465953 | -0.714550744 |
| Slco4a1 | 65.52497335 | 0.599072395 | 0.152654865 | 3.924358355 |
| Ntsr1 | 81.28326963 | -0.510711211 | 0.131667375 | -3.878798468 |
| Mrgbp | 164.3286957 | 0.053467408 | 0.090420293 | 0.591320894 |
| Ogfr | 1008.536141 | 1.676286334 | 0.132649406 | 12.63696824 |
| Col9a3 | 196.0069601 | -0.114687309 | 0.116860629 | -0.981402464 |
| Tcfl5 | 5.628819471 | -0.375168805 | 0.443330024 | -0.846251742 |
| Dido1 | 1582.103518 | -0.117258536 | 0.089590378 | -1.308829569 |
| Gm22502 | 0 | NA | NA | NA |
| Gid8 | 550.3352372 | 0.358500079 | 0.058897971 | 6.086798427 |
| Slc17a9 | 7.930149013 | 1.545228463 | 0.545627867 | 2.832018956 |
| Gm14343 | 0 | NA | NA | NA |
| Bhlhe23 | 1.454032994 | 0.486815487 | 0.811670191 | 0.599770069 |
| Gm14340 | 2.077246534 | -0.305328237 | 0.680182478 | -0.448891653 |
| Gm14342 | 4.321485189 | 0.472099451 | 0.462573749 | 1.02059283 |
| Gm14344 | 0.429573641 | -0.219755113 | 1.477387813 | -0.14874572 |
| Gm22842 | 0.112288909 | -0.687373951 | 3.724945049 | -0.184532642 |
| Gm14339 | 0.202674043 | -0.206490498 | 2.623803806 | -0.078698909 |
| Mir124a-3 | 0.458109599 | 0.905608692 | 1.772562591 | 0.510903647 |
| Gm27032 | 102.6108423 | -0.461967008 | 0.190687102 | -2.422644228 |
| Ythdf1 | 817.9793704 | 0.266028315 | 0.072759527 | 3.656267765 |
| Birc7 | 1.578816077 | -2.861135653 | 0.931051679 | -3.073014869 |
| Nkain4 | 83.76982434 | -0.947134037 | 0.242149025 | -3.911368374 |

| | | | | |
|---|---|---|---|---|
| Gm14341 | 1.152403953 | -0.436635286 | 0.876612835 | -0.498093649 |
| Arfgap1 | 898.465573 | 0.14422451 | 0.094924913 | 1.519353621 |
| 9230112E08Rik | 145.4154449 | -0.349102229 | 0.195183045 | -1.788588912 |
| Col20a1 | 78.79971225 | 0.386691847 | 0.227929382 | 1.696542341 |
| Chrna4 | 653.8085059 | 0.542132668 | 0.123413181 | 4.392826313 |
| Kcnq2 | 4581.893317 | -0.230658069 | 0.127839502 | -1.804278535 |
| Eef1a2 | 7678.32573 | -0.189324001 | 0.07419173 | -2.551820807 |
| Gm24892 | 0 | NA | NA | NA |
| Ppdpf | 209.3303033 | -0.223029416 | 0.096501159 | -2.311157888 |
| Ptk6 | 1.572767014 | -0.632479782 | 1.176687789 | -0.53750858 |
| Srms | 1.881316667 | 0.80042594 | 0.768516824 | 1.041520387 |
| Gm14398 | 0.631396267 | 2.200634523 | 1.25957881 | 1.747119359 |
| BC051628 | 0.900425091 | 1.38471502 | 1.163881633 | 1.189738699 |
| Helz2 | 1488.152321 | 4.634867168 | 0.279005358 | 16.61210812 |
| Gmeb2 | 338.4791566 | 0.366333629 | 0.117851068 | 3.108445569 |
| Gm7645 | 4.486974988 | -0.793531689 | 0.568298446 | -1.396329156 |
| Stmn3 | 2848.78216 | -0.167702595 | 0.106257906 | -1.578259935 |
| Rtel1 | 189.1322798 | -0.012039545 | 0.151881488 | -0.07926934 |
| Arfrp1 | 263.798925 | 0.040719795 | 0.098758854 | 0.412315385 |
| Zgpat | 398.0436084 | -0.392957739 | 0.122464587 | -3.208745892 |
| Lime1 | 3.160497059 | -0.345906131 | 0.645464982 | -0.535902242 |
| Slc2a4rg-ps | 70.84153722 | -0.893998722 | 0.353726289 | -2.527374272 |
| Zbtb46 | 247.6912476 | -0.265438465 | 0.134878434 | -1.967982994 |
| Gm22708 | 0.139376635 | 0.740973539 | 2.922770099 | 0.253517558 |
| Abhd16b | 0 | NA | NA | NA |
| Tpd52l2 | 603.1097156 | 0.066865454 | 0.062562404 | 1.068780129 |
| Gm25184 | 2.339668925 | -0.018670474 | 0.68337397 | -0.02732102 |
| Dnajc5 | 4139.797779 | -0.052190372 | 0.042605925 | -1.224955731 |
| Uckl1 | 187.8382027 | -0.166322395 | 0.129233465 | -1.286991693 |
| Uckl1os | 3.735935155 | -0.889117121 | 0.574341509 | -1.54806349 |
| Znf512b | 722.5500756 | 0.070128286 | 0.114087468 | 0.614688772 |
| Mir7006 | 0 | NA | NA | NA |
| Prpf6 | 758.9138183 | -0.006205307 | 0.082468555 | -0.075244519 |
| Samd10 | 162.7504342 | -0.322847898 | 0.100662299 | -3.207237478 |
| Gm25879 | 0 | NA | NA | NA |
| Sox18 | 27.0834614 | 0.420883675 | 0.239906634 | 1.754364474 |
| Tcea2 | 231.4654535 | -0.069214344 | 0.145559862 | -0.475504322 |
| Rgs19 | 143.057256 | -0.10626173 | 0.119491187 | -0.889285083 |
| Lkaaear1 | 0 | NA | NA | NA |
| Oprl1 | 406.903062 | 0.021361432 | 0.084762028 | 0.252016523 |
| Gm14489 | 0 | NA | NA | NA |
| Myt1 | 485.0253362 | -0.321815292 | 0.135285042 | -2.378794332 |
| Pcmtd2 | 953.1475173 | -0.370884202 | 0.088508392 | -4.190384572 |
| Polr3k | 406.1158035 | -0.325175752 | 0.061014814 | -5.329455729 |
| Gm5466 | 17.7294434 | -0.378858026 | 0.270684261 | -1.399630786 |
| Gm14496 | 0.046345709 | 0.274391604 | 3.724945049 | 0.073663262 |
| Gm28002 | 0 | NA | NA | NA |

| | | | | |
|---|---|---|---|---|
| Gm27460 | 0 | NA | NA | NA |
| Gm37338 | 0 | NA | NA | NA |
| Gm8747 | 0 | NA | NA | NA |
| Hnf4g | 1.172273398 | 2.921733577 | 1.100327852 | 2.655330019 |
| Gm37000 | 0.047948698 | -0.206487837 | 3.724945049 | -0.055433794 |
| Gm23237 | 0 | NA | NA | NA |
| Rps24-ps2 | 0.229688899 | -0.703282757 | 2.548865832 | -0.275919881 |
| Gm2071 | 0 | NA | NA | NA |
| Gm2077 | 0.293065915 | -0.206481216 | 1.787583577 | -0.115508566 |
| Gm8775 | 6.987551743 | 0.436210454 | 0.400057356 | 1.090369788 |
| Gm23976 | 0.841305426 | -1.237712707 | 1.2703514 | -0.974307351 |
| n-R5s194 | 0 | NA | NA | NA |
| Gm34023 | 0 | NA | NA | NA |
| Gm38131 | 0 | NA | NA | NA |
| Gm37029 | 0 | NA | NA | NA |
| Gm22944 | 0 | NA | NA | NA |
| Gm37763 | 0 | NA | NA | NA |
| Gm37801 | 0 | NA | NA | NA |
| 1110015O18Rik | 13.63005333 | -0.294897426 | 0.312598509 | -0.943374384 |
| Gm6140 | 0 | NA | NA | NA |
| Gm17308 | 1.32255022 | 0.095370422 | 0.998028961 | 0.095558772 |
| 2700069I18Rik | 10.07409941 | 0.051565446 | 0.332577778 | 0.15504778 |
| Gm10748 | 0.529004033 | 0.346509529 | 1.456512127 | 0.237903635 |
| Zfhx4 | 931.4280682 | 0.002717398 | 0.073125754 | 0.037160609 |
| 4930555M17Rik | 0 | NA | NA | NA |
| Pex2 | 255.6278662 | 0.17839085 | 0.105446974 | 1.691758834 |
| Gm8797 | 20.70001869 | 0.67511076 | 0.334080323 | 2.020803718 |
| Gm37350 | 0.062426873 | 0.755288839 | 3.724945049 | 0.202765096 |
| Gm25352 | 0 | NA | NA | NA |
| Gm25144 | 0 | NA | NA | NA |
| Gm17934 | 0.363515313 | 1.200675952 | 1.938902347 | 0.619255505 |
| Gm6162 | 0.35319577 | 0.681736677 | 1.551223188 | 0.439483295 |
| Gm38298 | 0 | NA | NA | NA |
| Gm23147 | 0.044544648 | 0.755286614 | 3.724945049 | 0.202764498 |
| Gm18658 | 0 | NA | NA | NA |
| Gm5272 | 0 | NA | NA | NA |
| Gm38291 | 0 | NA | NA | NA |
| 1700008P02Rik | 0.048949532 | -0.206487837 | 3.724945049 | -0.055433794 |
| Olfr289-ps1 | 0.152964131 | 0.274406991 | 2.843489581 | 0.096503604 |
| Gm22074 | 0 | NA | NA | NA |
| Gm8823 | 0 | NA | NA | NA |
| Pkia | 2228.089358 | 0.353517979 | 0.08297221 | 4.26067933 |
| Gm37918 | 0 | NA | NA | NA |
| Gm8826 | 0.880107612 | 0.958218243 | 1.096273399 | 0.87406868 |
| Gm23592 | 0 | NA | NA | NA |
| Zc2hc1a | 623.5639214 | -0.140082286 | 0.071178207 | -1.968050223 |
| Il7 | 17.75567427 | 3.289305549 | 0.460912069 | 7.136514257 |

| | | | | |
|---|---|---|---|---|
| Gm16685 | 14.13537915 | 5.209158891 | 0.802889293 | 6.488016386 |
| Gm29747 | 0.364508673 | 0.755307948 | 1.662390991 | 0.454350362 |
| 1700010I02Rik | 0.136120942 | 0.274393632 | 3.599844381 | 0.076223748 |
| Gm7103 | 0.4787828 | 2.527782887 | 1.439390185 | 1.756148481 |
| Gm5841 | 5.450194582 | 0.82402861 | 0.490898661 | 1.678612462 |
| Gm24614 | 0 | NA | NA | NA |
| 1700003N22Rik | 1.26814241 | -0.86406162 | 0.93998482 | -0.919229334 |
| Stmn2 | 2333.139224 | -0.225151915 | 0.068187376 | -3.301958917 |
| Gm18822 | 4.048703541 | 1.070133556 | 0.50652721 | 2.112687204 |
| Gm32496 | 1.115586983 | 3.618335776 | 1.222505762 | 2.959769916 |
| Hey1 | 408.0829732 | -0.105104379 | 0.069265052 | -1.51742293 |
| Gm38007 | 0.33971137 | -1.313579295 | 1.64630152 | -0.797897152 |
| Gm23670 | 0 | NA | NA | NA |
| C230057A21Rik | 0.383304402 | -0.687379107 | 1.716208523 | -0.400521905 |
| Gm37647 | 0.199458489 | -0.206483616 | 2.665489557 | -0.07746555 |
| Mrps28 | 80.2242516 | 0.460429179 | 0.15925042 | 2.891227409 |
| Gm15467 | 0 | NA | NA | NA |
| Gm15466 | 6.37553239 | -0.24989267 | 0.429743509 | -0.581492598 |
| Tpd52 | 690.3315485 | 0.249939194 | 0.071590971 | 3.49121114 |
| Gm37616 | 0 | NA | NA | NA |
| 4930539M17Rik | 0.310536081 | -0.664841845 | 1.598359638 | -0.415952599 |
| Gm8860 | 0.282898957 | 1.21531106 | 2.051177957 | 0.592494208 |
| Zbtb10 | 267.8821127 | -0.016993251 | 0.100441227 | -0.169186015 |
| Gm22604 | 0.061920613 | 0.274393828 | 3.724945049 | 0.073663859 |
| C030034L19Rik | 1.859233056 | 0.571809835 | 0.749363395 | 0.763060804 |
| Zfp704 | 1430.007501 | -0.499089934 | 0.091472655 | -5.456165395 |
| Gm38001 | 0 | NA | NA | NA |
| Gm17806 | 0 | NA | NA | NA |
| Pag1 | 670.6925854 | -0.267587426 | 0.112891174 | -2.370313084 |
| Gm16337 | 0 | NA | NA | NA |
| Gm24786 | 3.953076191 | -0.73677584 | 0.522491767 | -1.41011952 |
| Gm6236 | 0 | NA | NA | NA |
| Gm17877 | 0.154612032 | 0.274389336 | 3.501099905 | 0.078372324 |
| Fabp5 | 368.7894691 | 0.276167252 | 0.129722294 | 2.128911252 |
| Gm38335 | 0.498917054 | 0.252346892 | 1.47785457 | 0.170752181 |
| Gm9833 | 92.22302499 | -0.216970894 | 0.167022439 | -1.299052361 |
| Gm37308 | 0 | NA | NA | NA |
| Pmp2 | 0 | NA | NA | NA |
| Gm37389 | 0 | NA | NA | NA |
| Fabp9 | 0 | NA | NA | NA |
| Fabp4 | 3.144700407 | 3.006384623 | 0.992384252 | 3.029456197 |
| Gm38310 | 0.547572158 | 0.274410397 | 1.639286571 | 0.167396233 |
| Gm37421 | 0 | NA | NA | NA |
| Gm37384 | 0 | NA | NA | NA |
| Fabp12 | 1.277907819 | 1.085613415 | 1.149040503 | 0.944799954 |
| Gm37316 | 0.044544648 | 0.755286614 | 3.724945049 | 0.202764498 |
| Gm38364 | 0 | NA | NA | NA |

| | | | | |
|---|---|---|---|---|
| Impa1 | 639.4989438 | 0.04373312 | 0.056132351 | 0.77910722 |
| Gm38303 | 0 | NA | NA | NA |
| Slc10a5 | 0.840861136 | -1.159398828 | 1.203065632 | -0.963703723 |
| Zfand1 | 152.3432722 | -0.14824473 | 0.134723562 | -1.100362314 |
| Chmp4c | 3.570328421 | 0.460610739 | 0.554247465 | 0.831056105 |
| Snx16 | 391.7704708 | -0.284021585 | 0.108172668 | -2.625631685 |
| Gm37831 | 0.049553047 | 0.274391604 | 3.724945049 | 0.073663262 |
| Gm18692 | 0 | NA | NA | NA |
| Gm37244 | 0.092140247 | -0.687373951 | 3.724945049 | -0.184532642 |
| Gm22795 | 0 | NA | NA | NA |
| Gm21631 | 0 | NA | NA | NA |
| Gm37832 | 0 | NA | NA | NA |
| Gm22547 | 0.047948698 | -0.206487837 | 3.724945049 | -0.055433794 |
| Gm8919 | 0 | NA | NA | NA |
| Gm10745 | 0 | NA | NA | NA |
| Gm37996 | 0 | NA | NA | NA |
| Gm2429 | 0 | NA | NA | NA |
| Gm8935 | 0.243796007 | -0.206481298 | 1.85392735 | -0.111375075 |
| Gm8939 | 0 | NA | NA | NA |
| Gm38317 | 9.626198242 | -0.403169502 | 0.35846922 | -1.124697686 |
| Gm37813 | 0.0885994 | 0.274402725 | 3.724945049 | 0.073666248 |
| Gm37197 | 0.044544648 | 0.755286614 | 3.724945049 | 0.202764498 |
| Gm2464 | 10.69073118 | -0.870618784 | 0.338027661 | -2.575584436 |
| Gm38112 | 362.4382461 | -0.355277511 | 0.142988399 | -2.484659688 |
| Gm37164 | 1.862656823 | 0.026815551 | 0.756906276 | 0.035427835 |
| Gm2474 | 11.24932943 | -0.527126312 | 0.330797259 | -1.593502656 |
| Gm38400 | 64.08375772 | 0.00748826 | 0.169481009 | 0.044183474 |
| Gm19114 | 0.044544648 | 0.755286614 | 3.724945049 | 0.202764498 |
| Gm25981 | 24.11327504 | -0.408293554 | 0.294098765 | -1.388287209 |
| Ralyl | 450.4153493 | -0.073545194 | 0.075905702 | -0.968902088 |
| Gm22026 | 0.132101749 | 0.274387056 | 3.724945049 | 0.073662041 |
| Gm23112 | 0 | NA | NA | NA |
| Gm38311 | 0.286827043 | 1.231356598 | 2.03959799 | 0.603725148 |
| Gm6300 | 1.130984576 | 0.825931495 | 0.883488956 | 0.934852088 |
| Gm37358 | 0 | NA | NA | NA |
| Gm8955 | 0.038981395 | 0.274393828 | 3.724945049 | 0.073663859 |
| Slc7a12 | 0 | NA | NA | NA |
| Gm8957 | 0.226376398 | 0.274411419 | 2.151097738 | 0.127568085 |
| Lrrcc1 | 419.00625 | 0.024708581 | 0.090628017 | 0.272637329 |
| Gm17119 | 0 | NA | NA | NA |
| E2f5 | 109.6456114 | -0.120942565 | 0.123989279 | -0.9754276 |
| 1810022K09Rik | 264.8506355 | 0.303498888 | 0.117456208 | 2.583932277 |
| Gm38151 | 0.061920613 | 0.274393828 | 3.724945049 | 0.073663859 |
| Car13 | 144.2580462 | 1.78215704 | 0.245536058 | 7.258229413 |
| Gm5843 | 0 | NA | NA | NA |
| Car1 | 0.04393175 | -0.206490061 | 3.724945049 | -0.055434391 |
| Gm16399 | 0.953016895 | 1.327582973 | 1.136083445 | 1.168561146 |

| | | | | |
|---|---|---|---|---|
| Car3 | 3.829701666 | -0.941302002 | 0.518252815 | -1.81629887 |
| Car2 | 1336.217829 | -0.752774438 | 0.1824664 | -4.125551 |
| A930001A20Rik | 0.572612294 | -0.65320632 | 1.433781745 | -0.455582812 |
| Gm7132 | 0 | NA | NA | NA |
| Gm33819 | 0.263469446 | 0.755307631 | 2.106063317 | 0.358634817 |
| Gm9733 | 1.473887978 | 0.755310333 | 1.358130536 | 0.556139718 |
| Sirpb1a | 36.66926605 | 6.569095277 | 0.915030624 | 7.179098824 |
| Sirpb1b | 7.832067487 | 5.146856764 | 0.970734508 | 5.302023077 |
| Gm37468 | 0 | NA | NA | NA |
| Sirpb1c | 25.99497885 | 6.183048488 | 1.029807518 | 6.004081712 |
| Gm6321 | 0.066050875 | 0.274391604 | 3.724945049 | 0.073663262 |
| Gm7343 | 0.430172323 | 0.274410444 | 1.667287828 | 0.164584926 |
| Gm5150 | 10.40898798 | 4.998834926 | 0.952609792 | 5.247515793 |
| Gm24694 | 0 | NA | NA | NA |
| Gm37836 | 0 | NA | NA | NA |
| Gm25863 | 0 | NA | NA | NA |
| Ythdf3 | 1623.468186 | 0.039926984 | 0.048967662 | 0.815374539 |
| Gm26485 | 0 | NA | NA | NA |
| Gm5283 | 0.46900631 | 0.68120694 | 1.34585534 | 0.506151679 |
| Gm24963 | 0.253979914 | -0.216040925 | 1.828496474 | -0.118152224 |
| Gm6350 | 0.222549579 | 0.263136663 | 1.897660857 | 0.138663694 |
| Gm30340 | 13.55485612 | -0.229370848 | 0.310236066 | -0.73934295 |
| Gm19694 | 0 | NA | NA | NA |
| Gm38154 | 0 | NA | NA | NA |
| Mir124-2hg | 730.7247633 | 0.022798288 | 0.091933673 | 0.247986265 |
| Mir124a-2 | 2.558881719 | -0.115772996 | 0.602278712 | -0.192224951 |
| Gm23441 | 0 | NA | NA | NA |
| RP24-84E6.5 | 0.192405858 | -0.206487548 | 2.648345937 | -0.077968495 |
| Cypt12 | 0.109982625 | -0.20649451 | 3.724945049 | -0.055435586 |
| Gm5844 | 33.81437118 | 0.022054581 | 0.199652812 | 0.110464665 |
| Bhlhe22 | 439.9718104 | 0.364713397 | 0.157643485 | 2.31353295 |
| Gm23726 | 0 | NA | NA | NA |
| Cyp7b1 | 183.0659675 | 0.586212581 | 0.133543881 | 4.389662602 |
| Gm23686 | 3.150048424 | 0.473200594 | 0.530803811 | 0.891479269 |
| RP23-77D22.2 | 2.298046648 | 0.754475488 | 0.691457645 | 1.091137676 |
| Gm30667 | 0 | NA | NA | NA |
| 4930433B08Rik | 0 | NA | NA | NA |
| Gm6369 | 0 | NA | NA | NA |
| RP23-427B4.3 | 0 | NA | NA | NA |
| Gm37317 | 2.158753835 | 0.885215636 | 0.668388399 | 1.324403052 |
| RP23-427B4.2 | 0.137548931 | -0.206480418 | 2.938932991 | -0.070256933 |
| Gm30341 | 0 | NA | NA | NA |
| Armc1 | 876.5532839 | -0.089319044 | 0.079557524 | -1.122697629 |
| Gm37979 | 14.15384062 | -0.504219738 | 0.305825696 | -1.648716065 |
| Mtfr1 | 148.0536149 | -0.05147757 | 0.135766056 | -0.379163775 |
| Pde7a | 422.8626837 | -0.305097588 | 0.106691192 | -2.859632401 |
| Gm16093 | 2.00382768 | -0.20448586 | 0.699216622 | -0.292449941 |

| | | | | |
|---|---|---|---|---|
| Gm22282 | 0.538570099 | -0.131221752 | 1.254267692 | -0.104620212 |
| Dnajc5b | 1.101603561 | 0.753033575 | 1.2689277 | 0.593440883 |
| Gm23330 | 0.225375977 | -0.206480584 | 2.14758428 | -0.096145509 |
| Gm6141 | 0.044191549 | -0.206490061 | 3.724945049 | -0.055434391 |
| 1700064H15Rik | 1.062875418 | 0.82170041 | 0.834067772 | 0.985172234 |
| Trim55 | 5.56456899 | 5.145140228 | 0.991258346 | 5.190513905 |
| BC002189 | 0 | NA | NA | NA |
| Crh | 42.16080477 | 0.489768559 | 0.283682713 | 1.726465999 |
| Gm18864 | 0.120349074 | 1.264824188 | 3.723011833 | 0.339731445 |
| RP23-105K2.2 | 0 | NA | NA | NA |
| 4632415L05Rik | 336.8514353 | -0.057488432 | 0.073086098 | -0.786585052 |
| Gm37523 | 0.176667124 | -0.206481646 | 2.304379988 | -0.089603992 |
| Gm17771 | 0.368202395 | 1.251922108 | 1.542973488 | 0.811369812 |
| Cp | 857.3323739 | 2.736030214 | 0.25181029 | 10.86544247 |
| Hps3 | 339.45544 | 0.16349827 | 0.084696789 | 1.930395153 |
| Hltf | 534.1407693 | 0.049746707 | 0.071773623 | 0.6931057 |
| Gyg | 294.265161 | 0.085285548 | 0.128665739 | 0.662845822 |
| Cpa3 | 1.364377633 | -0.93380464 | 0.981533051 | -0.951373608 |
| Mir7007 | 0 | NA | NA | NA |
| Cpb1 | 1.12517466 | 0.255228024 | 1.140071343 | 0.223870222 |
| RP24-190F20.3 | 0.04393175 | -0.206490061 | 3.724945049 | -0.055434391 |
| Agtr1b | 1.030384379 | 1.244222695 | 0.966330117 | 1.287575201 |
| Gm18426 | 0.046345709 | 0.274391604 | 3.724945049 | 0.073663262 |
| RP24-190F20.4 | 0 | NA | NA | NA |
| Gm31466 | 0 | NA | NA | NA |
| Gm18491 | 0 | NA | NA | NA |
| Gm38323 | 0 | NA | NA | NA |
| Gm18493 | 0 | NA | NA | NA |
| Gm37480 | 0.122601409 | 1.236179401 | 3.724945049 | 0.331865138 |
| Gm7488 | 13.25965715 | 0.172439028 | 0.295025712 | 0.584488133 |
| RP23-105J10.1 | 0.180386268 | -1.298894529 | 3.333134146 | -0.389691645 |
| Gm29137 | 0.309067702 | -0.198718916 | 3.72036706 | -0.053413793 |
| RP23-458I4.1 | 0 | NA | NA | NA |
| RP23-458I4.2 | 0 | NA | NA | NA |
| RP23-458I4.3 | 0 | NA | NA | NA |
| 7530428D23Rik | 1.976191175 | 0.121780089 | 0.650799772 | 0.187123743 |
| RP23-105M18.2 | 4.557716346 | -0.093568991 | 0.486739345 | -0.192236341 |
| RP23-105M18.1 | 0 | NA | NA | NA |
| RP24-487F9.4 | 40.33606232 | -0.452178548 | 0.164549761 | -2.7479745 |
| Tbl1xr1 | 1839.527346 | 0.017601773 | 0.07137701 | 0.246602831 |
| Gm5842 | 2.255157929 | 0.344168456 | 0.644904243 | 0.533673736 |
| Rprl2 | 0.403845505 | -0.700268678 | 1.584937462 | -0.441827325 |
| Gm38324 | 0 | NA | NA | NA |
| Gm37636 | 0 | NA | NA | NA |
| Gm26354 | 0 | NA | NA | NA |
| Spin3-ps | 2.893792528 | 1.560094243 | 0.855920784 | 1.822708681 |
| Gm37889 | 0.097139547 | 0.274393828 | 3.724945049 | 0.073663859 |

| | | | | |
|---|---|---|---|---|
| Gm36983 | 1.90140223 | 3.430177892 | 1.037731417 | 3.305458267 |
| Gm37021 | 1.244068253 | 3.02554803 | 1.12646062 | 2.685888858 |
| Gm37492 | 0 | NA | NA | NA |
| 4930412E21Rik | 0.700626182 | 2.853159037 | 1.181211019 | 2.415452439 |
| Naaladl2 | 18.35136831 | 0.271881431 | 0.275054011 | 0.988465612 |
| Gm7536 | 51.1862964 | 0.554456293 | 0.19897549 | 2.78655573 |
| Gm24704 | 0 | NA | NA | NA |
| Gm24062 | 0 | NA | NA | NA |
| RP23-152K14.2 | 3.653026081 | -0.340583357 | 0.704155893 | -0.483676073 |
| Gm37136 | 0.341775432 | -0.05757732 | 2.012920634 | -0.02860387 |
| Gm38002 | 0 | NA | NA | NA |
| Gm10259 | 3.08375672 | 1.059094384 | 0.574108063 | 1.844764868 |
| Nlgn1 | 2154.133087 | 0.238813129 | 0.094165571 | 2.536098133 |
| Gm24250 | 0 | NA | NA | NA |
| Gm37573 | 125.0600198 | -0.255040211 | 0.192509504 | -1.324818804 |
| Gm6197 | 0.660619988 | 1.918538232 | 1.223418379 | 1.568178364 |
| A830092H15Rik | 35.52709241 | -0.769175685 | 0.272249252 | -2.825262802 |
| Gm4855 | 1.087037337 | -1.073252713 | 1.182698503 | -0.907460955 |
| Gm25773 | 0.04393175 | -0.206490061 | 3.724945049 | -0.055434391 |
| Spata16 | 0.312179078 | 1.030398814 | 1.703516234 | 0.604865861 |
| Gm37659 | 0 | NA | NA | NA |
| AC108829.1 | 0 | NA | NA | NA |
| Gm7558 | 0 | NA | NA | NA |
| Ect2 | 26.23418059 | 1.122257537 | 0.419653049 | 2.674250883 |
| Gm25152 | 0 | NA | NA | NA |
| 1700125G22Rik | 0.384795782 | -0.142140664 | 1.410805416 | -0.100751431 |
| Nceh1 | 1766.692954 | 0.177192976 | 0.074966008 | 2.363644272 |
| Gm37191 | 0.357797561 | 0.986114956 | 1.914033512 | 0.515202555 |
| RP23-131B7.3 | 0.060174537 | 0.755286614 | 3.724945049 | 0.202764498 |
| Tnfsf10 | 460.2181555 | 5.676025375 | 0.368624726 | 15.39784224 |
| Ghsr | 15.57801817 | 0.350687271 | 0.327446338 | 1.070976311 |
| Fndc3b | 605.684463 | 0.099917008 | 0.080671659 | 1.238563948 |
| RP24-62O16.3 | 0 | NA | NA | NA |
| Mir3092 | 1.750169107 | -0.432702722 | 0.767719716 | -0.563620698 |
| Tmem212 | 11.59725055 | -0.376217884 | 0.327490846 | -1.14878901 |
| Gm26040 | 0 | NA | NA | NA |
| Pld1 | 212.9629629 | -0.052227265 | 0.11902683 | -0.438785654 |
| Gm37098 | 0 | NA | NA | NA |
| Tnik | 3513.493683 | -0.202561814 | 0.129949498 | -1.558773347 |
| Gm37537 | 40.89549279 | -0.474353908 | 0.302267122 | -1.569320228 |
| Mir466q | 6.638222057 | -0.408032262 | 0.400851483 | -1.017913815 |
| Slc2a2 | 4.994744966 | 1.080976602 | 0.62176638 | 1.738557498 |
| 1700112D23Rik | 5.161670123 | 0.255316383 | 0.447825718 | 0.570124432 |
| Gm6505 | 17.67248122 | 0.195999318 | 0.255387872 | 0.76745742 |
| Eif5a2 | 1024.993172 | -0.116840267 | 0.121847978 | -0.95890198 |
| Gm32950 | 1.23849881 | 0.049677045 | 0.913793496 | 0.054363535 |
| Rpl22l1 | 357.7709375 | 0.066578255 | 0.145664411 | 0.457066035 |

| | | | | |
|---|---|---|---|---|
| Gm33051 | 0.1140246 | 0.75528439 | 3.724945049 | 0.202763901 |
| Gm37867 | 0 | NA | NA | NA |
| Gm37924 | 0 | NA | NA | NA |
| Gm1527 | 0.240368072 | 0.274401446 | 2.538652261 | 0.108089418 |
| Egfem1 | 145.805512 | 0.243081524 | 0.138488144 | 1.755251507 |
| Gm38029 | 61.7718118 | -0.343413772 | 0.361889796 | -0.948945716 |
| Gm37965 | 30.75954022 | -0.448748251 | 0.246207893 | -1.822639578 |
| Mir551b | 1.726387458 | 0.033098689 | 0.718677531 | 0.046054993 |
| Mir21b | 0.755943838 | -1.049795774 | 1.067996263 | -0.982958285 |
| Gm37557 | 0.247197661 | 1.797028564 | 2.096223027 | 0.857269738 |
| Gm33206 | 0.045882455 | 0.755288839 | 3.724945049 | 0.202765096 |
| Mannr | 0.312486075 | 1.630339157 | 1.749008516 | 0.932150497 |
| Gm37281 | 0.306751849 | -0.16289549 | 1.598072856 | -0.101932455 |
| Mecom | 32.25235124 | 0.960601245 | 0.269162837 | 3.568847971 |
| Mecomos | 0.404625785 | 0.274413984 | 1.808769711 | 0.151713058 |
| Gm38066 | 1.468378907 | 0.61963744 | 1.129291105 | 0.548695936 |
| Gm38197 | 1.573498709 | 0.313391328 | 0.922643104 | 0.339666905 |
| Gm10258 | 0.786098488 | -0.095556593 | 1.221967018 | -0.078198995 |
| Gm38362 | 0.449676329 | -1.067443396 | 1.442287211 | -0.740104598 |
| Gm37024 | 0 | NA | NA | NA |
| Gm37652 | 2.839259381 | 1.106091234 | 0.806320878 | 1.371775509 |
| Gm38258 | 0.044191549 | -0.206490061 | 3.724945049 | -0.055434391 |
| Gm38153 | 0.060174537 | 0.755286614 | 3.724945049 | 0.202764498 |
| 1600017P15Rik | 2.396959378 | 0.36805691 | 0.618293508 | 0.595278625 |
| Actrt3 | 12.26637337 | 0.55121224 | 0.310222782 | 1.776827084 |
| Mynn | 359.7065454 | 0.121295068 | 0.07097617 | 1.708954823 |
| Lrrc34 | 2.945986073 | -0.71594767 | 0.594955196 | -1.20336401 |
| Lrriq4 | 0.139037128 | 0.274384718 | 3.724945049 | 0.073661414 |
| Lrrc31 | 0.163642604 | 0.75530398 | 2.783881602 | 0.271313255 |
| Gm15462 | 0.843690133 | -0.924208779 | 1.030369075 | -0.896968669 |
| Samd7 | 0.989999884 | 1.382997436 | 1.196248026 | 1.156112617 |
| 4933429H19Rik | 2.000591198 | 1.236118043 | 0.700055918 | 1.765741866 |
| Sec62 | 1867.564255 | -0.032084219 | 0.058732174 | -0.54628011 |
| Gm38080 | 0.150086409 | 1.264824188 | 3.723011833 | 0.339731445 |
| Gpr160 | 12.95102122 | 1.023060634 | 0.368360724 | 2.777333652 |
| Phc3 | 2283.871593 | 0.116339607 | 0.072811803 | 1.59781247 |
| 9530022L04Rik | 9.381402519 | -0.392153022 | 0.338637083 | -1.15803331 |
| Gm2979 | 0 | NA | NA | NA |
| Prkci | 784.3797283 | -0.185747459 | 0.056400354 | -3.293374023 |
| Mir7008 | 0.429612315 | -1.960836368 | 1.405393489 | -1.395222323 |
| Skil | 1820.158226 | 0.284989173 | 0.095325461 | 2.989643796 |
| Cldn11 | 3469.821532 | 0.760183696 | 0.142360717 | 5.339841711 |
| Slc7a14 | 2555.251066 | 0.177878344 | 0.061774845 | 2.879462408 |
| Gm15496 | 15.67699621 | 0.754255815 | 0.257168204 | 2.932927951 |
| Gm38025 | 1.47797824 | 0.00774651 | 0.843795388 | 0.009180555 |
| Gm6558 | 0 | NA | NA | NA |
| Kcnmb2 | 45.73165487 | -0.351609555 | 0.176551245 | -1.991543894 |

| | | | | |
|---|---|---|---|---|
| Gm37834 | 14.36667755 | -0.594254176 | 0.294480507 | -2.017974579 |
| Gm37770 | 0 | NA | NA | NA |
| Zmat3 | 1732.179252 | -0.539836644 | 0.061507734 | -8.776727904 |
| Gm36990 | 0.044544648 | 0.755286614 | 3.724945049 | 0.202764498 |
| Pik3ca | 1418.733806 | -0.093289871 | 0.041628557 | -2.241006601 |
| Gm37123 | 26.09608814 | -0.323796803 | 0.322783484 | -1.003139315 |
| Kcnmb3 | 0.414197093 | -2.22753641 | 1.673890626 | -1.330753859 |
| Zfp639 | 349.0890839 | 0.21386122 | 0.089643609 | 2.38568285 |
| Gm37592 | 0 | NA | NA | NA |
| Mfn1 | 1164.365532 | -0.063135173 | 0.069743747 | -0.905244925 |
| Gnb4 | 421.880452 | 0.242931466 | 0.106844898 | 2.273683357 |
| Gm37640 | 0.609267175 | -0.152345194 | 1.272771763 | -0.119695611 |
| Actl6a | 159.1766196 | 0.137444221 | 0.15135573 | 0.908087329 |
| Gm37353 | 0.579969052 | -1.615140653 | 1.245738257 | -1.296532915 |
| Mrpl47 | 199.0986341 | 0.132661212 | 0.097108712 | 1.366110299 |
| Ndufb5 | 763.5212508 | -0.140752529 | 0.076766378 | -1.833517899 |
| Usp13 | 553.4368463 | -0.29113591 | 0.090294422 | -3.224295618 |
| Gm15574 | 0 | NA | NA | NA |
| Gm37459 | 3.866617621 | -0.393985331 | 0.601530196 | -0.654971826 |
| Pex5l | 2341.167297 | -0.141318839 | 0.080888417 | -1.747083744 |
| Gm23214 | 0 | NA | NA | NA |
| 4930419G24Rik | 1.068758569 | 0.264835911 | 1.043260113 | 0.253854152 |
| Gm37443 | 87.11784037 | -0.375413918 | 0.311332534 | -1.205829386 |
| Gm19445 | 130.8739924 | -0.492731392 | 0.29316961 | -1.680704193 |
| Gm37128 | 0.066050875 | 0.274391604 | 3.724945049 | 0.073663262 |
| Gm27974 | 0 | NA | NA | NA |
| Gm37031 | 0.110870145 | -0.206490066 | 3.724945049 | -0.055434393 |
| Gm24780 | 0 | NA | NA | NA |
| Gm37398 | 0.221866436 | 0.353169112 | 2.151656398 | 0.164138248 |
| Gm5845 | 1.420437351 | 0.905616826 | 0.925426125 | 0.978594403 |
| RP23-373G12.3 | 0.234927675 | -0.719608697 | 1.872171324 | -0.384371178 |
| RP23-373G12.1 | 1.69071356 | -1.20006368 | 0.911975535 | -1.315894598 |
| RP23-373G12.4 | 0.110242424 | -0.20649451 | 3.724945049 | -0.055435586 |
| Ttc14 | 2035.534113 | -0.42521665 | 0.13165307 | -3.229827075 |
| Ccdc39 | 244.5153087 | -0.512182365 | 0.185039095 | -2.767968383 |
| RP23-373G12.5 | 10.01486383 | 1.02782763 | 0.400205854 | 2.568247365 |
| RP23-373G12.2 | 0 | NA | NA | NA |
| RP24-457B22.2 | 0 | NA | NA | NA |
| Gm23175 | 0 | NA | NA | NA |
| Gm9791 | 24.4382261 | 1.021832594 | 0.276446711 | 3.696309459 |
| RP24-457B22.3 | 1.01049493 | 0.75669357 | 0.860531542 | 0.879332754 |
| Fxr1 | 1205.298799 | 0.064260693 | 0.053604459 | 1.1987938 |
| RP23-418O21.3 | 27.80606419 | -1.012210849 | 0.241016786 | -4.199752499 |
| Dnajc19 | 148.9300614 | -0.298757092 | 0.137994109 | -2.164998887 |
| RP23-418O21.5 | 200.8974819 | 0.121429712 | 0.163795717 | 0.741348521 |
| Sox2ot | 356.2706365 | -0.642500577 | 0.10359118 | -6.202271067 |
| RP23-418O21.1 | 0.658880446 | -1.722718889 | 1.33144812 | -1.293868581 |

| | | | | |
|---|---|---|---|---|
| Gm20446 | 0 | NA | NA | NA |
| RP24-396D16.3 | 2.524695083 | -1.248363054 | 0.699849578 | -1.783759102 |
| Gm20515 | 12.14791881 | -0.636743279 | 0.376022625 | -1.69336427 |
| Gm29135 | 0 | NA | NA | NA |
| Gm27403 | 0 | NA | NA | NA |
| RP24-277H8.2 | 0.177597096 | -0.206487109 | 2.279327799 | -0.09059123 |
| RP24-277H8.3 | 0.133275802 | -0.206488775 | 2.961732282 | -0.06971892 |
| Mir1897 | 0 | NA | NA | NA |
| mmu-mir-1897 | 0 | NA | NA | NA |
| RP24-277H8.1 | 54.03438621 | -1.284552468 | 0.233163699 | -5.509230109 |
| Sox2 | 458.0693419 | -0.091865914 | 0.127614499 | -0.719870508 |
| Gm27611 | 0 | NA | NA | NA |
| RP23-423J10.6 | 3.912046527 | 1.461255442 | 0.611908482 | 2.388029396 |
| RP23-423J10.7 | 0.80598721 | -1.938589894 | 1.116549523 | -1.736232791 |
| RP23-423J10.4 | 0 | NA | NA | NA |
| RP23-423J10.2 | 0.090665643 | -0.687373951 | 3.724945049 | -0.184532642 |
| Mir6378 | 0 | NA | NA | NA |
| RP23-477L20.1 | 1.570552818 | 0.662405386 | 0.902769802 | 0.733747833 |
| Gm25442 | 0.241676147 | -1.609528964 | 1.82625298 | -0.88132859 |
| RP24-141D20.1 | 2.674854522 | 0.193276251 | 0.619705363 | 0.311884103 |
| RP23-112A11.3 | 2.926346155 | -0.279908363 | 0.610499194 | -0.458490962 |
| Ube2l3-ps1 | 0 | NA | NA | NA |
| RP23-112A11.1 | 0.159330469 | 1.236180454 | 3.465376669 | 0.356723258 |
| Gm25696 | 0 | NA | NA | NA |
| Atp11b | 1725.242478 | 0.20281932 | 0.106733118 | 1.9002473 |
| RP23-331L5.1 | 12.60277648 | 0.699553271 | 0.431197997 | 1.622348144 |
| Gm7741 | 2.653371563 | -0.42941621 | 0.72492311 | -0.592361044 |
| Dcun1d1 | 831.5482228 | 0.07152411 | 0.063465726 | 1.126972215 |
| Gm15952 | 9.495619415 | 0.479702895 | 0.390356756 | 1.228883293 |
| RP23-111M12.3 | 21.60654816 | -0.448226194 | 0.396179747 | -1.131370791 |
| Mccc1 | 363.2980275 | -0.194130642 | 0.146954457 | -1.321025887 |
| RP23-111M12.2 | 53.110291 | -0.316205573 | 0.284203769 | -1.112601617 |
| AC116677.4 | 0.320207964 | -0.083238815 | 1.710825826 | -0.048654173 |
| Mccc1os | 32.77909455 | 0.06624078 | 0.191229061 | 0.346394946 |
| Ccdc144b | 0.1140246 | 0.75528439 | 3.724945049 | 0.202763901 |
| Acad9 | 366.3759439 | -0.080570562 | 0.064900744 | -1.241442813 |
| RP23-218L17.3 | 0 | NA | NA | NA |
| RP23-218L17.2 | 12.80704659 | -0.490082808 | 0.281718457 | -1.739619098 |
| D3Ertd254e | 555.8188512 | 0.114429173 | 0.092780491 | 1.233332261 |
| Qrfpr | 20.7155208 | -0.347039753 | 0.242357241 | -1.431934739 |
| RP24-389H4.1 | 1.921125148 | 0.919988299 | 0.681424519 | 1.35009568 |
| RP24-389H4.2 | 1.29860622 | -0.485430981 | 0.818353508 | -0.593180057 |
| RP24-363L5.1 | 0.700674929 | 1.062024474 | 1.30271734 | 0.815237843 |
| RP23-165D1.3 | 0.038981395 | 0.274393828 | 3.724945049 | 0.073663859 |
| RP23-165D1.2 | 0 | NA | NA | NA |
| Anxa5 | 799.2156265 | 0.314544937 | 0.14543342 | 2.162810559 |
| Mir7009 | 0 | NA | NA | NA |

| | | | | |
|---|---|---|---|---|
| 1810062G17Rik | 1.725215873 | -0.126741022 | 0.728830739 | -0.173896373 |
| Gm11548 | 0.199639456 | 1.293005945 | 3.220815344 | 0.401452988 |
| Gm11549 | 318.3834355 | 1.530124086 | 0.283052575 | 5.405794608 |
| RP23-151D16.2 | 14.17505885 | 2.388648499 | 0.435224905 | 5.488308395 |
| Gm11550 | 0 | NA | NA | NA |
| Exosc9 | 304.3832438 | 0.439269812 | 0.101657969 | 4.321056344 |
| Ccna2 | 45.411053 | 1.417400852 | 0.293153352 | 4.835014991 |
| Bbs7 | 206.6797326 | -0.305525479 | 0.133567279 | -2.287427597 |
| Trpc3 | 351.384491 | -0.15530345 | 0.113634042 | -1.366698288 |
| Gm12377 | 0 | NA | NA | NA |
| Gm12376 | 0 | NA | NA | NA |
| 4932438A13Rik | 5270.27695 | 0.060250369 | 0.067522981 | 0.892294263 |
| Gm12531 | 0.264018088 | 0.274411668 | 2.121866723 | 0.129325591 |
| Adad1 | 0.098502579 | -0.206492286 | 3.724945049 | -0.055434988 |
| Gm12582 | 0 | NA | NA | NA |
| Gm12583 | 0.046345709 | 0.274391604 | 3.724945049 | 0.073663262 |
| Gm12584 | 0 | NA | NA | NA |
| Il2 | 0.52475134 | 0.723859608 | 1.340919226 | 0.539823425 |
| Gm12532 | 1.446123796 | 0.094680669 | 0.785913157 | 0.120472177 |
| Il21 | 4.872996577 | 3.994522312 | 0.921158997 | 4.336409157 |
| Gm12534 | 11.60045621 | 5.99866469 | 0.975550212 | 6.149006596 |
| RP23-243E17.13 | 4.833786608 | 5.64245834 | 1.022653742 | 5.517467063 |
| Gm24169 | 0.075043205 | 0.755286614 | 3.724945049 | 0.202764498 |
| Cetn4 | 69.0187925 | 0.107882413 | 0.157320119 | 0.685750899 |
| RP23-328P19.3 | 0 | NA | NA | NA |
| Bbs12 | 67.61927239 | -0.141701764 | 0.207610182 | -0.68253764 |
| Gm12540 | 6.941958254 | -0.086329311 | 0.443902961 | -0.194477891 |
| Fgf2os | 0.170190954 | 0.755297539 | 2.753251924 | 0.274329251 |
| RP23-137F6.5 | 13.65974216 | -0.28855069 | 0.3554897 | -0.811699157 |
| Fgf2 | 254.8186985 | 1.197857319 | 0.225244594 | 5.318029149 |
| RP23-328P19.4 | 0.559063306 | 0.239954961 | 1.207668723 | 0.198692701 |
| Nudt6 | 28.00862341 | -0.110154597 | 0.216543046 | -0.508696073 |
| Spata5 | 377.9700528 | 0.387602372 | 0.0910645 | 4.256349878 |
| RP23-465A17.7 | 107.3750371 | 0.253470686 | 0.251661938 | 1.007187215 |
| Gm12564 | 0 | NA | NA | NA |
| Gm12565 | 0.230175392 | -0.206486422 | 2.133664997 | -0.096775465 |
| 4930594O21Rik | 0.042716945 | -0.206490061 | 3.724945049 | -0.055434391 |
| RP23-382L1.6 | 4.839488669 | 0.48952011 | 0.499124607 | 0.980757315 |
| Spry1 | 135.2416231 | 0.579889565 | 0.198665477 | 2.918924691 |
| RP23-382L1.4 | 8.157939626 | 0.029531362 | 0.37940202 | 0.077836595 |
| RP23-382L1.3 | 1.313092963 | -0.378524996 | 0.850204335 | -0.445216497 |
| Gm5148 | 59.44844248 | 0.024866864 | 0.211089782 | 0.117802313 |
| Rps23-ps1 | 35.46711007 | 0.814717037 | 0.189062229 | 4.309253305 |
| Rpl31-ps15 | 0.453368568 | 1.09741276 | 1.504788356 | 0.72928047 |
| RP23-382L1.7 | 0.0878635 | -0.687376175 | 3.724945049 | -0.184533239 |
| Gm26404 | 0 | NA | NA | NA |
| RP23-175A22.1 | 0.138375801 | 0.740964019 | 2.929302995 | 0.252948917 |

| | | | | |
|---|---|---|---|---|
| RP23-175A22.2 | 0.167536551 | 0.755304179 | 2.763312762 | 0.273332859 |
| RP23-112C14.1 | 0.320788412 | 0.806811804 | 1.733431724 | 0.465441928 |
| Gm7799 | 0 | NA | NA | NA |
| RP23-112C14.2 | 0 | NA | NA | NA |
| Gm22899 | 0 | NA | NA | NA |
| RP24-537K9.3 | 0.194322771 | 0.274408701 | 2.636226981 | 0.104091454 |
| RP24-537K9.1 | 6.837594151 | -0.218709635 | 0.423353356 | -0.516612497 |
| RP24-537K9.2 | 0 | NA | NA | NA |
| RP24-392E13.1 | 0 | NA | NA | NA |
| Gm25574 | 0 | NA | NA | NA |
| RP23-334N1.5 | 0 | NA | NA | NA |
| RP23-334N1.3 | 0.614372065 | 0.595996025 | 1.330855946 | 0.447829103 |
| RP23-334N1.1 | 2.007620227 | -1.151921603 | 0.760837527 | -1.514017859 |
| RP23-334N1.2 | 0.353731249 | -1.567469906 | 1.646120903 | -0.952220401 |
| Ankrd50 | 1197.125447 | -0.099125168 | 0.066694208 | -1.486263522 |
| Gm24462 | 0 | NA | NA | NA |
| RP23-33I24.2 | 0 | NA | NA | NA |
| RP23-33I24.1 | 0 | NA | NA | NA |
| C230034O21Rik | 45.21372464 | -0.397476438 | 0.171810771 | -2.313454717 |
| Fat4 | 722.5535298 | -0.095294662 | 0.100235051 | -0.950711955 |
| RP23-194E5.1 | 0 | NA | NA | NA |
| RP23-382L1.5 | 0.075043205 | 0.755286614 | 3.724945049 | 0.202764498 |
| RP23-49O22.2 | 0.068097359 | -0.206487837 | 3.724945049 | -0.055433794 |
| Gm9845 | 9.009532912 | 1.396965167 | 0.421969498 | 3.310583281 |
| n-R5s195 | 0 | NA | NA | NA |
| RP24-85O24.1 | 0.162691105 | -0.206480295 | 2.788998794 | -0.074033842 |
| Gm23796 | 0 | NA | NA | NA |
| Gm23831 | 0 | NA | NA | NA |
| RP23-256D7.1 | 0 | NA | NA | NA |
| RP23-256D7.2 | 0.608405602 | 0.009726226 | 1.497096417 | 0.006496726 |
| 1700017G19Rik | 0.863538605 | -0.964817417 | 1.0743286 | -0.898065468 |
| Gm23810 | 0 | NA | NA | NA |
| Intu | 226.3559068 | -0.07774612 | 0.089927315 | -0.864543995 |
| Gm16114 | 0 | NA | NA | NA |
| Slc25a31 | 2.917089469 | 0.617308759 | 0.763696786 | 0.80831656 |
| Hspa4l | 1914.656164 | -0.319210704 | 0.072806347 | -4.384380142 |
| Plk4 | 7.648973505 | 0.192313121 | 0.376565998 | 0.510702298 |
| AC146980.1 | 0 | NA | NA | NA |
| Mfsd8 | 158.5183234 | -0.310539315 | 0.103848723 | -2.990304603 |
| 3110057O12Rik | 161.8450865 | -0.109169926 | 0.191095579 | -0.571284416 |
| Gm10731 | 0.47056225 | 1.603636523 | 1.500453569 | 1.068767842 |
| 6430590A07Rik | 35.09747937 | -0.929948714 | 0.323740688 | -2.872511085 |
| 1700034I23Rik | 0.704088048 | 0.131619512 | 1.164892483 | 0.112988549 |
| Larp1b | 264.3748089 | 0.201713767 | 0.160258749 | 1.258675534 |
| Gm22043 | 4.896223194 | -0.068651794 | 0.452627939 | -0.151673787 |
| RP23-235G13.3 | 37.44091296 | 0.292788737 | 0.338234683 | 0.865637829 |
| RP23-235G13.4 | 1.041424966 | 2.258797966 | 1.089476821 | 2.073286849 |

| | | | | |
|---|---|---|---|---|
| Pgrmc2 | 739.9745203 | 0.024707188 | 0.07983972 | 0.309459857 |
| RP23-144N19.1 | 0.207481443 | -0.206489116 | 2.618749975 | -0.07885026 |
| RP23-149I5.1 | 0 | NA | NA | NA |
| RP23-325M18.3 | 0 | NA | NA | NA |
| Gm25487 | 0 | NA | NA | NA |
| RP23-325M18.5 | 0.106618422 | 0.274402725 | 3.724945049 | 0.073666248 |
| Platr4 | 0.110814304 | -0.687373951 | 3.724945049 | -0.184532642 |
| RP23-325M18.2 | 0.374922842 | 2.181357776 | 1.684535973 | 1.294930955 |
| RP23-325M18.1 | 0.284631292 | -2.02071992 | 2.351506599 | -0.859329896 |
| Jade1 | 693.3829749 | -0.101218809 | 0.067547059 | -1.498493206 |
| Gm31266 | 2.246458349 | 0.413189889 | 0.672439787 | 0.614463773 |
| Sclt1 | 316.1906754 | -0.40208803 | 0.122291181 | -3.287956071 |
| Gm38047 | 3.02071253 | -0.927097542 | 0.575468962 | -1.611029616 |
| D3Ertd751e | 153.9838688 | -0.415036006 | 0.152838037 | -2.715528233 |
| Gm37268 | 18.97111997 | -0.645725182 | 0.318138624 | -2.029697541 |
| Gm7899 | 12.35707777 | -0.184645261 | 0.347303457 | -0.531653969 |
| Gm37348 | 0.897478403 | 0.55365379 | 1.172496111 | 0.472200961 |
| Gm37846 | 1.825474869 | -1.476377173 | 0.738023403 | -2.000447638 |
| Gm38274 | 0 | NA | NA | NA |
| Gm37533 | 0.066050875 | 0.274391604 | 3.724945049 | 0.073663262 |
| Gm38044 | 0 | NA | NA | NA |
| Gm9723 | 0 | NA | NA | NA |
| Gm18865 | 0 | NA | NA | NA |
| Gm36976 | 0.179149492 | 0.808021847 | 2.712320448 | 0.297907958 |
| Gm29739 | 0 | NA | NA | NA |
| Gm25714 | 0 | NA | NA | NA |
| Gm32175 | 0 | NA | NA | NA |
| Gm37746 | 0 | NA | NA | NA |
| Gm37365 | 0 | NA | NA | NA |
| Gm38128 | 0 | NA | NA | NA |
| Gm37597 | 0 | NA | NA | NA |
| Gm24997 | 0 | NA | NA | NA |
| Gm27989 | 0 | NA | NA | NA |
| Gm38124 | 15.15835603 | -0.836147442 | 0.290175155 | -2.881526651 |
| 5930409G06Rik | 29.06761066 | -0.770358106 | 0.228061777 | -3.377848386 |
| 2610316D01Rik | 48.48921872 | -0.904783012 | 0.192952087 | -4.689158993 |
| Gm36996 | 0.744898608 | -1.39024253 | 1.118540583 | -1.242907544 |
| Pcdh10 | 3146.497889 | -0.288010092 | 0.080363584 | -3.583838341 |
| 1700027H10Rik | 0.202816535 | 1.265180537 | 3.204096853 | 0.394863387 |
| Gm2136 | 79.20150034 | -0.099194401 | 0.125089873 | -0.792985063 |
| Gm37345 | 48.63821402 | 0.580058878 | 0.206128358 | 2.814066368 |
| Gm37835 | 0 | NA | NA | NA |
| Gm38166 | 0.402076344 | 1.251057973 | 1.708351054 | 0.732319022 |
| Gm32340 | 0.086930093 | -0.206490058 | 3.724945049 | -0.05543439 |
| Gm25796 | 0 | NA | NA | NA |
| Gm23337 | 0 | NA | NA | NA |
| Gm26081 | 0 | NA | NA | NA |

| | | | | |
|---|---|---|---|---|
| Gm18843 | 0 | NA | NA | NA |
| Pabpc4l | 7.77852774 | -0.319361616 | 0.473250872 | -0.674825203 |
| Gm37445 | 0 | NA | NA | NA |
| Mir6379 | 0 | NA | NA | NA |
| Gm10356 | 11.08110355 | 0.330328992 | 0.310995979 | 1.062164832 |
| Gm5274 | 0 | NA | NA | NA |
| Gm23891 | 0 | NA | NA | NA |
| Rpl23a-ps5 | 0.044544648 | 0.755286614 | 3.724945049 | 0.202764498 |
| Gm10730 | 0 | NA | NA | NA |
| Gm2229 | 0 | NA | NA | NA |
| Gm7977 | 1.207653455 | 0.194686622 | 0.945352777 | 0.20594071 |
| Gm37295 | 0 | NA | NA | NA |
| 1700018B24Rik | 0 | NA | NA | NA |
| Gm37190 | 0.042716945 | -0.206490061 | 3.724945049 | -0.055434391 |
| Gm23511 | 0 | NA | NA | NA |
| Gm31415 | 0 | NA | NA | NA |
| Gm37819 | 0.126225412 | 0.755282661 | 3.698754989 | 0.20419916 |
| Gm37684 | 0 | NA | NA | NA |
| Gm20557 | 0.243142889 | 1.210289991 | 2.135956462 | 0.566626714 |
| Gm5846 | 20.89105308 | 0.269750544 | 0.25779433 | 1.046378886 |
| Gm22480 | 0 | NA | NA | NA |
| Gm37424 | 0 | NA | NA | NA |
| Gm38261 | 0.543563254 | 1.286010008 | 1.448824753 | 0.887622886 |
| Gm38088 | 0 | NA | NA | NA |
| Gm2328 | 0 | NA | NA | NA |
| Gm29741 | 0 | NA | NA | NA |
| Gm23096 | 0 | NA | NA | NA |
| Pcdh18 | 119.7553779 | -0.362110527 | 0.163724561 | -2.211705585 |
| Gm37550 | 0.066050875 | 0.274391604 | 3.724945049 | 0.073663262 |
| Slc7a11 | 1346.770682 | 0.49672271 | 0.147703864 | 3.362963546 |
| Gm37854 | 0 | NA | NA | NA |
| Gm37826 | 0 | NA | NA | NA |
| Gm37692 | 0 | NA | NA | NA |
| Gm2345 | 0 | NA | NA | NA |
| Gm37498 | 31.49812525 | -0.04285629 | 0.308289128 | -0.139013302 |
| Gm6209 | 0.762494802 | 0.54433306 | 1.137086764 | 0.478708465 |
| 5830415G21Rik | 0.198884431 | 0.755290731 | 3.229783255 | 0.233851832 |
| Gm38237 | 0.062426873 | 0.755288839 | 3.724945049 | 0.202765096 |
| Gm37199 | 2.566445083 | 1.429486139 | 0.9300808 | 1.536948338 |
| Gm37461 | 0.509388139 | 1.203976285 | 1.535815322 | 0.783932982 |
| Gm24503 | 0.088534442 | 0.274389469 | 3.724945049 | 0.073662689 |
| Gm37548 | 1.788766466 | 0.813359476 | 0.979069917 | 0.830747081 |
| Gm37843 | 13.03457715 | 3.671327569 | 0.660253629 | 5.560480712 |
| Gm37209 | 0.2407585 | -0.206484594 | 2.09221951 | -0.098691649 |
| Gm29230 | 0.044191549 | -0.206490061 | 3.724945049 | -0.055434391 |
| Gm38246 | 0 | NA | NA | NA |
| Ccrn4l | 783.7619654 | 0.157080265 | 0.128300917 | 1.224311324 |

| | | | | |
|---|---|---|---|---|
| Gm38357 | 107.4530042 | -0.359495995 | 0.23780546 | -1.511723048 |
| Elf2 | 758.9211253 | 0.166421479 | 0.07849626 | 2.120119862 |
| 4930577N17Rik | 6.225444594 | -0.270518826 | 0.435664683 | -0.620933569 |
| Gm24282 | 0 | NA | NA | NA |
| Gm37399 | 7.483535934 | 0.595208593 | 0.382770262 | 1.555002184 |
| Gm37203 | 0.04393175 | -0.206490061 | 3.724945049 | -0.055434391 |
| Gm5103 | 0.703642298 | 0.289376537 | 1.336441221 | 0.216527695 |
| Mgarp | 3.540390614 | 0.19763891 | 0.622896251 | 0.317290254 |
| Gm9442 | 4.775535441 | -1.033409641 | 0.578810684 | -1.785401806 |
| Ndufc1 | 324.928662 | -0.311887509 | 0.100268865 | -3.110512026 |
| Naa15 | 1426.402046 | 0.174663098 | 0.070230977 | 2.486980903 |
| Gm37646 | 5.007836009 | -0.460073841 | 0.565463781 | -0.81362212 |
| Gm38160 | 2.044538346 | -0.335615442 | 0.821185746 | -0.408696137 |
| Rab33b | 363.4884665 | 0.057623333 | 0.094793781 | 0.607880945 |
| Gm37968 | 16.21162866 | -0.485220856 | 0.33278983 | -1.458039917 |
| Setd7 | 3529.863564 | -0.217317155 | 0.085917758 | -2.529362496 |
| Gm38200 | 38.03688965 | 0.019613303 | 0.29535444 | 0.066405987 |
| 5031434O11Rik | 65.52831371 | -0.370341912 | 0.157804637 | -2.346837953 |
| Gm38334 | 0 | NA | NA | NA |
| 5430433H01Rik | 0.38231627 | 1.759262492 | 3.698171395 | 0.475711454 |
| Gm38247 | 0 | NA | NA | NA |
| Mgst2 | 4.6105502 | 2.772190416 | 0.686148736 | 4.040217915 |
| Gm37261 | 0.485780372 | 1.222491544 | 1.340286889 | 0.912111843 |
| Maml3 | 325.2409999 | -0.196218947 | 0.0941464 | -2.084189589 |
| Gm10729 | 0.120349074 | 1.264824188 | 3.723011833 | 0.339731445 |
| Gm37342 | 22.56178576 | -1.29869513 | 0.266226702 | -4.878155043 |
| Gm37229 | 0.104366086 | 0.274400501 | 3.724945049 | 0.073665651 |
| n-R5s196 | 0.213343419 | 1.548423363 | 2.562041716 | 0.604370863 |
| 3110080O07Rik | 13.48253032 | -1.37537525 | 0.434564744 | -3.164948996 |
| Gm38000 | 2.827666343 | -2.115798754 | 0.814120641 | -2.598876194 |
| Gm37675 | 6.636235977 | -0.893857938 | 0.466393807 | -1.916530463 |
| Gm38252 | 9.035061384 | -0.717641275 | 0.433359081 | -1.655996856 |
| Gm10728 | 9.423739219 | -0.771563081 | 0.414208463 | -1.862740987 |
| Gm20089 | 0.225870551 | -0.68737813 | 2.583824798 | -0.266031246 |
| Gm38055 | 26.67308699 | -0.726496504 | 0.30806738 | -2.35823898 |
| Gm37526 | 8.63944635 | -0.582043048 | 0.373883212 | -1.556750956 |
| Gm37465 | 35.32081551 | -1.000621888 | 0.317741898 | -3.149165702 |
| C130089K02Rik | 104.1019996 | -1.068963035 | 0.313995383 | -3.404390932 |
| 5430420F09Rik | 30.46786594 | -0.862343608 | 0.289670512 | -2.97698099 |
| Gm37202 | 0 | NA | NA | NA |
| Gm30074 | 0.368444795 | -0.20648182 | 1.486453026 | -0.138909078 |
| Gm38132 | 0 | NA | NA | NA |
| Gm37170 | 1.831335546 | -0.525484521 | 0.815338505 | -0.644498595 |
| Olfr1398-ps1 | 0 | NA | NA | NA |
| Foxo1 | 708.0426262 | 0.521598136 | 0.105843097 | 4.92803169 |
| Gm20402 | 0.491186624 | 1.999934744 | 3.296379369 | 0.606706486 |
| Gm38034 | 38.7415979 | -0.05999046 | 0.317785171 | -0.188776776 |

| | | | | |
|---|---|---|---|---|
| Gm38098 | 1.419478831 | -0.03550667 | 0.848890254 | -0.041827161 |
| Gm30173 | 1.249461252 | -1.303952406 | 0.978724782 | -1.332297321 |
| Gm10293 | 11.59886104 | 0.318685846 | 0.305444247 | 1.043351935 |
| Gm22798 | 7.790214668 | 0.080165182 | 0.429306004 | 0.18673203 |
| Gm2447 | 1.231911519 | 1.798198127 | 1.064517815 | 1.689213747 |
| Gm30292 | 0.646295659 | 1.034812942 | 1.420727018 | 0.728368595 |
| Gm20750 | 0.229084632 | -0.176823868 | 2.573849386 | -0.068700161 |
| Gm37007 | 0 | NA | NA | NA |
| Cog6 | 435.3182281 | -0.343380391 | 0.069299092 | -4.955048923 |
| RP23-256O22.3 | 10.07518621 | 0.356052371 | 0.328383914 | 1.084256434 |
| Lhfp | 547.2260578 | 0.083098776 | 0.102043886 | 0.814343509 |
| RP23-372C7.5 | 0.218578798 | -1.085583153 | 2.573659259 | -0.421805314 |
| RP23-372C7.4 | 86.73692889 | -0.539268655 | 0.290473822 | -1.856513783 |
| RP23-372C7.2 | 134.5349933 | -0.685480386 | 0.361328125 | -1.897113285 |
| RP23-372C7.7 | 21.66084825 | -0.476791937 | 0.328170321 | -1.452879514 |
| RP23-372C7.6 | 0 | NA | NA | NA |
| RP23-372C7.8 | 30.07124974 | -0.643773392 | 0.376340801 | -1.710612801 |
| RP23-335E20.1 | 0 | NA | NA | NA |
| RP23-335E20.2 | 0.902201473 | 0.581004843 | 1.119717386 | 0.518885256 |
| Nhlrc3 | 292.4384838 | 0.836944378 | 0.108958947 | 7.681281808 |
| Gm16206 | 0.175945213 | 0.755298819 | 2.714629575 | 0.278232738 |
| Proser1 | 377.6447841 | 0.059416952 | 0.114784825 | 0.517637694 |
| Stoml3 | 10.96884363 | -1.31642112 | 0.79614113 | -1.653502211 |
| RP23-335E20.3 | 0.440404983 | -0.057066903 | 1.525582538 | -0.037406631 |
| Gm23675 | 0.061920613 | 0.274393828 | 3.724945049 | 0.073663859 |
| Frem2 | 82.40792598 | -0.258400679 | 0.133147822 | -1.940705255 |
| RP24-291E8.1 | 0 | NA | NA | NA |
| RP24-291E8.2 | 16.32404086 | 0.876550254 | 0.285640832 | 3.068714817 |
| RP24-291E8.4 | 0.325009107 | 0.817959518 | 1.778946331 | 0.459799997 |
| RP24-291E8.3 | 0.344979474 | 1.358852864 | 1.759419237 | 0.772330344 |
| RP24-211P15.1 | 0.713939428 | -0.327985821 | 1.145286915 | -0.286378737 |
| Gm24851 | 0 | NA | NA | NA |
| Gm10985 | 0 | NA | NA | NA |
| Ufm1 | 702.309027 | 0.028806002 | 0.061712683 | 0.466776037 |
| RP23-391P1.1 | 0 | NA | NA | NA |
| Gm8109 | 0.115603922 | 0.274387056 | 3.724945049 | 0.073662041 |
| RP23-234G15.2 | 0 | NA | NA | NA |
| Trpc4 | 278.9395309 | -0.112456077 | 0.100268882 | -1.121545131 |
| Postn | 86.52036441 | 0.440079363 | 0.276376188 | 1.592320109 |
| Gm5641 | 12.39303311 | 0.23991238 | 0.439979889 | 0.545280331 |
| Gm26678 | 0 | NA | NA | NA |
| RP23-382K4.1 | 0 | NA | NA | NA |
| Supt20 | 511.0808974 | -0.056747975 | 0.100567852 | -0.564275498 |
| Gm21958 | 0.229069447 | 1.813222309 | 2.53328607 | 0.715759002 |
| Exosc8 | 123.3892681 | 0.023665217 | 0.115908834 | 0.204170954 |
| Alg5 | 141.422727 | 0.249889749 | 0.129821852 | 1.924866624 |
| Smad9 | 371.890028 | -0.185497222 | 0.101151471 | -1.833855897 |

| | | | | |
|---|---|---|---|---|
| Rfxap | 213.8530842 | 0.095428766 | 0.113714653 | 0.839194977 |
| Gm10254 | 54.85742217 | 0.137688408 | 0.191051548 | 0.720687215 |
| Sertm1 | 270.0278678 | 0.032080201 | 0.16804321 | 0.190904479 |
| Ccna1 | 5.128124108 | 0.936355901 | 0.561323959 | 1.668120317 |
| RP23-137P16.2 | 0.532719758 | 1.653633888 | 1.478108055 | 1.11875034 |
| RP23-137P16.1 | 8.16035312 | 0.487949273 | 0.365569435 | 1.33476496 |
| Spg20 | 505.3869147 | -0.207966147 | 0.079519807 | -2.615274797 |
| Ccdc169 | 8.621555654 | 0.009538271 | 0.392498261 | 0.024301436 |
| Gm22029 | 0.119587853 | 1.218392466 | 3.723618458 | 0.327206581 |
| Sohlh2 | 0.439952846 | -0.740422266 | 3.259019726 | -0.22719171 |
| Dclk1 | 10037.27044 | -0.281100097 | 0.119371393 | -2.354836364 |
| Gm25132 | 4.787591675 | -0.835948848 | 0.477099558 | -1.752147606 |
| RP24-372J8.2 | 44.9034577 | -0.888122609 | 0.314788535 | -2.821330866 |
| RP24-372J8.3 | 0.907267504 | -0.440907539 | 1.146863565 | -0.384446374 |
| Gm21954 | 0.194279094 | 0.274391173 | 3.724945049 | 0.073663147 |
| RP24-555P13.2 | 122.1621789 | -0.660432657 | 0.239333209 | -2.759469363 |
| RP24-555P13.4 | 139.4117742 | -0.723466632 | 0.259975642 | -2.782824679 |
| Gm25404 | 0.125889889 | -0.687378102 | 3.019818789 | -0.227622301 |
| WI1-49P9.1 | 0.657052355 | 1.613387144 | 1.284557288 | 1.255986914 |
| WI1-49P9.2 | 2.567514962 | 2.740951538 | 0.912938319 | 3.002340335 |
| Nbea | 6573.173349 | -0.251253903 | 0.087606229 | -2.867991307 |
| Mab21l1 | 155.2590693 | -0.122063161 | 0.139979219 | -0.872009153 |
| RP24-549I13.1 | 0.075043205 | 0.755286614 | 3.724945049 | 0.202764498 |
| RP24-549I13.2 | 0.126225412 | 0.755282661 | 3.698754989 | 0.20419916 |
| Gm22784 | 14.91209993 | -0.57482153 | 0.277553882 | -2.071026803 |
| RP24-293C5.5 | 80.47609759 | -0.665321045 | 0.272506174 | -2.441489804 |
| RP24-80L24.3 | 126.7577974 | -0.113501005 | 0.38762441 | -0.292811808 |
| RP24-80L24.2 | 91.10912563 | -0.236661071 | 0.418563852 | -0.565412111 |
| RP24-80L24.1 | 244.6441849 | -0.282582562 | 0.296007499 | -0.95464663 |
| Gm25727 | 0 | NA | NA | NA |
| RP24-111L5.2 | 0.108266322 | 0.274389235 | 3.724945049 | 0.073662626 |
| Gm22269 | 0.061920613 | 0.274393828 | 3.724945049 | 0.073663859 |
| Gm8177 | 1.764273044 | 1.102908477 | 0.80608726 | 1.368224671 |
| RP24-218M19.4 | 0.066050875 | 0.274391604 | 3.724945049 | 0.073663262 |
| RP24-218M19.5 | 0 | NA | NA | NA |
| Tm4sf1 | 103.6301723 | 0.994239397 | 0.243568119 | 4.081976749 |
| RP23-234F9.2 | 0.925167547 | 0.200178119 | 1.032864707 | 0.193808655 |
| Tm4sf4 | 0.133813382 | 0.274408832 | 2.960524421 | 0.092689265 |
| Wwtr1 | 465.1289583 | 1.006229022 | 0.147809562 | 6.807604391 |
| Gm16016 | 0 | NA | NA | NA |
| Gm26671 | 2.020290962 | 0.372704846 | 0.686648998 | 0.542788015 |
| Mir6377 | 0 | NA | NA | NA |
| RP23-132D7.2 | 73.00516792 | -0.036678043 | 0.170469395 | -0.215159111 |
| Commd2 | 168.5798793 | -0.108890561 | 0.110276648 | -0.98743082 |
| RP23-132D7.4 | 6.836736274 | -0.130136056 | 0.409274679 | -0.317967523 |
| Ankub1 | 37.54081804 | -0.265614945 | 0.265057853 | -1.002101774 |
| RP23-132D7.5 | 0 | NA | NA | NA |

| | | | | |
|---|---|---|---|---|
| RP24-352B14.1 | 0.588070563 | 1.480033204 | 1.384841253 | 1.068738529 |
| Rnf13 | 1345.661625 | 0.044677808 | 0.108335513 | 0.412402238 |
| RP24-352B14.2 | 0.154696864 | 0.274407077 | 2.833481316 | 0.096844498 |
| RP24-352B14.3 | 0.210855403 | -0.66139321 | 2.60734478 | -0.253665421 |
| Pfn2 | 3248.658378 | -0.062440486 | 0.079813882 | -0.782326144 |
| Gm24531 | 0.132790949 | -0.206481859 | 2.970952992 | -0.069500211 |
| RP23-1B20.1 | 0.115957212 | 0.274402725 | 3.724945049 | 0.073666248 |
| RP23-1B20.2 | 6.805132405 | -0.183467685 | 0.415593264 | -0.441459718 |
| RP23-1B20.3 | 0.497476243 | 0.277958567 | 1.603821426 | 0.173310172 |
| RP23-1B20.4 | 0.383276703 | 1.262964975 | 1.721022337 | 0.733845777 |
| Gm5537 | 1.183520696 | -0.041476908 | 0.828371177 | -0.050070439 |
| RP23-375H15.2 | 20.27877935 | -0.431799974 | 0.281207984 | -1.535518191 |
| 1700007F19Rik | 6.275202345 | -0.844809971 | 0.476227831 | -1.77396178 |
| Gm26166 | 0 | NA | NA | NA |
| RP24-376N4.1 | 0 | NA | NA | NA |
| RP23-413C3.1 | 0 | NA | NA | NA |
| Tsc22d2 | 1435.051753 | -0.044085178 | 0.070217155 | -0.627840555 |
| Serp1 | 878.7409205 | 0.701840864 | 0.131419027 | 5.340481372 |
| Eif2a | 501.9842971 | 0.377864049 | 0.071052432 | 5.318101575 |
| RP23-413C3.4 | 13.02020907 | -0.416865152 | 0.31671936 | -1.316197256 |
| Selt | 2800.757624 | 0.350806059 | 0.095103864 | 3.68866251 |
| Erich6 | 23.0190923 | -0.148200916 | 0.244005385 | -0.607367399 |
| RP24-275F18.1 | 4.381802891 | 0.604458071 | 0.518164709 | 1.166536547 |
| RP24-275F18.2 | 3.555894033 | -2.442324751 | 0.681254016 | -3.585042722 |
| Siah2 | 196.4969752 | -0.153329202 | 0.135951037 | -1.127826642 |
| RP24-275F18.4 | 3.656147857 | -0.626720849 | 0.505262357 | -1.240386981 |
| 4930593A02Rik | 1.120771396 | -0.020375211 | 0.878384641 | -0.023196229 |
| Fam188b2 | 11.97822246 | 0.682921928 | 0.345303489 | 1.977744072 |
| Clrn1 | 6.849902671 | 0.249377728 | 0.38399787 | 0.649424769 |
| Gm22491 | 0 | NA | NA | NA |
| Gm25572 | 0.089089296 | 1.194485492 | 3.723939576 | 0.320758559 |
| Gm10071 | 7.465476509 | 0.35311939 | 0.402709764 | 0.876858278 |
| Gm27793 | 0.08169834 | -0.206492286 | 3.724945049 | -0.055434988 |
| RP24-493L18.4 | 21.6695156 | -0.466812397 | 0.249939746 | -1.867699732 |
| Med12l | 1187.74355 | -0.392599665 | 0.093625286 | -4.193308054 |
| Gpr171 | 49.42350281 | 7.3068188 | 0.907592055 | 8.050774311 |
| P2ry14 | 82.16983007 | 2.340737937 | 0.265545731 | 8.814820451 |
| Gpr87 | 0.049553047 | 0.274391604 | 3.724945049 | 0.073663262 |
| P2ry13 | 94.23721791 | 0.601892131 | 0.189321009 | 3.179214678 |
| P2ry12 | 174.5045895 | -1.039173262 | 0.224438832 | -4.630095654 |
| RP24-427G24.5 | 16.25700884 | -0.271204979 | 0.38760352 | -0.699696892 |
| Igsf10 | 258.1667321 | 1.186875385 | 0.185390457 | 6.402030615 |
| Gm38186 | 1.889693139 | 0.467632542 | 0.889080111 | 0.525973459 |
| Gm37715 | 0.038981395 | 0.274393828 | 3.724945049 | 0.073663859 |
| Gm8276 | 1.405995652 | 1.68055704 | 0.979037912 | 1.716539288 |
| Gm38288 | 0 | NA | NA | NA |
| Gm37771 | 0 | NA | NA | NA |

| | | | | |
|---|---|---|---|---|
| Gm4856 | 0.075043205 | 0.755286614 | 3.724945049 | 0.202764498 |
| Gm5709 | 0 | NA | NA | NA |
| Gm37926 | 0 | NA | NA | NA |
| Gm5538 | 0 | NA | NA | NA |
| Gm38208 | 0 | NA | NA | NA |
| Gm37579 | 0 | NA | NA | NA |
| Gm5539 | 2.248417857 | 0.401483101 | 0.70693188 | 0.567923321 |
| 4930449A18Rik | 0.066050875 | 0.274391604 | 3.724945049 | 0.073663262 |
| Gm2756 | 0 | NA | NA | NA |
| Gm8298 | 0.066050875 | 0.274391604 | 3.724945049 | 0.073663262 |
| C130079G13Rik | 0.049553047 | 0.274391604 | 3.724945049 | 0.073663262 |
| Gm9696 | 0.049553047 | 0.274391604 | 3.724945049 | 0.073663262 |
| Aadacl2 | 0 | NA | NA | NA |
| Gm18427 | 0 | NA | NA | NA |
| Aadac | 0.048949532 | -0.206487837 | 3.724945049 | -0.055433794 |
| Sucnr1 | 0.697362497 | 2.395253923 | 1.508718377 | 1.58760837 |
| Gm24382 | 0 | NA | NA | NA |
| RP23-254F12.1 | 1.05607488 | 1.230684153 | 1.02339989 | 1.202544738 |
| Mbnl1 | 2752.745771 | 0.283239593 | 0.049900747 | 5.67605915 |
| Gm37589 | 184.5056286 | -0.178527274 | 0.199978456 | -0.892732534 |
| Gm37488 | 59.18839802 | 0.003573263 | 0.207711019 | 0.017203049 |
| Gm38280 | 0.072025462 | 0.755288839 | 3.724945049 | 0.202765096 |
| Gm33707 | 0.263895486 | -1.483256155 | 3.704486237 | -0.400394565 |
| Gm38326 | 0 | NA | NA | NA |
| Gm36985 | 0.325677113 | 1.22863582 | 1.586674913 | 0.774346282 |
| Gm8325 | 112.612681 | 0.17416792 | 0.289644914 | 0.601315304 |
| Gm37035 | 1.517989225 | -0.933624175 | 0.859939361 | -1.085686059 |
| P2ry1 | 94.52974778 | -0.52315105 | 0.133223063 | -3.926880523 |
| Gm37719 | 0.066050875 | 0.274391604 | 3.724945049 | 0.073663262 |
| Gm19901 | 0.038981395 | 0.274393828 | 3.724945049 | 0.073663859 |
| Gm37696 | 26.28877038 | 0.233421764 | 0.219501044 | 1.063419835 |
| Rap2b | 869.3150327 | 0.146613525 | 0.081442761 | 1.800203275 |
| B430305J03Rik | 35.36276701 | 0.140312223 | 0.195179001 | 0.718889955 |
| Gm8349 | 2.822290259 | -0.582595158 | 0.57125714 | -1.019847484 |
| Gm9700 | 0.470561891 | 0.267112728 | 1.477932635 | 0.180734034 |
| Gm37404 | 0.15100991 | 1.236179172 | 3.527597384 | 0.350430913 |
| 9330121J05Rik | 22.77484088 | 1.006192427 | 0.254994657 | 3.945935334 |
| Arhgef26 | 647.6477621 | -0.05648371 | 0.110458484 | -0.511356921 |
| Dhx36 | 1564.510516 | 0.062137226 | 0.063417783 | 0.979807596 |
| Gpr149 | 48.93600836 | 0.375599168 | 0.178083253 | 2.109121215 |
| Gm18382 | 0.246177344 | -1.231984836 | 2.49861285 | -0.493067518 |
| Gm9701 | 0.309747539 | 0.755285347 | 3.724945049 | 0.202764158 |
| Gm22433 | 0 | NA | NA | NA |
| Gm8388 | 2.905967886 | 0.79026213 | 0.729711558 | 1.082978776 |
| Mme | 252.6871563 | 0.017624578 | 0.088595973 | 0.198932046 |
| Gm34240 | 0 | NA | NA | NA |
| Gm34302 | 0.608629602 | -0.770546265 | 1.259608802 | -0.611734583 |

| | | | | |
|---|---|---|---|---|
| Gm37279 | 0 | NA | NA | NA |
| Gm24946 | 0 | NA | NA | NA |
| Plch1 | 346.9457733 | 0.356608291 | 0.092789919 | 3.843179242 |
| Gm23500 | 0 | NA | NA | NA |
| Gm16162 | 0 | NA | NA | NA |
| Gm16129 | 0 | NA | NA | NA |
| Gm34379 | 0.085327104 | 0.274389469 | 3.724945049 | 0.073662689 |
| E130311K13Rik | 62.02924702 | 0.127474605 | 0.188408122 | 0.676587632 |
| Slc33a1 | 460.1328625 | 0.075565176 | 0.085071979 | 0.888249894 |
| Gm26850 | 5.346930916 | 0.586810584 | 0.429747126 | 1.365478785 |
| Gmps | 2065.915186 | 0.057039037 | 0.054272214 | 1.050980466 |
| Vmn2r-ps3 | 0.09176491 | 1.15872478 | 3.72306111 | 0.311229052 |
| Vmn2r1 | 7.920741964 | 1.180896089 | 0.390303366 | 3.025585207 |
| Gm22165 | 0.115603922 | 0.274387056 | 3.724945049 | 0.073662041 |
| Vmn2r2 | 2.02639444 | 0.040915329 | 0.727783231 | 0.056219114 |
| Gm5139 | 0.217472118 | 1.27529724 | 2.167249336 | 0.58844048 |
| Gm18294 | 0.049553047 | 0.274391604 | 3.724945049 | 0.073663262 |
| Vmn2r3 | 0.204914957 | 1.221428882 | 2.200579739 | 0.555048681 |
| Gm23897 | 0.049553047 | 0.274391604 | 3.724945049 | 0.073663262 |
| Vmn2r-ps4 | 0.124853745 | 1.290908285 | 3.721447108 | 0.346883416 |
| Vmn2r-ps5 | 0.174149299 | 0.755285184 | 3.362794904 | 0.224600431 |
| Vmn2r4 | 0.145501641 | 0.755294927 | 2.8880725 | 0.261522149 |
| Vmn2r-ps6 | 0 | NA | NA | NA |
| Vmn2r-ps7 | 0 | NA | NA | NA |
| Gm20656 | 0.439965521 | 0.696263225 | 1.516901372 | 0.459003623 |
| Vmn2r-ps8 | 0 | NA | NA | NA |
| Vmn2r5 | 0.344244685 | 0.258595363 | 1.855973152 | 0.139331414 |
| Vmn2r6 | 0.549943695 | 0.590785635 | 1.487925653 | 0.397053196 |
| Vmn2r-ps9 | 0.898522788 | 0.305758987 | 1.089954914 | 0.280524436 |
| Gm20657 | 0.48442959 | -0.262016383 | 1.314234626 | -0.199368041 |
| Vmn2r-ps10 | 0 | NA | NA | NA |
| Vmn2r-ps11 | 0.791552024 | 0.251855425 | 1.092666457 | 0.230496162 |
| Gm26939 | 0 | NA | NA | NA |
| Vmn2r7 | 3.934381616 | 0.131906868 | 0.537363671 | 0.245470386 |
| Vmn2r-ps12 | 0.044544648 | 0.755286614 | 3.724945049 | 0.202764498 |
| Gm20698 | 0 | NA | NA | NA |
| Gm25222 | 0.066050875 | 0.274391604 | 3.724945049 | 0.073663262 |
| Vmn2r-ps13 | 0 | NA | NA | NA |
| Gm38048 | 50.77510117 | -0.071247575 | 0.217393466 | -0.327735586 |
| Kcnab1 | 1328.585016 | -0.120672057 | 0.093296864 | -1.293420289 |
| A330015K06Rik | 13.82607597 | -0.153493262 | 0.333916422 | -0.459675692 |
| A730090N16Rik | 2.24970321 | 0.988880797 | 0.775698639 | 1.274826005 |
| Ssr3 | 1935.252712 | 0.520407994 | 0.092616885 | 5.618932135 |
| Gm5540 | 6.757178092 | -0.413254731 | 0.417223694 | -0.990487206 |
| Gm34780 | 0.183309527 | 0.755297975 | 2.685861896 | 0.281212514 |
| Gm37009 | 0.786503428 | 0.173866035 | 1.046847438 | 0.166085362 |
| 4931440P22Rik | 18.83552398 | 0.363030669 | 0.307437603 | 1.18082715 |

| | | | | |
|---|---|---|---|---|
| Gm22279 | 0 | NA | NA | NA |
| Tiparp | 657.7234625 | 0.57898797 | 0.234275789 | 2.471394813 |
| Gm5847 | 0.982102764 | 0.536235176 | 1.259699113 | 0.425685127 |
| Gm38233 | 0 | NA | NA | NA |
| Mir8120 | 0.138110619 | 1.201070057 | 3.622723897 | 0.33153784 |
| Lekr1 | 106.5364298 | -0.558249188 | 0.194479637 | -2.870476291 |
| Gm37776 | 35.53555177 | -0.890952953 | 0.314000595 | -2.837424409 |
| Gm25755 | 0 | NA | NA | NA |
| Gm37359 | 0.166952883 | 0.274401811 | 2.756036668 | 0.099563919 |
| Gm37131 | 0.161076545 | 0.755297744 | 2.79165938 | 0.270555122 |
| Gm37037 | 0.659599365 | -0.584754559 | 1.362956216 | -0.429034002 |
| Ccnl1 | 947.3859406 | 0.495678989 | 0.136030131 | 3.643891128 |
| Gm37305 | 112.0234715 | -0.057902435 | 0.106683486 | -0.542749747 |
| Gm35106 | 0.709932405 | 0.385711453 | 1.372795769 | 0.280967834 |
| Gm37078 | 4.562323805 | -1.534931274 | 0.499344295 | -3.07389368 |
| Gm19024 | 0 | NA | NA | NA |
| Gm6546 | 0 | NA | NA | NA |
| Veph1 | 5.036914537 | -1.556029985 | 0.576592501 | -2.698664969 |
| Gm37822 | 0 | NA | NA | NA |
| Gm26442 | 0 | NA | NA | NA |
| Gm36973 | 0.066050875 | 0.274391604 | 3.724945049 | 0.073663262 |
| Tpi-rs2 | 0 | NA | NA | NA |
| Gm37330 | 0 | NA | NA | NA |
| Ptx3 | 208.6062493 | 5.920594402 | 0.652991043 | 9.066884551 |
| Gm17952 | 0.276493613 | 0.712261948 | 1.780142907 | 0.400115039 |
| Gm6555 | 0 | NA | NA | NA |
| Gm38356 | 0.174174109 | 0.274407947 | 2.729571827 | 0.100531499 |
| RP23-103D17.4 | 0 | NA | NA | NA |
| Shox2 | 134.0978986 | -0.21854084 | 0.14516002 | -1.505516737 |
| Rsrc1 | 525.7525617 | -0.057992597 | 0.062235727 | -0.931821628 |
| Gm22295 | 1.86515265 | -0.024116799 | 0.726055368 | -0.033216198 |
| Gm17402 | 0 | NA | NA | NA |
| Mlf1 | 15.4417281 | 0.399593918 | 0.324579283 | 1.23111344 |
| Gfm1 | 616.4629507 | -0.047851806 | 0.050218867 | -0.952865116 |
| Lxn | 276.0893367 | -0.42341402 | 0.158827452 | -2.665874289 |
| Gm37453 | 18.12819328 | -0.412473995 | 0.275480589 | -1.497288782 |
| Rarres1 | 12.46569881 | -0.457579228 | 0.315652702 | -1.449628737 |
| Gm35299 | 0.137416547 | -0.684638423 | 3.63730774 | -0.188226697 |
| 4930402C01Rik | 0 | NA | NA | NA |
| Gm25919 | 0 | NA | NA | NA |
| Mfsd1 | 388.2294903 | 0.407632199 | 0.113871202 | 3.579765477 |
| Gm8515 | 6.537093261 | -0.371483096 | 0.439112855 | -0.845985474 |
| Gm412 | 0 | NA | NA | NA |
| Gm37601 | 0.128477747 | 0.755285235 | 3.677131942 | 0.205400635 |
| Gm18717 | 0.045882455 | 0.755288839 | 3.724945049 | 0.202765096 |
| Iqcj | 18.38164937 | -0.965734792 | 0.339257616 | -2.84661197 |
| Schip1 | 1612.457602 | 0.08729538 | 0.087787904 | 0.994389622 |

| | | | | |
|---|---|---|---|---|
| Gm10292 | 11.84136725 | -0.136173755 | 0.458474971 | -0.297014589 |
| Gm10723 | 64.31874068 | -0.265962712 | 0.34528415 | -0.770271999 |
| Gm22073 | 0 | NA | NA | NA |
| Il12a | 10.96521039 | 0.055278857 | 0.366530875 | 0.150816373 |
| Gm10040 | 0.437154627 | -0.604560402 | 1.33550964 | -0.452681421 |
| 4930535E02Rik | 0 | NA | NA | NA |
| Gm35584 | 0.044191549 | -0.206490061 | 3.724945049 | -0.055434391 |
| Gm7270 | 0 | NA | NA | NA |
| Gm17641 | 6.199809403 | 0.352128744 | 0.476502585 | 0.73898601 |
| 1110032F04Rik | 89.81938694 | -0.022358646 | 0.183466098 | -0.121867998 |
| Ift80 | 422.7464304 | -0.43088633 | 0.125210471 | -3.441296297 |
| Gm20689 | 16.19484603 | 0.791243451 | 0.297909167 | 2.655988937 |
| Gm20690 | 0.088383099 | -0.687376175 | 3.724945049 | -0.184533239 |
| Gm18588 | 78.97133165 | 0.209189733 | 0.167873354 | 1.246116367 |
| Gm20691 | 11.60463875 | -0.841355262 | 0.310797046 | -2.707088996 |
| Smc4 | 201.8629177 | 0.753238126 | 0.140660535 | 5.355006827 |
| Mir15b | 0.132199999 | 1.236180999 | 3.642365845 | 0.339389576 |
| Mir16-2 | 0.358973896 | -0.647159268 | 1.45819836 | -0.443807431 |
| Trim59 | 149.282414 | -0.940251852 | 0.144251465 | -6.518144225 |
| Kpna4 | 1744.547512 | 0.152425681 | 0.058176253 | 2.620066981 |
| Gm22009 | 385.5335551 | -0.213539423 | 0.277696481 | -0.768966975 |
| Gm37558 | 126.3325027 | 0.169179233 | 0.215607322 | 0.78466367 |
| Gm1647 | 0.255219096 | -1.190091351 | 1.832121749 | -0.649570015 |
| Arl14 | 0.146692594 | 0.274392389 | 3.52332538 | 0.07787881 |
| Ppm1l | 3121.106828 | -0.158953507 | 0.076546518 | -2.076560908 |
| Gm17214 | 0 | NA | NA | NA |
| Gm17212 | 0 | NA | NA | NA |
| Gm17213 | 0 | NA | NA | NA |
| Gm8565 | 1.479948724 | -0.845164404 | 0.886047891 | -0.953858603 |
| B3galnt1 | 333.6927738 | -0.157438647 | 0.089343269 | -1.762176925 |
| Gm37380 | 0.038981395 | 0.274393828 | 3.724945049 | 0.073663859 |
| Gm25621 | 0 | NA | NA | NA |
| Rpl32-ps | 0.038981395 | 0.274393828 | 3.724945049 | 0.073663859 |
| Nmd3 | 566.0817292 | 0.381097899 | 0.101435235 | 3.757056402 |
| Sptssb | 37.86867121 | -0.558021459 | 0.189258011 | -2.94846943 |
| Gm17895 | 0 | NA | NA | NA |
| Gm23484 | 0 | NA | NA | NA |
| Gm37784 | 14.73966199 | 0.312696275 | 0.297353687 | 1.051597099 |
| Otol1 | 0.139883403 | 0.274393101 | 3.559413475 | 0.077089415 |
| Gm37681 | 0.143485256 | 0.274391988 | 3.549605256 | 0.077302113 |
| Gm37585 | 0 | NA | NA | NA |
| Gm23477 | 0 | NA | NA | NA |
| Gm6631 | 0 | NA | NA | NA |
| Gm37213 | 0 | NA | NA | NA |
| Gm10780 | 0 | NA | NA | NA |
| Gm6634 | 0.207012398 | -0.206483515 | 2.635059287 | -0.078360102 |
| Gm8599 | 0 | NA | NA | NA |

| | | | | |
|---|---|---|---|---|
| Gm38305 | 0 | NA | NA | NA |
| Gm19194 | 0 | NA | NA | NA |
| Gm8604 | 0 | NA | NA | NA |
| Gm37617 | 0 | NA | NA | NA |
| Gm3351 | 0 | NA | NA | NA |
| Gm37603 | 0 | NA | NA | NA |
| Gm27465 | 0 | NA | NA | NA |
| Gm37901 | 0 | NA | NA | NA |
| Gm22451 | 0 | NA | NA | NA |
| Sis | 3.661596079 | 1.343793953 | 0.559825787 | 2.400378805 |
| Gm37056 | 46.70151825 | -0.084147785 | 0.149129893 | -0.564258336 |
| Slitrk3 | 736.758703 | -0.193052692 | 0.083443491 | -2.313574008 |
| Gm20754 | 35.42307591 | 0.244136037 | 0.197796253 | 1.234280392 |
| Gm24544 | 0.953051642 | -0.794790224 | 1.071273156 | -0.741911827 |
| Gm37725 | 9.175969031 | 0.115144933 | 0.329205335 | 0.349766305 |
| Gm22538 | 0 | NA | NA | NA |
| Gm38353 | 21.39725853 | 0.227815142 | 0.405856263 | 0.561319763 |
| 4930509J09Rik | 0 | NA | NA | NA |
| Bche | 53.21720014 | -0.424013323 | 0.168875493 | -2.51080436 |
| Gm8625 | 0 | NA | NA | NA |
| Gm20356 | 0 | NA | NA | NA |
| Gm6098 | 0 | NA | NA | NA |
| Gm37050 | 0 | NA | NA | NA |
| Gm8643 | 0 | NA | NA | NA |
| Gm37464 | 5.385716294 | -1.157361137 | 0.451770155 | -2.561836201 |
| Zbbx | 20.8812694 | -0.625750781 | 0.261672517 | -2.391350794 |
| Serpini2 | 1.184533449 | 3.04760551 | 1.062128202 | 2.869338657 |
| Wdr49 | 44.34530745 | 1.420535863 | 0.3078457 | 4.614441141 |
| Gm17836 | 0 | NA | NA | NA |
| Pdcd10 | 402.9200995 | 0.341016751 | 0.130070781 | 2.621778296 |
| Serpini1 | 2225.756587 | 0.098376084 | 0.094828351 | 1.037412155 |
| Gm37672 | 32.21841053 | -0.289700354 | 0.248561281 | -1.165508776 |
| Platr10 | 0.223571151 | 1.387207844 | 2.53625993 | 0.546950187 |
| Gm37193 | 0 | NA | NA | NA |
| Gm25846 | 0.062426873 | 0.755288839 | 3.724945049 | 0.202765096 |
| Gm29133 | 0 | NA | NA | NA |
| Golim4 | 319.1997903 | -0.261247139 | 0.094297542 | -2.770455459 |
| Gm37685 | 0 | NA | NA | NA |
| Gm37256 | 0.048949532 | -0.206487837 | 3.724945049 | -0.055433794 |
| Gm18428 | 0 | NA | NA | NA |
| Fstl5 | 496.3711575 | -0.185269308 | 0.099861302 | -1.855266296 |
| Gm37897 | 17.02018816 | -0.511738479 | 0.30948524 | -1.653514978 |
| Gm37290 | 0.044544648 | 0.755286614 | 3.724945049 | 0.202764498 |
| Gm19231 | 0.236516182 | 0.728748314 | 2.107367726 | 0.345809754 |
| Gm23644 | 0 | NA | NA | NA |
| Dnmt3a-ps1 | 0.044544648 | 0.755286614 | 3.724945049 | 0.202764498 |
| Gm37806 | 0.072025462 | 0.755288839 | 3.724945049 | 0.202765096 |

| | | | | |
|---|---|---|---|---|
| Gm38195 | 0 | NA | NA | NA |
| Gm37576 | 0 | NA | NA | NA |
| RP23-257I21.2 | 0 | NA | NA | NA |
| Gm6677 | 0 | NA | NA | NA |
| Gm8684 | 0.072025462 | 0.755288839 | 3.724945049 | 0.202765096 |
| Gm6680 | 0 | NA | NA | NA |
| RP23-399J17.1 | 4.954402907 | 0.461008448 | 0.534913696 | 0.861837063 |
| Gm15442 | 3.380166461 | 0.208395505 | 0.529335143 | 0.393692932 |
| Gm18952 | 0 | NA | NA | NA |
| Gm24749 | 0 | NA | NA | NA |
| Gm17953 | 0.338292759 | 0.274382847 | 3.724945049 | 0.073660911 |
| Gm5277 | 0.066050875 | 0.274391604 | 3.724945049 | 0.073663262 |
| Gm10291 | 12.97480729 | 0.05911522 | 0.339778681 | 0.173981544 |
| Gm9762 | 88.84803059 | 0.404202005 | 0.180220474 | 2.242819562 |
| Gm23440 | 0 | NA | NA | NA |
| Rapgef2 | 2966.610995 | 0.358018585 | 0.076734662 | 4.665669647 |
| Gm24150 | 0 | NA | NA | NA |
| RP24-253I2.4 | 0.238666214 | 1.455558153 | 2.531266373 | 0.5750316 |
| 6430573P05Rik | 8.415433566 | 0.075572591 | 0.368074379 | 0.205318803 |
| Gm17359 | 0.372745468 | 1.611891597 | 1.431830031 | 1.125756244 |
| Fnip2 | 1106.231662 | 0.635092341 | 0.147472065 | 4.306526402 |
| Gm3513 | 13.83725982 | -0.026661862 | 0.33519227 | -0.079541997 |
| 4930589L23Rik | 0.532739632 | -0.134870827 | 1.268495912 | -0.106323423 |
| Gm35067 | 0.044544648 | 0.755286614 | 3.724945049 | 0.202764498 |
| Ppid | 668.4459743 | -0.211376257 | 0.07480814 | -2.825578293 |
| Etfdh | 350.8915493 | -0.258036024 | 0.104460624 | -2.470175009 |
| Gm37973 | 5.608718336 | -0.594605307 | 0.45263415 | -1.313655425 |
| 4930579G24Rik | 15.15182425 | -0.203590994 | 0.277556372 | -0.733512233 |
| Rxfp1 | 93.03486081 | -0.640061796 | 0.242741258 | -2.636806784 |
| Gm26984 | 4.172577245 | -1.165104084 | 0.566489695 | -2.056708345 |
| Tmem144 | 92.24781446 | -0.607842889 | 0.175391494 | -3.465634932 |
| Gm26420 | 0 | NA | NA | NA |
| Gm36569 | 0.140714442 | 0.638522383 | 3.579135419 | 0.178401292 |
| Fam198b | 79.79527804 | -0.814034081 | 0.170897511 | -4.763288091 |
| A330069K06Rik | 1.631297931 | 1.153137167 | 0.884131531 | 1.304259746 |
| Gm19066 | 0.045882455 | 0.755288839 | 3.724945049 | 0.202765096 |
| RP23-121N19.4 | 0.091880448 | -0.687373951 | 3.724945049 | -0.184532642 |
| Gm38295 | 0.117760149 | 0.274400501 | 3.724945049 | 0.073665651 |
| Gm37892 | 0.469035539 | 1.113505333 | 1.467518825 | 0.758767325 |
| Gm37971 | 2.078197752 | -0.276154508 | 0.731383814 | -0.377578097 |
| Gm6706 | 0.213298592 | 0.274406908 | 2.175755888 | 0.126120264 |
| Gria2 | 8463.13547 | 0.084723226 | 0.11410176 | 0.742523396 |
| Gm25749 | 0 | NA | NA | NA |
| Glrb | 2044.481699 | -0.132079258 | 0.066640215 | -1.981975265 |
| RP24-445D5.2 | 0 | NA | NA | NA |
| Gm29808 | 0.687463866 | 0.480373967 | 1.306843088 | 0.367583509 |
| RP23-187I18.1 | 0.148659141 | 0.274384287 | 3.724945049 | 0.073661298 |

| | | | | |
|---|---|---|---|---|
| Pdgfc | 119.0878968 | 0.295574809 | 0.117500893 | 2.515511161 |
| Gm16000 | 0.137483974 | -0.206488593 | 2.933430427 | -0.070391508 |
| Gm37300 | 0 | NA | NA | NA |
| Gm4857 | 0 | NA | NA | NA |
| RP23-144D4.1 | 0 | NA | NA | NA |
| A830029E22Rik | 4.685490163 | 0.413622978 | 0.508604082 | 0.813251392 |
| Gm22531 | 0 | NA | NA | NA |
| Gm9989 | 1.473237501 | -0.349994156 | 0.779466977 | -0.449017298 |
| Ctso | 423.7109783 | 0.833116431 | 0.114035132 | 7.305787394 |
| Tdo2 | 9.666145708 | -0.090102348 | 0.358195645 | -0.251545067 |
| Asic5 | 0.518949687 | 1.469643862 | 1.350534436 | 1.08819429 |
| Gucy1b3 | 1608.78633 | -0.158858357 | 0.090622163 | -1.752974674 |
| Gucy1a3 | 860.8261742 | -0.288867336 | 0.108231096 | -2.668986512 |
| Mir7010 | 0.40751274 | 0.688957393 | 1.347184141 | 0.51140551 |
| Map9 | 1296.963444 | -0.100102784 | 0.070025215 | -1.429524842 |
| Npy2r | 123.6004455 | 0.288061954 | 0.208097802 | 1.384262354 |
| Gm35323 | 0 | NA | NA | NA |
| Gm29999 | 0.156071933 | -0.206480922 | 3.489245271 | -0.059176385 |
| RP23-469J3.5 | 0.062426873 | 0.755288839 | 3.724945049 | 0.202765096 |
| RP23-223I2.3 | 0 | NA | NA | NA |
| Gm22038 | 0 | NA | NA | NA |
| RP23-223I2.2 | 0 | NA | NA | NA |
| Rbm46 | 23.88977253 | 0.16449612 | 0.246735495 | 0.666690133 |
| Rbm46os | 0.56520525 | -1.049009107 | 1.395837864 | -0.751526473 |
| Lrat | 10.45023781 | -0.49483243 | 0.48783271 | -1.014348606 |
| Gm30097 | 0.374039128 | 0.176345954 | 1.593058957 | 0.11069644 |
| Fgg | 0.874079576 | 0.94041492 | 1.267481665 | 0.741955443 |
| Fga | 10.18503337 | 5.097158926 | 0.995416193 | 5.120630911 |
| Fgb | 1.174237097 | 1.333051868 | 1.484349021 | 0.898071713 |
| Plrg1 | 372.6327839 | 0.132787597 | 0.09691265 | 1.370178163 |
| Gm10710 | 19.45997954 | -0.13771765 | 0.253511909 | -0.543239373 |
| Dchs2 | 240.9884021 | -0.188906036 | 0.111975777 | -1.687025897 |
| Gm38096 | 1.042898341 | 2.375035182 | 1.196119795 | 1.985616484 |
| 1700028M03Rik | 0.205907367 | -0.206487659 | 2.588140555 | -0.079782243 |
| Gm22342 | 0.083172944 | -0.206492286 | 3.724945049 | -0.055434988 |
| Rpl28-ps2 | 0 | NA | NA | NA |
| Gm24584 | 0 | NA | NA | NA |
| Sfrp2 | 19.78329196 | -0.47624263 | 0.26444824 | -1.800891664 |
| Gm26771 | 5.06408741 | 0.333834216 | 0.451363368 | 0.739613004 |
| Gm7115 | 0 | NA | NA | NA |
| Tlr2 | 267.1921433 | 4.506830524 | 0.246036158 | 18.31775685 |
| D930015E06Rik | 251.151783 | -0.379763125 | 0.149340152 | -2.542940529 |
| Gm36981 | 2.682934602 | 1.718256416 | 0.835818799 | 2.055776225 |
| Mnd1 | 6.972706414 | -0.022240985 | 0.590180922 | -0.037685028 |
| Trim2 | 5153.10411 | -0.075458486 | 0.06124819 | -1.232011696 |
| Gm6525 | 0.265820157 | -2.008731134 | 2.43893295 | -0.823610643 |
| 4930565D16Rik | 0.075043205 | 0.755286614 | 3.724945049 | 0.202764498 |

| | | | | |
|---|---|---|---|---|
| Fhdc1 | 188.3832 | -0.117286966 | 0.106776528 | -1.09843397 |
| Arfip1 | 76.98979055 | 0.086341707 | 0.125760759 | 0.686555234 |
| Gm37240 | 3.253759892 | -0.289592078 | 0.525889747 | -0.550670705 |
| Gm8813 | 4.188906716 | -0.67491597 | 0.53098049 | -1.271074894 |
| Gm38241 | 5.062387905 | -0.183042234 | 0.443830492 | -0.412414734 |
| Tigd4 | 1.127018203 | 0.422396428 | 0.960210436 | 0.439899851 |
| Tmem154 | 14.23682184 | 2.581491348 | 0.515170148 | 5.010949019 |
| Fbxw7 | 2169.618475 | -0.10984701 | 0.067098575 | -1.637099012 |
| Gm37726 | 37.63354527 | 0.023098874 | 0.281012152 | 0.082198842 |
| AC124496.1 | 0 | NA | NA | NA |
| Gm38041 | 0 | NA | NA | NA |
| Gm38313 | 0 | NA | NA | NA |
| 1700036G14Rik | 0.202694752 | -1.615763465 | 2.186041579 | -0.739127508 |
| RP23-94M23.3 | 2.853198453 | -0.646770557 | 0.546889725 | -1.182634319 |
| Mir466k | 0 | NA | NA | NA |
| RP24-389G6.1 | 1.736188672 | -2.342084505 | 0.964541241 | -2.42818493 |
| Gm15535 | 15.003366 | -1.146409597 | 0.504585391 | -2.271983331 |
| Gatb | 159.5508096 | -0.009482069 | 0.108271637 | -0.087576668 |
| Fam160a1 | 54.14219974 | 0.249319577 | 0.138917561 | 1.794730458 |
| Glt28d2 | 6.204872336 | 0.155040796 | 0.395971304 | 0.391545535 |
| RP23-329L9.3 | 0.488911873 | -0.16243441 | 1.414077864 | -0.114869495 |
| Gm26204 | 0 | NA | NA | NA |
| Gm38030 | 0 | NA | NA | NA |
| Gm9790 | 12.80588666 | -0.2911016 | 0.34690996 | -0.839127249 |
| Gm25876 | 0 | NA | NA | NA |
| Gm24501 | 0 | NA | NA | NA |
| Gm27811 | 0 | NA | NA | NA |
| Sh3d19 | 475.5047945 | -0.323682612 | 0.11337764 | -2.854906943 |
| Prss48 | 0.087261593 | 0.274400501 | 3.724945049 | 0.073665651 |
| Gm25188 | 1.524225208 | -0.243036366 | 0.902682551 | -0.269237913 |
| Rps3a1 | 1477.132456 | 0.380919249 | 0.095995066 | 3.96811277 |
| Snord73a | 1.006316758 | 0.335138654 | 0.930803716 | 0.360052983 |
| Rnu73b | 4.371881307 | -0.368264268 | 0.53353838 | -0.690230135 |
| Gm37933 | 3.2042328 | 0.82263083 | 0.635110904 | 1.295255403 |
| Gm37493 | 0 | NA | NA | NA |
| Gm23025 | 0 | NA | NA | NA |
| Lrba | 1713.427863 | -0.331685441 | 0.07180145 | -4.619481085 |
| Gm37288 | 0 | NA | NA | NA |
| Gm26343 | 0 | NA | NA | NA |
| Gm23850 | 3.443253566 | -0.649143744 | 0.657449859 | -0.987366161 |
| Gm25039 | 5.355852112 | 0.228975476 | 0.402039974 | 0.569534104 |
| Mab21l2 | 76.13297546 | 0.350401219 | 0.435144332 | 0.805252862 |
| Gm3788 | 95.68068835 | 1.058012392 | 0.287653033 | 3.678085298 |
| Gm37876 | 2.813292308 | -0.169766077 | 0.720715232 | -0.235552226 |
| Dclk2 | 1006.44227 | -0.158194015 | 0.076297951 | -2.073371742 |
| Gm37025 | 0.066050875 | 0.274391604 | 3.724945049 | 0.073663262 |
| Gm23376 | 0 | NA | NA | NA |

| | | | | |
|---|---|---|---|---|
| Gm37207 | 0.045882455 | 0.755288839 | 3.724945049 | 0.202765096 |
| Cd1d2 | 0.436249931 | 0.755309718 | 1.499543575 | 0.503693077 |
| Cd1d1 | 24.21988694 | 1.479424291 | 0.256489216 | 5.767978522 |
| Gm8869 | 0 | NA | NA | NA |
| Gm18403 | 0 | NA | NA | NA |
| Gm37575 | 0 | NA | NA | NA |
| Gm24472 | 0 | NA | NA | NA |
| Kirrel | 258.042677 | 0.474034209 | 0.151068382 | 3.137878368 |
| Gm37855 | 0.445528452 | 1.145913467 | 1.450999948 | 0.789740529 |
| Gm10705 | 72.15716491 | 0.806440873 | 0.152069099 | 5.303121273 |
| Fcrls | 15.92947883 | -3.01407794 | 0.408002664 | -7.387397703 |
| Cd5l | 3.379428938 | 3.562495599 | 1.097112123 | 3.247157263 |
| Fcrl1 | 1.476596612 | 2.610255114 | 1.126920511 | 2.31627261 |
| Fcrl5 | 1.398885825 | 0.204251374 | 1.11436466 | 0.183289529 |
| Etv3 | 674.3690235 | 0.42775782 | 0.114968265 | 3.720659942 |
| Etv3l | 0.263217861 | 0.714171662 | 2.116797803 | 0.337383033 |
| Gm25167 | 0 | NA | NA | NA |
| Gm24109 | 0 | NA | NA | NA |
| Arhgef11 | 1362.693994 | -0.114953773 | 0.040025292 | -2.872028324 |
| Lrrc71 | 7.798021523 | 0.430883903 | 0.437584066 | 0.984688283 |
| Pear1 | 78.16134709 | 0.645460814 | 0.150017842 | 4.302560316 |
| Ntrk1 | 25.65422587 | 0.637801343 | 0.263758721 | 2.418124185 |
| Insrr | 23.94499803 | 1.196766972 | 0.258393643 | 4.631565068 |
| Sh2d2a | 48.27836269 | 7.690807265 | 0.890647946 | 8.635069896 |
| Prcc | 536.6201964 | 0.071401907 | 0.062296446 | 1.146163415 |
| Hdgf | 1352.852318 | 0.116797059 | 0.058735652 | 1.98852069 |
| Mrpl24 | 277.5276893 | -0.098565173 | 0.092951731 | -1.060390938 |
| Rrnad1 | 232.3898889 | -0.26884917 | 0.123678344 | -2.173777254 |
| Isg20l2 | 217.9624788 | 0.153688932 | 0.094382524 | 1.628362177 |
| Crabp2 | 45.98201362 | 0.49206276 | 0.289376372 | 1.700424799 |
| Gm3745 | 0 | NA | NA | NA |
| Nes | 844.5112463 | 2.159597642 | 0.408778546 | 5.283050352 |
| Bcan | 1604.188113 | -0.502052301 | 0.147699256 | -3.399152546 |
| Hapln2 | 312.1547389 | 0.508634127 | 0.168736757 | 3.014364715 |
| RP23-240E15.3 | 115.0424175 | 0.646259634 | 0.196756781 | 3.28456092 |
| Gpatch4 | 115.8497267 | 0.405314566 | 0.185576135 | 2.184087765 |
| Apoa1bp | 265.3851978 | 0.028732573 | 0.102885527 | 0.27926739 |
| Ttc24 | 0.042716945 | -0.206490061 | 3.724945049 | -0.055434391 |
| Iqgap3 | 18.40657348 | 1.923918268 | 0.483883928 | 3.975991263 |
| Mef2d | 2901.836846 | -0.086820682 | 0.077119535 | -1.125793645 |
| 1700113A16Rik | 8.541957373 | -0.929575265 | 0.384237055 | -2.41927543 |
| AI849053 | 5.915945368 | -0.615610085 | 0.397962785 | -1.546903648 |
| AW047730 | 113.9278857 | -1.173612997 | 0.197777001 | -5.934021614 |
| Gm3764 | 811.9285128 | -0.58298356 | 0.180941977 | -3.2219365 |
| Mir3093 | 13.97251529 | -1.290257277 | 0.336882206 | -3.829995334 |
| Gm25641 | 0.574683629 | -1.912629029 | 1.262915079 | -1.514455771 |
| mmu-mir-9-1 | 11.97633615 | -1.031747739 | 0.347122796 | -2.972284597 |

| | | | | |
|---|---|---|---|---|
| Rhbg | 2.720079777 | 3.576381322 | 1.041760959 | 3.433015311 |
| Gm38392 | 2.080370332 | -0.302057406 | 0.724364853 | -0.41699622 |
| Tsacc | 7.84413125 | -0.485797522 | 0.404979602 | -1.199560472 |
| Gm37116 | 2.788796254 | -0.12798215 | 0.729339292 | -0.175476834 |
| Cct3 | 1136.589819 | 0.151476191 | 0.084093558 | 1.801281739 |
| Glmp | 457.6296344 | 0.334780576 | 0.126876675 | 2.638629801 |
| Tmem79 | 9.024460536 | 0.279929802 | 0.33935586 | 0.824885718 |
| Smg5 | 1241.608425 | 0.32223023 | 0.069895174 | 4.610192844 |
| Paqr6 | 85.91657556 | -0.592549576 | 0.167417409 | -3.539354593 |
| Bglap3 | 2.927464552 | -1.434063863 | 0.731250327 | -1.961112099 |
| Gm21956 | 0.097139547 | 0.274393828 | 3.724945049 | 0.073663859 |
| Bglap2 | 0.555158567 | -0.151723493 | 1.314000127 | -0.115466879 |
| Gm6821 | 0.068097359 | -0.206487837 | 3.724945049 | -0.055433794 |
| Bglap | 0.801792645 | 0.787909645 | 1.364367108 | 0.577490941 |
| Pmf1 | 65.8586388 | 0.66704551 | 0.185258296 | 3.60062423 |
| Slc25a44 | 1273.147583 | 0.360205275 | 0.064324788 | 5.599789535 |
| RP24-298F19.6 | 2.520287764 | 0.939787768 | 0.731249118 | 1.285181404 |
| Sema4a | 442.3120329 | -0.245762419 | 0.101268305 | -2.426844407 |
| Mir7011 | 0 | NA | NA | NA |
| Gm5979 | 0 | NA | NA | NA |
| Lmna | 660.5220702 | 0.408669258 | 0.169391066 | 2.412578588 |
| Gm37584 | 6.065917723 | -1.234786314 | 0.429311421 | -2.876201877 |
| Mex3a | 75.1280634 | -0.475119975 | 0.16376654 | -2.901202987 |
| Mir1905 | 0.318342637 | -2.080184072 | 3.682092106 | -0.564946235 |
| Rab25 | 1.64905476 | 0.649727045 | 0.742850032 | 0.874640933 |
| Lamtor2 | 189.4900279 | 0.015043776 | 0.113680608 | 0.132333707 |
| Ubqln4 | 1064.387744 | -0.184841906 | 0.062841515 | -2.941397987 |
| Ssr2 | 493.374432 | 0.327123684 | 0.086921563 | 3.763435373 |
| Gm10704 | 39.754764 | 0.882812674 | 0.232512453 | 3.796840399 |
| Arhgef2 | 1402.732759 | -0.180810236 | 0.064590519 | -2.799330921 |
| Gm20652 | 1.311025493 | 0.589287951 | 0.826925847 | 0.712624902 |
| Rxfp4 | 0 | NA | NA | NA |
| RP23-199B2.8 | 2.30646632 | 0.667563641 | 0.667054223 | 1.000763684 |
| 2810403A07Rik | 753.6655769 | -0.015470158 | 0.097094827 | -0.159330408 |
| Gm25945 | 6.725632374 | 0.212350786 | 0.406357408 | 0.522571465 |
| Rit1 | 282.0700927 | 0.007902019 | 0.10944447 | 0.07220117 |
| Gm17146 | 5.547323716 | -1.314208312 | 0.472383728 | -2.782077864 |
| Gm10253 | 5.431051021 | 1.063269486 | 0.567079679 | 1.874991337 |
| Syt11 | 7534.20102 | -0.033931985 | 0.051208819 | -0.662619944 |
| 5830417I10Rik | 63.92099027 | -0.548072938 | 0.153253198 | -3.576257755 |
| 1500004A13Rik | 4.959186419 | -0.307875945 | 0.47717251 | -0.645208888 |
| Gon4l | 808.0428841 | -0.111858673 | 0.073187447 | -1.528386044 |
| RP23-51N3.3 | 0.176220594 | -0.20648024 | 2.719488399 | -0.075926134 |
| Gm23054 | 2.17116889 | -1.290327505 | 0.77911514 | -1.656144823 |
| Msto1 | 187.4319594 | 0.182960131 | 0.086188108 | 2.12280018 |
| n-R5s197 | 0.163066018 | 0.813382652 | 3.445567682 | 0.236066369 |
| Dap3 | 627.5926466 | -0.106267502 | 0.054954281 | -1.933743831 |

| | | | | |
|---|---|---|---|---|
| Ash1l | 4532.149695 | -0.141986549 | 0.061444844 | -2.310796795 |
| Gm26465 | 0 | NA | NA | NA |
| Gm24403 | 0 | NA | NA | NA |
| RP24-394C15.7 | 0 | NA | NA | NA |
| Rusc1 | 959.1680519 | -0.069868279 | 0.063767142 | -1.095678369 |
| Fdps | 34.50620784 | -0.139259759 | 0.205640774 | -0.677199156 |
| Gm22935 | 1.101484168 | 1.269585104 | 1.103872848 | 1.150118971 |
| RP24-394C15.4 | 0 | NA | NA | NA |
| Pklr | 0.541363579 | -0.892662017 | 1.192681934 | -0.748449349 |
| Hcn3 | 139.9590133 | -0.452298694 | 0.131152674 | -3.44864258 |
| Clk2 | 311.1271416 | -0.236523543 | 0.144967554 | -1.63156193 |
| Scamp3 | 397.6184147 | 0.128344719 | 0.094362682 | 1.360121572 |
| Gm16069 | 1.160335885 | -0.453651619 | 0.913041722 | -0.496857491 |
| Fam189b | 352.3484391 | -0.063918796 | 0.068152587 | -0.937877758 |
| Gm23269 | 0 | NA | NA | NA |
| Gba | 494.4063209 | 0.537632092 | 0.118026921 | 4.555164939 |
| Mtx1 | 203.9215083 | 0.102087498 | 0.085023241 | 1.20070109 |
| RP23-350K3.2 | 104.7257883 | 0.071729538 | 0.114499965 | 0.626459035 |
| Thbs3 | 91.591488 | 0.219800663 | 0.192132537 | 1.144005416 |
| Mir92b | 0.280267168 | -0.206482567 | 1.81575733 | -0.11371705 |
| Muc1 | 17.55319324 | 2.810737116 | 0.399621167 | 7.033504108 |
| Trim46 | 576.4645691 | -0.252889501 | 0.126953365 | -1.991987383 |
| Krtcap2 | 231.132102 | 0.167223005 | 0.108819232 | 1.536704511 |
| Dpm3 | 102.4945791 | -0.057540685 | 0.138304871 | -0.416042363 |
| Slc50a1 | 178.718743 | 0.339452446 | 0.100613824 | 3.373815168 |
| Efna1 | 37.54655563 | 0.415297766 | 0.204128507 | 2.034491764 |
| Gm22642 | 0 | NA | NA | NA |
| Efna3 | 366.6839132 | -0.245418915 | 0.14652822 | -1.674891812 |
| Gm15998 | 1.818279517 | 1.843898422 | 0.999262856 | 1.845258644 |
| Efna4 | 3.626077548 | 0.048127431 | 0.594461135 | 0.08095976 |
| 4731419I09Rik | 0.772156528 | -0.121479851 | 1.159941528 | -0.104729288 |
| Adam15 | 928.1765902 | 0.398352427 | 0.075730675 | 5.260119874 |
| Dcst1 | 1.444845127 | 1.104094633 | 0.838284807 | 1.31708773 |
| Gm10702 | 17.15096642 | -0.037949515 | 0.27295432 | -0.139032476 |
| Zbtb7b | 489.1333668 | 0.69236565 | 0.134671711 | 5.141136502 |
| Gm15417 | 31.01182027 | 0.173064888 | 0.187058779 | 0.925189873 |
| Lenep | 0.241023442 | -0.604982388 | 1.829356696 | -0.330707723 |
| Flad1 | 214.0339016 | -0.143680497 | 0.088332866 | -1.626580273 |
| Cks1b | 21.556343 | 0.583920558 | 0.371678466 | 1.571036828 |
| Shc1 | 291.0704285 | 0.583115761 | 0.124502431 | 4.683569269 |
| Pygo2 | 447.2134043 | -0.212326545 | 0.097185432 | -2.184756915 |
| Pbxip1 | 1167.4111 | 0.732796308 | 0.128739805 | 5.692072532 |
| Pmvk | 242.2372548 | -0.038183745 | 0.121180548 | -0.315097971 |
| Kcnn3 | 548.2658225 | -0.105248973 | 0.094886542 | -1.109208645 |
| Adar | 3619.123941 | 0.828999023 | 0.091026875 | 9.107189757 |
| Chrnb2 | 737.8398015 | -0.258891887 | 0.069487261 | -3.725746015 |
| 4632404H12Rik | 215.8736812 | -0.315749339 | 0.13499596 | -2.338953985 |

| | | | | |
|---|---|---|---|---|
| Ube2q1 | 886.7801256 | -0.024974148 | 0.054801235 | -0.455722364 |
| RP24-122I22.4 | 0 | NA | NA | NA |
| She | 25.99300754 | 0.811448382 | 0.304145676 | 2.667959621 |
| Il6ra | 238.4815568 | 1.429323269 | 0.13414645 | 10.65494669 |
| RP24-122I22.6 | 1.56556854 | 0.773258362 | 0.987077147 | 0.783381891 |
| Atp8b2 | 1163.874269 | -0.207100974 | 0.097634447 | -2.121187561 |
| Aqp10-ps | 0.173853943 | 0.755298194 | 2.74376612 | 0.275277907 |
| Gm24046 | 0.066050875 | 0.274391604 | 3.724945049 | 0.073663262 |
| Hax1 | 320.365027 | -0.031771617 | 0.118325707 | -0.268509842 |
| RP23-114P22.5 | 1.940516017 | -0.777348912 | 0.74548845 | -1.042737701 |
| Ubap2l | 1956.985286 | 0.040317378 | 0.068418826 | 0.589273161 |
| Gm24608 | 5.024933871 | -0.074864004 | 0.503093786 | -0.148807252 |
| Mir7669 | 4.045153519 | -0.363589868 | 0.553280295 | -0.657153114 |
| 4933434E20Rik | 645.1823872 | 0.130487809 | 0.089378251 | 1.45995034 |
| Gm16540 | 6.216095886 | -0.306272618 | 0.479156466 | -0.639191244 |
| Mir190b | 0.194279094 | 0.274391173 | 3.724945049 | 0.073663147 |
| Tpm3 | 824.8814748 | 0.532504947 | 0.106548651 | 4.997763363 |
| Nup210l | 14.08624278 | -0.432621553 | 0.352173993 | -1.228431291 |
| Gm23723 | 0.35561486 | 0.276321738 | 1.597340044 | 0.172988675 |
| RP24-257C17.2 | 1.986149933 | 0.035630078 | 0.75381982 | 0.047266041 |
| Gm24185 | 0 | NA | NA | NA |
| Rps27 | 1242.923509 | 0.510157303 | 0.143344311 | 3.558964409 |
| Rab13 | 74.74244885 | 1.338123911 | 0.208218971 | 6.426522559 |
| Jtb | 232.6321241 | 0.12399797 | 0.098932158 | 1.253363647 |
| Creb3l4 | 5.167911745 | 0.238128157 | 0.513022638 | 0.464166958 |
| Slc39a1 | 919.2059152 | 1.013677528 | 0.134848028 | 7.517184678 |
| Crtc2 | 364.8494084 | 0.292286651 | 0.115809508 | 2.523857126 |
| Dennd4b | 682.1714222 | -0.050178655 | 0.064125727 | -0.782504263 |
| Mir7012 | 0 | NA | NA | NA |
| RP23-358O22.1 | 2.468095164 | -0.142967128 | 0.656058212 | -0.217918357 |
| Gatad2b | 2548.376701 | 0.170408446 | 0.06316914 | 2.697653412 |
| Slc27a3 | 56.99060069 | 0.111695832 | 0.165639637 | 0.674330336 |
| Ints3 | 947.6918643 | -0.113467087 | 0.083690736 | -1.355790292 |
| Npr1 | 184.6399887 | 1.146314874 | 0.208696578 | 5.492734392 |
| Gm16048 | 1.106761898 | -1.327197901 | 1.075501944 | -1.2340265 |
| RP24-333B11.3 | 0.066050875 | 0.274391604 | 3.724945049 | 0.073663262 |
| Ilf2 | 670.5056944 | 0.164048139 | 0.083079271 | 1.974597716 |
| Snapin | 374.92744 | -0.00252305 | 0.077136299 | -0.03270899 |
| Chtop | 1458.724535 | 0.074805416 | 0.075881737 | 0.985815808 |
| S100a1 | 401.8878366 | 0.199510548 | 0.151777896 | 1.314490138 |
| S100a13 | 63.775755 | 0.40931183 | 0.171620377 | 2.384983867 |
| S100a14 | 1.045271382 | 1.741039282 | 1.118844441 | 1.556104869 |
| S100a16 | 533.8051986 | 0.883742728 | 0.170417979 | 5.185736476 |
| Gm24593 | 0 | NA | NA | NA |
| Gm23464 | 0 | NA | NA | NA |
| S100a3 | 9.213372739 | 1.931923774 | 0.436187778 | 4.429110287 |
| S100a2 | 0.475453923 | -0.190481559 | 1.364853321 | -0.139561926 |

| | | | | |
|---|---|---|---|---|
| RP23-326D6.5 | 0 | NA | NA | NA |
| S100a4 | 14.92865454 | 2.098584181 | 0.519882768 | 4.036648853 |
| S100a5 | 23.65689645 | -1.927738395 | 1.465377334 | -1.315523552 |
| S100a6 | 362.9019992 | 2.395495825 | 0.275423067 | 8.697513439 |
| RP24-174D10.3 | 0 | NA | NA | NA |
| S100a7a | 5.924278019 | 0.456219772 | 0.515840039 | 0.884421018 |
| S100a8 | 13.16848317 | 3.285556264 | 0.835264126 | 3.933553666 |
| S100a9 | 10.76590429 | 2.712813261 | 0.598944059 | 4.529326604 |
| RP24-174D10.4 | 0 | NA | NA | NA |
| RP24-174D10.5 | 0 | NA | NA | NA |
| Pglyrp4 | 0.060174537 | 0.755286614 | 3.724945049 | 0.202764498 |
| RP24-174D10.6 | 2.240082864 | -0.731785446 | 0.721568756 | -1.014158998 |
| Gm5849 | 0 | NA | NA | NA |
| RP23-111A11.1 | 0 | NA | NA | NA |
| RP23-309N14.1 | 0 | NA | NA | NA |
| RP24-296O9.1 | 0 | NA | NA | NA |
| 4930529C04Rik | 0.183030605 | -0.20648243 | 2.280962878 | -0.090524239 |
| 9130204L05Rik | 0 | NA | NA | NA |
| Gm23697 | 0 | NA | NA | NA |
| Pglyrp3 | 2.336438979 | 3.820990521 | 1.077865504 | 3.544960392 |
| Mif-ps3 | 0.176744298 | -0.206480793 | 2.303963344 | -0.089619825 |
| Gm25145 | 0 | NA | NA | NA |
| Lor | 15.81042869 | -0.069122418 | 0.303456768 | -0.227783413 |
| RP23-381I16.5 | 0 | NA | NA | NA |
| RP23-381I16.3 | 0 | NA | NA | NA |
| RP23-381I16.4 | 0 | NA | NA | NA |
| Prr9 | 0 | NA | NA | NA |
| Lelp1 | 0 | NA | NA | NA |
| RP23-381I16.6 | 0.291617713 | 0.968593514 | 1.731084856 | 0.559529771 |
| Sprr2a1 | 0 | NA | NA | NA |
| Sprr2a2 | 0 | NA | NA | NA |
| Gm5850 | 0.099106094 | 0.274387056 | 3.724945049 | 0.073662041 |
| Gm37087 | 0.047948698 | -0.206487837 | 3.724945049 | -0.055433794 |
| Gm5851 | 0.36081128 | -0.206484761 | 1.750571986 | -0.117952739 |
| Gm37752 | 0 | NA | NA | NA |
| Sprr2a3 | 0 | NA | NA | NA |
| Sprr2b | 0.207368832 | 1.236194129 | 2.640009706 | 0.46825363 |
| Gm37428 | 0 | NA | NA | NA |
| Sprr2c-ps | 0 | NA | NA | NA |
| Sprr2d | 0 | NA | NA | NA |
| Sprr2e | 0.186228844 | 1.236183955 | 3.289587508 | 0.375786919 |
| Sprr2f | 0 | NA | NA | NA |
| Sprr2g | 0.097139547 | 0.274393828 | 3.724945049 | 0.073663859 |
| Sprr2h | 0 | NA | NA | NA |
| Sprr2i | 0 | NA | NA | NA |
| Sprr2j-ps | 0 | NA | NA | NA |
| Gm9774 | 28.33359312 | -0.104397374 | 0.240516288 | -0.43405532 |

| | | | | |
|---|---|---|---|---|
| Sprr2k | 0 | NA | NA | NA |
| Sprr1b | 0.278361807 | 1.296114431 | 3.705122054 | 0.349816932 |
| Sprr3 | 0 | NA | NA | NA |
| Gm37377 | 0 | NA | NA | NA |
| Sprr1a | 21.48993395 | 2.042037172 | 0.850924792 | 2.399785729 |
| Sprr4 | 0 | NA | NA | NA |
| 2310046K23Rik | 0.227721369 | 1.190314394 | 3.108233493 | 0.382955269 |
| Ivl | 0 | NA | NA | NA |
| Smcp | 0 | NA | NA | NA |
| 2210017I01Rik | 0 | NA | NA | NA |
| Lce6a | 0.09176491 | 1.15872478 | 3.72306111 | 0.311229052 |
| Lce1a1 | 0.475368968 | 2.48242336 | 3.690247613 | 0.672698317 |
| Lce1b | 0.18352982 | 1.625373558 | 3.713044335 | 0.437746876 |
| Rpl13-ps4 | 0 | NA | NA | NA |
| Lce1a2 | 0 | NA | NA | NA |
| Lce1c | 0.27529473 | 1.966307691 | 3.703391457 | 0.530947839 |
| Lce1d | 0.45492314 | 1.472211785 | 3.690964141 | 0.398869165 |
| Gm4202 | 299.4188111 | 0.023300051 | 0.17348118 | 0.134308809 |
| Lce1e | 0 | NA | NA | NA |
| Lce1f | 0.373862045 | 0.755307876 | 1.711842211 | 0.441225173 |
| Gm38119 | 0 | NA | NA | NA |
| Lce1g | 0.655362467 | 1.688316195 | 2.489219004 | 0.678251368 |
| Lce1h | 0.27529473 | 1.966307691 | 3.703391457 | 0.530947839 |
| Lce1i | 0.17962841 | 0.274411151 | 3.374012434 | 0.081330806 |
| Lce1j | 0.09176491 | 1.15872478 | 3.72306111 | 0.311229052 |
| Lce1k | 0 | NA | NA | NA |
| Kprp | 0.596471915 | 2.722327997 | 3.68431128 | 0.738897392 |
| Gm38201 | 0 | NA | NA | NA |
| Lce1l | 0.09176491 | 1.15872478 | 3.72306111 | 0.311229052 |
| 2310050C09Rik | 0.31922648 | 1.555688877 | 3.70283481 | 0.420134561 |
| Lce3a | 0 | NA | NA | NA |
| Lce3b | 0 | NA | NA | NA |
| Gm38078 | 0 | NA | NA | NA |
| Lce3c | 0.342841614 | 1.316489277 | 2.012057842 | 0.654299916 |
| Gm24481 | 0 | NA | NA | NA |
| Lce3d | 0 | NA | NA | NA |
| Gm22577 | 0 | NA | NA | NA |
| Lce3e | 0 | NA | NA | NA |
| Lce3f | 0.038981395 | 0.274393828 | 3.724945049 | 0.073663859 |
| Crct1 | 1.039606943 | 2.71326814 | 1.189061848 | 2.281856192 |
| Lce1m | 0.045882455 | 0.755288839 | 3.724945049 | 0.202765096 |
| Gm38040 | 0.139839889 | 0.274406293 | 2.92397156 | 0.093847114 |
| Gm4858 | 0 | NA | NA | NA |
| Gm37361 | 0 | NA | NA | NA |
| Crnn | 0 | NA | NA | NA |
| Gm37501 | 0 | NA | NA | NA |
| Flg2 | 0.73411928 | 2.954651731 | 3.678675968 | 0.803183471 |

| | | | | |
|---|---|---|---|---|
| Gm18432 | 0.092691418 | 0.274387304 | 3.724945049 | 0.073662108 |
| Flg | 1.695700131 | 3.810286967 | 3.64866258 | 1.044296885 |
| Rplp0-ps1 | 2.415407705 | 0.423912088 | 0.782085236 | 0.542027989 |
| Gm37922 | 0 | NA | NA | NA |
| Gm37245 | 0.045882455 | 0.755288839 | 3.724945049 | 0.202765096 |
| Gm37913 | 0.038981395 | 0.274393828 | 3.724945049 | 0.073663859 |
| Hrnr | 2.939544198 | 4.106278058 | 3.643600773 | 1.126983529 |
| AC123859.1 | 0 | NA | NA | NA |
| Gm37857 | 0 | NA | NA | NA |
| Rptn | 0.124596252 | 0.755282361 | 3.714681068 | 0.203323609 |
| Gm18296 | 0.119587853 | 1.218392466 | 3.723618458 | 0.327206581 |
| Gm22752 | 0.140442044 | -0.206481048 | 3.613090186 | -0.057148047 |
| Gm5278 | 0.680250709 | 0.193325462 | 1.495200854 | 0.129297319 |
| Tchh | 32.61220021 | 0.188420518 | 0.245717313 | 0.766818244 |
| Tchhl1 | 0.282572244 | 0.335418534 | 2.063456582 | 0.162551777 |
| Gm36953 | 0 | NA | NA | NA |
| S100a11 | 216.4239304 | 3.194025735 | 0.393029306 | 8.126685933 |
| Gm29950 | 0 | NA | NA | NA |
| S100a10 | 413.4047504 | 1.485301023 | 0.196996673 | 7.539726446 |
| Gm5541 | 0 | NA | NA | NA |
| Gm38207 | 0 | NA | NA | NA |
| Tdpoz2 | 0 | NA | NA | NA |
| Tdpoz1 | 0 | NA | NA | NA |
| Gm18433 | 0 | NA | NA | NA |
| Gm37596 | 0 | NA | NA | NA |
| Gm38086 | 0 | NA | NA | NA |
| Gm5852 | 0 | NA | NA | NA |
| Gm5773 | 0 | NA | NA | NA |
| Tdpoz4 | 0 | NA | NA | NA |
| Tdpoz3 | 0 | NA | NA | NA |
| Gm37430 | 0 | NA | NA | NA |
| Gm9113 | 0.044544648 | 0.755286614 | 3.724945049 | 0.202764498 |
| Gm9116 | 0 | NA | NA | NA |
| Gm37185 | 0 | NA | NA | NA |
| Gm9117 | 0 | NA | NA | NA |
| Gm37104 | 0 | NA | NA | NA |
| Tdpoz5 | 0 | NA | NA | NA |
| Gm37807 | 0 | NA | NA | NA |
| Gm37481 | 0 | NA | NA | NA |
| Gm9121 | 0.045882455 | 0.755288839 | 3.724945049 | 0.202765096 |
| Gm9124 | 0 | NA | NA | NA |
| Gm37085 | 0 | NA | NA | NA |
| Gm9125 | 0 | NA | NA | NA |
| Gm37081 | 0 | NA | NA | NA |
| Gm10697 | 0 | NA | NA | NA |
| Gm37778 | 0 | NA | NA | NA |
| Gm37155 | 0 | NA | NA | NA |

| | | | | |
|---|---|---|---|---|
| Gm5542 | 0 | NA | NA | NA |
| Gm9131 | 0 | NA | NA | NA |
| Gm37882 | 0 | NA | NA | NA |
| Gm5543 | 0 | NA | NA | NA |
| Gm37001 | 0 | NA | NA | NA |
| Gm16560 | 0 | NA | NA | NA |
| Gm10696 | 0.045882455 | 0.755288839 | 3.724945049 | 0.202765096 |
| Gm5286 | 0 | NA | NA | NA |
| Gm16584 | 0 | NA | NA | NA |
| Gm4982 | 0.038981395 | 0.274393828 | 3.724945049 | 0.073663859 |
| Gm9141 | 0 | NA | NA | NA |
| Gm4778 | 0 | NA | NA | NA |
| Gm16561 | 0 | NA | NA | NA |
| Them4 | 202.6068946 | -0.198509417 | 0.087800666 | -2.260910151 |
| RP23-148G12.2 | 0 | NA | NA | NA |
| Them5 | 0.249936274 | 0.461189648 | 2.48903363 | 0.185288637 |
| RP23-148G12.3 | 4.416068108 | -0.653621963 | 0.51314413 | -1.273759018 |
| C2cd4d | 4.518953556 | -0.5864338 | 0.508656779 | -1.152906683 |
| RP23-148G12.5 | 0.049553047 | 0.274391604 | 3.724945049 | 0.073663262 |
| Rorc | 43.04669428 | 0.133853539 | 0.234210979 | 0.571508388 |
| RP23-298M23.3 | 0.086648695 | -0.687376175 | 3.724945049 | -0.184533239 |
| Lingo4 | 68.20621523 | -0.359079285 | 0.244584433 | -1.468119944 |
| 1700040D17Rik | 7.265577632 | -1.334528634 | 0.400664055 | -3.330792016 |
| Tdrkh | 464.6804853 | -0.537696277 | 0.087428159 | -6.150149851 |
| RP23-298M23.7 | 44.92890297 | -1.245251153 | 0.261315285 | -4.765320756 |
| RP23-298M23.8 | 0.072025462 | 0.755288839 | 3.724945049 | 0.202765096 |
| Oaz3 | 2.190110068 | 3.056885078 | 1.149411149 | 2.65952273 |
| Mrpl9 | 334.7621041 | 0.401730934 | 0.088391249 | 4.544917519 |
| RP23-298M23.9 | 0.961922688 | -0.599531309 | 1.005488355 | -0.596258829 |
| Riiad1 | 12.40794134 | 0.237255464 | 0.316375961 | 0.749916218 |
| Celf3 | 927.0754436 | -0.443430931 | 0.141935975 | -3.124161668 |
| Snx27 | 1992.373795 | -0.132244061 | 0.057395799 | -2.304072121 |
| RP23-298M23.11 | 19.53939973 | -0.081284991 | 0.347898405 | -0.23364577 |
| RP23-298M23.12 | 7.754593511 | -0.11798827 | 0.482777946 | -0.244394491 |
| RP24-276A6.1 | 23.82234211 | 0.025661027 | 0.317548311 | 0.080809838 |
| AC127252.1 | 0.108123235 | 0.274402725 | 3.724945049 | 0.073666248 |
| Rpl31-ps11 | 1.676330775 | -0.100965759 | 0.859671815 | -0.117446865 |
| Tuft1 | 38.81743166 | 0.57285908 | 0.226310157 | 2.531300788 |
| Gm10972 | 0 | NA | NA | NA |
| Gm15234 | 0.403742549 | 1.125760615 | 1.436966351 | 0.783428655 |
| BC021767 | 1.492922606 | 1.10774839 | 0.824060977 | 1.344255366 |
| Selenbp2 | 2.718408089 | 0.278836632 | 0.696098755 | 0.400570507 |
| Gm15264 | 3.338245402 | 0.141257267 | 0.615955885 | 0.229330169 |
| RP23-449N19.1 | 0 | NA | NA | NA |
| Cgn | 50.96569556 | -0.691065562 | 0.173602721 | -3.980730009 |
| Gm15263 | 0.664861716 | -0.132756769 | 1.274154989 | -0.10419201 |
| Pogz | 1033.996817 | -0.17462088 | 0.062126534 | -2.810729466 |

| | | | | |
|---|---|---|---|---|
| Psmb4 | 959.4736378 | 0.385073015 | 0.115353992 | 3.338185431 |
| Gm26279 | 0.212513282 | 1.926509226 | 3.159662244 | 0.609719988 |
| Selenbp1 | 39.74177677 | 0.139578737 | 0.256366245 | 0.544450527 |
| Rfx5 | 386.701994 | 0.326737277 | 0.074465975 | 4.387739191 |
| B230398E01Rik | 27.22757332 | 4.126892211 | 0.48528465 | 8.504065009 |
| Pi4kb | 609.4035235 | -0.221586777 | 0.064293464 | -3.446489961 |
| A730011C13Rik | 177.9491982 | 3.564835795 | 0.213563457 | 16.69216186 |
| Gm15265 | 34.26348499 | 0.772180604 | 0.266499622 | 2.897492305 |
| Mir7013 | 0.881607526 | -0.251160202 | 0.981719462 | -0.25583704 |
| Zfp687 | 719.9016467 | 0.565852332 | 0.088699444 | 6.379434942 |
| 4930481B07Rik | 9.171066605 | 0.139850161 | 0.342609152 | 0.408191552 |
| Psmd4 | 779.7769061 | 0.294201471 | 0.097623429 | 3.013635894 |
| Pip5k1a | 758.4226424 | 0.262098545 | 0.086036867 | 3.046351567 |
| Gm24920 | 5.105149463 | -0.812666692 | 0.541848347 | -1.49980469 |
| Gm22301 | 0 | NA | NA | NA |
| Vps72 | 306.4355309 | 0.200464096 | 0.067424711 | 2.97315471 |
| Tmod4 | 6.657288501 | -1.57439816 | 0.471099975 | -3.341961884 |
| Scnm1 | 114.2649021 | 0.449105548 | 0.114732205 | 3.91438087 |
| Lysmd1 | 47.85101064 | -0.171727194 | 0.155115618 | -1.107091582 |
| Mir8099-2 | 0.049553047 | 0.274391604 | 3.724945049 | 0.073663262 |
| Tnfaip8l2 | 38.08952461 | 2.349777328 | 0.285496947 | 8.230481464 |
| Gm9169 | 2.545959807 | -0.247628279 | 0.713272335 | -0.347172135 |
| Sema6c | 68.04043718 | -0.406749033 | 0.1879883 | -2.163693337 |
| AC140190.1 | 0 | NA | NA | NA |
| Gabpb2 | 547.3167726 | -0.122419478 | 0.110258419 | -1.110295962 |
| Gm16740 | 8.225987562 | -0.167461702 | 0.366332914 | -0.457129828 |
| Mllt11 | 2083.533929 | -0.24796066 | 0.072465241 | -3.42178756 |
| Cdc42se1 | 925.5629768 | 0.521313721 | 0.111342513 | 4.682072535 |
| Gm128 | 1.475208495 | -2.081219385 | 0.916753463 | -2.270206188 |
| Bnipl | 3.731006451 | -2.241880705 | 0.640519474 | -3.500097651 |
| Prune | 303.723384 | -0.058269502 | 0.076362878 | -0.763060583 |
| Fam63a | 346.9866243 | 0.068487419 | 0.10119374 | 0.67679502 |
| Anxa9 | 8.904347822 | -0.980523279 | 0.405783607 | -2.416369862 |
| 6330562C20Rik | 7.360490999 | 0.27119544 | 0.400552 | 0.677054264 |
| Cers2 | 766.0749113 | 0.346494424 | 0.107609649 | 3.219919654 |
| Setdb1 | 649.1102505 | -0.187805728 | 0.092339621 | -2.033858563 |
| RP23-407B16.2 | 0 | NA | NA | NA |
| Gm37500 | 1.278257442 | 0.55322264 | 0.896774958 | 0.616902418 |
| RP23-407B16.3 | 0 | NA | NA | NA |
| Gm5070 | 0.368600158 | -0.433153198 | 1.477164956 | -0.293232788 |
| RP23-407B16.4 | 0.048949532 | -0.206487837 | 3.724945049 | -0.055433794 |
| Gm4349 | 3.988126414 | -1.317250559 | 0.50658934 | -2.600233475 |
| Arnt | 674.0580543 | -0.125265445 | 0.110156992 | -1.137153819 |
| RP23-422N18.1 | 29.38944538 | -0.29361262 | 0.258988946 | -1.133687843 |
| Ctsk | 101.6290436 | 1.318476887 | 0.200376084 | 6.580011247 |
| Ctss | 2705.512219 | 2.52149927 | 0.182781764 | 13.79513585 |
| Hmgb1-ps5 | 4.681085585 | 3.89372994 | 0.959030753 | 4.060067864 |

| | | | | |
|---|---|---|---|---|
| Hormad1 | 0.108266322 | 0.274389235 | 3.724945049 | 0.073662626 |
| Golph3l | 291.0130091 | 0.316812848 | 0.111604346 | 2.83871425 |
| RP23-422N18.3 | 2.848305637 | -0.137650087 | 0.704303504 | -0.195441434 |
| Rps10-ps1 | 80.90969664 | 0.264759133 | 0.190348781 | 1.39091583 |
| Ensa | 2238.110929 | -0.058504053 | 0.073505858 | -0.795910077 |
| RP23-231L15.4 | 47.75963671 | -0.468420512 | 0.261999047 | -1.787871053 |
| Mcl1 | 1783.072468 | 1.03328451 | 0.142477348 | 7.252272192 |
| Adamtsl4 | 317.4111096 | 0.567037329 | 0.144097961 | 3.935082251 |
| Ecm1 | 388.3138544 | 2.101987167 | 0.344759968 | 6.096958354 |
| Mir7014 | 0 | NA | NA | NA |
| Tars2 | 297.5330298 | -0.270853749 | 0.08938965 | -3.030034768 |
| Rprd2 | 1419.095113 | -0.131340518 | 0.05500125 | -2.387955132 |
| RP23-218K6.1 | 0.165206806 | 0.755296655 | 2.77541954 | 0.272137831 |
| Gm26361 | 3.436253382 | -0.979050645 | 0.5928808 | -1.651344832 |
| RP23-218K6.4 | 0.306564387 | 2.056164532 | 1.964695276 | 1.04655646 |
| Prpf3 | 300.0577214 | -0.036751891 | 0.115509521 | -0.318171963 |
| Gm26594 | 0.250472921 | 0.755298454 | 2.504502568 | 0.301576235 |
| Mrps21 | 159.8335192 | -0.287332753 | 0.122002578 | -2.355136738 |
| C920021L13Rik | 8.391545128 | 0.783198696 | 0.39191971 | 1.998365164 |
| Ciart | 209.0128952 | 0.203184293 | 0.191718463 | 1.059805559 |
| BC028528 | 36.09890605 | 1.072592115 | 0.205272714 | 5.225205495 |
| Aph1a | 300.5160635 | 0.335668102 | 0.103279893 | 3.250081801 |
| Car14 | 134.9063813 | 0.013353981 | 0.133862104 | 0.099759234 |
| Anp32e | 879.4154652 | -0.110198498 | 0.062083449 | -1.775006052 |
| RP23-49O17.4 | 9.354061232 | -0.807521776 | 0.390438944 | -2.068240858 |
| RP23-49O17.5 | 0 | NA | NA | NA |
| Plekho1 | 424.3660083 | 0.415591671 | 0.12354658 | 3.36384603 |
| Vps45 | 380.4768023 | -0.163210073 | 0.095476949 | -1.709418599 |
| RP23-261M19.2 | 0.092691418 | 0.274387304 | 3.724945049 | 0.073662108 |
| Otud7b | 1265.281876 | -0.066786516 | 0.061526018 | -1.085500382 |
| Mtmr11 | 145.3552089 | 0.657696156 | 0.160346725 | 4.101712437 |
| Gm17690 | 21.29907499 | -0.816169571 | 0.280564763 | -2.90902379 |
| Sf3b4 | 445.2184839 | 0.393514861 | 0.113902431 | 3.45484164 |
| Gm22027 | 1.636315489 | -0.243547969 | 0.73833187 | -0.329862463 |
| Sv2a | 5873.135983 | -0.17700626 | 0.040971618 | -4.32021654 |
| Bola1 | 50.41452498 | 0.017317574 | 0.181299977 | 0.095518899 |
| Gm15444 | 1.046383212 | -0.226836052 | 1.052175069 | -0.215587747 |
| Hist2h2ab | 34.08697731 | 0.124944157 | 0.297931442 | 0.419372175 |
| Hist2h2ac | 86.71087739 | 0.503260768 | 0.172031666 | 2.925396114 |
| AC125099.2 | 1.307878832 | 1.132172874 | 0.902061655 | 1.25509478 |
| Hist2h2be | 54.11456325 | -0.848962651 | 0.2124984 | -3.995148447 |
| Hist2h3c2 | 19.58774007 | 0.402968323 | 0.247293135 | 1.629516823 |
| Gm20632 | 14.19037399 | 0.888359437 | 0.286066747 | 3.105427126 |
| Hist2h2aa2 | 13.3101306 | 0.284582509 | 0.295611484 | 0.962690978 |
| AC125099.3 | 0 | NA | NA | NA |
| Gm20633 | 28.70123197 | 0.02610522 | 0.223447092 | 0.116829538 |
| Hist2h2aa1 | 0.135217741 | 1.264602945 | 3.618332936 | 0.349498779 |

| | | | | |
|---|---|---|---|---|
| Gm20634 | 1.134122097 | 1.309378124 | 1.192195548 | 1.098291406 |
| AC125099.1 | 0 | NA | NA | NA |
| Hist2h3c1 | 0 | NA | NA | NA |
| Gm20627 | 9.369992915 | -1.181052926 | 0.461523947 | -2.559028483 |
| Hist2h4 | 368.633472 | 0.050381547 | 0.155081733 | 0.32487093 |
| Gm38227 | 4.279540226 | 0.171993544 | 0.488409063 | 0.352150599 |
| Gm20628 | 12.09656833 | 0.189411616 | 0.329467945 | 0.5749015 |
| Hist2h3b | 50.50959981 | 0.809364044 | 0.289428273 | 2.796423568 |
| Hist2h2bb | 201.6384358 | 0.580594663 | 0.157047893 | 3.696927419 |
| Fcgr1 | 583.6067119 | 4.099562273 | 0.286691794 | 14.29954522 |
| Gm24830 | 0 | NA | NA | NA |
| Platr30 | 0 | NA | NA | NA |
| Gm26553 | 0 | NA | NA | NA |
| Rnu1b1 | 6.056136525 | 1.179821722 | 0.521965949 | 2.260342317 |
| Rnu1b2 | 0.421929236 | 1.11993233 | 1.431144884 | 0.782542943 |
| Gm17575 | 0 | NA | NA | NA |
| Gm26689 | 0 | NA | NA | NA |
| 4930477E14Rik | 0 | NA | NA | NA |
| Gm26654 | 2.969550979 | -2.623416683 | 0.998846976 | -2.626445037 |
| Pafah1b1-ps1 | 0.705160482 | 0.485019031 | 1.159820555 | 0.418184545 |
| Gm21960 | 0.551586351 | -0.502417049 | 1.33884183 | -0.375262438 |
| Terc | 7.890901767 | 0.513703473 | 0.501708063 | 1.023909145 |
| Gm24136 | 0 | NA | NA | NA |
| BC107364 | 0.047948698 | -0.206487837 | 3.724945049 | -0.055433794 |
| Gm26232 | 0 | NA | NA | NA |
| RP23-404J7.8 | 2.243072942 | 1.019451138 | 0.725714572 | 1.404754951 |
| RP23-404J7.10 | 0 | NA | NA | NA |
| Rnu1b6 | 0.046345709 | 0.274391604 | 3.724945049 | 0.073663262 |
| RP23-404J7.9 | 0 | NA | NA | NA |
| Gm28001 | 0 | NA | NA | NA |
| Gm25890 | 0 | NA | NA | NA |
| Gm22614 | 1.309217219 | 2.787176191 | 1.068208411 | 2.609206371 |
| Hfe2 | 3.500549162 | -1.194161586 | 0.531563469 | -2.246508002 |
| Gm15441 | 0.831325084 | 1.486422102 | 1.12134499 | 1.325570735 |
| Txnip | 698.8955186 | 2.351885351 | 0.18670065 | 12.59709248 |
| Gm16253 | 0.327232518 | 1.084434944 | 2.259148645 | 0.480019297 |
| Polr3gl | 127.7419174 | -0.235622463 | 0.124149111 | -1.897898912 |
| Ankrd34a | 497.5908006 | -0.540280296 | 0.107566686 | -5.022747443 |
| Lix1l | 406.2282066 | 0.20538773 | 0.151845722 | 1.352607942 |
| 6330549D23Rik | 6.560446392 | -0.110503498 | 0.45339828 | -0.2437228 |
| Rbm8a | 404.2810953 | 0.152164967 | 0.121891672 | 1.248362292 |
| Pex11b | 270.344613 | 0.055363943 | 0.076262656 | 0.725964005 |
| Itga10 | 23.71082201 | -0.597378051 | 0.281528599 | -2.121908936 |
| RP23-204O17.2 | 0.060174537 | 0.755286614 | 3.724945049 | 0.202764498 |
| Ankrd35 | 72.85459675 | -0.033393 | 0.170350035 | -0.196025787 |
| Pias3 | 236.294852 | 0.279759337 | 0.09902512 | 2.825135038 |
| Nudt17 | 21.37927619 | -0.210759094 | 0.21412332 | -0.984288373 |

| | | | | |
|---|---|---|---|---|
| Polr3c | 240.883843 | 0.17229405 | 0.095079003 | 1.812114615 |
| Gm22581 | 3.657530093 | 0.125517071 | 0.570206078 | 0.220125804 |
| RP23-204O17.4 | 9.840503469 | 0.366123839 | 0.353884395 | 1.034585996 |
| Rnf115 | 868.8846253 | 0.679532436 | 0.096871431 | 7.014786826 |
| RP23-395B13.2 | 10.83277785 | 0.044077868 | 0.418216399 | 0.105394883 |
| Cd160 | 2.898004674 | 3.607450242 | 0.982072362 | 3.673303906 |
| Pdzk1 | 2.857891658 | -0.273966925 | 0.743186274 | -0.368638301 |
| 4930442L01Rik | 0.150086409 | 1.264824188 | 3.723011833 | 0.339731445 |
| Gpr89 | 501.075764 | -0.183623054 | 0.060909235 | -3.014699699 |
| Gja5 | 5.28099507 | -0.540409431 | 0.521196054 | -1.036864011 |
| Gja8 | 0 | NA | NA | NA |
| RP24-86K23.1 | 0 | NA | NA | NA |
| RP23-97A19.1 | 0.188929682 | 0.755307772 | 2.260134566 | 0.334187081 |
| RP23-97A19.2 | 0.09349418 | 0.274402725 | 3.724945049 | 0.073666248 |
| Gm24435 | 0 | NA | NA | NA |
| Acp6 | 115.9459266 | -0.663192189 | 0.181897666 | -3.64596316 |
| Bcl9 | 929.9899206 | 0.180050388 | 0.060537103 | 2.974215452 |
| RP23-394K8.5 | 0.094831987 | 0.274404949 | 3.724945049 | 0.073666845 |
| RP23-394K8.3 | 12.87106279 | -0.677968615 | 0.342313663 | -1.980547925 |
| Olfr1402 | 0 | NA | NA | NA |
| RP23-290C4.1 | 0 | NA | NA | NA |
| Chd1l | 117.2581915 | 0.107077935 | 0.134466081 | 0.796319294 |
| RP24-316D12.2 | 2.928805847 | -1.778942497 | 0.718497483 | -2.475920291 |
| Fmo5 | 26.93201772 | -0.751642113 | 0.277400942 | -2.709587453 |
| Prkab2 | 567.66263 | -0.027904307 | 0.12676565 | -0.220125145 |
| Gm17651 | 0 | NA | NA | NA |
| Pde4dip | 3395.191701 | -0.343346997 | 0.093543581 | -3.670449589 |
| Mir7225 | 0 | NA | NA | NA |
| RP24-124P9.3 | 0.671139028 | -0.357374547 | 1.135685085 | -0.314677503 |
| Gm15999 | 0.08486385 | 0.755286617 | 3.724945049 | 0.202764499 |
| Sec22b | 837.8239496 | 0.203351708 | 0.06303513 | 3.226006024 |
| Gm5544 | 1.267397472 | -0.689274944 | 1.270463576 | -0.542538139 |
| Notch2 | 1217.583416 | 0.459775675 | 0.134053987 | 3.429779929 |
| RP23-97M20.1 | 25.75372241 | -0.224270333 | 0.274235169 | -0.817802961 |
| RP23-3C4.2 | 11.28696547 | -0.13556328 | 0.338338328 | -0.400673731 |
| Adam30 | 0.456191127 | 2.236343788 | 3.24747733 | 0.688640308 |
| RP23-3C4.5 | 0.046345709 | 0.274391604 | 3.724945049 | 0.073663262 |
| Reg4 | 0.197005158 | -0.687380037 | 3.279854003 | -0.209576413 |
| RP23-348H21.1 | 0 | NA | NA | NA |
| Hmgcs2 | 85.88327543 | 0.557110999 | 0.283692091 | 1.963787556 |
| RP23-348H21.2 | 2.607862967 | 0.660574065 | 0.73422457 | 0.899689403 |
| Phgdh | 330.7849704 | -0.276903014 | 0.183013318 | -1.513021113 |
| Zfp697 | 358.3939029 | -0.47959005 | 0.134913466 | -3.55479748 |
| Gm4450 | 0 | NA | NA | NA |
| RP23-381G13.3 | 0.131100043 | -1.037279138 | 3.663654906 | -0.283126868 |
| RP23-381G13.4 | 0 | NA | NA | NA |
| Hsd3b4 | 0 | NA | NA | NA |

| | | | | |
|---|---|---|---|---|
| RP23-381G13.6 | 0 | NA | NA | NA |
| Gm10681 | 0 | NA | NA | NA |
| RP24-285G1.2 | 0.109727584 | 0.755282166 | 3.724945049 | 0.202763304 |
| RP24-285G1.1 | 0 | NA | NA | NA |
| Hsd3b5 | 0 | NA | NA | NA |
| Gm12399 | 0 | NA | NA | NA |
| Gm12400 | 0 | NA | NA | NA |
| Hsd3b2 | 0.319559307 | -1.591777813 | 1.562192868 | -1.018938087 |
| Hsd3b3 | 0.048949532 | -0.206487837 | 3.724945049 | -0.055433794 |
| Hsd3b6 | 0 | NA | NA | NA |
| Gm12427 | 0 | NA | NA | NA |
| Gm12428 | 10.43905456 | -0.092437386 | 0.350046078 | -0.264072051 |
| Gm12431 | 0 | NA | NA | NA |
| Gm12432 | 0.046345709 | 0.274391604 | 3.724945049 | 0.073663262 |
| Gm12433 | 0.907786306 | 0.028836956 | 1.049885628 | 0.02746676 |
| Hsd3b1 | 0.244145947 | -0.677888742 | 1.840278149 | -0.36836211 |
| Hao2 | 0.110595523 | 0.755282166 | 3.724945049 | 0.202763304 |
| 5730437C11Rik | 0 | NA | NA | NA |
| Gm12440 | 0.266043621 | 1.404892413 | 2.452600267 | 0.572817524 |
| Gm12441 | 0 | NA | NA | NA |
| Gm22341 | 0 | NA | NA | NA |
| Gm24216 | 0 | NA | NA | NA |
| Wars2 | 117.2102137 | -0.416057714 | 0.154880062 | -2.68632198 |
| RP23-200E11.1 | 19.63971965 | -0.94596005 | 0.307863444 | -3.072661173 |
| RP23-200E11.3 | 0 | NA | NA | NA |
| Mir6481 | 0 | NA | NA | NA |
| RP23-200E11.2 | 15.98219274 | -0.696877428 | 0.331678583 | -2.101062488 |
| Tbx15 | 21.3154785 | 0.421545308 | 0.2937944 | 1.434830982 |
| RP23-200E11.4 | 0 | NA | NA | NA |
| Gm12449 | 0 | NA | NA | NA |
| M6pr-ps | 7.110781671 | 0.407483527 | 0.513857118 | 0.792989943 |
| Gm12448 | 0 | NA | NA | NA |
| Gm22505 | 0 | NA | NA | NA |
| Spag17 | 14.25668977 | 0.109644859 | 0.298852656 | 0.366886013 |
| Rpl36-ps7 | 0 | NA | NA | NA |
| Tra2b-ps | 0 | NA | NA | NA |
| Spag17os | 4.312461576 | -0.005241745 | 0.524069722 | -0.010002 |
| Wdr3 | 327.3352847 | -0.053235075 | 0.085039384 | -0.62600495 |
| Gdap2 | 404.7187721 | -0.064299227 | 0.058448668 | -1.100097375 |
| Gm25802 | 0 | NA | NA | NA |
| Gm12464 | 0 | NA | NA | NA |
| 4930406D18Rik | 0 | NA | NA | NA |
| RP23-335A19.3 | 0 | NA | NA | NA |
| Fam46c | 258.8118225 | 2.496926243 | 0.182045301 | 13.71596099 |
| Gm12474 | 1.5464145 | -0.091040358 | 0.911738254 | -0.099853612 |
| RP23-356K2.5 | 3.883316289 | 1.10894496 | 0.74559483 | 1.487329198 |
| Man1a2 | 2251.936784 | 0.066447804 | 0.066698637 | 0.996239311 |

| | | | | |
|---|---|---|---|---|
| RP23-356K2.6 | 73.44238526 | -0.296665371 | 0.210343675 | -1.410384083 |
| RP23-267O21.4 | 93.44924604 | 0.094746327 | 0.2090186 | 0.453291366 |
| RP23-267O21.3 | 152.3385727 | 0.074880868 | 0.241886163 | 0.309570698 |
| RP23-267O21.2 | 27.15683575 | -0.1466274 | 0.261804004 | -0.560065537 |
| Gm12485 | 0 | NA | NA | NA |
| RP23-267O21.5 | 2.75295346 | 2.915294954 | 1.019599863 | 2.859253968 |
| Gm23465 | 0 | NA | NA | NA |
| Vtcn1 | 9.162540417 | 1.973435512 | 0.675138884 | 2.923006743 |
| Trim45 | 89.1891917 | -0.247413999 | 0.128518524 | -1.925123258 |
| Ttf2 | 41.18065889 | 0.730788499 | 0.242339714 | 3.015554021 |
| Gm12488 | 0 | NA | NA | NA |
| Cd101 | 11.43243995 | 2.068817384 | 0.434928223 | 4.756686908 |
| Ptgfrn | 610.398054 | 0.192957254 | 0.091057112 | 2.119079438 |
| RP23-434L7.5 | 6.750937273 | 0.070394891 | 0.428090619 | 0.164439227 |
| Gm12486 | 0 | NA | NA | NA |
| Tpt1-ps1 | 0.068097359 | -0.206487837 | 3.724945049 | -0.055433794 |
| Gm12490 | 0.045882455 | 0.755288839 | 3.724945049 | 0.202765096 |
| Cd2 | 30.67976892 | 6.973959814 | 0.870206011 | 8.014148062 |
| Gm10355 | 2.030855555 | 0.841166481 | 0.72599795 | 1.158634789 |
| RP23-347G20.3 | 0 | NA | NA | NA |
| Igsf3 | 593.9020174 | -0.128932887 | 0.103137373 | -1.250108312 |
| RP23-347G20.4 | 4.3771345 | -1.108555292 | 0.551785844 | -2.009031772 |
| RP23-347G20.5 | 2.646385505 | -1.556898227 | 0.691691166 | -2.250857469 |
| RP23-347G20.6 | 0 | NA | NA | NA |
| Atp1a1 | 7569.239803 | -0.158051339 | 0.076554259 | -2.064566245 |
| RP23-103P11.2 | 13.65833158 | -1.019178373 | 0.346496855 | -2.941378424 |
| RP23-103P11.1 | 32.33019698 | -0.69812776 | 0.319301524 | -2.186421635 |
| RP24-115G23.1 | 0.049553047 | 0.274391604 | 3.724945049 | 0.073663262 |
| RP24-115G23.2 | 0.117046891 | -0.687371727 | 3.724945049 | -0.184532045 |
| Mab21l3 | 2.984303933 | 1.243358243 | 0.732021809 | 1.698526229 |
| Slc22a15 | 163.9002955 | -0.111363602 | 0.117113091 | -0.950906522 |
| Nhlh2 | 53.60793837 | -0.224951775 | 0.181291007 | -1.240832513 |
| RP23-226J22.2 | 0 | NA | NA | NA |
| RP23-226J22.3 | 0 | NA | NA | NA |
| Casq2 | 54.88658604 | -0.091126515 | 0.218434597 | -0.41717986 |
| RP24-89C11.1 | 0.931900496 | -1.303177688 | 1.018300481 | -1.27975751 |
| RP24-89C11.2 | 3.004992861 | -0.690310674 | 0.614799667 | -1.122822133 |
| Vangl1 | 127.7913766 | -0.172989436 | 0.11793097 | -1.466870289 |
| Gm16160 | 0.666323006 | -2.14402484 | 1.222842343 | -1.753312561 |
| RP23-191N23.1 | 0.144846928 | -0.990864828 | 3.528682026 | -0.280803093 |
| RP23-63A19.1 | 24.83821061 | -1.127013664 | 0.230282746 | -4.894043002 |
| RP24-409P22.1 | 3.653697379 | -0.571025043 | 0.529065954 | -1.07930786 |
| RP24-409P22.2 | 0 | NA | NA | NA |
| Ngf | 23.55789567 | 0.385447073 | 0.248624872 | 1.550315821 |
| RP23-63A19.3 | 5.480294069 | 0.119805113 | 0.508880859 | 0.23542861 |
| RP24-89D17.1 | 0.141094079 | 0.755285157 | 3.563850454 | 0.211929531 |
| RP24-89D17.3 | 0.352483205 | 0.752599922 | 1.732766114 | 0.434334395 |

| | | | | |
|---|---|---|---|---|
| RP24-89D17.2 | 0 | NA | NA | NA |
| RP23-183L19.1 | 0.184015763 | 0.755289308 | 3.314398464 | 0.227881263 |
| Tspan2os | 4.207560025 | 0.363805437 | 0.470662985 | 0.772963773 |
| Tspan2 | 1566.869316 | -0.428818806 | 0.0971256 | -4.415095559 |
| Tshb | 0.532135551 | -0.155209013 | 1.431607969 | -0.108415862 |
| RP23-183L19.3 | 0.133946075 | 0.755288219 | 3.626642455 | 0.20826101 |
| Sycp1 | 43.46231598 | 0.600903114 | 0.320790159 | 1.873196847 |
| RP23-183L19.5 | 0 | NA | NA | NA |
| RP23-183L19.6 | 0 | NA | NA | NA |
| RP23-119F18.1 | 0 | NA | NA | NA |
| Nr1h5 | 0.37220402 | 2.363008873 | 1.481180103 | 1.595355534 |
| Gm22826 | 0.042716945 | -0.206490061 | 3.724945049 | -0.055434391 |
| Sike1 | 543.965291 | -0.488169633 | 0.075005401 | -6.508459711 |
| Csde1 | 5133.717668 | 0.448349459 | 0.076548316 | 5.857078014 |
| Nras | 746.5356171 | 0.167797694 | 0.091858325 | 1.826700984 |
| RP23-106G24.2 | 13.15425346 | -0.375529229 | 0.310167746 | -1.210729464 |
| Ampd1 | 2.035201819 | 1.962684187 | 1.086583276 | 1.806289707 |
| Gm23820 | 0 | NA | NA | NA |
| RP24-409D4.3 | 0.140443404 | 0.755293356 | 2.913153115 | 0.259270051 |
| Dennd2c | 11.7142143 | 0.558646307 | 0.35054765 | 1.593638717 |
| Bcas2 | 429.1205886 | -0.115209433 | 0.06465368 | -1.781947021 |
| Gm27024 | 0 | NA | NA | NA |
| 2410024N13Rik | 0 | NA | NA | NA |
| RP23-70G12.2 | 0 | NA | NA | NA |
| Trim33 | 1476.330058 | -0.090988969 | 0.062456053 | -1.456847888 |
| RP23-70G12.3 | 47.1675988 | -0.81853328 | 0.348077792 | -2.351581457 |
| Gm25009 | 0 | NA | NA | NA |
| Gm25869 | 2.797214178 | 0.509157359 | 0.588924817 | 0.864554089 |
| Gm25199 | 0 | NA | NA | NA |
| Syt6 | 477.8102627 | -0.385916579 | 0.174937912 | -2.206020269 |
| RP23-184N6.1 | 10.82643969 | -1.504445565 | 0.461772456 | -3.257980298 |
| Atg4a-ps | 7.430379725 | 0.623813542 | 0.369270279 | 1.689314243 |
| Gm25325 | 0 | NA | NA | NA |
| Gm15986 | 0 | NA | NA | NA |
| Olfml3 | 289.0413337 | 1.05812567 | 0.163122786 | 6.486682185 |
| Gm10964 | 0 | NA | NA | NA |
| Hipk1 | 3690.970568 | 0.089221066 | 0.064111521 | 1.391654189 |
| RP24-559G13.2 | 18.01004 | -0.277922929 | 0.244417877 | -1.137081022 |
| Gm15886 | 0.872687401 | 0.757974793 | 1.155101968 | 0.656197301 |
| Gm16520 | 20.13453085 | -0.636004834 | 0.314261152 | -2.023809911 |
| Dclre1b | 139.879516 | 0.327572412 | 0.101334955 | 3.232570758 |
| Gm15471 | 0.37827039 | -0.557678123 | 1.563615609 | -0.356659348 |
| Ap4b1 | 32.06009994 | 0.066103785 | 0.206349694 | 0.320348355 |
| RP24-180F7.1 | 1.345434513 | -1.056287611 | 0.891306627 | -1.185100143 |
| Bcl2l15 | 59.73746658 | -0.135507333 | 0.20965846 | -0.646324181 |
| Ptpn22 | 140.1073571 | 2.854626056 | 0.352927643 | 8.088417317 |
| Gm15472 | 7.402758977 | 1.332525981 | 0.629802769 | 2.115782983 |

| | | | | |
|---|---|---|---|---|
| RP24-180F7.2 | 12.50366051 | 0.812223402 | 0.409294314 | 1.98444829 |
| Gm15602 | 0.044544648 | 0.755286614 | 3.724945049 | 0.202764498 |
| Rsbn1 | 982.5522449 | 0.100061678 | 0.055416493 | 1.805629905 |
| Phtf1os | 24.52540072 | -0.28740868 | 0.211618018 | -1.358148433 |
| Phtf1 | 538.6159203 | -0.402579824 | 0.097874163 | -4.113239004 |
| Magi3 | 898.9838019 | -0.251911767 | 0.086921654 | -2.898147411 |
| RP24-535M14.2 | 5.341278601 | -0.697581129 | 0.424307773 | -1.644045134 |
| RP24-535M14.1 | 11.88755228 | -0.359872884 | 0.303731301 | -1.184839636 |
| Gm23830 | 0 | NA | NA | NA |
| RP23-314M6.1 | 7.461541511 | -0.804409647 | 0.394290538 | -2.040144436 |
| RP23-333I5.9 | 6.272582582 | -0.366735319 | 0.5636325 | -0.650663897 |
| RP23-333I5.8 | 0.141094079 | 0.755285157 | 3.563850454 | 0.211929531 |
| RP23-333I5.7 | 34.16158652 | -0.417408511 | 0.310495976 | -1.344328246 |
| RP23-333I5.6 | 17.9477799 | -0.565358991 | 0.270210362 | -2.092292048 |
| RP23-333I5.5 | 22.39591524 | -0.619310511 | 0.352186843 | -1.758471458 |
| RP23-333I5.4 | 10.78090146 | -0.351618803 | 0.309038141 | -1.13778449 |
| RP23-333I5.3 | 57.87337 | -0.594720257 | 0.254419775 | -2.337555158 |
| RP23-333I5.2 | 44.8221039 | -0.431489137 | 0.29108209 | -1.482362368 |
| RP23-333I5.1 | 53.3220374 | -0.578009193 | 0.263425021 | -2.19420764 |
| Gm23077 | 0 | NA | NA | NA |
| Gm5546 | 0.706390846 | -0.55610551 | 1.126141754 | -0.49381484 |
| Lrig2 | 847.580948 | -0.12387764 | 0.064486468 | -1.920986596 |
| RP23-357D17.5 | 0.044191549 | -0.206490061 | 3.724945049 | -0.055434391 |
| RP23-357D17.6 | 0.315700266 | 0.782004898 | 1.784311128 | 0.43826712 |
| RP23-357D17.4 | 84.69226509 | -0.43355442 | 0.267750363 | -1.619248674 |
| RP23-357D17.3 | 51.88363519 | -0.281436228 | 0.250497328 | -1.123509901 |
| RP23-38L11.4 | 32.89618772 | -0.320395915 | 0.206710396 | -1.549974854 |
| Gm26091 | 0.643663771 | 0.056010453 | 1.217131845 | 0.046018394 |
| RP23-38L11.3 | 7.967952249 | -0.341516936 | 0.431194573 | -0.792025127 |
| Slc16a1 | 692.2362281 | 0.435989113 | 0.113469842 | 3.842334729 |
| RP23-38L11.1 | 26.81542658 | -0.167556156 | 0.279726513 | -0.598999908 |
| RP23-38L11.2 | 8.709900708 | -0.05828985 | 0.423760677 | -0.137553702 |
| Gm29561 | 0 | NA | NA | NA |
| Gm29560 | 0.214789954 | -0.206486577 | 2.566334377 | -0.080459732 |
| Gm24223 | 0 | NA | NA | NA |
| Gm29572 | 0 | NA | NA | NA |
| Gm6485 | 49.19972077 | 1.345114424 | 0.21660924 | 6.209866306 |
| RP24-503L4.2 | 0.134880151 | 0.274398319 | 2.937637886 | 0.093407809 |
| Fam19a3 | 6.79823111 | -0.758447821 | 0.50702646 | -1.495874241 |
| Ppm1j | 22.89084466 | 0.130575113 | 0.371002546 | 0.351952068 |
| Rhoc | 431.5423868 | 2.171752628 | 0.191761871 | 11.3252578 |
| Mov10 | 458.979696 | 1.884920226 | 0.149804203 | 12.58255904 |
| Capza1 | 545.1140594 | 0.358939935 | 0.093247355 | 3.849331013 |
| RP24-503L4.4 | 28.08267805 | 0.267701362 | 0.325339261 | 0.822837554 |
| RP24-503L4.3 | 42.80149299 | 0.417823757 | 0.335253533 | 1.246291883 |
| St7l | 574.3995767 | -0.186269855 | 0.067267814 | -2.769078449 |
| RP24-274H19.1 | 0.085327104 | 0.274389469 | 3.724945049 | 0.073662689 |

| | | | | |
|---|---|---|---|---|
| Wnt2b | 46.92012226 | 0.201805045 | 0.255941679 | 0.788480585 |
| RP24-274H19.3 | 0.479138389 | 0.542637634 | 1.312386877 | 0.413473834 |
| RP24-274H19.5 | 3.753741655 | -0.414786482 | 0.562435241 | -0.737483095 |
| Cttnbp2nl | 873.2904401 | 0.705577645 | 0.082376481 | 8.565280233 |
| 4930564D02Rik | 0.045882455 | 0.755288839 | 3.724945049 | 0.202765096 |
| Kcnd3os | 14.28524435 | -1.410808079 | 0.377043082 | -3.741768907 |
| Kcnd3 | 1457.767906 | -0.03254161 | 0.068453888 | -0.475380007 |
| RP23-443O1.1 | 42.61810582 | -0.906681841 | 0.248495172 | -3.648689972 |
| Ddx20 | 175.0365538 | 0.256252186 | 0.107758984 | 2.378012262 |
| Fam212b | 966.0315667 | 0.276254532 | 0.17370828 | 1.590336008 |
| RP24-500I24.6 | 43.0223173 | 0.157799331 | 0.202233168 | 0.780284127 |
| Rap1a | 552.0763281 | 0.15506437 | 0.10741429 | 1.443610249 |
| RP24-500I24.4 | 37.31668493 | -0.160716666 | 0.269640311 | -0.596040948 |
| RP24-500I24.3 | 48.64596038 | -0.319386518 | 0.253064986 | -1.262073124 |
| RP24-500I24.2 | 32.60373694 | -0.479399453 | 0.257729416 | -1.860088228 |
| RP24-500I24.5 | 33.83971767 | -0.361049047 | 0.268143328 | -1.346477832 |
| RP24-500I24.7 | 2.098595915 | 0.029100739 | 0.693603735 | 0.041955857 |
| Adora3 | 9.886871762 | 1.746437855 | 0.40630562 | 4.298335461 |
| Tmigd3 | 0.650450437 | 1.786601743 | 1.401654464 | 1.274637786 |
| I830077J02Rik | 45.37358814 | 1.805499337 | 0.299038752 | 6.037676796 |
| Atp5f1 | 2177.75384 | 0.07227625 | 0.084376694 | 0.856590214 |
| Wdr77 | 315.9051542 | -0.04065149 | 0.075388151 | -0.539229169 |
| RP23-169H9.3 | 8.021508243 | 0.57173237 | 0.416548995 | 1.372545312 |
| Ovgp1 | 35.01689641 | 0.13638927 | 0.295345467 | 0.461795711 |
| Pifo | 4.561976979 | -0.384118846 | 0.458911297 | -0.837021987 |
| RP23-169H9.6 | 8.449170836 | -0.536237692 | 0.503486159 | -1.06504952 |
| RP23-169H9.5 | 0 | NA | NA | NA |
| Chil5 | 6.28406316 | 0.659495508 | 0.476498781 | 1.384044482 |
| Gm4540 | 10.64549809 | -0.280752631 | 0.3134234 | -0.895761549 |
| RP23-169H9.8 | 0 | NA | NA | NA |
| RP23-62K10.1 | 0.969048608 | 0.603407697 | 0.934783127 | 0.64550555 |
| Chia1 | 8.605265352 | -0.198470112 | 0.393084314 | -0.504904687 |
| Chil3 | 390.2692537 | 9.659027794 | 1.321503263 | 7.309121413 |
| RP23-169H9.9 | 0 | NA | NA | NA |
| Chil4 | 0.092691418 | 0.274387304 | 3.724945049 | 0.073662108 |
| RP23-191E1.1 | 0.325695176 | 0.274407918 | 1.951119909 | 0.140641237 |
| RP23-191E1.2 | 0 | NA | NA | NA |
| Chil6 | 1.21043771 | 1.131593084 | 1.122950177 | 1.007696608 |
| Gm38244 | 0 | NA | NA | NA |
| 2010016I18Rik | 10.3751689 | 3.883991723 | 0.627817974 | 6.186493357 |
| Dennd2d | 26.9763545 | 2.739988774 | 0.315232907 | 8.691950336 |
| Cept1 | 592.7431419 | 0.314430391 | 0.116821398 | 2.69154791 |
| Dram2 | 391.8607469 | 0.534132597 | 0.113078271 | 4.723565305 |
| RP23-164N7.2 | 0 | NA | NA | NA |
| RP23-164N7.3 | 46.01383439 | 0.488843129 | 0.289410595 | 1.689098941 |
| Gm38271 | 1.062454588 | 1.856362338 | 1.065298624 | 1.742574614 |
| Lrif1 | 343.0611023 | -0.045621117 | 0.100262083 | -0.455018641 |

| | | | | |
|---|---|---|---|---|
| Gm38253 | 70.59424867 | -0.157645591 | 0.230881079 | -0.682799958 |
| Cd53 | 595.4541458 | 3.078287267 | 0.29222281 | 10.53404172 |
| Olfr266 | 0.317967701 | 0.764895615 | 1.78239561 | 0.429139081 |
| RP24-481E4.4 | 1.491118976 | 2.289331216 | 0.887414424 | 2.579776882 |
| Gm27008 | 4.962650854 | -0.470385165 | 0.472677572 | -0.995150168 |
| Kcna3 | 229.4345153 | -0.060039296 | 0.098007833 | -0.612596915 |
| AI504432 | 766.1594696 | 0.027431905 | 0.093679911 | 0.29282591 |
| A930002I21Rik | 0.066050875 | 0.274391604 | 3.724945049 | 0.073663262 |
| Kcna2 | 5020.598395 | -0.256263682 | 0.073878981 | -3.468695378 |
| Kcna10 | 0.10503227 | 0.274389469 | 3.724945049 | 0.073662689 |
| Cym | 1.145893073 | 2.101800676 | 1.134717027 | 1.852268562 |
| A630076J17Rik | 0 | NA | NA | NA |
| Prok1 | 4.595448637 | 0.511101352 | 0.656658014 | 0.778337188 |
| Lamtor5 | 258.6284024 | 0.15178627 | 0.13728271 | 1.105647388 |
| Slc16a4 | 15.80845376 | -0.514971515 | 0.333213318 | -1.545470982 |
| Rbm15 | 884.0175483 | 0.085120172 | 0.067059078 | 1.269331084 |
| RP23-332I1.7 | 0.048949532 | -0.206487837 | 3.724945049 | -0.055433794 |
| Gm24383 | 0.089089296 | 1.194485492 | 3.723939576 | 0.320758559 |
| Kcnc4 | 618.4655694 | -0.186437983 | 0.08082264 | -2.306754433 |
| Slc6a17 | 6021.526622 | -0.265432796 | 0.110215335 | -2.408310941 |
| Ubl4b | 0.42233865 | 0.18764662 | 1.574606141 | 0.119170512 |
| Alx3 | 13.73531081 | 0.780256864 | 0.489164446 | 1.595080898 |
| Strip1 | 672.0896654 | -0.149188618 | 0.073257059 | -2.036508431 |
| Gm10961 | 0.150451996 | 0.274405555 | 2.858243839 | 0.096004949 |
| Ahcyl1 | 4251.746979 | -0.131759829 | 0.076767782 | -1.716342793 |
| Csf1 | 1357.448556 | 2.2729115 | 0.229472381 | 9.904945837 |
| RP23-455E11.2 | 0.180126469 | -1.298925849 | 3.33473513 | -0.389513949 |
| RP23-455E11.1 | 0.049553047 | 0.274391604 | 3.724945049 | 0.073663262 |
| RP23-313F7.2 | 1.373329198 | 0.139898521 | 0.83861036 | 0.166821838 |
| RP23-313F7.8 | 0 | NA | NA | NA |
| Eps8l3 | 0.290796633 | -1.073074919 | 1.58261049 | -0.678041076 |
| RP23-313F7.4 | 0 | NA | NA | NA |
| 4933431E20Rik | 826.6688943 | -0.661969622 | 0.118365362 | -5.592595775 |
| Gstm5 | 640.7472393 | -0.178514672 | 0.130333527 | -1.369675751 |
| RP23-313F7.6 | 0.038981395 | 0.274393828 | 3.724945049 | 0.073663859 |
| Gm12489 | 0 | NA | NA | NA |
| Gstm7 | 114.4264593 | -0.143996917 | 0.184375633 | -0.780997546 |
| Gstm6 | 23.14962032 | -0.078455421 | 0.257724891 | -0.304415382 |
| Gm12494 | 1.001132476 | -2.433969194 | 1.190537912 | -2.044428128 |
| Gstm3 | 0.425236713 | -0.206687862 | 1.71706217 | -0.120372964 |
| Gm12497 | 0 | NA | NA | NA |
| Gstm2 | 20.7015664 | 0.039164108 | 0.317945395 | 0.123178723 |
| RP23-313F7.7 | 0.044544648 | 0.755286614 | 3.724945049 | 0.202764498 |
| Gstm1 | 1044.038813 | -0.218847979 | 0.180826453 | -1.210265288 |
| Gm12498 | 0.194528622 | 0.755289046 | 3.242107212 | 0.232962391 |
| Gm12502 | 2.704774846 | 1.020734851 | 0.647901045 | 1.575448687 |
| Gm12499 | 0 | NA | NA | NA |

| | | | | |
|---|---|---|---|---|
| Gm25592 | 0 | NA | NA | NA |
| Gstm4 | 56.39273402 | 0.20479014 | 0.175923663 | 1.164085241 |
| Ampd2 | 616.2955637 | -0.452257037 | 0.071541584 | -6.321596628 |
| Gm12500 | 6.670440749 | -0.522576653 | 0.383311325 | -1.363321715 |
| Gnat2 | 1.625657436 | -0.013788269 | 0.944470025 | -0.014598948 |
| Gnai3 | 600.7292191 | 0.281874406 | 0.133117621 | 2.117483804 |
| Gpr61 | 149.8257791 | -0.419212213 | 0.125975552 | -3.327726728 |
| Gm12524 | 0.213105667 | -0.206489796 | 2.612901063 | -0.079027024 |
| Amigo1 | 1089.407811 | -0.195815391 | 0.067942788 | -2.882062915 |
| Cyb561d1 | 232.812123 | -0.016226961 | 0.080870479 | -0.200653707 |
| Atxn7l2 | 171.8889948 | -0.307932308 | 0.164404514 | -1.873016134 |
| Sypl2 | 13.67443026 | -1.575455693 | 0.299844231 | -5.254247141 |
| RP23-89M15.15 | 1.384376546 | -1.019620348 | 0.876029609 | -1.163910829 |
| Gm12501 | 0.378789762 | -1.212144686 | 1.639481725 | -0.739346263 |
| 1700010K24Rik | 1.755101273 | -1.859629809 | 0.961674792 | -1.933740828 |
| Psma5 | 407.5269465 | 0.359276001 | 0.121273275 | 2.962532362 |
| Gm12523 | 1.090677807 | -0.107677684 | 0.943918466 | -0.114075196 |
| Sort1 | 4425.028415 | -0.325391479 | 0.103789648 | -3.135105332 |
| RP23-27B23.9 | 51.99739385 | -0.667340208 | 0.378198653 | -1.764522962 |
| Gm12525 | 7.569909558 | -0.435968198 | 0.465762937 | -0.936030248 |
| Mybphl | 0.67227865 | -2.460849731 | 1.196717736 | -2.056332632 |
| Gm12522 | 10.72785558 | -0.148396854 | 0.34871835 | -0.425549314 |
| Psrc1 | 41.89133445 | 0.397953242 | 0.223306282 | 1.782096046 |
| Celsr2 | 4197.124515 | -0.432910655 | 0.12160751 | -3.559900645 |
| Sars | 1527.302422 | 0.171639373 | 0.082973111 | 2.068614399 |
| Gm22942 | 0 | NA | NA | NA |
| RP23-37G19.3 | 0.129625439 | -1.028499994 | 3.675612039 | -0.279817343 |
| 5330417C22Rik | 475.2428445 | -0.308303212 | 0.129632104 | -2.378293665 |
| Gm23336 | 0 | NA | NA | NA |
| Gm37859 | 19.39367035 | -1.327943474 | 0.480193471 | -2.765434257 |
| 1700013F07Rik | 7.727506971 | -0.427510972 | 0.37529807 | -1.139123821 |
| Scarna2 | 566.7431568 | -0.018806265 | 0.235886025 | -0.079726069 |
| RP23-128G10.7 | 0 | NA | NA | NA |
| Tmem167b | 537.5116706 | 0.138035112 | 0.066961731 | 2.061402966 |
| Gm22860 | 0.047948698 | -0.206487837 | 3.724945049 | -0.055433794 |
| Taf13 | 217.8213017 | 0.048582678 | 0.119190135 | 0.407606532 |
| Wdr47 | 1240.073746 | -0.177777955 | 0.073363902 | -2.423234708 |
| Gm12517 | 0 | NA | NA | NA |
| Clcc1 | 305.3641183 | 0.413474938 | 0.081631139 | 5.065161971 |
| Gpsm2 | 151.0149954 | -0.202003351 | 0.164171217 | -1.230443158 |
| Gm27244 | 0 | NA | NA | NA |
| Aknad1 | 0.537146556 | 0.763452123 | 1.348551768 | 0.566127413 |
| Stxbp3 | 425.8909719 | 0.654629384 | 0.110849105 | 5.905590151 |
| Fndc7 | 3.245667884 | 1.759743517 | 0.820511538 | 2.144690763 |
| RP23-316M11.8 | 0.10503227 | 0.274389469 | 3.724945049 | 0.073662689 |
| Prpf38b | 771.447708 | -0.011475718 | 0.11556272 | -0.099302941 |
| Henmt1 | 3.246767699 | -0.103756623 | 0.603356 | -0.171965842 |

| | | | | |
|---|---|---|---|---|
| Gm9857 | 0.597180582 | 1.155938386 | 1.263754343 | 0.914685985 |
| Fam102b | 647.6814876 | -0.42037936 | 0.111809256 | -3.759790335 |
| RP23-316M11.10 | 0 | NA | NA | NA |
| AL671921.1 | 0 | NA | NA | NA |
| RP23-288B16.2 | 0 | NA | NA | NA |
| Slc25a54 | 0 | NA | NA | NA |
| RP23-288B16.4 | 8.294058351 | -0.40505009 | 0.390100783 | -1.038321654 |
| Slc25a24 | 72.40842471 | 0.918745294 | 0.237704418 | 3.865074538 |
| Gm13865 | 0 | NA | NA | NA |
| Vav3 | 309.6866449 | 0.07045793 | 0.079823431 | 0.882672288 |
| Ntng1 | 634.8071448 | -0.396638848 | 0.093099508 | -4.260375354 |
| Gm25681 | 0 | NA | NA | NA |
| Gm12535 | 0 | NA | NA | NA |
| RP23-327M1.2 | 2.031725428 | 2.06973701 | 0.837492726 | 2.471349238 |
| Prmt6 | 217.8360219 | 0.359597597 | 0.104263408 | 3.44893385 |
| RP23-327M1.1 | 30.13062775 | 0.33876879 | 0.222385799 | 1.523338234 |
| RP23-440G4.1 | 0 | NA | NA | NA |
| RP23-440G4.2 | 0.155610039 | 1.236192275 | 2.828260953 | 0.437085649 |
| RP23-43A4.1 | 0 | NA | NA | NA |
| Gm26076 | 0 | NA | NA | NA |
| RP24-366N1.1 | 0.538110248 | 0.739835099 | 1.629401912 | 0.454053167 |
| RP24-223F21.1 | 0.075043205 | 0.755286614 | 3.724945049 | 0.202764498 |
| RP24-230D18.1 | 0 | NA | NA | NA |
| RP23-14C8.1 | 0 | NA | NA | NA |
| RP23-14C8.2 | 0 | NA | NA | NA |
| RP24-93G19.1 | 0.12092566 | 1.236179401 | 3.724945049 | 0.331865138 |
| Gm25519 | 0 | NA | NA | NA |
| RP24-325G15.1 | 0 | NA | NA | NA |
| RP24-325G15.2 | 0.39879887 | 0.277529098 | 1.777004156 | 0.156178081 |
| RP23-117K10.1 | 0 | NA | NA | NA |
| RP24-221P14.1 | 0 | NA | NA | NA |
| RP23-430D14.1 | 5.464530712 | -0.063632014 | 0.483749662 | -0.13153914 |
| RP24-141M5.1 | 0 | NA | NA | NA |
| Gm5548 | 0.097139547 | 0.274393828 | 3.724945049 | 0.073663859 |
| RP23-470O13.1 | 0 | NA | NA | NA |
| Amy2b | 0.038981395 | 0.274393828 | 3.724945049 | 0.073663859 |
| Amy2-ps1 | 0 | NA | NA | NA |
| Amy2a5 | 0.143022002 | 0.755289479 | 3.547809333 | 0.212888971 |
| Amy2a4 | 0.089089296 | 1.194485492 | 3.723939576 | 0.320758559 |
| Amy2a3 | 0.060174537 | 0.755286614 | 3.724945049 | 0.202764498 |
| Amy2a2 | 0 | NA | NA | NA |
| Amy2a1 | 0.175778459 | 0.755296102 | 2.7274238 | 0.276926564 |
| Amy1 | 192.4654876 | 0.230999229 | 0.185123319 | 1.247812703 |
| Rnpc3 | 532.6768979 | 0.126317277 | 0.08199118 | 1.540620312 |
| RP24-237D17.3 | 0 | NA | NA | NA |
| RP24-182K13.1 | 0.088383099 | -0.687376175 | 3.724945049 | -0.184533239 |
| RP23-256J12.1 | 0.075043205 | 0.755286614 | 3.724945049 | 0.202764498 |

| | | | | |
|---|---|---|---|---|
| Col11a1 | 184.655142 | -0.368746786 | 0.151661095 | -2.431386814 |
| RP23-281B17.2 | 0 | NA | NA | NA |
| Gm23750 | 0 | NA | NA | NA |
| RP23-292P14.1 | 0 | NA | NA | NA |
| Olfm3 | 941.997379 | -0.37046177 | 0.146996895 | -2.520201338 |
| RP23-231J6.1 | 0.046345709 | 0.274391604 | 3.724945049 | 0.073663262 |
| Gm22337 | 0 | NA | NA | NA |
| Gm25347 | 0 | NA | NA | NA |
| RP23-238A12.1 | 0.12092566 | 1.236179401 | 3.724945049 | 0.331865138 |
| S1pr1 | 1328.879257 | 0.213360865 | 0.099093926 | 2.153117483 |
| Gm9889 | 3.625155807 | 1.511136511 | 0.554103197 | 2.727175226 |
| A930005H10Rik | 72.61233888 | -0.402844083 | 0.134440883 | -2.996440321 |
| Dph5 | 126.6011858 | 0.272748141 | 0.123145247 | 2.214849121 |
| RP24-155C13.8 | 0.199959441 | 0.755303988 | 2.66342588 | 0.283583633 |
| RP24-155C13.6 | 0.510666263 | -0.453115318 | 1.282058276 | -0.35342802 |
| RP24-155C13.7 | 0.595761752 | 1.69618938 | 1.279970611 | 1.32517838 |
| Slc30a7 | 561.2209546 | 0.232666945 | 0.088525455 | 2.628249074 |
| RP24-245I16.1 | 1.76977757 | 0.502594076 | 0.85267063 | 0.589435192 |
| RP24-155C13.3 | 4.122231334 | 0.774512114 | 0.521675502 | 1.484662614 |
| Mir669n | 0 | NA | NA | NA |
| Extl2 | 975.1969821 | 0.122255948 | 0.079228796 | 1.543074675 |
| Gm26344 | 0 | NA | NA | NA |
| Vcam1 | 1522.902751 | 2.324392717 | 0.215886324 | 10.76674371 |
| RP24-245I16.3 | 1.059433203 | -0.93034777 | 1.059725963 | -0.877913539 |
| RP23-93F11.2 | 1.275102148 | 2.712160226 | 1.084289072 | 2.501325795 |
| Gm26544 | 0.239854152 | 0.802056892 | 2.524148309 | 0.317753473 |
| RP23-93F11.4 | 0.265934997 | 0.296439582 | 2.091457959 | 0.141738246 |
| Gpr88 | 1159.493482 | -0.154884984 | 0.15388746 | -1.006482166 |
| Cdc14a | 108.7934714 | 0.201341888 | 0.132574178 | 1.518711186 |
| Gm29151 | 0 | NA | NA | NA |
| Gm22361 | 0 | NA | NA | NA |
| Gm26358 | 0 | NA | NA | NA |
| Rtca | 414.5859796 | 0.383889339 | 0.110664679 | 3.468941871 |
| Gm26530 | 0.207691825 | -0.206489716 | 2.623795641 | -0.078698856 |
| Dbt | 216.6118277 | 0.205096101 | 0.102593096 | 1.999121866 |
| Gm4596 | 16.93547575 | 0.355333035 | 0.277101651 | 1.282320166 |
| RP23-453M11.3 | 3.08271666 | 0.056065185 | 0.516412692 | 0.108566628 |
| Gm9761 | 55.52035785 | 0.231145813 | 0.160010719 | 1.444564554 |
| Lrrc39 | 5.281818806 | -0.601829875 | 0.538584708 | -1.117428448 |
| RP23-453M11.4 | 10.63896767 | 0.783628465 | 0.438862122 | 1.785591477 |
| Trmt13 | 134.1213703 | -0.029331497 | 0.133051695 | -0.220451885 |
| Sass6 | 170.9486267 | 0.047004311 | 0.117684008 | 0.399411199 |
| RP24-565B3.4 | 8.173995324 | 0.004569597 | 0.347577244 | 0.013146997 |
| Hiat1 | 617.2974484 | -0.10883837 | 0.066153375 | -1.645242892 |
| RP24-565B3.2 | 6.147691891 | -0.630261245 | 0.490922067 | -1.283831566 |
| RP24-565B3.5 | 0 | NA | NA | NA |
| RP24-565B3.3 | 0.186953752 | 0.274409582 | 2.668894796 | 0.102817684 |

| | | | | |
|---|---|---|---|---|
| Slc35a3 | 453.1222322 | 0.111759873 | 0.084069667 | 1.32937213 |
| Agl | 702.330067 | -0.492799022 | 0.075620831 | -6.516709956 |
| RP24-423B3.1 | 0.424532791 | 1.632513078 | 1.428094367 | 1.143140899 |
| RP24-423B3.2 | 0.346647033 | -0.221355722 | 1.612172145 | -0.137302783 |
| RP24-539I8.1 | 0 | NA | NA | NA |
| CH25-626J19.1 | 0.097139547 | 0.274393828 | 3.724945049 | 0.073663859 |
| Frrs1 | 66.10065566 | 1.541096519 | 0.231440643 | 6.658711719 |
| Palmd | 159.5277226 | -0.062761885 | 0.163071503 | -0.384873409 |
| RP24-370B22.3 | 26.73605718 | -0.450879477 | 0.355626833 | -1.267844367 |
| 4930455H04Rik | 0.11657011 | 1.236179401 | 3.724945049 | 0.331865138 |
| RP24-341P14.2 | 0 | NA | NA | NA |
| 1700061I17Rik | 0 | NA | NA | NA |
| RP24-83G17.1 | 0.038981395 | 0.274393828 | 3.724945049 | 0.073663859 |
| RP24-83G17.2 | 0 | NA | NA | NA |
| RP23-173L24.3 | 0.766169159 | 1.408074242 | 1.394785042 | 1.009527777 |
| RP23-173L24.2 | 1.114955615 | -0.18653141 | 1.387876008 | -0.13440063 |
| Lppr4 | 2571.916126 | -0.27772661 | 0.136383716 | -2.036361942 |
| Gm22408 | 0.596872081 | 0.298503516 | 1.496571583 | 0.199458228 |
| Lppr5 | 515.7492512 | -0.236868569 | 0.112730084 | -2.101201031 |
| Snx7 | 152.0630282 | 0.043322674 | 0.112653559 | 0.38456552 |
| Gm24811 | 0.346782673 | 0.168014165 | 1.536680854 | 0.109335757 |
| CH25-626J19.2 | 0 | NA | NA | NA |
| RP24-294N3.1 | 0 | NA | NA | NA |
| Gm26871 | 573.8481393 | 0.128899586 | 0.25741904 | 0.500738351 |
| Mir137 | 0 | NA | NA | NA |
| Gm24155 | 3.962088627 | 0.466938628 | 0.553802923 | 0.843149447 |
| Gm9916 | 100.2258943 | -0.043756595 | 0.387520173 | -0.112914367 |
| Gm37773 | 129.487562 | 0.214024001 | 0.169849088 | 1.260083308 |
| Gm37928 | 218.6973413 | 0.344011551 | 0.244465163 | 1.407200714 |
| Gm32444 | 100.5808754 | 0.025105205 | 0.190866997 | 0.131532455 |
| Dpyd | 164.1135791 | 0.764257714 | 0.117367871 | 6.511643333 |
| RP24-173A7.2 | 9.88657072 | 0.833360324 | 0.325887647 | 2.557201326 |
| Gm23432 | 0 | NA | NA | NA |
| RP23-235E15.1 | 152.0784296 | 0.518425448 | 0.147711083 | 3.509726141 |
| Ptbp2 | 2043.522665 | 0.298768636 | 0.117656981 | 2.539319239 |
| RP23-235E15.5 | 196.3079525 | 0.675176175 | 0.258245973 | 2.614469321 |
| RP23-235E15.6 | 474.5268742 | 0.631504365 | 0.210509747 | 2.999881835 |
| RP23-235E15.7 | 0 | NA | NA | NA |
| Gm24744 | 0 | NA | NA | NA |
| Gm26137 | 0 | NA | NA | NA |
| RP23-427K12.1 | 0 | NA | NA | NA |
| Gm23733 | 0 | NA | NA | NA |
| 6530403H02Rik | 55.37544644 | -0.147876273 | 0.190685799 | -0.775497041 |
| Gm25013 | 0.075043205 | 0.755286614 | 3.724945049 | 0.202764498 |
| RP23-324P1.1 | 0.575937436 | 0.609042184 | 1.18549148 | 0.513746572 |
| RP24-295P14.2 | 0.374268822 | -1.936076896 | 3.688737335 | -0.524861686 |
| RP24-295P14.1 | 0.066050875 | 0.274391604 | 3.724945049 | 0.073663262 |

| | | | | |
|---|---|---|---|---|
| Rwdd3 | 95.46200367 | -0.139720545 | 0.192514949 | -0.725764653 |
| RP24-210J6.4 | 0.060174537 | 0.755286614 | 3.724945049 | 0.202764498 |
| Gm5710 | 0 | NA | NA | NA |
| Tmem56 | 921.2771509 | -0.35410218 | 0.071960881 | -4.920759353 |
| RP24-291L17.1 | 8.847422939 | -0.619805172 | 0.454119398 | -1.364850686 |
| RP24-291L17.2 | 0.233612452 | 1.697742765 | 2.527723224 | 0.671648996 |
| Alg14 | 122.1469618 | 0.086987173 | 0.126813547 | 0.685945427 |
| Gm26027 | 1.314748965 | -0.491076628 | 0.995567471 | -0.49326303 |
| RP24-277J22.6 | 0 | NA | NA | NA |
| RP24-277J22.4 | 33.98944694 | 3.193072461 | 0.501474212 | 6.367371215 |
| Cnn3 | 2221.060378 | 1.427068073 | 0.170379297 | 8.375830285 |
| RP24-277J22.3 | 26.71658222 | 0.702252687 | 0.337362335 | 2.081597777 |
| Slc44a3 | 22.72182752 | 3.531421872 | 0.541833968 | 6.517535038 |
| A530020G20Rik | 0.979355264 | 0.638588591 | 1.199217978 | 0.532504184 |
| A730020M07Rik | 12.26765879 | 0.638886986 | 0.330434151 | 1.933477473 |
| 4930432M17Rik | 0 | NA | NA | NA |
| RP24-404P10.3 | 0.136120942 | 0.274393632 | 3.599844381 | 0.076223748 |
| RP24-404P10.4 | 0 | NA | NA | NA |
| RP24-404P10.2 | 5.861481922 | 0.539684595 | 0.399495262 | 1.350916134 |
| F3 | 702.2491587 | 0.769498695 | 0.150923348 | 5.098606062 |
| RP24-404P10.8 | 9.75452295 | -0.964941419 | 0.357464752 | -2.699402985 |
| Abcd3 | 888.7244778 | -0.178686932 | 0.085151334 | -2.098463096 |
| RP24-404P10.7 | 19.74428529 | -0.805542127 | 0.248315913 | -3.244021361 |
| RP24-404P10.6 | 21.98336171 | -0.408760481 | 0.306555387 | -1.333398458 |
| RP24-404P10.9 | 0.234491734 | 0.75530354 | 2.143011043 | 0.352449672 |
| RP23-54L16.1 | 0 | NA | NA | NA |
| Arhgap29 | 378.6264762 | 0.261335078 | 0.083417398 | 3.132860583 |
| RP23-54L16.2 | 5.712404952 | -0.553982006 | 0.378777009 | -1.462554466 |
| Abca4 | 76.95354951 | -0.188892526 | 0.183627195 | -1.028674029 |
| RP24-471B15.3 | 0.334549462 | 1.330665492 | 1.64163359 | 0.810573992 |
| RP24-471B15.2 | 0 | NA | NA | NA |
| RP24-471B15.1 | 0.04393175 | -0.206490061 | 3.724945049 | -0.055434391 |
| Gm4609 | 10.08219113 | 0.703431955 | 0.325136058 | 2.163500288 |
| Gm22903 | 0.085433889 | -0.687376175 | 3.724945049 | -0.184533239 |
| Gclm | 1174.677346 | 0.411127852 | 0.105433829 | 3.899392229 |
| Gm17494 | 67.68281203 | -0.409758446 | 0.274887708 | -1.49063939 |
| Gm6061 | 18.81787436 | 0.809104568 | 0.241948718 | 3.344115957 |
| Dnttip2 | 377.7972389 | 0.565076896 | 0.119540762 | 4.727064522 |
| RP23-71G16.7 | 1.463879734 | -1.593450618 | 0.941496916 | -1.692465042 |
| Mir760 | 0 | NA | NA | NA |
| Bcar3 | 197.0241314 | 0.420206247 | 0.122666001 | 3.425612996 |
| RP23-71G16.6 | 1.51534681 | 2.959049654 | 1.050995185 | 2.815474034 |
| RP23-192O12.4 | 0.11657011 | 1.236179401 | 3.724945049 | 0.331865138 |
| Gm25153 | 0.892441177 | -1.12465106 | 1.044672096 | -1.076558917 |
| Fnbp1l | 1523.630982 | -0.408689592 | 0.101325155 | -4.033446519 |
| mmu-mir-7657 | 0 | NA | NA | NA |
| RP23-1A12.4 | 0.048949532 | -0.206487837 | 3.724945049 | -0.055433794 |

| | | | | |
|---|---|---|---|---|
| RP23-1A12.2 | 29.85261811 | -0.621725173 | 0.274741846 | -2.262943127 |
| RP23-1A12.5 | 0.187986659 | -0.206495442 | 3.315974774 | -0.062272923 |
| RP23-1A12.1 | 0.044544648 | 0.755286614 | 3.724945049 | 0.202764498 |
| Pde5a | 353.2225523 | -0.439600543 | 0.165686857 | -2.653201052 |
| RP24-182A8.1 | 4.735351793 | -0.458116666 | 0.461196716 | -0.993321615 |
| RP24-182A8.2 | 0.332478248 | -0.821146155 | 1.940250675 | -0.423216528 |
| RP24-70L3.3 | 0.663651821 | 0.657882009 | 1.253157403 | 0.52497955 |
| 4933405D12Rik | 1.378174623 | -0.404068186 | 0.924248975 | -0.43718543 |
| Fabp2 | 0 | NA | NA | NA |
| Gm10959 | 0.205306308 | 1.089387126 | 2.603665196 | 0.418405226 |
| 1810037I17Rik | 393.3493054 | 0.226371706 | 0.135603607 | 1.669363451 |
| Usp53 | 810.7985606 | 0.389215877 | 0.114069016 | 3.412108666 |
| RP23-48L9.1 | 1.574470888 | -0.328656011 | 0.782774633 | -0.419860323 |
| Myoz2 | 0.617569467 | 1.71343114 | 1.297165853 | 1.320903674 |
| Gm22066 | 0.060174537 | 0.755286614 | 3.724945049 | 0.202764498 |
| RP23-48L9.2 | 0 | NA | NA | NA |
| Synpo2 | 278.5160077 | 0.010137349 | 0.131639209 | 0.077008582 |
| RP23-48L9.4 | 0.075043205 | 0.755286614 | 3.724945049 | 0.202764498 |
| RP23-48L9.3 | 2.800041622 | -0.550283953 | 0.570041413 | -0.965340307 |
| RP24-488I7.1 | 9.542608144 | -0.184526675 | 0.385735865 | -0.478375727 |
| RP24-488I7.2 | 12.44003729 | -0.317586857 | 0.361873093 | -0.877619428 |
| Sec24d | 210.6853567 | 0.702897825 | 0.133653963 | 5.259087049 |
| Mettl14 | 164.8105688 | -0.056828844 | 0.095661724 | -0.59406042 |
| Gm22713 | 0.066050875 | 0.274391604 | 3.724945049 | 0.073663262 |
| RP23-113O9.4 | 1.285708346 | 1.691479976 | 0.897622701 | 1.884399731 |
| Prss12 | 146.351687 | 0.007016015 | 0.15900452 | 0.044124626 |
| RP23-113O9.3 | 0.319644482 | 0.724122149 | 1.565815427 | 0.462456901 |
| Snhg8 | 70.39141616 | 0.310985816 | 0.340425145 | 0.913521873 |
| Snora24 | 0 | NA | NA | NA |
| RP24-149B7.2 | 0.066050875 | 0.274391604 | 3.724945049 | 0.073663262 |
| Ndst3 | 624.8561942 | -0.511079707 | 0.166354251 | -3.072237134 |
| RP24-374F22.1 | 20.67476562 | -0.910998326 | 0.290703655 | -3.133769768 |
| Gm24456 | 0 | NA | NA | NA |
| Gm15400 | 0.119587853 | 1.218392466 | 3.723618458 | 0.327206581 |
| Gm15399 | 0.060174537 | 0.755286614 | 3.724945049 | 0.202764498 |
| RP24-107N14.2 | 0 | NA | NA | NA |
| Gm9372 | 0.194279094 | 0.274391173 | 3.724945049 | 0.073663147 |
| RP24-107N14.4 | 0 | NA | NA | NA |
| Tram1l1 | 242.7416911 | -0.231230451 | 0.088957584 | -2.599333748 |
| Gm4617 | 19.97291991 | 1.211493707 | 0.357001388 | 3.393526606 |
| RP23-412L13.3 | 0 | NA | NA | NA |
| RP23-412L13.5 | 0 | NA | NA | NA |
| 1700006A11Rik | 0.255320825 | 2.173795215 | 2.379358213 | 0.913605696 |
| Gm22575 | 0.066050875 | 0.274391604 | 3.724945049 | 0.073663262 |
| RP23-412L13.4 | 0 | NA | NA | NA |
| Gm22900 | 0 | NA | NA | NA |
| 1700003H04Rik | 0 | NA | NA | NA |

| | | | | |
|---|---|---|---|---|
| RP24-332L11.2 | 0 | NA | NA | NA |
| RP23-24O21.1 | 0.192951104 | 0.802372537 | 3.260557261 | 0.24608448 |
| Gm23673 | 0.219094128 | 1.86792186 | 3.130988064 | 0.596591818 |
| Gm9381 | 0.088383099 | -0.687376175 | 3.724945049 | -0.184533239 |
| RP23-446O11.1 | 0 | NA | NA | NA |
| Ndst4 | 306.8135137 | -0.311462167 | 0.137478577 | -2.265532372 |
| Ugt8a | 1013.869147 | -0.969465427 | 0.224878802 | -4.311057411 |
| Gm26275 | 0.551648795 | -0.20424952 | 1.721431184 | -0.118650993 |
| RP24-297G3.1 | 0 | NA | NA | NA |
| RP23-301O12.1 | 0.159882028 | 0.274408585 | 2.804321407 | 0.097852045 |
| Arsj | 51.42050542 | 1.081912769 | 0.210228067 | 5.14637643 |
| Gm17225 | 0.384085174 | 1.177822688 | 3.287677935 | 0.358253671 |
| RP23-60D3.4 | 6.256257677 | 0.606544715 | 0.561320975 | 1.08056663 |
| RP23-60D3.3 | 21.83940584 | 0.736285349 | 0.33138663 | 2.221831792 |
| Camk2d | 2946.282952 | 0.162136838 | 0.049448853 | 3.278879653 |
| RP23-293K21.1 | 122.6972469 | -0.166386334 | 0.23709985 | -0.701756388 |
| RP23-293K21.2 | 86.00755271 | -0.057406162 | 0.21495312 | -0.267063635 |
| RP23-293K21.3 | 29.22029167 | -0.421329507 | 0.271482576 | -1.551957822 |
| RP23-293K21.4 | 16.36398547 | -0.020785043 | 0.400525664 | -0.051894411 |
| RP23-293K21.5 | 4.108293731 | -0.948770027 | 0.616000253 | -1.540210449 |
| Gm15551 | 0 | NA | NA | NA |
| RP23-293K21.7 | 52.63659607 | -0.248911882 | 0.275195603 | -0.904490766 |
| RP23-293K21.8 | 19.23585701 | 0.100896445 | 0.295749198 | 0.34115543 |
| Ank2 | 12996.91815 | -0.230024978 | 0.054657817 | -4.208455262 |
| RP24-423L4.5 | 54.25629816 | -0.071389587 | 0.285578387 | -0.249982457 |
| RP24-423L4.4 | 84.96617208 | -0.537554414 | 0.244349352 | -2.19994205 |
| RP24-423L4.3 | 17.18518708 | -0.361714389 | 0.297715625 | -1.214966092 |
| Gm25501 | 0 | NA | NA | NA |
| RP24-423L4.2 | 66.13758907 | -0.374091713 | 0.328016738 | -1.140465315 |
| RP24-423L4.1 | 97.87426476 | -0.650145363 | 0.276931124 | -2.347678919 |
| RP23-50E10.7 | 69.57910019 | -0.593234746 | 0.228000209 | -2.601904398 |
| Gm16958 | 3.547171809 | 0.573676026 | 0.546377269 | 1.049963201 |
| RP23-50E10.5 | 105.9269057 | -0.588901873 | 0.259813827 | -2.266630225 |
| RP23-50E10.4 | 141.5973348 | -0.535133467 | 0.211704289 | -2.527740319 |
| RP23-50E10.3 | 170.8359331 | -0.715677591 | 0.371314904 | -1.927414128 |
| RP23-50E10.2 | 42.82266199 | -0.958038172 | 0.341229513 | -2.807606419 |
| RP23-50E10.1 | 29.57400322 | -0.893771142 | 0.411142674 | -2.173871014 |
| RP24-163F11.1 | 44.61355707 | -0.116638676 | 0.312027092 | -0.373809451 |
| RP24-260H22.2 | 2.710738714 | -1.135837198 | 0.646738346 | -1.756254605 |
| RP24-260H22.1 | 2.472653692 | -1.009145393 | 0.727452594 | -1.38723183 |
| AC157817.1 | 0 | NA | NA | NA |
| RP24-166J20.2 | 0 | NA | NA | NA |
| Larp7 | 118.9071558 | 0.083265853 | 0.149198647 | 0.558087184 |
| Mir302b | 0 | NA | NA | NA |
| Mir302c | 0 | NA | NA | NA |
| Mir302a | 0 | NA | NA | NA |
| Mir302d | 0 | NA | NA | NA |

| | | | | |
|---|---|---|---|---|
| Mir367 | 0 | NA | NA | NA |
| Zgrf1 | 113.3311546 | 0.602670153 | 0.147779145 | 4.07818136 |
| RP23-182M12.2 | 0 | NA | NA | NA |
| Neurog2 | 4.696444918 | 0.085989751 | 0.465263041 | 0.184819646 |
| Alpk1 | 333.345234 | 1.220230032 | 0.162826919 | 7.494031338 |
| Gm16238 | 2.994207206 | 0.776683993 | 0.637733467 | 1.217881816 |
| RP23-182M12.4 | 9.869931754 | 1.142761693 | 0.462930373 | 2.46853903 |
| RP23-182M12.3 | 4.2109437 | 0.852597647 | 0.664250718 | 1.28354795 |
| Gm23279 | 0 | NA | NA | NA |
| Tifa | 352.3487698 | 2.235994334 | 0.163557958 | 13.67096024 |
| Ap1ar | 641.0884169 | 0.071933111 | 0.079741876 | 0.902074476 |
| RP23-173N16.4 | 70.08836606 | 0.19843139 | 0.349184513 | 0.56827088 |
| RP23-173N16.3 | 2.767049531 | -0.288544376 | 0.560508943 | -0.51478996 |
| RP23-173N16.5 | 0.803713496 | 0.396482565 | 1.282978151 | 0.309032983 |
| RP23-173N16.8 | 0.269164771 | 1.558512592 | 2.053565847 | 0.758929934 |
| 5730508B09Rik | 54.45665569 | 1.691416735 | 0.245487781 | 6.890024119 |
| RP24-285A14.1 | 0.588650639 | 2.498980238 | 1.338059729 | 1.867614864 |
| RP24-285A14.2 | 0.048949532 | -0.206487837 | 3.724945049 | -0.055433794 |
| Gm10650 | 0 | NA | NA | NA |
| RP24-380G1.1 | 1.298106709 | -0.176093574 | 0.901324091 | -0.195372093 |
| RP24-380G1.2 | 0 | NA | NA | NA |
| Gm22293 | 0 | NA | NA | NA |
| RP24-196G4.1 | 0 | NA | NA | NA |
| RP23-61G16.1 | 0.062426873 | 0.755288839 | 3.724945049 | 0.202765096 |
| RP24-261K17.1 | 0 | NA | NA | NA |
| Gm9387 | 8.054043807 | -0.65629422 | 0.37800584 | -1.73620127 |
| Gm28543 | 0 | NA | NA | NA |
| Pitx2 | 11.30362075 | -0.317434916 | 0.323072424 | -0.982550327 |
| Gm27195 | 0.265852463 | 0.791636136 | 1.827719615 | 0.433127778 |
| RP23-446B17.3 | 0.066050875 | 0.274391604 | 3.724945049 | 0.073663262 |
| Enpep | 24.04966124 | 0.079232389 | 0.273443048 | 0.289758285 |
| RP24-140K12.2 | 0 | NA | NA | NA |
| RP24-140K12.4 | 0 | NA | NA | NA |
| RP23-257P11.1 | 0 | NA | NA | NA |
| RP23-257P11.2 | 0.095897396 | -0.687371727 | 3.724945049 | -0.184532045 |
| Rpl7a-ps7 | 2.979957284 | 0.538538176 | 0.603936582 | 0.891713124 |
| RP23-257P11.4 | 0.174406792 | 1.259563417 | 3.365276104 | 0.374282341 |
| Elovl6 | 655.176359 | -0.744103863 | 0.077423421 | -9.610836763 |
| RP23-3L20.3 | 30.90056436 | -0.92267492 | 0.237399093 | -3.886598333 |
| Egf | 35.62468515 | 0.441736898 | 0.230092773 | 1.919820834 |
| Gm24515 | 0.098502579 | -0.206492286 | 3.724945049 | -0.055434988 |
| RP23-3L20.6 | 0.263299413 | -1.715741491 | 2.097131309 | -0.818137369 |
| RP23-3L20.8 | 0 | NA | NA | NA |
| RP23-3L20.5 | 1.405849262 | -0.477107713 | 0.814634259 | -0.585671063 |
| Lrit3 | 3.350603553 | 2.070868359 | 0.863203194 | 2.399050853 |
| Rrh | 25.69789624 | 1.223057206 | 0.463694919 | 2.637633402 |
| Gar1 | 114.8672286 | 0.205773215 | 0.203876181 | 1.009304831 |

| | | | | |
|---|---|---|---|---|
| Cfi | 0.976809389 | 1.372579706 | 1.345836694 | 1.019870919 |
| Pla2g12a | 138.6047462 | -0.109354588 | 0.111502145 | -0.98073977 |
| Casp6 | 39.32840354 | 0.222477742 | 0.19260296 | 1.15511071 |
| Ccdc109b | 33.80637658 | 0.551323607 | 0.269369282 | 2.046720404 |
| Sec24b | 993.3623414 | -0.049777386 | 0.049147746 | -1.012811166 |
| Gm9396 | 8.082634732 | 0.548590985 | 0.44206254 | 1.240980484 |
| RP23-123C15.1 | 0 | NA | NA | NA |
| Col25a1 | 767.761363 | 0.029786592 | 0.105499228 | 0.282339428 |
| RP23-123C15.2 | 0.147068667 | 1.236183405 | 3.514416608 | 0.351746404 |
| Gm22682 | 2.066582291 | -0.47054031 | 0.713912762 | -0.659100572 |
| RP23-105B8.2 | 34.01273389 | 0.303268535 | 0.305368998 | 0.99312156 |
| RP23-105B8.3 | 9.17532523 | 0.201227272 | 0.326245735 | 0.616796637 |
| RP23-105B8.4 | 6.910068968 | 0.113618929 | 0.381674621 | 0.297685313 |
| Etnppl | 214.5413129 | -0.142303009 | 0.184832998 | -0.769900455 |
| Ostc | 293.8716669 | 0.178788157 | 0.149461651 | 1.196214252 |
| RP23-246L14.2 | 8.0379673 | -0.922234433 | 0.461193826 | -1.999667776 |
| Rpl34 | 837.1533585 | 0.465532202 | 0.131741364 | 3.533682892 |
| RP23-246L14.3 | 2.763783036 | -0.345753105 | 0.548896851 | -0.629905426 |
| RP23-246L14.4 | 0.231616962 | 0.25980922 | 2.124643517 | 0.122283676 |
| RP23-246L14.5 | 0.678557026 | -0.116790015 | 1.215761809 | -0.096063237 |
| RP23-349J3.1 | 0 | NA | NA | NA |
| RP23-349J3.2 | 0.048949532 | -0.206487837 | 3.724945049 | -0.055433794 |
| RP23-349J3.3 | 0.086908494 | -0.687376175 | 3.724945049 | -0.184533239 |
| Gm26090 | 0 | NA | NA | NA |
| RP23-349J3.4 | 6.268689147 | 4.555677119 | 0.956187389 | 4.764418742 |
| RP23-349J3.5 | 0.049553047 | 0.274391604 | 3.724945049 | 0.073663262 |
| RP23-349J3.6 | 0 | NA | NA | NA |
| RP23-349J3.7 | 0.075043205 | 0.755286614 | 3.724945049 | 0.202764498 |
| RP23-349J3.8 | 18.61381697 | 0.067371348 | 0.241404037 | 0.27908128 |
| RP24-176P19.1 | 0.066050875 | 0.274391604 | 3.724945049 | 0.073663262 |
| RP24-176P19.3 | 3.678638758 | -2.115285531 | 0.672219327 | -3.1467193 |
| Lef1 | 120.0536891 | 0.026060608 | 0.145425381 | 0.179202614 |
| RP24-176P19.6 | 5.471034591 | -0.7732638 | 0.416174074 | -1.858029723 |
| Hadh | 222.3007882 | -0.105095602 | 0.145427858 | -0.722664855 |
| RP24-434E24.5 | 0.259720851 | -0.632263221 | 2.042091337 | -0.309615544 |
| Cyp2u1 | 190.2662416 | -0.509395002 | 0.096054861 | -5.303167327 |
| 4930534D22Rik | 5.431134866 | -0.670566549 | 0.429866422 | -1.55994168 |
| Sgms2 | 102.4694722 | 0.689170281 | 0.197580496 | 3.48804814 |
| RP23-86I19.3 | 0 | NA | NA | NA |
| Papss1 | 709.3146802 | -0.237807129 | 0.083613119 | -2.844136556 |
| RP24-188G23.3 | 0.068097359 | -0.206487837 | 3.724945049 | -0.055433794 |
| RP23-292C19.1 | 0.108772582 | 0.75528439 | 3.724945049 | 0.202763901 |
| Olfr375-ps1 | 0 | NA | NA | NA |
| Gm26103 | 0 | NA | NA | NA |
| Gm22421 | 0.047948698 | -0.206487837 | 3.724945049 | -0.055433794 |
| Dkk2 | 4.638032695 | -1.468465299 | 0.491704089 | -2.986481772 |
| RP23-435E17.1 | 0 | NA | NA | NA |

| | | | | |
|---|---|---|---|---|
| Gm25307 | 0 | NA | NA | NA |
| Gm5549 | 0 | NA | NA | NA |
| RP24-450N10.1 | 0.627611986 | 1.305665529 | 1.171304295 | 1.114710783 |
| Aimp1 | 305.8677058 | 0.107327344 | 0.118730246 | 0.90395959 |
| Tbck | 868.7732137 | -0.178116119 | 0.063574594 | -2.801687103 |
| RP24-450N10.3 | 11.22849639 | -0.306234255 | 0.311356172 | -0.983549653 |
| Gm22322 | 0 | NA | NA | NA |
| RP23-447N11.5 | 18.07808909 | -0.331748437 | 0.279192383 | -1.18824315 |
| RP23-447N11.2 | 0.788050755 | -0.83214835 | 1.031988714 | -0.806354119 |
| RP23-447N11.3 | 31.71702897 | -0.215336311 | 0.245926191 | -0.875613576 |
| Npnt | 229.9057745 | 0.031665749 | 0.134699512 | 0.235084361 |
| Gm23196 | 0 | NA | NA | NA |
| Gstcd | 129.9010644 | -0.186414379 | 0.13718202 | -1.358883465 |
| RP23-10D15.4 | 0 | NA | NA | NA |
| RP23-10D15.3 | 6.257617145 | -0.466460367 | 0.559837558 | -0.833206634 |
| RP23-10D15.2 | 9.313879906 | -0.46566607 | 0.422998483 | -1.100869361 |
| Ints12 | 209.4893335 | 0.207008761 | 0.103730939 | 1.995631805 |
| Arhgef38 | 3.684370567 | -0.8806667 | 0.580393315 | -1.517361896 |
| RP23-335C9.1 | 28.10162786 | -0.054959154 | 0.211626099 | -0.25969932 |
| Ppa2 | 314.4988618 | -0.329417923 | 0.075110992 | -4.38574855 |
| Gm25042 | 0.060174537 | 0.755286614 | 3.724945049 | 0.202764498 |
| Tet2 | 1445.087248 | 0.259021481 | 0.149901683 | 1.727942454 |
| RP24-333H9.2 | 2.368693662 | -0.725059236 | 0.68985053 | -1.051038167 |
| RP24-333H9.1 | 0 | NA | NA | NA |
| Gm23011 | 0 | NA | NA | NA |
| Gm37228 | 0 | NA | NA | NA |
| RP24-289J19.2 | 0 | NA | NA | NA |
| RP23-437C24.1 | 0.044544648 | 0.755286614 | 3.724945049 | 0.202764498 |
| RP23-437C24.2 | 0.116046057 | -0.687371727 | 3.724945049 | -0.184532045 |
| Gm26691 | 49.78358579 | -0.376084908 | 0.226431334 | -1.660922544 |
| AC123608.1 | 1.334504357 | 0.355697442 | 0.750268058 | 0.47409381 |
| RP23-304A10.3 | 0 | NA | NA | NA |
| RP23-304A10.1 | 0 | NA | NA | NA |
| RP23-304A10.2 | 4.293971641 | -0.471274861 | 0.464007593 | -1.01566196 |
| Cxxc4 | 699.6168967 | -0.308519146 | 0.080518665 | -3.831647553 |
| Mir1895 | 0 | NA | NA | NA |
| RP23-384A11.6 | 0.213180531 | 1.236194976 | 2.597654287 | 0.475888952 |
| RP23-384A11.5 | 0.844415878 | -0.712249137 | 0.98389074 | -0.723910805 |
| RP23-384A11.4 | 0.116217011 | 0.274402725 | 3.724945049 | 0.073666248 |
| RP23-424L10.1 | 0 | NA | NA | NA |
| Gm26820 | 0.046345709 | 0.274391604 | 3.724945049 | 0.073663262 |
| Tacr3 | 61.25689593 | 0.075935748 | 0.175898211 | 0.43170279 |
| RP23-465M13.2 | 0.165933562 | 1.204233133 | 2.762106933 | 0.435983531 |
| RP23-465M13.3 | 0.044544648 | 0.755286614 | 3.724945049 | 0.202764498 |
| RP23-465M13.4 | 0 | NA | NA | NA |
| RP23-93G15.1 | 0 | NA | NA | NA |
| Gm24048 | 0 | NA | NA | NA |

| | | | | |
|---|---|---|---|---|
| RP23-93G15.2 | 0 | NA | NA | NA |
| RP23-453I14.1 | 0 | NA | NA | NA |
| Cenpe | 78.90502966 | 2.205326939 | 0.197082293 | 11.18987862 |
| Bdh2 | 27.37528258 | 1.171364225 | 0.337588197 | 3.469802074 |
| Slc9b2 | 28.12143682 | -0.486511567 | 0.227745778 | -2.136204551 |
| Slc9b1 | 1.329624306 | -1.160517169 | 0.8219179 | -1.411962398 |
| Cisd2 | 673.1771875 | -0.061375144 | 0.058194885 | -1.054648428 |
| RP24-372J15.5 | 30.69028919 | -0.176494712 | 0.247556375 | -0.712947555 |
| RP24-372J15.4 | 100.7051577 | -0.028662998 | 0.19319692 | -0.148361567 |
| 4930539J05Rik | 36.18476307 | -0.032760832 | 0.256888896 | -0.127529187 |
| Ube2d3 | 3995.493788 | 0.321958817 | 0.079861985 | 4.031440207 |
| RP23-167C5.2 | 0 | NA | NA | NA |
| Manba | 173.3995577 | -0.351793476 | 0.103488778 | -3.399339368 |
| Gm16100 | 2.944243352 | -0.571592687 | 0.679300429 | -0.84144314 |
| Oaz2-ps | 15.79173318 | -0.226358153 | 0.258427959 | -0.875904271 |
| Nfkb1 | 567.2400552 | 0.681008782 | 0.124600277 | 5.465547889 |
| RP23-167C5.1 | 62.27141224 | 0.024391269 | 0.362570591 | 0.067273159 |
| Slc39a8 | 39.47806482 | 0.656554442 | 0.212180043 | 3.094327031 |
| RP24-349N15.3 | 0.185615945 | 0.274408391 | 2.674999822 | 0.102582583 |
| RP24-349N15.4 | 1.794573941 | 0.655867816 | 0.762239458 | 0.860448523 |
| RP24-349N15.5 | 1.687301136 | -0.950018865 | 0.782181067 | -1.214576657 |
| RP24-349N15.6 | 0 | NA | NA | NA |
| RP24-349N15.2 | 0.623481245 | 0.026134282 | 1.189875881 | 0.021963872 |
| RP24-349N15.1 | 0.362626786 | 0.209724098 | 1.771043275 | 0.118418392 |
| RP24-326F21.2 | 0.246385664 | 1.635357791 | 2.060318243 | 0.793740383 |
| Bank1 | 11.35624835 | 0.693285749 | 0.34129518 | 2.031337651 |
| RP24-335I21.1 | 0.047948698 | -0.206487837 | 3.724945049 | -0.055433794 |
| RP24-335I21.2 | 0.075043205 | 0.755286614 | 3.724945049 | 0.202764498 |
| 1700030L20Rik | 0.443470786 | -0.607714198 | 1.576041523 | -0.385595296 |
| Ppp3ca | 9465.004104 | -0.000298076 | 0.112270345 | -0.002654983 |
| RP24-164P24.2 | 107.1091577 | -0.471882185 | 0.380811032 | -1.239150513 |
| RP24-164P24.5 | 0.044191549 | -0.206490061 | 3.724945049 | -0.055434391 |
| AC124177.1 | 0 | NA | NA | NA |
| RP23-174N20.1 | 0.068097359 | -0.206487837 | 3.724945049 | -0.055433794 |
| RP23-174N20.3 | 0.099155932 | 0.75528439 | 3.724945049 | 0.202763901 |
| RP23-174N20.2 | 0 | NA | NA | NA |
| Gm26107 | 0 | NA | NA | NA |
| RP23-77E5.1 | 0.132101749 | 0.274387056 | 3.724945049 | 0.073662041 |
| Emcn | 39.45898547 | 1.501218051 | 0.276242805 | 5.434415034 |
| Gm4861 | 0.22850592 | -0.724857314 | 2.556398893 | -0.283546248 |
| Gm22468 | 0.075043205 | 0.755286614 | 3.724945049 | 0.202764498 |
| Ddit4l | 50.60904352 | -0.79802163 | 0.180704742 | -4.416163187 |
| RP24-134B9.2 | 0.132574648 | -1.045763943 | 3.713921268 | -0.28157946 |
| Gm21962 | 0.087261593 | 0.274400501 | 3.724945049 | 0.073665651 |
| RP24-134B9.3 | 0 | NA | NA | NA |
| Gm25080 | 0 | NA | NA | NA |
| Mir6380 | 0 | NA | NA | NA |

| | | | | |
|---|---|---|---|---|
| RP24-380C2.1 | 0 | NA | NA | NA |
| RP24-380C2.2 | 1.550909849 | 1.03450408 | 1.035644198 | 0.998899122 |
| H2afz | 499.3132004 | 0.178127285 | 0.096630625 | 1.843383336 |
| Dnajb14 | 1867.871122 | -0.27028634 | 0.049061665 | -5.509114687 |
| RP23-209N19.2 | 0 | NA | NA | NA |
| RP23-209N19.1 | 20.11453365 | -0.444259807 | 0.339824138 | -1.307322689 |
| Gm15460 | 2.001670347 | -0.304737042 | 0.708962068 | -0.429835467 |
| Gm25920 | 0 | NA | NA | NA |
| RP23-209N19.4 | 0.841319663 | 0.815491446 | 1.091028909 | 0.74745173 |
| Lamtor3 | 343.7921853 | -0.067728013 | 0.083342731 | -0.812644513 |
| Dapp1 | 62.05077585 | 1.007663192 | 0.224282573 | 4.49282874 |
| RP23-209N19.6 | 0.453752633 | 1.439393739 | 1.317980818 | 1.0921204 |
| Gm5105 | 1.256408573 | 0.113363903 | 0.877899295 | 0.129130873 |
| 1110002E22Rik | 61.32006928 | 0.279528535 | 0.184196065 | 1.517559754 |
| Mttp | 190.0120384 | -0.016230988 | 0.107195258 | -0.151415173 |
| RP24-553B2.3 | 0.388301457 | 0.274379295 | 3.724945049 | 0.073659958 |
| Trmt10a | 78.50456046 | -0.297686617 | 0.150746107 | -1.974754924 |
| 4930579F01Rik | 0.038981395 | 0.274393828 | 3.724945049 | 0.073663859 |
| 0610031O16Rik | 0 | NA | NA | NA |
| Adh7 | 0.650173001 | -0.100691649 | 1.309146325 | -0.076913976 |
| Adh1 | 2.324828864 | 0.407682613 | 0.826231082 | 0.493424445 |
| Gm16559 | 0.066050875 | 0.274391604 | 3.724945049 | 0.073663262 |
| Adh6a | 0.234786912 | 1.372501092 | 3.049930571 | 0.450010602 |
| Adh6b | 0.808183924 | -0.694554912 | 1.142887462 | -0.607719426 |
| Gm23467 | 0 | NA | NA | NA |
| Adh6-ps1 | 0 | NA | NA | NA |
| Adh4 | 0.195798127 | -1.22946224 | 3.71166609 | -0.331242685 |
| Adh5 | 412.48678 | 0.05536773 | 0.097432414 | 0.568268077 |
| Metap1 | 515.9695598 | -0.165646117 | 0.052523223 | -3.153769099 |
| RP24-129H9.2 | 0.047948698 | -0.206487837 | 3.724945049 | -0.055433794 |
| RP24-129H9.1 | 0.20323857 | 1.157552161 | 2.643759526 | 0.437843212 |
| Eif4e | 1261.786848 | 0.02344497 | 0.065129951 | 0.359972178 |
| Mir1956 | 0 | NA | NA | NA |
| RP24-129H9.7 | 0 | NA | NA | NA |
| RP24-129H9.6 | 0.597553932 | -2.319745429 | 1.347053396 | -1.722088698 |
| RP24-129H9.8 | 0 | NA | NA | NA |
| RP24-93M7.1 | 0 | NA | NA | NA |
| RP24-93M7.2 | 0.061920613 | 0.274393828 | 3.724945049 | 0.073663859 |
| Tspan5 | 1750.479894 | -0.196189073 | 0.080038816 | -2.451174109 |
| Gm16060 | 0.740177848 | 0.282555118 | 1.264369371 | 0.223475137 |
| Rap1gds1 | 3054.008212 | -0.129039278 | 0.049335959 | -2.615521848 |
| RP24-428N8.5 | 39.16231688 | -0.727905343 | 0.291958244 | -2.493183045 |
| Gm4862 | 0 | NA | NA | NA |
| Stpg2 | 4.136924382 | 0.377385237 | 0.486186063 | 0.776215663 |
| RP23-276I11.1 | 0.091666476 | -0.687373951 | 3.724945049 | -0.184532642 |
| Gm24260 | 1.280600893 | 1.834582928 | 0.97400531 | 1.883545098 |
| RP23-4F17.1 | 21.90285516 | 0.194047962 | 0.245577309 | 0.790170568 |

| | | | | |
|---|---|---|---|---|
| RP23-434M21.1 | 0.112396584 | 0.27438722 | 3.724945049 | 0.073662085 |
| RP23-434M21.2 | 0.115957212 | 0.274402725 | 3.724945049 | 0.073666248 |
| Mir7j | 0.266304556 | 0.759334413 | 3.724657033 | 0.203866935 |
| RP24-221D2.1 | 0 | NA | NA | NA |
| RP24-63C3.1 | 0.099155932 | 0.75528439 | 3.724945049 | 0.202763901 |
| RP23-201A9.1 | 0 | NA | NA | NA |
| RP23-460F15.1 | 0 | NA | NA | NA |
| RP23-205G9.1 | 0 | NA | NA | NA |
| Gm6214 | 0 | NA | NA | NA |
| RP23-148J20.2 | 0 | NA | NA | NA |
| RP23-148J20.3 | 0.226297694 | 1.663785031 | 2.123090118 | 0.783661992 |
| RP23-148J20.4 | 0.157356115 | 0.755296948 | 2.830742007 | 0.266819423 |
| Pdha2 | 0 | NA | NA | NA |
| Gm15688 | 15.75159018 | 0.11131123 | 0.285360651 | 0.390072105 |
| Unc5c | 1697.764115 | -0.233973192 | 0.089422223 | -2.616499387 |
| Gm15689 | 1.903171754 | 2.854004976 | 0.978216571 | 2.917559425 |
| RP23-97D22.3 | 43.75796597 | -0.04497852 | 0.278331623 | -0.161600467 |
| Bmpr1b | 318.2842301 | -0.223595851 | 0.155437059 | -1.438497694 |
| RP23-100B7.1 | 0.094097695 | 0.755282166 | 3.724945049 | 0.202763304 |
| Pdlim5 | 665.5298056 | 0.229711096 | 0.109068181 | 2.106123836 |
| Gbp5 | 1952.644448 | 8.085159565 | 0.427201621 | 18.9258635 |
| Gbp7 | 3441.7084 | 6.332700281 | 0.243098377 | 26.04994883 |
| RP23-100J23.5 | 0 | NA | NA | NA |
| Gbp3 | 3129.531274 | 7.289031623 | 0.25010702 | 29.14365073 |
| Gbp2b | 4052.460821 | 7.861075048 | 0.345172931 | 22.77430915 |
| Gbp2 | 2406.349028 | 6.881211474 | 0.287481513 | 23.93618774 |
| Mir6381 | 0.56675494 | 0.274414444 | 1.405535751 | 0.195238324 |
| RP24-314I8.2 | 0 | NA | NA | NA |
| RP24-314I8.3 | 0.091666476 | -0.687373951 | 3.724945049 | -0.184532642 |
| Ccbl2 | 63.67129629 | 0.573568311 | 0.140558162 | 4.080647493 |
| Gm15540 | 0.047948698 | -0.206487837 | 3.724945049 | -0.055433794 |
| Gtf2b | 311.3568098 | 0.365655055 | 0.077728779 | 4.704242904 |
| Gm24457 | 0 | NA | NA | NA |
| Pkn2 | 955.7411507 | 0.2715254 | 0.080966443 | 3.35355475 |
| Gm25965 | 0 | NA | NA | NA |
| RP24-445M20.2 | 0.255858694 | 0.274410542 | 2.153551622 | 0.127422319 |
| 9530052C20Rik | 8.046682377 | -0.309668344 | 0.362823061 | -0.853496861 |
| RP24-445M20.3 | 0 | NA | NA | NA |
| Gm2574 | 15.8626213 | -0.002236125 | 0.280225613 | -0.007979732 |
| RP23-49B23.1 | 0.306809375 | -0.206484225 | 1.770752026 | -0.11660821 |
| RP23-49B23.2 | 0.297888126 | 0.593412532 | 1.718899753 | 0.345228121 |
| RP24-377E5.1 | 10.93206221 | 0.602296311 | 0.367933076 | 1.636972456 |
| RP24-337A2.1 | 0.793655395 | 0.211834033 | 1.571906285 | 0.134762508 |
| RP24-337A2.2 | 1.407833646 | -0.738341689 | 0.927887037 | -0.795723682 |
| RP24-337A2.3 | 0.046345709 | 0.274391604 | 3.724945049 | 0.073663262 |
| RP23-402L6.2 | 0.617903777 | -0.002057134 | 1.074457731 | -0.001914579 |
| RP23-402L6.1 | 1.629102473 | -0.422521127 | 0.728226244 | -0.58020585 |

| | | | | |
|---|---|---|---|---|
| RP24-273H20.2 | 0.222190708 | -0.699692037 | 2.15133339 | -0.325236451 |
| Gm38006 | 0 | NA | NA | NA |
| RP24-273H20.3 | 0.068097359 | -0.206487837 | 3.724945049 | -0.055433794 |
| RP24-273H20.4 | 3.820552698 | -0.832446597 | 0.565744344 | -1.471418327 |
| Gm24728 | 0 | NA | NA | NA |
| RP23-229C5.1 | 0.29112196 | -0.1861353 | 1.610792651 | -0.115555096 |
| RP23-229C5.2 | 0.131251387 | -0.206490426 | 2.981475279 | -0.069257803 |
| RP24-224P21.1 | 0.177934686 | 0.808082304 | 2.718161593 | 0.297290016 |
| RP23-327I19.1 | 2.34560619 | -0.697181068 | 0.620665195 | -1.123280432 |
| RP24-224P21.2 | 0.044544648 | 0.755286614 | 3.724945049 | 0.202764498 |
| RP23-327I19.2 | 0.038981395 | 0.274393828 | 3.724945049 | 0.073663859 |
| Lmo4 | 1293.515492 | 0.18384246 | 0.075285769 | 2.441928447 |
| RP23-461P7.1 | 43.89163664 | -0.435902573 | 0.240086308 | -1.815607795 |
| RP23-461P7.2 | 0 | NA | NA | NA |
| RP24-232N19.1 | 0.421951652 | 2.782590745 | 1.668944233 | 1.667276047 |
| RP24-224M8.1 | 22.99821006 | 0.109404245 | 0.259603576 | 0.421428112 |
| Hs2st1 | 927.2177343 | -0.110071324 | 0.054069043 | -2.035754966 |
| RP23-225M12.2 | 46.28994543 | -0.660028774 | 0.247854651 | -2.662967067 |
| RP23-225M12.1 | 14.69975535 | -0.611220167 | 0.363192787 | -1.682908332 |
| 15-Sep | 1591.561157 | 0.324733952 | 0.124205293 | 2.614493661 |
| Gm24406 | 0 | NA | NA | NA |
| Rpl9-ps8 | 0.045882455 | 0.755288839 | 3.724945049 | 0.202765096 |
| Sh3glb1 | 1627.178665 | 0.489411324 | 0.084116454 | 5.818259139 |
| RP24-321M14.4 | 0.899213277 | -1.013437795 | 1.101691956 | -0.919892161 |
| Clca3a1 | 41.15834729 | 0.913332273 | 0.553876349 | 1.64898226 |
| RP24-321M14.5 | 0.600994565 | 0.299057214 | 1.363763613 | 0.219288161 |
| RP24-271G16.1 | 1.394083107 | -0.477986394 | 0.790038909 | -0.60501627 |
| Clca3a2 | 1.360869654 | 1.440623699 | 0.910080585 | 1.582962787 |
| RP24-271G16.2 | 0.93582689 | -0.954840887 | 1.03928677 | -0.918746311 |
| Clca3b | 0.093141081 | -0.687373951 | 3.724945049 | -0.184532642 |
| Gm16215 | 0.201808035 | 0.748603472 | 3.20149504 | 0.23382934 |
| Clca4c-ps | 0.221289295 | 2.003750107 | 2.520812838 | 0.794882538 |
| Clca4b | 0 | NA | NA | NA |
| Gm16233 | 0.233058144 | -0.648167255 | 1.863179717 | -0.347882305 |
| Clca4a | 0 | NA | NA | NA |
| RP24-277E17.1 | 0.140122821 | 0.274406091 | 3.572128793 | 0.076818644 |
| Clca1 | 0.274296751 | 0.755279098 | 3.724945049 | 0.202762481 |
| Clca2 | 0.311801716 | 1.583494709 | 3.701979767 | 0.42774267 |
| Odf2l | 262.3245237 | -0.095689547 | 0.107771969 | -0.887889009 |
| Col24a1 | 47.56753531 | -0.826768436 | 0.239959385 | -3.445451557 |
| RP23-307M9.1 | 0 | NA | NA | NA |
| RP23-307M9.2 | 0.046345709 | 0.274391604 | 3.724945049 | 0.073663262 |
| Znhit6 | 139.6637941 | -0.013203107 | 0.137401817 | -0.096091208 |
| Cyr61 | 545.855505 | 3.001998611 | 0.587093643 | 5.113321611 |
| Gm17501 | 9.319672861 | -0.578922248 | 0.385906661 | -1.500161326 |
| RP24-267M22.1 | 0 | NA | NA | NA |
| Ddah1 | 1181.961696 | -0.088101115 | 0.106165556 | -0.829846498 |

| | | | | |
|---|---|---|---|---|
| Rpl36a-ps2 | 93.94039235 | 0.499036141 | 0.164319179 | 3.036992656 |
| Bcl10 | 200.6606098 | 0.506805176 | 0.116207323 | 4.361215467 |
| 2410004B18Rik | 209.7691853 | 0.103897006 | 0.123235031 | 0.843080136 |
| Syde2 | 65.37702174 | -0.486994571 | 0.148589919 | -3.277440181 |
| RP24-267M11.2 | 0 | NA | NA | NA |
| Wdr63 | 17.44671922 | -0.627215832 | 0.303475313 | -2.066777114 |
| Gm25627 | 0 | NA | NA | NA |
| Mcoln3 | 1.139337436 | 0.958601324 | 1.030138496 | 0.930555772 |
| Mcoln2 | 13.29155609 | 5.113057285 | 0.895734924 | 5.708225892 |
| Lpar3 | 6.729774449 | -0.179699616 | 0.424276956 | -0.423543192 |
| RP24-455A5.3 | 0 | NA | NA | NA |
| RP24-364E17.1 | 0.399538249 | 1.267537438 | 1.691945445 | 0.749159757 |
| Gm22078 | 0 | NA | NA | NA |
| Gm22901 | 0 | NA | NA | NA |
| Ssx2ip | 795.7799406 | -0.093200304 | 0.079408515 | -1.173681487 |
| Ctbs | 130.2632904 | 0.379417189 | 0.145672659 | 2.604587521 |
| Spata1 | 56.80384134 | -0.3165509 | 0.183765153 | -1.722583928 |
| Gng5 | 243.1666755 | 0.617983536 | 0.158302162 | 3.90382247 |
| Rpf1 | 164.2090017 | 0.439326097 | 0.113370319 | 3.8751421 |
| RP24-100G24.2 | 0 | NA | NA | NA |
| Uox | 7.769118165 | -0.933348558 | 0.54462185 | -1.713755255 |
| Dnase2b | 0.061920613 | 0.274393828 | 3.724945049 | 0.073663859 |
| Gm16325 | 1.444443621 | 0.461427401 | 0.894492463 | 0.515853872 |
| 4930503B20Rik | 2.396082498 | 2.11247167 | 1.085439226 | 1.946190647 |
| Samd13 | 1.10620882 | -0.330229378 | 1.018109495 | -0.324355464 |
| RP24-86C15.1 | 0.038981395 | 0.274393828 | 3.724945049 | 0.073663859 |
| Prkacb | 5477.753168 | -0.094188807 | 0.047236334 | -1.993990605 |
| RP24-409N2.3 | 0.044544648 | 0.755286614 | 3.724945049 | 0.202764498 |
| Gm10288 | 30.25828668 | 0.219336174 | 0.255891018 | 0.857146824 |
| Ttll7 | 4265.777113 | -0.429768726 | 0.071831526 | -5.983009837 |
| Gm29771 | 2.774619275 | -0.629121208 | 0.588370419 | -1.069260432 |
| Gm6074 | 0 | NA | NA | NA |
| RP24-293H7.2 | 0 | NA | NA | NA |
| Gm37803 | 0 | NA | NA | NA |
| Gm37041 | 0.332175987 | 1.788505721 | 3.713642212 | 0.481604209 |
| Gm36888 | 0 | NA | NA | NA |
| RP24-184C18.1 | 0.044191549 | -0.206490061 | 3.724945049 | -0.055434391 |
| Gm23131 | 0 | NA | NA | NA |
| RP23-290J2.1 | 0 | NA | NA | NA |
| Gm38068 | 0 | NA | NA | NA |
| RP24-196A21.2 | 0 | NA | NA | NA |
| Adgrl2 | 1106.274129 | -0.131918915 | 0.126805858 | -1.040321936 |
| RP24-224A19.1 | 3.670148074 | -0.82688537 | 0.508801513 | -1.625162955 |
| RP24-224A19.2 | 2.247096996 | -1.285159759 | 0.668148936 | -1.923463002 |
| Gm25127 | 1.189027437 | -0.430784652 | 0.951949012 | -0.452529124 |
| RP23-7K18.3 | 0.083172944 | -0.206492286 | 3.724945049 | -0.055434988 |
| Gm10287 | 19.5831566 | -0.627319942 | 0.359152104 | -1.746669266 |

| | | | | |
|---|---|---|---|---|
| RP23-473G18.1 | 38.52654686 | -0.566112631 | 0.221439717 | -2.556509005 |
| RP24-512E12.1 | 0.471094982 | 0.742273034 | 1.459619804 | 0.508538615 |
| Gm26468 | 0.506521265 | -0.607888729 | 1.343271071 | -0.452543602 |
| RP24-248L23.1 | 0 | NA | NA | NA |
| Rpsa-ps10 | 128.4715849 | 0.221975926 | 0.131679906 | 1.6857236 |
| RP24-134A5.1 | 4.381038091 | 0.307827327 | 0.533819029 | 0.576651095 |
| RP24-257N6.1 | 0 | NA | NA | NA |
| Mir3963 | 0 | NA | NA | NA |
| RP24-232H5.1 | 0 | NA | NA | NA |
| RP24-232H5.2 | 0.216927137 | 0.728355557 | 2.161354775 | 0.336990283 |
| RP24-232H5.3 | 0.510381651 | -0.100710277 | 1.309855148 | -0.076886576 |
| RP23-239A13.1 | 0.433950198 | -0.687379246 | 1.454811854 | -0.472486696 |
| RP23-239A13.2 | 0.049553047 | 0.274391604 | 3.724945049 | 0.073663262 |
| Adgrl4 | 62.50733234 | 0.28939867 | 0.225484837 | 1.283450692 |
| Gm24458 | 0.493466105 | -2.114871963 | 1.486685207 | -1.422541876 |
| Ifi44 | 3535.414574 | 9.514928272 | 0.357834612 | 26.59029603 |
| Ifi44l | 509.0915789 | 8.613147072 | 0.613114671 | 14.04818297 |
| Ptgfr | 13.2398592 | -0.975599705 | 0.386057887 | -2.527081395 |
| RP24-267D17.1 | 0.162302921 | 0.274407436 | 2.79109695 | 0.098315265 |
| RP24-267D17.2 | 0 | NA | NA | NA |
| Gm25056 | 0 | NA | NA | NA |
| RP23-245O13.3 | 0.112396584 | 0.27438722 | 3.724945049 | 0.073662085 |
| RP23-245O13.1 | 0.13307909 | 0.755293646 | 2.968852744 | 0.254405897 |
| RP23-245O13.2 | 0.060174537 | 0.755286614 | 3.724945049 | 0.202764498 |
| RP24-499N1.1 | 0.129258854 | -0.206490532 | 2.995468109 | -0.068934312 |
| Gipc2 | 25.12877053 | -0.126181623 | 0.207196895 | -0.608993794 |
| RP24-499N1.2 | 2.888459128 | 0.125037933 | 0.678905402 | 0.184175781 |
| Gm16198 | 0.109869311 | -0.206490066 | 3.724945049 | -0.055434393 |
| D630002J18Rik | 0.719682426 | 0.92710113 | 1.124955644 | 0.824122386 |
| Dnajb4 | 820.1226806 | -0.237989703 | 0.078553097 | -3.029666699 |
| Fubp1 | 2019.104171 | -0.178306815 | 0.092189893 | -1.934125419 |
| Nexn | 60.15238185 | -0.625789155 | 0.187371965 | -3.339822776 |
| RP23-181C16.4 | 5.705262924 | -1.762522445 | 0.543903425 | -3.240506245 |
| Fam73a | 1902.314384 | -0.040781053 | 0.055207768 | -0.738683241 |
| Usp33 | 1820.049344 | -0.128766282 | 0.065580835 | -1.963474268 |
| RP24-511F2.1 | 7.968776044 | 0.035387984 | 0.33547758 | 0.105485392 |
| RP24-511F2.2 | 7.38870447 | -0.164399112 | 0.427860982 | -0.384234877 |
| RP24-511F2.3 | 58.00030583 | -0.481840525 | 0.371171329 | -1.298162028 |
| RP24-511F2.4 | 45.38790308 | -0.154031276 | 0.219891067 | -0.700489013 |
| Zzz3 | 889.1435612 | -0.149040154 | 0.067924153 | -2.194214383 |
| Ak5 | 942.1680984 | -0.196319045 | 0.12912464 | -1.520384063 |
| AI115009 | 13.76623626 | -0.553280235 | 0.27109665 | -2.04089661 |
| Pigk | 806.3801702 | 0.022559535 | 0.045487228 | 0.495953165 |
| Rpl29-ps1 | 0.333447913 | 2.380148287 | 1.65776529 | 1.435757101 |
| St6galnac5 | 598.912246 | -0.250400715 | 0.12468296 | -2.008299412 |
| Gm16232 | 3.327150153 | -0.614001123 | 0.564415452 | -1.087853143 |
| Gm16213 | 1.998664546 | 1.977034379 | 1.148843876 | 1.720890384 |

| | | | | |
|---|---|---|---|---|
| Gm16231 | 0.342796299 | 0.773411729 | 1.799978012 | 0.429678431 |
| RP23-394P9.1 | 0.038981395 | 0.274393828 | 3.724945049 | 0.073663859 |
| RP24-117O8.1 | 6.160904613 | 0.66901074 | 0.502904937 | 1.330292647 |
| St6galnac3 | 220.6741086 | -0.158324061 | 0.111961655 | -1.41409183 |
| RP24-303G10.2 | 9.238644217 | -0.347038471 | 0.533213615 | -0.650843228 |
| RP24-303G10.1 | 48.22872844 | -0.348689763 | 0.306474531 | -1.137744665 |
| Gm27405 | 0.046345709 | 0.274391604 | 3.724945049 | 0.073663262 |
| RP24-104D12.1 | 4.330890979 | -0.46047055 | 0.580862932 | -0.792735298 |
| RP24-104D12.2 | 7.42840581 | -0.624368449 | 0.507876475 | -1.229370683 |
| Gm22136 | 0 | NA | NA | NA |
| Gm22206 | 0 | NA | NA | NA |
| RP24-186A10.2 | 0 | NA | NA | NA |
| Gm10042 | 0 | NA | NA | NA |
| 5730460C07Rik | 0 | NA | NA | NA |
| Gm25757 | 0 | NA | NA | NA |
| Asb17os | 1.419315775 | 4.051029891 | 1.157018519 | 3.501266251 |
| Asb17 | 0.04393175 | -0.206490061 | 3.724945049 | -0.055434391 |
| Msh4 | 11.18809076 | -0.188955229 | 0.318348918 | -0.593547578 |
| Gm24395 | 0 | NA | NA | NA |
| Rabggtb | 426.99162 | 0.171930962 | 0.100710603 | 1.707178362 |
| Snord45b | 0 | NA | NA | NA |
| Gm24494 | 6.926426429 | 0.375735948 | 0.422055913 | 0.89025159 |
| Snord45c | 0 | NA | NA | NA |
| RP23-403C5.3 | 0.748230394 | -0.163738418 | 1.131704712 | -0.144682987 |
| Acadm | 231.7967069 | -0.657832021 | 0.129421099 | -5.082880826 |
| Slc44a5 | 191.2716275 | 0.199718416 | 0.149976943 | 1.331660803 |
| Gm3076 | 3.513132756 | -0.215903994 | 0.633905341 | -0.340593429 |
| RP24-334O8.1 | 5.951295928 | 0.018482419 | 0.523585812 | 0.035299694 |
| RP24-527P22.4 | 0 | NA | NA | NA |
| Lhx8 | 17.86896063 | -0.104594457 | 0.268651566 | -0.389331277 |
| AI606473 | 4.337481582 | 0.0708251 | 0.450701181 | 0.157144251 |
| Tyw3 | 99.42373595 | -0.036334927 | 0.127831849 | -0.284240016 |
| Cryz | 179.0070392 | -0.103080842 | 0.08844333 | -1.16550159 |
| Gm38139 | 1.950077631 | -1.820806782 | 0.794216466 | -2.292582515 |
| Erich3 | 339.8935089 | -0.176896109 | 0.151740301 | -1.165781985 |
| Gm20389 | 0.04393175 | -0.206490061 | 3.724945049 | -0.055434391 |
| Tnni3k | 1.221396067 | 1.62775608 | 1.130558624 | 1.43978034 |
| Gm37890 | 0 | NA | NA | NA |
| Fpgt | 339.8878321 | 0.115399953 | 0.091367218 | 1.263034542 |
| Gm26456 | 0 | NA | NA | NA |
| Lrriq3 | 12.459711 | -0.352774284 | 0.289991954 | -1.216496799 |
| Gm37482 | 1.621103067 | -0.932088493 | 0.819575448 | -1.137282108 |
| Gm5285 | 0 | NA | NA | NA |
| Scp2-ps1 | 0.068097359 | -0.206487837 | 3.724945049 | -0.055433794 |
| Gm37449 | 0.554117557 | 1.286296754 | 1.228986785 | 1.046631884 |
| Gm37324 | 0.156124879 | 0.274407146 | 2.825334012 | 0.09712379 |
| Gm23038 | 0 | NA | NA | NA |

| | | | | |
|---|---|---|---|---|
| Gm37102 | 0.042716945 | -0.206490061 | 3.724945049 | -0.055434391 |
| Gm37515 | 0.408926061 | -0.531529724 | 1.427666416 | -0.372306666 |
| RP23-322I5.1 | 0.044191549 | -0.206490061 | 3.724945049 | -0.055434391 |
| RP23-322I5.2 | 0 | NA | NA | NA |
| RP23-322I5.3 | 0 | NA | NA | NA |
| RP23-187E3.1 | 0 | NA | NA | NA |
| 4930570G19Rik | 328.3713293 | 0.67782137 | 0.168612924 | 4.019984676 |
| Negr1 | 3447.761138 | 0.230097567 | 0.091319811 | 2.519689485 |
| RP23-354G14.1 | 240.3499704 | -0.421391618 | 0.31329299 | -1.345040049 |
| RP23-354G14.2 | 93.63872821 | -0.523023834 | 0.257008841 | -2.035042187 |
| RP23-354G14.3 | 137.6961956 | -0.559530852 | 0.26833961 | -2.085159372 |
| RP23-354G14.4 | 101.0414782 | -0.498302572 | 0.281249114 | -1.77174806 |
| Gm15578 | 21.23400752 | -0.479477107 | 0.274015328 | -1.749818564 |
| Gm15577 | 12.27858454 | -0.031240881 | 0.42417359 | -0.07365117 |
| RP24-176M3.1 | 45.69791764 | -0.247459826 | 0.331620674 | -0.746213508 |
| RP23-103L12.1 | 51.84091573 | -0.27723329 | 0.253621956 | -1.093096568 |
| 4930566N20Rik | 0.044544648 | 0.755286614 | 3.724945049 | 0.202764498 |
| Gm24373 | 15.34325161 | -0.434324481 | 0.464775256 | -0.934482798 |
| Gm23725 | 0 | NA | NA | NA |
| Gm22458 | 0.088383099 | -0.687376175 | 3.724945049 | -0.184533239 |
| Gm23379 | 0 | NA | NA | NA |
| RP23-64P1.1 | 0.244387265 | 0.274408675 | 2.082774238 | 0.131751522 |
| RP23-210I6.1 | 1.128024331 | 1.255042883 | 1.114303807 | 1.126302249 |
| Zranb2 | 1982.94177 | -0.087092567 | 0.081396221 | -1.069982933 |
| Mir186 | 29.35376171 | -0.215146676 | 0.235166822 | -0.914868325 |
| Ptger3 | 53.06310274 | -0.312876455 | 0.185494083 | -1.686719328 |
| RP24-411A22.2 | 0.062426873 | 0.755288839 | 3.724945049 | 0.202765096 |
| RP23-287M3.1 | 0.332723394 | 0.257122135 | 1.552710658 | 0.165595653 |
| RP23-287M3.3 | 0 | NA | NA | NA |
| Cth | 54.84234108 | 0.385256664 | 0.202425045 | 1.903206514 |
| RP23-287M3.2 | 3.057191251 | -0.686057151 | 0.596061534 | -1.15098377 |
| 1810013D15Rik | 0.328075509 | 1.630385346 | 1.728074532 | 0.943469345 |
| Ankrd13c | 1568.866954 | -0.010196763 | 0.065212905 | -0.156361124 |
| RP24-68G17.2 | 78.17925785 | -0.435578781 | 0.298785358 | -1.457831747 |
| Srsf11 | 1647.601996 | 0.090659511 | 0.120904561 | 0.749843597 |
| RP24-68G17.3 | 48.08364435 | 0.435470411 | 0.263904231 | 1.650107729 |
| Lrrc40 | 326.5754133 | 0.200344882 | 0.072729843 | 2.754644784 |
| RP24-68G17.4 | 1897.008325 | -0.486842996 | 0.229243806 | -2.123690947 |
| Lrrc7 | 2770.36321 | -0.492402559 | 0.219522278 | -2.243064184 |
| RP24-213M16.2 | 0.56553622 | 2.54202168 | 3.452303624 | 0.736326221 |
| RP24-213M16.1 | 0.354487963 | 0.274416184 | 1.582259052 | 0.173433158 |
| RP24-481I21.3 | 0.26582355 | -1.012702949 | 2.467868246 | -0.410355355 |
| RP24-481I21.2 | 0 | NA | NA | NA |
| Gm9423 | 0 | NA | NA | NA |
| RP23-324D2.1 | 2.321564718 | -0.92221496 | 0.775605456 | -1.189025881 |
| RP24-209L5.1 | 0.316660866 | 1.133559987 | 1.98341358 | 0.571519727 |
| Depdc1a | 5.945976878 | 2.165613877 | 0.655189196 | 3.305325987 |

| | | | | |
|---|---|---|---|---|
| Rpe65 | 5.700803298 | -2.350566796 | 0.521484187 | -4.507455556 |
| Wls | 574.2412302 | -0.150043539 | 0.103862084 | -1.444642101 |
| Wl1-1749A12.1 | 12.79191494 | -0.942246867 | 0.401915928 | -2.344387971 |
| Gm23941 | 0 | NA | NA | NA |
| Gm27878 | 0 | NA | NA | NA |
| Vmn1r-ps2 | 0 | NA | NA | NA |
| Gm11812 | 0 | NA | NA | NA |
| Vmn1r2 | 0 | NA | NA | NA |
| Vmn1r3 | 0 | NA | NA | NA |
| AI838599 | 1.421121277 | -1.468579763 | 0.85102401 | -1.725661961 |
| Gm11783 | 0.04393175 | -0.206490061 | 3.724945049 | -0.055434391 |
| Gm11786 | 0 | NA | NA | NA |
| Gm11784 | 0 | NA | NA | NA |
| Gm11785 | 0.044544648 | 0.755286614 | 3.724945049 | 0.202764498 |
| Gm11787 | 2.521183062 | 1.712200802 | 0.780040781 | 2.195014469 |
| Mir3471-2 | 0 | NA | NA | NA |
| Tmem68 | 334.4267812 | -0.131546362 | 0.06933846 | -1.897163027 |
| Tgs1 | 548.3289277 | 0.332628716 | 0.090777003 | 3.664239878 |
| Gm22541 | 0.302788994 | -0.681185354 | 1.806339149 | -0.377108227 |
| 2210414B05Rik | 0.54041206 | 2.654199726 | 1.465860979 | 1.810676295 |
| Lyn | 538.8172436 | 1.722338996 | 0.138124671 | 12.46945233 |
| Gm26857 | 5.106476976 | -0.227523062 | 0.522594196 | -0.435372347 |
| Gm11805 | 0.72046455 | 1.022070783 | 1.175163747 | 0.86972627 |
| Gm22781 | 2.17130535 | 0.470603376 | 0.714152832 | 0.658967317 |
| 9430025C20Rik | 38.03548056 | -0.006480521 | 0.28968189 | -0.022371165 |
| Rps20 | 938.4544041 | 0.7054449 | 0.098156405 | 7.186947232 |
| Gm24016 | 0 | NA | NA | NA |
| Gm24125 | 0 | NA | NA | NA |
| Mos | 0.721375344 | 0.01903069 | 1.203016351 | 0.015819145 |
| Plag1 | 101.48303 | 0.050983231 | 0.152526564 | 0.334258045 |
| Chchd7 | 221.4013722 | -0.035671195 | 0.104709637 | -0.340667735 |
| Gm11808 | 144.4071686 | 0.360506412 | 0.211363717 | 1.705621081 |
| Rps10-ps3 | 0 | NA | NA | NA |
| Sdr16c5 | 0.092228164 | 0.75528439 | 3.724945049 | 0.202763901 |
| Sdr16c6 | 0.09176491 | 1.15872478 | 3.72306111 | 0.311229052 |
| Penk | 1553.820092 | -0.471435556 | 0.221934255 | -2.124212669 |
| A830012C17Rik | 4.915491126 | -0.172199262 | 0.457800263 | -0.376144962 |
| Gm11780 | 0.085433889 | -0.687376175 | 3.724945049 | -0.184533239 |
| Gm26436 | 0 | NA | NA | NA |
| Impad1 | 2842.589061 | 0.124957014 | 0.084837307 | 1.472901706 |
| Gm11779 | 0 | NA | NA | NA |
| Gm11782 | 0 | NA | NA | NA |
| Gm11781 | 0 | NA | NA | NA |
| 4930423M02Rik | 0 | NA | NA | NA |
| Fam110b | 526.7316921 | 0.057169014 | 0.078154257 | 0.731489447 |
| Gm24337 | 0 | NA | NA | NA |
| Gm11796 | 0 | NA | NA | NA |

| | | | | |
|---|---|---|---|---|
| Gm11797 | 0.624599851 | 0.218686315 | 3.583318143 | 0.061028998 |
| Ubxn2b | 545.8183538 | -0.44385427 | 0.124448806 | -3.566561104 |
| Gm11798 | 0 | NA | NA | NA |
| Cyp7a1 | 0.264608745 | -1.00250669 | 2.469844052 | -0.405898781 |
| 4930430E12Rik | 36.79472905 | 6.391946551 | 0.929747179 | 6.874929764 |
| Sdcbp | 2354.562233 | 0.549421709 | 0.126879721 | 4.330256289 |
| Gm37689 | 8.169802865 | 0.09954678 | 0.466892462 | 0.213211366 |
| Nsmaf | 844.8221621 | 0.332308271 | 0.112135021 | 2.963465558 |
| BX510318.1 | 42.81714402 | 0.052025634 | 0.278317092 | 0.186929353 |
| Gm11803 | 0 | NA | NA | NA |
| Gm11801 | 0 | NA | NA | NA |
| Gm22473 | 0 | NA | NA | NA |
| Tox | 323.9376703 | 0.079094065 | 0.122707972 | 0.644571526 |
| Gm11802 | 0.913351404 | 1.100698094 | 1.257886194 | 0.8750379 |
| Gm11804 | 0 | NA | NA | NA |
| 8430436N08Rik | 0.23059471 | 1.756319809 | 2.129271586 | 0.82484537 |
| Gm11795 | 0.166709681 | -0.206481506 | 2.767631179 | -0.074605861 |
| Gm11794 | 0.481267046 | 1.149123794 | 1.415800436 | 0.811642493 |
| Gm23860 | 0 | NA | NA | NA |
| Gm11800 | 0 | NA | NA | NA |
| Car8 | 1324.707954 | -0.272322295 | 0.304754004 | -0.893580698 |
| Gm11799 | 5.921281838 | -1.038472429 | 0.402392261 | -2.580746522 |
| Gm25355 | 0 | NA | NA | NA |
| Gm37386 | 0.066050875 | 0.274391604 | 3.724945049 | 0.073663262 |
| Gm11810 | 0.196054161 | -0.206480699 | 2.235905757 | -0.092347675 |
| Rab2a | 2207.111245 | -0.040764049 | 0.067471661 | -0.604165491 |
| Gm11809 | 0 | NA | NA | NA |
| Chd7 | 1522.223342 | 0.072920731 | 0.135969204 | 0.536303288 |
| Rps18-ps2 | 0.644285774 | 1.676308725 | 1.314892577 | 1.274863631 |
| Gm23423 | 0.075043205 | 0.755286614 | 3.724945049 | 0.202764498 |
| Gm26548 | 1.161740484 | 0.488841249 | 0.963320121 | 0.507454623 |
| Clvs1 | 941.1106066 | -0.100421004 | 0.07866838 | -1.27651039 |
| Gm11816 | 3.30561101 | 0.95186477 | 0.662288211 | 1.437236471 |
| Gm11817 | 0.206997213 | 1.992410212 | 2.574827493 | 0.773803378 |
| Asph | 1754.934329 | -0.207597803 | 0.104264437 | -1.991070099 |
| Gm25677 | 0.044191549 | -0.206490061 | 3.724945049 | -0.055434391 |
| Gm24152 | 0 | NA | NA | NA |
| 4930412C18Rik | 9.306510091 | -0.505974781 | 0.335848338 | -1.506557348 |
| Gdf6 | 2.455729561 | 0.222394832 | 0.801038579 | 0.277633111 |
| 4930448K20Rik | 0 | NA | NA | NA |
| Gm11834 | 0 | NA | NA | NA |
| Gm11835 | 0 | NA | NA | NA |
| Gm11814 | 0.850521408 | -0.101380309 | 1.099741604 | -0.092185572 |
| 1700123O12Rik | 0 | NA | NA | NA |
| Gm11815 | 0.853137171 | -0.964275914 | 0.981498737 | -0.982452526 |
| Gm11813 | 0 | NA | NA | NA |
| Gm12920 | 0.044191549 | -0.206490061 | 3.724945049 | -0.055434391 |

| | | | | |
|---|---|---|---|---|
| Mir3471-1 | 0 | NA | NA | NA |
| Gm12918 | 39.37875012 | 0.146517283 | 0.212468658 | 0.689594807 |
| Gm12919 | 1.164100891 | -1.139367582 | 1.125909019 | -1.011953509 |
| AL672160.1 | 0.087930927 | -0.206490058 | 3.724945049 | -0.05543439 |
| 2610301B20Rik | 374.8592291 | 0.098439871 | 0.108587568 | 0.906548261 |
| Rps11-ps3 | 0.170186936 | 0.27439223 | 3.410018679 | 0.080466489 |
| Plekhf2 | 240.1964193 | 1.172223834 | 0.14722184 | 7.962295782 |
| Ndufaf6 | 73.7669564 | 0.070879416 | 0.166372101 | 0.426029461 |
| Gm11827 | 1.479390131 | 1.293060575 | 0.813222363 | 1.59004552 |
| Gm11829 | 0 | NA | NA | NA |
| mmu-mir-684-2 | 0.974719181 | -1.648352141 | 1.338769443 | -1.231244222 |
| Gm11830 | 0.15906016 | 0.274395307 | 3.439462686 | 0.079778539 |
| Trp53inp1 | 416.2178088 | 0.539618506 | 0.125003123 | 4.316840185 |
| Ccne2 | 114.245643 | 0.392078922 | 0.149328231 | 2.625618206 |
| Ints8 | 627.572806 | -0.156449743 | 0.090545605 | -1.727855745 |
| Dpy19l4 | 659.7126594 | -0.102981511 | 0.070037534 | -1.47037602 |
| Gm22940 | 3.154500895 | 1.165614687 | 0.803041657 | 1.451499654 |
| Gm26663 | 0.326609485 | 0.274416089 | 1.607124276 | 0.170749763 |
| Esrp1 | 1.320956406 | -0.39518793 | 1.040842745 | -0.379680727 |
| Gm11821 | 0.370194986 | 1.647273607 | 1.548538694 | 1.063760055 |
| Gm11822 | 0 | NA | NA | NA |
| 1110037F02Rik | 957.3336285 | -0.020969016 | 0.053698908 | -0.390492416 |
| Gm11831 | 0 | NA | NA | NA |
| Rad54b | 15.9562118 | 0.445100043 | 0.328807446 | 1.35367994 |
| Fsbp | 18.80141561 | 0.509943449 | 0.240862499 | 2.117155856 |
| Gm11832 | 0 | NA | NA | NA |
| Gm11833 | 0 | NA | NA | NA |
| Gem | 164.20295 | 2.427223066 | 0.366463165 | 6.623375268 |
| Cdh17 | 2.087534489 | 1.04447954 | 1.000972861 | 1.043464394 |
| Gm22477 | 0.061920613 | 0.274393828 | 3.724945049 | 0.073663859 |
| Gm25002 | 0 | NA | NA | NA |
| Pdp1 | 1783.616816 | -0.061212116 | 0.126191485 | -0.485073263 |
| 1700123M08Rik | 21.10087427 | -0.463577647 | 0.238095205 | -1.947026389 |
| Gm10604 | 0.068097359 | -0.206487837 | 3.724945049 | -0.055433794 |
| Gm26895 | 2.384116323 | -1.393873971 | 0.7372389 | -1.890667966 |
| Gm11840 | 0.048949532 | -0.206487837 | 3.724945049 | -0.055433794 |
| Gm11842 | 0.297840172 | -0.691232368 | 1.763109119 | -0.392053084 |
| Tmem67 | 313.6983145 | 0.52021621 | 0.105092602 | 4.950074514 |
| Rbm12b2 | 274.5941422 | -0.426629614 | 0.08506078 | -5.015585511 |
| Gm11839 | 15.15531981 | -0.396708897 | 0.358851545 | -1.105495859 |
| Gm11841 | 7.772074058 | -0.289439726 | 0.439506434 | -0.65855629 |
| Gm11843 | 1.645711418 | -0.624504141 | 0.890691756 | -0.701145078 |
| Rbm12b1 | 103.5381567 | -0.372717032 | 0.172691538 | -2.158281963 |
| Fam92a | 431.8026252 | 0.047191305 | 0.072103059 | 0.654497968 |
| Gm11847 | 135.4062284 | -0.149458236 | 0.125239615 | -1.193378274 |
| Gm11846 | 0.044191549 | -0.206490061 | 3.724945049 | -0.055434391 |
| Gm37985 | 1.26802963 | 0.015757329 | 0.981374933 | 0.01605638 |

| | | | | |
|---|---|---|---|---|
| n-R5s182 | 0 | NA | NA | NA |
| Triqk | 145.0715167 | -0.059146214 | 0.133542012 | -0.442903418 |
| Gm11820 | 0.208450558 | 0.274415766 | 2.196616084 | 0.124926594 |
| Gm26250 | 0 | NA | NA | NA |
| Gm11819 | 0 | NA | NA | NA |
| Gm11826 | 3.374863659 | 0.193242302 | 0.578578658 | 0.333994867 |
| Gm11825 | 0.044544648 | 0.755286614 | 3.724945049 | 0.202764498 |
| Gm23162 | 0 | NA | NA | NA |
| Runx1t1 | 591.5908782 | -0.523050109 | 0.097269988 | -5.377302062 |
| Gm11823 | 0.253691411 | 0.274401845 | 2.475805474 | 0.110833362 |
| Gm24068 | 0 | NA | NA | NA |
| Gm11824 | 3.085586268 | 0.960814355 | 0.917105362 | 1.047659729 |
| 1700120G11Rik | 0 | NA | NA | NA |
| Gm24908 | 0 | NA | NA | NA |
| Gm11836 | 2.257326888 | 0.097670267 | 0.730155414 | 0.133766408 |
| Slc26a7 | 38.39735321 | -0.563936521 | 0.222972571 | -2.529174415 |
| Lrrc69 | 6.895850206 | -0.326930404 | 0.498367664 | -0.656002441 |
| Gm11838 | 0 | NA | NA | NA |
| Otud6b | 619.2074641 | 0.000307756 | 0.062634257 | 0.00491354 |
| Tmem55a | 534.2645095 | -0.161398273 | 0.092081619 | -1.752774059 |
| Gm11837 | 8.354147893 | 0.319263792 | 0.354040248 | 0.901772593 |
| Necab1 | 1188.342868 | -0.263528235 | 0.111887037 | -2.355306228 |
| Gm11844 | 0 | NA | NA | NA |
| Gm11845 | 0 | NA | NA | NA |
| Tmem64 | 564.4500706 | -0.526823225 | 0.085777766 | -6.141722382 |
| Gm24317 | 0 | NA | NA | NA |
| Gm11856 | 0 | NA | NA | NA |
| Gm11859 | 0.314457495 | 0.262217319 | 1.784424879 | 0.146947805 |
| Gm11857 | 0 | NA | NA | NA |
| Gm11858 | 0.914915251 | -0.415383511 | 1.046195489 | -0.397041963 |
| Gm11855 | 11.73213907 | -0.093337966 | 0.331351989 | -0.281688261 |
| Gm24567 | 0 | NA | NA | NA |
| Calb1 | 2166.896649 | -0.062019118 | 0.156094924 | -0.397316687 |
| Gm26148 | 0.061920613 | 0.274393828 | 3.724945049 | 0.073663859 |
| Decr1 | 158.7523432 | -0.167463984 | 0.134631006 | -1.243873826 |
| Mir6400 | 0.947067764 | -1.673214422 | 1.135348404 | -1.473745342 |
| Nbn | 245.1479976 | 0.307524846 | 0.081377027 | 3.77901305 |
| Gm11869 | 0.066050875 | 0.274391604 | 3.724945049 | 0.073663262 |
| Osgin2 | 252.2581438 | -0.135607403 | 0.099803352 | -1.358745969 |
| Gm11870 | 26.42157704 | -0.303721945 | 0.267332164 | -1.136121973 |
| Gm11874 | 4.610084374 | 0.949089323 | 0.474644029 | 1.999581299 |
| Ripk2 | 184.8884919 | 1.329117234 | 0.180673754 | 7.356448882 |
| Gm22690 | 1.641288957 | -0.321621743 | 0.804348616 | -0.399853666 |
| A530072M11Rik | 6.322029942 | -0.867946583 | 0.441104126 | -1.96766825 |
| Gm11865 | 0.577397264 | 0.490312463 | 1.374021466 | 0.356844835 |
| Gm37355 | 0.136963818 | 0.755286583 | 3.59954433 | 0.209828388 |
| Gm11864 | 0.066050875 | 0.274391604 | 3.724945049 | 0.073663262 |

| | | | | |
|---|---|---|---|---|
| Gm11861 | 0.117650406 | -0.206492286 | 3.724945049 | -0.055434988 |
| Gm24978 | 0.911472295 | 1.03951484 | 0.968403127 | 1.073431933 |
| Gm11862 | 0 | NA | NA | NA |
| Gm23975 | 0 | NA | NA | NA |
| Gm11848 | 0.044191549 | -0.206490061 | 3.724945049 | -0.055434391 |
| Gm11850 | 0 | NA | NA | NA |
| Gm11849 | 0.314045685 | -1.12199552 | 1.762884939 | -0.636454198 |
| Gm23384 | 1.375344072 | -0.176515267 | 1.214201041 | -0.145375651 |
| Gm11851 | 0.112396584 | 0.27438722 | 3.724945049 | 0.073662085 |
| Mmp16 | 803.4952648 | -0.149165412 | 0.093685557 | -1.59219219 |
| Gm11860 | 0 | NA | NA | NA |
| Gm11852 | 0.075043205 | 0.755286614 | 3.724945049 | 0.202764498 |
| Gm11867 | 0.108266322 | 0.274389235 | 3.724945049 | 0.073662626 |
| Gm11854 | 0 | NA | NA | NA |
| Gm11853 | 0 | NA | NA | NA |
| Gm25971 | 0 | NA | NA | NA |
| Gm24419 | 0 | NA | NA | NA |
| Gm11868 | 68.16477173 | -0.353146043 | 0.170663483 | -2.069253698 |
| Cnbd1 | 0 | NA | NA | NA |
| Gm12392 | 0 | NA | NA | NA |
| Gm12393 | 0 | NA | NA | NA |
| Cngb3 | 8.051515703 | 5.077056111 | 0.964991876 | 5.261242333 |
| Gm11877 | 0 | NA | NA | NA |
| Gm25655 | 0.263406493 | 1.167504843 | 2.094597371 | 0.55738867 |
| Cpne3 | 505.9295095 | 0.470047795 | 0.115987657 | 4.052567368 |
| Rmdn1 | 159.5744036 | -0.508751333 | 0.100308161 | -5.071883753 |
| Gm12353 | 18.78038428 | -0.336280627 | 0.274024356 | -1.227192474 |
| Wwp1 | 1103.425899 | -0.306035877 | 0.063390495 | -4.827788107 |
| Gm12354 | 0 | NA | NA | NA |
| Slc7a13 | 0.367515839 | 1.224755832 | 1.76320751 | 0.69461809 |
| Atp6v0d2 | 1.127083422 | 0.29859406 | 1.265153004 | 0.236014188 |
| 4930480G23Rik | 0.075043205 | 0.755286614 | 3.724945049 | 0.202764498 |
| Ttpa | 39.70977185 | -0.817050739 | 0.256024808 | -3.191295198 |
| 4930598A11Rik | 0.729378188 | 0.320590575 | 1.38558791 | 0.231375124 |
| Ggh | 127.1892972 | 0.237211443 | 0.138003677 | 1.718877694 |
| Nkain3 | 142.3883735 | -0.286194769 | 0.11357523 | -2.519869592 |
| Gm11872 | 3.989504448 | 0.66004954 | 0.51219112 | 1.288678218 |
| Gm11873 | 0.255186079 | -0.191032159 | 1.801104409 | -0.106063901 |
| Gm11871 | 0 | NA | NA | NA |
| Gm22817 | 0 | NA | NA | NA |
| Gm25977 | 0 | NA | NA | NA |
| Gm11875 | 0 | NA | NA | NA |
| Gm11876 | 0 | NA | NA | NA |
| Prdm13 | 1.264085207 | -1.603335524 | 1.079348023 | -1.48546668 |
| Ccnc | 534.707253 | 0.137967213 | 0.083350671 | 1.655262174 |
| Tstd3 | 155.9862153 | -0.076964988 | 0.099853781 | -0.770776908 |
| Usp45 | 1038.908871 | -0.100388601 | 0.055558894 | -1.806886238 |

| | | | | |
|---|---|---|---|---|
| Pnisr | 1930.889929 | -0.396403072 | 0.141318192 | -2.805039228 |
| Coq3 | 168.8413342 | 0.000915505 | 0.113524115 | 0.008064411 |
| Faxc | 940.4861022 | -0.183968501 | 0.08995486 | -2.045120205 |
| Gm11885 | 0.049553047 | 0.274391604 | 3.724945049 | 0.073663262 |
| Gm11880 | 0 | NA | NA | NA |
| Gm11882 | 0 | NA | NA | NA |
| Fbxl4 | 229.5017738 | 0.132750388 | 0.095521926 | 1.389737345 |
| Gm24607 | 2.356080661 | 1.32117529 | 0.756502768 | 1.746424924 |
| Gm37900 | 25.72877726 | 0.617701242 | 0.353723905 | 1.746280741 |
| Pou3f2 | 455.224007 | -0.586790998 | 0.128170706 | -4.578198987 |
| Gm11878 | 0.318934723 | -0.206481536 | 1.600598952 | -0.129002669 |
| Gm11881 | 0 | NA | NA | NA |
| Gm11879 | 0 | NA | NA | NA |
| Gm24078 | 0 | NA | NA | NA |
| 1700025O08Rik | 0 | NA | NA | NA |
| Gm11884 | 0.044544648 | 0.755286614 | 3.724945049 | 0.202764498 |
| Gm11888 | 0 | NA | NA | NA |
| Gm11889 | 2.706150538 | -0.809288485 | 0.638808391 | -1.266872033 |
| Gm25978 | 0.086648695 | -0.687376175 | 3.724945049 | -0.184533239 |
| Gm11890 | 1.145011917 | 1.837622449 | 1.211175868 | 1.517221815 |
| Gm28448 | 1.698407129 | -0.350482128 | 0.722300792 | -0.485230158 |
| Gm11892 | 0.26738533 | 0.274410359 | 2.109447118 | 0.130086389 |
| Gm11891 | 0 | NA | NA | NA |
| Gm27243 | 8.985200294 | -0.727201702 | 0.370742352 | -1.961474587 |
| Mms22l | 56.96292425 | 0.66668594 | 0.268579357 | 2.482267984 |
| Klhl32 | 321.673524 | -0.123131762 | 0.092532224 | -1.330690617 |
| Gm11899 | 0.512549389 | 0.587432289 | 1.373392811 | 0.427723434 |
| Ndufaf4 | 345.0672365 | -0.339476185 | 0.10316768 | -3.290528442 |
| C230012O17Rik | 12.79852409 | 0.086669116 | 0.307743686 | 0.2816276 |
| Gpr63 | 190.3210986 | -0.401724819 | 0.153350573 | -2.619649944 |
| Gm11906 | 2.251356483 | -1.275731919 | 0.684525346 | -1.863673751 |
| Fhl5 | 2.616435038 | -0.096629639 | 0.714696527 | -0.135203733 |
| Ufl1 | 454.4895708 | -0.14416198 | 0.074646905 | -1.931251936 |
| Prdx6-ps2 | 0.044191549 | -0.206490061 | 3.724945049 | -0.055434391 |
| Gm11897 | 0 | NA | NA | NA |
| Gm11894 | 0.044544648 | 0.755286614 | 3.724945049 | 0.202764498 |
| Gm25975 | 0.109124069 | 0.274402725 | 3.724945049 | 0.073666248 |
| Gm11896 | 0 | NA | NA | NA |
| Gm11893 | 3.313083698 | 0.016188485 | 0.601206442 | 0.026926666 |
| Gm24977 | 0 | NA | NA | NA |
| Fut9 | 4215.601891 | -0.201831783 | 0.102022609 | -1.978304461 |
| Gm24042 | 0 | NA | NA | NA |
| Gm11898 | 0 | NA | NA | NA |
| Gm26067 | 0 | NA | NA | NA |
| Gm11895 | 0 | NA | NA | NA |
| Gm24965 | 0 | NA | NA | NA |
| Manea | 627.7666335 | -0.021313089 | 0.111866954 | -0.190521757 |

| | | | | |
|---|---|---|---|---|
| Bhmt-ps1 | 0 | NA | NA | NA |
| Gm11903 | 0 | NA | NA | NA |
| Gng2-ps1 | 0.3346585 | -0.271326915 | 1.753976693 | -0.154692429 |
| Gm11905 | 0 | NA | NA | NA |
| 4930548K13Rik | 0 | NA | NA | NA |
| Gm11904 | 0 | NA | NA | NA |
| Gm11902 | 0 | NA | NA | NA |
| Gm11901 | 0.045882455 | 0.755288839 | 3.724945049 | 0.202765096 |
| Gm25173 | 0 | NA | NA | NA |
| Gm11914 | 3.923996896 | 0.788462072 | 0.530782056 | 1.485472358 |
| Tpm3-rs2 | 0 | NA | NA | NA |
| Gm11911 | 0 | NA | NA | NA |
| Gm11907 | 0.192582685 | 0.24569102 | 2.673283602 | 0.091906081 |
| Gm25378 | 0 | NA | NA | NA |
| Gm11908 | 0 | NA | NA | NA |
| Gm11909 | 0 | NA | NA | NA |
| Gm11910 | 0 | NA | NA | NA |
| Gm11912 | 0 | NA | NA | NA |
| Gm11913 | 0 | NA | NA | NA |
| Epha7 | 823.6749401 | -0.354457324 | 0.141179301 | -2.510689036 |
| Gm11915 | 4.082662799 | 0.122331556 | 0.469996781 | 0.26028169 |
| Gm11917 | 0.126225412 | 0.755282661 | 3.698754989 | 0.20419916 |
| Gm11916 | 0 | NA | NA | NA |
| Gm11918 | 0 | NA | NA | NA |
| Gm11919 | 0 | NA | NA | NA |
| Gm11923 | 0 | NA | NA | NA |
| Gm11925 | 0 | NA | NA | NA |
| Gm11924 | 0.272022412 | 1.286193694 | 2.446925674 | 0.525636601 |
| 4930556G01Rik | 0.046345709 | 0.274391604 | 3.724945049 | 0.073663262 |
| Gm22831 | 0 | NA | NA | NA |
| Gm11920 | 0.174512195 | -1.282572172 | 3.342629336 | -0.383701584 |
| Gm25705 | 0.044544648 | 0.755286614 | 3.724945049 | 0.202764498 |
| Gm11922 | 0.132199999 | 1.236180999 | 3.642365845 | 0.339389576 |
| Map3k7 | 821.1190469 | 0.152174973 | 0.054732551 | 2.780337664 |
| Gm11926 | 0 | NA | NA | NA |
| Gm26254 | 0 | NA | NA | NA |
| Gm11927 | 0.564158409 | 0.209931721 | 1.619513116 | 0.129626441 |
| Gm11928 | 0 | NA | NA | NA |
| Gm11929 | 0 | NA | NA | NA |
| Bach2 | 252.756904 | -0.266017338 | 0.082231759 | -3.234970784 |
| Bach2it1 | 12.10687002 | -0.908473025 | 0.390048259 | -2.329129804 |
| Gm24371 | 4.271398782 | -1.872513698 | 0.620866755 | -3.015967088 |
| Bach2os | 18.90185043 | 2.430379269 | 0.488122869 | 4.979031764 |
| Gja10 | 0.072025462 | 0.755288839 | 3.724945049 | 0.202765096 |
| Gm11933 | 0.233612452 | 1.697742765 | 2.527723224 | 0.671648996 |
| Casp8ap2 | 404.3248676 | -0.142124195 | 0.095865704 | -1.482534304 |
| Gm11936 | 0 | NA | NA | NA |

| | | | | |
|---|---|---|---|---|
| Mdn1 | 1768.721666 | -0.351045927 | 0.11811173 | -2.972151244 |
| Lyrm2 | 76.45699897 | 0.110767098 | 0.181677581 | 0.609690518 |
| Ankrd6 | 449.0953625 | -0.316613175 | 0.094703063 | -3.343220005 |
| Gm11941 | 0 | NA | NA | NA |
| Gm11942 | 24.81150096 | 0.160492481 | 0.276195407 | 0.581083093 |
| Rragd | 1716.34318 | -0.341296437 | 0.098092514 | -3.479332135 |
| Gm11943 | 0.062426873 | 0.755288839 | 3.724945049 | 0.202765096 |
| 4933421O10Rik | 104.7690149 | -0.37185346 | 0.126312635 | -2.943913405 |
| Ube2j1 | 1400.842074 | 0.119998515 | 0.065248999 | 1.839085919 |
| Gm11931 | 0 | NA | NA | NA |
| Gabrr2 | 12.63021313 | 0.418594764 | 0.303021299 | 1.381403764 |
| Gabrr1 | 1.88379093 | 0.573573829 | 0.895678072 | 0.640379447 |
| Pm20d2 | 35.06690641 | 0.060748636 | 0.180885294 | 0.335840656 |
| Gm11930 | 0 | NA | NA | NA |
| Srsf12 | 183.1024447 | -0.178103528 | 0.099761951 | -1.785285135 |
| Pnrc1 | 961.8622753 | 0.148875714 | 0.096853098 | 1.537129079 |
| Gm11934 | 0.906232762 | 0.55213044 | 1.034799669 | 0.533562636 |
| Rngtt | 426.6224941 | -0.146274509 | 0.062544468 | -2.338728164 |
| Mir8118 | 1.705950494 | 0.479889718 | 0.85969939 | 0.558206419 |
| Gm11935 | 0 | NA | NA | NA |
| Gm23304 | 0.144050924 | 1.290908285 | 3.721447108 | 0.346883416 |
| Gm24341 | 0 | NA | NA | NA |
| Cnr1 | 3191.885636 | -0.067054049 | 0.103153449 | -0.65004176 |
| Gm22010 | 0 | NA | NA | NA |
| Spaca1 | 0.461160426 | 0.287660681 | 1.340753553 | 0.214551496 |
| Gm25623 | 0 | NA | NA | NA |
| Gm12753 | 0.179340113 | -0.663437986 | 2.728510352 | -0.243150254 |
| Gm12751 | 94.08612612 | 0.319826661 | 0.129677649 | 2.466320624 |
| Gm12752 | 0.088534442 | 0.274389469 | 3.724945049 | 0.073662689 |
| Gm12754 | 0.046345709 | 0.274391604 | 3.724945049 | 0.073663262 |
| Gm23466 | 0 | NA | NA | NA |
| Gm12351 | 0 | NA | NA | NA |
| Gm12350 | 35.03170047 | -0.221558696 | 0.207825771 | -1.066079028 |
| Akirin2 | 531.2879269 | 0.000363391 | 0.071968558 | 0.005049306 |
| Orc3 | 556.144257 | 0.101867963 | 0.098409645 | 1.035142065 |
| Rars2 | 246.197839 | -0.155175538 | 0.098071999 | -1.58226139 |
| Slc35a1 | 169.2461004 | -0.269916639 | 0.107138268 | -2.519329886 |
| Gm22849 | 0.748256776 | 1.265378273 | 1.153450317 | 1.097037518 |
| 1700003M02Rik | 17.88018956 | -0.349380489 | 0.288979871 | -1.209013238 |
| Gm136 | 1.409557441 | -0.770929282 | 0.984310161 | -0.783217844 |
| Smim8 | 84.47012772 | 0.267662718 | 0.133113441 | 2.010786557 |
| Zfp292 | 1650.746143 | 0.182148264 | 0.094465064 | 1.928207692 |
| Gm12363 | 0 | NA | NA | NA |
| Cga | 2.158593695 | -2.514766186 | 1.680590136 | -1.496359007 |
| Platr9 | 0 | NA | NA | NA |
| Gm12361 | 0.042716945 | -0.206490061 | 3.724945049 | -0.055434391 |
| Gm12362 | 0.090537259 | -0.20649451 | 3.724945049 | -0.055435586 |

| | | | | |
|---|---|---|---|---|
| Mob3b | 188.1401066 | -0.473693167 | 0.160194146 | -2.956994243 |
| Gm12364 | 0 | NA | NA | NA |
| Ifnk | 0.038981395 | 0.274393828 | 3.724945049 | 0.073663859 |
| 3110043O21Rik | 349.8244095 | 0.034292175 | 0.076425596 | 0.44870014 |
| Gm12366 | 24.68870437 | 0.350976747 | 0.233708024 | 1.501774486 |
| Gm12367 | 4.278993874 | -0.857078158 | 0.535127975 | -1.601632129 |
| Gm12368 | 0 | NA | NA | NA |
| Gm12365 | 0 | NA | NA | NA |
| Lingo2 | 668.3004972 | -0.05309617 | 0.122286884 | -0.434193495 |
| Gm12369 | 25.01436732 | -0.089589896 | 0.249293937 | -0.35937455 |
| Gm12370 | 0 | NA | NA | NA |
| Mir876 | 9.32634141 | -0.521687803 | 0.32064432 | -1.626998425 |
| Mir873a | 8.772029568 | -0.310025961 | 0.352302596 | -0.879999083 |
| mmu-mir-873a | 0 | NA | NA | NA |
| Gm23314 | 0 | NA | NA | NA |
| AL844160.1 | 0 | NA | NA | NA |
| Gm12371 | 46.23436975 | 1.037525694 | 0.178217775 | 5.821673475 |
| Gm12372 | 1.336268985 | 0.462697911 | 0.90317757 | 0.512300047 |
| Rpl23a-ps6 | 0 | NA | NA | NA |
| Gm12374 | 2.339895364 | 0.085667107 | 0.664970828 | 0.128828369 |
| Gm22732 | 0.04393175 | -0.206490061 | 3.724945049 | -0.055434391 |
| Gm12375 | 0 | NA | NA | NA |
| Gm12378 | 0 | NA | NA | NA |
| Gm12379 | 0 | NA | NA | NA |
| Gm12380 | 0.504525611 | 0.795635765 | 1.301665041 | 0.611244629 |
| Gm12381 | 0 | NA | NA | NA |
| Gm12382 | 0 | NA | NA | NA |
| Gm25581 | 0 | NA | NA | NA |
| Gm12383 | 0 | NA | NA | NA |
| Gm23607 | 0 | NA | NA | NA |
| Gm17094 | 0.175174943 | 0.274407987 | 2.724622258 | 0.10071414 |
| 1700009N14Rik | 0 | NA | NA | NA |
| Gm12385 | 0.045882455 | 0.755288839 | 3.724945049 | 0.202765096 |
| Gm12384 | 0.119234754 | 0.274400501 | 3.724945049 | 0.073665651 |
| Gm12386 | 0 | NA | NA | NA |
| Gm22850 | 0 | NA | NA | NA |
| Gm12387 | 0.097139547 | 0.274393828 | 3.724945049 | 0.073663859 |
| Gm26087 | 0 | NA | NA | NA |
| Gm12388 | 0.062426873 | 0.755288839 | 3.724945049 | 0.202765096 |
| Gm12389 | 0 | NA | NA | NA |
| Aco1 | 373.8664707 | -0.280252832 | 0.09636499 | -2.908243243 |
| Ddx58 | 2204.242023 | 5.814961772 | 0.227902806 | 25.5150951 |
| Topors | 450.6972402 | 0.109092933 | 0.07251715 | 1.504374245 |
| Toporsos | 15.19562275 | 0.023471598 | 0.324117603 | 0.072416917 |
| Ndufb6 | 365.4765144 | -0.229196849 | 0.105811171 | -2.166093105 |
| 4930509K18Rik | 0 | NA | NA | NA |
| Gm23338 | 0.239936926 | 1.90922946 | 2.42746865 | 0.78651045 |

| | | | | |
|---|---|---|---|---|
| Tmem215 | 126.9200701 | 0.526818029 | 0.201619703 | 2.612929303 |
| 4933428C19Rik | 0.268438559 | 0.768540404 | 1.811148498 | 0.424338703 |
| Gm12391 | 0 | NA | NA | NA |
| Aptx | 267.7610802 | -0.092186971 | 0.090793775 | -1.015344622 |
| Gm12390 | 0.717871974 | 0.151738623 | 1.180149124 | 0.128575804 |
| Gm6297 | 15.05970898 | -0.465648821 | 0.277120678 | -1.680310631 |
| Dnaja1 | 2433.261954 | 0.130870983 | 0.102563471 | 1.275999941 |
| Mir207 | 0.128150834 | -1.019523341 | 3.714811905 | -0.274448173 |
| Smu1 | 643.1185477 | 0.149578748 | 0.078496455 | 1.905547825 |
| Gm12396 | 8.729157278 | 0.032150994 | 0.362817043 | 0.088614895 |
| B4galt1 | 382.463159 | 2.083699353 | 0.196381299 | 10.61047751 |
| Mir5123 | 0.217530652 | 1.2362011 | 2.176047851 | 0.568094631 |
| Gm24112 | 0.24398126 | -1.067272932 | 1.806934748 | -0.590653831 |
| Gm25931 | 0.653402772 | 2.483878687 | 1.43008079 | 1.736879976 |
| Gm12397 | 7.076756442 | -0.749916082 | 0.423857211 | -1.769265832 |
| Spink4 | 0.044544648 | 0.755286614 | 3.724945049 | 0.202764498 |
| Bag1 | 715.7527563 | 0.391584863 | 0.090850065 | 4.310232064 |
| Chmp5 | 658.9006136 | 0.351584111 | 0.114877788 | 3.060505588 |
| Nfx1 | 1354.303035 | 0.025047159 | 0.091452215 | 0.27388248 |
| Mir3094 | 0.798730366 | 0.576362704 | 1.118831428 | 0.515147046 |
| Aqp7 | 0.174839421 | -1.162972749 | 2.306333002 | -0.504251879 |
| Gm25637 | 0 | NA | NA | NA |
| Aqp3 | 1.154822103 | 1.682518883 | 1.126115444 | 1.49409094 |
| Nol6 | 1475.373276 | 0.143676535 | 0.046055757 | 3.119621613 |
| Gm12402 | 0.230106605 | -1.545253127 | 2.560976671 | -0.603384304 |
| Ube2r2 | 789.6006133 | 0.194607331 | 0.064654287 | 3.009967917 |
| Gm12403 | 11.82053707 | -0.995065631 | 0.419515602 | -2.371939509 |
| Gm24650 | 0 | NA | NA | NA |
| Ubap2 | 693.4431244 | -0.220409676 | 0.085393866 | -2.581094938 |
| Gm22888 | 2.630926952 | -0.243515239 | 0.645112453 | -0.377477195 |
| Gm24837 | 2.20124 | -0.113235349 | 0.71894247 | -0.157502657 |
| Gm23090 | 0.341439828 | 0.300364375 | 1.601077438 | 0.187601404 |
| Gm12401 | 0.164132501 | 1.537077023 | 3.390982371 | 0.453283696 |
| Dcaf12 | 631.7207173 | -0.070145488 | 0.086425533 | -0.811629226 |
| n-R5s183 | 0 | NA | NA | NA |
| Gm26049 | 4.518367824 | -0.298502569 | 0.604836044 | -0.493526422 |
| Gm26084 | 0.449030389 | -2.778002068 | 1.450219674 | -1.915573287 |
| Ubap1 | 662.4155343 | 0.146174669 | 0.071750864 | 2.037253086 |
| Kif24 | 28.6589064 | 0.563422454 | 0.20355861 | 2.76786353 |
| Nudt2 | 96.39035252 | 0.137937615 | 0.116358124 | 1.185457536 |
| AI464131 | 299.7303309 | -0.912866614 | 0.200062757 | -4.562901293 |
| 1110017D15Rik | 25.98366852 | -0.202692515 | 0.207919978 | -0.974858296 |
| Fam219aos | 0.808497328 | 0.447749366 | 1.122465571 | 0.398898084 |
| Fam219a | 1771.430822 | -0.063704487 | 0.051432854 | -1.238595209 |
| Gm12404 | 1.315597923 | 0.120513566 | 0.822350811 | 0.146547634 |
| Dnaic1 | 23.7871792 | -0.341718819 | 0.273303027 | -1.250329432 |
| Enho | 315.6348979 | -0.385345856 | 0.129910588 | -2.966239024 |

| | | | | |
|---|---|---|---|---|
| Gm12406 | 1.108658207 | -1.134131003 | 0.994956844 | -1.139879593 |
| Gm12405 | 0.625648376 | -2.087287211 | 1.521776276 | -1.371612401 |
| Cntfr | 660.2301978 | -0.154755473 | 0.10460534 | -1.479422307 |
| Il11ra1 | 183.2536711 | -0.041977759 | 0.104900581 | -0.40016708 |
| Rpp25l | 90.6293161 | 0.139022059 | 0.129748737 | 1.071471389 |
| Dctn3 | 483.0857412 | 0.130340065 | 0.069293831 | 1.880976476 |
| Arid3c | 1.591794818 | -0.441208269 | 1.036163177 | -0.42580964 |
| Sigmar1 | 271.0497585 | -0.382727905 | 0.087899602 | -4.354148331 |
| Galt | 238.390805 | -0.18224746 | 0.083888924 | -2.172485385 |
| Ccl27a | 57.35970255 | -1.196344883 | 0.153772922 | -7.77994505 |
| Gm12407 | 0.693596057 | 1.569732378 | 1.299238602 | 1.208194073 |
| Gm10601 | 0.106056992 | 1.236179401 | 3.724945049 | 0.331865138 |
| Gm13307 | 0 | NA | NA | NA |
| Gm13302 | 0 | NA | NA | NA |
| Gm21541 | 0 | NA | NA | NA |
| Ccl19-ps1 | 0 | NA | NA | NA |
| Gm20878 | 0 | NA | NA | NA |
| Gm20938 | 0 | NA | NA | NA |
| Gm13303 | 0 | NA | NA | NA |
| Gm21093 | 0.097139547 | 0.274393828 | 3.724945049 | 0.073663859 |
| Gm37530 | 0.382550573 | -0.794354517 | 1.377909336 | -0.576492586 |
| Gm3892 | 0 | NA | NA | NA |
| Gm3893 | 0 | NA | NA | NA |
| Gm13304 | 0.049553047 | 0.274391604 | 3.724945049 | 0.073663262 |
| Gm13299 | 0.097501745 | -0.206492286 | 3.724945049 | -0.055434988 |
| Ccl19-ps3 | 0.089089296 | 1.194485492 | 3.723939576 | 0.320758559 |
| Gm13306 | 1.067961214 | 0.644851161 | 1.055799092 | 0.61077071 |
| Gm13305 | 0.062426873 | 0.755288839 | 3.724945049 | 0.202765096 |
| 1700045I11Rik | 0 | NA | NA | NA |
| Gm17167 | 0.21074972 | -0.687379132 | 2.626897576 | -0.26166956 |
| Gm13301 | 0 | NA | NA | NA |
| Gm5859 | 0 | NA | NA | NA |
| Gm17081 | 0 | NA | NA | NA |
| Gm10600 | 0 | NA | NA | NA |
| Ccl21b | 0 | NA | NA | NA |
| Gm21980 | 0 | NA | NA | NA |
| Gm21953 | 8.239491219 | -0.901823116 | 0.359293321 | -2.509991316 |
| Gm21598 | 0 | NA | NA | NA |
| Gm21586 | 0.193603971 | 1.259282423 | 3.249745779 | 0.387501826 |
| Gm3883 | 0 | NA | NA | NA |
| Gm2163 | 0 | NA | NA | NA |
| Gm13298 | 0.089814205 | 0.274402725 | 3.724945049 | 0.073666248 |
| Gm10593 | 0.444623086 | 2.856481377 | 3.202684442 | 0.891902224 |
| Gm10591 | 0 | NA | NA | NA |
| Gm2564 | 0 | NA | NA | NA |
| Ccl27b | 1.785252223 | 0.839020168 | 0.710255104 | 1.181294106 |
| Il11ra2 | 0.424152111 | 0.044954669 | 1.663857999 | 0.027018333 |

| | | | | |
|---|---|---|---|---|
| Gm21955 | 83.35882431 | -0.796336496 | 0.209993265 | -3.792200165 |
| 4930578G10Rik | 26.07152981 | 0.201587916 | 0.350604926 | 0.574971716 |
| Gm12395 | 6.47297608 | -0.018005564 | 0.566659864 | -0.031774906 |
| Ccl19 | 26.89383427 | 6.521870944 | 0.86710172 | 7.521460048 |
| Ccl21a | 1.739286695 | 1.054244864 | 0.893734532 | 1.179595088 |
| Gm12394 | 0.61514822 | -0.369824864 | 1.163386972 | -0.317886372 |
| Gm12429 | 2.41835347 | -0.673700011 | 0.67293864 | -1.001131412 |
| BC049635 | 0.963622687 | 0.167257903 | 1.20332607 | 0.138996326 |
| N28178 | 2446.431538 | -0.448861676 | 0.150456642 | -2.98332908 |
| Dnajb5 | 1123.231011 | -0.588837601 | 0.165679406 | -3.554078424 |
| 1700022I11Rik | 5.07266936 | 1.721209468 | 0.905498278 | 1.900842343 |
| Vcp | 2874.0526 | 0.152190714 | 0.065565203 | 2.321211676 |
| Fancg | 91.45811176 | -0.402271775 | 0.170356802 | -2.361348479 |
| Pigo | 262.034385 | -0.081254149 | 0.082269511 | -0.9876581 |
| Stoml2 | 348.4296267 | 0.105626736 | 0.095660278 | 1.10418596 |
| Gm26643 | 0.357756446 | -2.169096518 | 1.352728745 | -1.603497025 |
| Fam214b | 535.3240349 | 0.2728207 | 0.082372011 | 3.312055841 |
| Gm26881 | 11.80662906 | -0.947775995 | 0.322373595 | -2.939992639 |
| Unc13b | 466.7629952 | -0.53951372 | 0.185568072 | -2.907362852 |
| Gm23709 | 0 | NA | NA | NA |
| Gm25010 | 0 | NA | NA | NA |
| Atp8b5 | 7.649160145 | -0.728310472 | 0.408363011 | -1.783487859 |
| Rusc2 | 1981.730262 | 0.019824416 | 0.07063009 | 0.280679471 |
| Fam166b | 3.554918289 | -0.148936498 | 0.549744815 | -0.270919333 |
| Tesk1 | 834.3782588 | 0.036870829 | 0.064026058 | 0.57587223 |
| Cd72 | 162.2518301 | 6.096992308 | 0.378045265 | 16.127678 |
| Sit1 | 2.004396688 | 3.812487378 | 0.999890299 | 3.812905659 |
| Rmrp | 17396.1871 | 0.025371921 | 0.169913545 | 0.149322531 |
| Ccdc107 | 211.9356722 | 0.075778545 | 0.151294795 | 0.500866835 |
| Arhgef39 | 4.559760072 | 0.381028005 | 0.497806335 | 0.765414134 |
| Gm12454 | 3.511636557 | 0.046590219 | 0.570090354 | 0.081724272 |
| Car9 | 11.98985167 | -0.195750129 | 0.349209176 | -0.560552651 |
| Tpm2 | 58.51175456 | -0.419452126 | 0.200921982 | -2.087636813 |
| Tln1 | 1972.671443 | 1.094651994 | 0.130792331 | 8.369389775 |
| Creb3 | 390.8704704 | 0.592542389 | 0.127691583 | 4.640418542 |
| Gba2 | 788.9852365 | -0.060887131 | 0.053086234 | -1.146947643 |
| Rgp1 | 609.9554589 | 0.112081504 | 0.056028818 | 2.000425977 |
| Msmp | 0.198131738 | -0.206481488 | 2.22921086 | -0.092625373 |
| Gm12472 | 0.580871419 | 2.054098034 | 1.37696402 | 1.491758684 |
| Gm25262 | 9.682546495 | -0.702312997 | 0.423773603 | -1.657283492 |
| Gm12473 | 1.944300039 | 0.579129272 | 0.782714097 | 0.739898865 |
| Npr2 | 447.7307936 | -0.252606306 | 0.096398151 | -2.620447617 |
| Spag8 | 14.00717955 | 0.198242696 | 0.319144378 | 0.621169317 |
| Hint2 | 86.66528165 | 0.27597813 | 0.153205045 | 1.801364502 |
| Fam221b | 17.22801957 | -0.910776429 | 0.279505543 | -3.25852725 |
| Gm12481 | 74.19163858 | 0.070151224 | 0.178910096 | 0.392103215 |
| Tmem8b | 757.5108737 | -0.116537001 | 0.079854731 | -1.459362504 |

| | | | | |
|---|---|---|---|---|
| Gm23257 | 4.230344492 | -1.282804456 | 0.542752352 | -2.363517084 |
| Olfr70 | 0 | NA | NA | NA |
| Olfr71 | 0 | NA | NA | NA |
| Hrct1 | 4.20546811 | 1.02789671 | 0.620024144 | 1.65783336 |
| 5430416O09Rik | 3.015746376 | 0.313390154 | 0.581286782 | 0.539131739 |
| A630077J23Rik | 0.321964759 | 0.997321839 | 1.992554776 | 0.500524177 |
| Olfr269-ps1 | 0 | NA | NA | NA |
| Olfr159 | 0 | NA | NA | NA |
| Olfr29-ps1 | 0 | NA | NA | NA |
| Gm12492 | 0 | NA | NA | NA |
| Olfr156 | 0.048949532 | -0.206487837 | 3.724945049 | -0.055433794 |
| Olfr157 | 0 | NA | NA | NA |
| Olfr155 | 0.08169834 | -0.206492286 | 3.724945049 | -0.055434988 |
| Reck | 121.3301511 | 0.218429722 | 0.185816414 | 1.175513607 |
| Gm27265 | 2.022028428 | -0.022236658 | 0.749481365 | -0.029669394 |
| Glipr2 | 555.7186848 | 4.334898061 | 0.355033651 | 12.20982308 |
| Ccin | 0.952506814 | 1.759563304 | 1.213725213 | 1.449721308 |
| Clta | 1302.037201 | 0.102905562 | 0.080750416 | 1.274365726 |
| Gne | 417.5634539 | -0.147713555 | 0.078492625 | -1.881878124 |
| Gm22706 | 0.185638728 | 1.204575731 | 2.665023689 | 0.451994381 |
| Gm12503 | 0.506185493 | 0.25834958 | 1.457389596 | 0.177268714 |
| Gm12504 | 0.323274704 | 0.803111663 | 1.598393851 | 0.50244917 |
| Rnf38 | 803.2801551 | -0.042264609 | 0.068724899 | -0.614982476 |
| Mir5106 | 2.834065591 | 0.387313746 | 0.619265851 | 0.62544018 |
| Melk | 12.45686811 | 2.332831203 | 0.439786124 | 5.304467499 |
| Gm12677 | 0 | NA | NA | NA |
| Gm12455 | 0 | NA | NA | NA |
| Gm24376 | 0 | NA | NA | NA |
| Pax5 | 62.51624943 | 1.129876484 | 0.197864051 | 5.710367676 |
| Mir5120 | 0.090665643 | -0.687373951 | 3.724945049 | -0.184532642 |
| Gm12462 | 5.015296868 | 0.63130613 | 0.512519782 | 1.231769296 |
| 4930517J16Rik | 1.109399417 | 0.940320552 | 1.235757514 | 0.760926429 |
| 5730488B01Rik | 1.238426347 | 0.637836159 | 0.903790576 | 0.705734466 |
| Gm12463 | 0.758386275 | -1.367594803 | 1.167688704 | -1.17119811 |
| Zcchc7 | 970.4223863 | -0.482244783 | 0.147499305 | -3.269471558 |
| Gm22639 | 3.577120001 | -1.400539425 | 0.587327565 | -2.384596788 |
| Gm12678 | 0 | NA | NA | NA |
| Gm12493 | 0 | NA | NA | NA |
| Gm12679 | 2.474621302 | 2.103130938 | 0.720838526 | 2.917617279 |
| Grhpr | 156.0778864 | -0.429907265 | 0.124256311 | -3.459842488 |
| Zbtb5 | 270.4928444 | 0.67228164 | 0.111567982 | 6.025757842 |
| 1700055D18Rik | 14.55774235 | 0.683407182 | 0.278949157 | 2.449934567 |
| Polr1e | 68.48577099 | -0.212753336 | 0.157084074 | -1.354391505 |
| Fbxo10 | 581.7003137 | 0.149180152 | 0.070146764 | 2.12668615 |
| Tomm5 | 134.9204341 | 0.226528709 | 0.147891363 | 1.531723727 |
| Gm12566 | 0 | NA | NA | NA |
| Frmpd1 | 273.0835449 | 0.669101649 | 0.138482355 | 4.831674393 |

| | | | | |
|---|---|---|---|---|
| Frmpd1os | 0.15456712 | -0.206480331 | 2.83422604 | -0.072852457 |
| Trmt10b | 109.9286799 | -0.066920846 | 0.104614251 | -0.639691487 |
| Exosc3 | 78.17585261 | 0.22706935 | 0.167891515 | 1.352476628 |
| Gm12585 | 0 | NA | NA | NA |
| Dcaf10 | 447.5914563 | 0.208053974 | 0.1070383 | 1.943733912 |
| Slc25a51 | 825.9827367 | 0.111603193 | 0.046184984 | 2.416438946 |
| Gm25692 | 0 | NA | NA | NA |
| Shb | 131.2113877 | 0.093761299 | 0.123379211 | 0.759944069 |
| Gm22518 | 0.171044682 | 0.274409026 | 2.745278379 | 0.099956721 |
| Gm22055 | 0 | NA | NA | NA |
| Gm12411 | 5.818705562 | -0.327300515 | 0.475281031 | -0.688646282 |
| Gm12408 | 2.792582105 | -0.246825093 | 0.54111846 | -0.456138742 |
| Gm829 | 0.154612032 | 0.274389336 | 3.501099905 | 0.078372324 |
| Gm12410 | 0.044191549 | -0.206490061 | 3.724945049 | -0.055434391 |
| Gm12409 | 0.36880298 | 1.221008595 | 1.569797421 | 0.777812843 |
| Gm12412 | 0.044544648 | 0.755286614 | 3.724945049 | 0.202764498 |
| Aldh1b1 | 125.3782894 | 1.421314665 | 0.180748075 | 7.863512041 |
| Igfbpl1 | 40.81830737 | 0.347785407 | 0.280283201 | 1.24083572 |
| Stra6l | 52.92012026 | 4.606347472 | 0.571985342 | 8.053261392 |
| E230008N13Rik | 12.72585994 | 0.01503293 | 0.361092056 | 0.041631848 |
| Tdrd7 | 1191.587 | 1.375617906 | 0.125688597 | 10.94465159 |
| Tmod1 | 407.2062115 | -0.637132762 | 0.12918348 | -4.93199874 |
| Tstd2 | 221.2458736 | -0.203290528 | 0.108416168 | -1.87509419 |
| Gm23250 | 0.397571503 | -0.134504525 | 3.261621106 | -0.04123855 |
| Ncbp1 | 647.6804023 | -0.003305551 | 0.07158034 | -0.046179594 |
| Xpa | 75.64072436 | 0.524410758 | 0.174806216 | 2.999954857 |
| Gm12444 | 2.312647122 | -0.094006805 | 0.644654832 | -0.145825022 |
| Gm12445 | 0 | NA | NA | NA |
| Gm12446 | 0.091666476 | -0.687373951 | 3.724945049 | -0.184532642 |
| Gm12447 | 2.955086367 | -0.006362985 | 0.55825527 | -0.011397984 |
| Foxe1 | 0.36004552 | 2.168174117 | 1.706854542 | 1.270274686 |
| 5830415F09Rik | 17.35131692 | -0.780959546 | 0.273943839 | -2.850801643 |
| Hemgn | 1.129829066 | 1.59466417 | 1.353326234 | 1.17832946 |
| Gm12443 | 0.104366086 | 0.274400501 | 3.724945049 | 0.073665651 |
| Anp32b | 407.7977659 | 0.278784442 | 0.122262244 | 2.280216956 |
| Nans | 132.4373756 | -0.049134419 | 0.147702012 | -0.332659107 |
| Trim14 | 69.63745902 | 4.040039539 | 0.347961928 | 11.61057925 |
| Gm22247 | 0.26079097 | -0.206481974 | 2.453645846 | -0.084153129 |
| Gm16731 | 0.468050163 | 1.754917252 | 1.437379514 | 1.220914334 |
| Coro2a | 311.7321339 | -0.065873583 | 0.080692084 | -0.816357435 |
| Tbc1d2 | 54.86226986 | 0.458611164 | 0.207226025 | 2.213096369 |
| Gabbr2 | 3104.900051 | -0.166624822 | 0.117769459 | -1.41483899 |
| Gm568 | 0.609243813 | -0.466779913 | 1.425212565 | -0.327515996 |
| Anks6 | 149.640635 | -0.363667858 | 0.195417904 | -1.86097512 |
| Gm12425 | 0 | NA | NA | NA |
| Gm12422 | 0.506756615 | 2.575331715 | 1.34617715 | 1.913070441 |
| Galnt12 | 14.31060596 | 0.486944228 | 0.315087931 | 1.545423295 |

| | | | | |
|---|---|---|---|---|
| Gm12424 | 0.961043033 | -0.899832534 | 1.050557557 | -0.856528544 |
| Gm12423 | 0 | NA | NA | NA |
| Gm12426 | 1.368259282 | -0.165745238 | 0.856039841 | -0.193618603 |
| Col15a1 | 61.55752535 | -0.557579659 | 0.180054823 | -3.096721592 |
| Tgfbr1 | 481.364332 | 0.12468766 | 0.144822046 | 0.860971541 |
| Gm12430 | 5.138410126 | 0.285454336 | 0.417406275 | 0.683876485 |
| Alg2 | 1382.70197 | 0.004074205 | 0.063740544 | 0.063918581 |
| Sec61b | 158.1254426 | 0.247852809 | 0.1604088 | 1.545132245 |
| Gm27601 | 0 | NA | NA | NA |
| Gm27668 | 0 | NA | NA | NA |
| Gm38343 | 0.066050875 | 0.274391604 | 3.724945049 | 0.073663262 |
| Gm22670 | 0 | NA | NA | NA |
| Gm12434 | 0 | NA | NA | NA |
| Nr4a3 | 542.7975049 | -0.818661003 | 0.309616303 | -2.644114652 |
| Stx17 | 355.1670537 | 0.11777415 | 0.073865566 | 1.594439154 |
| Erp44 | 524.436621 | 0.077678711 | 0.093517671 | 0.830631373 |
| Mir1958 | 0.694929783 | -2.083219981 | 1.231462812 | -1.691662925 |
| Invs | 182.2944049 | -0.400975634 | 0.115637237 | -3.467530392 |
| Gm12435 | 13.93294907 | 0.523080652 | 0.299642788 | 1.745680765 |
| Tex10 | 345.3321695 | -0.074886587 | 0.077590686 | -0.965149182 |
| Msantd3 | 126.8937499 | -0.03881078 | 0.113584375 | -0.341691191 |
| Tmeff1 | 650.557545 | -0.640148209 | 0.113365419 | -5.646767904 |
| Murc | 22.05245246 | -0.115152965 | 0.23440869 | -0.491248701 |
| Gm12439 | 2.996720251 | -1.653868026 | 0.705477204 | -2.344325253 |
| Gm12438 | 0 | NA | NA | NA |
| Gm24573 | 0 | NA | NA | NA |
| Gm23746 | 0 | NA | NA | NA |
| Gm12436 | 0.044544648 | 0.755286614 | 3.724945049 | 0.202764498 |
| Gm12442 | 0 | NA | NA | NA |
| Lppr1 | 119.1937067 | -0.108855868 | 0.144529107 | -0.753176091 |
| Gm22685 | 0.467201629 | -0.166819435 | 1.411050657 | -0.118223562 |
| Gm12453 | 0.624449418 | -0.93461921 | 1.273726911 | -0.733767342 |
| Rbpsuh-ps1 | 0.090074004 | 0.274402725 | 3.724945049 | 0.073666248 |
| Acnat2 | 0.218471152 | 0.755309375 | 1.911296337 | 0.395181721 |
| Gm12451 | 0.314054052 | 1.350329655 | 3.70375512 | 0.364583945 |
| Gm12452 | 0.075043205 | 0.755286614 | 3.724945049 | 0.202764498 |
| Acnat1 | 2.07728567 | 0.56414223 | 0.840665623 | 0.671066134 |
| Gm26424 | 0 | NA | NA | NA |
| Baat | 6.000090479 | -0.404220429 | 0.500819443 | -0.807118083 |
| Mrpl50 | 361.7674449 | 0.173009254 | 0.089846173 | 1.925616297 |
| Zfp189 | 205.1948466 | 0.454474458 | 0.097923056 | 4.64113844 |
| Aldob | 16.11872095 | 0.658656413 | 0.289769161 | 2.273038345 |
| Tmem246 | 593.6062876 | -0.142121851 | 0.074279139 | -1.913348133 |
| Rnf20 | 1042.085228 | -0.040801305 | 0.07103642 | -0.574371643 |
| Grin3a | 866.8310369 | 0.069747987 | 0.07499615 | 0.930020894 |
| Ppp3r2 | 15.47209461 | -0.25568962 | 0.33210459 | -0.769906914 |
| Gm12459 | 3.099509663 | -0.823062679 | 0.5139486 | -1.601449403 |

| | | | | |
|---|---|---|---|---|
| Gm12458 | 0.048949532 | -0.206487837 | 3.724945049 | -0.055433794 |
| Gm25485 | 0 | NA | NA | NA |
| Gm12460 | 0.083526043 | 0.75528439 | 3.724945049 | 0.202763901 |
| Cylc2 | 0 | NA | NA | NA |
| Amd-ps4 | 32.8894957 | 0.054196445 | 0.214796273 | 0.252315573 |
| Gm22563 | 0 | NA | NA | NA |
| Gm23043 | 0 | NA | NA | NA |
| Gm24943 | 0 | NA | NA | NA |
| Gm22042 | 0.482432794 | 1.167237562 | 1.456738958 | 0.801267485 |
| Gm12461 | 0 | NA | NA | NA |
| C630028M04Rik | 0.154174174 | -0.686749087 | 2.834967975 | -0.242242273 |
| Gm12465 | 0 | NA | NA | NA |
| Gm12466 | 0.144050924 | 1.290908285 | 3.721447108 | 0.346883416 |
| Gm12467 | 0 | NA | NA | NA |
| Gm12475 | 1.233559488 | 2.967868066 | 1.151600595 | 2.577167881 |
| Smc2os | 3.507361272 | -0.340088443 | 0.586827985 | -0.579536852 |
| Smc2 | 184.4916281 | -0.140027802 | 0.171646979 | -0.815789494 |
| Tmc4-ps | 0 | NA | NA | NA |
| Vma21-ps | 0.038981395 | 0.274393828 | 3.724945049 | 0.073663859 |
| Toporsl | 0 | NA | NA | NA |
| 4930552N02Rik | 0 | NA | NA | NA |
| Gm12482 | 0.399060758 | -0.687379237 | 1.728950558 | -0.39757021 |
| Olfr275 | 0 | NA | NA | NA |
| Olfr273 | 0.062426873 | 0.755288839 | 3.724945049 | 0.202765096 |
| Cct3-ps1 | 0.33824293 | -0.206486819 | 1.945198697 | -0.106152044 |
| Olfr272 | 0.046345709 | 0.274391604 | 3.724945049 | 0.073663262 |
| Olfr271-ps1 | 0.118974955 | 0.274400501 | 3.724945049 | 0.073665651 |
| Olfr270 | 0 | NA | NA | NA |
| Nipsnap3a | 0 | NA | NA | NA |
| Nipsnap3b | 127.01825 | 0.523838334 | 0.189242607 | 2.768078192 |
| Abca1 | 2154.257843 | 0.624007841 | 0.145757798 | 4.281128336 |
| 4930412L05Rik | 1.32543013 | 1.393910343 | 1.007625378 | 1.383361687 |
| 4930522O17Rik | 0.038981395 | 0.274393828 | 3.724945049 | 0.073663859 |
| AI427809 | 3.06421064 | 1.159138229 | 0.768098978 | 1.50910008 |
| Gm12495 | 0 | NA | NA | NA |
| Gm12496 | 0.503001268 | -1.36442152 | 1.480828439 | -0.921390679 |
| Slc44a1 | 1434.826822 | 0.01370279 | 0.086385953 | 0.158622899 |
| 1700060J05Rik | 0.251076704 | 0.279358265 | 2.507215301 | 0.111421729 |
| Fsd1l | 701.9750228 | -0.385358038 | 0.061405228 | -6.275655153 |
| Gm12470 | 15.34197034 | -0.32574106 | 0.351731327 | -0.926107614 |
| Fktn | 705.1094383 | -0.404981307 | 0.062419011 | -6.488108346 |
| Gm25445 | 0 | NA | NA | NA |
| Tal2 | 3.478949006 | 1.658694212 | 0.63686138 | 2.604482332 |
| Rps15a-ps8 | 0 | NA | NA | NA |
| Tmem38b | 167.1596389 | 0.165106264 | 0.113018068 | 1.46088379 |
| n-R5s185 | 0 | NA | NA | NA |
| Gm12468 | 0.068097359 | -0.206487837 | 3.724945049 | -0.055433794 |

| | | | | |
|---|---|---|---|---|
| Gm23745 | 0.047948698 | -0.206487837 | 3.724945049 | -0.055433794 |
| Gm24177 | 0 | NA | NA | NA |
| Gm12484 | 0 | NA | NA | NA |
| Gm12469 | 0.094831987 | 0.274404949 | 3.724945049 | 0.073666845 |
| Gm25687 | 0 | NA | NA | NA |
| Tpt1-ps2 | 0 | NA | NA | NA |
| Gm29067 | 0 | NA | NA | NA |
| Gm12478 | 0 | NA | NA | NA |
| Gm12480 | 0.086930093 | -0.206490058 | 3.724945049 | -0.05543439 |
| Gm12477 | 0 | NA | NA | NA |
| Gm24628 | 0 | NA | NA | NA |
| Gm12479 | 0 | NA | NA | NA |
| Zfp462 | 1321.864971 | -0.425496328 | 0.076389145 | -5.570115097 |
| Gm12515 | 0.097139547 | 0.274393828 | 3.724945049 | 0.073663859 |
| Gm12514 | 0.145451803 | 0.274387818 | 3.53800508 | 0.077554388 |
| Gm12509 | 0 | NA | NA | NA |
| Gm12508 | 0.167320703 | 0.274408884 | 2.764437523 | 0.099263912 |
| Gm12512 | 0 | NA | NA | NA |
| Gm25419 | 0 | NA | NA | NA |
| Rad23b | 2280.331224 | 0.184181321 | 0.07382847 | 2.494719467 |
| Gm12516 | 0 | NA | NA | NA |
| Gm12513 | 0 | NA | NA | NA |
| Gm12505 | 0.048949532 | -0.206487837 | 3.724945049 | -0.055433794 |
| Gm12510 | 0 | NA | NA | NA |
| Gm22150 | 0 | NA | NA | NA |
| Gm12507 | 0 | NA | NA | NA |
| Klf4 | 85.90030529 | 1.866271073 | 0.271468388 | 6.874727058 |
| Gm12511 | 0.772413307 | 0.59900329 | 1.235545483 | 0.484808774 |
| Gm12506 | 1.081462396 | -0.804584085 | 1.000205114 | -0.804419088 |
| Gm12519 | 0.384295504 | -0.670349841 | 1.59496592 | -0.420291012 |
| Gm12520 | 0 | NA | NA | NA |
| 2310081O03Rik | 1.122999083 | 2.337547794 | 1.232614963 | 1.896413612 |
| Gm12518 | 0 | NA | NA | NA |
| Gm26144 | 0.110595523 | 0.755282166 | 3.724945049 | 0.202763304 |
| Actl7b | 1.554701746 | -0.02202677 | 0.834234045 | -0.026403585 |
| Gm26657 | 0 | NA | NA | NA |
| Actl7a | 0.04393175 | -0.206490061 | 3.724945049 | -0.055434391 |
| Ikbkap | 924.710914 | -0.107683202 | 0.088267267 | -1.219967555 |
| Gm22952 | 4.197905476 | 0.135109644 | 0.456682508 | 0.29585027 |
| Fam206a | 278.4213948 | -0.220179134 | 0.091215644 | -2.413830841 |
| Ctnnal1 | 338.824844 | -0.114110738 | 0.131511441 | -0.867686776 |
| Tmem245 | 2131.110561 | -0.059723045 | 0.091566972 | -0.652233488 |
| Mir32 | 0.646815127 | -1.62454561 | 1.403548617 | -1.157455888 |
| Gm25053 | 0.048949532 | -0.206487837 | 3.724945049 | -0.055433794 |
| Gm22300 | 0.061920613 | 0.274393828 | 3.724945049 | 0.073663859 |
| Frrs1l | 3263.550938 | -0.435131523 | 0.103475578 | -4.205161574 |
| Epb4.1l4b | 724.2344794 | -0.345033298 | 0.083606427 | -4.126875309 |

| | | | | |
|---|---|---|---|---|
| Gm12530 | 6.592824524 | 2.156377781 | 0.717936598 | 3.003576902 |
| Ptpn3 | 752.3310739 | -0.261560656 | 0.116652929 | -2.242212507 |
| Gm12536 | 3.13007401 | -0.045886823 | 0.592234039 | -0.077480894 |
| 1700042G15Rik | 0 | NA | NA | NA |
| Gm12537 | 23.16944423 | 0.040673332 | 0.22358161 | 0.181917164 |
| Gm12538 | 1.143639422 | -0.587022853 | 1.302826257 | -0.450576467 |
| Palm2 | 230.289783 | -0.312349612 | 0.115196412 | -2.711452611 |
| Gm12539 | 0 | NA | NA | NA |
| Pakap | 9.557701205 | 0.276500622 | 0.295193232 | 0.936676699 |
| Akap2 | 22.84245767 | 0.218912991 | 0.251485719 | 0.870478816 |
| Gm12526 | 0.201955748 | 1.25779302 | 2.218326002 | 0.567000981 |
| Gm24277 | 1.91044629 | -0.814592601 | 0.804660149 | -1.012343662 |
| Gm26138 | 0 | NA | NA | NA |
| D630039A03Rik | 3.846205949 | -0.083366979 | 0.580751259 | -0.143550234 |
| Txn1 | 650.0859016 | 0.858973676 | 0.164807238 | 5.211989993 |
| Txndc8 | 0 | NA | NA | NA |
| Svep1 | 270.025923 | -0.686986557 | 0.217496934 | -3.158603404 |
| Gm26889 | 1.802555565 | -1.757044031 | 0.856920919 | -2.050415614 |
| Musk | 19.66503011 | 0.239434852 | 0.220377612 | 1.08647539 |
| Lpar1 | 579.1101207 | -0.226456556 | 0.137434224 | -1.647745006 |
| Mir3095 | 0.494865952 | -0.953131863 | 1.410115039 | -0.675924898 |
| Gm12577 | 4.644629704 | -1.119335932 | 0.634113706 | -1.765197506 |
| Gm12578 | 0 | NA | NA | NA |
| Gm12579 | 0.044191549 | -0.206490061 | 3.724945049 | -0.055434391 |
| Gm12580 | 0.526179982 | -1.590666009 | 1.305170163 | -1.218742241 |
| Olfr267 | 2.961215905 | -0.238158883 | 0.659236696 | -0.361264603 |
| AI314180 | 2945.683157 | 0.19588442 | 0.074295567 | 2.636555972 |
| Gm26400 | 0 | NA | NA | NA |
| Zkscan16 | 430.9535622 | -0.196354436 | 0.095802747 | -2.049569993 |
| Ptgr1 | 21.48653641 | 0.319261038 | 0.399875345 | 0.798401407 |
| Dnajc25 | 87.02722299 | 0.133975593 | 0.162689792 | 0.823503376 |
| Gm20503 | 0 | NA | NA | NA |
| Gng10 | 14.26907646 | -0.433056956 | 0.29278596 | -1.479090582 |
| AI481877 | 0.167924218 | 0.755296859 | 2.761499998 | 0.273509636 |
| Gm12593 | 0.308655657 | 2.071435376 | 3.704213912 | 0.559210517 |
| Gm12594 | 0 | NA | NA | NA |
| Rpsa-ps11 | 1.26617739 | -0.21332769 | 0.951474439 | -0.224207483 |
| Ugcg | 1300.167576 | -0.093643004 | 0.057654511 | -1.624209479 |
| Gm12596 | 0.346136167 | 1.21605331 | 1.693783257 | 0.717950957 |
| Susd1 | 107.3121581 | -0.174014681 | 0.16222945 | -1.072645452 |
| Gm12527 | 0.551374303 | -0.713189629 | 1.241421911 | -0.574494153 |
| Ptbp3 | 1030.313491 | 0.922201212 | 0.117624616 | 7.840205929 |
| Gm12529 | 0 | NA | NA | NA |
| 1700018M17Rik | 1.139841548 | 2.026583147 | 1.068549818 | 1.896573387 |
| Hsdl2 | 568.5576942 | 0.497454912 | 0.118024015 | 4.214861794 |
| E130308A19Rik | 348.4769703 | -0.200168133 | 0.068258767 | -2.932489733 |
| Inip | 231.4780127 | -0.043141134 | 0.086661299 | -0.497813143 |

| | | | | |
|---|---|---|---|---|
| Snx30 | 979.4902318 | -0.080263983 | 0.066856639 | -1.20053871 |
| Gm12543 | 7.392584082 | 0.397310901 | 0.428240491 | 0.927775184 |
| Gm12542 | 0.038981395 | 0.274393828 | 3.724945049 | 0.073663859 |
| Slc46a2 | 0 | NA | NA | NA |
| Mup4 | 0.187777844 | -1.407193087 | 3.298122832 | -0.426664851 |
| Mup6 | 0.095435502 | 0.75528439 | 3.724945049 | 0.202763901 |
| Mup7 | 0.045882455 | 0.755288839 | 3.724945049 | 0.202765096 |
| Mup-ps1 | 0 | NA | NA | NA |
| Mup2 | 0 | NA | NA | NA |
| Mup-ps2 | 0 | NA | NA | NA |
| Mup8 | 0 | NA | NA | NA |
| Mup-ps3 | 0 | NA | NA | NA |
| Mup-ps29 | 0 | NA | NA | NA |
| Mup-ps28 | 0 | NA | NA | NA |
| Mup9 | 0 | NA | NA | NA |
| Mup-ps4 | 0 | NA | NA | NA |
| Mup-ps26 | 0 | NA | NA | NA |
| Mup1 | 0 | NA | NA | NA |
| Gm13773 | 0 | NA | NA | NA |
| Mup-ps6 | 0 | NA | NA | NA |
| Mup-ps24 | 0 | NA | NA | NA |
| Mup10 | 0 | NA | NA | NA |
| Mup-ps5 | 0 | NA | NA | NA |
| Mup11 | 0 | NA | NA | NA |
| Mup-ps7 | 0 | NA | NA | NA |
| Mup12 | 0 | NA | NA | NA |
| Mup-ps8 | 0 | NA | NA | NA |
| Gm21286 | 0 | NA | NA | NA |
| Mup-ps9 | 0 | NA | NA | NA |
| Mup13 | 0 | NA | NA | NA |
| Mup-ps10 | 0 | NA | NA | NA |
| Mup14 | 0 | NA | NA | NA |
| Mup-ps11 | 0 | NA | NA | NA |
| Mup15 | 0 | NA | NA | NA |
| Mup-ps12 | 0 | NA | NA | NA |
| Mup16 | 0 | NA | NA | NA |
| Mup-ps13 | 0 | NA | NA | NA |
| Mup17 | 0 | NA | NA | NA |
| Mup-ps14 | 0 | NA | NA | NA |
| Mup18 | 0 | NA | NA | NA |
| Mup-ps15 | 0 | NA | NA | NA |
| Gm12573 | 0 | NA | NA | NA |
| Mup19 | 0 | NA | NA | NA |
| Gm13775 | 0 | NA | NA | NA |
| Mup-ps16 | 0 | NA | NA | NA |
| Mup5 | 0.135858026 | -1.143903624 | 2.941468349 | -0.388888639 |
| Mup-ps23 | 0 | NA | NA | NA |

| | | | | |
|---|---|---|---|---|
| Mup-ps17 | 0 | NA | NA | NA |
| Mup-ps18 | 0 | NA | NA | NA |
| Mup-ps19 | 0 | NA | NA | NA |
| Mup-ps20 | 0 | NA | NA | NA |
| Mup-ps21 | 0 | NA | NA | NA |
| Mup20 | 0 | NA | NA | NA |
| Mup3 | 1.273710888 | 0.309553666 | 0.92305392 | 0.335358162 |
| Gm12909 | 0 | NA | NA | NA |
| Gm12910 | 0 | NA | NA | NA |
| Mup21 | 0.088476398 | 0.274400501 | 3.724945049 | 0.073665651 |
| Zfp37 | 301.4106421 | 0.269400624 | 0.077764931 | 3.464294523 |
| Slc31a2 | 164.8023296 | 1.048029851 | 0.21492207 | 4.876324953 |
| Gm23148 | 0 | NA | NA | NA |
| Fkbp15 | 744.3892095 | 0.340160915 | 0.074291359 | 4.578741348 |
| Slc31a1 | 348.3933879 | 0.384259985 | 0.114161729 | 3.365926468 |
| Cdc26 | 169.15404 | 0.123042895 | 0.106183546 | 1.158775534 |
| Prpf4 | 294.0494031 | -0.014828825 | 0.076437677 | -0.193998894 |
| Rnf183 | 1.151952751 | 2.836954713 | 1.205994124 | 2.352378554 |
| Wdr31 | 53.77822627 | -0.215851392 | 0.152215961 | -1.418060178 |
| Bspry | 7.199256456 | 0.248211533 | 0.445371536 | 0.557313419 |
| Hdhd3 | 119.7174087 | -0.389427276 | 0.1431355 | -2.720689668 |
| Alad | 452.599456 | -0.081250973 | 0.084188797 | -0.965104337 |
| Pole3 | 241.0396122 | 0.375718604 | 0.098824279 | 3.801885621 |
| 4933430I17Rik | 8.456414667 | 0.730506533 | 0.387218876 | 1.886546806 |
| Rgs3 | 280.2490358 | 0.130460323 | 0.079736184 | 1.636149574 |
| Gm11209 | 0.195038611 | -0.672217163 | 3.240686736 | -0.207430467 |
| Gm11210 | 0.707441185 | 1.441827962 | 1.276613742 | 1.129415981 |
| Gm11211 | 0.066050875 | 0.274391604 | 3.724945049 | 0.073663262 |
| Gm24117 | 0 | NA | NA | NA |
| Gm25391 | 0 | NA | NA | NA |
| Gm11481 | 0 | NA | NA | NA |
| Gm11480 | 0 | NA | NA | NA |
| Zfp618 | 184.4983496 | -0.008775518 | 0.106876836 | -0.082108695 |
| Ambp | 0.720892307 | 1.116916381 | 1.470432987 | 0.759583327 |
| Kif12 | 0.229693479 | 0.776449392 | 2.138260945 | 0.363121907 |
| Col27a1 | 179.4777456 | -1.122421425 | 0.198771692 | -5.646787096 |
| Mir455 | 0.26766946 | -0.67667175 | 2.082674493 | -0.324905189 |
| Orm1 | 0.137647365 | 1.404948034 | 3.719469264 | 0.377728093 |
| Gm11212 | 0 | NA | NA | NA |
| Orm3 | 0.157314084 | 0.755289343 | 3.435173412 | 0.219869349 |
| Orm2 | 0.751984476 | 2.681272025 | 1.205659504 | 2.223904856 |
| Akna | 219.0544516 | 1.46306236 | 0.119684661 | 12.22430967 |
| Aknaos | 0.425011668 | 1.888748921 | 1.488734384 | 1.268694363 |
| Whrn | 563.7172746 | -0.32431044 | 0.082388707 | -3.936345806 |
| Atp6v1g1 | 617.2868754 | 0.309055798 | 0.113587639 | 2.720857669 |
| 6330416G13Rik | 190.6195337 | 0.53027389 | 0.115767142 | 4.580521552 |
| Gm11483 | 0 | NA | NA | NA |

| | | | | |
|---|---|---|---|---|
| Gm24448 | 0 | NA | NA | NA |
| Gm11213 | 0 | NA | NA | NA |
| 1700018C11Rik | 0 | NA | NA | NA |
| Gm24234 | 0 | NA | NA | NA |
| Gm11482 | 1.060662479 | 0.488054209 | 1.04443672 | 0.467289401 |
| Gm11214 | 0 | NA | NA | NA |
| Tnfsf15 | 4.678869644 | 1.785340655 | 0.732286248 | 2.438036573 |
| Tnfsf8 | 49.08655836 | 5.188384807 | 0.552775055 | 9.386068993 |
| Rpl17-ps4 | 3.619380876 | -0.271459149 | 0.546801352 | -0.496449302 |
| Tnc | 854.476176 | 1.052143784 | 0.320684896 | 3.280927156 |
| 8030451A03Rik | 1.099630792 | 1.142348643 | 1.043344342 | 1.094891301 |
| Gm11216 | 0.038981395 | 0.274393828 | 3.724945049 | 0.073663859 |
| Gm11217 | 0 | NA | NA | NA |
| Gm23950 | 0 | NA | NA | NA |
| Gm11218 | 0.060174537 | 0.755286614 | 3.724945049 | 0.202764498 |
| Gm11219 | 0 | NA | NA | NA |
| Pappa | 141.7302013 | 0.102734704 | 0.220003878 | 0.466967697 |
| Astn2 | 1043.79549 | -0.247587748 | 0.086285491 | -2.869401856 |
| Trim32 | 1762.038928 | 0.012126232 | 0.059500213 | 0.203801494 |
| Gm11484 | 0 | NA | NA | NA |
| Gm25480 | 0 | NA | NA | NA |
| Gm11220 | 0 | NA | NA | NA |
| Tlr4 | 178.7502272 | 1.399829375 | 0.242721051 | 5.76723513 |
| Gm11403 | 0 | NA | NA | NA |
| Hmgb1-rs18 | 0 | NA | NA | NA |
| AL589845.1 | 0 | NA | NA | NA |
| Gm11249 | 12.40046496 | 0.022549396 | 0.369839316 | 0.060970791 |
| Gm11751 | 0.126225412 | 0.755282661 | 3.698754989 | 0.20419916 |
| Gm23738 | 0 | NA | NA | NA |
| Tcp1-ps1 | 0.143022002 | 0.755289479 | 3.547809333 | 0.212888971 |
| Gm12911 | 0 | NA | NA | NA |
| Brinp1 | 2163.011251 | 0.145314338 | 0.116752129 | 1.244639724 |
| Gm24697 | 0 | NA | NA | NA |
| Gm11404 | 1.959659191 | -2.225685708 | 1.25163846 | -1.778217736 |
| Gm11221 | 11.46002724 | 0.986240527 | 0.347322591 | 2.839551907 |
| Gm11222 | 0.623048784 | -0.260910541 | 1.321215004 | -0.197477731 |
| Gm11223 | 93.55249628 | -0.175671747 | 0.153324947 | -1.145747968 |
| Gm11224 | 5.294292594 | 0.170716583 | 0.468546892 | 0.36435325 |
| Gm11225 | 0.044191549 | -0.206490061 | 3.724945049 | -0.055434391 |
| Cdk5rap2 | 668.7817866 | 0.149496445 | 0.09930025 | 1.505499182 |
| Gm11226 | 21.32553314 | 0.289130179 | 0.251081274 | 1.151540193 |
| Gm26320 | 0.207279166 | 0.755304285 | 2.634010914 | 0.286750628 |
| Megf9 | 2380.976153 | -0.071293766 | 0.082554993 | -0.863591212 |
| Gm11227 | 0 | NA | NA | NA |
| Gm11228 | 0 | NA | NA | NA |
| Gm11229 | 0 | NA | NA | NA |
| Rps18-ps1 | 0.478648982 | 2.094999381 | 1.389194328 | 1.508067906 |

| | | | | |
|---|---|---|---|---|
| Gm11231 | 0.345321076 | -0.204772951 | 1.748533964 | -0.117111223 |
| Gm11232 | 0.115000406 | -0.206492286 | 3.724945049 | -0.055434988 |
| Gm11233 | 0 | NA | NA | NA |
| Gm11234 | 0 | NA | NA | NA |
| Tle1 | 623.9206616 | -0.257350944 | 0.096480585 | -2.667385815 |
| Gm11250 | 0 | NA | NA | NA |
| C630043F03Rik | 39.86748014 | -0.05665825 | 0.189397253 | -0.299150328 |
| Gm11235 | 0.263040051 | 0.695664 | 1.811516539 | 0.384022991 |
| Aldoart1 | 2.66134452 | -0.168870948 | 0.60073776 | -0.281105933 |
| Gm5287 | 0 | NA | NA | NA |
| Gm25653 | 0 | NA | NA | NA |
| Gm11251 | 0 | NA | NA | NA |
| Gm25769 | 0 | NA | NA | NA |
| Gm11487 | 0 | NA | NA | NA |
| Gm11486 | 0 | NA | NA | NA |
| Gm11485 | 0.097139547 | 0.274393828 | 3.724945049 | 0.073663859 |
| Gm11488 | 14.4078922 | 0.363069556 | 0.319313301 | 1.13703236 |
| Gm24170 | 0 | NA | NA | NA |
| Gm11238 | 0 | NA | NA | NA |
| Gm11237 | 0 | NA | NA | NA |
| Gm11236 | 0 | NA | NA | NA |
| Gm11239 | 0 | NA | NA | NA |
| Gm428 | 0 | NA | NA | NA |
| Rasef | 2.980195284 | 0.317249971 | 0.608207464 | 0.521614729 |
| Gm11240 | 0.068097359 | -0.206487837 | 3.724945049 | -0.055433794 |
| Gm13867 | 0.251494677 | 1.835008761 | 2.49482048 | 0.735527376 |
| Gm11758 | 0 | NA | NA | NA |
| Gm11757 | 0 | NA | NA | NA |
| Gm13871 | 0 | NA | NA | NA |
| Gm11756 | 0 | NA | NA | NA |
| 2310002L09Rik | 0.092493346 | 0.274402725 | 3.724945049 | 0.073666248 |
| Frmd3 | 144.1270821 | -0.430831725 | 0.125836241 | -3.42374916 |
| Gm11760 | 4.795815343 | -1.088316307 | 0.548075651 | -1.985704537 |
| Gm11405 | 10.60243355 | -0.330996386 | 0.382510897 | -0.865325377 |
| Kdm4c | 648.3243287 | -0.033014141 | 0.085755682 | -0.384979053 |
| A230083N12Rik | 2.600875038 | 0.845787227 | 0.657544693 | 1.286280972 |
| Gm26379 | 0 | NA | NA | NA |
| Gm11254 | 0.066050875 | 0.274391604 | 3.724945049 | 0.073663262 |
| Gm11258 | 0 | NA | NA | NA |
| Gm11253 | 0 | NA | NA | NA |
| Gm11257 | 0 | NA | NA | NA |
| Gm12912 | 0.235437891 | -1.141966923 | 2.566528568 | -0.444946118 |
| Tmem261 | 148.8567893 | -0.112355289 | 0.114749479 | -0.979135502 |
| Gm22864 | 0 | NA | NA | NA |
| Gm11255 | 0 | NA | NA | NA |
| Gm11259 | 0 | NA | NA | NA |
| Gm11256 | 0 | NA | NA | NA |

| | | | | |
|---|---|---|---|---|
| Gm11241 | 0.044544648 | 0.755286614 | 3.724945049 | 0.202764498 |
| Ptprd | 4811.876378 | -0.384019467 | 0.099600277 | -3.855606422 |
| Gm11252 | 0 | NA | NA | NA |
| Gm11242 | 0.10503227 | 0.274389469 | 3.724945049 | 0.073662689 |
| Gm11243 | 0 | NA | NA | NA |
| Gm11244 | 7.743757494 | -0.347440684 | 0.35785561 | -0.97089629 |
| Gm11245 | 0 | NA | NA | NA |
| Gm11246 | 0.158325629 | 0.75529608 | 2.813153441 | 0.268487338 |
| Gm23159 | 10.5795326 | -0.285878462 | 0.394270032 | -0.725082909 |
| B230208B08Rik | 26.21483297 | -0.203506613 | 0.256978809 | -0.791919824 |
| Gm11260 | 0.431719086 | 0.291121966 | 1.463101164 | 0.198975965 |
| Gm11261 | 0.068097359 | -0.206487837 | 3.724945049 | -0.055433794 |
| Gm22262 | 0 | NA | NA | NA |
| Gm11262 | 0.136963818 | 0.755286583 | 3.59954433 | 0.209828388 |
| Gm25836 | 0 | NA | NA | NA |
| Gm11263 | 32.27155379 | 0.741349995 | 0.235045775 | 3.15406646 |
| Gm11409 | 0 | NA | NA | NA |
| Gm11407 | 1.154096829 | -1.490297246 | 0.867866617 | -1.717196188 |
| Gm11408 | 0.245813264 | 1.607879323 | 2.05284663 | 0.783243765 |
| Gm11410 | 0.40441216 | 0.134371366 | 1.698955119 | 0.079090592 |
| Gm11406 | 0 | NA | NA | NA |
| Tyrp1 | 7.568395809 | 0.516838138 | 0.459860508 | 1.123901985 |
| Lurap1l | 131.167045 | 0.11248431 | 0.111041138 | 1.012996728 |
| Gm27452 | 1.818514136 | 0.579925379 | 0.787709222 | 0.736217582 |
| Mpdz | 900.4834607 | -0.250213345 | 0.073242122 | -3.416249257 |
| Gm25803 | 3.471866242 | -0.798446049 | 0.545701846 | -1.463154386 |
| Gm11765 | 0 | NA | NA | NA |
| Gm11411 | 0.049553047 | 0.274391604 | 3.724945049 | 0.073663262 |
| Gm11412 | 0.477185738 | -1.755135289 | 1.316492543 | -1.333190452 |
| n-R5s187 | 0 | NA | NA | NA |
| Gm12914 | 0 | NA | NA | NA |
| Gm11264 | 0.922800452 | 0.425877331 | 1.108794825 | 0.384090294 |
| Gm22402 | 0 | NA | NA | NA |
| Gm5860 | 3.25720724 | 0.196315976 | 0.527711259 | 0.372013999 |
| Nfib | 2628.924338 | -0.248161358 | 0.097878866 | -2.535392671 |
| n-R5s188 | 0 | NA | NA | NA |
| Gm11266 | 18.26086893 | -0.708858732 | 0.270523418 | -2.620322994 |
| Gm11267 | 0.708875568 | -1.160333202 | 1.229323697 | -0.943879309 |
| Gm11268 | 0.189675868 | 0.274409669 | 2.656640733 | 0.103291975 |
| Gm11269 | 0.044544648 | 0.755286614 | 3.724945049 | 0.202764498 |
| Zdhhc21 | 1208.957435 | -0.218845533 | 0.052617414 | -4.159184535 |
| Cer1 | 0.481690985 | -0.058130084 | 1.324962883 | -0.043872991 |
| Frem1 | 91.57009446 | -0.287046006 | 0.171081408 | -1.67783285 |
| Gm24884 | 0 | NA | NA | NA |
| Gm11248 | 0 | NA | NA | NA |
| Gm11185 | 2.941272167 | -0.452348232 | 0.672919942 | -0.672217012 |
| Ttc39b | 1497.137622 | 0.464675905 | 0.099515112 | 4.669400427 |

| | | | | |
|---|---|---|---|---|
| Gm23412 | 0.392867355 | 1.241852422 | 1.968644856 | 0.630815872 |
| Gm11413 | 1.891350131 | 0.131551918 | 0.831758526 | 0.158161189 |
| Gm11184 | 0 | NA | NA | NA |
| Snapc3 | 534.9796752 | -0.2162536 | 0.098204227 | -2.202080369 |
| Psip1 | 2060.877521 | -0.005200927 | 0.076483216 | -0.068000889 |
| Gm11414 | 0.092493346 | 0.274402725 | 3.724945049 | 0.073666248 |
| Gm10154 | 11.65614726 | 0.602491329 | 0.503782056 | 1.195936459 |
| Ccdc171 | 44.96192503 | 0.047772843 | 0.174006832 | 0.274545787 |
| Gm25899 | 0 | NA | NA | NA |
| Gm27046 | 0.323557773 | -0.687379291 | 1.758682669 | -0.390848959 |
| Cbx3-ps3 | 0.066050875 | 0.274391604 | 3.724945049 | 0.073663262 |
| Gm26968 | 0.264809544 | -0.675096381 | 2.093082436 | -0.322536929 |
| Gm11415 | 0 | NA | NA | NA |
| 6030471H07Rik | 0 | NA | NA | NA |
| Gm12416 | 6.021123113 | 0.152048191 | 0.455395336 | 0.33388175 |
| Gm12414 | 0.092269992 | -0.20649451 | 3.724945049 | -0.055435586 |
| Bnc2 | 94.00517019 | -0.123226144 | 0.161502015 | -0.763000665 |
| Gm12421 | 0.473868817 | 0.703137735 | 1.39370251 | 0.504510633 |
| Gm12420 | 0 | NA | NA | NA |
| Gm12415 | 0.047948698 | -0.206487837 | 3.724945049 | -0.055433794 |
| Cntln | 269.0535505 | -0.225928713 | 0.089967242 | -2.511233081 |
| Gm12418 | 9.82225973 | -0.581192544 | 0.324801357 | -1.789378433 |
| Gm12419 | 0.575593679 | -0.911255149 | 1.308897669 | -0.696200452 |
| Gm12417 | 0 | NA | NA | NA |
| Sh3gl2 | 2324.421738 | -0.14670969 | 0.05944044 | -2.468179773 |
| Gm12413 | 0 | NA | NA | NA |
| Gm12680 | 8.074609009 | 0.549955666 | 0.421958978 | 1.303339175 |
| Gm26069 | 1.027412341 | -0.646530827 | 0.969280644 | -0.667021292 |
| Gm25811 | 0 | NA | NA | NA |
| Adamtsl1 | 125.2776745 | 0.37035848 | 0.137970295 | 2.684334914 |
| Gm12550 | 0.075043205 | 0.755286614 | 3.724945049 | 0.202764498 |
| Saxo1 | 0.416337256 | 0.590355097 | 1.650255329 | 0.35773561 |
| Gm26017 | 0 | NA | NA | NA |
| Saxo1os | 0 | NA | NA | NA |
| Rraga | 797.0060883 | 0.08153012 | 0.086368814 | 0.943976377 |
| Haus6 | 193.2480707 | 0.15842062 | 0.087442779 | 1.811706138 |
| Scarna8 | 0.537445765 | -0.968734783 | 1.216174849 | -0.796542359 |
| Gm12551 | 0.925310778 | 1.652251667 | 1.429262606 | 1.156016858 |
| Plin2 | 668.7166373 | 2.518969798 | 0.35568529 | 7.082018474 |
| Dennd4c | 657.3655427 | 0.226495796 | 0.073371031 | 3.08699214 |
| Gm12630 | 2.006602685 | 2.779148138 | 1.069648542 | 2.598188122 |
| Rps6 | 1228.330895 | 0.550955921 | 0.069427095 | 7.935747817 |
| Acer2 | 61.2692064 | 0.439762617 | 0.199401892 | 2.205408449 |
| Slc24a2 | 8155.610505 | -0.158786405 | 0.103544943 | -1.533502278 |
| Gm12600 | 0.561663198 | -0.111789918 | 1.257063616 | -0.088929404 |
| Gm23154 | 0 | NA | NA | NA |
| Gm12604 | 0.075043205 | 0.755286614 | 3.724945049 | 0.202764498 |

| | | | | |
|---|---|---|---|---|
| Mllt3 | 907.8146563 | 0.071153866 | 0.07160752 | 0.993664724 |
| Gm12631 | 3.345683444 | 0.483221718 | 0.521147542 | 0.92722632 |
| Gm12646 | 0.591515762 | 0.672312184 | 1.438326634 | 0.467426639 |
| Focad | 1053.950382 | -0.145526772 | 0.108478268 | -1.341529276 |
| Mir491 | 10.7430229 | -0.27631842 | 0.303131125 | -0.911547501 |
| Gm12645 | 3.761156174 | -0.15760165 | 0.530635477 | -0.297005491 |
| Hacd4 | 63.03965297 | 0.354335517 | 0.235774113 | 1.502860144 |
| Ifnb1 | 44.66037375 | 8.316976597 | 1.21508855 | 6.844749377 |
| Ifna15 | 0.599468885 | 2.817361754 | 1.483393798 | 1.899267583 |
| Ifna14 | 0.481692262 | 1.920848379 | 1.660891208 | 1.156516676 |
| Ifna-ps1 | 0 | NA | NA | NA |
| Ifna9 | 0.316561663 | 0.274407703 | 1.962550721 | 0.139821967 |
| Mrpl48-ps | 38.64384611 | 0.216389933 | 0.243912088 | 0.887163628 |
| Ifna12 | 0.176391548 | 0.755304606 | 2.718652661 | 0.277823135 |
| Gm12601 | 0.215725693 | -0.203102913 | 2.182795414 | -0.093047159 |
| C87499 | 0.779996687 | 1.45671341 | 1.239526786 | 1.175217371 |
| Ifna13 | 0.075043205 | 0.755286614 | 3.724945049 | 0.202764498 |
| Gm13281 | 0.046345709 | 0.274391604 | 3.724945049 | 0.073663262 |
| Ifna16 | 0.145446656 | 0.755296437 | 2.882544899 | 0.262024171 |
| Ifna2 | 8.251847137 | 5.922249076 | 1.063885342 | 5.566623433 |
| Ifnab | 1.397532748 | 3.574627179 | 1.07108271 | 3.337396024 |
| Gm13273 | 0.313664056 | 1.81576471 | 3.710734157 | 0.489327619 |
| Gm13282 | 2.459004477 | 0.734342843 | 0.709988482 | 1.034302472 |
| Klhl9 | 1871.695579 | -0.070287491 | 0.08737869 | -0.804400833 |
| Gm26566 | 4.232091902 | -0.18407034 | 0.508437212 | -0.362031604 |
| Gm13274 | 0.044191549 | -0.206490061 | 3.724945049 | -0.055434391 |
| Gm13271 | 0.08486385 | 0.755286617 | 3.724945049 | 0.202764499 |
| Gm16686 | 0 | NA | NA | NA |
| Gm13286 | 0 | NA | NA | NA |
| Gm10583 | 0.144244773 | -0.687376278 | 3.581436307 | -0.191927545 |
| Gm13283 | 0 | NA | NA | NA |
| Gm10582 | 0 | NA | NA | NA |
| Gm13284 | 0 | NA | NA | NA |
| Gm10581 | 0 | NA | NA | NA |
| Gm13290 | 0 | NA | NA | NA |
| Gm10580 | 0 | NA | NA | NA |
| Gm13289 | 0 | NA | NA | NA |
| Gm13272 | 0 | NA | NA | NA |
| Ifnz | 0 | NA | NA | NA |
| Gm26525 | 0.092269992 | -0.20649451 | 3.724945049 | -0.055435586 |
| Gm13276 | 0 | NA | NA | NA |
| Gm13277 | 0 | NA | NA | NA |
| Gm13278 | 0 | NA | NA | NA |
| Gm13275 | 0 | NA | NA | NA |
| Gm26867 | 0.045882455 | 0.755288839 | 3.724945049 | 0.202765096 |
| Gm13279 | 0 | NA | NA | NA |
| Gm13285 | 0 | NA | NA | NA |

| | | | | |
|---|---|---|---|---|
| Gm13287 | 0 | NA | NA | NA |
| Gm13288 | 0 | NA | NA | NA |
| Ifna7 | 0.305381988 | 2.030573186 | 2.416093053 | 0.840436664 |
| Ifna11 | 0.275757984 | 1.862467219 | 3.708580271 | 0.502204909 |
| Ifna6 | 0.418480079 | 2.78416884 | 3.678030391 | 0.756972766 |
| Ifna5 | 11.73825278 | 6.617175961 | 1.279797718 | 5.170485827 |
| Ifna4 | 14.65900129 | 7.236434174 | 1.401170483 | 5.16456367 |
| Ifna1 | 2.156732396 | 4.448978936 | 1.309175577 | 3.398305784 |
| 4930553M12Rik | 0.266535354 | -0.687379355 | 2.124587725 | -0.323535407 |
| Ifne | 0.083526043 | 0.75528439 | 3.724945049 | 0.202763901 |
| Mir31 | 1.252737907 | 1.616014364 | 0.898625134 | 1.79831868 |
| Gm12603 | 0.048949532 | -0.206487837 | 3.724945049 | -0.055433794 |
| Gm12602 | 0 | NA | NA | NA |
| Mtap | 151.2844411 | -0.105172166 | 0.151423096 | -0.694558283 |
| Gm12607 | 0 | NA | NA | NA |
| Gm26490 | 0 | NA | NA | NA |
| Gm12606 | 0.046345709 | 0.274391604 | 3.724945049 | 0.073663262 |
| Cdkn2a | 2.014914513 | 3.412047499 | 1.02429244 | 3.331126313 |
| Cdkn2b | 6.964193097 | 1.989668064 | 0.539691134 | 3.686679173 |
| Gm12609 | 0.942397274 | 0.568291324 | 1.165329172 | 0.487665921 |
| Gm12610 | 0.811625864 | 3.664786985 | 1.14951629 | 3.188112265 |
| Gm12608 | 0 | NA | NA | NA |
| Gm27580 | 0.090427103 | 1.236179401 | 3.724945049 | 0.331865138 |
| Dmrta1 | 4.432480129 | -0.163633115 | 0.449612566 | -0.363942486 |
| Gm12629 | 0 | NA | NA | NA |
| Zfp352 | 0 | NA | NA | NA |
| Gm12633 | 0.176993581 | -0.687378681 | 2.299612248 | -0.298910689 |
| Gm12634 | 0 | NA | NA | NA |
| Gm12635 | 0 | NA | NA | NA |
| Gm12636 | 0 | NA | NA | NA |
| Gm12632 | 0 | NA | NA | NA |
| Gm25244 | 0.236157963 | 2.079075828 | 2.481497198 | 0.837831221 |
| Gm12643 | 0 | NA | NA | NA |
| Gm12644 | 0.116217011 | 0.274402725 | 3.724945049 | 0.073666248 |
| Elavl2 | 1479.215199 | -0.02567637 | 0.122445145 | -0.20969692 |
| Gm12653 | 0 | NA | NA | NA |
| Mir6402 | 6.56355345 | -0.74649542 | 0.451752272 | -1.652444197 |
| Gm12668 | 0.048949532 | -0.206487837 | 3.724945049 | -0.055433794 |
| Gm12670 | 0.219222703 | 1.094295263 | 2.169844874 | 0.504319583 |
| Gm12667 | 0.171083144 | 0.27438982 | 3.398950499 | 0.080727807 |
| Gm12669 | 20.03569431 | 0.197027293 | 0.237088406 | 0.831028796 |
| Gm12671 | 13.22935111 | 0.171277291 | 0.291483235 | 0.587605977 |
| 4930577H14Rik | 0 | NA | NA | NA |
| Izumo3 | 0 | NA | NA | NA |
| Gm12666 | 1.628724105 | 0.225774862 | 0.854856528 | 0.264108485 |
| Gm12638 | 0 | NA | NA | NA |
| Gm12637 | 0 | NA | NA | NA |

| | | | | |
|---|---|---|---|---|
| Gm22605 | 0 | NA | NA | NA |
| Gm12639 | 0 | NA | NA | NA |
| Gm12641 | 0 | NA | NA | NA |
| Gm12642 | 0 | NA | NA | NA |
| Tusc1 | 50.66388116 | 0.194310168 | 0.169139049 | 1.148819092 |
| Gm12649 | 0.203196539 | 1.22244354 | 2.594324671 | 0.471199135 |
| Gm23443 | 0 | NA | NA | NA |
| Gm12640 | 0 | NA | NA | NA |
| Gm24796 | 0 | NA | NA | NA |
| Gm22619 | 0 | NA | NA | NA |
| Gm12650 | 0 | NA | NA | NA |
| Gm12647 | 0 | NA | NA | NA |
| Gm12648 | 0.255640523 | -0.687379848 | 2.125070327 | -0.323462165 |
| Gm23836 | 0 | NA | NA | NA |
| Gm12654 | 0 | NA | NA | NA |
| Gm26410 | 0.038981395 | 0.274393828 | 3.724945049 | 0.073663859 |
| Gm12655 | 23.30350443 | -0.095867991 | 0.227288773 | -0.421789382 |
| Gm22804 | 0 | NA | NA | NA |
| Gm12656 | 0 | NA | NA | NA |
| Caap1 | 152.6588818 | -0.390168599 | 0.117609368 | -3.317495908 |
| Gm10306 | 0.49643671 | -1.338775338 | 1.431072529 | -0.935504882 |
| Plaa | 900.5907788 | 0.303358192 | 0.084732078 | 3.580204779 |
| Gm12657 | 14.22801902 | 0.078154541 | 0.291908925 | 0.267736045 |
| Ift74 | 232.3827732 | 0.063859964 | 0.090777934 | 0.70347452 |
| Lrrc19 | 0.551940858 | 1.777113555 | 1.526911905 | 1.163861221 |
| Mir872 | 1.137124893 | 0.120675229 | 0.89794383 | 0.134390621 |
| Tek | 47.11216226 | 0.134311068 | 0.211215527 | 0.635895808 |
| Eqtn | 11.24146221 | -0.306463388 | 0.405572422 | -0.755631722 |
| Gm12693 | 29.98890503 | 0.047593669 | 0.194768838 | 0.244359774 |
| Mysm1 | 851.8413644 | 0.233603795 | 0.085121151 | 2.74436838 |
| Gm12694 | 0.203088869 | 0.716472593 | 2.636766587 | 0.271723935 |
| Gm12703 | 1.47600346 | -0.146402317 | 1.008282711 | -0.14519967 |
| Gm12704 | 0.126912322 | -0.206491353 | 3.697934765 | -0.055839642 |
| Jun | 1515.464005 | 1.137879236 | 0.231862514 | 4.907560155 |
| Junos | 29.43953063 | 0.197940512 | 0.231703369 | 0.854284135 |
| n-R5s189 | 0.792393139 | 0.68514648 | 0.964618125 | 0.710277428 |
| 4930551L18Rik | 0.046345709 | 0.274391604 | 3.724945049 | 0.073663262 |
| Gm28096 | 1.499901305 | -0.409452289 | 0.832176952 | -0.49202551 |
| Gm12708 | 1.926663799 | 0.681634469 | 0.961959518 | 0.708589557 |
| Gm29064 | 0 | NA | NA | NA |
| Gm830 | 0.19500349 | -0.687380054 | 3.29079171 | -0.208879842 |
| Gm12707 | 0 | NA | NA | NA |
| Gm25004 | 0 | NA | NA | NA |
| Fggy | 21.91263025 | 0.240800229 | 0.259222072 | 0.928934127 |
| Gm20651 | 0 | NA | NA | NA |
| 9530080O11Rik | 18.68130564 | 0.445132561 | 0.232912245 | 1.911159975 |
| Hook1 | 1088.16238 | -0.006355162 | 0.11095071 | -0.057279149 |

| | | | | |
|---|---|---|---|---|
| Cyp2j13 | 7.229154167 | 0.694616278 | 0.376136447 | 1.846713562 |
| Hspe1-ps6 | 0.042716945 | -0.206490061 | 3.724945049 | -0.055434391 |
| Cyp2j12 | 25.97268065 | -0.354624134 | 0.217121802 | -1.633295832 |
| Cyp2j7-ps3 | 0 | NA | NA | NA |
| Cyp2j7-ps2 | 0 | NA | NA | NA |
| Cyp2j7-ps1 | 0 | NA | NA | NA |
| Cyp2j7 | 0.068097359 | -0.206487837 | 3.724945049 | -0.055433794 |
| Cyp2j15-ps | 0 | NA | NA | NA |
| Cyp2j11 | 0.27158854 | -0.699252823 | 2.096046632 | -0.333605566 |
| Cyp2j14-ps | 0 | NA | NA | NA |
| Cyp2j8 | 2.219411473 | 0.971770966 | 0.623105234 | 1.559561551 |
| Cyp2j6 | 245.6472783 | -0.008820921 | 0.131318898 | -0.067171754 |
| Cyp2j9 | 418.0043846 | 0.503071097 | 0.14905452 | 3.375081132 |
| Cyp2j5 | 0.490888186 | -0.231081532 | 1.711754738 | -0.13499687 |
| Gm12695 | 0.642811059 | -0.539143146 | 1.09858844 | -0.490759893 |
| Gm27579 | 0 | NA | NA | NA |
| Gm27521 | 0 | NA | NA | NA |
| Gm10192 | 0 | NA | NA | NA |
| Gm12696 | 28.72011168 | -0.190720638 | 0.20858877 | -0.914337995 |
| Gm12697 | 3.534004104 | -1.130725364 | 0.713189745 | -1.585448151 |
| E130114P18Rik | 46.0958998 | -0.126645121 | 0.172071436 | -0.736003162 |
| Gm12676 | 1.650824257 | -0.533015912 | 1.18783612 | -0.448728493 |
| Nfia | 1906.233348 | 0.024674372 | 0.082172047 | 0.300276957 |
| Gm12691 | 1.556603733 | -0.378238063 | 0.808018279 | -0.468105825 |
| Gm12692 | 0.373707575 | -0.984243705 | 1.509383115 | -0.652083421 |
| Gm12787 | 0 | NA | NA | NA |
| Gm24203 | 0 | NA | NA | NA |
| 0610025J13Rik | 0.422542347 | -0.194144781 | 1.454642798 | -0.133465605 |
| Gm12788 | 0 | NA | NA | NA |
| Tm2d1 | 205.9564824 | -0.136407574 | 0.106568702 | -1.279996575 |
| Gm24960 | 0 | NA | NA | NA |
| Inadl | 702.0612203 | -0.287473079 | 0.128311349 | -2.24043377 |
| Gm22625 | 0.042716945 | -0.206490061 | 3.724945049 | -0.055434391 |
| Gm26154 | 4.026424441 | -0.055827787 | 0.601359061 | -0.092836028 |
| Gm12846 | 13.02039719 | -0.525021312 | 0.355592189 | -1.476470315 |
| L1td1 | 1.260641324 | -0.811933857 | 0.90107945 | -0.901068 |
| Gm17376 | 0.049553047 | 0.274391604 | 3.724945049 | 0.073663262 |
| Kank4 | 103.4907107 | -0.379673265 | 0.162668695 | -2.334027855 |
| Kank4os | 0.479560471 | 1.351298122 | 1.672452533 | 0.807973976 |
| I0C0044D17Rik | 0.072025462 | 0.755288839 | 3.724945049 | 0.202765096 |
| Gm12851 | 0 | NA | NA | NA |
| Usp1 | 392.4990393 | 0.519002243 | 0.104019305 | 4.989480028 |
| Dock7 | 964.8693113 | -0.149434526 | 0.08259234 | -1.809302491 |
| Angptl3 | 2.293039322 | 1.793945276 | 0.78488699 | 2.285609647 |
| Gm12852 | 6.635996975 | -0.531490655 | 0.516390675 | -1.029241388 |
| Gm12705 | 0.112642008 | 0.274402725 | 3.724945049 | 0.073666248 |
| Gm12681 | 0.364815146 | 0.258205302 | 1.649329638 | 0.156551666 |

| | | | | |
|---|---|---|---|---|
| Atg4c | 539.9595983 | -0.31169817 | 0.087977073 | -3.542947717 |
| Gm10305 | 27.16950442 | -0.373613488 | 0.275080467 | -1.358197086 |
| Gm12689 | 9.012307495 | -0.53913486 | 0.43656415 | -1.234949914 |
| Gm37556 | 0 | NA | NA | NA |
| Gm12685 | 0 | NA | NA | NA |
| Gm12686 | 0 | NA | NA | NA |
| Gm12684 | 0 | NA | NA | NA |
| Gm12683 | 0 | NA | NA | NA |
| Gm37117 | 0 | NA | NA | NA |
| Gm12690 | 0 | NA | NA | NA |
| Gm12687 | 0 | NA | NA | NA |
| Gm12688 | 2.558789337 | -0.728341658 | 0.837855854 | -0.869292319 |
| Foxd3 | 16.0637295 | 0.079472746 | 1.075202024 | 0.073914246 |
| Gm23366 | 0 | NA | NA | NA |
| Gm12682 | 0 | NA | NA | NA |
| Alg6 | 154.9068683 | -0.291209157 | 0.15903499 | -1.831101177 |
| Itgb3bp | 53.12261853 | 0.297841771 | 0.168238885 | 1.770350359 |
| Efcab7 | 42.69675528 | 0.366453663 | 0.195334798 | 1.876028578 |
| Gm26425 | 0.198326984 | 0.777338998 | 2.231313123 | 0.34837737 |
| Pgm2 | 636.1507098 | 0.020692825 | 0.082126013 | 0.251964315 |
| Gm12702 | 0 | NA | NA | NA |
| Ror1 | 118.724752 | 0.171542994 | 0.15179185 | 1.130119924 |
| Gm12701 | 11.03703998 | -1.12509695 | 0.416544315 | -2.701025818 |
| Gm12706 | 0.200296414 | 0.728012831 | 2.212362357 | 0.329065819 |
| Ube2u | 0.310183902 | 1.14413713 | 1.758379698 | 0.650676945 |
| Ube2uos | 0 | NA | NA | NA |
| Ccdc50-ps | 0.574153598 | -0.310084191 | 1.214654721 | -0.255285873 |
| Gm12700 | 0 | NA | NA | NA |
| Cachd1 | 422.3160165 | 0.012123773 | 0.075390518 | 0.160812968 |
| Gm10577 | 0 | NA | NA | NA |
| Gm10576 | 0 | NA | NA | NA |
| Raver2 | 272.6356933 | -0.59510515 | 0.187565047 | -3.172793445 |
| Gm22100 | 0.061920613 | 0.274393828 | 3.724945049 | 0.073663859 |
| Jak1 | 2519.077659 | 0.230084449 | 0.063050064 | 3.649234161 |
| Gm24468 | 0.165470308 | 1.646276139 | 2.753318586 | 0.597924318 |
| Gm12785 | 2.331474832 | -0.761993323 | 0.743634041 | -1.024688597 |
| Gm25124 | 0 | NA | NA | NA |
| Gm12801 | 0 | NA | NA | NA |
| Gm12796 | 0.250633908 | 0.74515247 | 1.836442681 | 0.405758632 |
| Gm12793 | 0.066050875 | 0.274391604 | 3.724945049 | 0.073663262 |
| E130102H24Rik | 106.6384263 | 0.592595199 | 0.211668896 | 2.799632874 |
| Mir101a | 0 | NA | NA | NA |
| 0610043K17Rik | 25.29227341 | -0.221080605 | 0.267299249 | -0.82709026 |
| Gm12795 | 0 | NA | NA | NA |
| Gm12798 | 2.114628594 | 0.137402962 | 0.73375392 | 0.187260277 |
| Ak4 | 432.3296171 | -0.708803056 | 0.147386224 | -4.809154043 |
| Dnajc6 | 4085.768198 | -0.129630922 | 0.057555299 | -2.252284727 |

| | | | | |
|---|---|---|---|---|
| Gm22533 | 11.8805187 | -0.740349762 | 0.356455376 | -2.076977404 |
| Leprot | 268.8974141 | 0.184653412 | 0.118008212 | 1.564750519 |
| Lepr | 95.56977508 | 0.452240144 | 0.146487986 | 3.087216623 |
| B020004J07Rik | 0.102891481 | 0.274400501 | 3.724945049 | 0.073665651 |
| Gm12791 | 3.064345978 | -0.076686387 | 0.580329546 | -0.132142827 |
| Gm12797 | 0.048949532 | -0.206487837 | 3.724945049 | -0.055433794 |
| C130073F10Rik | 0 | NA | NA | NA |
| Gm12792 | 0 | NA | NA | NA |
| Gm12800 | 0.092691418 | 0.274387304 | 3.724945049 | 0.073662108 |
| Gm12794 | 0.223364959 | 0.755300291 | 2.521604034 | 0.299531679 |
| Gm12790 | 0.047948698 | -0.206487837 | 3.724945049 | -0.055433794 |
| Gm12789 | 0.194279094 | 0.274391173 | 3.724945049 | 0.073663147 |
| Gm12799 | 4.949076095 | -0.727616993 | 0.473266715 | -1.537435382 |
| Pde4b | 1759.695653 | 0.552796513 | 0.094190022 | 5.868949821 |
| Gm24869 | 0 | NA | NA | NA |
| Sgip1 | 2873.427705 | -0.154231811 | 0.075698277 | -2.037454707 |
| Gm12709 | 0.447471404 | -0.335613996 | 1.532970868 | -0.218930446 |
| Tctex1d1 | 5.63377873 | -0.216135114 | 0.520104135 | -0.415561231 |
| BB031773 | 3.502581246 | -0.132396093 | 0.538808789 | -0.245719996 |
| Insl5 | 5.684128039 | -1.101505718 | 0.481837181 | -2.286053801 |
| Wdr78 | 212.7570655 | -0.715107477 | 0.161781976 | -4.42019249 |
| Gm12710 | 0.045882455 | 0.755288839 | 3.724945049 | 0.202765096 |
| Mier1 | 957.8111048 | 0.164201567 | 0.059912779 | 2.740676857 |
| Slc35d1 | 196.800066 | -0.352009611 | 0.104470535 | -3.36946309 |
| 4921539E11Rik | 0.165933562 | 1.204233133 | 2.762106933 | 0.435983531 |
| 4930456L15Rik | 0.038981395 | 0.274393828 | 3.724945049 | 0.073663859 |
| Oma1 | 95.40654656 | -0.173001849 | 0.175751542 | -0.984354655 |
| Gm12718 | 0.250798881 | 0.274411209 | 2.084063917 | 0.131671206 |
| Gm23064 | 0 | NA | NA | NA |
| Gm12720 | 0.060174537 | 0.755286614 | 3.724945049 | 0.202764498 |
| Gm12716 | 0 | NA | NA | NA |
| Gm12715 | 393.1600641 | 0.399498146 | 0.108262768 | 3.690078818 |
| Gm22801 | 0 | NA | NA | NA |
| Gm25877 | 0 | NA | NA | NA |
| Dab1 | 1235.718576 | -0.012731214 | 0.077258632 | -0.164786944 |
| Gm12719 | 0 | NA | NA | NA |
| Gm12717 | 3.781002861 | 0.089562613 | 0.529445513 | 0.169163041 |
| C8b | 6.711931076 | 2.685158815 | 0.982114366 | 2.734059198 |
| C8a | 0.626920446 | 0.257147831 | 1.279948851 | 0.200904771 |
| Gm17662 | 0.225129614 | 1.818209708 | 3.710727189 | 0.489987438 |
| 1700024P16Rik | 18.7794585 | 0.826471567 | 0.310184337 | 2.664452938 |
| Gm12721 | 0.110242424 | -0.20649451 | 3.724945049 | -0.055435586 |
| Prkaa2 | 803.30275 | -0.131718596 | 0.049900979 | -2.639599435 |
| Ppap2b | 1987.101825 | -0.361881574 | 0.131448299 | -2.753033534 |
| Gm12722 | 2.944400151 | 0.55058607 | 0.588795092 | 0.93510642 |
| Gm12723 | 0 | NA | NA | NA |
| Gm12726 | 0.106112162 | -0.206492286 | 3.724945049 | -0.055434988 |

| | | | | |
|---|---|---|---|---|
| Gm12728 | 0.174579622 | -1.161156722 | 2.306965692 | -0.503326393 |
| Gm24363 | 0 | NA | NA | NA |
| Gm12729 | 0.086930093 | -0.206490058 | 3.724945049 | -0.05543439 |
| Gm12727 | 0.640223289 | 1.205671051 | 1.369625105 | 0.880292751 |
| Gm26047 | 0 | NA | NA | NA |
| Hspd1-ps4 | 0.60453213 | 1.605687425 | 1.432695666 | 1.120745643 |
| Gm12724 | 0.146823232 | -0.194513984 | 2.876272087 | -0.067627115 |
| Gm12731 | 0 | NA | NA | NA |
| Gm12730 | 0.072025462 | 0.755288839 | 3.724945049 | 0.202765096 |
| Gm37253 | 0 | NA | NA | NA |
| Usp24 | 2364.206895 | 0.18222305 | 0.075462512 | 2.414749332 |
| Pcsk9 | 22.35786243 | -0.965713754 | 0.262971861 | -3.672308328 |
| Bsnd | 0.044191549 | -0.206490061 | 3.724945049 | -0.055434391 |
| Tmem61 | 0 | NA | NA | NA |
| 2700068H02Rik | 1.22619721 | 0.035925628 | 1.040240824 | 0.034535876 |
| Dhcr24 | 1131.153985 | -0.325508732 | 0.113524563 | -2.867297821 |
| Gm12744 | 0.280446515 | -1.571627912 | 1.798134543 | -0.874032434 |
| BC055111 | 2.227745575 | 0.539629718 | 0.658905749 | 0.818978617 |
| Ttc22 | 5.229776616 | -0.493374621 | 0.422984336 | -1.166413457 |
| Pars2 | 65.0940951 | 0.327791733 | 0.144417778 | 2.269746416 |
| Ttc4 | 363.5749445 | -0.083546596 | 0.060965374 | -1.370394218 |
| Mroh7 | 34.09442817 | -0.79519244 | 0.296841375 | -2.678846371 |
| Fam151a | 2.097884303 | -0.466289605 | 0.886709434 | -0.525865167 |
| Acot11 | 491.6282229 | -0.357009499 | 0.145631278 | -2.451461682 |
| Gm12745 | 0 | NA | NA | NA |
| Gm12746 | 0.048949532 | -0.206487837 | 3.724945049 | -0.055433794 |
| Ssbp3 | 1314.52194 | -0.262902569 | 0.0994853 | -2.642627294 |
| Gm12786 | 0.08486385 | 0.755286617 | 3.724945049 | 0.202764499 |
| Mrpl37 | 409.2304581 | 0.086322067 | 0.0789257 | 1.093713036 |
| Cyb5rl | 41.42110935 | -0.249694211 | 0.181883047 | -1.372828395 |
| Cdcp2 | 0.295634611 | 0.27441053 | 2.067073803 | 0.132753136 |
| Gm12802 | 0.337683083 | -0.687379668 | 1.776404758 | -0.386949913 |
| Tceanc2 | 245.3997247 | -0.114723481 | 0.097753441 | -1.173600435 |
| Tmem59 | 1671.351481 | 0.151383853 | 0.073660671 | 2.055151689 |
| Ldlrad1 | 1.155427534 | 0.308365586 | 1.097733034 | 0.280911275 |
| Lrrc42 | 182.0305453 | -0.412362636 | 0.112708459 | -3.658666244 |
| Hspb11 | 48.05584592 | -0.300021902 | 0.15662838 | -1.915501526 |
| Gm12850 | 2.64217102 | -0.998817767 | 0.640013098 | -1.560620821 |
| Dio1 | 0.200957741 | 0.274391329 | 3.253433744 | 0.084338994 |
| Yipf1 | 215.0974917 | -0.069399598 | 0.082916679 | -0.836979959 |
| Gm12870 | 0.418005171 | -0.594400898 | 1.431501855 | -0.415228871 |
| Gm12869 | 0 | NA | NA | NA |
| Ndc1 | 286.6300775 | -0.147598955 | 0.084050554 | -1.756073549 |
| Gm24804 | 0 | NA | NA | NA |
| Glis1 | 17.53901492 | 0.43436806 | 0.293820171 | 1.478346628 |
| Dmrtb1 | 26.37534565 | 1.041463801 | 0.237787945 | 4.379800673 |
| Gm12899 | 0 | NA | NA | NA |

| | | | | |
|---|---|---|---|---|
| 1700047F07Rik | 0.364322736 | -2.236138262 | 1.652447643 | -1.353227905 |
| Gm25123 | 0 | NA | NA | NA |
| Gm12906 | 0.718440912 | 0.790511237 | 1.461014216 | 0.541070188 |
| Gm12907 | 0 | NA | NA | NA |
| 4933424M12Rik | 0.383698231 | 0.274412545 | 1.755345608 | 0.156329639 |
| Lrp8os3 | 15.16761459 | -0.293552275 | 0.31006021 | -0.946758939 |
| Lrp8 | 1123.019012 | -0.312455117 | 0.09818859 | -3.182193747 |
| Lrp8os2 | 23.3868331 | 0.522507659 | 0.211601733 | 2.46929764 |
| Lrp8os1 | 0.049553047 | 0.274391604 | 3.724945049 | 0.073663262 |
| Magoh | 160.5650247 | 0.401157285 | 0.146144023 | 2.744944852 |
| 0610037L13Rik | 286.4495153 | -0.109194297 | 0.086271552 | -1.265704565 |
| Cpt2 | 89.15985385 | 0.306856788 | 0.194154369 | 1.580478411 |
| 4930407G08Rik | 0.217432726 | -0.707412018 | 2.167380711 | -0.32639029 |
| Slc1a7 | 1.139422845 | -0.506834826 | 1.135885811 | -0.446202269 |
| Podn | 28.84068978 | -0.45966105 | 0.281413825 | -1.633398967 |
| Gm23354 | 0.487093236 | 0.608554373 | 1.366737329 | 0.445260666 |
| Scp2 | 720.4369207 | -0.0563651 | 0.083780186 | -0.672773636 |
| Mir6397 | 0 | NA | NA | NA |
| Echdc2 | 50.39875141 | -0.840539487 | 0.198706843 | -4.230048021 |
| Zyg11a | 0.650364221 | 2.162440974 | 1.268616972 | 1.704565698 |
| Zyg11b | 3195.301796 | -0.267166607 | 0.047004934 | -5.683799198 |
| Gm12742 | 0 | NA | NA | NA |
| Gm12740 | 0 | NA | NA | NA |
| Coa7 | 154.9104418 | -0.415843978 | 0.116195075 | -3.578843411 |
| Fam159a | 1.112155548 | 0.663124466 | 0.980038017 | 0.67663137 |
| Gpx7 | 31.9655395 | 0.289428039 | 0.217998655 | 1.327659744 |
| Gm12738 | 0.049553047 | 0.274391604 | 3.724945049 | 0.073663262 |
| Zcchc11 | 1174.50692 | 0.229617851 | 0.09478429 | 2.422530702 |
| Gm24967 | 6.345655539 | 0.164575549 | 0.437286793 | 0.376356092 |
| Gm12739 | 0.090277459 | -0.20649451 | 3.724945049 | -0.055435586 |
| Gm23138 | 4.286477608 | 0.134581068 | 0.560554612 | 0.240085559 |
| Gm12737 | 23.64047355 | -0.626457414 | 0.361145841 | -1.734638317 |
| Gm25827 | 0.189729909 | 0.274410585 | 2.263759903 | 0.121218944 |
| Gm22345 | 0.183160349 | 0.274409134 | 2.268344043 | 0.120973331 |
| Prpf38a | 397.5696076 | 0.768242303 | 0.095643383 | 8.032362271 |
| Orc1 | 6.089138634 | 2.338654178 | 0.558683451 | 4.186009396 |
| Gm12743 | 18.25140362 | -0.136605273 | 0.291672524 | -0.468351531 |
| Cc2d1b | 307.3753207 | 0.121310807 | 0.10608186 | 1.143558441 |
| Gm20731 | 6.967362313 | 0.090714596 | 0.416821946 | 0.217633924 |
| Gm17354 | 78.27363649 | -0.516200448 | 0.190129406 | -2.714995327 |
| Zfyve9 | 2056.889909 | -0.034018404 | 0.0684767 | -0.496788013 |
| 3110021N24Rik | 19.00627441 | -0.331112965 | 0.248903089 | -1.330288695 |
| Gm12741 | 0 | NA | NA | NA |
| Btf3l4 | 356.4128181 | 0.197180865 | 0.085065305 | 2.317993983 |
| Txndc12 | 312.2245455 | 0.205239754 | 0.121660016 | 1.686994303 |
| A730015C16Rik | 2.046195117 | 0.291638686 | 0.718880702 | 0.4056844 |
| Kti12 | 133.8561621 | -0.005760492 | 0.103983252 | -0.055398266 |

| | | | | |
|---|---|---|---|---|
| Rab3b | 288.1114516 | -0.348131641 | 0.141217112 | -2.465222777 |
| Gm12736 | 0 | NA | NA | NA |
| 8030443G20Rik | 0.272094973 | 0.771219105 | 2.083206868 | 0.370207643 |
| Gm23589 | 0.08486385 | 0.755286617 | 3.724945049 | 0.202764499 |
| Nrd1 | 1613.530245 | -0.204052397 | 0.053076082 | -3.844526369 |
| Mir761 | 0.40746677 | -2.487795213 | 1.3973592 | -1.780354838 |
| Osbpl9 | 801.6924287 | -0.137701072 | 0.068774492 | -2.002211395 |
| Gm23743 | 1.374639076 | -1.256488657 | 0.810877918 | -1.549541096 |
| Calr4 | 3.300543358 | -1.546132106 | 0.588032991 | -2.629328845 |
| Eps15 | 2866.84323 | -0.332111927 | 0.091998223 | -3.609981969 |
| Gm12749 | 6.563129853 | 0.26359057 | 0.457370043 | 0.57631796 |
| Ttc39aos1 | 29.58635856 | 0.234582789 | 0.229776539 | 1.020917064 |
| Ttc39a | 61.40979687 | -0.266782916 | 0.180346092 | -1.479283046 |
| Rnf11 | 2222.627844 | -0.073642759 | 0.084206703 | -0.874547471 |
| 4930522H14Rik | 0 | NA | NA | NA |
| Gm22497 | 0 | NA | NA | NA |
| Gm12811 | 0.075043205 | 0.755286614 | 3.724945049 | 0.202764498 |
| Gm12803 | 0 | NA | NA | NA |
| 9630013D21Rik | 4.08627299 | -0.094339282 | 0.566322652 | -0.166582215 |
| Cdkn2c | 32.18597743 | 0.48410377 | 0.288495872 | 1.678026678 |
| Faf1 | 667.4892566 | 0.008936193 | 0.07084204 | 0.126142509 |
| Gm12808 | 0 | NA | NA | NA |
| Dmrta2os | 0.681178858 | -0.035426349 | 1.12532548 | -0.03148098 |
| Dmrta2 | 37.20715882 | 0.350024466 | 0.209431056 | 1.671311181 |
| Elavl4 | 1395.565762 | 0.011704014 | 0.079145885 | 0.147879002 |
| Agbl4 | 173.4577221 | -0.49231804 | 0.16738218 | -2.941281086 |
| Gm12806 | 4.671720804 | 0.35071393 | 0.463510128 | 0.756647825 |
| Gm12807 | 1.537250626 | 0.343209374 | 0.778549894 | 0.440831572 |
| Gm12804 | 7.93214406 | -0.419497732 | 0.400642166 | -1.047063358 |
| Gm12805 | 2.823428998 | 3.3936269 | 0.951709753 | 3.565821293 |
| Bend5 | 101.6608683 | 0.539273962 | 0.168841291 | 3.19396966 |
| n-R5s191 | 0 | NA | NA | NA |
| Spata6 | 105.6804718 | 0.854328251 | 0.144171705 | 5.925769198 |
| Gm12961 | 0.049553047 | 0.274391604 | 3.724945049 | 0.073663262 |
| Gm12960 | 11.3450459 | 0.291656023 | 0.384209125 | 0.759107486 |
| Slc5a9 | 0.45131048 | 0.755306699 | 1.734460635 | 0.435470649 |
| Skint8 | 0.165933562 | 1.204233133 | 2.762106933 | 0.435983531 |
| Skint7 | 0.891009806 | 0.723301546 | 1.141601324 | 0.633585062 |
| Gm25844 | 0 | NA | NA | NA |
| Skint1 | 0.404673549 | 0.755310243 | 1.612718078 | 0.468346113 |
| Skint4 | 0 | NA | NA | NA |
| Gm12820 | 0 | NA | NA | NA |
| Gm12815 | 0 | NA | NA | NA |
| Skint3 | 0 | NA | NA | NA |
| Skint9 | 0 | NA | NA | NA |
| Gm12819 | 0 | NA | NA | NA |
| Gm12821 | 0.143485256 | 0.274391988 | 3.549605256 | 0.077302113 |

| | | | | |
|---|---|---|---|---|
| Skint2 | 0.106112162 | -0.206492286 | 3.724945049 | -0.055434988 |
| Gm12814 | 0 | NA | NA | NA |
| Gm24099 | 0 | NA | NA | NA |
| Gm12817 | 0 | NA | NA | NA |
| Skint10 | 0 | NA | NA | NA |
| Skint6 | 0 | NA | NA | NA |
| Gm12823 | 0.15269957 | -0.675457068 | 2.841177074 | -0.237738462 |
| Gm12816 | 17.15975745 | 0.281002337 | 0.278035376 | 1.010671161 |
| Gm12818 | 0.762181831 | -0.206479216 | 1.169368285 | -0.176573299 |
| Gm12813 | 0 | NA | NA | NA |
| Skint5 | 0 | NA | NA | NA |
| Gm12825 | 0 | NA | NA | NA |
| Skint11 | 0 | NA | NA | NA |
| Gm15741 | 0 | NA | NA | NA |
| Trabd2b | 136.8658561 | 0.175641439 | 0.136110747 | 1.290430351 |
| Gm28864 | 0.044544648 | 0.755286614 | 3.724945049 | 0.202764498 |
| Gm12829 | 0.792430516 | 1.183278326 | 1.165893156 | 1.014911461 |
| 9130410C08Rik | 0.223616349 | 0.274411746 | 1.885690919 | 0.145523184 |
| Gm12830 | 0.072025462 | 0.755288839 | 3.724945049 | 0.202765096 |
| Gm23230 | 0 | NA | NA | NA |
| Foxd2 | 5.895850479 | -0.353220762 | 0.451719848 | -0.781946518 |
| Foxd2os | 3.321918732 | 1.272173386 | 0.653713626 | 1.946071391 |
| Foxe3 | 0.279606198 | 0.274384968 | 3.724945049 | 0.073661481 |
| Cmpk1 | 803.5483932 | 0.208539767 | 0.099572379 | 2.094353552 |
| Gm24045 | 1.058755838 | -0.079003788 | 1.156968514 | -0.068285167 |
| Stil | 19.48329208 | 0.636052224 | 0.300226143 | 2.118577073 |
| Tal1 | 51.3179016 | 0.225116166 | 0.167062969 | 1.347492908 |
| Pdzk1ip1 | 3.920075909 | -0.070713765 | 0.568005112 | -0.124494945 |
| Cyp4x1 | 182.705986 | -0.306373848 | 0.099364601 | -3.083329947 |
| Cyp4x1os | 7.5998123 | -0.116337995 | 0.381880271 | -0.30464521 |
| Cyp4a28-ps | 0 | NA | NA | NA |
| Cyp4a29-ps1 | 0 | NA | NA | NA |
| Gm12834 | 0.117650406 | -0.206492286 | 3.724945049 | -0.055434988 |
| Cyp4a29 | 0.128150834 | -1.019523341 | 3.714811905 | -0.274448173 |
| Cyp4a12a | 0.066050875 | 0.274391604 | 3.724945049 | 0.073663262 |
| Cyp4a30-ps | 0 | NA | NA | NA |
| Cyp4a12b | 2.349144555 | 4.141873214 | 1.007228861 | 4.112147076 |
| Gm12837 | 0 | NA | NA | NA |
| Cyp4a30b | 0 | NA | NA | NA |
| Gm12838 | 0 | NA | NA | NA |
| Gm12833 | 0 | NA | NA | NA |
| Cyp4a14 | 0.301400513 | 1.161833416 | 2.390943329 | 0.485930972 |
| Cyp4a10 | 0.11147366 | 0.274389235 | 3.724945049 | 0.073662626 |
| Cyp4b1-ps1 | 0 | NA | NA | NA |
| Cyp4a31 | 0 | NA | NA | NA |
| Cyp4b1-ps2 | 0 | NA | NA | NA |
| Cyp4a32 | 0 | NA | NA | NA |

| | | | | |
|---|---|---|---|---|
| Cyp4b1 | 3.175114547 | -0.353679112 | 0.54658874 | -0.647066224 |
| Gm12848 | 0 | NA | NA | NA |
| Gm12849 | 0 | NA | NA | NA |
| Efcab14 | 1217.01994 | 0.139491067 | 0.095687047 | 1.457784219 |
| Tex38 | 1.456205946 | 0.046461834 | 0.897521143 | 0.051766841 |
| Atpaf1 | 375.0164648 | 0.133157433 | 0.070690181 | 1.883676501 |
| Gm12847 | 4.394170468 | 0.239620222 | 0.595668739 | 0.402270938 |
| Mob3c | 80.03911941 | 1.076251784 | 0.173833673 | 6.191273319 |
| Mknk1 | 268.8412447 | 0.10149166 | 0.076249441 | 1.331047932 |
| Gm29434 | 0.203831703 | 1.08928842 | 2.609621464 | 0.417412423 |
| Kncn | 2.503085161 | 0.942745837 | 0.681562681 | 1.383212231 |
| 6430628N08Rik | 3.312938502 | -0.322616166 | 0.675342495 | -0.477707486 |
| Dmbx1 | 11.05645536 | -0.24374276 | 0.338873521 | -0.719273549 |
| Faah | 605.5629304 | -0.105233702 | 0.100426651 | -1.04786628 |
| Nsun4 | 130.3320412 | 0.180345919 | 0.132392103 | 1.362210548 |
| Uqcrh | 768.3040327 | -0.053978452 | 0.149883066 | -0.360137097 |
| Gm12854 | 9.36533067 | 2.777254784 | 0.64329715 | 4.317219161 |
| Lrrc41 | 643.5531935 | 0.418421319 | 0.090656523 | 4.615457372 |
| Rad54l | 35.66809566 | 0.944362538 | 0.301301184 | 3.134280873 |
| Pomgnt1 | 564.1232995 | -0.191047771 | 0.06157391 | -3.102738962 |
| Lurap1 | 633.8593911 | -0.387771642 | 0.154411674 | -2.511284494 |
| Tspan1 | 1.564275958 | -1.547086543 | 0.902148586 | -1.714891058 |
| 1700042G07Rik | 0.107078754 | -0.206490058 | 3.724945049 | -0.05543439 |
| Gm12951 | 1.402628926 | -1.603038239 | 1.021656991 | -1.569057182 |
| Llph-ps1 | 0.397229082 | -1.420371895 | 1.67100973 | -0.850008153 |
| Pik3r3 | 1226.032443 | -0.186542496 | 0.070249828 | -2.655415689 |
| Mast2 | 1537.82531 | -0.200241535 | 0.054592976 | -3.667899268 |
| Gm12950 | 0 | NA | NA | NA |
| Ipp | 171.4951214 | -0.096975834 | 0.096071042 | -1.009417943 |
| Tmem69 | 92.62743766 | -0.033024905 | 0.153236213 | -0.215516321 |
| Gm12953 | 2.041941669 | 1.167230863 | 0.751960859 | 1.552249495 |
| Gpbp1l1 | 655.1959422 | -0.023954723 | 0.068269159 | -0.350886456 |
| C530005A16Rik | 44.26883829 | -0.57145745 | 0.195527933 | -2.922638421 |
| Ccdc17 | 25.14108292 | 0.033840536 | 0.220766456 | 0.153286586 |
| Nasp | 221.6735445 | -0.040130655 | 0.129719028 | -0.309365984 |
| Akr1a1 | 2348.648558 | 0.412537938 | 0.097991206 | 4.209948568 |
| Gm25137 | 0 | NA | NA | NA |
| Prdx1 | 1494.062489 | 0.341717576 | 0.172834338 | 1.977139379 |
| Mmachc | 150.583502 | -0.251871955 | 0.107319581 | -2.346933836 |
| Ccdc163 | 28.56972285 | -0.478054772 | 0.258754638 | -1.847521556 |
| Tesk2 | 80.19330176 | -0.132291589 | 0.121902215 | -1.085227113 |
| Gm27688 | 0.669412454 | -0.297911353 | 1.12742568 | -0.264240347 |
| Toe1 | 130.6486763 | 0.238435832 | 0.106317967 | 2.242667347 |
| Mutyh | 16.21670338 | 0.142622172 | 0.265573082 | 0.537035495 |
| Hpdl | 10.50466602 | -0.120849279 | 0.357481678 | -0.338057268 |
| Gm12996 | 2.848165926 | -0.300074468 | 0.547578347 | -0.548002802 |
| Rpl36-ps8 | 0 | NA | NA | NA |

| | | | | |
|---|---|---|---|---|
| Gm12994 | 0.047948698 | -0.206487837 | 3.724945049 | -0.055433794 |
| Gm12993 | 8.790287878 | -0.372572169 | 0.330993199 | -1.125618802 |
| Zswim5 | 280.6732534 | -0.265311663 | 0.103766632 | -2.556810964 |
| Gm26355 | 2.145613104 | -0.637434379 | 0.669369089 | -0.952291328 |
| Gm25435 | 4.493034422 | -0.061291623 | 0.507769377 | -0.120707601 |
| Gm22092 | 0.088123299 | -0.687376175 | 3.724945049 | -0.184533239 |
| Gm12998 | 0.333118728 | 2.45285519 | 3.689232993 | 0.664868604 |
| Urod | 271.8023691 | -0.094531736 | 0.09259774 | -1.020885992 |
| Hectd3 | 678.4462804 | -0.039120932 | 0.054844124 | -0.713311285 |
| 1700021J08Rik | 0.083172944 | -0.206492286 | 3.724945049 | -0.055434988 |
| Gm12997 | 0 | NA | NA | NA |
| Eif2b3 | 158.1507724 | -0.015443315 | 0.085167944 | -0.181327784 |
| Gm22335 | 0.400019583 | -1.158955978 | 1.522966838 | -0.760985695 |
| Ptch2 | 34.03195546 | 0.316141283 | 0.225130356 | 1.404258798 |
| Btbd19 | 68.25176527 | 0.554599271 | 0.132684657 | 4.179829713 |
| Tctex1d4 | 5.254348095 | -0.04906544 | 0.43707644 | -0.112258257 |
| Plk3 | 168.7442851 | 0.216878651 | 0.135749109 | 1.597643273 |
| Best4-ps | 0.256651301 | -0.508397018 | 2.065791517 | -0.246102772 |
| Gm13015 | 0.494552988 | -0.688881198 | 1.315467051 | -0.523678033 |
| Rps8 | 1469.97404 | 0.313851232 | 0.112278126 | 2.795301667 |
| Gm22980 | 0.6528699 | 1.330112809 | 1.268416738 | 1.048640221 |
| Snord38a | 0 | NA | NA | NA |
| Gm26330 | 0 | NA | NA | NA |
| Snord55 | 5.259902018 | 1.153068546 | 0.481017038 | 2.397146995 |
| Gm22398 | 0 | NA | NA | NA |
| Kif2c | 10.61551923 | 1.580387167 | 0.45843367 | 3.447362772 |
| Gm25099 | 10.85160868 | 0.881151266 | 0.392159254 | 2.24692203 |
| Gm23143 | 1.628265577 | 1.301769151 | 0.769461212 | 1.691793076 |
| Gm23287 | 0.580091356 | 2.672635862 | 1.49447313 | 1.788346548 |
| Gm1661 | 2.47324152 | -0.420240169 | 0.671240024 | -0.626065422 |
| 1700012C08Rik | 0 | NA | NA | NA |
| Tmem53 | 15.28288558 | 0.498455647 | 0.277434748 | 1.796659038 |
| Rnf220 | 1260.662527 | 0.074727028 | 0.060651905 | 1.232064001 |
| Gm12843 | 2.662738786 | 1.039052069 | 0.62325144 | 1.667147483 |
| Gm12828 | 0.106520246 | 0.755282166 | 3.724945049 | 0.202763304 |
| Gm12827 | 1.000155967 | 0.296231474 | 1.045485793 | 0.283343376 |
| Gm12826 | 0.267394713 | 0.771241935 | 1.828905896 | 0.4216958 |
| Eri3 | 969.1178252 | -0.07943119 | 0.052561756 | -1.511197413 |
| Gm25559 | 3.182023478 | 0.917746282 | 0.563322796 | 1.629165886 |
| Dmap1 | 212.1899085 | 0.224095024 | 0.096366734 | 2.325439641 |
| Gm12840 | 0.644938119 | 2.404574976 | 1.231469027 | 1.952606946 |
| Gm12845 | 0.176646398 | 0.755284509 | 3.360121616 | 0.22477892 |
| Gm12844 | 4.659505038 | 0.07372131 | 0.494868403 | 0.148971545 |
| Klf17 | 0.667883529 | 0.497289258 | 1.339366123 | 0.371287021 |
| Gm12842 | 0.226488584 | -0.647112821 | 1.868523929 | -0.346323005 |
| Slc6a9 | 973.6643676 | -0.415213917 | 0.139214092 | -2.982556657 |
| Gm17114 | 0.046345709 | 0.274391604 | 3.724945049 | 0.073663262 |

| | | | | |
|---|---|---|---|---|
| Ccdc24 | 3.944649576 | -0.99822327 | 0.527608878 | -1.891975879 |
| B4galt2 | 391.215399 | -0.47802353 | 0.084435594 | -5.661398349 |
| Gm12841 | 2.986054906 | -1.119272693 | 0.608981393 | -1.837942351 |
| Atp6v0b | 1543.475499 | -0.061480233 | 0.066930135 | -0.918573268 |
| Dph2 | 90.95128011 | -0.054171801 | 0.120009099 | -0.451397443 |
| Ipo13 | 1183.676242 | 0.088062188 | 0.055635574 | 1.582839559 |
| Artn | 7.776272826 | -0.260610648 | 0.499010721 | -0.522254608 |
| St3gal3 | 444.5428053 | -0.111857112 | 0.062286828 | -1.795838954 |
| 9530034E10Rik | 0.241524581 | 0.274412266 | 2.124125594 | 0.129188343 |
| Kdm4a | 879.391759 | 0.26786358 | 0.067458085 | 3.970815046 |
| Gm23382 | 0 | NA | NA | NA |
| Ptprf | 1597.948165 | -0.219185168 | 0.046744947 | -4.688959595 |
| Mir7226 | 0 | NA | NA | NA |
| Hyi | 25.72014861 | 0.158905726 | 0.213280982 | 0.745053427 |
| Szt2 | 768.797515 | -0.194119894 | 0.123155614 | -1.576216367 |
| Med8 | 175.0860242 | 0.388932364 | 0.140958786 | 2.75919207 |
| Elovl1 | 262.3958902 | 0.558520647 | 0.140148119 | 3.985216864 |
| Cdc20 | 20.13486822 | 1.011865347 | 0.323234755 | 3.130434864 |
| Gm12858 | 0.124415284 | -0.687378399 | 3.724945049 | -0.184533836 |
| Mpl | 4.268644651 | 0.594706864 | 0.594618474 | 1.00014865 |
| Tie1 | 38.04113095 | 0.840285614 | 0.216762962 | 3.876518411 |
| Gm12859 | 0 | NA | NA | NA |
| Gm12857 | 0 | NA | NA | NA |
| 2610528J11Rik | 0.045882455 | 0.755288839 | 3.724945049 | 0.202765096 |
| Tmem125 | 85.91567833 | -0.277672256 | 0.190241949 | -1.459574281 |
| Cfap57 | 12.09672597 | -0.688424635 | 0.2956892 | -2.328203515 |
| Ebna1bp2 | 374.5410988 | 0.087728962 | 0.089097001 | 0.98464551 |
| Gm12853 | 0 | NA | NA | NA |
| Olfr62 | 0.260073237 | 0.274380235 | 3.724945049 | 0.07366021 |
| Olfr1343-ps1 | 0 | NA | NA | NA |
| Olfr1342 | 0.045882455 | 0.755288839 | 3.724945049 | 0.202765096 |
| Olfr1341 | 0.138076337 | 0.75528675 | 3.589802529 | 0.210397854 |
| Olfr1340 | 0.163066018 | 0.813382652 | 3.445567682 | 0.236066369 |
| Olfr1339 | 0.060174537 | 0.755286614 | 3.724945049 | 0.202764498 |
| Olfr1338 | 0 | NA | NA | NA |
| Olfr1337 | 0 | NA | NA | NA |
| Gm12855 | 0 | NA | NA | NA |
| Olfr1335 | 0 | NA | NA | NA |
| Olfr1333 | 0 | NA | NA | NA |
| Olfr1332-ps1 | 0 | NA | NA | NA |
| Olfr1331 | 0 | NA | NA | NA |
| Olfr1330 | 0 | NA | NA | NA |
| Olfr1329 | 0 | NA | NA | NA |
| Olfr1328 | 0.212007022 | 1.23618661 | 3.149720606 | 0.392475005 |
| Gm12856 | 0.270533821 | 0.274404422 | 2.445748984 | 0.112196478 |
| Gm12863 | 0.232395771 | 1.236201847 | 2.124734505 | 0.581814737 |
| Lao1 | 0.148650935 | 0.723036258 | 2.859657322 | 0.252840175 |

| | | | | |
|---|---|---|---|---|
| Gm12861 | 0 | NA | NA | NA |
| Gm24352 | 0 | NA | NA | NA |
| Gm12865 | 0.488817361 | 0.287326356 | 1.55684441 | 0.184556886 |
| Gm12864 | 0 | NA | NA | NA |
| Gm12862 | 0.177984204 | 0.755287302 | 3.339976663 | 0.226135503 |
| Gm12866 | 0.218085611 | 0.284900652 | 2.548594097 | 0.111787378 |
| Slc2a1 | 824.8283364 | 0.65290051 | 0.110273266 | 5.920750617 |
| Mir1957a | 1.374216805 | 0.351138992 | 1.009722483 | 0.347757922 |
| Gm12867 | 0.088534442 | 0.274389469 | 3.724945049 | 0.073662689 |
| Gm12868 | 0.130524232 | 0.274405456 | 3.657871338 | 0.075017799 |
| Zfp691 | 52.30840471 | -0.055333574 | 0.151997529 | -0.364042588 |
| Ermap | 7.770417513 | 0.119873913 | 0.416962225 | 0.287493461 |
| Ccdc23 | 99.45512681 | 1.047746681 | 0.184289293 | 5.685336707 |
| AU022252 | 374.3096476 | -0.546936908 | 0.108205863 | -5.054595861 |
| 4930538K18Rik | 9.22375745 | -0.750531034 | 0.332054109 | -2.260267265 |
| Gm12898 | 0.061920613 | 0.274393828 | 3.724945049 | 0.073663859 |
| P3h1 | 112.7927286 | -0.124577118 | 0.139244167 | -0.894666689 |
| Gm12927 | 0.131862677 | -0.687376665 | 2.97735462 | -0.230868255 |
| Cldn19 | 10.33246628 | 0.970455347 | 0.414325808 | 2.342251744 |
| Ybx1 | 865.8775829 | 0.510493 | 0.128210753 | 3.981670705 |
| Gm37817 | 0.046345709 | 0.274391604 | 3.724945049 | 0.073663262 |
| Ppih | 73.62859532 | 0.200710621 | 0.164552787 | 1.21973395 |
| Ccdc30 | 150.4461269 | -0.192481394 | 0.199546551 | -0.964593945 |
| Ldha-ps2 | 11.09854899 | -0.716426584 | 0.323788638 | -2.212636579 |
| Ppcs | 48.56475404 | 0.125076861 | 0.161786371 | 0.773098873 |
| Zmynd12 | 4.300457922 | -0.467846217 | 0.509785671 | -0.917731203 |
| Rimkla | 301.1007889 | -0.417017951 | 0.128300075 | -3.250332876 |
| Gm12954 | 0.144050924 | 1.290908285 | 3.721447108 | 0.346883416 |
| Gm12956 | 0.301349932 | 0.728294115 | 1.60592319 | 0.453504949 |
| Gm12955 | 1.414308396 | 0.037719023 | 0.823444454 | 0.045806396 |
| Gm12957 | 0.066050875 | 0.274391604 | 3.724945049 | 0.073663262 |
| Rnu6-ps2 | 0 | NA | NA | NA |
| AA415398 | 75.1633106 | -0.113798041 | 0.139352548 | -0.816619736 |
| Foxj3 | 1047.552468 | 0.06092882 | 0.076088484 | 0.800762705 |
| Gm12959 | 0 | NA | NA | NA |
| Guca2a | 0.577807171 | 1.187901014 | 1.232633321 | 0.963709965 |
| Guca2b | 0.198152624 | 0.27438216 | 3.724945049 | 0.073660727 |
| Hivep3 | 1525.201888 | -0.18128037 | 0.109022177 | -1.662784349 |
| Edn2 | 0.478833344 | 0.560042977 | 1.278911696 | 0.4379059 |
| Foxo6 | 102.9223404 | -0.546533513 | 0.13865941 | -3.941553722 |
| Foxo6os | 0.889815719 | 0.294348622 | 1.298190008 | 0.226737704 |
| Scmh1 | 665.7514625 | -0.175899143 | 0.092912673 | -1.893166322 |
| Slfnl1 | 5.927427543 | -0.639333999 | 0.472033894 | -1.354423925 |
| Ctps | 504.9997313 | 0.199298864 | 0.109170261 | 1.825578349 |
| Gm8439 | 0.147068667 | 1.236183405 | 3.514416608 | 0.351746404 |
| Gm12860 | 1.253221006 | -0.344175252 | 0.920671701 | -0.373830597 |
| Cited4 | 44.06063388 | 0.877499004 | 0.186832657 | 4.696711052 |

| | | | | |
|---|---|---|---|---|
| Kcnq4 | 121.7819068 | 0.397492885 | 0.158930186 | 2.50105342 |
| Nfyc | 366.6066488 | 0.074051582 | 0.08829641 | 0.838670352 |
| Mir30c-1 | 1.678046897 | -0.805953469 | 0.742403522 | -1.085600277 |
| mmu-mir-30e | 0 | NA | NA | NA |
| Mir30e | 2.389660156 | -0.601403898 | 0.762302008 | -0.788931279 |
| Gm23936 | 0.479339789 | -0.12436391 | 1.334528144 | -0.093189425 |
| Rims3 | 2593.580875 | -0.368880936 | 0.086037552 | -4.287441072 |
| Gm12871 | 0.804633503 | 1.462930758 | 1.148570733 | 1.273696705 |
| Gm24678 | 0.394656821 | 1.881885413 | 3.31226449 | 0.568156746 |
| Exo5 | 86.65934631 | 0.094004701 | 0.14945899 | 0.628966522 |
| Zfp69 | 17.8840471 | -0.256898345 | 0.27991901 | -0.917759551 |
| Smap2 | 1633.072005 | -0.058637942 | 0.077088493 | -0.760657522 |
| mmu-mir-7015 | 1.055430728 | -2.356541681 | 1.086823528 | -2.168283644 |
| Col9a2 | 128.0701482 | -0.384224172 | 0.133213313 | -2.884277569 |
| Zmpste24 | 545.8456237 | 0.136982567 | 0.078210312 | 1.751464269 |
| Gm26186 | 0 | NA | NA | NA |
| Gm23540 | 0.845517282 | 0.866220677 | 1.114469224 | 0.777249526 |
| Tmco2 | 0.224241997 | -0.687380109 | 2.572972695 | -0.267154063 |
| Gm12882 | 0 | NA | NA | NA |
| Gm12881 | 0.075043205 | 0.755286614 | 3.724945049 | 0.202764498 |
| Gm12879 | 0 | NA | NA | NA |
| Gm22982 | 0 | NA | NA | NA |
| Rlf | 853.6771386 | 0.20796737 | 0.115819475 | 1.795616588 |
| Gm12890 | 0 | NA | NA | NA |
| Gm12888 | 0 | NA | NA | NA |
| Gm12889 | 0 | NA | NA | NA |
| Gm12885 | 0 | NA | NA | NA |
| Gm12884 | 0 | NA | NA | NA |
| Gm12883 | 0.060174537 | 0.755286614 | 3.724945049 | 0.202764498 |
| Gm12886 | 0 | NA | NA | NA |
| Gm12880 | 0 | NA | NA | NA |
| Gm26162 | 0 | NA | NA | NA |
| Gm12887 | 0 | NA | NA | NA |
| Gm12878 | 0 | NA | NA | NA |
| Gm12892 | 61.92315877 | -0.262799264 | 0.172545893 | -1.523068786 |
| Gm12896 | 0 | NA | NA | NA |
| Gm12897 | 0 | NA | NA | NA |
| Gm12893 | 0 | NA | NA | NA |
| Gm12895 | 0 | NA | NA | NA |
| Gm12877 | 0.066050875 | 0.274391604 | 3.724945049 | 0.073663262 |
| Gm12874 | 0 | NA | NA | NA |
| Gm12875 | 0 | NA | NA | NA |
| Gm12876 | 0.038981395 | 0.274393828 | 3.724945049 | 0.073663859 |
| 9530002B09Rik | 0 | NA | NA | NA |
| Gm12872 | 0 | NA | NA | NA |
| Gm12873 | 0.136623169 | -0.206497534 | 3.634681883 | -0.056813097 |
| Ppt1 | 1536.560467 | 0.299136517 | 0.054280785 | 5.510909914 |

| | | | | |
|---|---|---|---|---|
| Cap1 | 2420.151979 | 0.220151359 | 0.065941047 | 3.338608772 |
| Gm12891 | 0 | NA | NA | NA |
| Mfsd2a | 324.2124731 | 0.736138713 | 0.153789378 | 4.786668108 |
| 4933421A08Rik | 2.139806348 | 0.856192416 | 0.743976564 | 1.15083251 |
| Mycl | 128.6415578 | -0.088832917 | 0.185600576 | -0.478624144 |
| Trit1 | 181.2275785 | 0.117425028 | 0.170473834 | 0.68881555 |
| Gm12923 | 0.164553454 | -0.206488957 | 2.77921914 | -0.074297472 |
| Bmp8b | 1.325872221 | 0.583799494 | 0.836749195 | 0.6976995 |
| Gm17244 | 11.29059836 | 0.267071283 | 0.286966533 | 0.93067049 |
| Oxct2b | 0.96273357 | 0.142742902 | 0.971170052 | 0.146980338 |
| Ppie | 115.1280594 | 0.34268405 | 0.139489061 | 2.456709138 |
| Hpcal4 | 5044.886741 | -0.2100671 | 0.163685646 | -1.283356878 |
| Nt5c1a | 53.31890087 | -0.399801553 | 0.163745097 | -2.4416093 |
| Heyl | 92.72271586 | 0.239148567 | 0.187832573 | 1.27320072 |
| Gm12901 | 0.083526043 | 0.75528439 | 3.724945049 | 0.202763901 |
| Pabpc4 | 351.2920336 | 0.004259457 | 0.104956871 | 0.04058293 |
| Gm12900 | 0 | NA | NA | NA |
| Gm25788 | 10.10737276 | 0.64186796 | 0.335343886 | 1.914058933 |
| Gm22154 | 9.014346252 | -0.48186284 | 0.357490557 | -1.347903687 |
| Bmp8a | 2.710969876 | 0.91700439 | 0.716559351 | 1.27973264 |
| Oxct2a | 0.793872245 | -0.03717835 | 1.052091797 | -0.035337554 |
| Macf1 | 11900.70528 | -0.195335722 | 0.061410983 | -3.180794571 |
| Gm12924 | 0.060174537 | 0.755286614 | 3.724945049 | 0.202764498 |
| Gm23548 | 0 | NA | NA | NA |
| Mir6398 | 0 | NA | NA | NA |
| Gm12926 | 0 | NA | NA | NA |
| Gm12925 | 2.717467516 | -0.624678857 | 0.658059303 | -0.949274411 |
| Gm12922 | 4.003874815 | 0.2214093 | 0.519054536 | 0.426562691 |
| Ndufs5 | 423.3196819 | -0.120214491 | 0.096560406 | -1.244966706 |
| Akirin1 | 729.4963505 | 0.26309286 | 0.098142138 | 2.68073291 |
| Rhbdl2 | 5.725445938 | 0.556786232 | 0.418370157 | 1.330845958 |
| 1700121C08Rik | 6.402408373 | 0.366704802 | 0.461538408 | 0.794527164 |
| Gm12904 | 0.192693611 | 0.274408644 | 2.64331282 | 0.103812399 |
| 4933427I04Rik | 0.332356096 | 2.075829886 | 3.704070232 | 0.560418609 |
| Gm26606 | 53.85320551 | -0.378741365 | 0.239017357 | -1.584576825 |
| D130007C19Rik | 26.17483582 | -0.657114879 | 0.296870912 | -2.213470072 |
| Gm12903 | 0 | NA | NA | NA |
| Mycbp | 64.1662213 | 0.128323149 | 0.136137606 | 0.942598839 |
| Gm12905 | 20.80298438 | -0.459105361 | 0.301664989 | -1.521904685 |
| Rragc | 863.0862859 | 0.274123149 | 0.088289476 | 3.104822484 |
| 4930535I16Rik | 1.505311134 | 1.084297411 | 0.847532758 | 1.279357524 |
| Gm12902 | 7.717556106 | -0.175080684 | 0.362794364 | -0.482589315 |
| Gm37667 | 0.274851136 | 1.260219029 | 3.723306324 | 0.338467727 |
| Gm12916 | 0.134481855 | 0.720143159 | 3.654633157 | 0.197049369 |
| 1700021L23Rik | 0.106971521 | 1.236179401 | 3.724945049 | 0.331865138 |
| 4933435F18Rik | 0.562550613 | -0.146819064 | 1.444629108 | -0.101630974 |
| Gm22983 | 0 | NA | NA | NA |

| | | | | |
|---|---|---|---|---|
| 1700057H15Rik | 1.747775439 | 0.185203615 | 0.779696074 | 0.237533087 |
| Gm27247 | 15.84162281 | -0.052942151 | 0.312495662 | -0.169417234 |
| 4933407E24Rik | 0 | NA | NA | NA |
| 3100002H09Rik | 4.323581818 | -0.270345987 | 0.627391447 | -0.430904801 |
| Pou3f1 | 370.5457929 | 0.439633077 | 0.130114776 | 3.378809774 |
| Utp11l | 181.9343619 | -0.129309067 | 0.087437936 | -1.478866881 |
| Gm24480 | 0 | NA | NA | NA |
| Fhl3 | 24.63805473 | 0.959637516 | 0.260293168 | 3.686756448 |
| Sf3a3 | 314.4333351 | 0.28029911 | 0.093129372 | 3.009782019 |
| Gm12917 | 6.099625106 | -0.473404934 | 0.596012914 | -0.794286369 |
| Mir697 | 1.448093177 | -0.180154686 | 0.824313253 | -0.218551243 |
| Gm12915 | 2.377012089 | 0.51219212 | 0.630071534 | 0.812911063 |
| Gm24721 | 0.117760149 | 0.274400501 | 3.724945049 | 0.073665651 |
| Inpp5b | 535.9436542 | 0.134984773 | 0.073624961 | 1.833410459 |
| Mir698 | 0.286901049 | 1.149108237 | 2.02294271 | 0.568037953 |
| Mtf1 | 507.9230992 | -0.275265306 | 0.092691237 | -2.969701504 |
| n-R5s192 | 1.205568873 | -0.649466045 | 0.876855973 | -0.740675852 |
| 1110065P20Rik | 188.1940773 | -0.221343859 | 0.110197961 | -2.008602129 |
| Yrdc | 152.5953946 | 0.380187461 | 0.195505155 | 1.944641615 |
| Maneal | 554.488513 | -0.094739294 | 0.081828788 | -1.157774616 |
| Epha10 | 353.1436211 | -0.402707187 | 0.170633696 | -2.360068357 |
| 1700125G02Rik | 0.77578221 | -0.908550843 | 1.148356427 | -0.791174954 |
| Cdca8 | 18.02129889 | 0.04858845 | 0.34602314 | 0.140419656 |
| 9930104L06Rik | 211.7189511 | 0.08730611 | 0.080951714 | 1.078496133 |
| Gm12930 | 0.650770932 | -1.517694692 | 1.224407746 | -1.239533724 |
| Rspo1 | 34.80671183 | -0.701436881 | 0.256753853 | -2.73194296 |
| Gnl2 | 487.0670719 | 0.316756938 | 0.103301116 | 3.066345735 |
| Gm12931 | 0 | NA | NA | NA |
| Dnali1 | 15.85509031 | -0.348669182 | 0.306767012 | -1.136592816 |
| Snip1 | 158.6932189 | 0.018671416 | 0.106561738 | 0.175216884 |
| Gm12932 | 3.658838888 | 0.059082464 | 0.554862201 | 0.106481328 |
| Meaf6 | 803.9924443 | -0.154171734 | 0.102883068 | -1.498514148 |
| Zc3h12a | 71.74715942 | 2.39879897 | 0.244201351 | 9.823037266 |
| 1700041M05Rik | 0 | NA | NA | NA |
| Grik3 | 1453.049993 | -0.365124567 | 0.081250872 | -4.493792601 |
| Ftl2 | 1.736617851 | 2.022288679 | 1.173843799 | 1.722791977 |
| Mir692-2 | 0.541671891 | 0.762998123 | 1.515795046 | 0.503364967 |
| 2610028E06Rik | 11.39828975 | 1.823881328 | 0.364954517 | 4.997557892 |
| Csf3r | 194.1912691 | 1.660889914 | 0.195531532 | 8.494230555 |
| Mrps15 | 201.7560872 | 0.193464593 | 0.105611827 | 1.831845913 |
| Oscp1 | 268.9501632 | -0.295529709 | 0.093222612 | -3.170150484 |
| Rpl28-ps3 | 19.78160788 | 0.898979329 | 0.266754325 | 3.370064675 |
| Lsm10 | 59.05863203 | -0.145433444 | 0.183026098 | -0.794604955 |
| Stk40 | 956.9610872 | 0.816931472 | 0.19018932 | 4.295359345 |
| Gm12946 | 0.363304151 | 0.275353194 | 1.728461886 | 0.159305332 |
| Eva1b | 26.65916579 | 0.679787939 | 0.267203339 | 2.544084747 |
| Sh3d21 | 14.96427342 | 0.766889526 | 0.294946676 | 2.600095503 |

| | | | | |
|---|---|---|---|---|
| Thrap3 | 1545.929057 | 0.116130652 | 0.059365334 | 1.956203127 |
| Gm25323 | 1.633037017 | -0.753152219 | 0.919607493 | -0.81899313 |
| Map7d1 | 3375.866482 | 0.072973832 | 0.076228799 | 0.957300037 |
| Trappc3 | 541.5073919 | 0.079423949 | 0.07343212 | 1.081596839 |
| Col8a2 | 71.84796312 | -0.094828648 | 0.216798406 | -0.437404729 |
| Adprhl2 | 208.9240687 | 0.304529584 | 0.106518783 | 2.858928503 |
| Tekt2 | 9.162665626 | 0.087952627 | 0.370592853 | 0.237329528 |
| Gm12945 | 0.40938346 | 0.61214737 | 1.44556878 | 0.423464714 |
| Ago3 | 1555.337107 | -0.718255458 | 0.10429462 | -6.886793017 |
| Ago1 | 1116.199882 | -0.262996079 | 0.072963368 | -3.604494753 |
| Gm12929 | 1.033962832 | 0.665619295 | 1.203001316 | 0.553298892 |
| Gm12928 | 17.68241393 | 0.292127118 | 0.268428442 | 1.088286754 |
| Ago4 | 190.9073592 | -0.350542045 | 0.12442452 | -2.817306787 |
| Gm12935 | 0.038981395 | 0.274393828 | 3.724945049 | 0.073663859 |
| Gm12934 | 4.064846038 | -0.424900125 | 0.605441542 | -0.701802066 |
| Clspn | 29.12900793 | 0.532499667 | 0.411053819 | 1.295449994 |
| Mir7119 | 0 | NA | NA | NA |
| Gm12936 | 0.066050875 | 0.274391604 | 3.724945049 | 0.073663262 |
| 5730409E04Rik | 892.9733379 | 0.038654929 | 0.062910176 | 0.614446362 |
| Gm12933 | 1.508740896 | -0.45950943 | 0.83110708 | -0.55288836 |
| Psmb2 | 628.3734742 | 0.707681602 | 0.109445779 | 6.466047463 |
| Tfap2e | 0.181508466 | 1.236194459 | 2.694094898 | 0.458853346 |
| Ncdn | 11330.12844 | -0.01346876 | 0.078460542 | -0.171662847 |
| AU040320 | 846.8326345 | 0.087004251 | 0.050099608 | 1.736625384 |
| Gm12937 | 0 | NA | NA | NA |
| Gm24161 | 0 | NA | NA | NA |
| Gm25604 | 3.572392357 | 0.654043451 | 0.50637248 | 1.291625191 |
| Zmym4 | 1413.511916 | -0.08965954 | 0.045361567 | -1.976552982 |
| Gm12939 | 0 | NA | NA | NA |
| Sfpq | 2483.364451 | -0.580209418 | 0.093955163 | -6.175386242 |
| Gm12940 | 393.2352481 | -1.054602763 | 0.254862821 | -4.137923141 |
| Gm12941 | 12.76876565 | -2.35402067 | 0.56183482 | -4.189880346 |
| Zmym1 | 160.873064 | -0.507848547 | 0.140020328 | -3.626962988 |
| Gm12944 | 1.541842 | -0.021090877 | 0.846061445 | -0.024928304 |
| Zmym6 | 362.4092715 | -0.386151524 | 0.118097142 | -3.269778747 |
| Gm25600 | 0 | NA | NA | NA |
| Gm12942 | 27.99737021 | -0.556452012 | 0.257457618 | -2.161334423 |
| Gm12943 | 0.338503588 | -1.360632367 | 1.518886239 | -0.895809266 |
| Dlgap3 | 2650.38369 | -0.336511198 | 0.122969289 | -2.736546663 |
| Smim12 | 221.697071 | -0.01942843 | 0.089014771 | -0.218260748 |
| Gm22221 | 0 | NA | NA | NA |
| Gm12948 | 0 | NA | NA | NA |
| Gja4 | 31.84417143 | 1.355996416 | 0.258139775 | 5.2529542 |
| Gjb3 | 2.20869273 | 1.264597604 | 0.903352666 | 1.399893586 |
| Gjb4 | 1.851903907 | 2.227253988 | 1.153315413 | 1.931175083 |
| Gjb5 | 0.85820172 | 1.178935887 | 1.481570396 | 0.795733966 |
| Gm12947 | 0 | NA | NA | NA |

| | | | | |
|---|---|---|---|---|
| 4930471C06Rik | 0.072025462 | 0.755288839 | 3.724945049 | 0.202765096 |
| 1700080G11Rik | 0.674986302 | 0.238840013 | 1.256913233 | 0.190021082 |
| A630031M04Rik | 0.919688069 | 0.986427324 | 1.151474894 | 0.856664204 |
| 1700112K13Rik | 0.099155932 | 0.75528439 | 3.724945049 | 0.202763901 |
| CK137956 | 8.83914943 | -0.220663838 | 0.339837128 | -0.649322337 |
| Csmd2 | 1037.327866 | -0.464943998 | 0.118935306 | -3.909217666 |
| Csmd2os | 0.11657011 | 1.236179401 | 3.724945049 | 0.331865138 |
| Hmgb4 | 0.309883016 | 0.274381181 | 3.724945049 | 0.073660464 |
| Hmgb4os | 3.44465028 | -1.070087948 | 0.654443588 | -1.635111059 |
| Zscan20 | 123.2244835 | -0.353852149 | 0.159318511 | -2.221036001 |
| Tlr12 | 8.189155563 | 0.662402891 | 0.515442166 | 1.285115837 |
| Gm12958 | 0 | NA | NA | NA |
| Phc2 | 1259.457039 | 0.406860138 | 0.094214534 | 4.318443442 |
| A3galt2 | 16.37679573 | 0.224866552 | 0.334376604 | 0.672494873 |
| Zfp362 | 336.5354883 | -0.140990581 | 0.091111454 | -1.547451775 |
| Gm12968 | 0.072025462 | 0.755288839 | 3.724945049 | 0.202765096 |
| Gm12969 | 0.192749185 | -0.537210589 | 3.283777405 | -0.163595312 |
| Trim62 | 671.0414608 | 0.238117044 | 0.155445034 | 1.531840795 |
| Azin2 | 295.6170531 | 0.059615866 | 0.087249435 | 0.68328083 |
| Gm15904 | 0.046345709 | 0.274391604 | 3.724945049 | 0.073663262 |
| Ak2 | 283.4084168 | 0.604787969 | 0.110137027 | 5.491232037 |
| 1700086P04Rik | 1.876204568 | 0.998202357 | 0.718079651 | 1.390099769 |
| Rnf19b | 1117.961967 | 1.173162334 | 0.16239872 | 7.223962934 |
| Tmem54 | 5.394435422 | 0.249918161 | 0.415523242 | 0.601454108 |
| Hpca | 2746.890777 | -0.071325401 | 0.156353111 | -0.456181527 |
| Gm15906 | 1.230936373 | -0.687769545 | 0.963972992 | -0.713473874 |
| Fndc5 | 973.7975932 | -0.318324949 | 0.09551238 | -3.332813495 |
| S100pbp | 678.3393157 | -0.22814354 | 0.068585433 | -3.326413955 |
| Yars | 969.2323716 | 0.200263569 | 0.084333524 | 2.374661464 |
| C77080 | 318.9680605 | -0.065834698 | 0.110017155 | -0.598403927 |
| Gm12976 | 2.076702053 | -1.112329819 | 0.652209331 | -1.705479769 |
| Gm22994 | 0 | NA | NA | NA |
| Sync | 16.23627725 | -0.413582116 | 0.262511041 | -1.575484653 |
| Rbbp4 | 1422.281671 | 0.145453021 | 0.071703088 | 2.028546119 |
| Gm26722 | 3.033581745 | 0.203348012 | 0.606028088 | 0.335542224 |
| Zbtb8os | 112.0160394 | -0.079718407 | 0.111596451 | -0.714345361 |
| Zbtb8a | 48.71610654 | 0.317447242 | 0.170606287 | 1.860700722 |
| Gm12980 | 0.075043205 | 0.755286614 | 3.724945049 | 0.202764498 |
| Zbtb8b | 38.62991341 | -0.492637086 | 0.233434137 | -2.110390077 |
| Gm24621 | 0 | NA | NA | NA |
| Bsdc1 | 654.7636716 | 0.18272171 | 0.065869893 | 2.773979156 |
| Tssk3 | 1.78535976 | 0.821182064 | 0.779819619 | 1.053041042 |
| Fam229a | 1.511769237 | -1.063727185 | 0.910787058 | -1.167920839 |
| Gm12979 | 0 | NA | NA | NA |
| Marcksl1 | 1179.090896 | 0.477034777 | 0.128544816 | 3.711038633 |
| Hdac1 | 211.8204638 | 0.595868529 | 0.117585063 | 5.067552934 |
| Lck | 123.7661852 | 3.663913511 | 0.269155951 | 13.61260452 |

| | | | | |
|---|---|---|---|---|
| Mir8119 | 0.156477978 | 1.236192357 | 2.823333319 | 0.437848535 |
| Fam167b | 2.785268692 | -0.362299372 | 0.762724469 | -0.475006881 |
| Eif3i | 596.7175796 | 0.175898971 | 0.086307232 | 2.038055994 |
| Tmem234 | 475.3810905 | 0.055871911 | 0.056526397 | 0.988421578 |
| Dcdc2b | 6.370853605 | -1.428177898 | 0.433817847 | -3.292114212 |
| Iqcc | 93.52464096 | -0.290810638 | 0.148315334 | -1.960759079 |
| Ccdc28b | 159.6210152 | -0.172741413 | 0.127408344 | -1.355809265 |
| Txlna | 633.3282999 | 0.183469269 | 0.067846817 | 2.704169143 |
| Kpna6 | 1687.26505 | 0.053989883 | 0.050697774 | 1.064935949 |
| Tmem39b | 67.54451293 | -0.241687997 | 0.171256826 | -1.411260518 |
| Mir7016 | 0.152586256 | -0.68737674 | 2.845690286 | -0.241550089 |
| Khdrbs1 | 1365.931483 | -0.047270389 | 0.062161936 | -0.760439455 |
| Gm12967 | 0 | NA | NA | NA |
| Gm28017 | 0 | NA | NA | NA |
| Ptp4a2 | 1712.805092 | 0.233720242 | 0.085379855 | 2.737416693 |
| Gm10571 | 11.18052223 | -0.58464719 | 0.346965179 | -1.685031308 |
| Gm23845 | 0.075043205 | 0.755286614 | 3.724945049 | 0.202764498 |
| Gm12966 | 92.3427904 | 0.980138052 | 0.191313172 | 5.123212583 |
| E330017L17Rik | 0.177020223 | 0.755302934 | 2.765325947 | 0.27313342 |
| Spocd1 | 2.098579804 | 0.240365311 | 0.728170033 | 0.330095032 |
| 1700003M07Rik | 24.71805353 | -0.418766807 | 0.27259415 | -1.536228153 |
| Adgrb2 | 3754.218321 | -0.22303356 | 0.128951031 | -1.729598881 |
| Gm12963 | 2.392410068 | -0.01435766 | 0.635474365 | -0.02259361 |
| Col16a1 | 419.3408002 | 1.207674597 | 0.15409409 | 7.837254468 |
| Pef1 | 447.0267585 | -0.046965663 | 0.096073357 | -0.488852111 |
| Hcrtr1 | 16.529312 | -0.997037042 | 0.323823457 | -3.078952496 |
| Tinagl1 | 47.82351247 | 1.401315333 | 0.255513589 | 5.484308438 |
| Gm853 | 1.50545653 | -0.633888915 | 0.757470854 | -0.836849248 |
| Serinc2 | 80.11973323 | -0.852501592 | 0.175759563 | -4.850385252 |
| Gm10570 | 4.998772346 | -0.340851465 | 0.460657969 | -0.739923085 |
| Fabp3 | 388.7351392 | -0.024836846 | 0.10324653 | -0.240558649 |
| Zcchc17 | 445.3923863 | 0.062934308 | 0.07544722 | 0.834150122 |
| Snrnp40 | 199.2041572 | 0.241358049 | 0.081269171 | 2.969859863 |
| Nkain1 | 233.0882025 | -0.382485902 | 0.126087769 | -3.033489319 |
| Pum1 | 1545.221209 | -0.141598084 | 0.05630732 | -2.514736696 |
| Snord85 | 3.381283184 | 0.106805841 | 0.560847371 | 0.190436555 |
| Gm12971 | 8.510907694 | -1.063363532 | 0.402309383 | -2.643148724 |
| Gm24091 | 2.986937453 | -1.412692851 | 0.628281088 | -2.248504496 |
| Gm26516 | 0.225317292 | -0.687379649 | 2.567000603 | -0.267775414 |
| Gm23711 | 1.081483968 | -2.19517459 | 1.0880178 | -2.017590696 |
| Gm12972 | 0.415245445 | -0.107669254 | 1.438543397 | -0.074846024 |
| Sdc3 | 6090.612101 | 1.200269321 | 0.122396573 | 9.806396473 |
| Gm12970 | 0.131862677 | -0.687376665 | 2.97735462 | -0.230868255 |
| Gm26716 | 0.436796958 | 1.21416304 | 1.676374679 | 0.724279038 |
| Gm22660 | 0 | NA | NA | NA |
| Laptm5 | 869.0779485 | 1.854190626 | 0.165512315 | 11.20273511 |
| Matn1 | 10.36744075 | 2.611882477 | 0.510299357 | 5.118333862 |

| | | | | |
|---|---|---|---|---|
| Gm12973 | 0.042716945 | -0.206490061 | 3.724945049 | -0.055434391 |
| A930031H19Rik | 0.159888962 | -0.206482815 | 3.461347753 | -0.059653878 |
| Gm25261 | 0 | NA | NA | NA |
| Gm831 | 0.963234904 | 1.902608866 | 1.073766028 | 1.771902647 |
| Gm12962 | 0.066050875 | 0.274391604 | 3.724945049 | 0.073663262 |
| Gm16080 | 0.066050875 | 0.274391604 | 3.724945049 | 0.073663262 |
| Ptpru | 445.2308485 | -0.236249519 | 0.088851556 | -2.658923808 |
| Mecr | 135.1425178 | -0.221759654 | 0.123506982 | -1.795523225 |
| Srsf4 | 621.5934249 | -0.020292876 | 0.075310512 | -0.269456089 |
| A930004J17Rik | 191.8201737 | -0.139537439 | 0.238735093 | -0.584486502 |
| Gm12992 | 46.74477129 | -0.287380963 | 0.200010609 | -1.436828596 |
| Gm12978 | 0 | NA | NA | NA |
| Tmem200b | 45.76488705 | -0.277245268 | 0.232164617 | -1.194175374 |
| Epb4.1 | 1119.732651 | -0.128189852 | 0.136951535 | -0.936023471 |
| Gm27445 | 1.407831039 | -0.354759062 | 0.747546051 | -0.474564827 |
| Gm13063 | 0.62973509 | 0.257204098 | 1.285565754 | 0.200070745 |
| Gm10300 | 1.165032807 | 0.915750793 | 0.982333033 | 0.932220298 |
| Gm13214 | 0 | NA | NA | NA |
| Gm37711 | 81.49722938 | -0.292446543 | 0.21821598 | -1.340170153 |
| Oprd1 | 249.9746901 | -0.033180428 | 0.154097814 | -0.215320565 |
| Gm13215 | 7.792582945 | 0.177409735 | 0.378222899 | 0.469061327 |
| Ythdf2 | 645.7985944 | 0.372050795 | 0.105049856 | 3.541659265 |
| Gm22033 | 0.373327567 | 0.274386425 | 3.724945049 | 0.073661872 |
| Rps15a-ps4 | 8.387413765 | 0.601742335 | 0.381422154 | 1.577628172 |
| Gmeb1 | 299.6588852 | 0.10212923 | 0.110685585 | 0.922696754 |
| Gm13252 | 3.851277915 | -0.683673534 | 0.520068077 | -1.314584695 |
| Gm28874 | 1.719071184 | -0.910537793 | 0.845682506 | -1.076689877 |
| Gm28872 | 0.044544648 | 0.755286614 | 3.724945049 | 0.202764498 |
| Rnu11 | 24.6768285 | 0.233932145 | 0.262037683 | 0.892742383 |
| Taf12 | 193.1512937 | 0.160689881 | 0.091598139 | 1.754291974 |
| Rab42 | 6.121133301 | -1.233662591 | 0.451803887 | -2.730526733 |
| Snhg12 | 177.2717645 | 0.897138051 | 0.162429548 | 5.523244147 |
| snopsi28S-3327 | 0 | NA | NA | NA |
| Snora16a | 0 | NA | NA | NA |
| Snora44 | 0 | NA | NA | NA |
| Snora61 | 0 | NA | NA | NA |
| Snord99 | 0 | NA | NA | NA |
| Trnau1ap | 131.2572649 | 0.04166769 | 0.108890076 | 0.382658285 |
| Rcc1 | 103.1344041 | -0.172797907 | 0.131973818 | -1.309334753 |
| Snhg3 | 63.93668591 | 0.361525317 | 0.148810031 | 2.429441852 |
| Gm17300 | 0.711576293 | 0.134030147 | 1.208769546 | 0.110881473 |
| Snora73b | 425.1513821 | -0.20966364 | 0.212082875 | -0.988592975 |
| Snora73a | 0.690105543 | -0.541614217 | 1.338717802 | -0.404576839 |
| Phactr4 | 351.1978573 | 0.868707276 | 0.142743242 | 6.085803191 |
| Gm25707 | 0 | NA | NA | NA |
| Gm23055 | 0 | NA | NA | NA |
| Med18 | 61.32883924 | 0.111486503 | 0.144446136 | 0.771820596 |

| | | | | |
|---|---|---|---|---|
| Sesn2 | 307.2550336 | 0.789968897 | 0.176589246 | 4.473482488 |
| Gm12981 | 4.426758316 | -0.477013694 | 0.521815007 | -0.914143303 |
| Atpif1 | 785.2790718 | 0.113276485 | 0.131719694 | 0.859981383 |
| Gm12999 | 2.573520391 | 0.256599434 | 0.602804876 | 0.425675778 |
| Dnajc8 | 648.8436146 | 0.261529345 | 0.088069382 | 2.969583064 |
| Gm24762 | 0 | NA | NA | NA |
| Ptafr | 215.886901 | 3.513396412 | 0.377006935 | 9.319182448 |
| Eya3 | 516.9045441 | -0.187539652 | 0.073178688 | -2.562763259 |
| Xkr8 | 74.57135612 | -0.256096809 | 0.14077387 | -1.819206988 |
| Smpdl3b | 33.7182008 | 0.823050144 | 0.212184805 | 3.87893066 |
| Rpa2 | 85.55301726 | 0.093578127 | 0.157727462 | 0.593290008 |
| Themis2 | 329.3702901 | 5.072919206 | 0.356011308 | 14.24932044 |
| Gm13022 | 0 | NA | NA | NA |
| Ppp1r8 | 368.0675459 | 0.174211733 | 0.080815633 | 2.155668734 |
| Gm22767 | 11.91181268 | 0.050266534 | 0.338654248 | 0.148430248 |
| Stx12 | 1297.871004 | 0.062563962 | 0.062698419 | 0.997855495 |
| Gm13033 | 2.338964691 | 1.054699692 | 0.716303008 | 1.47242114 |
| Fam76a | 504.2948798 | 0.17517657 | 0.074306944 | 2.357472403 |
| Gm24913 | 0 | NA | NA | NA |
| Fgr | 92.45807742 | 4.663401605 | 0.457127608 | 10.20153131 |
| Ahdc1 | 1531.214203 | -0.061698673 | 0.096904649 | -0.636694666 |
| Gm26615 | 5.273376998 | -0.403647333 | 0.541503601 | -0.745419481 |
| Wasf2 | 693.9247449 | 1.095237595 | 0.120306715 | 9.10371126 |
| Gm24636 | 0 | NA | NA | NA |
| Mir7017 | 0.269321163 | 1.295750407 | 3.708688725 | 0.349382357 |
| Gpr3 | 98.5239047 | -0.520830845 | 0.257570769 | -2.022088325 |
| Cd164l2 | 13.0383265 | -0.355629139 | 0.316541834 | -1.123482273 |
| Map3k6 | 67.01031991 | 2.737093534 | 0.377710174 | 7.24654436 |
| Sytl1 | 19.69897721 | -0.630060837 | 0.282274289 | -2.232087231 |
| Tmem222 | 442.6011969 | 0.211911892 | 0.089135447 | 2.377414363 |
| Gm13259 | 0.441858849 | -0.873970838 | 1.370796802 | -0.637564106 |
| Wdtc1 | 1171.200647 | 0.52953597 | 0.090219493 | 5.869418603 |
| Slc9a1 | 963.2912325 | 0.331541069 | 0.070236933 | 4.720323843 |
| Mir5122 | 0 | NA | NA | NA |
| Gm13257 | 0.719481786 | 1.882149088 | 1.336099343 | 1.40868948 |
| Fam46b | 4.767060263 | 1.538438489 | 0.62764023 | 2.451147037 |
| Trnp1 | 753.4664742 | -0.170598436 | 0.11602176 | -1.470400338 |
| Kdf1 | 1.792464502 | 0.059321786 | 0.835634547 | 0.070990107 |
| Nudc | 578.9021612 | 0.212501113 | 0.169329149 | 1.254958847 |
| Nr0b2 | 0.783927492 | -0.567012753 | 0.967672422 | -0.585955267 |
| Gm23158 | 0.049553047 | 0.274391604 | 3.724945049 | 0.073663262 |
| Gpatch3 | 56.4655971 | 0.652946203 | 0.174201066 | 3.748233115 |
| Gpn2 | 58.10242144 | 0.152484393 | 0.197253234 | 0.773038747 |
| Sfn | 14.04179457 | -0.300988622 | 0.412981641 | -0.728818407 |
| Zdhhc18 | 588.2128307 | 0.220071004 | 0.090743881 | 2.42518836 |
| Gm13213 | 0.097899064 | -0.687371727 | 3.724945049 | -0.184532045 |
| Gm22662 | 0 | NA | NA | NA |

| | | | | |
|---|---|---|---|---|
| Pigv | 97.35604423 | 0.526040534 | 0.175469744 | 2.997898799 |
| Arid1a | 2860.653318 | -0.030911565 | 0.064391349 | -0.480057724 |
| mmu-mir-7227 | 2.041220482 | -1.651094056 | 0.767944341 | -2.15001787 |
| Gm25270 | 0 | NA | NA | NA |
| Gm12974 | 0.15596086 | -1.15072961 | 3.483969583 | -0.330292668 |
| Rps6ka1 | 359.3082904 | -0.061767653 | 0.114889854 | -0.537624962 |
| Gm12977 | 0 | NA | NA | NA |
| Gm12985 | 2.189170689 | -1.712578936 | 0.862132664 | -1.986444786 |
| Gm22523 | 2.911568235 | -0.880536186 | 0.688330823 | -1.279233992 |
| Hmgn2 | 174.4388056 | 0.324150115 | 0.169000149 | 1.918046323 |
| Dhdds | 602.5039709 | -0.008753933 | 0.060254836 | -0.145281835 |
| Lin28a | 0.305648149 | 0.274414024 | 1.763148838 | 0.155638604 |
| Gm13061 | 0 | NA | NA | NA |
| Zfp683 | 0.393365814 | -0.206483808 | 1.544622786 | -0.133679115 |
| Aim1l | 11.08135394 | -0.115826864 | 0.319846428 | -0.362132742 |
| Cd52 | 180.6648992 | 5.139588114 | 0.379966288 | 13.5264319 |
| Ubxn11 | 97.44407537 | -0.185048517 | 0.201925045 | -0.916421819 |
| Sh3bgrl3 | 569.2295487 | -0.028095875 | 0.089034042 | -0.31556329 |
| Cep85 | 164.0264212 | -0.301113397 | 0.09931462 | -3.031914103 |
| Gm27619 | 0.047948698 | -0.206487837 | 3.724945049 | -0.055433794 |
| Gm7534 | 0 | NA | NA | NA |
| Gm13131 | 1.316625395 | -0.970781165 | 0.892008828 | -1.088308921 |
| Catsper4 | 0.735041978 | 1.734986748 | 1.195152196 | 1.451686867 |
| Cnksr1 | 18.73766402 | 2.400181528 | 0.432306881 | 5.552031742 |
| Zfp593 | 102.2427118 | 0.577006265 | 0.17287832 | 3.337643875 |
| E130218I03Rik | 0.926475344 | 1.547285973 | 1.301070632 | 1.189240565 |
| Grrp1 | 25.39142091 | 1.482331576 | 0.299184126 | 4.954579625 |
| Pdik1l | 224.1146395 | 0.150071258 | 0.086772959 | 1.72947033 |
| Trim63 | 3.249610149 | 1.44546114 | 0.719927296 | 2.007787659 |
| Gm13195 | 0 | NA | NA | NA |
| Slc30a2 | 1.413987658 | 0.635272592 | 0.908367916 | 0.699356044 |
| Extl1 | 547.7075812 | 0.276699758 | 0.185800446 | 1.489230858 |
| Pafah2 | 65.33746165 | -0.394151528 | 0.209933402 | -1.877507462 |
| 1700021N21Rik | 2.462361163 | 0.303708787 | 0.629953137 | 0.482113302 |
| Stmn1 | 1361.893308 | -0.24501912 | 0.091937141 | -2.665072225 |
| Gm23634 | 0.446933609 | 0.416920169 | 1.656891984 | 0.251627851 |
| Gm13250 | 0 | NA | NA | NA |
| Paqr7 | 388.1743221 | -0.660107218 | 0.100707165 | -6.554719517 |
| Aunip | 1.169101948 | 0.612334855 | 1.092320034 | 0.560581913 |
| Mtfr1l | 639.4674001 | 0.237210578 | 0.069626406 | 3.406905409 |
| Sepn1 | 472.582935 | -0.131890552 | 0.082257221 | -1.603391778 |
| Man1c1 | 640.4685701 | -0.145655857 | 0.114946837 | -1.267158465 |
| Mir6403 | 1.273845189 | -0.77241198 | 0.806348255 | -0.957913625 |
| Ldlrap1 | 252.7861691 | 0.771528242 | 0.164682151 | 4.684953645 |
| Tmem57 | 886.6149259 | -0.012158421 | 0.075220467 | -0.161637135 |
| Rhd | 1.579581474 | -1.250312242 | 0.82744924 | -1.511044039 |
| Tmem50a | 553.5327818 | 0.47809983 | 0.105405054 | 4.535834019 |

| | | | | |
|---|---|---|---|---|
| Rsrp1 | 2711.383038 | 0.291781084 | 0.091830118 | 3.177400741 |
| Syf2 | 379.4524818 | 0.420696083 | 0.135107872 | 3.113779217 |
| Gm16225 | 0 | NA | NA | NA |
| Runx3 | 97.20798678 | 7.328443688 | 0.714926777 | 10.25062136 |
| Gm16224 | 0 | NA | NA | NA |
| Clic4 | 3776.355182 | 1.562745813 | 0.23307736 | 6.704837463 |
| Gm12983 | 0.072025462 | 0.755288839 | 3.724945049 | 0.202765096 |
| Gm12984 | 0 | NA | NA | NA |
| Gm12982 | 0 | NA | NA | NA |
| Srrm1 | 1767.897914 | 0.170605153 | 0.094986658 | 1.796095955 |
| Ncmap | 11.39922601 | 0.511467875 | 0.298895519 | 1.71119285 |
| Gm12991 | 0 | NA | NA | NA |
| Gm12990 | 3.613492829 | -0.60448735 | 0.599404034 | -1.008480617 |
| Rcan3 | 591.8608699 | -0.283613435 | 0.095144417 | -2.98087313 |
| Mir700 | 0.366383937 | -0.03675816 | 1.695996489 | -0.021673488 |
| Gm15979 | 4.479156939 | -0.515328335 | 0.541617759 | -0.951461296 |
| Nipal3 | 1375.805614 | 0.130830109 | 0.064736925 | 2.020950308 |
| Gm37270 | 0.048949532 | -0.206487837 | 3.724945049 | -0.055433794 |
| Stpg1 | 13.32983798 | -0.149798102 | 0.275841656 | -0.543058304 |
| Gm25317 | 0 | NA | NA | NA |
| Grhl3 | 12.09498146 | 0.218339894 | 0.310425814 | 0.703356115 |
| Gm23106 | 0.133280846 | 0.671929825 | 3.652577664 | 0.183960448 |
| 1700029M20Rik | 0.369889 | 0.219977564 | 1.618736329 | 0.135894623 |
| Ifnlr1 | 9.250335689 | 1.150778733 | 0.379363198 | 3.033448527 |
| Gm12989 | 39.77014755 | 0.388234711 | 0.240676794 | 1.613095738 |
| Il22ra1 | 3.4240698 | 2.106464372 | 0.682835686 | 3.084877395 |
| Gm12988 | 0.337511725 | -0.632324038 | 1.579487666 | -0.400334901 |
| Myom3 | 10.58892132 | -0.507994129 | 0.312493544 | -1.625614794 |
| Gm13000 | 0 | NA | NA | NA |
| Srsf10 | 1164.791877 | 0.201185939 | 0.077959069 | 2.580661109 |
| Pnrc2 | 595.9273528 | 0.461240833 | 0.102844898 | 4.48481977 |
| Gm13006 | 0 | NA | NA | NA |
| Cnr2 | 15.77841223 | 2.858223739 | 0.517060335 | 5.527834078 |
| Fuca1 | 282.912463 | 0.325863225 | 0.094679665 | 3.441744582 |
| Hmgcl | 258.9557325 | -0.027072672 | 0.096072617 | -0.28179384 |
| Gale | 89.88904888 | -0.064794551 | 0.128035906 | -0.506065469 |
| Lypla2 | 893.641742 | 0.675016618 | 0.130860733 | 5.158282424 |
| Pithd1 | 571.8837247 | 0.286064456 | 0.080909407 | 3.535614296 |
| Gm13007 | 0.235668049 | 0.321145224 | 2.537749666 | 0.126547243 |
| Tceb3 | 686.4047522 | 0.412525582 | 0.078584861 | 5.249428181 |
| Gm13008 | 0.394976806 | 1.223707281 | 1.535887266 | 0.796742904 |
| Gm24362 | 2.378997451 | -0.907985201 | 0.854103885 | -1.063085201 |
| Rpl11 | 1068.809872 | 0.351954322 | 0.137627072 | 2.557304442 |
| Gm13009 | 0 | NA | NA | NA |
| Gm26001 | 0 | NA | NA | NA |
| Id3 | 323.3694017 | 0.633855048 | 0.162770883 | 3.894155002 |
| E2f2 | 28.44497551 | 0.842969682 | 0.287158093 | 2.935559551 |

| | | | | |
|---|---|---|---|---|
| Asap3 | 274.1575355 | 2.119136051 | 0.19287431 | 10.98713484 |
| Tcea3 | 4.626838346 | 0.006438391 | 0.467713189 | 0.013765683 |
| Zfp46 | 472.2796557 | 0.188658604 | 0.125379434 | 1.504701356 |
| Gm13013 | 0 | NA | NA | NA |
| Hnrnpr | 1542.097552 | -0.249905914 | 0.078319698 | -3.190843696 |
| Gm27022 | 0 | NA | NA | NA |
| Htr1d | 41.88636825 | 0.060109146 | 0.204787595 | 0.293519471 |
| 6030445D17Rik | 4.632584323 | -0.539893697 | 0.526589107 | -1.0252656 |
| Luzp1 | 1615.824644 | -0.037926903 | 0.061672441 | -0.614973278 |
| Kdm1a | 902.2554757 | 0.230078889 | 0.072028438 | 3.194278465 |
| Gm12987 | 8.499369334 | -0.459517113 | 0.510663449 | -0.899843359 |
| Gm12986 | 0 | NA | NA | NA |
| Gm22180 | 0 | NA | NA | NA |
| 4930549C01Rik | 0 | NA | NA | NA |
| Lactbl1 | 0.276289708 | 2.137939035 | 2.410215545 | 0.887032299 |
| Gm37583 | 0.251203948 | -0.564115414 | 2.087522995 | -0.270231952 |
| Ephb2 | 367.915314 | -0.386799672 | 0.10157545 | -3.808003537 |
| C1qb | 684.4735841 | 2.484305533 | 0.122138647 | 20.34004464 |
| C1qc | 767.5830084 | 2.428392265 | 0.153561509 | 15.81380829 |
| C1qa | 854.8487578 | 2.138463036 | 0.169863475 | 12.58930468 |
| Epha8 | 193.1729168 | -0.356753194 | 0.103044608 | -3.462123847 |
| Zbtb40 | 195.6296531 | -0.976303714 | 0.173179304 | -5.637531105 |
| Gm23834 | 0.094097695 | 0.755282166 | 3.724945049 | 0.202763304 |
| 1700037C06Rik | 0 | NA | NA | NA |
| Gm13001 | 0.262972053 | 1.178074198 | 1.799994087 | 0.654487815 |
| Gm13003 | 0.049553047 | 0.274391604 | 3.724945049 | 0.073663262 |
| Gm25399 | 0 | NA | NA | NA |
| Gm23263 | 0 | NA | NA | NA |
| Gm13005 | 0.48771898 | -2.000017577 | 1.420468066 | -1.407998973 |
| Gm13002 | 0.904140992 | -2.366174155 | 1.143446816 | -2.069334685 |
| Wnt4 | 161.8183686 | -0.096450013 | 0.140730815 | -0.685351062 |
| Cdc42 | 2845.278538 | 0.222613492 | 0.088298364 | 2.52115081 |
| Gm25772 | 0 | NA | NA | NA |
| 2810405F17Rik | 18.73872419 | -0.067085522 | 0.348376981 | -0.192565887 |
| Rpl31-ps10 | 1.205425868 | 1.056128273 | 0.858202509 | 1.230628274 |
| Cela3a | 0 | NA | NA | NA |
| Cela3b | 0 | NA | NA | NA |
| Gm13010 | 0.090890357 | 0.755282166 | 3.724945049 | 0.202763304 |
| 1700013G24Rik | 0 | NA | NA | NA |
| Hspg2 | 519.9526523 | 0.520490063 | 0.143362134 | 3.630596518 |
| Mir7018 | 0 | NA | NA | NA |
| Ldlrad2 | 0.178810541 | -1.089380053 | 2.733025976 | -0.398598499 |
| Usp48 | 1402.55254 | -0.172107671 | 0.124584005 | -1.381458811 |
| Rap1gap | 2225.616142 | -0.102633708 | 0.094153689 | -1.090065708 |
| Rap1gapos | 0.214365142 | 0.27441045 | 2.181068002 | 0.125814716 |
| Alpl | 62.50808463 | 0.109528021 | 0.175377761 | 0.624526285 |
| Ece1 | 1063.140229 | 0.118796682 | 0.089769296 | 1.323355399 |

| | | | | |
|---|---|---|---|---|
| Gm13012 | 0.199351704 | 0.755298188 | 2.622590944 | 0.287996948 |
| Eif4g3 | 3297.877854 | 0.044879287 | 0.072094543 | 0.622506012 |
| 1700095J12Rik | 0 | NA | NA | NA |
| Mir6399 | 2.27686558 | 0.783057521 | 0.756441657 | 1.035185614 |
| 2310026L22Rik | 1.951947418 | -0.030705585 | 0.815807748 | -0.037638262 |
| Hp1bp3 | 1903.997062 | 0.057287308 | 0.067414703 | 0.849774684 |
| Sh2d5 | 591.0024521 | 0.103886976 | 0.134644766 | 0.771563421 |
| Kif17 | 150.6944098 | 0.007785066 | 0.206503386 | 0.037699459 |
| Ddost | 796.5474421 | 0.334984485 | 0.087395489 | 3.832972241 |
| Pink1 | 2455.55194 | -0.032728737 | 0.094887376 | -0.344921935 |
| Mir7019 | 0 | NA | NA | NA |
| Gm27646 | 0 | NA | NA | NA |
| Cda | 12.16163641 | -0.018423075 | 0.341462944 | -0.053953366 |
| Fam43b | 205.4390261 | -0.511225494 | 0.142681109 | -3.582993553 |
| AB041806 | 16.25226339 | -0.112718442 | 0.275665342 | -0.408895951 |
| Rpl38-ps1 | 1.247502082 | 0.775659096 | 0.955560631 | 0.811731952 |
| Mul1 | 449.0477894 | 0.28292395 | 0.115607216 | 2.447286252 |
| Camk2n1 | 9074.818375 | 0.008090929 | 0.123033918 | 0.065761778 |
| Vwa5b1 | 72.56742202 | -0.515404622 | 0.182598737 | -2.822607819 |
| Gm22961 | 0.044191549 | -0.206490061 | 3.724945049 | -0.055434391 |
| Ubxn10 | 30.26165497 | 0.567685544 | 0.199935924 | 2.839337392 |
| Pla2g2c | 2.097114995 | -1.506297779 | 0.840457489 | -1.792235538 |
| Pla2g2f | 0.721011721 | 1.307073375 | 1.178973534 | 1.108653704 |
| Pla2g2d | 2.315173619 | 1.11458095 | 0.69593831 | 1.601551363 |
| Pla2g5 | 22.3998674 | 0.296252495 | 0.257787623 | 1.149211479 |
| Gm25280 | 0.126225412 | 0.755282661 | 3.698754989 | 0.20419916 |
| Pla2g2a | 0 | NA | NA | NA |
| Gm13030 | 0.123845245 | 0.755285145 | 3.724945049 | 0.202764104 |
| Pla2g2e | 6.503439099 | 5.002382827 | 0.948013319 | 5.276700999 |
| Otud3 | 118.1387416 | -0.245678236 | 0.136385586 | -1.801350442 |
| Rnf186 | 0.191999622 | 0.865561509 | 3.26411976 | 0.265174556 |
| Tmco4 | 88.13299696 | 1.581682326 | 0.220124797 | 7.185389147 |
| Htr6 | 39.32244043 | -0.187889861 | 0.19165437 | -0.980357823 |
| Nbl1 | 578.5377914 | 0.363565871 | 0.128992864 | 2.818496006 |
| Minos1 | 533.822241 | 0.139417784 | 0.101479361 | 1.373853586 |
| Hspe1-ps4 | 0 | NA | NA | NA |
| Gm16287 | 5.315121335 | 0.35707302 | 0.460298804 | 0.775741793 |
| Capzb | 1556.163037 | 0.309078319 | 0.064772704 | 4.771737156 |
| Pqlc2 | 134.0855149 | 0.071403011 | 0.120540643 | 0.592356315 |
| Akr7a5 | 131.3216444 | -0.355225307 | 0.126696018 | -2.803760619 |
| Gm25732 | 0 | NA | NA | NA |
| Gm24245 | 1.005478281 | 2.003409211 | 2.154219558 | 0.929993047 |
| Gm13019 | 0 | NA | NA | NA |
| Ccnd3-ps | 1.016942265 | -0.760379589 | 1.187463314 | -0.640339436 |
| Mrto4 | 197.6742609 | 0.350345591 | 0.10774526 | 3.251610235 |
| Emc1 | 1429.8396 | 0.090995904 | 0.071342909 | 1.275472303 |
| Ubr4 | 6917.676742 | 0.208774688 | 0.050801055 | 4.109652616 |

| | | | | |
|---|---|---|---|---|
| 2700016F22Rik | 3.508143679 | -0.547962568 | 0.589764839 | -0.929120441 |
| Iffo2 | 505.8284133 | 0.679148946 | 0.156328976 | 4.344357414 |
| Gm25571 | 0 | NA | NA | NA |
| Gm21969 | 1.469723789 | -0.351797038 | 0.87162471 | -0.403610676 |
| Aldh4a1 | 91.67888502 | 0.434291416 | 0.130490134 | 3.328155188 |
| Mir7020 | 0.163190422 | 0.274394252 | 3.398594894 | 0.080737558 |
| Tas1r2 | 0.187591875 | 0.716184432 | 2.702695168 | 0.264988979 |
| Pax7 | 25.98741359 | -0.09662625 | 0.232354924 | -0.41585626 |
| Gm1667 | 0.227050272 | 0.727506638 | 2.172346148 | 0.334894436 |
| Gm13028 | 0.115000406 | -0.206492286 | 3.724945049 | -0.055434988 |
| 4933427I22Rik | 0.61103801 | -0.957535271 | 1.444045641 | -0.66309211 |
| Klhdc7a | 224.24311 | -0.193212461 | 0.193313869 | -0.999475422 |
| Mir2139 | 0 | NA | NA | NA |
| Igsf21 | 330.4037649 | -0.298950947 | 0.082378009 | -3.62901397 |
| Gm13027 | 0.336994846 | 0.755310935 | 1.718134129 | 0.439611158 |
| Gm13029 | 0.895881185 | 0.540787772 | 1.11032997 | 0.487051405 |
| Gm9867 | 0 | NA | NA | NA |
| Gm13016 | 1.130346611 | 0.065283797 | 1.170772521 | 0.055761299 |
| Gm13017 | 0.248149725 | 0.274404813 | 2.424616978 | 0.113174499 |
| Gm13021 | 0 | NA | NA | NA |
| Gm13026 | 0.049553047 | 0.274391604 | 3.724945049 | 0.073663262 |
| Arhgef10l | 696.9675906 | 0.021901264 | 0.081110446 | 0.270017799 |
| Gm13025 | 0.470217137 | -0.794333173 | 1.308659494 | -0.606982317 |
| Gm23045 | 0 | NA | NA | NA |
| Rcc2 | 1056.188668 | -0.044286028 | 0.060495488 | -0.732055057 |
| Gm25951 | 0 | NA | NA | NA |
| Padi6 | 0.601311955 | -0.391990594 | 1.209476725 | -0.324099328 |
| Padi4 | 0.880251114 | 2.235657942 | 1.247396978 | 1.792258585 |
| Padi3 | 0.34999831 | -1.104824505 | 1.55487289 | -0.710556157 |
| Gm13032 | 0.111006857 | 0.755286617 | 3.724945049 | 0.202764499 |
| Padi1 | 3.269897503 | -0.124768699 | 0.593852353 | -0.210100538 |
| 4930515B02Rik | 0.062426873 | 0.755288839 | 3.724945049 | 0.202765096 |
| Padi2 | 199.5704136 | -1.212942784 | 0.231999157 | -5.228220663 |
| Gm26226 | 0.275809893 | -1.836141187 | 1.971958652 | -0.931125602 |
| Gm13031 | 7.163063276 | -1.378402665 | 0.429046551 | -3.21271121 |
| Sdhb | 803.5181823 | -0.021625146 | 0.07248087 | -0.298356605 |
| Atp13a2 | 1396.203774 | -0.063427732 | 0.065619919 | -0.966592657 |
| Mfap2 | 17.61408699 | -0.346213831 | 0.248668415 | -1.392271033 |
| Crocc | 384.0657555 | -0.388794523 | 0.176080725 | -2.208047036 |
| Necap2 | 429.1644651 | 0.660890235 | 0.122928176 | 5.376230727 |
| Gm13047 | 0 | NA | NA | NA |
| Spata21 | 3.445363304 | 0.15464508 | 0.568101674 | 0.272213738 |
| Szrd1 | 732.8062499 | 0.440240764 | 0.117321852 | 3.752419141 |
| 4921514A10Rik | 0.424480717 | 0.726213544 | 1.568465607 | 0.463008905 |
| Fbxo42 | 663.8042497 | 0.049059276 | 0.079440103 | 0.617563094 |
| Rsg1 | 15.89625249 | 0.17081067 | 0.28513759 | 0.599046481 |
| C630004L07Rik | 0.566706395 | -2.446730979 | 1.34586378 | -1.817963314 |

| | | | | |
|---|---|---|---|---|
| Arhgef19 | 124.7706795 | -0.792877566 | 0.200185461 | -3.960715038 |
| Gm13055 | 0.062426873 | 0.755288839 | 3.724945049 | 0.202765096 |
| Gm13056 | 0.112396584 | 0.27438722 | 3.724945049 | 0.073662085 |
| Epha2 | 56.26488158 | 2.052379655 | 0.273925209 | 7.492481831 |
| Gm25690 | 0 | NA | NA | NA |
| Gm13074 | 0 | NA | NA | NA |
| Gm13076 | 0 | NA | NA | NA |
| Fam131c | 33.32296123 | -0.2724188 | 0.208533357 | -1.306355991 |
| Clcnka | 0.814794334 | 1.987795035 | 1.181352315 | 1.682643704 |
| Gm13075 | 0.049553047 | 0.274391604 | 3.724945049 | 0.073663262 |
| Clcnkb | 0.117046891 | -0.687371727 | 3.724945049 | -0.184532045 |
| Hspb7 | 5.320519917 | 1.480895092 | 0.792507502 | 1.868619651 |
| Gm694 | 13.84823866 | 1.235331999 | 0.404114736 | 3.056884314 |
| Zbtb17 | 276.1721208 | 0.031439746 | 0.108249116 | 0.290438827 |
| Spen | 2269.995177 | -0.183662072 | 0.070367836 | -2.610028709 |
| B330016D10Rik | 28.20952649 | -0.078541013 | 0.245654849 | -0.319720994 |
| Fblim1 | 31.34586486 | 0.727794881 | 0.298957489 | 2.434442709 |
| Tmem82 | 7.023389443 | 0.597532049 | 0.406346516 | 1.470498762 |
| AI507597 | 0.761152721 | 1.550484939 | 1.401225809 | 1.106520397 |
| Slc25a34 | 67.64740712 | 0.039765674 | 0.284975304 | 0.139540771 |
| Plekhm2 | 1488.934527 | 0.251104741 | 0.097243074 | 2.582237795 |
| Ddi2 | 701.2059972 | -0.307482198 | 0.075807607 | -4.056086304 |
| Rsc1a1 | 0 | NA | NA | NA |
| Agmat | 14.27242461 | -0.036956563 | 0.35770832 | -0.103314798 |
| Dnajc16 | 741.3636435 | 0.049229112 | 0.059301573 | 0.83014851 |
| Casp9 | 369.2534085 | -0.000750533 | 0.067779312 | -0.011073192 |
| Cela2a | 0.706300188 | 0.076197115 | 1.514118461 | 0.050324408 |
| Ctrc | 0 | NA | NA | NA |
| Ctrcos | 0 | NA | NA | NA |
| Efhd2 | 2399.369846 | 0.584730135 | 0.096156071 | 6.081052688 |
| Gm29367 | 15.45718579 | -0.210805365 | 0.271218044 | -0.777254202 |
| Fhad1os2 | 0.207873529 | 1.275782664 | 2.195931359 | 0.580975657 |
| Fhad1 | 63.64227438 | 1.149252233 | 0.227617785 | 5.049044092 |
| Gm13059 | 0 | NA | NA | NA |
| Fhad1os1 | 0 | NA | NA | NA |
| 4930455G09Rik | 2.445960431 | 1.401769414 | 0.976425721 | 1.435612955 |
| Tmem51 | 80.10815602 | 0.487453543 | 0.202058983 | 2.412431934 |
| Gm13053 | 1.791442492 | 0.036139202 | 0.791383569 | 0.045665848 |
| Tmem51os1 | 3.504632306 | 0.44658821 | 0.568432926 | 0.785648032 |
| Kazn | 1222.293754 | 0.007559716 | 0.067238396 | 0.112431529 |
| Gm13062 | 0.107803068 | 0.75528661 | 3.724945049 | 0.202764497 |
| Gm13052 | 20.91063599 | -0.575297441 | 0.468377257 | -1.228277916 |
| Gm37624 | 0 | NA | NA | NA |
| Prdm2 | 1439.09533 | -0.058802499 | 0.051938469 | -1.132156959 |
| mmu-mir-7021 | 0.613445446 | 0.936768833 | 1.400817056 | 0.668730317 |
| Gm13038 | 0 | NA | NA | NA |
| Gm13039 | 0.205676178 | 1.260739251 | 2.642181475 | 0.477158463 |

| | | | | |
|---|---|---|---|---|
| Gm22039 | 0 | NA | NA | NA |
| Gm13050 | 32.91936983 | -0.055282062 | 0.196829852 | -0.280862181 |
| Pdpn | 594.7265377 | 2.004580076 | 0.257355399 | 7.789151035 |
| Gm26313 | 0 | NA | NA | NA |
| Lrrc38 | 24.86342066 | -0.257846618 | 0.219651428 | -1.173890017 |
| Anp32b-ps1 | 18.32970893 | 0.058766 | 0.334046614 | 0.175921555 |
| Pramel1 | 0.631666884 | 1.139745314 | 1.304504322 | 0.873699914 |
| Gm13045 | 0.09176491 | 1.15872478 | 3.72306111 | 0.311229052 |
| Pramef8 | 229.9222564 | 0.35837907 | 0.104278516 | 3.436748859 |
| Oog4 | 0 | NA | NA | NA |
| Gm13042 | 1.564646527 | -0.295910329 | 0.865714396 | -0.341810567 |
| Gm13046 | 0 | NA | NA | NA |
| Gm13041 | 0 | NA | NA | NA |
| Gm13043 | 0 | NA | NA | NA |
| Gm13040 | 0 | NA | NA | NA |
| BC080695 | 0 | NA | NA | NA |
| Gm13057 | 0 | NA | NA | NA |
| Gm13058 | 0 | NA | NA | NA |
| Gm13080 | 5.317189742 | -0.382600353 | 0.494101722 | -0.774335196 |
| Gm13083 | 0 | NA | NA | NA |
| Gm13088 | 0 | NA | NA | NA |
| Gm13085 | 0 | NA | NA | NA |
| Gm13082 | 2.119392185 | -0.396804668 | 0.72624149 | -0.546381161 |
| Gm13089 | 0 | NA | NA | NA |
| Gm13078 | 0 | NA | NA | NA |
| Gm13087 | 0 | NA | NA | NA |
| Gm13081 | 0 | NA | NA | NA |
| Gm13023 | 0 | NA | NA | NA |
| Gm13084 | 0 | NA | NA | NA |
| Gm13103 | 0 | NA | NA | NA |
| Pramef6 | 0 | NA | NA | NA |
| Gm13100 | 0 | NA | NA | NA |
| Gm13106 | 0 | NA | NA | NA |
| Pramef25 | 0 | NA | NA | NA |
| Gm13101 | 0 | NA | NA | NA |
| Pramef17 | 0 | NA | NA | NA |
| Gm13108 | 0 | NA | NA | NA |
| Gm13104 | 0 | NA | NA | NA |
| Pramel4 | 0 | NA | NA | NA |
| Gm13102 | 0 | NA | NA | NA |
| Oog3 | 0 | NA | NA | NA |
| Oog2 | 0 | NA | NA | NA |
| C87977 | 0 | NA | NA | NA |
| Gm13123 | 21.87672395 | 0.179652708 | 0.228448373 | 0.786403971 |
| Pramel5 | 0 | NA | NA | NA |
| Gm13120 | 0 | NA | NA | NA |
| Gm13130 | 0 | NA | NA | NA |

| | | | | |
|---|---|---|---|---|
| Gm13128 | 0 | NA | NA | NA |
| Gm13122 | 0 | NA | NA | NA |
| Gm13119 | 0 | NA | NA | NA |
| Gm13125 | 0 | NA | NA | NA |
| Pramef12 | 0 | NA | NA | NA |
| Pramef12os | 0 | NA | NA | NA |
| 1700012P22Rik | 0.273864066 | -1.268116147 | 2.403990248 | -0.527504697 |
| Gm9944 | 0.048949532 | -0.206487837 | 3.724945049 | -0.055433794 |
| Aadacl3 | 0 | NA | NA | NA |
| Gm13118 | 0 | NA | NA | NA |
| 9430007A20Rik | 0.068097359 | -0.206487837 | 3.724945049 | -0.055433794 |
| Gm13121 | 0.042716945 | -0.206490061 | 3.724945049 | -0.055434391 |
| Gm13126 | 0.046345709 | 0.274391604 | 3.724945049 | 0.073663262 |
| Gm13124 | 0 | NA | NA | NA |
| Gm13127 | 28.94697736 | -0.079924296 | 0.214568385 | -0.372488687 |
| Gm13177 | 0 | NA | NA | NA |
| Gm436 | 0 | NA | NA | NA |
| Gm13178 | 0.072025462 | 0.755288839 | 3.724945049 | 0.202765096 |
| Gm25938 | 0 | NA | NA | NA |
| Gm438 | 0.169902121 | 1.294024635 | 3.398864604 | 0.380722619 |
| Prdx2-rs3 | 0.044191549 | -0.206490061 | 3.724945049 | -0.055434391 |
| Dhrs3 | 254.6119046 | 0.292502476 | 0.130675453 | 2.23838884 |
| Gm38074 | 1.747383901 | -0.679938165 | 0.851787303 | -0.798248768 |
| Vps13d | 3201.77473 | -0.152256843 | 0.071723606 | -2.122827498 |
| Tnfrsf1b | 457.2834194 | 3.422808527 | 0.296927541 | 11.52742019 |
| Tnfrsf8 | 21.18745374 | 0.196422506 | 0.247380886 | 0.794008417 |
| Gm13024 | 0.288426457 | 0.182158831 | 2.066570972 | 0.088145451 |
| Gm13224 | 0 | NA | NA | NA |
| Platr16 | 0 | NA | NA | NA |
| Gm13230 | 0.097899064 | -0.687371727 | 3.724945049 | -0.184532045 |
| Gm13223 | 0.093141081 | -0.687373951 | 3.724945049 | -0.184532642 |
| Smarca5-ps | 7.147813683 | 0.307670499 | 0.3797118 | 0.810273737 |
| Gm13229 | 0 | NA | NA | NA |
| Gm13225 | 0.120679735 | -0.206494107 | 3.724945049 | -0.055435477 |
| Gm21978 | 0.060174537 | 0.755286614 | 3.724945049 | 0.202764498 |
| Gm13228 | 0 | NA | NA | NA |
| Gm13226 | 9.748555663 | 0.76967633 | 0.368617891 | 2.088005893 |
| Gm13222 | 0 | NA | NA | NA |
| Gm13212 | 8.306006014 | 2.685452112 | 0.624116927 | 4.302802884 |
| Gm26763 | 0 | NA | NA | NA |
| Gm13234 | 0.712482738 | -0.736704363 | 1.149990659 | -0.640617693 |
| Gm13231 | 1.074501331 | 1.084908064 | 1.118124194 | 0.970292987 |
| Gm13249 | 0 | NA | NA | NA |
| Gm13242 | 0.150086409 | 1.264824188 | 3.723011833 | 0.339731445 |
| Gm13239 | 0 | NA | NA | NA |
| Gm13232 | 0 | NA | NA | NA |
| Gm13244 | 0.205478627 | 0.27439469 | 3.197978754 | 0.085802537 |

| | | | | |
|---|---|---|---|---|
| Gm13236 | 0 | NA | NA | NA |
| Gm13243 | 1.675672734 | -0.059541506 | 0.823901711 | -0.07226773 |
| Vmn2r-ps15 | 0.062426873 | 0.755288839 | 3.724945049 | 0.202765096 |
| Znf41-ps | 66.78666031 | 0.62843525 | 0.213875219 | 2.938326626 |
| Gm13238 | 8.42982011 | -0.022123022 | 0.382497368 | -0.057838365 |
| Gm13246 | 0 | NA | NA | NA |
| Gm13235 | 2.295964185 | 2.013544438 | 0.824865733 | 2.441057202 |
| Gm13036 | 0 | NA | NA | NA |
| Gm13034 | 8.179144896 | 0.117883118 | 0.364665351 | 0.323263831 |
| Gm13035 | 0 | NA | NA | NA |
| Gm13051 | 0 | NA | NA | NA |
| Gm26573 | 0 | NA | NA | NA |
| Zfp600 | 0.463671963 | -0.098398828 | 1.246296943 | -0.078952956 |
| Gm13165 | 0 | NA | NA | NA |
| Gm13170 | 0 | NA | NA | NA |
| Gm13167 | 0 | NA | NA | NA |
| Gm13169 | 0.687288638 | -1.129012643 | 1.1673277 | -0.96717712 |
| Gm13166 | 0.061920613 | 0.274393828 | 3.724945049 | 0.073663859 |
| Gm13168 | 0.090665643 | -0.687373951 | 3.724945049 | -0.184532642 |
| Vmn2r-ps17 | 0 | NA | NA | NA |
| Gm13251 | 42.78458666 | 0.509314625 | 0.187632279 | 2.714429678 |
| Gm13241 | 5.328295998 | -0.056649329 | 0.469822552 | -0.120576011 |
| Gm13245 | 1.259358607 | 0.494858196 | 0.915550784 | 0.540503274 |
| Gm13247 | 3.086345102 | 1.974639718 | 0.763923361 | 2.584866257 |
| Gm26880 | 0 | NA | NA | NA |
| 1700095A21Rik | 0 | NA | NA | NA |
| Gm13237 | 0.2312517 | 0.244128917 | 2.172053336 | 0.112395452 |
| Gm13233 | 0.066050875 | 0.274391604 | 3.724945049 | 0.073663262 |
| Gm13240 | 0.060174537 | 0.755286614 | 3.724945049 | 0.202764498 |
| Vmn2r-ps16 | 0.194279094 | 0.274391173 | 3.724945049 | 0.073663147 |
| Gm13248 | 6.591240675 | 1.806523 | 0.499347432 | 3.617767679 |
| Gm21411 | 0.046345709 | 0.274391604 | 3.724945049 | 0.073663262 |
| Gm20760 | 0.333659237 | 0.796164155 | 1.748147859 | 0.45543296 |
| Gm13140 | 10.11168674 | -0.120813702 | 0.351874914 | -0.343342754 |
| Gm13149 | 0 | NA | NA | NA |
| Gm13150 | 0 | NA | NA | NA |
| C230088H06Rik | 0 | NA | NA | NA |
| Rex2 | 0.066050875 | 0.274391604 | 3.724945049 | 0.073663262 |
| Gm13153 | 0 | NA | NA | NA |
| 4933438K21Rik | 0 | NA | NA | NA |
| Gm13160 | 0.210629592 | 0.755303847 | 2.186357621 | 0.34546217 |
| Gm13142 | 2.102244023 | 0.237625528 | 0.733460744 | 0.323978523 |
| Gm13162 | 1.043892331 | 1.063750495 | 1.046617613 | 1.016369763 |
| Vmn2r-ps18 | 0.642632734 | 0.198311294 | 1.211088633 | 0.16374631 |
| Gm13139 | 85.76215223 | 0.675130138 | 0.135455905 | 4.9841322 |
| Gm26624 | 9.940525084 | -0.24957897 | 0.354913392 | -0.703210913 |
| Gm13146 | 6.463804552 | -0.576966749 | 0.407984311 | -1.414188571 |

| | | | | |
|---|---|---|---|---|
| Gm13163 | 0 | NA | NA | NA |
| Gm13147 | 1.544988629 | -0.750040055 | 0.828611345 | -0.905177149 |
| Gm13158 | 0.256692543 | -0.665809158 | 1.82158712 | -0.365510467 |
| Gm13135 | 13.09013645 | 0.259990328 | 0.366446443 | 0.709490657 |
| Gm13144 | 0 | NA | NA | NA |
| Gm13151 | 2.603243443 | -0.389405399 | 0.688928156 | -0.565233684 |
| Gm13145 | 0.352653913 | -0.687379209 | 1.980656247 | -0.347046192 |
| Gm8178 | 0 | NA | NA | NA |
| Gm13161 | 3.441294106 | -0.451324105 | 0.595532789 | -0.757849296 |
| Vmn2r-ps19 | 2.482748844 | -1.421343837 | 0.737892879 | -1.92621975 |
| Gm13152 | 33.96153438 | 0.144457502 | 0.236077373 | 0.611907445 |
| Gm15951 | 0 | NA | NA | NA |
| Vmn2r-ps20 | 4.516610571 | -0.917928373 | 0.509506944 | -1.801601302 |
| Gm13141 | 0 | NA | NA | NA |
| Gm16503 | 25.68570846 | -0.372497823 | 0.337854782 | -1.102538258 |
| Gm13154 | 0.973397315 | 0.296381947 | 1.053702942 | 0.281276568 |
| 2610305D13Rik | 21.91887679 | 0.224459188 | 0.282025417 | 0.795882832 |
| Gm13155 | 0.092140247 | -0.687373951 | 3.724945049 | -0.184532642 |
| Gm16211 | 0.547750696 | 2.089166707 | 1.50691096 | 1.38639028 |
| Zfp534 | 0.518969261 | 0.92011271 | 1.347226127 | 0.682968279 |
| Gm13137 | 0 | NA | NA | NA |
| Gm13136 | 14.06422963 | -0.078911608 | 0.28003363 | -0.281793326 |
| Gm13148 | 0.33580603 | 0.755307634 | 1.732797774 | 0.435889084 |
| Gm13157 | 104.6671989 | 1.107384204 | 0.14665208 | 7.551097837 |
| Gm20707 | 14.6133478 | 1.434434296 | 0.368083426 | 3.897035817 |
| Rps19-ps3 | 30.20379719 | 0.195634713 | 0.206706528 | 0.946437033 |
| Zfp933 | 375.230941 | -0.306241119 | 0.082836186 | -3.696948549 |
| Miip | 134.4730676 | 0.251225547 | 0.163474866 | 1.536783932 |
| Fv1 | 52.36557907 | -0.936528511 | 0.169625268 | -5.521161567 |
| Mfn2 | 2014.021368 | -0.089634788 | 0.037476034 | -2.391789564 |
| Gm13156 | 0.836555448 | -1.041726341 | 1.108570866 | -0.939702074 |
| Plod1 | 436.6411955 | 0.308099891 | 0.119186709 | 2.585018872 |
| 2510039O18Rik | 318.750318 | 0.411469314 | 0.103112409 | 3.990492699 |
| Nppb | 0.91071486 | 0.963619537 | 1.206975969 | 0.79837508 |
| Nppa | 2.986404308 | -0.378664634 | 0.771492635 | -0.490820803 |
| Gm13054 | 12.47324529 | -0.066744476 | 0.382157763 | -0.174651631 |
| Clcn6 | 1341.943323 | 0.008284927 | 0.047092943 | 0.175927152 |
| Gm13207 | 0.045882455 | 0.755288839 | 3.724945049 | 0.202765096 |
| Mthfr | 731.1208707 | 0.991072267 | 0.116583714 | 8.50094952 |
| Gm13202 | 5.417309824 | 0.501600559 | 0.446668748 | 1.122981094 |
| Gm13201 | 0.498362359 | 0.420497481 | 1.492631486 | 0.281715537 |
| Agtrap | 214.8520272 | 0.761803421 | 0.145417301 | 5.238739918 |
| Draxin | 11.61009598 | 0.168912861 | 0.33204742 | 0.508701018 |
| Mad2l2 | 105.7912503 | 0.882757368 | 0.163516036 | 5.398598157 |
| Fbxo6 | 181.4057765 | 0.391879073 | 0.117472287 | 3.33592784 |
| Mir7022 | 0.141872876 | 0.274406407 | 2.910933148 | 0.094267506 |
| Fbxo44 | 836.5623358 | 0.002876224 | 0.053625855 | 0.053635033 |

| | | | | |
|---|---|---|---|---|
| Fbxo2 | 551.7280945 | -0.238292058 | 0.114911887 | -2.073693715 |
| Ptchd2 | 379.3452522 | -0.386140699 | 0.125789834 | -3.06972897 |
| Gm13206 | 0.047948698 | -0.206487837 | 3.724945049 | -0.055433794 |
| Gm13200 | 2.555034347 | 0.388235499 | 0.604479324 | 0.642264315 |
| Ubiad1 | 175.5010009 | 0.188231823 | 0.122597034 | 1.53537012 |
| Gm23303 | 1.89929672 | -0.520944542 | 0.869041764 | -0.599447074 |
| Mtor | 2020.048309 | -0.094454994 | 0.035743694 | -2.642563901 |
| Angptl7 | 17.72480738 | 0.97383135 | 0.334693489 | 2.909621436 |
| Gm24002 | 0.589151384 | -1.924158433 | 1.253130649 | -1.535481104 |
| Gm23318 | 1.238646735 | -0.857215724 | 0.93157553 | -0.920178447 |
| Exosc10 | 484.0039437 | 0.270082823 | 0.080889846 | 3.338896481 |
| Gm13204 | 0 | NA | NA | NA |
| Gm13209 | 0.331673312 | 0.755307475 | 1.764708341 | 0.428006973 |
| Srm | 303.9037999 | -0.2583817 | 0.090574998 | -2.852682363 |
| Masp2 | 31.81998135 | -1.426106622 | 0.299010858 | -4.769414172 |
| Tardbp | 2054.279046 | -0.070099199 | 0.074369074 | -0.94258535 |
| Gm572 | 0.786597823 | 0.90101243 | 1.086328811 | 0.829410415 |
| Gm13203 | 0 | NA | NA | NA |
| Casz1 | 60.05755845 | 0.258547086 | 0.179264887 | 1.44226284 |
| Gm15969 | 0 | NA | NA | NA |
| Gm13205 | 27.32691937 | -0.515111922 | 0.216564569 | -2.378560461 |
| Gm17029 | 1.365296087 | 0.422080942 | 0.973964944 | 0.433363587 |
| Pex14 | 521.3504447 | 0.427918052 | 0.088105287 | 4.856894122 |
| Gm9506 | 0 | NA | NA | NA |
| Dffa | 268.8330888 | -0.44295958 | 0.073636318 | -6.015504177 |
| Cort | 13.59908539 | -0.213038128 | 0.282199152 | -0.754921219 |
| Apitd1 | 26.0334167 | -0.103715219 | 0.210951036 | -0.491655414 |
| Rpsa-ps12 | 20.84511033 | -0.052832791 | 0.339937236 | -0.155419253 |
| Pgd | 591.4610587 | 0.659068662 | 0.100076338 | 6.585659253 |
| Kif1b | 15537.60889 | -0.14138326 | 0.050588989 | -2.794743742 |
| Ube4b | 1942.055481 | 0.014882333 | 0.045546373 | 0.32675121 |
| Ube4bos3 | 0.106056992 | 1.236179401 | 3.724945049 | 0.331865138 |
| Ube4bos2 | 0.038981395 | 0.274393828 | 3.724945049 | 0.073663859 |
| Ube4bos1 | 0.415914162 | -0.60767789 | 3.3499784 | -0.181397555 |
| Rbp7 | 1.09427696 | 1.683234607 | 1.136178533 | 1.481487775 |
| Trmt112-ps2 | 31.55425196 | -0.006097487 | 0.233805581 | -0.026079306 |
| Nmnat1 | 53.5119218 | -0.714019067 | 0.174842631 | -4.083781312 |
| Gm13097 | 0.04393175 | -0.206490061 | 3.724945049 | -0.055434391 |
| Lzic | 141.3885089 | -0.100889403 | 0.116540344 | -0.865703661 |
| Gm13071 | 9.973492174 | 0.016330808 | 0.29788983 | 0.054821636 |
| Gm24381 | 0 | NA | NA | NA |
| Ctnnbip1 | 373.0545822 | -0.122590045 | 0.098805897 | -1.240715873 |
| Mir5616 | 0.908927132 | 1.870670976 | 1.209841262 | 1.546211917 |
| Gm13066 | 0.3242121 | 0.269992345 | 1.801239468 | 0.149892532 |
| Clstn1 | 10198.28505 | -0.17332901 | 0.081805906 | -2.118783569 |
| Pik3cd | 275.4671803 | 0.479440436 | 0.120210517 | 3.988340203 |
| Mir7023 | 0 | NA | NA | NA |

| | | | | |
|---|---|---|---|---|
| Tmem201 | 968.9539549 | -0.19043222 | 0.089619321 | -2.124901397 |
| Slc25a33 | 270.7812472 | -0.06663961 | 0.119759284 | -0.556446297 |
| Gm13068 | 0.185281328 | -1.530471326 | 2.686846338 | -0.569616247 |
| Gm16188 | 0.247276735 | 0.274407296 | 2.051440682 | 0.133763213 |
| Gm13073 | 0.234101041 | 0.755309654 | 1.878262897 | 0.402132021 |
| Gm13065 | 0.044544648 | 0.755286614 | 3.724945049 | 0.202764498 |
| Gm13064 | 0.150086409 | 1.264824188 | 3.723011833 | 0.339731445 |
| n-R5s193 | 0 | NA | NA | NA |
| AL626808.1 | 0 | NA | NA | NA |
| Spsb1 | 673.0771038 | 1.804095111 | 0.186573273 | 9.669633177 |
| Gm13070 | 0.062426873 | 0.755288839 | 3.724945049 | 0.202765096 |
| H6pd | 247.2321755 | 0.022873613 | 0.13855233 | 0.165090064 |
| Gm13067 | 8.956120355 | -0.870077186 | 0.38797679 | -2.242601126 |
| Mir34a | 0.164553454 | -0.206488957 | 2.77921914 | -0.074297472 |
| Gm37795 | 24.19206811 | -0.468228782 | 0.329789349 | -1.419781393 |
| Gpr157 | 52.57716929 | 0.870491901 | 0.193243997 | 4.504625838 |
| Gm13093 | 6.364908752 | -0.844043392 | 0.4309617 | -1.958511377 |
| Slc2a5 | 12.42258672 | -0.1386792 | 0.28871432 | -0.48033364 |
| Slc2a7 | 0 | NA | NA | NA |
| Car6 | 1.391048348 | 0.75530851 | 1.411103006 | 0.535261073 |
| Gm13094 | 0.173900533 | 0.755298197 | 2.743535063 | 0.275301091 |
| Eno1 | 2763.278328 | -0.202191954 | 0.0666621 | -3.033087057 |
| Rere | 3154.94565 | 0.157090594 | 0.060937729 | 2.577887259 |
| Gm13092 | 5.881560967 | 0.869513637 | 0.637314872 | 1.364339159 |
| Gm23209 | 6.850924464 | -1.088796355 | 0.525204677 | -2.0730896 |
| Gm13091 | 33.11693011 | 1.90275908 | 0.252250832 | 7.543123103 |
| Slc45a1 | 569.9271968 | -0.074895054 | 0.063207258 | -1.18491224 |
| Gm16079 | 0 | NA | NA | NA |
| Gm13049 | 0 | NA | NA | NA |
| 1700045H11Rik | 0.109982625 | -0.20649451 | 3.724945049 | -0.055435586 |
| Rps15a-ps3 | 6.113901122 | 0.256012472 | 0.409772641 | 0.624767118 |
| Errfi1 | 790.4060845 | 1.124383652 | 0.236281836 | 4.758654627 |
| Park7 | 548.339284 | 0.029380661 | 0.1021843 | 0.287526179 |
| Tnfrsf9 | 42.91515327 | 3.451598634 | 0.598270151 | 5.769297743 |
| Camta1 | 4644.702699 | -0.186454183 | 0.05348406 | -3.4861636 |
| Gm23405 | 0 | NA | NA | NA |
| Uts2 | 1.009670171 | 2.354867385 | 1.228512615 | 1.916844285 |
| Per3 | 705.3281863 | -0.129269805 | 0.236792139 | -0.545921014 |
| Vamp3 | 420.1066725 | 0.405370343 | 0.120344386 | 3.368419226 |
| Gm13090 | 0.329009797 | -0.180298043 | 1.557452601 | -0.115764707 |
| 4930589P08Rik | 0.062426873 | 0.755288839 | 3.724945049 | 0.202765096 |
| Gm37141 | 0.038981395 | 0.274393828 | 3.724945049 | 0.073663859 |
| Dnajc11 | 693.2058939 | 0.211410752 | 0.063051075 | 3.353007919 |
| Thap3 | 106.1287797 | 0.009798214 | 0.127575543 | 0.076803229 |
| Phf13 | 200.4894639 | -0.118364697 | 0.165790788 | -0.71394013 |
| Klhl21 | 246.8014045 | -0.280677964 | 0.112837973 | -2.487442446 |
| Zbtb48 | 85.35332904 | 0.079126652 | 0.161775822 | 0.489112967 |

| | | | | |
|---|---|---|---|---|
| Tas1r1 | 4.276683327 | 0.537488636 | 0.623943777 | 0.861437611 |
| Nol9 | 273.9583489 | -0.03290148 | 0.083522787 | -0.393922195 |
| Plekhg5 | 606.8905356 | -0.012086827 | 0.165458156 | -0.073050658 |
| Tnfrsf25 | 29.08316862 | -2.670096153 | 0.620055462 | -4.306221485 |
| Espn | 20.79395242 | 0.47675085 | 0.20362597 | 2.341306705 |
| Gm13077 | 2.185424001 | 0.866412959 | 0.697360115 | 1.242418287 |
| Hes2 | 0.293168959 | -0.186971471 | 2.049589042 | -0.091223883 |
| Acot7 | 1614.713023 | -0.230233049 | 0.072943147 | -3.156335558 |
| Gm27500 | 0 | NA | NA | NA |
| Gm13096 | 2.546102757 | -1.836809602 | 0.730496326 | -2.514467954 |
| Gpr153 | 338.048994 | -0.762888507 | 0.09217222 | -8.276772583 |
| Hes3 | 57.0044881 | 0.05992706 | 0.26467004 | 0.226421774 |
| Icmt | 1103.865666 | -0.367758898 | 0.11824446 | -3.110157541 |
| Rnf207 | 32.84951904 | -0.20704698 | 0.246322925 | -0.840550998 |
| Rpl22 | 661.306753 | 0.3085554 | 0.118177921 | 2.610939488 |
| Chd5 | 2858.157954 | -0.434785612 | 0.095360046 | -4.55941069 |
| Gm13645 | 1.241205481 | -0.926382549 | 0.936835852 | -0.988841905 |
| Kcnab2 | 3430.815311 | -0.002437319 | 0.079718949 | -0.030573899 |
| Gm16333 | 0.211164147 | 0.755300877 | 2.560729507 | 0.294955353 |
| Gm16334 | 14.13596111 | -0.408449741 | 0.335741366 | -1.216560668 |
| Nphp4 | 205.6488101 | -0.458180049 | 0.191623579 | -2.391042119 |
| Gm13172 | 0 | NA | NA | NA |
| Gm833 | 0.678447637 | 1.458727199 | 1.387413908 | 1.051400155 |
| Gm25779 | 0 | NA | NA | NA |
| Gm13173 | 0.184770788 | 1.288632945 | 2.685921747 | 0.479773078 |
| Gm13174 | 0.2360389 | 0.274409587 | 2.150752264 | 0.127587724 |
| Ajap1 | 418.755463 | -0.076048068 | 0.115065352 | -0.660911972 |
| Gm25880 | 0 | NA | NA | NA |
| Gm13115 | 0.044191549 | -0.206490061 | 3.724945049 | -0.055434391 |
| BC039966 | 5.183547717 | -0.412260403 | 0.449680853 | -0.916784427 |
| A430005L14Rik | 129.8028141 | -0.036430358 | 0.123310351 | -0.295436333 |
| Dffb | 64.42929542 | 0.386752317 | 0.198147499 | 1.951840522 |
| Cep104 | 594.1324898 | -0.090977924 | 0.070666672 | -1.287423356 |
| Lrrc47 | 577.7083269 | 0.231503387 | 0.09595977 | 2.412504598 |
| Smim1 | 121.5585543 | -0.158915669 | 0.144831349 | -1.097246351 |
| Ccdc27 | 0.478716937 | -0.77755428 | 1.477064548 | -0.52641862 |
| Gm13117 | 0 | NA | NA | NA |
| Gm13114 | 0 | NA | NA | NA |
| Trp73 | 20.9206457 | -0.198546461 | 0.290312839 | -0.683905204 |
| Trp73os | 0.141071297 | -0.206491178 | 3.564043836 | -0.057937328 |
| Gm26036 | 0 | NA | NA | NA |
| Wrap73 | 158.2566248 | -0.152111611 | 0.098563564 | -1.543284409 |
| Tprgl | 931.6745304 | 0.029130224 | 0.089698008 | 0.324758876 |
| Megf6 | 28.54359773 | -0.16393329 | 0.208768621 | -0.78523913 |
| Gm13132 | 0 | NA | NA | NA |
| Arhgef16 | 13.58180626 | -0.293927862 | 0.310215876 | -0.947494582 |
| Prdm16 | 164.5019658 | -0.275459425 | 0.126610581 | -2.175643004 |

| | | | | |
|---|---|---|---|---|
| B230104I21Rik | 0.989733885 | -0.673603235 | 1.090628519 | -0.617628481 |
| Gm13133 | 0.742042093 | -0.206482704 | 1.446310445 | -0.142765134 |
| Gm13134 | 0.593269415 | -0.228089768 | 1.317164719 | -0.173167232 |
| Gm13111 | 52.71521894 | -1.321321959 | 0.271163523 | -4.872786509 |
| Gm27202 | 0 | NA | NA | NA |
| Gm13110 | 1.937464243 | -0.440838234 | 0.788744676 | -0.558911201 |
| Actrt2 | 0 | NA | NA | NA |
| Gm13113 | 1.711017467 | 0.896836337 | 0.771734181 | 1.162105241 |
| Ttc34 | 32.00110063 | 0.088083749 | 0.182862138 | 0.481694844 |
| Gm13112 | 3.984283232 | -0.283865712 | 0.495917965 | -0.572404574 |
| Mmel1 | 12.70547386 | -0.048551505 | 0.315499211 | -0.153887879 |
| Fam213b | 258.5981968 | -0.106457283 | 0.134043821 | -0.794197612 |
| Tnfrsf14 | 53.00596033 | 3.108181911 | 0.287751855 | 10.80160512 |
| Gm20421 | 6.460871859 | -0.758737904 | 0.436908073 | -1.736607657 |
| Hes5 | 56.4435291 | -2.145942357 | 0.33612465 | -6.384364717 |
| Pank4 | 318.1852538 | -0.064949587 | 0.072996565 | -0.889762235 |
| Plch2 | 799.0502262 | -0.765839525 | 0.147921973 | -5.177320919 |
| Gm10564 | 2.918309484 | 1.078829448 | 0.643203889 | 1.677274448 |
| Pex10 | 129.4440069 | -0.153985793 | 0.117790576 | -1.307284492 |
| Rer1 | 559.9984354 | 0.212742477 | 0.090701434 | 2.345524963 |
| Morn1 | 92.30872075 | -0.267415471 | 0.229276752 | -1.166343594 |
| Ski | 2357.896731 | 0.03731376 | 0.090288907 | 0.413270703 |
| 2610002J02Rik | 142.9826346 | 0.061030568 | 0.120803755 | 0.505204233 |
| Prkcz | 1482.735323 | -0.365156992 | 0.106747066 | -3.420768427 |
| Gm27200 | 1.590510877 | 1.318518889 | 0.804418628 | 1.639095421 |
| Gabrd | 409.0842723 | -0.403879832 | 0.161534132 | -2.500275496 |
| 5830444B04Rik | 0.822034208 | -0.685063549 | 1.199356161 | -0.571192754 |
| Cfap74 | 161.3175935 | -0.391625059 | 0.18396553 | -2.12879586 |
| Tmem52 | 14.34551441 | 1.880338839 | 0.355002642 | 5.296689699 |
| Gnb1 | 6791.840275 | 0.020493827 | 0.056657034 | 0.361717249 |
| Gm13171 | 0.045882455 | 0.755288839 | 3.724945049 | 0.202765096 |
| Nadk | 615.6859532 | 0.064313457 | 0.093463512 | 0.688112997 |
| Slc35e2 | 942.3227332 | -0.143617777 | 0.060034893 | -2.392238414 |
| Gm16023 | 7.870775316 | -1.021639409 | 0.496740496 | -2.056686374 |
| Gm10563 | 0.110242424 | -0.20649451 | 3.724945049 | -0.055435586 |
| Gm16024 | 3.554695965 | 0.077607855 | 0.615131278 | 0.126164702 |
| Cdk11b | 500.6534106 | 0.266294664 | 0.131912776 | 2.018717757 |
| Mmp23 | 18.56308676 | 0.065483403 | 0.267586216 | 0.244718896 |
| Mib2 | 880.1110826 | -0.251383473 | 0.066940114 | -3.755348769 |
| AL670236.1 | 0 | NA | NA | NA |
| B930041F14Rik | 0 | NA | NA | NA |
| Ssu72 | 463.6570633 | 0.091929692 | 0.079934706 | 1.150059791 |
| Gm22573 | 0.048949532 | -0.206487837 | 3.724945049 | -0.055433794 |
| 1500002C15Rik | 18.92769097 | -0.47497859 | 0.249615082 | -1.902844116 |
| Tmem240 | 359.2263579 | -0.567804711 | 0.107902521 | -5.262200603 |
| Atad3a | 323.5868663 | -0.253620509 | 0.105404062 | -2.406173968 |
| Atad3aos | 2.955322402 | -1.690668066 | 0.680056984 | -2.486068233 |

| | | | | |
|---|---|---|---|---|
| Vwa1 | 254.228511 | 0.243701321 | 0.11267052 | 2.162955493 |
| Gm26840 | 23.01521584 | -0.038748486 | 0.238503743 | -0.162464895 |
| Tmem88b | 1074.722227 | -0.149426511 | 0.137936824 | -1.08329674 |
| Gm27743 | 0 | NA | NA | NA |
| Ankrd65 | 0.81397646 | -0.748150863 | 1.066278503 | -0.701646765 |
| Mrpl20 | 295.0369231 | 0.228077581 | 0.138628393 | 1.645244357 |
| Ccnl2 | 1143.506995 | 0.209314082 | 0.132729964 | 1.576991937 |
| Aurkaip1 | 435.2140168 | 0.171456073 | 0.11274459 | 1.520747681 |
| Mxra8os | 0.102891481 | 0.274400501 | 3.724945049 | 0.073665651 |
| Mxra8 | 155.5730168 | 1.032729638 | 0.182016941 | 5.67381054 |
| Dvl1 | 770.5694415 | -0.193541048 | 0.075848387 | -2.551683104 |
| Tas1r3 | 8.894659495 | -0.023752801 | 0.353743311 | -0.067146997 |
| Cptp | 215.1678806 | 0.074123293 | 0.101139341 | 0.732882893 |
| Cpsf3l | 232.6024494 | 0.141941812 | 0.078542038 | 1.807208172 |
| Pusl1 | 121.0895384 | -0.093473084 | 0.132319484 | -0.706419651 |
| Acap3 | 1132.056167 | 0.02560653 | 0.106951031 | 0.239422933 |
| Gm22991 | 0 | NA | NA | NA |
| Ube2j2 | 313.5792266 | 0.215402934 | 0.097625908 | 2.206411581 |
| Fam132a | 40.80633962 | -0.427248976 | 0.236499414 | -1.806554054 |
| B3galt6 | 79.13773638 | 0.278998502 | 0.166551028 | 1.675153283 |
| Sdf4 | 2638.964062 | 0.116703613 | 0.055083298 | 2.118675115 |
| Tnfrsf4 | 12.95165872 | 1.543023259 | 0.460637811 | 3.349753802 |
| Gm37090 | 21.13329923 | -1.026965405 | 0.292951444 | -3.505582322 |
| Gm16008 | 35.8777959 | -0.288418258 | 0.217611679 | -1.325380417 |
| Gm25982 | 1.203677616 | -0.854028976 | 1.099018736 | -0.777083181 |
| Tnfrsf18 | 8.73841548 | 5.931830944 | 0.971669658 | 6.104781491 |
| Ttll10 | 1.729677859 | -0.109929319 | 0.756874997 | -0.14524105 |
| Gm13648 | 0.840567365 | -1.420271806 | 1.189271167 | -1.194237147 |
| Mir429 | 0 | NA | NA | NA |
| Mir200a | 0.075043205 | 0.755286614 | 3.724945049 | 0.202764498 |
| Mir200b | 0.044191549 | -0.206490061 | 3.724945049 | -0.055434391 |
| 9430015G10Rik | 166.5177331 | -0.452903483 | 0.144730447 | -3.129289606 |
| Agrn | 2086.461589 | 0.389113695 | 0.094142286 | 4.133250989 |
| Isg15 | 2298.739811 | 9.14789139 | 0.440178002 | 20.78225478 |
| AW011738 | 181.0853581 | 2.329356954 | 0.218251472 | 10.67281209 |
| Perm1 | 7.861063657 | 1.096681023 | 0.413915615 | 2.649528027 |
| Plekhn1 | 36.96362493 | 0.116260403 | 0.206807558 | 0.562167089 |
| Klhl17 | 254.2446718 | -0.318218599 | 0.119776773 | -2.656763827 |
| Mir7658 | 0 | NA | NA | NA |
| Noc2l | 659.9224954 | 0.228588836 | 0.074195057 | 3.080917338 |
| Samd11 | 37.37872384 | -0.05518803 | 0.214800667 | -0.256926716 |
| Vmn2r-ps159 | 0.208220181 | -0.251440776 | 3.172055944 | -0.079267447 |
| Gm20782 | 0 | NA | NA | NA |
| Vmn1r-ps5 | 0 | NA | NA | NA |
| Gm3027 | 0 | NA | NA | NA |
| RP24-481N17.5 | 0 | NA | NA | NA |
| RP24-481N17.7 | 0 | NA | NA | NA |

| | | | | |
|---|---|---|---|---|
| RP24-481N17.8 | 0 | NA | NA | NA |
| Gm6486 | 0 | NA | NA | NA |
| Gm3053 | 0 | NA | NA | NA |
| RP23-128P4.1 | 0 | NA | NA | NA |
| Gm8715 | 0 | NA | NA | NA |
| RP23-135F23.1 | 0 | NA | NA | NA |
| Gm15772 | 876.5883103 | 0.449550797 | 0.165484606 | 2.716571705 |
| RP23-135F23.2 | 0 | NA | NA | NA |
| Cdk6 | 185.3456827 | 1.450168463 | 0.176850682 | 8.199959707 |
| RP23-454P8.1 | 0.082913145 | -0.206492286 | 3.724945049 | -0.055434988 |
| RP23-454P8.3 | 0.295949093 | 1.223293408 | 2.060243417 | 0.59376159 |
| Fam133b | 414.4885661 | -0.027356845 | 0.090030429 | -0.303862211 |
| RP23-434H17.4 | 7.609880518 | -0.374739994 | 0.392013946 | -0.955935366 |
| n-R5s169 | 0 | NA | NA | NA |
| 1700109H08Rik | 123.3221518 | 0.369202797 | 0.37530269 | 0.98374674 |
| Rbm48 | 131.5051051 | 0.425347225 | 0.135888703 | 3.130114672 |
| RP23-434H17.3 | 18.64070939 | -0.290040664 | 0.300135968 | -0.966364231 |
| Pex1 | 339.3891102 | -0.306762741 | 0.131747625 | -2.328411924 |
| RP23-434H17.5 | 1.254627161 | 0.607978539 | 0.83553839 | 0.727648839 |
| RP23-434H17.6 | 46.52130311 | -0.540195994 | 0.149731506 | -3.607764374 |
| Gatad1 | 720.1271276 | -0.17613979 | 0.085704133 | -2.055207662 |
| Tmbim7 | 0.149576918 | 0.755295313 | 2.863326609 | 0.263782452 |
| Gm26238 | 0.120349074 | 1.264824188 | 3.723011833 | 0.339731445 |
| Ankib1 | 1655.656652 | 0.105584048 | 0.0596774 | 1.769246773 |
| Gm22926 | 0 | NA | NA | NA |
| RP23-425I8.2 | 106.4878435 | -0.334664984 | 0.244355071 | -1.369584769 |
| Krit1 | 1289.563298 | -0.328111816 | 0.106982233 | -3.066974823 |
| Lrrd1 | 7.878715828 | -0.23464918 | 0.367456548 | -0.638576674 |
| Rps4x-ps | 7.918913376 | 0.372928684 | 0.383145647 | 0.973333997 |
| Mterf1a | 31.30236157 | -0.016382937 | 0.24160853 | -0.067807774 |
| Akap9 | 2881.578872 | -0.084802608 | 0.099950481 | -0.848446221 |
| Wdr46-ps | 30.12730886 | 0.01189335 | 0.265833029 | 0.044739927 |
| RP23-369O21.1 | 30.72591236 | -0.447262985 | 0.261985514 | -1.707205022 |
| Cyp51 | 835.8806558 | 0.034556012 | 0.137898733 | 0.250589775 |
| Mterf1b | 11.84616759 | -0.184884909 | 0.344423934 | -0.536794601 |
| Fzd1 | 479.4036875 | 0.272686852 | 0.140150886 | 1.945666269 |
| Gm15750 | 1.582330525 | -0.100348019 | 0.818388406 | -0.122616619 |
| RP23-114F4.1 | 8.620768254 | 0.402916633 | 0.43051477 | 0.935895028 |
| AC026231.1 | 0 | NA | NA | NA |
| Cdk14 | 2684.227082 | -0.098363371 | 0.081772804 | -1.20288612 |
| RP23-114F4.2 | 25.16791745 | -0.445664169 | 0.337572206 | -1.320203978 |
| RP23-114F4.3 | 17.31984568 | -0.296251361 | 0.28871593 | -1.026099808 |
| Gm25037 | 1.520499152 | 1.269763529 | 0.989169548 | 1.283666214 |
| RP23-114F4.4 | 36.06826517 | -0.336314146 | 0.40472126 | -0.830977217 |
| RP23-282L6.2 | 14.6407167 | -0.183352536 | 0.312472933 | -0.586778939 |
| RP23-282L6.1 | 20.90691181 | -0.487520946 | 0.299695542 | -1.626720714 |
| RP23-407F12.1 | 44.77564562 | -0.531347647 | 0.240140487 | -2.212653327 |

| | | | | |
|---|---|---|---|---|
| RP23-396K2.1 | 12.0291979 | -0.47932957 | 0.354775357 | -1.3510791 |
| RP23-396K2.2 | 12.35285954 | -0.029067967 | 0.36837688 | -0.078908228 |
| 1700015F17Rik | 1.819296732 | 0.680691806 | 0.751577533 | 0.905684079 |
| Cldn12 | 558.7465676 | -0.210531581 | 0.077298032 | -2.723634416 |
| Gtpbp10 | 131.1217465 | 0.294041537 | 0.16019657 | 1.83550457 |
| RP23-119M19.2 | 0 | NA | NA | NA |
| Gm8773 | 3.281282651 | 0.71155957 | 0.571586069 | 1.24488613 |
| Cfap69 | 276.8459991 | -0.56614502 | 0.116477441 | -4.86055512 |
| Steap2 | 544.5451271 | -0.324362061 | 0.11221131 | -2.890636065 |
| RP23-261D18.1 | 0.350305046 | 0.593730316 | 1.724903046 | 0.344210834 |
| Steap1 | 8.981601178 | 0.42264588 | 0.356247042 | 1.186384252 |
| Gm26825 | 3.449447184 | -0.150042597 | 0.604636285 | -0.248153478 |
| Gm15459 | 4780.466862 | -0.141842701 | 0.104283505 | -1.360164303 |
| RP23-261D18.2 | 0 | NA | NA | NA |
| Gm23838 | 0 | NA | NA | NA |
| RP24-323L24.1 | 0.269679299 | -0.962046401 | 1.590586128 | -0.604837666 |
| RP24-323L24.2 | 0 | NA | NA | NA |
| Gm24686 | 0 | NA | NA | NA |
| RP24-338C23.1 | 0.161949631 | 0.274399427 | 2.77761947 | 0.098789424 |
| RP24-338C23.2 | 0.322473324 | 1.168987837 | 1.75755816 | 0.665120429 |
| RP23-189N11.1 | 0.187072314 | -0.206481343 | 2.668356397 | -0.077381471 |
| RP23-74J16.1 | 1.533852754 | 0.167983003 | 0.774513611 | 0.216888381 |
| Zfp804b | 64.00592618 | -0.423437732 | 0.132993849 | -3.183889589 |
| Gm24525 | 5.266732454 | -1.340218142 | 0.500969063 | -2.675251307 |
| Tubb4b-ps1 | 53.69871365 | 0.095216279 | 0.187659076 | 0.507389682 |
| RP24-72E12.3 | 27.34862605 | -0.849274601 | 0.320725314 | -2.6479812 |
| RP24-72E12.2 | 24.54249589 | -0.935636753 | 0.286347182 | -3.267490697 |
| RP24-72E12.4 | 0 | NA | NA | NA |
| 4921511H03Rik | 0 | NA | NA | NA |
| Gm4960 | 0.287827876 | 1.231326611 | 2.037600285 | 0.604302336 |
| RP23-465J5.1 | 0.130017973 | -0.206489706 | 3.662785653 | -0.056375045 |
| Gm8802 | 0.060174537 | 0.755286614 | 3.724945049 | 0.202764498 |
| Gm22196 | 0 | NA | NA | NA |
| RP23-212F6.1 | 0 | NA | NA | NA |
| Steap4 | 41.56055496 | 4.38476496 | 0.693590014 | 6.321839806 |
| Gm15731 | 0.761917678 | 2.247237804 | 1.209006325 | 1.858747765 |
| Sri | 562.266319 | 0.103114805 | 0.08949303 | 1.152210464 |
| Gm15730 | 0.062426873 | 0.755288839 | 3.724945049 | 0.202765096 |
| Adam22 | 4541.002357 | -0.261773482 | 0.091997668 | -2.845436041 |
| Gm21759 | 0.083172944 | -0.206492286 | 3.724945049 | -0.055434988 |
| Gm24196 | 4.833961456 | -0.716918094 | 0.482206834 | -1.486743952 |
| Gm23993 | 0 | NA | NA | NA |
| RP23-201E13.1 | 54.61168029 | -0.812733069 | 0.27745635 | -2.929228576 |
| Dbf4 | 47.80200343 | 0.837979193 | 0.262149201 | 3.196573509 |
| Slc25a40 | 206.1233218 | -0.267877959 | 0.098670936 | -2.714861841 |
| Rundc3b | 435.3356137 | -0.095572146 | 0.094753767 | -1.008636901 |
| RP24-159A20.4 | 29.56068002 | -0.552712121 | 0.278589572 | -1.983965576 |

| | | | | |
|---|---|---|---|---|
| Abcb1a | 172.8364366 | 0.495726063 | 0.162022628 | 3.059610062 |
| RP24-223N5.1 | 5.526269997 | -0.198512168 | 0.521850504 | -0.380400452 |
| RP23-162P5.1 | 1.143206505 | -0.075831922 | 0.840292969 | -0.090244623 |
| Gm5106 | 1.599390384 | 1.052605538 | 0.847016179 | 1.242721879 |
| Abcb1b | 71.1266548 | 1.254160471 | 0.170603256 | 7.351327885 |
| RP23-338P12.2 | 0.602373977 | 1.831567401 | 1.373890204 | 1.333125016 |
| Abcb4 | 48.46738024 | -0.443072828 | 0.167081234 | -2.651840767 |
| RP23-338P12.3 | 11.71418499 | 0.972271902 | 0.393392388 | 2.471506646 |
| Crot | 573.9541497 | 0.600958621 | 0.122270061 | 4.915010383 |
| Gm15610 | 11.65246704 | 0.320809079 | 0.461317021 | 0.695419992 |
| Gm37153 | 0.09689823 | -0.687371727 | 3.724945049 | -0.184532045 |
| Gm27941 | 0 | NA | NA | NA |
| Gm15611 | 0.120349074 | 1.264824188 | 3.723011833 | 0.339731445 |
| Tmem243 | 183.1811946 | 0.86899134 | 0.129063755 | 6.733039358 |
| Dmtf1 | 881.7559076 | -0.379908625 | 0.099806595 | -3.806448106 |
| Gm24666 | 1.171812973 | -0.870433773 | 0.959205286 | -0.907453061 |
| Gm7332 | 10.24110484 | 0.426273535 | 0.384853055 | 1.10762674 |
| Gm15734 | 0.410151891 | 0.180597893 | 1.553919397 | 0.116220888 |
| 9330182L06Rik | 795.1077193 | 0.085884224 | 0.05702788 | 1.506004138 |
| Gm15733 | 3.241155436 | -1.293823912 | 0.613643916 | -2.108427835 |
| Mir879 | 0.599062483 | 0.572611049 | 1.23967795 | 0.461903069 |
| Grm3 | 803.8407714 | -0.746797809 | 0.127544742 | -5.855183033 |
| RP24-118C7.3 | 12.97903413 | -1.468573921 | 0.303005395 | -4.846692317 |
| Gm23312 | 0 | NA | NA | NA |
| RP23-193N17.1 | 0.49396662 | 1.460107799 | 1.384409513 | 1.054679114 |
| RP23-193C5.2 | 0.302344664 | 1.215460223 | 3.724447934 | 0.326346413 |
| Gm6586 | 0.313880683 | 1.65921614 | 1.791552852 | 0.926132956 |
| RP23-193C5.3 | 0.217113707 | 0.274411394 | 2.551113951 | 0.107565322 |
| Gm10482 | 0.386285301 | 1.43074663 | 1.561952644 | 0.915998725 |
| Gm28685 | 0.419836731 | -0.206481509 | 1.446630157 | -0.142732756 |
| Gm5152 | 0.268081508 | 0.755305095 | 2.038374038 | 0.370542933 |
| RP23-412L2.1 | 0.048949532 | -0.206487837 | 3.724945049 | -0.055433794 |
| Gm17091 | 0.90048758 | -0.047862592 | 0.886042053 | -0.05401842 |
| RP24-383C4.1 | 0.834920584 | -0.604361609 | 1.118408411 | -0.540376487 |
| RP23-281K10.3 | 0 | NA | NA | NA |
| RP23-281K10.1 | 0 | NA | NA | NA |
| RP23-281K10.2 | 0 | NA | NA | NA |
| Gm6455 | 0 | NA | NA | NA |
| RP24-113C21.2 | 0 | NA | NA | NA |
| RP23-307D8.3 | 0 | NA | NA | NA |
| Gm8871 | 0 | NA | NA | NA |
| Gm8857 | 0 | NA | NA | NA |
| RP24-79B1.2 | 0 | NA | NA | NA |
| RP23-307D8.4 | 0 | NA | NA | NA |
| RP24-79B1.3 | 0 | NA | NA | NA |
| RP23-307D8.5 | 0 | NA | NA | NA |
| Gm8879 | 0 | NA | NA | NA |

| | | | | |
|---|---|---|---|---|
| RP23-92D22.3 | 0 | NA | NA | NA |
| RP23-307D8.7 | 0 | NA | NA | NA |
| Gm5861 | 0 | NA | NA | NA |
| RP23-307D8.9 | 0 | NA | NA | NA |
| Gm8890 | 0 | NA | NA | NA |
| RP24-79B1.4 | 0 | NA | NA | NA |
| RP23-307D8.8 | 0 | NA | NA | NA |
| Speer1 | 0 | NA | NA | NA |
| RP24-390F5.2 | 0 | NA | NA | NA |
| RP23-307D8.10 | 0 | NA | NA | NA |
| Gm8897 | 0 | NA | NA | NA |
| RP23-299O17.3 | 0 | NA | NA | NA |
| RP23-299O17.4 | 0.739845808 | 0.556199438 | 1.405271508 | 0.395795001 |
| RP23-307D8.11 | 0 | NA | NA | NA |
| Gm8906 | 0 | NA | NA | NA |
| RP23-299O17.5 | 0 | NA | NA | NA |
| RP23-307D8.12 | 0 | NA | NA | NA |
| Gm6460 | 0 | NA | NA | NA |
| RP23-395E24.4 | 0 | NA | NA | NA |
| RP23-307D8.6 | 0 | NA | NA | NA |
| Gm8922 | 0 | NA | NA | NA |
| 1700003C15Rik | 0 | NA | NA | NA |
| RP23-395E24.3 | 0 | NA | NA | NA |
| Gm8926 | 0 | NA | NA | NA |
| RP23-408K8.3 | 0 | NA | NA | NA |
| RP23-307D8.13 | 0 | NA | NA | NA |
| Gm6465 | 0 | NA | NA | NA |
| RP24-341O14.4 | 0 | NA | NA | NA |
| RP23-307D8.14 | 0 | NA | NA | NA |
| 4933402N22Rik | 0 | NA | NA | NA |
| RP24-341O14.3 | 0.201351563 | -1.136224871 | 2.201968076 | -0.516004243 |
| Gm29066 | 0 | NA | NA | NA |
| Gm38242 | 0 | NA | NA | NA |
| RP24-235L14.3 | 0.152964131 | 0.274406991 | 2.843489581 | 0.096503604 |
| RP23-364E18.2 | 0.046345709 | 0.274391604 | 3.724945049 | 0.073663262 |
| Sema3d | 247.8894151 | 0.274668152 | 0.152921373 | 1.796139721 |
| RP23-237H9.3 | 0 | NA | NA | NA |
| RP23-237H9.2 | 4.98564735 | -0.279471797 | 0.443186667 | -0.63059613 |
| RP24-346C14.1 | 0 | NA | NA | NA |
| Gm23243 | 0 | NA | NA | NA |
| Gm10481 | 11.05583305 | 0.039028896 | 0.313006584 | 0.124690336 |
| RP23-30B3.2 | 0 | NA | NA | NA |
| Gm10108 | 1.450945253 | 0.918852808 | 0.913177347 | 1.00621507 |
| Sema3a | 401.2948366 | -0.05323755 | 0.090360164 | -0.589170581 |
| RP23-30B3.4 | 0 | NA | NA | NA |
| RP23-10F17.2 | 15.6478416 | -0.565987795 | 0.34985597 | -1.617773723 |
| RP23-10F17.3 | 7.940230631 | -0.513576909 | 0.376837088 | -1.362861898 |

| | | | | |
|---|---|---|---|---|
| RP23-10F17.4 | 8.41488845 | -0.764098913 | 0.434971367 | -1.756664853 |
| RP23-192D21.1 | 16.18937152 | -0.553224543 | 0.254430301 | -2.174365789 |
| RP23-192D21.2 | 0 | NA | NA | NA |
| RP24-405C17.3 | 0 | NA | NA | NA |
| RP24-405C17.4 | 0.101408268 | 0.755286617 | 3.724945049 | 0.202764499 |
| Gm6647 | 0.217530652 | 1.2362011 | 2.176047851 | 0.568094631 |
| Speer3 | 0 | NA | NA | NA |
| Gm6650 | 0 | NA | NA | NA |
| RP24-405C17.5 | 0 | NA | NA | NA |
| RP24-405C17.6 | 0 | NA | NA | NA |
| RP24-405C17.7 | 0 | NA | NA | NA |
| Gm24054 | 0 | NA | NA | NA |
| Sema3e | 550.5002642 | -0.136257606 | 0.116447555 | -1.170119942 |
| RP24-194J1.1 | 17.56400965 | -0.754184493 | 0.303186383 | -2.487527595 |
| Gm4128 | 0 | NA | NA | NA |
| RP24-481P22.1 | 12.20307049 | 0.399920409 | 0.320262207 | 1.248728075 |
| Pclo | 8266.158298 | -0.417187471 | 0.108054039 | -3.860915112 |
| Gm26918 | 1.104625153 | -0.758059923 | 1.003542217 | -0.755384188 |
| RP23-342M4.2 | 153.5905569 | -0.722662769 | 0.234942769 | -3.075909822 |
| RP23-342M4.3 | 100.8300785 | -0.679303671 | 0.231972863 | -2.928375597 |
| RP23-342M4.4 | 73.09923858 | -0.908206327 | 0.194051869 | -4.680224575 |
| RP23-342M4.5 | 112.0484156 | -0.896223886 | 0.25691267 | -3.488437864 |
| RP23-342M4.6 | 56.2336579 | -0.921257907 | 0.278107194 | -3.312600056 |
| RP23-342M4.7 | 8.648288359 | -0.487393283 | 0.351798266 | -1.38543401 |
| Gm26954 | 0.454340154 | -0.201321026 | 1.397855198 | -0.144021374 |
| RP23-451C1.2 | 88.75190297 | -0.141993448 | 0.212907182 | -0.666926531 |
| RP23-451C1.1 | 51.00169416 | -0.471964825 | 0.263619988 | -1.790322611 |
| Gm9758 | 0.061920613 | 0.274393828 | 3.724945049 | 0.073663859 |
| Speer4e | 0.072025462 | 0.755288839 | 3.724945049 | 0.202765096 |
| RP23-239L21.4 | 0.368800866 | -0.181817229 | 1.534007199 | -0.118524365 |
| Gm10354 | 0.066050875 | 0.274391604 | 3.724945049 | 0.073663262 |
| Gm17019 | 0.062426873 | 0.755288839 | 3.724945049 | 0.202765096 |
| RP24-152F23.2 | 0.163328881 | 0.755299253 | 2.776580496 | 0.27202498 |
| Gm8579 | 0.295956696 | 0.755307829 | 1.743917399 | 0.433109865 |
| Gm21847 | 0.231556444 | 0.755305291 | 2.129776049 | 0.354640711 |
| RP24-152F23.3 | 0.112396584 | 0.27438722 | 3.724945049 | 0.073662085 |
| RP23-465P18.1 | 0.409112724 | -0.206482056 | 1.715263135 | -0.12037923 |
| Gm21655 | 0 | NA | NA | NA |
| Speer4d | 0 | NA | NA | NA |
| 4930572O03Rik | 0 | NA | NA | NA |
| Speer4cos | 0.174149299 | 0.755285184 | 3.362794904 | 0.224600431 |
| Speer4c | 0.220272213 | 0.274409923 | 2.189171044 | 0.125348781 |
| RP23-77E18.1 | 0 | NA | NA | NA |
| RP23-77E18.2 | 0 | NA | NA | NA |
| RP23-77E18.3 | 0 | NA | NA | NA |
| RP23-77E18.5 | 0.109124069 | 0.274402725 | 3.724945049 | 0.073666248 |
| RP23-77E18.6 | 0 | NA | NA | NA |

| | | | | |
|---|---|---|---|---|
| Gm3510 | 30.33973269 | -0.480203373 | 0.211695854 | -2.268364566 |
| Cacna2d1 | 2942.12003 | -0.291276973 | 0.098266679 | -2.964147928 |
| RP23-123P10.5 | 0.04393175 | -0.206490061 | 3.724945049 | -0.055434391 |
| RP24-220P21.4 | 77.60211899 | -0.789779541 | 0.254105638 | -3.108075631 |
| RP24-372A2.2 | 0 | NA | NA | NA |
| Gm8984 | 0 | NA | NA | NA |
| Hgf | 116.1833121 | 0.864806924 | 0.160548138 | 5.386589576 |
| RP24-295A5.2 | 4.076536194 | 0.319493767 | 0.54727337 | 0.583791911 |
| RP24-295A5.3 | 3.112827446 | 0.358887324 | 0.680215108 | 0.527608574 |
| Gm8991 | 9.192051872 | 0.679438104 | 0.381638224 | 1.780319847 |
| Gm28710 | 0.299775621 | 0.720835012 | 1.783389855 | 0.404193738 |
| RP24-363D4.1 | 1.258311994 | 0.25413533 | 0.92438847 | 0.274922653 |
| Gm6673 | 0 | NA | NA | NA |
| Gm3495 | 0 | NA | NA | NA |
| Gm21999 | 0 | NA | NA | NA |
| Speer4f | 0 | NA | NA | NA |
| RP23-76L18.2 | 0 | NA | NA | NA |
| Sema3c | 248.8209663 | -0.241575642 | 0.125289523 | -1.928139215 |
| Cd36 | 46.54322315 | 1.779138647 | 0.553059074 | 3.216905262 |
| Gnat3 | 0.097139547 | 0.274393828 | 3.724945049 | 0.073663859 |
| Gnai1 | 1646.477261 | -0.251887366 | 0.074772983 | -3.368694894 |
| RP24-302O2.1 | 0 | NA | NA | NA |
| RP24-459D3.1 | 0.038981395 | 0.274393828 | 3.724945049 | 0.073663859 |
| RP24-332E15.1 | 0 | NA | NA | NA |
| Gm23822 | 0.060174537 | 0.755286614 | 3.724945049 | 0.202764498 |
| RP23-350E1.2 | 51.82168262 | -0.08083754 | 0.152032402 | -0.531712578 |
| 4921504A21Rik | 48.57949567 | -0.197012373 | 0.190564276 | -1.03383686 |
| RP23-350E1.3 | 0 | NA | NA | NA |
| Magi2 | 1985.599496 | -0.167904258 | 0.106502916 | -1.576522639 |
| RP24-388K6.6 | 0.090665643 | -0.687373951 | 3.724945049 | -0.184532642 |
| Gm25335 | 0 | NA | NA | NA |
| Gm23570 | 4.591749267 | -0.911406362 | 0.49392589 | -1.845228972 |
| Gm25761 | 0 | NA | NA | NA |
| RP24-531D1.4 | 0.048949532 | -0.206487837 | 3.724945049 | -0.055433794 |
| Gm22755 | 8.57913668 | -0.162450002 | 0.368124999 | -0.441290328 |
| RP23-281M20.1 | 6.048354155 | 0.009574423 | 0.421806689 | 0.022698604 |
| Gm29254 | 8.172606336 | -0.286599711 | 0.398387156 | -0.719399977 |
| RP23-343F16.2 | 0.689035189 | -0.086835866 | 1.228174284 | -0.070703212 |
| Phtf2 | 298.2117683 | -0.414501095 | 0.105616545 | -3.924584881 |
| Rpl31-ps5 | 0.299266739 | 0.75531055 | 1.76768002 | 0.427289182 |
| Tmem60 | 129.5434812 | -0.188753822 | 0.137305319 | -1.374701459 |
| Rsbn1l | 755.8552143 | 0.233616322 | 0.070092748 | 3.332959939 |
| A630072M18Rik | 116.1077269 | 0.711541318 | 0.134451064 | 5.2921955 |
| Ptpn12 | 569.2757863 | -0.117886602 | 0.092077845 | -1.280292796 |
| RP23-234P16.4 | 0.061920613 | 0.274393828 | 3.724945049 | 0.073663859 |
| Gm22490 | 0 | NA | NA | NA |
| Rpl31-ps21 | 0 | NA | NA | NA |

| | | | | |
|---|---|---|---|---|
| RP23-234P16.2 | 0.792202851 | -0.936023625 | 1.24112604 | -0.7541729 |
| RP23-234P16.3 | 0.272741579 | 1.227442778 | 1.795751354 | 0.683526021 |
| Gsap | 232.3497055 | 0.45208426 | 0.147726931 | 3.060269774 |
| RP23-250D22.2 | 3.212737209 | 2.014789279 | 0.760541475 | 2.649151091 |
| RP23-250D22.3 | 0 | NA | NA | NA |
| Ccdc146 | 10.59368012 | -0.808468221 | 0.354515838 | -2.280485477 |
| Fgl2 | 737.5148379 | 4.207538616 | 0.27495291 | 15.3027608 |
| Fam185a | 169.3008059 | -0.322262771 | 0.129516099 | -2.488206276 |
| Fbxl13 | 5.258800805 | -0.672511347 | 0.446646696 | -1.505689738 |
| Lrrc17 | 1.924082696 | 0.197276709 | 0.723988549 | 0.272485952 |
| RP23-6C10.3 | 0.285998089 | 0.274414007 | 2.09721176 | 0.130847067 |
| RP23-6C10.4 | 1.597009952 | 0.423603046 | 0.808927371 | 0.523660172 |
| Armc10 | 162.4313593 | 0.107520021 | 0.096098188 | 1.118855863 |
| Napepld | 190.2960083 | -0.184447942 | 0.110741675 | -1.665569364 |
| Gm15719 | 0.801310233 | -0.219836962 | 0.984139371 | -0.223379908 |
| Pmpcb | 597.1185698 | 0.370961372 | 0.076589644 | 4.84349255 |
| Gm15715 | 0.120974994 | 0.274404949 | 3.724945049 | 0.073666845 |
| Dnajc2 | 320.2722277 | 0.156594847 | 0.124667051 | 1.256104528 |
| Psmc2 | 881.0786388 | 0.243493616 | 0.08484172 | 2.869975009 |
| Slc26a5 | 5.142851575 | -0.401558732 | 0.413956187 | -0.970051287 |
| Reln | 2935.282946 | -0.438822646 | 0.127961517 | -3.429332933 |
| Gm16111 | 0 | NA | NA | NA |
| Gm10475 | 1.275826341 | 0.784279311 | 0.994510766 | 0.788608166 |
| Dnaja1-ps | 7.533972655 | 0.082971164 | 0.466892395 | 0.177709393 |
| RP23-205D3.1 | 11.79089672 | -1.089860933 | 0.390675155 | -2.789685802 |
| Gm16113 | 6.656687351 | -1.317831257 | 0.454783032 | -2.897714214 |
| Gm16112 | 0 | NA | NA | NA |
| Gm16110 | 0 | NA | NA | NA |
| Gm25459 | 0 | NA | NA | NA |
| RP23-143M9.2 | 0.121578509 | 0.75528439 | 3.724945049 | 0.202763901 |
| n-R5s170 | 0 | NA | NA | NA |
| RP24-86E4.1 | 1.834562507 | 0.522819855 | 0.93126765 | 0.561406654 |
| Orc5 | 252.3162393 | -0.012093757 | 0.098652546 | -0.12258941 |
| Gm15421 | 12.14808016 | -0.087244837 | 0.31956802 | -0.273008661 |
| Gm22887 | 3.183676125 | -0.488432023 | 0.657057248 | -0.743362963 |
| 6030443J06Rik | 71.11363277 | -0.581147137 | 0.225833167 | -2.573347145 |
| RP23-258C13.1 | 3.322405445 | -0.476182097 | 0.56343694 | -0.845138228 |
| Rpl17-ps5 | 55.4808862 | 0.578811512 | 0.16943756 | 3.416075594 |
| Gm24512 | 0 | NA | NA | NA |
| RP24-324L19.2 | 13.10249877 | 0.053585239 | 0.381616026 | 0.140416636 |
| RP24-324L19.3 | 161.9541949 | 0.074466895 | 0.116552878 | 0.638910823 |
| RP24-324L19.4 | 19.8545456 | 0.26921201 | 0.241962832 | 1.112617207 |
| AC164538.1 | 0.163066018 | 0.813382652 | 3.445567682 | 0.236066369 |
| Lhfpl3 | 375.6236598 | -0.16437548 | 0.098624177 | -1.666685444 |
| Gm21846 | 71.52268729 | -0.434017624 | 0.31667415 | -1.370549582 |
| RP23-211E12.6 | 0.049553047 | 0.274391604 | 3.724945049 | 0.073663262 |
| Gm6564 | 0 | NA | NA | NA |

| | | | | |
|---|---|---|---|---|
| RP23-240A11.2 | 0.172588697 | 0.274409367 | 2.318606165 | 0.118351004 |
| Gm28022 | 0 | NA | NA | NA |
| RP23-443N7.4 | 0.181654796 | 0.274390172 | 3.327599044 | 0.082458904 |
| RP23-443N7.5 | 0 | NA | NA | NA |
| 5031425E22Rik | 328.5648936 | -0.709827963 | 0.165916603 | -4.278221412 |
| RP23-443N7.6 | 0 | NA | NA | NA |
| RP23-443N7.10 | 6.257156534 | -0.555803527 | 0.566189614 | -0.981656167 |
| RP23-443N7.7 | 0.066050875 | 0.274391604 | 3.724945049 | 0.073663262 |
| Kmt2e | 2575.672994 | -0.140398794 | 0.078329302 | -1.792417269 |
| Gm25219 | 2.780058051 | 0.764051322 | 0.555944017 | 1.374331404 |
| Srpk2 | 2903.573116 | -0.132957715 | 0.046671396 | -2.848805182 |
| Gm24009 | 0.699061148 | -1.259317107 | 1.389195471 | -0.906508215 |
| RP23-110C17.1 | 13.01527202 | -0.336535873 | 0.287333365 | -1.171238407 |
| RP23-110C17.2 | 20.4004732 | -0.042034132 | 0.282842624 | -0.148613146 |
| Mir3096 | 0.548185132 | 0.347888563 | 1.453311203 | 0.23937651 |
| Pus7 | 222.5424342 | -0.34713638 | 0.13969281 | -2.484998195 |
| Rint1 | 243.1730144 | -0.148519947 | 0.098483224 | -1.508073566 |
| RP23-354E8.1 | 9.469473219 | -0.196193191 | 0.373438448 | -0.525369554 |
| 4933427G23Rik | 0.868720842 | 0.278797809 | 1.254308728 | 0.222272079 |
| Gm22054 | 0.593565611 | 0.334626826 | 1.209692152 | 0.276621473 |
| Tomm7 | 416.4948252 | 0.089478886 | 0.108722151 | 0.823005116 |
| 2700038G22Rik | 16.39405462 | -0.196782248 | 0.309592908 | -0.635616134 |
| Snord93 | 2.473235781 | -1.076210104 | 0.680739646 | -1.580942303 |
| Fam126a | 374.7933423 | -0.023417276 | 0.093506105 | -0.250435799 |
| RP23-170E15.2 | 3.709199715 | -0.49282891 | 0.543045158 | -0.907528413 |
| RP23-451E8.3 | 0 | NA | NA | NA |
| Klhl7 | 1038.582517 | -0.093369442 | 0.062875319 | -1.48499352 |
| RP23-451E8.2 | 13.50599551 | 0.024417516 | 0.286424836 | 0.085249296 |
| RP23-451E8.7 | 63.10412136 | -0.03638158 | 0.257374281 | -0.141356704 |
| Nupl2 | 116.6198412 | 0.106266666 | 0.132145588 | 0.804163556 |
| Gm5575 | 2.4388784 | 0.303841307 | 0.711772147 | 0.426880017 |
| RP23-451E8.8 | 0 | NA | NA | NA |
| RP23-451E8.9 | 0.083526043 | 0.75528439 | 3.724945049 | 0.202763901 |
| Gm9836 | 12.95280587 | -0.022700145 | 0.272072159 | -0.083434281 |
| Kcnh2 | 620.4579014 | -0.35450398 | 0.108964162 | -3.253399784 |
| Gm15589 | 0 | NA | NA | NA |
| Nos3 | 66.34634559 | 1.29145696 | 0.194514093 | 6.639400478 |
| Gm15587 | 5.162842253 | -2.085625209 | 0.606059643 | -3.441287066 |
| Atg9b | 48.1694312 | -0.178437742 | 0.175469412 | -1.016916512 |
| Abcb8 | 361.5327256 | -0.188149991 | 0.084756876 | -2.219878786 |
| Asic3 | 3.0483297 | 1.565552668 | 0.746458738 | 2.097306372 |
| Cdk5 | 656.490936 | -0.0810543 | 0.053946557 | -1.502492552 |
| Slc4a2 | 638.8880438 | 0.409309544 | 0.117416311 | 3.48596834 |
| Fastk | 605.3488211 | -0.039215067 | 0.061385281 | -0.638835012 |
| Gm22333 | 0 | NA | NA | NA |
| Tmub1 | 167.7441116 | 0.056764305 | 0.11155425 | 0.508849329 |
| RP24-239E2.2 | 2.781790008 | -1.324612433 | 0.631897293 | -2.096246412 |

| | | | | |
|---|---|---|---|---|
| Gm10472 | 2.635270342 | 0.412101668 | 0.673027019 | 0.612310734 |
| Agap3 | 2182.413688 | 0.013457195 | 0.064672423 | 0.208082429 |
| Gbx1 | 12.22572069 | -0.280081797 | 0.294896682 | -0.949762459 |
| Asb10 | 0.545953117 | -1.474586256 | 1.255121377 | -1.174855502 |
| 4931409K22Rik | 0.666655512 | -0.941929499 | 1.212606795 | -0.776780654 |
| Abcf2 | 862.9471066 | 0.167877129 | 0.069356738 | 2.420487661 |
| Gm10221 | 170.6058012 | -0.045138808 | 0.151327704 | -0.298285157 |
| Chpf2 | 822.9892482 | 0.389469501 | 0.081461845 | 4.781005164 |
| Smarcd3 | 501.6330187 | -0.235756244 | 0.082798839 | -2.84733757 |
| Mir671 | 0 | NA | NA | NA |
| 1700022A21Rik | 0 | NA | NA | NA |
| Nub1 | 1661.721015 | 0.97660511 | 0.082191354 | 11.88209056 |
| Gm26648 | 1.867746029 | -0.719712088 | 0.799769278 | -0.899899644 |
| Wdr86 | 10.62884738 | -0.10751806 | 0.416669392 | -0.258041655 |
| Crygn | 1.666884929 | 0.280959564 | 0.772139097 | 0.363871697 |
| Rheb | 967.3691609 | -0.067705519 | 0.103984253 | -0.651113199 |
| Gm25363 | 7.580824313 | -1.083837058 | 0.397834043 | -2.724344681 |
| Prkag2 | 719.0261586 | -0.475406247 | 0.112414 | -4.229066192 |
| Prkag2os2 | 1.088852924 | 0.708137971 | 1.017863555 | 0.695710115 |
| Prkag2os1 | 0 | NA | NA | NA |
| 1500035N22Rik | 3.98331357 | -0.498187123 | 0.474259095 | -1.050453493 |
| 2900005J15Rik | 14.08399665 | -0.324940408 | 0.318860208 | -1.019068545 |
| RP24-84G9.2 | 1.333500337 | -1.254669156 | 0.870303326 | -1.441645823 |
| Gm26051 | 0 | NA | NA | NA |
| Galntl5 | 0.110595523 | 0.755282166 | 3.724945049 | 0.202763304 |
| Gm26356 | 0 | NA | NA | NA |
| Galnt11 | 598.673327 | -0.256792252 | 0.071448561 | -3.594085722 |
| E130116L18Rik | 0 | NA | NA | NA |
| Kmt2c | 3673.91328 | -0.180862664 | 0.078993269 | -2.28959589 |
| RP23-171F16.5 | 0.112029109 | -0.687373951 | 3.724945049 | -0.184532642 |
| Hspe1-ps3 | 0.038981395 | 0.274393828 | 3.724945049 | 0.073663859 |
| Gm20432 | 5.780307253 | -0.127963311 | 0.525365669 | -0.24356999 |
| Gm18343 | 0 | NA | NA | NA |
| 4831440E17Rik | 12.8027681 | -0.701435422 | 0.300467333 | -2.33448147 |
| Gm22656 | 0.372618932 | -1.143449709 | 1.597875454 | -0.71560628 |
| Cct8l1 | 0.82877443 | -0.179139904 | 1.135145778 | -0.157812245 |
| 1700096K18Rik | 15.86316258 | 0.155326452 | 0.271175456 | 0.572789491 |
| RP23-405O11.3 | 0.04393175 | -0.206490061 | 3.724945049 | -0.055434391 |
| Gm10062 | 3.869150979 | -0.366926136 | 0.483170198 | -0.759413841 |
| RP24-121I11.1 | 0 | NA | NA | NA |
| RP24-121I11.4 | 17.85167491 | -0.748646124 | 0.270815381 | -2.764415087 |
| Xrcc2 | 97.35748713 | -0.446568696 | 0.145577168 | -3.067573728 |
| RP24-121I11.5 | 0.245367999 | -0.206481886 | 1.856229481 | -0.111237262 |
| Actr3b | 678.1794346 | 0.06778042 | 0.073270639 | 0.925069316 |
| RP23-56M18.4 | 0 | NA | NA | NA |
| RP23-56M18.5 | 0.232282403 | 1.713621339 | 2.537884778 | 0.675216367 |
| Gm21671 | 0.275686258 | 0.253313562 | 1.811319311 | 0.139850307 |

| | | | | |
|---|---|---|---|---|
| RP23-56M18.8 | 24.28531059 | -0.441616963 | 0.326489075 | -1.352624015 |
| RP23-56M18.6 | 0.144833758 | -0.206488293 | 2.88634391 | -0.07153974 |
| Gm21698 | 0 | NA | NA | NA |
| Gm1979 | 0.35614329 | 1.148814528 | 1.724612174 | 0.6661292 |
| Gm5067 | 39.82603458 | -0.826526158 | 0.24149858 | -3.42248868 |
| Gm5862 | 0 | NA | NA | NA |
| Speer4a | 0.739238851 | 2.000774609 | 1.248138553 | 1.603006816 |
| Gm7347 | 0.182828849 | -0.206486331 | 2.266107739 | -0.09111938 |
| RP24-162F23.9 | 8.999765542 | -1.224440424 | 0.408185082 | -2.999718704 |
| Gm10471 | 0.138573873 | 0.75528185 | 3.628901543 | 0.208129606 |
| 5031410I06Rik | 0 | NA | NA | NA |
| Gm10220 | 0.108266322 | 0.274389235 | 3.724945049 | 0.073662626 |
| Gm7361 | 6.292791096 | -0.200764041 | 0.380464931 | -0.527680805 |
| RP24-362M8.2 | 0.097139547 | 0.274393828 | 3.724945049 | 0.073663859 |
| 4930584F24Rik | 0 | NA | NA | NA |
| RP24-165O21.1 | 0 | NA | NA | NA |
| Dpp6 | 3300.38036 | 0.009912465 | 0.063133831 | 0.157007189 |
| RP23-474A22.1 | 0 | NA | NA | NA |
| Gm16056 | 0 | NA | NA | NA |
| Gm16057 | 0 | NA | NA | NA |
| RP23-478E6.1 | 84.06891195 | -0.631172497 | 0.308224532 | -2.047768529 |
| RP23-478E6.3 | 79.08630011 | -0.452586932 | 0.305021019 | -1.483789327 |
| Speer4b | 0.378318939 | 0.732159874 | 1.698155759 | 0.431150011 |
| Gm16058 | 0.092228164 | 0.75528439 | 3.724945049 | 0.202763901 |
| Paxip1 | 503.8218817 | 0.069486164 | 0.079082642 | 0.878652537 |
| RP23-397C5.2 | 0.110870145 | -0.206490066 | 3.724945049 | -0.055434393 |
| RP23-397C5.1 | 0 | NA | NA | NA |
| Htr5a | 258.8451686 | 0.079924124 | 0.1228396 | 0.650638094 |
| RP23-397C5.5 | 4.181737927 | -0.035015657 | 0.522946064 | -0.066958448 |
| RP23-397C5.4 | 1.516892051 | -0.161186361 | 0.873279339 | -0.184575947 |
| RP23-256A5.2 | 3.457092485 | -0.648773892 | 0.607580891 | -1.067798382 |
| Gm26608 | 2.045375739 | 0.28684158 | 0.6378088 | 0.44972973 |
| RP23-256A5.4 | 1.504629277 | -1.146388462 | 0.859163325 | -1.334307959 |
| Insig1 | 653.7555455 | 0.204642462 | 0.138315585 | 1.479532927 |
| En2 | 252.7745475 | -0.56109422 | 0.282357853 | -1.987174123 |
| RP23-358F24.3 | 0.075043205 | 0.755286614 | 3.724945049 | 0.202764498 |
| Cnpy1 | 429.1835849 | -0.775344831 | 0.295781776 | -2.621340779 |
| Rbm33 | 1626.312609 | 0.015260344 | 0.12508972 | 0.121995187 |
| Shh | 57.43406997 | 0.322385971 | 0.158851537 | 2.029479708 |
| 9530036O11Rik | 6.185525992 | -0.113680198 | 0.409439316 | -0.277648465 |
| Gm26894 | 16.72981752 | -0.534906588 | 0.25651604 | -2.085275402 |
| RP23-94F17.1 | 1.563650339 | -0.400144668 | 0.814408813 | -0.491331456 |
| RP23-94F17.2 | 0.559314221 | -0.942168525 | 1.479590001 | -0.636776759 |
| Rnf32 | 99.31368546 | -0.513365829 | 0.152685584 | -3.36224165 |
| Gm7420 | 6.011270768 | -0.782470295 | 0.60047396 | -1.303087806 |
| Lmbr1 | 450.4254583 | -0.298588917 | 0.077628534 | -3.846381004 |
| RP23-396H16.2 | 45.39463601 | -0.51291264 | 0.249839632 | -2.052967484 |

| | | | | |
|---|---|---|---|---|
| Gm23801 | 1.101786575 | -2.200937694 | 1.051941929 | -2.092261591 |
| 4632411P08Rik | 8.069396325 | -0.049434391 | 0.367050425 | -0.134680109 |
| Nom1 | 466.9739484 | 0.255851937 | 0.074078907 | 3.453775813 |
| Mnx1 | 0.180050447 | -0.206493189 | 3.327115185 | -0.062063733 |
| Gm24538 | 0 | NA | NA | NA |
| Gm6083 | 0 | NA | NA | NA |
| Ube3c | 1443.177322 | -0.040463044 | 0.046958029 | -0.861685323 |
| RP24-224N3.5 | 0.401137426 | -0.099114552 | 1.483114185 | -0.066828672 |
| Gm26346 | 0.288530209 | 1.105518389 | 1.732201355 | 0.638215866 |
| RP24-224N3.4 | 0 | NA | NA | NA |
| Gm5129 | 0.619575296 | -1.633108753 | 1.181603581 | -1.382112224 |
| Dnajb6 | 825.5467343 | 0.328049383 | 0.05721344 | 5.733781811 |
| RP24-224N3.8 | 0.381425259 | 0.762927648 | 1.587471198 | 0.480593064 |
| RP24-224N3.9 | 0 | NA | NA | NA |
| RP24-176O2.1 | 0 | NA | NA | NA |
| RP24-176O2.2 | 0.121388914 | 0.755282166 | 3.724945049 | 0.202763304 |
| RP24-176O2.4 | 0.21891683 | 0.755297267 | 2.560922434 | 0.294931723 |
| RP24-418J2.4 | 0.04393175 | -0.206490061 | 3.724945049 | -0.055434391 |
| Il6 | 43.66316143 | 8.23358321 | 1.157673218 | 7.112182505 |
| Tyms | 45.1820026 | 0.066772905 | 0.205649144 | 0.32469333 |
| RP24-418J2.5 | 0.550684032 | 1.04183166 | 1.23038856 | 0.846750119 |
| 3110082J24Rik | 7.185718025 | -1.321651327 | 0.440446164 | -3.000710266 |
| Gareml | 196.41537 | -0.921054458 | 0.156090876 | -5.900757831 |
| Hadha | 925.8447827 | 0.058717154 | 0.088220206 | 0.665574896 |
| Hadhb | 496.8674058 | -0.270452018 | 0.08458743 | -3.197307421 |
| Mir1960 | 2.414716579 | -1.149805634 | 0.8326927 | -1.380828287 |
| Adgrf3 | 4.779796961 | 1.975536763 | 0.538321357 | 3.669809374 |
| Ept1 | 1152.189136 | -0.376317563 | 0.050373939 | -7.470481103 |
| Gm16288 | 0.621982117 | -1.061842909 | 1.453553317 | -0.73051528 |
| Drc1 | 103.3566621 | -0.551025957 | 0.18994325 | -2.901003098 |
| WI1-2080J1.2 | 1.385170032 | -2.211322761 | 0.95573025 | -2.313751983 |
| Otof | 183.3860885 | -0.35356913 | 0.204368316 | -1.730058434 |
| 1700001C02Rik | 2.869726659 | -0.508684206 | 0.595029358 | -0.854889257 |
| Cib4 | 0.038981395 | 0.274393828 | 3.724945049 | 0.073663859 |
| Gm25666 | 0 | NA | NA | NA |
| RP23-446J15.1 | 0.136419714 | 0.274406094 | 2.946414804 | 0.093132201 |
| Gm9899 | 195.7601427 | -0.399250237 | 0.175963913 | -2.268932471 |
| Kcnk3 | 462.3649798 | -0.356464883 | 0.120025987 | -2.96989754 |
| Slc35f6 | 587.4667661 | 0.605885998 | 0.097184931 | 6.234361547 |
| Mir5625 | 0 | NA | NA | NA |
| Cenpa | 31.10520093 | 0.885311999 | 0.264703582 | 3.344541057 |
| Dpysl5 | 464.621931 | -0.440052751 | 0.091270145 | -4.821431501 |
| RP23-342N8.1 | 3.407592442 | -1.208363253 | 0.509300538 | -2.372593713 |
| Mapre3 | 1596.309188 | -0.036083728 | 0.064637025 | -0.558251682 |
| RP23-342N8.2 | 49.78286253 | -0.821027144 | 0.327020195 | -2.510631322 |
| Gm15461 | 0 | NA | NA | NA |
| Tmem214 | 593.0310336 | -0.004334598 | 0.061994316 | -0.069919282 |

| | | | | |
|---|---|---|---|---|
| RP23-113M20.5 | 2.578624624 | 1.011812457 | 0.783904698 | 1.290734014 |
| Agbl5 | 130.4690619 | -0.299598394 | 0.142029071 | -2.109415997 |
| RP23-113M20.4 | 16.2135406 | -1.061192978 | 0.277899729 | -3.818618252 |
| Ost4 | 231.3485647 | -0.00861883 | 0.149290355 | -0.057731992 |
| Emilin1 | 69.27241042 | 1.011296159 | 0.197417054 | 5.122638277 |
| Khk | 59.85891304 | -0.057070454 | 0.163542909 | -0.348963184 |
| Cgref1 | 143.7238615 | -0.177360324 | 0.19516538 | -0.908769394 |
| Abhd1 | 97.11033058 | -0.816056996 | 0.138403922 | -5.896198506 |
| Preb | 614.623067 | -0.114162266 | 0.066076176 | -1.727737181 |
| Tcf23 | 3.89979361 | 1.251229748 | 0.569202404 | 2.198215851 |
| Slc5a6 | 215.3588507 | -0.405455883 | 0.090081081 | -4.501010391 |
| Atraid | 300.310359 | 0.085908569 | 0.108437543 | 0.792240094 |
| Cad | 344.3964736 | -0.216712647 | 0.079371431 | -2.730360846 |
| Slc30a3 | 365.9223408 | -0.308323494 | 0.147313062 | -2.092981372 |
| Gm9924 | 4.140852306 | -0.23337992 | 0.469720808 | -0.496848162 |
| Dnajc5g | 0.288733551 | -0.20648635 | 2.054349134 | -0.10051181 |
| Trim54 | 0.834511584 | -0.757260652 | 0.997427876 | -0.759213443 |
| Gm15469 | 0 | NA | NA | NA |
| Ucn | 1.659798835 | -0.682010749 | 1.17191532 | -0.581962482 |
| Mpv17 | 189.2554659 | -0.211361613 | 0.103544806 | -2.041257513 |
| Gm29609 | 14.60460443 | -0.624880283 | 0.29970983 | -2.084950912 |
| Gtf3c2 | 322.248693 | -0.055504706 | 0.098308592 | -0.564596696 |
| Eif2b4 | 322.6252131 | 0.121664011 | 0.079589901 | 1.52863628 |
| Snx17 | 719.475255 | 0.21016148 | 0.072381449 | 2.903526848 |
| Zfp513 | 310.9180716 | 0.570183207 | 0.098633994 | 5.780798121 |
| Ppm1g | 655.6864786 | 0.117775155 | 0.077845051 | 1.512943399 |
| RP24-86I23.4 | 34.84001459 | 0.153742042 | 0.361936339 | 0.424776475 |
| Gm9970 | 1.14535829 | -0.185001296 | 1.014398999 | -0.182375274 |
| Nrbp1 | 774.3138472 | 0.255445849 | 0.062239899 | 4.10421377 |
| Krtcap3 | 6.967827923 | -0.858039495 | 0.439792041 | -1.951011878 |
| Ift172 | 621.7871443 | -0.477437998 | 0.082457369 | -5.790119262 |
| Fndc4 | 773.732385 | -0.086271947 | 0.088248596 | -0.977601363 |
| Gckr | 6.742714648 | -0.091263381 | 0.375003687 | -0.243366625 |
| RP23-383E21.3 | 0 | NA | NA | NA |
| RP23-383E21.5 | 0 | NA | NA | NA |
| RP23-383E21.4 | 0.38059087 | 0.785746584 | 1.73721072 | 0.452303554 |
| Zfp512 | 587.0033922 | -0.225759628 | 0.065135924 | -3.465977189 |
| RP23-383E21.1 | 18.84380616 | -1.009391938 | 0.323429389 | -3.120903576 |
| 4930548H24Rik | 1.368337452 | 0.577471518 | 0.972658958 | 0.593704004 |
| 4930566F21Rik | 0.183097766 | -0.687376827 | 2.686642813 | -0.255849726 |
| Gpn1 | 250.7486316 | 0.041191414 | 0.077967353 | 0.528316177 |
| Supt7l | 184.2270805 | 0.144472198 | 0.098585099 | 1.465456741 |
| Slc4a1ap | 405.404987 | -0.132865867 | 0.070008185 | -1.897861908 |
| RP24-80F7.3 | 2.627266017 | 0.580494162 | 1.019188755 | 0.569564921 |
| RP24-80F7.5 | 6.412813486 | -0.249930686 | 0.571358264 | -0.437432521 |
| RP24-80F7.6 | 3.22349573 | 1.754751765 | 0.611325836 | 2.870403413 |
| Mrpl33 | 188.1679828 | -0.011173107 | 0.119208592 | -0.093727363 |

| | | | | |
|---|---|---|---|---|
| RP24-80F7.7 | 0 | NA | NA | NA |
| Rbks | 30.53545577 | 0.443082095 | 0.220791933 | 2.006785705 |
| Mir3473e | 1.131450813 | -0.363609185 | 0.984659475 | -0.369274043 |
| Bre | 293.3718634 | 0.134954879 | 0.07017047 | 1.923243204 |
| RP24-80F7.8 | 3.022895243 | -0.158688294 | 0.672260468 | -0.2360518 |
| Gm24436 | 2.112795873 | -0.089730077 | 0.81937235 | -0.109510746 |
| RP23-181I1.4 | 6.442610278 | -0.114529365 | 0.391942376 | -0.292209702 |
| Gm17130 | 0.230862969 | 0.274410817 | 2.134669009 | 0.128549586 |
| Gm10463 | 2.064454091 | 0.349286998 | 0.764790767 | 0.456709224 |
| Fosl2 | 1087.482308 | 0.247195558 | 0.310262783 | 0.796729649 |
| Gm26335 | 0 | NA | NA | NA |
| RP23-415E17.3 | 0 | NA | NA | NA |
| Plb1 | 16.55472255 | 0.599899488 | 0.28985547 | 2.069650395 |
| Gm15615 | 0 | NA | NA | NA |
| RP23-262J12.5 | 0.060174537 | 0.755286614 | 3.724945049 | 0.202764498 |
| RP23-262J12.4 | 0 | NA | NA | NA |
| Gm5864 | 1.456934103 | 0.505336517 | 0.835795221 | 0.604617619 |
| Ppp1cb | 4028.577039 | 0.065723788 | 0.08744385 | 0.751611325 |
| RP23-262J12.3 | 12.26687181 | -0.256187373 | 0.309217804 | -0.828501366 |
| RP23-262J12.2 | 50.45321847 | -0.152587919 | 0.279402391 | -0.546122455 |
| Gm15613 | 0 | NA | NA | NA |
| Gm15614 | 0.985784616 | 1.823293197 | 1.157888383 | 1.574670948 |
| RP23-149A1.7 | 0.075043205 | 0.755286614 | 3.724945049 | 0.202764498 |
| RP23-149A1.5 | 0.156588133 | -0.206490485 | 2.822767146 | -0.073151796 |
| RP23-149A1.6 | 21.7617041 | -0.129674858 | 0.243938768 | -0.531587739 |
| Yes1 | 334.4835077 | -0.292254836 | 0.094822863 | -3.082113622 |
| RP23-149A1.2 | 1.316494506 | 0.46577723 | 0.908044073 | 0.51294562 |
| RP23-353M5.2 | 7.746880428 | -0.825724337 | 0.427779746 | -1.930255805 |
| Gm2420 | 4.384803307 | -1.168389007 | 0.515149592 | -2.268057715 |
| Pisd | 289.4875556 | 0.168546709 | 0.077629196 | 2.17117679 |
| Gm20671 | 0 | NA | NA | NA |
| RP23-353M5.1 | 0 | NA | NA | NA |
| Prr14l | 1555.512628 | -0.292608498 | 0.054910381 | -5.328837471 |
| RP23-118L1.4 | 11.68546513 | -0.600785601 | 0.375788648 | -1.598732704 |
| RP23-118L1.5 | 0 | NA | NA | NA |
| Depdc5 | 821.0906113 | -0.214847377 | 0.074841008 | -2.87071732 |
| Gm24003 | 1.689516722 | -1.681995322 | 0.857571254 | -1.961347602 |
| RP23-69J7.1 | 15.99339509 | -0.497895367 | 0.385976159 | -1.289964043 |
| Ywhah | 4942.475359 | -0.063554182 | 0.074669992 | -0.851134178 |
| RP23-69J7.4 | 0 | NA | NA | NA |
| Rpl35a-ps5 | 4.14321256 | 0.958543114 | 0.509154412 | 1.882617711 |
| Slc5a1 | 30.58515113 | 0.779934124 | 0.309247366 | 2.522039666 |
| Gm15936 | 0.038981395 | 0.274393828 | 3.724945049 | 0.073663859 |
| 5033423K11Rik | 0.310320078 | 1.245181267 | 1.967869152 | 0.632756129 |
| Spon2 | 4.327007151 | 3.061544031 | 0.83460249 | 3.668266113 |
| Ctbp1 | 1818.873768 | -0.00580413 | 0.074204218 | -0.078218329 |
| RP24-406N3.3 | 0 | NA | NA | NA |

| | | | | |
|---|---|---|---|---|
| Gm20465 | 0 | NA | NA | NA |
| Maea | 757.2802516 | 0.103680234 | 0.077320453 | 1.340916023 |
| Uvssa | 220.4651291 | 0.388550481 | 0.129949545 | 2.99001032 |
| Nkx1-1 | 0.498903533 | -1.547701026 | 1.566753802 | -0.98783933 |
| RP24-94L5.3 | 0.038981395 | 0.274393828 | 3.724945049 | 0.073663859 |
| Fam53a | 291.3243653 | 0.286654265 | 0.07829606 | 3.661158244 |
| Gm9903 | 6.764630232 | -0.333692291 | 0.41529436 | -0.803507879 |
| Slbp | 223.0429402 | 0.276271257 | 0.095605887 | 2.889688748 |
| RP23-391P21.2 | 0.046345709 | 0.274391604 | 3.724945049 | 0.073663262 |
| Tmem129 | 242.2710187 | -0.332127272 | 0.099333796 | -3.343547564 |
| Tacc3 | 66.98387769 | 0.772555848 | 0.211161705 | 3.658598271 |
| RP23-391P21.5 | 0 | NA | NA | NA |
| RP23-391P21.4 | 17.89361774 | -0.045008316 | 0.255900172 | -0.175882319 |
| Fgfr3 | 1284.08441 | -0.152369457 | 0.134493439 | -1.13291368 |
| Letm1 | 945.73928 | -0.04336741 | 0.049020711 | -0.884675262 |
| Gm10059 | 2.485764843 | 0.165052081 | 0.70279386 | 0.234851342 |
| RP23-391P21.3 | 0 | NA | NA | NA |
| Whsc1 | 2442.311656 | -0.144825795 | 0.045296838 | -3.197260606 |
| Nelfa | 291.9774525 | 0.177655112 | 0.090131115 | 1.971074167 |
| Mir7024 | 0 | NA | NA | NA |
| RP24-294K15.5 | 2.887331315 | 0.613549462 | 0.599839269 | 1.022856443 |
| RP24-294K15.4 | 13.77895346 | 1.360851111 | 0.359978894 | 3.780363608 |
| Gm1673 | 134.0090932 | -0.256955787 | 0.165576046 | -1.551889861 |
| RP24-294K15.7 | 56.16396208 | 0.019830919 | 0.154645025 | 0.128235093 |
| Nat8l | 3474.475194 | -0.427568555 | 0.069984597 | -6.109466588 |
| Poln | 11.42523575 | -0.600361296 | 0.298189514 | -2.013354823 |
| RP23-60E7.2 | 0.517722191 | 1.820076085 | 1.301366011 | 1.39858892 |
| RP23-60E7.3 | 0.042716945 | -0.206490061 | 3.724945049 | -0.055434391 |
| RP23-60E7.1 | 0.122601409 | 1.236179401 | 3.724945049 | 0.331865138 |
| Haus3 | 129.8840804 | 0.026922075 | 0.109429694 | 0.246021663 |
| Gm25918 | 0 | NA | NA | NA |
| Mxd4 | 934.5782134 | -0.191430916 | 0.116269018 | -1.646448211 |
| RP24-230P10.6 | 0.165470308 | 1.646276139 | 2.753318586 | 0.597924318 |
| Zfyve28 | 544.0876334 | -0.516408406 | 0.128611106 | -4.015270701 |
| Gm15513 | 73.95229317 | -0.655735057 | 0.303969618 | -2.157238811 |
| RP23-201D22.6 | 0 | NA | NA | NA |
| RP23-201D22.4 | 0.535802326 | 1.635610565 | 1.407387414 | 1.162160858 |
| RP23-201D22.5 | 0.419184927 | 1.401106156 | 3.327918146 | 0.421015811 |
| Rnf4 | 1197.186287 | 0.318751913 | 0.08353469 | 3.815802918 |
| Fam193a | 1067.198729 | -0.085297995 | 0.069742669 | -1.223038878 |
| RP23-429E19.4 | 44.14393845 | -0.707768557 | 0.283798322 | -2.493913822 |
| Gm26931 | 0.110870145 | -0.206490066 | 3.724945049 | -0.055434393 |
| RP23-429E19.5 | 37.84079439 | -0.716428082 | 0.245522189 | -2.917976924 |
| Tnip2 | 144.594261 | 0.712524148 | 0.164983352 | 4.318763926 |
| Sh3bp2 | 240.9989662 | 2.161143865 | 0.178507926 | 12.10671094 |
| Gm22847 | 1.623325927 | 1.1463131 | 1.044083893 | 1.097912829 |
| Add1 | 3389.415062 | -0.176232698 | 0.051201166 | -3.441966481 |

| | | | | |
|---|---|---|---|---|
| Mir7036b | 0 | NA | NA | NA |
| RP24-229D18.2 | 13.74406399 | 0.079675982 | 0.443570325 | 0.179624238 |
| Mfsd10 | 128.2944897 | 0.207390819 | 0.155962941 | 1.32974422 |
| Nop14 | 276.7616417 | 0.578455427 | 0.1411545 | 4.098030369 |
| Gm15522 | 0.123992736 | 0.274402763 | 3.720643035 | 0.073751435 |
| Grk4 | 52.31774275 | -0.556272393 | 0.193877913 | -2.869189091 |
| RP23-44L13.1 | 0.042716945 | -0.206490061 | 3.724945049 | -0.055434391 |
| Mir467g | 0 | NA | NA | NA |
| Htt | 3899.683035 | -0.026070244 | 0.065682078 | -0.396915642 |
| Msantd1 | 17.05411229 | 0.673169067 | 0.258132541 | 2.607842721 |
| Rgs12 | 414.5610989 | 0.018100373 | 0.092186396 | 0.196345376 |
| Mir3097 | 0.476896902 | 0.584307893 | 1.438227252 | 0.406269518 |
| RP23-352G19.2 | 0.30060319 | 1.184382532 | 1.60253836 | 0.739066572 |
| Hgfac | 2.316699699 | -0.213657184 | 0.659575392 | -0.323931406 |
| Dok7 | 162.3460107 | 0.193832866 | 0.136773733 | 1.41717903 |
| Lrpap1 | 1373.379439 | 0.286743982 | 0.101762341 | 2.817780924 |
| Adra2c | 169.5646234 | -0.18782783 | 0.116230349 | -1.615996435 |
| RP24-276N7.5 | 0.097139547 | 0.274393828 | 3.724945049 | 0.073663859 |
| E130018O15Rik | 0 | NA | NA | NA |
| Hmx1 | 0.038981395 | 0.274393828 | 3.724945049 | 0.073663859 |
| 4930442P19Rik | 0 | NA | NA | NA |
| RP24-276N7.7 | 0 | NA | NA | NA |
| Cpz | 3.532213938 | -1.202959851 | 0.709777025 | -1.694841912 |
| RP23-26P10.1 | 28.70925171 | -0.209871899 | 0.317203781 | -0.661631141 |
| Plk-ps1 | 0 | NA | NA | NA |
| RP23-26P10.2 | 15.37615298 | -0.458912539 | 0.321303477 | -1.428283763 |
| Trmt44 | 83.81309991 | -0.209014636 | 0.16606723 | -1.258614568 |
| 4931431C16Rik | 1.224874833 | 1.314313982 | 1.024941772 | 1.28233039 |
| Acox3 | 452.6412082 | 0.100290705 | 0.083156812 | 1.206043174 |
| Htra3 | 58.84703746 | 0.343188765 | 0.210560855 | 1.629879231 |
| Sh3tc1 | 32.64252681 | 1.869800933 | 0.249392531 | 7.497421527 |
| Gm15650 | 0.369202626 | 0.274414581 | 1.76321871 | 0.155632753 |
| Ablim2 | 1153.537665 | -0.155022713 | 0.129969599 | -1.192761344 |
| Afap1 | 482.3908662 | -0.138193937 | 0.069337619 | -1.993058588 |
| H2af-ps | 0.473803596 | -0.08440464 | 1.421344518 | -0.059383661 |
| Sorcs2 | 714.7892678 | -0.209816414 | 0.077501308 | -2.707262896 |
| Psapl1 | 0.670061835 | 0.059424657 | 1.275757829 | 0.046579888 |
| Mir7689 | 0.235888945 | -0.687380079 | 2.17290611 | -0.316341362 |
| 2210406O10Rik | 10.33515927 | -0.349790697 | 0.322374706 | -1.085043866 |
| Grpel1 | 361.9003875 | 0.337391728 | 0.113693662 | 2.967550897 |
| Tada2b | 496.4167252 | 0.345662595 | 0.107150549 | 3.225952626 |
| Ccdc96 | 19.96485888 | 0.091956115 | 0.266963633 | 0.34445184 |
| Tbc1d14 | 875.8556229 | 0.056985592 | 0.053044172 | 1.074304494 |
| RP23-463H10.2 | 0 | NA | NA | NA |
| D5Ertd579e | 1861.527523 | -0.175351068 | 0.055532631 | -3.157622187 |
| RP23-463H10.1 | 0 | NA | NA | NA |
| Gm24878 | 6.521421862 | 0.483162697 | 0.449673693 | 1.074474012 |

| | | | | |
|---|---|---|---|---|
| Gm25767 | 0 | NA | NA | NA |
| Bloc1s4 | 95.95579344 | 0.055483834 | 0.161695515 | 0.343137746 |
| RP23-382O15.2 | 0.348252234 | -0.660419358 | 1.484675969 | -0.444823903 |
| Mrfap1 | 2250.019853 | 0.115346785 | 0.077245304 | 1.493253051 |
| Man2b2 | 514.8886766 | 0.417793394 | 0.109168855 | 3.827038335 |
| Ppp2r2c | 4810.614684 | -0.214918434 | 0.099398004 | -2.162200697 |
| Ppp2r2cos | 0.150455055 | 0.274400633 | 2.848105408 | 0.096344971 |
| RP23-16N23.2 | 5.207155389 | -1.127014555 | 0.516524566 | -2.181918595 |
| Wfs1 | 1615.855355 | 0.314692351 | 0.134563745 | 2.338611717 |
| Jakmip1 | 619.5416328 | -0.328639402 | 0.186511807 | -1.762030015 |
| Gm26647 | 0.176363682 | -0.206487563 | 2.731427642 | -0.075596937 |
| Gm25551 | 0 | NA | NA | NA |
| RP23-106D12.1 | 30.55419668 | -0.708865223 | 0.284419831 | -2.492319962 |
| Gm1043 | 54.30076238 | -0.321751783 | 0.225225869 | -1.428573833 |
| Crmp1 | 1273.345192 | -0.311247844 | 0.073308118 | -4.245748649 |
| Evc | 77.51767331 | -0.38292026 | 0.204207015 | -1.875157227 |
| RP23-339H11.3 | 0 | NA | NA | NA |
| Evc2 | 92.04870115 | -0.225603882 | 0.135322409 | -1.667158333 |
| Gm15732 | 0 | NA | NA | NA |
| Stk32b | 51.08340483 | -0.390086905 | 0.181540466 | -2.148760072 |
| Cytl1 | 1.87157638 | -1.022930006 | 0.716915071 | -1.426849633 |
| RP24-199D21.1 | 0 | NA | NA | NA |
| Msx1 | 35.29830261 | -0.417675507 | 0.261380563 | -1.597959319 |
| RP24-292O13.1 | 0.159287055 | 0.755303748 | 2.807608812 | 0.269020294 |
| RP23-453O15.1 | 0.961470315 | -1.599465657 | 1.105234793 | -1.447172734 |
| RP23-453O15.4 | 0.066050875 | 0.274391604 | 3.724945049 | 0.073663262 |
| Stx18 | 243.7354772 | 0.120161274 | 0.085745146 | 1.401376984 |
| RP24-535K22.2 | 24.02896761 | -0.452340586 | 0.228867691 | -1.976428322 |
| Nsg1 | 2866.052638 | -0.130455411 | 0.085307273 | -1.529241362 |
| Zbtb49 | 139.0035971 | -0.036959168 | 0.135398124 | -0.272966618 |
| Lyar | 86.46303146 | 0.359789574 | 0.218112864 | 1.649556879 |
| Tmem128 | 186.1267044 | 0.267121197 | 0.128857035 | 2.073004376 |
| Otop1 | 1.049274302 | 0.224776405 | 0.969711861 | 0.231797108 |
| Gm26560 | 1.795106397 | 0.289627229 | 0.932181562 | 0.310698302 |
| Drd5 | 35.97084852 | 0.110068622 | 0.195382053 | 0.563350732 |
| RP23-426E16.4 | 0.045882455 | 0.755288839 | 3.724945049 | 0.202765096 |
| Slc2a9 | 46.88057285 | 2.373084159 | 0.254750513 | 9.315326327 |
| Gm15796 | 0.265425963 | 0.274410648 | 2.093237601 | 0.131093884 |
| Wdr1 | 2375.805282 | 0.445049967 | 0.104101832 | 4.275140608 |
| Gm22249 | 0 | NA | NA | NA |
| 4930421P07Rik | 0 | NA | NA | NA |
| Zfp518b | 296.1034572 | -0.052292443 | 0.076750687 | -0.681328669 |
| Gm22779 | 0 | NA | NA | NA |
| Clnk | 4.40624827 | 1.6926361 | 0.691268437 | 2.448594509 |
| Clnkos | 0.326535978 | -1.61380852 | 1.757043134 | -0.91847974 |
| RP24-258B22.1 | 0.156480458 | -1.154805639 | 3.480826907 | -0.331761869 |
| RP24-263A3.3 | 0 | NA | NA | NA |

| | | | | |
|---|---|---|---|---|
| RP24-263A3.2 | 0.133021046 | 0.671922749 | 3.654933084 | 0.183839959 |
| RP24-263A3.1 | 0 | NA | NA | NA |
| Gm25505 | 0.048949532 | -0.206487837 | 3.724945049 | -0.055433794 |
| RP23-477D20.1 | 0.049553047 | 0.274391604 | 3.724945049 | 0.073663262 |
| RP23-477D20.4 | 0.771325962 | 0.933593266 | 1.11531761 | 0.837064938 |
| RP23-477D20.3 | 0.202816535 | 1.265180537 | 3.204096853 | 0.394863387 |
| RP23-477D20.2 | 2.015324298 | -0.601115858 | 0.778731993 | -0.771916222 |
| Gm25377 | 0 | NA | NA | NA |
| RP23-2C16.2 | 158.46767 | 0.01212767 | 0.117645018 | 0.10308698 |
| Hs3st1 | 277.9990456 | -0.128828891 | 0.102489775 | -1.256992619 |
| Gm7816 | 17.85736311 | 0.081389832 | 0.269921126 | 0.301531906 |
| RP23-318O24.1 | 0.038981395 | 0.274393828 | 3.724945049 | 0.073663859 |
| Gm2810 | 23.07760637 | 0.06197995 | 0.251500065 | 0.246441089 |
| Gm23022 | 0 | NA | NA | NA |
| RP23-477O15.2 | 0 | NA | NA | NA |
| RP23-477O15.1 | 3.151818414 | 0.495918746 | 0.852938438 | 0.581423844 |
| RP24-377B20.1 | 0 | NA | NA | NA |
| Rab28 | 662.797046 | 0.037231318 | 0.056986993 | 0.653330108 |
| Nkx3-2 | 0.153659334 | 0.274405733 | 2.839457797 | 0.096640187 |
| Bod1l | 1615.891892 | 0.035389757 | 0.11275907 | 0.313852863 |
| RP24-188B2.2 | 1.139214769 | 1.315393618 | 0.964348975 | 1.364022415 |
| Mir6414 | 0 | NA | NA | NA |
| RP24-188B2.1 | 0.100902008 | 0.274391828 | 3.724945049 | 0.073663322 |
| Gm16223 | 2.960191199 | -0.499601453 | 0.612654596 | -0.815470015 |
| RP23-146B9.1 | 0.099155932 | 0.75528439 | 3.724945049 | 0.202763901 |
| RP24-167K2.2 | 0.184373794 | -0.673752294 | 2.724223369 | -0.247319035 |
| RP24-560A18.4 | 0.659798846 | -1.203096789 | 1.167139912 | -1.030807683 |
| RP24-560A18.1 | 1.041030589 | -1.633020531 | 0.970869413 | -1.682018725 |
| RP24-560A18.3 | 0.172203646 | -0.206481459 | 2.739420422 | -0.07537414 |
| RP24-560A18.2 | 14.93512307 | -1.280572191 | 0.42062776 | -3.044430996 |
| Gm7854 | 28.25369218 | -0.546532054 | 0.267934071 | -2.03980051 |
| Cpeb2 | 1599.741892 | -0.175888148 | 0.064512893 | -2.726403032 |
| RP24-230L5.2 | 66.96472975 | -0.219161665 | 0.215093901 | -1.018911574 |
| RP24-230L5.3 | 79.78188433 | -0.416834125 | 0.207686931 | -2.007031081 |
| RP24-230L5.4 | 0.343955217 | 0.289950119 | 1.753967529 | 0.165310996 |
| RP24-230L5.5 | 0.150451996 | 0.274405555 | 2.858243839 | 0.096004949 |
| RP23-15G1.1 | 0.062426873 | 0.755288839 | 3.724945049 | 0.202765096 |
| C1qtnf7 | 8.52901898 | -0.32001932 | 0.335737174 | -0.953184052 |
| RP23-15G1.2 | 2.619111478 | -0.435762681 | 0.567230437 | -0.768228665 |
| Gm15866 | 0.088534442 | 0.274389469 | 3.724945049 | 0.073662689 |
| RP23-15G1.3 | 0.396976673 | 0.525528726 | 1.527541516 | 0.344035642 |
| Cc2d2a | 374.1500514 | -0.142149335 | 0.120508618 | -1.179578172 |
| Fbxl5 | 1545.557561 | -0.154655304 | 0.07193777 | -2.149848442 |
| RP23-336O18.1 | 33.11233713 | 0.634155262 | 0.311935857 | 2.032966866 |
| RP23-336O18.2 | 3.500141698 | 0.047807234 | 0.505839075 | 0.094510758 |
| RP23-336O18.3 | 11.15943119 | 6.014371801 | 0.992820121 | 6.057866549 |
| RP23-336O18.4 | 0.062426873 | 0.755288839 | 3.724945049 | 0.202765096 |

| | | | | |
|---|---|---|---|---|
| Bst1 | 59.92677726 | 3.708191531 | 0.507038929 | 7.313425693 |
| Cd38 | 164.7434097 | 0.871177335 | 0.212451435 | 4.10059521 |
| Gm16014 | 2.851290958 | 0.884830928 | 0.778856745 | 1.136063768 |
| Gm16015 | 0 | NA | NA | NA |
| Gm7879 | 31.12801235 | 0.179476477 | 0.213536734 | 0.840494623 |
| Gm7882 | 0 | NA | NA | NA |
| Gm5292 | 0 | NA | NA | NA |
| RP23-58C20.3 | 0 | NA | NA | NA |
| RP23-58C20.4 | 0 | NA | NA | NA |
| Fgfbp1 | 20.48618477 | 1.00477339 | 0.284611943 | 3.53032757 |
| Prom1 | 116.7320341 | 0.225044109 | 0.196552027 | 1.144959493 |
| RP24-230J14.2 | 3.662822882 | 0.633936056 | 0.568137775 | 1.115813953 |
| Gm16401 | 1.371463738 | 1.07764896 | 0.805238491 | 1.338297873 |
| RP24-230J14.1 | 0.385783664 | 2.016721824 | 1.747280391 | 1.154206179 |
| RP24-230J14.3 | 0.273584676 | 0.755310238 | 1.806933275 | 0.418006712 |
| RP24-230J14.4 | 0.127971488 | 0.274389832 | 3.687653711 | 0.0744077 |
| RP24-230J14.5 | 0.770607605 | 1.421415791 | 1.089812413 | 1.304275648 |
| Tapt1 | 762.3494051 | 0.409852667 | 0.074540521 | 5.49838747 |
| RP24-230J14.8 | 46.64492421 | 0.157772485 | 0.277791976 | 0.567951915 |
| RP24-230J14.7 | 0 | NA | NA | NA |
| RP24-230J14.9 | 0 | NA | NA | NA |
| RP24-230J14.10 | 12.44662364 | -0.656519881 | 0.298349934 | -2.200502851 |
| RP24-137F10.3 | 0 | NA | NA | NA |
| RP24-137F10.4 | 0.038981395 | 0.274393828 | 3.724945049 | 0.073663859 |
| RP24-137F10.1 | 0.921219545 | -0.728078328 | 1.129030357 | -0.644870463 |
| RP24-137F10.2 | 0.094097695 | 0.755282166 | 3.724945049 | 0.202763304 |
| Ldb2 | 272.9695285 | -0.401997885 | 0.164239966 | -2.447625235 |
| RP24-188E19.2 | 17.17285275 | -0.998817429 | 0.489005308 | -2.042549257 |
| RP24-188E19.3 | 7.245259879 | -1.329175875 | 0.500233788 | -2.657109347 |
| RP24-188E19.4 | 5.313268904 | -0.176499683 | 0.682435347 | -0.258632095 |
| RP24-188E19.5 | 0 | NA | NA | NA |
| RP24-247A21.1 | 40.44753983 | -1.06520297 | 0.375587653 | -2.836096874 |
| RP24-496C22.1 | 0.159422358 | -0.206488113 | 2.819252093 | -0.07324216 |
| RP24-496C22.2 | 19.70483131 | -1.155085672 | 0.352272868 | -3.278951566 |
| RP24-496C22.3 | 8.581378291 | -1.061752141 | 0.446509704 | -2.377892644 |
| RP24-496C22.4 | 7.00608203 | -1.255238218 | 0.442040315 | -2.839646464 |
| RP24-496C22.5 | 14.99026705 | -1.304595758 | 0.407663808 | -3.200175565 |
| RP24-496C22.6 | 21.60948068 | -1.08344919 | 0.316237335 | -3.426063496 |
| RP23-133N3.1 | 0 | NA | NA | NA |
| 4930431F12Rik | 0.1567526 | 0.274409612 | 2.828090123 | 0.09703001 |
| Gm10205 | 0 | NA | NA | NA |
| 4930435H24Rik | 4.37057135 | 1.887965581 | 0.650607353 | 2.901850973 |
| RP23-194K16.1 | 0.718177591 | 0.636875385 | 1.309662877 | 0.48628956 |
| RP23-33C9.1 | 1.272457209 | 0.282203286 | 0.952988001 | 0.2961247 |
| Qdpr | 1058.229875 | -0.66584974 | 0.118679265 | -5.610497657 |
| AC092711.4 | 2.958230677 | -1.641081766 | 0.640460272 | -2.562347481 |
| Clrn2 | 1.123782183 | -1.079622414 | 0.922627622 | -1.170160515 |

| | | | | |
|---|---|---|---|---|
| Lap3 | 1007.429224 | 1.194659184 | 0.149539633 | 7.988913413 |
| Med28 | 563.4186552 | 0.354335024 | 0.086479231 | 4.097342449 |
| Fam184b | 121.4810835 | -0.629759199 | 0.205409302 | -3.065874776 |
| RP23-7D3.3 | 16.87252161 | 0.683341396 | 0.406820532 | 1.679712161 |
| RP23-118K12.1 | 0.502057302 | 1.09731767 | 1.358541962 | 0.80771717 |
| 9630001P10Rik | 17.90177705 | -0.491107622 | 0.2718646 | -1.806441966 |
| RP23-7E4.8 | 0.664660227 | -1.167430427 | 1.311415292 | -0.890206507 |
| RP23-7E4.7 | 0.154585317 | 0.274399357 | 2.835007735 | 0.096789632 |
| RP23-7E4.9 | 3.048203817 | -0.636169783 | 0.664922852 | -0.956757285 |
| Ncapg | 19.54545852 | 2.040541403 | 0.389663819 | 5.236671469 |
| Lcorl | 806.219256 | 0.156100962 | 0.067493958 | 2.312813882 |
| Gm36401 | 0.944820375 | 0.06257183 | 1.113868583 | 0.056175236 |
| RP23-7E4.3 | 66.02500977 | 0.029325111 | 0.174252622 | 0.168290786 |
| Gm29036 | 24.05605521 | 0.284071982 | 0.267237316 | 1.062995193 |
| RP23-7E4.4 | 46.18654444 | 0.226172617 | 0.210812568 | 1.072861162 |
| AC158786.1 | 0.746606991 | 1.909781642 | 1.181139259 | 1.616897946 |
| RP23-228M24.1 | 0 | NA | NA | NA |
| Gm7931 | 9.771161355 | 1.005950328 | 0.355640882 | 2.828556499 |
| RP24-400K16.1 | 0 | NA | NA | NA |
| RP23-277H1.1 | 0 | NA | NA | NA |
| RP23-287A8.1 | 0.858277593 | -0.943331836 | 1.126227872 | -0.837602993 |
| RP23-319E24.2 | 0 | NA | NA | NA |
| RP23-319E24.1 | 0 | NA | NA | NA |
| RP23-195A18.4 | 0 | NA | NA | NA |
| Slit2 | 695.3750179 | -0.412817234 | 0.088875163 | -4.644911153 |
| 4930518C09Rik | 0 | NA | NA | NA |
| Gm20475 | 5.876575912 | -0.995626156 | 0.410116433 | -2.427667059 |
| RP23-195A18.3 | 20.06045718 | -0.903160376 | 0.336534848 | -2.683705366 |
| RP23-219K16.2 | 8.550972595 | -1.371653054 | 0.395751602 | -3.465944418 |
| RP23-219K16.1 | 10.624249 | -0.22755321 | 0.385390905 | -0.590447794 |
| Mir218-1 | 2.27602632 | -1.258166339 | 0.61619325 | -2.041837263 |
| RP23-79E5.4 | 4.747594266 | 0.001080068 | 0.461851657 | 0.002338561 |
| RP23-79E5.5 | 0 | NA | NA | NA |
| Pacrgl | 220.5840451 | -0.136644407 | 0.143528771 | -0.952034955 |
| Kcnip4 | 719.2995326 | -0.310150408 | 0.08435974 | -3.676521616 |
| RP23-14C20.2 | 34.79027562 | -0.723290155 | 0.30914855 | -2.339620079 |
| Gm20700 | 0 | NA | NA | NA |
| RP24-174D16.1 | 87.81495278 | -0.50925464 | 0.227858523 | -2.234959804 |
| Gm19031 | 0 | NA | NA | NA |
| RP24-170C14.2 | 202.1229481 | -0.566885259 | 0.236891653 | -2.393014914 |
| RP23-309G12.4 | 37.2300334 | -0.553746741 | 0.33964319 | -1.630377872 |
| RP23-309G12.3 | 161.4099445 | -0.560187242 | 0.27697516 | -2.022517984 |
| RP23-309G12.2 | 70.38172967 | -0.568052194 | 0.265711026 | -2.137857062 |
| RP23-79E5.7 | 138.4994086 | -0.565306882 | 0.284398111 | -1.987730788 |
| RP23-309G12.1 | 115.2856778 | -0.539117746 | 0.260389311 | -2.070429638 |
| Gm10025 | 0 | NA | NA | NA |
| Gm10048 | 309.3515181 | -0.625387573 | 0.333272784 | -1.87650358 |

| | | | | |
|---|---|---|---|---|
| RP23-187D22.9 | 187.5255152 | -0.544330727 | 0.37128442 | -1.466074785 |
| RP23-187D22.8 | 194.8703213 | -0.476332766 | 0.21834893 | -2.181520954 |
| RP23-187D22.7 | 88.63220688 | -0.537133191 | 0.303371182 | -1.770547842 |
| RP23-187D22.6 | 375.4133243 | -0.520772857 | 0.291868474 | -1.784272377 |
| RP23-187D22.5 | 181.2108272 | -0.417250203 | 0.240981756 | -1.731459717 |
| RP23-187D22.4 | 188.0951161 | -0.581233292 | 0.356331821 | -1.631157417 |
| RP23-187D22.10 | 319.4630086 | -0.538823483 | 0.293990877 | -1.832789806 |
| RP23-187D22.11 | 0.476944705 | -1.135296052 | 1.431645984 | -0.79300055 |
| RP23-187D22.3 | 106.6585838 | -0.491490419 | 0.426338896 | -1.152816277 |
| RP23-187D22.1 | 388.4487423 | -0.546749148 | 0.296917956 | -1.841414897 |
| RP23-140B23.2 | 209.5621235 | -0.769468087 | 0.343745743 | -2.23848034 |
| RP23-140B23.1 | 603.2557397 | -0.500543173 | 0.25308786 | -1.977744695 |
| RP23-394G5.2 | 44.14745288 | -0.538279409 | 0.368136025 | -1.462175318 |
| RP23-394G5.1 | 396.9657275 | -0.581566909 | 0.308794069 | -1.883348699 |
| RP24-158C2.3 | 251.5456547 | -0.948648853 | 0.295477852 | -3.210558239 |
| Gm19932 | 0 | NA | NA | NA |
| RP24-158C2.4 | 0.083526043 | 0.75528439 | 3.724945049 | 0.202763901 |
| RP24-253A20.1 | 476.3543519 | -0.946355719 | 0.306838205 | -3.084217361 |
| Gm5555 | 28.60310479 | 0.083240962 | 0.214611501 | 0.387868132 |
| RP24-79O7.3 | 0.322897824 | 1.592261212 | 1.54687858 | 1.029338199 |
| RP24-79O7.1 | 1.170209419 | -1.12738606 | 0.878613712 | -1.283141891 |
| Adgra3 | 255.2158286 | -0.263137111 | 0.106649505 | -2.467307391 |
| RP24-315J23.1 | 0.596915583 | 1.346332604 | 1.24055724 | 1.085264396 |
| RP24-315J23.2 | 0 | NA | NA | NA |
| RP24-478I4.1 | 0.047948698 | -0.206487837 | 3.724945049 | -0.055433794 |
| Mir8117 | 0 | NA | NA | NA |
| Gm22618 | 0.389682156 | -1.078529122 | 1.531391888 | -0.70428029 |
| RP23-21O1.1 | 0.042716945 | -0.206490061 | 3.724945049 | -0.055434391 |
| RP23-21O1.3 | 0.04393175 | -0.206490061 | 3.724945049 | -0.055434391 |
| RP23-21O1.2 | 0.038981395 | 0.274393828 | 3.724945049 | 0.073663859 |
| RP23-125B15.1 | 0 | NA | NA | NA |
| RP24-363A3.1 | 0.195174816 | -0.687379431 | 2.237604007 | -0.307194405 |
| RP24-312A3.1 | 0 | NA | NA | NA |
| RP24-441G10.1 | 0 | NA | NA | NA |
| AC119939.1 | 0 | NA | NA | NA |
| Ppargc1a | 1065.622016 | 0.150060804 | 0.061357734 | 2.445670579 |
| RP23-385G21.4 | 13.87657411 | 0.011743208 | 0.317782328 | 0.036953622 |
| RP23-385G21.3 | 24.14107729 | -0.011651184 | 0.246425911 | -0.047280678 |
| RP23-385G21.2 | 0 | NA | NA | NA |
| RP23-385G21.1 | 28.94227565 | -0.197250639 | 0.24632916 | -0.800760411 |
| RP24-296L6.1 | 199.7747531 | -0.267941524 | 0.365742958 | -0.732595168 |
| RP24-296L6.2 | 266.7259097 | -0.266987871 | 0.232716167 | -1.147268255 |
| RP24-296L6.3 | 71.01041919 | -0.555177029 | 0.341134478 | -1.627443325 |
| RP24-296L6.4 | 241.104923 | -0.296824024 | 0.305611843 | -0.971245164 |
| RP24-299M10.1 | 94.15749016 | -0.67201225 | 0.458366361 | -1.466102896 |
| RP24-299M10.2 | 0.645477656 | -1.496965609 | 1.298601095 | -1.152752462 |
| Mir6417 | 1.584881738 | -1.432809762 | 0.935817829 | -1.531077649 |

| | | | | |
|---|---|---|---|---|
| RP24-309H3.1 | 30.5102578 | -0.311579537 | 0.278000428 | -1.12078798 |
| Dhx15 | 1662.063983 | 0.253121405 | 0.098520028 | 2.569238056 |
| RP24-309H3.3 | 27.29790927 | -0.31915052 | 0.238479165 | -1.338274226 |
| 9230114K14Rik | 31.71347572 | -0.613416283 | 0.208234648 | -2.945793555 |
| RP24-309H3.5 | 0.319554274 | 0.82507794 | 1.602441015 | 0.514888181 |
| RP24-309H3.6 | 0.802626749 | -0.422844144 | 1.229750937 | -0.343845352 |
| RP24-237I4.4 | 0 | NA | NA | NA |
| Gm25601 | 0.045882455 | 0.755288839 | 3.724945049 | 0.202765096 |
| Sod3 | 146.4300929 | 0.257008121 | 0.152116874 | 1.689543803 |
| Ccdc149 | 659.1518796 | 0.062743514 | 0.074726431 | 0.839642854 |
| RP24-237I4.3 | 0.09689823 | -0.687371727 | 3.724945049 | -0.184532045 |
| Lgi2 | 1350.453356 | -0.52102358 | 0.075232357 | -6.925525127 |
| RP23-181F22.4 | 0 | NA | NA | NA |
| RP23-181F22.3 | 4.780205594 | -0.46739585 | 0.512138481 | -0.912635678 |
| Gm5866 | 0.405467889 | -0.084723552 | 1.543932048 | -0.054875182 |
| 8030423F21Rik | 2.177346141 | -0.218236643 | 0.776251468 | -0.281141681 |
| Sepsecs | 236.6757126 | 0.11894535 | 0.100040774 | 1.188968711 |
| Pi4k2b | 58.07000918 | 0.789738214 | 0.218575468 | 3.613114601 |
| Zcchc4 | 173.3575901 | 0.026954383 | 0.16560299 | 0.162765077 |
| Anapc4 | 711.191152 | 0.228514541 | 0.051981676 | 4.396059478 |
| RP24-373A18.2 | 0 | NA | NA | NA |
| Gm23532 | 0 | NA | NA | NA |
| RP24-373A18.1 | 0 | NA | NA | NA |
| RP24-373A18.3 | 0.457291341 | 0.274409786 | 1.797948663 | 0.152623816 |
| Slc34a2 | 1.437183928 | 1.423420097 | 1.2116324 | 1.174795339 |
| Gm17182 | 0 | NA | NA | NA |
| Sel1l3 | 908.2236204 | -0.501151677 | 0.126653427 | -3.956874195 |
| n-R5s171 | 0 | NA | NA | NA |
| Smim20 | 129.4033044 | 0.214585539 | 0.142312418 | 1.507848307 |
| RP23-425N12.2 | 79.50346624 | -0.257254335 | 0.153085239 | -1.680464658 |
| n-R5s172 | 0.379461129 | -1.43116078 | 1.725433787 | -0.829449841 |
| RP23-425N12.1 | 0.485522898 | -0.340287412 | 1.513189088 | -0.224880958 |
| Gm26486 | 0 | NA | NA | NA |
| Rbpj | 346.6721826 | 0.047962692 | 0.101846249 | 0.470932335 |
| RP24-540G23.3 | 0.236145685 | -0.495795032 | 2.112236085 | -0.234725198 |
| RP23-137N6.2 | 0.29199547 | 0.790163704 | 1.785850368 | 0.442457956 |
| Cckar | 6.464590576 | -0.104852573 | 0.513362877 | -0.204246505 |
| Tbc1d19 | 456.9575057 | -0.117105371 | 0.103621616 | -1.130124924 |
| RP24-196M24.2 | 141.7804511 | 0.106578848 | 0.213004068 | 0.500360622 |
| RP24-196M24.3 | 10.28263209 | 0.248604695 | 0.375094124 | 0.662779499 |
| Stim2 | 905.9735999 | 0.008397897 | 0.061186129 | 0.137251655 |
| Gm15820 | 0 | NA | NA | NA |
| RP24-482F14.1 | 1.962603442 | 0.855154035 | 0.679760307 | 1.258022903 |
| RP23-148F6.1 | 0 | NA | NA | NA |
| RP23-187K3.1 | 6.300906923 | 0.009155772 | 0.420815485 | 0.021757213 |
| RP24-201P24.1 | 0.044544648 | 0.755286614 | 3.724945049 | 0.202764498 |
| Gm24235 | 0 | NA | NA | NA |

| | | | | |
|---|---|---|---|---|
| RP23-267P6.1 | 0 | NA | NA | NA |
| Gm25513 | 0 | NA | NA | NA |
| RP24-133H21.1 | 0 | NA | NA | NA |
| RP23-317I8.3 | 0 | NA | NA | NA |
| RP23-317I8.2 | 0 | NA | NA | NA |
| Gm15975 | 34.36835703 | 0.454163003 | 0.242193885 | 1.875204251 |
| RP23-317I8.1 | 0.181740481 | -0.682097809 | 2.280596453 | -0.299087464 |
| RP23-162F3.1 | 0.072025462 | 0.755288839 | 3.724945049 | 0.202765096 |
| RP23-162F3.2 | 1.035655884 | -0.458392015 | 1.024106654 | -0.447601832 |
| RP24-301M6.1 | 0.124853745 | 1.290908285 | 3.721447108 | 0.346883416 |
| RP24-394H13.1 | 0.707701802 | 0.954048796 | 1.379435718 | 0.691622512 |
| Gm23896 | 0 | NA | NA | NA |
| Gm24902 | 0 | NA | NA | NA |
| Gm23015 | 0 | NA | NA | NA |
| RP23-17A11.1 | 0 | NA | NA | NA |
| RP24-499N22.1 | 0 | NA | NA | NA |
| RP24-499N22.2 | 0 | NA | NA | NA |
| RP23-297D16.1 | 0 | NA | NA | NA |
| RP23-297D16.2 | 0.038981395 | 0.274393828 | 3.724945049 | 0.073663859 |
| 4932441J04Rik | 57.67652023 | -0.283687483 | 0.149969147 | -1.891638972 |
| Gm8121 | 18.9482281 | 0.079824598 | 0.255178175 | 0.312819065 |
| RP24-189M4.2 | 0 | NA | NA | NA |
| RP24-189M4.1 | 0.956833923 | -1.780472868 | 1.128849756 | -1.577245209 |
| Pcdh7 | 3274.629757 | -0.20222991 | 0.092555338 | -2.184962141 |
| RP23-458G12.1 | 132.1876841 | -0.504697076 | 0.130729186 | -3.860630447 |
| RP23-458G12.4 | 95.53520872 | -0.322701361 | 0.207728959 | -1.553473151 |
| RP23-458G12.3 | 115.8575128 | -0.460915889 | 0.176298708 | -2.614403093 |
| RP23-458G12.2 | 78.02550846 | -0.703878032 | 0.221808454 | -3.173359796 |
| RP23-358D12.4 | 15.63086198 | 0.07156286 | 0.391782705 | 0.182659568 |
| RP23-358D12.1 | 30.61318993 | 0.233460937 | 0.268831256 | 0.868429293 |
| RP23-358D12.3 | 67.73431069 | -0.621566511 | 0.175489715 | -3.541897091 |
| RP23-358D12.2 | 74.52970784 | -0.745376719 | 0.230644128 | -3.231717742 |
| RP24-231P10.2 | 87.86336388 | -0.691828966 | 0.20256366 | -3.415365647 |
| RP24-231P10.3 | 68.50219268 | -0.583113891 | 0.257950175 | -2.260567921 |
| RP24-231P10.4 | 91.67305281 | -0.81884551 | 0.236039831 | -3.469098866 |
| Gm37720 | 43.33085034 | -0.566638281 | 0.188405539 | -3.007545768 |
| RP24-231P10.5 | 54.31474321 | -0.404746727 | 0.203867647 | -1.985340656 |
| RP24-340K5.2 | 33.48457548 | -0.10332142 | 0.226394662 | -0.456377458 |
| RP24-340K5.4 | 0 | NA | NA | NA |
| RP24-340K5.3 | 0 | NA | NA | NA |
| RP24-502C10.1 | 0 | NA | NA | NA |
| Gm22536 | 0 | NA | NA | NA |
| RP24-375D24.2 | 0 | NA | NA | NA |
| 1700029E06Rik | 0 | NA | NA | NA |
| RP24-353N4.4 | 0 | NA | NA | NA |
| Gm6632 | 0 | NA | NA | NA |
| RP24-353N4.3 | 0 | NA | NA | NA |

| | | | | |
|---|---|---|---|---|
| Gm22308 | 0 | NA | NA | NA |
| RP23-324H16.1 | 0 | NA | NA | NA |
| RP23-324H16.2 | 3.023846459 | 0.428428631 | 0.560473579 | 0.764404687 |
| RP23-340N4.1 | 0 | NA | NA | NA |
| RP24-267O6.1 | 1.051957033 | 0.477138008 | 0.92018426 | 0.518524418 |
| RP24-543J12.1 | 0 | NA | NA | NA |
| RP24-543J12.2 | 0.062426873 | 0.755288839 | 3.724945049 | 0.202765096 |
| RP24-543J12.3 | 0 | NA | NA | NA |
| RP24-543J12.4 | 0.075043205 | 0.755286614 | 3.724945049 | 0.202764498 |
| RP23-183O3.1 | 0.136194719 | -0.687371727 | 3.724945049 | -0.184532045 |
| Cbfa2t2-ps1 | 30.26364068 | -0.123529393 | 0.191868721 | -0.643822465 |
| Gm23331 | 0 | NA | NA | NA |
| RP23-419P12.1 | 0 | NA | NA | NA |
| RP23-84N24.1 | 0 | NA | NA | NA |
| Gm23708 | 0 | NA | NA | NA |
| RP24-390N7.1 | 2.35460177 | -0.650089398 | 0.65441053 | -0.99339691 |
| RP24-390N7.2 | 0 | NA | NA | NA |
| RP24-390N7.3 | 0 | NA | NA | NA |
| RP24-300K9.1 | 0 | NA | NA | NA |
| RP23-210A16.1 | 0 | NA | NA | NA |
| RP23-86N20.1 | 0 | NA | NA | NA |
| G6pd2 | 0.044191549 | -0.206490061 | 3.724945049 | -0.055434391 |
| RP23-480E8.1 | 0 | NA | NA | NA |
| Gm22273 | 0 | NA | NA | NA |
| RP24-447I18.1 | 0.236971717 | 0.247865331 | 2.106041586 | 0.117692515 |
| Gm24079 | 0 | NA | NA | NA |
| Arap2 | 2567.739427 | 0.411810918 | 0.072630658 | 5.669932341 |
| RP23-387F14.1 | 16.56268708 | -0.152811681 | 0.289167704 | -0.528453487 |
| RP23-387F14.2 | 57.3555258 | 0.13287126 | 0.188422062 | 0.705178891 |
| Dthd1 | 4.062000091 | 1.813033638 | 0.781383716 | 2.320285924 |
| RP23-37I24.1 | 105.4712107 | -0.170008672 | 0.162783853 | -1.044382898 |
| Nwd2 | 487.7321807 | -0.486520969 | 0.160232835 | -3.036337517 |
| Gm3687 | 4.714620213 | -0.996258333 | 0.504983567 | -1.972852975 |
| 0610040J01Rik | 36.24267069 | 0.189069639 | 0.188640935 | 1.002272594 |
| Rell1 | 246.5447793 | 0.548275679 | 0.121829895 | 4.500337772 |
| Gm15819 | 0.356802427 | 1.785153219 | 3.709291204 | 0.48126532 |
| RP23-192C21.2 | 0.04393175 | -0.206490061 | 3.724945049 | -0.055434391 |
| Gm25608 | 0 | NA | NA | NA |
| RP23-192C21.3 | 1.013052095 | 0.312752136 | 1.248260355 | 0.250550404 |
| Pgm1 | 176.2902017 | 0.500364757 | 0.142187952 | 3.519037666 |
| Gm23967 | 0 | NA | NA | NA |
| Gm22395 | 0 | NA | NA | NA |
| Gm22559 | 0 | NA | NA | NA |
| Tbc1d1 | 746.5571268 | 0.674284921 | 0.099719742 | 6.761799687 |
| Gm15824 | 0.060174537 | 0.755286614 | 3.724945049 | 0.202764498 |
| RP23-339I6.1 | 13.2226886 | 0.148612411 | 0.331465985 | 0.448348903 |
| Gm15823 | 0 | NA | NA | NA |

| | | | | |
|---|---|---|---|---|
| Gm6044 | 4.011017284 | 0.531389643 | 0.5236523 | 1.014775727 |
| RP23-339I6.2 | 0.245047857 | 1.738500082 | 2.092747643 | 0.830726097 |
| RP23-339I6.3 | 0.541557618 | 1.762255569 | 1.491512381 | 1.181522588 |
| RP23-195B3.4 | 0.062426873 | 0.755288839 | 3.724945049 | 0.202765096 |
| 1700027F09Rik | 0 | NA | NA | NA |
| RP23-195B3.3 | 0 | NA | NA | NA |
| RP23-195B3.5 | 10.29678862 | -2.578554073 | 0.407864991 | -6.322077475 |
| RP24-448C16.4 | 0.540262476 | 0.359952582 | 1.27769919 | 0.281719348 |
| RP24-448C16.6 | 0.192501058 | -1.073580721 | 2.663543715 | -0.403064802 |
| RP24-448C16.5 | 1.291588001 | 0.432906793 | 0.875432454 | 0.49450622 |
| RP24-448C16.7 | 3.668071862 | -0.395257262 | 0.533199063 | -0.741293992 |
| RP24-448C16.9 | 0.116217011 | 0.274402725 | 3.724945049 | 0.073666248 |
| RP24-448C16.8 | 0 | NA | NA | NA |
| Gm25306 | 0.049553047 | 0.274391604 | 3.724945049 | 0.073663262 |
| Gm26761 | 0.503421469 | -1.101853852 | 1.376753589 | -0.800327568 |
| Gm20033 | 20.30300836 | -0.341801263 | 0.234844855 | -1.455434323 |
| Klf3 | 642.6073925 | 0.283051718 | 0.111386013 | 2.541178298 |
| RP23-326O2.2 | 26.4782624 | -0.236390297 | 0.226716421 | -1.042669497 |
| Tlr1 | 85.37836272 | 1.400585078 | 0.213829627 | 6.550004779 |
| Tlr6 | 42.58363877 | 1.739689245 | 0.278641585 | 6.243465934 |
| Fam114a1 | 72.52929912 | 0.501851969 | 0.185433015 | 2.706378733 |
| Mir574 | 0.038981395 | 0.274393828 | 3.724945049 | 0.073663859 |
| RP24-87P2.3 | 1.276825249 | 0.820147767 | 0.935121181 | 0.877049718 |
| Tmem156 | 6.703461548 | 1.772247769 | 0.600651408 | 2.950542936 |
| Klhl5 | 474.7177188 | -0.212236114 | 0.108340519 | -1.958972657 |
| Gm25154 | 2.287715794 | 1.227487733 | 0.841933899 | 1.457938366 |
| Wdr19 | 272.9715081 | -0.431847453 | 0.117283888 | -3.682069718 |
| Rfc1 | 611.6308179 | 0.374674377 | 0.077251496 | 4.850059798 |
| RP23-344O11.3 | 13.50952572 | -0.237845176 | 0.3844265 | -0.6187013 |
| Klb | 1.57634325 | -0.850268349 | 0.81583802 | -1.042202408 |
| Rpl9 | 1617.17554 | 0.284670978 | 0.087963553 | 3.236237824 |
| Lias | 326.7940019 | 0.202548071 | 0.085179682 | 2.377891828 |
| RP23-477N9.1 | 7.915089792 | 0.180197115 | 0.479232127 | 0.376012176 |
| RP23-477N9.5 | 5.012299285 | -0.657796479 | 0.468291338 | -1.404673599 |
| Ugdh | 393.579355 | 0.111519904 | 0.092606614 | 1.204232611 |
| AC136513.6 | 0.044544648 | 0.755286614 | 3.724945049 | 0.202764498 |
| RP23-477N9.2 | 5.168412192 | 0.079892115 | 0.460778604 | 0.173385037 |
| Smim14 | 1287.523276 | -0.133419648 | 0.054308298 | -2.456708307 |
| RP23-477N9.4 | 25.33295597 | -0.540944843 | 0.333301553 | -1.622989269 |
| RP23-477N9.6 | 5.250217998 | -1.041068291 | 0.454244028 | -2.291870065 |
| RP23-477N9.3 | 33.76352782 | -1.68059298 | 0.425909319 | -3.945893889 |
| Ube2k | 1596.82342 | 0.067290102 | 0.061781831 | 1.089156801 |
| RP23-403P24.1 | 88.56705528 | -0.523678882 | 0.289167259 | -1.81098954 |
| RP23-403P24.3 | 0 | NA | NA | NA |
| Pds5a | 1522.438719 | -0.131573222 | 0.065366253 | -2.012861633 |
| RP23-403P24.5 | 11.79713159 | -0.694800328 | 0.337573475 | -2.058219554 |
| RP23-403P24.6 | 4.819094623 | -0.010070657 | 0.483577425 | -0.020825324 |

| | | | | |
|---|---|---|---|---|
| Gm24399 | 0.0878635 | -0.687376175 | 3.724945049 | -0.184533239 |
| RP23-403P24.7 | 16.35270107 | -0.214388489 | 0.356308221 | -0.601693916 |
| RP23-403P24.8 | 29.36603598 | -0.488772145 | 0.270926024 | -1.80407972 |
| N4bp2os | 32.02296958 | 0.292119758 | 0.19609721 | 1.489668101 |
| N4bp2 | 387.2265267 | 0.080441805 | 0.116019285 | 0.693348565 |
| Gm24346 | 4.621381795 | -0.738345142 | 0.518733074 | -1.423362381 |
| RP23-42O8.7 | 30.0274551 | -0.403483735 | 0.336250279 | -1.199950632 |
| RP23-42O8.6 | 0.112396584 | 0.27438722 | 3.724945049 | 0.073662085 |
| RP23-42O8.5 | 0.165156969 | 0.274389239 | 3.414282971 | 0.080365114 |
| RP23-42O8.4 | 0 | NA | NA | NA |
| Rhoh | 32.07602456 | 1.975870854 | 0.246422977 | 8.018208693 |
| RP23-42O8.9 | 0.263966226 | -0.206481789 | 1.822725234 | -0.113281906 |
| Chrna9 | 0.40746857 | 1.627668997 | 1.536003361 | 1.059678018 |
| RP23-92N2.1 | 0 | NA | NA | NA |
| RP23-92N2.2 | 0 | NA | NA | NA |
| 9130230L23Rik | 1.235688375 | 2.102626545 | 1.217166218 | 1.727476917 |
| Gm25952 | 0 | NA | NA | NA |
| Rbm47 | 123.9754224 | 1.57388539 | 0.212991807 | 7.389417526 |
| RP24-351O18.5 | 7.439759207 | 1.657569893 | 0.586795245 | 2.824784126 |
| RP24-351O18.4 | 9.722078793 | 0.987755864 | 0.501420712 | 1.969914367 |
| Gm26725 | 0.204218301 | -0.206485372 | 2.585377521 | -0.079866623 |
| Gm22929 | 0 | NA | NA | NA |
| 1700126H18Rik | 0 | NA | NA | NA |
| RP24-351O18.2 | 0 | NA | NA | NA |
| Gm15794 | 0 | NA | NA | NA |
| RP24-351O18.7 | 0.049553047 | 0.274391604 | 3.724945049 | 0.073663262 |
| Nsun7 | 120.513026 | 0.087368061 | 0.178561119 | 0.489289388 |
| Apbb2 | 1193.808036 | 0.009595876 | 0.072255887 | 0.13280407 |
| RP23-21D6.2 | 18.43062459 | 0.364040047 | 0.272207013 | 1.337364688 |
| Gm23183 | 8.321784305 | 0.612747808 | 0.426286966 | 1.437406858 |
| RP23-21D6.1 | 24.68721265 | 0.030851435 | 0.329111405 | 0.093741617 |
| Gm16273 | 0.062426873 | 0.755288839 | 3.724945049 | 0.202765096 |
| RP23-157G2.2 | 24.13982844 | -0.055427428 | 0.265023778 | -0.20914134 |
| RP23-157G2.1 | 9.142804473 | -0.285655936 | 0.373669064 | -0.764462363 |
| RP24-360B3.2 | 21.21818474 | 0.081595878 | 0.354106933 | 0.230427224 |
| RP24-360B3.3 | 23.13608884 | -0.452432551 | 0.320358338 | -1.412270253 |
| RP24-360B3.1 | 33.09598643 | -0.096072153 | 0.362258418 | -0.265203369 |
| Uchl1os | 3.562613421 | 0.492017777 | 0.507239674 | 0.96999072 |
| Uchl1 | 2844.203176 | -0.071491647 | 0.077059289 | -0.927748591 |
| RP23-472J13.3 | 20.85386141 | 1.217942968 | 0.50736675 | 2.400517903 |
| RP23-352L15.1 | 0.061920613 | 0.274393828 | 3.724945049 | 0.073663859 |
| Limch1 | 1533.433457 | -0.254449506 | 0.055401895 | -4.592794301 |
| Gm6517 | 9.113272248 | -0.537471612 | 0.35377888 | -1.519230351 |
| RP24-79P8.1 | 29.56015194 | -0.460217978 | 0.27765405 | -1.657523015 |
| RP24-383P24.2 | 11.74058244 | -0.616085982 | 0.344482317 | -1.78844008 |
| Gm15949 | 0 | NA | NA | NA |
| Gm15948 | 0.038981395 | 0.274393828 | 3.724945049 | 0.073663859 |

| | | | | |
|---|---|---|---|---|
| RP24-383P24.3 | 0.808719243 | -0.32692543 | 1.090152108 | -0.299889738 |
| Gm23841 | 0 | NA | NA | NA |
| Phox2b | 13.13116622 | 0.182611445 | 0.560616329 | 0.325733369 |
| RP24-95M11.1 | 1.073023658 | -0.865115529 | 1.23733108 | -0.699178695 |
| RP24-353A8.1 | 1.943439641 | 0.615645873 | 0.864588913 | 0.71206774 |
| Tmem33 | 1537.582746 | -0.060141546 | 0.065106146 | -0.923746051 |
| Slc30a9 | 1760.179162 | -0.264029082 | 0.056276594 | -4.691632209 |
| RP24-353A8.2 | 0.708786783 | -0.011628838 | 1.165039956 | -0.009981493 |
| Bend4 | 204.8953353 | -0.372909598 | 0.111775298 | -3.336243366 |
| Gm26756 | 4.883969858 | -0.444062717 | 0.445401429 | -0.99699437 |
| RP24-103L16.3 | 7.272572362 | -0.771995622 | 0.519553088 | -1.485884001 |
| RP24-103L16.4 | 0 | NA | NA | NA |
| RP24-103L16.5 | 15.55852251 | -1.196046629 | 0.28457084 | -4.202983798 |
| RP23-213N8.1 | 1.075240245 | -0.325869333 | 0.947563841 | -0.343902246 |
| Shisa3 | 72.51937258 | -0.910945767 | 0.54090125 | -1.684125829 |
| Atp8a1 | 4334.315932 | 0.073347586 | 0.060607937 | 1.210197697 |
| Gm23617 | 2.388208885 | -1.154748908 | 0.730620059 | -1.580505344 |
| RP23-213N8.2 | 52.3886502 | -0.154299819 | 0.276895998 | -0.557248281 |
| Gm15478 | 2.676788521 | -0.799665206 | 0.68380528 | -1.16943409 |
| RP24-274I18.1 | 56.96681602 | -0.151611245 | 0.299580555 | -0.50607839 |
| RP23-130C18.1 | 59.36905083 | -0.159166499 | 0.273739863 | -0.581451665 |
| RP23-130C18.2 | 29.63687175 | 0.151899482 | 0.291509185 | 0.521079573 |
| Gm15477 | 9.54473322 | 0.514744257 | 0.349586438 | 1.47243772 |
| RP23-130C18.3 | 0.310726904 | -1.174596199 | 2.089583724 | -0.562119711 |
| Gm21905 | 0.178178593 | 1.650851562 | 3.716108103 | 0.444242072 |
| RP23-130C18.4 | 1.374506326 | 3.619059137 | 1.266678637 | 2.857124949 |
| Gm5108 | 0.225572436 | 0.755306741 | 2.199105652 | 0.34346087 |
| RP23-387L12.2 | 0.395257705 | 0.736289764 | 1.692863463 | 0.434937477 |
| RP23-387L12.3 | 1.482312138 | 0.477769631 | 0.75599783 | 0.631972225 |
| Grxcr1 | 0.175796035 | 0.27440939 | 2.309857975 | 0.118799248 |
| RP23-387L12.4 | 0.038981395 | 0.274393828 | 3.724945049 | 0.073663859 |
| RP24-224K15.1 | 0 | NA | NA | NA |
| Gm23605 | 0 | NA | NA | NA |
| RP24-441E22.1 | 0.086648695 | -0.687376175 | 3.724945049 | -0.184533239 |
| RP23-72G21.1 | 2.775994898 | -0.902968479 | 0.687894381 | -1.312655699 |
| Kctd8 | 186.6518599 | -0.154472425 | 0.095176733 | -1.623006175 |
| RP23-319P12.6 | 23.8423184 | -0.592988553 | 0.241352988 | -2.456934795 |
| RP23-319P12.5 | 17.40396653 | -0.66559952 | 0.273108954 | -2.437120829 |
| RP23-319P12.4 | 57.79128105 | -0.580874052 | 0.25318334 | -2.294282284 |
| RP23-319P12.3 | 56.51122244 | -0.227937932 | 0.257025681 | -0.886829404 |
| RP23-319P12.2 | 47.23969796 | -0.255700267 | 0.259717875 | -0.984530878 |
| RP23-319P12.1 | 45.02692642 | -0.812875167 | 0.300787461 | -2.702490201 |
| RP23-114M24.1 | 151.2873273 | -0.546047837 | 0.225206836 | -2.424650362 |
| Gm24368 | 0.334883495 | 0.677912909 | 1.558610813 | 0.434946879 |
| Gm23932 | 0 | NA | NA | NA |
| Gm5867 | 0.309731683 | 0.264138752 | 1.747949639 | 0.15111348 |
| RP24-351E22.1 | 0.956841293 | 0.646169632 | 1.095953592 | 0.589595797 |

| | | | | |
|---|---|---|---|---|
| Yipf7 | 2.071883479 | 0.953149884 | 0.753096255 | 1.265641515 |
| RP24-351E22.2 | 0.331853251 | -0.676488758 | 1.768799928 | -0.382456346 |
| Guf1 | 544.3541319 | 0.068959223 | 0.108494067 | 0.635603633 |
| Gnpda2 | 249.4753375 | -0.281480393 | 0.13026633 | -2.160806965 |
| Gm26044 | 0 | NA | NA | NA |
| Gm23661 | 0 | NA | NA | NA |
| RP23-236G19.1 | 0 | NA | NA | NA |
| Gm23067 | 0 | NA | NA | NA |
| Gm26072 | 0 | NA | NA | NA |
| RP23-332O18.1 | 0 | NA | NA | NA |
| RP23-332O18.2 | 0.480717083 | -1.123357289 | 1.524650005 | -0.736796829 |
| Gabrg1 | 384.6081965 | -0.034372534 | 0.135300811 | -0.254045289 |
| Gabra2 | 1383.339494 | -0.321108544 | 0.082438495 | -3.895128661 |
| Cox7b2 | 1.615076682 | -1.521194526 | 0.911595972 | -1.668715718 |
| Gabra4 | 1311.463824 | -0.154216962 | 0.109420805 | -1.409393405 |
| Gabrb1 | 2188.449478 | -0.016867146 | 0.080624003 | -0.209207495 |
| RP24-156O12.1 | 9.874941751 | -0.786773317 | 0.325827101 | -2.414695755 |
| Gm15617 | 0.150086409 | 1.264824188 | 3.723011833 | 0.339731445 |
| Gm15616 | 12.70121426 | -0.572186442 | 0.279607066 | -2.046394784 |
| Commd8 | 452.1455523 | 0.231716568 | 0.072970972 | 3.175462255 |
| RP23-27G7.2 | 0 | NA | NA | NA |
| Atp10d | 139.6844161 | 0.540737496 | 0.143772626 | 3.761060151 |
| Corin | 18.43000873 | -0.588224715 | 0.31927652 | -1.842367595 |
| Gm25768 | 0.088383099 | -0.687376175 | 3.724945049 | -0.184533239 |
| Gm20647 | 0 | NA | NA | NA |
| Gm19560 | 0 | NA | NA | NA |
| Gm23712 | 0.116046057 | -0.687371727 | 3.724945049 | -0.184532045 |
| AU023070 | 0.14021869 | -0.687378066 | 2.921525367 | -0.23528054 |
| Nfxl1 | 313.9107044 | -0.019707816 | 0.0745988 | -0.264184086 |
| RP24-316L21.3 | 23.03118454 | -0.137932273 | 0.228890744 | -0.602611842 |
| RP24-506B15.1 | 0.413335106 | 0.715301464 | 1.467426561 | 0.487452989 |
| Zar1 | 3.771757384 | -0.332716411 | 0.573397009 | -0.580254877 |
| Gm5868 | 0.655263372 | 2.242871773 | 1.406907149 | 1.594186066 |
| Cnga1 | 0.310549208 | 0.274378369 | 3.724945049 | 0.073659709 |
| RP24-506B15.2 | 5.852374332 | 0.56613045 | 0.467764272 | 1.210290064 |
| RP24-506B15.3 | 0 | NA | NA | NA |
| Nipal1 | 7.542068972 | 3.219503208 | 0.874903778 | 3.679836901 |
| Txk | 24.20729952 | 2.905593443 | 0.515213925 | 5.639586397 |
| RP23-415O9.1 | 0 | NA | NA | NA |
| Tec | 81.72636874 | 1.260360791 | 0.29860469 | 4.220833878 |
| RP23-204A8.1 | 6.319016411 | 0.50118492 | 0.461505593 | 1.085977998 |
| Gm9870 | 6.265701809 | -0.295580544 | 0.425105606 | -0.695310859 |
| Slain2 | 498.0036851 | -0.162940925 | 0.076174484 | -2.139048609 |
| Gm10135 | 0.045882455 | 0.755288839 | 3.724945049 | 0.202765096 |
| Gm24776 | 0 | NA | NA | NA |
| Slc10a4 | 54.26526042 | 0.108658229 | 0.18495771 | 0.587476074 |
| Gm15847 | 1.549834172 | -0.555417133 | 1.193280446 | -0.46545398 |

| | | | | |
|---|---|---|---|---|
| AC108848.6 | 1.313571363 | 0.349761602 | 0.812072324 | 0.430702527 |
| Fryl | 1628.389783 | -0.134733344 | 0.061422702 | -2.193543082 |
| RP23-300K5.1 | 35.04837016 | -0.374043753 | 0.256783325 | -1.456651255 |
| 1700071G01Rik | 0 | NA | NA | NA |
| RP23-318I19.2 | 17.43330629 | -0.723470976 | 0.330963075 | -2.185956775 |
| Gm17207 | 5.90520374 | -0.759933006 | 0.416483517 | -1.824641254 |
| RP23-153H17.2 | 0.097139547 | 0.274393828 | 3.724945049 | 0.073663859 |
| 1700025M24Rik | 0.068097359 | -0.206487837 | 3.724945049 | -0.055433794 |
| RP23-153H17.3 | 0 | NA | NA | NA |
| Ociad1 | 3089.545592 | -0.042430806 | 0.04654677 | -0.911573603 |
| RP23-153H17.6 | 0.3552573 | 0.734157771 | 2.06700577 | 0.355179352 |
| RP23-153H17.5 | 470.3152913 | -0.243844289 | 0.085316638 | -2.858109458 |
| Ociad2 | 527.2792456 | 0.14458437 | 0.149812179 | 0.965104247 |
| RP23-153H17.7 | 29.63938725 | -0.32613858 | 0.277640427 | -1.174679723 |
| RP23-153H17.9 | 0.315779812 | -0.206494968 | 3.724945049 | -0.055435708 |
| Cwh43 | 14.92583306 | 0.261772776 | 0.305393141 | 0.85716652 |
| RP24-274B10.1 | 0.149988742 | 0.75530322 | 2.860996817 | 0.264000021 |
| Gm15653 | 0 | NA | NA | NA |
| Dcun1d4 | 1162.429234 | -0.15687932 | 0.066871222 | -2.345991539 |
| Lrrc66 | 2.285793377 | 0.903193857 | 0.743723174 | 1.214422097 |
| Sgcb | 773.7364474 | 0.604395724 | 0.092526879 | 6.532109678 |
| RP23-105O12.2 | 2.772617781 | -0.296501747 | 0.600467839 | -0.493784559 |
| Spata18 | 12.87588817 | -0.397958268 | 0.355461851 | -1.11955268 |
| RP23-276J7.1 | 0 | NA | NA | NA |
| RP23-276J7.2 | 0.066050875 | 0.274391604 | 3.724945049 | 0.073663262 |
| Usp46 | 1284.997462 | -0.364077563 | 0.073295267 | -4.967272487 |
| Usp46os1 | 0.374638785 | 0.343322582 | 1.563198432 | 0.219628279 |
| Usp46os2 | 0.045882455 | 0.755288839 | 3.724945049 | 0.202765096 |
| CH25-194I22.2 | 46.11457756 | -0.574313374 | 0.301578182 | -1.904359824 |
| CH25-194I22.3 | 0 | NA | NA | NA |
| CH25-194I22.4 | 0.285862058 | -1.106953133 | 1.797152541 | -0.615948344 |
| Snora26 | 6.265788682 | -0.061425219 | 0.408967554 | -0.150195824 |
| Dancr | 0 | NA | NA | NA |
| RP23-422P4.2 | 0.766635478 | -0.864710717 | 1.201950665 | -0.719422803 |
| Rasl11b | 390.2960293 | -0.218509629 | 0.137336088 | -1.591057619 |
| Scfd2 | 235.5014393 | -0.057491464 | 0.078522156 | -0.73216868 |
| RP23-422P4.3 | 0.472351266 | 0.566161274 | 1.561168026 | 0.362652363 |
| Gm15981 | 4.746611391 | 0.158815321 | 0.465927017 | 0.340858793 |
| Gm15982 | 6.659240228 | -0.062733918 | 0.425236475 | -0.147527133 |
| RP24-465E5.2 | 15.61428043 | -0.407409075 | 0.394960889 | -1.031517515 |
| RP24-465E5.1 | 9.836186905 | 0.013335229 | 0.43629756 | 0.030564527 |
| Fip1l1 | 1364.710559 | -0.017895117 | 0.069746281 | -0.256574491 |
| RP24-245O20.1 | 42.86441423 | -0.367947596 | 0.205689974 | -1.788845555 |
| Lnx1 | 208.7467131 | -0.230190463 | 0.088408983 | -2.603699923 |
| Gm15984 | 0.100902008 | 0.274391828 | 3.724945049 | 0.073663322 |
| Gm15985 | 0 | NA | NA | NA |
| RP23-88L6.1 | 0 | NA | NA | NA |

| | | | | |
|---|---|---|---|---|
| RP24-245O20.2 | 0 | NA | NA | NA |
| Gm6116 | 0.042716945 | -0.206490061 | 3.724945049 | -0.055434391 |
| RP23-382I10.2 | 10.57589096 | 0.150563156 | 0.380988601 | 0.395190709 |
| Chic2 | 373.5242777 | 0.260104008 | 0.104612221 | 2.486363497 |
| RP23-382I10.1 | 56.62750935 | -0.30475332 | 0.272106501 | -1.11997809 |
| RP23-382I10.3 | 0.095897396 | -0.687371727 | 3.724945049 | -0.184532045 |
| Gsx2 | 1.492637219 | 0.17576613 | 0.85182762 | 0.206340022 |
| RP23-55P22.2 | 0 | NA | NA | NA |
| RP23-55P22.3 | 0.097139547 | 0.274393828 | 3.724945049 | 0.073663859 |
| Olfr1401-ps1 | 0 | NA | NA | NA |
| Pdgfra | 784.1705611 | 0.098661984 | 0.108824125 | 0.906618673 |
| RP23-55P22.6 | 0.141219523 | -0.687378064 | 2.915099341 | -0.23579919 |
| RP23-55P22.5 | 0 | NA | NA | NA |
| Mir7025 | 0.116046057 | -0.687371727 | 3.724945049 | -0.184532045 |
| RP23-55P22.7 | 0 | NA | NA | NA |
| RP23-149H20.2 | 0 | NA | NA | NA |
| Gm22084 | 0 | NA | NA | NA |
| RP23-149H20.3 | 0 | NA | NA | NA |
| Gm24502 | 0 | NA | NA | NA |
| Kit | 814.734425 | -0.090198269 | 0.113834911 | -0.792360341 |
| RP24-273B9.2 | 0.092228164 | 0.75528439 | 3.724945049 | 0.202763901 |
| RP24-273B9.1 | 0.222682676 | -0.206485734 | 2.151237311 | -0.095984638 |
| RP24-273B9.3 | 0 | NA | NA | NA |
| RP23-329A21.3 | 0.435363789 | -0.882657371 | 1.38440396 | -0.637572122 |
| Gm38180 | 0.072025462 | 0.755288839 | 3.724945049 | 0.202765096 |
| Gm28865 | 0.066050875 | 0.274391604 | 3.724945049 | 0.073663262 |
| Gm25374 | 0 | NA | NA | NA |
| Kdr | 411.3836948 | 2.179510529 | 0.296858799 | 7.341909802 |
| Gm24738 | 0 | NA | NA | NA |
| RP24-445F15.2 | 0 | NA | NA | NA |
| RP24-445F15.1 | 2.139707088 | 0.581159175 | 0.634562574 | 0.915842186 |
| Srd5a3 | 137.1421468 | 0.372557362 | 0.113656159 | 3.277933764 |
| Tmem165 | 351.682121 | 0.002438498 | 0.096839329 | 0.025180864 |
| Apc-ps1 | 7.578865732 | 0.309579275 | 0.493565882 | 0.627229893 |
| Clock | 2177.681423 | 0.094844147 | 0.060651958 | 1.563744192 |
| RP23-386C10.8 | 47.58777522 | 0.190347523 | 0.205295286 | 0.92718896 |
| RP23-386C10.5 | 89.03088876 | 0.129650095 | 0.28353714 | 0.457259657 |
| Gm7467 | 7.824312757 | -0.488416273 | 0.376968948 | -1.295640597 |
| Pdcl2 | 0.114443069 | -0.206492286 | 3.724945049 | -0.055434988 |
| 4930432L08Rik | 0.99774422 | -0.522239718 | 1.284669787 | -0.406516697 |
| Nmu | 1.605616479 | -0.022815335 | 0.907712344 | -0.025134984 |
| RP24-186A17.1 | 0 | NA | NA | NA |
| RP24-186A17.2 | 0.068097359 | -0.206487837 | 3.724945049 | -0.055433794 |
| Gm7271 | 1.031658297 | 0.5616182 | 1.117190066 | 0.502706046 |
| Exoc1 | 794.8721072 | 0.144706207 | 0.097822506 | 1.479273157 |
| Cep135 | 200.4137152 | 0.030988225 | 0.113387533 | 0.27329482 |
| A730089K16Rik | 11.31091771 | -0.230239772 | 0.330789148 | -0.696031819 |

| | | | | |
|---|---|---|---|---|
| C530008M17Rik | 974.9986135 | -0.341895918 | 0.07225635 | -4.731707559 |
| Gm7494 | 0.044191549 | -0.206490061 | 3.724945049 | -0.055434391 |
| Aasdh | 129.6955173 | -0.60401803 | 0.170474347 | -3.543160847 |
| Ppat | 241.3568791 | -0.002705733 | 0.083775807 | -0.032297306 |
| 2310040G07Rik | 1.39740727 | 0.522609502 | 1.00598931 | 0.519498068 |
| Paics | 931.5296296 | 0.033744917 | 0.091954697 | 0.366973287 |
| Srp72 | 1194.911166 | 0.2128348 | 0.063404093 | 3.356799086 |
| Arl9 | 3.563487532 | 0.595205182 | 0.610439203 | 0.975044164 |
| 1700112J05Rik | 0 | NA | NA | NA |
| 1700023E05Rik | 5.201511052 | -0.086260761 | 0.458433042 | -0.188164362 |
| Hopx | 352.4399641 | 0.657118724 | 0.140141782 | 4.688956541 |
| Hopxos | 6.015739958 | 1.741228418 | 0.440076903 | 3.956645776 |
| Gm15831 | 0.13681464 | 0.274404728 | 2.94378999 | 0.093214777 |
| RP23-3L23.1 | 0 | NA | NA | NA |
| Spink2 | 0.769070381 | -0.20657024 | 1.411268061 | -0.146372079 |
| RP24-439P5.1 | 0 | NA | NA | NA |
| Rest | 302.1062253 | 0.799482244 | 0.137312014 | 5.822376502 |
| Mir5098 | 0.366116714 | 0.250088675 | 1.557448018 | 0.160575937 |
| Noa1 | 216.3475351 | 0.132398966 | 0.085088016 | 1.556023667 |
| Polr2b | 926.3349238 | 0.099902574 | 0.056451282 | 1.7697131 |
| RP24-439P5.3 | 1.692103223 | -0.290102095 | 0.753110421 | -0.385205259 |
| Igfbp7 | 165.7570609 | 1.110342518 | 0.159702033 | 6.952588503 |
| RP24-89O12.2 | 0 | NA | NA | NA |
| RP24-89O12.4 | 0.126225412 | 0.755282661 | 3.698754989 | 0.20419916 |
| RP24-89O12.3 | 0 | NA | NA | NA |
| RP24-89O12.5 | 0 | NA | NA | NA |
| Pea15b | 0.138038828 | 0.755294464 | 3.633333205 | 0.207879218 |
| Gm22011 | 0 | NA | NA | NA |
| RP24-358F13.1 | 0.17692102 | -0.206487546 | 2.728717236 | -0.07567202 |
| RP23-465P13.1 | 0 | NA | NA | NA |
| Mthfr-ps1 | 0.438166815 | -0.045321096 | 1.436457111 | -0.031550608 |
| RP23-135B1.1 | 0 | NA | NA | NA |
| Gm25756 | 0 | NA | NA | NA |
| RP24-318J14.1 | 0 | NA | NA | NA |
| RP24-187F3.3 | 0 | NA | NA | NA |
| Gm22553 | 0 | NA | NA | NA |
| Gm23429 | 0 | NA | NA | NA |
| Gm24304 | 0 | NA | NA | NA |
| Gm22974 | 0 | NA | NA | NA |
| Gm2199 | 0.805896106 | -1.8623996 | 1.294518539 | -1.438681289 |
| Adgrl3 | 3423.719416 | -0.044820082 | 0.083022902 | -0.539852032 |
| RP24-248M2.5 | 137.1470277 | -0.387167601 | 0.143349186 | -2.700870588 |
| RP24-74E24.5 | 20.74210032 | -0.078759351 | 0.315449901 | -0.24967309 |
| AC104196.1 | 2.384345449 | 0.893943614 | 0.603235247 | 1.48191542 |
| Gm16054 | 1.328319734 | 0.442932321 | 1.089825108 | 0.406425139 |
| RP24-82B9.1 | 52.62095602 | 0.019321216 | 0.161721591 | 0.11947209 |
| RP23-393A24.1 | 0.106465261 | 0.75528439 | 3.724945049 | 0.202763901 |

| | | | | |
|---|---|---|---|---|
| 1700031L13Rik | 0.049553047 | 0.274391604 | 3.724945049 | 0.073663262 |
| RP23-151G22.1 | 9.983061651 | 0.707473109 | 0.359653031 | 1.967098975 |
| Mir1187 | 0 | NA | NA | NA |
| Gm24129 | 0 | NA | NA | NA |
| RP24-558F5.1 | 0 | NA | NA | NA |
| Tecrl | 1.615357459 | -1.037867421 | 0.849392786 | -1.221893379 |
| Gm15626 | 0.088383099 | -0.687376175 | 3.724945049 | -0.184533239 |
| AC120867.1 | 0 | NA | NA | NA |
| Gm25765 | 0 | NA | NA | NA |
| Gm25806 | 0 | NA | NA | NA |
| Gm24792 | 0 | NA | NA | NA |
| RP23-189P23.1 | 0 | NA | NA | NA |
| Epha5 | 1720.415688 | -0.276637103 | 0.094579105 | -2.924928319 |
| Hmgn2-ps1 | 6.552657376 | 0.66497291 | 0.408134908 | 1.629296826 |
| RP24-166L23.1 | 2.846346598 | 0.155222984 | 0.748695134 | 0.207324686 |
| RP24-138J15.1 | 0 | NA | NA | NA |
| RP24-273G7.1 | 0 | NA | NA | NA |
| RP23-227I15.1 | 0.342219405 | -1.159667875 | 1.718645153 | -0.674757016 |
| Gm24626 | 0 | NA | NA | NA |
| RP23-258M22.2 | 1.514822285 | -1.485582205 | 0.844216343 | -1.759717421 |
| Cenpc1 | 463.6409797 | -0.115086004 | 0.06158538 | -1.868722811 |
| RP23-19G21.1 | 0.150451996 | 0.274405555 | 2.858243839 | 0.096004949 |
| Stap1 | 5.819746149 | 3.021396233 | 0.819688392 | 3.686030277 |
| Gm24524 | 0.085327104 | 0.274389469 | 3.724945049 | 0.073662689 |
| Uba6 | 668.4127751 | -0.327442444 | 0.065419744 | -5.005254169 |
| Gnrhr | 0.310823584 | 1.021444524 | 1.716239197 | 0.59516443 |
| Tmprss11c | 0.04393175 | -0.206490061 | 3.724945049 | -0.055434391 |
| Gm25473 | 0 | NA | NA | NA |
| Tmprss11d | 1.632697194 | -0.730719848 | 0.86632277 | -0.84347298 |
| Tmprss11a | 3.616981247 | -1.389577328 | 0.527260464 | -2.635466575 |
| Tmprss11g | 0.130746305 | 1.120259715 | 3.664569624 | 0.305700213 |
| Tmprss11f | 0.18352982 | 1.625373558 | 3.713044335 | 0.437746876 |
| Tmprss11b | 2.079765874 | 2.896724449 | 1.038398734 | 2.789607069 |
| RP23-314N3.1 | 0.090074004 | 0.274402725 | 3.724945049 | 0.073666248 |
| Tmprss11e | 0.166952883 | 0.274401811 | 2.756036668 | 0.099563919 |
| Gm16579 | 0.090277459 | -0.20649451 | 3.724945049 | -0.055435586 |
| RP24-567O8.4 | 0 | NA | NA | NA |
| RP24-567O8.3 | 0.115000406 | -0.206492286 | 3.724945049 | -0.055434988 |
| Ythdc1 | 1204.211079 | 0.241198303 | 0.107793952 | 2.237586609 |
| RP24-158N23.1 | 0 | NA | NA | NA |
| Ugt2b34 | 1.309248412 | 3.305737097 | 1.161100041 | 2.847073448 |
| Gm25211 | 0 | NA | NA | NA |
| Ugt2b1 | 0 | NA | NA | NA |
| Gm24121 | 0 | NA | NA | NA |
| RP23-59B8.1 | 0 | NA | NA | NA |
| RP23-227D7.1 | 0.092228164 | 0.75528439 | 3.724945049 | 0.202763901 |
| RP23-59B8.2 | 33.07073811 | 0.336760019 | 0.242618752 | 1.388021397 |

| | | | | |
|---|---|---|---|---|
| Ugt2b35 | 0.394550875 | 1.554533879 | 1.432115503 | 1.085480798 |
| RP23-59B8.3 | 0 | NA | NA | NA |
| RP23-227D7.2 | 0 | NA | NA | NA |
| Ugt2b36 | 0.256447508 | 1.236196794 | 2.47295682 | 0.499886122 |
| RP23-227D7.3 | 0 | NA | NA | NA |
| RP23-67B9.4 | 0 | NA | NA | NA |
| Ugt2b5 | 0.491974008 | 1.199105169 | 1.384189143 | 0.866287079 |
| RP23-67B9.2 | 0.045882455 | 0.755288839 | 3.724945049 | 0.202765096 |
| RP23-227D7.4 | 0 | NA | NA | NA |
| RP23-67B9.1 | 0.072025462 | 0.755288839 | 3.724945049 | 0.202765096 |
| Ugt2b37 | 0.046345709 | 0.274391604 | 3.724945049 | 0.073663262 |
| RP23-67B9.3 | 0.075043205 | 0.755286614 | 3.724945049 | 0.202764498 |
| Ugt2a3 | 0.072025462 | 0.755288839 | 3.724945049 | 0.202765096 |
| RP23-227D7.7 | 0 | NA | NA | NA |
| RP23-67B9.5 | 0 | NA | NA | NA |
| RP23-227D7.8 | 0.093141081 | -0.687373951 | 3.724945049 | -0.184532642 |
| RP23-227D7.6 | 0 | NA | NA | NA |
| RP23-227D7.5 | 0 | NA | NA | NA |
| Ugt2b38 | 0.123841226 | 0.274391173 | 3.724945049 | 0.073663147 |
| Ugt2a2 | 0.062426873 | 0.755288839 | 3.724945049 | 0.202765096 |
| Ugt2a1 | 0 | NA | NA | NA |
| RP23-227D7.10 | 0 | NA | NA | NA |
| Sult1b1 | 0.042716945 | -0.206490061 | 3.724945049 | -0.055434391 |
| Sult1d1 | 1.935656465 | -1.310163844 | 1.172747699 | -1.117174517 |
| Sult1e1 | 0.061920613 | 0.274393828 | 3.724945049 | 0.073663859 |
| mmu-mir-3969 | 0 | NA | NA | NA |
| RP24-571N6.2 | 0 | NA | NA | NA |
| RP24-571N6.3 | 0 | NA | NA | NA |
| Csn1s1 | 0.044544648 | 0.755286614 | 3.724945049 | 0.202764498 |
| Csn2 | 0 | NA | NA | NA |
| Gm25066 | 0 | NA | NA | NA |
| RP23-440C1.3 | 0.266205562 | 1.63692817 | 1.791280146 | 0.91383147 |
| Gm24314 | 0 | NA | NA | NA |
| Csn1s2a | 0.075043205 | 0.755286614 | 3.724945049 | 0.202764498 |
| Csn1s2b | 0 | NA | NA | NA |
| Prr27 | 0 | NA | NA | NA |
| RP23-440C1.5 | 0 | NA | NA | NA |
| Gm7337 | 0 | NA | NA | NA |
| Odam | 0 | NA | NA | NA |
| 1700066N21Rik | 0 | NA | NA | NA |
| RP23-423C8.4 | 0 | NA | NA | NA |
| Csn3 | 0.488468934 | 0.75531038 | 1.432048631 | 0.527433471 |
| BC051076 | 0.204292078 | -1.244198191 | 3.71120435 | -0.335254562 |
| 2310003L06Rik | 0.63240349 | 0.977080035 | 1.264067691 | 0.772964962 |
| Gm28434 | 0 | NA | NA | NA |
| Cabs1 | 0.109869311 | -0.206490066 | 3.724945049 | -0.055434393 |
| Smr3a | 0 | NA | NA | NA |

| | | | | |
|---|---|---|---|---|
| RP24-324D14.5 | 0 | NA | NA | NA |
| RP24-324D14.6 | 0 | NA | NA | NA |
| Smr2 | 0 | NA | NA | NA |
| Gm11116 | 0 | NA | NA | NA |
| RP23-50G20.2 | 0 | NA | NA | NA |
| Gm11115 | 0 | NA | NA | NA |
| RP23-50G20.3 | 0 | NA | NA | NA |
| RP23-50G20.5 | 0 | NA | NA | NA |
| RP23-50G20.4 | 0 | NA | NA | NA |
| Gm7714 | 0 | NA | NA | NA |
| Prol1 | 0 | NA | NA | NA |
| Amtn | 0 | NA | NA | NA |
| RP23-265M4.3 | 1.532002984 | -1.063232753 | 0.973199125 | -1.092513059 |
| RP23-265M4.4 | 0 | NA | NA | NA |
| Ambn | 0 | NA | NA | NA |
| Enam | 0 | NA | NA | NA |
| Jchain | 15.78987055 | 0.685829954 | 0.593354767 | 1.155851427 |
| RP24-132B11.1 | 0.854816446 | -0.711751577 | 1.088205574 | -0.654059852 |
| Utp3 | 218.2673712 | 0.502454089 | 0.184001794 | 2.73070212 |
| Rufy3 | 2816.589347 | 0.043814559 | 0.050764246 | 0.863098794 |
| Grsf1 | 1026.519618 | 0.1779792 | 0.061115556 | 2.912175089 |
| Mob1b | 844.5626815 | 0.229196033 | 0.120045349 | 1.909245417 |
| Gm15682 | 3.374206779 | 0.278151608 | 0.669195857 | 0.415650523 |
| Dck | 405.0431763 | 0.499858488 | 0.112389692 | 4.44754746 |
| RP24-242L17.1 | 22.06035809 | -0.93249508 | 0.375853295 | -2.481008127 |
| RP24-242L17.2 | 0 | NA | NA | NA |
| Mkrn1-ps1 | 1.85792581 | -0.046555964 | 0.721181953 | -0.064555087 |
| Slc4a4 | 4189.981023 | -0.507521167 | 0.132868352 | -3.819729539 |
| Gc | 1.073168736 | -0.268666568 | 1.245841595 | -0.215650665 |
| RP24-365I5.1 | 0.045882455 | 0.755288839 | 3.724945049 | 0.202765096 |
| RP24-365I5.2 | 0 | NA | NA | NA |
| Npffr2 | 7.733873509 | -1.365812673 | 0.486806263 | -2.805659617 |
| Adamts3 | 373.7060012 | -0.171798889 | 0.210220823 | -0.8172306 |
| Gm25320 | 0.186976534 | 1.236194859 | 2.668791307 | 0.463204019 |
| Gm25758 | 0 | NA | NA | NA |
| Eif5al3-ps | 2.402331814 | 1.630568816 | 0.77733298 | 2.097645229 |
| Cox18 | 236.2894485 | 0.69639227 | 0.110084568 | 6.325975427 |
| Ankrd17 | 4578.10987 | 0.177083362 | 0.062590126 | 2.829253944 |
| Gm9958 | 14.24586082 | -0.835194935 | 0.293779331 | -2.84293293 |
| RP23-279P6.5 | 0 | NA | NA | NA |
| Alb | 7.938965767 | 0.23171093 | 0.593485953 | 0.390423614 |
| Afp | 3.204684021 | 0.943562599 | 0.695424434 | 1.356815425 |
| RP23-279P6.3 | 0.060174537 | 0.755286614 | 3.724945049 | 0.202764498 |
| Afm | 0.397459543 | 0.274411973 | 1.762534705 | 0.155691671 |
| RP24-63O24.1 | 0.290146865 | -0.635210674 | 2.027255783 | -0.313335238 |
| RP24-63O24.4 | 0 | NA | NA | NA |
| 5830473C10Rik | 0.606733601 | 1.008253226 | 1.244258167 | 0.81032478 |

| | | | | |
|---|---|---|---|---|
| Rassf6 | 13.93087578 | -0.555439172 | 0.295562345 | -1.879262296 |
| RP24-168A7.1 | 0.237346126 | 0.776241243 | 2.118027027 | 0.366492605 |
| Cxcl5 | 7.088183414 | 0.007330192 | 0.528217895 | 0.013877212 |
| Ppbp | 1.785008614 | 0.408282748 | 0.88026527 | 0.463817854 |
| Pf4 | 3.421604734 | -1.402651077 | 0.664947445 | -2.109416448 |
| Cxcl3 | 3.225786153 | 2.092786394 | 1.218951268 | 1.716874537 |
| Cxcl15 | 0.068097359 | -0.206487837 | 3.724945049 | -0.055433794 |
| Gm24322 | 0 | NA | NA | NA |
| RP24-168A7.7 | 0 | NA | NA | NA |
| RP23-307P18.1 | 0 | NA | NA | NA |
| Cxcl1 | 39.73533034 | 6.491330382 | 1.042238218 | 6.228259785 |
| Gm22816 | 0.235449268 | 1.731321027 | 2.531872866 | 0.683810412 |
| Cxcl2 | 19.17032838 | 6.684058095 | 1.168675849 | 5.719343052 |
| RP23-307P18.3 | 0 | NA | NA | NA |
| Mthfd2l | 111.8726162 | -0.217743864 | 0.119477775 | -1.822463325 |
| Epgn | 0.633611781 | -1.093111607 | 1.340036223 | -0.815732879 |
| RP23-42K10.4 | 0.186036187 | -0.206489511 | 2.713779053 | -0.076089286 |
| Ereg | 4.701452776 | 3.613866585 | 1.113018577 | 3.2469059 |
| Areg | 0.714414789 | 2.361791682 | 1.328045861 | 1.778396177 |
| Gm23092 | 5.083391071 | 0.072952576 | 0.520802438 | 0.140077255 |
| RP23-258C6.2 | 6.099374268 | 1.031998118 | 0.41560652 | 2.483113399 |
| Btc | 22.83547764 | 1.378602962 | 0.321761757 | 4.284545732 |
| Parm1 | 754.069321 | 0.039903393 | 0.057730974 | 0.69119556 |
| RP23-118H9.3 | 3.970608911 | 0.495939645 | 0.543724307 | 0.912116009 |
| RP23-17D18.3 | 4.213956554 | -0.227683841 | 0.481220259 | -0.473138519 |
| Gm28271 | 0.521165882 | -0.189976517 | 1.565288523 | -0.121368371 |
| 1700063O14Rik | 0.323348989 | -0.577982469 | 1.734225183 | -0.333279943 |
| RP24-266G11.2 | 0.541747793 | 1.682172984 | 1.545730755 | 1.088270372 |
| RP24-266G11.4 | 0 | NA | NA | NA |
| Gm5558 | 5.850622695 | -0.558550064 | 0.483939333 | -1.154173729 |
| RP23-442L7.2 | 0.329047954 | 2.077623354 | 1.72251734 | 1.206155263 |
| Rchy1 | 427.1403147 | 0.286951595 | 0.088060941 | 3.258557 |
| RP23-442L7.3 | 26.15808167 | -0.278743222 | 0.315137083 | -0.884514191 |
| Thap6 | 30.14759305 | 0.353301665 | 0.198890653 | 1.776361334 |
| Gm24113 | 0.102891481 | 0.274400501 | 3.724945049 | 0.073665651 |
| Gm1045 | 3.240137224 | 1.367946705 | 0.65055086 | 2.102751358 |
| Cdkl2 | 400.6574654 | -0.340456385 | 0.092016975 | -3.699930216 |
| Gm26582 | 7.944179279 | -0.480764929 | 0.361832496 | -1.328694727 |
| G3bp2 | 4286.859093 | 0.523327941 | 0.102619131 | 5.099711274 |
| Gm22728 | 6.520177888 | 1.452823162 | 0.476403993 | 3.04956126 |
| Gm24931 | 1.274348854 | -0.210117488 | 0.810126505 | -0.259363799 |
| Uso1 | 1247.998467 | -0.010066079 | 0.05554596 | -0.181220726 |
| Gm15710 | 147.9457653 | 0.205785394 | 0.158103453 | 1.301586971 |
| RP23-321C24.2 | 0 | NA | NA | NA |
| Gm25290 | 1.111053339 | -1.362385637 | 0.990833916 | -1.374988901 |
| U90926 | 0.097139547 | 0.274393828 | 3.724945049 | 0.073663859 |
| Ppef2 | 8.480238973 | -1.692438351 | 0.415492335 | -4.073332303 |

| | | | | |
|---|---|---|---|---|
| Naaa | 229.011174 | 0.610359761 | 0.144771524 | 4.216020832 |
| Sdad1 | 341.6351534 | 0.529670591 | 0.108074172 | 4.900991435 |
| RP23-447A7.2 | 0.049553047 | 0.274391604 | 3.724945049 | 0.073663262 |
| Gm23031 | 0.045882455 | 0.755288839 | 3.724945049 | 0.202765096 |
| Cxcl9 | 597.6617564 | 12.04803288 | 0.936288099 | 12.86786929 |
| Art3 | 40.01616527 | 1.21907582 | 0.195895317 | 6.223098325 |
| Cxcl10 | 3859.437308 | 11.86627287 | 0.713110118 | 16.64016898 |
| Cxcl11 | 885.873838 | 8.640438616 | 0.606426309 | 14.24812625 |
| Gm2673 | 0.689365755 | 1.236208226 | 1.340343768 | 0.92230684 |
| Nup54 | 228.7748306 | 0.169229819 | 0.123442543 | 1.370919745 |
| Gm26039 | 0 | NA | NA | NA |
| Scarb2 | 2989.989026 | 0.508099191 | 0.085587884 | 5.936578475 |
| Mir6415 | 0 | NA | NA | NA |
| Fam47e | 1.965355678 | -0.561410502 | 0.688617547 | -0.815271851 |
| Stbd1 | 13.78635077 | -0.029248909 | 0.387210893 | -0.075537413 |
| Ccdc158 | 6.083259113 | -0.92244215 | 0.436308008 | -2.114199448 |
| Shroom3 | 115.9678624 | 0.334911651 | 0.187314521 | 1.787964162 |
| Gm20500 | 0.399554685 | -0.69557456 | 1.542124333 | -0.451049598 |
| Mir1961 | 0 | NA | NA | NA |
| 4932430I15Rik | 0 | NA | NA | NA |
| RP23-9D24.2 | 1.173754143 | -1.400894765 | 1.019787807 | -1.373712016 |
| Gm25521 | 0.066050875 | 0.274391604 | 3.724945049 | 0.073663262 |
| Sowahb | 261.7368774 | -0.052632159 | 0.131390741 | -0.400577386 |
| Gm22915 | 1.677596521 | -1.196844602 | 0.837162363 | -1.42964454 |
| RP24-470E5.2 | 19.84950728 | -0.109379357 | 0.370529881 | -0.29519713 |
| RP24-470E5.3 | 3.144773293 | -0.742018345 | 0.602053516 | -1.232479049 |
| 11-Sep | 1923.005761 | -0.383195991 | 0.047159176 | -8.125587081 |
| RP24-470E5.4 | 138.2382822 | -0.235975754 | 0.150685701 | -1.566012919 |
| Ccni | 2707.469122 | -0.017336319 | 0.087337579 | -0.198497826 |
| RP24-470E5.9 | 57.10845697 | -0.63403949 | 0.236428496 | -2.681738884 |
| Gm2986 | 3.888599492 | 0.123603327 | 0.582058532 | 0.212355493 |
| 2010109A12Rik | 6.375499996 | -1.096415388 | 0.559018202 | -1.961323234 |
| Ccng2 | 404.8129049 | -0.475214117 | 0.104601318 | -4.543098739 |
| Kat2b-ps | 1.175757053 | 0.644570619 | 1.140339133 | 0.565244672 |
| RP23-388L12.2 | 0 | NA | NA | NA |
| RP23-173B14.3 | 0 | NA | NA | NA |
| Gm16427 | 0 | NA | NA | NA |
| C87414 | 0 | NA | NA | NA |
| RP23-388L12.3 | 0 | NA | NA | NA |
| RP23-388L12.1 | 0 | NA | NA | NA |
| RP23-57N5.1 | 0 | NA | NA | NA |
| RP23-173B14.4 | 0 | NA | NA | NA |
| RP23-57N5.3 | 0 | NA | NA | NA |
| Gm16429 | 0 | NA | NA | NA |
| RP23-173B14.2 | 0 | NA | NA | NA |
| RP23-173B14.1 | 0 | NA | NA | NA |
| BC061212 | 0 | NA | NA | NA |

| | | | | |
|---|---|---|---|---|
| Gm6351 | 0 | NA | NA | NA |
| Gm3106 | 0 | NA | NA | NA |
| Gm16522 | 0 | NA | NA | NA |
| Gm6502 | 0 | NA | NA | NA |
| AA792892 | 0 | NA | NA | NA |
| RP23-432M17.3 | 0 | NA | NA | NA |
| Gm7682 | 0 | NA | NA | NA |
| Gm3139 | 0 | NA | NA | NA |
| Gm3147 | 0 | NA | NA | NA |
| Gm6509 | 0 | NA | NA | NA |
| Gm6205 | 0 | NA | NA | NA |
| RP23-133M13.3 | 0 | NA | NA | NA |
| RP23-432M17.5 | 0 | NA | NA | NA |
| Gm3183 | 0 | NA | NA | NA |
| RP24-85B14.2 | 0 | NA | NA | NA |
| Gm7647 | 0 | NA | NA | NA |
| Gm6367 | 0 | NA | NA | NA |
| RP23-365E3.3 | 0 | NA | NA | NA |
| RP24-85B14.3 | 0 | NA | NA | NA |
| Gm7942 | 0 | NA | NA | NA |
| Gm16513 | 0 | NA | NA | NA |
| Gm10424 | 0 | NA | NA | NA |
| RP23-320H13.5 | 0 | NA | NA | NA |
| Gm3259 | 0 | NA | NA | NA |
| RP23-365E3.2 | 0 | NA | NA | NA |
| RP23-320H13.4 | 0 | NA | NA | NA |
| RP23-320H13.3 | 0 | NA | NA | NA |
| D5Ertd577e | 0 | NA | NA | NA |
| Gm3286 | 0 | NA | NA | NA |
| Gm3176 | 0 | NA | NA | NA |
| RP23-23P3.2 | 0 | NA | NA | NA |
| RP23-23P3.1 | 0 | NA | NA | NA |
| Gm7978 | 0 | NA | NA | NA |
| Gm7982 | 0 | NA | NA | NA |
| E330014E10Rik | 0 | NA | NA | NA |
| RP24-322E6.3 | 0 | NA | NA | NA |
| Gm5559 | 314.382447 | 0.201271281 | 0.129324524 | 1.556327248 |
| Gm25937 | 0 | NA | NA | NA |
| Cxcl13 | 78.29238741 | 7.377640548 | 0.656693915 | 11.23451943 |
| RP23-57A17.2 | 106.8115065 | 0.911696375 | 0.212711755 | 4.286064837 |
| Cnot6l | 1535.89927 | 0.27412765 | 0.072550852 | 3.77842084 |
| Gm2840 | 2.621041726 | -1.509097496 | 0.769294602 | -1.961663961 |
| Mrpl1 | 300.8320522 | 0.198215782 | 0.077458247 | 2.55900166 |
| Fras1 | 497.6484057 | -0.822400297 | 0.172331386 | -4.772202662 |
| RP23-381C14.2 | 0.206671645 | -0.206481294 | 2.63660209 | -0.078313407 |
| RP23-381C14.5 | 0.680615379 | -0.46004518 | 1.289957068 | -0.356636039 |
| RP23-480P21.2 | 0.07796279 | 0.274392189 | 3.724945049 | 0.073663419 |

| | | | | |
|---|---|---|---|---|
| Anxa3 | 185.5288032 | 2.093316334 | 0.240054461 | 8.720172601 |
| RP23-480P21.3 | 7.010785502 | 0.614681219 | 0.581822355 | 1.056475766 |
| RP24-142B15.2 | 0.147290723 | 1.192974065 | 2.861512785 | 0.41690328 |
| RP24-142B15.3 | 0.270258959 | 1.236204576 | 1.81862319 | 0.679747505 |
| RP24-142B15.4 | 0.176646398 | 0.755284509 | 3.360121616 | 0.22477892 |
| Gm24623 | 0 | NA | NA | NA |
| RP24-142B15.5 | 2.750814746 | 3.123629764 | 0.991867863 | 3.149239814 |
| RP24-142B15.6 | 0.520715546 | 1.249664313 | 1.580806796 | 0.790523115 |
| RP24-142B15.7 | 1.115544105 | 0.103959532 | 0.907230648 | 0.114589969 |
| RP24-142B15.8 | 0.895791331 | -0.24444838 | 0.997788809 | -0.2449901 |
| RP24-142B15.9 | 0.252544957 | -0.206485684 | 2.109116732 | -0.097901496 |
| RP24-142B15.11 | 5.79381536 | 0.511511956 | 0.473896419 | 1.079375019 |
| Bmp2k | 887.8164416 | 1.025394665 | 0.107218921 | 9.563560784 |
| RP24-142B15.10 | 11.84488042 | 0.940021938 | 0.429746389 | 2.187387638 |
| Paqr3 | 238.9609811 | -0.013844383 | 0.148429161 | -0.09327266 |
| RP23-45P1.1 | 1.824918576 | -0.082596052 | 0.637564496 | -0.129549329 |
| RP23-28F1.3 | 1.733684669 | 0.418796386 | 0.939119846 | 0.445945624 |
| Gm5560 | 26.0495091 | 0.002189705 | 0.260471483 | 0.008406698 |
| RP23-28F1.4 | 0 | NA | NA | NA |
| Naa11 | 0.23191841 | -0.691815593 | 2.580425347 | -0.268101379 |
| 4930467D21Rik | 11.74611447 | 0.15058272 | 0.304349488 | 0.494769092 |
| Gk2 | 0.04393175 | -0.206490061 | 3.724945049 | -0.055434391 |
| Antxr2 | 68.57005387 | 0.480454317 | 0.210196333 | 2.285740721 |
| RP23-178N8.7 | 1.361532721 | 1.491744555 | 0.991660862 | 1.504289028 |
| RP23-178N8.3 | 0.042716945 | -0.206490061 | 3.724945049 | -0.055434391 |
| RP23-178N8.4 | 0 | NA | NA | NA |
| RP23-178N8.5 | 0.202865515 | 0.274409289 | 2.223190743 | 0.123430385 |
| Gm8048 | 0.123841226 | 0.274391173 | 3.724945049 | 0.073663147 |
| Prdm8 | 199.5149206 | 0.018204929 | 0.134169294 | 0.135686255 |
| A730035I17Rik | 9.47161779 | 0.651883 | 0.359831126 | 1.811635941 |
| Fgf5 | 19.39898948 | -0.242752385 | 0.271780711 | -0.893192103 |
| 1700007G11Rik | 3.95741196 | -0.174268762 | 0.524150892 | -0.332478232 |
| RP23-425F8.1 | 1.376239274 | -1.791665582 | 1.139310217 | -1.572588006 |
| RP24-240P8.1 | 0.208122295 | 1.22105634 | 2.194123754 | 0.556512064 |
| mmu-mir-703 | 13.41816306 | -0.381760043 | 0.305371895 | -1.250147933 |
| Gm24279 | 0.141779851 | -0.180471455 | 3.595043549 | -0.050200075 |
| Gm11111 | 0.419266243 | 1.557044813 | 1.346466999 | 1.156392851 |
| 1700010H22Rik | 0 | NA | NA | NA |
| Bmp3 | 131.1088595 | -0.706251855 | 0.157239671 | -4.491562787 |
| Prkg2 | 141.6564417 | -0.374402312 | 0.134588201 | -2.781836073 |
| Gm3470 | 1.546431532 | 0.205055228 | 0.775904679 | 0.26427889 |
| RP24-548B7.1 | 0.424196832 | 1.256586902 | 1.6847285 | 0.745869083 |
| Rasgef1b | 664.9745574 | 0.368761378 | 0.073018832 | 5.050222935 |
| RP23-253I14.3 | 18.40400765 | 0.136108174 | 0.276242597 | 0.492712478 |
| A930011G23Rik | 101.3913818 | -0.263746702 | 0.291050595 | -0.906188501 |
| RP23-380D12.1 | 0.13861434 | -1.111919598 | 3.614496194 | -0.307627824 |
| Gm16227 | 6.434094977 | -0.591213932 | 0.448453407 | -1.318339704 |

| | | | | |
|---|---|---|---|---|
| Gm16228 | 3.733916757 | -0.57651554 | 0.541748102 | -1.06417639 |
| 4930405H06Rik | 0 | NA | NA | NA |
| Gm25969 | 0 | NA | NA | NA |
| Gm16226 | 0 | NA | NA | NA |
| 4930522N08Rik | 0 | NA | NA | NA |
| RP23-438O22.1 | 33.98607118 | -0.229240119 | 0.350697373 | -0.653669335 |
| RP23-57P23.3 | 0.038981395 | 0.274393828 | 3.724945049 | 0.073663859 |
| Gm26252 | 0 | NA | NA | NA |
| RP23-57P23.4 | 0.879409403 | 0.403240157 | 1.248759779 | 0.322912512 |
| Gm22838 | 0.115603922 | 0.274387056 | 3.724945049 | 0.073662041 |
| RP23-57P23.5 | 0.082913145 | -0.206492286 | 3.724945049 | -0.055434988 |
| Hnrnpd | 2150.330006 | 0.171836071 | 0.079518036 | 2.160969746 |
| Gm17092 | 3.730625592 | 0.709438495 | 0.548366 | 1.293731731 |
| 4930524J08Rik | 4.763368001 | 0.299742216 | 0.516025481 | 0.580867084 |
| Hnrnpdl | 2079.483625 | 0.323112343 | 0.084068385 | 3.843446533 |
| Enoph1 | 302.3695605 | -0.163931498 | 0.094216801 | -1.739939118 |
| Gm26363 | 0 | NA | NA | NA |
| Tmem150c | 388.2756464 | -0.161726737 | 0.071413393 | -2.264655534 |
| Tmem150cos | 0.555352798 | 2.510261636 | 1.342039688 | 1.870482414 |
| Gm9932 | 0.452014659 | -0.212632675 | 1.537988461 | -0.138253752 |
| 2310034O05Rik | 0.531171913 | 0.39832854 | 1.507899212 | 0.264161249 |
| Gm8091 | 0.417821482 | -0.916140844 | 1.334430183 | -0.686540859 |
| Sec31a | 1485.020543 | 0.015443751 | 0.040317334 | 0.383054878 |
| 5430416N02Rik | 76.01050414 | 0.275470777 | 0.176175104 | 1.563619209 |
| Gm27553 | 0.170081438 | -1.075631546 | 2.769839891 | -0.388337084 |
| Gm27997 | 0.038981395 | 0.274393828 | 3.724945049 | 0.073663859 |
| Gm28030 | 0 | NA | NA | NA |
| Gm27286 | 0.243228626 | -0.206488331 | 3.033562892 | -0.068067925 |
| RP23-325M7.4 | 4.625370955 | -0.069524066 | 0.550339994 | -0.126329299 |
| Lin54 | 315.1002167 | 0.099148052 | 0.097559703 | 1.01628079 |
| Gm23222 | 0.640027454 | -0.75553723 | 1.454623055 | -0.519404135 |
| RP23-313F5.1 | 0 | NA | NA | NA |
| Gm23173 | 0 | NA | NA | NA |
| Cops4 | 680.1477187 | 0.306243508 | 0.063257477 | 4.841222299 |
| Plac8 | 627.7400569 | 9.96179759 | 0.920496808 | 10.82219678 |
| RP23-313F5.2 | 0 | NA | NA | NA |
| Coq2 | 258.0378681 | 0.217528784 | 0.082754695 | 2.62859748 |
| Gm15777 | 0.230076657 | 1.606735026 | 3.108237438 | 0.516928021 |
| Hpse | 70.70159234 | 2.392817503 | 0.249746272 | 9.580993874 |
| Helq | 242.9045317 | -0.009212401 | 0.151304801 | -0.060886379 |
| Mrps18c | 124.8778561 | 0.53285529 | 0.14878756 | 3.58131613 |
| RP23-7M16.1 | 22.40968198 | -0.070679266 | 0.273579206 | -0.258350285 |
| Fam175a | 72.7916956 | -0.233516009 | 0.145459172 | -1.605371494 |
| RP23-7M16.2 | 5.191824483 | -0.968462581 | 0.550615092 | -1.758874021 |
| RP23-7M16.3 | 1.089839937 | 0.804437844 | 1.135593862 | 0.708385163 |
| Gm26286 | 0.158374819 | -0.687378027 | 2.812678226 | -0.24438559 |
| 4930458D05Rik | 1.568767244 | 0.459080383 | 0.831903637 | 0.551843221 |

| | | | | |
|---|---|---|---|---|
| Agpat9 | 94.53062151 | 0.317339503 | 0.220463115 | 1.439422206 |
| RP23-99A9.1 | 2.620592534 | -0.1630257 | 0.667385725 | -0.244275078 |
| RP23-99A9.2 | 0.075043205 | 0.755286614 | 3.724945049 | 0.202764498 |
| RP23-302F24.1 | 0.045882455 | 0.755288839 | 3.724945049 | 0.202765096 |
| RP23-336N16.1 | 0 | NA | NA | NA |
| Gm20484 | 0 | NA | NA | NA |
| Nkx6-1 | 10.48168021 | -0.364574484 | 0.550736935 | -0.661975729 |
| RP24-292F8.2 | 0 | NA | NA | NA |
| RP24-292F8.1 | 1.593328872 | -0.485569594 | 0.898854235 | -0.540209497 |
| Cds1 | 1169.603167 | 0.23982067 | 0.072627825 | 3.30204944 |
| RP24-292F8.3 | 17.79350931 | -0.655569826 | 0.330437691 | -1.983943857 |
| WI1-1010L24.1 | 36.41421054 | -0.063751983 | 0.27816659 | -0.229186341 |
| Wdfy3 | 5687.595008 | -0.247220506 | 0.060424588 | -4.091389205 |
| Gm20548 | 3.946887117 | 0.739554515 | 0.53393704 | 1.385096855 |
| RP24-491C20.4 | 0.127971488 | 0.274389832 | 3.687653711 | 0.0744077 |
| RP23-409I5.1 | 28.85031586 | -0.06165909 | 0.328976496 | -0.187427039 |
| RP23-409I5.2 | 1.952620281 | -0.981769488 | 0.738104854 | -1.330121978 |
| RP23-409I5.3 | 0.494178821 | 1.419347463 | 1.377346699 | 1.030493966 |
| RP23-409I5.4 | 2.040397007 | 0.087203171 | 0.67966407 | 0.128303341 |
| RP23-260N17.2 | 0.650505631 | -1.61334948 | 1.233921224 | -1.307497957 |
| RP23-260N17.1 | 0.199035941 | 0.755299442 | 3.217008196 | 0.234783189 |
| Arhgap24 | 215.5874552 | 0.562667486 | 0.109717964 | 5.128307747 |
| Mapk10 | 3784.726759 | -0.076109332 | 0.075967644 | -1.0018651 |
| RP23-286M22.2 | 10.7259886 | -0.20281284 | 0.337645642 | -0.600667724 |
| RP23-286M22.1 | 125.4642517 | 0.254277743 | 0.257019526 | 0.989332396 |
| RP24-212G3.4 | 0 | NA | NA | NA |
| RP24-212G3.3 | 0 | NA | NA | NA |
| RP24-270K7.2 | 0 | NA | NA | NA |
| RP24-270K7.1 | 0 | NA | NA | NA |
| RP24-270K7.3 | 2.638949413 | 0.939339468 | 0.789706617 | 1.189479039 |
| Ptpn13 | 508.0720606 | 0.292187628 | 0.094920143 | 3.078246814 |
| RP24-102B13.3 | 0 | NA | NA | NA |
| Gm23085 | 0.072025462 | 0.755288839 | 3.724945049 | 0.202765096 |
| Slc10a6 | 36.57358822 | 8.044986935 | 1.1688342 | 6.882915419 |
| Gm15844 | 0 | NA | NA | NA |
| RP24-102B13.5 | 0.358939267 | -0.692281286 | 1.475661706 | -0.469132785 |
| 1700016H13Rik | 1.060754565 | 0.663611468 | 1.240414967 | 0.534991503 |
| Gm25721 | 0 | NA | NA | NA |
| Aff1 | 1183.051492 | 1.266215398 | 0.154390849 | 8.20136301 |
| Klhl8 | 320.9854992 | -0.439822709 | 0.084355915 | -5.213892944 |
| RP23-337H1.2 | 4.609961798 | -1.124343329 | 0.598363662 | -1.879030097 |
| Gm24897 | 0 | NA | NA | NA |
| Hsd17b13 | 1.339240761 | 0.80079085 | 0.869629298 | 0.920841619 |
| Gm8200 | 0 | NA | NA | NA |
| Hsd17b11 | 306.1707364 | 0.117481489 | 0.095948575 | 1.224421404 |
| RP23-102H7.7 | 6.166014486 | -0.504789624 | 0.40472835 | -1.247230701 |
| Nudt9 | 452.8860333 | 0.354312297 | 0.103863073 | 3.411340393 |

| | | | | |
|---|---|---|---|---|
| Mir5619 | 0.706746895 | 0.048260141 | 1.137924126 | 0.042410685 |
| Gm17660 | 2.469786297 | 0.557071364 | 0.656720337 | 0.848262696 |
| RP23-102H7.9 | 2.342408071 | -0.147359598 | 0.663954434 | -0.221942336 |
| Sparcl1 | 15907.62479 | -0.153209814 | 0.098453492 | -1.556164349 |
| Gm26703 | 13.10610984 | -0.322498277 | 0.344670015 | -0.935672565 |
| Dspp | 1.924149456 | -0.663965696 | 0.833842964 | -0.79627187 |
| Dmp1 | 39.86314599 | -0.166655212 | 0.23199873 | -0.718345363 |
| RP24-191K3.1 | 0.046345709 | 0.274391604 | 3.724945049 | 0.073663262 |
| Gm23616 | 0 | NA | NA | NA |
| Ibsp | 1.789549342 | 0.056730968 | 0.854239688 | 0.066411065 |
| Mepe | 0 | NA | NA | NA |
| Spp1 | 1206.760665 | 2.808636335 | 0.321404508 | 8.738633911 |
| RP24-299A7.2 | 2.453753307 | -0.699001976 | 1.003651635 | -0.696458763 |
| Pkd2 | 529.8119712 | -0.213506144 | 0.113194381 | -1.886190315 |
| BC005561 | 622.8318008 | -0.251270082 | 0.116315055 | -2.160254158 |
| D930016D06Rik | 256.987905 | -0.687840908 | 0.154378818 | -4.455539424 |
| Gm8258 | 4.524171958 | 0.582620497 | 0.495643599 | 1.175482742 |
| Zfp951 | 41.98758719 | -0.322985483 | 0.206715954 | -1.562460358 |
| Abcg3 | 1.390608906 | 1.461438383 | 1.107009083 | 1.320168376 |
| Gm24097 | 0 | NA | NA | NA |
| RP23-370D2.2 | 0 | NA | NA | NA |
| RP23-370D2.5 | 0 | NA | NA | NA |
| RP23-370D2.4 | 0.124853745 | 1.290908285 | 3.721447108 | 0.346883416 |
| Gbp8 | 237.2535119 | 7.824300964 | 0.601083999 | 13.01698427 |
| Gbp9 | 1809.026709 | 6.678667544 | 0.252504181 | 26.4497305 |
| Gbp4 | 4374.309881 | 9.517562929 | 0.383326363 | 24.8288765 |
| Gbp6 | 1754.483529 | 7.698606612 | 0.28091392 | 27.40557182 |
| Gbp10 | 886.6277886 | 9.237791466 | 0.557948649 | 16.55670551 |
| Gbp11 | 1003.541595 | 5.936680926 | 0.310530122 | 19.11789071 |
| Gm26519 | 1.456550475 | 1.681195212 | 0.965007201 | 1.742158205 |
| RP24-63G23.2 | 1.81533978 | 2.559560113 | 0.96687561 | 2.64724861 |
| Lrrc8b | 1659.203879 | -0.087614708 | 0.066626311 | -1.315016636 |
| RP23-41J13.2 | 3.634356604 | 1.330600083 | 0.611083312 | 2.17744464 |
| Lrrc8c | 697.388411 | 0.259253098 | 0.108235767 | 2.395262717 |
| C230066G23Rik | 8.141609263 | -0.090811444 | 0.435596106 | -0.208476252 |
| Lrrc8d | 580.2514977 | 0.216273562 | 0.079256468 | 2.728781225 |
| Rps15a-ps5 | 17.05466322 | 0.616435193 | 0.287364239 | 2.145135368 |
| Lrrc8dos | 0.047948698 | -0.206487837 | 3.724945049 | -0.055433794 |
| Gm10359 | 23.82378434 | -0.08603833 | 0.254974379 | -0.337439121 |
| RP24-354P12.2 | 0.324285546 | 2.124971439 | 3.702490672 | 0.573930261 |
| Zfp326 | 429.4665804 | -0.259258059 | 0.09650828 | -2.686381511 |
| RP24-354P12.3 | 23.94047403 | -0.208745149 | 0.304915073 | -0.684600952 |
| RP24-400H4.1 | 0 | NA | NA | NA |
| Gm25672 | 0 | NA | NA | NA |
| RP24-476K21.1 | 0.34546946 | -0.206485004 | 1.741738153 | -0.118551117 |
| RP23-424O22.1 | 0.204914957 | 1.221428882 | 2.200579739 | 0.555048681 |
| RP23-424O22.2 | 0.075043205 | 0.755286614 | 3.724945049 | 0.202764498 |

| | | | | |
|---|---|---|---|---|
| RP23-424O22.3 | 0 | NA | NA | NA |
| RP24-289G19.1 | 0.04393175 | -0.206490061 | 3.724945049 | -0.055434391 |
| RP24-289G19.3 | 0.147068667 | 1.236183405 | 3.514416608 | 0.351746404 |
| RP24-289G19.2 | 0.089089296 | 1.194485492 | 3.723939576 | 0.320758559 |
| RP24-289G19.4 | 0 | NA | NA | NA |
| Barhl2 | 85.11172079 | -0.054718416 | 0.238914115 | -0.229029651 |
| Gm26872 | 0.927992339 | 1.167742807 | 1.065600788 | 1.095853926 |
| Gm28050 | 8.739896192 | 0.521413469 | 0.3530521 | 1.476874006 |
| Gm28039 | 0.075043205 | 0.755286614 | 3.724945049 | 0.202764498 |
| Gm29464 | 0.340386612 | 1.275931744 | 1.692864844 | 0.753711526 |
| Gm17304 | 0 | NA | NA | NA |
| RP23-167H23.3 | 0.042716945 | -0.206490061 | 3.724945049 | -0.055434391 |
| Zfp644 | 2759.468483 | -0.174533724 | 0.075312014 | -2.317475173 |
| RP23-167H23.2 | 69.78784834 | -0.37642557 | 0.316866193 | -1.187963808 |
| Gm24191 | 0 | NA | NA | NA |
| RP24-512F21.1 | 0 | NA | NA | NA |
| Hfm1 | 28.3914873 | 0.247518892 | 0.19991833 | 1.238100041 |
| Gm27527 | 0.338550938 | 1.591023006 | 1.531337526 | 1.038976045 |
| Olfr719-ps | 0 | NA | NA | NA |
| Cdc7 | 158.8450571 | -0.768034676 | 0.193451789 | -3.970160633 |
| RP23-24K16.1 | 0.066050875 | 0.274391604 | 3.724945049 | 0.073663262 |
| Tgfbr3 | 375.510489 | 0.383453295 | 0.11833292 | 3.24046171 |
| RP23-24K16.3 | 17.70335292 | 0.061587311 | 0.447006731 | 0.137777144 |
| RP23-24K16.2 | 5.270225733 | 0.196151848 | 0.529800732 | 0.370237027 |
| Gm8145 | 0 | NA | NA | NA |
| RP24-357G2.2 | 11.97838418 | -0.341857353 | 0.380636366 | -0.898120577 |
| Gm22557 | 0 | NA | NA | NA |
| Brdt | 172.1838994 | 0.331995441 | 0.106332702 | 3.122232722 |
| Gm8152 | 0.483474337 | -0.936044584 | 1.487466451 | -0.629287863 |
| RP24-140D11.1 | 19.5481605 | -0.671174951 | 0.238986394 | -2.808423278 |
| n-R5s173 | 0.044191549 | -0.206490061 | 3.724945049 | -0.055434391 |
| Ephx4 | 172.3772472 | -0.299091152 | 0.121388239 | -2.463921993 |
| Gm17202 | 1.45035625 | 0.900260741 | 0.78754096 | 1.143128784 |
| Lpcat2b | 2.065408989 | -1.091786367 | 0.672421055 | -1.623664753 |
| Btbd8 | 41.15341184 | -0.39213994 | 0.157014345 | -2.497478426 |
| Gm9727 | 10.68833693 | 0.30183943 | 0.352711488 | 0.855768639 |
| A830010M20Rik | 1635.382598 | 0.186742062 | 0.089553405 | 2.085259195 |
| RP23-32E20.1 | 2.258596541 | 1.325252611 | 0.794352767 | 1.668342664 |
| 1700028K03Rik | 21.85631515 | 0.724280553 | 0.302560366 | 2.393838169 |
| Glmn | 205.8721023 | -0.085333256 | 0.089014805 | -0.958641164 |
| Rpap2 | 173.555107 | 0.206533561 | 0.092912342 | 2.222886177 |
| A930041C12Rik | 0.201858732 | 1.23619586 | 2.604406118 | 0.474655566 |
| Gm26692 | 1.12480443 | -0.813179253 | 1.011167263 | -0.804198556 |
| RP23-32E20.3 | 0.156258025 | 0.711829273 | 2.812062138 | 0.253134262 |
| RP24-140D11.4 | 0.388935867 | 1.274168812 | 1.681052733 | 0.757958859 |
| Gfi1 | 6.525819315 | 4.841699842 | 0.936732214 | 5.168712862 |
| A430072P03Rik | 0.11147366 | 0.274389235 | 3.724945049 | 0.073662626 |

| | | | | |
|---|---|---|---|---|
| Evi5 | 1027.982555 | -0.328125511 | 0.061428414 | -5.3415918 |
| RP24-140D11.6 | 51.7496199 | -0.564067406 | 0.201826321 | -2.794815875 |
| Ube2d2b | 0 | NA | NA | NA |
| Gm16007 | 0 | NA | NA | NA |
| RP24-140D11.5 | 1.387064387 | -0.318591608 | 0.988120167 | -0.322421927 |
| Rpl5 | 1390.886114 | 0.324200058 | 0.095666894 | 3.388842727 |
| Snord21 | 0 | NA | NA | NA |
| Gm22270 | 2.785641484 | -0.761597118 | 0.600774005 | -1.267693195 |
| Gm26387 | 2.437479595 | 0.273467675 | 0.643088969 | 0.425240811 |
| Fam69a | 302.41774 | -0.007633172 | 0.087225125 | -0.087511163 |
| Gm9850 | 0.572539054 | -0.771190009 | 1.268842588 | -0.607790136 |
| Rps19-ps8 | 0 | NA | NA | NA |
| Mtf2 | 757.8828192 | -0.080269753 | 0.086879271 | -0.923922959 |
| Tmed5 | 501.0379422 | 0.545339733 | 0.114677755 | 4.75540992 |
| Ccdc18 | 25.54476345 | 0.112211136 | 0.239362251 | 0.468792118 |
| RP23-46J11.4 | 0 | NA | NA | NA |
| RP23-46J11.5 | 0 | NA | NA | NA |
| RP23-46J11.6 | 0 | NA | NA | NA |
| Dr1 | 585.9554827 | 0.066817108 | 0.09776698 | 0.683432263 |
| RP24-317M4.1 | 6.093084322 | -0.746540986 | 0.51185601 | -1.458498036 |
| Pigg | 279.9406767 | -0.354995992 | 0.08947533 | -3.967529262 |
| RP24-317M4.2 | 2.428646146 | -0.913178048 | 0.610705189 | -1.495284574 |
| RP24-317M4.3 | 9.503775188 | -1.014474707 | 0.367773023 | -2.758426103 |
| RP24-317M4.6 | 137.5888963 | -0.267760705 | 0.169918819 | -1.575815477 |
| Gm10419 | 867.0367941 | 0.03822537 | 0.121066627 | 0.315738291 |
| Pde6b | 10.02136167 | 0.809133848 | 0.659056081 | 1.227716232 |
| Atp5k | 305.7650685 | -0.175422763 | 0.181844812 | -0.9646839 |
| Mfsd7a | 18.35894452 | 1.339977361 | 0.301055531 | 4.450930884 |
| Pcgf3 | 699.3355955 | -0.133871439 | 0.069619324 | -1.922906326 |
| Gm10086 | 0.202977511 | -0.20648591 | 2.197917809 | -0.093946147 |
| Cplx1 | 5792.979381 | -0.096671368 | 0.079768342 | -1.211901432 |
| RP24-529E15.1 | 10.79973995 | 1.402870024 | 0.55550077 | 2.525415087 |
| Gak | 1581.201267 | 0.156375988 | 0.057244852 | 2.73170394 |
| RP24-529E15.3 | 21.02397872 | -0.195623766 | 0.286069144 | -0.683833856 |
| Tmem175 | 304.8091594 | -0.226063231 | 0.100568112 | -2.247861936 |
| Dgkq | 523.7346149 | -0.456978246 | 0.100859659 | -4.530832731 |
| Idua | 131.045456 | 0.300846671 | 0.130611554 | 2.303369504 |
| Slc26a1 | 12.6609391 | -0.273511921 | 0.324191081 | -0.843675033 |
| Fgfrl1 | 247.8383437 | 0.629999256 | 0.152497315 | 4.131215388 |
| Rnf212 | 0.048949532 | -0.206487837 | 3.724945049 | -0.055433794 |
| 1700047L14Rik | 0.220636191 | 0.274401165 | 2.573805194 | 0.106613028 |
| Tmed11 | 0 | NA | NA | NA |
| Vmn2r8 | 0 | NA | NA | NA |
| Gm18594 | 0 | NA | NA | NA |
| Vmn2r9 | 0 | NA | NA | NA |
| Gm20629 | 0.046345709 | 0.274391604 | 3.724945049 | 0.073663262 |
| Gm10417 | 1.121165163 | -0.698969939 | 0.994455281 | -0.70286714 |

| | | | | |
|---|---|---|---|---|
| Vmn2r-ps21 | 0 | NA | NA | NA |
| RP23-37G13.2 | 0.042716945 | -0.206490061 | 3.724945049 | -0.055434391 |
| RP23-37G13.3 | 0 | NA | NA | NA |
| Gm18595 | 0 | NA | NA | NA |
| Vmn2r10 | 0 | NA | NA | NA |
| Gm5871 | 0.397993176 | 0.755309163 | 1.767559089 | 0.427317631 |
| Vmn2r11 | 0 | NA | NA | NA |
| Vmn2r12 | 0 | NA | NA | NA |
| Vmn2r13 | 0 | NA | NA | NA |
| Vmn2r14 | 0 | NA | NA | NA |
| Gm8473 | 0 | NA | NA | NA |
| Vmn2r15 | 0 | NA | NA | NA |
| Vmn2r16 | 0 | NA | NA | NA |
| Gm7171 | 0 | NA | NA | NA |
| Vmn2r17 | 0.044191549 | -0.206490061 | 3.724945049 | -0.055434391 |
| RP23-410M21.3 | 0 | NA | NA | NA |
| Gm8488 | 0.787704992 | 0.675757162 | 1.179742064 | 0.572800769 |
| RP23-203O7.3 | 2.482507492 | -2.432098737 | 0.857428824 | -2.83650219 |
| Crlf2 | 173.2274311 | 0.633417277 | 0.124696041 | 5.079690361 |
| Vmn2r-ps23 | 0.148650935 | 0.723036258 | 2.859657322 | 0.252840175 |
| Gm28000 | 0 | NA | NA | NA |
| 5430403G16Rik | 36.45249734 | -0.204514162 | 0.258298943 | -0.791773126 |
| 4930522L14Rik | 39.79845386 | -0.199543843 | 0.193037852 | -1.03370319 |
| Gm8508 | 4.624103064 | -0.180427508 | 0.474336937 | -0.380378363 |
| RP23-396K22.3 | 1.154557212 | -0.019413985 | 0.822736487 | -0.023596844 |
| RP23-396K22.4 | 4.399392508 | 0.63170895 | 0.544544199 | 1.160069193 |
| Gm26779 | 37.79669302 | -0.465677723 | 0.189657751 | -2.455358251 |
| Gm26808 | 0.439601053 | -0.607461448 | 1.598094519 | -0.380116095 |
| Gm15446 | 190.6866757 | -0.273737567 | 0.131235469 | -2.085850486 |
| 1010001B22Rik | 0.952639941 | -1.098120613 | 1.059828144 | -1.036130829 |
| Zfp932 | 186.9135822 | -0.353211834 | 0.127287509 | -2.774913554 |
| Gm17655 | 4.402479275 | -0.646490016 | 0.518451081 | -1.246964353 |
| Plcxd1 | 122.7290074 | -0.269932751 | 0.146374877 | -1.844119412 |
| Gtpbp6 | 242.7828586 | 0.191404917 | 0.0732084 | 2.614521246 |
| Zfp605 | 230.5230773 | 0.138594631 | 0.08182699 | 1.693752041 |
| Gm15484 | 2.365693163 | 0.090458606 | 0.668061062 | 0.135404698 |
| Chfr | 523.8452094 | -0.285205711 | 0.112710858 | -2.530419115 |
| RP24-374B14.4 | 1.731816556 | 0.761560198 | 0.777701369 | 0.979245027 |
| Gm15787 | 10.57349902 | 1.239416549 | 0.395474228 | 3.134000807 |
| Golga3 | 1104.05704 | 0.168496158 | 0.065865051 | 2.558202802 |
| RP24-374B14.6 | 9.781179111 | 0.190515579 | 0.348821076 | 0.546169918 |
| RP24-374B14.8 | 3.53955661 | 1.607432663 | 0.534269567 | 3.008654737 |
| Ankle2 | 833.7400944 | 0.272000891 | 0.065779498 | 4.135040529 |
| Pgam5 | 509.9939935 | 0.083207578 | 0.067816336 | 1.22695479 |
| Gm26718 | 13.31072805 | -0.55403904 | 0.31156886 | -1.77822341 |
| Gm15788 | 2.377542658 | -0.460310885 | 0.684639081 | -0.672340943 |
| Pxmp2 | 26.9648925 | 0.112922489 | 0.262529887 | 0.430131937 |

| | | | | |
|---|---|---|---|---|
| Pole | 51.3992184 | 1.416336433 | 0.327566011 | 4.323819886 |
| Gm26277 | 0 | NA | NA | NA |
| Gm15791 | 0.092493346 | 0.274402725 | 3.724945049 | 0.073666248 |
| Gm15792 | 0.061920613 | 0.274393828 | 3.724945049 | 0.073663859 |
| P2rx2 | 1.822501977 | 0.115096069 | 0.765528925 | 0.150348426 |
| Lrcol1 | 0.046345709 | 0.274391604 | 3.724945049 | 0.073663262 |
| Fbrsl1 | 654.9798202 | -0.082075715 | 0.081325156 | -1.009229103 |
| Gm26711 | 1.656227335 | 0.599395866 | 0.83326495 | 0.719334067 |
| RP23-231C16.1 | 2.276862309 | 0.809768412 | 0.645203346 | 1.255059226 |
| RP23-231C16.2 | 0 | NA | NA | NA |
| A630023P12Rik | 2.678264972 | 0.744039998 | 0.642436785 | 1.158152857 |
| Galnt9 | 582.7762908 | -0.130215328 | 0.094549844 | -1.377213565 |
| Noc4l | 266.4007122 | 0.896077038 | 0.155245407 | 5.77200353 |
| Ddx51 | 235.8120389 | 0.135134514 | 0.101357571 | 1.333245386 |
| RP23-393E10.2 | 29.91262942 | 0.682127086 | 0.218352121 | 3.123977381 |
| Ep400 | 2474.898703 | 0.136725205 | 0.039860605 | 3.430083541 |
| Mir7026 | 1.550196443 | 0.562840383 | 0.760308145 | 0.740279302 |
| Gm22716 | 3.382263423 | -0.364542577 | 0.533613855 | -0.683158006 |
| Gm26079 | 0.044544648 | 0.755286614 | 3.724945049 | 0.202764498 |
| Gm22583 | 0 | NA | NA | NA |
| Gm15559 | 13.41128988 | 0.55653089 | 0.267708514 | 2.078868845 |
| Pus1 | 142.3904629 | 0.173638322 | 0.102553974 | 1.693140854 |
| Gm26515 | 2.087315227 | -0.269167207 | 0.682785408 | -0.394219331 |
| Ulk1 | 1100.014041 | -0.076810812 | 0.054569228 | -1.407584724 |
| Hscb | 47.36481145 | 0.265487502 | 0.194774541 | 1.363050331 |
| Chek2 | 15.72251656 | -0.18824797 | 0.292475768 | -0.643636126 |
| RP24-207A18.2 | 0.348192234 | 1.217001664 | 1.587692736 | 0.766522159 |
| Ttc28 | 598.1288249 | 0.069406652 | 0.05696326 | 1.218445919 |
| Gm15989 | 3.684955975 | -0.125687698 | 0.523419224 | -0.240128165 |
| RP23-326H18.1 | 27.43115472 | -0.231572647 | 0.220001549 | -1.052595526 |
| Mir701 | 2.293891663 | -0.077323633 | 0.596696149 | -0.129586277 |
| Gm15988 | 0 | NA | NA | NA |
| RP24-230N16.2 | 6.359331415 | -0.817996872 | 0.462231689 | -1.769668528 |
| RP23-72O23.3 | 4.724619101 | -0.640381916 | 0.513055955 | -1.248171684 |
| RP23-72O23.4 | 1.873630482 | -0.301071075 | 0.823633206 | -0.365540235 |
| Gm27602 | 0 | NA | NA | NA |
| RP23-131F7.2 | 0.716273392 | -1.60364709 | 1.140429497 | -1.406178194 |
| Gm27680 | 0 | NA | NA | NA |
| Pitpnb | 878.8123659 | 0.105799462 | 0.065057013 | 1.626257604 |
| RP23-131F7.5 | 0.255202296 | 0.745200604 | 1.828147006 | 0.407626193 |
| Mn1 | 649.1153886 | -0.414397292 | 0.143249738 | -2.892831069 |
| RP23-131F7.4 | 14.92077947 | -1.453890166 | 0.325666121 | -4.464358038 |
| C130026L21Rik | 34.50140241 | -0.568729074 | 0.266619652 | -2.133110102 |
| RP24-572L22.2 | 3.151250669 | -1.768244204 | 0.676951079 | -2.612070887 |
| Gm26897 | 0.18833836 | 0.285088417 | 2.670116479 | 0.10677003 |
| E130006D01Rik | 3.354483374 | -0.334339799 | 0.534715829 | -0.625266321 |
| Gm16019 | 0 | NA | NA | NA |

| | | | | |
|---|---|---|---|---|
| RP24-340B11.1 | 0 | NA | NA | NA |
| RP23-427D2.1 | 0.576279854 | -0.444261015 | 1.246414254 | -0.356431269 |
| RP24-317K11.4 | 0 | NA | NA | NA |
| 1700028D13Rik | 0 | NA | NA | NA |
| Miat | 1661.236509 | -0.682887783 | 0.490718507 | -1.391607964 |
| Gm27998 | 0 | NA | NA | NA |
| Gm27882 | 0 | NA | NA | NA |
| Gm26953 | 0.344925633 | 0.741812487 | 1.791155966 | 0.414152928 |
| Gm27627 | 0 | NA | NA | NA |
| Gm27357 | 0 | NA | NA | NA |
| Cryba4 | 4.194020461 | 1.161911357 | 0.614978057 | 1.889354172 |
| Crybb1 | 14.29264834 | 0.091404483 | 0.494256998 | 0.184933108 |
| Tpst2 | 280.8629906 | 0.60300726 | 0.095758269 | 6.297182097 |
| Tfip11 | 581.9359828 | 0.240130415 | 0.06416627 | 3.742315318 |
| Srrd | 100.6125371 | 0.028123411 | 0.118524334 | 0.237279636 |
| Hps4 | 178.5209335 | -0.407806472 | 0.134628752 | -3.0291187 |
| Gm6583 | 0.632098713 | 1.090536959 | 1.455765266 | 0.749115935 |
| RP23-459I1.3 | 20.2973942 | -1.197603778 | 0.435191046 | -2.751903535 |
| RP23-459I1.5 | 0 | NA | NA | NA |
| Gm20636 | 0.231217418 | -0.20648129 | 2.131487463 | -0.096871923 |
| Asphd2 | 618.5894786 | -0.04461496 | 0.081260493 | -0.549036297 |
| Sez6l | 3346.420789 | -0.000402803 | 0.089412822 | -0.004504985 |
| Gm6588 | 0 | NA | NA | NA |
| Gm27004 | 3.168227339 | -0.328556019 | 0.662241939 | -0.496126868 |
| 1700034G24Rik | 0.106465261 | 0.75528439 | 3.724945049 | 0.202763901 |
| RP23-156K23.3 | 0.158818843 | -0.666974881 | 2.81011023 | -0.237348298 |
| RP23-156K23.4 | 7.736953527 | -0.436110693 | 0.413020603 | -1.05590542 |
| RP24-304N8.2 | 2.801170497 | 0.588056835 | 0.556294129 | 1.057096964 |
| Myo18b | 28.0640786 | 0.311840402 | 0.236014467 | 1.321276642 |
| 1700095B10Rik | 0 | NA | NA | NA |
| Adrbk2 | 1355.458738 | -0.097052998 | 0.081415323 | -1.192072872 |
| Gm22740 | 0 | NA | NA | NA |
| Crybb2 | 0.278057897 | 1.260206682 | 2.411131497 | 0.522661947 |
| RP23-4P1.1 | 0.92508897 | 1.036259717 | 1.014923459 | 1.021022529 |
| Crybb3 | 5.690583048 | -0.80183103 | 0.464531519 | -1.726106836 |
| 2900026A02Rik | 622.1930781 | -0.048552903 | 0.09276796 | -0.523380086 |
| RP23-4P1.6 | 0.061920613 | 0.274393828 | 3.724945049 | 0.073663859 |
| RP23-4P1.5 | 5.731549506 | -0.763900887 | 0.462014952 | -1.653411611 |
| RP23-4P1.3 | 0.48780114 | -2.37547249 | 1.322596799 | -1.79606702 |
| RP23-4P1.7 | 0.066050875 | 0.274391604 | 3.724945049 | 0.073663262 |
| RP23-4P1.2 | 1.950711236 | -0.463008879 | 0.871168011 | -0.53148058 |
| Tmem211 | 0.590173444 | -1.606714454 | 1.31440439 | -1.22238975 |
| Sgsm1 | 1205.098046 | -0.51422007 | 0.081860419 | -6.281669186 |
| RP24-456F12.1 | 1.004172246 | 1.903820403 | 1.158180502 | 1.643802844 |
| RP24-456F12.2 | 0.236337014 | 1.235355044 | 2.166364679 | 0.570243346 |
| mmu-mir-7229 | 0 | NA | NA | NA |
| mmu-mir-7230 | 0 | NA | NA | NA |

| | | | | |
|---|---|---|---|---|
| Aym1 | 0.330879691 | 1.348825569 | 1.992275863 | 0.677027511 |
| Wscd2 | 877.7252593 | -0.477777818 | 0.129962674 | -3.676269534 |
| Cmklr1 | 152.364205 | 2.508823573 | 0.194717537 | 12.88442533 |
| Gm15736 | 0.075043205 | 0.755286614 | 3.724945049 | 0.202764498 |
| 1700069L16Rik | 1.680905595 | 0.660544664 | 0.958692767 | 0.689005578 |
| Ficd | 177.6330377 | 0.080877929 | 0.10650187 | 0.759403845 |
| Sart3 | 480.3375866 | 0.13808754 | 0.096007996 | 1.438292074 |
| Gm3511 | 0.172107867 | 1.236193725 | 2.739902689 | 0.451181617 |
| Iscu | 446.2315743 | -0.008769746 | 0.091123332 | -0.096240399 |
| Tmem119 | 184.4126201 | 0.471288647 | 0.217670007 | 2.165151985 |
| RP24-555M20.7 | 0 NA | NA | NA | |
| Selplg | 268.6005576 | 0.839589625 | 0.161476466 | 5.199455034 |
| Coro1c | 1185.306564 | 0.229753818 | 0.097107863 | 2.365965124 |
| Gm17122 | 0.291024678 | -0.206481897 | 2.061133605 | -0.100178803 |
| RP24-555M20.4 | 0.261813223 | -1.615559152 | 2.497637454 | -0.646834932 |
| RP24-555M20.3 | 33.32994184 | -0.246380548 | 0.359093506 | -0.686118083 |
| RP24-555M20.5 | 0.345648411 | -0.206482209 | 1.604093846 | -0.128722025 |
| Gm22056 | 0 NA | NA | NA | |
| Ssh1 | 648.036513 | -0.01157503 | 0.062470459 | -0.185288057 |
| Dao | 29.66175715 | -0.526069326 | 0.320730096 | -1.640224389 |
| Svop | 1130.54196 | -0.192745066 | 0.078349783 | -2.460058705 |
| Usp30 | 451.0626031 | 0.147905717 | 0.10642123 | 1.389814007 |
| RP24-226L20.2 | 8.375339354 | -0.220411581 | 0.394082113 | -0.559303694 |
| RP24-226L20.1 | 11.87863406 | 0.128883986 | 0.543054937 | 0.237331395 |
| Alkbh2 | 30.67206188 | -0.104252191 | 0.201636002 | -0.51703163 |
| Ung | 32.04545787 | 0.37775402 | 0.243466171 | 1.551566766 |
| Acacb | 113.2622031 | 0.338498541 | 0.134553649 | 2.515714308 |
| Gm13784 | 0 NA | NA | NA | |
| Foxn4 | 0.07796279 | 0.274392189 | 3.724945049 | 0.073663419 |
| Myo1h | 18.55275259 | 1.034409037 | 0.311192394 | 3.324017735 |
| Kctd10 | 678.4952583 | 0.502674754 | 0.120450193 | 4.173299688 |
| Ube3b | 1199.562226 | -0.107180339 | 0.075515692 | -1.419312149 |
| Mir7027 | 0.775147523 | -0.511935489 | 1.135642743 | -0.450789205 |
| Mmab | 164.6481927 | -0.111621355 | 0.088173458 | -1.265929195 |
| Mvk | 173.1109817 | -0.038883714 | 0.130257715 | -0.298513714 |
| Gm13790 | 0.110814304 | -0.687373951 | 3.724945049 | -0.184532642 |
| Fam222a | 196.5946661 | -0.301401029 | 0.127550208 | -2.362999109 |
| Trpv4 | 24.56884945 | 0.658514066 | 0.25279189 | 2.60496516 |
| Gltp | 531.1234222 | 0.353812022 | 0.123763414 | 2.858777167 |
| Tchp | 125.3804559 | -0.087200976 | 0.180847827 | -0.48217873 |
| RP23-3M10.11 | 0.104637558 | -0.206492286 | 3.724945049 | -0.055434988 |
| Mir7028 | 0.496352305 | -2.400550056 | 1.397691526 | -1.717510632 |
| Git2 | 916.5459927 | 0.103269633 | 0.099154073 | 1.041506723 |
| 4930515G01Rik | 7.627541774 | -0.730572237 | 0.406640672 | -1.796603947 |
| Ankrd13a | 914.9811008 | 0.695964528 | 0.103664578 | 6.713619451 |
| 1500011B03Rik | 398.2607319 | 0.285033607 | 0.085119485 | 3.348629376 |
| Gm20499 | 1.052193156 | 1.552641243 | 1.067849553 | 1.453988756 |

| | | | | |
|---|---|---|---|---|
| 2610524H06Rik | 87.22289935 | 0.461332969 | 0.130901314 | 3.524280653 |
| Ywhaq-ps2 | 0.12209515 | 0.75528439 | 3.724945049 | 0.202763901 |
| RP24-297N9.13 | 1.537880876 | 0.739508626 | 0.825345676 | 0.895998668 |
| Mir6240 | 0 | NA | NA | NA |
| 4930519G04Rik | 0.461107361 | 0.134083794 | 1.430046178 | 0.093761863 |
| Gm9936 | 3.314927513 | 1.233733356 | 0.623265963 | 1.979465316 |
| Gm13823 | 0.097139547 | 0.274393828 | 3.724945049 | 0.073663859 |
| Oasl2 | 5567.265836 | 8.525453234 | 0.170994676 | 49.85800393 |
| Gm13822 | 5.72139037 | 4.20357122 | 1.00922664 | 4.165140963 |
| Oasl1 | 994.9398756 | 9.189111743 | 0.611197052 | 15.03461397 |
| 2210016L21Rik | 838.2899106 | 0.10448141 | 0.06994711 | 1.493720175 |
| Hnf1a | 0.04393175 | -0.206490061 | 3.724945049 | -0.055434391 |
| Hnf1aos1 | 0.088476398 | 0.274400501 | 3.724945049 | 0.073665651 |
| Hnf1aos2 | 0.044544648 | 0.755286614 | 3.724945049 | 0.202764498 |
| RP24-556H22.2 | 0.075043205 | 0.755286614 | 3.724945049 | 0.202764498 |
| Sppl3 | 907.0167297 | -0.001884957 | 0.063284965 | -0.02978523 |
| RP24-556H22.3 | 18.34444885 | -0.061639938 | 0.321112796 | -0.191957278 |
| Gm13827 | 0.049553047 | 0.274391604 | 3.724945049 | 0.073663262 |
| Gm10401 | 0.066050875 | 0.274391604 | 3.724945049 | 0.073663262 |
| Rpl37rt | 126.6981566 | 0.494407804 | 0.162762491 | 3.037602842 |
| Acads | 60.73886381 | -0.260386142 | 0.198334918 | -1.312860812 |
| Unc119b | 384.4903987 | 0.120949525 | 0.123986558 | 0.975505142 |
| Mlec | 1723.019123 | -0.188795273 | 0.112673222 | -1.675600201 |
| Cabp1 | 800.6070211 | -0.43758008 | 0.124494149 | -3.514864635 |
| Gm13828 | 0.405693472 | -1.916392788 | 1.609780954 | -1.190468046 |
| Pop5 | 169.6778782 | -0.048573962 | 0.108267204 | -0.448648902 |
| Rnf10 | 2262.437902 | 0.058284652 | 0.066337322 | 0.87861027 |
| Gm13831 | 0.044544648 | 0.755286614 | 3.724945049 | 0.202764498 |
| Coq5 | 405.2808921 | 0.019000979 | 0.078963042 | 0.240631291 |
| Gm13829 | 0 | NA | NA | NA |
| Dynll1 | 1360.661757 | -0.378710702 | 0.115097749 | -3.290339775 |
| Gm13830 | 22.05592487 | -0.995918111 | 0.292625931 | -3.403382972 |
| Srsf9 | 360.4762658 | -0.148810459 | 0.084375775 | -1.763663317 |
| Gatc | 179.8265356 | -0.135434493 | 0.102403719 | -1.322554443 |
| Triap1 | 96.42262351 | -0.029025796 | 0.128881191 | -0.225213591 |
| Cox6a1 | 2038.657447 | 0.095144117 | 0.111770971 | 0.851241753 |
| RP24-132L16.11 | 0.287284012 | 0.768171323 | 1.781054835 | 0.431301332 |
| 4930401G09Rik | 0.097899064 | -0.687371727 | 3.724945049 | -0.184532045 |
| Gm13832 | 4.797855873 | 0.607756945 | 0.479689398 | 1.266980149 |
| RP24-132L16.12 | 3.340097371 | -1.455418758 | 0.553330289 | -2.630289335 |
| RP24-132L16.13 | 6.416277231 | -0.796847943 | 0.463420779 | -1.719491181 |
| Msi1 | 624.614844 | -0.176745944 | 0.097079677 | -1.820627643 |
| RP24-132L16.14 | 0.383028346 | -0.596107528 | 1.45449421 | -0.409838365 |
| Pla2g1b | 0.110595523 | 0.755282166 | 3.724945049 | 0.202763304 |
| Sirt4 | 90.54918992 | -0.630272282 | 0.197525885 | -3.190833853 |
| Gm24265 | 261.4064373 | 0.042846045 | 0.214783588 | 0.199484724 |
| Gm24407 | 234.2932087 | -0.178113343 | 0.179811408 | -0.990556411 |

| | | | | |
|---|---|---|---|---|
| Gm13836 | 0.227885852 | 1.096645348 | 2.508748414 | 0.437128467 |
| Pxn | 603.300534 | 0.250296255 | 0.125117434 | 2.000490634 |
| Gm13840 | 1.630127001 | -0.202430652 | 0.8303711 | -0.243783354 |
| Rplp0 | 924.8935804 | 0.730126606 | 0.13121575 | 5.564321408 |
| Gcn1l1 | 1454.726854 | -0.024480609 | 0.072938905 | -0.3356317 |
| Mir7029 | 0.140122821 | 0.274406091 | 3.572128793 | 0.076818644 |
| RP23-228F6.9 | 3.230897769 | -0.542681631 | 0.631912229 | -0.858792734 |
| Rab35 | 708.4090342 | 0.131289455 | 0.067592368 | 1.942370992 |
| Mir7030 | 0.697123821 | -0.682496956 | 1.245306786 | -0.548055278 |
| Ccdc64 | 496.8343264 | -0.266031596 | 0.101839654 | -2.612259431 |
| Gm13841 | 108.5809223 | 0.28580759 | 0.167065095 | 1.710755855 |
| Cit | 2100.160842 | -0.453319903 | 0.134634914 | -3.3670308 |
| RP23-265I19.3 | 0.244575677 | 0.740861492 | 2.091207798 | 0.35427445 |
| RP23-265I19.2 | 0.484822013 | -0.129493058 | 1.53669922 | -0.084267016 |
| Gm14508 | 2.937257927 | -0.358641084 | 0.548113869 | -0.654318572 |
| Prkab1 | 201.4925811 | 0.642269435 | 0.128667663 | 4.99169272 |
| Gm13842 | 0 | NA | NA | NA |
| Gm17420 | 0 | NA | NA | NA |
| Tmem233 | 0.047948698 | -0.206487837 | 3.724945049 | -0.055433794 |
| C330018A13Rik | 4.719781364 | -0.232174389 | 0.500692272 | -0.463706757 |
| Ccdc60 | 12.74564928 | 0.384878355 | 0.337943462 | 1.138883862 |
| Gm14507 | 0.354637581 | -2.056055302 | 1.63157277 | -1.260167698 |
| Gm13837 | 0.143618953 | -0.206488341 | 2.893930267 | -0.071352217 |
| Gm13839 | 0.393020699 | 0.718536371 | 1.723493579 | 0.416906904 |
| Gm13838 | 0.066050875 | 0.274391604 | 3.724945049 | 0.073663262 |
| B230112J18Rik | 0.528863779 | 0.380898224 | 1.753776758 | 0.217187406 |
| RP24-79A6.9 | 0.038981395 | 0.274393828 | 3.724945049 | 0.073663859 |
| 4930562A09Rik | 0.146089079 | -0.206488916 | 3.522538524 | -0.058619349 |
| Hspb8 | 433.011455 | 1.691658467 | 0.209783672 | 8.063823331 |
| Srrm4os | 0.683645101 | -1.180869154 | 1.115774323 | -1.058340499 |
| Srrm4 | 850.8466253 | -0.34818986 | 0.09139861 | -3.809575001 |
| RP24-79A6.8 | 32.11370528 | -0.865908153 | 0.317783016 | -2.724840883 |
| RP23-312N12.2 | 10.26602459 | -0.890708916 | 0.441562579 | -2.017174819 |
| RP23-312N12.1 | 15.57120856 | -1.367073058 | 0.402794978 | -3.393967482 |
| Gstm2-ps1 | 32.88268672 | -0.21780523 | 0.246112823 | -0.884981234 |
| RP23-418H8.3 | 0.072025462 | 0.755288839 | 3.724945049 | 0.202765096 |
| RP23-280O16.1 | 0.060174537 | 0.755286614 | 3.724945049 | 0.202764498 |
| 4930569F06Rik | 0 | NA | NA | NA |
| RP23-73P16.2 | 0.598047226 | 1.730276293 | 1.337163029 | 1.293990528 |
| Suds3 | 810.9948176 | -0.112477604 | 0.058308673 | -1.929002972 |
| 9530046B11Rik | 1.655761812 | -0.525214877 | 0.779410222 | -0.673861932 |
| Taok3 | 787.4816357 | -0.078031281 | 0.068906949 | -1.132415266 |
| RP23-376H8.2 | 15.63613524 | -0.408571256 | 0.357453043 | -1.143006793 |
| n-R5s174 | 1.199099511 | -0.307389632 | 0.988790705 | -0.310874314 |
| Pebp1 | 1757.043143 | 0.053955446 | 0.072544999 | 0.743751415 |
| RP23-76J15.3 | 0 | NA | NA | NA |
| RP23-76J15.4 | 0.215166026 | 0.775847198 | 2.554583147 | 0.303707945 |

| | | | | |
|---|---|---|---|---|
| BC051077 | 1.218454842 | 0.359378006 | 0.966736867 | 0.371743354 |
| Gm10399 | 0.929291888 | 0.204985932 | 1.164580181 | 0.176017019 |
| Vsig10 | 55.95498491 | 0.605945834 | 0.234663135 | 2.582194403 |
| Gm15727 | 0 | NA | NA | NA |
| Wsb2 | 1799.690181 | 0.310871185 | 0.066282747 | 4.690076938 |
| Rfc5 | 71.308591 | 0.368792536 | 0.165707207 | 2.225567274 |
| Gm15728 | 37.43701224 | 0.088745073 | 0.214200984 | 0.414307492 |
| Ksr2 | 1532.422263 | -0.396269622 | 0.077724234 | -5.098404962 |
| RP23-429P17.1 | 4.731857296 | -1.344459507 | 0.515673716 | -2.607190292 |
| Nos1 | 1164.626249 | -0.385303965 | 0.082099184 | -4.693152162 |
| Gm37028 | 0.69250253 | 0.631126509 | 1.268467344 | 0.497550459 |
| Gm26995 | 0.399888529 | -0.641728434 | 1.447690164 | -0.443277471 |
| Gm25343 | 0.046345709 | 0.274391604 | 3.724945049 | 0.073663262 |
| Fbxo21 | 1283.047223 | -0.000580026 | 0.06227437 | -0.009314037 |
| RP23-42D10.2 | 0.853574741 | 0.254004033 | 1.23282182 | 0.206034667 |
| Tesc | 196.9276744 | -0.691583421 | 0.166961954 | -4.142161756 |
| Fbxw8 | 652.4874552 | 0.090219565 | 0.097977168 | 0.920822332 |
| Gm9754 | 24.46512519 | -0.948412669 | 0.435026175 | -2.180127827 |
| Gm26411 | 1.298260455 | 0.874977116 | 0.872726851 | 1.002578431 |
| Hrk | 446.6006905 | 0.689275925 | 0.191270241 | 3.603675723 |
| Rnft2 | 775.8299915 | -0.16605973 | 0.081163841 | -2.045981658 |
| Gm24050 | 1.923134663 | -2.011544649 | 0.898216272 | -2.239488096 |
| 2410131K14Rik | 480.8992711 | 0.376904475 | 0.099753175 | 3.778370698 |
| Gm25076 | 4.628642042 | -0.760935793 | 0.490539491 | -1.551222291 |
| Gm28563 | 0.587040768 | -0.991932028 | 1.10715561 | -0.895928286 |
| Gm26455 | 0.133946075 | 0.755288219 | 3.626642455 | 0.20826101 |
| Gm15754 | 0.066050875 | 0.274391604 | 3.724945049 | 0.073663262 |
| Med13l | 2276.940129 | 0.061871358 | 0.066393967 | 0.931882224 |
| RP24-122N24.1 | 0 | NA | NA | NA |
| RP24-122N24.2 | 76.64565332 | -0.373681322 | 0.298825113 | -1.250501735 |
| RP24-122N24.3 | 27.41095622 | -0.477002851 | 0.299232354 | -1.59408849 |
| RP24-122N24.4 | 64.02437309 | -0.280054657 | 0.317230623 | -0.882810915 |
| RP23-155F13.2 | 26.31192174 | -0.228530738 | 0.330428943 | -0.691618403 |
| RP23-155F13.3 | 28.57824406 | -0.230912728 | 0.352387091 | -0.655281461 |
| RP24-372H3.1 | 0 | NA | NA | NA |
| RP24-372H3.2 | 0 | NA | NA | NA |
| RP24-372H3.4 | 0 | NA | NA | NA |
| RP24-372H3.3 | 0 | NA | NA | NA |
| 1700081H04Rik | 0.049553047 | 0.274391604 | 3.724945049 | 0.073663262 |
| RP23-316P20.1 | 0.048949532 | -0.206487837 | 3.724945049 | -0.055433794 |
| n-R5s175 | 0 | NA | NA | NA |
| RP23-73O10.1 | 0 | NA | NA | NA |
| Tbx3os1 | 2.722982093 | -0.591499606 | 0.72665161 | -0.81400715 |
| RP23-44J10.1 | 0.387852398 | -0.482832641 | 1.480012823 | -0.326235444 |
| Gm16063 | 0 | NA | NA | NA |
| Gm16064 | 0 | NA | NA | NA |
| Tbx3 | 28.477325 | 0.153821076 | 0.235024163 | 0.654490473 |

| | | | | |
|---|---|---|---|---|
| Tbx3os2 | 0.046345709 | 0.274391604 | 3.724945049 | 0.073663262 |
| RP24-150D8.1 | 0 | NA | NA | NA |
| RP24-150D8.2 | 0 | NA | NA | NA |
| Tbx5 | 0.360567166 | 1.590263686 | 1.986725544 | 0.800444576 |
| RP24-150D8.3 | 0 | NA | NA | NA |
| RP24-69A21.1 | 0 | NA | NA | NA |
| n-R5s176 | 0 | NA | NA | NA |
| Rbm19 | 227.3684793 | 0.14609426 | 0.080801841 | 1.808056071 |
| Gm10390 | 1.46581892 | 0.044422758 | 1.026826487 | 0.043262186 |
| RP23-213H20.1 | 1.459375539 | 2.629299511 | 1.059616845 | 2.481368168 |
| Gm26474 | 0 | NA | NA | NA |
| RP23-213H20.2 | 0.303117673 | 2.477041507 | 3.68805292 | 0.671639361 |
| Gm27199 | 3.757776822 | -0.799059909 | 0.546004501 | -1.463467623 |
| Lhx5 | 49.17951797 | -0.264445863 | 0.208142691 | -1.270502755 |
| Sdsl | 34.31511951 | -0.229387253 | 0.205564006 | -1.115892113 |
| Sds | 0.913087876 | 0.421946649 | 1.224406974 | 0.344613072 |
| Gm15690 | 5.787489139 | 0.277892997 | 0.432446982 | 0.642605935 |
| Plbd2 | 910.6907376 | 0.117304878 | 0.080412599 | 1.458787297 |
| Slc8b1 | 149.2518986 | 1.145347267 | 0.174806646 | 6.552080802 |
| Tpcn1 | 461.0027236 | -0.712474324 | 0.150216296 | -4.742989579 |
| RP23-297M4.7 | 22.51234608 | -0.277279896 | 0.258817985 | -1.071331637 |
| Iqcd | 5.452284118 | -0.971329589 | 0.490196777 | -1.981509538 |
| Rita1 | 142.064277 | -0.509551071 | 0.134662954 | -3.783899413 |
| Ddx54 | 525.9668418 | 0.330431569 | 0.091323751 | 3.618243519 |
| RP23-297M4.4 | 3.794821076 | -0.205284036 | 0.587277451 | -0.349552049 |
| Ccdc42b | 3.366237703 | 0.366659655 | 0.507553862 | 0.722405408 |
| RP23-297M4.1 | 0 | NA | NA | NA |
| Gm15749 | 0.639699895 | -1.170791662 | 1.603018996 | -0.73036668 |
| Rasal1 | 506.8043477 | -0.03900196 | 0.130152117 | -0.29966443 |
| Dtx1 | 1090.783309 | 0.119077143 | 0.074030521 | 1.608487172 |
| Oas2 | 477.0806149 | 7.646438713 | 0.462329591 | 16.53893425 |
| RP24-547J21.3 | 0.066050875 | 0.274391604 | 3.724945049 | 0.073663262 |
| Oas3 | 836.3377262 | 10.38333901 | 0.874897827 | 11.86805897 |
| RP24-547J21.4 | 1.429586369 | 3.464575993 | 1.10207664 | 3.14367973 |
| Oas1e | 0.736605981 | 0.183744895 | 1.466366467 | 0.125306258 |
| Oas1c | 32.60080593 | 1.930495865 | 0.270502828 | 7.136693829 |
| Oas1b | 318.3948597 | 7.150761008 | 0.437039772 | 16.3618084 |
| Oas1f | 0.049553047 | 0.274391604 | 3.724945049 | 0.073663262 |
| Gm22907 | 0 | NA | NA | NA |
| Oas1h | 0.100902008 | 0.274391828 | 3.724945049 | 0.073663322 |
| Oas1g | 272.4663882 | 8.813993112 | 0.771405844 | 11.42588325 |
| Oas1a | 611.2994765 | 8.943771573 | 0.584254136 | 15.30801585 |
| Oas1d | 0.674169638 | 0.755310476 | 1.384462508 | 0.545562246 |
| Rph3a | 3838.839031 | -0.282784966 | 0.088860604 | -3.182343493 |
| RP23-39M18.2 | 36.68051303 | -1.390386631 | 0.324203931 | -4.288617432 |
| Ptpn11 | 2663.867196 | 0.127358664 | 0.074534484 | 1.708721355 |
| RP23-257E17.2 | 45.93026392 | -0.756641885 | 0.33562955 | -2.254395909 |

| | | | | |
|---|---|---|---|---|
| Rpl6 | 1338.143399 | 0.435775854 | 0.084959976 | 5.129189952 |
| Gm26205 | 0 | NA | NA | NA |
| Gm15800 | 6892.751409 | -0.314951599 | 0.090100694 | -3.495551301 |
| Gm24671 | 2.5200291 | -0.188391951 | 0.632258123 | -0.297966833 |
| Trafd1 | 1684.87789 | 2.18122586 | 0.152856981 | 14.26971697 |
| RP23-50I16.1 | 5.493864583 | 1.413059385 | 0.465939469 | 3.032710214 |
| RP23-50I16.2 | 19.99381265 | 0.708789298 | 0.245876228 | 2.882707712 |
| Naa25 | 948.0018726 | 0.191951232 | 0.087906195 | 2.183591633 |
| RP23-50I16.3 | 14.51075867 | -0.27854291 | 0.368406936 | -0.756074011 |
| Erp29 | 574.7376244 | 0.140198245 | 0.100677432 | 1.392548878 |
| Tmem116 | 26.92021731 | -0.40956558 | 0.218471138 | -1.874689644 |
| Gm15546 | 0 | NA | NA | NA |
| Adam1b | 0.227186609 | -0.206489173 | 2.571337767 | -0.080304181 |
| RP24-435H23.1 | 2.953870943 | -0.592722823 | 0.705243818 | -0.840450931 |
| Adam1a | 36.9795004 | -0.793157022 | 0.286516564 | -2.768276333 |
| Mapkapk5 | 369.1571913 | 0.005310129 | 0.093732623 | 0.056651877 |
| Gm15547 | 4.040079771 | 0.165644922 | 0.657121146 | 0.252076688 |
| Aldh2 | 693.3685642 | -0.136010793 | 0.117172554 | -1.160773474 |
| RP23-181P9.3 | 1.724244418 | 0.090799703 | 0.789386973 | 0.115025591 |
| Acad12 | 28.57697775 | -0.948683022 | 0.244233306 | -3.88433108 |
| Acad10 | 41.29551714 | -0.509889673 | 0.201400804 | -2.531716174 |
| Brap | 823.207007 | 0.175089527 | 0.079501175 | 2.202351432 |
| Atxn2 | 1833.502759 | -0.228530054 | 0.069642665 | -3.281466268 |
| RP23-18K20.2 | 63.62259654 | -0.467961571 | 0.360909438 | -1.296617713 |
| Gm16552 | 0 | NA | NA | NA |
| RP23-18K20.3 | 37.1706614 | -0.874765749 | 0.329681915 | -2.65336286 |
| Sh2b3 | 323.628142 | 0.322404594 | 0.10525222 | 3.063161951 |
| Mir7031 | 0.877654538 | -1.536784616 | 1.207747228 | -1.272438951 |
| RP23-431M7.2 | 0.523632775 | 2.121114503 | 1.446475337 | 1.466402122 |
| RP23-431M7.4 | 0.25563639 | -0.675060942 | 2.113943058 | -0.319337335 |
| Fam109a | 123.4761975 | -0.208191187 | 0.111207497 | -1.872096692 |
| Cux2 | 1234.512826 | -0.335584937 | 0.128945861 | -2.602525845 |
| AC113302.1 | 0 | NA | NA | NA |
| RP23-254C8.1 | 0 | NA | NA | NA |
| Gm15637 | 0.291530733 | -0.632965994 | 1.795426472 | -0.352543534 |
| Myl2 | 1.403419482 | 0.182672071 | 0.790416211 | 0.23110871 |
| Ccdc63 | 13.49297597 | -0.00856598 | 0.361699218 | -0.023682607 |
| Ppp1cc | 926.7286209 | 0.026069289 | 0.069316893 | 0.376088544 |
| RP24-347B12.1 | 50.60541787 | -0.406658584 | 0.321184901 | -1.266119864 |
| Hvcn1 | 77.26580927 | 2.499037515 | 0.231276643 | 10.8054038 |
| Tctn1 | 251.1741939 | -0.245826825 | 0.100964385 | -2.434787527 |
| Gm15481 | 0 | NA | NA | NA |
| Pptc7 | 900.5757088 | -0.152345464 | 0.061630966 | -2.471898033 |
| Gm15842 | 8.407355864 | 0.42741124 | 0.447976005 | 0.954094048 |
| Rad9b | 39.0160455 | -0.356386621 | 0.200637106 | -1.776274728 |
| Vps29 | 780.350795 | -0.155338516 | 0.086791251 | -1.789794638 |
| Fam216a | 407.2222889 | 0.000721658 | 0.067854666 | 0.010635352 |

| | | | | |
|---|---|---|---|---|
| Gpn3 | 109.4263101 | 0.075764425 | 0.131309763 | 0.576990041 |
| RP23-407I6.1 | 0.395669761 | -0.968786143 | 1.661861216 | -0.582952495 |
| Arpc3 | 620.2922037 | 0.311367091 | 0.092500083 | 3.366127692 |
| RP23-407I6.2 | 78.71060985 | -0.21830955 | 0.151087574 | -1.444920616 |
| Anapc7 | 355.9121168 | 0.167726249 | 0.083322153 | 2.012985052 |
| Gm15846 | 0 | NA | NA | NA |
| Atp2a2 | 11857.11526 | 0.016596593 | 0.060937988 | 0.272352158 |
| RP23-407I6.6 | 93.54004702 | -0.471723229 | 0.406610855 | -1.160134373 |
| RP23-407I6.5 | 69.66663032 | -0.496156319 | 0.302619294 | -1.639539612 |
| RP23-407I6.4 | 0.863835993 | 1.816121529 | 1.313027728 | 1.383155504 |
| Gm22965 | 1.295159943 | 2.318883821 | 0.993063812 | 2.335080377 |
| Ift81 | 401.1137358 | -0.235857062 | 0.082885412 | -2.84557991 |
| RP23-407I6.7 | 1.063558902 | 0.306721928 | 0.821516005 | 0.373360867 |
| RP24-114E20.2 | 0.51473784 | -0.948887008 | 1.31285952 | -0.722763551 |
| P2rx7 | 217.2536391 | 1.047077625 | 0.167536485 | 6.249848347 |
| Mir8115 | 0 | NA | NA | NA |
| Gm10064 | 0 | NA | NA | NA |
| P2rx4 | 293.0467081 | 0.677834397 | 0.148556547 | 4.562803918 |
| Camkk2 | 1870.397667 | -0.815797605 | 0.14104946 | -5.783769796 |
| RP23-31J24.2 | 7.791943915 | -0.48782262 | 0.392942433 | -1.241460781 |
| Anapc5 | 1309.787107 | -0.138840708 | 0.054512393 | -2.546956735 |
| Rnf34 | 645.0189091 | 0.119303685 | 0.06719072 | 1.775597663 |
| Kdm2b | 295.4828908 | -0.258283074 | 0.092776499 | -2.783927776 |
| RP23-214I6.2 | 6.36596584 | -0.254895517 | 0.53576279 | -0.475761889 |
| RP23-214I6.5 | 0 | NA | NA | NA |
| A930024E05Rik | 27.86517634 | -0.263024984 | 0.207232858 | -1.269224321 |
| RP23-214I6.3 | 0.145501641 | 0.755294927 | 2.8880725 | 0.261522149 |
| Orai1 | 101.2846242 | 0.613281582 | 0.15587654 | 3.934405919 |
| Morn3 | 1.434593181 | -0.132367674 | 0.856525677 | -0.154540229 |
| RP23-214I6.4 | 5.158449836 | 1.493901132 | 0.576400325 | 2.591777046 |
| Gm26745 | 0.042716945 | -0.206490061 | 3.724945049 | -0.055434391 |
| Gm6444 | 0.573335652 | 0.526435529 | 1.161953298 | 0.453060833 |
| Tmem120b | 108.3890549 | -0.266098147 | 0.143868999 | -1.849586427 |
| Rhof | 259.5705259 | -0.279157992 | 0.143250853 | -1.948735283 |
| RP23-371A16.4 | 14.62755803 | -1.252423002 | 0.381146075 | -3.285939658 |
| AI480526 | 273.446315 | -0.331321275 | 0.220517109 | -1.50247424 |
| Gm38102 | 1.104086423 | 2.274095994 | 1.159673 | 1.960980373 |
| RP23-371A16.3 | 8.618846306 | -0.035244512 | 0.390065292 | -0.090355415 |
| Setd1b | 1298.15226 | -0.127344179 | 0.103020801 | -1.236101616 |
| Psmd9 | 209.0408427 | 0.271074439 | 0.089935731 | 3.014090564 |
| Hpd | 0.5402512 | 1.986071107 | 1.405077854 | 1.413495417 |
| RP23-297I15.1 | 0 | NA | NA | NA |
| Wdr66 | 48.81208703 | -0.312649676 | 0.154804733 | -2.019638992 |
| Rpl35a-ps6 | 0.817924277 | 0.37844458 | 1.172652282 | 0.322725318 |
| Gm15857 | 0.981461204 | 0.26996537 | 0.962438692 | 0.280501368 |
| Gm15860 | 13.45882266 | -0.327461956 | 0.264117378 | -1.239834951 |
| Bcl7a | 835.7958974 | -0.207160393 | 0.106700157 | -1.941519106 |

| | | | | |
|---|---|---|---|---|
| Gm23458 | 4.576786753 | -1.0241505 | 0.498927302 | -2.052704865 |
| RP23-370G17.1 | 16.44557487 | -0.728044574 | 0.346153131 | -2.103244226 |
| Gm15747 | 1.272170519 | 1.98547832 | 1.084855068 | 1.830178407 |
| Mlxip | 530.2039296 | 0.10995984 | 0.102309858 | 1.07477268 |
| RP23-370G17.2 | 23.88538577 | -0.197320624 | 0.294985475 | -0.668916408 |
| RP23-370G17.3 | 10.05076668 | -0.343758318 | 0.420439918 | -0.817615795 |
| RP23-370G17.4 | 55.71562266 | -0.270024582 | 0.25272456 | -1.068454061 |
| Rpl31-ps6 | 0.955200391 | -2.443247957 | 1.092589522 | -2.236199329 |
| Il31 | 0.27099349 | 0.755305772 | 2.070494665 | 0.364794841 |
| Lrrc43 | 5.510592113 | -1.012517283 | 0.578924693 | -1.748961991 |
| Gm15751 | 2.413806088 | -1.616318023 | 0.665314475 | -2.429404565 |
| Diablo | 222.7606092 | -0.116805081 | 0.099350346 | -1.175688717 |
| B3gnt4 | 5.682107971 | -1.114674533 | 0.451210742 | -2.470407789 |
| Gm24682 | 0.386251008 | -0.522314686 | 1.707422264 | -0.305908326 |
| Mir7647 | 1.111319392 | -0.481914064 | 0.933088472 | -0.516471995 |
| Vps33a | 1223.976526 | 0.157392622 | 0.052809543 | 2.980382227 |
| RP23-121M13.2 | 3.773545612 | -0.444273257 | 0.563397682 | -0.788560677 |
| Clip1 | 1856.591368 | -0.148607195 | 0.088653243 | -1.676274779 |
| Gm24840 | 0 | NA | NA | NA |
| RP23-121M13.3 | 21.46809472 | -0.371018507 | 0.307060812 | -1.208289993 |
| RP23-121M13.4 | 2.51877741 | 1.370617597 | 0.630653737 | 2.173328273 |
| Zcchc8 | 358.9731995 | -0.341169607 | 0.101833747 | -3.350260776 |
| Rsrc2 | 977.6377703 | 0.012213797 | 0.078909241 | 0.154782847 |
| Kntc1 | 15.85629698 | 0.899903075 | 0.412969331 | 2.179103891 |
| Hcar2 | 55.90105025 | 8.286108294 | 0.888125768 | 9.329881641 |
| Hcar1 | 26.46370296 | 0.562119228 | 0.266427216 | 2.109841616 |
| Denr | 363.9918925 | 0.206253626 | 0.107508355 | 1.918489269 |
| RP24-492O4.5 | 0.261825437 | -0.044072162 | 2.067944014 | -0.021312067 |
| Ccdc62 | 24.55556878 | -0.897276271 | 0.254443382 | -3.526428017 |
| Hip1r | 946.4062031 | -0.046003045 | 0.068119623 | -0.675327352 |
| Mir7032 | 0 | NA | NA | NA |
| Vps37b | 406.419246 | 0.392755663 | 0.104754762 | 3.749286956 |
| Gm37939 | 0.497466187 | -0.564165027 | 1.528256801 | -0.369155908 |
| RP24-492O4.9 | 0.226827357 | -0.206480579 | 2.143585549 | -0.09632486 |
| RP24-492O4.8 | 0.200931026 | 0.274406434 | 2.184849737 | 0.125595106 |
| Abcb9 | 408.729264 | -0.328769008 | 0.074544002 | -4.410401924 |
| Gm16001 | 32.41362582 | -0.540735178 | 0.18889027 | -2.86269472 |
| Ogfod2 | 183.0496373 | -0.022869482 | 0.085230333 | -0.268325614 |
| Arl6ip4 | 210.5407291 | 0.252491501 | 0.137818865 | 1.832053255 |
| Pitpnm2 | 2668.846213 | -0.15427665 | 0.141700514 | -1.08875152 |
| Pitpnm2os2 | 1.19134753 | -0.736210196 | 0.916987718 | -0.802857205 |
| Pitpnm2os1 | 0.048949532 | -0.206487837 | 3.724945049 | -0.055433794 |
| RP24-257M14.2 | 0 | NA | NA | NA |
| Mphosph9 | 491.0917071 | -0.287564067 | 0.169136046 | -1.700193861 |
| Gm23171 | 0 | NA | NA | NA |
| RP24-257M14.3 | 50.23293639 | -0.490887492 | 0.344635696 | -1.424366359 |
| RP23-180G16.2 | 6.931404989 | -0.315763148 | 0.429792308 | -0.73468776 |

| | | | | |
|---|---|---|---|---|
| 2810006K23Rik | 85.50115535 | -0.787014976 | 0.214594064 | -3.6674592 |
| Cdk2ap1 | 265.3510822 | -0.266482529 | 0.099540607 | -2.677123816 |
| Sbno1 | 3030.840915 | -0.297831442 | 0.052910714 | -5.628943936 |
| Gm37415 | 6.225233596 | -0.243529676 | 0.447583162 | -0.544099279 |
| Gm27486 | 0 | NA | NA | NA |
| Setd8 | 525.0620955 | 0.046695351 | 0.102471124 | 0.455692776 |
| Rilpl2 | 121.8049509 | 0.052515137 | 0.121400197 | 0.432578683 |
| Gm15621 | 0.143485256 | 0.274391988 | 3.549605256 | 0.077302113 |
| Snrnp35 | 68.78650209 | 0.189168235 | 0.214203257 | 0.883124926 |
| Rilpl1 | 373.8418609 | -0.262326603 | 0.093650005 | -2.801138166 |
| RP23-311A21.2 | 18.59305175 | 0.019298762 | 0.263073224 | 0.0733589 |
| Tmed2 | 550.4698307 | 0.378881664 | 0.090372329 | 4.192452123 |
| Ddx55 | 213.2561564 | -0.243623117 | 0.136670635 | -1.782556407 |
| Eif2b1 | 210.495533 | 0.186261452 | 0.098796776 | 1.885298885 |
| Gtf2h3 | 153.3355758 | -0.434725127 | 0.137835697 | -3.153937154 |
| Tctn2 | 238.196889 | -0.143355833 | 0.125519118 | -1.142103571 |
| Atp6v0a2 | 1102.977081 | 0.100960964 | 0.102287197 | 0.987034218 |
| Gm24854 | 0 | NA | NA | NA |
| WI1-2065I1.2 | 0.075043205 | 0.755286614 | 3.724945049 | 0.202764498 |
| Dnah10 | 37.63473061 | -0.71212783 | 0.226263532 | -3.147338073 |
| RP23-437D10.1 | 3.569206524 | 1.685397574 | 0.628025439 | 2.683645388 |
| Ccdc92 | 896.6833926 | 0.024634466 | 0.078417935 | 0.314143266 |
| Zfp664 | 1184.705232 | -0.215241802 | 0.06267208 | -3.434412924 |
| Fam101a | 19.01723143 | 1.425497217 | 0.59239786 | 2.406317295 |
| Ncor2 | 4028.896925 | 0.035869464 | 0.074317632 | 0.482650792 |
| RP24-291G12.3 | 0.185374628 | -0.680385936 | 2.266154646 | -0.300238087 |
| RP24-291G12.2 | 0.590920558 | 1.684844589 | 1.232882846 | 1.366589368 |
| RP23-363M4.1 | 0.852297073 | 1.200604896 | 1.160334877 | 1.034705515 |
| Scarb1 | 198.4544971 | 0.025711492 | 0.096034741 | 0.267731158 |
| RP23-363M4.2 | 6.758329482 | -0.842538792 | 0.456508959 | -1.84561283 |
| RP24-105J12.1 | 0 | NA | NA | NA |
| Ubc | 623.813315 | 0.543968491 | 0.147986505 | 3.675797939 |
| Gm10382 | 3.55196556 | -0.569188588 | 0.566600545 | -1.004567668 |
| Dhx37 | 317.3772995 | 0.264126722 | 0.084742897 | 3.11680072 |
| RP24-105J12.3 | 0.046345709 | 0.274391604 | 3.724945049 | 0.073663262 |
| Bri3bp | 1674.728707 | -0.384578313 | 0.066412939 | -5.790713684 |
| RP24-324M10.2 | 0.427937583 | -2.16950163 | 1.346249829 | -1.611514878 |
| Aacs | 461.1003154 | -0.004454863 | 0.092087704 | -0.048376302 |
| Gm27551 | 0.589769752 | -1.537189636 | 1.315765838 | -1.168285109 |
| Tmem132b | 1925.698646 | -0.167241882 | 0.119880931 | -1.395066599 |
| Gm4868 | 0 | NA | NA | NA |
| RP24-332C17.1 | 1.392497718 | 0.463277529 | 1.074299384 | 0.431236894 |
| RP24-332C17.2 | 0 | NA | NA | NA |
| RP24-332C17.3 | 0.274664082 | 0.274407626 | 2.080459595 | 0.1318976 |
| RP24-332C17.4 | 5.259892424 | 1.026460236 | 0.455828943 | 2.251854016 |
| Gm23151 | 0 | NA | NA | NA |
| Gm37700 | 0 | NA | NA | NA |

| | | | | |
|---|---|---|---|---|
| RP23-414E14.1 | 0.0878635 | -0.687376175 | 3.724945049 | -0.184533239 |
| Gm24839 | 0 | NA | NA | NA |
| RP23-74L20.1 | 1.146741161 | 0.214267271 | 0.983004043 | 0.217971912 |
| RP23-74L20.2 | 0.044191549 | -0.206490061 | 3.724945049 | -0.055434391 |
| RP23-145H8.1 | 0 | NA | NA | NA |
| RP23-145H8.2 | 0.512350924 | 2.875373745 | 1.398211366 | 2.05646572 |
| RP23-282A16.1 | 0.230183376 | 1.531095668 | 2.495000664 | 0.613665435 |
| RP23-37N15.1 | 0.075043205 | 0.755286614 | 3.724945049 | 0.202764498 |
| RP23-37N15.3 | 0 | NA | NA | NA |
| RP23-37N15.4 | 0.066050875 | 0.274391604 | 3.724945049 | 0.073663262 |
| RP23-37N15.2 | 0 | NA | NA | NA |
| Tmem132c | 179.9200518 | -0.010703414 | 0.113178412 | -0.094571162 |
| Tmem132cos | 0 | NA | NA | NA |
| RP23-71M14.1 | 1.788302827 | -1.643003394 | 1.01427878 | -1.619873576 |
| WI1-2285D16.1 | 0.302970058 | 2.023065343 | 1.960954654 | 1.0316737 |
| Slc15a4 | 300.9485309 | -0.067017587 | 0.11011613 | -0.608608266 |
| WI1-1024F3.1 | 0.545152564 | 1.359172118 | 1.272037203 | 1.068500289 |
| Glt1d1 | 207.4632902 | -0.23407332 | 0.122810433 | -1.905972606 |
| Tmem132d | 473.83412 | -0.202371706 | 0.144153546 | -1.403862147 |
| Gm17132 | 0.461821276 | -0.169588861 | 1.48254375 | -0.11439046 |
| Gm17131 | 0.522940275 | 0.173520831 | 1.506088038 | 0.11521294 |
| RP23-378C15.2 | 8.098685304 | -0.51099693 | 0.504866804 | -1.012142067 |
| RP23-378C15.1 | 6.802413532 | -1.645355307 | 0.489577232 | -3.36076762 |
| WI1-2255B14.1 | 0.141177696 | 0.274407731 | 2.91536683 | 0.094124598 |
| CH36-12I15.2 | 13.22303069 | 1.42782911 | 0.374121694 | 3.816483072 |
| CH36-12I15.3 | 7.173227082 | 3.145171263 | 0.598926985 | 5.251343393 |
| CH36-12I15.1 | 0 | NA | NA | NA |
| CH36-12I15.4 | 0.075043205 | 0.755286614 | 3.724945049 | 0.202764498 |
| 5930412G12Rik | 23.11388567 | -1.012716214 | 0.229369645 | -4.415214639 |
| RP24-258A20.1 | 1.894888719 | -1.154217386 | 0.784577418 | -1.47113256 |
| Fzd10 | 39.99764841 | -0.544730606 | 0.176554537 | -3.085339038 |
| Piwil1 | 0.809884792 | -0.379140174 | 1.096094053 | -0.345901132 |
| RP23-156F20.2 | 0 | NA | NA | NA |
| Rimbp2 | 1419.490251 | -0.532363972 | 0.223272797 | -2.384365582 |
| RP23-156F20.4 | 0.134481855 | 0.720143159 | 3.654633157 | 0.197049369 |
| RP23-156F20.5 | 1.664368785 | -3.070691848 | 1.010939474 | -3.037463595 |
| Stx2 | 147.9774137 | 0.056146234 | 0.122593622 | 0.457986583 |
| Ran | 1531.557904 | 0.143624139 | 0.090075933 | 1.594478509 |
| Adgrd1 | 48.98301064 | 0.370865723 | 0.178117004 | 2.082146648 |
| Rps16-ps2 | 75.19513727 | 0.371919366 | 0.163581252 | 2.273606303 |
| RP24-158G19.1 | 0 | NA | NA | NA |
| RP24-81J14.1 | 2.60471737 | 0.467804166 | 0.797602949 | 0.586512584 |
| Sfswap | 880.7266452 | -0.104037451 | 0.136945612 | -0.75969905 |
| RP24-161C20.4 | 61.43406924 | -0.939888939 | 0.35709307 | -2.632055949 |
| Mmp17 | 1263.227049 | -0.19510719 | 0.134342267 | -1.452314259 |
| Gm6139 | 8.767336692 | 0.419828398 | 0.412774935 | 1.017087915 |
| Zfp11 | 110.3957661 | 0.018170938 | 0.134101615 | 0.135501261 |

| | | | | |
|---|---|---|---|---|
| 14-Sep | 0.250988417 | 1.236196787 | 2.479133789 | 0.498640611 |
| Mrps17 | 416.7636034 | -0.212563118 | 0.062748326 | -3.387550409 |
| Gbas | 690.6986552 | -0.049573039 | 0.068654597 | -0.722064377 |
| Gm15903 | 27.46139881 | -0.466224957 | 0.327031509 | -1.425627025 |
| Psph | 139.9382442 | -0.106715459 | 0.150617614 | -0.708519115 |
| Cct6a | 1504.12042 | 0.156407933 | 0.088122327 | 1.774895626 |
| Gm23245 | 2.474741284 | -0.260243429 | 0.639305524 | -0.407072079 |
| Snora15 | 0 | NA | NA | NA |
| Sumf2 | 86.25597758 | 0.036337944 | 0.125597562 | 0.289320458 |
| Phkg1 | 43.76477553 | -1.67503992 | 0.26730822 | -6.266324009 |
| WI1-535J9.1 | 0.523518094 | -2.220378644 | 1.282366332 | -1.731469852 |
| Chchd2 | 843.0146962 | 0.125535634 | 0.117623645 | 1.067265295 |
| Zbed5 | 99.19524039 | -0.123121695 | 0.159268995 | -0.773042456 |
| Nupr1l | 8.851478164 | -0.515872934 | 0.328430038 | -1.57072397 |
| Vkorc1l1 | 758.1542901 | -0.230904471 | 0.063012893 | -3.664400423 |
| RP23-189N6.2 | 51.40024564 | -0.467014525 | 0.313710657 | -1.488679182 |
| RP23-189N6.3 | 26.74839225 | -0.316393877 | 0.365268264 | -0.866195911 |
| Gusb | 191.859854 | 0.931699758 | 0.139805128 | 6.664274554 |
| Asl | 102.1205115 | 0.322312165 | 0.134663125 | 2.393470108 |
| Crcp | 187.2081582 | 0.648084483 | 0.111880577 | 5.792645163 |
| Tpst1 | 399.6196954 | 0.267243431 | 0.09173616 | 2.913174382 |
| RP23-353L10.2 | 7.510072057 | -0.240555015 | 0.499334639 | -0.481751107 |
| Kctd7 | 165.5730604 | -0.594757289 | 0.10388479 | -5.72516238 |
| Rabgef1 | 458.137125 | -0.061432636 | 0.071449051 | -0.859810383 |
| Gm15920 | 252.9729714 | 0.151986798 | 0.106249804 | 1.430466618 |
| Gm15921 | 0.048949532 | -0.206487837 | 3.724945049 | -0.055433794 |
| Tmem248 | 679.0584884 | 0.327869953 | 0.104259411 | 3.144751627 |
| AC113029.6 | 6.876401235 | 0.317866946 | 0.407716788 | 0.779626829 |
| RP23-206J17.2 | 14.12832924 | 0.564045293 | 0.420617754 | 1.340992594 |
| Sbds | 553.098637 | 0.413122374 | 0.119648462 | 3.452801382 |
| Tyw1 | 213.1214519 | -0.298874077 | 0.132956809 | -2.247903507 |
| Caln1 | 4353.134745 | 0.177478821 | 0.095226382 | 1.863756828 |
| A330070K13Rik | 2.26626686 | -0.702047731 | 0.753164391 | -0.932130807 |
| RP24-571K14.1 | 0 | NA | NA | NA |
| Gm23761 | 4.136581771 | -0.327734511 | 0.534419497 | -0.613253282 |
| RP24-559P3.1 | 12.49495635 | -0.726408446 | 0.373102114 | -1.946942724 |
| Wbscr17 | 856.0949554 | -0.11687831 | 0.082571901 | -1.415473175 |
| RP23-323D23.1 | 6.092562558 | -0.709691646 | 0.453615334 | -1.56452305 |
| RP23-323D23.2 | 13.43160181 | -0.405696431 | 0.395738784 | -1.02516217 |
| RP23-323D23.3 | 19.70298692 | -0.192398308 | 0.408326962 | -0.471186882 |
| RP23-445H19.1 | 18.79099024 | -0.667889658 | 0.269192182 | -2.48108862 |
| RP23-114L5.4 | 5.83927098 | 0.044794555 | 0.440449096 | 0.101702002 |
| Auts2 | 2085.675582 | -0.256866234 | 0.126298472 | -2.033803188 |
| Gm27266 | 0 | NA | NA | NA |
| RP23-120C8.7 | 31.89874726 | -0.618502442 | 0.391563334 | -1.579571907 |
| RP23-120C8.6 | 19.56297173 | -0.498333755 | 0.337100719 | -1.47829336 |
| RP23-120C8.9 | 0.108767819 | -0.20649451 | 3.724945049 | -0.055435586 |

| | | | | |
|---|---|---|---|---|
| RP23-120C8.5 | 11.30893109 | -1.150676298 | 0.476989618 | -2.412371788 |
| RP23-120C8.4 | 44.54237424 | -0.434552869 | 0.34741972 | -1.250800817 |
| RP23-120C8.3 | 22.2103361 | -0.598656879 | 0.352810521 | -1.696822639 |
| RP23-120C8.2 | 22.94359978 | -0.676287863 | 0.291942173 | -2.316513084 |
| RP24-95O4.7 | 0 | NA | NA | NA |
| RP24-95O4.6 | 73.13605096 | -0.692163288 | 0.295095236 | -2.345559 |
| RP24-95O4.5 | 20.70201435 | -0.817912973 | 0.499119846 | -1.638710581 |
| RP24-95O4.4 | 58.37291526 | -0.629230506 | 0.337139828 | -1.866378439 |
| RP24-95O4.9 | 0 | NA | NA | NA |
| RP24-95O4.3 | 42.64488512 | -0.532634434 | 0.250587109 | -2.125546025 |
| RP24-95O4.2 | 132.5110994 | -0.750319001 | 0.368154459 | -2.038054905 |
| RP24-95O4.1 | 237.0691924 | -0.79962097 | 0.320290336 | -2.496550407 |
| RP23-378O9.2 | 72.35690436 | -0.85197789 | 0.320618226 | -2.657297123 |
| RP23-378O9.1 | 291.0443093 | -0.731824247 | 0.289267165 | -2.529925053 |
| RP24-147H20.3 | 37.48355553 | -0.915686828 | 0.314258978 | -2.913796877 |
| RP24-147H20.2 | 50.07029278 | -0.973341371 | 0.243505611 | -3.997203041 |
| RP23-276C23.3 | 53.66935453 | -0.982327855 | 0.325706751 | -3.015988625 |
| RP23-276C23.2 | 105.0397251 | -0.914817648 | 0.22938801 | -3.988079627 |
| RP23-276C23.1 | 58.80184518 | -1.120462363 | 0.337952507 | -3.315443266 |
| RP24-399A15.2 | 87.32223591 | -1.14477937 | 0.30661308 | -3.733628619 |
| RP24-399A15.3 | 100.1496488 | -0.977081566 | 0.283864072 | -3.442075495 |
| RP24-399A15.5 | 46.61621938 | -0.792158535 | 0.378380801 | -2.093548437 |
| Gm28616 | 0.118974955 | 0.274400501 | 3.724945049 | 0.073665651 |
| RP24-399A15.6 | 0.098502579 | -0.206492286 | 3.724945049 | -0.055434988 |
| Gm38373 | 0 | NA | NA | NA |
| Gm25722 | 0 | NA | NA | NA |
| RP23-455A10.3 | 0 | NA | NA | NA |
| Gm10051 | 4.533521042 | 0.563302779 | 0.613322313 | 0.918444947 |
| RP23-455A10.2 | 0 | NA | NA | NA |
| RP23-455A10.1 | 0.061920613 | 0.274393828 | 3.724945049 | 0.073663859 |
| RP23-409D19.1 | 5.435068931 | -0.356151038 | 0.407676333 | -0.873612249 |
| RP23-409D19.2 | 0 | NA | NA | NA |
| RP24-199E1.1 | 0.733529363 | -1.109988443 | 1.246669997 | -0.890362682 |
| Gm15627 | 4.57824934 | -0.185144577 | 0.461334467 | -0.401323963 |
| Gatsl2 | 1933.057981 | -0.262120671 | 0.076454207 | -3.428466292 |
| Wbscr16 | 129.6578973 | 0.220800593 | 0.121720274 | 1.814000128 |
| B230377A18Rik | 10.36497119 | 0.486595999 | 0.387808064 | 1.254734091 |
| Gtf2ird2 | 207.0338271 | 0.095228111 | 0.099151432 | 0.960431017 |
| Gm24139 | 2.685875012 | 0.429846535 | 0.591505535 | 0.726699092 |
| Ncf1 | 204.7455163 | 1.868272994 | 0.199483543 | 9.365549497 |
| Gtf2i | 2503.932034 | -0.473120267 | 0.103431248 | -4.574248825 |
| RP23-15K13.2 | 110.8580539 | -0.979639606 | 0.370399006 | -2.644822445 |
| RP23-15K13.3 | 5.451269312 | -1.419177396 | 0.5911889 | -2.400548111 |
| Mir3965 | 0 | NA | NA | NA |
| Gtf2ird1 | 384.0610061 | -0.143753515 | 0.13862725 | -1.036978764 |
| n-R5s178 | 0 | NA | NA | NA |
| Gm26340 | 0 | NA | NA | NA |

| | | | | |
|---|---|---|---|---|
| RP24-359H20.2 | 0.585862523 | -0.031464091 | 1.200452664 | -0.026210189 |
| Gm16020 | 0 | NA | NA | NA |
| RP24-359H20.3 | 3.016053333 | -3.244221697 | 0.878019418 | -3.694931605 |
| Clip2 | 1985.813075 | 0.097800295 | 0.051221516 | 1.909359646 |
| RP23-85H24.1 | 29.77219559 | -0.695836866 | 0.319494566 | -2.177930207 |
| RP23-85H24.2 | 8.735753554 | -0.30026547 | 0.333269796 | -0.900968146 |
| Syna | 7.492197562 | 0.54250292 | 0.402772267 | 1.346922228 |
| mmu-mir-7228 | 0.285083685 | -0.690265434 | 1.774655491 | -0.388957428 |
| Rfc2 | 247.4000499 | 0.004292289 | 0.0863546 | 0.049705391 |
| Lat2 | 113.3855065 | 2.466005311 | 0.250364823 | 9.849647745 |
| Eif4h | 3070.987643 | 0.096214503 | 0.049093781 | 1.959810419 |
| Limk1 | 829.834934 | -0.321721287 | 0.121331662 | -2.65158559 |
| Gm10369 | 1.019349322 | 2.339887886 | 1.252984006 | 1.867452318 |
| Eln | 72.96738645 | 0.028801025 | 0.188445059 | 0.152835128 |
| RP23-315E2.1 | 8.223192194 | -0.047491565 | 0.343469403 | -0.138270146 |
| Wbscr28 | 0.045882455 | 0.755288839 | 3.724945049 | 0.202765096 |
| Cldn13 | 0 | NA | NA | NA |
| RP23-201C9.3 | 0 | NA | NA | NA |
| RP23-201C9.2 | 0.038981395 | 0.274393828 | 3.724945049 | 0.073663859 |
| Wbscr27 | 62.20310215 | -0.127242505 | 0.159123157 | -0.799647938 |
| Cldn4 | 0.565252138 | 1.199700022 | 3.096598863 | 0.387425067 |
| Cldn3 | 2.938076242 | 0.457613133 | 0.632007866 | 0.724062402 |
| Wbscr25 | 2.760772158 | 0.345137602 | 0.54557425 | 0.632613438 |
| Rps29-ps | 0 | NA | NA | NA |
| RP23-38B15.1 | 4.804143474 | -0.923607565 | 0.561186384 | -1.645812501 |
| Abhd11 | 157.6817239 | 0.121959438 | 0.116723788 | 1.044855039 |
| Abhd11os | 7.140605062 | -1.025345108 | 0.537873104 | -1.906295555 |
| Stx1a | 708.4200384 | -0.588141592 | 0.159357458 | -3.690706407 |
| Wbscr22 | 194.0156334 | -0.187819899 | 0.082328937 | -2.281335165 |
| Gm25492 | 29.10926416 | -0.061529786 | 0.203818512 | -0.301885168 |
| Dnajc30 | 252.3546012 | -0.015162794 | 0.086418992 | -0.175456735 |
| Vps37d | 164.5301101 | 0.353259878 | 0.119803787 | 2.948653694 |
| Mlxipl | 13.0557667 | -0.320796872 | 0.348852615 | -0.919577087 |
| RP23-240H6.1 | 0 | NA | NA | NA |
| Tbl2 | 380.9054715 | -0.04284143 | 0.065892741 | -0.650169186 |
| Bcl7b | 442.8768574 | 0.210081285 | 0.057662606 | 3.643284597 |
| Baz1b | 2277.625964 | -0.242474957 | 0.087299166 | -2.777517442 |
| Gm25315 | 0 | NA | NA | NA |
| Fzd9 | 41.32503692 | -0.309450727 | 0.181880985 | -1.701391311 |
| RP23-240H6.2 | 0.095295241 | -0.206492286 | 3.724945049 | -0.055434988 |
| Fkbp6 | 0.371024948 | 1.347915677 | 1.67069175 | 0.806800942 |
| Trim50 | 0.160978641 | -1.177170116 | 2.79661493 | -0.42092678 |
| Nsun5 | 116.6234585 | 0.256099425 | 0.174852491 | 1.464659856 |
| Pom121 | 1177.537542 | 0.009634335 | 0.07311349 | 0.131772326 |
| Mir7033 | 0 | NA | NA | NA |
| Hip1 | 1047.747718 | 0.041767703 | 0.115219309 | 0.362506106 |
| RP23-333I24.3 | 8.728400981 | -0.866029649 | 0.384854266 | -2.250279457 |

| | | | | |
|---|---|---|---|---|
| Ccl26 | 0 | NA | NA | NA |
| Ccl24 | 1.573001602 | -3.853295396 | 0.99711974 | -3.864425947 |
| RP23-423A22.3 | 0.045882455 | 0.755288839 | 3.724945049 | 0.202765096 |
| Rhbdd2 | 635.3255966 | -0.088125539 | 0.055033877 | -1.601296203 |
| Gm15607 | 0.101408268 | 0.755286617 | 3.724945049 | 0.202764499 |
| Gm21839 | 0.540306067 | -0.611452699 | 1.337273188 | -0.457238434 |
| RP23-67P7.1 | 0.422515209 | 0.756815089 | 1.398437706 | 0.541186129 |
| Por | 973.1641497 | 0.06835762 | 0.101810846 | 0.671417863 |
| Gm16061 | 0.898502897 | 0.019217057 | 1.14244163 | 0.016821041 |
| Tmem120a | 119.540289 | 0.085613808 | 0.124241164 | 0.689093735 |
| Mir7034 | 0 | NA | NA | NA |
| Styxl1 | 3.717676903 | -0.483752313 | 0.552640591 | -0.875347053 |
| Mdh2 | 1871.37566 | -0.120228873 | 0.059216572 | -2.03032477 |
| Srrm3 | 617.0965085 | -0.533552485 | 0.144655424 | -3.688437464 |
| Gm23268 | 0 | NA | NA | NA |
| Srrm3os | 0.79881978 | 3.576715367 | 3.584673752 | 0.997779886 |
| Gm22450 | 0.295673938 | -0.677079412 | 1.623238414 | -0.41711643 |
| Hspb1 | 217.0812965 | 2.232831221 | 0.442255312 | 5.048738048 |
| RP23-11P12.2 | 1.422147357 | 3.771986353 | 1.22619379 | 3.076174732 |
| Ywhag | 12717.10653 | 0.035032756 | 0.068640536 | 0.51037999 |
| Ssc4d | 19.88563498 | 0.289081615 | 0.266227474 | 1.08584441 |
| Zp3 | 0.564341384 | -0.453600921 | 1.3659231 | -0.332083791 |
| Dtx2 | 165.5707377 | 0.386648762 | 0.136239516 | 2.838007455 |
| Upk3b | 0.665037802 | 1.844313356 | 1.346186403 | 1.370028217 |
| Upk3bl | 6.950616643 | 1.028098552 | 0.589176599 | 1.744975198 |
| Rasa4 | 403.5442416 | 2.038160797 | 0.137328985 | 14.84144656 |
| Gm22214 | 0 | NA | NA | NA |
| Gm15701 | 0.196892959 | -0.206480695 | 2.233140408 | -0.09246203 |
| Mir7035 | 0 | NA | NA | NA |
| Polr2j | 169.450244 | -0.016243356 | 0.102719665 | -0.158132873 |
| RP23-314O1.1 | 1.911784405 | -0.289329351 | 0.732619343 | -0.394924531 |
| Lrwd1 | 120.9503839 | -0.246489492 | 0.112679034 | -2.187536443 |
| Alkbh4 | 76.72992867 | 0.471829615 | 0.124430335 | 3.791917902 |
| Orai2 | 675.4374974 | -0.029206832 | 0.085671328 | -0.340917235 |
| Prkrip1 | 213.2039114 | 0.371486147 | 0.131046528 | 2.834765261 |
| RP23-284P20.2 | 0 | NA | NA | NA |
| Sh2b2 | 93.36965878 | 0.40736072 | 0.162009446 | 2.514425731 |
| Cux1 | 1881.15298 | -0.262522629 | 0.070844877 | -3.705597935 |
| Gm16599 | 1.050839697 | -1.982425684 | 1.113698261 | -1.780038412 |
| A430110C17Rik | 51.6013035 | -0.664601903 | 0.304361447 | -2.183594239 |
| mmu-mir-721 | 2.23838086 | -1.001613919 | 0.732713814 | -1.366991996 |
| Gm38082 | 75.94798503 | -0.899531065 | 0.283732599 | -3.170347956 |
| Myl10 | 0.304278192 | 2.496778883 | 3.686936961 | 0.677195978 |
| Col26a1 | 25.30413262 | -0.635385358 | 0.211398502 | -3.005628481 |
| Gm20485 | 0.218750444 | -0.659016028 | 2.603325808 | -0.253143892 |
| RP24-456B4.2 | 0 | NA | NA | NA |
| Ift22 | 390.4508874 | -0.126845713 | 0.08625465 | -1.470595657 |

| | | | | |
|---|---|---|---|---|
| 4933404O12Rik | 126.181175 | -0.302146722 | 0.108053989 | -2.796256998 |
| Fis1 | 968.7424033 | 0.211918387 | 0.080297513 | 2.63916501 |
| Cldn15 | 7.096013574 | -0.19853633 | 0.553403092 | -0.358755368 |
| Znhit1 | 217.8120474 | 0.122474544 | 0.101224565 | 1.209929066 |
| Plod3 | 660.8862868 | 0.794887029 | 0.112279795 | 7.079519774 |
| Mir702 | 0 | NA | NA | NA |
| Gm7284 | 0.113979814 | 0.274404949 | 3.724945049 | 0.073666845 |
| Gm26989 | 0 | NA | NA | NA |
| Vgf | 2915.612919 | -0.313122224 | 0.163835251 | -1.911201783 |
| Ap1s1 | 1188.779745 | -0.00663903 | 0.070678025 | -0.093933439 |
| Serpine1 | 1051.265589 | 4.107129364 | 0.511444878 | 8.030443821 |
| Trim56 | 1329.516906 | 3.394674857 | 0.166998756 | 20.32754582 |
| Muc3 | 0.044544648 | 0.755286614 | 3.724945049 | 0.202764498 |
| Gm3054 | 0.061920613 | 0.274393828 | 3.724945049 | 0.073663859 |
| CAAA01077340.1 | 0.185747766 | -0.687376472 | 3.292273734 | -0.208784727 |
| CAAA01154573.1 | 3.791680059 | 1.376928266 | 0.571197092 | 2.410600975 |
| Muc3a | 21.091838 | 0.135364906 | 0.456179721 | 0.296735913 |
| WI1-166E14.1 | 15.34798663 | -0.781753733 | 0.302555168 | -2.583838633 |
| Ache | 1111.230768 | 0.196247476 | 0.105288835 | 1.863896351 |
| Mir8116 | 1.629432974 | -0.224809916 | 0.911191553 | -0.246720808 |
| Ufsp1 | 91.40840618 | 0.345992114 | 0.160622076 | 2.154075722 |
| Srrt | 944.0836702 | -0.025239859 | 0.074926635 | -0.336860969 |
| Mir7036 | 0 | NA | NA | NA |
| RP24-210N1.1 | 0.752770407 | 1.140684808 | 1.299273293 | 0.877940625 |
| Trip6 | 103.7948532 | 0.959381104 | 0.216929175 | 4.422554522 |
| Slc12a9 | 407.3671883 | 0.484382281 | 0.085142039 | 5.689108297 |
| RP24-210N1.3 | 0.151841013 | -0.206481646 | 2.850049718 | -0.072448437 |
| Ephb4 | 94.20008279 | 0.611899245 | 0.153271717 | 3.992251521 |
| Zan | 8.749737854 | 0.412323539 | 0.372559602 | 1.106731746 |
| RP24-210N1.4 | 0 | NA | NA | NA |
| Epo | 0.229076001 | -1.613542751 | 2.528990758 | -0.638018445 |
| Pop7 | 101.0875345 | 0.15574833 | 0.135285186 | 1.151259305 |
| RP23-45G4.2 | 6.33503719 | -0.242471998 | 0.426740152 | -0.568195885 |
| RP23-45G4.3 | 32.98247416 | -0.293448517 | 0.211893227 | -1.384888613 |
| Gigyf1 | 841.5272401 | -0.142156478 | 0.121785911 | -1.167265384 |
| Gnb2 | 2014.783637 | 0.072160184 | 0.062587321 | 1.152952112 |
| Mir6418 | 0 | NA | NA | NA |
| Gm17112 | 0.842992085 | -1.410044477 | 1.157023019 | -1.218683167 |
| Actl6b | 409.7728831 | -0.345271167 | 0.108767896 | -3.174384903 |
| Tfr2 | 33.22771086 | 0.305299065 | 0.21785176 | 1.401407387 |
| Mospd3 | 114.8382337 | -0.070946565 | 0.110433105 | -0.642439284 |
| RP23-286M3.4 | 4.875430065 | -0.374973668 | 0.512994307 | -0.730950935 |
| Pcolce | 87.87249262 | 0.08494203 | 0.19360168 | 0.438746349 |
| Fbxo24 | 2.595989607 | -0.263975015 | 0.61512571 | -0.429139947 |
| RP23-286M3.5 | 4.924586763 | -1.422572013 | 0.532532575 | -2.671333322 |
| Gm16089 | 9.122415521 | -0.252799153 | 0.36835383 | -0.686294352 |
| Lrch4 | 4.483355963 | -0.598280788 | 0.591808967 | -1.01093566 |

| | | | | |
|---|---|---|---|---|
| Gm20605 | 6.061787305 | -0.47410195 | 0.486802636 | -0.973909989 |
| Sap25 | 0.075043205 | 0.755286614 | 3.724945049 | 0.202764498 |
| Irs3 | 1.938977309 | -1.046009904 | 0.751525223 | -1.391849364 |
| Agfg2 | 460.5243141 | 0.001774892 | 0.078507767 | 0.022607851 |
| Gm15498 | 0.332718546 | 1.663163316 | 1.997923232 | 0.832446057 |
| Gm15753 | 9.170347486 | 2.055910455 | 0.470751158 | 4.367297712 |
| Nyap1 | 848.2957187 | -0.1668133 | 0.077028331 | -2.16560969 |
| Tsc22d4 | 1384.328407 | 0.670858731 | 0.14585522 | 4.599483872 |
| Ppp1r35 | 80.27309234 | 0.067131594 | 0.154960988 | 0.433216095 |
| Mepce | 500.0316796 | 0.059440765 | 0.071640322 | 0.829711024 |
| Zcwpw1 | 45.4661166 | -0.548078028 | 0.191911693 | -2.855886578 |
| Pilra | 8.121710499 | 2.27065037 | 0.484500007 | 4.686584801 |
| Pilrb1 | 1.522009338 | 1.689763836 | 1.258946486 | 1.342204657 |
| RP23-425N17.3 | 0.609942678 | 1.728889023 | 1.471822173 | 1.174658906 |
| Pilrb2 | 1.815499725 | 2.276298954 | 0.953630356 | 2.386982483 |
| RP23-425N17.5 | 0.143022002 | 0.755289479 | 3.547809333 | 0.212888971 |
| RP23-425N17.6 | 0 | NA | NA | NA |
| RP23-131D6.3 | 0 | NA | NA | NA |
| Cyp3a13 | 18.8351161 | 0.623113108 | 0.321472708 | 1.938307958 |
| Gm15502 | 0.499480636 | -0.303818416 | 1.442242896 | -0.210656899 |
| RP23-131D6.4 | 4.460458762 | 0.515600006 | 0.510621386 | 1.009750119 |
| Gjc3 | 574.2169864 | -0.318443057 | 0.130290088 | -2.444108078 |
| Azgp1 | 0.218824723 | 0.245255888 | 2.580964285 | 0.095024906 |
| Gm4963 | 0.586951709 | 0.244321232 | 1.242801626 | 0.196589083 |
| Gm29127 | 0 | NA | NA | NA |
| Smok3a | 0.044544648 | 0.755286614 | 3.724945049 | 0.202764498 |
| Smok3b | 0 | NA | NA | NA |
| Smok3c | 0 | NA | NA | NA |
| Zkscan1 | 1601.280236 | 0.102860839 | 0.047695411 | 2.156619188 |
| RP23-147H4.1 | 0 | NA | NA | NA |
| Zscan21 | 197.9428209 | 0.047038029 | 0.092972807 | 0.505933186 |
| Zfp113 | 154.778503 | -0.493540816 | 0.169631633 | -2.909485729 |
| Cops6 | 678.0499843 | 0.062705015 | 0.089180961 | 0.703121099 |
| Mcm7 | 172.4442146 | 0.4115148 | 0.146248231 | 2.813810445 |
| Mir25 | 0 | NA | NA | NA |
| Mir93 | 0 | NA | NA | NA |
| Mir106b | 0.072025462 | 0.755288839 | 3.724945049 | 0.202765096 |
| Ap4m1 | 196.8680612 | 0.068395713 | 0.111971207 | 0.610833041 |
| Taf6 | 295.6377165 | -0.154559077 | 0.106293792 | -1.454074355 |
| Cnpy4 | 86.00518766 | -0.031928846 | 0.141964148 | -0.22490781 |
| Mblac1 | 28.62003678 | -0.409535783 | 0.225550074 | -1.815720008 |
| Gm454 | 0 | NA | NA | NA |
| RP24-86H20.4 | 0 | NA | NA | NA |
| Gm22956 | 0 | NA | NA | NA |
| Nxpe5 | 0.64031546 | -1.166470805 | 1.394655668 | -0.836386236 |
| Lamtor4 | 172.5096594 | -0.142143186 | 0.117602621 | -1.208673622 |
| BC037034 | 543.4878643 | -0.008999954 | 0.057998903 | -0.155174557 |

| | | | | |
|---|---|---|---|---|
| 6330418K02Rik | 28.7450452 | 0.120209046 | 0.223600693 | 0.537605875 |
| Gal3st4 | 0 | NA | NA | NA |
| Gpc2 | 31.36245639 | -0.771632201 | 0.221613625 | -3.481880687 |
| Stag3 | 27.52049443 | 0.852094802 | 0.2616353 | 3.256803651 |
| RP24-95J4.1 | 0.060174537 | 0.755286614 | 3.724945049 | 0.202764498 |
| Gm10874 | 0.888510269 | -0.206142893 | 1.299384196 | -0.158646606 |
| Got2-ps1 | 146.5220016 | 0.110985148 | 0.102240565 | 1.085529487 |
| Gm25525 | 0.061920613 | 0.274393828 | 3.724945049 | 0.073663859 |
| Zfp157 | 295.429195 | -0.064785703 | 0.096030744 | -0.674635016 |
| Gm5564 | 8.712984693 | -0.134438116 | 0.388401871 | -0.346131484 |
| RP24-255F19.2 | 1.435448476 | -0.492355586 | 0.92344405 | -0.533173164 |
| 1700123K08Rik | 2.221046269 | 1.377342144 | 0.789218687 | 1.745197074 |
| Zfp68 | 529.1369275 | 0.074558769 | 0.066778402 | 1.116510222 |
| Gm24082 | 0 | NA | NA | NA |
| Gm15497 | 0.148708979 | 0.755295232 | 2.868529146 | 0.263304012 |
| RP24-255F19.5 | 0.796574019 | 1.962795533 | 1.447638133 | 1.355860618 |
| A430033K04Rik | 179.5683617 | -0.161603943 | 0.114079972 | -1.416584703 |
| Gm38232 | 0.132101749 | 0.274387056 | 3.724945049 | 0.073662041 |
| Fam20c | 570.0411155 | -0.099647793 | 0.096933667 | -1.027999829 |
| Gm5294 | 0.042716945 | -0.206490061 | 3.724945049 | -0.055434391 |
| Pdgfa | 337.774443 | 0.007869014 | 0.083871574 | 0.093822187 |
| 6330403L08Rik | 206.3643988 | -0.712881437 | 0.135037426 | -5.279139705 |
| Prkar1b | 4853.393054 | -0.128624388 | 0.083395233 | -1.542347011 |
| Gm15672 | 16.59824677 | -0.277788779 | 0.306725298 | -0.905659825 |
| Dnaaf5 | 128.0720811 | 0.293038951 | 0.1234559 | 2.373632611 |
| Sun1 | 919.0838976 | -0.214688219 | 0.108095304 | -1.986101258 |
| RP23-430G6.3 | 0.066050875 | 0.274391604 | 3.724945049 | 0.073663262 |
| Get4 | 323.1872879 | 0.463136984 | 0.114320699 | 4.051208481 |
| Adap1 | 1303.85996 | -0.088163013 | 0.06866026 | -1.284047175 |
| Gm27316 | 0 | NA | NA | NA |
| Cox19 | 240.9171869 | -0.094172563 | 0.076499115 | -1.231028143 |
| Cyp2w1 | 0 | NA | NA | NA |
| 3110082I17Rik | 68.15138838 | 0.160068677 | 0.127354215 | 1.256877733 |
| Mir339 | 1.20342686 | -1.085985934 | 1.124603823 | -0.965660896 |
| Gpr146 | 293.4623801 | 0.579995936 | 0.097047878 | 5.976389696 |
| C130050O18Rik | 9.082211208 | 1.650781771 | 0.438403143 | 3.765442376 |
| Gper1 | 14.62948997 | 0.086503625 | 0.315068508 | 0.274554971 |
| Gm26285 | 0.142244466 | 0.27438755 | 3.560510919 | 0.077064094 |
| Zfand2a | 664.2777979 | 0.366743347 | 0.100200664 | 3.660088981 |
| RP23-400D9.3 | 0.170157162 | 0.274406568 | 2.749797554 | 0.099791553 |
| Uncx | 53.72924227 | -0.604573061 | 0.198420286 | -3.04693171 |
| RP23-92L4.4 | 0 | NA | NA | NA |
| Micall2 | 117.2123074 | 2.054257653 | 0.292282504 | 7.028329175 |
| Ints1 | 1337.189342 | -0.040323526 | 0.055721425 | -0.723662858 |
| Mir7037 | 0 | NA | NA | NA |
| Mafk | 342.8849901 | 2.020137027 | 0.206275048 | 9.793414425 |
| Tmem184a | 2.905166453 | 0.504574046 | 0.713344593 | 0.707335629 |

| | | | | |
|---|---|---|---|---|
| Gm26938 | 0.061920613 | 0.274393828 | 3.724945049 | 0.073663859 |
| Psmg3 | 54.0547785 | -0.009633948 | 0.166136589 | -0.057988119 |
| Gm24146 | 2.128603593 | 1.224952309 | 0.76837613 | 1.594209218 |
| RP23-168E11.1 | 0.493577707 | -0.944285126 | 1.411512464 | -0.66898816 |
| Elfn1 | 266.0295 | -0.450252458 | 0.13295712 | -3.38644865 |
| RP23-31L10.2 | 0 | NA | NA | NA |
| Mad1l1 | 200.5910728 | 0.328279585 | 0.0972775 | 3.374671279 |
| RP23-31L10.4 | 9.073372328 | -0.997432552 | 0.346012653 | -2.882647626 |
| Gm16122 | 0.099106094 | 0.274387056 | 3.724945049 | 0.073662041 |
| Gm16121 | 0.641775973 | -0.020895769 | 1.517713188 | -0.01376793 |
| RP24-97G6.1 | 10.02733677 | -0.706029102 | 0.347300338 | -2.032906466 |
| Gm16120 | 0.194043089 | -0.687377773 | 2.234735546 | -0.307587971 |
| RP23-307I19.1 | 0.437638936 | 1.487865958 | 1.854216344 | 0.802423063 |
| Nudt1 | 23.04760792 | 1.47E-05 | 0.264143371 | 5.55E-05 |
| Ftsj2 | 69.98359942 | -0.062475599 | 0.150886059 | -0.414058132 |
| Snx8 | 233.8617382 | 0.305188387 | 0.129352438 | 2.359355516 |
| Eif3b | 1148.304761 | 0.283043431 | 0.084171293 | 3.362707408 |
| Mir7038 | 0.519085648 | -1.374535779 | 1.428934234 | -0.96193075 |
| RP23-307I19.6 | 0 | NA | NA | NA |
| RP23-307I19.5 | 0.947788321 | -2.19908022 | 1.271222545 | -1.729893974 |
| Chst12 | 156.329386 | 0.092387932 | 0.104415298 | 0.884812231 |
| RP23-462C22.2 | 1.646628032 | 1.575858721 | 1.093417 | 1.441223908 |
| RP23-462C22.1 | 0.174406792 | 1.259563417 | 3.365276104 | 0.374282341 |
| Grifin | 1.541974883 | -0.96672386 | 0.97416864 | -0.992357811 |
| RP23-462C22.5 | 1.34756889 | 0.87449662 | 0.898115238 | 0.973702018 |
| Lfng | 314.5144813 | 0.310008184 | 0.175248945 | 1.768958914 |
| Ttyh3 | 2824.785642 | -0.180196698 | 0.092769141 | -1.942420683 |
| Iqce | 386.6329369 | -0.322076283 | 0.1414845 | -2.276406828 |
| Brat1 | 216.0259619 | -0.142721915 | 0.11091403 | -1.286779632 |
| Amz1 | 137.7511797 | -0.667857924 | 0.158394479 | -4.216421736 |
| Gna12 | 984.1548664 | -0.201830668 | 0.118322486 | -1.705767649 |
| Card11 | 40.90469028 | 4.434445048 | 0.447351031 | 9.912674263 |
| AC161570.1 | 0 | NA | NA | NA |
| Gm26113 | 0 | NA | NA | NA |
| Sdk1 | 231.3078279 | -0.033625845 | 0.097581056 | -0.344593979 |
| Gm6433 | 1.643128883 | 1.047826817 | 0.818334162 | 1.280438805 |
| RP23-49G7.2 | 0 | NA | NA | NA |
| RP23-286C5.1 | 0 | NA | NA | NA |
| RP23-286C5.2 | 0.061920613 | 0.274393828 | 3.724945049 | 0.073663859 |
| Gm16035 | 0 | NA | NA | NA |
| Gm24131 | 0 | NA | NA | NA |
| Gm16036 | 0 | NA | NA | NA |
| Gm26970 | 0 | NA | NA | NA |
| Foxk1 | 1176.476401 | 0.180787405 | 0.075677344 | 2.388923775 |
| Ap5z1 | 229.1563663 | -0.096602472 | 0.115574532 | -0.835845665 |
| Radil | 157.9503125 | -0.661800692 | 0.129669116 | -5.103764969 |
| Papolb | 13.6158541 | -0.781380729 | 0.319574032 | -2.445069534 |

| | | | | |
|---|---|---|---|---|
| RP23-106N8.2 | 0 | NA | NA | NA |
| Mmd2 | 1010.130574 | -0.309218491 | 0.156510505 | -1.975704377 |
| RP23-106N8.4 | 0 | NA | NA | NA |
| Wipi2 | 1453.522254 | -0.089251773 | 0.058353233 | -1.529508606 |
| Slc29a4 | 200.9815486 | -0.647522262 | 0.119623707 | -5.412992773 |
| Tnrc18 | 2698.838952 | -0.016226015 | 0.066862446 | -0.24267756 |
| Gm15770 | 0.972120366 | 3.348976454 | 1.239705539 | 2.701428968 |
| Fbxl18 | 720.5681582 | -0.102011494 | 0.070636695 | -1.44417139 |
| D430018E03Rik | 19.83389332 | -0.661801097 | 0.306028683 | -2.162545974 |
| Actb | 25647.77243 | 0.671783703 | 0.113825389 | 5.901879243 |
| Fscn1 | 1883.804362 | 0.113423131 | 0.087129524 | 1.301776082 |
| Rnf216 | 737.7813341 | -0.140008985 | 0.047029089 | -2.977072036 |
| RP23-393G8.5 | 4.643209367 | -1.103824488 | 0.506830376 | -2.177897262 |
| RP23-393G8.10 | 0.927313021 | 0.294345102 | 1.041654098 | 0.282574707 |
| Olfr718-ps1 | 0 | NA | NA | NA |
| RP23-301L9.8 | 0.114443069 | -0.206492286 | 3.724945049 | -0.055434988 |
| AC123686.1 | 0 | NA | NA | NA |
| Rbakdn | 0.061920613 | 0.274393828 | 3.724945049 | 0.073663859 |
| Rbak | 178.0893661 | 0.256442309 | 0.140374135 | 1.826848719 |
| 4933411G11Rik | 0.556792772 | 0.002491781 | 1.305090978 | 0.001909278 |
| Spdyb | 0.075043205 | 0.755286614 | 3.724945049 | 0.202764498 |
| 4930448H16Rik | 0 | NA | NA | NA |
| Zfp12 | 444.4098317 | 0.156265102 | 0.092954922 | 1.681084757 |
| Zfp316 | 378.1393857 | -0.127756753 | 0.099624308 | -1.282385351 |
| RP23-301L9.3 | 37.38971463 | -0.534598685 | 0.285533545 | -1.872279788 |
| Zfp853 | 32.64981703 | -0.167166706 | 0.195224701 | -0.856278458 |
| E130309D02Rik | 214.9676355 | 0.146004901 | 0.106762484 | 1.367567485 |
| RP23-301L9.6 | 1.497168834 | 1.188979259 | 0.798849463 | 1.488364596 |
| Zdhhc4 | 177.8279622 | 0.340294915 | 0.120141325 | 2.832455159 |
| 0610040B10Rik | 20.77576342 | 0.774080265 | 0.294939714 | 2.624537243 |
| RP23-301L9.4 | 0.904055358 | 1.239540784 | 1.136038159 | 1.091108405 |
| Grid2ip | 247.2010809 | -0.327154569 | 0.221400293 | -1.47766096 |
| Gm20635 | 4.77646696 | 0.05803773 | 0.463903832 | 0.125107243 |
| RP23-372N19.4 | 0.76879839 | -0.917664022 | 1.105141061 | -0.830359177 |
| RP23-372N19.2 | 152.8195759 | -0.469950207 | 0.215816118 | -2.17754916 |
| 9530056K15Rik | 3.981355527 | -0.345350403 | 0.489610189 | -0.705357876 |
| Kdelr2 | 375.7129238 | 0.170112478 | 0.129689774 | 1.311687675 |
| RP23-158A20.2 | 0 | NA | NA | NA |
| RP23-158A20.4 | 0 | NA | NA | NA |
| Daglb | 512.0781547 | -0.168151216 | 0.079076914 | -2.126426113 |
| Rac1 | 2571.473266 | 0.216088706 | 0.076909465 | 2.80965034 |
| RP23-158A20.7 | 0.049553047 | 0.274391604 | 3.724945049 | 0.073663262 |
| Fam220a | 378.2842679 | -0.009517604 | 0.081752582 | -0.116419615 |
| Cyth3 | 478.6545393 | 0.027610312 | 0.073015854 | 0.378141331 |
| Usp42 | 463.5942125 | 0.140953046 | 0.070726153 | 1.992940938 |
| D130017N08Rik | 140.3508044 | -0.585875993 | 0.152422739 | -3.843757154 |
| RP23-151C5.2 | 0 | NA | NA | NA |

| | | | | |
|---|---|---|---|---|
| RP23-151C5.3 | 4.718276104 | 0.019291264 | 0.463726258 | 0.041600544 |
| Eif2ak1 | 746.7429788 | 0.194424893 | 0.061533354 | 3.159666754 |
| Ankrd61 | 4.124592769 | -0.583454667 | 0.484253751 | -1.204853168 |
| Aimp2 | 134.5489307 | -0.130874648 | 0.125312126 | -1.044389336 |
| Pms2 | 217.1478713 | -0.036500446 | 0.100874156 | -0.361841401 |
| Gm17135 | 0.441552168 | 1.035749205 | 1.371014466 | 0.755461909 |
| PMS2 | 0.78876463 | 1.447373564 | 1.100749732 | 1.314897948 |
| Rsph10b | 55.58878597 | -0.270532108 | 0.186600266 | -1.449794871 |
| Ccz1 | 450.6103287 | 0.521195664 | 0.067168404 | 7.759536209 |
| Gm24311 | 0 | NA | NA | NA |
| Ocm | 1.753517396 | 1.329010703 | 0.768067024 | 1.730331679 |
| RP24-194F23.5 | 0.296409788 | -0.233348574 | 3.719653481 | -0.062733955 |
| Lmtk2 | 3277.074586 | 0.013641644 | 0.067530824 | 0.202006177 |
| Gm26925 | 0 | NA | NA | NA |
| Bhlha15 | 4.240558544 | -0.089837969 | 0.688952634 | -0.130397889 |
| Tecpr1 | 1209.265373 | -0.24120558 | 0.066695366 | -3.61652685 |
| RP24-82B23.5 | 8.247112194 | -1.443236401 | 0.384045968 | -3.757978268 |
| 2900089D17Rik | 4.905083075 | -1.351863992 | 0.465721343 | -2.902731458 |
| Bri3 | 199.49671 | 0.306117924 | 0.128440672 | 2.383341032 |
| Baiap2l1 | 221.2967602 | -0.285692141 | 0.123282335 | -2.317381004 |
| Gm15708 | 7.40733758 | 0.085656929 | 0.444893909 | 0.192533381 |
| RP24-182D2.1 | 0.265735694 | 1.353480912 | 3.7166316 | 0.364168704 |
| Gm15709 | 0 | NA | NA | NA |
| Nptx2 | 612.3205461 | -0.091880288 | 0.318432478 | -0.288539312 |
| Gm6272 | 0.315729261 | 0.263766816 | 1.607378837 | 0.16409748 |
| Tmem130 | 1967.351054 | -0.130095559 | 0.08805837 | -1.477378678 |
| Trrap | 3060.283518 | -0.213205382 | 0.076587961 | -2.783797587 |
| Gm23995 | 1.477838792 | -0.832797251 | 0.827109959 | -1.006876102 |
| Smurf1 | 700.542789 | 0.099138135 | 0.068940593 | 1.438022652 |
| Mir7039 | 0.22969359 | -0.206491565 | 3.142040024 | -0.065718948 |
| Kpna7 | 0.7084488 | 0.149851641 | 1.41662888 | 0.10578045 |
| RP23-424M12.7 | 0 | NA | NA | NA |
| Gm4871 | 0 | NA | NA | NA |
| RP23-424M12.4 | 0 | NA | NA | NA |
| RP23-424M12.5 | 0.075043205 | 0.755286614 | 3.724945049 | 0.202764498 |
| 1700018F24Rik | 0.068097359 | -0.206487837 | 3.724945049 | -0.055433794 |
| Arpc1a | 1136.742611 | 0.158849849 | 0.068536476 | 2.317741718 |
| Arpc1b | 585.2155934 | 2.351691763 | 0.167150393 | 14.06931641 |
| Pdap1 | 676.5616892 | 0.359744349 | 0.13151207 | 2.735447387 |
| Bud31 | 300.4083312 | 0.300170604 | 0.090635684 | 3.311836911 |
| Ptcd1 | 337.0823285 | 0.514233984 | 0.079843692 | 6.440508612 |
| Cpsf4 | 256.7055373 | 0.03477311 | 0.091503857 | 0.380017968 |
| Atp5j2 | 504.0996134 | -0.029924377 | 0.242922097 | -0.123185075 |
| Zkscan14 | 94.39392942 | 0.201228093 | 0.120951816 | 1.663704598 |
| Zkscan5 | 178.3410506 | 0.18841864 | 0.123519883 | 1.525411418 |
| Zfp655 | 776.8726219 | 0.300898888 | 0.099535937 | 3.023017596 |
| RP23-60E24.5 | 0 | NA | NA | NA |

| | | | | |
|---|---|---|---|---|
| Zscan25 | 198.2716498 | 0.661911767 | 0.162770627 | 4.066530784 |
| RP23-60E24.3 | 23.00508971 | 1.348237525 | 0.421494316 | 3.198708672 |
| Cyp3a57 | 13.25184546 | 3.921375349 | 0.599470338 | 6.541400137 |
| RP24-443E20.1 | 0.066050875 | 0.274391604 | 3.724945049 | 0.073663262 |
| RP24-443E20.2 | 0.066050875 | 0.274391604 | 3.724945049 | 0.073663262 |
| Cyp3a16 | 0 | NA | NA | NA |
| RP24-127J3.1 | 0.120974994 | 0.274404949 | 3.724945049 | 0.073666845 |
| Cyp3a41b | 0 | NA | NA | NA |
| RP23-379D24.1 | 0 | NA | NA | NA |
| RP23-379D24.2 | 0 | NA | NA | NA |
| RP24-443E20.4 | 0 | NA | NA | NA |
| Cyp3a41a | 0 | NA | NA | NA |
| RP23-102C14.1 | 0 | NA | NA | NA |
| Cyp3a44 | 0 | NA | NA | NA |
| RP23-102C14.2 | 0.060174537 | 0.755286614 | 3.724945049 | 0.202764498 |
| Cyp3a11 | 0.492793603 | -0.094217207 | 1.450388439 | -0.064959982 |
| RP23-161K8.1 | 0 | NA | NA | NA |
| RP23-161K8.2 | 0 | NA | NA | NA |
| RP23-161K8.3 | 0 | NA | NA | NA |
| Cyp3a25 | 0.261272317 | 0.75530529 | 2.041273979 | 0.370016616 |
| RP23-53G23.1 | 0 | NA | NA | NA |
| RP23-53G23.2 | 0 | NA | NA | NA |
| RP23-53G23.3 | 0 | NA | NA | NA |
| Cyp3a59 | 0 | NA | NA | NA |
| RP23-441I24.9 | 0.282778275 | 1.221993057 | 2.007159203 | 0.608817205 |
| Gm6309 | 0.038981395 | 0.274393828 | 3.724945049 | 0.073663859 |
| 1700001J03Rik | 1.161484436 | 1.731907026 | 1.001465153 | 1.72937323 |
| RP23-441I24.5 | 22.34345423 | 1.248938896 | 0.489162684 | 2.553217847 |
| RP23-441I24.4 | 2.948678684 | 0.875497717 | 0.677144447 | 1.292926082 |
| RP23-441I24.3 | 9.915980733 | 0.695766549 | 0.409013393 | 1.701085004 |
| Rnf6 | 700.4219294 | -0.075361273 | 0.069147032 | -1.089869964 |
| Cdk8 | 479.7397194 | -0.225973032 | 0.10469337 | -2.158427336 |
| Gm15739 | 20.56479697 | -0.231457919 | 0.267050696 | -0.866719025 |
| Wasf3 | 1567.401421 | -0.202366558 | 0.073863704 | -2.739729342 |
| Gm6408 | 0.779579221 | 1.360143335 | 1.217934955 | 1.116761884 |
| Gm6370 | 0.038981395 | 0.274393828 | 3.724945049 | 0.073663859 |
| 4930449I24Rik | 0 | NA | NA | NA |
| Gm3402 | 0 | NA | NA | NA |
| Gm3404 | 0 | NA | NA | NA |
| Gm3409 | 0 | NA | NA | NA |
| Gm3415 | 0 | NA | NA | NA |
| Gm8494 | 20.48103304 | 0.114526676 | 0.244568623 | 0.468280333 |
| Gpr12 | 122.0110685 | -0.62108005 | 0.159721712 | -3.888513615 |
| RP23-134K6.2 | 0.089089296 | 1.194485492 | 3.723939576 | 0.320758559 |
| RP23-134K6.3 | 0 | NA | NA | NA |
| Usp12 | 869.2011401 | 0.424825273 | 0.070830141 | 5.99780355 |
| RP24-126A19.2 | 0.044191549 | -0.206490061 | 3.724945049 | -0.055434391 |

| | | | | |
|---|---|---|---|---|
| Rpl21 | 82.06727034 | 0.031905064 | 0.144088941 | 0.221426183 |
| Gm25091 | 0 | NA | NA | NA |
| Gm23202 | 0.066050875 | 0.274391604 | 3.724945049 | 0.073663262 |
| Rasl11a | 102.228073 | 1.135560075 | 0.310853346 | 3.653041182 |
| Gtf3a | 110.5136172 | -0.168678679 | 0.127846642 | -1.319382948 |
| Mtif3 | 139.0778404 | 0.074903609 | 0.097332867 | 0.769561312 |
| Lnx2 | 154.2635579 | 0.028903171 | 0.128632282 | 0.224696088 |
| Polr1d | 179.4983994 | -0.122878967 | 0.125053384 | -0.982612092 |
| Rps10-ps4 | 0.048949532 | -0.206487837 | 3.724945049 | -0.055433794 |
| Gsx1 | 3.152922583 | 0.037753518 | 0.543387577 | 0.069478066 |
| Gm24556 | 0 | NA | NA | NA |
| 2210019I11Rik | 0 | NA | NA | NA |
| Pdx1 | 0.046345709 | 0.274391604 | 3.724945049 | 0.073663262 |
| Gm27033 | 0 | NA | NA | NA |
| Mir7k | 0 | NA | NA | NA |
| Cdx2 | 0 | NA | NA | NA |
| Urad | 0 | NA | NA | NA |
| RP24-510G5.6 | 0 | NA | NA | NA |
| Flt3 | 155.671954 | -0.545062496 | 0.152158019 | -3.582213416 |
| Gm26597 | 1.472170236 | -0.211586935 | 0.805423827 | -0.262702602 |
| Gm6054 | 0.773509025 | 1.646347838 | 1.206321389 | 1.364767178 |
| RP24-544J6.3 | 0.075043205 | 0.755286614 | 3.724945049 | 0.202764498 |
| RP24-544J6.4 | 1.088238703 | -1.135948297 | 1.023474276 | -1.109894331 |
| RP24-544J6.5 | 2.007455813 | -0.577768438 | 0.699844905 | -0.825566399 |
| RP24-544J6.6 | 0.135435006 | 1.15287194 | 3.633258134 | 0.317310771 |
| Pan3 | 1329.249898 | -0.163843149 | 0.100273462 | -1.633963229 |
| RP24-544J6.7 | 8.762375666 | -0.240589993 | 0.42632832 | -0.56433031 |
| Flt1 | 324.459667 | 0.298775635 | 0.200463317 | 1.49042548 |
| RP23-270L23.1 | 0.088534442 | 0.274389469 | 3.724945049 | 0.073662689 |
| RP23-270L23.2 | 0 | NA | NA | NA |
| Pomp | 809.6499027 | 0.588626083 | 0.095142722 | 6.18676941 |
| RP23-270L23.4 | 0.636753058 | -0.765983554 | 1.220549441 | -0.627572737 |
| Slc46a3 | 67.21071847 | 0.245949684 | 0.14446272 | 1.702513179 |
| Gm15721 | 25.84402214 | -0.247432821 | 0.216318588 | -1.143835228 |
| Mtus2 | 777.696009 | -0.249956654 | 0.107765119 | -2.319457868 |
| Gm10167 | 12.31383881 | 0.630886119 | 0.278053859 | 2.268934948 |
| RP24-71D19.1 | 4.780781177 | -0.264877969 | 0.537802406 | -0.492519123 |
| Slc7a1 | 875.5929286 | 0.572194657 | 0.162576371 | 3.519543788 |
| RP24-307I24.1 | 0.318558491 | -1.140364924 | 1.75933877 | -0.648178136 |
| Ubl3 | 1852.018664 | 0.037827407 | 0.05928683 | 0.638040643 |
| RP24-61O18.3 | 0 | NA | NA | NA |
| RP24-61O18.1 | 0.068097359 | -0.206487837 | 3.724945049 | -0.055433794 |
| RP24-61O18.2 | 0 | NA | NA | NA |
| 2210417A02Rik | 0 | NA | NA | NA |
| n-R5s179 | 0 | NA | NA | NA |
| 4930505K14Rik | 0 | NA | NA | NA |
| Gm15407 | 0.046345709 | 0.274391604 | 3.724945049 | 0.073663262 |

| | | | | |
|---|---|---|---|---|
| Katnal1 | 1032.752 | -0.298534993 | 0.068053283 | -4.386783113 |
| RP23-188F20.1 | 0 | NA | NA | NA |
| Gm15410 | 0.390771115 | -1.15834713 | 1.591054829 | -0.728037217 |
| Gm15406 | 5.088172819 | 0.776432835 | 0.475043971 | 1.634444142 |
| 8430423G03Rik | 0.143485256 | 0.274391988 | 3.549605256 | 0.077302113 |
| Gm15409 | 0.354937333 | 1.460480183 | 1.648061711 | 0.886180519 |
| RP23-134H19.3 | 52.59786348 | -0.044548198 | 0.201370523 | -0.22122502 |
| Gm15411 | 7.066285288 | -0.319970394 | 0.453411266 | -0.705695728 |
| Gm15408 | 2.719793829 | 1.419259851 | 0.75130204 | 1.889066949 |
| Hmgb1 | 340.808951 | 0.20204479 | 0.101504211 | 1.990506472 |
| 5730422E09Rik | 12.70416494 | -1.044790389 | 0.38251362 | -2.731380881 |
| RP23-408A1.6 | 1.653178163 | 0.543044135 | 0.719884508 | 0.754348967 |
| RP23-408A1.4 | 3.467320147 | 2.960490841 | 0.870775753 | 3.399831508 |
| Gm23362 | 0 | NA | NA | NA |
| Gm8615 | 17.8882258 | -0.486608139 | 0.255895102 | -1.901592234 |
| RP23-408A1.8 | 0.278466489 | 1.25349613 | 1.792053111 | 0.699474877 |
| Uspl1 | 565.005295 | 0.004295881 | 0.09426862 | 0.045570631 |
| 2310047D07Rik | 2.953893651 | -0.217706518 | 0.556465821 | -0.39123071 |
| Gm29264 | 0.433555451 | 0.272298176 | 1.443104116 | 0.18868921 |
| AC122546.1 | 0 | NA | NA | NA |
| Gm24105 | 0 | NA | NA | NA |
| Alox5ap | 91.57756619 | 2.237779369 | 0.248318982 | 9.011712895 |
| n-R5s180 | 0 | NA | NA | NA |
| WI1-1993I8.1 | 5.494432944 | 0.346929231 | 0.481194103 | 0.72097565 |
| BC028471 | 1.239154484 | -0.723735061 | 1.004033245 | -0.720827786 |
| Gm20488 | 0.478946765 | -0.592050305 | 1.563359871 | -0.378703788 |
| RP23-375E13.2 | 0.173926504 | 0.274410294 | 2.311249723 | 0.118728103 |
| Medag | 75.60539664 | 0.226403914 | 0.215548581 | 1.050361422 |
| RP23-375E13.4 | 0 | NA | NA | NA |
| RP23-375E13.3 | 0.045882455 | 0.755288839 | 3.724945049 | 0.202765096 |
| Tex26 | 5.173350472 | 0.183873517 | 0.552970486 | 0.332519586 |
| RP23-375E13.6 | 1.443591694 | 1.653974217 | 0.935670262 | 1.7676892 |
| Gm15997 | 2.38686755 | -1.572074959 | 0.703443115 | -2.234828838 |
| Gm2566 | 0.852004391 | -1.820068123 | 1.119012774 | -1.626494501 |
| Wdr95 | 6.751900483 | 0.585744027 | 0.403778668 | 1.450656195 |
| Hsph1 | 3279.88901 | -0.301755709 | 0.228834469 | -1.318663706 |
| RP23-256B18.1 | 1.058337859 | 0.630816098 | 1.305711397 | 0.483120619 |
| Gm5566 | 1.331862033 | 0.933303824 | 1.130319992 | 0.825698767 |
| B3glct | 236.8677375 | 0.183894723 | 0.108100157 | 1.701151298 |
| RP23-346P15.1 | 0 | NA | NA | NA |
| RP23-346P15.3 | 0.483795822 | 1.091516808 | 1.647699717 | 0.662448866 |
| RP23-346P15.4 | 0 | NA | NA | NA |
| RP23-337K7.2 | 0 | NA | NA | NA |
| RP23-337K7.1 | 0.29963536 | -0.2262938 | 1.606832618 | -0.140832217 |
| RP23-337K7.3 | 0 | NA | NA | NA |
| Rxfp2 | 8.210362686 | -0.642207533 | 0.377046295 | -1.703259101 |
| RP23-301H20.1 | 0 | NA | NA | NA |

| | | | | |
|---|---|---|---|---|
| RP23-301H20.2 | 0 | NA | NA | NA |
| Fry | 3743.235337 | -0.224115787 | 0.078926393 | -2.8395544 |
| RP23-327P24.3 | 2.277130459 | -0.467726098 | 0.60977135 | -0.767051613 |
| RP23-327P24.2 | 0 | NA | NA | NA |
| RP24-333I1.2 | 4.567880012 | -1.016906466 | 0.498190821 | -2.041198719 |
| RP24-333I1.3 | 1.611190043 | -0.89778835 | 0.789725984 | -1.136835268 |
| Zar1l | 2.866525085 | -0.370675352 | 0.540904073 | -0.685288522 |
| Gm15933 | 0.686705509 | -0.709678207 | 1.132711676 | -0.626530319 |
| Brca2 | 165.9286918 | 0.349216599 | 0.167356444 | 2.086663592 |
| Gm25284 | 0.075043205 | 0.755286614 | 3.724945049 | 0.202764498 |
| Gm23641 | 0.146089079 | -0.206488916 | 3.522538524 | -0.058619349 |
| RP24-333I1.6 | 1.253964362 | -1.55684659 | 0.965928951 | -1.611760978 |
| RP24-333I1.5 | 4.525610808 | -0.359964576 | 0.424705324 | -0.847563135 |
| RP24-333I1.7 | 13.83704783 | 0.73863218 | 0.262694439 | 2.811754149 |
| N4bp2l1 | 393.7981301 | -0.148780914 | 0.09573913 | -1.554024097 |
| RP23-34P11.1 | 29.55790751 | -0.084202829 | 0.381525258 | -0.220700536 |
| RP23-34P11.2 | 251.9245356 | -0.603049548 | 0.171935878 | -3.507409592 |
| N4bp2l2 | 1453.675956 | 0.032140696 | 0.08096057 | 0.396991966 |
| Pds5b | 2134.751181 | -0.176439678 | 0.070229394 | -2.512333754 |
| Gm24630 | 4.821230019 | -0.316761453 | 0.456141111 | -0.694437413 |
| Gm26013 | 0.049553047 | 0.274391604 | 3.724945049 | 0.073663262 |
| RP23-336F4.1 | 29.09809687 | -0.306383934 | 0.264221406 | -1.159572718 |
| RP23-336F4.4 | 4.80270818 | 0.595553743 | 0.458957617 | 1.297622528 |
| RP23-336F4.5 | 0 | NA | NA | NA |
| Kl | 209.3353533 | -0.672258946 | 0.215732137 | -3.116174319 |
| RP23-412C8.2 | 0.322046281 | 1.236204743 | 1.775749961 | 0.69615924 |
| Stard13 | 272.414666 | -0.074326979 | 0.085336596 | -0.870985984 |
| RP24-155K17.1 | 0.139883403 | 0.274393101 | 3.559413475 | 0.077089415 |
| RP24-343A12.4 | 0 | NA | NA | NA |
| RP24-343A12.3 | 0 | NA | NA | NA |
| D730045B01Rik | 4.778829002 | -0.488245208 | 0.519032519 | -0.940683272 |
| 1700028E10Rik | 23.33760472 | 0.122332822 | 0.209910611 | 0.582785318 |
| Vmn2r-ps25 | 0.141684195 | 0.755287278 | 3.558854591 | 0.212227631 |
| Vmn2r-ps26 | 0.233449105 | 1.549739465 | 2.493543112 | 0.621500971 |
| Vmn2r18 | 0.476067163 | 1.236204712 | 1.689764339 | 0.731584094 |
| RP24-211L21.1 | 0.060174537 | 0.755286614 | 3.724945049 | 0.202764498 |
| Rfc3 | 87.13724546 | 1.110761813 | 0.200296747 | 5.545580891 |
| RP24-211L21.2 | 0 | NA | NA | NA |
| RP24-211L21.3 | 0.154433973 | -0.688825965 | 3.500771551 | -0.196764043 |
| Gm24242 | 0.09689823 | -0.687371727 | 3.724945049 | -0.184532045 |
| Gm26463 | 0 | NA | NA | NA |
| Gm23551 | 0 | NA | NA | NA |
| Vmn1r-ps6 | 0 | NA | NA | NA |
| Vmn1r-ps7 | 0.046345709 | 0.274391604 | 3.724945049 | 0.073663262 |
| RP24-328P2.7 | 0.256207099 | -0.126256599 | 2.104666069 | -0.059988898 |
| RP24-328P2.8 | 0.219080626 | -0.68737862 | 2.171582349 | -0.316533527 |
| RP23-368D12.2 | 0 | NA | NA | NA |

| | | | | |
|---|---|---|---|---|
| RP23-368D12.3 | 0 | NA | NA | NA |
| RP23-258F1.1 | 0.066050875 | 0.274391604 | 3.724945049 | 0.073663262 |
| Gm18222 | 0 | NA | NA | NA |
| RP23-253G12.8 | 136.7192892 | 5.587174628 | 0.339901917 | 16.43760846 |
| RP24-328P2.5 | 839.9270636 | 5.741603529 | 0.200305965 | 28.66416651 |
| RP23-253G12.10 | 0.128477747 | 0.755285235 | 3.677131942 | 0.205400635 |
| RP23-253G12.9 | 311.8525769 | 5.391060068 | 0.252732184 | 21.3311181 |
| Samd9l | 3774.434653 | 6.413008456 | 0.206735061 | 31.02042024 |
| Hepacam2 | 3.625877892 | -0.748533501 | 0.54874514 | -1.364082243 |
| Ccdc132 | 896.7331518 | 0.038002979 | 0.075124125 | 0.505869169 |
| RP23-318M11.2 | 26.8210992 | 0.369418422 | 0.341170229 | 1.082797942 |
| Calcr | 25.19477301 | 0.031249444 | 0.257016264 | 0.121585474 |
| Mir653 | 0 | NA | NA | NA |
| Mir489 | 0.044191549 | -0.206490061 | 3.724945049 | -0.055434391 |
| Tfpi2 | 9.186610503 | 4.413762092 | 0.939278779 | 4.699096997 |
| Gngt1 | 0 | NA | NA | NA |
| Gng11 | 162.5242613 | 0.811734849 | 0.180559887 | 4.495654393 |
| Bet1 | 205.6506794 | -0.112381554 | 0.132427697 | -0.848625753 |
| Gm8602 | 0 | NA | NA | NA |
| Gm24442 | 0 | NA | NA | NA |
| Gm37883 | 0 | NA | NA | NA |
| Col1a2 | 642.3682694 | 0.467833015 | 0.13601109 | 3.439668154 |
| Gm26696 | 3.592646241 | -0.554573155 | 0.541540843 | -1.024065245 |
| Casd1 | 867.3601419 | -0.071197495 | 0.085752261 | -0.830269598 |
| Sgce | 237.5338859 | -0.233349302 | 0.115817885 | -2.014795055 |
| Peg10 | 1587.476363 | 0.662802863 | 0.220833776 | 3.001365438 |
| Gm20714 | 0 | NA | NA | NA |
| Gm26828 | 0 | NA | NA | NA |
| Gm9835 | 0.856801049 | 1.753847044 | 1.251092331 | 1.401852605 |
| Ppp1r9a | 4137.715075 | -0.228778042 | 0.104255579 | -2.194396161 |
| 2210017G18Rik | 30.57888805 | -0.620262953 | 0.335643613 | -1.847980802 |
| Gm26814 | 84.80524745 | -0.604442071 | 0.274152271 | -2.204767698 |
| Gm26736 | 22.89958731 | -0.837306456 | 0.338008178 | -2.477178098 |
| Gm26556 | 14.27493484 | -0.544878774 | 0.292773759 | -1.86109157 |
| Pon1 | 0.759396657 | -1.297334663 | 1.107903232 | -1.170981928 |
| Pon3 | 56.36395702 | 0.134309461 | 0.232278546 | 0.578225855 |
| Pon2 | 309.9460515 | 0.279591784 | 0.109601087 | 2.550994641 |
| Asb4 | 38.68988062 | 0.407631069 | 0.291453836 | 1.398612811 |
| Pdk4 | 314.1644295 | 0.581864553 | 0.190154375 | 3.059958804 |
| Dync1i1 | 1162.518359 | -0.111370284 | 0.060773951 | -1.832533219 |
| 1700019G24Rik | 1.274692649 | 0.424028227 | 1.005291018 | 0.421796495 |
| Slc25a13 | 101.7915475 | 0.403517689 | 0.173861428 | 2.320915538 |
| Gm20619 | 0.313425824 | 1.970880596 | 2.018860922 | 0.976233961 |
| Gm8649 | 0 | NA | NA | NA |
| Gm27018 | 0 | NA | NA | NA |
| Gm8652 | 0.294205253 | -0.206481239 | 1.635811928 | -0.126225537 |
| Gm20617 | 0.379053103 | 2.078785812 | 1.666341871 | 1.247514599 |

| | | | | |
|---|---|---|---|---|
| Gm20685 | 0 | NA | NA | NA |
| Shfm1 | 204.0907547 | 0.380595482 | 0.249489427 | 1.525497442 |
| Gm20618 | 0.47603017 | -0.921218692 | 1.311628668 | -0.702347177 |
| Gm18006 | 0 | NA | NA | NA |
| Gm5873 | 4.742073203 | 0.104645085 | 0.464029501 | 0.225513863 |
| Dlx6os1 | 337.2803174 | -1.137472381 | 0.262242849 | -4.337477201 |
| Gm27699 | 0.095295241 | -0.206492286 | 3.724945049 | -0.055434988 |
| Gm27905 | 0 | NA | NA | NA |
| Dlx6os3 | 6.490635638 | -2.004305682 | 0.510416004 | -3.926808069 |
| Dlx6 | 64.91335236 | -0.636564573 | 0.203934524 | -3.121416432 |
| Dlx6os2 | 8.021264403 | -1.73629299 | 0.538973483 | -3.221481286 |
| Dlx5 | 52.02918498 | -0.852636828 | 0.216295781 | -3.941994736 |
| Gm22155 | 0 | NA | NA | NA |
| Gm26774 | 0.048949532 | -0.206487837 | 3.724945049 | -0.055433794 |
| Sdhaf3 | 139.5460954 | -0.407437772 | 0.112437734 | -3.623674701 |
| Gm27190 | 0 | NA | NA | NA |
| Gm8669 | 0 | NA | NA | NA |
| Gm19272 | 0.25372768 | -0.206486202 | 2.095178707 | -0.098553026 |
| Tac1 | 258.728839 | -0.123568812 | 0.149977522 | -0.823915547 |
| Gm8676 | 0 | NA | NA | NA |
| Asns | 800.622713 | 0.121621124 | 0.090204503 | 1.348282185 |
| C1galt1 | 215.3562405 | 0.007901207 | 0.123468634 | 0.063993634 |
| Gm22276 | 0.301804762 | -1.119561048 | 2.021323482 | -0.55387525 |
| Gm9825 | 372.9565206 | 0.106276745 | 0.085762732 | 1.239194967 |
| Col28a1 | 8.368185802 | -0.195051646 | 0.364688852 | -0.534844004 |
| Mios | 475.6118824 | -0.013077194 | 0.06896081 | -0.189632252 |
| Rpa3 | 69.13831145 | 0.039249237 | 0.143169222 | 0.274145775 |
| Umad1 | 363.7124902 | -0.033434595 | 0.078325863 | -0.426865326 |
| Gm16055 | 0.775287707 | 1.077264388 | 1.254507918 | 0.858714698 |
| Gm8428 | 0 | NA | NA | NA |
| Gm16043 | 0.136120942 | 0.274393632 | 3.599844381 | 0.076223748 |
| Gm24059 | 4.143222737 | -0.002493479 | 0.481175224 | -0.00518206 |
| A430035B10Rik | 32.4009088 | -0.09883369 | 0.245234262 | -0.403017461 |
| Gm26075 | 0.250670796 | -1.178145318 | 2.480260474 | -0.475008706 |
| Glcci1 | 81.25189137 | -0.373852059 | 0.127483694 | -2.93254805 |
| Gm16042 | 0.476187739 | 0.274411212 | 1.759262335 | 0.155980837 |
| Ica1 | 575.9737311 | 0.018608406 | 0.122148246 | 0.152342802 |
| Nxph1 | 529.823522 | -0.136360022 | 0.075484735 | -1.806458257 |
| Gm26058 | 0.921802213 | -2.663243957 | 1.071648954 | -2.485183181 |
| Gm27894 | 0 | NA | NA | NA |
| Ndufa4 | 1081.345019 | -0.120554982 | 0.140672006 | -0.856993404 |
| Phf14 | 900.5267977 | -0.107597004 | 0.063688371 | -1.689429345 |
| Gm6578 | 0.045882455 | 0.755288839 | 3.724945049 | 0.202765096 |
| Thsd7a | 1278.572731 | -0.175597828 | 0.056980849 | -3.081699066 |
| Gm16123 | 0 | NA | NA | NA |
| n-R5s160 | 0 | NA | NA | NA |
| Gm15529 | 0 | NA | NA | NA |

| | | | | |
|---|---|---|---|---|
| Gm5300 | 0.044544648 | 0.755286614 | 3.724945049 | 0.202764498 |
| Tmem106b | 2433.505759 | -0.095353807 | 0.076507698 | -1.24632958 |
| Gm24985 | 0 | NA | NA | NA |
| Vwde | 1.568186543 | 3.833104164 | 0.984112148 | 3.894987143 |
| 1700016P04Rik | 0.132574648 | -1.045763943 | 3.713921268 | -0.28157946 |
| Gm25737 | 0.154696864 | 0.274407077 | 2.833481316 | 0.096844498 |
| Tmem168 | 343.227461 | 0.226132015 | 0.096708314 | 2.338289291 |
| B630005N14Rik | 527.4785247 | -0.076415836 | 0.091168021 | -0.838186846 |
| Gpr85 | 431.2033423 | 0.192154633 | 0.08908447 | 2.156993621 |
| 2610001J05Rik | 197.506035 | -0.150906743 | 0.108707222 | -1.388194275 |
| 1110019D14Rik | 33.23543665 | -0.034827949 | 0.204030258 | -0.17069992 |
| Gm23760 | 0.097899064 | -0.687371727 | 3.724945049 | -0.184532045 |
| Gm23013 | 0 | NA | NA | NA |
| Ppp1r3a | 6.253343815 | 0.197861427 | 0.380312537 | 0.520260069 |
| Foxp2 | 446.2894329 | -0.199454955 | 0.089399864 | -2.231043163 |
| Gm25470 | 1.281282525 | 1.052444901 | 1.009840915 | 1.04218881 |
| Mdfic | 320.1052487 | 2.171680125 | 0.234003631 | 9.280540287 |
| Gm29589 | 0 | NA | NA | NA |
| RP23-244M15.1 | 0.212513282 | 1.926509226 | 3.159662244 | 0.609719988 |
| Tfec | 15.16942446 | 2.390486039 | 0.755499819 | 3.164111996 |
| Gm5721 | 0 | NA | NA | NA |
| Gm15473 | 0.646370819 | 1.309323783 | 1.55688384 | 0.840990027 |
| Gm4876 | 26.61523984 | 0.26937283 | 0.225477248 | 1.194678544 |
| Tes | 60.86189865 | 1.72412771 | 0.260122747 | 6.628131253 |
| D830026I12Rik | 0 | NA | NA | NA |
| Gm15581 | 0.736762124 | -0.832008529 | 1.380193411 | -0.602820244 |
| Cav2 | 214.4350625 | 0.007114803 | 0.142846982 | 0.049807165 |
| Cav1 | 153.6111711 | -0.125697903 | 0.175063143 | -0.718014657 |
| Gm29591 | 0 | NA | NA | NA |
| Gm25021 | 0 | NA | NA | NA |
| Met | 239.4772955 | 0.749762998 | 0.174023879 | 4.308391493 |
| Capza2 | 2658.372187 | 0.494427549 | 0.079702212 | 6.203435729 |
| St7 | 410.6774712 | -0.262738748 | 0.111644468 | -2.353352139 |
| Gm27755 | 0 | NA | NA | NA |
| Gm27762 | 0 | NA | NA | NA |
| Gm27960 | 2.550499828 | -0.070447371 | 0.637699266 | -0.110471149 |
| Gm27365 | 1.826179369 | -0.636294427 | 0.715478625 | -0.889326955 |
| Gm27542 | 1.845381307 | -0.557227699 | 0.746092967 | -0.746860946 |
| Gm28008 | 1.623521978 | -0.562427678 | 0.804854603 | -0.698794137 |
| Gm13898 | 0.097139547 | 0.274393828 | 3.724945049 | 0.073663859 |
| Gm13899 | 0 | NA | NA | NA |
| RP23-479D21.3 | 13.97715435 | -0.301818751 | 0.341729714 | -0.883208977 |
| RP23-479D21.4 | 0.045882455 | 0.755288839 | 3.724945049 | 0.202765096 |
| Gm26738 | 0.238158741 | 1.261036919 | 2.528669932 | 0.498695739 |
| Gm27444 | 0 | NA | NA | NA |
| Gm27402 | 3.526582245 | -0.052895313 | 0.555251309 | -0.095263734 |
| RP23-86H7.3 | 81.37030569 | 0.004456309 | 0.285053096 | 0.01563326 |

| | | | | |
|---|---|---|---|---|
| Wnt2 | 29.40825172 | -0.361468596 | 0.316535644 | -1.141952267 |
| Asz1 | 0.284445263 | 1.574900939 | 2.007700686 | 0.784430145 |
| RP23-275N13.1 | 0 | NA | NA | NA |
| Gm15593 | 0.075043205 | 0.755286614 | 3.724945049 | 0.202764498 |
| Gm15596 | 0.396988867 | 0.758407754 | 1.468047995 | 0.516609645 |
| Cftr | 19.0807837 | -0.293300958 | 0.301808434 | -0.971811668 |
| Gm15592 | 0 | NA | NA | NA |
| Gm15595 | 0.243349506 | -1.285363256 | 2.447492637 | -0.525175536 |
| Cttnbp2 | 1527.7495 | -0.595195025 | 0.139489193 | -4.266961554 |
| Gm15594 | 1.482657567 | 2.322454992 | 1.040654515 | 2.231725283 |
| RP23-16J8.6 | 154.7156367 | -0.874629638 | 0.348429276 | -2.510207087 |
| Gm26233 | 3.766699628 | -0.387423878 | 0.565830738 | -0.684699243 |
| Gm23648 | 0 | NA | NA | NA |
| RP23-148L8.1 | 0 | NA | NA | NA |
| Gm5874 | 1.63591125 | -0.321413265 | 0.939901477 | -0.341964847 |
| Gm26809 | 2.591679042 | 0.790781093 | 0.635309332 | 1.244718208 |
| RP23-148L8.3 | 35.09407007 | 0.019713148 | 0.190502195 | 0.103479897 |
| Lsm8 | 195.2002428 | 0.089930388 | 0.089893787 | 1.000407151 |
| Ankrd7 | 2.04319782 | 1.063720673 | 0.749632186 | 1.418990131 |
| RP24-230L6.3 | 0.39809594 | -0.206483481 | 1.828500763 | -0.112925018 |
| Gm22718 | 0 | NA | NA | NA |
| RP23-18G14.1 | 0.045882455 | 0.755288839 | 3.724945049 | 0.202765096 |
| RP23-179A12.2 | 0 | NA | NA | NA |
| RP23-179A12.1 | 0 | NA | NA | NA |
| Gm24213 | 0 | NA | NA | NA |
| Mir6370 | 0 | NA | NA | NA |
| Gm22828 | 0 | NA | NA | NA |
| RP23-302D18.3 | 0 | NA | NA | NA |
| RP23-302D18.1 | 0 | NA | NA | NA |
| RP23-302D18.2 | 0.044191549 | -0.206490061 | 3.724945049 | -0.055434391 |
| RP23-441A23.1 | 0 | NA | NA | NA |
| Gm23960 | 0 | NA | NA | NA |
| Gm26373 | 0 | NA | NA | NA |
| Kcnd2 | 2260.738386 | -0.461399931 | 0.104784854 | -4.403307462 |
| RP23-307L24.2 | 1439.777736 | -0.860696589 | 0.330464049 | -2.604508997 |
| RP23-94H23.1 | 58.13546571 | -0.810789128 | 0.30196248 | -2.685065799 |
| RP23-301C7.2 | 101.8565513 | -0.494659354 | 0.241601829 | -2.0474156 |
| Tspan12 | 200.3279207 | -0.143600249 | 0.132465553 | -1.084057293 |
| RP24-169J24.2 | 0.152959369 | -0.677445784 | 2.840073126 | -0.238531106 |
| RP24-169J24.1 | 0.164540623 | 0.813330268 | 3.435283845 | 0.236757806 |
| Ing3 | 280.4244847 | -0.074585093 | 0.074320811 | -1.003555959 |
| Gm26719 | 0 | NA | NA | NA |
| Cped1 | 74.68292297 | -0.526349997 | 0.154984295 | -3.396150536 |
| RP24-279C2.3 | 0 | NA | NA | NA |
| Wnt16 | 10.34745373 | -0.168208589 | 0.351840186 | -0.478082366 |
| Fam3c | 626.0288257 | -0.187154467 | 0.093067767 | -2.010948282 |
| RP23-369P2.2 | 0 | NA | NA | NA |

| | | | | |
|---|---|---|---|---|
| Gm8927 | 0.687801143 | 2.673443984 | 1.333419832 | 2.004952919 |
| RP24-241H23.2 | 1.331472478 | 0.246465619 | 0.88237057 | 0.279322121 |
| RP24-241H23.1 | 0.086648695 | -0.687376175 | 3.724945049 | -0.184533239 |
| Gm23961 | 0 | NA | NA | NA |
| Gm25942 | 0 | NA | NA | NA |
| RP23-369P2.3 | 0.047948698 | -0.206487837 | 3.724945049 | -0.055433794 |
| Ptprz1 | 4928.564296 | -0.266577947 | 0.134710526 | -1.978894705 |
| Aass | 39.94585083 | -0.548028367 | 0.230105279 | -2.381641868 |
| RP24-301O17.2 | 0 | NA | NA | NA |
| RP24-301O17.3 | 0 | NA | NA | NA |
| RP24-301O17.1 | 0 | NA | NA | NA |
| Fezf1 | 7.883156094 | 0.581208937 | 0.452283986 | 1.285053098 |
| RP23-124H2.4 | 3.80210492 | -0.891364671 | 0.525722656 | -1.695503628 |
| Cadps2 | 1566.7278 | -0.07866715 | 0.128013903 | -0.61452036 |
| Gm27959 | 0 | NA | NA | NA |
| Gm24217 | 2.108983506 | 0.376141243 | 0.641454314 | 0.586388204 |
| Rnf133 | 16.93650597 | -0.07983713 | 0.408370545 | -0.195501687 |
| Rnf148 | 13.88597797 | -0.73659257 | 0.404954718 | -1.818950459 |
| Tas2r118 | 0 | NA | NA | NA |
| Slc13a1 | 10.97794222 | 0.043864479 | 0.383373409 | 0.114417113 |
| Gm20706 | 0 | NA | NA | NA |
| Gm26386 | 0.072025462 | 0.755288839 | 3.724945049 | 0.202765096 |
| Iqub | 42.66293401 | -0.306844197 | 0.175918011 | -1.744245495 |
| Ndufa5 | 270.339159 | 0.008291784 | 0.218600687 | 0.037931188 |
| Asb15 | 1.031776346 | -0.013037326 | 1.049729881 | -0.012419696 |
| RP23-392I2.4 | 0 | NA | NA | NA |
| Lmod2 | 0.965679557 | -0.0738598 | 0.944043048 | -0.078237746 |
| Wasl | 1854.805742 | 0.021584627 | 0.054887029 | 0.393255528 |
| RP23-392I2.3 | 73.97553726 | -0.018359343 | 0.180570638 | -0.101674023 |
| Hyal6 | 0.086908494 | -0.687376175 | 3.724945049 | -0.184533239 |
| Hyal4 | 0 | NA | NA | NA |
| Spam1 | 0 | NA | NA | NA |
| RP24-255M7.1 | 0 | NA | NA | NA |
| Hyal5 | 0 | NA | NA | NA |
| Gm22267 | 0 | NA | NA | NA |
| Tmem229a | 986.0139867 | -0.988233518 | 0.16149611 | -6.119240398 |
| RP23-339M5.2 | 0.278190828 | -0.206488454 | 2.443116518 | -0.084518463 |
| Gm24443 | 0 | NA | NA | NA |
| Gm22529 | 0 | NA | NA | NA |
| Gpr37 | 655.1826324 | -0.60823442 | 0.151255332 | -4.021242844 |
| Pot1a | 253.1759545 | -0.603183258 | 0.155191996 | -3.886690516 |
| RP23-302O24.1 | 0.342793059 | 0.240244137 | 1.652219547 | 0.145406909 |
| RP23-302O24.2 | 0 | NA | NA | NA |
| Gm24691 | 0 | NA | NA | NA |
| RP23-308F24.1 | 0 | NA | NA | NA |
| Grm8 | 291.4031537 | -0.607303658 | 0.129015784 | -4.707204355 |
| RP23-414I9.1 | 5.120409944 | -0.076087408 | 0.46588819 | -0.163316886 |

| | | | | |
|---|---|---|---|---|
| Gm26310 | 0 | NA | NA | NA |
| Mir592 | 0 | NA | NA | NA |
| RP23-23C4.6 | 185.154323 | -0.671991198 | 0.140163201 | -4.794348272 |
| 6530409C15Rik | 8.058619829 | 0.161977796 | 0.362976001 | 0.446249326 |
| RP23-23C4.5 | 118.802725 | -0.412374996 | 0.253102246 | -1.629282246 |
| RP23-23C4.4 | 75.79663319 | -0.61572709 | 0.227965294 | -2.700968548 |
| Zfp800 | 452.6169934 | -0.014722973 | 0.069391989 | -0.212171075 |
| Gm5303 | 1.49686415 | -0.384009107 | 0.768632524 | -0.499600387 |
| Gcc1 | 343.3632238 | 0.138707983 | 0.090741905 | 1.528598981 |
| Arf5 | 1016.757531 | -0.052480714 | 0.075444237 | -0.695622567 |
| Fscn3 | 1.623741216 | -0.247366543 | 0.785336785 | -0.314981481 |
| Pax4 | 0.249106506 | 0.729655795 | 1.83835317 | 0.396907301 |
| RP23-434O14.5 | 0 | NA | NA | NA |
| Snd1 | 1136.070214 | 0.166319659 | 0.074615529 | 2.229022049 |
| Gm37321 | 29.66800731 | -0.899166469 | 0.311064757 | -2.890608623 |
| Gm37978 | 39.62155469 | -0.717962686 | 0.310984591 | -2.308676083 |
| Lrrc4 | 977.950627 | 0.029026919 | 0.103561735 | 0.280286136 |
| Gm37940 | 70.33009473 | -0.214462174 | 0.273708594 | -0.783541982 |
| Gm3294 | 93.79479471 | -1.147755439 | 0.277918982 | -4.129820254 |
| Mir129-1 | 0 | NA | NA | NA |
| Lep | 0.088534442 | 0.274389469 | 3.724945049 | 0.073662689 |
| Gm25589 | 0 | NA | NA | NA |
| Rbm28 | 422.2309319 | 0.015608204 | 0.114924477 | 0.135812706 |
| Gm24303 | 1.400035301 | -0.907207213 | 1.026359103 | -0.883908186 |
| Prrt4 | 24.29690244 | -0.143076938 | 0.217706992 | -0.657199553 |
| Impdh1 | 348.4065021 | -0.553359073 | 0.077242103 | -7.163956616 |
| Gm26021 | 0 | NA | NA | NA |
| Hilpda | 32.36285512 | 0.356276033 | 0.217026566 | 1.641624068 |
| Fam71f2 | 3.490009053 | 1.032774239 | 0.577736719 | 1.78762091 |
| Fam71f1 | 0.2407585 | -0.206484594 | 2.09221951 | -0.098691649 |
| Gm17172 | 0.045882455 | 0.755288839 | 3.724945049 | 0.202765096 |
| Calu | 1240.173328 | 0.383033247 | 0.116588739 | 3.285336552 |
| Opn1sw | 0.679991162 | -2.844767615 | 1.365216906 | -2.083747719 |
| Ccdc136 | 845.6284117 | -0.069976377 | 0.099938 | -0.700197892 |
| Gm26220 | 0.160408561 | -0.206482807 | 3.457615423 | -0.059718269 |
| AC044807.1 | 0.581411716 | -0.33201065 | 1.18355494 | -0.280519847 |
| Flnc | 1393.252646 | 2.809679317 | 0.377330905 | 7.446194516 |
| Atp6v1f | 586.7322244 | 0.031655792 | 0.09281758 | 0.341053836 |
| Gm9047 | 2.295347566 | -0.308199387 | 0.621004905 | -0.49629139 |
| Kcp | 10.30552492 | -0.339540247 | 0.314536724 | -1.079493177 |
| Gm26484 | 0 | NA | NA | NA |
| Gm26627 | 0.072025462 | 0.755288839 | 3.724945049 | 0.202765096 |
| Irf5 | 262.9012068 | 3.040040343 | 0.220347275 | 13.79658695 |
| Tnpo3 | 1004.219965 | 0.035471246 | 0.075538899 | 0.469575895 |
| RP23-446B13.7 | 31.58542221 | -0.187029384 | 0.266300616 | -0.702324264 |
| Tspan33 | 165.1105126 | -0.125645889 | 0.124440608 | -1.009685592 |
| Smo | 219.4811244 | -0.146486023 | 0.143553432 | -1.020428569 |

| | | | | |
|---|---|---|---|---|
| Gm13717 | 1.21856855 | -1.041413357 | 0.862256551 | -1.207776683 |
| Ahcyl2 | 2068.520444 | -0.2159514 | 0.044930354 | -4.806358804 |
| RP23-353E8.7 | 0.12092566 | 1.236179401 | 3.724945049 | 0.331865138 |
| Strip2 | 859.1687429 | -0.337242471 | 0.076059329 | -4.433939596 |
| Gm13724 | 2.453329902 | -0.186043744 | 0.630959818 | -0.294858307 |
| Gm24536 | 0.309201626 | -1.608697426 | 1.572143558 | -1.023250973 |
| Gm13725 | 0.309323177 | -0.206482368 | 2.014677331 | -0.102489051 |
| Smkr-ps | 0.763164319 | 0.126822201 | 1.17953698 | 0.107518631 |
| Gm23049 | 0 | NA | NA | NA |
| Nrf1 | 255.4170055 | -0.021382709 | 0.094366073 | -0.226593188 |
| Gm25580 | 0 | NA | NA | NA |
| Mir182 | 0 | NA | NA | NA |
| Mir96 | 0 | NA | NA | NA |
| Mir183 | 0 | NA | NA | NA |
| Gm37046 | 0.075043205 | 0.755286614 | 3.724945049 | 0.202764498 |
| Ube2h | 1345.992283 | 0.000221195 | 0.058438503 | 0.003785084 |
| 1700095J07Rik | 0 | NA | NA | NA |
| Gm25625 | 0.044191549 | -0.206490061 | 3.724945049 | -0.055434391 |
| Zc3hc1 | 183.2490365 | 0.327213793 | 0.097456614 | 3.357532966 |
| n-R5s161 | 0 | NA | NA | NA |
| Klhdc10 | 1743.124813 | -0.003662825 | 0.042154601 | -0.086890276 |
| 1700012C14Rik | 0.900590653 | -1.347494321 | 1.085211396 | -1.241688326 |
| Tmem209 | 379.778664 | 0.356799745 | 0.092455229 | 3.859162399 |
| Ssmem1 | 1.951291546 | -0.839872142 | 0.69501969 | -1.20841489 |
| Cpa2 | 12.99980896 | 0.549834777 | 0.302719518 | 1.816317562 |
| Cpa4 | 11.88556507 | 1.623163189 | 0.580894227 | 2.794249133 |
| Gm13781 | 0.062426873 | 0.755288839 | 3.724945049 | 0.202765096 |
| Cpa5 | 0.592694123 | -0.641158177 | 1.335100601 | -0.480232108 |
| Cpa1 | 0.176646398 | 0.755284509 | 3.360121616 | 0.22477892 |
| Cep41 | 83.12317599 | 0.054668067 | 0.140965112 | 0.387812744 |
| Gm13782 | 0.049553047 | 0.274391604 | 3.724945049 | 0.073663262 |
| Gm14528 | 0 | NA | NA | NA |
| Mest | 610.9026533 | -1.052008696 | 0.118715408 | -8.86160202 |
| Gm27522 | 0 | NA | NA | NA |
| Gm27568 | 0 | NA | NA | NA |
| Mir335 | 5.501266716 | -0.904701535 | 0.544725988 | -1.660837844 |
| Copg2 | 853.6229713 | -0.022507338 | 0.05913165 | -0.380630979 |
| 4930412F09Rik | 0.271012069 | 2.190632929 | 3.700163284 | 0.592036827 |
| Tsga13 | 1.195665945 | 2.625947974 | 1.106761013 | 2.372642282 |
| Klf14 | 3.978034112 | -0.02170377 | 0.561466446 | -0.038655506 |
| Mir29a | 6.497893077 | 0.87060909 | 0.490230069 | 1.775919401 |
| Mir29b-1 | 0.108266322 | 0.274389235 | 3.724945049 | 0.073662626 |
| Gm27019 | 20.56482463 | -0.060651125 | 0.379963703 | -0.159623471 |
| Gm13834 | 35.42125562 | 0.032279307 | 0.336192585 | 0.096014333 |
| Lincpint | 345.1173544 | -0.199385167 | 0.216528112 | -0.920828086 |
| Gm13833 | 0.042716945 | -0.206490061 | 3.724945049 | -0.055434391 |
| 5330406M23Rik | 159.3696281 | -0.547835882 | 0.378772089 | -1.44634702 |

| | | | | |
|---|---|---|---|---|
| Gm13835 | 176.8303318 | 0.616756131 | 0.132078118 | 4.669631413 |
| Gm37728 | 114.4100306 | -0.017168308 | 0.417824649 | -0.041089745 |
| 2210408F21Rik | 141.5258413 | -0.282479727 | 0.161811401 | -1.745734385 |
| RP23-400L4.7 | 44.53988991 | -1.185467357 | 0.422092636 | -2.808547828 |
| Gm14532 | 12.88187382 | -0.63233565 | 0.327459764 | -1.931033122 |
| Mkln1os | 23.92253865 | -0.656718369 | 0.230753772 | -2.845970249 |
| Gm13844 | 13.1985606 | -0.482316241 | 0.469916897 | -1.026386249 |
| Mkln1 | 2442.640936 | 0.061189029 | 0.086847785 | 0.704554851 |
| Podxl | 278.0855086 | 1.085247706 | 0.130290352 | 8.329455653 |
| Gm6117 | 0.646697505 | -0.666108591 | 1.226500306 | -0.543096963 |
| RP24-414G20.1 | 3.468425366 | -0.377731409 | 0.595482251 | -0.634328577 |
| Gm13846 | 0.441802452 | 1.727412222 | 1.628652465 | 1.060638939 |
| Gm13848 | 0.595326517 | 0.760662492 | 1.63020701 | 0.466604847 |
| RP24-414G20.4 | 0.125911488 | -0.206491416 | 3.707612284 | -0.055693908 |
| Gm13849 | 0.554297541 | -1.656724625 | 1.34422316 | -1.232477371 |
| 1700012A03Rik | 0.048949532 | -0.206487837 | 3.724945049 | -0.055433794 |
| Gm13850 | 0 | NA | NA | NA |
| Plxna4 | 2374.903231 | -0.316744416 | 0.101082002 | -3.133539208 |
| Plxna4os3 | 0 | NA | NA | NA |
| Plxna4os2 | 0.093141081 | -0.687373951 | 3.724945049 | -0.184532642 |
| Plxna4os1 | 3.620095065 | 1.959425571 | 0.764717835 | 2.562285698 |
| Gm13852 | 0.049553047 | 0.274391604 | 3.724945049 | 0.073663262 |
| RP23-225G5.2 | 0.573159608 | -0.44427032 | 1.482239695 | -0.299729067 |
| Chchd3 | 512.9631055 | 0.115111876 | 0.097277024 | 1.183340851 |
| Gm14540 | 0 | NA | NA | NA |
| RP24-246G10.3 | 30.7240258 | -0.332771393 | 0.35756063 | -0.930671236 |
| Gm22321 | 0 | NA | NA | NA |
| Gm1401 | 0 | NA | NA | NA |
| Exoc4 | 1197.152527 | -0.088103033 | 0.067708551 | -1.301209843 |
| Gm13854 | 0 | NA | NA | NA |
| RP23-4H14.1 | 15.19819234 | -0.357296359 | 0.416252817 | -0.858363823 |
| RP24-365C10.2 | 0 | NA | NA | NA |
| Gm13853 | 0.045882455 | 0.755288839 | 3.724945049 | 0.202765096 |
| Lrguk | 29.47287266 | 0.039479481 | 0.209615805 | 0.188342101 |
| Gm13856 | 0.085327104 | 0.274389469 | 3.724945049 | 0.073662689 |
| Slc35b4 | 730.9802596 | -0.181313053 | 0.054472946 | -3.328497264 |
| Gm13855 | 0 | NA | NA | NA |
| Gm14546 | 0.956739943 | 0.254435161 | 1.010524792 | 0.251785175 |
| Akr1b3 | 294.7538566 | 0.665755116 | 0.088687142 | 7.506782853 |
| Akr1b8 | 26.07990468 | 2.134799296 | 0.383520867 | 5.566318494 |
| Gm13859 | 0.259720851 | -0.632263221 | 2.042091337 | -0.309615544 |
| Gm13858 | 0.289603658 | 1.586945312 | 2.047000082 | 0.775254152 |
| Akr1b10 | 145.9333712 | 0.160946586 | 0.100955609 | 1.594231236 |
| Akr1b7 | 1.910658871 | 0.470249242 | 0.759727489 | 0.618970945 |
| Gm13857 | 0.038981395 | 0.274393828 | 3.724945049 | 0.073663859 |
| Bpgm | 500.8527338 | 0.090706754 | 0.080587292 | 1.125571447 |
| Gm13860 | 0 | NA | NA | NA |

| | | | | |
|---|---|---|---|---|
| Cald1 | 388.6951145 | 0.127358697 | 0.12187806 | 1.044968193 |
| Gm13861 | 2.144383846 | -0.43790958 | 0.811898523 | -0.539364917 |
| Npn2 | 0 | NA | NA | NA |
| Gm14547 | 0 | NA | NA | NA |
| Agbl3 | 107.7106235 | 0.085221576 | 0.155102693 | 0.549452586 |
| Gm13864 | 2.929070206 | -0.103112648 | 0.648891645 | -0.158905804 |
| Gm13862 | 0.066050875 | 0.274391604 | 3.724945049 | 0.073663262 |
| Gm13863 | 0 | NA | NA | NA |
| RP23-386A17.7 | 1.526274188 | 1.247241118 | 0.912262579 | 1.367195309 |
| Tmem140 | 313.938631 | 4.966729158 | 0.271284926 | 18.30816492 |
| 3110062M04Rik | 36.44138757 | -1.046432564 | 0.247172487 | -4.233612629 |
| Wdr91 | 325.7975239 | 0.121725534 | 0.079735387 | 1.526618703 |
| Stra8 | 0.124347486 | 0.755286736 | 3.717316019 | 0.203180664 |
| 2010107G12Rik | 6.64236274 | 1.473708208 | 0.456970611 | 3.224951831 |
| Cnot4 | 879.3333849 | -0.133882579 | 0.069268144 | -1.932816017 |
| Nup205 | 597.9465125 | 0.029724294 | 0.086541997 | 0.343466697 |
| Gm23435 | 0.60633683 | -1.193166268 | 1.17494226 | -1.015510556 |
| 1810058I24Rik | 158.214639 | -0.035073802 | 0.13449893 | -0.260773834 |
| Slc13a4 | 187.4589476 | -0.011202074 | 0.204404071 | -0.054803576 |
| Fam180a | 13.25868466 | -0.703757354 | 0.31630885 | -2.224905674 |
| RP23-338E5.5 | 0.361225529 | 0.7553092 | 1.931570213 | 0.391033779 |
| Mtpn | 3362.617321 | 0.068030348 | 0.059249083 | 1.148209309 |
| Gm23053 | 1.826002588 | 0.192973958 | 0.729069828 | 0.264685152 |
| RP23-20C5.1 | 0 | NA | NA | NA |
| RP23-13B24.1 | 13.51087949 | -0.079509279 | 0.293197995 | -0.271179478 |
| Gm23273 | 0 | NA | NA | NA |
| RP23-352D5.1 | 6.269730523 | -0.044549067 | 0.454853853 | -0.097941497 |
| 9330158H04Rik | 22.23221858 | 0.097751651 | 0.228171252 | 0.428413526 |
| Chrm2 | 536.4163306 | 0.26255061 | 0.121079566 | 2.168413874 |
| Mir490 | 3.003392332 | 0.285613937 | 0.58292462 | 0.489967188 |
| RP23-81D20.2 | 0 | NA | NA | NA |
| RP23-5K15.1 | 0 | NA | NA | NA |
| Gm25111 | 0.298614356 | 1.783612334 | 1.958621935 | 0.910646564 |
| Ptn | 1821.983259 | -0.113484219 | 0.134480092 | -0.843873741 |
| Dgki | 1466.320099 | -0.236419362 | 0.114610752 | -2.062802639 |
| 1700111E14Rik | 0.088736197 | 0.274400501 | 3.724945049 | 0.073665651 |
| RP23-151F20.5 | 2.105924665 | -1.085905235 | 0.93171154 | -1.1654951 |
| Creb3l2 | 335.1776498 | -0.190823087 | 0.102003435 | -1.870751575 |
| Akr1d1 | 0.049553047 | 0.274391604 | 3.724945049 | 0.073663262 |
| RP24-549J13.2 | 0 | NA | NA | NA |
| RP23-245J24.1 | 9.865094166 | 0.066529431 | 0.369115053 | 0.180240362 |
| Gm15487 | 1555.447171 | 0.060299795 | 0.07088664 | 0.850651063 |
| RP23-459F14.2 | 0.11197992 | 0.75528439 | 3.724945049 | 0.202763901 |
| Trim24 | 495.6674005 | 0.065072882 | 0.10910679 | 0.596414591 |
| Svopl | 1.88997817 | 0.937280195 | 0.915424271 | 1.023875184 |
| Atp6v0a4 | 3.783295768 | 0.594705281 | 0.582649744 | 1.020690881 |
| Tmem213 | 0 | NA | NA | NA |

| | | | | |
|---|---|---|---|---|
| Gm23231 | 0 | NA | NA | NA |
| D630045J12Rik | 1061.614314 | -0.4414118 | 0.133349179 | -3.310195108 |
| RP23-387J8.3 | 0 | NA | NA | NA |
| Zc3hav1l | 80.97671505 | -0.765125616 | 0.186495012 | -4.102659935 |
| Zc3hav1 | 1446.894863 | 4.506751842 | 0.174092304 | 25.88713993 |
| RP24-142K18.4 | 0.189025308 | -0.687378639 | 2.256242799 | -0.304656325 |
| Rpl30-ps1 | 9.381830078 | -0.192522746 | 0.373405602 | -0.515586122 |
| Ttc26 | 133.6908354 | -0.427299782 | 0.119487005 | -3.576119266 |
| Gm16563 | 1.549622433 | -2.445376512 | 1.03302927 | -2.367189956 |
| Ubn2 | 2325.889383 | -0.22153693 | 0.101411175 | -2.184541586 |
| Mir7670 | 0 | NA | NA | NA |
| 1110001J03Rik | 118.1303116 | -0.382826369 | 0.14944218 | -2.56170225 |
| Gm15697 | 2.902282796 | -0.171967877 | 0.702061142 | -0.244947152 |
| Luc7l2 | 4182.925321 | -0.061603968 | 0.112037881 | -0.549849453 |
| Gm26699 | 1.935704857 | 0.304642493 | 0.729498907 | 0.417605139 |
| Klrg2 | 32.43324731 | 2.662539482 | 0.312390887 | 8.523102291 |
| Clec2l | 411.0156647 | 0.009701853 | 0.077692878 | 0.124874416 |
| Hipk2 | 3020.872203 | 0.042755337 | 0.099847966 | 0.428204382 |
| Tbxas1 | 29.76369512 | 0.845109996 | 0.28755081 | 2.938993622 |
| RP23-346B17.4 | 0.320565116 | 2.156562473 | 3.700650391 | 0.582752285 |
| Gm23010 | 0.282660793 | -2.0566163 | 3.682238457 | -0.558523388 |
| 1700025N23Rik | 9.679120195 | 6.731591496 | 0.922993468 | 7.293216834 |
| Parp12 | 1518.309522 | 4.246389286 | 0.156425596 | 27.14638399 |
| 4930599N23Rik | 26.63738886 | 3.468522864 | 0.438481984 | 7.91029733 |
| RP23-345F4.3 | 0.255942976 | 0.274394014 | 3.054170292 | 0.089842408 |
| Kdm7a | 1749.004419 | -0.45121371 | 0.074388378 | -6.065647921 |
| RP24-173B18.1 | 0.413211764 | 2.074553263 | 1.494565016 | 1.388064916 |
| Gm26008 | 0.222111871 | 1.942901157 | 3.112888699 | 0.624147326 |
| Slc37a3 | 628.5897833 | -0.030413026 | 0.070693053 | -0.430212376 |
| Rab19 | 12.00148128 | 5.221790825 | 0.891566166 | 5.856874141 |
| Mkrn1 | 1161.354 | -0.003540688 | 0.049828213 | -0.07105789 |
| Gm10244 | 4.940723394 | 0.66069956 | 0.450552363 | 1.466421251 |
| Gm25402 | 0.04393175 | -0.206490061 | 3.724945049 | -0.055434391 |
| Dennd2a | 408.9845452 | 0.085165569 | 0.094517888 | 0.901052395 |
| 8030453O22Rik | 22.33756815 | -0.093103603 | 0.296938972 | -0.313544574 |
| Adck2 | 176.9877061 | -0.097293458 | 0.095189384 | -1.022104091 |
| Gm16272 | 1.598725798 | 0.273687142 | 0.834836585 | 0.327833191 |
| RP23-366P9.8 | 0.091666476 | -0.687373951 | 3.724945049 | -0.184532642 |
| Ndufb2 | 352.2685807 | -0.352000404 | 0.093489277 | -3.765142003 |
| Braf | 3301.479618 | -0.067773112 | 0.077398544 | -0.87563807 |
| Fth-ps2 | 26.73179484 | -0.01441328 | 0.387386873 | -0.037206425 |
| Mrps33 | 137.537649 | 0.141007933 | 0.134169656 | 1.050967389 |
| RP23-380O16.6 | 0 | NA | NA | NA |
| RP23-380O16.7 | 0.107078754 | -0.206490058 | 3.724945049 | -0.05543439 |
| Gm26833 | 7.290008566 | 0.458519684 | 0.431701886 | 1.062121103 |
| Tmem178b | 2878.275615 | -0.234428144 | 0.084182765 | -2.784752259 |
| Gm37995 | 95.94723699 | -0.484404873 | 0.308909216 | -1.56811402 |

| | | | | |
|---|---|---|---|---|
| Gm37673 | 72.41137234 | -0.595824607 | 0.280824854 | -2.121694708 |
| Gm26638 | 42.96957209 | -0.831468477 | 0.400481119 | -2.076173975 |
| Gm26807 | 1.057496005 | 2.259620231 | 1.159906218 | 1.948105973 |
| Gm37003 | 29.74312774 | -0.675332767 | 0.370547396 | -1.822527355 |
| Agk | 341.3868807 | -0.077180535 | 0.082345287 | -0.937279325 |
| E330009J07Rik | 1236.299548 | -0.878385381 | 0.140552907 | -6.249499915 |
| Wee2 | 3.520209586 | 0.13829917 | 0.586279685 | 0.235892824 |
| Ssbp1 | 232.2371204 | 0.256287146 | 0.136208967 | 1.88157323 |
| Tas2r137 | 4.276107861 | -0.289646125 | 0.528901689 | -0.547636983 |
| Tas2r108 | 4.194868342 | 0.922288709 | 0.570183398 | 1.617529923 |
| Prss37 | 0 | NA | NA | NA |
| RP24-503P15.7 | 0 | NA | NA | NA |
| Olfr461 | 0 | NA | NA | NA |
| Olfr460 | 0.474871048 | 1.544168186 | 1.393189573 | 1.108369037 |
| Clec5a | 38.15951351 | 2.806056966 | 0.268073427 | 10.46749392 |
| Tas2r138 | 0 | NA | NA | NA |
| Mgam | 2.716524655 | 0.563619453 | 0.781254088 | 0.721429125 |
| Mgam2-ps | 0.962702212 | 3.039760081 | 1.126852518 | 2.697566923 |
| Moxd2 | 0 | NA | NA | NA |
| Gm38276 | 0 | NA | NA | NA |
| Trbv1 | 1.116438897 | 2.73752121 | 1.156164071 | 2.367761876 |
| Prss58 | 0 | NA | NA | NA |
| 1700074P13Rik | 0 | NA | NA | NA |
| Gm4744 | 0 | NA | NA | NA |
| Gm29588 | 0 | NA | NA | NA |
| 1810009J06Rik | 0 | NA | NA | NA |
| Gm2663 | 0 | NA | NA | NA |
| Gm38015 | 0 | NA | NA | NA |
| Gm37878 | 0 | NA | NA | NA |
| Gm38294 | 0 | NA | NA | NA |
| 2210010C04Rik | 0.16656219 | 0.274407806 | 2.783958904 | 0.098567477 |
| Trbv2 | 1.408278968 | 3.598575106 | 1.119079873 | 3.215655283 |
| Trbv3 | 0.461584485 | 2.390155163 | 1.62624763 | 1.469736293 |
| 5830405F06Rik | 0.95870128 | 2.892121538 | 1.094847981 | 2.641573613 |
| Trbv5 | 0.150914725 | 0.755296917 | 2.849470602 | 0.265065699 |
| Trbv6 | 0 | NA | NA | NA |
| Trbv7 | 0 | NA | NA | NA |
| Gm37610 | 0 | NA | NA | NA |
| Trbv8 | 0 | NA | NA | NA |
| Trbv9 | 0 | NA | NA | NA |
| Trbv10 | 0 | NA | NA | NA |
| Trbv11 | 0 | NA | NA | NA |
| Trbv12-1 | 0 | NA | NA | NA |
| Trbv13-1 | 2.762763099 | 3.264010098 | 1.077057331 | 3.030488726 |
| Trbv12-2 | 0.921345027 | 2.859402005 | 1.126117939 | 2.539167441 |
| Trbv13-2 | 3.038727124 | 4.684421195 | 0.991924912 | 4.722556252 |
| Trbv12-3 | 0 | NA | NA | NA |

| | | | | |
|---|---|---|---|---|
| Trbv13-3 | 1.194558986 | 2.544348656 | 1.211402832 | 2.100332432 |
| Trbv14 | 1.5023977 | 3.65187283 | 1.799493491 | 2.029389297 |
| Trbv15 | 0.168389893 | 1.210042036 | 2.766164025 | 0.437444065 |
| Trbv16 | 0.74889667 | 3.433089127 | 1.353544675 | 2.536369275 |
| Trbv17 | 1.696056422 | 3.756904395 | 1.102269284 | 3.408336285 |
| Trbv18 | 0 | NA | NA | NA |
| Trbv19 | 0.811168495 | 2.151742081 | 1.278509032 | 1.683008901 |
| Trbv20 | 0.334925437 | 1.202982073 | 2.020974135 | 0.595248624 |
| Trbv21 | 0 | NA | NA | NA |
| Trbv22 | 0.097139547 | 0.274393828 | 3.724945049 | 0.073663859 |
| Trbv23 | 0 | NA | NA | NA |
| Trbv24 | 0.038981395 | 0.274393828 | 3.724945049 | 0.073663859 |
| Trbv25 | 0 | NA | NA | NA |
| Trbv26 | 0.038981395 | 0.274393828 | 3.724945049 | 0.073663859 |
| Gm2719 | 0.3093278 | -0.668778914 | 1.788434983 | -0.37394645 |
| Trbv27 | 0 | NA | NA | NA |
| Trbv28 | 0 | NA | NA | NA |
| Trbv29 | 1.593432551 | 4.016264109 | 1.153907698 | 3.480576579 |
| Trbv30 | 0.049553047 | 0.274391604 | 3.724945049 | 0.073663262 |
| Try4 | 0.044544648 | 0.755286614 | 3.724945049 | 0.202764498 |
| Gm18964 | 0 | NA | NA | NA |
| Try5 | 0 | NA | NA | NA |
| Try10 | 0 | NA | NA | NA |
| Prss3 | 0 | NA | NA | NA |
| Gm5771 | 0 | NA | NA | NA |
| Gm38306 | 0 | NA | NA | NA |
| Gm5409 | 0 | NA | NA | NA |
| Gm10334 | 0.980132196 | 0.427680831 | 0.906022154 | 0.472042354 |
| Prss1 | 0 | NA | NA | NA |
| Gm37301 | 0 | NA | NA | NA |
| Gm29813 | 0.146823232 | -0.194513984 | 2.876272087 | -0.067627115 |
| Gm37841 | 0 | NA | NA | NA |
| Gm38018 | 0 | NA | NA | NA |
| Prss2 | 0.046345709 | 0.274391604 | 3.724945049 | 0.073663262 |
| Trbd1 | 0.960611118 | 3.085724222 | 1.116853953 | 2.762871737 |
| Trbj1-1 | 0.739650571 | 1.198115301 | 1.308591736 | 0.915576087 |
| Trbj1-2 | 0.363286374 | 0.274380111 | 3.724945049 | 0.073660177 |
| Trbj1-3 | 0.347715489 | 0.755309651 | 1.551958826 | 0.486681501 |
| Trbj1-4 | 0.471163529 | 1.236206525 | 1.533419397 | 0.806176398 |
| Trbj1-5 | 0.450331108 | 2.374707882 | 1.361749946 | 1.743864862 |
| Trbj1-6 | 0.583857846 | 1.228930376 | 1.44974606 | 0.847686646 |
| Trbj1-7 | 0.161076545 | 0.755297744 | 2.79165938 | 0.270555122 |
| Trbc1 | 24.37840666 | 7.070084673 | 0.897378869 | 7.878595002 |
| Trbd2 | 0.440524946 | 2.0560679 | 1.677762629 | 1.225481999 |
| Trbj2-1 | 0.112396584 | 0.27438722 | 3.724945049 | 0.073662085 |
| Trbj2-2 | 0.218053095 | -0.206486751 | 2.161793264 | -0.095516419 |
| Trbj2-3 | 0.696513874 | -1.264448787 | 1.158312448 | -1.091630147 |

| | | | | |
|---|---|---|---|---|
| Trbj2-4 | 0.088736197 | 0.274400501 | 3.724945049 | 0.073665651 |
| Trbj2-5 | 0.202417642 | -0.206482447 | 2.653394741 | -0.077818217 |
| Trbj2-6 | 0.354669972 | 0.755305511 | 1.918554246 | 0.39368473 |
| Trbj2-7 | 1.180533128 | -0.370382437 | 0.868229078 | -0.426595292 |
| Trbc2 | 69.31285373 | 1.178468993 | 0.212265285 | 5.551868708 |
| Trbv31 | 1.290124216 | 4.070298984 | 1.087674879 | 3.742201885 |
| Gm27549 | 0 | NA | NA | NA |
| Ephb6 | 693.8526302 | -0.35609921 | 0.13327241 | -2.67196496 |
| Trpv6 | 9.753117248 | 0.780535089 | 0.362296061 | 2.1544123 |
| Trpv5 | 0 | NA | NA | NA |
| 1700034O15Rik | 0.14896402 | -0.20648958 | 2.866996645 | -0.072022958 |
| Kel | 0.227427828 | 0.755305022 | 2.559553268 | 0.295092519 |
| Gm37527 | 0 | NA | NA | NA |
| Olfr459 | 0 | NA | NA | NA |
| Pip | 0.038981395 | 0.274393828 | 3.724945049 | 0.073663859 |
| Sval2 | 0.163190422 | 0.274394252 | 3.398594894 | 0.080737558 |
| Sval1 | 0.173394273 | 0.274392352 | 3.390823156 | 0.080922047 |
| Sval3 | 0.066050875 | 0.274391604 | 3.724945049 | 0.073663262 |
| Sva | 0.066050875 | 0.274391604 | 3.724945049 | 0.073663262 |
| Tas2r139 | 0 | NA | NA | NA |
| Tas2r144 | 0.049553047 | 0.274391604 | 3.724945049 | 0.073663262 |
| Gstk1 | 46.48690246 | -0.188926125 | 0.189242462 | -0.998328405 |
| Tmem139 | 0.247571123 | -0.629448325 | 2.47861698 | -0.253951429 |
| Casp2 | 320.4543346 | 0.420683728 | 0.127433693 | 3.301197009 |
| Gm25108 | 3.821797685 | 0.817738667 | 0.538364538 | 1.51893115 |
| Clcn1 | 102.7118201 | 0.448715221 | 0.282446052 | 1.588675848 |
| Fam131b | 1482.538637 | -0.104795703 | 0.087918792 | -1.191960223 |
| Gm24611 | 0 | NA | NA | NA |
| Zyx | 868.7508899 | 0.775869194 | 0.16936407 | 4.58107316 |
| Epha1 | 4.796176745 | -0.318092414 | 0.515342333 | -0.617244874 |
| 2010310C07Rik | 14.52291164 | 0.585035369 | 0.391154789 | 1.495662037 |
| Tas2r143 | 4.118545241 | 0.455911026 | 0.515614686 | 0.884208766 |
| Tas2r135 | 5.600410292 | 1.110936494 | 0.526578764 | 2.109725209 |
| Tas2r126 | 2.194003788 | 0.109984494 | 0.705466804 | 0.155903145 |
| Olfr458 | 0.062426873 | 0.755288839 | 3.724945049 | 0.202765096 |
| Olfr457 | 0 | NA | NA | NA |
| Olfr456 | 0 | NA | NA | NA |
| Gm4877 | 0.146848595 | -0.999135224 | 3.718157501 | -0.268717832 |
| Olfr455 | 0 | NA | NA | NA |
| Tcaf3 | 0.038981395 | 0.274393828 | 3.724945049 | 0.073663859 |
| Tcaf2 | 17.75907395 | 0.631430673 | 0.278056591 | 2.270871085 |
| Tcaf1 | 3402.596378 | -0.345944547 | 0.081490971 | -4.245188662 |
| Olfr453 | 0.410820671 | 2.222255248 | 1.505843078 | 1.475754865 |
| Olfr38 | 0.225129614 | 1.818209708 | 3.710727189 | 0.489987438 |
| Gm22363 | 0.272364125 | -0.206486215 | 2.046244987 | -0.100909821 |
| Olfr452 | 0.180050447 | -0.206493189 | 3.327115185 | -0.062063733 |
| Olfr450 | 0.211502678 | 0.274401415 | 2.552509456 | 0.107502605 |

| | | | | |
|---|---|---|---|---|
| Gm23982 | 0.194279094 | 0.274391173 | 3.724945049 | 0.073663147 |
| Olfr449 | 0.134148234 | -0.20649159 | 3.624632408 | -0.056968974 |
| Olfr448 | 0.07796279 | 0.274392189 | 3.724945049 | 0.073663419 |
| Olfr447 | 0 | NA | NA | NA |
| Olfr446 | 0.048949532 | -0.206487837 | 3.724945049 | -0.055433794 |
| Olfr444 | 0 | NA | NA | NA |
| Gm17472 | 0 | NA | NA | NA |
| Gm20730 | 0 | NA | NA | NA |
| Olfr443-ps1 | 0.336338362 | 1.249206505 | 1.741934795 | 0.717137351 |
| Olfr441 | 0 | NA | NA | NA |
| Olfr237-ps1 | 0 | NA | NA | NA |
| Olfr437 | 0 | NA | NA | NA |
| Olfr13 | 0 | NA | NA | NA |
| Olfr435 | 0.038981395 | 0.274393828 | 3.724945049 | 0.073663859 |
| Gm21827 | 0 | NA | NA | NA |
| Olfr434 | 0.061920613 | 0.274393828 | 3.724945049 | 0.073663859 |
| Olfr47 | 0 | NA | NA | NA |
| Arhgef5 | 73.22831166 | 0.607195859 | 0.183791864 | 3.303714573 |
| Nobox | 0.405536581 | 1.274615155 | 1.583212494 | 0.805081541 |
| Tpk1 | 61.31573283 | 0.084758719 | 0.174091488 | 0.486863085 |
| Gm15550 | 0.332944463 | -0.543195537 | 1.566878693 | -0.346673638 |
| Gm22939 | 0 | NA | NA | NA |
| Cntnap2 | 2123.978884 | 0.052551436 | 0.062196528 | 0.844925558 |
| Gm22839 | 8.429271064 | 0.279260906 | 0.348811669 | 0.800606548 |
| Gm25161 | 0 | NA | NA | NA |
| Rbpsuh-rs3 | 97.81620039 | 0.34020362 | 0.137035408 | 2.482596462 |
| Cul1 | 1508.243995 | 0.207266984 | 0.059500053 | 3.483475609 |
| A930035D04Rik | 12.16101713 | -0.046548776 | 0.303090803 | -0.1535803 |
| Ezh2 | 210.0637519 | -0.021112298 | 0.106474453 | -0.198285099 |
| Gm26625 | 0.182019239 | -0.206490577 | 2.725623728 | -0.075759018 |
| Gm25283 | 0 | NA | NA | NA |
| Rny3 | 804.362109 | -0.105037867 | 0.320334916 | -0.32790015 |
| Rpl31-ps7 | 0 | NA | NA | NA |
| Rny1 | 909.9952825 | 0.030673131 | 0.417755368 | 0.073423666 |
| Pdia4 | 818.8143374 | 0.452157014 | 0.125959529 | 3.589700732 |
| Mir704 | 0.123992736 | 0.274402763 | 3.720643035 | 0.073751435 |
| Rpl35a-ps7 | 0.314214724 | 2.510626975 | 1.667198491 | 1.505895661 |
| Zfp786 | 50.22326823 | 0.023272814 | 0.170110776 | 0.136809755 |
| Zfp398 | 324.873339 | -0.042150767 | 0.144431363 | -0.291839435 |
| Zfp282 | 307.4764343 | -0.221612897 | 0.109592111 | -2.022161044 |
| Zfp212 | 200.032767 | 0.222134311 | 0.133691979 | 1.661538052 |
| Zfp783 | 130.5839755 | 0.270811953 | 0.158045064 | 1.713510988 |
| Zfp956 | 173.5071749 | 0.155375384 | 0.141649489 | 1.09690042 |
| Zfp777 | 460.5844228 | 0.498732365 | 0.100572576 | 4.958930004 |
| Gm24563 | 1.500420242 | 0.006992228 | 0.836121372 | 0.008362695 |
| Zfp746 | 384.3199641 | 0.090402357 | 0.087545646 | 1.032631094 |
| Gm16630 | 7.347387118 | 0.52851179 | 0.37959722 | 1.392296262 |

| | | | | |
|---|---|---|---|---|
| Gm22122 | 0 | NA | NA | NA |
| Krba1 | 549.5583407 | -0.364009185 | 0.134585686 | -2.704664939 |
| Gm24325 | 1.916854426 | -0.878438489 | 0.684912763 | -1.282555292 |
| Zfp467 | 330.9730189 | -0.352087055 | 0.088145938 | -3.994365057 |
| Sspo | 38.70866699 | -0.570967777 | 0.348351656 | -1.639055726 |
| Gm10243 | 43.3226042 | 0.602547417 | 0.207937574 | 2.897732263 |
| Gm7887 | 75.72156547 | 0.210290159 | 0.157027505 | 1.339193147 |
| 1700026J14Rik | 0.552687697 | -1.158809557 | 1.364597848 | -0.849194917 |
| Atp6v0e2 | 1764.72768 | 0.011261582 | 0.067244962 | 0.167471012 |
| Lrrc61 | 226.76103 | 0.119084673 | 0.109715642 | 1.085393752 |
| Rarres2 | 195.195546 | 2.261427308 | 0.32289588 | 7.003580564 |
| Gm5111 | 0.049553047 | 0.274391604 | 3.724945049 | 0.073663262 |
| Repin1 | 624.3197349 | 0.181672327 | 0.089788657 | 2.023332723 |
| Zfp775 | 56.68412911 | -0.371532234 | 0.160011969 | -2.321902777 |
| AI854703 | 220.6699541 | -0.281748523 | 0.115726542 | -2.434605902 |
| Gimap8 | 51.49836854 | 1.858298359 | 0.287116526 | 6.472279332 |
| Gimap9 | 17.15776395 | 2.65114692 | 0.402229295 | 6.59113335 |
| Gimap4 | 77.7956503 | 6.17124677 | 0.575362562 | 10.72583998 |
| Gimap6 | 85.26288905 | 2.732326652 | 0.261085417 | 10.46525956 |
| Gm28053 | 0.321006207 | 1.193018826 | 2.381111246 | 0.501034476 |
| Gimap7 | 21.5292461 | 5.834774779 | 0.918305492 | 6.353849381 |
| Gimap1os | 2.672780519 | 2.695917214 | 0.976006545 | 2.762191737 |
| Gimap1 | 14.27405217 | 3.359211379 | 0.524462744 | 6.40505244 |
| Gimap5 | 14.86559326 | 3.507007654 | 0.523287492 | 6.701875564 |
| Gimap3 | 25.51805368 | 6.431897953 | 0.897712083 | 7.164767048 |
| Tmem176b | 421.451915 | 0.994042203 | 0.186399184 | 5.332867775 |
| Tmem176a | 202.5557947 | 1.20186964 | 0.190497675 | 6.309103982 |
| Gm7932 | 3.063139515 | 1.634507424 | 0.875804008 | 1.86629361 |
| Aoc1 | 3.736083012 | 2.760424205 | 0.893219926 | 3.090419418 |
| 1600015I10Rik | 0 | NA | NA | NA |
| Gm24348 | 0 | NA | NA | NA |
| Gm10242 | 0.061920613 | 0.274393828 | 3.724945049 | 0.073663859 |
| Doxl2 | 0.106112162 | -0.206492286 | 3.724945049 | -0.055434988 |
| Svs1 | 0.044191549 | -0.206490061 | 3.724945049 | -0.055434391 |
| Gpnmb | 840.8388242 | 3.961102434 | 0.439033097 | 9.022332175 |
| Malsu1 | 255.5915214 | 0.064457785 | 0.09598218 | 0.671559924 |
| Igf2bp3 | 6.6227068 | 4.33269821 | 0.849261091 | 5.101726969 |
| Tra2a | 624.172519 | -0.278496279 | 0.089736716 | -3.103481943 |
| Ccdc126 | 154.9854233 | 0.15429897 | 0.114865411 | 1.343302301 |
| Fam221a | 18.46478838 | 0.405256815 | 0.26602824 | 1.523360129 |
| Stk31 | 5.37787014 | 0.394407593 | 0.430453517 | 0.916260589 |
| Gm17060 | 0.066050875 | 0.274391604 | 3.724945049 | 0.073663262 |
| Npy | 399.7067098 | 0.063247098 | 0.185979752 | 0.340075182 |
| Gm21786 | 0.284351963 | 0.892102451 | 3.705884352 | 0.240725928 |
| Mpp6 | 900.1739491 | 0.027166767 | 0.055742908 | 0.487358271 |
| Dfna5 | 75.89232883 | -0.339516756 | 0.141794585 | -2.394426815 |
| Osbpl3 | 392.5077654 | -0.352139594 | 0.11134344 | -3.16264339 |

| | | | | |
|---|---|---|---|---|
| Gm26289 | 0 | NA | NA | NA |
| Cycs | 696.5473769 | 0.131809678 | 0.14698245 | 0.89677154 |
| 5430402O13Rik | 7.75721647 | -1.279948402 | 0.399582743 | -3.20321241 |
| Mir6371 | 0.990815542 | -0.881675603 | 1.347936373 | -0.654092894 |
| 4921507P07Rik | 15.81567947 | 0.551377587 | 0.239682593 | 2.30044902 |
| Gm4782 | 0.19378329 | -0.687377773 | 2.235607224 | -0.307468041 |
| Gm8038 | 0.04393175 | -0.206490061 | 3.724945049 | -0.055434391 |
| Npvf | 3.377670371 | 0.606143283 | 0.550454664 | 1.101168401 |
| G930045G22Rik | 0 | NA | NA | NA |
| Gm29004 | 0.141094079 | 0.755285157 | 3.563850454 | 0.211929531 |
| Mir148a | 0.208883516 | 1.261687113 | 2.63299814 | 0.479182683 |
| Gm26677 | 0.297812842 | -0.20648164 | 1.769499057 | -0.116689319 |
| Nfe2l3 | 68.43159579 | 0.240504963 | 0.166535558 | 1.444165833 |
| Hnrnpa2b1 | 3001.421392 | -0.163175925 | 0.091575498 | -1.781873191 |
| Cbx3 | 448.3183731 | 0.042582024 | 0.104937709 | 0.405783817 |
| Snx10 | 1492.510951 | 0.541930132 | 0.104534757 | 5.184210014 |
| Gm22914 | 0.310734401 | -0.185984945 | 1.768913445 | -0.105140783 |
| Skap2 | 251.8896519 | 0.989437605 | 0.148443113 | 6.665432891 |
| Halr1 | 0.212468866 | 1.221524374 | 2.177464967 | 0.560984628 |
| Rps8-ps3 | 0 | NA | NA | NA |
| Rps19-ps14 | 0 | NA | NA | NA |
| Hoxa1 | 1.953182484 | -0.650829656 | 0.914181014 | -0.711926463 |
| Hotairm1 | 2.721387227 | -1.757996115 | 0.735400758 | -2.390528017 |
| Gm27867 | 0 | NA | NA | NA |
| Gm27861 | 0.467025382 | -1.856884484 | 1.489880025 | -1.246331552 |
| Gm27868 | 0 | NA | NA | NA |
| Gm27477 | 0.460648582 | -2.313561222 | 1.310982862 | -1.764753216 |
| Hoxa2 | 5.976910094 | 0.292302968 | 0.672576537 | 0.434601791 |
| Gm28308 | 0 | NA | NA | NA |
| Hoxaas2 | 3.211323041 | -0.458374466 | 0.732130014 | -0.626083424 |
| Hoxa3 | 12.50440182 | -0.654547074 | 0.791982604 | -0.826466479 |
| Gm15050 | 0.11193333 | 0.75528439 | 3.724945049 | 0.202763901 |
| Hoxaas3 | 0.798143448 | 0.911596346 | 1.510394222 | 0.603548619 |
| 5730596B20Rik | 0.683443395 | -1.407109554 | 1.318345127 | -1.067330189 |
| Gm29430 | 0.397176565 | 1.891159384 | 1.835656463 | 1.030236007 |
| Hoxa4 | 7.87519976 | -0.215390524 | 0.66671581 | -0.323061972 |
| Hoxa5 | 22.4157456 | -0.784470709 | 0.92566415 | -0.84746796 |
| Hoxa6 | 2.164192679 | -0.864723046 | 1.688982837 | -0.511978586 |
| Hoxa7 | 1.260199239 | -1.703379789 | 1.19102273 | -1.430182436 |
| Hoxa9 | 0.137647365 | 1.404948034 | 3.719469264 | 0.377728093 |
| AC015583.1 | 0 | NA | NA | NA |
| Mir196b | 0 | NA | NA | NA |
| Hoxa10 | 0 | NA | NA | NA |
| Hoxa11 | 0 | NA | NA | NA |
| Hoxa11os | 0 | NA | NA | NA |
| Gm28032 | 0 | NA | NA | NA |
| Gm27492 | 0 | NA | NA | NA |

| | | | | |
|---|---|---|---|---|
| Gm27695 | 0 | NA | NA | NA |
| Gm27817 | 0 | NA | NA | NA |
| Gm27454 | 0 | NA | NA | NA |
| Gm27543 | 0 | NA | NA | NA |
| 9530018H14Rik | 0 | NA | NA | NA |
| Hoxa13 | 0 | NA | NA | NA |
| Gm27280 | 0 | NA | NA | NA |
| Gm27985 | 0 | NA | NA | NA |
| Hottip | 0 | NA | NA | NA |
| Gm27383 | 0 | NA | NA | NA |
| Gm27866 | 0 | NA | NA | NA |
| 5730457N03Rik | 3.810690039 | -0.190175178 | 0.553179774 | -0.343785487 |
| Evx1 | 6.587797916 | -0.234063533 | 0.561308421 | -0.416996297 |
| Gm8129 | 27.45075818 | 0.358364893 | 0.351414388 | 1.019778657 |
| Gm15054 | 0 | NA | NA | NA |
| Gm37351 | 0.106358623 | 0.274402725 | 3.724945049 | 0.073666248 |
| Hibadh | 577.34931 | 0.038344428 | 0.102065437 | 0.375684749 |
| Gm24355 | 3.634340466 | 0.040207828 | 0.571625228 | 0.070339492 |
| Tax1bp1 | 2381.399223 | 0.318566267 | 0.070520491 | 4.517357451 |
| Jazf1 | 226.345128 | -0.28955743 | 0.106127451 | -2.728393335 |
| Gm15572 | 3.883676966 | -0.245963647 | 0.521987854 | -0.47120569 |
| Gm26215 | 6.191745285 | -0.235969509 | 0.434367804 | -0.543248157 |
| Gm15573 | 0.191192637 | -1.042666354 | 2.65601198 | -0.392568393 |
| Gm25818 | 0.106618422 | 0.274402725 | 3.724945049 | 0.073666248 |
| Creb5 | 153.7701789 | 0.913703418 | 0.204913336 | 4.458974882 |
| Tril | 1025.612528 | -0.161134617 | 0.16746414 | -0.962203712 |
| Gm16499 | 0.523988269 | 1.022236501 | 1.439015017 | 0.710372365 |
| Gm22910 | 0 | NA | NA | NA |
| Cpvl | 0.864647981 | 0.263090767 | 1.246138928 | 0.211124748 |
| Chn2 | 1528.702195 | -0.248413195 | 0.220037576 | -1.128958059 |
| 9530036M11Rik | 0.430808853 | 1.521333417 | 1.540035961 | 0.987855775 |
| Gm15526 | 0.194037777 | -0.667437513 | 2.637297208 | -0.253076336 |
| Gm15527 | 0.824096613 | 0.776795868 | 1.196672049 | 0.649130117 |
| 9130019P16Rik | 5.722762008 | 0.290259525 | 0.424979249 | 0.682996936 |
| Prr15 | 15.96290937 | 1.356911436 | 0.321576166 | 4.219564696 |
| Wipf3 | 1686.195845 | 0.147155048 | 0.150851489 | 0.97549616 |
| Scrn1 | 2778.202593 | -0.150770748 | 0.052658688 | -2.863169495 |
| Fkbp14 | 126.0321638 | -0.08909282 | 0.16084315 | -0.553911184 |
| Plekha8 | 312.5409318 | -0.607235094 | 0.076290595 | -7.959501404 |
| Gm15863 | 0.061920613 | 0.274393828 | 3.724945049 | 0.073663859 |
| Gm28402 | 0.435961187 | -0.190795323 | 1.440599746 | -0.132441591 |
| Mturn | 1138.29535 | -0.465686369 | 0.101278806 | -4.598063387 |
| Znrf2 | 429.2306728 | 0.148958765 | 0.072072842 | 2.066780771 |
| Nod1 | 393.3170085 | 2.367430576 | 0.181550227 | 13.040086 |
| Gm24230 | 0 | NA | NA | NA |
| Ggct | 96.47110576 | 0.471669547 | 0.213261964 | 2.21169091 |
| Gars | 1477.498831 | 0.349009594 | 0.093463437 | 3.734183192 |

| | | | | |
|---|---|---|---|---|
| Crhr2 | 14.43709135 | -0.124790931 | 0.280638706 | -0.444667568 |
| Inmt | 2.148441887 | -0.097706048 | 0.755224496 | -0.129373515 |
| Fam188b | 89.47598892 | -0.19590705 | 0.150063123 | -1.305497625 |
| Gm25458 | 0 | NA | NA | NA |
| Aqp1 | 74.91150286 | 0.494981643 | 0.560178758 | 0.883613732 |
| Ghrhr | 0.426601203 | 0.755280747 | 3.26857073 | 0.231073705 |
| Adcyap1r1 | 2280.821534 | 0.028012555 | 0.049360368 | 0.567511067 |
| AC069309.1 | 0.075043205 | 0.755286614 | 3.724945049 | 0.202764498 |
| Neurod6 | 272.3533864 | -0.476386417 | 0.18596946 | -2.561637897 |
| Ccdc129 | 12.62732475 | 0.115158978 | 0.295599351 | 0.389577912 |
| Ppp1r17 | 219.4288715 | -0.157499694 | 0.298207198 | -0.528155239 |
| Pde1c | 512.5279336 | -0.755427831 | 0.109989623 | -6.868173639 |
| AC079956.1 | 1.707215572 | -1.120609628 | 0.771094032 | -1.45327234 |
| Gm24709 | 0 | NA | NA | NA |
| Gm22493 | 0.217296566 | -1.501333772 | 2.166387933 | -0.693012433 |
| Lsm5 | 26.01436472 | 0.37841972 | 0.209584786 | 1.805568651 |
| Avl9 | 1503.371839 | 0.469623871 | 0.083891593 | 5.597984876 |
| Kbtbd2 | 937.4819014 | 0.036269686 | 0.064121463 | 0.565640334 |
| Fkbp9 | 377.8815204 | 0.021469783 | 0.121722169 | 0.176383506 |
| Nt5c3 | 889.5108846 | 1.49951352 | 0.139932093 | 10.71600868 |
| Vmn1r4 | 1.941638245 | 1.248093179 | 0.682151707 | 1.829641655 |
| Vmn1r5 | 0.356992291 | 0.922333094 | 3.305960079 | 0.278990996 |
| Vmn1r6 | 0.118371171 | 0.75528439 | 3.724945049 | 0.202763901 |
| Vmn1r7 | 0 | NA | NA | NA |
| Vmn1r8 | 0 | NA | NA | NA |
| Vmn1r9 | 0 | NA | NA | NA |
| Vmn1r10 | 0 | NA | NA | NA |
| Vmn1r11 | 0 | NA | NA | NA |
| Vmn1r12 | 0 | NA | NA | NA |
| Vmn1r-ps8 | 0 | NA | NA | NA |
| Vmn1r13 | 0 | NA | NA | NA |
| Vmn1r14 | 0 | NA | NA | NA |
| Vmn1r15 | 0 | NA | NA | NA |
| Vmn1r-ps9 | 0 | NA | NA | NA |
| Vmn1r-ps10 | 0 | NA | NA | NA |
| Vmn1r16 | 0 | NA | NA | NA |
| Vmn1r17 | 0 | NA | NA | NA |
| Vmn1r18 | 0 | NA | NA | NA |
| Vmn1r19 | 0 | NA | NA | NA |
| Vmn1r20 | 0 | NA | NA | NA |
| Vmn1r-ps11 | 0 | NA | NA | NA |
| 4930533I22Rik | 0.062426873 | 0.755288839 | 3.724945049 | 0.202765096 |
| Ppm1k | 1833.8061 | 0.577439197 | 0.106594516 | 5.417156699 |
| Gm27896 | 0 | NA | NA | NA |
| Herc6 | 2713.472928 | 4.389463738 | 0.168642317 | 26.02824612 |
| Gm37397 | 18.0739852 | -0.724537894 | 0.260455327 | -2.781812536 |
| Pyurf | 116.2739024 | -0.453271712 | 0.113427092 | -3.996150336 |

| | | | | |
|---|---|---|---|---|
| Gm26712 | 0 | NA | NA | NA |
| Lancl2 | 881.9561078 | -0.008188169 | 0.062383004 | -0.131256415 |
| Vopp1 | 778.6664814 | -0.255078471 | 0.082010679 | -3.110307998 |
| Vmn1r21 | 0.060174537 | 0.755286614 | 3.724945049 | 0.202764498 |
| Gm25726 | 0 | NA | NA | NA |
| Vmn1r-ps12 | 0 | NA | NA | NA |
| Vmn1r22 | 0 | NA | NA | NA |
| Vmn1r23 | 0 | NA | NA | NA |
| Gm20660 | 0 | NA | NA | NA |
| Vmn1r24 | 0 | NA | NA | NA |
| Vmn1r25 | 0 | NA | NA | NA |
| Vmn1r26 | 0 | NA | NA | NA |
| Rps15-ps2 | 2.860642787 | 0.286608463 | 0.7209475 | 0.397544153 |
| Vmn1r-ps14 | 0 | NA | NA | NA |
| Vmn1r-ps15 | 0.09176491 | 1.15872478 | 3.72306111 | 0.311229052 |
| Gm6695 | 0.119587853 | 1.218392466 | 3.723618458 | 0.327206581 |
| Vmn1r-ps17 | 0 | NA | NA | NA |
| Gm20718 | 0.110595523 | 0.755282166 | 3.724945049 | 0.202763304 |
| Vmn1r-ps18 | 0 | NA | NA | NA |
| Vmn1r27 | 0 | NA | NA | NA |
| Vmn1r-ps19 | 0 | NA | NA | NA |
| Vmn1r28 | 0.068097359 | -0.206487837 | 3.724945049 | -0.055433794 |
| Vmn1r29 | 0.139856492 | -0.206491248 | 3.574411025 | -0.057769307 |
| Gm20674 | 0 | NA | NA | NA |
| Vmn1r-ps20 | 0 | NA | NA | NA |
| Vmn1r30 | 0.075043205 | 0.755286614 | 3.724945049 | 0.202764498 |
| Vmn1r-ps21 | 0 | NA | NA | NA |
| Gm20673 | 0 | NA | NA | NA |
| Vmn1r31 | 0 | NA | NA | NA |
| Vmn1r-ps22 | 0.047948698 | -0.206487837 | 3.724945049 | -0.055433794 |
| Vmn1r-ps23 | 0.066050875 | 0.274391604 | 3.724945049 | 0.073663262 |
| Gm20672 | 0 | NA | NA | NA |
| Abcg2 | 150.9866917 | 0.326813826 | 0.115685348 | 2.825023499 |
| Herc3 | 2335.927333 | 0.036878842 | 0.086687608 | 0.425422304 |
| Nap1l5 | 4015.777592 | 0.3366187 | 0.128653477 | 2.616475717 |
| Fam13a | 397.9236056 | 0.059582708 | 0.069124069 | 0.861967603 |
| BB365896 | 2.654833662 | 0.929112437 | 0.744403101 | 1.248130799 |
| Gm22367 | 0 | NA | NA | NA |
| Tigd2 | 330.9779313 | 0.544050104 | 0.110885672 | 4.906405799 |
| Gprin3 | 236.7557592 | -0.286371745 | 0.122576144 | -2.33627634 |
| Gm22622 | 0 | NA | NA | NA |
| Gm5570 | 0.128070691 | 1.236174986 | 3.723419581 | 0.331999915 |
| Gm22723 | 0 | NA | NA | NA |
| n-R5s162 | 0 | NA | NA | NA |
| A530053G22Rik | 0.106618422 | 0.274402725 | 3.724945049 | 0.073666248 |
| Snca | 1013.675009 | -0.128981362 | 0.12583754 | -1.02498318 |
| Mmrn1 | 0.717105192 | 1.627497879 | 1.305675698 | 1.246479414 |

| | | | | |
|---|---|---|---|---|
| A730020E08Rik | 23.88982311 | -0.021802484 | 0.271716744 | -0.08023975 |
| Ccser1 | 527.4917736 | -0.161047221 | 0.104852812 | -1.535936114 |
| Gm22212 | 0 | NA | NA | NA |
| Grid2 | 1329.763725 | -0.238116816 | 0.117926139 | -2.01920302 |
| Gm25205 | 1.741084885 | -0.618621363 | 0.972169719 | -0.636330623 |
| Atoh1 | 0.260468428 | 0.766818633 | 2.044308664 | 0.375099243 |
| Gm15534 | 0.038981395 | 0.274393828 | 3.724945049 | 0.073663859 |
| Smarcad1 | 613.7764423 | -0.133057857 | 0.067162868 | -1.981122313 |
| Hpgds | 46.4750562 | 0.363744904 | 0.262653503 | 1.384885029 |
| Gm17147 | 2.881706403 | 0.05548544 | 0.724023189 | 0.076634893 |
| C130060K24Rik | 9.901210978 | 0.151695602 | 0.376230647 | 0.403198419 |
| Tnip3 | 4.68769279 | 3.924668574 | 1.023614665 | 3.834126951 |
| Ndnf | 237.9647372 | -0.5956965 | 0.163979236 | -3.632755678 |
| Gm15490 | 0.045882455 | 0.755288839 | 3.724945049 | 0.202765096 |
| Gm16838 | 1.376958878 | 0.556757708 | 0.782210983 | 0.711774342 |
| Prdm5 | 99.54165155 | 0.156165037 | 0.113729259 | 1.373129811 |
| Gm23543 | 0 | NA | NA | NA |
| Gm20445 | 3.495574212 | -0.255719048 | 0.491556145 | -0.520223478 |
| 4930544G11Rik | 0 | NA | NA | NA |
| Gm23625 | 0 | NA | NA | NA |
| Mad2l1 | 71.55826495 | 0.045937047 | 0.173740523 | 0.264400305 |
| Vmn1r32 | 0 | NA | NA | NA |
| Vmn1r33 | 0.066050875 | 0.274391604 | 3.724945049 | 0.073663262 |
| Vmn1r34 | 0.225129614 | 1.818209708 | 3.710727189 | 0.489987438 |
| Vmn1r35 | 0.038981395 | 0.274393828 | 3.724945049 | 0.073663859 |
| Vmn1r36 | 0 | NA | NA | NA |
| Vmn1r37 | 0 | NA | NA | NA |
| Vmn1r-ps24 | 0.066050875 | 0.274391604 | 3.724945049 | 0.073663262 |
| Vmn1r38 | 0 | NA | NA | NA |
| Vmn1r39 | 0.047948698 | -0.206487837 | 3.724945049 | -0.055433794 |
| RP23-263B18.4 | 379.7491701 | 0.506817073 | 0.15629615 | 3.242671518 |
| 4930597O21Rik | 2.946845261 | 0.666851044 | 0.605143118 | 1.10197245 |
| Gng12 | 1315.427679 | 0.836663941 | 0.118777169 | 7.043979495 |
| Gm15644 | 4.54732579 | -0.245115487 | 0.538271848 | -0.455374896 |
| Gm25260 | 0.532344628 | 0.320089245 | 1.568864505 | 0.204026061 |
| Gadd45a | 98.08999083 | 0.840975366 | 0.276525653 | 3.041220072 |
| E230016M11Rik | 92.89630384 | -0.205979386 | 0.18649183 | -1.104495495 |
| A430010J10Rik | 11.54166846 | -0.501973883 | 0.354751178 | -1.415002721 |
| Gm27015 | 0.824269312 | 1.800992521 | 1.247790456 | 1.443345325 |
| Gm8566 | 29.8603066 | 0.439351863 | 0.217273083 | 2.022118232 |
| Serbp1 | 2115.70978 | -0.107989362 | 0.1113497 | -0.969821762 |
| Il12rb2 | 27.958596 | 3.614725477 | 0.595428107 | 6.070800883 |
| Gm4761 | 0 | NA | NA | NA |
| Il23r | 2.075698228 | 1.74292559 | 0.910437577 | 1.914382308 |
| RP23-123I10.3 | 0.062426873 | 0.755288839 | 3.724945049 | 0.202765096 |
| Tacstd2 | 4.381073283 | 1.530976673 | 0.608918217 | 2.514256646 |
| Igkv2-137 | 0.061920613 | 0.274393828 | 3.724945049 | 0.073663859 |

| | | | | |
|---|---|---|---|---|
| RP23-123I10.4 | 0 | NA | NA | NA |
| Igkv1-136 | 0 | NA | NA | NA |
| Igkv1-135 | 0.427013396 | 0.755308186 | 1.643620422 | 0.459539305 |
| RP24-322E13.1 | 0 | NA | NA | NA |
| RP24-322E13.2 | 0 | NA | NA | NA |
| Igkv14-134-1 | 0 | NA | NA | NA |
| Igkv17-134 | 0 | NA | NA | NA |
| Igkv1-133 | 0.216076386 | 1.736121978 | 3.709588981 | 0.468009256 |
| RP23-255N3.3 | 0 | NA | NA | NA |
| RP23-255N3.4 | 0 | NA | NA | NA |
| RP23-255N3.6 | 0 | NA | NA | NA |
| Igkv1-132 | 0.132101749 | 0.274387056 | 3.724945049 | 0.073662041 |
| Igkv1-131 | 0 | NA | NA | NA |
| RP23-255N3.8 | 0 | NA | NA | NA |
| Igkv14-130 | 0 | NA | NA | NA |
| Gm22425 | 0 | NA | NA | NA |
| Gm24911 | 0 | NA | NA | NA |
| Igkv9-129 | 0 | NA | NA | NA |
| Igkv9-128 | 0 | NA | NA | NA |
| Igkv17-127 | 0.143022002 | 0.755289479 | 3.547809333 | 0.212888971 |
| Igkv14-126-1 | 0 | NA | NA | NA |
| RP23-234A12.5 | 0 | NA | NA | NA |
| Igkv14-126 | 0 | NA | NA | NA |
| Gm25910 | 0 | NA | NA | NA |
| Igkv11-125 | 0.062426873 | 0.755288839 | 3.724945049 | 0.202765096 |
| Igkv9-124 | 0 | NA | NA | NA |
| Igkv9-123 | 0 | NA | NA | NA |
| Igkv1-122 | 0 | NA | NA | NA |
| Igkv17-121 | 0.097139547 | 0.274393828 | 3.724945049 | 0.073663859 |
| Igkv9-120 | 0.291418641 | 0.274388235 | 3.724945049 | 0.073662358 |
| Igkv9-119 | 0 | NA | NA | NA |
| Igkv14-118-2 | 0 | NA | NA | NA |
| Igkv14-118-1 | 0 | NA | NA | NA |
| RP23-185M15.10 | 0 | NA | NA | NA |
| RP23-185M15.7 | 0 | NA | NA | NA |
| Igkv11-118 | 0 | NA | NA | NA |
| Igkv1-117 | 0.45022207 | 0.274378393 | 3.724945049 | 0.073659716 |
| RP24-76K4.8 | 0 | NA | NA | NA |
| RP24-76K4.7 | 0.141094079 | 0.755285157 | 3.563850454 | 0.211929531 |
| Igkv2-116 | 0.060174537 | 0.755286614 | 3.724945049 | 0.202764498 |
| Igkv1-115 | 0 | NA | NA | NA |
| Igkv11-114 | 0 | NA | NA | NA |
| Igkv2-113 | 0 | NA | NA | NA |
| RP24-76K4.1 | 0 | NA | NA | NA |
| Igkv2-112 | 0.157314084 | 0.755289343 | 3.435173412 | 0.219869349 |
| Igkv14-111 | 0.94930219 | 2.327431739 | 1.214072798 | 1.917044631 |
| Gm25436 | 0 | NA | NA | NA |

| | | | | |
|---|---|---|---|---|
| Igkv1-110 | 0.099106094 | 0.274387056 | 3.724945049 | 0.073662041 |
| RP24-76K4.12 | 0 | NA | NA | NA |
| RP24-76K4.11 | 0 | NA | NA | NA |
| Igkv2-109 | 0.106971521 | 1.236179401 | 3.724945049 | 0.331865138 |
| Igkv1-108 | 0 | NA | NA | NA |
| Igkv2-107 | 0 | NA | NA | NA |
| Igkv19-93 | 0 | NA | NA | NA |
| Igkv2-105 | 0 | NA | NA | NA |
| RP23-29F15.2 | 0 | NA | NA | NA |
| Igkv16-104 | 0.862499085 | 3.432708692 | 3.660802972 | 0.937692828 |
| Igkv15-103 | 0.066050875 | 0.274391604 | 3.724945049 | 0.073663262 |
| Igkv15-102 | 0 | NA | NA | NA |
| Igkv20-101-2 | 0 | NA | NA | NA |
| Igkv15-101-1 | 0 | NA | NA | NA |
| Igkv15-101 | 0 | NA | NA | NA |
| Igkv14-100 | 0 | NA | NA | NA |
| Gm25833 | 0 | NA | NA | NA |
| Igkv1-99 | 0 | NA | NA | NA |
| Igkv12-98 | 0 | NA | NA | NA |
| Igkv15-97 | 0 | NA | NA | NA |
| Igkv10-96 | 0.163190422 | 0.274394252 | 3.398594894 | 0.080737558 |
| Igkv2-95-2 | 0 | NA | NA | NA |
| Igkv2-95-1 | 0 | NA | NA | NA |
| Igkv10-95 | 0 | NA | NA | NA |
| Igkv10-94 | 0.066050875 | 0.274391604 | 3.724945049 | 0.073663262 |
| Igkv2-93-1 | 0 | NA | NA | NA |
| RP24-405O23.8 | 0 | NA | NA | NA |
| AC120404.1 | 0 | NA | NA | NA |
| Igkv4-92 | 0 | NA | NA | NA |
| RP23-211L5.2 | 0 | NA | NA | NA |
| Igkv4-91 | 0 | NA | NA | NA |
| Igkv4-90 | 0 | NA | NA | NA |
| Igkv13-89-1 | 0 | NA | NA | NA |
| Igkv12-89 | 0.175102337 | 0.27439644 | 3.315835544 | 0.082753332 |
| Igkv1-88 | 0.108772582 | 0.75528439 | 3.724945049 | 0.202763901 |
| RP23-211L5.7 | 0 | NA | NA | NA |
| IGKV1D-42 | 0 | NA | NA | NA |
| RP23-211L5.9 | 0.045882455 | 0.755288839 | 3.724945049 | 0.202765096 |
| Igkv13-87 | 0.049553047 | 0.274391604 | 3.724945049 | 0.073663262 |
| RP23-211L5.12 | 0.088534442 | 0.274389469 | 3.724945049 | 0.073662689 |
| Igkv4-86 | 0 | NA | NA | NA |
| Igkv12-47 | 0.097139547 | 0.274393828 | 3.724945049 | 0.073663859 |
| Igkv13-84 | 0.060174537 | 0.755286614 | 3.724945049 | 0.202764498 |
| Igkv4-56 | 0 | NA | NA | NA |
| Igkv13-82 | 0 | NA | NA | NA |
| Igkv4-81 | 0 | NA | NA | NA |
| Igkv13-80-1 | 0 | NA | NA | NA |

| | | | | |
|---|---|---|---|---|
| Igkv4-80 | 0.097139547 | 0.274393828 | 3.724945049 | 0.073663859 |
| Igkv4-79 | 0 | NA | NA | NA |
| Igkv13-78-1 | 0 | NA | NA | NA |
| Igkv4-78 | 0 | NA | NA | NA |
| RP23-344N8.12 | 0 | NA | NA | NA |
| Igkv4-77 | 0 | NA | NA | NA |
| Igkv13-76 | 0 | NA | NA | NA |
| Igkv4-75 | 0 | NA | NA | NA |
| Igkv13-74-1 | 0 | NA | NA | NA |
| Igkv4-74 | 0.062426873 | 0.755288839 | 3.724945049 | 0.202765096 |
| Igkv13-73-1 | 0 | NA | NA | NA |
| Igkv4-73 | 0.046345709 | 0.274391604 | 3.724945049 | 0.073663262 |
| Igkv4-72 | 0.182573437 | -0.687378218 | 3.311478753 | -0.207574401 |
| Igkv13-71-1 | 0 | NA | NA | NA |
| Igkv4-71 | 0 | NA | NA | NA |
| Igkv4-70 | 0.046345709 | 0.274391604 | 3.724945049 | 0.073663262 |
| Igkv4-69 | 0 | NA | NA | NA |
| Igkv4-68 | 0.060174537 | 0.755286614 | 3.724945049 | 0.202764498 |
| Igkv12-67 | 0 | NA | NA | NA |
| Rprl1 | 0 | NA | NA | NA |
| Igkv12-66 | 0 | NA | NA | NA |
| Igkv4-65 | 0 | NA | NA | NA |
| Igkv13-64 | 0 | NA | NA | NA |
| Igkv4-63 | 0 | NA | NA | NA |
| Igkv13-62-1 | 0 | NA | NA | NA |
| Igkv4-62 | 0 | NA | NA | NA |
| Igkv13-61-1 | 0 | NA | NA | NA |
| Igkv4-61 | 0.266304556 | 0.759334413 | 3.724657033 | 0.203866935 |
| Igkv4-59 | 0 | NA | NA | NA |
| Igkv4-60 | 0 | NA | NA | NA |
| Igkv4-58 | 0 | NA | NA | NA |
| Rn7s6 | 476.0174306 | -0.159186283 | 0.334718912 | -0.47558198 |
| AC156953.1 | 0 | NA | NA | NA |
| Igkv13-57-2 | 0 | NA | NA | NA |
| Igkv4-57-1 | 0.097139547 | 0.274393828 | 3.724945049 | 0.073663859 |
| Igkv13-57-1 | 0 | NA | NA | NA |
| Igkv4-57 | 0.102891481 | 0.274400501 | 3.724945049 | 0.073665651 |
| Igkv13-56-1 | 0 | NA | NA | NA |
| Igkv13-55-1 | 0 | NA | NA | NA |
| Igkv4-55 | 0.097139547 | 0.274393828 | 3.724945049 | 0.073663859 |
| Igkv13-54-1 | 0 | NA | NA | NA |
| Igkv4-54 | 0 | NA | NA | NA |
| Igkv4-53 | 0 | NA | NA | NA |
| Gm11143 | 0 | NA | NA | NA |
| RP23-24I14.7 | 0 | NA | NA | NA |
| Igkv4-50 | 0 | NA | NA | NA |
| Igkv12-49 | 0 | NA | NA | NA |

| | | | | |
|---|---|---|---|---|
| Igkv5-48 | 0.183032319 | 1.343421106 | 3.313584731 | 0.405428325 |
| Igkv12-46 | 0.771993125 | 1.357873032 | 3.56639775 | 0.380740772 |
| Igkv5-45 | 0.049553047 | 0.274391604 | 3.724945049 | 0.073663262 |
| Igkv12-44 | 0.240624804 | 0.274397013 | 3.054169286 | 0.08984342 |
| Igkv5-43 | 0.240624804 | 0.274397013 | 3.054169286 | 0.08984342 |
| Igkv12-42 | 0 | NA | NA | NA |
| Igkv12-41 | 0.194279094 | 0.274391173 | 3.724945049 | 0.073663147 |
| Igkv5-40-1 | 0 | NA | NA | NA |
| Igkv12-40 | 0 | NA | NA | NA |
| Igkv5-39 | 0.975696798 | 2.486335721 | 2.655818481 | 0.936184359 |
| RP24-475M8.10 | 0 | NA | NA | NA |
| Igkv12-38 | 0.099106094 | 0.274387056 | 3.724945049 | 0.073662041 |
| Igkv5-37 | 0 | NA | NA | NA |
| Igkv18-36 | 0 | NA | NA | NA |
| Igkv1-35 | 0 | NA | NA | NA |
| Gm23694 | 0 | NA | NA | NA |
| RP24-314D8.6 | 0 | NA | NA | NA |
| Igkv8-34 | 0.066050875 | 0.274391604 | 3.724945049 | 0.073663262 |
| Igkv7-33 | 0.093831153 | 0.274404949 | 3.724945049 | 0.073666845 |
| Igkv6-32 | 0.998588786 | 1.68894227 | 3.672532198 | 0.459884946 |
| Igkv8-31 | 0 | NA | NA | NA |
| Igkv8-30 | 0.203286941 | 0.755296304 | 2.621314208 | 0.288136501 |
| Amd-ps2 | 0.04393175 | -0.206490061 | 3.724945049 | -0.055434391 |
| Igkv6-29 | 0 | NA | NA | NA |
| Igkv8-28 | 0 | NA | NA | NA |
| Gm24096 | 0 | NA | NA | NA |
| Igkv8-27 | 0.112396584 | 0.27438722 | 3.724945049 | 0.073662085 |
| Igkv8-26 | 0 | NA | NA | NA |
| RP23-64J23.3 | 0.049553047 | 0.274391604 | 3.724945049 | 0.073663262 |
| Igkv6-25 | 0.097139547 | 0.274393828 | 3.724945049 | 0.073663859 |
| Igkv8-24 | 0 | NA | NA | NA |
| Igkv8-23-1 | 0 | NA | NA | NA |
| Igkv6-23 | 1.206331356 | 0.274370149 | 3.724945049 | 0.073657503 |
| RP23-64J23.7 | 0 | NA | NA | NA |
| Gm11065 | 0 | NA | NA | NA |
| Igkv8-22 | 0 | NA | NA | NA |
| Igkv8-21 | 0.277388482 | 1.157005455 | 1.767937437 | 0.654438008 |
| Amd-ps5 | 0.048949532 | -0.206487837 | 3.724945049 | -0.055433794 |
| Igkv6-20 | 0.038981395 | 0.274393828 | 3.724945049 | 0.073663859 |
| Igkv8-19 | 0.269481931 | -1.32768445 | 2.410497298 | -0.550792756 |
| Igkv8-18 | 0 | NA | NA | NA |
| RP23-64J23.14 | 0.089814205 | 0.274402725 | 3.724945049 | 0.073666248 |
| Igkv6-17 | 0.659092009 | 0.274379374 | 3.724945049 | 0.073659979 |
| Igkv8-16 | 0.097139547 | 0.274393828 | 3.724945049 | 0.073663859 |
| RP23-175B24.3 | 0 | NA | NA | NA |
| Igkv6-15 | 0.421418085 | 0.274412545 | 1.736823192 | 0.157996822 |
| RP23-175B24.4 | 0.066050875 | 0.274391604 | 3.724945049 | 0.073663262 |

| | | | | |
|---|---|---|---|---|
| Gm10360 | 0 | NA | NA | NA |
| RP23-175B24.5 | 5.844746325 | -0.93980037 | 0.417510849 | -2.25096036 |
| Igkv6-14 | 0 | NA | NA | NA |
| RP23-175B24.8 | 0 | NA | NA | NA |
| Igkv6-13 | 0.044544648 | 0.755286614 | 3.724945049 | 0.202764498 |
| RP23-175B24.10 | 0 | NA | NA | NA |
| RP23-175B24.13 | 0.042716945 | -0.206490061 | 3.724945049 | -0.055434391 |
| RP23-175B24.14 | 0 | NA | NA | NA |
| Igkv3-12-1 | 0 | NA | NA | NA |
| Igkv3-12 | 0.232585366 | 1.280397609 | 2.540139519 | 0.504065859 |
| Igkv3-11 | 0 | NA | NA | NA |
| Igkv3-10 | 0.300744356 | 0.755279374 | 3.724945049 | 0.202762555 |
| Igkv3-9 | 0 | NA | NA | NA |
| Igkv3-8 | 0 | NA | NA | NA |
| Igkv3-7 | 4.685552657 | 2.051347108 | 1.683031371 | 1.218840685 |
| Igkv3-6 | 0 | NA | NA | NA |
| Igkv3-5 | 0.169165009 | 0.755292496 | 3.351424224 | 0.225364635 |
| Igkv3-4 | 0.266304556 | 0.759334413 | 3.724657033 | 0.203866935 |
| Igkv3-3 | 0 | NA | NA | NA |
| Igkv3-2 | 0.309883016 | 0.274381181 | 3.724945049 | 0.073660464 |
| Igkv3-1 | 0 | NA | NA | NA |
| RP23-435I4.11 | 0.318875346 | 0.755276743 | 3.724945049 | 0.202761848 |
| Igkj1 | 0.169165009 | 0.755292496 | 3.351424224 | 0.225364635 |
| Igkj2 | 0 | NA | NA | NA |
| Igkj3 | 0.062426873 | 0.755288839 | 3.724945049 | 0.202765096 |
| Igkj4 | 0.254453631 | 0.755294773 | 2.976423281 | 0.253759194 |
| Igkj5 | 0.189830966 | 0.274393147 | 3.269587874 | 0.083922854 |
| Igkc | 42.12345985 | 3.751857295 | 1.060434477 | 3.538037828 |
| Rpia | 130.920887 | -0.047277258 | 0.103308753 | -0.457630709 |
| Eif2ak3 | 463.4739881 | 0.008287796 | 0.07130626 | 0.116228168 |
| Gm23346 | 0.975898478 | -0.039844222 | 0.995184078 | -0.040037037 |
| RP24-122N23.3 | 0.09349418 | 0.274402725 | 3.724945049 | 0.073666248 |
| Tex37 | 0 | NA | NA | NA |
| Foxi3 | 0.042716945 | -0.206490061 | 3.724945049 | -0.055434391 |
| Gm23485 | 0 | NA | NA | NA |
| Rpl34-ps1 | 54.38793798 | 0.278780639 | 0.25585864 | 1.089588527 |
| Thnsl2 | 33.95158729 | 0.139246152 | 0.21345042 | 0.652358296 |
| Fabp1 | 0.161948271 | -0.68737637 | 3.446645093 | -0.199433464 |
| Smyd1 | 16.55316792 | -1.029040865 | 0.302695273 | -3.399593448 |
| Mir8112 | 5.618825651 | 0.432889784 | 0.519163104 | 0.833822322 |
| Krcc1 | 217.6208105 | 0.314635211 | 0.176282024 | 1.784840022 |
| Gm1070 | 1.970056041 | 1.173495343 | 0.832863328 | 1.40898909 |
| Cd8b1 | 33.23125073 | 7.967513733 | 0.894362881 | 8.908591693 |
| Cd8a | 74.76319009 | 8.338456033 | 0.890133912 | 9.367642239 |
| Rmnd5a | 2145.404065 | -0.264414183 | 0.061951901 | -4.268055985 |
| Rnf103 | 778.7289262 | -0.082397621 | 0.062630982 | -1.315604796 |
| Chmp3 | 861.0926457 | -0.041466103 | 0.073416394 | -0.564807133 |

| | | | | |
|---|---|---|---|---|
| Kdm3a | 752.2610037 | 0.08726833 | 0.076588638 | 1.139442238 |
| Reep1 | 1109.512451 | -0.329351992 | 0.07364554 | -4.472124077 |
| Mrpl35 | 321.4157534 | -0.167047701 | 0.075062613 | -2.225444788 |
| Immt | 1466.512924 | 0.068086856 | 0.059452601 | 1.145229226 |
| Ptcd3 | 534.5440444 | 0.046242391 | 0.075036276 | 0.616267141 |
| Gm22486 | 336.2636012 | -0.285763365 | 0.329688324 | -0.866768229 |
| Polr1a | 650.2299052 | 0.173439705 | 0.055876183 | 3.1040006 |
| Gm26628 | 0.134276636 | -0.206488731 | 2.954908363 | -0.06987991 |
| St3gal5 | 1041.7803 | 0.066570847 | 0.054398957 | 1.223752258 |
| Gm29438 | 74.24082104 | -0.672884385 | 0.306067796 | -2.198481494 |
| Atoh8 | 37.89726758 | 0.663418217 | 0.252165875 | 2.63088024 |
| Sftpb | 0.285479113 | 0.755310389 | 1.788126767 | 0.422403155 |
| Usp39 | 316.7202377 | 0.213032974 | 0.076536409 | 2.783420035 |
| 0610030E20Rik | 238.8730958 | -0.235577244 | 0.088130552 | -2.673048533 |
| Tmem150a | 75.99670554 | -0.461650889 | 0.163501144 | -2.823533082 |
| Rnf181 | 551.102173 | 0.162490087 | 0.094281142 | 1.723463288 |
| Vamp5 | 21.09757441 | 0.362103151 | 0.331641185 | 1.091852182 |
| Vamp8 | 157.0775941 | 1.338681598 | 0.179237286 | 7.46876738 |
| Ggcx | 228.9491815 | -0.225959294 | 0.093752662 | -2.410164028 |
| Mat2a | 2291.964425 | 0.205541822 | 0.123500804 | 1.664295419 |
| Sh2d6 | 8.896846792 | 0.855461618 | 0.381574399 | 2.241926135 |
| Capg | 296.8370465 | 4.050905451 | 0.431901092 | 9.379243356 |
| Elmod3 | 206.9098534 | -0.009012422 | 0.090832454 | -0.099220282 |
| Retsat | 77.16422045 | -0.098500853 | 0.175794405 | -0.560318475 |
| Tgoln1 | 2850.702626 | 0.247291437 | 0.076158755 | 3.247052011 |
| Gm20536 | 0.787340167 | 0.508093033 | 1.033459364 | 0.491642972 |
| Tcf7l1 | 89.55502137 | 0.080620224 | 0.144266853 | 0.558827077 |
| Gm15401 | 1.324332992 | 0.042564451 | 1.095989458 | 0.038836551 |
| Gm37969 | 4.217504979 | -0.963083016 | 0.507942771 | -1.896046309 |
| Gm38114 | 1.798572027 | -1.058669573 | 0.814041009 | -1.300511353 |
| Gm38320 | 0.230331698 | 0.792116967 | 2.557603658 | 0.309710601 |
| Gm15402 | 0.090074004 | 0.274402725 | 3.724945049 | 0.073666248 |
| Gm37736 | 8.749811602 | -0.908639897 | 0.403862054 | -2.249876878 |
| Kcmf1 | 755.6815892 | 0.200439408 | 0.079539495 | 2.519998502 |
| Gm26640 | 0.344365504 | 1.236172584 | 3.724945049 | 0.331863307 |
| Tmsb10 | 1562.811079 | 0.956471373 | 0.218423672 | 4.378973044 |
| Dnah6 | 180.1945128 | -0.340529172 | 0.171350419 | -1.98732617 |
| Gm24533 | 0.239208914 | 0.274416027 | 2.110573275 | 0.130019663 |
| Suclg1 | 851.7330975 | -0.105516804 | 0.057328681 | -1.840558735 |
| Gm24370 | 0 | NA | NA | NA |
| 4931417E11Rik | 0 | NA | NA | NA |
| Gm26179 | 0 | NA | NA | NA |
| Gm23924 | 0 | NA | NA | NA |
| Gm22254 | 0 | NA | NA | NA |
| Gm8993 | 0 | NA | NA | NA |
| Gm9001 | 4.797213872 | 0.368230414 | 0.517546203 | 0.711492832 |
| AC161363.1 | 0 | NA | NA | NA |

| | | | | |
|---|---|---|---|---|
| Gm22283 | 0 | NA | NA | NA |
| Gm9008 | 0.868283866 | -1.979690683 | 1.243702005 | -1.591772526 |
| Ctnna2 | 1519.198385 | -0.052657668 | 0.058986028 | -0.89271425 |
| Lrrtm1 | 446.8085958 | -0.069508115 | 0.109030465 | -0.637510951 |
| Gm21045 | 0 | NA | NA | NA |
| Gm5576 | 0.929768241 | -1.215202368 | 1.13363945 | -1.071947848 |
| Reg3b | 0.091880448 | -0.687373951 | 3.724945049 | -0.184532642 |
| Reg3d | 0 | NA | NA | NA |
| Reg3a | 0 | NA | NA | NA |
| Reg2 | 0 | NA | NA | NA |
| Reg1 | 0 | NA | NA | NA |
| Reg3g | 0.501377686 | 0.2744132 | 1.532128255 | 0.179105894 |
| Gm21392 | 0 | NA | NA | NA |
| Gm23727 | 0 | NA | NA | NA |
| Gm29005 | 0 | NA | NA | NA |
| Gm20594 | 0 | NA | NA | NA |
| Gm4409 | 0.396083486 | -0.813329391 | 1.522222894 | -0.534303744 |
| Lrrtm4 | 669.3441651 | -0.183012754 | 0.12051572 | -1.518579929 |
| Gm26264 | 0 | NA | NA | NA |
| Gm22530 | 0 | NA | NA | NA |
| mmu-mir-7232 | 0.163166504 | -0.205853182 | 2.784882282 | -0.073918091 |
| mmu-mir-468 | 0 | NA | NA | NA |
| 1700009C05Rik | 0.362568279 | -0.198087516 | 1.526543695 | -0.1297621 |
| Gcfc2 | 109.5450915 | -0.014384424 | 0.181986962 | -0.079040958 |
| Mrpl19 | 289.1779516 | -0.070952138 | 0.06932741 | -1.023435581 |
| Eva1a | 89.53357422 | -0.285302723 | 0.16823498 | -1.695858511 |
| Gm15864 | 0 | NA | NA | NA |
| Tacr1 | 94.67186864 | -0.269709952 | 0.127801041 | -2.110389319 |
| Pole4 | 302.9621574 | 0.011541759 | 0.115427796 | 0.099991154 |
| Hk2 | 352.4423473 | 1.394955544 | 0.199882133 | 6.978890615 |
| Gm17034 | 3.741498389 | 3.756837622 | 1.015643864 | 3.698971416 |
| 2310069B03Rik | 0.29717078 | 1.13011842 | 2.037892517 | 0.554552515 |
| Sema4f | 443.9451415 | -0.194831683 | 0.097227347 | -2.00387739 |
| M1ap | 2.42047183 | 1.967005808 | 0.866544435 | 2.269942232 |
| Dok1 | 94.45580167 | 2.624407726 | 0.297431539 | 8.823569053 |
| Loxl3 | 99.54709547 | 1.128595629 | 0.203913436 | 5.534680053 |
| Mir7040 | 0 | NA | NA | NA |
| Htra2 | 198.8763925 | 0.132124193 | 0.085554669 | 1.544324753 |
| Aup1 | 620.1822418 | 0.302202089 | 0.057228896 | 5.280585633 |
| Dqx1 | 10.03751516 | -0.195724655 | 0.41077518 | -0.476476342 |
| Tlx2 | 0.090665643 | -0.687373951 | 3.724945049 | -0.184532642 |
| Gm37092 | 0.240333628 | 0.745042294 | 1.855932626 | 0.401438222 |
| Pcgf1 | 96.34584019 | 0.183936528 | 0.11128495 | 1.652842795 |
| Lbx2 | 0.810336573 | 0.632231993 | 1.144737364 | 0.552294363 |
| Mir3470a | 0 | NA | NA | NA |
| Ccdc142 | 74.38601428 | -0.234064369 | 0.178491354 | -1.311348499 |
| Ccdc142os | 12.6558528 | -0.469651197 | 0.323087284 | -1.453635656 |

| | | | | |
|---|---|---|---|---|
| Mrpl53 | 0 | NA | NA | NA |
| Mogs | 256.0958102 | -0.355525565 | 0.105919228 | -3.356572481 |
| Wbp1 | 156.9620067 | -0.4219394 | 0.126948126 | -3.323715058 |
| Ino80b | 119.6851225 | 0.077594979 | 0.166786464 | 0.46523547 |
| Rtkn | 581.9726703 | 0.338272514 | 0.081333857 | 4.159061512 |
| Wdr54 | 115.1061269 | -0.297939123 | 0.11465844 | -2.598492731 |
| 1700003E16Rik | 151.7522411 | -0.226017082 | 0.105825953 | -2.135743401 |
| Dctn1 | 3696.021488 | -0.175337739 | 0.058662308 | -2.988933512 |
| Gm15624 | 21.19334217 | 0.233612431 | 0.359604538 | 0.64963705 |
| Slc4a5 | 42.25336671 | -0.227016171 | 0.240304759 | -0.944701106 |
| Mthfd2 | 159.228204 | 0.82658584 | 0.174725491 | 4.73076846 |
| Mob1a | 606.7587231 | 0.633423061 | 0.130998033 | 4.83536314 |
| Bola3 | 181.7266574 | 0.151940318 | 0.094436323 | 1.608918187 |
| Tet3 | 1814.49476 | -0.240481759 | 0.105785439 | -2.27329736 |
| Mir6374 | 7.73301019 | -0.705458794 | 0.495540525 | -1.423614735 |
| B230319C09Rik | 9.836627767 | 0.161402097 | 0.338601942 | 0.476672094 |
| Dguok | 113.566417 | -0.436930716 | 0.169062468 | -2.584433567 |
| Actg2 | 1.148058676 | -0.067764418 | 1.130462282 | -0.059943988 |
| Stambp | 312.4465512 | 0.201921321 | 0.082902684 | 2.435642743 |
| Gm24990 | 0 | NA | NA | NA |
| Clec4f | 0.297646291 | 0.768376032 | 1.816326233 | 0.423038559 |
| Cd207 | 0.357781856 | 1.236206851 | 1.588262199 | 0.778339277 |
| Vax2os | 1.069186981 | -0.159238215 | 0.954232567 | -0.166875687 |
| Gm27772 | 0 | NA | NA | NA |
| Gm27631 | 0 | NA | NA | NA |
| Gm27893 | 0 | NA | NA | NA |
| Vax2 | 0.095898756 | 0.27438722 | 3.724945049 | 0.073662085 |
| Gm25054 | 0 | NA | NA | NA |
| Atp6v1b1 | 0.540009155 | -0.884079097 | 1.543915517 | -0.572621421 |
| 1700124L16Rik | 10.03157174 | -0.380889816 | 0.372628754 | -1.022169684 |
| Ankrd53 | 0 | NA | NA | NA |
| Tex261 | 481.6030204 | 0.13625222 | 0.107592799 | 1.266369321 |
| Nagk | 216.7968119 | -0.036964672 | 0.101760998 | -0.363249897 |
| Paip2b | 685.0019845 | 0.172794287 | 0.075057737 | 2.302151577 |
| Gm23014 | 0 | NA | NA | NA |
| Gm5138 | 12.43742249 | 0.276119406 | 0.28754831 | 0.96025397 |
| Zfp638 | 2492.303382 | -0.162890108 | 0.079388349 | -2.051813773 |
| Dysf | 208.5045205 | 0.690888424 | 0.241450452 | 2.861408697 |
| Gm15475 | 0.044544648 | 0.755286614 | 3.724945049 | 0.202764498 |
| Cyp26b1 | 126.4459904 | -0.393861857 | 0.171478379 | -2.296860158 |
| Exoc6b | 1761.121593 | 0.190211044 | 0.046913462 | 4.054508808 |
| Gm7507 | 0.706207703 | 1.054839687 | 1.092990724 | 0.965094821 |
| Gm26353 | 0.038981395 | 0.274393828 | 3.724945049 | 0.073663859 |
| Npm3-ps1 | 4.293674138 | -0.549501277 | 0.647743529 | -0.848331559 |
| Gm5878 | 0.534626289 | 0.640237854 | 1.275865472 | 0.501806709 |
| Spr | 209.6465649 | -0.094081805 | 0.094269489 | -0.998009071 |
| Gm10444 | 0.134148234 | -0.20649159 | 3.624632408 | -0.056968974 |

| | | | | |
|---|---|---|---|---|
| Gm22893 | 0 | NA | NA | NA |
| Spr-ps1 | 0.674932609 | -0.56319257 | 1.404858706 | -0.40088912 |
| Emx1 | 100.6339851 | 0.173736139 | 0.170128884 | 1.021203074 |
| Sfxn5 | 2038.190871 | -0.555613147 | 0.114684654 | -4.844703535 |
| Rab11fip5 | 1169.630522 | 0.099092115 | 0.081127398 | 1.221438344 |
| Mir705 | 0 | NA | NA | NA |
| Noto | 0 | NA | NA | NA |
| Smyd5 | 326.4950938 | 0.146416244 | 0.075566529 | 1.93758064 |
| Pradc1 | 68.96069805 | -0.26767308 | 0.143401911 | -1.866593535 |
| Cct7 | 1480.602166 | 0.043982876 | 0.076383408 | 0.575817139 |
| Fbxo41 | 1325.807268 | -0.323820263 | 0.118893693 | -2.723611773 |
| Egr4 | 333.4082471 | -0.563093935 | 0.534387407 | -1.053718571 |
| Alms1 | 761.6584805 | -0.447969638 | 0.130428527 | -3.434598624 |
| RP23-27D8.2 | 34.95142274 | -1.178033739 | 0.251111031 | -4.691286305 |
| RP24-478I22.8 | 43.9950079 | -0.342850957 | 0.24004212 | -1.428294988 |
| Gm4477 | 0.754021959 | -0.692196564 | 1.106369639 | -0.625646745 |
| Alms1-ps1 | 10.38534064 | -0.907738228 | 0.319636978 | -2.839903671 |
| Cml3 | 8.402099551 | -0.813847026 | 0.382506691 | -2.127667424 |
| Gm27317 | 0.061920613 | 0.274393828 | 3.724945049 | 0.073663859 |
| Alms1-ps2 | 11.88847144 | -0.198914047 | 0.320572564 | -0.620496166 |
| mmu-mir-6375 | 0.167183452 | -0.206481502 | 2.765153652 | -0.074672705 |
| Gm11128 | 2.945191291 | -2.54946198 | 0.695013941 | -3.668217036 |
| Cml5 | 13.56449452 | -0.329892839 | 0.357369142 | -0.923115067 |
| Nat8 | 14.92973809 | -0.807654444 | 0.31693955 | -2.548291759 |
| Gm9769 | 437.2277849 | 0.188997928 | 0.085668089 | 2.20616486 |
| Cml2 | 0.711965957 | -0.491092332 | 1.152865387 | -0.425975433 |
| RP23-448G18.6 | 0 | NA | NA | NA |
| 1700019G17Rik | 48.23925206 | -0.037545707 | 0.203760139 | -0.184264238 |
| Cml1 | 59.08754969 | -0.130053766 | 0.183951914 | -0.706998709 |
| Tprkb | 600.9053318 | -0.016508646 | 0.138702176 | -0.119022256 |
| RP23-345O17.4 | 0 | NA | NA | NA |
| Nat8b | 0.082913145 | -0.206492286 | 3.724945049 | -0.055434988 |
| RP23-345O17.6 | 0.298750603 | 1.252756021 | 1.761006018 | 0.711386564 |
| Dusp11 | 1203.32624 | -0.442823458 | 0.122487169 | -3.615264049 |
| RP23-345O17.5 | 10.75741223 | -0.524027454 | 0.344960318 | -1.519094883 |
| RP23-345O17.7 | 10.85736135 | 0.05458898 | 0.33124299 | 0.16480041 |
| Figla | 0.579661142 | 1.311029642 | 1.358618942 | 0.964972297 |
| Add2 | 1996.526052 | 0.145683452 | 0.115053167 | 1.266227215 |
| Tgfa | 554.9848714 | 0.252734743 | 0.092548274 | 2.73084232 |
| Gm7545 | 0.358406704 | -0.206481624 | 1.75964664 | -0.117342664 |
| Gm10443 | 50.80569197 | 0.436241708 | 0.232603874 | 1.875470522 |
| Fam136a | 140.0252418 | -0.204500768 | 0.156657506 | -1.30540038 |
| Snrpg | 133.4774109 | 0.278398581 | 0.105540803 | 2.637828902 |
| Pcyox1 | 1206.150805 | 0.090843802 | 0.057256927 | 1.586599327 |
| Tia1 | 1452.959773 | -0.293784226 | 0.126282757 | -2.326400157 |
| C87436 | 200.9232409 | -0.109683558 | 0.115933008 | -0.946094294 |
| A430078I02Rik | 3.729996922 | -0.652988526 | 0.555618595 | -1.175245992 |

| | | | | |
|---|---|---|---|---|
| 2310040G24Rik | 5.093180958 | -0.150703817 | 0.438685093 | -0.343535305 |
| Gm28720 | 0.502304703 | 1.177429721 | 1.352133174 | 0.870794197 |
| Gm28719 | 0.796860523 | -1.176620386 | 1.499186438 | -0.784839267 |
| Pcbp1 | 1630.30914 | 0.129172243 | 0.083129295 | 1.553871508 |
| 1600020E01Rik | 80.68127648 | -0.033960694 | 0.15649514 | -0.217007977 |
| Gm22184 | 0 | NA | NA | NA |
| Asprv1 | 17.06775386 | 0.708419117 | 0.29540825 | 2.398102002 |
| Mxd1 | 914.3853629 | 0.662790295 | 0.190205392 | 3.484603087 |
| Snrnp27 | 334.3660018 | 0.233898655 | 0.111049627 | 2.106253401 |
| Gmcl1 | 364.6549906 | -0.218271298 | 0.0922086 | -2.36714686 |
| Anxa4 | 269.2860603 | 2.106482046 | 0.235981935 | 8.92645468 |
| 2610306M01Rik | 36.58485778 | -0.097969121 | 0.184975191 | -0.529633841 |
| Aak1 | 5511.101378 | -0.135081004 | 0.077341718 | -1.746547744 |
| Nfu1 | 197.0210827 | 0.066543502 | 0.091330989 | 0.728597191 |
| Gfpt1 | 1366.619225 | -0.176476945 | 0.054661439 | -3.228545571 |
| D6Ertd527e | 0.429069343 | 1.977321864 | 1.492671344 | 1.324686691 |
| Antxr1 | 334.4251876 | 0.030090883 | 0.122003639 | 0.246639228 |
| n-R5s164 | 0 | NA | NA | NA |
| Gkn1 | 0.539210579 | 0.274413737 | 1.509237773 | 0.181822733 |
| Gkn2 | 0 | NA | NA | NA |
| Gkn3 | 2.877452801 | -5.081836144 | 0.970984226 | -5.233695883 |
| Bmp10 | 1.435473298 | 2.630454513 | 1.119391787 | 2.349896205 |
| Arhgap25 | 146.1837619 | 1.476454303 | 0.156904594 | 9.409885777 |
| Prokr1 | 11.83858572 | 2.041822032 | 0.555427042 | 3.676130033 |
| Aplf | 74.79263325 | -0.660362028 | 0.15398848 | -4.288385901 |
| E230015B07Rik | 2.634345682 | -0.424043215 | 0.592945049 | -0.715147577 |
| 1810020O05Rik | 22.22824979 | -0.05595691 | 0.228248961 | -0.24515735 |
| Efcc1 | 88.67690409 | -0.241036831 | 0.182426403 | -1.321282598 |
| Gm5879 | 12.16014049 | 0.754871713 | 0.314328347 | 2.401538772 |
| Gp9 | 0.777280298 | -0.074095075 | 1.154374017 | -0.064186368 |
| Rab43 | 434.7988563 | 0.678135936 | 0.085387668 | 7.941848673 |
| Isy1 | 190.4889355 | 0.152929545 | 0.093636202 | 1.633230972 |
| Cnbp | 2353.702497 | 0.535286259 | 0.106156861 | 5.042408517 |
| Copg1 | 2888.377927 | -0.03109465 | 0.0317761 | -0.978554649 |
| Gm26636 | 0.2969049 | 2.281492459 | 1.906817059 | 1.196492578 |
| Hmces | 147.1776525 | -0.318459075 | 0.138934317 | -2.292155609 |
| Rps19-ps9 | 0 | NA | NA | NA |
| H1fx | 161.3903661 | -0.18077587 | 0.1191083 | -1.517743682 |
| Gm5577 | 38.22855657 | -0.458694797 | 0.271199454 | -1.691355898 |
| AC153872.1 | 0.355821277 | 0.250116633 | 1.477275319 | 0.169309423 |
| Rab7 | 2929.399181 | 0.249017269 | 0.077080565 | 3.230610309 |
| Rpn1 | 1428.832432 | 0.207238271 | 0.070234672 | 2.9506548 |
| Mir6376 | 0 | NA | NA | NA |
| Gata2 | 41.94893332 | 1.515784666 | 0.246770353 | 6.142490974 |
| Dnajb8 | 0 | NA | NA | NA |
| 1700031F10Rik | 0.520495925 | 2.334334435 | 1.563154307 | 1.493348688 |
| Eefsec | 187.7376768 | -0.137182378 | 0.103737258 | -1.322402197 |

| | | | | |
|---|---|---|---|---|
| Gm25178 | 0 | NA | NA | NA |
| Ruvbl1 | 250.3254379 | 0.011856605 | 0.098414963 | 0.12047563 |
| Sec61a1 | 1211.826051 | 0.406744507 | 0.102123599 | 3.982864999 |
| Kbtbd12 | 4.967738157 | 0.362801056 | 0.581022483 | 0.624418274 |
| Gm26588 | 16.08933923 | -0.423693933 | 0.365006425 | -1.160784863 |
| Mgll | 1645.494617 | 0.017539376 | 0.090206932 | 0.194434897 |
| Abtb1 | 225.610541 | 0.004115112 | 0.08827714 | 0.046615829 |
| Podxl2 | 1494.724341 | -0.145652237 | 0.082825928 | -1.758534305 |
| Gm15612 | 6.463977267 | -0.401994077 | 0.47650273 | -0.843634362 |
| Mcm2 | 112.1005328 | 0.762021016 | 0.226311539 | 3.367132845 |
| Tpra1 | 240.1461408 | -0.324660811 | 0.118631407 | -2.736718879 |
| 4933427D06Rik | 0.766530834 | 2.661502549 | 1.218291567 | 2.184618707 |
| Gm1965 | 0.219739983 | 0.274401974 | 2.588615737 | 0.106003363 |
| Gm6507 | 0 | NA | NA | NA |
| Gm839 | 0.209172349 | 0.755299119 | 2.602036237 | 0.290272329 |
| Gm25961 | 0.088534442 | 0.274389469 | 3.724945049 | 0.073662689 |
| Gm26811 | 0.627488786 | -0.410561857 | 1.322965021 | -0.310334628 |
| Plxna1 | 1720.837352 | -0.176459307 | 0.070526618 | -2.502024229 |
| Chchd6 | 271.9148547 | -0.17497593 | 0.100948764 | -1.733314248 |
| Txnrd3 | 60.18168024 | 0.189614293 | 0.18251086 | 1.038920602 |
| Vmn1r-ps25 | 0 | NA | NA | NA |
| Vmn1r40 | 0.046345709 | 0.274391604 | 3.724945049 | 0.073663262 |
| Vmn1r41 | 0 | NA | NA | NA |
| Vmn1r-ps26 | 0 | NA | NA | NA |
| Vmn1r-ps27 | 0 | NA | NA | NA |
| Vmn1r42 | 0 | NA | NA | NA |
| Vmn1r43 | 0.049553047 | 0.274391604 | 3.724945049 | 0.073663262 |
| Vmn1r44 | 0 | NA | NA | NA |
| Vmn1r-ps28 | 0 | NA | NA | NA |
| Vmn1r45 | 0 | NA | NA | NA |
| Vmn1r46 | 0 | NA | NA | NA |
| Vmn1r-ps29 | 0 | NA | NA | NA |
| Vmn1r47 | 0 | NA | NA | NA |
| Vmn1r48 | 0 | NA | NA | NA |
| Vmn1r-ps31 | 0 | NA | NA | NA |
| Gm20534 | 0 | NA | NA | NA |
| Vmn1r49 | 0 | NA | NA | NA |
| Vmn1r50 | 0 | NA | NA | NA |
| Vmn1r51 | 0.060174537 | 0.755286614 | 3.724945049 | 0.202764498 |
| Vmn1r-ps32 | 0 | NA | NA | NA |
| Vmn1r52 | 0 | NA | NA | NA |
| V1ra8 | 0 | NA | NA | NA |
| Vmn1r53 | 0.265601376 | -0.648208672 | 1.793603868 | -0.36140013 |
| Gm20426 | 1.074962736 | 1.417887818 | 0.993099748 | 1.42773958 |
| Gm4287 | 0.044544648 | 0.755286614 | 3.724945049 | 0.202764498 |
| Vmn1r-ps34 | 0 | NA | NA | NA |
| Vmn1r54 | 0 | NA | NA | NA |

| | | | | |
|---|---|---|---|---|
| BC048671 | 0 | NA | NA | NA |
| Chst13 | 0.497746636 | 0.705566261 | 1.39810028 | 0.504660696 |
| Uroc1 | 0.105143817 | 0.274402725 | 3.724945049 | 0.073666248 |
| Zxdc | 516.6253474 | -0.049186916 | 0.086743729 | -0.567037134 |
| Ccdc37 | 25.13015435 | -0.500411653 | 0.21479496 | -2.32971785 |
| Klf15 | 358.4959565 | 0.643282629 | 0.135090945 | 4.761848613 |
| Aldh1l1 | 1458.014872 | 0.909667168 | 0.111120228 | 8.186332809 |
| Gm15756 | 0.143022002 | 0.755289479 | 3.547809333 | 0.212888971 |
| Slc41a3 | 95.87477886 | 0.042110732 | 0.142449116 | 0.29561947 |
| Iqsec1 | 4998.243591 | 0.044333386 | 0.064212083 | 0.69042124 |
| Gm24816 | 4.887144399 | -0.560267461 | 0.646675362 | -0.866381331 |
| Rpl21-ps12 | 7.418190298 | -0.713310224 | 0.478700363 | -1.490097521 |
| Gm25185 | 0 | NA | NA | NA |
| Ppp1r2-ps2 | 19.04760772 | 0.489503902 | 0.29386047 | 1.66576982 |
| Nup210 | 552.2987281 | 0.077941675 | 0.087110015 | 0.894749867 |
| Hdac11 | 1873.279407 | 0.074224197 | 0.067030101 | 1.107326349 |
| Gm14573 | 0.08169834 | -0.206492286 | 3.724945049 | -0.055434988 |
| Fbln2 | 117.7950445 | 0.027436708 | 0.248033996 | 0.110616723 |
| Wnt7a | 199.3147353 | -0.5473928 | 0.187081922 | -2.925952401 |
| Gm4575 | 4.420079801 | -0.389733029 | 0.472440759 | -0.824935236 |
| 4930471M09Rik | 1.669552124 | 0.086421906 | 0.788365743 | 0.10962159 |
| 1810044D09Rik | 5.670729865 | 0.14809077 | 0.408189168 | 0.362799362 |
| Chchd4 | 184.8478106 | -0.019696095 | 0.097920423 | -0.201143895 |
| Tmem43 | 388.1633489 | 0.519985668 | 0.090909325 | 5.719827595 |
| Xpc | 207.6063262 | 0.009233705 | 0.10789502 | 0.085580453 |
| Lsm3 | 147.4136989 | 0.255117448 | 0.108722678 | 2.346497089 |
| Slc6a6 | 1683.442733 | 0.051418381 | 0.074119742 | 0.693720456 |
| Grip2 | 311.1559795 | -0.820000504 | 0.110712101 | -7.40660229 |
| Ccdc174 | 303.9255113 | -0.021751887 | 0.110616294 | -0.196642706 |
| 4930590J08Rik | 9.858077027 | 0.42849009 | 0.404880285 | 1.058313052 |
| Fgd5 | 79.82046982 | 0.093642106 | 0.175695278 | 0.532980207 |
| 4930466I24Rik | 0.042716945 | -0.206490061 | 3.724945049 | -0.055434391 |
| Nr2c2 | 2086.98739 | -0.011863196 | 0.056970848 | -0.208232752 |
| Mrps25 | 334.8028407 | -0.300212272 | 0.072908293 | -4.117669758 |
| Rbsn | 899.6634951 | 0.303887897 | 0.091170061 | 3.333198355 |
| Trh | 83.76368859 | 0.000731814 | 0.412515053 | 0.001774029 |
| Prickle2 | 2847.184781 | -0.025053824 | 0.106660129 | -0.234893999 |
| Gm17797 | 0.118974955 | 0.274400501 | 3.724945049 | 0.073665651 |
| Adamts9 | 272.0878833 | 1.340775644 | 0.199308819 | 6.727126546 |
| A730049H05Rik | 9.582230866 | 2.699791493 | 0.598208376 | 4.513128871 |
| Gm15737 | 0 | NA | NA | NA |
| 9530026P05Rik | 37.55723895 | 0.286615684 | 0.204752183 | 1.399817477 |
| Gm5313 | 0 | NA | NA | NA |
| Gm25094 | 0 | NA | NA | NA |
| Gm22312 | 0 | NA | NA | NA |
| Magi1 | 1204.640072 | -0.293011732 | 0.070149012 | -4.176990156 |
| Gm23035 | 0 | NA | NA | NA |

| | | | | |
|---|---|---|---|---|
| Gm22840 | 0 | NA | NA | NA |
| Slc25a26 | 101.4816024 | -0.033162513 | 0.119959711 | -0.276447088 |
| Lrig1 | 1206.313659 | 0.184899674 | 0.124869875 | 1.480738835 |
| Mir7041 | 0.248387756 | -0.648706618 | 1.83325957 | -0.353854211 |
| Gm7833 | 0.121388914 | 0.755282166 | 3.724945049 | 0.202763304 |
| Gm23364 | 0 | NA | NA | NA |
| Gm24601 | 0 | NA | NA | NA |
| Gm26501 | 0 | NA | NA | NA |
| Kbtbd8os | 0.511579953 | -0.02022514 | 1.161966137 | -0.017405963 |
| Kbtbd8 | 179.6004661 | -0.349400061 | 0.124744091 | -2.800934768 |
| AY512915 | 0.311830279 | 2.111551842 | 1.959330473 | 1.077690502 |
| Gm10234 | 0 | NA | NA | NA |
| Suclg2 | 141.9514173 | -0.02832078 | 0.148352639 | -0.190901762 |
| Fam19a1 | 272.9938384 | 0.025033743 | 0.141093331 | 0.177426835 |
| 1700123L14Rik | 0.257669488 | -1.300912567 | 2.092497368 | -0.621703323 |
| Gm22971 | 0 | NA | NA | NA |
| Gm26011 | 0 | NA | NA | NA |
| Fam19a4 | 11.35413066 | 0.076428859 | 0.308911418 | 0.247413513 |
| Eogt | 164.467491 | 0.170756323 | 0.130812966 | 1.305347076 |
| Tmf1 | 950.0693584 | 0.235066945 | 0.068134844 | 3.450025425 |
| Gm26175 | 2.7824929 | -0.360895426 | 0.668369453 | -0.539963974 |
| Uba3 | 755.4137509 | -0.013583478 | 0.087905321 | -0.154523951 |
| Arl6ip5 | 1177.848838 | 0.282514745 | 0.063730985 | 4.432926099 |
| Lmod3 | 0.289573199 | 0.301107727 | 2.055176004 | 0.146511893 |
| Frmd4b | 362.2741639 | 0.26211062 | 0.088914548 | 2.947893509 |
| Gm25852 | 1.201575708 | -0.510240964 | 0.875986414 | -0.582475887 |
| Gm23510 | 0.586937805 | -0.147367957 | 1.547393956 | -0.095236224 |
| Gm24565 | 0 | NA | NA | NA |
| Gm23703 | 0 | NA | NA | NA |
| Mitf | 247.2340933 | 0.166829739 | 0.114908089 | 1.451853738 |
| Gm15531 | 0.049553047 | 0.274391604 | 3.724945049 | 0.073663262 |
| Gm765 | 8.927308765 | -0.659810678 | 0.329840747 | -2.000391652 |
| Foxp1 | 1512.202388 | 0.029798401 | 0.123319081 | 0.241636578 |
| AC155839.1 | 0.110870145 | -0.206490066 | 3.724945049 | -0.055434393 |
| Gm20696 | 52.11740566 | -0.499056785 | 0.265071777 | -1.882723204 |
| Gm20705 | 26.36886051 | -0.664004591 | 0.379759067 | -1.748489102 |
| Gm22328 | 2.778696684 | -0.504725737 | 0.649379297 | -0.777243345 |
| Gm24936 | 0 | NA | NA | NA |
| Eif4e3 | 223.9200148 | 0.011605311 | 0.090394006 | 0.128385845 |
| Gpr27 | 69.76717276 | -0.136875222 | 0.188800755 | -0.724971793 |
| Prok2 | 9.10031652 | -0.379717348 | 0.43360061 | -0.875730661 |
| Gm26748 | 2.205804803 | -0.660179357 | 0.607792481 | -1.086192045 |
| Tpt1-ps3 | 354.4684848 | 0.529613073 | 0.168123461 | 3.150143768 |
| RP23-191I2.1 | 0 | NA | NA | NA |
| Gm24248 | 0 | NA | NA | NA |
| Rybp | 419.1411725 | 0.00987119 | 0.082246866 | 0.120019041 |
| Gm23234 | 0.066050875 | 0.274391604 | 3.724945049 | 0.073663262 |

| | | | | |
|---|---|---|---|---|
| 1700049E22Rik | 0 | NA | NA | NA |
| Shq1 | 52.12462995 | -0.207435167 | 0.14593425 | -1.421428942 |
| Gm6565 | 46.92634586 | 0.205359378 | 0.180930739 | 1.135016525 |
| Gxylt2 | 11.76095793 | -1.042618539 | 0.348279542 | -2.993625564 |
| Gm15576 | 0.092881282 | -0.687373951 | 3.724945049 | -0.184532642 |
| Ppp4r2 | 799.647895 | 0.019776628 | 0.080160396 | 0.246713203 |
| Pdzrn3 | 253.9254666 | 0.15211288 | 0.128825273 | 1.180768937 |
| Gm26911 | 0.672570739 | -0.425338078 | 1.19375506 | -0.356302639 |
| Gm9871 | 0.521928645 | 1.616166967 | 1.475886446 | 1.095048316 |
| Cntn3 | 401.4213994 | 0.048340125 | 0.14095258 | 0.342953101 |
| Mir6373 | 0 | NA | NA | NA |
| Gm26022 | 0 | NA | NA | NA |
| Chl1 | 3209.185577 | 0.027862119 | 0.075776503 | 0.367688101 |
| Gm24784 | 4.963107029 | -0.29279879 | 0.564973249 | -0.518252484 |
| Gm25656 | 0 | NA | NA | NA |
| RP24-151A23.1 | 0 | NA | NA | NA |
| Cntn6 | 392.3293275 | -0.294207479 | 0.110289623 | -2.667589861 |
| Gm23339 | 0 | NA | NA | NA |
| Gm25570 | 0 | NA | NA | NA |
| Gm15631 | 17.2682059 | -0.98998964 | 0.358110733 | -2.764479106 |
| Cntn4 | 819.1636369 | -0.084697436 | 0.072392303 | -1.169978473 |
| Il5ra | 0.538620109 | 1.647546394 | 1.408236353 | 1.169935992 |
| Trnt1 | 529.4811257 | 0.338316639 | 0.076807601 | 4.404728643 |
| Crbn | 1050.434189 | 0.420090424 | 0.089565244 | 4.690328572 |
| Gm22418 | 0 | NA | NA | NA |
| Gm27690 | 0 | NA | NA | NA |
| Lrrn1 | 1083.285535 | -0.133064945 | 0.075495409 | -1.762556782 |
| Setmar | 60.36555304 | 0.488274372 | 0.137612778 | 3.548176104 |
| Sumf1 | 349.8000208 | 0.47063888 | 0.090212553 | 5.216999873 |
| Gm4604 | 20.23814071 | 0.288093902 | 0.249527791 | 1.154556377 |
| Itpr1 | 13527.53245 | -0.043122227 | 0.165134297 | -0.261134286 |
| Gm27813 | 4.625054459 | -0.818267034 | 0.434654103 | -1.882570594 |
| mmu-mir-7661 | 0.67439107 | -2.115278996 | 1.192528074 | -1.773777105 |
| Gm16433 | 10.96864868 | 0.211860052 | 0.34341626 | 0.616919106 |
| 0610040F04Rik | 4.191962974 | -0.387421994 | 0.4547341 | -0.851974799 |
| Gm38167 | 1.527506968 | -1.155198024 | 0.937694665 | -1.231955419 |
| Gm17055 | 0.461373441 | -1.722955637 | 1.671751003 | -1.030629343 |
| Bhlhe40 | 1442.70058 | 0.178743377 | 0.13000319 | 1.374915312 |
| Arl8b | 2174.485167 | 0.019248862 | 0.055215778 | 0.348611619 |
| Edem1 | 637.9057285 | 0.989458718 | 0.136697486 | 7.238309554 |
| Gm20387 | 22.36903969 | 0.272124891 | 0.245100098 | 1.110260227 |
| Grm7 | 1035.879049 | -0.182713444 | 0.14385539 | -1.270118862 |
| Gm20386 | 13.98619218 | -0.795576662 | 0.319978903 | -2.486340984 |
| Gm22093 | 0.0878635 | -0.687376175 | 3.724945049 | -0.184533239 |
| Gm24005 | 0 | NA | NA | NA |
| Gm26362 | 0 | NA | NA | NA |
| Gm25454 | 0 | NA | NA | NA |

| | | | | |
|---|---|---|---|---|
| 1700054K19Rik | 0 | NA | NA | NA |
| n-R5s165 | 0 | NA | NA | NA |
| Lmcd1 | 50.45369243 | 0.470797564 | 0.220233795 | 2.13771716 |
| 5031434C07Rik | 0.668421634 | -1.222413159 | 1.159077615 | -1.054643057 |
| Ssu2 | 1.179082066 | 1.883098157 | 0.940808674 | 2.001573975 |
| Cav3 | 2.338446252 | 0.973307897 | 0.884087261 | 1.10091836 |
| Oxtr | 56.02683163 | -0.125378209 | 0.14261436 | -0.879141546 |
| Gm27944 | 0 | NA | NA | NA |
| Gm15519 | 37.45729361 | 0.633187538 | 0.180317894 | 3.51150695 |
| Gm23002 | 0.075043205 | 0.755286614 | 3.724945049 | 0.202764498 |
| Gm5578 | 12.94208794 | -0.006217199 | 0.34784319 | -0.017873568 |
| Rad18 | 111.1939575 | 0.207107572 | 0.171307709 | 1.20897987 |
| Gm26799 | 2.930040241 | 1.416876567 | 0.634848253 | 2.231835028 |
| Srgap3 | 4773.252643 | -0.271731242 | 0.07184984 | -3.781932458 |
| Gm6134 | 0.068097359 | -0.206487837 | 3.724945049 | -0.055433794 |
| Gm23244 | 0 | NA | NA | NA |
| Thumpd3 | 249.1100717 | 0.140847538 | 0.087600019 | 1.607848272 |
| Gm22591 | 0.068097359 | -0.206487837 | 3.724945049 | -0.055433794 |
| Gt(ROSA)26Sor | 121.8807241 | 0.101415309 | 0.141965502 | 0.714365869 |
| Setd5 | 2417.574525 | -0.067882388 | 0.072724448 | -0.933419089 |
| Lhfpl4 | 1265.24945 | -0.23608108 | 0.079418247 | -2.972630187 |
| Mtmr14 | 178.5916463 | 0.244510586 | 0.08630529 | 2.833089206 |
| Gm16161 | 0.217535639 | -0.648203836 | 3.180597456 | -0.203799395 |
| Cpne9 | 367.3936013 | -0.502771053 | 0.12969171 | -3.876662993 |
| Brpf1 | 657.1446642 | 0.261473341 | 0.11602045 | 2.253683211 |
| Gm15492 | 1.140702544 | -0.100710145 | 0.993677993 | -0.101350886 |
| Ogg1 | 60.44940537 | 0.111806773 | 0.180710318 | 0.618707188 |
| Camk1 | 512.5663802 | -0.460040271 | 0.091860432 | -5.008035122 |
| Tada3 | 319.0164571 | -0.178837354 | 0.073342207 | -2.438396124 |
| Arpc4 | 1219.326062 | 0.144569538 | 0.082646073 | 1.749260819 |
| Ttll3 | 119.1716164 | -0.272191306 | 0.197539321 | -1.377909495 |
| Rpusd3 | 80.45315769 | -0.196960149 | 0.14767337 | -1.33375536 |
| Gm24387 | 0.061920613 | 0.274393828 | 3.724945049 | 0.073663859 |
| Cidec | 0.392922208 | -0.16061338 | 1.704667365 | -0.094219778 |
| Jagn1 | 177.5032672 | 0.087131604 | 0.143000501 | 0.609309783 |
| Il17re | 7.25222539 | -0.59960317 | 0.468732204 | -1.279201993 |
| Il17rc | 69.78172622 | 1.425588432 | 0.23383322 | 6.096603508 |
| Creld1 | 601.1655518 | -0.230664329 | 0.069450625 | -3.321270754 |
| Prrt3 | 474.5330712 | -0.152592876 | 0.116102057 | -1.314299507 |
| Emc3 | 603.6486607 | 0.198310035 | 0.081545764 | 2.431886398 |
| Gm23365 | 0.654623538 | 0.379731112 | 1.290238807 | 0.29431072 |
| Fancd2 | 38.49030515 | 0.522900504 | 0.177565245 | 2.944835879 |
| Fancd2os | 0.484250087 | -0.701309447 | 1.450034459 | -0.483650194 |
| Brk1 | 853.6369896 | -0.110314445 | 0.063282929 | -1.743194358 |
| Vhl | 414.2383988 | 0.068475611 | 0.086824258 | 0.788669112 |
| 1700015O11Rik | 0.140088945 | -1.111948817 | 3.601861891 | -0.308715006 |
| Irak2 | 312.5390426 | 0.900221088 | 0.14787728 | 6.087622703 |

| | | | | |
|---|---|---|---|---|
| Tatdn2 | 686.0036907 | 0.322539963 | 0.102122996 | 3.158348035 |
| mmu-mir-7042 | 1.150390536 | -0.67215209 | 1.04259938 | -0.644688749 |
| Gm26982 | 73.25146826 | 0.001479962 | 0.215198801 | 0.006877185 |
| Ghrl | 1.379464875 | -0.065734465 | 0.845733932 | -0.077724758 |
| Sec13 | 453.3719348 | 0.2548995 | 0.085203974 | 2.991638639 |
| Atp2b2 | 10812.54371 | -0.424720429 | 0.114648748 | -3.704536134 |
| Gm15082 | 0 | NA | NA | NA |
| Gm15083 | 4.119074061 | -0.083099111 | 0.464814269 | -0.178779174 |
| Slc6a11 | 2799.545472 | -0.737045028 | 0.160107564 | -4.603436651 |
| Slc6a1 | 4290.118131 | -0.200379667 | 0.085562516 | -2.34190947 |
| Hrh1 | 148.4312325 | -0.099539651 | 0.128917883 | -0.772116708 |
| Atg7 | 275.6563131 | 0.135785162 | 0.096481586 | 1.407368684 |
| Vgll4 | 362.9008528 | -0.161455479 | 0.095674682 | -1.687546548 |
| Tamm41 | 94.56214301 | -0.021372257 | 0.149095098 | -0.143346477 |
| Gm24767 | 0 | NA | NA | NA |
| Gm17733 | 5.222249821 | -0.786015071 | 0.460956613 | -1.705182329 |
| Syn2 | 4179.711114 | 0.03255365 | 0.087775266 | 0.370874978 |
| Gm17482 | 8.270587696 | -0.16917649 | 0.329669938 | -0.513169296 |
| Timp4 | 415.701942 | 0.412641759 | 0.201696542 | 2.045854403 |
| Pparg | 54.34159874 | 0.169327921 | 0.215225605 | 0.786746171 |
| Tsen2 | 155.5268573 | -0.551077125 | 0.112906082 | -4.880845341 |
| Mkrn2os | 0.394336369 | 1.907183949 | 1.501374778 | 1.27029172 |
| Mkrn2 | 337.3089779 | 0.244729389 | 0.103635378 | 2.361446401 |
| Raf1 | 1017.633135 | 0.067448854 | 0.090927026 | 0.741791049 |
| Gm14335 | 18.11564472 | 0.150076071 | 0.360633234 | 0.41614598 |
| Gm24008 | 0 | NA | NA | NA |
| Tmem40 | 16.46128129 | -1.117060727 | 0.472597887 | -2.363660014 |
| Gm27853 | 0 | NA | NA | NA |
| Cand2 | 86.32620824 | -0.207520409 | 0.167011632 | -1.242550636 |
| Mir7660 | 0.118974955 | 0.274400501 | 3.724945049 | 0.073665651 |
| Rpl32 | 874.900214 | 0.464982237 | 0.142958533 | 3.252567215 |
| Snora7a | 4.44476562 | 0.61185139 | 0.580521172 | 1.053969122 |
| Efcab12 | 50.83190883 | -0.415150549 | 0.238131949 | -1.743363507 |
| Mbd4 | 111.5763811 | -0.098365433 | 0.174974046 | -0.562171567 |
| Ift122 | 349.0430765 | -0.108569712 | 0.1124566 | -0.965436553 |
| Rho | 4.082856454 | -1.707015657 | 0.659720387 | -2.587483563 |
| H1foo | 0.271840384 | 1.594335423 | 2.040867021 | 0.781204952 |
| Plxnd1 | 1058.405309 | 0.148365512 | 0.093172365 | 1.592376796 |
| Tmcc1 | 1022.727567 | -0.324819785 | 0.054880561 | -5.918667384 |
| Gm20404 | 1.065143116 | 1.11336376 | 1.083801007 | 1.027276919 |
| Gm8203 | 110.3300294 | 0.244185224 | 0.136470805 | 1.789285434 |
| Gm8213 | 4.570295035 | 0.535512966 | 0.486462053 | 1.100831941 |
| Fam21 | 1941.589181 | 0.151634893 | 0.082914633 | 1.828807385 |
| Zfand4 | 120.8501776 | 0.39798186 | 0.119888826 | 3.319590927 |
| 8-Mar | 943.7508335 | 0.158775016 | 0.080087987 | 1.982507251 |
| Gm23527 | 0 | NA | NA | NA |
| Alox5 | 9.062762248 | 0.093270446 | 0.347779594 | 0.268188381 |

| | | | | |
|---|---|---|---|---|
| Olfr211 | 0.287786674 | 0.755304973 | 2.026686947 | 0.372679646 |
| Olfr212 | 0.265357793 | 0.274413213 | 1.814365877 | 0.151244695 |
| Olfr213 | 0.046345709 | 0.274391604 | 3.724945049 | 0.073663262 |
| Gm5580 | 0.092493346 | 0.274402725 | 3.724945049 | 0.073666248 |
| Olfr214 | 0 | NA | NA | NA |
| Olfr215 | 0 | NA | NA | NA |
| Zfp422 | 293.8653895 | 0.082794939 | 0.070285538 | 1.177979732 |
| Rassf4 | 281.0708732 | -0.217225026 | 0.153406797 | -1.416006524 |
| Mir7043 | 0 | NA | NA | NA |
| 8430408G22Rik | 5.938959585 | 1.393364578 | 0.591406774 | 2.356017278 |
| Tmem72 | 34.03285026 | -0.3867987 | 0.234169552 | -1.651789041 |
| Gm22882 | 0 | NA | NA | NA |
| Cxcl12 | 287.0988006 | 0.545748084 | 0.12270301 | 4.447715544 |
| Rpl28-ps4 | 0 | NA | NA | NA |
| Gm9946 | 0.160978641 | -1.177170116 | 2.79661493 | -0.42092678 |
| 1700030F04Rik | 1.580617114 | -1.588029731 | 0.879165928 | -1.80629126 |
| Gm7292 | 1492.722712 | -0.09309239 | 0.075084309 | -1.23983814 |
| Gm6253 | 0.669418093 | -1.160850066 | 1.406039589 | -0.825616914 |
| Zfp637 | 220.273934 | -0.072847418 | 0.086037973 | -0.846689137 |
| Zfp239 | 252.8739187 | -0.430845522 | 0.110967852 | -3.882615693 |
| Gm29509 | 0.066050875 | 0.274391604 | 3.724945049 | 0.073663262 |
| Hnrnpf | 625.6970897 | 0.433928863 | 0.117420899 | 3.695499423 |
| 4933440N22Rik | 4.749668983 | 2.076582954 | 0.550767903 | 3.770341269 |
| Fxyd4 | 0.432778942 | -0.642780812 | 1.439065737 | -0.446665358 |
| Gm25535 | 0.060174537 | 0.755286614 | 3.724945049 | 0.202764498 |
| Gm4875 | 25.05432516 | -0.3532072 | 0.210944884 | -1.674405149 |
| Rasgef1a | 1668.685123 | -0.153370409 | 0.107332291 | -1.428930733 |
| Mir7044 | 0.497274054 | 0.599649106 | 1.315492932 | 0.455836053 |
| Csgalnact2 | 271.8412438 | -0.044299683 | 0.074999408 | -0.590667104 |
| Gm23908 | 2.493933935 | -0.522173455 | 0.609739207 | -0.856388188 |
| Ret | 333.8009791 | -0.345561777 | 0.148960232 | -2.319825716 |
| Bms1 | 452.1647011 | 0.012933762 | 0.119952295 | 0.107824217 |
| Zfp248 | 186.5369703 | -0.254131141 | 0.102973639 | -2.467924266 |
| Gm25953 | 0 | NA | NA | NA |
| Zfp9 | 323.1489127 | 0.012259171 | 0.079741082 | 0.153737208 |
| Ankrd26 | 319.7692467 | -0.750211332 | 0.175297894 | -4.279636885 |
| Cacna1c | 2127.985368 | -0.3840351 | 0.124110665 | -3.094295731 |
| Gm18981 | 18.01703932 | -0.979774443 | 0.396454272 | -2.471342882 |
| Gm25905 | 0 | NA | NA | NA |
| Dcp1b | 230.6682507 | -0.306211951 | 0.118292508 | -2.588599703 |
| Cacna2d4 | 45.15394803 | -0.37825475 | 0.154912536 | -2.441731045 |
| Lrtm2 | 701.9863131 | -0.057348864 | 0.071574654 | -0.801245428 |
| Adipor2 | 1326.364367 | 0.284390638 | 0.110713368 | 2.568710919 |
| Wnt5b | 23.55176808 | 0.315313137 | 0.254829835 | 1.237347804 |
| Fbxl14 | 457.0418174 | 0.352401364 | 0.09183985 | 3.837129144 |
| Erc1 | 1392.520042 | -0.371640509 | 0.083777393 | -4.436047651 |
| 3110021A11Rik | 3.049669005 | -0.636871708 | 0.593054664 | -1.073883651 |

| | | | | |
|---|---|---|---|---|
| Gm15532 | 0 | NA | NA | NA |
| 4930540M05Rik | 1.752268337 | 0.887499741 | 0.798054271 | 1.112079433 |
| Rad52 | 228.0315541 | 0.000656384 | 0.093920419 | 0.006988726 |
| Wnk1 | 6814.715123 | 0.211686794 | 0.062978565 | 3.361251479 |
| Gm16199 | 3.97190282 | 0.872548867 | 0.524771816 | 1.662720521 |
| Mir706 | 1.782601153 | -0.108444845 | 0.715436785 | -0.151578514 |
| Ninj2 | 7.372786058 | -0.462959895 | 0.375426362 | -1.233157661 |
| Gm25327 | 0 | NA | NA | NA |
| B4galnt3 | 7.654890081 | -0.983557616 | 0.356045041 | -2.762452783 |
| D330020A13Rik | 0 | NA | NA | NA |
| Ccdc77 | 82.48745712 | -0.444242673 | 0.177291435 | -2.505719875 |
| Kdm5a | 1440.026259 | 0.272521264 | 0.084335692 | 3.231387039 |
| Gm24592 | 0 | NA | NA | NA |
| Il17ra | 446.1211285 | 1.509647372 | 0.188095066 | 8.025980712 |
| Cecr6 | 417.0501185 | -0.597208397 | 0.167004846 | -3.575994428 |
| Cecr5 | 216.0841903 | -0.363113012 | 0.1450097 | -2.504060165 |
| Gm20692 | 2.176719344 | -0.435761959 | 0.628936231 | -0.692855551 |
| Gm24391 | 0.075043205 | 0.755286614 | 3.724945049 | 0.202764498 |
| Cecr2 | 227.0044623 | 0.036710362 | 0.155158445 | 0.236599185 |
| Slc25a18 | 412.4554696 | -0.412265457 | 0.142343917 | -2.896263253 |
| Atp6v1e1 | 2143.722746 | 0.082329785 | 0.066369039 | 1.240484809 |
| Gm25193 | 1.884175004 | -0.446708539 | 0.815062909 | -0.548066333 |
| Bcl2l13 | 678.3631799 | -0.149422966 | 0.071801856 | -2.081046026 |
| Bid | 179.7781668 | 1.820829518 | 0.172707734 | 10.5428372 |
| Mical3 | 1830.192808 | -0.374910157 | 0.123068926 | -3.046342955 |
| Gm4651 | 0.182574094 | 0.638973862 | 2.702311969 | 0.236454513 |
| Pex26 | 225.9169415 | 0.392111931 | 0.108754948 | 3.6054629 |
| Tuba8 | 200.697007 | 0.561513452 | 0.164281868 | 3.417988 |
| Gm15856 | 6.178163136 | 5.595472182 | 0.97604603 | 5.73279539 |
| Usp18 | 2014.636858 | 9.079844236 | 0.403187519 | 22.52015204 |
| Slc6a13 | 90.14607231 | -0.098958439 | 0.252190554 | -0.392395499 |
| Gm24855 | 0 | NA | NA | NA |
| Slc6a12 | 25.52923883 | 0.389684116 | 0.300906798 | 1.295032611 |
| Iqsec3 | 2651.332656 | -0.30410186 | 0.097293864 | -3.125601617 |
| A2m | 460.9649802 | 1.410895139 | 0.262441904 | 5.376028427 |
| Mug1 | 1.274577395 | 1.931069824 | 1.118612867 | 1.726307538 |
| Mug2 | 0.087930927 | -0.206490058 | 3.724945049 | -0.05543439 |
| Gm10319 | 0 | NA | NA | NA |
| Mug-ps1 | 0.15956642 | 0.75529148 | 3.418807262 | 0.22092251 |
| Gm27703 | 0 | NA | NA | NA |
| Klrg1 | 4.976324902 | 2.389093742 | 0.629161 | 3.79726929 |
| Gm27826 | 0.044544648 | 0.755286614 | 3.724945049 | 0.202764498 |
| M6pr | 1445.217025 | 0.75966612 | 0.106618247 | 7.125104233 |
| Phc1 | 481.579932 | -0.139975504 | 0.08362334 | -1.673880793 |
| 1700063H04Rik | 0 | NA | NA | NA |
| Rimklb | 372.2860085 | -0.198797108 | 0.061731899 | -3.22033035 |
| Gm8430 | 34.0404713 | 0.561194589 | 0.287091078 | 1.95476151 |

| | | | | |
|---|---|---|---|---|
| Mfap5 | 3.28087796 | 1.383385027 | 0.755634406 | 1.830759711 |
| Aicda | 0 | NA | NA | NA |
| Gm16556 | 1.225054178 | 2.365559169 | 1.202849951 | 1.966628645 |
| Gm16557 | 0.258216092 | 0.755306254 | 2.049883088 | 0.368463089 |
| Gm8459 | 0.839467773 | 1.676317352 | 2.371813909 | 0.706765967 |
| Apobec1 | 484.8295906 | 4.303602383 | 0.237515508 | 18.11924792 |
| Gdf3 | 0.810901572 | 1.768583478 | 1.20595251 | 1.466544879 |
| Gm26168 | 0 | NA | NA | NA |
| Dppa3 | 0.038981395 | 0.274393828 | 3.724945049 | 0.073663859 |
| Gm10420 | 14.33523795 | 0.160710298 | 0.29396735 | 0.54669438 |
| Gm10224 | 1.550258836 | 0.867199789 | 0.805201471 | 1.076997274 |
| Nanog | 0.725797938 | -0.154149483 | 1.66328341 | -0.092677822 |
| Slc2a3 | 1520.555383 | -0.175790356 | 0.082498635 | -2.130827447 |
| AC163108.1 | 10.5610363 | -0.307564232 | 0.350572393 | -0.877320172 |
| Gm26826 | 2.946188958 | 0.564813496 | 0.53642275 | 1.052926067 |
| Foxj2 | 640.279476 | -0.058352038 | 0.064837603 | -0.899972171 |
| mmu-mir-7231 | 1.544805485 | -1.650191597 | 0.982499454 | -1.679585255 |
| C3ar1 | 498.6296836 | 3.302226409 | 0.300863541 | 10.97582776 |
| Necap1 | 1570.429647 | -0.009752742 | 0.063794287 | -0.152877979 |
| Gm5316 | 0 | NA | NA | NA |
| Clec4a1 | 94.94432573 | 4.473638604 | 0.422228962 | 10.59529073 |
| Clec4a3 | 123.1859983 | 3.670563811 | 0.469514113 | 7.817792289 |
| Clec4a4 | 1.253602477 | 2.07105391 | 1.22329057 | 1.693018782 |
| Clec4b1 | 0.237601816 | 0.755308985 | 2.114704841 | 0.357169932 |
| Clec4a2 | 17.55388936 | 1.506176007 | 0.410746253 | 3.666925734 |
| Clec4b2 | 0.180312491 | -0.206487119 | 2.693521358 | -0.076660658 |
| Gm22946 | 0 | NA | NA | NA |
| Gm5883 | 0.359325923 | -0.178869105 | 1.983709163 | -0.090169017 |
| Clec4n | 6.544164894 | 1.391819531 | 0.519191916 | 2.680741915 |
| Clec4d | 34.33899703 | 6.731908295 | 1.171828261 | 5.744790872 |
| Clec4e | 72.26271789 | 8.991474946 | 1.107155752 | 8.121237621 |
| Vmn2r19 | 0 | NA | NA | NA |
| Vmn2r20 | 0 | NA | NA | NA |
| Gm22833 | 0 | NA | NA | NA |
| Vmn2r-ps31 | 0 | NA | NA | NA |
| Vmn2r21 | 0 | NA | NA | NA |
| Gm25891 | 0 | NA | NA | NA |
| Gm22722 | 0 | NA | NA | NA |
| Vmn2r-ps32 | 0 | NA | NA | NA |
| Vmn2r22 | 0 | NA | NA | NA |
| Gm25065 | 0 | NA | NA | NA |
| Vmn2r23 | 0.060174537 | 0.755286614 | 3.724945049 | 0.202764498 |
| Gm22724 | 0 | NA | NA | NA |
| Vmn2r24 | 0 | NA | NA | NA |
| Gm23079 | 0 | NA | NA | NA |
| Vmn2r25 | 0 | NA | NA | NA |
| A330044P14Rik | 0.221456285 | -0.206486871 | 2.183616399 | -0.09456188 |

| | | | | |
|---|---|---|---|---|
| Gm22763 | 0 | NA | NA | NA |
| Vmn2r26 | 0 | NA | NA | NA |
| Gm24843 | 0.072025462 | 0.755288839 | 3.724945049 | 0.202765096 |
| Vmn2r-ps33 | 0 | NA | NA | NA |
| Gm22916 | 0 | NA | NA | NA |
| Vmn2r-ps34 | 0 | NA | NA | NA |
| Gm23554 | 0 | NA | NA | NA |
| Vmn2r27 | 0.851816096 | 0.990616185 | 1.111600398 | 0.891162136 |
| Gm25311 | 0.355991514 | -1.247951177 | 1.631500949 | -0.764909869 |
| Cd163 | 12.17190331 | -0.150248091 | 0.294603625 | -0.510000823 |
| 1700013D24Rik | 0.329404011 | -0.206487284 | 2.014022334 | -0.102524823 |
| Pex5 | 534.6967276 | -0.066131311 | 0.073067301 | -0.905073952 |
| Clstn3 | 4659.131254 | -0.264240973 | 0.060731484 | -4.350971769 |
| C1rl | 16.99094215 | 2.345559941 | 0.350757818 | 6.687120921 |
| C1ra | 434.4635122 | 5.920939146 | 0.326453275 | 18.13717182 |
| C1s1 | 241.4617169 | 5.413592064 | 0.43477497 | 12.45148051 |
| Gm16567 | 0 | NA | NA | NA |
| C1rb | 42.44241018 | 5.048087303 | 0.535069437 | 9.434452713 |
| C1s2 | 22.96225106 | 2.938229442 | 0.441601774 | 6.653572548 |
| Lpcat3 | 223.387876 | 0.491511484 | 0.10011827 | 4.909308612 |
| Emg1 | 187.4943886 | 0.012568071 | 0.098295114 | 0.127860584 |
| Phb2 | 583.7592251 | 0.213654793 | 0.076020904 | 2.810474264 |
| Gm23547 | 0.819923432 | -1.615070874 | 1.146281332 | -1.40896552 |
| Gm15884 | 0.260697633 | -0.206482684 | 2.112272612 | -0.097753804 |
| Gm29008 | 0.068097359 | -0.206487837 | 3.724945049 | -0.055433794 |
| Mir141 | 0 | NA | NA | NA |
| Mir200c | 0 | NA | NA | NA |
| Ptpn6 | 244.4682648 | 3.041456795 | 0.227763544 | 13.35357164 |
| Gm20531 | 0.660431168 | 1.130394566 | 1.257752497 | 0.898741659 |
| Grcc10 | 122.9669071 | 0.154639197 | 0.115076394 | 1.343795987 |
| Atn1 | 2476.146781 | -0.124892446 | 0.074991353 | -1.665424633 |
| Rnu7 | 0 | NA | NA | NA |
| Eno2 | 8072.573921 | -0.257903813 | 0.057390091 | -4.493873559 |
| Lrrc23 | 22.31127345 | -0.780211771 | 0.266097047 | -2.932057238 |
| Spsb2 | 56.84758583 | 0.427133533 | 0.221226707 | 1.930750307 |
| Tpi1 | 3615.771022 | -0.137305259 | 0.059347173 | -2.313593923 |
| Usp5 | 1925.798522 | 0.019623953 | 0.045743806 | 0.428996934 |
| Cdca3 | 13.51901529 | 1.009211664 | 0.449451548 | 2.245429278 |
| Gnb3 | 5.506721047 | 1.276988408 | 0.625468924 | 2.041649646 |
| P3h3 | 139.641216 | -0.452692766 | 0.181714874 | -2.491225709 |
| Gpr162 | 1473.024775 | -0.133933655 | 0.072626025 | -1.844155105 |
| Cd4 | 50.03247416 | -0.147543716 | 0.215521653 | -0.684588829 |
| Lag3 | 118.9770841 | 2.306446544 | 0.227765344 | 10.12641564 |
| Ptms | 3060.359574 | -0.055535223 | 0.079110034 | -0.701999733 |
| A230083G16Rik | 12.40694247 | -0.002990446 | 0.288505694 | -0.010365294 |
| Mlf2 | 3556.102244 | 0.001435452 | 0.061382739 | 0.02338527 |
| Cops7a | 930.3160726 | -0.046244424 | 0.067730786 | -0.682768153 |

| | | | | |
|---|---|---|---|---|
| Gm23851 | 0 | NA | NA | NA |
| Pianp | 1373.810968 | -0.110373849 | 0.063618705 | -1.734927614 |
| Zfp384 | 433.8857635 | -0.033274192 | 0.073333481 | -0.453738072 |
| Ing4 | 294.8789085 | -0.284479557 | 0.080762765 | -3.522409815 |
| Acrbp | 30.74231455 | -1.104773501 | 0.288353377 | -3.831318064 |
| Lpar5 | 6.566982277 | 1.275700458 | 0.46003654 | 2.773041592 |
| Chd4 | 3499.470299 | 0.027751294 | 0.057924425 | 0.479094864 |
| Mir7045 | 0 | NA | NA | NA |
| Gm23751 | 1.854332816 | -0.81035049 | 0.785763996 | -1.031289922 |
| Nop2 | 279.2923541 | 0.105220438 | 0.138660734 | 0.758833699 |
| Iffo1 | 569.9560514 | -0.16351115 | 0.106950158 | -1.528853743 |
| Gapdh | 740.2095212 | 0.087702567 | 0.076229182 | 1.150511709 |
| Ncapd2 | 168.4681777 | -0.289491283 | 0.155717703 | -1.85907753 |
| Scarna10 | 0.038981395 | 0.274393828 | 3.724945049 | 0.073663859 |
| Gm28967 | 0 | NA | NA | NA |
| Mrpl51 | 347.6131604 | 0.049556722 | 0.083406593 | 0.594158333 |
| Mir3098 | 14.8885832 | -0.356348983 | 0.340334408 | -1.047055408 |
| Vamp1 | 2358.40077 | -0.193185086 | 0.164164058 | -1.176780642 |
| Tapbpl | 330.3516113 | 2.859325398 | 0.175807006 | 16.26400148 |
| Cd27 | 20.08776085 | 6.673381288 | 0.894605244 | 7.459582124 |
| Mir8113 | 0 | NA | NA | NA |
| Gm24175 | 0 | NA | NA | NA |
| Tuba3a | 0.099106094 | 0.274387056 | 3.724945049 | 0.073662041 |
| Ltbr | 255.90804 | 1.637647128 | 0.200533814 | 8.166438847 |
| Scnn1a | 38.07952501 | 0.006454208 | 0.191370762 | 0.033726196 |
| Tnfrsf1a | 469.1495194 | 2.183521129 | 0.189165006 | 11.54294431 |
| Plekhg6 | 3.530240572 | 0.453590599 | 0.489331251 | 0.926960209 |
| Cd9 | 682.5462011 | 0.472310215 | 0.133198707 | 3.545906905 |
| Gm28809 | 1.776315563 | 2.020491874 | 0.940705843 | 2.147846628 |
| Gm26728 | 0.409864138 | 1.396844932 | 1.65012201 | 0.84651009 |
| Vwf | 172.1922304 | 1.363212667 | 0.174950889 | 7.79197337 |
| Ano2 | 81.25229014 | -0.421718524 | 0.13934384 | -3.026459754 |
| Gm16303 | 0.512522738 | 0.274415395 | 1.489222201 | 0.184267596 |
| Gm24947 | 0.271265366 | -0.195215215 | 2.086420432 | -0.093564659 |
| Ntf3 | 14.00482386 | -0.580946905 | 0.29244714 | -1.98650226 |
| Gm29009 | 0 | NA | NA | NA |
| Kcna5 | 77.10056723 | 0.319128859 | 0.158916219 | 2.008157888 |
| Gm33320 | 17.12675634 | -0.301221337 | 0.308058478 | -0.977805705 |
| Kcna1 | 5474.951842 | 0.058706505 | 0.094761777 | 0.619516715 |
| Kcna6 | 1135.188091 | -0.537053915 | 0.09433598 | -5.692991307 |
| Gm26673 | 76.86612292 | 0.068775992 | 0.228853696 | 0.300523843 |
| Ndufa9 | 671.0675705 | -0.139537553 | 0.07051542 | -1.978823255 |
| RP23-222G22.5 | 0.062426873 | 0.755288839 | 3.724945049 | 0.202765096 |
| Akap3 | 1.281806 | -0.557520311 | 0.869760998 | -0.641004037 |
| Dyrk4 | 3.739156486 | 1.690161259 | 0.615898624 | 2.744219897 |
| Rad51ap1 | 20.13187148 | 0.432230777 | 0.329970442 | 1.30990756 |
| RP23-222G22.9 | 0 | NA | NA | NA |

| | | | | |
|---|---|---|---|---|
| D6Wsu163e | 358.0575039 | 0.054680356 | 0.079300761 | 0.689531293 |
| RP23-8L20.9 | 0 | NA | NA | NA |
| RP23-8L20.8 | 0.141823039 | -0.206497533 | 3.6001346 | -0.057358281 |
| Fgf6 | 0.246418769 | 0.274412349 | 1.860767557 | 0.147472664 |
| Fgf23 | 4.436253295 | 0.755374009 | 0.485366308 | 1.556296752 |
| Tigar | 254.7556436 | -0.233239918 | 0.092579053 | -2.519359497 |
| Platr31 | 1.918110815 | 0.2447122 | 0.686834949 | 0.356289674 |
| RP23-8L20.6 | 0.061920613 | 0.274393828 | 3.724945049 | 0.073663859 |
| Ccnd2 | 1274.440512 | -0.09262038 | 0.13517434 | -0.685192027 |
| 9330179D12Rik | 23.50546626 | 0.186741837 | 0.216936991 | 0.860811406 |
| RP23-387P23.5 | 2.073082813 | 0.189784416 | 0.712643748 | 0.266310365 |
| RP23-403P6.1 | 26.03534595 | -0.17094675 | 0.209838487 | -0.814658707 |
| Gm4968 | 0.813324618 | -0.356156379 | 1.123764298 | -0.316931566 |
| RP23-387P23.7 | 0 | NA | NA | NA |
| Gm7308 | 60.57587847 | -0.048847197 | 0.175518772 | -0.278301838 |
| RP23-387P23.3 | 0.11147366 | 0.274389235 | 3.724945049 | 0.073662626 |
| RP23-387P23.4 | 2.402022068 | 0.25213044 | 0.585617662 | 0.430537629 |
| RP23-387P23.9 | 0.145088245 | -0.206488961 | 3.53071253 | -0.058483651 |
| RP23-387P23.8 | 0 | NA | NA | NA |
| RP23-223J11.1 | 0.049553047 | 0.274391604 | 3.724945049 | 0.073663262 |
| Parp11 | 870.7509703 | 1.49862729 | 0.1832917 | 8.176187418 |
| RP23-422D12.3 | 30.48963593 | 0.520729658 | 0.335674288 | 1.551294445 |
| Cracr2a | 10.36781213 | 0.004400898 | 0.30021844 | 0.014658986 |
| RP23-42M8.2 | 6.20278278 | -0.36981265 | 0.465812156 | -0.793909401 |
| Prmt8 | 579.4218534 | -0.589475863 | 0.149072103 | -3.954300302 |
| RP23-42M8.9 | 0 | NA | NA | NA |
| RP23-42M8.10 | 14.99285838 | -0.731227972 | 0.313299088 | -2.333961384 |
| RP23-42M8.5 | 0.045882455 | 0.755288839 | 3.724945049 | 0.202765096 |
| RP23-42M8.7 | 0.12092566 | 1.236179401 | 3.724945049 | 0.331865138 |
| mmu-mir-7233 | 0 | NA | NA | NA |
| RP24-473H20.4 | 0.306891002 | 1.233679798 | 1.770354731 | 0.696854577 |
| Gm23098 | 0.762875166 | -1.418176128 | 1.228204807 | -1.154673976 |
| RP24-473H20.3 | 0.711610403 | -1.123176471 | 1.380804431 | -0.813421833 |
| Tspan11 | 99.16132127 | -0.059415929 | 0.16087477 | -0.369330315 |
| Gm15870 | 0.825566729 | -0.071914005 | 1.327930994 | -0.054154926 |
| Tspan9 | 1288.655582 | 0.063239306 | 0.134565627 | 0.469951407 |
| Gm26338 | 0.326202429 | 0.635700147 | 1.80659232 | 0.35187803 |
| Gm10010 | 121.3586711 | -0.674180968 | 0.179894925 | -3.747637513 |
| Gm4691 | 17.793641 | -0.258308025 | 0.284620666 | -0.907551895 |
| Tead4 | 29.15459458 | 0.211956001 | 0.26320568 | 0.805286576 |
| Gm21893 | 0 | NA | NA | NA |
| Gm26951 | 0.061920613 | 0.274393828 | 3.724945049 | 0.073663859 |
| Rpl18-ps2 | 5.774417445 | 0.561618244 | 0.474714245 | 1.183065918 |
| Tulp3 | 551.9556277 | 0.61979561 | 0.110223954 | 5.623057327 |
| Gm26770 | 0.55622196 | -1.41391917 | 1.321646787 | -1.06981622 |
| Rhno1 | 92.43520098 | 0.061017074 | 0.11976694 | 0.50946508 |
| Foxm1 | 45.1673141 | 0.844689624 | 0.371394038 | 2.274375831 |

| | | | | |
|---|---|---|---|---|
| 4933413G19Rik | 17.62518862 | -0.987352884 | 0.35862374 | -2.75317212 |
| Gm15862 | 1.729221609 | 0.177665364 | 0.779946774 | 0.227791652 |
| Nrip2 | 216.9234121 | 0.020152674 | 0.091029461 | 0.221386286 |
| Itfg2 | 280.4983282 | 0.251653783 | 0.102993912 | 2.443385028 |
| Fkbp4 | 1131.070024 | -0.334813759 | 0.082533221 | -4.056715032 |
| Gm10069 | 14.56716468 | -0.511258751 | 0.326839464 | -1.564250365 |
| Pzp | 1.32803898 | 1.238653001 | 0.971655128 | 1.274786666 |
| Gm8724 | 0.141071297 | -0.206491178 | 3.564043836 | -0.057937328 |
| BC048546 | 30.70108197 | 0.103558458 | 0.190647163 | 0.543194328 |
| Klrb1a | 2.241070021 | 3.518747779 | 1.038862115 | 3.387117239 |
| Gm5884 | 150.0674224 | 0.19087896 | 0.122343655 | 1.56018684 |
| Clec2j | 0 | NA | NA | NA |
| Clec2h | 0.169629266 | -0.687377704 | 2.768139436 | -0.248317587 |
| Klrb1 | 0 | NA | NA | NA |
| Klrb1c | 5.966816195 | 4.91355573 | 1.000914022 | 4.909068734 |
| Klrb1b | 4.841237096 | 3.896530263 | 0.887065153 | 4.392608873 |
| Gm26656 | 0.322709507 | 2.27651505 | 1.916398067 | 1.187913455 |
| BC035044 | 6.979976624 | 1.000974441 | 0.460671285 | 2.172860506 |
| Clec2i | 12.44756223 | 4.426214645 | 0.862629706 | 5.131071439 |
| Gm27336 | 0.319483352 | -0.206484901 | 2.014202452 | -0.102514472 |
| Clec2g | 0.594455495 | 0.946171812 | 1.397672847 | 0.676962291 |
| Gm15987 | 6.297883917 | 3.797801711 | 0.890382227 | 4.265361097 |
| Mir7649 | 0 | NA | NA | NA |
| Gm8778 | 0.092691418 | 0.274387304 | 3.724945049 | 0.073662108 |
| BC064078 | 21.92509358 | 1.209844139 | 0.311025179 | 3.889859149 |
| Gm16187 | 0.11197992 | 0.75528439 | 3.724945049 | 0.202763901 |
| Clec2f | 1.610080848 | 2.029343998 | 0.868113591 | 2.337647997 |
| Gm16186 | 0 | NA | NA | NA |
| Klrb1f | 4.97974666 | 4.03487847 | 0.934729556 | 4.316626604 |
| Clec2e | 0.752785761 | 1.070999571 | 1.411201968 | 0.758927209 |
| Klrb1-ps1 | 0 | NA | NA | NA |
| Gm26160 | 0.088534442 | 0.274389469 | 3.724945049 | 0.073662689 |
| Clec2d | 395.2909725 | 2.928757116 | 0.263757434 | 11.10397939 |
| Gm27514 | 0.12209515 | 0.75528439 | 3.724945049 | 0.202763901 |
| 2310001H17Rik | 41.45082234 | 3.231839028 | 0.370044248 | 8.733655627 |
| Cd69 | 49.45403992 | 8.295460321 | 0.892925269 | 9.290206703 |
| 4922502D21Rik | 0.201858732 | 1.23619586 | 2.604406118 | 0.474655566 |
| Clec12a | 44.34900859 | 2.591411612 | 0.388158704 | 6.676165142 |
| Clec12b | 2.028096924 | 3.147589389 | 1.140570985 | 2.759661109 |
| Clec1b | 1.612175364 | 0.972935047 | 0.895931456 | 1.085948082 |
| Clec9a | 4.424159844 | 2.620585337 | 0.765867757 | 3.421720411 |
| Clec1a | 25.40223335 | 1.322608727 | 0.299570043 | 4.415023319 |
| Clec7a | 77.62764879 | 3.504230255 | 0.310570778 | 11.28319373 |
| Olr1 | 7.563519298 | 4.281459703 | 0.939922498 | 4.555119928 |
| Tmem52b | 0.046345709 | 0.274391604 | 3.724945049 | 0.073663262 |
| 1700101I11Rik | 14.17165844 | 0.746609805 | 0.300303191 | 2.486186715 |
| Gabarapl1 | 4493.649597 | 0.052138935 | 0.059327471 | 0.878832921 |

| | | | | |
|---|---|---|---|---|
| Klre1 | 4.71579917 | 3.199534896 | 0.969652893 | 3.299670347 |
| Klrd1 | 23.39137982 | 5.999419002 | 0.88029277 | 6.815254203 |
| Klrk1 | 186.6532185 | 8.480902918 | 0.760520066 | 11.15145188 |
| Klrc3 | 5.985488163 | 5.107767716 | 0.946073041 | 5.398914769 |
| Klrc2 | 17.16294782 | 5.976201659 | 0.91627094 | 6.522308412 |
| Klrc1 | 33.05721631 | 7.893085794 | 0.87948263 | 8.974692075 |
| Gm7309 | 0 | NA | NA | NA |
| Mir680-1 | 0.066050875 | 0.274391604 | 3.724945049 | 0.073663262 |
| Klri1 | 2.598038951 | 2.919458143 | 1.096108791 | 2.6634748 |
| Klri2 | 8.227621471 | 3.758129039 | 1.023437616 | 3.6720646 |
| Gm156 | 0.097139547 | 0.274393828 | 3.724945049 | 0.073663859 |
| Gm16242 | 0.044544648 | 0.755286614 | 3.724945049 | 0.202764498 |
| Klra17 | 6.058344776 | 5.507688895 | 1.000747741 | 5.503573648 |
| Gm17631 | 0 | NA | NA | NA |
| Gm23977 | 0 | NA | NA | NA |
| Klra5 | 0.677147933 | 1.763415148 | 1.529765522 | 1.152735581 |
| Gm15854 | 0 | NA | NA | NA |
| Gm22555 | 0 | NA | NA | NA |
| Klra6 | 0 | NA | NA | NA |
| Klra4 | 0.378042144 | 1.236204876 | 1.729535435 | 0.714761231 |
| Gm24676 | 0 | NA | NA | NA |
| Klra11-ps | 0 | NA | NA | NA |
| Gm22621 | 0 | NA | NA | NA |
| Klra8 | 0.110595523 | 0.755282166 | 3.724945049 | 0.202763304 |
| Gm24712 | 0 | NA | NA | NA |
| Klra14-ps | 0 | NA | NA | NA |
| Gm23552 | 0 | NA | NA | NA |
| Klra9 | 0.604622052 | 1.675668373 | 1.361669298 | 1.230598631 |
| Klra7 | 3.947549917 | 3.799951039 | 0.997697867 | 3.808719218 |
| Klra10 | 0.285107744 | 0.755298724 | 2.430606519 | 0.310744959 |
| Klra13-ps | 0.52061208 | 0.755309804 | 1.474308514 | 0.512314619 |
| Gm24072 | 0 | NA | NA | NA |
| Klra3 | 12.4313916 | 4.870021968 | 0.921540212 | 5.284654868 |
| Klra1 | 0.328294527 | 1.246124403 | 1.724352023 | 0.722662418 |
| Gm6584 | 0 | NA | NA | NA |
| Gm15861 | 0 | NA | NA | NA |
| Gm20997 | 0 | NA | NA | NA |
| Gm27013 | 0.044544648 | 0.755286614 | 3.724945049 | 0.202764498 |
| Gm8816 | 0 | NA | NA | NA |
| Gm26996 | 0.113979814 | 0.274404949 | 3.724945049 | 0.073666845 |
| Gm26914 | 0 | NA | NA | NA |
| Gm27047 | 0 | NA | NA | NA |
| Gm26919 | 0.161486377 | 0.755296344 | 2.795731986 | 0.270160498 |
| Gm6600 | 0 | NA | NA | NA |
| Gm8837 | 0 | NA | NA | NA |
| Gm5581 | 0.915657815 | 1.895589319 | 1.501080897 | 1.26281623 |
| Klra2 | 81.98899445 | 4.793529134 | 0.611950751 | 7.833194295 |

| | | | | |
|---|---|---|---|---|
| Gm5582 | 1.579793576 | 0.169076725 | 0.73247923 | 0.230828012 |
| Magohb | 84.69062322 | 0.219245609 | 0.158323742 | 1.384792995 |
| Styk1 | 21.45282092 | 1.398169778 | 0.309049705 | 4.524093553 |
| Ybx3 | 331.3395029 | 0.352359124 | 0.105510962 | 3.3395499 |
| Gm22362 | 7.679914887 | 0.365467141 | 0.401540101 | 0.910163493 |
| 5430401F13Rik | 0.172107867 | 1.236193725 | 2.739902689 | 0.451181617 |
| Gm6619 | 0 | NA | NA | NA |
| Gm22190 | 0 | NA | NA | NA |
| Gm16571 | 0 | NA | NA | NA |
| Hmgb1l | 0 | NA | NA | NA |
| Tas2r130 | 0 | NA | NA | NA |
| Tas2r107 | 0 | NA | NA | NA |
| Tas2r106 | 0 | NA | NA | NA |
| Tas2r104 | 0 | NA | NA | NA |
| Tas2r105 | 0 | NA | NA | NA |
| Tas2r114 | 0 | NA | NA | NA |
| Gm4736 | 0 | NA | NA | NA |
| Prpmp5 | 0 | NA | NA | NA |
| Gm8882 | 0.046345709 | 0.274391604 | 3.724945049 | 0.073663262 |
| n-R5s166 | 0 | NA | NA | NA |
| Prh1 | 0 | NA | NA | NA |
| n-R5s167 | 0 | NA | NA | NA |
| Prp2 | 0 | NA | NA | NA |
| A630073D07Rik | 0 | NA | NA | NA |
| Tas2r120 | 0 | NA | NA | NA |
| Tas2r121 | 0 | NA | NA | NA |
| Tas2r122 | 0 | NA | NA | NA |
| Tas2r115 | 0 | NA | NA | NA |
| Tas2r124 | 0 | NA | NA | NA |
| Tas2r102 | 0 | NA | NA | NA |
| Tas2r136 | 0.061920613 | 0.274393828 | 3.724945049 | 0.073663859 |
| Tas2r117 | 0 | NA | NA | NA |
| Tas2r123 | 0 | NA | NA | NA |
| Tas2r116 | 0.061920613 | 0.274393828 | 3.724945049 | 0.073663859 |
| Tas2r110 | 0 | NA | NA | NA |
| Tas2r113 | 0 | NA | NA | NA |
| Tas2r125 | 0 | NA | NA | NA |
| Tas2r129 | 0 | NA | NA | NA |
| Tas2r131 | 0 | NA | NA | NA |
| Tas2r109 | 0 | NA | NA | NA |
| Tas2r103 | 0 | NA | NA | NA |
| Tas2r140 | 0 | NA | NA | NA |
| Smim10l1 | 1843.017742 | 0.083088347 | 0.060081596 | 1.382925091 |
| Gm23403 | 0 | NA | NA | NA |
| Gm25019 | 0.075043205 | 0.755286614 | 3.724945049 | 0.202764498 |
| 5530400C23Rik | 0 | NA | NA | NA |
| Gm5885 | 0 | NA | NA | NA |

| | | | | |
|---|---|---|---|---|
| Gm23375 | 0.502960085 | 1.588798003 | 1.489994876 | 1.066311051 |
| Kap | 0.042716945 | -0.206490061 | 3.724945049 | -0.055434391 |
| Etv6 | 690.3323098 | 1.303972604 | 0.128626603 | 10.1376587 |
| Gm17088 | 0.202171817 | 0.274403705 | 2.59796949 | 0.105622374 |
| Gm17089 | 0.375570109 | 0.755308261 | 1.712796302 | 0.440979619 |
| Bcl2l14 | 0.379111656 | 1.201446451 | 1.460777817 | 0.822470356 |
| Lrp6 | 1132.17589 | -0.286862167 | 0.063858686 | -4.492140121 |
| Mansc1 | 117.9240851 | 0.096668925 | 0.127068957 | 0.760759573 |
| Loh12cr1 | 180.7292652 | 0.267278725 | 0.091181952 | 2.931267841 |
| Dusp16 | 411.5721016 | 0.319612869 | 0.126025766 | 2.536091457 |
| Crebl2 | 520.8784886 | -0.136352858 | 0.111946524 | -1.218017793 |
| Gpr19 | 174.2410689 | -0.040650438 | 0.131629661 | -0.308824298 |
| 2810454H06Rik | 8.20098561 | -0.571027974 | 0.387094167 | -1.475165533 |
| Cdkn1b | 860.406846 | 0.059396342 | 0.100738948 | 0.589606529 |
| 1190002F15Rik | 1.018292001 | 2.201603613 | 1.160587415 | 1.896973537 |
| Apold1 | 80.86165162 | 0.957811689 | 0.257587208 | 3.718397736 |
| Ddx47 | 401.4348954 | 0.069108612 | 0.069695842 | 0.991574401 |
| Gprc5a | 6.993744504 | 2.424664552 | 0.614695849 | 3.944494751 |
| Gprc5d | 0.543589123 | -0.686446967 | 1.501951922 | -0.457036578 |
| Hebp1 | 102.192585 | -0.273933192 | 0.17224112 | -1.590405311 |
| Gm22881 | 0 | NA | NA | NA |
| 8430419L09Rik | 1464.187594 | -0.137429033 | 0.054883642 | -2.504007149 |
| Gsg1 | 0.811303554 | 0.83445208 | 1.252341324 | 0.666313619 |
| Pbp2 | 0 | NA | NA | NA |
| Dynlt1-ps1 | 35.59685521 | 0.309211753 | 0.272202955 | 1.13596031 |
| Emp1 | 465.3937255 | 3.252234762 | 0.419027891 | 7.76138016 |
| Gm25136 | 0.672405421 | -0.977619826 | 1.363002287 | -0.717254722 |
| Grin2b | 9373.268553 | -0.26704046 | 0.189501008 | -1.409176992 |
| Gm14329 | 0.341881936 | -0.539154002 | 1.560660397 | -0.345465294 |
| Gm22892 | 0.112396584 | 0.27438722 | 3.724945049 | 0.073662085 |
| Gm38202 | 40.04288314 | -0.632628654 | 0.335626384 | -1.884919315 |
| Gm37182 | 48.96267409 | -0.793571215 | 0.401923422 | -1.974433862 |
| Gm14330 | 0.038981395 | 0.274393828 | 3.724945049 | 0.073663859 |
| Gm26653 | 7.145952665 | -0.526830228 | 0.425540437 | -1.238026243 |
| Gm26105 | 0.110595523 | 0.755282166 | 3.724945049 | 0.202763304 |
| Gm8994 | 0.107803068 | 0.75528661 | 3.724945049 | 0.202764497 |
| Gm15476 | 0.04393175 | -0.206490061 | 3.724945049 | -0.055434391 |
| E330021D16Rik | 0 | NA | NA | NA |
| Gm25882 | 0 | NA | NA | NA |
| Gm6728 | 11.37417031 | -0.048868881 | 0.31962617 | -0.152893868 |
| Rpl36a-ps3 | 8.907631286 | 0.06492499 | 0.375346698 | 0.172973388 |
| Atf7ip | 2145.155086 | 0.117425163 | 0.075937244 | 1.546344802 |
| 5830415B17Rik | 15.21012998 | 0.17404438 | 0.312128912 | 0.557604162 |
| Plbd1 | 20.58495972 | 2.497803733 | 0.459529348 | 5.435569549 |
| Gucy2c | 13.73199816 | 0.281399513 | 0.326794498 | 0.861090119 |
| Hist4h4 | 361.153536 | -0.330141677 | 0.15867592 | -2.080603513 |
| H2afj | 164.4299334 | -0.125907898 | 0.130672974 | -0.963534349 |

| | | | | |
|---|---|---|---|---|
| AC122804.1 | 2.891209199 | 0.235649693 | 0.6430276 | 0.366469017 |
| Wbp11 | 918.8633835 | 0.23414828 | 0.085913946 | 2.725381516 |
| BC049715 | 12.24740255 | 0.334154981 | 0.316115151 | 1.057067272 |
| Smco3 | 28.46706308 | -0.230675822 | 0.192471364 | -1.198494244 |
| Art4 | 1.936338333 | -1.218253968 | 0.859321531 | -1.417692824 |
| Mgp | 118.7015475 | 0.631440358 | 0.184460376 | 3.423176149 |
| Erp27 | 1.418067754 | 2.398441511 | 1.029862614 | 2.328894629 |
| Arhgdib | 195.7130349 | 2.07691666 | 0.173985507 | 11.93729695 |
| Pde6h | 2.208047999 | 0.983394777 | 0.973833342 | 1.009818348 |
| Rerg | 82.27360041 | -0.350874304 | 0.137353619 | -2.554532648 |
| Ptpro | 585.5542697 | -0.449029393 | 0.162394953 | -2.765045246 |
| Eps8 | 778.7034068 | -0.014299391 | 0.122866398 | -0.116381626 |
| Strap | 1329.576707 | 0.149839968 | 0.083547499 | 1.79347042 |
| Dera | 50.46973726 | 0.905596638 | 0.234150927 | 3.867576559 |
| Gm24308 | 0 | NA | NA | NA |
| Slc15a5 | 0.807800175 | -0.041619665 | 1.081262259 | -0.03849174 |
| Mgst1 | 196.4190543 | 1.083819824 | 0.223772907 | 4.843391618 |
| Lmo3 | 872.8499117 | 0.052134941 | 0.122206011 | 0.426615191 |
| A830011K09Rik | 12.61465114 | 0.261479563 | 0.339050214 | 0.771211909 |
| B230110G15Rik | 1.657582323 | 0.118983098 | 0.718426554 | 0.165616231 |
| Igbp1b | 1.236116855 | 0.199710854 | 0.792344197 | 0.25205063 |
| Gm25434 | 0 | NA | NA | NA |
| Rergl | 0.244560259 | -1.117405499 | 2.108054029 | -0.530064924 |
| Pik3c2g | 12.46784403 | 2.135999724 | 0.391930245 | 5.449948682 |
| Gm37269 | 0.211089261 | 1.236201041 | 2.185019532 | 0.56576201 |
| Plcz1 | 6.588952628 | 0.424119054 | 0.373694748 | 1.134934477 |
| Capza3 | 0.060174537 | 0.755286614 | 3.724945049 | 0.202764498 |
| Gm24174 | 0 | NA | NA | NA |
| n-R5s168 | 0 | NA | NA | NA |
| Plekha5 | 1362.818818 | -0.0543444 | 0.107694971 | -0.504614091 |
| Gm25780 | 21.43207649 | 0.357137615 | 0.29921293 | 1.19359018 |
| Aebp2 | 1104.796083 | -0.060699604 | 0.063203729 | -0.960380107 |
| Gm11077 | 0.333293459 | -0.687379038 | 1.743021552 | -0.394360607 |
| Rpl38-ps2 | 307.4974746 | 0.295469828 | 0.090969558 | 3.248007728 |
| Gm28524 | 1.176308512 | -0.346990312 | 0.972914295 | -0.356650441 |
| Gm28523 | 12.22868002 | -0.196118468 | 0.305864849 | -0.641193222 |
| Pde3a | 315.7616379 | 0.548488557 | 0.104880067 | 5.229673996 |
| Slco1c1 | 209.1116899 | -0.565377677 | 0.228706133 | -2.472070464 |
| Slco1b2 | 3.268651982 | -0.948602738 | 0.595468386 | -1.593036272 |
| Gm5724 | 0.34217064 | -0.617238092 | 1.56032536 | -0.395582939 |
| Gm23069 | 0 | NA | NA | NA |
| Slco1a4 | 155.850488 | -0.480962398 | 0.343830614 | -1.398835296 |
| Slco1a1 | 0.113999573 | -0.206492286 | 3.724945049 | -0.055434988 |
| Gm22036 | 0 | NA | NA | NA |
| Gm6614 | 0.602494748 | 2.720468736 | 1.335207706 | 2.037487294 |
| Gm23775 | 0 | NA | NA | NA |
| Slco1a6 | 0.857960599 | 1.72429594 | 1.203419464 | 1.432830357 |

| | | | | |
|---|---|---|---|---|
| Gm18159 | 0.046345709 | 0.274391604 | 3.724945049 | 0.073663262 |
| Gm20400 | 0.439621597 | 2.190603649 | 1.5132265 | 1.447637646 |
| Slco1a5 | 10.05483302 | 0.988081208 | 0.458015602 | 2.15730906 |
| Gm25430 | 0 | NA | NA | NA |
| Iapp | 0.092228164 | 0.75528439 | 3.724945049 | 0.202763901 |
| Pyroxd1 | 173.2075479 | 0.094510244 | 0.085234485 | 1.108826357 |
| Recql | 183.7198923 | -0.321741113 | 0.14611536 | -2.201966396 |
| Golt1b | 467.436909 | 0.143584038 | 0.064387029 | 2.230014954 |
| B230216G23Rik | 18.91388396 | -1.064863503 | 0.278784429 | -3.819666355 |
| Gys2 | 1.458192719 | 2.758941732 | 1.122729431 | 2.457352285 |
| Ldhb | 2909.672582 | -0.22676403 | 0.072272135 | -3.137641226 |
| Kcnj8 | 29.13393712 | 0.822519947 | 0.218762075 | 3.759883636 |
| Abcc9 | 80.16356092 | -0.682440653 | 0.173140736 | -3.941537202 |
| 5330439B14Rik | 0 | NA | NA | NA |
| Cmas | 639.4281021 | -0.238274689 | 0.065904133 | -3.615474155 |
| Gm766 | 1.893318208 | 0.190711372 | 0.751179295 | 0.253882626 |
| Gm7457 | 1.3052917 | -1.163364918 | 1.030915285 | -1.128477709 |
| St8sia1 | 1060.037098 | -0.432970808 | 0.062533395 | -6.923833444 |
| C2cd5 | 774.2859654 | -0.571957818 | 0.121505106 | -4.707273935 |
| Etnk1 | 6574.447317 | 0.577970448 | 0.090900579 | 6.358270228 |
| D6Ertd474e | 2.829004364 | 0.574777575 | 0.549265706 | 1.046447229 |
| Gm23272 | 0 | NA | NA | NA |
| 4930579D09Rik | 0 | NA | NA | NA |
| 1700060C16Rik | 0.044191549 | -0.206490061 | 3.724945049 | -0.055434391 |
| Sox5 | 636.5935809 | -0.292992735 | 0.096374012 | -3.040163323 |
| Sox5os1 | 0.085433889 | -0.687376175 | 3.724945049 | -0.184533239 |
| Sox5os2 | 0 | NA | NA | NA |
| Sox5it | 4.538093024 | 0.795035093 | 0.461889554 | 1.721266665 |
| Sox5os3 | 0.137647365 | 1.404948034 | 3.719469264 | 0.377728093 |
| Sox5os4 | 0 | NA | NA | NA |
| Sox5os5 | 0 | NA | NA | NA |
| Gm22792 | 0 | NA | NA | NA |
| Bcat1 | 1548.257675 | 0.136940316 | 0.103392384 | 1.324471991 |
| Gm23071 | 1.299875216 | -0.209553529 | 0.826075039 | -0.253673721 |
| Gm26666 | 7.30636117 | -0.107704241 | 0.37670386 | -0.285912232 |
| Gm15687 | 0.219710209 | 0.274409735 | 2.172592196 | 0.12630522 |
| Lrmp | 42.64411814 | 1.613089645 | 0.221126958 | 7.294857484 |
| Casc1 | 19.07366442 | -0.104999584 | 0.258503403 | -0.4061826 |
| Gm15543 | 0.049553047 | 0.274391604 | 3.724945049 | 0.073663262 |
| Lyrm5 | 128.9520887 | -0.075431404 | 0.12380819 | -0.609260211 |
| Kras | 1419.910694 | 0.05471183 | 0.071916196 | 0.760772026 |
| Gm15706 | 24.72915098 | -0.127134594 | 0.217303246 | -0.58505612 |
| Rps25-ps1 | 12.76755632 | 0.86270596 | 0.327577313 | 2.633594956 |
| Gm22793 | 0 | NA | NA | NA |
| 2010013B24Rik | 0.275148976 | 0.274409807 | 2.093147516 | 0.131099125 |
| Gm15707 | 0.08486385 | 0.755286617 | 3.724945049 | 0.202764499 |
| Lmntd1 | 13.42183498 | 0.144205322 | 0.321195604 | 0.448964183 |

| | | | | |
|---|---|---|---|---|
| Gm23498 | 0 | NA | NA | NA |
| 1700073E17Rik | 0.253253743 | 1.537834452 | 2.465631522 | 0.623708141 |
| Gm15499 | 0 | NA | NA | NA |
| Gm25373 | 0 | NA | NA | NA |
| Tuba3b | 0.095898756 | 0.27438722 | 3.724945049 | 0.073662085 |
| Gm15704 | 2.261169167 | -0.848408649 | 0.659090058 | -1.28724237 |
| Rassf8 | 399.0644075 | -0.2127539 | 0.109868818 | -1.936435686 |
| Bhlhe41 | 914.9264729 | -0.182331583 | 0.062377618 | -2.923028968 |
| Sspn | 140.0205592 | -0.536594432 | 0.19022156 | -2.820891758 |
| Sspnos | 0.044191549 | -0.206490061 | 3.724945049 | -0.055434391 |
| Itpr2 | 644.2115075 | 0.193305233 | 0.098178236 | 1.968921434 |
| Gm15720 | 1.860207608 | -0.485718517 | 0.781996342 | -0.621126329 |
| Asun | 359.4234607 | 0.013251672 | 0.105206275 | 0.125958953 |
| Fgfr1op2 | 883.1771317 | 0.152808471 | 0.052500418 | 2.910614374 |
| Tm7sf3 | 427.7210704 | -0.108917032 | 0.079725987 | -1.366142163 |
| Med21 | 211.1955998 | 0.2463206 | 0.158784735 | 1.551286399 |
| Gm6654 | 0.926048393 | -0.702754253 | 1.281007735 | -0.548594856 |
| 4930479D17Rik | 2.935438062 | -0.209047127 | 0.560533253 | -0.372943311 |
| Stk38l | 426.5635259 | -0.038050355 | 0.083720719 | -0.4544915 |
| Arntl2 | 48.93835431 | -0.276003151 | 0.216928575 | -1.272322706 |
| Smco2 | 1.092136027 | 1.356987302 | 1.241000786 | 1.093462081 |
| Ppfibp1 | 368.3784692 | -0.071948534 | 0.085600511 | -0.84051524 |
| 1700034J05Rik | 0.062426873 | 0.755288839 | 3.724945049 | 0.202765096 |
| Rep15 | 3.085232335 | -0.892180851 | 0.572966807 | -1.557124845 |
| Mrps35 | 175.5694965 | -0.136878401 | 0.103221748 | -1.326061646 |
| Mansc4 | 2.80060554 | 0.044293616 | 0.665354902 | 0.066571414 |
| Klhl42 | 434.0929622 | -0.282379901 | 0.071886631 | -3.928128162 |
| Gm7571 | 0.356315666 | 1.253069309 | 1.567983235 | 0.79915989 |
| E330012B07Rik | 0.576204374 | 1.811634555 | 1.514372361 | 1.196293991 |
| Gm15767 | 0.045882455 | 0.755288839 | 3.724945049 | 0.202765096 |
| Pthlh | 19.04363101 | -0.202395519 | 0.295986375 | -0.683800121 |
| Gm15766 | 2.701807514 | 0.193128179 | 0.613369633 | 0.314864266 |
| Gm6288 | 5.185232385 | 0.266553273 | 0.474865498 | 0.56132373 |
| Ccdc91 | 322.5136679 | 0.044053946 | 0.101023694 | 0.436075383 |
| 1700049E15Rik | 0.04393175 | -0.206490061 | 3.724945049 | -0.055434391 |
| Gm15762 | 0.295243939 | -0.973264164 | 1.749139951 | -0.556424409 |
| Far2os2 | 3.668828368 | 0.780327453 | 0.714662704 | 1.091882154 |
| Far2 | 187.1712709 | -0.543251201 | 0.119938222 | -4.529425159 |
| Far2os1 | 2.690587419 | -0.580464209 | 0.67590287 | -0.85879826 |
| Ergic2 | 794.7085187 | 0.121941997 | 0.07608712 | 1.60266279 |
| Gm17216 | 1.982138501 | -0.878341215 | 0.848896444 | -1.034685939 |
| Tmtc1 | 1255.055725 | -0.280479731 | 0.085185204 | -3.292587408 |
| Gm15763 | 0.045882455 | 0.755288839 | 3.724945049 | 0.202765096 |
| Rps4l | 60.78732413 | 0.261051971 | 0.263778611 | 0.989663151 |
| Gm6313 | 0.072025462 | 0.755288839 | 3.724945049 | 0.202765096 |
| Gm23456 | 0.106358623 | 0.274402725 | 3.724945049 | 0.073666248 |
| 4930528G23Rik | 0.514450355 | -0.580760335 | 1.452972018 | -0.399705106 |

| | | | | |
|---|---|---|---|---|
| Gm15778 | 0 | NA | NA | NA |
| Ipo8 | 948.9724006 | -0.202504546 | 0.05301986 | -3.81940929 |
| Caprin2 | 102.1573326 | -0.529205088 | 0.156148856 | -3.389106398 |
| Gm7618 | 0.320079734 | 1.202742884 | 1.728819142 | 0.695701971 |
| Fam60a | 49.2104772 | 0.959842838 | 0.178232669 | 5.385336176 |
| 3010003L21Rik | 2.014334011 | 2.414986032 | 0.895258327 | 2.697529819 |
| Gm25539 | 0 | NA | NA | NA |
| Dennd5b | 2078.167419 | -0.392464627 | 0.052006591 | -7.546440199 |
| Gm26540 | 0.259164831 | -0.196120913 | 1.836486513 | -0.106791372 |
| Gm15779 | 0 | NA | NA | NA |
| Gm15781 | 0 | NA | NA | NA |
| Gm22578 | 4.906737206 | -0.396415068 | 0.450283645 | -0.880367458 |
| Gm15780 | 0 | NA | NA | NA |
| Gm23462 | 2.97460163 | -0.606507165 | 0.646125432 | -0.938683319 |
| Gm10203 | 0.141071297 | -0.206491178 | 3.564043836 | -0.057937328 |
| Mettl20 | 22.19794556 | -0.007423633 | 0.335246858 | -0.022143781 |
| Gm10011 | 0.371635155 | 1.148526589 | 1.549312983 | 0.741313474 |
| Amn1 | 149.0049197 | -0.172093328 | 0.113517814 | -1.516002833 |
| 2810474O19Rik | 949.1809365 | 1.196364457 | 0.146736547 | 8.153145789 |
| Gm15782 | 0.129871752 | 0.755293287 | 2.991137803 | 0.252510361 |
| Gm15784 | 6.345417981 | -0.242400849 | 0.39156067 | -0.619063322 |
| Gm15786 | 0.200503209 | 1.257028057 | 2.612813754 | 0.481101286 |
| Gm15783 | 0.047948698 | -0.206487837 | 3.724945049 | -0.055433794 |
| Bicd1 | 1658.556008 | -0.225796066 | 0.050165479 | -4.501024776 |
| Gm15785 | 0.570614426 | -0.486125527 | 1.197607719 | -0.405913822 |
| Gm24307 | 1.150914894 | 0.312528839 | 0.922944191 | 0.338621601 |
| Gm21814 | 6.725415989 | -0.93006862 | 0.424174622 | -2.192655034 |
| Gm3104 | 0 | NA | NA | NA |
| Gm7353 | 46.36268977 | -0.097194984 | 0.168125784 | -0.578108736 |
| AU018091 | 0.640879238 | 1.236313428 | 1.381462034 | 0.894931165 |
| Mir290a | 0 | NA | NA | NA |
| mmu-mir-290b | 0 | NA | NA | NA |
| Mir291a | 0 | NA | NA | NA |
| Mir292 | 0 | NA | NA | NA |
| mmu-mir-292b | 0 | NA | NA | NA |
| Mir291b | 0 | NA | NA | NA |
| Mir293 | 0 | NA | NA | NA |
| Mir294 | 0 | NA | NA | NA |
| Mir295 | 0 | NA | NA | NA |
| Nlrp12 | 0.605827841 | 0.33941185 | 1.19816085 | 0.283277366 |
| Myadm | 554.9737066 | -0.291140403 | 0.0903102 | -3.223782053 |
| Prkcg | 1983.083602 | -0.304068354 | 0.158808673 | -1.914683555 |
| Cacng7 | 1812.960324 | 0.186681518 | 0.07805434 | 2.391686577 |
| Cacng8 | 593.3089663 | -0.297504135 | 0.101860768 | -2.920694013 |
| Gm23450 | 0 | NA | NA | NA |
| Cacng6 | 9.245691091 | 0.460241136 | 0.407928122 | 1.128240765 |
| Tarm1 | 15.12490142 | 4.287462695 | 1.203529502 | 3.562407641 |

| | | | | |
|---|---|---|---|---|
| Gm25100 | 0 | NA | NA | NA |
| Oscar | 1.834747314 | 2.274618958 | 0.913235193 | 2.49072635 |
| Ndufa3 | 436.48222 | -0.08801157 | 0.118588801 | -0.742157516 |
| Tfpt | 142.0055102 | 0.178689316 | 0.130105565 | 1.373417926 |
| Prpf31 | 410.4576804 | 0.455389032 | 0.073446207 | 6.200307021 |
| Gm15927 | 8.516043887 | -0.102548792 | 0.339508365 | -0.302050854 |
| Cnot3 | 493.1599569 | -0.140025556 | 0.077183996 | -1.814178636 |
| Mir3572 | 0 | NA | NA | NA |
| Leng1 | 69.24448306 | 0.122745301 | 0.18018992 | 0.681199596 |
| Tmc4 | 26.68951419 | -0.068959825 | 0.283991903 | -0.24282321 |
| Gm15929 | 0.377779179 | 0.554117121 | 3.171857807 | 0.17469797 |
| Mboat7 | 1337.109882 | -0.025555124 | 0.051794159 | -0.493397788 |
| Tsen34 | 403.0156256 | -0.076078745 | 0.122219154 | -0.622478088 |
| Rps9 | 1343.617836 | 0.447153751 | 0.141301653 | 3.164533057 |
| Pirb | 100.508228 | 3.670177172 | 0.350751429 | 10.46375543 |
| Gm15922 | 4.272688704 | 4.54812219 | 0.935879524 | 4.859730418 |
| Gm15930 | 0 | NA | NA | NA |
| Gm15925 | 2.457332247 | 3.773946765 | 1.154606983 | 3.268598596 |
| Gm15448 | 3.00180885 | 3.392290772 | 1.118899618 | 3.031809751 |
| Pira2 | 10.48735136 | 5.717560609 | 0.908533662 | 6.293174207 |
| Gm10693 | 0.291730981 | 1.227259542 | 1.769789387 | 0.693449487 |
| Gm14548 | 29.20745553 | 5.104148108 | 0.857550361 | 5.952009744 |
| Lilra6 | 22.5183229 | 4.748884501 | 0.7877738 | 6.028233614 |
| Gm7363 | 4.253190973 | 0.545761161 | 0.498780643 | 1.094190741 |
| Gm15928 | 0.849955171 | -1.195855631 | 1.100724018 | -1.086426399 |
| Lair1 | 165.5340521 | 1.089816826 | 0.213374025 | 5.107542147 |
| 9430041J12Rik | 62.38036442 | -0.168953548 | 0.154251178 | -1.09531448 |
| Gm23741 | 0.042716945 | -0.206490061 | 3.724945049 | -0.055434391 |
| Ttyh1 | 4104.837726 | -0.261786878 | 0.116083795 | -2.255154384 |
| D030047H15Rik | 2.31889725 | -0.549428839 | 0.602945364 | -0.911241502 |
| Leng8 | 1590.8448 | -0.255149479 | 0.285544752 | -0.893553383 |
| Leng9 | 55.71701833 | -0.158354709 | 0.148809908 | -1.064140899 |
| Cdc42ep5 | 10.88113561 | 0.207457709 | 0.362499778 | 0.572297478 |
| Gm5319 | 0 | NA | NA | NA |
| Gm15926 | 0 | NA | NA | NA |
| Lilra5 | 0.951293484 | -2.365876862 | 1.276629976 | -1.853220515 |
| Gm15923 | 0 | NA | NA | NA |
| Gm15931 | 19.55735248 | 3.732483916 | 0.655133502 | 5.697287508 |
| Ncr1 | 7.234741561 | 4.199789423 | 0.943118808 | 4.45308628 |
| Gp6 | 0.386555381 | 0.755270813 | 3.724945049 | 0.202760256 |
| Gm25349 | 0.061920613 | 0.274393828 | 3.724945049 | 0.073663859 |
| Gm24752 | 0.04393175 | -0.206490061 | 3.724945049 | -0.055434391 |
| Rdh13 | 296.6943263 | -0.562661126 | 0.113488696 | -4.957860525 |
| Gm24605 | 0 | NA | NA | NA |
| Gm15494 | 14.28857811 | -0.869342525 | 0.284352913 | -3.057266113 |
| Eps8l1 | 3.37593979 | 0.452246008 | 0.48332294 | 0.935701515 |
| Ppp1r12c | 945.1906909 | 0.040528268 | 0.079091014 | 0.512425695 |

| | | | | |
|---|---|---|---|---|
| Tnnt1 | 15.65154636 | -0.187969543 | 0.244648535 | -0.768324824 |
| Tnni3 | 10.79028321 | 0.208289969 | 0.444503593 | 0.468590069 |
| Dnaaf3 | 15.48723177 | 0.785323586 | 0.326634132 | 2.404291253 |
| Gm15873 | 0.239904818 | 0.755298172 | 2.554358129 | 0.295690007 |
| Syt5 | 799.4033589 | -0.112626059 | 0.116413945 | -0.96746192 |
| Ptprh | 6.302761948 | 1.04136493 | 0.471187507 | 2.210086036 |
| Gm23577 | 0 | NA | NA | NA |
| Tmem86b | 45.98200731 | 0.05834903 | 0.25293013 | 0.230692289 |
| Ppp6r1 | 1889.707859 | 0.429553849 | 0.062638894 | 6.857621876 |
| Hspbp1 | 559.9998375 | 0.20563243 | 0.081564627 | 2.521098144 |
| Brsk1 | 2659.133262 | -0.209716974 | 0.083693944 | -2.505760438 |
| Tmem150b | 4.110811422 | -0.375525374 | 0.502800565 | -0.746867447 |
| Gm24537 | 0 | NA | NA | NA |
| Suv420h2 | 136.5439044 | -0.577858944 | 0.169455971 | -3.410083115 |
| Cox6b2 | 14.20000006 | -0.643633718 | 0.325765052 | -1.975760488 |
| Fam71e2 | 0.747386132 | 1.196324253 | 1.356744558 | 0.881760863 |
| Il11 | 12.48613724 | 0.077406425 | 0.607587906 | 0.127399548 |
| Tmem190 | 0.273106308 | -0.687379814 | 1.801284754 | -0.381605303 |
| Tmem238 | 19.96996773 | -0.039201629 | 0.250691989 | -0.156373679 |
| Rpl28 | 949.3495926 | 0.454136587 | 0.099892745 | 4.546241945 |
| Ube2s | 303.8563974 | 0.171449914 | 0.115545751 | 1.483827075 |
| Shisa7 | 1255.260455 | 0.017286742 | 0.236710146 | 0.073029156 |
| Gm15525 | 0 | NA | NA | NA |
| Isoc2b | 8.838166061 | -0.444459924 | 0.381349111 | -1.165493537 |
| Isoc2a | 117.0217193 | -0.196172725 | 0.121717378 | -1.611706794 |
| Zfp628 | 234.1479967 | 0.81317568 | 0.139936932 | 5.811015477 |
| Nat14 | 284.6791148 | -0.157867176 | 0.114132475 | -1.383192431 |
| Ssc5d | 87.37008735 | 1.323601003 | 0.179835891 | 7.360049184 |
| Sbk2 | 2.02355919 | 0.630969712 | 0.695330295 | 0.907438834 |
| Sbk3 | 14.88936924 | -0.263880851 | 0.352244055 | -0.749142102 |
| Zfp579 | 184.6131844 | -0.397714389 | 0.106569016 | -3.731988946 |
| Fiz1 | 274.5806166 | 0.347996753 | 0.082550476 | 4.215563243 |
| Zfp524 | 43.21558028 | -0.194172689 | 0.204748129 | -0.948349025 |
| Zfp865 | 368.9050981 | 0.129745588 | 0.076140047 | 1.704038715 |
| 4632433K11Rik | 17.57441645 | 0.643965837 | 0.265741828 | 2.423276159 |
| Zfp784 | 56.52918824 | -0.207332505 | 0.167867954 | -1.235092821 |
| Gm15510 | 3.40753209 | -0.554759235 | 0.532464381 | -1.041871072 |
| Zfp580 | 125.3343534 | 0.194330904 | 0.1347096 | 1.442591353 |
| Ccdc106 | 206.5384023 | -0.061229271 | 0.110000589 | -0.556626757 |
| U2af2 | 1303.244093 | 0.120816882 | 0.055396506 | 2.180947669 |
| Epn1 | 2656.292776 | 0.022619283 | 0.072677197 | 0.311229431 |
| Rfpl4 | 0 | NA | NA | NA |
| Rasl2-9 | 0.99583754 | 0.011234352 | 1.032725952 | 0.010878347 |
| Gm26217 | 0 | NA | NA | NA |
| Vmn1r55 | 0.048949532 | -0.206487837 | 3.724945049 | -0.055433794 |
| Vmn1r56 | 0 | NA | NA | NA |
| Vmn1r57 | 0.085327104 | 0.274389469 | 3.724945049 | 0.073662689 |

| | | | | |
|---|---|---|---|---|
| Gm23942 | 0 | NA | NA | NA |
| Nlrp2 | 0 | NA | NA | NA |
| Gm5065 | 0.07796279 | 0.274392189 | 3.724945049 | 0.073663419 |
| Vmn1r58 | 0 | NA | NA | NA |
| Vmn1r59 | 0 | NA | NA | NA |
| Vmn2r28 | 0 | NA | NA | NA |
| Vmn1r60 | 0 | NA | NA | NA |
| Vmn1r-ps40 | 0 | NA | NA | NA |
| Vmn1r61 | 0 | NA | NA | NA |
| AC161762.1 | 0 | NA | NA | NA |
| Vmn1r62 | 0 | NA | NA | NA |
| Vmn1r63 | 0 | NA | NA | NA |
| Vmn1r64 | 0 | NA | NA | NA |
| Gm20399 | 0 | NA | NA | NA |
| Vmn1r-ps45 | 0 | NA | NA | NA |
| Vmn1r-ps46 | 0 | NA | NA | NA |
| Vmn1r65 | 0.115603922 | 0.274387056 | 3.724945049 | 0.073662041 |
| Gm5321 | 0.234623521 | -0.206484815 | 2.121654474 | -0.097322546 |
| Nlrp4c | 0 | NA | NA | NA |
| Zfp787 | 156.8006384 | 0.080094124 | 0.102239821 | 0.783394607 |
| Zfp444 | 319.2682102 | -0.053073216 | 0.061593621 | -0.861667412 |
| Galp | 0.278580731 | 1.222132892 | 1.794338423 | 0.681105011 |
| Zscan5b | 0.398992304 | 1.274373616 | 1.677130507 | 0.759853577 |
| Gm6792 | 0.123841226 | 0.274391173 | 3.724945049 | 0.073663147 |
| Zfp667 | 214.0871925 | -0.177921222 | 0.106596808 | -1.669104589 |
| Zfp583 | 81.78065755 | -0.306239526 | 0.1358619 | -2.254050082 |
| Zfp78 | 26.1290204 | -0.362045295 | 0.224755911 | -1.6108377 |
| Zfp28 | 117.9628295 | -0.0033491 | 0.098132084 | -0.034128489 |
| Smim17 | 178.4907971 | -0.583900507 | 0.169718499 | -3.440405797 |
| Gm16306 | 2.355736216 | -0.257901571 | 0.686807212 | -0.375507954 |
| Gm16307 | 0 | NA | NA | NA |
| Gm20715 | 6.848240459 | -1.118860846 | 0.43333279 | -2.581989805 |
| Olfr1344 | 0 | NA | NA | NA |
| Olfr1336 | 0 | NA | NA | NA |
| Olfr1346 | 0.047948698 | -0.206487837 | 3.724945049 | -0.055433794 |
| Olfr5 | 0.046345709 | 0.274391604 | 3.724945049 | 0.073663262 |
| Olfr1347 | 0.072025462 | 0.755288839 | 3.724945049 | 0.202765096 |
| Olfr1348 | 0 | NA | NA | NA |
| n-R5s150 | 0 | NA | NA | NA |
| Olfr1349 | 0.115957212 | 0.274402725 | 3.724945049 | 0.073666248 |
| Olfr1350 | 0 | NA | NA | NA |
| Zim1 | 102.3587915 | 0.031049646 | 0.187724474 | 0.165400094 |
| Peg3 | 8284.965525 | -0.17184506 | 0.077332068 | -2.222170749 |
| Usp29 | 1064.211711 | -0.178333878 | 0.099800142 | -1.786910072 |
| Gm27675 | 0 | NA | NA | NA |
| Mir3099 | 1.920827858 | -0.627818417 | 0.7790998 | -0.805825412 |
| Gm22649 | 0 | NA | NA | NA |

| | | | | |
|---|---|---|---|---|
| Aurkc | 1.107598099 | -0.201581094 | 0.88634613 | -0.227429315 |
| Zfp954 | 110.343462 | 0.131652756 | 0.145687999 | 0.903662324 |
| Zfp773 | 25.68791501 | -0.877132343 | 0.250629966 | -3.499710574 |
| Zfp418 | 91.99907945 | -0.419695905 | 0.157465207 | -2.665324688 |
| Zfp772 | 42.36616452 | -0.706839934 | 0.170442445 | -4.147088684 |
| Vmn2r29 | 3.485300844 | -1.066760007 | 0.562124801 | -1.897728058 |
| Gm18190 | 0.639457996 | -0.523217148 | 1.293597472 | -0.404466737 |
| Clcn4-2 | 1846.754651 | -0.062463672 | 0.05668784 | -1.101888383 |
| Mir5620 | 0.831589853 | 0.619480551 | 1.048677247 | 0.590725653 |
| Vmn2r30 | 0.25221175 | 0.274414507 | 2.131649039 | 0.128733437 |
| Gm3912 | 1.415212796 | -0.230258772 | 0.845413415 | -0.272362335 |
| Vmn2r31 | 0.094294407 | -0.206492286 | 3.724945049 | -0.055434988 |
| Vmn2r-ps36 | 0 | NA | NA | NA |
| Vmn2r32 | 0 | NA | NA | NA |
| Gm3563 | 0 | NA | NA | NA |
| Vmn2r33 | 0 | NA | NA | NA |
| Gm7648 | 0 | NA | NA | NA |
| Gm7651 | 0 | NA | NA | NA |
| Vmn2r-ps37 | 0.072025462 | 0.755288839 | 3.724945049 | 0.202765096 |
| Vmn2r34 | 0 | NA | NA | NA |
| Vmn2r-ps38 | 0 | NA | NA | NA |
| Vmn2r35 | 0.093141081 | -0.687373951 | 3.724945049 | -0.184532642 |
| Vmn2r-ps40 | 0.038981395 | 0.274393828 | 3.724945049 | 0.073663859 |
| Vmn2r36 | 0 | NA | NA | NA |
| Gm7230 | 0 | NA | NA | NA |
| Gm3978 | 0.160408561 | -0.206482807 | 3.457615423 | -0.059718269 |
| Gm20438 | 0.038981395 | 0.274393828 | 3.724945049 | 0.073663859 |
| Gm18200 | 0.398333997 | -1.094552346 | 1.534246888 | -0.713413437 |
| Gm4073 | 0 | NA | NA | NA |
| Gm3675 | 0 | NA | NA | NA |
| 2810047C21Rik1 | 0.042716945 | -0.206490061 | 3.724945049 | -0.055434391 |
| Gm3665 | 0 | NA | NA | NA |
| Vmn2r41 | 0.042716945 | -0.206490061 | 3.724945049 | -0.055434391 |
| Vmn2r42 | 0 | NA | NA | NA |
| Vmn2r43 | 0 | NA | NA | NA |
| Gm3654 | 0 | NA | NA | NA |
| Vmn2r-ps45 | 0.124853745 | 1.290908285 | 3.721447108 | 0.346883416 |
| Gm7696 | 0.676743848 | 0.289436329 | 1.107382836 | 0.261369708 |
| Vmn2r44 | 0.263039019 | 0.274414615 | 2.106825953 | 0.130250254 |
| Vmn2r-ps46 | 0 | NA | NA | NA |
| Vmn2r45 | 0.068097359 | -0.206487837 | 3.724945049 | -0.055433794 |
| Vmn2r-ps47 | 0 | NA | NA | NA |
| Vmn2r-ps49 | 0 | NA | NA | NA |
| Gm3584 | 0.047948698 | -0.206487837 | 3.724945049 | -0.055433794 |
| Gm3956 | 0 | NA | NA | NA |
| Gm3585 | 0 | NA | NA | NA |
| Gm3962 | 0.333123609 | -0.949162823 | 1.68046324 | -0.564822128 |

| | | | | |
|---|---|---|---|---|
| Gm18191 | 0.264294608 | -0.690304661 | 1.807677817 | -0.381873724 |
| Vmn2r-ps44 | 0 | NA | NA | NA |
| Gm3644 | 0 | NA | NA | NA |
| Gm20482 | 0 | NA | NA | NA |
| Vmn2r-ps43 | 0 | NA | NA | NA |
| Vmn2r40 | 0 | NA | NA | NA |
| Vmn2r-ps42 | 0 | NA | NA | NA |
| Vmn2r39 | 0 | NA | NA | NA |
| Zfp419 | 0.038981395 | 0.274393828 | 3.724945049 | 0.073663859 |
| Vmn2r38 | 0.068097359 | -0.206487837 | 3.724945049 | -0.055433794 |
| Gm5583 | 0 | NA | NA | NA |
| Gm18201 | 0 | NA | NA | NA |
| Vmn2r-ps41 | 0 | NA | NA | NA |
| Vmn2r37 | 0 | NA | NA | NA |
| Gm4011 | 0 | NA | NA | NA |
| Gm3623 | 0 | NA | NA | NA |
| Gm3616 | 0 | NA | NA | NA |
| Gm20287 | 0 | NA | NA | NA |
| Gm3611 | 0 | NA | NA | NA |
| Gm3972 | 0.099106094 | 0.274387056 | 3.724945049 | 0.073662041 |
| Gm18199 | 0.40494807 | 1.310587367 | 1.70050719 | 0.770703808 |
| Gm10302 | 0 | NA | NA | NA |
| Gm3683 | 0 | NA | NA | NA |
| Gm20345 | 0 | NA | NA | NA |
| Vmn2r46 | 0.048949532 | -0.206487837 | 3.724945049 | -0.055433794 |
| Vmn2r47 | 0 | NA | NA | NA |
| Vmn2r-ps51 | 0 | NA | NA | NA |
| Vmn2r-ps52 | 0 | NA | NA | NA |
| Vmn2r48 | 0 | NA | NA | NA |
| Vmn2r49 | 0 | NA | NA | NA |
| Vmn2r50 | 0.038981395 | 0.274393828 | 3.724945049 | 0.073663859 |
| Vmn2r51 | 0 | NA | NA | NA |
| Vmn2r52 | 0 | NA | NA | NA |
| Gm20679 | 0 | NA | NA | NA |
| Vmn1r66 | 0 | NA | NA | NA |
| Vmn1r-ps47 | 0 | NA | NA | NA |
| Vmn1r-ps48 | 0 | NA | NA | NA |
| Nlrp4d | 0.062426873 | 0.755288839 | 3.724945049 | 0.202765096 |
| Vmn1r67 | 0 | NA | NA | NA |
| Zik1 | 251.415385 | 0.062214977 | 0.091542549 | 0.679629057 |
| Vmn1r68 | 0 | NA | NA | NA |
| Vmn1r-ps49 | 0 | NA | NA | NA |
| Vmn1r69 | 0 | NA | NA | NA |
| Vmn1r-ps50 | 0 | NA | NA | NA |
| Vmn1r70 | 0 | NA | NA | NA |
| Vmn1r-ps51 | 0 | NA | NA | NA |
| Gm6900 | 2.818881342 | 0.380675686 | 0.643499776 | 0.591570813 |

| | | | | |
|---|---|---|---|---|
| Vmn1r-ps52 | 0 | NA | NA | NA |
| Nlrp4b | 0 | NA | NA | NA |
| Vmn1r71 | 0 | NA | NA | NA |
| Zscan4b | 0 | NA | NA | NA |
| Mir297-1 | 0 | NA | NA | NA |
| Zscan4c | 0 | NA | NA | NA |
| Zscan4d | 0 | NA | NA | NA |
| Gm26248 | 0 | NA | NA | NA |
| Zscan4e | 0 | NA | NA | NA |
| Gm23087 | 0 | NA | NA | NA |
| Zscan4f | 0 | NA | NA | NA |
| Vmn1r-ps53 | 0 | NA | NA | NA |
| Vmn1r72 | 0 | NA | NA | NA |
| Vmn1r73 | 0 | NA | NA | NA |
| Vmn1r-ps54 | 0 | NA | NA | NA |
| Vmn1r74 | 0 | NA | NA | NA |
| Vmn1r-ps55 | 0 | NA | NA | NA |
| Vmn1r75 | 0 | NA | NA | NA |
| Vmn1r-ps56 | 0 | NA | NA | NA |
| Vmn1r76 | 0 | NA | NA | NA |
| Vmn1r-ps57 | 0 | NA | NA | NA |
| Gm24414 | 1.280135606 | -0.600038873 | 0.955283606 | -0.628126422 |
| Vmn1r-ps58 | 0 | NA | NA | NA |
| Vmn1r-ps59 | 0 | NA | NA | NA |
| Vmn1r-ps60 | 0 | NA | NA | NA |
| Vmn1r77 | 0 | NA | NA | NA |
| Gm10179 | 23.18691661 | -0.149574996 | 0.240097188 | -0.622976876 |
| Vmn1r-ps61 | 0 | NA | NA | NA |
| Vmn1r-ps62 | 0 | NA | NA | NA |
| Vmn1r-ps63 | 0 | NA | NA | NA |
| Vmn1r78 | 0 | NA | NA | NA |
| Vmn1r79 | 0 | NA | NA | NA |
| Vmn1r80 | 0 | NA | NA | NA |
| Vmn1r-ps64 | 0 | NA | NA | NA |
| Vmn1r-ps65 | 0 | NA | NA | NA |
| Vmn1r81 | 0 | NA | NA | NA |
| Vmn1r82 | 0 | NA | NA | NA |
| Vmn1r83 | 0 | NA | NA | NA |
| Gm23521 | 0 | NA | NA | NA |
| Vmn1r84 | 0.047948698 | -0.206487837 | 3.724945049 | -0.055433794 |
| Vmn1r-ps66 | 0 | NA | NA | NA |
| Vmn1r-ps67 | 0 | NA | NA | NA |
| Vmn1r-ps68 | 0 | NA | NA | NA |
| Zfp551 | 75.69228729 | -0.218402331 | 0.121729654 | -1.794158817 |
| Vmn2r-ps53 | 0 | NA | NA | NA |
| Zfp606 | 312.4159778 | -0.107194281 | 0.071807556 | -1.492799478 |
| 2900092C05Rik | 0.326351934 | 0.275584418 | 1.708746549 | 0.161278698 |

| | | | | |
|---|---|---|---|---|
| Gm15877 | 0.593305459 | 1.334749967 | 1.246247932 | 1.071014789 |
| Vmn2r53 | 2.096357237 | 0.262107964 | 0.654649097 | 0.400379326 |
| Gm15773 | 0.596424136 | -0.345062667 | 1.316857274 | -0.262034978 |
| Vmn2r54 | 1.895285101 | 0.959832741 | 0.839699929 | 1.14306636 |
| Vmn2r55 | 1.524911348 | 0.810716567 | 0.834819862 | 0.97112755 |
| Vmn2r56 | 2.422070769 | 1.923486143 | 0.923390631 | 2.083068724 |
| Gm6946 | 0 | NA | NA | NA |
| Zscan18 | 180.8411845 | 0.161250715 | 0.145370127 | 1.109242441 |
| Zfp329 | 519.818062 | -0.585326462 | 0.125095265 | -4.679045691 |
| Zfp110 | 270.5868238 | 0.634304303 | 0.116662809 | 5.437073793 |
| Zfp128 | 118.8763104 | -0.274360602 | 0.135502176 | -2.024768974 |
| Zscan22 | 156.5051589 | -0.351402863 | 0.160508113 | -2.189315277 |
| Rps5 | 1620.953915 | 1.048348617 | 0.129679439 | 8.084154505 |
| Rnf225 | 104.6098484 | 2.745966013 | 0.225710233 | 12.16589066 |
| Gm26325 | 0 | NA | NA | NA |
| 1810019N24Rik | 6.340267528 | 1.040725593 | 0.491056655 | 2.119359512 |
| Gm29638 | 0.046345709 | 0.274391604 | 3.724945049 | 0.073663262 |
| Zfp324 | 270.4925259 | 0.745466224 | 0.110300711 | 6.758489736 |
| Zfp446 | 119.2751431 | 0.398434843 | 0.144995215 | 2.747917189 |
| Slc27a5 | 10.7500977 | -0.577556945 | 0.349449574 | -1.652761906 |
| Zbtb45 | 249.6021384 | 0.23250783 | 0.099818642 | 2.329302669 |
| Trim28 | 1232.544845 | 0.291956267 | 0.072902131 | 4.004770006 |
| Chmp2a | 510.643161 | 0.307073657 | 0.108922175 | 2.819202409 |
| Ube2m | 1024.417635 | 0.18212251 | 0.082674909 | 2.202875239 |
| Mzf1 | 76.8160005 | -0.313335507 | 0.186054456 | -1.684106436 |
| Vmn1r85 | 0 | NA | NA | NA |
| Vmn1r86 | 0 | NA | NA | NA |
| Vmn1r87 | 0 | NA | NA | NA |
| Vmn1r88 | 0.289463044 | 1.635748776 | 2.011704171 | 0.813115964 |
| Gm4879 | 6.771121092 | 0.473890241 | 0.472500281 | 1.002941712 |
| Vmn1r89 | 0.099106094 | 0.274387056 | 3.724945049 | 0.073662041 |
| Vmn1r-ps69 | 0 | NA | NA | NA |
| 6330408A02Rik | 197.7921584 | -0.487365502 | 0.182631503 | -2.668573019 |
| Lig1 | 167.3687248 | 0.244036036 | 0.126581526 | 1.92789615 |
| Pla2g4c | 22.31652208 | 0.213558866 | 0.327407641 | 0.65227209 |
| Rps8-ps4 | 2.214157104 | 0.583131763 | 0.679330799 | 0.85839147 |
| Gm24909 | 0.201760502 | 0.274410885 | 2.223929814 | 0.123390083 |
| Gm23499 | 0 | NA | NA | NA |
| Cabp5 | 0.609661125 | -1.147705648 | 1.302557364 | -0.881117162 |
| Bsph1 | 0.222333927 | 1.636331882 | 2.128347805 | 0.768827293 |
| Bsph2-ps | 0 | NA | NA | NA |
| Bsph2 | 0 | NA | NA | NA |
| Sult2a5 | 0 | NA | NA | NA |
| Zbed4-ps1 | 0 | NA | NA | NA |
| Kat6b-ps1 | 0 | NA | NA | NA |
| Tmem167-ps1 | 0 | NA | NA | NA |
| Sult2a-ps1 | 0 | NA | NA | NA |

| | | | | |
|---|---|---|---|---|
| Sult2a2 | 0 | NA | NA | NA |
| Zbed4-ps2 | 0.047948698 | -0.206487837 | 3.724945049 | -0.055433794 |
| Sult2a1 | 0.169165009 | 0.755292496 | 3.351424224 | 0.225364635 |
| Kat6b-ps2 | 0 | NA | NA | NA |
| Sult2a-ps2 | 0 | NA | NA | NA |
| Tmem167-ps2 | 0 | NA | NA | NA |
| Sult2a4 | 0 | NA | NA | NA |
| Zbed4-ps3 | 0.112642008 | 0.274402725 | 3.724945049 | 0.073666248 |
| Tmem167-ps3 | 0 | NA | NA | NA |
| Sult2a3 | 0 | NA | NA | NA |
| Sult2a-ps3 | 0 | NA | NA | NA |
| Sult2a-ps4 | 0 | NA | NA | NA |
| Rpap3-ps1 | 0 | NA | NA | NA |
| Sult2a6 | 0 | NA | NA | NA |
| Rpap3-ps2 | 0 | NA | NA | NA |
| Phf20-ps | 17.46056627 | 0.144544991 | 0.250616473 | 0.576757742 |
| Obox8 | 0 | NA | NA | NA |
| Mrps36-ps3 | 0 | NA | NA | NA |
| Crs-ps | 0 | NA | NA | NA |
| Sult2a8 | 0.047948698 | -0.206487837 | 3.724945049 | -0.055433794 |
| Sult2a7 | 0.119587853 | 1.218392466 | 3.723618458 | 0.327206581 |
| Obox4-ps1 | 0.155414793 | 0.274408395 | 2.82937401 | 0.09698555 |
| Obox4-ps2 | 0.183158741 | -1.142233453 | 2.72428501 | -0.419278251 |
| Nlrp5-ps | 183.9697913 | 0.067142201 | 0.174082298 | 0.385692294 |
| Obox4-ps3 | 0 | NA | NA | NA |
| Vmn1r90 | 3.490537078 | -0.01744011 | 0.57226596 | -0.030475533 |
| Zfp-ps | 0 | NA | NA | NA |
| Ranbp2-ps1 | 0 | NA | NA | NA |
| Obox3-ps7 | 4.106584469 | -0.566983548 | 0.518971772 | -1.09251327 |
| Obox7 | 0.129647038 | -0.687376656 | 2.992845844 | -0.229673258 |
| Obox3-ps1 | 0 | NA | NA | NA |
| RP24-194K24.10 | 0 | NA | NA | NA |
| Gtpbp4-ps1 | 0 | NA | NA | NA |
| Ranbp2-ps2 | 0 | NA | NA | NA |
| Obox4-ps36 | 0 | NA | NA | NA |
| Obox4-ps37 | 0 | NA | NA | NA |
| Gm26056 | 0 | NA | NA | NA |
| Obox3-ps2 | 0 | NA | NA | NA |
| RP23-349I13.7 | 0 | NA | NA | NA |
| Gtpbp4-ps2 | 0 | NA | NA | NA |
| Ranbp2-ps3 | 0 | NA | NA | NA |
| Obox4-ps38 | 0 | NA | NA | NA |
| Obox4-ps39 | 0 | NA | NA | NA |
| Ranbp2-ps4 | 0 | NA | NA | NA |
| Obox4-ps6 | 0 | NA | NA | NA |
| Gm24701 | 0 | NA | NA | NA |
| Obox3-ps3 | 0 | NA | NA | NA |

| | | | | |
|---|---|---|---|---|
| RP23-263E7.8 | 0 | NA | NA | NA |
| Gtpbp4-ps3 | 0 | NA | NA | NA |
| Ranbp2-ps5 | 0 | NA | NA | NA |
| Obox4-ps8 | 0 | NA | NA | NA |
| Obox4-ps9 | 0 | NA | NA | NA |
| Obox3-ps4 | 0 | NA | NA | NA |
| RP23-263E7.7 | 0 | NA | NA | NA |
| Gtpbp4-ps4 | 0 | NA | NA | NA |
| Ranbp2-ps6 | 0 | NA | NA | NA |
| Obox4-ps11 | 0 | NA | NA | NA |
| Obox4-ps12 | 0 | NA | NA | NA |
| Obox4-ps13 | 0 | NA | NA | NA |
| Obox4-ps14 | 0 | NA | NA | NA |
| Obox4-ps15 | 0 | NA | NA | NA |
| Obox4-ps16 | 0 | NA | NA | NA |
| Obox4-ps17 | 0 | NA | NA | NA |
| Obox4-ps18 | 0 | NA | NA | NA |
| Obox4-ps19 | 0 | NA | NA | NA |
| Obox4-ps20 | 0 | NA | NA | NA |
| Obox4-ps21 | 0 | NA | NA | NA |
| Obox4-ps22 | 0 | NA | NA | NA |
| Obox4-ps23 | 0 | NA | NA | NA |
| Mrip-ps | 35.07510409 | 0.849805203 | 0.352954593 | 2.407689885 |
| Obox3-ps5 | 0 | NA | NA | NA |
| RP24-84K14.12 | 0 | NA | NA | NA |
| Gtpbp4-ps5 | 0 | NA | NA | NA |
| RP24-105I10.4 | 0 | NA | NA | NA |
| Obox4-ps26 | 0 | NA | NA | NA |
| Ranbp2-ps7 | 0 | NA | NA | NA |
| Obox2 | 0 | NA | NA | NA |
| Obox1 | 0 | NA | NA | NA |
| Ranbp2-ps8 | 0 | NA | NA | NA |
| Obox4-ps27 | 0 | NA | NA | NA |
| Obox4-ps28 | 0 | NA | NA | NA |
| Obox4-ps29 | 0 | NA | NA | NA |
| Obox3 | 0 | NA | NA | NA |
| Obox3-ps6 | 0 | NA | NA | NA |
| Obox4-ps30 | 0 | NA | NA | NA |
| Ranbp2-ps9 | 0 | NA | NA | NA |
| Ranbp2-ps10 | 0 | NA | NA | NA |
| Obox4-ps31 | 0 | NA | NA | NA |
| Gm25354 | 0 | NA | NA | NA |
| RP23-140B24.1 | 0 | NA | NA | NA |
| RP23-140B24.2 | 0 | NA | NA | NA |
| Obox4-ps33 | 0 | NA | NA | NA |
| Gm23714 | 0 | NA | NA | NA |
| Obox4-ps34 | 0 | NA | NA | NA |

| | | | | |
|---|---|---|---|---|
| RP24-370G23.12 | 0 | NA | NA | NA |
| Obox5 | 0 | NA | NA | NA |
| RP24-370G23.13 | 0 | NA | NA | NA |
| RP24-370G23.14 | 0 | NA | NA | NA |
| Obox4-ps35 | 0.220980774 | 0.274398268 | 3.11985971 | 0.087952118 |
| Obox6 | 0.123845245 | 0.755285145 | 3.724945049 | 0.202764104 |
| Crx | 0.603215896 | 2.044749381 | 2.55126862 | 0.801463776 |
| Crxos | 0.501748666 | -2.131596082 | 1.283847841 | -1.660318313 |
| RP24-466N11.6 | 0 | NA | NA | NA |
| Sepw1 | 5104.264334 | 0.964794346 | 0.096652969 | 9.982045604 |
| Gltscr2 | 737.8364871 | 0.603353651 | 0.093335733 | 6.464337182 |
| Snord23 | 0.926190085 | 1.049539159 | 1.20557344 | 0.870572562 |
| Gm24576 | 0.169505627 | 0.274410251 | 2.749549019 | 0.099801913 |
| Ehd2 | 172.4468587 | 1.501574602 | 0.194798955 | 7.708329871 |
| Gltscr1 | 438.3260183 | 0.101454163 | 0.077356863 | 1.31150824 |
| Zfp541 | 3.78555653 | -0.427769524 | 0.503188478 | -0.850117884 |
| Napa | 1603.381598 | -0.047368317 | 0.056174946 | -0.843228518 |
| Kptn | 155.1087845 | -0.451467228 | 0.103337446 | -4.368863817 |
| Slc8a2 | 2081.402497 | -0.468290495 | 0.139779474 | -3.350209303 |
| Meis3 | 261.2098945 | 0.202626716 | 0.102427213 | 1.978250806 |
| Dhx34 | 160.4271404 | 0.252636493 | 0.089946494 | 2.808741971 |
| C5ar2 | 2.109974762 | 1.536249894 | 0.942865441 | 1.629341609 |
| Gm26473 | 0 | NA | NA | NA |
| C5ar1 | 78.4729806 | 4.182367096 | 0.403459638 | 10.36625898 |
| Inafm1 | 160.8892379 | -0.227422926 | 0.110923443 | -2.050269272 |
| Ccdc9 | 366.0675831 | 0.016483661 | 0.102110848 | 0.161429087 |
| Bbc3 | 74.98142202 | 0.329949598 | 0.14725075 | 2.240732895 |
| Sae1 | 546.3005839 | -0.170525973 | 0.062572953 | -2.725234535 |
| Zc3h4 | 1014.843218 | -0.054125361 | 0.076727535 | -0.705422909 |
| Tmem160 | 153.6281965 | 0.031033543 | 0.149248044 | 0.20793266 |
| Npas1 | 76.47707439 | 0.083074183 | 0.143267039 | 0.579855514 |
| n-R5s151 | 0 | NA | NA | NA |
| Arhgap35 | 4302.533817 | 0.04203654 | 0.051514847 | 0.816008257 |
| Gm17826 | 0.07796279 | 0.274392189 | 3.724945049 | 0.073663419 |
| Gm17981 | 12.95166994 | -0.349192406 | 0.381571043 | -0.915143883 |
| Gm29443 | 4.384705177 | -0.373594946 | 0.457254247 | -0.817039859 |
| Gm29442 | 0.132101749 | 0.274387056 | 3.724945049 | 0.073662041 |
| Gm24708 | 0 | NA | NA | NA |
| Ceacam15 | 0.205464717 | 0.27440636 | 2.643378516 | 0.103808954 |
| Ceacam9 | 1.215943724 | -0.354964911 | 0.805843687 | -0.440488542 |
| Ap2s1 | 780.4435061 | 0.029655694 | 0.097456907 | 0.304295457 |
| Slc1a5 | 49.69459424 | 1.212395445 | 0.240413565 | 5.042957719 |
| Fkrp | 486.4658568 | 0.072493639 | 0.076372864 | 0.94920677 |
| Strn4 | 1924.374574 | 0.187954519 | 0.078474216 | 2.395111787 |
| Prkd2 | 82.10841754 | 0.261522206 | 0.185197161 | 1.412128592 |
| 9330104G04Rik | 30.06220494 | -0.368289017 | 0.189760644 | -1.940808215 |
| Dact3 | 1250.275623 | -0.136531491 | 0.098674975 | -1.383648594 |

| | | | | |
|---|---|---|---|---|
| Gng8 | 8.57217951 | -0.774237242 | 0.359676159 | -2.152595387 |
| Ptgir | 4.875732241 | 2.755168559 | 0.939174158 | 2.93360772 |
| Calm3 | 11988.4177 | -0.035867975 | 0.069338935 | -0.517284775 |
| Pnmal2 | 4095.934161 | -0.179912817 | 0.074347069 | -2.419904637 |
| Gm25815 | 0 | NA | NA | NA |
| Pnmal1 | 642.282333 | -0.249160466 | 0.104327707 | -2.388248265 |
| Ccdc8 | 31.97129462 | -0.717083174 | 0.253586414 | -2.827766523 |
| Ppp5c | 936.7635144 | -0.021296759 | 0.052915477 | -0.402467476 |
| Hif3a | 169.2274098 | 0.293350802 | 0.197954613 | 1.481909401 |
| Psg16 | 99.26004857 | 0.270376194 | 0.187510458 | 1.441925946 |
| Ceacam3 | 2.261397335 | -0.533074093 | 0.725501072 | -0.734766789 |
| Psg29 | 2.218822718 | 0.44769726 | 0.740994173 | 0.604184589 |
| Gm24520 | 0 | NA | NA | NA |
| Gm25068 | 0 | NA | NA | NA |
| Ceacam5 | 0 | NA | NA | NA |
| Gm6065 | 0.197744397 | -1.17479164 | 2.224521995 | -0.528109698 |
| Ceacam14 | 0 | NA | NA | NA |
| Gm5155 | 0 | NA | NA | NA |
| Ceacam11 | 0 | NA | NA | NA |
| Ceacam13 | 0 | NA | NA | NA |
| Gm22414 | 0 | NA | NA | NA |
| Ceacam12 | 0 | NA | NA | NA |
| Igfl3 | 0 | NA | NA | NA |
| Mill1 | 0 | NA | NA | NA |
| Psg18 | 0 | NA | NA | NA |
| Gm24715 | 0 | NA | NA | NA |
| Psg28 | 0.047948698 | -0.206487837 | 3.724945049 | -0.055433794 |
| Psg26 | 0 | NA | NA | NA |
| Psg25 | 0.049553047 | 0.274391604 | 3.724945049 | 0.073663262 |
| Gm26062 | 0 | NA | NA | NA |
| Psg27 | 0 | NA | NA | NA |
| Psg23 | 0.066050875 | 0.274391604 | 3.724945049 | 0.073663262 |
| Psg21 | 0 | NA | NA | NA |
| Psg20 | 0.124347486 | 0.755286736 | 3.717316019 | 0.203180664 |
| Psg22 | 0 | NA | NA | NA |
| Gm24464 | 0 | NA | NA | NA |
| Psg19 | 0 | NA | NA | NA |
| Psg17 | 1.789359587 | 0.978833471 | 1.051559174 | 0.930840123 |
| Mill2 | 14.66817284 | 3.375263139 | 0.485419139 | 6.95329637 |
| Pglyrp1 | 28.41791276 | 1.674730553 | 0.282478557 | 5.928699761 |
| Ccdc61 | 68.71592428 | -0.360054503 | 0.177356977 | -2.030111857 |
| Nova2 | 1854.132093 | -0.33865706 | 0.105920658 | -3.197271102 |
| Gm26821 | 0.975505809 | -0.799371728 | 1.212131333 | -0.659476169 |
| Nanos2 | 10.4316941 | 0.505555304 | 0.393651385 | 1.284271625 |
| Gm10676 | 6.752775585 | -0.32501489 | 0.378870532 | -0.857852122 |
| Mypop | 377.759842 | -0.425345532 | 0.087053078 | -4.886048145 |
| Mypopos | 3.140367151 | 0.461123077 | 0.674037656 | 0.684120647 |

| | | | | |
|---|---|---|---|---|
| Irf2bp1 | 595.038875 | 0.018060937 | 0.076832854 | 0.235067896 |
| Foxa3 | 0.267510113 | -0.206490028 | 2.425014883 | -0.085150005 |
| Sympk | 1388.597211 | 0.004858867 | 0.059083683 | 0.082237041 |
| Rsph6a | 1.359012152 | -1.025551284 | 0.910268083 | -1.126647527 |
| Dmwd | 973.7957556 | 0.238342073 | 0.078291186 | 3.044302754 |
| Dmpk | 178.6755614 | -0.291458632 | 0.203801651 | -1.430109272 |
| Mir3100 | 1.552752546 | -0.724882165 | 0.877446513 | -0.826126897 |
| Six5 | 17.71455458 | -0.758592412 | 0.281946764 | -2.69055194 |
| Gm4969 | 2.587362481 | 0.024474598 | 0.79690352 | 0.030712122 |
| Fbxo46 | 157.8854747 | -0.325477733 | 0.149243931 | -2.180844001 |
| Qpctl | 177.5083971 | 0.545333931 | 0.126605845 | 4.307336126 |
| Snrpd2 | 249.1707862 | 0.135329846 | 0.124110645 | 1.09039677 |
| Gipr | 9.076618891 | -0.568768947 | 0.447257722 | -1.271680552 |
| Eml2 | 1469.733199 | -0.069364915 | 0.071271463 | -0.973249489 |
| Mir330 | 0 | NA | NA | NA |
| Gpr4 | 26.0798917 | 0.273740605 | 0.221654135 | 1.234989838 |
| Opa3 | 585.1491548 | -0.050733736 | 0.064169908 | -0.790615681 |
| Gm25134 | 1.373839161 | -0.848032959 | 0.840803502 | -1.008598272 |
| Vasp | 344.2888316 | 1.331988598 | 0.144944038 | 9.189674958 |
| 1700058P15Rik | 0 | NA | NA | NA |
| Ppm1n | 0.85008157 | 1.515531229 | 1.298916864 | 1.16676538 |
| Rtn2 | 574.5812076 | 0.179163301 | 0.100428054 | 1.783996542 |
| Gm26802 | 0.561122788 | 1.835594781 | 1.648664857 | 1.11338261 |
| Fosb | 461.8672414 | 0.425326184 | 0.613238793 | 0.693573513 |
| Ercc1 | 157.0107263 | 0.174195487 | 0.12430881 | 1.401312478 |
| Cd3eap | 115.4671044 | 0.121882344 | 0.121949553 | 0.999448881 |
| Ppp1r13l | 62.86165383 | 0.000642224 | 0.175751048 | 0.003654167 |
| Ercc2 | 283.3150772 | -0.016639966 | 0.083067908 | -0.200317623 |
| Gm26852 | 0.604815377 | -0.148349325 | 1.275576264 | -0.116299847 |
| Mir343 | 1.738846634 | -0.811291136 | 0.748241618 | -1.08426358 |
| Klc3 | 30.34969473 | 0.875871425 | 0.217077467 | 4.034833454 |
| Ckm | 1.573181334 | 0.075169961 | 0.899003011 | 0.083614804 |
| A930016O22Rik | 2.771915612 | -0.797391709 | 0.574516886 | -1.38793433 |
| Mark4 | 736.3036542 | 0.06796612 | 0.06861545 | 0.990536686 |
| Gm23099 | 0 | NA | NA | NA |
| Exoc3l2 | 2.258282039 | -0.502003484 | 0.677261798 | -0.74122516 |
| Bloc1s3 | 84.9281982 | -0.255792008 | 0.124318492 | -2.057553989 |
| Trappc6a | 46.60324668 | -0.125397338 | 0.179433316 | -0.698852038 |
| Nkpd1 | 9.973239763 | 0.002359385 | 0.316834435 | 0.007446743 |
| Ppp1r37 | 1313.266029 | -0.030666689 | 0.07325888 | -0.418607127 |
| Gemin7 | 188.0515057 | 0.407419351 | 0.104777349 | 3.888429658 |
| Zfp296 | 23.1978082 | 1.400823223 | 0.282586317 | 4.95715164 |
| Clasrp | 352.6239218 | 0.054588248 | 0.175756032 | 0.310591032 |
| Gm26890 | 51.60349904 | -0.560068138 | 0.349430363 | -1.602803298 |
| Relb | 145.3464235 | 1.75482225 | 0.239276398 | 7.333871045 |
| Clptm1 | 1928.605293 | 0.170465257 | 0.064521631 | 2.641986184 |
| Apoc2 | 34.46812154 | 4.628521634 | 0.633221528 | 7.309482426 |

| | | | | |
|---|---|---|---|---|
| Apoc4 | 0.250169628 | 0.274414486 | 2.136490353 | 0.128441715 |
| Apoc1 | 11.42902807 | 0.915424491 | 0.371258819 | 2.4657313 |
| Apoe | 8648.195503 | 0.012830336 | 0.177167723 | 0.072419148 |
| Tomm40 | 595.2610474 | -0.055083389 | 0.065647277 | -0.839081101 |
| Gm20512 | 1.197749286 | -0.532103321 | 0.911767665 | -0.583595297 |
| Pvrl2 | 109.3428072 | 0.574704289 | 0.166265034 | 3.456555338 |
| Bcam | 112.1762543 | -0.217512738 | 0.15005303 | -1.449572451 |
| Gm26600 | 0 | NA | NA | NA |
| Cblc | 0.325981489 | -0.188726483 | 1.503910654 | -0.125490489 |
| Bcl3 | 136.661948 | 4.850031723 | 0.43973197 | 11.0295181 |
| Gm16175 | 0.542127486 | -1.122143301 | 1.202675103 | -0.933039437 |
| Gm16174 | 1.534203801 | 4.176259445 | 1.019196507 | 4.09759984 |
| Gm19345 | 14.35855545 | -0.432674857 | 0.287131825 | -1.506885758 |
| Ceacam16 | 1.967062318 | 1.20738023 | 0.784269899 | 1.539495819 |
| Ceacam19 | 1.492916405 | 2.401008842 | 1.173986088 | 2.045176571 |
| Pvr | 193.8546322 | 0.460302309 | 0.183655884 | 2.506330314 |
| Gm16184 | 0 | NA | NA | NA |
| Gm16251 | 0.990102919 | -0.655777023 | 0.958320303 | -0.684298372 |
| Igsf23 | 0.400433843 | 0.755307721 | 1.688546295 | 0.447312415 |
| Rpl7a-ps8 | 0.1140246 | 0.75528439 | 3.724945049 | 0.202763901 |
| Ceacam20 | 0.152347716 | 1.221984416 | 2.842792129 | 0.429853595 |
| Nlrp9b | 0 | NA | NA | NA |
| n-R5s152 | 0 | NA | NA | NA |
| Vmn1r91 | 0 | NA | NA | NA |
| Gm16442 | 0 | NA | NA | NA |
| Vmn1r93 | 0 | NA | NA | NA |
| Vmn1r94 | 0 | NA | NA | NA |
| Vmn1r95 | 0 | NA | NA | NA |
| Gm6164 | 0 | NA | NA | NA |
| Gm4498 | 0 | NA | NA | NA |
| Gm4133 | 0 | NA | NA | NA |
| Vmn1r-ps79 | 0 | NA | NA | NA |
| Gm5728 | 0 | NA | NA | NA |
| Vmn1r100 | 0 | NA | NA | NA |
| Vmn1r101 | 0 | NA | NA | NA |
| Gm10670 | 0 | NA | NA | NA |
| Vmn1r103 | 0 | NA | NA | NA |
| Vmn1r104 | 0 | NA | NA | NA |
| Gm5726 | 0 | NA | NA | NA |
| Gm4141 | 0 | NA | NA | NA |
| Vmn1r107 | 0 | NA | NA | NA |
| Vmn1r-ps71 | 0 | NA | NA | NA |
| Vmn1r111 | 0 | NA | NA | NA |
| Gm10668 | 0 | NA | NA | NA |
| Vmn1r112 | 0 | NA | NA | NA |
| Vmn1r113 | 0 | NA | NA | NA |
| Vmn1r114 | 0 | NA | NA | NA |

| | | | | |
|---|---|---|---|---|
| Vmn1r115 | 0 | NA | NA | NA |
| Vmn1r116 | 0 | NA | NA | NA |
| Vmn1r117 | 0 | NA | NA | NA |
| Vmn1r118 | 0 | NA | NA | NA |
| Gm4526 | 0 | NA | NA | NA |
| Vmn1r119 | 0 | NA | NA | NA |
| Vmn1r-ps73 | 0 | NA | NA | NA |
| Vmn1r120 | 0 | NA | NA | NA |
| Vmn1r-ps74 | 0 | NA | NA | NA |
| Vmn1r121 | 0 | NA | NA | NA |
| Vmn1r122 | 0 | NA | NA | NA |
| Vmn1r123 | 0 | NA | NA | NA |
| Gm5157 | 0 | NA | NA | NA |
| Vmn1r124 | 0 | NA | NA | NA |
| Vmn1r125 | 0 | NA | NA | NA |
| Vmn1r126 | 0 | NA | NA | NA |
| Vmn1r127 | 0 | NA | NA | NA |
| Vmn1r128 | 0 | NA | NA | NA |
| Vmn1r129 | 0 | NA | NA | NA |
| Vmn1r-ps75 | 0 | NA | NA | NA |
| Gm6882 | 0 | NA | NA | NA |
| Vmn1r130 | 0 | NA | NA | NA |
| Vmn1r-ps76 | 0 | NA | NA | NA |
| Vmn1r131 | 0 | NA | NA | NA |
| Gm4545 | 0 | NA | NA | NA |
| Vmn1r132 | 0 | NA | NA | NA |
| Gm8600 | 0 | NA | NA | NA |
| Gm4175 | 0 | NA | NA | NA |
| Gm4177 | 0 | NA | NA | NA |
| Vmn1r135 | 0 | NA | NA | NA |
| Gm5725 | 0 | NA | NA | NA |
| Vmn1r-ps77 | 0 | NA | NA | NA |
| Gm5891 | 0 | NA | NA | NA |
| Vmn1r137 | 0 | NA | NA | NA |
| Vmn1r-ps78 | 0 | NA | NA | NA |
| Vmn1r138 | 0 | NA | NA | NA |
| Gm6176 | 0 | NA | NA | NA |
| Vmn1r139 | 0 | NA | NA | NA |
| AC127227.1 | 0 | NA | NA | NA |
| Gm16451 | 0 | NA | NA | NA |
| Gm10666 | 0 | NA | NA | NA |
| Vmn1r142 | 0 | NA | NA | NA |
| Vmn1r143 | 0 | NA | NA | NA |
| Gm8653 | 0 | NA | NA | NA |
| Gm8453 | 0 | NA | NA | NA |
| Gm4557 | 0 | NA | NA | NA |
| Gm4558 | 0 | NA | NA | NA |

| | | | | |
|---|---|---|---|---|
| Gm4187 | 0 | NA | NA | NA |
| Vmn1r-ps70 | 0 | NA | NA | NA |
| Gm8660 | 0 | NA | NA | NA |
| Vmn1r148 | 0 | NA | NA | NA |
| Vmn1r149 | 0 | NA | NA | NA |
| Gm10665 | 0 | NA | NA | NA |
| Vmn1r151 | 0 | NA | NA | NA |
| Vmn1r152 | 0 | NA | NA | NA |
| Gm8677 | 0 | NA | NA | NA |
| Gm4201 | 0 | NA | NA | NA |
| Gm4565 | 0 | NA | NA | NA |
| Vmn1r155 | 0 | NA | NA | NA |
| Vmn1r-ps80 | 0 | NA | NA | NA |
| Gm8693 | 0 | NA | NA | NA |
| Gm4567 | 0 | NA | NA | NA |
| Vmn1r157 | 0 | NA | NA | NA |
| Vmn1r158 | 0 | NA | NA | NA |
| Gm4207 | 0 | NA | NA | NA |
| Vmn1r159 | 0 | NA | NA | NA |
| Vmn1r160 | 0 | NA | NA | NA |
| Gm8708 | 0 | NA | NA | NA |
| Gm4214 | 0 | NA | NA | NA |
| Gm4216 | 0 | NA | NA | NA |
| Vmn1r163 | 0 | NA | NA | NA |
| Gm8720 | 0 | NA | NA | NA |
| Gm16456 | 0 | NA | NA | NA |
| Gm10662 | 0 | NA | NA | NA |
| Vmn1r165 | 0 | NA | NA | NA |
| Vmn1r-ps83 | 0 | NA | NA | NA |
| Vmn1r166 | 0 | NA | NA | NA |
| Gm6902 | 0 | NA | NA | NA |
| Nlrp4e | 0 | NA | NA | NA |
| Nlrp5 | 1.484051332 | -2.584554619 | 1.132839763 | -2.281482962 |
| Gm25196 | 0.066050875 | 0.274391604 | 3.724945049 | 0.073663262 |
| Vmn1r-ps84 | 0 | NA | NA | NA |
| Vmn1r167 | 0 | NA | NA | NA |
| Vmn1r168 | 0.047948698 | -0.206487837 | 3.724945049 | -0.055433794 |
| Vmn1r169 | 0 | NA | NA | NA |
| Vmn1r170 | 0 | NA | NA | NA |
| Vmn1r171 | 0 | NA | NA | NA |
| Vmn1r172 | 0 | NA | NA | NA |
| Vmn1r173 | 0 | NA | NA | NA |
| Vmn1r174 | 0 | NA | NA | NA |
| Vmn1r-ps85 | 0 | NA | NA | NA |
| Vmn1r175 | 0 | NA | NA | NA |
| Vmn1r176 | 0 | NA | NA | NA |
| Vmn1r177 | 0 | NA | NA | NA |

| | | | | |
|---|---|---|---|---|
| Vmn1r178 | 0 | NA | NA | NA |
| Vmn1r179 | 0 | NA | NA | NA |
| Gm10175 | 13.75409139 | -0.035410822 | 0.341948687 | -0.103555953 |
| Vmn1r180 | 0 | NA | NA | NA |
| Vmn1r181 | 0 | NA | NA | NA |
| V1rd19 | 0 | NA | NA | NA |
| Vmn1r183 | 0 | NA | NA | NA |
| Ptma-ps1 | 0 | NA | NA | NA |
| Zfp180 | 620.834939 | -0.074637342 | 0.092019113 | -0.811106955 |
| Zfp112 | 130.5214199 | 0.019860561 | 0.163575347 | 0.12141537 |
| Zfp235 | 143.0420723 | -0.416649016 | 0.124014327 | -3.359684541 |
| Zfp114 | 11.47062778 | -0.229851525 | 0.349878107 | -0.656947436 |
| Zfp111 | 197.4067323 | -0.421509812 | 0.160778683 | -2.621677232 |
| Zfp109 | 90.1039288 | -0.550436324 | 0.15067987 | -3.653018324 |
| Zfp108 | 66.9010371 | -0.538725723 | 0.212264141 | -2.537996856 |
| Zfp93 | 102.2549038 | -0.418706345 | 0.139652559 | -2.998200296 |
| Zfp61 | 139.4922126 | 0.140601733 | 0.128901966 | 1.090764845 |
| Zfp94 | 74.342771 | -0.280755291 | 0.150770299 | -1.862139248 |
| Gm15465 | 0 | NA | NA | NA |
| Tescl | 0 | NA | NA | NA |
| Gm27592 | 0.061920613 | 0.274393828 | 3.724945049 | 0.073663859 |
| Lypd5 | 0.324847777 | 2.034049876 | 3.702850821 | 0.549319963 |
| Kcnn4 | 8.685732437 | 1.905200849 | 0.53401434 | 3.567696046 |
| Smg9 | 288.8311134 | 0.18031311 | 0.077695498 | 2.320766522 |
| Irgc1 | 0.04393175 | -0.206490061 | 3.724945049 | -0.055434391 |
| Plaur | 88.94456294 | 4.072120741 | 0.488865842 | 8.329730547 |
| Cadm4 | 2550.068439 | -0.074750219 | 0.088834366 | -0.841456096 |
| Zfp428 | 224.1959431 | -0.235745658 | 0.10110622 | -2.331663259 |
| Irgq | 1232.318714 | -0.347127709 | 0.049684021 | -6.986707235 |
| Pinlyp | 0.104366086 | 0.274400501 | 3.724945049 | 0.073665651 |
| Xrcc1 | 220.1632784 | -0.135285676 | 0.098064629 | -1.37955629 |
| Zfp575 | 72.63402446 | -0.475667624 | 0.122819878 | -3.87288792 |
| Ethe1 | 83.5924299 | -0.103437063 | 0.159923576 | -0.646790581 |
| Phldb3 | 6.837825038 | 2.992773958 | 0.668643619 | 4.475888012 |
| Gm26550 | 0 | NA | NA | NA |
| Lypd3 | 0.412745228 | -0.035740767 | 3.68668308 | -0.009694559 |
| Tex101 | 0 | NA | NA | NA |
| BC049730 | 0 | NA | NA | NA |
| Gm4763 | 0 | NA | NA | NA |
| Cd177 | 15.47993706 | 5.026630162 | 0.993332055 | 5.060372449 |
| Ceacam10 | 1.306976768 | 2.222843321 | 0.987315141 | 2.251402038 |
| Gm5893 | 0 | NA | NA | NA |
| Gm9844 | 66.59610774 | 1.062733231 | 0.19954171 | 5.325870125 |
| Lypd4 | 0 | NA | NA | NA |
| Dmrtc2 | 0.042716945 | -0.206490061 | 3.724945049 | -0.055434391 |
| Rps19 | 1020.693303 | 0.44446433 | 0.087789788 | 5.062824982 |
| Cd79a | 3.200457292 | 0.027587021 | 0.609622581 | 0.045252624 |

| | | | | |
|---|---|---|---|---|
| Arhgef1 | 583.8558966 | 0.194327005 | 0.120045009 | 1.618784551 |
| Rabac1 | 736.5228606 | 0.077766773 | 0.120888584 | 0.643292947 |
| Mir7046 | 0 | NA | NA | NA |
| Atp1a3 | 26298.87918 | -0.12401982 | 0.06268639 | -1.978417018 |
| Mir7047 | 0.284641305 | -1.778680843 | 1.71003639 | -1.040142101 |
| Grik5 | 1931.46398 | -0.302754091 | 0.10806816 | -2.801510539 |
| Zfp574 | 425.7976918 | 0.182752873 | 0.084049853 | 2.17433898 |
| Pou2f2 | 170.0044398 | 0.109505892 | 0.159777916 | 0.685363123 |
| Mir6537 | 0 | NA | NA | NA |
| D930028M14Rik | 1.834100151 | -1.033372573 | 0.884190188 | -1.168722056 |
| Dedd2 | 172.9180783 | -0.129005646 | 0.132894218 | -0.970739344 |
| Mir7048 | 0.228760906 | -1.40412778 | 2.111290149 | -0.665056757 |
| Zfp526 | 147.8424307 | -0.145827503 | 0.11557967 | -1.261705487 |
| Gsk3a | 1893.258904 | 0.135759478 | 0.047560564 | 2.854454759 |
| 9130221H12Rik | 75.48946259 | -0.259646612 | 0.13106443 | -1.981060852 |
| Erf | 724.7532469 | 0.325814675 | 0.126336868 | 2.578935826 |
| Cic | 3271.904299 | 0.004461788 | 0.058421981 | 0.076371731 |
| Pafah1b3 | 85.99027182 | -0.215550479 | 0.144153813 | -1.495281149 |
| Prr19 | 0.750844096 | -0.650943053 | 1.294243966 | -0.502952357 |
| Tmem145 | 351.2431402 | -0.408922335 | 0.145449595 | -2.811436735 |
| Megf8 | 2644.487032 | -0.392865168 | 0.082815108 | -4.743882811 |
| Cnfn | 0.044191549 | -0.206490061 | 3.724945049 | -0.055434391 |
| Lipe | 172.463792 | 0.764349528 | 0.135093024 | 5.657949668 |
| 4732471J01Rik | 9.165417789 | -0.260611923 | 0.38501426 | -0.676889015 |
| Cxcl17 | 0.234440768 | 1.229343644 | 2.125569239 | 0.578359727 |
| Gm15495 | 0.148650935 | 0.723036258 | 2.859657322 | 0.252840175 |
| Ceacam1 | 24.75964592 | 1.217132622 | 0.278782921 | 4.36587943 |
| Ceacam2 | 1.575759893 | -0.118595337 | 1.049165367 | -0.113037792 |
| Erich4 | 0.961427984 | -0.242970409 | 1.039902828 | -0.233647224 |
| Atp5sl | 334.5795391 | 0.017506264 | 0.066525984 | 0.263149261 |
| B3gnt8 | 13.86552076 | 1.403761421 | 0.465889726 | 3.013076578 |
| Bckdha | 305.8692003 | 0.194416939 | 0.122071884 | 1.59264306 |
| Exosc5 | 96.23120801 | -0.111240777 | 0.133057816 | -0.83603339 |
| Tmem91 | 111.1486534 | -0.782532669 | 0.146531624 | -5.340367125 |
| B9d2 | 96.55601536 | 0.03876632 | 0.140098206 | 0.276708182 |
| Tgfb1 | 180.5550002 | 1.658836953 | 0.196106105 | 8.458874634 |
| Ccdc97 | 599.3683796 | 0.036196096 | 0.07256701 | 0.498795473 |
| Hnrnpul1 | 1763.970679 | 0.278031477 | 0.063121117 | 4.404730012 |
| Axl | 1705.534697 | 1.624583831 | 0.131474605 | 12.35663594 |
| Cyp2s1 | 45.56318474 | -0.586772374 | 0.204545684 | -2.86866172 |
| Gm26707 | 1.31640046 | -1.575556125 | 0.888772496 | -1.772732766 |
| Gm4613 | 2.523943987 | 1.010904004 | 0.67024841 | 1.508252745 |
| Gm24865 | 0 | NA | NA | NA |
| Gm23921 | 0 | NA | NA | NA |
| Gm6434 | 0 | NA | NA | NA |
| Cyp2b10 | 0.605952604 | 1.270119647 | 1.371882093 | 0.925822746 |
| Cyp2b13 | 0.046345709 | 0.274391604 | 3.724945049 | 0.073663262 |

| | | | | |
|---|---|---|---|---|
| Rnf170-ps | 17.15945834 | -0.138818363 | 0.236374159 | -0.587282314 |
| Cyp2b9 | 0 | NA | NA | NA |
| Gm8902 | 0.060174537 | 0.755286614 | 3.724945049 | 0.202764498 |
| Vmn1r-ps86 | 0 | NA | NA | NA |
| Vmn1r184 | 0 | NA | NA | NA |
| Cyp2a4 | 0.118974955 | 0.274400501 | 3.724945049 | 0.073665651 |
| Nlrp9c | 0.044544648 | 0.755286614 | 3.724945049 | 0.202764498 |
| Nlrp4a | 0 | NA | NA | NA |
| Gm25947 | 0 | NA | NA | NA |
| Nlrp9a | 0 | NA | NA | NA |
| Vmn1r185 | 0 | NA | NA | NA |
| Cyp2b23 | 0 | NA | NA | NA |
| Cyp2b19 | 0.186596897 | 0.890994395 | 3.282617118 | 0.27142806 |
| Cyp2g1 | 8.028728541 | -2.283176093 | 1.652545336 | -1.381611774 |
| Cyp2a5 | 11.27708144 | -1.212436798 | 1.167583119 | -1.038415833 |
| Cyp2a22 | 0.038981395 | 0.274393828 | 3.724945049 | 0.073663859 |
| Cyp2a12 | 0.191971534 | 0.27441076 | 2.653024697 | 0.103433172 |
| Cyp2f2 | 13.0302823 | -0.223362045 | 0.459825872 | -0.48575354 |
| Gm15883 | 0.044544648 | 0.755286614 | 3.724945049 | 0.202764498 |
| Cyp2t4 | 0.092269992 | -0.20649451 | 3.724945049 | -0.055435586 |
| Egln2 | 888.7193016 | 0.401657318 | 0.076030188 | 5.282866313 |
| Rab4b | 72.36944043 | 0.226537702 | 0.164613892 | 1.376176088 |
| Gm21983 | 0 | NA | NA | NA |
| Mir3101 | 0 | NA | NA | NA |
| Mia | 1.133325952 | -2.455587973 | 1.100690069 | -2.230953146 |
| Snrpa | 438.9269378 | 0.119440802 | 0.083805476 | 1.425214769 |
| BC024978 | 169.2292599 | -0.033670969 | 0.097740836 | -0.34449234 |
| Mir1191 | 8.542722226 | -0.192403597 | 0.360478565 | -0.533744905 |
| Itpkc | 197.4348915 | 0.494715409 | 0.217839437 | 2.271009396 |
| Adck4 | 262.1854901 | 0.01083241 | 0.083595662 | 0.129581004 |
| Gm15567 | 1.049380316 | -1.086704494 | 1.025076892 | -1.06011998 |
| Numbl | 1053.281901 | -0.196225518 | 0.095695693 | -2.050515662 |
| Gm15570 | 0 | NA | NA | NA |
| Ltbp4 | 207.6280549 | -0.682070155 | 0.148467365 | -4.594074637 |
| Shkbp1 | 92.29759634 | -0.259101781 | 0.178374501 | -1.452571861 |
| Gm20479 | 0 | NA | NA | NA |
| Sptbn4 | 1935.821563 | -0.319550004 | 0.132735079 | -2.407426933 |
| Blvrb | 91.00347656 | 0.120443868 | 0.123380439 | 0.976199057 |
| Pgam1-ps2 | 38.75767613 | -0.032642626 | 0.174750925 | -0.186795153 |
| Sertad3 | 55.81009826 | 1.016284636 | 0.201639733 | 5.040101079 |
| Sertad1 | 143.2528654 | 0.659685187 | 0.235183023 | 2.80498643 |
| Prx | 13.03989321 | 0.938593295 | 0.361361552 | 2.597380072 |
| Gm15541 | 1.986167605 | 2.703514005 | 0.967764731 | 2.793565334 |
| Hipk4 | 1.250864467 | -0.739712554 | 1.014405686 | -0.729207816 |
| Pld3 | 3221.190747 | -0.157595784 | 0.074559623 | -2.113688045 |
| 2310022A10Rik | 301.4687265 | 0.131300559 | 0.105318907 | 1.24669505 |
| Akt2 | 915.7419638 | 0.101924045 | 0.07633077 | 1.33529433 |

| | | | | |
|---|---|---|---|---|
| Ttc9b | 367.5397197 | -0.09374095 | 0.090894499 | -1.031315985 |
| Map3k10 | 1510.75745 | -0.120741653 | 0.078187091 | -1.544265834 |
| C030039L03Rik | 80.29137703 | -0.85131216 | 0.169769211 | -5.014526225 |
| Zfp60 | 578.4836708 | -0.349958031 | 0.175083798 | -1.998803061 |
| Zfp626 | 57.40632837 | -0.017356347 | 0.154964064 | -0.112002397 |
| Zfp59 | 63.21053847 | 0.013623529 | 0.1711383 | 0.079605377 |
| Zfp607 | 29.83338546 | 0.215597893 | 0.243095006 | 0.886887381 |
| 1700049G17Rik | 131.7655322 | -0.184057551 | 0.101991992 | -1.804627484 |
| Gm26891 | 2.658590328 | 1.20283047 | 0.80276385 | 1.498361529 |
| Gm10651 | 26.89522568 | -0.190524492 | 0.233895092 | -0.814572425 |
| Zfp780b | 147.3540878 | -0.352382124 | 0.141487683 | -2.490549817 |
| Zfp850 | 137.3735265 | -0.107753921 | 0.109122358 | -0.987459606 |
| Psmc4 | 769.5473873 | 0.269346238 | 0.086920323 | 3.09877172 |
| Fcgbp | 5.334611605 | 0.225596785 | 0.396781016 | 0.568567487 |
| Gm10935 | 0.257434735 | -1.207303886 | 2.117967881 | -0.570029365 |
| Gm24356 | 0 | NA | NA | NA |
| 9530053A07Rik | 2.222860927 | 0.790049376 | 0.696345031 | 1.13456597 |
| Fbl | 235.6745308 | 0.444006218 | 0.157436986 | 2.82021543 |
| Dyrk1b | 419.6037486 | -0.326339285 | 0.102479254 | -3.184442431 |
| Eid2 | 336.2126705 | 0.125027489 | 0.094057651 | 1.329264421 |
| Eid2b | 286.8257227 | 0.231145322 | 0.09810894 | 2.356006724 |
| BC089491 | 5.091535971 | -0.300507251 | 0.414385859 | -0.725187031 |
| Dll3 | 12.77550445 | -0.678446405 | 0.341154694 | -1.988676744 |
| Timm50 | 228.1838604 | 0.031855647 | 0.10496165 | 0.303497968 |
| Supt5 | 1576.058628 | 0.101612388 | 0.057930752 | 1.75403193 |
| Rps16 | 917.1611222 | 0.435765721 | 0.097606986 | 4.464493175 |
| AF357399 | 3.515687709 | 0.150946215 | 0.608862412 | 0.247915148 |
| Plekhg2 | 230.709035 | 1.437618048 | 0.202459437 | 7.100770735 |
| Zfp36 | 478.3772212 | 3.562442677 | 0.332721699 | 10.7069743 |
| Med29 | 93.91206282 | 0.226210209 | 0.128887007 | 1.755104828 |
| Paf1 | 477.8677575 | 0.099381772 | 0.07940176 | 1.251631858 |
| Pak4 | 1507.799238 | 0.229430777 | 0.095865306 | 2.393261831 |
| Gmfg | 19.76928172 | 2.682260608 | 0.394517866 | 6.79883179 |
| Lrfn1 | 754.1318081 | -0.056747783 | 0.099895831 | -0.568069581 |
| 1700028B04Rik | 0.752450377 | 1.847851543 | 1.208661914 | 1.528840714 |
| Ifnl2 | 0.046345709 | 0.274391604 | 3.724945049 | 0.073663262 |
| Ifnl3 | 0.046345709 | 0.274391604 | 3.724945049 | 0.073663262 |
| Sycn | 0 | NA | NA | NA |
| Nccrp1 | 4.109619456 | -0.502880344 | 0.496728153 | -1.012385429 |
| Mir7049 | 0.300003454 | -1.152920221 | 2.020832197 | -0.570517544 |
| Papl | 10.97390351 | -0.195418916 | 0.308096622 | -0.634278023 |
| Fbxo27 | 104.6688318 | -0.033664413 | 0.117635004 | -0.286176832 |
| Fbxo17 | 17.81551388 | 0.221194715 | 0.254374569 | 0.869563004 |
| Mrps12 | 129.792039 | -0.295053132 | 0.126778496 | -2.327312133 |
| Sars2 | 103.3753545 | 0.162635748 | 0.109835942 | 1.480715189 |
| Gm6537 | 10.517279 | -0.009907328 | 0.361585124 | -0.027399711 |
| Nfkbib | 264.1176441 | 0.608692849 | 0.159994554 | 3.804459801 |

| | | | | |
|---|---|---|---|---|
| Sirt2 | 1431.100394 | 0.239144988 | 0.06451536 | 3.706791475 |
| Rinl | 62.25372522 | 1.572709443 | 0.256733447 | 6.125845548 |
| Hnrnpl | 2413.616906 | -0.071718758 | 0.059690982 | -1.201500714 |
| Ech1 | 280.1649813 | -0.130190825 | 0.101928465 | -1.277276422 |
| Lgals4 | 13.26484918 | -0.07895519 | 0.315465098 | -0.250281854 |
| Lgals7 | 1.097858384 | 0.977559251 | 1.25253058 | 0.780467372 |
| Capn12 | 3.372768181 | -0.452236578 | 0.55087812 | -0.82093763 |
| Actn4 | 1770.860706 | 0.13621332 | 0.090405105 | 1.506699429 |
| Eif3k | 526.8853122 | 0.227686885 | 0.113536965 | 2.005398722 |
| Map4k1 | 43.16772284 | 1.978256271 | 0.244532394 | 8.089955855 |
| Ryr1 | 194.7438043 | -1.144568506 | 0.222594378 | -5.141947053 |
| AC165142.1 | 0.214156338 | 0.274411302 | 2.175868539 | 0.126115754 |
| Gm26604 | 0.400939804 | -1.612643092 | 1.463263636 | -1.102086495 |
| Mir1963 | 0 | NA | NA | NA |
| Rasgrp4 | 17.50432713 | 0.044745407 | 0.339463577 | 0.131812101 |
| Gm16282 | 0 | NA | NA | NA |
| Fam98c | 181.7263762 | -0.230301463 | 0.102278684 | -2.251705377 |
| 1110035H17Rik | 2.173831266 | -0.091731185 | 0.639607959 | -0.14341783 |
| Spred3 | 426.8502981 | -0.509905184 | 0.112984036 | -4.513072855 |
| Ggn | 9.283367035 | -0.355615459 | 0.363609394 | -0.978015049 |
| Psmd8 | 837.7336857 | 0.443960014 | 0.104808482 | 4.235916856 |
| Catsperg1 | 14.9802426 | -0.518127416 | 0.33208083 | -1.560244884 |
| Gm17113 | 0 | NA | NA | NA |
| Kcnk6 | 62.8823065 | 1.168359382 | 0.190500144 | 6.133115492 |
| Yif1b | 285.5672007 | -0.020343468 | 0.107282528 | -0.189625177 |
| 2200002D01Rik | 2.362804395 | -0.354915069 | 0.727307328 | -0.487985004 |
| Spint2 | 203.5653366 | 0.168250962 | 0.10610637 | 1.585682002 |
| Ppp1r14a | 171.4189243 | -0.393761991 | 0.146642573 | -2.685181961 |
| Dpf1 | 418.7210972 | -0.439418961 | 0.115075954 | -3.818512421 |
| Sipa1l3 | 1249.288852 | -0.19581831 | 0.074987586 | -2.61134304 |
| 4932431P20Rik | 0.682519493 | 0.20160516 | 1.184744084 | 0.170167686 |
| 4930432E11Rik | 0.998548739 | -3.26548499 | 1.123589286 | -2.906297728 |
| Gm29326 | 0.495715849 | -0.937690223 | 1.43982952 | -0.651250867 |
| A330087D11Rik | 0.362932359 | -2.422644535 | 1.580125934 | -1.533197122 |
| n-R5s153 | 0 | NA | NA | NA |
| Catsperg2 | 7.493041037 | -0.701576235 | 0.394933315 | -1.776442272 |
| Gm23103 | 0 | NA | NA | NA |
| Zfp84 | 275.1985705 | -0.283570514 | 0.08762435 | -3.236206756 |
| Mir1964 | 1.817730626 | -1.085697087 | 0.751625741 | -1.444465014 |
| Zfp30 | 158.5141845 | -0.178288036 | 0.099225769 | -1.796791676 |
| Zfp790 | 215.9787552 | -0.421229297 | 0.117738351 | -3.5776728 |
| Zfp940 | 138.395624 | -0.255472102 | 0.106990315 | -2.387805859 |
| Gm26554 | 0 | NA | NA | NA |
| Gm17199 | 0 | NA | NA | NA |
| Zfp420 | 147.2350297 | -0.719622378 | 0.119995818 | -5.997062146 |
| Zfp27 | 203.8758977 | -0.323504863 | 0.081219619 | -3.983087671 |
| Mir7668 | 0.042716945 | -0.206490061 | 3.724945049 | -0.055434391 |

| | | | | |
|---|---|---|---|---|
| Zfp383 | 132.1721319 | -0.304656933 | 0.10204849 | -2.98541344 |
| Zfp74 | 213.3163599 | -0.196357038 | 0.092591409 | -2.120683116 |
| C230062I16Rik | 7.143933271 | 0.085753675 | 0.439517657 | 0.1951086 |
| Gm10169 | 7.433721481 | 0.849500949 | 0.40214492 | 2.112424919 |
| Zfp568 | 112.9697705 | 0.00484518 | 0.130626259 | 0.037091932 |
| Zfp14 | 53.58896386 | -0.351433198 | 0.174068262 | -2.018938966 |
| Zfp82 | 6.950672932 | -0.552539989 | 0.359996988 | -1.534846147 |
| Gm26920 | 0.228152142 | 1.718362781 | 2.550451095 | 0.673748571 |
| Zfp566 | 31.53288666 | -0.204803286 | 0.188647073 | -1.085642533 |
| Zfp260 | 833.8589085 | 0.156447612 | 0.06864562 | 2.279061839 |
| Gm23368 | 0 | NA | NA | NA |
| Zfp382 | 241.7030479 | 0.153392095 | 0.092391842 | 1.660234194 |
| Zfp146 | 303.8588219 | -0.083073199 | 0.07499085 | -1.107777803 |
| Gm5113 | 98.06848606 | -0.432049778 | 0.138608101 | -3.117060073 |
| Cox7a1 | 8.620735387 | -0.618038698 | 0.38699786 | -1.597008051 |
| Capns1 | 1143.806175 | 0.251774188 | 0.113864278 | 2.211178006 |
| Gm26810 | 0.29029136 | 1.096220824 | 1.775122021 | 0.617546744 |
| Tbcb | 367.5355379 | 0.04664724 | 0.086661492 | 0.538269525 |
| Polr2i | 115.9016702 | 0.213302213 | 0.135377553 | 1.575609902 |
| Ovol3 | 0.088736197 | 0.274400501 | 3.724945049 | 0.073665651 |
| Wdr62 | 63.98026938 | 2.276857205 | 0.248258149 | 9.171329167 |
| Rps12-ps5 | 0.252860697 | -0.709302944 | 1.836154608 | -0.38629805 |
| Thap8 | 1.268210335 | -0.28014777 | 0.943588243 | -0.296896206 |
| Clip3 | 6878.216135 | 0.103466501 | 0.079576325 | 1.300217126 |
| Alkbh6 | 168.5814911 | -0.256239982 | 0.095194302 | -2.691757561 |
| Gm24757 | 0 | NA | NA | NA |
| Syne4 | 17.72551575 | -0.282971076 | 0.260782981 | -1.085082605 |
| Sdhaf1 | 75.1581409 | 0.428690829 | 0.125982343 | 3.402785007 |
| E130208F15Rik | 4.251243736 | 0.971326998 | 0.530883778 | 1.82964151 |
| Lrfn3 | 312.5931955 | 0.133500217 | 0.088526281 | 1.508029202 |
| Gm25259 | 0.679915734 | -0.387123341 | 1.214468086 | -0.318759583 |
| Tyrobp | 264.1135694 | 1.890744171 | 0.200945104 | 9.409257204 |
| Hcst | 5.467031529 | 3.896865852 | 0.910774736 | 4.278627521 |
| Nfkbid | 33.79332328 | 0.28532494 | 0.265160846 | 1.076044764 |
| Aplp1 | 8728.071125 | -0.0616638 | 0.065890892 | -0.935847098 |
| Kirrel2 | 45.1116197 | -0.340176894 | 0.24914876 | -1.36535656 |
| Nphs1 | 8.109329026 | -0.049936553 | 0.353227161 | -0.141372349 |
| Nphs1os | 0.917433238 | -1.08649345 | 1.034568712 | -1.050189743 |
| Prodh2 | 0.146848595 | -0.999135224 | 3.718157501 | -0.268717832 |
| Arhgap33os | 1.185039339 | -0.847033599 | 0.920872924 | -0.919815945 |
| Arhgap33 | 1004.610349 | -0.54926961 | 0.158921107 | -3.456240768 |
| Proser3 | 27.82645344 | -0.814045052 | 0.270196107 | -3.012793417 |
| Hspb6 | 57.72339347 | 0.782700012 | 0.176723397 | 4.428955231 |
| mmu-mir-9769 | 0 | NA | NA | NA |
| Lin37 | 148.1215379 | -0.075792125 | 0.12692284 | -0.597151191 |
| Psenen | 209.1857129 | 0.272364204 | 0.099248433 | 2.744267047 |
| U2af1l4 | 195.8717764 | 0.245969563 | 0.077873139 | 3.158593148 |

| | | | | |
|---|---|---|---|---|
| Gm17104 | 0.139856492 | -0.206491248 | 3.574411025 | -0.057769307 |
| Igflr1 | 0 | NA | NA | NA |
| Kmt2b | 1103.022709 | 0.002170397 | 0.137025608 | 0.015839353 |
| Mir3569 | 0.55981669 | -2.376318568 | 1.199047339 | -1.981838824 |
| Zbtb32 | 35.12199024 | 7.693336103 | 0.901638177 | 8.532620181 |
| Upk1a | 5.454212958 | -0.261160803 | 0.513355994 | -0.508732354 |
| Cox6b1 | 1385.504029 | -0.052569103 | 0.070289065 | -0.74789873 |
| Etv2 | 0.288531392 | 0.755308018 | 1.779727156 | 0.4243954 |
| Rbm42 | 1.341866388 | 0.277673232 | 0.816854678 | 0.339929781 |
| Gm21982 | 0.195772763 | -0.653567824 | 2.219885219 | -0.294415143 |
| Gm26610 | 2.644816022 | -0.31025745 | 0.583367846 | -0.531838448 |
| Haus5 | 8.540474638 | 0.107877763 | 0.35810699 | 0.301244506 |
| Pmis2 | 0 | NA | NA | NA |
| 2200002J24Rik | 0.066050875 | 0.274391604 | 3.724945049 | 0.073663262 |
| Gm4883 | 0 | NA | NA | NA |
| Atp4a | 2.461184311 | 0.121723272 | 0.567115257 | 0.214635862 |
| Tmem147 | 361.1132233 | -0.103364109 | 0.083183214 | -1.242607776 |
| Tmem147os | 88.86610623 | -0.418545319 | 0.183425582 | -2.281826311 |
| Gapdhs | 0.221390349 | -0.142971524 | 2.608989397 | -0.05479958 |
| Sbsn | 30.54518145 | 0.492025013 | 0.241005384 | 2.041551953 |
| Gm26935 | 15.53492519 | -0.473130588 | 0.337264272 | -1.402848235 |
| Dmkn | 16.49677372 | 0.595900477 | 0.508680935 | 1.17146218 |
| Krtdap | 1.379211929 | 3.296850706 | 3.675597772 | 0.896956335 |
| Ffar2 | 3.090575541 | 2.842316031 | 1.129272573 | 2.516944182 |
| Ffar3 | 0.43344934 | -2.722692095 | 1.534030007 | -1.774862345 |
| Ffar1 | 0.265062405 | -0.131804489 | 2.438331845 | -0.054055189 |
| Cd22 | 18.32792618 | 2.386495231 | 0.477731364 | 4.995475309 |
| Gm29439 | 0.095898756 | 0.27438722 | 3.724945049 | 0.073662085 |
| Gm29440 | 0.110242424 | -0.20649451 | 3.724945049 | -0.055435586 |
| Mag | 2336.670031 | 0.069306089 | 0.129003249 | 0.53724297 |
| Hamp2 | 0.159795471 | -0.20649052 | 2.810463622 | -0.073472049 |
| Hamp | 0.755877001 | 1.341472651 | 1.163329455 | 1.153132197 |
| Usf2 | 555.4235933 | -0.08275234 | 0.060271221 | -1.372999237 |
| Gm17077 | 7.799693549 | 0.31851048 | 0.434958676 | 0.732277564 |
| Gm4673 | 75.79538166 | -0.142888141 | 0.23191211 | -0.61613057 |
| Lsr | 56.83401911 | 1.674075701 | 0.24058037 | 6.95848835 |
| Fam187b | 1.876867459 | 0.305315535 | 0.740788808 | 0.412149228 |
| Fxyd5 | 213.0889357 | 1.629759888 | 0.224461819 | 7.260744382 |
| Mir7050 | 0 | NA | NA | NA |
| Fxyd7 | 269.8762191 | -0.429015745 | 0.136441956 | -3.144309552 |
| Fxyd1 | 309.5145642 | 0.426081908 | 0.17500039 | 2.43474834 |
| Lgi4 | 451.055952 | 0.272786465 | 0.126617851 | 2.154407631 |
| Fxyd3 | 1.806277867 | 0.184126666 | 0.940691781 | 0.195735383 |
| Hpn | 35.43406477 | -0.509178552 | 0.28184832 | -1.806569405 |
| Scn1b | 2784.178309 | -0.176667815 | 0.076379542 | -2.313025325 |
| Gramd1a | 871.9470282 | 0.07056314 | 0.096388053 | 0.732073507 |
| G630030J09Rik | 3.656513056 | -0.404701419 | 0.584832175 | -0.691995818 |

| | | | | |
|---|---|---|---|---|
| Gm10640 | 1.251626804 | -1.18453995 | 0.996996422 | -1.188108527 |
| Gm9569 | 0 | NA | NA | NA |
| Gm37292 | 9.829030887 | -0.945122191 | 0.372575506 | -2.536726584 |
| Scgb1b1-ps | 0 | NA | NA | NA |
| Scgb2b1 | 0 | NA | NA | NA |
| Scgb1b2 | 0 | NA | NA | NA |
| Scgb2b2 | 0 | NA | NA | NA |
| Scgb2b3 | 0 | NA | NA | NA |
| Scgb1b3 | 0 | NA | NA | NA |
| Scgb2b4-ps | 0 | NA | NA | NA |
| Scgb1b4-ps | 0 | NA | NA | NA |
| Scgb1b28-ps | 0 | NA | NA | NA |
| Gm29492 | 0 | NA | NA | NA |
| Scgb2b5-ps | 0 | NA | NA | NA |
| Scgb1b5-ps | 0 | NA | NA | NA |
| Scgb2b28-ps | 0 | NA | NA | NA |
| Rpl23a-ps8 | 0 | NA | NA | NA |
| Scgb2b6 | 0 | NA | NA | NA |
| Scgb1b6-ps | 0 | NA | NA | NA |
| Gm4373 | 0 | NA | NA | NA |
| Rpl23a-ps9 | 0 | NA | NA | NA |
| Scgb2b7 | 0 | NA | NA | NA |
| Scgb1b7 | 0 | NA | NA | NA |
| Gm9114 | 0 | NA | NA | NA |
| Gm29627 | 0 | NA | NA | NA |
| Scgb2b8-ps | 0 | NA | NA | NA |
| Scgb1b8-ps | 0 | NA | NA | NA |
| Gm6669 | 0 | NA | NA | NA |
| Gm9120 | 0 | NA | NA | NA |
| Gm21673 | 0 | NA | NA | NA |
| Scgb2b9-ps | 0 | NA | NA | NA |
| Scgb1b9-ps | 0 | NA | NA | NA |
| Gm5329 | 0 | NA | NA | NA |
| Scgb2b29-ps | 0 | NA | NA | NA |
| Gm9127 | 0 | NA | NA | NA |
| Gm29262 | 0 | NA | NA | NA |
| Gm29263 | 0 | NA | NA | NA |
| Scgb2b10 | 0 | NA | NA | NA |
| Scgb1b10 | 0 | NA | NA | NA |
| Rpl23a-ps10 | 0 | NA | NA | NA |
| Scgb2b30 | 0 | NA | NA | NA |
| Gm9135 | 0 | NA | NA | NA |
| Gm28256 | 0 | NA | NA | NA |
| Gm28257 | 0 | NA | NA | NA |
| Scgb2b11 | 0 | NA | NA | NA |
| Scgb1b11 | 0 | NA | NA | NA |
| Scgb2b12 | 0 | NA | NA | NA |

| | | | | |
|---|---|---|---|---|
| Scgb1b12 | 0 | NA | NA | NA |
| Gm9142 | 0 | NA | NA | NA |
| Gm28862 | 0 | NA | NA | NA |
| Rpl23a-ps11 | 0 | NA | NA | NA |
| Scgb1b29 | 0 | NA | NA | NA |
| Gm9146 | 0 | NA | NA | NA |
| Gm5116 | 0 | NA | NA | NA |
| Scgb2b13-ps | 0 | NA | NA | NA |
| Scgb1b13-ps | 0 | NA | NA | NA |
| Gm5327 | 0 | NA | NA | NA |
| Gm9149 | 0 | NA | NA | NA |
| Gm21135 | 0 | NA | NA | NA |
| Scgb2b14-ps | 0 | NA | NA | NA |
| Scgb1b14-ps | 0 | NA | NA | NA |
| Gm24766 | 0 | NA | NA | NA |
| Gm5058 | 0 | NA | NA | NA |
| Gm29429 | 0 | NA | NA | NA |
| Scgb2b31-ps | 0 | NA | NA | NA |
| Gm4398 | 0 | NA | NA | NA |
| Gm29432 | 0 | NA | NA | NA |
| Gm28408 | 0 | NA | NA | NA |
| Scgb2b15 | 0 | NA | NA | NA |
| Scgb1b15 | 0 | NA | NA | NA |
| Scgb2b16-ps | 0 | NA | NA | NA |
| Scgb1b16-ps | 0 | NA | NA | NA |
| Gm25817 | 0 | NA | NA | NA |
| Gm9156 | 0 | NA | NA | NA |
| Gm28409 | 0 | NA | NA | NA |
| Scgb2b32-ps | 0 | NA | NA | NA |
| Gm4405 | 0 | NA | NA | NA |
| Gm28383 | 0 | NA | NA | NA |
| Gm28384 | 0 | NA | NA | NA |
| Scgb2b17 | 0 | NA | NA | NA |
| Scgb1b17 | 0 | NA | NA | NA |
| Gm28992 | 0 | NA | NA | NA |
| Rpl23a-ps7 | 0 | NA | NA | NA |
| Scgb2b33 | 0 | NA | NA | NA |
| Gm9162 | 0 | NA | NA | NA |
| Gm28559 | 0 | NA | NA | NA |
| Gm6003 | 0 | NA | NA | NA |
| Scgb2b18 | 0 | NA | NA | NA |
| Scgb1b18 | 0 | NA | NA | NA |
| Scgb2b19 | 0 | NA | NA | NA |
| Scgb1b19 | 0 | NA | NA | NA |
| Gm9167 | 0 | NA | NA | NA |
| Scgb2b20 | 0 | NA | NA | NA |
| Scgb1b20 | 0 | NA | NA | NA |

| | | | | |
|---|---|---|---|---|
| Gm5732 | 0 | NA | NA | NA |
| Gm28794 | 0 | NA | NA | NA |
| Gm5731 | 0 | NA | NA | NA |
| Scgb1b34-ps | 0 | NA | NA | NA |
| Scgb2b21 | 0 | NA | NA | NA |
| Scgb1b21 | 0 | NA | NA | NA |
| Gm12770 | 0 | NA | NA | NA |
| Gm12771 | 0 | NA | NA | NA |
| Scgb2b22-ps | 0 | NA | NA | NA |
| Scgb1b22-ps | 0 | NA | NA | NA |
| Scgb2b23-ps | 0 | NA | NA | NA |
| Scgb1b23-ps | 0 | NA | NA | NA |
| Gm12763 | 0.048949532 | -0.206487837 | 3.724945049 | -0.055433794 |
| Rpl23a-ps12 | 0 | NA | NA | NA |
| Scgb2b24 | 0 | NA | NA | NA |
| Scgb1b24 | 0 | NA | NA | NA |
| Gm12782 | 0 | NA | NA | NA |
| Gm28614 | 0 | NA | NA | NA |
| Gm26443 | 0 | NA | NA | NA |
| Gm12768 | 0 | NA | NA | NA |
| Scgb2b25-ps | 0 | NA | NA | NA |
| Scgb1b25-ps | 0 | NA | NA | NA |
| Gm12757 | 0 | NA | NA | NA |
| Gm28615 | 0 | NA | NA | NA |
| Gm12765 | 0 | NA | NA | NA |
| Scgb2b26 | 0 | NA | NA | NA |
| Scgb1b26-ps | 0 | NA | NA | NA |
| Gm28184 | 0 | NA | NA | NA |
| Gm12773 | 0 | NA | NA | NA |
| Gm12755 | 0 | NA | NA | NA |
| Scgb2b27 | 0.046345709 | 0.274391604 | 3.724945049 | 0.073663262 |
| Scgb1b27 | 0 | NA | NA | NA |
| Gm12760 | 0 | NA | NA | NA |
| Gm12779 | 0.044544648 | 0.755286614 | 3.724945049 | 0.202764498 |
| Scgb1b30 | 0 | NA | NA | NA |
| Wtip | 46.54900976 | 0.748521851 | 0.173664039 | 4.310171844 |
| Uba2 | 890.9651073 | -0.037937142 | 0.059534925 | -0.637224983 |
| Gm12778 | 0.649479285 | 0.25993217 | 1.440183897 | 0.180485402 |
| Gm12783 | 0 | NA | NA | NA |
| Pdcd2l | 117.7724749 | 0.161978606 | 0.142102397 | 1.139872438 |
| Gpi1 | 4331.119929 | -0.12741454 | 0.049367031 | -2.58096418 |
| Gm26762 | 2.694167648 | -0.183399079 | 0.711872602 | -0.257629074 |
| Gm12762 | 6.006325422 | -0.406078884 | 0.406229741 | -0.999628639 |
| Gm12758 | 0.954952387 | -1.53174525 | 1.091171412 | -1.403762262 |
| 4931406P16Rik | 686.0018774 | 0.016648907 | 0.060609939 | 0.274689392 |
| Gm6096 | 0.066050875 | 0.274391604 | 3.724945049 | 0.073663262 |
| Lsm14a | 923.1856796 | 0.161962432 | 0.058376096 | 2.774464935 |

| | | | | |
|---|---|---|---|---|
| Gm12774 | 0 | NA | NA | NA |
| Gm12764 | 10.54452331 | 3.473868252 | 0.557988372 | 6.225700081 |
| Gm12766 | 0.212513282 | 1.926509226 | 3.159662244 | 0.609719988 |
| Gm12780 | 0 | NA | NA | NA |
| Gm12784 | 0.530195705 | 0.185116747 | 1.322232033 | 0.140003224 |
| Kctd15 | 124.850857 | -0.188630809 | 0.116271062 | -1.622336677 |
| Gm12781 | 0.50824631 | -0.528953848 | 1.313944217 | -0.402569486 |
| Gm12756 | 2.131963076 | -1.747235309 | 0.74041199 | -2.359814984 |
| Chst8 | 122.9302308 | -0.178635205 | 0.124337584 | -1.436695165 |
| Pepd | 269.5084426 | 0.111843197 | 0.095505867 | 1.171061015 |
| Gm25922 | 0.089089296 | 1.194485492 | 3.723939576 | 0.320758559 |
| Cebpg | 578.773901 | 0.085591 | 0.095414028 | 0.897048392 |
| Gm25033 | 0 | NA | NA | NA |
| Gm22535 | 0 | NA | NA | NA |
| Cebpa | 259.903502 | 0.917043549 | 0.200211902 | 4.580364792 |
| Slc7a10 | 228.5227835 | -1.079301235 | 0.179331064 | -6.018484538 |
| Lrp3 | 882.6663157 | -0.146640048 | 0.076319129 | -1.921406222 |
| Gm24505 | 0.219162254 | 0.740546191 | 2.159165238 | 0.342978007 |
| Gpatch1 | 413.9756821 | 0.098958648 | 0.095931055 | 1.031560095 |
| Rhpn2 | 165.8571245 | 0.294207716 | 0.163312253 | 1.801504232 |
| C230052I12Rik | 49.69667937 | 0.515110281 | 0.199214825 | 2.585702545 |
| Cep89 | 116.2122036 | -0.129029106 | 0.139078298 | -0.92774436 |
| Gm18254 | 0 | NA | NA | NA |
| Slc7a9 | 0.108767819 | -0.20649451 | 3.724945049 | -0.055435586 |
| Tdrd12 | 4.458748399 | -0.369501542 | 0.525516543 | -0.703120668 |
| Gm18558 | 0.451259361 | -0.059095937 | 1.401176894 | -0.042175929 |
| Nudt19 | 464.3454165 | 0.419679297 | 0.105193087 | 3.989609098 |
| B230322F03Rik | 17.22929159 | 0.561955964 | 0.298525772 | 1.882437021 |
| Rgs9bp | 10.39343307 | 0.28294565 | 0.386563168 | 0.731951913 |
| Ankrd27 | 324.283844 | -0.463014715 | 0.14842595 | -3.119499746 |
| Pdcd5 | 124.9127266 | 0.650761154 | 0.219696042 | 2.962097764 |
| Dpy19l3 | 892.6871314 | -0.075711522 | 0.097206693 | -0.778871492 |
| Zfp507 | 543.6392318 | 0.019165936 | 0.071300844 | 0.26880378 |
| Gm26790 | 0.186516865 | 0.755298189 | 2.671088948 | 0.282767891 |
| E130304I02Rik | 2.861485906 | -0.575821256 | 0.570341573 | -1.009607723 |
| Gm28514 | 1.65996821 | -0.859055723 | 0.840935408 | -1.021547809 |
| Rpl17-ps9 | 11.99013794 | 0.436131839 | 0.304684155 | 1.431422777 |
| Gm9211 | 0.12092566 | 1.236179401 | 3.724945049 | 0.331865138 |
| Gm23292 | 2.206361005 | -0.512729422 | 0.618213615 | -0.829372581 |
| Gm25247 | 0 | NA | NA | NA |
| Gm29129 | 0 | NA | NA | NA |
| Gm38285 | 5.376801637 | 0.286005893 | 0.465514043 | 0.614387251 |
| Tshz3 | 524.5764167 | 0.021415365 | 0.134160768 | 0.159624644 |
| Gm37452 | 12.71943701 | -1.209278096 | 0.418041524 | -2.892722432 |
| Gm38340 | 32.55178755 | -0.663640115 | 0.368077892 | -1.80298825 |
| E230020A03Rik | 24.88402293 | -0.292997989 | 0.315733197 | -0.927992344 |
| Gm37827 | 13.69016405 | -0.847667619 | 0.447541401 | -1.894054085 |

| | | | | |
|---|---|---|---|---|
| Gm24550 | 0 | NA | NA | NA |
| Gm28078 | 0 | NA | NA | NA |
| Gm28076 | 0.144050924 | 1.290908285 | 3.721447108 | 0.346883416 |
| Gm28075 | 0 | NA | NA | NA |
| Gm28077 | 0.313601826 | 1.778002839 | 1.732157472 | 1.026467205 |
| Zfp536 | 718.5400894 | -0.113600899 | 0.076750408 | -1.480134149 |
| 4930505M18Rik | 0 | NA | NA | NA |
| Gm22302 | 0 | NA | NA | NA |
| Uri1 | 620.2418666 | 0.250705896 | 0.074241298 | 3.376906165 |
| Ccne1 | 105.1069268 | -0.093983076 | 0.123036572 | -0.763862928 |
| Gm22203 | 0 | NA | NA | NA |
| 1600014C10Rik | 402.5956352 | 0.584136924 | 0.134577301 | 4.340530833 |
| Plekhf1 | 106.4439667 | 0.364209646 | 0.157051873 | 2.31904045 |
| Pop4 | 57.53508151 | 0.630353052 | 0.189327604 | 3.32943024 |
| Gm28581 | 0 | NA | NA | NA |
| Gm9220 | 0 | NA | NA | NA |
| Gm4451 | 0 | NA | NA | NA |
| Gm28453 | 0.075043205 | 0.755286614 | 3.724945049 | 0.202764498 |
| Gm22638 | 0 | NA | NA | NA |
| Gm6818 | 0 | NA | NA | NA |
| Gm28583 | 0 | NA | NA | NA |
| Gm6605 | 0 | NA | NA | NA |
| Gm6819 | 0 | NA | NA | NA |
| Gm29087 | 0.10503227 | 0.274389469 | 3.724945049 | 0.073662689 |
| Gm5591 | 0 | NA | NA | NA |
| Gm4454 | 0 | NA | NA | NA |
| Gm20749 | 0 | NA | NA | NA |
| Gm29086 | 0 | NA | NA | NA |
| Gm25286 | 0 | NA | NA | NA |
| Gm21115 | 0 | NA | NA | NA |
| Gm21276 | 0 | NA | NA | NA |
| Gm21122 | 0 | NA | NA | NA |
| Gm29085 | 0 | NA | NA | NA |
| 1700010N08Rik | 0.146060149 | -0.206490268 | 3.528689472 | -0.058517552 |
| Gm24797 | 0 | NA | NA | NA |
| Gm21129 | 0 | NA | NA | NA |
| Gm21136 | 0 | NA | NA | NA |
| Gm21142 | 0 | NA | NA | NA |
| Gm21152 | 0 | NA | NA | NA |
| Gm25671 | 0 | NA | NA | NA |
| Gm28451 | 0 | NA | NA | NA |
| Gm1982 | 0 | NA | NA | NA |
| Gm1988 | 0 | NA | NA | NA |
| Gm6124 | 0 | NA | NA | NA |
| Gm16387 | 0 | NA | NA | NA |
| Gm9226 | 0 | NA | NA | NA |
| Gm23307 | 0 | NA | NA | NA |

| | | | | |
|---|---|---|---|---|
| Gm29258 | 0 | NA | NA | NA |
| Gm9271 | 0 | NA | NA | NA |
| Gm5114 | 0 | NA | NA | NA |
| Gm27437 | 0 | NA | NA | NA |
| Gm5590 | 0.783781752 | -0.474906348 | 1.227565203 | -0.386868532 |
| Zfp939 | 78.05214499 | 0.424740824 | 0.155870741 | 2.724955442 |
| Gm28455 | 4.111696988 | 0.374452108 | 0.516201525 | 0.725399073 |
| Zfp619 | 83.85373399 | -0.069280117 | 0.142361154 | -0.486650428 |
| Gm33989 | 0.779068608 | -1.506222223 | 1.187953939 | -1.267912984 |
| Gm37494 | 103.4889581 | -0.527723916 | 0.149677306 | -3.525744361 |
| Gm4790 | 216.6323139 | -0.473772892 | 0.150676235 | -3.144310657 |
| 6230415J03Rik | 105.8274719 | -0.400799239 | 0.189374867 | -2.116433115 |
| Gm28582 | 30.69961378 | -0.041068737 | 0.21686959 | -0.18937066 |
| Gm17983 | 0 | NA | NA | NA |
| Gm26526 | 0.0878635 | -0.687376175 | 3.724945049 | -0.184533239 |
| Gm2058 | 91.57356217 | 0.024201253 | 0.243931325 | 0.099213387 |
| Gm26327 | 0.206849446 | 1.048730835 | 2.582154035 | 0.40614573 |
| Gm2021 | 0.044544648 | 0.755286614 | 3.724945049 | 0.202764498 |
| Gm9230 | 0 | NA | NA | NA |
| Gm37302 | 0.167459142 | 0.755298223 | 2.757565248 | 0.2739004 |
| 4930558N11Rik | 0 | NA | NA | NA |
| 4930435C17Rik | 0 | NA | NA | NA |
| Gm38199 | 0 | NA | NA | NA |
| Gm6851 | 4.478289484 | 0.715899616 | 0.503159737 | 1.422807834 |
| 4933404I11Rik | 0 | NA | NA | NA |
| Gm24910 | 0 | NA | NA | NA |
| Gm28807 | 0.144050924 | 1.290908285 | 3.721447108 | 0.346883416 |
| 4921507H08Rik | 0 | NA | NA | NA |
| 4933402C06Rik | 0.04393175 | -0.206490061 | 3.724945049 | -0.055434391 |
| A230077H06Rik | 28.4887614 | -0.927043174 | 0.286809929 | -3.232256206 |
| Gm28307 | 0.072025462 | 0.755288839 | 3.724945049 | 0.202765096 |
| Gm25784 | 0 | NA | NA | NA |
| Vstm2b | 175.071028 | -0.224509074 | 0.135605804 | -1.655600771 |
| Gm29293 | 0.161114758 | 1.626894704 | 3.426332624 | 0.474821006 |
| 4930433I11Rik | 0.134148234 | -0.20649159 | 3.624632408 | -0.056968974 |
| Gm4884 | 0 | NA | NA | NA |
| 1700110I07Rik | 0.044191549 | -0.206490061 | 3.724945049 | -0.055434391 |
| Gm28061 | 0 | NA | NA | NA |
| Gm9239 | 0 | NA | NA | NA |
| Gm6820 | 0 | NA | NA | NA |
| Gm2128 | 0 | NA | NA | NA |
| Gm28060 | 0 | NA | NA | NA |
| Gm28667 | 0 | NA | NA | NA |
| Gm5896 | 0.091880448 | -0.687373951 | 3.724945049 | -0.184532642 |
| Gm6866 | 0 | NA | NA | NA |
| Gm9244 | 0 | NA | NA | NA |
| Gm5592 | 0 | NA | NA | NA |

| | | | | |
|---|---|---|---|---|
| Gm28059 | 0 | NA | NA | NA |
| Gm2108 | 0 | NA | NA | NA |
| Gm17791 | 0.292366188 | -0.772262631 | 2.05189631 | -0.37636533 |
| Gm9246 | 0 | NA | NA | NA |
| AI987944 | 114.8081952 | -0.137085678 | 0.15233507 | -0.899895725 |
| Gm17102 | 8.180380778 | -0.249575767 | 0.366139749 | -0.681640732 |
| Gm20449 | 4.437436714 | -1.267453714 | 0.577885338 | -2.19326159 |
| Vmn2r57 | 0.538693099 | -0.32816572 | 1.330865733 | -0.246580637 |
| AW146154 | 60.96357997 | -0.268088671 | 0.18464539 | -1.45191099 |
| Gm15470 | 0.332094496 | 1.348706996 | 1.990295162 | 0.677641699 |
| Gm6871 | 0 | NA | NA | NA |
| Gm17217 | 0 | NA | NA | NA |
| 2610021A01Rik | 362.0305132 | -0.041844061 | 0.072020871 | -0.580999101 |
| Zfp788 | 649.6253934 | 0.229819318 | 0.072434192 | 3.172801573 |
| Gm9252 | 0 | NA | NA | NA |
| Vmn2r-ps54 | 6.472406225 | 0.177412873 | 0.558390974 | 0.317721598 |
| Gm6872 | 0.060174537 | 0.755286614 | 3.724945049 | 0.202764498 |
| Cd9-ps | 0.178447459 | 0.27439001 | 3.343252531 | 0.082072774 |
| Vmn2r-ps55 | 0.046345709 | 0.274391604 | 3.724945049 | 0.073663262 |
| Vmn2r-ps56 | 0.066050875 | 0.274391604 | 3.724945049 | 0.073663262 |
| Vmn2r58 | 0 | NA | NA | NA |
| Vmn2r-ps57 | 0 | NA | NA | NA |
| Vmn2r59 | 0 | NA | NA | NA |
| Hmgb1-ps7 | 2.510100352 | -0.246452624 | 0.656665538 | -0.375309209 |
| Gm7224 | 23.28790071 | 0.047979711 | 0.217071531 | 0.221031797 |
| Gm5593 | 0.933648904 | 0.618241408 | 1.309959176 | 0.471954713 |
| Vmn2r60 | 0 | NA | NA | NA |
| Gm2308 | 2.058388305 | 0.430507987 | 0.68817102 | 0.625582849 |
| Vmn2r61 | 0 | NA | NA | NA |
| Vmn2r-ps58 | 0 | NA | NA | NA |
| Gm5594 | 0 | NA | NA | NA |
| Vmn2r-ps59 | 0 | NA | NA | NA |
| Vmn2r-ps60 | 0 | NA | NA | NA |
| 4933421I07Rik | 0 | NA | NA | NA |
| Gm6887 | 0.072025462 | 0.755288839 | 3.724945049 | 0.202765096 |
| Zfp141 | 173.0700645 | 0.274781872 | 0.093967164 | 2.924232888 |
| Gm19246 | 0.428879182 | -1.913947939 | 1.380425463 | -1.386491332 |
| Gm21028 | 1.10003083 | 0.917305329 | 1.013858416 | 0.904766696 |
| Gm7221 | 16.48036244 | -0.947561641 | 0.275171285 | -3.443533877 |
| Gm24938 | 1.200098617 | -0.328238596 | 1.079587858 | -0.304040652 |
| 9830147E19Rik | 96.87745312 | -0.109211367 | 0.182011486 | -0.600024591 |
| Gm5595 | 96.44755503 | -0.411440856 | 0.128351233 | -3.205585541 |
| Gm17067 | 0 | NA | NA | NA |
| Gm18128 | 0 | NA | NA | NA |
| Gm6004 | 0 | NA | NA | NA |
| Vmn2r62 | 0.088383099 | -0.687376175 | 3.724945049 | -0.184533239 |
| Gm2381 | 0.372704184 | 1.104510983 | 1.538562415 | 0.717885067 |

| | | | | |
|---|---|---|---|---|
| Gm7211 | 0 | NA | NA | NA |
| Vmn2r63 | 0 | NA | NA | NA |
| Gm9266 | 0 | NA | NA | NA |
| Gm9294 | 4.49822579 | 1.033810268 | 0.599409036 | 1.724715855 |
| Gm23256 | 0 | NA | NA | NA |
| Gm9268 | 0 | NA | NA | NA |
| RP23-367H8.3 | 0 | NA | NA | NA |
| Gm10351 | 0 | NA | NA | NA |
| Zfp936 | 0.092140247 | -0.687373951 | 3.724945049 | -0.184532642 |
| Gm22707 | 0 | NA | NA | NA |
| EU599041 | 0.135598227 | -1.141928459 | 2.942835517 | -0.388036794 |
| Zfp715 | 456.2127529 | 0.030153079 | 0.0815968 | 0.369537517 |
| Siglecf | 5.415502606 | 1.503842465 | 0.531256142 | 2.830729561 |
| Gm10298 | 0 | NA | NA | NA |
| Siglecg | 1.312064872 | 3.410833588 | 1.122329097 | 3.039067238 |
| Lim2 | 0.046345709 | 0.274391604 | 3.724945049 | 0.073663262 |
| Nkg7 | 96.73496261 | 9.305182707 | 0.946248522 | 9.83376195 |
| Cldnd2 | 1.387261314 | 2.391522752 | 1.268863102 | 1.884776024 |
| Etfb | 181.9463623 | -0.031616794 | 0.132056221 | -0.239419193 |
| Mir7051 | 0.307454254 | 0.755310641 | 1.756171436 | 0.430089355 |
| Vsig10l | 33.09847067 | 0.573759428 | 0.189635145 | 3.025596491 |
| Iglon5 | 396.8950678 | -0.518735978 | 0.101223523 | -5.124658407 |
| 4931406B18Rik | 0.154787072 | 0.274405795 | 2.832963972 | 0.096861731 |
| Cd33 | 51.25040222 | 0.540645247 | 0.327831208 | 1.649157353 |
| Zfp658 | 98.3557729 | -0.330828727 | 0.173596528 | -1.905733539 |
| Zfp719 | 304.4942731 | -0.265512738 | 0.158683049 | -1.673226853 |
| Zfp819 | 3.834671984 | 1.176160953 | 0.65904473 | 1.784645108 |
| Ceacam18 | 0.539507586 | 1.964570042 | 1.355733495 | 1.449082765 |
| Siglece | 5.430287587 | 0.762492971 | 0.538787389 | 1.415201965 |
| Ctu1 | 87.06700889 | -0.01324001 | 0.151031236 | -0.087664051 |
| Klk14 | 0 | NA | NA | NA |
| 2310002F09Rik | 9.427952603 | -0.281103798 | 0.334282302 | -0.840917381 |
| Klk13 | 0.134945109 | 0.274406006 | 2.95629408 | 0.092820944 |
| Klk12 | 0 | NA | NA | NA |
| Klk11 | 0.225356219 | 0.208529641 | 2.121598072 | 0.098288947 |
| Klk10 | 1.594085209 | -0.959163743 | 0.925522616 | -1.03634825 |
| Klk9 | 11.12317148 | 4.719458986 | 1.026487124 | 4.597679673 |
| Klk8 | 23.43242316 | 1.195052307 | 0.335030981 | 3.566990441 |
| Klk7 | 6.053054786 | 4.088882039 | 1.09569782 | 3.731760677 |
| Klk6 | 383.1545088 | 1.206364599 | 0.332526639 | 3.627873557 |
| Klk5 | 0.178695003 | 0.610518405 | 2.710791628 | 0.225217755 |
| Klk4 | 0.195922627 | 0.755302383 | 2.238503775 | 0.337413942 |
| Klk15 | 0 | NA | NA | NA |
| Klk1b7-ps | 0.097139547 | 0.274393828 | 3.724945049 | 0.073663859 |
| Klk1b8 | 0.138087489 | 0.274387362 | 3.620422247 | 0.075788774 |
| Klk1b1 | 0.049553047 | 0.274391604 | 3.724945049 | 0.073663262 |
| Klk1b9 | 0.07796279 | 0.274392189 | 3.724945049 | 0.073663419 |

| | | | | |
|---|---|---|---|---|
| Klk1b11 | 0.276660002 | -0.206481814 | 1.849077419 | -0.111667479 |
| Klk1b26 | 0.046345709 | 0.274391604 | 3.724945049 | 0.073663262 |
| Klk1b27 | 1.612931274 | 3.142531618 | 1.089976405 | 2.883118941 |
| Klk1b21 | 0 | NA | NA | NA |
| Klk1b22 | 0 | NA | NA | NA |
| Klk1b16 | 0.166231529 | 1.727084934 | 3.417706679 | 0.505334453 |
| Klk1b24 | 0.038981395 | 0.274393828 | 3.724945049 | 0.073663859 |
| Klk1b3 | 0 | NA | NA | NA |
| Klk1b4 | 0 | NA | NA | NA |
| Klk1b5 | 0 | NA | NA | NA |
| Gm10109 | 2.250427361 | 0.605722933 | 0.675170252 | 0.897141027 |
| Klk1 | 11.42971091 | 5.009781377 | 0.923493978 | 5.424812178 |
| 1700028J19Rik | 6.192777873 | 0.528197727 | 0.451262705 | 1.170488323 |
| Gm25386 | 0 | NA | NA | NA |
| 2410002F23Rik | 502.3878778 | 0.219636594 | 0.064631354 | 3.398297888 |
| Gm28496 | 4.360017817 | 1.024976561 | 0.630595527 | 1.625410453 |
| Gm26247 | 0 | NA | NA | NA |
| Snord88c | 1.895772021 | -1.442087723 | 0.94758507 | -1.521855682 |
| Snord88a | 0 | NA | NA | NA |
| Gm23991 | 0 | NA | NA | NA |
| Acpt | 0 | NA | NA | NA |
| Gm15517 | 0 | NA | NA | NA |
| Clec11a | 34.26572508 | -0.434673228 | 0.185102319 | -2.348286232 |
| Shank1 | 9409.0605 | -0.503344479 | 0.132459185 | -3.799996811 |
| 1700008O03Rik | 12.39141503 | 0.43969834 | 0.312888004 | 1.405289862 |
| Syt3 | 466.7523524 | -0.293808465 | 0.099092477 | -2.964992629 |
| Lrrc4b | 2121.923621 | -0.225360477 | 0.079664599 | -2.828866036 |
| Aspdh | 27.56526144 | -0.689940567 | 0.239197046 | -2.884402542 |
| Josd2 | 428.3256145 | -0.5126801 | 0.10781385 | -4.755234152 |
| Mir7052 | 5.276354033 | -0.844857842 | 0.425521818 | -1.985463039 |
| Emc10 | 1263.340695 | 0.008491966 | 0.075844555 | 0.111965403 |
| Fam71e1 | 39.06557393 | 0.172148841 | 0.179405951 | 0.959549224 |
| Mybpc2 | 3.531971173 | 0.510621255 | 0.562196942 | 0.908260463 |
| Spib | 15.69420495 | 3.527918934 | 0.624857229 | 5.645960021 |
| Pold1 | 95.92363125 | 0.075819292 | 0.193778981 | 0.391266855 |
| Mir7053 | 0.189266655 | 0.682551539 | 2.684314295 | 0.254274077 |
| Nr1h2 | 489.1957701 | 0.337205803 | 0.102306751 | 3.296026903 |
| Napsa | 53.67902564 | 5.506837234 | 0.67089292 | 8.208220815 |
| Gm15396 | 3.09078215 | -0.325163057 | 0.664857056 | -0.48907213 |
| Kcnc3 | 3773.256925 | -0.293938366 | 0.117880766 | -2.493522698 |
| Myh14 | 523.8678887 | -0.344077521 | 0.157970104 | -2.178117968 |
| 2310016G11Rik | 0.744214146 | -0.46723541 | 1.157088666 | -0.403802599 |
| Izumo2 | 0 | NA | NA | NA |
| Zfp473 | 10.07782813 | -0.043128653 | 0.303667908 | -0.142025721 |
| Vrk3 | 217.5211013 | 0.10441756 | 0.12860696 | 0.811912201 |
| Atf5 | 717.0131176 | 1.313557371 | 0.176470189 | 7.443508605 |
| Nup62 | 303.2451805 | 0.327729591 | 0.116467202 | 2.813921744 |

| | | | | |
|---|---|---|---|---|
| Il4i1 | 16.51100415 | -0.275291606 | 0.294729176 | -0.934049386 |
| Tbc1d17 | 424.275241 | 0.14329429 | 0.080812262 | 1.773175094 |
| Akt1s1 | 507.5705319 | 0.137854796 | 0.067194884 | 2.051566829 |
| Mir707 | 0 | NA | NA | NA |
| Pnkp | 197.282501 | -0.020856608 | 0.088897646 | -0.234613723 |
| Ptov1 | 973.5278823 | -0.060443066 | 0.054846682 | -1.102036879 |
| Med25 | 617.720589 | 0.233917821 | 0.073432354 | 3.185487177 |
| Fuz | 99.72731422 | -0.621279747 | 0.14075746 | -4.413831743 |
| Ap2a1 | 2601.373962 | -0.062542137 | 0.050608855 | -1.235794364 |
| Tsks | 3.95301002 | 0.533713747 | 0.458125953 | 1.164993477 |
| Cpt1c | 718.0796799 | -0.064829947 | 0.07276901 | -0.890900491 |
| Prmt1 | 509.9239553 | 0.090337575 | 0.095833259 | 0.942653681 |
| Gm15545 | 9.287909809 | -0.647224116 | 0.370765051 | -1.745644889 |
| Bcl2l12 | 25.31009217 | 0.515818712 | 0.287107715 | 1.796603452 |
| Irf3 | 149.8132699 | -0.394959921 | 0.172273033 | -2.292639268 |
| Scaf1 | 2253.505891 | 0.070912046 | 0.056301782 | 1.259499142 |
| Mir7054 | 0.567171362 | -1.227228405 | 1.206289111 | -1.017358438 |
| Rras | 111.4151899 | 1.289319112 | 0.19252062 | 6.697044254 |
| Prr12 | 1649.371309 | -0.119385052 | 0.065073583 | -1.834616232 |
| Prrg2 | 52.66422563 | 0.351569629 | 0.15722393 | 2.236107628 |
| Nosip | 442.5713574 | 0.506757702 | 0.089390758 | 5.669016752 |
| Rcn3 | 57.62462159 | 0.439958355 | 0.214159509 | 2.054348921 |
| Fcgrt | 111.4072777 | -0.143691349 | 0.191331343 | -0.751007895 |
| Mir150 | 0 | NA | NA | NA |
| Rps11 | 1103.617122 | 0.300322193 | 0.09885116 | 3.038125124 |
| Snord35b | 0.134597393 | -1.14191529 | 2.949594298 | -0.387143171 |
| Rpl13a | 2064.540611 | 0.467290675 | 0.144641528 | 3.230681273 |
| Snord35a | 0.656197145 | -1.523479092 | 1.246875292 | -1.221837582 |
| Snord34 | 0 | NA | NA | NA |
| Snord33 | 0 | NA | NA | NA |
| mmu-mir-5121 | 0 | NA | NA | NA |
| Snord32a | 0 | NA | NA | NA |
| Flt3l | 50.76228596 | 2.012609133 | 0.229942671 | 8.752656131 |
| Aldh16a1 | 113.740617 | 0.458120304 | 0.132738479 | 3.451299924 |
| Pih1d1 | 223.2642417 | 0.001170211 | 0.091739346 | 0.012755822 |
| Slc17a7 | 6082.783442 | -0.116353153 | 0.121591002 | -0.956922397 |
| mmu-mir-7055 | 0.339810401 | -0.7958245 | 1.704706025 | -0.46683973 |
| Gfy | 10.55721659 | 0.061128456 | 0.64395377 | 0.094926778 |
| Pth2 | 1.157525169 | -2.049999304 | 1.029495717 | -1.9912655 |
| Ccdc155 | 18.99181208 | -0.285894925 | 0.279544506 | -1.022717022 |
| Dkkl1 | 52.36991393 | -0.447110235 | 0.204740398 | -2.183790983 |
| Tead2 | 38.54700613 | -0.215741119 | 0.238035504 | -0.906340085 |
| Cd37 | 67.79623972 | 1.370431358 | 0.24422445 | 5.611360187 |
| Slc6a16 | 3.556493767 | -0.073304627 | 0.524827822 | -0.139673669 |
| Gm23585 | 0 | NA | NA | NA |
| 1700039E15Rik | 0.62192417 | 1.046411985 | 1.279569423 | 0.817784456 |
| Trpm4 | 109.1332828 | 0.224379255 | 0.172071194 | 1.303990804 |

| | | | | |
|---|---|---|---|---|
| Rpl14-ps1 | 281.4107973 | 0.615355013 | 0.13398667 | 4.592658463 |
| Hrc | 0.548344614 | -0.933740589 | 1.374039704 | -0.679558666 |
| Ppfia3 | 1660.177301 | -0.187026213 | 0.103804057 | -1.801723547 |
| Mtag2 | 16.72535298 | -0.453138533 | 0.336944527 | -1.344846101 |
| Lin7b | 211.863161 | -0.661090378 | 0.188289562 | -3.511030415 |
| Snrnp70 | 1046.054046 | -0.196488892 | 0.178750859 | -1.099233276 |
| Gm22478 | 0.046345709 | 0.274391604 | 3.724945049 | 0.073663262 |
| Kcna7 | 0.228985044 | 0.274409018 | 2.57304005 | 0.106647783 |
| Ntf5 | 1.628131443 | 1.355992856 | 0.896941643 | 1.511796075 |
| Lhb | 9.980967548 | 1.153879023 | 0.571226998 | 2.020000855 |
| Ruvbl2 | 254.9844317 | -0.20845477 | 0.11029562 | -1.889964168 |
| Gys1 | 207.1643521 | -0.563031194 | 0.126097565 | -4.465044149 |
| Ftl1 | 3794.633164 | 1.02317325 | 0.193641995 | 5.283839642 |
| Gm22133 | 0 | NA | NA | NA |
| Bax | 270.9969915 | 0.310008559 | 0.100468092 | 3.085641955 |
| Dhdh | 156.4792992 | -0.111893796 | 0.148472439 | -0.753633449 |
| Tulp2 | 0.658348016 | -0.659596323 | 1.316968802 | -0.500844304 |
| Nucb1 | 959.7799318 | 0.232754792 | 0.068512577 | 3.397256402 |
| Ppp1r15a | 417.9443 | 1.652681915 | 0.354711482 | 4.659228693 |
| Plekha4 | 287.7791967 | 3.573106786 | 0.319179244 | 11.19467149 |
| Gm16022 | 0.072025462 | 0.755288839 | 3.724945049 | 0.202765096 |
| Hsd17b14 | 2.84382331 | 0.012090406 | 0.647725423 | 0.018665944 |
| 0610005C13Rik | 1.694978974 | -1.606675676 | 0.909480747 | -1.766585694 |
| Bcat2 | 107.8855267 | -0.236345669 | 0.154254297 | -1.532182079 |
| Fgf21 | 1.959085883 | 2.070985504 | 1.188979159 | 1.74181817 |
| Fut1 | 0.956979021 | 0.770444318 | 1.062908511 | 0.724845374 |
| Izumo1 | 0.702218406 | -0.625953248 | 1.480798049 | -0.422713447 |
| Rasip1 | 78.58288542 | 1.441142199 | 0.222840528 | 6.467145866 |
| Gm16047 | 2.391421094 | 0.39336585 | 0.734605193 | 0.535479266 |
| Mamstr | 31.70904901 | 0.385538095 | 0.256599481 | 1.502489769 |
| Fut2 | 5.566764691 | 0.691565845 | 0.474327086 | 1.457993577 |
| Sec1 | 5.465091639 | 0.524307279 | 0.451453801 | 1.161375268 |
| Ntn5 | 29.63111439 | 0.02875497 | 0.309403287 | 0.092936861 |
| Car11 | 810.7850525 | -0.212316274 | 0.137566773 | -1.543368856 |
| Dbp | 869.3346952 | -0.61747556 | 0.152161689 | -4.058022511 |
| Sphk2 | 380.1522572 | 0.298488928 | 0.111243794 | 2.683196219 |
| Rpl18 | 1024.378528 | 0.516827217 | 0.125766024 | 4.10943433 |
| Fam83e | 2.171234967 | -1.667988273 | 0.772508658 | -2.159183921 |
| Spaca4 | 0 | NA | NA | NA |
| Sult2b1 | 46.17871227 | 0.343084615 | 0.263797722 | 1.300559431 |
| Lmtk3 | 2315.447368 | -0.259166101 | 0.109782407 | -2.360725249 |
| Cyth2 | 643.2901348 | -0.093727176 | 0.077752452 | -1.205456204 |
| Kcnj14 | 17.48480922 | 0.082589138 | 0.409646903 | 0.201610553 |
| Grwd1 | 158.3457012 | 0.425094925 | 0.155316729 | 2.736955177 |
| Grin2d | 305.5974088 | -0.099143434 | 0.092594839 | -1.07072311 |
| Kdelr1 | 538.7074619 | 0.059872169 | 0.074372948 | 0.805026155 |
| Gm22098 | 0 | NA | NA | NA |

| | | | | |
|---|---|---|---|---|
| Syngr4 | 0.321511422 | 1.235183055 | 1.76492991 | 0.699848219 |
| Tmem143 | 157.483315 | -0.234243224 | 0.116146608 | -2.016789196 |
| Emp3 | 69.37531286 | 1.569690142 | 0.294842314 | 5.323829272 |
| Ccdc114 | 21.307866 | -0.047837133 | 0.25041476 | -0.191031602 |
| Abcc6 | 6.348614184 | -1.05312359 | 0.470806044 | -2.236852316 |
| Nomo1 | 1371.14784 | -0.233301941 | 0.065522847 | -3.560619718 |
| Kcnj11 | 116.36677 | -0.301153455 | 0.125080421 | -2.407678622 |
| Abcc8 | 310.3174424 | -0.624174872 | 0.120563863 | -5.177130678 |
| Ush1c | 7.85454541 | 0.540629713 | 0.352558067 | 1.533448708 |
| Otog | 2.40295838 | -0.040659184 | 0.610900171 | -0.066556183 |
| Gm38176 | 0 | NA | NA | NA |
| Myod1 | 0.509027769 | 0.352150278 | 1.372280514 | 0.256616832 |
| Kcnc1 | 3456.691312 | -0.348477581 | 0.075470545 | -4.617398517 |
| Sergef | 366.9267911 | -0.168739617 | 0.104830769 | -1.609638262 |
| Gm22969 | 1.792854059 | 0.362349568 | 0.668056821 | 0.542393336 |
| Gm15700 | 0 | NA | NA | NA |
| Tph1 | 2.962511132 | 2.000020041 | 0.854314099 | 2.341082798 |
| Saal1 | 161.7785274 | 0.104028231 | 0.094132496 | 1.105125603 |
| Saa3 | 441.1512112 | 11.3626607 | 1.116772691 | 10.174551 |
| Saa4 | 0.385868569 | 1.240909231 | 1.692971908 | 0.732976859 |
| Saa1 | 4.201556626 | 3.006561268 | 0.789817979 | 3.806650834 |
| Saa2 | 1.450828853 | 4.157836 | 1.074643715 | 3.869036726 |
| Hps5 | 293.9049292 | -0.052137095 | 0.116823008 | -0.446291319 |
| Gtf2h1 | 540.0561687 | 0.377810653 | 0.086413215 | 4.372139767 |
| Gm9392 | 24.65709606 | 0.342403985 | 0.232368495 | 1.473538766 |
| Ldha | 1608.193775 | 0.125052807 | 0.081454172 | 1.535253551 |
| Ldhc | 0.061920613 | 0.274393828 | 3.724945049 | 0.073663859 |
| Tsg101 | 606.9962445 | 0.124431632 | 0.073794429 | 1.68619277 |
| Uevld | 206.960226 | -0.285621703 | 0.114157839 | -2.5019894 |
| Gm9999 | 0.062426873 | 0.755288839 | 3.724945049 | 0.202765096 |
| Spty2d1 | 614.2646612 | 0.673305071 | 0.120497488 | 5.587710419 |
| Tmem86a | 239.3545487 | 0.921284961 | 0.141498012 | 6.510939243 |
| Ptpn5 | 1663.361268 | -0.282945318 | 0.133010551 | -2.127239648 |
| Mir7056 | 0.65404267 | -0.430029663 | 1.206455282 | -0.356440615 |
| Gm14377 | 1.857966922 | 2.311169076 | 1.08819538 | 2.123854887 |
| Mrgpra6 | 0 | NA | NA | NA |
| Mrgpra9 | 0.097139547 | 0.274393828 | 3.724945049 | 0.073663859 |
| Mrgprx3-ps | 0 | NA | NA | NA |
| Mrgpra1 | 0 | NA | NA | NA |
| Gm29479 | 0 | NA | NA | NA |
| Gm25326 | 0.128477747 | 0.755285235 | 3.677131942 | 0.205400635 |
| Mrgpra2a | 0.092691418 | 0.274387304 | 3.724945049 | 0.073662108 |
| Mrgpra2b | 0.175945213 | 0.755298819 | 2.714629575 | 0.278232738 |
| Gm7255 | 0 | NA | NA | NA |
| Mrgprc4-ps | 0 | NA | NA | NA |
| Mrgpra14 | 0 | NA | NA | NA |
| Gm9322 | 0 | NA | NA | NA |

| | | | | |
|---|---|---|---|---|
| Mrgpra3 | 0 | NA | NA | NA |
| Mrgprc1-ps | 0 | NA | NA | NA |
| Gm2707 | 0 | NA | NA | NA |
| Gm28288 | 0 | NA | NA | NA |
| Mrgprc3-ps | 0 | NA | NA | NA |
| Gm5733 | 0 | NA | NA | NA |
| Gm22427 | 0 | NA | NA | NA |
| Gm18348 | 0 | NA | NA | NA |
| Gm29563 | 0 | NA | NA | NA |
| Mrgprc7-ps | 0 | NA | NA | NA |
| Gm18349 | 0 | NA | NA | NA |
| Mrgpra18-ps | 0 | NA | NA | NA |
| Gm18350 | 0 | NA | NA | NA |
| Gm5734 | 0 | NA | NA | NA |
| Mrgprc2-ps | 0 | NA | NA | NA |
| Mrgpra4 | 0 | NA | NA | NA |
| Mrgprx1 | 0 | NA | NA | NA |
| Mrgprb7-ps | 0 | NA | NA | NA |
| Gm28666 | 0 | NA | NA | NA |
| Mrgprb12-ps | 0 | NA | NA | NA |
| Gm18351 | 0 | NA | NA | NA |
| Gm28665 | 0 | NA | NA | NA |
| Mrgprb5 | 0 | NA | NA | NA |
| Gm18352 | 0 | NA | NA | NA |
| Mrgprb4 | 0 | NA | NA | NA |
| Gm26950 | 0 | NA | NA | NA |
| Gm18356 | 0 | NA | NA | NA |
| Mrgprb13 | 0 | NA | NA | NA |
| Gm18357 | 0 | NA | NA | NA |
| Mrgprb8 | 0.066050875 | 0.274391604 | 3.724945049 | 0.073663262 |
| Mrgprb1 | 0.19039241 | -0.566201134 | 2.660990838 | -0.21277831 |
| Mrgprx2 | 0.10503227 | 0.274389469 | 3.724945049 | 0.073662689 |
| Mrgprb2 | 0.044544648 | 0.755286614 | 3.724945049 | 0.202764498 |
| Mrgprb9-ps | 0 | NA | NA | NA |
| Gm18353 | 0 | NA | NA | NA |
| Mrgprb3 | 0 | NA | NA | NA |
| Mrgprb11 | 0.045882455 | 0.755288839 | 3.724945049 | 0.202765096 |
| Zdhhc13 | 253.9183394 | 0.063407659 | 0.09266679 | 0.6842544 |
| Csrp3 | 0.531680062 | 0.75530798 | 1.623046209 | 0.465364434 |
| E2f8 | 18.28425174 | 1.27447816 | 0.337860764 | 3.772199371 |
| Gm2788 | 0.049553047 | 0.274391604 | 3.724945049 | 0.073663262 |
| Nav2 | 2595.787216 | -0.126946453 | 0.102667841 | -1.236477275 |
| Gm37613 | 24.19551003 | -1.056098636 | 0.396645665 | -2.662574509 |
| Gm38059 | 24.62141553 | -0.744543407 | 0.43791694 | -1.700193211 |
| Dbx1 | 0.097139547 | 0.274393828 | 3.724945049 | 0.073663859 |
| Htatip2 | 67.03302061 | 0.277023574 | 0.205305962 | 1.349320652 |
| Prmt3 | 302.2229905 | -0.240633901 | 0.08948134 | -2.68920762 |

| | | | | |
|---|---|---|---|---|
| Gm26856 | 0 | NA | NA | NA |
| Slc6a5 | 485.8678254 | -0.614926458 | 0.562169709 | -1.093844879 |
| Nell1 | 633.6668325 | -0.106076594 | 0.101568971 | -1.044379915 |
| Nell1os | 0.048949532 | -0.206487837 | 3.724945049 | -0.055433794 |
| 4933405O20Rik | 9.680996758 | -0.342687162 | 0.377468753 | -0.907855708 |
| Ano5 | 52.81759069 | -0.066648588 | 0.192060511 | -0.347018695 |
| Slc17a6 | 1077.091468 | -0.189013635 | 0.154343535 | -1.224629431 |
| Gm7336 | 39.57013241 | 0.005384395 | 0.181818896 | 0.029614057 |
| Gm29296 | 0.142694128 | -0.68737806 | 2.905728216 | -0.236559654 |
| Fancf | 37.44894291 | -0.00721289 | 0.238790915 | -0.030205882 |
| Gas2 | 177.6414515 | 0.05871452 | 0.129289622 | 0.454131732 |
| Svip | 477.7009006 | -0.315178128 | 0.095074489 | -3.315065181 |
| 1700015G11Rik | 0.161582804 | 1.257837896 | 2.797750917 | 0.44958895 |
| Gm22211 | 0 | NA | NA | NA |
| Gm6181 | 1.4136546 | -1.746493054 | 1.064156867 | -1.641198876 |
| Gm22444 | 0 | NA | NA | NA |
| Mir6238 | 0 | NA | NA | NA |
| Gm27921 | 0 | NA | NA | NA |
| Gm25956 | 0 | NA | NA | NA |
| Gm5776 | 44.67075326 | 0.104569704 | 0.188260701 | 0.55545158 |
| Luzp2 | 603.6111141 | -0.350860274 | 0.107502317 | -3.26374616 |
| Gm24116 | 0 | NA | NA | NA |
| Gm26482 | 0 | NA | NA | NA |
| Siglech | 79.78094538 | -0.09513936 | 0.300048053 | -0.317080411 |
| Tubgcp5 | 553.9336148 | 0.091633264 | 0.067976515 | 1.348013574 |
| Cyfip1 | 1268.126837 | 0.228753735 | 0.084162101 | 2.718013583 |
| Gm17907 | 0.149576918 | 0.755295313 | 2.863326609 | 0.263782452 |
| Nipa2 | 555.4063321 | 0.274055865 | 0.079092334 | 3.465011743 |
| A230056P14Rik | 25.86046502 | -0.799066735 | 0.241903813 | -3.303241584 |
| Nipa1 | 678.816936 | -0.480618009 | 0.106935953 | -4.494447347 |
| Gm15888 | 0 | NA | NA | NA |
| Rps12l1 | 1.168735097 | 3.453140262 | 1.110586739 | 3.10929362 |
| Herc2 | 5466.518983 | -0.376303975 | 0.083718922 | -4.494849716 |
| Oca2 | 2.497417949 | -0.100273482 | 0.650017659 | -0.154262705 |
| Gabrg3 | 564.6589 | -0.044899708 | 0.097533727 | -0.460350583 |
| Gm9962 | 23.2644548 | -0.113263956 | 0.217799687 | -0.520037277 |
| Gabra5 | 375.6717185 | -0.149813936 | 0.106372371 | -1.408391441 |
| Gabrb3 | 2705.952044 | -0.090739148 | 0.077084807 | -1.17713402 |
| Mir6389 | 0 | NA | NA | NA |
| Gm23911 | 0.328919656 | 0.196303562 | 1.747593439 | 0.112327935 |
| Gm26288 | 0 | NA | NA | NA |
| Atp10a | 221.527901 | 1.239110005 | 0.205476549 | 6.03042055 |
| Gm6226 | 2.39103255 | -0.42331616 | 0.822037587 | -0.514959615 |
| Ube3a | 3913.118757 | 0.073164384 | 0.073998569 | 0.988727017 |
| Snhg14 | 6561.535641 | -0.154105835 | 0.124930866 | -1.233528915 |
| Gm25499 | 5.73832025 | 0.995027181 | 0.628359325 | 1.583532132 |
| Gm24966 | 65.59095034 | -0.332787919 | 0.232801097 | -1.429494635 |

| | | | | |
|---|---|---|---|---|
| Gm22496 | 12.77173823 | -0.137125798 | 0.38503658 | -0.356137067 |
| Gm25648 | 15.8722383 | -0.205251642 | 0.459792484 | -0.446400602 |
| Gm24306 | 4.90255571 | 0.500804795 | 0.454644732 | 1.101529964 |
| Gm25934 | 2.147180949 | 0.336504775 | 0.646042136 | 0.52087125 |
| Gm25156 | 8.000524403 | 0.368590668 | 0.407154495 | 0.905284536 |
| Gm26488 | 4.065985139 | -0.062205093 | 0.514943078 | -0.12079994 |
| Gm24220 | 8.515480277 | 0.023151662 | 0.335955184 | 0.068912947 |
| Gm24653 | 7.944433476 | 0.002958904 | 0.343082408 | 0.00862447 |
| Gm24969 | 3.545771104 | -0.816597776 | 0.604798508 | -1.350198066 |
| Gm24417 | 0.567759171 | -1.636571678 | 1.304282754 | -1.254767552 |
| Gm25121 | 85.00767218 | -0.299713559 | 0.267819046 | -1.119089787 |
| Gm24657 | 0.475242527 | -1.528454584 | 1.655822505 | -0.92307876 |
| Gm26230 | 0.075043205 | 0.755286614 | 3.724945049 | 0.202764498 |
| Gm23922 | 0.156071933 | -0.206480922 | 3.489245271 | -0.059176385 |
| Gm25314 | 2.261769448 | -1.254923681 | 0.876490124 | -1.431760207 |
| Gm26096 | 0.262111692 | 1.865729859 | 2.447786356 | 0.762211071 |
| Gm22996 | 6.119234212 | -0.1383247 | 0.557427151 | -0.24814848 |
| Gm26499 | 0.650021907 | -0.69125395 | 1.369387417 | -0.504790639 |
| Gm25646 | 2.157091062 | 0.188506315 | 0.664942023 | 0.283492859 |
| Gm26498 | 21.92470555 | 0.075151415 | 0.328311895 | 0.228902504 |
| Gm23660 | 2.095785588 | -0.521061267 | 0.65962244 | -0.789938661 |
| Gm23933 | 2.60823589 | -0.275762979 | 0.580953294 | -0.474673234 |
| Gm24862 | 0.637223532 | -0.19646543 | 1.139312922 | -0.172442028 |
| Gm24866 | 6.740698188 | -0.523726079 | 0.416297976 | -1.258055789 |
| Gm22348 | 0 | NA | NA | NA |
| Gm26466 | 4.89789051 | -0.281241449 | 0.57568888 | -0.488530279 |
| Gm24799 | 3.843037622 | -0.774289032 | 0.607084146 | -1.275422916 |
| Gm24088 | 1.480005993 | -0.040636711 | 0.736792693 | -0.055153521 |
| Gm25463 | 3.198438298 | -0.296568771 | 0.686426473 | -0.432047398 |
| Gm25229 | 2.607201442 | -0.744718492 | 0.576193754 | -1.292479288 |
| Gm24528 | 0.50461959 | -0.566740554 | 1.30093537 | -0.435640822 |
| Gm23356 | 0.770457139 | 1.153013582 | 1.293877808 | 0.891130194 |
| Gm25449 | 0.090074004 | 0.274402725 | 3.724945049 | 0.073666248 |
| Gm24872 | 0.860548126 | -0.804418997 | 0.919027947 | -0.875293292 |
| Gm26337 | 0.156716517 | -0.68737803 | 2.821984814 | -0.243579635 |
| Gm23145 | 0.371627904 | 0.690013262 | 1.571428355 | 0.439099409 |
| Gm22130 | 0.047948698 | -0.206487837 | 3.724945049 | -0.055433794 |
| Gm25988 | 0 | NA | NA | NA |
| Gm25221 | 0 | NA | NA | NA |
| Gm26389 | 0.044191549 | -0.206490061 | 3.724945049 | -0.055434391 |
| Gm23471 | 2.996745481 | 0.105590286 | 0.576013543 | 0.183312159 |
| Gm25120 | 0.150248541 | 0.755303235 | 2.859451661 | 0.264142684 |
| Gm24418 | 0 | NA | NA | NA |
| Gm23687 | 0 | NA | NA | NA |
| Gm22393 | 0 | NA | NA | NA |
| Gm24639 | 0.168184286 | -0.206481493 | 2.759948137 | -0.074813541 |
| Gm24566 | 0 | NA | NA | NA |

| | | | | |
|---|---|---|---|---|
| Gm22111 | 0 | NA | NA | NA |
| Gm23076 | 0.042716945 | -0.206490061 | 3.724945049 | -0.055434391 |
| Gm25874 | 0 | NA | NA | NA |
| Gm25088 | 0.11657011 | 1.236179401 | 3.724945049 | 0.331865138 |
| Gm22510 | 5.686408051 | 0.346504569 | 0.485529271 | 0.713663602 |
| Gm24983 | 1.441787318 | 0.44143366 | 0.77912172 | 0.566578557 |
| Gm24027 | 3.476611664 | 0.315747228 | 0.560830862 | 0.562999025 |
| Gm22524 | 0.172163037 | -0.206488682 | 2.739828592 | -0.075365547 |
| Gm22630 | 0.258404708 | 1.983258179 | 2.047008745 | 0.96885672 |
| Gm25741 | 0.657611471 | -0.647688593 | 1.173093181 | -0.552120329 |
| Gm26059 | 0.559779376 | -0.979995036 | 1.343397473 | -0.72949001 |
| Gm25175 | 0.444383067 | 0.246761201 | 1.5225835 | 0.162067434 |
| Gm22511 | 1.321027671 | 0.706305025 | 0.910615801 | 0.775634494 |
| Gm25077 | 0.739785724 | 1.374756041 | 1.371175895 | 1.002611005 |
| Gm22834 | 0.106971521 | 1.236179401 | 3.724945049 | 0.331865138 |
| Gm24926 | 1.408313197 | 0.092346294 | 0.958258834 | 0.096368842 |
| Gm26200 | 0.042716945 | -0.206490061 | 3.724945049 | -0.055434391 |
| Gm22632 | 0.060174537 | 0.755286614 | 3.724945049 | 0.202764498 |
| Gm23254 | 4.913973528 | 0.606057671 | 0.443258857 | 1.367277071 |
| Gm26390 | 0.52711509 | 1.698751913 | 1.306491323 | 1.300239721 |
| Gm26366 | 1.250668422 | 1.379623426 | 0.818252511 | 1.686060731 |
| Gm26336 | 0.636553247 | -0.376057636 | 1.155200859 | -0.325534415 |
| Gm26283 | 0.137685729 | -0.206481801 | 2.938031474 | -0.070278961 |
| Gm25098 | 0.174523186 | 0.189228852 | 3.364914203 | 0.056235862 |
| Gm24702 | 0.239175706 | 1.894637782 | 2.99364251 | 0.632887119 |
| Gm26432 | 0.437372945 | 1.58035512 | 1.399462779 | 1.129258415 |
| Gm25462 | 6.852308181 | 0.109484631 | 0.498660335 | 0.219557529 |
| Gm24495 | 12.03897181 | 0.111099087 | 0.32015701 | 0.347014381 |
| Gm25523 | 2.19067646 | -0.269441867 | 0.764629362 | -0.352382317 |
| Gm22255 | 1.065694062 | 1.306720557 | 0.978994188 | 1.334758237 |
| Gm23449 | 8.027250868 | 0.202598273 | 0.366780286 | 0.552369581 |
| Gm23310 | 0.295362062 | -2.172043576 | 1.951579377 | -1.112967067 |
| Gm25461 | 0.060174537 | 0.755286614 | 3.724945049 | 0.202764498 |
| Gm24310 | 0.210314706 | 0.74043346 | 2.185265531 | 0.338829973 |
| Gm22909 | 0.331212083 | -0.118676429 | 1.969685487 | -0.060251461 |
| Gm25823 | 1.425569217 | -1.995474803 | 0.941789177 | -2.118812632 |
| Gm25647 | 0.32719657 | 0.250651004 | 1.768822889 | 0.141704975 |
| Gm22252 | 2.966653922 | 0.086881566 | 0.604769427 | 0.143660645 |
| Gm26392 | 0.252618675 | -0.51469678 | 2.50787895 | -0.205231907 |
| Gm24219 | 2.893504144 | -0.360197963 | 0.555234867 | -0.648730807 |
| Gm25230 | 1.071417476 | 1.960863413 | 1.052717728 | 1.862667799 |
| Gm22627 | 0.53540973 | -0.409285359 | 1.210181395 | -0.338201662 |
| Gm24585 | 0.104637558 | -0.206492286 | 3.724945049 | -0.055434988 |
| Gm23305 | 0.469128522 | 0.47940981 | 1.566732143 | 0.305993473 |
| Gm24206 | 0.186417525 | 0.740120901 | 2.263344322 | 0.327003229 |
| Gm25085 | 2.147377591 | -0.099522221 | 0.656619561 | -0.151567554 |
| Gm25209 | 0.361503534 | -0.601067264 | 1.554689118 | -0.386615727 |

| | | | | |
|---|---|---|---|---|
| Gm23682 | 0.137685729 | -0.206481801 | 2.938031474 | -0.070278961 |
| Gm24654 | 0.322713358 | 0.369400663 | 1.703105916 | 0.216898233 |
| Gm26334 | 0.362177786 | 0.628319054 | 1.510603707 | 0.415939039 |
| Gm25742 | 0.531759696 | 1.090627061 | 1.326642598 | 0.822095614 |
| Gm25585 | 0 | NA | NA | NA |
| Gm25840 | 0 | NA | NA | NA |
| Gm25089 | 0.347269729 | 1.245764272 | 3.262018299 | 0.381899842 |
| Gm24040 | 0.042716945 | -0.206490061 | 3.724945049 | -0.055434391 |
| Gm22050 | 0.315224781 | -0.562196571 | 1.537812998 | -0.365581883 |
| Gm23286 | 0 | NA | NA | NA |
| Gm24021 | 0 | NA | NA | NA |
| Gm22274 | 0 | NA | NA | NA |
| Gm24652 | 0.045882455 | 0.755288839 | 3.724945049 | 0.202765096 |
| Gm23618 | 0 | NA | NA | NA |
| Gm23688 | 0 | NA | NA | NA |
| Gm24570 | 0 | NA | NA | NA |
| Gm26467 | 0 | NA | NA | NA |
| Gm25194 | 0.147068667 | 1.236183405 | 3.514416608 | 0.351746404 |
| Gm22640 | 0 | NA | NA | NA |
| Gm24952 | 0 | NA | NA | NA |
| Gm25087 | 0 | NA | NA | NA |
| Gm23944 | 0 | NA | NA | NA |
| Gm22629 | 0.292758571 | 1.222756596 | 2.033299417 | 0.601365734 |
| Gm26374 | 0.198152624 | 0.27438216 | 3.724945049 | 0.073660727 |
| Gm26284 | 0 | NA | NA | NA |
| Gm25452 | 0 | NA | NA | NA |
| Gm25984 | 0 | NA | NA | NA |
| Gm25146 | 0.176667124 | -0.206481646 | 2.304379988 | -0.089603992 |
| Gm24759 | 1.076799075 | -0.053790666 | 0.972025333 | -0.055338749 |
| Gm24100 | 1.19629966 | -0.803480677 | 0.895257332 | -0.897485726 |
| Gm22253 | 0.244208214 | 1.256183507 | 2.453152552 | 0.512069054 |
| Gm25017 | 0.060174537 | 0.755286614 | 3.724945049 | 0.202764498 |
| Gm23575 | 0.210511418 | -0.206481408 | 2.190351522 | -0.094268617 |
| Gm22391 | 0.323669746 | -1.403101863 | 1.503900664 | -0.932975094 |
| Gm22449 | 1.252922778 | 0.551615349 | 0.871616309 | 0.632864878 |
| Gm24137 | 0.11657011 | 1.236179401 | 3.724945049 | 0.331865138 |
| Gm22912 | 0.442601433 | 0.785091088 | 1.465775171 | 0.535614945 |
| Gm25710 | 0.427556563 | -0.36735285 | 1.484283752 | -0.247495029 |
| Gm26434 | 7.062431358 | -0.713045962 | 0.410883082 | -1.73539869 |
| Gm23560 | 8.541564187 | 0.049139333 | 0.406964358 | 0.120746036 |
| Gm25870 | 11.09216655 | -0.121634794 | 0.32563633 | -0.373529557 |
| Gm23862 | 2.015428526 | -1.13394019 | 0.692691669 | -1.63700567 |
| Gm26504 | 0.522929342 | 0.919033715 | 1.298577884 | 0.707723215 |
| Gm22285 | 0 | NA | NA | NA |
| Gm22131 | 0.271448477 | -1.225204947 | 2.428973351 | -0.504412676 |
| Gm24153 | 0.725964322 | -0.723523379 | 1.182897462 | -0.611653505 |
| Gm26365 | 0 | NA | NA | NA |

| | | | | |
|---|---|---|---|---|
| Gm24609 | 0.334054859 | 0.206145258 | 3.697742571 | 0.055748948 |
| Gm24264 | 0.41844861 | -0.258897083 | 3.288898524 | -0.078718477 |
| Gm25155 | 0.246032045 | 0.295116139 | 2.0960735 | 0.140794748 |
| Gm25816 | 0.401944266 | 0.887270117 | 1.429669844 | 0.620611899 |
| Gm23141 | 0.141693294 | -0.687378062 | 2.912076028 | -0.236043996 |
| Gm23767 | 0.37340313 | -0.033560602 | 1.712819267 | -0.019593779 |
| Gm22417 | 0.221310845 | -0.781324857 | 3.162078526 | -0.247092174 |
| Gm22584 | 1.21151113 | -0.779008776 | 0.93008839 | -0.83756424 |
| Gm26270 | 0.134148234 | -0.20649159 | 3.624632408 | -0.056968974 |
| Gm23089 | 0 | NA | NA | NA |
| Gm24711 | 0.104637558 | -0.206492286 | 3.724945049 | -0.055434988 |
| Gm26188 | 0 | NA | NA | NA |
| Gm23265 | 0.938116572 | 2.118993815 | 1.03018062 | 2.056914849 |
| Gm24518 | 0.272816007 | -0.152921808 | 2.071773138 | -0.073812043 |
| Gm25944 | 0.061920613 | 0.274393828 | 3.724945049 | 0.073663859 |
| Gm23047 | 0.652779004 | -0.533767148 | 1.218258492 | -0.438139485 |
| Gm23953 | 0 | NA | NA | NA |
| Gm26332 | 0 | NA | NA | NA |
| Gm26032 | 0 | NA | NA | NA |
| Gm22863 | 0 | NA | NA | NA |
| Gm24618 | 0 | NA | NA | NA |
| Gm22046 | 0 | NA | NA | NA |
| Gm24760 | 0.092140247 | -0.687373951 | 3.724945049 | -0.184532642 |
| Gm25350 | 0.472357773 | 0.378382703 | 1.379197557 | 0.274349894 |
| Gm26502 | 0 | NA | NA | NA |
| Gm23313 | 0 | NA | NA | NA |
| Gm25471 | 0.129625439 | -1.028499994 | 3.675612039 | -0.279817343 |
| Gm22776 | 0 | NA | NA | NA |
| Gm23446 | 0.198993444 | -1.171393095 | 2.222930108 | -0.526959031 |
| Gm25210 | 0 | NA | NA | NA |
| Gm22128 | 0 | NA | NA | NA |
| Gm22851 | 0.048949532 | -0.206487837 | 3.724945049 | -0.055433794 |
| Gm25074 | 0.045882455 | 0.755288839 | 3.724945049 | 0.202765096 |
| Gm25157 | 0.086648695 | -0.687376175 | 3.724945049 | -0.184533239 |
| Gm22631 | 0.573189294 | -1.330705712 | 1.187569728 | -1.120528488 |
| Gm25474 | 0.37594025 | -0.629289027 | 1.423623346 | -0.442033372 |
| Gm22941 | 0.150983267 | -0.206489499 | 2.855298352 | -0.072318011 |
| Gm26246 | 0.123992736 | 0.274402763 | 3.720643035 | 0.073751435 |
| Gm22047 | 0 | NA | NA | NA |
| Gm25597 | 0.783837108 | -1.024355846 | 1.049350114 | -0.976181193 |
| Gm26097 | 0.21090123 | -0.655403003 | 2.172089159 | -0.301738536 |
| Gm22258 | 0.307376478 | -1.060739279 | 1.74928669 | -0.606383896 |
| Gm26201 | 0 | NA | NA | NA |
| Gm23619 | 0 | NA | NA | NA |
| Gm22812 | 0.249740077 | -1.002029643 | 2.513367833 | -0.398680062 |
| Gm26433 | 0.126225412 | 0.755282661 | 3.698754989 | 0.20419916 |
| Gm23357 | 0.67569888 | 1.283683966 | 1.189114273 | 1.079529524 |

| | | | | |
|---|---|---|---|---|
| Gm25615 | 0.379570916 | 0.283136193 | 1.585128645 | 0.178620324 |
| Gm22110 | 0.872838561 | -0.715679005 | 1.09729383 | -0.652221843 |
| Gm23724 | 0 NA | NA | NA | |
| Gm22188 | 0.093141081 | -0.687373951 | 3.724945049 | -0.184532642 |
| Gm23549 | 0 NA | NA | NA | |
| Snord116l2 | 0.075043205 | 0.755286614 | 3.724945049 | 0.202764498 |
| Gm26223 | 0 NA | NA | NA | |
| Gm23524 | 0.112288909 | -0.687373951 | 3.724945049 | -0.184532642 |
| Gm26136 | 0 NA | NA | NA | |
| Gm24658 | 0.115000406 | -0.206492286 | 3.724945049 | -0.055434988 |
| Gm23359 | 0.04393175 | -0.206490061 | 3.724945049 | -0.055434391 |
| Gm24742 | 0.049553047 | 0.274391604 | 3.724945049 | 0.073663262 |
| Snord116l1 | 0.083526043 | 0.75528439 | 3.724945049 | 0.202763901 |
| Gm22173 | 0.200209858 | 0.274410098 | 2.224689937 | 0.12334757 |
| Gm23696 | 0.675101473 | 0.646130667 | 1.282779835 | 0.503695685 |
| Gm26094 | 1.506060395 | 0.449173244 | 0.801324126 | 0.560538775 |
| Gm22289 | 0.897799824 | -0.81331316 | 1.032720298 | -0.78754447 |
| Gm22373 | 29.41355203 | -0.241668083 | 0.283244586 | -0.853213426 |
| Gm38393 | 1891.788183 | -0.0692635 | 0.150331322 | -0.460738979 |
| Snord64 | 0 NA | NA | NA | |
| Snord107 | 0 NA | NA | NA | |
| Snrpn | 3523.734492 | 0.026509698 | 0.054541078 | 0.486050127 |
| Snurf | 0 NA | NA | NA | |
| Gm7367 | 30.85968858 | 1.349978577 | 0.2426318 | 5.563897953 |
| A230006K03Rik | 161.3268707 | -0.214143841 | 0.17026277 | -1.257725582 |
| Ube2nl | 15.82980231 | 0.32761364 | 0.401390134 | 0.81619754 |
| Gm25533 | 0.514202509 | 1.585232241 | 1.586976182 | 0.998901092 |
| Gm23962 | 0.605864455 | -1.278573338 | 1.351362441 | -0.946136505 |
| Mir344d-1 | 0.868248373 | -0.371574596 | 1.096690351 | -0.338814503 |
| Mir344d-2 | 0.131862677 | -0.687376665 | 2.97735462 | -0.230868255 |
| Gm23490 | 9.098233414 | 0.129086424 | 0.333329292 | 0.387263967 |
| Gm22466 | 0.385656404 | 0.794969414 | 1.464949675 | 0.54265988 |
| Mir344d-3 | 1.914600676 | 0.338680874 | 0.677205805 | 0.500115137 |
| Gm23917 | 0.903269115 | -0.389097486 | 1.11038651 | -0.350416258 |
| Mir344e | 0.296439348 | -1.893307113 | 1.538312073 | -1.230769196 |
| Mir344h-1 | 0.876086486 | -0.053050901 | 1.15361497 | -0.045986662 |
| Mir344h-2 | 2.928954034 | -0.26061109 | 0.6465463 | -0.403081868 |
| Gm23873 | 4.451125399 | 0.00796807 | 0.490082042 | 0.016258645 |
| Mir344b | 5.840177946 | -0.348824451 | 0.432802053 | -0.805967644 |
| Mir344c | 9.878672328 | -0.77614267 | 0.318769591 | -2.434807748 |
| Mir344 | 15.06924853 | -0.461984236 | 0.334652348 | -1.380490049 |
| Mir344-2 | 22.73106971 | -0.567612428 | 0.260016275 | -2.182988078 |
| Mir344g | 49.18631136 | 0.15492107 | 0.169755007 | 0.912615618 |
| Gm23431 | 7.186751093 | 0.097472993 | 0.394413602 | 0.247133955 |
| Mir344f | 6.212148508 | -0.016878183 | 0.384844467 | -0.043857154 |
| Gm24923 | 13.62120263 | -0.583464436 | 0.336715144 | -1.732813171 |
| Mir344i | 13.24250092 | -1.059636246 | 0.416304146 | -2.545341564 |

| | | | | |
|---|---|---|---|---|
| Gm9801 | 35.25565351 | -0.052525611 | 0.286920347 | -0.183066874 |
| Ndn | 2072.502296 | 0.111674681 | 0.10474858 | 1.066121198 |
| Magel2 | 65.47304857 | -0.375382179 | 0.186133142 | -2.016740134 |
| Mkrn3 | 19.82500287 | -0.560680773 | 0.25545708 | -2.194813992 |
| Peg12 | 5.656883354 | 0.460524085 | 0.465733377 | 0.988814862 |
| Atp5l-ps1 | 1.396333172 | -0.530227594 | 0.828864792 | -0.639703362 |
| Gm22494 | 0 | NA | NA | NA |
| Chrna7 | 175.1769349 | 0.087788869 | 0.084794874 | 1.035308677 |
| 4930554H23Rik | 0.967147681 | -1.955031241 | 1.219531378 | -1.603100401 |
| Otud7a | 560.9803126 | -0.06415461 | 0.126995744 | -0.505171337 |
| Gm5898 | 84.2128684 | 0.148824963 | 0.132160129 | 1.126095774 |
| Gm22228 | 3.911342548 | -0.234665152 | 0.464849466 | -0.504819667 |
| Gm3307 | 7.681223622 | -0.480733639 | 0.420851331 | -1.142288508 |
| Klf13 | 2862.082524 | 0.128344909 | 0.07476806 | 1.716574026 |
| Gm27252 | 12.51525409 | 1.949264766 | 0.352688973 | 5.526866213 |
| E030018B13Rik | 0.724732643 | 1.632288414 | 1.431113243 | 1.140572504 |
| Gm20670 | 0.072025462 | 0.755288839 | 3.724945049 | 0.202765096 |
| Trpm1 | 3.26130212 | 0.476582261 | 0.66697055 | 0.714547682 |
| Mir211 | 0 | NA | NA | NA |
| Mir6416 | 0 | NA | NA | NA |
| Mtmr10 | 533.9198899 | 0.205264278 | 0.092680257 | 2.214757333 |
| Gm20457 | 18.61572913 | 0.20151773 | 0.24672619 | 0.816766678 |
| Fan1 | 158.3085833 | 0.266092842 | 0.109765128 | 2.424201997 |
| Mphosph10 | 195.4413562 | 0.132345391 | 0.106631108 | 1.241151792 |
| Mcee | 82.01444615 | -0.162195853 | 0.148233513 | -1.094191522 |
| Apba2 | 1821.774854 | -0.328573948 | 0.083441668 | -3.937768217 |
| Fam189a1 | 495.1526885 | -0.187026681 | 0.110899306 | -1.686454926 |
| Ndnl2 | 270.3594914 | 0.131641898 | 0.121570183 | 1.082846913 |
| Gm25249 | 1.017734822 | -0.883443169 | 1.055804405 | -0.836748895 |
| Tjp1 | 2187.437935 | 0.264950454 | 0.050392869 | 5.257697365 |
| Gm7546 | 0 | NA | NA | NA |
| BC046251 | 8.000213675 | -0.212262147 | 0.381628646 | -0.55620077 |
| Tarsl2 | 377.8388073 | -0.018612321 | 0.074106067 | -0.251157853 |
| Tm2d3 | 199.3078344 | 0.034324233 | 0.090821516 | 0.377930636 |
| Gm7551 | 1.00142042 | -0.711787927 | 1.167022802 | -0.609917755 |
| 1810008I18Rik | 0.117907917 | 1.204448679 | 3.72248341 | 0.323560523 |
| Gm20684 | 1.680189541 | 0.49199756 | 0.748576772 | 0.657243958 |
| Pcsk6 | 300.8982086 | -0.394734784 | 0.135452533 | -2.914192711 |
| Snrpa1 | 232.9850694 | 0.506361405 | 0.134871383 | 3.754402115 |
| Vimp | 381.3181123 | 0.639994852 | 0.103170959 | 6.203246153 |
| Gm26827 | 0.677567072 | -0.192102283 | 1.302163916 | -0.147525424 |
| Chsy1 | 358.5432631 | 0.248117463 | 0.096295223 | 2.576633133 |
| Gm23042 | 0 | NA | NA | NA |
| Lrrk1 | 209.5656891 | 1.30331499 | 0.148074932 | 8.801726078 |
| Gm10974 | 0 | NA | NA | NA |
| Gm23957 | 0.352640022 | 1.695952166 | 1.731813185 | 0.97929279 |
| Mir7057 | 0.172107867 | 1.236193725 | 2.739902689 | 0.451181617 |

| | | | | |
|---|---|---|---|---|
| Aldh1a3 | 20.34501332 | 0.153175201 | 0.343042876 | 0.446519114 |
| Asb7 | 386.1973764 | 0.180401704 | 0.077258203 | 2.335049196 |
| Lins | 133.4462556 | -0.107090609 | 0.100196091 | -1.068810252 |
| Cers3 | 5.123955725 | 0.86216984 | 0.529056424 | 1.629636843 |
| Adamts17 | 97.65468606 | -0.704159891 | 0.196783694 | -3.578344709 |
| 1700112J16Rik | 8.55332328 | -0.522876539 | 0.382307702 | -1.367685076 |
| Lysmd4 | 213.3922501 | -0.21817789 | 0.090393634 | -2.413642204 |
| Mef2a | 2662.223246 | 0.213602206 | 0.061436168 | 3.476815275 |
| Lrrc28 | 238.448027 | -0.10679294 | 0.097505352 | -1.095252083 |
| Gm23233 | 1.460122852 | -2.040183699 | 0.999466024 | -2.041273691 |
| Ttc23 | 41.9836744 | 0.109939508 | 0.202200068 | 0.543716474 |
| Synm | 824.232063 | 0.336549501 | 0.126123828 | 2.668405376 |
| Gm23844 | 0 | NA | NA | NA |
| Gm7583 | 0.042716945 | -0.206490061 | 3.724945049 | -0.055434391 |
| 4833412C05Rik | 2.535954356 | -0.737946543 | 0.610756167 | -1.208250663 |
| Igf1r | 783.90039 | 0.034134802 | 0.078008462 | 0.437578201 |
| Pgpep1l | 5.087731363 | 0.306542007 | 0.45658968 | 0.671373054 |
| Gm16157 | 0.108309328 | 1.204448679 | 3.72248341 | 0.323560523 |
| Fam169b | 17.53390412 | 2.835323315 | 0.375070066 | 7.559449738 |
| Gm16158 | 0 | NA | NA | NA |
| Gm5334 | 0 | NA | NA | NA |
| Arrdc4 | 222.7753849 | 1.52069129 | 0.240521251 | 6.322482051 |
| Gm23795 | 0 | NA | NA | NA |
| Gm5342 | 0 | NA | NA | NA |
| Gm23815 | 0.0885994 | 0.274402725 | 3.724945049 | 0.073666248 |
| Gm7627 | 0 | NA | NA | NA |
| Gm24120 | 0 | NA | NA | NA |
| Gm29327 | 0 | NA | NA | NA |
| Nr2f2 | 456.5175412 | -0.015571898 | 0.113538311 | -0.137151045 |
| B130024G19Rik | 28.43045277 | -0.238264299 | 0.283577046 | -0.840210103 |
| Gm28258 | 0.072025462 | 0.755288839 | 3.724945049 | 0.202765096 |
| Gm7656 | 3.484008338 | -0.493968112 | 0.544364351 | -0.90742186 |
| Gm24880 | 0.045882455 | 0.755288839 | 3.724945049 | 0.202765096 |
| 6030442E23Rik | 0.042716945 | -0.206490061 | 3.724945049 | -0.055434391 |
| Gm10172 | 0 | NA | NA | NA |
| Gm10295 | 0.372348714 | -0.144535575 | 3.279307958 | -0.044075024 |
| Gm29295 | 0.243433062 | 0.255570114 | 1.844782334 | 0.138536731 |
| Gm29328 | 0.046345709 | 0.274391604 | 3.724945049 | 0.073663262 |
| Mctp2 | 119.8191935 | 1.169813926 | 0.159645944 | 7.32755183 |
| Gm37620 | 0 | NA | NA | NA |
| Gm28745 | 0 | NA | NA | NA |
| Rpl17-ps10 | 146.566573 | 0.295403339 | 0.114936192 | 2.570150727 |
| Gm28744 | 0 | NA | NA | NA |
| Kansl2-ps | 68.64857961 | 0.592208047 | 0.145032442 | 4.083279835 |
| Gm7693 | 0 | NA | NA | NA |
| Gm5335 | 0.044544648 | 0.755286614 | 3.724945049 | 0.202764498 |
| Gm28746 | 0.114742407 | 0.274402725 | 3.724945049 | 0.073666248 |

| | | | | |
|---|---|---|---|---|
| A730056A06Rik | 25.12540099 | -1.025355227 | 0.25386307 | -4.039009016 |
| Rgma | 807.0540894 | -0.044216629 | 0.111545583 | -0.396399641 |
| Chd2 | 1727.684492 | 0.099440338 | 0.105835724 | 0.939572527 |
| 1810026B05Rik | 562.1011433 | -0.073273349 | 0.109694976 | -0.667973611 |
| Gm26176 | 0 | NA | NA | NA |
| Gm23020 | 0.068097359 | -0.206487837 | 3.724945049 | -0.055433794 |
| A830073O21Rik | 30.67817576 | 0.136104338 | 0.282017409 | 0.482609704 |
| Fam174b | 232.8015944 | -0.218417002 | 0.124223931 | -1.758252223 |
| Gm10619 | 0 | NA | NA | NA |
| mmu-mir-7234 | 0 | NA | NA | NA |
| St8sia2 | 68.7914267 | 0.467465189 | 0.21098925 | 2.215587708 |
| Gm38143 | 0.044191549 | -0.206490061 | 3.724945049 | -0.055434391 |
| Gm27459 | 0 | NA | NA | NA |
| Slco3a1 | 783.7089768 | 0.089712492 | 0.083368074 | 1.076101295 |
| Gm7580 | 0 | NA | NA | NA |
| Gm22326 | 0 | NA | NA | NA |
| Sv2b | 4509.045788 | 0.025781094 | 0.114637026 | 0.224893253 |
| Gm24858 | 0.075043205 | 0.755286614 | 3.724945049 | 0.202764498 |
| 1500012K07Rik | 0.790435591 | -0.151923171 | 1.036824806 | -0.14652733 |
| Gm10161 | 0 | NA | NA | NA |
| Akap13 | 1197.678966 | 0.751504728 | 0.082555113 | 9.103067069 |
| Gm25282 | 0.642535093 | 1.740545474 | 1.19034885 | 1.462214606 |
| Gm23251 | 0.141684195 | 0.755287278 | 3.558854591 | 0.212227631 |
| AU020206 | 84.50291907 | 2.90694163 | 0.252454857 | 11.51469875 |
| Gm26729 | 0 | NA | NA | NA |
| Klhl25 | 139.2928838 | 0.556453447 | 0.152741556 | 3.643104474 |
| Gm5597 | 1.352145875 | 0.477642853 | 0.891500779 | 0.535773905 |
| Agbl1 | 4.71615247 | -0.07596444 | 0.458966475 | -0.165511959 |
| Gm23239 | 0.182574094 | 0.638973862 | 2.702311969 | 0.236454513 |
| AC134536.1 | 0 | NA | NA | NA |
| Ntrk3 | 1530.084009 | -0.067275371 | 0.112350932 | -0.598796732 |
| Gm9885 | 28.00701669 | 0.310491394 | 0.245758937 | 1.263398182 |
| Mrpl46 | 163.3248249 | 0.210805111 | 0.12323576 | 1.710583929 |
| Mrps11 | 173.3279118 | -0.063286364 | 0.097804925 | -0.647067258 |
| Det1 | 52.30670483 | -0.319818057 | 0.186313087 | -1.716562492 |
| Mir7-2 | 0.175313191 | -1.166555201 | 2.770426814 | -0.421074181 |
| Aen | 320.6797765 | 0.650358712 | 0.124936572 | 5.205511094 |
| Isg20 | 337.9177596 | 7.499034517 | 0.54948236 | 13.64745269 |
| Gm26633 | 15.61936475 | -0.753934526 | 0.264784608 | -2.847350272 |
| Acan | 162.5228664 | -0.639439038 | 0.23024826 | -2.777172076 |
| Hapln3 | 8.674975699 | 0.012566255 | 0.3227562 | 0.038934203 |
| Mfge8 | 724.7290691 | -1.054370104 | 0.19543697 | -5.394936808 |
| Abhd2 | 3069.461115 | 0.412981111 | 0.072823838 | 5.670960536 |
| Gm24224 | 0 | NA | NA | NA |
| Rlbp1 | 85.04677146 | -0.717871292 | 0.165742932 | -4.331233227 |
| Fanci | 15.02295619 | 0.576303727 | 0.352866548 | 1.633205897 |
| Gm16017 | 0 | NA | NA | NA |

| | | | | |
|---|---|---|---|---|
| Polg | 722.9823603 | -0.299332587 | 0.096466318 | -3.102975154 |
| AI854517 | 340.1156097 | 0.296634237 | 0.129837749 | 2.28465327 |
| Mir9-3 | 6.978027697 | -1.249946675 | 0.463026776 | -2.699512728 |
| A330074H02Rik | 52.62118314 | -0.696873271 | 0.30762083 | -2.265364382 |
| 2900037B21Rik | 15.38072168 | -1.178745082 | 0.3769188 | -3.127318354 |
| Gm37608 | 67.71388935 | -0.763249385 | 0.32528367 | -2.34641163 |
| Rhcg | 17.15633722 | -0.252002308 | 0.446572584 | -0.564303132 |
| Ticrr | 11.11653513 | 2.185862256 | 0.43195535 | 5.060389358 |
| Kif7 | 33.31114625 | -0.047081152 | 0.181499311 | -0.25940127 |
| 9330171B17Rik | 15.94460044 | -0.093284854 | 0.290374585 | -0.321256952 |
| Plin1 | 0.616538614 | -0.200904451 | 1.408923863 | -0.142594256 |
| Pex11a | 89.1075553 | 0.052577046 | 0.142701439 | 0.368440892 |
| Wdr93 | 5.435520937 | -1.173644357 | 0.501165619 | -2.34182935 |
| Mesp1 | 0.291110193 | -0.699915581 | 1.630984298 | -0.429136922 |
| Mesp2 | 2.119758866 | 0.858401678 | 0.705176866 | 1.217285647 |
| Anpep | 165.5971786 | 1.17950207 | 0.219004401 | 5.38574596 |
| Ap3s2 | 947.0018652 | 0.008707771 | 0.058096402 | 0.149884857 |
| Gm26646 | 0.14985154 | -0.206486716 | 2.85297593 | -0.072375905 |
| 5430400D12Rik | 13.24663712 | -0.009048935 | 0.309772998 | -0.029211504 |
| 2610034B18Rik | 234.4523726 | -0.033898917 | 0.113581229 | -0.298455275 |
| Gm24541 | 0 | NA | NA | NA |
| Zfp710 | 414.4888839 | 0.488060843 | 0.093791598 | 5.203673398 |
| Idh2 | 278.3167953 | 0.009516926 | 0.135783349 | 0.070089053 |
| Sema4b | 443.5342128 | 0.4024611 | 0.113768399 | 3.537547355 |
| Cib1 | 109.0301844 | 0.428821969 | 0.157437905 | 2.723753014 |
| Gdpgp1 | 63.61861299 | -0.357208749 | 0.168938543 | -2.114430146 |
| Gm15504 | 0 | NA | NA | NA |
| Ttll13 | 6.876810528 | 0.69909696 | 0.462692111 | 1.510933391 |
| Ngrn | 276.3196493 | 0.03549462 | 0.100011028 | 0.354907064 |
| Vps33b | 168.9997345 | -0.33193276 | 0.112031036 | -2.962864335 |
| Prc1 | 39.35090745 | 1.324516812 | 0.443122299 | 2.989054745 |
| 6330403N20Rik | 1.079531651 | 0.495635481 | 0.995562636 | 0.497844598 |
| Rccd1 | 59.77890577 | -0.17933674 | 0.155431041 | -1.153802602 |
| Unc45a | 445.147613 | 0.170444315 | 0.112381207 | 1.516662076 |
| Hddc3 | 174.8477751 | -0.125188618 | 0.1486852 | -0.841970941 |
| Man2a2 | 2375.928341 | -0.224053491 | 0.07220655 | -3.102952435 |
| Fes | 119.6055901 | 2.194181195 | 0.198008134 | 11.08126799 |
| Furin | 1024.487266 | 0.676391097 | 0.137768445 | 4.909622771 |
| Gm15544 | 0.049553047 | 0.274391604 | 3.724945049 | 0.073663262 |
| Blm | 98.86775975 | 0.44520853 | 0.129857425 | 3.428441084 |
| Gm17257 | 13.53931913 | 0.122163524 | 0.30627764 | 0.398865304 |
| Crtc3 | 380.3991371 | 0.053938832 | 0.111481243 | 0.483837733 |
| Gm15880 | 1.230449842 | 1.689740095 | 1.073876737 | 1.573495389 |
| Iqgap1 | 1218.575044 | 1.038628212 | 0.18115159 | 5.733475549 |
| Zscan2 | 88.28572028 | -0.002481405 | 0.131630861 | -0.018851242 |
| Wdr73 | 209.5163078 | 0.189308476 | 0.095813242 | 1.975807019 |
| Nmb | 36.76065315 | -0.04235484 | 0.316888363 | -0.133658553 |

| | | | | |
|---|---|---|---|---|
| Sec11a | 259.8445855 | -0.015204721 | 0.096376721 | -0.157763417 |
| Gm16171 | 0 | NA | NA | NA |
| Zfp592 | 595.8256381 | 0.042921729 | 0.072831048 | 0.589332852 |
| Alpk3 | 21.69556852 | 1.311079163 | 0.483321542 | 2.712643756 |
| Platr32 | 1.295334173 | 1.249212521 | 0.899866049 | 1.388220527 |
| Slc28a1 | 1.668031982 | -1.608369551 | 1.017129047 | -1.581283668 |
| Gm6415 | 6.092582732 | -1.305535793 | 0.448730326 | -2.909399512 |
| Pde8a | 237.4167335 | -0.588675204 | 0.122447984 | -4.807553252 |
| Gm7180 | 6.599784284 | 0.058391688 | 0.455332658 | 0.128239621 |
| Rps17 | 912.2411316 | 0.329431572 | 0.115039648 | 2.863635088 |
| Cpeb1 | 643.0184324 | -0.106021323 | 0.117136327 | -0.90511053 |
| Rpl7a-ps9 | 0 | NA | NA | NA |
| Ap3b2 | 1542.537639 | -0.174806254 | 0.05593806 | -3.124996697 |
| BC048679 | 0.097899064 | -0.687371727 | 3.724945049 | -0.184532045 |
| 2900076A07Rik | 53.12144243 | -0.276666559 | 0.179296745 | -1.543065156 |
| Mir1839 | 0 | NA | NA | NA |
| mmu-mir-1839 | 0 | NA | NA | NA |
| Gm37829 | 0.980535863 | -0.629149056 | 1.052471463 | -0.597782532 |
| Fsd2 | 3.637652488 | -1.188928238 | 0.524627377 | -2.266233694 |
| Whamm | 307.5779694 | 1.099106097 | 0.115736816 | 9.496598688 |
| Homer2 | 622.3176929 | -0.950732975 | 0.160449592 | -5.925430937 |
| Gm26149 | 0 | NA | NA | NA |
| Fam103a1 | 318.4598396 | 0.069444212 | 0.082785128 | 0.838848888 |
| 3110040N11Rik | 45.08078921 | -0.114216552 | 0.173591164 | -0.657962935 |
| Btbd1 | 1030.704126 | -0.257618267 | 0.101773663 | -2.531286186 |
| Gm10160 | 0.442186748 | -1.061537588 | 1.567789114 | -0.677092078 |
| Tm6sf1 | 282.4071646 | 0.312989549 | 0.098591968 | 3.174594816 |
| Hdgfrp3 | 1850.998309 | 0.010241204 | 0.064759584 | 0.158141902 |
| Gm25907 | 0 | NA | NA | NA |
| Bnc1 | 0.212279271 | 1.193729487 | 2.584506884 | 0.461879012 |
| 4833418N17Rik | 0 | NA | NA | NA |
| Sh3gl3 | 330.4796501 | -0.049487218 | 0.078333721 | -0.631748588 |
| Adamtsl3 | 118.3270371 | 0.695336957 | 0.137331556 | 5.063198711 |
| Saxo2 | 36.73756128 | -0.604984006 | 0.230808104 | -2.621155828 |
| Eftud1 | 227.2937603 | 0.088257332 | 0.081947408 | 1.076999681 |
| 4933406J10Rik | 0.25679625 | 1.79603842 | 2.073194179 | 0.866314617 |
| n-R5s154 | 0 | NA | NA | NA |
| 1700010L04Rik | 0.351906323 | 1.186793621 | 1.548524457 | 0.766402891 |
| Mex3b | 220.6262723 | -0.283505456 | 0.130605738 | -2.170696796 |
| Gm22934 | 0 | NA | NA | NA |
| A530021J07Rik | 0.162691105 | -0.206480295 | 2.788998794 | -0.074033842 |
| Gm10610 | 1.084114685 | 2.955241595 | 1.163337678 | 2.540312802 |
| Tmc3 | 14.78000229 | 1.287464245 | 0.339003584 | 3.797789483 |
| Gm16638 | 33.35336936 | 0.263405212 | 0.245806441 | 1.071596052 |
| Stard5 | 261.421678 | 1.284244974 | 0.137434048 | 9.344445486 |
| Il16 | 509.4594508 | -0.121199996 | 0.192726707 | -0.628869753 |
| 2310044K18Rik | 1.104418013 | 1.363373085 | 1.037620409 | 1.313942048 |

| | | | | |
|---|---|---|---|---|
| Gm7964 | 331.670112 | 0.173936773 | 0.094392468 | 1.842697587 |
| 1700026D08Rik | 9.589069244 | -0.301510053 | 0.409724319 | -0.735885177 |
| Mesdc1 | 387.3285222 | -0.076991632 | 0.098208696 | -0.78395942 |
| Mesdc2 | 527.7692794 | 0.451299424 | 0.098897789 | 4.563291344 |
| Cemip | 295.0602538 | -0.036365197 | 0.080937257 | -0.449301076 |
| Abhd17c | 249.4441419 | -0.052501757 | 0.082900894 | -0.633307484 |
| Gm26708 | 1.414184224 | 0.332489797 | 0.929451662 | 0.357726831 |
| Gm22177 | 0 | NA | NA | NA |
| Arnt2 | 3090.055291 | -0.31586807 | 0.066147156 | -4.775232802 |
| Gm25790 | 1.94490296 | 0.550295317 | 0.730353208 | 0.75346464 |
| Gm2115 | 123.9262083 | -0.643682446 | 0.152875717 | -4.210495036 |
| Fah | 224.6003258 | -0.263065102 | 0.140255095 | -1.875618865 |
| Zfand6 | 399.9024983 | -0.006021416 | 0.118964354 | -0.050615293 |
| 2610206C17Rik | 0.927099338 | 0.765786761 | 1.058069785 | 0.723758274 |
| Gm18958 | 0.18866986 | 0.755307765 | 2.26103487 | 0.33405401 |
| Vmn2r-ps62 | 0 | NA | NA | NA |
| Gm6155 | 2.358756262 | 0.297504662 | 0.741613421 | 0.401158682 |
| Olfr291 | 0.046345709 | 0.274391604 | 3.724945049 | 0.073663262 |
| Gm8018 | 0.038981395 | 0.274393828 | 3.724945049 | 0.073663859 |
| Olfr290 | 0.092881282 | -0.687373951 | 3.724945049 | -0.184532642 |
| Gm18560 | 0 | NA | NA | NA |
| Vmn2r65 | 0 | NA | NA | NA |
| Vmn2r66 | 0 | NA | NA | NA |
| Vmn2r-ps64 | 0 | NA | NA | NA |
| Vmn2r-ps65 | 0 | NA | NA | NA |
| Vmn2r-ps66 | 0 | NA | NA | NA |
| Vmn2r-ps67 | 0 | NA | NA | NA |
| Gm18358 | 0 | NA | NA | NA |
| Vmn2r67 | 0 | NA | NA | NA |
| Vmn2r-ps68 | 0 | NA | NA | NA |
| Vmn2r68 | 0 | NA | NA | NA |
| Gm20414 | 0 | NA | NA | NA |
| Gm20415 | 0 | NA | NA | NA |
| Vmn2r-ps69 | 0 | NA | NA | NA |
| Vmn2r69 | 0 | NA | NA | NA |
| Gm18359 | 0 | NA | NA | NA |
| Vmn2r-ps70 | 0 | NA | NA | NA |
| Gm18360 | 0 | NA | NA | NA |
| Vmn2r70 | 0 | NA | NA | NA |
| Vmn2r-ps71 | 0 | NA | NA | NA |
| Vmn2r-ps72 | 0 | NA | NA | NA |
| Vmn2r71 | 0 | NA | NA | NA |
| Vmn2r-ps73 | 0 | NA | NA | NA |
| Vmn2r-ps74 | 0 | NA | NA | NA |
| Vmn2r72 | 0 | NA | NA | NA |
| Gm18779 | 0 | NA | NA | NA |
| Gm18354 | 0 | NA | NA | NA |

| | | | | |
|---|---|---|---|---|
| Vmn2r-ps75 | 0 | NA | NA | NA |
| Vmn2r-ps76 | 0 | NA | NA | NA |
| Vmn2r-ps77 | 0 | NA | NA | NA |
| Vmn2r73 | 0.08486385 | 0.755286617 | 3.724945049 | 0.202764499 |
| Vmn2r-ps78 | 0 | NA | NA | NA |
| Vmn2r74 | 0 | NA | NA | NA |
| Vmn2r-ps79 | 0 | NA | NA | NA |
| Vmn2r-ps80 | 0 | NA | NA | NA |
| Vmn2r-ps81 | 0 | NA | NA | NA |
| Vmn2r-ps82 | 0 | NA | NA | NA |
| Vmn2r-ps83 | 0 | NA | NA | NA |
| Vmn2r-ps84 | 0 | NA | NA | NA |
| Vmn2r75 | 0 | NA | NA | NA |
| Vmn2r-ps85 | 0 | NA | NA | NA |
| Vmn2r-ps86 | 0 | NA | NA | NA |
| Vmn2r76 | 0 | NA | NA | NA |
| Olfr310 | 0 | NA | NA | NA |
| Gm20480 | 0 | NA | NA | NA |
| Gm23192 | 0 | NA | NA | NA |
| Olfr309 | 0.297212607 | 0.274413159 | 1.798759863 | 0.152556862 |
| Olfr308 | 0 | NA | NA | NA |
| Olfr307 | 0 | NA | NA | NA |
| Olfr306-ps1 | 0 | NA | NA | NA |
| Olfr305 | 0 | NA | NA | NA |
| Olfr304 | 0 | NA | NA | NA |
| Olfr303 | 0 | NA | NA | NA |
| Olfr301 | 0 | NA | NA | NA |
| Olfr300-ps1 | 0 | NA | NA | NA |
| Olfr299 | 0 | NA | NA | NA |
| Olfr298 | 0.066050875 | 0.274391604 | 3.724945049 | 0.073663262 |
| Olfr297 | 0 | NA | NA | NA |
| Gm17919 | 0 | NA | NA | NA |
| Olfr296-ps1 | 0 | NA | NA | NA |
| Olfr295 | 0 | NA | NA | NA |
| Olfr294 | 0 | NA | NA | NA |
| Olfr293 | 0 | NA | NA | NA |
| Olfr292 | 0 | NA | NA | NA |
| Folh1 | 228.0677052 | 0.651950059 | 0.17658636 | 3.691961586 |
| Vmn2r77 | 0.046345709 | 0.274391604 | 3.724945049 | 0.073663262 |
| Vmn2r-ps87 | 0.08486385 | 0.755286617 | 3.724945049 | 0.202764499 |
| Gm19203 | 0 | NA | NA | NA |
| Vmn2r78 | 0.360847685 | 0.037515718 | 1.968993482 | 0.019053247 |
| Vmn2r79 | 0.061920613 | 0.274393828 | 3.724945049 | 0.073663859 |
| Nox4 | 23.51386303 | 0.152019723 | 0.36110225 | 0.420988024 |
| Tyr | 0.679694067 | 0.638498851 | 1.09336393 | 0.583976509 |
| Grm5 | 3152.369327 | -0.046057695 | 0.105105482 | -0.438204492 |
| Gm24249 | 0.097139547 | 0.274393828 | 3.724945049 | 0.073663859 |

| | | | | |
|---|---|---|---|---|
| Ctsc | 748.4761849 | 2.770810527 | 0.182583749 | 15.17555942 |
| Gm26522 | 0.075043205 | 0.755286614 | 3.724945049 | 0.202764498 |
| Gm25345 | 0.066050875 | 0.274391604 | 3.724945049 | 0.073663262 |
| Rab38 | 4.431782484 | -0.474884664 | 0.575813197 | -0.824720007 |
| Gm15661 | 0 | NA | NA | NA |
| Rps13-ps2 | 13.83618781 | 0.221749053 | 0.366438732 | 0.605146328 |
| Gm8183 | 0.155358952 | -0.206481611 | 2.829692658 | -0.072969625 |
| Tmem135 | 828.7159243 | 0.00496695 | 0.057344908 | 0.086615364 |
| Gm24659 | 0.092269992 | -0.20649451 | 3.724945049 | -0.055435586 |
| Gm22171 | 0.104637558 | -0.206492286 | 3.724945049 | -0.055434988 |
| Gm28748 | 6.9950682 | -0.285280983 | 0.408558049 | -0.698263034 |
| Fzd4 | 116.5445208 | 0.375620558 | 0.183890619 | 2.042630347 |
| Prss23 | 214.4898237 | 0.4483548 | 0.192495369 | 2.329171883 |
| Prss23os | 0.31866137 | -0.714197177 | 2.003531189 | -0.356469208 |
| Gm2546 | 6.356979297 | -0.487953042 | 0.420744218 | -1.159737962 |
| A230065N10Rik | 4.255075643 | 0.678940152 | 0.473780693 | 1.433026214 |
| Me3 | 830.2853921 | 0.072340302 | 0.122358526 | 0.591215862 |
| Gm15744 | 0.161532967 | 0.755285283 | 3.44977679 | 0.218937435 |
| Ccdc81 | 14.73968261 | -1.141430924 | 0.289391377 | -3.944246496 |
| l7Rn6 | 302.6286143 | -0.348376053 | 0.115840289 | -3.007382466 |
| Eed | 245.5434659 | 0.18086312 | 0.090331465 | 2.002216176 |
| Gm26529 | 7.028223412 | -0.351727199 | 0.367530503 | -0.957001382 |
| Gm5341 | 0.068097359 | -0.206487837 | 3.724945049 | -0.055433794 |
| 2310010J17Rik | 13.38140942 | 0.156839787 | 0.282972641 | 0.554257779 |
| Picalm | 2942.135727 | 0.299002541 | 0.06438808 | 4.643756119 |
| Ccdc83 | 1.068251668 | 2.434458002 | 1.067898516 | 2.279671678 |
| n-R5s155 | 0 | NA | NA | NA |
| Sytl2 | 471.4003508 | -0.373978462 | 0.098137311 | -3.810767353 |
| Ccdc89 | 10.74032172 | -0.309116518 | 0.330737566 | -0.934627781 |
| Crebzf | 1021.356941 | -0.088004368 | 0.056086666 | -1.569078254 |
| Tmem126a | 179.1913439 | 0.052994244 | 0.146240363 | 0.362377684 |
| Tmem126b | 260.3332693 | -0.081003018 | 0.127027855 | -0.637679179 |
| Dlg2 | 4221.663311 | -0.072264877 | 0.10156445 | -0.711517439 |
| Gm27470 | 0.068097359 | -0.206487837 | 3.724945049 | -0.055433794 |
| Gm27660 | 0 | NA | NA | NA |
| Gm24552 | 6.758948741 | -0.47367872 | 0.363971395 | -1.301417436 |
| Gm22080 | 0 | NA | NA | NA |
| Gm23928 | 0 | NA | NA | NA |
| Ccdc90b | 140.9402912 | -0.050310838 | 0.150931957 | -0.333334565 |
| Ankrd42 | 228.9912909 | -0.254067503 | 0.124803545 | -2.03573948 |
| Gm37008 | 21.96798092 | -0.467106779 | 0.259486727 | -1.800118196 |
| Gm26944 | 56.56169118 | -0.50494206 | 0.274117734 | -1.842062724 |
| 6430511E19Rik | 167.4764364 | -0.729443322 | 0.229901729 | -3.172848348 |
| Pcf11 | 1375.530546 | 0.005857289 | 0.079289103 | 0.073872564 |
| Gm26981 | 0 | NA | NA | NA |
| 4632427E13Rik | 515.8795252 | -0.46690527 | 0.179636847 | -2.599162015 |
| Rab30 | 322.4076359 | -0.135743507 | 0.075202697 | -1.805035093 |

| | | | | |
|---|---|---|---|---|
| Ddias | 20.7592005 | 1.485882768 | 0.348683864 | 4.261403874 |
| Prcp | 124.058315 | -0.313933497 | 0.121698297 | -2.579604683 |
| Gm26862 | 2.576837399 | 0.270255701 | 0.661944698 | 0.408275347 |
| Gm9934 | 11.67004998 | -0.351540135 | 0.321513786 | -1.093390548 |
| Fam181b | 200.5250975 | -0.640525835 | 0.169688137 | -3.774723718 |
| Gm25860 | 0 | NA | NA | NA |
| Gm15501 | 19.75181214 | -0.264153822 | 0.278693713 | -0.947828423 |
| Gm5899 | 1.074124899 | 0.232936078 | 0.96830007 | 0.240561873 |
| Gm23884 | 0.681297893 | 0.806142422 | 1.122888137 | 0.717918727 |
| Gm25684 | 0 | NA | NA | NA |
| Gm9966 | 3.042324184 | -0.691054714 | 0.513870486 | -1.344803277 |
| Tenm4 | 3151.27742 | -0.134834971 | 0.125806611 | -1.071763799 |
| Mir708 | 5.493531071 | -0.927365633 | 0.455417802 | -2.036296407 |
| Mir6394 | 3.129376264 | -2.074432 | 0.692658048 | -2.994886157 |
| Gm15412 | 0 | NA | NA | NA |
| Rps11-ps5 | 0 | NA | NA | NA |
| Gm15414 | 0.304874617 | 0.755275357 | 3.724945049 | 0.202761476 |
| Gm15413 | 2.589705772 | 4.036378253 | 1.086596953 | 3.714696827 |
| Gm15416 | 0.568378176 | 0.692130538 | 1.337580599 | 0.517449594 |
| Gm25712 | 0 | NA | NA | NA |
| Gm15415 | 0.44530367 | 1.220530637 | 1.548310974 | 0.788298125 |
| Gm22226 | 0 | NA | NA | NA |
| C230038L03Rik | 4.597572509 | 0.613885756 | 0.562465471 | 1.091419452 |
| Nars2 | 182.0977735 | 0.002059497 | 0.090893256 | 0.022658414 |
| Gab2 | 640.0412359 | 1.013985412 | 0.123339038 | 8.22112306 |
| Usp35 | 246.8318203 | -0.099061589 | 0.086941085 | -1.139410543 |
| Kctd21 | 137.2874877 | -0.921265767 | 0.228112337 | -4.038649462 |
| Alg8 | 69.82562007 | -0.217173284 | 0.161160783 | -1.347556646 |
| Ndufc2 | 551.0014102 | 0.230138731 | 0.13946414 | 1.650164196 |
| Gm16053 | 0.824460844 | -2.749003264 | 1.098940074 | -2.501504249 |
| Thrsp | 209.116264 | 0.451966261 | 0.138648243 | 3.259805169 |
| Kctd14 | 10.62226215 | -0.980529694 | 0.326394882 | -3.00412092 |
| Ints4 | 375.7085532 | -0.172696437 | 0.098138218 | -1.759726644 |
| Gm24412 | 1.785420374 | -1.038000177 | 0.848211139 | -1.223752117 |
| Aamdc | 120.3980682 | -0.212814991 | 0.102644542 | -2.0733201 |
| Rsf1 | 1617.979498 | -0.107812679 | 0.058017303 | -1.858284897 |
| Rsf1os1 | 0 | NA | NA | NA |
| Rsf1os2 | 1.842452978 | -1.294366065 | 0.80140758 | -1.615115824 |
| Clns1a | 571.5874786 | 0.020646167 | 0.072085211 | 0.286413357 |
| Gm9990 | 8.809819601 | -1.066232537 | 0.460262253 | -2.316576104 |
| Aqp11 | 58.08426219 | -0.495738165 | 0.164359495 | -3.016182089 |
| Pak1 | 1544.109605 | 0.080366288 | 0.07572881 | 1.061237956 |
| Gdpd4 | 2.28172379 | -2.28238533 | 0.987989799 | -2.310130462 |
| Myo7a | 84.81893064 | 0.541067877 | 0.123184533 | 4.392336164 |
| Capn5 | 668.0892524 | 0.71345759 | 0.098895852 | 7.214231704 |
| Omp | 79.7450718 | -1.028932062 | 0.739113258 | -1.392116907 |
| Gm16938 | 29.29590882 | 0.330417526 | 0.189633022 | 1.742405001 |

| | | | | |
|---|---|---|---|---|
| B3gnt6 | 6.185203872 | 0.378922341 | 0.420895028 | 0.90027754 |
| Acer3 | 413.2037177 | 0.434249605 | 0.095081455 | 4.567132535 |
| Tsku | 55.8336496 | 0.597630134 | 0.224801585 | 2.658478296 |
| Gucy2d | 0.893137065 | 1.704962167 | 1.011943974 | 1.6848385 |
| Lrrc32 | 74.30774312 | 1.122194532 | 0.271531779 | 4.132829453 |
| Gm23040 | 0.075043205 | 0.755286614 | 3.724945049 | 0.202764498 |
| A630091E08Rik | 0 | NA | NA | NA |
| 2210018M11Rik | 690.3005469 | -0.446118203 | 0.104008842 | -4.289233422 |
| Gm15506 | 8.670677481 | -0.618281811 | 0.359185297 | -1.72134499 |
| Prkrir | 815.4013897 | 0.071463241 | 0.072896218 | 0.980342229 |
| Wnt11 | 8.862266067 | 0.832852445 | 0.352407193 | 2.363324196 |
| Gm8398 | 0.117760149 | 0.274400501 | 3.724945049 | 0.073665651 |
| Uvrag | 562.8884481 | 0.381129937 | 0.088362534 | 4.313252666 |
| Gm23479 | 0 | NA | NA | NA |
| Dgat2 | 363.7510314 | 0.01166811 | 0.110944517 | 0.105170681 |
| Mogat2 | 0.144988549 | 0.755282891 | 3.575547253 | 0.211235606 |
| Gm26705 | 2.196663442 | 0.069736491 | 0.741414977 | 0.094058649 |
| Map6 | 1290.520615 | -0.339483669 | 0.074440443 | -4.560473519 |
| Serpinh1 | 377.8077043 | 0.721375608 | 0.208951739 | 3.452355127 |
| Gdpd5 | 476.3039452 | -0.202169441 | 0.121353858 | -1.665949837 |
| Klhl35 | 5.303271472 | 0.426887329 | 0.422750729 | 1.009784964 |
| Rps3 | 2214.045192 | 0.456020646 | 0.110427841 | 4.129580403 |
| Snord15b | 0 | NA | NA | NA |
| Snord15a | 45.47276232 | 0.17927574 | 0.21588098 | 0.830437862 |
| n-R5s156 | 0 | NA | NA | NA |
| Arrb1 | 3455.425929 | -0.23391803 | 0.07340238 | -3.186790812 |
| Mir326 | 2.043157693 | -0.078793814 | 0.65072901 | -0.121085449 |
| Gm4980 | 50.35484282 | -0.337887714 | 0.196308033 | -1.721211859 |
| Gm10605 | 21.76352257 | -0.502906231 | 0.24196641 | -2.078413412 |
| Gm25409 | 0 | NA | NA | NA |
| Slco2b1 | 87.4821162 | -0.787719816 | 0.32250067 | -2.44253699 |
| Gm15635 | 0.464401238 | 1.078447224 | 1.546001059 | 0.697572112 |
| Olfr520 | 0.044544648 | 0.755286614 | 3.724945049 | 0.202764498 |
| Olfr521 | 0 | NA | NA | NA |
| F730035P03Rik | 0.645028293 | 0.339700807 | 1.335648966 | 0.254333897 |
| Neu3 | 60.27959619 | 0.129104958 | 0.145822302 | 0.885358111 |
| Spcs2 | 666.5559383 | 0.270737867 | 0.108456933 | 2.496270727 |
| Xrra1 | 7.264420559 | -0.161968761 | 0.365683759 | -0.442920302 |
| Rnf169 | 651.1755701 | -0.000996599 | 0.078631578 | -0.012674283 |
| Chrdl2 | 5.288126676 | -0.610845464 | 0.463452114 | -1.318033614 |
| Pold3 | 335.8617301 | 0.325757244 | 0.074454296 | 4.375264591 |
| Lipt2 | 20.2504011 | 0.184502355 | 0.229425181 | 0.804194008 |
| Kcne3 | 1.196981447 | 0.409137564 | 0.892140347 | 0.458602243 |
| Pgm2l1 | 3618.346017 | -0.350757265 | 0.11035195 | -3.17853256 |
| Gpx2-ps1 | 5.150345274 | -0.240228288 | 0.53428064 | -0.449629409 |
| P4ha3 | 123.5044264 | 0.624015982 | 0.192955831 | 3.233983543 |
| Ppme1 | 1499.055743 | 0.036981094 | 0.05899295 | 0.626873108 |

| | | | | |
|---|---|---|---|---|
| C2cd3 | 640.0813776 | -0.200840176 | 0.138135461 | -1.453936412 |
| Gm10603 | 1.405100988 | -0.701651645 | 0.82516368 | -0.850318139 |
| Gm37716 | 3.584490054 | -0.614292284 | 0.572793195 | -1.072450389 |
| Ucp3 | 15.93899765 | -0.421441526 | 0.304101649 | -1.385857418 |
| Ucp2 | 1246.309197 | 2.09169114 | 0.15950111 | 13.11395978 |
| Dnajb13 | 7.766840755 | 0.168493945 | 0.423854601 | 0.397527701 |
| Coa4 | 46.56779514 | -0.112851107 | 0.216108287 | -0.522197035 |
| Mrpl48 | 372.182467 | -0.035444877 | 0.072831004 | -0.48667292 |
| Cox20-ps | 0.105143817 | 0.274402725 | 3.724945049 | 0.073666248 |
| Rab6a | 4454.768753 | -0.017575653 | 0.051707677 | -0.339904134 |
| Gm14382 | 10.15270512 | -1.036111276 | 0.429437315 | -2.412718319 |
| Plekhb1 | 2340.080905 | -0.147573797 | 0.243511111 | -0.606024901 |
| Fam168a | 4198.767291 | -0.059865122 | 0.046512249 | -1.287082931 |
| Gm3200 | 5.569043595 | 0.382669915 | 0.447699931 | 0.854746423 |
| Relt | 198.9215136 | 0.400461415 | 0.122505394 | 3.268928828 |
| Arhgef17 | 2322.379047 | -0.092024685 | 0.063012345 | -1.460423097 |
| Mir3102 | 0 | NA | NA | NA |
| P2ry6 | 83.22755977 | 2.341703791 | 0.245554423 | 9.536394253 |
| P2ry2 | 41.59914068 | 3.436186426 | 0.438202541 | 7.841548382 |
| Fchsd2 | 883.8523491 | -0.106871239 | 0.064804514 | -1.649132647 |
| Gm6341 | 2.08700676 | -0.460772085 | 0.624651544 | -0.737646596 |
| Gm15673 | 11.22830619 | -0.44048847 | 0.341486792 | -1.289913636 |
| Atg16l2 | 153.6349008 | -0.238293644 | 0.159920097 | -1.490079411 |
| Stard10 | 401.7502125 | -0.200229072 | 0.091019743 | -2.199842203 |
| Gm20476 | 0.089814205 | 0.274402725 | 3.724945049 | 0.073666248 |
| Arap1 | 603.6362813 | 0.574427406 | 0.098239994 | 5.847184845 |
| Pde2a | 2506.890893 | -0.126718959 | 0.124115553 | -1.020975659 |
| Mir139 | 0.438507067 | 0.755033697 | 1.480256753 | 0.510069416 |
| Art2a-ps | 2.02887955 | 4.076508072 | 1.089501256 | 3.741627693 |
| Gm7027 | 32.67737545 | 0.252515051 | 0.240318584 | 1.050751246 |
| Art2b | 0.341803366 | 0.743875166 | 3.72410795 | 0.199745865 |
| Clpb | 506.8075599 | -0.247637817 | 0.072251666 | -3.427434011 |
| Phox2a | 2.085398638 | -0.183553779 | 0.780177739 | -0.235271746 |
| Inppl1 | 617.7038875 | -0.182121552 | 0.113029733 | -1.611271184 |
| Gm10602 | 1.695248615 | 0.406652194 | 0.787760476 | 0.516212994 |
| Folr2 | 8.630795641 | 0.690332556 | 0.50103289 | 1.377818841 |
| Folr1 | 27.39965009 | -0.244240042 | 0.24059842 | -1.015135683 |
| Gm22655 | 0.092269992 | -0.20649451 | 3.724945049 | -0.055435586 |
| Anapc15 | 157.008238 | 0.205049911 | 0.092825132 | 2.208991331 |
| Tomt | 1.048945322 | -1.147650336 | 1.078992555 | -1.063631377 |
| Lamtor1 | 591.6362557 | 0.13839548 | 0.099334351 | 1.393228815 |
| Lrrc51 | 20.02074397 | -0.300643211 | 0.26806048 | -1.121549923 |
| Numa1 | 1753.65198 | -0.038274007 | 0.083365162 | -0.459112725 |
| Il18bp | 580.6338207 | 2.931458608 | 0.197615823 | 14.834129 |
| Rnf121 | 426.15743 | 0.054231339 | 0.094785369 | 0.572148845 |
| Xndc1 | 64.93653681 | -0.159326614 | 0.209032784 | -0.762208734 |
| Xntrpc | 2.817151494 | 0.722755908 | 0.693049447 | 1.042863409 |

| | | | | |
|---|---|---|---|---|
| Trpc2 | 0 | NA | NA | NA |
| Art5 | 0.775564919 | -0.919120287 | 1.198088674 | -0.767155476 |
| Art1 | 0.377906432 | -1.066397819 | 1.581050245 | -0.674486989 |
| Chrna10 | 6.942902888 | 0.919095811 | 0.420482185 | 2.185813914 |
| Nup98 | 911.2468387 | 0.335462781 | 0.095051901 | 3.529259037 |
| Pgap2 | 160.2875351 | 0.237910669 | 0.151345742 | 1.571968029 |
| Rhog | 475.1594366 | 0.934297563 | 0.150055577 | 6.226343492 |
| Stim1 | 903.1380357 | 0.087756707 | 0.063775849 | 1.376017845 |
| Rrm1 | 306.720304 | 0.178100321 | 0.123654571 | 1.440305201 |
| Olfr543 | 0 | NA | NA | NA |
| Olfr544 | 0 | NA | NA | NA |
| Olfr545 | 0 | NA | NA | NA |
| Olfr547 | 0 | NA | NA | NA |
| Olfr548-ps1 | 0 | NA | NA | NA |
| Olfr549 | 0 | NA | NA | NA |
| Trim21 | 602.064124 | 5.150572968 | 0.187621237 | 27.45197211 |
| Olfr550 | 0.446685102 | 1.660314459 | 1.627896094 | 1.019914272 |
| Olfr551 | 0 | NA | NA | NA |
| Olfr552 | 0.044544648 | 0.755286614 | 3.724945049 | 0.202764498 |
| Olfr553 | 0 | NA | NA | NA |
| Olfr554 | 0.152980968 | -0.206487987 | 2.83734303 | -0.072775123 |
| Olfr555 | 0 | NA | NA | NA |
| Olfr556 | 0.166709681 | -0.206481506 | 2.767631179 | -0.074605861 |
| Trim68 | 38.7507144 | 0.109809845 | 0.184849104 | 0.594051274 |
| Olfr557 | 0.253342923 | 1.988542029 | 2.063655748 | 0.963601623 |
| Olfr558 | 0.693780966 | -0.344670917 | 1.083690743 | -0.318052839 |
| Olfr33 | 0 | NA | NA | NA |
| Olfr559 | 0 | NA | NA | NA |
| Gm24210 | 0 | NA | NA | NA |
| Olfr78 | 4.264554404 | -1.184306707 | 0.60147565 | -1.969001915 |
| Olfr561 | 0 | NA | NA | NA |
| Olfr564 | 0 | NA | NA | NA |
| Olfr566 | 0.049553047 | 0.274391604 | 3.724945049 | 0.073663262 |
| Olfr568 | 0.044544648 | 0.755286614 | 3.724945049 | 0.202764498 |
| Olfr569 | 0 | NA | NA | NA |
| Olfr570 | 0 | NA | NA | NA |
| Olfr571 | 0 | NA | NA | NA |
| Olfr572 | 0 | NA | NA | NA |
| Olfr573-ps1 | 0 | NA | NA | NA |
| Olfr574 | 0.224720342 | 0.274412326 | 2.141112919 | 0.128163407 |
| Olfr575 | 0 | NA | NA | NA |
| Olfr576 | 0 | NA | NA | NA |
| Olfr577 | 0 | NA | NA | NA |
| Olfr578 | 0 | NA | NA | NA |
| Olfr582 | 0 | NA | NA | NA |
| Olfr583 | 0 | NA | NA | NA |
| Olfr584 | 0.046345709 | 0.274391604 | 3.724945049 | 0.073663262 |

| | | | | |
|---|---|---|---|---|
| Olfr585 | 0.062426873 | 0.755288839 | 3.724945049 | 0.202765096 |
| Olfr586 | 0 | NA | NA | NA |
| Olfr587-ps1 | 0.046345709 | 0.274391604 | 3.724945049 | 0.073663262 |
| Gm15120 | 0.07796279 | 0.274392189 | 3.724945049 | 0.073663419 |
| Olfr588-ps1 | 0 | NA | NA | NA |
| Gm15116 | 0 | NA | NA | NA |
| Olfr589 | 0 | NA | NA | NA |
| Olfr590-ps1 | 0 | NA | NA | NA |
| Olfr591 | 0 | NA | NA | NA |
| Olfr592 | 0 | NA | NA | NA |
| Olfr593 | 0.044191549 | -0.206490061 | 3.724945049 | -0.055434391 |
| Olfr594 | 0 | NA | NA | NA |
| Olfr595-ps1 | 0 | NA | NA | NA |
| Usp17ld | 0 | NA | NA | NA |
| Gm25171 | 0 | NA | NA | NA |
| Gm23550 | 0 | NA | NA | NA |
| Gm23973 | 0 | NA | NA | NA |
| Olfr596 | 0 | NA | NA | NA |
| Olfr597 | 0 | NA | NA | NA |
| Olfr598 | 0 | NA | NA | NA |
| Olfr599 | 0.097139547 | 0.274393828 | 3.724945049 | 0.073663859 |
| Olfr600 | 0 | NA | NA | NA |
| Olfr601 | 0 | NA | NA | NA |
| Olfr602-ps1 | 0 | NA | NA | NA |
| Olfr603 | 0 | NA | NA | NA |
| Usp17lc | 0 | NA | NA | NA |
| Gm25774 | 0.116217011 | 0.274402725 | 3.724945049 | 0.073666248 |
| Olfr605 | 0 | NA | NA | NA |
| Olfr606 | 0 | NA | NA | NA |
| Gm15121 | 0 | NA | NA | NA |
| Olfr607 | 0 | NA | NA | NA |
| Olfr608 | 0 | NA | NA | NA |
| Olfr609 | 0 | NA | NA | NA |
| Olfr610 | 0 | NA | NA | NA |
| Olfr611 | 0 | NA | NA | NA |
| Olfr612 | 0 | NA | NA | NA |
| Olfr613 | 0 | NA | NA | NA |
| Olfr615 | 0 | NA | NA | NA |
| Olfr616 | 0.04393175 | -0.206490061 | 3.724945049 | -0.055434391 |
| Gm26506 | 0.044544648 | 0.755286614 | 3.724945049 | 0.202764498 |
| Olfr617 | 0 | NA | NA | NA |
| Olfr618 | 0 | NA | NA | NA |
| Olfr619 | 0 | NA | NA | NA |
| Olfr620 | 0 | NA | NA | NA |
| Olfr621-ps1 | 0 | NA | NA | NA |
| Olfr622 | 0 | NA | NA | NA |
| Olfr623 | 0 | NA | NA | NA |

| | | | | |
|---|---|---|---|---|
| Olfr624 | 0 | NA | NA | NA |
| Olfr625-ps1 | 0 | NA | NA | NA |
| Olfr626-ps1 | 0 | NA | NA | NA |
| Olfr260-ps1 | 0 | NA | NA | NA |
| Olfr243 | 0 | NA | NA | NA |
| Olfr628 | 0.061920613 | 0.274393828 | 3.724945049 | 0.073663859 |
| Olfr629 | 0 | NA | NA | NA |
| Olfr630 | 0 | NA | NA | NA |
| Olfr69 | 0 | NA | NA | NA |
| Olfr68 | 0 | NA | NA | NA |
| Olfr67 | 0.045882455 | 0.755288839 | 3.724945049 | 0.202765096 |
| Hbb-bt | 10.74344224 | 3.148425826 | 0.645527155 | 4.877294163 |
| Hbb-bs | 223.1020574 | 2.645114838 | 0.692381586 | 3.82031367 |
| Gm15115 | 0 | NA | NA | NA |
| Hbb-bh3 | 0 | NA | NA | NA |
| Hbb-bh2 | 0 | NA | NA | NA |
| Hbb-bh1 | 0 | NA | NA | NA |
| Hbb-bh0 | 0 | NA | NA | NA |
| Hbb-y | 0 | NA | NA | NA |
| Olfr66 | 0 | NA | NA | NA |
| Dnajc19-ps | 11.1957499 | 0.212282923 | 0.341685417 | 0.621281777 |
| Olfr64 | 0 | NA | NA | NA |
| Olfr65 | 0.042716945 | -0.206490061 | 3.724945049 | -0.055434391 |
| Olfr631 | 0 | NA | NA | NA |
| Olfr632 | 0.042716945 | -0.206490061 | 3.724945049 | -0.055434391 |
| Olfr633 | 0 | NA | NA | NA |
| Olfr634-ps1 | 0 | NA | NA | NA |
| Gm4886 | 0 | NA | NA | NA |
| Olfr635 | 0.092691418 | 0.274387304 | 3.724945049 | 0.073662108 |
| Olfr636-ps1 | 0.038981395 | 0.274393828 | 3.724945049 | 0.073663859 |
| Olfr637-ps1 | 0 | NA | NA | NA |
| 4930516K23Rik | 0 | NA | NA | NA |
| Olfr638 | 0 | NA | NA | NA |
| Olfr639 | 0 | NA | NA | NA |
| Olfr640 | 0 | NA | NA | NA |
| Olfr641 | 0 | NA | NA | NA |
| Olfr642 | 0 | NA | NA | NA |
| Olfr643 | 0.062426873 | 0.755288839 | 3.724945049 | 0.202765096 |
| Olfr644 | 0 | NA | NA | NA |
| Olfr645 | 0 | NA | NA | NA |
| Olfr646 | 0 | NA | NA | NA |
| 4931431F19Rik | 0.042716945 | -0.206490061 | 3.724945049 | -0.055434391 |
| Ubqln3 | 0 | NA | NA | NA |
| Ubqlnl | 0 | NA | NA | NA |
| E030002O03Rik | 0.160237607 | -1.169544531 | 2.799265637 | -0.417804054 |
| Olfr647-ps1 | 0 | NA | NA | NA |
| Olfr648 | 0 | NA | NA | NA |

| | | | | |
|---|---|---|---|---|
| Olfr649 | 0.060174537 | 0.755286614 | 3.724945049 | 0.202764498 |
| Olfr650-ps1 | 0.524522254 | 1.259894479 | 1.62439136 | 0.775610183 |
| Trim6 | 12.76726646 | 2.77625965 | 0.449732308 | 6.17313811 |
| Trim34a | 130.1339841 | 4.659616832 | 0.259892071 | 17.92904573 |
| Gm15135 | 0 | NA | NA | NA |
| Trim5 | 3.073902706 | 1.691310818 | 0.9271508 | 1.824202511 |
| Gm24846 | 0 | NA | NA | NA |
| Trim12a | 0.470008961 | 2.13190257 | 1.640653554 | 1.29942276 |
| Trim34b | 43.17410611 | 5.366017161 | 0.509966001 | 10.52230373 |
| Gm15133 | 0.044544648 | 0.755286614 | 3.724945049 | 0.202764498 |
| Trim12c | 411.4065939 | 3.423787165 | 0.27539696 | 12.4321894 |
| Gm27995 | 0 | NA | NA | NA |
| Trim30b | 23.44767613 | 6.582811819 | 0.952157077 | 6.913577577 |
| Trim30c | 106.4359115 | 5.244867584 | 0.787707798 | 6.658392361 |
| Trim30a | 1972.4724 | 7.136599129 | 0.293941408 | 24.27898535 |
| Gm25405 | 0.545822006 | -0.755254691 | 1.173745273 | -0.643457067 |
| Trim30d | 381.5932247 | 7.320150151 | 0.495473656 | 14.77404513 |
| Gm16464 | 7.703381671 | 0.38253533 | 0.422171728 | 0.90611309 |
| Gm6574 | 0 | NA | NA | NA |
| Trim30e-ps1 | 0 | NA | NA | NA |
| Olfr651 | 0.060174537 | 0.755286614 | 3.724945049 | 0.202764498 |
| Olfr652 | 0.038981395 | 0.274393828 | 3.724945049 | 0.073663859 |
| Olfr653 | 0 | NA | NA | NA |
| Olfr654 | 0.212522253 | 1.651753349 | 2.159845708 | 0.764755252 |
| Olfr655 | 0.733075257 | 0.878239253 | 1.25656858 | 0.69891868 |
| Olfr656 | 0 | NA | NA | NA |
| Olfr657 | 0 | NA | NA | NA |
| Olfr658 | 7.112035581 | 2.059713132 | 0.515159536 | 3.998204418 |
| Usp-ps | 0 | NA | NA | NA |
| Olfr659 | 0 | NA | NA | NA |
| Olfr660-ps1 | 0 | NA | NA | NA |
| Olfr661 | 0 | NA | NA | NA |
| Olfr663 | 0 | NA | NA | NA |
| Olfr664 | 0 | NA | NA | NA |
| Usp17le | 0 | NA | NA | NA |
| Gm4887 | 2.328839425 | -0.341710485 | 0.786636573 | -0.434394353 |
| Usp17lb | 0 | NA | NA | NA |
| Usp17la | 0 | NA | NA | NA |
| Olfr665 | 0 | NA | NA | NA |
| Olfr666 | 0.045882455 | 0.755288839 | 3.724945049 | 0.202765096 |
| Olfr667 | 0.038981395 | 0.274393828 | 3.724945049 | 0.073663859 |
| Olfr668 | 0.044544648 | 0.755286614 | 3.724945049 | 0.202764498 |
| Olfr669 | 0 | NA | NA | NA |
| Gm5900 | 87.04897631 | -0.028974705 | 0.138784839 | -0.20877428 |
| Olfr670 | 0 | NA | NA | NA |
| Olfr671 | 0.038981395 | 0.274393828 | 3.724945049 | 0.073663859 |
| Olfr672 | 0.101408268 | 0.755286617 | 3.724945049 | 0.202764499 |

| | | | | |
|---|---|---|---|---|
| Olfr675 | 0 | NA | NA | NA |
| Olfr676 | 0 | NA | NA | NA |
| Olfr677 | 0 | NA | NA | NA |
| Olfr678 | 0 | NA | NA | NA |
| Olfr679 | 0 | NA | NA | NA |
| Olfr680-ps1 | 0.149335367 | -0.206481672 | 2.864896933 | -0.072072984 |
| Olfr681 | 0 | NA | NA | NA |
| Olfr682-ps1 | 0.201268616 | 1.287832049 | 2.613944264 | 0.492677701 |
| Olfr683 | 0.362463993 | 0.755307759 | 1.691902168 | 0.446425197 |
| Olfr684 | 0.256199707 | 0.274399549 | 2.992454766 | 0.091697142 |
| Olfr685 | 0.357595806 | 1.293112424 | 3.306128559 | 0.391125874 |
| Olfr686 | 0.169924525 | -0.193116811 | 3.387745846 | -0.057004516 |
| Olfr687 | 0 | NA | NA | NA |
| Olfr688 | 0 | NA | NA | NA |
| Olfr689 | 0 | NA | NA | NA |
| Olfr690 | 5.292052314 | -0.157338995 | 0.479821779 | -0.327911325 |
| Olfr691 | 5.347759076 | -0.349694147 | 0.607176859 | -0.575934576 |
| Olfr692 | 0.049553047 | 0.274391604 | 3.724945049 | 0.073663262 |
| Fam160a2 | 1185.683241 | -0.656874624 | 0.100816325 | -6.515558106 |
| Gm5901 | 0.044544648 | 0.755286614 | 3.724945049 | 0.202764498 |
| Cnga4 | 8.850792061 | -0.387110206 | 0.361194159 | -1.071751014 |
| Cckbr | 263.3216545 | -0.579545772 | 0.167961991 | -3.450457869 |
| Prkcdbp | 205.5345052 | 0.481177687 | 0.203219491 | 2.367773306 |
| Smpd1 | 1173.492348 | 0.067429787 | 0.069326811 | 0.972636505 |
| Apbb1 | 2731.609977 | -0.111607833 | 0.062369869 | -1.789451149 |
| Hpx | 16.70994554 | 3.481594084 | 0.504065961 | 6.907020813 |
| Trim3 | 1334.924561 | 0.038364824 | 0.083778356 | 0.457932408 |
| Arfip2 | 801.3768636 | 0.24323899 | 0.069636214 | 3.492995636 |
| Timm10b | 130.0419406 | 0.003485105 | 0.103054218 | 0.033818168 |
| Dnhd1 | 163.944155 | 0.010504352 | 0.1980065 | 0.05305054 |
| Gm22504 | 0.072025462 | 0.755288839 | 3.724945049 | 0.202765096 |
| Mrps36-ps2 | 0.9406005 | 1.470801164 | 1.205323367 | 1.220254418 |
| Rrp8 | 163.2337327 | 0.116377641 | 0.12864782 | 0.904621945 |
| Ilk | 832.9278343 | 0.481577158 | 0.105695504 | 4.556269072 |
| Taf10 | 390.2000004 | 0.009434596 | 0.113371361 | 0.083218511 |
| Tpp1 | 810.0873032 | 0.067012212 | 0.118288638 | 0.566514357 |
| Dchs1 | 413.6250274 | -0.246114928 | 0.104295289 | -2.359789495 |
| Gm15645 | 4.914941411 | 0.112499475 | 0.454685096 | 0.247422833 |
| Mrpl17 | 334.4169704 | 0.250289514 | 0.115360494 | 2.169629351 |
| 9130208D14Rik | 37.17479923 | 6.744039664 | 0.855942193 | 7.879083095 |
| Gm4070 | 48.02494048 | 5.135965134 | 0.569909734 | 9.011892284 |
| Gm20663 | 0.150903271 | 0.755303274 | 2.855571776 | 0.26450159 |
| Gm18852 | 4.812241163 | 4.050615638 | 0.942327158 | 4.298523719 |
| Gm27228 | 0 | NA | NA | NA |
| Gm8979 | 1.384150549 | 2.347157263 | 1.154673857 | 2.032744786 |
| Gm27149 | 0.066050875 | 0.274391604 | 3.724945049 | 0.073663262 |
| Gm8982 | 0 | NA | NA | NA |

| | | | | |
|---|---|---|---|---|
| Gvin1 | 6.195143366 | 4.852478548 | 0.986349384 | 4.919634591 |
| Gm18853 | 1.133034309 | 1.7149466 | 1.306430489 | 1.312696401 |
| Gm27150 | 0 | NA | NA | NA |
| Gm8989 | 0.700564593 | 2.02260672 | 1.368638192 | 1.477824257 |
| Gm4759 | 67.3121842 | 5.977034462 | 0.776241568 | 7.699967007 |
| Gm22372 | 0 | NA | NA | NA |
| Gm8995 | 675.8586595 | 4.85446378 | 0.22050488 | 22.01522157 |
| Gm1966 | 19.76971248 | 5.101458126 | 0.906379618 | 5.628390162 |
| Olfr693 | 0.090277459 | -0.20649451 | 3.724945049 | -0.055435586 |
| Olfr694 | 0 | NA | NA | NA |
| Olfr695 | 0 | NA | NA | NA |
| Olfr697 | 0 | NA | NA | NA |
| Olfr698 | 0 | NA | NA | NA |
| Olfr699 | 0.049553047 | 0.274391604 | 3.724945049 | 0.073663262 |
| Olfr700 | 0 | NA | NA | NA |
| Olfr701 | 0.038981395 | 0.274393828 | 3.724945049 | 0.073663859 |
| Olfr702 | 0 | NA | NA | NA |
| Olfr703 | 0.085433889 | -0.687376175 | 3.724945049 | -0.184533239 |
| Olfr704 | 0 | NA | NA | NA |
| Olfr705 | 0.136194719 | -0.687371727 | 3.724945049 | -0.184532045 |
| Olfr706 | 0 | NA | NA | NA |
| Olfr707 | 0.068097359 | -0.206487837 | 3.724945049 | -0.055433794 |
| Olfr1532-ps1 | 0 | NA | NA | NA |
| Olfr709-ps1 | 0 | NA | NA | NA |
| Olfr710 | 0.04393175 | -0.206490061 | 3.724945049 | -0.055434391 |
| Olfr6 | 0 | NA | NA | NA |
| Olfr711 | 0 | NA | NA | NA |
| Olfr2 | 0 | NA | NA | NA |
| Olfr713 | 0 | NA | NA | NA |
| Olfr714 | 0 | NA | NA | NA |
| Olfr17 | 0.048949532 | -0.206487837 | 3.724945049 | -0.055433794 |
| Gm10081 | 0 | NA | NA | NA |
| Olfr715 | 0.097139547 | 0.274393828 | 3.724945049 | 0.073663859 |
| Olfr716 | 0.092691418 | 0.274387304 | 3.724945049 | 0.073662108 |
| Nlrp14 | 0 | NA | NA | NA |
| Gm3765 | 0.678124035 | 1.294566597 | 1.469156235 | 0.881163328 |
| Rbmxl2 | 0 | NA | NA | NA |
| Platr28 | 0.049553047 | 0.274391604 | 3.724945049 | 0.073663262 |
| Gm24932 | 0.060174537 | 0.755286614 | 3.724945049 | 0.202764498 |
| Syt9 | 336.6996381 | -0.026292163 | 0.085580086 | -0.307222915 |
| Olfml1 | 105.9701042 | 0.066706831 | 0.17477262 | 0.381677812 |
| Ppfibp2 | 154.4178595 | 0.095485503 | 0.127893268 | 0.746603044 |
| 5330417H12Rik | 0 | NA | NA | NA |
| Cyb5r2 | 9.463141173 | 1.348680793 | 0.450612218 | 2.992996501 |
| Ovch2 | 0 | NA | NA | NA |
| Olfr467 | 0 | NA | NA | NA |
| Olfr469 | 0 | NA | NA | NA |

| | | | | |
|---|---|---|---|---|
| Olfr470 | 0 | NA | NA | NA |
| Gm10156 | 22.50808326 | 1.536196056 | 0.346575768 | 4.432497005 |
| Olfr472 | 0.092881282 | -0.687373951 | 3.724945049 | -0.184532642 |
| Olfr473 | 0.038981395 | 0.274393828 | 3.724945049 | 0.073663859 |
| Olfr474 | 0 | NA | NA | NA |
| Olfr476 | 0 | NA | NA | NA |
| Olfr477 | 0.165236907 | -0.206488136 | 3.378398333 | -0.061120127 |
| Olfr478 | 0 | NA | NA | NA |
| Olfr479 | 0 | NA | NA | NA |
| Olfr480 | 0 | NA | NA | NA |
| Olfr481 | 0 | NA | NA | NA |
| Olfr482 | 0 | NA | NA | NA |
| Olfr483 | 0.062426873 | 0.755288839 | 3.724945049 | 0.202765096 |
| Gm25424 | 0 | NA | NA | NA |
| Olfr484 | 0 | NA | NA | NA |
| Olfr485 | 0 | NA | NA | NA |
| Olfr486 | 0 | NA | NA | NA |
| Olfr487 | 0.123841226 | 0.274391173 | 3.724945049 | 0.073663147 |
| Olfr488 | 0 | NA | NA | NA |
| Olfr490 | 0 | NA | NA | NA |
| Olfr491 | 0 | NA | NA | NA |
| Olfr492 | 0 | NA | NA | NA |
| Olfr493 | 0 | NA | NA | NA |
| Olfr494 | 0 | NA | NA | NA |
| Olfr495 | 0 | NA | NA | NA |
| Olfr497 | 0.049553047 | 0.274391604 | 3.724945049 | 0.073663262 |
| Olfr498 | 0 | NA | NA | NA |
| Olfr502 | 0 | NA | NA | NA |
| Olfr503 | 0 | NA | NA | NA |
| Olfr504 | 0 | NA | NA | NA |
| Olfr506 | 0 | NA | NA | NA |
| Olfr507 | 0 | NA | NA | NA |
| Olfr508 | 0 | NA | NA | NA |
| Olfr509 | 0 | NA | NA | NA |
| Olfr510 | 0.046345709 | 0.274391604 | 3.724945049 | 0.073663262 |
| Olfr512 | 0 | NA | NA | NA |
| Olfr513 | 0 | NA | NA | NA |
| Olfr514 | 0 | NA | NA | NA |
| Olfr516 | 0 | NA | NA | NA |
| Olfr517 | 0 | NA | NA | NA |
| Olfr518 | 0 | NA | NA | NA |
| Olfr519 | 0 | NA | NA | NA |
| Nlrp10 | 10.06809784 | 0.844897044 | 0.328550353 | 2.571590736 |
| Eif3f | 1059.444892 | 0.4020273 | 0.076354655 | 5.265262513 |
| Gm26599 | 4.712628857 | 3.16969431 | 0.770634886 | 4.113094757 |
| Tub | 1801.451592 | 0.064515891 | 0.111024729 | 0.581094789 |
| Ric3 | 1280.403782 | 0.375413567 | 0.07203773 | 5.211346427 |

| | | | | |
|---|---|---|---|---|
| Gm25450 | 3.817416342 | -0.291014881 | 0.556328845 | -0.523098674 |
| Lmo1 | 71.3151019 | 0.377844641 | 0.137366138 | 2.750638885 |
| Stk33 | 14.06609556 | -0.370013483 | 0.291758107 | -1.268220055 |
| Gm9105 | 5.082316306 | 0.530706286 | 0.452913538 | 1.171760702 |
| 1700095J03Rik | 3.157210742 | 0.678783873 | 0.621728012 | 1.09176981 |
| Trim66 | 632.3889432 | 0.088719022 | 0.105373231 | 0.841950288 |
| Rpl27a | 1290.635761 | 0.32648001 | 0.102450437 | 3.18671174 |
| Snora3 | 0 | NA | NA | NA |
| AC124457.1 | 0.875090066 | 0.469265552 | 1.049523574 | 0.44712245 |
| Gm24888 | 1.185922749 | -1.619411773 | 1.014351122 | -1.596500204 |
| St5 | 362.0690004 | 0.101365705 | 0.085718489 | 1.182541904 |
| Akip1 | 31.41517011 | 0.50672077 | 0.237176392 | 2.136472211 |
| BC051019 | 3.117097637 | 0.98263882 | 0.695895796 | 1.412048795 |
| Ascl3 | 1.052193792 | 0.502122346 | 0.984905299 | 0.509817895 |
| Tmem9b | 679.663783 | 0.041150246 | 0.073902689 | 0.556816629 |
| Nrip3 | 1443.710783 | -0.523062983 | 0.111939958 | -4.672710189 |
| Gm22822 | 1.435595059 | 0.321817862 | 0.798936208 | 0.402807957 |
| Scube2 | 22.45075123 | 0.185576399 | 0.300728763 | 0.617088957 |
| Dennd5a | 3399.692085 | -0.22843545 | 0.068672018 | -3.326470617 |
| Tmem41b | 489.61422 | -0.089241575 | 0.100680262 | -0.886386004 |
| Gm24842 | 0.126912322 | -0.206491353 | 3.697934765 | -0.055839642 |
| 1600010M07Rik | 2.455553585 | 1.838527425 | 0.901585074 | 2.039216795 |
| Gm23100 | 0 | NA | NA | NA |
| Ipo7 | 1735.407786 | -0.026561559 | 0.076096852 | -0.349049384 |
| Gm25636 | 6.97985599 | 0.004276235 | 0.401274418 | 0.010656635 |
| Snora23 | 31.16499087 | -0.366420827 | 0.234707555 | -1.561180371 |
| AA474408 | 22.01653641 | -1.077084725 | 0.270125418 | -3.98735051 |
| Zfp143 | 153.6133789 | 0.166383577 | 0.096754549 | 1.719646036 |
| Wee1 | 194.9507975 | -0.13948806 | 0.119102065 | -1.171164079 |
| Swap70 | 161.8502858 | 0.1787737 | 0.112700334 | 1.586274798 |
| Sbf2 | 1399.071276 | 0.069116997 | 0.072727011 | 0.950362138 |
| Gm9064 | 0 | NA | NA | NA |
| Gm17219 | 0.046345709 | 0.274391604 | 3.724945049 | 0.073663262 |
| Gm10087 | 0 | NA | NA | NA |
| Gm28863 | 0.430307537 | 0.281239605 | 1.456347697 | 0.193112953 |
| Adm | 37.32539147 | 1.094586843 | 0.243191511 | 4.500925378 |
| Ampd3 | 786.3935543 | 1.201264128 | 0.16283651 | 7.377117881 |
| Rnf141 | 819.8137337 | -0.015769826 | 0.108430978 | -0.145436534 |
| AC122844.1 | 0.956902809 | 0.204964459 | 1.211522873 | 0.16917919 |
| Lyve1 | 8.973244537 | -0.199235156 | 0.391216996 | -0.509270195 |
| Mrvi1 | 43.82669296 | -0.155683045 | 0.179201305 | -0.868760666 |
| Gm16336 | 0.075043205 | 0.755286614 | 3.724945049 | 0.202764498 |
| Ctr9 | 691.8584163 | -0.137813483 | 0.059890559 | -2.301088604 |
| Eif4g2 | 8499.079975 | 0.112125842 | 0.082386862 | 1.36096751 |
| Gm23262 | 0 | NA | NA | NA |
| Galnt18 | 280.6267264 | -0.473746305 | 0.129290823 | -3.664191283 |
| Gm25787 | 0 | NA | NA | NA |

| | | | | |
|---|---|---|---|---|
| Gm23345 | 0.066050875 | 0.274391604 | 3.724945049 | 0.073663262 |
| Usp47 | 1697.490691 | 0.211007401 | 0.062903266 | 3.354474518 |
| Dkk3 | 1672.01163 | 0.368726516 | 0.122007195 | 3.022170254 |
| Mical2 | 2673.306519 | -0.34590447 | 0.141290392 | -2.448181119 |
| Micalcl | 33.92286921 | -0.977137821 | 0.260774696 | -3.747057653 |
| Parva | 887.2698577 | -0.053461036 | 0.086722427 | -0.616461489 |
| Parvaos | 2.062454624 | 0.963094972 | 0.740528915 | 1.300550124 |
| 2310014F06Rik | 4.100390628 | 0.126108126 | 0.51532875 | 0.244713934 |
| Gm24154 | 0.181894773 | 0.274410467 | 2.698759046 | 0.10168024 |
| Tead1 | 787.7264761 | 0.548577579 | 0.122663658 | 4.472209517 |
| AC126052.1 | 1.51359592 | -0.896918982 | 0.841447182 | -1.06592428 |
| Rassf10 | 10.84181486 | -0.010445973 | 0.292337923 | -0.035732529 |
| AC104930.1 | 0 | NA | NA | NA |
| Gm23662 | 0 | NA | NA | NA |
| Arntl | 352.8719286 | 0.101618878 | 0.128010669 | 0.793831317 |
| Btbd10 | 732.5892807 | -0.072366475 | 0.077757055 | -0.930674075 |
| Gm15994 | 0.277092755 | 0.755303684 | 2.076192735 | 0.363792663 |
| Pth | 0 | NA | NA | NA |
| Far1 | 1848.391941 | -0.180841448 | 0.064048071 | -2.823526834 |
| Far1os | 0.286281128 | 0.274409398 | 3.724945049 | 0.073668039 |
| Gm23700 | 0 | NA | NA | NA |
| Gm5600 | 0.213133906 | -0.184302513 | 2.170898092 | -0.084896898 |
| Gm29507 | 0.046345709 | 0.274391604 | 3.724945049 | 0.073663262 |
| Spon1 | 1904.191878 | 0.695382088 | 0.107858751 | 6.447155045 |
| Rras2 | 177.862388 | 0.171269814 | 0.119909846 | 1.428321521 |
| Copb1 | 1041.818964 | 0.259692943 | 0.086728053 | 2.994336149 |
| Psma1 | 653.5961011 | 0.356294354 | 0.107300169 | 3.320538592 |
| 4933406I18Rik | 91.43073966 | -0.296192048 | 0.124993421 | -2.369661108 |
| Pde3b | 366.815521 | 0.108755631 | 0.096233905 | 1.130117613 |
| Cyp2r1 | 3.701896043 | 0.9538234 | 0.587816604 | 1.622654741 |
| Calca | 61.65923974 | 1.226103057 | 0.258231792 | 4.748071679 |
| Gm15500 | 180.1198848 | 0.554586404 | 0.165179703 | 3.357473065 |
| Calcb | 5.761881877 | 1.574172998 | 0.59933419 | 2.626536286 |
| Insc | 59.233201 | -0.737204102 | 0.260028228 | -2.835092585 |
| Gm24982 | 0 | NA | NA | NA |
| Rps4l-ps | 0.061920613 | 0.274393828 | 3.724945049 | 0.073663859 |
| Gm22669 | 0 | NA | NA | NA |
| Gm27684 | 0.263733274 | 1.244013933 | 2.101824633 | 0.591873325 |
| Sox6 | 489.52169 | -0.039053727 | 0.087103352 | -0.448360782 |
| Sox6os | 0.611953183 | -1.073858442 | 1.209943253 | -0.887527939 |
| 1700003G18Rik | 0.77082754 | -1.092139366 | 1.100387714 | -0.992504143 |
| Gm4353 | 19.37501129 | -0.13387294 | 0.300865249 | -0.444959796 |
| 1110004F10Rik | 588.1908301 | 0.286388312 | 0.092538135 | 3.094813959 |
| Plekha7 | 339.2168702 | -0.232857872 | 0.100881673 | -2.308227708 |
| Rps13 | 763.6410431 | 0.447164846 | 0.102423619 | 4.365837207 |
| Snord14a | 3.97447326 | 0.756447585 | 0.549228652 | 1.377290829 |
| Pik3c2a | 1083.338475 | 0.03408319 | 0.066718255 | 0.510852545 |

| | | | | |
|---|---|---|---|---|
| Nucb2 | 157.5399997 | 0.542086524 | 0.170659932 | 3.176413578 |
| n-R5s157 | 0 | NA | NA | NA |
| Gm25423 | 0 | NA | NA | NA |
| RP24-246N21.1 | 393.097115 | 0.310709916 | 0.121998529 | 2.546833285 |
| Gm26054 | 0 | NA | NA | NA |
| Xylt1 | 443.4184742 | -0.132288819 | 0.065056897 | -2.033432654 |
| Gm24063 | 0 | NA | NA | NA |
| Rps15a | 1296.320441 | 0.295000875 | 0.075084217 | 3.928933244 |
| Arl6ip1 | 2631.753197 | 0.715841499 | 0.118380459 | 6.046956599 |
| Smg1 | 4666.36924 | 0.032964124 | 0.103780396 | 0.317633437 |
| Gm25683 | 0 | NA | NA | NA |
| 4930583K01Rik | 0.241683003 | 1.664107227 | 2.082187402 | 0.799211073 |
| Gm23229 | 0.044544648 | 0.755286614 | 3.724945049 | 0.202764498 |
| Syt17 | 179.5473353 | -0.038857521 | 0.152742026 | -0.254399672 |
| Itpripl2 | 154.921355 | 1.667534616 | 0.171075979 | 9.747333451 |
| Coq7 | 126.5384451 | 0.126121856 | 0.15033642 | 0.838930817 |
| Tmc7 | 194.611435 | -0.906732782 | 0.119927809 | -7.560654973 |
| Tmc5 | 2.207900702 | 0.213477732 | 0.750209167 | 0.284557616 |
| Gde1 | 1198.036269 | 0.011085794 | 0.058761844 | 0.188656333 |
| Ccp110 | 802.7760504 | -0.651973139 | 0.080928848 | -8.056127786 |
| 9030624J02Rik | 685.4574909 | -0.014463612 | 0.069944582 | -0.206786745 |
| Gm26147 | 1.083234348 | -1.901939203 | 1.029917452 | -1.846690916 |
| Knop1 | 359.1550831 | 0.006092091 | 0.07106425 | 0.085726522 |
| Iqck | 21.26371758 | 0.533134114 | 0.245606253 | 2.170686244 |
| Gprc5b | 1912.548548 | -0.525020966 | 0.094291926 | -5.568037342 |
| Gpr139 | 17.82217913 | -0.837636803 | 0.303352948 | -2.761261461 |
| Gm23321 | 0 | NA | NA | NA |
| Gp2 | 0.244487964 | 0.263406501 | 1.853142167 | 0.142140471 |
| Umod | 0 | NA | NA | NA |
| Pdilt | 0.092881282 | -0.687373951 | 3.724945049 | -0.184532642 |
| Gm37518 | 0.130524232 | 0.274405456 | 3.657871338 | 0.075017799 |
| Gm26967 | 0.180739367 | -0.20648076 | 3.322505087 | -0.062146108 |
| Acsm5 | 5.692917749 | 0.481276096 | 0.464345066 | 1.036462171 |
| Acsm2 | 0.062426873 | 0.755288839 | 3.724945049 | 0.202765096 |
| Acsm1 | 0.038981395 | 0.274393828 | 3.724945049 | 0.073663859 |
| Acsm4 | 1.49381288 | -0.823614289 | 0.80098935 | -1.028246243 |
| Thumpd1 | 343.5482368 | 0.390493058 | 0.105930291 | 3.686321009 |
| Gm5601 | 215.6914571 | 0.025083866 | 0.094709696 | 0.264850035 |
| Acsm3 | 0.623781035 | 1.422877869 | 1.305190822 | 1.09016846 |
| Eri2 | 94.72177517 | -0.108587576 | 0.152378562 | -0.712617144 |
| 2610020H08Rik | 22.09157058 | 0.060428045 | 0.284031975 | 0.212750854 |
| Dcun1d3 | 429.5927485 | -0.207745755 | 0.092722865 | -2.240501901 |
| Lyrm1 | 64.24767704 | 0.113076432 | 0.145318636 | 0.778127536 |
| Dnah3 | 28.90499188 | -0.886316419 | 0.230842503 | -3.839485403 |
| Gm25217 | 0.280224789 | -0.2064822 | 1.81573473 | -0.113718263 |
| Tmem159 | 54.2618588 | 0.623181492 | 0.166214487 | 3.749260984 |
| Zp2 | 1.125736089 | 2.089583461 | 1.091674552 | 1.914108429 |

| | | | | |
|---|---|---|---|---|
| Anks4b | 4.77461678 | 2.45523132 | 0.59001893 | 4.1612755 |
| Crym | 493.688269 | 1.01187933 | 0.125026556 | 8.093315205 |
| Abca14 | 0.666404725 | 0.727814711 | 1.296427967 | 0.561400039 |
| Abca15 | 0.612941558 | 0.283424426 | 1.440789195 | 0.196714708 |
| Abca16 | 0.254958917 | 0.274407753 | 2.095474205 | 0.13095258 |
| E130201H02Rik | 1.649955311 | 0.921258844 | 0.824711372 | 1.117068196 |
| Uqcrc2 | 1539.021604 | 0.111711475 | 0.078490785 | 1.423243181 |
| Pdzd9 | 1.255624142 | 0.588820718 | 1.127938972 | 0.522032426 |
| 9030407P20Rik | 49.63574211 | -0.127835345 | 0.173658895 | -0.736128979 |
| BC030336 | 1048.447558 | 0.062532853 | 0.065122437 | 0.960235144 |
| Vwa3a | 56.37564545 | -0.286634529 | 0.166889709 | -1.717508714 |
| Eef2k | 574.705251 | 0.208161266 | 0.104889261 | 1.984581299 |
| Gm15774 | 0.046345709 | 0.274391604 | 3.724945049 | 0.073663262 |
| Polr3e | 380.9951259 | 0.138238578 | 0.10674263 | 1.295064377 |
| Cdr2 | 189.8166021 | -0.302821107 | 0.177542873 | -1.70562243 |
| 4933427G17Rik | 3.26880693 | 0.598941863 | 0.562077292 | 1.065586303 |
| Gm9234 | 7.527412146 | -0.026340735 | 0.389483872 | -0.067629847 |
| Gm25924 | 0 | NA | NA | NA |
| Gm9905 | 21.38643968 | 0.534943566 | 0.264083001 | 2.025664527 |
| Mettl9 | 576.9325256 | 0.082117664 | 0.112222518 | 0.731739635 |
| Gm27991 | 0 | NA | NA | NA |
| Igsf6 | 72.32097277 | 4.171894814 | 0.409718189 | 10.18235199 |
| Otoa | 4.95839639 | 0.482265843 | 0.476704042 | 1.011667198 |
| Hs3st2 | 198.3223821 | -1.018157434 | 0.313930444 | -3.243258033 |
| Usp31 | 2536.972672 | -0.372622573 | 0.078852618 | -4.725557383 |
| 1700069B07Rik | 1.411393777 | 0.703074371 | 0.847006081 | 0.83007004 |
| Scnn1g | 16.54571043 | 1.857288854 | 0.31812513 | 5.838233698 |
| Scnn1b | 3.3334194 | 0.800687156 | 0.867278409 | 0.923218136 |
| Cog7 | 494.7223238 | -0.013893205 | 0.065596944 | -0.211796536 |
| Gga2 | 393.4606856 | 0.033100333 | 0.096885727 | 0.341643023 |
| Ears2 | 110.4529885 | -0.112951416 | 0.124578187 | -0.906670897 |
| Ubfd1 | 1136.037908 | -0.14177022 | 0.063228102 | -2.242202677 |
| Ndufab1 | 625.6134519 | -0.20414511 | 0.080332883 | -2.541239662 |
| Palb2 | 22.09319308 | 0.217929026 | 0.262212052 | 0.831117501 |
| Gm16326 | 0.136194719 | -0.687371727 | 3.724945049 | -0.184532045 |
| Dctn5 | 636.704053 | 0.009604793 | 0.055444464 | 0.173232673 |
| Gm15489 | 0.496441948 | -0.69367545 | 1.572154792 | -0.441225924 |
| Plk1 | 22.12491667 | 0.726862806 | 0.310848059 | 2.338321841 |
| Ern2 | 3.478004289 | -0.274850402 | 0.529657994 | -0.518920521 |
| Chp2 | 6.555594633 | -0.783580389 | 0.405293986 | -1.933362981 |
| Gm23614 | 0 | NA | NA | NA |
| Prkcb | 4097.896829 | -0.339528551 | 0.119819575 | -2.833665119 |
| Gm14388 | 23.59541605 | 0.037039103 | 0.249899366 | 0.148216075 |
| Gm26414 | 4.07614265 | -0.921544583 | 0.585084316 | -1.575062872 |
| Gm14389 | 0 | NA | NA | NA |
| Cacng3 | 479.8047837 | -0.381720079 | 0.164672272 | -2.318059221 |
| Rbbp6 | 1656.438353 | 0.194961399 | 0.072847159 | 2.676307525 |

| | | | | |
|---|---|---|---|---|
| Tnrc6a | 1721.479868 | -0.157037135 | 0.094319443 | -1.664949773 |
| Slc5a11 | 15.52195762 | -0.138662696 | 0.277534421 | -0.499623418 |
| Arhgap17 | 309.4984358 | 0.328691866 | 0.141462287 | 2.323529987 |
| Lcmt1 | 343.5567309 | -0.146607782 | 0.068685136 | -2.134490658 |
| Gm23559 | 0 | NA | NA | NA |
| Aqp8 | 0.525182132 | 1.515632015 | 1.456755741 | 1.040416023 |
| Zkscan2 | 230.1962317 | 0.149785565 | 0.158460451 | 0.945255193 |
| Gm23788 | 0.167772976 | -0.206492322 | 3.417713432 | -0.060418267 |
| Gm26974 | 9.009306514 | 0.250771143 | 0.335334897 | 0.747822983 |
| Gm27040 | 8.361238006 | 0.297351751 | 0.398253077 | 0.746640185 |
| Hs3st4 | 521.3318301 | 0.29404003 | 0.086610278 | 3.394978479 |
| Gm25759 | 6.10632452 | -0.037697517 | 0.505266508 | -0.074609175 |
| Gm15338 | 14.24273356 | -0.860744597 | 0.422270004 | -2.038374948 |
| Gm15339 | 0 | NA | NA | NA |
| Gm23168 | 0.097139547 | 0.274393828 | 3.724945049 | 0.073663859 |
| 3100003L05Rik | 1.759517327 | 1.920481938 | 0.910322502 | 2.109672048 |
| Gm21957 | 0 | NA | NA | NA |
| 4930533L02Rik | 0 | NA | NA | NA |
| 4933440M02Rik | 0.422952957 | -0.135342054 | 3.695029225 | -0.036628142 |
| 4930571K23Rik | 0 | NA | NA | NA |
| Kdm8 | 38.64691656 | 0.603556812 | 0.24359605 | 2.4776954 |
| Nsmce1 | 119.9047304 | 0.612005437 | 0.171264968 | 3.573442039 |
| Il4ra | 356.9425988 | 2.433100522 | 0.195803673 | 12.42622513 |
| Il21r | 78.43361135 | 3.523029637 | 0.263641318 | 13.36296475 |
| Gtf3c1 | 2212.631377 | -0.027697791 | 0.062666555 | -0.441986813 |
| D430042O09Rik | 320.9984056 | -0.185572046 | 0.07579936 | -2.448200718 |
| Gsg1l | 413.6031104 | -0.41617407 | 0.159613106 | -2.607392843 |
| Xpo6 | 1490.157562 | 0.240967053 | 0.066880137 | 3.60296887 |
| Gm17137 | 0.048949532 | -0.206487837 | 3.724945049 | -0.055433794 |
| Gm17136 | 0.044544648 | 0.755286614 | 3.724945049 | 0.202764498 |
| Sbk1 | 1551.156001 | -0.382685226 | 0.143689108 | -2.663286245 |
| Gm10155 | 8.990861825 | 0.280793739 | 0.366379388 | 0.766401572 |
| Lat | 33.16436472 | 3.167411223 | 0.361025213 | 8.773379547 |
| Mir7058 | 0 | NA | NA | NA |
| Spns1 | 414.4773662 | -0.079014063 | 0.080535821 | -0.981104574 |
| Nfatc2ip | 98.35474023 | 0.279265193 | 0.141857646 | 1.968629835 |
| Cd19 | 0.620398479 | 0.755308267 | 1.617683556 | 0.466907303 |
| Rabep2 | 121.2584971 | -0.091171653 | 0.108509745 | -0.840216264 |
| Atp2a1 | 15.93822915 | 0.157255748 | 0.265670656 | 0.591919899 |
| Sh2b1 | 913.839647 | -0.079008695 | 0.058491924 | -1.350762459 |
| Tufm | 383.5821207 | -0.310830518 | 0.1066793 | -2.913691017 |
| Atxn2l | 1896.105242 | 0.046176033 | 0.084393652 | 0.547150543 |
| Eif3c | 2058.465012 | 0.246364263 | 0.084805585 | 2.905047638 |
| Gm24596 | 0.884909101 | -0.609946062 | 1.084014298 | -0.562673447 |
| G730046D07Rik | 2.910496466 | 2.396187205 | 1.024586984 | 2.338685969 |
| Gm25579 | 0 | NA | NA | NA |
| Cln3 | 171.9378215 | 0.191709369 | 0.109267513 | 1.754495587 |

| | | | | |
|---|---|---|---|---|
| Mir7059 | 0.228481116 | -0.640270733 | 2.168157963 | -0.295306312 |
| Apobr | 49.34344806 | 2.764207222 | 0.404840199 | 6.827897106 |
| Il27 | 13.85132785 | 5.663008644 | 0.903941838 | 6.264793158 |
| Nupr1 | 177.6816824 | 2.23631966 | 0.283472026 | 7.889031196 |
| 2510046G10Rik | 4.55499157 | -0.9663302 | 0.480329237 | -2.011807995 |
| Ccdc101 | 124.2951638 | -0.180406871 | 0.100309546 | -1.798501524 |
| Sult1a1 | 88.10259738 | 1.238787703 | 0.250343966 | 4.948342572 |
| Gm6939 | 1.140676537 | 1.109711964 | 1.01095634 | 1.09768535 |
| Slx1b | 109.1217166 | -0.010286994 | 0.121750015 | -0.084492751 |
| Bola2 | 141.4468845 | 0.058968611 | 0.148175077 | 0.397965787 |
| Coro1a | 641.2179666 | 0.435521864 | 0.087833908 | 4.958470748 |
| Mapk3 | 1099.213407 | -0.093193435 | 0.092993746 | -1.002147338 |
| Gdpd3 | 17.57179256 | -0.620255829 | 0.281994454 | -2.199532017 |
| Gm9967 | 7.556454724 | 0.156271387 | 0.416519119 | 0.375184188 |
| Ypel3 | 1100.394795 | -0.095608004 | 0.071648641 | -1.334400803 |
| Tbx6 | 9.328334444 | -0.832057756 | 0.420575299 | -1.978379991 |
| Ppp4c | 390.6480823 | 0.463272822 | 0.123726017 | 3.744344426 |
| Aldoa | 12436.05844 | -0.035017249 | 0.049795685 | -0.703218537 |
| Gm15676 | 14.57694585 | 0.369159911 | 0.279520426 | 1.320690283 |
| Fam57b | 780.1775602 | -0.309253051 | 0.067702508 | -4.567822648 |
| 4930451I11Rik | 0.843446786 | -2.212797075 | 1.059826989 | -2.087885191 |
| Doc2a | 204.7161822 | -0.77512017 | 0.163015518 | -4.754885773 |
| Ino80e | 331.245082 | 0.091789615 | 0.073810063 | 1.243592146 |
| Hirip3 | 134.4899788 | -0.145679132 | 0.139755801 | -1.042383434 |
| Taok2 | 2099.502443 | -0.086433633 | 0.047743687 | -1.810367809 |
| Tmem219 | 154.4928956 | 0.546169208 | 0.159564672 | 3.422870496 |
| Gm20650 | 0.099106094 | 0.274387056 | 3.724945049 | 0.073662041 |
| Gm23445 | 0.097899064 | -0.687371727 | 3.724945049 | -0.184532045 |
| Kctd13 | 647.9350141 | -0.042357879 | 0.085897147 | -0.493123239 |
| Gm21984 | 106.5590713 | 0.339797316 | 0.197727808 | 1.718510516 |
| Asphd1 | 163.222168 | -0.39276498 | 0.111973279 | -3.507667027 |
| Sez6l2 | 4188.453118 | 0.070917695 | 0.072319343 | 0.980618626 |
| D830044I16Rik | 2.339863245 | 0.30397764 | 0.724958006 | 0.419303791 |
| Cdipt | 964.396228 | 0.155945805 | 0.044663291 | 3.491587908 |
| Mvp | 599.1504578 | 2.879046525 | 0.267060361 | 10.78050861 |
| Pagr1b | 0.194631057 | 1.693379269 | 3.232136674 | 0.52391945 |
| Pagr1a | 269.1231763 | -0.198650975 | 0.145577715 | -1.364569953 |
| Prrt2 | 12.36009762 | -0.314589079 | 0.38930413 | -0.808080509 |
| Maz | 1509.464906 | -0.105361603 | 0.073126547 | -1.440811954 |
| Kif22 | 26.64433543 | 0.761398637 | 0.261832246 | 2.907963581 |
| Gm25333 | 0 | NA | NA | NA |
| AC122863.1 | 0 | NA | NA | NA |
| Zg16 | 0.046345709 | 0.274391604 | 3.724945049 | 0.073663262 |
| Gm22747 | 0 | NA | NA | NA |
| AI467606 | 37.52684094 | 1.346581011 | 0.267693132 | 5.030315882 |
| Qprt | 3.756627285 | 1.035199619 | 0.535945611 | 1.93153857 |
| Spn | 82.89016059 | 5.164473196 | 0.426907202 | 12.09741408 |

| | | | | |
|---|---|---|---|---|
| Gm25735 | 0 | NA | NA | NA |
| Cd2bp2 | 785.0302282 | 0.214049924 | 0.059816102 | 3.578466646 |
| Tbc1d10b | 882.4067378 | 0.025578677 | 0.067862392 | 0.376919768 |
| Mylpf | 4.837556069 | 0.260593916 | 0.51055171 | 0.510416301 |
| 1-Sep | 35.87249771 | 3.098491626 | 0.331970385 | 9.333638692 |
| Zfp553 | 281.7915856 | 0.049528976 | 0.081669701 | 0.606454723 |
| Zfp771 | 217.3863754 | -0.000644926 | 0.134254703 | -0.004803748 |
| Dctpp1 | 50.21803819 | 0.109927715 | 0.163678184 | 0.671608841 |
| Sephs2 | 242.8260877 | 0.300130868 | 0.104065068 | 2.884069295 |
| Gm17511 | 0.08486385 | 0.755286617 | 3.724945049 | 0.202764499 |
| Itgal | 262.0288419 | 3.377465428 | 0.220320664 | 15.32977144 |
| Zfp768 | 163.5362928 | 0.208961044 | 0.101929185 | 2.050060965 |
| Zfp747 | 52.03124678 | 0.144446252 | 0.158195486 | 0.913087069 |
| 9130019O22Rik | 50.28528677 | -0.36595475 | 0.179401972 | -2.039859132 |
| E430018J23Rik | 67.77893546 | 0.101266987 | 0.146869826 | 0.689501648 |
| Smt3h2-ps2 | 0 | NA | NA | NA |
| Zfp764 | 37.42744171 | 0.021792984 | 0.190546706 | 0.114370826 |
| Zfp688 | 66.55993122 | 0.366699161 | 0.143175505 | 2.561186426 |
| Zfp689 | 64.00074658 | 0.047464069 | 0.142961653 | 0.332005598 |
| B130055M24Rik | 80.27483843 | -0.897374606 | 0.189624823 | -4.732368863 |
| Prr14 | 312.8086992 | 0.102675054 | 0.073581023 | 1.395401273 |
| Fbrs | 544.6972203 | 0.241503443 | 0.097912674 | 2.466518729 |
| Mir7060 | 0.754347455 | -0.494388193 | 1.16360674 | -0.42487567 |
| AC136146.1 | 0 | NA | NA | NA |
| 1700008J07Rik | 27.76791627 | -0.01672723 | 0.214348303 | -0.078037615 |
| Srcap | 3720.548078 | -0.063549547 | 0.059211764 | -1.073258814 |
| Snora30 | 11.72211257 | -0.032857187 | 0.301054986 | -0.109140152 |
| Gm28198 | 0.198870357 | -0.206482483 | 2.667920877 | -0.07739453 |
| Tmem265 | 0 | NA | NA | NA |
| Phkg2 | 467.0988482 | 0.107902143 | 0.071545085 | 1.508169892 |
| Gm166 | 30.45493143 | -0.304823619 | 0.198736257 | -1.533809805 |
| Rnf40 | 789.9745989 | 0.114217862 | 0.066362079 | 1.72113146 |
| 1700120K04Rik | 1.43115425 | 1.06801658 | 0.75839222 | 1.408264158 |
| Zfp629 | 457.8583134 | -0.240697507 | 0.064800472 | -3.71444063 |
| Bcl7c | 293.7258915 | 0.093022557 | 0.079782322 | 1.165954502 |
| Mir762 | 1.435335593 | -0.753480268 | 0.965908561 | -0.780074117 |
| Ctf1 | 11.31378371 | -0.224738255 | 0.311068422 | -0.722472096 |
| Ctf2 | 0 | NA | NA | NA |
| Fbxl19 | 610.6258965 | -0.629657777 | 0.098112937 | -6.417683532 |
| Orai3 | 179.2087718 | 0.401039293 | 0.148051227 | 2.708787365 |
| Setd1a | 391.9559766 | 0.117000006 | 0.126103318 | 0.927810682 |
| Hsd3b7 | 112.3828733 | 0.398809212 | 0.144192222 | 2.765816406 |
| Stx1b | 6333.347927 | -0.305312715 | 0.074153995 | -4.117279384 |
| Stx4a | 301.4532526 | 0.161697655 | 0.144696006 | 1.117499084 |
| Zfp668 | 218.1486246 | -0.174511135 | 0.106672034 | -1.63595957 |
| Zfp646 | 586.462955 | 0.075867778 | 0.073499666 | 1.032219351 |
| Prss53 | 5.985399039 | -1.239207222 | 0.427628683 | -2.897858048 |

| | | | | |
|---|---|---|---|---|
| Gm21974 | 0.049553047 | 0.274391604 | 3.724945049 | 0.073663262 |
| Vkorc1 | 34.89639478 | -0.55406122 | 0.262073749 | -2.114142384 |
| Bckdk | 313.1884233 | 0.111910866 | 0.086109675 | 1.299631724 |
| Kat8 | 146.4017955 | -0.07657356 | 0.097372251 | -0.786400227 |
| Prss8 | 2.192463596 | -0.441263588 | 0.626957476 | -0.703817413 |
| Prss36 | 34.47699352 | -0.417592735 | 0.185869981 | -2.246692727 |
| Fus | 2383.412294 | 0.040990012 | 0.118576076 | 0.345685344 |
| Gm26690 | 0.089089296 | 1.194485492 | 3.723939576 | 0.320758559 |
| B230325K18Rik | 116.590507 | 0.62528472 | 0.274809657 | 2.275337508 |
| Pycard | 48.5114067 | 4.01154757 | 0.430624533 | 9.315650313 |
| Gm15533 | 0 | NA | NA | NA |
| Trim72 | 39.26137666 | 2.169629061 | 0.388482682 | 5.584879737 |
| Gm22809 | 0 | NA | NA | NA |
| Itgam | 625.9394924 | 1.780469128 | 0.153317655 | 11.61294262 |
| Itgax | 181.6100497 | 4.553395728 | 0.373032176 | 12.20644229 |
| Itgad | 28.13752969 | 2.050607846 | 0.302806865 | 6.771999193 |
| Cox6a2 | 20.40538094 | 0.954305195 | 0.228083172 | 4.184022822 |
| 9130023H24Rik | 70.01739671 | 0.115302059 | 0.155610698 | 0.740964858 |
| Armc5 | 244.6664953 | 0.750495564 | 0.097777209 | 7.675567431 |
| Tgfb1i1 | 119.9274198 | 0.258575618 | 0.173205531 | 1.49288315 |
| Slc5a2 | 1.211082793 | -0.747789677 | 1.190077273 | -0.628353884 |
| BC017158 | 206.5939799 | -0.153687926 | 0.083743904 | -1.8352133 |
| Mir3103 | 0 | NA | NA | NA |
| Rgs10 | 178.444261 | 0.07942513 | 0.125083972 | 0.634974482 |
| Gm15503 | 2.556904923 | 0.22177437 | 0.593891817 | 0.373425536 |
| Tial1 | 1499.062975 | -0.162205585 | 0.070545801 | -2.299294665 |
| Gm24365 | 0 | NA | NA | NA |
| Bag3 | 352.4742549 | 0.686674583 | 0.159516539 | 4.304723431 |
| Gm7258 | 0 | NA | NA | NA |
| Inpp5f | 1743.240792 | -0.141520274 | 0.073665187 | -1.921128282 |
| RP24-143L4.2 | 0 | NA | NA | NA |
| Gm16044 | 15.72144794 | 0.022998756 | 0.361449863 | 0.063629173 |
| Mcmbp | 481.2577654 | 0.124827878 | 0.069138498 | 1.805475693 |
| Gm23783 | 0 | NA | NA | NA |
| Sec23ip | 696.8616121 | 0.020909081 | 0.074344887 | 0.281244372 |
| n-R5s158 | 0 | NA | NA | NA |
| Gm25778 | 0.04393175 | -0.206490061 | 3.724945049 | -0.055434391 |
| Ppapdc1a | 43.3421754 | -0.417457743 | 0.246486176 | -1.693635517 |
| Wdr11 | 633.8080906 | -0.018650282 | 0.064927497 | -0.287247817 |
| Gm23847 | 0 | NA | NA | NA |
| Fgfr2 | 819.5441903 | -0.284007014 | 0.09685985 | -2.932143868 |
| Gm5903 | 24.16642656 | 0.211356937 | 0.225647776 | 0.936667493 |
| Ate1 | 1165.905196 | 0.016969607 | 0.056970123 | 0.297868532 |
| Gm4275 | 0.372589196 | 0.755310342 | 1.629310831 | 0.46357658 |
| Gm27884 | 0.4630892 | 1.079103809 | 1.476336708 | 0.730933399 |
| Nsmce4a | 299.1185009 | 0.457675223 | 0.116818183 | 3.91784233 |
| Tacc2 | 858.9102761 | -0.151080282 | 0.074625133 | -2.024522768 |

| | | | | |
|---|---|---|---|---|
| Etos1 | 0 | NA | NA | NA |
| Btbd16 | 23.45110209 | 0.736340818 | 0.293409824 | 2.509598378 |
| Gm5602 | 13.63298114 | 0.284696083 | 0.294548381 | 0.966551174 |
| Plekha1 | 1609.865415 | 0.047264447 | 0.071314978 | 0.66275625 |
| mmu-mir-7061 | 2.359287033 | 0.837418259 | 0.756600132 | 1.106817491 |
| Htra1 | 1343.622786 | 0.798283293 | 0.147813364 | 5.400616504 |
| Dmbt1 | 0.462783582 | 1.720370782 | 3.401619965 | 0.505750437 |
| 4933402N03Rik | 0.061920613 | 0.274393828 | 3.724945049 | 0.073663859 |
| 1700029B22Rik | 0 | NA | NA | NA |
| 5430419D17Rik | 1.767964381 | -1.171278074 | 0.75935938 | -1.542455529 |
| Cuzd1 | 2.321753028 | 0.061350746 | 0.708153985 | 0.086634753 |
| 1700007K09Rik | 3.665048522 | 1.846782883 | 0.603459274 | 3.060327286 |
| Fam24a | 0 | NA | NA | NA |
| 2310057M21Rik | 207.8650457 | 0.113953871 | 0.129320064 | 0.881177041 |
| Pstk | 158.2756739 | -0.464558322 | 0.111240104 | -4.176176637 |
| Ikzf5 | 441.795929 | 0.240635455 | 0.082654445 | 2.91134319 |
| Acadsb | 717.8726284 | -0.093813908 | 0.085450256 | -1.097877429 |
| Hmx3 | 4.229700681 | 1.090088693 | 0.683627047 | 1.594566361 |
| Hmx2 | 1.376538491 | -0.308119642 | 0.822290212 | -0.37470912 |
| Bub3 | 556.6642844 | 0.189401937 | 0.08205499 | 2.308231783 |
| Gm16477 | 27.01530151 | 0.262084061 | 0.227938901 | 1.149799618 |
| Gpr26 | 2155.295409 | -0.690620489 | 0.177812039 | -3.883991737 |
| Cpxm2 | 34.2720618 | 0.21895515 | 0.205255101 | 1.066746448 |
| n-R5s159 | 0.155551505 | 1.257968755 | 2.831330457 | 0.444303049 |
| Gm15677 | 0.494008362 | -0.1384385 | 1.564203338 | -0.088504159 |
| Chst15 | 526.7601878 | 0.016264658 | 0.089085536 | 0.182573501 |
| Oat | 670.5788141 | -0.150980324 | 0.098012285 | -1.540422458 |
| Nkx1-2 | 0.886311283 | -1.517807647 | 1.143992489 | -1.326763647 |
| Gm16764 | 0 | NA | NA | NA |
| Lhpp | 114.0965916 | -0.236978014 | 0.122656534 | -1.932045582 |
| Gm15582 | 0.853586773 | 1.424474579 | 1.183977725 | 1.203126164 |
| Fam53b | 689.2067909 | -0.35591842 | 0.083987266 | -4.237766492 |
| Mettl10 | 199.6988259 | 0.007864924 | 0.095217745 | 0.082599356 |
| Fam175b | 351.9015338 | 0.103400052 | 0.074073043 | 1.395920127 |
| 1500002F19Rik | 8.219991273 | 0.057874757 | 0.345580703 | 0.167471032 |
| Zranb1 | 1066.949998 | 0.211512865 | 0.076498822 | 2.764916611 |
| Gm15718 | 1.397387052 | 1.568261736 | 0.80717681 | 1.942897413 |
| Ctbp2 | 328.6048763 | -0.296782259 | 0.091320911 | -3.24988283 |
| Tex36 | 0.64913551 | 0.35466711 | 1.199496129 | 0.295680079 |
| Gm28747 | 0.061920613 | 0.274393828 | 3.724945049 | 0.073663859 |
| Edrf1 | 475.154792 | -0.193391406 | 0.123856205 | -1.561418784 |
| Mmp21 | 0.96439918 | 0.399763171 | 1.067959422 | 0.374324308 |
| Uros | 150.8894574 | -0.264531207 | 0.114270201 | -2.314962289 |
| Bccip | 345.0185436 | 0.191738562 | 0.082407355 | 2.326716602 |
| Dhx32 | 391.5252559 | -0.02380176 | 0.074737107 | -0.31847313 |
| Gm15483 | 5.172158321 | -1.152500707 | 0.468540658 | -2.459766699 |
| Fank1 | 22.86474296 | -0.647190386 | 0.225912754 | -2.86478021 |

| | | | | |
|---|---|---|---|---|
| Adam12 | 174.7221536 | 0.350803195 | 0.193723014 | 1.810849357 |
| Gm23631 | 0.152314784 | -0.206481641 | 2.847275348 | -0.072519028 |
| A130023I24Rik | 1.325949706 | 0.287590487 | 1.16202569 | 0.247490644 |
| D7Ertd443e | 78.15056778 | 0.125950465 | 0.189371355 | 0.665097765 |
| Dock1 | 717.8821439 | -0.15493488 | 0.080181592 | -1.932299877 |
| Fam196a | 171.3787744 | -0.470183693 | 0.12628198 | -3.723284122 |
| Gm24581 | 0 | NA | NA | NA |
| 1700120G07Rik | 0.35282422 | 0.27437713 | 3.724945049 | 0.073659377 |
| Nps | 2.222986028 | 1.173603315 | 0.727179933 | 1.613910482 |
| 4930544L04Rik | 0.117650406 | -0.206492286 | 3.724945049 | -0.055434988 |
| Foxi2 | 0.120349074 | 1.264824188 | 3.723011833 | 0.339731445 |
| Clrn3 | 0 | NA | NA | NA |
| Ptpre | 396.1280751 | -0.237462132 | 0.09634733 | -2.464646716 |
| Mir1962 | 1.121758522 | -0.642918134 | 0.978361761 | -0.657137431 |
| 5830432E09Rik | 5.577809048 | 2.931167225 | 0.743896045 | 3.940291446 |
| Mki67 | 177.6667962 | 2.636819994 | 0.256985845 | 10.26056512 |
| Gm9341 | 0 | NA | NA | NA |
| Gm24444 | 0 | NA | NA | NA |
| Mgmt | 8.081835135 | 0.27341742 | 0.353647604 | 0.773135225 |
| Ebf3 | 140.1280894 | -0.173694388 | 0.164929093 | -1.053145837 |
| 9430038I01Rik | 31.06055986 | 0.061840237 | 0.242038592 | 0.255497424 |
| Glrx3 | 666.4075028 | 0.038094795 | 0.10603842 | 0.359254645 |
| Gm25798 | 0 | NA | NA | NA |
| Tcerg1l | 114.589252 | -0.218553001 | 0.134617318 | -1.623513264 |
| Mapk1ip1 | 278.2027142 | -0.222268597 | 0.109228931 | -2.034887596 |
| Ppp2r2d | 650.2346655 | 0.25414066 | 0.116295184 | 2.185306832 |
| Bnip3 | 859.8363811 | 0.230352602 | 0.086134122 | 2.674347822 |
| Gm9358 | 0.765055553 | -1.301956969 | 1.147248995 | -1.134851261 |
| Jakmip3 | 408.0351068 | -0.420624451 | 0.139594281 | -3.013192578 |
| Dpysl4 | 522.9033322 | 0.098794626 | 0.106003087 | 0.931997631 |
| Stk32c | 865.5787329 | 0.079233906 | 0.075266036 | 1.052717934 |
| Gm27975 | 0 | NA | NA | NA |
| Lrrc27 | 79.21060963 | -0.101325213 | 0.139489737 | -0.726399053 |
| Pwwp2b | 187.2201883 | 0.470593662 | 0.102383147 | 4.596397697 |
| Inpp5a | 914.9916766 | -0.11326274 | 0.104882571 | -1.079900491 |
| Nkx6-2 | 248.995156 | 0.835830551 | 0.172223039 | 4.853186642 |
| Cfap46 | 121.7543508 | -0.515647583 | 0.165739039 | -3.111201712 |
| Gm4459 | 68.82999537 | 0.000656836 | 0.160300006 | 0.00409754 |
| Rpl21-ps13 | 0 | NA | NA | NA |
| Mir6401 | 0 | NA | NA | NA |
| Adgra1 | 707.4917962 | -0.073123429 | 0.070636523 | -1.035207088 |
| Kndc1 | 2147.099969 | 0.061440795 | 0.07113472 | 0.863724425 |
| Utf1 | 1.176348267 | 0.678548967 | 1.036700158 | 0.654527698 |
| Gm16201 | 8.156929821 | 0.559766128 | 0.39057645 | 1.433179415 |
| Gm2044 | 0.046345709 | 0.274391604 | 3.724945049 | 0.073663262 |
| mmu-mir-7686 | 0 | NA | NA | NA |
| Mir202 | 0 | NA | NA | NA |

| | | | | |
|---|---|---|---|---|
| 6430531B16Rik | 3.025363873 | 0.627736964 | 0.548028169 | 1.14544653 |
| Adam8 | 325.8439531 | 2.610406372 | 0.491436937 | 5.311783008 |
| Mir7062 | 0 | NA | NA | NA |
| Tubgcp2 | 416.699423 | 0.223540371 | 0.073215768 | 3.053172513 |
| Zfp511 | 95.06049784 | 0.240794049 | 0.122217244 | 1.970213375 |
| Msx3 | 0.762104533 | -2.240647913 | 1.198515898 | -1.869518725 |
| Caly | 767.6017605 | -0.042494372 | 0.072787434 | -0.583814675 |
| Prap1 | 0 | NA | NA | NA |
| Fuom | 141.6570441 | -0.176873674 | 0.113292256 | -1.561215926 |
| Echs1 | 365.0393526 | 0.212915429 | 0.098998349 | 2.150696761 |
| Paox | 52.9831637 | 0.516757811 | 0.156251579 | 3.307216567 |
| Mtg1 | 103.7910862 | -0.113070389 | 0.123956922 | -0.912174863 |
| Sprn | 1168.69905 | 0.06622142 | 0.14310334 | 0.462752443 |
| Olfr522 | 0 | NA | NA | NA |
| Olfr523 | 0.038981395 | 0.274393828 | 3.724945049 | 0.073663859 |
| Olfr524 | 0.042716945 | -0.206490061 | 3.724945049 | -0.055434391 |
| Cd163l1 | 0.514333851 | 1.27045668 | 3.190229018 | 0.398233692 |
| 5830411N06Rik | 0.560014729 | 0.256115433 | 1.329432025 | 0.192650266 |
| Olfr525 | 0.068097359 | -0.206487837 | 3.724945049 | -0.055433794 |
| Olfr527 | 0.1567526 | 0.274409612 | 2.828090123 | 0.09703001 |
| Olfr60 | 0.072025462 | 0.755288839 | 3.724945049 | 0.202765096 |
| Olfr530 | 0.095435502 | 0.75528439 | 3.724945049 | 0.202763901 |
| Olfr531 | 0.095898756 | 0.27438722 | 3.724945049 | 0.073662085 |
| Olfr532 | 0.042716945 | -0.206490061 | 3.724945049 | -0.055434391 |
| Olfr533 | 0.120974994 | 0.274404949 | 3.724945049 | 0.073666845 |
| Olfr535 | 0 | NA | NA | NA |
| Olfr536 | 0.638507312 | 0.238318394 | 1.397478148 | 0.170534612 |
| Olfr538 | 0 | NA | NA | NA |
| Olfr46 | 0.111376404 | 0.274404949 | 3.724945049 | 0.073666845 |
| Olfr61 | 0.224318018 | -1.73519907 | 2.570571478 | -0.675024634 |
| Olfr53 | 0.178643884 | 0.776586404 | 2.710504808 | 0.286509879 |
| Olfr539 | 0.207883488 | -0.682179401 | 2.193075849 | -0.31106056 |
| Olfr45 | 0.066050875 | 0.274391604 | 3.724945049 | 0.073663262 |
| Olfr541 | 0 | NA | NA | NA |
| Cyp2e1 | 1.071126076 | 1.223860097 | 1.158951483 | 1.056006326 |
| Syce1 | 4.365775959 | -0.392807473 | 0.644764335 | -0.609226428 |
| Zfp941 | 465.5985334 | 0.655934265 | 0.102431103 | 6.403662977 |
| Urah | 8.499830305 | 1.020811668 | 0.391370351 | 2.608300977 |
| C330022C24Rik | 1.303324972 | 2.468930392 | 1.198066702 | 2.06076205 |
| Scgb1c1 | 1.058089029 | -1.617135776 | 1.11580412 | -1.449300775 |
| Odf3 | 0.418471408 | -2.413390192 | 1.630918139 | -1.47977396 |
| Bet1l | 215.3921604 | 0.196937956 | 0.099584457 | 1.977597323 |
| Ric8 | 517.4466835 | 0.342692921 | 0.074304858 | 4.611985386 |
| Sirt3 | 296.3129455 | -0.129634972 | 0.111125553 | -1.166563123 |
| Psmd13 | 560.5188265 | 0.302449094 | 0.097873203 | 3.090213513 |
| Gm15542 | 83.64766115 | 0.335791444 | 0.174697786 | 1.922127646 |
| Cox8b | 4.291398722 | -0.862417513 | 0.479136829 | -1.799939935 |

| | | | | |
|---|---|---|---|---|
| Nlrp6 | 8.710158728 | 1.046997471 | 0.375487621 | 2.78836748 |
| Athl1 | 114.0507651 | 1.111357937 | 0.163396961 | 6.801582677 |
| Ifitm5 | 0.496272132 | 2.152233653 | 1.333303395 | 1.614211485 |
| Ifitm2 | 214.7202724 | 2.16083748 | 0.23452989 | 9.213484374 |
| Ifitm1 | 100.7816241 | 2.672423801 | 0.27507383 | 9.715296452 |
| Ifitm3 | 2374.688831 | 5.988698122 | 0.270685699 | 22.12417632 |
| Ifitm6 | 40.88379849 | 4.289360732 | 0.692115748 | 6.197461545 |
| Gm17387 | 0.276680467 | 0.75530721 | 1.805778928 | 0.418272247 |
| B4galnt4 | 713.1671909 | 0.181545226 | 0.107730578 | 1.685178242 |
| Pkp3 | 31.38673572 | 0.701972287 | 0.262272264 | 2.676502183 |
| Sigirr | 8.814058262 | 1.592600062 | 0.532811027 | 2.989052365 |
| Ano9 | 0.759896141 | 1.130396059 | 1.283914314 | 0.880429517 |
| Gm24962 | 0 | NA | NA | NA |
| Ptdss2 | 397.5875048 | -0.157586846 | 0.0815865 | -1.931530914 |
| Rnh1 | 785.7956867 | 1.347698275 | 0.193551525 | 6.962994856 |
| Gm22019 | 0.061920613 | 0.274393828 | 3.724945049 | 0.073663859 |
| Hras | 652.7768577 | -0.175614142 | 0.081157561 | -2.163866688 |
| Lrrc56 | 44.78247708 | -0.042859659 | 0.21271812 | -0.201485697 |
| Lmntd2 | 1.504148252 | 0.395664251 | 0.819128298 | 0.483030866 |
| Rassf7 | 15.32468764 | 0.152370236 | 0.291279317 | 0.523106952 |
| Mir210 | 0.042716945 | -0.206490061 | 3.724945049 | -0.055434391 |
| Phrf1 | 827.2765508 | 0.00588788 | 0.05650857 | 0.104194463 |
| Irf7 | 4351.088714 | 9.50412825 | 0.317598896 | 29.92494107 |
| Cdhr5 | 0.766219994 | 1.203889179 | 1.279748469 | 0.940723281 |
| Sct | 31.79202062 | 6.337717343 | 0.818146931 | 7.746429283 |
| Drd4 | 5.233666748 | 2.769349869 | 0.785039459 | 3.527656907 |
| Deaf1 | 397.411066 | 0.048332596 | 0.080417619 | 0.60101999 |
| Tmem80 | 167.3778283 | -0.2996992 | 0.107729456 | -2.781961509 |
| Eps8l2 | 83.71994531 | 0.272863505 | 0.147947981 | 1.844320568 |
| B230206H07Rik | 17.64932275 | -1.005892305 | 0.375481678 | -2.678938452 |
| Gm25725 | 0 | NA | NA | NA |
| Taldo1 | 760.0746265 | 0.020407309 | 0.115385495 | 0.176861994 |
| Gm4535 | 4.201928798 | -0.513988921 | 0.546304821 | -0.940846394 |
| Pddc1 | 201.5225707 | -0.080187475 | 0.108844815 | -0.736713784 |
| Cend1 | 1948.915175 | -0.063148605 | 0.105413232 | -0.599057663 |
| Slc25a22 | 2538.937697 | 0.87620079 | 0.125566214 | 6.977997999 |
| Pidd1 | 23.70766093 | 0.358083553 | 0.267381122 | 1.339225262 |
| Rplp2 | 1141.878145 | 0.41665348 | 0.093907336 | 4.436857633 |
| Snora52 | 0 | NA | NA | NA |
| Pnpla2 | 482.1622727 | 1.791354085 | 0.195219327 | 9.176110335 |
| Cracr2b | 10.14241748 | -0.191501135 | 0.329048601 | -0.581984346 |
| Gm10575 | 13.15716283 | -0.358949663 | 0.330923054 | -1.084692221 |
| Cd151 | 360.834311 | 0.883246953 | 0.197661304 | 4.468486927 |
| Polr2l | 159.7689273 | 0.317880663 | 0.1596564 | 1.991029886 |
| Tspan4 | 266.6277742 | 0.929984225 | 0.221237531 | 4.203555425 |
| Chid1 | 475.4024328 | -0.38266312 | 0.119427135 | -3.204155566 |
| Ap2a2 | 2741.755715 | -0.05165619 | 0.057319519 | -0.901197207 |

| | | | | |
|---|---|---|---|---|
| Gm16982 | 0.330744509 | 1.245163583 | 1.770900871 | 0.703124384 |
| Mir7063 | 2.38820333 | 0.48165782 | 0.607634354 | 0.792677071 |
| Muc6 | 0.236992836 | 0.755303948 | 2.091273358 | 0.361169402 |
| Muc2 | 8.969982396 | 0.543842189 | 0.56634401 | 0.960268282 |
| Muc5ac | 0.957274486 | 0.706470537 | 1.408997067 | 0.50139958 |
| Muc5b | 2.416483532 | 0.405706845 | 1.024350157 | 0.396062657 |
| Tollip | 1722.402167 | 0.104370412 | 0.063309717 | 1.648568615 |
| Gm20501 | 7.216052579 | -0.486604069 | 0.383358967 | -1.269317039 |
| Brsk2 | 1833.13049 | -0.149498992 | 0.085975203 | -1.738861755 |
| Mir3104 | 0.562632007 | -2.149843823 | 1.203344014 | -1.786557956 |
| Mob2 | 221.1798062 | 0.051978778 | 0.080642354 | 0.64455929 |
| Gm25416 | 0.290864751 | -1.195151484 | 2.045403122 | -0.584310971 |
| Dusp8 | 1691.653108 | -0.311826738 | 0.079088248 | -3.94276958 |
| Gm25370 | 0.286235171 | -1.138164625 | 1.622826626 | -0.701347024 |
| Gm4553 | 0 | NA | NA | NA |
| Krtap5-2 | 0 | NA | NA | NA |
| Gm10153 | 0 | NA | NA | NA |
| Krtap5-3 | 0.115000406 | -0.206492286 | 3.724945049 | -0.055434988 |
| Gm7579 | 0 | NA | NA | NA |
| Krtap5-5 | 0 | NA | NA | NA |
| Gm2431 | 0.097139547 | 0.274393828 | 3.724945049 | 0.073663859 |
| Gm4559 | 0 | NA | NA | NA |
| Gm10013 | 0 | NA | NA | NA |
| Krtap5-1 | 0.044544648 | 0.755286614 | 3.724945049 | 0.202764498 |
| Krtap5-4 | 0 | NA | NA | NA |
| Gm5054 | 0.042716945 | -0.206490061 | 3.724945049 | -0.055434391 |
| Ifitm10 | 182.7874866 | 0.132711009 | 0.118587041 | 1.119102119 |
| Ctsd | 5752.611281 | 0.672606516 | 0.170363633 | 3.948063933 |
| Gm26143 | 0.044544648 | 0.755286614 | 3.724945049 | 0.202764498 |
| Syt8 | 0.543884849 | -0.184224814 | 1.317042325 | -0.139877672 |
| Tnni2 | 2.708583151 | 2.128541207 | 0.892818046 | 2.384070546 |
| Lsp1 | 166.316609 | 2.569950315 | 0.193474865 | 13.28312241 |
| Prr33 | 0.642141737 | -1.605707864 | 1.17731244 | -1.363875731 |
| Tnnt3 | 1.402656864 | 0.67432223 | 0.915219819 | 0.73678718 |
| Mrpl23 | 104.8815584 | -0.138750394 | 0.122461961 | -1.133008106 |
| Nctc1 | 0 | NA | NA | NA |
| H19 | 64.97075824 | 2.871152474 | 0.354200412 | 8.106011101 |
| Gm27483 | 0.184422046 | 0.75529805 | 2.68070113 | 0.281753919 |
| Mir675 | 0 | NA | NA | NA |
| Gm27786 | 0.110595523 | 0.755282166 | 3.724945049 | 0.202763304 |
| Igf2 | 848.6181956 | -0.060319466 | 0.17986871 | -0.335352749 |
| Mir483 | 0 | NA | NA | NA |
| Igf2os | 4.725164502 | 1.338561847 | 0.493674565 | 2.711425587 |
| Ins2 | 0.386429078 | 0.731613451 | 1.444513245 | 0.506477496 |
| Gm7290 | 0.08486385 | 0.755286617 | 3.724945049 | 0.202764499 |
| Gm6471 | 0 | NA | NA | NA |
| Th | 127.7817552 | -0.472930582 | 0.262212732 | -1.803614104 |

| | | | | |
|---|---|---|---|---|
| Ascl2 | 2.369287799 | 0.989526042 | 0.925135199 | 1.069601549 |
| Tspan32os | 6.17140379 | -0.898443642 | 0.446931716 | -2.010248122 |
| Tspan32 | 1.376815522 | 1.428757495 | 1.062718205 | 1.344436831 |
| R74862 | 40.83695903 | 0.014532344 | 0.189286048 | 0.076774511 |
| Cd81 | 3734.509123 | 0.334264481 | 0.106430314 | 3.140688676 |
| Trpm5 | 0.502574849 | -0.210157341 | 3.430093716 | -0.061268688 |
| Tssc4 | 336.1484664 | 0.44724969 | 0.086740044 | 5.156207773 |
| Kcnq1 | 3.259092159 | 0.254323874 | 0.597142375 | 0.425901569 |
| Gm38321 | 0 | NA | NA | NA |
| Gm37235 | 0.046345709 | 0.274391604 | 3.724945049 | 0.073663262 |
| Gm28821 | 0 | NA | NA | NA |
| Kcnq1ot1 | 4980.818381 | -0.572760611 | 0.250609552 | -2.285469993 |
| Gm27395 | 0.113999573 | -0.206492286 | 3.724945049 | -0.055434988 |
| Gm27389 | 0.179789776 | -1.47888837 | 2.234383842 | -0.661877491 |
| Gm27803 | 0 | NA | NA | NA |
| Gm27539 | 0.072025462 | 0.755288839 | 3.724945049 | 0.202765096 |
| Gm37364 | 1.783218599 | 1.51652281 | 0.885495571 | 1.712626081 |
| 4933417O13Rik | 0.382744327 | -2.553475356 | 1.361539479 | -1.875432476 |
| Gm38095 | 0.038981395 | 0.274393828 | 3.724945049 | 0.073663859 |
| Gm38065 | 0 | NA | NA | NA |
| Gm27901 | 0.136963818 | 0.755286583 | 3.59954433 | 0.209828388 |
| Cdkn1c | 70.29508147 | 0.257617327 | 0.142069397 | 1.813320343 |
| Slc22a18 | 7.798117052 | 1.095249706 | 0.507701123 | 2.15727257 |
| Phlda2 | 0.124308499 | 0.274388155 | 3.719824689 | 0.073763733 |
| Nap1l4 | 877.3075028 | -0.037301649 | 0.083315298 | -0.447716685 |
| Gm23297 | 5.403433764 | -0.830716347 | 0.515228141 | -1.612327204 |
| Cars | 570.4979151 | 0.41351567 | 0.114668713 | 3.606176942 |
| Mir7064 | 0.434125078 | -1.878862743 | 1.254300128 | -1.497937137 |
| Gm15580 | 0 | NA | NA | NA |
| Gm15579 | 0.072025462 | 0.755288839 | 3.724945049 | 0.202765096 |
| Tnfrsf26 | 16.36698174 | 3.609347922 | 0.781537675 | 4.618264785 |
| Tnfrsf22 | 17.56781374 | 0.186968033 | 0.280634561 | 0.666233097 |
| Tnfrsf23 | 21.91112602 | 0.122754748 | 0.324739909 | 0.378009429 |
| Osbpl5 | 343.6236498 | -0.049874676 | 0.074859552 | -0.666243309 |
| Mrgprg | 0.045882455 | 0.755288839 | 3.724945049 | 0.202765096 |
| Mrgpre | 90.26427259 | 0.134039833 | 0.139195992 | 0.962957564 |
| Gm22064 | 0.123841226 | 0.274391173 | 3.724945049 | 0.073663147 |
| Nadsyn1 | 56.75566904 | -0.512681307 | 0.193557244 | -2.648732209 |
| Dhcr7 | 321.2309411 | -0.383249897 | 0.105230879 | -3.641990854 |
| Gm498 | 0.049553047 | 0.274391604 | 3.724945049 | 0.073663262 |
| Mir6404 | 0 | NA | NA | NA |
| Shank2 | 2746.162572 | -0.117035764 | 0.116950858 | -1.000725994 |
| Mir3105 | 0.102891481 | 0.274400501 | 3.724945049 | 0.073665651 |
| Gm14372 | 0.097899064 | -0.687371727 | 3.724945049 | -0.184532045 |
| Cttn | 1215.2151 | 0.120478266 | 0.075815607 | 1.58909585 |
| Gm14376 | 6.097514919 | -0.287661673 | 0.582898478 | -0.493502186 |
| Ppfia1 | 744.1652023 | -0.072414748 | 0.082075508 | -0.882294247 |

| | | | | |
|---|---|---|---|---|
| Faddos | 0.049553047 | 0.274391604 | 3.724945049 | 0.073663262 |
| Fadd | 59.54792491 | 0.406131686 | 0.188178983 | 2.158220215 |
| Ano1 | 32.22433977 | -0.653896922 | 0.229443591 | -2.849924548 |
| Gm10152 | 0 | NA | NA | NA |
| Fgf3 | 4.236074083 | -0.991649998 | 0.572375877 | -1.732515359 |
| Fgf4 | 0 | NA | NA | NA |
| Gm26793 | 3.941161089 | 0.526983391 | 0.483525879 | 1.08987629 |
| Fgf15 | 0.844302062 | 1.667541888 | 1.215912028 | 1.371433007 |
| Oraov1 | 196.1845682 | -0.051723089 | 0.080804179 | -0.640104127 |
| Ccnd1 | 896.4298962 | 0.923802018 | 0.24419983 | 3.782975679 |
| Tpcn2 | 87.43800306 | 0.007048396 | 0.202954317 | 0.034728978 |
| Mrgprf | 52.34489822 | -0.437109547 | 0.257022234 | -1.700668227 |
| Mrgprd | 0.203212381 | 0.274405474 | 2.643427671 | 0.103806689 |
| Gm7346 | 0 | NA | NA | NA |
| Gm25775 | 0 | NA | NA | NA |
| Insr | 1190.786105 | -0.064682552 | 0.056237366 | -1.150170365 |
| Gm16180 | 0 | NA | NA | NA |
| A430078G23Rik | 4.533747209 | 3.171889909 | 0.795220271 | 3.988693478 |
| Arhgef18 | 750.8432807 | -0.181728992 | 0.072755125 | -2.497817052 |
| Pex11g | 11.94680139 | 0.23612033 | 0.394091224 | 0.599151454 |
| 1700019B03Rik | 3.816282625 | 1.70006711 | 0.613552132 | 2.770860084 |
| Zfp358 | 245.1757097 | 0.565635338 | 0.110872162 | 5.101689433 |
| Mcoln1 | 510.2888465 | -0.127570356 | 0.058251699 | -2.189985142 |
| Pnpla6 | 618.1933405 | -0.183814122 | 0.090845283 | -2.023375526 |
| C330021F23Rik | 0.687005404 | -1.507326087 | 1.139827428 | -1.322416052 |
| Rps23-ps2 | 34.30525878 | 0.578336776 | 0.19495735 | 2.966478444 |
| Camsap3 | 321.1724317 | -0.292761023 | 0.09592162 | -3.052085909 |
| Xab2 | 460.0137195 | 0.197217577 | 0.070962846 | 2.779166659 |
| Gm22204 | 0.066050875 | 0.274391604 | 3.724945049 | 0.073663262 |
| Pet100 | 155.4280518 | 0.0879723 | 0.113858297 | 0.772647248 |
| Pcp2 | 337.4770539 | 0.015085045 | 0.413262252 | 0.036502355 |
| Stxbp2 | 289.3398245 | 0.106977255 | 0.141501607 | 0.756014421 |
| Retn | 3.230460487 | -0.663755795 | 0.772811471 | -0.8588845 |
| Mcemp1 | 19.01625202 | 3.491994993 | 0.634189931 | 5.506229007 |
| Fcor | 19.48329625 | -0.393035257 | 0.310414012 | -1.266164677 |
| Trappc5 | 363.4451168 | 0.353409751 | 0.087587382 | 4.034939079 |
| Fcer2a | 1.327801032 | 0.12000938 | 0.853995697 | 0.140526914 |
| Clec4g | 0.624219342 | -0.213830405 | 1.40517147 | -0.152173887 |
| Gm16553 | 0.238000031 | 1.220560956 | 2.464171736 | 0.495323008 |
| Gm23886 | 0 | NA | NA | NA |
| Cd209a | 1.799118211 | -0.435482375 | 0.928205887 | -0.469165711 |
| Gm24934 | 0 | NA | NA | NA |
| Rprl3 | 2178.394221 | 0.181424477 | 0.183413727 | 0.989154304 |
| Gm7381 | 0.061920613 | 0.274393828 | 3.724945049 | 0.073663859 |
| Cd209e | 0 | NA | NA | NA |
| Cd209d | 0.062426873 | 0.755288839 | 3.724945049 | 0.202765096 |
| Cd209b | 0.371584567 | -2.49487354 | 1.596794342 | -1.562426341 |

| | | | | |
|---|---|---|---|---|
| Cd209c | 0.190986542 | -0.687380379 | 2.703604247 | -0.254245931 |
| Gm7389 | 8.537660985 | 0.211517483 | 0.41972795 | 0.503939475 |
| Gm16589 | 85.35553407 | 1.774519372 | 0.2442046 | 7.266527206 |
| Gm26750 | 0 | NA | NA | NA |
| Cd209f | 0.631752858 | -1.710778682 | 1.174299698 | -1.456850143 |
| Gm23888 | 0 | NA | NA | NA |
| Cd209g | 0.34920187 | -1.72212294 | 1.676297403 | -1.027337355 |
| 4932443L11Rik | 1.299081245 | 1.142206032 | 0.874135712 | 1.306668994 |
| Evi5l | 1072.773425 | -0.29646441 | 0.07475275 | -3.965933192 |
| Prr36 | 298.1056952 | -0.49828438 | 0.119870856 | -4.156843437 |
| Lrrc8e | 1.436219828 | 0.263376914 | 1.162053383 | 0.226647861 |
| Map2k7 | 1090.223813 | -0.002860964 | 0.058640447 | -0.04878823 |
| Tgfbr3l | 14.69813157 | -0.438165024 | 0.394668199 | -1.110211121 |
| Snapc2 | 202.0178337 | 0.324582908 | 0.119912822 | 2.706824032 |
| Ctxn1 | 1372.549575 | -0.331738222 | 0.123177552 | -2.693171086 |
| Timm44 | 376.9137615 | -0.108226945 | 0.103127167 | -1.049451354 |
| Gm26102 | 0 | NA | NA | NA |
| Elavl1 | 1614.161581 | -0.082422817 | 0.053322387 | -1.545745054 |
| Ccl25 | 51.47610217 | -0.268164263 | 0.191334185 | -1.401549147 |
| Cers4 | 639.9032004 | -0.119756909 | 0.123702607 | -0.968103356 |
| Gm36956 | 0.066050875 | 0.274391604 | 3.724945049 | 0.073663262 |
| Zfp958 | 156.442319 | -0.045221886 | 0.114604994 | -0.394589137 |
| Gm7461 | 2.204253537 | -0.470855101 | 0.726000397 | -0.64856039 |
| Gm6410 | 0.983694817 | 0.778496809 | 0.963374436 | 0.808093696 |
| Shcbp1 | 11.59037295 | 0.911703779 | 0.458535318 | 1.988295649 |
| Gm9457 | 0.283539495 | 1.260097264 | 2.396768941 | 0.525748328 |
| Rpl21-ps14 | 0.417309641 | -0.014136185 | 1.731202608 | -0.008165529 |
| n-R5s93 | 0 | NA | NA | NA |
| Slc10a2 | 0 | NA | NA | NA |
| n-R5s94 | 0 | NA | NA | NA |
| Gm1840 | 36.14079938 | 0.111468088 | 0.176661099 | 0.630971325 |
| Gm22565 | 0 | NA | NA | NA |
| Gm24288 | 0 | NA | NA | NA |
| Gm17215 | 0.061920613 | 0.274393828 | 3.724945049 | 0.073663859 |
| Gm26427 | 0.673557475 | 0.657297963 | 1.157875103 | 0.567676049 |
| Gm27585 | 0 | NA | NA | NA |
| Gm27545 | 0 | NA | NA | NA |
| Gm25169 | 0 | NA | NA | NA |
| Gm28262 | 0 | NA | NA | NA |
| Efnb2 | 602.2040592 | -0.191041248 | 0.07237607 | -2.639563692 |
| 4921522P10Rik | 0.133946075 | 0.755288219 | 3.626642455 | 0.20826101 |
| Arglu1 | 500.6182152 | -0.071268265 | 0.069654749 | -1.023164482 |
| Mir7654 | 0 | NA | NA | NA |
| Fam155a | 942.9312957 | -0.052474024 | 0.067006887 | -0.783113886 |
| n-R5s95 | 14.75670795 | -0.504136568 | 0.524512609 | -0.961152428 |
| AC099760.1 | 0 | NA | NA | NA |
| Gm10217 | 0.045882455 | 0.755288839 | 3.724945049 | 0.202765096 |

| | | | | |
|---|---|---|---|---|
| Gm10067 | 0 | NA | NA | NA |
| Lig4 | 157.1064013 | -0.616288675 | 0.137735628 | -4.474431808 |
| Abhd13 | 722.6949606 | 0.248521975 | 0.084080457 | 2.955763825 |
| Tnfsf13b | 20.67774298 | 1.098208376 | 0.254512432 | 4.31494983 |
| Myo16 | 882.108668 | -0.107743934 | 0.112658798 | -0.956373904 |
| Rps16-ps3 | 0 | NA | NA | NA |
| 3930402G23Rik | 10.57984639 | 2.553023819 | 0.561552452 | 4.546367503 |
| Irs2 | 1258.437026 | 0.388182391 | 0.130513729 | 2.974264813 |
| 9530052E02Rik | 11.90754286 | -0.095508556 | 0.338236846 | -0.282371825 |
| Gm15418 | 2.963241567 | -0.288461995 | 0.62079989 | -0.4646618 |
| Col4a1 | 1092.321152 | 1.53904807 | 0.16411426 | 9.377905781 |
| Col4a2 | 772.5542793 | 0.818031785 | 0.15986485 | 5.117020951 |
| Gm15419 | 0.932741312 | -0.52898777 | 1.04909942 | -0.504230352 |
| Rab20 | 71.74611719 | 4.564525548 | 0.446798144 | 10.21607991 |
| E230013L22Rik | 6.9128239 | 1.516421583 | 0.51781818 | 2.928482702 |
| Carkd | 279.2439548 | -0.046469849 | 0.072346984 | -0.642319095 |
| Cars2 | 87.5486135 | -0.268982555 | 0.12909414 | -2.083615536 |
| Ing1 | 228.1766961 | -0.205495172 | 0.101762941 | -2.019351748 |
| Ankrd10 | 320.9466642 | -0.379885421 | 0.11441267 | -3.320309019 |
| mmu-mir-7238 | 0 | NA | NA | NA |
| 1700016D06Rik | 0.194626871 | 1.808438157 | 2.64426296 | 0.683910104 |
| Gm27854 | 0 | NA | NA | NA |
| 1700128E19Rik | 0.068097359 | -0.206487837 | 3.724945049 | -0.055433794 |
| Arhgef7 | 1845.245703 | -0.325217791 | 0.067385002 | -4.826263734 |
| Gm15875 | 0 | NA | NA | NA |
| Tex29 | 5.082148535 | -2.316239861 | 0.593718533 | -3.901242308 |
| A230072I06Rik | 0.719797104 | -3.13800976 | 1.282825679 | -2.446170054 |
| Gm29175 | 0.569306118 | 0.014070491 | 1.487479605 | 0.009459283 |
| Gm22207 | 0.197462999 | -1.510588839 | 2.627104102 | -0.575001515 |
| Gm5607 | 118.2838747 | -0.460038257 | 0.131869572 | -3.488585351 |
| Sox1 | 235.5645892 | -0.431128804 | 0.156328791 | -2.757833669 |
| Gm25239 | 0 | NA | NA | NA |
| D630011A20Rik | 0 | NA | NA | NA |
| Spaca7 | 0 | NA | NA | NA |
| Tubgcp3 | 535.0497003 | 0.005442727 | 0.082878694 | 0.065671001 |
| Gm26483 | 0 | NA | NA | NA |
| 4931415C17Rik | 18.23396489 | -0.167210916 | 0.283164421 | -0.590508212 |
| Gm15348 | 0.194797293 | -1.231370682 | 3.207102014 | -0.383951205 |
| Atp11a | 1406.61443 | 0.048430599 | 0.071521009 | 0.67715206 |
| Gm15350 | 0.251121349 | -0.206484597 | 2.080245602 | -0.099259721 |
| Gm15347 | 0.159763836 | -1.177185737 | 2.803296028 | -0.419929157 |
| Mcf2l | 2158.382309 | -0.30323965 | 0.071466882 | -4.243079323 |
| Gm15353 | 0.04393175 | -0.206490061 | 3.724945049 | -0.055434391 |
| Gm15351 | 6.732411598 | -1.142647168 | 0.457983485 | -2.494952779 |
| mmu-mir-9768 | 0 | NA | NA | NA |
| Gm15349 | 0 | NA | NA | NA |
| Gm15352 | 1.728865685 | -0.572508881 | 1.000975489 | -0.571950949 |

| | | | | |
|---|---|---|---|---|
| F7 | 20.47438508 | 2.905428981 | 0.609159151 | 4.769572906 |
| F10 | 60.6296221 | 6.182719773 | 0.902047576 | 6.854094993 |
| Proz | 0.405129103 | -0.20648241 | 1.605794875 | -0.128585795 |
| Gm17023 | 0.048949532 | -0.206487837 | 3.724945049 | -0.055433794 |
| Pcid2 | 338.1239794 | 0.030186453 | 0.093184499 | 0.323942861 |
| Gm17022 | 0 | NA | NA | NA |
| Cul4a | 779.7913125 | -0.055987472 | 0.053927973 | -1.038189806 |
| Mir7065 | 0.26402779 | -0.967112707 | 3.699374841 | -0.261425984 |
| Lamp1 | 3957.92154 | 0.359037752 | 0.124547752 | 2.882731682 |
| Grtp1 | 32.55675301 | 0.012019002 | 0.222467393 | 0.054025906 |
| Mir1968 | 0.779555773 | -1.570957557 | 1.15849046 | -1.356038407 |
| 2810030D12Rik | 0.867078918 | -1.648463989 | 1.10628542 | -1.490089229 |
| Adprhl1 | 5.011785424 | 0.294687132 | 0.48978803 | 0.601662585 |
| Dcun1d2 | 379.2848758 | -0.419760046 | 0.096177565 | -4.364427896 |
| Gm25014 | 4.915973584 | -1.092473585 | 0.46810666 | -2.333813379 |
| Tmco3 | 370.903991 | -0.393883909 | 0.091954608 | -4.283460279 |
| Tfdp1 | 412.9010695 | 0.138268217 | 0.084997676 | 1.626729371 |
| Atp4b | 0 | NA | NA | NA |
| Grk1 | 1.091106207 | 3.42078141 | 1.182116879 | 2.893775963 |
| Tmem255b | 12.33555406 | -0.283729585 | 0.343472019 | -0.82606317 |
| Gas6 | 898.867002 | -0.171153296 | 0.078007027 | -2.194075374 |
| 1700029H14Rik | 5.973066547 | 0.824839487 | 0.582652255 | 1.415663426 |
| Rasa3 | 1272.496422 | -0.004205201 | 0.075650243 | -0.055587413 |
| 4932443I19Rik | 8.590972356 | -1.123981912 | 0.417307922 | -2.693411392 |
| Cdc16 | 437.6639685 | -0.023529154 | 0.083098943 | -0.283146251 |
| Upf3a | 175.6020734 | 0.229566049 | 0.16752786 | 1.370315657 |
| AF366264 | 0.110595523 | 0.755282166 | 3.724945049 | 0.202763304 |
| Champ1 | 447.8929496 | 0.092535626 | 0.068738298 | 1.346201894 |
| AC139186.1 | 0 | NA | NA | NA |
| Gm24698 | 0.093141081 | -0.687373951 | 3.724945049 | -0.184532642 |
| Coprs | 191.5374814 | -0.278344268 | 0.122470092 | -2.272752998 |
| Gm7676 | 0.12936564 | -1.026948348 | 3.677739829 | -0.27923355 |
| Fbxo25 | 406.2625769 | -0.215546906 | 0.070867857 | -3.041532739 |
| Tdrp | 156.0986126 | -0.246032853 | 0.11269021 | -2.183267317 |
| Erich1 | 48.89166254 | -0.006456323 | 0.165264207 | -0.039066675 |
| Gm10699 | 61.82645713 | -0.136664747 | 0.169651692 | -0.805560764 |
| Dlgap2 | 896.9164479 | -0.258849086 | 0.18292489 | -1.415056673 |
| Gm3160 | 8.920971897 | -1.185178889 | 0.526472707 | -2.251168718 |
| Gm26184 | 0 | NA | NA | NA |
| AC130213.1 | 3.509323303 | -1.336080276 | 0.51790421 | -2.579782614 |
| C030037F17Rik | 0 | NA | NA | NA |
| Cln8 | 468.4529787 | 0.054538343 | 0.098841995 | 0.551772994 |
| Arhgef10 | 845.0217701 | 0.257786185 | 0.111491541 | 2.31215913 |
| Gm16350 | 0.676711058 | 0.718990108 | 1.4687504 | 0.489525047 |
| Kbtbd11 | 1707.888606 | 0.091161849 | 0.093148484 | 0.978672379 |
| 5830468F06Rik | 0.645565975 | 1.242285105 | 1.48643535 | 0.835747821 |
| Gm37844 | 22.02823964 | -0.377419292 | 0.230551714 | -1.637026614 |

| | | | | |
|---|---|---|---|---|
| BB014433 | 0 | NA | NA | NA |
| Myom2 | 7.622785529 | 0.60521448 | 0.421774881 | 1.434923007 |
| Gm15678 | 0 | NA | NA | NA |
| Gm24231 | 0 | NA | NA | NA |
| Gm24222 | 0 | NA | NA | NA |
| Gm25871 | 1.401691129 | 0.299998514 | 0.8233955 | 0.364343155 |
| Gm25255 | 0 | NA | NA | NA |
| Csmd1 | 2110.968194 | -0.205019049 | 0.239415266 | -0.856332398 |
| Mir3106 | 1.89102523 | -0.057435179 | 0.787431667 | -0.07293989 |
| Gm16346 | 0 | NA | NA | NA |
| Gm16347 | 0 | NA | NA | NA |
| Gm25278 | 0 | NA | NA | NA |
| Gm25665 | 0 | NA | NA | NA |
| Gm24469 | 0 | NA | NA | NA |
| Mcph1 | 236.5966961 | -0.230973286 | 0.099988148 | -2.310006629 |
| Gm25763 | 0 | NA | NA | NA |
| Gm16725 | 0 | NA | NA | NA |
| Gm17149 | 0.323327586 | 0.895281678 | 1.997767721 | 0.448141027 |
| Angpt2 | 48.84955761 | 1.524048335 | 0.316377395 | 4.817184659 |
| Agpat5 | 782.015753 | -0.285342572 | 0.068608408 | -4.159002992 |
| Xkr5 | 3.737801717 | 0.145616945 | 0.864459259 | 0.168448592 |
| Gm22551 | 0.097139547 | 0.274393828 | 3.724945049 | 0.073663859 |
| Defb40 | 0 | NA | NA | NA |
| Defb37 | 0 | NA | NA | NA |
| Defb38 | 0 | NA | NA | NA |
| Gm15366 | 0.11193333 | 0.75528439 | 3.724945049 | 0.202763901 |
| Defb39 | 0 | NA | NA | NA |
| Defb12 | 0 | NA | NA | NA |
| Defb34 | 0 | NA | NA | NA |
| Spag11b | 0.25135723 | -0.675044178 | 2.124018352 | -0.317814664 |
| Spag11a | 0.093744596 | -0.20649451 | 3.724945049 | -0.055435586 |
| Defb14 | 0 | NA | NA | NA |
| Defb4 | 0 | NA | NA | NA |
| Defa-ps12 | 0 | NA | NA | NA |
| Defb6 | 0.199888145 | 1.066503726 | 2.620047295 | 0.407055143 |
| Defb46 | 0 | NA | NA | NA |
| Defb5 | 0 | NA | NA | NA |
| Gm15357 | 0 | NA | NA | NA |
| Defb3 | 0 | NA | NA | NA |
| Defa-ps13 | 0 | NA | NA | NA |
| Defa-ps14 | 0 | NA | NA | NA |
| Defb8 | 0 | NA | NA | NA |
| Defa-ps15 | 0 | NA | NA | NA |
| Rpl19-ps11 | 69.27011738 | 0.450363286 | 0.218019334 | 2.065703431 |
| Gm27288 | 0 | NA | NA | NA |
| Defb7 | 0 | NA | NA | NA |
| Gm15358 | 0 | NA | NA | NA |

| | | | | |
|---|---|---|---|---|
| Gm27852 | 0 | NA | NA | NA |
| Gm16425 | 10.9074589 | 0.080982107 | 0.342478137 | 0.236459202 |
| Gm15359 | 1.904847302 | 0.979890682 | 0.759609945 | 1.289991908 |
| Gm6483 | 149.5723312 | 0.099057842 | 0.124075239 | 0.798369137 |
| 4930467E23Rik | 2.550781979 | 0.980998748 | 0.649989289 | 1.509253714 |
| Gm26853 | 52.27869009 | 0.031183307 | 0.227693786 | 0.136952824 |
| Gm21769 | 152.8336227 | -0.12649305 | 0.16680557 | -0.758326297 |
| Gm21811 | 150.012751 | 0.058593073 | 0.215396455 | 0.272024314 |
| 6820431F20Rik | 191.9164182 | 0.324351602 | 0.117689742 | 2.755988723 |
| Gm26804 | 0 | NA | NA | NA |
| Gm15319 | 17.21563751 | 0.745226338 | 0.363108735 | 2.052350347 |
| 2610005L07Rik | 198.0277139 | -0.105463359 | 0.213060414 | -0.494992742 |
| Gm21092 | 212.4801284 | 0.125000839 | 0.098432188 | 1.269918313 |
| Gm21182 | 0.374378724 | -1.548217744 | 1.71944972 | -0.900414665 |
| Gm21119 | 1.839005393 | -0.476053012 | 0.831671049 | -0.572405415 |
| Gm20946 | 1.715564851 | -0.189205962 | 0.778928501 | -0.24290543 |
| Gm20797 | 0.04393175 | -0.206490061 | 3.724945049 | -0.055434391 |
| Gm20945 | 0.044191549 | -0.206490061 | 3.724945049 | -0.055434391 |
| Gm20796 | 3.51668492 | 0.613889146 | 0.723990014 | 0.847924881 |
| Defb33 | 0.682587184 | 1.584953546 | 1.532217617 | 1.034418041 |
| Gm15056 | 287.1653067 | 10.68096449 | 1.090896305 | 9.790998864 |
| Gm6040 | 1.098746849 | 2.669055172 | 1.159589902 | 2.301723366 |
| AY761185 | 1.275844484 | 3.16913323 | 1.112753037 | 2.848011305 |
| Defa21 | 0 | NA | NA | NA |
| Gm23142 | 0 | NA | NA | NA |
| Defa23 | 0 | NA | NA | NA |
| Gm10104 | 0 | NA | NA | NA |
| Defa-ps3 | 0 | NA | NA | NA |
| Defa25 | 0 | NA | NA | NA |
| Gm6689 | 0 | NA | NA | NA |
| Gm14851 | 0 | NA | NA | NA |
| Gm24568 | 0 | NA | NA | NA |
| Gm2869 | 0.109727584 | 0.755282166 | 3.724945049 | 0.202763304 |
| Defa-ps4 | 0 | NA | NA | NA |
| Gm15284 | 0.075043205 | 0.755286614 | 3.724945049 | 0.202764498 |
| Defa22 | 0 | NA | NA | NA |
| Gm26240 | 0 | NA | NA | NA |
| Defa-rs7 | 0 | NA | NA | NA |
| Gm15293 | 0 | NA | NA | NA |
| Defa-ps5 | 0 | NA | NA | NA |
| Defa-ps6 | 0 | NA | NA | NA |
| Defa-ps7 | 0 | NA | NA | NA |
| Gm15303 | 0 | NA | NA | NA |
| Gm15292 | 0 | NA | NA | NA |
| Defa-ps8 | 0 | NA | NA | NA |
| Defa3 | 0 | NA | NA | NA |
| Defa5 | 0 | NA | NA | NA |

| | | | | |
|---|---|---|---|---|
| Gm15300 | 0 | NA | NA | NA |
| Gm15299 | 0 | NA | NA | NA |
| Defa-rs1 | 0 | NA | NA | NA |
| Gm23561 | 0 | NA | NA | NA |
| Gm15302 | 0 | NA | NA | NA |
| Defa-ps16 | 0 | NA | NA | NA |
| Defa2 | 0 | NA | NA | NA |
| Gm21002 | 0 | NA | NA | NA |
| Gm21498 | 0 | NA | NA | NA |
| Gm15305 | 0 | NA | NA | NA |
| Defa-ps18 | 0 | NA | NA | NA |
| Gm15309 | 0 | NA | NA | NA |
| Gm7849 | 0 | NA | NA | NA |
| Gm23568 | 0 | NA | NA | NA |
| Gm15304 | 0 | NA | NA | NA |
| Defa-ps17 | 0 | NA | NA | NA |
| Defa20 | 0 | NA | NA | NA |
| Gm15308 | 0 | NA | NA | NA |
| Gm24423 | 0 | NA | NA | NA |
| Gm14850 | 0 | NA | NA | NA |
| Gm15311 | 0 | NA | NA | NA |
| Gm6696 | 0 | NA | NA | NA |
| Gm15312 | 0 | NA | NA | NA |
| Gm7861 | 0 | NA | NA | NA |
| Gm24829 | 0 | NA | NA | NA |
| Gm15314 | 0 | NA | NA | NA |
| Defa26 | 0 | NA | NA | NA |
| Gm15313 | 0 | NA | NA | NA |
| Defa17 | 0 | NA | NA | NA |
| Gm15315 | 0 | NA | NA | NA |
| Defa-ps9 | 0 | NA | NA | NA |
| Defa-ps1 | 0 | NA | NA | NA |
| AY761184 | 0.101408268 | 0.755286617 | 3.724945049 | 0.202764499 |
| Gm22447 | 0 | NA | NA | NA |
| Gm7869 | 0.109869311 | -0.206490066 | 3.724945049 | -0.055434393 |
| Defa-ps10 | 0 | NA | NA | NA |
| Defa24 | 0.046345709 | 0.274391604 | 3.724945049 | 0.073663262 |
| Defa-ps11 | 0 | NA | NA | NA |
| Defb1 | 2.378549416 | 0.340724211 | 0.699302696 | 0.48723423 |
| Gm15316 | 0.049553047 | 0.274391604 | 3.724945049 | 0.073663262 |
| Defb50 | 0 | NA | NA | NA |
| Defb2 | 0.062426873 | 0.755288839 | 3.724945049 | 0.202765096 |
| Defb10 | 0.392953962 | -1.064591663 | 1.528670434 | -0.696416729 |
| Defb9 | 0.27226464 | -0.570104691 | 2.040937322 | -0.279334738 |
| Defb11 | 0.092881282 | -0.687373951 | 3.724945049 | -0.184532642 |
| Defb15 | 0 | NA | NA | NA |
| Defb35 | 0 | NA | NA | NA |

| | | | | |
|---|---|---|---|---|
| Defb13 | 0 | NA | NA | NA |
| C86695 | 0 | NA | NA | NA |
| Ccdc70 | 0 | NA | NA | NA |
| Atp7b | 61.90632721 | -0.078724014 | 0.220936247 | -0.356320046 |
| Alg11 | 609.1938378 | 0.179189795 | 0.07691505 | 2.329710452 |
| Nek5 | 10.77738955 | 0.349761609 | 0.34831968 | 1.004139671 |
| Nek3 | 31.0393675 | -0.372321795 | 0.268501268 | -1.386666805 |
| Ckap2 | 17.47088672 | 1.177316412 | 0.528761624 | 2.226554195 |
| Vps36 | 310.0223612 | 0.175423273 | 0.088828153 | 1.974861201 |
| Thsd1 | 30.08606779 | 0.18970527 | 0.266350206 | 0.712239998 |
| Rpl30-ps2 | 0 | NA | NA | NA |
| Tpte | 0.493569458 | -0.312699806 | 1.333193335 | -0.234549482 |
| Slc25a15 | 143.2256164 | -0.365793764 | 0.118865191 | -3.07738338 |
| 1810012K16Rik | 2.758229591 | 0.132825494 | 0.593457536 | 0.22381634 |
| Mrps31 | 170.4153543 | 0.119181373 | 0.089922208 | 1.325383075 |
| Smim19 | 208.6627446 | 0.128547314 | 0.103725234 | 1.23930609 |
| Slc20a2 | 881.8761026 | 0.469796975 | 0.109391825 | 4.294625975 |
| Gm17491 | 71.03296188 | -0.356189848 | 0.25465398 | -1.398720914 |
| Vdac3 | 786.9911644 | 0.107245688 | 0.087136426 | 1.230779049 |
| Dkk4 | 0.121388914 | 0.755282166 | 3.724945049 | 0.202763304 |
| Polb | 409.3894603 | -0.049178244 | 0.072467963 | -0.678620488 |
| Gm26909 | 0.178821284 | 0.274410435 | 2.292793971 | 0.119683861 |
| Ikbkb | 547.644006 | -0.144617476 | 0.097435164 | -1.484243164 |
| Gm15346 | 0 | NA | NA | NA |
| Plat | 1098.82342 | 1.491705885 | 0.18530989 | 8.049791002 |
| Ap3m2 | 1163.088985 | 0.262361663 | 0.076802067 | 3.416075533 |
| 1700041G16Rik | 3.442441086 | 1.417440285 | 0.640303833 | 2.213699517 |
| Gm24735 | 0.372097677 | -0.220847528 | 1.551161773 | -0.142375561 |
| Kat6a | 2202.941112 | 0.08527512 | 0.070940982 | 1.202057226 |
| Ank1 | 2360.60079 | -0.401849366 | 0.091364729 | -4.398298687 |
| AC126445.1 | 0 | NA | NA | NA |
| Gm15815 | 3.535656746 | -0.597230129 | 0.558177588 | -1.069964367 |
| Gm15816 | 4.564354496 | -0.54546384 | 0.51800829 | -1.053002143 |
| Mir486 | 1.840863337 | -0.573058383 | 0.679058552 | -0.843901282 |
| mmu-mir-486b | 0 | NA | NA | NA |
| Nkx6-3 | 1.853806923 | 0.887669315 | 0.793175888 | 1.119133005 |
| Agpat6 | 875.7847179 | -0.048058389 | 0.064713117 | -0.742637525 |
| B230112G18Rik | 0.914342362 | -1.524744686 | 1.111269728 | -1.372074346 |
| Gins4 | 158.5957338 | -0.036182386 | 0.094180245 | -0.384182331 |
| Golga7 | 765.1713379 | 0.063607508 | 0.104600184 | 0.608101301 |
| Gm24335 | 0 | NA | NA | NA |
| Sfrp1 | 332.0038061 | 1.324642477 | 0.189302875 | 6.997476822 |
| Zmat4 | 554.6012764 | -0.065369445 | 0.098232327 | -0.665457564 |
| A730045E13Rik | 1.027651671 | -0.12399796 | 1.147462708 | -0.108062737 |
| Gm26714 | 0.777682211 | 1.848980019 | 1.264176814 | 1.462596053 |
| 1810011O10Rik | 28.96321082 | 0.966411539 | 0.307049812 | 3.147409641 |
| Ido2 | 21.23577249 | 0.576323277 | 0.244169367 | 2.360342269 |

| | | | | |
|---|---|---|---|---|
| Ido1 | 30.3065116 | 1.145868065 | 0.24425859 | 4.69120888 |
| Adam18 | 1.533690469 | 1.115044742 | 1.042671649 | 1.069411202 |
| Adam3 | 2.60343246 | 0.492267216 | 0.6961597 | 0.707118232 |
| Gm26208 | 0 | NA | NA | NA |
| Adam5 | 3.408984752 | 0.830086372 | 0.579551515 | 1.43229092 |
| Adam32 | 15.27834067 | 0.331061087 | 0.28283402 | 1.170513668 |
| Gm23898 | 0 | NA | NA | NA |
| Adam9 | 978.4277452 | 0.383917737 | 0.089051156 | 4.31120441 |
| Tm2d2 | 316.9476182 | -0.048273252 | 0.079746621 | -0.60533288 |
| Htra4 | 16.56292703 | 2.018464303 | 0.495557235 | 4.073120444 |
| Gm16933 | 1.72427518 | -0.45679599 | 0.892324052 | -0.511917155 |
| Plekha2 | 504.2642388 | 1.303356282 | 0.111986357 | 11.63852739 |
| Tacc1 | 2873.630983 | -0.256416538 | 0.089447763 | -2.866662385 |
| Mir8108 | 0 | NA | NA | NA |
| Gm10689 | 1.990043323 | 0.688464707 | 0.749339081 | 0.91876258 |
| Fgfr1 | 1712.090942 | 0.275988006 | 0.111974413 | 2.464741702 |
| Gm16159 | 0.092228164 | 0.75528439 | 3.724945049 | 0.202763901 |
| Letm2 | 106.132572 | -0.325272058 | 0.141683511 | -2.295765093 |
| D830025C05Rik | 3.021300136 | -0.475603855 | 0.568584221 | -0.836470373 |
| Whsc1l1 | 2360.967649 | 0.22774145 | 0.080263663 | 2.837416597 |
| Gm23184 | 0 | NA | NA | NA |
| Gm24992 | 3.122516278 | 0.31274273 | 0.548359741 | 0.570324017 |
| Gm23094 | 7.166830998 | 0.673198355 | 0.559448332 | 1.203325341 |
| Ppapdc1b | 162.0488265 | -0.425721128 | 0.132674209 | -3.208770813 |
| Ddhd2 | 814.1538713 | -0.156111647 | 0.078333874 | -1.992900884 |
| Gm17484 | 32.71518712 | -1.161942941 | 0.308045981 | -3.771978899 |
| Bag4 | 936.8776659 | 0.076588089 | 0.063304744 | 1.209831752 |
| Lsm1 | 143.0297487 | 0.1260763 | 0.118435842 | 1.064511369 |
| Star | 22.62371583 | 0.036230081 | 0.290696539 | 0.124631967 |
| Ash2l | 496.3350927 | -0.134324424 | 0.06094862 | -2.20389609 |
| Kcnu1 | 15.88801623 | -0.244119423 | 0.265289238 | -0.920201002 |
| Hgsnat | 490.0921398 | -0.218987322 | 0.071272752 | -3.072525147 |
| Pomk | 265.4581434 | -0.189329628 | 0.079741675 | -2.374287064 |
| Fnta | 475.5451638 | -0.181553357 | 0.070832511 | -2.56313597 |
| Hook3 | 2198.855902 | -0.295873427 | 0.041966395 | -7.050246362 |
| 1700047A11Rik | 0.066050875 | 0.274391604 | 3.724945049 | 0.073663262 |
| Rnf170 | 300.8605897 | -0.268317292 | 0.101344167 | -2.647584951 |
| Thap1 | 73.51708713 | -0.089000404 | 0.133434918 | -0.666994856 |
| AC154102.1 | 0 | NA | NA | NA |
| Gm22227 | 0 | NA | NA | NA |
| 2310008N11Rik | 0.04393175 | -0.206490061 | 3.724945049 | -0.055434391 |
| Zfp703 | 588.8755344 | 0.359617186 | 0.091234344 | 3.941686547 |
| Erlin2 | 723.3406109 | 0.21675583 | 0.081473433 | 2.660447992 |
| Proscos | 5.115524656 | 0.873627869 | 0.444503703 | 1.965400654 |
| Prosc | 626.0658266 | -0.233331034 | 0.066509708 | -3.508225185 |
| Adgra2 | 197.3133489 | 0.069959056 | 0.125916509 | 0.555598757 |
| Brf2 | 100.3354801 | 0.239626221 | 0.137007202 | 1.749004563 |

| | | | | |
|---|---|---|---|---|
| Rab11fip1 | 53.92462846 | 0.721939524 | 0.217890016 | 3.313320814 |
| 4933416M07Rik | 0.124596252 | 0.755282361 | 3.714681068 | 0.203323609 |
| Got1l1 | 2.891827676 | 0.145256027 | 0.597947243 | 0.242924486 |
| Adrb3 | 6.872911924 | -0.657880139 | 0.41750449 | -1.575743864 |
| Eif4ebp1 | 90.4572564 | 2.461969891 | 0.282307793 | 8.720871172 |
| Tex24 | 0.451016279 | 0.728860985 | 1.566621838 | 0.465243729 |
| Chrnb3 | 25.45046697 | -0.502146643 | 0.22752835 | -2.206962976 |
| Chrna6 | 16.6337043 | 0.100103116 | 0.303849558 | 0.329449601 |
| Gm24410 | 0 | NA | NA | NA |
| Poteg | 0 | NA | NA | NA |
| Gm1698 | 3.756750159 | 0.154938542 | 0.529082864 | 0.292843622 |
| AC111034.1 | 0 | NA | NA | NA |
| Gm24202 | 0 | NA | NA | NA |
| Gm8100 | 1.189584463 | -1.115529323 | 1.000924092 | -1.114499423 |
| Gm26795 | 1.493046057 | 2.216353877 | 1.006303269 | 2.202471111 |
| Unc5d | 798.2940023 | 0.076803728 | 0.077401114 | 0.992281935 |
| 4933433F19Rik | 0 | NA | NA | NA |
| Dusp26 | 500.6484086 | 0.290625868 | 0.109601052 | 2.651670413 |
| Rnf122 | 133.8921804 | 0.165816107 | 0.135457075 | 1.224122897 |
| Snord13 | 7.690137191 | 0.61830627 | 0.501682937 | 1.232464221 |
| Tti2 | 237.0865655 | 0.29692437 | 0.098993397 | 2.9994361 |
| Mak16 | 240.9495093 | 0.233780538 | 0.097225267 | 2.404524515 |
| Fut10 | 112.6731054 | 1.211958234 | 0.134573817 | 9.005899203 |
| Gm8051 | 0 | NA | NA | NA |
| Gm23539 | 0.319751656 | 0.209753709 | 1.831687699 | 0.114513904 |
| Gm24184 | 0.141071297 | -0.206491178 | 3.564043836 | -0.057937328 |
| Gm5117 | 3.857935733 | 0.818447178 | 0.527714727 | 1.550927302 |
| Nrg1 | 264.3991212 | 0.540757031 | 0.144957766 | 3.730445388 |
| Gm26578 | 5.176536363 | -1.127696843 | 0.548063806 | -2.057601378 |
| Gm22140 | 0 | NA | NA | NA |
| Gm5908 | 9.444744683 | 0.253076824 | 0.361556148 | 0.699965482 |
| Gm3985 | 0.861087095 | 0.356521102 | 1.117300399 | 0.319091537 |
| Gm6877 | 13.36504497 | 0.724949067 | 0.314764624 | 2.303146578 |
| Wrn | 282.207455 | 0.118354011 | 0.1088766 | 1.087047269 |
| Purg | 493.6010051 | -0.527203798 | 0.08874619 | -5.940579533 |
| 5930422O12Rik | 8.011746571 | -0.960177066 | 0.387992039 | -2.474733937 |
| Tex15 | 53.01275634 | -0.473038616 | 0.253333795 | -1.867254289 |
| Ppp2cb | 866.8510396 | 0.057176673 | 0.083252664 | 0.686784902 |
| Ubxn8 | 210.3585288 | 0.342291043 | 0.08452766 | 4.049456045 |
| Gsr | 438.4005393 | 0.5138816 | 0.089562114 | 5.737711811 |
| Gm24727 | 28.80332305 | 0.325838313 | 0.33129001 | 0.983544036 |
| 1700104B16Rik | 0.424137699 | 0.146650666 | 1.567293812 | 0.093569352 |
| Gtf2e2 | 144.048718 | -0.282948399 | 0.102529422 | -2.759680033 |
| Smim18 | 19.2016788 | -0.450870658 | 0.266916931 | -1.689179688 |
| Rbpms | 30.315901 | 0.937942125 | 0.265166292 | 3.537184601 |
| Gm26632 | 0 | NA | NA | NA |
| Gm26978 | 0.712265815 | 0.650365169 | 1.274750247 | 0.510190267 |

| | | | | |
|---|---|---|---|---|
| Gm9951 | 0.166952883 | 0.274401811 | 2.756036668 | 0.099563919 |
| Dctn6 | 437.7144974 | 0.048210909 | 0.073144959 | 0.659114585 |
| Gm26768 | 0.185862605 | 1.585182872 | 2.679501876 | 0.591596104 |
| Gm10131 | 44.98302574 | 0.391450242 | 0.205872701 | 1.90141889 |
| Mboat4 | 2.274616701 | 0.180262193 | 0.719887148 | 0.250403405 |
| Leprotl1 | 852.1230496 | 0.27906393 | 0.088545843 | 3.151632187 |
| Saraf | 1822.362344 | 0.268551774 | 0.095892189 | 2.800559442 |
| Mir6395 | 0 | NA | NA | NA |
| Gm29243 | 0.741879394 | 0.260038307 | 1.352185323 | 0.192309665 |
| Gm23177 | 0.047948698 | -0.206487837 | 3.724945049 | -0.055433794 |
| Gm6100 | 0 | NA | NA | NA |
| Gm8254 | 0 | NA | NA | NA |
| Gm4889 | 0 | NA | NA | NA |
| Gm29174 | 0.095435502 | 0.75528439 | 3.724945049 | 0.202763901 |
| Dusp4 | 233.7081911 | -0.366527399 | 0.208646575 | -1.756690221 |
| Tnks | 1939.237959 | 0.105154709 | 0.055453613 | 1.896264348 |
| Gm9939 | 2.193369661 | -1.101538327 | 0.74369234 | -1.481174766 |
| Rpl31-ps23 | 0 | NA | NA | NA |
| Ppp1r3b | 56.58455372 | 0.406169629 | 0.262883329 | 1.545056621 |
| Eri1 | 199.30024 | -0.303002897 | 0.129989086 | -2.330987218 |
| Gm16793 | 8.177702489 | 0.349940763 | 0.371600663 | 0.941711891 |
| Mfhas1 | 794.683006 | -0.008902117 | 0.084722056 | -0.105074378 |
| Gm22216 | 0.304980231 | 0.732736221 | 1.742700509 | 0.420460209 |
| AC129211.1 | 0 | NA | NA | NA |
| Cldn23 | 10.064079 | 1.282454777 | 0.436792127 | 2.936075762 |
| Gm22030 | 0 | NA | NA | NA |
| Rps12l2 | 3.933964785 | 2.071865686 | 0.882660039 | 2.34729748 |
| D8Ertd82e | 662.5180749 | -0.552012425 | 0.094592359 | -5.835697833 |
| Lonrf1 | 282.0614182 | -0.020707676 | 0.118230565 | -0.175146556 |
| Gm5787 | 0.203967006 | 0.274410937 | 2.216948771 | 0.123778655 |
| Gm26150 | 0.099155932 | 0.75528439 | 3.724945049 | 0.202763901 |
| 6430573F11Rik | 301.8566133 | 0.019956946 | 0.183547566 | 0.108729016 |
| Dlc1 | 1005.579271 | -0.321608011 | 0.066688521 | -4.822539256 |
| AI429214 | 46.9821826 | 0.161792297 | 0.162924714 | 0.993049445 |
| Gm25126 | 0 | NA | NA | NA |
| Sgcz | 92.85252959 | -0.211191127 | 0.157142462 | -1.343946915 |
| Gm16205 | 2.502004683 | 0.151409642 | 0.659727942 | 0.229503151 |
| Gm16204 | 0 | NA | NA | NA |
| Mir383 | 2.557294913 | 0.115158256 | 0.584349843 | 0.19707074 |
| Gm23128 | 0 | NA | NA | NA |
| Tusc3 | 992.2313355 | 0.042100372 | 0.076698684 | 0.548906059 |
| Msr1 | 385.467979 | 6.648465654 | 0.468734335 | 14.18386741 |
| Fgf20 | 6.705059015 | -0.21445696 | 0.375706998 | -0.570809064 |
| Micu3 | 1360.461354 | -0.041564584 | 0.065409302 | -0.635453718 |
| Mir7666 | 6.340121748 | 0.169190274 | 0.410865599 | 0.411789827 |
| Zdhhc2 | 665.2779763 | -0.442913791 | 0.099681485 | -4.443290441 |
| Cnot7 | 1005.736534 | -0.058705554 | 0.057198722 | -1.026343802 |

| | | | | |
|---|---|---|---|---|
| Vps37a | 864.8152951 | -0.105139909 | 0.071753893 | -1.465285067 |
| Mtmr7 | 611.4343167 | 0.031392952 | 0.063331801 | 0.495690178 |
| Adam24 | 0.11193333 | 0.75528439 | 3.724945049 | 0.202763901 |
| Adam25 | 0 | NA | NA | NA |
| Adam20 | 0 | NA | NA | NA |
| Adam39 | 0 | NA | NA | NA |
| Gm5345 | 0.200503209 | 1.257028057 | 2.612813754 | 0.481101286 |
| Gm26584 | 0.132101749 | 0.274387056 | 3.724945049 | 0.073662041 |
| Slc7a2 | 630.8632614 | 0.340831227 | 0.135887501 | 2.508186728 |
| Pdgfrl | 37.56964625 | 1.091608695 | 0.270768045 | 4.031527035 |
| Mtus1 | 1041.21851 | -0.257521322 | 0.080377735 | -3.203888753 |
| B430010I23Rik | 0.577003193 | 1.063847243 | 1.365395387 | 0.77914958 |
| Gm16192 | 0.177745132 | -0.206480788 | 2.300214322 | -0.089765891 |
| Gm16193 | 1.147365236 | 0.68107547 | 0.99035761 | 0.687706605 |
| Fgl1 | 0.121388914 | 0.755282166 | 3.724945049 | 0.202763304 |
| Gm16348 | 0.652886586 | 0.755263999 | 3.724945049 | 0.202758427 |
| Pcm1 | 2899.225793 | 0.019535312 | 0.060681408 | 0.321932407 |
| Asah1 | 694.5391705 | 0.145255637 | 0.092468885 | 1.570859625 |
| Frg1 | 216.5532014 | 0.101271061 | 0.107782215 | 0.939589721 |
| Gapdh-ps14 | 0.166952883 | 0.274401811 | 2.756036668 | 0.099563919 |
| Smarce1-ps1 | 16.22242323 | -0.163958191 | 0.278489323 | -0.588741391 |
| 2810404M03Rik | 0.402944817 | -1.060025824 | 1.727988835 | -0.613444834 |
| Zfp353-ps | 0 | NA | NA | NA |
| Gm6180 | 51.37583421 | 0.267581819 | 0.149305607 | 1.792175285 |
| Gm23531 | 0 | NA | NA | NA |
| Gm26181 | 0 | NA | NA | NA |
| Triml1 | 0.169875191 | 0.741257059 | 2.746838206 | 0.269858289 |
| Triml2 | 0.045882455 | 0.755288839 | 3.724945049 | 0.202765096 |
| Zfp42 | 0.483255338 | 0.274413736 | 1.443321439 | 0.190126557 |
| Gm22986 | 0 | NA | NA | NA |
| Adam26b | 0 | NA | NA | NA |
| Adam26a | 0 | NA | NA | NA |
| Gm5346 | 0.068097359 | -0.206487837 | 3.724945049 | -0.055433794 |
| Adam34 | 0.137685729 | -0.206481801 | 2.938031474 | -0.070278961 |
| Gm24591 | 0 | NA | NA | NA |
| Gm37972 | 0 | NA | NA | NA |
| Gm26241 | 0 | NA | NA | NA |
| Gm26089 | 0 | NA | NA | NA |
| Fat1 | 3170.327197 | -0.482224967 | 0.08019409 | -6.013223262 |
| Gm2366 | 0.106971521 | 1.236179401 | 3.724945049 | 0.331865138 |
| AY512931 | 0.183958792 | 1.222273706 | 2.67815496 | 0.456386477 |
| Mtnr1a | 1.009506168 | -0.469622529 | 1.013756559 | -0.463249806 |
| F11 | 1.357931586 | 2.97185272 | 1.020753683 | 2.911429827 |
| Klkb1 | 0.521565614 | 2.036002159 | 1.413014023 | 1.440893102 |
| Cyp4v3 | 176.7598727 | 1.043989275 | 0.152085193 | 6.864503085 |
| Fam149a | 309.9705733 | -0.324110101 | 0.102414256 | -3.164697124 |
| Tlr3 | 1099.900133 | 3.087621077 | 0.193145261 | 15.98600485 |

| | | | | |
|---|---|---|---|---|
| Sorbs2 | 2803.02876 | -0.402784846 | 0.131084737 | -3.072705904 |
| Gm23604 | 0 | NA | NA | NA |
| Gm16351 | 0.27308476 | 0.755306614 | 2.016745306 | 0.374517601 |
| Sorbs2os | 1.437005151 | -1.236101928 | 0.901299795 | -1.371465893 |
| Gm16352 | 7.186401275 | -0.485319457 | 0.370518427 | -1.309838921 |
| Gm25309 | 0 | NA | NA | NA |
| Pdlim3 | 13.56838278 | -0.450737108 | 0.364584538 | -1.23630341 |
| Ccdc110 | 5.316494593 | -0.962337021 | 0.488674671 | -1.969279519 |
| Gm15878 | 0.075043205 | 0.755286614 | 3.724945049 | 0.202764498 |
| 1700029J07Rik | 30.24045965 | -0.356492127 | 0.269577926 | -1.322408449 |
| Ufsp2 | 315.566229 | -0.207092796 | 0.119236522 | -1.736823516 |
| Ankrd37 | 31.55324993 | -0.437099984 | 0.194957893 | -2.242022508 |
| Lrp2bp | 13.15399262 | 1.283686972 | 0.305459524 | 4.202478136 |
| Snx25 | 564.8783107 | -0.186458231 | 0.054439095 | -3.425079534 |
| Gm23030 | 0 | NA | NA | NA |
| Cfap97 | 360.4521644 | -0.074542899 | 0.074376103 | -1.002242605 |
| Gm15634 | 3.396395574 | 0.007347822 | 0.571079219 | 0.012866555 |
| Slc25a4 | 4403.300027 | -0.021369973 | 0.084171025 | -0.25388752 |
| Actg-ps1 | 1.057130107 | 1.771593548 | 1.018335628 | 1.739695144 |
| Gm23812 | 0 | NA | NA | NA |
| Gm10313 | 0.79470004 | -0.588442747 | 1.219133052 | -0.482673114 |
| Gm24669 | 0 | NA | NA | NA |
| Helt | 1.906358267 | 0.383683825 | 1.141316623 | 0.336176497 |
| Acsl1 | 635.4438796 | 0.162481144 | 0.089654638 | 1.812300486 |
| Cenpu | 23.99853257 | -0.445443339 | 0.223420328 | -1.993745796 |
| Primpol | 71.59947947 | 0.046725019 | 0.15397049 | 0.303467364 |
| Casp3 | 141.296778 | 0.475221226 | 0.151682825 | 3.132992971 |
| Gm16675 | 21.12817838 | 4.287510069 | 0.527890515 | 8.12196837 |
| Irf2 | 579.6276244 | 1.2800013 | 0.151244436 | 8.463129832 |
| Enpp6 | 117.2534975 | -0.039214843 | 0.124738512 | -0.31437639 |
| 4930579M01Rik | 0.29175089 | 0.729183191 | 1.778011001 | 0.410111743 |
| Stox2 | 2896.450262 | -0.171228149 | 0.063267936 | -2.706396968 |
| 4930448N21Rik | 21.17633594 | -1.184702073 | 0.336045387 | -3.525422811 |
| Gm10083 | 4.406530871 | -1.048539871 | 0.495619839 | -2.115613196 |
| Gm22857 | 0 | NA | NA | NA |
| Trappc11 | 834.2127417 | 0.02068178 | 0.057297453 | 0.360954612 |
| Rwdd4a | 348.2097546 | -0.301064568 | 0.085077386 | -3.538714342 |
| Gm27409 | 0 | NA | NA | NA |
| Gm8623 | 3.204879861 | -0.190201439 | 0.563724769 | -0.337401245 |
| Ing2 | 227.0032211 | -0.198794737 | 0.104579615 | -1.900893747 |
| AA386476 | 7.480509175 | 0.15005129 | 0.376298528 | 0.398755983 |
| Cdkn2aip | 219.8046481 | 0.042250818 | 0.105691022 | 0.399757875 |
| E030037K01Rik | 3.293407775 | -0.708103796 | 0.573908289 | -1.233827441 |
| n-R5s96 | 0.212081199 | 1.089879014 | 2.576983506 | 0.422928207 |
| Cldn24 | 0.048949532 | -0.206487837 | 3.724945049 | -0.055433794 |
| Cldn22 | 1.967947156 | 1.013537519 | 0.683583161 | 1.482683566 |
| Wwc2 | 675.3607201 | 0.085799677 | 0.064219343 | 1.336041012 |

| | | | | |
|---|---|---|---|---|
| n-R5s97 | 0 | NA | NA | NA |
| Dctd | 53.07648255 | -0.671544773 | 0.1905785 | -3.523717386 |
| Tenm3 | 1960.745608 | -0.280426595 | 0.112820161 | -2.485607131 |
| Gm19744 | 64.58441548 | 0.737346826 | 0.263036078 | 2.80321556 |
| Gm25830 | 0.249706104 | 0.274415662 | 1.848218408 | 0.148475776 |
| 4930555F03Rik | 2.112369234 | -1.319806325 | 0.713257871 | -1.850391533 |
| Gm23986 | 0.048949532 | -0.206487837 | 3.724945049 | -0.055433794 |
| AC140437.1 | 0.038981395 | 0.274393828 | 3.724945049 | 0.073663859 |
| Gm24929 | 0 | NA | NA | NA |
| Gm9892 | 89.20159694 | 0.685914592 | 0.167227971 | 4.101673827 |
| Gm6463 | 0 | NA | NA | NA |
| Gm25813 | 1.893990199 | 0.158112769 | 0.858522448 | 0.184168474 |
| Gm23666 | 0 | NA | NA | NA |
| Gm27964 | 0 | NA | NA | NA |
| Gm22841 | 0 | NA | NA | NA |
| Aga | 79.49866868 | 0.492113716 | 0.157118676 | 3.132114705 |
| Neil3 | 7.05498645 | 1.853539305 | 0.700340327 | 2.646626552 |
| Vegfc | 39.65073296 | 0.044536288 | 0.167600516 | 0.265728825 |
| Spcs3 | 857.5164723 | -0.179544589 | 0.078373872 | -2.290873032 |
| Asb5 | 7.025391348 | 0.409467214 | 0.414159402 | 0.988670576 |
| 5330439A09Rik | 2.898308479 | -0.859099044 | 0.757453128 | -1.134194331 |
| Spata4 | 0.398944595 | 1.127471471 | 1.634046274 | 0.68998748 |
| Wdr17 | 405.2573486 | -0.294120372 | 0.13127622 | -2.240469548 |
| Gm15758 | 0.38984224 | 0.432808796 | 1.418553034 | 0.305105826 |
| n-R5s98 | 6.806331161 | -1.102331471 | 0.419633958 | -2.626888148 |
| Gpm6a | 9377.570103 | -0.11537728 | 0.084237851 | -1.369660777 |
| Adam29 | 0.15906016 | 0.274395307 | 3.439462686 | 0.079778539 |
| Glra3 | 117.655716 | 0.157541618 | 0.130277337 | 1.209278769 |
| Hpgd | 129.4286017 | 1.430326994 | 0.254663729 | 5.616532039 |
| Gm24375 | 0.075043205 | 0.755286614 | 3.724945049 | 0.202764498 |
| Cep44 | 122.059907 | 0.221516855 | 0.148033497 | 1.496396828 |
| Fbxo8 | 283.448663 | 0.236902082 | 0.117499285 | 2.016200196 |
| Gm25992 | 6.166697826 | 0.501675699 | 0.383908079 | 1.30675994 |
| Gm29173 | 0.088736197 | 0.274400501 | 3.724945049 | 0.073665651 |
| AV026068 | 0.061920613 | 0.274393828 | 3.724945049 | 0.073663859 |
| Hand2 | 0.062426873 | 0.755288839 | 3.724945049 | 0.202765096 |
| Gm28063 | 0 | NA | NA | NA |
| 5033428I22Rik | 0 | NA | NA | NA |
| Scrg1 | 178.3231663 | 0.226541913 | 0.210918214 | 1.074074681 |
| Sap30 | 157.6272062 | 1.137647661 | 0.154001591 | 7.387246169 |
| 2500002B13Rik | 14.59964785 | -0.530636815 | 0.319336258 | -1.661686706 |
| Hmgb2 | 86.7028358 | 2.367541662 | 0.290991006 | 8.136133474 |
| Galnt7 | 420.2620518 | -0.128263568 | 0.073109042 | -1.754414557 |
| Gm25379 | 0 | NA | NA | NA |
| Gm10284 | 11.17616707 | -0.020237795 | 0.278577543 | -0.072646899 |
| Galntl6 | 211.6445591 | -0.38248445 | 0.105712599 | -3.618153878 |
| Gm26905 | 1.736582332 | -0.725981446 | 0.73467558 | -0.988166023 |

| | | | | |
|---|---|---|---|---|
| Gm15882 | 0 | NA | NA | NA |
| Gm15881 | 19.66782202 | -0.701778965 | 0.282749233 | -2.481983619 |
| BC030500 | 65.02402097 | -0.488043894 | 0.163521328 | -2.984588614 |
| Gm24847 | 0 | NA | NA | NA |
| Gm26164 | 0 | NA | NA | NA |
| Gm10283 | 0 | NA | NA | NA |
| Aadat | 2.615189709 | -0.410166899 | 0.641228081 | -0.639658354 |
| Mfap3l | 976.8064727 | -0.505135911 | 0.082700166 | -6.108039838 |
| Gm23329 | 0.066050875 | 0.274391604 | 3.724945049 | 0.073663262 |
| 2700029M09Rik | 270.2446424 | 0.103533211 | 0.118316969 | 0.875049555 |
| Clcn3 | 3034.91923 | -0.117870905 | 0.043049691 | -2.738019736 |
| B230317F23Rik | 9.640489829 | -1.083853475 | 0.404723788 | -2.678007838 |
| Nek1 | 677.7359105 | -0.55063825 | 0.15122789 | -3.641115724 |
| Gm16179 | 0 | NA | NA | NA |
| Gm16178 | 0.095898756 | 0.27438722 | 3.724945049 | 0.073662085 |
| Sh3rf1 | 531.7020943 | -0.37369048 | 0.101034246 | -3.698651641 |
| 1700001D01Rik | 0 | NA | NA | NA |
| Gm10015 | 0.445602023 | 1.144871059 | 1.43502621 | 0.797804982 |
| Cbr4 | 83.07961058 | 0.077845185 | 0.145589847 | 0.534688282 |
| Palld | 202.5425458 | 0.941480665 | 0.206253951 | 4.564667286 |
| 4930512H18Rik | 0.155551505 | 1.257968755 | 2.831330457 | 0.444303049 |
| Ddx60 | 1763.240931 | 6.90516961 | 0.223816381 | 30.85194027 |
| Anxa10 | 0.200514362 | 0.755297509 | 2.651420135 | 0.284865269 |
| Gm22324 | 0 | NA | NA | NA |
| Spock3 | 1216.4572 | -0.236656176 | 0.098282297 | -2.407922729 |
| Gm24236 | 0 | NA | NA | NA |
| Gm5350 | 0 | NA | NA | NA |
| Gm24580 | 0 | NA | NA | NA |
| Sgol2b | 2.142608394 | -1.392277476 | 0.730452169 | -1.906048795 |
| Tll1 | 166.2460114 | -0.767769571 | 0.193689379 | -3.963921906 |
| Gm22586 | 0 | NA | NA | NA |
| Mir710 | 0.327710148 | 1.913840423 | 1.945250294 | 0.983853044 |
| Cpe | 12511.93844 | 0.014767576 | 0.093897937 | 0.157272636 |
| Msmo1 | 801.7074196 | -0.113953511 | 0.142588991 | -0.79917468 |
| Klhl2 | 1600.801606 | -0.056336346 | 0.091496493 | -0.615721364 |
| Gm26210 | 0 | NA | NA | NA |
| Tmem192 | 214.6384359 | 0.887309474 | 0.13908536 | 6.379603652 |
| Trim75 | 0 | NA | NA | NA |
| Trim60 | 0 | NA | NA | NA |
| Apela | 0.083526043 | 0.75528439 | 3.724945049 | 0.202763901 |
| Gm10663 | 0 | NA | NA | NA |
| BC030870 | 0 | NA | NA | NA |
| Gm24333 | 0 | NA | NA | NA |
| Gm25403 | 0 | NA | NA | NA |
| 1-Mar | 492.5184607 | 0.39128585 | 0.067161323 | 5.826059249 |
| Gm23515 | 0 | NA | NA | NA |
| Gm15355 | 0.044191549 | -0.206490061 | 3.724945049 | -0.055434391 |

| | | | | |
|---|---|---|---|---|
| Gm26123 | 1.969797064 | 0.026891252 | 0.675942411 | 0.039783347 |
| Gm24205 | 4.278889304 | 0.083951704 | 0.525288984 | 0.159820035 |
| Gm15354 | 0.20829534 | 0.274401349 | 2.561195153 | 0.107138009 |
| Gm15356 | 0.306626524 | 0.274412571 | 1.784120459 | 0.15380832 |
| Tma16 | 219.0930399 | 1.183288971 | 0.183640821 | 6.443496399 |
| Tktl2 | 2.437974107 | 0.943629614 | 0.580938449 | 1.624319437 |
| Gm16330 | 0 | NA | NA | NA |
| Npy5r | 36.55808565 | -0.810694526 | 0.195002704 | -4.157350172 |
| Npy1r | 157.9670103 | -0.561969441 | 0.144029589 | -3.901763827 |
| Naf1 | 250.5788205 | -0.284883978 | 0.116641853 | -2.442382141 |
| Gm25275 | 0 | NA | NA | NA |
| Gm24274 | 0 | NA | NA | NA |
| Gm24677 | 0 | NA | NA | NA |
| Gm10660 | 0 | NA | NA | NA |
| Nat1 | 7.474725557 | -1.495944795 | 0.437215049 | -3.421530886 |
| Nat2 | 24.83237878 | 1.294852389 | 0.283176447 | 4.572599183 |
| Gm9755 | 12.27831489 | -0.044625085 | 0.314388253 | -0.141942598 |
| Nat3 | 0.254414262 | 1.28862501 | 2.466219951 | 0.522510172 |
| AC141868.1 | 0 | NA | NA | NA |
| Psd3 | 7016.150575 | -0.056663648 | 0.104799116 | -0.540688225 |
| Gm15992 | 24.57258576 | -0.29363051 | 0.293863096 | -0.999208524 |
| Gm15991 | 12.26391693 | -0.42830348 | 0.28282786 | -1.514360998 |
| Gm22018 | 0.176406687 | 0.712240329 | 2.705670667 | 0.263239846 |
| Gm21807 | 5.900355672 | -0.85511655 | 0.443791391 | -1.926843484 |
| Gm26659 | 0.106520246 | 0.755282166 | 3.724945049 | 0.202763304 |
| Sh2d4a | 4.03121477 | 0.097631901 | 0.745817534 | 0.130905881 |
| Csgalnact1 | 201.9711671 | -0.258308658 | 0.115638313 | -2.233763625 |
| 1700125H03Rik | 0.107078754 | -0.206490058 | 3.724945049 | -0.05543439 |
| Gm15656 | 0 | NA | NA | NA |
| Gm15654 | 0 | NA | NA | NA |
| Ints10 | 560.1978323 | -0.528691822 | 0.11958846 | -4.420926739 |
| Gm15549 | 5.491253898 | -0.285401354 | 0.566309812 | -0.503966818 |
| Lpl | 272.7124532 | -0.021857349 | 0.128898424 | -0.169570335 |
| Slc18a1 | 6.521337389 | 0.679135314 | 0.40491411 | 1.677233017 |
| Gm15716 | 0 | NA | NA | NA |
| Atp6v1b2 | 5572.862093 | 0.118695156 | 0.048797305 | 2.432412127 |
| Gm15717 | 14.06246178 | -0.351486719 | 0.397450918 | -0.884352515 |
| Lzts1 | 982.0473404 | -0.285878932 | 0.162445075 | -1.759849788 |
| Zfp930 | 119.9920175 | 0.247765151 | 0.135671952 | 1.826207612 |
| D130040H23Rik | 96.03445661 | -0.420838416 | 0.140189738 | -3.00192027 |
| Gm10033 | 37.55209856 | -0.528302411 | 0.239596755 | -2.204964804 |
| Gm10311 | 21.9424117 | -0.407052097 | 0.252832056 | -1.609970282 |
| Gm25661 | 0.130524232 | 0.274405456 | 3.657871338 | 0.075017799 |
| RP23-468H22.1 | 1.045653151 | 2.715278346 | 1.253198122 | 2.166679233 |
| RP23-468H22.3 | 0.584793824 | 1.435284207 | 1.22363254 | 1.172969956 |
| RP23-468H22.2 | 0.794752679 | 1.995963273 | 1.337187035 | 1.49265826 |
| Zfp868 | 254.8017484 | 0.098748253 | 0.085451821 | 1.155601512 |

| | | | | |
|---|---|---|---|---|
| Zfp964 | 24.06935087 | -0.767039146 | 0.228687867 | -3.354087638 |
| Zfp869 | 250.7494571 | 0.280905018 | 0.088413122 | 3.177186944 |
| Zfp963 | 46.44832303 | 0.15785177 | 0.15449408 | 1.021733455 |
| Gm20422 | 0 | NA | NA | NA |
| Zfp866 | 322.7321497 | -0.269685211 | 0.113320123 | -2.379852792 |
| Atp13a1 | 689.8934044 | 0.56682192 | 0.086792141 | 6.530797753 |
| Gmip | 150.8857527 | 0.404874968 | 0.143901667 | 2.813553012 |
| Lpar2 | 13.32151877 | 0.333213959 | 0.292993806 | 1.137273047 |
| Pbx4 | 59.57232151 | -0.041155503 | 0.138183658 | -0.29783191 |
| Cilp2 | 10.39023466 | -0.265737226 | 0.353624068 | -0.751468155 |
| Yjefn3 | 43.79218575 | 0.001428379 | 0.183987557 | 0.007763454 |
| Ndufa13 | 488.3269015 | 0.015429141 | 0.343313363 | 0.044941861 |
| Tssk6 | 2.109268871 | -0.680159025 | 0.80921877 | -0.84051316 |
| Gatad2a | 755.2849878 | 0.566166688 | 0.102814335 | 5.506690154 |
| Mau2 | 1031.702982 | 0.141924694 | 0.065459301 | 2.168136435 |
| Sugp1 | 554.3624155 | 0.249959604 | 0.060326591 | 4.143439865 |
| Tm6sf2 | 17.28368832 | -2.069372594 | 0.495548895 | -4.175920101 |
| Hapln4 | 1233.584627 | -0.21775582 | 0.074707371 | -2.914783612 |
| Ncan | 3935.767652 | -0.306518599 | 0.169874476 | -1.804382893 |
| Mir7066 | 0.192965486 | 0.296605227 | 2.249093267 | 0.131877691 |
| Gm25906 | 0 | NA | NA | NA |
| Rfxank | 136.35992 | -0.257110513 | 0.114464344 | -2.246206154 |
| Nr2c2ap | 176.3814956 | 0.438404 | 0.156133676 | 2.807875993 |
| 2310045N01Rik | 134.2125858 | 0.162829908 | 0.103792223 | 1.568806436 |
| Mef2b | 0.295756179 | -0.302280633 | 2.074058164 | -0.145743566 |
| Tmem161a | 299.8727589 | -0.114738802 | 0.095651776 | -1.199547014 |
| Slc25a42 | 283.822492 | -0.631223871 | 0.162606397 | -3.88191291 |
| Armc6 | 187.6904792 | -0.229970518 | 0.102681041 | -2.239659 |
| Sugp2 | 1115.657998 | -0.135723009 | 0.094588227 | -1.434882688 |
| Homer3 | 488.6342012 | -0.112092502 | 0.201469502 | -0.55637454 |
| Ddx49 | 235.0236882 | 0.401952438 | 0.088746901 | 4.529199713 |
| Cope | 659.244897 | 0.028642438 | 0.075350812 | 0.380121153 |
| Cers1 | 345.2784473 | -0.214029783 | 0.091739345 | -2.333020619 |
| Gdf1 | 0 | NA | NA | NA |
| Upf1 | 1210.834756 | 0.142588 | 0.06074259 | 2.34741391 |
| Comp | 10.12622545 | -0.90782078 | 0.423043432 | -2.14592808 |
| Crtc1 | 1987.206553 | 0.092077589 | 0.099974538 | 0.921010394 |
| Klhl26 | 487.3158746 | 0.127235656 | 0.074248837 | 1.713638366 |
| Tmem59l | 1446.494139 | 0.117691481 | 0.055343075 | 2.126580064 |
| Crlf1 | 37.87736275 | 0.296924798 | 0.234027168 | 1.268762084 |
| 2810428I15Rik | 262.175928 | 0.216413861 | 0.106349996 | 2.034921192 |
| Uba52 | 19.53769797 | -0.088317503 | 0.273439347 | -0.322987544 |
| Kxd1 | 196.5815617 | 0.196389715 | 0.084169393 | 2.33326757 |
| Fkbp8 | 2500.961044 | 0.225098575 | 0.076406031 | 2.946083852 |
| Ell | 288.2178684 | 0.125126048 | 0.121397149 | 1.030716529 |
| Isyna1 | 251.2867603 | -0.215889211 | 0.143759028 | -1.501743673 |
| Ssbp4 | 876.1567567 | -0.098344692 | 0.063732981 | -1.54307379 |

| | | | | |
|---|---|---|---|---|
| Lrrc25 | 65.01020252 | 4.104929786 | 0.441197372 | 9.304066724 |
| Gdf15 | 12.20491556 | 3.930348645 | 0.941783413 | 4.17330417 |
| Pgpep1 | 166.5791501 | -0.224817406 | 0.170784191 | -1.316383005 |
| Lsm4 | 218.808746 | 0.212631442 | 0.097510669 | 2.180596702 |
| Jund | 1476.72973 | 0.337327989 | 0.137950638 | 2.445280383 |
| Gm11175 | 17.16041749 | -0.384487029 | 0.288020834 | -1.334927837 |
| Gm3336 | 0.36285418 | 0.093710473 | 1.720168599 | 0.054477493 |
| Pde4c | 6.152153976 | -0.228728772 | 0.412735147 | -0.55417808 |
| Rab3a | 3789.848654 | -0.100466493 | 0.055480285 | -1.81085034 |
| Mpv17l2 | 124.5250829 | 0.145944783 | 0.146137077 | 0.998684151 |
| Ifi30 | 138.4088315 | 1.834145648 | 0.216046226 | 8.489598164 |
| Pik3r2 | 1054.806414 | -0.10664129 | 0.075433972 | -1.413703763 |
| 2010320M18Rik | 39.80095612 | 0.139596639 | 0.193825777 | 0.720217101 |
| Mast3 | 1971.583413 | -0.276502032 | 0.117443078 | -2.354349337 |
| Mir7240 | 0.07796279 | 0.274392189 | 3.724945049 | 0.073663419 |
| Il12rb1 | 265.2067133 | 4.34022525 | 0.240325029 | 18.05981366 |
| Arrdc2 | 123.0827081 | 1.238226172 | 0.180652603 | 6.854183953 |
| Kcnn1 | 121.7377044 | -0.522425643 | 0.13051284 | -4.002867782 |
| A230052G05Rik | 2.786635864 | -1.352830741 | 0.701024649 | -1.929790547 |
| Ccdc124 | 301.1606489 | -0.111756429 | 0.113810494 | -0.981951887 |
| Slc5a5 | 42.07648303 | -0.393003319 | 0.181352321 | -2.167070802 |
| Rpl18a | 1403.238223 | 0.649056586 | 0.133733971 | 4.853341173 |
| Mir7067 | 0 | NA | NA | NA |
| Snora68 | 19.80479055 | -0.042304481 | 0.290725189 | -0.145513642 |
| Map1s | 619.2559287 | 0.077827864 | 0.069048004 | 1.12715588 |
| Mir1969 | 0.386072962 | 0.274403512 | 3.26857073 | 0.083952141 |
| Gm10654 | 0.749724794 | 0.312042605 | 1.260930621 | 0.247470083 |
| 1700026F02Rik | 0.167183452 | -0.206481502 | 2.765153652 | -0.074672705 |
| Gm17576 | 0.045882455 | 0.755288839 | 3.724945049 | 0.202765096 |
| Gm27628 | 0.044544648 | 0.755286614 | 3.724945049 | 0.202764498 |
| Haus8 | 58.62861715 | -0.033954092 | 0.145287593 | -0.233702628 |
| Myo9b | 1047.53723 | 0.295352682 | 0.094447697 | 3.127156005 |
| Use1 | 300.4227806 | 0.253793267 | 0.096899178 | 2.619147782 |
| Ocel1 | 128.1701314 | -0.103018453 | 0.140442372 | -0.733528289 |
| Nr2f6 | 250.0485222 | -0.135707515 | 0.120934071 | -1.12216114 |
| Ushbp1 | 11.95745292 | -0.162171225 | 0.393936679 | -0.411668255 |
| Babam1 | 547.0408888 | 0.06545169 | 0.075225645 | 0.870071509 |
| Ankle1 | 4.029650605 | 1.097397904 | 0.516494897 | 2.124702318 |
| Abhd8 | 1731.379142 | 0.034953975 | 0.082969307 | 0.421288018 |
| Mrpl34 | 112.3878888 | -0.440528522 | 0.119032682 | -3.700903945 |
| Dda1 | 548.7093606 | 0.089476754 | 0.06867147 | 1.302968381 |
| Ano8 | 471.5065914 | -0.282820572 | 0.097780243 | -2.892410206 |
| Gtpbp3 | 220.5550811 | -0.225329964 | 0.082719121 | -2.724037224 |
| Plvap | 140.3657839 | 0.281931444 | 0.123424221 | 2.284247301 |
| Bst2 | 1856.786934 | 7.787799617 | 0.312006903 | 24.96034396 |
| Mvb12a | 194.6062646 | 0.844445499 | 0.099441635 | 8.491870614 |
| Tmem221 | 4.789566606 | 2.924766787 | 0.902500427 | 3.240737287 |

| | | | | |
|---|---|---|---|---|
| Nxnl1 | 1.098494234 | 0.48509436 | 1.00418003 | 0.483075091 |
| Slc27a1 | 744.5497192 | -0.056970037 | 0.11121827 | -0.512236315 |
| Pgls | 165.2612445 | 0.045706568 | 0.111569487 | 0.40966907 |
| Fam129c | 18.63060221 | -0.874352491 | 0.295750779 | -2.956382713 |
| Colgalt1 | 425.2055677 | -0.410443299 | 0.108127748 | -3.795910927 |
| Unc13a | 2849.774368 | -0.617194276 | 0.134500906 | -4.588774133 |
| Mir6769b | 0.250814404 | -1.030920997 | 1.784277208 | -0.577780735 |
| Jak3 | 179.3964878 | 2.046693327 | 0.12799578 | 15.99031876 |
| Insl3 | 0.723819272 | 1.85068607 | 1.291867025 | 1.432567001 |
| B3gnt3 | 7.857962099 | 0.773010162 | 0.484152925 | 1.596623963 |
| Fcho1 | 729.7161296 | -0.353120006 | 0.100280336 | -3.521328503 |
| Zfp709 | 191.7633267 | -0.129992058 | 0.143963617 | -0.902950769 |
| Zfp882 | 121.9521505 | -0.641423239 | 0.152769039 | -4.198646826 |
| Zfp617 | 199.055627 | -0.031844381 | 0.125779354 | -0.253176538 |
| Ccnb2-ps | 2.364122105 | -1.024516748 | 0.871851917 | -1.175104084 |
| Zfp961 | 151.8196963 | -0.276153068 | 0.144049725 | -1.917067653 |
| Cyp4f18 | 43.06846795 | 5.435762736 | 0.892028394 | 6.093710437 |
| Gm25052 | 0 | NA | NA | NA |
| Olfr372 | 0 | NA | NA | NA |
| Olfr373 | 0 | NA | NA | NA |
| Olfr374 | 0 | NA | NA | NA |
| Gm24747 | 0 | NA | NA | NA |
| Tpm4 | 485.8967539 | 1.009773651 | 0.177098372 | 5.701767002 |
| Gm26586 | 0.363579626 | -0.809082327 | 1.701614559 | -0.475479199 |
| Gm16091 | 1.601615687 | -1.836486092 | 0.92359218 | -1.988416675 |
| Rab8a | 470.4426857 | 0.420263961 | 0.085523898 | 4.913994449 |
| Hsh2d | 15.58486664 | 5.598480298 | 0.881422744 | 6.351640389 |
| Cib3 | 0.321217965 | 1.870116373 | 3.708947011 | 0.504217603 |
| Gm11034 | 0 | NA | NA | NA |
| Fam32a | 524.8981796 | 0.244760467 | 0.119752244 | 2.043890447 |
| Ap1m1 | 552.8521804 | -0.040460191 | 0.058687944 | -0.689412305 |
| Gm10282 | 57.72901061 | 0.329906647 | 0.162258813 | 2.033212502 |
| Klf2 | 122.111689 | 1.842661287 | 0.215028092 | 8.569397934 |
| Eps15l1 | 756.2958603 | 0.046406227 | 0.079662902 | 0.582532472 |
| Calr3 | 22.05166792 | 0.474178529 | 0.256353316 | 1.849707021 |
| Gm3835 | 0.038981395 | 0.274393828 | 3.724945049 | 0.073663859 |
| 1700030K09Rik | 65.49681279 | -0.145226748 | 0.1677573 | -0.865695548 |
| Cherp | 754.5952941 | 0.215925836 | 0.070910957 | 3.045027811 |
| Mir7068 | 0 | NA | NA | NA |
| Slc35e1 | 803.4949084 | -0.016057016 | 0.065362247 | -0.245661937 |
| Gm17435 | 24.35316493 | 0.808335309 | 0.287449696 | 2.812093104 |
| Med26 | 154.280037 | 0.195855239 | 0.124511756 | 1.572985928 |
| Gm28019 | 1.785542434 | 0.457541902 | 0.692547255 | 0.660665245 |
| Smim7 | 545.215953 | -0.060370566 | 0.106423782 | -0.567265745 |
| Tmem38a | 713.1160059 | -0.150073853 | 0.082787486 | -1.81276013 |
| Nwd1 | 810.0404047 | -0.595695757 | 0.192158755 | -3.100018822 |
| Gm26237 | 0.846376972 | -1.545392184 | 1.151746367 | -1.341781688 |

| | | | | |
|---|---|---|---|---|
| Sin3b | 1202.221482 | -0.046516323 | 0.089371146 | -0.520484796 |
| F2rl3 | 3.0074759 | -0.253453847 | 0.680577916 | -0.372409743 |
| Large | 1767.595343 | 0.106001432 | 0.07335633 | 1.445020936 |
| Mir28b | 0 | NA | NA | NA |
| Gm22532 | 8.167798302 | 0.974536637 | 0.414128334 | 2.353223762 |
| Gm23255 | 0 | NA | NA | NA |
| Crry-ps | 1.276549371 | 0.371588044 | 0.99864228 | 0.372093242 |
| Gm27864 | 0 | NA | NA | NA |
| Gm11033 | 0 | NA | NA | NA |
| Isx | 0.612261413 | -0.153149716 | 1.564666409 | -0.097880108 |
| Hmgxb4 | 272.0156998 | -0.223589256 | 0.133112383 | -1.679702901 |
| Tom1 | 187.7576606 | 0.069089322 | 0.09835892 | 0.7024205 |
| Hmox1 | 1627.230529 | 3.557215074 | 0.4571007 | 7.782125634 |
| Mcm5 | 102.0398356 | 0.936824078 | 0.216024086 | 4.336664931 |
| Rasd2 | 1068.740531 | 0.099741841 | 0.144967459 | 0.688029172 |
| Gm7984 | 22.42806742 | 0.35779446 | 0.26645889 | 1.342775465 |
| Gm10358 | 5.720807359 | -0.206279512 | 0.443467556 | -0.465151305 |
| 1700007B14Rik | 4.206508402 | -0.149102293 | 0.564253974 | -0.264246776 |
| Gm38270 | 0 | NA | NA | NA |
| Gm7901 | 51.21551474 | -0.148521498 | 0.260008684 | -0.571217454 |
| Gm22544 | 0 | NA | NA | NA |
| Gm22509 | 0 | NA | NA | NA |
| Gm7069 | 0.097139547 | 0.274393828 | 3.724945049 | 0.073663859 |
| Gm10649 | 41.3073234 | -0.34883224 | 0.208313658 | -1.674552903 |
| Gm27355 | 0 | NA | NA | NA |
| Gm5353 | 7.21028118 | -0.48055281 | 0.464150899 | -1.035337454 |
| Nr3c2 | 828.7814193 | 0.088440602 | 0.063033974 | 1.403062451 |
| Gm24838 | 0 | NA | NA | NA |
| Gm23260 | 3.058609062 | -0.119519572 | 0.536279123 | -0.222868217 |
| Arhgap10 | 375.9267285 | -0.02803487 | 0.075053795 | -0.373530346 |
| 0610038B21Rik | 3.891239497 | -0.252378416 | 0.56387629 | -0.447577634 |
| Prmt10 | 268.6822526 | 0.104585034 | 0.077851333 | 1.343394265 |
| Gm23468 | 0 | NA | NA | NA |
| Tmem184c | 547.9950958 | -0.415268761 | 0.058383504 | -7.112775507 |
| 4933431K23Rik | 0.309777098 | -0.200474303 | 1.61058947 | -0.124472628 |
| 1700092C02Rik | 0 | NA | NA | NA |
| Ednra | 46.53120417 | -0.262718639 | 0.168663399 | -1.557650571 |
| Gm23981 | 0 | NA | NA | NA |
| Ttc29 | 3.962620848 | -0.249528017 | 0.582134611 | -0.428643156 |
| Pou4f2 | 9.952480325 | 0.08109542 | 0.345940727 | 0.234419984 |
| Rbmxl1 | 229.4117921 | 0.067916842 | 0.091627448 | 0.741228123 |
| Slc10a7 | 127.4608844 | 0.100276644 | 0.126891627 | 0.790254225 |
| Lsm6 | 350.1356414 | 0.198336735 | 0.093826398 | 2.113869215 |
| Gm8054 | 0.763271679 | 1.928924234 | 1.186679498 | 1.625480375 |
| Gm22044 | 0 | NA | NA | NA |
| Zfp827 | 1018.675046 | -0.156454326 | 0.067926756 | -2.303279811 |
| 1700011L22Rik | 0.586370721 | 0.60492827 | 1.155521617 | 0.523510994 |

| | | | | |
|---|---|---|---|---|
| Mmaa | 175.5439761 | -0.35016768 | 0.11182929 | -3.13126982 |
| Gm4890 | 7.177265976 | 0.271977144 | 0.436115967 | 0.623634915 |
| Smad1 | 552.5127933 | 0.345143837 | 0.099153689 | 3.480897576 |
| Gm8079 | 0.101408268 | 0.755286617 | 3.724945049 | 0.202764499 |
| Otud4 | 1259.849304 | 0.00726624 | 0.085060622 | 0.085424254 |
| Tpd52-ps | 12.01690052 | 0.297966056 | 0.319947181 | 0.931297645 |
| Abce1 | 1227.066757 | 0.083230263 | 0.078551751 | 1.059559613 |
| Gm27006 | 0.04393175 | -0.206490061 | 3.724945049 | -0.055434391 |
| Anapc10 | 137.3995736 | -0.22542117 | 0.111512426 | -2.021489235 |
| Hhip | 182.8551312 | 0.266614199 | 0.109262695 | 2.440121024 |
| Gm27820 | 0 | NA | NA | NA |
| Gm5909 | 4.376335103 | 0.079162524 | 0.470535684 | 0.168239152 |
| Gypa | 2.705694399 | 1.88859127 | 0.810274296 | 2.330804863 |
| 4930505O20Rik | 0.533083002 | -0.141851062 | 1.390155149 | -0.102039734 |
| Frem3 | 59.61918439 | 1.158135315 | 0.264605053 | 4.37684504 |
| Smarca5 | 1563.50047 | 0.203081312 | 0.07472835 | 2.717593938 |
| Gab1 | 641.4208176 | -0.153629586 | 0.101273253 | -1.516980851 |
| Gm27048 | 0 | NA | NA | NA |
| Gm4891 | 0.083526043 | 0.75528439 | 3.724945049 | 0.202763901 |
| Usp38 | 593.7032144 | 0.322725749 | 0.09427759 | 3.423143814 |
| Gm9725 | 0.99585445 | 0.5869904 | 1.16136475 | 0.505431563 |
| Gm4899 | 0.173022395 | 1.236193799 | 2.735311368 | 0.451938969 |
| Gm5354 | 0 | NA | NA | NA |
| Inpp4b | 528.8835549 | 0.415133363 | 0.085086684 | 4.878946296 |
| 4930579O11Rik | 0 | NA | NA | NA |
| Gm17169 | 0 | NA | NA | NA |
| Gm17072 | 10.76767044 | 0.774291235 | 0.351514891 | 2.202726698 |
| Gm25267 | 0 | NA | NA | NA |
| Il15 | 30.29114704 | 2.111205983 | 0.352293963 | 5.992739599 |
| Gm25138 | 0 | NA | NA | NA |
| Gm8167 | 0 | NA | NA | NA |
| Zfp330 | 179.4385362 | 0.136711527 | 0.118725809 | 1.151489539 |
| Rnf150 | 857.5833068 | -0.315015483 | 0.121584487 | -2.590918375 |
| Gm10645 | 5.503045566 | -0.667061211 | 0.439706663 | -1.517059594 |
| Tbc1d9 | 1881.448684 | 0.131370118 | 0.073182487 | 1.79510321 |
| Ucp1 | 2.960476708 | 2.002428703 | 0.759030657 | 2.638139427 |
| Elmod2 | 319.9756207 | -0.234607355 | 0.075043749 | -3.126274471 |
| Mgat4d | 0.674779921 | 1.45847527 | 1.386541698 | 1.051879847 |
| Gm15660 | 0 | NA | NA | NA |
| Clgn | 175.9442896 | -0.420983015 | 0.118844362 | -3.542305304 |
| Scoc | 643.4231515 | -0.127452875 | 0.100215834 | -1.271783812 |
| Olfr370 | 0.049553047 | 0.274391604 | 3.724945049 | 0.073663262 |
| Ndufb7 | 290.6417338 | 0.210895182 | 0.207304994 | 1.017318385 |
| Tecr | 2387.710836 | -0.136661001 | 0.066997561 | -2.039790684 |
| Dnajb1 | 549.0730292 | -0.091640906 | 0.173066085 | -0.529513948 |
| Gipc1 | 557.0990177 | 0.154374015 | 0.065234989 | 2.366429698 |
| Ptger1 | 17.78461194 | 0.461329853 | 0.327192349 | 1.40996528 |

| | | | | |
|---|---|---|---|---|
| Pkn1 | 501.3936003 | 0.225407897 | 0.097791952 | 2.304973902 |
| Ddx39 | 165.8939195 | 0.734288512 | 0.147100583 | 4.991744413 |
| Adgre5 | 155.9202038 | 1.19232953 | 0.142867514 | 8.345700857 |
| Mir1668 | 0 | NA | NA | NA |
| Gm22024 | 0 | NA | NA | NA |
| Gm26721 | 9.148635249 | -0.765179346 | 0.440700084 | -1.73628137 |
| Adgrl1 | 7570.249026 | -0.383468536 | 0.102498047 | -3.74122774 |
| AC156028.1 | 0.257982025 | 0.288963106 | 1.834375612 | 0.157526683 |
| AC156028.2 | 0.523560496 | -2.261940331 | 1.348929092 | -1.676841536 |
| Gm10644 | 10.70655375 | 0.228827373 | 0.435352304 | 0.525614246 |
| Asf1b | 15.09267688 | 1.732003187 | 0.405532231 | 4.270938417 |
| Prkaca | 1912.644388 | -0.260083558 | 0.051574949 | -5.04282724 |
| Samd1 | 227.0946354 | -0.346406511 | 0.104674108 | -3.309381077 |
| 1700067K01Rik | 1.769403265 | -1.583710226 | 0.796912362 | -1.987307892 |
| Mir8111 | 0 | NA | NA | NA |
| 2210011C24Rik | 4.04060031 | -0.677865626 | 0.48613359 | -1.394401949 |
| Mir1199 | 0 | NA | NA | NA |
| Palm3 | 9.849211078 | -0.920992736 | 0.364981745 | -2.523393973 |
| Il27ra | 17.46056049 | 2.511957917 | 0.382152812 | 6.573176581 |
| Rln3 | 9.200686208 | -0.310181643 | 0.868664576 | -0.357078729 |
| Gm10643 | 25.86145265 | 0.186036243 | 0.220882772 | 0.842239713 |
| C330011M18Rik | 25.69760837 | -0.112205334 | 0.198432359 | -0.565458853 |
| Rfx1 | 271.1011036 | -0.335911472 | 0.12016016 | -2.795531162 |
| Mir709 | 0.78198097 | 0.141681916 | 1.224834862 | 0.1156743 |
| Dcaf15 | 175.8113705 | 0.178566279 | 0.119583955 | 1.493229418 |
| Podnl1 | 7.966600571 | 2.598041395 | 0.567875141 | 4.575022234 |
| Cc2d1a | 577.8296229 | 0.047478515 | 0.049255291 | 0.96392722 |
| 4930432K21Rik | 28.89408785 | 0.272935894 | 0.198494196 | 1.375032114 |
| Gm25110 | 0 | NA | NA | NA |
| Nanos3 | 8.912972197 | 0.064350688 | 0.349709126 | 0.184012036 |
| Mir181d | 2.185676227 | -0.777890871 | 0.614102719 | -1.266711328 |
| Mir181c | 0.986137052 | -0.479589607 | 0.964884166 | -0.497043712 |
| Gm26887 | 0.282313603 | -0.207975173 | 1.794502079 | -0.115895755 |
| Gm37352 | 38.80286291 | -0.202394883 | 0.228727261 | -0.884874335 |
| Mir23a | 1.248589509 | 0.9296904 | 1.227189456 | 0.757576913 |
| Mir27a | 0.444870878 | -0.206484152 | 1.495043404 | -0.13811248 |
| Mir24-2 | 0 | NA | NA | NA |
| mmu-mir-3074-2 | 0.551097043 | -1.296099407 | 1.597822695 | -0.811165977 |
| Gm26532 | 9.112776141 | -0.275853108 | 0.426055898 | -0.64745755 |
| Zswim4 | 345.6336222 | 0.193331272 | 0.154513773 | 1.251223555 |
| D8Ertd738e | 305.0049009 | 0.152174914 | 0.123362807 | 1.233555867 |
| Mri1 | 76.67094331 | 0.034620365 | 0.147499545 | 0.234715064 |
| Ccdc130 | 95.20490353 | -0.002991348 | 0.127133028 | -0.023529274 |
| Gm23626 | 0 | NA | NA | NA |
| Cacna1a | 3274.65666 | -0.363095457 | 0.126244259 | -2.876134404 |
| Gm16183 | 0.17506888 | 0.776661459 | 2.727910138 | 0.284709327 |
| Ier2 | 148.7573661 | 0.567350281 | 0.27660463 | 2.051123585 |

| | | | | |
|---|---|---|---|---|
| Gm26664 | 2.727852729 | -1.905022707 | 0.672440879 | -2.832996575 |
| Nacc1 | 1589.037921 | 0.205150769 | 0.062117547 | 3.30262186 |
| Trmt1 | 439.8140685 | 0.095722109 | 0.094054985 | 1.017724995 |
| Nfix | 4316.174095 | 0.226064685 | 0.089910119 | 2.51434084 |
| Lyl1 | 28.85883109 | 1.019401526 | 0.321643152 | 3.169355605 |
| G430095P16Rik | 2.042153182 | 1.524596345 | 0.872079495 | 1.748230928 |
| Dand5 | 53.8269615 | -0.302429031 | 0.144308369 | -2.095713741 |
| Gadd45gip1 | 231.7276792 | -0.066647369 | 0.108294788 | -0.615425452 |
| Rad23a | 572.6689497 | -0.044881548 | 0.061474463 | -0.730084426 |
| Calr | 4607.859221 | 0.324016889 | 0.108142624 | 2.996199608 |
| 1700122E12Rik | 0.913679305 | -2.15655891 | 1.341976093 | -1.60700248 |
| Farsa | 355.8983591 | 0.434360731 | 0.107767568 | 4.030532911 |
| Mir7069 | 0 | NA | NA | NA |
| Syce2 | 15.66997929 | 0.369065799 | 0.303940369 | 1.21427042 |
| Gcdh | 224.1995156 | -0.123877591 | 0.080926125 | -1.530749084 |
| Klf1 | 0.981329651 | 0.589398421 | 1.062624654 | 0.554662852 |
| Dnase2a | 91.10452649 | 0.49119684 | 0.183890858 | 2.671132459 |
| Mast1 | 1321.949102 | -0.288557628 | 0.08070129 | -3.575625969 |
| Gm24197 | 0 | NA | NA | NA |
| Rtbdn | 11.67166441 | -0.515933826 | 0.321839323 | -1.603078895 |
| Rnaseh2a | 132.5774289 | -0.187322279 | 0.093936407 | -1.994139286 |
| Prdx2 | 1409.970294 | -0.005648401 | 0.073454114 | -0.076897002 |
| Junb | 1137.476461 | 0.872025109 | 0.331710887 | 2.628870935 |
| Hook2 | 101.5563122 | 0.346036966 | 0.147310404 | 2.349032764 |
| Best2 | 0.602809315 | 1.151744264 | 1.293272366 | 0.890565897 |
| Asna1 | 647.8422268 | 0.106154966 | 0.081871344 | 1.296607095 |
| 2310036O22Rik | 524.574075 | 0.128216695 | 0.106837931 | 1.20010462 |
| Tnpo2 | 1895.024094 | -0.165901014 | 0.053712328 | -3.088695278 |
| Gm25506 | 0.455189762 | -0.74418196 | 1.323329326 | -0.562355829 |
| A230103J11Rik | 33.92778836 | -0.174953453 | 0.193675072 | -0.903334906 |
| Fbxw9 | 120.3613756 | 0.05226397 | 0.11060749 | 0.472517457 |
| Mir7070 | 0 | NA | NA | NA |
| Gm5741 | 3.225606799 | 0.579098745 | 0.561087764 | 1.032100114 |
| Dhps | 207.0919154 | 0.147576877 | 0.095777923 | 1.540823521 |
| Wdr83 | 132.9258923 | -0.022511071 | 0.108547536 | -0.20738445 |
| Wdr83os | 100.509984 | 0.005568964 | 0.142550032 | 0.039066735 |
| Man2b1 | 495.4554757 | 0.220463947 | 0.143464243 | 1.536717037 |
| Zfp791 | 32.04972151 | 0.139370765 | 0.198703416 | 0.701400949 |
| Gm23388 | 0 | NA | NA | NA |
| Cks1brt | 0 | NA | NA | NA |
| Olfr371 | 0 | NA | NA | NA |
| Vps35 | 2002.604974 | 0.155140998 | 0.087991461 | 1.76313696 |
| Orc6 | 141.8270405 | 0.305924036 | 0.122949339 | 2.488212129 |
| Gm22336 | 0.044544648 | 0.755286614 | 3.724945049 | 0.202764498 |
| Mylk3 | 3.257485277 | -1.0546791 | 0.601709246 | -1.752805209 |
| AA672651 | 0 | NA | NA | NA |
| 4921524J17Rik | 80.96275031 | -0.08498165 | 0.143921039 | -0.590474128 |

| | | | | |
|---|---|---|---|---|
| 1700051O22Rik | 0.111006857 | 0.755286617 | 3.724945049 | 0.202764499 |
| Gpt2 | 699.6609475 | -0.27508296 | 0.150338231 | -1.829760517 |
| Dnaja2 | 1599.493447 | 0.244877538 | 0.078607585 | 3.115189694 |
| Neto2 | 1594.195169 | -0.024719442 | 0.092202576 | -0.268099256 |
| Mir8109 | 0.42925394 | 2.078020933 | 1.683782758 | 1.234138385 |
| Itfg1 | 2429.683998 | -0.007758224 | 0.080957424 | -0.095830915 |
| Phkb | 740.2327041 | -0.114020039 | 0.102791392 | -1.109237233 |
| Gm16167 | 0 | NA | NA | NA |
| Hm629797 | 14.11744488 | -0.412428597 | 0.350988871 | -1.175047504 |
| Gm23904 | 0 | NA | NA | NA |
| Gm24490 | 0.060174537 | 0.755286614 | 3.724945049 | 0.202764498 |
| Abcc12 | 27.6985328 | 0.270522869 | 0.25296058 | 1.069426979 |
| AC132147.1 | 0 | NA | NA | NA |
| Lonp2 | 859.6725007 | 0.037992261 | 0.08045487 | 0.472218289 |
| Gm16177 | 0.509424666 | 0.680240593 | 1.228345013 | 0.553786262 |
| Gm22305 | 0.045882455 | 0.755288839 | 3.724945049 | 0.202765096 |
| Siah1a | 351.8844398 | 0.409306493 | 0.1128309 | 3.627609942 |
| Gm10638 | 6.090970552 | 0.078661047 | 0.409914992 | 0.191895998 |
| N4bp1 | 1537.795251 | 0.615227942 | 0.093858316 | 6.554858105 |
| 2010110E17Rik | 5.305919521 | 2.360943255 | 0.611396294 | 3.861559643 |
| Gm27167 | 3.969393191 | 1.36221976 | 0.573239787 | 2.37635243 |
| Gm27168 | 0.090665643 | -0.687373951 | 3.724945049 | -0.184532642 |
| Gm24781 | 0 | NA | NA | NA |
| Rps13-ps1 | 58.84124857 | 0.561270963 | 0.187803989 | 2.988599804 |
| Gm5118 | 1.461301642 | 0.903432419 | 0.818471915 | 1.103803811 |
| Gm27246 | 0.76513295 | 0.015668176 | 1.137377613 | 0.013775703 |
| Gm10637 | 0.117650406 | -0.206492286 | 3.724945049 | -0.055434988 |
| Gm27169 | 0.24523014 | 0.755299097 | 2.525599258 | 0.29905738 |
| Gm27207 | 0.273479446 | -0.568869916 | 2.461054195 | -0.231148878 |
| Gm27208 | 0 | NA | NA | NA |
| 4930535O05Rik | 0 | NA | NA | NA |
| Gm19872 | 0 | NA | NA | NA |
| Cbln1 | 1077.308355 | -0.354164707 | 0.238423388 | -1.485444487 |
| Gm2694 | 162.596044 | -0.769293586 | 0.252148511 | -3.050954311 |
| Gm27016 | 0.046345709 | 0.274391604 | 3.724945049 | 0.073663262 |
| 1700120C18Rik | 0 | NA | NA | NA |
| 4933402J07Rik | 1.368974137 | 2.417913339 | 1.054283584 | 2.293418371 |
| Gm24841 | 0.060174537 | 0.755286614 | 3.724945049 | 0.202764498 |
| Zfp423 | 479.0759597 | -0.010419158 | 0.085454783 | -0.121925982 |
| Gm2716 | 0.511053402 | 0.27441385 | 1.450807755 | 0.189145563 |
| Cnep1r1 | 300.1426235 | 0.071756347 | 0.084614421 | 0.848039218 |
| Heatr3 | 504.4058116 | -0.184156827 | 0.061712123 | -2.984127224 |
| Gm9988 | 0.95664278 | -0.475487314 | 0.926152702 | -0.513400558 |
| Mir7071 | 0 | NA | NA | NA |
| 9430002A10Rik | 3.207016025 | -0.826985074 | 0.598057596 | -1.382785002 |
| Papd5 | 693.6320357 | -0.165462195 | 0.072098335 | -2.294951671 |
| Adcy7 | 236.1657784 | 0.536305297 | 0.125727499 | 4.265616508 |

| | | | | |
|---|---|---|---|---|
| Brd7 | 454.0573306 | -0.030692554 | 0.092886183 | -0.330431858 |
| Nkd1 | 108.148034 | -0.461824622 | 0.139047598 | -3.321341961 |
| Snx20 | 40.8375519 | 2.292082983 | 0.288567976 | 7.942956867 |
| Nod2 | 50.86176278 | 2.241079796 | 0.258454201 | 8.671090628 |
| Cyld | 2133.072429 | 0.03311346 | 0.063078401 | 0.52495719 |
| Rps6-ps2 | 0.516489969 | -0.430111681 | 1.407445884 | -0.305597313 |
| Mir8110 | 0 | NA | NA | NA |
| Sall1 | 352.191393 | -0.446897712 | 0.154681528 | -2.889147254 |
| Gm6625 | 0 | NA | NA | NA |
| Gm5356 | 0.530154252 | 0.257013137 | 1.37953302 | 0.186304448 |
| Gm26331 | 1.106943244 | 0.031752458 | 1.008816615 | 0.031474956 |
| Gm24212 | 0 | NA | NA | NA |
| Tox3 | 156.4269144 | 0.133719355 | 0.109464761 | 1.22157445 |
| AC129321.1 | 0 | NA | NA | NA |
| Chd9 | 2318.475297 | -0.319870765 | 0.071631155 | -4.465525739 |
| Gm6658 | 1.730564267 | 0.141064877 | 0.915958425 | 0.154007947 |
| Gm22428 | 0.075043205 | 0.755286614 | 3.724945049 | 0.202764498 |
| Rbl2 | 619.2188564 | -0.020533909 | 0.07382171 | -0.278155423 |
| Aktip | 891.7499773 | -0.089865382 | 0.062326161 | -1.44185653 |
| Gm23408 | 0 | NA | NA | NA |
| Rpgrip1l | 501.0190372 | -0.575480505 | 0.148866496 | -3.865748985 |
| Fto | 1709.590573 | -0.124146447 | 0.047581062 | -2.609156705 |
| Gm3235 | 0.204751453 | 0.274411094 | 2.204478766 | 0.124478901 |
| Irx3 | 54.79817873 | 0.009679793 | 0.229674155 | 0.042145767 |
| Irx3os | 5.187223665 | -0.796575645 | 0.427514174 | -1.863273065 |
| Gm28511 | 0.044544648 | 0.755286614 | 3.724945049 | 0.202764498 |
| Gm28515 | 0.165933562 | 1.204233133 | 2.762106933 | 0.435983531 |
| Gm28516 | 0.048949532 | -0.206487837 | 3.724945049 | -0.055433794 |
| Gm24254 | 0 | NA | NA | NA |
| Crnde | 9.401908757 | 0.776440784 | 0.386616791 | 2.008295557 |
| Gm21817 | 2.549468396 | -0.869922065 | 0.788153545 | -1.103746941 |
| Irx5 | 30.17646328 | 0.210121643 | 0.223592133 | 0.939754188 |
| Gm29177 | 0 | NA | NA | NA |
| Irx6 | 3.738898168 | -0.29500322 | 0.561620609 | -0.525271358 |
| Gm3272 | 14.76791741 | 0.413582509 | 0.303843584 | 1.361169137 |
| Mmp2 | 39.1239749 | 0.40630871 | 0.17899689 | 2.269920498 |
| Lpcat2 | 68.00775929 | 0.262706002 | 0.206139946 | 1.274406088 |
| Capns2 | 4.884769305 | 0.308359084 | 0.551771625 | 0.558852739 |
| Gm22151 | 0 | NA | NA | NA |
| Slc6a2 | 13.59669148 | -0.581542392 | 0.272111937 | -2.137143995 |
| Ces1a | 0.156716517 | -0.68737803 | 2.821984814 | -0.243579635 |
| Ces1b | 0 | NA | NA | NA |
| Ces1c | 0.230645793 | 0.27441552 | 1.885418553 | 0.145546208 |
| Ces1d | 1.445763283 | -1.684040892 | 1.011818629 | -1.664370317 |
| Ces1e | 0 | NA | NA | NA |
| Ces1f | 0.742894727 | -0.81999536 | 1.111676819 | -0.737620274 |
| Ces1g | 0.379165397 | -0.687379409 | 1.572924405 | -0.437007276 |

| | | | | |
|---|---|---|---|---|
| Ces1h | 0 | NA | NA | NA |
| Ces5a | 9.541882804 | 0.723728852 | 0.363315584 | 1.992011581 |
| Gm26843 | 19.49403685 | -0.747306814 | 0.29481551 | -2.534828694 |
| Gnao1 | 5592.419096 | -0.221306216 | 0.063176997 | -3.502955614 |
| Gm24159 | 0.183166439 | 0.755304911 | 2.686322637 | 0.281166864 |
| Amfr | 1992.652925 | 0.249213632 | 0.070294892 | 3.545259473 |
| Nudt21 | 505.4581536 | 0.037142362 | 0.078844595 | 0.471083172 |
| Ogfod1 | 1369.192929 | -0.06148475 | 0.054262044 | -1.133107884 |
| Bbs2 | 229.083479 | -0.300524311 | 0.102935006 | -2.919554029 |
| Mt4 | 0.564343484 | 2.693115611 | 1.479413859 | 1.820393661 |
| Mt3 | 1914.641927 | 0.33115161 | 0.124597757 | 2.657765428 |
| Mt2 | 2941.273043 | 2.475561247 | 0.319381518 | 7.751109905 |
| Mt1 | 3127.581112 | 1.575682153 | 0.231842008 | 6.796361746 |
| Nup93 | 580.310508 | 0.061520294 | 0.129015772 | 0.476843202 |
| Mir138-2 | 5.437333865 | 0.945743028 | 0.561442781 | 1.684486931 |
| Slc12a3 | 11.0278551 | -0.120523363 | 0.440181156 | -0.273804004 |
| Gm15889 | 0.042716945 | -0.206490061 | 3.724945049 | -0.055434391 |
| Gm15890 | 0.82545695 | 1.427882846 | 1.237768099 | 1.153594803 |
| Herpud1 | 583.7203636 | 0.188566785 | 0.095077326 | 1.983299197 |
| Ap3s1-ps2 | 8.423014622 | 0.21907862 | 0.36491857 | 0.600349333 |
| 9330175E14Rik | 54.28032715 | 6.902480795 | 0.745953666 | 9.253229941 |
| Nlrc5 | 1448.823358 | 8.234492125 | 0.252248155 | 32.64440971 |
| Mir7072 | 0.934615587 | 3.130311725 | 1.365326735 | 2.292719864 |
| Cpne2 | 360.9580537 | 0.353072028 | 0.147826773 | 2.38841734 |
| Fam192a | 433.1676574 | 0.326826155 | 0.078747889 | 4.150284635 |
| Rspry1 | 440.5534813 | -0.36065348 | 0.091984198 | -3.920819951 |
| Arl2bp | 939.3975919 | -0.138761507 | 0.07429198 | -1.867785819 |
| Pllp | 348.7112223 | -0.71681951 | 0.173495875 | -4.13162279 |
| Gm22846 | 0 | NA | NA | NA |
| Ccl22 | 3.029817251 | -0.130245327 | 0.578304475 | -0.225219296 |
| Cx3cl1 | 4526.862607 | -0.152433977 | 0.094212493 | -1.6179805 |
| Ccl17 | 8.845676815 | 0.838113847 | 0.430265024 | 1.94790141 |
| Ciapin1 | 565.2127423 | -0.135771905 | 0.084569436 | -1.605448862 |
| Gm7418 | 0.075043205 | 0.755286614 | 3.724945049 | 0.202764498 |
| Coq9 | 367.4866646 | -0.211508041 | 0.073985756 | -2.858767029 |
| Polr2c | 334.1966378 | 0.399471974 | 0.10687462 | 3.737762772 |
| Dok4 | 119.6154433 | -0.591585682 | 0.128357236 | -4.608900142 |
| Ccdc102a | 31.06208967 | 1.321281122 | 0.29960614 | 4.410060234 |
| Adgrg5 | 33.49054987 | 4.366223097 | 0.417117482 | 10.46760992 |
| Gm10286 | 0.154272232 | 1.204026025 | 2.826489749 | 0.425979265 |
| Adgrg1 | 915.8714062 | 0.088442618 | 0.137730439 | 0.642142858 |
| Adgrg3 | 4.584657955 | 2.202674562 | 0.68806427 | 3.201262814 |
| Drc7 | 36.74700655 | 0.361318795 | 0.225579033 | 1.601739266 |
| Katnb1 | 374.5637383 | -0.009375341 | 0.084974912 | -0.110330695 |
| Kifc3 | 519.7752177 | -0.309964666 | 0.093662056 | -3.309394206 |
| Cngb1 | 16.48700249 | -0.15525673 | 0.289052844 | -0.537122306 |
| Tepp | 2.391530947 | 0.133192748 | 0.6773372 | 0.196641714 |

| | | | | |
|---|---|---|---|---|
| Zfp319 | 384.2111316 | 0.050606413 | 0.083968992 | 0.602679769 |
| Usb1 | 184.5297308 | 0.853462892 | 0.151766258 | 5.62353518 |
| Mmp15 | 359.2212613 | -0.542726778 | 0.111196543 | -4.88078824 |
| Cfap20 | 234.0850033 | -0.145217294 | 0.087271003 | -1.663981025 |
| Csnk2a2 | 377.4618607 | 0.173146336 | 0.101652424 | 1.703317335 |
| Ccdc113 | 14.24627225 | 0.383058111 | 0.287706909 | 1.331417839 |
| Prss54 | 0.189486273 | 0.274408531 | 2.657487022 | 0.103258653 |
| Gm5912 | 6.84788527 | 0.039515265 | 0.405038043 | 0.097559392 |
| Gins3 | 30.02162402 | -0.004933807 | 0.235186074 | -0.020978311 |
| Ndrg4 | 14045.15733 | -0.29763629 | 0.062184492 | -4.786342704 |
| Setd6 | 77.58215835 | -0.17125761 | 0.163815452 | -1.045430138 |
| Cnot1 | 2334.865287 | 0.029687294 | 0.045371177 | 0.654320566 |
| Gm26493 | 9.543263168 | -0.307429842 | 0.370840451 | -0.829008381 |
| mmu-mir-7073 | 0.68581662 | -2.164056651 | 1.165067874 | -1.857451141 |
| SNORA76 | 0 | NA | NA | NA |
| Gm26265 | 0 | NA | NA | NA |
| 4930513N10Rik | 11.81172818 | 0.267435408 | 0.319780213 | 0.836310058 |
| Gm10094 | 48.65651831 | 0.747408991 | 0.239391544 | 3.122119428 |
| Slc38a7 | 365.4018507 | 0.061350649 | 0.07510113 | 0.816907141 |
| Got2 | 2359.910276 | 0.161186853 | 0.052759374 | 3.055132011 |
| Gm22780 | 0 | NA | NA | NA |
| Gm24132 | 0 | NA | NA | NA |
| Gm25160 | 0 | NA | NA | NA |
| Gm26399 | 0.072025462 | 0.755288839 | 3.724945049 | 0.202765096 |
| Gm5913 | 8.128038844 | 0.493429555 | 0.371956614 | 1.326578253 |
| Gm7191 | 0 | NA | NA | NA |
| Gm23494 | 0 | NA | NA | NA |
| Gm15679 | 0.097899064 | -0.687371727 | 3.724945049 | -0.184532045 |
| Cdh8 | 694.165117 | -0.313290123 | 0.095346378 | -3.28581043 |
| Gm24693 | 0.415711858 | 0.300583249 | 1.589101878 | 0.189152913 |
| Gm15681 | 35.39573601 | -0.731436636 | 0.319037046 | -2.292638562 |
| Gm15680 | 7.35783165 | -0.999500934 | 0.420213191 | -2.37855678 |
| A330008L17Rik | 88.53438246 | -0.46036808 | 0.216123279 | -2.13011797 |
| Gm8688 | 2.373068043 | 1.097241067 | 0.680252812 | 1.612990122 |
| Gm36942 | 0 | NA | NA | NA |
| Gm15210 | 118.5129343 | 0.131987221 | 0.164341018 | 0.803130117 |
| Gm5742 | 0.038981395 | 0.274393828 | 3.724945049 | 0.073663859 |
| Gm22223 | 0 | NA | NA | NA |
| Gm24602 | 0 | NA | NA | NA |
| Cdh11 | 712.29886 | -0.499867621 | 0.085899122 | -5.819240173 |
| Gm26301 | 4.260983332 | -0.647936238 | 0.564090823 | -1.148638148 |
| Gm8730 | 715.9366467 | 0.730187817 | 0.129374165 | 5.644000241 |
| Cdh5 | 215.0038206 | 1.917159918 | 0.162455055 | 11.8011712 |
| Bean1 | 97.56300219 | -0.623159945 | 0.113756342 | -5.478023783 |
| Gm10631 | 0.136963818 | 0.755286583 | 3.59954433 | 0.209828388 |
| Tk2 | 352.362705 | 0.36743168 | 0.081496253 | 4.508571463 |
| Cklf | 22.85197058 | -0.175753508 | 0.256521202 | -0.685142229 |

| | | | | |
|---|---|---|---|---|
| Cmtm2a | 0.047948698 | -0.206487837 | 3.724945049 | -0.055433794 |
| Cmtm1 | 0.461873605 | -0.659116081 | 1.573679518 | -0.418837554 |
| Gm11020 | 0 | NA | NA | NA |
| Cmtm2b | 0.088534442 | 0.274389469 | 3.724945049 | 0.073662689 |
| Cmtm3 | 162.243504 | 0.561993956 | 0.186693344 | 3.010251702 |
| Cmtm4 | 1562.719887 | -0.42840519 | 0.076795658 | -5.578508001 |
| Dync1li2 | 3022.19108 | -0.174569035 | 0.050712024 | -3.442359827 |
| Ccdc79 | 18.23009573 | -1.042976705 | 0.284866957 | -3.661276537 |
| Gm22148 | 0 | NA | NA | NA |
| Nae1 | 544.9477949 | -0.04007737 | 0.076586086 | -0.523298313 |
| Car7 | 137.94085 | -0.633182727 | 0.12509235 | -5.061722207 |
| Pdp2 | 131.2365206 | -0.118377121 | 0.113548521 | -1.042524553 |
| Cdh16 | 0 | NA | NA | NA |
| Rrad | 32.51512905 | 0.554966951 | 0.308480879 | 1.799031933 |
| Fam96b | 134.4805842 | 0.028541515 | 0.142170758 | 0.200755171 |
| 1700082M22Rik | 0.890885745 | 0.856300745 | 0.941871186 | 0.909148468 |
| AC156564.1 | 0 | NA | NA | NA |
| Ces2a | 0.086648695 | -0.687376175 | 3.724945049 | -0.184533239 |
| 4932416K20Rik | 0 | NA | NA | NA |
| Ces2b | 0.087930927 | -0.206490058 | 3.724945049 | -0.05543439 |
| Matr3-ps2 | 0 | NA | NA | NA |
| Ces2c | 0 | NA | NA | NA |
| Ces2d-ps | 0 | NA | NA | NA |
| Ces2e | 0.208319007 | 0.274411174 | 2.193432295 | 0.125105833 |
| Ces2f | 0 | NA | NA | NA |
| Ces2g | 3.472075296 | 0.363490335 | 0.539607652 | 0.673619682 |
| Ces2h | 0 | NA | NA | NA |
| Ces3a | 0.090890357 | 0.755282166 | 3.724945049 | 0.202763304 |
| Ces3b | 0.093831153 | 0.274404949 | 3.724945049 | 0.073666845 |
| Ces4a | 0.088534442 | 0.274389469 | 3.724945049 | 0.073662689 |
| Gm3830 | 0 | NA | NA | NA |
| Cbfb | 387.4469817 | 0.358887634 | 0.133558752 | 2.68711431 |
| Gm22063 | 3.344407031 | 0.259376932 | 0.516852418 | 0.501839448 |
| Gm24629 | 1.944273076 | 0.516010938 | 0.763073307 | 0.676227216 |
| D230025D16Rik | 379.8537354 | -0.111619237 | 0.085016685 | -1.31290977 |
| B3gnt9 | 40.27416652 | 0.085178584 | 0.183775146 | 0.463493493 |
| Tradd | 49.19104427 | -0.040905767 | 0.220345547 | -0.185643722 |
| Fbxl8 | 12.45441639 | -0.403103602 | 0.372773041 | -1.081364685 |
| Hsf4 | 105.9523221 | -0.098631043 | 0.133774403 | -0.737293834 |
| Nol3 | 174.8745832 | -0.134731816 | 0.11882732 | -1.13384545 |
| 4931428F04Rik | 216.122387 | -0.587937906 | 0.101334865 | -5.80193113 |
| Exoc3l | 19.59593397 | 0.210592 | 0.259189651 | 0.812501576 |
| E2f4 | 317.7432688 | 0.168808493 | 0.075114028 | 2.247363062 |
| Elmo3 | 30.0446705 | 0.465574352 | 0.202855018 | 2.295108874 |
| Gm27320 | 4.749146687 | 0.148651102 | 0.589565643 | 0.252136643 |
| Mir328 | 3.399306995 | 0.133989778 | 0.630640863 | 0.212466058 |
| Lrrc29 | 48.99782354 | 0.161115223 | 0.230107023 | 0.700175168 |

| | | | | |
|---|---|---|---|---|
| Tmem208 | 199.915556 | 0.084097733 | 0.112292297 | 0.748918094 |
| Fhod1 | 83.53890257 | 0.974338165 | 0.221123813 | 4.406301386 |
| Slc9a5 | 310.0134594 | -0.245532414 | 0.152849815 | -1.606363825 |
| Plekhg4 | 37.98167613 | -0.456033002 | 0.364022605 | -1.252760119 |
| Kctd19 | 6.042255962 | 4.974384932 | 0.922684878 | 5.391206736 |
| Lrrc36 | 11.57992328 | 0.102556392 | 0.29136928 | 0.351980799 |
| Tppp3 | 487.5931413 | 0.292309951 | 0.157906958 | 1.851153079 |
| Zdhhc1 | 228.3370938 | -0.190701978 | 0.094830663 | -2.010973785 |
| Hsd11b2 | 0.385063269 | -1.353685643 | 1.402233338 | -0.965378305 |
| Atp6v0d1 | 1618.375995 | -0.107189336 | 0.049079441 | -2.183996667 |
| Gm38250 | 12.77996492 | -0.553334029 | 0.366976677 | -1.50781797 |
| Agrp | 1.967041363 | -0.862885427 | 0.927382312 | -0.930452755 |
| Fam65a | 1008.594529 | -0.045183897 | 0.09473218 | -0.476964611 |
| Mir1966 | 0 NA | NA | NA | |
| Gm5914 | 15.31483393 | 0.0143263 | 0.303731593 | 0.047167633 |
| Gm38384 | 0 NA | NA | NA | |
| Ctcf | 822.3735919 | -0.022070497 | 0.066801768 | -0.330387911 |
| Gm5915 | 1.873087375 | -0.850619132 | 0.802072549 | -1.060526424 |
| Rltpr | 568.0764324 | -0.002577766 | 0.084263211 | -0.030591834 |
| Acd | 255.1203144 | 0.032097452 | 0.084681146 | 0.379038948 |
| Pard6a | 178.3545304 | -0.230320988 | 0.133805582 | -1.72131076 |
| Enkd1 | 47.5336699 | 0.002647897 | 0.173065376 | 0.015299981 |
| 4933405L10Rik | 1.137565759 | -1.349428203 | 0.997202308 | -1.353214079 |
| Gfod2 | 210.1736066 | -0.072041693 | 0.099607503 | -0.723255685 |
| Ranbp10 | 548.0493242 | -0.190482468 | 0.063758567 | -2.987558798 |
| Tsnaxip1 | 3.0628478 | -0.591895972 | 0.523511148 | -1.130627253 |
| Cenpt | 114.9710339 | -0.070426811 | 0.119428501 | -0.589698522 |
| Thap11 | 254.668566 | -0.038288402 | 0.08321634 | -0.46010678 |
| Nutf2 | 250.6042831 | 0.052080982 | 0.109681747 | 0.474837275 |
| Edc4 | 794.3036961 | -0.029200872 | 0.106110942 | -0.275191905 |
| Nrn1l | 0.575698224 | -0.426330628 | 1.315584775 | -0.324061692 |
| Pskh1 | 139.9944564 | 0.28436318 | 0.107838144 | 2.63694431 |
| Ctrl | 5.684503014 | 0.336914078 | 0.473303897 | 0.711834575 |
| Gm16156 | 0.885244197 | -0.252565367 | 1.095725879 | -0.230500503 |
| Psmb10 | 1108.11009 | 3.287474967 | 0.185158123 | 17.75495943 |
| Lcat | 338.7334527 | 0.619912826 | 0.181481044 | 3.41585443 |
| Slc12a4 | 501.7697974 | 0.494012593 | 0.119285117 | 4.141443663 |
| Mir7074 | 0.042716945 | -0.206490061 | 3.724945049 | -0.055434391 |
| Dpep3 | 0.127971488 | 0.274389832 | 3.687653711 | 0.0744077 |
| Dpep2 | 27.40815167 | 5.312485207 | 0.924636025 | 5.745488024 |
| Gm22972 | 0 NA | NA | NA | |
| Dus2 | 119.8929111 | -0.45025834 | 0.132400809 | -3.400721976 |
| Ddx28 | 135.9796744 | 0.447204378 | 0.124026718 | 3.605710018 |
| Nfatc3 | 441.1062357 | 0.256226622 | 0.076331302 | 3.356769976 |
| Gm27956 | 0 NA | NA | NA | |
| Esrp2 | 1.734636384 | 1.400600141 | 0.86181302 | 1.625178675 |
| 1810019D21Rik | 1.035937776 | 1.729983049 | 1.251158954 | 1.382704447 |

| | | | | |
|---|---|---|---|---|
| Pla2g15 | 252.7616184 | 0.034478068 | 0.114852067 | 0.300195452 |
| Slc7a6 | 332.464804 | -0.080308091 | 0.130778152 | -0.614078803 |
| Slc7a6os | 176.7031225 | 0.362435906 | 0.130106108 | 2.785694773 |
| Prmt7 | 349.6739077 | -0.105556202 | 0.073644748 | -1.433316091 |
| Gm26786 | 0.213709657 | 1.256766036 | 2.560135076 | 0.490898331 |
| Smpd3 | 843.1766415 | -0.121620305 | 0.146486204 | -0.830250913 |
| Gm10629 | 2.127429481 | -1.996793256 | 0.807541005 | -2.472683422 |
| Zfp90 | 174.4767697 | 0.52543941 | 0.094166881 | 5.579874855 |
| 6030452D12Rik | 0.119587853 | 1.218392466 | 3.723618458 | 0.327206581 |
| Cdh3 | 29.60771109 | 0.948302333 | 0.268734957 | 3.528764347 |
| Gm10073 | 77.24332829 | -0.052572441 | 0.20683739 | -0.254172811 |
| 1110028F18Rik | 0.166497232 | 0.27439225 | 3.399989659 | 0.080703848 |
| Cdh1 | 87.82624501 | 0.382442146 | 0.178542669 | 2.142021005 |
| Tango6 | 76.05254567 | -0.243361277 | 0.176582114 | -1.378176259 |
| Has3 | 26.28930354 | -0.814250966 | 0.197172785 | -4.129631608 |
| Chtf8 | 670.8970226 | -0.234580864 | 0.108767595 | -2.156716478 |
| Cirh1a | 288.9954583 | 0.143191978 | 0.084254012 | 1.699527116 |
| Rpl10-ps2 | 1.60060964 | 0.956216189 | 0.979007948 | 0.976719537 |
| Sntb2 | 331.1315353 | 0.269121254 | 0.103471936 | 2.600910588 |
| Gm22085 | 0 | NA | NA | NA |
| Gm16208 | 0.439742618 | -0.11549741 | 1.723090336 | -0.067029225 |
| Vps4a | 870.3055482 | 0.071337142 | 0.053567357 | 1.331727855 |
| Gm16209 | 1.711796903 | 0.404173379 | 0.736555161 | 0.548734705 |
| Pdf | 11.11514947 | 0.535248768 | 0.305262685 | 1.753403853 |
| Cog8 | 288.9214874 | 0.170379504 | 0.097303118 | 1.751017932 |
| Nip7 | 304.7822817 | 0.425956195 | 0.095445141 | 4.462837947 |
| Tmed6 | 0.075043205 | 0.755286614 | 3.724945049 | 0.202764498 |
| Terf2 | 412.9827996 | 0.083696512 | 0.0678102 | 1.234276131 |
| Mir7075 | 0 | NA | NA | NA |
| C630050I24Rik | 0 | NA | NA | NA |
| Cyb5b | 1556.047896 | 0.016728778 | 0.051064315 | 0.327602126 |
| Rps18-ps3 | 31.42787796 | 0.494366645 | 0.212730658 | 2.323908791 |
| Nfat5 | 2785.630128 | -0.02023638 | 0.081797695 | -0.247395482 |
| Nqo1 | 48.27701717 | -0.037030526 | 0.218705938 | -0.16931651 |
| Nob1 | 223.6934552 | -0.016649275 | 0.09825771 | -0.16944497 |
| Wwp2 | 543.3421393 | 0.390913451 | 0.08163678 | 4.78844767 |
| Rps26-ps1 | 30.55227829 | 0.435722808 | 0.20815395 | 2.093271872 |
| Mir140 | 0.882355932 | 1.127271749 | 1.004361412 | 1.122376602 |
| Psmd7 | 734.7468457 | 0.39592337 | 0.095145242 | 4.161252445 |
| Gm23646 | 0.066050875 | 0.274391604 | 3.724945049 | 0.073663262 |
| Gm22825 | 0.0885994 | 0.274402725 | 3.724945049 | 0.073666248 |
| Zfhx3 | 1136.418799 | -0.065552157 | 0.090899818 | -0.721147286 |
| Gm38042 | 32.44911505 | -0.57242478 | 0.347153775 | -1.648908411 |
| Gm38318 | 5.263687666 | -0.627542674 | 0.497575125 | -1.261201861 |
| Gm20686 | 0 | NA | NA | NA |
| Mir3108 | 0.47188914 | 0.302111186 | 1.354892444 | 0.222977984 |
| D030068K23Rik | 6.705867478 | 0.053879683 | 0.379282186 | 0.142056983 |

| | | | | |
|---|---|---|---|---|
| Gm23163 | 0 | NA | NA | NA |
| Pmfbp1 | 9.435034997 | -0.145277536 | 0.457633587 | -0.317453833 |
| Dhx38 | 483.563221 | 0.393611404 | 0.098041249 | 4.014753037 |
| Txnl4b | 126.0385784 | -0.06017118 | 0.107166505 | -0.561473757 |
| Hp | 55.13014879 | 6.62384452 | 1.045901318 | 6.333144826 |
| Dhodh | 150.7597164 | -0.132377135 | 0.115384016 | -1.14727446 |
| Pkd1l3 | 15.56061734 | 0.354435011 | 0.307390853 | 1.153043454 |
| Ist1 | 750.1340784 | 0.090296091 | 0.078246731 | 1.153991869 |
| Zfp821 | 241.0196306 | 0.109340863 | 0.100843942 | 1.084258118 |
| Atxn1l | 914.3276386 | 0.13518875 | 0.088641952 | 1.52511026 |
| Ap1g1 | 1809.349129 | -0.019189116 | 0.047339782 | -0.405348629 |
| Snord71 | 7.711840907 | -0.099437281 | 0.370186894 | -0.268613726 |
| Gm25321 | 0 | NA | NA | NA |
| Gm17344 | 3.933162639 | -2.193862771 | 0.676978077 | -3.240670335 |
| Phlpp2 | 846.0254909 | -0.342577134 | 0.073468412 | -4.662917371 |
| Gm16349 | 3.039183673 | -0.957123129 | 0.710339436 | -1.347416573 |
| Marveld3 | 4.627756683 | 0.314534764 | 0.512191253 | 0.614096319 |
| Tat | 1.816527333 | -0.204474839 | 0.720577079 | -0.283765394 |
| Chst4 | 0.843638514 | -1.413506024 | 1.239137666 | -1.140717502 |
| Zfp612 | 420.3106796 | -0.275362788 | 0.091606035 | -3.00594591 |
| Gm17720 | 0 | NA | NA | NA |
| Gm21964 | 0.038981395 | 0.274393828 | 3.724945049 | 0.073663859 |
| Calb2 | 675.5295577 | -0.339244212 | 0.134809136 | -2.516477897 |
| Cmtr2 | 86.97394878 | -0.005493061 | 0.125630411 | -0.043723973 |
| Gm26832 | 0 | NA | NA | NA |
| Hydin | 49.41383929 | -0.754157813 | 0.204868599 | -3.681178165 |
| Gm23627 | 0 | NA | NA | NA |
| Gm22701 | 0 | NA | NA | NA |
| AC069308.1 | 0 | NA | NA | NA |
| AC069308.2 | 0 | NA | NA | NA |
| Vac14 | 425.0913073 | 0.167876395 | 0.072568613 | 2.31334715 |
| Mtss1l | 3340.251647 | -0.45164727 | 0.106365033 | -4.246200632 |
| Il34 | 286.7182791 | -0.107473932 | 0.110388866 | -0.97359395 |
| Gm15894 | 3.21488869 | 0.570960622 | 0.655168798 | 0.871471023 |
| Gm15895 | 1.066917243 | 1.803238402 | 0.9908748 | 1.819844849 |
| Sf3b3 | 1283.257394 | 0.191672843 | 0.069882003 | 2.742806934 |
| Snord111 | 4.735066032 | 0.903328741 | 0.521174854 | 1.733254653 |
| Gm22193 | 2.867962316 | 0.66010062 | 0.578432599 | 1.141188484 |
| Cog4 | 549.572635 | 0.018944166 | 0.076153735 | 0.248762139 |
| Gm26816 | 0.327007429 | 0.755308741 | 1.984282364 | 0.380645796 |
| Fuk | 171.2700549 | -0.304851457 | 0.130035994 | -2.344362096 |
| St3gal2 | 1026.922896 | -0.265600017 | 0.065014546 | -4.085239916 |
| Gm26132 | 2.688917129 | -1.52749039 | 0.754674984 | -2.024037397 |
| Ddx19a | 222.5239366 | 0.126341868 | 0.09747531 | 1.296142257 |
| Ddx19b | 400.244091 | 0.068200625 | 0.093566483 | 0.728900168 |
| Mir3473d | 0.67306248 | 0.259901308 | 1.126025794 | 0.230812926 |
| Aars | 2524.454962 | 0.365135191 | 0.100987323 | 3.615653732 |

| | | | | |
|---|---|---|---|---|
| Gm24103 | 0 | NA | NA | NA |
| Clec18a | 9.750147804 | -0.248856242 | 0.328500256 | -0.75755266 |
| Pdpr | 569.8637507 | 0.024212195 | 0.061925407 | 0.390989675 |
| 9430091E24Rik | 22.80186376 | -0.870423155 | 0.249454084 | -3.489312107 |
| Glg1 | 3651.900435 | -0.1088341 | 0.060158034 | -1.809136593 |
| Gm23193 | 0 | NA | NA | NA |
| Rfwd3 | 350.9863053 | -0.023101919 | 0.090423554 | -0.255485636 |
| Mlkl | 209.3845513 | 7.203131786 | 0.617631216 | 11.66251251 |
| Fa2h | 504.9529093 | -0.247339207 | 0.168415579 | -1.468624271 |
| Gm24793 | 0 | NA | NA | NA |
| Wdr59 | 307.9043311 | -0.268494173 | 0.071943031 | -3.732038635 |
| Mir1957b | 0.047948698 | -0.206487837 | 3.724945049 | -0.055433794 |
| Znrf1 | 1233.211656 | -0.000885388 | 0.079550013 | -0.011129959 |
| Ldhd | 179.3875016 | -0.122840716 | 0.096372892 | -1.274639715 |
| Zfp1 | 213.1490368 | 0.596363866 | 0.111655202 | 5.341120307 |
| Ctrb1 | 0.649033156 | 1.121714775 | 1.456131543 | 0.770338903 |
| Mir7076 | 0 | NA | NA | NA |
| Bcar1 | 823.9151968 | 0.196329239 | 0.105985882 | 1.852409344 |
| Cfdp1 | 449.2752954 | -0.065486245 | 0.130767447 | -0.500784 |
| Tmem170 | 50.89811066 | -0.640114835 | 0.166196653 | -3.85155069 |
| Gm23853 | 0 | NA | NA | NA |
| Chst5 | 6.122444543 | -0.340385372 | 0.492144329 | -0.691637294 |
| Tmem231 | 128.7512586 | -0.535640927 | 0.108754069 | -4.92524952 |
| Gabarapl2 | 905.821491 | -0.075705938 | 0.081280712 | -0.931413328 |
| Adat1 | 132.6219706 | -1.053955503 | 0.214065003 | -4.923530185 |
| Kars | 1033.948333 | 0.966759149 | 0.120400278 | 8.029542484 |
| Terf2ip | 508.2144975 | 0.243610076 | 0.081279064 | 2.997205745 |
| Gm6793 | 13.31818528 | 0.477340326 | 0.428742447 | 1.11334982 |
| Gm22827 | 0 | NA | NA | NA |
| Gm26994 | 1.945104103 | 0.298848412 | 0.679978954 | 0.43949656 |
| Cntnap4 | 605.8576847 | -0.098648278 | 0.076968936 | -1.281663532 |
| Gm22664 | 1.761579563 | -1.137388074 | 0.878507423 | -1.294682373 |
| Gm25766 | 0 | NA | NA | NA |
| Gm24291 | 0 | NA | NA | NA |
| Mon1b | 489.7840867 | 0.041732913 | 0.073481958 | 0.56793415 |
| Syce1l | 2.596219419 | 1.312406806 | 0.67922929 | 1.932199956 |
| Adamts18 | 110.6812341 | -0.148194018 | 0.204463786 | -0.724793472 |
| Nudt7 | 34.66451103 | -0.319506233 | 0.243458475 | -1.312364392 |
| Vat1l | 1139.282187 | -0.21177018 | 0.109043239 | -1.942075284 |
| 4933408N05Rik | 0.707695957 | 1.014059948 | 1.446106429 | 0.701234659 |
| Clec3a | 0.137470077 | 1.236181898 | 3.595099416 | 0.343851937 |
| Wwox | 259.2877687 | 0.311486413 | 0.071400012 | 4.362554039 |
| Gm16116 | 0 | NA | NA | NA |
| Gm16117 | 0.066050875 | 0.274391604 | 3.724945049 | 0.073663262 |
| Gm16118 | 0.567921522 | 1.700056587 | 1.354429519 | 1.255182764 |
| Gm22556 | 0 | NA | NA | NA |
| Gm23677 | 1.922491501 | -1.000715653 | 0.713719313 | -1.402113736 |

| | | | | |
|---|---|---|---|---|
| Maf | 623.4348012 | 0.0071569 | 0.105862846 | 0.067605402 |
| Gm15655 | 1.118566861 | -0.96657388 | 1.097087113 | -0.881036582 |
| Dynlrb2 | 11.99619067 | 0.242383365 | 0.289966514 | 0.835901229 |
| Cdyl2 | 447.4685583 | -0.186624461 | 0.090135605 | -2.070485479 |
| Gm10620 | 0 | NA | NA | NA |
| Cmc2 | 85.51804082 | -0.62278683 | 0.125740563 | -4.952950865 |
| Cenpn | 11.50529104 | 0.019733389 | 0.390424927 | 0.050543363 |
| Atmin | 1056.148415 | 0.097162522 | 0.076110015 | 1.276606268 |
| 1700030J22Rik | 85.33078503 | -0.672078146 | 0.201086476 | -3.342234441 |
| Gcsh | 322.5962855 | 0.008290732 | 0.112743424 | 0.073536276 |
| Pkd1l2 | 5.568506049 | 1.139015162 | 0.520052537 | 2.190192491 |
| Gm15395 | 0 | NA | NA | NA |
| Bco1 | 1.165159028 | 1.72685226 | 1.05053779 | 1.643779287 |
| Gm20694 | 0 | NA | NA | NA |
| Gan | 644.4361812 | -0.097172311 | 0.08344207 | -1.164548183 |
| Cmip | 3352.599209 | 0.010071025 | 0.074423796 | 0.135319953 |
| Mir7077 | 0.151026272 | 0.741083323 | 2.850563012 | 0.259977878 |
| Mir7078 | 0.769887329 | -0.534870618 | 1.371817846 | -0.389899155 |
| Plcg2 | 185.0113621 | 1.538009365 | 0.177205705 | 8.679231649 |
| Gm27361 | 0 | NA | NA | NA |
| Sdr42e1 | 58.48922315 | 0.036318941 | 0.177175557 | 0.204988437 |
| Hsd17b2 | 4.082108608 | -1.183718627 | 0.57964979 | -2.042127241 |
| Mphosph6 | 123.4766612 | 0.57316876 | 0.117560035 | 4.875540926 |
| Gm10617 | 0.340027382 | -2.382804059 | 1.424142586 | -1.673149924 |
| Gm25200 | 0 | NA | NA | NA |
| Cdh13 | 744.3496955 | -0.0904577 | 0.064391143 | -1.404815875 |
| Gm24035 | 0 | NA | NA | NA |
| Gm22999 | 0.35553296 | -1.488290995 | 1.712411637 | -0.869119879 |
| Hsbp1 | 1254.7106 | 0.180259347 | 0.053752393 | 3.353512975 |
| Mlycd | 201.1556644 | 0.08163465 | 0.098609748 | 0.827855779 |
| Osgin1 | 24.0618504 | 1.89064732 | 0.387292176 | 4.881708016 |
| Necab2 | 468.9419142 | -0.20310553 | 0.148501864 | -1.367696839 |
| Slc38a8 | 4.05865859 | -0.195730296 | 0.571581081 | -0.342436624 |
| Mbtps1 | 1032.591658 | 0.106168977 | 0.068541298 | 1.54897821 |
| Hsdl1 | 556.8575977 | 0.119654411 | 0.080740541 | 1.481961968 |
| Dnaaf1 | 4.031485904 | -0.250588379 | 0.558657267 | -0.448554765 |
| Taf1c | 112.5897251 | -0.069214828 | 0.113420365 | -0.610250444 |
| Rps13-ps4 | 4.918628032 | 0.619080468 | 0.463943124 | 1.334388715 |
| Adad2 | 0.577193238 | 0.226017278 | 1.262156033 | 0.179072374 |
| Kcng4 | 392.3468948 | 0.125468915 | 0.174307256 | 0.71981464 |
| Wfdc1 | 23.0470312 | 0.245398106 | 0.297195936 | 0.825711516 |
| Atp2c2 | 14.74911433 | 0.891453432 | 0.34615459 | 2.575304378 |
| Tldc1 | 91.32362537 | -0.397373268 | 0.118701946 | -3.347655873 |
| Cotl1 | 860.7212699 | 0.771053396 | 0.112505853 | 6.853451408 |
| Klhl36 | 75.98809112 | 0.245540355 | 0.129611789 | 1.894429182 |
| Usp10 | 645.8447964 | -0.217646134 | 0.05761872 | -3.777351089 |
| Gm20388 | 0 | NA | NA | NA |

| | | | | |
|---|---|---|---|---|
| Crispld2 | 132.7086816 | 1.762338109 | 0.294524552 | 5.98367131 |
| Gm15684 | 0.208379788 | 1.721655425 | 2.623716312 | 0.656189626 |
| Zdhhc7 | 282.094203 | 0.170886017 | 0.102639056 | 1.664921949 |
| Gm15898 | 0 | NA | NA | NA |
| A130014A01Rik | 33.99075599 | -0.310791078 | 0.171207757 | -1.815286201 |
| Gm38379 | 3.088015912 | -1.924038383 | 0.609119811 | -3.158719104 |
| 6430548M08Rik | 3271.759844 | -0.329120446 | 0.09545869 | -3.447778791 |
| Fam92b | 3.336165937 | -0.431526075 | 0.573786149 | -0.752067778 |
| A330074K22Rik | 139.4201663 | 0.041234876 | 0.149512364 | 0.275795762 |
| Gse1 | 1501.615146 | -0.000475932 | 0.093809677 | -0.00507338 |
| Gm22715 | 0 | NA | NA | NA |
| Gm26971 | 9.453064617 | -0.632053725 | 0.509333592 | -1.240942548 |
| 1700016A09Rik | 5.586702209 | -0.456324439 | 0.478745276 | -0.9531675 |
| Mir7687 | 1.495561828 | -0.678126741 | 0.871682456 | -0.777951577 |
| Gins2 | 20.25107559 | 0.744093811 | 0.378468586 | 1.966064922 |
| Gm10614 | 0.913620126 | -0.454469968 | 1.022791724 | -0.444342634 |
| Gm26739 | 0.232719486 | 0.776366685 | 2.130156226 | 0.36446467 |
| 1190005I06Rik | 8.331890292 | -0.421646443 | 0.350894232 | -1.201634009 |
| Gm27021 | 0.176744647 | 1.71275451 | 2.709446337 | 0.632141883 |
| Gm26537 | 0 | NA | NA | NA |
| Emc8 | 634.0511647 | 0.091708293 | 0.071923263 | 1.275085272 |
| Cox4i1 | 2787.69123 | -0.047690043 | 0.091892789 | -0.518974818 |
| Gm33142 | 0.833731302 | 2.781178173 | 1.113263204 | 2.498221591 |
| Gm24239 | 0 | NA | NA | NA |
| Gm18709 | 0.119587853 | 1.218392466 | 3.723618458 | 0.327206581 |
| Irf8 | 490.9940863 | 3.49208041 | 0.191452276 | 18.23995241 |
| Gm26878 | 0.075043205 | 0.755286614 | 3.724945049 | 0.202764498 |
| Gm28324 | 0 | NA | NA | NA |
| Fendrr | 2.814059987 | 1.628612951 | 0.671369793 | 2.425806117 |
| Foxf1 | 17.00115138 | 2.855497646 | 0.381159972 | 7.491598947 |
| Gm27530 | 0.136963818 | 0.755286583 | 3.59954433 | 0.209828388 |
| Mthfsd | 357.8554343 | -0.342744294 | 0.113754217 | -3.013024963 |
| Foxc2 | 30.92999064 | -0.077377251 | 0.262767752 | -0.29447012 |
| AC127554.1 | 0 | NA | NA | NA |
| Foxl1 | 1.257454774 | -0.868230935 | 1.201930293 | -0.722363801 |
| Gm37609 | 0.320352182 | -1.666105614 | 1.955047831 | -0.852207086 |
| 5033426O07Rik | 1.128616957 | 0.532565994 | 1.013109718 | 0.525674548 |
| Gm26815 | 0 | NA | NA | NA |
| Gm26747 | 6.573009254 | -0.962724641 | 0.405634089 | -2.373381986 |
| Gm26784 | 0.109869311 | -0.206490066 | 3.724945049 | -0.055434393 |
| 1700018B08Rik | 0.229212118 | 1.217882219 | 2.536134947 | 0.480211915 |
| Gm27045 | 0.361384634 | -0.637006127 | 1.772192819 | -0.359445158 |
| 1700030M09Rik | 0.55198573 | -0.20648228 | 1.423206267 | -0.145082469 |
| Fbxo31 | 1046.295583 | 0.097461241 | 0.07383941 | 1.319908188 |
| Gm17786 | 1.168706595 | -1.633111217 | 1.115428396 | -1.464111209 |
| Map1lc3b | 1461.563508 | 0.246958377 | 0.077209724 | 3.198539821 |
| Zcchc14 | 1352.76978 | -0.113063364 | 0.096810534 | -1.167882867 |

| | | | | |
|---|---|---|---|---|
| Gm27030 | 11.36160826 | -0.171508202 | 0.290598744 | -0.590189068 |
| Gm23299 | 0.157818009 | -0.659606625 | 3.475069758 | -0.189811046 |
| Jph3 | 2592.754649 | -0.117616121 | 0.080305285 | -1.46461246 |
| Klhdc4 | 215.8889794 | 0.268748865 | 0.143151811 | 1.877369643 |
| Gm20406 | 0 | NA | NA | NA |
| Gm24283 | 0.323040168 | 1.117437258 | 1.744042526 | 0.64071675 |
| Gm26812 | 0.227390921 | 1.652392026 | 2.11704536 | 0.780518008 |
| 9530085P06Rik | 0.295432452 | 1.223944372 | 1.796332956 | 0.681357188 |
| Slc7a5 | 545.5188095 | 0.80426156 | 0.137738884 | 5.839030628 |
| BC048644 | 2.632535955 | -0.241439471 | 0.631916159 | -0.382075165 |
| Car5a | 0.305395298 | -0.429796355 | 1.779303981 | -0.241553079 |
| Gm26999 | 0.075043205 | 0.755286614 | 3.724945049 | 0.202764498 |
| Gm37497 | 0 | NA | NA | NA |
| Gm27011 | 0.490536136 | 1.909484346 | 1.515952629 | 1.259593677 |
| Banp | 309.1391948 | -1.228721424 | 0.17456982 | -7.038567301 |
| Gm17709 | 4.900990937 | 0.966573406 | 0.588790292 | 1.641625922 |
| Mir7237 | 0.780673905 | -1.808999962 | 1.151755289 | -1.570646107 |
| Gm22 | 32.71914304 | -0.037860605 | 0.218199583 | -0.173513646 |
| Zfpm1 | 722.8421094 | 1.364828574 | 0.174101524 | 7.839268412 |
| Trhr2 | 33.68978649 | -0.495279295 | 0.268565322 | -1.844166965 |
| Gm20681 | 0.36185533 | 0.78259765 | 1.555566708 | 0.503094882 |
| Zc3h18 | 664.5062256 | 0.002772429 | 0.086785209 | 0.031945868 |
| Il17c | 0.855647594 | 2.728282418 | 1.376974071 | 1.981360779 |
| Cyba | 233.8961265 | 2.943620312 | 0.234489073 | 12.55333684 |
| Mvd | 137.2979585 | -0.543613841 | 0.120723339 | -4.502972209 |
| 9330133O14Rik | 42.62923582 | 0.239458385 | 0.167995502 | 1.425385691 |
| Snai3 | 6.952624755 | -0.647176882 | 0.554272386 | -1.167615235 |
| Rnf166 | 241.1004881 | -0.110114701 | 0.091665147 | -1.201271207 |
| Gm24366 | 2.393340753 | -0.872347786 | 0.616725426 | -1.41448325 |
| Ctu2 | 170.7851 | -0.152717211 | 0.100518356 | -1.519296744 |
| Piezo1 | 341.3561795 | 1.237107795 | 0.162344092 | 7.620282191 |
| Gm26497 | 0.280709254 | -0.707978847 | 1.788608539 | -0.395826606 |
| Cdt1 | 76.18453122 | 1.655810978 | 0.293379387 | 5.643924058 |
| Aprt | 172.1584049 | 0.462196281 | 0.140987047 | 3.278288979 |
| Galns | 112.6437773 | 0.89380788 | 0.157379479 | 5.679316536 |
| Trappc2l | 257.9503243 | -0.012798037 | 0.078549867 | -0.162928816 |
| Pabpn1l | 0.259153158 | 0.233010239 | 2.078289693 | 0.112116342 |
| Mir6396 | 0 | NA | NA | NA |
| Cbfa2t3 | 329.1625399 | -0.660969251 | 0.164919872 | -4.007820543 |
| Gm15899 | 22.41855246 | 0.414582306 | 0.236675343 | 1.751692008 |
| Gm10612 | 0.141684195 | 0.755287278 | 3.558854591 | 0.212227631 |
| Acsf3 | 163.1806245 | -0.103128921 | 0.113160293 | -0.911352543 |
| Gm16378 | 9.858353745 | -0.409741331 | 0.322991182 | -1.268583647 |
| Cdh15 | 97.85341112 | 0.039104316 | 0.237419093 | 0.164705859 |
| Ankrd11 | 2871.409642 | -0.020584601 | 0.0796738 | -0.258360972 |
| Gm24445 | 2.473368846 | -0.060282388 | 0.706469329 | -0.085329094 |
| 2810013P06Rik | 123.5382447 | -0.228230314 | 0.111223741 | -2.051992787 |

| | | | | |
|---|---|---|---|---|
| Spg7 | 394.1502427 | -0.115513993 | 0.086495551 | -1.335490581 |
| Rpl13 | 1506.813565 | 0.241356275 | 0.113446681 | 2.127486428 |
| Snord68 | 0 | NA | NA | NA |
| Mir7079 | 0 | NA | NA | NA |
| Cpne7 | 380.383942 | 0.056987779 | 0.170763111 | 0.333724177 |
| Mir7080 | 0 | NA | NA | NA |
| Sult5a1 | 0.948467964 | 0.281836266 | 1.030442448 | 0.273509954 |
| Dpep1 | 2.460632423 | 2.23304611 | 0.828370588 | 2.695709074 |
| Chmp1a | 416.7948105 | 0.319233137 | 0.090323132 | 3.534345313 |
| Rps12-ps9 | 5.02878481 | 0.952011974 | 0.559317827 | 1.702094818 |
| Spata33 | 6.260054198 | -0.244364006 | 0.41645356 | -0.586773723 |
| Cdk10 | 231.2422657 | -0.124780239 | 0.091471436 | -1.364144311 |
| Spata2l | 264.1685552 | -0.442933089 | 0.158368617 | -2.796848874 |
| 4933417D19Rik | 2.606279457 | -0.763077853 | 0.645214669 | -1.182672822 |
| Vps9d1 | 418.2806154 | 0.216664984 | 0.098200555 | 2.206351927 |
| Zfp276 | 246.8177323 | -0.089147434 | 0.115598052 | -0.77118457 |
| Fanca | 30.7735149 | 1.398940336 | 0.301863338 | 4.634349925 |
| Spire2 | 197.4572324 | -0.107268259 | 0.106633345 | -1.005954178 |
| Tcf25 | 4948.09245 | -0.010391988 | 0.056353494 | -0.184407162 |
| Mc1r | 2.754460718 | 1.120356533 | 0.788577866 | 1.420730381 |
| Gm38092 | 0.099155932 | 0.75528439 | 3.724945049 | 0.202763901 |
| Tubb3 | 2462.821606 | -0.132772483 | 0.097778642 | -1.357888394 |
| Def8 | 517.338147 | -0.207209718 | 0.060639022 | -3.417101933 |
| Gm25428 | 0 | NA | NA | NA |
| Afg3l1 | 423.4960045 | 0.117442428 | 0.064592493 | 1.818205537 |
| Dbndd1 | 119.4149113 | -0.650545505 | 0.109421141 | -5.945336525 |
| Gas8 | 123.9867013 | 0.117361621 | 0.181636519 | 0.6461345 |
| Gm22314 | 2.308872772 | 0.790040328 | 0.651746381 | 1.212189821 |
| n-R5s100 | 0 | NA | NA | NA |
| n-R5s101 | 0 | NA | NA | NA |
| n-R5s102 | 0 | NA | NA | NA |
| n-R5s103 | 0 | NA | NA | NA |
| n-R5s104 | 0 | NA | NA | NA |
| n-R5s105 | 0 | NA | NA | NA |
| n-R5s106 | 0 | NA | NA | NA |
| n-R5s107 | 0 | NA | NA | NA |
| n-R5s108 | 0 | NA | NA | NA |
| n-R5s109 | 0 | NA | NA | NA |
| n-R5s110 | 0 | NA | NA | NA |
| n-R5s111 | 0 | NA | NA | NA |
| n-R5s112 | 0 | NA | NA | NA |
| n-R5s113 | 0 | NA | NA | NA |
| n-R5s114 | 0 | NA | NA | NA |
| n-R5s115 | 0 | NA | NA | NA |
| Gm25018 | 0 | NA | NA | NA |
| n-R5s117 | 0 | NA | NA | NA |
| n-R5s118 | 0 | NA | NA | NA |

| | | | | |
|---|---|---|---|---|
| n-R5s119 | 0 | NA | NA | NA |
| n-R5s120 | 0 | NA | NA | NA |
| n-R5s121 | 0 | NA | NA | NA |
| n-R5s122 | 0 | NA | NA | NA |
| n-R5s123 | 0 | NA | NA | NA |
| n-R5s124 | 0 | NA | NA | NA |
| n-R5s125 | 0 | NA | NA | NA |
| Gm24312 | 0 | NA | NA | NA |
| n-R5s127 | 0 | NA | NA | NA |
| n-R5s128 | 0 | NA | NA | NA |
| n-R5s129 | 0 | NA | NA | NA |
| n-R5s130 | 0 | NA | NA | NA |
| n-R5s131 | 0 | NA | NA | NA |
| Gm26391 | 0 | NA | NA | NA |
| n-R5s133 | 0 | NA | NA | NA |
| n-R5s134 | 0 | NA | NA | NA |
| Rn5s | 0 | NA | NA | NA |
| n-R5s136 | 0 | NA | NA | NA |
| Gm24089 | 0 | NA | NA | NA |
| n-R5s138 | 0 | NA | NA | NA |
| n-R5s139 | 0 | NA | NA | NA |
| Gm22109 | 0 | NA | NA | NA |
| n-R5s141 | 0 | NA | NA | NA |
| n-R5s142 | 0 | NA | NA | NA |
| n-R5s143 | 0 | NA | NA | NA |
| n-R5s144 | 0 | NA | NA | NA |
| Gm25212 | 0 | NA | NA | NA |
| n-R5s146 | 0 | NA | NA | NA |
| Gm22291 | 0 | NA | NA | NA |
| Gm23284 | 0 | NA | NA | NA |
| n-R5s149 | 0 | NA | NA | NA |
| Gm23647 | 0 | NA | NA | NA |
| Rhou | 659.2759545 | 0.615447745 | 0.125208945 | 4.915365628 |
| 4732419C18Rik | 5.763193032 | -0.037846368 | 0.558468748 | -0.067768104 |
| Rab4a | 299.8235972 | 0.002101427 | 0.076156117 | 0.02759368 |
| 2810455O05Rik | 58.83641319 | -0.608498355 | 0.259131556 | -2.348221743 |
| Ccsap | 351.2028742 | 0.039091754 | 0.147805971 | 0.264480206 |
| Gm4342 | 0.738388108 | 0.573522213 | 1.317874973 | 0.435187119 |
| AC147266.1 | 3.758891091 | 0.133307324 | 0.534659085 | 0.249331449 |
| Acta1 | 38.57868714 | 0.035622695 | 0.175375662 | 0.20312223 |
| Nup133 | 281.5212268 | 0.022634944 | 0.069787649 | 0.32434025 |
| Abcb10 | 231.5059475 | -0.081143995 | 0.110708564 | -0.73295138 |
| Taf5l | 237.4112607 | 0.324645824 | 0.112136694 | 2.89509003 |
| Urb2 | 291.7584138 | -0.258353217 | 0.089366914 | -2.890926915 |
| Mir1967 | 0.881693066 | 0.825230341 | 1.071238187 | 0.770351871 |
| Galnt2 | 11.54263694 | -0.432359107 | 0.319741407 | -1.352214939 |
| 1810008B01Rik | 0.299839612 | -1.62779111 | 1.581002964 | -1.029593964 |

| | | | | |
|---|---|---|---|---|
| Gm15775 | 0.11147366 | 0.274389235 | 3.724945049 | 0.073662626 |
| Gm24459 | 0 | NA | NA | NA |
| Pgbd5 | 1529.511435 | 0.245389418 | 0.126022255 | 1.947191142 |
| Cog2 | 399.9630909 | -0.018868512 | 0.081646874 | -0.231099012 |
| Agt | 480.964286 | 0.083667572 | 0.258758242 | 0.323342636 |
| Capn9 | 3.44454021 | 1.687696848 | 0.604592819 | 2.791460293 |
| 2310022B05Rik | 690.1050064 | -0.511834409 | 0.185458524 | -2.759832221 |
| Ttc13 | 501.1809921 | -0.128143915 | 0.132407151 | -0.967802076 |
| Arv1 | 50.9902722 | -0.010528986 | 0.170863132 | -0.061622341 |
| Fam89a | 4.128209337 | 0.931103068 | 0.545408585 | 1.707166137 |
| Trim67 | 432.3924619 | 0.195249226 | 0.133295494 | 1.464784897 |
| Gm16163 | 3.256114358 | -0.334449358 | 0.647126612 | -0.516822136 |
| 2810004N23Rik | 135.240311 | -0.015577176 | 0.116862102 | -0.133295359 |
| Gnpat | 451.9781084 | -0.324408643 | 0.076118819 | -4.261871701 |
| Exoc8 | 423.6774759 | -0.281079359 | 0.070484844 | -3.987798568 |
| Sprtn | 158.0541678 | 0.210384918 | 0.11868966 | 1.77256315 |
| Egln1 | 684.0866431 | -0.139000197 | 0.084981078 | -1.635660551 |
| Tsnax | 775.1924979 | 0.000398471 | 0.079750097 | 0.0049965 |
| Disc1 | 23.96939849 | -0.257952952 | 0.21733021 | -1.186917146 |
| Gm22197 | 0.208494586 | -0.206482388 | 2.629256435 | -0.078532617 |
| Gm16237 | 0.184167273 | 0.755304955 | 2.681673463 | 0.281654335 |
| Sipa1l2 | 1159.737547 | 0.12908883 | 0.084668723 | 1.524634192 |
| Map10 | 80.06219714 | 0.500247853 | 0.133467393 | 3.748090381 |
| Ntpcr | 128.5931328 | 0.128938189 | 0.163058644 | 0.790747341 |
| Pcnxl2 | 1292.263944 | -0.194362828 | 0.12161093 | -1.598234869 |
| A730098A19Rik | 4.29871648 | -0.300467605 | 0.497994091 | -0.603355765 |
| BC021891 | 32.34379101 | 0.256105799 | 0.285464306 | 0.897155242 |
| Kcnk1 | 670.0428367 | -0.03881801 | 0.092942139 | -0.417657806 |
| Slc35f3 | 269.0447631 | -0.52901084 | 0.169823934 | -3.115054674 |
| Coa6 | 108.6067078 | -0.086826238 | 0.154623699 | -0.56153254 |
| Gm17296 | 160.9155127 | -0.781793907 | 0.18547244 | -4.215148654 |
| Gm6091 | 0 | NA | NA | NA |
| Gm26759 | 0.554140925 | 0.084211656 | 1.483122944 | 0.056779956 |
| Irf2bp2 | 1675.480812 | 0.405430133 | 0.116584909 | 3.477552432 |
| Tomm20 | 2682.24209 | -0.026811292 | 0.04643051 | -0.577449876 |
| Gm26397 | 173.2653863 | -0.023235017 | 0.174929217 | -0.132825248 |
| Rbm34 | 225.3370361 | 0.18460264 | 0.116701229 | 1.581839729 |
| Gm22730 | 0 | NA | NA | NA |
| Pard3 | 532.7837827 | -0.139417098 | 0.069243411 | -2.013434881 |
| Gm6921 | 2.797109942 | -0.706462939 | 0.598777011 | -1.179843124 |
| Gm22968 | 0.822547242 | -0.463083042 | 1.080189458 | -0.428705389 |
| Mir21c | 0.121578509 | 0.75528439 | 3.724945049 | 0.202763901 |
| Nrp1 | 639.8097654 | -0.137839953 | 0.104763488 | -1.315725125 |
| Mir1903 | 0 | NA | NA | NA |
| Gm26002 | 0 | NA | NA | NA |
| Itgb1 | 1279.915761 | 0.415771244 | 0.101381348 | 4.101062501 |
| Gm3952 | 0.143618953 | -0.206488341 | 2.893930267 | -0.071352217 |

| | | | | |
|---|---|---|---|---|
| Gm26100 | 0.368403345 | 1.716149512 | 1.718767127 | 0.998477039 |
| Ccdc7 | 0.294450523 | -0.199878538 | 1.752272367 | -0.114068191 |
| Gm22545 | 0 | NA | NA | NA |
| 1700008F21Rik | 0.249972158 | 1.1127079 | 2.089027157 | 0.532644057 |
| Gm22374 | 0.159888962 | -0.206482815 | 3.461347753 | -0.059653878 |
| Gm10999 | 0.583900451 | 0.274414601 | 1.495875204 | 0.183447523 |
| 2610044O15Rik8 | 322.1898291 | -0.397616604 | 0.131263433 | -3.029149816 |
| Gm21399 | 59.65666709 | 0.294080152 | 0.173541762 | 1.694578571 |
| Gm26870 | 1.94439448 | 2.329305312 | 0.991892601 | 2.348344275 |
| Gm10722 | 0.242739364 | 0.274412626 | 2.093495053 | 0.131078708 |
| Gm11168 | 0.318248637 | -1.088437297 | 2.059036832 | -0.528614777 |
| Gm10721 | 0.187213551 | 0.27440959 | 2.667715587 | 0.102863136 |
| Gm10720 | 0 | NA | NA | NA |
| Gm10719 | 0.044544648 | 0.755286614 | 3.724945049 | 0.202764498 |
| Gm10718 | 0.690752925 | 0.111690315 | 1.186293044 | 0.094150696 |
| Gm10717 | 0.158171364 | -0.206480315 | 2.813813679 | -0.073380948 |
| Gm17535 | 0 | NA | NA | NA |
| Gm10715 | 0 | NA | NA | NA |
| mmu-mir-101c | 0 | NA | NA | NA |
| 4930433N12Rik | 0 | NA | NA | NA |
| Gm27679 | 0 | NA | NA | NA |
| Alkbh8 | 460.6917347 | -0.12817546 | 0.058264841 | -2.199876587 |
| Gm24918 | 2.468298956 | 0.039194282 | 0.768358516 | 0.051010408 |
| Gm23275 | 1.098824885 | 1.349425184 | 1.067345569 | 1.264281432 |
| Cwf19l2 | 426.6056693 | 0.707092164 | 0.114898627 | 6.154052324 |
| Gucy1a2 | 1402.939796 | -0.239821731 | 0.095786021 | -2.503723693 |
| Gm22195 | 0.192501058 | -1.073580721 | 2.663543715 | -0.403064802 |
| Aasdhppt | 467.0250061 | -0.124555423 | 0.088645745 | -1.405091965 |
| Kbtbd3 | 165.3891345 | -0.181703961 | 0.134773598 | -1.348216299 |
| Gm23992 | 0.044191549 | -0.206490061 | 3.724945049 | -0.055434391 |
| Msantd4 | 720.9495236 | -0.044337716 | 0.07045597 | -0.62929679 |
| Gria4 | 2416.785679 | -0.376128483 | 0.085205986 | -4.414343419 |
| Gm23811 | 8.724062623 | -0.575207802 | 0.345516304 | -1.664777597 |
| Gm24896 | 0 | NA | NA | NA |
| Casp1 | 190.837744 | 3.242541516 | 0.24732635 | 13.11037629 |
| Casp4 | 185.281841 | 5.94030992 | 0.469692449 | 12.64723317 |
| Casp12 | 173.1181621 | 3.02784812 | 0.23307293 | 12.99099005 |
| Pdgfd | 56.87235133 | -0.243186674 | 0.234245669 | -1.038169349 |
| Ddi1 | 0 | NA | NA | NA |
| Gm23097 | 0 | NA | NA | NA |
| Dync2h1 | 1480.682915 | -0.284537868 | 0.100768465 | -2.823679704 |
| Dcun1d5 | 410.911556 | 0.150891849 | 0.090776576 | 1.662233312 |
| Mmp13 | 18.19977551 | 3.792148809 | 0.625420627 | 6.063357438 |
| Mmp12 | 7.267951943 | 1.07688813 | 0.643683285 | 1.673009313 |
| Mmp1b | 0 | NA | NA | NA |
| Mmp3 | 11.53096597 | 6.770634949 | 0.952623902 | 7.107353632 |
| Mmp1a | 0.181654796 | 0.274390172 | 3.327599044 | 0.082458904 |

| | | | | |
|---|---|---|---|---|
| Mmp10 | 0.099106094 | 0.274387056 | 3.724945049 | 0.073662041 |
| Mmp8 | 40.56306267 | 6.427797656 | 1.19551136 | 5.376609433 |
| Mmp27 | 4.159708068 | 2.554627442 | 1.159199484 | 2.20378587 |
| Mmp20 | 0.336931769 | -0.20648347 | 1.822656745 | -0.113287085 |
| AV064505 | 0 | NA | NA | NA |
| Mmp7 | 0.107803068 | 0.75528661 | 3.724945049 | 0.202764497 |
| Gm10709 | 0.713745664 | -1.155647712 | 1.220672934 | -0.94673002 |
| Tmem123 | 239.3847869 | 0.700583955 | 0.135703912 | 5.162592194 |
| Mir5618 | 0.424774107 | -0.367947408 | 1.379343672 | -0.266755425 |
| Birc2 | 772.896626 | 0.592116617 | 0.107155548 | 5.525767232 |
| Birc3 | 154.0540683 | 3.417444098 | 0.275811205 | 12.39051946 |
| Yap1 | 390.0541373 | 0.121056961 | 0.103489982 | 1.169745699 |
| Gm20416 | 0.109982625 | -0.20649451 | 3.724945049 | -0.055435586 |
| 9230110C19Rik | 76.10268791 | -0.410151055 | 0.141161817 | -2.905538234 |
| Cep126 | 298.1066087 | -0.208793502 | 0.085913676 | -2.430270846 |
| 1700128F08Rik | 0.650208918 | -0.725572321 | 1.248462079 | -0.581172895 |
| Mir1899 | 0.060174537 | 0.755286614 | 3.724945049 | 0.202764498 |
| Trpc6 | 122.6815528 | -0.174164667 | 0.15093175 | -1.153929954 |
| Pgr | 476.6556311 | -0.352945104 | 0.088308255 | -3.996739662 |
| Gm16485 | 192.3635005 | -0.35319655 | 0.135183808 | -2.612713427 |
| Arhgap42 | 263.2227951 | -0.16792904 | 0.080223295 | -2.093270281 |
| Gm16833 | 0 | NA | NA | NA |
| Gm22815 | 0 | NA | NA | NA |
| Cntn5 | 249.9584538 | -0.141427385 | 0.123316709 | -1.146863112 |
| Gm26195 | 0.87738752 | 1.212339595 | 0.982377578 | 1.234087201 |
| Mir6237 | 0 | NA | NA | NA |
| Gm24496 | 0.646963711 | -0.137384208 | 1.127086115 | -0.121893267 |
| Gm25086 | 0.47668666 | 1.405400122 | 1.624946562 | 0.864890055 |
| Gm25365 | 0 | NA | NA | NA |
| Gm21070 | 0 | NA | NA | NA |
| Gm29176 | 0 | NA | NA | NA |
| Jrkl | 160.5671298 | -0.105360306 | 0.139014571 | -0.757908363 |
| Ccdc82 | 240.4966967 | -0.231605895 | 0.095590795 | -2.422889099 |
| Gm26189 | 1.402298525 | 1.432111334 | 0.932054467 | 1.536510349 |
| Phxr4 | 43.51084174 | -0.055922502 | 0.330349944 | -0.169282613 |
| Gm17571 | 0 | NA | NA | NA |
| Maml2 | 397.5331643 | 0.40625505 | 0.087839773 | 4.624955602 |
| Gm22643 | 0 | NA | NA | NA |
| Mtmr2 | 890.4922701 | -0.031358629 | 0.103374028 | -0.303351139 |
| Cep57 | 215.6600884 | 0.390674171 | 0.119943956 | 3.257139283 |
| Fam76b | 356.5380025 | 0.397953504 | 0.142982402 | 2.783234161 |
| Sesn3 | 970.3136834 | 0.235496227 | 0.082794088 | 2.844360425 |
| Endod1 | 2711.618671 | 0.919714365 | 0.179778774 | 5.115811754 |
| Kdm4dl | 4.695898157 | -0.135468171 | 0.571666781 | -0.236970513 |
| Kdm4d | 25.20782128 | 0.031818133 | 0.221197824 | 0.143844691 |
| Cwc15 | 455.3458575 | 0.375882027 | 0.131272862 | 2.863364307 |
| Amotl1 | 1488.185642 | 0.399181711 | 0.112944308 | 3.534323415 |

| | | | | |
|---|---|---|---|---|
| Piwil4 | 14.13799992 | 4.795621961 | 0.753933631 | 6.360801225 |
| Gm10706 | 3.101177844 | -1.138885278 | 0.63470885 | -1.794342836 |
| Gm15586 | 0 | NA | NA | NA |
| Fut4 | 18.43119656 | 1.0346733 | 0.310892874 | 3.328070174 |
| 1700012B09Rik | 5.943573362 | -0.186757117 | 0.448215105 | -0.416668503 |
| Ankrd49 | 112.3845942 | 0.530951453 | 0.128125393 | 4.143998641 |
| Mre11a | 230.8957433 | 0.009052055 | 0.091265725 | 0.099183513 |
| Gm16379 | 0.870448589 | 0.802246928 | 1.110940768 | 0.722132944 |
| Gm16302 | 2.36738784 | -0.105399388 | 0.708391871 | -0.148786841 |
| Gpr83 | 316.7124723 | -0.21369583 | 0.126610493 | -1.687820852 |
| Izumo1r | 3.609677315 | 4.458315239 | 0.951293245 | 4.686583517 |
| Panx1 | 176.4686585 | 0.060439309 | 0.116813452 | 0.517400254 |
| Hephl1 | 1.803729066 | -1.164397087 | 0.980174907 | -1.187948272 |
| Gm22658 | 0.042716945 | -0.206490061 | 3.724945049 | -0.055434391 |
| Vstm5 | 75.69295347 | 0.139184034 | 0.172041587 | 0.809013891 |
| Med17 | 334.3072139 | 0.285289551 | 0.075992476 | 3.75418154 |
| 4931406C07Rik | 230.1835539 | 0.214406591 | 0.11836944 | 1.811334007 |
| Taf1d | 188.0883961 | 0.612957909 | 0.151015174 | 4.058916027 |
| Gm25500 | 1.755293934 | 0.091077273 | 0.781235325 | 0.116581099 |
| mmu-mir-7650 | 0 | NA | NA | NA |
| Gm24455 | 11.03006032 | 0.605175746 | 0.316662481 | 1.911106564 |
| Gm24299 | 7.628192187 | 0.136892819 | 0.359100595 | 0.381210225 |
| AC154296.1 | 0.044191549 | -0.206490061 | 3.724945049 | -0.055434391 |
| Gm25791 | 5.041681076 | 1.223311241 | 0.47538518 | 2.57330538 |
| Gm22620 | 6.972143724 | 0.608740368 | 0.387256148 | 1.571932096 |
| Gm24357 | 17.62472295 | 0.931888636 | 0.292133218 | 3.189944102 |
| Gm23455 | 26.8443768 | 0.924369087 | 0.273847154 | 3.375492766 |
| Gm22579 | 6.383805412 | -0.018857238 | 0.456552236 | -0.041303573 |
| Cep295 | 648.5597578 | -0.104061083 | 0.112863551 | -0.922007874 |
| Scarna9 | 8.758301848 | -0.385512181 | 0.337095212 | -1.143629953 |
| Gm24339 | 46.69042643 | -0.194613008 | 0.272598657 | -0.713917707 |
| Gm16568 | 0.41713571 | -0.151973124 | 1.326794376 | -0.11454158 |
| Smco4 | 17.55463808 | -0.215683578 | 0.301643375 | -0.715028394 |
| Ccdc67 | 5.175431821 | 0.737546774 | 0.448410074 | 1.644804202 |
| Slc36a4 | 909.8448179 | -0.014665508 | 0.073414209 | -0.199763896 |
| Mtnr1b | 0.066050875 | 0.274391604 | 3.724945049 | 0.073663262 |
| Fat3 | 3920.904863 | -0.468389585 | 0.132376823 | -3.538305075 |
| Gm5611 | 1.991326778 | -0.126693592 | 0.748501404 | -0.169262999 |
| CT943659.1 | 0 | NA | NA | NA |
| Chordc1 | 659.9625695 | -0.562093534 | 0.111300944 | -5.05021352 |
| Naalad2 | 9.142264245 | -0.682742973 | 0.455535188 | -1.498771096 |
| Ubtfl1 | 0.110595523 | 0.755282166 | 3.724945049 | 0.202763304 |
| Gm5612 | 0.507208347 | -0.215578386 | 1.466885971 | -0.146963288 |
| Mbd3l2 | 0.241190471 | 1.348328565 | 3.031116258 | 0.44482905 |
| Zfp558 | 31.78569487 | -0.218737723 | 0.207589853 | -1.053701424 |
| Gm22854 | 0 | NA | NA | NA |
| Mbd3l1 | 0.181654796 | 0.274390172 | 3.327599044 | 0.082458904 |

| | | | | |
|---|---|---|---|---|
| Muc16 | 0.130524232 | 0.274405456 | 3.657871338 | 0.075017799 |
| Gm25295 | 0 | NA | NA | NA |
| Gm24546 | 0 | NA | NA | NA |
| Olfr24 | 0 | NA | NA | NA |
| Olfr828 | 0 | NA | NA | NA |
| Olfr829 | 0.068097359 | -0.206487837 | 3.724945049 | -0.055433794 |
| Olfr830 | 0 | NA | NA | NA |
| Olfr831-ps1 | 0 | NA | NA | NA |
| Olfr832 | 0 | NA | NA | NA |
| Olfr833-ps1 | 0 | NA | NA | NA |
| Olfr834 | 0 | NA | NA | NA |
| Olfr835 | 0.046345709 | 0.274391604 | 3.724945049 | 0.073663262 |
| Olfr836 | 0 | NA | NA | NA |
| Olfr837 | 0 | NA | NA | NA |
| Olfr843 | 0 | NA | NA | NA |
| Olfr844 | 0 | NA | NA | NA |
| Olfr845 | 0 | NA | NA | NA |
| Olfr846 | 0.049553047 | 0.274391604 | 3.724945049 | 0.073663262 |
| Olfr847 | 0 | NA | NA | NA |
| Gm23048 | 0 | NA | NA | NA |
| Olfr849 | 0 | NA | NA | NA |
| Olfr850 | 0 | NA | NA | NA |
| Olfr851 | 0 | NA | NA | NA |
| Olfr853 | 0 | NA | NA | NA |
| Olfr854 | 0 | NA | NA | NA |
| Olfr855 | 0 | NA | NA | NA |
| Zfp317 | 272.3857193 | 0.127302985 | 0.08531916 | 1.49207968 |
| Olfr857 | 1.183877471 | 1.157036269 | 0.954666537 | 1.211979497 |
| Olfr58 | 0.474412771 | 2.120703276 | 1.509223595 | 1.405161755 |
| Olfr859 | 0.318756133 | 0.368438406 | 1.776793051 | 0.207361463 |
| Olfr860 | 0 | NA | NA | NA |
| Olfr862 | 0 | NA | NA | NA |
| Olfr77 | 0 | NA | NA | NA |
| Gm7808 | 2.442408819 | -0.579811379 | 0.661922981 | -0.875949914 |
| Olfr866 | 0 | NA | NA | NA |
| AC121953.1 | 0 | NA | NA | NA |
| Gm26231 | 0 | NA | NA | NA |
| Olfr867 | 0.092691418 | 0.274387304 | 3.724945049 | 0.073662108 |
| Olfr868 | 0.089062654 | -0.20649451 | 3.724945049 | -0.055435586 |
| Olfr869 | 0 | NA | NA | NA |
| Olfr870 | 0 | NA | NA | NA |
| Gm25914 | 0 | NA | NA | NA |
| Gm26290 | 0 | NA | NA | NA |
| Olfr871 | 0 | NA | NA | NA |
| Olfr872 | 0 | NA | NA | NA |
| Gm25425 | 0.049553047 | 0.274391604 | 3.724945049 | 0.073663262 |
| Olfr39 | 0 | NA | NA | NA |

| | | | | |
|---|---|---|---|---|
| Olfr873 | 0 | NA | NA | NA |
| Olfr18 | 0.954195627 | -0.92048598 | 1.449999762 | -0.634818021 |
| Zfp560 | 108.4598925 | -0.29130827 | 0.143060359 | -2.036261274 |
| Gm26733 | 0 | NA | NA | NA |
| Zfp26 | 822.179251 | 0.189333658 | 0.089535163 | 2.114629048 |
| Zfp426 | 384.7268472 | -0.349179601 | 0.117946663 | -2.960487323 |
| Gm23316 | 0 | NA | NA | NA |
| Zfp266 | 1050.28284 | -0.175638958 | 0.069780416 | -2.517023661 |
| Zfp846 | 184.0291169 | -0.396098222 | 0.120672224 | -3.282430765 |
| Fbxl12os | 22.46068426 | -0.771901013 | 0.263390443 | -2.9306341 |
| Gm26521 | 12.01662456 | 0.176257366 | 0.380684186 | 0.463001544 |
| Fbxl12 | 213.0329372 | -0.078351917 | 0.161662763 | -0.484662736 |
| Ubl5 | 521.6630355 | 0.241876805 | 0.111502777 | 2.16924467 |
| Pin1 | 657.0847573 | 0.051832016 | 0.068144066 | 0.760624051 |
| Olfm2 | 557.2935379 | -0.385406871 | 0.086594131 | -4.45072738 |
| Col5a3 | 212.1757937 | 2.131618417 | 0.238303853 | 8.94495992 |
| Gm26274 | 0 | NA | NA | NA |
| Rdh8 | 0.705140734 | -0.206485821 | 1.594113792 | -0.129530164 |
| A230050P20Rik | 616.357269 | 0.591597371 | 0.125458864 | 4.715468912 |
| Angptl6 | 38.80084079 | -0.780206776 | 0.182589508 | -4.273009894 |
| Ppan | 173.1887125 | 0.529792714 | 0.147993614 | 3.579834956 |
| Gm24067 | 0 | NA | NA | NA |
| Gm23008 | 1.246341358 | 0.838112065 | 0.898254251 | 0.933045475 |
| Eif3g | 495.9541041 | 0.213450694 | 0.106332456 | 2.007389858 |
| Dnmt1 | 868.320093 | -0.077698183 | 0.097317113 | -0.79840206 |
| Mir7081 | 0.867462339 | 0.347450026 | 0.995683866 | 0.348956167 |
| S1pr2 | 119.737217 | 1.131025145 | 0.23940262 | 4.724364117 |
| Mrpl4 | 356.3497304 | 0.092721768 | 0.067813913 | 1.367297111 |
| Icam1 | 316.3122786 | 5.02054545 | 0.376622618 | 13.33044063 |
| Icam4 | 2.361925876 | 0.022845202 | 0.595630187 | 0.038354675 |
| Icam5 | 1112.549133 | -0.109265792 | 0.230315436 | -0.474418017 |
| Mir1900 | 0.298647883 | 0.806503888 | 2.086679815 | 0.386501025 |
| Gm26592 | 23.65836997 | 0.069603345 | 0.218857275 | 0.318030759 |
| Zglp1 | 9.660883422 | -1.528234097 | 0.428200565 | -3.56896796 |
| Fdx1l | 166.5463135 | 0.1488629 | 0.115051052 | 1.293885598 |
| Raver1 | 630.7586862 | 0.233959565 | 0.074353576 | 3.146581216 |
| Mir7082 | 0 | NA | NA | NA |
| 1700084C06Rik | 2.554977676 | -0.490473425 | 0.624393119 | -0.785520228 |
| Tyk2 | 239.0143666 | -0.000588837 | 0.157345287 | -0.003742322 |
| Cdc37 | 916.0704561 | 0.029415171 | 0.092986728 | 0.316337309 |
| Pde4a | 1129.44041 | -0.393864723 | 0.100664851 | -3.912634056 |
| Gm16754 | 3.260813172 | -0.447841684 | 0.626575872 | -0.714744541 |
| Keap1 | 857.7412683 | 0.459342601 | 0.066221562 | 6.936450679 |
| S1pr5 | 222.4130938 | 0.137606096 | 0.142494137 | 0.965696546 |
| Atg4d | 243.847421 | -0.329652653 | 0.08538729 | -3.860675905 |
| Kri1 | 243.0174812 | 0.033511322 | 0.111659474 | 0.300120722 |
| Gm37379 | 20.17791531 | -0.62093422 | 0.292082575 | -2.125885873 |

| | | | | |
|---|---|---|---|---|
| Cdkn2d | 273.9573276 | 0.164159191 | 0.086792256 | 1.891403657 |
| Ap1m2 | 9.647799368 | 0.072779272 | 0.373035563 | 0.195100091 |
| Slc44a2 | 583.7110967 | 0.397551739 | 0.120821137 | 3.290415465 |
| Ilf3 | 1100.802251 | 0.055511839 | 0.053717644 | 1.033400479 |
| Gm16853 | 1.441198087 | -0.346372003 | 0.850408434 | -0.407300762 |
| Qtrt1 | 87.06778592 | 0.18315992 | 0.152071555 | 1.204432481 |
| Dnm2 | 1057.753246 | 0.544515027 | 0.104122308 | 5.229571212 |
| Mir199a-1 | 0.092691418 | 0.274387304 | 3.724945049 | 0.073662108 |
| Tmed1 | 82.01844689 | 0.255189983 | 0.150173517 | 1.699300834 |
| AB124611 | 105.0329745 | 5.266493057 | 0.440252397 | 11.96244038 |
| Gm20519 | 0.907036939 | -1.532047921 | 1.186107197 | -1.291660589 |
| Carm1 | 707.8332732 | -0.022342074 | 0.061300802 | -0.364466262 |
| Yipf2 | 186.1268908 | -0.145173117 | 0.095337081 | -1.522735074 |
| 1810026J23Rik | 718.2922187 | 0.473831886 | 0.113608174 | 4.170755246 |
| Mir1946b | 0.611133411 | -1.06827471 | 1.32539484 | -0.806004881 |
| Smarca4 | 2424.773633 | -0.021837588 | 0.051673603 | -0.422606248 |
| Gm27373 | 1.398170896 | 0.300228574 | 0.859932328 | 0.349130466 |
| Gm26511 | 0.192512644 | -1.179165976 | 2.691637281 | -0.438085022 |
| Ldlr | 610.6285346 | 0.484553832 | 0.181742846 | 2.666150783 |
| Spc24 | 9.830363938 | 0.519057658 | 0.386385106 | 1.343368702 |
| Kank2 | 390.6615536 | 0.164251636 | 0.144827808 | 1.1341167 |
| Dock6 | 147.3513096 | -0.330325531 | 0.13887473 | -2.378586303 |
| Mir7083 | 0 NA | NA | NA | |
| Gm6484 | 0.140389662 | 0.755287681 | 3.575868592 | 0.211217963 |
| Rab3d | 110.9505493 | -0.132964447 | 0.164913552 | -0.806267564 |
| Tmem205 | 101.5694773 | -0.166774136 | 0.192840436 | -0.864829698 |
| Ccdc159 | 62.58867534 | -0.526752165 | 0.196965632 | -2.67433541 |
| Lppr2 | 1319.490486 | -0.249588098 | 0.095798684 | -2.605339542 |
| Swsap1 | 101.2171663 | 0.167380161 | 0.124107895 | 1.3486665 |
| Epor | 34.63035605 | -0.336863586 | 0.242035721 | -1.391792848 |
| Rgl3 | 100.740861 | -0.528575937 | 0.214954882 | -2.459008757 |
| Ccdc151 | 32.36726646 | -0.159433126 | 0.228012213 | -0.699230639 |
| Prkcsh | 759.8445769 | 0.083804112 | 0.077297543 | 1.084175622 |
| Elavl3 | 2565.303786 | -0.388783997 | 0.07069234 | -5.499662274 |
| Zfp653 | 159.2068104 | 0.046690012 | 0.104114825 | 0.448447295 |
| Gm16845 | 10.87916454 | 0.086763463 | 0.315765846 | 0.274771523 |
| Ecsit | 303.4651564 | -0.209560071 | 0.069761799 | -3.00393733 |
| Cnn1 | 3.417651458 | -1.314919845 | 0.610100798 | -2.155250165 |
| Elof1 | 264.1837638 | -0.001691472 | 0.11405252 | -0.014830645 |
| Mir7084 | 0 NA | NA | NA | |
| Acp5 | 37.28268211 | 3.807143414 | 0.404599828 | 9.409651597 |
| Pigyl | 121.7874717 | -0.148058945 | 0.148150849 | -0.999379659 |
| Rpl15-ps3 | 5.54092659 | 0.774392092 | 0.441206336 | 1.755169925 |
| Zfp872 | 0.324168737 | -0.206484192 | 1.746873753 | -0.118202127 |
| 1810064F22Rik | 0.089062654 | -0.20649451 | 3.724945049 | -0.055435586 |
| Zfp809 | 131.6943064 | -0.157944765 | 0.117030848 | -1.349599423 |
| Zfp599 | 85.70332244 | 0.285872161 | 0.168004458 | 1.701574852 |

| | | | | |
|---|---|---|---|---|
| Zfp810 | 217.69277 | 0.463088792 | 0.112267596 | 4.124866005 |
| Gm10182 | 10.2804556 | 0.61038738 | 0.319988294 | 1.907530342 |
| Anln | 682.8292644 | 0.315935103 | 0.182159724 | 1.734385052 |
| 9530077C05Rik | 75.06799202 | -0.230100947 | 0.172386198 | -1.334799131 |
| Rp9 | 229.1720738 | 0.051266076 | 0.076099974 | 0.673667451 |
| Gm17545 | 0 | NA | NA | NA |
| Bbs9 | 315.6451564 | -0.125818813 | 0.088299915 | -1.424902992 |
| Gm2976 | 0 | NA | NA | NA |
| Gm16165 | 1.13143437 | -3.265494306 | 1.060709673 | -3.078593879 |
| Gm16164 | 0 | NA | NA | NA |
| Gm27639 | 0 | NA | NA | NA |
| Bmper | 91.07485983 | -0.148070426 | 0.114625768 | -1.291772595 |
| Npsr1 | 33.2953994 | 0.403588406 | 0.183785447 | 2.19597586 |
| Gm15600 | 0.84374619 | 2.668190894 | 1.156994042 | 2.306140566 |
| Gm10701 | 1.895990101 | 2.291500937 | 1.022300636 | 2.241513754 |
| Gm22380 | 0.229628521 | 0.755303593 | 2.143139772 | 0.352428527 |
| Dpy19l1 | 977.1122373 | -0.306335728 | 0.05956499 | -5.142882186 |
| Dpy19l2 | 3.565532606 | 0.228068073 | 0.561077871 | 0.406482031 |
| Cypt4 | 0 | NA | NA | NA |
| Gm38179 | 0.167183452 | -0.206481502 | 2.765153652 | -0.074672705 |
| Tbx20 | 7.101997061 | 0.88494549 | 0.530204816 | 1.669063469 |
| Gm23893 | 0 | NA | NA | NA |
| Gm10181 | 0 | NA | NA | NA |
| Gm10180 | 2.742928344 | -0.901166254 | 0.752399732 | -1.19772272 |
| Herpud2 | 587.9118386 | 0.011609381 | 0.085128682 | 0.136374493 |
| Gm29642 | 3.768175155 | -0.873810732 | 0.545634241 | -1.601458755 |
| Gm29375 | 1.122258044 | -0.634072587 | 1.019732776 | -0.621802693 |
| 7-Sep | 3678.656398 | -0.024673798 | 0.076181175 | -0.323883143 |
| Eepd1 | 188.7348402 | 0.027051206 | 0.094219446 | 0.287108525 |
| Gm25861 | 0 | NA | NA | NA |
| Gm25346 | 0 | NA | NA | NA |
| Gm26190 | 0 | NA | NA | NA |
| B3gat1 | 1268.762376 | -0.272868683 | 0.112859963 | -2.417763354 |
| Glb1l2 | 10.70062985 | -0.048618567 | 0.377800426 | -0.128688491 |
| Glb1l3 | 0.449952257 | -0.98746806 | 1.361278391 | -0.725397587 |
| Gm1110 | 2.025517295 | 0.277584583 | 0.822731061 | 0.337394072 |
| Acad8 | 225.253486 | -0.022407498 | 0.101611707 | -0.220520834 |
| Thyn1 | 253.8863181 | 0.184001937 | 0.124669497 | 1.475917855 |
| Vps26b | 1272.753477 | 0.063441416 | 0.059539336 | 1.065537852 |
| Ncapd3 | 405.8601585 | -0.160671817 | 0.167261897 | -0.960600234 |
| Jam3 | 390.9541926 | 0.01212186 | 0.132860137 | 0.09123775 |
| Gm29172 | 0.038981395 | 0.274393828 | 3.724945049 | 0.073663859 |
| Igsf9b | 818.461646 | -0.455576712 | 0.239161669 | -1.904890171 |
| Spata19 | 4.643680173 | 0.589535702 | 0.584002377 | 1.009474832 |
| Opcml | 3000.095646 | -0.212127295 | 0.082743932 | -2.563659823 |
| Gm15606 | 43.86298618 | -0.618473066 | 0.515024335 | -1.200861832 |
| AC157511.1 | 4.057004922 | -0.266384181 | 0.488400182 | -0.545421953 |

| | | | | |
|---|---|---|---|---|
| Ntm | 1996.252357 | -0.310246553 | 0.079576545 | -3.898718552 |
| Gm15520 | 1.48836363 | 1.401226593 | 1.035330773 | 1.353409586 |
| Gm15521 | 42.12449345 | -0.60706916 | 0.305879541 | -1.984667425 |
| Gm37605 | 0 | NA | NA | NA |
| Gm26435 | 0 | NA | NA | NA |
| Snx19 | 1050.451712 | 0.067690433 | 0.048799754 | 1.387106027 |
| Adamts15 | 127.7747721 | -0.327209265 | 0.119364392 | -2.741263609 |
| Adamts8 | 29.93733091 | -0.346638562 | 0.240936351 | -1.438714251 |
| Zbtb44 | 1100.679016 | -0.324779248 | 0.075138188 | -4.322425863 |
| Gm17508 | 0 | NA | NA | NA |
| St14 | 17.50112768 | 1.191327652 | 0.27521073 | 4.328783445 |
| Gm16071 | 0.620902317 | 2.442811405 | 1.31255998 | 1.861104592 |
| Aplp2 | 5929.023008 | -0.123529974 | 0.063836762 | -1.935091475 |
| Gm7244 | 0.710933469 | -2.533760218 | 1.184057601 | -2.139896079 |
| Prdm10 | 216.453307 | -0.535831841 | 0.134409386 | -3.986565643 |
| Nfrkb | 530.6418161 | 0.211324233 | 0.141182562 | 1.496815401 |
| Tmem45b | 0.191334258 | 0.755296462 | 2.673199576 | 0.282543985 |
| Barx2 | 29.27701895 | 0.199421783 | 0.317303914 | 0.628488254 |
| Gm3756 | 26.83663404 | -0.0620505 | 0.257884892 | -0.240613165 |
| AC160635.1 | 2.239755095 | -0.245138982 | 0.609008955 | -0.402521145 |
| Arhgap32 | 5811.857867 | -0.00292957 | 0.097967094 | -0.029903611 |
| Kcnj5 | 21.9094618 | -0.01474544 | 0.28251579 | -0.052193329 |
| Kcnj1 | 0.551016524 | 2.041315764 | 1.526647497 | 1.337123185 |
| Fli1 | 127.9880946 | 1.308001854 | 0.205440716 | 6.366809258 |
| Gm22060 | 0.733464214 | 1.756689048 | 1.167607517 | 1.50452016 |
| Gm38292 | 5.098635657 | -0.337005419 | 0.503834586 | -0.668881074 |
| Gm38346 | 11.05146628 | 0.899648159 | 0.447460955 | 2.010562371 |
| Gm27201 | 0.169165009 | 0.755292496 | 3.351424224 | 0.225364635 |
| Gm37474 | 11.13391284 | 0.592153086 | 0.527060624 | 1.123500902 |
| Gm37082 | 13.07759707 | 0.616550613 | 0.470156379 | 1.311373495 |
| Gm27163 | 0 | NA | NA | NA |
| Ets1 | 209.3380529 | 1.568209894 | 0.210477434 | 7.450726974 |
| Gm27240 | 0 | NA | NA | NA |
| Gm3331 | 0 | NA | NA | NA |
| Gm37167 | 0.620529807 | -2.783781475 | 1.30963355 | -2.125618632 |
| Gm27162 | 0 | NA | NA | NA |
| Gm37192 | 0.061920613 | 0.274393828 | 3.724945049 | 0.073663859 |
| Gm27166 | 0 | NA | NA | NA |
| Gm37758 | 2.309104371 | 1.105251995 | 0.65892763 | 1.677349597 |
| Gm10698 | 69.88234991 | 0.560177994 | 0.168282064 | 3.328803915 |
| Gm25439 | 0 | NA | NA | NA |
| Gm27165 | 0 | NA | NA | NA |
| Gm27164 | 0 | NA | NA | NA |
| 7630403G23Rik | 0.458133289 | -0.712515452 | 1.545771679 | -0.460944823 |
| Gm27161 | 0 | NA | NA | NA |
| Gm38354 | 0 | NA | NA | NA |
| Kirrel3 | 816.5410815 | -0.066684854 | 0.068007362 | -0.980553468 |

| | | | | |
|---|---|---|---|---|
| Gm38217 | 71.70747919 | -0.686778888 | 0.293412285 | -2.340661666 |
| Gm37038 | 54.63741103 | -0.729332342 | 0.272395631 | -2.677474446 |
| Gm26662 | 0.042716945 | -0.206490061 | 3.724945049 | -0.055434391 |
| Kirrel3os | 3.153914063 | -0.278568034 | 0.680192837 | -0.409542734 |
| Gm23702 | 2.111071361 | -0.046192334 | 0.629566992 | -0.073371595 |
| St3gal4 | 323.0715017 | 0.266277028 | 0.097236989 | 2.738433509 |
| 4930581F22Rik | 31.67819825 | -0.262658146 | 0.213587913 | -1.229742556 |
| Dcps | 130.3926271 | 0.03861166 | 0.131551416 | 0.293510026 |
| Gm24262 | 0 | NA | NA | NA |
| Tirap | 88.4573936 | 0.763207548 | 0.198526779 | 3.844355663 |
| Srpr | 878.7068951 | 0.260740237 | 0.072488844 | 3.596970556 |
| Foxred1 | 265.905042 | 0.331523191 | 0.095958022 | 3.454877277 |
| Fam118b | 211.7892874 | -0.273169947 | 0.103923801 | -2.628560017 |
| Rpusd4 | 78.94005876 | -0.153128737 | 0.159443653 | -0.960394061 |
| Gm5614 | 0 | NA | NA | NA |
| 2610105M22Rik | 0.044191549 | -0.206490061 | 3.724945049 | -0.055434391 |
| Cdon | 259.9464149 | -0.177128405 | 0.14523395 | -1.219607433 |
| Gm25630 | 1.242022914 | -1.184321304 | 0.852935009 | -1.388524672 |
| Gm24188 | 0 | NA | NA | NA |
| Gm1113 | 0 | NA | NA | NA |
| Ddx25 | 512.2202113 | -0.096093331 | 0.054129768 | -1.775240035 |
| Pus3 | 127.6005767 | 0.108001132 | 0.107775252 | 1.00209584 |
| Hyls1 | 86.77207718 | -0.144120368 | 0.146897715 | -0.981093329 |
| Pate2 | 15.21999782 | 0.053280532 | 0.314505947 | 0.169410253 |
| Gm6762 | 0 | NA | NA | NA |
| Gm10105 | 0 | NA | NA | NA |
| Pate4 | 0 | NA | NA | NA |
| Gm17365 | 0 | NA | NA | NA |
| Gm17252 | 0 | NA | NA | NA |
| Gm17677 | 0.297465678 | 1.227042182 | 1.757296257 | 0.698255731 |
| Gm27235 | 0 | NA | NA | NA |
| Gm17727 | 0 | NA | NA | NA |
| D730048I06Rik | 0 | NA | NA | NA |
| Gm36974 | 1.090715414 | -0.588568154 | 0.912199364 | -0.645218773 |
| 9230110F15Rik | 0 | NA | NA | NA |
| Gm3428 | 0.042716945 | -0.206490061 | 3.724945049 | -0.055434391 |
| 9230113P08Rik | 1.958122126 | -0.388506527 | 0.71254957 | -0.545234386 |
| Gm27187 | 0 | NA | NA | NA |
| Gm5916 | 0 | NA | NA | NA |
| Gm3434 | 0 | NA | NA | NA |
| Gm3867 | 0.061920613 | 0.274393828 | 3.724945049 | 0.073663859 |
| Gm7257 | 0 | NA | NA | NA |
| Gm9513 | 0 | NA | NA | NA |
| Gm27160 | 0 | NA | NA | NA |
| Gm5615 | 0 | NA | NA | NA |
| Gm27218 | 0 | NA | NA | NA |
| Gm17689 | 0.194279094 | 0.274391173 | 3.724945049 | 0.073663147 |

| | | | | |
|---|---|---|---|---|
| A630095E13Rik | 0.217428854 | 0.274401121 | 2.581304093 | 0.106303291 |
| 1700027I24Rik | 0.357600009 | 0.75530742 | 1.728441349 | 0.436987591 |
| Acrv1 | 0.095898756 | 0.27438722 | 3.724945049 | 0.073662085 |
| Chek1 | 39.39245048 | -0.028932231 | 0.205287632 | -0.140935093 |
| Stt3a | 958.0649057 | 0.157244138 | 0.067572877 | 2.327030368 |
| Gm26787 | 0.495189146 | 1.006500435 | 1.24207491 | 0.810337949 |
| Ei24 | 869.5859806 | -0.048509906 | 0.070001479 | -0.692984015 |
| Gm27219 | 6.744631513 | 0.481694412 | 0.430520474 | 1.118865282 |
| Fez1 | 1689.638317 | -0.264571422 | 0.060878288 | -4.345907702 |
| Pknox2 | 477.999823 | -0.3024567 | 0.106157767 | -2.849124551 |
| Gm3896 | 0.132101749 | 0.274387056 | 3.724945049 | 0.073662041 |
| n-R5s82 | 0 | NA | NA | NA |
| Gm25273 | 0 | NA | NA | NA |
| Tmem218 | 64.43814503 | -0.602194839 | 0.136691339 | -4.405508381 |
| Slc37a2 | 140.0947805 | 0.955981484 | 0.266313204 | 3.589688647 |
| Gm10177 | 68.34373515 | 0.134927472 | 0.206003388 | 0.654976958 |
| Ccdc15 | 56.68857134 | -0.228084692 | 0.187029795 | -1.219509932 |
| Hepacam | 1455.687833 | 0.405214293 | 0.127636941 | 3.174741485 |
| Robo4 | 52.91085951 | 1.709018692 | 0.256238035 | 6.669652667 |
| Robo3 | 261.3456688 | 1.381321275 | 0.424252884 | 3.255891302 |
| Msantd2 | 153.0368494 | -0.303754559 | 0.104221159 | -2.914519111 |
| Esam | 83.18252856 | 0.973175068 | 0.167310375 | 5.816585289 |
| Vsig2 | 19.39362468 | 1.461104796 | 0.349482823 | 4.180762834 |
| Nrgn | 5315.665183 | -0.121394472 | 0.205480347 | -0.59078386 |
| Spa17 | 8.072628227 | -0.533763855 | 0.420366124 | -1.269759443 |
| Siae | 198.8133421 | -0.062590221 | 0.133429367 | -0.469088795 |
| Tbrg1 | 650.3765193 | 1.005192428 | 0.124201589 | 8.09323321 |
| Panx3 | 0.252375374 | 0.262029633 | 2.126445668 | 0.123224231 |
| Gm8343 | 0 | NA | NA | NA |
| Tpt1-ps5 | 7.622765194 | 0.325536616 | 0.44540908 | 0.730871082 |
| Olfr160 | 0 | NA | NA | NA |
| Olfr874 | 0 | NA | NA | NA |
| Olfr875 | 0.072025462 | 0.755288839 | 3.724945049 | 0.202765096 |
| Olfr876 | 0.066050875 | 0.274391604 | 3.724945049 | 0.073663262 |
| Gm24764 | 0 | NA | NA | NA |
| Olfr877 | 0.162452565 | 1.662034851 | 2.773995847 | 0.599148284 |
| Olfr145 | 0.104637558 | -0.206492286 | 3.724945049 | -0.055434988 |
| Olfr878 | 0.170434537 | -0.206482566 | 2.328336882 | -0.088682427 |
| Olfr881 | 0 | NA | NA | NA |
| Olfr883 | 0 | NA | NA | NA |
| Olfr884 | 0.092140247 | -0.687373951 | 3.724945049 | -0.184532642 |
| Olfr885 | 0 | NA | NA | NA |
| Olfr887 | 0.072025462 | 0.755288839 | 3.724945049 | 0.202765096 |
| Olfr888 | 0 | NA | NA | NA |
| Olfr889 | 0 | NA | NA | NA |
| Olfr890 | 0 | NA | NA | NA |
| Olfr891 | 0 | NA | NA | NA |

| | | | | |
|---|---|---|---|---|
| Gm17798 | 0 | NA | NA | NA |
| Olfr892-ps1 | 0 | NA | NA | NA |
| Olfr893 | 0 | NA | NA | NA |
| Olfr894 | 0 | NA | NA | NA |
| Olfr143 | 0 | NA | NA | NA |
| Olfr895 | 0 | NA | NA | NA |
| Olfr25 | 0.111376404 | 0.274404949 | 3.724945049 | 0.073666845 |
| Olfr898 | 0 | NA | NA | NA |
| Olfr250 | 0.072025462 | 0.755288839 | 3.724945049 | 0.202765096 |
| Olfr251 | 0 | NA | NA | NA |
| Olfr147 | 0.068097359 | -0.206487837 | 3.724945049 | -0.055433794 |
| Olfr901 | 0.114443069 | -0.206492286 | 3.724945049 | -0.055434988 |
| Olfr902 | 0 | NA | NA | NA |
| Olfr904 | 0 | NA | NA | NA |
| Olfr905 | 0 | NA | NA | NA |
| Olfr906 | 0 | NA | NA | NA |
| Olfr907 | 0 | NA | NA | NA |
| Gm23271 | 0 | NA | NA | NA |
| Olfr908 | 0 | NA | NA | NA |
| Olfr911-ps1 | 0 | NA | NA | NA |
| Olfr910 | 0 | NA | NA | NA |
| Olfr912 | 0 | NA | NA | NA |
| Olfr913 | 0 | NA | NA | NA |
| Olfr914 | 0 | NA | NA | NA |
| Olfr916 | 0 | NA | NA | NA |
| Olfr917 | 0 | NA | NA | NA |
| Olfr918 | 0 | NA | NA | NA |
| Olfr919 | 0 | NA | NA | NA |
| Vwa5a | 1119.696084 | 2.600102808 | 0.165807085 | 15.68149402 |
| Olfr920 | 22.44487698 | 3.569526773 | 0.494514438 | 7.218245814 |
| Olfr921 | 0.704586541 | 2.652381506 | 1.502308975 | 1.765536618 |
| Olfr922 | 0.673070397 | -1.134889593 | 1.206335594 | -0.940774357 |
| Olfr923 | 0.176917541 | -0.687380517 | 2.767967824 | -0.248333998 |
| Olfr924 | 0.326455024 | -2.11858144 | 1.663865731 | -1.273288704 |
| Olfr26 | 0.232584034 | 0.776370376 | 2.13051607 | 0.364404844 |
| Olfr926 | 0.106056992 | 1.236179401 | 3.724945049 | 0.331865138 |
| Olfr930 | 0 | NA | NA | NA |
| Olfr933 | 0 | NA | NA | NA |
| Olfr934 | 0 | NA | NA | NA |
| Olfr935 | 0 | NA | NA | NA |
| Olfr146 | 0 | NA | NA | NA |
| Olfr936 | 0 | NA | NA | NA |
| Gm25529 | 0 | NA | NA | NA |
| Olfr937 | 0 | NA | NA | NA |
| Gm25032 | 0 | NA | NA | NA |
| Olfr938 | 0 | NA | NA | NA |
| Olfr27 | 0 | NA | NA | NA |

| | | | | |
|---|---|---|---|---|
| Olfr943 | 0 | NA | NA | NA |
| Olfr944 | 0 | NA | NA | NA |
| Olfr1537 | 0 | NA | NA | NA |
| Olfr945 | 0 | NA | NA | NA |
| Olfr948 | 0 | NA | NA | NA |
| Olfr951 | 0 | NA | NA | NA |
| Olfr952 | 0 | NA | NA | NA |
| Olfr954 | 0 | NA | NA | NA |
| Olfr955 | 0 | NA | NA | NA |
| Olfr44 | 0.046345709 | 0.274391604 | 3.724945049 | 0.073663262 |
| Olfr957 | 0 | NA | NA | NA |
| Gm24278 | 0 | NA | NA | NA |
| Olfr958 | 0 | NA | NA | NA |
| Olfr959 | 0.090427103 | 1.236179401 | 3.724945049 | 0.331865138 |
| AW551984 | 246.7130081 | -0.137458029 | 0.155531802 | -0.883793719 |
| Olfr148 | 0 | NA | NA | NA |
| Olfr960 | 0 | NA | NA | NA |
| Olfr961 | 0 | NA | NA | NA |
| Olfr963 | 0 | NA | NA | NA |
| Olfr964-ps1 | 0.048949532 | -0.206487837 | 3.724945049 | -0.055433794 |
| Olfr149 | 0 | NA | NA | NA |
| Olfr965 | 0 | NA | NA | NA |
| Olfr150 | 0 | NA | NA | NA |
| Olfr967 | 0 | NA | NA | NA |
| Olfr968 | 0.046345709 | 0.274391604 | 3.724945049 | 0.073663262 |
| Olfr969 | 0 | NA | NA | NA |
| Olfr970 | 0 | NA | NA | NA |
| Olfr971 | 0 | NA | NA | NA |
| Olfr972 | 0 | NA | NA | NA |
| Olfr229 | 0 | NA | NA | NA |
| Olfr974 | 0.049553047 | 0.274391604 | 3.724945049 | 0.073663262 |
| Olfr975 | 0 | NA | NA | NA |
| Olfr976 | 0.111006857 | 0.755286617 | 3.724945049 | 0.202764499 |
| Olfr978 | 0 | NA | NA | NA |
| Olfr979 | 0.046345709 | 0.274391604 | 3.724945049 | 0.073663262 |
| Olfr980 | 0.09176491 | 1.15872478 | 3.72306111 | 0.311229052 |
| Olfr981 | 0 | NA | NA | NA |
| 1700001J11Rik | 0.049553047 | 0.274391604 | 3.724945049 | 0.073663262 |
| Olfr982 | 0 | NA | NA | NA |
| Olfr983 | 0 | NA | NA | NA |
| Olfr984 | 0.072025462 | 0.755288839 | 3.724945049 | 0.202765096 |
| Olfr985 | 0.09176491 | 1.15872478 | 3.72306111 | 0.311229052 |
| Tmem225 | 0 | NA | NA | NA |
| Olfr986 | 0 | NA | NA | NA |
| Zfp202 | 72.13492 | 0.003176867 | 0.148250142 | 0.021429102 |
| Scn3b | 970.5865091 | -0.290027317 | 0.104684828 | -2.770480896 |
| Gramd1b | 3247.828285 | -0.431683096 | 0.080860803 | -5.338595213 |

| | | | | |
|---|---|---|---|---|
| 1700110K17Rik | 1.343476617 | -0.264692834 | 0.937573195 | -0.282316981 |
| Gm17540 | 0 | NA | NA | NA |
| Gm21915 | 0 | NA | NA | NA |
| Clmp | 308.4342746 | -0.419920008 | 0.128100314 | -3.27805604 |
| Gm16095 | 0.148850706 | -0.206486747 | 2.85897618 | -0.072224018 |
| Gm16096 | 0.692884898 | 1.195682668 | 1.405202515 | 0.850897045 |
| Hspa8 | 7490.665187 | -0.154750096 | 0.080094205 | -1.932101023 |
| Snord14c | 0 | NA | NA | NA |
| Snord14d | 0 | NA | NA | NA |
| Snord14e | 0 | NA | NA | NA |
| Bsx | 2.789926683 | 0.531460654 | 0.596081822 | 0.891590103 |
| 4931429I11Rik | 11.72828212 | 0.471167536 | 0.326509781 | 1.44304264 |
| Crtam | 123.3542397 | 0.469779533 | 0.248075039 | 1.893699317 |
| Ubash3b | 620.5872832 | -0.160238118 | 0.093380705 | -1.715966028 |
| 1700063D05Rik | 6.117005653 | -0.459573305 | 0.388143899 | -1.184028154 |
| Gm25401 | 0 | NA | NA | NA |
| 2610203C20Rik | 984.4296156 | 0.047978727 | 0.145309566 | 0.330182853 |
| Mir100 | 2.750782513 | -0.223511972 | 0.678119077 | -0.329605787 |
| Mirlet7a-2 | 2.112107905 | 0.634080388 | 0.698904312 | 0.907249215 |
| Gm28119 | 36.18489172 | 0.25987481 | 0.237435391 | 1.094507473 |
| Mir125b-1 | 0 | NA | NA | NA |
| Gm8707 | 28.79399415 | 0.362098836 | 0.205019833 | 1.766164912 |
| Sorl1 | 7353.79371 | -0.013089763 | 0.075592051 | -0.17316322 |
| Gm16214 | 0 | NA | NA | NA |
| Sc5d | 491.2699012 | -0.200164327 | 0.105126577 | -1.904031628 |
| Gm26272 | 0 | NA | NA | NA |
| Tecta | 9.746868474 | 0.771105532 | 0.463516498 | 1.66359889 |
| Tbcel | 691.6961961 | -0.141610543 | 0.058817414 | -2.407629522 |
| AC136903.1 | 0 | NA | NA | NA |
| Gm16322 | 0.291180489 | 1.776633033 | 3.708688794 | 0.479046135 |
| Grik4 | 287.3585913 | -0.25716567 | 0.09744688 | -2.639034415 |
| Gm24420 | 0 | NA | NA | NA |
| Gm27509 | 1.432503634 | -0.976700129 | 0.825800836 | -1.182730855 |
| Arhgef12 | 4563.170901 | -0.094760827 | 0.038429454 | -2.465838485 |
| Tmem136 | 181.8275258 | -0.263279491 | 0.105242386 | -2.501648811 |
| Pou2f3 | 2.121136815 | 1.492192902 | 0.787713086 | 1.894335549 |
| Oaf | 220.6241672 | 0.826427657 | 0.167748402 | 4.926590355 |
| D630033O11Rik | 2.615308024 | 0.153787709 | 0.625669117 | 0.245797187 |
| Trim29 | 4.261450001 | 0.279937093 | 0.626256549 | 0.447000663 |
| Gm28215 | 0 | NA | NA | NA |
| Pvrl1 | 1033.013847 | -0.131534474 | 0.079014663 | -1.664684355 |
| Gm3898 | 0.044544648 | 0.755286614 | 3.724945049 | 0.202764498 |
| Gm23326 | 0.048949532 | -0.206487837 | 3.724945049 | -0.055433794 |
| Thy1 | 4661.495848 | -0.02630174 | 0.077846239 | -0.337867831 |
| Rnf26 | 340.15711 | 0.08015092 | 0.077215695 | 1.038013321 |
| Usp2 | 947.8979254 | 0.084463659 | 0.062081665 | 1.360525041 |
| Mfrp | 54.45536832 | 0.103427631 | 0.251694319 | 0.41092557 |

| | | | | |
|---|---|---|---|---|
| C1qtnf5 | 0 | NA | NA | NA |
| Gm20444 | 3.364322678 | -1.410966957 | 0.614806032 | -2.294979039 |
| Gm26737 | 7.342448606 | 0.506349715 | 0.401293375 | 1.261794352 |
| Gm10687 | 8.115134709 | -0.509466004 | 0.369116797 | -1.380229803 |
| Mcam | 257.1599992 | 0.032234492 | 0.154828217 | 0.2081952 |
| Cbl | 967.2924709 | -0.021244359 | 0.073037203 | -0.29087038 |
| Ccdc153 | 11.25998896 | -1.10077005 | 0.368465637 | -2.987442893 |
| Pdzd3 | 3.822296293 | -0.659584414 | 0.620490074 | -1.063005586 |
| Nlrx1 | 97.08268411 | 0.612083964 | 0.143776495 | 4.257190744 |
| Abcg4 | 564.6710316 | -0.344634825 | 0.084371976 | -4.084707255 |
| Hinfp | 148.4591195 | -0.11787186 | 0.135345406 | -0.870896645 |
| C2cd2l | 1532.300989 | -0.243067525 | 0.109253686 | -2.224799301 |
| Dpagt1 | 146.9363667 | -0.042319232 | 0.116114823 | -0.364460201 |
| H2afx | 329.2993814 | 0.153272997 | 0.099434399 | 1.541448423 |
| AC122428.1 | 1.088551472 | 0.054038761 | 1.027487255 | 0.05259312 |
| Hmbs | 132.3982402 | 0.099738866 | 0.128700127 | 0.774970999 |
| Vps11 | 770.3516232 | 0.247433655 | 0.0663159 | 3.731136178 |
| Gm22141 | 0.240440413 | -0.174660559 | 2.14647057 | -0.081371047 |
| Gm10080 | 110.1720267 | 0.125406691 | 0.167687183 | 0.747860922 |
| Hyou1 | 2036.023468 | 0.107495752 | 0.08130666 | 1.322102668 |
| Gm26306 | 0 | NA | NA | NA |
| Slc37a4 | 109.665464 | -0.271516555 | 0.13206573 | -2.055919849 |
| Mir7085 | 0.487060322 | 0.379353938 | 1.36692295 | 0.27752401 |
| Trappc4 | 251.412174 | 0.055731519 | 0.12736009 | 0.437590136 |
| Rps25 | 1102.353914 | 0.44566865 | 0.131807103 | 3.381218767 |
| Ccdc84 | 64.93599903 | -0.432468985 | 0.164337096 | -2.63159686 |
| Foxr1 | 0.310849204 | 1.297135236 | 3.72109831 | 0.348589349 |
| Upk2 | 0.044191549 | -0.206490061 | 3.724945049 | -0.055434391 |
| Gm9830 | 5.961436335 | -0.252817828 | 0.410942539 | -0.615214549 |
| Gm22540 | 5.014127157 | -0.781983828 | 0.440597707 | -1.774825006 |
| C030014I23Rik | 47.83992227 | -0.25032693 | 0.160082478 | -1.563737224 |
| Bcl9l | 785.1067627 | -0.052590155 | 0.081705024 | -0.643658771 |
| Cxcr5 | 2.247268683 | 4.440139822 | 1.007765543 | 4.405925419 |
| Ddx6 | 4111.857025 | 0.080856648 | 0.053521797 | 1.510723711 |
| Treh | 0.042716945 | -0.206490061 | 3.724945049 | -0.055434391 |
| Phldb1 | 1542.489038 | -0.224178341 | 0.071175516 | -3.149655293 |
| Gm24166 | 0 | NA | NA | NA |
| Arcn1 | 1826.393337 | 0.240957593 | 0.081384926 | 2.960715281 |
| Ift46 | 313.3153656 | -0.106402839 | 0.074681913 | -1.424747093 |
| Tmem25 | 356.7404894 | -0.493372395 | 0.094944512 | -5.196428763 |
| Ttc36 | 3.625487916 | -0.233240317 | 0.611099336 | -0.381673327 |
| Kmt2a | 3944.583612 | -0.286685246 | 0.093737994 | -3.058367628 |
| Gm26249 | 0 | NA | NA | NA |
| Atp5l | 692.9294976 | -0.071024375 | 0.138330763 | -0.51343876 |
| Ube4a | 1166.455959 | -0.033311253 | 0.057919226 | -0.575132902 |
| Cd3g | 29.68427331 | 6.85164152 | 0.907385327 | 7.550972353 |
| Cd3d | 21.88467821 | 7.47897194 | 0.876930216 | 8.528582782 |

| | | | | |
|---|---|---|---|---|
| Cd3e | 72.02395476 | 8.616257932 | 0.881605131 | 9.773375434 |
| Mpzl2 | 55.99719323 | 1.032644821 | 0.236695016 | 4.362765373 |
| Mpzl3 | 77.05801845 | 0.573284958 | 0.147028894 | 3.899131281 |
| Amica1 | 164.8869608 | 3.71123045 | 0.347144901 | 10.69072435 |
| Gm10684 | 5.682893444 | -0.077710439 | 0.457036049 | -0.170031312 |
| Scn2b | 3837.721104 | -0.265507352 | 0.097809412 | -2.714537856 |
| Scn4b | 2000.069508 | -0.33610606 | 0.125834421 | -2.671018451 |
| Tmprss4 | 3.058856891 | 0.186244409 | 0.592498902 | 0.314337139 |
| Gm17099 | 0.153914375 | -0.684655047 | 3.503921201 | -0.195396816 |
| BC049352 | 7.258084432 | -0.256352205 | 0.472737913 | -0.542271305 |
| Il10ra | 224.5975288 | 2.120925498 | 0.152684649 | 13.89088888 |
| Gm20612 | 0.047948698 | -0.206487837 | 3.724945049 | -0.055433794 |
| Mir7086 | 0.58926593 | 1.474488551 | 1.311575849 | 1.124211422 |
| 1700003G13Rik | 0 | NA | NA | NA |
| Tmprss13 | 0.459044381 | 1.981327464 | 1.531773097 | 1.293486266 |
| Fxyd6 | 526.6237023 | -0.052408428 | 0.109925183 | -0.476764529 |
| Fxyd2 | 9.473000343 | 0.278499743 | 0.346860823 | 0.802914959 |
| 4833428L15Rik | 0.565850957 | -2.261821324 | 1.147947007 | -1.970318586 |
| Dscaml1 | 663.8123879 | -0.138704828 | 0.113801247 | -1.218833996 |
| Gm22069 | 0 | NA | NA | NA |
| Cep164 | 262.2380998 | -0.241717129 | 0.1337845 | -1.806764826 |
| Gm7286 | 0.119587853 | 1.218392466 | 3.723618458 | 0.327206581 |
| Bace1 | 1452.453558 | -0.142523028 | 0.044334418 | -3.214726505 |
| Gm16536 | 127.1699055 | -0.137244915 | 0.106310194 | -1.290985465 |
| Rnf214 | 859.5718938 | -0.198393104 | 0.04781822 | -4.148901934 |
| Gm24646 | 2.996628699 | 0.557632184 | 0.893722119 | 0.623943587 |
| Pcsk7 | 359.8709397 | 0.362705582 | 0.085954915 | 4.21971895 |
| Tagln | 130.3382176 | 1.219709096 | 0.446022887 | 2.734633428 |
| Sidt2 | 826.3607848 | 0.138748439 | 0.08484745 | 1.635269396 |
| Mir7087 | 1.036669434 | 1.432113576 | 1.014133679 | 1.412154636 |
| Pafah1b2 | 2591.382288 | 0.058162749 | 0.066877974 | 0.869684674 |
| Sik3 | 2827.253407 | 0.310630535 | 0.081657211 | 3.804079644 |
| Apoa1 | 4.148602219 | 0.901796351 | 0.525421535 | 1.716329252 |
| Apoc3 | 5.512392282 | 2.336401616 | 0.519316808 | 4.498990943 |
| Apoa4 | 0.183366429 | 0.274411468 | 2.271808962 | 0.120789852 |
| Gm10680 | 0 | NA | NA | NA |
| Apoa5 | 0.33367842 | -0.206486564 | 2.011886159 | -0.102633324 |
| Zpr1 | 322.0139224 | 0.239669239 | 0.074143646 | 3.232498701 |
| Bud13 | 129.6764734 | 0.109774698 | 0.103620102 | 1.05939578 |
| 4931429L15Rik | 0.137762903 | -0.206480416 | 2.937523348 | -0.070290647 |
| 4930448E22Rik | 1.471695147 | 0.643622192 | 0.990653454 | 0.649694592 |
| Gm22659 | 0.246177344 | -1.231984836 | 2.49861285 | -0.493067518 |
| Gm22805 | 0 | NA | NA | NA |
| 2900052N01Rik | 275.6261956 | 0.647465373 | 0.15952958 | 4.058591361 |
| Gm4791 | 0.045882455 | 0.755288839 | 3.724945049 | 0.202765096 |
| Gm22286 | 0.075043205 | 0.755286614 | 3.724945049 | 0.202764498 |
| Cadm1 | 2092.339405 | 0.093393909 | 0.066450065 | 1.40547505 |

| | | | | |
|---|---|---|---|---|
| Nxpe2 | 2.03192395 | 2.629695766 | 0.989345411 | 2.658015831 |
| Nxpe4 | 71.31690674 | 0.407596386 | 0.135393341 | 3.010461091 |
| Gm5616 | 4.81606905 | -0.329176073 | 0.489165514 | -0.672933933 |
| Rexo2 | 393.6205041 | 0.002795876 | 0.103855765 | 0.026920762 |
| Rbm7 | 414.2773601 | 0.818111833 | 0.112441598 | 7.275882313 |
| Gm5617 | 29.90601096 | 0.201006141 | 0.235757477 | 0.852597098 |
| Gm23653 | 0 | NA | NA | NA |
| Nnmt | 5.720995077 | 2.526549668 | 0.656136981 | 3.850643604 |
| Zbtb16 | 1098.476404 | 0.469761228 | 0.175856676 | 2.67127321 |
| Htr3a | 33.96213922 | 0.301670673 | 0.220519128 | 1.368002293 |
| Htr3b | 0.066050875 | 0.274391604 | 3.724945049 | 0.073663262 |
| Usp28 | 333.1237609 | -0.69009384 | 0.11942676 | -5.778385366 |
| Cldn21 | 0 | NA | NA | NA |
| RP23-136A1.2 | 0 | NA | NA | NA |
| Zw10 | 188.7291143 | -0.173419153 | 0.094024054 | -1.844412627 |
| Tmprss5 | 18.34493739 | -1.106957793 | 0.277413063 | -3.990287198 |
| Gm4894 | 0.044544648 | 0.755286614 | 3.724945049 | 0.202764498 |
| Drd2 | 144.2458334 | -0.373738195 | 0.222648072 | -1.678605124 |
| Ankk1 | 0.421296087 | -0.67748766 | 1.475295045 | -0.459221809 |
| Ttc12 | 60.41204789 | -0.357704608 | 0.177822267 | -2.011585016 |
| Ncam1 | 4558.689867 | -0.221461187 | 0.05658628 | -3.913690543 |
| Gm11149 | 1.147320409 | -0.83012754 | 0.980373894 | -0.846745864 |
| Gm25633 | 0 | NA | NA | NA |
| Gm8907 | 0.242574275 | -0.687379871 | 2.536878224 | -0.270955013 |
| Rpl10-ps3 | 4.878838789 | 0.43231654 | 0.460608061 | 0.938577884 |
| Plet1os | 0.85456564 | -2.012818785 | 1.147910482 | -1.753463198 |
| Plet1 | 4.418087894 | 1.011647184 | 0.502356047 | 2.013805128 |
| Pts | 145.2821715 | -0.120744758 | 0.133031312 | -0.907641636 |
| Bco2 | 15.34273135 | -0.187588872 | 0.273341765 | -0.686279581 |
| Tex12 | 0.048949532 | -0.206487837 | 3.724945049 | -0.055433794 |
| Il18 | 136.7701011 | 0.519513184 | 0.138212239 | 3.75880739 |
| Sdhd | 647.1493877 | 0.116569425 | 0.073087668 | 1.594926035 |
| Timm8b | 499.1712965 | 3.94E-05 | 0.148935735 | 0.000264826 |
| AU019823 | 218.1162646 | 0.269427584 | 0.117530068 | 2.292414096 |
| Pih1d2 | 29.01466556 | 0.038577433 | 0.190745041 | 0.202246059 |
| Dlat | 1112.83225 | -0.009770239 | 0.058877841 | -0.165940855 |
| Dixdc1 | 1202.736387 | -0.505651131 | 0.074771803 | -6.762591124 |
| 2310030G06Rik | 16.94180656 | 0.518994321 | 0.245535919 | 2.11372056 |
| Hspb2 | 5.620585044 | 1.789821932 | 0.522141676 | 3.427847294 |
| Cryab | 871.6678754 | 0.031158823 | 0.200445599 | 0.155447779 |
| 1110032A03Rik | 144.5010807 | 0.242697236 | 0.110748221 | 2.191432369 |
| Fdxacb1 | 52.09799359 | -0.007442441 | 0.152128883 | -0.048921944 |
| Alg9 | 194.0505543 | -0.033844861 | 0.086055103 | -0.3932929 |
| Ppp2r1b | 243.7699036 | 0.252494791 | 0.104881538 | 2.40742838 |
| Gm25558 | 0.177937505 | -0.600440611 | 2.732390334 | -0.219749208 |
| Sik2 | 790.3337553 | -0.313125348 | 0.105105465 | -2.979153835 |
| Layn | 62.49321233 | 0.761954148 | 0.255174236 | 2.986015206 |

| | | | | |
|---|---|---|---|---|
| 4833427G06Rik | 5.495872342 | 0.713766827 | 0.485665288 | 1.469668194 |
| Mir34c | 0 | NA | NA | NA |
| Mir34b | 0 | NA | NA | NA |
| Btg4 | 0.260468428 | 0.766818633 | 2.044308664 | 0.375099243 |
| Pou2af1 | 4.531271601 | 1.855493463 | 0.839878748 | 2.209239688 |
| Gm684 | 7.066067338 | -0.963957451 | 0.429818777 | -2.242706701 |
| 2010007H06Rik | 9.722359338 | -0.604801243 | 0.352757384 | -1.714496337 |
| 1810046K07Rik | 0.216202823 | -0.206480616 | 2.17332255 | -0.09500689 |
| Gm7293 | 4.644993497 | -0.44471931 | 0.535135502 | -0.831040566 |
| Arhgap20 | 1285.799934 | -0.186453374 | 0.087845135 | -2.122523628 |
| Arhgap20os | 1.805117362 | -0.154173824 | 0.949429819 | -0.162385698 |
| Fdx1 | 101.2393919 | 0.14135457 | 0.152264449 | 0.928349137 |
| Gm6981 | 16.53353035 | 0.25249435 | 0.288314224 | 0.87576099 |
| Rdx | 1573.201998 | 0.235132252 | 0.091433757 | 2.571613146 |
| Gm27686 | 0 | NA | NA | NA |
| Zc3h12c | 704.8556827 | 0.334848771 | 0.121675158 | 2.751989609 |
| Mir6244 | 0.506103766 | 1.353694486 | 3.320024167 | 0.407736335 |
| Gm25562 | 0 | NA | NA | NA |
| AI593442 | 2277.537907 | -0.310375341 | 0.100573749 | -3.086047252 |
| Gm22242 | 0 | NA | NA | NA |
| Ddx10 | 340.9961152 | 0.049257253 | 0.08690697 | 0.566781395 |
| 4930510E17Rik | 0 | NA | NA | NA |
| Exph5 | 843.2123267 | -0.354535434 | 0.18514241 | -1.914933659 |
| Kdelc2 | 102.2218059 | -0.021973924 | 0.123565186 | -0.177832646 |
| Mir130c | 0.049553047 | 0.274391604 | 3.724945049 | 0.073663262 |
| 4930550C14Rik | 3.372172578 | 0.344385566 | 0.572845763 | 0.601183754 |
| Atm | 883.8572749 | 0.665697451 | 0.097577872 | 6.822217308 |
| Npat | 649.966786 | 0.087382074 | 0.094828215 | 0.921477578 |
| Acat1 | 830.5020792 | 0.201496034 | 0.064298199 | 3.133774166 |
| Cul5 | 1233.233245 | -0.144166749 | 0.081469043 | -1.76958933 |
| Gm22304 | 0 | NA | NA | NA |
| Gm16124 | 9.053711666 | 0.113131789 | 0.362244479 | 0.312307835 |
| Rab39 | 22.16151509 | 0.06513858 | 0.242244572 | 0.268895933 |
| Rps6-ps3 | 1.738215179 | 1.059019213 | 0.737193255 | 1.436555755 |
| Slc35f2 | 23.18056578 | 0.172231909 | 0.227781572 | 0.756127494 |
| Sln | 3.22564905 | 0.296747618 | 0.616419969 | 0.481404939 |
| Gm16380 | 0.31602048 | 1.140372985 | 1.560088186 | 0.730967002 |
| Elmod1 | 1587.467376 | -0.211494241 | 0.062844165 | -3.365375927 |
| Tnfaip8l3 | 95.49101603 | 0.491914558 | 0.14262047 | 3.449116087 |
| Cyp19a1 | 1.568920191 | -0.150277461 | 0.799841726 | -0.187883998 |
| Gldnos | 0.038981395 | 0.274393828 | 3.724945049 | 0.073663859 |
| Gldn | 61.66008331 | 1.286046416 | 0.26686548 | 4.819081195 |
| Dmxl2 | 4593.55088 | -0.301213456 | 0.067770568 | -4.444605743 |
| Gm25980 | 0 | NA | NA | NA |
| Sh2d7 | 0.788119377 | 2.392776924 | 1.407829815 | 1.699620862 |
| Cib2 | 155.6054035 | -0.304461927 | 0.134675492 | -2.260707729 |
| Idh3a | 1642.060698 | -0.191335291 | 0.066056413 | -2.896543764 |

| | | | | |
|---|---|---|---|---|
| Acsbg1 | 1565.688373 | -0.151639421 | 0.147353697 | -1.0290846 |
| Dnaja4 | 456.041311 | -0.226574017 | 0.119163036 | -1.901378357 |
| Mir5710 | 0.417007765 | -0.049512218 | 1.381909977 | -0.03582883 |
| Wdr61 | 268.6245129 | -0.096722347 | 0.093986613 | -1.029107695 |
| Crabp1 | 9.084845767 | 0.083036301 | 0.349775691 | 0.237398719 |
| Ireb2 | 1512.434034 | -0.059432251 | 0.068992817 | -0.861426648 |
| Hykk | 65.99315051 | -0.569473695 | 0.175988936 | -3.235849404 |
| AY074887 | 0.108309328 | 1.204448679 | 3.72248341 | 0.323560523 |
| Psma4 | 828.6826824 | 0.656696775 | 0.096245138 | 6.823168268 |
| Chrna5 | 21.98151757 | 0.107008688 | 0.228260273 | 0.468801193 |
| Chrna3 | 28.52374879 | -0.517217547 | 0.247943692 | -2.086028256 |
| Chrnb4 | 16.87880904 | 0.561102111 | 0.32993621 | 1.700638168 |
| Ube2q2 | 800.5171951 | -0.078760715 | 0.069741278 | -1.129327107 |
| Fbxo22 | 362.6766692 | 0.039227672 | 0.068884526 | 0.56947002 |
| Nrg4 | 36.91795082 | 0.365823975 | 0.298356603 | 1.226129979 |
| Gm15625 | 0 | NA | NA | NA |
| AI118078 | 183.6671056 | -0.431872642 | 0.140594847 | -3.071753001 |
| Etfa | 264.841253 | -0.015485376 | 0.138398738 | -0.111889571 |
| Isl2 | 0.471858061 | 0.975124582 | 1.356433113 | 0.718888807 |
| Gm17226 | 0 | NA | NA | NA |
| Scaper | 701.5265708 | -0.129286092 | 0.080281008 | -1.610419389 |
| Gm23407 | 0.627732484 | -0.698705574 | 1.165951854 | -0.599257655 |
| 4930563M21Rik | 0.038981395 | 0.274393828 | 3.724945049 | 0.073663859 |
| Rcn2 | 1128.177669 | -0.157649429 | 0.088790311 | -1.775525122 |
| Pstpip1 | 94.70433045 | 2.784717558 | 0.2662226 | 10.46010954 |
| Tspan3 | 2389.846793 | -0.022212034 | 0.089306141 | -0.248717874 |
| Peak1 | 1406.181793 | -0.118841545 | 0.072324498 | -1.643171389 |
| Gm24270 | 405.7586766 | 1.002427585 | 0.8600356 | 1.165565222 |
| Gm29322 | 6.151976361 | -1.020496145 | 0.439977649 | -2.319427243 |
| Peak1os | 8.730896443 | 0.795935843 | 0.57778841 | 1.377555917 |
| Gm37842 | 2.222650432 | 1.570948274 | 1.253081611 | 1.253667966 |
| Gm36940 | 30.43733313 | -0.777179317 | 0.293738648 | -2.645819067 |
| Hmg20a | 798.0476359 | 0.010499389 | 0.060558165 | 0.173376933 |
| Gm26868 | 0.201261165 | -0.206482256 | 2.21910279 | -0.09304763 |
| Lingo1 | 2194.009453 | -0.339644255 | 0.187607022 | -1.810402679 |
| Odf3l1 | 0.839566529 | 0.854394009 | 1.268877388 | 0.67334639 |
| Cspg4 | 408.2241713 | 0.332260147 | 0.092524491 | 3.591050802 |
| Snx33 | 323.7784099 | 0.532764903 | 0.145932185 | 3.650770412 |
| Imp3 | 174.2311039 | 0.178441291 | 0.14372411 | 1.241554322 |
| Snupn | 97.8417329 | -0.32922303 | 0.130735451 | -2.518238386 |
| Ptpn9 | 554.0054497 | -0.024667774 | 0.092066543 | -0.267934188 |
| Gm10658 | 5.404131218 | 0.316147941 | 0.488541976 | 0.64712544 |
| Sin3a | 642.6567267 | 0.09324125 | 0.060119435 | 1.550933579 |
| 2700012I20Rik | 1.975685927 | 0.009525169 | 0.787432313 | 0.012096493 |
| Man2c1os | 25.71371313 | -0.352023207 | 0.267003711 | -1.318420651 |
| Man2c1 | 469.3784322 | -0.068962216 | 0.100707181 | -0.684779528 |
| Neil1 | 55.964215 | -0.224050977 | 0.194950882 | -1.149268853 |

| | | | | |
|---|---|---|---|---|
| Commd4 | 212.8717519 | -0.014511376 | 0.087259105 | -0.166302138 |
| 1700041C23Rik | 0.044544648 | 0.755286614 | 3.724945049 | 0.202764498 |
| Trcg1 | 0.119587853 | 1.218392466 | 3.723618458 | 0.327206581 |
| 1700017B05Rik | 450.6317575 | 1.108851359 | 0.155418875 | 7.134599054 |
| Gm5121 | 0.778482992 | 0.255729843 | 1.305104839 | 0.195945824 |
| Ppcdc | 133.777483 | 0.18752464 | 0.136718714 | 1.371609157 |
| Scamp5 | 3375.801194 | -0.099108128 | 0.05013525 | -1.976815277 |
| Gm6166 | 226.0029141 | 0.212224362 | 0.123627145 | 1.716648577 |
| Rpp25 | 85.87965751 | -0.266764463 | 0.161608514 | -1.650683229 |
| Cox5a | 985.6266964 | -0.123953217 | 0.101683959 | -1.219004631 |
| Gm26631 | 4.769631826 | 0.569389945 | 0.646572108 | 0.880628683 |
| Fam219b | 230.0016874 | -0.13418634 | 0.087043255 | -1.541605276 |
| Mpi | 358.5215663 | 0.027684045 | 0.070294955 | 0.393826908 |
| Scamp2 | 323.5294745 | 0.771529743 | 0.121230249 | 6.364168591 |
| Ulk3 | 153.3484004 | -0.097588138 | 0.146992271 | -0.663899789 |
| Cplx3 | 72.27239396 | -0.271508812 | 0.223728591 | -1.213563323 |
| Lman1l | 1.399662294 | 1.665020417 | 1.25116775 | 1.330773125 |
| Csk | 512.8013575 | 0.215140676 | 0.116107628 | 1.852941779 |
| Cyp1a2 | 0.530050215 | -1.64207987 | 1.521828789 | -1.079017483 |
| Cyp1a1 | 0.692555501 | -0.126999124 | 1.162090847 | -0.109285022 |
| Edc3 | 308.5919229 | -0.105307734 | 0.102556295 | -1.026828573 |
| Clk3 | 437.4151506 | -0.108305835 | 0.102796387 | -1.053595735 |
| Gm17231 | 0.756247029 | 2.86059713 | 3.160237386 | 0.905184257 |
| Arid3b | 117.0600692 | -0.73452201 | 0.216817267 | -3.387746829 |
| Ubl7 | 525.5396743 | -0.004572061 | 0.078369361 | -0.058339905 |
| Sema7a | 1597.276098 | -0.271307864 | 0.058330806 | -4.651193472 |
| Gm17322 | 37.01146921 | -0.325676394 | 0.231125365 | -1.409089799 |
| Cyp11a1 | 16.97152848 | -1.010689856 | 0.288193391 | -3.506984856 |
| Ccdc33 | 7.670793784 | -0.605020912 | 0.44180121 | -1.3694415 |
| Gm16130 | 0.257325455 | -0.504010529 | 2.062921351 | -0.244318829 |
| Stra6 | 207.0557773 | 1.927086527 | 0.264049884 | 7.298191139 |
| Gm16131 | 0 | NA | NA | NA |
| Islr | 283.8085694 | 0.720309515 | 0.176571173 | 4.079428715 |
| Islr2 | 662.2018967 | 0.546830719 | 0.179159416 | 3.052201945 |
| 1600029O15Rik | 9.217140603 | 0.768891816 | 0.363529819 | 2.115072204 |
| Pml | 1126.656421 | 2.913791379 | 0.176229341 | 16.53408774 |
| Mir6385 | 0.090427103 | 1.236179401 | 3.724945049 | 0.331865138 |
| Stoml1 | 580.7474093 | 1.030107793 | 0.111194431 | 9.264023259 |
| Loxl1 | 78.19031371 | 0.783183755 | 0.220120779 | 3.557972846 |
| Tbc1d21 | 0 | NA | NA | NA |
| 1700072B07Rik | 0.112288909 | -0.687373951 | 3.724945049 | -0.184532642 |
| 4930461G14Rik | 0.112396584 | 0.27438722 | 3.724945049 | 0.073662085 |
| 6030419C18Rik | 1394.926258 | -0.370598168 | 0.084243597 | -4.399125646 |
| Cd276 | 125.0400392 | -0.012935116 | 0.112049465 | -0.115441122 |
| Gm10657 | 8.005046079 | 0.615136996 | 0.34940523 | 1.760526013 |
| Nptn | 4736.976647 | -0.115190751 | 0.046486034 | -2.47796469 |
| Rec114 | 2.938016276 | 0.935629341 | 0.626672207 | 1.493012344 |

| | | | | |
|---|---|---|---|---|
| Gm20620 | 0 | NA | NA | NA |
| Hcn4 | 140.1825926 | -0.44115633 | 0.121170397 | -3.640792969 |
| Neo1 | 1678.641993 | -0.048244813 | 0.044191069 | -1.091732198 |
| Gm7589 | 1.467463761 | 0.112475383 | 0.768261523 | 0.146402468 |
| Adpgk | 228.1330696 | 0.156530352 | 0.082493025 | 1.897498014 |
| Bbs4 | 465.2733797 | 0.153076922 | 0.062438913 | 2.451626955 |
| Arih1 | 1854.328923 | 0.082222951 | 0.04744888 | 1.732874457 |
| Rps11-ps1 | 39.22874844 | -0.067455483 | 0.232453259 | -0.290189449 |
| Tmem202 | 11.87611272 | 0.064760281 | 0.344764975 | 0.187838922 |
| Hexa | 567.217733 | 0.152878518 | 0.149918975 | 1.019740949 |
| Celf6 | 249.5287256 | -0.210478145 | 0.099318863 | -2.119216208 |
| Parp6 | 1004.151157 | -0.096281687 | 0.077900737 | -1.235953478 |
| Pkm | 10594.62968 | 0.135376793 | 0.062125338 | 2.17909144 |
| Gramd2 | 45.41966732 | -1.019494618 | 0.248801668 | -4.097619713 |
| Senp8 | 100.069545 | -0.719299796 | 0.199364078 | -3.607970917 |
| Myo9a | 3366.723381 | -0.224049287 | 0.071807356 | -3.120143952 |
| Gm7613 | 0 | NA | NA | NA |
| Gm2735 | 0.060174537 | 0.755286614 | 3.724945049 | 0.202764498 |
| Nr2e3 | 0.25324161 | -1.440693827 | 2.98720403 | -0.482288392 |
| Gm7616 | 0.135217741 | 1.264602945 | 3.618332936 | 0.349498779 |
| Gm15507 | 0.044544648 | 0.755286614 | 3.724945049 | 0.202764498 |
| Thsd4 | 209.2641097 | 0.108503831 | 0.156347621 | 0.693990931 |
| Gm23668 | 0.21890733 | -0.725235155 | 2.590555603 | -0.279953518 |
| Gm22099 | 0 | NA | NA | NA |
| Lrrc49 | 843.7713801 | 0.028123843 | 0.06611078 | 0.425404803 |
| B930082K07Rik | 9.514292225 | 0.567199786 | 0.312095632 | 1.817390979 |
| Larp6 | 390.5730594 | 0.131293552 | 0.092109877 | 1.425401459 |
| 1700036A12Rik | 5.346398063 | 0.765097728 | 0.456859978 | 1.674687573 |
| Uaca | 260.6470978 | 1.848360527 | 0.239677322 | 7.71187075 |
| Gm9869 | 0.554399409 | 1.577906307 | 1.61086395 | 0.979540393 |
| Gm10655 | 0.545210175 | 0.504635662 | 1.332570917 | 0.378693288 |
| Tle3 | 1433.600019 | 0.527095047 | 0.130528246 | 4.038168481 |
| Rplp1 | 2533.538003 | 0.393285813 | 0.127525018 | 3.08398947 |
| Kif23 | 18.40282969 | 1.103724659 | 0.381923547 | 2.889909951 |
| Paqr5 | 21.19486437 | 0.606571135 | 0.290574991 | 2.087485685 |
| Gm23221 | 0 | NA | NA | NA |
| Glce | 558.6811353 | -0.190488787 | 0.066228863 | -2.876220109 |
| Mir5133 | 0 | NA | NA | NA |
| Gm25603 | 0.463251208 | 0.274413527 | 1.533391716 | 0.17895853 |
| Spesp1 | 1.257510835 | 1.238863654 | 1.314680832 | 0.942330354 |
| Anp32a | 1220.827071 | -0.063446352 | 0.091709609 | -0.691817931 |
| Coro2b | 2114.073909 | -0.108714028 | 0.083943672 | -1.295083058 |
| n-R5s84 | 0 | NA | NA | NA |
| Itga11 | 34.17670704 | 0.85722792 | 0.240828337 | 3.55949773 |
| Fem1b | 2918.112869 | -0.042337009 | 0.072234263 | -0.586107029 |
| Gm26609 | 0.048949532 | -0.206487837 | 3.724945049 | -0.055433794 |
| Cln6 | 139.097147 | 0.500668833 | 0.097958177 | 5.111046878 |

| | | | | |
|---|---|---|---|---|
| Gm10653 | 0.777406778 | 0.584986451 | 1.168246789 | 0.500738762 |
| Calml4 | 9.674253487 | -0.594775197 | 0.379481528 | -1.567336362 |
| Pias1 | 838.3302677 | 0.214435351 | 0.06912469 | 3.10215282 |
| Gm24526 | 0 | NA | NA | NA |
| Gm25064 | 0.490308497 | 0.274414377 | 1.442846927 | 0.190189529 |
| Skor1 | 43.41229914 | 0.615209523 | 0.232051543 | 2.651176179 |
| Map2k5 | 366.7989628 | -0.176518026 | 0.073578711 | -2.399036686 |
| 2300009A05Rik | 131.9294766 | -0.282878513 | 0.099793013 | -2.834652484 |
| Gm16759 | 25.37684091 | -0.847937474 | 0.367235417 | -2.308975214 |
| Iqch | 1.370360641 | 0.230927959 | 0.804178489 | 0.287160079 |
| Gm25995 | 0.525324978 | -1.210057061 | 1.69512699 | -0.713844489 |
| Gm6257 | 8.599489497 | 0.201112891 | 0.422677238 | 0.475807242 |
| Aagab | 585.5754181 | 0.056558637 | 0.065725194 | 0.86053207 |
| Smad3 | 973.5426315 | 0.220411672 | 0.09409699 | 2.342388135 |
| Smad6 | 31.70011363 | 0.356727606 | 0.245633501 | 1.452275868 |
| 1700055C04Rik | 0 | NA | NA | NA |
| Gm25606 | 0 | NA | NA | NA |
| 1110036E04Rik | 0.062426873 | 0.755288839 | 3.724945049 | 0.202765096 |
| Scarletltr | 0 | NA | NA | NA |
| Lctl | 0.650092409 | 0.287304299 | 1.125175149 | 0.255341846 |
| Zwilch | 32.2726367 | 0.289301836 | 0.247455915 | 1.169104549 |
| Rpl4 | 3612.185091 | 0.307301914 | 0.085036092 | 3.613782161 |
| Gm23344 | 0.928033173 | -1.108187184 | 1.071569448 | -1.034172061 |
| Snord16a | 3.328548066 | 1.120421712 | 0.583051105 | 1.921652669 |
| Gm22571 | 2.155635849 | -0.580420659 | 0.672568634 | -0.862990971 |
| Gm23136 | 2.475450049 | 0.097811817 | 0.693182335 | 0.141105467 |
| Gm22455 | 1.074104802 | 0.387455994 | 0.950046642 | 0.407828392 |
| Snapc5 | 90.39216346 | -0.346355372 | 0.134178573 | -2.581301649 |
| Map2k1 | 2292.231272 | 0.324536305 | 0.089437398 | 3.628642057 |
| Gm24106 | 0 | NA | NA | NA |
| Uchl4 | 10.29624034 | -0.553481337 | 0.359051806 | -1.541508295 |
| Tipin | 120.2139083 | 0.309277133 | 0.132020586 | 2.342643228 |
| Gm24289 | 1.279592076 | 0.104412847 | 0.881864092 | 0.118400157 |
| Dis3l | 359.2069684 | -0.182317092 | 0.088376417 | -2.062960897 |
| Megf11 | 632.3694475 | -0.420834848 | 0.117917365 | -3.568896299 |
| Rab11a | 1359.033138 | -0.090099631 | 0.087254164 | -1.032611241 |
| AC147560.1 | 0 | NA | NA | NA |
| Dennd4a | 1982.107441 | 0.132617278 | 0.071954211 | 1.84307877 |
| Gm24347 | 0 | NA | NA | NA |
| Slc24a1 | 2.426791053 | 0.039461404 | 0.714625027 | 0.055219734 |
| Vwa9 | 254.7088933 | 0.205243252 | 0.083446783 | 2.45957057 |
| Gm25640 | 1.475874304 | -1.181401467 | 1.012804751 | -1.166465171 |
| Hacd3 | 1635.485628 | -0.160136442 | 0.06731011 | -2.379084554 |
| Dpp8 | 2380.590348 | -0.206324491 | 0.054805993 | -3.76463378 |
| Igdcc4 | 562.3809832 | -1.03839063 | 0.149562802 | -6.942840168 |
| Igdcc3 | 23.91639695 | -0.57340771 | 0.224555467 | -2.553523715 |
| Gm25313 | 3.277280862 | 1.536380413 | 0.650456768 | 2.362002347 |

| | | | | |
|---|---|---|---|---|
| Rnu5g | 4.132676428 | 0.383333811 | 0.57549375 | 0.666095524 |
| Parp16 | 35.8728912 | -0.108458547 | 0.185702344 | -0.58404512 |
| Cilp | 2.578207035 | -0.590014056 | 0.853118385 | -0.69159693 |
| Clpx | 458.2865452 | 0.272279554 | 0.08419673 | 3.23384949 |
| Gm16218 | 0 | NA | NA | NA |
| Pdcd7 | 276.9287299 | 0.044445933 | 0.07891932 | 0.563181905 |
| Ubap1l | 57.59547211 | -0.450004161 | 0.333464495 | -1.349481482 |
| Kbtbd13 | 1.752461279 | 1.360159826 | 0.753974772 | 1.803985858 |
| Rasl12 | 22.15572106 | -0.296908157 | 0.228759101 | -1.297907518 |
| Slc51b | 0.090890357 | 0.755282166 | 3.724945049 | 0.202763304 |
| Mtfmt | 124.7687732 | -0.0317467 | 0.125990952 | -0.251976027 |
| Spg21 | 367.5189745 | 0.595772866 | 0.138408145 | 4.304463935 |
| Ankdd1a | 14.47040722 | -0.928722819 | 0.322909968 | -2.876104518 |
| Plekho2 | 544.0730231 | 0.417214986 | 0.150358203 | 2.774806948 |
| Pif1 | 5.794504373 | -0.000567985 | 0.47045049 | -0.001207321 |
| Rbpms2 | 39.33216484 | 0.511504847 | 0.224362082 | 2.279818591 |
| Oaz2 | 881.559079 | -0.13989725 | 0.080910599 | -1.72903491 |
| Zfp609 | 1323.683813 | -0.204475144 | 0.057721812 | -3.542424192 |
| Trip4 | 581.8480211 | 0.060152156 | 0.057269605 | 1.050333001 |
| n-R5s85 | 0 | NA | NA | NA |
| Gm16073 | 0.045882455 | 0.755288839 | 3.724945049 | 0.202765096 |
| 2810417H13Rik | 14.58011455 | 3.250022641 | 0.557946469 | 5.824972145 |
| Gm23248 | 0 | NA | NA | NA |
| Csnk1g1 | 818.8766249 | -0.192369576 | 0.093599547 | -2.055240469 |
| Gm22657 | 0 | NA | NA | NA |
| Ppib | 745.1195212 | 0.509839514 | 0.111095379 | 4.589205426 |
| Snx22 | 48.15551827 | -0.090294974 | 0.188406497 | -0.479256156 |
| Snx1 | 927.1821691 | 0.453264803 | 0.083883655 | 5.4034937 |
| Fam96a | 202.4742243 | 0.166069618 | 0.110767267 | 1.499266181 |
| Dapk2 | 23.66058682 | 0.630748637 | 0.236309316 | 2.669165349 |
| Herc1 | 7489.340024 | 0.061306496 | 0.095421017 | 0.6424842 |
| Gm23428 | 0 | NA | NA | NA |
| Gm28379 | 0.527778125 | 0.689726289 | 1.42643203 | 0.483532531 |
| Fbxl22 | 15.26684527 | -1.209994903 | 0.322383821 | -3.75327428 |
| Usp3 | 603.065053 | -0.172918766 | 0.077319598 | -2.236415753 |
| Gm15563 | 0 | NA | NA | NA |
| Car12 | 301.1637606 | 0.255518023 | 0.151913712 | 1.681994475 |
| Aph1b | 309.2262609 | -0.302312055 | 0.084089926 | -3.595104313 |
| Gm10647 | 6.118905949 | -1.100714809 | 0.532695026 | -2.066313285 |
| BC050972 | 1.031670571 | -0.357011748 | 0.926428412 | -0.385363557 |
| Ppp1r2-ps4 | 7.934796858 | 0.192985752 | 0.391811049 | 0.492548009 |
| Aph1c | 99.73584743 | 0.234192331 | 0.122891559 | 1.905682795 |
| Gm17098 | 1.018413619 | -0.272340114 | 1.027651347 | -0.26501217 |
| Rab8b | 840.6987422 | 0.656907461 | 0.114707251 | 5.726817227 |
| Rps27l | 178.3460037 | 0.916439244 | 0.177742998 | 5.155979452 |
| Lactb | 129.5314519 | -0.020245652 | 0.108299987 | -0.186940483 |
| Gm24225 | 0 | NA | NA | NA |

| | | | | |
|---|---|---|---|---|
| Tpm1 | 1521.522177 | -0.208391051 | 0.118131048 | -1.764066721 |
| Gm22145 | 0 | NA | NA | NA |
| Tln2 | 1967.481299 | -0.413255186 | 0.091787521 | -4.502302495 |
| Mir190a | 0.401901456 | 0.642089074 | 1.525198241 | 0.42098729 |
| Mir7241 | 0.17692102 | -0.206487546 | 2.728717236 | -0.07567202 |
| Mir7242 | 0.162691105 | -0.206480295 | 2.788998794 | -0.074033842 |
| M5C1000I18Rik | 30.04157137 | -0.193645483 | 0.201606004 | -0.960514462 |
| C2cd4b | 11.79896043 | 0.348524882 | 0.431717159 | 0.80729912 |
| C2cd4a | 1.024692053 | 3.125942648 | 1.087228697 | 2.875147296 |
| Vps13c | 2530.722944 | -0.355095137 | 0.096611069 | -3.675511944 |
| Gm28195 | 0 | NA | NA | NA |
| Gm22382 | 0 | NA | NA | NA |
| Gm22169 | 0 | NA | NA | NA |
| Gm22962 | 0.044191549 | -0.206490061 | 3.724945049 | -0.055434391 |
| Rora | 3475.748826 | -0.010989863 | 0.071418018 | -0.153880815 |
| AC127262.1 | 19.3378465 | -0.594597999 | 0.345512219 | -1.720917427 |
| Gm15511 | 0.358080405 | 0.268900704 | 1.577643814 | 0.170444496 |
| Ice2 | 215.338252 | -0.392637789 | 0.120644954 | -3.254489929 |
| Gm4978 | 0.090277459 | -0.20649451 | 3.724945049 | -0.055435586 |
| Anxa2 | 640.8248345 | 2.22812338 | 0.345531939 | 6.448386177 |
| Mir3109 | 0.728690676 | -0.166716616 | 1.295621705 | -0.128676924 |
| Gm16144 | 0 | NA | NA | NA |
| Foxb1 | 12.10284569 | -0.93027511 | 0.318037681 | -2.925046823 |
| B230323A14Rik | 7.674232682 | -0.887799852 | 0.445180705 | -1.994246028 |
| Gm28731 | 0.659754175 | 0.863317897 | 1.252344127 | 0.689361557 |
| Bnip2 | 412.8066768 | 0.502960601 | 0.11601255 | 4.335398218 |
| Gtf2a2 | 313.4260604 | -0.029942335 | 0.099840082 | -0.299902946 |
| Gcnt3 | 0.134276636 | -0.206488731 | 2.954908363 | -0.06987991 |
| Fam81a | 750.583868 | -0.345213869 | 0.144264538 | -2.392922571 |
| Gm24257 | 0 | NA | NA | NA |
| Gm26849 | 0.04393175 | -0.206490061 | 3.724945049 | -0.055434391 |
| Myo1e | 393.0843238 | 0.705872485 | 0.139756991 | 5.050713222 |
| Mir5626 | 0.091666476 | -0.687373951 | 3.724945049 | -0.184532642 |
| Ccnb2 | 19.87963883 | 0.345690873 | 0.334356382 | 1.033899429 |
| Rnf111 | 823.3856292 | -0.067712978 | 0.066052632 | -1.025136711 |
| Gm24615 | 0 | NA | NA | NA |
| Gm23588 | 0 | NA | NA | NA |
| Gm23730 | 0.04393175 | -0.206490061 | 3.724945049 | -0.055434391 |
| Gm22229 | 0 | NA | NA | NA |
| Sltm | 725.1236 | -0.11605258 | 0.115896737 | -1.001344675 |
| Fam63b | 1775.64688 | -0.273376368 | 0.092473587 | -2.956264344 |
| Gm10642 | 11.00079997 | 0.544244225 | 0.321145862 | 1.694694805 |
| Adam10 | 1465.376635 | -0.015592453 | 0.092888838 | -0.167861423 |
| Lipc | 1.303781876 | -0.093319099 | 1.106663846 | -0.084324702 |
| Gm25149 | 0 | NA | NA | NA |
| Gm3436 | 6.640611746 | 0.454238168 | 0.41979529 | 1.082046843 |
| Aqp9 | 16.64346168 | 0.575704247 | 0.337491443 | 1.705833613 |

| | | | | |
|---|---|---|---|---|
| Aldh1a2 | 316.9562088 | 1.668516973 | 0.202136407 | 8.254410961 |
| Polr2m | 1314.180415 | 0.229579947 | 0.081509475 | 2.81660443 |
| Gcom1 | 0 | NA | NA | NA |
| Myzap | 9.380444241 | 0.343369072 | 0.3452499 | 0.99455227 |
| Cgnl1 | 362.6537632 | -0.14358083 | 0.125042557 | -1.148255715 |
| Gm27220 | 0 | NA | NA | NA |
| Gm37660 | 16.37804138 | 0.014031758 | 0.336313516 | 0.041722253 |
| Tcf12 | 1303.134971 | 0.005513096 | 0.068525076 | 0.080453704 |
| Gm38057 | 46.80930091 | -1.070874295 | 0.304116429 | -3.521264198 |
| Gm38111 | 32.02976615 | -0.728150189 | 0.316555976 | -2.300225696 |
| Gm18821 | 0 | NA | NA | NA |
| Gm37879 | 56.51000279 | -0.064392405 | 0.24896809 | -0.258637182 |
| Gm27152 | 0 | NA | NA | NA |
| Gm37663 | 62.49855959 | -0.408625359 | 0.180135267 | -2.268436184 |
| Gm37023 | 32.44644427 | 0.381928692 | 0.324520706 | 1.176900842 |
| A430027C01Rik | 29.11976123 | 0.363281812 | 0.27046635 | 1.343168244 |
| Gm27450 | 0 | NA | NA | NA |
| Gm37137 | 44.47354403 | -0.115233691 | 0.275961019 | -0.41757235 |
| Rpl15-ps2 | 4.494967837 | 0.554159249 | 0.614650553 | 0.901584236 |
| Gm25045 | 0.207243203 | 1.833032826 | 2.596246052 | 0.706032013 |
| Gm7863 | 0 | NA | NA | NA |
| Gm7866 | 2.795202431 | -0.079891516 | 0.692133262 | -0.115427939 |
| Gm27159 | 0.158742293 | 0.27438884 | 3.451672432 | 0.079494461 |
| Gm27188 | 5.84902935 | -0.347175112 | 0.519496205 | -0.668291913 |
| Gm7870 | 0.124347486 | 0.755286736 | 3.717316019 | 0.203180664 |
| Gm25163 | 0.730530645 | 0.103545578 | 1.104982703 | 0.093707872 |
| Gm27253 | 3.522083342 | -0.507887153 | 0.536157144 | -0.947272939 |
| Zfp280d | 1238.243116 | -0.077022837 | 0.107067469 | -0.719385987 |
| Gm19353 | 21.12517525 | -0.715960793 | 0.307158185 | -2.330918814 |
| Gm17828 | 1.657074577 | -0.015531972 | 0.786386615 | -0.019751063 |
| Gm37983 | 73.73482861 | -0.076075213 | 0.235344435 | -0.323250529 |
| Gm37303 | 69.68360701 | -0.334074105 | 0.236104624 | -1.414940965 |
| Gm37581 | 38.67009967 | -0.494780727 | 0.30046296 | -1.646727861 |
| BC065403 | 0.430924493 | 0.693517881 | 1.404254072 | 0.493869233 |
| Gm37150 | 0.370677815 | 0.769961722 | 1.749138892 | 0.44019473 |
| Gm27230 | 0.066050875 | 0.274391604 | 3.724945049 | 0.073663262 |
| Gm27255 | 0.075043205 | 0.755286614 | 3.724945049 | 0.202764498 |
| Gm27254 | 0.194277958 | 0.812300801 | 3.250214206 | 0.249922236 |
| Mns1 | 77.00500091 | 0.970069354 | 0.154659316 | 6.27229822 |
| Gm36948 | 4.487847835 | 1.402567685 | 0.607833553 | 2.307486446 |
| Gm27231 | 0.460076145 | 0.437469354 | 1.543787014 | 0.283374164 |
| Tex9 | 177.6432418 | 0.155720274 | 0.126209846 | 1.23382033 |
| Gm28010 | 0.172182752 | 0.755291024 | 3.331044598 | 0.226742994 |
| 4930509E16Rik | 0.362648911 | -0.206482793 | 1.76179073 | -0.117200522 |
| Rfx7 | 1180.076257 | -0.330888524 | 0.076768085 | -4.310235492 |
| Gm19427 | 12.05712943 | -0.069359652 | 0.321685199 | -0.215613438 |
| Gm37105 | 45.12253491 | -0.418262833 | 0.335249522 | -1.247616493 |

| | | | | |
|---|---|---|---|---|
| Gm18014 | 0 | NA | NA | NA |
| Gm27241 | 0.220148986 | 0.802838658 | 2.591704594 | 0.309772441 |
| Gm27232 | 0.138573873 | 0.75528185 | 3.628901543 | 0.208129606 |
| Gm23057 | 0.072025462 | 0.755288839 | 3.724945049 | 0.202765096 |
| Nedd4 | 6315.797856 | -0.185362617 | 0.048826638 | -3.796342001 |
| A530064N14Rik | 50.37861278 | -0.49652881 | 0.329488746 | -1.506967433 |
| Gm7265 | 195.1401439 | -0.210948515 | 0.101949255 | -2.069152089 |
| Gm37955 | 4.498464866 | -1.089295554 | 0.536062021 | -2.032032697 |
| Gm36936 | 103.529168 | -0.656967232 | 0.285532822 | -2.300846633 |
| Gm27204 | 0 | NA | NA | NA |
| Prtg | 50.92406102 | -0.60640914 | 0.205087621 | -2.956829559 |
| Gm27203 | 0 | NA | NA | NA |
| A730062M13Rik | 2.892325926 | -1.360977124 | 0.593960999 | -2.291357726 |
| Gm37234 | 1.821448544 | 0.132276533 | 0.75290636 | 0.175687894 |
| Pygo1 | 611.8774595 | -0.525646836 | 0.098930574 | -5.313290051 |
| Dyx1c1 | 5.660739534 | 0.156072774 | 0.417686135 | 0.373660415 |
| Dyx1c1Ccpg1 | 0 | NA | NA | NA |
| Gm23567 | 0.220111735 | -0.687378617 | 2.168634177 | -0.31696384 |
| Gm37710 | 0.046345709 | 0.274391604 | 3.724945049 | 0.073663262 |
| Ccpg1os | 38.04226344 | -0.383559437 | 0.215862315 | -1.776870764 |
| Ccpg1 | 158.4319805 | -0.469661099 | 0.112500576 | -4.174743963 |
| Gm27353 | 16.47684853 | -0.963054651 | 0.329756728 | -2.920500386 |
| Pigb | 210.9882647 | -0.11318949 | 0.09027323 | -1.253854442 |
| Gm38192 | 17.09336393 | -0.837407161 | 0.276529266 | -3.028276803 |
| 2310009A05Rik | 48.02464468 | -0.471547634 | 0.226716463 | -2.079900281 |
| Rab27a | 175.4812393 | 0.348186563 | 0.120683428 | 2.885123232 |
| Gm27186 | 0.89763465 | -0.536689518 | 1.347507475 | -0.398283148 |
| Khdc3 | 3.338837625 | -0.937289091 | 0.594420537 | -1.57681142 |
| Gm20509 | 0 | NA | NA | NA |
| Rsl24d1 | 296.3744691 | 0.351471128 | 0.081053472 | 4.336287126 |
| Gm38215 | 2.049787317 | -2.013007401 | 0.92140605 | -2.184712594 |
| Gm5745 | 0.038981395 | 0.274393828 | 3.724945049 | 0.073663859 |
| Gm27211 | 0.328170597 | -0.206503948 | 3.724945049 | -0.055438119 |
| Unc13c | 1536.822218 | -0.627554705 | 0.097647197 | -6.426755952 |
| Gm37611 | 8.958307541 | -0.53260905 | 0.410648596 | -1.296994693 |
| Gm27148 | 1.654347319 | -0.936804683 | 0.792631752 | -1.181891441 |
| Gm25105 | 4.734499335 | -0.226731624 | 0.444873673 | -0.509653948 |
| Gm37326 | 31.32772203 | -1.377268847 | 0.324988312 | -4.237902709 |
| n-R5s86 | 0 | NA | NA | NA |
| Wdr72 | 4.387779186 | 1.002086813 | 0.558354255 | 1.7947151 |
| Gm24141 | 0 | NA | NA | NA |
| Nr1h2-ps | 0 | NA | NA | NA |
| Gm28622 | 0.044544648 | 0.755286614 | 3.724945049 | 0.202764498 |
| Gm27189 | 0 | NA | NA | NA |
| Gm27233 | 0 | NA | NA | NA |
| Gm22315 | 0 | NA | NA | NA |
| Gm16551 | 2.498792093 | 0.541307998 | 0.698361921 | 0.775110987 |

| | | | | |
|---|---|---|---|---|
| Gm20649 | 0 | NA | NA | NA |
| Gm28182 | 0.458414965 | -1.883093545 | 1.383496238 | -1.361112154 |
| Onecut1 | 8.345816899 | -0.203857218 | 0.372663245 | -0.547027968 |
| Fam214a | 701.9545256 | 0.235492713 | 0.084558955 | 2.784952971 |
| Arpp19 | 2304.068667 | -0.373766311 | 0.122922045 | -3.040677618 |
| Myo5a | 9406.867657 | -0.11891081 | 0.065363115 | -1.819234132 |
| Gm9719 | 0.038981395 | 0.274393828 | 3.724945049 | 0.073663859 |
| Gm16353 | 0 | NA | NA | NA |
| Myo5c | 38.56207624 | 0.012377659 | 0.341308185 | 0.036265345 |
| Gnb5 | 588.9032945 | -0.42509835 | 0.076780615 | -5.536532238 |
| Bcl2l10 | 2.578686406 | -0.672826004 | 0.636387608 | -1.057258179 |
| Mapk6 | 1348.202811 | 0.240709518 | 0.095831186 | 2.511807774 |
| 4933433G15Rik | 7.660142694 | 0.079644733 | 0.427187593 | 0.186439714 |
| A130057D12Rik | 0.39127578 | 0.274415358 | 1.594577459 | 0.172092836 |
| Leo1 | 251.7819994 | 0.396184844 | 0.180291311 | 2.197470535 |
| Tmod3 | 491.2591432 | 0.532590366 | 0.095633485 | 5.569078332 |
| Tmod2 | 9543.757063 | -0.075895977 | 0.06112342 | -1.241684064 |
| Lysmd2 | 376.8025865 | 0.095611848 | 0.107589663 | 0.888671322 |
| Scg3 | 2648.530245 | 0.178351135 | 0.079223533 | 2.251239358 |
| Gm37871 | 0.090665643 | -0.687373951 | 3.724945049 | -0.184532642 |
| Gm3671 | 0.843494912 | 0.852212495 | 1.105877409 | 0.770621127 |
| Bmp5 | 56.01127082 | 0.543122767 | 0.219333574 | 2.476240902 |
| Gm26903 | 0.668743806 | 0.357852935 | 1.377225573 | 0.25983611 |
| AC079245.1 | 0 | NA | NA | NA |
| Gm25300 | 0 | NA | NA | NA |
| Hmgcll1 | 224.7683895 | -0.445275243 | 0.117749101 | -3.781559601 |
| Gm38083 | 27.55755119 | -0.762636223 | 0.262114091 | -2.909558274 |
| Gfral | 1.244634631 | -1.030162757 | 0.978913868 | -1.052352808 |
| Hcrtr2 | 58.13001265 | -0.599225752 | 0.235653872 | -2.542821586 |
| Gm27234 | 0.038981395 | 0.274393828 | 3.724945049 | 0.073663859 |
| Fam83b | 2.476850474 | 0.782753639 | 0.638661466 | 1.225615887 |
| Gm27156 | 0 | NA | NA | NA |
| Gm27245 | 0 | NA | NA | NA |
| Gm22938 | 0 | NA | NA | NA |
| Gm37408 | 0 | NA | NA | NA |
| Gm27155 | 0 | NA | NA | NA |
| Gm23791 | 0 | NA | NA | NA |
| Gm18486 | 0 | NA | NA | NA |
| Tinag | 0.568818845 | -1.132997589 | 1.304343606 | -0.868634295 |
| Gm27157 | 0 | NA | NA | NA |
| Gm27158 | 0 | NA | NA | NA |
| Gm19541 | 0.139037128 | 0.274384718 | 3.724945049 | 0.073661414 |
| Gm5746 | 0.044544648 | 0.755286614 | 3.724945049 | 0.202764498 |
| Mlip | 470.9327951 | -0.086987195 | 0.10602702 | -0.820424781 |
| Gm27210 | 1.42879052 | 0.387190514 | 0.854780797 | 0.452970534 |
| Gm38254 | 2.89654368 | 0.22017959 | 0.588923746 | 0.37386774 |
| 4930590H14Rik | 0 | NA | NA | NA |

| | | | | |
|---|---|---|---|---|
| 5730403I07Rik | 4.072737608 | -0.420445416 | 0.589754445 | -0.71291606 |
| Lrrc1 | 172.5600245 | -0.272651735 | 0.122325191 | -2.22890912 |
| Gm19569 | 3.396866737 | 1.155265907 | 0.574267 | 2.011722609 |
| Gm21013 | 0.284055481 | -1.227323997 | 2.435667199 | -0.503896426 |
| Gm25250 | 0 | NA | NA | NA |
| Klhl31 | 1.14819629 | 0.005376842 | 0.970419205 | 0.005540741 |
| Gclc | 631.4586054 | 0.099550628 | 0.054484115 | 1.827149575 |
| Elovl5 | 1099.939201 | 0.021761111 | 0.101779538 | 0.213806347 |
| Gm23609 | 0 | NA | NA | NA |
| Gcm1 | 0.062426873 | 0.755288839 | 3.724945049 | 0.202765096 |
| Fbxo9 | 758.0697948 | -0.168932307 | 0.090956679 | -1.857283149 |
| Ick | 452.7014818 | -0.118079228 | 0.087439425 | -1.350411762 |
| Rn7sk | 139206.5501 | 0.199463444 | 0.172334177 | 1.157422441 |
| C920006O11Rik | 1.914692995 | 0.93076162 | 0.75097597 | 1.239402668 |
| Gsta4 | 217.5686189 | -0.1802256 | 0.156037283 | -1.15501627 |
| Gsta1 | 0.047948698 | -0.206487837 | 3.724945049 | -0.055433794 |
| Gm10639 | 0.087930927 | -0.206490058 | 3.724945049 | -0.05543439 |
| Omt2a | 0 | NA | NA | NA |
| Gm8074 | 0 | NA | NA | NA |
| Omt2b | 0 | NA | NA | NA |
| Gsta2 | 0.44449992 | 1.696470899 | 1.710014367 | 0.99207991 |
| Dppa5a | 0.045882455 | 0.755288839 | 3.724945049 | 0.202765096 |
| Ooep | 0.251515219 | 1.17754919 | 1.820460073 | 0.646841536 |
| Gm8087 | 0.090074004 | 0.274402725 | 3.724945049 | 0.073666248 |
| 4930542C12Rik | 0 | NA | NA | NA |
| Ddx43 | 4.317706999 | 1.029268904 | 0.463881592 | 2.218818169 |
| Gm8093 | 1.913696796 | 2.610985073 | 0.943597429 | 2.767054034 |
| Mb21d1 | 40.10846614 | 2.022823142 | 0.272643207 | 7.419305118 |
| Gm17324 | 2.766932339 | 0.42250862 | 0.664670286 | 0.635666479 |
| Mto1 | 264.8336796 | -0.102445853 | 0.077134042 | -1.328153565 |
| Eef1a1 | 14473.51984 | 0.444193789 | 0.094550866 | 4.697934656 |
| Gm26377 | 0 | NA | NA | NA |
| Slc17a5 | 273.2142388 | 0.008487376 | 0.102331724 | 0.082939835 |
| Cd109 | 135.8248537 | 1.973712189 | 0.267253459 | 7.385169845 |
| 4921509A06Rik | 0.07796279 | 0.274392189 | 3.724945049 | 0.073663419 |
| Gm8116 | 61.44867496 | -0.254338936 | 0.147251753 | -1.727238759 |
| Gm28620 | 0 | NA | NA | NA |
| Col12a1 | 350.8884372 | -0.138153925 | 0.160824574 | -0.859034917 |
| 4930562D21Rik | 0 | NA | NA | NA |
| Cox7a2 | 993.8036579 | -0.024406932 | 0.13239114 | -0.184354727 |
| Tmem30a | 5117.156797 | 0.162912814 | 0.090072659 | 1.808682179 |
| 4930429F24Rik | 2.804294664 | 0.212396159 | 0.616246992 | 0.344660763 |
| Filip1 | 118.2003987 | -0.0458414 | 0.123404672 | -0.37147216 |
| Gm20477 | 0.402709178 | -0.974239542 | 1.325726304 | -0.734872303 |
| Gm3211 | 0 | NA | NA | NA |
| Gm25433 | 0 | NA | NA | NA |
| Senp6 | 1335.192463 | -0.022740089 | 0.055717311 | -0.408133277 |

| | | | | |
|---|---|---|---|---|
| Gm20648 | 0 | NA | NA | NA |
| Gm17477 | 0 | NA | NA | NA |
| Myo6 | 1685.660187 | -0.281783162 | 0.080056037 | -3.519824007 |
| Gm27227 | 0 | NA | NA | NA |
| Impg1 | 1.92279121 | -0.364754608 | 0.70031944 | -0.520840329 |
| Gm27224 | 0.046345709 | 0.274391604 | 3.724945049 | 0.073663262 |
| Gm27225 | 0.281850154 | -0.162234664 | 1.751686006 | -0.092616293 |
| Gm8151 | 0.792246753 | 1.267706509 | 1.089655553 | 1.163401136 |
| Gm26713 | 0.150086409 | 1.264824188 | 3.723011833 | 0.339731445 |
| Gm27226 | 0 | NA | NA | NA |
| Htr1b | 292.4427632 | -0.054656089 | 0.130729599 | -0.418085036 |
| D430036J16Rik | 57.68366793 | 0.313333185 | 0.160509067 | 1.952121407 |
| Gm9531 | 81.22717314 | 0.056373012 | 0.132637579 | 0.425015392 |
| Gm26907 | 0.36944391 | 1.519121794 | 3.253720289 | 0.466887642 |
| Mei4 | 18.14348477 | -0.250491756 | 0.239912709 | -1.044095398 |
| Gm37837 | 2.253692409 | 0.247243401 | 0.693976583 | 0.356270524 |
| Gm15896 | 1.074470558 | 0.867522649 | 1.02343225 | 0.847660066 |
| Gm23748 | 0.232637982 | 0.298792896 | 2.544854123 | 0.117410618 |
| 1700063H06Rik | 1.312371618 | 0.809413369 | 0.86601389 | 0.934642478 |
| Gm22836 | 0 | NA | NA | NA |
| Gm24393 | 0 | NA | NA | NA |
| Gm27249 | 0.180969744 | 0.235169692 | 2.733013994 | 0.086047745 |
| Irak1bp1 | 163.4822182 | 0.130336635 | 0.100925436 | 1.291415125 |
| Phip | 3482.802041 | 0.208285322 | 0.086559318 | 2.406272672 |
| Gm27248 | 0.875572801 | 1.157839841 | 1.267565539 | 0.913435878 |
| Hmgn3 | 287.8474579 | -0.005730723 | 0.075860833 | -0.07554258 |
| Gm2065 | 1.354044546 | 0.712585001 | 1.144108557 | 0.622829885 |
| Gm29054 | 6.179723687 | -0.578373752 | 0.372822611 | -1.551337644 |
| Gm38279 | 0.200264842 | 0.27440964 | 2.210050429 | 0.124164425 |
| Gm27216 | 0 | NA | NA | NA |
| Lca5 | 153.3277019 | -0.326070383 | 0.11313933 | -2.882025037 |
| Rps27a-ps2 | 369.3263099 | 0.563617182 | 0.141556049 | 3.981583168 |
| Gm2087 | 1.000715493 | 2.774971453 | 1.137344962 | 2.43986789 |
| Gm37307 | 1.831656741 | 1.122770394 | 0.759759513 | 1.477797085 |
| Gm28552 | 0.11657011 | 1.236179401 | 3.724945049 | 0.331865138 |
| Gm28477 | 0.334445592 | 0.290608087 | 1.973878812 | 0.147226914 |
| Sh3bgrl2 | 558.0448667 | 0.005709506 | 0.085664611 | 0.066649524 |
| Gm2109 | 2.113095361 | 0.078711176 | 0.683305798 | 0.115191729 |
| Elovl4 | 771.8863038 | 0.243624124 | 0.10090225 | 2.414456802 |
| Ttk | 15.5422173 | 1.038635632 | 0.411848526 | 2.521887454 |
| Bckdhb | 244.9808363 | -0.175540753 | 0.075313531 | -2.33079966 |
| Gm29163 | 0 | NA | NA | NA |
| Gm29164 | 0 | NA | NA | NA |
| Gm26146 | 0.204782255 | -0.699723755 | 2.205163063 | -0.317311571 |
| Gm8226 | 4.794882048 | 0.652284015 | 0.492733006 | 1.323808245 |
| Gm26126 | 0 | NA | NA | NA |
| 4930554C24Rik | 0.203235021 | 1.210233894 | 2.195041552 | 0.551348968 |

| | | | | |
|---|---|---|---|---|
| Gm28625 | 0 | NA | NA | NA |
| Gm8228 | 8.163884718 | 0.059238499 | 0.382057008 | 0.155051464 |
| Gm28070 | 0 | NA | NA | NA |
| Gm11114 | 0 | NA | NA | NA |
| Fam46a | 425.1094954 | 1.344524548 | 0.185132613 | 7.262494322 |
| Gm36927 | 0 | NA | NA | NA |
| Gm25125 | 0 | NA | NA | NA |
| Ibtk | 540.332865 | -0.034000358 | 0.054843752 | -0.619949523 |
| 4933431K14Rik | 24.95481328 | -0.244500807 | 0.222634647 | -1.098215441 |
| Gm22830 | 0.143846094 | -0.985158148 | 3.718482058 | -0.264935566 |
| Gm5619 | 19.25933968 | 0.974290832 | 0.384133896 | 2.536331323 |
| Gm18766 | 0.416300838 | 0.869061371 | 1.467983613 | 0.592010267 |
| 9330154J02Rik | 12.67333215 | 0.213235206 | 0.277389917 | 0.768720105 |
| Tpbg | 242.3331958 | -0.111425145 | 0.119588445 | -0.931738392 |
| Ube2cbp | 42.39662063 | 0.015289064 | 0.207218463 | 0.073782343 |
| Gm16202 | 0.329441611 | 1.155265223 | 1.695197882 | 0.681492843 |
| Dopey1 | 1570.3345 | 0.021762234 | 0.109568963 | 0.198616776 |
| Gm17530 | 6.903802255 | -0.979715891 | 0.416056119 | -2.354768612 |
| Pgm3 | 195.6218196 | 0.033223533 | 0.090934636 | 0.365356194 |
| Rwdd2a | 134.250853 | 0.173105584 | 0.118419844 | 1.4617954 |
| Me1 | 717.0322805 | -0.101298174 | 0.103968836 | -0.974312861 |
| Gm38230 | 5.852527318 | -0.382721189 | 0.414420765 | -0.923508717 |
| A330041J22Rik | 15.09013069 | -0.485801467 | 0.261140412 | -1.860307497 |
| Gm25949 | 0 | NA | NA | NA |
| Gm22138 | 0.271189253 | 0.684853454 | 3.714955618 | 0.184350373 |
| Gm37484 | 1.311065665 | -1.996615247 | 0.953677874 | -2.09359502 |
| Prss35 | 55.61444896 | 0.244705486 | 0.174599756 | 1.401522493 |
| Snap91 | 4128.306371 | -0.268891768 | 0.063428244 | -4.239306484 |
| Gm37941 | 94.46575361 | -0.339360777 | 0.269884425 | -1.257430017 |
| Ripply2 | 33.98125597 | -0.265266413 | 0.205043909 | -1.293705401 |
| Cyb5r4 | 351.6163489 | -0.298112761 | 0.079922741 | -3.730011735 |
| Gm8282 | 3.540589861 | -0.228067212 | 0.520083816 | -0.43852011 |
| Mrap2 | 94.81455933 | 0.574596611 | 0.206590332 | 2.781333499 |
| Gm37012 | 4.127795301 | 0.045916883 | 0.506516424 | 0.090652309 |
| Cep162 | 341.6360393 | -0.255748322 | 0.106600501 | -2.399128701 |
| Gm24649 | 0.157258859 | 0.711849947 | 2.806391105 | 0.253653151 |
| Gm9540 | 0 | NA | NA | NA |
| Gm37375 | 0.154920219 | 0.777094091 | 2.834936002 | 0.274113451 |
| Tbx18 | 38.37151821 | 0.026711511 | 0.183998799 | 0.145172203 |
| D030062O11Rik | 0.22856443 | 1.662033007 | 2.147910842 | 0.773790501 |
| Gm37374 | 0.748906342 | -0.215161821 | 1.120310638 | -0.192055501 |
| Gm9541 | 0 | NA | NA | NA |
| Gm25528 | 0 | NA | NA | NA |
| Gm23176 | 0 | NA | NA | NA |
| Gm28877 | 0.115603922 | 0.274387056 | 3.724945049 | 0.073662041 |
| Gm2324 | 0 | NA | NA | NA |
| Gm5368 | 0 | NA | NA | NA |

| | | | | |
|---|---|---|---|---|
| Gm28231 | 0 | NA | NA | NA |
| Gm5066 | 2.101151607 | 0.833755482 | 0.786171442 | 1.060526289 |
| Nt5e | 129.2471619 | 0.682322709 | 0.155504939 | 4.387788019 |
| Snx14 | 532.7999784 | -0.405333504 | 0.069943918 | -5.795121489 |
| Gm10163 | 3.877733872 | 0.403341056 | 0.464966628 | 0.867462376 |
| Gm20537 | 1.59229893 | -0.271522199 | 0.775228135 | -0.350248123 |
| Gm28229 | 64.56740978 | -0.40274225 | 0.17320041 | -2.325296173 |
| Syncrip | 1823.979902 | 0.016568463 | 0.066073327 | 0.250758725 |
| Snhg5 | 101.6600101 | 0.574614241 | 0.157150991 | 3.656446809 |
| Snord50a | 0 | NA | NA | NA |
| Snord50b | 0 | NA | NA | NA |
| Gm22985 | 0 | NA | NA | NA |
| Zfp949 | 311.469506 | -0.01781264 | 0.094820909 | -0.187855614 |
| Gm26461 | 289.3437332 | -0.242176283 | 0.199311788 | -1.21506252 |
| Trim43a | 11.4592027 | 0.782789543 | 0.357815489 | 2.187690489 |
| 9430037G07Rik | 40.90789348 | -0.006226382 | 0.214914991 | -0.028971371 |
| Gm29603 | 0.044544648 | 0.755286614 | 3.724945049 | 0.202764498 |
| Gm23017 | 0 | NA | NA | NA |
| Gm26567 | 0 | NA | NA | NA |
| Gm29415 | 1.336334664 | 0.882776702 | 0.896294601 | 0.984918018 |
| Gm38214 | 23.29150933 | -0.986302366 | 0.317056385 | -3.110810603 |
| Gm37226 | 40.31519145 | -1.130868764 | 0.343684001 | -3.29043179 |
| Mthfsl | 134.439119 | -0.282181167 | 0.158069128 | -1.785175702 |
| Gm10634 | 2.499553143 | -0.961559272 | 0.641863057 | -1.498075425 |
| Bcl2a1d | 28.6456985 | 5.937933023 | 0.713085916 | 8.327093395 |
| Gm37560 | 0.141331096 | -0.206491164 | 3.56184083 | -0.057973159 |
| Gm26082 | 0.238313115 | 0.679411292 | 2.150603872 | 0.31591652 |
| Gm37614 | 0.29572682 | -0.685773704 | 1.764980694 | -0.388544592 |
| Gm29562 | 2.519137971 | 0.996803304 | 0.639485904 | 1.558757272 |
| Trim43c | 0 | NA | NA | NA |
| 9330159M07Rik | 56.07294295 | -0.134198416 | 0.199695134 | -0.672016452 |
| Gm9775 | 0 | NA | NA | NA |
| Gm26135 | 0 | NA | NA | NA |
| Gm26611 | 0 | NA | NA | NA |
| Gm2396 | 3.793477816 | 2.556883255 | 0.708349124 | 3.609637067 |
| Gm23707 | 0 | NA | NA | NA |
| Bcl2a1a | 13.58752781 | 7.062053695 | 0.891789205 | 7.918971942 |
| Gm37833 | 3.739270958 | -1.467678125 | 0.650312997 | -2.256879581 |
| Gm37993 | 0.55378357 | 1.637218456 | 1.407100389 | 1.163540618 |
| Gm37471 | 0.168528002 | 0.274406491 | 2.758169185 | 0.099488636 |
| Gm24463 | 0.710913347 | 0.73603076 | 1.223279209 | 0.601686642 |
| Gm19498 | 0 | NA | NA | NA |
| Trim43b | 0.095295241 | -0.206492286 | 3.724945049 | -0.055434988 |
| 4930579C12Rik | 0.595794717 | 1.453669919 | 1.418282984 | 1.024950546 |
| Gm26423 | 0 | NA | NA | NA |
| Bcl2a1b | 8.616394077 | 4.990139772 | 0.889133811 | 5.612360831 |
| Gm29094 | 0.444881751 | 0.755308145 | 1.680175443 | 0.449541236 |

| | | | | |
|---|---|---|---|---|
| Mthfs | 70.61009405 | -0.32010338 | 0.2489229 | -1.285953924 |
| Gm38091 | 0.04393175 | -0.206490061 | 3.724945049 | -0.055434391 |
| AF529169 | 248.5876439 | 0.086205234 | 0.175497697 | 0.491204359 |
| Tmed3 | 223.4678697 | 0.042162216 | 0.121258933 | 0.347703996 |
| Ankrd34c | 157.1958196 | -0.68524872 | 0.126471877 | -5.41819049 |
| Gm29608 | 0.210489921 | -0.687379134 | 2.627895457 | -0.261570197 |
| Mir184 | 0 | NA | NA | NA |
| 1700017I07Rik | 0 | NA | NA | NA |
| 4930524O08Rik | 0.357611442 | 1.375385776 | 1.943831689 | 0.707564232 |
| Rasgrf1 | 5020.678698 | 0.080554824 | 0.076286393 | 1.05595271 |
| Gm23759 | 0.203875117 | 1.640672188 | 2.619671844 | 0.626289202 |
| Gm17944 | 0.117650406 | -0.206492286 | 3.724945049 | -0.055434988 |
| Ctsh | 415.3048284 | 2.604723671 | 0.20844775 | 12.4958109 |
| Morf4l1 | 2238.38623 | 0.203604615 | 0.065528981 | 3.107092654 |
| Gm26882 | 0.372996712 | 1.632283201 | 1.923135069 | 0.848761601 |
| Gm6223 | 7.567416792 | -0.098913219 | 0.374544361 | -0.264089461 |
| 9430062P05Rik | 1.427283908 | 0.480445226 | 0.92711336 | 0.518216269 |
| Adamts7 | 9.932952153 | -0.043281832 | 0.342874279 | -0.126232368 |
| Tbc1d2b | 421.9794416 | 0.137657531 | 0.088354276 | 1.558017767 |
| Gm16200 | 6.214365972 | -0.689503712 | 0.396774108 | -1.737773961 |
| Gm22717 | 0 | NA | NA | NA |
| Gm22866 | 0.060174537 | 0.755286614 | 3.724945049 | 0.202764498 |
| Gm2497 | 0 | NA | NA | NA |
| Gm29605 | 0 | NA | NA | NA |
| Gm29604 | 0.044544648 | 0.755286614 | 3.724945049 | 0.202764498 |
| Gm28703 | 3.32610108 | 0.165082558 | 0.560992098 | 0.294268955 |
| Gm37665 | 0.317495958 | -0.583499069 | 1.763147803 | -0.330941665 |
| Gm20561 | 0.047948698 | -0.206487837 | 3.724945049 | -0.055433794 |
| Gm29408 | 0.136071997 | -1.15312687 | 2.941811982 | -0.39197844 |
| Gm5620 | 30.93367375 | -0.018642811 | 0.226424882 | -0.082335524 |
| Gm36960 | 3.033497949 | 0.568867027 | 0.663569999 | 0.857282621 |
| Gm29602 | 0 | NA | NA | NA |
| Gm37812 | 6.137587838 | -0.577238054 | 0.410622278 | -1.405764093 |
| Gm37457 | 5.572788009 | -0.527327383 | 0.448071973 | -1.176880982 |
| Gm37040 | 5.798168318 | 0.698586556 | 0.488974291 | 1.428677478 |
| Gm28652 | 0 | NA | NA | NA |
| A730094K22Rik | 3.673889323 | -1.044705464 | 0.755390508 | -1.383000519 |
| Zic1 | 2091.234844 | -0.501744314 | 0.184265974 | -2.722935238 |
| Zic4 | 124.4589485 | -0.175608576 | 0.139548612 | -1.25840432 |
| Gm37477 | 5.296492507 | 0.036386409 | 0.45742137 | 0.079546805 |
| Gm29478 | 1.174654913 | -0.14327328 | 0.819741889 | -0.174778527 |
| Gm37987 | 0.060174537 | 0.755286614 | 3.724945049 | 0.202764498 |
| Gm26101 | 0.062426873 | 0.755288839 | 3.724945049 | 0.202765096 |
| Plscr5 | 1.323501177 | -0.802585173 | 0.850607917 | -0.943543032 |
| Plscr1 | 60.43889054 | 1.568891521 | 0.285285124 | 5.499380754 |
| Mir6386 | 0 | NA | NA | NA |
| Plscr2 | 121.3806642 | 2.661597563 | 0.240913986 | 11.04791634 |

| | | | | |
|---|---|---|---|---|
| Gm29139 | 0 | NA | NA | NA |
| 1700057G04Rik | 0.707934375 | -1.073888547 | 1.345802407 | -0.797954099 |
| Gm15493 | 0 | NA | NA | NA |
| Plscr4 | 44.4848455 | 0.021986785 | 0.183146103 | 0.120050522 |
| Gm25417 | 0.083526043 | 0.75528439 | 3.724945049 | 0.202763901 |
| Gm38391 | 19.9576536 | 0.328187023 | 0.236718602 | 1.386401494 |
| Gm37545 | 0 | NA | NA | NA |
| Plod2 | 158.0764622 | 0.025461414 | 0.111420449 | 0.228516527 |
| Gm2495 | 0 | NA | NA | NA |
| Gm29396 | 0 | NA | NA | NA |
| 4933400C23Rik | 0.454057983 | -0.326977431 | 1.449936159 | -0.225511605 |
| Gm28054 | 0.110242424 | -0.20649451 | 3.724945049 | -0.055435586 |
| Gm29397 | 3.384086637 | -0.125722658 | 0.603800295 | -0.208218942 |
| 1700034K08Rik | 0 | NA | NA | NA |
| Gm9621 | 0 | NA | NA | NA |
| Gm24200 | 0 | NA | NA | NA |
| Gm5370 | 0 | NA | NA | NA |
| Gm28934 | 0 | NA | NA | NA |
| 1190002N15Rik | 756.6875991 | -0.215283755 | 0.139439811 | -1.543918869 |
| Gm24178 | 0.049553047 | 0.274391604 | 3.724945049 | 0.073663262 |
| Gm29395 | 0.832501709 | -0.137232451 | 1.014130603 | -0.135320293 |
| Slc9a9 | 216.5651415 | -0.22075421 | 0.121771153 | -1.812861293 |
| Gm16262 | 0.060174537 | 0.755286614 | 3.724945049 | 0.202764498 |
| Mir7656 | 0.276333946 | -0.196378714 | 1.796789905 | -0.109294199 |
| Chst2 | 1637.467997 | 0.275667812 | 0.107414686 | 2.566388475 |
| Gm37805 | 7.969835414 | 0.417110384 | 0.358792529 | 1.162539212 |
| Gm8449 | 2.212778765 | 2.234010558 | 0.825796599 | 2.705279435 |
| Gm19193 | 0 | NA | NA | NA |
| U2surp | 1949.775422 | 0.258162119 | 0.084884679 | 3.041327607 |
| Gm28424 | 12.26829614 | -0.64382053 | 0.274200191 | -2.347994463 |
| Paqr9 | 964.539708 | -0.188867467 | 0.098575339 | -1.915970763 |
| Gm16126 | 0.766553141 | 1.714815736 | 1.292268819 | 1.326980664 |
| Gm28164 | 0.74026972 | 1.293876301 | 1.162020019 | 1.113471609 |
| Gm25582 | 0.857373094 | 0.024688676 | 0.932545627 | 0.026474496 |
| Gm16127 | 0.660063185 | 2.244886071 | 1.3474877 | 1.665978896 |
| Pcolce2 | 60.65372678 | -0.049822868 | 0.165218377 | -0.301557665 |
| Itpa-ps2 | 0.066050875 | 0.274391604 | 3.724945049 | 0.073663262 |
| Trpc1 | 585.9086551 | -0.137494133 | 0.064746104 | -2.123589284 |
| Pls1 | 183.634994 | -0.817091202 | 0.119864746 | -6.816776647 |
| 1700065D16Rik | 3.065206492 | -0.208929282 | 0.574969639 | -0.36337446 |
| Atr | 564.2015883 | -0.143596879 | 0.097127193 | -1.478441561 |
| Gm24004 | 0 | NA | NA | NA |
| Gm16794 | 29.63475761 | 0.047926895 | 0.201896448 | 0.237383547 |
| Xrn1 | 1619.277144 | 0.522504952 | 0.083350491 | 6.268768735 |
| C78334 | 0.042716945 | -0.206490061 | 3.724945049 | -0.055434391 |
| Gm28085 | 0 | NA | NA | NA |
| Gm38375 | 61.21253001 | 0.269870135 | 0.263224527 | 1.025246918 |

| | | | | |
|---|---|---|---|---|
| Gk5 | 59.46593611 | 0.097832637 | 0.15373736 | 0.636362155 |
| Tfdp2 | 637.9968549 | -0.247066443 | 0.08522752 | -2.898904522 |
| Gm37195 | 34.22903916 | -1.062107102 | 0.308325942 | -3.444754257 |
| Gm28924 | 0.049553047 | 0.274391604 | 3.724945049 | 0.073663262 |
| Atp1b3 | 1332.613662 | 0.321382161 | 0.080447417 | 3.994934481 |
| BC043934 | 5.01349092 | -0.288446501 | 0.519297954 | -0.555454723 |
| Rnf7 | 496.7963806 | -0.063068267 | 0.09707333 | -0.649697167 |
| Gm27289 | 0 | NA | NA | NA |
| Gm28729 | 2.066931119 | 0.151363028 | 0.738919349 | 0.204843774 |
| Gm8495 | 0.930891567 | -2.41643491 | 1.036600171 | -2.331115677 |
| Rasa2 | 837.5261912 | -0.251783014 | 0.084462669 | -2.980997592 |
| n-R5s87 | 2.429268922 | -0.624548564 | 0.648147669 | -0.963589925 |
| Gm19280 | 0 | NA | NA | NA |
| Zbtb38 | 1623.281997 | 0.062816159 | 0.039946418 | 1.572510416 |
| A930006L05Rik | 0.095435502 | 0.75528439 | 3.724945049 | 0.202763901 |
| C430002N11Rik | 6.432897433 | 2.525569687 | 1.048741509 | 2.408190833 |
| Gm28688 | 0.279349467 | 0.274407326 | 2.04879268 | 0.133936112 |
| Gm8520 | 7.409839226 | 0.271336768 | 0.428224605 | 0.633631895 |
| Pxylp1 | 198.9385142 | -0.355951657 | 0.120913401 | -2.943856137 |
| Gm10123 | 609.9706294 | 0.094209735 | 0.094340045 | 0.998618716 |
| Gm8524 | 2.21064614 | -0.033654951 | 0.684825858 | -0.049143808 |
| Spsb4 | 74.46297335 | -0.330679465 | 0.135185133 | -2.446123012 |
| Gm16010 | 1.553703116 | -1.656101312 | 0.898354706 | -1.843482648 |
| Gm18291 | 0.371128506 | 1.240096368 | 1.992730596 | 0.622310096 |
| Gm19325 | 18.24486811 | -0.920031734 | 0.341672653 | -2.692728623 |
| Gm26767 | 0.08169834 | -0.206492286 | 3.724945049 | -0.055434988 |
| Slc25a36 | 1229.888798 | -0.044046411 | 0.060744482 | -0.725109676 |
| Gm16185 | 0.038981395 | 0.274393828 | 3.724945049 | 0.073663859 |
| Rpl7a-ps10 | 2.493931419 | 1.526686087 | 0.827783609 | 1.844305771 |
| Gm5371 | 0 | NA | NA | NA |
| Gm29241 | 0 | NA | NA | NA |
| Gm29240 | 0.044544648 | 0.755286614 | 3.724945049 | 0.202764498 |
| A030012G06Rik | 0.136880458 | -0.675115491 | 3.636764192 | -0.185636312 |
| Trim42 | 0.23239932 | 1.236201655 | 2.134821991 | 0.579065449 |
| Gm15891 | 0.046345709 | 0.274391604 | 3.724945049 | 0.073663262 |
| Gm25607 | 0.495675163 | 1.326784964 | 1.638457513 | 0.80977685 |
| Clstn2 | 1155.089609 | -0.010065227 | 0.08516651 | -0.118182921 |
| Gm37661 | 32.39089962 | -0.660280612 | 0.287336197 | -2.29793746 |
| 4921534H16Rik | 0.120974994 | 0.274404949 | 3.724945049 | 0.073666845 |
| 2610303G11Rik | 0 | NA | NA | NA |
| Nmnat3 | 20.17326344 | 0.123257252 | 0.228792249 | 0.538730015 |
| Gm28530 | 3.500655289 | 0.783847443 | 0.61583959 | 1.272811063 |
| Rbp1 | 140.0093858 | 0.846168316 | 0.190114065 | 4.450845423 |
| Rbp2 | 0.656451805 | 0.614313468 | 1.248003064 | 0.492237147 |
| Gm37113 | 0 | NA | NA | NA |
| 4930579K19Rik | 5.920659574 | -0.003826237 | 0.462010662 | -0.008281706 |
| Copb2 | 1299.911645 | 0.306836877 | 0.077445863 | 3.961953107 |

| | | | | |
|---|---|---|---|---|
| Mrps22 | 205.9148466 | 0.073431914 | 0.090908113 | 0.807759734 |
| Gm37637 | 13.96030438 | -0.510119282 | 0.288690837 | -1.767008912 |
| Gm27919 | 0 | NA | NA | NA |
| Pisrt1 | 0 | NA | NA | NA |
| E330023G01Rik | 0.110595523 | 0.755282166 | 3.724945049 | 0.202763304 |
| Gm37282 | 0 | NA | NA | NA |
| Gm37917 | 0.58604645 | -0.150050656 | 1.443061333 | -0.103980789 |
| Gm38226 | 0.474407165 | -0.145847839 | 1.33136851 | -0.10954731 |
| Prr23a | 0.228844266 | -0.206481303 | 2.137880943 | -0.096582227 |
| Gm6406 | 0 | NA | NA | NA |
| 2410012M07Rik | 0 | NA | NA | NA |
| Gm5920 | 2.082893185 | -0.220724946 | 0.76959713 | -0.286805834 |
| 7420426K07Rik | 0 | NA | NA | NA |
| Gm28266 | 0 | NA | NA | NA |
| Foxl2os | 2.17543179 | -0.050497194 | 0.787286345 | -0.064140823 |
| Foxl2 | 2.235327434 | -0.731698416 | 0.700105659 | -1.045125698 |
| Faim | 196.4395141 | 0.135041376 | 0.105345844 | 1.281886126 |
| Gm1123 | 1.879715592 | 1.102269893 | 0.817578722 | 1.348212549 |
| Pik3cb | 544.1368007 | 0.094590731 | 0.065710817 | 1.439500145 |
| Gm2950 | 0.061920613 | 0.274393828 | 3.724945049 | 0.073663859 |
| Gm6432 | 0.11197992 | 0.75528439 | 3.724945049 | 0.202763901 |
| Gm38327 | 0.266212691 | -1.203653844 | 3.708343979 | -0.32457988 |
| Cep70 | 260.9692196 | -0.253205079 | 0.157502091 | -1.607629954 |
| 4930422M22Rik | 0.158279039 | 0.755296076 | 2.813413384 | 0.26846253 |
| Gm37249 | 36.58656883 | -1.136374513 | 0.307275359 | -3.698228576 |
| Esyt3 | 26.59934933 | -0.256884936 | 0.227592171 | -1.128707261 |
| 1700008A23Rik | 0.963690934 | 0.042607644 | 1.031776371 | 0.041295425 |
| Gm28087 | 0 | NA | NA | NA |
| Mras | 2060.728902 | 0.190552805 | 0.063743117 | 2.989386384 |
| Nme9 | 2.199217757 | -1.034736271 | 0.747093465 | -1.385015825 |
| Armc8 | 1137.205985 | 0.32531486 | 0.078622159 | 4.137699399 |
| Dbr1 | 168.1373829 | 0.539215266 | 0.14178119 | 3.803150934 |
| A4gnt | 0.979608328 | -0.967255976 | 1.001052353 | -0.966239152 |
| Dzip1l | 286.0060264 | -0.439516376 | 0.153910636 | -2.85565955 |
| Cldn18 | 0.262323822 | 2.122512014 | 3.701790295 | 0.573374461 |
| Gm16004 | 0 | NA | NA | NA |
| Gm23949 | 0 | NA | NA | NA |
| Sox14 | 8.749592595 | -0.718131782 | 0.38792403 | -1.851217576 |
| Gm23594 | 0 | NA | NA | NA |
| Gm8641 | 0.504639468 | 0.788974917 | 1.385457922 | 0.569468696 |
| 4930519F24Rik | 0.061920613 | 0.274393828 | 3.724945049 | 0.073663859 |
| 4930540E01Rik | 0 | NA | NA | NA |
| Gm28167 | 0 | NA | NA | NA |
| Gm28267 | 0 | NA | NA | NA |
| Gm28166 | 0.08486385 | 0.755286617 | 3.724945049 | 0.202764499 |
| Il20rb | 65.86335234 | 0.357096447 | 0.263308303 | 1.356191366 |
| Nck1 | 358.9705822 | 0.017862153 | 0.078794348 | 0.226693323 |

| | | | | |
|---|---|---|---|---|
| Gm28586 | 8.399474289 | -0.716260732 | 0.5007555 | -1.430360189 |
| Slc35g2 | 192.5839218 | -0.319891956 | 0.154296431 | -2.073229777 |
| Stag1 | 1284.878169 | -0.0018474 | 0.069189676 | -0.026700521 |
| Gm8661 | 0.119234754 | 0.274400501 | 3.724945049 | 0.073665651 |
| Gm37204 | 117.4552302 | -0.499010643 | 0.267171969 | -1.867750737 |
| Gm38297 | 15.65520555 | -0.510009123 | 0.389881922 | -1.308111748 |
| Gm3081 | 62.39629019 | 0.03289306 | 0.23478945 | 0.140095988 |
| Pccb | 410.7052558 | -0.047123473 | 0.074063264 | -0.636259742 |
| Gm3096 | 0 | NA | NA | NA |
| Msl2 | 1119.25172 | 0.106686814 | 0.067144942 | 1.588903208 |
| Gm37553 | 42.26828888 | -0.486228098 | 0.257043001 | -1.891621617 |
| Ppp2r3a | 1068.136217 | 0.241275192 | 0.070219197 | 3.436028913 |
| Gm37962 | 38.30761735 | -0.472978842 | 0.268354427 | -1.762515517 |
| Gm37621 | 78.87355947 | -0.51184532 | 0.222271632 | -2.302791933 |
| Gm37953 | 20.22364659 | -0.711848024 | 0.299126046 | -2.379759417 |
| Gm38344 | 18.31202231 | 0.251336408 | 0.313144682 | 0.802620712 |
| Gm37314 | 38.77458494 | 0.305193854 | 0.231245822 | 1.319781049 |
| Gm24338 | 0.290020514 | 0.263458215 | 1.63464772 | 0.161171249 |
| Gm3121 | 0 | NA | NA | NA |
| Gm28660 | 0 | NA | NA | NA |
| Gm28979 | 0.404019895 | 0.693061014 | 1.54095927 | 0.449759464 |
| Gm24638 | 0 | NA | NA | NA |
| Gm29385 | 0 | NA | NA | NA |
| Gm5161 | 0.106056992 | 1.236179401 | 3.724945049 | 0.331865138 |
| Gm29521 | 6.846922448 | -0.44506682 | 0.449081345 | -0.991060583 |
| Gm37460 | 0.652148592 | 1.537509336 | 1.345686517 | 1.142546438 |
| Ephb1 | 568.496847 | -0.247407351 | 0.071143381 | -3.477587741 |
| 9630041A04Rik | 1.37032714 | 0.857468963 | 0.943032733 | 0.909267444 |
| Gm37683 | 0.138485956 | -0.68737807 | 2.932778697 | -0.234377749 |
| Gm37562 | 15.3341162 | -0.265645982 | 0.360669314 | -0.736536134 |
| n-R5s88 | 0 | NA | NA | NA |
| Gm29388 | 6.105350049 | -0.761196816 | 0.549884801 | -1.384284154 |
| Gm37945 | 0.049553047 | 0.274391604 | 3.724945049 | 0.073663262 |
| Gm37260 | 0.11197992 | 0.75528439 | 3.724945049 | 0.202763901 |
| Mir7243 | 1.350235793 | -1.932303722 | 0.907385968 | -2.129527885 |
| Gm26846 | 1.767979395 | -0.49617021 | 0.794777171 | -0.624288452 |
| Gm22894 | 0.570356529 | -1.06468568 | 1.351520867 | -0.787768584 |
| Ky | 45.80996683 | -0.370160322 | 0.220963353 | -1.675211374 |
| Gm25669 | 0.261093266 | 1.950434934 | 3.710027945 | 0.525719742 |
| Gm7364 | 0.312630927 | 2.551532587 | 3.684752935 | 0.692456898 |
| Gm29143 | 0 | NA | NA | NA |
| Gm29387 | 0.088534442 | 0.274389469 | 3.724945049 | 0.073662689 |
| Cep63 | 192.8660548 | -0.22991141 | 0.098161046 | -2.342185826 |
| Anapc13 | 138.6441374 | -0.026777337 | 0.154962799 | -0.172798488 |
| 4930533D04Rik | 0.164893722 | 1.582203397 | 3.387385449 | 0.467086908 |
| Gm36957 | 0.137470077 | 1.236181898 | 3.595099416 | 0.343851937 |
| Amotl2 | 701.5681414 | 0.863534952 | 0.182272536 | 4.73760322 |

| | | | | |
|---|---|---|---|---|
| Gm5627 | 0.038981395 | 0.274393828 | 3.724945049 | 0.073663859 |
| Gm36958 | 0.813964436 | 0.258216264 | 1.023904778 | 0.252187771 |
| Gm28621 | 0 | NA | NA | NA |
| Ryk | 428.6544998 | -0.188203968 | 0.074600007 | -2.522841158 |
| Slco2a1 | 62.94329372 | 0.663117153 | 0.181196252 | 3.65966264 |
| Gm25842 | 0.189042777 | 0.274408515 | 2.65947082 | 0.103181623 |
| n-R5s89 | 0 | NA | NA | NA |
| Gm22595 | 0 | NA | NA | NA |
| Rab6b | 7282.57726 | -0.188505717 | 0.087051021 | -2.165462452 |
| Gm37166 | 45.91226253 | -0.859742192 | 0.392590002 | -2.189923808 |
| Srprb | 192.082496 | -0.166802523 | 0.104879464 | -1.590421203 |
| Gm20425 | 0 | NA | NA | NA |
| Trf | 281.060478 | 0.844067305 | 0.210771957 | 4.00464709 |
| 1300017J02Rik | 2.855215039 | -0.036137081 | 0.650008843 | -0.055594752 |
| Topbp1 | 318.7285085 | 0.124796181 | 0.092084406 | 1.355236857 |
| Cdv3 | 1213.725262 | 0.347811818 | 0.094969756 | 3.662342979 |
| Gm29600 | 0.038981395 | 0.274393828 | 3.724945049 | 0.073663859 |
| 5830418P13Rik | 5.81979985 | 0.135222363 | 0.405812801 | 0.333213646 |
| Bfsp2 | 26.48732468 | 0.278154283 | 0.249938534 | 1.112890752 |
| Gm37938 | 2.10811511 | -0.005694046 | 0.925464797 | -0.006152633 |
| Gm37546 | 0.280931382 | 0.456318254 | 3.704888373 | 0.123166532 |
| Gm16252 | 0.115957212 | 0.274402725 | 3.724945049 | 0.073666248 |
| Tmem108 | 366.5574461 | -0.08460102 | 0.067321162 | -1.256677956 |
| Gm26360 | 2.674678528 | 0.121598762 | 0.643609231 | 0.188932595 |
| Gm37649 | 24.6588879 | -0.839957829 | 0.405943714 | -2.069148505 |
| 4932413F04Rik | 0.450128165 | -1.067086471 | 1.42814079 | -0.747185767 |
| Gm38032 | 30.93777638 | -1.111569668 | 0.317314399 | -3.503054605 |
| Gm5372 | 0 | NA | NA | NA |
| Gm29154 | 0.341479653 | 0.362364348 | 3.713618932 | 0.097577149 |
| Nphp3 | 143.1369062 | 0.058962531 | 0.139153482 | 0.423723 |
| Gm28305 | 0 | NA | NA | NA |
| Gm37200 | 6.440447106 | -0.202415272 | 0.404247972 | -0.500720563 |
| Uba5 | 589.0417371 | -0.101084075 | 0.065525383 | -1.542670488 |
| Gm37750 | 0.049553047 | 0.274391604 | 3.724945049 | 0.073663262 |
| Acad11 | 231.7177883 | -0.469734315 | 0.121562523 | -3.864137601 |
| Ackr4 | 2.634801621 | -0.341359336 | 0.645948585 | -0.528462085 |
| Dnajc13 | 1701.736079 | 0.277029603 | 0.069110503 | 4.008502188 |
| Hmgb1-rs16 | 11.38856345 | 0.616623012 | 0.420032925 | 1.468034944 |
| Gm37563 | 1.931745907 | 2.778922719 | 1.013119531 | 2.742936677 |
| Acpp | 6.437954073 | -0.120514968 | 0.423700974 | -0.284434012 |
| Gm37188 | 0.048949532 | -0.206487837 | 3.724945049 | -0.055433794 |
| Gm28996 | 0 | NA | NA | NA |
| Gm28548 | 0 | NA | NA | NA |
| Mir2136 | 0 | NA | NA | NA |
| Gm25904 | 0 | NA | NA | NA |
| Cpne4 | 536.3195774 | -0.402545645 | 0.155466537 | -2.589275176 |
| Gm23391 | 1.103801871 | -0.942585934 | 0.908329633 | -1.037713513 |

| | | | | |
|---|---|---|---|---|
| Mrpl3 | 481.1686626 | 0.066978784 | 0.059103974 | 1.133236563 |
| Gm38077 | 99.57854452 | -0.040832198 | 0.307945788 | -0.13259541 |
| Nudt16 | 104.3868709 | -0.034468133 | 0.124526894 | -0.276792683 |
| 1700080E11Rik | 0.3422147 | 0.755304834 | 1.9602451 | 0.385311425 |
| Nek11 | 22.44919175 | 0.123326037 | 0.27830181 | 0.443137748 |
| Gm37582 | 0.092140247 | -0.687373951 | 3.724945049 | -0.184532642 |
| Gm20643 | 0 | NA | NA | NA |
| Gm15619 | 0 | NA | NA | NA |
| Aste1 | 73.3563129 | 0.414109476 | 0.138189549 | 2.996677237 |
| Atp2c1 | 2093.602576 | -0.240567834 | 0.058834483 | -4.08889182 |
| Gm19667 | 0.638749378 | 3.056387392 | 1.517419959 | 2.014200073 |
| Gm7436 | 0.368385343 | 0.833618245 | 1.778331166 | 0.468764345 |
| Gm5621 | 1.423724791 | -0.793788083 | 0.976687444 | -0.812735014 |
| Gm28555 | 0 | NA | NA | NA |
| Pik3r4 | 491.4021009 | -0.152414811 | 0.07290477 | -2.090601362 |
| Gm22720 | 0 | NA | NA | NA |
| Col6a6 | 14.261268 | 0.673407286 | 0.333441017 | 2.019569433 |
| Gm38314 | 0 | NA | NA | NA |
| Col6a5 | 28.81713568 | -0.1914582 | 0.265248955 | -0.721805669 |
| Col6a4 | 6.820926848 | 0.046157776 | 0.378172511 | 0.122054814 |
| Gm24631 | 0 | NA | NA | NA |
| Gm36993 | 0 | NA | NA | NA |
| Gm9451 | 0.136331796 | -1.154745002 | 3.64305592 | -0.316971528 |
| Gm29208 | 0 | NA | NA | NA |
| D030055H07Rik | 7.771284628 | -2.331040021 | 0.447389686 | -5.210312379 |
| Glyctk | 39.68295307 | -0.038129917 | 0.218346609 | -0.174630225 |
| Mir135a-1 | 0.193036943 | -0.660002567 | 3.248873595 | -0.203148121 |
| Wdr82 | 1508.867638 | -0.210681633 | 0.058089448 | -3.626848618 |
| Mirlet7g | 2.956734474 | -0.269462337 | 0.748778079 | -0.359869426 |
| Ppm1m | 99.17848393 | 0.203732874 | 0.199505414 | 1.021189704 |
| Twf2 | 271.214745 | -0.038854796 | 0.099424523 | -0.390796907 |
| Tlr9 | 362.3896947 | 4.968508619 | 0.38189593 | 13.01011147 |
| Alas1 | 600.2556519 | 0.09738652 | 0.064650423 | 1.506355497 |
| 4930500F10Rik | 7.519553513 | -0.451270549 | 0.401827275 | -1.123046086 |
| Poc1a | 35.43149234 | 0.141400478 | 0.220335979 | 0.64174938 |
| Gm29123 | 0.049553047 | 0.274391604 | 3.724945049 | 0.073663262 |
| Dusp7 | 640.9294056 | -0.098989812 | 0.084366489 | -1.173330944 |
| Rpl29 | 8.095074919 | 0.276120194 | 0.382523857 | 0.721837839 |
| Acy1 | 102.5946641 | -0.239131577 | 0.139083447 | -1.719338874 |
| Abhd14a | 66.26385201 | -0.811177158 | 0.170262181 | -4.764282677 |
| Abhd14b | 60.06866735 | -0.447384189 | 0.214385904 | -2.086817185 |
| Gm28959 | 0.650039631 | -1.542024282 | 1.213815349 | -1.270394449 |
| Pcbp4 | 1192.577122 | -0.216100795 | 0.063985465 | -3.377341946 |
| Gpr62 | 124.4874142 | 0.219988806 | 0.147499139 | 1.491458235 |
| Parp3 | 396.144577 | 3.076656766 | 0.15279627 | 20.13567978 |
| Rrp9 | 173.3168163 | 0.121368532 | 0.100562735 | 1.2068937 |
| Iqcf1 | 0.403414764 | -0.206482556 | 1.560726172 | -0.132299029 |

| | | | | |
|---|---|---|---|---|
| Iqcf3 | 0.731923588 | -2.639429817 | 1.18398739 | -2.229271899 |
| Iqcf5 | 0.458372996 | -2.632115839 | 1.612005388 | -1.632820745 |
| Iqcf4 | 0 | NA | NA | NA |
| Iqcf6 | 0 | NA | NA | NA |
| Gm26185 | 0 | NA | NA | NA |
| Gm28111 | 0 | NA | NA | NA |
| Grm2 | 419.4914956 | -0.36754566 | 0.131534181 | -2.79429771 |
| Tex264 | 547.2087676 | 0.115597037 | 0.080277197 | 1.439973503 |
| Rad54l2 | 692.1787233 | 0.02968171 | 0.094957899 | 0.312577578 |
| Gm22384 | 0.206359549 | 0.299355767 | 2.640765838 | 0.113359451 |
| Vprbp | 629.97184 | -0.275866683 | 0.085973063 | -3.208757185 |
| Manf | 616.8659586 | 0.096365065 | 0.10983979 | 0.877323829 |
| Gm37612 | 1.882200288 | -1.030235018 | 0.771263947 | -1.335774895 |
| Rbm15b | 823.0495333 | -0.08529628 | 0.068101974 | -1.252478822 |
| 4930524O07Rik | 4.402341252 | 0.053626086 | 0.610428788 | 0.087849864 |
| Dock3 | 4718.325882 | -0.253072216 | 0.112615868 | -2.247216317 |
| Gm17205 | 0.109124069 | 0.274402725 | 3.724945049 | 0.073666248 |
| Gm22454 | 0 | NA | NA | NA |
| Gm17117 | 0 | NA | NA | NA |
| Gm23595 | 0 | NA | NA | NA |
| AA543401 | 0.072025462 | 0.755288839 | 3.724945049 | 0.202765096 |
| Gm22238 | 0 | NA | NA | NA |
| 1700039M15Rik | 0.049553047 | 0.274391604 | 3.724945049 | 0.073663262 |
| Mapkapk3 | 63.6118118 | 0.804024415 | 0.229071671 | 3.509925137 |
| Gm17040 | 0.094097695 | 0.755282166 | 3.724945049 | 0.202763304 |
| Cish | 89.05641637 | 0.226977461 | 0.174365561 | 1.301733322 |
| Gm17041 | 0.049553047 | 0.274391604 | 3.724945049 | 0.073663262 |
| Hemk1 | 115.6953326 | -0.227482312 | 0.115564403 | -1.96844622 |
| 6430571L13Rik | 122.123982 | -0.16769499 | 0.166768912 | -1.005553066 |
| 4930429P21Rik | 0.791934515 | -1.276335882 | 1.207989179 | -1.056578903 |
| Cacna2d2 | 1015.043882 | -0.161100816 | 0.073317236 | -2.19731165 |
| Gm17141 | 0.093831153 | 0.274404949 | 3.724945049 | 0.073666845 |
| Gm17118 | 0 | NA | NA | NA |
| Tmem115 | 377.5379067 | 0.257099867 | 0.095493959 | 2.692315501 |
| Cyb561d2 | 55.18192089 | -0.024524982 | 0.208383316 | -0.117691678 |
| Nprl2 | 184.5566623 | -0.356456767 | 0.087054899 | -4.09462042 |
| Zmynd10 | 22.83363798 | -0.239783189 | 0.257334909 | -0.931794251 |
| Gm34106 | 0.182655267 | 1.584997757 | 2.694309109 | 0.588276138 |
| Rassf1 | 176.3853711 | 0.485810407 | 0.136113645 | 3.569152867 |
| Gm9917 | 9.204737304 | -0.535953356 | 0.471194302 | -1.137435986 |
| Tusc2 | 617.3101563 | -0.086743409 | 0.071063239 | -1.220650946 |
| Hyal2 | 95.99414798 | -0.071825544 | 0.135571552 | -0.529798053 |
| Gm16343 | 0.186399948 | 1.293453504 | 3.295513693 | 0.392489191 |
| Hyal1 | 60.01938957 | 0.242647684 | 0.199104492 | 1.218695177 |
| Nat6 | 87.56719576 | 0.397291049 | 0.162895842 | 2.438926894 |
| Hyal3 | 5.312334351 | 0.212044299 | 0.55489996 | 0.382130681 |
| Ifrd2 | 67.19084161 | -0.236340704 | 0.217454163 | -1.086852977 |

| | | | | |
|---|---|---|---|---|
| Lsmem2 | 2.036564983 | 0.892863408 | 0.900773046 | 0.991219056 |
| Gm38150 | 3.547776317 | -0.14164192 | 0.464444335 | -0.304970713 |
| Sema3b | 58.08865837 | 0.05141829 | 0.197776316 | 0.259982038 |
| Gnai2 | 2404.56764 | 0.627931163 | 0.114897758 | 5.465129808 |
| Gm19721 | 19.767715 | -0.069379069 | 0.411349605 | -0.168662053 |
| Slc38a3 | 598.0789621 | -0.298329881 | 0.180440192 | -1.653344959 |
| Gnat1 | 1.440190137 | 0.270662597 | 0.769794778 | 0.351603576 |
| A930036K24Rik | 0.18066559 | 0.755299131 | 2.691842631 | 0.280588145 |
| Sema3f | 139.5279565 | -0.274309749 | 0.109512696 | -2.504821441 |
| Rbm5 | 1611.009065 | -0.447743756 | 0.158467908 | -2.825453822 |
| Gm37850 | 0 | NA | NA | NA |
| Rbm6 | 1030.666851 | -0.27578478 | 0.123403196 | -2.234826891 |
| Gm37436 | 36.96383849 | -0.144995166 | 0.242943279 | -0.59682724 |
| Mon1a | 192.9155728 | -0.130926947 | 0.099632621 | -1.314097186 |
| Gm23856 | 0.487068913 | -1.394360371 | 1.538812223 | -0.906127694 |
| Mst1r | 16.57438961 | 0.156793576 | 0.262704517 | 0.596843855 |
| Actl11 | 0.123841226 | 0.274391173 | 3.724945049 | 0.073663147 |
| Gm37974 | 0.217247303 | 0.274414419 | 2.596047673 | 0.105704692 |
| Camkv | 2202.703855 | -0.178799685 | 0.15608249 | -1.145546088 |
| Traip | 114.0875043 | -0.170418974 | 0.24379368 | -0.699029499 |
| Uba7 | 688.1155965 | 6.068435165 | 0.197827387 | 30.67540469 |
| Gm20661 | 4.55440648 | 0.989907002 | 0.599518938 | 1.651168862 |
| Fam212a | 16.26337642 | 0.516200861 | 0.256145382 | 2.01526515 |
| Gm24944 | 1.177026929 | 3.745877107 | 1.093324143 | 3.426135911 |
| 4921523L03Rik | 0.198550338 | 1.17024797 | 2.203705519 | 0.531036457 |
| Cdhr4 | 13.4482997 | 0.917034211 | 0.367174491 | 2.497543359 |
| Gm20662 | 0.066050875 | 0.274391604 | 3.724945049 | 0.073663262 |
| 4930535L15Rik | 1.219811481 | -0.334004998 | 0.920154079 | -0.362988119 |
| Ip6k1 | 2129.145135 | 0.227466331 | 0.052985556 | 4.292987522 |
| Gm38134 | 0.127971488 | 0.274389832 | 3.687653711 | 0.0744077 |
| Gmppb | 158.7886363 | 0.343438293 | 0.150880121 | 2.27623289 |
| Rnf123 | 811.9292402 | -0.188813966 | 0.071049739 | -2.65748993 |
| Gm20529 | 0.04393175 | -0.206490061 | 3.724945049 | -0.055434391 |
| Amigo3 | 37.0261778 | 1.378674173 | 0.307569744 | 4.482476578 |
| Mst1 | 0.439905485 | 1.191569858 | 1.45064971 | 0.821404265 |
| Mir7088 | 0 | NA | NA | NA |
| Apeh | 282.2906194 | -0.105772964 | 0.083816548 | -1.261958015 |
| Bsn | 11059.17198 | -0.218125302 | 0.121386858 | -1.796943304 |
| AC137678.1 | 2.386774017 | 0.512731775 | 0.834346941 | 0.614530659 |
| 4930447F24Rik | 14.97539824 | 1.647189057 | 0.410558634 | 4.012067755 |
| Gm37201 | 2.168403122 | 0.454428867 | 0.862538164 | 0.526850736 |
| Gm37247 | 41.41364108 | -0.400903381 | 0.353223071 | -1.134986398 |
| Gm37230 | 0.567387455 | -0.416637532 | 1.269814927 | -0.328108863 |
| Dag1 | 1797.643256 | 0.29416621 | 0.082641689 | 3.559537735 |
| Nicn1 | 612.9850749 | -0.15238538 | 0.071412182 | -2.133884935 |
| Amt | 190.2692267 | -0.657774566 | 0.139147259 | -4.727183051 |
| Tcta | 207.060682 | 0.189168046 | 0.110640279 | 1.709757491 |

| | | | | |
|---|---|---|---|---|
| Rhoa | 1859.627478 | 0.34083423 | 0.106817744 | 3.190801618 |
| Gm37568 | 3.725366168 | -0.434904398 | 0.561926446 | -0.773952536 |
| Gm37401 | 0.049553047 | 0.274391604 | 3.724945049 | 0.073663262 |
| Gpx1 | 610.2857368 | 0.71689289 | 0.146182989 | 4.904078751 |
| Usp4 | 1105.767034 | -0.12832633 | 0.054072546 | -2.373225207 |
| Rps2-ps11 | 0.578661668 | 0.762240534 | 1.326255149 | 0.574731442 |
| Gm37963 | 32.08673899 | -0.734897731 | 0.339160834 | -2.166811897 |
| 1700102P08Rik | 15.20131833 | -1.048009001 | 0.300817302 | -3.483872087 |
| Ccdc36 | 5.280767892 | 0.077731869 | 0.430135564 | 0.180714816 |
| Gm25456 | 0.042716945 | -0.206490061 | 3.724945049 | -0.055434391 |
| BC048562 | 0 | NA | NA | NA |
| Klhdc8b | 159.0275307 | -0.735571666 | 0.125192139 | -5.875541979 |
| Ccdc71 | 275.4617965 | 0.242176775 | 0.075280731 | 3.216982243 |
| Gm37475 | 3.601477103 | -1.486622125 | 0.643998028 | -2.308426516 |
| Lamb2 | 362.809454 | -0.042794418 | 0.148958541 | -0.287290795 |
| Usp19 | 1729.388652 | 0.084367187 | 0.050465402 | 1.671782714 |
| Qars | 489.2628777 | 0.259638766 | 0.063539342 | 4.086267775 |
| Qrich1 | 1514.838001 | 0.221715187 | 0.069058692 | 3.210532674 |
| Gm38366 | 45.47164434 | -0.05121918 | 0.334324794 | -0.153201859 |
| Gm37678 | 6.993644233 | -0.403771427 | 0.448920226 | -0.899428013 |
| Impdh2 | 294.2527551 | -0.02986634 | 0.099146132 | -0.301235554 |
| Ndufaf3 | 182.2571026 | -0.117214067 | 0.08192809 | -1.430694484 |
| 4833445I07Rik | 30.38428203 | 0.015406331 | 0.194521547 | 0.079201152 |
| Gm24083 | 0 | NA | NA | NA |
| mmu-mir-191 | 0 | NA | NA | NA |
| Mir425 | 0 | NA | NA | NA |
| Dalrd3 | 336.6293249 | -0.165605055 | 0.077614984 | -2.133673776 |
| Wdr6 | 1862.952255 | 0.053222252 | 0.089628858 | 0.593807093 |
| P4htm | 622.680632 | 0.085706112 | 0.0778808 | 1.100478072 |
| Gm38163 | 3.058745156 | -1.379608684 | 0.648309921 | -2.128007977 |
| Arih2 | 618.9684878 | 0.252868401 | 0.082767742 | 3.055156447 |
| Gm26298 | 0 | NA | NA | NA |
| Slc25a20 | 138.8267023 | 0.309982932 | 0.157177509 | 1.972183772 |
| Prkar2a | 906.3458615 | -0.156143701 | 0.068546658 | -2.277918525 |
| Gm24259 | 0.258327161 | 0.755287902 | 3.043292301 | 0.248181189 |
| Ip6k2 | 843.8946416 | 0.323951404 | 0.085976079 | 3.767924838 |
| Gm23034 | 0.062426873 | 0.755288839 | 3.724945049 | 0.202765096 |
| Nckipsd | 578.0213764 | -0.114129264 | 0.080188238 | -1.423266894 |
| Gm35025 | 6.065554424 | 0.003866319 | 0.437544278 | 0.008836407 |
| Gm37714 | 0.267221877 | 0.274413853 | 1.803924478 | 0.152120477 |
| Celsr3 | 1375.868327 | -0.335818908 | 0.161766337 | -2.075950499 |
| Gm23156 | 0.616331352 | 1.463056173 | 1.545495385 | 0.94665839 |
| Slc26a6 | 70.00938289 | -0.31618737 | 0.161125433 | -1.962367853 |
| Tmem89 | 0.562660809 | 1.294748716 | 1.244373103 | 1.040482724 |
| Uqcrc1 | 1892.13114 | 0.02363199 | 0.067344107 | 0.350914008 |
| Gm37284 | 0 | NA | NA | NA |
| Col7a1 | 41.89192254 | -0.409356336 | 0.265091471 | -1.544207869 |

| | | | | |
|---|---|---|---|---|
| Mir711 | 0 | NA | NA | NA |
| Ucn2 | 10.62182747 | 4.817298229 | 1.151360155 | 4.184006375 |
| Gm38004 | 0.533662682 | 1.041500647 | 1.351579704 | 0.770580265 |
| Pfkfb4 | 187.9455917 | 0.31563024 | 0.107718142 | 2.930149315 |
| Shisa5 | 1519.355689 | 2.320679506 | 0.14770752 | 15.71131593 |
| Trex1 | 520.4087808 | 2.340149022 | 0.195766701 | 11.95376442 |
| Atrip | 28.91987549 | -0.209506394 | 0.39064838 | -0.536304269 |
| Tma7 | 371.0569712 | 0.218103249 | 0.089418213 | 2.439136758 |
| Ccdc51 | 52.13264183 | -0.240871691 | 0.267352034 | -0.900953274 |
| Gm7628 | 6.615983682 | 0.710571295 | 0.4320196 | 1.644766339 |
| Plxnb1 | 1318.803774 | -0.403095576 | 0.12397682 | -3.251378582 |
| Fbxw21 | 0 | NA | NA | NA |
| Fbxw13 | 0.127364494 | -0.687378399 | 3.724945049 | -0.184533836 |
| Fbxw20 | 0.066050875 | 0.274391604 | 3.724945049 | 0.073663262 |
| Fbxw14 | 0.088383099 | -0.687376175 | 3.724945049 | -0.184533239 |
| Gm22821 | 0 | NA | NA | NA |
| Gm25484 | 0 | NA | NA | NA |
| Gm24641 | 0 | NA | NA | NA |
| Fbxw22 | 0.038981395 | 0.274393828 | 3.724945049 | 0.073663859 |
| Fbxw16 | 0.066050875 | 0.274391604 | 3.724945049 | 0.073663262 |
| RP23-62L19.6 | 0 | NA | NA | NA |
| RP23-62L19.5 | 0 | NA | NA | NA |
| Fbxw19 | 0.038981395 | 0.274393828 | 3.724945049 | 0.073663859 |
| RP24-187P11.4 | 0.450885768 | 0.788898848 | 1.647052474 | 0.478976147 |
| RP24-187P11.2 | 9.209079983 | 2.056593627 | 0.568304087 | 3.618826038 |
| RP24-187P11.5 | 0.238621995 | 0.755297565 | 2.519154582 | 0.299821841 |
| Fbxw15 | 0.204053819 | 0.202155946 | 2.621632496 | 0.077110711 |
| RP24-187P11.3 | 0.046345709 | 0.274391604 | 3.724945049 | 0.073663262 |
| Fbxw24 | 0.124596252 | 0.755282361 | 3.714681068 | 0.203323609 |
| Fbxw25 | 0.109982625 | -0.20649451 | 3.724945049 | -0.055435586 |
| Fbxw18 | 0.15906016 | 0.274395307 | 3.439462686 | 0.079778539 |
| RP24-344B20.5 | 0 | NA | NA | NA |
| Fbxw26 | 0.166952883 | 0.274401811 | 2.756036668 | 0.099563919 |
| RP23-357H5.5 | 0.394548002 | 0.119015251 | 1.748490956 | 0.06806741 |
| Spink8 | 16.32657011 | 1.022693817 | 0.286562087 | 3.568838526 |
| 3000002C10Rik | 56.32280498 | -0.123856348 | 0.179037442 | -0.691790197 |
| Nme6 | 38.63028386 | -0.141767118 | 0.216858939 | -0.653729647 |
| Camp | 2.243591458 | 3.902242651 | 1.052079292 | 3.709076571 |
| Cdc25a | 165.9711309 | 0.336014307 | 0.109057802 | 3.081066189 |
| Mir7089 | 0.220081591 | 1.715799776 | 2.153657697 | 0.796691034 |
| RP24-421E18.9 | 0 | NA | NA | NA |
| Map4 | 7553.036783 | 0.193786259 | 0.080664194 | 2.402382644 |
| RP24-421E18.8 | 10.03429169 | -0.137925591 | 0.32168557 | -0.428759025 |
| RP24-421E18.10 | 0 | NA | NA | NA |
| RP24-421E18.5 | 78.64983153 | -0.218203002 | 0.123828936 | -1.762132568 |
| AC161246.1 | 0 | NA | NA | NA |
| RP24-421E18.7 | 15.20321581 | -0.355045441 | 0.374676947 | -0.94760418 |

| | | | | |
|---|---|---|---|---|
| Dhx30 | 1202.629578 | -0.023933646 | 0.050491478 | -0.474013573 |
| Gm27855 | 0.367828087 | -0.264099482 | 1.509438653 | -0.174965363 |
| Smarcc1 | 1075.459101 | -0.066733971 | 0.050684996 | -1.316641531 |
| Gm23377 | 0 | NA | NA | NA |
| Cspg5 | 2416.656234 | 0.291139482 | 0.078192094 | 3.723387725 |
| RP24-285I4.2 | 0.569412722 | -0.669571285 | 1.200116727 | -0.557921801 |
| Mir6236 | 2.610552123 | -0.505200273 | 1.12175129 | -0.450367454 |
| RP24-330A11.9 | 0.916112882 | -0.08655024 | 0.954265007 | -0.090698328 |
| Elp6 | 89.70397602 | 0.569863319 | 0.119201523 | 4.780671467 |
| RP24-330A11.12 | 0.042716945 | -0.206490061 | 3.724945049 | -0.055434391 |
| Scap | 1121.694434 | 0.055058697 | 0.047990086 | 1.14729315 |
| 5830462I19Rik | 0.695310992 | -0.707755574 | 1.573162243 | -0.449893568 |
| Ptpn23 | 992.3070621 | 0.224435765 | 0.092777453 | 2.419076587 |
| Ngp | 7.348605377 | 1.653927088 | 0.971835299 | 1.701859452 |
| Klhl18 | 377.384243 | -0.123621499 | 0.076284971 | -1.620522335 |
| Kif9 | 65.23232709 | -0.047011761 | 0.160753602 | -0.292446084 |
| Setd2 | 2113.532458 | -0.047357723 | 0.070764574 | -0.669229263 |
| RP24-363K20.7 | 41.47908056 | -0.041959289 | 0.228068412 | -0.183976765 |
| Nradd | 21.00591379 | 0.067462756 | 0.34928191 | 0.193147007 |
| Nbeal2 | 229.3785941 | 0.057028403 | 0.150863276 | 0.37801382 |
| Mir8107 | 0 | NA | NA | NA |
| RP24-179L19.4 | 0.417968807 | 1.062368998 | 1.496387965 | 0.709955588 |
| Ccdc12 | 189.7858103 | 0.352330877 | 0.120996911 | 2.911899781 |
| RP24-179L19.2 | 0.211345377 | -1.168306176 | 2.640924559 | -0.44238529 |
| Pth1r | 56.35433901 | 0.120792731 | 0.155658499 | 0.776011152 |
| Myl3 | 1.171543132 | 0.462679645 | 1.050562308 | 0.440411427 |
| Prss42 | 0.145501641 | 0.755294927 | 2.8880725 | 0.261522149 |
| Prss44 | 0.225957929 | 1.236201616 | 2.142169805 | 0.57707919 |
| Prss43 | 1.895398054 | 1.893919918 | 0.838048413 | 2.259917075 |
| Prss45 | 3.856439033 | 2.228235153 | 0.685468903 | 3.250672852 |
| Prss46 | 14.47717373 | 4.842807871 | 0.978369154 | 4.949877918 |
| Prss50 | 6.552872895 | 0.731359125 | 0.455531467 | 1.605507366 |
| Tmie | 70.89016146 | 0.232653581 | 0.172575132 | 1.348129235 |
| Als2cl | 108.0650459 | -0.23868321 | 0.140417408 | -1.699812109 |
| RP23-265F20.14 | 0.087930927 | -0.206490058 | 3.724945049 | -0.05543439 |
| Gm590 | 0.446134451 | 0.75530802 | 1.629100739 | 0.463634938 |
| Gm24269 | 0 | NA | NA | NA |
| Tdgf1 | 2.307514072 | 0.34916071 | 0.643736802 | 0.542396688 |
| Lrrc2 | 11.32380672 | -0.566744408 | 0.373710745 | -1.51653228 |
| RP24-166N8.8 | 0.109982625 | -0.20649451 | 3.724945049 | -0.055435586 |
| Rtp3 | 1.735450405 | 1.375740773 | 1.1529761 | 1.193208405 |
| 1700061E17Rik | 0.467110691 | 0.740934752 | 1.580798278 | 0.468709235 |
| Gm10030 | 2.141449619 | -0.410448548 | 0.692001594 | -0.593132373 |
| Ltf | 6.616994115 | 2.0149147 | 0.695854739 | 2.895596722 |
| RP24-166N8.9 | 4.949110328 | 5.506117994 | 0.914039828 | 6.023936624 |
| Ccrl2 | 163.4421883 | 5.776066894 | 0.456838677 | 12.64355929 |
| Rp31-ps19 | 0 | NA | NA | NA |

| | | | | |
|---|---|---|---|---|
| Gm26409 | 0.044544648 | 0.755286614 | 3.724945049 | 0.202764498 |
| Lrrfip2 | 487.0826255 | 0.110254829 | 0.091692712 | 1.202438308 |
| Mlh1 | 123.8432082 | -0.164276872 | 0.131924995 | -1.245229324 |
| Gm24860 | 1.095639389 | -0.789017012 | 1.090661967 | -0.723429473 |
| Epm2aip1 | 2946.703591 | -0.093581743 | 0.067063809 | -1.395413478 |
| Trank1 | 3806.887753 | -0.189824017 | 0.277240958 | -0.684689659 |
| RP23-361O11.4 | 10.70607882 | -1.072021189 | 0.405847265 | -2.641439977 |
| Dclk3 | 344.3535978 | -0.360681768 | 0.147155316 | -2.451027781 |
| RP24-362N9.3 | 0.122601409 | 1.236179401 | 3.724945049 | 0.331865138 |
| RP24-362N9.4 | 0 | NA | NA | NA |
| Stac | 39.84967809 | -0.67906338 | 0.196007886 | -3.464469696 |
| n-R5s90 | 0.892725456 | 1.079207755 | 1.047562839 | 1.030208132 |
| Arpp21 | 4354.712088 | -0.411592144 | 0.150099787 | -2.74212344 |
| Mir128-2 | 2.542567827 | -0.112959238 | 0.606549487 | -0.186232517 |
| 2310075C17Rik | 2.414579911 | 1.642240466 | 0.702890816 | 2.336409053 |
| 2900079G21Rik | 54.14502989 | 0.06157094 | 0.166254519 | 0.370341454 |
| Gm27847 | 0 | NA | NA | NA |
| Gm24957 | 0 | NA | NA | NA |
| Gm22492 | 0 | NA | NA | NA |
| Pdcd6ip | 1431.871971 | 0.006875883 | 0.061192273 | 0.112365222 |
| Gm25059 | 0 | NA | NA | NA |
| Clasp2 | 4322.484509 | -0.294935788 | 0.054593283 | -5.402418961 |
| Ubp1 | 888.9733183 | -0.053618487 | 0.116148116 | -0.461638886 |
| Fbxl2 | 321.4966191 | -0.282715527 | 0.097166869 | -2.909587714 |
| Susd5 | 65.24506384 | -0.213756651 | 0.169675237 | -1.259798749 |
| 4921528I07Rik | 0.068097359 | -0.206487837 | 3.724945049 | -0.055433794 |
| Bcl2a1c | 0.061920613 | 0.274393828 | 3.724945049 | 0.073663859 |
| 4930520O04Rik | 10.94558613 | 0.472518394 | 0.299743053 | 1.576411492 |
| Crtap | 52.46044785 | 0.546152816 | 0.242719135 | 2.250143221 |
| Glb1 | 239.9047864 | 0.143419552 | 0.088576516 | 1.619159997 |
| Tmppe | 289.5667612 | -0.037206023 | 0.075207185 | -0.494713668 |
| Ccr4 | 0.577241561 | 2.776952469 | 1.401822692 | 1.980958423 |
| Trim71 | 15.97479315 | -0.365929067 | 0.379606784 | -0.963968723 |
| C130032M10Rik | 0.045882455 | 0.755288839 | 3.724945049 | 0.202765096 |
| Cnot10 | 446.9158187 | -0.031264877 | 0.072340946 | -0.432187836 |
| Gm23232 | 2.256171834 | 0.209856442 | 0.771809616 | 0.271901823 |
| Dync1li1 | 1024.902468 | -0.007442239 | 0.05319658 | -0.139900711 |
| Cmtm6 | 352.6464989 | 0.633550815 | 0.097308782 | 6.51072598 |
| Cmtm7 | 42.27447401 | 1.420980402 | 0.290383422 | 4.893462547 |
| Gm9888 | 1.350285312 | 1.625703758 | 0.969919157 | 1.676122949 |
| Cmtm8 | 7.699545734 | 1.349579708 | 0.399350305 | 3.379438281 |
| Gm23889 | 0 | NA | NA | NA |
| Gpd1l | 938.6073185 | -0.11959966 | 0.07878625 | -1.518027071 |
| Osbpl10 | 236.991532 | -0.168709516 | 0.137335608 | -1.228446999 |
| Rps27rt | 78.90118146 | 0.275147857 | 0.228461388 | 1.204351682 |
| Gm4665 | 0.066050875 | 0.274391604 | 3.724945049 | 0.073663262 |
| Gm26962 | 0.061920613 | 0.274393828 | 3.724945049 | 0.073663859 |

| | | | | |
|---|---|---|---|---|
| Gm27002 | 10.03766149 | -1.579968312 | 0.373086539 | -4.234857455 |
| Stt3b | 1311.019778 | -0.07245087 | 0.0656313 | -1.103907291 |
| Mir467h | 0 | NA | NA | NA |
| Gm26303 | 0 | NA | NA | NA |
| Gm25510 | 0 | NA | NA | NA |
| Gm22243 | 0 | NA | NA | NA |
| Gm5921 | 6.413797967 | 0.385589556 | 0.444977632 | 0.86653694 |
| n-R5s92 | 0 | NA | NA | NA |
| Gadl1 | 3.709958051 | -0.968374218 | 0.570792065 | -1.696544639 |
| Gm16142 | 0.404993149 | -0.180240451 | 1.360660576 | -0.132465403 |
| Gm9385 | 43.14016071 | 0.558222208 | 0.251503769 | 2.219538142 |
| Tgfbr2 | 420.264414 | 1.658884235 | 0.111158573 | 14.92358333 |
| Rbms3 | 651.4525007 | 0.127616911 | 0.096690447 | 1.319850263 |
| Gm18489 | 0.042716945 | -0.206490061 | 3.724945049 | -0.055434391 |
| Gm20397 | 0 | NA | NA | NA |
| Gm17396 | 26.24111455 | 0.322446801 | 0.255256124 | 1.263228463 |
| Gm20396 | 0 | NA | NA | NA |
| Gm26614 | 3.2216295 | 0.222759958 | 0.634377982 | 0.351147051 |
| Zcwpw2 | 38.02155974 | -0.198364657 | 0.198917584 | -0.997220318 |
| Gm22008 | 0 | NA | NA | NA |
| Azi2 | 1280.08819 | 0.642034957 | 0.096779094 | 6.634025253 |
| Cmc1 | 102.3559624 | 0.073057568 | 0.114228334 | 0.639574833 |
| Gm17399 | 4.930845136 | -0.48881654 | 0.490635386 | -0.996292877 |
| Gm22198 | 0 | NA | NA | NA |
| 2610509F24Rik | 0.046345709 | 0.274391604 | 3.724945049 | 0.073663262 |
| Platr11 | 0.154746054 | -1.142176591 | 2.823763514 | -0.404487339 |
| Gm22479 | 0.124853745 | 1.290908285 | 3.721447108 | 0.346883416 |
| Eomes | 133.6882393 | -0.706791687 | 0.419111312 | -1.686405655 |
| Golga4 | 1496.655227 | -0.04060228 | 0.129642238 | -0.313187124 |
| Itga9 | 160.656506 | -0.366252903 | 0.103899521 | -3.525068253 |
| Gm10157 | 51.89187838 | -0.042070389 | 0.155361594 | -0.270790149 |
| Gm2415 | 124.2848123 | -0.707243963 | 0.153623698 | -4.603742608 |
| Gm16295 | 0.648502457 | -0.328598341 | 1.124746919 | -0.292153137 |
| Ctdspl | 341.7777662 | -0.220262085 | 0.132866588 | -1.657768803 |
| Mir26a-1 | 1.512289855 | 1.541730995 | 0.994955586 | 1.549547555 |
| Vill | 6.901825543 | 1.065892317 | 0.421595852 | 2.52823246 |
| Plcd1 | 118.7634359 | 0.279467107 | 0.132095842 | 2.11563893 |
| Dlec1 | 29.71330161 | -1.285077763 | 0.236574193 | -5.432028516 |
| Acaa1b | 17.68007261 | -0.760474655 | 0.288304448 | -2.637748606 |
| Gm10608 | 0.106465261 | 0.75528439 | 3.724945049 | 0.202763901 |
| Slc22a14 | 0.191792366 | 0.755303231 | 2.240923694 | 0.337049956 |
| Slc22a13 | 1.365498172 | -1.718433828 | 1.014401277 | -1.694037524 |
| Slc22a13b-ps | 14.19279457 | -1.333923605 | 0.365503586 | -3.649549984 |
| Oxsr1 | 850.5060215 | -0.114318707 | 0.068992341 | -1.656976776 |
| Gm22729 | 1.308834292 | -0.680287012 | 0.834563925 | -0.815140688 |
| Myd88 | 293.492393 | 3.188397536 | 0.233269797 | 13.66828271 |
| Acaa1a | 291.2994478 | 0.295617278 | 0.103761883 | 2.84899686 |

| | | | | |
|---|---|---|---|---|
| Xylb | 75.68413251 | 0.131330594 | 0.149034966 | 0.881206592 |
| Acvr2b | 67.3176329 | -0.319570141 | 0.133744465 | -2.389408337 |
| 4930516B21Rik | 0 | NA | NA | NA |
| Exog | 250.3105798 | -0.07831785 | 0.08803447 | -0.889627103 |
| Scn5a | 99.30242439 | -0.393446932 | 0.132821007 | -2.962234226 |
| Scn10a | 7.212801384 | 2.894070376 | 0.484321151 | 5.975519286 |
| Scn11a | 3.528093251 | -0.084546143 | 0.611209297 | -0.13832601 |
| Gm2449 | 4.853795414 | 0.552261335 | 0.45447834 | 1.215154356 |
| Wdr48 | 797.9104278 | -0.089888986 | 0.050057762 | -1.795705245 |
| Cx3cr1 | 232.9636481 | -0.556078757 | 0.481734575 | -1.154326024 |
| Gorasp1 | 319.6341586 | -0.262316776 | 0.132129632 | -1.985298625 |
| Ttc21a | 29.13279244 | -0.423965566 | 0.27765112 | -1.526972291 |
| Csrnp1 | 561.5797647 | 2.127331078 | 0.34831888 | 6.107423971 |
| Xirp1 | 5.455029321 | 2.870026163 | 0.933376658 | 3.074885298 |
| Ccr8 | 0.637648044 | 2.19543532 | 1.260414319 | 1.741836226 |
| Slc25a38 | 245.895353 | 0.311854681 | 0.103062926 | 3.025866746 |
| Rpsa | 2894.845799 | 0.317256596 | 0.097150325 | 3.265625673 |
| Gm24044 | 12.71611818 | 0.083378554 | 0.350878493 | 0.237627999 |
| Gm26448 | 99.87717075 | -0.231448769 | 0.187131911 | -1.236821487 |
| Snora62 | 1.810539642 | 0.829633724 | 0.77748114 | 1.067078906 |
| Mobp | 12397.77805 | 0.51235843 | 0.158718737 | 3.228090397 |
| Gm37111 | 46.80149452 | -0.298321573 | 0.190231076 | -1.568206303 |
| Myrip | 677.5660742 | -0.377479588 | 0.146024123 | -2.585049513 |
| Gm15566 | 4.5585197 | -0.325500056 | 0.564567261 | -0.576547876 |
| Gm15565 | 3.283342549 | -1.313314296 | 0.705942125 | -1.860371054 |
| Eif1b | 736.4044258 | 0.157238395 | 0.094727824 | 1.659896616 |
| Entpd3 | 153.1227425 | -0.041386468 | 0.113313504 | -0.365238621 |
| Rpl14 | 1081.536881 | 0.34847799 | 0.11277559 | 3.090012572 |
| 5830454E08Rik | 6.315262053 | -0.109359496 | 0.381518951 | -0.286642369 |
| Gm23624 | 0 | NA | NA | NA |
| 1700020M21Rik | 0.275587006 | 0.274402717 | 2.437434975 | 0.112578477 |
| 4930593C16Rik | 1.144931742 | 0.404812707 | 0.971867428 | 0.41653079 |
| Ctnnb1 | 5088.011416 | 0.248454891 | 0.090632741 | 2.741337039 |
| mmu-mir-7090 | 0.464416953 | -0.935358924 | 1.268640178 | -0.737292528 |
| Ulk4 | 49.64989384 | -0.614274097 | 0.19263225 | -3.188843484 |
| Gm22987 | 0 | NA | NA | NA |
| Gm17157 | 0 | NA | NA | NA |
| Trak1 | 1974.275711 | -0.202583473 | 0.072015665 | -2.813047318 |
| Cck | 1108.958497 | -0.057129784 | 0.121310316 | -0.470939204 |
| Lyzl4 | 13.61068275 | -0.650084348 | 0.381234117 | -1.705210311 |
| Lyzl4os | 0.219234697 | 0.755300834 | 2.549697479 | 0.296231549 |
| Vipr1 | 229.8541341 | -0.418493502 | 0.220102976 | -1.901353216 |
| Sec22c | 369.7565343 | -0.248723916 | 0.071829083 | -3.462718798 |
| Deb1 | 212.2253457 | -0.142840351 | 0.085359847 | -1.673390425 |
| Nktr | 2807.808864 | -0.077960098 | 0.170898307 | -0.456178292 |
| E530011L22Rik | 71.35472267 | -0.408559344 | 0.147438509 | -2.771049081 |
| Zfp651 | 995.5133322 | 0.055425834 | 0.083017744 | 0.667638406 |

| | | | | |
|---|---|---|---|---|
| Klhl40 | 81.7128723 | 0.582172124 | 0.232681048 | 2.502017799 |
| Hhatl | 134.4694746 | -0.223503952 | 0.144508815 | -1.546645801 |
| Ccdc13 | 49.28911322 | 0.265360675 | 0.176390259 | 1.504395289 |
| Gm17163 | 0.831375036 | 1.049657492 | 0.990945314 | 1.059248656 |
| Higd1a | 526.1847965 | -0.064303418 | 0.079422714 | -0.809635113 |
| Ackr2 | 29.87282185 | 0.434474579 | 0.304659339 | 1.426099657 |
| Cyp8b1 | 0.044191549 | -0.206490061 | 3.724945049 | -0.055434391 |
| 1700048O20Rik | 89.81562415 | 0.366591889 | 0.172662464 | 2.123170719 |
| Fam198a | 45.49996011 | -0.509232749 | 0.250155884 | -2.035661688 |
| Pomgnt2 | 324.6867476 | -0.141169259 | 0.087872265 | -1.606528067 |
| Snrk | 1696.951144 | -0.293115834 | 0.107048742 | -2.738152994 |
| Ano10 | 333.863595 | 0.617461559 | 0.07919112 | 7.797106037 |
| Mir8106 | 0.421183792 | 1.807673262 | 1.59683345 | 1.132036195 |
| Gm26797 | 96.16516166 | 4.012498703 | 0.308150266 | 13.02124042 |
| Abhd5 | 237.0914825 | 0.124600081 | 0.121854844 | 1.022528745 |
| A730085K08Rik | 3.727958404 | 0.24750069 | 0.488632126 | 0.506517433 |
| 9530059O14Rik | 357.5582499 | -1.072122125 | 0.342744274 | -3.128052618 |
| Mir138-1 | 4.764516023 | -0.469028783 | 0.458589156 | -1.022764662 |
| Topaz1 | 1.867918734 | -0.473031514 | 0.978712072 | -0.483320404 |
| Tcaim | 210.4247602 | -0.176195166 | 0.102780155 | -1.714291698 |
| Gm25341 | 1.808478971 | -0.001364945 | 0.710361059 | -0.00192148 |
| Zfp445 | 1117.035592 | -0.149551584 | 0.107427962 | -1.392110399 |
| A530083I20Rik | 14.49954602 | 0.087980402 | 0.318926767 | 0.275863962 |
| Zkscan7 | 42.10827637 | -0.463162236 | 0.201278633 | -2.301099865 |
| Zfp105 | 88.67340489 | 0.289783645 | 0.126823769 | 2.284931654 |
| 1110059G10Rik | 126.7733861 | 0.4425384 | 0.110678486 | 3.998413911 |
| Kif15 | 30.3785014 | 1.492449664 | 0.338742961 | 4.405847012 |
| Tmem42 | 63.48058778 | -0.221470001 | 0.131356196 | -1.686026306 |
| Tgm4 | 6.449499428 | -0.496211849 | 0.406521828 | -1.220627811 |
| Zdhhc3 | 963.4259801 | 0.141660173 | 0.051751599 | 2.737310048 |
| Exosc7 | 112.7568782 | -0.485316355 | 0.10127946 | -4.791853706 |
| Clec3b | 48.60893226 | -0.936639204 | 0.208543992 | -4.491326714 |
| Gm23323 | 0.148038037 | -0.203753848 | 3.549417237 | -0.057404874 |
| Cdcp1 | 2.585897959 | -0.577709781 | 0.645390657 | -0.895131926 |
| Tmem158 | 284.2320044 | -0.078140144 | 0.105690178 | -0.739332125 |
| Scp2-ps2 | 91.28446377 | 0.034629572 | 0.162843061 | 0.212656113 |
| Lars2 | 5438.700325 | 0.572591094 | 0.817352628 | 0.700543528 |
| Limd1 | 249.5361358 | 0.498247064 | 0.125919986 | 3.956854517 |
| Sacm1l | 1325.845621 | -0.047656914 | 0.067341899 | -0.707685932 |
| Slc6a20b | 4.677638664 | 1.074433788 | 0.470526514 | 2.283471295 |
| Gm17120 | 4.624289095 | 2.489614196 | 0.85759481 | 2.903019196 |
| Slc6a20a | 203.7172933 | 0.475065479 | 0.200414346 | 2.37041653 |
| Gm17200 | 1.771524358 | 0.657246609 | 0.916597737 | 0.71705022 |
| Ccr9 | 18.10541206 | 2.157298763 | 0.494649888 | 4.36126403 |
| Gm10052 | 130.6631812 | 0.231256854 | 0.128886793 | 1.794263395 |
| Lztfl1 | 833.9950152 | -0.137389603 | 0.06300202 | -2.180717449 |
| Gm17020 | 0.049553047 | 0.274391604 | 3.724945049 | 0.073663262 |

| | | | | |
|---|---|---|---|---|
| Gm17021 | 0.832525973 | -0.308523167 | 1.137127433 | -0.271318023 |
| Fyco1 | 809.1842188 | 0.145907763 | 0.085104279 | 1.714458597 |
| Cxcr6 | 20.53480371 | 6.898899059 | 0.90378952 | 7.633302785 |
| Xcr1 | 0.667406772 | 2.013034409 | 2.581269237 | 0.77986224 |
| Ccr1 | 181.3036799 | 4.753546007 | 0.656189827 | 7.244162911 |
| Ccr1l1 | 0.461281393 | 1.760878427 | 1.52781741 | 1.152545072 |
| Ccr3 | 1.785934789 | 4.467991655 | 1.068244848 | 4.182553901 |
| Ccr2 | 315.4917145 | 5.307697235 | 0.448783203 | 11.8268625 |
| Ccr5 | 468.5098695 | 2.772331262 | 0.304619187 | 9.100973875 |
| 2010315B03Rik | 245.0633303 | -0.103547466 | 0.121486618 | -0.852336398 |
| Nlrp4g | 0.042716945 | -0.206490061 | 3.724945049 | -0.055434391 |
| Ppp2r3d | 158.3671903 | 0.159780168 | 0.115809399 | 1.379682213 |
| Gm21836 | 0 | NA | NA | NA |
| SNORA28 | 0 | NA | NA | NA |
| RP24-571H12.4 | 0 | NA | NA | NA |
| RP24-571H12.6 | 0 | NA | NA | NA |
| AC168054.1 | 0 | NA | NA | NA |
| WI1-2477B13.3 | 0 | NA | NA | NA |
| CAAA01180111.1 | 0 | NA | NA | NA |
| CAAA01202134.1 | 0 | NA | NA | NA |
| CAAA01125382.1 | 0 | NA | NA | NA |
| AC109138.1 | 0 | NA | NA | NA |
| AC109138.3 | 0 | NA | NA | NA |
| AC109138.2 | 0 | NA | NA | NA |
| AC116115.1 | 0 | NA | NA | NA |
| mmu-mir-505 | 0 | NA | NA | NA |
| CAAA01000182.1 | 0 | NA | NA | NA |
| RP23-6C3.5 | 0 | NA | NA | NA |
| RP23-6C3.4 | 0 | NA | NA | NA |
| Hspa9-ps1 | 7.785073125 | -0.001471098 | 0.417346652 | -0.003524882 |
| RP23-299A23.1 | 0 | NA | NA | NA |
| RP23-6C3.2 | 0 | NA | NA | NA |
| RP23-6C3.3 | 0 | NA | NA | NA |
| RP23-445P15.4 | 0 | NA | NA | NA |
| RP23-445P15.5 | 0 | NA | NA | NA |
| RP23-445P15.2 | 0 | NA | NA | NA |
| RP23-163F11.2 | 0 | NA | NA | NA |
| RP23-346N8.3 | 0 | NA | NA | NA |
| CU041226.1 | 0 | NA | NA | NA |
| RP23-402O17.1 | 0 | NA | NA | NA |
| RP23-16C4.1 | 0 | NA | NA | NA |
| AL593853.1 | 0 | NA | NA | NA |
| RP23-199A2.7 | 0 | NA | NA | NA |
| SNORA77 | 0 | NA | NA | NA |
| mmu-mir-1306 | 0 | NA | NA | NA |
| mmu-mir-3618 | 0 | NA | NA | NA |
| mmu-mir-185 | 0 | NA | NA | NA |

| | | | | |
|---|---|---|---|---|
| AC084822.2 | 0 | NA | NA | NA |
| U1 | 0 | NA | NA | NA |
| AC084822.1 | 0 | NA | NA | NA |
| RP23-285E22.4 | 0 | NA | NA | NA |
| RP23-348F1.1 | 0 | NA | NA | NA |
| 7SK | 0 | NA | NA | NA |
| RP23-58H5.2 | 0 | NA | NA | NA |
| RP23-58H5.5 | 0 | NA | NA | NA |
| RP23-58H5.6 | 0 | NA | NA | NA |
| RP23-58H5.8 | 0 | NA | NA | NA |
| OTTMUSG000000101 | 0 | NA | NA | NA |
| RP24-364D22.1 | 0 | NA | NA | NA |
| AC132586.1 | 0 | NA | NA | NA |
| AC245272.1 | 0 | NA | NA | NA |
| AC245272.3 | 0 | NA | NA | NA |
| mmu-mir-290a | 0 | NA | NA | NA |
| mmu-mir-291a | 0 | NA | NA | NA |
| mmu-mir-292a | 0 | NA | NA | NA |
| mmu-mir-291b | 0 | NA | NA | NA |
| mmu-mir-293 | 0 | NA | NA | NA |
| mmu-mir-294 | 0 | NA | NA | NA |
| mmu-mir-295 | 0 | NA | NA | NA |
| AC245272.2 | 0 | NA | NA | NA |
| mmu-mir-935 | 0 | NA | NA | NA |
| Mico1 | 0 | NA | NA | NA |
| MEG3_1 | 0 | NA | NA | NA |
| MEG3_2 | 0 | NA | NA | NA |
| mmu-mir-1906-1 | 0 | NA | NA | NA |
| mmu-mir-770 | 0 | NA | NA | NA |
| AC152063.1 | 0 | NA | NA | NA |
| mmu-mir-673 | 0 | NA | NA | NA |
| mmu-mir-493 | 0 | NA | NA | NA |
| mmu-mir-337 | 0 | NA | NA | NA |
| mmu-mir-540 | 0 | NA | NA | NA |
| mmu-mir-665 | 0 | NA | NA | NA |
| mmu-mir-3070-1 | 0 | NA | NA | NA |
| mmu-mir-3070-2 | 0 | NA | NA | NA |
| mmu-mir-431 | 0 | NA | NA | NA |
| mmu-mir-433 | 0 | NA | NA | NA |
| mmu-mir-127 | 0 | NA | NA | NA |
| mmu-mir-434 | 0 | NA | NA | NA |
| mmu-mir-432 | 0 | NA | NA | NA |
| mmu-mir-3071 | 0 | NA | NA | NA |
| MEG8_1 | 0 | NA | NA | NA |
| mmu-mir-341 | 0 | NA | NA | NA |
| mmu-mir-1188 | 0 | NA | NA | NA |
| AC152063.2 | 0 | NA | NA | NA |

| | | | | |
|---|---|---|---|---|
| mmu-mir-370 | 0 | NA | NA | NA |
| SNORD113 | 0 | NA | NA | NA |
| RP23-204I16.2 | 0 | NA | NA | NA |
| RP23-204I16.3 | 0 | NA | NA | NA |
| mmu-mir-882 | 0 | NA | NA | NA |
| CH36-472P4.1 | 0 | NA | NA | NA |
| AC233742.1 | 0 | NA | NA | NA |
| CH36-472P4.2 | 0 | NA | NA | NA |
| CH36-472P4.3 | 0 | NA | NA | NA |
| CH36-169F23.1 | 0 | NA | NA | NA |
| CH36-169F23.2 | 0 | NA | NA | NA |
| AC240971.1 | 0 | NA | NA | NA |
| CH36-169F23.4 | 0 | NA | NA | NA |
| CH36-169F23.3 | 0 | NA | NA | NA |
| CH36-169F23.5 | 0 | NA | NA | NA |
| RP24-152F23.1 | 0 | NA | NA | NA |
| RP24-152F23.4 | 0 | NA | NA | NA |
| RP24-152F23.5 | 0 | NA | NA | NA |
| AC244694.1 | 0 | NA | NA | NA |
| RP24-505E20.1 | 0 | NA | NA | NA |
| RP24-410B18.1 | 0 | NA | NA | NA |
| RP24-410B18.2 | 0 | NA | NA | NA |
| RP24-410B18.3 | 0 | NA | NA | NA |
| RP24-410B18.5 | 0 | NA | NA | NA |
| AC202135.1 | 0 | NA | NA | NA |
| RP24-410B18.4 | 0 | NA | NA | NA |
| Rhox5 | 0.049553047 | 0.274391604 | 3.724945049 | 0.073663262 |
| Rhox6 | 0 | NA | NA | NA |
| Rhox7a | 0 | NA | NA | NA |
| Rhox8 | 0.068097359 | -0.206487837 | 3.724945049 | -0.055433794 |
| Rhox7b | 0 | NA | NA | NA |
| Rhox9 | 0.095435502 | 0.75528439 | 3.724945049 | 0.202763901 |
| RP23-202F23.7 | 0 | NA | NA | NA |
| Rhox11-ps | 0 | NA | NA | NA |
| AC154679.1 | 0 | NA | NA | NA |
| AC182748.2 | 0 | NA | NA | NA |
| AC182748.1 | 0 | NA | NA | NA |
| U6 | 0 | NA | NA | NA |
| RP24-176A1.1 | 0 | NA | NA | NA |
| RP24-176A1.2 | 0 | NA | NA | NA |
| AC127556.1 | 0 | NA | NA | NA |
| mmu-mir-194-2 | 0 | NA | NA | NA |
| mmu-mir-192 | 0 | NA | NA | NA |
| AC127556.5 | 0 | NA | NA | NA |
| mmu-mir-6989 | 0 | NA | NA | NA |
| AC127556.10 | 0 | NA | NA | NA |
| AC167245.6 | 0 | NA | NA | NA |

| | | | | |
|---|---|---|---|---|
| AC167245.5 | 0 | NA | NA | NA |
| AC124394.1 | 0 | NA | NA | NA |
| Gm14423 | 0 | NA | NA | NA |
| AC124394.3 | 0 | NA | NA | NA |
| AC124394.4 | 0 | NA | NA | NA |
| RP23-188E3.2 | 0 | NA | NA | NA |
| RP23-188E3.1 | 0 | NA | NA | NA |
| BX890553.1 | 0 | NA | NA | NA |
| RP23-324J18.1 | 0 | NA | NA | NA |
| RP23-324J18.2 | 0 | NA | NA | NA |
| AC164883.2 | 0 | NA | NA | NA |
| AC164883.1 | 0 | NA | NA | NA |
| CH25-484K21.2 | 0 | NA | NA | NA |
| CH25-484K21.3 | 0 | NA | NA | NA |
| CH25-484K21.5 | 0 | NA | NA | NA |
| CH25-484K21.6 | 0 | NA | NA | NA |
| Gm9919 | 0 | NA | NA | NA |
| FO907130.1 | 0 | NA | NA | NA |
| SNORD103 | 0 | NA | NA | NA |
| RP23-157N11.1 | 0 | NA | NA | NA |
| RP23-157N11.4 | 0 | NA | NA | NA |
| RP23-157N11.7 | 0 | NA | NA | NA |
| RP23-157N11.6 | 0 | NA | NA | NA |
| RP23-146J17.3 | 0 | NA | NA | NA |
| AC125149.4 | 0 | NA | NA | NA |
| AC125149.1 | 0 | NA | NA | NA |
| AC125149.2 | 0 | NA | NA | NA |
| AC125149.3 | 0 | NA | NA | NA |
| AC133103.4 | 0 | NA | NA | NA |
| AC133103.6 | 0 | NA | NA | NA |
| AC133103.7 | 0 | NA | NA | NA |
| AC133103.2 | 0 | NA | NA | NA |
| AC133103.5 | 0 | NA | NA | NA |
| AC133103.1 | 0 | NA | NA | NA |
| AC133103.3 | 0 | NA | NA | NA |
| AC168977.2 | 0 | NA | NA | NA |
| AC168977.1 | 0 | NA | NA | NA |
| DHRSX | 0 | NA | NA | NA |
| U2 | 0 | NA | NA | NA |
| Vmn1r186 | 0 | NA | NA | NA |
| AC102264.1 | 0 | NA | NA | NA |
| AC132444.1 | 0 | NA | NA | NA |
| AC132444.3 | 0 | NA | NA | NA |
| AC132444.5 | 0 | NA | NA | NA |
| AC132444.4 | 0 | NA | NA | NA |
| AC132444.2 | 0 | NA | NA | NA |
| Csprs | 0 | NA | NA | NA |

| | | | | |
|---|---|---|---|---|
| AC132444.6 | 0 | NA | NA | NA |
| Vamp7 | 0 | NA | NA | NA |
| CAAA01165726.1 | 0 | NA | NA | NA |
| Spry3 | 0 | NA | NA | NA |
| Tmlhe | 0 | NA | NA | NA |
| CR524822.1 | 0 | NA | NA | NA |
| AC087559.4 | 0 | NA | NA | NA |
| AC087559.5 | 0 | NA | NA | NA |
| AC087559.1 | 0 | NA | NA | NA |
| AC087559.3 | 0 | NA | NA | NA |
| AC087559.2 | 0 | NA | NA | NA |
| Ccl21c | 0 | NA | NA | NA |
| AC140325.2 | 0 | NA | NA | NA |
| AC140325.1 | 0 | NA | NA | NA |
| AC140325.3 | 0 | NA | NA | NA |
| AC140325.4 | 0 | NA | NA | NA |
| CAAA01098150.1 | 0 | NA | NA | NA |
| CAAA01064564.1 | 0 | NA | NA | NA |
| CAAA01205117.2 | 0 | NA | NA | NA |
| CAAA01205117.1 | 0 | NA | NA | NA |
| Vmn2r122 | 0 | NA | NA | NA |
| Gm2002 | 0 | NA | NA | NA |
| CR974586.2 | 0 | NA | NA | NA |
| CR974586.4 | 0 | NA | NA | NA |
| CR974586.8 | 0 | NA | NA | NA |
| CR974586.5 | 0 | NA | NA | NA |
| CR974586.7 | 0 | NA | NA | NA |
| CR974586.6 | 0 | NA | NA | NA |
| CR974586.3 | 0 | NA | NA | NA |
| CR974586.1 | 0 | NA | NA | NA |
| Scya19 | 0 | NA | NA | NA |
| CT868723.1 | 0 | NA | NA | NA |
| Gm10931 | 0 | NA | NA | NA |
| CT868723.2 | 0 | NA | NA | NA |
| CAAA01147332.1 | 0 | NA | NA | NA |
| AC164084.2 | 0 | NA | NA | NA |
| AC164084.3 | 0 | NA | NA | NA |
| AC164084.1 | 0 | NA | NA | NA |
| AC165294.1 | 0 | NA | NA | NA |
| AC165294.2 | 0 | NA | NA | NA |
| AC165294.3 | 0 | NA | NA | NA |
| AC125178.1 | 0 | NA | NA | NA |
| AC125178.3 | 0 | NA | NA | NA |
| AC125178.2 | 0 | NA | NA | NA |
| AC123873.3 | 0 | NA | NA | NA |
| AC123873.1 | 0 | NA | NA | NA |
| AC123873.2 | 0 | NA | NA | NA |

| | | | | |
|---|---|---|---|---|
| AC126035.1 | 0 | NA | NA | NA |
| Gm16367 | 0 | NA | NA | NA |
| AC163611.1 | 0 | NA | NA | NA |
| AC140365.1 | 0 | NA | NA | NA |
| AC124606.2 | 0 | NA | NA | NA |
| AC124606.1 | 0 | NA | NA | NA |
| AC133095.2 | 0 | NA | NA | NA |
| AC133095.1 | 0 | NA | NA | NA |
| AC234645.1 | 0 | NA | NA | NA |
| PISD | 0 | NA | NA | NA |
| mt-Tf | 0 | NA | NA | NA |
| mt-Rnr1 | 119593.2604 | 0.724996047 | 0.923923368 | 0.784692835 |
| mt-Tv | 0 | NA | NA | NA |
| mt-Rnr2 | 65951.93075 | 0.948974582 | 0.794777654 | 1.194012662 |
| mt-Tl1 | 208.0973662 | 1.099849769 | 0.21797199 | 5.04583075 |
| mt-Nd1 | 34373.33605 | 0.087457414 | 0.157385347 | 0.555689685 |
| mt-Ti | 143.1190915 | -0.741967576 | 0.337013743 | -2.201594422 |
| mt-Tq | 55.05421077 | 0.26111969 | 0.187409345 | 1.393312003 |
| mt-Tm | 149.670499 | -0.725277463 | 0.245281672 | -2.956916661 |
| mt-Nd2 | 20368.9686 | 0.044810681 | 0.224871032 | 0.199272805 |
| mt-Tw | 186.9799014 | 0.716350599 | 0.184918931 | 3.873862972 |
| mt-Ta | 11.02334367 | 0.302450394 | 0.343733976 | 0.879896709 |
| mt-Tn | 0.72655531 | 1.304367176 | 1.175198243 | 1.109912463 |
| mt-Tc | 1.449755302 | 1.533100344 | 0.965540228 | 1.587816125 |
| mt-Ty | 0.807376506 | 0.25997956 | 1.280283385 | 0.203064074 |
| mt-Co1 | 112876.7948 | 0.139734745 | 0.120039697 | 1.164071124 |
| mt-Ts1 | 1.308396679 | 2.070569912 | 0.994135335 | 2.082784748 |
| mt-Td | 0.616049689 | -0.100071765 | 1.387394026 | -0.072129304 |
| mt-Co2 | 1688.977157 | 0.189755855 | 0.128016366 | 1.482278089 |
| mt-Tk | 0.097139547 | 0.274393828 | 3.724945049 | 0.073663859 |
| mt-Atp8 | 110.3540989 | 0.415511337 | 0.427308224 | 0.97239256 |
| mt-Atp6 | 624.2525603 | 0.233205123 | 0.196843394 | 1.184724147 |
| mt-Co3 | 1101.033194 | 0.2730378 | 0.146752188 | 1.860536478 |
| mt-Tg | 0 | NA | NA | NA |
| mt-Nd3 | 119.756487 | -0.118637912 | 0.172828191 | -0.68645 |
| mt-Tr | 0 | NA | NA | NA |
| mt-Nd4l | 129.7203189 | 0.029821767 | 0.214498155 | 0.139030412 |
| mt-Nd4 | 21602.70211 | -0.028905502 | 0.143048378 | -0.202068015 |
| mt-Th | 0.134005754 | -0.206481845 | 2.962652837 | -0.069694917 |
| mt-Ts2 | 0.119587853 | 1.218392466 | 3.723618458 | 0.327206581 |
| mt-Tl2 | 0.731462286 | 0.767785233 | 1.225115087 | 0.626704578 |
| mt-Nd5 | 23114.44776 | 0.12110918 | 0.210317472 | 0.575839846 |
| mt-Nd6 | 1598.689532 | 0.501727491 | 0.154952281 | 3.23794841 |
| mt-Te | 5.226567033 | 0.066292493 | 0.478195728 | 0.138630459 |
| mt-Cytb | 37159.01855 | -0.120675408 | 0.1406862 | -0.857762936 |
| mt-Tt | 18.73040197 | 1.247073294 | 0.341900977 | 3.647469235 |
| mt-Tp | 56.15899216 | 0.68837653 | 0.210774636 | 3.26593627 |

| | | | | |
|---|---|---|---|---|
| Gm21950 | 0 | NA | NA | NA |
| Gm21364 | 0 | NA | NA | NA |
| Gm14331 | 0 | NA | NA | NA |
| Gm14333 | 0 | NA | NA | NA |
| Gm14332 | 0 | NA | NA | NA |
| Gm14334 | 0 | NA | NA | NA |
| Gm14352 | 0 | NA | NA | NA |
| Gm14346 | 0 | NA | NA | NA |
| Gm14353 | 0 | NA | NA | NA |
| Gm14360 | 0 | NA | NA | NA |
| Gm14361 | 0 | NA | NA | NA |
| Gm14345 | 0 | NA | NA | NA |
| Gm14351 | 0 | NA | NA | NA |
| Gm14357 | 0 | NA | NA | NA |
| Gm14359 | 0 | NA | NA | NA |
| Gm14355 | 0 | NA | NA | NA |
| Spin2-ps1 | 0 | NA | NA | NA |
| Gm3701 | 0 | NA | NA | NA |
| Gm15251 | 0 | NA | NA | NA |
| Gm15257 | 0 | NA | NA | NA |
| Gm3706 | 0 | NA | NA | NA |
| Gm15258 | 0 | NA | NA | NA |
| Gm14350 | 0 | NA | NA | NA |
| Gm14349 | 0 | NA | NA | NA |
| Gm14358 | 0 | NA | NA | NA |
| Gm14362 | 0 | NA | NA | NA |
| Gm14356 | 0 | NA | NA | NA |
| Gm14347 | 0 | NA | NA | NA |
| Gm10921 | 0 | NA | NA | NA |
| Gm14348 | 0 | NA | NA | NA |
| Gm14364 | 0 | NA | NA | NA |
| Gm14368 | 0 | NA | NA | NA |
| Gm15260 | 0 | NA | NA | NA |
| Gm10922 | 0 | NA | NA | NA |
| Gm6019 | 0 | NA | NA | NA |
| Gm9427 | 0 | NA | NA | NA |
| Gm15256 | 0 | NA | NA | NA |
| Gm5925 | 0 | NA | NA | NA |
| Gm4725 | 0 | NA | NA | NA |
| Gm9498 | 0 | NA | NA | NA |
| Gm15252 | 0 | NA | NA | NA |
| Gm3750 | 0 | NA | NA | NA |
| Gm4726 | 0 | NA | NA | NA |
| Gm3757 | 0 | NA | NA | NA |
| Gm15259 | 0 | NA | NA | NA |
| Gm15254 | 0 | NA | NA | NA |
| Gm3763 | 0 | NA | NA | NA |

| | | | | |
|---|---|---|---|---|
| Gm14370 | 0 | NA | NA | NA |
| Gm14367 | 0 | NA | NA | NA |
| Gm14365 | 0 | NA | NA | NA |
| Gm9428 | 0 | NA | NA | NA |
| Gm5924 | 0 | NA | NA | NA |
| Gm14369 | 0 | NA | NA | NA |
| Gm14363 | 0 | NA | NA | NA |
| Mycs | 0 | NA | NA | NA |
| Gm14366 | 0 | NA | NA | NA |
| Gm14375 | 0 | NA | NA | NA |
| Gm14374 | 0 | NA | NA | NA |
| Gm14373 | 4.562379813 | 0.23806724 | 0.540484836 | 0.440469786 |
| Gm14452 | 0.11997416 | 0.274404949 | 3.724945049 | 0.073666845 |
| Gm14451 | 0.112642008 | 0.274402725 | 3.724945049 | 0.073666248 |
| Gm14448 | 2.617244783 | -2.00326962 | 0.682631938 | -2.934626271 |
| Gm14450 | 0 | NA | NA | NA |
| Gm26618 | 11.11537949 | -0.405253872 | 0.305973349 | -1.324474413 |
| Nudt11 | 271.955026 | 0.078446299 | 0.102671751 | 0.764049492 |
| Rpl19-ps12 | 2.084829622 | 0.879942813 | 0.812075614 | 1.083572513 |
| Gm27360 | 0 | NA | NA | NA |
| AU022751 | 17.60759048 | -0.479964253 | 0.365853966 | -1.311901188 |
| n-R5s2 | 0 | NA | NA | NA |
| Nudt10 | 171.3843856 | -0.191772599 | 0.099652701 | -1.924409444 |
| Bmp15 | 3.216309252 | 0.360565687 | 0.689275969 | 0.523107874 |
| Shroom4 | 90.11915289 | -0.120285226 | 0.157047377 | -0.765916809 |
| Dgkk | 132.1695717 | -0.228762834 | 0.216369047 | -1.057280779 |
| Ccnb3 | 0.629652017 | 0.274411239 | 1.529884859 | 0.179367249 |
| Akap4 | 0 | NA | NA | NA |
| Gm14380 | 0.089089296 | 1.194485492 | 3.723939576 | 0.320758559 |
| Clcn5 | 513.7578379 | -0.169349443 | 0.095286846 | -1.777259402 |
| Gm26401 | 0 | NA | NA | NA |
| Mir500 | 0.181074517 | -0.676080656 | 3.345017058 | -0.202115757 |
| Mir501 | 0.726672012 | -1.395145233 | 1.156513791 | -1.206336876 |
| Mir362 | 0.060174537 | 0.755286614 | 3.724945049 | 0.202764498 |
| Mir188 | 0.206496347 | 0.254321523 | 2.177473318 | 0.11679662 |
| Mir532 | 0.516362146 | -1.741109473 | 1.253884867 | -1.388572045 |
| Usp27x | 183.7584339 | 0.207829085 | 0.129245998 | 1.608011758 |
| Gm14379 | 5.282615923 | 0.355490622 | 0.469300112 | 0.757491024 |
| 2010204K13Rik | 54.17774226 | 0.025185287 | 0.152159799 | 0.165518665 |
| Gm14381 | 0.045882455 | 0.755288839 | 3.724945049 | 0.202765096 |
| Gm14395 | 0 | NA | NA | NA |
| Gm14394 | 0.287498921 | 0.274408133 | 2.023842827 | 0.13558767 |
| Gm23082 | 0.046345709 | 0.274391604 | 3.724945049 | 0.073663262 |
| Ppp1r3f | 110.0662308 | -0.124674389 | 0.110055443 | -1.132832554 |
| Gm36995 | 5.062940487 | 0.092777906 | 0.45980461 | 0.201776806 |
| Ppp1r3fos | 7.493797218 | -0.12337885 | 0.377951556 | -0.32644091 |
| Foxp3 | 2.783566837 | 2.544116399 | 0.81791834 | 3.110477264 |

| | | | | |
|---|---|---|---|---|
| Ccdc22 | 22.02494333 | -0.127470984 | 0.227059513 | -0.561399004 |
| Cacna1f | 2.348133422 | 1.499312923 | 0.760980992 | 1.970237022 |
| Syp | 9951.334057 | -0.058341455 | 0.06868002 | -0.849467655 |
| Gm14703 | 9.786129742 | -0.117280674 | 0.533053941 | -0.220016521 |
| Prickle3 | 15.66947633 | 0.007557199 | 0.262386907 | 0.028801738 |
| Plp2 | 46.01570745 | 1.110781433 | 0.235999386 | 4.706713242 |
| Magix | 1.802146662 | -0.581934004 | 0.778513292 | -0.747493986 |
| Gm6079 | 0 | NA | NA | NA |
| Gpkow | 510.2906317 | 0.270904714 | 0.079437823 | 3.41027366 |
| Gm14720 | 0 | NA | NA | NA |
| Wdr45 | 250.0876241 | 0.035245596 | 0.069914167 | 0.504126668 |
| Praf2 | 530.2469731 | -0.205556166 | 0.064314932 | -3.196087755 |
| Ccdc120 | 144.7907492 | -0.326852604 | 0.139542187 | -2.342321072 |
| Smt3h2-ps | 1.463749607 | 0.375807356 | 0.937750925 | 0.400753917 |
| Tfe3 | 940.6091959 | 0.166458597 | 0.061290665 | 2.715888263 |
| Gripap1 | 833.0662841 | 0.072265408 | 0.116001814 | 0.62296791 |
| Mir1198 | 4.250207662 | -0.864130218 | 0.510255415 | -1.693524837 |
| Kcnd1 | 104.6643679 | -0.612127171 | 0.138762808 | -4.411320158 |
| Otud5 | 1030.482185 | 0.21635705 | 0.047654661 | 4.540102564 |
| Pim2 | 309.3189051 | -0.440569933 | 0.090306096 | -4.878628904 |
| Slc35a2 | 203.1775785 | -0.12176519 | 0.105180553 | -1.15767779 |
| Pqbp1 | 363.0226281 | 0.005495069 | 0.072388005 | 0.075911313 |
| Timm17b | 118.9161447 | -0.302770385 | 0.152202982 | -1.989253955 |
| Gm10491 | 0.046345709 | 0.274391604 | 3.724945049 | 0.073663262 |
| Gm10490 | 0.314722824 | -1.174104552 | 1.73897752 | -0.675169482 |
| Pcsk1n | 5132.141306 | 0.198089677 | 0.101676933 | 1.948226318 |
| Eras | 0.25849572 | 0.274412531 | 2.083255822 | 0.131722916 |
| Hdac6 | 414.7095717 | -0.12555538 | 0.083008782 | -1.512555394 |
| Psmb6-ps | 0 | NA | NA | NA |
| Gata1 | 0.556661178 | 1.645409231 | 1.58684576 | 1.036905585 |
| Glod5 | 2.482897828 | 0.750569852 | 0.568219395 | 1.320915582 |
| Gm9429 | 0.112029109 | -0.687373951 | 3.724945049 | -0.184532642 |
| Rpl22-ps1 | 1.865863811 | 0.843285002 | 0.746864781 | 1.12909997 |
| Gm14799 | 1.762166695 | -0.699412841 | 0.885359841 | -0.789975791 |
| Gm14820 | 0.044544648 | 0.755286614 | 3.724945049 | 0.202764498 |
| Gm7079 | 50.37483144 | -0.416449117 | 0.302146718 | -1.37830098 |
| Suv39h1 | 269.5752616 | 0.115631165 | 0.093975152 | 1.230444032 |
| Was | 67.79222777 | 2.976662807 | 0.273921136 | 10.86686062 |
| Gm22083 | 0 | NA | NA | NA |
| Gm6787 | 1.653410964 | -1.037261265 | 0.82084324 | -1.263653294 |
| Psmb7-ps2 | 0.216874018 | -0.68658363 | 2.593170119 | -0.264766135 |
| Wdr13 | 1616.98844 | -0.056812707 | 0.077827059 | -0.729986559 |
| Rbm3 | 813.7615764 | 0.859247969 | 0.132256949 | 6.496807715 |
| Rbm3os | 4.093244608 | -0.605320146 | 0.590018239 | -1.025934634 |
| Tbc1d25 | 154.8391368 | 0.257168776 | 0.103900252 | 2.475150636 |
| Ebp | 209.4144051 | 0.08783884 | 0.098414773 | 0.892537145 |
| Porcn | 473.2143967 | -0.266283167 | 0.113247131 | -2.351345801 |

| | | | | |
|---|---|---|---|---|
| Ftsj1 | 170.553967 | -0.208945379 | 0.086626944 | -2.412013753 |
| Slc38a5 | 13.85939591 | 0.113613403 | 0.349141088 | 0.325408287 |
| Ssxb10 | 0 | NA | NA | NA |
| Ssxb9 | 0 | NA | NA | NA |
| Ssxb1 | 0 | NA | NA | NA |
| Ssxb2 | 0 | NA | NA | NA |
| Gm14459 | 0 | NA | NA | NA |
| Ssxb6 | 0 | NA | NA | NA |
| Ssxb3 | 0.117907917 | 1.204448679 | 3.72248341 | 0.323560523 |
| Gm14500 | 0 | NA | NA | NA |
| Gm6797 | 0 | NA | NA | NA |
| Gm6798 | 0 | NA | NA | NA |
| Gm9430 | 0.048949532 | -0.206487837 | 3.724945049 | -0.055433794 |
| Ssxb8 | 0.176646398 | 0.755284509 | 3.360121616 | 0.22477892 |
| Ssx9 | 0 | NA | NA | NA |
| Ssxb5 | 0 | NA | NA | NA |
| Gm14502 | 0.848591665 | -1.409551782 | 1.163414026 | -1.211565058 |
| Gm6592 | 0 | NA | NA | NA |
| Gm5751 | 0.339985237 | -0.656877091 | 1.741400421 | -0.377211975 |
| B630019K06Rik | 470.2877301 | 0.327024378 | 0.133611709 | 2.447572742 |
| Gm14504 | 0.188390742 | -0.551732295 | 3.306242977 | -0.166875907 |
| Gm5752 | 0 | NA | NA | NA |
| Gm16479 | 0 | NA | NA | NA |
| Gm14510 | 0 | NA | NA | NA |
| Gm5634 | 0 | NA | NA | NA |
| Gm14509 | 0 | NA | NA | NA |
| Gm14511 | 0 | NA | NA | NA |
| Gm14457 | 0 | NA | NA | NA |
| Gm14458 | 0 | NA | NA | NA |
| Gm9431 | 0.150711795 | 0.27440557 | 2.856704362 | 0.096056692 |
| Gm14865 | 0 | NA | NA | NA |
| Fthl17 | 0 | NA | NA | NA |
| Gm14503 | 0 | NA | NA | NA |
| Gm14499 | 0 | NA | NA | NA |
| Gm14864 | 0.066050875 | 0.274391604 | 3.724945049 | 0.073663262 |
| Gm5379 | 0 | NA | NA | NA |
| Gm5635 | 0 | NA | NA | NA |
| Gm16480 | 0 | NA | NA | NA |
| Gm6826 | 0 | NA | NA | NA |
| 4930402K13Rik | 0 | NA | NA | NA |
| Gm5753 | 0 | NA | NA | NA |
| Lancl3 | 181.9738499 | -0.4106696 | 0.099075619 | -4.145011694 |
| Gm14862 | 0.045882455 | 0.755288839 | 3.724945049 | 0.202765096 |
| Xk | 302.6762856 | -0.374065649 | 0.092306329 | -4.052437702 |
| 1700012L04Rik | 1.468848822 | -1.522824774 | 0.841124581 | -1.810462813 |
| Gm25481 | 0.305717649 | -0.961050068 | 1.732081029 | -0.554852834 |
| Gm6829 | 5.233925965 | -0.085310159 | 0.508973679 | -0.167612124 |

| | | | | |
|---|---|---|---|---|
| Gm14863 | 0.303434138 | 0.274413009 | 1.762680052 | 0.15567942 |
| Gm14501 | 0.106056992 | 1.236179401 | 3.724945049 | 0.331865138 |
| Cybb | 408.2087086 | 5.709718727 | 0.248443207 | 22.98198769 |
| Gm25728 | 0.044191549 | -0.206490061 | 3.724945049 | -0.055434391 |
| Gm5132 | 0.273271296 | 0.755309648 | 2.030826423 | 0.371922307 |
| Dynlt3 | 1172.505651 | 0.102194735 | 0.098537533 | 1.03711481 |
| Hypm | 1.020378237 | -0.424899717 | 0.978810218 | -0.434098162 |
| 4930557A04Rik | 0.934106423 | -1.492173907 | 1.154610287 | -1.292361521 |
| Sytl5 | 63.53451429 | -0.678895813 | 0.16704019 | -4.064266288 |
| Gm5754 | 0.681090533 | 0.952325309 | 1.343310917 | 0.708938859 |
| Srpx | 4.924570278 | -0.571560021 | 0.52666175 | -1.085250677 |
| Rpgr | 331.1611277 | -0.140455585 | 0.143144116 | -0.981218013 |
| Otc | 0.279231709 | -1.570346286 | 2.08289701 | -0.753924116 |
| 4933416E14Rik | 0.989536815 | -1.367057273 | 1.21517834 | -1.124984891 |
| Tspan7 | 5607.471727 | -0.283095527 | 0.086307433 | -3.280082783 |
| Gm25885 | 9.492997069 | -0.771100962 | 0.334714908 | -2.303754463 |
| Gm5931 | 6.023005853 | 0.195993051 | 0.442494348 | 0.442927807 |
| Gm14470 | 0.083526043 | 0.75528439 | 3.724945049 | 0.202763901 |
| Gm14480 | 0.946116905 | -0.135548571 | 0.928408615 | -0.146000984 |
| Gm10489 | 4.50082467 | 0.442462543 | 0.426369971 | 1.03774321 |
| Mid1ip1 | 964.1006633 | 0.015118163 | 0.10823046 | 0.139684918 |
| Gm14493 | 2.463509541 | -0.589826415 | 0.650765629 | -0.906357662 |
| Gm14473 | 0 | NA | NA | NA |
| Gm14485 | 0.048949532 | -0.206487837 | 3.724945049 | -0.055433794 |
| Gm14481 | 0.306362211 | 2.153768497 | 1.985737501 | 1.084618937 |
| Gm14483 | 0.04393175 | -0.206490061 | 3.724945049 | -0.055434391 |
| Gm14474 | 0 | NA | NA | NA |
| Gm14477 | 0 | NA | NA | NA |
| Gm14476 | 0 | NA | NA | NA |
| Gm14484 | 0 | NA | NA | NA |
| Gm14479 | 0 | NA | NA | NA |
| Gm14482 | 0 | NA | NA | NA |
| Gm14478 | 0 | NA | NA | NA |
| Gm14475 | 0 | NA | NA | NA |
| Gm4906 | 0 | NA | NA | NA |
| Gm14515 | 0 | NA | NA | NA |
| Gm14513 | 0 | NA | NA | NA |
| Gm26314 | 0 | NA | NA | NA |
| Rpl23a-ps13 | 0 | NA | NA | NA |
| Gm14491 | 1.107727442 | 0.171235585 | 0.822086839 | 0.208293791 |
| Gm14512 | 0 | NA | NA | NA |
| Bcor | 487.1376583 | -0.127954777 | 0.123673811 | -1.03461497 |
| Gm14521 | 0 | NA | NA | NA |
| Gm14635 | 0.042716945 | -0.206490061 | 3.724945049 | -0.055434391 |
| Rpl30-ps11 | 0.439488226 | 0.274413988 | 1.574258081 | 0.174313216 |
| Atp6ap2 | 2034.077482 | 0.07990697 | 0.06435671 | 1.241626089 |
| Gm9432 | 2.504601602 | 0.763405583 | 0.737590718 | 1.034998901 |

| | | | | |
|---|---|---|---|---|
| Gm4984 | 0.075043205 | 0.755286614 | 3.724945049 | 0.202764498 |
| 1810030O07Rik | 260.8927199 | 0.043449278 | 0.074144606 | 0.586007264 |
| Gm16265 | 0.351984013 | 0.75530905 | 1.808336542 | 0.417681683 |
| Med14 | 1262.541672 | -0.249756533 | 0.042518628 | -5.874049715 |
| Gm14634 | 22.48773761 | 0.036492977 | 0.23248271 | 0.156970716 |
| Gm25063 | 0.044544648 | 0.755286614 | 3.724945049 | 0.202764498 |
| Gm14524 | 0 | NA | NA | NA |
| AA414768 | 46.27663079 | 0.40444141 | 0.196158782 | 2.061806291 |
| Gm14534 | 0.163190422 | 0.274394252 | 3.398594894 | 0.080737558 |
| 5730405O15Rik | 64.71272437 | -0.388827079 | 0.173052563 | -2.246872702 |
| Usp9x | 6600.196581 | -0.014270276 | 0.04593936 | -0.310632898 |
| Gm15027 | 0.158569248 | 1.236192552 | 2.811594796 | 0.43967664 |
| Gm14536 | 0 | NA | NA | NA |
| Rpl3-ps1 | 956.7057751 | 0.324670466 | 0.106525353 | 3.047823429 |
| n-R5s3 | 0 | NA | NA | NA |
| Llph-ps2 | 8.542226199 | 0.600448233 | 0.418036836 | 1.436352449 |
| 2010308F09Rik | 1.66644364 | 0.415380462 | 0.732111206 | 0.567373453 |
| Gm16481 | 0 | NA | NA | NA |
| Ddx3x | 4505.87406 | 0.479759935 | 0.100352884 | 4.780728897 |
| Gm5380 | 0.854198329 | -0.9464486 | 1.058468946 | -0.894167565 |
| Gm7129 | 0.097899064 | -0.687371727 | 3.724945049 | -0.184532045 |
| Gm9434 | 0.22969359 | -0.206491565 | 3.142040024 | -0.065718948 |
| Nyx | 9.174112265 | 0.059528472 | 0.409181379 | 0.145481871 |
| Gm14506 | 0 | NA | NA | NA |
| Cask | 1849.363507 | 0.32544588 | 0.067976642 | 4.787613353 |
| Gpr34 | 42.59505678 | -2.139156558 | 0.652786688 | -3.27696106 |
| Gm25246 | 0 | NA | NA | NA |
| Gpr82 | 1.132182296 | 1.609010864 | 1.179031305 | 1.36468884 |
| Gm5381 | 0 | NA | NA | NA |
| Gm25552 | 0 | NA | NA | NA |
| Gm5382 | 0 | NA | NA | NA |
| Gm14867 | 0 | NA | NA | NA |
| Gm25202 | 0 | NA | NA | NA |
| Gm14505 | 0.222175754 | 0.274406964 | 2.58033673 | 0.106345409 |
| Gm9435 | 0 | NA | NA | NA |
| Ppp1r2-ps9 | 0 | NA | NA | NA |
| Gm14519 | 0.126242988 | 0.274405458 | 3.017245153 | 0.090945695 |
| Gm14518 | 0 | NA | NA | NA |
| Gm14517 | 0 | NA | NA | NA |
| Gm25207 | 0 | NA | NA | NA |
| Gm22464 | 0 | NA | NA | NA |
| Gm23516 | 0 | NA | NA | NA |
| Drr1 | 0.510920245 | -0.501820419 | 1.38167543 | -0.363197035 |
| Gm14520 | 0 | NA | NA | NA |
| Gm14516 | 0.07796279 | 0.274392189 | 3.724945049 | 0.073663419 |
| Cypt1 | 0.038981395 | 0.274393828 | 3.724945049 | 0.073663859 |
| Csnk2a1-ps | 0.061920613 | 0.274393828 | 3.724945049 | 0.073663859 |

| | | | | |
|---|---|---|---|---|
| Maoa | 471.3466681 | -0.12574772 | 0.09912135 | -1.268623968 |
| Maob | 199.4044433 | 0.715573107 | 0.139325348 | 5.135986516 |
| Ndp | 53.73532709 | -1.170027188 | 0.290993008 | -4.020808591 |
| Efhc2 | 28.39597078 | 0.454353 | 0.233668187 | 1.94443671 |
| Gm14514 | 0 | NA | NA | NA |
| Gm22294 | 0.044191549 | -0.206490061 | 3.724945049 | -0.055434391 |
| Fundc1 | 517.0185679 | 0.069552718 | 0.137091461 | 0.507345371 |
| Gm5073 | 0.100902008 | 0.274391828 | 3.724945049 | 0.073663322 |
| Gm9436 | 0 | NA | NA | NA |
| Gm5755 | 0 | NA | NA | NA |
| Gm5123 | 0 | NA | NA | NA |
| Dusp21 | 0 | NA | NA | NA |
| Kdm6a | 813.7693481 | 0.039289873 | 0.083315683 | 0.471578356 |
| n-R5s5 | 0 | NA | NA | NA |
| Gm14883 | 0 | NA | NA | NA |
| 4930578C19Rik | 7.057036865 | 0.794524507 | 0.521137103 | 1.524597849 |
| Gm6923 | 0 | NA | NA | NA |
| Gm5383 | 0 | NA | NA | NA |
| Mir221 | 8.635094361 | -0.140836846 | 0.432114664 | -0.325924709 |
| Mir222 | 2.111772912 | 0.002956241 | 0.772434018 | 0.003827176 |
| Gm14636 | 161.0641105 | -0.332573414 | 0.185894042 | -1.789048264 |
| Gm26652 | 38.71122557 | -0.579744226 | 0.277700723 | -2.087658323 |
| Gm22783 | 0.140090306 | -0.206491113 | 2.91543191 | -0.070826937 |
| Gm14885 | 0 | NA | NA | NA |
| BC049702 | 0 | NA | NA | NA |
| Gm14531 | 0 | NA | NA | NA |
| Gm23678 | 0 | NA | NA | NA |
| Gm14530 | 0.578381701 | 1.489079397 | 1.483788889 | 1.00356554 |
| Rpl36-ps10 | 0.038981395 | 0.274393828 | 3.724945049 | 0.073663859 |
| Gm5384 | 0.309182497 | 0.383879924 | 1.760532012 | 0.218047682 |
| Chst7 | 15.98671876 | -0.079333249 | 0.293978817 | -0.269860427 |
| Slc9a7 | 557.1947993 | -0.24381729 | 0.068358133 | -3.566763451 |
| Gm9083 | 2.845117016 | -0.680259862 | 0.635797567 | -1.069931527 |
| Gm14529 | 0 | NA | NA | NA |
| Rp2h | 271.8265236 | -0.093039139 | 0.135218115 | -0.688067122 |
| Gm9085 | 0.090427103 | 1.236179401 | 3.724945049 | 0.331865138 |
| Gm14537 | 3.667719452 | 0.009169685 | 0.547169818 | 0.01675839 |
| Jade3 | 162.3751509 | -0.251733922 | 0.124117087 | -2.028197152 |
| Gm14884 | 0.062426873 | 0.755288839 | 3.724945049 | 0.202765096 |
| Gm14538 | 0 | NA | NA | NA |
| Rgn | 3.195312812 | 1.297293937 | 0.606034921 | 2.140625716 |
| Gm14539 | 108.3056856 | 0.13021147 | 0.165575612 | 0.786416963 |
| Gm23628 | 1.296764287 | 0.724574807 | 0.901433742 | 0.803802624 |
| Ndufb11 | 499.4740233 | -0.022373516 | 0.137857131 | -0.162294949 |
| Rbm10 | 598.4934086 | -0.335690696 | 0.072951962 | -4.601530757 |
| Uba1 | 4405.966393 | 0.279898127 | 0.072925129 | 3.838157434 |
| Cdk16 | 2633.067354 | 0.013339792 | 0.058059511 | 0.229760662 |

| | | | | |
|---|---|---|---|---|
| Usp11 | 1238.553824 | -0.327217799 | 0.059846668 | -5.467602597 |
| Araf | 1405.997007 | -0.075234707 | 0.051513377 | -1.460488732 |
| Gm26065 | 0 | NA | NA | NA |
| Gm24824 | 0 | NA | NA | NA |
| Gm15028 | 0.262683741 | 0.755297982 | 2.479720904 | 0.304589916 |
| Syn1 | 5420.391119 | -0.198882226 | 0.093780015 | -2.12073144 |
| Mir5617 | 0 | NA | NA | NA |
| Timp1 | 225.2635421 | 5.34801455 | 0.526645111 | 10.15487363 |
| Gm15029 | 0 | NA | NA | NA |
| Cfp | 118.3901945 | -0.263315559 | 0.212559062 | -1.238787734 |
| Elk1 | 876.8854966 | 0.545446836 | 0.131916852 | 4.134777528 |
| Uxt | 53.85454816 | 0.21402557 | 0.164380943 | 1.302009619 |
| A230072C01Rik | 65.61270515 | -0.356178543 | 0.139727712 | -2.549090213 |
| Zfp182 | 201.9227118 | -0.015797202 | 0.115670827 | -0.136570324 |
| Spaca5 | 0.708400758 | 1.557778977 | 1.315950387 | 1.183767255 |
| Zfp300 | 69.84368967 | -0.241087627 | 0.163187177 | -1.477368696 |
| Ssxa1 | 0 | NA | NA | NA |
| Gm1848 | 2.174616668 | -0.875115678 | 0.810244524 | -1.080063675 |
| Gm21876 | 0 | NA | NA | NA |
| 4930453H23Rik | 0.062426873 | 0.755288839 | 3.724945049 | 0.202765096 |
| Gm6938 | 0.624139293 | 1.526790651 | 1.254110594 | 1.217429035 |
| Gm5124 | 229.7670475 | -0.178520149 | 0.117030188 | -1.525419654 |
| Gm26593 | 1.567651568 | 0.512456052 | 1.007212908 | 0.508786224 |
| Agtr2 | 19.19641764 | 0.023381974 | 0.299739659 | 0.078007607 |
| Gm22592 | 0 | NA | NA | NA |
| Gm9439 | 0.758270043 | 1.050768879 | 1.079774109 | 0.973137687 |
| Gm14558 | 0 | NA | NA | NA |
| Slc6a14 | 1.053279316 | 1.61415967 | 0.992400234 | 1.626520847 |
| Gm25553 | 0.068097359 | -0.206487837 | 3.724945049 | -0.055433794 |
| Gm28269 | 0 | NA | NA | NA |
| Gm28268 | 0.530320922 | -0.51729248 | 1.417967015 | -0.364812774 |
| Gm24535 | 0 | NA | NA | NA |
| Gm14563 | 0 | NA | NA | NA |
| Gm14561 | 0 | NA | NA | NA |
| Gm6071 | 0 | NA | NA | NA |
| Gm25812 | 0 | NA | NA | NA |
| Gm14562 | 0 | NA | NA | NA |
| Gm15030 | 0 | NA | NA | NA |
| Ppp1r2-ps7 | 0 | NA | NA | NA |
| Gm26131 | 0 | NA | NA | NA |
| Klhl13 | 551.8407492 | -0.141723791 | 0.096395425 | -1.470233587 |
| Gm14523 | 0.533370488 | -0.92767471 | 1.174400542 | -0.789913388 |
| Gm7164 | 0.047948698 | -0.206487837 | 3.724945049 | -0.055433794 |
| Wdr44 | 445.6653234 | -0.018781681 | 0.069113025 | -0.271753124 |
| Gm4789 | 0 | NA | NA | NA |
| Gm4907 | 0.266401621 | -0.681967052 | 2.506477774 | -0.272081827 |
| Gm4985 | 0 | NA | NA | NA |

| | | | | | |
|---|---|---|---|---|---|
| Gm25299 | 0 | NA | NA | NA | |
| Gm4732 | 0 | NA | NA | NA | |
| Gm9440 | 0 | NA | NA | NA | |
| Gm6956 | 0 | NA | NA | NA | |
| Gm27192 | 0 | NA | NA | NA | |
| Gm5934 | 0.046345709 | | 0.274391604 | 3.724945049 | 0.073663262 |
| Gm4297 | 0 | NA | NA | NA | |
| Gm27910 | 0 | NA | NA | NA | |
| Gm27839 | 0 | NA | NA | NA | |
| Gm5935 | 0 | NA | NA | NA | |
| Gm2759 | 0 | NA | NA | NA | |
| CR938728.1 | 0 | NA | NA | NA | |
| Gm2768 | 0 | NA | NA | NA | |
| Gm5169 | 0 | NA | NA | NA | |
| Gm1989 | 0 | NA | NA | NA | |
| Gm1993 | 0 | NA | NA | NA | |
| Gm2003 | 0 | NA | NA | NA | |
| Gm27892 | 0 | NA | NA | NA | |
| Gm7350 | 0 | NA | NA | NA | |
| E330010L02Rik | 0 | NA | NA | NA | |
| Gm14533 | 0 | NA | NA | NA | |
| Gm5168 | 0 | NA | NA | NA | |
| Gm2012 | 0 | NA | NA | NA | |
| Gm26302 | 0 | NA | NA | NA | |
| Gm2030 | 0 | NA | NA | NA | |
| Gm2005 | 0 | NA | NA | NA | |
| Slx | 0 | NA | NA | NA | |
| Gm27341 | 0 | NA | NA | NA | |
| Gm14525 | 0 | NA | NA | NA | |
| Gm6113 | 0 | NA | NA | NA | |
| Gm6121 | 0 | NA | NA | NA | |
| Gm7365 | 0 | NA | NA | NA | |
| Gm7375 | 0 | NA | NA | NA | |
| Gm10230 | 0 | NA | NA | NA | |
| BX813306.1 | 0 | NA | NA | NA | |
| Gm2092 | 0 | NA | NA | NA | |
| CR589930.1 | 0 | NA | NA | NA | |
| Gm2101 | 0 | NA | NA | NA | |
| Gm7386 | 0 | NA | NA | NA | |
| Gm10058 | 0 | NA | NA | NA | |
| Gm2117 | 0 | NA | NA | NA | |
| Gm7391 | 0 | NA | NA | NA | |
| Gm4836 | 0 | NA | NA | NA | |
| Gm7398 | 0 | NA | NA | NA | |
| Gm10147 | 0 | NA | NA | NA | |
| BX813332.1 | 0 | NA | NA | NA | |
| Gm2165 | 0 | NA | NA | NA | |

| | | | | |
|---|---|---|---|---|
| Gm7408 | 0 | NA | NA | NA |
| Gm10096 | 0 | NA | NA | NA |
| Gm2200 | 0 | NA | NA | NA |
| Gm7415 | 0 | NA | NA | NA |
| Gm26818 | 0 | NA | NA | NA |
| Gm2066 | 0 | NA | NA | NA |
| Gm7421 | 0 | NA | NA | NA |
| Gm15432 | 1.975319427 | 0.62241796 | 0.844994127 | 0.736594421 |
| Gm14527 | 0 | NA | NA | NA |
| BX294196.1 | 0 | NA | NA | NA |
| Gm3657 | 0 | NA | NA | NA |
| Gm3669 | 0 | NA | NA | NA |
| Gm10488 | 0 | NA | NA | NA |
| BX294192.1 | 0 | NA | NA | NA |
| E330016L19Rik | 0 | NA | NA | NA |
| Gm3681 | 0 | NA | NA | NA |
| Gm14632 | 0 | NA | NA | NA |
| Gm7437 | 0 | NA | NA | NA |
| Gm14974 | 0 | NA | NA | NA |
| Gm7438 | 0 | NA | NA | NA |
| Gm10487 | 0 | NA | NA | NA |
| Gm21447 | 0 | NA | NA | NA |
| Spin2f | 0 | NA | NA | NA |
| Gm2784 | 0 | NA | NA | NA |
| Gm15280 | 0 | NA | NA | NA |
| Gm15276 | 0 | NA | NA | NA |
| Gm2777 | 0 | NA | NA | NA |
| Gm21883 | 0 | NA | NA | NA |
| Spin2e | 0 | NA | NA | NA |
| Gm21608 | 0 | NA | NA | NA |
| Gm21637 | 0 | NA | NA | NA |
| Gm21645 | 0 | NA | NA | NA |
| Gm2799 | 0 | NA | NA | NA |
| Gmcl1l | 0 | NA | NA | NA |
| Gm5926 | 0 | NA | NA | NA |
| Gm21951 | 0 | NA | NA | NA |
| Gm21657 | 0 | NA | NA | NA |
| Gm21789 | 0 | NA | NA | NA |
| Gm2825 | 0 | NA | NA | NA |
| Spin2-ps6 | 0 | NA | NA | NA |
| Gm15275 | 0 | NA | NA | NA |
| Gm15274 | 0 | NA | NA | NA |
| Gm2863 | 0 | NA | NA | NA |
| Gm2182 | 0 | NA | NA | NA |
| Gm2854 | 0 | NA | NA | NA |
| Gm21979 | 0 | NA | NA | NA |
| Gm21990 | 0 | NA | NA | NA |

| | | | | |
|---|---|---|---|---|
| Gm2892 | 0 | NA | NA | NA |
| Gm2223 | 89.01610747 | 0.214313289 | 0.150383739 | 1.425109453 |
| Gm2231 | 0 | NA | NA | NA |
| Gm2913 | 0 | NA | NA | NA |
| Gm15273 | 0 | NA | NA | NA |
| Gm15281 | 0 | NA | NA | NA |
| Gm15279 | 0 | NA | NA | NA |
| Gm7023 | 0 | NA | NA | NA |
| Gm2927 | 0 | NA | NA | NA |
| Gm2933 | 0 | NA | NA | NA |
| Gm2964 | 0 | NA | NA | NA |
| Gm21870 | 0 | NA | NA | NA |
| Gm21681 | 0 | NA | NA | NA |
| Spin2g | 0 | NA | NA | NA |
| Gm21699 | 0 | NA | NA | NA |
| Gm2288 | 0 | NA | NA | NA |
| Gm14552 | 0 | NA | NA | NA |
| Gm10486 | 0 | NA | NA | NA |
| Gm2309 | 0 | NA | NA | NA |
| Gm2318 | 0 | NA | NA | NA |
| Gm14553 | 0 | NA | NA | NA |
| Gm14819 | 0 | NA | NA | NA |
| Gm4764 | 0 | NA | NA | NA |
| Gm14554 | 0.066050875 | 0.274391604 | 3.724945049 | 0.073663262 |
| Gm25624 | 0 | NA | NA | NA |
| Dock11 | 371.5272742 | 0.180411499 | 0.084819821 | 2.12699693 |
| Il13ra1 | 318.8934833 | 1.719214991 | 0.193283519 | 8.894783162 |
| Zcchc12 | 687.811863 | -0.079379302 | 0.131525778 | -0.603526573 |
| Gm14560 | 0 | NA | NA | NA |
| Gm6023 | 0.589180756 | 2.05249377 | 1.551884901 | 1.322581184 |
| Gm14559 | 0 | NA | NA | NA |
| Lonrf3 | 281.4852912 | 0.104224737 | 0.134244238 | 0.776381455 |
| Gm25047 | 24.14149978 | 0.21577238 | 0.252035072 | 0.856120451 |
| Gm14568 | 0.274682661 | 1.776633033 | 3.708688794 | 0.479046135 |
| AL450395.1 | 0 | NA | NA | NA |
| Gm6268 | 0.075043205 | 0.755286614 | 3.724945049 | 0.202764498 |
| Gm22364 | 0 | NA | NA | NA |
| Gm14569 | 4.657189074 | 1.55524053 | 0.493617172 | 3.150701834 |
| Pgrmc1 | 2005.442762 | 0.064453055 | 0.07873592 | 0.818597856 |
| Akap17b | 352.8596161 | -0.225485735 | 0.07106793 | -3.172819807 |
| Gm6274 | 10.81761153 | 0.229628209 | 0.39140541 | 0.586676123 |
| Slc25a43 | 7.740311858 | 1.244055455 | 0.545918785 | 2.278828807 |
| Gm23613 | 0.350819806 | 0.755304775 | 1.994187518 | 0.378753135 |
| Slc25a5 | 1945.709052 | 0.011963753 | 0.07440054 | 0.160801961 |
| Gm14549 | 0 | NA | NA | NA |
| 2310010G23Rik | 9.327370753 | 1.009875325 | 0.377434078 | 2.675633666 |
| C330007P06Rik | 639.7924073 | 0.15083281 | 0.075461468 | 1.998805676 |

| | | | | | |
|---|---|---|---|---|---|
| Gm14541 | | 2.652685225 | 1.324930509 | 0.724850013 | 1.827868503 |
| Ube2a | | 329.5173645 | -0.146940894 | 0.100080188 | -1.468231601 |
| Nkrf | | 449.5931371 | -0.289751976 | 0.079229825 | -3.657107376 |
| Gm15008 | | 0.250168298 | -0.074760622 | 2.061797522 | -0.036259924 |
| | 6-Sep | 721.875096 | -0.078850396 | 0.064344578 | -1.22543963 |
| Gm14542 | | 0 | NA | NA | NA |
| Sowahd | | 0.087930927 | -0.206490058 | 3.724945049 | -0.05543439 |
| Gm26099 | | 0 | NA | NA | NA |
| Gm24809 | | 0 | NA | NA | NA |
| Rpl39 | | 722.6800404 | 0.486705411 | 0.127558055 | 3.815559991 |
| Snora69 | | 1.244281449 | -0.451297425 | 0.907717588 | -0.497178231 |
| Upf3b | | 273.0236683 | -0.0878306 | 0.118734929 | -0.73971999 |
| Nkap | | 229.4254487 | 0.179286759 | 0.084287138 | 2.127095118 |
| Akap14 | | 1.404017228 | -0.896828381 | 1.276678871 | -0.702469824 |
| Ndufa1 | | 279.6891087 | -0.119224113 | 0.133349966 | -0.894069321 |
| Rnf113a1 | | 34.43701755 | 0.225964634 | 0.196751899 | 1.148474981 |
| Gm9 | | 0 | NA | NA | NA |
| Rhox1 | | 0 | NA | NA | NA |
| Rhox2a | | 0 | NA | NA | NA |
| Rhox3a | | 0 | NA | NA | NA |
| Rhox4a | | 0 | NA | NA | NA |
| Rhox3a2 | | 0 | NA | NA | NA |
| Rhox4a2 | | 0 | NA | NA | NA |
| Rhox2b | | 0 | NA | NA | NA |
| Rhox3b-ps | | 0 | NA | NA | NA |
| Rhox4b | | 0 | NA | NA | NA |
| Rhox2c | | 0 | NA | NA | NA |
| Rhox3c | | 0 | NA | NA | NA |
| Rhox4c | | 0 | NA | NA | NA |
| Rhox2d | | 0 | NA | NA | NA |
| Gm26494 | | 0 | NA | NA | NA |
| Rhox3d-ps | | 0 | NA | NA | NA |
| Rhox4d | | 0 | NA | NA | NA |
| Rhox2e | | 0 | NA | NA | NA |
| Rhox3e | | 0 | NA | NA | NA |
| Rhox4e | | 0 | NA | NA | NA |
| Rhox2f | | 0 | NA | NA | NA |
| Rhox3f | | 0 | NA | NA | NA |
| Rhox4f | | 0 | NA | NA | NA |
| Rhox3g | | 0 | NA | NA | NA |
| Dppa3-ps | | 0 | NA | NA | NA |
| Gm24493 | | 0 | NA | NA | NA |
| Rhox2g | | 0 | NA | NA | NA |
| Rhox4g | | 0 | NA | NA | NA |
| Rhox3h | | 0 | NA | NA | NA |
| Gm24147 | | 0 | NA | NA | NA |
| Rhox2h | | 0 | NA | NA | NA |

| | | | | |
|---|---|---|---|---|
| RP23-457N24.12 | 0 | NA | NA | NA |
| Rhox2-ps | 0 | NA | NA | NA |
| Rhox11-ps1 | 0 | NA | NA | NA |
| Rhox11-ps2 | 0 | NA | NA | NA |
| Btg1-ps1 | 0.062426873 | 0.755288839 | 3.724945049 | 0.202765096 |
| Btg1-ps2 | 0 | NA | NA | NA |
| Rhox10 | 0 | NA | NA | NA |
| Rhox11 | 0 | NA | NA | NA |
| Rhox11-ps3 | 0 | NA | NA | NA |
| Rhox12 | 0.066050875 | 0.274391604 | 3.724945049 | 0.073663262 |
| Rhox13 | 0 | NA | NA | NA |
| Zfp36l1-ps | 0.671662539 | 2.368378756 | 1.257332779 | 1.883653075 |
| Zbtb33 | 539.0021361 | 0.285967476 | 0.101243862 | 2.824541365 |
| Tmem255a | 487.0062224 | -0.045400604 | 0.111653409 | -0.406620849 |
| Mir3110 | 0 | NA | NA | NA |
| Rhox7-ps1 | 0.075043205 | 0.755286614 | 3.724945049 | 0.202764498 |
| Atp1b4 | 0.134880151 | 0.274398319 | 2.937637886 | 0.093407809 |
| Gm40 | 0 | NA | NA | NA |
| Gm4853 | 2.301314035 | -0.20311737 | 0.716876265 | -0.28333672 |
| Lamp2 | 2710.25613 | 1.135140323 | 0.126882201 | 8.94641103 |
| Gm7598 | 7.758147147 | -0.139654268 | 0.387258989 | -0.360622405 |
| Cul4b | 778.5921849 | 0.184449608 | 0.059694769 | 3.089878914 |
| Mcts1 | 310.2787056 | -0.126235341 | 0.095181664 | -1.326256922 |
| C1galt1c1 | 195.6813856 | -0.080408429 | 0.191757709 | -0.419323058 |
| Ap3s1-ps1 | 43.23259583 | 0.052388464 | 0.169857643 | 0.308425712 |
| Gm14565 | 0.060174537 | 0.755286614 | 3.724945049 | 0.202764498 |
| Gm25642 | 0 | NA | NA | NA |
| Gm14564 | 0 | NA | NA | NA |
| 6030498E09Rik | 0.374199036 | 0.250390819 | 1.464129066 | 0.171016903 |
| Gm6024 | 0 | NA | NA | NA |
| Gm14567 | 9.440869076 | 0.054577434 | 0.362458544 | 0.150575659 |
| Gm16423 | 0 | NA | NA | NA |
| Gm14551 | 0 | NA | NA | NA |
| Gm14566 | 0 | NA | NA | NA |
| Gm26437 | 0 | NA | NA | NA |
| Cypt15 | 0.048949532 | -0.206487837 | 3.724945049 | -0.055433794 |
| Gm14550 | 0 | NA | NA | NA |
| Gm14570 | 0 | NA | NA | NA |
| Gm14571 | 0.04393175 | -0.206490061 | 3.724945049 | -0.055434391 |
| Cypt14 | 0 | NA | NA | NA |
| Gm14643 | 0.048949532 | -0.206487837 | 3.724945049 | -0.055433794 |
| Gm26194 | 0 | NA | NA | NA |
| Gm7189 | 0 | NA | NA | NA |
| Gm16383 | 0 | NA | NA | NA |
| Gm7190 | 0 | NA | NA | NA |
| Gm4986 | 0.708087976 | 2.540563515 | 1.197498408 | 2.121558991 |
| Gm14575 | 0 | NA | NA | NA |

| | | | | |
|---|---|---|---|---|
| Gm14574 | 0 | NA | NA | NA |
| Gm14576 | 0 | NA | NA | NA |
| Gm5385 | 0 | NA | NA | NA |
| Gm14577 | 0 | NA | NA | NA |
| Gria3 | 2075.810427 | -0.184265548 | 0.099936768 | -1.843821366 |
| Gm22413 | 0 | NA | NA | NA |
| Gm14614 | 0 | NA | NA | NA |
| Gm14613 | 0 | NA | NA | NA |
| Thoc2 | 1453.981551 | 0.144897962 | 0.062380901 | 2.322793685 |
| Gm7657 | 0 | NA | NA | NA |
| Gm5135 | 0 | NA | NA | NA |
| AL954355.1 | 0 | NA | NA | NA |
| Xiap | 1927.944407 | 0.139597974 | 0.077445787 | 1.802525093 |
| Gm14615 | 0.068097359 | -0.206487837 | 3.724945049 | -0.055433794 |
| Gm37564 | 0.632215448 | -0.838251607 | 1.114146967 | -0.752370765 |
| Stag2 | 1734.40788 | 0.098935211 | 0.081464087 | 1.214464118 |
| Gm14619 | 0.123841226 | 0.274391173 | 3.724945049 | 0.073663147 |
| RP23-280E8.2 | 0.589937099 | 1.637794284 | 1.360104837 | 1.204167678 |
| Sh2d1a | 7.357494345 | 4.089708289 | 0.785835188 | 5.204282463 |
| Tenm1 | 2192.810765 | -0.210113733 | 0.102045954 | -2.059010919 |
| Gm23705 | 2.363489277 | -0.351715994 | 0.622702803 | -0.5648216 |
| Gm14578 | 7.50102364 | -0.312389459 | 0.519590529 | -0.601222389 |
| Gm24763 | 0 | NA | NA | NA |
| Gm14587 | 8.400665238 | -1.695759188 | 0.53131704 | -3.191614537 |
| Gm14588 | 0 | NA | NA | NA |
| Gm14579 | 0 | NA | NA | NA |
| Gm362 | 0 | NA | NA | NA |
| Gm14580 | 0 | NA | NA | NA |
| Gm14628 | 0 | NA | NA | NA |
| Gm14603 | 0 | NA | NA | NA |
| Dcaf12l2 | 11.20309874 | -0.285739 | 0.299942572 | -0.952645695 |
| Itpa-ps3 | 0 | NA | NA | NA |
| Gm14602 | 0 | NA | NA | NA |
| Dcaf12l1 | 439.0135136 | -0.281777603 | 0.074600035 | -3.777177909 |
| Gm7700 | 0 | NA | NA | NA |
| Prr32 | 4.737155157 | -1.165239873 | 0.537830925 | -2.166554242 |
| Gm14606 | 0 | NA | NA | NA |
| Gm14605 | 0 | NA | NA | NA |
| Gm5386 | 0 | NA | NA | NA |
| Gm14652 | 0 | NA | NA | NA |
| Gm25078 | 0 | NA | NA | NA |
| Gm14653 | 0 | NA | NA | NA |
| 4930515L19Rik | 0 | NA | NA | NA |
| Actrt1 | 0 | NA | NA | NA |
| Gm14654 | 0.049553047 | 0.274391604 | 3.724945049 | 0.073663262 |
| Gm4987 | 0 | NA | NA | NA |
| Gm29242 | 0 | NA | NA | NA |

| | | | | |
|---|---|---|---|---|
| Gm7722 | 0.112396584 | 0.27438722 | 3.724945049 | 0.073662085 |
| Gm14609 | 0.13648314 | -0.206488635 | 2.940069555 | -0.070232568 |
| Gm14610 | 0.187229393 | -0.673484163 | 2.704274152 | -0.249044337 |
| Gm4908 | 0 | NA | NA | NA |
| Gm14608 | 0.401923326 | -0.624020236 | 1.43056146 | -0.43620652 |
| Gm6480 | 0.088476398 | 0.274400501 | 3.724945049 | 0.073665651 |
| Smarca1 | 656.7356124 | -0.232201951 | 0.085807856 | -2.706068683 |
| Gm14648 | 4.891543041 | 0.838555714 | 0.508311997 | 1.649687039 |
| Ocrl | 1059.362513 | -0.158461815 | 0.048537849 | -3.264706163 |
| Apln | 136.796465 | -1.264741254 | 0.180460245 | -7.008420348 |
| Xpnpep2 | 2.216825317 | -0.89656996 | 0.59746961 | -1.500611823 |
| Sash3 | 86.39783744 | 2.825845277 | 0.246649636 | 11.45692051 |
| Zdhhc9 | 642.249467 | -0.306466233 | 0.072040768 | -4.254066721 |
| Gm24038 | 0 | NA | NA | NA |
| Utp14a | 243.5098185 | 0.214786929 | 0.098299886 | 2.185017079 |
| 9530027J09Rik | 0.791221454 | -2.510751779 | 1.145769279 | -2.191324052 |
| Gm22528 | 0 | NA | NA | NA |
| Gm7212 | 0 | NA | NA | NA |
| Gm22612 | 0 | NA | NA | NA |
| Bcorl1 | 414.200989 | -0.709187413 | 0.108930181 | -6.510476748 |
| Elf4 | 210.3602389 | 3.286150201 | 0.203530434 | 16.14574356 |
| Aifm1 | 270.3649458 | -0.359183555 | 0.091605207 | -3.920994953 |
| Rab33a | 124.0995529 | 0.354813043 | 0.106235237 | 3.339880934 |
| Zfp280c | 445.3017861 | 0.004498772 | 0.099900999 | 0.045032306 |
| Slc25a14 | 322.987207 | -0.151630802 | 0.077074803 | -1.967320011 |
| Gm27828 | 1.991846454 | -0.140194163 | 0.668720144 | -0.209645491 |
| Gpr119 | 0.257827269 | -0.206481351 | 1.84476804 | -0.111928083 |
| Rbmx2 | 38.78282912 | 0.388181162 | 0.230743739 | 1.682304202 |
| Gm595 | 0.555008095 | 1.883729619 | 1.553339673 | 1.212696522 |
| Gm7219 | 0 | NA | NA | NA |
| Enox2 | 126.1449739 | -0.002901062 | 0.132471299 | -0.021899548 |
| Gm15034 | 0 | NA | NA | NA |
| Gm15036 | 0 | NA | NA | NA |
| Gm14696 | 0.319846959 | 1.260738242 | 1.776010005 | 0.709871137 |
| Gm14697 | 0.131474741 | 0.274407218 | 2.980045311 | 0.092081559 |
| Arhgap36 | 111.713745 | 0.105503547 | 0.231805922 | 0.455137409 |
| Gm15037 | 0 | NA | NA | NA |
| Gm14719 | 6.72125269 | 0.12274947 | 0.361020349 | 0.340007067 |
| Olfr1320 | 0 | NA | NA | NA |
| Olfr1321 | 0 | NA | NA | NA |
| Igsf1 | 120.6751455 | -0.987375866 | 0.158200654 | -6.241288144 |
| n-R5s6 | 0 | NA | NA | NA |
| Olfr1322 | 0.062426873 | 0.755288839 | 3.724945049 | 0.202765096 |
| Gm15033 | 0 | NA | NA | NA |
| Olfr1323 | 0.233510017 | 0.75529888 | 2.541434124 | 0.297193963 |
| Gm23385 | 0 | NA | NA | NA |
| Gm7803 | 2.841226851 | 0.822608738 | 0.662025983 | 1.242562618 |

| | | | | |
|---|---|---|---|---|
| Gm14612 | 0.038981395 | 0.274393828 | 3.724945049 | 0.073663859 |
| Gm22288 | 0.45637343 | -0.210066999 | 1.412305523 | -0.148740478 |
| Olfr1324 | 0.046345709 | 0.274391604 | 3.724945049 | 0.073663262 |
| Gm14620 | 0 | NA | NA | NA |
| Firre | 150.6157184 | -0.582866539 | 0.213243415 | -2.733338981 |
| Stk26 | 85.23139706 | 0.325580337 | 0.135342836 | 2.405597123 |
| Frmd7 | 56.57679548 | -0.909630045 | 0.656297261 | -1.386003111 |
| Rap2c | 496.5610328 | 0.259495109 | 0.112058192 | 2.315717439 |
| Gm5387 | 0 | NA | NA | NA |
| Mbnl3 | 8.934417775 | 0.714778358 | 0.408480821 | 1.749845577 |
| Gm7834 | 0 | NA | NA | NA |
| Gm5388 | 0.810280881 | 0.876127175 | 1.123411092 | 0.779881186 |
| Gm23417 | 0 | NA | NA | NA |
| Gm14621 | 0 | NA | NA | NA |
| Hs6st2 | 613.6901847 | -0.246538465 | 0.084943271 | -2.902389569 |
| Gm14581 | 4.970320586 | 0.034146191 | 0.49651275 | 0.068772033 |
| Usp26 | 0 | NA | NA | NA |
| Gm7840 | 0 | NA | NA | NA |
| Gm22539 | 0.072025462 | 0.755288839 | 3.724945049 | 0.202765096 |
| 1700080O16Rik | 0 | NA | NA | NA |
| Gpc4 | 223.3113565 | 0.425292682 | 0.093161631 | 4.565105602 |
| Gpc3 | 54.37477559 | -0.17596761 | 0.15831855 | -1.111478153 |
| Gm14582 | 0 | NA | NA | NA |
| mmu-mir-6384 | 0.085433889 | -0.687376175 | 3.724945049 | -0.184533239 |
| Mir717 | 0 | NA | NA | NA |
| A630012P03Rik | 15.99725564 | 6.204448615 | 0.928249112 | 6.684033988 |
| Mir363 | 0 | NA | NA | NA |
| Mir92-2 | 0 | NA | NA | NA |
| Mir19b-2 | 0 | NA | NA | NA |
| Mir20b | 0 | NA | NA | NA |
| Mir18b | 0 | NA | NA | NA |
| Mir106a | 0 | NA | NA | NA |
| Kis2 | 0.066050875 | 0.274391604 | 3.724945049 | 0.073663262 |
| Gm14586 | 42.19938222 | 0.378301301 | 0.225193472 | 1.679894618 |
| Gm7851 | 0 | NA | NA | NA |
| Ccdc160 | 20.72630267 | -0.02555649 | 0.252920227 | -0.101045657 |
| Gm14607 | 0.060174537 | 0.755286614 | 3.724945049 | 0.202764498 |
| Gm6539 | 0 | NA | NA | NA |
| Rps2-ps13 | 156.9088428 | 0.410198875 | 0.153399182 | 2.674061676 |
| Phf6 | 556.50205 | 0.260988926 | 0.082575445 | 3.160611809 |
| Hprt | 957.8882762 | 0.001884081 | 0.076660025 | 0.024577099 |
| Mir450b | 0 | NA | NA | NA |
| Mir450-1 | 0 | NA | NA | NA |
| Mir450-2 | 0 | NA | NA | NA |
| Mir542 | 0 | NA | NA | NA |
| Gm28730 | 0.097899064 | -0.687371727 | 3.724945049 | -0.184532045 |
| Mir351 | 0 | NA | NA | NA |

| | | | | |
|---|---|---|---|---|
| Mir503 | 0.047948698 | -0.206487837 | 3.724945049 | -0.055433794 |
| Mir322 | 0 | NA | NA | NA |
| C430049B03Rik | 3.09373086 | -0.942621605 | 0.588201704 | -1.602548239 |
| Plac1 | 0 | NA | NA | NA |
| Fam122b | 69.8401722 | -0.288717082 | 0.213816252 | -1.350304661 |
| Fam122c | 0 | NA | NA | NA |
| Gm23718 | 0 | NA | NA | NA |
| Gm14585 | 0 | NA | NA | NA |
| Mospd1 | 150.5973759 | 0.308640031 | 0.110787684 | 2.785869501 |
| Gm14584 | 111.7814324 | 0.118017285 | 0.117582892 | 1.003694356 |
| Etd | 0 | NA | NA | NA |
| Gm14597 | 0.48133134 | 0.2645355 | 1.583325445 | 0.167075885 |
| Cxx1c | 578.0914899 | -0.115768262 | 0.106857558 | -1.083388615 |
| Gm14592 | 0.289161111 | -1.255096502 | 1.71370776 | -0.732386543 |
| Gm14600 | 0 | NA | NA | NA |
| Cxx1a | 186.6998316 | -0.112949845 | 0.092886773 | -1.215994925 |
| Cxx1b | 332.64126 | -0.163714311 | 0.077958998 | -2.100005332 |
| 4933416I08Rik | 0.129625439 | -1.028499994 | 3.675612039 | -0.279817343 |
| AW822252 | 37.35059821 | -0.333062118 | 0.243540443 | -1.367584427 |
| 4930502E18Rik | 2.580172256 | -0.562363724 | 0.62443666 | -0.900593703 |
| 1700013H16Rik | 0.066050875 | 0.274391604 | 3.724945049 | 0.073663262 |
| Gm6025 | 0 | NA | NA | NA |
| Gm14593 | 0 | NA | NA | NA |
| Zfp36l3 | 0.434206664 | -1.38066973 | 1.431457366 | -0.96452033 |
| Xlr | 8.912034157 | 1.70400471 | 0.408431795 | 4.172066751 |
| Gm14599 | 0.044544648 | 0.755286614 | 3.724945049 | 0.202764498 |
| Gm14589 | 0 | NA | NA | NA |
| Gm1862 | 2.065160235 | -0.523698923 | 0.70140138 | -0.746646554 |
| Gm14591 | 0.119587853 | 1.218392466 | 3.723618458 | 0.327206581 |
| Gm15482 | 0.887798776 | 1.349175634 | 1.006432025 | 1.340553162 |
| Gm14601 | 0 | NA | NA | NA |
| Gm14644 | 0 | NA | NA | NA |
| Gm22550 | 0 | NA | NA | NA |
| Gm16428 | 0 | NA | NA | NA |
| Gm7916 | 1.902020358 | 0.0516074 | 0.774006525 | 0.066675666 |
| Gm14598 | 0.066050875 | 0.274391604 | 3.724945049 | 0.073663262 |
| Gm6591 | 0 | NA | NA | NA |
| Gm14596 | 0 | NA | NA | NA |
| Gm14594 | 0 | NA | NA | NA |
| Gm16405 | 0 | NA | NA | NA |
| Gm16430 | 0 | NA | NA | NA |
| Gm14595 | 0 | NA | NA | NA |
| Gm14590 | 0 | NA | NA | NA |
| Gm16431 | 0 | NA | NA | NA |
| Gm7950 | 0 | NA | NA | NA |
| Gm7958 | 0 | NA | NA | NA |
| Gm7974 | 0 | NA | NA | NA |

| | | | | |
|---|---|---|---|---|
| Slxl1 | 0 | NA | NA | NA |
| Gm6660 | 0 | NA | NA | NA |
| Gm14625 | 0 | NA | NA | NA |
| Gm16404 | 0 | NA | NA | NA |
| Gm14622 | 0 | NA | NA | NA |
| Gm6664 | 0 | NA | NA | NA |
| Gm14626 | 0 | NA | NA | NA |
| Gm7927 | 0 | NA | NA | NA |
| Gm14623 | 0 | NA | NA | NA |
| Gm14624 | 0 | NA | NA | NA |
| Gm22103 | 0.038981395 | 0.274393828 | 3.724945049 | 0.073663859 |
| Gm14638 | 0 | NA | NA | NA |
| 3830403N18Rik | 1.591056397 | 2.390641498 | 1.038840916 | 2.301258509 |
| Gm14639 | 1.176771261 | 0.543799843 | 1.168258326 | 0.465479107 |
| Gm773 | 0 | NA | NA | NA |
| Gm14627 | 0 | NA | NA | NA |
| 1600025M17Rik | 0 | NA | NA | NA |
| Zfp449 | 65.59676692 | -0.369048572 | 0.138125398 | -2.671837174 |
| Gm2155 | 3.901369078 | 0.405519774 | 0.472694299 | 0.857890131 |
| Smim10l2a | 178.3604988 | -0.173533681 | 0.10841185 | -1.600689236 |
| Gm14633 | 2.099444623 | 0.480619203 | 0.677207264 | 0.709707689 |
| Gm2174 | 0.09349418 | 0.274402725 | 3.724945049 | 0.073666248 |
| Ddx26b | 661.1565071 | -0.226065889 | 0.131695932 | -1.716574577 |
| Gm10477 | 0 | NA | NA | NA |
| Gm648 | 0.240593192 | -1.495001879 | 2.543642901 | -0.587740472 |
| Mmgt1 | 503.4834972 | -0.176188657 | 0.063894661 | -2.757486362 |
| Mir5116 | 0 | NA | NA | NA |
| Slc9a6 | 1377.918565 | -0.213837084 | 0.057140262 | -3.742318942 |
| Gm26312 | 0 | NA | NA | NA |
| Fhl1 | 841.3587556 | -0.111110468 | 0.05924656 | -1.875391049 |
| Gm23258 | 1.567065625 | -1.993580552 | 0.917008704 | -2.174003959 |
| Mtap7d3 | 3.192641839 | 0.674882174 | 0.653208265 | 1.033180702 |
| Adgrg4 | 0.589632284 | 1.040253107 | 1.425045451 | 0.729978897 |
| Brs3 | 2.988176321 | 0.428632822 | 0.692551607 | 0.618918241 |
| Htatsf1 | 484.4912738 | -0.059438938 | 0.128794175 | -0.461503309 |
| Vgll1 | 0 | NA | NA | NA |
| Gm14718 | 0.17796667 | -0.684257857 | 2.715150213 | -0.252014733 |
| Cd40lg | 0.63650561 | 2.699771766 | 1.293413841 | 2.087322464 |
| Arhgef6 | 229.7557389 | 0.087767814 | 0.085236039 | 1.029703112 |
| Rbmx | 637.3674608 | -0.314167994 | 0.094805617 | -3.313812041 |
| Snord61 | 0.989940444 | 0.400563084 | 1.020281882 | 0.392600409 |
| Gm364 | 0.352737941 | 0.274377118 | 3.724945049 | 0.073659373 |
| Gpr101 | 120.5546104 | -0.559030886 | 0.151384553 | -3.692786855 |
| Gm14630 | 0.097139547 | 0.274393828 | 3.724945049 | 0.073663859 |
| Gm5389 | 0 | NA | NA | NA |
| Gm14629 | 0 | NA | NA | NA |
| Gm8107 | 0.038981395 | 0.274393828 | 3.724945049 | 0.073663859 |

| | | | | |
|---|---|---|---|---|
| Gm8111 | 0 | NA | NA | NA |
| Gm14631 | 0 | NA | NA | NA |
| Gm25023 | 0 | NA | NA | NA |
| Zic3 | 63.76716955 | -0.522215063 | 0.159280905 | -3.278579204 |
| Gm14649 | 0 | NA | NA | NA |
| Gm14647 | 0 | NA | NA | NA |
| Gm25851 | 0 | NA | NA | NA |
| Gm14645 | 0 | NA | NA | NA |
| Gm16436 | 0.046345709 | 0.274391604 | 3.724945049 | 0.073663262 |
| Gm14650 | 0.045882455 | 0.755288839 | 3.724945049 | 0.202765096 |
| Gm7071 | 0 | NA | NA | NA |
| AL713977.1 | 0 | NA | NA | NA |
| 4930550L24Rik | 0.32299001 | 0.27572125 | 1.956846722 | 0.14090079 |
| Fgf13 | 698.6388779 | -0.382773957 | 0.102707073 | -3.726850986 |
| Gm26351 | 1.389516516 | -1.039446784 | 1.006823071 | -1.032402628 |
| Mir504 | 1.452681833 | -0.292050385 | 0.78875794 | -0.370266174 |
| Gm26487 | 0 | NA | NA | NA |
| Gm14890 | 0.104106287 | 0.274400501 | 3.724945049 | 0.073665651 |
| AL670091.1 | 0 | NA | NA | NA |
| AL672036.1 | 0 | NA | NA | NA |
| Gm14887 | 0 | NA | NA | NA |
| F9 | 3.199446176 | 3.3065024 | 0.894028437 | 3.698430904 |
| Gm5637 | 10.86361093 | 1.43925515 | 0.487117391 | 2.954637172 |
| Gm14891 | 0.066050875 | 0.274391604 | 3.724945049 | 0.073663262 |
| Mcf2 | 115.7996196 | -0.400659598 | 0.184822914 | -2.167802621 |
| Atp11c | 543.5317974 | 0.13210781 | 0.082649504 | 1.598410194 |
| Gm22287 | 0 | NA | NA | NA |
| Mir505 | 2.930256519 | -0.940856924 | 0.657947487 | -1.429987869 |
| Gm7073 | 0 | NA | NA | NA |
| Gm715 | 17.31431649 | -0.260235758 | 0.230075921 | -1.131086456 |
| Gm14661 | 0.27928303 | 0.750201004 | 2.056224415 | 0.364843934 |
| Gm14651 | 0 | NA | NA | NA |
| Gm4909 | 0 | NA | NA | NA |
| Gm8163 | 0.296485438 | 0.274413924 | 2.095974795 | 0.130924248 |
| Gm14660 | 0 | NA | NA | NA |
| Sox3 | 13.97423569 | -0.154080981 | 0.301581385 | -0.510910119 |
| Gm14662 | 1.313401919 | -0.093446682 | 0.83691767 | -0.111655765 |
| Gm7076 | 0 | NA | NA | NA |
| Rpl7l1-ps1 | 0 | NA | NA | NA |
| Gm14664 | 0.089814205 | 0.274402725 | 3.724945049 | 0.073666248 |
| Gm14663 | 0.061920613 | 0.274393828 | 3.724945049 | 0.073663859 |
| C230004F18Rik | 1115.002913 | -0.238985661 | 0.161393468 | -1.480764143 |
| Gm24396 | 21.15961452 | 0.266704741 | 0.453381235 | 0.588257122 |
| Cdr1 | 62.47634879 | 0.131583345 | 0.484952501 | 0.27133244 |
| Gm14665 | 0.120974994 | 0.274404949 | 3.724945049 | 0.073666845 |
| Gm14666 | 0.43984171 | 1.631062683 | 1.417785242 | 1.150430005 |
| Gm14821 | 0.106112162 | -0.206492286 | 3.724945049 | -0.055434988 |

| | | | | |
|---|---|---|---|---|
| Gm14655 | 0.060174537 | 0.755286614 | 3.724945049 | 0.202764498 |
| Ldoc1 | 14.5236453 | -0.454134128 | 0.273405214 | -1.661029508 |
| Gm7327 | 29.50965119 | 0.065921791 | 0.303757296 | 0.217021261 |
| Gm6743 | 0.441426087 | -0.532207073 | 1.529284385 | -0.348010532 |
| Gm25412 | 0 | NA | NA | NA |
| Gm8215 | 0 | NA | NA | NA |
| Gm5390 | 0 | NA | NA | NA |
| Gm14656 | 0 | NA | NA | NA |
| Gm16411 | 0.044191549 | -0.206490061 | 3.724945049 | -0.055434391 |
| 4933402E13Rik | 0 | NA | NA | NA |
| Gm14679 | 0 | NA | NA | NA |
| 4931400O07Rik | 0 | NA | NA | NA |
| 1700019B21Rik | 0 | NA | NA | NA |
| Gm14678 | 0 | NA | NA | NA |
| Gm14677 | 0 | NA | NA | NA |
| Gm14680 | 169.5303447 | -0.184098136 | 0.093329576 | -1.972559428 |
| Gm8244 | 0 | NA | NA | NA |
| Gm6754 | 0 | NA | NA | NA |
| Gm14659 | 0 | NA | NA | NA |
| Gm14657 | 0 | NA | NA | NA |
| Gm8260 | 0 | NA | NA | NA |
| Gm6760 | 0 | NA | NA | NA |
| 3830417A13Rik | 0 | NA | NA | NA |
| Slitrk4 | 1198.015236 | -0.10545946 | 0.098479879 | -1.070873164 |
| Gm24598 | 1.413287166 | -0.837220299 | 0.792957818 | -1.055819464 |
| Gm14676 | 0 | NA | NA | NA |
| Gm26111 | 0 | NA | NA | NA |
| Gm14667 | 0 | NA | NA | NA |
| Gm14673 | 0 | NA | NA | NA |
| Ctag2 | 0 | NA | NA | NA |
| Gm14669 | 0 | NA | NA | NA |
| Gm14670 | 0 | NA | NA | NA |
| Gm14668 | 0 | NA | NA | NA |
| Gm14671 | 0 | NA | NA | NA |
| Gm14674 | 0 | NA | NA | NA |
| 4930447F04Rik | 0.792081435 | 0.387119028 | 1.168619303 | 0.331261881 |
| Gm23028 | 0 | NA | NA | NA |
| Slitrk2 | 714.9978112 | -0.021025454 | 0.096410562 | -0.218082477 |
| Gm23000 | 0.301026439 | 1.990480505 | 2.015938029 | 0.987371872 |
| Gm8310 | 0.044544648 | 0.755286614 | 3.724945049 | 0.202764498 |
| Mir743 | 0 | NA | NA | NA |
| Mir743b | 0 | NA | NA | NA |
| Gm14681 | 55.50849165 | 0.552041852 | 0.147577183 | 3.740699204 |
| Mir742 | 0 | NA | NA | NA |
| Mir883a | 0 | NA | NA | NA |
| Mir883b | 0 | NA | NA | NA |
| Mir471 | 0 | NA | NA | NA |

| | | | | |
|---|---|---|---|---|
| Mir741 | 0 | NA | NA | NA |
| Mir463 | 0 | NA | NA | NA |
| Mir880 | 0 | NA | NA | NA |
| Mir878 | 0 | NA | NA | NA |
| Mir881 | 0 | NA | NA | NA |
| Mir871 | 0 | NA | NA | NA |
| Mir470 | 0 | NA | NA | NA |
| mmu-mir-465d | 0 | NA | NA | NA |
| Mir465c-1 | 0 | NA | NA | NA |
| Mir465b-1 | 0 | NA | NA | NA |
| Mir465c-2 | 0 | NA | NA | NA |
| Mir465b-2 | 0 | NA | NA | NA |
| Mir465 | 0 | NA | NA | NA |
| Gm6783 | 0 | NA | NA | NA |
| Gm14682 | 0.089089296 | 1.194485492 | 3.723939576 | 0.320758559 |
| Gm8312 | 0 | NA | NA | NA |
| 1700036O09Rik | 0 | NA | NA | NA |
| Gm26276 | 0 | NA | NA | NA |
| Gm14687 | 0 | NA | NA | NA |
| Gm1140 | 0 | NA | NA | NA |
| Gm14692 | 0 | NA | NA | NA |
| Rpl7a-ps13 | 0 | NA | NA | NA |
| 4933436I01Rik | 0 | NA | NA | NA |
| Mir201 | 0 | NA | NA | NA |
| Mir547 | 0 | NA | NA | NA |
| Gm14688 | 0 | NA | NA | NA |
| mmu-mir-509 | 0 | NA | NA | NA |
| Gm4910 | 0 | NA | NA | NA |
| Gm5391 | 0 | NA | NA | NA |
| Gm6806 | 0.624564003 | 2.347912452 | 1.386757705 | 1.693094939 |
| Gm22332 | 0 | NA | NA | NA |
| Gm14701 | 0 | NA | NA | NA |
| Gm5638 | 0.108772582 | 0.75528439 | 3.724945049 | 0.202763901 |
| Fmr1os | 1.106201151 | 2.095258203 | 1.01188706 | 2.07064433 |
| Gm27979 | 0 | NA | NA | NA |
| Gm27838 | 0 | NA | NA | NA |
| Fmr1 | 1356.393382 | 0.390165212 | 0.076316518 | 5.112460845 |
| Gm22435 | 0 | NA | NA | NA |
| Fmr1nb | 0.234997993 | 1.256887608 | 2.122840449 | 0.592078227 |
| Gm14699 | 0 | NA | NA | NA |
| AC055766.1 | 0 | NA | NA | NA |
| Gm14698 | 5.615076243 | -0.288480287 | 0.415892818 | -0.693640944 |
| Gm6812 | 0 | NA | NA | NA |
| Gm8344 | 0 | NA | NA | NA |
| n-R5s7 | 0 | NA | NA | NA |
| Gm14705 | 0.713101696 | -1.295478977 | 1.242380414 | -1.042739376 |
| Aff2 | 361.4846116 | -0.067703063 | 0.08703107 | -0.777918317 |

| | | | | |
|---|---|---|---|---|
| Gm8172 | 2.258517424 | 0.353478903 | 0.739359589 | 0.47808794 |
| 1700111N16Rik | 0.153668717 | 0.755303435 | 2.839403532 | 0.266007782 |
| 1700020N15Rik | 0.116217011 | 0.274402725 | 3.724945049 | 0.073666248 |
| Ids | 5255.302793 | -0.041945102 | 0.070500098 | -0.594965151 |
| 1110012L19Rik | 65.12692351 | 0.08899798 | 0.18847654 | 0.472196593 |
| 4930567H17Rik | 0 | NA | NA | NA |
| Gm16407 | 0 | NA | NA | NA |
| Gm14704 | 0 | NA | NA | NA |
| Gm6823 | 0.187945415 | -0.206489201 | 2.684043509 | -0.076932136 |
| Gm16441 | 0 | NA | NA | NA |
| Gm14722 | 0 | NA | NA | NA |
| Gm14721 | 0 | NA | NA | NA |
| BC023829 | 354.2798382 | -0.160270781 | 0.095382752 | -1.680291023 |
| Gm14727 | 0 | NA | NA | NA |
| Gm14729 | 0 | NA | NA | NA |
| Gm6038 | 0 | NA | NA | NA |
| Gm8401 | 0.88643884 | 0.64811995 | 1.143654378 | 0.566709631 |
| Gm14726 | 0 | NA | NA | NA |
| Gm14724 | 0 | NA | NA | NA |
| Gm14725 | 0 | NA | NA | NA |
| Gm14728 | 0.144050924 | 1.290908285 | 3.721447108 | 0.346883416 |
| Gm14732 | 0 | NA | NA | NA |
| Gm14731 | 0 | NA | NA | NA |
| Rplp1-ps1 | 0.046345709 | 0.274391604 | 3.724945049 | 0.073663262 |
| Gm22785 | 0 | NA | NA | NA |
| Gm8410 | 0.503546025 | 0.43404144 | 1.436888125 | 0.302070448 |
| Gm6062 | 0.075043205 | 0.755286614 | 3.724945049 | 0.202764498 |
| Mamld1 | 657.3894617 | -0.172075173 | 0.095070677 | -1.80997104 |
| n-R5s8 | 8.139267235 | -0.055954054 | 0.365454269 | -0.153108223 |
| Gm14730 | 1.043158539 | -0.175145344 | 1.099174994 | -0.159342548 |
| Mtm1 | 330.8783525 | -0.119924452 | 0.135678477 | -0.88388707 |
| Mtmr1 | 659.908289 | 0.156458472 | 0.074461265 | 2.101206196 |
| Cd99l2 | 1057.193581 | -0.082180047 | 0.055596168 | -1.478160278 |
| Gm16189 | 0 | NA | NA | NA |
| Gm8421 | 0 | NA | NA | NA |
| Gm23945 | 0 | NA | NA | NA |
| Hmgb3 | 158.3977876 | 0.054636261 | 0.17054106 | 0.320370129 |
| Gpr50 | 5.432542524 | -1.063905001 | 0.449433328 | -2.3672143 |
| Gm14723 | 0 | NA | NA | NA |
| Rpl30-ps10 | 27.35008078 | 0.409570446 | 0.218905929 | 1.870988362 |
| Vma21 | 809.8468664 | 0.050717973 | 0.101558996 | 0.499394197 |
| Gm1141 | 6.002430894 | -1.000085641 | 0.482197311 | -2.074017458 |
| Prrg3 | 424.0651492 | -0.443333485 | 0.107640688 | -4.118642246 |
| Fate1 | 0.616736355 | -0.938780998 | 1.259559721 | -0.745324722 |
| Cnga2 | 3.751482543 | -0.963294012 | 0.785394748 | -1.226509363 |
| Magea4 | 0 | NA | NA | NA |
| Gabre | 35.49517603 | -0.105152378 | 0.255456799 | -0.411624894 |

| | | | | |
|---|---|---|---|---|
| Mir224 | 0.097139547 | 0.274393828 | 3.724945049 | 0.073663859 |
| Mir452 | 0.132574648 | -1.045763943 | 3.713921268 | -0.28157946 |
| Magea10 | 0 | NA | NA | NA |
| Gabra3 | 880.1056003 | -0.171267453 | 0.107017757 | -1.600364813 |
| Gm14735 | 0 | NA | NA | NA |
| Mir767 | 0.267172041 | -1.926887225 | 1.977856494 | -0.974230047 |
| Mir105 | 0.580314088 | -0.223063109 | 1.445770165 | -0.154286701 |
| Gm23461 | 0 | NA | NA | NA |
| Gm14736 | 0 | NA | NA | NA |
| Gabrq | 110.4557477 | -0.161382905 | 0.177140839 | -0.91104291 |
| Magea9-ps | 0 | NA | NA | NA |
| Cetn2 | 224.1480122 | -0.076285687 | 0.087262917 | -0.874205109 |
| Nsdhl | 258.6958972 | -0.126545173 | 0.0949609 | -1.332602933 |
| Gm14684 | 0.411174732 | -1.720048605 | 1.632728895 | -1.053480838 |
| Zfp185 | 38.8421771 | 0.823259153 | 0.204856437 | 4.018712647 |
| Mir2137 | 0.131193343 | -0.194387882 | 2.978303909 | -0.065267981 |
| Pnma5 | 2.351169846 | -0.243741614 | 0.665421745 | -0.366296436 |
| Pnma3 | 162.7565536 | -0.297762121 | 0.101724909 | -2.927130854 |
| Xlr4a | 1.613791847 | 0.676656314 | 1.218151561 | 0.555477935 |
| Xlr3a | 10.70218797 | 0.056081083 | 0.353274112 | 0.158746653 |
| Gm20455 | 0 | NA | NA | NA |
| Xlr5a | 0 | NA | NA | NA |
| Gm14685 | 1.153352726 | 0.071609237 | 1.009785223 | 0.070915314 |
| DXBay18 | 1.088618379 | -0.05043964 | 1.143441539 | -0.044112128 |
| Xlr5b | 0.044544648 | 0.755286614 | 3.724945049 | 0.202764498 |
| Spin2d | 0 | NA | NA | NA |
| Xlr3b | 10.24179207 | -0.083286251 | 0.871783004 | -0.09553553 |
| Xlr4b | 4.01574399 | 3.737196705 | 0.980056764 | 3.813245151 |
| F8a | 148.8077908 | 0.085039552 | 0.147315501 | 0.577261401 |
| Xlr4c | 1.800570423 | 3.906032017 | 1.071030458 | 3.646985001 |
| Xlr3c | 1.249216686 | 1.404128187 | 1.254011174 | 1.11970947 |
| Xlr5c | 0 | NA | NA | NA |
| Gm5639 | 0.465319851 | 1.192878003 | 1.324367224 | 0.900715437 |
| Gm8501 | 0 | NA | NA | NA |
| Zfp275 | 599.4099204 | -0.020867259 | 0.072727391 | -0.286924343 |
| Gm18336 | 1.728761539 | 1.854469334 | 0.933318187 | 1.986963673 |
| Gm26726 | 0.597994088 | 2.614510087 | 1.471758403 | 1.776453311 |
| Gm37741 | 0.240513512 | 2.072179727 | 2.473954268 | 0.837598234 |
| Zfp92 | 61.40182776 | 0.067559566 | 0.169170838 | 0.399357044 |
| Trex2 | 0.567095806 | 0.287506005 | 1.342144854 | 0.214213842 |
| Haus7 | 77.87449152 | 0.100852866 | 0.119946173 | 0.840817709 |
| Bgn | 275.5973356 | 0.807880764 | 0.183853493 | 4.3941551 |
| Atp2b3 | 1878.820538 | -0.202322504 | 0.092438092 | -2.188735189 |
| Dusp9 | 7.63352917 | 0.19097436 | 0.424466472 | 0.449916243 |
| Pnck | 377.2175805 | -0.22496391 | 0.073349607 | -3.067009068 |
| Slc6a8 | 1645.774035 | 0.073363096 | 0.064702006 | 1.133861217 |
| Bcap31 | 456.597487 | 0.076467394 | 0.106751118 | 0.716314693 |

| | | | | |
|---|---|---|---|---|
| Abcd1 | 224.9704374 | 0.430669668 | 0.119837665 | 3.593775514 |
| Gm25226 | 0 | NA | NA | NA |
| Plxnb3 | 370.4438468 | -0.352628266 | 0.120160402 | -2.934646189 |
| Srpk3 | 30.04135515 | -0.702194492 | 0.23808903 | -2.94929377 |
| Idh3g | 816.0853577 | 0.005513837 | 0.049097553 | 0.112303698 |
| Ssr4 | 382.7763301 | 0.333330347 | 0.101196257 | 3.293899949 |
| Pdzd4 | 1049.261182 | -0.150933894 | 0.083750776 | -1.802179055 |
| Gm14817 | 2.398130883 | 0.636477389 | 0.722560959 | 0.880863242 |
| Gm23615 | 0.046345709 | 0.274391604 | 3.724945049 | 0.073663262 |
| L1cam | 1414.37122 | -0.377484702 | 0.065665356 | -5.748612694 |
| Arhgap4 | 118.1046268 | 0.638556261 | 0.174480444 | 3.659758343 |
| Avpr2 | 0.170941961 | 0.75529616 | 2.738626201 | 0.275793812 |
| Naa10 | 112.0862865 | -0.314831948 | 0.114445807 | -2.75092601 |
| Renbp | 87.59544553 | 0.988070263 | 0.188534983 | 5.240779428 |
| Hcfc1 | 2331.747261 | 0.137292411 | 0.054204921 | 2.532840347 |
| Gm8545 | 0.428564514 | -0.677260946 | 1.550245609 | -0.43687332 |
| Irak1 | 580.922968 | 0.147031264 | 0.068018712 | 2.161629633 |
| Mir5132 | 0 | NA | NA | NA |
| Mir718 | 0.042716945 | -0.206490061 | 3.724945049 | -0.055434391 |
| Mecp2 | 3093.872843 | 0.127520167 | 0.050558152 | 2.522247393 |
| Opn1mw | 3.912128538 | -0.106074508 | 0.552653652 | -0.191936681 |
| Gm6758 | 1.061458158 | 0.075734409 | 0.956652406 | 0.079166068 |
| Tex28 | 0.045882455 | 0.755288839 | 3.724945049 | 0.202765096 |
| Rpl3-ps2 | 16.62506886 | 0.717401385 | 0.272233128 | 2.63524645 |
| Tktl1 | 1.027298454 | 3.37297736 | 1.135351732 | 2.970865561 |
| Flna | 2140.441872 | 0.940267907 | 0.140921471 | 6.672282796 |
| Emd | 326.2260813 | 0.315843792 | 0.117249344 | 2.693778752 |
| Gm14686 | 0 | NA | NA | NA |
| Rpl10 | 1410.009698 | 0.348855201 | 0.096583721 | 3.611946171 |
| Snora70 | 0 | NA | NA | NA |
| Dnase1l1 | 38.9238987 | 1.29348376 | 0.278066246 | 4.651710805 |
| Mir7091 | 0 | NA | NA | NA |
| Gm14846 | 0 | NA | NA | NA |
| Taz | 323.2193373 | -0.215484408 | 0.127594223 | -1.688825741 |
| Atp6ap1 | 2828.059804 | 0.039247838 | 0.055707767 | 0.704530814 |
| Gdi1 | 4514.466658 | -0.133580431 | 0.044762591 | -2.984197947 |
| Fam50a | 152.1449703 | 0.026733716 | 0.106196492 | 0.251738217 |
| mmu-mir-7092 | 0.044191549 | -0.206490061 | 3.724945049 | -0.055434391 |
| Plxna3 | 360.6800962 | -0.428692482 | 0.091658274 | -4.677073482 |
| Lage3 | 80.66907545 | -0.111256897 | 0.152208804 | -0.73094916 |
| Ubl4 | 481.5895494 | -0.105480672 | 0.079049877 | -1.334355924 |
| Slc10a3 | 39.38722088 | 0.506165335 | 0.237563433 | 2.130653395 |
| Fam3a | 178.954657 | -0.168008426 | 0.130669481 | -1.285751081 |
| Ikbkg | 603.2886534 | 0.245789176 | 0.079555386 | 3.089535325 |
| G6pdx | 686.6621862 | 0.621818524 | 0.150722114 | 4.125595827 |
| Gm6880 | 0.066050875 | 0.274391604 | 3.724945049 | 0.073663262 |
| Gm6881 | 0 | NA | NA | NA |

| | | | | |
|---|---|---|---|---|
| Gm15361 | 20.55718455 | -0.316055936 | 0.26051346 | -1.213203863 |
| Gm15362 | 0 | NA | NA | NA |
| Gm15364 | 0 | NA | NA | NA |
| Gm26125 | 0 | NA | NA | NA |
| Gm4989 | 0 | NA | NA | NA |
| Olfr1326-ps1 | 0 | NA | NA | NA |
| Olfr1325 | 0 | NA | NA | NA |
| Gm26417 | 0 | NA | NA | NA |
| Gm5640 | 1.316030826 | 0.744462654 | 0.871612729 | 0.854120907 |
| Xlr5d-ps | 0.066050875 | 0.274391604 | 3.724945049 | 0.073663262 |
| Spin2-ps2 | 0 | NA | NA | NA |
| Xlr3d-ps | 0.16282664 | 0.755289343 | 3.41893252 | 0.220913791 |
| Gm15365 | 0.152853976 | 1.641852012 | 2.820813903 | 0.582049036 |
| Gm22351 | 0 | NA | NA | NA |
| Xlr4d-ps | 0 | NA | NA | NA |
| Gm6890 | 0 | NA | NA | NA |
| Gm14755 | 0 | NA | NA | NA |
| Olfr1327-ps1 | 0 | NA | NA | NA |
| Gm14758 | 0 | NA | NA | NA |
| Gm14695 | 0 | NA | NA | NA |
| Gm8617 | 0 | NA | NA | NA |
| Spin2-ps3 | 0 | NA | NA | NA |
| Xlr5e-ps | 0 | NA | NA | NA |
| Gm5936 | 1.015210553 | 0.799948227 | 1.079823044 | 0.740814184 |
| Gm14694 | 0 | NA | NA | NA |
| Gm6897 | 0 | NA | NA | NA |
| Gm14756 | 0 | NA | NA | NA |
| Spin2-ps4 | 0 | NA | NA | NA |
| Xlr3e-ps | 0.045882455 | 0.755288839 | 3.724945049 | 0.202765096 |
| Xlr4e-ps | 0.114443069 | -0.206492286 | 3.724945049 | -0.055434988 |
| Gm15363 | 0 | NA | NA | NA |
| Gab3 | 34.85498717 | 0.367254575 | 0.198321581 | 1.851813476 |
| Gm25629 | 0.361882898 | 1.493731208 | 1.854648342 | 0.805398616 |
| Gm15367 | 0 | NA | NA | NA |
| Dkc1 | 249.3852457 | -0.18047981 | 0.089624652 | -2.01372954 |
| Gm25520 | 1.152914642 | -0.938764327 | 0.961171428 | -0.976687716 |
| Mpp1 | 513.8380435 | 0.590300275 | 0.097307447 | 6.06634223 |
| Gm15378 | 0.117046891 | -0.687371727 | 3.724945049 | -0.184532045 |
| Fam220-ps | 0.313388536 | 1.354077336 | 2.016859421 | 0.671379136 |
| Smim9 | 2.883149428 | 1.555157985 | 0.724164123 | 2.147521445 |
| Gm8503 | 0.097139547 | 0.274393828 | 3.724945049 | 0.073663859 |
| F8 | 100.3533705 | -0.017007849 | 0.151420184 | -0.112322208 |
| Gm6039 | 5.682746135 | 0.317725998 | 0.418839303 | 0.758586875 |
| Gm8522 | 1.810659408 | -0.927150667 | 0.744092485 | -1.246015362 |
| Fundc2 | 533.8816821 | 0.189908847 | 0.104899656 | 1.810385789 |
| Mtcp1 | 186.8893314 | -0.22378514 | 0.130350088 | -1.716800843 |
| Brcc3 | 516.7646188 | 0.233784093 | 0.073935505 | 3.162000335 |

| | | | | |
|---|---|---|---|---|
| Gm15379 | 11.17773374 | -0.089962859 | 0.323409439 | -0.278170172 |
| Gm15385 | 2.75264805 | 1.088838688 | 0.742003714 | 1.46743024 |
| Gm8666 | 2.699454663 | 0.410960658 | 0.626798429 | 0.655650427 |
| Vbp1 | 703.5174424 | 0.082228377 | 0.094301453 | 0.871973595 |
| Gm15384 | 1.213939271 | -0.912381801 | 1.021079078 | -0.893546662 |
| Rab39b | 757.2087334 | 0.098413536 | 0.088987871 | 1.105920778 |
| Gm15063 | 0.98952171 | 2.907751462 | 1.236398069 | 2.351792302 |
| Gm25421 | 0 | NA | NA | NA |
| Gm15065 | 1.438276361 | -1.55745702 | 1.005258619 | -1.54930979 |
| 4933407K13Rik | 98.06458889 | 0.540134492 | 0.254781713 | 2.119989249 |
| Gm7153 | 0.141331096 | -0.206491164 | 3.56184083 | -0.057973159 |
| Pls3 | 1303.00074 | -0.017098401 | 0.061831158 | -0.276533728 |
| Gm23320 | 0 | NA | NA | NA |
| Oat-rs1 | 0 | NA | NA | NA |
| Gm14715 | 0 | NA | NA | NA |
| Gm7155 | 0 | NA | NA | NA |
| Gm8692 | 1.34351349 | 0.176139225 | 0.89827417 | 0.196086263 |
| Gm14710 | 0 | NA | NA | NA |
| Gm5397 | 0 | NA | NA | NA |
| Gm14711 | 0.062426873 | 0.755288839 | 3.724945049 | 0.202765096 |
| Gm14707 | 0 | NA | NA | NA |
| Gm14717 | 0 | NA | NA | NA |
| Cldn34b3 | 0.209261529 | 1.220404626 | 2.616217222 | 0.466476796 |
| Gm14712 | 0 | NA | NA | NA |
| Magea7-ps | 0.046345709 | 0.274391604 | 3.724945049 | 0.073663262 |
| Gm14709 | 0 | NA | NA | NA |
| Cldn34b4 | 0 | NA | NA | NA |
| Gm5793 | 0.038981395 | 0.274393828 | 3.724945049 | 0.073663859 |
| Cldn34d | 0.044191549 | -0.206490061 | 3.724945049 | -0.055434391 |
| Gm14716 | 0 | NA | NA | NA |
| Gm14713 | 0 | NA | NA | NA |
| Gm26041 | 0.046345709 | 0.274391604 | 3.724945049 | 0.073663262 |
| Gm23280 | 0 | NA | NA | NA |
| Gm14706 | 0.190396633 | 1.25932864 | 3.267946179 | 0.385357827 |
| Gm14714 | 0 | NA | NA | NA |
| Gm5398 | 2.350384682 | -0.242978922 | 0.717618542 | -0.33859064 |
| Gm5757 | 0 | NA | NA | NA |
| Gm5937 | 0.810948363 | 0.539261829 | 1.121960731 | 0.480642338 |
| Tbl1x | 1255.905752 | 0.324202433 | 0.053157986 | 6.098847176 |
| Gm23121 | 6.016926621 | 0.034452401 | 0.402615762 | 0.085571417 |
| Cldn34-ps | 0 | NA | NA | NA |
| Gm14746 | 0.224947322 | -0.239275989 | 2.154193279 | -0.111074522 |
| Gm14748 | 0 | NA | NA | NA |
| Gm8722 | 2.37024928 | -1.381854659 | 0.768470027 | -1.798189403 |
| Prkx | 384.5278709 | 0.37093679 | 0.140565125 | 2.638896318 |
| Gm14742 | 0.093141081 | -0.687373951 | 3.724945049 | -0.184532642 |
| Pbsn | 0 | NA | NA | NA |

| | | | | |
|---|---|---|---|---|
| Gm14744 | 0.130840244 | -1.039642477 | 2.947200097 | -0.352755986 |
| 5430402E10Rik | 0.090665643 | -0.687373951 | 3.724945049 | -0.184532642 |
| Gm14750 | 0 | NA | NA | NA |
| Gm14753 | 0 | NA | NA | NA |
| Gm16458 | 0 | NA | NA | NA |
| Obp1a | 0.143846094 | -0.985158148 | 3.718482058 | -0.264935566 |
| Gm5938 | 0 | NA | NA | NA |
| Obp1b | 0.091880448 | -0.687373951 | 3.724945049 | -0.184532642 |
| Gm14743 | 0.116046057 | -0.687371727 | 3.724945049 | -0.184532045 |
| Gm8749 | 0 | NA | NA | NA |
| 4930480E11Rik | 0.066050875 | 0.274391604 | 3.724945049 | 0.073663262 |
| Gm14749 | 0 | NA | NA | NA |
| Gm8754 | 0 | NA | NA | NA |
| Gm14752 | 0 | NA | NA | NA |
| Gm14745 | 0 | NA | NA | NA |
| Prrg1 | 108.3946446 | -1.070096963 | 0.155218712 | -6.89412346 |
| Gm22843 | 1.022987396 | 0.006939435 | 0.900043543 | 0.007710111 |
| Gm8757 | 0 | NA | NA | NA |
| Gm8761 | 0 | NA | NA | NA |
| Gm14747 | 0 | NA | NA | NA |
| Fam47c | 0 | NA | NA | NA |
| Gm8770 | 0.097139547 | 0.274393828 | 3.724945049 | 0.073663859 |
| Gm5939 | 0.62385056 | 1.4439606 | 1.253257078 | 1.152166323 |
| Gm23556 | 0 | NA | NA | NA |
| Gm14751 | 0 | NA | NA | NA |
| Rps24-ps3 | 11.66239001 | 0.781857189 | 0.294508419 | 2.654787233 |
| Gm7173 | 57.33673308 | -0.018960274 | 0.159898414 | -0.118576996 |
| Mageb16 | 1.972874096 | 0.930474535 | 0.776799258 | 1.19783139 |
| Gm4772 | 0.201649101 | -1.13452089 | 2.200631607 | -0.515543304 |
| Gm4773 | 0.396881693 | 1.229632864 | 1.543636742 | 0.796581755 |
| Gm14760 | 16.61937256 | 0.558336147 | 0.257798367 | 2.165786207 |
| Gm6952 | 0 | NA | NA | NA |
| Gm14768 | 0 | NA | NA | NA |
| Gm5940 | 3.69570757 | 0.342022105 | 0.570047604 | 0.599988672 |
| Gm8806 | 5.085933841 | 0.524274541 | 0.510840065 | 1.026298789 |
| Gm6954 | 0 | NA | NA | NA |
| Gm14733 | 0 | NA | NA | NA |
| Gm26775 | 0.462969528 | -0.583213135 | 1.45232732 | -0.401571414 |
| Tmem47 | 1344.966627 | -0.297726076 | 0.128403299 | -2.318679343 |
| Gm14754 | 0 | NA | NA | NA |
| 4930595M18Rik | 0.072025462 | 0.755288839 | 3.724945049 | 0.202765096 |
| Gm8822 | 164.0860792 | 0.457334892 | 0.132266203 | 3.457685195 |
| Gm26174 | 0 | NA | NA | NA |
| Gm22686 | 0 | NA | NA | NA |
| Rpl10-ps4 | 0 | NA | NA | NA |
| Gm24706 | 0 | NA | NA | NA |
| Gm14740 | 12.83618259 | -0.570610002 | 0.773617335 | -0.737586887 |

| | | | | |
|---|---|---|---|---|
| Gm14739 | 0 | NA | NA | NA |
| Gm24470 | 0 | NA | NA | NA |
| Gm14741 | 0 | NA | NA | NA |
| Gm23671 | 0 | NA | NA | NA |
| Dmd | 2142.472415 | -0.068520196 | 0.100484058 | -0.681901161 |
| n-R5s9 | 0.908782567 | 1.435256812 | 1.064259213 | 1.348597029 |
| Tsga8 | 0 | NA | NA | NA |
| Gm22580 | 0.283050572 | 0.27441596 | 1.646635187 | 0.166652555 |
| Gm14769 | 8.178611007 | -0.032378762 | 0.438909091 | -0.073770999 |
| Gm14770 | 2.258554499 | -0.509392552 | 0.593636429 | -0.858088431 |
| Gm14771 | 0 | NA | NA | NA |
| Gm16463 | 0.470240119 | 0.739793491 | 1.593993689 | 0.464113187 |
| 1600014K23Rik | 0 | NA | NA | NA |
| Gm14765 | 0 | NA | NA | NA |
| Tab3 | 521.751977 | 0.273613971 | 0.090699806 | 3.016698529 |
| AL645586.1 | 2.461055123 | 0.187417184 | 0.728088472 | 0.257409905 |
| Gyk | 504.0204661 | 0.027180574 | 0.056831617 | 0.47826501 |
| Gm14767 | 0.044544648 | 0.755286614 | 3.724945049 | 0.202764498 |
| Gm14764 | 0 | NA | NA | NA |
| Gm14766 | 0.576946983 | -0.022989126 | 1.14045153 | -0.020157916 |
| Gm6964 | 0 | NA | NA | NA |
| Gm14761 | 0.106520246 | 0.755282166 | 3.724945049 | 0.202763304 |
| Gm14762 | 0.702946434 | -0.49613646 | 1.291283241 | -0.384219701 |
| 5430427O19Rik | 37.85834025 | 3.059543084 | 0.336988132 | 9.079082602 |
| Gm14772 | 0 | NA | NA | NA |
| Samt3 | 0.047948698 | -0.206487837 | 3.724945049 | -0.055433794 |
| Nr0b1 | 0.363180662 | 1.142807947 | 1.530348998 | 0.74676296 |
| Mageb4 | 0.097139547 | 0.274393828 | 3.724945049 | 0.073663859 |
| Gm8839 | 0 | NA | NA | NA |
| Mageb10-ps | 0 | NA | NA | NA |
| Gm8844 | 0.25781401 | 0.239224385 | 1.821347396 | 0.131344732 |
| Mageb8-ps | 0 | NA | NA | NA |
| Gm41 | 0 | NA | NA | NA |
| Mageb6-ps | 0 | NA | NA | NA |
| Mageb7-ps | 0 | NA | NA | NA |
| n-R5s10 | 0 | NA | NA | NA |
| Il1rapl1 | 848.3376803 | -0.062435716 | 0.110653117 | -0.564247244 |
| Gm14775 | 0 | NA | NA | NA |
| Gm8855 | 54.40271974 | -0.626142379 | 0.251500081 | -2.489630926 |
| Gm14776 | 77.87756624 | -0.538889919 | 0.294301225 | -1.831082828 |
| Gm24812 | 2.806226544 | -0.726217676 | 0.667155367 | -1.088528567 |
| Gm14774 | 28.40504812 | -0.679473141 | 0.284252338 | -2.390387167 |
| Gm25006 | 0 | NA | NA | NA |
| Gm8858 | 0 | NA | NA | NA |
| Gm14793 | 0 | NA | NA | NA |
| Gm8864 | 0.154920219 | 0.777094091 | 2.834936002 | 0.274113451 |
| Gm14795 | 0 | NA | NA | NA |

| | | | | |
|---|---|---|---|---|
| Gm14792 | 0 | NA | NA | NA |
| Smek3-ps | 0 | NA | NA | NA |
| Gm14794 | 0.838406277 | -0.984378205 | 1.172986262 | -0.839206934 |
| Gm8876 | 0 | NA | NA | NA |
| Gm27000 | 21.60956527 | -0.683467715 | 0.385252355 | -1.774077967 |
| Mir1906-2 | 0 | NA | NA | NA |
| Gm27335 | 0 | NA | NA | NA |
| Gm5759 | 0 | NA | NA | NA |
| Gm8880 | 0.146451277 | -0.68737672 | 2.882358805 | -0.238477152 |
| Gm16412 | 0.045882455 | 0.755288839 | 3.724945049 | 0.202765096 |
| Gm14791 | 0 | NA | NA | NA |
| Gm14796 | 0 | NA | NA | NA |
| Gm14797 | 0 | NA | NA | NA |
| Gm649 | 0 | NA | NA | NA |
| Gm14805 | 0 | NA | NA | NA |
| Pet2 | 0 | NA | NA | NA |
| Gm14804 | 0 | NA | NA | NA |
| Gm6973 | 0 | NA | NA | NA |
| 4932429P05Rik | 0 | NA | NA | NA |
| Gm7108 | 0.816803644 | 1.674560019 | 1.120512104 | 1.494459555 |
| Gm24954 | 0 | NA | NA | NA |
| Nanog-ps1 | 0 | NA | NA | NA |
| 4930415L06Rik | 0 | NA | NA | NA |
| Gm6027 | 0 | NA | NA | NA |
| Gm4746 | 0 | NA | NA | NA |
| Gm6977 | 10.13847199 | 1.001739338 | 0.408515519 | 2.452145125 |
| Gm44 | 0 | NA | NA | NA |
| Gm14773 | 0 | NA | NA | NA |
| Gm24835 | 0 | NA | NA | NA |
| Gm14786 | 0.042716945 | -0.206490061 | 3.724945049 | -0.055434391 |
| Gm8908 | 0 | NA | NA | NA |
| Gm14788 | 0 | NA | NA | NA |
| Mageb2 | 0.045882455 | 0.755288839 | 3.724945049 | 0.202765096 |
| Gm14783 | 0.291614092 | 0.755310462 | 1.778857155 | 0.424604337 |
| Gm5072 | 0 | NA | NA | NA |
| Gm8914 | 0 | NA | NA | NA |
| Gm14787 | 0 | NA | NA | NA |
| Gm8916 | 0 | NA | NA | NA |
| Gm14785 | 0 | NA | NA | NA |
| 1700084M14Rik | 0 | NA | NA | NA |
| Mageb17-ps | 0 | NA | NA | NA |
| Gm14781 | 0 | NA | NA | NA |
| Mageb5 | 0 | NA | NA | NA |
| Gm14782 | 0 | NA | NA | NA |
| Gm7113 | 0 | NA | NA | NA |
| Gm16465 | 0.297592784 | 0.703271246 | 1.608072868 | 0.437337922 |
| Gm5071 | 0 | NA | NA | NA |

| | | | | |
|---|---|---|---|---|
| Gm14784 | 0 | NA | NA | NA |
| Mageb1 | 0.062426873 | 0.755288839 | 3.724945049 | 0.202765096 |
| Gm4911 | 0 | NA | NA | NA |
| Gm4912 | 0.306841164 | 0.755304244 | 2.018021183 | 0.374279641 |
| Mageb18 | 0.274198742 | 0.626796938 | 1.808329691 | 0.346616516 |
| Gm14777 | 0 | NA | NA | NA |
| Gm5941 | 0 | NA | NA | NA |
| Gm6982 | 0 | NA | NA | NA |
| Gm25332 | 0 | NA | NA | NA |
| Gm24973 | 0.33162218 | -1.410046247 | 1.720545792 | -0.819534274 |
| 1700003E24Rik | 0 | NA | NA | NA |
| Gm24123 | 0 | NA | NA | NA |
| BC061195 | 0 | NA | NA | NA |
| Gm23454 | 0 | NA | NA | NA |
| Arx | 155.2305208 | -0.336905551 | 0.170811975 | -1.972376652 |
| Pola1 | 222.1831865 | 0.054277176 | 0.125643257 | 0.431994344 |
| Gm14780 | 0 | NA | NA | NA |
| Gm23557 | 1.675207292 | -1.758004463 | 0.851450748 | -2.064716563 |
| Gm6985 | 0 | NA | NA | NA |
| Pcyt1b | 616.1736765 | -0.075045828 | 0.067880892 | -1.105551596 |
| Pdk3 | 429.9123672 | 0.115706118 | 0.065452353 | 1.767791582 |
| Supt20-ps | 0 | NA | NA | NA |
| Gm15164 | 0 | NA | NA | NA |
| AU015836 | 0.093831153 | 0.274404949 | 3.724945049 | 0.073666845 |
| Gm14790 | 0 | NA | NA | NA |
| Gm15165 | 0.044191549 | -0.206490061 | 3.724945049 | -0.055434391 |
| Gm6987 | 0 | NA | NA | NA |
| Gm14798 | 0.269031123 | 1.213956081 | 1.798199927 | 0.675095168 |
| Zfx | 737.8114658 | 0.090345111 | 0.093404451 | 0.967246323 |
| Eif2s3x | 977.9631196 | 0.245318815 | 0.074043729 | 3.313161241 |
| Klhl15 | 561.3122414 | -0.037473852 | 0.063900287 | -0.586442613 |
| Mir7673 | 0 | NA | NA | NA |
| Gm5400 | 0.277195307 | -0.206487002 | 2.101880076 | -0.098239193 |
| Gm14826 | 0 | NA | NA | NA |
| Gm14825 | 0.060174537 | 0.755286614 | 3.724945049 | 0.202764498 |
| 4932442L08Rik | 2.026572498 | 0.801734091 | 0.774164982 | 1.035611413 |
| Apoo | 208.2198411 | -0.259143306 | 0.10989803 | -2.358034133 |
| AL627302.1 | 0 | NA | NA | NA |
| Gm9003 | 0 | NA | NA | NA |
| Gm14827 | 54.27178075 | -1.142028034 | 0.396842655 | -2.877785489 |
| Gm24595 | 0 | NA | NA | NA |
| Maged1 | 4444.520797 | -0.049406707 | 0.055113567 | -0.896452729 |
| Gm9005 | 0 | NA | NA | NA |
| Gm14831 | 0.038981395 | 0.274393828 | 3.724945049 | 0.073663859 |
| Gm5401 | 0.15966113 | -0.206492938 | 3.449727875 | -0.059857747 |
| Gm9009 | 0.046345709 | 0.274391604 | 3.724945049 | 0.073663262 |
| Gspt2 | 293.6766047 | -0.137187295 | 0.083963064 | -1.633900536 |

| | | | | |
|---|---|---|---|---|
| Gm5402 | 0.046345709 | 0.274391604 | 3.724945049 | 0.073663262 |
| Zxdb | 233.650166 | 0.093959092 | 0.098407393 | 0.954797089 |
| Gm7061 | 8.775414701 | -0.544067376 | 0.450542322 | -1.207583284 |
| Gm14830 | 3.497870509 | -0.220942019 | 0.491714215 | -0.449330145 |
| Rpl13-ps5 | 0.088383099 | -0.687376175 | 3.724945049 | -0.184533239 |
| Gm26617 | 0.179489757 | 0.740028 | 2.288344906 | 0.323390061 |
| Zxda | 5.296744549 | 0.284845585 | 0.428132114 | 0.665321697 |
| Gm371 | 0.049553047 | 0.274391604 | 3.724945049 | 0.073663262 |
| Spin4 | 40.24504316 | 1.028409828 | 0.24277598 | 4.236044399 |
| Arhgef9 | 4283.077243 | -0.150027955 | 0.10168076 | -1.475480269 |
| Gm6092 | 0.047948698 | -0.206487837 | 3.724945049 | -0.055433794 |
| Gm14807 | 0.075043205 | 0.755286614 | 3.724945049 | 0.202764498 |
| Gm3712 | 0.119587853 | 1.218392466 | 3.723618458 | 0.327206581 |
| Gm14806 | 0.045882455 | 0.755288839 | 3.724945049 | 0.202765096 |
| Amer1 | 313.2545163 | 0.227036693 | 0.099994435 | 2.270493282 |
| Asb12 | 0.245805814 | 0.204722109 | 2.118679232 | 0.096627232 |
| Gm14818 | 0 | NA | NA | NA |
| Gm9043 | 0 | NA | NA | NA |
| Zc4h2 | 203.86252 | 0.006460752 | 0.099134257 | 0.06517174 |
| Zc3h12b | 477.9337351 | -0.376814566 | 0.117028336 | -3.219857502 |
| 1700010D01Rik | 0 | NA | NA | NA |
| Las1l | 521.4809968 | 0.059363827 | 0.117677542 | 0.50446182 |
| Msn | 2032.338517 | 2.552564068 | 0.188818176 | 13.51863532 |
| Gm24718 | 0 | NA | NA | NA |
| Gm14925 | 0.293430937 | -0.19105372 | 2.033018266 | -0.093975407 |
| Gm9051 | 0 | NA | NA | NA |
| F630028O10Rik | 26.3940196 | 1.376415256 | 0.464707132 | 2.961898285 |
| Mir223 | 0 | NA | NA | NA |
| Vsig4 | 0.106465261 | 0.75528439 | 3.724945049 | 0.202763901 |
| Hsf3 | 1.054375976 | 0.91860646 | 0.946570355 | 0.970457669 |
| Gm14933 | 0.042716945 | -0.206490061 | 3.724945049 | -0.055434391 |
| Heph | 82.76888646 | 0.474331293 | 0.188790595 | 2.512473111 |
| Rpl17-ps8 | 13.95711591 | 0.560131898 | 0.279074107 | 2.007108089 |
| Gm14935 | 0.695213882 | -0.349363923 | 1.13305378 | -0.308338341 |
| Gpr165 | 370.7971054 | 0.560856555 | 0.130004478 | 4.314132573 |
| Gm16394 | 0 | NA | NA | NA |
| Gm14803 | 0 | NA | NA | NA |
| Gm9059 | 0 | NA | NA | NA |
| Gm9060 | 0 | NA | NA | NA |
| Pgr15l | 28.47859877 | 0.367238051 | 0.288600939 | 1.272476978 |
| Gm14800 | 0 | NA | NA | NA |
| Eda2r | 27.60206714 | 1.543881589 | 0.419231337 | 3.682648347 |
| Gm9062 | 0 | NA | NA | NA |
| Gm14802 | 0.124347486 | 0.755286736 | 3.717316019 | 0.203180664 |
| Ar | 593.9010294 | 0.656009668 | 0.122065315 | 5.37425122 |
| Gm14811 | 0 | NA | NA | NA |
| Gm14815 | 0.75340593 | 1.831758011 | 1.209968363 | 1.513889178 |

| | | | | |
|---|---|---|---|---|
| Gm14810 | 2.604223114 | 1.958569068 | 0.816192848 | 2.399640076 |
| Ophn1 | 626.2746632 | -0.194420553 | 0.089327383 | -2.176494446 |
| Gm27554 | 3.06361838 | -1.016711302 | 0.689077087 | -1.47546816 |
| Gm26000 | 0.038981395 | 0.274393828 | 3.724945049 | 0.073663859 |
| Yipf6 | 767.4798748 | 0.213782511 | 0.070516919 | 3.031648476 |
| Gm14813 | 0.038981395 | 0.274393828 | 3.724945049 | 0.073663859 |
| Stard8 | 229.0060904 | -0.501122016 | 0.160707054 | -3.118232853 |
| Gm5760 | 0 | NA | NA | NA |
| Efnb1 | 173.9851955 | -0.078788572 | 0.128219459 | -0.614482177 |
| Gm14812 | 1.182411106 | 3.079798263 | 3.660292437 | 0.841407706 |
| Gm14809 | 0.233612452 | 1.697742765 | 2.527723224 | 0.671648996 |
| Gm14808 | 0.179754939 | 0.235146722 | 2.738727615 | 0.085859843 |
| Pja1 | 1595.105195 | -0.021195684 | 0.082682246 | -0.25635109 |
| Gm14816 | 0.266066471 | -0.206481877 | 2.094083908 | -0.098602485 |
| Pcna-ps1 | 0.075043205 | 0.755286614 | 3.724945049 | 0.202764498 |
| Tmem28 | 262.17291 | -0.18017612 | 0.104581308 | -1.722832925 |
| Eda | 61.73840336 | 0.110370078 | 0.158703961 | 0.695446272 |
| Gm14832 | 1.244840331 | -0.386253996 | 0.924476714 | -0.417808248 |
| Gm14829 | 1.581404248 | -0.506504034 | 0.72775035 | -0.695985971 |
| Gm14828 | 1.203205628 | -0.446866166 | 0.909176594 | -0.491506457 |
| Mir676 | 0 | NA | NA | NA |
| Awat2 | 2.157021785 | 0.501092384 | 0.778515779 | 0.643650902 |
| Otud6a | 0 | NA | NA | NA |
| Igbp1 | 392.0809371 | 0.32244439 | 0.074731248 | 4.314719709 |
| Dgat2l6 | 5.685626011 | -1.051509755 | 0.562362144 | -1.869808923 |
| Awat1 | 0.271018715 | 0.274412376 | 1.835610563 | 0.149493788 |
| Gm6175 | 0 | NA | NA | NA |
| P2ry4 | 1.586926032 | 2.923568894 | 1.064121518 | 2.74740135 |
| Arr3 | 2.257394224 | -1.97717686 | 0.740838878 | -2.668835179 |
| Pdzd11 | 261.6832867 | 0.312872148 | 0.08537607 | 3.66463518 |
| Kif4 | 26.59334305 | 1.154286712 | 0.355369793 | 3.248128384 |
| Gdpd2 | 50.75686142 | -0.910061038 | 0.274279782 | -3.318002634 |
| Gm14866 | 1.936928019 | 0.231776878 | 0.842551645 | 0.275089224 |
| Gm14902 | 0.875624081 | 2.490723311 | 1.162375879 | 2.142786473 |
| Gm22160 | 0.092691418 | 0.274387304 | 3.724945049 | 0.073662108 |
| Dlg3 | 1696.534007 | -0.169879751 | 0.088838155 | -1.912238616 |
| Tex11 | 4.291613721 | 0.87982076 | 0.519753351 | 1.692765921 |
| Gm24577 | 0 | NA | NA | NA |
| Gm27510 | 0 | NA | NA | NA |
| Gm14901 | 0 | NA | NA | NA |
| Gm9078 | 0 | NA | NA | NA |
| Slc7a3 | 112.9406466 | 0.301882953 | 0.168387025 | 1.792792247 |
| Snx12 | 585.2370213 | -0.244099839 | 0.068170542 | -3.580723176 |
| Gm24061 | 0.522088832 | -0.154411356 | 1.328131498 | -0.116262099 |
| Gm14842 | 0 | NA | NA | NA |
| Gm22472 | 0.04393175 | -0.206490061 | 3.724945049 | -0.055434391 |
| Foxo4 | 217.5431392 | -0.224622651 | 0.150198917 | -1.495501137 |

| | | | | |
|---|---|---|---|---|
| Gm614 | 0.87838318 | 3.167366798 | 1.164324891 | 2.720346205 |
| Gm20489 | 0 | NA | NA | NA |
| Il2rg | 109.8692796 | 5.369598899 | 0.397739763 | 13.50028185 |
| Med12 | 591.9911033 | 0.02302449 | 0.120444274 | 0.19116301 |
| Gm21986 | 6.701345333 | -0.305404674 | 0.455052154 | -0.671142136 |
| Nlgn3 | 1054.489164 | -0.195566031 | 0.092224117 | -2.120551947 |
| Gjb1 | 235.1220583 | 0.050036528 | 0.284670932 | 0.175769714 |
| Zmym3 | 1188.784781 | -0.16534024 | 0.059689592 | -2.770001188 |
| Nono | 1296.303821 | 0.348256481 | 0.056382863 | 6.176637059 |
| Itgb1bp2 | 18.97263099 | 0.551478258 | 0.419157443 | 1.315682847 |
| Gm14843 | 1.315890906 | -0.07679161 | 0.986712944 | -0.077825684 |
| Gm6184 | 0.809862658 | -0.797531493 | 1.283523396 | -0.621361087 |
| Taf1 | 1383.236683 | -0.113374392 | 0.045777532 | -2.476638374 |
| Ogt | 3519.482034 | -0.369415196 | 0.12531529 | -2.947886049 |
| Gm14845 | 0.98816996 | 1.354761497 | 0.906354834 | 1.494736328 |
| Gm9089 | 0.045882455 | 0.755288839 | 3.724945049 | 0.202765096 |
| Cxcr3 | 9.848512992 | 6.691801054 | 0.890607843 | 7.513745925 |
| Tpt1-ps6 | 6.439785805 | 0.133834804 | 0.444303048 | 0.301224141 |
| Gm9095 | 0 | NA | NA | NA |
| Gm4779 | 0.781258755 | -1.98232769 | 1.107387217 | -1.790094431 |
| 8030474K03Rik | 0 | NA | NA | NA |
| Gm9097 | 0 | NA | NA | NA |
| H3f3a-ps1 | 4.580768557 | -0.041408064 | 0.592058616 | -0.069939129 |
| Nhsl2 | 1474.969298 | 0.030025605 | 0.084511978 | 0.355282242 |
| Gm25828 | 0 | NA | NA | NA |
| Gm27843 | 0 | NA | NA | NA |
| Rgag4 | 516.2445288 | 0.469736581 | 0.135016829 | 3.47909652 |
| Gm5125 | 0 | NA | NA | NA |
| Pin4 | 43.34765536 | 0.386697495 | 0.298658856 | 1.294779935 |
| Gm16471 | 0.089089296 | 1.194485492 | 3.723939576 | 0.320758559 |
| Ercc6l | 15.19884806 | 1.642777585 | 0.48473602 | 3.389014879 |
| Gm9103 | 0.062426873 | 0.755288839 | 3.724945049 | 0.202765096 |
| Rps4x | 1364.482336 | 0.408710341 | 0.115452503 | 3.540073455 |
| Gm14859 | 0 | NA | NA | NA |
| Cited1 | 20.65218563 | 0.415715307 | 0.280082018 | 1.484262755 |
| Gm14858 | 2.504509112 | -0.180175302 | 0.571456042 | -0.315291621 |
| Gm26318 | 0.128271896 | 0.274403127 | 3.679089475 | 0.074584521 |
| Hdac8 | 196.3053901 | -0.373238863 | 0.115803423 | -3.223038257 |
| Phka1 | 482.7901274 | -0.310106999 | 0.08211566 | -3.776466001 |
| Gm9109 | 0 | NA | NA | NA |
| Gm14840 | 0 | NA | NA | NA |
| Gm3880 | 0 | NA | NA | NA |
| Gm9112 | 0 | NA | NA | NA |
| Dmrtc1b | 0 | NA | NA | NA |
| Gm9115 | 4.289267201 | 0.633474341 | 0.597875522 | 1.059542191 |
| Gm14839 | 0 | NA | NA | NA |
| Gm4234 | 0 | NA | NA | NA |

| | | | | |
|---|---|---|---|---|
| Gm9119 | 0 | NA | NA | NA |
| Gm5166 | 0 | NA | NA | NA |
| Gm25358 | 0 | NA | NA | NA |
| Dmrtc1c1 | 0 | NA | NA | NA |
| Gm9126 | 0 | NA | NA | NA |
| Gm16511 | 0 | NA | NA | NA |
| Gm24509 | 0 | NA | NA | NA |
| Dmrtc1c2 | 0 | NA | NA | NA |
| Gm26152 | 0 | NA | NA | NA |
| 1700031F05Rik | 0 | NA | NA | NA |
| Gm9133 | 0 | NA | NA | NA |
| Dmrtc1a | 21.57087845 | 0.188355657 | 0.215846769 | 0.872635982 |
| 1700011M02Rik | 0.046345709 | 0.274391604 | 3.724945049 | 0.073663262 |
| Gm14841 | 0 | NA | NA | NA |
| Gm14837 | 0.193036943 | -0.660002567 | 3.248873595 | -0.203148121 |
| Gm21998 | 0.296810931 | 0.288111613 | 1.771981385 | 0.162592911 |
| Pabpc1l2b-ps | 6.125206409 | -0.166295912 | 0.427895816 | -0.388636453 |
| Gm9143 | 0.044191549 | -0.206490061 | 3.724945049 | -0.055434391 |
| Pabpc1l2a-ps | 3.182304534 | -0.933036353 | 0.592986278 | -1.573453531 |
| Gm3928 | 0.473680182 | 0.213006603 | 1.34593145 | 0.158259623 |
| Gm14838 | 0.200176296 | -0.574396872 | 3.251120522 | -0.176676585 |
| Gm6206 | 202.4849468 | -0.07288652 | 0.123674416 | -0.58934194 |
| 4930519F16Rik | 7.945419585 | -0.706736832 | 0.380192985 | -1.858889723 |
| Nap1l2 | 1056.694576 | -0.072927633 | 0.067022713 | -1.088103266 |
| Gm9050 | 0 | NA | NA | NA |
| Gm14847 | 0 | NA | NA | NA |
| Gm19200 | 0.512704799 | -2.299002311 | 1.492776705 | -1.540084531 |
| Gm14848 | 4.393179665 | 1.857578916 | 1.335561323 | 1.390860071 |
| Gm3931 | 0.286297534 | 0.72736331 | 2.039346529 | 0.356664892 |
| Cdx4 | 2.430558831 | 1.753015469 | 1.055813545 | 1.660345689 |
| Gm9516 | 0 | NA | NA | NA |
| Chic1 | 1013.690449 | 0.101301609 | 0.064845517 | 1.562199121 |
| Gm26952 | 0 | NA | NA | NA |
| Tsx | 2.523036181 | 0.977714304 | 0.643629522 | 1.519063794 |
| Gm26992 | 0 | NA | NA | NA |
| Tsix | 4.036643334 | 0.899032643 | 0.592188125 | 1.518153785 |
| Xist | 7370.143953 | -0.286764971 | 0.215986395 | -1.327699234 |
| Gm27733 | 0 | NA | NA | NA |
| Gm27927 | 0 | NA | NA | NA |
| Gm27520 | 0.255832186 | -1.050251068 | 1.783000394 | -0.589035803 |
| Gm14861 | 0.384616848 | 0.713032966 | 1.413164143 | 0.504564858 |
| Rps12-ps1 | 0.281032441 | -0.736785947 | 2.031605795 | -0.362661865 |
| Jpx | 90.00391533 | -0.457469556 | 0.216502367 | -2.113000255 |
| Gm27875 | 0 | NA | NA | NA |
| Gm9670 | 11.86610392 | -0.963572168 | 0.367140524 | -2.624532313 |
| Gm14860 | 4.132034165 | -1.610793589 | 0.562999885 | -2.861090442 |
| Gm9785 | 0.044544648 | 0.755286614 | 3.724945049 | 0.202764498 |

| | | | | |
|---|---|---|---|---|
| Gm9157 | 5.137711797 | -0.262042332 | 0.498184138 | -0.525994933 |
| Ftx | 498.1238306 | -0.436962764 | 0.205604344 | -2.125260372 |
| Mir421 | 14.61544255 | -0.499402505 | 0.367607009 | -1.358522806 |
| Mir374b | 9.615598131 | -0.533592576 | 0.45105225 | -1.182995043 |
| mmu-mir-374c | 0 | NA | NA | NA |
| Gm38020 | 450.5061 | -0.54271257 | 0.276454234 | -1.963119042 |
| Gm27798 | 0.19090462 | 1.25932167 | 3.265182271 | 0.38568189 |
| Gm9159 | 0.296935004 | -0.110262002 | 3.720347245 | -0.029637557 |
| Gm27987 | 0.094097695 | 0.755282166 | 3.724945049 | 0.202763304 |
| Gm27285 | 0 | NA | NA | NA |
| Gm27321 | 0.436808164 | 0.475225925 | 1.528657177 | 0.310878026 |
| Zcchc13 | 0.559888142 | 1.10144216 | 1.274983993 | 0.86388705 |
| Gm14874 | 0.823502631 | -1.087370239 | 1.017128456 | -1.06905891 |
| Gm14875 | 0 | NA | NA | NA |
| Gm3943 | 0.062426873 | 0.755288839 | 3.724945049 | 0.202765096 |
| Slc16a2 | 413.5769034 | 0.264350265 | 0.088008214 | 3.00369992 |
| Gm6222 | 51.51232969 | 0.117736298 | 0.198446884 | 0.593288721 |
| Gm37956 | 12.17933112 | -0.317114529 | 0.301050736 | -1.053359089 |
| Gm14893 | 0 | NA | NA | NA |
| Gm9164 | 0.119587853 | 1.218392466 | 3.723618458 | 0.327206581 |
| Gm9166 | 0.719693239 | 3.336472801 | 1.466606758 | 2.274960743 |
| Rlim | 2303.194065 | 0.417551262 | 0.098182617 | 4.252802327 |
| Gm14892 | 0 | NA | NA | NA |
| Gm23378 | 0 | NA | NA | NA |
| Gm22266 | 0.086648695 | -0.687376175 | 3.724945049 | -0.184533239 |
| Gm24993 | 0 | NA | NA | NA |
| C77370 | 997.2398713 | -0.299809999 | 0.08285497 | -3.618491405 |
| Mir672 | 0.913602969 | -0.992036032 | 0.980809406 | -1.011446287 |
| Abcb7 | 785.2393539 | -0.123827833 | 0.081184907 | -1.525256819 |
| Vcp-rs | 650.7603801 | 0.147143595 | 0.07998112 | 1.839729128 |
| Gm6238 | 4.907369357 | -0.89323497 | 0.503997726 | -1.772299605 |
| Uprt | 137.0040339 | -0.16906846 | 0.133133239 | -1.269919226 |
| Zdhhc15 | 117.8220007 | -0.319490734 | 0.128689943 | -2.482639498 |
| Gm22430 | 0 | NA | NA | NA |
| Gm14822 | 2.347436586 | -0.089338958 | 0.697369128 | -0.128108565 |
| 1700121L16Rik | 0.182267886 | 0.274408265 | 2.690523896 | 0.101990644 |
| Magee2 | 43.93040372 | -0.08893225 | 0.160889592 | -0.552753283 |
| Gm6285 | 2.055059636 | 1.106639435 | 0.753332513 | 1.468991999 |
| Gm22090 | 0.110870145 | -0.206490066 | 3.724945049 | -0.055434393 |
| Gm9673 | 0 | NA | NA | NA |
| 5530601H04Rik | 228.1153272 | -0.050544722 | 0.103986845 | -0.486068421 |
| Pbdc1 | 243.3133995 | -0.149710506 | 0.104401192 | -1.433992296 |
| Magee1 | 1870.496208 | -0.089210224 | 0.058915311 | -1.514211187 |
| Gm6292 | 2.268236772 | 0.029903742 | 0.655006042 | 0.045654147 |
| E530001F21Rik | 15.1906836 | -0.473590409 | 0.332359594 | -1.42493377 |
| Rpl30-ps9 | 19.84181199 | -0.561798317 | 0.283963901 | -1.978414558 |
| Gm14824 | 57.64436276 | -0.725098232 | 0.225846005 | -3.210586934 |

| | | | | |
|---|---|---|---|---|
| Mir384 | 25.25469777 | -0.735266754 | 0.23214632 | -3.167255701 |
| 5330434G04Rik | 854.7814095 | -0.370324161 | 0.162012026 | -2.28578193 |
| Mir325 | 9.585432933 | 0.039869409 | 0.483469378 | 0.08246522 |
| Rab9b-ps1 | 0.047948698 | -0.206487837 | 3.724945049 | -0.055433794 |
| Cypt2 | 0 | NA | NA | NA |
| Gm14823 | 0 | NA | NA | NA |
| Gm14855 | 0 | NA | NA | NA |
| Gm14854 | 0 | NA | NA | NA |
| Gm7117 | 0.290452866 | 1.213756166 | 1.763948201 | 0.688090594 |
| Fgf16 | 4.769070217 | -0.376207905 | 0.446783383 | -0.842036475 |
| Atrx | 4307.043563 | -0.073220296 | 0.085268846 | -0.858699279 |
| Gm26020 | 0 | NA | NA | NA |
| Gm14853 | 16.35826218 | -0.168332264 | 0.372642813 | -0.451725508 |
| Gm23656 | 0 | NA | NA | NA |
| Magt1 | 450.0307998 | -0.062251296 | 0.120692246 | -0.515785377 |
| Cox7b | 1087.530571 | 0.023728599 | 0.118120624 | 0.200884468 |
| Atp7a | 328.8971482 | 0.266251216 | 0.105211572 | 2.530626719 |
| Tlr13 | 410.2003387 | 3.507389992 | 0.30219184 | 11.60650132 |
| Gm14856 | 0 | NA | NA | NA |
| Pgk1 | 2368.453568 | 0.099363861 | 0.065363461 | 1.520174408 |
| Taf9b | 306.5549042 | 0.256884559 | 0.119520655 | 2.149290096 |
| Fnd3c2 | 0.045882455 | 0.755288839 | 3.724945049 | 0.202765096 |
| Gm14852 | 0 | NA | NA | NA |
| Gm14857 | 0.631966691 | -0.187777741 | 1.346900268 | -0.139414733 |
| Gm5133 | 0 | NA | NA | NA |
| Mir6382 | 0 | NA | NA | NA |
| Gm14876 | 0 | NA | NA | NA |
| Gm6325 | 0.676409176 | 0.721514676 | 1.365584272 | 0.528356024 |
| Fndc3c1 | 1.066871938 | -0.535846603 | 1.045613924 | -0.512470799 |
| Gm27428 | 0.038981395 | 0.274393828 | 3.724945049 | 0.073663859 |
| Cysltr1 | 35.52481597 | 1.066704079 | 0.33988815 | 3.138397376 |
| Gm5127 | 6.92021651 | 2.094934886 | 0.597315152 | 3.507252208 |
| Gm24622 | 0.231961655 | 0.274410909 | 2.128142741 | 0.128943846 |
| Gm14877 | 0.75733687 | 0.436788575 | 1.466122851 | 0.297920856 |
| Gm14878 | 0.046345709 | 0.274391604 | 3.724945049 | 0.073663262 |
| Gm14879 | 0 | NA | NA | NA |
| Gm9200 | 1.650574175 | 0.431869818 | 0.737979019 | 0.585206092 |
| Gm24491 | 0 | NA | NA | NA |
| Gm22102 | 0 | NA | NA | NA |
| Gm14882 | 0 | NA | NA | NA |
| Gm14881 | 0 | NA | NA | NA |
| Gm14880 | 0 | NA | NA | NA |
| Zcchc5 | 3.290042265 | -0.221810421 | 0.54470278 | -0.407213676 |
| Lpar4 | 34.6581693 | -0.55179022 | 0.242607075 | -2.274419329 |
| P2ry10 | 28.76235556 | 7.279978401 | 0.87287459 | 8.340234075 |
| A630033H20Rik | 13.48921018 | 2.902979467 | 0.589246906 | 4.926592632 |
| Gm14834 | 0 | NA | NA | NA |

| | | | | |
|---|---|---|---|---|
| Gm14836 | 0 | NA | NA | NA |
| Gpr174 | 17.71929983 | 4.034051133 | 0.581123699 | 6.941811423 |
| Gm9208 | 0.060174537 | 0.755286614 | 3.724945049 | 0.202764498 |
| Gm14833 | 0 | NA | NA | NA |
| Gm5392 | 0 | NA | NA | NA |
| Itm2a | 152.8149106 | -0.903792455 | 0.24799784 | -3.644356151 |
| Gm14835 | 0 | NA | NA | NA |
| 4933401B06Rik | 0 | NA | NA | NA |
| Tbx22 | 1.078738829 | -1.53715752 | 0.959159004 | -1.602609695 |
| 2610002M06Rik | 1022.333748 | 0.216002149 | 0.092398566 | 2.337721879 |
| Fam46d | 2.160810062 | 2.462773509 | 0.847867265 | 2.904668701 |
| Gm732 | 0 | NA | NA | NA |
| Gm14849 | 0 | NA | NA | NA |
| Gm378 | 0 | NA | NA | NA |
| Gm9218 | 0 | NA | NA | NA |
| Gm379 | 0.046345709 | 0.274391604 | 3.724945049 | 0.073663262 |
| Brwd3 | 697.4322742 | -0.207475994 | 0.080121061 | -2.589531281 |
| 2810403D21Rik | 30.21590391 | 0.067597631 | 0.193739479 | 0.348909943 |
| Gm14870 | 0.48610956 | 0.138790432 | 1.482554498 | 0.093615737 |
| 5730416F02Rik | 0.253264426 | 0.274411906 | 2.082453259 | 0.131773381 |
| Hmgn5 | 36.85669999 | 0.007325385 | 0.254210244 | 0.028816245 |
| Gm23322 | 0.212513282 | 1.926509226 | 3.159662244 | 0.609719988 |
| Gm25797 | 0 | NA | NA | NA |
| Sh3bgrl | 620.2824578 | 0.077515475 | 0.133162203 | 0.582113188 |
| Gm26415 | 0 | NA | NA | NA |
| Gm14869 | 0 | NA | NA | NA |
| Gm6377 | 31.64666742 | 4.530240056 | 0.688498875 | 6.579880115 |
| Gm14868 | 0 | NA | NA | NA |
| Gm4784 | 0.759591101 | -0.206482632 | 1.357652062 | -0.152088033 |
| Gm4991 | 0 | NA | NA | NA |
| Gm7134 | 0 | NA | NA | NA |
| Gm9225 | 0 | NA | NA | NA |
| Gm14886 | 0 | NA | NA | NA |
| Gm26108 | 0.089814205 | 0.274402725 | 3.724945049 | 0.073666248 |
| Pou3f4 | 86.02711921 | -0.461138554 | 0.143632194 | -3.210551482 |
| Cylc1 | 0 | NA | NA | NA |
| Gm10112 | 0 | NA | NA | NA |
| Rps6ka6 | 115.5535552 | -0.252429577 | 0.139506133 | -1.809451465 |
| Hdx | 271.1100276 | -0.328456043 | 0.106010396 | -3.098338029 |
| Gm7340 | 0 | NA | NA | NA |
| Gm14898 | 0 | NA | NA | NA |
| Gm14899 | 0 | NA | NA | NA |
| Tex16 | 4.590268127 | -0.167606178 | 0.508655713 | -0.329508101 |
| Gm14900 | 0 | NA | NA | NA |
| Gm6335 | 3.683185528 | 0.482399255 | 0.624165532 | 0.772870704 |
| 4933403O08Rik | 4.019211381 | 0.069310963 | 0.528057692 | 0.131256422 |
| Gm5393 | 0 | NA | NA | NA |

| | | | | |
|---|---|---|---|---|
| Apool | 63.23518737 | 0.589529282 | 0.183455926 | 3.213465454 |
| Satl1 | 0 | NA | NA | NA |
| Gm6418 | 0.152980968 | -0.206487987 | 2.83734303 | -0.072775123 |
| Gm14908 | 0.684617987 | 1.806159003 | 1.391367025 | 1.298118305 |
| 2010106E10Rik | 0.828534694 | 1.1457272 | 1.054459859 | 1.086553641 |
| Zfp711 | 267.1925573 | 0.047025588 | 0.096558451 | 0.487016798 |
| Pof1b | 1.108407561 | 1.57769999 | 1.394720545 | 1.131194343 |
| Gm14921 | 0 | NA | NA | NA |
| Gm14922 | 0 | NA | NA | NA |
| Gm14923 | 0 | NA | NA | NA |
| Gm6427 | 0 | NA | NA | NA |
| Gm6429 | 0 | NA | NA | NA |
| Gm14937 | 0 | NA | NA | NA |
| Gm14938 | 0 | NA | NA | NA |
| Gm14936 | 0.124596252 | 0.755282361 | 3.714681068 | 0.203323609 |
| Gm14939 | 1.913926814 | 0.738053267 | 0.795254153 | 0.928072195 |
| Chm | 978.6370897 | 0.111262408 | 0.066435475 | 1.674743936 |
| Mir361 | 1.21006551 | 0.89685332 | 0.898806631 | 0.997826772 |
| Gm7405 | 0 | NA | NA | NA |
| Dach2 | 22.20089386 | 0.275601757 | 0.241708516 | 1.140223615 |
| Gm24831 | 0 | NA | NA | NA |
| Gm23985 | 0 | NA | NA | NA |
| Gm4915 | 0 | NA | NA | NA |
| Gm14896 | 0 | NA | NA | NA |
| Gm7416 | 0 | NA | NA | NA |
| Gm7146 | 0.194279094 | 0.274391173 | 3.724945049 | 0.073663147 |
| Klhl4 | 212.9303465 | -0.456546932 | 0.136776133 | -3.337913727 |
| Gm14895 | 3.529479033 | -0.822175099 | 0.59335786 | -1.385631091 |
| Gm1866 | 7.003298379 | 0.118880106 | 0.387198548 | 0.307026219 |
| Gm14894 | 0.088476398 | 0.274400501 | 3.724945049 | 0.073665651 |
| Gm5942 | 0 | NA | NA | NA |
| Gm14897 | 0.045882455 | 0.755288839 | 3.724945049 | 0.202765096 |
| Ube2dnl1 | 0 | NA | NA | NA |
| Ube2dnl2 | 0 | NA | NA | NA |
| 4930555B12Rik | 0 | NA | NA | NA |
| Gm4914 | 0 | NA | NA | NA |
| Gm7429 | 0.963943105 | -1.89209576 | 1.165741209 | -1.623083875 |
| Gm5943 | 0.089062654 | -0.20649451 | 3.724945049 | -0.055435586 |
| Gm14905 | 0 | NA | NA | NA |
| Gm25366 | 0 | NA | NA | NA |
| Gm27414 | 0 | NA | NA | NA |
| Gm6461 | 0 | NA | NA | NA |
| Gm5944 | 0.066050875 | 0.274391604 | 3.724945049 | 0.073663262 |
| Gm14903 | 0 | NA | NA | NA |
| Gm14906 | 0 | NA | NA | NA |
| Gm14907 | 0 | NA | NA | NA |
| Gm24683 | 0 | NA | NA | NA |

| | | | | |
|---|---|---|---|---|
| Gm14904 | 0 | NA | NA | NA |
| Gm24624 | 0 | NA | NA | NA |
| Cpxcr1 | 0 | NA | NA | NA |
| H2afb2 | 0 | NA | NA | NA |
| Gm24438 | 0 | NA | NA | NA |
| Gm16373 | 0.048949532 | -0.206487837 | 3.724945049 | -0.055433794 |
| Gm14919 | 0 | NA | NA | NA |
| Gm14920 | 0 | NA | NA | NA |
| Gm14918 | 0 | NA | NA | NA |
| Gm14913 | 0 | NA | NA | NA |
| Gm6491 | 0 | NA | NA | NA |
| Gm28579 | 0 | NA | NA | NA |
| Gm14911 | 0 | NA | NA | NA |
| Gm14914 | 0.265235714 | 0.755309511 | 2.048392787 | 0.368732753 |
| Gm14917 | 0 | NA | NA | NA |
| Gm6500 | 0 | NA | NA | NA |
| Gm14916 | 0 | NA | NA | NA |
| Gm14912 | 0 | NA | NA | NA |
| Gm14915 | 0 | NA | NA | NA |
| Gm14909 | 0 | NA | NA | NA |
| Gm14910 | 0 | NA | NA | NA |
| Tgif2lx2 | 0 | NA | NA | NA |
| Tgif2lx1 | 0 | NA | NA | NA |
| Gm7485 | 0 | NA | NA | NA |
| Gm14926 | 0 | NA | NA | NA |
| Gm26182 | 0 | NA | NA | NA |
| Gm14929 | 0 | NA | NA | NA |
| Pabpc5 | 33.05411456 | -0.054292635 | 0.200915417 | -0.270226324 |
| Gm4913 | 0.123841226 | 0.274391173 | 3.724945049 | 0.073663147 |
| Gm16420 | 0 | NA | NA | NA |
| Gm5863 | 0.479236126 | 0.580990946 | 1.284857698 | 0.452183107 |
| Gm14928 | 0.622864522 | -0.094325729 | 1.255375254 | -0.075137477 |
| Pcdh11x | 761.4248623 | 0.03305705 | 0.097734663 | 0.338232613 |
| H2afb3 | 0 | NA | NA | NA |
| Gm14931 | 0 | NA | NA | NA |
| Gm14927 | 1.075015726 | -1.193652981 | 1.014056105 | -1.177107436 |
| Gm14930 | 0 | NA | NA | NA |
| Gm4993 | 0.279479172 | -0.206485447 | 2.498164641 | -0.082654859 |
| Gm22590 | 0.042716945 | -0.206490061 | 3.724945049 | -0.055434391 |
| Gm5394 | 0 | NA | NA | NA |
| Gm25825 | 0 | NA | NA | NA |
| Gm5648 | 1.049721047 | 1.91605405 | 1.228225526 | 1.560018098 |
| Gm14932 | 0.047948698 | -0.206487837 | 3.724945049 | -0.055433794 |
| Gm23277 | 0 | NA | NA | NA |
| Gm14943 | 0 | NA | NA | NA |
| Gm14945 | 0 | NA | NA | NA |
| Gm4992 | 0.108772582 | 0.75528439 | 3.724945049 | 0.202763901 |

| | | | | |
|---|---|---|---|---|
| Gm24783 | 0 | NA | NA | NA |
| Gm16421 | 0.491147404 | 1.448982157 | 1.306015761 | 1.109467588 |
| Gm14946 | 0.141684195 | 0.755287278 | 3.558854591 | 0.212227631 |
| Gm14942 | 0.160761659 | 0.755303827 | 2.799488298 | 0.269800673 |
| Nap1l3 | 532.8242286 | -0.31605649 | 0.123015084 | -2.569249888 |
| Gm14947 | 0 | NA | NA | NA |
| Gm14940 | 0 | NA | NA | NA |
| Gm17521 | 0 | NA | NA | NA |
| Cldn34c1 | 0.062426873 | 0.755288839 | 3.724945049 | 0.202765096 |
| Astx6 | 0 | NA | NA | NA |
| Srsx | 0 | NA | NA | NA |
| Gm17577 | 0 | NA | NA | NA |
| Gm14951 | 0 | NA | NA | NA |
| Rps12-ps21 | 0 | NA | NA | NA |
| Astx2 | 0 | NA | NA | NA |
| Gm17412 | 0 | NA | NA | NA |
| Rps12-ps22 | 0.049553047 | 0.274391604 | 3.724945049 | 0.073663262 |
| Cldn34c2 | 0 | NA | NA | NA |
| Gm14950 | 0 | NA | NA | NA |
| Gm17467 | 0 | NA | NA | NA |
| Rps12-ps23 | 0 | NA | NA | NA |
| Cldn34c3 | 0 | NA | NA | NA |
| Astx5 | 0 | NA | NA | NA |
| Rps12-ps20 | 0 | NA | NA | NA |
| Vmn2r121 | 0 | NA | NA | NA |
| Rps12-ps6 | 0.072025462 | 0.755288839 | 3.724945049 | 0.202765096 |
| Astx1a | 0 | NA | NA | NA |
| Rps12-ps11 | 0 | NA | NA | NA |
| Gm17584 | 0 | NA | NA | NA |
| Astx4a | 0 | NA | NA | NA |
| Rps12-ps12 | 0 | NA | NA | NA |
| Gm17469 | 0 | NA | NA | NA |
| Astx4b | 0 | NA | NA | NA |
| Rps12-ps18 | 0 | NA | NA | NA |
| Astx1b | 0 | NA | NA | NA |
| Rps12-ps15 | 0 | NA | NA | NA |
| Gm17361 | 0 | NA | NA | NA |
| Gm21616 | 0 | NA | NA | NA |
| Rps12-ps17 | 0.094831987 | 0.274404949 | 3.724945049 | 0.073666845 |
| Astx4c | 0 | NA | NA | NA |
| Rps12-ps13 | 0 | NA | NA | NA |
| Gm17693 | 0 | NA | NA | NA |
| Astx1c | 0 | NA | NA | NA |
| Rps12-ps14 | 0 | NA | NA | NA |
| Gm17522 | 0 | NA | NA | NA |
| Astx4d | 0 | NA | NA | NA |
| Rps12-ps16 | 0 | NA | NA | NA |

| | | | | |
|---|---|---|---|---|
| Gm17267 | 0 | NA | NA | NA |
| Astx3 | 0 | NA | NA | NA |
| Gm14957 | 0.04393175 | -0.206490061 | 3.724945049 | -0.055434391 |
| Gm14971 | 0 | NA | NA | NA |
| Gm7746 | 0 | NA | NA | NA |
| 4932411N23Rik | 0 | NA | NA | NA |
| Gm23586 | 0 | NA | NA | NA |
| Gm381 | 0 | NA | NA | NA |
| Gm14973 | 0 | NA | NA | NA |
| Gm14972 | 0 | NA | NA | NA |
| Gm382 | 0 | NA | NA | NA |
| Gm22139 | 0 | NA | NA | NA |
| 4921511C20Rik | 0 | NA | NA | NA |
| Gm22636 | 0 | NA | NA | NA |
| Gm14970 | 0.307002753 | 1.92983037 | 1.72380604 | 1.119517118 |
| Cldn34c4 | 0.191938586 | -0.206489615 | 3.27925298 | -0.062968492 |
| Gm25061 | 0 | NA | NA | NA |
| Gm5403 | 0.038981395 | 0.274393828 | 3.724945049 | 0.073663859 |
| Mdm4-ps | 2.28277985 | -0.750925736 | 0.729563305 | -1.029281119 |
| Gm14975 | 0 | NA | NA | NA |
| Gm4916 | 0 | NA | NA | NA |
| Gm38390 | 0 | NA | NA | NA |
| Gm26475 | 0 | NA | NA | NA |
| Gm24522 | 0 | NA | NA | NA |
| 4930558G05Rik | 0 | NA | NA | NA |
| Gm26029 | 0 | NA | NA | NA |
| Gm22225 | 0 | NA | NA | NA |
| Gm14986 | 0 | NA | NA | NA |
| Gm14985 | 0 | NA | NA | NA |
| Gm14983 | 0 | NA | NA | NA |
| Diap2 | 874.9384135 | 0.085170088 | 0.074808372 | 1.138510107 |
| Gm14984 | 0 | NA | NA | NA |
| Gm14987 | 0 | NA | NA | NA |
| Gm7781 | 0.389466168 | -0.599992701 | 1.447629093 | -0.414465766 |
| Gm14982 | 0.044191549 | -0.206490061 | 3.724945049 | -0.055434391 |
| Gm26209 | 0 | NA | NA | NA |
| Gm25170 | 0 | NA | NA | NA |
| Gm14976 | 0 | NA | NA | NA |
| Gm7790 | 0 | NA | NA | NA |
| Gm14977 | 0 | NA | NA | NA |
| Gm4994 | 0 | NA | NA | NA |
| Gm7058 | 0 | NA | NA | NA |
| Gm14981 | 0 | NA | NA | NA |
| Gm26406 | 0 | NA | NA | NA |
| Gm23768 | 0 | NA | NA | NA |
| Gm22143 | 0 | NA | NA | NA |
| Gm7820 | 0 | NA | NA | NA |

| | | | | |
|---|---|---|---|---|
| Gm7821 | 0 | NA | NA | NA |
| Gm7823 | 0 | NA | NA | NA |
| Gm14979 | 0 | NA | NA | NA |
| Gm22552 | 0 | NA | NA | NA |
| Gm14980 | 0.189785611 | 0.755305193 | 2.656151365 | 0.284360749 |
| Pcdh19 | 966.848733 | -0.086632014 | 0.060891287 | -1.422732518 |
| Gm26851 | 2.30478967 | -1.471768671 | 0.711069253 | -2.069796529 |
| Tnmd | 3.21833566 | 0.158808867 | 0.553281873 | 0.28703067 |
| Tspan6 | 133.5839573 | -0.259975321 | 0.141422895 | -1.838283125 |
| 4930570D08Rik | 0 | NA | NA | NA |
| Srpx2 | 13.98697368 | -0.111615538 | 0.28175277 | -0.396147084 |
| Sytl4 | 25.13300853 | -0.121969361 | 0.283064772 | -0.43088852 |
| Gm7841 | 0.106465261 | 0.75528439 | 3.724945049 | 0.202763901 |
| Cstf2 | 1299.774295 | 0.01537885 | 0.075244094 | 0.204386139 |
| Mir3112 | 0 | NA | NA | NA |
| Nox1 | 3.798596485 | 0.506799373 | 0.487041967 | 1.040566126 |
| Xkrx | 14.27994078 | -0.126955371 | 0.255317319 | -0.497245433 |
| Arl13a | 0.299311364 | 0.274383636 | 3.724945049 | 0.073661123 |
| Trmt2b | 186.6964886 | -0.307961713 | 0.119870219 | -2.56912615 |
| Tmem35 | 346.804519 | -0.382593084 | 0.103591407 | -3.69328976 |
| Cenpi | 15.04443857 | 0.976604056 | 0.349323298 | 2.795702608 |
| Drp2 | 836.9736946 | -0.578821045 | 0.109562366 | -5.283027985 |
| Taf7l | 0.599737706 | 2.481464355 | 1.437489114 | 1.726249145 |
| Gm16410 | 0.92854447 | 1.793922339 | 1.114353339 | 1.609832606 |
| Gm23124 | 0.094831987 | 0.274404949 | 3.724945049 | 0.073666845 |
| Timm8a1 | 33.17250175 | 0.359985454 | 0.240146709 | 1.499023054 |
| Btk | 35.98129176 | 1.784405262 | 0.251718106 | 7.088903097 |
| Gm15154 | 0 | NA | NA | NA |
| Rpl36a | 325.2517739 | 0.315332691 | 0.204401552 | 1.542711822 |
| Gla | 403.723937 | 0.884957848 | 0.167821892 | 5.273196713 |
| Hnrnph2 | 1054.6625 | 0.23103157 | 0.092252937 | 2.504327528 |
| Gm7855 | 0.152402701 | 1.236191965 | 2.846761649 | 0.434244984 |
| Gm7857 | 0.072025462 | 0.755288839 | 3.724945049 | 0.202765096 |
| B230119M05Rik | 0.783798686 | 1.703507311 | 1.126965966 | 1.511587184 |
| Armcx4 | 981.8027628 | -0.267257835 | 0.100605894 | -2.656482889 |
| Armcx1 | 541.5903477 | -0.002999286 | 0.08026656 | -0.037366569 |
| Gm16397 | 0.442616626 | 0.968297869 | 1.452054494 | 0.666846784 |
| Gm10344 | 1.729071564 | 0.915690417 | 0.811666221 | 1.128161298 |
| Armcx6 | 82.02743705 | 1.194817596 | 0.209429326 | 5.705111227 |
| Armcx3 | 414.6905456 | -0.030554345 | 0.077310231 | -0.395217355 |
| Mir7093 | 0.745433557 | -0.919693544 | 1.352925937 | -0.679781146 |
| Armcx2 | 966.547524 | -0.066142112 | 0.075409726 | -0.877103192 |
| Gm7222 | 0 | NA | NA | NA |
| Gm15018 | 0 | NA | NA | NA |
| Nxf2 | 1.760497888 | 1.349301024 | 0.810386281 | 1.665009707 |
| Zmat1 | 230.6846648 | -0.460133134 | 0.155182376 | -2.965112049 |
| 3632454L22Rik | 43.94593524 | -0.495128429 | 0.184042783 | -2.690289835 |

| | | | | |
|---|---|---|---|---|
| Gm6207 | 0 | NA | NA | NA |
| Gm15020 | 0 | NA | NA | NA |
| Gm25868 | 0 | NA | NA | NA |
| Gm15023 | 0.072025462 | 0.755288839 | 3.724945049 | 0.202765096 |
| Tceal6 | 310.5221001 | -0.055588478 | 0.126184386 | -0.44053373 |
| Gm15022 | 0 | NA | NA | NA |
| Pramel | 0 | NA | NA | NA |
| Pramel3 | 0.045882455 | 0.755288839 | 3.724945049 | 0.202765096 |
| Gm25936 | 0 | NA | NA | NA |
| Gm15017 | 0 | NA | NA | NA |
| Gm6215 | 0 | NA | NA | NA |
| Gm5128 | 0 | NA | NA | NA |
| Gm22006 | 0 | NA | NA | NA |
| Gm6221 | 0 | NA | NA | NA |
| Gm7903 | 0 | NA | NA | NA |
| Gm24416 | 0 | NA | NA | NA |
| Gm7905 | 0 | NA | NA | NA |
| AV320801 | 0 | NA | NA | NA |
| Gm22233 | 0 | NA | NA | NA |
| Gm15015 | 0 | NA | NA | NA |
| Gm6228 | 0 | NA | NA | NA |
| Nxf7 | 11.01788405 | 0.061159223 | 0.383017144 | 0.159677508 |
| Prame | 0 | NA | NA | NA |
| Gm15021 | 0.141317957 | 1.201108798 | 3.595358929 | 0.334072014 |
| Tcp11x2 | 0.28975322 | 0.758890736 | 2.39051243 | 0.317459439 |
| Tmsb15a | 0.072025462 | 0.755288839 | 3.724945049 | 0.202765096 |
| Gm15019 | 0 | NA | NA | NA |
| Gprasp1 | 6518.488455 | -0.189418739 | 0.063923543 | -2.963207775 |
| Armcx5 | 396.6884651 | 0.798250756 | 0.170440869 | 4.683446887 |
| mmu-mir-7646 | 0 | NA | NA | NA |
| Mir1970 | 0 | NA | NA | NA |
| Gprasp2 | 1597.674091 | 0.100536386 | 0.111141772 | 0.90457786 |
| Bhlhb9 | 800.6765123 | 0.297595954 | 0.090576669 | 3.285569653 |
| Gm15016 | 2.223121331 | 0.809181151 | 0.707486286 | 1.14374111 |
| Arxes2 | 329.4047032 | -0.340951532 | 0.099016062 | -3.443396209 |
| Arxes1 | 156.3845472 | -0.249848985 | 0.142984246 | -1.747388207 |
| Bex2 | 883.5005139 | 0.182924597 | 0.166998452 | 1.095367023 |
| Nxf3 | 2.723995911 | 0.797513245 | 0.579669864 | 1.375805945 |
| 5730412P04Rik | 0.591118678 | 2.017916393 | 1.259383556 | 1.60230486 |
| Bex4 | 82.98433973 | 0.239523587 | 0.149531889 | 1.601822785 |
| Tceal8 | 272.1310813 | 0.298214892 | 0.142289139 | 2.095837349 |
| Mir6383 | 0 | NA | NA | NA |
| Tceal5 | 256.9790592 | -0.02708197 | 0.231942344 | -0.116761645 |
| Bex1 | 254.7050612 | 0.22795735 | 0.142362511 | 1.601245639 |
| Tceal7 | 6.33875668 | 0.22603758 | 0.427365012 | 0.528909887 |
| Wbp5 | 588.6657451 | 0.308103819 | 0.101133293 | 3.046512277 |
| Ngfrap1 | 809.4390861 | -0.021694814 | 0.155587372 | -0.139438145 |

| | | | | |
|---|---|---|---|---|
| n-R5s12 | 0.061920613 | 0.274393828 | 3.724945049 | 0.073663859 |
| Nsa2-ps2 | 0 | NA | NA | NA |
| Gm14997 | 0 | NA | NA | NA |
| Gm4917 | 0.046345709 | 0.274391604 | 3.724945049 | 0.073663262 |
| Gm7973 | 0.061920613 | 0.274393828 | 3.724945049 | 0.073663859 |
| Gm14994 | 0.090427103 | 1.236179401 | 3.724945049 | 0.331865138 |
| Gm14999 | 0 | NA | NA | NA |
| Gm14998 | 0 | NA | NA | NA |
| Gm14993 | 0 | NA | NA | NA |
| Gm14992 | 0 | NA | NA | NA |
| Gm14996 | 0 | NA | NA | NA |
| Gm14990 | 0 | NA | NA | NA |
| Gm14995 | 0.106112162 | -0.206492286 | 3.724945049 | -0.055434988 |
| Kir3dl2 | 0.620158706 | 1.178175727 | 1.178689139 | 0.999564421 |
| Gm14991 | 0.047948698 | -0.206487837 | 3.724945049 | -0.055433794 |
| Kir3dl1 | 0 | NA | NA | NA |
| Gm15006 | 0 | NA | NA | NA |
| Gm15003 | 0 | NA | NA | NA |
| Gm15005 | 0 | NA | NA | NA |
| Gm15004 | 0 | NA | NA | NA |
| Tceal3 | 620.6626587 | 0.085987568 | 0.119864788 | 0.717371381 |
| Gm15010 | 0.784324088 | 1.335563806 | 1.105307765 | 1.208318487 |
| Gm23068 | 0 | NA | NA | NA |
| Tceal1 | 253.1864694 | 0.224667495 | 0.122591229 | 1.832655543 |
| Morf4l2 | 2139.902612 | 0.264056518 | 0.084277129 | 3.133193065 |
| BC065397 | 34.23957864 | -1.361255413 | 0.257583211 | -5.284721036 |
| Glra4 | 6.335166068 | -0.886575346 | 0.496309245 | -1.786336556 |
| Gm15009 | 0 | NA | NA | NA |
| Plp1 | 16662.56891 | -0.209760067 | 0.142926071 | -1.467612348 |
| Rab9b | 322.6084878 | -0.327618431 | 0.109676623 | -2.987130922 |
| Gm15026 | 0 | NA | NA | NA |
| Gm15025 | 0 | NA | NA | NA |
| H2bfm | 0 | NA | NA | NA |
| Gm6275 | 0.048949532 | -0.206487837 | 3.724945049 | -0.055433794 |
| Tmsb15l | 3.858449061 | 0.156940155 | 0.592561094 | 0.26485059 |
| Tmsb15b2 | 0.857879072 | -0.753444297 | 1.032688166 | -0.729595169 |
| Tmsb15b1 | 3.641358966 | -1.098348036 | 0.547876691 | -2.004735836 |
| Slc25a53 | 178.3064943 | -0.055615325 | 0.129379404 | -0.429862272 |
| Zcchc18 | 1940.436389 | 0.007894074 | 0.068656937 | 0.114978531 |
| Gm15038 | 0 | NA | NA | NA |
| Fam199x | 561.8101836 | -0.066743381 | 0.089342262 | -0.747052729 |
| Gm25770 | 0 | NA | NA | NA |
| Gm8019 | 0 | NA | NA | NA |
| Gm5761 | 0.368947036 | 1.17106188 | 1.554455726 | 0.753358143 |
| Gm15039 | 0 | NA | NA | NA |
| Gm15042 | 0 | NA | NA | NA |
| Gm15045 | 0 | NA | NA | NA |

| | | | | |
|---|---|---|---|---|
| Gm15043 | 0 | NA | NA | NA |
| Gm15044 | 0 | NA | NA | NA |
| Esx1 | 0 | NA | NA | NA |
| Gm15040 | 0.045882455 | 0.755288839 | 3.724945049 | 0.202765096 |
| Gm15041 | 0 | NA | NA | NA |
| Gm23679 | 0 | NA | NA | NA |
| Gm8036 | 0.810529001 | -2.124404308 | 1.168062244 | -1.818742382 |
| Il1rapl2 | 78.53886856 | 0.051295602 | 0.145959047 | 0.351438322 |
| Gm14988 | 0 | NA | NA | NA |
| Gm25393 | 0 | NA | NA | NA |
| Gm14989 | 0 | NA | NA | NA |
| Gm24675 | 0 | NA | NA | NA |
| Tex13a | 0 | NA | NA | NA |
| Gm22057 | 0 | NA | NA | NA |
| Gm23958 | 0 | NA | NA | NA |
| Gm15000 | 0 | NA | NA | NA |
| Gm15001 | 0 | NA | NA | NA |
| Nrk | 62.1717717 | -0.046871839 | 0.258961559 | -0.180999215 |
| Serpina7 | 0.108772582 | 0.75528439 | 3.724945049 | 0.202763901 |
| 4930513O06Rik | 0 | NA | NA | NA |
| 4933428M09Rik | 0 | NA | NA | NA |
| Gm15007 | 0 | NA | NA | NA |
| Mum1l1 | 62.92430637 | -0.473377188 | 0.144636456 | -3.272876014 |
| Platr21 | 0.194631057 | 1.693379269 | 3.232136674 | 0.52391945 |
| Gm15002 | 0 | NA | NA | NA |
| Trap1a | 0 | NA | NA | NA |
| Gm8061 | 0 | NA | NA | NA |
| D330045A20Rik | 6.283406809 | 0.079973589 | 0.411550365 | 0.194322726 |
| Rnf128 | 220.8248196 | 0.078962425 | 0.109261375 | 0.72269295 |
| Gm24380 | 0 | NA | NA | NA |
| Gm6322 | 0 | NA | NA | NA |
| Tbc1d8b | 118.5380032 | -0.457883693 | 0.138248033 | -3.312044906 |
| Gm15013 | 2.788702176 | 0.30917507 | 0.610690396 | 0.50627138 |
| Ripply1 | 0.677889948 | 0.242229825 | 1.414009617 | 0.171307056 |
| Gm15012 | 0.066050875 | 0.274391604 | 3.724945049 | 0.073663262 |
| Cldn2 | 23.20106824 | -0.643230412 | 0.277114229 | -2.321174244 |
| Morc4 | 90.84373692 | -0.231521932 | 0.169671578 | -1.364529848 |
| Rbm41 | 210.9730729 | -0.221987693 | 0.10299332 | -2.155360103 |
| Gm15011 | 0 | NA | NA | NA |
| Nup62cl | 3.861479097 | 1.068540536 | 0.533475955 | 2.002977876 |
| Pih1h3b | 0.733953745 | -0.631318028 | 1.068950461 | -0.590596151 |
| Gm6298 | 0.042716945 | -0.206490061 | 3.724945049 | -0.055434391 |
| Gm15046 | 0.104637558 | -0.206492286 | 3.724945049 | -0.055434988 |
| Mir3475 | 0.844812178 | -0.441133944 | 1.327186379 | -0.332382815 |
| Frmpd3 | 67.41637008 | -0.328040569 | 0.151858586 | -2.160171369 |
| Gm8097 | 0.313170507 | 0.70469739 | 2.070888775 | 0.340287416 |
| Prps1 | 548.0358506 | 0.019940335 | 0.062617587 | 0.318446233 |

| | | | | |
|---|---|---|---|---|
| Tsc22d3 | 1422.675399 | 0.774535188 | 0.154602052 | 5.009863565 |
| Gm15047 | 0 | NA | NA | NA |
| Gm22025 | 0 | NA | NA | NA |
| Mid2 | 552.0542967 | 0.025385578 | 0.07531454 | 0.33706079 |
| Gm23199 | 2.503876714 | -0.780156683 | 0.786956533 | -0.991359307 |
| Eif2c5 | 0 | NA | NA | NA |
| Gm6088 | 0.360750703 | -0.687379379 | 1.589640975 | -0.432411714 |
| Tex13 | 0 | NA | NA | NA |
| Gm7091 | 0 | NA | NA | NA |
| Vsig1 | 0.30513904 | 1.364035085 | 3.711809654 | 0.367485193 |
| Psmd10 | 105.8155227 | 0.185527316 | 0.153970956 | 1.204950087 |
| Atg4a | 98.38075259 | 0.581357286 | 0.170727114 | 3.405184296 |
| Col4a6 | 3.411580873 | 0.297227811 | 0.635434992 | 0.467754868 |
| Gm15077 | 0 | NA | NA | NA |
| Gm15078 | 0 | NA | NA | NA |
| Gm15079 | 0 | NA | NA | NA |
| Col4a5 | 123.2002736 | 0.153573105 | 0.208373231 | 0.737009758 |
| Gm15268 | 0.88858162 | -0.993983898 | 1.328956909 | -0.74794291 |
| Gm15282 | 0 | NA | NA | NA |
| Irs4 | 86.40347737 | -0.300803506 | 0.238716806 | -1.260085166 |
| Gm15295 | 0.642988239 | 0.51836731 | 1.319757493 | 0.392774667 |
| Gm15297 | 0 | NA | NA | NA |
| Hmgn2l6 | 0 | NA | NA | NA |
| Gm15294 | 0.045882455 | 0.755288839 | 3.724945049 | 0.202765096 |
| Gm15298 | 0.198041555 | 0.274404852 | 2.604148426 | 0.105372201 |
| Gm6368 | 8.824265348 | 0.040819355 | 0.374779551 | 0.10891564 |
| Gm5762 | 0 | NA | NA | NA |
| Gm6373 | 0 | NA | NA | NA |
| Gucy2f | 26.45992321 | -0.166402358 | 0.228206999 | -0.729172896 |
| Gm6382 | 25.01211358 | 0.442450894 | 0.229664152 | 1.926512647 |
| Gm15014 | 0 | NA | NA | NA |
| Nxt2 | 433.560056 | 0.094064164 | 0.090287647 | 1.041827624 |
| Gm5643 | 2.51713653 | -0.373618607 | 0.626783657 | -0.596088622 |
| Kcne1l | 125.6411286 | 0.624373481 | 0.247775314 | 2.519918032 |
| Acsl4 | 1433.382608 | 0.220275568 | 0.07208094 | 3.055947505 |
| Gm6067 | 0 | NA | NA | NA |
| Gm7123 | 0 | NA | NA | NA |
| Gm15031 | 0 | NA | NA | NA |
| Gm25915 | 0.060174537 | 0.755286614 | 3.724945049 | 0.202764498 |
| Gm4995 | 0.090537259 | -0.20649451 | 3.724945049 | -0.055435586 |
| Gm15032 | 0 | NA | NA | NA |
| Gm5763 | 8.930037149 | 0.463853827 | 0.340073458 | 1.363981269 |
| Tmem164 | 493.7209216 | -0.158032453 | 0.080154367 | -1.971601271 |
| Mir652 | 2.416261644 | -0.928966727 | 0.77488515 | -1.198844407 |
| Gm15049 | 0.046345709 | 0.274391604 | 3.724945049 | 0.073663262 |
| Ammecr1 | 116.0557192 | 0.434013986 | 0.115574855 | 3.755263083 |
| AL731548.1 | 1.863015902 | -0.318643222 | 0.710504955 | -0.448474313 |

| | | | | |
|---|---|---|---|---|
| Rgag1 | 33.41144712 | 0.387017069 | 0.216427501 | 1.788206518 |
| Gm4760 | 0.046345709 | 0.274391604 | 3.724945049 | 0.073663262 |
| Chrdl1 | 139.3962317 | -0.101807675 | 0.179940869 | -0.565784058 |
| Gm23325 | 0 | NA | NA | NA |
| Pak3 | 2356.012352 | -0.015975583 | 0.102071091 | -0.15651428 |
| Capn6 | 31.3985991 | 0.517982538 | 0.218490747 | 2.370729858 |
| BX530055.1 | 0 | NA | NA | NA |
| Dcx | 707.4354374 | -0.09396051 | 0.160237528 | -0.586382676 |
| Gm15048 | 0.045882455 | 0.755288839 | 3.724945049 | 0.202765096 |
| A730046J19Rik | 10.6199982 | -0.249354628 | 0.360692059 | -0.691322756 |
| Gm8199 | 0.207515118 | 0.274411166 | 2.588142969 | 0.106026278 |
| Gm8202 | 0 | NA | NA | NA |
| Gm15067 | 0 | NA | NA | NA |
| Gm15066 | 0.044544648 | 0.755286614 | 3.724945049 | 0.202764498 |
| Mir680-2 | 0 | NA | NA | NA |
| Alg13 | 168.7701936 | -0.182316437 | 0.110311752 | -1.652738111 |
| Gm2214 | 0.090665643 | -0.687373951 | 3.724945049 | -0.184532642 |
| Trpc5 | 161.8271037 | -0.234416284 | 0.091755276 | -2.554798977 |
| Trpc5os | 0.750003179 | 2.858014483 | 2.514059431 | 1.136812618 |
| Gm15071 | 0 | NA | NA | NA |
| Mageb16-ps1 | 3.715911654 | -1.866868632 | 0.62620397 | -2.981246878 |
| Zcchc16 | 32.00016923 | -0.26752206 | 0.194739064 | -1.373746256 |
| Gm26024 | 2.396750964 | -0.348116113 | 0.628229991 | -0.554122086 |
| Gm8216 | 4.713227208 | -0.262433583 | 0.448222298 | -0.585498724 |
| Mir3472 | 0.066050875 | 0.274391604 | 3.724945049 | 0.073663262 |
| Gm25097 | 0 | NA | NA | NA |
| Gm15073 | 0.066050875 | 0.274391604 | 3.724945049 | 0.073663262 |
| Gm6441 | 0.044191549 | -0.206490061 | 3.724945049 | -0.055434391 |
| Lhfpl1 | 2.080192563 | 0.59686327 | 0.738897117 | 0.807775882 |
| Amot | 561.2978999 | -0.30316647 | 0.105017183 | -2.886827282 |
| Gm5644 | 2.782059835 | 0.338745527 | 0.633959799 | 0.53433282 |
| Gm15060 | 0 | NA | NA | NA |
| Gm27659 | 0.042716945 | -0.206490061 | 3.724945049 | -0.055434391 |
| Gm15058 | 0.106618422 | 0.274402725 | 3.724945049 | 0.073666248 |
| Gm6446 | 0.038981395 | 0.274393828 | 3.724945049 | 0.073663859 |
| Gm15057 | 0.867677903 | -0.486557285 | 1.066822175 | -0.456080963 |
| Gm6447 | 0 | NA | NA | NA |
| Gm15062 | 0 | NA | NA | NA |
| Gm4918 | 0 | NA | NA | NA |
| Hmgb1-ps2 | 15.03640688 | 0.314153953 | 0.258085064 | 1.217249647 |
| Gm15061 | 0 | NA | NA | NA |
| Gm26151 | 0.325583935 | -1.824082651 | 1.991831669 | -0.915781529 |
| Gm15076 | 0 | NA | NA | NA |
| Gm8261 | 2.625663271 | 0.053977403 | 0.735352149 | 0.073403476 |
| Gm15075 | 1.268895825 | -1.102730786 | 1.024539875 | -1.076318075 |
| Gm15074 | 0 | NA | NA | NA |
| Gm4996 | 0 | NA | NA | NA |

| | | | | |
|---|---|---|---|---|
| Htr2c | 718.8775726 | -0.464077744 | 0.110944517 | -4.182971436 |
| Snora35 | 4.50470804 | -0.311492552 | 0.52835986 | -0.589546207 |
| Gm22650 | 4.136795173 | -0.499001021 | 0.56580565 | -0.881930077 |
| Mir764 | 3.164642609 | -1.45221755 | 0.628290751 | -2.311378207 |
| Mir1912 | 2.763607753 | -1.301432101 | 0.677362243 | -1.921323657 |
| Mir1264 | 5.146865135 | -0.746989204 | 0.518087403 | -1.44182082 |
| Mir1298 | 1.82841141 | -1.190150408 | 1.02530484 | -1.160777128 |
| Gm24627 | 1.916860434 | -0.420575975 | 0.811612004 | -0.518198318 |
| Mir448 | 0.601491876 | -1.945846321 | 1.379183114 | -1.410868724 |
| Il13ra2 | 7.235467095 | 0.578942905 | 0.375776835 | 1.540656183 |
| Vmn1r239-ps | 0 | NA | NA | NA |
| Lrch2 | 586.4938171 | -0.024401225 | 0.087999659 | -0.277287728 |
| Gm25107 | 111.0918953 | -0.22427606 | 0.210089548 | -1.067526024 |
| Gm16400 | 0 | NA | NA | NA |
| Gm15167 | 0 | NA | NA | NA |
| Gm8289 | 0 | NA | NA | NA |
| Gm15128 | 0 | NA | NA | NA |
| Gm15081 | 0 | NA | NA | NA |
| Gm15080 | 0 | NA | NA | NA |
| Gm15108 | 0 | NA | NA | NA |
| Gm15110 | 0 | NA | NA | NA |
| Gm8303 | 0 | NA | NA | NA |
| Gm15107 | 0 | NA | NA | NA |
| Gm15109 | 0 | NA | NA | NA |
| Gm15111 | 0 | NA | NA | NA |
| Gm15113 | 0 | NA | NA | NA |
| Gm8320 | 0 | NA | NA | NA |
| Gm15114 | 0 | NA | NA | NA |
| Gm15106 | 0 | NA | NA | NA |
| Gm15112 | 0 | NA | NA | NA |
| Gm8331 | 0 | NA | NA | NA |
| Gm8334 | 0 | NA | NA | NA |
| Gm15127 | 0 | NA | NA | NA |
| Gm15125 | 0 | NA | NA | NA |
| Gm15126 | 0 | NA | NA | NA |
| Kars-ps1 | 0 | NA | NA | NA |
| Luzp4 | 0.048949532 | -0.206487837 | 3.724945049 | -0.055433794 |
| Gm15099 | 0 | NA | NA | NA |
| Gm15084 | 0 | NA | NA | NA |
| Gm15094 | 0 | NA | NA | NA |
| Gm15096 | 0 | NA | NA | NA |
| Ott | 0 | NA | NA | NA |
| Gm15092 | 0 | NA | NA | NA |
| Gm15088 | 0 | NA | NA | NA |
| Gm15090 | 0 | NA | NA | NA |
| Gm8346 | 0 | NA | NA | NA |
| Gm15093 | 0 | NA | NA | NA |

| | | | | |
|---|---|---|---|---|
| Gm15098 | 0 | NA | NA | NA |
| Gm15089 | 0 | NA | NA | NA |
| Gm8366 | 0 | NA | NA | NA |
| Gm15100 | 0 | NA | NA | NA |
| Gm15085 | 0 | NA | NA | NA |
| Gm15086 | 0 | NA | NA | NA |
| Gm10439 | 0 | NA | NA | NA |
| Gm15101 | 0 | NA | NA | NA |
| Gm8383 | 0 | NA | NA | NA |
| Gm15097 | 0 | NA | NA | NA |
| Gm15095 | 0 | NA | NA | NA |
| Gm15087 | 0 | NA | NA | NA |
| Gm15091 | 0 | NA | NA | NA |
| Gm15102 | 0 | NA | NA | NA |
| Gm15103 | 0 | NA | NA | NA |
| Gm8402 | 0 | NA | NA | NA |
| Gm5946 | 2.587997239 | 0.799651523 | 0.729054016 | 1.096834397 |
| Gm15104 | 1.895634264 | 0.130723627 | 0.670503825 | 0.194963284 |
| Gm15105 | 0.604613898 | 0.177248036 | 1.541629027 | 0.114974506 |
| Tmem29 | 269.2719752 | -0.088311385 | 0.108007302 | -0.817642729 |
| Gm15129 | 0 | NA | NA | NA |
| 3110067C02Rik | 4.62715315 | -1.257331649 | 0.541122456 | -2.323562133 |
| Apex2 | 300.6728437 | -0.202015792 | 0.182891767 | -1.10456471 |
| Alas2 | 56.13115503 | 2.703985294 | 0.801902453 | 3.371962866 |
| Mir3620 | 3.543203576 | 0.195471354 | 0.477938189 | 0.408988775 |
| Pfkfb1 | 11.63733137 | 0.533666565 | 0.338958497 | 1.574430406 |
| Tro | 2279.729976 | 0.155216324 | 0.092651923 | 1.675262841 |
| Gm15132 | 0 | NA | NA | NA |
| Gm15131 | 0 | NA | NA | NA |
| Maged2 | 434.343474 | 0.034752679 | 0.115739876 | 0.300265392 |
| Gm24907 | 0.178529143 | -1.465050858 | 2.705292286 | -0.541549933 |
| Itih5l-ps | 0.139037128 | 0.274384718 | 3.724945049 | 0.073661414 |
| Gm27191 | 0 | NA | NA | NA |
| Gm8424 | 0.517778811 | 0.259648617 | 1.484244299 | 0.174936577 |
| Gnl3l | 3189.542655 | -0.151029932 | 0.055039402 | -2.744032957 |
| Gm15152 | 0 | NA | NA | NA |
| Fgd1 | 278.5392714 | -0.285095699 | 0.081537371 | -3.496503427 |
| Tsr2 | 790.8463341 | 0.172478091 | 0.062570913 | 2.756521888 |
| Gm15149 | 0 | NA | NA | NA |
| Rps12-ps7 | 0 | NA | NA | NA |
| Gm15138 | 7.688685255 | -0.209829745 | 0.452181057 | -0.464039219 |
| Wnk3 | 948.2720462 | -0.27732787 | 0.067025469 | -4.137649065 |
| Rpl21-ps15 | 49.38234276 | 0.420990565 | 0.210181298 | 2.002987744 |
| Gm26441 | 0.12209515 | 0.75528439 | 3.724945049 | 0.202763901 |
| A230072E10Rik | 43.55964103 | 0.977564579 | 0.177472116 | 5.508271397 |
| Fam120c | 1332.354665 | 0.138440099 | 0.071368119 | 1.93980309 |
| Gm22248 | 2.415551224 | -0.20411574 | 0.634173527 | -0.321861023 |

| | | | | |
|---|---|---|---|---|
| Gm6451 | 2.057373881 | 0.820275615 | 0.64972819 | 1.262490418 |
| Gm15148 | 0.123992736 | 0.274402763 | 3.720643035 | 0.073751435 |
| Phf8 | 763.8243954 | 0.012922696 | 0.092444876 | 0.139788129 |
| Gm24460 | 0 | NA | NA | NA |
| Gm5395 | 1.549805528 | 2.035780306 | 0.995975998 | 2.044005387 |
| Gm15151 | 0.220031754 | 1.26143515 | 2.591130944 | 0.486828021 |
| Gm4919 | 0 | NA | NA | NA |
| n-R5s13 | 0 | NA | NA | NA |
| Gm15161 | 0 | NA | NA | NA |
| Huwe1 | 8349.182081 | -0.269721307 | 0.052096104 | -5.177379594 |
| Mir3113 | 0 | NA | NA | NA |
| Mirlet7f-2 | 0.623206311 | 0.594822543 | 1.238247457 | 0.480374532 |
| Mir98 | 2.834035009 | -0.052112893 | 0.607013549 | -0.085851285 |
| Gm15191 | 3.302051428 | -0.630704332 | 0.57956341 | -1.088240426 |
| n-R5s14 | 0 | NA | NA | NA |
| Hsd17b10 | 170.8953565 | 0.321618462 | 0.12337841 | 2.60676451 |
| Ribc1 | 4.147795559 | -0.577282271 | 0.565440314 | -1.020942894 |
| Smc1a | 891.0140031 | 0.136642331 | 0.104041924 | 1.313339142 |
| Gm15267 | 3.127372856 | 0.185378636 | 0.567568422 | 0.326619009 |
| Gm15266 | 0 | NA | NA | NA |
| Iqsec2 | 2091.297293 | -0.023168159 | 0.099173764 | -0.233611773 |
| Kdm5c | 1433.211927 | -0.11961167 | 0.057221072 | -2.09034304 |
| Kantr | 325.3470963 | -0.155896568 | 0.086233831 | -1.807835357 |
| n-R5s15 | 0 | NA | NA | NA |
| Tspyl2 | 1239.902915 | 0.35884157 | 0.107820664 | 3.328133547 |
| Gpr173 | 222.5226893 | 0.075775441 | 0.085151238 | 0.889892417 |
| 3010001F23Rik | 25.65934517 | -0.551183034 | 0.357782009 | -1.540555478 |
| Gm4750 | 1.060705812 | 1.401357196 | 1.140438816 | 1.228787705 |
| Gm8464 | 0.320494736 | -0.625308466 | 1.752236684 | -0.356863015 |
| Gm7158 | 0 | NA | NA | NA |
| Cldn34a | 0.10503227 | 0.274389469 | 3.724945049 | 0.073662689 |
| Gm8475 | 0.097139547 | 0.274393828 | 3.724945049 | 0.073663859 |
| Shroom2 | 979.6660779 | -0.359578096 | 0.090350027 | -3.979833865 |
| Gpr143 | 0.64528552 | 1.62659489 | 1.49599326 | 1.087300948 |
| Gm24687 | 0 | NA | NA | NA |
| Rpl7a-ps12 | 0.049553047 | 0.274391604 | 3.724945049 | 0.073663262 |
| Gm6472 | 317.0452057 | 0.411223114 | 0.16424366 | 2.50373813 |
| Gm26035 | 140.1127517 | 0.265486057 | 0.248746051 | 1.067297578 |
| Usp51 | 28.56090422 | 0.271137856 | 0.21276337 | 1.274363425 |
| Mageh1 | 200.6419644 | -0.167388406 | 0.137791318 | -1.214796468 |
| 9530051G07Rik | 22.2217983 | -0.230887968 | 0.262751919 | -0.878729903 |
| Gm5396 | 0.044544648 | 0.755286614 | 3.724945049 | 0.202764498 |
| Gm26960 | 1.043034268 | 0.119384285 | 0.950076045 | 0.12565761 |
| Foxr2 | 13.36216574 | 0.338235971 | 0.290842166 | 1.162953694 |
| Rragb | 646.2289516 | -0.149472846 | 0.071837244 | -2.080715197 |
| Gm15153 | 0.202306569 | -1.142272835 | 2.641239443 | -0.432476063 |
| Klf8 | 93.85799211 | 0.139128637 | 0.130054802 | 1.069769314 |

| | | | | |
|---|---|---|---|---|
| Gm7150 | 0 | NA | NA | NA |
| Ubqln2 | 2353.389334 | -0.140532628 | 0.05338531 | -2.632421336 |
| Gm15166 | 0 | NA | NA | NA |
| Cypt3 | 0 | NA | NA | NA |
| Gm25793 | 0 | NA | NA | NA |
| Gm8534 | 0 | NA | NA | NA |
| Kctd12b | 83.33022565 | 0.190551568 | 0.153270262 | 1.243239006 |
| 2210013O21Rik | 216.6134074 | 0.095357064 | 0.099031739 | 0.962893971 |
| Smt3h1-ps | 0.284401297 | -0.194272558 | 2.051614158 | -0.094692541 |
| Spin2c | 37.12736702 | 0.131546689 | 0.205748794 | 0.639355817 |
| Gm25895 | 0 | NA | NA | NA |
| Samt1 | 0 | NA | NA | NA |
| 4921511M17Rik | 0 | NA | NA | NA |
| Gm10057 | 0 | NA | NA | NA |
| Gm15143 | 0 | NA | NA | NA |
| Gm5947 | 0 | NA | NA | NA |
| Gm15140 | 0 | NA | NA | NA |
| Gm5645 | 0 | NA | NA | NA |
| Gm16445 | 0 | NA | NA | NA |
| Gm6620 | 0 | NA | NA | NA |
| Gm15147 | 0 | NA | NA | NA |
| Gm5060 | 0 | NA | NA | NA |
| Gm15141 | 0 | NA | NA | NA |
| 4930524N10Rik | 0 | NA | NA | NA |
| Gm5646 | 0 | NA | NA | NA |
| Gm8569 | 0 | NA | NA | NA |
| Samt4 | 0 | NA | NA | NA |
| Samt2 | 0 | NA | NA | NA |
| Gm15142 | 0 | NA | NA | NA |
| Gm22359 | 0 | NA | NA | NA |
| Gm8577 | 0.253240508 | -0.055957178 | 2.091809948 | -0.026750603 |
| Cldn34b1 | 0 | NA | NA | NA |
| Magea6 | 0 | NA | NA | NA |
| Magea3 | 0 | NA | NA | NA |
| Gm6645 | 1.418926095 | -0.286026432 | 0.802372201 | -0.356475999 |
| Gm25257 | 0 | NA | NA | NA |
| Magea8 | 0 | NA | NA | NA |
| Magea2 | 0 | NA | NA | NA |
| Gm24268 | 0 | NA | NA | NA |
| Magea5 | 0 | NA | NA | NA |
| Magea1 | 0 | NA | NA | NA |
| Cldn34b2 | 0 | NA | NA | NA |
| Gm25429 | 0.066050875 | 0.274391604 | 3.724945049 | 0.073663262 |
| Gm4997 | 0 | NA | NA | NA |
| Amd-ps1 | 0.844289582 | 0.134465708 | 1.074325016 | 0.125162968 |
| Gm8595 | 0.04393175 | -0.206490061 | 3.724945049 | -0.055434391 |
| Gm15157 | 0.250163086 | 0.801658619 | 2.491669664 | 0.321735514 |

| | | | | |
|---|---|---|---|---|
| Sat1 | 845.2085452 | 1.458558813 | 0.158343974 | 9.211331359 |
| Gm15159 | 0 | NA | NA | NA |
| Acot9 | 212.0243755 | 0.458504479 | 0.122850474 | 3.732215786 |
| Gm650 | 0 | NA | NA | NA |
| Prdx4 | 114.043658 | 0.649763223 | 0.216706954 | 2.998349669 |
| Gm15158 | 0 | NA | NA | NA |
| Gm8606 | 0.172182752 | 0.755291024 | 3.331044598 | 0.226742994 |
| n-R5s16 | 0 | NA | NA | NA |
| Ptchd1 | 278.1680322 | -0.221369641 | 0.091770521 | -2.412208628 |
| Gm15156 | 5.651205916 | 1.055926658 | 0.436133025 | 2.421111442 |
| Gm15155 | 12.34800111 | -0.643020837 | 0.291225849 | -2.207979952 |
| Gm15160 | 1.808882573 | 0.474474153 | 0.67169093 | 0.706387614 |
| Gm23404 | 0 | NA | NA | NA |
| 4930503H13Rik | 1.186325201 | -0.445474828 | 0.780233136 | -0.570950921 |
| Gm7312 | 0 | NA | NA | NA |
| Gm15162 | 0.389425067 | 1.738931155 | 3.266995524 | 0.53227228 |
| Gm15163 | 28.67019377 | 0.241938094 | 0.19873757 | 1.217374722 |
| Gm22666 | 0 | NA | NA | NA |
| Phex | 65.77163163 | 0.22973691 | 0.196039836 | 1.171888908 |
| Gm23798 | 0 | NA | NA | NA |
| Gm25795 | 0.325008436 | 1.129972727 | 2.056638555 | 0.549426988 |
| Gm15172 | 0.412251148 | -1.842152033 | 1.672423068 | -1.10148686 |
| Gm1947 | 0.704217346 | -0.687380412 | 1.484203069 | -0.463130974 |
| Gm24281 | 0 | NA | NA | NA |
| Sms | 958.8198068 | -0.139569191 | 0.056836762 | -2.455614762 |
| Mbtps2 | 467.2768864 | 0.08910793 | 0.063493504 | 1.403418062 |
| Gm6568 | 0.666747433 | -0.18033779 | 1.403536071 | -0.128488176 |
| Yy2 | 0 | NA | NA | NA |
| Gm15176 | 0 | NA | NA | NA |
| Gm15175 | 0.320922609 | 1.693538175 | 1.73160525 | 0.978016309 |
| Gm15173 | 0.066050875 | 0.274391604 | 3.724945049 | 0.073663262 |
| Gm23381 | 0 | NA | NA | NA |
| Gm8644 | 0.329848065 | -0.206485379 | 1.936344079 | -0.106636719 |
| Gm15171 | 0 | NA | NA | NA |
| Gm8648 | 0 | NA | NA | NA |
| Smpx | 10.49823735 | -0.139830528 | 0.307607231 | -0.454574904 |
| Gm15169 | 2.861039839 | 2.484656775 | 0.81323121 | 3.0552895 |
| Klhl34 | 1.755040647 | 0.652700243 | 0.722133933 | 0.903849291 |
| Cnksr2 | 3647.587889 | -0.235787859 | 0.12625107 | -1.867610772 |
| Gm8659 | 0.100902008 | 0.274391828 | 3.724945049 | 0.073663322 |
| Gm15170 | 0.046345709 | 0.274391604 | 3.724945049 | 0.073663262 |
| Gm8662 | 0 | NA | NA | NA |
| Gm15177 | 0 | NA | NA | NA |
| Gm22522 | 0 | NA | NA | NA |
| Gm15168 | 0 | NA | NA | NA |
| Gm5764 | 0.044544648 | 0.755286614 | 3.724945049 | 0.202764498 |
| Gm15185 | 0 | NA | NA | NA |

| | | | | |
|---|---|---|---|---|
| Gm15188 | 0 | NA | NA | NA |
| Gm15189 | 0 | NA | NA | NA |
| Rps6ka3 | 1265.248659 | -0.134118563 | 0.07306466 | -1.835614695 |
| Eif1ax | 1820.91249 | -0.183396799 | 0.069370598 | -2.643725216 |
| Gm15187 | 0 | NA | NA | NA |
| Map7d2 | 1767.738311 | -0.030309703 | 0.089947747 | -0.336970119 |
| Gm23806 | 1.294438978 | 0.033709795 | 0.854930909 | 0.039429847 |
| A830080D01Rik | 225.115431 | -0.015621178 | 0.090656466 | -0.172311787 |
| Gm15190 | 9.37392129 | 0.034117322 | 0.357825646 | 0.095346218 |
| Gm15186 | 6.883933299 | -0.270720425 | 0.464190531 | -0.583209711 |
| Sh3kbp1 | 688.8231907 | -0.0033031 | 0.059380246 | -0.055626245 |
| Map3k15 | 25.20550808 | 0.408898997 | 0.255610688 | 1.599694441 |
| Gm15194 | 0 | NA | NA | NA |
| Gm8682 | 0 | NA | NA | NA |
| Pdha1 | 1985.030852 | -0.128011521 | 0.052246723 | -2.450134934 |
| Gm15195 | 0 | NA | NA | NA |
| Gm15192 | 0 | NA | NA | NA |
| Gm15196 | 0.075043205 | 0.755286614 | 3.724945049 | 0.202764498 |
| Gm15193 | 0 | NA | NA | NA |
| Adgrg2 | 34.96479159 | 0.595095405 | 0.249692007 | 2.383317804 |
| Gm15241 | 0.873806335 | 2.008638072 | 1.135470065 | 1.768992538 |
| Phka2 | 591.6410219 | 0.559811953 | 0.130309952 | 4.296003052 |
| Gm15243 | 0.261996957 | 0.755309454 | 2.055677216 | 0.367426096 |
| Ppef1 | 10.0328463 | 1.258973357 | 0.380756711 | 3.306503392 |
| Gm15242 | 0.08486385 | 0.755286617 | 3.724945049 | 0.202764499 |
| Gm15244 | 0.061920613 | 0.274393828 | 3.724945049 | 0.073663859 |
| Rs1 | 1.55354348 | -0.287138331 | 0.825393625 | -0.347880481 |
| Cdkl5 | 3363.729212 | -0.068499066 | 0.103677848 | -0.660691436 |
| Gja6 | 33.35128538 | -0.39768653 | 0.283614868 | -1.402206215 |
| Scml2 | 17.41690607 | -0.436281154 | 0.264047376 | -1.652283617 |
| Gm5419 | 0 | NA | NA | NA |
| Gm15262 | 0 | NA | NA | NA |
| Prkaca-ps1 | 0.150961315 | 0.755296921 | 2.849194963 | 0.265091344 |
| Rai2 | 127.3185232 | 0.05628884 | 0.097909847 | 0.574904787 |
| Scml1 | 0.926668879 | 0.054313939 | 1.060632666 | 0.051209001 |
| Gm15205 | 1.789402658 | -1.420445024 | 0.797867868 | -1.780301077 |
| Nhs | 212.8785834 | -0.205428027 | 0.093089737 | -2.206774167 |
| Gm15202 | 0.888501202 | -1.103008353 | 1.218192897 | -0.905446383 |
| Gm26317 | 0.531781488 | 0.79247626 | 1.471226279 | 0.538650153 |
| Gm23794 | 0.04393175 | -0.206490061 | 3.724945049 | -0.055434391 |
| Gm27490 | 0 | NA | NA | NA |
| Gm23867 | 0.167772976 | -0.206492322 | 3.417713432 | -0.060418267 |
| Gm15204 | 0.395626157 | -0.573775266 | 1.56248642 | -0.367219362 |
| Reps2 | 2262.978211 | -0.172725594 | 0.078143667 | -2.210359479 |
| Gm7331 | 62.14126764 | 0.209684947 | 0.176165117 | 1.190275067 |
| Gm15206 | 2.290494867 | -0.435517822 | 0.60729198 | -0.71714733 |
| Gm15203 | 0 | NA | NA | NA |

| | | | | |
|---|---|---|---|---|
| Gm22257 | 0 | NA | NA | NA |
| Gm25557 | 0 | NA | NA | NA |
| Gm8737 | 0 | NA | NA | NA |
| Gm15201 | 8.230899355 | 0.238250314 | 0.355112757 | 0.670914545 |
| Gm23786 | 0 | NA | NA | NA |
| Rbbp7 | 1339.028074 | 0.417867317 | 0.107006624 | 3.905060265 |
| Txlng | 376.7131286 | -0.046389443 | 0.097685842 | -0.474883997 |
| Gm16459 | 0 | NA | NA | NA |
| Gm15207 | 0 | NA | NA | NA |
| Gm23901 | 0.060174537 | 0.755286614 | 3.724945049 | 0.202764498 |
| Syap1 | 318.0290325 | 0.150273204 | 0.086077525 | 1.745789089 |
| Mir3473a | 0.568149222 | -2.14702507 | 1.414135204 | -1.51826011 |
| Gm15209 | 0 | NA | NA | NA |
| Ctps2 | 433.4122222 | -0.064471142 | 0.062908281 | -1.024843488 |
| S100g | 0.048949532 | -0.206487837 | 3.724945049 | -0.055433794 |
| Gm23501 | 0 | NA | NA | NA |
| Gm22746 | 0 | NA | NA | NA |
| Gm15215 | 0 | NA | NA | NA |
| Gm15212 | 0 | NA | NA | NA |
| Gm15214 | 0 | NA | NA | NA |
| Gm26007 | 0 | NA | NA | NA |
| Gm7199 | 0 | NA | NA | NA |
| Grpr | 12.17739777 | -0.208629959 | 0.313044614 | -0.666454395 |
| Rnf138rt1 | 7.321455992 | 0.239177028 | 0.380481443 | 0.628616802 |
| Gm15213 | 0 | NA | NA | NA |
| Ap1s2 | 658.6125027 | 0.024996123 | 0.106498126 | 0.234709514 |
| Zrsr2 | 430.8776927 | 0.095744214 | 0.085213841 | 1.123575858 |
| Car5b | 61.00096164 | 0.064758013 | 0.189718759 | 0.341336897 |
| Siah1b | 50.98963704 | 0.308015729 | 0.156573433 | 1.967228554 |
| Tmem27 | 1.459315314 | 0.960699128 | 0.918435983 | 1.04601643 |
| n-R5s18 | 0.046345709 | 0.274391604 | 3.724945049 | 0.073663262 |
| Ace2 | 16.8793861 | 0.682129953 | 0.34894205 | 1.954851678 |
| Bmx | 4.886164753 | -0.179645183 | 0.492522889 | -0.364744841 |
| Gm25367 | 0 | NA | NA | NA |
| Pir | 44.18337336 | -0.208991705 | 0.176760035 | -1.182347042 |
| Gm22023 | 0.437539382 | -1.349186383 | 1.661222204 | -0.812164911 |
| Figf | 22.19425402 | 0.344570192 | 0.290790157 | 1.18494448 |
| Piga | 134.018528 | -0.45041555 | 0.116511728 | -3.865838715 |
| Asb11 | 36.5181848 | 0.938026024 | 0.231353 | 4.054522837 |
| Gm15211 | 0.30015424 | 0.755305567 | 2.014987115 | 0.374843869 |
| Asb9 | 0.209460102 | 1.236200975 | 2.189949797 | 0.564488271 |
| Gm15216 | 0 | NA | NA | NA |
| Mospd2 | 329.2346432 | -0.043033722 | 0.096485966 | -0.446010174 |
| Fancb | 25.83413697 | -0.205566305 | 0.231100642 | -0.889509885 |
| Gm17604 | 0 | NA | NA | NA |
| Glra2 | 135.3189216 | -0.04741988 | 0.143636728 | -0.330137567 |
| Gm15236 | 0.167734623 | 0.75528487 | 3.394819337 | 0.22248161 |

| | | | | |
|---|---|---|---|---|
| Gm22062 | 0 | NA | NA | NA |
| Gm15224 | 0 | NA | NA | NA |
| Gm15225 | 0 | NA | NA | NA |
| Gemin8 | 60.76505143 | -0.473328962 | 0.171451436 | -2.760717395 |
| Gm15227 | 0 | NA | NA | NA |
| Gpm6b | 8313.7276 | -0.290783266 | 0.111732895 | -2.602485732 |
| Ofd1 | 183.0566898 | 0.277321538 | 0.109399235 | 2.534949513 |
| Trappc2 | 241.3367123 | -0.239762516 | 0.102514539 | -2.33881474 |
| Rab9 | 264.7892433 | 0.273907589 | 0.096124541 | 2.849507375 |
| Gm15229 | 2.091380692 | 1.700747698 | 1.292254662 | 1.316108773 |
| Gm15223 | 0.211816017 | 1.670237526 | 2.167791704 | 0.770478789 |
| Tceanc | 44.56431763 | 0.666226033 | 0.20745168 | 3.211475728 |
| Egfl6 | 12.75317809 | -0.498415498 | 0.331624346 | -1.502952072 |
| Gm15226 | 0.151064894 | 1.203968472 | 2.845233505 | 0.423152782 |
| Gm1720 | 0.332179105 | -1.048680196 | 1.573683691 | -0.666385629 |
| Rpl7a-ps11 | 9.849636779 | 0.807899745 | 0.346232692 | 2.333401105 |
| Gm15230 | 0.199910846 | 0.274411844 | 2.215429956 | 0.123863922 |
| Gm15231 | 0 | NA | NA | NA |
| AL671888.1 | 0.061920613 | 0.274393828 | 3.724945049 | 0.073663859 |
| Gm22368 | 0 | NA | NA | NA |
| Gm8814 | 0.110595523 | 0.755282166 | 3.724945049 | 0.202763304 |
| Gm8817 | 0.926280627 | 2.584686314 | 1.145570986 | 2.256242823 |
| Gm15232 | 1.683504739 | 0.023097685 | 0.928488761 | 0.024876645 |
| Gm15228 | 2.037836299 | 3.678812751 | 0.981722058 | 3.74730579 |
| Tmsb4x | 4681.12954 | 0.688413957 | 0.106178834 | 6.483532827 |
| Tlr8 | 77.00154128 | 3.901988042 | 0.590500935 | 6.60792864 |
| Tlr7 | 273.9150659 | 2.829084287 | 0.208260637 | 13.58434475 |
| Prps2 | 362.3824202 | -0.285023268 | 0.099508499 | -2.864310788 |
| Gm15239 | 0.195968864 | 1.791179267 | 3.234710441 | 0.553737127 |
| Gm26368 | 0.238567011 | 0.7552969 | 2.57003757 | 0.293885548 |
| Gm6744 | 0.523620968 | 2.291425766 | 3.095881838 | 0.74015285 |
| Frmpd4 | 1476.981339 | -0.028259162 | 0.07034257 | -0.401736284 |
| Gm15238 | 0.110870145 | -0.206490066 | 3.724945049 | -0.055434393 |
| Gm24772 | 0 | NA | NA | NA |
| Gm15240 | 0 | NA | NA | NA |
| Gm15237 | 0 | NA | NA | NA |
| Gm8832 | 0.092228164 | 0.75528439 | 3.724945049 | 0.202763901 |
| Gm7209 | 0 | NA | NA | NA |
| Gm23197 | 0 | NA | NA | NA |
| Msl3 | 245.0695052 | -0.025951028 | 0.075203022 | -0.345079589 |
| Gm15233 | 0 | NA | NA | NA |
| Arhgap6 | 136.4378488 | -0.066493131 | 0.126870338 | -0.524103051 |
| Gm15261 | 0.130746305 | 1.120259715 | 3.664569624 | 0.305700213 |
| Amelx | 0.092691418 | 0.274387304 | 3.724945049 | 0.073662108 |
| Hccs | 283.4725314 | -0.091137392 | 0.102417194 | -0.889864179 |
| Gm27980 | 0.850639233 | 0.596364251 | 1.17353977 | 0.508175578 |
| Gm15246 | 24.67077973 | -0.418720574 | 0.294868206 | -1.420026186 |

| | | | | |
|---|---|---|---|---|
| BC022960 | 47.81857404 | -0.296347324 | 0.165813069 | -1.787237434 |
| Gm15245 | 0.062426873 | 0.755288839 | 3.724945049 | 0.202765096 |
| Mid1 | 46.97430459 | 0.07121724 | 0.208152327 | 0.342140014 |
| Gm25651 | 0 | NA | NA | NA |
| 4933400A11Rik | 0 | NA | NA | NA |
| G530011O06Rik | 0.809500859 | 2.750630419 | 1.387884272 | 1.981887449 |
| Gm15726 | 1.372866643 | -0.072884688 | 0.834839955 | -0.087303785 |
| Gm15247 | 0.097139547 | 0.274393828 | 3.724945049 | 0.073663859 |
| Gm21887 | 20.27296318 | -0.098668047 | 0.371607925 | -0.26551653 |
| Asmt | 1.358880654 | 0.421075976 | 0.909990217 | 0.462725827 |
| Gm15069 | 0 | NA | NA | NA |
| Gm15068 | 0 | NA | NA | NA |
| Gm29277 | 0 | NA | NA | NA |
| Gm29089 | 0 | NA | NA | NA |
| Gm2098 | 0.060174537 | 0.755286614 | 3.724945049 | 0.202764498 |
| Zfy1 | 0.044191549 | -0.206490061 | 3.724945049 | -0.055434391 |
| Uba1y | 4.871321793 | 0.212076604 | 0.553441363 | 0.383196159 |
| Gm28588 | 0 | NA | NA | NA |
| Gm28587 | 0.444799355 | 0.789694816 | 3.072797206 | 0.256995422 |
| Rhoay-ps3 | 0.088383099 | -0.687376175 | 3.724945049 | -0.184533239 |
| Kdm5d | 320.9905738 | -0.107633952 | 0.186646748 | -0.576671992 |
| n-R5s1 | 0 | NA | NA | NA |
| AC145392.1 | 0 | NA | NA | NA |
| Eif2s3y | 295.8023547 | -0.222597663 | 0.161380612 | -1.37933337 |
| Gm29650 | 9.742440598 | -1.585176214 | 0.602198472 | -2.632315237 |
| Tspy-ps | 2.062938699 | 0.691620293 | 0.950327059 | 0.72777081 |
| Uty | 395.1440632 | 0.103752721 | 0.125984722 | 0.82353415 |
| Ddx3y | 555.3045698 | 0.449552019 | 0.146108458 | 3.07683775 |
| Gm4017 | 4.097680658 | 0.646890346 | 0.605892197 | 1.067665749 |
| Gm18665 | 3.611403245 | 0.698781203 | 0.580194021 | 1.204392286 |
| Usp9y | 1.14946738 | -2.752938855 | 3.67318078 | -0.749470015 |
| Gm25565 | 0 | NA | NA | NA |
| Gm17790 | 0 | NA | NA | NA |
| Gm29167 | 0 | NA | NA | NA |
| Gm8446 | 0.334381441 | -1.215657078 | 1.743550105 | -0.69723094 |
| Gm28534 | 0 | NA | NA | NA |
| Gm28975 | 0 | NA | NA | NA |
| Gm28974 | 0 | NA | NA | NA |
| Gm18797 | 0 | NA | NA | NA |
| Rhoay-ps1 | 0 | NA | NA | NA |
| Zfy2 | 0 | NA | NA | NA |
| Uba1y-ps2 | 0 | NA | NA | NA |
| Gm18796 | 0 | NA | NA | NA |
| Gm29552 | 0 | NA | NA | NA |
| Gm29550 | 0 | NA | NA | NA |
| Gm29551 | 0 | NA | NA | NA |
| Gm8480 | 0 | NA | NA | NA |

| | | | | |
|---|---|---|---|---|
| Gm29553 | 0 | NA | NA | NA |
| Gm22119 | 0 | NA | NA | NA |
| Gm28630 | 0 | NA | NA | NA |
| Gm28629 | 0 | NA | NA | NA |
| Gm6026 | 0 | NA | NA | NA |
| Uba1y-ps1 | 0 | NA | NA | NA |
| Sry | 0 | NA | NA | NA |
| Gm18798 | 0 | NA | NA | NA |
| Gm16501 | 0 | NA | NA | NA |
| Gm28628 | 0 | NA | NA | NA |
| Gm4064 | 0 | NA | NA | NA |
| Gm6137 | 0 | NA | NA | NA |
| Gm8498 | 0 | NA | NA | NA |
| Rbmy | 0 | NA | NA | NA |
| Gm10256 | 0 | NA | NA | NA |
| Gm8502 | 0 | NA | NA | NA |
| Gm8506 | 0 | NA | NA | NA |
| Gm10352 | 0 | NA | NA | NA |
| Gm8510 | 0 | NA | NA | NA |
| Gm29289 | 0 | NA | NA | NA |
| Gm21677 | 0 | NA | NA | NA |
| Gm29288 | 0 | NA | NA | NA |
| Gm21693 | 0 | NA | NA | NA |
| Gm21139 | 0 | NA | NA | NA |
| Gm21704 | 0 | NA | NA | NA |
| Gm21147 | 0 | NA | NA | NA |
| Gm21708 | 0 | NA | NA | NA |
| Gm21156 | 0 | NA | NA | NA |
| Rhoay-ps2 | 0 | NA | NA | NA |
| Gm2316 | 0 | NA | NA | NA |
| Gm3376 | 0 | NA | NA | NA |
| Gm2325 | 0 | NA | NA | NA |
| Gm28649 | 0 | NA | NA | NA |
| Gm8521 | 0 | NA | NA | NA |
| Gm18177 | 0 | NA | NA | NA |
| Gm28429 | 0 | NA | NA | NA |
| Gm28413 | 0 | NA | NA | NA |
| Gm28412 | 0 | NA | NA | NA |
| Gm8525 | 0 | NA | NA | NA |
| Gm28414 | 0 | NA | NA | NA |
| Gm28648 | 0 | NA | NA | NA |
| Gm28647 | 0 | NA | NA | NA |
| Gm29435 | 0 | NA | NA | NA |
| Vmn2r-ps139 | 0.046345709 | 0.274391604 | 3.724945049 | 0.073663262 |
| Gm29038 | 0 | NA | NA | NA |
| Gm28191 | 0 | NA | NA | NA |
| Gm28186 | 0 | NA | NA | NA |

| | | | | |
|---|---|---|---|---|
| Gm21064 | 0 | NA | NA | NA |
| Gm28241 | 0 | NA | NA | NA |
| Gm28242 | 0 | NA | NA | NA |
| Gm37222 | 0 | NA | NA | NA |
| Gm28356 | 0 | NA | NA | NA |
| Gm28357 | 0 | NA | NA | NA |
| Gm29352 | 0 | NA | NA | NA |
| Gm29351 | 0 | NA | NA | NA |
| Gm29350 | 0 | NA | NA | NA |
| Gm29349 | 0 | NA | NA | NA |
| Gm29355 | 0 | NA | NA | NA |
| Gm20918 | 0 | NA | NA | NA |
| Gm29354 | 0 | NA | NA | NA |
| Gm29353 | 0 | NA | NA | NA |
| Gm21820 | 0 | NA | NA | NA |
| Gm29357 | 0 | NA | NA | NA |
| Gm29356 | 0 | NA | NA | NA |
| Gm29161 | 0 | NA | NA | NA |
| Gm28767 | 0 | NA | NA | NA |
| Gm29160 | 0 | NA | NA | NA |
| Gm20915 | 0 | NA | NA | NA |
| Gm28766 | 0 | NA | NA | NA |
| Gm29159 | 0 | NA | NA | NA |
| Gm27663 | 0 | NA | NA | NA |
| Gm29158 | 0 | NA | NA | NA |
| Gm21854 | 0 | NA | NA | NA |
| Gm28768 | 0 | NA | NA | NA |
| Gm28769 | 0 | NA | NA | NA |
| Gm20914 | 0 | NA | NA | NA |
| Gm28171 | 0 | NA | NA | NA |
| Gm28173 | 0 | NA | NA | NA |
| Gm37236 | 0 | NA | NA | NA |
| Gm29194 | 0.066050875 | 0.274391604 | 3.724945049 | 0.073663262 |
| Gm21778 | 0 | NA | NA | NA |
| Gm29195 | 0 | NA | NA | NA |
| Gm29196 | 0 | NA | NA | NA |
| Gm28435 | 0 | NA | NA | NA |
| Gm28436 | 0 | NA | NA | NA |
| Gm29197 | 0 | NA | NA | NA |
| Gm29198 | 0 | NA | NA | NA |
| Gm29193 | 0 | NA | NA | NA |
| Gm28430 | 0 | NA | NA | NA |
| Gm21746 | 0 | NA | NA | NA |
| Gm28593 | 0 | NA | NA | NA |
| Gm20873 | 0 | NA | NA | NA |
| Gm28916 | 0 | NA | NA | NA |
| Gm28915 | 0 | NA | NA | NA |

| | | | | |
|---|---|---|---|---|
| Gm28918 | 0.060174537 | 0.755286614 | 3.724945049 | 0.202764498 |
| Gm28917 | 0 | NA | NA | NA |
| Gm28919 | 0 | NA | NA | NA |
| Gm21719 | 0 | NA | NA | NA |
| Gm28921 | 0 | NA | NA | NA |
| Gm28920 | 0 | NA | NA | NA |
| Gm28444 | 0 | NA | NA | NA |
| Gm28443 | 0 | NA | NA | NA |
| Gm20830 | 0 | NA | NA | NA |
| Gm28442 | 0 | NA | NA | NA |
| Gm28445 | 0 | NA | NA | NA |
| Gm20788 | 0 | NA | NA | NA |
| Gm28999 | 0 | NA | NA | NA |
| Gm21244 | 0.139376635 | 0.740973539 | 2.922770099 | 0.253517558 |
| Gm28997 | 0 | NA | NA | NA |
| Gm28998 | 0 | NA | NA | NA |
| Gm28398 | 0 | NA | NA | NA |
| Gm28397 | 0 | NA | NA | NA |
| Gm28400 | 0 | NA | NA | NA |
| Gm28399 | 0 | NA | NA | NA |
| Gm28395 | 0 | NA | NA | NA |
| Gm21788 | 0 | NA | NA | NA |
| Gm28394 | 0 | NA | NA | NA |
| Gm28396 | 0 | NA | NA | NA |
| Gm28393 | 0 | NA | NA | NA |
| Gm20826 | 0.204257686 | -0.620360393 | 2.624259596 | -0.236394446 |
| Vmn2r-ps149 | 0 | NA | NA | NA |
| Gm28576 | 0 | NA | NA | NA |
| Gm28575 | 0 | NA | NA | NA |
| Gm28574 | 0 | NA | NA | NA |
| Gm28573 | 0 | NA | NA | NA |
| Gm28572 | 0 | NA | NA | NA |
| Gm28571 | 0 | NA | NA | NA |
| Gm28570 | 0 | NA | NA | NA |
| Gm29049 | 0 | NA | NA | NA |
| Gm20825 | 0 | NA | NA | NA |
| Gm29047 | 0 | NA | NA | NA |
| Gm29048 | 0 | NA | NA | NA |
| Gm29045 | 0 | NA | NA | NA |
| Gm29046 | 0 | NA | NA | NA |
| Gm29043 | 0 | NA | NA | NA |
| Gm29044 | 0 | NA | NA | NA |
| Gm37951 | 0 | NA | NA | NA |
| Gm20824 | 0 | NA | NA | NA |
| Gm28146 | 0 | NA | NA | NA |
| Gm28145 | 0 | NA | NA | NA |
| Gm28148 | 0 | NA | NA | NA |

| | | | | |
|---|---|---|---|---|
| Gm28147 | 0 | NA | NA | NA |
| Gm28142 | 0 | NA | NA | NA |
| Gm28141 | 0 | NA | NA | NA |
| Gm28144 | 0 | NA | NA | NA |
| Gm28143 | 0 | NA | NA | NA |
| Gm28150 | 0 | NA | NA | NA |
| Gm28149 | 0 | NA | NA | NA |
| Gm28098 | 0 | NA | NA | NA |
| Gm28330 | 0 | NA | NA | NA |
| Gm28331 | 0 | NA | NA | NA |
| Gm28099 | 0 | NA | NA | NA |
| Gm20815 | 0 | NA | NA | NA |
| Gm28328 | 0 | NA | NA | NA |
| Gm28329 | 0 | NA | NA | NA |
| Gm28270 | 0 | NA | NA | NA |
| Gm28327 | 0 | NA | NA | NA |
| Gm28097 | 0 | NA | NA | NA |
| Gm29365 | 0 | NA | NA | NA |
| Gm29364 | 0 | NA | NA | NA |
| Gm29363 | 0 | NA | NA | NA |
| Gm21292 | 0 | NA | NA | NA |
| Gm21721 | 0 | NA | NA | NA |
| Gm29366 | 0 | NA | NA | NA |
| Gm29281 | 0 | NA | NA | NA |
| Gm29330 | 0 | NA | NA | NA |
| Gm29329 | 0 | NA | NA | NA |
| Gm21812 | 0 | NA | NA | NA |
| Gm29535 | 0 | NA | NA | NA |
| Gm29533 | 0 | NA | NA | NA |
| Gm29534 | 0 | NA | NA | NA |
| Gm21874 | 0 | NA | NA | NA |
| Gm28849 | 0 | NA | NA | NA |
| Gm28847 | 0 | NA | NA | NA |
| Gm28848 | 0 | NA | NA | NA |
| Gm28846 | 0 | NA | NA | NA |
| Gm29471 | 0 | NA | NA | NA |
| Gm29474 | 0 | NA | NA | NA |
| Gm29475 | 0 | NA | NA | NA |
| Gm20821 | 0 | NA | NA | NA |
| Gm28506 | 0 | NA | NA | NA |
| Gm28507 | 0 | NA | NA | NA |
| Gm21301 | 0 | NA | NA | NA |
| Gm29142 | 0 | NA | NA | NA |
| Gm21310 | 0 | NA | NA | NA |
| Gm28509 | 0 | NA | NA | NA |
| Gm28508 | 0 | NA | NA | NA |
| Gm20834 | 0 | NA | NA | NA |

| | | | | |
|---|---|---|---|---|
| Gm28510 | 0 | NA | NA | NA |
| Gm29525 | 0 | NA | NA | NA |
| Gm29526 | 0 | NA | NA | NA |
| Gm20737 | 0.044544648 | 0.755286614 | 3.724945049 | 0.202764498 |
| Gm21775 | 0 | NA | NA | NA |
| Gm21900 | 0 | NA | NA | NA |
| Gm29527 | 0 | NA | NA | NA |
| Gm20775 | 1.978445198 | 0.922487958 | 0.791662649 | 1.165253861 |
| Gm29522 | 0 | NA | NA | NA |
| Gm29523 | 0 | NA | NA | NA |
| Gm29524 | 0 | NA | NA | NA |
| Gm29528 | 0 | NA | NA | NA |
| Gm20777 | 0 | NA | NA | NA |
| Gm28955 | 0 | NA | NA | NA |
| Gm28954 | 0 | NA | NA | NA |
| Gm28953 | 0 | NA | NA | NA |
| Gm20776 | 0 | NA | NA | NA |
| Gm28958 | 0 | NA | NA | NA |
| Gm20828 | 0 | NA | NA | NA |
| Gm28957 | 0 | NA | NA | NA |
| Gm28956 | 0 | NA | NA | NA |
| Gm28952 | 0 | NA | NA | NA |
| Gm28951 | 0 | NA | NA | NA |
| Gm28390 | 0 | NA | NA | NA |
| Gm28391 | 0 | NA | NA | NA |
| Gm28389 | 0 | NA | NA | NA |
| Gm28655 | 0 | NA | NA | NA |
| Gm28654 | 0 | NA | NA | NA |
| Gm28657 | 0 | NA | NA | NA |
| Gm28656 | 0 | NA | NA | NA |
| Gm29554 | 0 | NA | NA | NA |
| Gm29556 | 0 | NA | NA | NA |
| Gm20812 | 0 | NA | NA | NA |
| Gm35108 | 0 | NA | NA | NA |
| Gm37739 | 0 | NA | NA | NA |
| Gm21736 | 0 | NA | NA | NA |
| Gm21379 | 0 | NA | NA | NA |
| Gm36928 | 0 | NA | NA | NA |
| Gm20807 | 0 | NA | NA | NA |
| Gm38084 | 0 | NA | NA | NA |
| Gm37231 | 0 | NA | NA | NA |
| Gm21891 | 0 | NA | NA | NA |
| Gm21425 | 0 | NA | NA | NA |
| Gm37252 | 0 | NA | NA | NA |
| Gm21828 | 0 | NA | NA | NA |
| Gm21440 | 0 | NA | NA | NA |
| Gm21454 | 0 | NA | NA | NA |

| | | | | |
|---|---|---|---|---|
| Gm37147 | 0 | NA | NA | NA |
| Gm38159 | 0 | NA | NA | NA |
| Gm21725 | 0 | NA | NA | NA |
| Gm37870 | 0 | NA | NA | NA |
| Gm37378 | 0 | NA | NA | NA |
| Gm21866 | 0 | NA | NA | NA |
| Gm20773 | 0 | NA | NA | NA |
| Gm36398 | 0 | NA | NA | NA |
| Gm37441 | 0 | NA | NA | NA |
| Gm37561 | 0 | NA | NA | NA |
| Gm36728 | 0 | NA | NA | NA |
| Gm32295 | 0 | NA | NA | NA |
| Gm20836 | 0 | NA | NA | NA |
| Gm37263 | 0 | NA | NA | NA |
| Gm21767 | 0 | NA | NA | NA |
| Gm37130 | 0 | NA | NA | NA |
| Gm37467 | 0 | NA | NA | NA |
| Gm20928 | 0 | NA | NA | NA |
| Gm38363 | 0 | NA | NA | NA |
| Gm30174 | 0 | NA | NA | NA |
| Gm30353 | 0 | NA | NA | NA |
| Gm30686 | 0 | NA | NA | NA |
| Gm30737 | 0 | NA | NA | NA |
| Gm36987 | 0 | NA | NA | NA |
| Gm31186 | 0 | NA | NA | NA |
| Gm37654 | 0 | NA | NA | NA |
| Gm37044 | 0 | NA | NA | NA |
| Gm31511 | 0 | NA | NA | NA |
| Gm37572 | 0 | NA | NA | NA |
| Gm37344 | 0 | NA | NA | NA |
| Gm37059 | 0 | NA | NA | NA |
| Gm31942 | 0 | NA | NA | NA |
| Gm37440 | 0 | NA | NA | NA |
| Gm38185 | 0 | NA | NA | NA |
| Gm37395 | 0 | NA | NA | NA |
| Gm37577 | 0 | NA | NA | NA |
| Gm20822 | 0 | NA | NA | NA |
| Gm38183 | 0 | NA | NA | NA |
| Gm38149 | 0 | NA | NA | NA |
| Gm38145 | 0 | NA | NA | NA |
| Gm32033 | 0 | NA | NA | NA |
| Gm32447 | 0 | NA | NA | NA |
| Gm20770 | 0 | NA | NA | NA |
| Gm38016 | 0 | NA | NA | NA |
| Gm37158 | 0.04393175 | -0.206490061 | 3.724945049 | -0.055434391 |
| Gm36998 | 0 | NA | NA | NA |
| Gm33112 | 0 | NA | NA | NA |

| | | | | |
|---|---|---|---|---|
| Gm21809 | 0 | NA | NA | NA |
| Gm34015 | 0 | NA | NA | NA |
| Gm20877 | 0 | NA | NA | NA |
| Gm33191 | 0 | NA | NA | NA |
| Gm38003 | 0 | NA | NA | NA |
| Gm20781 | 0 | NA | NA | NA |
| Rbm31y | 0 | NA | NA | NA |
| Gm37325 | 0 | NA | NA | NA |
| Gm20833 | 0 | NA | NA | NA |
| Gm37865 | 0 | NA | NA | NA |
| Gm36941 | 0 | NA | NA | NA |
| Gm21904 | 0 | NA | NA | NA |
| Gm37937 | 0 | NA | NA | NA |
| Gm20772 | 0 | NA | NA | NA |
| Gm20831 | 0 | NA | NA | NA |
| Gm37286 | 0 | NA | NA | NA |
| Srsy | 0 | NA | NA | NA |
| Gm38028 | 0 | NA | NA | NA |
| Gm21764 | 0 | NA | NA | NA |
| Ssty1 | 0 | NA | NA | NA |
| Gm38072 | 0 | NA | NA | NA |
| Gm20851 | 0 | NA | NA | NA |
| Gm30312 | 0 | NA | NA | NA |
| Gm37433 | 0 | NA | NA | NA |
| Gm37473 | 0 | NA | NA | NA |
| Gm37656 | 0 | NA | NA | NA |
| Gm38127 | 0 | NA | NA | NA |
| Gm30638 | 0 | NA | NA | NA |
| Gm38361 | 0 | NA | NA | NA |
| Gm31260 | 0 | NA | NA | NA |
| Gm21660 | 0 | NA | NA | NA |
| Gm38209 | 0 | NA | NA | NA |
| Gm21822 | 0 | NA | NA | NA |
| Gm37734 | 0 | NA | NA | NA |
| Gm38013 | 0 | NA | NA | NA |
| Gm21678 | 0 | NA | NA | NA |
| Gm37948 | 0 | NA | NA | NA |
| Gm37442 | 0 | NA | NA | NA |
| Gm34550 | 0 | NA | NA | NA |
| Gm37544 | 0 | NA | NA | NA |
| Gm34716 | 0 | NA | NA | NA |
| Gm37635 | 0 | NA | NA | NA |
| Gm35070 | 0 | NA | NA | NA |
| Gm35134 | 0 | NA | NA | NA |
| Gm37998 | 0 | NA | NA | NA |
| Gm38094 | 0 | NA | NA | NA |
| Gm21763 | 0 | NA | NA | NA |

| | | | | |
|---|---|---|---|---|
| Gm38369 | 0 | NA | NA | NA |
| Gm35843 | 0 | NA | NA | NA |
| Gm38168 | 0 | NA | NA | NA |
| Gm37740 | 0 | NA | NA | NA |
| Gm27701 | 0 | NA | NA | NA |
| Gm37723 | 0 | NA | NA | NA |
| Gm36261 | 0 | NA | NA | NA |
| Gm37898 | 0 | NA | NA | NA |
| Gm36467 | 0 | NA | NA | NA |
| Gm37927 | 0 | NA | NA | NA |
| Gm36891 | 0 | NA | NA | NA |
| Gm36929 | 0 | NA | NA | NA |
| Gm38174 | 0 | NA | NA | NA |
| Gm29866 | 0 | NA | NA | NA |
| Gm36950 | 0 | NA | NA | NA |
| Gm20910 | 0 | NA | NA | NA |
| Gm30598 | 0 | NA | NA | NA |
| Gm20909 | 0 | NA | NA | NA |
| Gm20865 | 0 | NA | NA | NA |
| Gm37462 | 0 | NA | NA | NA |
| Gm37658 | 0 | NA | NA | NA |
| Gm32347 | 0 | NA | NA | NA |
| Gm32419 | 0 | NA | NA | NA |
| Gm38370 | 0 | NA | NA | NA |
| Gm38054 | 0 | NA | NA | NA |
| Gm37476 | 0 | NA | NA | NA |
| Gm37369 | 0 | NA | NA | NA |
| Gm38181 | 0 | NA | NA | NA |
| Gm31335 | 0 | NA | NA | NA |
| Gm37690 | 0 | NA | NA | NA |
| Gm38107 | 0 | NA | NA | NA |
| Gm31571 | 0 | NA | NA | NA |
| Gm32114 | 0 | NA | NA | NA |
| Gm37075 | 0 | NA | NA | NA |
| Gm37921 | 0 | NA | NA | NA |
| Gm32180 | 0 | NA | NA | NA |
| Gm38371 | 0 | NA | NA | NA |
| Gm37657 | 0 | NA | NA | NA |
| Gm37448 | 0 | NA | NA | NA |
| Gm33163 | 0 | NA | NA | NA |
| Gm21065 | 0.044191549 | -0.206490061 | 3.724945049 | -0.055434391 |
| Gm36978 | 0 | NA | NA | NA |
| Gm37721 | 0 | NA | NA | NA |
| Gm37129 | 0 | NA | NA | NA |
| Gm37990 | 0 | NA | NA | NA |
| Gm21366 | 0 | NA | NA | NA |
| Gm33954 | 0 | NA | NA | NA |

| | | | | |
|---|---|---|---|---|
| Gm37538 | 0 | NA | NA | NA |
| Gm21871 | 0 | NA | NA | NA |
| Gm34274 | 0 | NA | NA | NA |
| Gm34217 | 0 | NA | NA | NA |
| Gm21469 | 0 | NA | NA | NA |
| Gm21661 | 0 | NA | NA | NA |
| Gm37952 | 0 | NA | NA | NA |
| Gm21907 | 0 | NA | NA | NA |
| Gm37627 | 0 | NA | NA | NA |
| Gm20925 | 0 | NA | NA | NA |
| Gm20926 | 0 | NA | NA | NA |
| Gm37454 | 0 | NA | NA | NA |
| Gm38138 | 0 | NA | NA | NA |
| Gm29740 | 0 | NA | NA | NA |
| Gm37840 | 0 | NA | NA | NA |
| Gm37574 | 0 | NA | NA | NA |
| Gm21768 | 0 | NA | NA | NA |
| Gm37172 | 0 | NA | NA | NA |
| Gm35670 | 0 | NA | NA | NA |
| Gm20840 | 0 | NA | NA | NA |
| Gm35955 | 0 | NA | NA | NA |
| Gm37687 | 0 | NA | NA | NA |
| Gm38296 | 0 | NA | NA | NA |
| Gm38290 | 0 | NA | NA | NA |
| Gm38121 | 0 | NA | NA | NA |
| Gm36047 | 0 | NA | NA | NA |
| Gm36345 | 0 | NA | NA | NA |
| Gm37157 | 0 | NA | NA | NA |
| Gm38017 | 0 | NA | NA | NA |
| Gm36468 | 0 | NA | NA | NA |
| Gm21845 | 0 | NA | NA | NA |
| Gm20809 | 0 | NA | NA | NA |
| Gm37434 | 0 | NA | NA | NA |
| Gm37765 | 0 | NA | NA | NA |
| Gm38229 | 0 | NA | NA | NA |
| Gm36866 | 0 | NA | NA | NA |
| Gm30045 | 0 | NA | NA | NA |
| Gm21921 | 0 | NA | NA | NA |
| Gm37112 | 0 | NA | NA | NA |
| Gm37287 | 0 | NA | NA | NA |
| Gm30175 | 0 | NA | NA | NA |
| Gm21350 | 0 | NA | NA | NA |
| Gm21412 | 0 | NA | NA | NA |
| Gm37071 | 0 | NA | NA | NA |
| Gm37547 | 0 | NA | NA | NA |
| Gm36968 | 0 | NA | NA | NA |
| Gm38035 | 0 | NA | NA | NA |

| | | | | |
|---|---|---|---|---|
| Gm20857 | 0 | NA | NA | NA |
| Gm30705 | 0 | NA | NA | NA |
| Gm21773 | 0 | NA | NA | NA |
| Gm37451 | 0 | NA | NA | NA |
| Gm37793 | 0 | NA | NA | NA |
| Gm30858 | 0 | NA | NA | NA |
| Gm21791 | 0 | NA | NA | NA |
| Gm21427 | 0 | NA | NA | NA |
| Gm37875 | 0 | NA | NA | NA |
| Gm37237 | 0 | NA | NA | NA |
| Gm38350 | 0 | NA | NA | NA |
| Gm20858 | 0 | NA | NA | NA |
| Gm31422 | 0 | NA | NA | NA |
| Gm37798 | 0 | NA | NA | NA |
| Gm37318 | 0 | NA | NA | NA |
| Gm31572 | 0 | NA | NA | NA |
| Gm21890 | 0 | NA | NA | NA |
| Gm21443 | 0 | NA | NA | NA |
| Gm37346 | 0 | NA | NA | NA |
| Gm38136 | 0 | NA | NA | NA |
| Gm37991 | 0 | NA | NA | NA |
| Gm37808 | 0 | NA | NA | NA |
| Gm33571 | 0 | NA | NA | NA |
| Gm32181 | 0 | NA | NA | NA |
| Gm28662 | 0 | NA | NA | NA |
| Gm28780 | 0 | NA | NA | NA |
| Gm21755 | 0 | NA | NA | NA |
| Gm20895 | 0 | NA | NA | NA |
| Gm28853 | 0 | NA | NA | NA |
| Gm28852 | 0 | NA | NA | NA |
| Gm28855 | 0 | NA | NA | NA |
| Gm28854 | 0 | NA | NA | NA |
| Gm28858 | 0 | NA | NA | NA |
| Gm20890 | 0 | NA | NA | NA |
| Gm28245 | 0 | NA | NA | NA |
| Gm28246 | 0 | NA | NA | NA |
| Gm28247 | 0 | NA | NA | NA |
| Gm28248 | 0 | NA | NA | NA |
| Gm20892 | 0 | NA | NA | NA |
| Gm28249 | 0 | NA | NA | NA |
| Gm28250 | 0 | NA | NA | NA |
| Gm28251 | 0 | NA | NA | NA |
| Gm28243 | 0 | NA | NA | NA |
| Gm28244 | 0 | NA | NA | NA |
| Gm21771 | 0 | NA | NA | NA |
| Gm21476 | 0 | NA | NA | NA |
| Gm29021 | 0 | NA | NA | NA |

| | | | | |
|---|---|---|---|---|
| Gm28880 | 0.044191549 | -0.206490061 | 3.724945049 | -0.055434391 |
| Gm28452 | 0 | NA | NA | NA |
| Gm21488 | 0 | NA | NA | NA |
| Gm28454 | 0 | NA | NA | NA |
| Gm28878 | 0 | NA | NA | NA |
| Gm28879 | 0 | NA | NA | NA |
| Gm29369 | 0 | NA | NA | NA |
| Gm21506 | 0 | NA | NA | NA |
| Gm29399 | 0 | NA | NA | NA |
| Gm29400 | 0 | NA | NA | NA |
| Gm29401 | 0 | NA | NA | NA |
| Gm20891 | 0 | NA | NA | NA |
| Gm20893 | 0 | NA | NA | NA |
| Gm28799 | 0 | NA | NA | NA |
| Gm28798 | 0 | NA | NA | NA |
| Gm28797 | 0 | NA | NA | NA |
| Gm28796 | 0 | NA | NA | NA |
| Gm20894 | 0 | NA | NA | NA |
| Gm28938 | 0 | NA | NA | NA |
| Gm28937 | 0 | NA | NA | NA |
| Gm28936 | 0 | NA | NA | NA |
| Gm21642 | 0 | NA | NA | NA |
| Gm21804 | 0 | NA | NA | NA |
| Gm29122 | 0 | NA | NA | NA |
| Gm28939 | 0 | NA | NA | NA |
| Gm28983 | 0 | NA | NA | NA |
| Gm28985 | 0 | NA | NA | NA |
| Gm20810 | 0 | NA | NA | NA |
| Gm20811 | 0 | NA | NA | NA |
| Gm28725 | 0 | NA | NA | NA |
| Gm28984 | 0 | NA | NA | NA |
| Gm28987 | 0 | NA | NA | NA |
| Gm28986 | 0 | NA | NA | NA |
| Gm29171 | 0 | NA | NA | NA |
| Gm28129 | 0 | NA | NA | NA |
| Gm28127 | 0 | NA | NA | NA |
| Gm28128 | 0 | NA | NA | NA |
| Gm21853 | 0 | NA | NA | NA |
| Gm28132 | 0 | NA | NA | NA |
| Gm28130 | 0 | NA | NA | NA |
| Gm28131 | 0 | NA | NA | NA |
| Gm21679 | 0 | NA | NA | NA |
| Gm29317 | 0 | NA | NA | NA |
| Gm29316 | 0 | NA | NA | NA |
| Gm29315 | 0 | NA | NA | NA |
| Gm28216 | 0 | NA | NA | NA |
| Gm28217 | 0 | NA | NA | NA |

| | | | | |
|---|---|---|---|---|
| Gm28093 | 0 | NA | NA | NA |
| Gm28092 | 0 | NA | NA | NA |
| Gm28091 | 0 | NA | NA | NA |
| Gm28090 | 0 | NA | NA | NA |
| Gm29580 | 0 | NA | NA | NA |
| Gm28089 | 0 | NA | NA | NA |
| Gm28088 | 0 | NA | NA | NA |
| Gm29579 | 0 | NA | NA | NA |
| Gm29582 | 0 | NA | NA | NA |
| Gm21529 | 0 | NA | NA | NA |
| Gm29581 | 0 | NA | NA | NA |
| Gm21539 | 0 | NA | NA | NA |
| Gm29578 | 0 | NA | NA | NA |
| Gm21908 | 0 | NA | NA | NA |
| Gm21780 | 0 | NA | NA | NA |
| Gm28486 | 0 | NA | NA | NA |
| Gm21737 | 0 | NA | NA | NA |
| Gm28487 | 0 | NA | NA | NA |
| Gm28489 | 0 | NA | NA | NA |
| Gm28488 | 0 | NA | NA | NA |
| Gm28491 | 0 | NA | NA | NA |
| Gm28490 | 0 | NA | NA | NA |
| Gm21562 | 0 | NA | NA | NA |
| Gm28298 | 0 | NA | NA | NA |
| Gm21572 | 0 | NA | NA | NA |
| Gm28297 | 0 | NA | NA | NA |
| Gm21753 | 0 | NA | NA | NA |
| Gm21723 | 0 | NA | NA | NA |
| Gm28296 | 0 | NA | NA | NA |
| Gm21821 | 0 | NA | NA | NA |
| Gm28295 | 0 | NA | NA | NA |
| Gm21842 | 0 | NA | NA | NA |
| Gm28170 | 0 | NA | NA | NA |
| Gm21744 | 0 | NA | NA | NA |
| Gm28764 | 0 | NA | NA | NA |
| Gm28762 | 0 | NA | NA | NA |
| Gm28763 | 0 | NA | NA | NA |
| Gm21094 | 0 | NA | NA | NA |
| Gm21588 | 0 | NA | NA | NA |
| Gm28761 | 0 | NA | NA | NA |
| Gm21599 | 0 | NA | NA | NA |
| Gm28765 | 0 | NA | NA | NA |
| Gm21810 | 0 | NA | NA | NA |
| Gm21916 | 0 | NA | NA | NA |
| Gm29191 | 0 | NA | NA | NA |
| Gm21756 | 0 | NA | NA | NA |
| Gm28210 | 0 | NA | NA | NA |

| | | | | |
|---|---|---|---|---|
| Gm29074 | 0 | NA | NA | NA |
| Gm28211 | 0 | NA | NA | NA |
| Gm28518 | 0 | NA | NA | NA |
| Gm28519 | 0 | NA | NA | NA |
| Gm28520 | 0 | NA | NA | NA |
| Gm20838 | 0 | NA | NA | NA |
| Gm21626 | 0 | NA | NA | NA |
| Gm28521 | 0 | NA | NA | NA |
| Gm21633 | 0 | NA | NA | NA |
| Gm28522 | 0 | NA | NA | NA |
| Gm21745 | 0 | NA | NA | NA |
| Gm28517 | 0 | NA | NA | NA |
| Gm29532 | 0 | NA | NA | NA |
| Gm29531 | 0 | NA | NA | NA |
| Gm28464 | 0 | NA | NA | NA |
| Gm29656 | 0 | NA | NA | NA |
| Gm29655 | 0 | NA | NA | NA |
| Gm21647 | 0 | NA | NA | NA |
| Gm29653 | 0 | NA | NA | NA |
| Gm29654 | 0 | NA | NA | NA |
| Gm29651 | 0 | NA | NA | NA |
| Gm29652 | 0 | NA | NA | NA |
| Gm29080 | 0 | NA | NA | NA |
| Gm21184 | 0 | NA | NA | NA |
| Gm21201 | 0 | NA | NA | NA |
| Gm29081 | 0 | NA | NA | NA |
| Gm29077 | 0 | NA | NA | NA |
| Gm29076 | 0 | NA | NA | NA |
| Gm29079 | 0 | NA | NA | NA |
| Gm29078 | 0 | NA | NA | NA |
| Gm20853 | 0 | NA | NA | NA |
| Gm29075 | 0 | NA | NA | NA |
| Gm29225 | 0 | NA | NA | NA |
| Gm29226 | 0 | NA | NA | NA |
| Gm29227 | 0 | NA | NA | NA |
| Gm28339 | 0 | NA | NA | NA |
| Gm20855 | 0 | NA | NA | NA |
| Gm29386 | 0 | NA | NA | NA |
| Gm29384 | 0 | NA | NA | NA |
| Gm21896 | 0 | NA | NA | NA |
| Gm29280 | 0 | NA | NA | NA |
| Gm20799 | 0 | NA | NA | NA |
| Gm29279 | 0 | NA | NA | NA |
| Gm28260 | 0 | NA | NA | NA |
| Gm21795 | 0 | NA | NA | NA |
| Gm28261 | 0 | NA | NA | NA |
| Gm28264 | 0 | NA | NA | NA |

| | | | | |
|---|---|---|---|---|
| Gm21872 | 0 | NA | NA | NA |
| Gm20798 | 0 | NA | NA | NA |
| Gm28463 | 0 | NA | NA | NA |
| Gm28584 | 0 | NA | NA | NA |
| Gm28585 | 0 | NA | NA | NA |
| Gm20896 | 0 | NA | NA | NA |
| Gm28470 | 0 | NA | NA | NA |
| Gm28469 | 0 | NA | NA | NA |
| Gm21898 | 0 | NA | NA | NA |
| Gm28468 | 0 | NA | NA | NA |
| Gm20898 | 0 | NA | NA | NA |
| Gm29333 | 0 | NA | NA | NA |
| Gm28332 | 0 | NA | NA | NA |
| Gm28333 | 0 | NA | NA | NA |
| Gm28334 | 0 | NA | NA | NA |
| Gm28335 | 0 | NA | NA | NA |
| Gm28336 | 0 | NA | NA | NA |
| Gm28337 | 0 | NA | NA | NA |
| Gm28338 | 0 | NA | NA | NA |
| Gm28812 | 0 | NA | NA | NA |
| Gm28811 | 0 | NA | NA | NA |
| Gm28810 | 0 | NA | NA | NA |
| Gm20835 | 0 | NA | NA | NA |
| Gm29027 | 0 | NA | NA | NA |
| Gm29028 | 0 | NA | NA | NA |
| Gm29029 | 0 | NA | NA | NA |
| Gm29030 | 0 | NA | NA | NA |
| Gm29023 | 0 | NA | NA | NA |
| Gm29024 | 0 | NA | NA | NA |
| Gm29025 | 0 | NA | NA | NA |
| Gm29026 | 0 | NA | NA | NA |
| Gm28213 | 0 | NA | NA | NA |
| Gm28212 | 0 | NA | NA | NA |
| Gm21799 | 0 | NA | NA | NA |
| Gm28227 | 0 | NA | NA | NA |
| Gm20905 | 0 | NA | NA | NA |
| Gm29411 | 0 | NA | NA | NA |
| Gm28222 | 0 | NA | NA | NA |
| Gm21835 | 0 | NA | NA | NA |
| Gm28820 | 0 | NA | NA | NA |
| Gm28223 | 0 | NA | NA | NA |
| Gm29412 | 0 | NA | NA | NA |
| Gm29413 | 0 | NA | NA | NA |
| Gm29671 | 0 | NA | NA | NA |
| Gm21878 | 0 | NA | NA | NA |
| Gm28162 | 0 | NA | NA | NA |
| Gm20922 | 0 | NA | NA | NA |

| | | | | |
|---|---|---|---|---|
| Gm20738 | 0 | NA | NA | NA |
| Gm28545 | 0 | NA | NA | NA |
| Gm28547 | 0 | NA | NA | NA |
| Gm28546 | 0 | NA | NA | NA |
| Gm21924 | 0 | NA | NA | NA |
| Gm21735 | 0 | NA | NA | NA |
| Gm28771 | 0 | NA | NA | NA |
| Gm28772 | 0 | NA | NA | NA |
| Gm28773 | 0 | NA | NA | NA |
| Gm20897 | 0 | NA | NA | NA |
| Gm28774 | 0 | NA | NA | NA |
| Gm21732 | 0 | NA | NA | NA |
| Gm28775 | 0 | NA | NA | NA |
| Gm21765 | 0 | NA | NA | NA |
| Gm20899 | 12.70112357 | 0.018517305 | 0.328852488 | 0.056308848 |
| Gm28776 | 0 | NA | NA | NA |
| Gm21787 | 0 | NA | NA | NA |
| Gm28770 | 0 | NA | NA | NA |
| Gm20901 | 0 | NA | NA | NA |
| Gm28176 | 0 | NA | NA | NA |
| Gm28786 | 0 | NA | NA | NA |
| Gm29360 | 0 | NA | NA | NA |
| Gm29496 | 0 | NA | NA | NA |
| Gm29497 | 0 | NA | NA | NA |
| Gm29499 | 0 | NA | NA | NA |
| Gm29498 | 0 | NA | NA | NA |
| Gm36782 | 0 | NA | NA | NA |
| Gm29839 | 0 | NA | NA | NA |
| Gm29495 | 0 | NA | NA | NA |
| Gm28371 | 0.044191549 | -0.206490061 | 3.724945049 | -0.055434391 |
| Gm28595 | 0 | NA | NA | NA |
| Gm37032 | 0 | NA | NA | NA |
| Gm21865 | 0 | NA | NA | NA |
| Gm29433 | 0 | NA | NA | NA |
| Gm29215 | 0 | NA | NA | NA |
| Gm21882 | 0 | NA | NA | NA |
| Gm21801 | 0 | NA | NA | NA |
| Gm28850 | 0 | NA | NA | NA |
| Gm21914 | 0 | NA | NA | NA |
| Gm28617 | 0 | NA | NA | NA |
| Gm28618 | 0 | NA | NA | NA |
| Gm28619 | 0 | NA | NA | NA |
| Gm21852 | 0 | NA | NA | NA |
| Gm20793 | 0 | NA | NA | NA |
| Gm27466 | 0 | NA | NA | NA |
| Gm28018 | 0 | NA | NA | NA |
| Gm28993 | 0 | NA | NA | NA |

| | | | | |
|---|---|---|---|---|
| Gm28994 | 0 | NA | NA | NA |
| Gm21797 | 0 | NA | NA | NA |
| Gm20795 | 0 | NA | NA | NA |
| Gm28596 | 0 | NA | NA | NA |
| Gm20832 | 0 | NA | NA | NA |
| Gm28597 | 0 | NA | NA | NA |
| Gm28428 | 0 | NA | NA | NA |
| Gm20827 | 0 | NA | NA | NA |
| Gm28427 | 0 | NA | NA | NA |
| Gm21800 | 0 | NA | NA | NA |
| Gm28426 | 0 | NA | NA | NA |
| Gm28425 | 0 | NA | NA | NA |
| Gm21729 | 0 | NA | NA | NA |
| Gm28834 | 0 | NA | NA | NA |
| Gm28833 | 0 | NA | NA | NA |
| Gm21774 | 0 | NA | NA | NA |
| Gm29537 | 0 | NA | NA | NA |
| Gm29274 | 0 | NA | NA | NA |
| Gm29275 | 0.04393175 | -0.206490061 | 3.724945049 | -0.055434391 |
| Gm29276 | 0 | NA | NA | NA |
| Gm21805 | 0 | NA | NA | NA |
| Gm29645 | 0 | NA | NA | NA |
| Gm28312 | 0 | NA | NA | NA |
| Gm29165 | 0 | NA | NA | NA |
| Gm29042 | 0 | NA | NA | NA |
| Gm28311 | 0 | NA | NA | NA |
| Gm28310 | 0 | NA | NA | NA |
| Gm28504 | 0 | NA | NA | NA |
| Gm28567 | 0 | NA | NA | NA |
| Gm28568 | 0 | NA | NA | NA |
| Gm29286 | 0 | NA | NA | NA |
| Gm29285 | 0 | NA | NA | NA |
| Gm28280 | 0 | NA | NA | NA |
| Gm21241 | 0 | NA | NA | NA |
| Gm28278 | 0 | NA | NA | NA |
| Gm28279 | 0 | NA | NA | NA |
| Gm21739 | 0 | NA | NA | NA |
| Gm28886 | 0 | NA | NA | NA |
| Gm29426 | 0 | NA | NA | NA |
| Gm28885 | 0 | NA | NA | NA |
| Gm21856 | 0 | NA | NA | NA |
| Gm28887 | 0 | NA | NA | NA |
| Gm29444 | 0 | NA | NA | NA |
| Gm21783 | 0 | NA | NA | NA |
| Gm29445 | 0 | NA | NA | NA |
| Gm28457 | 0 | NA | NA | NA |
| Gm21779 | 0 | NA | NA | NA |

| | | | | |
|---|---|---|---|---|
| Gm28458 | 0 | NA | NA | NA |
| Gm28459 | 0 | NA | NA | NA |
| Gm28461 | 0 | NA | NA | NA |
| Gm28460 | 0 | NA | NA | NA |
| Gm28851 | 0 | NA | NA | NA |
| Gm28456 | 0 | NA | NA | NA |
| Gm29073 | 0 | NA | NA | NA |
| Gm29648 | 0 | NA | NA | NA |
| Gm29647 | 0 | NA | NA | NA |
| Gm29646 | 0 | NA | NA | NA |
| Gm28549 | 0 | NA | NA | NA |
| Gm29211 | 0 | NA | NA | NA |
| Gm29212 | 0 | NA | NA | NA |
| Gm21247 | 0 | NA | NA | NA |
| Gm29210 | 0 | NA | NA | NA |
| Gm29213 | 0 | NA | NA | NA |
| Gm28550 | 0 | NA | NA | NA |
| Gm28604 | 0 | NA | NA | NA |
| Gm28603 | 0 | NA | NA | NA |
| Gm28565 | 0 | NA | NA | NA |
| Gm28566 | 0 | NA | NA | NA |
| Gm28697 | 0 | NA | NA | NA |
| Gm28554 | 0 | NA | NA | NA |
| Gm21256 | 0 | NA | NA | NA |
| Gm29056 | 0 | NA | NA | NA |
| Gm29098 | 0 | NA | NA | NA |
| Gm21170 | 0 | NA | NA | NA |
| Gm29373 | 0 | NA | NA | NA |
| Gm29370 | 0 | NA | NA | NA |
| Gm21751 | 0 | NA | NA | NA |
| Gm29166 | 0 | NA | NA | NA |
| Gm28194 | 0 | NA | NA | NA |
| Gm20792 | 0 | NA | NA | NA |
| Gm28754 | 0 | NA | NA | NA |
| Gm28753 | 0 | NA | NA | NA |
| Gm21191 | 0 | NA | NA | NA |
| Gm21271 | 0 | NA | NA | NA |
| Gm21281 | 0 | NA | NA | NA |
| Gm28193 | 0 | NA | NA | NA |
| Gm20802 | 0 | NA | NA | NA |
| Gm29070 | 0 | NA | NA | NA |
| Gm28553 | 0 | NA | NA | NA |
| Gm28751 | 0 | NA | NA | NA |
| Gm28752 | 0 | NA | NA | NA |
| Gm28726 | 0 | NA | NA | NA |
| Gm21308 | 0 | NA | NA | NA |
| Gm29267 | 0 | NA | NA | NA |

| | | | | |
|---|---|---|---|---|
| Gm29268 | 0 | NA | NA | NA |
| Gm29269 | 0 | NA | NA | NA |
| Gm29270 | 0 | NA | NA | NA |
| Gm29271 | 0 | NA | NA | NA |
| Gm21209 | 0 | NA | NA | NA |
| Gm28682 | 0 | NA | NA | NA |
| Gm28569 | 0 | NA | NA | NA |
| Gm28681 | 0 | NA | NA | NA |
| Gm28684 | 0 | NA | NA | NA |
| Gm21316 | 0 | NA | NA | NA |
| Gm21923 | 0 | NA | NA | NA |
| Gm28683 | 0 | NA | NA | NA |
| Gm28680 | 0 | NA | NA | NA |
| Gm28679 | 0 | NA | NA | NA |
| Gm28870 | 0 | NA | NA | NA |
| Gm28871 | 0 | NA | NA | NA |
| Gm29512 | 0 | NA | NA | NA |
| Gm29511 | 0 | NA | NA | NA |
| Gm20880 | 0 | NA | NA | NA |
| Gm20881 | 0 | NA | NA | NA |
| Gm29121 | 0 | NA | NA | NA |
| Gm29120 | 0 | NA | NA | NA |
| Gm29117 | 0 | NA | NA | NA |
| Gm20883 | 0 | NA | NA | NA |
| Gm29119 | 0 | NA | NA | NA |
| Gm29118 | 0 | NA | NA | NA |
| Gm29116 | 0 | NA | NA | NA |
| Gm20889 | 0 | NA | NA | NA |
| Gm20887 | 0 | NA | NA | NA |
| Gm29320 | 0 | NA | NA | NA |
| Gm21849 | 0 | NA | NA | NA |
| Gm29321 | 0 | NA | NA | NA |
| Gm21117 | 0 | NA | NA | NA |
| Gm29319 | 0 | NA | NA | NA |
| Gm28317 | 0 | NA | NA | NA |
| Gm28318 | 0 | NA | NA | NA |
| Gm20884 | 0 | NA | NA | NA |
| Gm28313 | 0 | NA | NA | NA |
| Gm28314 | 0 | NA | NA | NA |
| Gm28315 | 0 | NA | NA | NA |
| Gm28316 | 0 | NA | NA | NA |
| Gm21387 | 0 | NA | NA | NA |
| Gm21396 | 0 | NA | NA | NA |
| Gm28944 | 0 | NA | NA | NA |
| Gm28943 | 0 | NA | NA | NA |
| Gm28946 | 0 | NA | NA | NA |
| Gm28945 | 0 | NA | NA | NA |

| | | | | |
|---|---|---|---|---|
| Gm28947 | 0 | NA | NA | NA |
| Gm20920 | 0 | NA | NA | NA |
| Gm28949 | 0 | NA | NA | NA |
| Gm28948 | 0 | NA | NA | NA |
| Gm28950 | 0 | NA | NA | NA |
| Gm21245 | 0 | NA | NA | NA |
| Gm29169 | 0 | NA | NA | NA |
| Gm21899 | 0 | NA | NA | NA |
| Gm29063 | 0 | NA | NA | NA |
| Gm28737 | 0 | NA | NA | NA |
| Gm28738 | 0 | NA | NA | NA |
| Gm20923 | 0 | NA | NA | NA |
| Gm28197 | 0 | NA | NA | NA |
| Gm21249 | 0 | NA | NA | NA |
| Gm28736 | 0 | NA | NA | NA |
| Gm21840 | 0 | NA | NA | NA |
| Gm28735 | 0 | NA | NA | NA |
| Gm21258 | 0 | NA | NA | NA |
| Gm28734 | 0 | NA | NA | NA |
| Gm28232 | 0 | NA | NA | NA |
| Gm28732 | 0 | NA | NA | NA |
| Gm28733 | 0 | NA | NA | NA |
| Gm21268 | 0 | NA | NA | NA |
| Gm21275 | 0 | NA | NA | NA |
| Gm28233 | 0 | NA | NA | NA |
| Gm28235 | 0 | NA | NA | NA |
| Gm28234 | 0 | NA | NA | NA |
| Gm21902 | 0 | NA | NA | NA |
| Gm28236 | 0 | NA | NA | NA |
| Gm21285 | 0 | NA | NA | NA |
| Gm20747 | 0 | NA | NA | NA |
| Gm29250 | 0 | NA | NA | NA |
| Gm29251 | 0 | NA | NA | NA |
| Gm21906 | 0 | NA | NA | NA |
| Gm29252 | 0 | NA | NA | NA |
| Gm29248 | 0 | NA | NA | NA |
| Gm29249 | 0 | NA | NA | NA |
| Gm29484 | 0 | NA | NA | NA |
| Gm21920 | 0 | NA | NA | NA |
| Gm28908 | 0 | NA | NA | NA |
| Gm28907 | 0 | NA | NA | NA |
| Gm21749 | 0 | NA | NA | NA |
| Gm20929 | 0 | NA | NA | NA |
| Gm28910 | 0 | NA | NA | NA |
| Gm28909 | 0 | NA | NA | NA |
| Gm28276 | 0 | NA | NA | NA |
| Gm28274 | 0 | NA | NA | NA |

| | | | | |
|---|---|---|---|---|
| Gm28275 | 0 | NA | NA | NA |
| Gm21302 | 0 | NA | NA | NA |
| Gm28290 | 0 | NA | NA | NA |
| Gm28292 | 0 | NA | NA | NA |
| Gm28291 | 0 | NA | NA | NA |
| Gm28293 | 0 | NA | NA | NA |
| Gm21740 | 0 | NA | NA | NA |
| Gm21912 | 0 | NA | NA | NA |
| Gm28698 | 0 | NA | NA | NA |
| Gm29304 | 0 | NA | NA | NA |
| Gm29305 | 0 | NA | NA | NA |
| Gm29303 | 0 | NA | NA | NA |
| Gm21894 | 0 | NA | NA | NA |
| Gm29307 | 0 | NA | NA | NA |
| Gm29557 | 0 | NA | NA | NA |
| Gm23052 | 0 | NA | NA | NA |
| Gm20927 | 0 | NA | NA | NA |
| Gm20930 | 0 | NA | NA | NA |
| Gm29306 | 0 | NA | NA | NA |
| Gm29300 | 0 | NA | NA | NA |
| Gm29301 | 0 | NA | NA | NA |
| Gm20931 | 0 | NA | NA | NA |
| Gm29559 | 0 | NA | NA | NA |
| Gm29558 | 0 | NA | NA | NA |
| Gm29555 | 0 | NA | NA | NA |
| Gm21733 | 0 | NA | NA | NA |
| Gm21330 | 0 | NA | NA | NA |
| Gm28355 | 0 | NA | NA | NA |
| Gm28354 | 0 | NA | NA | NA |
| Gm28353 | 0 | NA | NA | NA |
| Gm28352 | 0 | NA | NA | NA |
| Gm21858 | 0 | NA | NA | NA |
| Gm29244 | 0 | NA | NA | NA |
| Gm28200 | 0 | NA | NA | NA |
| Gm28201 | 0 | NA | NA | NA |
| Gm28202 | 0 | NA | NA | NA |
| Gm28203 | 0 | NA | NA | NA |
| Gm21340 | 0 | NA | NA | NA |
| Gm28204 | 0 | NA | NA | NA |
| Gm28205 | 0 | NA | NA | NA |
| Gm28206 | 0 | NA | NA | NA |
| Gm28207 | 0 | NA | NA | NA |
| Gm28208 | 0 | NA | NA | NA |
| Gm21851 | 0 | NA | NA | NA |
| Gm21819 | 0 | NA | NA | NA |
| Gm29224 | 0 | NA | NA | NA |
| Gm29223 | 0 | NA | NA | NA |

| | | | | |
|---|---|---|---|---|
| Gm29222 | 0 | NA | NA | NA |
| Gm29221 | 0 | NA | NA | NA |
| Gm29220 | 0 | NA | NA | NA |
| Gm21815 | 0 | NA | NA | NA |
| Gm29407 | 0 | NA | NA | NA |
| Gm29405 | 0 | NA | NA | NA |
| Gm29406 | 0 | NA | NA | NA |
| Gm20913 | 0 | NA | NA | NA |
| Gm29404 | 0 | NA | NA | NA |
| Gm29403 | 0 | NA | NA | NA |
| Gm28350 | 0 | NA | NA | NA |
| Gm28964 | 0 | NA | NA | NA |
| Sly | 0 | NA | NA | NA |
| Gm28165 | 0 | NA | NA | NA |
| Gm28709 | 0 | NA | NA | NA |
| Gm28965 | 0 | NA | NA | NA |
| Gm21868 | 0 | NA | NA | NA |
| Gm21943 | 0 | NA | NA | NA |
| Gm28532 | 0 | NA | NA | NA |
| Gm28533 | 0 | NA | NA | NA |
| Gm28966 | 0 | NA | NA | NA |
| Gm28961 | 0 | NA | NA | NA |
| Gm28962 | 0 | NA | NA | NA |
| Gm28963 | 0 | NA | NA | NA |
| Gm29466 | 0 | NA | NA | NA |
| Gm29467 | 0 | NA | NA | NA |
| Gm21724 | 0 | NA | NA | NA |
| Gm20963 | 0 | NA | NA | NA |
| Gm28326 | 0 | NA | NA | NA |
| Gm28325 | 0 | NA | NA | NA |
| Gm21405 | 0 | NA | NA | NA |
| Gm28284 | 0 | NA | NA | NA |
| Gm29368 | 0 | NA | NA | NA |
| Gm21861 | 0 | NA | NA | NA |
| Gm21413 | 0 | NA | NA | NA |
| Gm28073 | 0 | NA | NA | NA |
| Gm21419 | 0 | NA | NA | NA |
| Gm28074 | 0 | NA | NA | NA |
| Gm21428 | 0 | NA | NA | NA |
| Gm28072 | 0 | NA | NA | NA |
| Gm21758 | 0 | NA | NA | NA |
| Gm29672 | 0 | NA | NA | NA |
| Gm29131 | 0 | NA | NA | NA |
| Gm20871 | 0 | NA | NA | NA |
| Gm29130 | 0 | NA | NA | NA |
| Gm20978 | 0 | NA | NA | NA |
| Gm29132 | 0 | NA | NA | NA |

| | | | | |
|---|---|---|---|---|
| Gm21450 | 0 | NA | NA | NA |
| Gm21843 | 0 | NA | NA | NA |
| Gm28889 | 0 | NA | NA | NA |
| Gm28890 | 0 | NA | NA | NA |
| Gm28891 | 0 | NA | NA | NA |
| Gm28691 | 0 | NA | NA | NA |
| Gm28692 | 0 | NA | NA | NA |
| Gm20794 | 0 | NA | NA | NA |
| Gm28690 | 0 | NA | NA | NA |
| Gm28689 | 0 | NA | NA | NA |
| Gm28687 | 0 | NA | NA | NA |
| Gm20987 | 0 | NA | NA | NA |
| Gm21918 | 0 | NA | NA | NA |
| 1700040F15Rik | 0 | NA | NA | NA |
| Gm29207 | 0 | NA | NA | NA |
| Gm29204 | 0 | NA | NA | NA |
| Gm21489 | 0 | NA | NA | NA |
| Gm21832 | 0 | NA | NA | NA |
| Gm21573 | 0 | NA | NA | NA |
| Gm28179 | 0 | NA | NA | NA |
| Gm21577 | 0 | NA | NA | NA |
| Gm29206 | 0 | NA | NA | NA |
| Gm29203 | 0 | NA | NA | NA |
| Gm21497 | 0 | NA | NA | NA |
| Gm29614 | 0 | NA | NA | NA |
| Gm29613 | 0 | NA | NA | NA |
| Gm29612 | 0 | NA | NA | NA |
| Gm29343 | 0 | NA | NA | NA |
| Gm29342 | 0 | NA | NA | NA |
| Gm20803 | 0 | NA | NA | NA |
| Gm20804 | 0 | NA | NA | NA |
| Gm28598 | 0 | NA | NA | NA |
| Gm28599 | 0 | NA | NA | NA |
| Gm28600 | 0 | NA | NA | NA |
| Gm21518 | 0 | NA | NA | NA |
| Gm28185 | 0 | NA | NA | NA |
| Gm29037 | 0 | NA | NA | NA |
| Gm20823 | 0 | NA | NA | NA |
| Gm21877 | 0 | NA | NA | NA |
| Gm28835 | 0 | NA | NA | NA |
| Gm29548 | 0 | NA | NA | NA |
| Gm29547 | 0 | NA | NA | NA |
| Gm29546 | 0 | NA | NA | NA |
| Gm29545 | 0 | NA | NA | NA |
| Gm29544 | 0 | NA | NA | NA |
| Gm29209 | 0 | NA | NA | NA |
| Gm29549 | 0 | NA | NA | NA |

| | | | | |
|---|---|---|---|---|
| Gm21530 | 0 | NA | NA | NA |
| Gm21617 | 0 | NA | NA | NA |
| Gm28589 | 0 | NA | NA | NA |
| Gm29420 | 0 | NA | NA | NA |
| Gm29421 | 0 | NA | NA | NA |
| Gm21627 | 0 | NA | NA | NA |
| Gm29422 | 0 | NA | NA | NA |
| Gm28808 | 0 | NA | NA | NA |
| Gm21634 | 0 | NA | NA | NA |
| Gm28446 | 0 | NA | NA | NA |
| Gm29060 | 0 | NA | NA | NA |
| Gm29061 | 0 | NA | NA | NA |
| Gm20885 | 0 | NA | NA | NA |
| Gm29062 | 0 | NA | NA | NA |
| Gm29057 | 0 | NA | NA | NA |
| Gm29625 | 0 | NA | NA | NA |
| Gm21654 | 0 | NA | NA | NA |
| Gm29058 | 0 | NA | NA | NA |
| Gm29059 | 0 | NA | NA | NA |
| Gm29628 | 0 | NA | NA | NA |
| Gm20882 | 0 | NA | NA | NA |
| Gm20907 | 0 | NA | NA | NA |
| Gm29624 | 0 | NA | NA | NA |
| Gm29623 | 0 | NA | NA | NA |
| Gm29622 | 0 | NA | NA | NA |
| Gm29265 | 0 | NA | NA | NA |
| Gm20908 | 0 | NA | NA | NA |
| Gm29259 | 0 | NA | NA | NA |
| Gm28860 | 0 | NA | NA | NA |
| Gm28252 | 0 | NA | NA | NA |
| Gm28859 | 0 | NA | NA | NA |
| Gm20921 | 0 | NA | NA | NA |
| Gm28253 | 0 | NA | NA | NA |
| Gm21826 | 0 | NA | NA | NA |
| Gm28254 | 0 | NA | NA | NA |
| Gm28861 | 0 | NA | NA | NA |
| Gm28255 | 0 | NA | NA | NA |
| Gm21171 | 0 | NA | NA | NA |
| Gm20924 | 0 | NA | NA | NA |
| Gm28990 | 0 | NA | NA | NA |
| Gm21848 | 0 | NA | NA | NA |
| Gm29457 | 0 | NA | NA | NA |
| Gm20736 | 0 | NA | NA | NA |
| Gm28558 | 0 | NA | NA | NA |
| Gm28560 | 0 | NA | NA | NA |
| Gm28561 | 0 | NA | NA | NA |
| Gm29584 | 0 | NA | NA | NA |

| | | | | |
|---|---|---|---|---|
| Gm20936 | 0 | NA | NA | NA |
| Gm20934 | 0 | NA | NA | NA |
| Gm29586 | 0 | NA | NA | NA |
| Gm21895 | 0 | NA | NA | NA |
| Gm28134 | 0 | NA | NA | NA |
| Gm28135 | 0 | NA | NA | NA |
| Gm20933 | 0 | NA | NA | NA |
| Gm28133 | 0 | NA | NA | NA |
| Gm28137 | 0 | NA | NA | NA |
| Gm28138 | 0 | NA | NA | NA |
| Gm21837 | 0 | NA | NA | NA |
| Gm28136 | 0 | NA | NA | NA |
| Gm21910 | 0 | NA | NA | NA |
| Gm29377 | 0 | NA | NA | NA |
| Gm21889 | 0 | NA | NA | NA |
| Gm29378 | 0 | NA | NA | NA |
| Gm29379 | 0 | NA | NA | NA |
| Gm29380 | 0 | NA | NA | NA |
| Gm29381 | 0 | NA | NA | NA |
| Gm28795 | 0 | NA | NA | NA |
| Gm20852 | 0 | NA | NA | NA |
| Gm29003 | 0 | NA | NA | NA |
| Gm28749 | 0 | NA | NA | NA |
| Gm29001 | 0 | NA | NA | NA |
| Gm29002 | 0 | NA | NA | NA |
| Ssty2 | 0 | NA | NA | NA |
| Gm29000 | 0 | NA | NA | NA |
| Gm28646 | 0 | NA | NA | NA |
| Gm28612 | 0 | NA | NA | NA |
| Gm28613 | 0 | NA | NA | NA |
| Gm28611 | 0 | NA | NA | NA |
| Gm21114 | 0 | NA | NA | NA |
| Gm29633 | 0 | NA | NA | NA |
| Gm21794 | 0 | NA | NA | NA |
| Gm29219 | 0 | NA | NA | NA |
| Gm29632 | 0 | NA | NA | NA |
| Gm29631 | 0 | NA | NA | NA |
| Gm29637 | 0 | NA | NA | NA |
| Gm20874 | 0 | NA | NA | NA |
| Gm29082 | 0 | NA | NA | NA |
| Gm29217 | 0 | NA | NA | NA |
| Gm29636 | 0 | NA | NA | NA |
| Gm29635 | 0 | NA | NA | NA |
| Gm29634 | 0 | NA | NA | NA |
| Gm29639 | 0 | NA | NA | NA |
| Gm29218 | 0 | NA | NA | NA |
| Gm21806 | 0 | NA | NA | NA |

| | | | | |
|---|---|---|---|---|
| Gm20935 | 0 | NA | NA | NA |
| Gm29662 | 0 | NA | NA | NA |
| Gm29660 | 0 | NA | NA | NA |
| Gm29661 | 0 | NA | NA | NA |
| Gm29659 | 0 | NA | NA | NA |
| Gm20937 | 0 | NA | NA | NA |
| Gm29450 | 0 | NA | NA | NA |
| Gm29449 | 0 | NA | NA | NA |
| Gm21867 | 0 | NA | NA | NA |
| Gm29448 | 0 | NA | NA | NA |
| Gm21127 | 0 | NA | NA | NA |
| Gm20816 | 0 | NA | NA | NA |
| Gm29447 | 0 | NA | NA | NA |
| Gm29446 | 0 | NA | NA | NA |
| Gm29452 | 0 | NA | NA | NA |
| Gm29451 | 0.044191549 | -0.206490061 | 3.724945049 | -0.055434391 |
| Gm20817 | 0 | NA | NA | NA |
| Gm28633 | 0 | NA | NA | NA |
| Gm29410 | 0 | NA | NA | NA |
| Gm28819 | 0 | NA | NA | NA |
| Gm28462 | 0 | NA | NA | NA |
| Gm20818 | 0 | NA | NA | NA |
| Gm21151 | 0 | NA | NA | NA |
| Gm29409 | 0 | NA | NA | NA |
| Gm28632 | 0 | NA | NA | NA |
| Gm28101 | 0 | NA | NA | NA |
| Gm33815 | 0 | NA | NA | NA |
| Gm21160 | 0 | NA | NA | NA |
| Gm28637 | 0 | NA | NA | NA |
| Gm28636 | 0 | NA | NA | NA |
| Gm20932 | 0 | NA | NA | NA |
| Gm29134 | 0 | NA | NA | NA |
| Gm28303 | 0 | NA | NA | NA |
| Gm28304 | 0 | NA | NA | NA |
| Gm28824 | 0 | NA | NA | NA |
| Gm28081 | 0 | NA | NA | NA |
| Gm28080 | 0 | NA | NA | NA |
| Gm21524 | 0 | NA | NA | NA |
| Gm28079 | 0 | NA | NA | NA |
| Gm28302 | 0 | NA | NA | NA |
| Gm28082 | 0 | NA | NA | NA |
| Gm21672 | 0 | NA | NA | NA |
| Gm29644 | 0 | NA | NA | NA |
| Gm21118 | 0 | NA | NA | NA |
| Gm29423 | 0 | NA | NA | NA |
| Gm29424 | 0 | NA | NA | NA |
| 4932431L22Rik | 0 | NA | NA | NA |

| | | | | |
|---|---|---|---|---|
| Gm29425 | 0 | NA | NA | NA |
| Gm20888 | 0 | NA | NA | NA |
| Gm28238 | 0 | NA | NA | NA |
| Gm28433 | 0 | NA | NA | NA |
| Gm28431 | 0 | NA | NA | NA |
| Gm28432 | 0 | NA | NA | NA |
| Gm20886 | 0 | NA | NA | NA |
| Gm21808 | 0 | NA | NA | NA |
| Gm28840 | 0 | NA | NA | NA |
| Gm28841 | 0 | NA | NA | NA |
| Gm28842 | 0 | NA | NA | NA |
| Gm28843 | 0 | NA | NA | NA |
| Gm28837 | 0 | NA | NA | NA |
| Gm28838 | 0 | NA | NA | NA |
| Gm28839 | 0 | NA | NA | NA |
| Gm20866 | 0 | NA | NA | NA |
| Gm20868 | 0 | NA | NA | NA |
| Gm28844 | 0 | NA | NA | NA |
| Gm21841 | 0 | NA | NA | NA |
| Gm28668 | 0 | NA | NA | NA |
| Gm20869 | 0 | NA | NA | NA |
| Gm28481 | 0 | NA | NA | NA |
| Gm28482 | 0 | NA | NA | NA |
| Gm28485 | 0 | NA | NA | NA |
| Gm20870 | 0 | NA | NA | NA |
| Gm28483 | 0 | NA | NA | NA |
| Gm28484 | 0 | NA | NA | NA |
| Gm28480 | 0 | NA | NA | NA |
| Gm20842 | 0 | NA | NA | NA |
| Gm29179 | 0 | NA | NA | NA |
| Gm21880 | 0 | NA | NA | NA |
| Gm29180 | 0 | NA | NA | NA |
| Gm29181 | 0 | NA | NA | NA |
| Gm20843 | 0 | NA | NA | NA |
| Gm29182 | 0 | NA | NA | NA |
| Gm20844 | 0 | NA | NA | NA |
| Gm29178 | 0 | NA | NA | NA |
| Gm21757 | 0 | NA | NA | NA |
| Gm28968 | 0 | NA | NA | NA |
| Gm28495 | 0 | NA | NA | NA |
| Gm20846 | 0 | NA | NA | NA |
| Gm20847 | 0 | NA | NA | NA |
| Gm20848 | 0 | NA | NA | NA |
| Gm28969 | 0 | NA | NA | NA |
| Gm21766 | 0 | NA | NA | NA |
| Gm20800 | 0 | NA | NA | NA |
| Gm21683 | 0 | NA | NA | NA |

| | | | | |
|---|---|---|---|---|
| Gm28677 | 0 | NA | NA | NA |
| Gm21855 | 0 | NA | NA | NA |
| Gm29255 | 0 | NA | NA | NA |
| Gm20902 | 0 | NA | NA | NA |
| Gm28678 | 0 | NA | NA | NA |
| Gm20903 | 0 | NA | NA | NA |
| Gm28272 | 0 | NA | NA | NA |
| Gm29256 | 0 | NA | NA | NA |
| Gm28259 | 0 | NA | NA | NA |
| Gm28273 | 0 | NA | NA | NA |
| Gm21715 | 0 | NA | NA | NA |
| Gm29617 | 0 | NA | NA | NA |
| Gm29615 | 0 | NA | NA | NA |
| Gm29616 | 0 | NA | NA | NA |
| Gm21784 | 0 | NA | NA | NA |
| Gm20805 | 0 | NA | NA | NA |
| Gm29618 | 0 | NA | NA | NA |
| Gm29109 | 0 | NA | NA | NA |
| Gm29108 | 0 | NA | NA | NA |
| Gm29110 | 0 | NA | NA | NA |
| Gm28781 | 0 | NA | NA | NA |
| Gm21913 | 0 | NA | NA | NA |
| Gm28174 | 0 | NA | NA | NA |
| Gm22832 | 0 | NA | NA | NA |
| Gm21076 | 0 | NA | NA | NA |
| Gm29416 | 0 | NA | NA | NA |
| Gm25359 | 0 | NA | NA | NA |
| Gm21869 | 0 | NA | NA | NA |
| Gm20813 | 0 | NA | NA | NA |
| Gm28790 | 0 | NA | NA | NA |
| Gm29272 | 0 | NA | NA | NA |
| Gm28789 | 0 | NA | NA | NA |
| Gm20814 | 0 | NA | NA | NA |
| Gm28787 | 0 | NA | NA | NA |
| Gm28788 | 0 | NA | NA | NA |
| Gm29278 | 0 | NA | NA | NA |
| Gm21922 | 0 | NA | NA | NA |
| Gm21111 | 0 | NA | NA | NA |
| Gm28608 | 0 | NA | NA | NA |
| Gm28607 | 0 | NA | NA | NA |
| Gm28606 | 0 | NA | NA | NA |
| Gm20850 | 0 | NA | NA | NA |
| Gm29574 | 0 | NA | NA | NA |
| Gm29575 | 0 | NA | NA | NA |
| Gm21875 | 0 | NA | NA | NA |
| Gm20904 | 0 | NA | NA | NA |
| Gm29576 | 0 | NA | NA | NA |

| | | | | |
|---|---|---|---|---|
| Gm29577 | 0 | NA | NA | NA |
| Gm29573 | 0 | NA | NA | NA |
| Gm28377 | 0 | NA | NA | NA |
| Gm21776 | 0 | NA | NA | NA |
| Gm21155 | 0 | NA | NA | NA |
| Gm21163 | 0 | NA | NA | NA |
| Gm28122 | 0 | NA | NA | NA |
| Gm28121 | 0 | NA | NA | NA |
| Gm21173 | 0 | NA | NA | NA |
| Gm28123 | 0 | NA | NA | NA |
| Gm28120 | 0 | NA | NA | NA |
| Gm29313 | 0 | NA | NA | NA |
| Gm21180 | 0 | NA | NA | NA |
| Gm20859 | 0 | NA | NA | NA |
| Gm29314 | 0 | NA | NA | NA |
| Gm29310 | 0 | NA | NA | NA |
| Gm29311 | 0 | NA | NA | NA |
| Gm21638 | 0 | NA | NA | NA |
| Gm29312 | 0 | NA | NA | NA |
| Gm29308 | 0 | NA | NA | NA |
| Gm29309 | 0 | NA | NA | NA |
| Gm28702 | 0 | NA | NA | NA |
| Gm28701 | 0 | NA | NA | NA |
| Gm28700 | 0 | NA | NA | NA |
| Gm28699 | 0 | NA | NA | NA |
| Gm28705 | 0 | NA | NA | NA |
| Gm28704 | 0 | NA | NA | NA |
| Gm28540 | 0 | NA | NA | NA |
| Gm21650 | 0 | NA | NA | NA |
| Gm28541 | 0 | NA | NA | NA |
| Gm28542 | 0 | NA | NA | NA |
| Gm28537 | 0 | NA | NA | NA |
| Gm28538 | 0 | NA | NA | NA |
| Gm29569 | 0 | NA | NA | NA |
| Gm20867 | 0 | NA | NA | NA |
| Gm21658 | 0 | NA | NA | NA |
| Gm29568 | 0 | NA | NA | NA |
| Gm29567 | 0 | NA | NA | NA |
| Gm21198 | 0 | NA | NA | NA |
| Gm29566 | 0 | NA | NA | NA |
| Gm29565 | 0 | NA | NA | NA |
| Gm29564 | 0 | NA | NA | NA |
| Gm29147 | 0 | NA | NA | NA |
| Gm29146 | 0 | NA | NA | NA |
| Gm28664 | 0 | NA | NA | NA |
| Gm28663 | 0 | NA | NA | NA |
| Gm29149 | 0 | NA | NA | NA |

| | | | | |
|---|---|---|---|---|
| Gm29148 | 0 | NA | NA | NA |
| Gm29145 | 0 | NA | NA | NA |
| Gm29144 | 0 | NA | NA | NA |
| Gm28817 | 0 | NA | NA | NA |
| Gm28816 | 0 | NA | NA | NA |
| Gm20808 | 0 | NA | NA | NA |
| Gm20806 | 0 | NA | NA | NA |
| Gm28815 | 0 | NA | NA | NA |
| Gm28814 | 0 | NA | NA | NA |
| Gm28813 | 0 | NA | NA | NA |
| Gm20919 | 0 | NA | NA | NA |
| Gm20917 | 0 | NA | NA | NA |
| Gm28421 | 0 | NA | NA | NA |
| Gm28420 | 0 | NA | NA | NA |
| Gm28423 | 0 | NA | NA | NA |
| Gm28422 | 0 | NA | NA | NA |
| Gm20916 | 0 | NA | NA | NA |
| Gm29033 | 0 | NA | NA | NA |
| Gm29034 | 0 | NA | NA | NA |
| Gm21823 | 0 | NA | NA | NA |
| Gm21242 | 0 | NA | NA | NA |
| Gm29031 | 0 | NA | NA | NA |
| Gm29032 | 0 | NA | NA | NA |
| Gm29606 | 0 | NA | NA | NA |
| Gm29607 | 0 | NA | NA | NA |
| Gm21248 | 0 | NA | NA | NA |
| Gm29189 | 0 | NA | NA | NA |
| Gm21873 | 0 | NA | NA | NA |
| Gm29190 | 0 | NA | NA | NA |
| Gm21760 | 0 | NA | NA | NA |
| Gm28159 | 0 | NA | NA | NA |
| Gm28160 | 0 | NA | NA | NA |
| Gm28157 | 0 | NA | NA | NA |
| Gm28158 | 0 | NA | NA | NA |
| Gm21257 | 0 | NA | NA | NA |
| Gm28161 | 0 | NA | NA | NA |
| Gm28695 | 0 | NA | NA | NA |
| Gm28696 | 0 | NA | NA | NA |
| Gm20791 | 0 | NA | NA | NA |
| Gm20841 | 0 | NA | NA | NA |
| Gm28899 | 0 | NA | NA | NA |
| Gm21722 | 0 | NA | NA | NA |
| Gm28898 | 0 | NA | NA | NA |
| Gm28897 | 0 | NA | NA | NA |
| Gm28896 | 0 | NA | NA | NA |
| Gm21282 | 0 | NA | NA | NA |
| Gm28895 | 0 | NA | NA | NA |

| | | | | |
|---|---|---|---|---|
| Gm28894 | 0 | NA | NA | NA |
| Gm21829 | 0 | NA | NA | NA |
| Gm29324 | 0 | NA | NA | NA |
| Gm20911 | 0 | NA | NA | NA |
| Gm28739 | 0 | NA | NA | NA |
| Gm28741 | 0 | NA | NA | NA |
| Gm20912 | 0 | NA | NA | NA |
| Gm20801 | 0 | NA | NA | NA |
| Gm28742 | 0 | NA | NA | NA |
| Gm21919 | 0 | NA | NA | NA |
| Gm28743 | 0 | NA | NA | NA |
| Gm28124 | 0 | NA | NA | NA |
| Gm21790 | 0 | NA | NA | NA |
| Gm21728 | 0 | NA | NA | NA |
| Gm28126 | 0 | NA | NA | NA |
| Gm28125 | 0 | NA | NA | NA |
| Gm21825 | 0 | NA | NA | NA |
| Gm21317 | 0 | NA | NA | NA |
| Gm29468 | 0 | NA | NA | NA |
| Gm29469 | 0 | NA | NA | NA |
| Gm29472 | 0 | NA | NA | NA |
| Gm29473 | 0 | NA | NA | NA |
| Gm29470 | 0 | NA | NA | NA |
| Gm21333 | 0 | NA | NA | NA |
| Gm29382 | 0 | NA | NA | NA |
| Gm28224 | 0 | NA | NA | NA |
| Gm28226 | 0 | NA | NA | NA |
| Gm28225 | 0 | NA | NA | NA |
| Gm21344 | 0 | NA | NA | NA |
| Gm20849 | 0 | NA | NA | NA |
| Gm28829 | 0 | NA | NA | NA |
| Gm21901 | 0 | NA | NA | NA |
| Gm28827 | 0 | NA | NA | NA |
| Gm28828 | 0 | NA | NA | NA |
| Gm28831 | 0 | NA | NA | NA |
| Gm28832 | 0 | NA | NA | NA |
| Gm21864 | 0 | NA | NA | NA |
| Gm21802 | 0 | NA | NA | NA |
| Gm28658 | 0 | NA | NA | NA |
| Gm29238 | 0 | NA | NA | NA |
| Gm21088 | 0 | NA | NA | NA |
| Gm29239 | 0 | NA | NA | NA |
| Gm25729 | 0 | NA | NA | NA |
| Gm21741 | 0 | NA | NA | NA |
| Gm20862 | 0 | NA | NA | NA |
| Gm22624 | 0 | NA | NA | NA |
| Gm20863 | 0 | NA | NA | NA |

| | | | | |
|---|---|---|---|---|
| Gm20864 | 0 | NA | NA | NA |
| Gm29476 | 0 | NA | NA | NA |
| Gm21717 | 0 | NA | NA | NA |
| Gm28472 | 0 | NA | NA | NA |
| Gm21095 | 0 | NA | NA | NA |
| Gm28473 | 0 | NA | NA | NA |
| Gm28474 | 0 | NA | NA | NA |
| Gm28475 | 0 | NA | NA | NA |
| Gm28476 | 0 | NA | NA | NA |
| Gm21380 | 0 | NA | NA | NA |
| Gm21782 | 0 | NA | NA | NA |
| Gm21394 | 0 | NA | NA | NA |
| Gm28471 | 0 | NA | NA | NA |
| Gm21824 | 0 | NA | NA | NA |
| Gm29338 | 0 | NA | NA | NA |
| Gm21409 | 0 | NA | NA | NA |
| Gm29339 | 0 | NA | NA | NA |
| Gm28759 | 0 | NA | NA | NA |
| Gm28758 | 0 | NA | NA | NA |
| Gm29071 | 0 | NA | NA | NA |
| Gm21879 | 0 | NA | NA | NA |
| Gm28757 | 0 | NA | NA | NA |
| Gm20856 | 0 | NA | NA | NA |
| Gm28348 | 0 | NA | NA | NA |
| Gm28973 | 0 | NA | NA | NA |
| Gm21734 | 0 | NA | NA | NA |
| Gm28972 | 0 | NA | NA | NA |
| Gm28970 | 0 | NA | NA | NA |
| Gm28971 | 0 | NA | NA | NA |
| Gm24828 | 0 | NA | NA | NA |
| Gm21785 | 0 | NA | NA | NA |
| Gm20854 | 0 | NA | NA | NA |
| Gm28349 | 0 | NA | NA | NA |
| Gm21892 | 0 | NA | NA | NA |
| Gm28346 | 0 | NA | NA | NA |
| Gm28345 | 0 | NA | NA | NA |
| Gm29090 | 0 | NA | NA | NA |
| Gm28152 | 0 | NA | NA | NA |
| Gm23770 | 0 | NA | NA | NA |
| Gm21881 | 0 | NA | NA | NA |
| Gm21435 | 0 | NA | NA | NA |
| Gm29505 | 0 | NA | NA | NA |
| Gm21862 | 0 | NA | NA | NA |
| Gm28823 | 0 | NA | NA | NA |
| Gm28220 | 0 | NA | NA | NA |
| Gm20820 | 0 | NA | NA | NA |
| Gm28219 | 0 | NA | NA | NA |

| | | | | |
|---|---|---|---|---|
| Gm28218 | 0 | NA | NA | NA |
| Gm23863 | 0 | NA | NA | NA |
| Gm20861 | 0 | NA | NA | NA |
| Gm20860 | 0 | NA | NA | NA |
| Gm28057 | 0 | NA | NA | NA |
| Gm21796 | 0 | NA | NA | NA |
| Gm29302 | 0 | NA | NA | NA |
| Gm28440 | 0 | NA | NA | NA |
| Gm29299 | 0 | NA | NA | NA |
| Gm29298 | 0 | NA | NA | NA |
| Gm29092 | 0 | NA | NA | NA |
| Gm29091 | 0 | NA | NA | NA |
| Gm29297 | 0 | NA | NA | NA |
| Gm21462 | 0 | NA | NA | NA |
| Gm21470 | 0 | NA | NA | NA |
| Gm28366 | 0 | NA | NA | NA |
| Gm21752 | 0 | NA | NA | NA |
| Gm28365 | 0 | NA | NA | NA |
| Gm21477 | 0 | NA | NA | NA |
| Gm28368 | 0 | NA | NA | NA |
| Gm28369 | 0 | NA | NA | NA |
| Gm28367 | 0 | NA | NA | NA |
| Gm29115 | 0 | NA | NA | NA |
| Gm21813 | 0 | NA | NA | NA |
| Gm20906 | 0 | NA | NA | NA |
| Gm29111 | 0 | NA | NA | NA |
| Gm28110 | 0 | NA | NA | NA |
| Gm28109 | 0 | NA | NA | NA |
| Gm28108 | 0 | NA | NA | NA |
| Gm28107 | 0 | NA | NA | NA |
| Gm28106 | 0 | NA | NA | NA |
| Gm28105 | 0 | NA | NA | NA |
| Gm28104 | 0 | NA | NA | NA |
| Gm28103 | 0 | NA | NA | NA |
| Gm28102 | 0 | NA | NA | NA |
| Gm28716 | 0 | NA | NA | NA |
| Gm28717 | 0 | NA | NA | NA |
| Gm28718 | 0 | NA | NA | NA |
| Gm29162 | 0.046345709 | 0.274391604 | 3.724945049 | 0.073663262 |
| Gm21777 | 0 | NA | NA | NA |
| Gm28866 | 0 | NA | NA | NA |
| Gm28405 | 0 | NA | NA | NA |
| Gm21792 | 0 | NA | NA | NA |
| Gm28406 | 0 | NA | NA | NA |
| Gm28978 | 0 | NA | NA | NA |
| Gm29436 | 0 | NA | NA | NA |
| Gm20819 | 0 | NA | NA | NA |

| | | | | |
|---|---|---|---|---|
| Gm29437 | 0 | NA | NA | NA |
| Gm28977 | 0 | NA | NA | NA |
| Gm28407 | 0 | NA | NA | NA |
| Gm21287 | 0 | NA | NA | NA |
| Gm29390 | 0 | NA | NA | NA |
| Gm29391 | 0 | NA | NA | NA |
| Gm21720 | 0 | NA | NA | NA |
| Gm29392 | 0 | NA | NA | NA |
| Gm29393 | 0 | NA | NA | NA |
| Gm21294 | 0 | NA | NA | NA |
| Gm28672 | 0 | NA | NA | NA |
| Gm21838 | 0 | NA | NA | NA |
| Gm28670 | 0 | NA | NA | NA |
| Gm28671 | 0 | NA | NA | NA |
| Gm28673 | 0 | NA | NA | NA |
| Gm28674 | 0 | NA | NA | NA |
| Gm28531 | 0 | NA | NA | NA |
| Gm21996 | 0 | NA | NA | NA |
| Gm21803 | 0 | NA | NA | NA |
| Gm28930 | 0 | NA | NA | NA |
| Gm20879 | 0 | NA | NA | NA |
| Gm29504 | 0 | NA | NA | NA |
| Gm20837 | 0 | NA | NA | NA |
| Gm29503 | 0 | NA | NA | NA |
| Gm28300 | 0 | NA | NA | NA |
| Gm28301 | 0 | NA | NA | NA |
| Gm21857 | 2.299789819 | 1.284534418 | 0.676305313 | 1.899341013 |
| Gm21860 | 0.344895107 | 1.980383029 | 1.530800511 | 1.293691121 |
| Erdr1 | 44.24775491 | -0.278982493 | 0.221144497 | -1.261539385 |
| Gm21742 | 0 | NA | NA | NA |
| Gm21748 | 0 | NA | NA | NA |

| pvalue | padj |
|---|---|
| NA | NA |
| NA | NA |
| 0.000236547 | 0.001480487 |
| 0.764353414 | NA |
| 0.405880477 | 0.567920367 |
| 0.652269273 | 0.773105252 |
| 0.867076241 | 0.920555406 |
| NA | NA |
| 0.946562919 | 0.967538031 |
| 0.856530291 | NA |
| 0.528001489 | 0.677127772 |
| 0.538285192 | 0.685954821 |
| 3.47E-14 | 9.90E-13 |
| NA | NA |
| NA | NA |
| 0.955792398 | NA |
| 0.941277857 | NA |
| 0.281036127 | 0.441212447 |
| NA | NA |
| 0.366880244 | 0.5300767 |
| 0.417322004 | NA |
| 0.687978645 | NA |
| 0.937023989 | NA |
| NA | NA |
| 0.200280695 | NA |
| 0.839318641 | NA |
| 0.691676967 | 0.802187554 |
| 0.738860418 | 0.835678947 |
| 0.955792398 | NA |
| 0.151575773 | NA |
| 0.474174132 | 0.631613 |
| 0.343306613 | 0.506878233 |
| 0.02788224 | 0.078327696 |
| 0.247978518 | 0.405188761 |
| 0.941279303 | NA |
| 1.73E-06 | 1.85E-05 |
| 0.356607408 | 0.520255564 |
| 1.36E-05 | 0.000119655 |
| 0.809074484 | 0.882967072 |
| 0.568632138 | 0.70980607 |
| 0.188266142 | 0.332624325 |
| 0.492512807 | NA |
| 0.584145947 | 0.721303603 |
| 0.079508476 | 0.177047501 |
| NA | NA |
| 0.950067215 | 0.969454679 |

| | | |
|---|---|---|
| | 0.661823912 | NA |
| | 0.108726874 | 0.223192501 |
| NA | | NA |
| NA | | NA |
| NA | | NA |
| | 0.233413017 | 0.388021835 |
| | 0.941278332 | NA |
| | 0.445281304 | 0.60591045 |
| | 1.70E-05 | 0.000145084 |
| | 0.451068288 | NA |
| | 0.46635575 | NA |
| NA | | NA |
| | 0.025974908 | 0.073915904 |
| | 3.16E-05 | 0.000252513 |
| NA | | NA |
| | 0.941275956 | NA |
| | 0.778655412 | 0.863592382 |
| NA | | NA |
| NA | | NA |
| NA | | NA |
| NA | | NA |
| | 0.359248579 | 0.522773255 |
| NA | | NA |
| | 0.000515118 | 0.002910933 |
| | 0.447346309 | 0.607678167 |
| NA | | NA |
| NA | | NA |
| | 0.941278788 | NA |
| | 0.047015947 | 0.11797603 |
| NA | | NA |
| NA | | NA |
| NA | | NA |
| | 0.941278332 | NA |
| NA | | NA |
| NA | | NA |
| | 0.841948209 | 0.904742721 |
| | 0.576979418 | 0.715572327 |
| | 0.31203317 | 0.474015448 |
| | 0.929351239 | 0.958004957 |
| | 0.585012353 | 0.721736589 |
| | 0.485609072 | 0.640961965 |
| | 0.593657013 | 0.728800997 |
| | 0.941279268 | NA |
| | 0.586653289 | NA |
| | 0.773912456 | NA |
| | 0.808552995 | NA |
| | 0.226228745 | 0.379230691 |

| | |
|---|---|
| 0.016245139 | 0.050823711 |
| 0.084316511 | 0.185019046 |
| NA | NA |
| 0.95136208 | NA |
| NA | NA |
| NA | NA |
| NA | NA |
| 0.349637347 | 0.5134855 |
| 0.084688665 | 0.185647413 |
| 3.39E-08 | 4.90E-07 |
| 0.307894863 | 0.46941542 |
| 0.142684797 | 0.273049006 |
| 0.228254794 | 0.3817753 |
| NA | NA |
| 0.351112002 | 0.51504794 |
| 4.55E-07 | 5.40E-06 |
| 0.493410022 | 0.646989056 |
| 6.00E-09 | 9.69E-08 |
| 0.7943415 | 0.873834799 |
| NA | NA |
| 0.511739958 | 0.663169515 |
| 0.03846761 | 0.100958801 |
| 0.922447883 | 0.953783863 |
| 0.462067633 | 0.620802314 |
| 0.721826942 | NA |
| 0.510547907 | 0.662075483 |
| NA | NA |
| NA | NA |
| 0.035310066 | 0.09435839 |
| 0.000101368 | 0.000708799 |
| NA | NA |
| 0.385658782 | 0.548452551 |
| NA | NA |
| 0.447115013 | 0.607433308 |
| 0.015984766 | 0.05018723 |
| 0.000225726 | 0.00142286 |
| 0.031215964 | 0.085589492 |
| 0.617780997 | NA |
| 0.955792398 | NA |
| NA | NA |
| 0.941278332 | NA |
| NA | NA |
| NA | NA |
| 9.73E-05 | 0.000684709 |
| NA | NA |
| NA | NA |
| 0.839319108 | NA |

| | | | |
|---|---|---|---|
| | 0.013427875 | | 0.04371175 |
| | 0.208434858 | | 0.35776783 |
| NA | | NA | |
| | 0.993067811 | | 0.995116407 |
| | 0.311281777 | NA | |
| NA | | NA | |
| NA | | NA | |
| | 0.85565056 | NA | |
| NA | | NA | |
| | 0.839319574 | NA | |
| NA | | NA | |
| | 0.08086659 | | 0.179130044 |
| | 0.062659799 | | 0.147944503 |
| NA | | NA | |
| NA | | NA | |
| NA | | NA | |
| | 0.939052821 | NA | |
| | 0.026627201 | | 0.075504009 |
| | 0.522407829 | | 0.672390001 |
| | 0.065518359 | | 0.153219491 |
| NA | | NA | |
| | 0.004224771 | | 0.01697468 |
| | 0.00132349 | | 0.006461891 |
| | 0.184266019 | | 0.327227407 |
| | 0.008468529 | | 0.030026504 |
| | 0.050733523 | | 0.125307392 |
| | 0.610512907 | | 0.742043008 |
| | 0.941277857 | NA | |
| | 0.001410532 | | 0.006818338 |
| | 0.005430704 | | 0.020841925 |
| | 0.613085286 | | 0.743802232 |
| | 0.084405617 | NA | |
| NA | | NA | |
| | 0.841773098 | NA | |
| | 0.853596091 | NA | |
| NA | | NA | |
| NA | | NA | |
| | 0.788698113 | | 0.869975354 |
| NA | | NA | |
| | 0.625757024 | | 0.753436846 |
| | 0.51732044 | | 0.668165833 |
| | 0.76518108 | | 0.854255406 |
| | 0.779953808 | | 0.864527319 |
| | 0.213089263 | | 0.363789316 |
| | 0.340463199 | | 0.50405529 |
| | 0.888648697 | | 0.93342843 |
| NA | | NA | |

| | | |
|---|---|---|
| | 0.396709452 | 0.559322259 |
| | 0.003200021 | 0.013535971 |
| | 0.000168602 | 0.001109192 |
| | 0.034607973 | 0.092805201 |
| NA | | NA |
| | 0.685230358 | NA |
| | 0.941278332 | NA |
| | 0.839319574 | NA |
| | 0.633849255 | 0.759386128 |
| | 0.955424541 | 0.972877644 |
| | 0.071361868 | 0.163416893 |
| | 0.941277857 | NA |
| NA | | NA |
| | 0.593677949 | 0.728800997 |
| | 6.80E-07 | 7.82E-06 |
| | 0.404465926 | 0.566499821 |
| | 0.29397874 | 0.455166979 |
| NA | | NA |
| | 0.362940198 | 0.526621714 |
| | 0.629770122 | NA |
| | 0.95804761 | 0.974181957 |
| | 0.013131032 | 0.042890739 |
| | 0.541098223 | 0.688286656 |
| | 0.761140935 | NA |
| | 0.695950266 | NA |
| | 0.171281745 | 0.311423733 |
| | 0.072337204 | 0.165062806 |
| NA | | NA |
| | 0.196781258 | 0.343537297 |
| | 0.929794865 | NA |
| NA | | NA |
| | 0.492716375 | 0.646542541 |
| | 0.398724708 | NA |
| | 0.038490365 | 0.100973664 |
| | 0.133178346 | 0.259255009 |
| | 0.941278332 | NA |
| | 0.82515277 | NA |
| | 0.989858493 | 0.993351574 |
| | 0.941277857 | NA |
| NA | | NA |
| | 0.940116961 | 0.963971218 |
| | 0.399399343 | 0.562049377 |
| | 0.065087062 | 0.152525575 |
| | 0.344044307 | 0.507652507 |
| | 0.807923066 | 0.882239978 |
| NA | | NA |
| | 0.011957887 | 0.039773845 |

| | |
|---|---|
| NA | NA |
| 0.789329655 | 0.870361551 |
| 0.188128925 | 0.332495519 |
| NA | NA |
| 0.955792873 | NA |
| NA | NA |
| NA | NA |
| 0.744938588 | NA |
| NA | NA |
| NA | NA |
| NA | NA |
| NA | NA |
| NA | NA |
| 0.190392409 | 0.335161609 |
| 0.240852842 | 0.396943619 |
| 0.432020485 | 0.593287695 |
| 0.341076967 | 0.504595534 |
| 0.839320041 | NA |
| NA | NA |
| 0.314973131 | 0.477302506 |
| NA | NA |
| 0.853595154 | NA |
| 0.494122303 | 0.647310858 |
| NA | NA |
| 0.652228611 | NA |
| 0.705253805 | NA |
| NA | NA |
| NA | NA |
| 0.148776139 | 0.281648745 |
| NA | NA |
| NA | NA |
| 0.462543667 | NA |
| 2.01E-09 | 3.43E-08 |
| NA | NA |
| 0.955792873 | NA |
| NA | NA |
| 0.471788704 | 0.629386544 |
| 0.162275709 | 0.299603353 |
| NA | NA |
| 0.705906915 | 0.812318525 |
| 0.731788199 | 0.830893609 |
| 5.11E-17 | 1.79E-15 |
| 0.924082144 | NA |
| 0.542094607 | 0.689112146 |
| 0.330255745 | 0.493206501 |
| 0.068604685 | 0.158721686 |
| 0.501529412 | 0.653874786 |

| | | |
|---|---|---|
| | 0.520036752 | 0.670380999 |
| | 0.379695132 | 0.542422487 |
| | 0.895320695 | NA |
| | 0.941278332 | NA |
| | 0.839319108 | NA |
| NA | | NA |
| NA | | NA |
| | 0.941277857 | NA |
| NA | | NA |
| NA | | NA |
| NA | | NA |
| NA | | NA |
| | 0.939568003 | 0.963757575 |
| | 0.990341018 | 0.993561791 |
| | 0.002785449 | 0.012056601 |
| | 0.018291146 | 0.055851939 |
| | 0.230874088 | 0.384795536 |
| NA | | NA |
| NA | | NA |
| | 9.81E-05 | 0.000689247 |
| | 0.955792873 | NA |
| | 0.688807755 | NA |
| | 0.338226044 | NA |
| | 0.33811345 | NA |
| NA | | NA |
| | 0.955792398 | NA |
| | 0.532034988 | 0.680613608 |
| | 0.005200522 | 0.020165948 |
| NA | | NA |
| | 0.065996332 | 0.154140387 |
| | 0.723035965 | NA |
| | 0.743186025 | 0.83854495 |
| | 0.941278332 | NA |
| | 0.636165355 | 0.760828449 |
| | 0.003502656 | 0.014572666 |
| | 6.01E-07 | 6.97E-06 |
| | 0.078795326 | 0.175837836 |
| | 0.941276431 | NA |
| | 0.18028656 | 0.322305896 |
| | 0.319320788 | 0.481866062 |
| NA | | NA |
| | 0.000570808 | 0.003175868 |
| | 0.188197457 | 0.332573175 |
| | 0.560415076 | NA |
| | 0.519726342 | 0.670198635 |
| | 0.570652967 | 0.7112545 |
| | 0.925672843 | 0.955348429 |

| | | |
|---|---|---|
| | 0.478675392 | 0.63543958 |
| | 0.441721764 | 0.602275552 |
| | 0.160363597 | 0.297110169 |
| | 0.308059295 | 0.469578047 |
| | 0.428697445 | 0.589988817 |
| | 0.032978003 | 0.089463228 |
| NA | | NA |
| | 0.934281796 | NA |
| | 0.941278332 | NA |
| | 0.657518622 | NA |
| | 0.042394578 | 0.108971463 |
| | 0.919265124 | NA |
| | 0.00924207 | 0.032264557 |
| | 0.090278922 | 0.194745232 |
| | 0.107459214 | 0.221201858 |
| | 0.002980741 | 0.012751833 |
| | 0.00534264 | 0.020586939 |
| | 0.010914022 | 0.03698399 |
| | 0.011254123 | 0.037899072 |
| | 0.016290916 | 0.050940146 |
| NA | | NA |
| | 0.000128152 | 0.000872002 |
| | 0.014692172 | 0.046891133 |
| | 0.002430804 | 0.010776838 |
| | 0.002862866 | 0.012331149 |
| | 0.006965436 | 0.025581131 |
| | 0.002406043 | 0.010688434 |
| | 0.008841657 | 0.031103921 |
| | 6.19E-06 | 5.91E-05 |
| | 0.000333931 | 0.002002072 |
| | 0.000453312 | 0.002605588 |
| | 8.07E-05 | 0.000580075 |
| | 0.000570763 | 0.003175868 |
| | 0.001729069 | 0.008103443 |
| | 0.003897519 | 0.015874185 |
| NA | | NA |
| NA | | NA |
| NA | | NA |
| NA | | NA |
| NA | | NA |
| NA | | NA |
| | 0.955792398 | NA |
| NA | | NA |
| | 0.728678906 | NA |
| NA | | NA |
| | 0.521295104 | 0.671493996 |
| | 0.955792873 | NA |

| | | |
|---|---|---|
| | 0.809735213 | NA |
| NA | | NA |
| NA | | NA |
| NA | | NA |
| NA | | NA |
| NA | | NA |
| | 0.955792398 | NA |
| | 0.085346179 | 0.186676141 |
| NA | | NA |
| NA | | NA |
| | 0.853596091 | NA |
| | 0.880060338 | NA |
| | 0.87203901 | 0.923527142 |
| | 0.237315046 | NA |
| | 0.034695725 | 0.0929672 |
| | 0.003663994 | 0.015096137 |
| | 0.147678155 | 0.280238814 |
| | 0.035349111 | 0.094452126 |
| NA | | NA |
| NA | | NA |
| | 0.941277857 | NA |
| NA | | NA |
| NA | | NA |
| NA | | NA |
| | 0.955792873 | NA |
| NA | | NA |
| NA | | NA |
| | 0.621801627 | NA |
| NA | | NA |
| NA | | NA |
| NA | | NA |
| | 0.941278332 | NA |
| NA | | NA |
| NA | | NA |
| | 0.103038077 | 0.214511705 |
| NA | | NA |
| NA | | NA |
| | 0.300482317 | 0.462050527 |
| | 0.955792873 | NA |
| | 0.644618225 | 0.767313612 |
| | 0.479980117 | 0.636603519 |
| | 0.490838422 | 0.645047538 |
| | 0.833801616 | NA |
| | 0.227048772 | 0.380257168 |
| | 0.608886309 | NA |
| NA | | NA |
| | 0.566229837 | 0.707921135 |

| | | |
|---|---|---|
| | 0.691921803 | NA |
| | 0.158184106 | 0.294240238 |
| | 0.845832418 | NA |
| | 0.764114553 | 0.853590585 |
| | 0.460562726 | 0.619619829 |
| | 0.613563113 | NA |
| NA | | NA |
| | 0.029405037 | NA |
| NA | | NA |
| NA | | NA |
| | 0.703178671 | 0.810589826 |
| | 0.853595154 | NA |
| NA | | NA |
| | 0.354798665 | 0.518602175 |
| NA | | NA |
| | 0.000256406 | 0.001585191 |
| NA | | NA |
| | 0.091554248 | 0.196605395 |
| | 0.093515839 | NA |
| | 0.001395865 | 0.006754296 |
| | 0.099129362 | 0.208176908 |
| | 0.370012529 | NA |
| | 0.016109861 | 0.050486756 |
| | 0.135772683 | 0.263100335 |
| | 0.110468716 | 0.225833237 |
| | 0.002120736 | 0.009600407 |
| | 0.439115682 | 0.600059219 |
| | 0.005448707 | 0.02089754 |
| NA | | NA |
| NA | | NA |
| | 0.152401098 | 0.28650588 |
| | 0.445612705 | 0.606118842 |
| | 0.112062459 | 0.228131255 |
| | 0.022851664 | 0.066705335 |
| | 0.805468719 | 0.880933991 |
| NA | | NA |
| | 0.628514554 | NA |
| NA | | NA |
| | 0.005265069 | 0.02035063 |
| | 0.139856244 | 0.268890009 |
| NA | | NA |
| | 0.033100794 | 0.089751245 |
| NA | | NA |
| | 0.909389044 | 0.945084837 |
| | 5.21E-10 | 9.62E-09 |
| | 0.08046023 | 0.178582095 |
| | 0.186828908 | 0.330673831 |

| | | | |
|---|---|---|---|
| | 0.906584818 | NA | |
| NA | | NA | |
| | 0.911696613 | NA | |
| NA | | NA | |
| NA | | NA | |
| | 0.843811749 | NA | |
| | 0.011665748 | | 0.039064174 |
| | 0.839318641 | NA | |
| | 0.682860478 | NA | |
| | 0.006696438 | NA | |
| | 7.68E-08 | | 1.05E-06 |
| | 0.943837575 | | 0.966121445 |
| NA | | NA | |
| | 0.646183886 | | 0.768447107 |
| | 0.47708805 | | 0.633898055 |
| | 0.011700131 | | 0.039124266 |
| | 0.323655626 | | 0.486310916 |
| | 0.774196026 | | 0.860571326 |
| | 0.328630442 | | 0.491551674 |
| | 0.704491558 | NA | |
| | 0.750331332 | NA | |
| | 0.839318641 | NA | |
| NA | | NA | |
| NA | | NA | |
| | 0.44766333 | NA | |
| NA | | NA | |
| NA | | NA | |
| | 0.032074578 | | 0.087420027 |
| NA | | NA | |
| NA | | NA | |
| | 0.687496744 | NA | |
| NA | | NA | |
| | 0.955792873 | NA | |
| | 0.420322013 | | 0.581889993 |
| | 0.717544793 | NA | |
| | 0.888611307 | NA | |
| | 0.00081802 | | 0.004309544 |
| | 0.545668023 | NA | |
| | 8.14E-24 | | 4.29E-22 |
| | 1.76E-15 | | 5.57E-14 |
| | 0.172321256 | | 0.312562496 |
| | 0.000179665 | | 0.001169368 |
| | 0.955792398 | NA | |
| | 0.123205726 | | 0.244994933 |
| | 9.52E-05 | | 0.000671793 |
| | 0.445348439 | | 0.605912779 |
| | 0.056480983 | | 0.136483285 |

| | |
|---|---|
| NA | NA |
| 0.66789474 | 0.784735757 |
| 0.659927484 | NA |
| 0.137284395 | 0.265381561 |
| 0.491433146 | 0.645465283 |
| 0.000385471 | 0.00226319 |
| 9.48E-06 | 8.63E-05 |
| 0.605455221 | 0.73815731 |
| 0.305388167 | 0.466954365 |
| NA | NA |
| 0.688995107 | 0.800559509 |
| 0.000100641 | 0.000705377 |
| 0.934418963 | 0.960973711 |
| 0.231645276 | 0.385810711 |
| 0.443475623 | 0.604029687 |
| 5.57E-11 | 1.16E-09 |
| NA | NA |
| 0.169000244 | 0.30854827 |
| 0.360148433 | 0.523848265 |
| 0.157172824 | 0.29295439 |
| 0.18727937 | 0.331277929 |
| 0.499633634 | NA |
| NA | NA |
| 0.355971526 | 0.519582649 |
| 0.422517306 | 0.583812662 |
| 0.195476834 | 0.341836721 |
| NA | NA |
| 0.000194495 | 0.001251182 |
| 0.941275481 | NA |
| NA | NA |
| 0.762190332 | 0.852321085 |
| 0.177852427 | 0.319321374 |
| 0.385542451 | 0.54841154 |
| 0.389715674 | 0.552275094 |
| 0.915545493 | 0.949079475 |
| 0.941278332 | NA |
| 0.938650974 | NA |
| 0.046078318 | 0.116180046 |
| 5.16E-07 | 6.07E-06 |
| 0.318630183 | 0.481071469 |
| 0.023443606 | 0.068115809 |
| 1.26E-25 | 7.24E-24 |
| 1.22E-08 | 1.88E-07 |
| 0.302015871 | NA |
| 0.899436455 | NA |
| 0.939241089 | NA |
| NA | NA |

| | |
|---|---|
| 7.01E-06 | 6.58E-05 |
| 0.033495909 | 0.090567947 |
| 0.706651361 | 0.812901033 |
| 0.882295097 | 0.929590895 |
| 0.832162011 | NA |
| 0.084601737 | 0.185473937 |
| 0.555926592 | 0.700231223 |
| NA | NA |
| 0.171770776 | 0.311912279 |
| NA | NA |
| NA | NA |
| NA | NA |
| 5.89E-07 | 6.84E-06 |
| 4.26E-12 | 1.00E-10 |
| NA | NA |
| 0.612697083 | NA |
| 0.648790618 | 0.770392331 |
| 0.361346847 | 0.525029113 |
| 0.023240975 | 0.067659208 |
| 0.045907121 | NA |
| 0.764646056 | NA |
| NA | NA |
| 0.542269551 | 0.689224107 |
| 0.519350754 | NA |
| 0.042270242 | 0.108745818 |
| 1.33E-05 | 0.000117108 |
| 0.011826784 | 0.039448455 |
| 0.03171171 | 0.086676751 |
| NA | NA |
| 0.152525895 | 0.28664993 |
| 0.013517916 | 0.043901411 |
| 0.96753686 | 0.980513149 |
| 0.636233408 | 0.760856295 |
| 2.74E-05 | 0.000222904 |
| 0.055080825 | 0.133847248 |
| 0.538391315 | NA |
| 0.828904408 | 0.896188177 |
| 0.382225119 | 0.544806412 |
| 0.468016401 | 0.626214311 |
| 0.955792398 | NA |
| 0.735799551 | NA |
| 0.026030635 | 0.07403288 |
| 0.849056857 | 0.909220827 |
| 1.64E-09 | 2.83E-08 |
| 0.082132421 | 0.181228876 |
| 0.000234808 | 0.001471659 |
| 1.66E-12 | 4.07E-11 |

| | | |
|---|---|---|
| | 0.388308615 | 0.551167134 |
| | 0.55111917 | 0.696459896 |
| | 0.554635497 | NA |
| | 0.296064065 | 0.457532585 |
| | 1.11E-10 | 2.21E-09 |
| | 8.67E-18 | 3.19E-16 |
| | 0.839318641 | NA |
| | 0.277923439 | 0.43753778 |
| | 0.941279303 | NA |
| | 0.018875535 | 0.057290931 |
| NA | | NA |
| | 0.86296073 | NA |
| | 0.549674733 | 0.695259294 |
| | 0.12867066 | 0.252996404 |
| NA | | NA |
| | 0.839319108 | NA |
| | 0.831413565 | NA |
| | 0.749443074 | 0.842618992 |
| NA | | NA |
| NA | | NA |
| | 0.896304533 | NA |
| NA | | NA |
| | 0.485731422 | 0.641087907 |
| | 0.941278284 | NA |
| | 0.569297955 | NA |
| | 0.09157549 | 0.196633272 |
| NA | | NA |
| | 0.003179854 | 0.013477011 |
| | 0.00249734 | 0.011026098 |
| | 0.009696558 | 0.033601108 |
| | 0.476528681 | 0.633524321 |
| | 0.40269048 | 0.564744214 |
| NA | | NA |
| | 0.010413785 | 0.035572462 |
| | 0.701106418 | NA |
| | 0.454276599 | 0.614113228 |
| | 0.014420655 | 0.046221444 |
| | 0.656802162 | 0.776226666 |
| | 0.000177757 | 0.001160124 |
| | 3.75E-06 | 3.75E-05 |
| | 0.02554241 | 0.072879458 |
| | 0.955792873 | NA |
| NA | | NA |
| NA | | NA |
| | 0.4493659 | 0.609482429 |
| | 0.068983398 | 0.159396452 |
| | 0.020090622 | 0.060082547 |

| | | |
|---|---|---|
| | 0.056767737 | 0.13696763 |
| | 0.701455261 | 0.809387149 |
| | 0.455001291 | NA |
| | 0.7683391 | 0.856460791 |
| | 0.349070104 | 0.513064507 |
| NA | | NA |
| | 0.166803331 | 0.305662706 |
| NA | | NA |
| NA | | NA |
| | 0.923929221 | NA |
| | 0.016718696 | 0.051990908 |
| | 0.208752972 | 0.358164611 |
| NA | | NA |
| | 0.443679493 | 0.604248973 |
| | 0.081761209 | 0.18059394 |
| NA | | NA |
| | 0.011775228 | 0.039303599 |
| | 0.015800527 | 0.049733422 |
| | 0.83029655 | 0.897000658 |
| | 0.170980999 | NA |
| | 0.947374177 | NA |
| NA | | NA |
| NA | | NA |
| | 0.07496296 | 0.169381846 |
| | 0.885515126 | 0.931457974 |
| | 0.180057174 | 0.322007571 |
| | 0.072457595 | 0.165210814 |
| | 0.934281796 | NA |
| | 0.92675519 | 0.95599493 |
| | 0.739991096 | NA |
| NA | | NA |
| NA | | NA |
| NA | | NA |
| | 2.07E-06 | 2.18E-05 |
| NA | | NA |
| | 0.012172889 | 0.040348015 |
| | 0.403992851 | 0.566137053 |
| | 0.955792873 | NA |
| | 0.897156917 | NA |
| NA | | NA |
| NA | | NA |
| NA | | NA |
| | 0.792316268 | NA |
| NA | | NA |
| | 0.02129376 | 0.062911162 |
| NA | | NA |
| | 0.932084447 | 0.959371849 |

| | | |
|---|---|---|
| | 0.922811894 | NA |
| | 0.818573585 | NA |
| | 6.21E-07 | 7.19E-06 |
| NA | | NA |
| NA | | NA |
| | 0.900085681 | NA |
| NA | | NA |
| NA | | NA |
| | 4.36E-05 | 0.000335463 |
| | 0.955792398 | NA |
| | 0.321008603 | 0.483821296 |
| | 0.08729199 | 0.189850508 |
| | 0.300819817 | 0.462211174 |
| | 0.860538198 | 0.916609361 |
| | 0.117050174 | 0.235624851 |
| | 0.432676357 | 0.59382618 |
| | 0.311165759 | NA |
| | 0.011932033 | NA |
| | 0.254226378 | 0.412243084 |
| NA | | NA |
| NA | | NA |
| | 0.032323845 | 0.088018781 |
| | 0.612508824 | 0.743447088 |
| | 0.030166329 | 0.083345207 |
| NA | | NA |
| NA | | NA |
| NA | | NA |
| NA | | NA |
| NA | | NA |
| NA | | NA |
| NA | | NA |
| NA | | NA |
| NA | | NA |
| NA | | NA |
| NA | | NA |
| NA | | NA |
| NA | | NA |
| NA | | NA |
| NA | | NA |
| NA | | NA |
| NA | | NA |
| | 0.955792873 | NA |
| NA | | NA |
| NA | | NA |
| NA | | NA |
| NA | | NA |
| NA | | NA |

| | | |
|---|---|---|
| | 0.94127925 | NA |
| | 0.000700336 | 0.003781622 |
| | 0.01873445 | 0.056960062 |
| | 0.001936625 | 0.008915969 |
| | 0.786035489 | NA |
| | 0.001120338 | 0.005626987 |
| | 0.293737178 | 0.454911336 |
| | 0.716613369 | 0.819843341 |
| | 0.000249316 | 0.001548997 |
| | 0.091657687 | 0.196721034 |
| | 0.82229013 | NA |
| | 0.680821291 | 0.79455603 |
| | 0.023316829 | 0.067805396 |
| | 0.757538328 | NA |
| | 0.398364818 | NA |
| | 0.808547564 | 0.882557513 |
| | 0.461268178 | 0.620186255 |
| | 0.001456487 | 0.006996427 |
| NA | | NA |
| NA | | NA |
| | 0.93278691 | NA |
| | 8.81E-11 | 1.78E-09 |
| | 0.000624872 | 0.003423062 |
| | 0.61475642 | NA |
| | 0 | 0 |
| | 0.839319574 | NA |
| | 4.35E-33 | 3.45E-31 |
| | 0.955792398 | NA |
| NA | | NA |
| | 0.783172287 | 0.866703867 |
| | 0.941277857 | NA |
| | 0.494291425 | 0.647467091 |
| | 0.971958683 | 0.983068907 |
| | 0.848371998 | NA |
| | 0.309399324 | 0.471075463 |
| | 0.456908199 | 0.616585154 |
| | 0.209220693 | 0.358754479 |
| | 0.781987388 | 0.865974863 |
| | 0.06705257 | 0.155964554 |
| | 0.767651348 | NA |
| | 8.56E-13 | 2.17E-11 |
| | 0.000178599 | 0.001163913 |
| | 0.003110068 | 0.013211823 |
| | 0.772289837 | 0.859375535 |
| | 0.089707327 | 0.193740476 |
| NA | | NA |
| | 0.197041046 | 0.343802457 |

| | |
|---|---|
| 0.098617346 | 0.207431078 |
| NA | NA |
| 0.267889282 | 0.427611267 |
| 0.955792398 | NA |
| 0.243793878 | 0.400288755 |
| 0.725155089 | 0.82635809 |
| 0.272268678 | 0.432127426 |
| 0.069232901 | 0.159895354 |
| 0.941277857 | NA |
| NA | NA |
| NA | NA |
| 0.458863333 | 0.618277105 |
| 0.941275956 | NA |
| 1.78E-09 | 3.06E-08 |
| NA | NA |
| 4.92E-05 | 0.000374153 |
| 0.955792873 | NA |
| 0.364605548 | 0.527948116 |
| 0.015683279 | 0.049423197 |
| 0.922668352 | NA |
| 0.16967175 | 0.309252392 |
| 0.303051149 | 0.464557615 |
| 4.65E-08 | 6.54E-07 |
| 1.24E-05 | 0.000109995 |
| 0.839319574 | NA |
| 0.000337031 | 0.002019132 |
| 0.269295965 | NA |
| 0.031923277 | 0.087127375 |
| NA | NA |
| 0.647979795 | 0.769890435 |
| 7.33E-05 | 0.000531771 |
| 0.446636501 | 0.607095084 |
| 0.063360632 | 0.149288235 |
| 0.134362123 | 0.261047036 |
| 0.402536934 | 0.564710866 |
| 0.195549398 | 0.341938493 |
| 0.555383355 | NA |
| 0.955180557 | 0.972692863 |
| 0.986196307 | 0.991444691 |
| 0.892669506 | NA |
| 0.436745476 | 0.597746871 |
| 0.080644096 | 0.17887343 |
| 0.563705343 | 0.705986435 |
| 0.213843969 | 0.364711532 |
| 0.070007661 | 0.161262204 |
| NA | NA |
| 0.955792873 | NA |

| | | | |
|---|---|---|---|
| | 0.016875504 | | 0.052389653 |
| | 0.862209327 | | 0.917648997 |
| | 0.046996024 | | 0.117945946 |
| | 0.749795858 | | 0.84281796 |
| | 0.294141159 | | 0.455359211 |
| NA | | NA | |
| NA | | NA | |
| | 0.666972928 | NA | |
| NA | | NA | |
| NA | | NA | |
| | 0.348589586 | | 0.512525091 |
| | 0.536924101 | | 0.684880356 |
| | 0.371495109 | | 0.534828923 |
| NA | | NA | |
| | 0.839319108 | NA | |
| | 0.762957868 | | 0.852875853 |
| NA | | NA | |
| | 0.560397267 | | 0.703585597 |
| | 0.609595592 | | 0.741490928 |
| | 1.70E-06 | | 1.83E-05 |
| | 0.000173525 | | 0.001136554 |
| NA | | NA | |
| NA | | NA | |
| NA | | NA | |
| | 0.628128348 | NA | |
| | 0.839319108 | NA | |
| | 1.43E-24 | | 7.88E-23 |
| | 0.839319107 | NA | |
| | 0.043980534 | | 0.112007776 |
| | 0.003679072 | | 0.015139916 |
| | 9.55E-05 | | 0.000673829 |
| | 0.39639024 | | 0.559037772 |
| NA | | NA | |
| | 0.540370857 | | 0.687508399 |
| | 0.72601533 | NA | |
| | 0.098185531 | | 0.206870144 |
| | 0.056331591 | | 0.136232932 |
| | 0.021512631 | | 0.063442404 |
| | 0.003783181 | | 0.015493337 |
| | 0.826630988 | | 0.894768681 |
| | 0.032768002 | | 0.089063737 |
| | 0.556576182 | | 0.700747225 |
| | 0.536616371 | | 0.684634587 |
| | 0.78594389 | | 0.868438628 |
| | 0.031164521 | | 0.085487833 |
| | 0.849111524 | | 0.909238461 |
| NA | | NA | |

| | |
|---|---|
| NA | NA |
| 0.73753467 | 0.834591624 |
| 0.211733265 | NA |
| 1.91E-06 | 2.03E-05 |
| NA | NA |
| 6.41E-31 | 4.62E-29 |
| NA | NA |
| 4.14E-34 | 3.44E-32 |
| 1.64E-07 | 2.12E-06 |
| 0.431811769 | 0.593074652 |
| 0.86646369 | 0.920243129 |
| 0.002769358 | 0.012008792 |
| 0.839319574 | NA |
| 0.724546952 | NA |
| 0.919271727 | NA |
| NA | NA |
| 0.831497906 | 0.897834783 |
| 0.70531761 | 0.812065262 |
| 0.001767124 | 0.008252124 |
| 0.839319108 | NA |
| NA | NA |
| 0.026227132 | 0.074493819 |
| 0.002791389 | 0.012073529 |
| NA | NA |
| 0.950425349 | NA |
| 0.562504662 | 0.705152787 |
| 0.955792398 | NA |
| 0.07955573 | 0.177103019 |
| 0.839319574 | NA |
| 0.939241089 | NA |
| 0.000109415 | 0.000757942 |
| 0.724596388 | NA |
| 0.596407264 | 0.73113338 |
| 0.176819192 | 0.318424986 |
| 0.002385345 | 0.010612295 |
| 0.671109562 | 0.787184599 |
| NA | NA |
| 0.90873405 | 0.944606605 |
| 0.628135633 | 0.755171448 |
| 0.425316073 | 0.58672616 |
| 0.000765761 | 0.004075723 |
| 0.723678964 | NA |
| 0.102920762 | 0.214361215 |
| 0.427150293 | 0.588567314 |
| NA | NA |
| 0.13844459 | 0.266908968 |
| 0.194548876 | 0.340714617 |

| | |
|---|---|
| 0.088942033 | 0.192541907 |
| 0.577748861 | 0.716153442 |
| 1.74E-06 | 1.86E-05 |
| NA | NA |
| 0.020045206 | 0.059969331 |
| 0.611642181 | 0.742780616 |
| 0.10687401 | 0.220490135 |
| 0.641756959 | 0.764985934 |
| 0.184639059 | 0.327715849 |
| 0.276740028 | NA |
| 0.001471044 | 0.007054967 |
| 0.7132227 | NA |
| NA | NA |
| NA | NA |
| 0.0004974 | 0.002824893 |
| 1.21E-29 | 8.33E-28 |
| 0.941278788 | NA |
| 0.75956874 | NA |
| 0.399676379 | NA |
| 0.941278332 | NA |
| 0.811511034 | NA |
| 0.241657277 | 0.397919509 |
| 4.36E-13 | 1.13E-11 |
| 0.081594244 | 0.180326746 |
| 0.955792873 | NA |
| 3.68E-18 | 1.37E-16 |
| 0.000365861 | 0.002159775 |
| 0.073959032 | 0.167734433 |
| NA | NA |
| NA | NA |
| NA | NA |
| 0.940685982 | NA |
| NA | NA |
| NA | NA |
| NA | NA |
| 0.941278838 | NA |
| NA | NA |
| 0.941278332 | NA |
| NA | NA |
| 0.00254561 | 0.011212178 |
| 0.099742145 | 0.209186935 |
| 0.236495501 | 0.391859792 |
| 0.123602075 | 0.245553508 |
| NA | NA |
| 0.924171713 | 0.954592539 |
| NA | NA |
| 0.955792398 | NA |

| | | |
|---|---|---|
| | 0.42931079 | NA |
| | 0.123165491 | 0.244994933 |
| | 0.699397208 | NA |
| | 0.260026987 | 0.418571387 |
| NA | | NA |
| | 0.284443096 | NA |
| | 0.165075722 | 0.303622909 |
| | 0.282529005 | 0.442942654 |
| | 0.015799022 | 0.049733422 |
| | 0.892970784 | NA |
| | 0.323033313 | 0.48574382 |
| NA | | NA |
| NA | | NA |
| NA | | NA |
| NA | | NA |
| | 0.044972768 | 0.113949523 |
| | 0.002881146 | 0.012391948 |
| | 0.043716759 | 0.111494136 |
| | 0.532530836 | NA |
| | 0.043063246 | 0.110297147 |
| NA | | NA |
| | 0.825972908 | NA |
| NA | | NA |
| | 0.510309061 | 0.661855725 |
| | 0.105890402 | 0.218875761 |
| | 0.941278332 | NA |
| | 0.941278332 | NA |
| | 0.853596091 | NA |
| | 0.853595622 | NA |
| | 0.941275956 | NA |
| | 6.79E-05 | 0.000497055 |
| | 0.779703156 | NA |
| NA | | NA |
| | 0.332035115 | 0.49514351 |
| | 3.05E-05 | 0.000244552 |
| NA | | NA |
| | 0.959890376 | 0.975380411 |
| NA | | NA |
| | 0.165570237 | 0.304246335 |
| | 0.288022055 | 0.448658439 |
| | 0.698290703 | 0.806977488 |
| | 0.862183265 | NA |
| | 0.795452505 | 0.874520116 |
| | 0.008351527 | 0.029686834 |
| | 0.8393708 | 0.902910642 |
| | 0.270030872 | NA |
| | 0.798526191 | 0.87680585 |

| | |
|---|---|
| 0.898451625 | NA |
| 0.63070114 | 0.756917689 |
| 0.261125013 | NA |
| NA | NA |
| NA | NA |
| 7.81E-05 | 0.000563119 |
| 0.436421497 | 0.597464541 |
| 0.701901976 | 0.809706336 |
| NA | NA |
| 0.744702092 | 0.839594856 |
| 0.007515792 | 0.027247735 |
| 0.00021338 | 0.001357262 |
| 0.704328242 | 0.811325597 |
| 0.176124578 | 0.317608698 |
| NA | NA |
| 0.022705168 | 0.066342794 |
| NA | NA |
| 0.631076715 | NA |
| NA | NA |
| NA | NA |
| NA | NA |
| NA | NA |
| 0.734058779 | NA |
| 0.917711286 | 0.950457225 |
| 0.449440157 | 0.609529407 |
| 0.602288948 | NA |
| 5.44E-05 | 0.000408166 |
| 0.487111032 | 0.64208987 |
| 5.55E-06 | 5.35E-05 |
| 0.137397557 | 0.265547692 |
| NA | NA |
| NA | NA |
| NA | NA |
| NA | NA |
| NA | NA |
| NA | NA |
| NA | NA |
| NA | NA |
| NA | NA |
| NA | NA |
| NA | NA |
| NA | NA |
| 0.037940335 | 0.099884511 |
| 0.806855712 | 0.88170566 |
| 0.481003843 | 0.637487957 |
| 0.69165438 | 0.802187554 |
| 0.000564552 | 0.003146209 |

| | | |
|---|---|---|
| | 0.520748741 | NA |
| | 0.026083374 | 0.07415629 |
| | 0.598076238 | 0.732537983 |
| | 0.222327029 | 0.37474881 |
| | 0.70397249 | 0.811033553 |
| | 0.001564879 | 0.007439035 |
| | 0.000310548 | 0.00187701 |
| | 0.746088573 | 0.840278884 |
| | 0.971643449 | 0.98291714 |
| | 0.000345431 | 0.00205773 |
| | 0.004262585 | 0.017100631 |
| | 0.000648478 | 0.00353529 |
| | 0.003319296 | 0.013958638 |
| | 0.839319108 | NA |
| | 0.739991096 | NA |
| | 0.00091761 | 0.004740821 |
| | 0.644591548 | 0.767313612 |
| | 0.033397663 | NA |
| | 0.168858796 | 0.30842651 |
| | 0.08769611 | 0.19052052 |
| | 0.941278788 | NA |
| NA | | NA |
| NA | | NA |
| NA | | NA |
| NA | | NA |
| NA | | NA |
| NA | | NA |
| NA | | NA |
| | 5.41E-05 | 0.000406831 |
| | 0.244695703 | 0.40122948 |
| | 0.51128908 | 0.66276564 |
| | 0.341410008 | 0.504899283 |
| | 0.170558558 | 0.310376457 |
| | 0.002051219 | 0.009340564 |
| | 0.039061936 | 0.102193032 |
| | 0.358597751 | 0.522231013 |
| | 0.002202357 | 0.009916858 |
| NA | | NA |
| NA | | NA |
| | 0.955792873 | NA |
| | 0.114006352 | 0.231060899 |
| NA | | NA |
| | 0.002001293 | 0.009171974 |
| | 0.360407241 | 0.5240966 |
| | 0.888752522 | 0.93342843 |
| | 0.955792398 | NA |
| | 0.513784057 | 0.664877297 |

| | | | |
|---|---|---|---|
| NA | | NA | |
| NA | | NA | |
| NA | | NA | |
| | 0.955792398 | NA | |
| | 0.328485026 | NA | |
| NA | | NA | |
| | 0.291344723 | | 0.452558919 |
| | 3.68E-09 | | 6.10E-08 |
| | 0.658031568 | | 0.777255093 |
| NA | | NA | |
| | 0.09098143 | | 0.195869725 |
| NA | | NA | |
| | 0.043153881 | | 0.110468375 |
| | 0.89683367 | | 0.937779975 |
| | 0.987743959 | | 0.992246211 |
| | 0.316703589 | | 0.479162143 |
| | 0.456343938 | | 0.616068199 |
| NA | | NA | |
| NA | | NA | |
| | 0.741994575 | NA | |
| | 0.697628254 | NA | |
| NA | | NA | |
| NA | | NA | |
| | 0.799654241 | NA | |
| | 0.842722675 | NA | |
| | 0.866626138 | | 0.920374591 |
| | 0.524437077 | | 0.67400925 |
| | 0.009028711 | | 0.031670714 |
| | 0.876015401 | | 0.925856434 |
| | 0.858138287 | | 0.915075826 |
| | 0.63296532 | | 0.758793279 |
| | 0.933804048 | | 0.960684818 |
| | 0.308070801 | | 0.469578047 |
| | 0.083630958 | | 0.183817714 |
| | 0.000195424 | | 0.001255461 |
| | 0.640874253 | NA | |
| | 0.128503364 | | 0.252759034 |
| | 0.743467922 | NA | |
| | 0.031063776 | | 0.085260609 |
| | 0.049080829 | | 0.121905655 |
| | 9.58E-07 | | 1.07E-05 |
| | 0.21857877 | | 0.369897752 |
| NA | | NA | |
| NA | | NA | |
| | 0.839319108 | NA | |
| NA | | NA | |
| | 0.914564281 | NA | |

| | |
|---|---|
| NA | NA |
| NA | NA |
| 0.941278332 | NA |
| 0.575597695 | 0.714417079 |
| NA | NA |
| 0.024154842 | 0.069807375 |
| 0.173886985 | 0.314622426 |
| NA | NA |
| 0.425256555 | NA |
| 0.000835279 | 0.004383971 |
| 0.570687826 | 0.7112545 |
| 0.001902986 | 0.008779777 |
| 0.927397734 | NA |
| NA | NA |
| NA | NA |
| 2.72E-08 | 3.98E-07 |
| 0.839319574 | NA |
| 0.17318386 | 0.313636327 |
| 0.696058876 | NA |
| 0.00070249 | 0.003788098 |
| 0.311328313 | 0.473412747 |
| 9.06E-05 | 0.000643297 |
| 0.012704173 | 0.041754313 |
| 0.237064643 | 0.392480571 |
| 0.094668747 | 0.201367786 |
| 1.33E-36 | 1.21E-34 |
| 1.43E-05 | 0.000124767 |
| 0.745670336 | NA |
| 0.856207614 | 0.913590067 |
| 0.048422124 | 0.120647764 |
| 0.079973353 | 0.177733496 |
| 0.340387555 | 0.504006507 |
| 0.001624732 | 0.007686715 |
| 0.458148773 | 0.617733931 |
| 0.751758557 | 0.844145736 |
| 3.23E-06 | 3.27E-05 |
| 0.119398214 | 0.239399907 |
| 0.002526371 | 0.011137744 |
| 0.000771787 | 0.00410156 |
| 0.074079341 | 0.167939025 |
| 0.047114067 | 0.118117656 |
| NA | NA |
| 0.086949877 | 0.18931403 |
| 0.955791922 | NA |
| 0.632489899 | 0.758414336 |
| 0.170524494 | 0.310355101 |
| 0.065102563 | 0.15254688 |

| | | |
|---|---|---|
| | 0.977441967 | 0.986100695 |
| | 0.129434039 | 0.253771747 |
| NA | | NA |
| | 0.002756957 | 0.011963734 |
| | 0.955792873 | NA |
| | 0.38014309 | 0.542889364 |
| | 0.839319108 | NA |
| | 0.441928715 | 0.602450589 |
| | 0.440208456 | 0.600913741 |
| | 0.000694329 | 0.003752591 |
| | 0.132886176 | 0.258834432 |
| | 0.140411959 | 0.269762325 |
| | 0.949974795 | 0.969401932 |
| | 0.762636936 | NA |
| | 0.213117066 | 0.363810687 |
| | 0.005860697 | 0.022223405 |
| | 0.625806505 | 0.753436846 |
| NA | | NA |
| | 0.84636739 | 0.907402217 |
| | 3.00E-06 | 3.05E-05 |
| | 0.6450162 | 0.767595399 |
| | 7.00E-05 | 0.000510714 |
| | 0.363090192 | 0.526804426 |
| | 0.746307949 | 0.840457525 |
| | 0.003503326 | 0.014572908 |
| | 0.022496402 | NA |
| | 0.022398018 | 0.065590259 |
| | 0.034385049 | 0.092332234 |
| | 0.250883374 | 0.408484061 |
| | 0.009492567 | 0.032989061 |
| NA | | NA |
| | 0.113164249 | 0.229941562 |
| | 3.48E-06 | 3.49E-05 |
| | 0.268769431 | 0.428597748 |
| | 0.347838434 | 0.511759822 |
| | 0.004330677 | 0.017324166 |
| | 0.005049382 | 0.019686387 |
| | 0.028036392 | 0.078647285 |
| NA | | NA |
| | 0.31850285 | 0.48096719 |
| | 0.955791922 | NA |
| | 3.15E-08 | 4.57E-07 |
| | 0.593100719 | NA |
| | 0.941279303 | NA |
| | 0.955791922 | NA |
| NA | | NA |
| NA | | NA |

| | | |
|---|---|---|
| | 0.697358527 | NA |
| | 0.155834254 | 0.291166255 |
| NA | | NA |
| | 0.359195003 | NA |
| NA | | NA |
| NA | | NA |
| | 0.174052239 | 0.314897491 |
| | 0.229500839 | 0.383238849 |
| | 0.632530573 | 0.758421641 |
| | 0.634471986 | NA |
| NA | | NA |
| | 0.839319108 | NA |
| | 0.103308276 | 0.214848718 |
| | 1.47E-05 | 0.000128098 |
| | 0.113743935 | NA |
| | 0.120086621 | 0.240456107 |
| | 0.868368412 | 0.921216246 |
| NA | | NA |
| | 0.124850159 | 0.247454699 |
| | 0.006463816 | 0.024039335 |
| | 0.804255216 | 0.880415888 |
| | 0.548664777 | NA |
| NA | | NA |
| | 0.949946287 | 0.969401932 |
| | 0.551999018 | 0.697308175 |
| | 0.956422879 | 0.973311273 |
| | 0.486147188 | NA |
| | 0.177923455 | 0.319413243 |
| | 0.720442805 | NA |
| | 0.837025118 | 0.901225435 |
| | 0.937861084 | 0.963153577 |
| | 0.941277857 | NA |
| NA | | NA |
| | 1.42E-09 | 2.46E-08 |
| | 0.579073769 | 0.71712351 |
| | 0.002611102 | 0.011443462 |
| | 0.655548261 | NA |
| | 0.107914655 | 0.221949584 |
| | 0.66059706 | 0.779473093 |
| | 0.515612568 | 0.666627395 |
| | 0.064336814 | 0.151124528 |
| NA | | NA |
| | 0.666691291 | NA |
| | 0.72169607 | NA |
| NA | | NA |
| | 0.00051149 | 0.002893178 |
| NA | | NA |

| | |
|---|---|
| 0.014111576 | 0.045432669 |
| 0.105346176 | 0.218058064 |
| 0.890024473 | 0.933769151 |
| 0.178540499 | 0.320146222 |
| 0.562613757 | 0.705194168 |
| 0.286089414 | 0.446729662 |
| 0.955791922 | NA |
| 0.066011067 | 0.154159677 |
| 0.464103174 | 0.622666014 |
| 0.41636328 | 0.578104111 |
| 0.852519494 | 0.911411651 |
| NA | NA |
| NA | NA |
| 0.388890313 | 0.551561603 |
| NA | NA |
| NA | NA |
| 0.009013603 | 0.031629336 |
| NA | NA |
| NA | NA |
| NA | NA |
| 1.75E-06 | 1.87E-05 |
| 0.940375891 | NA |
| 0.005439904 | 0.02086714 |
| 0.132037777 | 0.257624654 |
| 0.851951877 | NA |
| 0.839319109 | NA |
| 0.906874546 | 0.943625266 |
| 0.572949665 | 0.712654051 |
| NA | NA |
| 0.8882061 | 0.933255773 |
| 0.486805667 | 0.641900641 |
| 0.633654147 | 0.759236647 |
| 0.955791446 | NA |
| NA | NA |
| 0.941278284 | NA |
| 0.501720419 | NA |
| NA | NA |
| 0.839106964 | NA |
| 0.837198879 | 0.901225435 |
| 0.931958979 | 0.959371849 |
| 0.745812696 | 0.840243944 |
| NA | NA |
| NA | NA |
| NA | NA |
| 0.955792398 | NA |
| 0.941278332 | NA |
| 0.935677476 | NA |

| | | | |
|---|---|---|---|
| NA | | NA | |
| | 0.669963072 | | 0.786469769 |
| NA | | NA | |
| NA | | NA | |
| | 0.327324278 | | 0.49010728 |
| | 0.955792873 | NA | |
| | 0.021355508 | | 0.063049344 |
| NA | | NA | |
| | 0.884793625 | | 0.93106936 |
| | 0.734058779 | NA | |
| NA | | NA | |
| | 0.405639188 | | 0.567708822 |
| | 0.019367197 | | 0.058396 |
| NA | | NA | |
| | 0.257803405 | | 0.416016044 |
| | 0.156562406 | | 0.292159811 |
| | 0.206665898 | | 0.355629099 |
| | 0.019163666 | | 0.057938115 |
| | 0.996916857 | NA | |
| | 9.20E-08 | | 1.25E-06 |
| | 0.203957339 | | 0.3524461 |
| | 0.178174014 | | 0.319705834 |
| | 0.695951391 | | 0.805261534 |
| | 0.805571242 | | 0.881005638 |
| | 0.005213247 | | 0.020199529 |
| | 0.120897166 | NA | |
| | 4.03E-13 | | 1.05E-11 |
| | 0.815379411 | | 0.88708569 |
| NA | | NA | |
| | 9.17E-06 | | 8.39E-05 |
| NA | | NA | |
| | 1.22E-06 | | 1.34E-05 |
| | 1.44E-11 | | 3.20E-10 |
| | 0.941279303 | NA | |
| | 1.09E-11 | | 2.46E-10 |
| | 0.941277857 | NA | |
| | 0.941278788 | NA | |
| | 0.379293375 | | 0.542132974 |
| | 2.68E-09 | | 4.52E-08 |
| | 0.941279303 | NA | |
| | 3.21E-08 | | 4.66E-07 |
| | 2.39E-07 | | 3.00E-06 |
| | 0.907103498 | | 0.943657232 |
| | 0.490278258 | NA | |
| | 0.371675193 | NA | |
| | 0.046473543 | | 0.116866668 |
| | 0.681553592 | | 0.795134722 |

| | |
|---|---|
| 0.091200024 | 0.196090073 |
| NA | NA |
| 2.01E-85 | 5.63E-83 |
| 4.24E-06 | 4.20E-05 |
| 2.61E-66 | 5.40E-64 |
| 0.039990549 | 0.104175884 |
| 0.145490891 | 0.277049486 |
| 2.56E-76 | 6.22E-74 |
| NA | NA |
| 4.00E-11 | 8.48E-10 |
| NA | NA |
| NA | NA |
| 0.905655164 | NA |
| NA | NA |
| 0.195113918 | 0.341418618 |
| 0.000312696 | 0.001887508 |
| 0.001277803 | 0.006277455 |
| 0.459394601 | NA |
| 4.95E-06 | 4.83E-05 |
| 0.911291597 | NA |
| 0.303199599 | 0.464637964 |
| 0.243881764 | 0.400299586 |
| 0.873578682 | 0.924581786 |
| 0.006157817 | 0.023126007 |
| 0.312732059 | 0.474684561 |
| 0.000116116 | 0.00079832 |
| 0.126952301 | 0.25043251 |
| 0.336177639 | NA |
| 0.256880482 | 0.414976872 |
| 0.446837499 | NA |
| 0.003573479 | 0.014802634 |
| 0.008298646 | 0.02956509 |
| NA | NA |
| NA | NA |
| 9.48E-05 | 0.000669661 |
| NA | NA |
| 0.406958217 | NA |
| NA | NA |
| 0.35124831 | 0.515109595 |
| 0.2255983 | 0.378493744 |
| 0.258498892 | 0.416811967 |
| 0.955792398 | NA |
| NA | NA |
| 5.74E-06 | 5.51E-05 |
| 0.019958227 | 0.059746663 |
| 0.794361236 | NA |
| 0.934268043 | 0.960921951 |

| | |
|---|---|
| NA | NA |
| 0.057722487 | 0.138680794 |
| 0.036449197 | 0.09671837 |
| 0.542716111 | 0.689533942 |
| 0.955792398 | NA |
| 0.574001298 | 0.71353814 |
| 0.176523948 | 0.318151226 |
| 0.041684706 | 0.10764667 |
| 0.010945393 | 0.037074461 |
| 0.908462926 | 0.944452264 |
| 0.233296804 | 0.387882393 |
| 0.074868549 | 0.169192188 |
| 0.028539662 | 0.079795728 |
| 0.839318641 | NA |
| 0.928896475 | 0.957705705 |
| 0.06031187 | 0.143597391 |
| 0.915196604 | NA |
| NA | NA |
| NA | NA |
| NA | NA |
| NA | NA |
| 0.704491558 | NA |
| 0.119286 | 0.23925553 |
| 0.264615756 | 0.423591649 |
| 0.273391098 | 0.433292143 |
| NA | NA |
| 0.823387982 | NA |
| 0.189410287 | 0.333973899 |
| 0.003584636 | 0.014838521 |
| 2.57E-06 | 2.66E-05 |
| 0.873237998 | NA |
| 0.000143904 | 0.000963219 |
| NA | NA |
| 0.199752172 | 0.347183534 |
| NA | NA |
| 0.725402998 | NA |
| 9.25E-06 | 8.45E-05 |
| 0.001079889 | 0.005454905 |
| 0.678981292 | NA |
| 0.248174296 | 0.405397337 |
| 2.35E-39 | 2.37E-37 |
| 0.338137036 | 0.50161033 |
| 0.464435914 | 0.622942626 |
| 0.402469887 | 0.564667742 |
| 0.088302976 | 0.191541733 |
| 0.016248016 | 0.050826031 |
| 0.264044513 | 0.423096 |

| | | |
|---|---|---|
| | 0.574963913 | 0.714074899 |
| | 0.562489473 | 0.705152787 |
| | 0.178628416 | 0.320255616 |
| NA | | NA |
| | 0.955791922 | NA |
| | 0.941278332 | NA |
| | 0.921305466 | NA |
| | 0.941277857 | NA |
| | 0.941277857 | NA |
| NA | | NA |
| | 0.000120873 | 0.000828156 |
| | 0.955792398 | NA |
| | 0.107474481 | 0.221201858 |
| | 0.463072223 | 0.621906283 |
| | 0.937794426 | NA |
| | 0.14783934 | 0.280405674 |
| | 0.398439404 | NA |
| | 0.418464573 | 0.580098646 |
| | 0.033026479 | NA |
| | 0.136737018 | 0.264516784 |
| | 0.438430911 | NA |
| | 0.311891534 | NA |
| | 0.755202218 | 0.846654571 |
| | 0.748833093 | 0.842331062 |
| | 0.87254855 | NA |
| | 0.926158431 | 0.955568264 |
| | 0.177653782 | 0.319205926 |
| | 0.038101542 | 0.100242389 |
| | 0.839320041 | NA |
| | 0.104370687 | 0.216471862 |
| | 0.274126508 | NA |
| NA | | NA |
| | 0.0003234 | 0.001945798 |
| | 0.941277857 | NA |
| | 0.941277857 | NA |
| | 0.853595154 | NA |
| | 0.013346494 | 0.043481424 |
| NA | | NA |
| | 0.955792873 | NA |
| NA | | NA |
| NA | | NA |
| NA | | NA |
| NA | | NA |
| | 0.238919157 | 0.394776381 |
| | 0.098129839 | 0.206789413 |
| | 0.839318641 | NA |
| | 0.941278332 | NA |

| | | |
|---|---|---|
| | 0.875897797 | NA |
| | 0.415206554 | 0.577060983 |
| | 0.108096607 | 0.222137953 |
| | 0.73093217 | 0.830396998 |
| | 0.613970292 | 0.74423105 |
| | 0.308099737 | NA |
| | 0.625432393 | 0.75325314 |
| | 0.286808668 | 0.447353861 |
| | 0.591030347 | 0.726862887 |
| | 0.804681543 | 0.880599085 |
| | 0.673757418 | 0.789413075 |
| | 0.076415575 | 0.171783482 |
| | 0.839319574 | NA |
| | 0.001295483 | 0.006347281 |
| | 0.955792398 | NA |
| | 0.955792873 | NA |
| | 0.107149663 | 0.220850041 |
| NA | | NA |
| | 0.921487119 | NA |
| | 0.091510452 | 0.196546813 |
| | 0.044763459 | 0.113503765 |
| | 0.941277857 | NA |
| | 0.081811638 | 0.180671794 |
| NA | | NA |
| NA | | NA |
| | 0.923999972 | 0.954592539 |
| | 0.669296973 | 0.786030809 |
| | 0.45578918 | NA |
| NA | | NA |
| | 0.643608344 | 0.76634146 |
| | 0.942433339 | 0.965107777 |
| | 0.3473542 | NA |
| | 0.022567548 | 0.066021727 |
| | 0.053961476 | 0.131651398 |
| | 0.019070893 | 0.057729289 |
| | 0.941275956 | NA |
| | 0.00868126 | 0.030630203 |
| | 0.955792398 | NA |
| | 0.681427421 | 0.795026492 |
| | 0.13219849 | 0.257832551 |
| | 0.052629878 | 0.129130642 |
| | 0.85964713 | 0.916299154 |
| | 0.061580717 | 0.14599046 |
| | 0.052420263 | 0.128659235 |
| | 0.001957348 | 0.009002667 |
| NA | | NA |
| NA | | NA |

| | |
|---|---|
| 0.749589203 | NA |
| 0.000544913 | 0.003058967 |
| 1.00E-05 | 9.08E-05 |
| 0.940163126 | NA |
| 0.089730755 | 0.193755912 |
| 0.268531884 | 0.428276326 |
| 0.403422049 | NA |
| NA | NA |
| 0.254056419 | 0.412120963 |
| 0.343746273 | NA |
| NA | NA |
| 0.09229744 | 0.197666337 |
| NA | NA |
| 0.833137156 | NA |
| 0.331352223 | 0.494473079 |
| 0.327882359 | NA |
| NA | NA |
| NA | NA |
| NA | NA |
| NA | NA |
| NA | NA |
| NA | NA |
| NA | NA |
| NA | NA |
| NA | NA |
| NA | NA |
| NA | NA |
| 0.111437162 | 0.227201246 |
| 0.413755774 | 0.575952683 |
| 0.971243032 | 0.982637365 |
| NA | NA |
| 0.790233216 | NA |
| 0.000659425 | 0.003590037 |
| 0.031648733 | 0.086536833 |
| 0.884748968 | 0.93106353 |
| 0.002282229 | 0.010224284 |
| 1.70E-12 | 4.16E-11 |
| 0.181564778 | 0.323958385 |
| 5.51E-08 | 7.68E-07 |
| 0.399096271 | 0.561795122 |
| 0.043128952 | 0.110429789 |
| 4.96E-22 | 2.39E-20 |
| 0.473792954 | NA |
| 0.934047677 | NA |
| 0.328509933 | 0.491402302 |
| 0.153147319 | NA |
| 0.839319574 | NA |

| | | |
|---|---|---|
| | 0.244227161 | NA |
| | 0.36292948 | 0.526621714 |
| | 6.69E-10 | 1.21E-08 |
| NA | | NA |
| | 0.159179526 | 0.295537526 |
| | 0.813753049 | 0.886358628 |
| | 0.854869393 | 0.912653119 |
| | 0.038641444 | 0.101271046 |
| | 0.948780888 | 0.968760795 |
| | 0.806818975 | NA |
| | 0.500321246 | 0.65307792 |
| | 0.404141552 | 0.566178765 |
| | 0.198163323 | 0.3452799 |
| | 0.307205413 | 0.468844783 |
| | 0.437312651 | 0.598213547 |
| | 5.58E-06 | 5.38E-05 |
| | 0.091987923 | 0.197145376 |
| | 0.000981728 | 0.005026261 |
| | 0.773502714 | 0.860363176 |
| | 0.023996739 | 0.069426325 |
| | 0.043673213 | 0.111438144 |
| | 0.053960186 | 0.131651398 |
| | 0.928473254 | NA |
| | 0.00253454 | 0.011171691 |
| | 0.001433841 | 0.006905763 |
| | 0.048339677 | 0.120517544 |
| NA | | NA |
| | 0.011682019 | 0.039096657 |
| | 0.50544411 | NA |
| | 0.988947876 | 0.992997975 |
| | 0.157344598 | 0.293090946 |
| | 0.214229126 | 0.365159099 |
| | 0.625314499 | 0.753149261 |
| | 0.072258759 | 0.16493124 |
| | 1.91E-12 | 4.66E-11 |
| | 0.605825006 | NA |
| | 0.954699433 | 0.972513734 |
| NA | | NA |
| | 0.929914733 | NA |
| NA | | NA |
| | 0.853595154 | NA |
| | 0.623918681 | 0.752000833 |
| | 0.376619684 | 0.539647272 |
| | 2.84E-05 | 0.000230039 |
| | 0.401656586 | 0.563925563 |
| | 0.386556518 | 0.549433789 |
| NA | | NA |

| | | |
|---|---|---|
| NA | | NA |
| | 0.326811132 | 0.48963321 |
| NA | | NA |
| NA | | NA |
| NA | | NA |
| NA | | NA |
| | 0.839319108 | NA |
| NA | | NA |
| NA | | NA |
| | 0.921305466 | NA |
| NA | | NA |
| | 0.111799418 | 0.227751604 |
| | 0.543835411 | 0.690513733 |
| | 0.936856978 | NA |
| NA | | NA |
| NA | | NA |
| | 0.005339034 | 0.020583039 |
| | 0.798033642 | 0.876467182 |
| | 0.706920711 | NA |
| | 0.832162011 | NA |
| | 0.152570588 | 0.286677459 |
| | 0.888175782 | 0.933255773 |
| | 0.015849776 | 0.049842299 |
| | 0.482919309 | 0.63889975 |
| | 0.597155917 | 0.731822498 |
| | 0.616311401 | 0.746152947 |
| NA | | NA |
| | 0.00202372 | 0.009245412 |
| | 0.129569726 | 0.253954103 |
| | 0.941278332 | NA |
| | 0.432769997 | NA |
| | 0.59638709 | NA |
| NA | | NA |
| | 0.123107715 | NA |
| | 0.665311229 | NA |
| | 0.647084319 | 0.769172041 |
| NA | | NA |
| | 0.08057761 | 0.178792604 |
| | 0.195032749 | 0.341335992 |
| | 0.839319108 | NA |
| | 0.839319108 | NA |
| | 0.051728697 | 0.127352756 |
| | 0.076396103 | 0.171772157 |
| | 0.038863766 | 0.101741682 |
| | 0.889617413 | 0.933545263 |
| NA | | NA |
| | 0.382343913 | 0.544910473 |

| | |
|---|---|
| 0.6664498 | 0.783775786 |
| NA | NA |
| NA | NA |
| NA | NA |
| NA | NA |
| NA | NA |
| NA | NA |
| 0.926627763 | NA |
| NA | NA |
| 0.839318641 | NA |
| 0.973203611 | 0.983784606 |
| NA | NA |
| NA | NA |
| NA | NA |
| NA | NA |
| NA | NA |
| NA | NA |
| NA | NA |
| NA | NA |
| 0.000133949 | 0.000906012 |
| 0.791947557 | NA |
| NA | NA |
| 0.955792398 | NA |
| NA | NA |
| NA | NA |
| NA | NA |
| 0.929914733 | NA |
| 0.553242705 | 0.698212241 |
| 0.005947532 | 0.022486074 |
| NA | NA |
| 0.33353112 | 0.496800273 |
| NA | NA |
| 0.748393366 | NA |
| 0.941278332 | NA |
| 0.007527003 | 0.027277363 |
| NA | NA |
| 0.347769271 | 0.511721326 |
| 0.47730569 | 0.63415183 |
| 0.627788416 | 0.755020963 |
| 0.938986848 | 0.963601145 |
| 0.257076982 | 0.415237943 |
| 0.293949893 | 0.455166979 |
| NA | NA |
| NA | NA |
| 0.264354206 | 0.423343794 |
| 0.586763803 | 0.723374155 |
| 0.272155161 | 0.432086255 |

| | | |
|---|---|---|
| | 1.27E-10 | 2.52E-09 |
| | 0.643986189 | NA |
| NA | | NA |
| | 0.017579761 | 0.054088279 |
| | 0.000135222 | 0.000913586 |
| | 4.14E-07 | 4.98E-06 |
| | 0.941279268 | NA |
| | 0.951557634 | 0.970393093 |
| | 0.888821685 | 0.93342843 |
| | 0.000366678 | 0.002163525 |
| | 0.00816257 | 0.02915011 |
| | 0.77058979 | NA |
| | 0.227628114 | NA |
| | 0.467936518 | 0.626177827 |
| | 0.085854323 | NA |
| | 0.880266462 | NA |
| | 0.204679066 | 0.353278652 |
| | 0.311775791 | 0.473765214 |
| | 3.18E-05 | 0.000253635 |
| | 0.292573728 | 0.453611248 |
| NA | | NA |
| | 0.639104957 | NA |
| NA | | NA |
| | 0.955792873 | NA |
| | 0.941275956 | NA |
| | 0.839319108 | NA |
| NA | | NA |
| | 0.571497609 | NA |
| | 0.839318641 | NA |
| NA | | NA |
| | 0.617155265 | NA |
| | 0.941278788 | NA |
| | 0.322550852 | NA |
| NA | | NA |
| | 0.873630445 | NA |
| NA | | NA |
| | 0.01508086 | 0.047911725 |
| | 0.436003781 | NA |
| | 0.388531194 | 0.551363288 |
| NA | | NA |
| NA | | NA |
| | 0.839319108 | NA |
| NA | | NA |
| | 0.941278838 | NA |
| NA | | NA |
| NA | | NA |
| | 0.955792398 | NA |

| | | |
|---|---|---|
| NA | NA | |
| NA | NA | |
| NA | NA | |
| NA | NA | |
| 0.955792873 | NA | |
| NA | NA | |
| 0.665906302 | NA | |
| 0.050120311 | | 0.124034481 |
| NA | NA | |
| NA | NA | |
| NA | NA | |
| 0.091232832 | | 0.196109805 |
| 0.805637616 | | 0.88102772 |
| 0.637814711 | | 0.761920489 |
| 0.583879395 | | 0.721086357 |
| 0.955792873 | NA | |
| 0.00665632 | | 0.024632109 |
| NA | NA | |
| 0.01365493 | | 0.044231715 |
| 0.923505151 | NA | |
| NA | NA | |
| NA | NA | |
| NA | NA | |
| 0.18919599 | | 0.333820827 |
| NA | NA | |
| NA | NA | |
| NA | NA | |
| NA | NA | |
| NA | NA | |
| NA | NA | |
| NA | NA | |
| 0.208289372 | | 0.357621267 |
| NA | NA | |
| 0.21906846 | | 0.370594703 |
| 0.393773799 | | 0.556270637 |
| NA | NA | |
| 0.663521599 | NA | |
| 0.009957749 | | 0.034280352 |
| NA | NA | |
| NA | NA | |
| NA | NA | |
| NA | NA | |
| NA | NA | |
| NA | NA | |
| NA | NA | |
| NA | NA | |
| NA | NA | |

| | |
|---|---|
| NA | NA |
| NA | NA |
| 0.561982861 | NA |
| NA | NA |
| NA | NA |
| NA | NA |
| NA | NA |
| NA | NA |
| 0.955792398 | NA |
| NA | NA |
| 0.941278332 | NA |
| NA | NA |
| 0.795902187 | 0.874871471 |
| 0.000907371 | 0.004699142 |
| 0.950512078 | NA |
| 5.31E-05 | 0.000400539 |
| NA | NA |
| 0.604097715 | 0.737106362 |
| NA | NA |
| 1.05E-05 | 9.47E-05 |
| NA | NA |
| NA | NA |
| NA | NA |
| NA | NA |
| 0.9305767 | 0.958607575 |
| NA | NA |
| 0.189931383 | NA |
| NA | NA |
| NA | NA |
| NA | NA |
| 0.297871584 | 0.4592986 |
| 0.79325905 | 0.872939996 |
| 5.00E-09 | 8.16E-08 |
| NA | NA |
| 0.32960172 | 0.49258328 |
| 0.543463336 | 0.690225406 |
| NA | NA |
| 0.649480029 | 0.770749546 |
| 0.741178832 | NA |
| 0.359590576 | 0.523164724 |
| 0.641001468 | 0.76453783 |
| 0.034216646 | 0.091968843 |
| 0.514441657 | NA |
| 0.240864494 | 0.396943619 |
| NA | NA |
| NA | NA |
| 0.839319108 | NA |

| | | |
|---|---|---|
| | 0.91768099 | NA |
| NA | | NA |
| | 3.07E-07 | 3.77E-06 |
| | 0.960266783 | 0.975605716 |
| NA | | NA |
| | 0.815443614 | 0.887114991 |
| | 0.008837471 | 0.031098388 |
| | 0.002550958 | 0.011233654 |
| | 0.140108522 | 0.269266293 |
| | 0.015982042 | 0.050186729 |
| | 0.391854286 | 0.554183941 |
| | 0.839319108 | NA |
| | 0.922194178 | NA |
| | 0.252955316 | 0.410782727 |
| | 0.297429299 | NA |
| NA | | NA |
| | 0.839319574 | NA |
| | 0.839319108 | NA |
| NA | | NA |
| | 0.698899688 | NA |
| | 0.839318641 | NA |
| NA | | NA |
| | 0.019869709 | 0.059520654 |
| NA | | NA |
| | 0.409674554 | 0.571575248 |
| | 0.021434716 | 0.063230421 |
| | 0.627041031 | 0.754362457 |
| | 0.941279303 | NA |
| | 0.001009599 | 0.005152333 |
| NA | | NA |
| | 0.941277857 | NA |
| | 0.9459869 | 0.967272298 |
| NA | | NA |
| | 0.001505262 | 0.007195871 |
| NA | | NA |
| NA | | NA |
| NA | | NA |
| NA | | NA |
| NA | | NA |
| NA | | NA |
| NA | | NA |
| NA | | NA |
| NA | | NA |
| NA | | NA |
| | 0.800391721 | NA |
| | 0.941278788 | NA |
| | 0.941276431 | NA |

| | |
|---|---|
| 0.839320041 | NA |
| NA | NA |
| 0.097913483 | 0.206370027 |
| 0.952262799 | 0.970945817 |
| 0.460261041 | 0.619248957 |
| 0.0147505 | 0.047025368 |
| 0.003207886 | 0.013566827 |
| 0.205152309 | 0.353783609 |
| 0.024261623 | 0.070047932 |
| 0.028502573 | 0.079738862 |
| NA | NA |
| 0.007333613 | 0.026698478 |
| 0.000818717 | 0.004312262 |
| 0.8141273 | 0.886443528 |
| NA | NA |
| 0.951059547 | 0.970148286 |
| 0.325588043 | 0.488470883 |
| 0.284819993 | 0.445417002 |
| 0.941277857 | NA |
| 7.02E-06 | 6.58E-05 |
| NA | NA |
| 0.009696917 | 0.033601108 |
| NA | NA |
| 0.015843339 | 0.049841812 |
| 0.340420003 | 0.504023197 |
| 0.014642096 | 0.046780022 |
| 0.633517302 | 0.759182999 |
| 0.108421969 | 0.222624198 |
| 0.902118556 | 0.940856448 |
| NA | NA |
| NA | NA |
| 0.182171007 | 0.324553466 |
| 0.967232536 | 0.980329961 |
| 0.074063141 | 0.167922546 |
| 0.541879523 | 0.688949116 |
| 0.839319574 | NA |
| NA | NA |
| 0.230325562 | 0.384177226 |
| 0.174258551 | 0.315055225 |
| 0.339811869 | 0.503498637 |
| 0.253482672 | 0.41138651 |
| 0.529342902 | 0.678445397 |
| 0.98910091 | 0.993064489 |
| 0.030921032 | 0.084947184 |
| 0.201852753 | 0.349860741 |
| 0.836827905 | 0.901121952 |
| 0.045323305 | 0.114672998 |

| | |
|---|---|
| 0.853289006 | 0.911742904 |
| 0.198942288 | 0.34626869 |
| 0.270208093 | 0.430289039 |
| 0.004731225 | 0.018672387 |
| 2.04E-05 | 0.000171109 |
| NA | NA |
| 0.01706277 | 0.052874373 |
| 0.253666181 | 0.411628201 |
| 0.044628094 | 0.113257052 |
| 0.273654096 | 0.433454102 |
| 0.093620898 | 0.19971004 |
| 0.00661619 | 0.024532423 |
| 4.06E-10 | 7.59E-09 |
| 0.424172168 | 0.585578803 |
| NA | NA |
| NA | NA |
| NA | NA |
| NA | NA |
| 0.162722984 | 0.300335955 |
| 9.13E-15 | 2.69E-13 |
| 0.839318641 | NA |
| 0.670847576 | NA |
| NA | NA |
| 6.05E-05 | 0.000449637 |
| 0.025402537 | 0.072584775 |
| 0.681267123 | 0.794897637 |
| 0.374671925 | 0.537784746 |
| 0.704866819 | 0.811749565 |
| 0.220073673 | 0.371740371 |
| 0.05804931 | 0.139283234 |
| 0.177758794 | 0.319321374 |
| 0.111408017 | 0.227187597 |
| 0.724091948 | 0.825581364 |
| NA | NA |
| 0.82580308 | 0.894113212 |
| 0.955792873 | NA |
| 0.892359954 | 0.935040806 |
| 0.261914828 | 0.420711077 |
| 0.25473129 | 0.412742327 |
| 0.176671155 | 0.318280868 |
| 0.061305239 | 0.145511299 |
| 0.650880393 | 0.771974087 |
| 0.839319108 | NA |
| 0.324777489 | 0.487719482 |
| 0.312283959 | 0.474291683 |
| 0.595951042 | 0.730837586 |
| 0.081255375 | 0.179809615 |

| | | |
|---|---|---|
| | 0.506424034 | 0.65810793 |
| | 0.941278332 | NA |
| | 0.955705192 | 0.972955303 |
| | 0.941278332 | NA |
| | 0.920159482 | NA |
| | 0.860633211 | NA |
| | 0.955792398 | NA |
| | 0.062088425 | NA |
| | 0.055162908 | 0.134017205 |
| | 0.941276431 | NA |
| | 0.941279268 | NA |
| | 0.009183738 | 0.03209858 |
| | 1.83E-13 | 4.92E-12 |
| | 3.55E-08 | 5.11E-07 |
| | 0.55807198 | 0.701696417 |
| | 0.948501871 | 0.968548661 |
| | 0.941277857 | NA |
| | 3.70E-08 | 5.30E-07 |
| | 0.006134782 | 0.023057014 |
| | 0.050046159 | 0.123876735 |
| | 0.549890433 | NA |
| | 8.80E-20 | 3.67E-18 |
| | 0.452437263 | 0.612426737 |
| | 0.922618691 | 0.953877599 |
| | 0.129599409 | 0.253974067 |
| | 0.213770104 | 0.364654304 |
| | 0.239146971 | 0.395052456 |
| | 0.24056854 | 0.396593184 |
| | 0.707597021 | 0.813437541 |
| | 0.108403842 | 0.222623085 |
| | 0.022589971 | 0.066071085 |
| | 0.499201163 | 0.652068164 |
| | 0.259478539 | 0.417925848 |
| | 7.56E-24 | 4.00E-22 |
| | 0.027574962 | 0.07770087 |
| | 0.218062402 | 0.369339023 |
| | 0.900406269 | 0.940018234 |
| | 6.08E-06 | 5.81E-05 |
| | 5.99E-11 | 1.24E-09 |
| NA | | NA |
| | 0.005923455 | 0.022411393 |
| | 0.900678806 | 0.940137567 |
| | 0.839318641 | NA |
| NA | | NA |
| | 0.010170368 | 0.034876112 |
| | 2.40E-06 | 2.50E-05 |
| | 0.804092407 | 0.880359129 |

| | | | |
|---|---|---|---|
| | 0.110443347 | | 0.225820166 |
| | 0.839319574 | NA | |
| | 0.002711214 | | 0.011808294 |
| | 0.010595965 | | 0.036080851 |
| | 0.357654478 | | 0.521229189 |
| | 3.86E-10 | | 7.23E-09 |
| | 0.252195454 | NA | |
| NA | | NA | |
| | 0.839319108 | NA | |
| NA | | NA | |
| | 0.690791855 | NA | |
| | 0.881721874 | | 0.929274887 |
| | 1.66E-27 | | 1.05E-25 |
| | 0.001926569 | | 0.008873103 |
| | 0.020391595 | | 0.060803013 |
| | 0.955792873 | NA | |
| | 0.547879723 | | 0.693835231 |
| | 0.007315574 | | 0.026649109 |
| | 0.562428469 | | 0.705131547 |
| | 0.458650827 | | 0.618146885 |
| | 0.745156098 | | 0.839941534 |
| | 0.572741986 | | 0.712520944 |
| | 0.542634801 | NA | |
| | 0.748393366 | NA | |
| | 0.048413127 | | 0.120647764 |
| | 0.12147167 | | 0.242576449 |
| | 0.075530465 | | 0.17045379 |
| | 0.171658796 | | 0.311817556 |
| | 0.441724406 | | 0.602275552 |
| | 1.77E-11 | | 3.90E-10 |
| | 0.385583864 | | 0.54841154 |
| | 0.07967772 | | 0.177291679 |
| | 2.69E-06 | | 2.77E-05 |
| | 0.000157475 | | 0.001044072 |
| | 0.323887256 | | 0.486567474 |
| | 0.784759155 | NA | |
| | 0.941278332 | NA | |
| | 0.500353267 | NA | |
| | 0.497480216 | NA | |
| | 0.023781637 | | 0.068904507 |
| | 0.209586058 | | 0.359199838 |
| NA | | NA | |
| NA | | NA | |
| | 0.011014116 | | 0.037248934 |
| | 0.838284155 | | 0.901986323 |
| | 0.305431383 | | 0.466954365 |
| | 0.511714315 | | 0.663169515 |

| | |
|---|---|
| NA | NA |
| 8.44E-18 | 3.11E-16 |
| 4.08E-05 | 0.000316658 |
| 0.941279895 | NA |
| 0.149849604 | 0.283028113 |
| 0.82922232 | NA |
| 0.941277857 | NA |
| 0.481037291 | 0.637487957 |
| 0.899148902 | NA |
| 0.065628564 | 0.153401854 |
| 0.69942835 | NA |
| 0.941275956 | NA |
| 0.077507424 | 0.17361488 |
| 0.651041961 | 0.772064209 |
| 0.684155959 | NA |
| 0.955791922 | NA |
| 0.273611993 | 0.433416256 |
| 3.30E-09 | 5.50E-08 |
| 0.000319967 | 0.001926598 |
| 0.363745253 | 0.527404028 |
| 0.839319108 | NA |
| 0.897733123 | NA |
| 0.000855648 | 0.00447214 |
| 0.263414038 | 0.422378128 |
| NA | NA |
| 0.310017324 | 0.471774972 |
| 0.058472658 | 0.140101288 |
| 0.764761771 | 0.853946599 |
| 0.363520621 | 0.527174668 |
| 0.64867813 | 0.770297188 |
| 0.016637145 | 0.051825328 |
| 0.619999167 | NA |
| 0.881556902 | NA |
| 0.337632113 | NA |
| 0.955792398 | NA |
| 0.000375528 | 0.002209722 |
| 0.483120549 | 0.639094942 |
| 0.123861067 | 0.245985909 |
| 5.93E-06 | 5.68E-05 |
| 0.010299841 | 0.035243935 |
| 0.955792398 | NA |
| 0.000505868 | 0.002865607 |
| 0.887311837 | 0.932811993 |
| 0.089518017 | 0.19340182 |
| 0.702895817 | 0.810420763 |
| 0.592200811 | 0.727775904 |
| 0.835025174 | NA |

| | | |
|---|---|---|
| | 4.35E-07 | 5.20E-06 |
| | 0.172258717 | 0.312555419 |
| | 0.955792873 | NA |
| | 0.907578776 | 0.943904135 |
| | 0.180186215 | 0.322157527 |
| | 0.71551127 | 0.819176213 |
| | 0.940074473 | NA |
| | 0.33532103 | 0.498520372 |
| | 0.001116357 | 0.005611725 |
| | 0.015470914 | 0.048883406 |
| | 0.918775095 | 0.951267337 |
| | 0.955792398 | NA |
| | 0.123541401 | 0.245454505 |
| | 0.007461609 | 0.027089876 |
| | 0.363832889 | 0.527434705 |
| | 2.84E-06 | 2.90E-05 |
| NA | | NA |
| | 0.255865006 | 0.413898251 |
| | 0.005865615 | 0.022231427 |
| | 0.086113828 | 0.187861408 |
| | 4.77E-05 | 0.000364005 |
| | 3.28E-07 | 4.00E-06 |
| | 0.012472006 | 0.041156294 |
| | 0.027784578 | 0.078152746 |
| | 0.975515323 | 0.985178051 |
| | 0.388179437 | 0.551049499 |
| | 0.633944391 | 0.759431497 |
| | 0.489108219 | 0.643583465 |
| | 0.941278207 | NA |
| | 7.49E-08 | 1.03E-06 |
| | 8.92E-08 | 1.21E-06 |
| | 0.801195789 | 0.878492596 |
| | 0.004181403 | 0.016837379 |
| NA | | NA |
| | 0.532581653 | NA |
| | 0.005249399 | 0.020309848 |
| | 3.92E-06 | 3.90E-05 |
| | 0.204812385 | 0.353410496 |
| | 0.000877169 | 0.004563578 |
| | 0.941277857 | NA |
| NA | | NA |
| | 0.427049586 | NA |
| | 0.470381419 | 0.628186331 |
| | 0.148002153 | 0.280674627 |
| | 0.932478619 | 0.95969456 |
| | 0.125493054 | 0.248439297 |
| | 0.018224799 | 0.055706519 |

| | |
|---|---|
| 0.000615835 | 0.003386024 |
| 0.941277857 | NA |
| 0.635102114 | NA |
| 0.94127925 | NA |
| 0.108032746 | 0.222093231 |
| 0.726674946 | NA |
| 0.097759126 | 0.206154222 |
| 0.00029081 | 0.001774838 |
| 0.006157997 | 0.023126007 |
| 0.06146731 | 0.145794212 |
| 0.941278332 | NA |
| NA | NA |
| NA | NA |
| 0.361892595 | 0.525582253 |
| NA | NA |
| 0.681283581 | 0.794897637 |
| NA | NA |
| 0.941278332 | NA |
| 0.533449556 | NA |
| NA | NA |
| NA | NA |
| 0.359864322 | NA |
| NA | NA |
| 0.292155841 | NA |
| NA | NA |
| 0.00230324 | 0.010312589 |
| 6.63E-06 | 6.28E-05 |
| NA | NA |
| 0.000113269 | 0.000781185 |
| 0.000298121 | 0.001813874 |
| NA | NA |
| 0.001347902 | 0.006562236 |
| 9.41E-06 | 8.58E-05 |
| 3.13E-06 | 3.18E-05 |
| 0.302160795 | NA |
| 0.000312983 | 0.001888368 |
| NA | NA |
| 2.19E-05 | 0.000182134 |
| NA | NA |
| 1.02E-95 | 3.31E-93 |
| 3.44E-29 | 2.33E-27 |
| NA | NA |
| NA | NA |
| NA | NA |
| 0.94127873 | NA |
| 0.817844966 | NA |
| 1.49E-05 | 0.000129433 |

| | | |
|---|---|---|
| | 0.761350405 | NA |
| | 0.131555572 | 0.256978723 |
| | 0.026968996 | 0.076345867 |
| | 0.955792398 | NA |
| NA | | NA |
| | 0.341871868 | 0.505261359 |
| | 0.372580561 | 0.535808261 |
| NA | | NA |
| | 0.839319108 | NA |
| NA | | NA |
| | 0.703547947 | 0.810844101 |
| | 2.19E-10 | 4.23E-09 |
| | 0.482409506 | 0.638554587 |
| | 0.976294661 | 0.985480846 |
| | 0.839318641 | NA |
| NA | | NA |
| | 0.145821265 | 0.277511613 |
| | 0.356948324 | 0.520593382 |
| | 0.955792398 | NA |
| NA | | NA |
| | 0.140168504 | 0.269338074 |
| | 0.126465816 | 0.2497212 |
| | 0.853595622 | NA |
| NA | | NA |
| | 5.69E-06 | 5.47E-05 |
| | 0.139413744 | 0.268234252 |
| | 0.656767342 | NA |
| | 0.955791922 | NA |
| NA | | NA |
| NA | | NA |
| | 0.709251061 | NA |
| NA | | NA |
| | 0.761638124 | NA |
| | 0.191112594 | 0.33618695 |
| | 0.414046347 | 0.576109104 |
| | 0.890314379 | 0.934029569 |
| NA | | NA |
| NA | | NA |
| | 0.916478513 | 0.949674427 |
| | 8.58E-06 | 7.89E-05 |
| | 0.543789309 | 0.690492032 |
| | 0.091405026 | 0.19639595 |
| NA | | NA |
| | 0.500613143 | NA |
| NA | | NA |
| NA | | NA |
| NA | | NA |

| | | | |
|---|---|---|---|
| | 0.586844692 | NA | |
| NA | | NA | |
| NA | | NA | |
| | 0.839319108 | NA | |
| NA | | NA | |
| NA | | NA | |
| | 0.612543954 | | 0.743448635 |
| | 0.204203307 | | 0.352666312 |
| | 0.052062998 | | 0.128030115 |
| | 0.206152282 | | 0.35492489 |
| | 0.000219611 | | 0.001390578 |
| | 0.440012034 | | 0.600846627 |
| | 0.802738672 | | 0.879402857 |
| | 0.144104043 | | 0.275057292 |
| NA | | NA | |
| | 0.635191309 | | 0.760094348 |
| | 0.082667445 | | 0.182044258 |
| | 0.755880571 | NA | |
| | 4.64E-28 | | 3.00E-26 |
| | 0.444770669 | NA | |
| NA | | NA | |
| NA | | NA | |
| | 0.008303793 | | 0.029570149 |
| NA | | NA | |
| NA | | NA | |
| NA | | NA | |
| NA | | NA | |
| NA | | NA | |
| NA | | NA | |
| NA | | NA | |
| | 0.888175583 | NA | |
| | 0.000813126 | | 0.004287556 |
| | 0.955792873 | NA | |
| | 0.0002268 | | 0.001428874 |
| | 0.009326521 | | 0.032502174 |
| | 0.295687421 | | 0.457158297 |
| | 0.514353663 | NA | |
| | 0.940729964 | NA | |
| | 0.839319574 | NA | |
| NA | | NA | |
| | 0.833801616 | NA | |
| | 0.839319108 | NA | |
| NA | | NA | |
| NA | | NA | |
| NA | | NA | |
| NA | | NA | |
| | 0.955792398 | NA | |

| | |
|---|---|
| NA | NA |
| NA | NA |
| NA | NA |
| 0.675678903 | 0.790917147 |
| NA | NA |
| NA | NA |
| NA | NA |
| 0.839318641 | NA |
| 0.766500887 | NA |
| 0.742335473 | NA |
| 0.853595154 | NA |
| 0.839319108 | NA |
| NA | NA |
| NA | NA |
| 1.99E-06 | 2.10E-05 |
| NA | NA |
| 0.667446504 | NA |
| 0.941276431 | NA |
| 0.764679347 | 0.853900906 |
| 0.000831561 | 0.004367348 |
| 0.047630063 | NA |
| 0.111309307 | 0.227046619 |
| 5.74E-14 | 1.60E-12 |
| 0.34242253 | NA |
| 0.941276431 | NA |
| 0.066634601 | NA |
| NA | NA |
| 0.001350602 | 0.006571808 |
| 0.094892555 | 0.201753693 |
| 2.45E-07 | 3.07E-06 |
| 0.524534879 | NA |
| 6.75E-06 | 6.38E-05 |
| 0.941277857 | NA |
| NA | NA |
| NA | NA |
| 0.941275956 | NA |
| NA | NA |
| 0.839319108 | NA |
| 0.053926496 | NA |
| NA | NA |
| 5.62E-07 | 6.56E-06 |
| 0.053940988 | 0.131628398 |
| 2.70E-06 | 2.78E-05 |
| 0.014974295 | 0.047643117 |
| 0.581847675 | 0.71952505 |
| 0.000455257 | 0.002615502 |
| 4.05E-08 | 5.76E-07 |

| | |
|---|---|
| 0.953634578 | 0.971969778 |
| NA | NA |
| 0.783102737 | NA |
| 0.941278332 | NA |
| 3.91E-14 | 1.11E-12 |
| 0.479223762 | 0.635954727 |
| 0.565098911 | 0.706989976 |
| 3.48E-13 | 9.11E-12 |
| 0.859346214 | NA |
| 0.839319574 | NA |
| NA | NA |
| 0.955792873 | NA |
| 0.831370221 | 0.897753108 |
| 0.03083138 | 0.08477852 |
| 0.925579587 | NA |
| 0.011880138 | 0.039587155 |
| 0.03645552 | 0.096724366 |
| 0.054389517 | 0.132451329 |
| 0.261839165 | 0.420646315 |
| 0.222057497 | NA |
| 2.85E-08 | 4.16E-07 |
| 3.82E-20 | 1.65E-18 |
| 1.21E-07 | 1.61E-06 |
| 0.939641213 | 0.963757575 |
| 0.751955238 | NA |
| 0.277202993 | NA |
| 0.006069206 | 0.022857475 |
| NA | NA |
| 0.953751371 | 0.971970805 |
| 4.55E-07 | 5.41E-06 |
| 0.570215281 | 0.710959587 |
| 0.379604511 | 0.542381878 |
| 0.292021957 | 0.453257687 |
| 0.004280409 | 0.01715768 |
| 0.19852485 | 0.345753261 |
| 0.038328477 | 0.100739224 |
| 0.000576743 | 0.003204395 |
| 0.180865615 | 0.322928766 |
| 0.03181436 | 0.08687992 |
| 0.081263946 | 0.179809615 |
| 0.745673167 | NA |
| 0.761580242 | 0.851817685 |
| 0.79583063 | 0.874871471 |
| 0.36515923 | 0.528456253 |
| 0.214705555 | NA |
| 3.72E-19 | 1.50E-17 |
| 0.711157924 | 0.815773823 |

| | | | |
|---|---|---|---|
| NA | 0.675440788 | | 0.790858573 |
| | 0.908636247 | | 0.944549939 |
| | 0.955792873 | NA | |
| | 0.028928257 | | 0.080626507 |
| | 0.941277857 | NA | |
| | 0.784318202 | NA | |
| | 0.974083195 | | 0.984381107 |
| | 0.793501896 | NA | |
| | 0.183746509 | NA | |
| | 0.223075025 | | 0.375504367 |
| | 0.610727417 | | 0.742190032 |
| NA | | NA | |
| | 0.519677273 | | 0.670188807 |
| | 0.823759658 | NA | |
| | 0.878734945 | NA | |
| | 0.631549638 | NA | |
| | 0.530332228 | NA | |
| | 0.038680255 | | 0.101350444 |
| | 0.764254325 | | 0.853630988 |
| | 0.839319108 | NA | |
| | 0.414856817 | | 0.576711191 |
| | 0.678339339 | | 0.792783705 |
| | 0.015051939 | | 0.047844923 |
| | 0.35682457 | | 0.520444783 |
| | 0.990272689 | | 0.993561791 |
| | 0.584546753 | | 0.721611007 |
| | 0.489295711 | NA | |
| | 0.000185729 | | 0.001202266 |
| NA | | NA | |
| | 0.006288813 | | 0.023528186 |
| | 0.003622081 | | 0.014946735 |
| | 0.53292253 | | 0.681273004 |
| NA | | NA | |
| | 0.003870678 | | 0.015781854 |
| NA | | NA | |
| | 0.548319409 | NA | |
| | 0.421922544 | | 0.583193984 |
| NA | | NA | |
| | 0.74627078 | NA | |
| | 0.066537679 | | 0.155161147 |
| | 0.941277857 | NA | |
| | 0.004822878 | | 0.018955441 |
| | 8.45E-24 | | 4.45E-22 |
| | 1.93E-05 | | 0.000162306 |
| | 1.28E-21 | | 6.03E-20 |
| | 0.900446002 | | 0.940018234 |
| NA | | NA | |

| | | |
|---|---|---|
| | 0.097182929 | 0.205248276 |
| | 0.577388158 | 0.715818165 |
| | 0.863897378 | NA |
| | 0.667655877 | 0.784651419 |
| | 0.079593777 | 0.177171147 |
| | 6.50E-06 | 6.17E-05 |
| | 0.501069745 | 0.653604528 |
| | 0.953932387 | NA |
| | 0.33994626 | 0.503648372 |
| NA | | NA |
| | 0.023699437 | 0.06871653 |
| | 0.293772836 | 0.454936958 |
| | 2.53E-65 | 5.10E-63 |
| | 0.158560814 | 0.294800886 |
| NA | | NA |
| | 0.7906082 | NA |
| | 1.07E-05 | 9.57E-05 |
| | 0.036943182 | 0.097756712 |
| | 0.69734637 | 0.806139125 |
| | 0.19085028 | 0.335793788 |
| | 0.022236228 | 0.065224676 |
| | 0.304567697 | 0.466194233 |
| NA | | NA |
| | 0.941278332 | NA |
| | 0.000159642 | 0.001055498 |
| | 0.306377355 | 0.46806122 |
| | 0.008815021 | 0.031023972 |
| | 0.001348575 | 0.00656417 |
| | 0.701387195 | 0.809347846 |
| | 0.025996677 | 0.073953974 |
| | 0.312990257 | 0.474932685 |
| | 0.526480282 | 0.675514301 |
| | 0.850698945 | NA |
| | 0.097641622 | 0.205961174 |
| NA | | NA |
| NA | | NA |
| NA | | NA |
| NA | | NA |
| | 0.941276431 | NA |
| | 0.561045037 | 0.70404203 |
| | 0.920092708 | 0.952258746 |
| | 0.405083333 | 0.567097606 |
| | 0.922943376 | 0.954130395 |
| | 0.921508665 | 0.953114487 |
| | 0.052156986 | 0.12815532 |
| | 0.000175607 | 0.001147979 |
| | 0.223516307 | 0.375954713 |

| | |
|---|---|
| 0.941278332 | NA |
| NA | NA |
| NA | NA |
| 0.941277857 | NA |
| 0.901607713 | 0.940488508 |
| 0.04702288 | 0.11797603 |
| 0.839319108 | NA |
| 0.003619039 | 0.014943441 |
| NA | NA |
| 0.138395656 | 0.266836243 |
| 0.941278332 | NA |
| 0.571538436 | 0.711794109 |
| 0.001418828 | 0.006844535 |
| 0.038395802 | 0.100815221 |
| 0.880073666 | NA |
| NA | NA |
| NA | NA |
| 3.92E-06 | 3.90E-05 |
| 0.879403857 | NA |
| 0.544365915 | 0.690985353 |
| 0.989783539 | 0.993351574 |
| 0.000279597 | 0.001714313 |
| 0.896741927 | 0.937779975 |
| 0.047818893 | 0.119569837 |
| 0.941278332 | NA |
| 0.13046296 | 0.255410377 |
| 0.00025344 | 0.001570934 |
| 0.07601315 | 0.171137437 |
| 0.068235626 | 0.158113744 |
| 0.10664838 | 0.220159995 |
| 0.024240935 | 0.069996691 |
| 0.013479614 | 0.04381887 |
| NA | NA |
| 0.146967915 | 0.279158113 |
| 0.079951211 | 0.177703143 |
| 0.801418659 | 0.878603792 |
| 0.839319574 | NA |
| 0.486914676 | NA |
| 0.941278424 | NA |
| NA | NA |
| 0.955792873 | NA |
| 0.42226494 | 0.583497829 |
| 0.157748589 | 0.293659621 |
| 0.047339204 | 0.118532268 |
| 0.053682477 | 0.131105101 |
| 0.122258989 | 0.243760132 |
| 0.955792873 | NA |

| | |
|---|---|
| 0.507309252 | 0.65900643 |
| 0.000348537 | 0.002072998 |
| 0.724546952 | NA |
| 1.33E-15 | 4.24E-14 |
| NA | NA |
| NA | NA |
| NA | NA |
| NA | NA |
| NA | NA |
| NA | NA |
| NA | NA |
| NA | NA |
| NA | NA |
| NA | NA |
| 0.001243458 | 0.006141687 |
| 0.07357688 | 0.167122522 |
| 0.024194152 | 0.06988704 |
| 0.169599889 | 0.309229312 |
| 0.397134618 | 0.559725832 |
| NA | NA |
| 0.930055273 | 0.958402269 |
| 0.059767309 | 0.142565352 |
| NA | NA |
| 0.634530591 | NA |
| 0.003291422 | 0.013867299 |
| NA | NA |
| NA | NA |
| 0.291011846 | 0.452102601 |
| NA | NA |
| 2.35E-11 | 5.11E-10 |
| 0.572927917 | 0.712654051 |
| 3.18E-05 | 0.00025388 |
| NA | NA |
| 0.941278332 | NA |
| 0.352731004 | 0.516378095 |
| 0.054775099 | 0.133199521 |
| 0.825159559 | NA |
| 0.924799522 | 0.95491588 |
| 0.91372523 | NA |
| 0.487015006 | 0.642037206 |
| 0.268378759 | 0.428146864 |
| NA | NA |
| NA | NA |
| NA | NA |
| 0.385432473 | 0.54832728 |
| 0.30959305 | 0.471249841 |
| 0.006775964 | 0.025016519 |

| | |
|---|---|
| 0.941277857 | NA |
| 0.021697736 | 0.063901201 |
| 0.556227914 | NA |
| 0.01691815 | 0.052501407 |
| 0.662290727 | 0.780542874 |
| 0.320150113 | 0.482838753 |
| 0.326873484 | 0.489693017 |
| 0.538029118 | 0.68577536 |
| 0.320112731 | NA |
| 0.909324538 | NA |
| 0.955792398 | NA |
| 0.955792398 | NA |
| NA | NA |
| 0.94054529 | NA |
| NA | NA |
| 0.27063742 | 0.430624498 |
| 0.724561778 | NA |
| 0.614815318 | 0.744800194 |
| 0.002783912 | 0.012052141 |
| 5.87E-06 | 5.63E-05 |
| NA | NA |
| 0.941277857 | NA |
| 0.317836229 | 0.480276311 |
| 0.404534177 | 0.566562077 |
| NA | NA |
| 0.002641742 | 0.011558592 |
| 0.000337944 | 0.002023621 |
| 0.2392298 | 0.395092085 |
| 0.557800463 | 0.701476257 |
| NA | NA |
| 3.57E-09 | 5.92E-08 |
| 9.52E-24 | 5.00E-22 |
| 5.19E-15 | 1.57E-13 |
| 0.679842222 | NA |
| 0.883551838 | NA |
| 0.033189919 | 0.089936503 |
| 0.839319574 | NA |
| 0.028053568 | 0.078686191 |
| 0.644581885 | 0.767313612 |
| 0.089085692 | 0.192686977 |
| 0.19254797 | 0.338081055 |
| 0.208548641 | 0.357885707 |
| 0.061431354 | 0.145723449 |
| 0.189646271 | 0.334264369 |
| NA | NA |
| 0.020678999 | 0.061479899 |
| 0.267862657 | NA |

| | | |
|---|---|---|
| | 0.679956778 | NA |
| NA | | NA |
| | 0.000675622 | 0.003666474 |
| NA | | NA |
| | 0.254666536 | 0.412688786 |
| | 0.915449404 | 0.949021199 |
| | 0.300388845 | 0.461936638 |
| | 0.025003823 | 0.071660567 |
| | 0.010601224 | 0.036093597 |
| | 0.839318641 | NA |
| NA | | NA |
| | 0.801165519 | 0.878492596 |
| | 0.278188025 | NA |
| | 0.945232475 | 0.966925536 |
| | 0.353256021 | 0.516950495 |
| | 0.550670985 | 0.696221756 |
| | 0.034090399 | 0.091699109 |
| | 0.001105512 | 0.005565438 |
| | 0.044852026 | 0.113682714 |
| | 0.941277857 | NA |
| | 0.955792398 | NA |
| | 0.034665089 | 0.092916492 |
| | 8.18E-06 | 7.56E-05 |
| | 0.545186151 | 0.691601598 |
| | 0.039483036 | 0.103113367 |
| | 0.709265415 | NA |
| | 0.342338765 | 0.505825768 |
| | 0.978607153 | NA |
| | 0.930062302 | NA |
| | 0.587988688 | NA |
| | 0.062201247 | 0.147153769 |
| | 0.294634376 | 0.455944829 |
| | 0.929178944 | NA |
| | 2.40E-06 | 2.50E-05 |
| NA | | NA |
| | 0.941277857 | NA |
| | 0.526393539 | NA |
| | 0.447786185 | 0.607987046 |
| NA | | NA |
| | 0.036123686 | 0.096101037 |
| NA | | NA |
| | 3.85E-19 | 1.55E-17 |
| | 0.941276431 | NA |
| | 0.955791922 | NA |
| | 0.784161164 | NA |
| | 0.955792398 | NA |
| | 0.141068882 | 0.270783995 |

| | |
|---|---|
| NA | NA |
| NA | NA |
| NA | NA |
| 0.955792873 | NA |
| 0.021884429 | 0.064347521 |
| 0.453165822 | NA |
| 0.231836137 | 0.386063949 |
| 0.151743199 | 0.285753935 |
| 0.045865544 | 0.115755666 |
| 0.794351884 | 0.873834799 |
| 5.81E-05 | 0.000433238 |
| 0.058989667 | 0.14108441 |
| 0.028732885 | 0.080241713 |
| NA | NA |
| NA | NA |
| 0.955792398 | NA |
| 0.346528632 | NA |
| 0.862364208 | NA |
| NA | NA |
| 0.839318641 | NA |
| 0.000638736 | 0.003488569 |
| 0.013428296 | 0.04371175 |
| 9.20E-05 | 0.000651305 |
| 0.700136044 | NA |
| 0.839319108 | NA |
| 0.583141297 | 0.720526464 |
| 0.955792398 | NA |
| 0.473761994 | 0.631205316 |
| NA | NA |
| 0.941277857 | NA |
| 0.163888444 | 0.301996227 |
| 0.331060608 | 0.494130846 |
| 0.599278245 | 0.73325558 |
| 0.484980824 | 0.640530022 |
| 0.29011481 | 0.451150706 |
| 0.651316995 | NA |
| 2.50E-05 | 0.000205551 |
| 0.588740061 | 0.724945212 |
| 0.827823515 | NA |
| 0.074345427 | 0.16837011 |
| 0.01227102 | 0.040616718 |
| 0.001970261 | 0.009049819 |
| 0.839319108 | NA |
| 0.738325218 | NA |
| NA | NA |
| NA | NA |
| NA | NA |

| | |
|---|---|
| 0.229955253 | 0.383747804 |
| 0.839319108 | NA |
| NA | NA |
| 0.000552913 | 0.003095114 |
| 0.030000557 | 0.083016655 |
| NA | NA |
| NA | NA |
| 1.02E-05 | 9.24E-05 |
| 0.794684693 | 0.873922535 |
| 0.009896436 | 0.034088997 |
| 0.775523927 | 0.86152434 |
| 8.87E-07 | 9.96E-06 |
| 0.00962327 | 0.03336623 |
| 0.080661391 | 0.178895121 |
| 0.940528311 | NA |
| 0.000734358 | 0.003934984 |
| 0.732599086 | NA |
| 0.010581892 | 0.036038088 |
| 0.526393642 | 0.675466142 |
| 4.41E-05 | 0.000338579 |
| 0.861394394 | 0.917109919 |
| 0.455171899 | 0.614869554 |
| NA | NA |
| 0.177084881 | NA |
| 0.000342579 | 0.002045554 |
| 1.04E-05 | 9.41E-05 |
| 0.1994091 | 0.346827578 |
| 0.419308926 | 0.580930795 |
| 0.105147463 | 0.217722481 |
| 0.212014105 | 0.362447729 |
| 0.011392117 | 0.038300494 |
| 0.000558182 | 0.003119478 |
| 0.011096185 | 0.037489198 |
| 0.279231763 | 0.438996785 |
| 0.839318641 | NA |
| 6.01E-69 | 1.26E-66 |
| 0.839318641 | NA |
| 0.839319574 | NA |
| 4.82E-30 | 3.36E-28 |
| 0.848659217 | NA |
| 1.06E-20 | 4.71E-19 |
| 0.383920331 | NA |
| 0.941278332 | NA |
| NA | NA |
| 0.191266748 | 0.336389747 |
| 0.257639163 | 0.415835584 |
| 0.580400251 | 0.718208103 |

| | | |
|---|---|---|
| | 0.175379666 | 0.316553182 |
| | 0.922199077 | 0.953631614 |
| | 0.00033795 | 0.002023621 |
| | 0.818181825 | NA |
| | 4.73E-46 | 5.89E-44 |
| | 0.280457929 | 0.440391848 |
| | 0.545049159 | 0.691560783 |
| | 0.077311728 | 0.173258051 |
| | 0.006661638 | 0.024647957 |
| | 0.000917271 | 0.004740821 |
| | 0.002316568 | 0.010354734 |
| | 0.275307808 | 0.43517575 |
| NA | | NA |
| | 0.000201398 | 0.001288621 |
| | 0.000548879 | 0.003077949 |
| | 0.000423888 | 0.002457227 |
| | 0.316514686 | 0.478937225 |
| | 0.318380049 | 0.480812253 |
| | 0.005979348 | 0.022594134 |
| | 0.740923169 | NA |
| | 4.06E-14 | 1.15E-12 |
| | 5.81E-53 | 8.65E-51 |
| | 1.93E-18 | 7.37E-17 |
| | 0.409630359 | 0.57154709 |
| | 0.787938448 | NA |
| | 0.000866607 | 0.004521475 |
| | 0.190007479 | 0.334657246 |
| NA | | NA |
| | 0.287452635 | 0.448064677 |
| | 0.941275481 | NA |
| | 0.031022459 | NA |
| NA | | NA |
| | 1.38E-15 | 4.39E-14 |
| | 8.27E-28 | 5.28E-26 |
| | 1.63E-25 | 9.24E-24 |
| | 0.086073172 | NA |
| | 0.46907235 | NA |
| | 0.260233254 | NA |
| | 0.941277857 | NA |
| | 3.41E-23 | 1.76E-21 |
| | 0.058857947 | 0.140825982 |
| | 0.110912653 | 0.226417477 |
| NA | | NA |
| | 0.818872766 | NA |
| NA | | NA |
| NA | | NA |
| | 4.69E-07 | 5.55E-06 |

| | |
|---|---|
| 1.47E-38 | 1.41E-36 |
| 2.06E-07 | 2.62E-06 |
| 0.094540372 | 0.201184616 |
| 1.11E-19 | 4.57E-18 |
| 0.116736218 | 0.235171855 |
| 6.29E-09 | 1.01E-07 |
| 8.50E-11 | 1.72E-09 |
| 9.84E-06 | 8.94E-05 |
| 0.415696879 | 0.577472692 |
| NA | NA |
| 0.055888039 | 0.135380329 |
| 0.275252795 | 0.435117687 |
| 0.001438006 | 0.006921622 |
| 0.748874096 | 0.842337377 |
| 0.000229632 | 0.001442521 |
| 0.271422114 | 0.431296425 |
| 0.057692644 | 0.138679087 |
| 0.051365074 | 0.126693419 |
| 4.11E-11 | 8.68E-10 |
| 0.030148597 | 0.083325234 |
| 0.010499093 | 0.035807336 |
| 0.512860246 | 0.664175099 |
| 0.112867692 | 0.229476108 |
| 0.06223454 | 0.14720327 |
| 0.635548107 | NA |
| 0.700372618 | NA |
| 4.50E-32 | 3.43E-30 |
| 0.982898814 | 0.989178714 |
| 4.80E-31 | 3.49E-29 |
| 0.897270553 | NA |
| 0.441534361 | 0.602154407 |
| 0.807276539 | 0.881922851 |
| 0.680476641 | 0.794365168 |
| 5.93E-22 | 2.84E-20 |
| 0.43458159 | NA |
| 0.827498797 | 0.895237779 |
| 0.064237142 | 0.150920191 |
| NA | NA |
| 0.798492012 | NA |
| NA | NA |
| NA | NA |
| NA | NA |
| 0.941278332 | NA |
| 0.955792398 | NA |
| NA | NA |
| 0.941278332 | NA |
| NA | NA |

| | | |
|---|---|---|
| NA | | NA |
| | 0.955792398 | NA |
| NA | | NA |
| | 0.808663822 | NA |
| NA | | NA |
| NA | | NA |
| | 0.835127771 | NA |
| NA | | NA |
| | 0.941278434 | NA |
| | 8.37E-06 | 7.72E-05 |
| | 0.957889065 | 0.974159522 |
| | 5.65E-29 | 3.77E-27 |
| | 1.25E-23 | 6.54E-22 |
| | 5.92E-21 | 2.71E-19 |
| NA | | NA |
| | 1.86E-17 | 6.68E-16 |
| | 1.73E-26 | 1.03E-24 |
| NA | | NA |
| | 4.19E-22 | 2.03E-20 |
| | 4.03E-08 | 5.73E-07 |
| | 0.413940145 | NA |
| | 4.13E-55 | 6.34E-53 |
| | 7.40E-41 | 7.83E-39 |
| | 6.06E-11 | 1.25E-09 |
| | 0.572197637 | 0.712174354 |
| | 3.64E-21 | 1.68E-19 |
| | 9.68E-53 | 1.41E-50 |
| | 1.16E-24 | 6.44E-23 |
| | 1.40E-58 | 2.31E-56 |
| | 3.50E-63 | 6.77E-61 |
| | 0.000113873 | 0.000784264 |
| | 8.88E-22 | 4.23E-20 |
| | 0.029331587 | 0.081512018 |
| | 0.658672295 | NA |
| | 0.616170843 | NA |
| NA | | NA |
| | 0.940729964 | NA |
| | 0.941278332 | NA |
| NA | | NA |
| NA | | NA |
| NA | | NA |
| NA | | NA |
| NA | | NA |
| | 0.941277857 | NA |
| NA | | NA |
| NA | | NA |
| | 0.000213225 | 0.001356639 |

| | | |
|---|---|---|
| NA | | NA |
| | 0.941278332 | NA |
| NA | | NA |
| NA | | NA |
| NA | | NA |
| | 0.853596091 | NA |
| | 0.785212979 | NA |
| | 2.72E-08 | 3.98E-07 |
| | 0.992827072 | 0.994981334 |
| | 0.491580661 | 0.645586123 |
| | 0.098593917 | 0.207400124 |
| | 0.917168494 | NA |
| | 0.001733328 | 0.008116601 |
| | 5.50E-20 | 2.33E-18 |
| | 0.006515279 | 0.024204255 |
| | 0.000346492 | 0.002062316 |
| | 0.261784535 | NA |
| | 0.955792398 | NA |
| | 0.009805798 | 0.033877154 |
| NA | | NA |
| | 0.839320041 | NA |
| | 0.426537031 | 0.587967476 |
| | 6.01E-07 | 6.96E-06 |
| | 0.019190786 | 0.057981607 |
| | 0.022962112 | 0.066978455 |
| | 0.955792873 | NA |
| | 0.164127471 | 0.302271902 |
| | 0.955792398 | NA |
| | 0.941278424 | NA |
| | 0.63827362 | NA |
| | 0.786733216 | NA |
| NA | | NA |
| | 0.01369334 | 0.044332002 |
| | 0.262471705 | 0.421378088 |
| | 0.015004408 | 0.047720007 |
| | 0.557166012 | NA |
| | 0.555323195 | 0.699910124 |
| NA | | NA |
| | 0.101271273 | 0.211708504 |
| | 0.736184774 | 0.833647157 |
| | 0.955792873 | NA |
| | 0.194147898 | 0.340212638 |
| | 0.050601187 | 0.125029528 |
| | 0.002218905 | 0.009978154 |
| | 0.074613829 | 0.16880129 |
| | 0.038482101 | 0.100972468 |
| | 0.68587595 | 0.798345944 |

| | | |
|---|---|---|
| | 0.199937283 | 0.347390847 |
| | 0.059313837 | 0.141760011 |
| | 0.255755725 | 0.4137837 |
| | 0.759573459 | 0.850193626 |
| | 0.38742082 | 0.550235119 |
| | 0.198048652 | 0.345167607 |
| | 0.113710548 | 0.230690211 |
| | 0.178373708 | 0.319967668 |
| | 0.377695226 | 0.540550885 |
| | 0.000608202 | 0.003350249 |
| | 0.068677195 | 0.158858561 |
| | 0.810636471 | NA |
| | 0.771840744 | NA |
| | 0.702700529 | NA |
| | 0.831413565 | NA |
| | 0.770458155 | 0.857898812 |
| NA | | NA |
| | 0.816900108 | 0.888212342 |
| | 0.004286153 | 0.017177811 |
| | 0.955043037 | NA |
| NA | | NA |
| | 0.341528134 | 0.504972832 |
| | 0.000386209 | 0.002266969 |
| | 0.921493168 | NA |
| | 0.305761115 | NA |
| | 2.46E-11 | 5.35E-10 |
| | 1.70E-05 | 0.000145061 |
| | 0.433810874 | 0.594996337 |
| | 0.315475568 | 0.477911753 |
| | 0.958031554 | 0.974181957 |
| | 4.27E-05 | 0.000329081 |
| | 0.486664075 | 0.64182213 |
| | 0.955792398 | NA |
| | 0.839319108 | NA |
| | 0.561857094 | 0.704693643 |
| | 0.005219783 | 0.020211693 |
| | 0.089024747 | 0.192633366 |
| | 0.671628797 | 0.787609431 |
| | 0.381198026 | 0.543825835 |
| | 0.217666871 | 0.369036746 |
| | 0.921403047 | NA |
| | 0.877020671 | 0.926329976 |
| NA | | NA |
| | 0.000261986 | 0.001616325 |
| | 0.003926287 | 0.015964326 |
| | 0.255902903 | NA |
| | 0.033890589 | 0.091334217 |

| | | |
|---|---|---|
| | 0.986430049 | 0.991595913 |
| | 0.122397906 | 0.243934937 |
| | 0.000740147 | 0.003961543 |
| | 0.027805273 | 0.078201715 |
| | 0.906098388 | NA |
| | 0.80729918 | 0.881922851 |
| | 0.678477789 | 0.79290658 |
| | 0.006570723 | 0.024387394 |
| | 0.550820136 | 0.696290294 |
| | 0.429467207 | 0.59074078 |
| | 2.47E-09 | 4.18E-08 |
| | 0.016411337 | 0.051235919 |
| | 0.808055709 | 0.882288452 |
| | 0.491044378 | 0.645132982 |
| | 0.932049627 | 0.959371849 |
| | 0.002109705 | 0.009559338 |
| | 0.678451645 | NA |
| | 0.095388124 | 0.202499834 |
| | 0.034075196 | 0.091686392 |
| | 0.97882581 | 0.986743972 |
| | 0.433512993 | NA |
| NA | | NA |
| | 0.989811182 | 0.993351574 |
| | 0.941275956 | NA |
| | 0.823033496 | NA |
| | 0.329800132 | 0.492805747 |
| | 0.667909329 | NA |
| | 0.479321996 | 0.636049626 |
| NA | | NA |
| | 0.767158023 | 0.855464758 |
| | 0.445085976 | 0.605713916 |
| NA | | NA |
| | 0.903529766 | 0.941652848 |
| | 0.128923231 | 0.25318582 |
| | 0.159889416 | 0.296601034 |
| | 0.182718746 | 0.325310165 |
| | 0.955792873 | NA |
| NA | | NA |
| | 0.749023973 | 0.842445421 |
| | 0.941278332 | NA |
| NA | | NA |
| | 0.839319108 | NA |
| | 0.91776262 | NA |
| NA | | NA |
| | 0.941277857 | NA |
| | 0.036222035 | 0.096293768 |
| NA | | NA |

| | | | |
|---|---|---|---|
| | 0.366530841 | | 0.529666785 |
| | 0.001244025 | | 0.006142873 |
| | 0.016653124 | | 0.05184118 |
| | 0.956108727 | | 0.973116391 |
| | 0.91975729 | | 0.951952995 |
| | 0.046203845 | | 0.116411974 |
| | 0.036653204 | | 0.097151399 |
| NA | | NA | |
| NA | | NA | |
| | 0.609496594 | | 0.741490928 |
| | 0.001566412 | | 0.007444834 |
| | 0.755626499 | NA | |
| | 0.789048417 | | 0.870172377 |
| | 0.000158742 | | 0.001050134 |
| | 0.613404983 | NA | |
| | 0.941278332 | NA | |
| | 0.231845983 | | 0.386063949 |
| | 0.327180096 | | 0.48996883 |
| | 0.252849842 | | 0.410667482 |
| | 0.296710126 | | 0.458174024 |
| | 0.052329074 | | 0.128511826 |
| | 0.592984036 | NA | |
| | 0.002863929 | | 0.012331267 |
| | 0.839319108 | NA | |
| NA | | NA | |
| | 0.006297391 | | 0.023552876 |
| | 7.30E-07 | | 8.33E-06 |
| | 0.240050497 | | 0.396068344 |
| | 0.955792398 | NA | |
| | 0.286803405 | | 0.447353861 |
| | 0.006300652 | | 0.023558122 |
| | 0.426494014 | | 0.587942229 |
| | 0.661074709 | | 0.779701886 |
| | 0.15308024 | | 0.28732875 |
| NA | | NA | |
| | 0.957324029 | NA | |
| | 0.839319108 | NA | |
| NA | | NA | |
| | 0.829290316 | NA | |
| | 0.051462409 | | 0.126846113 |
| | 0.941599939 | NA | |
| | 0.182759534 | | 0.32532898 |
| | 0.896843949 | NA | |
| | 0.796379111 | NA | |
| | 0.000224383 | | 0.001416648 |
| | 0.860398971 | | 0.916609361 |
| | 0.164298034 | | 0.302515805 |

| | | |
|---|---|---|
| | 6.83E-32 | 5.18E-30 |
| | 0.791138588 | 0.871710058 |
| | 0.821936329 | 0.891651634 |
| | 0.357980955 | 0.521556142 |
| | 0.010258308 | 0.035127067 |
| | 0.109975054 | 0.2251468 |
| | 0.112873507 | NA |
| | 0.790651661 | NA |
| | 0.955792398 | NA |
| | 0.643239334 | NA |
| | 0.839319108 | NA |
| | 0.748393366 | NA |
| | 0.136908707 | 0.264778419 |
| | 0.468274072 | 0.626453432 |
| | 0.941278332 | NA |
| NA | | NA |
| | 0.955792873 | NA |
| | 0.839319574 | NA |
| NA | | NA |
| | 0.572530033 | NA |
| | 0.084095614 | 0.184636454 |
| | 0.882079476 | NA |
| NA | | NA |
| | 5.05E-19 | 2.01E-17 |
| | 0.839322136 | NA |
| | 0.794688654 | 0.873922535 |
| | 0.000699211 | 0.003776407 |
| NA | | NA |
| NA | | NA |
| | 0.018796834 | 0.05710303 |
| | 7.39E-05 | 0.000535342 |
| | 0.928729626 | NA |
| | 0.829234815 | NA |
| NA | | NA |
| NA | | NA |
| | 2.68E-05 | 0.00021816 |
| | 0.464354203 | NA |
| NA | | NA |
| NA | | NA |
| | 0.127238597 | 0.250815802 |
| NA | | NA |
| NA | | NA |
| NA | | NA |
| | 0.25928589 | 0.417700355 |
| | 0.000104196 | 0.000725377 |
| | 0.575013392 | 0.714074899 |
| | 0.000190961 | 0.001231773 |

| | | |
|---|---|---|
| | 0.752446484 | 0.844680199 |
| | 0.103148545 | 0.21468535 |
| | 1.71E-06 | 1.84E-05 |
| | 0.620504254 | 0.74932736 |
| NA | | NA |
| | 0.708636092 | NA |
| | 0.149224821 | 0.282257672 |
| | 0.002376593 | 0.010581305 |
| NA | | NA |
| NA | | NA |
| NA | | NA |
| NA | | NA |
| | 0.839320041 | NA |
| | 0.000281147 | 0.001722487 |
| | 0.941277857 | NA |
| | 0.359037687 | 0.522583931 |
| | 0.108852587 | 0.223392764 |
| | 0.338195836 | 0.50166629 |
| | 0.198930978 | 0.34626869 |
| NA | | NA |
| | 0.722225122 | 0.823887005 |
| | 0.947074013 | NA |
| | 0.705632596 | NA |
| | 0.955792397 | NA |
| | 0.00752683 | 0.027277363 |
| | 0.940547608 | NA |
| | 0.229381964 | 0.383104282 |
| | 0.005150256 | 0.020014104 |
| | 0.235189529 | NA |
| | 0.519604705 | 0.670150701 |
| | 0.830665589 | 0.897271536 |
| | 0.654536157 | 0.774433047 |
| | 0.686017752 | NA |
| | 0.001992504 | 0.009141398 |
| | 0.238187116 | 0.393785492 |
| | 0.067991351 | 0.157639796 |
| NA | | NA |
| | 0.839318641 | NA |
| | 0.465942026 | 0.624105351 |
| | 0.839318641 | NA |
| | 0.539497451 | 0.686727544 |
| NA | | NA |
| NA | | NA |
| | 0.007377627 | 0.026838191 |
| | 0.388208547 | 0.551057958 |
| | 0.900182131 | NA |
| | 0.056405021 | 0.136341253 |

| | | |
|---|---|---|
| | 0.654474796 | 0.77439894 |
| | 0.106912429 | 0.220513833 |
| | 0.012069598 | 0.040061437 |
| | 0.955792398 | NA |
| | 0.839318641 | NA |
| | 0.005319145 | 0.020522983 |
| | 0.853595622 | NA |
| | 0.001363736 | 0.006625772 |
| | 0.582577391 | 0.720188379 |
| | 0.927548224 | NA |
| | 0.817371544 | 0.888562571 |
| | 0.011218256 | 0.03779968 |
| | 0.006120311 | 0.023009997 |
| | 0.355127801 | 0.518882667 |
| | 0.679873317 | NA |
| | 0.754194652 | 0.846066498 |
| | 0.919861069 | NA |
| | 0.941278424 | NA |
| | 0.839318641 | NA |
| NA | | NA |
| NA | | NA |
| | 0.040858619 | 0.105961741 |
| | 0.655346245 | 0.775083296 |
| | 0.01984934 | 0.059489089 |
| | 0.941277857 | NA |
| | 0.370245202 | 0.533460847 |
| | 0.606563713 | 0.739151523 |
| | 0.56726518 | 0.708880443 |
| | 0.661067802 | NA |
| | 0.720864842 | NA |
| | 0.107005721 | 0.220665888 |
| | 0.004026594 | 0.016296718 |
| NA | | NA |
| | 3.11E-06 | 3.15E-05 |
| NA | | NA |
| | 0.262079941 | NA |
| | 1.28E-18 | 4.97E-17 |
| NA | | NA |
| | 0.000177077 | 0.001156638 |
| | 0.001173575 | 0.00584871 |
| | 0.11332037 | 0.230180189 |
| | 0.245030725 | 0.401640474 |
| | 0.002035482 | 0.009284889 |
| | 0.05083609 | 0.125518735 |
| NA | | NA |
| | 0.480991462 | 0.637487957 |
| | 0.839319574 | NA |

| | |
|---|---|
| 0.735704024 | NA |
| 0.055766051 | 0.135151999 |
| 8.68E-05 | 0.000618441 |
| 0.028909014 | 0.080603732 |
| 0.279021662 | 0.438802351 |
| 0.002783504 | 0.012052141 |
| NA | NA |
| NA | NA |
| NA | NA |
| 0.676927949 | 0.791715346 |
| 0.007015841 | 0.025714666 |
| 0.065785851 | 0.153739304 |
| 8.74E-05 | 0.000622337 |
| 0.00103547 | 0.005260675 |
| 0.163389467 | 0.301332411 |
| 0.689429516 | NA |
| 0.002894832 | 0.012446314 |
| NA | NA |
| 0.778548485 | 0.863592382 |
| NA | NA |
| 0.271888118 | 0.431777488 |
| 0.941275956 | NA |
| 0.62493479 | 0.752858299 |
| 0.29219074 | 0.453357998 |
| 0.002031868 | 0.009271957 |
| 0.873207054 | NA |
| NA | NA |
| 0.074871915 | 0.169192188 |
| 0.898523458 | NA |
| 0.618482378 | 0.74787226 |
| 0.503042694 | 0.655164129 |
| 0.935434275 | NA |
| 0.152581438 | 0.286677459 |
| 0.775089149 | 0.861161925 |
| 0.559218163 | 0.702666895 |
| NA | NA |
| 0.589317643 | 0.725393728 |
| 0.246002307 | 0.402900084 |
| 0.77157138 | 0.858857129 |
| 3.60E-26 | 2.13E-24 |
| 0.941275956 | NA |
| 0.839318641 | NA |
| 0.003268781 | 0.01378029 |
| 0.475839865 | 0.633023196 |
| 0.154882158 | 0.289723172 |
| 0.305053515 | 0.466704268 |
| 0.387108155 | 0.549955133 |

| | | |
|---|---|---|
| NA | | NA |
| | 0.017426187 | 0.05372672 |
| NA | | NA |
| NA | | NA |
| | 0.839320041 | NA |
| NA | | NA |
| NA | | NA |
| NA | | NA |
| | 0.902532926 | NA |
| | 0.046059513 | 0.116153186 |
| | 0.237036468 | 0.392480571 |
| | 0.105897541 | 0.218875761 |
| | 0.184224295 | 0.327207422 |
| | 0.452855774 | 0.612784146 |
| | 0.955792873 | NA |
| | 0.840135214 | NA |
| | 0.941280349 | NA |
| | 0.220693925 | 0.372629411 |
| | 0.529352589 | 0.678445397 |
| | 0.121852528 | 0.243194187 |
| | 0.839318641 | NA |
| | 0.094867686 | 0.201718838 |
| | 0.24384677 | 0.400299586 |
| | 0.114949621 | 0.232537045 |
| | 0.627619356 | 0.75486182 |
| NA | | NA |
| NA | | NA |
| | 0.415929485 | 0.577694673 |
| | 0.279066305 | 0.438843564 |
| | 0.888101699 | 0.933255773 |
| | 0.43553078 | 0.596667643 |
| | 0.941278332 | NA |
| | 0.955792398 | NA |
| NA | | NA |
| NA | | NA |
| | 0.452711094 | NA |
| NA | | NA |
| NA | | NA |
| | 0.74102849 | 0.83729508 |
| | 0.011914683 | 0.039680047 |
| | 0.941275956 | NA |
| | 0.602873218 | 0.736145071 |
| | 0.000166648 | 0.001097854 |
| | 0.839318641 | NA |
| NA | | NA |
| | 0.941278332 | NA |
| | 0.689529376 | 0.800908677 |

| | | |
|---|---|---|
| | 0.375163194 | 0.538264876 |
| | 0.298281152 | NA |
| | 0.916477071 | NA |
| | 0.003910182 | 0.015920311 |
| | 0.009353127 | 0.032571049 |
| | 0.561429734 | 0.704287474 |
| | 0.000405311 | 0.002361616 |
| | 0.000899919 | 0.004667656 |
| | 0.620784019 | 0.749468748 |
| | 0.360884195 | 0.524597875 |
| | 0.941276431 | NA |
| | 4.29E-09 | 7.05E-08 |
| | 0.038333423 | 0.1007411 |
| | 0.004848298 | 0.01903021 |
| | 0.615176539 | NA |
| NA | | NA |
| | 0.872701145 | 0.924063911 |
| | 0.000100672 | 0.000705382 |
| | 0.00036247 | 0.002142951 |
| | 0.42404278 | 0.585444678 |
| | 0.834845185 | 0.899970585 |
| | 0.839319108 | NA |
| | 0.067602545 | 0.156930802 |
| | 0.073619066 | 0.167154536 |
| | 0.040892101 | 0.106037019 |
| | 0.037995987 | 0.100019961 |
| | 3.02E-05 | 0.000242568 |
| | 0.285010365 | 0.44554162 |
| | 0.649708963 | NA |
| | 0.472613189 | 0.630097974 |
| | 0.511285889 | 0.66276564 |
| | 0.008149271 | 0.029107229 |
| | 3.77E-09 | 6.23E-08 |
| | 0.94239861 | NA |
| | 0.071473608 | 0.163625547 |
| | 0.266999795 | 0.426633265 |
| | 0.316802775 | 0.479251281 |
| | 0.790651661 | NA |
| | 5.47E-18 | 2.03E-16 |
| | 0.759538946 | NA |
| NA | | NA |
| | 0.000704139 | 0.00379613 |
| | 0.009586428 | 0.033266681 |
| NA | | NA |
| | 0.549676052 | 0.695259294 |
| | 0.378234114 | 0.541061866 |
| | 0.898005787 | NA |

| | |
|---|---|
| 0.157744217 | 0.293659621 |
| 0.071505548 | 0.163682925 |
| 1.94E-09 | 3.32E-08 |
| 0.696152597 | 0.805326553 |
| 0.19974378 | 0.347183534 |
| 0.492597616 | 0.646457985 |
| 0.327735749 | 0.49050002 |
| 0.323169137 | 0.485855976 |
| 0.839319108 | NA |
| 1.37E-17 | 4.96E-16 |
| 0.405309809 | NA |
| NA | NA |
| NA | NA |
| 0.034958419 | 0.093523692 |
| 0.044762739 | 0.113503765 |
| 0.977221983 | 0.985962343 |
| 0.01850542 | 0.056396652 |
| 0.45141919 | 0.611361568 |
| 0.955792398 | NA |
| 0.664833656 | 0.782560827 |
| NA | NA |
| 0.72784503 | 0.828590368 |
| 0.941277857 | NA |
| NA | NA |
| NA | NA |
| NA | NA |
| NA | NA |
| 0.442898313 | 0.603495857 |
| 0.270373252 | 0.430463153 |
| 0.113730509 | 0.230698276 |
| 7.65E-17 | 2.64E-15 |
| 0.155668207 | 0.290930361 |
| 0.595372016 | 0.730240375 |
| 0.839318641 | NA |
| 0.843230766 | NA |
| 0.424703651 | 0.586048211 |
| 0.050443421 | 0.124680492 |
| 0.053339491 | 0.130516861 |
| 0.132033327 | 0.257624654 |
| 0.796063389 | 0.874908486 |
| 0.013985629 | 0.045100386 |
| NA | NA |
| 0.917447524 | NA |
| 0.008288892 | 0.029534761 |
| 0.91807123 | NA |
| 0.853595154 | NA |
| 0.94752471 | NA |

| | | |
|---|---|---|
| | 0.674461232 | NA |
| | 0.000834075 | 0.004378616 |
| | 0.077111604 | 0.172958979 |
| | 1.30E-05 | 0.000114147 |
| | 0.69498483 | NA |
| | 0.322094415 | 0.484852724 |
| | 0.955792398 | NA |
| | 0.4850496 | 0.640543197 |
| | 0.892137165 | NA |
| NA | | NA |
| | 0.894657804 | 0.936563533 |
| NA | | NA |
| | 0.010853925 | 0.036811786 |
| | 0.974189511 | 0.984409087 |
| | 0.002924001 | 0.01255132 |
| | 0.739991096 | NA |
| | 0.004276626 | 0.017148291 |
| | 4.69E-07 | 5.54E-06 |
| | 0.955791446 | NA |
| | 0.219304736 | 0.370809926 |
| | 0.777353288 | 0.862914199 |
| | 0.001384854 | 0.006714666 |
| | 0.500330891 | 0.65307792 |
| | 0.001153704 | 0.00576639 |
| | 0.792598653 | 0.872430315 |
| | 0.017833337 | 0.054720779 |
| NA | | NA |
| | 0.671574274 | 0.787584296 |
| NA | | NA |
| | 1.68E-14 | 4.87E-13 |
| | 0.273178745 | NA |
| | 5.28E-13 | 1.36E-11 |
| | 0.921188999 | NA |
| NA | | NA |
| | 0.860281579 | 0.916609361 |
| | 0.25072215 | NA |
| | 4.86E-05 | 0.000370073 |
| | 0.330959657 | 0.49404213 |
| | 0.088775564 | 0.192282299 |
| | 0.106891676 | 0.220490135 |
| NA | | NA |
| | 0.149387676 | 0.282411771 |
| | 6.62E-06 | 6.28E-05 |
| | 0.659743598 | 0.778620444 |
| | 0.986640112 | 0.991639555 |
| | 0.012039255 | 0.039999769 |
| | 0.653656977 | 0.774003044 |

| | |
|---|---|
| 0.829727987 | 0.896589892 |
| 8.11E-06 | 7.50E-05 |
| 0.084783418 | 0.185838006 |
| 0.110721165 | 0.226248291 |
| NA | NA |
| 0.810054404 | 0.88359447 |
| 0.275338102 | 0.435194734 |
| 0.933274841 | NA |
| 0.434265324 | NA |
| 0.941275956 | NA |
| NA | NA |
| 0.982269016 | 0.988869696 |
| 0.611816556 | NA |
| 3.91E-17 | 1.37E-15 |
| 0.941278332 | NA |
| 0.276148805 | 0.435955021 |
| NA | NA |
| NA | NA |
| 0.224780535 | NA |
| 0.935866248 | NA |
| 0.270600127 | NA |
| NA | NA |
| 0.903954661 | 0.941698566 |
| 0.346320375 | 0.510123058 |
| 0.353372911 | 0.517057972 |
| 0.003164335 | 0.013418406 |
| 0.941280925 | NA |
| 0.915258534 | NA |
| 0.604606515 | NA |
| 0.007174133 | 0.026202044 |
| 1.16E-08 | 1.81E-07 |
| 0.563477564 | 0.705811384 |
| 0.112299887 | 0.228536414 |
| 0.59901969 | NA |
| 0.941278284 | NA |
| 0.941278332 | NA |
| 0.029611198 | 0.082128877 |
| 0.817872151 | 0.888822616 |
| 0.413523767 | 0.575764417 |
| 0.941277857 | NA |
| 1.98E-06 | 2.09E-05 |
| 0.646427217 | NA |
| 4.03E-09 | 6.64E-08 |
| 0.060660727 | 0.144298082 |
| 0.057420317 | 0.138177978 |
| 0.246494437 | 0.403483983 |
| 0.000103201 | 0.000719717 |

| | | | |
|---|---|---|---|
| | 0.000186628 | | 0.001207751 |
| | 4.51E-06 | | 4.43E-05 |
| | 0.665840783 | NA | |
| NA | | NA | |
| | 0.941277857 | NA | |
| | 0.941278207 | NA | |
| NA | | NA | |
| NA | | NA | |
| NA | | NA | |
| | 0.941278332 | NA | |
| NA | | NA | |
| NA | | NA | |
| NA | | NA | |
| NA | | NA | |
| NA | | NA | |
| NA | | NA | |
| NA | | NA | |
| NA | | NA | |
| NA | | NA | |
| NA | | NA | |
| NA | | NA | |
| | 0.000316615 | | 0.001908347 |
| | 0.207512419 | | 0.356762964 |
| | 0.746205156 | NA | |
| | 0.562036317 | NA | |
| | 0.011314725 | | 0.038081602 |
| | 0.164787956 | NA | |
| | 2.63E-05 | | 0.00021528 |
| | 0.001137923 | | 0.005699673 |
| | 0.942338624 | | 0.965107777 |
| | 0.174844279 | | 0.315850298 |
| NA | | NA | |
| | 0.908720435 | NA | |
| | 0.002901834 | | 0.012474165 |
| | 0.858392524 | NA | |
| | 0.532810455 | | 0.681202904 |
| | 0.667816868 | NA | |
| | 0.639003685 | | 0.762946033 |
| | 0.020857658 | | 0.061864832 |
| | 0.922180658 | NA | |
| | 0.072792453 | | 0.165720321 |
| | 0.122982313 | | 0.244744678 |
| | 0.164550986 | | 0.302911258 |
| | 0.936446456 | | 0.962372303 |
| | 0.108843747 | | 0.223392764 |
| | 0.131773148 | | 0.257298152 |
| NA | | NA | |

| | |
|---|---|
| NA | NA |
| 0.000733219 | 0.003931535 |
| 0.124952524 | 0.247575126 |
| 0.486599889 | NA |
| 0.955792398 | NA |
| 0.023676857 | 0.068667791 |
| 0.039360112 | 0.102826187 |
| 0.003935562 | 0.015996285 |
| NA | NA |
| 1.83E-18 | 7.01E-17 |
| 0.709683894 | NA |
| NA | NA |
| 0.941278332 | NA |
| 0.788231337 | 0.869831931 |
| 0.099928988 | 0.209486505 |
| 0.111614514 | 0.227472272 |
| 0.009847285 | 0.033978828 |
| 0.011790718 | 0.039349786 |
| 0.000979226 | 0.005015608 |
| 0.688724443 | 0.800376082 |
| NA | NA |
| 0.016639337 | 0.051825373 |
| 0.000689413 | 0.003730257 |
| 0.238053155 | 0.39368904 |
| 0.62020941 | NA |
| 0.101719602 | 0.212435725 |
| 0.006033982 | 0.022775268 |
| 1.38E-05 | 0.00012088 |
| 0.845515797 | 0.906897706 |
| 0.513025364 | 0.664282495 |
| NA | NA |
| NA | NA |
| 0.290684995 | 0.451801243 |
| 0.039984485 | 0.104171487 |
| NA | NA |
| 2.98E-08 | 4.34E-07 |
| 0.000905907 | 0.0046936 |
| 0.317256805 | 0.479724698 |
| 0.874676852 | NA |
| 0.085211766 | 0.186486321 |
| 0.051508338 | 0.126923853 |
| 0.409043025 | 0.571072235 |
| 0.533302863 | 0.681616287 |
| NA | NA |
| 0.085458665 | 0.186870662 |
| 0.517176137 | NA |
| 1.06E-08 | 1.66E-07 |

| | |
|---|---|
| 2.47E-34 | 2.08E-32 |
| 0.136879078 | 0.264748563 |
| 0.074162087 | 0.168058408 |
| 0.521370754 | 0.671552762 |
| 0.899507177 | NA |
| 0.005198252 | 0.020161118 |
| 2.82E-14 | 8.10E-13 |
| 0.00187945 | 0.008682971 |
| 0.138822853 | 0.267486566 |
| 1.98E-08 | 2.96E-07 |
| 0.217080172 | 0.368495675 |
| 0.263694965 | 0.42271468 |
| 0.985687955 | 0.991027527 |
| 0.773592768 | 0.860423133 |
| NA | NA |
| 0.007645389 | 0.027613893 |
| 2.88E-05 | 0.000232892 |
| NA | NA |
| 1.82E-05 | 0.000154406 |
| NA | NA |
| 0.085699172 | 0.187285679 |
| 0.084990527 | 0.186171961 |
| 0.690904438 | NA |
| NA | NA |
| NA | NA |
| NA | NA |
| NA | NA |
| 0.650613611 | NA |
| 0.019430553 | 0.058535126 |
| 0.007510032 | 0.027232448 |
| 0.582670395 | 0.720221546 |
| 0.963496083 | 0.977841478 |
| 0.901510694 | 0.940439722 |
| NA | NA |
| 0.006701501 | 0.024772343 |
| 0.346554581 | 0.510375704 |
| 8.28E-12 | 1.89E-10 |
| 0.000698118 | 0.003771358 |
| 0.610838256 | 0.742227061 |
| 0.609210691 | 0.741368826 |
| NA | NA |
| 0.955792873 | NA |
| 0.015532779 | 0.04904435 |
| 0.817775131 | 0.888757758 |
| 0.613130135 | NA |
| 0.080264982 | 0.178298375 |
| 0.046397908 | 0.116777308 |

| | |
|---|---|
| 0.606406325 | 0.739051887 |
| 0.63823723 | 0.7623487 |
| 2.76E-07 | 3.42E-06 |
| NA | NA |
| 0.003311383 | 0.01393521 |
| 0.879599661 | 0.927901038 |
| 0.002771672 | 0.012012259 |
| 0.839319574 | NA |
| NA | NA |
| 2.63E-08 | 3.87E-07 |
| 0.216958988 | 0.36834062 |
| 0.01525599 | 0.04835194 |
| NA | NA |
| 0.050428284 | 0.124679964 |
| 0.929145386 | 0.957879352 |
| 0.005179304 | 0.020110569 |
| 0.068311864 | 0.158244184 |
| 0.06624373 | 0.154627171 |
| 0.037579118 | 0.099076003 |
| 0.739308425 | 0.83608206 |
| NA | NA |
| 0.147383853 | 0.279747247 |
| 0.921939591 | NA |
| NA | NA |
| 0.822484197 | 0.892002043 |
| 2.41E-12 | 5.82E-11 |
| 0.726781288 | 0.827696086 |
| 0.086116957 | 0.187861408 |
| 2.15E-05 | 0.00017937 |
| 0.14859301 | 0.281503567 |
| 0.796941232 | NA |
| 0.27499212 | NA |
| 0.39523242 | NA |
| 0.784704236 | 0.867673057 |
| 0.102435402 | 0.213593294 |
| NA | NA |
| 0.39419447 | 0.556732734 |
| NA | NA |
| 0.876432709 | 0.926140201 |
| 0.016243822 | 0.050823711 |
| 0.011425868 | 0.03837806 |
| 0.530085946 | 0.678924703 |
| NA | NA |
| 0.0036353 | 0.014990891 |
| 0.084504816 | 0.185312661 |
| NA | NA |
| NA | NA |

| | | |
|---|---|---|
| | 0.403674787 | 0.565791266 |
| | 0.015290595 | 0.048435819 |
| | 0.000357262 | 0.002118999 |
| | 0.897532873 | NA |
| | 0.941278332 | NA |
| | 0.023694271 | 0.068709923 |
| | 0.887924633 | NA |
| | 0.941278332 | NA |
| | 1.00E-13 | 2.73E-12 |
| | 0.00036261 | 0.002143242 |
| NA | | NA |
| | 0.197300576 | 0.344141017 |
| | 0.056137376 | 0.13585987 |
| | 0.032953496 | 0.089416225 |
| | 0.742832758 | 0.838474193 |
| NA | | NA |
| | 0.281082073 | 0.441255474 |
| | 7.16E-05 | 0.000520933 |
| | 0.15232907 | 0.28639309 |
| | 0.528337249 | 0.677412368 |
| | 0.000451987 | 0.00260087 |
| | 0.663931658 | NA |
| | 0.839318641 | NA |
| | 0.005726423 | 0.021797588 |
| | 0.003642961 | 0.015017279 |
| | 2.37E-21 | 1.10E-19 |
| | 0.41683321 | 0.578440607 |
| | 0.941278332 | NA |
| | 0.319001565 | 0.481475919 |
| | 0.159964563 | 0.296671071 |
| | 0.000301573 | 0.001830214 |
| NA | | NA |
| NA | | NA |
| NA | | NA |
| NA | | NA |
| | 0.626917375 | NA |
| | 0.715334491 | 0.819127853 |
| NA | | NA |
| | 0.520934974 | 0.671156554 |
| NA | | NA |
| NA | | NA |
| | 0.941278332 | NA |
| | 0.004953528 | 0.019379334 |
| | 0.709067061 | 0.814301563 |
| NA | | NA |
| | 0.022063272 | 0.064761375 |
| NA | | NA |

| | |
|---|---|
| NA | NA |
| NA | NA |
| NA | NA |
| NA | NA |
| NA | NA |
| 0.596229743 | 0.731120237 |
| 0.898764838 | 0.939004507 |
| 1.89E-33 | 1.53E-31 |
| NA | NA |
| 0.051255952 | 0.126463582 |
| NA | NA |
| 7.56E-32 | 5.71E-30 |
| 8.59E-44 | 1.00E-41 |
| NA | NA |
| 0.914158046 | NA |
| 0.520746927 | NA |
| NA | NA |
| NA | NA |
| 0.439889844 | 0.60073774 |
| 0.955792398 | NA |
| NA | NA |
| NA | NA |
| NA | NA |
| 0.167499292 | 0.306654731 |
| 0.183171254 | 0.325826878 |
| NA | NA |
| NA | NA |
| 0.392111609 | NA |
| 0.372329746 | 0.535674119 |
| 0.244517905 | 0.401020821 |
| 0.097203125 | 0.205267396 |
| 0.600166032 | 0.734115416 |
| 0.664832697 | 0.782560827 |
| 0.007346816 | 0.026734278 |
| 0.814102131 | NA |
| 4.54E-15 | 1.39E-13 |
| 0.706239142 | 0.812504425 |
| 0.000589211 | 0.003260731 |
| 0.042624927 | 0.109445364 |
| 0.270785973 | 0.430745688 |
| NA | NA |
| NA | NA |
| 0.121012356 | 0.241878854 |
| 8.22E-07 | 9.30E-06 |
| 0.334998774 | 0.498259128 |
| 0.018769312 | 0.057041261 |
| 0.818055147 | 0.888899731 |

| | | |
|---|---|---|
| | 0.726165958 | NA |
| NA | | NA |
| | 0.91823575 | NA |
| | 0.484961563 | 0.640530022 |
| | 0.180017071 | 0.321993412 |
| NA | | NA |
| | 0.563191042 | 0.705530167 |
| | 0.951797818 | NA |
| | 0.711077766 | 0.815773823 |
| | 0.001191513 | 0.005928175 |
| | 0.743511643 | NA |
| | 0.748393366 | NA |
| | 0.941277857 | NA |
| | 0.618441186 | NA |
| | 0.901547656 | NA |
| | 0.305320797 | 0.4669266 |
| | 0.000799114 | 0.004223962 |
| NA | | NA |
| | 0.030283819 | 0.083601886 |
| | 0.041840958 | 0.107933075 |
| | 0.889442731 | 0.933455095 |
| NA | | NA |
| NA | | NA |
| NA | | NA |
| | 0.86734997 | 0.920732494 |
| | 0.033436224 | 0.090468229 |
| | 0.083679964 | 0.183876524 |
| | 0.696023042 | 0.805305291 |
| | 0.941277857 | NA |
| NA | | NA |
| NA | | NA |
| | 0.066296784 | 0.154720663 |
| | 0.374230921 | 0.537369956 |
| | 0.94020055 | NA |
| | 0.008516889 | 0.030157543 |
| | 0.001505194 | 0.007195871 |
| | 0.918357387 | NA |
| | 0.010605712 | 0.036098548 |
| | 0.340134913 | 0.50383243 |
| | 0.839319108 | NA |
| | 0.007720177 | 0.027845986 |
| NA | | NA |
| | 0.020724379 | 0.061537134 |
| | 3.42E-08 | 4.94E-07 |
| | 0.007829959 | 0.028173647 |
| | 0.578349748 | 0.716599724 |
| | 0.089247042 | 0.192938973 |

| | | |
|---|---|---|
| | 0.772041469 | 0.859139326 |
| | 0.836894448 | 0.901142802 |
| | 0.955792398 | NA |
| | 0.000419252 | 0.002434524 |
| | 0.265024932 | 0.42416107 |
| | 0.983301731 | 0.989370601 |
| | 0.63727439 | 0.761580812 |
| | 5.67E-08 | 7.89E-07 |
| | 8.06E-05 | 0.000579318 |
| NA | | NA |
| | 0.888421782 | NA |
| | 0.154686355 | 0.289475952 |
| | 0.853595154 | NA |
| | 0.859274891 | NA |
| | 0.941278332 | NA |
| NA | | NA |
| | 0.874080268 | 0.924824793 |
| | 4.67E-06 | 4.58E-05 |
| | 0.459058641 | 0.618451782 |
| | 0.064030501 | 0.150553588 |
| | 2.39E-16 | 8.04E-15 |
| | 1.66E-19 | 6.83E-18 |
| | 0.698569821 | NA |
| | 0.267828565 | 0.427555701 |
| | 0.114531589 | 0.231947664 |
| | 0.005463399 | 0.020947137 |
| | 0.622859524 | 0.751218768 |
| | 0.204196015 | 0.352666312 |
| | 0.2434202 | NA |
| | 0.356659629 | 0.520299859 |
| | 2.80E-10 | 5.34E-09 |
| | 0.000113404 | 0.000781709 |
| | 0.307088932 | 0.468757183 |
| | 0.126769773 | 0.250176116 |
| | 0.352594019 | 0.516235643 |
| NA | | NA |
| | 0.019870226 | 0.059520654 |
| | 0.017925566 | 0.054947884 |
| | 0.001448609 | 0.006965614 |
| | 0.941275956 | NA |
| | 0.916526481 | 0.949682787 |
| | 0.258506775 | 0.416811967 |
| | 0.769070129 | 0.857023917 |
| | 0.005002018 | 0.019527316 |
| | 0.61059758 | 0.742070136 |
| | 0.028014198 | 0.078594289 |
| | 0.028580011 | 0.079889773 |

| | |
|---|---|
| 0.138909981 | 0.267611119 |
| 3.02E-10 | 5.73E-09 |
| 5.60E-29 | 3.74E-27 |
| 0.01601419 | 0.050266353 |
| 0.009420473 | 0.032772011 |
| 2.29E-07 | 2.89E-06 |
| 0.738662407 | 0.83551027 |
| 0.647970274 | 0.769890435 |
| NA | NA |
| 0.585691632 | 0.722349978 |
| 0.206799836 | 0.355756699 |
| 0.016365084 | 0.051145186 |
| NA | NA |
| 0.941275956 | NA |
| 0.308888043 | 0.470417379 |
| NA | NA |
| 0.08168421 | 0.180490862 |
| 0.00273326 | 0.011882228 |
| 0.551396716 | 0.696685493 |
| 0.799735322 | 0.877620713 |
| 0.523794811 | 0.673511106 |
| 0.379310795 | 0.542132974 |
| 0.728678906 | NA |
| 0.625829043 | NA |
| 0.83481082 | 0.899970585 |
| 7.15E-05 | 0.000520799 |
| 0.342950694 | 0.506417195 |
| 0.219403589 | 0.370898027 |
| 0.853595622 | NA |
| 0.189683891 | 0.334264369 |
| 0.843117477 | NA |
| 0.917207309 | 0.950263189 |
| 0.839319108 | NA |
| 9.17E-07 | 1.02E-05 |
| 0.056118002 | 0.135854424 |
| 4.54E-10 | 8.44E-09 |
| 0.057098197 | 0.137625448 |
| NA | NA |
| NA | NA |
| 0.940294516 | NA |
| NA | NA |
| NA | NA |
| NA | NA |
| 0.197696371 | 0.344755588 |
| 0.359015776 | 0.522583931 |
| 0.663492045 | 0.781571694 |
| 0.166989684 | 0.305980636 |

| | |
|---|---|
| NA | NA |
| 0.012400218 | 0.040953237 |
| 0.063501173 | 0.149560155 |
| 0.048422321 | 0.120647764 |
| 0.967610821 | 0.980522415 |
| 0.623202006 | 0.751385543 |
| 7.24E-18 | 2.68E-16 |
| 0.839319108 | NA |
| 0.148139386 | 0.280823006 |
| 0.05242153 | 0.128659235 |
| 0.689806998 | 0.801033623 |
| NA | NA |
| 0.705710115 | 0.812209632 |
| 0.045279369 | 0.114585871 |
| 4.92E-05 | 0.000374153 |
| 0.000181875 | 0.001182135 |
| 9.14E-09 | 1.45E-07 |
| 0.00575834 | 0.021891946 |
| 8.15E-07 | 9.23E-06 |
| 0.941277857 | NA |
| 0.839319574 | NA |
| 1.05E-09 | 1.87E-08 |
| 0.785573783 | NA |
| 0.013540089 | 0.043943441 |
| 0.365069126 | NA |
| 0.898829255 | NA |
| 0.000190952 | 0.001231773 |
| NA | NA |
| 0.207933855 | 0.357217006 |
| 0.000791961 | 0.004193607 |
| 0.818207482 | NA |
| 0.038815868 | 0.101638645 |
| 0.000104916 | 0.000729756 |
| NA | NA |
| NA | NA |
| 0.126532607 | 0.249811639 |
| NA | NA |
| 0.064614436 | 0.151671874 |
| 0.200737329 | 0.348348636 |
| NA | NA |
| NA | NA |
| 0.812386596 | NA |
| 0.177827182 | 0.319321374 |
| 0.117542644 | 0.236476208 |
| NA | NA |
| 0.776514509 | 0.862302806 |
| 0.092952654 | 0.198747671 |

| | | |
|---|---|---|
| | 0.035088237 | 0.093828809 |
| | 0.05827472 | 0.139725484 |
| | 0.412830267 | 0.575067948 |
| | 0.596820033 | 0.731564034 |
| | 0.015359347 | 0.0486146 |
| | 0.269740159 | 0.429685134 |
| | 0.247285257 | 0.404416908 |
| | 0.839319108 | NA |
| | 0.010828601 | 0.036736369 |
| | 1.92E-05 | 0.000161972 |
| | 0.906843833 | 0.943625266 |
| | 6.15E-09 | 9.91E-08 |
| | 0.69863054 | NA |
| NA | | NA |
| | 0.000716645 | 0.003854874 |
| | 0.879964369 | 0.928121235 |
| | 0.102507022 | 0.213690337 |
| NA | | NA |
| | 0.143376735 | 0.273954648 |
| | 0.000869293 | 0.004532511 |
| | 0.928181296 | NA |
| | 0.315733955 | 0.478211874 |
| | 4.39E-08 | 6.20E-07 |
| | 0.050150235 | 0.124069836 |
| | 0.952799628 | 0.971368344 |
| | 0.322659151 | 0.485470987 |
| | 0.008454757 | 0.029989646 |
| | 0.002237148 | 0.010050633 |
| | 0.593489266 | 0.728755032 |
| | 0.955791922 | NA |
| | 0.685401233 | 0.797955725 |
| NA | | NA |
| NA | | NA |
| NA | | NA |
| | 0.002395974 | 0.010647383 |
| | 0.926974691 | NA |
| | 0.191809764 | NA |
| | 0.669025802 | NA |
| | 0.000749957 | 0.004005941 |
| NA | | NA |
| | 0.000828697 | 0.004355186 |
| | 1.84E-13 | 4.93E-12 |
| | 0.955792398 | NA |
| | 0.002315183 | 0.010354376 |
| | 0.522067772 | NA |
| | 0.428170964 | NA |
| | 0.154035145 | 0.288753878 |

| | |
|---|---|
| 0.031610931 | 0.086484461 |
| 0.062079185 | 0.146879598 |
| NA | NA |
| 0.212560637 | NA |
| NA | NA |
| 0.197381292 | NA |
| 0.641100192 | 0.764578954 |
| 0.58298883 | 0.720435083 |
| 0.01272454 | 0.041809706 |
| 0.815291998 | NA |
| 0.839319108 | NA |
| 0.51241028 | 0.66382134 |
| 0.148505628 | 0.28140522 |
| 0.226132783 | 0.379176648 |
| 0.600848019 | 0.734592824 |
| 1.37E-08 | 2.11E-07 |
| 0.98698539 | 0.991860936 |
| 0.662071668 | 0.78032334 |
| 2.75E-77 | 6.88E-75 |
| 6.17E-05 | 0.000457948 |
| 6.20E-06 | 5.91E-05 |
| 0.002875105 | 0.012370437 |
| NA | NA |
| NA | NA |
| 0.637581518 | NA |
| 0.760958431 | NA |
| NA | NA |
| 0.773557461 | NA |
| 0.899267487 | NA |
| NA | NA |
| NA | NA |
| NA | NA |
| 0.839318641 | NA |
| NA | NA |
| NA | NA |
| 0.839319108 | NA |
| NA | NA |
| NA | NA |
| NA | NA |
| 0.225268683 | 0.37807398 |
| 0.81942151 | NA |
| 0.335835872 | NA |
| 0.60208899 | 0.73552259 |
| 0.098370184 | 0.207105344 |
| 0.941278332 | NA |
| 0.839319108 | NA |
| 0.435122187 | 0.59635245 |

| | |
|---|---|
| 0.128494508 | 0.252759034 |
| 4.59E-07 | 5.44E-06 |
| 0.103883747 | 0.215762734 |
| 3.03E-40 | 3.13E-38 |
| 0.059255185 | 0.141662506 |
| 0.987253915 | 0.991921388 |
| 0.313403781 | 0.475408629 |
| 0.685731546 | 0.798248264 |
| NA | NA |
| 0.033834568 | 0.091234891 |
| NA | NA |
| 0.631758097 | 0.757871772 |
| 0.790219486 | 0.871100603 |
| 0.626929147 | 0.754295434 |
| 1.16E-07 | 1.54E-06 |
| NA | NA |
| 0.510431017 | 0.661977827 |
| 0.514889439 | NA |
| 0.939477865 | NA |
| NA | NA |
| 0.955792398 | NA |
| 0.874892397 | 0.925190548 |
| 1.77E-09 | 3.05E-08 |
| 0.449858303 | NA |
| NA | NA |
| 0.755626499 | NA |
| 0.432041661 | 0.593287695 |
| 0.025408666 | 0.072593574 |
| NA | NA |
| 0.941275956 | NA |
| 0.941278838 | NA |
| NA | NA |
| 0.941277857 | NA |
| NA | NA |
| NA | NA |
| 0.953932387 | NA |
| 0.002364725 | 0.010534381 |
| NA | NA |
| NA | NA |
| 0.941278332 | NA |
| NA | NA |
| 0.191889082 | 0.337222676 |
| 0.955792873 | NA |
| 0.139012599 | 0.267765468 |
| 0.818799758 | 0.889505789 |
| 0.025762037 | 0.073400527 |
| 2.16E-05 | 0.000180484 |

| | |
|---|---|
| 0.85099766 | 0.91048553 |
| 0.001408225 | 0.006808564 |
| 0.029036781 | 0.08082095 |
| 4.78E-05 | 0.00036417 |
| 0.458673787 | 0.618146885 |
| NA | NA |
| 0.783393506 | 0.866827774 |
| 0.11869143 | 0.238285025 |
| 0.083530557 | 0.183666912 |
| 0.15457783 | 0.289409384 |
| 0.144655015 | 0.275865608 |
| 0.002503629 | 0.011049766 |
| 0.748028893 | 0.841684363 |
| 0.051380182 | 0.126710301 |
| 1.23E-08 | 1.90E-07 |
| 0.090496809 | 0.194998386 |
| 1.20E-13 | 3.26E-12 |
| 4.39E-07 | 5.24E-06 |
| NA | NA |
| 0.697988805 | NA |
| 0.09720923 | 0.205267396 |
| 0.663475436 | 0.781571694 |
| 0.839319107 | NA |
| 0.077659966 | 0.173891111 |
| 0.21252427 | 0.363033062 |
| 0.833618019 | 0.899282818 |
| 0.9278563 | NA |
| 0.043720034 | 0.111494136 |
| 0.786261696 | 0.868628494 |
| 0.298995906 | 0.460687503 |
| 0.001011229 | 0.005159544 |
| 0.975306314 | 0.985157056 |
| 0.067340224 | 0.156465444 |
| 1.95E-17 | 6.95E-16 |
| 2.20E-08 | 3.27E-07 |
| 2.45E-42 | 2.77E-40 |
| 0.373964356 | 0.537090133 |
| 0.219502415 | 0.371038739 |
| 2.66E-21 | 1.23E-19 |
| 9.86E-05 | 0.00069243 |
| 0.018752715 | 0.057005379 |
| 0.088546983 | 0.19194857 |
| 0.875032236 | 0.92525621 |
| 2.54E-10 | 4.87E-09 |
| 0.827693041 | 0.895308202 |
| 9.02E-07 | 1.01E-05 |
| 0.014423805 | 0.046225314 |

| | |
|---|---|
| 0.870308767 | 0.922433507 |
| 4.35E-06 | 4.29E-05 |
| 0.245954606 | 0.402877402 |
| 0.354749914 | NA |
| NA | NA |
| 0.002478465 | 0.010956986 |
| 0.00145039 | 0.006971364 |
| 0.000297195 | 0.001808705 |
| 0.734894545 | 0.832977211 |
| 0.001428651 | 0.006882157 |
| 0.048905157 | 0.121608915 |
| 1.64E-05 | 0.000140752 |
| NA | NA |
| 0.160149192 | NA |
| 0.051802018 | 0.127520077 |
| NA | NA |
| 0.067399865 | 0.156573452 |
| 0.031203996 | 0.085578113 |
| 0.95259943 | 0.971205844 |
| 0.955791922 | NA |
| 0.011640519 | 0.039003574 |
| 0.583186547 | NA |
| 0.223407318 | 0.375877641 |
| 0.034008332 | 0.091537522 |
| 0.00091568 | 0.004738046 |
| 0.486370132 | 0.641664161 |
| 0.695607795 | 0.805081541 |
| 0.098344443 | 0.207105344 |
| 0.131481174 | 0.256934691 |
| 0.494832434 | 0.647854874 |
| 0.143782465 | 0.274597628 |
| 0.985176375 | 0.990712243 |
| 0.365027964 | 0.528362661 |
| 0.001649802 | 0.007786752 |
| 0.631475822 | NA |
| 0.06007802 | 0.143183842 |
| 0.101684271 | 0.212380573 |
| 0.902504014 | 0.941134744 |
| 0.048293138 | 0.120459038 |
| 0.748393366 | NA |
| 0.501300487 | 0.653734067 |
| 0.002417651 | 0.01073047 |
| 0.134080196 | 0.260584374 |
| 0.923943097 | 0.954592539 |
| 0.689483513 | 0.800908677 |
| NA | NA |
| 0.859256872 | 0.916063427 |

| | | |
|---|---|---|
| | 0.371004606 | 0.534252019 |
| | 0.684266352 | 0.797279344 |
| | 0.100746027 | 0.210920655 |
| | 0.749692375 | 0.84281796 |
| | 6.82E-09 | 1.09E-07 |
| | 1.96E-10 | 3.82E-09 |
| | 0.022262764 | 0.065273033 |
| | 0.091714384 | 0.196778847 |
| | 3.65E-06 | 3.66E-05 |
| | 0.515104363 | 0.666223661 |
| | 0.098138643 | 0.206789659 |
| | 0.061671079 | 0.14617208 |
| | 0.012630992 | 0.041565446 |
| | 0.923140031 | 0.954250805 |
| | 0.941278284 | NA |
| NA | | NA |
| | 0.355119705 | 0.518882667 |
| | 0.40085242 | 0.563228413 |
| | 0.811781384 | 0.884991549 |
| | 0.091783625 | 0.196846835 |
| | 0.024619635 | 0.070866676 |
| | 0.728678906 | NA |
| | 0.713354003 | 0.817686106 |
| | 0.423006049 | 0.584352291 |
| | 0.56848707 | 0.70973665 |
| | 2.15E-07 | 2.72E-06 |
| | 0.003143551 | 0.013339775 |
| | 0.041873665 | 0.107970043 |
| | 0.004337783 | 0.017343847 |
| | 0.016137677 | 0.050547308 |
| | 0.911933701 | NA |
| | 0.082540747 | 0.181843287 |
| | 0.115469797 | 0.233252322 |
| | 0.760107969 | 0.850711895 |
| | 0.066404013 | 0.15492534 |
| | 0.131821098 | 0.257349555 |
| | 0.520948548 | 0.671156554 |
| | 0.795155502 | 0.874314744 |
| | 0.203845455 | 0.352355085 |
| | 0.172072883 | 0.31233728 |
| | 0.017365566 | 0.053615044 |
| | 0.226714203 | 0.379883947 |
| | 0.952950987 | NA |
| | 0.179982097 | 0.321993412 |
| | 0.913035137 | 0.947550135 |
| | 0.883855033 | 0.930740057 |
| | 0.965458583 | 0.979108402 |

| | | |
|---|---|---|
| | 0.021526256 | 0.063474721 |
| | 0.458530302 | NA |
| | 2.31E-05 | 0.000191695 |
| | 4.42E-08 | 6.25E-07 |
| | 1.23E-05 | 0.000109021 |
| | 0.853595154 | NA |
| NA | | NA |
| | 0.654012956 | 0.774121854 |
| | 0.675525259 | 0.790858573 |
| | 0.582810471 | NA |
| | 0.00628527 | 0.023518628 |
| | 0.209793016 | 0.359476881 |
| | 0.281840773 | 0.442038355 |
| | 0.955791922 | NA |
| | 0.814941777 | 0.886893337 |
| | 0.00430311 | 0.017225466 |
| | 0.274311037 | 0.43419697 |
| | 0.941277857 | NA |
| | 0.656107022 | 0.775636204 |
| | 0.291494171 | 0.45273372 |
| | 0.189939591 | 0.334636725 |
| | 0.474960562 | 0.632341722 |
| | 0.007015765 | 0.025714666 |
| | 0.100950209 | 0.211272613 |
| | 0.021950947 | 0.064495301 |
| | 0.814074659 | 0.886443528 |
| | 0.941278332 | NA |
| | 0.231313119 | 0.385392332 |
| NA | | NA |
| | 0.027223783 | 0.076929892 |
| | 0.215317884 | 0.36625429 |
| | 0.611330928 | 0.742468556 |
| | 0.891532448 | 0.934565988 |
| | 1.64E-08 | 2.48E-07 |
| | 0.339423717 | 0.503080099 |
| | 0.777388411 | NA |
| | 0.008810841 | 0.031023025 |
| | 0.333150194 | 0.496409223 |
| | 0.351622764 | 0.515282306 |
| | 0.839319108 | NA |
| NA | | NA |
| | 5.25E-05 | 0.000395918 |
| | 0.014398223 | 0.046167569 |
| | 0.124741811 | 0.247301732 |
| | 2.49E-15 | 7.78E-14 |
| | 4.80E-06 | 4.70E-05 |
| | 0.208451864 | 0.35777122 |

| | |
|---|---|
| NA | NA |
| 0.915432081 | 0.949021199 |
| 0.941277857 | NA |
| 0.001996836 | 0.009157744 |
| 0.70907796 | NA |
| 0.804607847 | 0.880558921 |
| 0.779189193 | NA |
| 0.000319895 | 0.001926598 |
| 0.007775411 | 0.02801125 |
| 0.126722212 | 0.250123732 |
| 0.839319108 | NA |
| 1.04E-05 | 9.36E-05 |
| 0.800313954 | 0.87808024 |
| 0.006345457 | 0.023687983 |
| 0.614088469 | 0.744311876 |
| 0.418173757 | 0.579763034 |
| NA | NA |
| 0.001825031 | 0.008469366 |
| NA | NA |
| 0.86971473 | 0.921968112 |
| 0.025980805 | 0.073917657 |
| 0.32560621 | 0.488470883 |
| 0.702673468 | 0.810252828 |
| 3.30E-05 | 0.000262013 |
| 0.037724532 | 0.099426345 |
| 0.217513359 | 0.368835107 |
| 3.78E-10 | 7.09E-09 |
| 0.849641404 | NA |
| 0.045581542 | 0.115197692 |
| 0.941278332 | NA |
| 6.35E-05 | 0.000469692 |
| 1.81E-07 | 2.33E-06 |
| NA | NA |
| 0.079754777 | 0.177413384 |
| 0.55593099 | 0.700231223 |
| 0.01443234 | 0.046246439 |
| 0.003534934 | 0.014676174 |
| NA | NA |
| 0.328467107 | 0.491369121 |
| 0.165437652 | 0.304073127 |
| 0.067348416 | 0.156469205 |
| 0.527168646 | 0.676241881 |
| 0.147784776 | 0.280405674 |
| NA | NA |
| 0.055610043 | 0.13485042 |
| 0.609470037 | NA |
| NA | NA |

| | | |
|---|---|---|
| | 0.424304047 | NA |
| | 0.941275481 | NA |
| NA | | NA |
| | 0.001625117 | 0.007687012 |
| | 0.620039831 | 0.748968897 |
| | 0.631388758 | NA |
| | 0.078254367 | 0.174860162 |
| | 0.408276385 | 0.570360048 |
| | 0.052110824 | 0.12808156 |
| | 0.662883024 | 0.781086225 |
| NA | | NA |
| | 0.145894772 | 0.277629306 |
| | 2.14E-06 | 2.25E-05 |
| | 0.400138711 | 0.562723875 |
| | 0.435144451 | 0.59635245 |
| | 0.882808626 | 0.929843832 |
| | 9.03E-07 | 1.01E-05 |
| | 0.174153525 | 0.31498497 |
| | 0.035761407 | 0.095350398 |
| | 0.002998154 | 0.012821065 |
| NA | | NA |
| | 0.544495894 | 0.691042149 |
| | 0.001889525 | 0.008722743 |
| | 0.673589745 | 0.789286817 |
| | 0.001013732 | 0.00516899 |
| | 0.012187018 | 0.040389223 |
| | 0.012867341 | 0.042191548 |
| | 7.60E-05 | 0.000549234 |
| | 0.584793116 | 0.721733424 |
| | 0.421888507 | 0.583193984 |
| | 3.41E-07 | 4.14E-06 |
| | 0.59853755 | 0.732876731 |
| | 0.105854247 | 0.21883815 |
| | 0.839318641 | NA |
| | 0.191944261 | 0.337269843 |
| | 0.542126333 | 0.689115672 |
| | 0.000124716 | 0.000851305 |
| NA | | NA |
| | 0.009795984 | 0.033850756 |
| | 0.310978709 | 0.473026281 |
| | 0.052339924 | 0.128525212 |
| | 0.0040116 | 0.016252614 |
| | 0.593019324 | NA |
| | 0.42876819 | 0.590052062 |
| | 0.263037384 | 0.422001605 |
| | 0.723106317 | 0.824615588 |
| NA | | NA |

| | | | |
|---|---|---|---|
| NA | | NA | |
| | 0.539235523 | | 0.686499168 |
| NA | | NA | |
| | 0.955792398 | NA | |
| | 0.941279303 | NA | |
| NA | | NA | |
| | 0.451219312 | | 0.611195201 |
| | 0.611527083 | NA | |
| | 0.938609606 | NA | |
| NA | | NA | |
| NA | | NA | |
| | 0.166060323 | | 0.304887983 |
| | 0.470302444 | NA | |
| | 0.006247028 | | 0.023401284 |
| | 0.469455235 | | 0.627469312 |
| | 0.018369074 | | 0.05604675 |
| | 2.92E-12 | | 7.00E-11 |
| | 0.700582578 | NA | |
| | 0.591414637 | | 0.727147641 |
| | 2.23E-10 | | 4.29E-09 |
| | 0.039512224 | | 0.103155639 |
| | 0.980766005 | | 0.988113966 |
| | 0.579271026 | | 0.717293151 |
| | 0.944591465 | NA | |
| | 0.000727928 | | 0.003908852 |
| | 0.6547772 | NA | |
| | 0.147823187 | NA | |
| | 0.740666951 | | 0.837021877 |
| | 0.058975595 | | 0.141064929 |
| NA | | NA | |
| | 0.582614617 | NA | |
| | 0.029691461 | | 0.082310292 |
| | 0.955792398 | NA | |
| NA | | NA | |
| | 0.003801483 | | 0.015552235 |
| NA | | NA | |
| | 2.15E-05 | | 0.000179654 |
| | 0.530446272 | | 0.679274878 |
| NA | | NA | |
| | 0.237522979 | | 0.392973805 |
| | 0.955792873 | NA | |
| | 0.000535685 | | 0.003011427 |
| | 0.787935297 | | 0.86971044 |
| | 0.006510127 | | 0.024196447 |
| NA | | NA | |
| | 0.072839815 | | 0.165812287 |
| | 0.06389718 | | 0.150314357 |

| | |
|---|---|
| 0.295900041 | 0.457397894 |
| 0.455007015 | 0.614819783 |
| 0.904458512 | NA |
| 0.011947617 | 0.039750795 |
| NA | NA |
| 0.575354264 | 0.714283149 |
| 0.555817378 | 0.700162224 |
| 0.287521176 | 0.448112823 |
| 0.183312885 | 0.325977832 |
| 0.567661155 | 0.709187479 |
| 0.809376489 | 0.883178802 |
| NA | NA |
| 0.154340739 | 0.289033695 |
| 0.901447884 | NA |
| NA | NA |
| 0.780839874 | 0.865187191 |
| NA | NA |
| NA | NA |
| 0.799654241 | NA |
| NA | NA |
| 5.31E-05 | 0.000400393 |
| NA | NA |
| NA | NA |
| 1.37E-07 | 1.80E-06 |
| 0.807905191 | 0.882239978 |
| 0.839319108 | NA |
| 0.051483258 | 0.126879738 |
| 0.095801028 | 0.203158952 |
| NA | NA |
| 0.184578037 | 0.327655821 |
| 0.010200252 | 0.034948379 |
| 0.941278332 | NA |
| 0.925588805 | 0.955329983 |
| 0.739812114 | 0.836336778 |
| 0.135826418 | 0.263183034 |
| NA | NA |
| 0.941278332 | NA |
| 0.464445838 | 0.622942626 |
| 0.168285203 | 0.307738507 |
| 0.507364649 | 0.659042424 |
| 0.083220628 | 0.183019254 |
| 0.84326491 | 0.905661115 |
| NA | NA |
| NA | NA |
| NA | NA |
| NA | NA |
| NA | NA |

| | | | |
|---|---|---|---|
| NA | | NA | |
| NA | | NA | |
| NA | | NA | |
| NA | | NA | |
| NA | | NA | |
| NA | | NA | |
| NA | | NA | |
| NA | | NA | |
| NA | | NA | |
| | 3.56E-05 | | 0.000280371 |
| | 0.116890308 | | 0.235375222 |
| | 0.043342118 | | 0.110796809 |
| | 0.604518398 | NA | |
| NA | | NA | |
| | 0.853596091 | NA | |
| | 0.024700688 | | 0.070996954 |
| | 0.505864417 | | 0.657596115 |
| | 0.955791446 | NA | |
| | 0.003978183 | | 0.016136452 |
| | 0.155111875 | | 0.290044217 |
| | 0.665955935 | | 0.783351568 |
| NA | | NA | |
| | 0.091026954 | | 0.195916388 |
| NA | | NA | |
| | 0.012735959 | | 0.041835673 |
| NA | | NA | |
| NA | | NA | |
| NA | | NA | |
| NA | | NA | |
| NA | | NA | |
| NA | | NA | |
| NA | | NA | |
| NA | | NA | |
| NA | | NA | |
| NA | | NA | |
| NA | | NA | |
| NA | | NA | |
| NA | | NA | |
| NA | | NA | |
| | 0.454596458 | NA | |
| | 0.861833744 | NA | |
| | 0.009166363 | | 0.032056682 |
| NA | | NA | |
| | 0.143999043 | | 0.274923014 |
| NA | | NA | |
| | 3.96E-05 | | 0.000307906 |
| | 0.284667466 | | 0.44526878 |

| | | | |
|---|---|---|---|
| | 0.017727682 | | 0.05445977 |
| | 0.001794447 | | 0.00835513 |
| NA | | NA | |
| NA | | NA | |
| NA | | NA | |
| | 0.01455593 | | 0.04656098 |
| | 0.30435238 | | 0.465933994 |
| | 0.892378243 | | 0.935040806 |
| | 0.955792873 | NA | |
| | 0.004190981 | | 0.01686166 |
| NA | | NA | |
| | 0.793759147 | NA | |
| | 0.922893457 | NA | |
| | 0.839319108 | NA | |
| | 0.865603351 | | 0.919767844 |
| | 0.939734005 | | 0.963757575 |
| | 0.084265641 | | 0.184941521 |
| | 0.000447856 | | 0.00258046 |
| | 0.773242253 | NA | |
| NA | | NA | |
| | 0.584836065 | | 0.721733424 |
| | 0.650344471 | NA | |
| | 0.941275956 | NA | |
| | 0.873518507 | NA | |
| | 0.612091074 | | 0.74305053 |
| | 0.189423058 | | 0.333973899 |
| | 0.000157524 | | 0.001044104 |
| NA | | NA | |
| NA | | NA | |
| NA | | NA | |
| NA | | NA | |
| | 0.769668652 | | 0.857314759 |
| | 0.107418741 | | 0.221201858 |
| | 0.07536305 | | 0.170168046 |
| NA | | NA | |
| | 0.000382607 | | 0.002246932 |
| | 0.126421127 | | 0.249653668 |
| | 0.434859332 | | 0.596228215 |
| | 0.607662797 | | 0.740244637 |
| | 0.332515995 | | 0.495712496 |
| | 0.871775126 | | 0.92337093 |
| NA | | NA | |
| | 0.839320041 | NA | |
| | 0.32953093 | | 0.492538266 |
| | 9.61E-15 | | 2.83E-13 |
| | 0.814797224 | NA | |
| | 0.93827938 | NA | |

| | |
|---|---|
| NA | NA |
| NA | NA |
| 0.747404088 | 0.841359824 |
| 0.032813872 | 0.089149159 |
| 0.702218908 | NA |
| 0.355944444 | 0.519582649 |
| 0.381644675 | 0.544142 |
| 0.837259124 | NA |
| NA | NA |
| NA | NA |
| 0.000380342 | 0.00223473 |
| NA | NA |
| 0.199855473 | 0.347299406 |
| NA | NA |
| NA | NA |
| 0.689749924 | 0.801006425 |
| 0.002083836 | 0.009461935 |
| 0.839319108 | NA |
| 3.12E-10 | 5.90E-09 |
| 0.237717682 | 0.393200575 |
| 0.000138438 | 0.000932663 |
| 0.526219138 | 0.675351446 |
| 0.000112283 | 0.000775777 |
| 0.002014843 | 0.009213695 |
| 0.297457717 | 0.45887334 |
| 0.033831125 | 0.091234891 |
| 0.839318641 | NA |
| 0.366933049 | 0.530095138 |
| 0.955792398 | NA |
| 0.630671701 | 0.756917689 |
| 0.871209835 | NA |
| 0.92403559 | 0.954592539 |
| 0.271391631 | 0.431276779 |
| 0.724931882 | 0.826238017 |
| 0.731130588 | 0.830477641 |
| 0.955792398 | NA |
| 0.941278332 | NA |
| 0.023991179 | 0.069418678 |
| 0.01066781 | 0.036278783 |
| NA | NA |
| 0.000343154 | 0.002048084 |
| 0.002012899 | 0.009206574 |
| 0.941430486 | NA |
| 0.034676295 | 0.092936062 |
| 0.160399776 | 0.29712966 |
| NA | NA |
| 0.718880549 | 0.821055414 |

| | |
|---|---|
| 0.833357167 | NA |
| NA | NA |
| 6.72E-05 | 0.000492572 |
| 0.730900578 | 0.830396998 |
| 0.147825768 | 0.280405674 |
| NA | NA |
| NA | NA |
| 0.043854149 | 0.111781586 |
| 0.087590747 | 0.190326361 |
| 4.00E-11 | 8.47E-10 |
| 0.027856398 | 0.078280753 |
| 0.84547085 | 0.906890364 |
| 2.28E-13 | 6.05E-12 |
| NA | NA |
| 0.584339232 | 0.721417285 |
| 0.014679345 | 0.046870857 |
| 0.205581635 | NA |
| NA | NA |
| NA | NA |
| 0.955792873 | NA |
| NA | NA |
| NA | NA |
| 0.689941883 | NA |
| 1.11E-11 | 2.49E-10 |
| 0.991386729 | 0.994143365 |
| 0.351772839 | NA |
| NA | NA |
| NA | NA |
| 0.18130863 | 0.323646919 |
| 0.788543491 | 0.869909616 |
| 0.030500309 | 0.084040488 |
| 0.082506089 | 0.181817418 |
| 0.002153499 | 0.00972814 |
| NA | NA |
| 0.839318641 | NA |
| 3.59E-08 | 5.17E-07 |
| 1.61E-12 | 3.95E-11 |
| 0.066848815 | 0.155608688 |
| 0.233523006 | 0.388123357 |
| NA | NA |
| 5.59E-06 | 5.39E-05 |
| 0.50717989 | NA |
| NA | NA |
| 0.562580242 | NA |
| NA | NA |
| 0.657782955 | NA |
| 0.941277857 | NA |

| | |
|---|---|
| NA | NA |
| 0.342782611 | 0.506230173 |
| 0.744245682 | 0.839341252 |
| 0.587643687 | 0.723857322 |
| 0.182029502 | NA |
| 0.085097926 | 0.186321482 |
| 0.728878986 | NA |
| 0.360948306 | NA |
| 0.223318905 | 0.375782014 |
| 2.71E-10 | 5.16E-09 |
| 3.58E-14 | 1.02E-12 |
| 6.26E-06 | 5.96E-05 |
| 2.77E-08 | 4.06E-07 |
| 0.037568894 | 0.09906002 |
| 0.911801817 | NA |
| 0.928613358 | 0.95745528 |
| 0.23747256 | 0.392953742 |
| 0.941278332 | NA |
| 0.07730633 | 0.173258051 |
| 1.86E-06 | 1.98E-05 |
| 0.967357373 | 0.980414741 |
| 0.028453735 | 0.07961159 |
| NA | NA |
| 0.826131618 | 0.894368452 |
| NA | NA |
| 0.004498604 | 0.017896669 |
| 1.28E-05 | 0.000112844 |
| 0.304798482 | 0.466427509 |
| NA | NA |
| 0.784609422 | 0.867627515 |
| 0.431344212 | 0.592774534 |
| NA | NA |
| 6.78E-06 | 6.39E-05 |
| 0.345330216 | 0.509013488 |
| NA | NA |
| 0.756098927 | NA |
| 0.407826015 | NA |
| 0.17653236 | 0.318151226 |
| 0.001117724 | 0.005617414 |
| 0.769681501 | 0.857314759 |
| NA | NA |
| 0.097435111 | 0.20563491 |
| 0.476342216 | 0.633402091 |
| 0.396185893 | 0.558887141 |
| 0.135429634 | 0.262614991 |
| 0.019330502 | 0.058329692 |
| 0.415587815 | 0.577422286 |

| | |
|---|---|
| 0.652949583 | 0.773665923 |
| 5.85E-05 | 0.000435924 |
| 0.128231884 | 0.252391904 |
| 7.71E-09 | 1.23E-07 |
| 0.839318641 | NA |
| 0.385053853 | 0.547854135 |
| 0.446140805 | 0.606629159 |
| 0.06789362 | 0.157459217 |
| 0.038508176 | 0.100999626 |
| 0.934140779 | 0.960895291 |
| 0.035034294 | 0.093705614 |
| 0.704826286 | 0.811742162 |
| 0.844635451 | 0.90649091 |
| 0.030861853 | 0.084823768 |
| 0.603844203 | 0.736872583 |
| 0.882014033 | 0.929459335 |
| 1.41E-09 | 2.46E-08 |
| 0.406134021 | 0.568033962 |
| 0.943708389 | 0.966072302 |
| 9.19E-22 | 4.37E-20 |
| 0.180594803 | 0.322663077 |
| 0.266818297 | 0.426543697 |
| 0.839318641 | NA |
| 0.000390581 | 0.002289807 |
| NA | NA |
| 4.11E-05 | 0.000318276 |
| 0.870935168 | 0.922814656 |
| 0.001600914 | 0.007584583 |
| 0.686705874 | 0.7989917 |
| 0.806931413 | 0.88170566 |
| 0.136608825 | 0.264333246 |
| 2.84E-11 | 6.13E-10 |
| 0.172922256 | 0.313271336 |
| 0.667378337 | NA |
| 0.427121 | 0.588567314 |
| 0.169664613 | 0.309252392 |
| 0.21725511 | 0.368604775 |
| 0.018662166 | 0.056780886 |
| 0.166188148 | 0.305028556 |
| 0.000138851 | 0.000934651 |
| 0.335405238 | 0.498588213 |
| NA | NA |
| 0.095137256 | 0.202111475 |
| 0.158766623 | 0.295047093 |
| 0.457118699 | NA |
| 0.941277857 | NA |
| 0.955792873 | NA |

| | | |
|---|---|---|
| NA | | NA |
| | 0.6720955 | NA |
| NA | | NA |
| | 0.389478266 | 0.55206213 |
| NA | | NA |
| | 0.955792873 | NA |
| | 0.394568594 | 0.55709584 |
| | 0.003040031 | 0.012967531 |
| | 0.042342224 | 0.108891503 |
| | 0.159537699 | 0.296063957 |
| | 0.110381154 | 0.225712391 |
| | 0.839319108 | NA |
| NA | | NA |
| | 0.301937219 | 0.46327014 |
| | 0.00080249 | 0.004238045 |
| | 2.90E-10 | 5.51E-09 |
| | 0.308000962 | 0.469547102 |
| NA | | NA |
| | 0.154428789 | 0.289175834 |
| | 1.68E-14 | 4.86E-13 |
| | 0.584969408 | NA |
| | 0.007004805 | 0.025682127 |
| | 2.15E-212 | 3.65E-209 |
| | 0.004831489 | 0.018979883 |
| NA | | NA |
| | 0.250926656 | 0.408514581 |
| | 0.697062943 | 0.805881687 |
| | 0.220517238 | 0.372383916 |
| | 0.947005058 | 0.967625072 |
| | 0.510093707 | 0.661684584 |
| | 0.00478017 | 0.018831117 |
| NA | | NA |
| | 0.349837246 | NA |
| | 0.818345676 | NA |
| | 0.886603457 | NA |
| | 0.203403574 | 0.351718987 |
| | 0.256268715 | 0.414385156 |
| NA | | NA |
| | 0.70223435 | 0.810011243 |
| | 0.172303084 | 0.312562496 |
| | 0.936826522 | NA |
| | 0.074515317 | 0.168610473 |
| | 0.24630715 | 0.403260613 |
| | 0.005216525 | 0.020208939 |
| | 6.83E-06 | 6.44E-05 |
| | 1.90E-14 | 5.51E-13 |
| NA | | NA |

| | | | |
|---|---|---|---|
| | 0.524339107 | NA | |
| | 0.037283016 | | 0.098448007 |
| | 0.010407314 | | 0.035560564 |
| | 0.004184557 | | 0.016844379 |
| NA | | NA | |
| | 0.020385085 | | 0.060795114 |
| | 0.955792398 | NA | |
| | 5.22E-08 | | 7.29E-07 |
| | 0.690999566 | | 0.801792603 |
| | 0.808884665 | | 0.882884982 |
| | 0.163403942 | | 0.301335758 |
| | 0.791812046 | | 0.871788609 |
| | 1.49E-07 | | 1.94E-06 |
| | 0.64853733 | | 0.770264094 |
| | 0.042607527 | | 0.109412488 |
| | 2.59E-05 | | 0.000212045 |
| | 0.016335373 | | 0.051059035 |
| | 2.91E-10 | | 5.53E-09 |
| | 0.219521949 | | 0.371045407 |
| | 0.217312763 | | 0.368646453 |
| | 0.04058004 | | 0.105372254 |
| | 0.049275862 | | 0.122262489 |
| | 0.214688693 | | 0.365549772 |
| | 8.18E-08 | | 1.11E-06 |
| | 0.978402328 | | 0.986609566 |
| | 0.964186428 | | 0.978200646 |
| | 0.00197371 | | 0.009060412 |
| | 0.606434341 | NA | |
| | 0.941277857 | NA | |
| NA | | NA | |
| | 0.955792398 | NA | |
| NA | | NA | |
| NA | | NA | |
| NA | | NA | |
| NA | | NA | |
| NA | | NA | |
| NA | | NA | |
| NA | | NA | |
| NA | | NA | |
| NA | | NA | |
| NA | | NA | |
| | 0.839318641 | NA | |
| NA | | NA | |
| NA | | NA | |
| NA | | NA | |
| NA | | NA | |
| NA | | NA | |

| | | |
|---|---|---|
| NA | | NA |
| NA | | NA |
| | 0.839319108 | NA |
| NA | | NA |
| NA | | NA |
| NA | | NA |
| NA | | NA |
| | 0.955791922 | NA |
| NA | | NA |
| NA | | NA |
| NA | | NA |
| NA | | NA |
| NA | | NA |
| NA | | NA |
| | 0.941278332 | NA |
| NA | | NA |
| NA | | NA |
| NA | | NA |
| NA | | NA |
| NA | | NA |
| NA | | NA |
| NA | | NA |
| NA | | NA |
| NA | | NA |
| NA | | NA |
| NA | | NA |
| | 0.941278332 | NA |
| | 0.941278332 | NA |
| NA | | NA |
| NA | | NA |
| NA | | NA |
| NA | | NA |
| NA | | NA |
| | 0.839318641 | NA |
| NA | | NA |
| NA | | NA |
| NA | | NA |
| NA | | NA |
| NA | | NA |
| NA | | NA |
| NA | | NA |
| | 0.839318641 | NA |
| NA | | NA |
| NA | | NA |
| | 0.839318641 | NA |
| NA | | NA |
| | 0.839318641 | NA |

| | |
|---|---|
| 0.023514381 | 0.068254792 |
| 2.98E-06 | 3.03E-05 |
| 0.667803686 | 0.784735757 |
| 0.009271421 | 0.032348045 |
| 0.348608883 | 0.512525091 |
| 0.016984903 | 0.052660499 |
| 0.101329956 | 0.211789218 |
| NA | NA |
| 0.939052821 | NA |
| 0.567219657 | NA |
| 0.125510149 | 0.248452477 |
| 0.376633076 | 0.539647272 |
| 0.040204358 | 0.104606924 |
| 0.992126026 | 0.994465748 |
| 0.098555407 | 0.207355767 |
| 0.017566071 | 0.05407117 |
| 0.204894188 | 0.353497802 |
| 0.134998 | 0.261983074 |
| 0.840751682 | 0.903905046 |
| 1.62E-05 | 0.000139161 |
| 0.009623526 | 0.03336623 |
| 2.14E-22 | 1.06E-20 |
| 0.000849902 | 0.00444537 |
| 0.988095283 | 0.992515358 |
| 0.009053033 | 0.031736453 |
| 0.018277908 | 0.055833005 |
| 0.869409418 | 0.921726558 |
| 0.913964304 | NA |
| 5.99E-16 | 1.96E-14 |
| 4.76E-06 | 4.66E-05 |
| 0.000280348 | 0.001718417 |
| 0.119379394 | 0.239382337 |
| 0.459572986 | 0.618750054 |
| NA | NA |
| NA | NA |
| NA | NA |
| 0.372331545 | NA |
| 0.048362337 | 0.120560924 |
| NA | NA |
| NA | NA |
| NA | NA |
| NA | NA |
| 0.363110386 | 0.526804426 |
| 0.832162011 | NA |
| 0.576147374 | 0.71487566 |
| 0.0005034 | 0.002853521 |
| 0.839319574 | NA |

| | |
|---|---|
| NA | NA |
| 0.140790317 | 0.270336489 |
| 0.116652255 | 0.235062392 |
| 0.00013901 | 0.000935459 |
| 3.29E-05 | 0.000261191 |
| 0.715122903 | 0.818964364 |
| 0.476544784 | 0.633524321 |
| 0.003379733 | 0.014154287 |
| 0.042905475 | 0.109975891 |
| 0.124889695 | 0.247512451 |
| 0.381291517 | 0.543833934 |
| 0.390129147 | 0.552745344 |
| 0.082661033 | 0.182044258 |
| 0.494418693 | 0.647540921 |
| NA | NA |
| 0.397762721 | 0.560442093 |
| 0.554797209 | 0.699580587 |
| 0.991205924 | 0.994087701 |
| 0.416947144 | 0.578490542 |
| 0.11505332 | 0.232687494 |
| 0.014726244 | 0.046979502 |
| 0.003191089 | 0.013512415 |
| 0.839318641 | NA |
| 1.56E-09 | 2.70E-08 |
| 1.48E-12 | 3.64E-11 |
| NA | NA |
| NA | NA |
| 0.782683574 | 0.866267226 |
| 0.128290677 | 0.252467072 |
| 0.001037729 | 0.00527009 |
| 0.50887713 | 0.660466407 |
| NA | NA |
| 0.47903379 | NA |
| 0.135958446 | 0.263374536 |
| 0.989860654 | 0.993351574 |
| 0.876779417 | 0.926202931 |
| 0.930494535 | 0.958605909 |
| 0.043382124 | 0.110878265 |
| 0.274323756 | 0.43419697 |
| 0.919159156 | NA |
| 0.431791798 | 0.593074652 |
| 0.698601548 | 0.807190003 |
| 0.070838798 | 0.162618687 |
| 0.15072635 | 0.284345612 |
| 0.725028456 | NA |
| NA | NA |
| 0.601688556 | 0.735222086 |

| | |
|---|---|
| 0.001532174 | 0.007303987 |
| 0.070862254 | 0.162626193 |
| 0.989197874 | 0.993078063 |
| 0.003443145 | 0.01436273 |
| 4.16E-05 | 0.000321739 |
| NA | NA |
| 0.43515038 | 0.59635245 |
| NA | NA |
| NA | NA |
| 0.063037629 | 0.148671756 |
| 3.28E-05 | 0.000260343 |
| NA | NA |
| 0.955792873 | NA |
| 0.775439144 | 0.861470362 |
| 0.838197868 | NA |
| 0.835561971 | 0.900129066 |
| 0.000224279 | 0.001416369 |
| 0.71749638 | 0.820340515 |
| 9.46E-13 | 2.38E-11 |
| 0.053365922 | 0.13051949 |
| 0.264088295 | 0.423114913 |
| 0.050133664 | 0.124053309 |
| 0.566927662 | 0.708644691 |
| 0.679554793 | 0.793747937 |
| 0.499134737 | NA |
| 0.645572798 | 0.767952477 |
| 0.784943973 | NA |
| 1.20E-08 | 1.87E-07 |
| 0.011179526 | 0.037696759 |
| 0.149322938 | 0.282370094 |
| 0.554992115 | 0.699641032 |
| 0.736143523 | 0.833647157 |
| 0.270876216 | 0.430802866 |
| 0.344538879 | 0.508098786 |
| 0.819307323 | NA |
| 0.003434715 | 0.014332588 |
| 0.009715704 | 0.03365151 |
| NA | NA |
| 3.42E-05 | 0.000270458 |
| 0.013796469 | 0.044593096 |
| 0.986695861 | 0.991653713 |
| 0.185782085 | 0.329212715 |
| 0.012275654 | 0.040626406 |
| 0.794886897 | NA |
| 0.526176698 | NA |
| 0.783578429 | NA |
| 0.123615457 | 0.245559598 |

| | |
|---|---|
| NA | NA |
| 0.15732585 | 0.293090946 |
| 2.68E-33 | 2.15E-31 |
| 1.27E-05 | 0.000112206 |
| NA | NA |
| 0.839318641 | NA |
| 0.18216927 | 0.324553466 |
| NA | NA |
| 0.004709994 | 0.018610107 |
| 0.116292149 | 0.234594943 |
| 0.300218389 | 0.461793853 |
| 0.642207747 | NA |
| 0.941278332 | NA |
| 0.932196201 | NA |
| 0.062030994 | 0.14678017 |
| 0.941278332 | NA |
| NA | NA |
| NA | NA |
| 0.678173923 | NA |
| NA | NA |
| NA | NA |
| 0.933274841 | NA |
| NA | NA |
| 6.13E-07 | 7.09E-06 |
| NA | NA |
| 0.000548941 | 0.003077949 |
| 8.99E-08 | 1.22E-06 |
| 0.326901704 | NA |
| NA | NA |
| NA | NA |
| NA | NA |
| NA | NA |
| 0.165869885 | NA |
| 0.22159652 | 0.373755696 |
| NA | NA |
| 0.295006216 | 0.456401557 |
| NA | NA |
| 0.039305986 | 0.102727959 |
| 0.590753046 | 0.726709598 |
| 0.093471863 | 0.199504256 |
| 4.60E-12 | 1.08E-10 |
| 0.933393074 | 0.960386545 |
| 0.525453154 | 0.674768598 |
| NA | NA |
| 0.134776344 | 0.261673875 |
| NA | NA |
| NA | NA |

| | | |
|---|---|---|
| | 0.000743774 | 0.003977378 |
| | 0.762555078 | 0.852608792 |
| | 0.244777948 | 0.401336689 |
| | 0.292887673 | NA |
| | 0.723490139 | NA |
| | 0.362530103 | 0.526251236 |
| | 0.069572146 | 0.160491978 |
| | 0.787036781 | NA |
| | 0.941275956 | NA |
| | 0.772910771 | NA |
| | 9.38E-36 | 8.30E-34 |
| | 0.839319108 | NA |
| | 0.411626023 | 0.573726217 |
| | 0.128682123 | 0.252996404 |
| | 0.556897593 | NA |
| NA | | NA |
| | 0.185423639 | 0.328699893 |
| NA | | NA |
| | 0.941277857 | NA |
| | 0.010296039 | 0.035235993 |
| | 0.202329427 | 0.350447171 |
| NA | | NA |
| NA | | NA |
| | 0.654823247 | NA |
| NA | | NA |
| | 0.917747111 | NA |
| NA | | NA |
| NA | | NA |
| NA | | NA |
| NA | | NA |
| NA | | NA |
| NA | | NA |
| | 0.561086181 | 0.70404203 |
| NA | | NA |
| NA | | NA |
| NA | | NA |
| NA | | NA |
| NA | | NA |
| | 0.015476823 | 0.048889097 |
| | 0.598478652 | 0.732842313 |
| | 0.003027126 | 0.012924062 |
| | 0.445927242 | 0.606402228 |
| | 0.26910788 | 0.428965026 |
| NA | | NA |
| NA | | NA |
| NA | | NA |
| | 0.001762611 | 0.008235898 |

| | |
|---|---|
| 0.994951302 | 0.996332656 |
| 0.243117403 | 0.399577436 |
| 0.789718901 | 0.870710083 |
| 4.48E-54 | 6.79E-52 |
| NA | NA |
| 0.350841328 | 0.51471433 |
| 0.557612777 | 0.701317451 |
| 0.865292103 | 0.919767844 |
| 0.008840644 | 0.031103921 |
| 0.002911128 | 0.01250734 |
| 0.041014551 | 0.106285066 |
| NA | NA |
| NA | NA |
| NA | NA |
| 0.597181074 | 0.731822498 |
| 0.306272243 | 0.467960709 |
| 0.948400634 | 0.968501454 |
| 0.955791922 | NA |
| NA | NA |
| 0.900024947 | NA |
| NA | NA |
| 0.140836786 | NA |
| 0.533682294 | 0.681914642 |
| 0.694227704 | NA |
| 0.020419997 | 0.060861104 |
| NA | NA |
| 0.040676204 | 0.105580699 |
| NA | NA |
| 0.000476755 | 0.002725196 |
| 0.022992009 | 0.067041013 |
| 0.295741011 | 0.457211453 |
| 0.670532819 | 0.786722102 |
| 0.007811819 | 0.028116877 |
| 0.312520877 | NA |
| 0.690542601 | 0.801512694 |
| 0.689776726 | NA |
| 0.625153932 | NA |
| 0.955792398 | NA |
| 0.941278788 | NA |
| 0.941278424 | NA |
| 0.839319108 | NA |
| 0.042729979 | 0.109626259 |
| NA | NA |
| NA | NA |
| 2.70E-07 | 3.36E-06 |
| 2.32E-06 | 2.42E-05 |
| NA | NA |

| | |
|---|---|
| 0.702523972 | 0.810227522 |
| 0.457440119 | NA |
| 0.010676947 | 0.036299486 |
| 0.153901951 | 0.288575251 |
| NA | NA |
| 1.05E-10 | 2.09E-09 |
| 0.708744856 | 0.814125883 |
| 0.725402955 | 0.826561402 |
| 0.065239274 | 0.152716877 |
| NA | NA |
| 3.35E-07 | 4.08E-06 |
| NA | NA |
| 0.158826585 | 0.295088312 |
| 0.000126389 | 0.000861734 |
| 0.955792398 | NA |
| NA | NA |
| NA | NA |
| 1.46E-11 | 3.25E-10 |
| 0.801069536 | 0.878492596 |
| NA | NA |
| 0.698470709 | NA |
| 0.6364747 | NA |
| 5.19E-10 | 9.59E-09 |
| NA | NA |
| NA | NA |
| 0.86843227 | 0.921216246 |
| 0.910280269 | 0.945786154 |
| 0.027541593 | 0.07761603 |
| 0.503477285 | 0.655567086 |
| NA | NA |
| NA | NA |
| NA | NA |
| NA | NA |
| 0.523056034 | 0.672888321 |
| 0.611721269 | 0.74282892 |
| 0.556353716 | 0.700585869 |
| 0.212376344 | NA |
| 0.942552478 | NA |
| 0.955792873 | NA |
| 0.019267294 | 0.058161081 |
| 0.61127205 | 0.742468556 |
| NA | NA |
| 0.818573585 | NA |
| NA | NA |
| 0.496563397 | 0.649564155 |
| 0.739991096 | NA |
| 0.835087402 | 0.899970585 |

| | | |
|---|---|---|
| | 0.941444399 | NA |
| | 0.718448802 | NA |
| NA | | NA |
| | 0.876977509 | 0.926329976 |
| | 0.867912408 | 0.921060969 |
| | 0.839319108 | NA |
| | 0.0278401 | 0.078250391 |
| | 0.941279303 | NA |
| NA | | NA |
| | 0.995839335 | 0.996917253 |
| NA | | NA |
| | 0.197193254 | 0.343979027 |
| | 0.06689451 | 0.155672969 |
| | 0.770803126 | 0.858202639 |
| | 0.116483738 | 0.234874018 |
| | 0.923082389 | NA |
| | 0.006070835 | 0.022859992 |
| | 0.941278332 | NA |
| NA | | NA |
| | 0.153868776 | 0.288535775 |
| | 0.002620369 | 0.011477736 |
| | 0.043962779 | 0.11197454 |
| | 4.38E-07 | 5.23E-06 |
| | 0.940617272 | NA |
| | 0.757667044 | 0.848573462 |
| | 0.00608917 | 0.02291843 |
| | 0.910227504 | 0.945786154 |
| | 0.758628764 | 0.849415836 |
| | 0.33262103 | 0.495806974 |
| | 0.013625884 | 0.044161668 |
| | 0.003669408 | 0.01511321 |
| | 1.43E-07 | 1.88E-06 |
| | 0.941277857 | NA |
| | 0.029363362 | 0.081590794 |
| | 0.438524701 | NA |
| NA | | NA |
| NA | | NA |
| | 0.791357188 | 0.871750204 |
| | 0.562626555 | 0.705194168 |
| | 0.169777818 | 0.309374645 |
| NA | | NA |
| | 0.361370782 | NA |
| NA | | NA |
| | 0.748393366 | NA |
| | 0.251338441 | NA |
| | 0.941277857 | NA |
| NA | | NA |

| | |
|---|---|
| NA | NA |
| NA | NA |
| NA | NA |
| NA | NA |
| 0.177847182 | NA |
| NA | NA |
| 0.943962327 | NA |
| 0.308080447 | 0.469578047 |
| NA | NA |
| NA | NA |
| NA | NA |
| 0.194540823 | 0.340714617 |
| 0.697507184 | 0.806238635 |
| 6.59E-05 | 0.000484881 |
| 0.839319574 | NA |
| NA | NA |
| 0.614526115 | 0.744487738 |
| 0.951172953 | NA |
| NA | NA |
| NA | NA |
| NA | NA |
| NA | NA |
| NA | NA |
| NA | NA |
| NA | NA |
| NA | NA |
| 0.55909211 | 0.702614065 |
| NA | NA |
| 0.709653002 | 0.814699308 |
| NA | NA |
| 0.635690741 | 0.760375447 |
| 0.214168237 | NA |
| NA | NA |
| NA | NA |
| 0.745010854 | 0.839817587 |
| 0.941278332 | NA |
| NA | NA |
| 0.515699945 | 0.666639079 |
| 0.114875066 | NA |
| 1.60E-37 | 1.51E-35 |
| 0.042726322 | 0.109626259 |
| 1.17E-05 | 0.000104125 |
| 0.327696404 | 0.49050002 |
| NA | NA |
| 0.839319574 | NA |
| NA | NA |
| 0.838882104 | NA |

| | |
|---|---|
| 0.807105069 | 0.88179174 |
| 0.067199047 | 0.156244184 |
| 0.367474604 | 0.530690553 |
| 0.221671505 | 0.373829189 |
| 0.072914755 | 0.165951137 |
| 0.205233143 | 0.353854348 |
| NA | NA |
| 0.893925643 | NA |
| 0.564205093 | 0.70628009 |
| 0.345337947 | NA |
| 0.865828316 | NA |
| 0.70030162 | 0.808638955 |
| 0.038690891 | 0.101367154 |
| 0.080310015 | 0.178348478 |
| 0.573428653 | NA |
| NA | NA |
| NA | NA |
| 0.445359384 | 0.605912779 |
| 0.839319108 | NA |
| 0.989271752 | 0.993110344 |
| 0.178422382 | 0.320006748 |
| 0.200951962 | 0.348594023 |
| 0.745861135 | NA |
| 0.486418123 | 0.641664161 |
| 0.641475628 | 0.764873405 |
| NA | NA |
| 0.653833302 | NA |
| 0.663776053 | NA |
| 0.01756711 | 0.05407117 |
| 0.839318641 | NA |
| 1.98E-06 | 2.09E-05 |
| 0.034596015 | 0.092794044 |
| 1.08E-05 | 9.67E-05 |
| 0.653575439 | 0.774003044 |
| 0.465453889 | 0.62380256 |
| NA | NA |
| 0.083189205 | 0.182967053 |
| NA | NA |
| NA | NA |
| 0.177104109 | 0.318650843 |
| 0.403700081 | 0.565793401 |
| 0.955792873 | NA |
| 0.797317158 | 0.875841932 |
| NA | NA |
| 0.019582427 | 0.058910634 |
| 0.020884986 | 0.06192133 |
| 0.814008435 | 0.886443528 |

| | | |
|---|---|---|
| | 0.856311252 | NA |
| | 0.839319108 | NA |
| | 0.641955948 | NA |
| | 0.681030079 | 0.794757695 |
| | 6.31E-06 | 6.01E-05 |
| | 0.97777821 | 0.986308467 |
| | 0.107066157 | 0.22071615 |
| | 0.941278332 | NA |
| | 0.027974609 | 0.07852024 |
| | 0.820371618 | NA |
| NA | | NA |
| | 0.732568743 | 0.831241253 |
| | 0.000521434 | 0.002943828 |
| | 0.685055705 | 0.797704755 |
| | 0.127733924 | 0.251613962 |
| | 0.07478784 | 0.169082463 |
| | 1.98E-09 | 3.39E-08 |
| NA | | NA |
| NA | | NA |
| | 0.941278332 | NA |
| NA | | NA |
| NA | | NA |
| NA | | NA |
| | 0.737678111 | NA |
| | 0.000256018 | 0.001583613 |
| NA | | NA |
| | 0.490268408 | 0.644680882 |
| | 0.955792398 | NA |
| | 0.00447752 | 0.017818749 |
| | 2.70E-07 | 3.37E-06 |
| | 0.431702431 | 0.592992917 |
| | 0.080616366 | 0.178842888 |
| | 0.079952229 | 0.177703143 |
| | 0.432583249 | 0.593825855 |
| | 0.426321112 | 0.587737916 |
| | 0.844382166 | 0.90637613 |
| | 0.941278207 | NA |
| | 0.862076964 | NA |
| NA | | NA |
| | 0.59375042 | 0.728852372 |
| | 0.955792398 | NA |
| | 0.839318641 | NA |
| | 0.117400268 | 0.236249677 |
| | 0.345161473 | 0.508826117 |
| | 1.80E-63 | 3.51E-61 |
| NA | | NA |
| | 0.691969511 | NA |

| | |
|---|---|
| 1.24E-71 | 2.72E-69 |
| 0.839318641 | NA |
| 0.206148585 | 0.35492489 |
| 7.63E-05 | 0.000550882 |
| NA | NA |
| NA | NA |
| NA | NA |
| 0.020692637 | 0.061483366 |
| 0.472354261 | 0.629823322 |
| NA | NA |
| 0.00244744 | 0.010839962 |
| 0.631162277 | 0.757347253 |
| 0.000648265 | 0.003534936 |
| 0.652736227 | NA |
| 0.924381304 | 0.954663244 |
| 0.575369751 | 0.714283149 |
| 2.53E-07 | 3.16E-06 |
| 0.001083021 | 0.005467247 |
| NA | NA |
| 0.457788003 | 0.617514572 |
| NA | NA |
| NA | NA |
| NA | NA |
| 0.955792873 | NA |
| 0.955792873 | NA |
| 0.941278332 | NA |
| 0.165025299 | 0.303620018 |
| 0.955791922 | NA |
| 0.0507347 | 0.125307392 |
| NA | NA |
| NA | NA |
| NA | NA |
| NA | NA |
| 0.93700822 | NA |
| NA | NA |
| NA | NA |
| NA | NA |
| 0.168650358 | 0.308169493 |
| 0.132314618 | 0.257992071 |
| 0.001062295 | 0.005374015 |
| 0.279577968 | 0.439357903 |
| 0.017300156 | 0.053442999 |
| NA | NA |
| 0.11205981 | 0.228131255 |
| NA | NA |
| NA | NA |
| 0.048870335 | 0.121547724 |

| | |
|---|---|
| NA | NA |
| 0.005825664 | 0.022101125 |
| NA | NA |
| 0.052098607 | 0.128077984 |
| 0.571716917 | 0.711883249 |
| 0.074007661 | 0.16782873 |
| 0.052712331 | 0.129292962 |
| NA | NA |
| 0.074165533 | NA |
| NA | NA |
| 2.74E-07 | 3.40E-06 |
| 0.042785912 | 0.109740331 |
| 0.635681956 | 0.760375447 |
| NA | NA |
| 0.941277857 | NA |
| NA | NA |
| 3.09E-22 | 1.52E-20 |
| 8.07E-06 | 7.47E-05 |
| NA | NA |
| 0.635235384 | 0.760094348 |
| 0.74456844 | 0.839557441 |
| NA | NA |
| 0.327704611 | 0.49050002 |
| NA | NA |
| NA | NA |
| NA | NA |
| 0.006184238 | 0.023206249 |
| 0.008912466 | 0.031329864 |
| 0.007024315 | 0.025737799 |
| 0.710868217 | NA |
| NA | NA |
| 0.941277857 | NA |
| 4.71E-15 | 1.43E-13 |
| 0.301061578 | 0.462373819 |
| NA | NA |
| 0.267116505 | NA |
| 0.182475379 | 0.324949795 |
| 0.101559758 | NA |
| 0.995959687 | 0.996917253 |
| NA | NA |
| 0.26446107 | 0.423469553 |
| NA | NA |
| NA | NA |
| NA | NA |
| NA | NA |
| NA | NA |
| 0.025056332 | 0.071777667 |

| | |
|---|---|
| NA | NA |
| NA | NA |
| 0.839318641 | NA |
| 0.955792398 | NA |
| 0.086495015 | 0.188530618 |
| NA | NA |
| 0.003406461 | 0.014237155 |
| 6.36E-09 | 1.02E-07 |
| 0.17677702 | 0.318375199 |
| 0.420136459 | 0.581772528 |
| NA | NA |
| 0.005337163 | 0.020582494 |
| 0.001646524 | 0.007775905 |
| 0.231631918 | NA |
| 0.80585951 | NA |
| 0.938171476 | NA |
| 0.344192627 | 0.5078084 |
| 0.790771535 | NA |
| 0.414017431 | 0.576109104 |
| 0.168107496 | 0.307513317 |
| 0.062603005 | 0.147839738 |
| 1.74E-31 | 1.30E-29 |
| 0.838993826 | NA |
| 0.123299047 | NA |
| 0.000974572 | 0.004992846 |
| 2.00E-09 | 3.42E-08 |
| 0.718082559 | 0.820734745 |
| 1.76E-27 | 1.11E-25 |
| 0.941198385 | NA |
| NA | NA |
| 0.675193969 | NA |
| NA | NA |
| 0.686695938 | NA |
| 0.831413565 | NA |
| 0.790166171 | NA |
| NA | NA |
| 0.839319574 | NA |
| 0.04581274 | 0.115646922 |
| 0.035522397 | 0.094819345 |
| 1.97E-07 | 2.51E-06 |
| NA | NA |
| 0.027580792 | 0.077708097 |
| 0.581435825 | 0.719157093 |
| 0.094727088 | 0.201455875 |
| 0.790609348 | 0.87134363 |
| 0.955791922 | NA |
| NA | NA |

| | |
|---|---|
| 0.955791922 | NA |
| NA | NA |
| 0.884606772 | NA |
| 0.006253278 | 0.023413641 |
| 0.000138761 | 0.000934309 |
| NA | NA |
| NA | NA |
| NA | NA |
| 5.84E-06 | 5.61E-05 |
| 0.147074781 | 0.27931652 |
| 0.484676724 | 0.64037579 |
| 1.51E-09 | 2.61E-08 |
| 1.71E-261 | 1.02E-257 |
| 4.13E-49 | 5.52E-47 |
| 1.18E-23 | 6.21E-22 |
| 6.39E-40 | 6.52E-38 |
| 1.29E-36 | 1.18E-34 |
| 9.76E-14 | 2.67E-12 |
| 3.89E-48 | 5.14E-46 |
| 0.917052906 | NA |
| 2.81E-79 | 7.36E-77 |
| 3.25E-44 | 3.85E-42 |
| 0.406654676 | NA |
| 9.79E-80 | 2.59E-77 |
| 0.481878042 | NA |
| NA | NA |
| 0.148207304 | NA |
| 0.941278332 | NA |
| 0.000708104 | 0.003812323 |
| 0.262469387 | 0.421378088 |
| 3.19E-05 | 0.000254706 |
| 0.948316757 | 0.96845735 |
| 0.15692713 | 0.29268479 |
| 0.886571234 | NA |
| 0.838882104 | NA |
| 0.582239028 | NA |
| NA | NA |
| 0.941277857 | NA |
| NA | NA |
| 0.941278332 | NA |
| NA | NA |
| NA | NA |
| 0.839318641 | NA |
| NA | NA |
| NA | NA |
| NA | NA |
| NA | NA |

| | |
|---|---|
| 0.941278332 | NA |
| NA | NA |
| 1.89E-12 | 4.62E-11 |
| 2.36E-06 | 2.46E-05 |
| 0.953559598 | 0.971934975 |
| 0.004008575 | 0.016243121 |
| NA | NA |
| NA | NA |
| 0.839319108 | NA |
| 0.06939049 | 0.160166063 |
| 0.133405046 | 0.259611389 |
| 0.955792398 | NA |
| 0.334298887 | 0.497591276 |
| 0.002770489 | 0.012011506 |
| 0.440026435 | 0.600846627 |
| 0.941278207 | NA |
| 0.634442885 | 0.75971383 |
| NA | NA |
| 0.059865087 | 0.142733527 |
| 1.01E-06 | 1.12E-05 |
| 0.390813777 | 0.553337072 |
| 0.941276431 | NA |
| 0.173871382 | 0.314618112 |
| 0.130237038 | 0.255114968 |
| 0.013840868 | 0.044718385 |
| 0.217896378 | 0.36921608 |
| 0.955792873 | NA |
| 0.661086158 | 0.779701886 |
| 0.937031162 | NA |
| 0.000194904 | 0.001253137 |
| 0.344523425 | 0.508098786 |
| NA | NA |
| 0.31470708 | 0.476929703 |
| 0.169054116 | 0.308622953 |
| NA | NA |
| 0.839318641 | NA |
| NA | NA |
| 0.54761397 | 0.693703524 |
| 0.990652875 | 0.993700524 |
| NA | NA |
| NA | NA |
| 0.000105483 | 0.000732837 |
| 0.119724982 | 0.239832859 |
| 1.43E-06 | 1.56E-05 |
| 0.839319108 | NA |
| 0.262182696 | NA |
| 0.955792873 | NA |

| | | |
|---|---|---|
| | 0.941275481 | NA |
| NA | | NA |
| | 0.941278332 | NA |
| NA | | NA |
| NA | | NA |
| NA | | NA |
| NA | | NA |
| | 0.023664905 | 0.068641491 |
| | 0.732083377 | 0.830990916 |
| NA | | NA |
| | 0.955791922 | NA |
| NA | | NA |
| NA | | NA |
| | 0.941278332 | NA |
| | 0.542047311 | NA |
| | 0.005486999 | 0.021013927 |
| | 0.082607163 | 0.181972762 |
| | 0.096625624 | 0.204325081 |
| | 0.002496393 | 0.011026004 |
| | 0.001849192 | 0.008559796 |
| | 0.000181183 | 0.001178344 |
| | 0.017439963 | 0.053749319 |
| | 0.054451908 | 0.132549019 |
| | 0.873979071 | 0.924785321 |
| | 0.221332514 | 0.373389786 |
| | 0.215684398 | 0.366720483 |
| | 0.984654639 | 0.990354943 |
| | 0.665272121 | 0.782932362 |
| | 0.031697294 | 0.086649706 |
| | 3.32E-14 | 9.48E-13 |
| | 0.157019402 | 0.292806001 |
| NA | | NA |
| | 0.68643273 | 0.798840218 |
| | 0.019449487 | 0.058580925 |
| | 0.396530553 | NA |
| | 0.042234272 | 0.108665025 |
| NA | | NA |
| | 0.648368663 | 0.770121809 |
| | 0.718482907 | 0.820995207 |
| | 0.194324632 | NA |
| NA | | NA |
| NA | | NA |
| | 0.845958754 | 0.907209293 |
| | 0.941278332 | NA |
| NA | | NA |
| | 0.729235808 | 0.829416758 |
| | 0.110819816 | 0.226354281 |

| | | | |
|---|---|---|---|
| | 0.273538784 | | 0.433386816 |
| | 0.395127569 | | 0.557715742 |
| | 0.020667636 | | 0.061460653 |
| | 2.89E-10 | | 5.50E-09 |
| | 0.988787462 | | 0.992959209 |
| NA | | NA | |
| NA | | NA | |
| | 0.613180976 | | 0.743842495 |
| | 0.884297813 | NA | |
| | 0.819592629 | | 0.890082756 |
| | 0.012451768 | | 0.041103127 |
| | 0.419591037 | | 0.581230426 |
| | 0.839318641 | NA | |
| | 0.470952974 | | 0.628518233 |
| | 0.513408279 | | 0.664499394 |
| | 0.650461686 | | 0.771683649 |
| | 0.712464925 | NA | |
| | 0.775321454 | | 0.861379819 |
| | 0.088611227 | | 0.192070342 |
| | 0.303223373 | | 0.464644481 |
| | 0.461287318 | | 0.620186255 |
| | 0.971483088 | | 0.982809378 |
| | 0.447829881 | NA | |
| | 0.097454322 | | 0.205655554 |
| | 0.024641941 | | 0.070896589 |
| | 0.290494808 | | 0.451561444 |
| | 6.92E-11 | | 1.42E-09 |
| | 0.162253117 | | 0.29958488 |
| | 4.89E-05 | | 0.000372003 |
| | 0.783831607 | NA | |
| NA | | NA | |
| | 0.689161697 | NA | |
| | 0.043413672 | | 0.110943978 |
| NA | | NA | |
| | 0.78056066 | | 0.864979973 |
| | 0.014189031 | | 0.045669684 |
| NA | | NA | |
| | 0.251913173 | | 0.409649338 |
| | 5.90E-89 | | 1.73E-86 |
| | 4.28E-06 | | 4.22E-05 |
| | 9.86E-18 | | 3.61E-16 |
| | 0.758443687 | | 0.849248559 |
| NA | | NA | |
| NA | | NA | |
| NA | | NA | |
| | 0.021102778 | | 0.062450373 |
| | 0.212176598 | | 0.362595303 |

| | | |
|---|---|---|
| | 0.850833465 | 0.910467928 |
| | 4.79E-12 | 1.12E-10 |
| | 0.005884797 | 0.022293481 |
| | 0.902966024 | NA |
| NA | | NA |
| NA | | NA |
| | 0.360515352 | NA |
| | 0.525622658 | 0.674913443 |
| | 0.955792873 | NA |
| | 8.15E-12 | 1.86E-10 |
| | 0.209021404 | NA |
| | 4.72E-05 | 0.000360198 |
| | 0.136827856 | 0.264670998 |
| NA | | NA |
| | 9.11E-05 | 0.000645771 |
| | 0.096411783 | 0.204065997 |
| | 0.00535593 | 0.020628129 |
| NA | | NA |
| | 0.339252917 | 0.502876626 |
| | 0.677837662 | NA |
| | 0.97870325 | 0.986662209 |
| | 0.748393366 | NA |
| | 0.001163144 | 0.005805233 |
| NA | | NA |
| | 0.019101203 | 0.05779164 |
| | 0.097083571 | 0.205074827 |
| | 3.50E-12 | 8.33E-11 |
| | 0.839319574 | NA |
| | 5.20E-05 | 0.000392708 |
| | 0.000421662 | 0.002446708 |
| | 0.01545868 | 0.048851234 |
| | 0.164103445 | 0.30225104 |
| NA | | NA |
| | 0.597596214 | NA |
| | 0.027536454 | NA |
| | 0.005951548 | 0.022496229 |
| | 0.941276431 | NA |
| | 0.052138915 | 0.128124144 |
| | 0.062936125 | 0.148479101 |
| | 0.008812312 | 0.031023613 |
| | 0.870718518 | 0.92266237 |
| NA | | NA |
| | 0.409295435 | 0.571381229 |
| | 0.434428787 | NA |
| NA | | NA |
| NA | | NA |
| | 0.0330618 | 0.08965899 |

| | | |
|---|---|---|
| | 0.839319107 | NA |
| | 0.941278332 | NA |
| | 0.364360893 | 0.527750136 |
| | 0.924265429 | 0.954626415 |
| | 0.941198385 | NA |
| NA | | NA |
| | 0.633030082 | NA |
| NA | | NA |
| | 0.032730807 | 0.088984428 |
| | 0.955792873 | NA |
| | 9.52E-05 | 0.000672041 |
| | 0.003557884 | 0.014756013 |
| | 0.04381085 | 0.11168318 |
| NA | | NA |
| | 0.156420433 | 0.291959612 |
| NA | | NA |
| | 0.940051175 | NA |
| | 0.938328926 | NA |
| | 0.303160989 | 0.46460871 |
| | 0.93765294 | 0.963105753 |
| | 0.941276431 | NA |
| NA | | NA |
| | 0.839319107 | NA |
| NA | | NA |
| NA | | NA |
| | 8.98E-05 | 0.000637888 |
| | 0.721324319 | NA |
| | 0.689864321 | NA |
| | 0.002635084 | 0.011531581 |
| | 0.172274101 | 0.312559506 |
| | 0.299475691 | 0.461008951 |
| | 0.000471964 | 0.002701056 |
| | 8.97E-78 | 2.27E-75 |
| | 4.26E-200 | 5.33E-197 |
| | 2.93E-196 | 3.17E-193 |
| | 0.695640024 | NA |
| | 0.005057865 | 0.019706548 |
| | 0.000475454 | 0.002718415 |
| | 0.023809617 | 0.068952001 |
| | 0.00678289 | 0.025038204 |
| | 0.440364905 | NA |
| | 0.333218967 | 0.496480616 |
| NA | | NA |
| | 0.941279268 | NA |
| | 2.14E-09 | 3.65E-08 |
| NA | | NA |
| NA | | NA |

| | | |
|---|---|---|
| NA | | NA |
| | 0.839318641 | NA |
| NA | | NA |
| | 0.02074357 | 0.061578754 |
| | 0.839318641 | NA |
| NA | | NA |
| | 0.853595154 | NA |
| | 0.955792873 | NA |
| NA | | NA |
| NA | | NA |
| | 0.150513601 | NA |
| | 0.955791922 | NA |
| | 0.941275956 | NA |
| | 0.21537009 | 0.366316913 |
| NA | | NA |
| NA | | NA |
| NA | | NA |
| NA | | NA |
| NA | | NA |
| NA | | NA |
| NA | | NA |
| NA | | NA |
| | 0.941278332 | NA |
| NA | | NA |
| | 0.913662933 | 0.947953669 |
| NA | | NA |
| NA | | NA |
| NA | | NA |
| | 0.00778482 | 0.028036659 |
| | 0.178081403 | 0.319587843 |
| | 0.041292603 | 0.106796321 |
| | 3.77E-05 | 0.000294167 |
| | 0.811511034 | NA |
| | 0.955792873 | NA |
| | 0.647371279 | 0.769436292 |
| | 0.73994852 | 0.836368823 |
| | 0.695530082 | 0.805081541 |
| | 0.36135732 | 0.525029113 |
| | 0.000229077 | 0.001439789 |
| | 0.221666183 | 0.373829189 |
| | 0.177426498 | 0.31903747 |
| | 0.086896939 | 0.189216078 |
| | 0.001104652 | 0.005562287 |
| | 0.640110287 | 0.763742797 |
| | 0.250887677 | 0.408484061 |
| | 0.69814447 | NA |
| | 0.598886136 | 0.733026998 |

| | | |
|---|---|---|
| | 2.56E-05 | 0.000209687 |
| | 0.523034808 | 0.672888321 |
| | 0.326145387 | 0.48909479 |
| | 0.962586234 | 0.977452113 |
| | 0.645420781 | 0.767868888 |
| | 0.016585825 | 0.051699294 |
| | 0.007074778 | 0.025906749 |
| | 0.641418153 | NA |
| | 4.97E-08 | 6.96E-07 |
| NA | | NA |
| | 0.241118171 | 0.397280706 |
| NA | | NA |
| | 0.160133573 | 0.29679951 |
| | 0.012223863 | 0.040483147 |
| | 0.000228434 | 0.001436886 |
| | 0.146302592 | 0.27818296 |
| NA | | NA |
| | 8.91E-10 | 1.59E-08 |
| | 4.61E-05 | 0.000352608 |
| | 1.86E-23 | 9.68E-22 |
| | 0.894522667 | NA |
| | 0.860297619 | NA |
| NA | | NA |
| NA | | NA |
| | 0.152107838 | 0.286135363 |
| | 0.14642157 | 0.278304386 |
| | 0.551936962 | NA |
| | 0.31769664 | 0.480145751 |
| | 0.410856956 | 0.572822005 |
| | 0.039191491 | 0.102475653 |
| | 0.729079852 | NA |
| | 1.50E-05 | 0.000130342 |
| | 0.98174139 | 0.988636341 |
| | 0.955792398 | NA |
| | 0.841923686 | NA |
| | 0.074699347 | 0.168962639 |
| | 0.731584008 | 0.830741022 |
| | 2.69E-09 | 4.53E-08 |
| | 0.907184928 | 0.943666829 |
| NA | | NA |
| | 0.000115553 | 0.000794911 |
| | 0.123468724 | 0.245370507 |
| | 3.00E-05 | 0.000241275 |
| NA | | NA |
| NA | | NA |
| | 0.114628651 | 0.232124481 |
| | 0.860885092 | NA |

| | |
|---|---|
| 0.14633222 | 0.278214703 |
| NA | NA |
| 0.277612792 | 0.437434077 |
| 0.068348062 | 0.158274032 |
| 0.015869987 | 0.049886085 |
| 0.060169709 | 0.143372899 |
| 0.570403058 | NA |
| 0.100112016 | 0.209796284 |
| 0.300559278 | 0.46205492 |
| 0.302044291 | 0.463344819 |
| NA | NA |
| NA | NA |
| NA | NA |
| 0.955792398 | NA |
| NA | NA |
| NA | NA |
| NA | NA |
| NA | NA |
| NA | NA |
| NA | NA |
| NA | NA |
| NA | NA |
| NA | NA |
| NA | NA |
| NA | NA |
| NA | NA |
| NA | NA |
| 0.399851872 | 0.562486658 |
| 0.178490212 | 0.320104282 |
| 0.040026137 | 0.104245773 |
| 0.482395317 | NA |
| 0.955792873 | NA |
| 1.57E-07 | 2.04E-06 |
| 4.63E-06 | 4.55E-05 |
| 4.94E-07 | 5.82E-06 |
| 0.955792873 | NA |
| 0.941277857 | NA |
| 0.941279303 | NA |
| 0.141896758 | 0.271978309 |
| 2.99E-05 | 0.000241051 |
| 0.056523562 | 0.136558448 |
| 0.824718718 | 0.893568108 |
| 0.857080091 | 0.914316078 |
| 0.001350697 | 0.006571808 |
| 0.004507258 | 0.017928101 |
| 9.47E-05 | 0.000668871 |
| NA | NA |

| | | | |
|---|---|---|---|
| | 0.022270494 | | 0.065273033 |
| | 0.626324182 | | 0.753830727 |
| | 0.911350558 | | 0.946338276 |
| | 0.15876026 | | 0.295047093 |
| | 0.106421607 | | 0.219825315 |
| NA | | NA | |
| | 0.018383054 | | 0.056075029 |
| | 0.503133183 | NA | |
| | 0.029932199 | | 0.082871489 |
| | 0.123541928 | | 0.245454505 |
| | 0.151777052 | | 0.285753935 |
| | 0.1439929 | | 0.274923014 |
| | 0.48090814 | | 0.637443646 |
| | 8.90E-07 | | 9.98E-06 |
| | 0.202552023 | | 0.350623963 |
| | 1.09E-12 | | 2.74E-11 |
| | 0.05227591 | | 0.128421013 |
| | 3.69E-05 | | 0.000288953 |
| | 0.995588208 | | 0.996802697 |
| | 0.002767433 | | 0.012002629 |
| | 0.001466252 | | 0.007036234 |
| | 0.718996938 | NA | |
| | 0.007214328 | | 0.026340763 |
| NA | | NA | |
| | 0.894684093 | NA | |
| | 0.473931185 | | 0.631360053 |
| | 0.941278332 | NA | |
| | 0.453903339 | | 0.613717579 |
| | 0.006969929 | | 0.025593683 |
| | 0.003551918 | | 0.014736404 |
| | 0.741345279 | | 0.837390866 |
| | 0.320168223 | | 0.482838753 |
| | 0.000599133 | | 0.003309483 |
| | 0.045515935 | | 0.115068539 |
| | 0.081107535 | NA | |
| NA | | NA | |
| | 0.820178735 | | 0.890475494 |
| | 0.478966798 | | 0.635685872 |
| | 4.34E-05 | | 0.000334427 |
| NA | | NA | |
| | 0.0277609 | | 0.078104603 |
| | 7.30E-13 | | 1.86E-11 |
| | 0.459379058 | | 0.618621801 |
| | 0.571169008 | | 0.711537167 |
| | 0.70089322 | | 0.808877387 |
| | 0.005717387 | | 0.021770157 |
| | 0.61440924 | | 0.744430877 |

| | |
|---|---|
| NA | NA |
| 7.41E-06 | 6.93E-05 |
| 0.734862468 | NA |
| 0.830212607 | 0.896950682 |
| 0.613880434 | NA |
| 0.241739577 | 0.398027491 |
| 0.955792398 | NA |
| 0.400619171 | 0.563101117 |
| 0.4928841 | NA |
| NA | NA |
| NA | NA |
| 0.000473057 | 0.00270551 |
| 0.478570816 | 0.635382784 |
| 0.025930863 | 0.073819686 |
| 0.73062559 | 0.830207203 |
| NA | NA |
| 0.178530814 | 0.320146222 |
| NA | NA |
| 0.760793432 | NA |
| 0.408571564 | 0.570638445 |
| 0.098994271 | NA |
| NA | NA |
| 0.641594348 | 0.764914359 |
| 0.119486931 | 0.239537433 |
| 0.839319108 | NA |
| NA | NA |
| 0.393356143 | 0.555845578 |
| 1.77E-05 | 0.000150243 |
| 0.47676245 | 0.633602986 |
| 0.853596091 | NA |
| NA | NA |
| 0.564678424 | 0.706620985 |
| 0.264322172 | 0.423320975 |
| 0.844080117 | 0.906163487 |
| 0.298840064 | 0.460509471 |
| NA | NA |
| 0.910342827 | NA |
| 0.637962718 | NA |
| 0.087797637 | 0.190671474 |
| 0.104317624 | 0.21639952 |
| 0.322688702 | 0.485470987 |
| 0.254953835 | 0.412985653 |
| 0.320194222 | 0.482847369 |
| 0.941278332 | NA |
| 0.103793529 | 0.215675475 |
| 0.677282567 | 0.791961605 |
| NA | NA |

| | |
|---|---|
| 0.017483273 | 0.053866439 |
| 0.005903532 | 0.022350224 |
| 0.616045947 | 0.745981842 |
| 0.175469065 | 0.316690528 |
| 0.942761152 | 0.965351729 |
| 0.835001975 | 0.899970585 |
| 0.467744953 | 0.625999994 |
| NA | NA |
| 0.574020179 | 0.71353814 |
| NA | NA |
| 0.927302814 | NA |
| 0.959170745 | 0.975065434 |
| 0.234756861 | 0.389602762 |
| 1.19E-13 | 3.24E-12 |
| 0.071355042 | 0.163416893 |
| NA | NA |
| 0.919596446 | NA |
| 0.939503242 | 0.963757575 |
| 0.839318641 | NA |
| 0.713981777 | 0.818089346 |
| 0.233204353 | 0.387864581 |
| 0.590308782 | 0.72631324 |
| 0.195890981 | 0.342384868 |
| 0.892003351 | 0.934936094 |
| NA | NA |
| 0.591546603 | 0.727272327 |
| 0.922101645 | NA |
| 7.14E-12 | 1.65E-10 |
| 0.487232351 | 0.642178659 |
| 0.911129419 | NA |
| 0.479205849 | NA |
| 0.107978333 | 0.22201972 |
| 0.043571155 | 0.111236014 |
| 0.613535657 | 0.74408314 |
| 0.149524553 | 0.282551092 |
| 0.839319108 | NA |
| 0.420266537 | 0.5818511 |
| NA | NA |
| 0.66787841 | 0.784735757 |
| 4.15E-06 | 4.11E-05 |
| 0.839319574 | NA |
| 0.046616263 | 0.117190567 |
| 1.38E-14 | 4.01E-13 |
| 0.002929585 | 0.012573022 |
| 0.839973075 | 0.903321135 |
| NA | NA |
| 0.171668652 | 0.311817556 |

| | | |
|---|---|---|
| | 0.000181473 | 0.001179848 |
| | 0.161127631 | 0.298084865 |
| | 0.553929807 | 0.698931152 |
| | 0.007858318 | 0.028250067 |
| | 0.953798549 | NA |
| | 0.507588337 | 0.659153123 |
| | 0.488893655 | 0.643513066 |
| | 0.003072019 | 0.013073518 |
| | 1.51E-05 | 0.000130637 |
| | 7.15E-06 | 6.70E-05 |
| NA | | NA |
| NA | | NA |
| | 0.084356557 | 0.185072796 |
| | 7.12E-07 | 8.15E-06 |
| | 0.298932852 | 0.460620169 |
| | 0.947971791 | 0.968205894 |
| | 0.000105391 | 0.000732412 |
| | 0.207526999 | 0.356762964 |
| | 0.675061907 | NA |
| | 0.880951048 | NA |
| | 0.258270244 | 0.416599916 |
| | 0.130623121 | 0.255660845 |
| | 0.748640618 | 0.842225958 |
| | 8.46E-05 | 0.000604357 |
| | 0.02429373 | 0.070132125 |
| | 3.76E-06 | 3.76E-05 |
| | 0.459850255 | 0.618941176 |
| | 0.104937508 | 0.217382294 |
| NA | | NA |
| NA | | NA |
| | 0.029781912 | 0.082497406 |
| | 0.176478605 | 0.318102518 |
| | 0.06744855 | 0.156640685 |
| | 0.170840428 | 0.310716273 |
| | 0.800638551 | 0.878395889 |
| | 2.29E-05 | 0.000190188 |
| | 0.963574058 | 0.977841478 |
| | 0.935923044 | 0.962074726 |
| | 0.075980349 | 0.171095957 |
| | 0.688681653 | 0.800376082 |
| | 0.063069504 | 0.148705333 |
| | 0.573406653 | 0.71303484 |
| | 0.766912144 | NA |
| | 5.80E-05 | 0.000433033 |
| | 0.093607784 | 0.19971004 |
| | 0.941277857 | NA |
| | 0.003706293 | 0.015233496 |

| | |
|---|---|
| 0.886769079 | 0.932447323 |
| 4.41E-05 | 0.000339053 |
| 0.572870451 | 0.712630003 |
| 0.019064111 | 0.057723442 |
| 0.920985196 | 0.952651559 |
| NA | NA |
| NA | NA |
| 0.057812178 | 0.138828162 |
| NA | NA |
| 0.735497356 | 0.833355932 |
| 0.002781559 | 0.012047725 |
| 0.84156035 | NA |
| 2.35E-12 | 5.68E-11 |
| 0.000454796 | 0.002613484 |
| NA | NA |
| 0.11765297 | 0.236658158 |
| NA | NA |
| 0.448311887 | 0.608468924 |
| 0.035943776 | 0.09575084 |
| 0.26354609 | 0.42253294 |
| 0.813167211 | 0.885973907 |
| 0.559783313 | 0.703111473 |
| 0.839319108 | NA |
| 1.69E-06 | 1.82E-05 |
| 0.303850751 | NA |
| 0.003714966 | 0.015263873 |
| 0.004886286 | 0.01916036 |
| 4.57E-15 | 1.39E-13 |
| 0.571214713 | 0.711537188 |
| NA | NA |
| 0.660896634 | 0.779633329 |
| 0.000469756 | 0.002689712 |
| 0.019202534 | 0.058009737 |
| 0.004700226 | 0.018574593 |
| 0.772683327 | 0.85973301 |
| 0.077107138 | 0.172958979 |
| 0.173768491 | 0.314520754 |
| 1.92E-13 | 5.13E-12 |
| NA | NA |
| 0.539604193 | 0.686789957 |
| 0.000556521 | 0.003111654 |
| 0.004812953 | 0.018922681 |
| 1.06E-06 | 1.18E-05 |
| 0.001191962 | 0.005929165 |
| 0.719499564 | 0.821486479 |
| NA | NA |
| 0.699470372 | NA |

| | |
|---|---|
| 0.665413608 | 0.783021441 |
| 0.404100387 | 0.566154418 |
| 0.292899264 | 0.454056811 |
| 0.005764223 | 0.021909937 |
| 0.043448921 | 0.110990869 |
| 0.009573562 | 0.033226876 |
| 0.850821017 | 0.910467928 |
| NA | NA |
| 0.095708454 | 0.203034993 |
| 0.355800575 | 0.519524279 |
| 0.626348326 | NA |
| NA | NA |
| 0.003182147 | 0.013484326 |
| 0.106459288 | 0.219864987 |
| 0.009281545 | 0.032378621 |
| 0.172188028 | 0.312450976 |
| 0.031526833 | 0.086292742 |
| 0.839318641 | NA |
| 0.520186446 | NA |
| 0.211781063 | 0.362179397 |
| 0.204061106 | 0.352574219 |
| 0.025950388 | 0.073857605 |
| 0.87602776 | 0.925856434 |
| 0.506423926 | 0.65810793 |
| 0.86854781 | NA |
| 0.059647631 | 0.142385053 |
| 0.324877357 | 0.487746364 |
| 0.170133727 | 0.30980973 |
| NA | NA |
| 0.941278332 | NA |
| 4.94E-14 | 1.38E-12 |
| NA | NA |
| 0.360099489 | 0.523833247 |
| 1.94E-84 | 5.37E-82 |
| 3.31E-18 | 1.24E-16 |
| 0.433044123 | 0.59424579 |
| 0.693933354 | 0.803747042 |
| 0.955791532 | NA |
| 0.702195248 | NA |
| 3.03E-05 | 0.00024319 |
| 0.224530925 | 0.377261406 |
| 0.129326025 | 0.253622646 |
| 0.113695764 | 0.230687329 |
| 0.812667916 | NA |
| 0.470465567 | 0.628208244 |
| 0.896771737 | 0.937779975 |
| 0.589297746 | NA |

| | | |
|---|---|---|
| | 0.783831607 | NA |
| | 0.21459162 | 0.365515225 |
| | 0.001142856 | 0.005721972 |
| | 6.80E-07 | 7.82E-06 |
| | 0.485009367 | 0.640530022 |
| | 0.296013774 | 0.457514276 |
| | 0.23469585 | 0.389555831 |
| | 0.498903401 | 0.651822305 |
| | 0.484289188 | 0.640178344 |
| | 0.546234503 | 0.692488609 |
| NA | | NA |
| | 0.843230766 | NA |
| NA | | NA |
| NA | | NA |
| | 0.181397714 | 0.323757378 |
| | 3.62E-17 | 1.28E-15 |
| | 0.16960712 | 0.309229312 |
| | 0.011953856 | 0.039765993 |
| | 0.289866643 | 0.450853155 |
| | 0.766245103 | NA |
| | 0.00142237 | 0.006857454 |
| | 0.075484769 | 0.170382974 |
| | 0.077141288 | 0.172973804 |
| | 0.003007096 | 0.012854685 |
| | 0.417112415 | 0.578628611 |
| | 0.808342493 | 0.882455076 |
| | 0.001303061 | 0.006373911 |
| | 0.131604211 | 0.257010456 |
| NA | | NA |
| NA | | NA |
| | 0.920537114 | NA |
| | 4.79E-09 | 7.82E-08 |
| NA | | NA |
| NA | | NA |
| NA | | NA |
| | 0.898269265 | NA |
| | 0.921305466 | NA |
| | 0.001245352 | 0.006145938 |
| NA | | NA |
| | 0.048918774 | 0.121622904 |
| | 0.000701937 | 0.003786831 |
| NA | | NA |
| | 0.941278788 | NA |
| NA | | NA |
| | 0.418145175 | 0.579763034 |
| | 0.079165207 | 0.176514309 |
| | 6.55E-05 | 0.000482014 |

| | | | |
|---|---|---|---|
| | 0.036097191 | | 0.096062761 |
| | 0.061821771 | | 0.146424602 |
| | 0.043325342 | | 0.110778889 |
| | 0.167441095 | | 0.306571765 |
| | 0.829165904 | | 0.896348753 |
| | 0.33764948 | NA | |
| NA | | NA | |
| | 1.47E-237 | | 3.90E-234 |
| | 0.000587844 | | 0.003255435 |
| NA | | NA | |
| | 0.383516044 | | 0.546253898 |
| NA | | NA | |
| | 0.19966611 | | 0.347183534 |
| | 0.488548381 | NA | |
| | 0.000260469 | | 0.001608217 |
| | 0.000245287 | | 0.001527159 |
| | 0.080375303 | | 0.17845255 |
| | 0.459843482 | | 0.618941176 |
| | 0.490868801 | | 0.645047538 |
| NA | | NA | |
| | 0.902594122 | | 0.941187475 |
| | 4.43E-41 | | 4.73E-39 |
| | 0.132546549 | | 0.258320885 |
| | 0.080608745 | | 0.178842888 |
| NA | | NA | |
| | 0.893747456 | | 0.935954744 |
| | 0.502127489 | | 0.654242278 |
| NA | | NA | |
| | 0.679630933 | | 0.793786519 |
| | 0.06368509 | | 0.149919149 |
| | 0.089489607 | | 0.19335799 |
| | 0.235174405 | | 0.390159699 |
| | 5.48E-05 | | 0.000411185 |
| | 0.036847445 | | 0.097535909 |
| | 0.00076958 | | 0.004092392 |
| | 0.540212977 | | 0.687381016 |
| | 0.589965786 | | 0.726003807 |
| | 0.000170176 | | 0.001118005 |
| | 0.125853749 | | 0.248863581 |
| | 0.273401746 | | 0.433292143 |
| NA | | NA | |
| | 0.1458199 | | 0.277511613 |
| | 0.625492824 | | 0.753287806 |
| | 0.014325764 | | 0.046010231 |
| | 0.416298854 | NA | |
| NA | | NA | |
| NA | | NA | |

| | | |
|---|---|---|
| NA | | NA |
| | 0.74627078 | NA |
| NA | | NA |
| NA | | NA |
| NA | | NA |
| NA | | NA |
| NA | | NA |
| NA | | NA |
| NA | | NA |
| NA | | NA |
| NA | | NA |
| NA | | NA |
| NA | | NA |
| NA | | NA |
| NA | | NA |
| NA | | NA |
| | 0.941278284 | NA |
| NA | | NA |
| | 0.941277857 | NA |
| NA | | NA |
| | 0.941278332 | NA |
| NA | | NA |
| NA | | NA |
| NA | | NA |
| NA | | NA |
| NA | | NA |
| NA | | NA |
| NA | | NA |
| NA | | NA |
| | 0.955792398 | NA |
| NA | | NA |
| NA | | NA |
| NA | | NA |
| | 0.876610858 | NA |
| NA | | NA |
| NA | | NA |
| | 0.955792398 | NA |
| NA | | NA |
| NA | | NA |
| NA | | NA |
| NA | | NA |
| NA | | NA |
| | 0.941278332 | NA |
| | 0.839318641 | NA |
| NA | | NA |
| NA | | NA |
| NA | | NA |

| | | |
|---|---|---|
| NA | | NA |
| | 0.941278332 | NA |
| | 0.941278332 | NA |
| | 0.955792873 | NA |
| | 0.138321747 | 0.266836243 |
| | 0.264194597 | NA |
| | 0.673324997 | 0.789093136 |
| | 0.092064058 | 0.197259115 |
| | 0.301499561 | 0.462837314 |
| NA | | NA |
| | 0.839319108 | NA |
| | 8.48E-09 | 1.35E-07 |
| | 0.231014596 | 0.384966158 |
| | 3.72E-05 | 0.000290937 |
| NA | | NA |
| | 0.005432754 | 0.02084643 |
| | 0.469848707 | 0.627798041 |
| | 0.903093988 | 0.941378781 |
| | 0.018845801 | 0.057237177 |
| | 0.10930319 | 0.224009531 |
| NA | | NA |
| NA | | NA |
| NA | | NA |
| | 0.901457499 | 0.940439722 |
| | 0.095335187 | 0.20240981 |
| | 0.854273887 | 0.912262867 |
| | 0.012003447 | 0.039903078 |
| | 0.390528734 | 0.553065087 |
| | 0.042180314 | 0.108573141 |
| | 0.762476208 | 0.852560656 |
| | 0.286638621 | 0.447176539 |
| | 0.605098914 | 0.737911894 |
| | 0.082426413 | 0.181742797 |
| | 0.955792873 | NA |
| | 0.005185966 | 0.020126586 |
| | 0.19820455 | 0.345312688 |
| | 0.955791446 | NA |
| | 0.614036818 | NA |
| | 0.001123894 | 0.005641275 |
| | 0.004175105 | 0.016820558 |
| | 0.000144536 | 0.000967176 |
| | 0.969811485 | 0.981982089 |
| NA | | NA |
| | 0.000660708 | 0.003596199 |
| | 0.306718516 | 0.468301888 |
| | 1.84E-05 | 0.000156007 |
| | 0.114138792 | 0.231309623 |

| | |
|---|---|
| 0.757684509 | 0.848573462 |
| 0.94162623 | 0.964770674 |
| NA | NA |
| 0.402995742 | 0.565005869 |
| 0.002317761 | 0.010358122 |
| 2.39E-06 | 2.48E-05 |
| 0.699258896 | 0.807871092 |
| 0.477625377 | 0.63450509 |
| 0.955792873 | NA |
| 0.001012787 | 0.005165279 |
| 0.175609003 | 0.316851507 |
| 0.703624059 | 0.810844101 |
| 0.934012062 | 0.960815762 |
| 0.070337029 | 0.161786043 |
| 0.088519094 | 0.191906668 |
| 0.988951223 | 0.992997975 |
| 0.887970624 | 0.933174833 |
| NA | NA |
| 0.000402469 | 0.002348507 |
| 0.52928279 | 0.678441971 |
| 0.202561763 | 0.350623963 |
| 0.508599786 | 0.660250455 |
| 0.341150015 | 0.504641667 |
| 0.022814977 | 0.06663093 |
| 0.736955521 | 0.834253557 |
| 0.179384753 | NA |
| 0.128973103 | 0.25318582 |
| 0.933540812 | NA |
| 0.852144747 | 0.91117472 |
| 0.064658476 | 0.151745321 |
| 5.51E-05 | 0.000413263 |
| 0.832527038 | 0.898431654 |
| 0.659794719 | NA |
| 0.244188041 | 0.400668251 |
| 0.690126483 | 0.801248259 |
| 0.734321918 | 0.83253931 |
| NA | NA |
| 0.061970732 | 0.14669591 |
| NA | NA |
| 0.229583913 | NA |
| 0.361194245 | 0.524891938 |
| 0.658217398 | 0.777381584 |
| 0.007729181 | 0.027874236 |
| 0.061515012 | 0.145863749 |
| NA | NA |
| 0.049208516 | 0.122133587 |
| 0.030153862 | 0.083330107 |

| | | |
|---|---|---|
| | 0.351304523 | 0.515109595 |
| | 0.341777099 | 0.505170493 |
| | 0.154119375 | 0.288812524 |
| | 0.148245069 | 0.281000965 |
| | 7.97E-14 | 2.20E-12 |
| | 0.415139905 | NA |
| | 0.032880347 | 0.089268623 |
| | 0.102158856 | 0.213161486 |
| NA | | NA |
| NA | | NA |
| | 0.94388205 | NA |
| | 0.654625801 | 0.774462116 |
| | 0.940475126 | NA |
| | 0.281672003 | NA |
| | 0.000323992 | 0.001948372 |
| | 0.673866117 | NA |
| | 0.000617343 | 0.003393532 |
| | 0.087039366 | 0.189441633 |
| | 0.00048716 | 0.002773352 |
| | 0.104551439 | 0.216752316 |
| | 0.151687393 | 0.285723529 |
| | 0.026141363 | 0.074294533 |
| | 0.224917621 | 0.377671196 |
| NA | | NA |
| NA | | NA |
| | 0.940773688 | NA |
| | 0.609653704 | 0.741490928 |
| | 0.092871284 | 0.19866292 |
| | 0.905108875 | 0.94244724 |
| | 0.283889963 | 0.444450387 |
| | 0.971184779 | 0.982620196 |
| | 6.13E-06 | 5.85E-05 |
| NA | | NA |
| | 0.000154415 | 0.001026934 |
| | 0.813203594 | 0.885973907 |
| | 0.480793522 | 0.637327212 |
| | 0.786788685 | 0.869049342 |
| | 0.238244265 | 0.393852618 |
| | 0.021263122 | 0.062854549 |
| | 0.00357594 | 0.014810254 |
| | 0.175202926 | 0.316330126 |
| | 4.09E-08 | 5.79E-07 |
| | 0.84615939 | NA |
| | 0.270532131 | 0.430545107 |
| | 0.955792398 | NA |
| | 0.601947285 | 0.735387223 |
| | 0.79328871 | 0.872939996 |

| | |
|---|---|
| 0.029254973 | 0.081327594 |
| 0.000460968 | 0.002644486 |
| 0.812422254 | NA |
| NA | NA |
| NA | NA |
| 0.839318641 | NA |
| 0.062311548 | 0.147370773 |
| 0.000356791 | 0.002116733 |
| 0.630299417 | 0.756552074 |
| 0.001113716 | 0.005600817 |
| 0.575881222 | 0.714694449 |
| 0.48613854 | 0.641486957 |
| 0.107160224 | 0.220852683 |
| 0.040587044 | 0.105372254 |
| 0.347900984 | 0.511788579 |
| 0.947989176 | 0.968205894 |
| 0.011482751 | NA |
| 8.16E-06 | 7.55E-05 |
| NA | NA |
| 0.944287042 | 0.966373729 |
| NA | NA |
| 0.033850217 | 0.091246078 |
| NA | NA |
| NA | NA |
| 0.006646406 | 0.024606901 |
| 2.47E-09 | 4.18E-08 |
| 8.53E-98 | 2.86E-95 |
| 0.769965383 | 0.857510576 |
| 0.69596767 | NA |
| NA | NA |
| 0.964970465 | 0.978745771 |
| NA | NA |
| 1.80E-17 | 6.46E-16 |
| NA | NA |
| NA | NA |
| 0.403671677 | 0.565791266 |
| 0.688831147 | 0.800447149 |
| 0.012077022 | 0.040074901 |
| NA | NA |
| 0.018986505 | 0.057533049 |
| 4.50E-05 | 0.000344947 |
| 0.584938448 | 0.721733424 |
| 0.941275956 | NA |
| NA | NA |
| NA | NA |
| 0.839318641 | NA |
| 0.571281864 | NA |

| | | |
|---|---|---|
| | 0.955792398 | NA |
| | 0.730177973 | NA |
| | 0.473658321 | NA |
| | 0.698470709 | NA |
| NA | | NA |
| | 0.925313117 | NA |
| | 0.618028203 | 0.747460621 |
| | 0.000218952 | 0.001387145 |
| | 0.097869602 | 0.206317718 |
| | 0.217514326 | 0.368835107 |
| | 0.01372288 | 0.044403477 |
| | 0.057712049 | 0.138680794 |
| | 2.09E-23 | 1.08E-21 |
| | 0.947074013 | NA |
| | 2.73E-08 | 4.00E-07 |
| | 0.031973031 | 0.087223161 |
| | 0.159192067 | 0.295537756 |
| | 0.4422202 | 0.602778903 |
| | 0.042207357 | 0.108619257 |
| | 0.002756288 | 0.011963734 |
| | 9.61E-05 | 0.000677247 |
| | 0.159135648 | 0.295479108 |
| | 0.154674114 | 0.289475803 |
| | 0.142911181 | 0.273356632 |
| NA | | NA |
| | 0.389751402 | NA |
| | 0.839318641 | NA |
| | 0.759558452 | 0.850193626 |
| NA | | NA |
| | 0.955792398 | NA |
| NA | | NA |
| | 0.170587729 | 0.310376457 |
| NA | | NA |
| | 0.478158004 | NA |
| | 0.676213523 | 0.791324721 |
| | 0.941277857 | NA |
| | 1.92E-34 | 1.63E-32 |
| | 6.55E-23 | 3.34E-21 |
| | 1.08E-08 | 1.69E-07 |
| | 2.22E-37 | 2.09E-35 |
| | 2.65E-11 | 5.75E-10 |
| | 0.694213201 | NA |
| | 3.63E-05 | 0.000284644 |
| NA | | NA |
| NA | | NA |
| | 0.032306682 | 0.087982111 |
| NA | | NA |

| | | |
|---|---|---|
| NA | | NA |
| | 0.008120701 | NA |
| | 0.336342165 | 0.49960161 |
| | 0.805761743 | 0.881052049 |
| | 0.165294602 | 0.303951028 |
| | 0.05762682 | 0.138548844 |
| | 0.376682241 | 0.539685228 |
| | 0.852666549 | NA |
| | 0.264104624 | 0.423114913 |
| | 0.000446799 | 0.002574989 |
| | 0.205734174 | 0.354384484 |
| | 0.607166797 | NA |
| | 1.30E-60 | 2.30E-58 |
| | 0.933089962 | NA |
| | 1.75E-85 | 4.96E-83 |
| | 0.08491191 | 0.186082318 |
| | 4.65E-127 | 2.05E-124 |
| | 4.60E-07 | 5.44E-06 |
| | 1.27E-18 | 4.93E-17 |
| | 0.694980698 | NA |
| | 6.84E-64 | 1.34E-61 |
| | 1.66E-30 | 1.17E-28 |
| | 4.27E-39 | 4.25E-37 |
| | 4.70E-14 | 1.32E-12 |
| | 2.28E-18 | 8.66E-17 |
| | 0.000689669 | 0.003730796 |
| | 0.547048294 | 0.69326182 |
| | 0.609116469 | NA |
| | 0.228906777 | 0.382588239 |
| | 0.0425634 | 0.109322763 |
| | 0.068478706 | 0.158491848 |
| | 0.424853008 | 0.586140534 |
| NA | | NA |
| | 0.002511344 | 0.011079706 |
| | 0.196721063 | 0.343482596 |
| | 0.275635268 | 0.435490915 |
| | 0.001386229 | 0.006718981 |
| | 0.094552719 | 0.201192903 |
| | 6.49E-11 | 1.34E-09 |
| | 0.105074497 | 0.217628191 |
| | 2.39E-33 | 1.92E-31 |
| | 3.82E-16 | 1.27E-14 |
| | 0.335492832 | 0.498651204 |
| | 5.85E-13 | 1.50E-11 |
| NA | | NA |
| | 1.75E-17 | 6.31E-16 |
| | 2.14E-20 | 9.35E-19 |

| | | |
|---|---|---|
| | 0.941278332 | NA |
| NA | | NA |
| | 0.941277857 | NA |
| NA | | NA |
| NA | | NA |
| | 2.99E-11 | 6.40E-10 |
| | 0.700438931 | 0.808645653 |
| | 0.544953841 | NA |
| | 0.489139968 | 0.643583465 |
| NA | | NA |
| | 0.955792398 | NA |
| | 0.267036021 | 0.426647517 |
| NA | | NA |
| | 0.743802896 | 0.839091736 |
| | 9.16E-06 | 8.38E-05 |
| | 0.030242079 | 0.083525406 |
| | 0.025426315 | 0.072607281 |
| | 0.269069453 | 0.428961226 |
| | 0.846313479 | 0.907398403 |
| | 0.739991096 | NA |
| | 0.547326993 | 0.693541156 |
| | 0.001204214 | 0.005980119 |
| | 0.025366308 | 0.072507369 |
| | 0.060377222 | 0.143724236 |
| NA | | NA |
| NA | | NA |
| | 0.009738611 | 0.033701428 |
| | 0.097500824 | 0.205718873 |
| | 0.733565361 | NA |
| | 0.014731988 | 0.046985232 |
| | 0.745597105 | 0.840134719 |
| | 0.018326185 | 0.055944575 |
| | 0.861806363 | 0.917466458 |
| | 0.00358942 | 0.014850577 |
| | 6.64E-05 | 0.000487692 |
| | 1.32E-08 | 2.03E-07 |
| | 0.277716051 | 0.437434077 |
| | 0.530666745 | 0.679484096 |
| | 0.301181992 | 0.462490658 |
| NA | | NA |
| | 0.35290716 | NA |
| | 0.654609042 | 0.774462116 |
| | 0.91911295 | NA |
| | 0.105523963 | 0.218274213 |
| NA | | NA |
| | 0.132323637 | 0.257992071 |
| | 0.396263951 | 0.5589259 |

| | |
|---|---|
| 0.875029313 | NA |
| 0.955792873 | NA |
| NA | NA |
| NA | NA |
| 0.718015804 | 0.820697857 |
| 0.894945802 | NA |
| 0.010857176 | 0.036817568 |
| NA | NA |
| 0.65645341 | NA |
| NA | NA |
| 6.19E-22 | 2.96E-20 |
| 0.564066266 | 0.70628009 |
| 5.83E-08 | 8.10E-07 |
| 0.935180275 | 0.96151883 |
| 0.031101096 | 0.085353197 |
| NA | NA |
| NA | NA |
| NA | NA |
| 0.136364431 | 0.263967647 |
| 0.892633265 | 0.935102195 |
| 0.021105699 | 0.06245125 |
| 0.003498243 | 0.014556848 |
| NA | NA |
| 0.47504001 | 0.632341722 |
| 0.781476852 | 0.86563254 |
| 0.484892784 | 0.640530022 |
| 0.111519484 | 0.227317527 |
| 2.33E-06 | 2.43E-05 |
| NA | NA |
| 0.000141607 | 0.000949709 |
| 0.790651661 | NA |
| 0.004684537 | 0.018521818 |
| 0.955792398 | NA |
| 5.86E-05 | 0.0004361 |
| 0.764316556 | 0.853630988 |
| 0.754322207 | 0.846066498 |
| 0.622755046 | NA |
| 0.000662744 | 0.003604808 |
| 0.067668079 | 0.157058568 |
| NA | NA |
| 0.517546942 | 0.668311565 |
| 1.87E-06 | 1.99E-05 |
| 0.026866018 | 0.076099603 |
| 0.220033918 | 0.371699596 |
| 0.937028989 | 0.962629548 |
| 0.939241089 | NA |
| NA | NA |

| | | |
|---|---|---|
| | 0.839318641 | NA |
| NA | | NA |
| | 0.621251189 | NA |
| NA | | NA |
| | 0.451772834 | NA |
| | 1.48E-05 | 0.000128489 |
| | 0.009320483 | 0.032485888 |
| | 0.628008473 | 0.75505671 |
| | 0.113029302 | 0.22974581 |
| NA | | NA |
| | 0.759036704 | 0.849712709 |
| | 0.009847936 | 0.033978828 |
| | 9.15E-09 | 1.45E-07 |
| | 0.336989848 | NA |
| | 0.739991096 | NA |
| | 0.002008187 | 0.009188556 |
| | 0.0178257 | 0.054704397 |
| | 0.954298825 | NA |
| | 0.000208615 | 0.001329795 |
| | 0.437239558 | 0.598147937 |
| | 0.651878225 | NA |
| | 0.797830044 | NA |
| | 0.941277857 | NA |
| | 0.206483595 | NA |
| NA | | NA |
| | 0.941278332 | NA |
| | 0.003531441 | 0.014664227 |
| | 0.241958623 | 0.39822287 |
| | 0.694723588 | 0.804427458 |
| | 0.521453158 | 0.671588554 |
| | 0.600926681 | 0.734592824 |
| | 0.26691773 | 0.426544813 |
| | 0.00090906 | 0.004705838 |
| | 0.097570771 | 0.205829966 |
| | 0.359661529 | NA |
| NA | | NA |
| | 0.01003969 | 0.034502556 |
| NA | | NA |
| | 0.433189914 | NA |
| | 0.345796066 | 0.509542374 |
| | 7.01E-06 | 6.58E-05 |
| NA | | NA |
| | 2.24E-07 | 2.83E-06 |
| | 0.853595154 | NA |
| NA | | NA |
| NA | | NA |
| | 5.37E-05 | 0.000404111 |

| | |
|---|---|
| 0.955792873 | NA |
| 0.003937757 | 0.015999743 |
| 0.955792398 | NA |
| 0.429405153 | NA |
| 3.60E-95 | 1.16E-92 |
| 0.044517995 | 0.11308616 |
| 0.238488268 | 0.394201233 |
| NA | NA |
| 0.941277857 | NA |
| NA | NA |
| NA | NA |
| 0.007672122 | 0.027703503 |
| 0.861526366 | NA |
| 0.282345673 | 0.442713551 |
| 0.091804454 | 0.196858524 |
| 0.26299433 | 0.421960973 |
| 0.180830802 | 0.322890828 |
| NA | NA |
| NA | NA |
| 0.837561094 | 0.901433656 |
| 0.543413938 | 0.690199496 |
| 0.22862721 | 0.382228339 |
| 0.230100561 | 0.383936457 |
| 0.362743723 | 0.526432906 |
| 0.88038897 | 0.928322255 |
| 0.000828998 | 0.004355807 |
| NA | NA |
| 0.794888697 | NA |
| NA | NA |
| NA | NA |
| NA | NA |
| NA | NA |
| NA | NA |
| NA | NA |
| NA | NA |
| 3.42E-07 | 4.15E-06 |
| NA | NA |
| 0.839319108 | NA |
| NA | NA |
| 0.08592622 | 0.187689071 |
| 0.066737818 | 0.155453807 |
| 0.687682291 | 0.799596632 |
| 0.941278332 | NA |
| 0.877409759 | 0.926494259 |
| 1.07E-08 | 1.68E-07 |
| 0.012487868 | 0.041191273 |
| 0.086106026 | 0.187861408 |

| | | |
|---|---|---|
| | 0.031446492 | 0.086102541 |
| | 0.462419281 | 0.621169576 |
| | 0.955792873 | NA |
| | 0.111083496 | 0.226681188 |
| | 0.177313076 | 0.318906289 |
| | 0.108063936 | 0.222118981 |
| | 8.61E-06 | 7.91E-05 |
| | 0.360288905 | 0.523988544 |
| | 0.000287325 | 0.001754465 |
| | 0.010158834 | 0.034846603 |
| | 3.04E-05 | 0.000243859 |
| | 0.487291935 | 0.642221629 |
| | 0.860522469 | 0.916609361 |
| NA | | NA |
| | 5.08E-05 | 0.000384294 |
| | 0.004511043 | 0.017938294 |
| | 4.38E-11 | 9.23E-10 |
| | 0.031946327 | 0.087170295 |
| | 0.635039462 | 0.760020207 |
| | 0.924819347 | NA |
| | 0.983959136 | 0.989822719 |
| | 0.938182526 | NA |
| | 0.839319108 | NA |
| NA | | NA |
| NA | | NA |
| NA | | NA |
| NA | | NA |
| NA | | NA |
| NA | | NA |
| | 0.17683142 | 0.318424986 |
| NA | | NA |
| | 0.078192048 | 0.17475372 |
| NA | | NA |
| | 0.961700941 | 0.976719825 |
| | 0.054112885 | 0.131953171 |
| | 0.42076515 | 0.582304326 |
| | 0.491766886 | 0.645723805 |
| | 0.232489881 | 0.38689255 |
| | 0.658236637 | 0.777381584 |
| | 0.947799563 | 0.968179709 |
| | 0.005099668 | 0.019833708 |
| | 0.789316206 | 0.870361551 |
| | 0.463050011 | NA |
| | 0.929003981 | NA |
| | 0.000750695 | 0.004008983 |
| | 0.505499883 | 0.657409475 |
| | 0.608445944 | NA |

| | |
|---|---|
| 0.001247264 | 0.006152823 |
| 0.797449741 | NA |
| 0.000187382 | 0.001212303 |
| 0.005666169 | 0.021606589 |
| NA | NA |
| 0.01450129 | 0.046411148 |
| 0.602961428 | 0.73616871 |
| NA | NA |
| 0.531887641 | 0.680464813 |
| 0.133232154 | 0.259317338 |
| 0.010621403 | 0.036146788 |
| 0.673911263 | 0.789546952 |
| 9.57E-07 | 1.07E-05 |
| 0.137813347 | 0.266097037 |
| 0.852708464 | 0.911466704 |
| 0.007779503 | 0.028021753 |
| 2.32E-19 | 9.45E-18 |
| 9.00E-11 | 1.81E-09 |
| 0.473516435 | 0.631019438 |
| 0.976235747 | 0.985471179 |
| 0.859782921 | NA |
| NA | NA |
| NA | NA |
| 0.074084521 | 0.167939025 |
| NA | NA |
| NA | NA |
| NA | NA |
| NA | NA |
| NA | NA |
| 0.695759609 | 0.805170278 |
| 0.255415369 | 0.413494241 |
| 0.010640435 | 0.036196033 |
| NA | NA |
| 0.067432355 | NA |
| 0.002061818 | 0.00937925 |
| NA | NA |
| NA | NA |
| NA | NA |
| NA | NA |
| 0.372088967 | 0.535415634 |
| 0.955791446 | NA |
| NA | NA |
| 0.839318641 | NA |
| 0.738484193 | 0.835388089 |
| 0.902934338 | 0.941349627 |
| 0.835908301 | NA |
| 0.676588443 | 0.791584276 |

| | | |
|---|---|---|
| | 0.975559893 | 0.985178051 |
| | 0.798410691 | NA |
| | 0.557048593 | 0.701045346 |
| | 0.670399876 | 0.786710932 |
| | 0.173217978 | NA |
| | 0.829569353 | 0.896499875 |
| | 0.920721722 | 0.952578389 |
| | 0.941277857 | NA |
| | 0.839318641 | NA |
| | 0.890452366 | NA |
| | 8.12E-08 | 1.11E-06 |
| | 0.566219642 | NA |
| | 0.067618973 | NA |
| | 0.001205738 | 0.005986434 |
| | 0.013306535 | 0.043392539 |
| | 0.028770524 | 0.080337402 |
| | 0.000441736 | 0.002548896 |
| | 0.329439646 | 0.492452581 |
| | 0.853595622 | NA |
| | 0.853386821 | NA |
| NA | | NA |
| | 0.176731042 | 0.318340567 |
| | 0.797021383 | 0.875600533 |
| | 0.202877785 | 0.351013813 |
| | 0.096214211 | 0.203744719 |
| | 0.367872587 | 0.530994743 |
| | 1.55E-07 | 2.01E-06 |
| | 0.537648596 | 0.685452502 |
| | 0.506416779 | 0.65810793 |
| | 0.080883324 | 0.179136957 |
| | 0.534811446 | 0.68295413 |
| | 0.879007908 | NA |
| | 0.177600617 | 0.319182263 |
| | 0.963039644 | 0.977662271 |
| | 0.094203288 | 0.200610777 |
| | 0.102544423 | 0.213708289 |
| NA | | NA |
| | 0.035372077 | 0.094502883 |
| | 0.839319108 | NA |
| | 0.252933051 | 0.410774596 |
| NA | | NA |
| | 0.291569564 | 0.452787934 |
| | 0.68082389 | 0.79455603 |
| | 0.015928502 | 0.050056801 |
| | 0.577671745 | 0.716095145 |
| | 0.206118997 | 0.35492489 |
| | 0.941278332 | NA |

| | | | |
|---|---|---|---|
| | 8.04E-07 | | 9.12E-06 |
| | 0.735841202 | | 0.833507461 |
| | 0.005390896 | | 0.020722562 |
| | 0.88358665 | | 0.930516537 |
| | 0.000282253 | | 0.001728374 |
| | 0.00302299 | | 0.012913355 |
| | 0.926473738 | NA | |
| | 0.897760934 | NA | |
| | 0.216962251 | | 0.36834062 |
| | 0.003116417 | | 0.01323171 |
| NA | | NA | |
| | 0.79607812 | | 0.874908486 |
| NA | | NA | |
| | 0.774440642 | | 0.860707236 |
| | 0.245006844 | | 0.401640474 |
| | 0.000101941 | | 0.000711973 |
| | 0.024893692 | | 0.071456784 |
| NA | | NA | |
| | 0.475080177 | NA | |
| | 0.011108235 | | 0.037513934 |
| | 0.393241241 | | 0.555716204 |
| | 0.046194465 | | 0.116400663 |
| | 0.000117889 | | 0.000809112 |
| | 0.293684743 | | 0.454889325 |
| | 0.742243177 | | 0.837967657 |
| | 0.038344173 | | 0.100744508 |
| | 0.002306455 | | 0.010321159 |
| | 0.727370069 | | 0.828101606 |
| | 0.70091215 | NA | |
| | 0.1613001 | | 0.29828801 |
| | 0.835323277 | NA | |
| | 0.839319574 | NA | |
| | 9.24E-06 | | 8.45E-05 |
| | 0.001520678 | | 0.007254995 |
| | 0.882521351 | | 0.929664669 |
| | 0.002495969 | | 0.011026004 |
| NA | | NA | |
| | 0.198907799 | | 0.346259302 |
| | 0.839318641 | NA | |
| | 0.180084912 | | 0.32201796 |
| NA | | NA | |
| | 0.697353356 | | 0.806139125 |
| | 0.080867552 | | 0.179130044 |
| | 0.009482042 | | 0.032957296 |
| | 0.888126719 | | 0.933255773 |
| | 0.27749209 | | 0.437366025 |
| NA | | NA | |

| | | |
|---|---|---|
| | 0.011828745 | 0.039449038 |
| | 0.148036636 | 0.280698833 |
| | 0.662659873 | 0.780900599 |
| | 0.920147381 | 0.952273924 |
| | 0.000403301 | 0.002352213 |
| | 0.003249741 | 0.0137146 |
| | 0.299059359 | 0.460705725 |
| | 0.329229525 | 0.492262149 |
| | 1.35E-05 | 0.00011889 |
| NA | | NA |
| | 0.021634204 | 0.063737754 |
| | 0.098727593 | 0.207552921 |
| | 0.100380979 | 0.210248862 |
| | 2.23E-31 | 1.65E-29 |
| | 0.839319108 | NA |
| | 0.77505655 | NA |
| | 0.00075605 | 0.004031253 |
| NA | | NA |
| | 0.766987831 | 0.855355121 |
| | 0.586844692 | NA |
| NA | | NA |
| | 0.564192635 | 0.70628009 |
| | 0.000625603 | 0.003426277 |
| | 0.642887391 | 0.765727445 |
| | 0.002961769 | 0.012683706 |
| | 0.148838149 | 0.281728267 |
| | 0.654074986 | NA |
| | 0.54310988 | 0.689900157 |
| | 0.050157948 | 0.124076022 |
| | 7.19E-05 | 0.000522958 |
| | 0.482835358 | 0.63882419 |
| | 0.876704245 | 0.926202931 |
| | 0.003854463 | 0.015734315 |
| | 0.853415982 | 0.911796714 |
| NA | | NA |
| | 0.354014258 | 0.517646356 |
| | 0.183709338 | 0.326463316 |
| | 0.271030251 | NA |
| | 0.072034142 | 0.164513207 |
| | 2.66E-08 | 3.91E-07 |
| NA | | NA |
| NA | | NA |
| | 0.004297357 | 0.017205331 |
| | 0.051464285 | 0.126846113 |
| | 1.09E-08 | 1.70E-07 |
| | 0.352434583 | 0.516033953 |
| | 0.496505379 | 0.649544962 |

| | | | |
|---|---|---|---|
| | 0.29345261 | NA | |
| NA | | NA | |
| | 0.43635475 | | 0.597439452 |
| | 0.901868174 | NA | |
| | 0.287453145 | NA | |
| | 0.145423941 | | 0.276954764 |
| | 0.825843457 | | 0.894113212 |
| | 0.088713601 | | 0.192239214 |
| | 0.057199793 | | 0.137814503 |
| | 0.009415682 | | 0.032760132 |
| | 0.911498357 | NA | |
| | 0.735089012 | NA | |
| NA | | NA | |
| NA | | NA | |
| NA | | NA | |
| NA | | NA | |
| NA | | NA | |
| NA | | NA | |
| NA | | NA | |
| NA | | NA | |
| NA | | NA | |
| NA | | NA | |
| NA | | NA | |
| NA | | NA | |
| NA | | NA | |
| NA | | NA | |
| NA | | NA | |
| NA | | NA | |
| NA | | NA | |
| NA | | NA | |
| | 0.839318641 | NA | |
| NA | | NA | |
| NA | | NA | |
| | 0.915881972 | NA | |
| NA | | NA | |
| NA | | NA | |
| NA | | NA | |
| NA | | NA | |
| NA | | NA | |
| NA | | NA | |
| NA | | NA | |
| | 0.941278332 | NA | |
| NA | | NA | |
| NA | | NA | |
| NA | | NA | |
| NA | | NA | |
| NA | | NA | |

| | | |
|---|---|---|
| NA | NA | |
| NA | NA | |
| NA | NA | |
| 0.839319108 | NA | |
| NA | NA | |
| NA | NA | |
| NA | NA | |
| NA | NA | |
| NA | NA | |
| NA | NA | |
| 0.941277857 | NA | |
| NA | NA | |
| 0.274648864 | | 0.434566321 |
| NA | NA | |
| NA | NA | |
| NA | NA | |
| NA | NA | |
| 0.839318641 | NA | |
| 0.795327666 | NA | |
| NA | NA | |
| 0.710678838 | NA | |
| NA | NA | |
| NA | NA | |
| NA | NA | |
| NA | NA | |
| NA | NA | |
| NA | NA | |
| NA | NA | |
| 0.82515277 | NA | |
| NA | NA | |
| 0.20501326 | NA | |
| 0.022236277 | | 0.065224676 |
| 0.359698944 | NA | |
| 0.708801934 | | 0.814125883 |
| 0.082462765 | | 0.1817556 |
| NA | NA | |
| 0.5519202 | NA | |
| 0.582829116 | | 0.72031251 |
| 0.487591632 | | 0.64243875 |
| NA | NA | |
| 0.125119334 | NA | |
| 8.76E-05 | | 0.000623589 |
| 0.066347348 | | 0.154823487 |
| 0.29748297 | NA | |
| 4.91E-20 | | 2.09E-18 |
| 0.013002826 | | 0.042553728 |
| 0.083628113 | | 0.183817714 |

| | | |
|---|---|---|
| | 0.475401002 | 0.632634449 |
| | 0.125660789 | NA |
| | 0.219223409 | 0.370777771 |
| | 0.002938738 | 0.012603215 |
| | 0.015134497 | 0.048037244 |
| | 0.41736893 | 0.578916973 |
| | 0.953689491 | 0.971970805 |
| | 0.240117094 | 0.396105389 |
| | 0.047770873 | 0.119462314 |
| | 0.773720255 | NA |
| | 0.066754316 | 0.155458183 |
| | 0.009472101 | 0.032932363 |
| | 0.078450296 | 0.175194981 |
| | 4.38E-05 | 0.0003366 |
| | 0.440954136 | 0.601673374 |
| | 0.405225795 | 0.567254505 |
| | 1.96E-104 | 7.08E-102 |
| | 3.11E-11 | 6.66E-10 |
| | 0.933394574 | NA |
| | 1.12E-09 | 1.98E-08 |
| | 0.010799438 | 0.036651265 |
| | 0.514030132 | 0.665123414 |
| | 0.482571961 | 0.638617692 |
| | 0.955792873 | NA |
| | 0.548471567 | 0.694416527 |
| | 1.01E-07 | 1.35E-06 |
| | 7.69E-34 | 6.29E-32 |
| | 1.28E-08 | 1.98E-07 |
| | 0.270730386 | 0.430686048 |
| | 0.45868481 | 0.618146885 |
| | 5.90E-05 | 0.000438654 |
| | 0.001747437 | 0.008174624 |
| | 0.181803712 | 0.324214576 |
| | 0.740262023 | 0.836603992 |
| | 0.570243235 | 0.710959587 |
| | 0.321073239 | 0.483835722 |
| | 0.020164358 | 0.060265201 |
| NA | | NA |
| | 0.027865508 | 0.07829711 |
| | 0.063215675 | 0.148976189 |
| | 0.000779278 | 0.004134727 |
| | 0.003293173 | 0.013868384 |
| | 0.980175233 | 0.987685995 |
| | 2.32E-05 | 0.000192154 |
| | 0.452634974 | 0.612572516 |
| | 0.681951729 | 0.795482237 |
| | 0.788864934 | NA |

| | |
|---|---|
| 0.212099332 | 0.362525595 |
| 0.102869536 | 0.214273274 |
| 0.005252315 | 0.020317248 |
| NA | NA |
| 0.000477998 | 0.002730335 |
| 0.074831775 | 0.169133596 |
| 0.002162594 | 0.009758116 |
| NA | NA |
| NA | NA |
| NA | NA |
| NA | NA |
| 0.923191794 | NA |
| NA | NA |
| 0.013278691 | 0.043313607 |
| 0.146904192 | 0.279059343 |
| 0.835336407 | 0.899994439 |
| 0.839318641 | NA |
| 0.070034223 | 0.161288038 |
| 0.000210436 | 0.001340324 |
| 2.61E-98 | 8.86E-96 |
| 0.414618629 | 0.576583207 |
| 0.28141111 | 0.441655495 |
| 0.096528902 | 0.204229416 |
| 0.941278788 | NA |
| 0.000856943 | 0.004477929 |
| 0.552168475 | 0.697448209 |
| NA | NA |
| 5.90E-27 | 3.63E-25 |
| NA | NA |
| 0.148134504 | 0.280823006 |
| 0.941278332 | NA |
| 0.017156 | 0.053084602 |
| 0.474620228 | 0.632101089 |
| 0.519911847 | 0.670292589 |
| 0.148722073 | 0.281568785 |
| 0.60364849 | 0.736822633 |
| 0.351186547 | 0.515109595 |
| 0.088400455 | 0.191700768 |
| 0.00101644 | 0.005179472 |
| 0.196935634 | 0.343731169 |
| 0.814684374 | NA |
| 0.598783733 | NA |
| 0.941278424 | NA |
| 0.004832546 | 0.018980901 |
| 0.609311996 | NA |
| 6.85E-06 | 6.45E-05 |
| 0.924198761 | NA |

| | | |
|---|---|---|
| | 0.002605282 | 0.011424267 |
| | 0.925970516 | 0.955475364 |
| | 0.017958114 | 0.055033193 |
| | 1.53E-06 | 1.66E-05 |
| | 0.941277857 | NA |
| | 0.714421554 | NA |
| | 0.941278332 | NA |
| | 0.688368799 | NA |
| | 0.391208931 | NA |
| NA | | NA |
| | 0.12640096 | 0.249653668 |
| | 0.27181054 | 0.431683089 |
| | 0.722999792 | 0.824533614 |
| | 2.50E-05 | 0.000205551 |
| NA | | NA |
| | 0.178369422 | 0.319967668 |
| | 0.908214309 | 0.944400043 |
| | 0.174665282 | 0.315622764 |
| | 0.034174643 | 0.091870888 |
| | 0.12587161 | 0.248878223 |
| | 0.008337611 | 0.029663948 |
| | 0.941276431 | NA |
| | 0.345118493 | 0.508795935 |
| | 0.83756937 | 0.901433656 |
| NA | | NA |
| | 9.64E-61 | 1.75E-58 |
| | 1.75E-05 | 0.000149027 |
| | 0.040335251 | 0.104840458 |
| | 0.739991096 | NA |
| | 0.44472451 | 0.605360447 |
| | 0.166207474 | 0.305040504 |
| NA | | NA |
| | 0.726207708 | 0.827161596 |
| | 1.94E-05 | 0.000163147 |
| | 0.000973672 | 0.004989311 |
| | 0.614370516 | NA |
| | 0.130645359 | 0.255683345 |
| NA | | NA |
| NA | | NA |
| | 0.329904767 | NA |
| | 0.038808306 | 0.101630024 |
| | 0.109999469 | 0.2251468 |
| | 0.672109911 | NA |
| | 0.073528645 | 0.167076736 |
| | 3.63E-05 | 0.000284874 |
| NA | | NA |
| | 0.086799419 | 0.189055616 |

| | | |
|---|---|---|
| | 0.076061865 | 0.171214726 |
| | 0.878834859 | NA |
| | 0.730414265 | 0.830111521 |
| | 0.224543481 | NA |
| | 1.71E-45 | 2.08E-43 |
| | 0.000647073 | 0.003529247 |
| | 0.013175089 | 0.043016936 |
| | 0.653046503 | 0.773708953 |
| | 0.38879798 | 0.551561603 |
| | 0.733409253 | NA |
| | 1.42E-09 | 2.46E-08 |
| | 0.421650391 | 0.583069609 |
| | 0.208860529 | 0.358237437 |
| | 0.000514729 | 0.002910115 |
| | 2.72E-05 | 0.000220897 |
| | 0.132242873 | 0.25788805 |
| | 0.102030252 | 0.212935034 |
| | 0.82256539 | 0.892041855 |
| | 0.147316236 | 0.279641204 |
| | 0.000323492 | 0.001945858 |
| | 0.855489631 | 0.913040498 |
| | 6.10E-10 | 1.11E-08 |
| | 3.98E-06 | 3.95E-05 |
| | 0.323387359 | 0.486083105 |
| | 0.730066049 | 0.830046911 |
| | 0.608104376 | 0.740539236 |
| | 0.613229071 | NA |
| | 0.002296119 | 0.010282639 |
| | 0.941277857 | NA |
| | 0.838882104 | NA |
| | 0.366036564 | 0.529243245 |
| | 0.00180285 | 0.0083817 |
| | 0.286258911 | 0.44693569 |
| | 0.507293758 | 0.65900643 |
| | 0.569205859 | 0.710298724 |
| | 0.022237291 | 0.065224676 |
| | 0.017140604 | 0.0530534 |
| | 0.623226852 | 0.751385543 |
| | 0.011665391 | 0.039064174 |
| | 7.82E-16 | 2.54E-14 |
| | 0.955791922 | NA |
| | 1.33E-11 | 2.97E-10 |
| | 0.941276431 | NA |
| NA | | NA |
| NA | | NA |
| | 0.025699945 | 0.07324115 |
| | 0.418280796 | NA |

| | |
|---|---|
| NA | NA |
| 0.270709284 | 0.430681264 |
| 0.00444332 | 0.017697439 |
| NA | NA |
| 0.879710995 | NA |
| NA | NA |
| 0.644348794 | NA |
| 0.580297798 | 0.718190941 |
| 0.955792398 | NA |
| 0.305057768 | 0.466704268 |
| 0.839320041 | NA |
| 0.853596091 | NA |
| 0.941275956 | NA |
| 0.955792873 | NA |
| 0.376967007 | 0.539833265 |
| NA | NA |
| 0.477504224 | 0.634380193 |
| 0.966687636 | NA |
| 0.908945009 | NA |
| 0.05186049 | 0.127624419 |
| 0.003321348 | 0.013962333 |
| 3.46E-07 | 4.20E-06 |
| 0.157066755 | 0.292827238 |
| 0.242019626 | 0.398268193 |
| 0.018092431 | 0.055370261 |
| 0.87510137 | NA |
| 0.015134031 | 0.048037244 |
| 0.169201848 | 0.308743683 |
| 0.194116843 | 0.340194574 |
| 0.00256571 | 0.011281919 |
| 0.272184261 | 0.432103634 |
| 0.804053432 | 0.880356952 |
| 0.808771762 | NA |
| 0.002041405 | 0.009304773 |
| 0.748393366 | NA |
| 0.007058953 | 0.025860736 |
| 0.86997928 | 0.922166413 |
| 0.552870784 | NA |
| 0.452838174 | 0.612784146 |
| 0.143441194 | 0.274033815 |
| 0.06258488 | 0.147811602 |
| 0.6404377 | NA |
| 1.74E-07 | 2.24E-06 |
| 0.912887466 | 0.947479504 |
| 0.009440273 | 0.032835218 |
| NA | NA |
| 0.511466606 | 0.662959656 |

| | |
|---|---|
| 0.622732467 | 0.751103586 |
| 0.132705811 | 0.258588924 |
| 0.029497562 | 0.081868118 |
| 0.532212028 | 0.680766912 |
| 0.006798962 | 0.025070316 |
| 0.028407616 | 0.079501244 |
| NA | NA |
| 0.000563937 | 0.003143714 |
| 0.006457683 | 0.02402028 |
| 0.859970729 | 0.916519447 |
| 1.66E-05 | 0.00014213 |
| 0.633999902 | 0.759459772 |
| 0.759462165 | 0.850149013 |
| 0.931001941 | NA |
| NA | NA |
| 2.68E-05 | 0.0002183 |
| 1.89E-21 | 8.88E-20 |
| 2.13E-15 | 6.69E-14 |
| 0.905963008 | NA |
| 0.034903543 | 0.093422154 |
| 0.170330715 | 0.310073551 |
| NA | NA |
| NA | NA |
| 0.024850775 | 0.071350802 |
| 0.588307973 | 0.724563095 |
| 0.085848952 | 0.187569006 |
| 0.839319108 | NA |
| 0.240152793 | NA |
| 0.705462454 | 0.812065262 |
| NA | NA |
| 0.769030771 | 0.857023917 |
| 0.617649467 | 0.747168765 |
| 0.020866195 | 0.061881034 |
| 0.953932387 | NA |
| 0.00511431 | 0.019884158 |
| 0.24268726 | 0.399118508 |
| 0.941277857 | NA |
| 8.56E-14 | 2.35E-12 |
| NA | NA |
| 0.565802428 | NA |
| 0.839318641 | NA |
| 0.955792873 | NA |
| 0.000586117 | 0.003247384 |
| 0.002734107 | 0.011882228 |
| 0.47692163 | 0.633714903 |
| 0.950097919 | NA |
| 0.611044785 | NA |

| | |
|---|---|
| NA | NA |
| 7.41E-07 | 8.45E-06 |
| 0.680234895 | NA |
| 0.175596598 | 0.316851507 |
| 1.24E-09 | 2.17E-08 |
| 0.043873351 | 0.111815564 |
| NA | NA |
| NA | NA |
| 0.420005747 | 0.581659207 |
| 0.062230263 | 0.14720327 |
| NA | NA |
| NA | NA |
| 0.955792873 | NA |
| 0.044075343 | 0.112159698 |
| 0.839318641 | NA |
| NA | NA |
| 0.103256211 | 0.214777971 |
| NA | NA |
| 0.955792873 | NA |
| 0.227871049 | 0.381279199 |
| 0.949717216 | 0.96926375 |
| 0.020448506 | 0.060918964 |
| 0.246787645 | 0.40382025 |
| 0.343592009 | NA |
| 0.839319108 | NA |
| 0.789126276 | NA |
| 0.839319108 | NA |
| 0.003362462 | 0.014100716 |
| NA | NA |
| 0.141234701 | 0.271014861 |
| 6.80E-17 | 2.37E-15 |
| 0.018776112 | 0.057054643 |
| 0.000487927 | 0.002777054 |
| NA | NA |
| 0.001670668 | 0.007863404 |
| 0.95013655 | 0.96946979 |
| 0.000492574 | 0.002800155 |
| NA | NA |
| 0.276592128 | 0.436307631 |
| 0.839319108 | NA |
| 0.842463131 | 0.905132592 |
| 0.523880694 | 0.673572118 |
| NA | NA |
| 0.941278788 | NA |
| NA | NA |
| NA | NA |
| 0.99994737 | 0.999955691 |

| | |
|---|---|
| NA | NA |
| 1.87E-12 | 4.58E-11 |
| 0.00052268 | 0.002948691 |
| 0.000739861 | 0.0039609 |
| 0.010455688 | 0.035689986 |
| 3.17E-05 | 0.000253095 |
| 0.774063949 | NA |
| 0.402385271 | 0.564648875 |
| 0.497008777 | 0.649989171 |
| 0.036239568 | 0.096312439 |
| NA | NA |
| 0.955792397 | NA |
| 0.839318641 | NA |
| NA | NA |
| NA | NA |
| 0.004177027 | 0.016825452 |
| 1.31E-07 | 1.73E-06 |
| 0.113612597 | 0.23059666 |
| NA | NA |
| 0.888717793 | 0.93342843 |
| 0.647818075 | NA |
| 0.088154618 | 0.19126738 |
| 0.013051465 | 0.042677699 |
| 0.000824791 | 0.004339449 |
| 0.125307887 | 0.248155282 |
| 0.040025031 | 0.104245773 |
| 0.894527985 | NA |
| 0.15912254 | 0.295477819 |
| 0.894961891 | 0.936758264 |
| 0.090790323 | 0.195511741 |
| 0.901960009 | NA |
| 0.051247317 | 0.126455385 |
| 0.941275956 | NA |
| 0.74185626 | NA |
| 0.992625137 | NA |
| 0.955792398 | NA |
| 0.955792873 | NA |
| NA | NA |
| 0.311505368 | 0.473511091 |
| NA | NA |
| 0.101367025 | 0.211848094 |
| 0.256194689 | 0.414318928 |
| 0.889868706 | 0.933726633 |
| 0.743430676 | 0.838711582 |
| 6.65E-09 | 1.07E-07 |
| 0.925509841 | NA |
| 0.053468152 | 0.130675526 |

| | |
|---|---|
| 0.003557305 | 0.014756013 |
| 0.180469309 | 0.322511487 |
| 0.804840741 | 0.880692319 |
| NA | NA |
| 0.113716841 | 0.230690211 |
| 2.07E-06 | 2.18E-05 |
| 1.30E-17 | 4.73E-16 |
| 0.23329198 | 0.387882393 |
| 0.246412304 | 0.403405025 |
| 0.603620142 | 0.736822633 |
| 0.055428238 | 0.134527474 |
| 0.013332831 | 0.043450859 |
| 0.659037444 | 0.778095696 |
| 7.61E-48 | 9.95E-46 |
| 1.85E-05 | 0.000156394 |
| 0.918150685 | 0.950786279 |
| 2.58E-05 | 0.000211121 |
| 0.000299565 | 0.001820954 |
| 0.748393366 | NA |
| 0.00084629 | 0.004429107 |
| 3.39E-06 | 3.42E-05 |
| 0.157961219 | 0.293894547 |
| 0.360758375 | 0.524479039 |
| 0.701386364 | 0.809347846 |
| 0.025393828 | 0.072568602 |
| 0.675318099 | 0.790767029 |
| 0.284397518 | 0.445022006 |
| 0.210972228 | NA |
| 0.266813003 | 0.426543697 |
| 0.200480108 | 0.348079913 |
| 0.470867757 | 0.628474981 |
| NA | NA |
| 0.918879521 | 0.951334074 |
| 0.697762208 | NA |
| 0.003422313 | 0.014288351 |
| 0.104418861 | 0.216552908 |
| 0.613740817 | 0.744153273 |
| 0.357769148 | 0.521343288 |
| 0.098112365 | 0.206770897 |
| 7.29E-05 | 0.000529142 |
| 9.40E-06 | 8.58E-05 |
| NA | NA |
| 0.076881945 | 0.172613279 |
| 0.028199297 | 0.079029751 |
| 3.84E-10 | 7.20E-09 |
| 0.955792398 | NA |
| 0.05725803 | 0.137871081 |

| | |
|---|---|
| 0.437068759 | 0.59798302 |
| 0.001131219 | 0.005672059 |
| NA | NA |
| NA | NA |
| 8.96E-05 | 0.000636179 |
| 0.80695542 | NA |
| 0.123130637 | 0.244964513 |
| 0.155911597 | 0.291242179 |
| 3.15E-06 | 3.19E-05 |
| 0.435383311 | 0.596499947 |
| 0.128960253 | 0.25318582 |
| 0.042668983 | 0.109528331 |
| 0.559158638 | NA |
| 0.839318641 | NA |
| 0.029464098 | 0.081784778 |
| 3.74E-05 | 0.000292017 |
| 0.22247744 | 0.374869605 |
| 0.791832533 | 0.871788609 |
| 0.3870726 | 0.549937445 |
| 5.24E-13 | 1.35E-11 |
| 8.19E-05 | 0.000587222 |
| 2.51E-06 | 2.60E-05 |
| 0.027680313 | 0.077896289 |
| 6.72E-13 | 1.71E-11 |
| 0.02483186 | 0.071305094 |
| 0.184781142 | 0.32785395 |
| 0.640556491 | 0.764160264 |
| 0.064963154 | 0.152340196 |
| 0.24415678 | 0.40064941 |
| 0.8876752 | 0.933029151 |
| 0.046289589 | 0.116566313 |
| 0.082268627 | 0.181478949 |
| 0.65182562 | 0.772647082 |
| 0.531357915 | 0.679966725 |
| 0.53438836 | 0.682660906 |
| NA | NA |
| 0.00165463 | 0.007803348 |
| 0.075384741 | 0.170189472 |
| 0.135213545 | 0.262294441 |
| 5.20E-11 | 1.09E-09 |
| 0.438047879 | NA |
| NA | NA |
| 0.028935937 | 0.08063847 |
| NA | NA |
| 1.17E-05 | 0.000103913 |
| 2.40E-05 | 0.000198376 |
| 0.016735899 | 0.052037605 |

| | |
|---|---|
| NA | NA |
| 0.171315928 | 0.311438305 |
| 2.60E-41 | 2.81E-39 |
| 0.001127947 | 0.005658038 |
| 0.175116857 | 0.316246695 |
| 0.010472923 | 0.035743692 |
| NA | NA |
| NA | NA |
| NA | NA |
| 0.630376237 | 0.756604064 |
| 0.941277857 | NA |
| NA | NA |
| 0.937574928 | NA |
| 0.000341627 | 0.002042041 |
| 0.825482462 | 0.893990126 |
| NA | NA |
| 0.078464919 | 0.175199063 |
| 0.179055523 | 0.320659059 |
| 0.466283921 | 0.624387617 |
| 6.59E-05 | 0.000484881 |
| 0.644996973 | NA |
| 0.955792873 | NA |
| 0.848917688 | 0.909153602 |
| NA | NA |
| NA | NA |
| 0.84401219 | NA |
| 0.452548143 | 0.61254199 |
| 3.85E-08 | 5.49E-07 |
| 0.030799114 | 0.084700232 |
| 0.119346136 | 0.239335809 |
| 2.60E-06 | 2.68E-05 |
| 0.340221686 | 0.50383243 |
| 0.386045269 | NA |
| 0.934411206 | NA |
| 0.097415864 | 0.205612521 |
| 0.085942029 | 0.187689071 |
| 0.640071162 | 0.7637344 |
| 0.923684082 | 0.954564454 |
| 0.045252672 | 0.114560415 |
| 0.000174243 | 0.001140318 |
| 1.84E-07 | 2.37E-06 |
| 0.941275956 | NA |
| 0.839319108 | NA |
| 0.839318641 | NA |
| 0.839319108 | NA |
| 1.14E-26 | 6.87E-25 |
| 6.32E-05 | 0.000467743 |

| | | |
|---|---|---|
| | 0.011851329 | 0.039507749 |
| | 0.001285107 | 0.006304228 |
| | 0.433499004 | NA |
| | 0.384774206 | 0.547554446 |
| | 0.835006512 | 0.899970585 |
| | 0.941277857 | NA |
| | 6.05E-30 | 4.20E-28 |
| | 0.001632751 | 0.007718519 |
| | 0.694696796 | NA |
| | 0.97027729 | 0.982229107 |
| | 0.000149673 | 0.000997903 |
| | 1.55E-07 | 2.01E-06 |
| | 0.943630623 | 0.966034241 |
| | 0.650936214 | 0.771977268 |
| | 0.032050059 | 0.087363204 |
| | 0.206377408 | 0.355261088 |
| | 0.142272588 | 0.272479252 |
| | 7.84E-217 | 1.43E-213 |
| | 0.000108432 | 0.00075157 |
| | 6.82E-10 | 1.24E-08 |
| | 0.671016509 | NA |
| | 0.52792109 | 0.677061145 |
| | 0.612511444 | 0.743447088 |
| | 0.009883483 | 0.034069277 |
| | 0.466794169 | NA |
| | 0.17382346 | 0.31455531 |
| | 0.003383569 | 0.01416386 |
| | 0.941277857 | NA |
| | 0.176628528 | 0.318266022 |
| | 0.000873715 | 0.004551579 |
| | 0.574144835 | 0.713655823 |
| | 3.64E-08 | 5.22E-07 |
| | 0.839319107 | NA |
| | 0.190594866 | 0.335443581 |
| | 0.000244055 | 0.001521474 |
| | 0.368802711 | NA |
| NA | | NA |
| | 0.21781929 | 0.369163681 |
| | 0.955792398 | NA |
| | 0.539217174 | 0.686499168 |
| | 0.00389253 | 0.015862011 |
| | 0.015665743 | 0.049394094 |
| | 0.321983959 | 0.484747766 |
| | 0.049586582 | 0.122916501 |
| | 0.002542963 | 0.011206741 |
| | 4.60E-07 | 5.44E-06 |
| | 5.51E-231 | 1.19E-227 |

| | |
|---|---|
| 6.65E-07 | 7.67E-06 |
| 0.747726593 | 0.841497435 |
| 0.076944374 | 0.172694954 |
| 0.095816107 | 0.203172826 |
| NA | NA |
| 0.563034309 | 0.705445208 |
| 0.453682263 | NA |
| 0.067223034 | 0.156269964 |
| 5.73E-05 | 0.00042829 |
| 0.201698564 | 0.349685281 |
| 0.695784594 | 0.805170278 |
| 0.000486844 | 0.002772691 |
| 0.9031781 | 0.941425231 |
| 0.954665737 | NA |
| 0.072543754 | 0.165328076 |
| 0.995649069 | NA |
| 0.008455614 | 0.029989646 |
| 0.950163485 | 0.96946979 |
| 0.351240853 | 0.515109595 |
| 0.934514019 | NA |
| 0.011676048 | 0.039082167 |
| 0.194355983 | NA |
| 0.225098025 | 0.377840846 |
| 0.532531113 | 0.680918912 |
| 1.28E-18 | 4.96E-17 |
| NA | NA |
| 0.008040618 | 0.028779368 |
| NA | NA |
| NA | NA |
| 0.182783987 | 0.32532898 |
| 0.406497045 | 0.568277552 |
| 0.000741252 | 0.003966151 |
| 3.96E-06 | 3.94E-05 |
| 0.219937269 | 0.371589074 |
| 0.078091764 | 0.174562376 |
| 0.945374456 | 0.967029215 |
| 0.094624163 | 0.201308932 |
| 0.03239239 | 0.088195342 |
| 0.037809698 | 0.099606688 |
| 0.330011747 | 0.492998155 |
| 0.4914076 | NA |
| 0.922849885 | NA |
| 0.069106973 | 0.159635497 |
| 0.941277857 | NA |
| 0.984825125 | 0.990484561 |
| 0.351326879 | 0.515109595 |
| 0.734058779 | NA |

| | |
|---|---|
| 2.70E-05 | 0.000219773 |
| 2.74E-06 | 2.81E-05 |
| 0.270569436 | 0.430545107 |
| 0.012682298 | 0.04169969 |
| 0.731092135 | 0.830477641 |
| 0.095131931 | 0.202111475 |
| 0.050721083 | 0.125299763 |
| 0.198746454 | 0.346038369 |
| NA | NA |
| 2.62E-05 | 0.000214386 |
| NA | NA |
| 0.014114557 | 0.045436122 |
| 0.005727352 | 0.021797637 |
| 0.685063561 | 0.797704755 |
| 0.898544353 | 0.938856571 |
| 0.003321165 | 0.013962333 |
| 0.788084881 | 0.869754548 |
| 0.004147691 | 0.016729365 |
| 0.957954501 | NA |
| 0.600892241 | 0.734592824 |
| 0.013706787 | 0.044369501 |
| 6.66E-05 | 0.000488886 |
| 0.093382474 | 0.199380126 |
| 0.037259278 | 0.098407162 |
| 3.25E-06 | 3.29E-05 |
| NA | NA |
| 0.024336881 | 0.070222626 |
| 0.41068674 | 0.572618229 |
| 0.36989981 | 0.533112642 |
| 0.151227082 | 0.285064308 |
| 0.505747963 | 0.65752944 |
| 0.639777971 | NA |
| 3.07E-06 | 3.12E-05 |
| 0.074385331 | 0.16842843 |
| NA | NA |
| 0.941278332 | NA |
| 0.955791446 | NA |
| 0.955792398 | NA |
| 3.30E-07 | 4.03E-06 |
| 0.417132119 | NA |
| 5.81E-11 | 1.20E-09 |
| NA | NA |
| NA | NA |
| 0.260968176 | 0.419728666 |
| NA | NA |
| 0.941277857 | NA |
| 0.024474513 | 0.070508638 |

| | |
|---|---|
| 0.008244219 | NA |
| 0.105521092 | 0.218274213 |
| 0.112031937 | 0.228122022 |
| 1.69E-07 | 2.19E-06 |
| 8.52E-08 | 1.16E-06 |
| 0.000215224 | 0.001367527 |
| 0.009182549 | 0.03209858 |
| 0.07331444 | 0.16671733 |
| 0.7604942 | 0.851024116 |
| NA | NA |
| 0.000586473 | 0.003248601 |
| 6.24E-05 | 0.000462387 |
| 0.67853317 | 0.79293237 |
| 0.000100784 | 0.000705963 |
| 0.566053354 | 0.707849213 |
| 0.011461564 | 0.038477877 |
| 0.45151442 | NA |
| NA | NA |
| NA | NA |
| NA | NA |
| NA | NA |
| NA | NA |
| NA | NA |
| NA | NA |
| NA | NA |
| NA | NA |
| 0.078439145 | 0.175194981 |
| 0.013434165 | 0.043718894 |
| 0.367907501 | 0.531012937 |
| 0.002083465 | 0.009461935 |
| 0.024348343 | 0.070238668 |
| 0.336873043 | 0.500078095 |
| 0.076235792 | 0.171512054 |
| 0.172328276 | 0.312562496 |
| 0.852104236 | 0.911172335 |
| 5.06E-06 | 4.92E-05 |
| 0.034266497 | 0.092076222 |
| 0.657518966 | 0.776985372 |
| 0.562563345 | 0.705189211 |
| 0.006596448 | 0.02446761 |
| 1.15E-09 | 2.02E-08 |
| 0.750778411 | 0.843499437 |
| 0.340227655 | 0.50383243 |
| 0.00352186 | 0.014639769 |
| NA | NA |
| NA | NA |
| 0.45260725 | 0.612572516 |

| | |
|---|---|
| 0.075574595 | 0.170521045 |
| 0.839319574 | NA |
| 0.000929737 | 0.004793068 |
| 0.000236025 | 0.001477994 |
| NA | NA |
| NA | NA |
| 0.12202426 | 0.243455275 |
| 0.946009346 | 0.967272298 |
| 0.014196186 | 0.045686536 |
| 0.017384825 | 0.053634945 |
| 3.73E-06 | 3.73E-05 |
| 0.176610112 | 0.318266022 |
| 0.010088928 | 0.034651755 |
| 0.134503922 | 0.261258556 |
| NA | NA |
| 0.009740296 | 0.03370224 |
| 5.33E-06 | 5.15E-05 |
| 0.955792873 | NA |
| 0.396464283 | NA |
| 0.718549988 | NA |
| 0.081732616 | 0.180547537 |
| 2.58E-05 | 0.000211724 |
| 0.000406559 | 0.002368309 |
| 0.447011221 | 0.607326964 |
| 0.729887658 | 0.829923376 |
| 2.54E-08 | 3.75E-07 |
| 0.941277857 | NA |
| 0.513230828 | 0.664414238 |
| 0.93946514 | NA |
| 0.008491463 | 0.030080933 |
| 0.790344476 | NA |
| 4.44E-15 | 1.36E-13 |
| 0.680129346 | 0.794122482 |
| 0.000338433 | 0.002025999 |
| 0.237531053 | 0.392973805 |
| 0.418363116 | 0.579991781 |
| NA | NA |
| 0.519449501 | 0.669986833 |
| NA | NA |
| NA | NA |
| 0.675096287 | 0.790546188 |
| 0.732358088 | 0.831176115 |
| 0.202726038 | 0.350812801 |
| 0.838518173 | 0.902115782 |
| NA | NA |
| 0.941276431 | NA |
| 0.23869367 | 0.394458566 |

| | |
|---|---|
| 0.889223388 | 0.933455095 |
| 0.79974672 | 0.877620713 |
| 0.061136935 | 0.145198696 |
| 0.707848148 | 0.813678645 |
| 0.957527636 | 0.973935849 |
| NA | NA |
| 0.916760998 | NA |
| 0.00286347 | 0.012331267 |
| NA | NA |
| 0.640962192 | 0.764529295 |
| 0.941278332 | NA |
| NA | NA |
| NA | NA |
| NA | NA |
| 0.839319108 | NA |
| 0.887414837 | NA |
| 5.68E-10 | 1.04E-08 |
| 0.955792398 | NA |
| 0.21615112 | 0.367386629 |
| 0.764827897 | 0.853946599 |
| 0.084369455 | 0.185084035 |
| NA | NA |
| 0.937902578 | NA |
| 0.381362447 | NA |
| 0.001340669 | 0.006532365 |
| 0.640856509 | NA |
| 0.955792398 | NA |
| NA | NA |
| 0.941278332 | NA |
| 0.31466388 | 0.476894599 |
| 1.81E-11 | 3.99E-10 |
| 0.200138306 | 0.347613249 |
| 0.286233093 | 0.446924698 |
| 0.126289607 | 0.249601031 |
| 0.000100156 | 0.000702595 |
| 0.233753159 | 0.388343176 |
| NA | NA |
| 0.000112394 | 0.000776323 |
| 0.495610283 | 0.6485522 |
| 0.427774037 | 0.589041703 |
| 1.25E-07 | 1.65E-06 |
| 0.380222592 | 0.542960052 |
| 0.006147091 | 0.023099632 |
| 0.106434527 | 0.219832923 |
| 0.194325015 | 0.340397705 |
| 1.90E-06 | 2.02E-05 |
| 0.207583813 | 0.356770158 |

| | |
|---|---|
| 1.57E-06 | 1.70E-05 |
| 0.443252545 | 0.603805704 |
| NA | NA |
| 0.490633082 | 0.64498225 |
| 0.036343298 | 0.096515142 |
| 0.346706142 | 0.510567319 |
| 0.505214143 | 0.65720799 |
| 0.680090566 | NA |
| 0.828316384 | 0.895633788 |
| NA | NA |
| 0.839319108 | NA |
| NA | NA |
| 0.955792873 | NA |
| 0.839319108 | NA |
| 0.895630503 | NA |
| 0.69896501 | 0.807570757 |
| NA | NA |
| 0.825063007 | 0.893704414 |
| 0.031623893 | 0.086506386 |
| 0.270286819 | 0.430354329 |
| 0.305417703 | 0.466954365 |
| 0.035950169 | 0.095757154 |
| 0.02154429 | 0.063520028 |
| 0.283867828 | 0.444450387 |
| 0.917107353 | 0.950201942 |
| 0.522834451 | 0.672772166 |
| 0.957142932 | 0.973669377 |
| 0.251564388 | 0.409227276 |
| 0.003997106 | 0.016202167 |
| NA | NA |
| 0.091191783 | 0.196090073 |
| 0.824927724 | 0.893634157 |
| 0.149278737 | 0.282330752 |
| NA | NA |
| 0.833205059 | 0.898918811 |
| NA | NA |
| 0.839318641 | NA |
| 0.735978364 | 0.833623157 |
| 0.953798549 | NA |
| 0.671993917 | NA |
| NA | NA |
| 0.853520029 | NA |
| NA | NA |
| 0.699962715 | NA |
| NA | NA |
| 0.479038878 | NA |
| 0.955792398 | NA |

| | |
|---|---|
| NA | NA |
| 0.922535448 | NA |
| 0.941278332 | NA |
| 0.844737756 | NA |
| 0.868770136 | 0.921377123 |
| 0.093668833 | 0.199794387 |
| 0.914516818 | 0.94855017 |
| 0.006794717 | 0.025058545 |
| 0.001091131 | 0.005502352 |
| 0.00474438 | 0.018711802 |
| NA | NA |
| 0.00592132 | 0.022410439 |
| 0.056674719 | 0.136831766 |
| 6.83E-10 | 1.24E-08 |
| NA | NA |
| NA | NA |
| 0.000129168 | 0.000877345 |
| 0.189677951 | 0.334264369 |
| 4.02E-89 | 1.19E-86 |
| 4.35E-47 | 5.59E-45 |
| 0.546489996 | 0.692701826 |
| 0.342780861 | 0.506230173 |
| NA | NA |
| NA | NA |
| NA | NA |
| 0.985430733 | 0.990884302 |
| 0.945883454 | NA |
| 0.169578725 | 0.309224914 |
| 0.103958977 | 0.21588131 |
| 9.68E-05 | 0.000681469 |
| 0.639130484 | 0.763032742 |
| 0.218167749 | 0.369464871 |
| NA | NA |
| 0.000463399 | 0.002657149 |
| 0.372395848 | 0.535736837 |
| NA | NA |
| 0.306580787 | 0.468161666 |
| 0.09105113 | 0.195949276 |
| 0.477651551 | 0.63450509 |
| 0.002475644 | 0.010948583 |
| 0.823641682 | 0.89285054 |
| 0.895318201 | NA |
| NA | NA |
| NA | NA |
| 0.941277857 | NA |
| 0.043284041 | 0.110710016 |
| 0.326554328 | 0.489405996 |

| | | | |
|---|---|---|---|
| | 0.001839101 | | 0.008516431 |
| | 0.039906389 | | 0.10399079 |
| NA | | NA | |
| | 0.251700444 | | 0.409415292 |
| | 0.955792873 | NA | |
| | 0.705171568 | NA | |
| | 0.557171066 | | 0.701125335 |
| | 0.247326614 | | 0.404456756 |
| | 0.282255435 | | 0.442601216 |
| NA | | NA | |
| | 0.073618695 | | 0.167154536 |
| NA | | NA | |
| NA | | NA | |
| | 0.37248546 | | 0.535768856 |
| | 0.502511562 | | 0.654560291 |
| | 0.744137954 | | 0.839270829 |
| | 0.0724455 | | 0.16519906 |
| | 0.000158648 | | 0.001049805 |
| | 0.391825678 | | 0.554183941 |
| | 0.008383311 | | 0.029777582 |
| | 2.08E-09 | | 3.55E-08 |
| | 2.53E-36 | | 2.27E-34 |
| NA | | NA | |
| | 0.077103988 | | 0.172958979 |
| | 1.01E-55 | | 1.57E-53 |
| | 3.10E-12 | | 7.42E-11 |
| | 0.299123867 | | 0.460705725 |
| | 0.193322064 | | 0.339090034 |
| NA | | NA | |
| | 0.073769031 | | 0.167415185 |
| | 0.839319108 | NA | |
| | 0.003244329 | | 0.013694186 |
| NA | | NA | |
| | 0.000309798 | | 0.001873429 |
| | 0.941275956 | NA | |
| NA | | NA | |
| | 0.983381527 | | 0.989370601 |
| | 0.001724357 | | 0.008089633 |
| NA | | NA | |
| | 0.189666966 | NA | |
| | 0.535904425 | | 0.684056263 |
| | 0.103997565 | | 0.215926776 |
| | 2.34E-06 | | 2.43E-05 |
| | 0.307363037 | | 0.468904948 |
| NA | | NA | |
| NA | | NA | |
| NA | | NA | |

| | |
|---|---|
| 0.839319108 | NA |
| 0.010100808 | 0.034682551 |
| NA | NA |
| NA | NA |
| NA | NA |
| 0.820501087 | 0.890678686 |
| 0.955791922 | NA |
| 0.74718063 | NA |
| 0.121669499 | 0.242898774 |
| 0.562840953 | 0.705330475 |
| 0.569695969 | 0.710736881 |
| 0.951068588 | 0.970148286 |
| 0.668868666 | NA |
| 0.000819312 | 0.004314257 |
| 0.941278332 | NA |
| 0.128120195 | 0.252213785 |
| 9.78E-06 | 8.90E-05 |
| NA | NA |
| NA | NA |
| NA | NA |
| 0.193256277 | NA |
| 0.609322887 | 0.741429546 |
| 0.000140434 | 0.00094291 |
| 0.000133878 | 0.000905789 |
| NA | NA |
| NA | NA |
| 3.99E-12 | 9.42E-11 |
| 0.459085027 | 0.618451782 |
| 0.017499796 | 0.053905804 |
| 0.005037025 | 0.019647863 |
| 0.911032953 | NA |
| 1.36E-09 | 2.37E-08 |
| 0.705620799 | 0.812185611 |
| 0.015414261 | 0.048730274 |
| 0.591618408 | 0.727323038 |
| 0.786931931 | 0.86916723 |
| 0.36942513 | 0.532718826 |
| 0.174117159 | 0.314943128 |
| 0.725028456 | NA |
| 0.796035895 | 0.874908486 |
| NA | NA |
| NA | NA |
| 0.001823243 | 0.008462722 |
| 0.955792398 | NA |
| 0.705728919 | 0.812209632 |
| 0.455429171 | 0.615042561 |
| 0.000268761 | 0.001653413 |

| | | |
|---|---|---|
| | 0.941279303 | NA |
| | 0.067856236 | 0.157387852 |
| | 0.855970579 | NA |
| NA | | NA |
| | 0.955792873 | NA |
| NA | | NA |
| | 0.00049241 | 0.002799892 |
| NA | | NA |
| NA | | NA |
| | 0.956095558 | 0.973116391 |
| NA | | NA |
| NA | | NA |
| | 0.81598515 | 0.887480947 |
| | 0.741391406 | 0.837403229 |
| | 0.590622342 | 0.726595333 |
| | 0.529917992 | 0.678894592 |
| | 0.685162694 | NA |
| NA | | NA |
| | 0.955792398 | NA |
| | 0.000619866 | 0.003401904 |
| NA | | NA |
| | 0.839319108 | NA |
| | 0.000478508 | 0.002732594 |
| | 9.82E-07 | 1.09E-05 |
| | 1.76E-07 | 2.27E-06 |
| | 0.551151945 | 0.696459896 |
| | 0.827997411 | 0.895532984 |
| | 0.574837547 | NA |
| | 0.142925942 | 0.273356632 |
| | 0.914652032 | 0.948649077 |
| | 0.546271299 | NA |
| | 0.024042912 | 0.069534554 |
| | 0.790921156 | NA |
| | 0.586374685 | 0.723080005 |
| | 0.000965379 | 0.004951076 |
| | 0.941279303 | NA |
| | 0.449339837 | 0.60948181 |
| | 0.706716776 | 0.812936048 |
| | 0.10251274 | 0.213690337 |
| NA | | NA |
| | 0.844641434 | 0.90649091 |
| | 0.941278207 | NA |
| | 0.019578848 | 0.058907313 |
| NA | | NA |
| NA | | NA |
| | 0.046476722 | 0.116866668 |
| NA | | NA |

| | |
|---|---|
| 0.77558268 | NA |
| 0.963368296 | 0.977829091 |
| 0.408789539 | 0.570842394 |
| 0.17479446 | 0.315808238 |
| 0.014928515 | 0.047510163 |
| 0.65754207 | 0.776985372 |
| 0.93859322 | 0.963595218 |
| 0.69344886 | 0.803616036 |
| 0.056061181 | 0.13574448 |
| 0.183647903 | 0.326378511 |
| 4.28E-05 | 0.000330088 |
| 0.073123984 | 0.16630012 |
| 2.12E-08 | 3.16E-07 |
| NA | NA |
| NA | NA |
| NA | NA |
| 0.00016939 | 0.001113762 |
| 0.403738614 | 0.565814089 |
| 0.395235855 | 0.557773261 |
| 0.218727587 | 0.370123277 |
| 0.288328375 | 0.44898868 |
| 0.350564917 | 0.514435645 |
| 0.018051964 | 0.055263387 |
| 0.019534663 | 0.058804104 |
| 0.597681265 | 0.732253475 |
| 5.54E-14 | 1.55E-12 |
| 0.739865916 | NA |
| 0.749851532 | 0.842839354 |
| 0.000704996 | 0.003799031 |
| 0.311516266 | 0.473511091 |
| NA | NA |
| 0.002846593 | 0.012267718 |
| NA | NA |
| NA | NA |
| 0.839319108 | NA |
| 0.863068525 | NA |
| 0.044675241 | 0.113340447 |
| 0.875647365 | 0.925643364 |
| 0.043669963 | 0.111438144 |
| 0.68849763 | 0.800333101 |
| NA | NA |
| 0.304259211 | NA |
| 0.225030917 | 0.377808131 |
| 0.000344927 | 0.002056085 |
| NA | NA |
| 0.897690844 | NA |
| 0.737805714 | 0.834858645 |

| | | |
|---|---|---|
| | 0.718805497 | 0.821055414 |
| | 0.067235095 | 0.1562822 |
| NA | | NA |
| | 0.839319108 | NA |
| NA | | NA |
| NA | | NA |
| | 0.424853819 | 0.586140534 |
| | 0.831861374 | 0.897998296 |
| | 0.941277857 | NA |
| NA | | NA |
| | 0.001757981 | 0.008219105 |
| | 2.61E-07 | 3.26E-06 |
| | 0.839318641 | NA |
| | 0.138306512 | 0.266836243 |
| | 0.578801235 | 0.7168925 |
| | 0.009045296 | 0.031717169 |
| | 0.393077536 | 0.555550828 |
| | 0.964634459 | 0.978571768 |
| | 0.080684438 | 0.178919034 |
| | 0.052764948 | 0.129408686 |
| | 0.829551407 | 0.896499875 |
| | 0.063391673 | 0.149317031 |
| | 0.601787059 | 0.735298604 |
| | 0.450926164 | 0.61093719 |
| NA | | NA |
| | 0.955792398 | NA |
| NA | | NA |
| NA | | NA |
| NA | | NA |
| NA | | NA |
| | 0.506132846 | 0.657873198 |
| NA | | NA |
| | 0.220799628 | 0.37270213 |
| | 0.642317966 | 0.76534102 |
| NA | | NA |
| NA | | NA |
| NA | | NA |
| NA | | NA |
| NA | | NA |
| NA | | NA |
| | 0.831413565 | NA |
| NA | | NA |
| | 0.050814433 | 0.125478278 |
| | 0.027634186 | 0.077821687 |
| | 2.95E-05 | 0.000238067 |
| | 0.060271157 | 0.143522077 |
| | 0.465204845 | 0.623644366 |

| | |
|---|---|
| 0.067728692 | 0.15713796 |
| 0.649868679 | 0.77113387 |
| 0.027883514 | 0.078327696 |
| 2.75E-05 | 0.000223595 |
| 0.833801616 | NA |
| NA | NA |
| NA | NA |
| NA | NA |
| NA | NA |
| 0.941277857 | NA |
| 2.34E-05 | 0.000193882 |
| 0.634271698 | 0.759670657 |
| NA | NA |
| NA | NA |
| NA | NA |
| 0.833801616 | NA |
| 0.925732935 | NA |
| 0.013905292 | 0.044889972 |
| 0.468263314 | 0.626453432 |
| NA | NA |
| NA | NA |
| 0.874004109 | 0.924785321 |
| 0.001030532 | 0.005240062 |
| 0.000301575 | 0.001830214 |
| 0.191943919 | 0.337269843 |
| 0.203952095 | 0.3524461 |
| 0.023289441 | 0.067758866 |
| 0.003295962 | 0.013877675 |
| NA | NA |
| 0.266276205 | 0.425762879 |
| 0.655494093 | 0.775104098 |
| NA | NA |
| 0.156724412 | 0.292416277 |
| NA | NA |
| 0.940604701 | 0.964388245 |
| 2.16E-05 | 0.000180484 |
| 0.941277857 | NA |
| 9.02E-09 | 1.43E-07 |
| 3.85E-06 | 3.84E-05 |
| 0.218881233 | 0.370330615 |
| 4.30E-05 | 0.00033129 |
| 0.04116357 | 0.106612234 |
| 0.005360517 | 0.020639119 |
| 0.106763828 | 0.220303461 |
| 0.752330038 | 0.844632517 |
| 0.108082575 | 0.222137953 |
| 0.941278207 | NA |

| | |
|---|---|
| 0.000679867 | 0.003683761 |
| 0.906987752 | 0.943641717 |
| 0.765228187 | 0.854255406 |
| 0.355856155 | 0.519573561 |
| 7.89E-10 | 1.42E-08 |
| NA | NA |
| 0.542485792 | 0.68931668 |
| 0.751475366 | 0.844026834 |
| 0.068207394 | 0.158063712 |
| 0.865474973 | NA |
| 0.108556629 | 0.22288147 |
| 0.010359561 | 0.035433004 |
| 0.839319108 | NA |
| 0.053700277 | 0.131135117 |
| NA | NA |
| 0.409627129 | 0.57154709 |
| 4.88E-08 | 6.84E-07 |
| 0.010212879 | 0.034981572 |
| 0.970564834 | 0.98224347 |
| 0.937924455 | NA |
| 0.327259993 | 0.490056141 |
| 0.204721135 | 0.353278652 |
| 0.561871942 | 0.704693643 |
| 0.449496249 | NA |
| NA | NA |
| NA | NA |
| 0.598224204 | 0.732643813 |
| 0.864172833 | 0.918999275 |
| 0.326185015 | 0.4891234 |
| 0.09160441 | 0.196642231 |
| 0.027429533 | 0.077373519 |
| 0.054208588 | 0.132131906 |
| NA | NA |
| 0.043956393 | 0.111970255 |
| 0.248203381 | 0.405417023 |
| NA | NA |
| 0.004133879 | 0.016682704 |
| 0.356217581 | 0.519878035 |
| 0.930028384 | 0.958402269 |
| 0.289920051 | 0.450877298 |
| 0.944569603 | 0.966578634 |
| NA | NA |
| NA | NA |
| 0.241033435 | 0.397194532 |
| 0.012719894 | 0.041800211 |
| 0.375342433 | 0.538446275 |
| NA | NA |

| | |
|---|---|
| 0.264735706 | 0.423755163 |
| 0.48863287 | 0.643347692 |
| NA | NA |
| 0.80447946 | 0.880558921 |
| 8.51E-05 | 0.000607885 |
| 0.002388355 | 0.010621755 |
| 0.234100795 | 0.388812165 |
| NA | NA |
| 0.544512877 | 0.691042149 |
| 0.701509449 | 0.809410438 |
| NA | NA |
| 0.251835446 | 0.409578906 |
| 0.361218987 | 0.524892206 |
| 0.404017768 | 0.566138639 |
| 0.00251487 | 0.011093204 |
| 1.20E-19 | 4.95E-18 |
| 0.031249024 | 0.085650537 |
| 0.001405795 | 0.006798197 |
| 0.504864325 | NA |
| 0.00111002 | 0.005583816 |
| 3.18E-07 | 3.89E-06 |
| 0.006941911 | 0.025514418 |
| 0.955792398 | NA |
| 0.537162198 | 0.685091592 |
| 0.534865146 | 0.682986062 |
| 0.472728571 | NA |
| 0.00454499 | 0.018042003 |
| 0.00359471 | 0.014864711 |
| 0.008945263 | 0.031417318 |
| 0.880078911 | 0.928159788 |
| 0.416753354 | 0.578400194 |
| 0.142638097 | 0.272981587 |
| 0.747738457 | 0.841497435 |
| NA | NA |
| 0.706072751 | 0.812470081 |
| 0.955792873 | NA |
| 0.114802039 | 0.232337227 |
| 7.36E-05 | 0.000533464 |
| NA | NA |
| 0.657949796 | 0.777197074 |
| 0.318759483 | 0.481152762 |
| 0.903852968 | 0.941698566 |
| 0.186067379 | 0.329620107 |
| 1.10E-11 | 2.48E-10 |
| NA | NA |
| 0.311971382 | 0.473960735 |
| 0.212105346 | 0.362525595 |

| | |
|---|---|
| 0.485407235 | 0.640802163 |
| NA | NA |
| NA | NA |
| NA | NA |
| 0.755176059 | 0.846654571 |
| 0.096861 | 0.204672581 |
| 0.161375868 | 0.298358584 |
| 0.941277857 | NA |
| 9.12E-08 | 1.24E-06 |
| 0.000710493 | 0.003824322 |
| 9.89E-05 | 0.000694332 |
| 0.002007586 | 0.009187572 |
| 1.44E-09 | 2.51E-08 |
| 0.167556317 | 0.306711948 |
| NA | NA |
| 0.004183318 | 0.016842243 |
| 0.004288801 | 0.017181743 |
| 0.002076211 | 0.009438117 |
| NA | NA |
| 0.743272628 | 0.838573024 |
| 0.790622989 | 0.87134363 |
| 0.048386315 | 0.120595924 |
| 0.380493526 | 0.543151427 |
| NA | NA |
| 2.59E-08 | 3.82E-07 |
| 0.230613989 | NA |
| 0.075755878 | 0.1707682 |
| 0.945837653 | 0.967272298 |
| 0.133920626 | 0.260401825 |
| NA | NA |
| 0.289202856 | 0.450026567 |
| 0.003526412 | 0.014653571 |
| 0.402445449 | 0.56466674 |
| 4.53E-08 | 6.38E-07 |
| 0.941278207 | NA |
| 0.149202011 | 0.282252923 |
| 0.068351112 | 0.158274032 |
| 0.792089576 | 0.871990938 |
| 0.077961966 | 0.174403264 |
| 8.53E-06 | 7.85E-05 |
| 0.180755498 | 0.322829021 |
| 7.36E-10 | 1.33E-08 |
| NA | NA |
| 0.940685982 | NA |
| NA | NA |
| 0.038494035 | 0.100973664 |
| 0.941278332 | NA |

| | |
|---|---|
| 0.050026724 | 0.123841505 |
| 0.304678213 | NA |
| 7.98E-05 | 0.00057416 |
| 0.000318235 | 0.00191714 |
| 0.525843921 | 0.675044348 |
| 0.122099199 | 0.243523139 |
| 0.400097684 | 0.562723875 |
| 0.001414677 | 0.006834802 |
| NA | NA |
| 0.018051461 | 0.055263387 |
| 0.557131185 | 0.701112216 |
| 3.34E-06 | 3.37E-05 |
| 0.941277857 | NA |
| 0.3501828 | 0.514001666 |
| 0.014399968 | 0.046167569 |
| 0.192131454 | 0.337522961 |
| 0.005298473 | 0.020455249 |
| 0.260070129 | NA |
| 0.592135906 | 0.727761843 |
| 0.00605564 | 0.022828054 |
| 0.118665682 | 0.23827236 |
| 0.001788875 | 0.008334076 |
| NA | NA |
| 0.349528172 | 0.513389049 |
| 0.361661122 | 0.525332776 |
| NA | NA |
| NA | NA |
| 0.001153908 | 0.00576639 |
| NA | NA |
| NA | NA |
| 0.727611677 | NA |
| 0.125759265 | 0.248738742 |
| 0.839319108 | NA |
| NA | NA |
| 0.144282797 | 0.275332249 |
| 0.000727036 | 0.003905421 |
| 0.973583965 | 0.984043719 |
| 0.001002815 | 0.005122104 |
| 0.955792873 | NA |
| 0.345017522 | 0.508678585 |
| 0.204700053 | 0.353278652 |
| 0.0042893 | 0.017181743 |
| 0.010099103 | 0.034681699 |
| 0.001279855 | 0.006286236 |
| 0.56498106 | 0.706879688 |
| 0.25454392 | 0.412546261 |
| 0.290772694 | 0.451878536 |

| | |
|---|---|
| 3.67E-05 | 0.00028725 |
| 6.77E-06 | 6.39E-05 |
| 0.012613302 | 0.041518712 |
| 0.235015819 | 0.389923779 |
| 0.142921018 | 0.273356632 |
| 0.001719682 | 0.008073346 |
| NA | NA |
| 0.042382986 | 0.108964505 |
| 0.000191822 | 0.001235992 |
| 0.097809636 | 0.206224197 |
| 3.02E-07 | 3.71E-06 |
| 0.216198797 | 0.367437613 |
| 0.002967507 | 0.012705991 |
| 0.056676934 | 0.136831766 |
| 0.000638894 | 0.003488636 |
| 0.789899811 | 0.87082887 |
| 2.34E-06 | 2.43E-05 |
| 5.59E-06 | 5.39E-05 |
| NA | NA |
| NA | NA |
| 0.300659442 | 0.462099415 |
| 0.522860441 | 0.672772166 |
| 0.84771872 | 0.908395855 |
| 0.059565916 | 0.142276763 |
| 0.180687457 | 0.322731717 |
| 1.90E-08 | 2.85E-07 |
| 9.89E-21 | 4.43E-19 |
| 0.597869194 | 0.732322075 |
| 0.8997227 | 0.939592826 |
| 0.756437498 | 0.847480599 |
| 0.415880978 | NA |
| 2.75E-11 | 5.94E-10 |
| 0.046018295 | 0.116091941 |
| 4.52E-05 | 0.000345741 |
| 0.001182497 | 0.005885778 |
| 0.000500552 | 0.002839404 |
| 0.740209347 | 0.836584183 |
| 0.634467335 | 0.75971383 |
| 0.144765657 | 0.276032377 |
| 0.2172327 | 0.368604775 |
| 6.61E-05 | 0.000485751 |
| 1.02E-05 | 9.19E-05 |
| 0.001268788 | 0.006239603 |
| 0.730202571 | NA |
| 0.066291019 | 0.154720663 |
| 0.979957962 | 0.987592489 |
| 8.40E-06 | 7.75E-05 |

| | | | |
|---|---|---|---|
| | 1.14E-05 | | 0.000101325 |
| | 0.312522139 | | 0.474585424 |
| | 0.633153176 | | 0.758942031 |
| | 1.12E-10 | | 2.24E-09 |
| NA | | NA | |
| NA | | NA | |
| | 0.824032901 | | 0.893071264 |
| | 2.66E-05 | | 0.000217031 |
| | 2.64E-06 | | 2.72E-05 |
| | 0.726714887 | NA | |
| | 0.036498433 | | 0.096816638 |
| | 0.311784678 | | 0.473765214 |
| | 0.297057205 | | 0.45855092 |
| | 0.839810269 | NA | |
| | 0.31768047 | | 0.480145751 |
| | 0.739991096 | NA | |
| | 0.901521432 | | 0.940439722 |
| NA | | NA | |
| NA | | NA | |
| | 0.000131203 | | 0.000890107 |
| | 0.126703192 | | 0.25010693 |
| NA | | NA | |
| | 0.643986189 | NA | |
| NA | | NA | |
| | 0.956562856 | | 0.97337139 |
| | 0.525327403 | | 0.674679914 |
| | 3.63E-07 | | 4.39E-06 |
| | 0.059727977 | | 0.142535123 |
| | 0.11996117 | | 0.240265549 |
| | 0.373310814 | | 0.536562863 |
| | 0.874583107 | | 0.925009245 |
| | 0.445638282 | | 0.606118994 |
| | 0.012544216 | | 0.041331318 |
| | 0.441528235 | | 0.602154407 |
| | 0.201597873 | | 0.349561636 |
| | 0.866842833 | | 0.92052258 |
| | 0.000252392 | | 0.001565631 |
| | 0.707252171 | | 0.813272637 |
| | 0.50654283 | | 0.658190438 |
| | 0.941277857 | NA | |
| | 0.37146893 | | 0.534823583 |
| | 1.10E-05 | | 9.84E-05 |
| | 0.655009216 | NA | |
| | 0.005048419 | | 0.019685858 |
| | 0.955792398 | NA | |
| | 0.839319108 | NA | |
| NA | | NA | |

| | |
|---|---|
| 0.001342927 | 0.006542029 |
| 8.46E-10 | 1.52E-08 |
| NA | NA |
| NA | NA |
| NA | NA |
| NA | NA |
| NA | NA |
| NA | NA |
| NA | NA |
| NA | NA |
| NA | NA |
| NA | NA |
| NA | NA |
| NA | NA |
| NA | NA |
| NA | NA |
| NA | NA |
| NA | NA |
| NA | NA |
| NA | NA |
| 0.942848144 | NA |
| NA | NA |
| NA | NA |
| NA | NA |
| NA | NA |
| 0.94054529 | NA |
| NA | NA |
| 0.557587101 | 0.701317451 |
| 0.137833391 | 0.266097037 |
| 0.734307777 | NA |
| NA | NA |
| 0.724660557 | 0.826150522 |
| 0.032699774 | 0.08891174 |
| 0.217893551 | 0.36921608 |
| 0.147640004 | NA |
| NA | NA |
| NA | NA |
| NA | NA |
| 0.78833772 | 0.869831931 |
| NA | NA |
| NA | NA |
| 0.799668177 | 0.877620713 |
| 0.941278838 | NA |
| 0.538993393 | 0.68637958 |
| 0.011805528 | 0.039388166 |
| 0.985406089 | 0.990884302 |
| 0.151894355 | 0.285846653 |

| | | |
|---|---|---|
| | 0.941278788 | NA |
| NA | | NA |
| | 0.007403347 | 0.026915299 |
| | 0.743511643 | NA |
| NA | | NA |
| NA | | NA |
| | 0.704737765 | 0.811679487 |
| | 0.941278332 | NA |
| | 0.839319108 | NA |
| | 0.752447507 | NA |
| | 0.000202282 | 0.001293928 |
| NA | | NA |
| NA | | NA |
| | 0.061331727 | 0.145545141 |
| NA | | NA |
| | 0.005388286 | 0.020717819 |
| | 5.37E-31 | 3.90E-29 |
| | 0.003086534 | 0.013128251 |
| NA | | NA |
| | 0.185212638 | 0.328448161 |
| | 0.817998496 | 0.888899731 |
| | 0.017243407 | 0.05328844 |
| | 0.021620168 | 0.063712175 |
| | 0.000340486 | 0.002036242 |
| | 0.936257905 | 0.962255791 |
| | 0.73980791 | NA |
| | 0.719061629 | 0.821183423 |
| | 0.7996816 | NA |
| | 0.417542195 | 0.57905607 |
| | 0.431860796 | 0.593107764 |
| | 0.467443775 | 0.625694339 |
| | 0.225696692 | 0.378605445 |
| | 0.269459104 | 0.429381106 |
| | 0.000121295 | 0.000830492 |
| | 0.808410289 | 0.882488612 |
| | 0.839319108 | NA |
| | 0.009782222 | 0.033808111 |
| | 0.000686198 | 0.00371532 |
| | 0.394091658 | 0.556620558 |
| | 0.029514227 | 0.08189527 |
| | 0.125631186 | 0.248549359 |
| | 0.084404577 | 0.185127569 |
| | 0.900939253 | 0.940203267 |
| | 0.624142788 | NA |
| | 0.668049022 | 0.784842192 |
| | 2.68E-05 | 0.000218303 |
| | 0.151695222 | 0.285723529 |

| | |
|---|---|
| 0.48384647 | 0.639699699 |
| 0.509057355 | NA |
| 0.189659604 | NA |
| 0.89143918 | 0.934565988 |
| 0.772883918 | 0.859898977 |
| NA | NA |
| NA | NA |
| NA | NA |
| 0.955792873 | NA |
| NA | NA |
| 0.661151297 | 0.779701886 |
| NA | NA |
| 2.05E-11 | 4.50E-10 |
| 0.075729105 | 0.170740187 |
| NA | NA |
| NA | NA |
| 0.897090734 | 0.937872959 |
| 0.126783332 | 0.250182132 |
| 4.28E-07 | 5.12E-06 |
| 0.025067959 | 0.071783873 |
| 0.072011063 | 0.164482143 |
| NA | NA |
| 9.50E-27 | 5.77E-25 |
| 1.42E-24 | 7.86E-23 |
| 0.216339541 | 0.367608712 |
| 0.154175871 | 0.288856135 |
| 2.10E-07 | 2.67E-06 |
| 0.056514821 | 0.136551189 |
| 0.082817278 | NA |
| 0.388036553 | 0.550945242 |
| 0.823894593 | 0.893002691 |
| 0.839319108 | NA |
| NA | NA |
| 0.275101859 | 0.434965753 |
| 0.584796088 | 0.721733424 |
| 8.34E-05 | 0.000596932 |
| 0.086152276 | 0.18792123 |
| 0.071025638 | 0.162850615 |
| 0.07652669 | 0.172000782 |
| NA | NA |
| 0.00085006 | 0.00444537 |
| 0.061825361 | 0.146424602 |
| 0.214975312 | 0.365918068 |
| 0.796857977 | 0.875503962 |
| 0.616339513 | 0.746152947 |
| 0.017565184 | 0.05407117 |
| 0.012645677 | 0.041602269 |

| | |
|---|---|
| 0.26898047 | NA |
| 0.001135566 | 0.00569026 |
| 0.259193878 | 0.417636927 |
| 3.77E-17 | 1.33E-15 |
| 0.00912188 | 0.031943356 |
| 2.29E-104 | 8.12E-102 |
| 0.955792398 | NA |
| 0.438747946 | 0.599659984 |
| 0.004243603 | 0.017041715 |
| NA | NA |
| 0.811511034 | NA |
| NA | NA |
| 0.941278332 | NA |
| NA | NA |
| NA | NA |
| 0.026773923 | 0.075873764 |
| NA | NA |
| NA | NA |
| NA | NA |
| 0.147908866 | NA |
| NA | NA |
| 0.050138343 | 0.124053309 |
| 0.94127925 | NA |
| 0.549810138 | NA |
| 0.645665705 | NA |
| NA | NA |
| 0.957006653 | NA |
| 0.001762266 | 0.008235898 |
| NA | NA |
| 0.078051593 | 0.174521749 |
| 0.853595154 | NA |
| 0.686468028 | NA |
| 0.941278788 | NA |
| 0.941278332 | NA |
| NA | NA |
| 0.912976495 | NA |
| 0.941279303 | NA |
| 0.014680391 | 0.046870857 |
| NA | NA |
| 3.06E-05 | 0.000244976 |
| 8.58E-12 | 1.96E-10 |
| NA | NA |
| 1.63E-22 | 8.08E-21 |
| 1.08E-19 | 4.47E-18 |
| 1.21E-12 | 3.01E-11 |
| NA | NA |
| 8.62E-10 | 1.54E-08 |

| | |
|---|---|
| 0.742081037 | 0.837824345 |
| NA | NA |
| 2.57E-21 | 1.19E-19 |
| NA | NA |
| 0.86007986 | NA |
| 0.739991096 | NA |
| 0.629286565 | 0.755982376 |
| 0.019210911 | 0.058027679 |
| 0.247419722 | 0.404581219 |
| 0.56581895 | 0.707593267 |
| 0.031646721 | 0.086536833 |
| 0.263312868 | 0.42224435 |
| NA | NA |
| 0.834309892 | 0.899651662 |
| 0.684781302 | NA |
| 0.739991096 | NA |
| NA | NA |
| NA | NA |
| NA | NA |
| NA | NA |
| 0.000234827 | 0.001471659 |
| 0.046465144 | 0.116866668 |
| 7.29E-05 | 0.000528735 |
| 5.73E-05 | 0.000428472 |
| 0.000239817 | 0.001497802 |
| NA | NA |
| 0.161143789 | 0.298091589 |
| 0.210937675 | 0.361022402 |
| 0.010604111 | 0.036098263 |
| NA | NA |
| 0.797416071 | 0.875910163 |
| NA | NA |
| 0.729947671 | NA |
| NA | NA |
| 0.839322351 | NA |
| 0.941278284 | NA |
| 0.839319574 | NA |
| NA | NA |
| NA | NA |
| NA | NA |
| NA | NA |
| 0.25192919 | NA |
| 0.755445343 | NA |
| NA | NA |
| NA | NA |
| 0.837539384 | NA |
| NA | NA |

| | | |
|---|---|---|
| | 0.756794958 | NA |
| NA | | NA |
| NA | | NA |
| | 0.810681736 | NA |
| | 0.741178832 | NA |
| | 0.07035778 | 0.161818134 |
| | 0.682619303 | 0.796065882 |
| | 0.047709377 | 0.119333605 |
| | 1.13E-08 | 1.76E-07 |
| | 0.081237422 | 0.179787366 |
| | 0.2525687 | 0.410434915 |
| | 0.254189071 | 0.412243084 |
| | 7.38E-06 | 6.90E-05 |
| NA | | NA |
| | 0.86523519 | 0.919767844 |
| | 0.642899641 | 0.765727445 |
| | 0.677488175 | 0.792053855 |
| | 0.2158718 | 0.366986685 |
| NA | | NA |
| | 0.479449372 | 0.63614772 |
| | 0.008482273 | 0.030057327 |
| | 0.794510888 | 0.873888547 |
| | 0.749990477 | 0.842955722 |
| | 0.003329245 | 0.013989051 |
| | 0.516635816 | 0.667442775 |
| | 0.471935466 | 0.629405948 |
| | 0.05756399 | 0.138439739 |
| | 0.951046422 | 0.970148286 |
| NA | | NA |
| NA | | NA |
| | 0.803218817 | 0.879726407 |
| NA | | NA |
| | 0.225903874 | 0.378907288 |
| | 0.236106397 | 0.391296788 |
| NA | | NA |
| NA | | NA |
| NA | | NA |
| | 0.717894476 | 0.820606787 |
| | 0.850346079 | 0.910250071 |
| | 0.491863855 | 0.645815505 |
| | 0.027282595 | 0.07705949 |
| NA | | NA |
| | 0.781439423 | 0.865631365 |
| | 0.612906852 | 0.743661563 |
| | 0.16033555 | 0.297104457 |
| | 0.060428871 | 0.143832798 |
| | 8.60E-05 | 0.000613807 |

| | |
|---|---|
| 0.100738232 | 0.210920655 |
| 0.076761429 | 0.172463237 |
| 0.210168647 | 0.360016851 |
| 0.070842435 | 0.162618687 |
| 0.081594784 | 0.180326746 |
| NA | NA |
| 0.901149433 | 0.940279321 |
| 0.2947006 | 0.455988018 |
| 6.69E-07 | 7.71E-06 |
| 5.22E-05 | 0.00039399 |
| 0.003870151 | 0.015781854 |
| 0.081351367 | 0.179939154 |
| 0.319366172 | 0.481903995 |
| 0.000820882 | 0.004320792 |
| NA | NA |
| NA | NA |
| 5.01E-08 | 7.01E-07 |
| NA | NA |
| 0.971110122 | 0.982586427 |
| NA | NA |
| NA | NA |
| 0.233671502 | 0.388288822 |
| NA | NA |
| NA | NA |
| 0.0062461 | 0.023401284 |
| 0.00347478 | 0.014469334 |
| NA | NA |
| 0.20611659 | 0.35492489 |
| NA | NA |
| 0.4822029 | 0.638438338 |
| 0.845429532 | 0.906886914 |
| 0.815511148 | 0.887147914 |
| 0.205966674 | 0.354733631 |
| 0.344431779 | 0.508039216 |
| 0.004399113 | 0.017554568 |
| 0.299038275 | 0.460705725 |
| 0.502074053 | 0.654241028 |
| 0.114748257 | 0.232287635 |
| 0.805224366 | 0.880868577 |
| 0.187135457 | 0.331093298 |
| 0.619475793 | 0.748579114 |
| 0.320433491 | 0.483116358 |
| 0.129489198 | 0.253858982 |
| NA | NA |
| NA | NA |
| 0.263942486 | 0.422997512 |
| 0.649366446 | 0.770749546 |

| | | |
|---|---|---|
| | 0.415658415 | 0.577452961 |
| | 0.24656583 | 0.40354534 |
| | 0.244398477 | 0.400880197 |
| | 0.354255996 | 0.517968008 |
| | 0.493562554 | 0.646993147 |
| | 0.337368978 | 0.500626958 |
| | 0.321820602 | 0.484685774 |
| | 0.22696196 | 0.380138524 |
| | 0.541352305 | 0.688462683 |
| | 0.106664543 | 0.220174265 |
| | 0.189068706 | 0.333659718 |
| | 0.275969979 | 0.435788326 |
| | 0.743070083 | 0.83852368 |
| | 0.955792873 | NA |
| | 0.038672473 | 0.101341209 |
| | 0.04566934 | 0.11533386 |
| | 0.231403915 | 0.385516622 |
| | 0.990476838 | 0.993604524 |
| NA | | NA |
| | 0.004105326 | 0.016587163 |
| | 0.104587467 | 0.216760263 |
| | 0.061396811 | 0.145656024 |
| | 0.008666879 | 0.030593067 |
| | 0.278742677 | 0.438508473 |
| NA | | NA |
| NA | | NA |
| NA | | NA |
| | 0.469857794 | 0.627798041 |
| | 0.057196655 | 0.137814503 |
| | 0.109315252 | 0.224009531 |
| NA | | NA |
| | 0.014613587 | 0.046714021 |
| | 0.268491247 | 0.42826358 |
| NA | | NA |
| | 0.260199334 | 0.418775072 |
| | 0.429587506 | 0.590769691 |
| NA | | NA |
| | 0.181969146 | 0.324363784 |
| | 0.953770009 | NA |
| | 0.63671547 | 0.76118881 |
| | 0.23559984 | 0.390756561 |
| | 0.115943654 | 0.23403077 |
| | 0.496079398 | NA |
| | 0.928413336 | NA |
| | 0.668847797 | NA |
| | 0.929499469 | 0.95803691 |
| | 0.224612394 | 0.377286014 |

| | | |
|---|---|---|
| | 9.94E-06 | 9.02E-05 |
| | 0.404529714 | NA |
| | 0.44743572 | 0.607730256 |
| | 0.06521495 | 0.152674952 |
| | 0.035853938 | 0.095564998 |
| | 0.196764177 | 0.343532674 |
| | 0.03783557 | 0.099663814 |
| | 0.987617721 | 0.992161271 |
| | 0.083637755 | 0.183817714 |
| | 0.005054835 | 0.019697968 |
| | 0.401316549 | 0.563741795 |
| | 0.869493434 | 0.921774573 |
| | 0.969810139 | 0.981982089 |
| | 0.005038844 | 0.019651737 |
| | 0.168917741 | 0.308444961 |
| | 0.249051982 | 0.406356956 |
| | 0.033442977 | 0.090474567 |
| | 0.001334761 | 0.006506244 |
| | 4.42E-28 | 2.86E-26 |
| | 0.584948319 | 0.721733424 |
| | 2.96E-06 | 3.02E-05 |
| | 0.627593828 | 0.75486182 |
| | 0.448236812 | 0.608427316 |
| | 0.106583663 | 0.220064569 |
| | 0.010442278 | 0.035649326 |
| | 0.335408557 | 0.498588213 |
| | 0.015113612 | 0.04800296 |
| | 0.154288375 | 0.289003849 |
| | 0.955792398 | NA |
| | 0.955792398 | NA |
| | 0.030368967 | 0.083749525 |
| | 0.01444988 | 0.046283951 |
| | 0.318606748 | 0.481071469 |
| | 0.105373108 | 0.218094844 |
| | 0.193599377 | 0.339426368 |
| | 0.541790131 | 0.688872259 |
| | 0.953902938 | 0.97203518 |
| | 1.65E-30 | 1.17E-28 |
| | 0.4484913 | NA |
| NA | | NA |
| NA | | NA |
| | 0.731194986 | 0.830477641 |
| | 0.634166387 | NA |
| | 0.882471426 | 0.929664669 |
| NA | | NA |
| | 0.300240116 | 0.461797431 |
| | 0.000113295 | 0.000781185 |

| | | |
|---|---|---|
| | 0.114204423 | 0.231384569 |
| | 0.254852035 | 0.41287695 |
| | 0.099628191 | 0.209003192 |
| | 0.470097205 | 0.627939484 |
| | 4.72E-11 | 9.88E-10 |
| | 0.103524165 | 0.215203683 |
| | 0.06930959 | 0.160044875 |
| | 0.82566599 | NA |
| | 1.21E-73 | 2.83E-71 |
| | 0.85946517 | 0.91620342 |
| | 0.013326057 | 0.043438346 |
| | 5.27E-05 | 0.000397719 |
| | 7.64E-10 | 1.38E-08 |
| | 0.232439362 | 0.386862576 |
| | 0.290667848 | NA |
| | 0.330275541 | 0.493206501 |
| | 0.021346915 | 0.063039611 |
| | 0.113696037 | 0.230687329 |
| | 0.261064335 | 0.419798224 |
| | 0.255275443 | 0.413381617 |
| | 0.051946695 | 0.127796923 |
| | 0.894355039 | NA |
| | 0.239265974 | 0.395102589 |
| | 0.560878081 | 0.703988552 |
| | 0.400907867 | 0.563273068 |
| | 0.00016848 | 0.001108695 |
| | 0.161174069 | 0.298101266 |
| | 0.839319108 | NA |
| | 0.939713734 | 0.963757575 |
| NA | | NA |
| | 0.064766851 | 0.151954716 |
| | 0.939241089 | NA |
| | 0.136437457 | 0.26405425 |
| | 0.006912408 | 0.025422379 |
| | 0.955792873 | NA |
| | 0.650540105 | 0.77173517 |
| | 0.017368386 | 0.053615044 |
| | 0.941277857 | NA |
| | 0.070726067 | 0.162516782 |
| NA | | NA |
| | 0.941277857 | NA |
| | 0.843562056 | 0.905863481 |
| | 0.865184175 | 0.919767844 |
| NA | | NA |
| NA | | NA |
| NA | | NA |
| NA | | NA |

| | | |
|---|---|---|
| | 0.680466704 | NA |
| | 0.941278207 | NA |
| | 0.562555139 | NA |
| | 0.99038934 | 0.993561791 |
| NA | | NA |
| NA | | NA |
| NA | | NA |
| NA | | NA |
| NA | | NA |
| NA | | NA |
| NA | | NA |
| NA | | NA |
| NA | | NA |
| NA | | NA |
| NA | | NA |
| NA | | NA |
| NA | | NA |
| NA | | NA |
| NA | | NA |
| NA | | NA |
| NA | | NA |
| NA | | NA |
| NA | | NA |
| | 0.820026198 | NA |
| NA | | NA |
| NA | | NA |
| NA | | NA |
| NA | | NA |
| NA | | NA |
| NA | | NA |
| NA | | NA |
| NA | | NA |
| NA | | NA |
| | 0.890156278 | NA |
| NA | | NA |
| | 0.839318641 | NA |
| NA | | NA |
| | 0.941278332 | NA |
| | 0.638357712 | 0.76241608 |
| NA | | NA |
| NA | | NA |
| NA | | NA |
| | 0.941277857 | NA |
| NA | | NA |
| NA | | NA |
| NA | | NA |

| | | |
|---|---|---|
| NA | | NA |
| NA | | NA |
| | 0.839319108 | NA |
| NA | | NA |
| NA | | NA |
| NA | | NA |
| NA | | NA |
| NA | | NA |
| NA | | NA |
| NA | | NA |
| NA | | NA |
| NA | | NA |
| | 0.941278332 | NA |
| NA | | NA |
| NA | | NA |
| NA | | NA |
| NA | | NA |
| NA | | NA |
| | 0.941279465 | NA |
| NA | | NA |
| NA | | NA |
| | 0.941281677 | NA |
| NA | | NA |
| NA | | NA |
| NA | | NA |
| | 0.6223855 | NA |
| | 0.839318641 | NA |
| NA | | NA |
| NA | | NA |
| NA | | NA |
| | 0.769784654 | NA |
| NA | | NA |
| NA | | NA |
| NA | | NA |
| NA | | NA |
| NA | | NA |
| NA | | NA |
| NA | | NA |
| NA | | NA |
| NA | | NA |
| NA | | NA |
| NA | | NA |
| NA | | NA |
| NA | | NA |
| NA | | NA |
| NA | | NA |
| NA | | NA |

| | | |
|---|---|---|
| NA | | NA |
| NA | | NA |
| NA | | NA |
| NA | | NA |
| NA | | NA |
| | 0.715465242 | NA |
| NA | | NA |
| NA | | NA |
| NA | | NA |
| | 0.941277857 | NA |
| NA | | NA |
| NA | | NA |
| NA | | NA |
| NA | | NA |
| NA | | NA |
| NA | | NA |
| NA | | NA |
| | 0.936176462 | NA |
| NA | | NA |
| NA | | NA |
| NA | | NA |
| NA | | NA |
| | 0.941278332 | NA |
| NA | | NA |
| NA | | NA |
| NA | | NA |
| NA | | NA |
| NA | | NA |
| NA | | NA |
| NA | | NA |
| NA | | NA |
| NA | | NA |
| NA | | NA |
| NA | | NA |
| | 0.709386887 | NA |
| NA | | NA |
| NA | | NA |
| NA | | NA |
| NA | | NA |
| NA | | NA |
| NA | | NA |
| NA | | NA |
| NA | | NA |
| NA | | NA |
| | 0.941278424 | NA |
| NA | | NA |
| NA | | NA |

| | |
|---|---|
| NA | NA |
| NA | NA |
| NA | NA |
| NA | NA |
| 0.941277857 | NA |
| NA | NA |
| NA | NA |
| NA | NA |
| NA | NA |
| NA | NA |
| NA | NA |
| NA | NA |
| NA | NA |
| NA | NA |
| 0.941277857 | NA |
| NA | NA |
| NA | NA |
| NA | NA |
| NA | NA |
| NA | NA |
| NA | NA |
| NA | NA |
| NA | NA |
| NA | NA |
| NA | NA |
| NA | NA |
| NA | NA |
| NA | NA |
| NA | NA |
| NA | NA |
| NA | NA |
| NA | NA |
| NA | NA |
| NA | NA |
| NA | NA |
| NA | NA |
| 0.839318641 | NA |
| NA | NA |
| NA | NA |
| NA | NA |
| NA | NA |
| NA | NA |
| NA | NA |
| NA | NA |
| 0.001383712 | 0.006710498 |
| NA | NA |
| NA | NA |

| | | |
|---|---|---|
| NA | | NA |
| | 0.728678906 | NA |
| NA | | NA |
| NA | | NA |
| NA | | NA |
| NA | | NA |
| NA | | NA |
| NA | | NA |
| NA | | NA |
| NA | | NA |
| NA | | NA |
| NA | | NA |
| NA | | NA |
| NA | | NA |
| NA | | NA |
| | 0.941278332 | NA |
| NA | | NA |
| NA | | NA |
| NA | | NA |
| NA | | NA |
| NA | | NA |
| | 0.924755761 | NA |
| | 0.941278332 | NA |
| NA | | NA |
| NA | | NA |
| | 0.839319108 | NA |
| NA | | NA |
| NA | | NA |
| NA | | NA |
| NA | | NA |
| NA | | NA |
| NA | | NA |
| NA | | NA |
| NA | | NA |
| NA | | NA |
| NA | | NA |
| NA | | NA |
| NA | | NA |
| NA | | NA |
| | 0.939241089 | NA |
| NA | | NA |
| NA | | NA |
| NA | | NA |
| NA | | NA |
| NA | | NA |
| NA | | NA |
| NA | | NA |

| | |
|---|---|
| 0.941278332 | NA |
| NA | NA |
| NA | NA |
| NA | NA |
| NA | NA |
| 0.760389963 | 0.850947478 |
| NA | NA |
| NA | NA |
| NA | NA |
| NA | NA |
| NA | NA |
| NA | NA |
| NA | NA |
| 0.116906389 | 0.235375222 |
| 0.839318641 | NA |
| NA | NA |
| 0.001372518 | 0.006661638 |
| 0.013736938 | 0.044442216 |
| 0.164191969 | NA |
| NA | NA |
| NA | NA |
| 0.01473848 | 0.046993342 |
| 0.853595154 | NA |
| NA | NA |
| 0.915240755 | 0.949011563 |
| 2.68E-05 | 0.00021816 |
| 0.687007173 | NA |
| 0.955792398 | NA |
| 0.785271835 | 0.868018401 |
| 0.832999916 | 0.898738226 |
| 0.009143885 | 0.03199597 |
| 0.654222994 | 0.774331959 |
| 0.649328197 | 0.770749546 |
| 0.698592845 | 0.807190003 |
| 0.034810497 | 0.093253736 |
| 0.505698002 | 0.657528461 |
| 0.575993664 | 0.714759459 |
| 0.941278332 | NA |
| 3.55E-09 | 5.90E-08 |
| 0.4453369 | 0.605912779 |
| 0.941278332 | NA |
| NA | NA |
| 0.106845027 | 0.220451225 |
| NA | NA |
| 0.949201544 | 0.968903643 |
| 0.12231707 | 0.243814664 |
| 0.741535438 | 0.83740696 |

| | |
|---|---|
| NA | NA |
| 0.029446839 | 0.081761507 |
| 0.207536305 | 0.356762964 |
| NA | NA |
| NA | NA |
| 0.967101623 | 0.980280761 |
| 0.029241803 | 0.081309975 |
| 0.053365922 | 0.13051949 |
| 4.56E-12 | 1.07E-10 |
| 0.000733429 | 0.003931775 |
| 0.915100837 | 0.948990507 |
| 0.471845042 | NA |
| NA | NA |
| 0.839318641 | NA |
| NA | NA |
| 6.78E-07 | 7.80E-06 |
| 0.941277857 | NA |
| 0.604575545 | 0.73746257 |
| NA | NA |
| 0.839813226 | 0.903304912 |
| 0.875550712 | 0.925639959 |
| 0.01204373 | 0.040009053 |
| 0.839319108 | NA |
| NA | NA |
| 0.244833927 | NA |
| 0.941278788 | NA |
| 0.151099203 | 0.284868374 |
| NA | NA |
| 0.012810066 | 0.042050088 |
| 0.450561346 | 0.610547176 |
| 0.95456991 | NA |
| NA | NA |
| NA | NA |
| 0.654933711 | NA |
| 1.77E-12 | 4.33E-11 |
| NA | NA |
| NA | NA |
| 0.50150175 | 0.653874786 |
| 0.0052308 | 0.020244472 |
| 0.009441401 | 0.032835218 |
| NA | NA |
| 0.896276044 | NA |
| 6.24E-06 | 5.95E-05 |
| 0.017720928 | 0.054446046 |
| 0.684398405 | 0.797279344 |
| 0.03000207 | 0.083016655 |
| 0.935658422 | NA |

| | | |
|---|---|---|
| | 0.204346225 | NA |
| | 0.955791922 | NA |
| | 0.941278332 | NA |
| | 0.941279303 | NA |
| | 0.655250979 | NA |
| | 0.941278332 | NA |
| | 0.006422097 | 0.023945957 |
| | 0.659103881 | 0.778096943 |
| | 0.748548076 | 0.842209469 |
| NA | | NA |
| | 0.286470078 | 0.44706011 |
| | 0.153352153 | 0.287657651 |
| | 0.955792873 | NA |
| | 0.162095912 | 0.299341058 |
| NA | | NA |
| | 0.247852469 | 0.405038411 |
| | 0.629505315 | 0.756092522 |
| NA | | NA |
| NA | | NA |
| | 0.053288799 | 0.130438091 |
| | 0.941278434 | NA |
| | 0.092920254 | 0.198696244 |
| | 0.093932858 | 0.200231943 |
| | 0.625312138 | 0.753149261 |
| | 0.039581969 | 0.10331506 |
| | 4.43E-20 | 1.90E-18 |
| | 0.737167795 | 0.834335134 |
| | 0.005807694 | 0.022044818 |
| | 0.134470624 | 0.261215195 |
| | 0.248106621 | 0.405314605 |
| NA | | NA |
| | 0.874847664 | NA |
| | 0.172900741 | 0.313271336 |
| | 0.326503584 | NA |
| | 0.941278332 | NA |
| | 0.8996056 | 0.93958188 |
| NA | | NA |
| NA | | NA |
| | 0.06954556 | 0.16046175 |
| NA | | NA |
| NA | | NA |
| | 0.001359661 | 0.006607321 |
| | 0.030398706 | 0.083802408 |
| | 0.001028838 | 0.0052348 |
| | 0.266926518 | 0.426544813 |
| | 0.048867027 | 0.121547724 |
| | 0.002577171 | 0.01132334 |

| | | |
|---|---|---|
| | 0.865781802 | 0.919806215 |
| | 0.064443935 | 0.151331348 |
| | 0.093042399 | 0.198800088 |
| | 0.355773132 | 0.51951608 |
| NA | | NA |
| | 0.001684771 | 0.007925081 |
| | 0.00331263 | 0.013937992 |
| | 1.11E-07 | 1.48E-06 |
| | 0.941278332 | NA |
| | 0.002140393 | 0.009674447 |
| | 0.001389546 | 0.006729193 |
| | 0.010150893 | 0.034824382 |
| | 0.021054317 | 0.062337959 |
| | 0.368570651 | 0.531712142 |
| | 0.732603905 | NA |
| | 0.478574485 | 0.635382784 |
| NA | | NA |
| NA | | NA |
| NA | | NA |
| | 0.731908309 | 0.830950707 |
| | 0.125112817 | 0.247851458 |
| | 0.879917634 | 0.92811307 |
| | 0.401451576 | NA |
| | 0.955792873 | NA |
| | 0.936508021 | NA |
| | 0.143806646 | 0.274599742 |
| | 0.043123943 | 0.110428843 |
| | 0.748393366 | NA |
| | 0.383643903 | 0.546403313 |
| | 0.847828498 | 0.908395855 |
| | 0.889348489 | 0.933455095 |
| | 0.941278332 | NA |
| | 0.118211082 | 0.237539906 |
| NA | | NA |
| | 0.839319109 | NA |
| | 0.941277857 | NA |
| | 0.839319108 | NA |
| | 0.817153331 | NA |
| NA | | NA |
| | 0.439497768 | NA |
| NA | | NA |
| NA | | NA |
| | 0.839319574 | NA |
| | 0.085239417 | NA |
| NA | | NA |
| | 0.941277857 | NA |
| NA | | NA |

| | |
|---|---|
| NA | NA |
| 0.425012839 | NA |
| 0.268505899 | 0.42826358 |
| 0.032226139 | 0.087782853 |
| 0.200456691 | NA |
| 0.007505677 | 0.027220803 |
| 0.064112019 | 0.150670841 |
| 0.234180297 | NA |
| 0.589088461 | NA |
| 8.92E-17 | 3.06E-15 |
| 0.070767558 | 0.162572089 |
| 0.395661152 | 0.558241064 |
| NA | NA |
| 9.22E-19 | 3.62E-17 |
| 0.053586003 | NA |
| NA | NA |
| NA | NA |
| 0.936273762 | NA |
| NA | NA |
| 0.651665587 | 0.772647082 |
| NA | NA |
| NA | NA |
| 0.220762397 | 0.372692146 |
| 0.941278332 | NA |
| 0.030367197 | 0.083749525 |
| 0.701341553 | NA |
| 1.77E-06 | 1.89E-05 |
| 0.017584031 | 0.054088279 |
| 7.09E-11 | 1.45E-09 |
| 0.453183035 | 0.613157264 |
| 0.002553157 | 0.011239292 |
| NA | NA |
| 0.589620556 | 0.725709409 |
| 0.868019256 | 0.921073262 |
| 0.826961652 | NA |
| NA | NA |
| 0.060150475 | 0.143342171 |
| NA | NA |
| NA | NA |
| NA | NA |
| 0.802085915 | 0.87905189 |
| 0.091905013 | 0.197020905 |
| 0.941278332 | NA |
| NA | NA |
| 0.853596091 | NA |
| NA | NA |
| 0.853595622 | NA |

| | |
|---|---|
| 0.941277857 | NA |
| NA | NA |
| 0.839318641 | NA |
| 0.017114216 | 0.052985506 |
| 0.067021836 | 0.155908303 |
| 0.739991096 | NA |
| 0.105555415 | 0.218320298 |
| NA | NA |
| 0.840631062 | NA |
| 6.56E-07 | 7.56E-06 |
| 0.03809661 | 0.100240494 |
| 0.091778196 | 0.196846835 |
| 0.000135486 | 0.000914845 |
| 0.941278207 | NA |
| 0.182781318 | 0.32532898 |
| 0.824597445 | 0.893479692 |
| 7.77E-06 | 7.23E-05 |
| 0.011704638 | 0.03913384 |
| 0.313171353 | 0.475116614 |
| NA | NA |
| 0.160789044 | 0.297713017 |
| 0.031907163 | 0.087093382 |
| 0.947904361 | 0.968205894 |
| 0.839318641 | NA |
| 0.480556466 | 0.637160636 |
| 0.539289196 | 0.686499168 |
| 0.28019364 | 0.44015107 |
| 0.839319108 | NA |
| 0.38849343 | 0.551363288 |
| NA | NA |
| 0.363203708 | 0.526883223 |
| 0.941277857 | NA |
| 0.900908777 | NA |
| 0.941278788 | NA |
| 0.004266561 | 0.017113698 |
| NA | NA |
| 0.955792873 | NA |
| 0.360578369 | NA |
| 0.003846563 | 0.015712354 |
| NA | NA |
| NA | NA |
| 0.839320041 | NA |
| 0.955792397 | NA |
| 0.273141857 | 0.433134088 |
| 0.234570203 | 0.389401588 |
| 0.061320854 | 0.145533849 |
| 0.071538711 | 0.163727346 |

| | | | |
|---|---|---|---|
| | 0.941277857 | NA | |
| | 0.002579072 | | 0.011328117 |
| NA | | NA | |
| NA | | NA | |
| NA | | NA | |
| NA | | NA | |
| NA | | NA | |
| | 0.554845657 | NA | |
| NA | | NA | |
| NA | | NA | |
| NA | | NA | |
| NA | | NA | |
| | 0.319026333 | NA | |
| NA | | NA | |
| NA | | NA | |
| NA | | NA | |
| NA | | NA | |
| | 0.839319108 | NA | |
| NA | | NA | |
| | 0.941278332 | NA | |
| NA | | NA | |
| | 0.981915534 | NA | |
| NA | | NA | |
| NA | | NA | |
| | 0.941278332 | NA | |
| NA | | NA | |
| | 0.941277857 | NA | |
| NA | | NA | |
| | 0.711618573 | NA | |
| NA | | NA | |
| NA | | NA | |
| | 0.839319108 | NA | |
| NA | | NA | |
| NA | | NA | |
| NA | | NA | |
| NA | | NA | |
| NA | | NA | |
| | 0.941277857 | NA | |
| NA | | NA | |
| NA | | NA | |
| | 0.955792398 | NA | |
| | 0.196378518 | | 0.34308584 |
| | 0.082732105 | NA | |
| | 0.478914724 | | 0.635685872 |
| NA | | NA | |
| | 0.941277857 | NA | |
| NA | | NA | |

| | |
|---|---|
| 0.414318759 | NA |
| NA | NA |
| 0.771629974 | 0.858863398 |
| NA | NA |
| NA | NA |
| NA | NA |
| 0.680284881 | NA |
| 0.925579587 | NA |
| NA | NA |
| NA | NA |
| NA | NA |
| NA | NA |
| 0.941277857 | NA |
| NA | NA |
| NA | NA |
| NA | NA |
| NA | NA |
| 0.839319108 | NA |
| NA | NA |
| 0.955791446 | NA |
| NA | NA |
| NA | NA |
| NA | NA |
| NA | NA |
| NA | NA |
| NA | NA |
| NA | NA |
| NA | NA |
| NA | NA |
| NA | NA |
| NA | NA |
| 0.941277857 | NA |
| NA | NA |
| NA | NA |
| NA | NA |
| NA | NA |
| NA | NA |
| NA | NA |
| 0.939241089 | NA |
| 0.839319108 | NA |
| NA | NA |
| 0.838882104 | NA |
| 2.34E-05 | 0.000193882 |
| NA | NA |
| 0.033561043 | 0.090713144 |
| 1.69E-07 | 2.19E-06 |
| 7.64E-12 | 1.75E-10 |

| | | |
|---|---|---|
| | 0.912727722 | 0.94739633 |
| NA | | NA |
| | 0.879100996 | NA |
| | 0.058717783 | 0.140561266 |
| | 1.99E-05 | 0.00016714 |
| | 0.941278788 | NA |
| | 0.00101203 | 0.005162522 |
| | 3.28E-06 | 3.32E-05 |
| | 2.49E-08 | 3.68E-07 |
| | 0.641331856 | 0.764778599 |
| | 5.60E-10 | 1.03E-08 |
| | 1.87E-10 | 3.64E-09 |
| | 0.374230751 | 0.537369956 |
| | 0.499894744 | NA |
| | 3.19E-10 | 6.02E-09 |
| | 0.002923873 | 0.01255132 |
| | 0.642107193 | 0.765281429 |
| | 0.955792873 | NA |
| | 0.419307133 | 0.580930795 |
| | 2.63E-06 | 2.71E-05 |
| | 2.78E-09 | 4.67E-08 |
| NA | | NA |
| | 5.15E-06 | 4.99E-05 |
| NA | | NA |
| | 0.839319574 | NA |
| | 0.693114219 | 0.803382559 |
| | 0.955792873 | NA |
| | 0.000363749 | 0.002148906 |
| | 0.853178115 | NA |
| | 0.236647945 | 0.39208509 |
| | 3.13E-14 | 8.97E-13 |
| NA | | NA |
| | 1.47E-24 | 8.12E-23 |
| | 2.37E-10 | 4.55E-09 |
| | 0.839320041 | NA |
| | 0.955792873 | NA |
| | 0.00210415 | 0.009539615 |
| | 1.60E-05 | 0.000137967 |
| | 0.941277857 | NA |
| | 3.69E-06 | 3.70E-05 |
| | 0.534578722 | NA |
| | 0.065838432 | 0.153816879 |
| | 0.027083279 | 0.076587407 |
| | 0.124769352 | 0.247335731 |
| | 0.000139953 | 0.000940207 |
| | 0.586681577 | 0.723346019 |
| NA | | NA |

| | |
|---|---|
| 0.85262745 | NA |
| 0.201186895 | 0.348976131 |
| 0.941278332 | NA |
| NA | NA |
| NA | NA |
| 0.671019701 | NA |
| NA | NA |
| NA | NA |
| 0.535168584 | 0.683300217 |
| NA | NA |
| 8.83E-25 | 4.96E-23 |
| NA | NA |
| 0.955792398 | NA |
| NA | NA |
| NA | NA |
| 0.034402555 | 0.092368824 |
| 0.839318641 | NA |
| 0.724271383 | NA |
| NA | NA |
| NA | NA |
| 0.000600589 | 0.00331545 |
| 0.162514583 | 0.299997836 |
| 0.95978906 | 0.975319097 |
| 0.94608657 | 0.967272298 |
| 0.672061656 | 0.78792294 |
| 4.12E-09 | 6.78E-08 |
| 0.741120567 | NA |
| 0.571451226 | 0.711757305 |
| 0.260497747 | 0.419113653 |
| 0.000153532 | 0.001022484 |
| 0.739785246 | 0.836336778 |
| 1.64E-26 | 9.88E-25 |
| 0.402146775 | 0.564423919 |
| 0.812553923 | 0.885468752 |
| 0.023723013 | 0.068768134 |
| 2.77E-05 | 0.000224567 |
| NA | NA |
| 0.531890087 | 0.680464813 |
| 0.941278332 | NA |
| 0.404650676 | 0.566691892 |
| NA | NA |
| 0.932442827 | NA |
| 0.839319108 | NA |
| NA | NA |
| 0.076152077 | 0.171385376 |
| NA | NA |
| NA | NA |

| | |
|---|---|
| NA | NA |
| 0.459578279 | 0.618750054 |
| 0.941278332 | NA |
| NA | NA |
| NA | NA |
| NA | NA |
| NA | NA |
| NA | NA |
| NA | NA |
| NA | NA |
| 0.941278332 | NA |
| NA | NA |
| NA | NA |
| NA | NA |
| NA | NA |
| NA | NA |
| NA | NA |
| NA | NA |
| NA | NA |
| NA | NA |
| NA | NA |
| NA | NA |
| 0.941278332 | NA |
| 0.839318641 | NA |
| 0.839318641 | NA |
| NA | NA |
| NA | NA |
| NA | NA |
| NA | NA |
| NA | NA |
| NA | NA |
| 0.020009547 | 0.059870173 |
| 0.238149302 | 0.393781288 |
| 0.403188254 | NA |
| 0.941208171 | NA |
| NA | NA |
| 0.838197868 | NA |
| 0.587351209 | 0.723721772 |
| 0.141698163 | 0.271641404 |
| 0.008519951 | 0.030163894 |
| 0.722068857 | NA |
| 0.745286722 | 0.839969438 |
| NA | NA |
| 0.505237872 | 0.65720799 |
| 0.955792398 | NA |
| 0.955791922 | NA |
| NA | NA |

| | |
|---|---|
| 0.002792595 | 0.012076238 |
| NA | NA |
| 0.757622602 | NA |
| 0.035753664 | 0.095340435 |
| 0.941278788 | NA |
| 0.770024608 | 0.857536412 |
| 0.955792398 | NA |
| 0.161672314 | 0.298744228 |
| NA | NA |
| 1.46E-12 | 3.62E-11 |
| 2.02E-05 | 0.000169385 |
| 0.161457542 | 0.29846322 |
| 0.848453389 | NA |
| 0.941278332 | NA |
| NA | NA |
| 0.9455074 | NA |
| NA | NA |
| 0.912223085 | NA |
| 0.60172312 | NA |
| 0.013786026 | 0.044565395 |
| NA | NA |
| 0.867181263 | 0.920594461 |
| NA | NA |
| 0.518770418 | 0.669465062 |
| 0.262551678 | 0.421478049 |
| NA | NA |
| 0.975029822 | NA |
| NA | NA |
| 0.006185302 | 0.023206584 |
| 0.839319108 | NA |
| 0.498585262 | NA |
| 0.834629079 | NA |
| 1.34E-05 | 0.000117658 |
| 0.376267794 | 0.539416131 |
| 0.839319108 | NA |
| NA | NA |
| 0.976461758 | 0.985529524 |
| NA | NA |
| 0.717885827 | 0.820606787 |
| 0.291889445 | 0.453182029 |
| 2.18E-16 | 7.35E-15 |
| 0.491654103 | NA |
| 0.122455799 | 0.244019293 |
| 0.889822732 | NA |
| 7.61E-14 | 2.10E-12 |
| 3.57E-23 | 1.84E-21 |
| 1.56E-07 | 2.03E-06 |

| | | | |
|---|---|---|---|
| | 0.013316998 | | 0.043418027 |
| NA | | NA | |
| NA | | NA | |
| | 0.763816898 | NA | |
| NA | | NA | |
| NA | | NA | |
| | 0.941277857 | NA | |
| NA | | NA | |
| NA | | NA | |
| NA | | NA | |
| NA | | NA | |
| NA | | NA | |
| NA | | NA | |
| NA | | NA | |
| NA | | NA | |
| NA | | NA | |
| NA | | NA | |
| NA | | NA | |
| NA | | NA | |
| NA | | NA | |
| NA | | NA | |
| NA | | NA | |
| NA | | NA | |
| | 0.663181692 | NA | |
| | 0.252533713 | | 0.41040608 |
| | 0.011723376 | | 0.039190983 |
| | 0.180356447 | | 0.322358201 |
| | 0.113222292 | | 0.230020229 |
| | 4.95E-16 | | 1.63E-14 |
| | 0.544863229 | NA | |
| | 0.094761619 | | 0.201511307 |
| | 0.468899634 | | 0.626973153 |
| | 0.256614534 | | 0.414744289 |
| | 1.57E-06 | | 1.70E-05 |
| | 4.15E-08 | | 5.87E-07 |
| | 0.070045981 | | 0.161288038 |
| | 0.193269211 | | 0.339022312 |
| | 0.625079463 | | 0.752980484 |
| | 0.940845277 | NA | |
| | 0.941277857 | NA | |
| | 0.06405514 | | 0.150568463 |
| | 0.332457704 | | 0.49565664 |
| | 0.853595622 | NA | |
| | 0.941278332 | NA | |
| | 0.707592684 | | 0.813437541 |
| | 0.401611031 | | 0.563925563 |
| | 0.142981023 | | 0.27341257 |

| | | |
|---|---|---|
| | 0.789608199 | NA |
| NA | | NA |
| | 0.327512106 | 0.490280829 |
| NA | | NA |
| | 0.00566626 | 0.021606589 |
| | 0.002251934 | 0.010101873 |
| | 0.196659639 | 0.343425732 |
| | 0.654374086 | 0.77439894 |
| | 0.839319108 | NA |
| | 0.031481502 | 0.086188487 |
| | 0.000416689 | 0.002422195 |
| | 1.29E-08 | 1.98E-07 |
| | 0.590110082 | NA |
| NA | | NA |
| | 0.706550721 | NA |
| NA | | NA |
| NA | | NA |
| | 0.324464556 | 0.487372546 |
| | 0.227864038 | 0.381279199 |
| | 0.036628193 | 0.097106734 |
| | 0.004806911 | 0.018902049 |
| NA | | NA |
| | 0.824978079 | NA |
| | 0.063379899 | 0.14930516 |
| | 0.01693343 | 0.05252828 |
| | 0.975207174 | 0.985098721 |
| | 0.036223781 | 0.096293768 |
| | 0.022645887 | 0.066193957 |
| | 0.174197097 | 0.31499197 |
| | 0.941278332 | NA |
| | 0.006650658 | 0.024614984 |
| | 0.370827012 | 0.534093219 |
| | 0.145325383 | 0.27678921 |
| NA | | NA |
| NA | | NA |
| | 0.955792398 | NA |
| | 0.853980414 | NA |
| | 0.923130627 | 0.954250805 |
| | 0.163678023 | 0.301771054 |
| NA | | NA |
| | 0.414801959 | NA |
| | 2.25E-06 | 2.36E-05 |
| | 0.217712905 | NA |
| | 0.463197287 | 0.621999042 |
| | 0.084972199 | 0.186148945 |
| | 0.736725738 | 0.834046731 |
| | 0.931484409 | 0.959252019 |

| | |
|---|---|
| 0.291534388 | NA |
| 0.57338242 | 0.71303484 |
| 0.740206831 | 0.836584183 |
| 0.000104095 | 0.000724888 |
| 0.939680037 | NA |
| 9.86E-08 | 1.33E-06 |
| 0.188546273 | 0.332970647 |
| 0.54007343 | NA |
| 0.054974885 | 0.133630742 |
| 0.805805278 | NA |
| 0.096781703 | 0.204546092 |
| 0.955791922 | NA |
| 0.941278332 | NA |
| NA | NA |
| 0.941278424 | NA |
| 0.958607487 | 0.974623149 |
| 0.103999015 | 0.215926776 |
| 1.29E-09 | 2.27E-08 |
| NA | NA |
| 0.353908388 | 0.517555139 |
| 0.839028726 | NA |
| NA | NA |
| 0.59636047 | 0.731120237 |
| NA | NA |
| 0.839319574 | NA |
| 0.48106085 | 0.637487957 |
| 3.30E-05 | 0.000262272 |
| 0.0007211 | 0.003877035 |
| 0.771194799 | 0.858558401 |
| 1.82E-05 | 0.000153884 |
| 0.352759641 | 0.51638284 |
| 1.57E-18 | 6.04E-17 |
| 8.46E-06 | 7.79E-05 |
| 0.000138329 | 0.000932193 |
| NA | NA |
| 0.051192608 | 0.126333485 |
| NA | NA |
| 0.941278332 | NA |
| NA | NA |
| 0.228166734 | 0.381699789 |
| 0.955792873 | NA |
| 0.00715252 | 0.026143162 |
| NA | NA |
| 0.142713785 | 0.273082524 |
| 0.668821413 | NA |
| 0.80028342 | NA |
| NA | NA |

| | |
|---|---|
| 0.4565437 | 0.616233024 |
| NA | NA |
| 0.000293743 | 0.001791355 |
| 0.231281746 | 0.385367037 |
| 0.00525659 | 0.02032776 |
| 0.941277857 | NA |
| 0.007268213 | 0.026496859 |
| 0.152588721 | 0.286677459 |
| 0.000102024 | 0.00071234 |
| 0.020218169 | 0.06040327 |
| 0.195653758 | 0.342095846 |
| 0.059967731 | 0.142949619 |
| 1.77E-06 | 1.89E-05 |
| 0.442341301 | 0.602854094 |
| NA | NA |
| NA | NA |
| 0.476786114 | 0.633602986 |
| 0.005635021 | 0.021516356 |
| 0.177955597 | 0.319425711 |
| 0.024316864 | 0.070181884 |
| 1.32E-12 | 3.29E-11 |
| 0.152748682 | 0.28684214 |
| 0.786557101 | NA |
| 0.298274275 | 0.459783971 |
| 6.23E-05 | 0.00046133 |
| 0.006627304 | 0.024560457 |
| 0.013489468 | 0.04383893 |
| 0.014830268 | 0.047241709 |
| 0.372828891 | 0.536022585 |
| NA | NA |
| NA | NA |
| 0.707277508 | 0.813272637 |
| 0.341630389 | 0.505061603 |
| 0.181185777 | 0.323451875 |
| NA | NA |
| 0.017903473 | 0.054907673 |
| NA | NA |
| 0.808473115 | 0.88251672 |
| 0.028213138 | 0.079040619 |
| 0.348616974 | 0.512525091 |
| 0.920589933 | 0.952524847 |
| 0.335985401 | 0.499154772 |
| 5.12E-07 | 6.02E-06 |
| 5.04E-06 | 4.91E-05 |
| 0.146717458 | 0.278771369 |
| 0.238636923 | 0.394419556 |
| 0.001655729 | 0.007806983 |

| | | |
|---|---|---|
| | 0.699688018 | 0.808249173 |
| | 0.366542003 | 0.529666785 |
| | 0.190056539 | 0.334718885 |
| | 1.65E-06 | 1.79E-05 |
| | 6.84E-05 | 0.000499826 |
| | 0.001542632 | 0.007349423 |
| | 0.588346562 | 0.72457313 |
| | 0.165007954 | 0.303611574 |
| | 0.485217038 | 0.640657676 |
| | 0.518734234 | 0.669465062 |
| | 0.013526658 | 0.043911827 |
| | 0.336179905 | NA |
| NA | | NA |
| NA | | NA |
| NA | | NA |
| NA | | NA |
| NA | | NA |
| NA | | NA |
| NA | | NA |
| | 2.85E-11 | 6.13E-10 |
| | 0.47492248 | NA |
| NA | | NA |
| | 0.853595154 | NA |
| | 0.941277857 | NA |
| NA | | NA |
| NA | | NA |
| NA | | NA |
| | 0.000285233 | 0.001743927 |
| | 0.118177279 | 0.237516703 |
| | 0.000504467 | 0.002858885 |
| | 0.736569455 | NA |
| | 3.46E-17 | 1.22E-15 |
| | 3.00E-13 | 7.89E-12 |
| | 0.73517979 | 0.833163949 |
| | 0.681577836 | NA |
| | 0.536567465 | 0.684608888 |
| | 0.096417368 | 0.204065997 |
| | 0.001897264 | 0.008756774 |
| NA | | NA |
| | 2.04E-07 | 2.60E-06 |
| | 0.941278788 | NA |
| NA | | NA |
| | 4.92E-08 | 6.89E-07 |
| | 8.96E-18 | 3.29E-16 |
| | 0.000333924 | 0.002002072 |
| | 0.595241544 | 0.730117971 |
| | 0.000191111 | 0.001232273 |

| | |
|---|---|
| 0.353786013 | 0.517470008 |
| 0.042929898 | 0.110014797 |
| 0.134050037 | 0.260547034 |
| NA | NA |
| 0.540481889 | 0.687576158 |
| 0.839319107 | NA |
| 0.01720228 | 0.053202789 |
| 0.311564619 | 0.473524109 |
| 0.251446641 | 0.409170229 |
| 0.839319109 | NA |
| 0.012536937 | 0.041318771 |
| NA | NA |
| NA | NA |
| 0.082766683 | 0.182206122 |
| 5.88E-05 | 0.000437562 |
| 0.002018766 | 0.00922809 |
| 0.673053947 | 0.78893086 |
| 0.00972478 | 0.033673148 |
| 0.862724885 | 0.91782853 |
| 0.696453196 | 0.805524481 |
| 8.26E-05 | 0.000591947 |
| 0.014219906 | 0.045742301 |
| 0.000591106 | 0.003269698 |
| 0.008442047 | 0.029954919 |
| 0.388910948 | 0.551561603 |
| 0.010006788 | 0.034410374 |
| 0.416743532 | 0.578400194 |
| NA | NA |
| NA | NA |
| 0.697421248 | 0.806178454 |
| NA | NA |
| 0.126068115 | NA |
| 0.323887701 | 0.486567474 |
| 0.60183193 | NA |
| 0.241418959 | 0.397637123 |
| 1.77E-29 | 1.21E-27 |
| 0.218318732 | 0.36958908 |
| 0.422960468 | 0.584352291 |
| 0.613387926 | 0.744003219 |
| NA | NA |
| 0.002164655 | 0.009764091 |
| 0.000170535 | 0.001119746 |
| 0.002588108 | 0.011357176 |
| 0.003080644 | 0.013105538 |
| 0.991848041 | 0.994361977 |
| 0.608797438 | 0.740979577 |
| 0.251554173 | 0.409227276 |

| | |
|---|---|
| 0.782175445 | NA |
| 0.962865378 | NA |
| 0.532268665 | 0.680783261 |
| 0.889102486 | 0.933455095 |
| NA | NA |
| 0.070307025 | 0.161732658 |
| 0.327006636 | 0.489769188 |
| NA | NA |
| 0.284474022 | 0.445083196 |
| NA | NA |
| 0.006335728 | 0.023661672 |
| 0.0887706 | 0.192282299 |
| 0.001461175 | 0.007016078 |
| 2.05E-12 | 4.98E-11 |
| 0.598192607 | 0.732642814 |
| 0.941276431 | NA |
| 3.63E-05 | 0.000284789 |
| 0.08913911 | 0.192758163 |
| 0.139724077 | 0.268679309 |
| 0.794434937 | 0.873845389 |
| 0.546691624 | 0.692920498 |
| NA | NA |
| 0.045622974 | 0.115258126 |
| 0.977346232 | 0.986045906 |
| 0.955792398 | NA |
| 3.34E-07 | 4.06E-06 |
| NA | NA |
| 2.36E-09 | 4.00E-08 |
| NA | NA |
| 0.325445159 | NA |
| 0.28928268 | 0.450091931 |
| NA | NA |
| NA | NA |
| 0.386693764 | 0.54949761 |
| NA | NA |
| 0.059770593 | 0.142565352 |
| 0.839318641 | NA |
| 5.86E-52 | 8.36E-50 |
| NA | NA |
| 0.174174987 | 0.314991348 |
| 0.941278332 | NA |
| 0.526127558 | NA |
| 0.044077885 | 0.112159698 |
| 2.29E-05 | 0.000189674 |
| NA | NA |
| 0.224359164 | 0.377079284 |
| 0.000198007 | 0.001269659 |

| | | |
|---|---|---|
| | 0.010032882 | 0.03448414 |
| | 0.185661761 | 0.329048492 |
| | 0.846376806 | NA |
| | 6.11E-24 | 3.26E-22 |
| | 0.690228974 | 0.801297308 |
| | 0.709958751 | 0.814893099 |
| | 0.465723048 | 0.623968899 |
| | 0.75299461 | 0.845095139 |
| | 0.955792398 | NA |
| NA | | NA |
| | 0.020549959 | 0.061156554 |
| | 0.000111027 | 0.000767966 |
| | 0.010974581 | 0.037146907 |
| | 0.008661275 | 0.030577821 |
| | 0.074830814 | 0.169133596 |
| | 4.18E-05 | 0.00032277 |
| | 0.004795181 | 0.018876301 |
| | 0.00066111 | 0.003597198 |
| | 2.80E-05 | 0.00022662 |
| | 0.791669975 | NA |
| | 0.885853159 | 0.931651465 |
| | 0.308153251 | 0.46960479 |
| | 0.606376941 | 0.739051887 |
| | 0.941275956 | NA |
| | 0.470568355 | 0.628216276 |
| | 0.014775554 | 0.047086323 |
| | 2.11E-07 | 2.68E-06 |
| | 7.75E-26 | 4.51E-24 |
| | 0.005275835 | 0.020384294 |
| | 0.093958523 | NA |
| NA | | NA |
| NA | | NA |
| | 0.026312697 | 0.074710104 |
| NA | | NA |
| | 0.955792873 | NA |
| NA | | NA |
| | 4.06E-06 | 4.03E-05 |
| | 9.14E-23 | 4.60E-21 |
| NA | | NA |
| NA | | NA |
| | 7.42E-55 | 1.13E-52 |
| NA | | NA |
| NA | | NA |
| NA | | NA |
| NA | | NA |
| NA | | NA |
| NA | | NA |

| | | |
|---|---|---|
| NA | | NA |
| NA | | NA |
| NA | | NA |
| NA | | NA |
| | 0.985736229 | 0.991027527 |
| NA | | NA |
| NA | | NA |
| | 0.389113289 | 0.551717073 |
| | 0.839319108 | NA |
| | 0.675551596 | 0.790858573 |
| | 0.379709414 | 0.542422487 |
| | 0.918276767 | NA |
| | 0.253381804 | 0.411306943 |
| | 0.022873053 | 0.066751397 |
| | 0.941277857 | NA |
| NA | | NA |
| | 0.487518746 | 0.642378277 |
| | 0.955791446 | NA |
| | 0.819098269 | 0.889708256 |
| | 0.559279003 | 0.70270053 |
| | 0.710184827 | NA |
| | 0.026720938 | 0.075742738 |
| | 0.017199269 | 0.053200389 |
| | 0.955792398 | NA |
| | 0.00537837 | 0.020689862 |
| NA | | NA |
| | 0.062008457 | 0.146741428 |
| NA | | NA |
| NA | | NA |
| NA | | NA |
| | 0.23324119 | 0.387871641 |
| | 0.733054088 | 0.831587399 |
| | 0.926622562 | 0.955899546 |
| | 0.110680068 | 0.226207023 |
| | 0.505701874 | 0.657528461 |
| | 0.257276359 | 0.41534777 |
| | 0.864515488 | NA |
| | 0.780941982 | 0.865201129 |
| | 0.09563491 | 0.202915147 |
| | 0.046193085 | NA |
| | 0.033190582 | 0.089936503 |
| NA | | NA |
| | 7.15E-05 | 0.000520509 |
| | 0.022595762 | 0.066071784 |
| | 0.140563521 | 0.269988131 |
| | 0.955792398 | NA |
| | 7.43E-06 | 6.94E-05 |

| | |
|---|---|
| 0.583299433 | 0.720557043 |
| 0.027246586 | 0.076966916 |
| 0.910302286 | 0.945786154 |
| 0.134109223 | 0.260606843 |
| 0.002544778 | 0.0112114 |
| 0.852586121 | 0.911441942 |
| 1.99E-06 | 2.10E-05 |
| 0.617448794 | 0.747039912 |
| 0.751007199 | NA |
| 0.85157531 | 0.910934139 |
| 0.00784914 | 0.028229926 |
| NA | NA |
| NA | NA |
| 0.839319107 | NA |
| 0.909650485 | 0.945315266 |
| NA | NA |
| NA | NA |
| 0.622095166 | NA |
| 0.319464757 | 0.482022196 |
| 0.000277209 | 0.001700108 |
| NA | NA |
| NA | NA |
| 0.89863617 | NA |
| NA | NA |
| NA | NA |
| NA | NA |
| 0.181679851 | 0.32409082 |
| NA | NA |
| NA | NA |
| NA | NA |
| NA | NA |
| NA | NA |
| NA | NA |
| NA | NA |
| NA | NA |
| NA | NA |
| NA | NA |
| NA | NA |
| NA | NA |
| NA | NA |
| 0.601279474 | NA |
| 0.287737879 | 0.448370958 |
| 0.031985874 | 0.087231669 |
| 0.425385239 | 0.586787567 |
| 0.859306226 | 0.916075012 |
| 0.638660233 | 0.762662596 |
| 0.032040189 | 0.087346305 |

| | | |
|---|---|---|
| | 0.20383864 | NA |
| | 0.010148039 | 0.034819607 |
| | 0.002690207 | 0.011738277 |
| | 0.534701848 | 0.682850809 |
| | 0.190755617 | 0.33567686 |
| | 0.099538917 | 0.208852724 |
| | 0.006419596 | 0.023942214 |
| | 0.167943276 | 0.307278488 |
| | 0.676846707 | 0.791715346 |
| | 0.922131067 | NA |
| | 0.299718809 | 0.461234052 |
| | 0.92300134 | NA |
| | 0.412614774 | 0.574868699 |
| | 0.248256619 | 0.405473683 |
| | 0.005074276 | 0.01976402 |
| | 0.874328214 | 0.924906832 |
| | 0.227745774 | 0.381183103 |
| | 0.67929368 | NA |
| NA | | NA |
| | 0.032214158 | 0.08776026 |
| | 0.023077862 | 0.067258391 |
| NA | | NA |
| | 0.028868924 | 0.080536583 |
| | 0.718245818 | 0.82084251 |
| | 0.938135643 | 0.963267529 |
| | 0.006430664 | 0.023962282 |
| | 0.991855393 | 0.994361977 |
| | 0.158263591 | 0.294365085 |
| NA | | NA |
| | 0.056732607 | 0.136910627 |
| NA | | NA |
| | 0.631317902 | 0.757431042 |
| | 0.000113874 | 0.000784264 |
| | 0.065579184 | 0.153316545 |
| NA | | NA |
| | 0.098363743 | 0.207105344 |
| | 0.297924513 | 0.459334062 |
| | 5.34E-09 | 8.68E-08 |
| | 0.826954775 | 0.894974654 |
| | 0.505557433 | 0.657448397 |
| | 0.599258661 | NA |
| | 0.025925988 | 0.073814639 |
| | 0.003561279 | 0.014764948 |
| | 0.338022401 | 0.501502786 |
| | 4.70E-07 | 5.55E-06 |
| NA | | NA |
| | 0.325432512 | 0.488271851 |

| | |
|---|---|
| NA | NA |
| NA | NA |
| NA | NA |
| 3.72E-15 | 1.15E-13 |
| 0.040384878 | 0.104927505 |
| 0.757514053 | NA |
| 0.721119883 | 0.822981178 |
| 0.233709473 | 0.388320446 |
| 0.002800839 | 0.012096821 |
| 0.45961135 | 0.618759578 |
| 0.839319574 | NA |
| 0.94127925 | NA |
| 0.896018124 | NA |
| NA | NA |
| NA | NA |
| 0.355269245 | 0.519003165 |
| 0.003400694 | 0.014218045 |
| 0.492426709 | 0.646344116 |
| 0.679490564 | NA |
| NA | NA |
| 0.797023827 | 0.875600533 |
| 0.662652279 | 0.780900599 |
| 0.571344423 | 0.711661519 |
| 0.132248615 | 0.25788805 |
| 0.420526488 | 0.582075568 |
| 0.515981125 | 0.666850342 |
| 0.609299942 | 0.741429546 |
| 0.027448784 | 0.077418645 |
| 0.005361959 | 0.02064133 |
| 0.847589041 | 0.908343735 |
| 0.042096049 | 0.108391407 |
| 0.009374021 | 0.032629491 |
| 0.642789706 | NA |
| 0.081973825 | 0.180946024 |
| 0.955792398 | NA |
| 1.11E-14 | 3.27E-13 |
| 0.29859743 | 0.460192699 |
| NA | NA |
| 0.001134091 | 0.005685267 |
| 0.741290097 | NA |
| 0.774145277 | 0.860571326 |
| 0.942552478 | NA |
| 0.769345625 | 0.857221448 |
| 0.230530091 | 0.384491431 |
| 8.48E-05 | 0.000606201 |
| 0.287633997 | NA |
| 0.228989981 | 0.382673561 |

| | |
|---|---|
| 0.079483668 | 0.17700882 |
| 0.852087452 | 0.911172335 |
| 0.481665764 | 0.638128049 |
| 0.005328036 | 0.020550628 |
| 0.300974 | 0.462358536 |
| NA | NA |
| 0.000201124 | 0.001287218 |
| 0.015732702 | 0.049572381 |
| 0.000145998 | 0.000975864 |
| 1.44E-05 | 0.000125798 |
| NA | NA |
| 0.448710679 | 0.60881295 |
| 3.03E-107 | 1.11E-104 |
| 1.45E-13 | 3.93E-12 |
| 0.702385047 | NA |
| 0.735303781 | 0.833234344 |
| 0.946961298 | 0.967625072 |
| NA | NA |
| NA | NA |
| 0.077125316 | 0.172970579 |
| 0.66423517 | 0.782114139 |
| 5.08E-06 | 4.94E-05 |
| 4.07E-08 | 5.77E-07 |
| NA | NA |
| 0.636284542 | 0.760856301 |
| 0.378308579 | 0.541103347 |
| 0.92443807 | NA |
| 0.01723337 | 0.053271257 |
| 0.338522878 | 0.502026264 |
| 0.429388101 | 0.590700241 |
| 0.000370033 | 0.0021817 |
| 0.489719399 | 0.644101295 |
| NA | NA |
| 0.01519675 | 0.048202697 |
| NA | NA |
| NA | NA |
| 0.222561676 | 0.374965311 |
| 0.035494846 | 0.094763503 |
| 0.130458594 | 0.255410377 |
| 0.105013685 | 0.217521167 |
| 0.955792398 | NA |
| 0.751530428 | 0.844048849 |
| 0.878693598 | 0.927232888 |
| 7.66E-05 | 0.000552758 |
| 2.04E-05 | 0.00017081 |
| NA | NA |
| NA | NA |

| | | |
|---|---|---|
| | 0.941278332 | NA |
| NA | | NA |
| | 0.876935462 | NA |
| | 0.955791446 | NA |
| NA | | NA |
| | 0.021566452 | 0.06356175 |
| | 0.894835952 | NA |
| | 0.991972551 | 0.99440104 |
| | 0.969233236 | 0.981563608 |
| | 0.106091388 | 0.219207916 |
| NA | | NA |
| | 0.000585094 | 0.003243223 |
| | 0.002923932 | 0.01255132 |
| | 0.177848871 | 0.319321374 |
| | 0.00329175 | 0.013867299 |
| | 0.011693299 | 0.039117908 |
| | 0.534153332 | 0.682443243 |
| NA | | NA |
| NA | | NA |
| | 0.758800944 | 0.849513895 |
| | 0.012826546 | 0.042098378 |
| | 0.554720525 | 0.699549914 |
| | 0.524317998 | 0.673899987 |
| NA | | NA |
| | 0.008062593 | 0.028840673 |
| | 0.118698333 | 0.238285025 |
| NA | | NA |
| | 0.000438285 | 0.002530825 |
| | 0.011115972 | 0.03752409 |
| | 0.939641795 | NA |
| | 0.955792398 | NA |
| | 0.76242508 | 0.852543539 |
| | 0.001651252 | 0.007790506 |
| | 0.000793274 | 0.004199625 |
| NA | | NA |
| | 0.749040939 | 0.842445421 |
| | 0.48256983 | 0.638617692 |
| NA | | NA |
| NA | | NA |
| NA | | NA |
| NA | | NA |
| NA | | NA |
| NA | | NA |
| | 0.022544403 | 0.065962124 |
| | 0.007763474 | 0.027972483 |
| | 0.93194745 | 0.959371849 |
| | 5.45E-13 | 1.40E-11 |

| | |
|---|---|
| 0.955792873 | NA |
| 5.64E-20 | 2.38E-18 |
| 0.835333057 | 0.899994439 |
| 0.090174229 | 0.19455466 |
| 0.382960662 | 0.545691432 |
| 0.696778986 | 0.805780803 |
| 0.884488497 | 0.930962684 |
| 0.097703593 | NA |
| NA | NA |
| 0.610596841 | NA |
| NA | NA |
| NA | NA |
| 0.308798294 | 0.470326584 |
| 1.30E-09 | 2.27E-08 |
| 0.37767334 | 0.540550885 |
| NA | NA |
| 6.47E-06 | 6.15E-05 |
| 0.97320208 | 0.983784606 |
| 0.483423922 | 0.639389652 |
| 0.01284284 | 0.042134428 |
| 0.944796895 | 0.966729244 |
| 0.423438993 | 0.584780673 |
| NA | NA |
| NA | NA |
| 0.494716908 | 0.647786973 |
| 0.798917656 | 0.87715476 |
| 0.402922314 | 0.564969479 |
| 0.414401867 | 0.576380135 |
| 0.00130258 | 0.006373911 |
| NA | NA |
| NA | NA |
| 5.21E-05 | 0.000393452 |
| 0.310246106 | 0.472032589 |
| 0.00710937 | 0.026005414 |
| 0.113447687 | 0.230360673 |
| 4.01E-05 | 0.000311391 |
| NA | NA |
| 0.000170911 | 0.001121597 |
| 0.5927265 | 0.728233942 |
| 0.000312692 | 0.001887508 |
| 0.131263284 | 0.256553487 |
| 0.001110081 | 0.005583816 |
| 0.175864247 | NA |
| 0.037197057 | 0.098264633 |
| 0.102930257 | 0.214362234 |
| 0.440542783 | 0.601284439 |
| 0.440287342 | NA |

| | | |
|---|---|---|
| | 0.741011381 | 0.83729508 |
| | 0.143207776 | 0.273675752 |
| | 0.55461425 | 0.699535179 |
| NA | | NA |
| | 0.501719135 | 0.653993365 |
| | 0.71911292 | NA |
| | 0.839318641 | NA |
| | 0.670330133 | NA |
| | 0.10573187 | NA |
| | 0.007938924 | 0.028463278 |
| | 0.404441063 | NA |
| | 0.000490381 | 0.00278902 |
| | 0.006350941 | 0.023704736 |
| | 0.248344437 | 0.405563931 |
| | 0.166708522 | 0.305559544 |
| | 0.482340778 | NA |
| | 0.013160689 | 0.042981712 |
| | 0.453491457 | 0.61343508 |
| | 0.544916578 | 0.691480748 |
| | 0.002958762 | 0.012673107 |
| | 0.839319108 | NA |
| | 0.000360346 | 0.002133573 |
| | 0.737044842 | 0.834275315 |
| | 0.657715521 | 0.777100741 |
| | 0.106645377 | 0.220159995 |
| | 0.816003719 | 0.887480947 |
| NA | | NA |
| | 0.199759621 | 0.347183534 |
| | 0.034793106 | 0.093217641 |
| | 1.52E-32 | 1.18E-30 |
| | 0.003093762 | 0.013154293 |
| | 0.739991096 | NA |
| | 0.036344224 | 0.096515142 |
| | 0.941275956 | NA |
| | 0.988441053 | 0.992695103 |
| | 0.764543749 | 0.853789543 |
| NA | | NA |
| | 0.67191961 | 0.787834017 |
| | 0.316198001 | 0.478683643 |
| | 0.113016758 | 0.229739933 |
| NA | | NA |
| | 0.577274569 | 0.71579985 |
| | 0.001085065 | 0.005475243 |
| NA | | NA |
| | 9.96E-09 | 1.56E-07 |
| NA | | NA |
| | 0.033174244 | 0.089912702 |

| | |
|---|---|
| 0.028947382 | 0.080660921 |
| 0.000400582 | 0.002339794 |
| 0.03662463 | 0.097106734 |
| 0.033659742 | 0.090907656 |
| 0.724945582 | 0.826238017 |
| 0.734058779 | NA |
| 0.025921448 | 0.07381054 |
| 0.596348285 | 0.731120237 |
| 0.005469473 | 0.020967047 |
| 0.738136366 | 0.835193087 |
| 2.57E-06 | 2.65E-05 |
| 0.678709093 | 0.793021149 |
| 0.212737311 | 0.363270923 |
| 0.421499988 | 0.582963638 |
| 0.610949833 | NA |
| 7.91E-08 | 1.08E-06 |
| 0.229103902 | 0.382783313 |
| 0.497831235 | 0.650850217 |
| 0.773458358 | 0.860354044 |
| NA | NA |
| NA | NA |
| 0.136771567 | NA |
| NA | NA |
| NA | NA |
| 0.838993826 | NA |
| 2.17E-08 | 3.23E-07 |
| 0.928528315 | 0.957409069 |
| 0.003425619 | 0.014299647 |
| 0.23063049 | 0.384551097 |
| NA | NA |
| 0.853595154 | NA |
| NA | NA |
| 0.009817215 | 0.033900245 |
| 0.055564752 | 0.13478926 |
| 0.667378035 | 0.784363618 |
| 0.799986307 | 0.877807949 |
| 0.848786497 | NA |
| 0.646477499 | 0.768719461 |
| 0.351478655 | 0.515109595 |
| 0.939413471 | NA |
| NA | NA |
| 0.126548887 | 0.249823058 |
| 0.684401321 | 0.797279344 |
| 0.867943192 | 0.921060969 |
| 0.170886519 | NA |
| 0.079425214 | 0.176911748 |
| 0.01301932 | 0.042601851 |

| | |
|---|---|
| 6.06E-51 | 8.44E-49 |
| 7.46E-17 | 2.58E-15 |
| 1.47E-11 | 3.25E-10 |
| 0.033055245 | 0.089651429 |
| 0.057546984 | 0.138412825 |
| 0.600019781 | 0.733974244 |
| 0.796437796 | 0.875158692 |
| 0.213273297 | 0.363920785 |
| 0.057977395 | 0.139124705 |
| NA | NA |
| 0.725722875 | 0.826767619 |
| 0.024115101 | 0.069717921 |
| 0.041901555 | 0.108008698 |
| 0.00216779 | 0.009772298 |
| 1.06E-05 | 9.53E-05 |
| 0.104256365 | 0.216329003 |
| 0.450624781 | 0.610583839 |
| 0.613380078 | NA |
| 0.071708832 | 0.163974793 |
| 0.013523776 | 0.043908459 |
| 0.26614522 | 0.42558203 |
| 0.923025346 | NA |
| 0.095246431 | 0.202291988 |
| NA | NA |
| 0.094929152 | 0.201813476 |
| 0.906259947 | NA |
| NA | NA |
| NA | NA |
| 1.71E-06 | 1.84E-05 |
| 0.460106832 | 0.619114165 |
| NA | NA |
| NA | NA |
| 3.39E-72 | 7.55E-70 |
| 0.115241115 | 0.23291063 |
| 0.292219914 | 0.453357998 |
| 0.427248177 | NA |
| 0.634613813 | 0.759812776 |
| 0.001034862 | 0.005258708 |
| NA | NA |
| 0.864761328 | NA |
| NA | NA |
| NA | NA |
| 0.609566993 | 0.741490928 |
| 9.79E-06 | 8.90E-05 |
| 0.499446647 | 0.652296322 |
| 0.012395929 | 0.040944755 |
| 0.905103057 | 0.94244724 |

| | | | |
|---|---|---|---|
| | 0.107899591 | | 0.221949584 |
| | 0.903555608 | | 0.941652848 |
| | 0.07217363 | | 0.164752733 |
| | 6.35E-24 | | 3.37E-22 |
| | 0.127333786 | | 0.250977044 |
| | 0.955792873 | NA | |
| | 0.057000585 | | 0.137437124 |
| | 0.145180993 | | 0.276601996 |
| | 0.439356535 | NA | |
| | 0.002195223 | | 0.009886603 |
| | 0.825665293 | NA | |
| | 0.12554833 | | 0.248467545 |
| | 0.578826992 | | 0.7168925 |
| | 0.412924937 | | 0.575132504 |
| NA | | NA | |
| | 0.976287078 | NA | |
| | 0.366492398 | NA | |
| | 0.03284872 | | 0.089232921 |
| | 0.940528311 | NA | |
| | 0.002797958 | | 0.012090853 |
| | 3.30E-08 | | 4.78E-07 |
| | 0.490956719 | | 0.645087045 |
| | 0.035402235 | | 0.09457284 |
| | 0.55292219 | | 0.698038772 |
| | 0.817257543 | NA | |
| | 0.137745232 | | 0.266034729 |
| | 0.547138144 | | 0.69333877 |
| | 0.879287201 | | 0.927735891 |
| | 0.305644929 | | 0.467212164 |
| | 0.886451512 | | 0.932154579 |
| | 0.002005592 | | 0.009183742 |
| | 0.921122847 | NA | |
| | 0.296421868 | | 0.457907145 |
| | 0.227449048 | | 0.38084715 |
| NA | | NA | |
| NA | | NA | |
| | 0.058758186 | | 0.140640627 |
| | 0.456644421 | | 0.616334024 |
| | 0.380939814 | | 0.54365808 |
| | 0.432164884 | | 0.593354209 |
| | 0.000293914 | | 0.001791942 |
| | 0.687541395 | NA | |
| | 0.011174517 | | 0.037689649 |
| | 0.055215645 | | 0.134120092 |
| | 0.003939664 | | 0.016004761 |
| | 0.840920899 | NA | |
| NA | | NA | |

| | |
|---|---|
| NA | NA |
| 0.001171588 | 0.005840031 |
| NA | NA |
| NA | NA |
| 0.839319108 | NA |
| NA | NA |
| 0.024187262 | 0.069884099 |
| 0.049482455 | 0.122711128 |
| 0.918525826 | NA |
| 0.923001222 | NA |
| 0.163244932 | 0.301112513 |
| 0.860676077 | 0.916679745 |
| NA | NA |
| NA | NA |
| 0.027886266 | 0.078327696 |
| 0.012557558 | 0.041363824 |
| 0.640393708 | NA |
| 0.957024918 | NA |
| NA | NA |
| 0.30365949 | 0.465043314 |
| 0.108226617 | NA |
| 0.218231201 | 0.369520313 |
| 0.203946656 | 0.3524461 |
| 0.046371687 | 0.116737903 |
| 1.63E-07 | 2.11E-06 |
| 0.212446819 | 0.362952856 |
| 0.225153393 | 0.377907133 |
| 0.761944111 | NA |
| 0.000514851 | 0.002910115 |
| 0.040256614 | 0.104708548 |
| 0.356435443 | 0.520164097 |
| 0.22764344 | 0.381038618 |
| 0.955792398 | NA |
| 0.941277857 | NA |
| NA | NA |
| 0.085031519 | 0.186227476 |
| 0.002043003 | 0.009308489 |
| 1.31E-22 | 6.49E-21 |
| NA | NA |
| 6.40E-15 | 1.93E-13 |
| 5.62E-07 | 6.55E-06 |
| 0.061708725 | 0.14623567 |
| NA | NA |
| 0.002358418 | 0.010512186 |
| 0.004559876 | 0.018089125 |
| 0.183538702 | 0.32625752 |
| 0.75237696 | NA |

| | |
|---|---|
| 0.046082506 | 0.116180046 |
| 0.036392339 | 0.096610579 |
| NA | NA |
| NA | NA |
| 1.80E-06 | 1.92E-05 |
| 0.000467877 | 0.00268024 |
| 0.852831423 | 0.911540416 |
| 0.555197568 | 0.699825903 |
| NA | NA |
| NA | NA |
| 0.364073825 | 0.52755777 |
| 0.067959989 | 0.157582431 |
| NA | NA |
| 1.77E-07 | 2.28E-06 |
| 0.077558786 | 0.173697235 |
| 0.165355051 | 0.303999222 |
| 0.450198539 | 0.610159755 |
| 0.677926304 | 0.7923788 |
| NA | NA |
| 0.839318641 | NA |
| 0.127759798 | 0.251629185 |
| 0.557474212 | 0.701317451 |
| NA | NA |
| 0.033845797 | 0.091244496 |
| 0.360693485 | 0.524416733 |
| NA | NA |
| 7.34E-05 | 0.000532351 |
| 0.11954608 | 0.239615647 |
| 0.072328205 | 0.165058095 |
| 0.726802277 | NA |
| 0.085311651 | 0.186617766 |
| 0.189400625 | 0.333973899 |
| 0.023510531 | 0.068254792 |
| 0.065566361 | 0.153301622 |
| NA | NA |
| 0.049273157 | 0.122262489 |
| 0.005876053 | 0.0222639 |
| 0.000689854 | 0.003730829 |
| NA | NA |
| NA | NA |
| 0.020255623 | 0.060484801 |
| NA | NA |
| 0.647122701 | NA |
| NA | NA |
| NA | NA |
| NA | NA |
| NA | NA |

| | | | |
|---|---|---|---|
| | 0.616104153 | | 0.745981842 |
| NA | | NA | |
| NA | | NA | |
| | 0.258913643 | NA | |
| NA | | NA | |
| | 0.343572854 | | 0.507114044 |
| NA | | NA | |
| | 0.941279268 | NA | |
| | 0.000125128 | | 0.000853626 |
| NA | | NA | |
| NA | | NA | |
| | 0.267624334 | | 0.427258327 |
| | 9.28E-13 | | 2.34E-11 |
| NA | | NA | |
| NA | | NA | |
| | 0.118637616 | | 0.238236104 |
| | 0.426505729 | NA | |
| NA | | NA | |
| | 0.853801216 | NA | |
| NA | | NA | |
| NA | | NA | |
| | 0.853595154 | NA | |
| | 0.113447938 | | 0.230360673 |
| | 0.239649995 | | 0.39565438 |
| NA | | NA | |
| | 0.071204835 | | 0.163151472 |
| | 0.285975095 | | 0.446609751 |
| NA | | NA | |
| NA | | NA | |
| | 0.867243025 | NA | |
| | 0.559084473 | | 0.702614065 |
| NA | | NA | |
| | 0.955792398 | NA | |
| NA | | NA | |
| NA | | NA | |
| NA | | NA | |
| NA | | NA | |
| | 0.839318641 | NA | |
| | 0.957911252 | | 0.974159522 |
| NA | | NA | |
| NA | | NA | |
| NA | | NA | |
| NA | | NA | |
| | 0.941277857 | NA | |
| | 0.827630909 | | 0.895299318 |
| NA | | NA | |
| NA | | NA | |

| | | |
|---|---|---|
| | 0.304346876 | NA |
| NA | | NA |
| | 0.955792398 | NA |
| NA | | NA |
| | 0.915941647 | 0.949448789 |
| NA | | NA |
| | 0.941279268 | NA |
| | 3.10E-05 | 0.000248568 |
| NA | | NA |
| | 0.839318641 | NA |
| | 0.744071634 | NA |
| NA | | NA |
| | 0.853595154 | NA |
| | 0.000206387 | 0.001317507 |
| NA | | NA |
| | 0.839319107 | NA |
| NA | | NA |
| | 0.839319108 | NA |
| | 5.92E-06 | 5.67E-05 |
| NA | | NA |
| | 0.839319108 | NA |
| | 0.161391849 | 0.298364954 |
| | 0.000301282 | 0.001829364 |
| NA | | NA |
| | 0.955792873 | NA |
| NA | | NA |
| | 0.941275481 | NA |
| NA | | NA |
| | 0.955792398 | NA |
| NA | | NA |
| | 0.853595622 | NA |
| NA | | NA |
| | 0.271504173 | 0.431360635 |
| NA | | NA |
| | 0.941278332 | NA |
| | 0.008251751 | 0.029417187 |
| NA | | NA |
| | 0.682756412 | NA |
| NA | | NA |
| NA | | NA |
| | 0.003871534 | 0.015781854 |
| NA | | NA |
| | 0.501012028 | NA |
| NA | | NA |
| NA | | NA |
| NA | | NA |
| | 0.449747864 | 0.609722542 |

| | |
|---|---|
| 0.941278788 | NA |
| 0.177839096 | 0.319321374 |
| NA | NA |
| 0.935650667 | NA |
| NA | NA |
| 0.037331937 | 0.09855532 |
| NA | NA |
| 0.955792873 | NA |
| NA | NA |
| 0.943962327 | NA |
| NA | NA |
| NA | NA |
| NA | NA |
| 0.306141282 | NA |
| NA | NA |
| 0.355734749 | NA |
| NA | NA |
| 0.955792398 | NA |
| 0.915332007 | 0.949020885 |
| NA | NA |
| NA | NA |
| NA | NA |
| NA | NA |
| NA | NA |
| 0.582335606 | 0.720005533 |
| NA | NA |
| NA | NA |
| 0.839319107 | NA |
| 0.787661003 | NA |
| NA | NA |
| NA | NA |
| 0.444873182 | NA |
| NA | NA |
| 0.322676557 | 0.485470987 |
| 0.941279803 | NA |
| NA | NA |
| NA | NA |
| 0.00984445 | 0.033978771 |
| NA | NA |
| 0.941278332 | NA |
| NA | NA |
| 0.568255961 | 0.709522551 |
| 0.063968501 | 0.150452397 |
| 0.910917927 | NA |
| NA | NA |
| NA | NA |
| 0.807962317 | NA |

| | | | |
|---|---|---|---|
| NA | | NA | |
| NA | | NA | |
| NA | | NA | |
| NA | | NA | |
| | 0.100254318 | | 0.210020532 |
| NA | | NA | |
| NA | | NA | |
| NA | | NA | |
| NA | | NA | |
| | 0.872047401 | NA | |
| | 0.091229012 | | 0.196109805 |
| NA | | NA | |
| | 0.839318641 | NA | |
| | 0.927474672 | | 0.956612738 |
| NA | | NA | |
| | 0.839319574 | NA | |
| NA | | NA | |
| NA | | NA | |
| | 0.699133929 | NA | |
| | 0.145678637 | | 0.277315078 |
| NA | | NA | |
| | 0.941278332 | NA | |
| | 0.010842013 | | 0.036776627 |
| | 0.839318641 | NA | |
| | 0.000184031 | | 0.00119227 |
| | 0.955791922 | NA | |
| | 0.000538134 | | 0.003023766 |
| | 0.955791446 | NA | |
| | 0.004607913 | | 0.018255251 |
| NA | | NA | |
| | 0.218058374 | | 0.369339023 |
| | 0.885719245 | NA | |
| | 0.955791922 | NA | |
| | 0.003904608 | | 0.015900338 |
| | 0.941278332 | NA | |
| | 0.941277857 | NA | |
| NA | | NA | |
| | 0.839319574 | NA | |
| | 0.941278332 | NA | |
| | 0.967409267 | | 0.980425588 |
| | 0.108364799 | | 0.222602826 |
| | 0.923246595 | NA | |
| | 0.320347986 | | 0.483048639 |
| | 0.853595622 | NA | |
| NA | | NA | |
| | 0.204564423 | NA | |
| | 0.207564639 | | 0.356762964 |

| | | |
|---|---|---|
| | 0.242029755 | NA |
| | 0.955792873 | NA |
| | 0.955791446 | NA |
| | 0.913133778 | 0.947569873 |
| | 1.02E-13 | 2.78E-12 |
| | 0.002773526 | 0.012018106 |
| | 0.011419591 | 0.03836944 |
| | 5.38E-05 | 0.000405045 |
| | 0.116835415 | 0.235331856 |
| | 0.588710781 | 0.724945212 |
| | 0.306024734 | 0.467672598 |
| | 1.17E-08 | 1.82E-07 |
| NA | | NA |
| | 0.815955091 | 0.887480947 |
| NA | | NA |
| | 5.87E-06 | 5.63E-05 |
| | 0.275944081 | 0.435776341 |
| NA | | NA |
| | 0.723056233 | NA |
| NA | | NA |
| NA | | NA |
| | 0.598016271 | NA |
| | 0.718757622 | NA |
| | 0.943347828 | NA |
| NA | | NA |
| NA | | NA |
| NA | | NA |
| | 0.272271868 | 0.432127426 |
| | 0.941276431 | NA |
| | 0.613120502 | 0.743807044 |
| | 0.839319108 | NA |
| | 0.349226625 | 0.513218471 |
| | 0.012478672 | 0.041172352 |
| | 0.959353542 | 0.975112026 |
| | 0.690850697 | 0.801737118 |
| | 0.041219059 | 0.106680051 |
| | 0.493646312 | 0.64701374 |
| NA | | NA |
| | 0.556705169 | 0.70087255 |
| | 0.956624997 | 0.97337139 |
| NA | | NA |
| | 0.611889436 | NA |
| NA | | NA |
| | 0.14747283 | 0.279893812 |
| | 0.425622301 | 0.587016513 |
| | 0.745631365 | 0.840134719 |
| | 0.001552821 | 0.007390571 |

| | |
|---|---|
| NA | NA |
| 0.941275956 | NA |
| 1.15E-09 | 2.03E-08 |
| NA | NA |
| 0.09067163 | 0.195326829 |
| 0.681804729 | 0.795349743 |
| 0.086896211 | 0.189216078 |
| 0.133289048 | 0.259406862 |
| 0.728920736 | 0.829338975 |
| 0.000869184 | 0.004532511 |
| 0.644926281 | 0.767565145 |
| 0.048367207 | 0.120560924 |
| 2.53E-05 | 0.000207846 |
| NA | NA |
| NA | NA |
| 0.831413565 | NA |
| NA | NA |
| 0.009950477 | 0.034260271 |
| 0.01521887 | 0.048259998 |
| 0.000115881 | 0.000796938 |
| 0.06491547 | 0.152258377 |
| 0.276139944 | NA |
| 0.007108775 | 0.026005414 |
| 0.471860395 | 0.629405948 |
| 0.501220825 | 0.653701813 |
| 0.129167535 | 0.25341065 |
| NA | NA |
| NA | NA |
| NA | NA |
| 0.600186687 | NA |
| 0.632862227 | NA |
| NA | NA |
| 0.570360632 | 0.711039416 |
| NA | NA |
| 0.916274145 | NA |
| 0.000377404 | 0.002219662 |
| 0.702554175 | NA |
| NA | NA |
| 0.955791922 | NA |
| 1.35E-05 | 0.000118328 |
| 1.63E-11 | 3.61E-10 |
| 0.001799043 | 0.008368346 |
| 0.006574525 | 0.024397699 |
| 0.01286586 | 0.042191548 |
| NA | NA |
| 0.941277857 | NA |
| 7.74E-06 | 7.20E-05 |

| | |
|---|---|
| 0.271609451 | 0.431421298 |
| 0.543742551 | 0.690469495 |
| 0.341278011 | 0.504791152 |
| 0.013467121 | 0.043788163 |
| 0.306514601 | 0.468115236 |
| NA | NA |
| 4.35E-10 | 8.10E-09 |
| 0.009174375 | 0.032079988 |
| 0.110738717 | 0.22624918 |
| 0.729118507 | 0.829416758 |
| 0.421795858 | 0.583142855 |
| 0.718606524 | 0.821015225 |
| 0.967929434 | 0.980741119 |
| 0.056111766 | 0.135853143 |
| 0.520467402 | 0.670806851 |
| 0.468723423 | 0.626807961 |
| 0.558543466 | NA |
| 5.51E-07 | 6.43E-06 |
| 0.490742553 | 0.645047538 |
| 0.000635148 | 0.003472229 |
| 0.063749359 | 0.150040763 |
| 0.000142049 | 0.000952411 |
| 0.003870997 | 0.015781854 |
| 0.000420746 | 0.002442586 |
| 0.916085589 | 0.949473947 |
| 0.921520196 | 0.953114487 |
| 0.284837081 | 0.445417002 |
| 0.284981311 | 0.445525463 |
| 0.580694081 | NA |
| 0.000572057 | 0.003181335 |
| 0.232380503 | NA |
| 0.794395917 | 0.873842851 |
| 0.234046694 | NA |
| 0.08249655 | 0.18181323 |
| 0.63385545 | NA |
| 0.06741181 | 0.156585918 |
| 0.666390826 | 0.783745155 |
| 0.578098685 | 0.716366394 |
| NA | NA |
| 0.116345614 | 0.234682907 |
| 0.959236667 | 0.975090803 |
| 0.00463642 | 0.018352915 |
| NA | NA |
| NA | NA |
| 0.092065856 | 0.197259115 |
| 0.014335277 | 0.046034575 |
| 0.056740641 | 0.136916132 |

| | |
|---|---|
| NA | NA |
| 0.691365965 | 0.801978899 |
| NA | NA |
| 0.000485938 | 0.002769045 |
| 0.398793041 | 0.561575367 |
| 0.631816231 | NA |
| 0.465448658 | 0.62380256 |
| 2.44E-14 | 7.01E-13 |
| 0.955792873 | NA |
| 0.713983126 | NA |
| 0.877657494 | NA |
| NA | NA |
| 0.044267102 | 0.112556945 |
| 0.024638638 | 0.070895656 |
| 0.188026381 | 0.332397796 |
| 0.250940748 | 0.408514581 |
| 0.6445622 | NA |
| 0.929323384 | 0.958004957 |
| 0.711084425 | NA |
| 0.623260168 | 0.751385543 |
| NA | NA |
| NA | NA |
| 0.000403725 | 0.002354108 |
| 0.01780771 | 0.054656238 |
| NA | NA |
| NA | NA |
| NA | NA |
| NA | NA |
| 0.955791922 | NA |
| 0.10543528 | 0.218147647 |
| 0.941277857 | NA |
| 0.843742925 | NA |
| NA | NA |
| NA | NA |
| NA | NA |
| NA | NA |
| 0.839319108 | NA |
| NA | NA |
| NA | NA |
| NA | NA |
| NA | NA |
| NA | NA |
| 0.660171324 | NA |
| NA | NA |
| 0.00016321 | 0.001077593 |
| NA | NA |
| NA | NA |

| | |
|---|---|
| 0.000375979 | 0.002211825 |
| 0.95507426 | 0.972692863 |
| 1.12E-11 | 2.51E-10 |
| 0.015066774 | 0.047873363 |
| NA | NA |
| 0.208327354 | 0.35766068 |
| 0.001517927 | 0.007244777 |
| 0.096745119 | 0.204488831 |
| 0.092628201 | 0.198232013 |
| NA | NA |
| 0.133808825 | 0.260226951 |
| 0.051876598 | 0.127650861 |
| NA | NA |
| 0.00397393 | 0.016121951 |
| 0.001386309 | 0.006718981 |
| 0.730174297 | 0.830051037 |
| NA | NA |
| 0.075123558 | 0.169680292 |
| 0.273079759 | 0.43309198 |
| 0.006203387 | 0.02325977 |
| 0.108257934 | 0.222402497 |
| 0.571720047 | 0.711883249 |
| 0.954389586 | NA |
| 0.365501085 | 0.52869382 |
| 0.03678604 | 0.097449229 |
| 0.706994061 | 0.813097828 |
| 0.49087245 | 0.645047538 |
| 0.21397869 | 0.364877858 |
| 0.00685871 | 0.025255928 |
| 0.08947788 | 0.193350205 |
| 0.129601253 | 0.253974067 |
| 0.598297466 | 0.73265814 |
| 0.725720549 | NA |
| 4.11E-06 | 4.07E-05 |
| 6.33E-06 | 6.02E-05 |
| 0.003951685 | 0.016037173 |
| 0.955792873 | NA |
| 0.15125119 | 0.285087174 |
| 0.08137747 | 0.179963443 |
| 0.016823232 | 0.052250624 |
| 4.76E-05 | 0.00036335 |
| 0.040053824 | 0.104306467 |
| NA | NA |
| 5.15E-05 | 0.000389339 |
| 0.048437127 | 0.120672021 |
| 0.005499832 | 0.02105291 |
| 0.000229145 | 0.001439838 |

| | | |
|---|---|---|
| | 0.281409381 | 0.441655495 |
| | 0.835747487 | 0.900202342 |
| | 0.039092339 | 0.102261332 |
| | 0.178667165 | 0.320290774 |
| | 0.466157345 | NA |
| | 0.453368746 | 0.613303944 |
| | 0.199346928 | 0.346770122 |
| | 0.014290415 | 0.045909092 |
| | 0.171620088 | 0.311790966 |
| | 0.007706288 | 0.027804314 |
| | 3.53E-07 | 4.27E-06 |
| | 0.068351597 | 0.158274032 |
| | 0.709750782 | 0.814772263 |
| | 0.171808733 | 0.311942805 |
| | 0.002006025 | 0.00918396 |
| | 0.821023302 | 0.890864179 |
| | 0.025664151 | 0.073174189 |
| | 0.076231411 | 0.171512054 |
| | 0.004882901 | 0.01915024 |
| | 0.940504279 | NA |
| | 0.011826913 | 0.039448455 |
| | 0.210374654 | 0.360291949 |
| | 0.044611789 | 0.113247766 |
| | 0.955792873 | NA |
| | 0.487587355 | NA |
| NA | | NA |
| | 1.40E-10 | 2.75E-09 |
| | 0.916453578 | 0.949674427 |
| | 1.29E-05 | 0.000113936 |
| NA | | NA |
| | 0.050742063 | 0.125312574 |
| | 0.011032526 | 0.037305894 |
| | 0.504998318 | 0.657044385 |
| | 0.517522827 | 0.668311565 |
| | 0.77119739 | NA |
| | 0.853596091 | NA |
| | 0.671134235 | 0.787184599 |
| | 0.604485095 | NA |
| | 0.56768965 | 0.709187479 |
| | 0.733406527 | 0.831782395 |
| | 0.31274459 | 0.474684561 |
| | 0.000372231 | 0.002191927 |
| | 0.59780782 | 0.732284583 |
| | 0.757375908 | 0.848372223 |
| | 0.519509776 | NA |
| | 0.889140612 | 0.933455095 |
| | 0.839318641 | NA |

| | |
|---|---|
| 0.322197843 | 0.484976247 |
| 0.964100011 | NA |
| 2.09E-25 | 1.18E-23 |
| 1.09E-05 | 9.75E-05 |
| NA | NA |
| 0.058120669 | 0.139440397 |
| 0.014236077 | 0.045777775 |
| 0.000195007 | 0.001253458 |
| 0.311877734 | 0.473848706 |
| NA | NA |
| 0.451483176 | 0.611413436 |
| 0.852753984 | NA |
| 0.076889799 | 0.172613279 |
| 0.798746984 | 0.877007828 |
| NA | NA |
| 0.94127925 | NA |
| NA | NA |
| 0.01974011 | 0.059280104 |
| NA | NA |
| 0.86290373 | 0.917895718 |
| 0.001016855 | 0.005179567 |
| 0.360320431 | 0.524002377 |
| 0.15857221 | 0.294800886 |
| 0.240166405 | 0.396149742 |
| 0.001993368 | 0.009143599 |
| 0.541506137 | NA |
| 0.975426346 | 0.985178051 |
| 4.23E-07 | 5.07E-06 |
| 0.77675006 | 0.862483902 |
| 0.859677326 | 0.916299154 |
| 0.839319108 | NA |
| 0.041990531 | 0.108189934 |
| 0.853206133 | 0.911742904 |
| 0.001642129 | 0.007756686 |
| 0.2013819 | NA |
| 0.007799938 | 0.028082608 |
| 0.826689272 | 0.894768681 |
| 0.793820284 | NA |
| NA | NA |
| NA | NA |
| NA | NA |
| NA | NA |
| NA | NA |
| 0.523292837 | NA |
| 0.873150626 | 0.924280679 |
| 0.921388903 | NA |
| 0.777319749 | NA |

| | | |
|---|---|---|
| | 0.9557893 | NA |
| | 0.110525208 | 0.22592932 |
| | 0.357302247 | 0.520886144 |
| | 0.071084199 | 0.162953491 |
| NA | | NA |
| NA | | NA |
| | 0.354408242 | 0.518126957 |
| NA | | NA |
| NA | | NA |
| | 0.180627432 | 0.322672931 |
| | 0.953932387 | NA |
| NA | | NA |
| | 0.072380486 | 0.165107922 |
| NA | | NA |
| | 0.955792398 | NA |
| | 0.941275481 | NA |
| NA | | NA |
| | 0.941275956 | NA |
| | 0.749117729 | 0.842491977 |
| | 0.010194333 | 0.034937243 |
| | 0.832936663 | 0.898710718 |
| | 0.387293388 | 0.550086957 |
| | 0.829975065 | NA |
| | 0.040420793 | 0.104997894 |
| | 0.839318641 | NA |
| NA | | NA |
| NA | | NA |
| NA | | NA |
| | 0.219997683 | NA |
| | 0.408726625 | NA |
| NA | | NA |
| NA | | NA |
| NA | | NA |
| | 0.839319108 | NA |
| | 0.955792873 | NA |
| NA | | NA |
| NA | | NA |
| NA | | NA |
| NA | | NA |
| NA | | NA |
| NA | | NA |
| NA | | NA |
| NA | | NA |
| NA | | NA |
| NA | | NA |
| NA | | NA |
| NA | | NA |

| | | |
|---|---|---|
| NA | | NA |
| NA | | NA |
| NA | | NA |
| NA | | NA |
| NA | | NA |
| NA | | NA |
| NA | | NA |
| NA | | NA |
| NA | | NA |
| NA | | NA |
| NA | | NA |
| NA | | NA |
| NA | | NA |
| NA | | NA |
| NA | | NA |
| NA | | NA |
| NA | | NA |
| NA | | NA |
| | 0.839318641 | NA |
| NA | | NA |
| NA | | NA |
| NA | | NA |
| NA | | NA |
| NA | | NA |
| NA | | NA |
| NA | | NA |
| NA | | NA |
| NA | | NA |
| NA | | NA |
| NA | | NA |
| NA | | NA |
| NA | | NA |
| NA | | NA |
| NA | | NA |
| | 0.955792873 | NA |
| NA | | NA |
| NA | | NA |
| NA | | NA |
| | 0.955792873 | NA |
| NA | | NA |
| NA | | NA |
| NA | | NA |
| NA | | NA |
| NA | | NA |
| NA | | NA |
| NA | | NA |
| NA | | NA |

| | |
|---|---|
| NA | NA |
| NA | NA |
| NA | NA |
| NA | NA |
| NA | NA |
| NA | NA |
| NA | NA |
| NA | NA |
| NA | NA |
| 0.955792398 | NA |
| 0.414702553 | 0.576663561 |
| 0.478967752 | 0.635685872 |
| NA | NA |
| 0.827840652 | NA |
| 0.609839695 | 0.741565568 |
| NA | NA |
| NA | NA |
| 0.399582031 | NA |
| NA | NA |
| NA | NA |
| 0.171741569 | 0.311902397 |
| NA | NA |
| NA | NA |
| 0.839319107 | NA |
| 0.839321471 | NA |
| NA | NA |
| 0.941278332 | NA |
| NA | NA |
| NA | NA |
| NA | NA |
| NA | NA |
| 0.496114211 | NA |
| NA | NA |
| NA | NA |
| NA | NA |
| 0.941279303 | NA |
| 0.974544494 | 0.984554671 |
| NA | NA |
| 0.474311796 | NA |
| NA | NA |
| NA | NA |
| 0.955791922 | NA |
| NA | NA |
| NA | NA |
| NA | NA |
| 0.839318641 | NA |
| NA | NA |

| | |
|---|---|
| NA | NA |
| 0.143161754 | 0.273613231 |
| NA | NA |
| NA | NA |
| NA | NA |
| NA | NA |
| NA | NA |
| NA | NA |
| NA | NA |
| NA | NA |
| 0.941278332 | NA |
| NA | NA |
| 0.243181005 | NA |
| NA | NA |
| NA | NA |
| 0.877379615 | NA |
| 0.955792873 | NA |
| 0.839318641 | NA |
| NA | NA |
| 0.941278332 | NA |
| NA | NA |
| NA | NA |
| NA | NA |
| 0.839318641 | NA |
| NA | NA |
| 0.12957094 | NA |
| 0.734058779 | NA |
| NA | NA |
| NA | NA |
| 0.939988807 | NA |
| 0.955792398 | NA |
| 0.594907112 | 0.729917873 |
| NA | NA |
| NA | NA |
| NA | NA |
| NA | NA |
| NA | NA |
| 0.839319108 | NA |
| NA | NA |
| NA | NA |
| 0.021985265 | 0.064572222 |
| 0.489046337 | 0.643583465 |
| NA | NA |
| 0.223165665 | 0.375603815 |
| 0.130999825 | 0.256250644 |
| 0.036386305 | 0.096605336 |
| 0.620825238 | 0.749468748 |

| | |
|---|---|
| NA | NA |
| 2.83E-08 | 4.13E-07 |
| 0.015053803 | 0.047844923 |
| 0.000816724 | 0.004303666 |
| 0.080869446 | 0.179130044 |
| 0.115400424 | 0.233151761 |
| NA | NA |
| 0.030977232 | 0.085089299 |
| 0.978205139 | 0.986494311 |
| 0.593080279 | NA |
| 9.19E-05 | 0.000650538 |
| NA | NA |
| 0.000157319 | 0.00104362 |
| 0.941276431 | NA |
| 0.276411938 | 0.436265361 |
| 0.748393366 | NA |
| 0.672355001 | NA |
| NA | NA |
| 0.953336382 | NA |
| NA | NA |
| 0.000260174 | 0.001606812 |
| 0.839318641 | NA |
| 0.036653087 | 0.097151399 |
| 0.654997175 | NA |
| 0.839318641 | NA |
| 0.941278332 | NA |
| 0.941278332 | NA |
| 0.054157326 | 0.132034484 |
| 0.410974651 | 0.572952535 |
| 0.388562633 | 0.551363288 |
| 0.891419176 | 0.934565988 |
| 2.38E-06 | 2.48E-05 |
| 0.219771046 | 0.371387323 |
| NA | NA |
| 0.17796422 | 0.319425711 |
| 0.743511643 | NA |
| 8.58E-12 | 1.96E-10 |
| 0.068928945 | 0.159286091 |
| 0.401350131 | 0.563741795 |
| 0.410359874 | 0.572330114 |
| 6.99E-09 | 1.12E-07 |
| 4.14E-28 | 2.70E-26 |
| 9.51E-06 | 8.66E-05 |
| 0.13665527 | 0.26438013 |
| 0.602989041 | 0.73616871 |
| 0.004469753 | 0.017791985 |
| 0.184724203 | 0.327826236 |

| | |
|---|---|
| 0.000104836 | 0.000729622 |
| 0.241898255 | 0.398206118 |
| 0.749401441 | 0.842611984 |
| NA | NA |
| 0.079727621 | 0.177386132 |
| 0.001437937 | 0.006921622 |
| NA | NA |
| 0.331946717 | 0.495080879 |
| 0.024770707 | 0.071155245 |
| NA | NA |
| NA | NA |
| 0.441461158 | NA |
| 0.161643756 | 0.298714649 |
| 0.000155185 | 0.001031476 |
| 0.021988429 | 0.064573545 |
| NA | NA |
| 0.465716876 | 0.623968899 |
| 0.081639222 | 0.180408203 |
| NA | NA |
| 6.08E-12 | 1.41E-10 |
| 0.109387545 | 0.224123469 |
| NA | NA |
| NA | NA |
| NA | NA |
| NA | NA |
| NA | NA |
| 0.458716569 | NA |
| NA | NA |
| NA | NA |
| 0.955792873 | NA |
| NA | NA |
| NA | NA |
| 0.941278332 | NA |
| NA | NA |
| 0.853595622 | NA |
| 0.955792397 | NA |
| 0.941278332 | NA |
| 0.395164864 | NA |
| NA | NA |
| NA | NA |
| NA | NA |
| NA | NA |
| 0.955792873 | NA |
| NA | NA |
| NA | NA |
| NA | NA |
| NA | NA |

| | |
|---|---|
| NA | NA |
| NA | NA |
| NA | NA |
| NA | NA |
| 0.782629377 | NA |
| 0.322072584 | 0.484850525 |
| 0.133205702 | 0.259287056 |
| 0.010032368 | 0.03448414 |
| 0.089363631 | 0.193155935 |
| 0.413145225 | 0.575304665 |
| 7.96E-07 | 9.03E-06 |
| 0.00914497 | 0.03199597 |
| 0.016179316 | 0.050664397 |
| 0.492793359 | 0.646572269 |
| 0.191154371 | 0.336229406 |
| NA | NA |
| 0.008472492 | 0.03003608 |
| 0.032124684 | 0.087546568 |
| 9.38E-10 | 1.67E-08 |
| 0.941277857 | NA |
| 0.013576124 | 0.044042375 |
| 0.098678108 | 0.207485542 |
| 0.941278332 | NA |
| 0.941277857 | NA |
| 0.839318641 | NA |
| 0.941275956 | NA |
| 0.140533106 | 0.269973283 |
| 0.290993771 | 0.452102601 |
| NA | NA |
| 1.54E-12 | 3.79E-11 |
| 0.544749467 | 0.691305545 |
| 0.941275956 | NA |
| 0.941278332 | NA |
| 0.941278332 | NA |
| 0.571723109 | NA |
| NA | NA |
| NA | NA |
| NA | NA |
| 0.941277857 | NA |
| NA | NA |
| NA | NA |
| NA | NA |
| NA | NA |
| NA | NA |
| 0.839319108 | NA |
| NA | NA |
| NA | NA |

| | |
|---|---|
| NA | NA |
| NA | NA |
| NA | NA |
| 0.751797079 | NA |
| NA | NA |
| 0.839319574 | NA |
| NA | NA |
| NA | NA |
| 0.918525826 | NA |
| NA | NA |
| NA | NA |
| NA | NA |
| 0.07176136 | 0.164047626 |
| 4.21E-05 | 0.000325288 |
| 0.031626694 | 0.086506386 |
| 0.660927207 | NA |
| 0.787419151 | NA |
| 0.765269649 | 0.854255406 |
| 0.052918801 | 0.129732547 |
| 0.035084591 | 0.093828809 |
| 0.970426842 | 0.982229107 |
| 0.941277857 | NA |
| 0.941276431 | NA |
| 0.572001533 | 0.71203287 |
| 0.731846995 | 0.83092073 |
| 0.003547085 | 0.01471892 |
| 0.38784236 | 0.550735226 |
| 0.939735321 | 0.963757575 |
| 0.555981466 | 0.700257745 |
| 0.460929774 | 0.619889027 |
| 0.871866359 | 0.923391864 |
| NA | NA |
| 2.56E-07 | 3.19E-06 |
| 0.000187573 | 0.001213209 |
| 0.000752293 | 0.004015718 |
| 0.942245039 | NA |
| NA | NA |
| 0.002721166 | 0.011842968 |
| NA | NA |
| NA | NA |
| NA | NA |
| 0.839318641 | NA |
| 0.941277857 | NA |
| 0.441958536 | NA |
| 0.385787934 | NA |
| 0.928113852 | NA |
| NA | NA |

| | | |
|---|---|---|
| 0.744930053 | | 0.839766275 |
| 0.839319108 | NA | |
| 0.782629377 | NA | |
| 1.47E-05 | | 0.00012782 |
| 0.203929813 | NA | |
| NA | NA | |
| 0.888191094 | NA | |
| 0.347484158 | NA | |
| NA | NA | |
| 0.571626004 | NA | |
| 0.171222799 | NA | |
| NA | NA | |
| NA | NA | |
| NA | NA | |
| 0.941280668 | NA | |
| 0.190241655 | NA | |
| 0.941277857 | NA | |
| NA | NA | |
| 0.455763687 | NA | |
| 0.839319108 | NA | |
| 0.839318641 | NA | |
| 0.839319107 | NA | |
| 0.941277857 | NA | |
| NA | NA | |
| NA | NA | |
| NA | NA | |
| 0.941277857 | NA | |
| NA | NA | |
| 0.786944116 | NA | |
| NA | NA | |
| NA | NA | |
| NA | NA | |
| NA | NA | |
| 0.476928417 | NA | |
| NA | NA | |
| 0.839319108 | NA | |
| 0.931036445 | NA | |
| NA | NA | |
| NA | NA | |
| NA | NA | |
| NA | NA | |
| NA | NA | |
| NA | NA | |
| NA | NA | |
| 0.941278838 | NA | |
| NA | NA | |
| NA | NA | |

| | | |
|---|---|---|
| NA | | NA |
| | 0.941278332 | NA |
| NA | | NA |
| | 0.495904397 | NA |
| NA | | NA |
| | 0.890864862 | NA |
| | 0.632001781 | NA |
| | 0.941278332 | NA |
| | 0.661065506 | NA |
| NA | | NA |
| NA | | NA |
| | 0.832162011 | NA |
| NA | | NA |
| NA | | NA |
| | 0.723896324 | NA |
| NA | | NA |
| NA | | NA |
| NA | | NA |
| | 0.941278332 | NA |
| NA | | NA |
| | 0.619594186 | NA |
| NA | | NA |
| NA | | NA |
| NA | | NA |
| NA | | NA |
| NA | | NA |
| NA | | NA |
| NA | | NA |
| NA | | NA |
| NA | | NA |
| NA | | NA |
| NA | | NA |
| NA | | NA |
| NA | | NA |
| NA | | NA |
| NA | | NA |
| | 0.839319107 | NA |
| | 0.739991096 | NA |
| NA | | NA |
| NA | | NA |
| NA | | NA |
| NA | | NA |
| NA | | NA |
| | 0.839318641 | NA |
| | 0.767053257 | NA |
| | 0.941278332 | NA |
| | 0.935650667 | NA |

| | | |
|---|---|---|
| NA | NA | |
| NA | NA | |
| 0.839318641 | NA | |
| 0.82515277 | NA | |
| 0.394086907 | NA | |
| 0.786168672 | NA | |
| 0.709202663 | NA | |
| 0.955792398 | NA | |
| NA | NA | |
| 0.766609892 | NA | |
| 0.838993826 | NA | |
| 0.839319108 | NA | |
| 0.795327666 | NA | |
| NA | NA | |
| NA | NA | |
| 0.941277857 | NA | |
| 0.23822416 | NA | |
| NA | NA | |
| 0.941279303 | NA | |
| NA | NA | |
| 0.725854934 | NA | |
| NA | NA | |
| NA | NA | |
| NA | NA | |
| 0.606757562 | NA | |
| 0.728678906 | NA | |
| NA | NA | |
| NA | NA | |
| NA | NA | |
| 0.229628136 | | 0.383390074 |
| 0.778547407 | NA | |
| 0.554428278 | NA | |
| NA | NA | |
| NA | NA | |
| 0.755626499 | NA | |
| NA | NA | |
| NA | NA | |
| NA | NA | |
| NA | NA | |
| 0.93838321 | NA | |
| NA | NA | |
| NA | NA | |
| 0.641893687 | NA | |
| 0.839318641 | NA | |
| 0.941278838 | NA | |
| 0.085925059 | NA | |
| NA | NA | |

| | | |
|---|---|---|
| NA | NA | |
| NA | NA | |
| 0.941277857 | NA | |
| 0.839321638 | NA | |
| 0.921083112 | NA | |
| NA | NA | |
| NA | NA | |
| 0.941278332 | NA | |
| 0.853595154 | NA | |
| NA | NA | |
| 0.855324909 | NA | |
| 0.164613452 | NA | |
| NA | NA | |
| NA | NA | |
| 0.63918867 | NA | |
| 0.740238263 | NA | |
| 0.809226989 | | 0.883056152 |
| 0.48521234 | NA | |
| 1.68E-06 | | 1.81E-05 |
| 0.741826584 | NA | |
| 0.622486081 | NA | |
| NA | NA | |
| 0.768034207 | NA | |
| 0.839319108 | NA | |
| 0.941278332 | NA | |
| 0.941278332 | NA | |
| 0.941277857 | NA | |
| NA | NA | |
| 0.572299172 | NA | |
| 0.941278332 | NA | |
| 0.938328926 | NA | |
| 0.941278332 | NA | |
| 0.839320041 | NA | |
| 0.900199862 | NA | |
| NA | NA | |
| 0.783102737 | NA | |
| 0.748393366 | NA | |
| 0.504836877 | NA | |
| 0.427248177 | NA | |
| 0.653185856 | NA | |
| 0.839318641 | NA | |
| 0.293459933 | NA | |
| 0.651601095 | NA | |
| 0.891951608 | NA | |
| 0.875819469 | NA | |
| 0.861781972 | | 0.917466458 |
| 0.941278838 | NA | |

| | | |
|---|---|---|
| | 0.653693256 | NA |
| | 0.612414261 | NA |
| | 0.941280862 | NA |
| | 0.443790957 | NA |
| | 0.155075433 | NA |
| | 0.607275292 | NA |
| | 0.541563517 | NA |
| | 0.1718865 | NA |
| | 0.941278332 | NA |
| | 0.427284748 | NA |
| | 0.089835991 | NA |
| | 0.196416147 | NA |
| | 0.214783334 | NA |
| | 0.035983981 | 0.095825767 |
| | 0.028117407 | NA |
| | 0.552804446 | NA |
| | 0.72965869 | NA |
| | 0.488190142 | NA |
| | 0.839319574 | NA |
| | 0.321689004 | NA |
| | 0.752173964 | NA |
| | 0.916208241 | NA |
| | 0.199160905 | NA |
| | 0.029057459 | NA |
| | 0.922893457 | NA |
| | 0.772306232 | NA |
| | 0.787036781 | NA |
| | 0.300975784 | NA |
| | 0.418726949 | NA |
| | 0.718362369 | NA |
| NA | | NA |
| | 0.486803806 | NA |
| | 0.934588843 | NA |
| | 7.29E-21 | 3.30E-19 |
| | 0.347495287 | 0.511413023 |
| | 0.009737559 | 0.033701428 |
| | 0.955792398 | NA |
| | 0.826045997 | NA |
| NA | | NA |
| NA | | NA |
| | 0.955791446 | NA |
| | 5.58E-08 | 7.76E-07 |
| | 5.10E-14 | 1.43E-12 |
| | 0.004150362 | 0.016735036 |
| | 3.80E-17 | 1.34E-15 |
| NA | | NA |
| | 2.17E-08 | 3.23E-07 |

| | |
|---|---|
| 0.820893994 | 0.890764479 |
| 0.525768382 | 0.675044348 |
| 0.006088269 | 0.02291843 |
| 0.000948019 | 0.004874589 |
| NA | NA |
| 0.039847021 | 0.103881577 |
| 0.000943629 | 0.004856274 |
| NA | NA |
| 0.013584772 | 0.044058421 |
| 8.33E-06 | 7.70E-05 |
| 0.941276431 | NA |
| 0.758155519 | 0.8489686 |
| 0.080819387 | 0.179128695 |
| 0.308457873 | NA |
| 0.486962766 | 0.642036657 |
| 0.066455798 | 0.155000578 |
| 0.066801246 | 0.155547179 |
| 0.089405308 | 0.193217795 |
| NA | NA |
| 0.009311301 | 0.032469192 |
| 0.25791741 | 0.416143586 |
| 0.041247953 | 0.106715627 |
| 0.208197406 | 0.357514946 |
| 0.530726226 | NA |
| 0.039752387 | 0.103668929 |
| 0.191303581 | 0.336392422 |
| 0.285086584 | 0.445602238 |
| 0.68588254 | 0.798345944 |
| 3.48E-05 | 0.000274783 |
| 0.000549232 | 0.003078856 |
| NA | NA |
| 0.243881769 | 0.400299586 |
| 2.39E-07 | 3.00E-06 |
| 0.794361236 | NA |
| NA | NA |
| 0.941277857 | NA |
| 0.01162236 | 0.038953271 |
| 0.550814622 | 0.696290294 |
| 0.941275481 | NA |
| 0.583186199 | 0.720526464 |
| 2.21E-05 | 0.000183743 |
| 0.038069658 | 0.10019173 |
| 0.001485745 | 0.007118161 |
| 0.000103766 | 0.000722809 |
| 0.001227854 | 0.006078492 |
| 0.133433097 | 0.259634466 |
| NA | NA |

| | | |
|---|---|---|
| | 0.004410418 | 0.017588902 |
| | 0.733834733 | 0.832164192 |
| | 0.704062397 | 0.811097874 |
| | 3.88E-05 | 0.000302635 |
| | 0.042329588 | 0.108874958 |
| | 0.078594276 | 0.175465318 |
| | 0.098656592 | 0.207476958 |
| | 0.294082358 | 0.455297794 |
| | 0.023410911 | 0.068042984 |
| | 0.713983597 | NA |
| | 0.935866248 | NA |
| | 3.76E-79 | 9.72E-77 |
| | 1.73E-05 | 0.000147678 |
| | 2.01E-50 | 2.78E-48 |
| | 3.76E-33 | 3.01E-31 |
| | 4.95E-140 | 2.51E-137 |
| | 0.063781436 | 0.150086578 |
| | 0.033975891 | 0.091491588 |
| | 0.576056424 | 0.714800073 |
| | 0.560134809 | 0.703441605 |
| | 0.411062064 | 0.572994741 |
| | 0.730893681 | 0.830396998 |
| | 0.148375514 | 0.281225831 |
| | 0.003277732 | 0.013815579 |
| | 0.743511643 | NA |
| | 8.44E-37 | 7.85E-35 |
| | 0.941413981 | 0.964703895 |
| | 1.06E-09 | 1.87E-08 |
| | 0.012161239 | 0.040320631 |
| | 0.001164662 | 0.005811589 |
| | 0.693905889 | 0.803747042 |
| | 1.31E-05 | 0.000114959 |
| | 2.11E-07 | 2.68E-06 |
| | 8.77E-15 | 2.59E-13 |
| | 0.687164306 | 0.799229101 |
| | 0.166359185 | 0.305201274 |
| | 0.107926746 | 0.221951997 |
| | 0.971721497 | 0.982929833 |
| | 0.001087712 | 0.005487438 |
| | 0.036761864 | 0.097406867 |
| | 0.530103151 | 0.678924703 |
| | 0.523403107 | NA |
| | 0.593452396 | 0.728755032 |
| | 0.631730384 | NA |
| NA | | NA |
| | 0.638274809 | NA |
| | 0.647924866 | NA |

| | |
|---|---|
| NA | NA |
| NA | NA |
| 0.777526424 | NA |
| 0.839318641 | NA |
| 0.941278332 | NA |
| NA | NA |
| NA | NA |
| 0.044833769 | 0.113657831 |
| 0.008077063 | 0.028883751 |
| NA | NA |
| NA | NA |
| 2.66E-34 | 2.23E-32 |
| 0.700051223 | 0.80855101 |
| 0.900129495 | 0.939852704 |
| 0.941278332 | NA |
| 7.51E-06 | 7.01E-05 |
| NA | NA |
| 7.45E-37 | 6.95E-35 |
| NA | NA |
| 0.141068416 | 0.270783995 |
| 0.896089585 | 0.937237962 |
| 0.011422319 | 0.038373188 |
| 0.157304072 | 0.293088871 |
| 0.955791922 | NA |
| 0.594440307 | 0.729511121 |
| 0.032929528 | 0.089361375 |
| NA | NA |
| NA | NA |
| NA | NA |
| 0.618277972 | 0.747701061 |
| 0.835112743 | NA |
| 0.195122992 | 0.341418618 |
| 2.04E-05 | 0.000170888 |
| 0.038327763 | 0.100739224 |
| 0.923989501 | 0.954592539 |
| 0.245797653 | 0.40264802 |
| 0.2412734 | 0.397479891 |
| 0.922801652 | 0.954025319 |
| 0.955792873 | NA |
| 0.22473099 | 0.377411065 |
| 0.005078896 | 0.019778776 |
| 0.003065803 | 0.013056404 |
| 0.123359997 | 0.245256819 |
| 0.039498343 | 0.10313071 |
| 0.010541221 | 0.035930422 |
| 0.941276431 | NA |
| 0.449178006 | NA |

| | |
|---|---|
| 0.079010158 | 0.17625115 |
| NA | NA |
| 0.042167707 | 0.10855243 |
| 0.439564579 | 0.60043134 |
| 0.61097617 | 0.742227061 |
| 0.081119165 | 0.179559042 |
| 1.58E-05 | 0.000136574 |
| 0.018337047 | 0.055963379 |
| 4.75E-09 | 7.77E-08 |
| NA | NA |
| NA | NA |
| 0.647122701 | NA |
| 0.073419517 | 0.166920634 |
| 0.941277857 | NA |
| NA | NA |
| 0.404758449 | 0.566776126 |
| 1.43E-08 | 2.19E-07 |
| 4.93E-22 | 2.38E-20 |
| 4.66E-48 | 6.13E-46 |
| 2.74E-56 | 4.32E-54 |
| 3.71E-20 | 1.60E-18 |
| 1.46E-25 | 8.33E-24 |
| 0.00213729 | 0.009662254 |
| 0.62852403 | 0.755481581 |
| 0.934851471 | 0.961237201 |
| 0.355185156 | 0.51891217 |
| 7.87E-06 | 7.31E-05 |
| 0.141159415 | 0.270914078 |
| 0.900732041 | NA |
| NA | NA |
| NA | NA |
| 0.667247529 | 0.784326421 |
| 0.805067502 | NA |
| 0.003933472 | 0.015990522 |
| 0.000129195 | 0.000877345 |
| 0.765925232 | 0.854626639 |
| 2.60E-25 | 1.47E-23 |
| 0.161704982 | 0.298781399 |
| 0.91391405 | NA |
| 0.718563264 | 0.821008199 |
| 0.021343566 | 0.063037542 |
| 0.000328621 | 0.001973716 |
| 0.564744601 | 0.706620985 |
| 0.250109568 | 0.407663684 |
| 7.16E-17 | 2.49E-15 |
| 0.145496957 | 0.277049486 |
| 0.931008106 | NA |

| | | |
|---|---|---|
| | 2.57E-06 | 2.65E-05 |
| | 0.001963824 | 0.009030708 |
| | 0.569135669 | 0.710248372 |
| | 0.120620783 | 0.241363251 |
| | 0.777409674 | 0.862934527 |
| | 0.098629293 | 0.207437876 |
| | 0.941275956 | NA |
| | 0.415880978 | NA |
| | 0.857660682 | NA |
| | 0.113091943 | 0.229814261 |
| | 0.739991096 | NA |
| NA | | NA |
| | 0.911283366 | 0.946309789 |
| | 0.955792873 | NA |
| | 0.026443431 | 0.075043674 |
| | 0.153204527 | 0.287414027 |
| | 0.157204264 | 0.292987364 |
| NA | | NA |
| | 0.97856308 | 0.986646267 |
| | 0.025227594 | 0.072180213 |
| | 0.005433636 | 0.020846455 |
| | 0.832068545 | 0.89814047 |
| NA | | NA |
| | 0.000391516 | 0.002294159 |
| NA | | NA |
| | 0.94952172 | NA |
| NA | | NA |
| | 0.002416012 | 0.010728716 |
| | 0.125447993 | 0.248391408 |
| | 0.765975552 | 0.854626639 |
| | 0.838498056 | 0.902115782 |
| NA | | NA |
| | 0.95308087 | 0.971571839 |
| | 0.97591174 | 0.985350438 |
| NA | | NA |
| | 0.981343513 | NA |
| NA | | NA |
| | 0.839319107 | NA |
| | 0.671527646 | 0.787568417 |
| NA | | NA |
| | 2.01E-11 | 4.42E-10 |
| | 0.082289137 | 0.181507371 |
| | 0.717207087 | 0.82021693 |
| | 0.954694081 | NA |
| | 0.13137268 | NA |
| | 0.839320041 | NA |
| | 5.98E-07 | 6.94E-06 |

| | | |
|---|---|---|
| | 0.36459969 | NA |
| | 0.568452065 | NA |
| | 0.896016071 | 0.937202256 |
| | 0.051382772 | 0.126710301 |
| | 1.00E-05 | 9.09E-05 |
| | 0.135149029 | 0.26220415 |
| | 0.508542311 | 0.660211852 |
| | 0.003350482 | 0.014064935 |
| | 0.72601533 | NA |
| | 0.258374746 | NA |
| | 0.009550669 | 0.033157092 |
| NA | | NA |
| | 0.957294836 | 0.97374067 |
| | 0.783663444 | NA |
| | 0.120890126 | 0.241699015 |
| | 0.00761543 | 0.027534935 |
| | 0.023718196 | 0.068762547 |
| NA | | NA |
| NA | | NA |
| | 0.004054818 | 0.01640258 |
| | 0.713461678 | 0.817770146 |
| | 0.001302802 | 0.006373911 |
| | 0.240769071 | 0.396841315 |
| NA | | NA |
| | 0.288810243 | 0.449650798 |
| | 0.598892477 | 0.733026998 |
| | 0.124125887 | 0.246450146 |
| | 0.881176883 | 0.928947302 |
| | 0.010998377 | 0.037206278 |
| | 0.822144812 | 0.891751494 |
| | 0.639829343 | 0.763522412 |
| | 0.625801876 | 0.753436846 |
| | 0.566607596 | 0.708356216 |
| | 4.88E-06 | 4.77E-05 |
| | 0.436438786 | 0.597464541 |
| | 0.024906848 | 0.071481913 |
| | 2.13E-09 | 3.64E-08 |
| | 3.78E-09 | 6.24E-08 |
| | 7.20E-07 | 8.24E-06 |
| | 0.25842378 | 0.416739267 |
| | 0.469401253 | NA |
| | 0.897624959 | NA |
| | 0.133057752 | 0.259083819 |
| | 0.491948199 | 0.645890619 |
| | 0.009891925 | 0.034078389 |
| | 0.13343872 | 0.259634466 |
| | 4.33E-05 | 0.000333766 |

| | |
|---|---|
| 0.811392644 | 0.884608268 |
| NA | NA |
| 8.28E-06 | 7.65E-05 |
| 0.939019179 | NA |
| 0.892460899 | 0.935045088 |
| 0.379801574 | 0.542521579 |
| 2.39E-05 | 0.000197719 |
| NA | NA |
| NA | NA |
| 0.066690791 | 0.155396397 |
| 0.000938655 | 0.004833812 |
| NA | NA |
| 0.937924455 | NA |
| 0.820220087 | 0.890479771 |
| NA | NA |
| 0.936737936 | NA |
| NA | NA |
| NA | NA |
| NA | NA |
| NA | NA |
| 0.325357561 | 0.488190169 |
| 0.007795524 | 0.028070961 |
| 0.100983815 | 0.211306969 |
| 0.201654465 | 0.349634292 |
| 0.017960009 | 0.055033193 |
| 0.429978793 | NA |
| 0.502643231 | 0.654695972 |
| 0.070399741 | 0.161877563 |
| NA | NA |
| 0.699845128 | NA |
| 0.486161474 | NA |
| 0.481796441 | 0.638194607 |
| 0.863481633 | 0.91838732 |
| 0.116489697 | 0.234874018 |
| NA | NA |
| NA | NA |
| 0.002396355 | 0.010647383 |
| 0.617155265 | NA |
| 0.012048301 | 0.040011729 |
| NA | NA |
| 0.941277857 | NA |
| NA | NA |
| NA | NA |
| NA | NA |
| NA | NA |
| 0.852729976 | NA |
| NA | NA |

| | |
|---|---|
| 0.955792398 | NA |
| NA | NA |
| NA | NA |
| NA | NA |
| 0.839319107 | NA |
| 0.941277857 | NA |
| NA | NA |
| NA | NA |
| 0.174099803 | 0.314935667 |
| NA | NA |
| 0.839319108 | NA |
| 0.000224843 | 0.001418801 |
| NA | NA |
| NA | NA |
| 0.45783042 | 0.617514572 |
| 0.729287796 | 0.829416758 |
| 0.007901217 | 0.028365728 |
| 3.78E-05 | 0.000294872 |
| 0.00361459 | 0.01493654 |
| 0.98934571 | NA |
| 0.023978108 | 0.069389292 |
| 5.36E-07 | 6.28E-06 |
| 0.340761834 | 0.504321235 |
| 0.593060764 | 0.728456617 |
| 0.053190844 | 0.130253502 |
| 0.007537551 | 0.027307274 |
| 0.773889083 | 0.860508485 |
| 0.500162636 | 0.652929911 |
| 0.667090471 | 0.784230075 |
| 9.80E-08 | 1.32E-06 |
| 0.848828812 | 0.909153602 |
| 0.0023339 | 0.010420464 |
| 0.839319574 | NA |
| 0.941276431 | NA |
| 0.802065762 | 0.87905189 |
| 0.317913062 | 0.480350836 |
| 0.60374608 | 0.736872583 |
| 0.003731101 | 0.015322235 |
| 0.494144958 | 0.647310858 |
| 0.042859281 | 0.109904827 |
| 0.016563522 | 0.051643301 |
| 0.052925405 | 0.129735374 |
| 0.070782147 | 0.162574222 |
| 0.019512148 | 0.058743758 |
| 0.004787028 | 0.018851894 |
| 0.488798609 | 0.643461309 |
| 0.738295709 | 0.835293967 |

| | | |
|---|---|---|
| | 0.001320142 | 0.006449509 |
| | 6.58E-10 | 1.20E-08 |
| | 0.941278332 | NA |
| | 1.44E-07 | 1.89E-06 |
| NA | | NA |
| NA | | NA |
| NA | | NA |
| | 1.08E-19 | 4.47E-18 |
| | 0.422110559 | NA |
| | 1.33E-09 | 2.33E-08 |
| | 0.513407061 | 0.664499394 |
| | 0.22424288 | 0.376990326 |
| | 0.039711998 | 0.103586439 |
| | 0.132824927 | 0.258778662 |
| | 0.022043601 | 0.064715 |
| | 4.85E-14 | 1.36E-12 |
| | 0.520379579 | 0.670786609 |
| | 0.428154846 | 0.589412473 |
| | 0.85604224 | 0.913536469 |
| | 0.357530888 | NA |
| | 0.592436978 | 0.727990962 |
| | 0.514260368 | 0.665240505 |
| NA | | NA |
| | 0.895514026 | 0.936820599 |
| | 0.158975762 | 0.295274373 |
| | 0.007260888 | 0.026478269 |
| | 0.004382293 | 0.017507107 |
| | 5.51E-19 | 2.19E-17 |
| | 0.813199652 | NA |
| | 7.54E-60 | 1.29E-57 |
| NA | | NA |
| | 0.653693256 | NA |
| NA | | NA |
| NA | | NA |
| | 0.816887035 | 0.888212342 |
| | 0.131120499 | 0.256444546 |
| | 0.941275956 | NA |
| | 0.941276431 | NA |
| NA | | NA |
| | 0.637944937 | 0.76203781 |
| | 0.585453205 | 0.722180687 |
| | 0.086393154 | 0.188377598 |
| | 0.035852883 | 0.095564998 |
| | 6.08E-06 | 5.81E-05 |
| | 0.175992054 | 0.317417813 |
| | 3.40E-05 | 0.000269356 |
| | 0.013884634 | 0.044847614 |

| | |
|---|---|
| 0.006821324 | 0.025137196 |
| 0.270084502 | 0.430147295 |
| 0.009514219 | 0.033041201 |
| 0.116032237 | 0.234150005 |
| 0.166692521 | 0.305553745 |
| 0.530017917 | 0.678924703 |
| 0.374759411 | 0.537869241 |
| 0.577283166 | 0.71579985 |
| 2.14E-18 | 8.13E-17 |
| 0.000499132 | 0.002833374 |
| 0.007595296 | 0.027478831 |
| 0.044614097 | 0.113247766 |
| NA | NA |
| 0.096555888 | 0.204268353 |
| NA | NA |
| 0.743511643 | NA |
| 0.115680953 | 0.233579747 |
| 1.59E-52 | 2.31E-50 |
| NA | NA |
| 0.142586518 | 0.272948714 |
| NA | NA |
| NA | NA |
| 0.475205236 | 0.632515602 |
| NA | NA |
| 0.981385528 | 0.988464598 |
| 0.706127211 | 0.812493468 |
| 0.839318641 | NA |
| 1.16E-06 | 1.28E-05 |
| NA | NA |
| 0.92960928 | 0.958108604 |
| 0.000173094 | 0.00113471 |
| 0.000326383 | 0.001960767 |
| 0.000276958 | 0.001699008 |
| 0.765163993 | 0.854255406 |
| NA | NA |
| 0.027213752 | 0.076910676 |
| 2.53E-21 | 1.18E-19 |
| NA | NA |
| 0.128291288 | 0.252467072 |
| 0.323756943 | 0.486432442 |
| NA | NA |
| 0.34033594 | 0.503961431 |
| 0.874483448 | 0.925004668 |
| 0.485890159 | 0.641261855 |
| 0.839318641 | NA |
| 0.26295349 | 0.421923889 |
| NA | NA |

| | |
|---|---|
| NA | NA |
| NA | NA |
| 0.610094329 | 0.741788737 |
| NA | NA |
| 0.839319108 | NA |
| 0.08655514 | 0.188644395 |
| NA | NA |
| NA | NA |
| NA | NA |
| 0.62095206 | 0.749468748 |
| 0.007126736 | 0.026060926 |
| 0.955792873 | NA |
| 0.955792873 | NA |
| 0.002465562 | 0.010908051 |
| NA | NA |
| NA | NA |
| 0.955792873 | NA |
| 0.394302349 | 0.556811936 |
| NA | NA |
| NA | NA |
| NA | NA |
| NA | NA |
| 6.48E-06 | 6.15E-05 |
| 0.664281528 | 0.782114609 |
| NA | NA |
| NA | NA |
| 0.762133242 | NA |
| 0.018008498 | 0.055151605 |
| 0.955792398 | NA |
| NA | NA |
| NA | NA |
| NA | NA |
| NA | NA |
| 0.91534172 | NA |
| 0.000146652 | 0.000979405 |
| 0.901015421 | 0.940241529 |
| NA | NA |
| NA | NA |
| NA | NA |
| NA | NA |
| 0.081720342 | 0.180537181 |
| 0.447995611 | 0.60817838 |
| 0.579393332 | 0.71730661 |
| 0.048053792 | 0.119993324 |
| 0.045994701 | 0.116044719 |
| NA | NA |
| 0.665670207 | 0.783218414 |

| | | | |
|---|---|---|---|
| NA | | NA | |
| NA | | NA | |
| | 0.672361601 | NA | |
| NA | | NA | |
| | 0.921495793 | NA | |
| | 0.604335971 | | 0.737245902 |
| | 0.478629287 | NA | |
| NA | | NA | |
| NA | | NA | |
| NA | | NA | |
| NA | | NA | |
| NA | | NA | |
| | 0.622185675 | NA | |
| NA | | NA | |
| | 5.11E-06 | | 4.97E-05 |
| | 0.189984596 | | 0.334641705 |
| | 0.169212934 | | 0.308743683 |
| | 0.6223372 | NA | |
| | 0.357328173 | | 0.520892036 |
| NA | | NA | |
| NA | | NA | |
| | 0.009588929 | | 0.033270509 |
| NA | | NA | |
| | 0.920510596 | | 0.952509904 |
| NA | | NA | |
| | 4.03E-21 | | 1.86E-19 |
| | 0.124903273 | | 0.247518749 |
| | 0.556558043 | | 0.700747225 |
| | 0.895398722 | | 0.936820599 |
| | 0.414742054 | | 0.576672972 |
| | 0.175242014 | | 0.3163767 |
| | 2.32E-08 | | 3.44E-07 |
| | 0.938767819 | NA | |
| | 0.941275956 | NA | |
| | 1.33E-08 | | 2.04E-07 |
| | 0.941278332 | NA | |
| | 0.37555794 | | 0.538690496 |
| | 0.41314334 | | 0.575304665 |
| NA | | NA | |
| NA | | NA | |
| NA | | NA | |
| | 0.030027479 | | 0.083077304 |
| | 0.000819459 | | 0.004314257 |
| NA | | NA | |
| | 0.000284459 | | 0.001740377 |
| NA | | NA | |
| | 0.941279303 | NA | |

| | |
|---|---|
| 0.653836094 | NA |
| 0.059245895 | 0.141654525 |
| 0.036207987 | 0.096293016 |
| NA | NA |
| 0.065909459 | 0.153952595 |
| NA | NA |
| NA | NA |
| NA | NA |
| NA | NA |
| NA | NA |
| NA | NA |
| NA | NA |
| NA | NA |
| NA | NA |
| 0.083042941 | 0.182729342 |
| 0.000256339 | 0.001585184 |
| NA | NA |
| NA | NA |
| NA | NA |
| NA | NA |
| NA | NA |
| NA | NA |
| 0.121673896 | 0.242898774 |
| NA | NA |
| 0.435758714 | 0.596911195 |
| NA | NA |
| NA | NA |
| 7.30E-15 | 2.18E-13 |
| NA | NA |
| NA | NA |
| NA | NA |
| 0.035924863 | 0.095721884 |
| NA | NA |
| NA | NA |
| NA | NA |
| NA | NA |
| NA | NA |
| NA | NA |
| 0.033344696 | 0.090302704 |
| NA | NA |
| NA | NA |
| 0.955792873 | NA |
| NA | NA |
| 0.319789983 | 0.482421167 |
| NA | NA |
| 0.981165621 | 0.988399698 |
| 0.839319108 | NA |

| | | |
|---|---|---|
| | 0.463271952 | 0.621999042 |
| | 0.039397393 | 0.102912285 |
| | 5.49E-07 | 6.42E-06 |
| | 0.664175906 | 0.782114139 |
| | 0.839319108 | NA |
| NA | | NA |
| NA | | NA |
| | 0.941277857 | NA |
| NA | | NA |
| | 0.595106824 | 0.729990343 |
| NA | | NA |
| | 0.388941054 | NA |
| NA | | NA |
| | 0.955792398 | NA |
| | 0.00014552 | 0.000973212 |
| NA | | NA |
| | 0.626598137 | 0.754008536 |
| | 0.030345906 | 0.083715029 |
| | 0.000290988 | 0.001775465 |
| | 0.423945906 | 0.58537884 |
| | 0.096050253 | 0.203494722 |
| | 0.370195825 | 0.533460847 |
| | 0.716488136 | 0.819778427 |
| | 0.026448467 | 0.075043674 |
| | 0.277128063 | 0.437008226 |
| | 0.192265592 | 0.337684889 |
| | 0.834314869 | NA |
| NA | | NA |
| | 0.014376945 | 0.046131039 |
| | 0.25562178 | 0.41364536 |
| | 0.017469689 | 0.053833965 |
| | 0.02246243 | 0.065762701 |
| | 0.595891908 | 0.73080272 |
| | 0.091407207 | 0.19639595 |
| | 0.667054526 | 0.784230075 |
| | 0.296578246 | 0.458057588 |
| | 1.70E-10 | 3.32E-09 |
| | 0.102526832 | 0.213690337 |
| | 0.006764657 | 0.024978649 |
| | 0.485192085 | 0.640657676 |
| | 0.473368753 | 0.630893278 |
| | 0.231913875 | 0.386149986 |
| | 0.540419784 | 0.687533898 |
| | 0.07723951 | 0.173161422 |
| | 0.853595154 | NA |
| | 0.033913235 | NA |
| | 0.017544117 | 0.054035335 |

| | | |
|---|---|---|
| | 0.055304263 | 0.134266837 |
| | 0.276540814 | 0.436298252 |
| | 0.707531809 | 0.813437541 |
| | 0.72930272 | 0.829416758 |
| | 0.839319108 | NA |
| | 0.607292906 | 0.739905085 |
| | 0.059305255 | 0.141753734 |
| | 0.784061025 | 0.867243855 |
| | 0.175749468 | 0.317076391 |
| | 0.122279082 | 0.243779772 |
| | 0.002156588 | 0.009738398 |
| | 0.804990865 | 0.880816097 |
| NA | | NA |
| | 0.955792873 | NA |
| | 0.252402617 | 0.41024905 |
| | 0.435139847 | 0.59635245 |
| | 0.877764985 | 0.926663806 |
| | 0.839319108 | NA |
| | 0.822350961 | NA |
| | 0.046037345 | 0.116127691 |
| | 0.676280056 | NA |
| | 0.000365954 | 0.002159791 |
| NA | | NA |
| | 0.941278332 | NA |
| NA | | NA |
| | 0.705388782 | 0.812065262 |
| | 0.246687651 | 0.403689203 |
| | 0.14952582 | 0.282551092 |
| | 0.027654511 | 0.077856499 |
| | 0.001966795 | 0.009039131 |
| NA | | NA |
| | 0.006423617 | 0.023945957 |
| | 1.58E-05 | 0.000136268 |
| | 0.911301254 | NA |
| | 0.006903616 | 0.025399429 |
| NA | | NA |
| | 0.955792398 | NA |
| | 1.66E-08 | 2.52E-07 |
| | 0.003893866 | 0.015864737 |
| | 0.777761052 | 0.863123301 |
| | 0.124443258 | 0.246874347 |
| | 0.007635648 | 0.027591268 |
| | 0.467751025 | 0.625999994 |
| | 0.014398209 | 0.046167569 |
| | 2.55E-12 | 6.14E-11 |
| | 0.832456097 | 0.898395829 |
| | 0.700016935 | NA |

| | |
|---|---|
| NA | NA |
| 3.84E-09 | 6.33E-08 |
| NA | NA |
| 0.819878119 | NA |
| NA | NA |
| 1.36E-40 | 1.42E-38 |
| 0.617726531 | NA |
| 0.174664661 | 0.315622764 |
| 0.001796111 | 0.008357365 |
| 4.26E-41 | 4.57E-39 |
| 0.721298992 | NA |
| 0.004997237 | 0.019511851 |
| 0.029038768 | 0.08082095 |
| NA | NA |
| 0.955792873 | NA |
| 0.941277857 | NA |
| 0.001711991 | 0.008041997 |
| 0.434516159 | 0.595826341 |
| 0.022022798 | 0.064666498 |
| 0.262785315 | 0.421739335 |
| 0.383246951 | 0.545935958 |
| 0.534438422 | 0.682660906 |
| 0.385712286 | 0.5484631 |
| NA | NA |
| 0.941276431 | NA |
| 0.000622041 | 0.003411066 |
| 0.000143397 | 0.000960363 |
| 0.46702223 | NA |
| NA | NA |
| 0.000917329 | 0.004740821 |
| 0.006530221 | 0.024252197 |
| 0.00988574 | 0.034069277 |
| 7.48E-06 | 6.98E-05 |
| 0.734058779 | NA |
| 0.647122701 | NA |
| 0.001056373 | 0.005346331 |
| 2.28E-10 | 4.40E-09 |
| 0.052486299 | 0.128804917 |
| NA | NA |
| 0.006505066 | 0.024181413 |
| NA | NA |
| 0.838882104 | NA |
| 0.007172477 | 0.026200013 |
| 0.045328973 | 0.114672998 |
| 7.21E-05 | 0.000523721 |
| 0.879990063 | NA |
| 0.941277857 | NA |

| | | |
|---|---|---|
| | 0.641177885 | NA |
| | 8.88E-12 | 2.02E-10 |
| | 0.194291352 | 0.340388838 |
| | 0.04092647 | 0.106103022 |
| | 0.011741225 | 0.039223108 |
| | 0.204096826 | 0.352588358 |
| | 0.234665904 | 0.38953329 |
| | 0.830647889 | 0.897271536 |
| NA | | NA |
| | 0.000198121 | 0.001270046 |
| | 0.250193759 | 0.407744735 |
| | 0.937924455 | NA |
| NA | | NA |
| | 0.000714257 | 0.003843713 |
| | 0.839319108 | NA |
| | 0.080912281 | 0.179184418 |
| NA | | NA |
| | 0.901209652 | 0.940279321 |
| | 0.676990415 | 0.791715346 |
| | 0.187263729 | 0.331277929 |
| | 0.061223431 | 0.145360609 |
| | 0.081560833 | 0.180285192 |
| | 0.263595637 | 0.422583913 |
| | 0.287982974 | 0.448626922 |
| | 0.167331867 | 0.306418919 |
| NA | | NA |
| | 0.01113668 | 0.037583334 |
| | 0.000213023 | 0.001355717 |
| NA | | NA |
| | 0.267281263 | NA |
| | 0.901349383 | 0.940383888 |
| NA | | NA |
| NA | | NA |
| NA | | NA |
| | 0.05625381 | 0.13605865 |
| NA | | NA |
| NA | | NA |
| | 0.839319108 | NA |
| NA | | NA |
| | 0.171816241 | 0.311942805 |
| | 0.156138385 | 0.291597163 |
| | 0.003618779 | 0.014943441 |
| NA | | NA |
| NA | | NA |
| | 0.947800832 | 0.968179709 |
| | 0.343734744 | 0.507321533 |
| | 0.061113114 | 0.145156605 |

| | |
|---|---|
| 0.583565264 | 0.720810608 |
| 0.043919423 | 0.111900033 |
| 0.039160276 | 0.102405282 |
| NA | NA |
| NA | NA |
| 0.00731861 | 0.026656092 |
| 0.118410152 | NA |
| 0.00873159 | 0.030803217 |
| 0.449168674 | 0.609319092 |
| NA | NA |
| NA | NA |
| 0.025607101 | 0.073043755 |
| 0.136027273 | NA |
| 0.576018057 | NA |
| 0.183521175 | 0.326250728 |
| 0.388872667 | 0.551561603 |
| 0.001390385 | 0.006731883 |
| 0.194574277 | 0.34073403 |
| 0.884668389 | 0.931019897 |
| 0.16841439 | 0.307871036 |
| 0.002592407 | 0.011371999 |
| 0.000117384 | 0.000806203 |
| 0.853234755 | 0.911742904 |
| 0.158130589 | 0.294186672 |
| NA | NA |
| 3.43E-05 | 0.000270844 |
| NA | NA |
| 4.47E-05 | 0.000342433 |
| 0.78876011 | 0.869975354 |
| 0.295386338 | 0.456752135 |
| 1.32E-11 | 2.95E-10 |
| NA | NA |
| NA | NA |
| NA | NA |
| NA | NA |
| NA | NA |
| 0.545095868 | 0.691560783 |
| 0.135662961 | 0.262951937 |
| 0.967689332 | 0.980542421 |
| 0.455304884 | NA |
| 0.003672035 | 0.015116766 |
| 0.955792873 | NA |
| 0.001690583 | 0.00794771 |
| 0.642188715 | 0.765284664 |
| 0.421404073 | 0.582917402 |
| 3.51E-05 | 0.000276521 |
| 0.053351148 | 0.130516861 |

| | |
|---|---|
| 0.151443823 | 0.28533728 |
| 0.001518853 | 0.00724774 |
| NA | NA |
| 0.09339116 | 0.199380787 |
| 0.793302807 | NA |
| 0.008051888 | 0.02881104 |
| 0.53990563 | 0.687090149 |
| 0.000631972 | 0.003458772 |
| 0.886432157 | NA |
| 0.941281114 | NA |
| 0.085808894 | 0.187498696 |
| 0.008853478 | 0.031140903 |
| NA | NA |
| 0.456491281 | 0.616197214 |
| NA | NA |
| 1.70E-05 | 0.000145061 |
| 0.649885232 | NA |
| 6.18E-06 | 5.90E-05 |
| NA | NA |
| 0.30645949 | 0.468073802 |
| 0.037164896 | 0.098201472 |
| 0.035480857 | 0.094750966 |
| 0.667924231 | 0.784735757 |
| 0.111319807 | 0.227046619 |
| 0.027842306 | 0.078250391 |
| 0.001213297 | 0.006016438 |
| 0.941275956 | NA |
| 0.252111 | 0.409836261 |
| 0.087958299 | 0.190968115 |
| NA | NA |
| NA | NA |
| 0.435023582 | 0.59635245 |
| 0.240015968 | 0.396038827 |
| NA | NA |
| 0.814533335 | 0.886610986 |
| 0.419881562 | 0.581554899 |
| NA | NA |
| 0.063132563 | 0.148809792 |
| 1.11E-05 | 9.95E-05 |
| 0.235929929 | NA |
| 0.150670128 | 0.284262078 |
| NA | NA |
| 0.629165156 | 0.755874674 |
| 0.016259823 | 0.050856282 |
| 0.728512078 | 0.829110414 |
| NA | NA |
| 1.32E-17 | 4.82E-16 |

| | |
|---|---|
| 0.445021833 | 0.605661253 |
| NA | NA |
| 0.093308993 | 0.199276865 |
| 0.000448695 | 0.002583415 |
| 0.69531065 | NA |
| 0.922876666 | NA |
| 0.35670989 | 0.520341288 |
| NA | NA |
| 0.381219811 | NA |
| 0.294605623 | 0.455929977 |
| 0.898402164 | NA |
| 0.628889531 | 0.755696114 |
| 0.135953166 | 0.263374536 |
| NA | NA |
| 0.955792873 | NA |
| NA | NA |
| 0.179941751 | NA |
| 0.576319571 | NA |
| 0.126099215 | 0.249307544 |
| 0.614460566 | 0.744446218 |
| 0.023562854 | 0.068370482 |
| 0.095574443 | 0.202810289 |
| 0.513846732 | 0.664922252 |
| NA | NA |
| 0.94054529 | NA |
| NA | NA |
| 0.344833561 | NA |
| 0.917617786 | 0.95044119 |
| 0.679724392 | NA |
| 0.040129678 | 0.104458289 |
| 0.477944328 | NA |
| 0.941278788 | NA |
| 0.65133367 | 0.772339667 |
| 0.653553867 | 0.774003044 |
| 8.09E-06 | 7.49E-05 |
| 0.141297395 | 0.271047759 |
| 0.755626499 | NA |
| 0.000425207 | 0.002463678 |
| 0.074438724 | 0.168517264 |
| 0.000202908 | 0.001297586 |
| 0.941278207 | NA |
| 0.825764651 | 0.894113212 |
| 0.005696174 | 0.021699796 |
| NA | NA |
| 0.205596348 | 0.354275268 |
| 0.69055629 | 0.801512694 |
| 3.68E-07 | 4.45E-06 |

| | |
|---|---|
| 0.095104022 | 0.202078602 |
| NA | NA |
| NA | NA |
| 0.102624927 | 0.213857337 |
| 0.275593684 | 0.43545412 |
| NA | NA |
| 0.000394879 | 0.002312153 |
| 0.73505188 | NA |
| 0.121731226 | 0.242992841 |
| 0.940685982 | NA |
| 3.00E-19 | 1.21E-17 |
| 1.36E-24 | 7.54E-23 |
| 0.809817973 | 0.883463483 |
| 0.1446247 | 0.275852001 |
| 0.905419891 | NA |
| 7.29E-08 | 1.00E-06 |
| 0.467439395 | 0.625694339 |
| 0.649135993 | 0.770648656 |
| 0.556470944 | 0.700688854 |
| 0.729132981 | 0.829416758 |
| NA | NA |
| NA | NA |
| 0.152217625 | 0.286251396 |
| 0.001584822 | 0.007519717 |
| 0.810206423 | 0.883701523 |
| 0.008002404 | 0.02867277 |
| 0.675900719 | 0.791098983 |
| 1.19E-09 | 2.09E-08 |
| 0.513137932 | 0.66433011 |
| 0.312304035 | 0.474291683 |
| 0.859135812 | NA |
| 0.460138025 | NA |
| 0.941275956 | NA |
| 3.09E-05 | 0.00024741 |
| 0.879244848 | NA |
| NA | NA |
| NA | NA |
| NA | NA |
| 0.941278332 | NA |
| NA | NA |
| 0.000195612 | 0.001256332 |
| 0.144398328 | 0.275503212 |
| 0.839319108 | NA |
| NA | NA |
| NA | NA |
| NA | NA |
| 0.04404767 | 0.112130763 |

| | | |
|---|---|---|
| | 0.273067845 | 0.43309198 |
| | 5.40E-05 | 0.000405754 |
| | 0.163380531 | 0.301332411 |
| | 0.071018495 | 0.162850615 |
| | 0.191273794 | 0.336389747 |
| | 0.275953244 | NA |
| | 0.180301468 | 0.32230834 |
| NA | | NA |
| NA | | NA |
| | 0.128815849 | 0.25310177 |
| | 0.356484876 | 0.520192878 |
| | 0.941278332 | NA |
| | 0.000524541 | 0.002957157 |
| | 0.004879 | 0.019138094 |
| | 0.279547025 | 0.439338282 |
| NA | | NA |
| NA | | NA |
| | 0.000563972 | 0.003143714 |
| NA | | NA |
| | 0.692348907 | NA |
| | 0.108000196 | 0.222045492 |
| | 0.608415806 | 0.740735178 |
| NA | | NA |
| | 0.000529034 | 0.00297896 |
| | 0.34747731 | 0.511413023 |
| | 1.83E-06 | 1.95E-05 |
| | 0.270223928 | 0.430289039 |
| | 0.240718638 | 0.396785667 |
| | 0.006971018 | 0.025593734 |
| NA | | NA |
| | 0.047184049 | 0.118243285 |
| | 0.839319108 | NA |
| | 0.117528831 | 0.236468405 |
| | 3.14E-06 | 3.18E-05 |
| | 0.184397659 | 0.327368769 |
| | 4.24E-09 | 6.97E-08 |
| | 0.243893143 | 0.400299586 |
| NA | | NA |
| | 0.46923105 | NA |
| | 0.00012902 | 0.000877162 |
| | 0.085402069 | 0.186764061 |
| | 0.551488293 | NA |
| | 0.000196841 | 0.001262518 |
| | 0.000675826 | 0.003666582 |
| | 0.255873647 | NA |
| | 0.206721598 | 0.355699231 |
| | 0.056243596 | 0.136047771 |

| | | |
|---|---|---|
| | 0.469083876 | NA |
| | 0.168500151 | 0.307965953 |
| | 0.287128911 | 0.447706672 |
| | 0.05644026 | 0.136412577 |
| NA | | NA |
| | 0.314488986 | 0.476720599 |
| | 0.000468337 | 0.002682234 |
| | 0.078115033 | 0.174597992 |
| | 0.973547161 | 0.984043719 |
| NA | | NA |
| | 0.488729998 | 0.643439981 |
| | 0.128933097 | 0.25318582 |
| NA | | NA |
| | 0.421491224 | 0.582963638 |
| | 0.015610946 | 0.049247414 |
| | 0.054441899 | 0.132549019 |
| NA | | NA |
| | 0.006868817 | 0.02528872 |
| | 9.10E-13 | 2.30E-11 |
| | 0.102639384 | 0.21386874 |
| | 0.645696389 | 0.768020862 |
| | 0.004157398 | 0.016760079 |
| | 0.125632188 | 0.248549359 |
| | 0.943163377 | NA |
| | 0.002915833 | 0.012523058 |
| | 0.396876545 | 0.559491561 |
| | 0.778829006 | 0.863664182 |
| | 0.006436909 | 0.023976731 |
| | 0.941277857 | NA |
| NA | | NA |
| NA | | NA |
| | 0.839319108 | NA |
| | 0.955792398 | NA |
| | 0.184266532 | 0.327227407 |
| | 0.003227842 | 0.013639109 |
| | 0.634914559 | NA |
| NA | | NA |
| | 0.941277857 | NA |
| | 1.05E-11 | 2.37E-10 |
| | 0.360933671 | 0.524637755 |
| NA | | NA |
| | 0.051645271 | 0.127189034 |
| NA | | NA |
| | 0.317671008 | NA |
| | 0.839319957 | NA |
| NA | | NA |
| | 0.696041823 | NA |

| | | |
|---|---|---|
| | 0.177330217 | 0.31891302 |
| | 0.955792398 | NA |
| NA | | NA |
| | 0.925586354 | 0.955329983 |
| | 0.941278332 | NA |
| NA | | NA |
| | 0.941279268 | NA |
| NA | | NA |
| | 0.89379076 | NA |
| NA | | NA |
| | 0.789230896 | 0.870333292 |
| NA | | NA |
| | 0.024145744 | 0.069789555 |
| NA | | NA |
| | 4.25E-07 | 5.09E-06 |
| | 0.087921402 | NA |
| | 5.30E-28 | 3.41E-26 |
| NA | | NA |
| NA | | NA |
| | 0.941277857 | NA |
| NA | | NA |
| | 0.297470106 | 0.45887334 |
| | 0.01245251 | 0.041103127 |
| | 0.106800296 | NA |
| | 0.299619522 | 0.46114089 |
| | 0.206759257 | 0.355735164 |
| | 0.087875226 | 0.190822564 |
| | 0.839319108 | NA |
| | 0.021681751 | 0.063869929 |
| | 0.382880085 | 0.545609279 |
| | 2.44E-08 | 3.62E-07 |
| | 0.549408124 | 0.695031207 |
| | 0.004491685 | 0.017872131 |
| NA | | NA |
| | 1.23E-36 | 1.13E-34 |
| | 0.002781879 | 0.012047725 |
| | 0.030083427 | 0.083203082 |
| | 0.576168639 | NA |
| | 0.465242698 | 0.623659985 |
| | 0.000966833 | 0.004957469 |
| | 0.728055127 | NA |
| | 0.018088105 | 0.055366908 |
| | 0.412866271 | 0.575084444 |
| | 0.00732543 | 0.026672769 |
| | 0.016241206 | 0.050823711 |
| | 0.257272618 | 0.41534777 |
| | 0.012531731 | 0.041307336 |

| | |
|---|---|
| 0.196458314 | 0.343174567 |
| NA | NA |
| NA | NA |
| 0.39926878 | 0.561898865 |
| 0.238382727 | 0.394054147 |
| NA | NA |
| NA | NA |
| 0.001099192 | 0.005537138 |
| NA | NA |
| 0.955792398 | NA |
| 0.029175244 | 0.081172314 |
| 0.74627078 | NA |
| 0.376965837 | 0.539833265 |
| NA | NA |
| 0.056852671 | 0.137130854 |
| 0.078841283 | 0.175923898 |
| 0.95948551 | 0.97513553 |
| 2.25E-09 | 3.82E-08 |
| 0.893522079 | NA |
| 2.99E-06 | 3.05E-05 |
| 0.000398556 | 0.002329675 |
| 0.560316924 | 0.703558948 |
| 0.270561616 | 0.430545107 |
| NA | NA |
| 7.85E-08 | 1.07E-06 |
| 0.000762023 | 0.004059461 |
| 0.563975414 | NA |
| 0.374540941 | 0.537715299 |
| 0.030311174 | 0.083648297 |
| 0.087570871 | 0.190300545 |
| 0.503994291 | 0.656096702 |
| 0.816624829 | 0.888075304 |
| 0.890901433 | 0.934367626 |
| 0.537206265 | 0.685093416 |
| 0.282791801 | 0.443208695 |
| 0.013934666 | 0.044960405 |
| NA | NA |
| 0.642789706 | NA |
| 0.001835293 | 0.00850538 |
| 0.719359067 | 0.82140487 |
| 0.714991074 | 0.818892192 |
| 0.014734888 | 0.046988184 |
| NA | NA |
| 0.008216919 | 0.029317808 |
| 0.05613073 | 0.135857599 |
| 0.342281612 | 0.505772717 |
| 0.094308113 | NA |

| | | |
|---|---|---|
| | 0.802858539 | 0.879453216 |
| | 3.95E-07 | 4.76E-06 |
| | 0.941278332 | NA |
| | 0.941277857 | NA |
| | 0.472371411 | NA |
| NA | | NA |
| NA | | NA |
| | 0.063195562 | NA |
| | 0.983710119 | 0.989655871 |
| | 0.397995795 | 0.560704109 |
| | 0.099654462 | 0.209039878 |
| | 0.212734153 | NA |
| | 9.18E-70 | 1.95E-67 |
| | 5.58E-11 | 1.16E-09 |
| | 0.000888169 | 0.004613262 |
| | 0.684644908 | 0.797412316 |
| | 3.19E-126 | 1.35E-123 |
| | 0.026189922 | 0.07441477 |
| | 0.839318641 | NA |
| | 2.25E-27 | 1.41E-25 |
| | 3.38E-18 | 1.27E-16 |
| | 0.780833705 | NA |
| | 3.61E-29 | 2.44E-27 |
| | 1.42E-08 | 2.17E-07 |
| | 0.852150132 | NA |
| NA | | NA |
| | 0.839319108 | NA |
| NA | | NA |
| NA | | NA |
| NA | | NA |
| NA | | NA |
| | 0.941278207 | NA |
| | 0.941277857 | NA |
| NA | | NA |
| | 0.829016785 | NA |
| NA | | NA |
| | 0.826698785 | NA |
| NA | | NA |
| | 0.002228721 | 0.01001851 |
| | 0.677846219 | NA |
| | 0.813998572 | 0.886443528 |
| | 0.101081017 | 0.21147318 |
| | 0.26167368 | 0.420465599 |
| | 0.710896617 | NA |
| | 0.687696103 | 0.799596632 |
| | 0.580666101 | 0.718421247 |
| | 0.574569078 | 0.713786167 |

| | |
|---|---|
| NA | NA |
| 0.839319574 | NA |
| 8.67E-59 | 1.45E-56 |
| 0.430064316 | 0.591322908 |
| 0.451549724 | 0.611468768 |
| 0.094143225 | 0.200518752 |
| 0.541685325 | 0.688789265 |
| NA | NA |
| 0.259649911 | 0.41817357 |
| 0.013937591 | 0.04496375 |
| 0.544280561 | 0.690931515 |
| 0.211168896 | 0.3613319 |
| 0.010116045 | 0.034724848 |
| 0.635042512 | 0.760020207 |
| 0.024381986 | 0.070293125 |
| 0.131188929 | 0.256536221 |
| 0.456770842 | NA |
| 0.008658932 | 0.030574085 |
| NA | NA |
| 0.228688436 | 0.382303845 |
| 0.2683206 | 0.428111471 |
| 0.494969066 | 0.647926838 |
| 0.523422396 | 0.6732824 |
| 0.005636571 | 0.021517509 |
| 0.077668081 | 0.173892924 |
| 7.51E-169 | 5.95E-166 |
| 7.11E-13 | 1.81E-11 |
| NA | NA |
| 0.581776013 | 0.719496656 |
| 0.833801616 | NA |
| NA | NA |
| 0.366432669 | 0.529609009 |
| 0.299113414 | 0.460705725 |
| 2.03E-05 | 0.00017035 |
| 0.325076596 | 0.487953153 |
| 0.245145065 | 0.401800223 |
| 0.272960087 | 0.432959882 |
| 0.008750447 | 0.03085602 |
| 0.000158017 | 0.0010465 |
| 0.120480046 | 0.241122176 |
| 0.042671045 | 0.109528331 |
| 0.40578245 | 0.567842547 |
| NA | NA |
| 5.69E-05 | 0.000425969 |
| 0.784245724 | 0.867327233 |
| 0.801122024 | 0.878492596 |
| 0.001347517 | 0.006562236 |

| | | |
|---|---|---|
| | 6.43E-08 | 8.89E-07 |
| | 0.895530027 | 0.936820599 |
| | 0.890560573 | 0.934198497 |
| | 0.013358329 | 0.043513746 |
| | 0.22975188 | 0.383542871 |
| | 0.001426544 | 0.006874794 |
| | 0.417507245 | 0.579041339 |
| | 0.000164587 | 0.001085935 |
| | 0.237456081 | 0.392953742 |
| | 0.664940897 | 0.78261995 |
| | 0.160080433 | 0.296772677 |
| | 1.93E-11 | 4.23E-10 |
| | 0.900772543 | 0.940152952 |
| | 0.539534234 | 0.686737639 |
| | 0.683919911 | 0.797113981 |
| | 0.113907125 | 0.230958122 |
| | 0.301910866 | 0.463259568 |
| | 0.839319108 | NA |
| | 0.000103335 | 0.000720255 |
| | 0.036023109 | 0.095919236 |
| | 0.062527157 | 0.147719245 |
| | 0.002029308 | 0.009265601 |
| | 0.832352077 | 0.898324301 |
| | 2.04E-10 | 3.95E-09 |
| | 0.451044035 | 0.611061625 |
| | 8.57E-32 | 6.44E-30 |
| | 0.302735719 | 0.464166167 |
| | 0.346275904 | 0.510091538 |
| | 0.955792398 | NA |
| | 0.046043598 | 0.116131156 |
| | 6.42E-13 | 1.64E-11 |
| | 0.416658832 | NA |
| | 5.38E-30 | 3.75E-28 |
| | 0.45869317 | 0.618146885 |
| | 0.728878986 | NA |
| | 0.413140175 | NA |
| | 0.72601533 | NA |
| NA | | NA |
| | 0.705817268 | NA |
| | 0.735197453 | NA |
| | 4.94E-12 | 1.15E-10 |
| | 0.842227873 | NA |
| | 1.90E-06 | 2.02E-05 |
| | 0.002030673 | 0.009270059 |
| | 1.44E-21 | 6.79E-20 |
| | 0.586527991 | 0.723231583 |
| | 2.59E-12 | 6.22E-11 |

| | |
|---|---|
| 0.21522196 | 0.366186989 |
| 0.569768534 | 0.710740037 |
| 0.211623082 | 0.361987252 |
| 0.000613665 | 0.003375656 |
| 0.046062092 | 0.116153186 |
| 0.001500909 | 0.00717939 |
| 0.000702476 | 0.003788098 |
| 0.752447452 | 0.844680199 |
| 0.755948455 | 0.84719517 |
| 5.12E-35 | 4.40E-33 |
| 2.20E-52 | 3.17E-50 |
| NA | NA |
| 1.08E-60 | 1.93E-58 |
| NA | NA |
| NA | NA |
| 0.128713621 | NA |
| 0.201361361 | 0.349202412 |
| 0.642273985 | 0.765326929 |
| 0.465724281 | 0.623968899 |
| 5.15E-06 | 5.00E-05 |
| 2.58E-32 | 1.99E-30 |
| 0.296381776 | 0.457874929 |
| 7.74E-05 | 0.000557857 |
| 0.904622105 | NA |
| 1.44E-60 | 2.54E-58 |
| 4.48E-34 | 3.69E-32 |
| 0.003213872 | 0.01358973 |
| 0.002062079 | 0.00937925 |
| 0.495206453 | 0.64816629 |
| 0.009620136 | 0.033364194 |
| 0.427424456 | 0.588713436 |
| 0.941278788 | NA |
| 0.954054724 | 0.972065003 |
| 0.127425757 | 0.251075155 |
| 0.884500336 | 0.930962684 |
| 0.384738812 | 0.547536814 |
| 9.16E-07 | 1.02E-05 |
| 0.001293242 | 0.006340217 |
| 8.38E-08 | 1.14E-06 |
| NA | NA |
| 0.000228865 | 0.001438837 |
| 0.89548397 | 0.936820599 |
| 0.335193578 | 0.498425638 |
| 0.00080113 | 0.004231799 |
| NA | NA |
| 0.005962397 | 0.022533656 |
| 0.073424882 | 0.166920634 |

| | | | |
|---|---|---|---|
| | 0.524012388 | | 0.673645307 |
| | 0.891963165 | | 0.934936094 |
| | 0.002942804 | | 0.012618379 |
| | 0.016042995 | | 0.050325409 |
| | 0.075858497 | | 0.170934767 |
| NA | | NA | |
| | 0.145683041 | | 0.277315078 |
| | 0.000465062 | | 0.002665547 |
| NA | | NA | |
| | 1.96E-13 | | 5.24E-12 |
| | 7.71E-21 | | 3.47E-19 |
| | 0.221465112 | | 0.373560527 |
| | 0.620967699 | | 0.749468748 |
| | 0.636775731 | NA | |
| NA | | NA | |
| | 0.001732555 | | 0.008114578 |
| | 0.157451197 | | 0.293243614 |
| | 1.19E-05 | | 0.000105331 |
| | 0.005308021 | | 0.020486706 |
| | 0.862977508 | NA | |
| | 1.54E-05 | | 0.000133317 |
| | 0.009230065 | | 0.032227375 |
| | 0.376188905 | | 0.539400501 |
| | 0.908526836 | | 0.944477453 |
| | 0.570919911 | | 0.711393348 |
| | 0.046249213 | | 0.116513946 |
| | 0.578433658 | | 0.716629082 |
| | 2.29E-07 | | 2.88E-06 |
| | 0.525364471 | NA | |
| | 0.203346701 | | 0.351646192 |
| | 0.007158428 | | 0.026160741 |
| | 0.023759689 | | 0.068866064 |
| | 0.181457222 | NA | |
| | 0.000716655 | | 0.003854874 |
| | 7.07E-46 | | 8.72E-44 |
| | 0.223864792 | | 0.37643446 |
| | 0.065475556 | | 0.153134439 |
| NA | | NA | |
| | 0.001368511 | | 0.006643543 |
| | 0.881637946 | | 0.929274887 |
| NA | | NA | |
| | 0.741510485 | | 0.83740696 |
| | 0.885782546 | NA | |
| | 8.06E-06 | | 7.47E-05 |
| | 0.107609118 | | 0.22145186 |
| | 0.001831178 | | 0.008489619 |
| | 0.093282836 | | 0.199238879 |

| | |
|---|---|
| 5.65E-07 | 6.59E-06 |
| 0.765347663 | 0.854255406 |
| 2.04E-06 | 2.15E-05 |
| 0.002003843 | 0.009179269 |
| NA | NA |
| 1.83E-18 | 7.03E-17 |
| NA | NA |
| 0.000512223 | 0.002896635 |
| 4.55E-05 | 0.000348199 |
| NA | NA |
| 0.013934128 | 0.044960405 |
| 0.157507432 | 0.293292374 |
| 0.000214495 | 0.001363987 |
| 0.003196686 | 0.013529077 |
| NA | NA |
| 0.777970902 | NA |
| 0.12154125 | 0.242695037 |
| 0.056404079 | 0.136341253 |
| 0.077710008 | 0.173937709 |
| 0.362841499 | 0.5265427 |
| 0.95456991 | NA |
| 0.330807178 | NA |
| 0.491612044 | 0.645589606 |
| 0.941277857 | NA |
| 0.582338733 | 0.720005533 |
| 0.272393539 | 0.432262885 |
| NA | NA |
| 0.000354191 | 0.002101836 |
| 0.983062248 | 0.989254962 |
| 0.974467673 | 0.984554671 |
| 0.628424978 | 0.755481152 |
| 0.001231346 | 0.006091978 |
| 0.045642659 | 0.115278712 |
| 0.096505197 | 0.204197412 |
| 0.857570418 | 0.914757152 |
| 0.854067199 | 0.91208307 |
| 0.087048279 | 0.189441633 |
| 0.001413724 | 0.006832379 |
| 0.091103685 | 0.196044653 |
| 0.072010318 | 0.164482143 |
| 7.48E-07 | 8.52E-06 |
| 0.566226497 | 0.707921135 |
| 0.000254062 | 0.001573556 |
| 0.839319108 | NA |
| 0.018624765 | 0.056681584 |
| 0.000276112 | 0.001694258 |
| 0.189076644 | 0.333659718 |

| | |
|---|---|
| 0.839318641 | NA |
| 0.080689843 | 0.178919034 |
| 0.021061269 | 0.062350786 |
| 3.98E-08 | 5.66E-07 |
| 0.783578429 | NA |
| 0.843878874 | 0.906040189 |
| 0.00863929 | 0.030522841 |
| 0.091027617 | 0.195916388 |
| 3.12E-12 | 7.46E-11 |
| 0.608560118 | 0.740804374 |
| 0.300582269 | NA |
| 1.10E-05 | 9.81E-05 |
| 0.561917676 | 0.704702602 |
| 0.439082134 | 0.600047824 |
| NA | NA |
| 0.231974446 | 0.386196808 |
| 0.897896696 | 0.938413841 |
| NA | NA |
| NA | NA |
| NA | NA |
| NA | NA |
| NA | NA |
| 0.941278332 | NA |
| NA | NA |
| NA | NA |
| 0.05740467 | 0.138154292 |
| NA | NA |
| 0.941278332 | NA |
| NA | NA |
| NA | NA |
| 0.776403287 | NA |
| 0.941278424 | NA |
| 0.941278332 | NA |
| 0.17879337 | NA |
| 0.667608755 | NA |
| NA | NA |
| 0.283312112 | 0.443878021 |
| 0.001790746 | 0.008341163 |
| 0.424002237 | 0.585422661 |
| 0.590629632 | 0.726595333 |
| 0.51689982 | 0.667747599 |
| 0.372508142 | 0.535768856 |
| 7.87E-07 | 8.94E-06 |
| 0.002134898 | 0.009653277 |
| 0.941278332 | NA |
| 0.020715908 | 0.061527333 |
| 0.0093116 | 0.032469192 |

| | |
|---|---|
| 0.436120081 | 0.597234346 |
| 0.002345798 | 0.010469658 |
| 0.00154416 | 0.007355232 |
| 4.99E-05 | 0.000378488 |
| 0.210611306 | 0.360645345 |
| 0.791559055 | 0.871750204 |
| 0.833914293 | 0.899431054 |
| 2.23E-07 | 2.81E-06 |
| NA | NA |
| 0.117218183 | NA |
| 1.22E-07 | 1.62E-06 |
| 2.10E-48 | 2.80E-46 |
| 0.069728121 | 0.160773877 |
| 7.10E-07 | 8.15E-06 |
| 0.004795605 | 0.018876301 |
| 0.164675357 | 0.303069886 |
| NA | NA |
| 0.00108226 | 0.005464564 |
| NA | NA |
| 0.00028209 | 0.001727822 |
| 0.242767328 | 0.399222602 |
| 0.905456649 | 0.942618177 |
| 0.065344685 | 0.15288845 |
| 0.941278332 | NA |
| 3.76E-14 | 1.07E-12 |
| 2.60E-05 | 0.000212737 |
| 0.123122222 | 0.244964513 |
| 0.021948186 | 0.06449515 |
| 1.68E-16 | 5.71E-15 |
| 0.003585818 | 0.014840835 |
| 0.509168235 | 0.660808196 |
| 0.037797607 | 0.099596883 |
| 0.830586818 | 0.897271536 |
| 4.67E-13 | 1.21E-11 |
| NA | NA |
| 0.513342915 | NA |
| 0.512244678 | 0.66367905 |
| 0.013772138 | 0.044538646 |
| 0.574254137 | 0.713717141 |
| 0.294195074 | 0.45538344 |
| 5.67E-06 | 5.46E-05 |
| 0.876763954 | 0.926202931 |
| NA | NA |
| NA | NA |
| 0.021770031 | 0.064082399 |
| 0.000924829 | 0.004771899 |
| 0.378643291 | 0.541443708 |

| | |
|---|---|
| 0.88743979 | 0.932905303 |
| 0.020755696 | 0.061607068 |
| 0.052108068 | 0.12808156 |
| 0.176010961 | 0.317427861 |
| 0.280450706 | 0.440391848 |
| 0.344030427 | 0.507652507 |
| 0.007911434 | 0.028393842 |
| 0.005277185 | 0.020384294 |
| 1.42E-05 | 0.000123854 |
| 0.842081017 | 0.904803737 |
| 0.006979081 | 0.025615434 |
| 4.04E-05 | 0.000313715 |
| NA | NA |
| 0.001049009 | 0.005315843 |
| NA | NA |
| 0.713184137 | 0.817609519 |
| 0.14417324 | NA |
| 0.000107251 | 0.000744039 |
| 0.839319108 | NA |
| NA | NA |
| 0.259412621 | 0.417847954 |
| NA | NA |
| NA | NA |
| 0.857977734 | 0.914986604 |
| 0.453802299 | 0.613681173 |
| 0.905471053 | 0.942618177 |
| 0.136359417 | 0.263967647 |
| NA | NA |
| 0.0005485 | 0.003076929 |
| 0.269923645 | 0.429948645 |
| 0.636545487 | 0.761067975 |
| 0.074447133 | 0.168520271 |
| 0.582439539 | NA |
| NA | NA |
| 0.000753015 | 0.004018476 |
| 0.147529006 | 0.279978105 |
| 0.404946464 | 0.56697269 |
| 0.338379646 | 0.501907668 |
| 0.164722223 | 0.3031327 |
| 0.177046104 | 0.318594644 |
| 0.334025696 | 0.497277935 |
| 0.000147478 | 0.000984096 |
| 0.039428601 | 0.102982505 |
| NA | NA |
| 0.050871728 | 0.1255937 |
| NA | NA |
| 0.851002719 | NA |

| | | |
|---|---|---|
| | 0.539146778 | 0.686485706 |
| | 0.004983843 | 0.019478724 |
| | 0.702940561 | 0.810433094 |
| | 0.215385777 | NA |
| | 0.839319574 | NA |
| | 0.079362738 | 0.176834121 |
| | 0.538376431 | 0.68603436 |
| | 0.719646338 | 0.821614646 |
| | 0.088727132 | 0.192239214 |
| | 0.210652393 | 0.360689754 |
| | 0.00018474 | 0.00119651 |
| NA | | NA |
| | 0.297306167 | 0.458735296 |
| | 0.850722131 | 0.910458133 |
| | 0.058362927 | 0.13990879 |
| | 0.252646871 | 0.410505893 |
| | 0.410999559 | 0.572953702 |
| | 0.00575857 | 0.021891946 |
| | 1.70E-05 | 0.000145061 |
| | 0.05612896 | 0.135857599 |
| | 0.008210688 | 0.029304362 |
| | 0.114686435 | 0.232201984 |
| | 0.000786706 | 0.004169487 |
| NA | | NA |
| NA | | NA |
| | 0.010181625 | 0.034909686 |
| NA | | NA |
| | 0.039309564 | 0.102727959 |
| NA | | NA |
| | 0.047170518 | 0.118234274 |
| | 1.83E-06 | 1.95E-05 |
| | 0.000164076 | 0.00108301 |
| | 0.002738473 | 0.011898712 |
| | 0.21499507 | 0.365918068 |
| | 0.140036545 | 0.269193171 |
| | 0.000446743 | 0.002574989 |
| | 0.001301785 | 0.0063716 |
| | 0.281588877 | 0.441788837 |
| | 0.839319108 | NA |
| NA | | NA |
| | 0.000195067 | 0.001253507 |
| | 0.015344362 | 0.048573813 |
| | 0.91270796 | 0.94739633 |
| | 0.463068662 | 0.621906283 |
| | 3.05E-07 | 3.74E-06 |
| | 0.820430545 | 0.890667632 |
| | 0.003376922 | 0.014145987 |

| | |
|---|---|
| 0.038399748 | 0.100815221 |
| 0.651732135 | 0.772647082 |
| 0.222393583 | 0.374807708 |
| 0.548842488 | 0.694694389 |
| 0.866019476 | 0.920017655 |
| 0.955792398 | NA |
| 0.834299218 | 0.899651662 |
| 0.259054162 | 0.417468325 |
| 0.03171447 | 0.086676751 |
| 0.682988369 | 0.796379227 |
| 0.891456772 | 0.934565988 |
| 1.42E-06 | 1.54E-05 |
| 0.005623796 | 0.021479075 |
| 0.054485939 | 0.132618295 |
| NA | NA |
| NA | NA |
| 0.303131436 | 0.46460871 |
| 0.936127164 | 0.962201441 |
| NA | NA |
| 0.870540667 | 0.922591889 |
| 0.020858133 | 0.061864832 |
| 0.099398273 | 0.20862693 |
| 2.36E-05 | 0.000195142 |
| 0.741550119 | NA |
| 0.839319108 | NA |
| 0.921134577 | NA |
| 0.003398544 | 0.014211549 |
| 0.279237514 | 0.438996785 |
| 6.71E-06 | 6.35E-05 |
| NA | NA |
| 3.40E-05 | 0.000269356 |
| 1.36E-12 | 3.36E-11 |
| NA | NA |
| 0.446449941 | 0.606910818 |
| 0.896747263 | NA |
| NA | NA |
| 0.18475187 | 0.327831314 |
| 9.28E-07 | 1.04E-05 |
| 8.20E-10 | 1.47E-08 |
| 0.004427634 | 0.017643821 |
| 3.21E-07 | 3.93E-06 |
| 0.723161097 | 0.824638549 |
| 0.839318641 | NA |
| 0.380103223 | 0.542887304 |
| 0.135608971 | 0.262885849 |
| 0.093043987 | 0.198800088 |
| 2.47E-06 | 2.57E-05 |

| | | |
|---|---|---|
| NA | | NA |
| | 0.065141529 | 0.152578102 |
| | 0.092462248 | 0.197965856 |
| | 1.03E-05 | 9.27E-05 |
| | 0.839319108 | NA |
| | 0.092912256 | 0.198696244 |
| | 0.769016229 | 0.857023917 |
| | 0.213243984 | 0.363920785 |
| | 7.49E-07 | 8.52E-06 |
| | 0.455119513 | 0.614869554 |
| | 0.092870566 | 0.19866292 |
| | 0.673653179 | NA |
| NA | | NA |
| | 0.000350322 | 0.002081471 |
| NA | | NA |
| NA | | NA |
| | 0.955791922 | NA |
| | 0.839318641 | NA |
| NA | | NA |
| | 0.941277857 | NA |
| | 0.416855379 | 0.578440607 |
| | 0.854044483 | NA |
| | 0.587198038 | 0.723682892 |
| | 0.193794493 | 0.339693388 |
| | 0.1789566 | 0.320594424 |
| | 0.026881551 | 0.07613454 |
| | 0.007850344 | 0.028229926 |
| | 0.831619968 | 0.89785986 |
| | 0.938062266 | NA |
| | 0.955791446 | NA |
| | 0.000302232 | 0.001833261 |
| | 0.006041586 | 0.022796738 |
| | 0.308397332 | 0.469850431 |
| | 0.205695227 | 0.35434304 |
| | 0.1645458 | 0.302911258 |
| | 0.042688624 | 0.109561638 |
| | 0.079030562 | 0.176263629 |
| | 0.002660549 | 0.011632328 |
| | 0.90405324 | NA |
| | 0.575075701 | 0.714074899 |
| | 0.278887402 | 0.438637044 |
| | 0.609038738 | 0.741235365 |
| | 0.38672673 | 0.549511648 |
| | 0.454725366 | 0.614545375 |
| | 0.001300487 | 0.006367866 |
| | 0.453571508 | 0.613508497 |
| | 0.745217213 | 0.839969438 |

| | |
|---|---|
| 0.569706283 | 0.710736881 |
| 0.141986151 | 0.272083919 |
| 0.839319108 | NA |
| 0.009963167 | 0.034284126 |
| 0.783984584 | 0.867199604 |
| 0.005224139 | 0.02022527 |
| NA | NA |
| 0.620707727 | 0.749468748 |
| 0.31834543 | 0.480812253 |
| 0.835303564 | 0.899994439 |
| NA | NA |
| NA | NA |
| 0.010049555 | 0.034531472 |
| 0.148917694 | 0.281815108 |
| 0.918473857 | 0.951079559 |
| NA | NA |
| 0.284418905 | 0.445026215 |
| 0.001263534 | 0.006217622 |
| 7.89E-06 | 7.33E-05 |
| 0.882620335 | NA |
| 2.38E-08 | 3.52E-07 |
| 0.648617899 | 0.770264094 |
| 0.853595622 | NA |
| 0.922154267 | 0.953631614 |
| 0.885509615 | 0.931457974 |
| 0.941278332 | NA |
| 0.896966548 | NA |
| 0.914812071 | 0.948773722 |
| 0.402548074 | 0.564710866 |
| 0.098470503 | 0.207226907 |
| 3.92E-08 | 5.58E-07 |
| 0.113764389 | 0.230727675 |
| 0.509840612 | 0.66146442 |
| 0.182106917 | 0.32448786 |
| 0.675041154 | 0.790546188 |
| 0.068284543 | 0.158196291 |
| 0.000946456 | 0.004868717 |
| 0.417195223 | NA |
| 6.33E-05 | 0.00046854 |
| 0.234343694 | NA |
| 0.537704182 | 0.685452502 |
| 0.941279303 | NA |
| 0.47917148 | NA |
| 0.18033578 | 0.322345466 |
| 0.894407662 | NA |
| NA | NA |
| 0.001807598 | 0.008400065 |

| | |
|---|---|
| 0.174759949 | 0.315769856 |
| 1.04E-09 | 1.85E-08 |
| 0.041771556 | 0.107790221 |
| 0.579692524 | NA |
| 0.020396771 | 0.060807106 |
| 0.245541658 | 0.402339429 |
| NA | NA |
| 0.941278332 | NA |
| NA | NA |
| 0.217599126 | 0.368948168 |
| 0.701649258 | NA |
| NA | NA |
| 0.037857184 | 0.099698682 |
| 0.177366304 | 0.318953821 |
| 0.110617925 | 0.22609943 |
| 0.000307079 | 0.001859346 |
| 0.368783743 | 0.531955072 |
| 0.907895244 | NA |
| 0.853595154 | NA |
| 0.161339112 | 0.298336976 |
| 0.000101012 | 0.000706935 |
| 0.011012528 | 0.037248854 |
| 8.74E-17 | 3.00E-15 |
| NA | NA |
| 2.97E-06 | 3.03E-05 |
| 0.014298157 | 0.045927764 |
| 0.941276431 | NA |
| 0.005517553 | 0.021113956 |
| 0.001757283 | 0.008217458 |
| 0.033591569 | 0.090773574 |
| 0.501144459 | 0.653638031 |
| 1.40E-20 | 6.15E-19 |
| 0.941277857 | NA |
| 0.286348959 | 0.447017638 |
| 0.242896131 | 0.399351641 |
| 0.024673184 | 0.0709436 |
| 0.404835295 | 0.566850385 |
| 0.025114922 | 0.071901056 |
| 0.518774989 | 0.669465062 |
| 0.945203115 | 0.966925536 |
| 0.063529962 | 0.149613154 |
| 0.941278332 | NA |
| 0.941674575 | 0.964778678 |
| 0.377945235 | NA |
| 0.955792398 | NA |
| NA | NA |
| 0.081033492 | NA |

| | |
|---|---|
| 0.853596091 | NA |
| NA | NA |
| NA | NA |
| 0.484695456 | NA |
| NA | NA |
| 0.832124654 | 0.898160303 |
| NA | NA |
| NA | NA |
| NA | NA |
| NA | NA |
| 0.463956783 | NA |
| NA | NA |
| 0.575513388 | 0.71434969 |
| NA | NA |
| 0.012441061 | 0.041076727 |
| NA | NA |
| 0.852438342 | NA |
| NA | NA |
| 0.592902214 | NA |
| 0.721320905 | 0.823168869 |
| NA | NA |
| 0.578131336 | 0.716366394 |
| 0.868332523 | 0.921216246 |
| 0.941279303 | NA |
| NA | NA |
| 0.615865114 | NA |
| 0.746832673 | NA |
| 0.959951701 | NA |
| 0.000507275 | 0.002872064 |
| 0.020598759 | 0.061286458 |
| 5.13E-06 | 4.98E-05 |
| 1.92E-07 | 2.45E-06 |
| 0.196230725 | 0.342880156 |
| 0.891315628 | 0.934565988 |
| 4.04E-12 | 9.51E-11 |
| 0.947283607 | 0.967817505 |
| 0.006045438 | 0.022807658 |
| 2.22E-13 | 5.91E-12 |
| 0.123029923 | 0.244805068 |
| 0.56973822 | 0.710739471 |
| 0.031344835 | 0.085883476 |
| 0.041410478 | 0.107043027 |
| 0.042658828 | 0.109520595 |
| 0.152505891 | 0.28664993 |
| 0.993246906 | 0.995209152 |
| 4.44E-05 | 0.000340971 |
| NA | NA |

| | |
|---|---|
| 0.006643844 | 0.024601241 |
| 0.068736766 | 0.158980907 |
| 0.177387913 | NA |
| 0.941277857 | NA |
| 0.007544628 | 0.027320436 |
| 0.000139125 | 0.000935965 |
| 0.037522271 | 0.09896998 |
| 4.74E-07 | 5.59E-06 |
| 0.364384305 | 0.52775193 |
| 0.599112029 | 0.733130419 |
| 0.292948266 | 0.454103199 |
| NA | NA |
| 0.181829146 | 0.324235641 |
| NA | NA |
| 0.331600611 | 0.49468867 |
| 0.38387533 | 0.546569397 |
| NA | NA |
| 0.562839861 | 0.705330475 |
| 0.788262679 | NA |
| NA | NA |
| NA | NA |
| NA | NA |
| NA | NA |
| NA | NA |
| NA | NA |
| NA | NA |
| NA | NA |
| 0.355143162 | 0.518882667 |
| NA | NA |
| 0.839319574 | NA |
| 0.941278284 | NA |
| 0.502419073 | 0.654475633 |
| NA | NA |
| 0.73204378 | 0.830985599 |
| 0.62795391 | NA |
| 0.351685744 | 0.515306516 |
| 0.638781254 | NA |
| 0.679670902 | NA |
| 0.536402599 | 0.684435223 |
| 0.059843029 | 0.142709526 |
| 0.001120655 | 0.005627393 |
| 0.297820552 | 0.459263024 |
| 0.049372862 | 0.122464867 |
| 0.003353438 | 0.01407486 |
| 0.579853176 | 0.717715304 |
| 7.49E-27 | 4.60E-25 |
| 0.845155857 | 0.906756804 |

| | | |
|---|---|---|
| | 3.14E-16 | 1.05E-14 |
| | 0.839319574 | NA |
| | 0.58741289 | NA |
| | 0.484712025 | NA |
| | 0.818858132 | NA |
| | 1.76E-39 | 1.78E-37 |
| | 0.010767164 | 0.036559207 |
| | 0.001615582 | 0.007647989 |
| | 0.836151551 | NA |
| | 0.955792398 | NA |
| NA | | NA |
| | 0.944114978 | NA |
| NA | | NA |
| NA | | NA |
| NA | | NA |
| NA | | NA |
| | 1.38E-15 | 4.39E-14 |
| | 0.617247231 | NA |
| | 0.630027639 | 0.756391124 |
| | 0.941277857 | NA |
| | 0.124074419 | 0.246389065 |
| | 0.405781209 | 0.567842547 |
| | 0.04349962 | 0.111080137 |
| | 0.022237553 | 0.065224676 |
| | 0.694607889 | 0.804340433 |
| | 0.21092032 | 0.361018658 |
| | 0.272534718 | 0.432400438 |
| NA | | NA |
| | 0.066622318 | 0.15529766 |
| | 0.951249806 | 0.970245452 |
| | 0.094405162 | 0.200932814 |
| | 0.615494011 | NA |
| | 0.048704765 | 0.121249956 |
| | 7.84E-07 | 8.91E-06 |
| | 0.000539085 | 0.003028392 |
| | 0.470272461 | NA |
| | 0.844913997 | 0.906676878 |
| | 0.19358002 | 0.339417431 |
| | 0.324840022 | 0.487742898 |
| | 0.013781414 | 0.044556536 |
| | 0.66972665 | NA |
| | 0.677781635 | 0.792248611 |
| | 0.744071595 | 0.839257403 |
| | 4.14E-06 | 4.11E-05 |
| NA | | NA |
| | 0.00235251 | 0.01049568 |
| | 0.499150804 | 0.652038166 |

| | | |
|---|---|---|
| | 0.549792709 | 0.695369895 |
| | 0.301061652 | 0.462373819 |
| | 0.010882118 | 0.036886383 |
| | 0.014959886 | 0.047603638 |
| | 0.132647743 | 0.258496936 |
| | 3.07E-07 | 3.77E-06 |
| | 0.245431673 | 0.402214589 |
| | 0.000588007 | 0.00325558 |
| | 0.204970759 | 0.35355562 |
| | 0.160640055 | NA |
| | 0.525194144 | 0.674545166 |
| | 0.358513151 | 0.522139755 |
| | 0.000303805 | 0.001841863 |
| | 0.145044927 | 0.276409877 |
| | 0.104122631 | 0.216099999 |
| | 0.003854583 | 0.015734315 |
| | 0.955792398 | NA |
| | 0.30800866 | NA |
| | 0.658105384 | NA |
| | 0.518044729 | 0.668822943 |
| | 0.004531028 | 0.017992579 |
| | 0.700404876 | 0.80864556 |
| NA | | NA |
| | 0.009710953 | 0.033639951 |
| | 0.017642089 | 0.054231824 |
| | 0.019450678 | 0.058580925 |
| | 0.455367657 | 0.614994381 |
| | 0.941278332 | NA |
| | 0.922661129 | NA |
| | 0.853596091 | NA |
| | 0.554445404 | 0.699359274 |
| | 0.818053899 | 0.888899731 |
| | 0.057248834 | 0.137871081 |
| NA | | NA |
| | 0.281780944 | 0.441980048 |
| | 0.494934457 | 0.647917168 |
| NA | | NA |
| NA | | NA |
| NA | | NA |
| | 0.31148654 | 0.473511091 |
| NA | | NA |
| | 0.026712414 | 0.075736615 |
| NA | | NA |
| | 0.051322089 | 0.126612742 |
| NA | | NA |
| | 0.977625341 | 0.986224845 |
| | 0.110116731 | 0.225299392 |

| | |
|---|---|
| 0.042202185 | 0.108617691 |
| 1.33E-07 | 1.75E-06 |
| 0.019002936 | NA |
| 0.340532852 | 0.504064859 |
| NA | NA |
| 0.186309636 | 0.330000145 |
| NA | NA |
| 0.276850781 | 0.436657762 |
| 0.179464775 | 0.321246941 |
| NA | NA |
| NA | NA |
| 0.020681857 | 0.061479899 |
| 0.15947192 | 0.295964963 |
| NA | NA |
| NA | NA |
| 0.216454548 | 0.367740981 |
| 0.001473687 | 0.007066218 |
| 0.710073848 | 0.814958195 |
| 0.013204052 | 0.043105589 |
| 2.66E-24 | 1.44E-22 |
| 0.522004298 | 0.672090432 |
| 0.641692976 | 0.764979276 |
| 0.48750182 | NA |
| 2.18E-28 | 1.43E-26 |
| 0.471774222 | NA |
| NA | NA |
| 0.798190692 | NA |
| 0.84317998 | 0.905616618 |
| 0.001098495 | 0.005534794 |
| 0.839319108 | NA |
| 0.102843858 | 0.214257287 |
| 0.000155693 | 0.001034275 |
| 0.658077826 | NA |
| 2.82E-10 | 5.37E-09 |
| NA | NA |
| NA | NA |
| 0.841312351 | NA |
| 0.147808514 | 0.280405674 |
| 0.000123431 | 0.00084374 |
| 2.28E-05 | 0.000189041 |
| 0.097907185 | 0.206370027 |
| 0.597690132 | 0.732253475 |
| 0.659075227 | 0.778096943 |
| 0.862585668 | 0.91780349 |
| 0.463588406 | 0.622318608 |
| 2.75E-07 | 3.41E-06 |
| 0.335479913 | 0.498651204 |

| | | |
|---|---|---|
| | 0.787205367 | 0.869227229 |
| | 0.888238183 | NA |
| | 0.94885658 | 0.968760795 |
| | 0.000249677 | 0.001550836 |
| | 0.748393366 | NA |
| | 0.72600519 | NA |
| | 0.006892501 | 0.025364147 |
| | 0.833801616 | NA |
| | 3.72E-08 | 5.33E-07 |
| | 0.955792398 | NA |
| | 0.181947811 | NA |
| | 0.723863988 | 0.825360989 |
| | 0.201328716 | 0.34917124 |
| | 0.758365546 | NA |
| | 0.410048876 | NA |
| | 0.019808437 | 0.059417822 |
| NA | | NA |
| | 0.977647896 | 0.986224845 |
| | 0.941275481 | NA |
| | 0.274699313 | 0.434566321 |
| | 0.000156738 | 0.001040347 |
| | 0.308239386 | 0.469648333 |
| NA | | NA |
| | 0.839320041 | NA |
| | 0.225908467 | 0.378907288 |
| | 0.939563728 | 0.963757575 |
| | 0.748046005 | 0.841684363 |
| | 0.028783553 | 0.080348909 |
| | 0.182683423 | 0.325271606 |
| | 0.546487763 | 0.692701826 |
| | 0.607166797 | NA |
| | 0.075477249 | 0.170382159 |
| | 0.21073338 | 0.36075057 |
| | 0.941277857 | NA |
| NA | | NA |
| | 0.035135239 | 0.093943939 |
| | 0.0036246 | 0.014954536 |
| NA | | NA |
| | 0.490379715 | NA |
| | 9.70E-11 | 1.95E-09 |
| | 0.040857255 | 0.105961741 |
| | 0.42195404 | 0.58320365 |
| NA | | NA |
| | 0.162474053 | 0.299946282 |
| | 0.007761468 | 0.027969487 |
| | 0.839319108 | NA |
| | 0.897976098 | NA |

| | |
|---|---|
| NA | NA |
| 0.488884454 | 0.643513066 |
| 0.99393866 | 0.995569919 |
| 0.277854413 | 0.437515927 |
| NA | NA |
| 0.80028342 | NA |
| NA | NA |
| 0.169839273 | 0.309439252 |
| 0.006252048 | 0.02341272 |
| 0.000679891 | 0.003683761 |
| 0.011771138 | 0.039300972 |
| 3.24E-22 | 1.58E-20 |
| 2.55E-07 | 3.19E-06 |
| 0.845990506 | NA |
| 0.002477159 | 0.010953248 |
| 0.33014518 | 0.493135587 |
| 0.041203956 | 0.106659767 |
| NA | NA |
| 0.001364177 | 0.006626561 |
| NA | NA |
| 0.948894218 | NA |
| 0.914113352 | 0.948236923 |
| 0.056842806 | 0.137120956 |
| 0.941279268 | NA |
| 0.839319108 | NA |
| 0.297282089 | 0.458735296 |
| 0.617860238 | 0.74738575 |
| 0.000274935 | 0.00168747 |
| 0.057614633 | 0.138533538 |
| NA | NA |
| 0.785038992 | 0.867881936 |
| 0.05661142 | 0.136744037 |
| 0.984537213 | 0.990320532 |
| 0.768501027 | 0.85656106 |
| 0.696167789 | 0.805326553 |
| 0.827850942 | 0.895415256 |
| 0.839319108 | NA |
| 0.941276431 | NA |
| 0.23356192 | 0.388160928 |
| 0.393464295 | 0.555932397 |
| NA | NA |
| 0.011162498 | 0.037654448 |
| 3.11E-05 | 0.000248714 |
| 0.40979795 | 0.571713897 |
| 0.955791446 | NA |
| 0.157227443 | 0.292987364 |
| 0.438622138 | 0.59955689 |

| | | | |
|---|---|---|---|
| | 0.751805331 | | 0.844158435 |
| | 0.081015649 | | 0.179363266 |
| | 0.2983805 | | 0.459917929 |
| | 1.37E-06 | | 1.49E-05 |
| | 0.952894722 | | 0.971423684 |
| | 0.551335734 | | 0.696655154 |
| | 0.00132338 | | 0.006461891 |
| NA | | NA | |
| | 0.007939152 | | 0.028463278 |
| | 0.000410452 | | 0.002388064 |
| | 0.074769617 | | 0.169057991 |
| | 0.027923569 | | 0.078404718 |
| | 0.142917456 | | 0.273356632 |
| | 0.761752229 | | 0.851911222 |
| | 0.741119769 | | 0.83729508 |
| | 0.637756582 | | 0.761889287 |
| | 0.000365842 | | 0.002159775 |
| NA | | NA | |
| | 0.406266801 | | 0.568152902 |
| | 0.03263335 | | 0.088770171 |
| | 0.267110503 | | 0.426724223 |
| | 0.739666725 | | 0.83628859 |
| | 0.990515425 | | 0.993604524 |
| | 0.771365799 | NA | |
| | 0.955792398 | NA | |
| | 0.002384672 | | 0.010611359 |
| | 0.620374041 | NA | |
| | 0.022831114 | | 0.066656327 |
| | 0.23259654 | | 0.387042984 |
| | 0.955792398 | NA | |
| | 0.254214881 | | 0.412243084 |
| | 0.728626463 | | 0.829123061 |
| | 0.018614044 | | 0.056656198 |
| | 0.005299011 | | 0.020455249 |
| NA | | NA | |
| | 0.853595622 | NA | |
| | 1.84E-07 | | 2.36E-06 |
| | 6.23E-08 | | 8.63E-07 |
| | 0.391093229 | | 0.553556313 |
| | 0.037385557 | | 0.098653108 |
| | 0.056380701 | | 0.13631684 |
| NA | | NA | |
| NA | | NA | |
| | 0.313741571 | | 0.475860376 |
| | 0.140696791 | | 0.270200501 |
| | 0.001817887 | | 0.00844115 |
| | 0.143036261 | | 0.273427363 |

| | |
|---|---|
| NA | NA |
| 0.497374853 | 0.65032958 |
| 0.924192525 | 0.954592539 |
| 0.989171322 | 0.993078063 |
| 0.525144121 | 0.674517315 |
| 0.138027345 | 0.266406654 |
| 0.684075033 | 0.797216702 |
| 0.774998739 | NA |
| 0.269252992 | 0.429158998 |
| 0.003776222 | 0.015468042 |
| 1.72E-24 | 9.38E-23 |
| 0.303069097 | 0.464557615 |
| 4.05E-11 | 8.56E-10 |
| 3.97E-05 | 0.000308517 |
| 0.955792873 | NA |
| 0.037001453 | 0.097878261 |
| 0.812515923 | 0.885467882 |
| 0.894007063 | 0.936129343 |
| NA | NA |
| NA | NA |
| 0.906888475 | NA |
| 0.468810502 | NA |
| 0.40119901 | NA |
| NA | NA |
| 0.003297388 | 0.013881223 |
| NA | NA |
| 0.839318641 | NA |
| 0.478762875 | NA |
| NA | NA |
| NA | NA |
| NA | NA |
| 0.43290567 | 0.594097138 |
| NA | NA |
| 0.941278332 | NA |
| 0.330553309 | NA |
| 0.941277857 | NA |
| 0.011521493 | 0.03864636 |
| 0.941277857 | NA |
| 0.789892712 | NA |
| 0.95458992 | NA |
| 0.941278332 | NA |
| NA | NA |
| 0.943087778 | NA |
| 0.565552975 | NA |
| NA | NA |
| NA | NA |
| 0.853595154 | NA |

| | |
|---|---|
| NA | NA |
| NA | NA |
| 0.317720524 | NA |
| 0.839318641 | NA |
| 5.87E-15 | 1.77E-13 |
| 0.028139732 | 0.078899978 |
| NA | NA |
| 2.64E-29 | 1.80E-27 |
| 0.075625998 | 0.170588515 |
| 0.006620103 | 0.024540056 |
| 0.096441126 | 0.204098131 |
| 7.90E-12 | 1.81E-10 |
| 0.000617561 | 0.003393576 |
| 0.00341415 | 0.014259272 |
| 0.001181488 | 0.005883099 |
| 0.77558268 | NA |
| NA | NA |
| 0.955792398 | NA |
| 0.207751573 | 0.356955385 |
| 0.561421954 | NA |
| 2.67E-05 | 0.00021816 |
| 0.267134138 | 0.42673334 |
| 0.93867433 | NA |
| 0.082415122 | 0.181741803 |
| 0.0730687 | 0.166237927 |
| NA | NA |
| 0.423675643 | 0.585073547 |
| 0.006050527 | 0.022816009 |
| 0.206846236 | 0.355810805 |
| 0.097117988 | 0.205129325 |
| 0.518925347 | 0.669580662 |
| 0.335999401 | 0.499154772 |
| 0.141971872 | 0.272078462 |
| 0.015383265 | 0.048664604 |
| 0.435252793 | 0.596391144 |
| 0.272451985 | 0.43232681 |
| 0.214997235 | 0.365918068 |
| 0.855500764 | 0.913040498 |
| 0.839319108 | NA |
| 0.002041008 | 0.009304743 |
| NA | NA |
| 0.839319108 | NA |
| NA | NA |
| 1.30E-09 | 2.27E-08 |
| 0.941275956 | NA |
| NA | NA |
| 0.456047309 | 0.615737594 |

| | |
|---|---|
| 0.339645382 | 0.503345955 |
| 0.63003993 | 0.756391124 |
| NA | NA |
| 0.000779606 | 0.004135541 |
| 0.009995622 | 0.034380894 |
| 7.04E-15 | 2.11E-13 |
| 0.880540847 | 0.928400143 |
| 0.928449386 | NA |
| 0.6347486 | 0.759935928 |
| 0.168913768 | 0.308444961 |
| 0.689531157 | 0.800908677 |
| 0.897269196 | 0.937977136 |
| 4.47E-05 | 0.00034283 |
| NA | NA |
| 0.905557324 | NA |
| NA | NA |
| 0.004993117 | 0.019498961 |
| 0.217764862 | 0.369117494 |
| 1.47E-07 | 1.92E-06 |
| 0.624819646 | 0.752819904 |
| 0.955791922 | NA |
| 0.00899472 | 0.031572382 |
| 0.275004861 | 0.434870164 |
| NA | NA |
| NA | NA |
| 0.622508214 | NA |
| 0.791700657 | 0.871750204 |
| 0.010480529 | 0.035759397 |
| 0.473717331 | 0.631205316 |
| 0.453659832 | 0.613593097 |
| 0.215564139 | NA |
| 0.1308078 | 0.255917085 |
| 0.111855159 | 0.227806666 |
| NA | NA |
| 0.876727271 | NA |
| 0.079290164 | 0.176726705 |
| NA | NA |
| NA | NA |
| NA | NA |
| NA | NA |
| NA | NA |
| 0.331510499 | 0.494616242 |
| 0.87416197 | 0.924870125 |
| 0.731769345 | NA |
| 0.702086231 | NA |
| 0.839318641 | NA |
| NA | NA |

| | |
|---|---|
| 0.964035877 | 0.978172988 |
| 0.660767965 | 0.779597458 |
| NA | NA |
| NA | NA |
| 0.182966488 | 0.325565658 |
| 4.70E-05 | 0.000358696 |
| 4.15E-186 | 3.65E-183 |
| 0.165780215 | NA |
| 0.432551489 | 0.593816504 |
| 0.145172039 | 0.276601996 |
| 0.000472582 | 0.002703944 |
| 0.246176167 | NA |
| 0.000119774 | 0.000821577 |
| 2.90E-05 | 0.000234446 |
| NA | NA |
| 0.616990099 | 0.746603528 |
| 0.836545299 | NA |
| 8.29E-07 | 9.38E-06 |
| 0.63960062 | NA |
| 0.193742867 | 0.339627907 |
| NA | NA |
| 0.92470012 | NA |
| 0.033677538 | 0.090935084 |
| 0.466004219 | 0.624153532 |
| 0.198218942 | 0.345312688 |
| 1.80E-07 | 2.32E-06 |
| NA | NA |
| 0.803238435 | NA |
| 1.41E-10 | 2.78E-09 |
| 0.73378892 | NA |
| 0.031840902 | 0.08694243 |
| 0.222630044 | 0.375020545 |
| NA | NA |
| 0.729234634 | 0.829416758 |
| 0.000101327 | 0.000708722 |
| 0.250369528 | 0.407947392 |
| 0.003451027 | 0.014388044 |
| 0.073474441 | 0.167001399 |
| 0.152794835 | 0.286906174 |
| 0.000554694 | 0.003104205 |
| 0.080423587 | NA |
| 2.64E-07 | 3.29E-06 |
| 0.178882091 | 0.320517279 |
| 0.939901802 | 0.963833643 |
| 0.662116105 | NA |
| 0.941281123 | NA |
| 0.960247567 | 0.975605716 |

| | |
|---|---|
| 0.001177473 | 0.00586568 |
| NA | NA |
| NA | NA |
| 0.941275956 | NA |
| 0.936265536 | NA |
| 0.137832022 | 0.266097037 |
| 0.021930956 | 0.064460437 |
| 0.646783226 | 0.768929341 |
| 9.67E-09 | 1.52E-07 |
| 0.010958374 | 0.037107871 |
| 0.04247042 | 0.109131053 |
| 0.554379721 | NA |
| NA | NA |
| 0.003030513 | 0.012933884 |
| 0.199833159 | 0.347285984 |
| 0.200338224 | 0.347909704 |
| 1.97E-05 | 0.00016602 |
| NA | NA |
| 7.14E-35 | 6.11E-33 |
| 0.952208693 | 0.970932242 |
| 0.99200425 | 0.99440104 |
| 0.036773192 | 0.097426037 |
| 0.282684117 | 0.443101592 |
| NA | NA |
| 0.00178071 | 0.008305794 |
| 0.893618279 | NA |
| 0.080184431 | 0.178152696 |
| 0.013653116 | 0.044231715 |
| NA | NA |
| 0.000301018 | 0.00182823 |
| 0.896841699 | 0.937779975 |
| 0.894609351 | 0.936554001 |
| 0.839318641 | NA |
| 0.724913094 | 0.826238017 |
| 0.865732619 | 0.919795018 |
| 0.081365776 | 0.179954302 |
| NA | NA |
| 0.619754249 | 0.748801555 |
| 0.069842543 | 0.160990918 |
| 0.527571807 | 0.676649646 |
| 3.52E-20 | 1.52E-18 |
| 0.186489123 | NA |
| 0.280303093 | 0.44029395 |
| 0.1079563 | 0.221993593 |
| 0.128973334 | 0.25318582 |
| NA | NA |
| 0.06242483 | 0.147550725 |

| | |
|---|---|
| 0.959408188 | 0.975112026 |
| 0.006250146 | 0.02340928 |
| 0.001072391 | 0.005421635 |
| 0.281572121 | 0.441788837 |
| 0.326558526 | 0.489405996 |
| 0.00549742 | 0.021047061 |
| NA | NA |
| 0.44760339 | NA |
| NA | NA |
| 0.669324949 | 0.786030809 |
| 0.358727085 | 0.522306941 |
| 0.677367608 | NA |
| NA | NA |
| 0.465633097 | NA |
| 0.177402328 | NA |
| 0.941277857 | NA |
| NA | NA |
| 0.000154047 | 0.001025485 |
| 0.134873851 | 0.261784832 |
| 0.085365179 | 0.186700542 |
| 0.120452826 | 0.241108248 |
| 0.941277857 | NA |
| 0.955043037 | NA |
| 0.982534179 | 0.988974438 |
| 3.83E-05 | 0.000298792 |
| 7.63E-05 | 0.000550882 |
| 0.421117485 | NA |
| 0.319451643 | NA |
| NA | NA |
| 0.623517136 | 0.751592974 |
| 3.92E-07 | 4.73E-06 |
| 0.380820554 | 0.543520467 |
| 0.162156031 | 0.299428848 |
| 0.27941463 | 0.43924406 |
| 0.00020655 | 0.001317819 |
| NA | NA |
| 0.941278424 | NA |
| 2.38E-05 | 0.000196898 |
| 0.530036848 | 0.678924703 |
| 0.16196486 | 0.299122254 |
| 0.594676495 | NA |
| 0.066991582 | 0.155853156 |
| 0.41234417 | NA |
| 0.287920951 | 0.448559659 |
| 7.50E-07 | 8.53E-06 |
| 0.650344471 | NA |
| 4.46E-08 | 6.28E-07 |

| | |
|---|---|
| 0.000717757 | 0.003859932 |
| NA | NA |
| 0.843838486 | 0.906037696 |
| NA | NA |
| 0.941277857 | NA |
| 0.090465653 | 0.19497134 |
| 0.911323478 | NA |
| 0.941278332 | NA |
| 3.74E-06 | 3.74E-05 |
| 0.405247267 | 0.567254505 |
| 0.735026293 | 0.833060132 |
| 0.326225775 | 0.489153704 |
| 3.42E-05 | 0.000270401 |
| NA | NA |
| 1.74E-250 | 5.93E-247 |
| 0.955792398 | NA |
| 9.82E-195 | 1.02E-191 |
| 1.24E-197 | 1.48E-194 |
| 0.594540316 | 0.729596236 |
| 0.693408296 | 0.803616036 |
| 0.464085975 | 0.622666014 |
| 0.002060051 | 0.009373608 |
| 0.441165269 | 0.601799195 |
| NA | NA |
| 0.839318641 | NA |
| NA | NA |
| 0.234177455 | NA |
| NA | NA |
| 0.166057761 | NA |
| NA | NA |
| 0.661466448 | NA |
| 0.839319574 | NA |
| NA | NA |
| 0.029694466 | NA |
| NA | NA |
| NA | NA |
| NA | NA |
| NA | NA |
| NA | NA |
| NA | NA |
| NA | NA |
| NA | NA |
| 0.839319108 | NA |
| 0.955791922 | NA |
| 0.839319108 | NA |
| NA | NA |
| NA | NA |

| | | |
|---|---|---|
| | 0.831413565 | NA |
| NA | | NA |
| NA | | NA |
| | 0.799654241 | NA |
| | 0.100892875 | 0.211190943 |
| NA | | NA |
| | 4.99E-46 | 6.18E-44 |
| | 2.59E-06 | 2.68E-05 |
| | 0.000864349 | 0.004512664 |
| | 0.918236882 | NA |
| NA | | NA |
| | 0.006675977 | 0.024693333 |
| | 0.916377354 | 0.949652289 |
| | 0.071968716 | 0.164438039 |
| | 0.763339271 | NA |
| | 1.40E-64 | 2.77E-62 |
| | 0.241119111 | 0.397280706 |
| | 0.711482928 | 0.816052267 |
| | 0.098811381 | 0.207674038 |
| | 0.300284112 | 0.461835257 |
| | 0.558320827 | 0.701904206 |
| | 0.000903376 | 0.004681506 |
| | 0.76134955 | 0.851700991 |
| | 0.491491949 | 0.645505235 |
| | 5.05E-05 | 0.000382342 |
| NA | | NA |
| | 0.01975057 | 0.059296552 |
| NA | | NA |
| | 0.168982443 | 0.308539438 |
| | 0.739991096 | NA |
| | 0.004410897 | 0.017588902 |
| NA | | NA |
| NA | | NA |
| NA | | NA |
| | 0.679276724 | NA |
| | 0.98140473 | 0.988464598 |
| | 0.839318641 | NA |
| | 0.082262431 | 0.181478949 |
| | 0.92212858 | NA |
| | 0.461981158 | 0.620756211 |
| | 6.59E-21 | 3.00E-19 |
| | 0.805753412 | 0.881052049 |
| | 0.370739595 | 0.533999627 |
| | 1.75E-06 | 1.87E-05 |
| | 0.008609763 | 0.030441115 |
| NA | | NA |
| | 0.165987081 | 0.304847569 |

| | |
|---|---|
| 0.641238093 | 0.764705099 |
| 0.721298992 | NA |
| 4.00E-05 | 0.000311061 |
| 0.262678734 | 0.421625142 |
| 0.628560472 | 0.755481581 |
| NA | NA |
| 0.000270247 | 0.001661263 |
| NA | NA |
| NA | NA |
| NA | NA |
| 0.173601504 | 0.314297026 |
| 0.107275539 | 0.221013796 |
| NA | NA |
| 0.784161164 | NA |
| 0.172929674 | 0.313271336 |
| 0.674715461 | 0.79037243 |
| 0.023456869 | 0.068146026 |
| 0.522001535 | 0.672090432 |
| NA | NA |
| NA | NA |
| 0.823705906 | 0.892879496 |
| NA | NA |
| NA | NA |
| NA | NA |
| 0.00476868 | 0.018788963 |
| 0.126480883 | 0.249730235 |
| 0.310544675 | 0.472396264 |
| 0.99174373 | 0.994333794 |
| 0.210723061 | 0.36075057 |
| 0.317840873 | 0.480276311 |
| 0.729614795 | 0.82965275 |
| NA | NA |
| 0.500993148 | 0.653548115 |
| 0.171143339 | 0.311219631 |
| 0.955792398 | NA |
| NA | NA |
| 1.12E-14 | 3.27E-13 |
| 0.179990609 | 0.321993412 |
| 0.952611725 | NA |
| NA | NA |
| 0.004519205 | 0.017954617 |
| 0.168731984 | 0.308294978 |
| 0.002094962 | 0.009505203 |
| 0.035708572 | 0.095241532 |
| 0.92635534 | 0.955706723 |
| NA | NA |
| 0.053272451 | 0.130413907 |

| | |
|---|---|
| 0.150978762 | 0.284663853 |
| 0.045835536 | 0.115692198 |
| 0.018964985 | 0.057496442 |
| 0.00398837 | 0.016169506 |
| 3.59E-06 | 3.61E-05 |
| 5.82E-05 | 0.000433734 |
| 0.004563003 | 0.0180954 |
| 0.040323153 | 0.104837712 |
| 0.569968322 | 0.71082719 |
| 0.093025207 | 0.198800088 |
| 9.45E-06 | 8.61E-05 |
| 7.28E-10 | 1.32E-08 |
| 0.252736063 | 0.410566733 |
| 0.607269599 | NA |
| 2.98E-07 | 3.68E-06 |
| 0.107478341 | 0.221201858 |
| 0.036798468 | 0.097460458 |
| 0.943520468 | 0.965963017 |
| 0.000816255 | 0.004302146 |
| 8.57E-15 | 2.54E-13 |
| 0.810924482 | 0.884219366 |
| 0.985991465 | NA |
| 0.801903069 | NA |
| 0.941278332 | NA |
| 0.941277857 | NA |
| 0.99337258 | NA |
| 0.127034324 | 0.250573546 |
| 0.006992587 | 0.025649183 |
| 0.006060917 | 0.022844331 |
| 0.015004476 | 0.047720007 |
| 0.839319108 | NA |
| 0.013834507 | 0.0447039 |
| 0.901177113 | NA |
| 0.941278838 | NA |
| 4.98E-15 | 1.51E-13 |
| NA | NA |
| 2.26E-05 | 0.000188095 |
| 0.099835723 | 0.209346302 |
| NA | NA |
| 1.89E-10 | 3.69E-09 |
| NA | NA |
| NA | NA |
| NA | NA |
| NA | NA |
| NA | NA |
| 0.586980103 | 0.723508833 |
| NA | NA |

| | | |
|---|---|---|
| NA | 0.653589117 | NA |
| NA | NA | |
| | 0.608126087 | 0.740539236 |
| | 0.001914275 | 0.00882502 |
| NA | NA | |
| | 0.839319108 | NA |
| | 0.213604669 | 0.364433971 |
| | 0.017580075 | 0.054088279 |
| | 0.4361606 | 0.597255471 |
| | 0.000433285 | 0.002504385 |
| | 0.389573665 | 0.552106721 |
| | 0.955792873 | NA |
| | 0.839319108 | NA |
| | 0.955792398 | NA |
| | 0.150971536 | 0.284663853 |
| | 0.895448395 | NA |
| | 0.188117058 | NA |
| | 1.49E-14 | 4.33E-13 |
| NA | NA | |
| | 0.955792873 | NA |
| | 8.95E-13 | 2.26E-11 |
| | 0.923558813 | NA |
| | 0.955792398 | NA |
| | 0.850721696 | 0.910458133 |
| | 1.30E-06 | 1.42E-05 |
| NA | NA | |
| NA | NA | |
| NA | NA | |
| | 0.377589961 | 0.540530237 |
| | 2.61E-09 | 4.41E-08 |
| NA | NA | |
| | 0.702716122 | 0.810252828 |
| NA | NA | |
| NA | NA | |
| NA | NA | |
| NA | NA | |
| NA | NA | |
| NA | NA | |
| | 0.409333908 | 0.571401424 |
| NA | NA | |
| NA | NA | |
| NA | NA | |
| | 2.46E-12 | 5.94E-11 |
| | 0.383014131 | 0.545734951 |
| | 8.37E-06 | 7.72E-05 |
| | 0.138389714 | 0.266836243 |
| NA | NA | |

| | |
|---|---|
| 0.010690709 | 0.036341082 |
| 0.440095423 | 0.600880586 |
| 0.000224783 | 0.0014188 |
| 0.003051267 | 0.013003805 |
| 0.780043219 | 0.864527319 |
| 0.015977564 | 0.05018448 |
| 0.941275956 | NA |
| 0.877758616 | 0.926663806 |
| 0.420244121 | 0.5818511 |
| 0.157952943 | 0.293894547 |
| 0.239428073 | NA |
| 0.955792398 | NA |
| 0.000128656 | 0.000875187 |
| 0.920236531 | NA |
| NA | NA |
| 0.003792682 | 0.015518895 |
| 0.005218961 | 0.020211693 |
| 0.90674772 | 0.943575798 |
| 0.932197423 | 0.959446643 |
| 0.248581058 | 0.405783384 |
| 0.872748495 | 0.92407294 |
| 0.600850951 | NA |
| 0.142989651 | 0.27341257 |
| 0.107441598 | 0.221201858 |
| 7.57E-05 | 0.00054746 |
| NA | NA |
| 0.839318641 | NA |
| NA | NA |
| 0.457221098 | 0.616937447 |
| 0.113586604 | 0.230570276 |
| 0.001966408 | 0.009039096 |
| 0.001156976 | 0.005779293 |
| NA | NA |
| 0.098804488 | 0.207674038 |
| 0.000259627 | 0.001603854 |
| 0.006150527 | 0.023105248 |
| NA | NA |
| NA | NA |
| NA | NA |
| NA | NA |
| 0.941278332 | NA |
| NA | NA |
| NA | NA |
| NA | NA |
| NA | NA |
| NA | NA |
| 0.782629377 | NA |

| | |
|---|---|
| 0.839320041 | NA |
| NA | NA |
| NA | NA |
| 0.941278332 | NA |
| NA | NA |
| NA | NA |
| 0.941277857 | NA |
| NA | NA |
| NA | NA |
| 0.761470755 | NA |
| NA | NA |
| NA | NA |
| NA | NA |
| NA | NA |
| 0.955792873 | NA |
| 0.941278332 | NA |
| 0.839319108 | NA |
| 0.955792873 | NA |
| 0.052771906 | 0.129412416 |
| 0.941277857 | NA |
| 0.938961853 | 0.963601145 |
| 0.363803323 | 0.527434705 |
| 0.791422408 | 0.871750204 |
| 0.866981407 | 0.920524624 |
| 0.001663538 | 0.007836046 |
| 0.885755859 | NA |
| 0.046463126 | 0.116866668 |
| 0.318747227 | 0.481152762 |
| 0.853596091 | NA |
| NA | NA |
| NA | NA |
| NA | NA |
| NA | NA |
| NA | NA |
| 0.609758473 | 0.741537274 |
| NA | NA |
| NA | NA |
| 0.431540789 | 0.592931833 |
| 1.70E-09 | 2.93E-08 |
| 0.103508427 | 0.215189762 |
| NA | NA |
| 0.839318641 | NA |
| 9.41E-16 | 3.03E-14 |
| 0.023913931 | 0.069212807 |
| 0.474742219 | 0.632190321 |
| 0.364495061 | 0.527880221 |
| 3.44E-11 | 7.33E-10 |

| | |
|---|---|
| 0.01523109 | 0.048279451 |
| 0.028991855 | 0.080729292 |
| 0.729473462 | NA |
| 0.699566997 | 0.808148595 |
| 0.194123296 | 0.340194574 |
| NA | NA |
| NA | NA |
| 0.955788948 | NA |
| 0.159673883 | 0.296270484 |
| 0.169876835 | 0.309467427 |
| 0.839318641 | NA |
| NA | NA |
| 0.574756967 | 0.713969278 |
| 0.157514344 | 0.293292374 |
| NA | NA |
| 0.11291171 | 0.229545997 |
| NA | NA |
| NA | NA |
| 0.941278332 | NA |
| 0.939787392 | NA |
| 5.01E-10 | 9.27E-09 |
| 0.784634927 | NA |
| NA | NA |
| 0.09529384 | 0.202353821 |
| NA | NA |
| 0.045693809 | 0.115371173 |
| NA | NA |
| 0.292201025 | 0.453357998 |
| 0.341155746 | 0.504641667 |
| NA | NA |
| 0.643380396 | NA |
| 0.628361553 | NA |
| 5.04E-11 | 1.05E-09 |
| NA | NA |
| 0.236058509 | 0.391286321 |
| NA | NA |
| 0.941275481 | NA |
| 9.11E-25 | 5.10E-23 |
| NA | NA |
| NA | NA |
| NA | NA |
| NA | NA |
| 0.000205963 | 0.00131571 |
| 0.06748786 | 0.156716686 |
| 0.839319108 | NA |
| 0.941277857 | NA |
| 3.41E-22 | 1.66E-20 |

| | | |
|---|---|---|
| | 0.406795049 | 0.568590977 |
| | 0.723368839 | NA |
| | 0.249421052 | 0.406875462 |
| | 0.029612219 | 0.082128877 |
| | 0.62960689 | 0.756176365 |
| | 0.03172307 | 0.086688194 |
| | 0.000515264 | 0.002911062 |
| | 0.885264869 | 0.931359371 |
| | 0.834976807 | 0.899970585 |
| | 0.602415714 | 0.735846188 |
| | 0.763809417 | 0.853490059 |
| | 0.955792398 | NA |
| | 0.641459596 | 0.764873405 |
| | 0.373042644 | 0.536245985 |
| | 0.930552689 | 0.958607575 |
| | 0.495874267 | NA |
| | 0.639304142 | 0.763125235 |
| | 0.127241442 | 0.250815802 |
| | 0.598879167 | 0.733026998 |
| | 0.642631011 | NA |
| | 0.941278332 | NA |
| NA | | NA |
| | 0.598997116 | 0.733062335 |
| | 0.000196042 | 0.001258409 |
| NA | | NA |
| | 0.264221347 | 0.423187975 |
| NA | | NA |
| | 0.894347231 | 0.936363116 |
| | 0.750107491 | NA |
| NA | | NA |
| NA | | NA |
| NA | | NA |
| NA | | NA |
| NA | | NA |
| NA | | NA |
| | 0.147053233 | 0.279297881 |
| | 0.473051832 | NA |
| NA | | NA |
| | 0.853596091 | NA |
| NA | | NA |
| NA | | NA |
| | 0.91776262 | NA |
| | 3.39E-10 | 6.40E-09 |
| NA | | NA |
| | 0.001248528 | 0.006157781 |
| | 0.839318641 | NA |
| | 3.53E-05 | 0.000277821 |

| | | |
|---|---|---|
| | 0.156308325 | NA |
| | 0.871584857 | 0.923330406 |
| | 0.854789616 | 0.91261431 |
| | 0.096597129 | 0.204300535 |
| NA | | NA |
| | 2.15E-15 | 6.75E-14 |
| | 0.782676663 | 0.866267226 |
| | 1.21E-09 | 2.13E-08 |
| | 0.032891779 | 0.089289475 |
| NA | | NA |
| | 0.97376774 | 0.984145885 |
| | 2.24E-06 | 2.35E-05 |
| | 0.179844439 | 0.321829738 |
| | 0.601427292 | 0.735091527 |
| | 0.071199885 | 0.163151472 |
| | 0.173097988 | 0.313552383 |
| | 0.4441259 | 0.604683942 |
| NA | | NA |
| | 0.839319574 | NA |
| | 0.262249989 | 0.421107349 |
| NA | | NA |
| | 9.34E-08 | 1.26E-06 |
| NA | | NA |
| | 0.047257848 | 0.118376563 |
| | 0.001662519 | 0.007832796 |
| | 0.955792398 | NA |
| | 0.955146572 | NA |
| NA | | NA |
| NA | | NA |
| NA | | NA |
| NA | | NA |
| NA | | NA |
| NA | | NA |
| | 0.941279303 | NA |
| NA | | NA |
| NA | | NA |
| NA | | NA |
| NA | | NA |
| NA | | NA |
| | 0.941277857 | NA |
| NA | | NA |
| NA | | NA |
| NA | | NA |
| NA | | NA |
| NA | | NA |
| NA | | NA |
| NA | | NA |

| | | |
|---|---|---|
| NA | NA | |
| NA | NA | |
| NA | NA | |
| NA | NA | |
| NA | NA | |
| NA | NA | |
| NA | NA | |
| NA | NA | |
| NA | NA | |
| NA | NA | |
| NA | NA | |
| NA | NA | |
| NA | NA | |
| NA | NA | |
| NA | NA | |
| NA | NA | |
| NA | NA | |
| NA | NA | |
| NA | NA | |
| 0.941278332 | NA | |
| NA | NA | |
| 0.941275956 | NA | |
| 0.839318641 | NA | |
| 0.470483159 | | 0.628208244 |
| 0.242601213 | | 0.399004566 |
| 0.690838319 | | 0.801737118 |
| 0.801495326 | | 0.878647372 |
| 0.853290739 | NA | |
| 0.941275956 | NA | |
| 0.273460788 | | 0.433293424 |
| 0.292081035 | | 0.453290265 |
| 6.82E-05 | | 0.000498239 |
| 0.443064724 | | 0.603618944 |
| 0.000606945 | | 0.003345117 |
| 0.13561782 | | 0.262885849 |
| 0.287334558 | | 0.447909958 |
| 0.895438704 | | 0.936820599 |
| NA | NA | |
| 0.839318641 | NA | |
| 0.591007616 | NA | |
| 0.563324819 | | 0.705660615 |
| 0.016544627 | | 0.051597905 |
| 0.995993632 | | 0.996917253 |
| 0.508296872 | | 0.659929206 |
| 0.230779186 | | 0.384664299 |
| 0.189149957 | | 0.333764347 |
| 0.748393366 | NA | |

| | |
|---|---|
| 0.874618418 | 0.925009245 |
| NA | NA |
| NA | NA |
| 1.30E-16 | 4.44E-15 |
| NA | NA |
| 1.11E-11 | 2.49E-10 |
| 0.917246314 | 0.950263189 |
| 0.998621397 | 0.998831218 |
| 0.58866046 | NA |
| NA | NA |
| 0.790545158 | 0.87134363 |
| 0.389326772 | 0.551909739 |
| 0.004805027 | 0.018900323 |
| 0.000371626 | 0.002188922 |
| 0.007371145 | 0.026818709 |
| 0.655046828 | 0.774844677 |
| 0.654443697 | 0.77439894 |
| 0.013617308 | 0.044145895 |
| 0.51705499 | 0.667911802 |
| 0.905338864 | 0.942563015 |
| 0.001028233 | 0.005232842 |
| 2.11E-11 | 4.63E-10 |
| 0.799158518 | NA |
| 0.85434824 | NA |
| 0.880365302 | 0.928322255 |
| NA | NA |
| NA | NA |
| 9.95E-05 | 0.000697907 |
| 0.700267162 | 0.808638955 |
| 0.001591413 | 0.007544077 |
| 0.057002743 | 0.137437124 |
| 0.037141228 | 0.098160359 |
| NA | NA |
| 0.903980219 | NA |
| 0.502389983 | NA |
| 0.008573851 | 0.030326232 |
| 0.414377527 | 0.576380135 |
| 0.094439043 | 0.200986956 |
| 4.18E-19 | 1.67E-17 |
| 0.010322447 | 0.035316213 |
| NA | NA |
| NA | NA |
| NA | NA |
| 0.941278332 | NA |
| 0.506258844 | 0.658001038 |
| 0.106061192 | 0.219175942 |
| 0.455744505 | 0.615398577 |

| | |
|---|---|
| NA | NA |
| 6.69E-05 | 0.000490695 |
| 0.951073023 | 0.970148286 |
| 0.000595612 | 0.003291562 |
| 2.96E-11 | 6.35E-10 |
| 9.49E-09 | 1.50E-07 |
| 0.010550904 | 0.035951081 |
| 0.770638186 | 0.858059133 |
| 0.034832803 | 0.093281996 |
| 0.2712247 | 0.43109785 |
| 0.714565978 | 0.818562875 |
| 0.966976561 | 0.980279233 |
| 0.955792873 | NA |
| 0.000446651 | 0.002574989 |
| 0.001307592 | 0.006393446 |
| 0.530960429 | 0.679713878 |
| 0.006017295 | 0.022715886 |
| 0.211644365 | 0.361997641 |
| 0.000132264 | 0.000896906 |
| NA | NA |
| 0.920689011 | NA |
| 0.029723191 | 0.082359882 |
| 0.00020657 | 0.001317819 |
| 0.006450242 | 0.024007608 |
| 0.941278332 | NA |
| 0.955791922 | NA |
| 0.233218372 | NA |
| NA | NA |
| 0.046391913 | 0.116774568 |
| 0.200596529 | 0.348231243 |
| NA | NA |
| 0.72037966 | 0.822294105 |
| 0.312308779 | 0.474291683 |
| 0.349611696 | NA |
| 0.34767385 | 0.511644181 |
| 1.94E-05 | 0.000163614 |
| 0.955792398 | NA |
| 0.124377693 | 0.246785415 |
| 0.07653882 | 0.172011802 |
| 0.246281954 | 0.403247099 |
| 0.531052338 | 0.679758416 |
| 0.918921406 | NA |
| 0.001300323 | 0.006367866 |
| 0.06955726 | 0.160473189 |
| 0.103237839 | 0.214777971 |
| 6.94E-94 | 2.20E-91 |
| 5.59E-05 | 0.000418263 |

| | |
|---|---|
| 0.036754904 | 0.097399269 |
| NA | NA |
| 8.17E-138 | 4.05E-135 |
| 0.410011193 | NA |
| 0.130394254 | 0.25531787 |
| 6.98E-11 | 1.43E-09 |
| 1.43E-05 | 0.000125202 |
| 0.955792873 | NA |
| 0.908365267 | 0.944452264 |
| 0.247795015 | 0.405000133 |
| 3.16E-05 | 0.000252513 |
| 0.087100522 | 0.189520672 |
| NA | NA |
| NA | NA |
| 0.488854721 | NA |
| 0.570858251 | 0.711393348 |
| 0.060254635 | 0.143504185 |
| 0.001686591 | 0.007930511 |
| 0.05963356 | 0.142381181 |
| 0.055771067 | 0.135151999 |
| 0.479708997 | NA |
| NA | NA |
| 0.021827812 | 0.064228655 |
| 0.270468742 | 0.430536601 |
| 0.941854745 | NA |
| 0.250756988 | 0.40840834 |
| 0.610141158 | 0.741788737 |
| 0.941277857 | NA |
| 0.90570436 | 0.942778588 |
| 0.053071555 | NA |
| 0.96152524 | 0.976583049 |
| 0.853596091 | NA |
| NA | NA |
| 0.924771793 | 0.95491588 |
| 7.59E-06 | 7.08E-05 |
| 2.71E-05 | 0.000220437 |
| 0.801927359 | 0.878999539 |
| 0.000265042 | 0.001632644 |
| 0.112097458 | 0.228163477 |
| 0.708016206 | 0.813747658 |
| 0.001176233 | 0.005860729 |
| 0.063054975 | 0.148685804 |
| 0.293560688 | 0.454763099 |
| 0.003982275 | 0.0161503 |
| 0.848903005 | 0.909153602 |
| 0.65131295 | NA |
| 0.542487188 | 0.68931668 |

| | | |
|---|---|---|
| | 0.258674178 | 0.417053633 |
| | 0.018685092 | 0.056836111 |
| | 0.333146275 | 0.496409223 |
| | 0.10382534 | 0.215699827 |
| | 0.051978156 | 0.127847895 |
| | 7.88E-15 | 2.35E-13 |
| | 3.45E-12 | 8.21E-11 |
| | 0.920706464 | 0.952578389 |
| | 0.955792873 | NA |
| NA | | NA |
| | 0.208829757 | 0.358237437 |
| | 0.116409187 | 0.234791244 |
| | 0.503395985 | 0.655497086 |
| | 0.543896911 | 0.690554983 |
| NA | | NA |
| | 0.006785444 | 0.025039866 |
| | 0.839320041 | NA |
| | 0.902279739 | NA |
| | 0.00155541 | 0.007399212 |
| NA | | NA |
| NA | | NA |
| | 0.233475505 | 0.388088566 |
| | 0.072930454 | 0.165970999 |
| | 5.29E-24 | 2.83E-22 |
| | 0.309676117 | 0.471346139 |
| | 0.939591567 | 0.963757575 |
| NA | | NA |
| | 0.02978322 | 0.082497406 |
| | 0.43056991 | 0.591812958 |
| | 3.17E-05 | 0.000253095 |
| | 0.020169811 | 0.060273927 |
| | 0.142550071 | 0.272922843 |
| | 0.376960262 | 0.539833265 |
| | 0.000852436 | 0.004456334 |
| | 0.831350787 | 0.897753108 |
| | 0.650646147 | 0.771787082 |
| | 0.37917532 | NA |
| | 0.003538737 | 0.014689401 |
| NA | | NA |
| | 0.853595154 | NA |
| | 0.955792873 | NA |
| | 0.568006021 | 0.709430128 |
| | 0.036432379 | 0.096684522 |
| | 0.390457315 | 0.553029754 |
| | 0.93976723 | 0.963757575 |
| NA | | NA |
| | 0.941278788 | NA |

| | | |
|---|---|---|
| | 0.007573973 | 0.027405853 |
| | 0.00119883 | 0.005955866 |
| | 0.056383467 | 0.13631684 |
| | 0.001290525 | 0.006328198 |
| | 0.043140373 | 0.110447152 |
| | 0.069311163 | 0.160044875 |
| | 0.447739103 | 0.607987046 |
| | 0.763316643 | 0.853019519 |
| | 0.941279268 | NA |
| | 0.924148601 | 0.954592539 |
| | 0.839321638 | NA |
| NA | | NA |
| | 0.608015819 | NA |
| | 0.341649214 | 0.505061603 |
| | 0.676631207 | 0.791584276 |
| | 0.115378066 | 0.233146169 |
| NA | | NA |
| | 0.359071466 | 0.522601139 |
| | 0.136442451 | 0.26405425 |
| | 0.027320871 | 0.07714014 |
| | 0.006050238 | 0.022816009 |
| | 0.889084175 | 0.933455095 |
| | 0.889401596 | 0.933455095 |
| | 0.000189861 | 0.001226675 |
| | 0.055743379 | 0.135112414 |
| | 0.003926358 | 0.015964326 |
| | 0.865222268 | NA |
| | 0.039686523 | 0.103553894 |
| NA | | NA |
| | 0.000734135 | 0.003934675 |
| | 0.032978336 | 0.089463228 |
| | 0.16605545 | 0.304887983 |
| | 0.05342724 | 0.130629189 |
| NA | | NA |
| | 0.333721263 | 0.497011232 |
| | 0.476298532 | 0.633402091 |
| | 0.001617897 | 0.007657426 |
| | 0.172177313 | 0.312450976 |
| | 0.018974046 | 0.057516587 |
| | 0.42082939 | 0.582325513 |
| | 5.56E-11 | 1.16E-09 |
| | 0.112546303 | 0.228979153 |
| | 5.78E-05 | 0.00043202 |
| NA | | NA |
| | 0.941278332 | NA |
| NA | | NA |
| | 0.895286056 | NA |

| | | |
|---|---|---|
| | 0.898565669 | NA |
| | 0.256785696 | 0.414908231 |
| | 0.839319108 | NA |
| | 0.668867796 | NA |
| | 0.224313455 | 0.37705571 |
| | 0.955792873 | NA |
| NA | | NA |
| NA | | NA |
| | 0.483712227 | 0.639628801 |
| NA | | NA |
| | 0.180013103 | 0.321993412 |
| | 0.014358357 | 0.046077608 |
| | 0.003226326 | 0.013635123 |
| | 0.010786343 | 0.03661388 |
| | 0.695263449 | NA |
| | 0.887587644 | NA |
| NA | | NA |
| NA | | NA |
| | 0.839318641 | NA |
| | 0.24436562 | 0.400880197 |
| | 0.008351058 | 0.029686834 |
| | 0.008440897 | 0.029954919 |
| | 0.719273501 | 0.821385975 |
| NA | | NA |
| | 0.198728842 | 0.346038369 |
| NA | | NA |
| | 0.010967522 | 0.037133566 |
| | 0.045140432 | 0.114313498 |
| | 0.850502193 | NA |
| | 5.36E-10 | 9.88E-09 |
| NA | | NA |
| | 6.34E-06 | 6.03E-05 |
| | 0.70365339 | 0.810844101 |
| NA | | NA |
| | 1.19E-06 | 1.31E-05 |
| | 2.94E-10 | 5.58E-09 |
| | 0.040330355 | 0.104840458 |
| | 1.25E-05 | 0.000110598 |
| | 0.631881107 | 0.75791343 |
| | 0.189960007 | 0.334641705 |
| | 0.358035914 | 0.521604286 |
| NA | | NA |
| | 0.644852306 | 0.767515481 |
| | 0.633520668 | 0.759182999 |
| NA | | NA |
| | 0.48596282 | 0.641322192 |
| | 0.373566993 | 0.536772808 |

| | | |
|---|---|---|
| | 0.562853838 | 0.705330475 |
| | 0.019096187 | 0.05778381 |
| | 0.543523214 | 0.690264622 |
| | 0.038370455 | 0.100771661 |
| | 0.147394875 | NA |
| | 0.001682459 | 0.007915773 |
| | 0.058845262 | 0.140809784 |
| | 0.941277857 | NA |
| NA | | NA |
| | 0.005493007 | 0.021033551 |
| | 0.941278332 | NA |
| | 0.175362236 | 0.316545726 |
| | 0.087343549 | 0.189910584 |
| NA | | NA |
| | 0.2758331 | 0.435658889 |
| | 3.89E-08 | 5.54E-07 |
| | 5.62E-09 | 9.12E-08 |
| | 0.866306409 | NA |
| | 0.481580386 | NA |
| | 0.941277857 | NA |
| | 0.148605594 | NA |
| | 0.800646606 | NA |
| | 0.790939096 | NA |
| NA | | NA |
| NA | | NA |
| | 0.941278332 | NA |
| NA | | NA |
| NA | | NA |
| NA | | NA |
| NA | | NA |
| NA | | NA |
| NA | | NA |
| NA | | NA |
| NA | | NA |
| NA | | NA |
| NA | | NA |
| | 0.839318641 | NA |
| NA | | NA |
| NA | | NA |
| NA | | NA |
| NA | | NA |
| NA | | NA |
| NA | | NA |
| NA | | NA |
| NA | | NA |
| NA | | NA |
| NA | | NA |

| | |
|---|---|
| NA | NA |
| NA | NA |
| NA | NA |
| NA | NA |
| NA | NA |
| NA | NA |
| NA | NA |
| NA | NA |
| NA | NA |
| NA | NA |
| NA | NA |
| NA | NA |
| NA | NA |
| 0.764982398 | NA |
| 0.576796754 | 0.715383061 |
| 0.835025174 | NA |
| NA | NA |
| NA | NA |
| NA | NA |
| 0.941278788 | NA |
| NA | NA |
| NA | NA |
| 0.955791446 | NA |
| 0.937796074 | NA |
| NA | NA |
| 0.054864783 | 0.133390354 |
| NA | NA |
| NA | NA |
| 0.021291602 | 0.062911162 |
| NA | NA |
| NA | NA |
| 0.853595622 | NA |
| 0.58965087 | NA |
| 0.941278332 | NA |
| NA | NA |
| NA | NA |
| NA | NA |
| NA | NA |
| NA | NA |
| NA | NA |
| 0.839319574 | NA |
| NA | NA |
| 0.251519045 | 0.409227276 |
| NA | NA |
| 0.941277857 | NA |
| NA | NA |
| 0.365491411 | 0.52869382 |

| | | |
|---|---|---|
| | 0.110096988 | 0.225286151 |
| | 0.010371965 | 0.035464165 |
| | 0.938317216 | 0.963370848 |
| | 0.924900956 | 0.95495131 |
| | 0.015285734 | 0.048426867 |
| | 0.255158551 | 0.413289135 |
| | 0.785746152 | 0.868300752 |
| | 0.013411742 | 0.043669805 |
| | 0.149911987 | 0.283070521 |
| | 0.182946152 | 0.325565658 |
| | 0.02554225 | 0.072879458 |
| | 0.493389634 | 0.646989056 |
| NA | | NA |
| | 0.941278332 | NA |
| | 0.453896807 | 0.613717579 |
| | 0.254750849 | 0.412742327 |
| | 0.458083674 | 0.617733931 |
| | 0.261889562 | 0.420698884 |
| NA | | NA |
| | 0.58676518 | 0.723374155 |
| NA | | NA |
| | 0.941277857 | NA |
| | 0.633003562 | 0.758800905 |
| | 0.013174845 | 0.043016936 |
| | 0.062628504 | 0.147885283 |
| | 0.632434833 | 0.758414336 |
| | 0.161170738 | 0.298101266 |
| | 0.644940217 | NA |
| NA | | NA |
| | 0.894406549 | NA |
| NA | | NA |
| | 0.895241625 | 0.936820599 |
| | 0.003021778 | 0.012910494 |
| | 2.96E-21 | 1.37E-19 |
| NA | | NA |
| | 0.839319108 | NA |
| NA | | NA |
| NA | | NA |
| NA | | NA |
| NA | | NA |
| NA | | NA |
| NA | | NA |
| NA | | NA |
| | 0.868358141 | NA |
| | 0.409458965 | 0.57144194 |
| NA | | NA |
| NA | | NA |

| | | |
|---|---|---|
| NA | | NA |
| NA | | NA |
| NA | | NA |
| | 0.661496085 | NA |
| | 0.721405302 | NA |
| | 0.243148358 | 0.399600719 |
| | 0.406132082 | 0.568033962 |
| | 0.009752384 | 0.033729475 |
| | 0.496595261 | NA |
| | 8.20E-10 | 1.47E-08 |
| | 0.324501999 | NA |
| | 0.066399058 | 0.15492534 |
| | 0.907039534 | 0.943641717 |
| | 7.26E-07 | 8.29E-06 |
| | 0.873495974 | NA |
| | 0.927040016 | 0.9562473 |
| | 0.619222136 | NA |
| | 0.606486227 | 0.739111421 |
| | 0.122090649 | 0.243523139 |
| | 0.967967834 | 0.980741119 |
| | 2.10E-47 | 2.73E-45 |
| | 0.892627295 | 0.935102195 |
| | 0.923256181 | NA |
| | 0.036122461 | 0.096101037 |
| | 0.000868704 | 0.004531427 |
| | 0.544381033 | 0.690985353 |
| | 0.126330311 | 0.249619299 |
| | 0.786333773 | 0.868667802 |
| | 0.716408829 | 0.819759372 |
| | 0.375281313 | 0.538391044 |
| | 0.114902796 | 0.232487119 |
| | 0.941278788 | NA |
| NA | | NA |
| NA | | NA |
| NA | | NA |
| NA | | NA |
| NA | | NA |
| NA | | NA |
| | 0.046454191 | 0.116866668 |
| | 0.059010299 | 0.141119576 |
| NA | | NA |
| NA | | NA |
| NA | | NA |
| | 0.924780815 | NA |
| | 0.068177819 | 0.158013689 |
| | 3.24E-05 | 0.000257737 |
| | 0.523898858 | 0.673572118 |

| | | |
|---|---|---|
| NA | | NA |
| | 0.522012261 | 0.672090432 |
| | 0.06377635 | 0.150086578 |
| | 0.814848042 | 0.886831874 |
| | 0.374677945 | 0.537784746 |
| | 0.732386214 | 0.831176115 |
| | 0.941276431 | NA |
| | 0.941278332 | NA |
| | 0.637505676 | 0.761744398 |
| NA | | NA |
| | 0.000100248 | 0.000703037 |
| NA | | NA |
| | 2.95E-07 | 3.64E-06 |
| | 0.441009633 | 0.601714604 |
| | 0.035497496 | 0.094763503 |
| | 0.001673519 | 0.007875266 |
| | 0.451382035 | 0.611346033 |
| | 0.027053504 | 0.076530484 |
| | 0.850144914 | 0.910141082 |
| | 0.171773219 | 0.311912279 |
| | 8.01E-15 | 2.38E-13 |
| | 0.620952107 | 0.749468748 |
| | 0.037127785 | 0.098146996 |
| | 0.587271041 | 0.723698747 |
| NA | | NA |
| | 0.087336406 | 0.189910584 |
| NA | | NA |
| | 0.837175376 | 0.901225435 |
| | 0.627894891 | 0.755021328 |
| | 0.815021277 | 0.886939303 |
| | 0.445695379 | 0.606162014 |
| | 0.636897475 | 0.761245039 |
| | 7.98E-05 | 0.00057416 |
| | 0.194260972 | 0.340360666 |
| | 0.015361331 | 0.0486146 |
| | 0.052829039 | 0.129525834 |
| | 0.019782482 | 0.059347445 |
| | 0.000591267 | 0.00326983 |
| | 0.955792398 | NA |
| | 0.126821349 | 0.250236405 |
| NA | | NA |
| | 0.087569183 | 0.190300545 |
| | 0.989863222 | 0.993351574 |
| | 3.85E-08 | 5.49E-07 |
| | 0.428932045 | 0.590243426 |
| | 0.14233912 | NA |
| | 0.002740963 | 0.011907355 |

| | | |
|---|---|---|
| | 0.091362557 | 0.19635319 |
| | 0.594162716 | NA |
| | 0.00185614 | 0.008588617 |
| | 0.008099892 | 0.028952339 |
| | 0.316210783 | 0.478683643 |
| | 0.806938299 | 0.88170566 |
| | 2.22E-07 | 2.81E-06 |
| | 0.941278332 | NA |
| | 2.60E-06 | 2.68E-05 |
| | 0.36325847 | 0.526910083 |
| | 0.00058602 | 0.003247384 |
| | 0.566810234 | 0.708535119 |
| | 0.020976523 | 0.062138544 |
| | 0.053530512 | 0.130787646 |
| | 0.012344774 | 0.040820586 |
| | 0.578974163 | 0.717037464 |
| | 0.240316583 | 0.396315063 |
| | 0.869256252 | 0.921646279 |
| | 0.357060242 | 0.5206928 |
| | 0.00075659 | 0.004033226 |
| | 0.050193015 | 0.124136964 |
| NA | | NA |
| | 0.000882154 | 0.004586505 |
| | 0.941277857 | NA |
| NA | | NA |
| NA | | NA |
| NA | | NA |
| | 0.772940871 | 0.859898977 |
| | 0.485534407 | 0.640898955 |
| | 0.660471271 | 0.779363303 |
| | 0.955792398 | NA |
| | 0.108922037 | 0.223477487 |
| | 0.01876738 | 0.057041261 |
| | 0.449665967 | 0.609646238 |
| | 0.001468828 | 0.007045756 |
| | 0.0234031 | 0.068031336 |
| | 0.628502169 | 0.755481581 |
| NA | | NA |
| | 0.317534977 | 0.480084325 |
| | 0.857646416 | 0.914797222 |
| | 0.131086266 | 0.256398662 |
| | 0.088985455 | 0.192618388 |
| NA | | NA |
| | 1.30E-06 | 1.42E-05 |
| | 0.264776506 | 0.423791971 |
| | 0.000124787 | 0.000851542 |
| NA | | NA |

| | |
|---|---|
| 0.220941779 | 0.37286275 |
| 0.005900069 | 0.022344222 |
| 0.049150221 | 0.122027075 |
| 0.048239253 | 0.120368036 |
| 0.102943928 | 0.214371949 |
| 0.341789161 | 0.505170493 |
| 0.235716064 | 0.390922085 |
| 0.226066686 | 0.379092523 |
| 0.634950636 | 0.760020207 |
| 0.000731077 | 0.003923582 |
| NA | NA |
| 0.135733935 | 0.263046663 |
| 0.033793445 | 0.091158231 |
| 0.613796013 | 0.744153273 |
| 0.000361424 | 0.002137828 |
| 0.057438056 | 0.138206694 |
| 0.899280055 | 0.939460317 |
| 0.718818345 | 0.821055414 |
| 0.640629492 | 0.764207288 |
| 0.106949109 | 0.220570375 |
| 0.355007813 | 0.518812303 |
| 0.001062529 | 0.005374058 |
| 0.333119936 | 0.496409223 |
| 0.000328807 | 0.001974336 |
| 0.815962245 | 0.887480947 |
| 0.002601174 | 0.011408356 |
| 0.03059746 | 0.08424771 |
| 0.661753735 | 0.78014028 |
| 0.474092888 | 0.631540123 |
| 0.378494311 | 0.541303946 |
| 0.351452804 | 0.515109595 |
| 0.273610369 | 0.433416256 |
| 0.955791446 | NA |
| 0.011967446 | 0.039800078 |
| 0.030600799 | 0.08424771 |
| 0.265837096 | 0.425203586 |
| 0.176636171 | 0.318266022 |
| 0.1550843 | 0.290038227 |
| 0.329622544 | 0.49258328 |
| 0.839320041 | NA |
| 9.47E-05 | 0.000668871 |
| 0.817176631 | 0.888457846 |
| 0.416412534 | NA |
| 0.460098437 | 0.619114165 |
| 0.442480366 | 0.602995401 |
| NA | NA |
| NA | NA |

| | |
|---|---|
| 0.011179793 | 0.037696759 |
| 0.000248545 | 0.00154461 |
| 0.380541636 | 0.543186333 |
| NA | NA |
| 0.003413255 | 0.014259272 |
| 0.643492597 | NA |
| 0.510066649 | 0.661684584 |
| NA | NA |
| 0.525797423 | 0.675044348 |
| 0.941277857 | NA |
| 0.574209337 | 0.713698727 |
| 0.645085969 | 0.767640048 |
| NA | NA |
| NA | NA |
| 0.101207153 | 0.211644056 |
| 6.59E-20 | 2.78E-18 |
| 2.85E-07 | 3.52E-06 |
| NA | NA |
| NA | NA |
| 0.986845644 | 0.991762372 |
| 0.001785314 | 0.008320745 |
| 0.564341013 | 0.706375935 |
| 0.941278332 | NA |
| 0.839318641 | NA |
| NA | NA |
| 0.252428965 | 0.410263859 |
| NA | NA |
| NA | NA |
| 9.62E-05 | 0.000677564 |
| NA | NA |
| 0.703339397 | 0.810735838 |
| 5.01E-05 | 0.000379459 |
| 0.143763455 | 0.274583354 |
| 0.867938234 | 0.921060969 |
| 0.01566837 | 0.049395832 |
| 0.649465216 | 0.770749546 |
| 0.000886103 | 0.004605026 |
| 0.00172451 | 0.008089633 |
| 0.922686566 | NA |
| 8.33E-07 | 9.41E-06 |
| 0.419487145 | NA |
| 3.79E-08 | 5.42E-07 |
| 0.231619793 | 0.385795264 |
| 0.19274186 | 0.33837157 |
| 0.000600631 | 0.00331545 |
| NA | NA |
| 0.885029945 | 0.931219182 |

| | |
|---|---|
| 0.914054361 | 0.948235787 |
| 0.801606314 | 0.878728572 |
| 0.089098095 | 0.192686977 |
| 0.002763486 | 0.011987698 |
| 4.90E-19 | 1.96E-17 |
| 0.00091751 | 0.004740821 |
| 0.014353252 | 0.046077608 |
| 3.44E-06 | 3.47E-05 |
| 0.000209441 | 0.001334702 |
| 0.003590497 | 0.014852451 |
| 1.79E-15 | 5.64E-14 |
| NA | NA |
| 0.209561747 | 0.359199838 |
| 0.065085158 | 0.152525575 |
| NA | NA |
| 0.48983611 | NA |
| NA | NA |
| 0.941278788 | NA |
| NA | NA |
| 0.054170796 | 0.132053799 |
| 2.70E-07 | 3.36E-06 |
| 0.059640535 | 0.142383553 |
| 0.6710142 | 0.787121384 |
| 0.945031075 | 0.966914213 |
| 0.70813986 | 0.813813099 |
| 0.776403287 | NA |
| 0.184157315 | 0.327112866 |
| 2.06E-07 | 2.62E-06 |
| 0.672719997 | NA |
| 0.941278332 | NA |
| 0.955792398 | NA |
| 0.000178598 | 0.001163913 |
| 0.009481767 | 0.032957296 |
| 0.842899965 | 0.90539758 |
| 0.652841662 | NA |
| 0.941277857 | NA |
| 0.030852257 | 0.084807186 |
| 0.010770956 | 0.036566867 |
| 0.955792398 | NA |
| 3.87E-14 | 1.10E-12 |
| 0.943209995 | NA |
| 0.124313458 | 0.246740237 |
| 0.109225297 | 0.223964552 |
| 0.130373798 | 0.25531787 |
| 0.62489624 | 0.752858299 |
| 4.86E-05 | 0.000370106 |
| 0.864121003 | 0.918985215 |

| | |
|---|---|
| 0.384981651 | 0.54778415 |
| 0.002623332 | 0.011488603 |
| 0.0325819 | 0.088650478 |
| 0.004315243 | 0.017265324 |
| 8.15E-14 | 2.24E-12 |
| NA | NA |
| 0.854128083 | NA |
| 0.574359935 | 0.713786167 |
| NA | NA |
| NA | NA |
| 0.727483389 | NA |
| 0.369507296 | 0.532772756 |
| 0.027916425 | 0.078393907 |
| 9.00E-07 | 1.01E-05 |
| 3.62E-12 | 8.59E-11 |
| 7.67E-07 | 8.72E-06 |
| 0.003467667 | 0.014444768 |
| NA | NA |
| 0.000769108 | 0.004090795 |
| 0.000105345 | 0.000732307 |
| NA | NA |
| 0.941277857 | NA |
| 0.032994944 | 0.08949808 |
| 0.452461206 | NA |
| 0.036226995 | 0.096293768 |
| NA | NA |
| 0.598874978 | 0.733026998 |
| 0.69024427 | 0.801297308 |
| NA | NA |
| 2.50E-05 | 0.000205688 |
| 0.461491948 | 0.620320853 |
| 0.625718289 | 0.753436846 |
| NA | NA |
| 0.72411748 | NA |
| 0.479047927 | NA |
| 0.019151412 | 0.057928822 |
| 5.33E-06 | 5.16E-05 |
| 0.500162176 | 0.652929911 |
| 0.012246586 | 0.04055276 |
| 0.322563675 | 0.485405613 |
| 0.766648079 | 0.855078721 |
| 0.216792744 | 0.368189616 |
| 0.459350844 | 0.618621801 |
| 0.853596091 | NA |
| 0.002305844 | 0.010320363 |
| 0.941275956 | NA |
| 0.51539977 | 0.666460875 |

| | |
|---|---|
| 0.002954649 | 0.012662326 |
| 0.456951286 | NA |
| 0.840598341 | 0.903781811 |
| 0.716437042 | 0.819759372 |
| NA | NA |
| NA | NA |
| 0.037145213 | 0.098160359 |
| 0.106528991 | 0.219989854 |
| 0.067606438 | 0.156930802 |
| 0.000255966 | 0.001583613 |
| 0.753420314 | 0.84541346 |
| 0.291920278 | 0.453188525 |
| 0.006116351 | 0.023005908 |
| 0.334715508 | 0.49796228 |
| 0.068538036 | 0.158598322 |
| 0.39727683 | 0.559856904 |
| 0.870086898 | NA |
| 0.955792398 | NA |
| 8.80E-16 | 2.84E-14 |
| 2.01E-10 | 3.90E-09 |
| NA | NA |
| 0.024759032 | 0.071130294 |
| 0.1903761 | 0.335157688 |
| 0.6145627 | NA |
| 0.839319574 | NA |
| 0.254466066 | 0.412504346 |
| 0.161815304 | 0.298962032 |
| 0.778243253 | 0.863416888 |
| 0.054842874 | 0.133350709 |
| 0.949001926 | 0.968816796 |
| 0.552092249 | 0.697388935 |
| 0.924944782 | NA |
| 2.07E-07 | 2.63E-06 |
| 0.006336421 | 0.023661672 |
| 0.941408783 | 0.964703895 |
| 0.408536545 | 0.570638445 |
| 0.639540791 | 0.76329288 |
| 0.725370641 | NA |
| 0.068853262 | 0.159203938 |
| 0.58754181 | 0.723813843 |
| 0.990167648 | 0.993506886 |
| 0.951159056 | NA |
| 0.001226083 | 0.006074776 |
| 0.013032316 | 0.042632652 |
| 0.839318641 | NA |
| 0.955792398 | NA |
| 0.000645091 | 0.003520049 |

| | |
|---|---|
| 0.139376818 | 0.268186826 |
| 0.875888922 | NA |
| 0.004511328 | 0.017938294 |
| 0.392486222 | 0.554912804 |
| 0.560720836 | 0.703843336 |
| 0.014404567 | 0.046176094 |
| NA | NA |
| NA | NA |
| 0.003789193 | 0.015512619 |
| 2.68E-24 | 1.45E-22 |
| 0.045965092 | 0.115994605 |
| 0.855216 | 0.912900316 |
| 0.336212134 | NA |
| 0.214001477 | 0.364877858 |
| 0.146199096 | 0.278052808 |
| NA | NA |
| 0.608155636 | 0.740539236 |
| 0.074571054 | 0.168720554 |
| 7.11E-11 | 1.46E-09 |
| 0.61217847 | NA |
| 0.000181192 | 0.001178344 |
| 0.002263267 | 0.010148887 |
| 0.374561787 | 0.537715299 |
| 0.181460715 | 0.323845539 |
| 0.941277857 | NA |
| 0.615359583 | 0.745383654 |
| 0.429577734 | 0.590769691 |
| NA | NA |
| 0.013949852 | 0.044996361 |
| 0.117483152 | 0.236396482 |
| 4.17E-34 | 3.44E-32 |
| 3.64E-142 | 1.89E-139 |
| 0.044221587 | 0.112489283 |
| 1.98E-13 | 5.28E-12 |
| 2.75E-11 | 5.94E-10 |
| 0.866210287 | 0.920118511 |
| 0.011101433 | 0.037501605 |
| 0.013500954 | 0.04386428 |
| 0.011121907 | 0.037538802 |
| 0.126383937 | 0.249642362 |
| 0.053008472 | 0.129912237 |
| 6.35E-19 | 2.51E-17 |
| NA | NA |
| 8.11E-245 | 2.41E-241 |
| 0.001148306 | 0.005745237 |
| 0.955792873 | NA |
| 0.955792873 | NA |

| | |
|---|---|
| 0.25423562 | 0.412243084 |
| 0.317045312 | 0.479496283 |
| 0.827139998 | 0.89505302 |
| 0.001250876 | 0.006166808 |
| 0.962869632 | 0.97753137 |
| 2.29E-30 | 1.61E-28 |
| 0.533310506 | NA |
| 0.12113309 | NA |
| NA | NA |
| 2.35E-05 | 0.000194007 |
| 5.71E-10 | 1.05E-08 |
| NA | NA |
| 1.06E-05 | 9.51E-05 |
| 5.33E-07 | 6.25E-06 |
| 2.70E-10 | 5.16E-09 |
| 4.97E-102 | 1.74E-99 |
| 2.10E-269 | 1.67E-265 |
| 6.93E-144 | 3.75E-141 |
| 0.132317205 | NA |
| 5.80E-155 | 3.84E-152 |
| 0.05713447 | 0.137698933 |
| 0.078347261 | 0.175034871 |
| 0.827417822 | NA |
| 5.49E-39 | 5.44E-37 |
| 7.65E-35 | 6.53E-33 |
| 0.583405454 | NA |
| 5.58E-22 | 2.68E-20 |
| 0.473414138 | 0.630918438 |
| 8.35E-32 | 6.29E-30 |
| 0.624843052 | NA |
| 0.001367393 | 0.006640827 |
| NA | NA |
| NA | NA |
| 0.223404061 | 0.375877641 |
| 0.256850094 | NA |
| NA | NA |
| NA | NA |
| NA | NA |
| NA | NA |
| NA | NA |
| 0.095394139 | NA |
| 0.006716536 | 0.024824067 |
| NA | NA |
| 1.25E-18 | 4.87E-17 |
| 0.00022592 | 0.00142371 |
| 0.008433359 | 0.029937482 |
| 0.00336562 | 0.014111059 |

| | |
|---|---|
| 0.941276431 | NA |
| 0.006327663 | 0.023636383 |
| 0.013346579 | 0.043481424 |
| NA | NA |
| 4.90E-06 | 4.79E-05 |
| 0.922220507 | 0.953631614 |
| 0.955792398 | NA |
| 0.003852015 | 0.015731924 |
| 0.67295593 | 0.788854817 |
| 0.009166161 | 0.032056682 |
| 0.955792873 | NA |
| 0.3313268 | 0.494466142 |
| 2.20E-152 | 1.38E-149 |
| 0.892231652 | NA |
| 0.955791446 | NA |
| NA | NA |
| 0.941278332 | NA |
| 0.941278434 | NA |
| 3.75E-27 | 2.32E-25 |
| 0.090933768 | 0.195785214 |
| 0.550701187 | 0.696222959 |
| 0.017580061 | 0.054088279 |
| NA | NA |
| NA | NA |
| 0.040995768 | 0.10624796 |
| 5.48E-24 | 2.93E-22 |
| NA | NA |
| NA | NA |
| 4.32E-33 | 3.44E-31 |
| 0.021558199 | 0.063553165 |
| 0.005094405 | 0.019822956 |
| 0.972315143 | 0.983304088 |
| 0.136921685 | 0.264778419 |
| 6.42E-05 | 0.000474214 |
| NA | NA |
| NA | NA |
| 0.630844315 | NA |
| 0.281444348 | 0.441678537 |
| 0.253945778 | 0.411997642 |
| 0.901054354 | NA |
| 0.013899643 | 0.044883909 |
| 0.006741536 | 0.024904864 |
| 0.347971076 | 0.511828424 |
| 9.26E-08 | 1.25E-06 |
| 0.011054805 | 0.037359999 |
| 0.024753769 | 0.071123757 |
| 0.392251079 | 0.554613294 |

| | |
|---|---|
| NA | NA |
| 5.62E-26 | 3.30E-24 |
| 0.002342063 | 0.010454947 |
| 0.321449123 | 0.484310167 |
| 0.861401243 | NA |
| 0.116480178 | 0.234874018 |
| 0.977190604 | 0.985962343 |
| 0.110750998 | 0.22625485 |
| 2.10E-05 | 0.00017539 |
| 0.839319574 | NA |
| 0.79141541 | 0.871750204 |
| 0.006041353 | 0.022796738 |
| 0.091686468 | 0.196765063 |
| 0.168876148 | 0.30842651 |
| 0.487204296 | 0.642177242 |
| 0.102427652 | NA |
| 0.777355107 | 0.862914199 |
| 0.042359291 | 0.108916047 |
| 0.9290454 | 0.957817759 |
| 0.450299652 | 0.6102273 |
| 8.77E-32 | 6.57E-30 |
| 0.24507433 | NA |
| 9.22E-11 | 1.85E-09 |
| 5.40E-15 | 1.63E-13 |
| 0.900888604 | NA |
| 9.35E-21 | 4.19E-19 |
| 0.000154235 | 0.001026306 |
| NA | NA |
| 2.68E-05 | 0.000218303 |
| 0.892875804 | NA |
| 0.050359126 | 0.124521911 |
| 0.285542395 | 0.446197485 |
| NA | NA |
| 0.674780796 | NA |
| 0.340808069 | 0.504346783 |
| 0.242572656 | 0.398998053 |
| 3.28E-148 | 1.91E-145 |
| 0.416132611 | 0.577875637 |
| 0.270913728 | 0.430805531 |
| 3.61E-38 | 3.46E-36 |
| 0.412038733 | 0.574166965 |
| 1.86E-19 | 7.62E-18 |
| 0.839319108 | NA |
| 3.00E-15 | 9.31E-14 |
| 4.59E-191 | 4.55E-188 |
| 8.27E-31 | 5.93E-29 |
| 0.858794724 | NA |

| | |
|---|---|
| 7.71E-77 | 1.91E-74 |
| 5.83E-47 | 7.46E-45 |
| 0.393496447 | NA |
| 6.29E-11 | 1.30E-09 |
| NA | NA |
| NA | NA |
| 0.013620222 | 0.04414933 |
| 1.37E-05 | 0.00012042 |
| 0.504259717 | NA |
| 0.205339949 | 0.353987214 |
| 0.941275956 | NA |
| NA | NA |
| 0.941275956 | NA |
| 0.941275956 | NA |
| 0.936418804 | NA |
| 0.595601101 | NA |
| 8.25E-05 | 0.000591537 |
| 0.004827625 | 0.018967834 |
| 0.0334104 | 0.090423733 |
| 0.730126526 | 0.830051037 |
| 0.0741337 | NA |
| 0.866989486 | NA |
| 0.635032494 | NA |
| 0.839319107 | NA |
| 2.47E-05 | 0.000203864 |
| 0.146252145 | 0.278118887 |
| 0.696996115 | 0.805881687 |
| 0.010883738 | 0.03688662 |
| 0.272586936 | 0.432454458 |
| 0.250126899 | 0.407663684 |
| 5.16E-23 | 2.63E-21 |
| 0.955791922 | NA |
| 0.49970428 | 0.652546292 |
| 0.000483727 | 0.002759746 |
| 0.767341402 | 0.855592455 |
| 3.01E-05 | 0.000241774 |
| 0.033470418 | 0.090509304 |
| 0.076237187 | 0.171512054 |
| NA | NA |
| 0.87187248 | 0.923391864 |
| 0.464468773 | NA |
| 0.669311246 | 0.786030809 |
| NA | NA |
| 0.260329902 | 0.418899367 |
| 0.020652574 | 0.061431212 |
| 0.116898739 | 0.235375222 |
| 5.42E-11 | 1.13E-09 |

| | |
|---|---|
| 1.81E-109 | 6.97E-107 |
| 1.10E-255 | 4.37E-252 |
| 3.17E-160 | 2.22E-157 |
| 2.58E-19 | 1.05E-17 |
| 4.34E-79 | 1.11E-76 |
| 4.18E-22 | 2.03E-20 |
| 2.45E-07 | 3.07E-06 |
| NA | NA |
| 0.009157498 | 0.032035094 |
| 8.08E-11 | 1.64E-09 |
| 1.69E-08 | 2.56E-07 |
| 0.615780011 | NA |
| 2.95E-61 | 5.40E-59 |
| 2.83E-11 | 6.10E-10 |
| 0.941279303 | NA |
| 4.42E-11 | 9.29E-10 |
| 0.948397483 | NA |
| 0.941277857 | NA |
| 0.838993826 | NA |
| 1.02E-06 | 1.13E-05 |
| 0.163909848 | 0.302011318 |
| 0.941981387 | NA |
| NA | NA |
| 0.588665496 | NA |
| 0.955792873 | NA |
| NA | NA |
| 0.846118829 | NA |
| 0.725854934 | NA |
| 0.415220865 | NA |
| 0.092235586 | 0.197551644 |
| 0.55979178 | NA |
| NA | NA |
| NA | NA |
| NA | NA |
| NA | NA |
| NA | NA |
| 0.941275956 | NA |
| 0.750685413 | NA |
| 0.638326593 | 0.76241608 |
| 0.045606507 | 0.115236314 |
| 0.955792873 | NA |
| NA | NA |
| NA | NA |
| NA | NA |
| NA | NA |
| NA | NA |
| NA | NA |

| | | |
|---|---|---|
| | 0.839318641 | NA |
| NA | | NA |
| | 0.941279803 | NA |
| | 0.839318641 | NA |
| NA | | NA |
| NA | | NA |
| | 0.941278332 | NA |
| NA | | NA |
| NA | | NA |
| | 0.605457358 | NA |
| | 0.935650667 | NA |
| | 0.936000936 | NA |
| NA | | NA |
| NA | | NA |
| NA | | NA |
| NA | | NA |
| NA | | NA |
| NA | | NA |
| NA | | NA |
| | 0.779478011 | NA |
| | 0.941278332 | NA |
| NA | | NA |
| | 3.38E-31 | 2.49E-29 |
| NA | | NA |
| | 0.013883034 | 0.044847614 |
| | 0.000151946 | 0.00101221 |
| | 0.652309587 | 0.773105252 |
| | 0.122952731 | NA |
| | 0.941278332 | NA |
| | 0.664504823 | 0.782300119 |
| | 0.016643467 | 0.051831453 |
| | 0.094711508 | 0.20144074 |
| | 0.902853906 | 0.941349627 |
| NA | | NA |
| | 0.435148188 | 0.59635245 |
| | 0.802372627 | 0.87912324 |
| | 0.18044022 | 0.322483715 |
| | 0.010113601 | 0.034721466 |
| | 0.23331284 | 0.387882393 |
| | 0.996128702 | NA |
| | 0.308466873 | 0.4699263 |
| | 0.941278332 | NA |
| | 0.839320041 | NA |
| | 0.743511643 | NA |
| | 0.941278332 | NA |
| | 0.661984724 | 0.780259502 |
| | 0.488027515 | NA |

| | | |
|---|---|---|
| | 6.54E-07 | 7.54E-06 |
| | 1.73E-10 | 3.38E-09 |
| | 2.28E-13 | 6.05E-12 |
| | 0.941277857 | NA |
| | 2.14E-14 | 6.17E-13 |
| | 0.360680628 | NA |
| | 0.955791922 | NA |
| | 0.941278332 | NA |
| NA | | NA |
| | 0.000175398 | 0.001146931 |
| | 0.009112621 | 0.031915628 |
| NA | | NA |
| | 0.839318641 | NA |
| | 6.49E-66 | 1.33E-63 |
| | 0.000101584 | 0.000710106 |
| NA | | NA |
| NA | | NA |
| | 0.955792397 | NA |
| | 0.955792873 | NA |
| NA | | NA |
| NA | | NA |
| NA | | NA |
| NA | | NA |
| NA | | NA |
| NA | | NA |
| NA | | NA |
| NA | | NA |
| NA | | NA |
| NA | | NA |
| NA | | NA |
| | 0.839319574 | NA |
| | 0.004198399 | 0.016882969 |
| | 0.454499864 | NA |
| | 0.103007803 | 0.214486198 |
| | 0.839318641 | NA |
| NA | | NA |
| NA | | NA |
| NA | | NA |
| | 0.941278332 | NA |
| NA | | NA |
| | 0.941278332 | NA |
| NA | | NA |
| | 0.941277857 | NA |
| NA | | NA |
| | 0.608127712 | 0.740539236 |
| | 0.828980389 | NA |
| NA | | NA |

| | | | |
|---|---|---|---|
| | 0.839319107 | NA | |
| NA | | NA | |
| | 0.955792398 | NA | |
| NA | | NA | |
| NA | | NA | |
| | 0.941279895 | NA | |
| NA | | NA | |
| | 0.926584778 | NA | |
| NA | | NA | |
| NA | | NA | |
| NA | | NA | |
| NA | | NA | |
| NA | | NA | |
| NA | | NA | |
| NA | | NA | |
| NA | | NA | |
| | 0.584146217 | | 0.721303603 |
| NA | | NA | |
| | 0.955792873 | NA | |
| NA | | NA | |
| | 0.955792873 | NA | |
| | 0.955792398 | NA | |
| | 0.941277857 | NA | |
| NA | | NA | |
| NA | | NA | |
| NA | | NA | |
| NA | | NA | |
| NA | | NA | |
| | 0.955791446 | NA | |
| NA | | NA | |
| NA | | NA | |
| NA | | NA | |
| NA | | NA | |
| | 0.955792398 | NA | |
| NA | | NA | |
| NA | | NA | |
| NA | | NA | |
| NA | | NA | |
| NA | | NA | |
| NA | | NA | |
| NA | | NA | |
| | 1.57E-05 | | 0.000135496 |
| | 0.255468929 | | 0.413518659 |
| | 0.180616418 | | 0.322672931 |
| NA | | NA | |
| NA | | NA | |
| | 0.847873162 | NA | |

| | | |
|---|---|---|
| NA | | NA |
| NA | | NA |
| | 0.661082449 | NA |
| NA | | NA |
| NA | | NA |
| NA | | NA |
| NA | | NA |
| NA | | NA |
| NA | | NA |
| | 0.659816134 | NA |
| | 0.58094093 | NA |
| NA | | NA |
| | 0.366445289 | 0.529609009 |
| | 0.670994104 | NA |
| | 0.973686558 | 0.984105625 |
| | 7.79E-07 | 8.85E-06 |
| | 0.068895678 | 0.159240137 |
| | 0.839318641 | NA |
| | 0.033963338 | 0.091478484 |
| | 0.613886747 | NA |
| | 0.419079254 | 0.580747812 |
| | 0.519181096 | 0.669754153 |
| | 0.004694666 | 0.0185557 |
| | 0.414052241 | 0.576109104 |
| | 0.181545295 | 0.323947905 |
| | 0.000192212 | 0.001238166 |
| | 0.941278332 | NA |
| | 0.506467142 | 0.658128018 |
| | 0.190372574 | 0.335157688 |
| | 0.968669488 | 0.981284949 |
| | 0.768977635 | NA |
| | 0.49683967 | 0.649803715 |
| | 0.011688087 | 0.039111465 |
| | 0.020655403 | 0.06143195 |
| NA | | NA |
| | 0.949627112 | 0.969213349 |
| | 0.839318641 | NA |
| | 0.787468404 | 0.869396677 |
| | 7.08E-12 | 1.64E-10 |
| | 4.21E-08 | 5.95E-07 |
| | 0.002408783 | 0.010698609 |
| NA | | NA |
| | 0.535829721 | NA |
| | 0.955791446 | NA |
| | 0.832575205 | 0.898442899 |
| | 0.009501266 | 0.033009653 |
| NA | | NA |

| | |
|---|---|
| 0.240221752 | 0.396213578 |
| 1.34E-07 | 1.77E-06 |
| 0.766975962 | 0.855355121 |
| 0.803586652 | 0.879967312 |
| 0.001509841 | 0.007209074 |
| 0.862230598 | 0.917648997 |
| 0.044614919 | 0.113247766 |
| 5.22E-20 | 2.21E-18 |
| 0.128412591 | 0.252646834 |
| 0.148038503 | 0.280698833 |
| 0.776392246 | 0.862247491 |
| 0.349476657 | 0.513345062 |
| NA | NA |
| NA | NA |
| 0.264122759 | 0.423115487 |
| 4.39E-07 | 5.24E-06 |
| 0.545267882 | NA |
| 0.661818021 | 0.78014028 |
| 0.610507208 | NA |
| 0.194748242 | 0.340919995 |
| 7.64E-15 | 2.28E-13 |
| 0.708388557 | 0.813993648 |
| 0.716404603 | 0.819759372 |
| 0.903876561 | 0.941698566 |
| 0.062419568 | 0.147550725 |
| 0.32345276 | 0.486083105 |
| 0.089556206 | 0.193466764 |
| 0.260120701 | NA |
| NA | NA |
| 0.196260953 | 0.342905617 |
| 0.037061781 | 0.097994282 |
| 0.251516498 | 0.409227276 |
| 0.23715576 | 0.392598511 |
| 0.136738107 | NA |
| 0.000389432 | 0.002284758 |
| 0.005053392 | 0.019695568 |
| 0.80339144 | 0.879874986 |
| 0.079952164 | 0.177703143 |
| 1.59E-08 | 2.42E-07 |
| 0.176913471 | 0.318509012 |
| 0.713793247 | 0.817953246 |
| 0.003101062 | 0.013182975 |
| 0.733565214 | NA |
| 0.882499196 | 0.929664669 |
| 0.037344095 | 0.098576482 |
| 0.600500223 | 0.734428947 |
| 0.163725521 | NA |

| | | |
|---|---|---|
| | 0.589073419 | 0.725205623 |
| | 0.000431454 | 0.002496226 |
| | 0.00143642 | 0.006916785 |
| | 0.029448845 | 0.081761507 |
| | 0.152139337 | 0.286172001 |
| | 0.131236144 | 0.256544242 |
| | 0.815614202 | 0.88721947 |
| | 0.364068462 | 0.52755777 |
| | 0.000940106 | 0.004840234 |
| | 0.001059553 | 0.005361283 |
| | 2.71E-07 | 3.37E-06 |
| | 0.818163141 | NA |
| | 0.138337797 | 0.266836243 |
| | 0.380269622 | 0.542991079 |
| | 0.023887373 | 0.06915194 |
| | 7.75E-06 | 7.21E-05 |
| | 0.160011718 | 0.296712287 |
| | 0.840790871 | NA |
| | 0.000129179 | 0.000877345 |
| | 0.960656747 | 0.975909171 |
| | 0.691851497 | 0.802311782 |
| NA | | NA |
| | 0.002732517 | 0.011882228 |
| | 0.242962888 | 0.39940622 |
| | 0.556839801 | 0.700930824 |
| | 0.862282396 | NA |
| | 0.003732816 | 0.015326631 |
| | 0.022718512 | 0.066373637 |
| | 0.870821571 | NA |
| | 0.003769993 | 0.015449961 |
| | 0.005641783 | 0.021526362 |
| | 9.36E-08 | 1.26E-06 |
| | 0.8195762 | 0.890082756 |
| | 0.678697985 | 0.793021149 |
| | 0.149931521 | 0.283070521 |
| | 0.917390342 | 0.950349045 |
| | 0.028806032 | 0.080398824 |
| | 0.460192175 | 0.619191302 |
| | 0.083909076 | NA |
| | 0.178947006 | 0.320594424 |
| | 0.630961561 | 0.757188207 |
| | 0.165570235 | 0.304246335 |
| | 0.991140406 | NA |
| | 0.198307955 | NA |
| | 0.742581523 | 0.838270117 |
| | 0.18405173 | 0.326974121 |
| | 0.955792873 | NA |

| | |
|---|---|
| NA | NA |
| 0.778242583 | 0.863416888 |
| 0.932576996 | NA |
| NA | NA |
| NA | NA |
| NA | NA |
| NA | NA |
| NA | NA |
| 1.12E-08 | 1.75E-07 |
| 1.64E-09 | 2.83E-08 |
| 0.000140616 | 0.000943866 |
| 0.01639416 | 0.051202441 |
| 3.20E-41 | 3.45E-39 |
| 6.46E-16 | 2.10E-14 |
| 0.445212954 | 0.60585208 |
| 0.642206364 | 0.765284664 |
| 0.009850439 | 0.033979731 |
| 0.679442995 | 0.793683852 |
| 0.690126481 | 0.801248259 |
| NA | NA |
| 0.155618006 | 0.290876376 |
| NA | NA |
| NA | NA |
| 0.915299134 | 0.949020885 |
| 0.697988805 | NA |
| NA | NA |
| 0.941278332 | NA |
| 0.892117846 | NA |
| 8.69E-07 | 9.77E-06 |
| 0.970513518 | 0.98224187 |
| 0.828903441 | 0.896188177 |
| NA | NA |
| 0.839319108 | NA |
| 0.336642901 | 0.49992353 |
| NA | NA |
| NA | NA |
| 0.581725998 | 0.719472186 |
| 0.109948885 | 0.225124368 |
| 0.015991696 | 0.050202368 |
| 0.480414705 | 0.637073419 |
| 0.686180884 | 0.7986151 |
| 0.027533085 | 0.077610433 |
| 0.237889077 | 0.393456731 |
| 0.233793619 | 0.388379701 |
| 0.470565387 | 0.628216276 |
| 0.496258325 | 0.649293132 |
| 0.524843946 | 0.6743142 |

| | | |
|---|---|---|
| | 0.086250184 | 0.188100319 |
| | 0.031987144 | 0.087231669 |
| | 1.78E-05 | 0.000151234 |
| | 0.01728255 | 0.053395542 |
| | 1.79E-05 | 0.000151662 |
| | 0.656836812 | NA |
| | 0.798151223 | 0.876515429 |
| | 1.06E-12 | 2.66E-11 |
| | 0.001796331 | 0.008357365 |
| | 0.210390448 | NA |
| | 1.92E-05 | 0.000161981 |
| | 0.839319574 | NA |
| | 0.002251133 | 0.010100184 |
| | 0.812893715 | NA |
| NA | | NA |
| | 0.64584516 | NA |
| NA | | NA |
| NA | | NA |
| NA | | NA |
| | 0.719952033 | NA |
| | 0.362323682 | 0.526015754 |
| NA | | NA |
| NA | | NA |
| NA | | NA |
| NA | | NA |
| NA | | NA |
| | 0.899690248 | 0.939592826 |
| NA | | NA |
| NA | | NA |
| | 0.691115045 | 0.80183126 |
| | 0.941278332 | NA |
| NA | | NA |
| NA | | NA |
| | 0.583506769 | 0.72077576 |
| | 0.000123871 | 0.000846508 |
| | 0.437616151 | NA |
| | 0.782471228 | 0.866267226 |
| | 0.620897668 | 0.749468748 |
| NA | | NA |
| | 0.016820306 | 0.052250624 |
| | 0.023495669 | 0.068233763 |
| | 0.014284474 | 0.045902395 |
| | 0.36685227 | 0.53006846 |
| | 0.002497279 | 0.011026098 |
| | 1.55E-05 | 0.000133957 |
| | 0.000199464 | 0.001277277 |
| | 0.841094678 | 0.904070427 |

| | | |
|---|---|---|
| | 0.042703606 | 0.109588273 |
| | 0.036255461 | 0.096343919 |
| | 8.59E-05 | 0.000613419 |
| | 0.006005509 | 0.022678584 |
| | 0.301719623 | 0.4631135 |
| | 0.520562884 | NA |
| | 0.383099995 | 0.545824619 |
| | 0.014663535 | 0.046832209 |
| | 0.811253261 | 0.884537339 |
| | 0.260705882 | 0.419381625 |
| | 2.13E-18 | 8.09E-17 |
| | 0.484765702 | 0.640452556 |
| | 1.74E-20 | 7.61E-19 |
| | 0.225022244 | 0.377808131 |
| | 0.021248745 | 0.062835462 |
| | 0.743511643 | NA |
| | 0.001578966 | 0.007498512 |
| | 3.74E-07 | 4.51E-06 |
| | 0.011039794 | 0.037325168 |
| | 0.002652656 | 0.011599947 |
| | 0.000237289 | 0.00148357 |
| | 0.137438587 | 0.265550226 |
| NA | | NA |
| | 4.55E-11 | 9.55E-10 |
| | 0.205650605 | 0.354291813 |
| | 0.000379237 | 0.002229196 |
| NA | | NA |
| | 0.94127925 | NA |
| | 0.571184801 | 0.711537167 |
| | 0.859708961 | 0.916299154 |
| | 0.651820815 | 0.772647082 |
| | 0.032458147 | 0.088354165 |
| | 0.7172559 | 0.82021693 |
| | 0.273296465 | 0.433292143 |
| | 0.276568382 | 0.436299087 |
| | 0.782649457 | 0.866267226 |
| | 0.367843209 | 0.530984539 |
| | 0.192035572 | 0.337405379 |
| | 0.935768685 | 0.961999147 |
| | 0.833717044 | 0.899348882 |
| | 0.048664492 | 0.121162367 |
| | 0.873486508 | NA |
| | 0.696621971 | 0.805645798 |
| | 0.35529317 | 0.519006263 |
| | 0.941278332 | NA |
| | 0.405347253 | 0.567333606 |
| | 0.043218477 | 0.110587637 |

| | |
|---|---|
| 0.954490871 | NA |
| 2.26E-07 | 2.84E-06 |
| 0.822774418 | 0.892194838 |
| 0.014205938 | 0.045705561 |
| 0.700795509 | NA |
| 0.003150551 | 0.013367097 |
| 0.0571626 | 0.13775278 |
| 1.14E-18 | 4.47E-17 |
| 0.001479084 | 0.007089239 |
| 0.36359975 | 0.527257295 |
| 0.000578456 | 0.003210167 |
| 0.968798515 | 0.981329578 |
| 8.22E-07 | 9.30E-06 |
| 7.38E-33 | 5.80E-31 |
| NA | NA |
| 0.006479987 | 0.024091947 |
| 8.31E-05 | 0.000595113 |
| 2.51E-60 | 4.35E-58 |
| 5.58E-09 | 9.07E-08 |
| NA | NA |
| 0.839319108 | NA |
| NA | NA |
| 0.754643971 | 0.846307756 |
| 0.066711248 | 0.155428848 |
| 0.686595173 | 0.798901951 |
| 6.61E-06 | 6.27E-05 |
| 0.941279303 | NA |
| 0.025231955 | 0.072184014 |
| NA | NA |
| NA | NA |
| NA | NA |
| NA | NA |
| NA | NA |
| NA | NA |
| 0.336441612 | 0.499686961 |
| 0.947960636 | NA |
| NA | NA |
| 0.011944987 | 0.039747599 |
| 0.160537412 | 0.297316486 |
| 0.478184378 | 0.635000255 |
| NA | NA |
| 0.104156109 | 0.216139818 |
| 0.03544395 | 0.094664299 |
| NA | NA |
| 0.646332965 | 0.768585992 |
| 0.755956642 | 0.84719517 |
| 0.147834224 | 0.280405674 |

| | |
|---|---|
| 0.550842208 | 0.696290294 |
| NA | NA |
| NA | NA |
| 0.10000853 | 0.209617732 |
| NA | NA |
| NA | NA |
| NA | NA |
| 0.912957345 | 0.947510715 |
| NA | NA |
| 0.016648496 | 0.051833553 |
| 0.096085656 | 0.203544926 |
| 0.001517056 | 0.00724207 |
| 0.664180348 | 0.782114139 |
| 0.15250512 | 0.28664993 |
| 5.06E-08 | 7.07E-07 |
| 0.602760757 | 0.736078782 |
| 0.291543475 | 0.45278075 |
| 0.927279492 | NA |
| 0.003914466 | 0.015932303 |
| 0.002677419 | 0.011693196 |
| 4.69E-08 | 6.59E-07 |
| 0.242429334 | 0.398832112 |
| 0.016818381 | 0.052250624 |
| 0.660175566 | NA |
| 0.292386705 | 0.453439429 |
| 0.70927587 | 0.814423477 |
| NA | NA |
| 0.955792873 | NA |
| 0.805346207 | 0.880868577 |
| 1.46E-07 | 1.91E-06 |
| 0.330848143 | NA |
| 0.325652984 | 0.488510261 |
| 0.839318641 | NA |
| 0.200707253 | 0.348339248 |
| 1.35E-05 | 0.000118536 |
| 0.923957337 | NA |
| 0.636352688 | 0.760899562 |
| 0.006879644 | 0.025324666 |
| 0.149801985 | 0.282983084 |
| 0.222961784 | 0.375371361 |
| NA | NA |
| 0.521204642 | 0.671413806 |
| 0.620046716 | 0.748968897 |
| 0.000177053 | 0.001156638 |
| 4.72E-26 | 2.78E-24 |
| 0.047338019 | 0.118532268 |
| NA | NA |

| | | |
|---|---|---|
| | 0.0086347 | 0.030511155 |
| | 0.185571305 | 0.328912667 |
| | 0.631905808 | NA |
| NA | | NA |
| | 0.593492202 | 0.728755032 |
| | 3.59E-11 | 7.63E-10 |
| | 0.339456379 | 0.503097182 |
| | 0.013802609 | 0.044606885 |
| | 8.22E-13 | 2.08E-11 |
| | 0.000800243 | 0.004228053 |
| | 0.996045637 | 0.996917253 |
| | 0.043268369 | 0.110691501 |
| | 0.915369359 | 0.949020885 |
| | 8.46E-07 | 9.54E-06 |
| | 0.839319108 | NA |
| | 0.244485241 | NA |
| | 0.703087085 | 0.810523504 |
| | 0.271553961 | 0.431361944 |
| | 5.76E-09 | 9.33E-08 |
| | 0.722760118 | 0.824299776 |
| | 0.186570514 | 0.330339312 |
| | 0.992008508 | NA |
| | 8.75E-05 | 0.0006227 |
| NA | | NA |
| | 0.725073091 | 0.826343774 |
| | 0.475223784 | 0.632515602 |
| | 0.057482711 | 0.138300161 |
| | 0.288241123 | 0.448882177 |
| NA | | NA |
| | 2.46E-08 | 3.64E-07 |
| | 0.002710923 | 0.011808294 |
| | 0.839319108 | NA |
| | 0.099301036 | 0.208445477 |
| | 0.748393366 | NA |
| | 0.00060713 | 0.003345117 |
| | 0.904887302 | NA |
| | 0.150866204 | 0.284519245 |
| | 0.349658557 | 0.5134855 |
| | 0.003039349 | 0.012966945 |
| | 1.61E-65 | 3.27E-63 |
| NA | | NA |
| | 0.667178247 | 0.784283714 |
| | 0.953450885 | 0.971893304 |
| | 0.582694997 | 0.720221546 |
| | 0.080484532 | 0.178619379 |
| | 0.994523722 | 0.995988288 |
| | 4.18E-05 | 0.00032277 |

| | | |
|---|---|---|
| | 0.123207058 | 0.244994933 |
| | 0.424624424 | 0.586019796 |
| | 0.682810693 | NA |
| | 0.005348806 | 0.020607363 |
| | 4.82E-22 | 2.33E-20 |
| | 0.955792397 | NA |
| | 0.1978659 | 0.34500067 |
| | 0.537707938 | 0.685452502 |
| NA | | NA |
| NA | | NA |
| NA | | NA |
| | 0.099435242 | 0.208671982 |
| | 0.839318641 | NA |
| | 4.59E-07 | 5.44E-06 |
| | 0.31039885 | NA |
| | 0.000550732 | 0.003085084 |
| | 5.90E-07 | 6.85E-06 |
| | 0.282853248 | 0.44327581 |
| | 0.941275956 | NA |
| | 2.27E-06 | 2.38E-05 |
| | 2.11E-09 | 3.60E-08 |
| | 7.64E-05 | 0.000551567 |
| | 1.36E-70 | 2.94E-68 |
| | 1.74E-186 | 1.59E-183 |
| | 0.833801616 | NA |
| | 1.17E-06 | 1.29E-05 |
| | 0.665745297 | 0.783218414 |
| | 0.294174616 | 0.455381388 |
| | 0.751160976 | 0.843777629 |
| | 2.99E-06 | 3.04E-05 |
| | 0.730437187 | 0.830111521 |
| NA | | NA |
| | 0.081930898 | 0.180868043 |
| | 0.036032587 | 0.095933739 |
| | 0.959421416 | 0.975112026 |
| | 0.661076623 | 0.779701886 |
| | 0.269265628 | 0.429158998 |
| | 4.81E-20 | 2.05E-18 |
| | 0.833801616 | NA |
| | 0.003517787 | 0.014625391 |
| | 0.613771515 | 0.744153273 |
| NA | | NA |
| NA | | NA |
| | 0.137228322 | 0.265294714 |
| | 0.048951498 | 0.121676717 |
| | 0.93962099 | 0.963757575 |
| | 0.165645098 | 0.304336906 |

| | | | |
|---|---|---|---|
| | 0.875719276 | | 0.925643364 |
| | 0.000509382 | | 0.002881936 |
| | 0.885340114 | | 0.931368125 |
| | 0.728407578 | | 0.829072167 |
| | 0.858549938 | NA | |
| | 0.000605197 | | 0.003337559 |
| | 3.33E-07 | | 4.06E-06 |
| | 0.901932562 | | 0.940738118 |
| | 0.525604672 | NA | |
| | 0.214460066 | NA | |
| | 0.925490591 | | 0.955311464 |
| | 0.018439388 | | 0.056232455 |
| | 0.264517459 | | 0.42349126 |
| | 0.4601755 | NA | |
| NA | | NA | |
| NA | | NA | |
| NA | | NA | |
| NA | | NA | |
| | 0.169570011 | | 0.309224914 |
| NA | | NA | |
| | 0.156866361 | | 0.292626697 |
| NA | | NA | |
| | 0.158941915 | | 0.295257587 |
| | 0.015017299 | | 0.047754409 |
| | 0.525958347 | | 0.675089548 |
| | 0.010979261 | | 0.037157452 |
| | 0.370252852 | | 0.533460847 |
| | 8.90E-06 | | 8.16E-05 |
| | 0.684834438 | | 0.797594035 |
| | 0.941277857 | NA | |
| | 0.161968327 | NA | |
| | 1.15E-08 | | 1.78E-07 |
| | 0.000560563 | | 0.003130577 |
| | 0.57175994 | | 0.711883249 |
| | 0.599188867 | NA | |
| | 0.001013957 | | 0.005169033 |
| | 0.77544211 | NA | |
| | 0.274703171 | | 0.434566321 |
| | 0.332894399 | NA | |
| | 0.941276431 | NA | |
| | 0.120085894 | | 0.240456107 |
| | 0.258829308 | | 0.417218961 |
| | 0.001713551 | | 0.008047739 |
| | 0.697001278 | | 0.805881687 |
| | 0.01660402 | | 0.051749232 |
| | 0.490819203 | | 0.645047538 |
| | 0.492127208 | | 0.646090006 |

| | | |
|---|---|---|
| NA | | NA |
| | 0.02798345 | 0.078535794 |
| | 0.955791446 | NA |
| | 0.563412163 | NA |
| | 0.003069697 | 0.013068312 |
| | 0.622718728 | NA |
| | 0.945798094 | NA |
| | 0.955792398 | NA |
| | 0.853595622 | NA |
| | 0.285675934 | NA |
| | 0.841310243 | 0.904179636 |
| | 0.297632801 | 0.459015008 |
| | 0.630616686 | NA |
| | 1.47E-05 | 0.000128263 |
| | 0.843323855 | 0.905661115 |
| NA | | NA |
| | 2.71E-09 | 4.57E-08 |
| NA | | NA |
| | 0.528608511 | 0.677723661 |
| | 0.734058779 | NA |
| NA | | NA |
| | 0.937429796 | NA |
| | 0.12874724 | 0.253008737 |
| | 0.032795306 | 0.089119063 |
| | 0.500804087 | 0.653373102 |
| NA | | NA |
| | 0.820672062 | 0.890678686 |
| | 5.79E-05 | 0.000432228 |
| | 0.955792398 | NA |
| | 0.011397582 | 0.038308663 |
| | 0.72890697 | 0.829338975 |
| | 0.259929019 | 0.4184815 |
| | 0.843743286 | NA |
| | 0.694710383 | NA |
| | 0.281224472 | 0.441449904 |
| | 0.705105239 | 0.811945569 |
| | 0.955792398 | NA |
| | 0.936856978 | NA |
| | 0.034689255 | 0.09296033 |
| | 0.137764442 | 0.266050243 |
| | 3.06E-05 | 0.000245404 |
| | 0.719399859 | NA |
| | 0.034639505 | 0.092868834 |
| | 0.861920861 | 0.917492965 |
| | 0.727482472 | NA |
| | 0.829040225 | NA |
| | 0.880685324 | NA |

| | | |
|---|---|---|
| NA | | NA |
| | 0.201858565 | 0.349860741 |
| | 0.004191209 | 0.01686166 |
| | 0.005981776 | 0.022599718 |
| | 0.000260625 | 0.001608764 |
| | 0.787756395 | 0.869553296 |
| | 0.370645992 | 0.533961742 |
| | 0.839318641 | NA |
| | 0.657762036 | 0.777100741 |
| | 0.0045599 | 0.018089125 |
| NA | | NA |
| | 0.336400493 | 0.49965707 |
| | 0.20021299 | 0.347717591 |
| | 0.076389748 | 0.171772157 |
| | 0.853596091 | NA |
| | 0.581820743 | NA |
| | 0.852634895 | 0.911453147 |
| NA | | NA |
| | 0.470509764 | 0.628208527 |
| | 0.748393366 | NA |
| NA | | NA |
| NA | | NA |
| NA | | NA |
| NA | | NA |
| | 0.305196929 | 0.466887159 |
| NA | | NA |
| | 0.902169966 | NA |
| | 0.020238177 | 0.060447871 |
| | 0.079908752 | 0.177689473 |
| | 0.42161586 | 0.583069609 |
| | 0.469489883 | 0.627480386 |
| | 0.953932387 | NA |
| | 0.694086701 | 0.803858336 |
| NA | | NA |
| | 0.614382688 | 0.744430877 |
| NA | | NA |
| | 0.94127925 | NA |
| | 0.839320041 | NA |
| | 1.49E-07 | 1.95E-06 |
| | 0.049633713 | 0.122996526 |
| | 0.001991288 | 0.009137581 |
| | 0.792316268 | NA |
| | 0.093615702 | 0.19971004 |
| NA | | NA |
| NA | | NA |
| NA | | NA |
| | 0.076891568 | 0.172613279 |

| | | |
|---|---|---|
| | 4.46E-20 | 1.91E-18 |
| | 0.634774166 | NA |
| | 0.087462153 | 0.190116361 |
| | 1.51E-06 | 1.64E-05 |
| | 0.872478966 | NA |
| | 0.280111723 | 0.440083782 |
| | 0.116613195 | 0.235043379 |
| | 0.449891103 | 0.609847259 |
| | 0.693919533 | 0.803747042 |
| | 0.712520586 | 0.816954935 |
| | 0.003742275 | 0.01536017 |
| | 0.012119418 | 0.040193171 |
| NA | | NA |
| | 0.024706781 | 0.071005893 |
| | 0.047079324 | 0.118080301 |
| | 0.411077145 | 0.572994741 |
| | 0.83521988 | 0.899994439 |
| | 0.592983709 | 0.728399558 |
| NA | | NA |
| | 0.417464175 | 0.579015343 |
| | 0.700372444 | 0.80864556 |
| NA | | NA |
| | 0.839318641 | NA |
| | 0.668601204 | 0.785297106 |
| | 0.095457689 | 0.20262944 |
| | 0.003090368 | 0.01314221 |
| | 0.148683939 | 0.281534875 |
| | 0.911838245 | NA |
| NA | | NA |
| | 3.16E-06 | 3.20E-05 |
| | 0.634466419 | NA |
| | 0.018952202 | 0.05747233 |
| NA | | NA |
| NA | | NA |
| | 0.094051783 | 0.200345758 |
| | 0.766815453 | NA |
| | 0.001772856 | 0.008275647 |
| NA | | NA |
| | 0.941277857 | NA |
| | 8.94E-06 | 8.20E-05 |
| NA | | NA |
| | 0.809080343 | 0.882967072 |
| NA | | NA |
| NA | | NA |
| NA | | NA |
| | 0.095940186 | 0.203345353 |
| | 0.865573144 | 0.919767844 |

| | |
|---|---|
| 0.006642524 | 0.02460018 |
| 0.289164794 | 0.449996759 |
| 0.041265927 | 0.106738925 |
| 0.005314984 | 0.020510254 |
| NA | NA |
| 0.462355688 | 0.621119205 |
| 0.218130906 | 0.369428763 |
| 0.433660978 | 0.594859301 |
| 0.104274222 | 0.216347197 |
| 0.003528747 | 0.014655601 |
| 0.231440329 | 0.385550302 |
| 5.39E-05 | 0.000405126 |
| 0.548660663 | 0.694528592 |
| 0.941278332 | NA |
| NA | NA |
| 0.680152127 | 0.794122482 |
| NA | NA |
| 0.839319574 | NA |
| 0.489664602 | 0.644064813 |
| 0.000166888 | 0.001098833 |
| 0.658208852 | 0.777381584 |
| 0.008784005 | 0.030949125 |
| 0.076995957 | 0.172794435 |
| 0.017000936 | 0.052696481 |
| 0.000139335 | 0.000936584 |
| 0.955791446 | NA |
| 0.383763314 | 0.546507981 |
| 0.070973707 | 0.162815609 |
| 0.09301854 | 0.198800088 |
| 6.07E-09 | 9.77E-08 |
| 0.209551596 | 0.359199838 |
| 0.014493122 | 0.046396663 |
| 0.323405865 | NA |
| NA | NA |
| 0.619100117 | NA |
| 0.941277857 | NA |
| 0.015328875 | 0.0485377 |
| 0.3988363 | 0.561575367 |
| 0.001045341 | 0.00530064 |
| 0.473761827 | 0.631205316 |
| 0.82762261 | 0.895299318 |
| 0.932871902 | NA |
| 0.000436693 | 0.002522855 |
| 0.229377671 | 0.383104282 |
| 0.947486646 | NA |
| NA | NA |
| 0.311010428 | 0.473044297 |

| | |
|---|---|
| 0.635774788 | 0.760437764 |
| 0.005672484 | 0.021626857 |
| 0.609519371 | 0.741490928 |
| 0.530707348 | 0.679499531 |
| 0.344294131 | 0.507895191 |
| 2.27E-06 | 2.38E-05 |
| 0.12685552 | 0.250262994 |
| 0.954335976 | 0.972268326 |
| 2.50E-10 | 4.80E-09 |
| 0.773077527 | 0.86001081 |
| 0.376798491 | 0.539734848 |
| 0.839318641 | NA |
| 0.887057937 | 0.9326293 |
| 0.679956555 | 0.794122482 |
| NA | NA |
| 6.80E-05 | 0.000497434 |
| 0.000530205 | 0.002984161 |
| NA | NA |
| 1.48E-05 | 0.000128742 |
| NA | NA |
| 0.941278332 | NA |
| NA | NA |
| NA | NA |
| 0.955792873 | NA |
| NA | NA |
| NA | NA |
| 0.139202288 | 0.268000716 |
| 0.956589265 | 0.97337139 |
| NA | NA |
| 0.870411946 | 0.922501787 |
| NA | NA |
| 0.202015337 | 0.350055988 |
| NA | NA |
| 0.12564295 | 0.248549991 |
| 0.011484417 | 0.03853829 |
| 0.790757221 | NA |
| 0.048807689 | 0.121445485 |
| 0.774462939 | 0.860707236 |
| 9.14E-05 | 0.000647916 |
| 0.237484766 | 0.392953742 |
| 0.277331619 | 0.437213353 |
| 7.33E-15 | 2.18E-13 |
| 0.274681986 | 0.434566321 |
| NA | NA |
| 0.922729691 | NA |
| 0.129132992 | 0.253390229 |
| 0.955792398 | NA |

| | |
|---|---|
| NA | NA |
| 0.008009369 | 0.028685129 |
| 0.536390138 | 0.684435223 |
| 0.166655803 | 0.305549823 |
| 0.7026159 | 0.810252828 |
| 9.23E-06 | 8.44E-05 |
| NA | NA |
| 3.03E-05 | 0.00024319 |
| 0.493547505 | 0.646993147 |
| 3.62E-05 | 0.000284502 |
| NA | NA |
| NA | NA |
| 1.32E-05 | 0.000116187 |
| 0.061874491 | 0.146497227 |
| 0.528309895 | 0.677412368 |
| 0.004801289 | 0.018889302 |
| 2.92E-07 | 3.60E-06 |
| 0.000345462 | 0.00205773 |
| 0.060175416 | 0.143372899 |
| 0.930675509 | NA |
| 0.007674755 | 0.027707344 |
| 0.36417985 | 0.52755777 |
| 0.941277857 | NA |
| 0.972178551 | 0.983249506 |
| 0.820670639 | 0.890678686 |
| 7.88E-25 | 4.43E-23 |
| 0.839318641 | NA |
| 0.918115069 | 0.950786279 |
| 0.168623537 | 0.308144141 |
| 0.09395526 | 0.200244258 |
| 3.48E-07 | 4.21E-06 |
| 0.762081253 | 0.852239146 |
| 0.610177202 | 0.741788737 |
| 0.747649392 | 0.841476774 |
| 0.003739182 | 0.015350121 |
| NA | NA |
| NA | NA |
| NA | NA |
| 0.534403946 | 0.682660906 |
| NA | NA |
| 0.36257595 | 0.526253598 |
| 0.385542064 | 0.54841154 |
| NA | NA |
| 0.626490016 | NA |
| 0.002351402 | 0.010492701 |
| NA | NA |
| 0.30435853 | 0.465933994 |

| | |
|---|---|
| 0.837221788 | 0.901225435 |
| NA | NA |
| 0.80296295 | 0.879527112 |
| 0.441172765 | 0.601799195 |
| 0.036225698 | 0.096293768 |
| 0.002675481 | 0.011686879 |
| 0.951761677 | 0.970559591 |
| 0.048280959 | 0.120454189 |
| 0.437517593 | 0.598364817 |
| 0.25627046 | 0.414385156 |
| 0.839318641 | NA |
| 0.209753276 | 0.359434664 |
| 0.656742065 | 0.776226666 |
| 0.000439158 | 0.00253525 |
| 0.285595573 | NA |
| 0.96517123 | NA |
| 0.839319108 | NA |
| 0.717240338 | 0.82021693 |
| 0.955792873 | NA |
| 0.941278332 | NA |
| 0.681779667 | 0.795349743 |
| 0.019069659 | 0.057729289 |
| NA | NA |
| 0.380990413 | 0.543697692 |
| 0.01258315 | 0.041442389 |
| 7.94E-18 | 2.93E-16 |
| NA | NA |
| 0.640282394 | NA |
| 0.295143527 | 0.456495304 |
| 0.839318641 | NA |
| 0.23577376 | 0.390990527 |
| NA | NA |
| 0.842596259 | 0.905193905 |
| 0.400667192 | 0.563101117 |
| NA | NA |
| 4.33E-09 | 7.12E-08 |
| 7.11E-07 | 8.15E-06 |
| 0.100907193 | 0.21120234 |
| 0.071685517 | 0.163937229 |
| 0.088730628 | 0.192239214 |
| 0.068599124 | 0.158721686 |
| NA | NA |
| NA | NA |
| NA | NA |
| 0.064638953 | NA |
| 0.000239808 | 0.001497802 |
| 0.025173701 | 0.072051999 |

| | | | |
|---|---|---|---|
| | 0.905021871 | | 0.942439103 |
| | 0.013515389 | | 0.043899193 |
| | 0.225073991 | | 0.377827149 |
| | 0.947395696 | NA | |
| | 0.925662219 | NA | |
| | 0.016686744 | | 0.051925465 |
| | 0.8053719 | | 0.880868577 |
| | 0.013558667 | | 0.043997735 |
| | 8.62E-05 | | 0.000614278 |
| | 0.017096983 | | 0.052949083 |
| NA | | NA | |
| NA | | NA | |
| | 0.045406021 | | 0.114851659 |
| | 0.936986733 | NA | |
| | 0.363867509 | NA | |
| | 0.690269725 | | 0.801297308 |
| | 0.71090035 | | 0.815698522 |
| | 0.873897584 | NA | |
| | 0.349348574 | | 0.513256858 |
| | 0.208098078 | | 0.357421738 |
| | 1.67E-05 | | 0.00014315 |
| | 0.000551863 | | 0.003090693 |
| | 0.443493102 | | 0.604029687 |
| | 0.412530957 | | 0.574785566 |
| | 0.00022557 | | 0.001422635 |
| | 0.012596109 | | 0.041477806 |
| | 0.005341636 | | 0.020586401 |
| | 0.940815805 | NA | |
| | 0.000695413 | | 0.003757597 |
| | 0.708307015 | | 0.813939244 |
| NA | | NA | |
| NA | | NA | |
| NA | | NA | |
| | 0.955792873 | NA | |
| | 0.308631232 | | 0.470146598 |
| | 1.32E-05 | | 0.000116338 |
| | 0.448709926 | | 0.60881295 |
| | 0.768031606 | NA | |
| | 0.459148871 | | 0.618451782 |
| | 0.000987397 | | 0.0050542 |
| | 0.48553152 | | 0.640898955 |
| | 0.876313213 | | 0.926075882 |
| | 0.84492899 | | 0.906676878 |
| | 1.18E-11 | | 2.63E-10 |
| | 2.15E-22 | | 1.06E-20 |
| NA | | NA | |
| | 0.550588572 | | 0.696197727 |

| | |
|---|---|
| 4.51E-10 | 8.39E-09 |
| 0.054241648 | 0.132156028 |
| NA | NA |
| 1.05E-07 | 1.41E-06 |
| 0.000337013 | 0.002019132 |
| 1.63E-08 | 2.48E-07 |
| 0.267258096 | 0.426846296 |
| 0.056150227 | 0.135877157 |
| 9.15E-05 | 0.00064806 |
| 0.718521799 | 0.821000233 |
| 0.273380721 | 0.433292143 |
| 0.000768217 | 0.004087884 |
| 0.168531422 | 0.307999456 |
| 0.630083709 | 0.75640553 |
| 0.031858468 | 0.086980415 |
| 0.436273175 | 0.597375258 |
| 2.60E-09 | 4.38E-08 |
| 0.032012788 | 0.087291601 |
| 0.271480439 | 0.431360308 |
| 0.061531612 | 0.145888577 |
| 0.651794398 | 0.772647082 |
| 0.032750715 | 0.089028383 |
| 0.158731443 | 0.295027784 |
| 0.341524637 | 0.504972832 |
| 2.66E-05 | 0.000216925 |
| 0.303411804 | 0.464796771 |
| 0.941278332 | NA |
| 0.761055315 | 0.851491896 |
| 0.004394018 | 0.017539229 |
| 0.524577643 | 0.674080713 |
| 0.000165605 | 0.001092196 |
| 4.34E-07 | 5.19E-06 |
| 0.419051365 | 0.580742974 |
| 0.567531825 | 0.709139193 |
| 0.000214759 | 0.001365301 |
| 0.015230571 | 0.048279451 |
| 2.68E-05 | 0.000218285 |
| NA | NA |
| 0.000679845 | 0.003683761 |
| NA | NA |
| 0.000599449 | 0.00331046 |
| 0.002524777 | 0.01113278 |
| 2.25E-07 | 2.84E-06 |
| 9.34E-07 | 1.04E-05 |
| 0.09152482 | 0.196559936 |
| 0.014000616 | 0.045130372 |
| 5.24E-07 | 6.16E-06 |

| | |
|---|---|
| 0.004911379 | 0.019233408 |
| 0.000668926 | 0.003634279 |
| 0.532433225 | 0.680836641 |
| 0.24028148 | 0.396284631 |
| 0.941278207 | NA |
| 0.523666716 | 0.673469863 |
| 0.003242054 | 0.013687011 |
| 0.004287776 | 0.017181423 |
| 0.003539363 | 0.014689438 |
| 4.07E-05 | 0.000315797 |
| 0.063561243 | 0.149642404 |
| 0.110888053 | 0.226401126 |
| 0.05302105 | 0.129929684 |
| 0.358635555 | 0.522254113 |
| 0.672462541 | 0.788354106 |
| 0.31616492 | 0.478683643 |
| 0.070748011 | 0.162542871 |
| 0.822447828 | NA |
| 0.675562411 | 0.790858573 |
| 0.918584643 | 0.951111522 |
| 0.504978666 | 0.657044385 |
| 0.131761088 | 0.257295711 |
| 0.020546637 | 0.061154316 |
| 0.001454062 | 0.006986191 |
| 0.042893684 | 0.109969125 |
| 0.054225444 | 0.132146414 |
| 0.113921125 | 0.230966832 |
| 0.011288379 | 0.038003678 |
| 0.255285118 | 0.413381617 |
| 0.032546071 | 0.088563117 |
| 0.839319108 | NA |
| 0.077138617 | 0.172973804 |
| 0.016657376 | 0.051847634 |
| 0.193066108 | 0.338765904 |
| 0.824493216 | 0.893407427 |
| 0.939356797 | 0.963757575 |
| 0.941278332 | NA |
| 0.326975601 | 0.489753525 |
| 0.235208693 | 0.390189385 |
| 0.072676727 | 0.165567712 |
| 0.126110162 | 0.249308478 |
| 0.126363636 | 0.249623478 |
| 0.062572203 | 0.147810994 |
| 0.75429539 | 0.846066498 |
| 0.050286161 | 0.124354412 |
| 0.617516953 | 0.747046431 |
| 0.002915838 | 0.012523058 |

| | |
|---|---|
| 0.000182645 | 0.001185523 |
| 0.018219858 | 0.05569857 |
| 0.01655565 | 0.051625517 |
| 0.000835536 | 0.004384355 |
| 0.008466061 | 0.030022227 |
| 0.581214112 | 0.718951164 |
| 0.642018719 | 0.765214299 |
| 0.024939247 | 0.071535787 |
| 0.003254766 | 0.013730936 |
| 0.071685429 | 0.163937229 |
| 0.001825522 | 0.008469468 |
| 0.263134232 | 0.422043194 |
| 0.347188594 | 0.511182909 |
| 0.000390379 | 0.002289186 |
| NA | NA |
| 0.158969276 | 0.295274373 |
| 0.00218901 | 0.009862355 |
| 0.626471365 | NA |
| 0.032771204 | 0.089063737 |
| 0.891066883 | 0.934449599 |
| 0.151785018 | 0.285753935 |
| 0.041625686 | 0.107505922 |
| NA | NA |
| NA | NA |
| 0.003743119 | 0.015360986 |
| 0.813353139 | 0.886089811 |
| 0.846525845 | 0.907490347 |
| 0.484534551 | 0.640324879 |
| 0.460108834 | 0.619114165 |
| 0.177817948 | 0.319321374 |
| 4.62E-06 | 4.54E-05 |
| 0.283576005 | 0.444143427 |
| 0.405266905 | 0.567254505 |
| NA | NA |
| 0.429152434 | 0.590466111 |
| 0.005789676 | 0.021985619 |
| NA | NA |
| NA | NA |
| 0.007253559 | 0.026455594 |
| 0.293099051 | 0.454277764 |
| NA | NA |
| 0.839319108 | NA |
| 0.027060336 | 0.076540714 |
| 0.004900918 | 0.019198758 |
| 0.941275956 | NA |
| NA | NA |
| 0.000456641 | 0.002622189 |

| | | |
|---|---|---|
| | 0.075895042 | 0.170972717 |
| | 0.941278332 | NA |
| | 0.038361419 | 0.100759047 |
| | 0.001258636 | 0.006199927 |
| | 0.002686863 | 0.011727988 |
| | 0.879293939 | NA |
| NA | | NA |
| | 0.941278332 | NA |
| NA | | NA |
| | 0.077486563 | 0.173584486 |
| | 0.618234337 | NA |
| | 0.0531333 | 0.130177953 |
| | 0.401329101 | NA |
| | 0.902022945 | NA |
| | 0.155103983 | 0.290044217 |
| | 0.787331887 | 0.869286278 |
| | 0.34248864 | 0.505952993 |
| NA | | NA |
| | 0.843335558 | 0.905661115 |
| | 0.454141095 | 0.613980893 |
| | 0.036186257 | 0.096245983 |
| | 0.887786452 | 0.93306386 |
| | 0.005196095 | 0.020156037 |
| | 2.63E-08 | 3.87E-07 |
| | 0.839319108 | NA |
| NA | | NA |
| | 0.847555821 | 0.908343735 |
| | 0.56300489 | 0.705445208 |
| | 0.128726704 | 0.253008737 |
| | 0.644739074 | 0.767419084 |
| | 0.223114029 | 0.375543463 |
| NA | | NA |
| | 0.484643539 | 0.64037579 |
| NA | | NA |
| NA | | NA |
| | 0.955792873 | NA |
| | 0.661569785 | NA |
| | 0.500700461 | 0.65330827 |
| NA | | NA |
| NA | | NA |
| NA | | NA |
| NA | | NA |
| NA | | NA |
| NA | | NA |
| | 0.955792873 | NA |
| NA | | NA |
| | 0.600629095 | NA |

| | |
|---|---|
| 0.649369671 | 0.770749546 |
| 5.89E-15 | 1.78E-13 |
| 0.227768514 | 0.381194358 |
| 0.081195914 | 0.179712214 |
| NA | NA |
| 0.091484739 | 0.196527053 |
| NA | NA |
| 1.26E-09 | 2.20E-08 |
| 0.312465428 | NA |
| 0.37802309 | 0.540922543 |
| NA | NA |
| 0.31503064 | NA |
| 0.000110617 | 0.000765823 |
| 0.891106079 | NA |
| 0.072066096 | 0.164570394 |
| 0.335810143 | 0.498998191 |
| 0.735774479 | NA |
| 0.779882339 | 0.864527319 |
| 0.022956141 | NA |
| 0.074488688 | 0.168582271 |
| 0.00541066 | 0.020781756 |
| 0.023378315 | 0.067967591 |
| 2.89E-11 | 6.21E-10 |
| 0.059630591 | 0.142381181 |
| 0.563734525 | 0.705986435 |
| 0.290440142 | 0.451561444 |
| 0.013482975 | 0.043823813 |
| NA | NA |
| 0.174086579 | 0.314935667 |
| 0.782278252 | 0.866117172 |
| NA | NA |
| NA | NA |
| 0.002080788 | 0.009453504 |
| 0.548884237 | 0.694694389 |
| NA | NA |
| NA | NA |
| NA | NA |
| 0.709395965 | 0.814522082 |
| 0.545496505 | 0.691847809 |
| NA | NA |
| 0.132035425 | 0.257624654 |
| 3.18E-06 | 3.21E-05 |
| 0.000103051 | 0.000718878 |
| 1.54E-05 | 0.000133317 |
| 1.01E-06 | 1.12E-05 |
| NA | NA |
| NA | NA |

| | |
|---|---|
| 0.955791922 | NA |
| 0.003672176 | 0.015116766 |
| NA | NA |
| NA | NA |
| 0.112526891 | 0.228959229 |
| NA | NA |
| NA | NA |
| 0.166607819 | 0.305509772 |
| NA | NA |
| 2.91E-09 | 4.88E-08 |
| 0.273718436 | 0.433520547 |
| 0.941278838 | NA |
| NA | NA |
| 0.202176302 | 0.350232922 |
| 4.51E-11 | 9.48E-10 |
| 0.105084096 | 0.217629135 |
| 0.645606632 | 0.767952477 |
| 0.163040059 | 0.300804548 |
| 1.50E-07 | 1.96E-06 |
| 0.10967291 | 0.224611478 |
| 0.556128084 | 0.700368288 |
| 0.116767067 | NA |
| 0.388546754 | 0.551363288 |
| 0.265418561 | NA |
| 0.031960569 | 0.08719916 |
| NA | NA |
| 0.885931856 | NA |
| NA | NA |
| NA | NA |
| 0.805262409 | NA |
| 0.176848935 | NA |
| NA | NA |
| 0.412098124 | NA |
| NA | NA |
| NA | NA |
| 0.955792873 | NA |
| NA | NA |
| NA | NA |
| 0.023501404 | 0.06824209 |
| 0.000228265 | 0.001436206 |
| NA | NA |
| 0.001835976 | 0.00850689 |
| 0.077822018 | 0.174122924 |
| 0.002104979 | 0.009541558 |
| 0.018022725 | 0.055188076 |
| 0.584335096 | 0.721417285 |
| 7.12E-05 | 0.000518741 |

| | |
|---|---|
| 0.288157457 | 0.448815917 |
| 0.375784309 | 0.538982714 |
| 0.114001929 | 0.231060899 |
| 0.22750421 | 0.380885577 |
| 0.396029459 | NA |
| 0.829128485 | 0.896348753 |
| NA | NA |
| 0.955792398 | NA |
| 7.57E-09 | 1.21E-07 |
| 0.000207821 | 0.001325444 |
| 0.013777049 | 0.044548473 |
| NA | NA |
| 0.81031009 | 0.883711375 |
| NA | NA |
| 1.19E-08 | 1.85E-07 |
| 0.261055127 | 0.419798224 |
| 1.21E-11 | 2.70E-10 |
| 0.002261184 | 0.010141457 |
| NA | NA |
| NA | NA |
| NA | NA |
| 6.53E-87 | 1.89E-84 |
| 1.38E-25 | 7.89E-24 |
| 2.31E-30 | 1.62E-28 |
| 1.56E-72 | 3.50E-70 |
| 1.23E-259 | 5.83E-256 |
| 0.004157981 | 0.016760079 |
| 7.56E-24 | 4.00E-22 |
| 2.06E-18 | 7.89E-17 |
| 5.69E-08 | 7.90E-07 |
| 0.000963945 | 0.004944787 |
| 0.839319108 | NA |
| 0.086709629 | 0.188911908 |
| 1.69E-08 | 2.56E-07 |
| 9.81E-06 | 8.91E-05 |
| 0.955792397 | NA |
| 2.88E-32 | 2.21E-30 |
| 0.839319108 | NA |
| 0.012674239 | 0.041678948 |
| 0.000137815 | 0.000928991 |
| 0.826525793 | 0.894713795 |
| 0.011398284 | 0.038308663 |
| 0.10720089 | 0.220898241 |
| 0.002154105 | 0.009729032 |
| 0.426918462 | NA |
| 0.749357084 | 0.842611984 |
| 0.642752594 | 0.765697809 |

| | |
|---|---|
| NA | NA |
| 0.04259865 | 0.109401497 |
| 0.973011095 | 0.983726508 |
| 0.013085135 | 0.042770171 |
| 0.383790518 | 0.546509909 |
| 0.00748483 | 0.027165892 |
| 1.65E-05 | 0.000141475 |
| NA | NA |
| 0.010531319 | 0.035906956 |
| 0.941277857 | NA |
| 0.064684421 | 0.151776281 |
| 0.086722573 | 0.188922817 |
| 0.044297579 | 0.112622408 |
| 0.600334566 | NA |
| 0.725028456 | NA |
| 0.708854511 | 0.814125883 |
| 0.328439144 | 0.491358172 |
| 0.464310462 | NA |
| 0.000798397 | 0.004221114 |
| 0.279795053 | 0.439670024 |
| 0.261246091 | 0.419948636 |
| 0.611826548 | 0.742854383 |
| 1.04E-12 | 2.61E-11 |
| NA | NA |
| 0.168810745 | 0.308377668 |
| 0.342488504 | 0.505952993 |
| 0.056196945 | 0.135976384 |
| 0.922173452 | 0.953631614 |
| 0.030618108 | 0.084280351 |
| 0.85183151 | 0.911044421 |
| 0.941279303 | NA |
| 0.322802181 | NA |
| 0.878913386 | 0.927385945 |
| 0.815192248 | 0.88702503 |
| 0.616022498 | 0.745981842 |
| 0.157733934 | 0.293659621 |
| 0.577193663 | 0.715763445 |
| NA | NA |
| NA | NA |
| NA | NA |
| NA | NA |
| NA | NA |
| 0.066756495 | 0.155458183 |
| 0.13936425 | 0.268186826 |
| NA | NA |
| 0.370672335 | 0.533967374 |
| 0.982880415 | 0.989178714 |

| | |
|---|---|
| 0.763159467 | 0.85297792 |
| 0.802699165 | 0.879400052 |
| 0.358969746 | 0.52254895 |
| 0.010536133 | 0.035918222 |
| 0.355731382 | 0.519502723 |
| NA | NA |
| NA | NA |
| 0.806287724 | 0.881424719 |
| 0.941278332 | NA |
| 0.559628327 | 0.702990999 |
| 0.941278332 | NA |
| 1.69E-06 | 1.82E-05 |
| 0.27085228 | 0.430793583 |
| 0.002015301 | 0.00921402 |
| 0.021979326 | 0.064562744 |
| 0.714118466 | 0.818158107 |
| 0.017176463 | 0.053136753 |
| NA | NA |
| 0.474041685 | NA |
| NA | NA |
| 0.702244941 | NA |
| 0.009405843 | 0.032730684 |
| 0.526163483 | 0.675316431 |
| 0.543946302 | NA |
| 0.71361624 | 0.817894557 |
| 7.80E-18 | 2.88E-16 |
| NA | NA |
| NA | NA |
| 0.014445659 | 0.046276659 |
| 0.000556046 | 0.003109732 |
| 0.591456882 | NA |
| 0.002006885 | 0.00918613 |
| 0.342715692 | 0.506194161 |
| 0.278331402 | 0.43798984 |
| 0.129527433 | 0.253892116 |
| 1.26E-18 | 4.90E-17 |
| 0.977813667 | 0.986308467 |
| NA | NA |
| 0.02477418 | 0.071156633 |
| 0.184370441 | 0.327344864 |
| 0.739991096 | NA |
| 0.404077035 | 0.566154418 |
| 0.341477558 | 0.504960787 |
| 0.033833875 | 0.091234891 |
| 0.630301135 | 0.756552074 |
| 1.30E-10 | 2.57E-09 |
| 0.497609531 | 0.650596103 |

| | |
|---|---|
| 0.157470282 | 0.293256212 |
| 4.12E-11 | 8.70E-10 |
| NA | NA |
| 0.531328796 | 0.67996602 |
| 0.007533827 | 0.027297936 |
| 0.016075375 | 0.050385314 |
| NA | NA |
| 0.786944116 | NA |
| NA | NA |
| 0.853510826 | NA |
| NA | NA |
| 0.551418284 | 0.696685493 |
| 0.538153298 | 0.685860184 |
| NA | NA |
| 0.628000254 | 0.75505671 |
| 0.839318641 | NA |
| 0.800086083 | NA |
| 0.151811051 | 0.28575774 |
| 0.839318641 | NA |
| 0.710607293 | 0.815440883 |
| 0.569249977 | 0.710316538 |
| 0.23682099 | 0.392262575 |
| 0.812759173 | NA |
| 1.81E-08 | 2.73E-07 |
| 0.002704018 | 0.011785576 |
| NA | NA |
| NA | NA |
| 0.590969165 | 0.726825198 |
| NA | NA |
| 0.955792398 | NA |
| 0.325138148 | 0.488014768 |
| 0.393524878 | 0.55597471 |
| 0.429886533 | 0.591125376 |
| 5.43E-05 | 0.000407343 |
| 0.399186339 | 0.561816062 |
| 0.604488298 | 0.737393935 |
| 0.323111354 | 0.485799789 |
| 0.001195304 | 0.00593959 |
| 0.009699838 | 0.033606338 |
| 0.006257321 | 0.023421413 |
| 0.004390962 | 0.017529971 |
| 0.028777598 | 0.080347728 |
| 0.11283246 | 0.229463279 |
| NA | NA |
| 0.532417967 | 0.680836641 |
| 0.805322933 | 0.880868577 |
| 0.000725555 | 0.003899224 |

| | |
|---|---|
| NA | NA |
| NA | NA |
| 5.96E-10 | 1.09E-08 |
| 4.21E-07 | 5.05E-06 |
| 0.974213595 | 0.984409087 |
| 0.552965257 | 0.698038772 |
| 0.200717284 | 0.348339248 |
| 0.684214243 | 0.797279344 |
| 0.000744333 | 0.003978825 |
| 0.733672831 | NA |
| NA | NA |
| NA | NA |
| 0.565440805 | 0.707194684 |
| 0.124591524 | 0.247106696 |
| 0.297882215 | 0.4592986 |
| 0.156045597 | 0.291469615 |
| NA | NA |
| 7.57E-06 | 7.06E-05 |
| NA | NA |
| 0.538725397 | 0.686332061 |
| 0.148853654 | 0.281728267 |
| 0.115138379 | 0.232770247 |
| 0.004801095 | 0.018889302 |
| 3.52E-05 | 0.000277467 |
| 0.298430504 | 0.459965218 |
| 0.026452249 | 0.075043674 |
| 0.001964775 | 0.009033334 |
| NA | NA |
| 0.048731302 | 0.121290645 |
| 0.000563455 | 0.003142303 |
| 0.090392061 | 0.194883319 |
| NA | NA |
| 0.143043338 | 0.273427363 |
| 0.006975948 | 0.025607885 |
| 0.696806414 | 0.805780803 |
| 0.947982043 | 0.968205894 |
| 0.792678571 | 0.872437591 |
| 0.761632803 | 0.851817685 |
| 0.955791922 | NA |
| 0.001260853 | 0.006207952 |
| 0.436905984 | NA |
| 6.83E-07 | 7.84E-06 |
| 0.244606819 | 0.401129014 |
| 0.60162034 | 0.735209972 |
| NA | NA |
| NA | NA |
| 0.0040227 | 0.01629481 |

| | |
|---|---|
| NA | NA |
| 0.941279303 | NA |
| 0.458024479 | 0.617706309 |
| 0.849602377 | NA |
| 0.685680319 | 0.79822767 |
| 0.941492992 | 0.964703895 |
| 3.39E-06 | 3.42E-05 |
| 0.001135233 | 0.005689791 |
| 0.686754807 | 0.799009576 |
| 0.29946988 | 0.461008951 |
| 0.000149891 | 0.000999075 |
| 0.576396977 | 0.715036266 |
| 0.00274473 | 0.011921544 |
| 0.006685104 | 0.024723253 |
| 0.085067481 | 0.186289096 |
| 1.32E-21 | 6.21E-20 |
| 0.027376027 | 0.077250052 |
| 0.328921265 | 0.49189394 |
| 6.31E-05 | 0.000467176 |
| 0.000364159 | 0.0021508 |
| 0.258728531 | 0.417113011 |
| NA | NA |
| 0.839318641 | NA |
| NA | NA |
| NA | NA |
| 0.664204707 | 0.782114139 |
| NA | NA |
| 0.196952175 | 0.343734834 |
| 0.74609083 | 0.840278884 |
| 0.787098947 | 0.869192807 |
| 2.85E-09 | 4.79E-08 |
| 0.25352829 | 0.411432492 |
| 0.542931073 | NA |
| 0.955792398 | NA |
| 0.911872328 | NA |
| 0.002419982 | 0.010738331 |
| NA | NA |
| NA | NA |
| 0.017092581 | 0.052949083 |
| 0.000294398 | 0.001794435 |
| 0.389092011 | 0.551717073 |
| 0.40699387 | 0.568768676 |
| 1.73E-08 | 2.61E-07 |
| 0.811295591 | 0.884542973 |
| 3.87E-10 | 7.25E-09 |
| 0.791922885 | 0.871847757 |
| 0.199991748 | 0.347434756 |

| | |
|---|---|
| NA | NA |
| NA | NA |
| NA | NA |
| 0.436834636 | 0.597834521 |
| 0.656443299 | NA |
| 0.749387171 | 0.842611984 |
| 0.064582598 | 0.151627046 |
| NA | NA |
| 0.487017131 | 0.642037206 |
| 5.14E-06 | 4.99E-05 |
| 0.923505151 | NA |
| NA | NA |
| 0.004227144 | 0.016981346 |
| 0.877248243 | 0.926447009 |
| 0.839319108 | NA |
| 5.53E-06 | 5.34E-05 |
| 0.463652145 | 0.622333974 |
| 0.247490133 | 0.404668554 |
| 0.730492118 | 0.830111521 |
| NA | NA |
| NA | NA |
| NA | NA |
| NA | NA |
| 0.839980114 | 0.903321135 |
| NA | NA |
| 0.237184896 | 0.392619438 |
| 0.935650667 | NA |
| NA | NA |
| 0.001953936 | 0.008990446 |
| 0.453713639 | 0.613631004 |
| 0.394320729 | 0.556811936 |
| 1.73E-05 | 0.000147234 |
| 0.234279193 | 0.389027023 |
| 0.694939119 | 0.80459874 |
| 0.268291101 | 0.4280931 |
| NA | NA |
| 0.002707135 | 0.011794841 |
| 0.154612391 | 0.289451325 |
| 0.001490567 | 0.007137092 |
| 0.663217698 | 0.781441892 |
| 0.867105798 | 0.920555406 |
| 3.27E-42 | 3.67E-40 |
| 0.472191185 | 0.629664543 |
| 0.035192573 | 0.094068275 |
| 5.29E-37 | 4.96E-35 |
| 0.503307628 | 0.655453749 |
| 0.121020724 | 0.241878854 |

| | |
|---|---|
| 0.055868016 | 0.135345599 |
| 0.876578686 | NA |
| 0.006932759 | 0.025484715 |
| 0.899925148 | NA |
| 0.096235224 | 0.203771089 |
| 0.025423714 | 0.072607281 |
| 0.392830424 | 0.555300496 |
| 0.036729413 | 0.097342554 |
| 0.000139186 | 0.000935983 |
| NA | NA |
| 0.243040524 | 0.399506253 |
| 0.944468596 | 0.966517973 |
| 0.397092253 | 0.559725832 |
| 0.249320826 | 0.406739843 |
| 0.011456445 | 0.038466116 |
| 0.000221973 | 0.001404039 |
| 0.32006316 | 0.482741482 |
| 5.49E-07 | 6.42E-06 |
| 3.66E-15 | 1.13E-13 |
| 0.192120706 | 0.337522961 |
| 2.99E-06 | 3.05E-05 |
| 0.43268328 | 0.59382618 |
| 3.05E-19 | 1.23E-17 |
| 0.587721326 | 0.723915494 |
| 0.062525925 | 0.147719245 |
| 1.09E-14 | 3.19E-13 |
| 0.853594685 | NA |
| 0.030103327 | 0.083248448 |
| 0.77863929 | 0.863592382 |
| NA | NA |
| 0.000114221 | 0.000786428 |
| 0.097233686 | 0.205300825 |
| 0.255779116 | 0.413787434 |
| 0.016492357 | 0.051455115 |
| 0.735650765 | 0.833490076 |
| 0.465045192 | 0.623500572 |
| 0.697340629 | NA |
| 0.645239751 | 0.767737465 |
| 0.023312727 | 0.067801752 |
| 0.954021193 | 0.972065003 |
| 0.955792398 | NA |
| 0.954654799 | 0.972509888 |
| 0.085934484 | 0.187689071 |
| 0.15995141 | 0.296669794 |
| 3.44E-09 | 5.72E-08 |
| 0.567654271 | 0.709187479 |
| 0.012799529 | 0.042021297 |

| | | |
|---|---|---|
| | 0.000124346 | 0.000849266 |
| | 0.894114519 | NA |
| | 0.463872564 | 0.622524513 |
| | 0.444652937 | 0.605297639 |
| | 0.145566632 | 0.277137815 |
| | 0.700236667 | 0.808638955 |
| | 0.087602046 | 0.190333537 |
| | 5.25E-07 | 6.16E-06 |
| NA | | NA |
| | 0.661149593 | 0.779701886 |
| | 0.002117845 | 0.009592568 |
| | 0.165371898 | 0.303999222 |
| | 0.065600452 | 0.153351203 |
| | 4.56E-07 | 5.41E-06 |
| | 0.042732226 | 0.109626259 |
| | 1.19E-07 | 1.58E-06 |
| | 0.337204445 | 0.500445205 |
| | 0.674787658 | 0.790418102 |
| | 0.446290393 | 0.606763234 |
| | 0.014664333 | 0.046832209 |
| | 0.251567663 | 0.409227276 |
| | 0.025451015 | 0.072662236 |
| | 0.522414431 | 0.672390001 |
| | 0.054984191 | 0.133639714 |
| | 0.102462365 | 0.2136308 |
| | 0.523622917 | 0.673469863 |
| | 0.676602037 | 0.791584276 |
| NA | | NA |
| | 0.868192527 | 0.921186627 |
| | 0.793264646 | 0.872939996 |
| | 0.002047444 | 0.009325155 |
| | 0.10959435 | 0.224489218 |
| | 0.49050307 | NA |
| | 0.006858849 | 0.025255928 |
| | 0.839318641 | NA |
| | 0.000411254 | 0.002392145 |
| | 1.73E-12 | 4.25E-11 |
| NA | | NA |
| | 8.58E-14 | 2.36E-12 |
| | 0.574473925 | 0.713786167 |
| | 0.212257273 | 0.36270713 |
| | 6.99E-23 | 3.55E-21 |
| | 0.000299464 | 0.001820954 |
| | 0.018986728 | 0.057533049 |
| | 0.000271353 | 0.001667201 |
| | 4.50E-06 | 4.43E-05 |
| | 0.058380726 | 0.139923274 |

| | |
|---|---|
| 0.101229365 | 0.211671909 |
| 0.113490887 | 0.230408572 |
| 0.000534581 | 0.00300664 |
| 0.058393517 | 0.139939841 |
| 1.90E-07 | 2.43E-06 |
| 6.40E-10 | 1.17E-08 |
| 0.169926866 | 0.309504076 |
| NA | NA |
| 0.001778844 | 0.008298714 |
| 0.445976538 | NA |
| 0.972443393 | 0.983356741 |
| 0.914708757 | NA |
| 0.764881077 | NA |
| 0.623272727 | 0.751385543 |
| 2.95E-05 | 0.000237781 |
| 0.717284909 | 0.82021693 |
| 0.443949861 | 0.60451342 |
| 0.686522685 | 0.798856658 |
| NA | NA |
| 0.565398967 | 0.707179517 |
| NA | NA |
| 1.32E-05 | 0.000116338 |
| NA | NA |
| NA | NA |
| NA | NA |
| 0.23068563 | 0.384589155 |
| 0.024211012 | 0.069927255 |
| 0.060274207 | 0.143522077 |
| 0.771490658 | 0.858807438 |
| 0.351439324 | 0.515109595 |
| 7.28E-05 | 0.000528311 |
| 9.88E-06 | 8.96E-05 |
| 0.953798549 | NA |
| 4.98E-05 | 0.000377733 |
| 0.486328409 | 0.64166235 |
| 0.18850393 | 0.332921245 |
| 7.27E-05 | 0.000527885 |
| NA | NA |
| 0.105707336 | 0.218585879 |
| 3.86E-06 | 3.85E-05 |
| 0.001570453 | 0.007461064 |
| 3.59E-05 | 0.000282073 |
| 0.832809068 | 0.898613783 |
| 0.944595199 | NA |
| 0.941278332 | NA |
| 0.00100468 | 0.005130529 |
| 0.002734181 | 0.011882228 |

| | |
|---|---|
| 0.167761397 | NA |
| 0.446346157 | 0.606804389 |
| 0.001416419 | 0.006838458 |
| 0.759906494 | 0.850526397 |
| 2.87E-32 | 2.20E-30 |
| NA | NA |
| 0.742752807 | 0.838423708 |
| 6.78E-07 | 7.80E-06 |
| 0.469731667 | NA |
| 0.614416628 | 0.744430877 |
| 0.128660904 | 0.252996404 |
| 0.941279303 | NA |
| 0.760026928 | NA |
| 0.384615451 | 0.547426716 |
| 0.00084592 | 0.004429079 |
| 0.294254888 | 0.455446406 |
| 0.001276225 | 0.006270995 |
| 0.005035637 | 0.019645669 |
| NA | NA |
| 0.16711569 | 0.306141868 |
| 0.005207779 | 0.020184913 |
| 0.000776963 | 0.004124283 |
| 0.000157061 | 0.0010422 |
| 0.087263532 | 0.189805958 |
| 0.006789865 | 0.025048413 |
| 0.48529829 | 0.640729416 |
| 0.24461758 | 0.401129014 |
| NA | NA |
| 0.270508818 | NA |
| 0.399510502 | NA |
| NA | NA |
| 0.941275481 | NA |
| 0.111478378 | 0.227253199 |
| NA | NA |
| 1.01E-39 | 1.03E-37 |
| 0.077703594 | 0.173937709 |
| 0.028603533 | 0.079946135 |
| 8.16E-08 | 1.11E-06 |
| NA | NA |
| 0.163212251 | 0.301075565 |
| 0.009218986 | 0.032198138 |
| 0.004136602 | 0.016690863 |
| 0.307461602 | 0.468965142 |
| 0.060243734 | 0.143492579 |
| 0.478116279 | 0.634945248 |
| 0.6851589 | 0.79777674 |
| 0.004204107 | 0.016900211 |

| | |
|---|---|
| 0.163761701 | 0.301878563 |
| 0.151954011 | 0.285891184 |
| 0.011572428 | 0.038800806 |
| 5.87E-58 | 9.56E-56 |
| 0.141270903 | 0.271040627 |
| 0.665834422 | 0.783284284 |
| 3.55E-08 | 5.11E-07 |
| 0.174603059 | 0.315558238 |
| 0.057957775 | 0.139091648 |
| 0.187408349 | 0.331428302 |
| NA | NA |
| NA | NA |
| 0.037764237 | 0.099519971 |
| 0.01544058 | 0.048800515 |
| 0.941278332 | NA |
| 0.067703782 | 0.157110794 |
| 0.501711785 | 0.653993365 |
| 0.15571648 | 0.290991886 |
| 1.06E-05 | 9.51E-05 |
| 0.81428631 | NA |
| 0.015534214 | 0.04904435 |
| 0.373624869 | 0.536791159 |
| 2.93E-09 | 4.90E-08 |
| 0.111721471 | 0.227670758 |
| 0.000917339 | 0.004740821 |
| 0.265218276 | 0.424384902 |
| NA | NA |
| NA | NA |
| NA | NA |
| 0.955792398 | NA |
| NA | NA |
| NA | NA |
| NA | NA |
| 0.314168915 | 0.476348818 |
| 0.839319574 | NA |
| 0.176975826 | 0.318533381 |
| 3.68E-05 | 0.000288391 |
| 0.004899628 | 0.019197266 |
| 0.026226576 | 0.074493819 |
| 0.941276431 | NA |
| 1.76E-08 | 2.65E-07 |
| 0.191628887 | 0.336864901 |
| 0.040362446 | 0.10488067 |
| 0.005097463 | 0.019828375 |
| 0.017098002 | 0.052949083 |
| 0.597458228 | 0.73204472 |
| 0.01289687 | 0.042259265 |

| | |
|---|---|
| 0.490037247 | 0.644428598 |
| 0.692080023 | 0.802420415 |
| 0.101150722 | 0.211581817 |
| 0.49706233 | 0.650023065 |
| 0.04373374 | 0.111510496 |
| NA | NA |
| 0.239857553 | 0.395887211 |
| 0.123733725 | 0.245753514 |
| 0.001288646 | 0.006320288 |
| 0.022857606 | 0.066714497 |
| 1.03E-07 | 1.38E-06 |
| 0.081715881 | 0.180537181 |
| 0.00339233 | 0.014195543 |
| 0.780040514 | 0.864527319 |
| 0.919073652 | 0.951369534 |
| 0.001419365 | 0.006845741 |
| 0.319534076 | 0.482096228 |
| 0.615826671 | 0.745835568 |
| 0.032861102 | 0.089246914 |
| 0.718850174 | NA |
| 0.511727952 | 0.663169515 |
| 0.038871043 | 0.101749541 |
| 0.677419616 | 0.792019928 |
| 0.715623464 | 0.819182883 |
| 0.366388442 | 0.529591164 |
| 0.211896872 | 0.362328017 |
| 0.003231663 | 0.013650406 |
| 0.541546158 | 0.688672418 |
| 0.058369196 | 0.139909728 |
| 9.66E-18 | 3.54E-16 |
| NA | NA |
| 8.86E-05 | 0.000629702 |
| 0.716857308 | 0.820043141 |
| NA | NA |
| 0.118281893 | 0.237622014 |
| 0.941276431 | NA |
| 0.03179328 | 0.086832318 |
| 0.856636058 | NA |
| 0.542187485 | 0.689156602 |
| 0.942188863 | NA |
| NA | NA |
| 0.465021917 | 0.623500572 |
| 0.941277857 | NA |
| 1.54E-06 | 1.66E-05 |
| 0.011389176 | 0.038299685 |
| 0.01992258 | 0.059654956 |
| 0.372678493 | 0.535851969 |

| | | |
|---|---|---|
| NA | 0.469183972 | 0.627247636 |
| | 0.144011039 | 0.274923864 |
| | 0.124610974 | 0.247124679 |
| | 0.687142604 | 0.799229101 |
| | 3.97E-09 | 6.55E-08 |
| | 0.599370768 | 0.733284819 |
| | 1.29E-06 | 1.41E-05 |
| | 0.334805153 | 0.498033389 |
| | 0.000273669 | 0.001680131 |
| | 0.183146478 | 0.325826878 |
| | 0.517114196 | 0.667952031 |
| | 0.341082159 | 0.504595534 |
| | 0.135091356 | 0.262142872 |
| | 0.308155565 | 0.46960479 |
| | 0.465143747 | 0.623597581 |
| | 0.854031865 | 0.91208307 |
| | 0.160089196 | 0.296772677 |
| | 0.559222712 | 0.702666895 |
| | 0.516095519 | 0.666925759 |
| | 0.57499573 | 0.714074899 |
| | 0.941278332 | NA |
| | 0.010759524 | 0.036538476 |
| | 3.79E-15 | 1.17E-13 |
| | 0.000182883 | 0.0011861 |
| | 4.61E-17 | 1.62E-15 |
| | 4.95E-44 | 5.83E-42 |
| | 0.148660012 | 0.281534875 |
| | 0.001545898 | 0.007362039 |
| | 0.009514511 | 0.033041201 |
| | 0.615229199 | NA |
| | 0.928502123 | 0.957409069 |
| | 3.28E-14 | 9.39E-13 |
| | 0.839319108 | NA |
| | 0.122890741 | 0.244649738 |
| | 0.909894411 | NA |
| | 0.000846298 | 0.004429107 |
| | 5.96E-41 | 6.33E-39 |
| | 7.33E-12 | 1.68E-10 |
| | 0.003791734 | 0.015517685 |
| | 0.362358416 | NA |
| | 0.114345414 | 0.231629748 |
| | 0.163255471 | NA |
| NA | | NA |
| | 0.855521257 | NA |
| | 0.648513854 | NA |
| | 0.015188059 | 0.048194397 |
| NA | | NA |

| | |
|---|---|
| 6.86E-07 | 7.87E-06 |
| 1.02E-22 | 5.10E-21 |
| 0.081184596 | NA |
| 7.25E-20 | 3.05E-18 |
| 0.678529608 | NA |
| 6.06E-39 | 5.98E-37 |
| 0.005253018 | 0.020317248 |
| 1.66E-27 | 1.05E-25 |
| 2.83E-07 | 3.50E-06 |
| 9.63E-70 | 2.03E-67 |
| 5.94E-10 | 1.09E-08 |
| 1.57E-17 | 5.68E-16 |
| 0.106574259 | NA |
| 1.79E-56 | 2.85E-54 |
| 3.55E-11 | 7.55E-10 |
| 0.545862296 | 0.692127337 |
| 0.829075137 | 0.89633205 |
| 3.59E-51 | 5.06E-49 |
| NA | NA |
| 0.131257156 | 0.256553487 |
| 0.128010737 | 0.25206085 |
| 0.158205982 | NA |
| 0.65737011 | NA |
| NA | NA |
| NA | NA |
| NA | NA |
| 0.695949126 | NA |
| 0.941278332 | NA |
| 0.180652514 | 0.322693519 |
| 0.064559278 | 0.151587249 |
| 0.839318641 | NA |
| NA | NA |
| 0.269923956 | NA |
| 0.012103721 | 0.040146708 |
| 0.839319574 | NA |
| 0.010711054 | 0.03640504 |
| 0.678318884 | NA |
| NA | NA |
| 0.043684611 | 0.111444922 |
| NA | NA |
| 0.941278332 | NA |
| 0.839318641 | NA |
| NA | NA |
| NA | NA |
| 0.955791446 | NA |
| 0.839318641 | NA |
| NA | NA |

| | | |
|---|---|---|
| NA | NA | |
| NA | NA | |
| NA | NA | |
| 0.941276431 | NA | |
| NA | NA | |
| NA | NA | |
| 0.941280538 | NA | |
| 0.839319108 | NA | |
| 0.955791922 | NA | |
| NA | NA | |
| 3.91E-86 | | 1.12E-83 |
| 0.022801852 | | 0.066600769 |
| 8.59E-07 | | 9.68E-06 |
| 0.41363169 | | 0.575813632 |
| 0.344724139 | | 0.508340497 |
| 0.017257725 | | 0.053325766 |
| 0.179032139 | | 0.320641308 |
| 0.941278332 | NA | |
| 0.894370951 | NA | |
| 0.94127925 | NA | |
| 0.941280876 | NA | |
| NA | NA | |
| 0.868389157 | | 0.921216246 |
| 1.71E-05 | | 0.000145635 |
| 6.25E-31 | | 4.52E-29 |
| 0.164820814 | NA | |
| 0.840100075 | NA | |
| 0.941278332 | NA | |
| NA | NA | |
| NA | NA | |
| 0.839319574 | NA | |
| 0.941278332 | NA | |
| NA | NA | |
| NA | NA | |
| NA | NA | |
| NA | NA | |
| NA | NA | |
| NA | NA | |
| NA | NA | |
| 0.941278332 | NA | |
| NA | NA | |
| NA | NA | |
| NA | NA | |
| NA | NA | |
| NA | NA | |
| NA | NA | |
| 0.839318641 | NA | |

| | | |
|---|---|---|
| NA | | NA |
| NA | | NA |
| NA | | NA |
| NA | | NA |
| NA | | NA |
| NA | | NA |
| NA | | NA |
| NA | | NA |
| | 0.955792873 | NA |
| NA | | NA |
| NA | | NA |
| NA | | NA |
| NA | | NA |
| NA | | NA |
| NA | | NA |
| NA | | NA |
| | 0.941277857 | NA |
| NA | | NA |
| NA | | NA |
| | 0.955792398 | NA |
| NA | | NA |
| NA | | NA |
| | 0.955792398 | NA |
| NA | | NA |
| NA | | NA |
| NA | | NA |
| NA | | NA |
| NA | | NA |
| | 1.34E-05 | 0.000117552 |
| NA | | NA |
| NA | | NA |
| | 0.279203229 | 0.438996785 |
| NA | | NA |
| | 0.601033226 | 0.734685336 |
| NA | | NA |
| | 0.042898216 | 0.109969125 |
| | 0.053934379 | 0.131625765 |
| | 0.807604926 | 0.882094918 |
| | 0.456737072 | 0.616424121 |
| | 0.568556378 | 0.709785949 |
| | 0.846130433 | 0.907311644 |
| NA | | NA |
| | 0.026014073 | 0.073994619 |
| | 0.232899874 | 0.38743939 |
| | 0.874720941 | NA |
| | 0.962629766 | 0.977454617 |
| | 0.005428327 | 0.020839523 |

| | |
|---|---|
| NA | NA |
| NA | NA |
| 0.089577931 | 0.193496135 |
| 0.810620976 | 0.884009918 |
| 0.582801678 | 0.72031251 |
| NA | NA |
| NA | NA |
| NA | NA |
| 7.84E-11 | 1.59E-09 |
| 0.839319108 | NA |
| 0.848090802 | 0.908595097 |
| 0.499051057 | 0.651943647 |
| 0.001238161 | 0.006120604 |
| 0.440159792 | 0.600905345 |
| 1.10E-08 | 1.72E-07 |
| NA | NA |
| 0.99284941 | 0.994981334 |
| NA | NA |
| NA | NA |
| 2.86E-08 | 4.17E-07 |
| 0.340176682 | 0.50383243 |
| NA | NA |
| 0.000483583 | 0.002759591 |
| 0.323257441 | 0.485958038 |
| 0.751411037 | 0.843994408 |
| 0.006552711 | 0.024328128 |
| 0.025940258 | 0.073837602 |
| 0.839318641 | NA |
| 0.334492821 | 0.4976932 |
| 0.046854517 | 0.117702503 |
| 0.015260585 | 0.048360065 |
| 0.877914847 | 0.926725968 |
| 0.001108809 | 0.005580859 |
| NA | NA |
| 0.922522153 | 0.953819221 |
| 0.747905452 | 0.841645576 |
| 0.114787446 | 0.232337227 |
| 0.530229078 | 0.679033278 |
| 8.87E-05 | 0.000630412 |
| 0.938855181 | 0.963601145 |
| 0.718428074 | 0.820971963 |
| 0.937924455 | NA |
| 0.072284386 | 0.164973915 |
| 0.945105437 | 0.96692025 |
| 0.030838267 | 0.08477852 |
| NA | NA |
| NA | NA |

| | | |
|---|---|---|
| | 0.001227095 | 0.006076155 |
| | 0.024499963 | 0.070564873 |
| | 0.030554566 | 0.084157375 |
| | 0.006049447 | 0.022816009 |
| | 0.975958796 | 0.985356151 |
| | 0.941278424 | NA |
| | 1.55E-08 | 2.35E-07 |
| | 0.742461047 | 0.838173868 |
| | 0.955792397 | NA |
| | 0.687178537 | 0.799229101 |
| | 0.603594117 | NA |
| | 0.288406072 | 0.449080291 |
| | 1.87E-26 | 1.12E-24 |
| | 0.016027681 | 0.050295433 |
| | 0.655245464 | NA |
| | 0.829902243 | 0.896696772 |
| | 0.127439478 | NA |
| | 0.001358196 | 0.006602896 |
| NA | | NA |
| | 0.853595154 | NA |
| | 2.34E-23 | 1.21E-21 |
| NA | | NA |
| | 0.007291178 | 0.0265681 |
| NA | | NA |
| NA | | NA |
| | 0.044928844 | 0.113850351 |
| | 0.801686363 | NA |
| | 0.581441018 | 0.719157093 |
| NA | | NA |
| | 0.941277857 | NA |
| | 0.02852667 | 0.079777422 |
| | 0.941277857 | NA |
| | 0.819650091 | 0.890104547 |
| | 0.955791922 | NA |
| | 3.60E-11 | 7.65E-10 |
| | 0.949147149 | 0.96888967 |
| | 0.830831238 | 0.897374649 |
| NA | | NA |
| | 8.01E-05 | 0.000576067 |
| | 0.728678906 | NA |
| NA | | NA |
| | 0.955792873 | NA |
| | 1.43E-17 | 5.19E-16 |
| | 0.994589076 | 0.996011831 |
| | 0.854582317 | NA |
| | 0.955792398 | NA |
| | 0.754154009 | 0.846066498 |

| | | |
|---|---|---|
| | 0.941277857 | NA |
| | 0.065040712 | 0.152461988 |
| | 0.498578038 | 0.651540269 |
| | 0.645159104 | 0.767688696 |
| | 0.839197272 | 0.902764778 |
| | 0.115674752 | 0.233579747 |
| | 0.321620902 | 0.484476976 |
| | 0.918558898 | 0.951111522 |
| | 0.02407681 | 0.069615675 |
| | 0.187704179 | 0.331902153 |
| | 0.281607103 | NA |
| | 0.424866928 | 0.586140534 |
| | 1.41E-12 | 3.50E-11 |
| | 0.001781763 | 0.008309078 |
| | 0.044036719 | 0.112117978 |
| | 1.19E-06 | 1.31E-05 |
| | 2.83E-129 | 1.27E-126 |
| | 2.05E-05 | 0.00017135 |
| | 0.058660094 | 0.14046555 |
| | 0.542905881 | 0.689701418 |
| | 0.003745947 | 0.015367291 |
| | 3.11E-05 | 0.000248949 |
| | 5.35E-05 | 0.00040294 |
| | 0.457827098 | 0.617514572 |
| | 0.022261072 | 0.065273033 |
| NA | | NA |
| NA | | NA |
| | 0.004990751 | 0.019494356 |
| | 0.001208841 | 0.00599934 |
| NA | | NA |
| | 0.959251204 | NA |
| NA | | NA |
| NA | | NA |
| | 0.316828028 | 0.479259023 |
| | 0.942148541 | NA |
| | 0.031367699 | 0.085916453 |
| NA | | NA |
| | 0.756057708 | 0.84719517 |
| | 0.116495317 | NA |
| | 0.018588702 | 0.05660802 |
| | 0.739827345 | 0.836336778 |
| | 0.487948913 | 0.642661519 |
| | 0.029669545 | 0.082278283 |
| | 6.67E-07 | 7.69E-06 |
| | 0.096336729 | 0.203967873 |
| | 8.47E-06 | 7.80E-05 |
| | 0.2764187 | 0.436265361 |

| | |
|---|---|
| 0.839318641 | NA |
| 0.512009113 | 0.663446073 |
| NA | NA |
| 0.940685982 | NA |
| NA | NA |
| 0.748395463 | NA |
| 0.001597637 | 0.007570626 |
| NA | NA |
| NA | NA |
| 0.955792398 | NA |
| 0.415552553 | 0.577406998 |
| 0.955791446 | NA |
| 0.537474019 | NA |
| NA | NA |
| 0.739991096 | NA |
| 0.941278332 | NA |
| 0.853595622 | NA |
| 0.721004972 | NA |
| NA | NA |
| 0.21777653 | 0.369117494 |
| 0.537487499 | NA |
| 0.001207617 | 0.005994517 |
| 0.386343171 | 0.54919614 |
| 4.87E-20 | 2.07E-18 |
| 0.039272875 | 0.10265462 |
| NA | NA |
| 3.86E-33 | 3.09E-31 |
| 0.085090468 | 0.186321482 |
| 0.446613121 | 0.607095084 |
| 3.45E-118 | 1.42E-115 |
| 2.35E-184 | 2.00E-181 |
| 3.61E-13 | 9.45E-12 |
| 1.62E-119 | 6.78E-117 |
| 3.13E-40 | 3.21E-38 |
| 7.46E-201 | 9.87E-198 |
| 0.015624825 | 0.049278134 |
| NA | NA |
| NA | NA |
| 9.92E-08 | 1.33E-06 |
| 0.620092261 | 0.748968897 |
| 0.000222207 | 0.001405148 |
| NA | NA |
| NA | NA |
| NA | NA |
| NA | NA |
| NA | NA |
| 0.839279346 | NA |

| | | |
|---|---|---|
| | 0.556760685 | 0.70090537 |
| | 0.017609451 | 0.054159473 |
| | 0.230546868 | 0.384492472 |
| | 0.158189044 | NA |
| | 6.04E-19 | 2.39E-17 |
| NA | | NA |
| | 0.000486928 | 0.002772691 |
| | 0.089016617 | 0.192633286 |
| | 0.001245194 | 0.006145938 |
| | 0.715092394 | 0.818964364 |
| | 0.006995307 | 0.025651388 |
| | 0.628596775 | 0.755481581 |
| | 0.000116507 | 0.000800777 |
| | 0.141515734 | 0.271335388 |
| | 0.793891254 | 0.873529939 |
| | 0.651339137 | 0.772339667 |
| | 2.81E-18 | 1.06E-16 |
| NA | | NA |
| | 0.080428612 | 0.178528567 |
| | 0.197735837 | 0.344799149 |
| | 0.003253248 | 0.013726966 |
| | 0.217272708 | 0.368604775 |
| | 0.005290061 | 0.020430639 |
| NA | | NA |
| | 0.782075196 | 0.866013204 |
| | 0.518133711 | 0.668869649 |
| | 0.267899303 | 0.427611267 |
| | 0.000919467 | 0.004749382 |
| | 0.105023001 | NA |
| | 7.04E-05 | 0.000513138 |
| NA | | NA |
| | 0.626780322 | NA |
| | 0.846128227 | 0.907311644 |
| | 0.95248084 | 0.971126534 |
| | 0.167142637 | 0.306141868 |
| | 0.758157993 | 0.8489686 |
| | 0.96846484 | NA |
| | 0.780292537 | 0.86472311 |
| NA | | NA |
| | 3.35E-09 | 5.58E-08 |
| NA | | NA |
| | 0.97931117 | 0.987066037 |
| | 0.066105567 | 0.154319803 |
| | 0.205063584 | 0.353690104 |
| | 0.872195017 | 0.923626646 |
| | 0.940913137 | 0.964662926 |
| NA | | NA |

| | | |
|---|---|---|
| | 0.18054638 | NA |
| | 0.941277857 | NA |
| | 0.941277857 | NA |
| | 0.936418804 | NA |
| | 0.839318641 | NA |
| | 0.795855186 | NA |
| NA | | NA |
| | 0.839318641 | NA |
| | 0.943837118 | NA |
| | 0.839319108 | NA |
| | 0.839318641 | NA |
| | 0.839318641 | NA |
| | 0.941932416 | NA |
| | 0.724742537 | NA |
| | 0.721661039 | 0.82332244 |
| | 0.981515839 | 0.988492849 |
| | 0.000129508 | 0.000879218 |
| | 0.755000438 | 0.846547975 |
| | 0.068533184 | 0.158598322 |
| | 0.618030501 | 0.747460621 |
| | 0.04142681 | 0.107066088 |
| | 2.32E-06 | 2.42E-05 |
| | 0.729036655 | 0.829402115 |
| | 0.047385268 | 0.118635126 |
| | 0.807857167 | 0.882239978 |
| | 0.274534269 | 0.434472384 |
| NA | | NA |
| | 0.00023064 | 0.001448085 |
| | 1.50E-13 | 4.06E-12 |
| | 0.118017063 | 0.237230144 |
| | 0.946094012 | 0.967272298 |
| | 0.877314982 | 0.926476383 |
| | 0.54612717 | 0.692426299 |
| | 2.80E-92 | 8.78E-90 |
| | 0.839318641 | NA |
| | 0.27559216 | 0.43545412 |
| | 0.009034249 | 0.031683099 |
| | 0.115441139 | 0.233214224 |
| | 0.177022399 | 0.318576074 |
| | 0.44169219 | 0.602275552 |
| | 0.853595154 | NA |
| | 0.461644799 | 0.62041928 |
| | 0.178982508 | 0.320600666 |
| | 0.44383793 | 0.604395584 |
| | 0.096737708 | 0.204488831 |
| | 0.077342058 | 0.173309703 |
| | 0.296596233 | 0.458057588 |

| | |
|---|---|
| 0.043332215 | 0.110783393 |
| 0.201889036 | 0.349888076 |
| NA | NA |
| 0.004733521 | 0.018678262 |
| 0.111962666 | 0.228006108 |
| 0.391108164 | 0.553556313 |
| 0.474970892 | 0.632341722 |
| 0.093418783 | 0.199421872 |
| 9.41E-11 | 1.89E-09 |
| 0.075959107 | 0.171064307 |
| 0.823985791 | 0.893060871 |
| 0.913988416 | 0.948208703 |
| 0.276082159 | 0.435882801 |
| 5.99E-06 | 5.73E-05 |
| 0.077265126 | 0.173202536 |
| 0.697358527 | NA |
| 0.002388887 | 0.010622134 |
| 0.496699109 | 0.649691279 |
| 0.481355454 | NA |
| 1.24E-05 | 0.000110011 |
| NA | NA |
| 0.05870599 | 0.140547171 |
| 0.061030099 | 0.145017311 |
| 0.525809999 | 0.675044348 |
| 0.045702334 | 0.115380457 |
| NA | NA |
| 0.031234427 | 0.08562039 |
| 0.001040902 | 0.005284886 |
| 0.906158045 | 0.943127106 |
| 0.941278207 | NA |
| 0.955792398 | NA |
| NA | NA |
| 1.46E-16 | 4.99E-15 |
| 0.449451753 | 0.609529407 |
| 1.89E-06 | 2.01E-05 |
| 1.16E-07 | 1.55E-06 |
| 0.246946346 | 0.403989506 |
| 0.39311215 | 0.555566762 |
| 0.085013779 | 0.186205758 |
| NA | NA |
| 0.306464252 | 0.468073802 |
| 0.003336017 | 0.014014098 |
| 0.80093399 | 0.878492596 |
| 0.149906888 | 0.283070521 |
| 0.219896126 | 0.371545935 |
| 0.165343137 | 0.303999222 |
| 0.18500795 | 0.328158524 |

| | | | |
|---|---|---|---|
| | 0.054291898 | | 0.132240663 |
| | 0.001558008 | | 0.007409331 |
| | 2.15E-07 | | 2.72E-06 |
| NA | | NA | |
| | 0.481297332 | | 0.637710927 |
| | 0.038882579 | | 0.101768544 |
| | 0.714560263 | | 0.818562875 |
| | 0.666168831 | | 0.7835615 |
| | 0.941277857 | NA | |
| | 0.616638929 | | 0.746287679 |
| | 2.09E-07 | | 2.66E-06 |
| | 0.541271123 | | 0.688462683 |
| | 0.80910826 | | 0.882967072 |
| NA | | NA | |
| | 0.955792398 | NA | |
| | 0.853596091 | NA | |
| | 0.416435452 | | 0.578127538 |
| | 0.373968355 | | 0.537090133 |
| | 0.008348085 | | 0.029685575 |
| | 0.060797893 | | 0.144494404 |
| | 5.21E-05 | | 0.000393431 |
| | 0.004028396 | | 0.016301238 |
| | 0.073865378 | | 0.167569931 |
| | 0.349560165 | | 0.51340436 |
| | 0.861074899 | | 0.91693382 |
| | 0.846388989 | NA | |
| NA | | NA | |
| | 0.933368057 | | 0.960386545 |
| | 0.823767788 | | 0.892905911 |
| | 0.106264852 | | 0.219539624 |
| | 0.634694337 | NA | |
| | 0.169073099 | | 0.308633937 |
| | 0.348588528 | | 0.512525091 |
| | 0.092539377 | | 0.198095354 |
| | 0.907883957 | | 0.944180274 |
| | 0.001816466 | | 0.008436199 |
| | 0.446232809 | | 0.6067196 |
| | 0.74432685 | NA | |
| | 0.155133989 | | 0.290062782 |
| | 0.691107379 | | 0.80183126 |
| | 0.055633262 | | 0.134872979 |
| | 0.036063642 | | 0.095994947 |
| | 0.071804887 | | 0.164131366 |
| | 0.082439973 | | 0.1817556 |
| | 0.372507124 | | 0.535768856 |
| | 0.792645162 | | 0.872437591 |
| | 0.995346713 | | 0.996644755 |

| | | |
|---|---|---|
| | 0.533457226 | 0.681736854 |
| | 0.842996373 | 0.905460274 |
| | 0.66861577 | NA |
| | 0.001054483 | 0.005337901 |
| | 0.928443192 | NA |
| | 0.899436455 | NA |
| | 1.60E-24 | 8.82E-23 |
| | 0.04957498 | 0.122902164 |
| | 0.78267199 | 0.866267226 |
| | 0.982347028 | 0.988869696 |
| | 0.127385229 | 0.251016079 |
| | 0.900678514 | 0.940137567 |
| | 0.690206221 | NA |
| | 0.023250203 | 0.067677795 |
| | 0.056359944 | 0.136287655 |
| | 0.27338167 | 0.433292143 |
| | 0.580485414 | 0.71823641 |
| | 0.073466688 | 0.166999725 |
| | 0.00944511 | 0.032843319 |
| | 5.67E-05 | 0.00042396 |
| | 6.84E-08 | 9.43E-07 |
| | 0.674190684 | NA |
| | 0.839318641 | NA |
| | 0.125549079 | 0.248467545 |
| | 0.350325977 | 0.514116709 |
| | 0.00034241 | 0.00204518 |
| | 0.487048405 | 0.642042874 |
| | 0.082351579 | 0.181628268 |
| | 6.20E-05 | 0.000459978 |
| | 0.955792398 | NA |
| | 0.598679533 | 0.733012873 |
| | 0.452142667 | 0.612132402 |
| | 0.17490059 | 0.315928044 |
| | 0.325212979 | 0.488096307 |
| NA | | NA |
| | 0.936054503 | NA |
| | 0.63354558 | 0.759182999 |
| | 0.00127159 | 0.006252092 |
| | 0.033794649 | 0.091158231 |
| | 0.568210142 | 0.70950256 |
| | 0.014591467 | 0.046655842 |
| | 0.442934969 | 0.603511256 |
| | 0.008250517 | 0.029417187 |
| | 0.921367846 | NA |
| | 0.035686243 | 0.095203312 |
| NA | | NA |
| NA | | NA |

| | |
|---|---|
| 0.122460737 | 0.244019293 |
| 1.49E-08 | 2.27E-07 |
| 0.941278332 | NA |
| 0.125834649 | 0.248846487 |
| 0.055523732 | 0.134730948 |
| NA | NA |
| 0.000132665 | 0.000899115 |
| 0.750793172 | 0.843499437 |
| NA | NA |
| NA | NA |
| 0.694449693 | NA |
| 0.713724286 | 0.817913599 |
| NA | NA |
| 0.853596091 | NA |
| NA | NA |
| NA | NA |
| NA | NA |
| NA | NA |
| 4.55E-05 | 0.000347678 |
| 0.579314459 | 0.71730661 |
| 0.94127925 | NA |
| 0.745592468 | 0.840134719 |
| NA | NA |
| 0.000184035 | 0.00119227 |
| 0.498065445 | NA |
| 0.000903073 | 0.004680953 |
| 0.504566491 | 0.656733833 |
| 0.033646814 | 0.090893369 |
| 0.693711379 | 0.803747042 |
| 0.409993542 | 0.571919725 |
| 0.787013578 | 0.869176744 |
| 0.00128356 | 0.006297939 |
| 0.941275481 | NA |
| 0.523706146 | 0.673469863 |
| 0.471079261 | 0.628611354 |
| 0.066558263 | 0.155188788 |
| 0.661764583 | 0.78014028 |
| 0.833891412 | 0.899431054 |
| 0.122646948 | 0.244288089 |
| 0.880010356 | 0.928128611 |
| NA | NA |
| 4.75E-07 | 5.60E-06 |
| 2.22E-15 | 6.97E-14 |
| 0.783946712 | 0.867199604 |
| 0.404044847 | 0.566143253 |
| 0.01128428 | 0.03799525 |
| 0.291362612 | 0.452558919 |

| | |
|---|---|
| 0.231950456 | 0.386183881 |
| NA | NA |
| 0.839319108 | NA |
| 5.50E-12 | 1.28E-10 |
| 0.883674194 | NA |
| 1.37E-34 | 1.17E-32 |
| 0.131211559 | 0.256544242 |
| 2.91E-05 | 0.000235225 |
| 0.144650492 | 0.275865608 |
| 0.005477704 | 0.02098846 |
| 0.168356216 | 0.307821067 |
| 0.254391756 | 0.412411973 |
| 0.381452753 | 0.543966116 |
| 0.454686434 | NA |
| 0.709147909 | 0.814355118 |
| 0.000485678 | 0.002768887 |
| 0.015637514 | 0.04931162 |
| 0.931313412 | 0.959158922 |
| 0.891451011 | 0.934565988 |
| NA | NA |
| 0.027968451 | 0.078512215 |
| 0.91515478 | 0.949005101 |
| 0.955791922 | NA |
| 3.67E-12 | 8.68E-11 |
| 0.950339281 | 0.969607594 |
| 0.887839888 | NA |
| 0.470627847 | 0.628248834 |
| NA | NA |
| 0.955791446 | NA |
| 0.925906778 | NA |
| 0.250258801 | 0.407794892 |
| NA | NA |
| NA | NA |
| 0.001825764 | 0.008469468 |
| 0.47692358 | 0.633714903 |
| 0.189355458 | 0.333953662 |
| 0.616209754 | 0.746071752 |
| 0.400839662 | 0.563228413 |
| 0.955792398 | NA |
| 0.689738514 | 0.801006425 |
| 0.899633268 | 0.93958188 |
| 0.491373528 | 0.645465283 |
| 0.152648589 | 0.286744671 |
| 0.089093815 | 0.192686977 |
| 0.292513493 | 0.453567428 |
| 0.763995555 | NA |
| 0.327335326 | 0.49010728 |

| | |
|---|---|
| 0.352162541 | 0.515730807 |
| 0.615923078 | 0.745914373 |
| 0.025309461 | 0.072379645 |
| 0.271244441 | 0.431100439 |
| 0.36207536 | NA |
| NA | NA |
| 0.375991959 | 0.53918307 |
| 0.005739374 | 0.021832922 |
| 0.536858637 | 0.684839603 |
| 0.658660816 | 0.777805355 |
| 0.115509452 | 0.233312625 |
| 0.762898218 | 0.852872264 |
| 0.016215035 | 0.05074954 |
| 0.000794181 | 0.004203492 |
| NA | NA |
| 0.007856445 | 0.028247599 |
| 0.566175962 | 0.707921135 |
| 0.003577841 | 0.014812973 |
| 0.801029676 | NA |
| NA | NA |
| 0.000181984 | 0.001182526 |
| NA | NA |
| 0.409539166 | 0.571486851 |
| NA | NA |
| 0.000744379 | 0.003978825 |
| NA | NA |
| NA | NA |
| NA | NA |
| 0.019616055 | 0.058974528 |
| 2.08E-05 | 0.000174082 |
| 0.839319108 | NA |
| 0.061252818 | 0.145415876 |
| 0.59294803 | 0.728393322 |
| 0.018082628 | NA |
| 0.941277857 | NA |
| 0.497089313 | 0.650023065 |
| 0.002091915 | 0.009493987 |
| 0.006995343 | 0.025651388 |
| 0.030300863 | 0.083638398 |
| 0.302824379 | 0.464272197 |
| 0.857838792 | 0.914920416 |
| 0.839319108 | NA |
| 0.75204143 | NA |
| 0.015190565 | 0.048195924 |
| NA | NA |
| 0.392195266 | NA |
| 0.955792398 | NA |

| | | |
|---|---|---|
| | 0.941276431 | NA |
| | 0.839318641 | NA |
| | 0.004985166 | 0.019480697 |
| | 0.894265368 | NA |
| NA | | NA |
| | 0.012165164 | 0.040328026 |
| | 0.075993585 | 0.171109575 |
| NA | | NA |
| | 0.713348974 | 0.817686106 |
| | 0.108101164 | 0.222137953 |
| | 0.067536345 | 0.156813976 |
| | 0.16653803 | 0.305482289 |
| | 0.366084276 | 0.529247931 |
| | 0.00012706 | 0.000865566 |
| NA | | NA |
| NA | | NA |
| | 0.116489995 | NA |
| | 0.114448945 | 0.231819741 |
| | 0.941278332 | NA |
| | 0.049094863 | 0.121927788 |
| NA | | NA |
| | 0.223967323 | 0.376580265 |
| | 2.12E-18 | 8.09E-17 |
| | 0.159626747 | 0.296206115 |
| NA | | NA |
| | 0.46228343 | 0.621057188 |
| | 0.075540456 | 0.170460175 |
| | 0.001314397 | 0.00642408 |
| | 0.000282557 | 0.001729788 |
| | 0.923246595 | NA |
| | 0.276518016 | 0.436298252 |
| | 0.025503477 | 0.072785822 |
| | 0.427036374 | 0.58851947 |
| NA | | NA |
| | 0.955792873 | NA |
| | 0.041924957 | 0.108056073 |
| | 0.955791922 | NA |
| | 0.313808621 | 0.475931745 |
| | 0.000314285 | 0.001895264 |
| | 0.21927293 | 0.370803078 |
| | 0.01688609 | 0.052415587 |
| | 0.869230086 | NA |
| | 0.996887411 | 0.997557982 |
| | 0.186623657 | 0.330384254 |
| | 0.01779507 | 0.054631529 |
| NA | | NA |
| | 0.464595526 | NA |

| | | |
|---|---|---|
| | 0.489553891 | 0.643954778 |
| | 0.950279069 | NA |
| | 0.893210198 | NA |
| | 0.520421767 | 0.670804662 |
| | 0.506584869 | NA |
| | 0.016311431 | 0.050997594 |
| | 0.404302029 | 0.566336917 |
| | 0.160908695 | 0.297865045 |
| | 0.68386772 | NA |
| | 0.119649142 | 0.23976165 |
| | 0.941278332 | NA |
| | 0.11652671 | 0.234928743 |
| | 0.362509862 | 0.526251236 |
| | 0.764141674 | NA |
| NA | | NA |
| | 0.664418005 | NA |
| | 0.236390589 | NA |
| | 0.749114657 | NA |
| | 0.839318641 | NA |
| NA | | NA |
| NA | | NA |
| | 0.901063737 | 0.940250726 |
| NA | | NA |
| NA | | NA |
| NA | | NA |
| NA | | NA |
| NA | | NA |
| NA | | NA |
| NA | | NA |
| | 0.04050097 | 0.105194684 |
| | 0.941278332 | NA |
| | 0.342759774 | NA |
| | 0.222200656 | NA |
| NA | | NA |
| | 0.713272423 | 0.817671349 |
| | 0.741030003 | 0.83729508 |
| | 0.766093558 | 0.854718237 |
| | 0.128894952 | 0.25318582 |
| | 0.477765167 | 0.634620599 |
| | 1.59E-07 | 2.07E-06 |
| | 0.064622215 | 0.151675176 |
| | 0.941277857 | NA |
| | 3.88E-08 | 5.54E-07 |
| | 0.678149665 | 0.792600949 |
| | 0.29662737 | NA |
| | 0.899252177 | 0.939460317 |
| | 0.754729442 | 0.846334728 |

| | | |
|---|---|---|
| | 0.899822829 | 0.939614942 |
| | 0.118862833 | 0.238508659 |
| | 0.625895829 | 0.753506273 |
| | 0.076316277 | 0.171641314 |
| | 0.699138859 | NA |
| | 0.613998741 | NA |
| | 0.35046443 | NA |
| | 0.966588632 | NA |
| | 0.214414996 | 0.365339168 |
| | 0.466176982 | NA |
| | 0.427694197 | 0.589041703 |
| | 0.435120531 | 0.59635245 |
| | 0.926703138 | NA |
| | 0.955792873 | NA |
| | 0.302445785 | 0.463781379 |
| | 0.340211345 | 0.50383243 |
| | 0.178413103 | 0.320006748 |
| | 0.713963613 | 0.818089346 |
| | 0.69783312 | NA |
| | 0.311332229 | 0.473412747 |
| | 0.576997204 | NA |
| | 0.848872661 | NA |
| | 0.093504703 | 0.199515816 |
| | 0.6585545 | NA |
| | 0.251703844 | NA |
| | 0.509297078 | 0.660939373 |
| | 0.913131503 | NA |
| | 0.532841603 | NA |
| | 0.716282505 | 0.819759372 |
| | 0.734544671 | 0.832752191 |
| | 0.373320109 | 0.536562863 |
| | 0.788433264 | 0.869873726 |
| NA | | NA |
| NA | | NA |
| | 0.186521411 | NA |
| NA | | NA |
| | 0.682470711 | NA |
| | 0.919670573 | NA |
| NA | | NA |
| NA | | NA |
| | 0.839318641 | NA |
| | 0.839318641 | NA |
| NA | | NA |
| NA | | NA |
| NA | | NA |
| | 0.941277857 | NA |
| NA | | NA |

| | | |
|---|---|---|
| | 0.839319108 | NA |
| NA | | NA |
| | 0.941275481 | NA |
| | 0.379698747 | NA |
| | 0.756130802 | NA |
| | 0.897402082 | NA |
| NA | | NA |
| NA | | NA |
| NA | | NA |
| | 0.941278838 | NA |
| | 0.941277857 | NA |
| NA | | NA |
| | 0.941599939 | NA |
| NA | | NA |
| | 0.853595154 | NA |
| | 0.929751076 | NA |
| | 0.839319108 | NA |
| | 0.955792873 | NA |
| | 0.923516642 | NA |
| | 0.035873597 | 0.095595986 |
| | 0.663388713 | 0.781527329 |
| | 0.358825499 | NA |
| | 0.806626069 | NA |
| | 0.875203635 | NA |
| | 0.449149741 | 0.609319092 |
| | 0.135684814 | NA |
| | 0.401917477 | NA |
| NA | | NA |
| | 0.951414318 | NA |
| | 0.476628786 | 0.633535398 |
| | 0.312355781 | NA |
| | 0.562227852 | 0.704917159 |
| | 0.700765382 | 0.808877387 |
| | 0.987378806 | NA |
| | 0.669098616 | NA |
| | 0.774308074 | NA |
| | 0.168384419 | 0.307848985 |
| | 0.613988252 | NA |
| | 0.260309652 | NA |
| | 0.376446961 | NA |
| | 0.629713959 | NA |
| | 0.959095933 | NA |
| | 0.282899038 | NA |
| | 0.940615927 | NA |
| | 0.350258065 | 0.514078611 |
| | 0.551683003 | NA |
| | 0.599275367 | NA |

| | | |
|---|---|---|
| | 0.199310718 | NA |
| | 0.320106465 | NA |
| | 0.845289779 | NA |
| | 0.73912886 | NA |
| | 0.550122886 | 0.695713562 |
| | 0.481762974 | NA |
| | 0.769066103 | NA |
| | 0.701939657 | NA |
| | 0.081726155 | NA |
| | 0.878796294 | NA |
| | 0.435724075 | NA |
| | 0.350542492 | NA |
| | 0.771662056 | NA |
| NA | | NA |
| | 0.399580211 | NA |
| | 0.059311491 | NA |
| NA | | NA |
| NA | | NA |
| | 0.641293676 | NA |
| NA | | NA |
| NA | | NA |
| | 0.284051471 | 0.444597429 |
| | 4.46E-05 | 0.000341816 |
| | 0.746320068 | 0.840457525 |
| | 0.920237177 | NA |
| | 0.501559889 | 0.653874786 |
| | 0.044853137 | 0.113682714 |
| NA | | NA |
| | 0.593337579 | 0.728721417 |
| | 0.000203601 | 0.001301306 |
| | 0.171363963 | NA |
| | 0.046922218 | NA |
| | 0.839318641 | NA |
| | 0.177930434 | 0.319413243 |
| NA | | NA |
| | 0.019161459 | 0.057938115 |
| | 0.200050134 | 0.347506808 |
| | 0.003413875 | 0.014259272 |
| | 0.041161049 | 0.106612234 |
| | 0.939587148 | NA |
| NA | | NA |
| | 5.79E-28 | 3.72E-26 |
| | 0.478417251 | 0.635274054 |
| | 3.91E-16 | 1.30E-14 |
| | 0.020950297 | 0.062084034 |
| | 0.980591934 | 0.988000796 |
| | 0.600334566 | NA |

| | | |
|---|---|---|
| | 0.839319108 | NA |
| | 0.140389811 | 0.269741546 |
| | 0.217896743 | 0.36921608 |
| | 0.860040617 | 0.916529517 |
| | 1.45E-06 | 1.58E-05 |
| | 0.01036968 | 0.035462517 |
| | 0.098933503 | 0.207857291 |
| | 0.547489608 | 0.693636432 |
| NA | | NA |
| | 0.932029703 | NA |
| | 0.08273171 | 0.182162827 |
| | 0.000652604 | 0.00355534 |
| | 0.611765021 | 0.74284415 |
| | 0.330602901 | 0.493602449 |
| | 0.954487334 | 0.972380908 |
| | 0.413009807 | 0.575191868 |
| | 0.017428116 | 0.05372672 |
| | 0.004805678 | 0.018900323 |
| | 0.000522791 | 0.002948691 |
| NA | | NA |
| | 0.001030077 | 0.005238865 |
| | 0.336028868 | 0.499167391 |
| | 0.028152773 | 0.078927245 |
| | 2.68E-05 | 0.00021816 |
| | 2.14E-15 | 6.73E-14 |
| | 0.871751571 | 0.92337093 |
| NA | | NA |
| | 0.016573327 | 0.051667104 |
| | 0.058173673 | 0.139539431 |
| | 0.151389333 | 0.285257196 |
| | 0.400579744 | 0.563101117 |
| | 0.105230704 | 0.217856937 |
| | 0.091848697 | 0.196918669 |
| | 0.075711423 | 0.170716491 |
| | 0.521979623 | NA |
| | 0.102523053 | 0.213690337 |
| | 0.839320041 | NA |
| | 0.673653179 | NA |
| | 0.000667058 | 0.003624957 |
| | 0.374934707 | 0.538088387 |
| | 0.014029629 | 0.04519329 |
| | 0.749787588 | 0.84281796 |
| | 0.321877377 | 0.484700548 |
| | 0.038340066 | 0.100744508 |
| | 0.433803971 | 0.594996337 |
| | 0.357217617 | 0.520794667 |
| | 0.938239498 | NA |

| | | |
|---|---|---|
| | 0.249402726 | NA |
| | 0.073510705 | 0.16705192 |
| | 0.496574631 | 0.649564155 |
| | 0.417698259 | 0.579238753 |
| | 0.75170613 | 0.844145736 |
| | 0.772189073 | NA |
| | 0.905301575 | NA |
| | 0.481733894 | 0.638181958 |
| | 0.001355443 | 0.006592205 |
| | 0.315375558 | 0.477790659 |
| | 0.422667092 | 0.583985727 |
| | 0.069368783 | 0.160131488 |
| | 0.107430126 | 0.221201858 |
| | 0.832029136 | NA |
| NA | | NA |
| NA | | NA |
| | 0.941275481 | NA |
| NA | | NA |
| | 0.005425853 | 0.020833385 |
| NA | | NA |
| NA | | NA |
| | 0.436893939 | 0.597881305 |
| | 0.353037758 | 0.516694627 |
| | 0.839318641 | NA |
| | 0.334016915 | 0.497277935 |
| NA | | NA |
| | 0.05862048 | 0.140384814 |
| | 0.085267003 | 0.186545368 |
| | 0.839318641 | NA |
| NA | | NA |
| NA | | NA |
| | 0.935946872 | NA |
| | 0.839319574 | NA |
| | 0.884415598 | 0.930962684 |
| | 0.680821984 | NA |
| | 0.76433004 | 0.853630988 |
| | 0.708037727 | 0.813747658 |
| | 0.005864403 | 0.022230376 |
| | 0.327806428 | NA |
| NA | | NA |
| | 0.038923109 | 0.101863424 |
| | 0.27345262 | 0.433293424 |
| | 0.013738586 | 0.044442216 |
| | 0.029511759 | 0.08189527 |
| | 0.009507571 | 0.033026736 |
| NA | | NA |
| | 0.941277857 | NA |

| | | |
|---|---|---|
| | 0.926049883 | NA |
| | 0.069645112 | 0.160625398 |
| | 0.156081994 | 0.291514721 |
| | 0.715379247 | 0.819139694 |
| | 0.181795366 | 0.324214576 |
| | 0.955792873 | NA |
| | 0.004674979 | 0.018490171 |
| | 0.837200141 | 0.901225435 |
| | 0.720087583 | 0.822000127 |
| NA | | NA |
| | 0.095648825 | 0.202926583 |
| | 0.535744361 | 0.683925303 |
| | 0.145260095 | 0.276687003 |
| NA | | NA |
| NA | | NA |
| | 0.784597358 | NA |
| | 0.038006232 | 0.100035867 |
| | 0.723917489 | NA |
| NA | | NA |
| | 0.953255299 | NA |
| | 0.935358748 | NA |
| | 0.057716844 | 0.138680794 |
| | 0.491576435 | NA |
| | 0.551710476 | NA |
| NA | | NA |
| | 0.941208171 | NA |
| | 0.000352728 | 0.002093674 |
| | 0.286541291 | 0.447112614 |
| | 0.941279303 | NA |
| | 0.040076502 | 0.104342692 |
| | 0.941278332 | NA |
| | 0.148688571 | 0.281534875 |
| | 0.437391285 | 0.598286728 |
| | 0.49294408 | 0.646663083 |
| | 0.772315678 | NA |
| | 0.066573125 | 0.155198191 |
| | 0.521414287 | NA |
| | 0.333537869 | 0.496800273 |
| | 0.182257231 | 0.324682779 |
| | 0.087885367 | 0.190827176 |
| | 0.780259584 | NA |
| | 0.839319108 | NA |
| NA | | NA |
| | 0.339955245 | 0.503648372 |
| | 0.817835508 | NA |
| | 0.002164739 | 0.009764091 |
| | 0.73486321 | 0.832977211 |

| | |
|---|---|
| 0.895552582 | 0.936820599 |
| 0.182378945 | 0.32485098 |
| NA | NA |
| NA | NA |
| 0.569516761 | 0.71061218 |
| 0.451979387 | 0.611980964 |
| 0.183630542 | 0.326372026 |
| 0.624267248 | 0.75230667 |
| 0.427250806 | 0.588567314 |
| 0.568119346 | 0.709438556 |
| 0.943764033 | 0.966087713 |
| 0.748618959 | 0.842225958 |
| 0.545464641 | 0.69184666 |
| 0.891675751 | NA |
| 0.839318641 | NA |
| 0.821695621 | NA |
| 9.20E-46 | 1.13E-43 |
| 0.100608662 | 0.210707203 |
| 0.355458975 | 0.519216602 |
| 0.723220731 | 0.824667042 |
| NA | NA |
| 0.000366873 | 0.002164139 |
| 0.303351053 | 0.464773666 |
| 0.073586651 | 0.167128765 |
| 0.081324448 | 0.179896331 |
| 0.000151447 | 0.001009165 |
| 0.885053371 | 0.931219182 |
| 0.008049415 | 0.028806522 |
| 0.146881837 | 0.279039147 |
| 0.45171412 | 0.611656587 |
| 0.791947557 | NA |
| 0.955126543 | 0.972692863 |
| 0.5480605 | 0.693991357 |
| 0.725143439 | NA |
| NA | NA |
| 0.540591317 | NA |
| 0.260019614 | 0.418571387 |
| NA | NA |
| NA | NA |
| 0.264143303 | 0.423119921 |
| 0.013522275 | 0.043908459 |
| NA | NA |
| NA | NA |
| 0.93838321 | NA |
| 0.632324948 | NA |
| NA | NA |
| 0.941277857 | NA |

| | |
|---|---|
| 2.68E-27 | 1.68E-25 |
| 7.66E-23 | 3.88E-21 |
| 0.07778824 | 0.174064696 |
| 0.357138769 | NA |
| NA | NA |
| NA | NA |
| 0.74884637 | NA |
| NA | NA |
| 0.026167429 | 0.074359734 |
| NA | NA |
| 0.096604038 | 0.204300535 |
| NA | NA |
| 0.09884305 | 0.207722256 |
| 0.098432882 | 0.207189553 |
| 0.219367658 | 0.370863627 |
| 0.002771094 | 0.012011942 |
| 0.839319108 | NA |
| 0.24649122 | 0.403483983 |
| 0.006753698 | 0.024942051 |
| 0.042509654 | 0.109208288 |
| 8.80E-16 | 2.84E-14 |
| 0.15193519 | 0.285878371 |
| 0.914215261 | NA |
| NA | NA |
| NA | NA |
| 0.385651245 | 0.548452551 |
| NA | NA |
| 0.362572837 | 0.526253598 |
| 0.395671306 | NA |
| NA | NA |
| 0.07123006 | 0.163170084 |
| 0.000154068 | 0.001025485 |
| 0.470797184 | NA |
| 1.44E-10 | 2.83E-09 |
| 0.627050719 | 0.754362457 |
| NA | NA |
| 0.955791922 | NA |
| 0.41711069 | NA |
| 0.650082722 | 0.771310948 |
| 0.616378724 | 0.746162466 |
| 0.839319108 | NA |
| 0.765031347 | NA |
| 0.000399341 | 0.002333691 |
| 0.554949093 | 0.699623852 |
| 0.113744153 | 0.230706292 |
| 0.004356953 | 0.017417566 |
| 0.226901332 | 0.380117222 |

| | | |
|---|---|---|
| | 0.152267752 | 0.28632304 |
| | 0.839321651 | NA |
| | 0.928797125 | NA |
| | 0.747521775 | 0.841412716 |
| | 0.71692576 | 0.820082033 |
| | 0.103143612 | 0.21468535 |
| | 0.14974212 | 0.282914902 |
| | 0.18482877 | 0.327889572 |
| | 0.773873981 | 0.860508485 |
| | 0.000562992 | 0.003141193 |
| | 0.267301596 | 0.426861948 |
| | 0.019322839 | 0.05831396 |
| | 0.483682074 | 0.639624465 |
| | 0.01981434 | 0.059426929 |
| | 0.731227753 | 0.830477641 |
| NA | | NA |
| | 9.84E-08 | 1.33E-06 |
| | 0.333604892 | 0.49686901 |
| | 0.855322399 | 0.912932007 |
| | 0.941278207 | NA |
| | 0.573616813 | 0.713222973 |
| | 0.176931609 | 0.318509012 |
| | 0.000133294 | 0.000902865 |
| NA | | NA |
| | 0.955792398 | NA |
| | 0.755671502 | 0.847100885 |
| | 0.000262627 | 0.00161986 |
| | 0.004854094 | 0.019046679 |
| | 0.001416037 | 0.006838458 |
| | 0.997911132 | 0.998288604 |
| | 0.054001101 | 0.13172107 |
| | 0.598858251 | 0.733026998 |
| | 0.526679943 | NA |
| | 0.35922194 | 0.522773255 |
| | 0.037533266 | 0.098980773 |
| | 0.853595622 | NA |
| | 0.750685413 | NA |
| | 0.900610105 | NA |
| | 0.941279268 | NA |
| | 0.920589357 | NA |
| | 0.962709292 | 0.977493669 |
| | 0.680262131 | 0.794173006 |
| | 0.431584621 | 0.592931833 |
| | 0.955792873 | NA |
| NA | | NA |
| | 0.941278332 | NA |
| | 0.941278332 | NA |

| | | |
|---|---|---|
| NA | NA | |
| | 0.955792398 | NA |
| | 0.813102129 | NA |
| | 0.47473503 | 0.632190321 |
| | 0.574815092 | 0.714004217 |
| | 0.487375264 | 0.642260328 |
| | 0.083015606 | 0.182703418 |
| | 0.057544424 | 0.138412825 |
| | 0.838882104 | NA |
| | 0.890869635 | 0.934367626 |
| | 0.148653939 | 0.281534875 |
| | 0.941277857 | NA |
| | 0.233616069 | 0.388223812 |
| | 0.822047081 | NA |
| | 0.841275891 | 0.904179636 |
| | 0.01827269 | 0.05582423 |
| | 0.063543084 | 0.149629253 |
| | 0.014494815 | 0.046396663 |
| | 0.113070794 | 0.229790901 |
| | 1.50E-08 | 2.29E-07 |
| | 0.016538175 | 0.051584541 |
| | 0.681265948 | 0.794897637 |
| | 0.0134021 | 0.043650355 |
| | 0.079479851 | 0.17700882 |
| | 0.475846186 | 0.633023196 |
| | 0.435253783 | 0.596391144 |
| | 0.2029588 | 0.35110294 |
| | 0.058885419 | 0.140863392 |
| | 0.987087149 | 0.991879441 |
| | 3.44E-07 | 4.17E-06 |
| | 0.899564145 | NA |
| NA | NA | |
| | 3.35E-05 | 0.000265951 |
| NA | NA | |
| NA | NA | |
| NA | NA | |
| | 9.36E-05 | 0.000661946 |
| | 0.11888066 | 0.238522926 |
| | 0.923272503 | NA |
| NA | NA | |
| | 0.129990546 | 0.254674045 |
| | 0.680412907 | NA |
| | 0.352688865 | NA |
| NA | NA | |
| | 0.64311933 | NA |
| | 0.077105836 | 0.172958979 |
| | 5.68E-06 | 5.46E-05 |

| | |
|---|---|
| 0.000170467 | 0.001119608 |
| 0.269956296 | 0.429971878 |
| NA | NA |
| NA | NA |
| 0.000199108 | 0.00127534 |
| 0.69140968 | 0.801978899 |
| 5.58E-06 | 5.38E-05 |
| 0.970784048 | 0.982410877 |
| 0.024678757 | 0.070951055 |
| 0.314105527 | 0.476290996 |
| 0.486184327 | 0.641507809 |
| 0.453163613 | 0.613157264 |
| 0.89366839 | NA |
| 0.250766503 | NA |
| 0.167154405 | 0.306141868 |
| 0.094945949 | 0.201831159 |
| 0.000126971 | 0.000865202 |
| 0.338483489 | 0.501999128 |
| 0.852882821 | 0.911554419 |
| 0.823289968 | 0.892591228 |
| NA | NA |
| 0.498011095 | 0.651013844 |
| NA | NA |
| NA | NA |
| 0.001439866 | 0.006926373 |
| 0.848319127 | NA |
| 0.513770604 | 0.664877297 |
| 0.29654903 | 0.458044128 |
| 0.20755424 | 0.356762964 |
| 2.61E-05 | 0.000213223 |
| 0.496427151 | 0.649478319 |
| 0.397856316 | 0.560540786 |
| NA | NA |
| 0.60601608 | NA |
| 0.879909693 | NA |
| 0.70505564 | 0.811927735 |
| 0.120467948 | 0.241118238 |
| 0.37930064 | 0.542132974 |
| 0.000342075 | 0.00204369 |
| 0.15640943 | 0.291959612 |
| 0.095337214 | 0.20240981 |
| 0.111443347 | 0.227201246 |
| 0.021164995 | 0.06261892 |
| 0.13837763 | 0.266836243 |
| 3.28E-05 | 0.000260687 |
| 0.202573096 | NA |
| 0.941278332 | NA |

| | |
|---|---|
| 0.240495528 | 0.396555217 |
| NA | NA |
| 0.839318641 | NA |
| 0.11767331 | NA |
| 0.009136147 | 0.031979203 |
| 0.773339775 | 0.860262341 |
| 0.889530478 | 0.933495192 |
| NA | NA |
| NA | NA |
| 0.941278332 | NA |
| 0.321814876 | 0.484685774 |
| 0.265718182 | 0.425070514 |
| 0.149182914 | 0.282239228 |
| 0.68944985 | 0.800908677 |
| 0.071973594 | 0.164438039 |
| 0.002121564 | 0.009600407 |
| 0.833357167 | NA |
| 0.734058779 | NA |
| 6.51E-05 | 0.000479455 |
| 0.001164929 | 0.005811702 |
| 0.000132552 | 0.000898603 |
| 0.558654328 | 0.702212205 |
| 0.944004221 | NA |
| 0.744707445 | 0.839594856 |
| NA | NA |
| 0.0417008 | 0.10766487 |
| 1.65E-05 | 0.000141908 |
| 0.393554593 | NA |
| 0.10452296 | 0.216712151 |
| 0.941278332 | NA |
| 0.00032274 | 0.001942318 |
| 0.16963467 | 0.309232168 |
| 0.000340394 | 0.002036207 |
| 8.44E-05 | 0.000603402 |
| 0.826682198 | 0.894768681 |
| 0.202576678 | 0.350623963 |
| 0.044464052 | 0.112961188 |
| 0.286514348 | 0.447099883 |
| 0.192505949 | 0.33805715 |
| NA | NA |
| 0.937862804 | 0.963153577 |
| 0.094955389 | 0.2018332 |
| 0.515594233 | 0.666627395 |
| 0.456424199 | 0.616141605 |
| 0.099498424 | 0.208786167 |
| 3.21E-05 | 0.000255413 |
| 0.000508105 | 0.002875394 |

| | | | |
|---|---|---|---|
| | 0.008260941 | | 0.029439576 |
| | 0.131234291 | | 0.256544242 |
| | 0.299546782 | | 0.461058749 |
| | 0.515705629 | | 0.666639079 |
| | 0.865209187 | | 0.919767844 |
| | 6.48E-05 | | 0.000478031 |
| | 0.149828693 | | 0.283011075 |
| | 1.30E-10 | | 2.57E-09 |
| | 0.862811936 | | 0.917839093 |
| | 0.007097357 | | 0.025973445 |
| NA | | NA | |
| | 0.631931881 | | 0.757936137 |
| | 0.568122068 | | 0.709438556 |
| NA | | NA | |
| | 0.372419598 | | 0.535738623 |
| | 0.156409448 | | 0.291959612 |
| NA | | NA | |
| | 0.19804414 | | 0.345167607 |
| | 0.270533862 | | 0.430545107 |
| | 0.699310731 | | 0.807891764 |
| | 0.693809724 | | 0.803747042 |
| | 0.488342284 | | 0.643036238 |
| | 0.023823041 | | 0.068982484 |
| | 0.151781174 | | 0.285753935 |
| | 0.338711051 | | 0.502180169 |
| | 0.004750624 | | 0.01872402 |
| | 0.04187797 | | 0.107970043 |
| | 0.342189291 | NA | |
| | 0.859180371 | | 0.916022897 |
| | 0.853594685 | NA | |
| | 0.010758364 | | 0.036538476 |
| | 0.002174641 | | 0.00979947 |
| | 0.09237461 | | 0.197813808 |
| | 1.05E-11 | | 2.37E-10 |
| | 0.402153821 | | 0.564423919 |
| | 0.680071606 | | 0.794122482 |
| | 0.015982498 | | 0.050186729 |
| | 0.679564552 | | 0.793747937 |
| | 0.015064711 | | 0.047873198 |
| | 4.51E-08 | | 6.36E-07 |
| | 0.032019738 | | 0.08730055 |
| | 0.036185819 | | 0.096245983 |
| | 0.669937696 | | 0.786469769 |
| | 0.853596091 | NA | |
| | 0.101614853 | | 0.212298853 |
| | 0.17825759 | | 0.319807579 |
| | 3.65E-07 | | 4.41E-06 |

| | |
|---|---|
| NA | NA |
| 0.01193063 | 0.039705377 |
| 0.824216337 | 0.893188729 |
| 0.168002472 | 0.307363169 |
| 0.464678929 | 0.623185028 |
| NA | NA |
| 0.007163908 | 0.026172732 |
| 0.931258788 | 0.959144166 |
| 0.346382771 | 0.510154242 |
| 0.416461317 | 0.578129726 |
| 0.003466783 | 0.014443616 |
| 0.739991096 | NA |
| 0.117307498 | 0.236122881 |
| 0.030160478 | 0.083338716 |
| 0.760325989 | NA |
| 0.009070056 | 0.031789939 |
| 0.322217224 | 0.484976247 |
| 0.718824888 | NA |
| 0.903830733 | 0.941698566 |
| 0.477531587 | NA |
| 0.852666549 | NA |
| 0.601812962 | 0.735298604 |
| 0.069806515 | 0.160923455 |
| 0.454152911 | 0.613980893 |
| 0.520805963 | NA |
| 0.634331999 | 0.759704652 |
| 0.136300528 | 0.263908162 |
| 0.005719605 | 0.021775118 |
| 0.706215269 | 0.812504425 |
| 0.015547759 | 0.049080605 |
| 0.30817691 | 0.46960479 |
| 0.456797075 | 0.616470147 |
| 0.292140355 | 0.453352767 |
| 0.839319108 | NA |
| 0.955792398 | NA |
| 0.006113264 | 0.02299793 |
| 5.76E-09 | 9.33E-08 |
| 6.16E-19 | 2.44E-17 |
| 0.257693225 | 0.41589464 |
| 0.90326073 | NA |
| 0.914297969 | 0.948364503 |
| 0.499485454 | 0.652296322 |
| NA | NA |
| NA | NA |
| 0.642399721 | 0.765400118 |
| 0.787101158 | 0.869192807 |
| 0.361661473 | 0.525332776 |

| | | |
|---|---|---|
| | 6.77E-06 | 6.39E-05 |
| | 0.0312698 | 0.085697615 |
| | 0.839319108 | NA |
| | 0.016208956 | 0.050737188 |
| | 0.020378329 | 0.06078258 |
| | 4.53E-23 | 2.32E-21 |
| | 0.949802528 | NA |
| NA | | NA |
| | 0.941275956 | NA |
| | 0.40198952 | 0.564293144 |
| NA | | NA |
| | 0.424461229 | 0.585920442 |
| NA | | NA |
| NA | | NA |
| | 0.545532949 | 0.691857164 |
| | 0.028918629 | 0.080618554 |
| | 0.190660457 | 0.335534211 |
| | 0.260920459 | 0.419680279 |
| | 0.771717895 | 0.858863398 |
| | 0.024479826 | 0.070515408 |
| | 0.545466533 | 0.69184666 |
| NA | | NA |
| | 0.848055616 | 0.908595097 |
| | 0.096931773 | 0.2047797 |
| | 4.37E-06 | 4.30E-05 |
| | 0.009766011 | 0.033766793 |
| | 3.30E-06 | 3.33E-05 |
| NA | | NA |
| | 0.00166068 | 0.00782723 |
| | 0.146072645 | 0.277856717 |
| | 0.222409212 | 0.374807708 |
| | 0.435868522 | 0.597027253 |
| | 0.851837912 | NA |
| | 0.046912945 | 0.117799547 |
| | 0.012880821 | 0.042229932 |
| | 0.941277857 | NA |
| | 0.388000727 | 0.550927239 |
| | 0.005473661 | 0.020979724 |
| | 0.044440484 | 0.112913364 |
| | 0.201286916 | 0.349124184 |
| | 0.367597327 | 0.530716839 |
| | 0.903026469 | 0.941349627 |
| NA | | NA |
| | 0.524129515 | 0.673723104 |
| | 0.839319574 | NA |
| | 0.941278332 | NA |
| | 0.720647675 | 0.822521145 |

| | |
|---|---|
| 0.806723297 | NA |
| 0.449610191 | 0.609640068 |
| 0.562701093 | 0.705250457 |
| 0.590254613 | 0.726284135 |
| 0.035786551 | 0.095406753 |
| 0.002790649 | 0.012072525 |
| 0.761817804 | NA |
| 0.028206368 | 0.079032563 |
| NA | NA |
| NA | NA |
| 0.086814743 | 0.189071692 |
| 0.22859715 | 0.382204929 |
| 0.014506861 | 0.046422735 |
| 0.020445567 | 0.060918964 |
| 0.526487933 | 0.675514301 |
| 0.027473324 | 0.077460324 |
| 0.839319108 | NA |
| 0.735533011 | NA |
| 0.172632495 | 0.312875859 |
| 0.663375488 | 0.781527329 |
| 0.212739828 | 0.363270923 |
| NA | NA |
| NA | NA |
| 1.54E-10 | 3.03E-09 |
| 0.839319108 | NA |
| 0.256704945 | 0.41484716 |
| 0.834791567 | NA |
| 0.158511326 | 0.294756764 |
| 4.05E-05 | 0.000314046 |
| 0.683654582 | 0.796921806 |
| NA | NA |
| 0.809878984 | 0.883463483 |
| 0.529816674 | 0.678833852 |
| 2.20E-07 | 2.78E-06 |
| 0.089925421 | 0.194105819 |
| NA | NA |
| 0.94127925 | NA |
| 6.75E-20 | 2.84E-18 |
| 0.779183709 | 0.863856288 |
| 0.955792873 | NA |
| NA | NA |
| 0.915881972 | NA |
| 0.000176399 | 0.00115284 |
| 0.001422369 | 0.006857454 |
| 0.117830596 | 0.23693536 |
| 0.01154048 | 0.038704593 |
| 0.955792873 | NA |

| | | | |
|---|---|---|---|
| | 0.636435736 | NA | |
| | 0.885248035 | | 0.931359371 |
| | 0.010262088 | | 0.035134956 |
| NA | | NA | |
| | 0.089607945 | | 0.193543403 |
| NA | | NA | |
| | 0.243618679 | | 0.400175894 |
| | 0.548908384 | | 0.694694389 |
| NA | | NA | |
| | 0.991306527 | | 0.994118841 |
| | 0.414327733 | NA | |
| | 0.049558797 | | 0.122874842 |
| | 0.955792873 | NA | |
| | 0.582345509 | NA | |
| | 0.853595622 | NA | |
| | 0.739991096 | NA | |
| | 0.941278424 | NA | |
| | 0.816636144 | NA | |
| | 0.672019408 | | 0.787912219 |
| NA | | NA | |
| NA | | NA | |
| NA | | NA | |
| | 0.941278424 | NA | |
| NA | | NA | |
| | 0.941278332 | NA | |
| NA | | NA | |
| NA | | NA | |
| NA | | NA | |
| NA | | NA | |
| NA | | NA | |
| NA | | NA | |
| | 0.009495088 | | 0.032993004 |
| NA | | NA | |
| NA | | NA | |
| NA | | NA | |
| | 0.941277857 | NA | |
| NA | | NA | |
| NA | | NA | |
| NA | | NA | |
| NA | | NA | |
| NA | | NA | |
| NA | | NA | |
| NA | | NA | |
| NA | | NA | |
| NA | | NA | |
| NA | | NA | |
| NA | | NA | |

| | |
|---|---|
| NA | NA |
| NA | NA |
| NA | NA |
| NA | NA |
| NA | NA |
| NA | NA |
| NA | NA |
| NA | NA |
| 0.955792873 | NA |
| NA | NA |
| NA | NA |
| NA | NA |
| 0.441844443 | 0.60240471 |
| 0.265128323 | 0.424269487 |
| 0.862681537 | 0.917823438 |
| 0.082111079 | 0.181202 |
| 0.000233838 | 0.001466231 |
| 0.021295489 | 0.062911162 |
| 7.08E-10 | 1.28E-08 |
| 0.326384952 | 0.489313939 |
| NA | NA |
| 0.002166231 | 0.009768973 |
| 0.236083642 | 0.391286321 |
| 5.68E-07 | 6.62E-06 |
| 0.279546964 | 0.439338282 |
| 0.955792398 | NA |
| 0.286439079 | 0.44706011 |
| 0.292290783 | 0.453408838 |
| NA | NA |
| 0.974400451 | 0.984534514 |
| 0.750685413 | NA |
| 0.839318641 | NA |
| 3.02E-11 | 6.48E-10 |
| 0.001968719 | 0.009046225 |
| 0.633714211 | NA |
| 0.914949096 | 0.948874488 |
| NA | NA |
| NA | NA |
| 0.414757777 | 0.576672972 |
| 0.743511643 | NA |
| 0.008809699 | 0.031023025 |
| NA | NA |
| 0.12816646 | 0.252283996 |
| 0.814391444 | NA |
| 0.873616258 | NA |
| 0.125358985 | 0.24823582 |
| NA | NA |

| | |
|---|---|
| 0.839320041 | NA |
| 0.142491619 | 0.272854828 |
| 0.466483497 | 0.624619722 |
| 1.92E-07 | 2.46E-06 |
| 0.00010824 | 0.000750461 |
| 0.000122692 | 0.000839168 |
| NA | NA |
| 0.281007617 | 0.441196787 |
| 0.033667539 | 0.090918398 |
| 0.051662909 | 0.127217108 |
| 0.233723162 | 0.388320446 |
| NA | NA |
| 0.941278332 | NA |
| NA | NA |
| NA | NA |
| NA | NA |
| 0.941278332 | NA |
| 0.719993123 | 0.821931718 |
| NA | NA |
| 0.941277857 | NA |
| NA | NA |
| NA | NA |
| 0.000166299 | 0.001096166 |
| 0.002028853 | 0.009265303 |
| 0.885727382 | NA |
| NA | NA |
| NA | NA |
| 0.85309909 | 0.911703688 |
| NA | NA |
| 0.611129382 | 0.742375296 |
| NA | NA |
| 0.488800266 | 0.643461309 |
| 0.001240122 | 0.006126479 |
| 8.57E-09 | 1.36E-07 |
| 0.787269278 | NA |
| 0.955792398 | NA |
| 0.678978277 | 0.793218855 |
| 0.428586609 | NA |
| 0.248999711 | 0.406299521 |
| 0.050566069 | 0.124968703 |
| 8.56E-10 | 1.54E-08 |
| NA | NA |
| 0.212395653 | NA |
| NA | NA |
| NA | NA |
| 0.951503881 | NA |
| 0.437478163 | 0.598364817 |

| | | | |
|---|---|---|---|
| | 0.955792398 | NA | |
| | 0.917409042 | | 0.950349045 |
| NA | | NA | |
| NA | | NA | |
| | 0.955792398 | NA | |
| NA | | NA | |
| NA | | NA | |
| | 0.839319108 | NA | |
| NA | | NA | |
| | 0.154341425 | NA | |
| NA | | NA | |
| | 0.101013287 | | 0.211350058 |
| | 0.262591433 | | 0.42151344 |
| NA | | NA | |
| NA | | NA | |
| | 0.839319574 | NA | |
| | 0.839318641 | NA | |
| | 0.828628445 | NA | |
| | 0.000743747 | | 0.003977378 |
| | 0.013988289 | | 0.045102852 |
| | 0.480585815 | | 0.637160636 |
| | 0.267186436 | | 0.426788239 |
| | 0.561938299 | | 0.704702602 |
| NA | | NA | |
| | 0.941278207 | NA | |
| | 2.53E-06 | | 2.62E-05 |
| | 0.388635633 | | 0.551434 |
| | 0.200775603 | | 0.348364259 |
| | 3.06E-08 | | 4.46E-07 |
| | 0.134276243 | NA | |
| NA | | NA | |
| | 0.507265838 | | 0.65900643 |
| | 0.3152337 | | 0.477636548 |
| | 0.114155161 | | 0.231323101 |
| | 0.819312393 | NA | |
| | 0.95456991 | NA | |
| | 0.949260615 | NA | |
| | 0.465761509 | | 0.623968899 |
| NA | | NA | |
| | 0.018601665 | | 0.056640251 |
| | 0.002553183 | | 0.011239292 |
| NA | | NA | |
| NA | | NA | |
| | 0.856133874 | NA | |
| | 0.955792873 | NA | |
| | 0.615234574 | NA | |
| | 0.941278332 | NA | |

| | |
|---|---|
| NA | NA |
| 0.839318641 | NA |
| NA | NA |
| 0.853595154 | NA |
| NA | NA |
| 0.539926651 | 0.687090149 |
| 7.16E-42 | 7.89E-40 |
| 0.000235874 | 0.001477442 |
| 3.46E-50 | 4.74E-48 |
| 0.331002453 | 0.494075027 |
| 0.000634232 | 0.003469543 |
| 0.253145779 | 0.410979868 |
| NA | NA |
| NA | NA |
| 0.915887873 | NA |
| 0.911737031 | 0.946698286 |
| 0.154891495 | 0.289723172 |
| 4.23E-06 | 4.18E-05 |
| NA | NA |
| 0.15182884 | 0.285768627 |
| 0.649280837 | 0.770749546 |
| 0.94199564 | 0.964941479 |
| 2.11E-10 | 4.08E-09 |
| 0.838457485 | NA |
| 1.27E-05 | 0.000112244 |
| 0.978661818 | 0.986662209 |
| NA | NA |
| NA | NA |
| NA | NA |
| 0.468761305 | 0.626823403 |
| NA | NA |
| 0.011115471 | 0.03752409 |
| 0.936169846 | NA |
| NA | NA |
| NA | NA |
| 0.570111267 | NA |
| 0.111082377 | 0.226681188 |
| NA | NA |
| 0.230287304 | 0.384177226 |
| 0.607085438 | NA |
| 0.221385632 | 0.373452929 |
| 0.611650285 | 0.742780616 |
| 5.15E-07 | 6.06E-06 |
| 0.8948617 | NA |
| NA | NA |
| 0.941277857 | NA |
| NA | NA |

| | |
|---|---|
| NA | NA |
| 0.006439039 | 0.023980912 |
| 0.834056753 | 0.899511501 |
| 0.955792398 | NA |
| NA | NA |
| 0.18004788 | 0.322007571 |
| 0.003187088 | 0.013498057 |
| NA | NA |
| NA | NA |
| 0.197486294 | 0.344439713 |
| NA | NA |
| NA | NA |
| 0.235171416 | NA |
| 0.027307694 | 0.07712123 |
| 0.107328444 | 0.221084519 |
| 1.12E-35 | 9.82E-34 |
| 0.049408971 | 0.122541662 |
| 2.17E-06 | 2.28E-05 |
| 0.040531294 | 0.105261961 |
| 0.941277857 | NA |
| 0.155911319 | 0.291242179 |
| NA | NA |
| 0.003491985 | 0.014538146 |
| 0.469782964 | 0.627798041 |
| 0.860204404 | 0.916609361 |
| 0.549091441 | 0.69480519 |
| 0.853596091 | NA |
| 0.839318641 | NA |
| 0.893761674 | 0.935954744 |
| 0.820043384 | 0.890409773 |
| 1.11E-05 | 9.95E-05 |
| 0.939397584 | NA |
| NA | NA |
| NA | NA |
| 0.275690199 | 0.435519885 |
| 0.885010317 | 0.931219182 |
| 0.016618191 | 0.051773059 |
| 0.09113234 | 0.196063556 |
| 0.216871391 | 0.368238897 |
| 0.378922642 | 0.541721228 |
| 0.178716675 | 0.320317342 |
| 0.465746973 | 0.623968899 |
| 0.042781286 | 0.109740293 |
| 0.912234101 | NA |
| 0.044996187 | 0.113984593 |
| 1.69E-07 | 2.19E-06 |
| 0.754265378 | 0.846066498 |

| | | |
|---|---|---|
| | 1.41E-05 | 0.000123024 |
| | 0.582079303 | 0.71975955 |
| | 0.935946872 | NA |
| | 0.033355357 | 0.090321297 |
| NA | | NA |
| NA | | NA |
| NA | | NA |
| | 0.000330687 | 0.00198462 |
| | 1.56E-06 | 1.68E-05 |
| | 0.766609892 | NA |
| NA | | NA |
| NA | | NA |
| NA | | NA |
| NA | | NA |
| | 1.79E-05 | 0.000152236 |
| | 0.850103266 | NA |
| | 0.000351094 | 0.002085538 |
| NA | | NA |
| | 0.377189711 | 0.540054711 |
| | 0.409445431 | 0.57144194 |
| NA | | NA |
| | 0.799198978 | 0.877382689 |
| | 2.97E-14 | 8.53E-13 |
| | 0.495149463 | 0.648127338 |
| NA | | NA |
| | 0.398683546 | NA |
| | 0.942661476 | 0.965291191 |
| | 1.71E-05 | 0.000146008 |
| | 1.85E-130 | 8.63E-128 |
| | 0.000242261 | 0.001511083 |
| | 0.917880069 | NA |
| | 0.012885294 | 0.042232961 |
| | 0.225576132 | 0.37848323 |
| | 0.955792873 | NA |
| NA | | NA |
| | 2.50E-05 | 0.000205688 |
| NA | | NA |
| | 0.653964645 | 0.774103167 |
| NA | | NA |
| | 0.90675032 | NA |
| | 0.154737616 | 0.289549114 |
| | 0.115004745 | 0.232609019 |
| | 0.348592734 | 0.512525091 |
| NA | | NA |
| NA | | NA |
| | 0.549093729 | 0.69480519 |
| NA | | NA |

| | | |
|---|---|---|
| | 0.734058779 | NA |
| | 0.011693291 | 0.039117908 |
| | 0.839318641 | NA |
| | 0.080332111 | 0.178380903 |
| | 0.063714976 | 0.149974669 |
| | 0.329062174 | 0.492042836 |
| | 8.47E-07 | 9.55E-06 |
| | 0.197031937 | 0.343802457 |
| | 0.941279303 | NA |
| NA | | NA |
| | 3.26E-05 | 0.000259061 |
| | 0.773378018 | NA |
| | 1.45E-07 | 1.90E-06 |
| | 0.695543117 | 0.805081541 |
| | 0.068403065 | 0.158332174 |
| | 0.8091288 | NA |
| | 0.177078294 | 0.318628481 |
| | 0.732211732 | NA |
| | 0.106641471 | NA |
| | 0.794228114 | 0.8738198 |
| | 0.955792873 | NA |
| | 0.955791446 | NA |
| NA | | NA |
| NA | | NA |
| NA | | NA |
| NA | | NA |
| NA | | NA |
| NA | | NA |
| | 0.685413192 | 0.797955725 |
| | 0.703671782 | 0.810844101 |
| | 3.42E-08 | 4.94E-07 |
| | 0.746060028 | 0.840278884 |
| | 0.839319574 | NA |
| | 0.941275956 | NA |
| | 0.985507561 | 0.990919693 |
| | 0.12793801 | 0.251938489 |
| | 0.223802442 | 0.376356206 |
| | 0.264597949 | 0.423591632 |
| | 0.061079058 | 0.145090192 |
| | 0.284689819 | NA |
| | 0.955792398 | NA |
| | 0.51021902 | 0.661775005 |
| | 0.734058779 | NA |
| | 0.636063985 | NA |
| | 0.808758998 | NA |
| | 0.005894292 | 0.022325896 |
| | 0.005228035 | 0.02023706 |

| | | |
|---|---|---|
| | 0.079022898 | 0.17626305 |
| | 0.08696329 | 0.189325915 |
| | 0.145192225 | 0.276601996 |
| | 0.347744802 | 0.511716957 |
| | 0.224595133 | 0.377286014 |
| | 1.79E-06 | 1.91E-05 |
| | 0.000230344 | 0.001446607 |
| | 5.91E-05 | NA |
| | 0.376503859 | 0.53956256 |
| NA | | NA |
| NA | | NA |
| | 0.020132631 | 0.060193051 |
| | 0.191492071 | 0.336649256 |
| | 0.18997269 | 0.334641705 |
| | 0.787531964 | 0.869426522 |
| | 0.000360805 | 0.00213576 |
| | 0.710323167 | 0.81515414 |
| | 0.202732126 | 0.350812801 |
| | 0.029973652 | 0.08295731 |
| | 0.811511034 | NA |
| | 0.356504426 | 0.520192878 |
| | 0.626330629 | NA |
| | 0.046957105 | 0.117885563 |
| | 0.000629025 | 0.003444227 |
| | 0.355951147 | 0.519582649 |
| | 0.486643875 | 0.64182213 |
| | 0.938938621 | 0.963601145 |
| | 0.137776446 | 0.266051839 |
| | 0.726110181 | NA |
| | 0.030502267 | 0.084040488 |
| | 0.707909309 | 0.813678645 |
| | 0.307720889 | 0.46930048 |
| | 0.962798624 | 0.977522084 |
| | 0.193118742 | 0.338808306 |
| | 0.782502018 | 0.866267226 |
| | 0.946624856 | NA |
| | 0.000746555 | 0.003988663 |
| | 0.941275956 | NA |
| | 0.379681089 | 0.542422487 |
| | 0.00077635 | 0.004121948 |
| | 0.460945126 | 0.619889027 |
| | 0.725389859 | NA |
| | 0.853595154 | NA |
| | 0.000195949 | 0.001258157 |
| | 0.023551901 | 0.068347029 |
| | 0.283522013 | 0.444089955 |
| | 0.141286205 | 0.271047759 |

| | | |
|---|---|---|
| | 0.002097458 | 0.009512898 |
| | 0.728678906 | NA |
| | 0.000193903 | 0.001247708 |
| | 0.069056874 | 0.159535253 |
| NA | | NA |
| | 0.017742304 | 0.054497654 |
| | 0.982590119 | NA |
| | 1.18E-06 | 1.30E-05 |
| | 0.08823494 | 0.191411596 |
| | 0.105419057 | 0.218133042 |
| | 0.839319107 | NA |
| | 0.592980962 | NA |
| | 0.421685583 | 0.583069609 |
| | 0.941277857 | NA |
| | 0.500550293 | 0.653188912 |
| NA | | NA |
| NA | | NA |
| | 0.012204923 | 0.040437302 |
| NA | | NA |
| | 0.57496721 | 0.714074899 |
| NA | | NA |
| | 0.920288337 | NA |
| | 0.903987238 | NA |
| | 0.03163536 | 0.086520147 |
| | 0.425625199 | 0.587016513 |
| | 0.769855935 | NA |
| | 0.881408354 | NA |
| | 0.005512935 | 0.021099676 |
| NA | | NA |
| NA | | NA |
| | 0.941277857 | NA |
| | 0.247753448 | 0.404960003 |
| | 0.553434523 | NA |
| | 0.107380384 | 0.221172369 |
| | 0.012622949 | 0.041544722 |
| | 0.378132653 | 0.541046793 |
| NA | | NA |
| | 0.415021475 | 0.576837439 |
| | 0.754739137 | 0.846334728 |
| | 0.344266432 | 0.507885808 |
| NA | | NA |
| | 0.525239582 | NA |
| NA | | NA |
| | 0.017993136 | 0.055111649 |
| | 0.102053379 | 0.212964627 |
| | 0.008651859 | 0.030553641 |
| | 1.49E-05 | 0.000129287 |

| | | |
|---|---|---|
| | 0.169307779 | NA |
| | 0.954214716 | NA |
| | 0.350707554 | 0.514581506 |
| | 0.696244425 | 0.805326553 |
| | 1.01E-11 | 2.29E-10 |
| NA | | NA |
| | 0.211587905 | 0.361953092 |
| | 0.741471171 | 0.83740696 |
| | 0.799992019 | 0.877807949 |
| | 0.671248618 | 0.787279964 |
| | 0.8699089 | 0.922132877 |
| NA | | NA |
| | 0.084591981 | 0.185469632 |
| NA | | NA |
| | 0.019818712 | 0.059426929 |
| | 0.090340807 | 0.194825759 |
| NA | | NA |
| | 0.941276431 | NA |
| NA | | NA |
| NA | | NA |
| | 0.955792398 | NA |
| | 0.122041385 | 0.243469034 |
| NA | | NA |
| | 0.210297015 | 0.3601849 |
| NA | | NA |
| NA | | NA |
| | 0.277647621 | 0.437434077 |
| | 0.459407417 | NA |
| NA | | NA |
| NA | | NA |
| NA | | NA |
| NA | | NA |
| NA | | NA |
| NA | | NA |
| NA | | NA |
| NA | | NA |
| | 0.926320375 | NA |
| | 0.723771192 | NA |
| | 0.239012883 | 0.394903835 |
| | 0.631319824 | 0.757431042 |
| | 0.955792398 | NA |
| | 0.955792873 | NA |
| | 0.034943064 | 0.093493123 |
| | 0.955792398 | NA |
| NA | | NA |
| | 0.865092243 | 0.919767844 |
| | 0.374376735 | 0.537546905 |

| | | | |
|---|---|---|---|
| | 0.641036688 | | 0.764541527 |
| NA | | NA | |
| NA | | NA | |
| | 0.447803584 | | 0.607987046 |
| | 0.189307881 | | 0.333894494 |
| | 0.082623147 | | 0.18199113 |
| | 0.002248646 | | 0.010094733 |
| | 0.856416059 | | 0.913730559 |
| NA | | NA | |
| | 0.839319108 | NA | |
| NA | | NA | |
| | 0.955792398 | NA | |
| | 0.955792398 | NA | |
| | 0.017557696 | | 0.054070162 |
| | 9.65E-08 | | 1.30E-06 |
| | 0.321841968 | | 0.484687284 |
| | 0.613793006 | | 0.744153273 |
| NA | | NA | |
| | 0.632086258 | NA | |
| | 0.017204916 | | 0.053204029 |
| | 0.023914214 | | 0.069212807 |
| | 0.524068167 | | 0.673680628 |
| | 0.213838178 | | 0.364711532 |
| | 0.20210759 | | 0.350190351 |
| | 0.249526927 | | 0.406970974 |
| | 0.686023046 | | 0.798470442 |
| NA | | NA | |
| NA | | NA | |
| | 0.48777245 | | 0.642605848 |
| | 6.03E-05 | | 0.000448267 |
| | 0.462065084 | | 0.620802314 |
| | 0.693776052 | | 0.803747042 |
| | 0.728923768 | NA | |
| | 0.027658143 | | 0.077856499 |
| | 0.918934822 | | 0.951349947 |
| | 0.367104262 | | 0.530271614 |
| | 0.68451196 | | 0.797335503 |
| | 0.421725804 | | 0.58309135 |
| | 0.88535145 | | 0.931368125 |
| | 0.01520499 | | 0.048222409 |
| | 0.002368214 | | 0.010547948 |
| | 0.388550262 | | 0.551363288 |
| NA | | NA | |
| | 0.839318641 | NA | |
| | 0.001584738 | | 0.007519717 |
| | 0.214531216 | | 0.365464644 |
| | 0.181610992 | | 0.324013442 |

| | | |
|---|---|---|
| | 0.563477158 | NA |
| | 0.000773559 | 0.004108962 |
| NA | | NA |
| | 0.740854781 | NA |
| | 0.839701948 | NA |
| NA | | NA |
| NA | | NA |
| | 0.26970656 | NA |
| | 0.853596091 | NA |
| NA | | NA |
| | 0.120051283 | 0.2404258 |
| | 0.7186384 | 0.821015225 |
| | 0.914708757 | NA |
| | 0.941278332 | NA |
| | 0.839319108 | NA |
| | 3.06E-09 | 5.11E-08 |
| | 0.771721347 | 0.858863398 |
| NA | | NA |
| | 0.941277857 | NA |
| | 0.032872718 | 0.089258093 |
| | 0.010478602 | 0.035757949 |
| | 1.03E-10 | 2.06E-09 |
| | 0.000135736 | 0.000916048 |
| | 0.98170931 | 0.988636341 |
| NA | | NA |
| NA | | NA |
| | 0.363344383 | 0.526983304 |
| NA | | NA |
| | 0.266897197 | 0.426544813 |
| NA | | NA |
| | 0.002522561 | 0.011125067 |
| | 0.290511738 | 0.451561444 |
| NA | | NA |
| | 0.391707535 | 0.554141025 |
| | 1.22E-06 | 1.33E-05 |
| | 0.011142995 | 0.037599315 |
| | 0.755790479 | 0.847194357 |
| | 0.536048111 | NA |
| | 0.731986345 | NA |
| | 0.355078112 | 0.518882667 |
| NA | | NA |
| NA | | NA |
| | 0.424706568 | NA |
| NA | | NA |
| NA | | NA |
| NA | | NA |
| | 0.011885766 | 0.039595573 |

| | | |
|---|---|---|
| NA | NA | |
| 0.955792873 | NA | |
| 0.754166792 | NA | |
| NA | NA | |
| NA | NA | |
| 0.198362517 | NA | |
| 0.839320041 | NA | |
| 0.156312131 | | 0.291852945 |
| NA | NA | |
| NA | NA | |
| NA | NA | |
| NA | NA | |
| NA | NA | |
| 0.281357927 | | 0.441630267 |
| NA | NA | |
| 0.144782917 | | 0.276043174 |
| 0.955791922 | NA | |
| 0.941277857 | NA | |
| 0.563014633 | | 0.705445208 |
| 0.46740271 | NA | |
| NA | NA | |
| NA | NA | |
| NA | NA | |
| NA | NA | |
| 0.955792873 | NA | |
| 0.941277857 | NA | |
| NA | NA | |
| NA | NA | |
| NA | NA | |
| 0.955792398 | NA | |
| 0.683118363 | NA | |
| 0.060366574 | | 0.143713261 |
| 0.000389757 | | 0.002286102 |
| 0.827033247 | | 0.894978193 |
| 0.775911599 | | 0.861874551 |
| NA | NA | |
| 0.941277857 | NA | |
| 0.676904027 | | 0.791715346 |
| 0.173481663 | | 0.314127846 |
| 2.57E-05 | | 0.000210515 |
| 0.756306129 | | 0.847373305 |
| 0.09321062 | | 0.199138245 |
| 0.839318641 | NA | |
| 0.539263245 | | 0.686499168 |
| 0.614338506 | | 0.744430877 |
| 0.761738057 | NA | |
| 0.693915177 | | 0.803747042 |

| | | |
|---|---|---|
| | 1.72E-59 | 2.93E-57 |
| | 1.96E-84 | 5.37E-82 |
| | 0.839319574 | NA |
| | 0.022312381 | 0.06537165 |
| | 0.026600495 | 0.075446257 |
| | 0.743121937 | 0.83852368 |
| | 0.939677211 | NA |
| | 1.40E-16 | 4.78E-15 |
| | 0.955790032 | NA |
| | 0.771513859 | NA |
| | 0.023614596 | 0.068512264 |
| | 0.000198931 | 0.001274548 |
| | 1.55E-12 | 3.81E-11 |
| | 0.955118169 | NA |
| | 0.823458367 | 0.892733139 |
| | 0.371229776 | NA |
| | 0.000173342 | 0.00113567 |
| | 0.017886623 | 0.054877211 |
| | 0.264057267 | 0.423096 |
| | 0.066099777 | 0.154319803 |
| | 0.955792873 | NA |
| | 0.941278332 | NA |
| | 0.955792398 | NA |
| NA | | NA |
| NA | | NA |
| NA | | NA |
| NA | | NA |
| | 0.955792398 | NA |
| NA | | NA |
| | 0.224624826 | 0.377286014 |
| NA | | NA |
| NA | | NA |
| | 0.941277857 | NA |
| NA | | NA |
| NA | | NA |
| | 0.941278332 | NA |
| NA | | NA |
| NA | | NA |
| NA | | NA |
| NA | | NA |
| | 0.955792873 | NA |
| NA | | NA |
| NA | | NA |
| | 0.941277857 | NA |
| | 0.955791446 | NA |
| NA | | NA |
| NA | | NA |

| | |
|---|---|
| NA | NA |
| 0.955792398 | NA |
| NA | NA |
| NA | NA |
| NA | NA |
| 0.839318641 | NA |
| NA | NA |
| NA | NA |
| NA | NA |
| NA | NA |
| NA | NA |
| NA | NA |
| NA | NA |
| NA | NA |
| NA | NA |
| NA | NA |
| 0.839319574 | NA |
| NA | NA |
| NA | NA |
| NA | NA |
| NA | NA |
| NA | NA |
| NA | NA |
| NA | NA |
| NA | NA |
| 0.944004221 | NA |
| NA | NA |
| NA | NA |
| NA | NA |
| NA | NA |
| NA | NA |
| NA | NA |
| 0.839319574 | NA |
| NA | NA |
| NA | NA |
| NA | NA |
| NA | NA |
| NA | NA |
| NA | NA |
| NA | NA |
| 0.839319108 | NA |
| NA | NA |
| 0.126320779 | 0.249619299 |
| NA | NA |
| NA | NA |
| NA | NA |
| NA | NA |

| | | |
|---|---|---|
| NA | | NA |
| NA | | NA |
| NA | | NA |
| NA | | NA |
| NA | | NA |
| NA | | NA |
| NA | | NA |
| NA | | NA |
| NA | | NA |
| NA | | NA |
| NA | | NA |
| NA | | NA |
| NA | | NA |
| NA | | NA |
| NA | | NA |
| NA | | NA |
| NA | | NA |
| NA | | NA |
| NA | | NA |
| NA | | NA |
| NA | | NA |
| NA | | NA |
| NA | | NA |
| NA | | NA |
| NA | | NA |
| NA | | NA |
| NA | | NA |
| NA | | NA |
| NA | | NA |
| NA | | NA |
| NA | | NA |
| NA | | NA |
| NA | | NA |
| | 0.955792873 | NA |
| NA | | NA |
| NA | | NA |
| NA | | NA |
| NA | | NA |
| NA | | NA |
| NA | | NA |
| NA | | NA |
| NA | | NA |
| | 0.941278332 | NA |
| NA | | NA |
| NA | | NA |
| NA | | NA |
| NA | | NA |

| | | |
|---|---|---|
| NA | | NA |
| | 0.89958943 | NA |
| NA | | NA |
| NA | | NA |
| NA | | NA |
| NA | | NA |
| NA | | NA |
| NA | | NA |
| NA | | NA |
| NA | | NA |
| | 0.955792398 | NA |
| NA | | NA |
| NA | | NA |
| NA | | NA |
| NA | | NA |
| NA | | NA |
| NA | | NA |
| NA | | NA |
| NA | | NA |
| | 0.116420427 | 0.234794018 |
| NA | | NA |
| NA | | NA |
| NA | | NA |
| NA | | NA |
| NA | | NA |
| NA | | NA |
| NA | | NA |
| | 0.084412634 | 0.185127569 |
| NA | | NA |
| NA | | NA |
| | 0.949019802 | NA |
| NA | | NA |
| | 0.230736037 | 0.384619312 |
| NA | | NA |
| NA | | NA |
| NA | | NA |
| NA | | NA |
| NA | | NA |
| NA | | NA |
| NA | | NA |
| NA | | NA |
| NA | | NA |
| NA | | NA |
| NA | | NA |
| NA | | NA |
| NA | | NA |
| NA | | NA |

| | | |
|---|---|---|
| NA | | NA |
| NA | | NA |
| NA | | NA |
| NA | | NA |
| NA | | NA |
| NA | | NA |
| NA | | NA |
| NA | | NA |
| NA | | NA |
| NA | | NA |
| NA | | NA |
| NA | | NA |
| NA | | NA |
| NA | | NA |
| NA | | NA |
| NA | | NA |
| NA | | NA |
| NA | | NA |
| NA | | NA |
| NA | | NA |
| NA | | NA |
| NA | | NA |
| NA | | NA |
| NA | | NA |
| NA | | NA |
| NA | | NA |
| NA | | NA |
| | 0.941278332 | NA |
| NA | | NA |
| NA | | NA |
| | 0.941278332 | NA |
| NA | | NA |
| NA | | NA |
| NA | | NA |
| NA | | NA |
| NA | | NA |
| NA | | NA |
| NA | | NA |
| NA | | NA |
| | 0.941277857 | NA |
| NA | | NA |
| NA | | NA |
| NA | | NA |
| NA | | NA |
| NA | | NA |
| NA | | NA |
| NA | | NA |

| | | |
|---|---|---|
| NA | | NA |
| NA | | NA |
| NA | | NA |
| NA | | NA |
| NA | | NA |
| NA | | NA |
| NA | | NA |
| NA | | NA |
| NA | | NA |
| | 0.839318641 | NA |
| NA | | NA |
| NA | | NA |
| | 0.955792398 | NA |
| | 0.817417153 | NA |
| NA | | NA |
| NA | | NA |
| NA | | NA |
| NA | | NA |
| NA | | NA |
| NA | | NA |
| NA | | NA |
| NA | | NA |
| NA | | NA |
| NA | | NA |
| NA | | NA |
| NA | | NA |
| NA | | NA |
| | 0.955792873 | NA |
| NA | | NA |
| NA | | NA |
| NA | | NA |
| NA | | NA |
| NA | | NA |
| NA | | NA |
| NA | | NA |
| NA | | NA |
| NA | | NA |
| | 0.839318641 | NA |
| | 0.839319574 | NA |
| NA | | NA |
| | 0.955792873 | NA |
| | 0.955792873 | NA |
| NA | | NA |
| NA | | NA |
| NA | | NA |
| NA | | NA |
| NA | | NA |

| | | | |
|---|---|---|---|
| | 0.839318641 | NA | |
| NA | | NA | |
| NA | | NA | |
| | 0.941278332 | NA | |
| NA | | NA | |
| | 0.955792873 | NA | |
| | 0.955792873 | NA | |
| NA | | NA | |
| NA | | NA | |
| NA | | NA | |
| NA | | NA | |
| NA | | NA | |
| NA | | NA | |
| | 0.941275956 | NA | |
| NA | | NA | |
| | 0.839318641 | NA | |
| NA | | NA | |
| NA | | NA | |
| NA | | NA | |
| | 0.955792398 | NA | |
| NA | | NA | |
| NA | | NA | |
| NA | | NA | |
| | 0.955792398 | NA | |
| NA | | NA | |
| NA | | NA | |
| | 0.682314696 | NA | |
| NA | | NA | |
| | 0.720046287 | NA | |
| NA | | NA | |
| | 0.955792398 | NA | |
| | 0.916080473 | NA | |
| | 0.839321695 | NA | |
| NA | | NA | |
| | 0.955792873 | NA | |
| | 0.941277857 | NA | |
| | 0.936155565 | NA | |
| | 0.941278332 | NA | |
| | 0.937272113 | NA | |
| | 0.839318641 | NA | |
| | 0.391499786 | NA | |
| NA | | NA | |
| | 0.941278332 | NA | |
| | 0.972945367 | | 0.983726508 |
| NA | | NA | |
| | 0.66962732 | | 0.78623601 |
| | 0.941278424 | NA | |

| | |
|---|---|
| NA | NA |
| NA | NA |
| 0.839319108 | NA |
| NA | NA |
| 0.636993064 | NA |
| 0.940836846 | NA |
| 0.068271408 | 0.158181256 |
| 0.131522896 | NA |
| 0.840934832 | 0.903976548 |
| 3.74E-06 | 3.75E-05 |
| 0.057292688 | 0.137940578 |
| 0.167637662 | 0.306837253 |
| 0.554370338 | 0.699308533 |
| 0.008426936 | 0.029919141 |
| 0.000242122 | 0.001510627 |
| 0.698315912 | 0.806977488 |
| 0.277735746 | 0.437434077 |
| 0.077679139 | 0.173901323 |
| 0.015778215 | 0.049689477 |
| 0.056637968 | 0.136793192 |
| 0.941278332 | NA |
| 0.350006639 | 0.513806465 |
| 0.285854633 | 0.446509514 |
| 0.058439666 | 0.140036337 |
| 0.091207176 | 0.196090073 |
| 3.36E-42 | 3.75E-40 |
| 0.595651136 | 0.730545078 |
| 0.000807488 | 0.004260749 |
| 0.000916997 | 0.004740821 |
| 0.059552246 | 0.142258386 |
| NA | NA |
| 1.23E-11 | 2.74E-10 |
| 0.763899107 | 0.853550209 |
| 0.06299298 | 0.148598506 |
| 0.015941527 | 0.050084506 |
| 6.78E-05 | 0.000496127 |
| 0.00109751 | 0.005531006 |
| 0.190278953 | 0.335036221 |
| 0.065799012 | 0.153754966 |
| 0.65787052 | NA |
| 0.607696 | 0.740244637 |
| 0.93919101 | 0.963727557 |
| 6.79E-05 | 0.000496795 |
| 0.701199638 | NA |
| 0.398465795 | 0.561233378 |
| 0.653627341 | 0.774003044 |
| 0.007434666 | 0.027008536 |

| | |
|---|---|
| 0.006732669 | 0.024879831 |
| 3.27E-05 | 0.000260067 |
| 0.019601803 | 0.058946571 |
| NA | NA |
| 0.34540562 | 0.509061583 |
| 0.776436168 | 0.862256038 |
| 0.236065521 | 0.391286321 |
| 0.533076553 | 0.681433304 |
| 0.012472076 | 0.041156294 |
| 0.131134917 | 0.256451671 |
| 0.85867375 | NA |
| 4.90E-13 | 1.27E-11 |
| NA | NA |
| 0.169887132 | 0.309467427 |
| 0.012494042 | 0.041205928 |
| 0.222763399 | 0.375192076 |
| 0.078201019 | 0.174757362 |
| 0.329860584 | 0.492865136 |
| 0.116898158 | 0.235375222 |
| 0.043574683 | 0.111236014 |
| 0.288974106 | 0.449788234 |
| 0.07482823 | 0.169133596 |
| 0.607196876 | 0.739825955 |
| 0.561080786 | 0.70404203 |
| 0.8312373 | 0.897731758 |
| 0.154304667 | 0.289011621 |
| 0.26745598 | NA |
| NA | NA |
| 0.941278332 | NA |
| NA | NA |
| NA | NA |
| 0.484905434 | 0.640530022 |
| 0.686050917 | NA |
| 0.299765469 | 0.461246215 |
| 0.813165824 | NA |
| 0.571100563 | 0.711506705 |
| 0.000621793 | 0.003410906 |
| 0.02826645 | 0.079139198 |
| NA | NA |
| 2.31E-05 | 0.000191779 |
| 0.024812645 | 0.071258516 |
| NA | NA |
| NA | NA |
| 0.000218717 | 0.001386272 |
| 0.554603737 | NA |
| 0.839319108 | NA |
| NA | NA |

| | | | |
|---|---|---|---|
| | 0.946758856 | | 0.967625072 |
| | 0.025392017 | | 0.072568602 |
| NA | | NA | |
| | 0.555298093 | | 0.699910124 |
| | 0.075894714 | | 0.170972717 |
| | 0.128936304 | | 0.25318582 |
| | 0.050444212 | | 0.124680492 |
| NA | | NA | |
| | 0.481893354 | | 0.638221925 |
| | 0.09432636 | | 0.200801003 |
| | 0.025887265 | | 0.073722025 |
| | 0.012702924 | | 0.041754313 |
| | 0.889426439 | NA | |
| | 0.753041079 | NA | |
| NA | | NA | |
| | 0.000101124 | | 0.00070751 |
| | 0.884502146 | NA | |
| | 0.769396625 | | 0.857238144 |
| | 0.006958355 | | 0.025563016 |
| | 0.208029087 | | 0.357354817 |
| | 0.065331155 | | 0.152871819 |
| NA | | NA | |
| NA | | NA | |
| NA | | NA | |
| NA | | NA | |
| NA | | NA | |
| NA | | NA | |
| NA | | NA | |
| NA | | NA | |
| NA | | NA | |
| NA | | NA | |
| NA | | NA | |
| | 0.29705154 | | 0.45855092 |
| NA | | NA | |
| | 0.410476548 | | 0.572425755 |
| | 0.398058198 | | 0.560725646 |
| | 0.098512816 | | 0.207284481 |
| | 0.482544757 | | 0.638617692 |
| NA | | NA | |
| NA | | NA | |
| | 0.229101704 | | 0.382783313 |
| NA | | NA | |
| NA | | NA | |
| | 0.92469464 | NA | |
| | 0.216653863 | | 0.368000803 |
| NA | | NA | |
| NA | | NA | |

| | | |
|---|---|---|
| NA | | NA |
| | 0.839320041 | NA |
| NA | | NA |
| | 0.955792873 | NA |
| NA | | NA |
| | 0.711945489 | 0.816425444 |
| NA | | NA |
| | 0.938552414 | NA |
| | 0.271535014 | 0.431360635 |
| | 0.003397816 | 0.014211549 |
| | 0.399643981 | 0.562340948 |
| | 0.839319574 | NA |
| | 0.43943054 | 0.60041129 |
| | 0.070880246 | 0.162626916 |
| | 0.116586903 | 0.235010286 |
| | 0.149481362 | 0.282534375 |
| | 0.381084409 | NA |
| | 0.806784551 | 0.88170566 |
| | 0.901186426 | 0.940279321 |
| | 0.867968974 | 0.921060969 |
| | 9.12E-06 | 8.36E-05 |
| | 0.00024711 | 0.001537705 |
| | 0.078077093 | 0.174545972 |
| | 0.986560852 | 0.991639555 |
| | 1.91E-24 | 1.04E-22 |
| NA | | NA |
| | 0.147819446 | 0.280405674 |
| | 2.81E-07 | 3.48E-06 |
| | 0.117388091 | 0.236245147 |
| | 0.297537158 | NA |
| | 0.92420459 | NA |
| NA | | NA |
| | 0.821760476 | 0.891558606 |
| NA | | NA |
| NA | | NA |
| | 0.338884898 | NA |
| | 0.064055804 | 0.150568463 |
| | 0.001239786 | 0.006126095 |
| NA | | NA |
| | 9.43E-05 | 0.000666652 |
| | 0.221935963 | 0.374169131 |
| | 0.004172531 | 0.016814873 |
| | 0.709902598 | NA |
| NA | | NA |
| | 0.75606819 | 0.84719517 |
| NA | | NA |
| NA | | NA |

| | |
|---|---|
| NA | NA |
| NA | NA |
| NA | NA |
| NA | NA |
| 0.853596091 | NA |
| 1.60E-29 | 1.10E-27 |
| 0.256053999 | NA |
| 0.151696842 | 0.285723529 |
| 0.817583941 | 0.888671704 |
| 0.003183948 | 0.013489558 |
| 0.853596091 | NA |
| 0.17317309 | 0.313636327 |
| 0.554099835 | 0.699108629 |
| 0.582399869 | 0.720034467 |
| 0.010292899 | 0.035230309 |
| 0.972850905 | 0.983678728 |
| 0.208902599 | 0.358260657 |
| NA | NA |
| 0.286278774 | 0.446937386 |
| 0.001791955 | 0.00834516 |
| NA | NA |
| NA | NA |
| 0.735073227 | NA |
| 0.85230442 | 0.911222648 |
| NA | NA |
| 0.767436123 | NA |
| 0.584720745 | 0.721733424 |
| 5.30E-07 | 6.22E-06 |
| 0.710735323 | NA |
| 0.116712299 | 0.23514357 |
| 0.174978584 | 0.316044943 |
| 0.239175619 | 0.395063017 |
| 0.043325799 | 0.110778889 |
| 0.560006216 | 0.703354333 |
| 0.73508158 | NA |
| 2.88E-05 | 0.000232892 |
| NA | NA |
| 0.782629377 | NA |
| NA | NA |
| NA | NA |
| NA | NA |
| NA | NA |
| NA | NA |
| 0.781946562 | 0.865974863 |
| 0.906653477 | NA |
| 5.19E-06 | 5.03E-05 |
| NA | NA |

| | | |
|---|---|---|
| | 0.656048262 | 0.775605262 |
| | 0.941275956 | NA |
| | 0.868468447 | 0.921216246 |
| | 0.781781668 | 0.865889588 |
| | 0.031405802 | 0.086010919 |
| | 0.356542585 | 0.520192878 |
| | 0.283982432 | NA |
| | 0.687328218 | 0.799299743 |
| | 0.482326307 | 0.638541118 |
| | 0.955791922 | NA |
| NA | | NA |
| | 0.002544962 | 0.0112114 |
| | 0.955792873 | NA |
| NA | | NA |
| | 0.422496847 | NA |
| | 0.910423936 | 0.94586711 |
| | 0.839318641 | NA |
| | 0.754263642 | 0.846066498 |
| | 0.000562102 | 0.003138436 |
| | 0.941278332 | NA |
| | 0.907318207 | NA |
| | 0.022149874 | 0.065007557 |
| | 0.839319107 | NA |
| | 0.989775754 | NA |
| | 0.92796026 | NA |
| NA | | NA |
| NA | | NA |
| | 0.007993616 | 0.028645596 |
| | 0.703928909 | 0.811026251 |
| NA | | NA |
| | 0.955791922 | NA |
| | 0.069878039 | 0.161025954 |
| | 0.839319416 | NA |
| | 0.941277857 | NA |
| NA | | NA |
| NA | | NA |
| | 0.619441729 | 0.748575957 |
| | 0.057642803 | 0.138573276 |
| NA | | NA |
| | 0.047483159 | 0.118842708 |
| | 0.662011802 | NA |
| | 0.495254432 | 0.648193445 |
| NA | | NA |
| NA | | NA |
| | 0.921188999 | NA |
| | 0.008382842 | 0.029777582 |
| | 0.645699248 | NA |

| | | | |
|---|---|---|---|
| | 0.028235227 | | 0.07906528 |
| NA | | NA | |
| | 0.01177314 | | 0.039302141 |
| | 0.031731809 | | 0.086694274 |
| NA | | NA | |
| | 0.200062423 | | 0.347506808 |
| | 0.941533104 | NA | |
| | 0.420368624 | NA | |
| | 0.000193673 | | 0.001246904 |
| | 0.003894598 | | 0.015865004 |
| | 0.955792398 | NA | |
| | 0.941278332 | NA | |
| | 0.703857386 | NA | |
| | 0.031429586 | | 0.086066154 |
| | 0.001110101 | | 0.005583816 |
| NA | | NA | |
| NA | | NA | |
| NA | | NA | |
| | 0.941278424 | NA | |
| | 0.158715032 | | 0.295027459 |
| NA | | NA | |
| NA | | NA | |
| | 0.839319108 | NA | |
| NA | | NA | |
| NA | | NA | |
| NA | | NA | |
| NA | | NA | |
| NA | | NA | |
| NA | | NA | |
| | 0.941277857 | NA | |
| NA | | NA | |
| NA | | NA | |
| NA | | NA | |
| NA | | NA | |
| | 0.839319574 | NA | |
| NA | | NA | |
| | 0.839318641 | NA | |
| NA | | NA | |
| NA | | NA | |
| NA | | NA | |
| | 0.855715846 | NA | |
| | 0.839319107 | NA | |
| NA | | NA | |
| NA | | NA | |
| NA | | NA | |
| NA | | NA | |
| NA | | NA | |

| | | |
|---|---|---|
| NA | NA | |
| NA | NA | |
| 0.941278332 | NA | |
| NA | NA | |
| 0.941278332 | NA | |
| NA | NA | |
| 0.955792873 | NA | |
| NA | NA | |
| NA | NA | |
| NA | NA | |
| NA | NA | |
| NA | NA | |
| NA | NA | |
| NA | NA | |
| 0.955792873 | NA | |
| NA | NA | |
| NA | NA | |
| 0.941278332 | NA | |
| NA | NA | |
| NA | NA | |
| 0.941278332 | NA | |
| NA | NA | |
| NA | NA | |
| 0.000192736 | | 0.001241209 |
| 0.859580522 | NA | |
| 0.445948433 | | 0.606402228 |
| 1.09E-05 | | 9.79E-05 |
| 0.689773905 | NA | |
| 0.025581118 | | 0.072981154 |
| 0.334881691 | | 0.498116111 |
| NA | NA | |
| 0.613843599 | | 0.744153273 |
| NA | NA | |
| 0.852252414 | | 0.911218679 |
| 0.055785657 | | 0.135173594 |
| 0.718926177 | NA | |
| 0.740230094 | NA | |
| 0.412260893 | | 0.57444291 |
| 3.40E-05 | | 0.000269356 |
| 0.633920942 | NA | |
| 0.1289147 | | 0.25318582 |
| 0.198041156 | | 0.345167607 |
| 0.609495728 | NA | |
| 0.482221849 | | 0.638438338 |
| 0.955792398 | NA | |
| NA | NA | |
| 0.839319108 | NA | |

| | | |
|---|---|---|
| | 0.839318641 | NA |
| | 0.365663357 | 0.528835528 |
| | 0.332667904 | 0.495845792 |
| | 0.001974384 | 0.009061761 |
| | 0.463746817 | 0.622425946 |
| | 0.012325922 | 0.04076443 |
| | 0.009332303 | 0.032517563 |
| | 0.369647157 | NA |
| | 0.101479792 | 0.212065145 |
| | 0.000218681 | 0.001386272 |
| | 0.756827946 | NA |
| | 0.033970526 | 0.091487494 |
| | 0.839319108 | NA |
| | 0.345740207 | 0.509491606 |
| | 0.161356227 | 0.298345444 |
| | 0.896825903 | NA |
| | 0.955792398 | NA |
| | 0.655689238 | 0.775219315 |
| NA | | NA |
| | 0.359685626 | NA |
| NA | | NA |
| | 0.714686589 | NA |
| | 0.896228543 | NA |
| | 0.839319108 | NA |
| | 0.882110255 | 0.929478431 |
| | 0.772061138 | NA |
| | 0.001532866 | 0.007305821 |
| | 0.645081937 | NA |
| | 0.019647321 | 0.059038698 |
| | 0.677026055 | 0.791715346 |
| | 0.110153711 | 0.225344102 |
| | 0.475445972 | 0.632634449 |
| | 0.949608511 | 0.969213349 |
| | 0.396352021 | NA |
| | 0.202691443 | 0.350793406 |
| | 0.000228651 | 0.001437874 |
| NA | | NA |
| | 0.940685982 | NA |
| | 0.048784197 | 0.121409604 |
| | 0.74627078 | NA |
| | 0.335872713 | 0.499060011 |
| | 0.358659308 | 0.522256751 |
| | 0.43607648 | 0.597208998 |
| | 0.920200528 | NA |
| | 0.575844711 | 0.714686402 |
| | 0.701885699 | 0.809706336 |
| | 0.824978079 | NA |

| | | |
|---|---|---|
| | 0.92069054 | NA |
| | 0.839320041 | NA |
| NA | | NA |
| | 0.299330603 | 0.46090484 |
| NA | | NA |
| | 0.955792873 | NA |
| | 2.55E-15 | 7.94E-14 |
| | 0.818573585 | NA |
| | 0.273461634 | 0.433293424 |
| NA | | NA |
| | 0.941278332 | NA |
| | 0.607439481 | NA |
| | 0.000117398 | 0.000806203 |
| | 0.89202369 | NA |
| NA | | NA |
| | 0.64752523 | 0.769542417 |
| | 0.722400472 | 0.824008053 |
| NA | | NA |
| | 0.000380594 | 0.002235661 |
| NA | | NA |
| | 2.81E-10 | 5.35E-09 |
| | 0.003357134 | 0.014087889 |
| | 0.03409148 | 0.091699109 |
| | 2.81E-07 | 3.48E-06 |
| | 0.820711782 | 0.890678686 |
| | 0.010974584 | 0.037146907 |
| | 0.611009185 | NA |
| | 0.019705459 | 0.059198453 |
| | 0.003924165 | 0.015963592 |
| | 0.125540431 | 0.248467545 |
| | 0.593392137 | 0.72875082 |
| | 0.025677872 | 0.073196908 |
| | 0.005084845 | 0.019795465 |
| | 0.530173712 | NA |
| | 0.953770009 | NA |
| | 4.54E-07 | 5.39E-06 |
| | 0.285898747 | 0.446549116 |
| | 2.28E-09 | 3.87E-08 |
| | 0.349010366 | 0.513008383 |
| | 5.95E-05 | 0.000442352 |
| | 0.047119579 | 0.118119032 |
| | 0.013951481 | 0.044996361 |
| | 0.048315295 | 0.120481996 |
| | 0.682978371 | 0.796379227 |
| | 0.030120238 | 0.083285538 |
| | 0.941275956 | NA |
| | 0.00980647 | 0.033877154 |

| | | | |
|---|---|---|---|
| | 6.33E-07 | | 7.32E-06 |
| NA | | NA | |
| | 0.110343628 | | 0.225686551 |
| | 0.233730456 | NA | |
| | 0.030629698 | | 0.084302505 |
| | 0.000877078 | | 0.004563578 |
| | 0.000967178 | | 0.004958166 |
| | 0.839319574 | NA | |
| | 0.002575352 | | 0.011318045 |
| | 0.007930508 | | 0.028445142 |
| | 1.62E-08 | | 2.47E-07 |
| | 0.318632539 | | 0.481071469 |
| | 3.27E-06 | | 3.30E-05 |
| NA | | NA | |
| | 0.429895818 | | 0.591125376 |
| | 0.40213852 | | 0.564423919 |
| | 0.873791205 | NA | |
| | 0.4469411 | | 0.607301026 |
| | 0.257249897 | | 0.41534777 |
| | 0.360831322 | | 0.524553052 |
| | 0.408562341 | | 0.570638445 |
| | 0.063977319 | | 0.150458269 |
| NA | | NA | |
| | 9.10E-12 | | 2.07E-10 |
| NA | | NA | |
| | 0.003369984 | | 0.014121895 |
| | 0.051976458 | | 0.127847895 |
| | 0.853595154 | NA | |
| NA | | NA | |
| | 0.970812799 | | 0.982410877 |
| | 0.941278788 | NA | |
| | 0.114315639 | | 0.231589144 |
| | 0.749797088 | | 0.84281796 |
| | 2.47E-20 | | 1.07E-18 |
| | 0.493977717 | | 0.647305528 |
| | 0.757698167 | | 0.848573462 |
| | 0.039330345 | | 0.1027597 |
| | 0.773811206 | | 0.860508485 |
| | 0.063077914 | | 0.14871043 |
| | 8.66E-07 | | 9.74E-06 |
| | 0.019722891 | | 0.059235867 |
| | 7.63E-05 | | 0.000550882 |
| | 0.125123849 | | 0.247852685 |
| | 0.983345437 | | 0.989370601 |
| | 0.732002055 | | 0.830985599 |
| | 0.007926173 | | 0.028433877 |
| | 0.000477031 | | 0.002726118 |

| | |
|---|---|
| 0.087060757 | 0.189451467 |
| NA | NA |
| 0.421812055 | 0.583142855 |
| 1.87E-07 | 2.39E-06 |
| 0.028621278 | 0.07997695 |
| 0.019095488 | 0.05778381 |
| 0.411858255 | 0.573982679 |
| 0.605287818 | 0.738028822 |
| 0.172458432 | 0.312683811 |
| 0.112653105 | 0.22915727 |
| 0.208055127 | 0.357373755 |
| 0.852357389 | NA |
| NA | NA |
| 0.108441786 | NA |
| 0.014707106 | 0.046931029 |
| 0.948745379 | NA |
| 0.043915461 | 0.111900033 |
| 0.104605474 | 0.216769936 |
| 6.28E-09 | 1.01E-07 |
| 1.70E-05 | 0.000144982 |
| 0.026617589 | 0.075485745 |
| 0.427766195 | 0.589041703 |
| 0.012894636 | 0.042257761 |
| 0.000310535 | 0.00187701 |
| 0.205845344 | 0.354550321 |
| 0.233807778 | 0.388379701 |
| 0.001461757 | 0.007016078 |
| NA | NA |
| 6.75E-05 | 0.000494563 |
| 0.663780225 | NA |
| 0.1178092 | 0.23691235 |
| 0.35199524 | 0.51562315 |
| 0.594029794 | 0.729103976 |
| 0.789840839 | 0.870804189 |
| 0.204950654 | 0.353546562 |
| 0.949948562 | 0.969401932 |
| 0.199320006 | 0.346770122 |
| 0.027178154 | 0.076819193 |
| 0.054241262 | 0.132156028 |
| 0.518937672 | 0.669580662 |
| 0.070567766 | 0.162207046 |
| 0.955792398 | NA |
| 0.905194605 | 0.942495275 |
| 0.499312689 | 0.652178051 |
| NA | NA |
| 0.377721751 | 0.540556345 |
| 0.066847973 | 0.155608688 |

| | |
|---|---|
| 0.255759423 | 0.4137837 |
| NA | NA |
| 0.019075941 | 0.057737228 |
| 0.618047748 | 0.747460621 |
| 7.45E-217 | 1.43E-213 |
| 0.001294631 | 0.00634572 |
| 0.941279803 | NA |
| NA | NA |
| 0.955792398 | NA |
| 0.258352829 | NA |
| 0.670348345 | 0.786689247 |
| 0.79167613 | 0.871750204 |
| 0.58290728 | NA |
| 0.107039874 | 0.220681081 |
| 0.507851321 | 0.659458655 |
| 0.594786425 | 0.729860622 |
| NA | NA |
| 2.44E-15 | 7.61E-14 |
| NA | NA |
| 0.017210725 | 0.053215078 |
| 1.97E-14 | 5.69E-13 |
| 0.11000051 | 0.2251468 |
| 0.837080309 | 0.901225435 |
| 1.44E-18 | 5.57E-17 |
| NA | NA |
| 0.008771019 | 0.030914822 |
| 0.485112555 | 0.640590792 |
| 0.259032413 | NA |
| 9.26E-11 | 1.86E-09 |
| 0.015391619 | 0.04868456 |
| 0.941278332 | NA |
| 1.75E-06 | 1.87E-05 |
| 0.666153715 | NA |
| 0.91951298 | NA |
| NA | NA |
| 0.977192456 | 0.985962343 |
| 0.090809634 | 0.195535635 |
| NA | NA |
| 0.826745498 | 0.894788847 |
| 0.381282956 | 0.543833934 |
| 0.580420543 | 0.718208103 |
| 0.872908827 | 0.924201588 |
| 0.295292106 | 0.456695432 |
| NA | NA |
| 0.941278434 | NA |
| 3.06E-17 | 1.08E-15 |
| 0.161274679 | 0.298264174 |

| | | |
|---|---|---|
| | 0.273916318 | 0.433725133 |
| | 0.739853959 | 0.836336778 |
| | 0.766230692 | 0.854831166 |
| | 0.001050252 | 0.005319876 |
| | 0.000178999 | 0.001165353 |
| | 0.882797128 | 0.929843832 |
| | 0.000570724 | 0.003175868 |
| | 0.020913905 | 0.061991628 |
| | 0.413331177 | NA |
| | 0.127073889 | 0.250630817 |
| NA | | NA |
| NA | | NA |
| | 1.70E-23 | 8.86E-22 |
| | 0.296655964 | 0.45812011 |
| | 5.73E-05 | 0.000428544 |
| | 4.85E-10 | 8.98E-09 |
| | 0.054268283 | 0.13219667 |
| | 0.635484616 | 0.760250222 |
| | 0.064087315 | 0.150627655 |
| | 0.289123383 | 0.449961734 |
| | 0.829885487 | 0.896696772 |
| NA | | NA |
| | 0.930327156 | 0.958516447 |
| NA | | NA |
| | 7.14E-17 | 2.48E-15 |
| | 0.142575574 | 0.272948714 |
| | 0.340848806 | 0.504375709 |
| NA | | NA |
| | 0.473155194 | 0.630714602 |
| | 1.86E-05 | 0.000157262 |
| | 3.55E-09 | 5.90E-08 |
| | 0.530115772 | 0.678924703 |
| | 0.000436679 | 0.002522855 |
| | 0.09439274 | 0.200924343 |
| | 0.09050278 | 0.194998386 |
| | 0.010729837 | 0.036453267 |
| | 0.59635769 | 0.731120237 |
| | 8.03E-20 | 3.37E-18 |
| | 0.045009402 | 0.114005937 |
| | 0.000418274 | 0.002430014 |
| | 0.392091841 | 0.554454016 |
| | 0.059715616 | 0.142519913 |
| | 0.415807039 | 0.577592013 |
| | 0.075212455 | 0.169864964 |
| | 0.431316146 | 0.592770189 |
| NA | | NA |
| | 0.002356324 | 0.010504818 |

| | |
|---|---|
| 0.904638855 | 0.942122688 |
| 0.612822035 | 0.743634467 |
| NA | NA |
| NA | NA |
| 0.955792398 | NA |
| 0.263535318 | 0.42253294 |
| 0.053433063 | 0.13063001 |
| 0.419926432 | 0.581583204 |
| 5.00E-05 | 0.000379447 |
| NA | NA |
| 0.955792398 | NA |
| NA | NA |
| 4.72E-08 | 6.63E-07 |
| 0.941275481 | NA |
| 4.02E-07 | 4.83E-06 |
| 0.679490564 | NA |
| 0.390279246 | NA |
| NA | NA |
| 0.839319108 | NA |
| 0.955792398 | NA |
| NA | NA |
| NA | NA |
| 0.722518141 | 0.824102779 |
| 0.017219941 | 0.05323666 |
| NA | NA |
| 0.910970337 | 0.946067279 |
| 0.019247469 | 0.058115048 |
| 0.018333301 | 0.055959122 |
| NA | NA |
| 0.019406203 | 0.058476573 |
| NA | NA |
| 0.804947949 | NA |
| 0.1662719 | 0.305099615 |
| 0.000999117 | 0.005108695 |
| 0.366937547 | 0.530095138 |
| 0.749297104 | 0.842611984 |
| 0.009987846 | 0.034359115 |
| 5.21E-09 | 8.49E-08 |
| 0.73639067 | 0.833733089 |
| 0.01542454 | 0.048756294 |
| 0.678744243 | 0.79302329 |
| 0.79461495 | 0.873922535 |
| 0.629859887 | 0.756365725 |
| 0.068866006 | 0.159217939 |
| 0.572752845 | 0.712520944 |
| 0.420045025 | NA |
| 0.375859494 | 0.539058067 |

| | |
|---|---|
| 0.939019179 | NA |
| 0.091393294 | 0.19639595 |
| 0.62005024 | 0.748968897 |
| 2.30E-12 | 5.58E-11 |
| NA | NA |
| NA | NA |
| 4.07E-11 | 8.60E-10 |
| 0.927839204 | NA |
| 0.113677584 | 0.230687329 |
| 0.463378211 | 0.622106615 |
| 0.595651783 | NA |
| 0.105159248 | 0.217727941 |
| 0.012096582 | 0.040128621 |
| 0.501885325 | 0.654102529 |
| 0.941278332 | NA |
| 0.675649535 | 0.790917147 |
| 0.867506714 | NA |
| 0.068179169 | 0.158013689 |
| NA | NA |
| 0.941278332 | NA |
| 0.941277857 | NA |
| 0.941278332 | NA |
| 0.569001703 | NA |
| 4.83E-05 | 0.000367972 |
| 0.794379235 | NA |
| 0.133767924 | 0.260189926 |
| 0.351241292 | NA |
| NA | NA |
| NA | NA |
| NA | NA |
| NA | NA |
| 0.048036359 | 0.119962376 |
| 4.04E-05 | 0.000313715 |
| 0.888907853 | 0.93342843 |
| 0.664473315 | 0.782300119 |
| 0.941278332 | NA |
| NA | NA |
| 0.605206406 | NA |
| 0.941278332 | NA |
| 0.197039558 | 0.343802457 |
| 0.562096936 | 0.704827271 |
| 0.055624744 | 0.134866065 |
| NA | NA |
| 0.480739303 | 0.637290832 |
| 0.364090009 | 0.52755777 |
| 0.926180987 | 0.955568264 |
| 0.850560949 | 0.910421896 |

| | |
|---|---|
| 0.660716216 | 0.779575044 |
| 0.001054284 | 0.005337901 |
| 0.401789777 | 0.56407929 |
| 0.000206376 | 0.001317507 |
| 0.036520119 | 0.096863369 |
| 4.38E-17 | 1.54E-15 |
| 0.807863526 | 0.882239978 |
| 2.26E-12 | 5.47E-11 |
| 0.560691476 | NA |
| 0.41418105 | 0.576207689 |
| 0.026316145 | 0.074710982 |
| 5.84E-05 | 0.000435356 |
| 0.398871681 | 0.561575367 |
| 0.394355095 | 0.556827429 |
| NA | NA |
| 0.181372564 | 0.323736765 |
| 5.36E-147 | 2.97E-144 |
| 0.001656116 | 0.007807262 |
| 0.940685982 | NA |
| 1.90E-05 | 0.000160344 |
| 0.335742887 | NA |
| 0.001851975 | 0.008571012 |
| 0.001252862 | 0.006175321 |
| 0.019759859 | 0.059299252 |
| 0.060459224 | 0.143876271 |
| 0.088718692 | 0.192239214 |
| 9.20E-05 | 0.000651286 |
| 0.142242898 | 0.272466237 |
| 0.457319522 | 0.617000299 |
| 0.05267471 | 0.129214 |
| 0.156484638 | 0.292037585 |
| NA | NA |
| NA | NA |
| 0.955792873 | NA |
| 0.000114313 | 0.000786835 |
| 0.001160446 | 0.005795411 |
| 0.560357214 | 0.703572422 |
| 0.287794463 | NA |
| 0.121265559 | 0.242266479 |
| NA | NA |
| 0.839318641 | NA |
| 0.035231076 | 0.094157879 |
| NA | NA |
| 6.67E-10 | 1.21E-08 |
| 0.493436726 | NA |
| 0.206269267 | NA |
| 0.262014768 | 0.420814812 |

| | | |
|---|---|---|
| | 1.17E-07 | 1.55E-06 |
| | 0.839318641 | NA |
| | 0.378314165 | NA |
| | 0.103462 | 0.215112031 |
| | 0.002054347 | 0.009353017 |
| | 0.027020353 | 0.07646382 |
| | 2.80E-05 | 0.000226674 |
| | 4.33E-05 | 0.000333599 |
| | 0.000344639 | 0.002054887 |
| NA | | NA |
| | 0.336899941 | NA |
| | 0.167803387 | 0.307046141 |
| | 0.925438299 | NA |
| NA | | NA |
| NA | | NA |
| | 0.839319108 | NA |
| | 0.192758446 | 0.33837573 |
| | 0.412164923 | NA |
| | 0.124086533 | 0.246392563 |
| | 0.343363548 | 0.506899408 |
| | 0.06503256 | 0.152457893 |
| | 0.816851017 | 0.888212342 |
| | 0.439683357 | 0.60052466 |
| | 0.120654416 | 0.241410256 |
| | 0.819510255 | 0.890074519 |
| | 0.579375092 | 0.71730661 |
| | 0.941276431 | NA |
| NA | | NA |
| NA | | NA |
| NA | | NA |
| | 0.941278332 | NA |
| NA | | NA |
| | 0.941275481 | NA |
| NA | | NA |
| | 0.189059785 | 0.333659718 |
| NA | | NA |
| NA | | NA |
| | 2.75E-07 | 3.41E-06 |
| | 0.604940096 | 0.737777095 |
| | 0.955792398 | NA |
| | 0.620357757 | 0.749188459 |
| | 0.000437999 | 0.00252979 |
| | 0.405057931 | 0.567095398 |
| | 0.405750656 | NA |
| | 0.450121917 | 0.610090647 |
| | 0.941279303 | NA |
| | 0.000308295 | 0.001866233 |

| | | |
|---|---|---|
| | 0.05758269 | 0.13847072 |
| | 0.941278332 | NA |
| NA | | NA |
| | 0.940528311 | NA |
| | 0.992832688 | 0.994981334 |
| | 7.23E-06 | 6.77E-05 |
| NA | | NA |
| | 0.863065563 | NA |
| | 0.118055975 | NA |
| | 0.354465641 | 0.518179043 |
| | 0.839319108 | NA |
| | 3.78E-05 | 0.000294872 |
| | 0.936413394 | NA |
| | 3.22E-05 | 0.000256369 |
| | 0.011356068 | 0.038209941 |
| | 4.86E-07 | 5.73E-06 |
| | 0.839320041 | NA |
| | 0.53949491 | NA |
| NA | | NA |
| | 1.59E-11 | 3.52E-10 |
| | 0.000223548 | 0.001412874 |
| | 0.86201527 | 0.917524705 |
| NA | | NA |
| | 0.93838321 | NA |
| NA | | NA |
| NA | | NA |
| | 0.479739064 | 0.636390192 |
| NA | | NA |
| | 0.602655685 | 0.736025994 |
| NA | | NA |
| | 0.352734705 | 0.516378095 |
| | 0.567939503 | 0.709387912 |
| NA | | NA |
| | 0.929820078 | 0.958284369 |
| | 0.941278332 | NA |
| | 0.004335067 | 0.017336402 |
| NA | | NA |
| | 0.630006596 | 0.756391124 |
| NA | | NA |
| | 0.618922969 | 0.748252971 |
| NA | | NA |
| | 0.816110485 | 0.88755651 |
| | 0.839319574 | NA |
| | 0.135155984 | 0.26220415 |
| NA | | NA |
| NA | | NA |
| | 0.693562443 | 0.803708533 |

| | |
|---|---|
| 0.249562298 | 0.406994301 |
| 0.534802699 | NA |
| NA | NA |
| 0.93827938 | NA |
| 0.001281805 | 0.006293219 |
| 0.748393366 | NA |
| 0.583956716 | NA |
| 5.27E-06 | 5.10E-05 |
| 0.889604348 | NA |
| NA | NA |
| NA | NA |
| NA | NA |
| 0.736113218 | 0.833647157 |
| 0.607886107 | 0.740400426 |
| 0.476014285 | 0.63314289 |
| 0.610237126 | NA |
| 0.793318421 | 0.872939996 |
| 0.941279303 | NA |
| NA | NA |
| 0.853595154 | NA |
| 0.784947319 | 0.867820897 |
| 6.21E-11 | 1.28E-09 |
| NA | NA |
| 0.995741188 | NA |
| 0.609651406 | 0.741490928 |
| 0.984587485 | 0.990329248 |
| 0.955792398 | NA |
| 7.36E-06 | 6.89E-05 |
| 0.300797066 | 0.462211174 |
| 0.57540795 | 0.714289314 |
| 0.347827706 | 0.511759822 |
| 0.001054239 | 0.005337901 |
| 0.153197918 | 0.287414027 |
| 0.090989478 | 0.195869725 |
| NA | NA |
| 0.835574813 | 0.900129066 |
| 0.572684795 | 0.712510767 |
| NA | NA |
| 3.43E-07 | 4.16E-06 |
| 5.26E-05 | 0.000397102 |
| 0.014810537 | 0.04718517 |
| 0.006445698 | 0.0239982 |
| 2.35E-05 | 0.000194389 |
| 0.390140399 | 0.552745344 |
| 0.415964646 | 0.577709797 |
| NA | NA |
| 0.615488861 | 0.745502309 |

| | | |
|---|---|---|
| | 0.116127767 | NA |
| NA | | NA |
| | 0.695600067 | 0.805081541 |
| | 0.177460176 | 0.31903747 |
| | 0.189551613 | 0.334154336 |
| NA | | NA |
| | 0.681272871 | NA |
| | 0.955792873 | NA |
| | 0.976022678 | 0.985378851 |
| | 0.941278332 | NA |
| | 9.01E-11 | 1.81E-09 |
| | 0.101735976 | 0.212451282 |
| NA | | NA |
| | 7.92E-08 | 1.08E-06 |
| | 0.253922696 | 0.411988275 |
| | 0.000518684 | 0.002929693 |
| | 0.003862871 | 0.015757378 |
| | 0.259829352 | 0.418377637 |
| | 0.941278838 | NA |
| | 0.427243147 | 0.588567314 |
| NA | | NA |
| | 0.501204517 | NA |
| | 0.809508728 | 0.883282605 |
| | 0.84376266 | 0.906007373 |
| | 9.60E-12 | 2.18E-10 |
| | 0.946093878 | 0.967272298 |
| | 0.743511643 | NA |
| | 0.394687717 | 0.557174909 |
| | 0.744270921 | 0.839341252 |
| | 0.645931921 | 0.768224234 |
| NA | | NA |
| | 0.955792873 | NA |
| | 0.837236233 | 0.901225435 |
| NA | | NA |
| NA | | NA |
| | 0.874187622 | NA |
| NA | | NA |
| NA | | NA |
| | 0.941278332 | NA |
| NA | | NA |
| NA | | NA |
| | 0.270724906 | NA |
| | 0.395207422 | 0.557766206 |
| NA | | NA |
| NA | | NA |
| NA | | NA |
| | 0.006648448 | 0.024610632 |

| | | |
|---|---|---|
| | 0.008546469 | 0.03024878 |
| | 0.839320041 | NA |
| | 1.72E-44 | 2.06E-42 |
| | 0.208874009 | 0.358237437 |
| NA | | NA |
| | 0.755971481 | 0.84719517 |
| | 0.066673992 | 0.155375718 |
| | 0.515067982 | 0.666212809 |
| | 0.199675582 | 0.347183534 |
| | 0.955792873 | NA |
| | 0.941278434 | NA |
| | 0.188807523 | NA |
| | 0.941279303 | NA |
| | 0.833357167 | NA |
| | 0.941277857 | NA |
| | 0.833562826 | NA |
| | 0.276045834 | 0.435879191 |
| | 0.839319108 | NA |
| NA | | NA |
| NA | | NA |
| NA | | NA |
| NA | | NA |
| | 0.459105875 | 0.618451782 |
| | 0.839319574 | NA |
| NA | | NA |
| | 0.853595154 | NA |
| | 0.646733543 | 0.768908681 |
| | 0.831318179 | 0.897753108 |
| | 0.00723377 | 0.026395552 |
| | 0.014022439 | 0.045176245 |
| | 0.741580204 | 0.837417783 |
| | 0.93147884 | 0.959252019 |
| NA | | NA |
| | 0.826372953 | NA |
| | 0.002237556 | 0.010050633 |
| | 0.441534886 | NA |
| | 0.390581419 | 0.553106791 |
| NA | | NA |
| | 0.193056734 | 0.338765904 |
| | 0.05327344 | 0.130413907 |
| | 0.518223389 | NA |
| | 0.112855601 | 0.229476108 |
| | 0.011665015 | 0.039064174 |
| | 0.699972323 | NA |
| | 0.395943244 | NA |
| | 0.169313674 | 0.308883579 |
| NA | | NA |

| | |
|---|---|
| 2.43E-26 | 1.44E-24 |
| 2.46E-06 | 2.55E-05 |
| 0.054658692 | 0.132984381 |
| 4.74E-05 | 0.00036158 |
| 0.951394244 | 0.970309615 |
| 0.085594692 | 0.187099353 |
| 0.88085999 | NA |
| NA | NA |
| 0.839322216 | NA |
| 0.113428337 | 0.230360673 |
| 0.113175943 | 0.229945694 |
| 0.003154401 | 0.013381045 |
| NA | NA |
| 1.06E-05 | 9.51E-05 |
| 0.839319574 | NA |
| 0.26543156 | 0.424697633 |
| 0.500727717 | NA |
| NA | NA |
| 0.600828719 | 0.734592824 |
| NA | NA |
| NA | NA |
| NA | NA |
| NA | NA |
| NA | NA |
| NA | NA |
| 0.68961623 | NA |
| NA | NA |
| NA | NA |
| NA | NA |
| NA | NA |
| NA | NA |
| 0.944289243 | NA |
| NA | NA |
| NA | NA |
| 0.652563176 | 0.773328788 |
| NA | NA |
| 0.837327665 | NA |
| 0.000618167 | 0.003395708 |
| 0.28719406 | 0.447778922 |
| 0.000134301 | 0.000907617 |
| 1.76E-08 | 2.65E-07 |
| 0.061506637 | 0.145863749 |
| 0.814603302 | 0.886646597 |
| 0.553221921 | 0.698212241 |
| 0.487865806 | 0.642657697 |
| 0.002215896 | 0.009968115 |
| 2.16E-08 | 3.22E-07 |

| | | |
|---|---|---|
| | 0.518783902 | NA |
| NA | | NA |
| NA | | NA |
| | 0.173681419 | 0.314393881 |
| | 0.595923859 | NA |
| | 0.052359019 | 0.128558837 |
| | 0.06998866 | 0.161234038 |
| | 0.839319574 | NA |
| | 0.767625476 | 0.855824691 |
| | 1.06E-08 | 1.66E-07 |
| | 0.052394151 | 0.12863183 |
| | 0.705401621 | 0.812065262 |
| | 0.009959359 | 0.034280935 |
| | 0.41046402 | 0.572425755 |
| | 0.041181363 | 0.106636074 |
| | 0.060716957 | 0.144402979 |
| | 0.004132201 | 0.016678762 |
| | 0.30953806 | 0.471196272 |
| | 3.84E-11 | 8.16E-10 |
| | 0.388177528 | NA |
| | 0.368771425 | 0.531955072 |
| | 0.294296867 | 0.455481763 |
| | 0.132749763 | 0.258653393 |
| | 0.377693926 | 0.540550885 |
| | 0.611278409 | 0.742468556 |
| | 0.088362859 | 0.191654162 |
| | 0.52524381 | NA |
| | 0.941277857 | NA |
| NA | | NA |
| | 0.392634368 | NA |
| | 0.941277857 | NA |
| NA | | NA |
| NA | | NA |
| NA | | NA |
| | 0.839319574 | NA |
| NA | | NA |
| | 0.955792398 | NA |
| NA | | NA |
| | 0.941278207 | NA |
| NA | | NA |
| NA | | NA |
| | 0.903961571 | NA |
| | 0.08420083 | 0.184833363 |
| | 0.640158568 | NA |
| | 0.267258648 | 0.426846296 |
| | 0.990037266 | 0.993459823 |
| | 0.344259614 | NA |

| | |
|---|---|
| 1.31E-10 | 2.58E-09 |
| 0.955792398 | NA |
| 0.090437984 | 0.194947011 |
| 0.633874615 | 0.759386128 |
| 1.57E-06 | 1.69E-05 |
| 1.26E-14 | 3.68E-13 |
| NA | NA |
| 4.43E-08 | 6.25E-07 |
| 0.020253423 | 0.060484801 |
| 1.93E-08 | 2.89E-07 |
| 0.933218787 | 0.960331758 |
| 0.713198238 | NA |
| 1.40E-05 | 0.000122944 |
| 1.16E-08 | 1.81E-07 |
| 0.229392581 | 0.383104282 |
| 1.07E-76 | 2.62E-74 |
| NA | NA |
| 0.622486252 | 0.750882717 |
| 0.547676096 | 0.693703741 |
| 1.01E-10 | 2.03E-09 |
| 0.925811825 | 0.955435877 |
| 0.400474491 | 0.56309628 |
| 0.00100684 | 0.005140458 |
| 0.11819159 | 0.237520789 |
| 0.024628102 | 0.070873907 |
| 0.605440737 | NA |
| 0.504718438 | 0.656895684 |
| 0.378175911 | 0.541061866 |
| NA | NA |
| 0.536196334 | 0.684245397 |
| 0.823103704 | 0.892429935 |
| 0.066633934 | 0.155309528 |
| 0.003380093 | 0.014154287 |
| 0.036484401 | 0.096790204 |
| 0.838882104 | NA |
| 0.563504484 | 0.705811384 |
| 0.981995551 | 0.98868314 |
| 0.237455808 | 0.392953742 |
| 0.982967197 | 0.989201134 |
| 0.687109015 | NA |
| NA | NA |
| NA | NA |
| 0.741178832 | NA |
| NA | NA |
| 0.007430679 | 0.02699817 |
| 0.945731563 | NA |
| 0.901343176 | NA |

| | | |
|---|---|---|
| | 0.615354465 | NA |
| | 0.751622423 | 0.844112339 |
| | 0.627884254 | 0.755021328 |
| | 0.34723652 | 0.51122185 |
| | 0.987195473 | 0.99190454 |
| | 0.009746781 | 0.033714994 |
| | 0.138495555 | 0.266963978 |
| | 5.52E-07 | 6.44E-06 |
| | 0.624063531 | 0.752137332 |
| | 0.913661969 | NA |
| | 0.151733236 | 0.285753935 |
| | 0.257748417 | 0.41595551 |
| | 0.592158794 | 0.727761843 |
| | 6.31E-05 | 0.000467039 |
| | 0.805230931 | NA |
| | 5.58E-08 | 7.76E-07 |
| | 0.291644048 | 0.452848302 |
| NA | | NA |
| | 0.839319574 | NA |
| NA | | NA |
| | 2.11E-06 | 2.22E-05 |
| | 0.954405206 | NA |
| | 3.91E-06 | 3.90E-05 |
| | 0.427742703 | NA |
| | 0.035654831 | 0.095130175 |
| | 0.955792398 | NA |
| | 0.006171331 | 0.023172428 |
| | 0.953798549 | NA |
| | 0.154755776 | 0.289560332 |
| | 0.037373258 | 0.098642524 |
| | 0.040318668 | 0.104837712 |
| NA | | NA |
| | 0.623456994 | 0.75155854 |
| | 0.106764155 | 0.220303461 |
| | 0.019695118 | 0.059174855 |
| | 0.25200439 | 0.409741682 |
| | 0.999498898 | 0.999624891 |
| | 0.354626813 | 0.518382817 |
| | 0.769537261 | 0.857314759 |
| | 0.766536545 | 0.855022354 |
| | 0.530947828 | NA |
| | 0.209714292 | 0.359393734 |
| | 0.736279747 | 0.833647157 |
| | 0.40747315 | 0.56933818 |
| | 0.941277857 | NA |
| | 0.203623577 | 0.352022689 |
| NA | | NA |

| | | | |
|---|---|---|---|
| | 0.108584608 | | 0.222919686 |
| | 0.301215451 | | 0.462490658 |
| | 0.154967003 | | 0.289841632 |
| NA | | NA | |
| | 2.01E-10 | | 3.90E-09 |
| | 0.025461629 | | 0.072683819 |
| | 0.835419619 | | 0.900043352 |
| | 0.000292358 | | 0.001783367 |
| | 0.233485768 | | 0.388088566 |
| | 0.804681148 | NA | |
| | 0.028717716 | | 0.080208764 |
| | 0.918822427 | NA | |
| | 0.000803403 | | 0.004241923 |
| | 0.49982603 | | 0.65263367 |
| | 0.000733037 | | 0.003931535 |
| | 0.044634155 | | 0.113260358 |
| | 0.437160638 | NA | |
| NA | | NA | |
| | 0.007868033 | | 0.028272181 |
| | 0.101240219 | | 0.21167601 |
| | 0.553826967 | NA | |
| | 0.000642958 | | 0.00351002 |
| | 0.161857129 | | 0.299002589 |
| | 0.507121492 | | 0.658870402 |
| | 0.079875517 | | 0.177632164 |
| | 0.002810575 | | 0.012134467 |
| | 0.929476525 | | 0.95803691 |
| | 0.055167657 | | 0.134017205 |
| | 0.568786958 | | 0.709962094 |
| | 0.350788725 | NA | |
| NA | | NA | |
| | 0.739991096 | NA | |
| | 0.000885346 | | 0.004602098 |
| | 0.807281316 | NA | |
| | 0.609719265 | NA | |
| | 0.35783729 | NA | |
| | 0.684336061 | | 0.797279344 |
| | 0.298547941 | NA | |
| NA | | NA | |
| | 1.53E-06 | | 1.66E-05 |
| NA | | NA | |
| | 0.941277857 | NA | |
| | 0.000129122 | | 0.000877345 |
| | 0.202578806 | | 0.350623963 |
| | 0.003315856 | | 0.013946634 |
| | 0.071725456 | | 0.163981299 |
| | 0.539255181 | NA | |

| | |
|---|---|
| NA | NA |
| 0.879829547 | 0.928061284 |
| 0.160162205 | 0.296829464 |
| 0.941278838 | NA |
| 0.018464404 | 0.056294317 |
| 3.22E-07 | 3.93E-06 |
| 0.696202495 | 0.805326553 |
| 0.631782695 | 0.757871772 |
| 0.07102233 | 0.162850615 |
| NA | NA |
| 0.34995473 | 0.513799278 |
| 0.000401107 | 0.002341708 |
| 0.247228296 | 0.404351536 |
| NA | NA |
| 0.390689522 | NA |
| 3.57E-06 | 3.59E-05 |
| 0.675071617 | 0.790546188 |
| 0.204670501 | 0.353278652 |
| 0.226029378 | 0.379056666 |
| 0.406961014 | 0.568756154 |
| 0.239392579 | 0.39525681 |
| 2.35E-60 | 4.12E-58 |
| 1.42E-28 | 9.42E-27 |
| 0.857927945 | 0.914974502 |
| 0.101296294 | 0.211737455 |
| 0.00045705 | 0.002623904 |
| 0.369039439 | 0.532291646 |
| 0.000577609 | 0.003206966 |
| 0.628858455 | 0.755696114 |
| 0.024373118 | 0.070281073 |
| 0.828207646 | 0.89562173 |
| 0.402639625 | 0.564739444 |
| 0.194629125 | 0.340786742 |
| 0.924826521 | 0.95491588 |
| 0.792589703 | 0.872430315 |
| 0.941278838 | NA |
| NA | NA |
| NA | NA |
| 0.129106138 | 0.253379313 |
| 0.650901099 | 0.771974087 |
| 0.000946677 | 0.004868799 |
| 0.034360826 | 0.092304992 |
| 0.62034145 | 0.749188459 |
| 0.839319108 | NA |
| 1.88E-44 | 2.24E-42 |
| 0.015941187 | 0.050084506 |
| 5.88E-06 | 5.64E-05 |

| | |
|---|---|
| 0.382385528 | NA |
| NA | NA |
| 0.839319108 | NA |
| 1.07E-07 | 1.44E-06 |
| 0.111825582 | 0.227765916 |
| NA | NA |
| NA | NA |
| NA | NA |
| 0.295128949 | 0.456495304 |
| NA | NA |
| 0.635657279 | 0.760375447 |
| NA | NA |
| NA | NA |
| 0.023513279 | 0.068254792 |
| 0.655162421 | NA |
| 0.396901366 | 0.559493414 |
| 0.015129517 | 0.048034249 |
| 0.230672637 | 0.384589155 |
| 9.67E-09 | 1.52E-07 |
| 0.000308826 | 0.00186815 |
| 0.673981444 | 0.78959031 |
| 0.460665238 | 0.619722714 |
| 0.916134243 | NA |
| 0.094053588 | 0.200345758 |
| 0.156423407 | 0.291959612 |
| 0.316322888 | 0.47870787 |
| 0.828229976 | 0.89562173 |
| 0.360924073 | NA |
| NA | NA |
| 0.853595622 | NA |
| NA | NA |
| NA | NA |
| NA | NA |
| NA | NA |
| NA | NA |
| NA | NA |
| 0.038347418 | 0.100744508 |
| 6.29E-16 | 2.05E-14 |
| 0.563049438 | NA |
| NA | NA |
| 0.839318641 | NA |
| 2.96E-08 | 4.31E-07 |
| 0.33325358 | 0.496501108 |
| 0.068831899 | 0.159170005 |
| 0.233050975 | 0.38763657 |
| 0.139377827 | 0.268186826 |
| 0.941275956 | NA |

| | | |
|---|---|---|
| | 9.92E-23 | 4.98E-21 |
| | 0.687825027 | 0.799707469 |
| | 0.494859177 | NA |
| | 0.002441227 | 0.010818049 |
| NA | | NA |
| | 0.955792398 | NA |
| NA | | NA |
| | 0.941275956 | NA |
| | 0.757062716 | NA |
| NA | | NA |
| | 0.30869946 | NA |
| | 0.486537955 | NA |
| | 1.04E-05 | 9.36E-05 |
| | 0.558695729 | NA |
| | 0.120820348 | 0.241625524 |
| | 0.736254635 | 0.833647157 |
| NA | | NA |
| NA | | NA |
| NA | | NA |
| | 0.955792398 | NA |
| | 0.461500784 | 0.620320853 |
| | 0.923475547 | NA |
| | 0.023259485 | 0.067696536 |
| | 0.000131224 | 0.000890107 |
| | 0.941277857 | NA |
| | 0.545858338 | 0.692127337 |
| | 0.912514484 | 0.947298925 |
| | 0.332360105 | 0.495542172 |
| | 0.941278332 | NA |
| | 0.597263383 | NA |
| NA | | NA |
| | 0.864445397 | 0.91922223 |
| NA | | NA |
| | 0.680529146 | 0.794367845 |
| | 0.271157558 | 0.431046513 |
| | 0.229907691 | 0.38372224 |
| | 0.016384454 | 0.051178842 |
| | 0.467228813 | 0.625512132 |
| | 5.92E-06 | 5.67E-05 |
| | 0.697148915 | NA |
| | 0.966703695 | NA |
| NA | | NA |
| | 0.967072719 | NA |
| NA | | NA |
| | 0.001788376 | 0.008333385 |
| | 2.51E-09 | 4.24E-08 |
| | 0.418032939 | NA |

| | |
|---|---|
| 0.221035159 | 0.372958886 |
| 0.000351694 | 0.002088579 |
| 7.76E-19 | 3.06E-17 |
| 0.000128817 | 0.00087603 |
| 0.018735414 | 0.056960062 |
| 2.57E-06 | 2.65E-05 |
| 0.673292067 | 0.789093136 |
| 0.004742336 | 0.018706842 |
| 0.72959231 | 0.82965275 |
| 0.566164895 | 0.707921135 |
| 0.194666763 | 0.340820764 |
| NA | NA |
| 0.049026919 | 0.121809888 |
| 0.955792398 | NA |
| 0.935817683 | NA |
| 0.897690844 | NA |
| 0.941278332 | NA |
| NA | NA |
| NA | NA |
| 0.953770009 | NA |
| NA | NA |
| NA | NA |
| 0.941277857 | NA |
| 0.737330128 | NA |
| 0.001250559 | 0.006166523 |
| 0.941277857 | NA |
| 8.79E-07 | 9.88E-06 |
| 0.003028104 | 0.012925921 |
| 3.89E-10 | 7.28E-09 |
| 0.118246988 | 0.237571943 |
| 0.657287669 | 0.776761883 |
| 0.661392917 | NA |
| 0.955791446 | NA |
| 0.16910015 | 0.308659645 |
| 0.003314197 | 0.013942119 |
| 0.606482586 | NA |
| 0.121438563 | 0.242530683 |
| NA | NA |
| 0.839318641 | NA |
| 0.839319108 | NA |
| 0.839319416 | NA |
| NA | NA |
| NA | NA |
| NA | NA |
| 0.941277857 | NA |
| NA | NA |
| 0.941278332 | NA |

| | | |
|---|---|---|
| NA | 0.823387982 | NA |
| NA | | NA |
| NA | | NA |
| NA | | NA |
| NA | | NA |
| | 0.161094304 | 0.298082621 |
| | 0.141091908 | 0.270806353 |
| | 0.96134632 | NA |
| | 0.978270291 | 0.986518216 |
| | 0.000952754 | 0.004895823 |
| | 0.473573944 | 0.631060745 |
| | 7.05E-05 | 0.000513506 |
| | 0.413889333 | 0.576060243 |
| | 9.74E-05 | 0.000684791 |
| | 0.124917424 | 0.247526185 |
| | 0.314512714 | 0.476726206 |
| | 0.610333763 | 0.741901038 |
| | 1.36E-05 | 0.00011928 |
| | 0.951249535 | 0.970245452 |
| | 0.00042883 | 0.002482854 |
| | 0.256462171 | 0.414554336 |
| | 0.247942123 | 0.405157107 |
| | 0.177261188 | 0.318837059 |
| | 8.45E-06 | 7.79E-05 |
| | 0.018454845 | 0.056272382 |
| | 1.17E-25 | 6.77E-24 |
| NA | | NA |
| | 0.013468328 | 0.043788163 |
| NA | | NA |
| NA | | NA |
| | 0.003692117 | 0.015185722 |
| | 0.86904209 | 0.921575098 |
| | 0.121844601 | NA |
| | 0.000955867 | 0.004908638 |
| | 0.005262116 | 0.020342519 |
| | 0.286723653 | 0.447279879 |
| NA | | NA |
| | 0.955090162 | 0.972692863 |
| | 0.07276507 | 0.165703494 |
| | 0.544209221 | 0.690882394 |
| NA | | NA |
| | 0.72258685 | NA |
| | 0.493319745 | 0.64694196 |
| | 0.960939642 | NA |
| | 0.700852965 | 0.808877387 |
| | 0.74627078 | NA |
| NA | | NA |

| | | | |
|---|---|---|---|
| NA | | NA | |
| NA | | NA | |
| | 0.622744509 | NA | |
| | 0.287002776 | | 0.447568636 |
| | 1.22E-10 | | 2.42E-09 |
| NA | | NA | |
| NA | | NA | |
| NA | | NA | |
| | 0.582378164 | NA | |
| | 0.871180627 | NA | |
| | 0.825431323 | | 0.893990126 |
| | 0.743123204 | | 0.83852368 |
| | 0.6706969 | NA | |
| | 0.002702387 | | 0.011780624 |
| | 0.619073124 | | 0.748320477 |
| NA | | NA | |
| NA | | NA | |
| | 0.005217977 | | 0.020211275 |
| | 0.045281687 | | 0.114585871 |
| | 0.618590021 | | 0.747919547 |
| | 0.552492141 | | 0.697708932 |
| | 0.081773843 | | 0.180605086 |
| | 0.542706964 | NA | |
| | 0.497935299 | | 0.650950513 |
| | 1.12E-51 | | 1.58E-49 |
| | 4.95E-22 | | 2.38E-20 |
| NA | | NA | |
| NA | | NA | |
| NA | | NA | |
| NA | | NA | |
| | 0.086998593 | | 0.18938545 |
| | 0.839318641 | NA | |
| | 0.00293713 | | 0.012599693 |
| | 0.003860039 | | 0.015751183 |
| | 0.048896992 | | 0.121601316 |
| | 0.9665525 | | 0.980058047 |
| | 0.528864728 | | 0.677979117 |
| | 0.734058779 | NA | |
| | 0.003685939 | | 0.015165553 |
| | 0.003394255 | | 0.014201101 |
| | 0.128634605 | | 0.252996271 |
| | 1.29E-38 | | 1.24E-36 |
| | 0.941278332 | NA | |
| | 0.008012911 | | 0.028688823 |
| | 0.581049165 | NA | |
| NA | | NA | |
| | 0.936260897 | | 0.962255791 |

| | |
|---|---|
| NA | NA |
| 0.004840489 | 0.019005827 |
| 0.691954794 | 0.802392478 |
| 0.062449315 | 0.147593943 |
| 4.06E-15 | 1.25E-13 |
| 0.711189516 | 0.815773823 |
| 0.101950631 | 0.212824849 |
| 0.892533482 | 0.935079975 |
| 0.005939568 | 0.022465216 |
| 0.955792873 | NA |
| 6.24E-07 | 7.21E-06 |
| 0.358802691 | 0.522337715 |
| 0.305821528 | NA |
| NA | NA |
| 0.726028365 | 0.827036474 |
| 0.897671564 | NA |
| 0.465532067 | 0.623872206 |
| 0.788281903 | 0.869831931 |
| 0.419798235 | 0.581473323 |
| 0.822198034 | 0.891751494 |
| 0.710977027 | 0.815747178 |
| 0.49285763 | 0.646601807 |
| 0.839319108 | NA |
| 0.000692088 | 0.003741332 |
| NA | NA |
| 0.941276431 | NA |
| 0.839319108 | NA |
| 0.145745957 | 0.277412657 |
| 0.635264132 | 0.760094348 |
| 0.836541617 | 0.900925912 |
| 0.301737844 | 0.4631135 |
| 0.853596091 | NA |
| 0.734058779 | NA |
| 0.031752023 | 0.08672959 |
| 0.955792398 | NA |
| NA | NA |
| 0.903633842 | 0.941652848 |
| 0.839319108 | NA |
| 0.650133202 | NA |
| NA | NA |
| NA | NA |
| NA | NA |
| 0.560533652 | NA |
| NA | NA |
| 0.3210973 | 0.483841348 |
| 0.946519504 | 0.967535204 |
| 0.854794699 | 0.91261431 |

| | |
|---|---|
| NA | NA |
| NA | NA |
| 8.14E-17 | 2.80E-15 |
| 0.941278284 | NA |
| 0.30379733 | 0.465224477 |
| 0.778970799 | 0.863740938 |
| 0.271524992 | 0.431360635 |
| 0.957802558 | 0.974132226 |
| 0.024016917 | 0.069467816 |
| 0.764267591 | 0.853630988 |
| 0.01092894 | 0.037024001 |
| 0.375114543 | 0.538264876 |
| 0.004637893 | 0.018355692 |
| 0.839319574 | NA |
| 0.532690624 | NA |
| 0.001261047 | 0.006207952 |
| NA | NA |
| 0.64880369 | NA |
| NA | NA |
| 0.941278788 | NA |
| NA | NA |
| NA | NA |
| NA | NA |
| NA | NA |
| 0.015586703 | 0.049183972 |
| 0.882201029 | NA |
| NA | NA |
| NA | NA |
| 0.265609727 | 0.424925576 |
| NA | NA |
| 0.896246089 | NA |
| 0.928424343 | 0.957384811 |
| 0.00739173 | 0.026877171 |
| 0.105860955 | 0.21883815 |
| 0.106042962 | 0.219157296 |
| 0.423090694 | 0.584435303 |
| 1.89E-06 | 2.01E-05 |
| 0.842530716 | NA |
| 0.505474915 | 0.657409475 |
| 0.261951426 | 0.42074147 |
| 0.074163462 | 0.168058408 |
| NA | NA |
| 0.72881242 | 0.82929502 |
| 0.509690494 | 0.661377809 |
| 7.07E-05 | 0.000514912 |
| 0.149404631 | 0.282411771 |
| NA | NA |

| | | |
|---|---|---|
| NA | NA | |
| | 0.85958895 | NA |
| | 0.039450831 | NA |
| | 0.717663493 | 0.820452759 |
| | 9.39E-65 | 1.88E-62 |
| | 0.500670908 | 0.653306779 |
| | 0.079919976 | 0.177697831 |
| NA | NA | |
| NA | NA | |
| | 0.66943952 | NA |
| | 0.033984349 | 0.091504013 |
| | 0.139487397 | 0.268332347 |
| NA | NA | |
| | 0.922345353 | NA |
| | 0.000902522 | 0.004679116 |
| NA | NA | |
| NA | NA | |
| | 0.988501572 | 0.992713995 |
| | 0.344434452 | 0.508039216 |
| | 0.061304596 | 0.145511299 |
| | 0.627869531 | 0.755021328 |
| | 0.031353596 | 0.085887707 |
| NA | NA | |
| | 0.916064961 | NA |
| | 0.733502242 | NA |
| | 0.895199992 | 0.936820599 |
| | 0.692162318 | 0.802476741 |
| | 0.532122271 | NA |
| | 0.891583055 | NA |
| | 0.943357016 | NA |
| | 0.438056317 | 0.598886643 |
| | 0.931578855 | 0.959272227 |
| | 3.26E-47 | 4.22E-45 |
| NA | NA | |
| | 0.660740905 | NA |
| | 0.005377696 | 0.020689862 |
| | 0.636252571 | 0.760856295 |
| NA | NA | |
| | 0.955792398 | NA |
| | 0.025820778 | 0.073541481 |
| | 0.000304575 | 0.001845596 |
| | 0.836568982 | NA |
| | 0.928771217 | NA |
| NA | NA | |
| | 0.427280465 | 0.588567314 |
| | 0.774527713 | 0.860736444 |
| | 0.777079586 | NA |

| | | |
|---|---|---|
| | 0.665065306 | 0.782727671 |
| | 0.940475126 | NA |
| | 0.055550928 | 0.134769462 |
| | 0.499025187 | 0.651943647 |
| | 0.000178631 | 0.001163913 |
| NA | | NA |
| | 0.000897037 | 0.00465575 |
| | 0.274915308 | 0.434813608 |
| | 0.926462868 | NA |
| | 0.692500545 | 0.802829768 |
| | 0.303474841 | 0.464820345 |
| NA | | NA |
| | 0.4024305 | 0.56466674 |
| | 0.339265308 | 0.502876626 |
| NA | | NA |
| | 0.270914734 | 0.430805531 |
| | 0.314900624 | NA |
| | 0.73569649 | 0.833502207 |
| NA | | NA |
| | 0.744418632 | 0.839468056 |
| | 0.459435058 | NA |
| NA | | NA |
| | 0.028103838 | 0.078808618 |
| | 0.247667057 | 0.404886179 |
| | 0.955792873 | NA |
| | 0.686441433 | 0.798840218 |
| NA | | NA |
| | 0.005862047 | 0.022224982 |
| | 0.029457313 | 0.081775479 |
| | 0.43323847 | NA |
| | 0.758365546 | NA |
| | 0.661661731 | 0.78014028 |
| | 0.911236573 | NA |
| | 0.004961857 | 0.019405541 |
| | 0.316148471 | 0.478683643 |
| | 0.872383611 | NA |
| | 0.750685413 | NA |
| | 0.551429294 | NA |
| NA | | NA |
| | 0.712154784 | 0.81662611 |
| | 0.556967407 | 0.701017303 |
| NA | | NA |
| | 0.658702622 | 0.777816134 |
| | 0.502206914 | 0.654270879 |
| | 0.103362934 | 0.21494361 |
| | 0.871323624 | NA |
| | 0.941278332 | NA |

| | | |
|---|---|---|
| | 0.416554439 | 0.578191554 |
| | 0.560090525 | NA |
| | 0.775856086 | 0.861853108 |
| | 0.904596468 | 0.942119766 |
| NA | | NA |
| | 0.55979178 | NA |
| | 0.983815119 | 0.989719673 |
| | 0.110463094 | 0.225833237 |
| | 0.121297754 | 0.242309129 |
| | 0.81419634 | NA |
| | 0.373353254 | 0.536562863 |
| | 0.003164035 | 0.013418406 |
| | 0.50305771 | 0.655164129 |
| | 0.530889725 | 0.679659921 |
| | 0.005260176 | 0.020338321 |
| | 0.017991064 | 0.055111649 |
| | 0.88771576 | NA |
| | 0.483300935 | 0.639262509 |
| | 0.557980395 | 0.701624458 |
| | 0.484806429 | 0.640470812 |
| | 0.839319574 | NA |
| | 0.774865444 | 0.860993758 |
| | 0.074113018 | 0.167987625 |
| | 0.006541447 | 0.024290096 |
| | 0.853596091 | NA |
| | 0.500489672 | 0.653188912 |
| | 0.955043037 | NA |
| | 0.350962545 | 0.514860431 |
| | 0.47809302 | 0.634945248 |
| | 0.732507364 | 0.83123434 |
| | 0.9771208 | 0.985962343 |
| | 0.302312231 | 0.463666198 |
| | 0.227583911 | 0.380965767 |
| | 0.227520004 | 0.380885577 |
| | 0.398985303 | NA |
| | 0.002324196 | 0.01038493 |
| | 0.024968964 | 0.071577898 |
| | 0.014084177 | 0.045350592 |
| | 0.256346336 | 0.414479452 |
| | 0.938708243 | 0.963601145 |
| | 0.283593472 | 0.444143427 |
| | 0.37634622 | 0.539463579 |
| | 0.004279345 | 0.017156301 |
| NA | | NA |
| | 0.032916936 | 0.089337391 |
| NA | | NA |
| NA | | NA |

| | |
|---|---|
| NA | NA |
| 0.013128025 | 0.042886804 |
| 0.955792398 | NA |
| 0.000826627 | 0.00434719 |
| 0.027038447 | 0.076496982 |
| 0.941278788 | NA |
| 6.40E-08 | 8.86E-07 |
| 5.22E-10 | 9.63E-09 |
| 0.008441988 | 0.029954919 |
| 3.44E-05 | 0.00027154 |
| 0.160411567 | 0.29712966 |
| 0.989800937 | NA |
| 0.154108677 | 0.288812524 |
| 6.73E-06 | 6.37E-05 |
| 4.41E-06 | 4.34E-05 |
| NA | NA |
| 0.176398764 | 0.317982685 |
| 0.925860344 | 0.955444522 |
| 0.069634846 | 0.160621048 |
| 0.955792398 | NA |
| 0.839319109 | NA |
| NA | NA |
| 0.006338889 | 0.023667173 |
| 0.839319108 | NA |
| 0.47488669 | NA |
| 8.70E-05 | 0.000619345 |
| 0.000104974 | 0.00072994 |
| 0.554305432 | 0.699293894 |
| 1.32E-36 | 1.20E-34 |
| 0.326394312 | 0.489313939 |
| 0.397412323 | 0.560014688 |
| 0.190592088 | 0.335443581 |
| NA | NA |
| 1.15E-09 | 2.03E-08 |
| 0.004625511 | 0.018312775 |
| NA | NA |
| 0.548659483 | 0.694528592 |
| 0.653509818 | 0.774003044 |
| 0.307447388 | 0.468965142 |
| 0.881754284 | NA |
| 0.853595622 | NA |
| 0.937272113 | NA |
| 0.609418528 | NA |
| 0.015408008 | 0.04872345 |
| 0.000255914 | 0.001583613 |
| 0.002119079 | 0.009596333 |
| 9.18E-05 | 0.000650125 |

| | |
|---|---|
| 0.618418037 | 0.74783245 |
| 0.128673509 | 0.252996404 |
| 0.073681045 | 0.16726335 |
| 0.089783219 | 0.193834029 |
| 1.12E-05 | 9.99E-05 |
| 0.071187653 | 0.163143512 |
| 0.010716162 | 0.036417202 |
| NA | NA |
| 0.020824135 | 0.061787095 |
| 0.59091636 | 0.726797809 |
| 0.297634097 | 0.459015008 |
| 0.080616636 | NA |
| 0.23414911 | 0.388865275 |
| 5.70E-62 | 1.08E-59 |
| 0.001880743 | 0.008687258 |
| 0.162615398 | 0.300160659 |
| 0.114505906 | 0.231915383 |
| 0.936818392 | 0.962579382 |
| 0.680108274 | 0.794122482 |
| 0.001333153 | 0.006499733 |
| 0.592026115 | 0.727711505 |
| 0.011491896 | 0.038557952 |
| 0.049069994 | 0.121891464 |
| 0.799868295 | NA |
| NA | NA |
| 0.285168755 | 0.445672141 |
| 0.978203691 | 0.986494311 |
| 0.2205919 | 0.372483571 |
| 0.198097184 | 0.345201637 |
| 0.121607011 | 0.24280598 |
| 0.538760276 | 0.686339761 |
| NA | NA |
| 0.940020164 | 0.963913491 |
| 0.001340163 | 0.006531237 |
| NA | NA |
| 0.079368075 | 0.176834121 |
| 0.634427554 | 0.75971383 |
| 0.373849886 | 0.536984792 |
| NA | NA |
| 0.801028292 | 0.878492596 |
| NA | NA |
| 0.017369364 | 0.053615044 |
| 2.78E-05 | 0.000225841 |
| 9.85E-08 | 1.33E-06 |
| 0.16162391 | 0.298701165 |
| 0.941278332 | NA |
| NA | NA |

| | |
|---|---|
| NA | NA |
| NA | NA |
| NA | NA |
| 0.007923085 | 0.028427084 |
| 0.955792873 | NA |
| NA | NA |
| 0.782609604 | NA |
| NA | NA |
| 0.908042031 | NA |
| 0.275550285 | 0.435443359 |
| 0.329903956 | 0.492868062 |
| NA | NA |
| NA | NA |
| NA | NA |
| NA | NA |
| NA | NA |
| NA | NA |
| NA | NA |
| 0.345489441 | 0.509153592 |
| NA | NA |
| 0.923871011 | 0.954592539 |
| 0.87678365 | 0.926202931 |
| 0.811955836 | NA |
| 0.970356947 | 0.982229107 |
| NA | NA |
| 0.090691966 | 0.195335279 |
| 0.043300088 | 0.110736939 |
| 0.839318641 | NA |
| NA | NA |
| NA | NA |
| 0.535748052 | NA |
| 0.660311384 | NA |
| NA | NA |
| 0.839319108 | NA |
| NA | NA |
| NA | NA |
| NA | NA |
| 0.955792873 | NA |
| 0.923120612 | NA |
| NA | NA |
| NA | NA |
| 2.04E-05 | 0.00017081 |
| NA | NA |
| 0.382080853 | 0.544633396 |
| NA | NA |
| 0.049062259 | 0.12188497 |
| 9.57E-13 | 2.41E-11 |

| | | |
|---|---|---|
| | 8.70E-11 | 1.76E-09 |
| | 0.649576673 | NA |
| | 0.939241089 | NA |
| | 0.079063059 | NA |
| | 0.093227596 | 0.19914182 |
| NA | | NA |
| | 0.357975631 | 0.521556142 |
| | 0.000960121 | 0.004927298 |
| | 0.034627549 | 0.092847238 |
| | 0.003078689 | 0.013099562 |
| | 0.129159938 | 0.25341065 |
| | 0.424930176 | NA |
| NA | | NA |
| | 0.688772153 | NA |
| | 0.938253196 | NA |
| | 0.003837403 | 0.015685707 |
| NA | | NA |
| | 0.560908506 | 0.703988552 |
| | 0.000480836 | 0.002744571 |
| NA | | NA |
| | 0.677444689 | NA |
| | 0.553519698 | NA |
| | 0.865650329 | 0.919767844 |
| | 0.941277857 | NA |
| | 0.445427134 | 0.605935863 |
| | 4.87E-08 | 6.82E-07 |
| NA | | NA |
| NA | | NA |
| | 0.017773028 | 0.0545709 |
| NA | | NA |
| | 0.158504394 | 0.294756764 |
| NA | | NA |
| | 0.937531888 | NA |
| | 0.033261604 | 0.090108433 |
| | 0.86441863 | NA |
| | 0.19392596 | 0.3398988 |
| NA | | NA |
| NA | | NA |
| NA | | NA |
| NA | | NA |
| | 0.002449944 | 0.010847019 |
| | 0.867058289 | NA |
| NA | | NA |
| NA | | NA |
| | 0.344761015 | 0.50836338 |
| | 0.839319108 | NA |
| NA | | NA |

| | |
|---|---|
| 0.435916557 | 0.597058689 |
| NA | NA |
| 0.335194487 | 0.498425638 |
| 0.271174292 | 0.431046513 |
| 0.405941936 | 0.567920367 |
| 0.008648835 | 0.030548704 |
| 0.941278332 | NA |
| NA | NA |
| 0.853595622 | NA |
| NA | NA |
| NA | NA |
| NA | NA |
| 0.955792873 | NA |
| NA | NA |
| NA | NA |
| NA | NA |
| NA | NA |
| 0.911318924 | NA |
| NA | NA |
| 0.260717163 | 0.419381625 |
| 0.941275956 | NA |
| 0.839319108 | NA |
| 0.010007084 | 0.034410374 |
| 0.012967533 | 0.042444063 |
| 0.971738589 | 0.982929833 |
| 0.111047473 | 0.22664654 |
| 0.964758155 | 0.978629815 |
| 0.839319108 | NA |
| 0.165049605 | 0.303622909 |
| 0.332594044 | 0.495797797 |
| 0.941279303 | NA |
| NA | NA |
| 0.546026392 | NA |
| 0.349864531 | 0.513724588 |
| NA | NA |
| 0.941277857 | NA |
| NA | NA |
| 0.898490789 | NA |
| 0.785132014 | 0.867944461 |
| NA | NA |
| 0.329348199 | 0.492346805 |
| 0.009768097 | 0.0337691 |
| 0.941277857 | NA |
| 3.92E-13 | 1.02E-11 |
| NA | NA |
| 0.955792398 | NA |
| 0.24258049 | 0.398998053 |

| | | |
|---|---|---|
| | 0.069324538 | 0.160044875 |
| | 3.70E-05 | 0.000289291 |
| | 0.648690005 | NA |
| NA | | NA |
| | 0.719868297 | NA |
| | 0.578115345 | 0.716366394 |
| | 7.02E-13 | 1.79E-11 |
| | 1.15E-07 | 1.53E-06 |
| NA | | NA |
| | 1.92E-09 | 3.30E-08 |
| | 0.941278332 | NA |
| | 0.869270697 | NA |
| | 1.54E-07 | 2.01E-06 |
| NA | | NA |
| NA | | NA |
| NA | | NA |
| | 0.414857954 | 0.576711191 |
| NA | | NA |
| | 0.612750174 | NA |
| | 0.905947044 | NA |
| | 0.889715909 | NA |
| | 0.459698775 | 0.618816122 |
| NA | | NA |
| NA | | NA |
| | 0.804145034 | 0.880376252 |
| | 0.847566002 | 0.908343735 |
| NA | | NA |
| | 0.937853114 | NA |
| | 0.955791446 | NA |
| | 0.912040872 | 0.946889856 |
| | 0.020693355 | 0.061483366 |
| NA | | NA |
| | 1.14E-05 | 0.000101242 |
| | 0.372672109 | 0.535851969 |
| | 0.275212307 | 0.435102939 |
| NA | | NA |
| NA | | NA |
| NA | | NA |
| NA | | NA |
| | 0.185369221 | 0.328627902 |
| | 0.94398916 | NA |
| NA | | NA |
| | 0.261565936 | 0.420320848 |
| | 0.099205817 | 0.208282338 |
| | 0.70456625 | 0.811560486 |
| | 0.004241323 | 0.017035433 |
| | 0.769942629 | 0.857510576 |

| | | |
|---|---|---|
| | 0.916677176 | NA |
| | 0.552886135 | 0.698038772 |
| | 0.923405008 | NA |
| | 0.955792398 | NA |
| | 0.324539477 | 0.487454321 |
| | 2.10E-07 | 2.66E-06 |
| NA | | NA |
| | 0.084263611 | 0.184941521 |
| | 0.734058779 | NA |
| NA | | NA |
| | 0.431524813 | 0.592931833 |
| | 0.928601912 | NA |
| | 0.41715333 | NA |
| | 1.68E-27 | 1.06E-25 |
| | 0.053557895 | 0.130841119 |
| | 0.488243221 | 0.642976936 |
| | 0.507429303 | 0.659090438 |
| | 0.341414753 | 0.504899283 |
| NA | | NA |
| | 0.822858287 | 0.892245135 |
| | 0.955792398 | NA |
| | 0.197893876 | 0.345024176 |
| | 0.941278332 | NA |
| NA | | NA |
| NA | | NA |
| NA | | NA |
| NA | | NA |
| NA | | NA |
| | 0.739991096 | NA |
| | 0.558891947 | 0.702438074 |
| | 0.696764575 | NA |
| | 0.957402216 | NA |
| NA | | NA |
| NA | | NA |
| NA | | NA |
| | 0.851563613 | 0.910934139 |
| | 0.847557081 | 0.908343735 |
| NA | | NA |
| | 0.005996467 | 0.022651629 |
| | 0.805215613 | 0.880868577 |
| | 0.593567283 | 0.728777906 |
| | 0.658614164 | NA |
| NA | | NA |
| NA | | NA |
| NA | | NA |
| | 0.068347525 | 0.158274032 |
| | 0.941277857 | NA |

| | |
|---|---|
| 0.000948212 | 0.004874589 |
| 0.007233715 | 0.026395552 |
| NA | NA |
| 0.015715675 | NA |
| 0.322924668 | 0.485641811 |
| 0.005327146 | 0.020550522 |
| NA | NA |
| NA | NA |
| 0.628615777 | 0.755481581 |
| 0.977180525 | NA |
| NA | NA |
| 0.06507175 | 0.152519724 |
| 0.011209532 | 0.03777635 |
| NA | NA |
| 0.185231257 | 0.328456706 |
| 0.116839512 | NA |
| 0.004724185 | 0.018656886 |
| 0.364163088 | 0.52755777 |
| 0.955792398 | NA |
| 0.545268128 | NA |
| NA | NA |
| NA | NA |
| NA | NA |
| 0.00748964 | 0.02717092 |
| NA | NA |
| 0.919747783 | NA |
| 0.018096179 | 0.055370261 |
| 0.606411484 | NA |
| 0.839319108 | NA |
| 1.69E-53 | 2.55E-51 |
| 0.284180081 | 0.444769483 |
| 0.215507026 | 0.366471261 |
| NA | NA |
| 0.573012297 | 0.712694712 |
| 0.250642992 | 0.408281173 |
| NA | NA |
| 0.660816856 | 0.779611457 |
| NA | NA |
| 0.119050035 | 0.238830132 |
| 0.116573341 | 0.235002852 |
| 0.308718916 | 0.470250072 |
| 0.082112628 | 0.181202 |
| 0.568593306 | 0.709794821 |
| 0.4428096 | 0.603409521 |
| 0.337608127 | 0.500950604 |
| 0.956645531 | 0.97337139 |
| 0.647623582 | 0.769582451 |

| | | |
|---|---|---|
| | 0.839319574 | NA |
| NA | | NA |
| NA | | NA |
| | 0.913924762 | NA |
| | 0.079216297 | 0.176611681 |
| | 0.342648221 | 0.506125913 |
| | 0.068357997 | 0.158274032 |
| | 0.963266418 | 0.977809079 |
| | 0.32562797 | NA |
| | 0.391295829 | NA |
| | 0.839318641 | NA |
| | 0.351258755 | NA |
| | 0.918810289 | NA |
| | 0.000358554 | 0.002124096 |
| | 0.879413258 | NA |
| | 0.583214136 | 0.720526464 |
| | 0.734107387 | 0.832335732 |
| | 0.937669761 | NA |
| | 0.459236529 | NA |
| NA | | NA |
| | 0.170133334 | 0.30980973 |
| | 0.955792398 | NA |
| | 0.839319108 | NA |
| | 0.551657241 | 0.696950406 |
| | 0.075596694 | 0.170538576 |
| | 0.087459318 | 0.190116361 |
| | 0.228835489 | 0.38252095 |
| | 0.941279803 | NA |
| | 0.786150112 | NA |
| | 0.369735635 | 0.532950055 |
| | 0.247635096 | 0.404877769 |
| | 0.077439166 | 0.173494636 |
| | 0.584873406 | 0.721733424 |
| | 0.734058779 | NA |
| | 0.005480689 | 0.020996516 |
| | 0.110084718 | 0.225280412 |
| | 0.246850443 | 0.403872302 |
| NA | | NA |
| | 0.000989927 | 0.00506497 |
| | 0.162948807 | NA |
| | 0.002793029 | 0.012076238 |
| | 9.30E-08 | 1.26E-06 |
| | 0.003983538 | 0.016152669 |
| | 0.003357818 | 0.014088275 |
| | 0.99267508 | 0.9949323 |
| NA | | NA |
| | 0.046421125 | 0.116823388 |

| | | |
|---|---|---|
| | 0.043593909 | 0.11127316 |
| NA | | NA |
| | 1.68E-18 | 6.48E-17 |
| | 0.839319108 | NA |
| | 0.025025647 | 0.071714479 |
| | 0.315793648 | 0.478271851 |
| | 0.183270015 | NA |
| | 0.017047445 | 0.052833762 |
| NA | | NA |
| | 0.365335646 | 0.52861514 |
| | 0.022985032 | 0.067037094 |
| | 0.904724277 | NA |
| | 0.363832085 | 0.527434705 |
| | 0.194791948 | NA |
| | 0.171904346 | 0.312055159 |
| | 0.066725589 | 0.155447047 |
| | 0.00126283 | 0.006215441 |
| NA | | NA |
| | 0.512485893 | 0.663847033 |
| | 0.080622813 | 0.178842888 |
| NA | | NA |
| | 0.799608236 | 0.877620713 |
| | 0.227883282 | 0.381279199 |
| | 0.092820386 | 0.198607589 |
| | 0.941278332 | NA |
| NA | | NA |
| | 0.955792397 | NA |
| NA | | NA |
| | 0.869622312 | NA |
| | 0.327780426 | 0.490525426 |
| | 0.70070336 | NA |
| | 0.188209422 | 0.332573175 |
| | 0.955791446 | NA |
| | 0.001238651 | 0.006121755 |
| | 0.005640693 | 0.021526338 |
| | 0.010221419 | 0.035005785 |
| NA | | NA |
| NA | | NA |
| NA | | NA |
| | 0.000218756 | 0.001386272 |
| | 0.379220881 | 0.54211504 |
| | 0.230608134 | 0.38454076 |
| | 2.67E-05 | 0.000218034 |
| | 0.030387771 | 0.083781966 |
| | 0.458482149 | 0.618109189 |
| | 5.57E-10 | 1.02E-08 |
| | 0.195710819 | NA |

| | |
|---|---|
| NA | NA |
| 0.074462818 | 0.168539748 |
| 0.090386152 | 0.194883319 |
| NA | NA |
| NA | NA |
| 0.927817401 | NA |
| 0.944417383 | NA |
| NA | NA |
| NA | NA |
| 3.60E-08 | 5.18E-07 |
| 0.471604728 | 0.629211644 |
| NA | NA |
| 0.016938987 | 0.052538671 |
| 0.082522681 | NA |
| NA | NA |
| 0.853595622 | NA |
| NA | NA |
| 0.46310244 | 0.621911773 |
| 0.378139996 | NA |
| 0.755128604 | 0.846651799 |
| 0.646599754 | 0.768788019 |
| NA | NA |
| 0.721298992 | NA |
| NA | NA |
| 0.057400674 | 0.138154292 |
| 0.104728826 | 0.216973968 |
| 0.553608847 | 0.698604238 |
| 0.259754229 | 0.418284971 |
| 0.219115566 | 0.37064805 |
| 0.25789906 | 0.416142189 |
| 0.186492733 | 0.330250727 |
| 0.265879565 | 0.425214367 |
| 0.961194897 | NA |
| 0.729045923 | 0.829402115 |
| 0.839319574 | NA |
| 0.214442211 | 0.365339168 |
| NA | NA |
| 0.081925924 | 0.180868043 |
| 0.217451826 | 0.368803504 |
| 0.152162499 | 0.286192952 |
| 0.176985294 | 0.318533381 |
| 0.553060658 | 0.698093534 |
| 0.41493618 | NA |
| 0.941277857 | NA |
| NA | NA |
| 0.030555753 | 0.084157375 |
| NA | NA |

| | | | |
|---|---|---|---|
| | 0.861946903 | | 0.917492965 |
| | 0.688086644 | NA | |
| | 6.45E-08 | | 8.92E-07 |
| | 4.06E-08 | | 5.76E-07 |
| NA | | NA | |
| | 1.55E-05 | | 0.000134451 |
| | 1.33E-06 | | 1.45E-05 |
| | 0.022170877 | | 0.065061178 |
| | 0.171719895 | | 0.311886832 |
| NA | | NA | |
| NA | | NA | |
| | 0.372235228 | | 0.535570507 |
| | 0.897100023 | NA | |
| | 0.955791922 | NA | |
| NA | | NA | |
| | 0.941278332 | NA | |
| NA | | NA | |
| | 0.589318811 | NA | |
| | 0.904109119 | | 0.941735821 |
| | 1.45E-05 | | 0.000126318 |
| | 7.80E-10 | | 1.40E-08 |
| | 3.44E-08 | | 4.96E-07 |
| | 0.839319108 | NA | |
| | 0.492870204 | | 0.646601807 |
| NA | | NA | |
| | 0.494899051 | | 0.647906452 |
| | 0.845801706 | | 0.907158717 |
| | 0.783831607 | NA | |
| | 0.416964278 | | 0.578490542 |
| | 1.05E-07 | | 1.41E-06 |
| | 0.842503134 | NA | |
| | 0.610965275 | | 0.742227061 |
| | 2.08E-05 | | 0.000173783 |
| | 0.313844816 | | 0.475956311 |
| NA | | NA | |
| | 0.922904709 | NA | |
| | 0.955792398 | NA | |
| | 0.326712432 | | 0.489587969 |
| | 0.003512411 | | 0.01460559 |
| | 0.937958036 | | 0.963177176 |
| | 0.656163306 | | 0.775664217 |
| | 0.906224297 | | 0.943154819 |
| | 1.64E-05 | | 0.000140752 |
| | 0.465830116 | NA | |
| | 0.853595154 | NA | |
| NA | | NA | |
| | 0.800307688 | NA | |

| | | |
|---|---|---|
| | 0.784597358 | NA |
| | 0.641615014 | NA |
| NA | | NA |
| NA | | NA |
| NA | | NA |
| | 0.917096772 | NA |
| | 0.60542669 | 0.73815731 |
| NA | | NA |
| NA | | NA |
| NA | | NA |
| NA | | NA |
| | 0.654276538 | NA |
| | 0.13002131 | 0.254713351 |
| | 0.340985217 | NA |
| | 0.137209431 | 0.265279741 |
| NA | | NA |
| NA | | NA |
| NA | | NA |
| | 0.020697652 | 0.061488456 |
| | 0.341750618 | 0.505170493 |
| NA | | NA |
| | 0.839319108 | NA |
| | 0.955792873 | NA |
| | 0.000931017 | 0.004798631 |
| NA | | NA |
| | 0.940983457 | NA |
| NA | | NA |
| NA | | NA |
| NA | | NA |
| | 0.994816399 | NA |
| | 0.369150648 | 0.532355269 |
| NA | | NA |
| | 0.387289123 | 0.550086957 |
| NA | | NA |
| | 0.418908372 | 0.58064623 |
| | 1.16E-05 | 0.000103498 |
| | 0.609559531 | 0.741490928 |
| NA | | NA |
| | 0.002786994 | 0.012061095 |
| | 0.567806864 | 0.709296682 |
| | 0.285174293 | NA |
| | 0.004072238 | 0.016470248 |
| | 0.910039633 | NA |
| | 0.208147552 | 0.357480917 |
| | 0.879444233 | 0.927860444 |
| | 0.386688804 | 0.54949761 |
| | 0.038145592 | 0.10031393 |

| | | | |
|---|---|---|---|
| | 0.756971744 | | 0.847999315 |
| | 0.937151727 | NA | |
| NA | | NA | |
| NA | | NA | |
| NA | | NA | |
| | 0.941275956 | NA | |
| | 0.853595622 | NA | |
| | 0.195344049 | NA | |
| | 0.390158535 | NA | |
| | 0.134005156 | | 0.260502347 |
| | 0.538908905 | | 0.68636008 |
| | 0.001009176 | | 0.005151277 |
| | 0.107173267 | | 0.220860443 |
| | 0.006617013 | | 0.024532423 |
| | 0.042387293 | | 0.108964505 |
| | 0.594965679 | | 0.729930059 |
| | 0.636783354 | | 0.76118881 |
| | 0.045451949 | | 0.114934253 |
| NA | | NA | |
| | 0.941278332 | NA | |
| NA | | NA | |
| NA | | NA | |
| NA | | NA | |
| | 0.765773414 | NA | |
| NA | | NA | |
| NA | | NA | |
| NA | | NA | |
| NA | | NA | |
| NA | | NA | |
| NA | | NA | |
| NA | | NA | |
| NA | | NA | |
| NA | | NA | |
| | 0.003957538 | | 0.016058185 |
| | 0.000730553 | | 0.003921659 |
| | 2.74E-06 | | 2.81E-05 |
| | 0.213901905 | NA | |
| | 0.000338582 | | 0.002026382 |
| | 0.692943712 | NA | |
| | 0.427786525 | | 0.589041703 |
| | 0.004891913 | | 0.019179263 |
| NA | | NA | |
| | 0.463973872 | NA | |
| | 0.955792398 | NA | |
| NA | | NA | |
| NA | | NA | |
| NA | | NA | |

| | | | |
|---|---|---|---|
| NA | | NA | |
| | 0.499786826 | | 0.652618281 |
| NA | | NA | |
| NA | | NA | |
| | 0.288160862 | | 0.448815917 |
| NA | | NA | |
| NA | | NA | |
| | 0.839319108 | NA | |
| NA | | NA | |
| NA | | NA | |
| | 0.836837225 | | 0.901121952 |
| NA | | NA | |
| NA | | NA | |
| | 0.955792398 | NA | |
| NA | | NA | |
| NA | | NA | |
| | 0.838197868 | NA | |
| NA | | NA | |
| | 0.570967822 | NA | |
| | 0.295386137 | | 0.456752135 |
| NA | | NA | |
| NA | | NA | |
| | 0.374743639 | NA | |
| NA | | NA | |
| NA | | NA | |
| NA | | NA | |
| NA | | NA | |
| | 0.026987014 | | 0.076381577 |
| | 0.941278332 | NA | |
| | 0.000771105 | | 0.004099587 |
| NA | | NA | |
| NA | | NA | |
| NA | | NA | |
| NA | | NA | |
| | 0.889439637 | | 0.933455095 |
| | 0.632146043 | NA | |
| | 0.815100009 | NA | |
| | 0.839318641 | NA | |
| | 0.124305961 | | 0.246740237 |
| | 0.433079436 | NA | |
| | 0.941278788 | NA | |
| | 0.406116523 | | 0.568033962 |
| | 2.69E-08 | | 3.95E-07 |
| | 0.921383099 | NA | |
| | 0.955792398 | NA | |
| NA | | NA | |
| | 0.220834807 | | 0.372735078 |

| | |
|---|---|
| 0.130604333 | 0.255645093 |
| 0.033995938 | 0.091524864 |
| 0.534643333 | 0.682812718 |
| NA | NA |
| 0.119945549 | 0.240254478 |
| 0.955792398 | NA |
| 0.828576439 | NA |
| 0.751023372 | 0.843678417 |
| 0.074196166 | 0.168112056 |
| 0.001867633 | 0.008636098 |
| 0.012883231 | 0.042232017 |
| 0.415861462 | 0.5776339 |
| 0.682762669 | 0.796194064 |
| 0.543266438 | 0.690048976 |
| 0.144829541 | 0.27610995 |
| 0.011426994 | 0.03837806 |
| 0.947054604 | 0.967625072 |
| 0.018933487 | 0.057430211 |
| NA | NA |
| 0.63427996 | NA |
| NA | NA |
| 5.34E-05 | 0.000402363 |
| 0.361709845 | NA |
| 0.037142925 | 0.098160359 |
| 0.734058779 | NA |
| 1.07E-06 | 1.18E-05 |
| 0.941276431 | NA |
| 0.545597103 | NA |
| 0.001551101 | 0.00738534 |
| 0.00935295 | 0.032571049 |
| 0.05529761 | 0.134264378 |
| 0.097722455 | 0.206095152 |
| 0.062498716 | 0.147696033 |
| 0.790215145 | NA |
| 0.018361868 | 0.056031946 |
| 0.119529625 | 0.239602842 |
| 0.001637373 | 0.00773576 |
| 0.000663118 | 0.003606015 |
| 0.002911021 | 0.01250734 |
| NA | NA |
| 0.889521997 | NA |
| NA | NA |
| 0.519252168 | 0.669768607 |
| NA | NA |
| 8.31E-07 | 9.39E-06 |
| 0.544045717 | NA |
| 0.850267775 | 0.910230888 |

| | | |
|---|---|---|
| | 0.966636482 | 0.980101451 |
| | 0.182762507 | 0.32532898 |
| | 0.296785326 | 0.458230691 |
| | 0.851870742 | 0.911045444 |
| | 0.091178476 | 0.196090073 |
| | 0.466387982 | NA |
| | 0.94522829 | NA |
| NA | | NA |
| | 7.23E-07 | 8.27E-06 |
| | 0.278251116 | 0.437937781 |
| | 0.415448191 | 0.577295689 |
| | 0.673167126 | NA |
| | 0.06338036 | 0.14930516 |
| | 0.057812988 | 0.138828162 |
| | 0.14625721 | 0.278118887 |
| NA | | NA |
| | 0.087152614 | 0.189593628 |
| NA | | NA |
| | 0.60384076 | 0.736872583 |
| | 1.58E-14 | 4.58E-13 |
| | 0.780833705 | NA |
| | 0.604711077 | 0.737590092 |
| | 0.098228696 | 0.206924456 |
| | 0.970160785 | NA |
| | 0.941277857 | NA |
| | 0.052294039 | 0.128439037 |
| NA | | NA |
| | 0.002149817 | 0.009713353 |
| | 0.645659785 | NA |
| | 0.439918798 | NA |
| | 0.774588058 | NA |
| NA | | NA |
| NA | | NA |
| | 0.640660122 | 0.764207288 |
| NA | | NA |
| | 0.941279303 | NA |
| NA | | NA |
| | 0.262055892 | 0.420824068 |
| | 0.111312797 | 0.227046619 |
| | 0.585560729 | 0.722263382 |
| NA | | NA |
| NA | | NA |
| | 0.572566654 | 0.71242494 |
| | 0.474140172 | NA |
| | 0.838219909 | 0.901957968 |
| | 0.054246046 | 0.132156028 |
| | 0.066675388 | 0.155375718 |

| | | |
|---|---|---|
| NA | 0.401359905 | 0.563741795 |
| | 0.47110198 | 0.628611354 |
| | 0.848600431 | 0.908979709 |
| | 0.095291845 | 0.202353821 |
| | 0.263246663 | NA |
| | 0.181953285 | 0.324363727 |
| | 0.008915569 | 0.031336146 |
| | 0.980612168 | 0.988000796 |
| | 0.743511643 | NA |
| | 0.820274673 | NA |
| | 0.018530864 | 0.056453555 |
| | 0.07974843 | 0.177413384 |
| | 0.004782485 | 0.018837118 |
| | 0.700647635 | 0.808808138 |
| | 0.941278424 | NA |
| | 0.005789532 | 0.021985619 |
| | 0.005388792 | 0.020717819 |
| | 0.819939881 | NA |
| | 0.209120715 | NA |
| | 0.002679125 | 0.011698501 |
| | 0.004130869 | 0.016676211 |
| | 0.383203384 | 0.545934203 |
| | 0.839319108 | NA |
| | 0.838197868 | NA |
| | 0.038356291 | 0.100756699 |
| | 0.014626801 | 0.046737431 |
| | 0.769666002 | 0.857314759 |
| | 0.571793489 | 0.711885586 |
| | 0.339756428 | 0.50344783 |
| NA | | NA |
| | 0.941278838 | NA |
| | 0.941277857 | NA |
| | 0.171241761 | 0.311374821 |
| | 0.941278332 | NA |
| NA | | NA |
| | 4.47E-05 | 0.000342417 |
| | 0.84632571 | 0.907398403 |
| | 0.926150426 | NA |
| | 9.92E-12 | 2.25E-10 |
| NA | | NA |
| | 0.587275766 | 0.723698747 |
| NA | | NA |
| | 0.829643269 | 0.896539049 |
| | 0.323431482 | 0.486083105 |
| | 0.750509572 | 0.843340045 |
| | 0.316294441 | 0.478699593 |
| NA | | NA |

| | |
|---|---|
| 0.285187505 | NA |
| 0.680044623 | 0.794122482 |
| 0.922849885 | NA |
| 0.799754051 | NA |
| 0.434022926 | 0.595218581 |
| 0.944591465 | NA |
| 0.941275956 | NA |
| 0.658880217 | 0.777987247 |
| 0.862407629 | NA |
| 0.463042737 | NA |
| 0.960066257 | 0.975517483 |
| 0.124656592 | 0.247194552 |
| 0.076069472 | 0.171215658 |
| NA | NA |
| NA | NA |
| NA | NA |
| 0.530108396 | 0.678924703 |
| 9.27E-08 | 1.26E-06 |
| 1.05E-07 | 1.41E-06 |
| 0.188107845 | 0.332492419 |
| 0.000225436 | 0.001422165 |
| 0.543607119 | 0.690334345 |
| 0.243397581 | 0.399927463 |
| 0.000337023 | 0.002019132 |
| 0.259393139 | 0.41784485 |
| 0.214832294 | 0.365715796 |
| 0.981493747 | 0.988492849 |
| 0.047957586 | 0.119803364 |
| 0.516063859 | 0.666921054 |
| NA | NA |
| 0.748393366 | NA |
| 0.38056362 | 0.543186333 |
| 0.955791922 | NA |
| 0.061803936 | 0.146417566 |
| 2.75E-05 | 0.000223253 |
| 8.23E-16 | 2.66E-14 |
| 1.20E-18 | 4.69E-17 |
| 0.941278332 | NA |
| 0.001476747 | 0.007079461 |
| 3.65E-06 | 3.66E-05 |
| 0.484116618 | 0.639985767 |
| 1.53E-10 | 3.01E-09 |
| 0.598906648 | 0.733026998 |
| 0.941277857 | NA |
| 0.086063384 | 0.187830612 |
| NA | NA |
| NA | NA |

| | | | |
|---|---|---|---|
| | 0.839319108 | NA | |
| NA | | NA | |
| NA | | NA | |
| NA | | NA | |
| NA | | NA | |
| NA | | NA | |
| | 0.570087036 | | 0.710839312 |
| | 0.941278332 | NA | |
| NA | | NA | |
| | 0.941278332 | NA | |
| | 0.941278332 | NA | |
| | 0.941278332 | NA | |
| NA | | NA | |
| NA | | NA | |
| | 0.955792873 | NA | |
| | 0.112374924 | NA | |
| NA | | NA | |
| | 0.229152543 | | 0.382837709 |
| | 1.38E-08 | | 2.11E-07 |
| | 0.372000423 | | 0.535343845 |
| | 0.986274629 | | 0.991481553 |
| | 0.839318641 | NA | |
| | 0.688865927 | NA | |
| NA | | NA | |
| | 0.438726041 | NA | |
| | 0.547629998 | | 0.693703524 |
| | 0.277617911 | | 0.437434077 |
| | 8.61E-05 | | 0.000613807 |
| | 0.941278332 | NA | |
| | 0.941277857 | NA | |
| | 0.287591602 | | 0.448193237 |
| | 0.071828547 | | 0.164169682 |
| | 0.472208726 | | 0.629664543 |
| | 0.307800799 | | 0.469362203 |
| | 0.856576343 | NA | |
| | 0.72601533 | NA | |
| | 7.95E-05 | | 0.000572565 |
| | 0.609101154 | | 0.741273426 |
| | 0.327181102 | | 0.48996883 |
| | 0.03493412 | | 0.093479699 |
| | 0.621964881 | NA | |
| | 0.839319374 | NA | |
| NA | | NA | |
| | 0.278817844 | | 0.438597734 |
| | 0.842315901 | | 0.905015261 |
| NA | | NA | |
| | 0.540713377 | NA | |

| | | | |
|---|---|---|---|
| NA | | NA | |
| NA | | NA | |
| | 0.000121451 | | 0.000831159 |
| NA | | NA | |
| NA | | NA | |
| NA | | NA | |
| | 0.941278788 | NA | |
| | 0.498667629 | | 0.651585798 |
| | 0.374406346 | | 0.537556994 |
| | 0.172102616 | | 0.31236743 |
| | 0.293267562 | | 0.45445947 |
| | 0.755626499 | NA | |
| | 0.002481525 | | 0.010964405 |
| | 0.941279303 | NA | |
| | 0.955167254 | | 0.972692863 |
| | 0.556236673 | NA | |
| | 0.941278332 | NA | |
| | 0.578861363 | NA | |
| | 0.941278332 | NA | |
| | 0.728678906 | NA | |
| | 0.82229013 | NA | |
| | 0.793689871 | NA | |
| NA | | NA | |
| NA | | NA | |
| | 0.646231565 | NA | |
| NA | | NA | |
| | 0.889188267 | NA | |
| | 0.691328233 | NA | |
| | 0.779075181 | | 0.863795819 |
| | 0.841974858 | NA | |
| NA | | NA | |
| | 0.817706245 | NA | |
| NA | | NA | |
| | 0.806092235 | | 0.881251487 |
| | 0.839319108 | NA | |
| NA | | NA | |
| | 0.941278332 | NA | |
| NA | | NA | |
| | 0.743111594 | | 0.83852368 |
| | 0.195865737 | | 0.342365886 |
| | 0.64574902 | | 0.768045082 |
| | 0.202370834 | | 0.350492004 |
| | 1.92E-08 | | 2.88E-07 |
| | 0.32193604 | | 0.484706282 |
| | 0.778547407 | NA | |
| | 0.868089774 | NA | |
| | 0.237671396 | | 0.393178661 |

| | |
|---|---|
| NA | NA |
| 0.013458714 | 0.04377522 |
| 0.670337301 | 0.786689247 |
| NA | NA |
| 0.740238263 | NA |
| 0.004098539 | 0.016568178 |
| 0.004547911 | 0.018048022 |
| NA | NA |
| 0.92069054 | NA |
| 0.786733216 | NA |
| 0.667898481 | NA |
| 0.000268547 | 0.001652522 |
| 0.587302118 | 0.723698747 |
| 0.778735071 | NA |
| 0.002112847 | 0.009571753 |
| NA | NA |
| NA | NA |
| 0.006961822 | 0.025571805 |
| NA | NA |
| NA | NA |
| 0.941278332 | NA |
| NA | NA |
| NA | NA |
| 1.22E-19 | 5.04E-18 |
| 0.689071788 | NA |
| NA | NA |
| 0.919922377 | NA |
| NA | NA |
| 0.132191277 | 0.257832551 |
| 0.351428715 | 0.515109595 |
| 0.973502181 | 0.984043719 |
| NA | NA |
| 0.21828044 | 0.369576826 |
| 0.340658417 | 0.504219366 |
| 0.00767884 | 0.027717888 |
| 0.134318133 | 0.260982873 |
| 0.147162078 | 0.279415426 |
| 0.850698945 | NA |
| NA | NA |
| NA | NA |
| 0.000343903 | 0.002051008 |
| 0.397560848 | 0.560190818 |
| NA | NA |
| 0.837259124 | NA |
| 0.839318641 | NA |
| 0.004418719 | 0.017614192 |
| 0.320033219 | 0.482726912 |

| | | |
|---|---|---|
| | 0.766455371 | 0.855009424 |
| | 0.441138563 | 0.601799195 |
| NA | | NA |
| | 0.88012057 | 0.928162597 |
| | 0.65077816 | NA |
| NA | | NA |
| | 0.955792398 | NA |
| NA | | NA |
| | 0.459915493 | 0.618993981 |
| | 0.90300356 | 0.941349627 |
| | 0.000578934 | 0.003211434 |
| | 0.007907621 | 0.028384435 |
| | 0.853595154 | NA |
| | 0.212721677 | 0.363270923 |
| | 0.006787606 | 0.025043961 |
| | 8.56E-08 | 1.16E-06 |
| | 0.73431627 | NA |
| | 0.657181815 | NA |
| | 7.12E-11 | 1.46E-09 |
| | 0.00879125 | 0.030967786 |
| | 0.441912916 | 0.602450589 |
| | 0.432650789 | 0.59382618 |
| | 0.515970007 | NA |
| | 0.937924455 | NA |
| | 0.037842104 | 0.099669997 |
| NA | | NA |
| NA | | NA |
| NA | | NA |
| | 0.34015522 | 0.50383243 |
| | 0.078039409 | 0.174521749 |
| | 0.941277857 | NA |
| NA | | NA |
| | 0.941277857 | NA |
| | 0.000171924 | 0.001127619 |
| | 0.003193517 | 0.013518068 |
| NA | | NA |
| NA | | NA |
| | 0.292984439 | 0.454129697 |
| | 0.938552414 | NA |
| | 0.93838321 | NA |
| NA | | NA |
| NA | | NA |
| NA | | NA |
| NA | | NA |
| NA | | NA |
| | 0.93754161 | NA |
| NA | | NA |

| | | | |
|---|---|---|---|
| NA | | NA | |
| NA | | NA | |
| NA | | NA | |
| NA | | NA | |
| NA | | NA | |
| NA | | NA | |
| NA | | NA | |
| NA | | NA | |
| NA | | NA | |
| | 0.016378113 | | 0.05116575 |
| | 0.572578332 | | 0.71242494 |
| | 0.020691101 | | 0.061483366 |
| | 0.21709844 | | 0.368495675 |
| | 0.45814076 | | 0.617733931 |
| | 0.726514089 | | 0.82747097 |
| NA | | NA | |
| | 0.574579571 | | 0.713786167 |
| NA | | NA | |
| | 0.012045643 | | 0.040009825 |
| NA | | NA | |
| NA | | NA | |
| NA | | NA | |
| NA | | NA | |
| NA | | NA | |
| | 0.010412041 | | 0.03557161 |
| | 0.016786505 | | 0.052167698 |
| | 0.004113311 | | 0.016613785 |
| | 3.94E-06 | | 3.92E-05 |
| NA | | NA | |
| | 0.008747231 | | 0.030853823 |
| | 0.299543816 | | 0.461058749 |
| | 0.24381311 | | 0.400288755 |
| | 0.584412955 | NA | |
| NA | | NA | |
| | 0.839318641 | NA | |
| NA | | NA | |
| | 0.005597796 | | 0.021390067 |
| NA | | NA | |
| | 0.955792873 | NA | |
| NA | | NA | |
| | 0.063558209 | | 0.149642404 |
| | 0.0982261 | | 0.206924456 |
| | 0.839319108 | NA | |
| | 0.729485695 | NA | |
| NA | | NA | |
| | 0.839319108 | NA | |
| | 0.839318641 | NA | |

| | |
|---|---|
| NA | NA |
| NA | NA |
| NA | NA |
| NA | NA |
| 0.839318641 | NA |
| NA | NA |
| 0.388777186 | 0.551561603 |
| 0.69380776 | 0.803747042 |
| NA | NA |
| NA | NA |
| 0.941280202 | NA |
| 0.941278332 | NA |
| 0.861879967 | 0.917492965 |
| 0.024908452 | 0.071481913 |
| NA | NA |
| 3.08E-06 | 3.12E-05 |
| NA | NA |
| 0.565269926 | NA |
| 0.837323055 | 0.901278133 |
| 0.26026871 | NA |
| 1.66E-05 | 0.000142142 |
| 0.936601529 | 0.9623981 |
| 0.915325748 | NA |
| 0.839319108 | NA |
| 0.004719535 | 0.01864471 |
| 0.013504697 | 0.04387045 |
| 0.188962188 | 0.33355666 |
| 0.463246069 | 0.621999042 |
| 0.008369049 | 0.029740235 |
| 0.039714289 | 0.103586439 |
| 0.000528981 | 0.00297896 |
| NA | NA |
| 0.858407835 | NA |
| 1.90E-06 | 2.02E-05 |
| 0.192145035 | 0.337522961 |
| 0.839318641 | NA |
| 0.853595622 | NA |
| 0.941276431 | NA |
| 0.447991755 | NA |
| 0.705744039 | 0.812209632 |
| 0.899636727 | NA |
| 0.457770282 | 0.617514572 |
| NA | NA |
| 0.047482009 | 0.118842708 |
| NA | NA |
| 0.713183813 | NA |
| 0.941279895 | NA |

| | | | |
|---|---|---|---|
| | 0.011885991 | | 0.039595573 |
| | 0.94388205 | NA | |
| NA | | NA | |
| NA | | NA | |
| NA | | NA | |
| | 0.41607394 | | 0.577827873 |
| NA | | NA | |
| | 0.653419178 | | 0.774003044 |
| | 2.76E-13 | | 7.27E-12 |
| | 0.801392723 | | 0.878603792 |
| | 0.276509345 | NA | |
| | 0.079606354 | | 0.177182574 |
| | 0.007608051 | | 0.027516611 |
| | 0.609067137 | NA | |
| | 0.15285344 | | 0.286964594 |
| | 0.166278137 | | 0.305099615 |
| NA | | NA | |
| | 0.952811619 | NA | |
| | 0.839318641 | NA | |
| NA | | NA | |
| NA | | NA | |
| NA | | NA | |
| | 0.504970083 | | 0.657044385 |
| | 0.452335875 | NA | |
| | 0.310416438 | | 0.472261557 |
| | 0.911857069 | NA | |
| | 0.458114332 | | 0.617733931 |
| | 3.05E-07 | | 3.74E-06 |
| | 0.369147319 | | 0.532355269 |
| | 0.170631293 | | 0.310430791 |
| | 0.586965001 | | 0.723508833 |
| | 0.091598372 | | 0.196642231 |
| | 0.047075925 | | 0.118080301 |
| | 0.936410447 | NA | |
| | 0.955791922 | NA | |
| NA | | NA | |
| NA | | NA | |
| | 0.071719957 | | 0.163981299 |
| | 0.459534849 | | 0.618750054 |
| NA | | NA | |
| | 5.97E-75 | | 1.44E-72 |
| | 0.010992394 | | 0.03719661 |
| | 0.039804091 | | 0.103781024 |
| | 0.969938813 | | 0.982023444 |
| | 0.217944715 | | 0.369244794 |
| | 0.410160827 | NA | |
| | 0.839319108 | NA | |

| | |
|---|---|
| 0.272015036 | 0.431950222 |
| 0.492363047 | 0.646328328 |
| 0.58185943 | 0.71952505 |
| 0.203702006 | 0.3521327 |
| 0.680035465 | 0.794122482 |
| 0.660009644 | 0.778895807 |
| 5.42E-07 | 6.34E-06 |
| 0.101609783 | 0.212298853 |
| 0.934488594 | 0.960973711 |
| NA | NA |
| NA | NA |
| NA | NA |
| 0.459829575 | NA |
| 0.236954105 | 0.392401142 |
| NA | NA |
| 0.015174604 | 0.048158122 |
| 0.023087517 | 0.067278291 |
| 0.930213147 | 0.958481962 |
| 0.072696655 | 0.165586992 |
| 0.695394038 | 0.805047125 |
| 0.908548565 | NA |
| NA | NA |
| NA | NA |
| 0.401397907 | 0.563761903 |
| NA | NA |
| NA | NA |
| NA | NA |
| 0.004304944 | 0.017229911 |
| 0.941276431 | NA |
| 0.787746606 | 0.869553296 |
| 7.24E-05 | 0.000526347 |
| 0.718807512 | 0.821055414 |
| 0.490049476 | 0.644428598 |
| 0.195232138 | 0.341559379 |
| NA | NA |
| NA | NA |
| 3.85E-06 | 3.84E-05 |
| NA | NA |
| NA | NA |
| 0.323463167 | 0.486083105 |
| 0.568993733 | 0.71014571 |
| 0.420673777 | 0.582211725 |
| 0.000234991 | 0.001472299 |
| 0.813780148 | 0.886358628 |
| 0.038137697 | 0.10031393 |
| 0.941278332 | NA |
| NA | NA |

| | |
|---|---|
| 0.839318641 | NA |
| 0.614477074 | NA |
| 8.02E-09 | 1.27E-07 |
| NA | NA |
| NA | NA |
| NA | NA |
| NA | NA |
| 0.001701755 | 0.007997072 |
| 0.429679317 | NA |
| 1.14E-07 | 1.52E-06 |
| 1.50E-13 | 4.05E-12 |
| 0.00116564 | 0.005814031 |
| 0.020543389 | 0.061152294 |
| 0.85457085 | 0.91249811 |
| 0.000198703 | 0.001273432 |
| 0.735828174 | NA |
| NA | NA |
| NA | NA |
| 0.004078464 | 0.016492627 |
| 0.32477721 | 0.487719482 |
| 1.69E-05 | 0.000144399 |
| 0.015600752 | 0.049221777 |
| 3.63E-06 | 3.64E-05 |
| 5.87E-18 | 2.18E-16 |
| 0.251727543 | 0.409431391 |
| 0.046754133 | 0.117499934 |
| 0.288966783 | 0.449788234 |
| 0.029721868 | 0.082359882 |
| 0.103448113 | 0.215101946 |
| 0.08905105 | 0.192672766 |
| NA | NA |
| 1.27E-07 | 1.68E-06 |
| 0.00067595 | 0.003666582 |
| 0.00257518 | 0.011318045 |
| 0.001021414 | 0.005199251 |
| 0.0289558 | 0.080674932 |
| 0.78003963 | 0.864527319 |
| 0.955792398 | NA |
| 7.01E-05 | 0.000511092 |
| 0.260252875 | 0.418832922 |
| 0.01555146 | 0.049085778 |
| 0.121886481 | 0.243241555 |
| 2.96E-09 | 4.95E-08 |
| 0.001273274 | 0.006259078 |
| 0.000128146 | 0.000872002 |
| 0.129910284 | NA |
| 0.002955925 | 0.012665515 |

| | | |
|---|---|---|
| | 0.000596908 | 0.00329796 |
| | 0.676681172 | 0.791584276 |
| | 0.230310086 | 0.384177226 |
| | 0.860704972 | 0.916679745 |
| | 0.071658485 | 0.163906905 |
| | 0.008324183 | 0.029625029 |
| | 0.409436475 | 0.57144194 |
| | 4.02E-06 | 4.00E-05 |
| | 0.000401107 | 0.002341708 |
| | 0.049865948 | 0.12349488 |
| | 0.941277857 | NA |
| | 0.908075072 | NA |
| | 0.955792873 | NA |
| | 0.563607856 | NA |
| | 0.000317454 | 0.001912922 |
| | 2.15E-08 | 3.20E-07 |
| | 0.198728901 | 0.346038369 |
| | 0.015230786 | 0.048279451 |
| NA | | NA |
| NA | | NA |
| | 0.015840122 | 0.049841812 |
| | 0.004024922 | 0.016296718 |
| | 0.00371733 | 0.01527095 |
| | 0.572110335 | NA |
| | 0.381769308 | 0.544287096 |
| | 0.894720347 | 0.936587813 |
| | 0.003267344 | 0.013779987 |
| | 0.000167595 | 0.00110318 |
| | 0.000146552 | 0.000979016 |
| | 0.005120863 | 0.019903131 |
| | 0.476077898 | 0.633163124 |
| NA | | NA |
| | 0.316941071 | 0.479396656 |
| | 0.873408565 | 0.924442843 |
| | 0.601272502 | 0.734940076 |
| | 0.942441814 | 0.965107777 |
| | 0.005401208 | 0.020758849 |
| | 0.060793915 | 0.144494404 |
| | 0.507573994 | 0.659153123 |
| | 0.000348548 | 0.002072998 |
| | 0.518791827 | 0.669465062 |
| | 0.126416718 | 0.249653668 |
| | 0.939477865 | NA |
| | 0.097692494 | 0.206050221 |
| | 0.0337706 | 0.091124341 |
| | 0.81338117 | NA |
| | 0.053144632 | 0.130179821 |

| | | | |
|---|---|---|---|
| | 0.020844082 | | 0.061838568 |
| NA | | NA | |
| NA | | NA | |
| NA | | NA | |
| | 0.273219552 | | 0.433227086 |
| | 0.498279603 | | 0.651329074 |
| | 0.250094874 | | 0.407663684 |
| NA | | NA | |
| | 0.454189166 | NA | |
| | 0.000563412 | | 0.003142303 |
| | 0.102771804 | | 0.214125916 |
| | 0.173791456 | | 0.314521307 |
| | 0.61928955 | | 0.748476006 |
| | 0.348307235 | | 0.51229122 |
| NA | | NA | |
| | 5.23E-06 | | 5.07E-05 |
| | 0.229867171 | | 0.383681515 |
| | 0.531013898 | | 0.679745767 |
| | 0.252621391 | | 0.410492514 |
| | 0.909462096 | NA | |
| | 2.01E-12 | | 4.90E-11 |
| | 0.046372442 | | 0.116737903 |
| | 0.124365686 | | 0.246782164 |
| | 0.677379005 | | 0.792011351 |
| | 0.000741341 | | 0.003966151 |
| | 0.041902038 | | 0.108008698 |
| NA | | NA | |
| | 0.09395546 | | 0.200244258 |
| | 0.064999923 | | 0.152406508 |
| | 0.935473954 | | 0.961779234 |
| | 0.916590621 | NA | |
| | 1.44E-07 | | 1.89E-06 |
| | 0.187809219 | | 0.332063218 |
| | 0.889424486 | | 0.933455095 |
| | 2.73E-07 | | 3.39E-06 |
| | 0.35486715 | | 0.518670427 |
| | 0.740865268 | NA | |
| | 0.103826272 | | 0.215699827 |
| | 0.116174095 | | 0.234416395 |
| | 2.82E-06 | | 2.88E-05 |
| | 0.028906675 | | 0.080603732 |
| | 1.26E-08 | | 1.94E-07 |
| | 0.752687269 | | 0.844869718 |
| | 0.267340184 | | 0.426861948 |
| | 8.45E-20 | | 3.54E-18 |
| | 0.000194738 | | 0.001252408 |
| | 0.019337815 | | 0.058344362 |

| | |
|---|---|
| 0.648589638 | 0.770264094 |
| NA | NA |
| 0.007631343 | 0.0275799 |
| 1.65E-26 | 9.91E-25 |
| 0.433402887 | 0.594608076 |
| 0.033906022 | 0.091365462 |
| 0.783102737 | NA |
| 0.941278332 | NA |
| 0.7883069 | 0.869831931 |
| 0.297069792 | 0.45855092 |
| 0.555678046 | 0.700060808 |
| 0.881705728 | 0.929274887 |
| 0.51108247 | 0.662619148 |
| 0.144303723 | 0.275350105 |
| 0.522698528 | 0.672646537 |
| 0.941278424 | NA |
| 5.80E-07 | 6.75E-06 |
| 0.219285107 | 0.370803078 |
| 0.862660329 | NA |
| 0.962301193 | 0.977246054 |
| NA | NA |
| 0.00037232 | 0.002191927 |
| 1.31E-10 | 2.58E-09 |
| 0.210073395 | 0.359879585 |
| 0.642528132 | 0.765514796 |
| 5.60E-14 | 1.56E-12 |
| 0.011607509 | 0.038912949 |
| 0.433918281 | 0.595109359 |
| NA | NA |
| 0.827492724 | 0.895237779 |
| 0.00698301 | 0.025625903 |
| 0.500101321 | 0.652921488 |
| 0.175165901 | 0.316287269 |
| 3.96E-08 | 5.63E-07 |
| 0.21719303 | 0.368600962 |
| 0.941278332 | NA |
| 0.048313818 | 0.120481996 |
| 0.973906655 | 0.984244489 |
| 0.324223507 | 0.487041206 |
| 0.188681315 | 0.333159693 |
| 0.01707987 | 0.052920473 |
| 0.119683157 | 0.239789437 |
| 2.15E-07 | 2.72E-06 |
| NA | NA |
| NA | NA |
| 9.46E-06 | 8.62E-05 |
| 0.889006124 | NA |

| | |
|---|---|
| NA | NA |
| 5.42E-05 | 0.000407241 |
| 0.188334007 | 0.332669836 |
| 3.39E-18 | 1.27E-16 |
| NA | NA |
| 0.376468981 | 0.53956256 |
| 8.37E-05 | 0.00059862 |
| 5.92E-06 | 5.67E-05 |
| NA | NA |
| NA | NA |
| 0.839319108 | NA |
| 0.31050689 | 0.472368976 |
| NA | NA |
| NA | NA |
| NA | NA |
| NA | NA |
| 0.927870633 | NA |
| NA | NA |
| NA | NA |
| 0.000392672 | 0.002299798 |
| 0.928589329 | NA |
| NA | NA |
| 0.819814622 | 0.890201991 |
| NA | NA |
| NA | NA |
| NA | NA |
| NA | NA |
| NA | NA |
| 0.575800219 | NA |
| NA | NA |
| NA | NA |
| 0.941279303 | NA |
| 0.955792873 | NA |
| 0.906105105 | NA |
| NA | NA |
| NA | NA |
| 0.639603224 | NA |
| NA | NA |
| NA | NA |
| NA | NA |
| 0.707075313 | NA |
| NA | NA |
| 0.941277857 | NA |
| NA | NA |
| NA | NA |
| NA | NA |
| 0.66424827 | 0.782114139 |

| | |
|---|---|
| NA | NA |
| 0.726476091 | NA |
| NA | NA |
| NA | NA |
| 0.016404671 | 0.051228547 |
| NA | NA |
| 0.70175293 | NA |
| NA | NA |
| NA | NA |
| NA | NA |
| 0.755626499 | NA |
| 0.501139238 | NA |
| 0.661569785 | NA |
| NA | NA |
| NA | NA |
| 0.595454926 | NA |
| 0.689989611 | NA |
| 0.893158387 | 0.93548761 |
| NA | NA |
| 0.659049991 | NA |
| NA | NA |
| 0.497612328 | NA |
| 0.595454926 | NA |
| 0.935178876 | NA |
| 0.755626499 | NA |
| NA | NA |
| 0.459969306 | NA |
| NA | NA |
| 0.755626499 | NA |
| 0.674387156 | NA |
| NA | NA |
| NA | NA |
| NA | NA |
| 0.5129186 | NA |
| NA | NA |
| NA | NA |
| NA | NA |
| NA | NA |
| 0.941277857 | NA |
| 0.022497832 | 0.065855313 |
| 0.839318641 | NA |
| 0.925230605 | NA |
| NA | NA |
| NA | NA |
| NA | NA |
| NA | NA |
| NA | NA |

| | | |
|---|---|---|
| 0.94127925 | NA | |
| NA | NA | |
| 0.587799225 | | 0.723973979 |
| NA | NA | |
| 0.839318641 | NA | |
| 0.941277857 | NA | |
| NA | NA | |
| NA | NA | |
| NA | NA | |
| 0.838882104 | NA | |
| 0.743511643 | NA | |
| 0.954427263 | NA | |
| 0.897122393 | NA | |
| 0.443189588 | | 0.603754497 |
| 0.870871358 | NA | |
| NA | NA | |
| 4.41E-16 | | 1.46E-14 |
| NA | NA | |
| 4.71E-14 | | 1.32E-12 |
| NA | NA | |
| NA | NA | |
| NA | NA | |
| NA | NA | |
| NA | NA | |
| NA | NA | |
| NA | NA | |
| NA | NA | |
| NA | NA | |
| NA | NA | |
| NA | NA | |
| NA | NA | |
| 0.839319108 | NA | |
| NA | NA | |
| NA | NA | |
| NA | NA | |
| NA | NA | |
| NA | NA | |
| NA | NA | |
| NA | NA | |
| 0.839318641 | NA | |
| NA | NA | |
| NA | NA | |
| NA | NA | |
| NA | NA | |
| NA | NA | |
| NA | NA | |
| NA | NA | |

| | | |
|---|---|---|
| NA | | NA |
| NA | | NA |
| NA | | NA |
| NA | | NA |
| NA | | NA |
| NA | | NA |
| | 0.839318641 | NA |
| NA | | NA |
| NA | | NA |
| | 0.941277857 | NA |
| NA | | NA |
| NA | | NA |
| NA | | NA |
| | 0.023764821 | 0.068872553 |
| NA | | NA |
| | 0.85300265 | NA |
| | 0.202748833 | 0.350816206 |
| | 0.248948689 | 0.40627197 |
| | 0.941278332 | NA |
| | 0.567655076 | 0.709187479 |
| | 0.853595154 | NA |
| | 0.142071645 | 0.27220392 |
| | 0.000865993 | 0.004519263 |
| | 7.74E-10 | 1.40E-08 |
| | 1.89E-06 | 2.01E-05 |
| | 0.839318641 | NA |
| | 0.007825145 | 0.028160584 |
| | 5.50E-06 | 5.31E-05 |
| | 0.55100233 | 0.696418727 |
| | 0.453305166 | 0.613287646 |
| | 0.001783125 | 0.008313796 |
| | 0.021218595 | 0.0627619 |
| | 0.815259992 | 0.88702503 |
| | 0.806925308 | 0.88170566 |
| | 0.935593184 | 0.96186027 |
| | 0.941275956 | NA |
| | 0.906505948 | 0.94336545 |
| | 0.011364035 | 0.038227533 |
| NA | | NA |
| | 0.433375434 | NA |
| | 0.178865823 | 0.320512259 |
| | 0.688736363 | 0.800376082 |
| | 0.818612307 | 0.889383336 |
| NA | | NA |
| | 6.87E-05 | 0.000501469 |
| | 0.917016975 | NA |
| | 0.004942933 | 0.019344243 |

| | |
|---|---|
| 0.000843275 | 0.004419115 |
| 0.542047311 | NA |
| 0.586131484 | 0.722855004 |
| 1.15E-05 | 0.000102027 |
| 1.83E-17 | 6.57E-16 |
| 0.00056792 | 0.003162757 |
| 1.49E-62 | 2.87E-60 |
| 0.00376159 | 0.015418178 |
| 0.798076671 | 0.876473997 |
| 1.78E-10 | 3.47E-09 |
| 0.68313305 | 0.796469893 |
| 0.002581374 | 0.011336137 |
| 0.002316368 | 0.010354734 |
| 0.133665002 | 0.260010982 |
| NA | NA |
| 0.002947558 | 0.012636489 |
| 0.000831885 | 0.004368084 |
| 9.06E-05 | 0.000643297 |
| 0.268254335 | 0.428063132 |
| 0.941278332 | NA |
| 1.86E-16 | 6.32E-15 |
| 0.728462004 | 0.829094468 |
| 0.030487893 | 0.084028812 |
| NA | NA |
| 0.266871513 | 0.426544813 |
| 0.647577732 | 0.769566388 |
| 0.000622109 | 0.003411066 |
| 2.84E-06 | 2.90E-05 |
| 0.023195076 | 0.067558638 |
| 0.000465088 | 0.002665547 |
| 0.445427266 | 0.605935863 |
| 0.498536018 | 0.651521128 |
| 0.015676129 | 0.049407206 |
| 0.498371526 | 0.651377687 |
| 0.001282265 | 0.00629418 |
| 0.041965858 | 0.108149778 |
| NA | NA |
| 0.537299092 | 0.685138379 |
| NA | NA |
| 0.769344224 | NA |
| 0.955792873 | NA |
| 0.009316035 | 0.0324799 |
| 0.255473989 | 0.413518659 |
| 0.256925511 | 0.415021446 |
| 4.70E-11 | 9.86E-10 |
| 2.73E-43 | 3.15E-41 |
| 4.91E-05 | 0.000373183 |

| | |
|---|---|
| 0.941278838 | NA |
| 0.00452957 | 0.017989792 |
| 0.845047382 | 0.906722165 |
| 0.164250952 | 0.302475912 |
| 0.426084304 | 0.587535369 |
| 0.073796814 | 0.167446308 |
| 4.10E-13 | 1.06E-11 |
| 8.32E-05 | 0.000595359 |
| 1.08E-09 | 1.92E-08 |
| NA | NA |
| 0.002445256 | 0.010832306 |
| 0.01694241 | 0.052542442 |
| 0.785516042 | NA |
| 0.098668183 | 0.207483001 |
| 0.295304192 | NA |
| 0.750354499 | 0.84324541 |
| 0.762975127 | NA |
| 0.018515891 | 0.056415161 |
| 0.045677086 | 0.115341183 |
| 0.289233067 | 0.450044157 |
| 1.74E-07 | 2.24E-06 |
| 0.001153718 | 0.00576639 |
| 0.920535473 | 0.952509904 |
| 0.075896888 | 0.170972717 |
| 0.03861738 | 0.101219126 |
| NA | NA |
| 0.000768644 | 0.004089242 |
| 0.087373438 | 0.189958219 |
| 0.94127925 | NA |
| 0.277700095 | 0.437434077 |
| 4.10E-05 | 0.000317803 |
| 0.003625593 | 0.014956041 |
| 0.000550616 | 0.003085084 |
| 0.741503888 | 0.83740696 |
| 1.56E-05 | 0.000134915 |
| 0.92390268 | 0.954592539 |
| 0.829309108 | 0.896381426 |
| 0.674944156 | 0.790543237 |
| 0.00344018 | 0.014352878 |
| 0.209444355 | 0.359108386 |
| 6.47E-05 | 0.000477027 |
| 0.103203656 | 0.214762496 |
| 0.001900044 | 0.008767905 |
| 0.335702626 | 0.49886957 |
| NA | NA |
| 0.906995136 | 0.943641717 |
| 0.726714887 | NA |

| | |
|---|---|
| 0.272077264 | 0.432020215 |
| NA | NA |
| NA | NA |
| 0.010496514 | 0.035806435 |
| 0.745278759 | 0.839969438 |
| 0.724725323 | 0.826184786 |
| 0.565357916 | 0.707165332 |
| 0.005167163 | 0.020073252 |
| 0.000218225 | 0.001384009 |
| 2.20E-46 | 2.80E-44 |
| NA | NA |
| NA | NA |
| NA | NA |
| 0.023800014 | 0.068940967 |
| 0.433895558 | NA |
| NA | NA |
| NA | NA |
| NA | NA |
| 0.008628193 | 0.030492686 |
| 0.675812195 | NA |
| 0.707465299 | NA |
| 0.30587819 | 0.46750318 |
| NA | NA |
| 0.955792873 | NA |
| NA | NA |
| 0.160094159 | 0.296772677 |
| NA | NA |
| 0.941278332 | NA |
| NA | NA |
| NA | NA |
| NA | NA |
| 0.009075249 | 0.031803392 |
| 0.024671488 | 0.0709436 |
| 0.184981925 | 0.328136815 |
| 2.19E-36 | 1.98E-34 |
| 0.631213672 | NA |
| 0.0577094 | 0.138680794 |
| 5.09E-07 | 6.00E-06 |
| 0.176180914 | 0.317662155 |
| 0.807445514 | 0.88200175 |
| 0.211898412 | 0.362328017 |
| 0.467860837 | 0.626111753 |
| 0.033845386 | 0.091244496 |
| 0.839319108 | NA |
| 0.844589983 | 0.90649091 |
| 0.004726069 | 0.018661231 |
| 0.324973745 | 0.487829532 |

| | |
|---|---|
| 0.069968489 | 0.161203171 |
| 0.825773179 | 0.894113212 |
| 0.300862297 | 0.462216768 |
| 2.30E-12 | 5.58E-11 |
| 0.916062476 | 0.949473947 |
| 0.000239434 | 0.001496198 |
| 0.712397343 | 0.816864899 |
| 0.734058779 | NA |
| 0.002572338 | 0.011308975 |
| 0.299799239 | 0.461268357 |
| NA | NA |
| NA | NA |
| 0.738238392 | NA |
| 0.941275956 | NA |
| NA | NA |
| 0.000266392 | 0.00164011 |
| 0.002937387 | 0.012599693 |
| 0.941275481 | NA |
| 0.047641994 | 0.119177587 |
| NA | NA |
| NA | NA |
| 0.425846474 | 0.587287663 |
| 0.013289324 | 0.043342354 |
| 0.006736694 | 0.024890841 |
| 0.825773692 | 0.894113212 |
| NA | NA |
| 0.000242124 | 0.001510627 |
| NA | NA |
| 0.753006527 | NA |
| 0.839319107 | NA |
| 0.001255307 | 0.006185965 |
| 0.587447841 | 0.723803371 |
| 0.000604071 | 0.003332892 |
| 0.413469707 | 0.575722825 |
| 0.688660354 | 0.800376082 |
| 0.491049649 | NA |
| 0.941278332 | NA |
| 0.833998291 | NA |
| NA | NA |
| 0.049554734 | 0.122874842 |
| 0.368285564 | 0.531365279 |
| 0.130274296 | 0.255166951 |
| 0.00037827 | 0.002224207 |
| NA | NA |
| 0.777079586 | NA |
| NA | NA |
| NA | NA |

| | | |
|---|---|---|
| NA | NA | |
| NA | NA | |
| 0.839320041 | NA | |
| NA | NA | |
| NA | NA | |
| NA | NA | |
| NA | NA | |
| 0.30823236 | NA | |
| 0.955792873 | NA | |
| NA | NA | |
| NA | NA | |
| 0.7917244 | | 0.871750204 |
| NA | NA | |
| 0.941278332 | NA | |
| 0.978087452 | | 0.986494311 |
| 0.712603246 | NA | |
| 0.839320041 | NA | |
| NA | NA | |
| 0.566768252 | NA | |
| NA | NA | |
| NA | NA | |
| NA | NA | |
| 0.007224343 | | 0.026369241 |
| 0.002121592 | | 0.009600407 |
| NA | NA | |
| NA | NA | |
| 0.035635481 | | 0.095089207 |
| 0.151335267 | | 0.285177897 |
| NA | NA | |
| NA | NA | |
| 0.427783682 | | 0.589041703 |
| NA | NA | |
| NA | NA | |
| 0.713704045 | | 0.817913599 |
| NA | NA | |
| NA | NA | |
| 0.992019692 | | 0.99440104 |
| 0.531311694 | | 0.67996602 |
| 0.271289697 | | 0.431143579 |
| NA | NA | |
| NA | NA | |
| NA | NA | |
| NA | NA | |
| 8.15E-43 | | 9.37E-41 |
| 0.920460545 | | 0.952509904 |
| 0.136927887 | | 0.264778419 |
| 0.319133883 | | 0.48161455 |

| | |
|---|---|
| 0.158426303 | 0.294644699 |
| 0.650338949 | 0.771576496 |
| 0.756887442 | 0.847944784 |
| 0.575434736 | 0.714289314 |
| NA | NA |
| 0.004246387 | 0.017050016 |
| NA | NA |
| 0.00346669 | 0.014443616 |
| 0.05421394 | 0.132131906 |
| 0.002565102 | 0.011281332 |
| NA | NA |
| 1.97E-06 | 2.08E-05 |
| 0.034083754 | 0.091699052 |
| 0.869385385 | 0.921726558 |
| NA | NA |
| 0.955792873 | NA |
| 0.839318641 | NA |
| 1.11E-15 | 3.54E-14 |
| 0.246605082 | 0.403581832 |
| NA | NA |
| 0.211259984 | 0.361418108 |
| 0.044533764 | 0.113114146 |
| 0.024394566 | 0.07031236 |
| NA | NA |
| 0.03896406 | 0.101948175 |
| 0.003267551 | 0.013779987 |
| 0.028784772 | 0.080348909 |
| 0.941278332 | NA |
| 0.853596091 | NA |
| 0.089408487 | 0.193217795 |
| 0.341651831 | 0.505061603 |
| 0.214667628 | 0.365540054 |
| NA | NA |
| NA | NA |
| 0.676546855 | 0.791584276 |
| 0.200630431 | 0.348254488 |
| 0.261513043 | 0.420264227 |
| 0.142411344 | 0.272723052 |
| 0.079548371 | NA |
| 0.778861431 | NA |
| 9.88E-07 | 1.10E-05 |
| 0.28045051 | 0.440391848 |
| NA | NA |
| 0.121065732 | 0.241948493 |
| 0.813876082 | 0.886422555 |
| 0.832162011 | NA |
| 0.66404563 | NA |

| | |
|---|---|
| NA | NA |
| 0.81973855 | NA |
| 0.439543821 | 0.60043134 |
| 1.01E-05 | 9.14E-05 |
| 0.913665819 | NA |
| 0.835025174 | NA |
| 0.061041208 | 0.14502923 |
| NA | NA |
| NA | NA |
| NA | NA |
| 0.110632754 | NA |
| 0.955792398 | NA |
| 7.59E-11 | 1.54E-09 |
| 4.71E-09 | 7.72E-08 |
| 0.06774474 | 0.157159874 |
| 0.225999106 | 0.379032605 |
| 0.070873099 | 0.162626193 |
| NA | NA |
| 0.795426886 | NA |
| 0.111016977 | 0.22660373 |
| 0.074757872 | 0.169057991 |
| NA | NA |
| NA | NA |
| NA | NA |
| 0.145158374 | 0.276601996 |
| 0.018693796 | 0.05685532 |
| NA | NA |
| 0.387283581 | 0.550086957 |
| NA | NA |
| 0.027382581 | 0.077259386 |
| 0.001122082 | 0.005633367 |
| 0.091159225 | 0.196090073 |
| NA | NA |
| NA | NA |
| 8.77E-11 | 1.77E-09 |
| NA | NA |
| 0.164027141 | 0.302157264 |
| 0.255504417 | 0.413539788 |
| 0.511697185 | 0.663169515 |
| 0.042989714 | 0.110144368 |
| 0.001226817 | 0.006076155 |
| 0.721346842 | NA |
| 0.748704271 | 0.842225958 |
| 0.235977854 | 0.391274461 |
| 0.518069431 | 0.668822943 |
| 6.04E-16 | 1.98E-14 |
| 0.034363269 | 0.092304992 |

| | | |
|---|---|---|
| | 0.047205887 | 0.118273108 |
| | 0.839319108 | NA |
| | 0.070976168 | 0.162815609 |
| | 0.174416589 | 0.315269112 |
| | 3.90E-05 | 0.000303969 |
| | 0.003753741 | 0.015396613 |
| | 0.10016688 | 0.209892779 |
| | 0.236080858 | 0.391286321 |
| NA | | NA |
| | 0.041335942 | 0.106896794 |
| | 0.515263473 | 0.666342374 |
| | 0.832162011 | NA |
| | 0.178842266 | 0.320494173 |
| | 0.036412399 | 0.096649501 |
| | 0.078667324 | 0.175585113 |
| | 0.255210483 | 0.413343937 |
| | 0.019410336 | 0.058481624 |
| | 0.138243901 | 0.266759736 |
| | 0.028220486 | 0.079042598 |
| NA | | NA |
| | 0.621436952 | NA |
| | 0.054733397 | 0.133111712 |
| | 0.955792398 | NA |
| | 0.661192658 | NA |
| | 0.105393769 | 0.218118641 |
| | 0.261220998 | 0.419948636 |
| | 0.121147552 | 0.242091682 |
| | 0.963295589 | NA |
| | 0.428346025 | 0.589607454 |
| | 0.000121869 | 0.000833785 |
| | 0.549172946 | 0.694807551 |
| | 0.890593148 | 0.934198497 |
| NA | | NA |
| | 0.935871621 | NA |
| NA | | NA |
| NA | | NA |
| | 5.30E-10 | 9.78E-09 |
| | 0.925579587 | NA |
| | 0.13468643 | 0.261549025 |
| | 0.724874209 | 0.826238017 |
| | 9.84E-30 | 6.81E-28 |
| | 2.63E-36 | 2.36E-34 |
| | 0.000118441 | 0.000812663 |
| | 0.410600377 | 0.572531352 |
| | 0.212657257 | 0.363208096 |
| | 0.00562151 | 0.021473788 |
| | 0.941278788 | NA |

| | | |
|---|---|---|
| | 0.43041565 | 0.591635095 |
| | 0.679259489 | NA |
| | 0.46082862 | 0.61983741 |
| | 1.08E-17 | 3.94E-16 |
| | 0.839318641 | NA |
| | 0.00018273 | 0.001185749 |
| | 0.634516142 | 0.759734051 |
| | 0.000263581 | 0.001624483 |
| | 0.017406247 | 0.053687119 |
| | 0.111759086 | 0.22768893 |
| | 0.435223654 | 0.596391144 |
| | 0.14884864 | 0.281728267 |
| | 0.551147872 | 0.696459896 |
| | 0.206922475 | 0.355916227 |
| | 0.062873044 | 0.148344985 |
| | 0.178148465 | 0.31968409 |
| | 0.966533889 | 0.980058047 |
| | 1.72E-05 | 0.000146704 |
| | 0.202437477 | NA |
| | 1.56E-09 | 2.71E-08 |
| | 0.391671396 | 0.554141025 |
| | 0.589728736 | 0.725787144 |
| | 0.169893743 | 0.309467427 |
| | 0.644227827 | 0.767040705 |
| | 0.402580209 | 0.56472252 |
| | 0.286853521 | 0.447394503 |
| NA | | NA |
| | 0.166344835 | 0.305198471 |
| | 0.37038013 | 0.533611032 |
| NA | | NA |
| | 0.518599623 | 0.669398493 |
| | 0.613625782 | 0.744153273 |
| | 2.69E-13 | 7.10E-12 |
| NA | | NA |
| | 0.94127925 | NA |
| | 0.888153369 | NA |
| NA | | NA |
| | 0.313600134 | 0.475676166 |
| NA | | NA |
| | 6.15E-10 | 1.12E-08 |
| | 3.56E-18 | 1.33E-16 |
| | 0.007112128 | 0.026011505 |
| | 2.32E-06 | 2.42E-05 |
| NA | | NA |
| | 0.091200471 | 0.196090073 |
| | 0.081407948 | 0.180014118 |
| | 0.649095828 | 0.770639411 |

| | | |
|---|---|---|
| NA | 0.494733256 | 0.647786973 |
| | 6.02E-26 | 3.52E-24 |
| | 0.667822013 | NA |
| | 0.009886417 | 0.034069277 |
| | 0.319663234 | 0.482260524 |
| | 0.540142899 | 0.68732859 |
| | 0.769655223 | 0.857314759 |
| | 0.941278332 | NA |
| | 0.000522992 | 0.002949125 |
| | 0.941278788 | NA |
| | 0.063987268 | 0.1504668 |
| NA | | NA |
| | 0.436370258 | 0.597439452 |
| | 0.26887917 | 0.428744022 |
| | 0.122232382 | 0.24374792 |
| | 0.204323004 | 0.35283048 |
| | 0.955792873 | NA |
| | 0.748393366 | NA |
| | 0.021068514 | 0.062364478 |
| | 0.016026525 | 0.050295433 |
| | 0.905140268 | NA |
| | 0.110694146 | 0.226216371 |
| | 0.041699327 | 0.10766487 |
| | 0.923516642 | NA |
| | 0.08609932 | 0.187861408 |
| | 3.96E-23 | 2.03E-21 |
| | 0.696895994 | NA |
| | 0.941278332 | NA |
| | 0.867510235 | 0.920820488 |
| NA | | NA |
| | 0.49774565 | NA |
| NA | | NA |
| | 2.24E-08 | 3.33E-07 |
| | 0.170788141 | 0.310677161 |
| | 0.941277857 | NA |
| NA | | NA |
| | 0.434803938 | 0.596186608 |
| | 0.76081145 | 0.851299085 |
| | 0.040911271 | 0.106075171 |
| | 0.904187708 | NA |
| NA | | NA |
| | 0.901965574 | 0.940738118 |
| | 0.839319108 | NA |
| | 0.226177113 | 0.379197552 |
| | 0.81579061 | NA |
| | 0.115152914 | 0.232770247 |
| NA | | NA |

| | |
|---|---|
| NA | NA |
| 0.244389469 | 0.400880197 |
| 2.59E-10 | 4.95E-09 |
| 0.172781141 | 0.31312142 |
| 0.988352138 | 0.99264769 |
| 0.034218808 | 0.091968843 |
| 0.000875577 | 0.00455729 |
| 0.937011128 | NA |
| 0.003950808 | 0.016037173 |
| 0.84096936 | 0.903976548 |
| 0.061066157 | 0.145074027 |
| 1.49E-07 | 1.94E-06 |
| 0.244460151 | 0.400953728 |
| 0.459696764 | NA |
| 0.053145001 | 0.130179821 |
| 0.003051197 | 0.013003805 |
| 0.909178185 | 0.944906958 |
| 0.001717924 | 0.008066686 |
| 0.077643973 | 0.173871657 |
| 0.349257614 | 0.513218471 |
| 0.039750465 | NA |
| 0.670436281 | 0.786714866 |
| 0.074733571 | 0.169023988 |
| 0.000370995 | 0.002186832 |
| 0.038582286 | 0.101138278 |
| NA | NA |
| 0.779617645 | NA |
| 0.017392968 | 0.053653114 |
| NA | NA |
| 0.005684706 | 0.021666514 |
| 0.254651513 | 0.412688786 |
| 0.936455126 | 0.962372303 |
| NA | NA |
| 0.039264612 | 0.102644294 |
| 0.955792873 | NA |
| 0.683562568 | 0.796853572 |
| 0.015382984 | 0.048664604 |
| NA | NA |
| 4.08E-07 | 4.91E-06 |
| 0.2185312 | 0.369843546 |
| NA | NA |
| 0.57130717 | NA |
| 3.51E-09 | 5.84E-08 |
| 0.03197758 | 0.087225574 |
| 0.941278788 | NA |
| 0.920897743 | 0.952651559 |
| 0.863464383 | 0.91838732 |

| | |
|---|---|
| 0.360356504 | NA |
| 0.000170056 | 0.001117523 |
| NA | NA |
| NA | NA |
| NA | NA |
| NA | NA |
| 0.299120331 | 0.460705725 |
| 0.000111055 | 0.000767966 |
| NA | NA |
| 0.377413359 | 0.540309923 |
| 2.04E-05 | 0.000170922 |
| NA | NA |
| NA | NA |
| 0.013460429 | 0.04377522 |
| 0.000562805 | 0.003140885 |
| 0.127674106 | 0.25152285 |
| NA | NA |
| 0.662049234 | NA |
| NA | NA |
| NA | NA |
| 0.64979056 | NA |
| 0.839319108 | NA |
| NA | NA |
| NA | NA |
| NA | NA |
| 0.739991096 | NA |
| NA | NA |
| NA | NA |
| 0.875892654 | NA |
| NA | NA |
| NA | NA |
| 0.895348826 | 0.936820599 |
| NA | NA |
| 0.941277857 | NA |
| NA | NA |
| 0.941277857 | NA |
| NA | NA |
| 0.831413565 | NA |
| 0.748393366 | NA |
| 0.839319108 | NA |
| NA | NA |
| 0.781836495 | NA |
| 0.212099656 | 0.362525595 |
| 0.123409221 | 0.245313704 |
| NA | NA |
| 0.853595154 | NA |
| 0.839319108 | NA |

| | | | |
|---|---|---|---|
| | 0.015041147 | | 0.047817468 |
| NA | | NA | |
| NA | | NA | |
| NA | | NA | |
| | 0.011728772 | | 0.039203517 |
| | 0.941278332 | NA | |
| NA | | NA | |
| NA | | NA | |
| | 0.739991096 | NA | |
| | 0.031309447 | | 0.08579639 |
| | 0.006387911 | | 0.023831513 |
| | 0.002731517 | | 0.0118815 |
| | 0.026770422 | | 0.075873764 |
| | 0.776729481 | NA | |
| | 0.723767582 | NA | |
| | 0.185111995 | NA | |
| | 0.008582565 | | 0.030349459 |
| | 0.555569375 | | 0.700035059 |
| | 0.13763322 | | 0.265883119 |
| NA | | NA | |
| | 0.122812659 | | 0.24452044 |
| NA | | NA | |
| | 4.94E-27 | | 3.05E-25 |
| | 0.379990641 | | 0.542759076 |
| | 0.01237293 | | 0.040880132 |
| | 0.750671946 | NA | |
| | 0.887286764 | NA | |
| | 0.314183686 | | 0.476348818 |
| | 0.12883521 | | 0.253118916 |
| NA | | NA | |
| NA | | NA | |
| NA | | NA | |
| | 0.000522512 | | 0.002948516 |
| | 0.937272155 | NA | |
| | 0.04559517 | | 0.115219899 |
| | 0.199730357 | | 0.347183534 |
| | 0.913546231 | | 0.947873905 |
| | 0.148580235 | | 0.281501771 |
| | 0.263811177 | | 0.422844016 |
| | 0.074165428 | | 0.168058408 |
| | 0.825519241 | | 0.893990126 |
| | 0.689590244 | | 0.800938222 |
| | 0.989510516 | | 0.99330814 |
| | 0.099919731 | | 0.20948555 |
| | 0.199200896 | | 0.346642765 |
| NA | | NA | |
| | 0.91810767 | NA | |

| | |
|---|---|
| 0.183725228 | 0.32646718 |
| 7.19E-11 | 1.47E-09 |
| 0.252980096 | NA |
| 0.890791471 | NA |
| NA | NA |
| 0.941277857 | NA |
| 2.76E-11 | 5.97E-10 |
| 0.700331208 | 0.808638955 |
| 0.204853534 | 0.353455877 |
| 0.739991096 | NA |
| NA | NA |
| NA | NA |
| 0.941277857 | NA |
| NA | NA |
| 0.312721587 | NA |
| 0.893085783 | 0.935461019 |
| 0.041714024 | 0.107687332 |
| 0.841904316 | NA |
| 0.035623323 | 0.095067422 |
| 0.700559344 | 0.808745441 |
| 0.912936186 | NA |
| NA | NA |
| NA | NA |
| 0.616555278 | 0.746239537 |
| NA | NA |
| 0.399144867 | 0.561816062 |
| 0.910098446 | 0.945698211 |
| 0.207639311 | 0.356814012 |
| 0.15936788 | 0.295794938 |
| 0.895354114 | 0.936820599 |
| 7.43E-11 | 1.52E-09 |
| 0.010551813 | 0.035951081 |
| NA | NA |
| 0.000448568 | 0.002583311 |
| 0.011106842 | 0.037513934 |
| 0.008936619 | 0.031396221 |
| 0.002700844 | 0.011778212 |
| NA | NA |
| NA | NA |
| NA | NA |
| NA | NA |
| NA | NA |
| 0.438046013 | 0.598886643 |
| 0.839319108 | NA |
| 0.607429187 | NA |
| 0.599679346 | NA |
| 0.941278332 | NA |

| | | |
|---|---|---|
| | 0.467983059 | 0.626204901 |
| | 0.839319108 | NA |
| NA | | NA |
| | 8.62E-07 | 9.71E-06 |
| | 0.172299984 | 0.312562496 |
| | 0.501807179 | NA |
| | 0.492747523 | 0.646547769 |
| | 0.621826747 | 0.750277297 |
| NA | | NA |
| | 1.92E-10 | 3.75E-09 |
| | 5.48E-17 | 1.92E-15 |
| | 0.037379225 | 0.098647334 |
| | 7.15E-11 | 1.46E-09 |
| | 0.594376844 | 0.729470849 |
| | 0.053177406 | 0.130245792 |
| NA | | NA |
| | 0.939241089 | NA |
| NA | | NA |
| | 0.176722299 | 0.318340567 |
| | 3.42E-07 | 4.15E-06 |
| | 0.006946401 | 0.025523036 |
| | 0.035864257 | 0.095581798 |
| | 0.001178549 | 0.005869811 |
| | 0.182401078 | 0.324866093 |
| | 0.724501056 | NA |
| NA | | NA |
| | 0.001731116 | 0.008109434 |
| | 0.143589331 | 0.274294803 |
| | 0.303632878 | 0.465032479 |
| | 0.417610358 | NA |
| NA | | NA |
| | 0.955792398 | NA |
| | 0.030502722 | 0.084040488 |
| | 0.853595154 | NA |
| | 9.64E-05 | 0.000679294 |
| | 0.136056197 | 0.263542448 |
| | 0.000825452 | 0.00434197 |
| | 2.28E-06 | 2.38E-05 |
| | 0.09055734 | 0.195098281 |
| NA | | NA |
| | 0.000613414 | 0.003375052 |
| | 0.004870533 | 0.019108032 |
| | 0.739991096 | NA |
| | 0.281677366 | 0.441869418 |
| | 5.50E-05 | 0.000412053 |
| NA | | NA |
| | 0.955792873 | NA |

| | | |
|---|---|---|
| | 0.023639195 | 0.068575275 |
| | 0.950345491 | NA |
| | 0.839319108 | NA |
| | 0.007973236 | 0.028576867 |
| | 0.320553247 | 0.483235695 |
| | 0.672137289 | NA |
| | 0.599597407 | NA |
| | 0.661976875 | 0.780259502 |
| NA | | NA |
| | 0.675650866 | NA |
| | 0.095045372 | 0.202006425 |
| | 0.000644624 | 0.003518308 |
| | 0.674587494 | 0.79026142 |
| | 0.186533485 | NA |
| | 0.839319108 | NA |
| NA | | NA |
| | 0.938616718 | 0.963595218 |
| | 0.839319108 | NA |
| | 0.334374389 | 0.497641419 |
| | 0.632382807 | 0.758400542 |
| | 0.380150282 | 0.542889364 |
| | 1.45E-07 | 1.90E-06 |
| | 0.552471702 | 0.697708932 |
| | 0.941278332 | NA |
| | 0.059510926 | 0.14217395 |
| | 0.964805063 | 0.978629815 |
| | 0.643753703 | NA |
| | 0.360968135 | 0.52465581 |
| NA | | NA |
| | 0.941278332 | NA |
| | 0.002124609 | 0.009610402 |
| | 0.001725762 | 0.008092314 |
| NA | | NA |
| | 0.743511643 | NA |
| | 0.839319108 | NA |
| NA | | NA |
| | 0.941278424 | NA |
| NA | | NA |
| | 0.009340491 | 0.03254133 |
| | 0.000689989 | 0.003730829 |
| NA | | NA |
| NA | | NA |
| | 0.360924073 | NA |
| | 0.941278332 | NA |
| NA | | NA |
| NA | | NA |
| NA | | NA |

| | | |
|---|---|---|
| NA | | NA |
| | 0.805616837 | NA |
| | 0.550779936 | NA |
| | 0.853595154 | NA |
| NA | | NA |
| | 0.02348003 | 0.068204988 |
| | 1.62E-05 | 0.000139814 |
| | 0.905551864 | NA |
| NA | | NA |
| | 0.92204978 | NA |
| | 2.66E-07 | 3.31E-06 |
| | 0.720153486 | NA |
| | 0.279889933 | 0.439790084 |
| | 0.026294675 | 0.074667843 |
| | 0.001042201 | 0.005289224 |
| | 0.482831101 | 0.63882419 |
| | 0.789420166 | 0.870421031 |
| | 0.120672316 | 0.241425777 |
| | 0.958612828 | 0.974623149 |
| | 0.123509064 | 0.245430183 |
| NA | | NA |
| | 0.365735229 | 0.528871805 |
| | 0.732986575 | 0.831579907 |
| | 2.57E-05 | 0.000210817 |
| | 0.802600916 | 0.879332888 |
| | 0.027811007 | 0.078208599 |
| | 0.224379024 | 0.377086037 |
| NA | | NA |
| | 0.254092495 | 0.412151395 |
| | 0.018890798 | 0.057329947 |
| | 0.009270769 | 0.032348045 |
| | 0.293735032 | 0.454911336 |
| | 0.023412824 | 0.068042984 |
| | 0.011479923 | 0.038528641 |
| | 0.05392804 | 0.131623792 |
| | 0.004991119 | 0.019494356 |
| | 0.029714823 | 0.082355872 |
| | 0.708546088 | 0.814056764 |
| | 0.079044945 | 0.176279188 |
| | 0.165371082 | 0.303999222 |
| NA | | NA |
| NA | | NA |
| | 0.576784851 | 0.715383061 |
| NA | | NA |
| NA | | NA |
| NA | | NA |
| NA | | NA |

| | |
|---|---|
| NA | NA |
| 4.54E-05 | 0.00034727 |
| NA | NA |
| 0.853370498 | 0.911789047 |
| 6.68E-14 | 1.85E-12 |
| 0.223268887 | 0.37572441 |
| 0.013566586 | 0.044017431 |
| 0.199300172 | 0.346764817 |
| NA | NA |
| 1.51E-42 | 1.73E-40 |
| 0.367017306 | 0.530178183 |
| 0.569851049 | 0.710770841 |
| 0.606699802 | 0.739258149 |
| 0.757296437 | NA |
| 0.447894472 | NA |
| 5.58E-12 | 1.30E-10 |
| 0.061815773 | NA |
| 0.955792873 | NA |
| NA | NA |
| 0.845101662 | 0.906739531 |
| NA | NA |
| NA | NA |
| NA | NA |
| 0.839318641 | NA |
| NA | NA |
| 0.082528253 | 0.18183259 |
| NA | NA |
| 0.325828802 | 0.488712404 |
| 0.664921943 | NA |
| 0.941278332 | NA |
| 0.772001163 | 0.85913464 |
| NA | NA |
| NA | NA |
| NA | NA |
| 0.853596091 | NA |
| 0.372546717 | 0.535791962 |
| 0.708194269 | NA |
| 7.20E-22 | 3.43E-20 |
| 0.000101659 | 0.000710209 |
| 0.054880534 | 0.133415022 |
| 0.955791922 | NA |
| 0.413278753 | NA |
| NA | NA |
| 0.558096583 | 0.701696417 |
| 0.016437632 | 0.05129981 |
| 0.008348678 | 0.029685575 |
| 0.312828464 | 0.474747711 |

| | |
|---|---|
| 0.307789683 | 0.469362203 |
| 0.326721087 | 0.489587969 |
| 0.248045093 | 0.405269722 |
| 0.040685545 | 0.105593429 |
| 0.311150366 | 0.473196666 |
| 0.214612959 | 0.365523127 |
| NA | NA |
| 0.777683256 | 0.863096307 |
| 0.725028456 | NA |
| 0.509831188 | 0.66146442 |
| 0.320650718 | 0.48335202 |
| 0.537368871 | 0.68519065 |
| 0.765943354 | 0.854626639 |
| 0.441358944 | 0.60198416 |
| 0.23161296 | 0.385795264 |
| 0.04553615 | 0.115107417 |
| 0.000409812 | 0.002384928 |
| 0.528756463 | 0.677876835 |
| 0.90267436 | NA |
| 0.923470349 | NA |
| NA | NA |
| 0.955792873 | NA |
| 0.853595154 | NA |
| NA | NA |
| 1.89E-06 | 2.01E-05 |
| 0.941278332 | NA |
| NA | NA |
| 0.839319108 | NA |
| 0.780182449 | 0.864641368 |
| 0.941278332 | NA |
| 0.001651134 | 0.007790506 |
| 0.857778612 | 0.914897227 |
| 0.063164793 | 0.148871018 |
| 0.469885812 | 0.627798041 |
| 0.756853334 | NA |
| 1.14E-07 | 1.52E-06 |
| 0.118773664 | 0.238368528 |
| 0.00048656 | 0.002771927 |
| NA | NA |
| 0.004453196 | 0.017730841 |
| 0.955792873 | NA |
| 0.839319574 | NA |
| NA | NA |
| NA | NA |
| 0.955792873 | NA |
| 0.002822077 | 0.012175289 |
| NA | NA |

| | | | |
|---|---|---|---|
| NA | | NA | |
| NA | | NA | |
| | 0.264974379 | NA | |
| | 0.366016829 | | 0.529243245 |
| | 0.005083615 | | 0.019793916 |
| | 0.32533699 | | 0.488190169 |
| NA | | NA | |
| | 0.234737628 | | 0.389598007 |
| | 0.420038694 | NA | |
| | 0.381240143 | | 0.543825835 |
| | 0.814143257 | | 0.886443528 |
| NA | | NA | |
| | 0.174183519 | | 0.314991348 |
| NA | | NA | |
| | 0.404728202 | | 0.566767115 |
| | 0.270953519 | | 0.430807785 |
| | 0.045974015 | | 0.116004824 |
| | 0.129175339 | | 0.25341065 |
| | 0.795095718 | | 0.874289396 |
| | 1.16E-05 | | 0.000102888 |
| | 0.839319108 | NA | |
| | 0.083998542 | | 0.1844914 |
| | 0.293241061 | | 0.45445947 |
| NA | | NA | |
| NA | | NA | |
| NA | | NA | |
| NA | | NA | |
| | 0.839319108 | NA | |
| | 0.853596091 | NA | |
| | 0.096729004 | | 0.204488831 |
| | 0.635433013 | | 0.760250222 |
| NA | | NA | |
| NA | | NA | |
| | 0.309790388 | | 0.471489918 |
| | 0.000127288 | | 0.00086687 |
| NA | | NA | |
| | 0.634153493 | NA | |
| | 0.469120496 | | 0.627198003 |
| | 0.941275956 | NA | |
| NA | | NA | |
| | 0.941278332 | NA | |
| | 0.665957439 | | 0.783351568 |
| | 0.662848679 | NA | |
| | 0.839319108 | NA | |
| NA | | NA | |
| NA | | NA | |
| NA | | NA | |

| | |
|---|---|
| NA | NA |
| NA | NA |
| 4.57E-29 | 3.06E-27 |
| 0.000520842 | 0.002941182 |
| 0.032662745 | 0.088829827 |
| 0.157961033 | 0.293894547 |
| 0.291586146 | 0.452787934 |
| 0.475878211 | 0.633023196 |
| 0.88205743 | 0.929463916 |
| 0.898521574 | 0.938856571 |
| 5.54E-05 | 0.000415195 |
| NA | NA |
| 0.000675488 | 0.003666474 |
| 0.400099726 | 0.562723875 |
| 0.381082077 | 0.5437633 |
| 4.61E-08 | 6.49E-07 |
| 0.946364244 | 0.967501149 |
| 0.001972599 | 0.009058017 |
| 0.918294267 | NA |
| 0.389541848 | 0.552094502 |
| 0.224527597 | 0.377261406 |
| NA | NA |
| 0.982476774 | NA |
| 0.905736153 | NA |
| 0.427346586 | NA |
| 0.042220755 | 0.108641989 |
| 0.955792873 | NA |
| 0.839319108 | NA |
| 0.699796417 | NA |
| 0.997881633 | 0.998288604 |
| 0.215289763 | 0.366232629 |
| 0.955792398 | NA |
| NA | NA |
| 0.955792873 | NA |
| 0.839319574 | NA |
| NA | NA |
| NA | NA |
| 0.941279303 | NA |
| 5.50E-08 | 7.67E-07 |
| 0.776758135 | NA |
| 0.839319108 | NA |
| 1.00E-05 | 9.10E-05 |
| 0.778265992 | NA |
| 0.941276431 | NA |
| NA | NA |
| NA | NA |
| NA | NA |

| | |
|---|---|
| NA | NA |
| 0.317843562 | 0.480276311 |
| 0.065273061 | 0.15276592 |
| 3.61E-08 | 5.18E-07 |
| NA | NA |
| 0.191103141 | 0.33618695 |
| 0.667315331 | 0.784328651 |
| NA | NA |
| 0.454790929 | 0.614564193 |
| 0.416421903 | 0.578127538 |
| 7.03E-06 | 6.59E-05 |
| 0.274780184 | NA |
| 0.897254093 | 0.937977136 |
| 0.129125419 | 0.253390229 |
| 0.879648222 | 0.92791114 |
| 0.941280961 | NA |
| 0.048295967 | 0.120459038 |
| 0.941277857 | NA |
| NA | NA |
| 0.938691979 | NA |
| 0.621712714 | 0.750200458 |
| 0.941278332 | NA |
| 0.652702796 | NA |
| 0.543373577 | NA |
| NA | NA |
| NA | NA |
| 0.740461181 | NA |
| 0.569852952 | 0.710770841 |
| 0.001611766 | 0.007631439 |
| 0.955792873 | NA |
| 0.661499944 | NA |
| 0.718867939 | 0.821055414 |
| NA | NA |
| NA | NA |
| 0.085053452 | NA |
| NA | NA |
| NA | NA |
| 0.941277857 | NA |
| 0.014239106 | 0.045778592 |
| 0.823165732 | NA |
| 0.008909122 | 0.031322736 |
| 0.012660356 | 0.041639049 |
| NA | NA |
| 0.43762166 | 0.598464277 |
| 0.853595622 | NA |
| 0.059626534 | 0.142381181 |
| 0.429428162 | 0.590721211 |

| | | |
|---|---|---|
| | 0.941279268 | NA |
| | 0.941275956 | NA |
| | 0.838457485 | NA |
| NA | | NA |
| | 0.839319574 | NA |
| NA | | NA |
| NA | | NA |
| NA | | NA |
| NA | | NA |
| NA | | NA |
| | 0.43323847 | NA |
| | 0.789608199 | NA |
| NA | | NA |
| | 0.696483229 | 0.805524481 |
| | 0.008883652 | 0.03123997 |
| | 0.003527824 | 0.014654323 |
| | 0.87162049 | 0.923330406 |
| | 0.150292891 | 0.283662786 |
| | 0.839320041 | NA |
| | 0.035193597 | 0.094068275 |
| | 6.98E-80 | 1.87E-77 |
| | 1.35E-149 | 8.22E-147 |
| NA | | NA |
| | 1.01E-186 | 9.57E-184 |
| | 8.24E-115 | 3.27E-112 |
| | 1.29E-126 | 5.57E-124 |
| | 0.845206378 | NA |
| NA | | NA |
| | 0.853595622 | NA |
| | 4.49E-05 | 0.000344048 |
| | 0.955792873 | NA |
| | 2.55E-06 | 2.64E-05 |
| NA | | NA |
| | 0.000797806 | 0.004219563 |
| NA | | NA |
| | 0.898606152 | NA |
| | 0.393383828 | 0.555851699 |
| NA | | NA |
| | 0.993633163 | 0.995389585 |
| | 0.907170531 | NA |
| | 0.72992288 | NA |
| | 0.101636225 | 0.212298853 |
| | 0.89279965 | NA |
| | 0.426192659 | 0.58759486 |
| | 0.941278332 | NA |
| | 0.998472388 | NA |
| | 0.561775809 | 0.704647334 |

| | | |
|---|---|---|
| | 0.745002123 | NA |
| NA | | NA |
| | 0.955792873 | NA |
| | 0.14117802 | 0.270927939 |
| NA | | NA |
| | 0.908005152 | NA |
| | 0.944784414 | NA |
| | 0.766245103 | NA |
| | 0.261318412 | 0.420036523 |
| | 0.839319108 | NA |
| | 0.941277857 | NA |
| | 0.014609042 | 0.046705764 |
| | 0.069430557 | 0.160227469 |
| NA | | NA |
| | 0.095459527 | NA |
| | 0.673442495 | 0.789191978 |
| | 0.041774967 | 0.107790221 |
| | 0.007745499 | 0.027920396 |
| | 0.092392838 | 0.197835043 |
| | 0.008935982 | 0.031396221 |
| NA | | NA |
| | 0.839318641 | NA |
| | 5.95E-09 | 9.60E-08 |
| | 0.357629115 | 0.521229189 |
| | 0.099151269 | 0.208204544 |
| | 0.826425584 | NA |
| | 0.545168186 | 0.691601598 |
| | 0.113429947 | 0.230360673 |
| | 0.35822828 | 0.521820657 |
| | 0.853595622 | NA |
| | 0.815117471 | NA |
| | 0.426681833 | NA |
| NA | | NA |
| | 0.727928575 | NA |
| NA | | NA |
| | 0.938767819 | NA |
| | 0.839320685 | NA |
| | 0.66883848 | NA |
| | 0.374600457 | 0.53773838 |
| | 0.000570106 | 0.003174191 |
| NA | | NA |
| | 0.941278332 | NA |
| | 0.923448134 | 0.954444938 |
| | 3.17E-07 | 3.88E-06 |
| | 0.133572619 | 0.259852511 |
| NA | | NA |
| | 0.406625578 | 0.568420861 |

| | |
|---|---|
| 0.002389513 | 0.010622933 |
| 1.29E-05 | 0.000114022 |
| 0.399183627 | 0.561816062 |
| 0.001047529 | 0.005309474 |
| NA | NA |
| 0.038755161 | 0.101513188 |
| NA | NA |
| 0.352083402 | 0.515678366 |
| 1.14E-08 | 1.78E-07 |
| 0.671898993 | 0.787834017 |
| NA | NA |
| 0.453760922 | NA |
| NA | NA |
| NA | NA |
| 0.240522628 | 0.39657243 |
| 0.009198492 | 0.03214071 |
| 0.084963793 | 0.186147663 |
| 9.47E-05 | 0.000668871 |
| 0.000106562 | 0.000739691 |
| NA | NA |
| 0.086573684 | 0.188667536 |
| 0.941277857 | NA |
| 0.605956473 | 0.73865492 |
| 0.051631843 | 0.127180081 |
| 0.745668933 | 0.840134719 |
| 0.941277857 | NA |
| 0.046153081 | 0.116321012 |
| 0.839319108 | NA |
| 0.391363746 | 0.553819256 |
| 2.19E-09 | 3.72E-08 |
| 0.284952335 | 0.445509424 |
| NA | NA |
| NA | NA |
| NA | NA |
| 0.630087135 | NA |
| NA | NA |
| 0.955792398 | NA |
| NA | NA |
| NA | NA |
| NA | NA |
| NA | NA |
| 0.298190356 | 0.459684382 |
| 0.104127842 | 0.216099999 |
| 0.054421929 | 0.1325167 |
| 0.650887845 | 0.771974087 |
| 0.955791922 | NA |
| 0.080694725 | 0.178919034 |

| | |
|---|---|
| 0.010572833 | 0.03601239 |
| 0.611075665 | NA |
| 0.650877417 | NA |
| NA | NA |
| 0.091849049 | 0.196918669 |
| 0.564175176 | 0.70628009 |
| NA | NA |
| NA | NA |
| NA | NA |
| 0.736124225 | NA |
| 0.938713777 | NA |
| 0.636579436 | NA |
| 0.941278332 | NA |
| 0.199334224 | 0.346770122 |
| 0.154869004 | NA |
| 8.79E-156 | 5.98E-153 |
| 7.90E-45 | 9.50E-43 |
| 0.011501484 | 0.038584684 |
| 0.921681957 | NA |
| NA | NA |
| NA | NA |
| 0.941279268 | NA |
| 0.799181995 | NA |
| 0.839319108 | NA |
| 0.945041907 | NA |
| 0.542528553 | 0.68933244 |
| 0.853875559 | 0.912048218 |
| 0.955792397 | NA |
| 0.409870021 | NA |
| 0.002448238 | 0.010841479 |
| 0.053097709 | 0.130104146 |
| 0.000838319 | 0.004397017 |
| 0.001193177 | 0.005931181 |
| 0.460099365 | 0.619114165 |
| 0.049591092 | 0.122916501 |
| 0.915990661 | 0.949458246 |
| 0.700804371 | 0.808877387 |
| 0.194231663 | 0.340334368 |
| 0.483621964 | 0.639580508 |
| 0.028220001 | 0.079042598 |
| 0.128414478 | 0.252646834 |
| 0.041261105 | 0.106738052 |
| 0.619927455 | 0.748934793 |
| 0.151071471 | NA |
| 0.044611484 | 0.113247766 |
| 0.276659945 | 0.436385686 |
| 0.085270717 | 0.186545368 |

| | | |
|---|---|---|
| | 0.667429575 | NA |
| | 0.941277857 | NA |
| | 0.183421869 | 0.326125219 |
| | 0.157334942 | 0.293090946 |
| | 0.515147692 | 0.6662435 |
| | 0.255227117 | 0.413343937 |
| | 0.941278332 | NA |
| | 0.42793206 | 0.589208012 |
| | 0.218932857 | 0.370391631 |
| NA | | NA |
| NA | | NA |
| NA | | NA |
| NA | | NA |
| NA | | NA |
| NA | | NA |
| | 0.000463053 | 0.002655804 |
| | 0.955792398 | NA |
| | 0.552814752 | 0.698005237 |
| NA | | NA |
| | 0.087788906 | 0.190671474 |
| NA | | NA |
| | 0.373330808 | 0.536562863 |
| NA | | NA |
| | 0.884961172 | NA |
| | 3.72E-07 | 4.49E-06 |
| | 0.182971675 | 0.325565658 |
| | 0.733409677 | 0.831782395 |
| | 0.971840767 | 0.982991414 |
| NA | | NA |
| | 0.697031102 | 0.805881687 |
| | 0.875131163 | 0.925319708 |
| | 0.776226451 | 0.862103588 |
| | 0.243816017 | 0.400288755 |
| | 0.021872052 | 0.064327021 |
| | 0.243702603 | 0.400262859 |
| | 0.955792398 | NA |
| | 0.149929555 | 0.283070521 |
| NA | | NA |
| | 0.206576766 | 0.355527129 |
| NA | | NA |
| | 0.22379573 | 0.376356206 |
| | 0.255420372 | 0.413494241 |
| NA | | NA |
| | 0.955792873 | NA |
| | 0.295269391 | NA |
| | 0.922628089 | NA |
| NA | | NA |

| | | |
|---|---|---|
| | 0.955792398 | NA |
| | 0.709664536 | NA |
| | 0.955792398 | NA |
| NA | | NA |
| NA | | NA |
| NA | | NA |
| | 5.82E-05 | 0.000433862 |
| | 0.01174584 | 0.039233018 |
| | 0.178612302 | 0.320250848 |
| | 0.041846628 | 0.107936004 |
| | 0.037054836 | 0.097986803 |
| | 0.076436388 | 0.171814043 |
| | 0.080149626 | 0.178091991 |
| | 0.941287954 | 0.964703895 |
| | 0.45553845 | 0.615155235 |
| | 0.274351403 | 0.434211855 |
| | 0.839319108 | NA |
| | 0.350054905 | 0.513845628 |
| NA | | NA |
| | 0.853595154 | NA |
| NA | | NA |
| | 0.895180832 | NA |
| | 0.260037606 | 0.418571387 |
| | 0.284626991 | 0.44526878 |
| | 0.360260761 | 0.523979629 |
| | 0.091657334 | 0.196721034 |
| | 0.839318641 | NA |
| | 0.868475169 | NA |
| NA | | NA |
| | 0.057013603 | 0.137449386 |
| | 0.249738913 | 0.407226526 |
| | 0.345440888 | NA |
| | 0.875748378 | 0.925643364 |
| | 0.144886925 | 0.276152995 |
| | 0.453348908 | 0.613303944 |
| | 0.098920904 | 0.207849167 |
| | 0.005875589 | 0.0222639 |
| | 0.033695995 | 0.090964282 |
| | 0.024892676 | 0.071456784 |
| | 0.461532193 | NA |
| | 0.862310964 | NA |
| | 0.681545291 | NA |
| NA | | NA |
| NA | | NA |
| | 0.234429483 | 0.389222276 |
| | 0.567647392 | NA |
| | 0.00094866 | 0.004875837 |

| | |
|---|---|
| 6.56E-06 | 6.22E-05 |
| 0.148558441 | 0.281482885 |
| 0.019058335 | 0.057723001 |
| NA | NA |
| NA | NA |
| NA | NA |
| NA | NA |
| NA | NA |
| NA | NA |
| 0.084408245 | 0.185127569 |
| 0.955792398 | NA |
| NA | NA |
| NA | NA |
| 0.839319108 | NA |
| 0.02816256 | 0.078945383 |
| NA | NA |
| 0.057806425 | 0.138828162 |
| 0.000248074 | 0.001542383 |
| 0.706093177 | NA |
| 0.070190974 | NA |
| 1.10E-35 | 9.66E-34 |
| 0.663292179 | 0.781490964 |
| 0.384450012 | NA |
| 0.509916756 | 0.661527151 |
| 0.982151882 | 0.988798709 |
| 6.63E-13 | 1.69E-11 |
| NA | NA |
| NA | NA |
| 0.987378675 | NA |
| 0.738184847 | 0.835208239 |
| 0.733353731 | 0.831782395 |
| 0.088078652 | 0.191159667 |
| NA | NA |
| 0.839319574 | NA |
| 0.755626499 | NA |
| 0.033652364 | 0.090898045 |
| 0.70680913 | 0.813002992 |
| 0.853595154 | NA |
| NA | NA |
| 0.140777542 | 0.270333768 |
| NA | NA |
| NA | NA |
| NA | NA |
| NA | NA |
| 0.464480248 | 0.622953675 |
| NA | NA |
| NA | NA |

| | |
|---|---|
| 0.951336115 | NA |
| 0.000361697 | 0.002138907 |
| NA | NA |
| 0.684816974 | NA |
| 6.20E-12 | 1.44E-10 |
| 1.49E-05 | 0.000129426 |
| 0.831162103 | 0.897691278 |
| 0.003041961 | 0.012973438 |
| 0.851716025 | 0.911002778 |
| NA | NA |
| NA | NA |
| NA | NA |
| 0.519204892 | 0.669754153 |
| 0.381553284 | 0.544044286 |
| NA | NA |
| 0.409459385 | NA |
| 0.940528311 | NA |
| 0.416996801 | NA |
| NA | NA |
| NA | NA |
| 0.371546278 | 0.534831768 |
| 0.009858694 | 0.033998353 |
| NA | NA |
| 0.941278332 | NA |
| 0.926421811 | NA |
| 0.545733622 | 0.692037915 |
| NA | NA |
| 0.591748959 | 0.727408393 |
| 0.202357514 | NA |
| 0.839319108 | NA |
| 0.611835867 | 0.742854383 |
| 0.201775156 | 0.349767116 |
| 0.150650813 | 0.28424817 |
| 0.439047075 | NA |
| 0.046473181 | 0.116866668 |
| 0.955792398 | NA |
| NA | NA |
| 0.131924157 | 0.257487395 |
| 0.781294012 | 0.865510569 |
| NA | NA |
| NA | NA |
| NA | NA |
| 0.926550601 | 0.955866739 |
| NA | NA |
| 0.32587696 | 0.488753838 |
| NA | NA |
| 0.955792398 | NA |

| | |
|---|---|
| NA | NA |
| 0.490449034 | 0.64488277 |
| 0.311560283 | 0.473524109 |
| 0.955792398 | NA |
| 0.364645733 | 0.527969933 |
| 0.935866248 | NA |
| 1.69E-15 | 5.36E-14 |
| 0.670086379 | 0.786575726 |
| 0.111824545 | 0.227765916 |
| NA | NA |
| 0.218231537 | 0.369520313 |
| 0.936413394 | NA |
| 1.58E-05 | 0.000136647 |
| 0.008649177 | 0.030548704 |
| 0.08401409 | 0.184508523 |
| 0.141459942 | 0.271272119 |
| 0.146640779 | 0.278647917 |
| 0.864420532 | NA |
| 0.704182428 | 0.811196891 |
| 0.287437411 | NA |
| NA | NA |
| 0.696172462 | 0.805326553 |
| NA | NA |
| 0.175838511 | 0.317212993 |
| 0.034246622 | 0.092033206 |
| NA | NA |
| NA | NA |
| 3.51E-11 | 7.48E-10 |
| 0.296733263 | 0.458180028 |
| 0.941277857 | NA |
| NA | NA |
| 0.627624377 | 0.75486182 |
| 0.051531571 | 0.126959368 |
| 0.955792873 | NA |
| 0.058668681 | 0.14047198 |
| 0.955792873 | NA |
| 0.695018992 | NA |
| 7.42E-07 | 8.46E-06 |
| 5.29E-07 | 6.21E-06 |
| 0.268944788 | 0.428791201 |
| 0.510180739 | 0.661761415 |
| 0.483212481 | 0.639181031 |
| 0.030905916 | 0.084925262 |
| 0.512791036 | 0.664140121 |
| 0.23272126 | 0.387196381 |
| 0.955792398 | NA |
| 0.987189413 | 0.99190454 |

| | | |
|---|---|---|
| NA | | NA |
| | 0.657835595 | 0.777100741 |
| | 0.900581664 | NA |
| NA | | NA |
| NA | | NA |
| | 0.738383432 | 0.835353508 |
| | 0.839319108 | NA |
| NA | | NA |
| | 7.56E-08 | 1.04E-06 |
| | 0.911748489 | NA |
| NA | | NA |
| | 0.294795408 | 0.456105063 |
| NA | | NA |
| NA | | NA |
| | 0.893587291 | 0.935895657 |
| | 0.01143312 | 0.038393216 |
| | 0.511822554 | 0.663240441 |
| NA | | NA |
| | 0.996079578 | 0.996917253 |
| | 0.079640798 | 0.17722609 |
| | 0.367177682 | 0.530345481 |
| | 0.018507447 | 0.056396652 |
| NA | | NA |
| NA | | NA |
| | 8.16E-10 | 1.47E-08 |
| NA | | NA |
| NA | | NA |
| | 0.883173219 | NA |
| NA | | NA |
| | 0.691336516 | 0.801978899 |
| | 0.778182557 | 0.863416888 |
| NA | | NA |
| | 0.691133944 | 0.80183126 |
| | 0.941277857 | NA |
| | 0.213546004 | 0.364359999 |
| | 0.14055017 | 0.269984274 |
| | 0.000157451 | 0.001044072 |
| | 0.941278332 | NA |
| | 0.1742271 | 0.315022291 |
| | 0.25590552 | 0.413935657 |
| | 0.045545495 | 0.115118814 |
| | 1.89E-13 | 5.06E-12 |
| | 0.689264301 | 0.800794108 |
| | 0.049106221 | 0.121943274 |
| | 0.72120797 | NA |
| | 0.833801616 | NA |
| | 0.941278332 | NA |

| | | |
|---|---|---|
| | 0.955791922 | NA |
| | 0.283077364 | 0.443568625 |
| NA | | NA |
| NA | | NA |
| | 0.955792398 | NA |
| NA | | NA |
| | 0.524480425 | NA |
| | 0.884414288 | 0.930962684 |
| | 0.941279268 | NA |
| | 0.111341528 | 0.227071462 |
| NA | | NA |
| | 0.839319108 | NA |
| | 0.941278838 | NA |
| NA | | NA |
| NA | | NA |
| NA | | NA |
| NA | | NA |
| | 0.038522286 | 0.101014372 |
| NA | | NA |
| NA | | NA |
| NA | | NA |
| | 1.43E-07 | 1.88E-06 |
| NA | | NA |
| | 0.577261905 | NA |
| | 5.07E-05 | 0.000383517 |
| | 3.94E-07 | 4.75E-06 |
| | 0.219750257 | 0.371378559 |
| | 1.38E-06 | 1.50E-05 |
| NA | | NA |
| | 0.487294672 | NA |
| | 0.81342166 | 0.886089811 |
| | 0.839319108 | NA |
| | 0.001416365 | 0.006838458 |
| | 0.817023389 | NA |
| | 0.085636645 | 0.187173867 |
| | 0.011739832 | 0.039223108 |
| | 0.197509978 | 0.344455781 |
| | 0.915531652 | NA |
| NA | | NA |
| NA | | NA |
| NA | | NA |
| NA | | NA |
| NA | | NA |
| | 0.137420244 | 0.265547692 |
| | 0.097871328 | 0.206317718 |
| | 0.440839177 | 0.601585488 |
| | 0.070780019 | 0.162574222 |

| | |
|---|---|
| 0.005031046 | 0.019634191 |
| 0.993565601 | 0.995363796 |
| 0.040843009 | 0.105944343 |
| 0.941278332 | NA |
| NA | NA |
| NA | NA |
| 0.16460865 | 0.302970545 |
| 0.080737142 | 0.178979739 |
| 0.080762181 | 0.179018572 |
| 4.69E-06 | 4.60E-05 |
| 0.897355537 | NA |
| NA | NA |
| NA | NA |
| NA | NA |
| NA | NA |
| 0.839319108 | NA |
| NA | NA |
| 0.20520105 | 0.353824645 |
| 0.853595154 | NA |
| 0.12921069 | 0.253459111 |
| 0.627513091 | 0.754842402 |
| 0.896498078 | NA |
| NA | NA |
| 0.049823687 | 0.123428746 |
| 0.013054905 | 0.042683085 |
| 0.183290832 | 0.325962969 |
| 0.668852487 | NA |
| 0.000999994 | 0.00511194 |
| 0.778229074 | 0.863416888 |
| 0.008802007 | 0.03100109 |
| 0.062367534 | 0.147459227 |
| 0.892450796 | 0.935045088 |
| 0.053451904 | 0.13066265 |
| 0.955792398 | NA |
| NA | NA |
| 0.839319108 | NA |
| 0.941275956 | NA |
| NA | NA |
| 0.978518225 | 0.986646267 |
| NA | NA |
| 0.04789437 | 0.119683129 |
| NA | NA |
| NA | NA |
| NA | NA |
| NA | NA |
| NA | NA |
| 0.848900296 | 0.909153602 |

| | | |
|---|---|---|
| NA | NA | |
| NA | NA | |
| 0.877063799 | NA | |
| NA | NA | |
| NA | NA | |
| NA | NA | |
| NA | NA | |
| 0.839318641 | NA | |
| NA | NA | |
| 0.137418741 | | 0.265547692 |
| NA | NA | |
| NA | NA | |
| 0.92677266 | NA | |
| NA | NA | |
| NA | NA | |
| NA | NA | |
| NA | NA | |
| NA | NA | |
| NA | NA | |
| 0.012049579 | | 0.040011729 |
| 0.794646495 | | 0.873922535 |
| 0.838197868 | NA | |
| NA | NA | |
| NA | NA | |
| NA | NA | |
| NA | NA | |
| NA | NA | |
| 0.599140721 | NA | |
| 0.941278332 | NA | |
| NA | NA | |
| 0.701199638 | NA | |
| 0.839319108 | NA | |
| 0.73431627 | NA | |
| 0.00543024 | | 0.020841925 |
| NA | NA | |
| NA | NA | |
| 0.896861983 | NA | |
| NA | NA | |
| NA | NA | |
| 0.001216552 | | 0.00603132 |
| 0.01985219 | | 0.059489089 |
| 0.002561611 | | 0.011268057 |
| 6.39E-07 | | 7.38E-06 |
| 0.839318641 | NA | |
| 0.501807179 | NA | |
| 0.138198183 | | 0.266693137 |
| NA | NA | |

| | |
|---|---|
| 0.00295721 | 0.012668737 |
| 0.542066837 | 0.689112146 |
| 0.000828122 | 0.004353128 |
| NA | NA |
| 0.561184453 | 0.704091105 |
| 0.000502665 | 0.002850712 |
| 0.839318641 | NA |
| 0.003240907 | 0.013687011 |
| 0.06590255 | 0.153951565 |
| NA | NA |
| 0.167154849 | 0.306141868 |
| 0.521925941 | 0.672090432 |
| 0.73699103 | 0.834254078 |
| NA | NA |
| 0.074215079 | 0.168138902 |
| 0.124261703 | 0.246678654 |
| 0.593644164 | 0.728800997 |
| 0.019349507 | 0.058364036 |
| 0.576703438 | 0.715341875 |
| NA | NA |
| 0.728678906 | NA |
| NA | NA |
| 0.515665248 | 0.666639079 |
| NA | NA |
| 0.830117013 | NA |
| NA | NA |
| 0.807889002 | NA |
| 0.013650908 | 0.044230726 |
| 0.941278788 | NA |
| 0.941278332 | NA |
| NA | NA |
| NA | NA |
| 0.286387919 | 0.447019821 |
| 0.995971254 | 0.996917253 |
| 0.30060257 | 0.462056337 |
| 0.113589911 | 0.230570276 |
| 0.011757844 | 0.039262094 |
| 0.272624992 | NA |
| 0.22665776 | 0.37983759 |
| 0.433499198 | 0.594705932 |
| 0.044348007 | 0.112714496 |
| 0.053829298 | 0.131423217 |
| NA | NA |
| 0.134560126 | 0.261325075 |
| NA | NA |
| 0.955792398 | NA |
| 0.955791446 | NA |

| | | | |
|---|---|---|---|
| | 0.003106539 | | 0.013203901 |
| NA | | NA | |
| | 0.941277857 | NA | |
| | 0.653647984 | | 0.774003044 |
| | 0.13315536 | | 0.259246961 |
| | 0.109236982 | | 0.223969217 |
| NA | | NA | |
| NA | | NA | |
| | 0.664147936 | | 0.782114139 |
| | 0.719314911 | | 0.821393855 |
| NA | | NA | |
| | 0.103737432 | | 0.215609358 |
| | 0.378859806 | | 0.541696469 |
| NA | | NA | |
| NA | | NA | |
| NA | | NA | |
| | 5.83E-09 | | 9.42E-08 |
| | 0.608441033 | | 0.740735178 |
| NA | | NA | |
| | 0.897493458 | | 0.938170368 |
| | 0.955792398 | NA | |
| NA | | NA | |
| NA | | NA | |
| NA | | NA | |
| | 0.541037642 | NA | |
| NA | | NA | |
| NA | | NA | |
| NA | | NA | |
| NA | | NA | |
| NA | | NA | |
| | 0.919777386 | NA | |
| NA | | NA | |
| | 0.839318641 | NA | |
| | 0.941276431 | NA | |
| NA | | NA | |
| NA | | NA | |
| | 0.941277857 | NA | |
| NA | | NA | |
| | 0.839318641 | NA | |
| NA | | NA | |
| | 0.003634655 | | 0.014990826 |
| | 1.34E-143 | | 7.09E-141 |
| | 0.132485032 | | 0.258243283 |
| | 0.942270122 | | 0.965098044 |
| | 0.030304073 | | 0.083638398 |
| NA | | NA | |
| | 0.431568499 | NA | |

| | | |
|---|---|---|
| | 0.008976987 | 0.031524084 |
| | 0.671318817 | NA |
| NA | | NA |
| | 0.309941581 | 0.471689866 |
| | 0.897693313 | NA |
| | 0.092896894 | 0.198690145 |
| | 0.20195554 | 0.349977849 |
| | 0.783740227 | NA |
| | 0.056708897 | 0.136867285 |
| | 0.92938797 | 0.958004957 |
| | 2.66E-26 | 1.58E-24 |
| | 0.569970713 | NA |
| | 0.554752388 | NA |
| | 0.082408359 | NA |
| | 0.076849523 | 0.172586849 |
| | 0.839319108 | NA |
| | 1.63E-05 | 0.000140223 |
| | 0.002209636 | 0.009947752 |
| | 0.784174936 | 0.867327233 |
| | 0.60645027 | NA |
| | 0.614084389 | NA |
| NA | | NA |
| | 0.135151852 | 0.26220415 |
| | 0.001810835 | 0.008413331 |
| | 0.546253061 | NA |
| | 0.002612753 | 0.011448591 |
| | 0.017694987 | 0.054373366 |
| NA | | NA |
| | 0.009848749 | 0.033978828 |
| | 0.705819009 | 0.812256638 |
| | 0.874848714 | 0.925185458 |
| | 0.851189126 | NA |
| | 0.650344471 | NA |
| | 0.417004416 | 0.578512509 |
| NA | | NA |
| | 0.621640675 | 0.75018753 |
| | 0.055419432 | NA |
| | 0.041624687 | 0.107505922 |
| | 0.005642508 | 0.021526362 |
| | 0.235836595 | 0.391067482 |
| | 5.05E-06 | 4.91E-05 |
| | 0.329630555 | 0.49258328 |
| | 0.689968301 | NA |
| | 0.215495443 | 0.366471261 |
| | 0.883489089 | 0.930478199 |
| | 0.211179231 | 0.3613319 |
| | 0.003014661 | 0.012882399 |

| | | |
|---|---|---|
| | 0.254336468 | 0.41235526 |
| | 0.170184131 | NA |
| | 0.139027482 | 0.267772465 |
| | 0.68903346 | 0.800564991 |
| | 0.283957529 | 0.444508852 |
| | 0.059975119 | 0.142952912 |
| | 0.670246562 | 0.786686161 |
| | 1.34E-05 | 0.000117458 |
| | 0.029819071 | 0.08257749 |
| | 7.26E-15 | 2.17E-13 |
| | 0.226972621 | NA |
| | 0.739991096 | NA |
| NA | | NA |
| NA | | NA |
| NA | | NA |
| NA | | NA |
| NA | | NA |
| NA | | NA |
| NA | | NA |
| | 0.941277857 | NA |
| | 0.56428228 | NA |
| NA | | NA |
| NA | | NA |
| | 0.941278332 | NA |
| | 0.955791922 | NA |
| | 0.748393366 | NA |
| | 0.541351386 | 0.688462683 |
| | 0.839318641 | NA |
| NA | | NA |
| | 0.793576211 | NA |
| NA | | NA |
| NA | | NA |
| NA | | NA |
| NA | | NA |
| NA | | NA |
| NA | | NA |
| | 0.012073413 | 0.040068513 |
| NA | | NA |
| | 0.698384734 | NA |
| NA | | NA |
| NA | | NA |
| | 0.941275956 | NA |
| | 0.372445343 | NA |
| NA | | NA |
| NA | | NA |
| | 0.237485908 | 0.392953742 |
| | 0.978445112 | NA |

| | | |
|---|---|---|
| | 0.000149318 | 0.000996098 |
| | 0.565310426 | 0.707165332 |
| | 0.974651559 | 0.984621043 |
| | 5.42E-14 | 1.52E-12 |
| | 0.238161297 | 0.393781288 |
| | 0.75057113 | NA |
| | 0.31676328 | 0.479221992 |
| | 0.889453052 | 0.933455095 |
| | 0.002851312 | 0.012285832 |
| | 0.000379306 | 0.002229196 |
| | 0.057322666 | 0.13798484 |
| | 0.020275422 | 0.060534763 |
| | 0.018208611 | 0.055671338 |
| | 0.323320122 | 0.48602157 |
| | 0.269512479 | 0.42943741 |
| | 0.108824965 | NA |
| | 0.000926131 | 0.004776733 |
| | 0.0032822 | 0.013831964 |
| | 0.003644901 | 0.015022674 |
| NA | | NA |
| NA | | NA |
| | 0.074506923 | 0.168607508 |
| | 0.778956256 | 0.863740938 |
| | 0.786453076 | 0.868759275 |
| | 0.564701547 | 0.706620985 |
| | 1.63E-58 | 2.68E-56 |
| | 0.000137343 | 0.00092607 |
| | 0.881299138 | 0.929035036 |
| | 0.616464845 | 0.746228766 |
| | 0.444024979 | 0.60458112 |
| | 0.934865977 | 0.961237201 |
| | 0.575102537 | 0.714074899 |
| | 0.036830609 | 0.097513031 |
| | 5.79E-17 | 2.02E-15 |
| | 3.48E-06 | 3.49E-05 |
| | 0.25140326 | 0.409155583 |
| | 0.045454286 | 0.114934253 |
| | 0.926201186 | NA |
| | 0.135762419 | NA |
| | 0.097462173 | 0.205655554 |
| | 0.459361363 | 0.618621801 |
| | 0.008781442 | 0.030946977 |
| | 0.534488225 | 0.682687374 |
| | 0.071645447 | 0.1638979 |
| | 0.001119921 | 0.005626079 |
| | 0.694981952 | 0.804609195 |
| | 0.14446536 | 0.275592251 |

| | | | |
|---|---|---|---|
| | 0.018102392 | | 0.055382151 |
| NA | | NA | |
| NA | | NA | |
| | 0.097351106 | | 0.205530515 |
| | 0.589795957 | | 0.725832344 |
| | 0.616706037 | NA | |
| NA | | NA | |
| NA | | NA | |
| NA | | NA | |
| NA | | NA | |
| | 0.955792873 | NA | |
| NA | | NA | |
| | 0.955791922 | NA | |
| | 0.239789297 | | 0.39582237 |
| | 0.976330722 | | 0.985480846 |
| | 2.75E-34 | | 2.30E-32 |
| | 0.147136251 | | 0.279388676 |
| | 0.202533833 | | 0.350623963 |
| | 0.059852565 | | 0.142717968 |
| | 0.651273031 | NA | |
| | 0.859297324 | NA | |
| | 0.615351599 | NA | |
| | 0.538566293 | | 0.68617383 |
| | 0.531682198 | | 0.680308551 |
| | 1.13E-07 | | 1.51E-06 |
| NA | | NA | |
| NA | | NA | |
| NA | | NA | |
| | 1.13E-08 | | 1.76E-07 |
| | 0.853595622 | NA | |
| | 0.218035276 | | 0.369339023 |
| | 0.446701012 | | 0.607148098 |
| | 0.480353283 | | 0.637059414 |
| | 0.241519155 | NA | |
| | 0.001077486 | | 0.005445077 |
| | 0.01709785 | | 0.052949083 |
| NA | | NA | |
| NA | | NA | |
| | 0.00352717 | | 0.014654162 |
| | 0.000540491 | | 0.003035578 |
| | 1.68E-09 | | 2.91E-08 |
| | 0.014288218 | | 0.045908229 |
| | 0.175611506 | | 0.316851507 |
| | 0.033446169 | | 0.090474567 |
| | 0.125590625 | | 0.248508442 |
| NA | | NA | |
| | 1.35E-06 | | 1.48E-05 |

| | | |
|---|---|---|
| | 0.941923527 | NA |
| | 0.522373191 | 0.672390001 |
| | 0.176222891 | 0.317713774 |
| NA | | NA |
| | 0.051927547 | 0.127763022 |
| | 0.015673154 | 0.049404373 |
| NA | | NA |
| | 0.447288018 | 0.607633663 |
| | 0.920378685 | NA |
| NA | | NA |
| | 0.491045888 | 0.645132982 |
| | 0.648290213 | 0.770105487 |
| | 0.937531888 | NA |
| | 0.955792398 | NA |
| | 0.436679355 | NA |
| | 0.839319108 | NA |
| | 3.74E-15 | 1.15E-13 |
| | 0.214666443 | 0.365540054 |
| | 8.06E-16 | 2.61E-14 |
| | 0.966792184 | 0.980159365 |
| | 7.05E-28 | 4.51E-26 |
| | 8.14E-07 | 9.23E-06 |
| | 0.060779767 | 0.144480176 |
| | 0.967105721 | NA |
| | 0.963167107 | 0.977749967 |
| | 0.002700196 | 0.011777546 |
| | 0.884059522 | 0.930873026 |
| NA | | NA |
| | 0.853595622 | NA |
| | 0.990905922 | 0.993912463 |
| | 0.203986802 | NA |
| | 0.004360916 | 0.017430484 |
| | 0.238665288 | 0.394439048 |
| | 0.941276431 | NA |
| | 0.022594824 | 0.066071784 |
| | 0.739391621 | 0.83613642 |
| | 3.64E-31 | 2.67E-29 |
| | 0.932934684 | NA |
| | 0.222118457 | NA |
| | 0.414295734 | 0.576333551 |
| | 0.026891002 | 0.076152247 |
| | 0.157115709 | 0.292870868 |
| | 0.743277646 | NA |
| | 0.06274769 | 0.148122635 |
| NA | | NA |
| | 0.055739035 | NA |
| | 0.122243909 | 0.243750484 |

| | | | |
|---|---|---|---|
| | 0.391705491 | | 0.554141025 |
| NA | | NA | |
| | 0.846474531 | | 0.907476209 |
| | 0.001956736 | | 0.009001589 |
| | 0.389253717 | | 0.551850436 |
| | 0.494053164 | | 0.647310858 |
| | 0.949035057 | | 0.968816796 |
| | 0.122314279 | | 0.243814664 |
| NA | | NA | |
| NA | | NA | |
| | 0.941278332 | NA | |
| NA | | NA | |
| NA | | NA | |
| | 0.008190491 | | 0.029241047 |
| | 0.110837709 | | 0.226354281 |
| | 0.406181906 | | 0.568067555 |
| | 0.090710261 | NA | |
| | 0.000525264 | | 0.002960536 |
| | 0.080866439 | | 0.179130044 |
| | 0.334470096 | | 0.4976932 |
| | 0.732583302 | | 0.831241253 |
| | 1.63E-08 | | 2.48E-07 |
| | 0.623250557 | | 0.751385543 |
| | 0.01906154 | | 0.057723001 |
| NA | | NA | |
| NA | | NA | |
| NA | | NA | |
| | 0.839319108 | NA | |
| NA | | NA | |
| | 0.451344105 | | 0.611329448 |
| | 0.905890521 | NA | |
| | 0.463090548 | NA | |
| | 0.941275956 | NA | |
| | 0.692708782 | NA | |
| | 0.715422001 | NA | |
| | 0.839319108 | NA | |
| | 0.5021784 | | 0.654269543 |
| NA | | NA | |
| | 0.419598455 | | 0.581230426 |
| | 0.054152302 | | 0.132034484 |
| | 3.46E-06 | | 3.48E-05 |
| | 0.023023869 | | 0.067117469 |
| | 0.0557035 | | 0.135029504 |
| | 0.565716338 | | 0.707502116 |
| | 0.352360267 | | 0.515977991 |
| | 0.441355112 | | 0.60198416 |
| | 0.109277418 | | 0.223994239 |

| | |
|---|---|
| 0.955792873 | NA |
| NA | NA |
| 0.839319574 | NA |
| NA | NA |
| 0.800797152 | 0.878492596 |
| NA | NA |
| NA | NA |
| NA | NA |
| 0.42297681 | NA |
| NA | NA |
| 0.808592441 | NA |
| NA | NA |
| 0.728678906 | NA |
| NA | NA |
| 0.009961356 | 0.03428285 |
| 0.562226988 | 0.704917159 |
| 0.414620543 | 0.576583207 |
| NA | NA |
| 0.941277857 | NA |
| NA | NA |
| NA | NA |
| 0.690947023 | NA |
| NA | NA |
| 0.839318641 | NA |
| 0.915461719 | NA |
| 0.941278332 | NA |
| 0.941276431 | NA |
| NA | NA |
| NA | NA |
| 0.005638793 | 0.021522537 |
| 1.86E-05 | 0.000157227 |
| 0.166553992 | 0.305488026 |
| 0.941277857 | NA |
| 0.131273209 | 0.256553487 |
| NA | NA |
| 0.356846493 | NA |
| 0.873965987 | 0.924785321 |
| 0.91128193 | NA |
| 3.48E-10 | 6.56E-09 |
| 0.354390058 | 0.518126957 |
| 8.69E-11 | 1.76E-09 |
| NA | NA |
| 0.009201316 | 0.032141141 |
| NA | NA |
| 0.144047336 | 0.274971103 |
| NA | NA |
| 0.955792873 | NA |

| | | |
|---|---|---|
| | 0.955792873 | NA |
| NA | | NA |
| NA | | NA |
| | 0.941275481 | NA |
| NA | | NA |
| NA | | NA |
| NA | | NA |
| NA | | NA |
| | 0.955792398 | NA |
| NA | | NA |
| NA | | NA |
| NA | | NA |
| | 2.55E-08 | 3.76E-07 |
| | 0.941277857 | NA |
| | 0.938182526 | NA |
| NA | | NA |
| | 0.920928731 | NA |
| NA | | NA |
| NA | | NA |
| | 0.012605675 | 0.041499345 |
| NA | | NA |
| NA | | NA |
| | 0.955792873 | NA |
| NA | | NA |
| NA | | NA |
| NA | | NA |
| | 6.21E-12 | 1.44E-10 |
| | 0.627811999 | NA |
| | 0.421154981 | 0.582722881 |
| | 0.674272875 | NA |
| NA | | NA |
| | 0.057905373 | 0.139007937 |
| NA | | NA |
| | 0.839320041 | NA |
| | 0.978935435 | 0.986770898 |
| NA | | NA |
| | 0.955792398 | NA |
| | 0.222477175 | 0.374869605 |
| | 0.76734443 | 0.855592455 |
| | 0.015785789 | 0.049706754 |
| | 0.385565827 | 0.54841154 |
| | 0.514250559 | 0.665240505 |
| | 0.247086152 | NA |
| | 0.955792873 | NA |
| | 0.941277857 | NA |
| | 2.61E-05 | 0.00021347 |
| | 3.68E-05 | 0.000288009 |

| | |
|---|---|
| 0.002668261 | 0.01165748 |
| 0.024947641 | 0.071544442 |
| 0.93824099 | 0.963334141 |
| NA | NA |
| 0.855647734 | 0.913156408 |
| 0.65229483 | 0.773105252 |
| 0.006698912 | 0.024768638 |
| NA | NA |
| 0.348924886 | 0.512921373 |
| 0.384038791 | 0.546736725 |
| 0.570713515 | NA |
| 0.311373765 | 0.473415418 |
| NA | NA |
| 0.885855641 | 0.931651465 |
| 1.87E-07 | 2.39E-06 |
| NA | NA |
| 0.001585271 | 0.007519717 |
| 0.040323891 | 0.104837712 |
| 0.277268723 | 0.437172108 |
| 0.099405008 | 0.20862693 |
| 0.499088329 | NA |
| 0.077530578 | 0.173650401 |
| NA | NA |
| 0.955792398 | NA |
| 0.22294204 | NA |
| 0.717901652 | 0.820606787 |
| 0.008375238 | 0.029757788 |
| NA | NA |
| 0.040406411 | 0.104971993 |
| 0.424637587 | 0.586019796 |
| 0.410221798 | 0.572171066 |
| NA | NA |
| 0.139116107 | 0.267878131 |
| 0.784461486 | NA |
| 0.576018057 | NA |
| NA | NA |
| 0.822595861 | 0.892041855 |
| 0.104331109 | 0.216408631 |
| 0.472787527 | NA |
| 0.283430223 | 0.444004619 |
| 0.565633456 | NA |
| 4.50E-15 | 1.38E-13 |
| NA | NA |
| 0.057884266 | 0.138971282 |
| 2.50E-05 | 0.000205551 |
| 0.00336256 | 0.014100716 |
| 0.618615752 | 0.747919547 |

| | | |
|---|---|---|
| | 0.229930188 | 0.38373288 |
| | 0.353524199 | 0.517143784 |
| | 0.941277857 | NA |
| NA | | NA |
| | 0.669623455 | NA |
| | 0.839319574 | NA |
| | 0.839318641 | NA |
| NA | | NA |
| NA | | NA |
| NA | | NA |
| NA | | NA |
| NA | | NA |
| NA | | NA |
| NA | | NA |
| NA | | NA |
| NA | | NA |
| NA | | NA |
| NA | | NA |
| NA | | NA |
| NA | | NA |
| NA | | NA |
| NA | | NA |
| NA | | NA |
| NA | | NA |
| NA | | NA |
| NA | | NA |
| NA | | NA |
| NA | | NA |
| NA | | NA |
| NA | | NA |
| NA | | NA |
| NA | | NA |
| NA | | NA |
| NA | | NA |
| NA | | NA |
| NA | | NA |
| NA | | NA |
| NA | | NA |
| NA | | NA |
| NA | | NA |
| NA | | NA |
| NA | | NA |
| NA | | NA |
| NA | | NA |
| | 0.697358527 | NA |
| NA | | NA |

| | |
|---|---|
| NA | NA |
| NA | NA |
| NA | NA |
| NA | NA |
| NA | NA |
| NA | NA |
| 0.737354924 | 0.834432515 |
| NA | NA |
| NA | NA |
| 0.941276431 | NA |
| 0.000531624 | 0.002990715 |
| 1.08E-06 | 1.19E-05 |
| NA | NA |
| 4.68E-06 | 4.59E-05 |
| 0.00076287 | 0.004062157 |
| 0.246547693 | 0.403543404 |
| 0.846176748 | 0.907320432 |
| 0.01865378 | 0.056762627 |
| 0.156173196 | 0.291639292 |
| 0.577313309 | 0.715799937 |
| 0.006514589 | 0.024204255 |
| 0.334492555 | 0.4976932 |
| 0.000143599 | 0.000961447 |
| 0.059221307 | 0.141609961 |
| 0.101808287 | 0.212583634 |
| 0.835673691 | NA |
| 0.258722395 | NA |
| 0.941278332 | NA |
| NA | NA |
| NA | NA |
| NA | NA |
| NA | NA |
| 0.934560279 | 0.961005908 |
| 0.447503688 | NA |
| 0.716513821 | NA |
| 1.63E-08 | 2.48E-07 |
| 0.745252829 | NA |
| 0.705632596 | NA |
| NA | NA |
| 0.825972908 | NA |
| 0.026154846 | NA |
| 2.31E-34 | 1.95E-32 |
| 0.204550094 | NA |
| 8.27E-05 | 0.000592591 |
| 0.006511279 | 0.02419695 |
| 4.64E-06 | 4.55E-05 |
| NA | NA |

| | |
|---|---|
| NA | NA |
| NA | NA |
| NA | NA |
| NA | NA |
| 0.640292839 | 0.763922313 |
| NA | NA |
| 0.01476728 | 0.047066255 |
| 6.23E-21 | 2.84E-19 |
| 0.619577448 | 0.748663947 |
| 0.001034665 | 0.005258708 |
| 0.273564261 | 0.433398332 |
| 0.941277857 | NA |
| NA | NA |
| NA | NA |
| 0.839319108 | NA |
| NA | NA |
| 0.64052299 | 0.764158599 |
| 0.004112489 | 0.016613284 |
| 0.838508625 | 0.902115782 |
| NA | NA |
| NA | NA |
| NA | NA |
| 8.06E-09 | 1.28E-07 |
| NA | NA |
| NA | NA |
| NA | NA |
| 0.951382471 | 0.970309615 |
| 0.838197868 | NA |
| NA | NA |
| 0.831413565 | NA |
| NA | NA |
| 0.213264212 | 0.363920785 |
| NA | NA |
| 0.075368101 | 0.170168046 |
| 0.004517694 | 0.017951613 |
| 0.843453706 | NA |
| 0.25189945 | 0.409649338 |
| 0.715594241 | 0.819182883 |
| 0.955792398 | NA |
| 0.132195787 | 0.257832551 |
| 0.24951007 | 0.406970974 |
| 0.774303269 | NA |
| 0.387812494 | 0.550725671 |
| NA | NA |
| NA | NA |
| NA | NA |
| 0.131537148 | NA |

| | | | |
|---|---|---|---|
| | 0.906771916 | NA | |
| | 0.955791922 | NA | |
| NA | | NA | |
| NA | | NA | |
| | 0.007644386 | | 0.027613893 |
| NA | | NA | |
| | 0.764825348 | | 0.853946599 |
| | 0.700961407 | NA | |
| | 0.778629151 | | 0.863592382 |
| NA | | NA | |
| NA | | NA | |
| NA | | NA | |
| NA | | NA | |
| NA | | NA | |
| NA | | NA | |
| | 0.941277857 | NA | |
| | 0.255524758 | | 0.413544591 |
| NA | | NA | |
| NA | | NA | |
| NA | | NA | |
| NA | | NA | |
| NA | | NA | |
| NA | | NA | |
| | 0.601938606 | | 0.735387223 |
| | 0.955792873 | NA | |
| | 0.462018374 | NA | |
| NA | | NA | |
| NA | | NA | |
| NA | | NA | |
| NA | | NA | |
| | 0.941275956 | NA | |
| | 0.000617636 | | 0.003393576 |
| | 0.047066141 | | 0.118072121 |
| | 0.386860226 | | 0.549668523 |
| | 0.700252935 | | 0.808638955 |
| | 0.198345021 | | 0.345507039 |
| NA | | NA | |
| | 0.941278332 | NA | |
| NA | | NA | |
| NA | | NA | |
| NA | | NA | |
| | 0.656358693 | NA | |
| | 0.327513032 | | 0.490280829 |
| NA | | NA | |
| NA | | NA | |
| NA | | NA | |
| NA | | NA | |

| | | |
|---|---|---|
| | 0.839319108 | NA |
| | 0.000115443 | 0.000794385 |
| NA | | NA |
| | 0.941278788 | NA |
| NA | | NA |
| | 0.331599926 | 0.49468867 |
| NA | | NA |
| | 0.78832422 | NA |
| | 0.468401169 | 0.62651782 |
| | 0.428407427 | 0.589657872 |
| | 0.842281546 | NA |
| | 0.955792873 | NA |
| NA | | NA |
| | 0.833801616 | NA |
| NA | | NA |
| | 0.001610124 | 0.007626701 |
| NA | | NA |
| | 0.085943332 | 0.187689071 |
| | 0.433483979 | NA |
| | 0.936960566 | NA |
| NA | | NA |
| | 0.261054609 | 0.419798224 |
| | 0.311061724 | 0.473092086 |
| | 0.461598294 | 0.620416884 |
| | 0.000634901 | 0.003472229 |
| | 0.143425139 | 0.274025137 |
| NA | | NA |
| | 0.941278332 | NA |
| | 0.182469312 | NA |
| NA | | NA |
| NA | | NA |
| | 0.700911525 | 0.808877387 |
| NA | | NA |
| | 0.709882426 | 0.814844787 |
| | 0.011232134 | 0.037841084 |
| NA | | NA |
| | 0.008784652 | 0.030949125 |
| | 0.345231326 | NA |
| | 0.917731244 | NA |
| | 0.839319108 | NA |
| | 3.19E-05 | 0.000254706 |
| | 0.965005645 | NA |
| | 0.093379735 | 0.199380126 |
| NA | | NA |
| NA | | NA |
| | 0.501445553 | 0.653851597 |
| | 3.02E-06 | 3.07E-05 |

| | | |
|---|---|---|
| | 0.528160924 | NA |
| | 0.874329785 | 0.924906832 |
| NA | | NA |
| | 0.027659632 | 0.077856499 |
| | 0.945784926 | 0.967272298 |
| | 0.941275956 | NA |
| | 0.231721356 | 0.38591042 |
| | 0.783665222 | 0.866967214 |
| NA | | NA |
| | 0.695908883 | NA |
| | 0.941278332 | NA |
| | 0.747045966 | NA |
| NA | | NA |
| NA | | NA |
| | 0.738468792 | 0.835388089 |
| | 0.955791446 | NA |
| | 0.445463 | 0.605949841 |
| | 0.613902592 | NA |
| NA | | NA |
| | 0.955792873 | NA |
| | 0.012031023 | 0.039978 |
| | 0.07355389 | 0.167102195 |
| | 0.486303305 | NA |
| NA | | NA |
| | 0.01358021 | 0.044049626 |
| NA | | NA |
| | 0.192458992 | 0.337999626 |
| | 0.504758533 | 0.656898006 |
| NA | | NA |
| | 0.007267426 | 0.026496859 |
| | 0.839319108 | NA |
| | 0.720541181 | NA |
| NA | | NA |
| NA | | NA |
| | 0.34518172 | 0.508826117 |
| | 0.070031615 | 0.161288038 |
| | 0.425716865 | NA |
| | 0.247674262 | 0.404886179 |
| | 1.42E-12 | 3.52E-11 |
| | 0.002021929 | 0.009239003 |
| | 0.009371714 | 0.032626229 |
| | 2.09E-15 | 6.59E-14 |
| | 0.027425446 | 0.077371158 |
| | 0.125152146 | 0.247888106 |
| | 0.929138015 | NA |
| NA | | NA |
| | 0.839319108 | NA |

| | | |
|---|---|---|
| | 0.320386122 | 0.48307554 |
| | 0.35380904 | 0.517470008 |
| | 0.640194667 | NA |
| | 0.179748671 | 0.321682546 |
| | 0.362006969 | 0.52568421 |
| | 0.766462317 | 0.855009424 |
| | 0.132875113 | 0.258834064 |
| | 7.66E-12 | 1.76E-10 |
| | 0.057529303 | NA |
| | 0.247469883 | NA |
| NA | | NA |
| | 0.888800656 | NA |
| | 0.374990812 | 0.538136458 |
| | 0.781148131 | NA |
| | 0.925866096 | NA |
| | 0.239907762 | NA |
| | 0.839319108 | NA |
| | 0.941278332 | NA |
| | 0.793302807 | NA |
| | 2.60E-08 | 3.83E-07 |
| | 0.000845674 | 0.004428762 |
| | 0.624609771 | NA |
| | 0.300994782 | 0.462360619 |
| | 0.42116552 | 0.582722881 |
| | 0.717328413 | 0.820227272 |
| | 0.955792398 | NA |
| | 0.839319107 | NA |
| NA | | NA |
| NA | | NA |
| | 0.847798961 | NA |
| NA | | NA |
| NA | | NA |
| NA | | NA |
| NA | | NA |
| NA | | NA |
| NA | | NA |
| NA | | NA |
| NA | | NA |
| NA | | NA |
| | 0.955791446 | NA |
| NA | | NA |
| NA | | NA |
| NA | | NA |
| NA | | NA |
| | 0.839318641 | NA |
| NA | | NA |
| NA | | NA |

| | |
|---|---|
| NA | NA |
| NA | NA |
| 0.400663599 | NA |
| 0.61552339 | NA |
| 0.449066179 | NA |
| 2.33E-07 | 2.94E-06 |
| 2.41E-07 | 3.02E-06 |
| 0.000678046 | 0.003676381 |
| 0.746289797 | NA |
| 0.839319574 | NA |
| 0.072126521 | 0.164676782 |
| 0.955792873 | NA |
| NA | NA |
| 0.487332163 | 0.642239087 |
| NA | NA |
| NA | NA |
| 0.941278332 | NA |
| 0.000864953 | 0.004514829 |
| 0.000227199 | 0.001430636 |
| 0.625786494 | 0.753436846 |
| 0.001432049 | NA |
| NA | NA |
| 0.739991096 | NA |
| 0.715900958 | 0.819421717 |
| NA | NA |
| NA | NA |
| 0.765008191 | NA |
| NA | NA |
| NA | NA |
| NA | NA |
| NA | NA |
| 0.402125501 | NA |
| NA | NA |
| 0.941275956 | NA |
| 0.833904229 | 0.899431054 |
| NA | NA |
| 0.098444032 | 0.207194699 |
| 0.955792873 | NA |
| 0.614036818 | NA |
| 0.935658422 | NA |
| 0.405957362 | 0.567920367 |
| 0.556796798 | 0.700913761 |
| NA | NA |
| NA | NA |
| 0.791696326 | 0.871750204 |
| NA | NA |
| NA | NA |

| | | | |
|---|---|---|---|
| NA | | NA | |
| NA | | NA | |
| NA | | NA | |
| NA | | NA | |
| | 0.250630585 | | 0.408281173 |
| | 0.637498537 | NA | |
| NA | | NA | |
| NA | | NA | |
| NA | | NA | |
| NA | | NA | |
| NA | | NA | |
| NA | | NA | |
| | 0.746345257 | NA | |
| NA | | NA | |
| NA | | NA | |
| | 0.941277857 | NA | |
| | 0.673178757 | | 0.788999447 |
| NA | | NA | |
| NA | | NA | |
| | 0.000908282 | | 0.004702836 |
| | 0.349528168 | NA | |
| | 0.000343325 | | 0.002048589 |
| | 0.788902502 | | 0.870055922 |
| | 0.48176008 | | 0.638181958 |
| | 0.244480257 | NA | |
| | 0.893093698 | | 0.935461019 |
| | 0.524844335 | | 0.6743142 |
| | 0.449870031 | | 0.609847259 |
| | 0.806952193 | | 0.88170566 |
| | 0.006062746 | | 0.022847607 |
| | 0.785834296 | NA | |
| | 0.884553227 | | 0.930962684 |
| | 0.955469555 | NA | |
| | 9.22E-07 | | 1.03E-05 |
| | 0.392947576 | | 0.555433113 |
| | 0.477532114 | NA | |
| | 0.941278332 | NA | |
| | 0.62270131 | | 0.751103586 |
| | 0.478579219 | | 0.635382784 |
| NA | | NA | |
| | 0.83454204 | NA | |
| NA | | NA | |
| NA | | NA | |
| | 0.352923226 | | 0.516558764 |
| NA | | NA | |
| | 0.055984026 | | 0.135585244 |
| | 0.95432283 | | 0.972268326 |

| | | |
|---|---|---|
| | 0.064788665 | 0.151983638 |
| | 0.955792398 | NA |
| | 0.102406793 | 0.213589773 |
| NA | | NA |
| NA | | NA |
| NA | | NA |
| | 0.955792873 | NA |
| NA | | NA |
| | 0.738677218 | NA |
| NA | | NA |
| | 0.118863529 | 0.238508659 |
| | 0.946444971 | 0.967520714 |
| | 0.000737939 | 0.003952392 |
| | 0.892614349 | NA |
| | 0.62359628 | NA |
| NA | | NA |
| NA | | NA |
| NA | | NA |
| | 0.360539272 | 0.524256567 |
| | 0.112864548 | 0.229476108 |
| | 0.461728774 | 0.620487143 |
| | 0.653627528 | 0.774003044 |
| | 0.763965908 | 0.853554567 |
| | 0.639708914 | 0.763416975 |
| | 0.514347359 | NA |
| NA | | NA |
| NA | | NA |
| | 0.893825164 | NA |
| NA | | NA |
| | 0.200546341 | 0.348169511 |
| NA | | NA |
| | 0.025062776 | 0.071777667 |
| | 0.955792398 | NA |
| | 0.926033828 | 0.955499271 |
| | 0.139817679 | 0.268837579 |
| | 0.367552166 | 0.530716839 |
| | 0.941278332 | NA |
| | 0.019594263 | 0.058938792 |
| | 0.419105564 | NA |
| | 0.839318641 | NA |
| NA | | NA |
| | 6.05E-07 | 7.01E-06 |
| | 0.070404022 | 0.161877563 |
| | 0.022277105 | 0.065284371 |
| | 0.303366258 | 0.464773666 |
| | 0.941275956 | NA |
| | 0.875598198 | NA |

| | | | |
|---|---|---|---|
| | 0.000395681 | | 0.002315712 |
| | 0.174401155 | | 0.315265155 |
| | 0.216849145 | | 0.368227392 |
| NA | | NA | |
| NA | | NA | |
| NA | | NA | |
| NA | | NA | |
| NA | | NA | |
| NA | | NA | |
| NA | | NA | |
| NA | | NA | |
| | 0.384687263 | | 0.547496187 |
| | 0.941078621 | | 0.964703895 |
| NA | | NA | |
| NA | | NA | |
| | 0.067085443 | | 0.156016892 |
| | 0.076668794 | | 0.172271371 |
| | 0.060651356 | | 0.144290212 |
| | 0.727556795 | NA | |
| | 0.801068647 | | 0.878492596 |
| NA | | NA | |
| | 0.258425695 | | 0.416739267 |
| | 0.006912597 | | 0.025422379 |
| | 0.742105939 | NA | |
| | 0.515255049 | NA | |
| NA | | NA | |
| | 0.79850231 | NA | |
| NA | | NA | |
| | 0.872240711 | | 0.923626646 |
| NA | | NA | |
| NA | | NA | |
| | 0.001509799 | | 0.007209074 |
| | 0.941277857 | NA | |
| | 0.000263023 | | 0.001621466 |
| | 0.549890433 | NA | |
| | 0.305510145 | | 0.467036125 |
| NA | | NA | |
| NA | | NA | |
| | 0.684919957 | NA | |
| | 0.941278332 | NA | |
| | 0.005116076 | | 0.019887772 |
| NA | | NA | |
| | 0.408185904 | | 0.570267118 |
| NA | | NA | |
| | 0.851456566 | | 0.910888992 |
| | 1.52E-06 | | 1.64E-05 |
| | 0.024304285 | | 0.070154085 |

| | | |
|---|---|---|
| | 0.037803645 | 0.099601766 |
| | 0.117641418 | 0.236654921 |
| | 0.002020403 | 0.009233801 |
| | 0.941276431 | NA |
| | 0.894871322 | 0.936704658 |
| | 0.955792873 | NA |
| NA | | NA |
| NA | | NA |
| | 0.94127925 | NA |
| | 0.764534404 | NA |
| | 0.955792873 | NA |
| | 0.941278424 | NA |
| | 0.124186728 | 0.246550379 |
| | 4.39E-09 | 7.19E-08 |
| NA | | NA |
| | 0.041604496 | 0.10747452 |
| | 0.826704229 | NA |
| | 0.677731101 | 0.792228448 |
| | 0.805898992 | 0.881121168 |
| | 0.022251111 | 0.065256401 |
| | 9.86E-06 | 8.95E-05 |
| | 0.839318641 | NA |
| | 0.006131277 | 0.023047484 |
| | 0.000753148 | 0.004018476 |
| | 0.662848679 | NA |
| | 0.941277857 | NA |
| | 0.324941165 | 0.487811391 |
| | 0.895244362 | NA |
| NA | | NA |
| | 0.839319108 | NA |
| NA | | NA |
| | 0.000224185 | 0.001416147 |
| NA | | NA |
| NA | | NA |
| | 0.869111681 | 0.921575098 |
| NA | | NA |
| | 0.8656684 | 0.919767844 |
| | 0.00625588 | 0.023419699 |
| | 0.840773037 | NA |
| | 0.624142788 | NA |
| | 0.007711366 | 0.027818418 |
| | 0.955791446 | NA |
| | 0.008300407 | 0.029565632 |
| | 0.005904585 | 0.022350656 |
| | 0.349733458 | 0.513563809 |
| NA | | NA |
| | 0.955791922 | NA |

| | | |
|---|---|---|
| | 0.614734813 | NA |
| NA | | NA |
| | 0.955792398 | NA |
| | 0.378700739 | NA |
| NA | | NA |
| | 0.262396148 | NA |
| | 0.94608247 | NA |
| NA | | NA |
| | 0.839318641 | NA |
| NA | | NA |
| | 0.015746042 | 0.049607848 |
| | 0.000240369 | 0.001500859 |
| | 0.955792398 | NA |
| NA | | NA |
| | 0.972449835 | 0.983356741 |
| | 0.004139932 | 0.016701469 |
| | 0.382100591 | NA |
| | 0.412798614 | 0.575057509 |
| | 0.24344732 | 0.39995397 |
| | 0.023222973 | 0.067615072 |
| | 0.170563876 | 0.310376457 |
| | 0.007387628 | 0.026866359 |
| | 0.598981892 | 0.733062335 |
| | 0.014227733 | 0.045757129 |
| NA | | NA |
| | 0.955792873 | NA |
| | 0.008226554 | 0.029347786 |
| | 0.839319107 | NA |
| | 0.274080855 | 0.433927931 |
| | 0.169805701 | 0.309401768 |
| | 0.894388618 | NA |
| | 0.698793288 | NA |
| | 0.240555107 | 0.396593184 |
| | 0.039864356 | 0.10389263 |
| | 0.778778453 | 0.863664182 |
| | 0.000253531 | 0.001571089 |
| | 0.055428575 | 0.134527474 |
| | 0.118613243 | 0.238207257 |
| | 0.93278691 | NA |
| | 0.402603833 | 0.56472252 |
| | 0.677974366 | NA |
| NA | | NA |
| | 0.079075851 | 0.176331591 |
| NA | | NA |
| | 0.139315023 | 0.268174378 |
| | 1.19E-05 | 0.000105539 |
| NA | | NA |

| | | |
|---|---|---|
| | 0.175982833 | NA |
| NA | | NA |
| | 0.588459205 | NA |
| NA | | NA |
| | 0.875773195 | NA |
| | 0.343761629 | 0.507329756 |
| | 0.00146164 | 0.007016078 |
| | 0.013537856 | 0.043942187 |
| | 0.941278332 | NA |
| | 0.006052107 | 0.02281835 |
| | 0.205618874 | 0.354288434 |
| | 0.113997347 | 0.231060899 |
| | 0.744129083 | NA |
| | 0.655451144 | 0.775091819 |
| | 0.102385114 | 0.213563271 |
| | 0.65613139 | NA |
| | 0.501091313 | 0.653604528 |
| NA | | NA |
| | 2.34E-05 | 0.000193431 |
| | 0.088275453 | NA |
| | 1.32E-08 | 2.03E-07 |
| NA | | NA |
| NA | | NA |
| | 0.000345118 | 0.002056711 |
| | 0.49863987 | 0.651585298 |
| | 0.184290538 | 0.327227407 |
| | 0.941278332 | NA |
| | 0.015412823 | 0.048730274 |
| | 0.706652184 | 0.812901033 |
| | 0.955791446 | NA |
| | 0.810263929 | 0.883701523 |
| | 0.082804888 | 0.18227337 |
| | 0.90351762 | NA |
| | 0.903712159 | NA |
| | 9.56E-16 | 3.08E-14 |
| | 2.84E-05 | 0.000229923 |
| | 0.639533223 | 0.76329288 |
| | 0.252806805 | 0.410625602 |
| | 0.827714351 | 0.895308202 |
| | 0.006627671 | 0.024560457 |
| | 0.619338549 | 0.748489268 |
| | 0.18342317 | 0.326125219 |
| NA | | NA |
| | 0.020449648 | 0.060918964 |
| | 0.091604447 | 0.196642231 |
| | 0.684974506 | 0.797679104 |
| | 0.955821178 | 0.973031764 |

| | | |
|---|---|---|
| | 0.013692809 | 0.044332002 |
| NA | | NA |
| | 0.711227782 | NA |
| | 0.839319107 | NA |
| | 0.000120785 | 0.000828035 |
| | 0.075017908 | NA |
| | 0.045262001 | 0.114560415 |
| | 0.1212517 | 0.242259127 |
| | 0.008555359 | 0.030275743 |
| | 0.000306218 | 0.00185508 |
| | 0.564400285 | 0.706412977 |
| | 0.307293734 | 0.468851733 |
| | 0.139064682 | 0.267805861 |
| | 0.38182018 | 0.54432702 |
| NA | | NA |
| NA | | NA |
| | 0.839319108 | NA |
| NA | | NA |
| | 0.867698789 | 0.920979557 |
| | 0.093341891 | 0.199329238 |
| | 0.899619119 | 0.93958188 |
| NA | | NA |
| | 0.97488596 | NA |
| | 0.094660229 | 0.201367662 |
| | 0.88243826 | 0.929664669 |
| | 0.003268578 | 0.01378029 |
| | 0.449260881 | 0.609409443 |
| | 0.659334935 | 0.778331108 |
| | 0.295070358 | 0.456447125 |
| | 0.000362718 | 0.002143353 |
| | 0.001403309 | 0.006787554 |
| NA | | NA |
| | 3.11E-09 | 5.19E-08 |
| | 0.941278332 | NA |
| | 0.447788262 | 0.607987046 |
| | 0.663220839 | NA |
| | 0.662848679 | NA |
| | 0.526351652 | 0.675448675 |
| NA | | NA |
| | 0.639537097 | NA |
| NA | | NA |
| NA | | NA |
| NA | | NA |
| NA | | NA |
| NA | | NA |
| NA | | NA |
| | 0.93838321 | NA |

| | | |
|---|---|---|
| | 0.955791922 | NA |
| NA | | NA |
| NA | | NA |
| NA | | NA |
| NA | | NA |
| NA | | NA |
| | 0.81208395 | NA |
| | 0.312173844 | 0.474177527 |
| | 0.859843569 | NA |
| NA | | NA |
| NA | | NA |
| NA | | NA |
| NA | | NA |
| NA | | NA |
| | 0.19690128 | 0.343696411 |
| | 0.839319108 | NA |
| | 0.310148034 | NA |
| | 0.884297813 | NA |
| | 0.839318641 | NA |
| NA | | NA |
| | 0.434246005 | 0.595490201 |
| | 0.051646165 | 0.127189034 |
| | 0.941279749 | NA |
| | 0.036228497 | 0.096293768 |
| | 0.945558632 | 0.967176044 |
| | 0.034126225 | 0.091751091 |
| | 0.177821536 | 0.319321374 |
| | 0.900923403 | 0.940203267 |
| | 0.002046981 | 0.009324829 |
| | 0.760636383 | 0.851143209 |
| NA | | NA |
| NA | | NA |
| | 0.955791922 | NA |
| | 0.783740227 | NA |
| | 0.941278332 | NA |
| NA | | NA |
| | 3.92E-05 | 0.000305211 |
| NA | | NA |
| NA | | NA |
| NA | | NA |
| NA | | NA |
| | 0.627016108 | NA |
| | 0.941278838 | NA |
| NA | | NA |
| NA | | NA |
| NA | | NA |
| NA | | NA |

| | |
|---|---|
| 0.517589084 | 0.668311565 |
| NA | NA |
| NA | NA |
| 0.144900029 | 0.276155857 |
| 0.958714477 | 0.974684862 |
| 0.059608747 | 0.14236478 |
| 0.687484641 | 0.799428864 |
| 5.97E-10 | 1.09E-08 |
| 0.18317324 | 0.325826878 |
| 0.676376769 | NA |
| 0.1665998 | 0.305509772 |
| 0.632858416 | 0.758703335 |
| 0.47197239 | 0.629419926 |
| 0.29470022 | 0.455988018 |
| 0.173131448 | 0.313589127 |
| 0.718744612 | 0.821055414 |
| 1.58E-05 | 0.000136511 |
| 3.92E-06 | 3.90E-05 |
| 0.001722759 | 0.008086197 |
| 0.001917387 | 0.008834232 |
| 0.012029271 | 0.039977758 |
| 0.086365202 | 0.188333903 |
| 0.955792398 | NA |
| 0.116634616 | 0.235046746 |
| 0.395320553 | NA |
| 0.007921073 | 0.028424149 |
| 0.000244551 | 0.001523375 |
| NA | NA |
| 0.312774237 | 0.474695682 |
| 0.829364788 | 0.896400903 |
| 0.120602537 | 0.241347032 |
| 0.725673533 | 0.826750978 |
| 0.003470793 | 0.014455261 |
| 0.878172278 | 0.926929338 |
| 0.757043143 | 0.848039387 |
| 2.55E-05 | 0.000209645 |
| NA | NA |
| 0.048025878 | 0.119948787 |
| 0.018928616 | 0.057422752 |
| 0.064671587 | 0.151761127 |
| 0.277821078 | 0.437492379 |
| 0.791594723 | NA |
| 0.024918274 | 0.071492858 |
| 0.591243096 | 0.727011839 |
| 0.735320032 | 0.833234344 |
| 0.583689979 | 0.720889834 |
| NA | NA |

| | | |
|---|---|---|
| | 0.955792873 | NA |
| | 0.260326906 | 0.418899367 |
| | 0.010563661 | 0.035986296 |
| | 0.340949254 | 0.504492985 |
| | 0.903922638 | 0.941698566 |
| | 0.853595154 | NA |
| | 0.506134538 | NA |
| | 0.307308457 | 0.468851733 |
| | 0.475653156 | 0.632839375 |
| | 0.955791922 | NA |
| NA | | NA |
| | 0.856110299 | 0.913568139 |
| | 0.446665612 | NA |
| | 0.160241799 | 0.296953853 |
| | 2.92E-05 | 0.00023538 |
| | 0.910618638 | 0.94586711 |
| | 0.11012239 | 0.225299392 |
| | 0.805602678 | NA |
| | 0.600502497 | NA |
| | 0.005185131 | 0.020126586 |
| | 0.294343737 | NA |
| NA | | NA |
| NA | | NA |
| | 0.016523294 | 0.05154488 |
| NA | | NA |
| | 0.000566088 | 0.003153294 |
| | 0.024645011 | 0.07089685 |
| | 0.090685434 | 0.195335279 |
| | 0.073720114 | NA |
| | 0.531272031 | 0.67996602 |
| NA | | NA |
| | 0.072389766 | 0.165107922 |
| | 0.217925177 | 0.369237975 |
| | 0.095485082 | 0.202651443 |
| | 0.839320041 | NA |
| | 0.77691363 | 0.862550475 |
| | 0.673247071 | NA |
| | 0.130738161 | 0.25584393 |
| | 0.103277906 | 0.214804328 |
| | 0.020048464 | 0.059971539 |
| | 0.050866187 | NA |
| | 0.822151266 | NA |
| | 0.881576087 | 0.929274887 |
| | 0.710423764 | NA |
| | 0.729099982 | NA |
| | 0.002858517 | 0.012314648 |
| | 0.941278332 | NA |

| | |
|---|---|
| 0.058494195 | 0.140138784 |
| 1.50E-08 | 2.29E-07 |
| 0.066070901 | 0.154269136 |
| 0.35831882 | 0.521920601 |
| 0.651703126 | 0.772647082 |
| 0.113458038 | 0.23036153 |
| 0.601493069 | 0.735134174 |
| 0.072520134 | 0.165305901 |
| 0.89720862 | NA |
| 7.16E-05 | 0.000521048 |
| NA | NA |
| 2.75E-06 | 2.81E-05 |
| NA | NA |
| 0.456239422 | 0.615962038 |
| 0.114975948 | 0.232570537 |
| 0.005794447 | 0.022000227 |
| 6.74E-05 | 0.000493904 |
| 0.001745477 | 0.008168668 |
| 0.853594685 | NA |
| 0.317238575 | 0.479724698 |
| 0.000105962 | 0.000735736 |
| NA | NA |
| NA | NA |
| 0.839318641 | NA |
| 0.144407112 | 0.275503212 |
| 0.0199013 | 0.059598733 |
| 0.324798227 | 0.487719853 |
| NA | NA |
| 0.941280761 | NA |
| NA | NA |
| 0.839318641 | NA |
| 0.833357167 | NA |
| 0.81338117 | NA |
| 0.839319108 | NA |
| NA | NA |
| NA | NA |
| NA | NA |
| NA | NA |
| NA | NA |
| NA | NA |
| NA | NA |
| NA | NA |
| NA | NA |
| 0.694707276 | NA |
| 0.910667622 | NA |
| 0.560691476 | NA |
| 0.800391721 | NA |

| | | |
|---|---|---|
| NA | | NA |
| NA | | NA |
| | 0.853576605 | NA |
| NA | | NA |
| | 0.821096032 | NA |
| | 0.910991996 | NA |
| | 3.20E-09 | 5.35E-08 |
| | 0.728021993 | 0.828712553 |
| | 0.941278788 | NA |
| | 0.94020055 | NA |
| | 0.715826207 | 0.81937556 |
| | 0.773734508 | 0.860500363 |
| | 1.31E-08 | 2.01E-07 |
| | 4.31E-07 | 5.16E-06 |
| | 0.023804668 | 0.068946059 |
| | 0.941277857 | NA |
| | 0.370965291 | 0.534227726 |
| | 0.817417153 | NA |
| | 0.019167781 | 0.05794152 |
| | 6.84E-05 | 0.000499826 |
| | 0.941278332 | NA |
| | 0.222565747 | 0.374965311 |
| | 0.334748233 | 0.49797984 |
| | 0.026922711 | 0.076223905 |
| | 0.439463868 | 0.60041129 |
| | 0.358759607 | 0.522306941 |
| | 0.0011527 | 0.005763984 |
| | 0.728678906 | NA |
| | 0.650185181 | NA |
| | 0.963464561 | 0.977841478 |
| | 0.941278332 | NA |
| NA | | NA |
| | 0.414145773 | 0.576192289 |
| | 0.423269034 | 0.58461563 |
| NA | | NA |
| | 0.335191357 | NA |
| | 0.941280349 | NA |
| | 0.096355611 | 0.203989704 |
| | 0.661454499 | NA |
| | 8.10E-05 | 0.000580957 |
| | 0.820627707 | 0.890678686 |
| | 0.058335755 | 0.139857739 |
| | 0.175601169 | 0.316851507 |
| | 0.0679138 | 0.157490672 |
| | 0.725028456 | NA |
| | 0.708530354 | 0.814056764 |
| | 2.64E-06 | 2.72E-05 |

| | |
|---|---|
| 0.01238245 | 0.040905909 |
| 0.40165432 | 0.563925563 |
| 0.277655877 | 0.437434077 |
| NA | NA |
| 0.430152173 | 0.591409544 |
| 0.925753076 | NA |
| 1.81E-05 | 0.000153261 |
| 0.202770925 | NA |
| 0.569928539 | NA |
| 0.529370974 | 0.678445397 |
| 0.358744763 | 0.522306941 |
| 0.446861653 | 0.607254912 |
| 0.030137112 | 0.083312839 |
| 0.003923128 | 0.015962101 |
| 0.079865971 | 0.177627531 |
| NA | NA |
| 0.437011568 | 0.597944132 |
| 0.789350543 | NA |
| NA | NA |
| 0.839319108 | NA |
| NA | NA |
| NA | NA |
| 0.072555516 | 0.165339051 |
| NA | NA |
| NA | NA |
| NA | NA |
| NA | NA |
| NA | NA |
| 0.839319108 | NA |
| NA | NA |
| NA | NA |
| NA | NA |
| NA | NA |
| NA | NA |
| 0.127741497 | 0.251613962 |
| NA | NA |
| NA | NA |
| NA | NA |
| NA | NA |
| 0.941278332 | NA |
| NA | NA |
| NA | NA |
| 0.941277857 | NA |
| NA | NA |
| NA | NA |
| 0.954694081 | NA |
| 3.57E-08 | 5.14E-07 |

| | | |
|---|---|---|
| | 0.000841991 | 0.004413358 |
| NA | NA | |
| | 1.70E-06 | 1.82E-05 |
| | 0.249801147 | 0.407300104 |
| | 0.632206039 | 0.758226753 |
| | 0.490939349 | 0.645087045 |
| | 0.940773688 | NA |
| | 0.485365141 | 0.640782134 |
| | 0.352024039 | 0.51562315 |
| | 0.88314754 | 0.930159642 |
| | 0.014021616 | 0.045176245 |
| | 0.199367074 | 0.346779823 |
| | 0.014621961 | 0.046728239 |
| | 0.202946825 | 0.35110294 |
| | 0.839319574 | NA |
| | 0.967628393 | 0.980522415 |
| NA | NA | |
| | 0.055612626 | 0.13485042 |
| | 0.177689362 | 0.319245335 |
| | 0.200639181 | 0.348254488 |
| | 0.971810578 | NA |
| | 0.001468717 | 0.007045756 |
| | 0.839319108 | NA |
| NA | NA | |
| NA | NA | |
| NA | NA | |
| | 0.342481065 | 0.505952993 |
| | 0.669697877 | 0.78623601 |
| | 0.213143989 | 0.363830552 |
| | 0.007346112 | 0.026734278 |
| | 0.183239702 | 0.325920744 |
| | 0.426888601 | 0.588383953 |
| | 0.917318227 | NA |
| | 0.575193942 | NA |
| | 0.113062514 | 0.229790901 |
| | 0.026865254 | 0.076099603 |
| NA | NA | |
| | 0.345886134 | 0.50959062 |
| | 0.128032965 | 0.252083765 |
| | 0.001903933 | 0.008782442 |
| | 0.200771184 | 0.348364259 |
| | 0.629387368 | 0.755989005 |
| | 0.735010743 | NA |
| | 0.843788904 | NA |
| | 0.739991096 | NA |
| | 0.919049593 | NA |
| NA | NA | |

| | |
|---|---|
| 0.812243254 | 0.885413965 |
| 0.865468469 | 0.919767844 |
| NA | NA |
| 0.666537593 | 0.783815501 |
| 0.000728004 | 0.003908852 |
| 0.139175897 | 0.267971581 |
| NA | NA |
| 0.000227131 | 0.00143058 |
| 0.002614352 | 0.011453491 |
| 0.427028737 | 0.58851947 |
| 0.826999637 | 0.894978193 |
| 0.416269051 | 0.578031384 |
| 0.941276431 | NA |
| 0.066741553 | 0.155453807 |
| 0.570009197 | NA |
| 0.002980892 | 0.012751833 |
| 0.458890004 | 0.618278042 |
| 0.044579345 | 0.113217838 |
| 0.051818127 | 0.127546542 |
| 0.246956025 | 0.403989506 |
| 0.018271568 | 0.05582423 |
| 0.428841905 | NA |
| 0.888328429 | 0.93326492 |
| 0.280812407 | 0.440919382 |
| 0.215147908 | NA |
| 0.006296204 | 0.023552137 |
| 0.002166926 | 0.009770254 |
| NA | NA |
| 0.255708545 | 0.413757633 |
| 0.860909218 | 0.916839428 |
| 0.915200469 | 0.949011134 |
| 0.133999719 | 0.260502347 |
| 8.96E-23 | 4.52E-21 |
| NA | NA |
| 7.00E-06 | 6.57E-05 |
| 0.08492615 | 0.186082318 |
| 0.614707698 | NA |
| 5.81E-07 | 6.75E-06 |
| 1.99E-17 | 7.11E-16 |
| 0.066974381 | 0.155828368 |
| 0.0015236 | 0.007266025 |
| 0.000751505 | 0.004012412 |
| 0.426843335 | 0.588355633 |
| 1.74E-05 | 0.000148474 |
| 0.87342832 | NA |
| 0.01095645 | 0.037106633 |
| 0.009319782 | 0.032485888 |

| | | |
|---|---|---|
| | 0.05044123 | 0.124680492 |
| | 0.412790333 | 0.575057509 |
| | 0.338415837 | 0.501930069 |
| | 0.27943171 | 0.43924406 |
| | 0.661817855 | 0.78014028 |
| | 0.004250746 | 0.01706464 |
| | 0.812401156 | 0.88542389 |
| | 0.671956238 | NA |
| | 5.71E-12 | 1.33E-10 |
| | 0.000312761 | 0.001887508 |
| | 0.58005876 | 0.717932432 |
| | 0.276468536 | 0.436286157 |
| | 0.004842824 | 0.019011859 |
| | 0.941277857 | NA |
| | 0.48280261 | 0.63882419 |
| | 0.195165041 | 0.341467091 |
| NA | | NA |
| | 0.941278332 | NA |
| | 0.538920407 | 0.68636008 |
| | 0.580339859 | 0.718205652 |
| | 1.01E-10 | 2.01E-09 |
| | 0.646339483 | NA |
| | 0.863702596 | 0.918581286 |
| | 0.082453316 | 0.1817556 |
| NA | | NA |
| NA | | NA |
| | 0.196486976 | 0.343174567 |
| | 0.048092181 | 0.120063992 |
| NA | | NA |
| | 6.60E-10 | 1.20E-08 |
| | 3.50E-05 | 0.000276491 |
| | 2.79E-05 | 0.000226266 |
| | 0.000286774 | 0.001752003 |
| | 0.980112151 | 0.987685995 |
| | 0.001076316 | 0.005440323 |
| NA | | NA |
| | 0.030669516 | 0.08439258 |
| | 0.37035464 | NA |
| | 0.006208779 | 0.023272654 |
| | 0.827225956 | 0.895105341 |
| NA | | NA |
| NA | | NA |
| | 1.50E-07 | 1.95E-06 |
| | 0.161545187 | 0.298602045 |
| | 0.053461404 | 0.130672453 |
| | 0.42618668 | 0.58759486 |
| NA | | NA |

| | | |
|---|---|---|
| | 0.839318641 | NA |
| | 0.84929261 | NA |
| | 0.391630492 | 0.554130838 |
| | 0.839319574 | NA |
| | 0.516130051 | 0.666934173 |
| | 9.26E-05 | 0.00065516 |
| | 0.739991096 | NA |
| | 0.941280558 | NA |
| | 0.102025768 | 0.212935034 |
| | 0.026348523 | 0.074793983 |
| | 0.198751808 | 0.346038369 |
| NA | | NA |
| | 1.57E-05 | 0.000135855 |
| | 0.501268703 | 0.653728435 |
| | 0.121754345 | 0.243018609 |
| | 0.839318641 | NA |
| | 0.870049735 | NA |
| | 0.125561726 | 0.248471916 |
| | 0.494429405 | 0.647540921 |
| | 0.941278332 | NA |
| | 3.99E-08 | 5.67E-07 |
| | 0.164498583 | 0.30286164 |
| | 5.05E-13 | 1.30E-11 |
| | 0.547537569 | 0.693660274 |
| | 0.648259448 | 0.770105487 |
| | 0.475552574 | 0.632740923 |
| | 0.000859726 | 0.00448951 |
| | 0.000879712 | 0.004574809 |
| | 0.017565052 | 0.05407117 |
| | 0.549570446 | 0.695199605 |
| | 0.088104982 | 0.191199379 |
| NA | | NA |
| | 0.115144618 | 0.232770247 |
| | 0.042504541 | 0.10920694 |
| | 0.737216099 | 0.83435013 |
| | 0.475013649 | 0.632341722 |
| | 0.062786452 | 0.148199438 |
| | 0.839319108 | NA |
| | 0.03482477 | 0.093271153 |
| NA | | NA |
| | 0.005537523 | 0.021176753 |
| | 0.292322182 | 0.453426099 |
| | 0.242838698 | 0.399309981 |
| NA | | NA |
| | 0.000206411 | 0.001317507 |
| | 4.03E-07 | 4.85E-06 |
| | 3.37E-42 | 3.75E-40 |

| | | |
|---|---|---|
| | 0.661496085 | NA |
| | 0.634782068 | 0.759937772 |
| | 0.041544333 | 0.107331034 |
| | 0.322946224 | 0.485643545 |
| | 0.000994372 | 0.005085528 |
| | 0.049907134 | 0.123571164 |
| | 0.175159863 | 0.316287269 |
| | 0.006847542 | 0.025222099 |
| | 0.286904916 | 0.447445341 |
| | 0.158167812 | 0.294232923 |
| | 0.8091288 | NA |
| | 0.446991947 | 0.607326964 |
| NA | | NA |
| NA | | NA |
| | 0.00619238 | 0.023229477 |
| | 0.091982531 | 0.197145376 |
| | 0.839319108 | NA |
| | 3.00E-07 | 3.70E-06 |
| | 0.784750658 | NA |
| | 0.741328157 | 0.837390866 |
| | 0.124482433 | 0.246931484 |
| | 0.083701966 | 0.183907892 |
| | 0.981974441 | 0.98868314 |
| | 4.61E-15 | 1.40E-13 |
| | 0.624946402 | 0.752858299 |
| | 0.002077298 | 0.009441254 |
| | 4.15E-08 | 5.87E-07 |
| | 0.402677311 | 0.564744214 |
| | 1.23E-06 | 1.35E-05 |
| | 0.459346666 | 0.618621801 |
| | 0.809897202 | 0.883463483 |
| | 0.404196394 | 0.566222269 |
| | 0.002979356 | 0.012749845 |
| | 0.002417433 | 0.01073047 |
| | 0.011912131 | 0.039677098 |
| | 0.848967055 | 0.909165565 |
| | 0.008213895 | 0.029311413 |
| | 0.024544039 | 0.070683268 |
| | 0.788872197 | NA |
| | 0.043633907 | 0.111363312 |
| | 0.940337222 | NA |
| | 1.06E-22 | 5.28E-21 |
| | 0.817417153 | NA |
| | 0.468894443 | NA |
| NA | | NA |
| | 3.95E-29 | 2.67E-27 |
| | 3.08E-07 | 3.78E-06 |

| | | |
|---|---|---|
| | 0.955792398 | NA |
| | 0.952431307 | NA |
| NA | | NA |
| | 0.076410719 | 0.171783482 |
| | 0.941278332 | NA |
| | 0.941278332 | NA |
| | 0.007839069 | 0.028202164 |
| | 0.072570376 | 0.165357083 |
| | 0.787578729 | 0.869437823 |
| | 0.558893044 | 0.702438074 |
| | 0.150766703 | 0.284399196 |
| NA | | NA |
| | 0.232409374 | 0.386839715 |
| | 0.349261104 | 0.513218471 |
| | 0.635097181 | 0.760047419 |
| | 0.841425253 | NA |
| | 0.351222687 | 0.515109595 |
| NA | | NA |
| | 0.180190033 | 0.322157527 |
| | 0.829517397 | 0.896499875 |
| | 0.639025801 | 0.762946033 |
| | 0.000397619 | 0.002324766 |
| | 0.941279438 | NA |
| | 0.114651056 | 0.232150102 |
| | 0.356165258 | 0.519833546 |
| | 0.188649516 | 0.333128258 |
| | 0.281618836 | 0.441806719 |
| | 0.839319108 | NA |
| | 0.371995148 | 0.535343845 |
| | 0.07938049 | 0.176845229 |
| | 0.00632332 | 0.023627578 |
| | 3.33E-08 | 4.82E-07 |
| NA | | NA |
| NA | | NA |
| NA | | NA |
| NA | | NA |
| NA | | NA |
| | 0.701973148 | 0.809749194 |
| | 0.190420982 | 0.335187119 |
| | 0.01512209 | 0.048022014 |
| | 0.911710338 | NA |
| | 0.322862325 | 0.485578734 |
| | 0.685788586 | NA |
| | 1.16E-09 | 2.04E-08 |
| NA | | NA |
| NA | | NA |
| | 0.440220692 | 0.600913741 |

| | |
|---|---|
| 7.70E-06 | 7.17E-05 |
| 0.360641551 | 0.524373256 |
| 0.389799305 | 0.552360723 |
| 0.670344114 | 0.786689247 |
| 0.002982042 | 0.012754458 |
| NA | NA |
| 1.17E-20 | 5.21E-19 |
| 0.010384283 | 0.035497157 |
| 0.068879857 | 0.159219031 |
| 0.000104917 | 0.000729756 |
| 0.552987085 | 0.698038772 |
| 4.53E-46 | 5.70E-44 |
| NA | NA |
| 0.031109541 | 0.085366532 |
| 0.88200323 | 0.929459335 |
| 0.318349436 | 0.480812253 |
| 0.140907188 | 0.270517252 |
| 0.018399823 | 0.056118988 |
| NA | NA |
| 1.95E-24 | 1.06E-22 |
| 0.524323749 | 0.673899987 |
| 0.456018171 | 0.61573318 |
| 8.73E-20 | 3.65E-18 |
| NA | NA |
| 0.726802277 | NA |
| 0.043167228 | 0.110468375 |
| 0.261232725 | 0.419948636 |
| 4.28E-13 | 1.11E-11 |
| 0.025609198 | 0.073043755 |
| 0.017434491 | 0.053739413 |
| 0.523757467 | NA |
| 4.37E-09 | 7.17E-08 |
| 2.35E-06 | 2.45E-05 |
| NA | NA |
| 0.158927007 | NA |
| 0.014240177 | 0.045778592 |
| 0.14145336 | 0.271272119 |
| 0.943405629 | 0.96592854 |
| 0.2094937 | 0.35914499 |
| 0.557905576 | NA |
| 0.941278332 | NA |
| 0.000178085 | 0.001161944 |
| 0.43949945 | 0.60041129 |
| 0.46611275 | 0.624228644 |
| 0.015300446 | 0.048460574 |
| 0.853596091 | NA |
| NA | NA |

| | |
|---|---|
| 0.002718479 | 0.011833439 |
| 0.631186348 | 0.757347253 |
| 0.031553801 | 0.086356626 |
| NA | NA |
| 0.741178832 | NA |
| 0.590835994 | 0.726752727 |
| NA | NA |
| 0.046983958 | 0.117928099 |
| 0.200814669 | 0.348400793 |
| 0.055105138 | 0.133892659 |
| 0.884488356 | 0.930962684 |
| 0.876317894 | NA |
| NA | NA |
| 0.89365632 | NA |
| 0.717252837 | 0.82021693 |
| 1.09E-41 | 1.20E-39 |
| 0.359445696 | 0.523017878 |
| 0.752334005 | 0.844632517 |
| 0.002430083 | 0.010776838 |
| 0.955792873 | NA |
| NA | NA |
| 0.276458753 | 0.436286157 |
| 0.146588693 | NA |
| 2.82E-08 | 4.13E-07 |
| 0.00084492 | 0.004425787 |
| 0.234345016 | 0.389109178 |
| 7.25E-07 | 8.28E-06 |
| 0.083724952 | 0.183941417 |
| 0.044665861 | 0.11332873 |
| NA | NA |
| 0.484329551 | 0.640196144 |
| 0.136426588 | 0.26405425 |
| 0.060448576 | 0.143865313 |
| 0.629725461 | 0.756242453 |
| 0.007697179 | 0.02777566 |
| 0.801328729 | NA |
| NA | NA |
| 5.57E-11 | 1.16E-09 |
| 0.575082584 | 0.714074899 |
| 0.000657039 | 0.003577863 |
| 0.108848185 | 0.223392764 |
| 0.205098634 | 0.353724924 |
| 0.338106316 | 0.501596019 |
| 2.80E-06 | 2.87E-05 |
| 0.871591612 | 0.923330406 |
| 0.130777231 | 0.255899347 |
| 5.74E-06 | 5.51E-05 |

| | |
|---|---|
| 0.001486015 | 0.007118161 |
| 0.001847076 | 0.008551666 |
| NA | NA |
| 1.18E-24 | 6.54E-23 |
| NA | NA |
| 2.02E-11 | 4.42E-10 |
| 0.839318641 | NA |
| NA | NA |
| NA | NA |
| 0.072479258 | 0.16522855 |
| 0.087045518 | 0.189441633 |
| NA | NA |
| 0.313223788 | 0.475165877 |
| 0.002874278 | 0.012369115 |
| 0.982708412 | NA |
| 0.341370257 | 0.50489622 |
| 0.043284909 | 0.110710016 |
| 0.955792873 | NA |
| 0.58708966 | 0.72358679 |
| NA | NA |
| 0.481833848 | 0.638208639 |
| 0.854044483 | NA |
| 0.891904586 | NA |
| 0.00241776 | 0.01073047 |
| 0.10672371 | 0.220277293 |
| 0.002036361 | 0.009287116 |
| 0.688909865 | NA |
| 0.104031626 | 0.215975645 |
| NA | NA |
| 0.009861133 | 0.034001839 |
| 7.30E-06 | 6.83E-05 |
| NA | NA |
| 3.24E-08 | 4.70E-07 |
| 0.000577976 | 0.00320825 |
| 0.778101595 | 0.863416888 |
| 0.61281069 | 0.743634467 |
| 2.49E-07 | 3.12E-06 |
| 0.000406828 | 0.002369299 |
| 0.899298755 | NA |
| 1.53E-07 | 1.99E-06 |
| 0.425600361 | NA |
| 0.287743315 | 0.448370958 |
| 0.010548685 | 0.035950716 |
| NA | NA |
| NA | NA |
| 9.85E-05 | 0.000692088 |
| 0.003329467 | 0.013989051 |

| | |
|---|---|
| 4.41E-28 | 2.86E-26 |
| 0.989016921 | 0.993022052 |
| 0.132400874 | 0.258121518 |
| NA | NA |
| 0.00141858 | 0.006844535 |
| NA | NA |
| 0.769125119 | 0.857023917 |
| 0.305237883 | 0.466903274 |
| 0.538572368 | 0.68617383 |
| 0.001401808 | 0.006781675 |
| 0.368203616 | 0.531279248 |
| NA | NA |
| NA | NA |
| NA | NA |
| 0.375061512 | NA |
| 0.786981812 | NA |
| 0.000140093 | 0.000940886 |
| 5.69E-92 | 1.76E-89 |
| 2.50E-56 | 3.97E-54 |
| 2.42E-36 | 2.18E-34 |
| 0.00053593 | 0.003012094 |
| 1.73E-08 | 2.60E-07 |
| 0.839320041 | NA |
| NA | NA |
| 0.512797575 | NA |
| 0.941278332 | NA |
| NA | NA |
| NA | NA |
| 0.159131374 | NA |
| 0.038514691 | 0.101005583 |
| 0.493122418 | 0.646825735 |
| 0.011697171 | 0.039124266 |
| NA | NA |
| 0.847298963 | 0.908223719 |
| 0.218461925 | 0.36977889 |
| NA | NA |
| NA | NA |
| 0.839320041 | NA |
| NA | NA |
| 0.000282767 | 0.00173063 |
| NA | NA |
| 0.690189067 | NA |
| 0.167137934 | 0.306141868 |
| 0.275684201 | 0.435519885 |
| 0.899878593 | NA |
| 0.532282014 | 0.680783261 |
| 0.185717217 | 0.329122269 |

| | | |
|---|---|---|
| | 0.773349075 | NA |
| | 0.533609192 | 0.681881296 |
| NA | | NA |
| | 0.300582236 | 0.46205492 |
| | 0.969976101 | 0.982023444 |
| | 0.395450367 | 0.558009819 |
| | 0.440373046 | 0.601087237 |
| | 0.969927307 | 0.982023444 |
| | 0.000126604 | 0.000862954 |
| | 0.73015306 | 0.830051037 |
| NA | | NA |
| NA | | NA |
| | 0.956972319 | 0.973537425 |
| | 0.000339679 | 0.002032438 |
| | 0.682616021 | 0.796065882 |
| | 0.416945456 | 0.578490542 |
| | 0.014393647 | 0.04616596 |
| | 0.947567487 | 0.968065985 |
| | 0.00476348 | 0.018771582 |
| | 0.955792398 | NA |
| | 0.004520733 | 0.017957691 |
| | 0.073095244 | 0.166266533 |
| | 0.267579601 | NA |
| | 0.109254853 | 0.223986565 |
| | 0.250468789 | 0.408053259 |
| | 0.838197868 | NA |
| NA | | NA |
| | 0.839319416 | NA |
| | 1.32E-07 | 1.73E-06 |
| | 0.071647662 | 0.1638979 |
| | 0.790874995 | NA |
| | 6.70E-13 | 1.71E-11 |
| | 0.326909521 | 0.48969689 |
| | 0.004824921 | 0.018960339 |
| | 0.169487141 | 0.309152651 |
| NA | | NA |
| | 0.437901458 | 0.598778107 |
| | 1.83E-06 | 1.95E-05 |
| | 0.553612013 | 0.698604238 |
| | 0.005051039 | 0.019689622 |
| NA | | NA |
| | 0.352374684 | 0.515977991 |
| NA | | NA |
| | 0.521951948 | 0.672090432 |
| | 0.001147533 | 0.005742972 |
| | 0.202142121 | 0.350224689 |
| | 3.96E-05 | 0.000308325 |

| | |
|---|---|
| 0.352826672 | 0.516449201 |
| 1.40E-05 | 0.000122279 |
| NA | NA |
| 0.686499034 | 0.798856658 |
| 0.000874232 | 0.004552631 |
| 0.935650667 | NA |
| 0.791017952 | NA |
| 0.677515187 | 0.792053855 |
| 0.737704719 | NA |
| 0.955791922 | NA |
| 0.507271563 | NA |
| 0.31756444 | 0.480087309 |
| NA | NA |
| 0.000284506 | 0.001740377 |
| 0.660218758 | NA |
| 0.626221902 | 0.753775851 |
| NA | NA |
| 0.955531966 | 0.972945408 |
| 0.909892212 | NA |
| NA | NA |
| 0.941278332 | NA |
| 0.787146561 | 0.869202618 |
| 0.543862647 | NA |
| NA | NA |
| 0.464134964 | 0.622666014 |
| NA | NA |
| 0.745862835 | NA |
| 0.073091554 | 0.166266533 |
| 0.47735932 | NA |
| 0.839319107 | NA |
| 0.833589206 | 0.899282818 |
| 0.839318641 | NA |
| 1.71E-07 | 2.21E-06 |
| 0.351788597 | NA |
| 0.001314884 | 0.006425144 |
| 0.765431005 | 0.85429939 |
| 0.333747707 | 0.497019314 |
| 0.16384034 | 0.301976751 |
| 0.027240994 | 0.076966916 |
| 7.61E-08 | 1.04E-06 |
| NA | NA |
| 0.785457679 | 0.868102883 |
| 0.000175136 | 0.001145533 |
| 0.643357984 | NA |
| 0.536863379 | 0.684839603 |
| 0.549141889 | 0.69480519 |
| 0.06906974 | NA |

| | | |
|---|---|---|
| | 7.47E-05 | 0.000540778 |
| | 0.839318641 | NA |
| | 0.941279268 | NA |
| | 6.76E-14 | 1.87E-12 |
| NA | | NA |
| NA | | NA |
| NA | | NA |
| | 0.191431525 | 0.336567672 |
| | 0.092444086 | NA |
| | 0.941278332 | NA |
| | 0.853596091 | NA |
| | 0.06167575 | 0.14617208 |
| | 0.002236506 | 0.010049709 |
| | 0.771480543 | 0.858807438 |
| | 0.009053462 | 0.031736453 |
| | 0.749179844 | 0.842522024 |
| | 0.014914741 | 0.047472674 |
| | 0.141426721 | 0.27125212 |
| | 0.268501341 | NA |
| | 0.88902284 | 0.933455095 |
| | 0.009816192 | 0.033900245 |
| | 4.99E-05 | 0.00037851 |
| NA | | NA |
| | 0.917713131 | 0.950457225 |
| | 0.406454823 | 0.568277552 |
| | 0.991165052 | 0.994087701 |
| | 0.959863874 | NA |
| NA | | NA |
| NA | | NA |
| | 1.19E-09 | 2.10E-08 |
| | 0.437008809 | 0.597944132 |
| | 0.56125686 | NA |
| | 4.44E-07 | 5.29E-06 |
| NA | | NA |
| NA | | NA |
| | 0.151112506 | 0.284870891 |
| | 0.015846496 | 0.049842299 |
| | 0.963576585 | 0.977841478 |
| | 0.432073717 | 0.593297485 |
| | 0.910481257 | 0.94586711 |
| | 0.839319109 | NA |
| | 0.219342658 | 0.370847702 |
| NA | | NA |
| | 0.257568452 | 0.415749647 |
| | 0.503667525 | NA |
| NA | | NA |
| | 0.633249287 | NA |

| | |
|---|---|
| NA | NA |
| 0.778816109 | 0.863664182 |
| 6.75E-15 | 2.02E-13 |
| NA | NA |
| 0.240439082 | 0.39648961 |
| 0.860355571 | 0.916609361 |
| 0.382281698 | NA |
| 0.755626499 | NA |
| 0.000588741 | 0.003258888 |
| NA | NA |
| 0.732493456 | 0.83123434 |
| NA | NA |
| NA | NA |
| NA | NA |
| NA | NA |
| NA | NA |
| NA | NA |
| NA | NA |
| 0.438732595 | 0.599659984 |
| NA | NA |
| NA | NA |
| NA | NA |
| 0.584803957 | 0.721733424 |
| NA | NA |
| NA | NA |
| NA | NA |
| NA | NA |
| NA | NA |
| NA | NA |
| NA | NA |
| NA | NA |
| NA | NA |
| NA | NA |
| NA | NA |
| NA | NA |
| NA | NA |
| NA | NA |
| NA | NA |
| NA | NA |
| NA | NA |
| NA | NA |
| NA | NA |
| NA | NA |
| 0.431630858 | 0.592931833 |
| NA | NA |
| NA | NA |
| NA | NA |

| | | |
|---|---|---|
| NA | | NA |
| NA | | NA |
| NA | | NA |
| NA | | NA |
| NA | | NA |
| NA | | NA |
| | 0.597843156 | NA |
| | 0.955792873 | NA |
| NA | | NA |
| NA | | NA |
| | 0.955792873 | NA |
| | 0.955792398 | NA |
| | 0.941278332 | NA |
| NA | | NA |
| | 0.709529034 | 0.814635573 |
| NA | | NA |
| NA | | NA |
| | 0.839318641 | NA |
| NA | | NA |
| | 0.703409083 | NA |
| | 0.955792398 | NA |
| | 0.025195709 | 0.072097652 |
| | 0.424726142 | 0.586048211 |
| | 0.03376831 | 0.091124341 |
| | 9.60E-31 | 6.86E-29 |
| | 0.427190531 | 0.588567314 |
| | 0.929761072 | NA |
| NA | | NA |
| NA | | NA |
| | 0.853596091 | NA |
| | 0.853595622 | NA |
| | 0.417782866 | 0.57932233 |
| NA | | NA |
| | 0.955791532 | NA |
| | 0.839319108 | NA |
| NA | | NA |
| | 0.036797303 | 0.097460458 |
| NA | | NA |
| | 1.69E-05 | 0.000144293 |
| NA | | NA |
| | 0.521771101 | NA |
| | 0.331900472 | 0.495042927 |
| NA | | NA |
| | 0.734058779 | NA |
| NA | | NA |
| NA | | NA |
| | 0.93162339 | NA |

| | | | |
|---|---|---|---|
| NA | | NA | |
| | 0.942388846 | | 0.965107777 |
| | 0.839318641 | NA | |
| | 0.003299892 | | 0.013889305 |
| | 0.953877378 | | 0.97203518 |
| NA | | NA | |
| | 0.014644335 | | 0.046780897 |
| NA | | NA | |
| | 0.746495443 | | 0.84054706 |
| NA | | NA | |
| NA | | NA | |
| NA | | NA | |
| | 0.937070042 | NA | |
| NA | | NA | |
| NA | | NA | |
| NA | | NA | |
| | 0.333455498 | NA | |
| | 0.941277857 | NA | |
| | 0.853595622 | NA | |
| NA | | NA | |
| | 0.006638999 | | 0.024590953 |
| | 0.90402687 | | 0.941732573 |
| | 0.588850004 | | 0.725043081 |
| | 0.009741678 | | 0.03370224 |
| NA | | NA | |
| NA | | NA | |
| | 0.910509861 | NA | |
| | 0.941278332 | NA | |
| | 0.839319108 | NA | |
| | 0.941278424 | NA | |
| | 0.000297155 | | 0.001808705 |
| | 0.941278332 | NA | |
| | 0.648797787 | NA | |
| | 0.731340621 | | 0.830543893 |
| NA | | NA | |
| NA | | NA | |
| NA | | NA | |
| | 0.941278332 | NA | |
| NA | | NA | |
| NA | | NA | |
| | 0.729746946 | NA | |
| | 0.745954294 | | 0.840278884 |
| | 0.309453339 | | 0.471127566 |
| | 0.86993086 | NA | |
| | 6.22E-07 | | 7.20E-06 |
| | 0.48192432 | | 0.638221925 |
| | 0.157306543 | | 0.293088871 |

| | |
|---|---|
| NA | NA |
| 0.365371546 | 0.528634948 |
| 0.714730395 | NA |
| 0.478020047 | 0.634888297 |
| NA | NA |
| 0.571914829 | 0.711962173 |
| 0.728556616 | NA |
| NA | NA |
| 0.448541207 | 0.608710749 |
| 0.054076941 | 0.131879031 |
| 0.540598995 | 0.687688381 |
| NA | NA |
| 0.071608156 | 0.163854772 |
| NA | NA |
| 0.270227736 | 0.430289039 |
| 0.778498281 | 0.863592382 |
| 0.426100141 | 0.587535369 |
| 0.853595622 | NA |
| 0.165627763 | NA |
| 0.494626887 | NA |
| NA | NA |
| 0.778101989 | 0.863416888 |
| 0.662917206 | NA |
| 4.32E-14 | 1.22E-12 |
| 9.74E-05 | 0.000684791 |
| 0.343925724 | 0.50754046 |
| 0.000218207 | 0.001384009 |
| 0.12434623 | 0.246782164 |
| 3.37E-08 | 4.87E-07 |
| 0.016766452 | 0.052118989 |
| 0.347370401 | 0.511355698 |
| 0.009737367 | 0.033701428 |
| 6.59E-05 | 0.000485135 |
| 0.42465286 | 0.586019796 |
| 0.623553195 | 0.751598377 |
| 0.861353377 | 0.917109919 |
| 0.860351174 | 0.916609361 |
| 0.839318641 | NA |
| 1.88E-17 | 6.73E-16 |
| 0.261445524 | 0.42018409 |
| 0.77816164 | NA |
| 1.62E-07 | 2.10E-06 |
| 0.610961809 | 0.742227061 |
| 6.72E-08 | 9.26E-07 |
| 0.000850152 | 0.00444537 |
| 0.924896662 | NA |
| 0.957225944 | 0.973712207 |

| | | |
|---|---|---|
| | 0.038107765 | 0.100247682 |
| | 0.002142531 | 0.009682271 |
| | 0.955792873 | NA |
| | 0.52070158 | 0.670947326 |
| | 0.124692939 | 0.247246029 |
| | 0.548874794 | 0.694694389 |
| | 0.008228094 | 0.029348882 |
| | 0.003618668 | 0.014943441 |
| | 0.124665695 | NA |
| | 0.357479516 | 0.521048832 |
| | 0.000841119 | 0.004409761 |
| NA | | NA |
| | 0.668646044 | NA |
| | 0.004335193 | 0.017336402 |
| | 1.85E-06 | 1.97E-05 |
| | 0.345893036 | 0.50959062 |
| | 0.40687221 | NA |
| NA | | NA |
| | 0.149228239 | 0.282257672 |
| NA | | NA |
| | 0.017380386 | 0.053628738 |
| | 0.66475065 | 0.782550817 |
| | 1.19E-06 | 1.31E-05 |
| NA | | NA |
| | 1.79E-09 | 3.08E-08 |
| | 0.450296253 | 0.6102273 |
| | 0.622962962 | 0.751305453 |
| | 0.876490807 | 0.926140201 |
| | 4.53E-11 | 9.51E-10 |
| | 0.005194087 | 0.020151533 |
| | 0.743856064 | 0.839111945 |
| | 0.739991096 | NA |
| | 0.941277857 | NA |
| | 0.856055538 | NA |
| | 0.13847664 | 0.266949136 |
| | 0.979194083 | 0.986989818 |
| | 4.43E-05 | 0.000340205 |
| | 0.955792398 | NA |
| | 0.386652692 | 0.54949761 |
| | 0.956280563 | 0.973222697 |
| NA | | NA |
| | 0.214710728 | 0.365561142 |
| | 0.122053396 | 0.24347259 |
| | 0.880849404 | NA |
| | 0.034108762 | 0.091724862 |
| | 6.65E-05 | 0.000488651 |
| NA | | NA |

| | | |
|---|---|---|
| | 0.033594841 | 0.090773574 |
| | 0.577905797 | 0.716273369 |
| | 0.568938007 | NA |
| | 0.893589818 | NA |
| | 0.687586868 | NA |
| | 0.839319108 | NA |
| | 0.734058779 | NA |
| NA | | NA |
| NA | | NA |
| | 4.06E-22 | 1.98E-20 |
| | 0.839318641 | NA |
| | 0.868873093 | 0.921445258 |
| | 0.024922548 | 0.071496502 |
| | 0.940773688 | NA |
| | 0.155671334 | 0.290930361 |
| | 6.65E-06 | 6.30E-05 |
| | 0.050170037 | 0.12409303 |
| | 0.630990172 | 0.757188207 |
| NA | | NA |
| | 0.59246934 | 0.727993146 |
| | 0.783084926 | NA |
| | 0.002420658 | 0.010739327 |
| | 0.009940643 | 0.034231362 |
| | 0.172460847 | 0.312683811 |
| | 0.038163942 | 0.100340013 |
| | 4.59E-14 | 1.29E-12 |
| | 0.236052147 | 0.391286321 |
| NA | | NA |
| NA | | NA |
| | 0.955791446 | NA |
| | 0.532123915 | 0.680690784 |
| | 1.95E-06 | 2.06E-05 |
| | 0.77370946 | 0.860500363 |
| | 7.96E-09 | 1.27E-07 |
| | 0.000490001 | 0.002788193 |
| NA | | NA |
| | 0.055257717 | 0.134181206 |
| | 0.585120233 | 0.721832268 |
| | 0.000756005 | 0.004031253 |
| | 0.907839022 | NA |
| | 0.839318641 | NA |
| | 0.941277857 | NA |
| | 0.000799384 | 0.004224454 |
| | 0.938780082 | 0.963601145 |
| | 0.475264201 | 0.632517113 |
| | 0.012866527 | 0.042191548 |
| | 0.624761722 | 0.752819904 |

| | | |
|---|---|---|
| | 0.388997069 | 0.551626565 |
| | 0.693638483 | 0.803747042 |
| | 0.941765806 | 0.964806618 |
| | 1.66E-05 | 0.000142287 |
| | 0.019216374 | 0.058036813 |
| | 0.214082272 | 0.364934917 |
| | 0.927314698 | NA |
| | 0.001597649 | 0.007570626 |
| NA | | NA |
| | 0.011921214 | 0.039695441 |
| | 1.27E-16 | 4.33E-15 |
| | 0.820873394 | 0.890764479 |
| | 0.001869876 | 0.008642094 |
| | 0.40059952 | 0.563101117 |
| | 0.009029387 | 0.031670714 |
| | 5.13E-06 | 4.98E-05 |
| | 0.322740498 | 0.485507446 |
| | 0.975609358 | 0.985178051 |
| | 0.768027986 | NA |
| | 0.223771415 | 0.376356206 |
| | 0.016800625 | 0.052204763 |
| NA | | NA |
| | 0.293074845 | NA |
| NA | | NA |
| | 0.631388758 | NA |
| | 0.898475246 | NA |
| | 0.508668768 | 0.660303993 |
| NA | | NA |
| | 0.955792398 | NA |
| | 0.359255615 | 0.522773255 |
| | 0.767660581 | 0.855824691 |
| | 0.050957142 | 0.12577848 |
| | 0.197946768 | 0.345091113 |
| | 0.015843338 | 0.049841812 |
| | 0.272533716 | 0.432400438 |
| | 0.598597381 | NA |
| NA | | NA |
| NA | | NA |
| | 0.494035027 | 0.647310858 |
| | 0.953798549 | NA |
| NA | | NA |
| | 0.122761785 | 0.244475904 |
| | 0.745363571 | 0.840016274 |
| | 0.432313377 | 0.593523851 |
| NA | | NA |
| | 0.343386816 | 0.506902319 |
| | 0.029581965 | 0.082064095 |

| | |
|---|---|
| 0.536820266 | 0.684839603 |
| 0.886475674 | NA |
| 0.86251998 | NA |
| 1.10E-06 | 1.21E-05 |
| NA | NA |
| 0.576222325 | 0.714928093 |
| NA | NA |
| 0.245192719 | 0.401850657 |
| 0.630022739 | 0.756391124 |
| 0.567047918 | 0.70872057 |
| 0.877698141 | 0.926663806 |
| 0.427080399 | 0.588546066 |
| 3.38E-27 | 2.10E-25 |
| 0.082456447 | 0.1817556 |
| 1.72E-10 | 3.37E-09 |
| 0.373593569 | 0.53677859 |
| 2.25E-07 | 2.84E-06 |
| 0.093488827 | 0.199515816 |
| 0.191116109 | 0.33618695 |
| 0.019000306 | 0.057559538 |
| 0.243475554 | 0.399972748 |
| 0.679408292 | 0.793682266 |
| 0.613415398 | 0.744003219 |
| 0.000624445 | 0.003421509 |
| 0.101193389 | 0.211644056 |
| 0.012409676 | 0.040978789 |
| 0.567868989 | NA |
| 0.033271154 | 0.090124047 |
| 1.18E-07 | 1.57E-06 |
| 0.717563336 | 0.820377661 |
| 0.839318641 | NA |
| 0.491381627 | 0.645465283 |
| 0.01674596 | 0.052062089 |
| 0.039716404 | 0.103586439 |
| 0.955791446 | NA |
| 0.899601552 | 0.93958188 |
| 0.043516563 | 0.111111482 |
| 0.806674096 | 0.881648833 |
| 0.0001731 | 0.00113471 |
| NA | NA |
| NA | NA |
| 0.250119246 | 0.407663684 |
| 0.955792873 | NA |
| 0.057060889 | 0.137549451 |
| 1.42E-07 | 1.87E-06 |
| 0.016120586 | 0.05050042 |
| 0.01291632 | 0.042299703 |

| | |
|---|---|
| 0.030544603 | 0.084146139 |
| 0.870939771 | 0.922814656 |
| 0.278676732 | 0.438433708 |
| NA | NA |
| 0.48289948 | NA |
| 0.099919429 | 0.20948555 |
| 0.11479738 | 0.232337227 |
| 0.128323168 | 0.252508933 |
| 0.941276431 | NA |
| 1.40E-08 | 2.14E-07 |
| 0.010720398 | 0.036426396 |
| 0.94646468 | 0.967520714 |
| 0.463629858 | 0.622333974 |
| 0.070729827 | 0.162516782 |
| 0.479927204 | 0.636595229 |
| 0.810777649 | 0.884102924 |
| NA | NA |
| 0.027355195 | 0.077218732 |
| 0.070831839 | 0.162618687 |
| 0.093904161 | 0.2001887 |
| 0.034117933 | 0.091739159 |
| 0.000808834 | 0.004266815 |
| 0.00045561 | 0.002616897 |
| 0.185045009 | 0.328175344 |
| 0.437109696 | 0.598004655 |
| 1.03E-09 | 1.83E-08 |
| 0.884520557 | 0.930962684 |
| 0.232385216 | 0.386826555 |
| NA | NA |
| 0.839319108 | NA |
| 0.955792398 | NA |
| 0.001752295 | 0.008195741 |
| 3.58E-05 | 0.000281335 |
| 6.26E-96 | 2.07E-93 |
| 1.36E-26 | 8.22E-25 |
| 0.008060429 | 0.028837265 |
| 0.574002183 | 0.71353814 |
| 0.007889469 | 0.028332098 |
| NA | NA |
| 0.002063639 | 0.009382759 |
| 0.797235346 | 0.875792482 |
| 0.936819898 | NA |
| NA | NA |
| NA | NA |
| NA | NA |
| NA | NA |
| NA | NA |

| | |
|---|---|
| NA | NA |
| NA | NA |
| NA | NA |
| NA | NA |
| NA | NA |
| NA | NA |
| 0.006596191 | 0.02446761 |
| NA | NA |
| 2.40E-16 | 8.08E-15 |
| 0.955791922 | NA |
| 0.552671582 | NA |
| 0.761232869 | 0.851610488 |
| 0.33910489 | 0.502701457 |
| NA | NA |
| 0.325240052 | 0.488106162 |
| 0.001747381 | 0.008174624 |
| 0.333861956 | 0.497127449 |
| 0.019890241 | 0.059573112 |
| 0.466828576 | 0.625011462 |
| 0.000308847 | 0.00186815 |
| 0.039858951 | 0.103889921 |
| 0.791947557 | NA |
| 0.734058779 | NA |
| 0.076852702 | 0.172586849 |
| NA | NA |
| 0.170816557 | 0.310696598 |
| 0.002162371 | 0.009758116 |
| 0.52309836 | 0.672906402 |
| 0.330387331 | 0.493342488 |
| 0.945938655 | 0.967272298 |
| 0.396189511 | 0.558887141 |
| 0.964314609 | 0.978288991 |
| 0.087783952 | 0.190671474 |
| 0.802131288 | 0.879054029 |
| 0.591409501 | 0.727147641 |
| 0.051694841 | 0.12728257 |
| 0.902410694 | 0.94110113 |
| 0.349327237 | 0.513256858 |
| NA | NA |
| 0.229020392 | 0.382697513 |
| 0.186766924 | 0.330609741 |
| 0.304844541 | 0.466468001 |
| 0.199258771 | 0.346718127 |
| 0.4059865 | 0.567927751 |
| 0.557352183 | 0.701283505 |
| 0.103796427 | 0.215675475 |
| 0.026921555 | 0.076223905 |

| | |
|---|---|
| 0.176670096 | 0.318280868 |
| 0.937105618 | 0.962666721 |
| 0.365103092 | 0.528425723 |
| 0.006456794 | 0.02402028 |
| 0.066430963 | 0.154957837 |
| NA | NA |
| 0.21317361 | 0.363855021 |
| 1.17E-06 | 1.29E-05 |
| 0.003844631 | 0.015708529 |
| 0.730687735 | NA |
| 0.235470588 | 0.390569403 |
| 0.80401566 | 0.880356952 |
| 0.173777936 | 0.314520754 |
| NA | NA |
| NA | NA |
| 0.545286864 | NA |
| NA | NA |
| NA | NA |
| 0.921305466 | NA |
| 0.505973469 | NA |
| 0.938320081 | NA |
| 0.828295333 | 0.895633788 |
| 0.001453104 | 0.006982998 |
| 0.007467325 | 0.027106493 |
| 0.611881424 | 0.742871648 |
| 0.008097403 | 0.028947789 |
| 0.001085054 | 0.005475243 |
| NA | NA |
| NA | NA |
| 0.545642654 | NA |
| 0.955043037 | NA |
| 0.839319108 | NA |
| NA | NA |
| NA | NA |
| 2.58E-10 | 4.95E-09 |
| 0.0630629 | NA |
| 0.249234599 | 0.406627043 |
| 0.839318641 | NA |
| 0.004435067 | 0.017667524 |
| 0.955791922 | NA |
| 0.137082449 | 0.265055765 |
| NA | NA |
| 0.003398044 | 0.014211549 |
| 0.001390704 | 0.006732059 |
| 0.006630342 | 0.024566533 |
| 0.313148803 | 0.475116614 |
| 0.047259678 | 0.118376563 |

| | | |
|---|---|---|
| | 0.002216253 | 0.009968115 |
| | 0.703648179 | 0.810844101 |
| | 0.928092825 | 0.957084417 |
| | 0.213970339 | 0.364877858 |
| | 1.96E-13 | 5.24E-12 |
| | 0.182490781 | NA |
| | 0.008005429 | 0.028679291 |
| | 0.013454505 | 0.043773118 |
| | 8.88E-07 | 9.96E-06 |
| | 0.486792135 | 0.641900641 |
| | 0.853596091 | NA |
| NA | | NA |
| | 0.734058779 | NA |
| | 1.66E-11 | 3.67E-10 |
| | 0.000140977 | 0.000945753 |
| | 0.364167261 | 0.52755777 |
| | 0.26802305 | 0.427751418 |
| | 0.907477482 | NA |
| | 0.132066111 | 0.257658818 |
| | 0.034993999 | 0.093608357 |
| | 0.644150833 | NA |
| | 4.76E-09 | 7.79E-08 |
| | 1.26E-06 | 1.38E-05 |
| NA | | NA |
| | 0.291572107 | NA |
| | 0.744162276 | NA |
| | 0.354376889 | NA |
| | 0.914340506 | NA |
| | 0.359667564 | NA |
| | 0.886501246 | NA |
| | 0.710977993 | NA |
| | 0.955792873 | NA |
| | 0.956920489 | 0.973526307 |
| | 0.58893742 | 0.725113204 |
| NA | | NA |
| NA | | NA |
| NA | | NA |
| NA | | NA |
| NA | | NA |
| NA | | NA |
| NA | | NA |
| NA | | NA |
| NA | | NA |
| NA | | NA |
| NA | | NA |
| NA | | NA |
| NA | | NA |

| | | | |
|---|---|---|---|
| NA | | NA | |
| NA | | NA | |
| NA | | NA | |
| NA | | NA | |
| NA | | NA | |
| NA | | NA | |
| NA | | NA | |
| NA | | NA | |
| NA | | NA | |
| NA | | NA | |
| NA | | NA | |
| NA | | NA | |
| | 0.692256265 | NA | |
| NA | | NA | |
| NA | | NA | |
| NA | | NA | |
| NA | | NA | |
| NA | | NA | |
| NA | | NA | |
| NA | | NA | |
| NA | | NA | |
| NA | | NA | |
| NA | | NA | |
| NA | | NA | |
| NA | | NA | |
| NA | | NA | |
| NA | | NA | |
| NA | | NA | |
| NA | | NA | |
| NA | | NA | |
| | 0.605851446 | NA | |
| NA | | NA | |
| NA | | NA | |
| | 0.923120612 | NA | |
| | 0.941278332 | NA | |
| | 0.072472299 | | 0.165228511 |
| NA | | NA | |
| | 0.52830463 | | 0.677412368 |
| NA | | NA | |
| NA | | NA | |
| | 0.900768708 | | 0.940152952 |
| NA | | NA | |
| | 0.314312124 | | 0.476513196 |
| | 0.55574685 | | 0.700110433 |
| NA | | NA | |
| | 0.105711371 | | 0.218585879 |
| | 0.172926039 | | 0.313271336 |

| | | | |
|---|---|---|---|
| | 0.078974952 | | 0.176189127 |
| | 0.029677675 | | 0.082287235 |
| NA | | NA | |
| NA | | NA | |
| | 0.839319107 | NA | |
| | 0.569970713 | NA | |
| NA | | NA | |
| NA | | NA | |
| NA | | NA | |
| NA | | NA | |
| NA | | NA | |
| NA | | NA | |
| | 0.241952704 | | 0.39822287 |
| | 0.012863447 | | 0.042191548 |
| NA | | NA | |
| | 0.211764549 | | 0.362177178 |
| | 0.000112963 | | 0.0007798 |
| | 0.450018498 | | 0.60998521 |
| | 0.002098613 | | 0.009516325 |
| | 0.003407382 | | 0.0142385 |
| | 2.87E-06 | | 2.93E-05 |
| | 0.000485852 | | 0.002769045 |
| | 0.000924331 | | 0.004770363 |
| | 0.165919798 | | 0.304747513 |
| | 0.885483592 | NA | |
| | 0.504819063 | | 0.656918886 |
| | 0.073402063 | | 0.166900641 |
| | 0.941277857 | NA | |
| | 0.839318641 | NA | |
| | 0.905652191 | NA | |
| | 0.941278332 | NA | |
| | 0.839318641 | NA | |
| | 0.785602812 | NA | |
| | 0.664934957 | NA | |
| | 0.722858773 | NA | |
| | 0.941279268 | NA | |
| | 0.904182744 | NA | |
| NA | | NA | |
| NA | | NA | |
| NA | | NA | |
| | 0.82229013 | NA | |
| | 0.900247435 | NA | |
| NA | | NA | |
| NA | | NA | |
| NA | | NA | |
| | 0.941275956 | NA | |
| NA | | NA | |

| | |
|---|---|
| 0.023306996 | 0.067793365 |
| 0.003035223 | 0.012951665 |
| 0.955792398 | NA |
| 0.001883099 | 0.008696452 |
| NA | NA |
| NA | NA |
| 7.18E-08 | 9.87E-07 |
| 0.559360321 | 0.702728511 |
| 0.59777104 | 0.732277216 |
| 0.075023631 | 0.169493692 |
| 0.686070258 | NA |
| 0.783375663 | 0.866827774 |
| NA | NA |
| NA | NA |
| NA | NA |
| NA | NA |
| NA | NA |
| 0.053837812 | 0.131430524 |
| 0.001295814 | 0.006347594 |
| 0.941277857 | NA |
| 0.00075525 | 0.004028789 |
| NA | NA |
| 0.941277857 | NA |
| NA | NA |
| 0.839319108 | NA |
| 0.594925079 | 0.729917873 |
| 0.301212436 | 0.462490658 |
| NA | NA |
| 0.114905406 | 0.232487119 |
| 0.853595622 | NA |
| NA | NA |
| 0.065004238 | 0.152406508 |
| NA | NA |
| 0.955792873 | NA |
| 0.659002827 | 0.778093422 |
| 0.981890689 | 0.98868314 |
| 0.471894511 | 0.629405948 |
| 0.943633965 | NA |
| 8.69E-05 | 0.000618948 |
| 0.669168711 | NA |
| 0.169224019 | 0.308743683 |
| 0.000859273 | 0.004488132 |
| 1.21E-07 | 1.61E-06 |
| 0.20044218 | 0.34803945 |
| 0.941277857 | NA |
| NA | NA |
| NA | NA |

| | | |
|---|---|---|
| | 0.450745408 | NA |
| | 0.494274514 | NA |
| | 0.002211377 | 0.009951824 |
| | 0.008069425 | 0.028860773 |
| NA | | NA |
| | 0.022578911 | 0.066046853 |
| | 7.33E-53 | 1.08E-50 |
| | 0.012838924 | 0.042127388 |
| | 0.13214684 | 0.257795187 |
| | 0.785248391 | 0.868018401 |
| | 0.895896289 | NA |
| | 0.600514922 | 0.734428947 |
| | 0.263201635 | 0.422094416 |
| | 0.095799217 | 0.203158952 |
| | 0.82323984 | NA |
| | 1.28E-06 | 1.40E-05 |
| | 0.941275481 | NA |
| | 0.209078075 | 0.358535759 |
| | 0.004105042 | 0.016587163 |
| | 0.332020929 | 0.49514351 |
| | 0.000605067 | 0.003337559 |
| NA | | NA |
| | 0.430341057 | 0.591600891 |
| | 0.858951208 | 0.915860621 |
| | 0.005275921 | 0.020384294 |
| | 0.003758929 | 0.015409927 |
| NA | | NA |
| NA | | NA |
| NA | | NA |
| | 0.839319574 | NA |
| NA | | NA |
| | 0.574520349 | 0.713786167 |
| | 0.902432241 | 0.94110113 |
| | 0.784846558 | 0.867785536 |
| | 0.457261961 | 0.61695761 |
| | 0.010072013 | 0.034602826 |
| | 0.398033667 | 0.560724274 |
| | 0.000635306 | 0.003472229 |
| NA | | NA |
| | 0.888330814 | 0.93326492 |
| | 0.522880947 | 0.672772166 |
| | 0.265872867 | 0.425214367 |
| | 0.81338117 | NA |
| | 0.095576969 | 0.202810289 |
| | 0.170515409 | 0.310355101 |
| | 0.941278332 | NA |
| NA | | NA |

| | | |
|---|---|---|
| | 0.905789546 | NA |
| NA | | NA |
| | 0.934281796 | NA |
| NA | | NA |
| | 1.88E-05 | 0.000159068 |
| NA | | NA |
| | 0.326269266 | 0.4891881 |
| | 0.941278332 | NA |
| | 0.073066151 | 0.166237927 |
| | 0.169338821 | 0.30890578 |
| | 0.004388374 | 0.017522578 |
| | 0.364666919 | NA |
| | 0.241502961 | 0.397747957 |
| | 0.881858899 | 0.92937815 |
| | 0.810813643 | NA |
| | 0.012955209 | 0.042409557 |
| | 0.1315357 | 0.256978723 |
| | 0.599326314 | 0.733276701 |
| | 0.824102086 | 0.89310558 |
| | 0.782070774 | NA |
| | 0.410505084 | 0.572432012 |
| | 0.525026651 | 0.674449691 |
| | 0.113891231 | 0.230945567 |
| | 0.802250348 | 0.879097643 |
| | 0.364127431 | 0.52755777 |
| NA | | NA |
| | 0.137545541 | 0.265735305 |
| | 0.932063201 | 0.959371849 |
| | 0.887588156 | 0.932978859 |
| | 0.421302523 | 0.582810801 |
| | 0.669466721 | 0.786132294 |
| NA | | NA |
| | 0.839319574 | NA |
| | 0.933506231 | 0.960461456 |
| | 0.001140329 | 0.005710524 |
| NA | | NA |
| | 3.15E-11 | 6.74E-10 |
| | 0.000578954 | 0.003211434 |
| | 0.309193148 | 0.470791665 |
| | 0.026426997 | 0.075007796 |
| | 0.035966463 | 0.095789835 |
| | 0.13296995 | 0.258943241 |
| | 0.000490359 | 0.00278902 |
| | 0.522930269 | 0.672799258 |
| NA | | NA |
| | 0.610857839 | 0.742227061 |
| | 0.036060338 | 0.095994947 |

| | | |
|---|---|---|
| | 0.540332189 | 0.687495951 |
| | 0.835164607 | 0.899994439 |
| | 0.342232965 | 0.505732231 |
| | 0.240052531 | NA |
| | 0.437288192 | NA |
| | 0.015499705 | 0.048954881 |
| | 0.765485532 | 0.854320187 |
| | 1.74E-06 | 1.87E-05 |
| | 0.004408659 | 0.017585871 |
| NA | | NA |
| NA | | NA |
| | 1.47E-32 | 1.14E-30 |
| | 0.368173659 | 0.531271239 |
| | 0.796374759 | 0.875129825 |
| | 0.71595382 | 0.819442817 |
| | 0.514973417 | 0.666126692 |
| | 0.006442923 | 0.023991622 |
| | 2.35E-05 | 0.000194007 |
| | 0.48661039 | 0.64182213 |
| NA | | NA |
| | 0.293509663 | 0.454763099 |
| | 0.308170426 | 0.46960479 |
| | 0.149402314 | 0.282411771 |
| NA | | NA |
| | 0.839320041 | NA |
| NA | | NA |
| | 0.000325533 | 0.001956649 |
| NA | | NA |
| | 0.022044753 | 0.064715 |
| | 0.853595622 | NA |
| | 0.941277857 | NA |
| | 0.807563873 | 0.882090556 |
| NA | | NA |
| | 0.019570525 | 0.058889714 |
| | 0.474234495 | NA |
| | 0.874604749 | 0.925009245 |
| | 0.566787235 | 0.708535119 |
| | 0.955792398 | NA |
| | 0.447605037 | 0.607898971 |
| NA | | NA |
| | 0.005702495 | 0.021716925 |
| | 0.002158042 | 0.009743118 |
| | 0.911428204 | NA |
| | 0.354929853 | 0.518730219 |
| NA | | NA |
| | 0.49953831 | NA |
| | 0.888778264 | NA |

| | | |
|---|---|---|
| | 0.176175777 | 0.317662155 |
| | 0.942968197 | NA |
| NA | | NA |
| | 0.505328522 | NA |
| | 0.000620507 | 0.003404634 |
| NA | | NA |
| | 0.108933149 | NA |
| | 0.927397734 | NA |
| | 0.00270229 | 0.011780624 |
| | 0.835127771 | NA |
| NA | | NA |
| | 0.941278838 | NA |
| | 0.597720898 | 0.732253478 |
| | 0.941277857 | NA |
| NA | | NA |
| NA | | NA |
| | 0.875239182 | 0.9253821 |
| NA | | NA |
| NA | | NA |
| NA | | NA |
| | 0.040582685 | 0.105372254 |
| | 0.137864819 | 0.266114543 |
| | 0.666359299 | NA |
| | 0.839319574 | NA |
| | 0.379589701 | 0.542381878 |
| | 0.955792397 | NA |
| NA | | NA |
| | 0.515280133 | 0.666342374 |
| | 0.946614783 | 0.967549494 |
| | 0.853561653 | 0.911911417 |
| | 0.285611468 | 0.446276122 |
| | 0.652905331 | 0.773657226 |
| | 0.18210295 | 0.32448786 |
| | 0.138997938 | 0.267758896 |
| | 0.046903104 | 0.117787264 |
| | 0.839319108 | NA |
| | 0.008758466 | 0.030879724 |
| | 0.90290283 | 0.941349627 |
| | 0.042409454 | 0.108997929 |
| | 0.781282224 | 0.865510569 |
| | 0.037044309 | 0.097973737 |
| | 0.623192035 | 0.751385543 |
| | 0.524270267 | NA |
| | 0.000773124 | 0.004107568 |
| | 0.192544786 | 0.338081055 |
| | 0.000119875 | 0.000822032 |
| | 0.040075731 | 0.104342692 |

| | | |
|---|---|---|
| | 0.036415122 | 0.096649501 |
| | 0.892864801 | 0.935303582 |
| | 0.000552797 | 0.003095114 |
| | 0.950512078 | NA |
| NA | | NA |
| NA | | NA |
| | 0.388860704 | 0.551561603 |
| | 0.946718097 | NA |
| | 0.52333317 | NA |
| NA | | NA |
| | 0.166937246 | NA |
| | 9.82E-09 | 1.55E-07 |
| | 0.630805746 | NA |
| NA | | NA |
| NA | | NA |
| | 0.839320041 | NA |
| | 0.768046038 | NA |
| | 0.955792398 | NA |
| | 1.14E-12 | 2.85E-11 |
| | 0.745413183 | 0.840032412 |
| | 0.397134416 | NA |
| | 0.002693507 | 0.011750524 |
| | 3.62E-09 | 5.99E-08 |
| | 0.505682857 | 0.657528461 |
| | 0.00138717 | 0.006720143 |
| | 0.167331764 | 0.306418919 |
| | 0.000242731 | 0.001513621 |
| | 7.99E-14 | 2.20E-12 |
| | 0.465075276 | NA |
| | 0.003719702 | 0.015278058 |
| | 0.020681331 | 0.061479899 |
| | 0.083619836 | 0.183817714 |
| | 0.392612458 | 0.555058311 |
| | 0.941277857 | NA |
| NA | | NA |
| | 0.925798536 | NA |
| | 0.023272433 | 0.067717658 |
| | 0.002978991 | 0.012749845 |
| | 4.54E-10 | 8.44E-09 |
| NA | | NA |
| | 0.000824188 | 0.004337239 |
| | 1.43E-06 | 1.55E-05 |
| | 0.017663682 | 0.054284177 |
| | 0.576672534 | 0.71534082 |
| | 0.012051549 | 0.040012688 |
| NA | | NA |
| | 0.944257906 | 0.966373729 |

| | |
|---|---|
| 0.196795926 | 0.343537706 |
| 0.03490869 | 0.093422154 |
| 0.000134201 | 0.00090746 |
| 0.95396211 | 0.972053861 |
| 3.01E-07 | 3.71E-06 |
| 0.727116949 | 0.827925808 |
| 0.363471865 | 0.52713608 |
| 3.72E-09 | 6.15E-08 |
| 0.084035354 | 0.184538196 |
| 0.027933728 | 0.07842399 |
| 6.76E-06 | 6.39E-05 |
| 0.428220695 | 0.589469028 |
| 0.006326504 | 0.023635762 |
| 0.036350818 | 0.096521883 |
| 0.619296129 | 0.748476006 |
| 0.919938007 | NA |
| 0.447724886 | 0.607987046 |
| NA | NA |
| 0.560591952 | 0.703718667 |
| 0.041225238 | 0.106680051 |
| 0.037073757 | 0.098015057 |
| 0.572348113 | 0.712315283 |
| 0.126354637 | 0.249623478 |
| 0.003689853 | 0.015179034 |
| 7.43E-09 | 1.19E-07 |
| 0.130294037 | 0.255184618 |
| 0.670999629 | 0.787121384 |
| 0.855288229 | 0.912932007 |
| 4.06E-05 | 0.000314845 |
| 0.051055633 | 0.126008521 |
| 7.03E-09 | 1.12E-07 |
| 0.328271523 | 0.491138273 |
| 0.807721396 | 0.882181649 |
| NA | NA |
| NA | NA |
| 0.651050316 | NA |
| 0.000528308 | 0.002976644 |
| 0.001802971 | 0.0083817 |
| 0.552710105 | 0.697910128 |
| 0.798066876 | NA |
| 0.597279906 | 0.731901581 |
| 0.142796346 | 0.27321854 |
| 0.057714275 | 0.138680794 |
| 0.568972827 | 0.71014571 |
| 0.661797704 | 0.78014028 |
| 0.004099484 | 0.016569184 |
| 0.925325733 | 0.955182718 |

| | |
|---|---|
| NA | NA |
| 0.044772486 | 0.113514561 |
| 0.711923471 | 0.816425444 |
| 0.054449514 | 0.132549019 |
| 0.813392475 | 0.886089811 |
| 0.912797399 | 0.947427337 |
| 0.770126294 | 0.857609528 |
| 0.89771406 | NA |
| 0.64788006 | 0.769848794 |
| 0.425608061 | 0.587016513 |
| NA | NA |
| NA | NA |
| 0.038485095 | 0.100972468 |
| NA | NA |
| 0.839319108 | NA |
| NA | NA |
| 0.545433097 | 0.69184666 |
| 0.452284831 | 0.612290043 |
| 0.407386622 | 0.569250697 |
| 0.584981766 | 0.721736263 |
| NA | NA |
| 0.115332417 | 0.233073713 |
| 0.839319108 | NA |
| 0.941685325 | NA |
| 0.595011558 | 0.729948722 |
| 0.002055364 | 0.009355855 |
| 0.607989361 | 0.740488296 |
| 0.05357515 | 0.13086984 |
| 0.023325689 | 0.067822875 |
| 0.029917811 | 0.082841291 |
| NA | NA |
| NA | NA |
| 9.88E-08 | 1.33E-06 |
| 0.109880008 | 0.225016256 |
| NA | NA |
| 0.004095416 | 0.016558364 |
| 0.049838488 | 0.123446754 |
| 0.197063143 | 0.343802457 |
| 0.394694821 | 0.557174909 |
| NA | NA |
| 0.059752194 | 0.142564328 |
| 0.011667655 | 0.039065062 |
| 0.941277857 | NA |
| 0.52689291 | NA |
| 0.000244201 | 0.001521881 |
| 0.937654382 | 0.963105753 |
| NA | NA |

| | | |
|---|---|---|
| NA | | NA |
| | 0.179947719 | 0.321966144 |
| | 0.00278968 | 0.012070527 |
| | 0.323231342 | NA |
| | 0.941277857 | NA |
| | 0.000251078 | 0.001558318 |
| | 0.421681244 | 0.583069609 |
| | 0.003856234 | 0.015738355 |
| | 0.941278332 | NA |
| | 0.000827145 | 0.004348953 |
| | 0.000253598 | 0.001571096 |
| NA | | NA |
| | 0.860386396 | 0.916609361 |
| | 0.257250505 | 0.41534777 |
| | 0.376331802 | 0.539463579 |
| | 0.814324117 | 0.886545425 |
| NA | | NA |
| | 0.001387395 | 0.006720143 |
| | 0.048715393 | 0.12126373 |
| NA | | NA |
| | 0.306375726 | 0.46806122 |
| | 0.000156599 | 0.001039716 |
| | 0.120688579 | 0.24143802 |
| | 0.897962929 | 0.938413841 |
| | 1.00E-09 | 1.78E-08 |
| | 0.044077315 | 0.112159698 |
| | 0.161936291 | NA |
| | 0.955792398 | NA |
| | 0.739991096 | NA |
| | 0.805665463 | 0.88102772 |
| NA | | NA |
| | 0.099671514 | 0.209057224 |
| | 0.549890433 | NA |
| | 5.94E-05 | 0.000441658 |
| | 0.030987062 | 0.085089299 |
| NA | | NA |
| | 0.245170131 | NA |
| | 0.673743537 | NA |
| | 0.000135741 | 0.000916048 |
| | 0.221315012 | 0.373386725 |
| | 0.012634322 | 0.041570655 |
| | 0.955792397 | NA |
| | 0.003523104 | 0.014642383 |
| | 1.57E-05 | 0.000135707 |
| | 9.73E-34 | 7.93E-32 |
| | 0.272242556 | 0.432127426 |
| | 0.000577502 | 0.003206966 |

| | |
|---|---|
| NA | NA |
| 0.857447575 | 0.914667108 |
| 0.18360256 | 0.326346661 |
| 4.17E-05 | 0.000322321 |
| 0.941208171 | NA |
| 0.004115257 | 0.016618824 |
| 0.955792398 | NA |
| NA | NA |
| 0.691429669 | 0.801978899 |
| 0.009111481 | 0.031915628 |
| 0.844339847 | 0.90637158 |
| 0.68454458 | NA |
| 0.459866574 | NA |
| 0.745989966 | 0.840278884 |
| 0.156430591 | 0.291959612 |
| 0.004835679 | 0.018990074 |
| 0.106095084 | 0.219207916 |
| 0.941277857 | NA |
| NA | NA |
| 0.941277857 | NA |
| NA | NA |
| NA | NA |
| 0.090105414 | 0.194459069 |
| 0.508207645 | 0.659849354 |
| NA | NA |
| 0.1532102 | 0.287414027 |
| 0.208169582 | 0.357492958 |
| 0.199726772 | 0.347183534 |
| 0.227800844 | 0.381221661 |
| 0.103127024 | 0.214678103 |
| 6.51E-14 | 1.81E-12 |
| 0.876322507 | NA |
| 0.232962853 | 0.387517075 |
| 0.046255032 | 0.116516274 |
| 0.952646527 | NA |
| 0.006784051 | 0.025038608 |
| 0.962848061 | NA |
| 0.751743419 | NA |
| 0.277902227 | 0.437533326 |
| 0.003001826 | 0.012834463 |
| 0.001255541 | 0.006185965 |
| 0.730506509 | 0.830111521 |
| 0.282686231 | 0.443101592 |
| NA | NA |
| 0.001590616 | 0.007541799 |
| NA | NA |
| 0.282610286 | 0.443040903 |

| | |
|---|---|
| NA | NA |
| 0.731494836 | 0.830679393 |
| 0.656447019 | NA |
| 0.13537096 | 0.262547607 |
| 0.000129694 | 0.000880235 |
| 0.03060271 | 0.08424771 |
| 0.923246595 | NA |
| 0.029115541 | 0.081025151 |
| 0.019355538 | 0.058368239 |
| 0.078064226 | 0.174533601 |
| 0.939739774 | NA |
| NA | NA |
| 0.012691165 | 0.041723082 |
| 0.15312676 | 0.287370743 |
| 2.18E-05 | 0.000181569 |
| 0.060771097 | 0.144473995 |
| NA | NA |
| 0.095482925 | 0.202651443 |
| NA | NA |
| 0.031653424 | 0.086539719 |
| 0.153623241 | 0.288120746 |
| NA | NA |
| 0.110052031 | 0.225232884 |
| 0.787914066 | NA |
| 0.147848551 | 0.280405674 |
| 0.941278332 | NA |
| 0.161101371 | 0.298082621 |
| 0.048106286 | 0.12008661 |
| 0.12620462 | 0.249453776 |
| 0.784878876 | 0.867785536 |
| 0.099033601 | 0.208030868 |
| 0.038171872 | 0.10034978 |
| 0.816695603 | 0.888111694 |
| 0.756029988 | 0.84719517 |
| 0.573196079 | 0.712886042 |
| 0.839318641 | NA |
| 1.22E-20 | 5.38E-19 |
| 0.895701039 | NA |
| 1.91E-05 | 0.000161169 |
| NA | NA |
| NA | NA |
| 0.495663547 | 0.648586242 |
| NA | NA |
| 0.01434148 | 0.046048281 |
| 0.358367761 | NA |
| 0.740069079 | NA |
| NA | NA |

| | | |
|---|---|---|
| | 0.854139007 | NA |
| NA | | NA |
| | 0.955792873 | NA |
| | 0.941278332 | NA |
| | 0.402556067 | NA |
| | 0.692943712 | NA |
| | 0.440164049 | 0.600905345 |
| NA | | NA |
| | 0.917893938 | 0.95060312 |
| | 0.208756308 | 0.358164611 |
| | 0.763008924 | 0.852875853 |
| | 0.941277857 | NA |
| | 0.805340803 | 0.880868577 |
| NA | | NA |
| NA | | NA |
| | 0.560954831 | 0.703988552 |
| NA | | NA |
| | 0.513543486 | 0.66463825 |
| | 0.923012141 | NA |
| | 0.753632792 | 0.845612017 |
| | 0.172560511 | 0.312792683 |
| NA | | NA |
| | 0.941278284 | NA |
| | 0.414803321 | 0.576702608 |
| | 0.839319574 | NA |
| | 0.804661329 | NA |
| | 0.302631015 | NA |
| | 0.092565209 | 0.198125081 |
| | 0.939917034 | NA |
| | 0.002331209 | 0.010412354 |
| | 0.041370199 | 0.106953055 |
| | 0.006402876 | 0.023883602 |
| | 0.308244949 | 0.469648333 |
| | 0.044746354 | 0.113484572 |
| | 0.868699204 | NA |
| | 0.923516642 | NA |
| | 0.839318641 | NA |
| | 0.340496825 | 0.50405529 |
| | 0.442351346 | 0.602854094 |
| | 0.941278788 | NA |
| | 0.730819482 | NA |
| | 0.238168029 | 0.393781288 |
| | 0.031567205 | 0.086383377 |
| | 0.042055871 | 0.108323107 |
| | 0.924703437 | 0.954913064 |
| | 1.38E-09 | 2.40E-08 |
| | 0.839318641 | NA |

| | | | |
|---|---|---|---|
| | 2.60E-13 | | 6.89E-12 |
| | 4.12E-05 | | 0.000318977 |
| | 0.255929877 | | 0.413946924 |
| NA | | NA | |
| | 0.400631115 | | 0.563101117 |
| NA | | NA | |
| NA | | NA | |
| NA | | NA | |
| NA | | NA | |
| | 0.000415045 | | 0.002413612 |
| | 0.252225937 | | 0.409989876 |
| | 0.26450178 | | 0.42349126 |
| | 0.180799355 | | 0.322872657 |
| | 0.24841565 | NA | |
| | 0.675942209 | NA | |
| | 0.940685982 | NA | |
| | 0.192139615 | NA | |
| | 3.83E-08 | | 5.47E-07 |
| | 0.570067619 | | 0.710839312 |
| NA | | NA | |
| NA | | NA | |
| | 0.027771238 | | 0.078124454 |
| NA | | NA | |
| | 0.941277857 | NA | |
| | 0.519011135 | | 0.669639149 |
| | 0.839320041 | NA | |
| | 0.014380114 | | 0.046134988 |
| | 0.041097079 | | 0.106475745 |
| | 0.007881387 | | 0.028307346 |
| | 0.795919118 | | 0.874871471 |
| NA | | NA | |
| | 0.004566859 | | 0.018107676 |
| | 0.941613418 | NA | |
| | 0.001041935 | | 0.005289004 |
| | 0.017411894 | | 0.053690741 |
| | 0.004516356 | | 0.017951613 |
| | 0.001373439 | | 0.006664749 |
| | 0.000612397 | | 0.003370238 |
| NA | | NA | |
| | 0.922702563 | NA | |
| NA | | NA | |
| | 0.00370965 | | 0.015244662 |
| | 0.62676188 | NA | |
| | 0.767134851 | | 0.855464758 |
| | 2.02E-08 | | 3.01E-07 |
| | 0.010396724 | | 0.035534582 |
| | 0.241936379 | | 0.39822287 |

| | |
|---|---|
| 1.36E-15 | 4.34E-14 |
| 4.18E-05 | 0.00032277 |
| 0.002170342 | 0.00978195 |
| 0.093013333 | 0.198800088 |
| 0.419253417 | NA |
| 0.070849331 | 0.162618687 |
| 0.373355011 | NA |
| 0.922893457 | NA |
| 0.338689777 | 0.502179908 |
| 1.63E-07 | 2.12E-06 |
| 0.020732874 | 0.06155468 |
| 0.955202209 | 0.972692863 |
| 0.866354522 | 0.920209306 |
| 0.287784131 | 0.448375852 |
| 0.283333414 | 0.443882177 |
| 0.105900312 | NA |
| NA | NA |
| 0.004675845 | 0.018490523 |
| NA | NA |
| NA | NA |
| 0.402253711 | 0.564528097 |
| NA | NA |
| NA | NA |
| NA | NA |
| 3.40E-06 | 3.43E-05 |
| NA | NA |
| 0.015196286 | 0.048202697 |
| 0.007281124 | 0.026539865 |
| 0.000528372 | 0.002976644 |
| 0.554890476 | 0.699587007 |
| 0.041167677 | 0.106612234 |
| 0.9981341 | 0.998469692 |
| NA | NA |
| 0.341079255 | 0.504595534 |
| 0.000236436 | 0.00148018 |
| 0.019303365 | 0.058262579 |
| NA | NA |
| 0.025419988 | 0.072607281 |
| NA | NA |
| 0.016710562 | 0.051979194 |
| 0.103021658 | 0.214496284 |
| 0.043122868 | 0.110428843 |
| 0.032528346 | 0.088535128 |
| 0.046841472 | 0.117682153 |
| 0.038412129 | 0.100825484 |
| NA | NA |
| 0.060586164 | 0.144149527 |

| | | |
|---|---|---|
| | 0.142627914 | 0.272981587 |
| | 0.029144906 | 0.081097389 |
| | 0.076635922 | 0.172213766 |
| | 0.074379417 | 0.16842843 |
| | 0.083369804 | 0.183330383 |
| | 0.102857108 | 0.214266137 |
| | 0.066833836 | 0.155607841 |
| | 0.427777502 | NA |
| | 0.248985802 | 0.406299521 |
| | 0.065560781 | 0.153301622 |
| | 0.025189748 | 0.072089261 |
| | 0.047957515 | 0.119803364 |
| | 0.143693173 | 0.274471143 |
| | 0.059653128 | 0.142385053 |
| | 0.001324774 | 0.006463431 |
| NA | | NA |
| | 0.839319574 | NA |
| | 0.002040884 | 0.009304743 |
| | 0.698113622 | 0.806861243 |
| | 0.303320779 | NA |
| | 0.19944237 | 0.346860097 |
| | 0.013613344 | 0.044139056 |
| | 0.27780457 | NA |
| NA | | NA |
| | 0.955792873 | NA |
| NA | | NA |
| | 0.481258238 | NA |
| | 0.955792398 | NA |
| | 0.955792398 | NA |
| | 0.941277857 | NA |
| NA | | NA |
| | 0.758695404 | NA |
| NA | | NA |
| NA | | NA |
| NA | | NA |
| | 0.014458304 | 0.046304702 |
| | 0.970521985 | 0.98224187 |
| | 0.962289528 | 0.977246054 |
| NA | | NA |
| | 0.423270362 | 0.58461563 |
| | 0.463805379 | 0.622469446 |
| | 0.25127077 | 0.40896792 |
| | 0.103642976 | 0.215431851 |
| | 0.331426209 | 0.49455248 |
| | 0.142620257 | 0.272981587 |
| | 0.249012002 | NA |
| | 0.125750203 | 0.248738742 |

| | |
|---|---|
| 0.262378122 | 0.421284678 |
| 0.010192241 | 0.034936021 |
| 0.18080706 | 0.322872657 |
| 0.003221273 | 0.013616186 |
| 0.60663116 | NA |
| 0.730962591 | NA |
| NA | NA |
| 0.839318641 | NA |
| 0.091115265 | 0.196051848 |
| 0.401108664 | 0.563521923 |
| 0.853596091 | NA |
| 4.34E-12 | 1.02E-10 |
| NA | NA |
| 0.361434183 | 0.525092019 |
| 0.956237904 | NA |
| 0.778601733 | 0.863592382 |
| 0.23445198 | 0.389232477 |
| 0.000302541 | 0.00183467 |
| 0.870703406 | 0.92266237 |
| 1.10E-05 | 9.86E-05 |
| NA | NA |
| NA | NA |
| NA | NA |
| 0.878694943 | NA |
| 0.240076605 | 0.396083965 |
| NA | NA |
| 7.59E-05 | 0.000549088 |
| NA | NA |
| 0.131593355 | 0.257010456 |
| 0.092866945 | 0.19866292 |
| 0.406849909 | NA |
| 0.822071883 | NA |
| NA | NA |
| 0.637689056 | 0.761846856 |
| 0.814422029 | NA |
| 0.658157848 | NA |
| 0.838160871 | 0.901935217 |
| 0.258423589 | 0.416739267 |
| 0.616821175 | 0.746470287 |
| 0.507471785 | 0.659093027 |
| 0.890831883 | 0.934367626 |
| NA | NA |
| 0.208383474 | 0.357705426 |
| NA | NA |
| 0.982641625 | 0.989040763 |
| 0.839319108 | NA |
| NA | NA |

| | | |
|---|---|---|
| NA | NA | |
| NA | NA | |
| NA | NA | |
| NA | NA | |
| NA | NA | |
| 0.06076463 | | 0.144473051 |
| 0.764873311 | NA | |
| 0.839318641 | NA | |
| 0.654440581 | | 0.77439894 |
| 0.728678906 | NA | |
| 0.489174421 | NA | |
| NA | NA | |
| NA | NA | |
| NA | NA | |
| NA | NA | |
| NA | NA | |
| NA | NA | |
| NA | NA | |
| 0.941277857 | NA | |
| 0.0585391 | | 0.140223682 |
| 0.75441813 | | 0.846134216 |
| NA | NA | |
| 0.114739114 | | 0.232287635 |
| 0.028891623 | | 0.080581017 |
| 0.000113095 | | 0.000780256 |
| 0.120310139 | | 0.240842887 |
| 0.008938351 | | 0.031397674 |
| 0.001506857 | | 0.007202048 |
| 0.855065142 | | 0.912821157 |
| 0.385159364 | | 0.5479715 |
| 0.00039726 | | 0.002323241 |
| 0.001230485 | | 0.006088984 |
| 0.000636964 | | 0.003479694 |
| 0.023786026 | | 0.068908836 |
| 0.000522207 | | 0.002947493 |
| 0.002633665 | | 0.011529609 |
| 0.047106584 | | 0.118117656 |
| 0.648118573 | | 0.770016885 |
| NA | NA | |
| NA | NA | |
| NA | NA | |
| NA | NA | |
| NA | NA | |
| NA | NA | |
| NA | NA | |
| NA | NA | |
| NA | NA | |

| | |
|---|---|
| NA | NA |
| NA | NA |
| 0.444626117 | 0.605295746 |
| NA | NA |
| 0.604092428 | 0.737106362 |
| NA | NA |
| 0.839318641 | NA |
| NA | NA |
| 0.839319108 | NA |
| 0.853596091 | NA |
| 0.519690565 | 0.670188807 |
| NA | NA |
| NA | NA |
| NA | NA |
| NA | NA |
| 0.320516567 | 0.483211004 |
| NA | NA |
| NA | NA |
| NA | NA |
| NA | NA |
| NA | NA |
| 0.955792398 | NA |
| NA | NA |
| NA | NA |
| 0.906311298 | NA |
| NA | NA |
| 1.43E-08 | 2.19E-07 |
| 0.597184622 | 0.731822498 |
| 0.480698916 | 0.637272786 |
| 0.020325415 | 0.060655157 |
| 0.296308273 | 0.45779322 |
| 0.002394711 | 0.010644056 |
| 0.048512316 | 0.120846691 |
| 0.316211955 | 0.478683643 |
| 6.78E-06 | 6.40E-05 |
| 0.630327941 | NA |
| 0.955792398 | NA |
| NA | NA |
| 0.802161731 | 0.879054029 |
| 0.000433115 | 0.00250401 |
| NA | NA |
| NA | NA |
| NA | NA |
| 1.36E-11 | 3.03E-10 |
| 0.839319108 | NA |
| 0.653901412 | 0.774066813 |
| NA | NA |

| | | |
|---|---|---|
| | 0.310212747 | 0.472012005 |
| | 0.40612838 | NA |
| | 0.237395185 | NA |
| | 0.839318641 | NA |
| NA | | NA |
| NA | | NA |
| | 2.58E-10 | 4.94E-09 |
| | 0.778158718 | NA |
| | 0.686900553 | NA |
| | 0.620948715 | 0.749468748 |
| | 0.458515204 | 0.618109189 |
| | 0.941275956 | NA |
| NA | | NA |
| | 0.941278332 | NA |
| | 0.423521035 | NA |
| | 0.14554906 | 0.277126528 |
| | 0.011047956 | 0.037342155 |
| | 0.29710139 | 0.458569955 |
| | 5.75E-11 | 1.19E-09 |
| | 4.28E-10 | 7.98E-09 |
| | 0.006802142 | 0.025074274 |
| | 0.941277857 | NA |
| | 0.380459634 | 0.543135621 |
| | 0.00317216 | 0.013446793 |
| | 0.050115989 | 0.124034481 |
| | 0.144857532 | 0.276119083 |
| | 0.000231348 | 0.001451766 |
| | 1.23E-06 | 1.35E-05 |
| | 0.536113153 | 0.68417593 |
| | 0.297317847 | 0.458735296 |
| | 0.001211164 | 0.006009018 |
| | 0.017411933 | 0.053690741 |
| | 0.706907844 | 0.813037956 |
| | 0.16011836 | 0.296794425 |
| | 0.228499685 | 0.382068809 |
| | 0.839319108 | NA |
| | 0.862348786 | 0.91768129 |
| | 0.014021649 | 0.045176245 |
| | 0.104591699 | 0.216760263 |
| | 0.021913142 | 0.064416032 |
| | 7.95E-05 | 0.000572565 |
| | 0.276084745 | 0.435882801 |
| | 0.070142471 | 0.161478972 |
| NA | | NA |
| | 0.044129188 | 0.112266239 |
| | 0.039569062 | 0.103292698 |
| | 0.983384996 | 0.989370601 |

| | | |
|---|---|---|
| | 0.853595154 | NA |
| | 0.5473779 | 0.69355825 |
| | 0.07121881 | 0.163167786 |
| | 0.136311526 | 0.263908162 |
| | 0.488090832 | 0.642811818 |
| | 0.154631132 | 0.289463642 |
| | 0.230158514 | 0.384006235 |
| | 0.941279268 | NA |
| | 0.935946872 | NA |
| NA | | NA |
| | 1.07E-15 | 3.43E-14 |
| | 0.909807062 | NA |
| | 0.289291106 | NA |
| NA | | NA |
| NA | | NA |
| | 0.084082047 | 0.184623698 |
| NA | | NA |
| | 1.47E-13 | 3.99E-12 |
| | 0.004731248 | 0.018672387 |
| | 0.048848186 | 0.12151803 |
| | 0.936343336 | NA |
| NA | | NA |
| NA | | NA |
| NA | | NA |
| NA | | NA |
| | 0.941278332 | NA |
| | 0.624636833 | 0.752713948 |
| | 0.894348336 | 0.936363116 |
| | 0.181103625 | 0.323329469 |
| | 0.150602421 | 0.284179391 |
| | 0.92531441 | 0.955182718 |
| | 0.839318641 | NA |
| | 0.834337904 | 0.899651662 |
| | 0.444591757 | 0.60528359 |
| | 0.817759809 | 0.888757758 |
| | 0.157870402 | 0.2938634 |
| | 0.7908528 | 0.871556549 |
| | 0.332051118 | 0.49514351 |
| | 0.353537997 | 0.517143784 |
| | 0.01637189 | 0.051153022 |
| | 0.941277857 | NA |
| | 4.37E-06 | 4.31E-05 |
| | 0.128704524 | 0.253008348 |
| | 0.09741378 | 0.205612521 |
| | 0.073705035 | 0.167301854 |
| NA | | NA |
| | 0.941277857 | NA |

| | | |
|---|---|---|
| | 0.764261262 | NA |
| NA | | NA |
| | 0.744626095 | 0.839582677 |
| | 0.484440363 | 0.640271505 |
| | 0.47642283 | 0.633473895 |
| | 0.355618549 | 0.519385943 |
| | 2.71E-06 | 2.79E-05 |
| | 0.992036053 | NA |
| | 0.000849188 | 0.004443256 |
| | 0.318767242 | 0.481152762 |
| | 0.137309806 | 0.265409128 |
| NA | | NA |
| | 2.63E-05 | 0.00021528 |
| | 0.730919803 | 0.830396998 |
| | 0.092157354 | 0.197409874 |
| | 0.226203042 | 0.379214312 |
| | 0.113991184 | 0.231060899 |
| | 0.577357807 | 0.71581782 |
| | 0.242228782 | 0.39855727 |
| | 0.61280162 | 0.743634467 |
| | 0.560936085 | 0.703988552 |
| | 0.602311341 | 0.735756455 |
| | 0.140902714 | 0.270517252 |
| | 0.574034459 | NA |
| | 0.656867578 | NA |
| | 0.004274975 | 0.017144559 |
| | 0.731251773 | NA |
| | 0.66360781 | NA |
| | 0.527405027 | 0.676508647 |
| | 0.905434405 | NA |
| | 0.941277857 | NA |
| NA | | NA |
| NA | | NA |
| | 0.853595154 | NA |
| | 0.189298992 | 0.333894494 |
| | 0.104588085 | 0.216760263 |
| | 0.014012812 | 0.045163568 |
| | 0.014804732 | 0.047172988 |
| | 0.02177429 | 0.064087009 |
| | 0.375170754 | 0.538264876 |
| | 0.324854557 | 0.487742898 |
| | 0.006882221 | 0.025330235 |
| | 0.015323136 | 0.048525984 |
| | 0.663600986 | NA |
| NA | | NA |
| | 0.879886199 | NA |
| | 0.55546167 | 0.699973458 |

| | | | |
|---|---|---|---|
| | 0.205641456 | | 0.354291813 |
| | 0.702122903 | NA | |
| | 0.525034801 | | 0.674449691 |
| | 0.030710254 | | 0.084485144 |
| NA | | NA | |
| NA | | NA | |
| NA | | NA | |
| NA | | NA | |
| NA | | NA | |
| NA | | NA | |
| | 0.461245912 | NA | |
| | 0.79946057 | | 0.877588917 |
| | 9.81E-05 | | 0.000689721 |
| | 0.095173733 | | 0.202170922 |
| | 0.158718873 | | 0.295027459 |
| | 0.834286262 | | 0.899651662 |
| | 0.015748358 | | 0.049608578 |
| | 0.734058779 | NA | |
| | 0.040717545 | | 0.105664959 |
| | 0.00149598 | | 0.007160127 |
| NA | | NA | |
| | 0.000169195 | | 0.001112786 |
| | 0.065421398 | | 0.15303785 |
| | 0.853595154 | NA | |
| NA | | NA | |
| NA | | NA | |
| | 0.853596091 | NA | |
| | 0.813990998 | NA | |
| | 0.791638073 | | 0.871750204 |
| | 0.546766941 | | 0.692942161 |
| | 0.62593735 | NA | |
| | 0.561742752 | | 0.704643003 |
| | 0.110894366 | NA | |
| | 0.941281159 | NA | |
| | 0.226167609 | | 0.379197552 |
| NA | | NA | |
| | 0.000233383 | | 0.001463765 |
| | 1.70E-08 | | 2.57E-07 |
| NA | | NA | |
| | 2.43E-05 | | 0.000200812 |
| | 0.277488726 | | 0.437366025 |
| | 0.486860511 | | 0.641937396 |
| | 0.032431732 | | 0.088292357 |
| | 0.839318641 | NA | |
| NA | | NA | |
| | 0.556884015 | | 0.700949409 |
| | 0.641606385 | | 0.764914359 |

| | | | |
|---|---|---|---|
| | 0.666684681 | | 0.783935816 |
| | 0.028268274 | | 0.079139198 |
| | 0.145212672 | | 0.276618808 |
| NA | | NA | |
| | 0.028818768 | | 0.08042494 |
| | 0.068055184 | | 0.157772425 |
| | 0.941277857 | NA | |
| | 0.955792873 | NA | |
| NA | | NA | |
| | 0.361993224 | | 0.52568421 |
| | 0.722455232 | NA | |
| | 0.004261734 | | 0.017100098 |
| | 0.3344926 | | 0.4976932 |
| | 0.240122875 | | 0.396105389 |
| | 0.955791348 | NA | |
| | 0.391352862 | | 0.553819256 |
| | 0.791779903 | NA | |
| NA | | NA | |
| | 0.018976538 | | 0.057516817 |
| | 0.224586582 | | 0.377286014 |
| | 6.48E-11 | | 1.34E-09 |
| | 0.621458339 | | 0.750022888 |
| | 0.26290443 | | 0.421873609 |
| NA | | NA | |
| | 0.941278332 | NA | |
| | 6.79E-07 | | 7.81E-06 |
| | 0.826160666 | NA | |
| | 0.839318641 | NA | |
| | 0.056863338 | | 0.137142686 |
| NA | | NA | |
| | 0.537928613 | NA | |
| | 0.880610121 | | 0.928432056 |
| NA | | NA | |
| | 0.471880451 | NA | |
| | 0.111596609 | | 0.227455257 |
| | 0.464065618 | | 0.622666014 |
| | 0.716864589 | NA | |
| | 0.733209887 | | 0.831674692 |
| | 0.882715965 | | 0.929828527 |
| | 0.302298199 | | 0.463666198 |
| | 0.975616832 | | 0.985178051 |
| | 0.797507269 | | 0.875969913 |
| | 0.073639692 | | 0.167185421 |
| | 0.009222346 | | 0.032205148 |
| | 0.941278284 | NA | |
| NA | | NA | |
| NA | | NA | |

| | | |
|---|---|---|
| NA | | NA |
| | 0.955792398 | NA |
| | 0.692702149 | 0.80298527 |
| | 0.012905607 | 0.042278928 |
| | 0.262723099 | 0.421667916 |
| | 0.853596091 | NA |
| | 0.836525318 | 0.900925912 |
| NA | | NA |
| | 0.941277857 | NA |
| NA | | NA |
| | 0.364608484 | 0.527948116 |
| | 0.813588497 | NA |
| NA | | NA |
| | 0.853596091 | NA |
| NA | | NA |
| NA | | NA |
| NA | | NA |
| NA | | NA |
| NA | | NA |
| | 0.428150597 | 0.589412473 |
| | 0.839319574 | NA |
| | 0.923532773 | NA |
| NA | | NA |
| | 0.523752248 | NA |
| | 0.839318641 | NA |
| | 0.941278332 | NA |
| NA | | NA |
| | 2.11E-13 | 5.61E-12 |
| NA | | NA |
| NA | | NA |
| | 0.359749674 | 0.523364203 |
| | 0.001045699 | 0.005301328 |
| | 0.979910701 | 0.987592489 |
| | 0.530508551 | 0.679318083 |
| | 0.117877655 | 0.236993235 |
| | 0.353828434 | 0.517470008 |
| | 0.647484423 | 0.769532343 |
| | 0.195099336 | 0.341418618 |
| | 0.955791922 | NA |
| | 0.684362991 | 0.797279344 |
| | 0.979947296 | 0.987592489 |
| NA | | NA |
| | 0.955792873 | NA |
| | 0.615170958 | 0.745193096 |
| | 0.139067324 | 0.267805861 |
| | 0.784626597 | 0.867627515 |
| | 0.486408903 | 0.641664161 |

| | |
|---|---|
| 2.23E-06 | 2.34E-05 |
| 0.955792398 | NA |
| 0.000395361 | 0.002314411 |
| 0.974234957 | 0.984409087 |
| 0.60341346 | 0.736611303 |
| 0.713638944 | 0.817894557 |
| 0.000788504 | 0.004177597 |
| 0.329538354 | 0.492538266 |
| NA | NA |
| 0.850747809 | 0.910458133 |
| 2.75E-06 | 2.81E-05 |
| 7.60E-05 | 0.000549234 |
| 0.925732935 | NA |
| NA | NA |
| 0.883627668 | NA |
| NA | NA |
| 5.80E-09 | 9.39E-08 |
| 0.872427408 | NA |
| 0.119702465 | 0.239807934 |
| 0.076774947 | 0.172477327 |
| 0.700085347 | 0.808551188 |
| 3.59E-12 | 8.51E-11 |
| NA | NA |
| 0.838197868 | NA |
| NA | NA |
| NA | NA |
| 0.835323277 | NA |
| NA | NA |
| 0.939680037 | NA |
| NA | NA |
| 0.974830433 | NA |
| NA | NA |
| NA | NA |
| NA | NA |
| NA | NA |
| NA | NA |
| NA | NA |
| NA | NA |
| NA | NA |
| 0.150240843 | NA |
| 0.58929908 | 0.725393728 |
| 0.006915824 | 0.02542639 |
| 0.802840171 | 0.879453216 |
| 0.138362802 | 0.266836243 |
| 0.684430251 | 0.797279344 |
| 0.90490135 | 0.942354828 |
| 0.839319574 | NA |

| | | |
|---|---|---|
| | 0.941278332 | NA |
| | 0.049171801 | 0.12205519 |
| NA | | NA |
| NA | | NA |
| NA | | NA |
| | 0.22174795 | 0.373931612 |
| | 0.853595154 | NA |
| NA | | NA |
| NA | | NA |
| NA | | NA |
| NA | | NA |
| NA | | NA |
| | 0.003445356 | 0.014366916 |
| | 0.103250198 | 0.214777971 |
| | 0.835756296 | 0.900202342 |
| NA | | NA |
| NA | | NA |
| | 0.499830153 | NA |
| NA | | NA |
| | 0.078455731 | 0.175194981 |
| | 0.061661389 | 0.146167152 |
| | 0.923516642 | NA |
| | 0.000227779 | 0.001433528 |
| | 0.941278788 | NA |
| | 5.58E-07 | 6.51E-06 |
| | 0.551733564 | NA |
| | 0.955792398 | NA |
| NA | | NA |
| | 0.398963971 | 0.561669157 |
| | 0.00840217 | 0.029840115 |
| | 0.759832911 | NA |
| | 0.661569785 | NA |
| | 0.005277204 | 0.020384294 |
| | 0.941275956 | NA |
| | 0.92069054 | NA |
| | 0.955791446 | NA |
| NA | | NA |
| | 0.955791922 | NA |
| | 0.025248026 | 0.072221308 |
| NA | | NA |
| | 0.004412318 | 0.017591624 |
| NA | | NA |
| NA | | NA |
| NA | | NA |
| NA | | NA |
| | 0.839319574 | NA |
| | 0.16513053 | 0.30369625 |

| | | | |
|---|---|---|---|
| | 0.277708764 | NA | |
| NA | | NA | |
| NA | | NA | |
| | 0.617155265 | NA | |
| NA | | NA | |
| NA | | NA | |
| | 0.38633275 | NA | |
| | 0.839318641 | NA | |
| NA | | NA | |
| | 0.839318641 | NA | |
| | 0.941278332 | NA | |
| | 0.839319108 | NA | |
| | 0.839318641 | NA | |
| NA | | NA | |
| NA | | NA | |
| | 0.853595622 | NA | |
| NA | | NA | |
| NA | | NA | |
| | 0.941278424 | NA | |
| | 0.839318641 | NA | |
| NA | | NA | |
| NA | | NA | |
| | 0.955792398 | NA | |
| | 0.263919714 | | 0.422989498 |
| | 0.941277857 | NA | |
| NA | | NA | |
| NA | | NA | |
| NA | | NA | |
| | 0.839319108 | NA | |
| NA | | NA | |
| NA | | NA | |
| | 0.360805407 | NA | |
| NA | | NA | |
| | 0.839319108 | NA | |
| NA | | NA | |
| NA | | NA | |
| NA | | NA | |
| NA | | NA | |
| NA | | NA | |
| NA | | NA | |
| NA | | NA | |
| | 0.597892605 | NA | |
| | 0.737433066 | NA | |
| | 0.439543117 | NA | |
| NA | | NA | |
| | 0.955792397 | NA | |
| NA | | NA | |

| | |
|---|---|
| NA | NA |
| NA | NA |
| NA | NA |
| NA | NA |
| NA | NA |
| NA | NA |
| NA | NA |
| NA | NA |
| NA | NA |
| NA | NA |
| NA | NA |
| NA | NA |
| 0.274607654 | 0.434530739 |
| NA | NA |
| NA | NA |
| NA | NA |
| 0.247741933 | 0.404960003 |
| 0.513073246 | 0.664282495 |
| 0.006319957 | 0.023618717 |
| 0.388083151 | 0.550978536 |
| 0.003589214 | 0.014850577 |
| 0.05623044 | 0.136029773 |
| 0.677665751 | 0.792190963 |
| 8.69E-06 | 7.97E-05 |
| 0.013101139 | 0.042804846 |
| NA | NA |
| 0.948528245 | 0.968548661 |
| 0.000133598 | 0.000904641 |
| 0.82926006 | 0.896381426 |
| 0.839318641 | NA |
| NA | NA |
| 0.00502137 | 0.019599649 |
| 0.413796658 | 0.57597591 |
| 0.643218136 | NA |
| NA | NA |
| 0.035936497 | 0.095742164 |
| 2.52E-10 | 4.83E-09 |
| 0.004665666 | 0.018459472 |
| 0.004470048 | 0.017791985 |
| NA | NA |
| 0.696223329 | 0.805326553 |
| 0.174839865 | 0.315850298 |
| 0.839319108 | NA |
| 0.876276063 | NA |
| 0.754025981 | NA |
| NA | NA |
| 0.417753537 | NA |

| | |
|---|---|
| 0.060208687 | 0.143437811 |
| 0.71399753 | NA |
| 0.988927942 | 0.992997975 |
| 0.642778251 | 0.765697809 |
| 0.034908651 | 0.093422154 |
| 0.086002098 | 0.187800178 |
| 0.955792873 | NA |
| NA | NA |
| NA | NA |
| NA | NA |
| 4.72E-10 | 8.76E-09 |
| 0.494094891 | NA |
| 1.07E-08 | 1.67E-07 |
| NA | NA |
| 0.068384714 | 0.158305092 |
| 0.41465293 | NA |
| 0.939348064 | NA |
| 0.00116667 | 0.00581673 |
| 0.07533881 | NA |
| 0.888598951 | 0.93342843 |
| 0.013023962 | 0.042611181 |
| 1.83E-05 | 0.00015505 |
| 0.489442654 | 0.643844039 |
| 0.361707652 | 0.525345712 |
| 0.636114357 | 0.760805684 |
| 0.903399268 | NA |
| 0.738922978 | NA |
| 0.276475766 | NA |
| NA | NA |
| 0.248428951 | 0.405646284 |
| 0.22775763 | NA |
| 0.001119804 | 0.005626079 |
| 0.376418706 | 0.539534989 |
| 0.07567338 | 0.170663046 |
| 0.941276431 | NA |
| 0.03548751 | 0.094758101 |
| 0.000215659 | 0.001369925 |
| 0.183948705 | 0.326815488 |
| 3.40E-07 | 4.14E-06 |
| 0.002291759 | 0.010265044 |
| 0.795354559 | 0.874452825 |
| 0.856194327 | 0.913590067 |
| 0.193057616 | 0.338765904 |
| NA | NA |
| 0.169134886 | 0.308675706 |
| 0.941277857 | NA |
| 4.63E-05 | 0.000354128 |

| | | |
|---|---|---|
| | 2.49E-05 | 0.000204788 |
| | 9.54E-07 | 1.06E-05 |
| | 0.941278332 | NA |
| | 0.839318641 | NA |
| | 6.83E-38 | 6.47E-36 |
| | 4.87E-10 | 9.02E-09 |
| | 3.57E-62 | 6.79E-60 |
| | 4.60E-46 | 5.77E-44 |
| | 0.356368547 | NA |
| | 0.170399976 | 0.310175911 |
| NA | | NA |
| | 2.91E-09 | 4.88E-08 |
| NA | | NA |
| | 0.414916718 | 0.576759196 |
| | 0.939787132 | 0.963757575 |
| | 0.034498236 | 0.092583946 |
| | 0.07378179 | 0.167428179 |
| | 0.651953803 | NA |
| NA | | NA |
| NA | | NA |
| | 0.169531105 | 0.309185454 |
| | 0.941278332 | NA |
| | 0.688731298 | 0.800376082 |
| | 0.152819065 | 0.286929036 |
| | 0.767843295 | 0.855988295 |
| | 0.217770188 | 0.369117494 |
| | 4.45E-16 | 1.48E-14 |
| | 0.117345593 | 0.236179587 |
| | 0.842655586 | 0.905216782 |
| | 0.00732406 | 0.026671861 |
| | 0.831829705 | 0.897998296 |
| | 0.049841329 | 0.123446754 |
| | 5.54E-06 | 5.35E-05 |
| | 0.571907356 | 0.711962173 |
| NA | | NA |
| NA | | NA |
| NA | | NA |
| NA | | NA |
| NA | | NA |
| NA | | NA |
| NA | | NA |
| NA | | NA |
| NA | | NA |
| NA | | NA |
| NA | | NA |
| NA | | NA |
| NA | | NA |

| | | | |
|---|---|---|---|
| NA | | NA | |
| NA | | NA | |
| NA | | NA | |
| NA | | NA | |
| NA | | NA | |
| NA | | NA | |
| NA | | NA | |
| NA | | NA | |
| NA | | NA | |
| NA | | NA | |
| NA | | NA | |
| NA | | NA | |
| NA | | NA | |
| NA | | NA | |
| NA | | NA | |
| NA | | NA | |
| NA | | NA | |
| NA | | NA | |
| NA | | NA | |
| NA | | NA | |
| NA | | NA | |
| NA | | NA | |
| NA | | NA | |
| NA | | NA | |
| NA | | NA | |
| NA | | NA | |
| NA | | NA | |
| NA | | NA | |
| NA | | NA | |
| NA | | NA | |
| NA | | NA | |
| NA | | NA | |
| NA | | NA | |
| NA | | NA | |
| NA | | NA | |
| NA | | NA | |
| | 0.119630295 | | 0.239758593 |
| NA | | NA | |
| | 2.76E-29 | | 1.88E-27 |
| | 1.82E-05 | | 0.000154104 |
| | 0.000157826 | | 0.001045736 |
| | 0.04980162 | | 0.123386923 |
| | 0.010497324 | | 0.035806435 |
| | 1.82E-06 | | 1.94E-05 |
| | 0.937578753 | NA | |
| | 0.721364294 | NA | |
| | 0.941278207 | NA | |

| | | |
|---|---|---|
| | 2.78E-18 | 1.05E-16 |
| | 0.290750907 | 0.451874181 |
| | 0.676749155 | NA |
| | 0.496664351 | NA |
| | 0.822151266 | NA |
| NA | | NA |
| | 0.001636958 | 0.007735334 |
| | 0.429222328 | NA |
| | 0.908770131 | 0.944606605 |
| | 0.806464097 | 0.881532419 |
| | 0.922010512 | NA |
| | 0.280420583 | 0.440391848 |
| | 1.14E-21 | 5.38E-20 |
| | 0.028714238 | 0.08020846 |
| | 0.925686951 | 0.955348429 |
| | 0.896922995 | 0.937779975 |
| | 0.655636524 | 0.775195497 |
| | 0.993292504 | 0.995209152 |
| NA | | NA |
| | 0.788621285 | NA |
| | 0.620763124 | 0.749468748 |
| | 0.955792398 | NA |
| | 0.022269431 | 0.065273033 |
| | 0.132506963 | 0.258264881 |
| | 0.955792398 | NA |
| NA | | NA |
| | 0.901766297 | NA |
| | 0.941278424 | NA |
| | 0.892069314 | 0.93496406 |
| | 0.070042468 | 0.161288038 |
| | 0.371754307 | 0.535072625 |
| | 0.739528174 | 0.836211382 |
| | 0.115814247 | 0.23380922 |
| | 0.577860849 | NA |
| | 0.211245513 | 0.361418108 |
| | 0.959962952 | NA |
| | 0.247520504 | NA |
| NA | | NA |
| | 7.07E-06 | 6.63E-05 |
| | 0.005405233 | 0.020765768 |
| | 0.791565025 | 0.871750204 |
| | 0.455746503 | NA |
| | 4.41E-07 | 5.26E-06 |
| | 0.62221576 | 0.750670562 |
| | 0.364836088 | 0.528149165 |
| | 0.758365546 | NA |
| | 0.187389955 | 0.331420398 |

| | |
|---|---|
| 0.2872488 | 0.447830664 |
| NA | NA |
| NA | NA |
| NA | NA |
| NA | NA |
| 0.513324863 | 0.664463694 |
| 0.941277857 | NA |
| NA | NA |
| 0.7467615 | 0.840715979 |
| 0.941279303 | NA |
| 0.955791922 | NA |
| 0.030697676 | 0.084460305 |
| 0.195758101 | 0.342218777 |
| 0.561330038 | 0.704199524 |
| 0.000121318 | 0.000830492 |
| 0.081869709 | 0.180783265 |
| NA | NA |
| 0.023533826 | 0.068302906 |
| 0.061416857 | NA |
| 0.890039875 | NA |
| 0.791655669 | NA |
| 0.492372108 | NA |
| 0.701679075 | 0.809527669 |
| 0.117907021 | 0.237028963 |
| 0.697766597 | NA |
| 0.941277857 | NA |
| NA | NA |
| 0.945731563 | NA |
| 0.899471264 | 0.93958188 |
| 0.309495694 | 0.471161912 |
| 0.603478948 | NA |
| NA | NA |
| NA | NA |
| 1.29E-06 | 1.41E-05 |
| 2.70E-27 | 1.69E-25 |
| NA | NA |
| 0.008573778 | 0.030326232 |
| 0.605206406 | NA |
| 9.61E-22 | 4.56E-20 |
| 0.951449697 | 0.970324594 |
| 0.000341868 | 0.002042966 |
| 0.796136581 | 0.874908486 |
| 0.108412074 | 0.222623085 |
| 0.078598909 | 0.175465318 |
| 0.478706104 | 0.63544491 |
| 0.806932201 | NA |
| 0.581055772 | 0.718830015 |

| | |
|---|---|
| 0.150030936 | 0.283213288 |
| 0.807017784 | 0.881736851 |
| 0.839319108 | NA |
| 0.839318641 | NA |
| NA | NA |
| NA | NA |
| 0.507986777 | 0.659598563 |
| NA | NA |
| 0.589052564 | 0.725205623 |
| 0.000959812 | 0.00492677 |
| 0.047262099 | 0.118376563 |
| 0.818724089 | 0.889464182 |
| 4.29E-05 | 0.000330619 |
| 0.166022853 | 0.304887983 |
| 0.940685982 | NA |
| 0.851325824 | 0.910790061 |
| 0.183478085 | 0.32619849 |
| 0.302778183 | NA |
| 0.897908921 | 0.938413841 |
| 0.191043648 | NA |
| 0.814377017 | NA |
| 2.92E-07 | 3.61E-06 |
| 0.31640875 | 0.478807367 |
| 0.54806132 | 0.693991357 |
| 0.322500539 | 0.485341289 |
| NA | NA |
| NA | NA |
| NA | NA |
| NA | NA |
| 0.234251214 | 0.389007703 |
| 0.002082224 | 0.009458224 |
| NA | NA |
| 0.839318641 | NA |
| 5.86E-12 | 1.36E-10 |
| NA | NA |
| 0.638974726 | NA |
| 0.592655733 | 0.728184583 |
| NA | NA |
| 2.38E-16 | 8.01E-15 |
| 1.85E-07 | 2.37E-06 |
| 0.060240384 | 0.143492579 |
| NA | NA |
| 0.357133122 | 0.520718058 |
| NA | NA |
| 0.220793299 | 0.37270213 |
| 0.212312919 | 0.362776173 |
| 0.000646443 | 0.00352662 |

| | | |
|---|---|---|
| | 0.966171311 | NA |
| | 0.396291688 | 0.558931901 |
| | 0.824358773 | 0.893302413 |
| | 0.119669016 | 0.239781289 |
| | 0.3494418 | 0.51332554 |
| | 0.425874032 | 0.58729164 |
| | 0.472544366 | 0.630041509 |
| | 0.941278332 | NA |
| NA | | NA |
| | 0.947050561 | 0.967625072 |
| NA | | NA |
| | 2.36E-18 | 8.93E-17 |
| | 0.486141574 | 0.641486957 |
| | 0.059269313 | 0.141682052 |
| | 0.030753 | 0.084592962 |
| | 8.37E-06 | 7.72E-05 |
| | 0.239801638 | 0.39582237 |
| | 0.118179578 | 0.237516703 |
| | 0.186778808 | 0.330609741 |
| NA | | NA |
| NA | | NA |
| NA | | NA |
| | 0.728678906 | NA |
| | 9.80E-39 | 9.56E-37 |
| | 3.67E-154 | 2.36E-151 |
| | 4.37E-136 | 2.08E-133 |
| | 2.35E-165 | 1.75E-162 |
| | 1.43E-61 | 2.68E-59 |
| | 1.79E-81 | 4.85E-79 |
| | 0.081480765 | 0.180141666 |
| | 0.008114967 | 0.028997515 |
| | 0.188504312 | 0.332921245 |
| | 0.029447408 | 0.081761507 |
| | 0.01660846 | 0.051756294 |
| | 0.834857123 | 0.899970585 |
| | 0.006356886 | 0.023723205 |
| | 0.031942025 | 0.087168549 |
| | 0.955792873 | NA |
| | 0.735785898 | 0.833507461 |
| | 0.566014994 | NA |
| | 0.007223056 | 0.026368586 |
| | 0.493595766 | 0.646993147 |
| NA | | NA |
| NA | | NA |
| | 0.905630995 | NA |
| | 0.578861363 | NA |
| | 0.839319108 | NA |

| | |
|---|---|
| NA | NA |
| 0.955792398 | NA |
| 0.725028456 | NA |
| 0.748393366 | NA |
| NA | NA |
| 0.81884587 | 0.889515285 |
| 0.273142705 | 0.433134088 |
| 0.139709414 | 0.268672818 |
| 0.839319108 | NA |
| 0.45102246 | NA |
| NA | NA |
| 0.955792398 | NA |
| 0.020477862 | 0.060987745 |
| 0.234847669 | 0.389682113 |
| NA | NA |
| NA | NA |
| 0.215678969 | 0.366720483 |
| 0.298815877 | NA |
| NA | NA |
| 7.18E-05 | 0.000522322 |
| 0.941278332 | NA |
| 0.001193363 | 0.005931181 |
| 0.890416548 | 0.934095583 |
| 0.71120589 | 0.815773823 |
| NA | NA |
| 0.36912127 | 0.532355269 |
| NA | NA |
| 0.00179485 | 0.008355372 |
| 0.529160616 | NA |
| 0.004978474 | 0.019460937 |
| 0.955792398 | NA |
| 0.013742602 | 0.044449165 |
| 0.252985125 | 0.410803107 |
| 0.104447377 | 0.216593176 |
| 0.012508008 | 0.041234847 |
| 0.392125765 | 0.554469046 |
| 0.037045779 | 0.097973737 |
| 0.095247723 | 0.202291988 |
| 0.016673103 | 0.051889802 |
| 0.337739548 | 0.501114369 |
| 0.026223476 | 0.074493819 |
| 0.635032494 | NA |
| 0.421282313 | 0.582810801 |
| 0.800164464 | NA |
| 0.448475612 | NA |
| 2.36E-07 | 2.96E-06 |
| 0.941278838 | NA |

| | |
|---|---|
| 9.21E-08 | 1.25E-06 |
| 0.005192928 | 0.020150322 |
| NA | NA |
| NA | NA |
| 0.747133075 | 0.841094521 |
| 0.000701883 | 0.003786831 |
| NA | NA |
| 0.204907536 | 0.353497802 |
| 0.670661136 | 0.786823559 |
| 0.930265213 | 0.958494117 |
| 0.543326672 | NA |
| NA | NA |
| 0.355526429 | 0.519283263 |
| 1.98E-06 | 2.09E-05 |
| 0.639218235 | 0.763060957 |
| NA | NA |
| NA | NA |
| NA | NA |
| 0.494333739 | 0.647486884 |
| 0.144703316 | 0.275935614 |
| 7.26E-05 | 0.000527476 |
| 0.134840189 | 0.261742575 |
| 0.005808044 | 0.022044818 |
| 0.115068334 | 0.232698088 |
| 0.75220116 | 0.844563046 |
| 0.219553511 | 0.371072404 |
| 0.334703162 | 0.49796228 |
| 8.55E-06 | 7.87E-05 |
| 0.054491815 | 0.132619037 |
| 0.925151936 | NA |
| 0.225550114 | 0.378466254 |
| 0.01155617 | 0.038751757 |
| 0.006300773 | 0.023558122 |
| 0.494080085 | 0.647310858 |
| 0.024584995 | 0.070784088 |
| 5.88E-06 | 5.64E-05 |
| 0.021258062 | 0.062854549 |
| 0.398851023 | 0.561575367 |
| 3.61E-05 | 0.000283464 |
| 0.955792873 | NA |
| 0.915095983 | NA |
| NA | NA |
| NA | NA |
| NA | NA |
| NA | NA |
| 0.941278332 | NA |
| 0.482138551 | 0.638399076 |

| | |
|---|---|
| NA | NA |
| 0.955792398 | NA |
| NA | NA |
| NA | NA |
| NA | NA |
| 0.669147992 | NA |
| NA | NA |
| NA | NA |
| NA | NA |
| NA | NA |
| NA | NA |
| NA | NA |
| NA | NA |
| NA | NA |
| 0.955792398 | NA |
| NA | NA |
| 0.566779598 | NA |
| 0.004561067 | 0.018090736 |
| 3.78E-07 | 4.56E-06 |
| 0.800391721 | NA |
| NA | NA |
| 0.428492975 | 0.589741517 |
| 0.301274941 | 0.462552161 |
| 0.703664573 | 0.810844101 |
| 0.98117419 | 0.988399698 |
| 0.246020636 | 0.402902379 |
| 0.014074425 | 0.045325322 |
| 0.703859239 | NA |
| 0.036992169 | 0.097864577 |
| 0.300141112 | 0.461704824 |
| 0.005521638 | 0.021126191 |
| 0.212410569 | 0.362916973 |
| 0.06516574 | 0.152589762 |
| 0.00893526 | 0.031396221 |
| 0.090312411 | 0.194799819 |
| 0.89229191 | 0.935040806 |
| 0.011392635 | 0.038300494 |
| 0.327458925 | 0.490261501 |
| 0.001724404 | 0.008089633 |
| 0.010521471 | 0.035878518 |
| 0.584949142 | 0.721733424 |
| 0.002624071 | 0.011489725 |
| 3.55E-05 | 0.000279429 |
| 0.219839584 | 0.371476769 |
| 0.07536717 | 0.170168046 |
| 0.50136667 | 0.653784555 |
| 0.66709967 | 0.784230075 |

| | | |
|---|---|---|
| | 1.53E-05 | 0.000133019 |
| NA | | NA |
| | 0.941275956 | NA |
| | 0.941277857 | NA |
| | 0.880489729 | 0.928387372 |
| | 0.941278332 | NA |
| | 0.312864772 | 0.474772541 |
| | 0.47193511 | 0.629405948 |
| | 0.209457261 | 0.359108386 |
| NA | | NA |
| | 0.246801663 | 0.40382025 |
| | 0.168446229 | 0.307891042 |
| | 7.83E-09 | 1.25E-07 |
| | 0.18245129 | 0.324931208 |
| | 0.001784242 | 0.008317376 |
| | 0.000603395 | 0.003329937 |
| | 0.459130537 | 0.618451782 |
| | 0.494507005 | 0.647606918 |
| | 0.839319108 | NA |
| NA | | NA |
| | 0.037629407 | 0.099197602 |
| | 0.09042866 | 0.194944566 |
| | 0.693419115 | 0.803616036 |
| | 0.159254073 | 0.295629812 |
| | 0.172866649 | 0.31325253 |
| | 0.519811419 | 0.670199414 |
| | 0.443365672 | NA |
| | 0.223054564 | 0.375496481 |
| | 0.8102309 | 0.883701523 |
| | 0.292526409 | 0.453567428 |
| | 0.896893761 | 0.937779975 |
| NA | | NA |
| | 0.076782376 | 0.172477737 |
| | 0.211968191 | 0.362420859 |
| | 0.714708179 | 0.818686369 |
| NA | | NA |
| | 0.159671218 | NA |
| NA | | NA |
| | 0.103894867 | 0.215767003 |
| | 0.683548132 | NA |
| | 0.003817866 | 0.01561389 |
| | 8.03E-06 | 7.45E-05 |
| | 0.032915695 | 0.089337391 |
| | 0.008999559 | 0.031584711 |
| | 0.914971424 | NA |
| | 0.53179628 | 0.680417947 |
| NA | | NA |

| | | |
|---|---|---|
| NA | | NA |
| | 0.721517616 | NA |
| NA | | NA |
| NA | | NA |
| | 0.164041146 | 0.302159678 |
| NA | | NA |
| NA | | NA |
| | 0.678762113 | NA |
| NA | | NA |
| NA | | NA |
| | 0.058844387 | 0.140809784 |
| | 0.853281503 | 0.911742904 |
| | 3.03E-10 | 5.74E-09 |
| | 0.000182333 | 0.001184142 |
| | 0.812439859 | 0.885425528 |
| | 0.002452683 | 0.010855106 |
| | 0.453787331 | NA |
| | 0.005924997 | 0.022413665 |
| NA | | NA |
| | 0.922828106 | NA |
| | 0.58298054 | 0.720435083 |
| | 0.996405554 | 0.997159645 |
| NA | | NA |
| | 0.619804904 | 0.748824746 |
| | 0.839319574 | NA |
| | 0.812386596 | NA |
| | 0.291011428 | 0.452102601 |
| | 0.290467337 | 0.451561444 |
| | 0.18640914 | 0.330127257 |
| NA | | NA |
| | 0.233232673 | 0.387871641 |
| NA | | NA |
| | 0.601209524 | NA |
| | 0.307243765 | 0.468851733 |
| | 0.084328194 | 0.185027625 |
| | 0.600709779 | 0.734592824 |
| | 0.941277857 | NA |
| | 0.098247121 | 0.206944954 |
| | 0.072483859 | NA |
| | 0.941278332 | NA |
| | 0.595085795 | 0.729990343 |
| | 0.221560273 | NA |
| | 3.35E-10 | 6.32E-09 |
| | 0.100216934 | 0.209970893 |
| | 0.56851266 | NA |
| NA | | NA |
| NA | | NA |

| | | |
|---|---|---|
| | 0.498388499 | NA |
| | 0.000236669 | 0.001480864 |
| | 5.51E-38 | 5.24E-36 |
| | 0.839319108 | NA |
| | 0.490819758 | 0.645047538 |
| | 0.447611014 | 0.607898971 |
| | 0.1503512 | 0.283750338 |
| | 0.651858658 | NA |
| | 0.923329646 | 0.954363915 |
| | 0.030376048 | 0.083759348 |
| NA | | NA |
| | 2.00E-07 | 2.55E-06 |
| | 0.017983138 | 0.055095207 |
| | 0.920202374 | NA |
| | 0.517738782 | NA |
| | 0.492638648 | 0.646476189 |
| | 0.897577607 | NA |
| NA | | NA |
| | 0.853003105 | 0.911642041 |
| | 0.10095852 | 0.211272613 |
| | 0.013891429 | 0.044863473 |
| | 0.164585364 | 0.302951111 |
| | 0.575954475 | 0.714748092 |
| | 0.812399708 | 0.88542389 |
| | 0.605134079 | 0.737916971 |
| | 0.12076592 | 0.241572438 |
| | 0.011879143 | 0.039587155 |
| NA | | NA |
| | 0.941278207 | NA |
| | 0.000887306 | 0.00461027 |
| | 3.00E-05 | 0.000241413 |
| | 0.155808032 | 0.291140114 |
| | 0.652141483 | NA |
| | 0.205538433 | 0.354201114 |
| | 0.765311111 | 0.854255406 |
| | 0.853595622 | NA |
| | 0.018127713 | 0.055445365 |
| | 0.00918836 | 0.032110021 |
| | 0.004252774 | 0.017067571 |
| | 0.629678985 | 0.756224794 |
| | 0.955791922 | NA |
| | 0.085885914 | NA |
| | 0.297640435 | 0.459015008 |
| | 0.072398517 | 0.165107922 |
| | 1.90E-11 | 4.18E-10 |
| | 0.000812124 | 0.004283218 |
| | 0.145949423 | 0.277688903 |

| | | |
|---|---|---|
| | 0.000424634 | 0.002460955 |
| | 0.839319574 | NA |
| | 0.370253475 | 0.533460847 |
| NA | | NA |
| | 0.925298327 | NA |
| | 0.047763641 | 0.119456781 |
| | 0.941277857 | NA |
| | 0 | 0 |
| | 3.11E-05 | 0.000248949 |
| | 4.36E-51 | 6.10E-49 |
| | 0.135248773 | 0.262341398 |
| | 0.955792398 | NA |
| | 0.941276431 | NA |
| | 0.839319108 | NA |
| | 0.839319108 | NA |
| | 0.976238338 | 0.985471179 |
| | 0.84777567 | 0.908395855 |
| | 0.941278332 | NA |
| | 0.941278332 | NA |
| | 0.00238468 | 0.010611359 |
| | 0.189229854 | 0.333831084 |
| | 0.329309753 | 0.492326909 |
| | 0.09381653 | 0.200055638 |
| | 0.000439978 | 0.00253937 |
| | 0.233862482 | NA |
| | 0.653684951 | 0.774003044 |
| | 0.379612625 | 0.542381878 |
| | 0.839319108 | NA |
| | 0.809840896 | 0.883463483 |
| NA | | NA |
| | 0.001000665 | 0.005112218 |
| | 0.000665569 | 0.003618519 |
| | 0.077788681 | 0.174064696 |
| | 0.185983593 | 0.32954526 |
| | 0.821813117 | 0.891558606 |
| | 0.394635074 | 0.557156654 |
| | 0.666249282 | NA |
| | 0.853596091 | NA |
| | 0.205162388 | 0.353783609 |
| | 0.008531223 | 0.03019931 |
| | 0.085524972 | 0.186981298 |
| | 0.068663478 | 0.158842269 |
| | 0.681924521 | NA |
| | 0.839320041 | NA |
| | 0.001418628 | 0.006844535 |
| | 0.841883592 | 0.904714129 |
| | 0.321902231 | 0.484700548 |

| | | |
|---|---|---|
| | 0.662018184 | NA |
| | 0.04544731 | 0.114934253 |
| | 0.807398613 | 0.881990995 |
| | 2.63E-08 | 3.87E-07 |
| | 0.737148616 | 0.834335134 |
| | 0.938767819 | NA |
| | 0.390454877 | 0.553029754 |
| | 0.052092204 | 0.128075472 |
| | 0.583653948 | NA |
| | 0.008994597 | 0.031572382 |
| | 0.087126192 | 0.189559198 |
| | 0.000759822 | 0.004048642 |
| | 0.723133214 | NA |
| | 0.932844136 | NA |
| | 0.512906583 | 0.664175099 |
| | 5.99E-07 | 6.94E-06 |
| NA | | NA |
| NA | | NA |
| | 0.955792873 | NA |
| | 0.642857856 | 0.765727445 |
| | 0.254751598 | 0.412742327 |
| | 0.207608873 | NA |
| | 0.943117438 | NA |
| | 0.676746504 | NA |
| | 0.941278332 | NA |
| | 0.828062299 | NA |
| | 0.941277857 | NA |
| | 0.953255299 | NA |
| | 7.39E-16 | 2.41E-14 |
| | 0.289900236 | NA |
| | 0.000139206 | 0.000935983 |
| | 0.006433248 | 0.023966845 |
| | 0.043677273 | 0.111438144 |
| | 0.000688879 | 0.003728215 |
| | 0.376166753 | 0.539400501 |
| | 0.839318641 | NA |
| | 0.839319108 | NA |
| NA | | NA |
| | 0.195668693 | NA |
| | 0.053730493 | 0.131195445 |
| | 0.500399097 | 0.653131131 |
| | 0.25745989 | 0.415602597 |
| | 0.253035771 | 0.410829292 |
| | 0.755896172 | 0.84719517 |
| | 0.457026878 | 0.616710343 |
| NA | | NA |
| | 0.761350405 | NA |

| | |
|---|---|
| 0.710083943 | 0.814958195 |
| 0.860280572 | 0.916609361 |
| 0.009817427 | 0.033900245 |
| NA | NA |
| 2.73E-06 | 2.80E-05 |
| 0.02604318 | 0.074059711 |
| 0.678648928 | 0.793021149 |
| 3.43E-07 | 4.16E-06 |
| 0.009128861 | 0.031958399 |
| 2.69E-06 | 2.77E-05 |
| 0.618800929 | NA |
| 0.657565049 | NA |
| 0.941278332 | NA |
| 0.992568581 | 0.994867451 |
| 0.836763832 | 0.901121952 |
| 3.44E-05 | 0.000271699 |
| 0.357143193 | 0.520718058 |
| 0.029247988 | 0.081317673 |
| 0.316064307 | 0.478620857 |
| 0.000313748 | 0.001892509 |
| 0.040758175 | 0.105758867 |
| 0.025124175 | 0.071918895 |
| 0.000157858 | 0.001045736 |
| 0.120848422 | 0.241635932 |
| 0.370291066 | NA |
| 0.835025174 | NA |
| 0.941278332 | NA |
| 0.351397395 | 0.515109595 |
| NA | NA |
| 0.211116322 | 0.361276202 |
| 0.110916217 | 0.226417477 |
| 0.377338442 | 0.540235164 |
| 0.489177002 | 0.643583465 |
| 0.512286555 | 0.663697179 |
| NA | NA |
| NA | NA |
| NA | NA |
| NA | NA |
| 0.941278332 | NA |
| 0.955792873 | NA |
| NA | NA |
| NA | NA |
| 0.415640855 | 0.577452961 |
| 0.744246226 | NA |
| NA | NA |
| NA | NA |
| 0.512795863 | 0.664140121 |

| | | |
|---|---|---|
| | 0.941278332 | NA |
| NA | | NA |
| NA | | NA |
| | 0.423453263 | NA |
| NA | | NA |
| NA | | NA |
| NA | | NA |
| | 0.070597774 | 0.162260354 |
| | 0.965492534 | 0.979108402 |
| | 0.013087911 | 0.042773372 |
| NA | | NA |
| | 0.501813314 | NA |
| | 0.143339467 | 0.273905426 |
| | 0.203905602 | 0.352433457 |
| | 0.264468318 | 0.423469553 |
| | 0.730385277 | 0.830111521 |
| | 0.520479831 | 0.670806851 |
| | 0.14462366 | 0.275852001 |
| | 5.67E-11 | 1.18E-09 |
| | 2.11E-06 | 2.22E-05 |
| | 0.284020339 | 0.444577938 |
| | 0.047534161 | 0.118957848 |
| | 0.00015439 | 0.001026934 |
| | 0.000296609 | 0.001806523 |
| | 0.726674903 | 0.827614529 |
| | 0.470045263 | 0.627939484 |
| NA | | NA |
| | 0.46516608 | NA |
| | 0.764433133 | 0.853706069 |
| | 0.107728524 | 0.22165926 |
| | 1.92E-61 | 3.58E-59 |
| | 0.941278332 | NA |
| | 1.73E-32 | 1.34E-30 |
| | 0.001668379 | 0.007854186 |
| | 0.900281098 | NA |
| | 9.56E-13 | 2.41E-11 |
| | 3.58E-60 | 6.18E-58 |
| | 0.941278332 | NA |
| NA | | NA |
| | 0.941278284 | NA |
| | 3.10E-30 | 2.17E-28 |
| | 6.76E-53 | 9.99E-51 |
| | 0.585366882 | NA |
| | 0.001460884 | 0.007016078 |
| | 1.80E-05 | 0.000152561 |
| | 0.087502576 | 0.190186862 |
| | 0.024171272 | 0.069846379 |

| | |
|---|---|
| 2.91E-07 | 3.59E-06 |
| NA | NA |
| 0.000473084 | 0.00270551 |
| 0.765728478 | 0.854551258 |
| 3.38E-46 | 4.28E-44 |
| 0.002423682 | 0.010750743 |
| 0.003942731 | 0.016011753 |
| 0.028992269 | 0.080729292 |
| 0.449604823 | 0.609640068 |
| 0.163756251 | 0.301878563 |
| 0.060835432 | 0.144569186 |
| NA | NA |
| 0.935995333 | NA |
| 0.400655603 | 0.563101117 |
| 0.005635365 | 0.021516356 |
| 0.954822509 | 0.972597482 |
| 0.800981788 | 0.878492596 |
| 0.245734033 | 0.402571509 |
| 0.908424838 | 0.944452264 |
| 0.000102612 | 0.000716238 |
| 0.011350583 | 0.038196885 |
| 0.027640494 | 0.077830242 |
| 0.001032689 | 0.005249906 |
| 0.194762755 | 0.340919995 |
| NA | NA |
| 0.007969415 | 0.028567471 |
| 0.002190115 | 0.009865465 |
| 0.203217202 | 0.351473321 |
| 0.142538769 | NA |
| 0.749470725 | NA |
| 0.061193228 | 0.145303392 |
| 0.009253984 | 0.032300454 |
| NA | NA |
| NA | NA |
| 0.724430676 | NA |
| 0.817230349 | 0.888457846 |
| 0.98110578 | 0.988399698 |
| 0.706851071 | 0.813011945 |
| 0.205470172 | 0.354109118 |
| 3.25E-27 | 2.02E-25 |
| 0.014900536 | 0.047445288 |
| NA | NA |
| 0.013439784 | 0.043731203 |
| 0.340036039 | 0.503736715 |
| 0.075687646 | 0.170679049 |
| 0.073486931 | 0.16701384 |
| 0.991514377 | 0.994187598 |

| | |
|---|---|
| 0.563946183 | 0.706178822 |
| 0.55992528 | NA |
| 0.000762314 | 0.004060103 |
| 0.148480185 | 0.281379408 |
| 0.044116202 | 0.112245199 |
| NA | NA |
| 0.785351255 | 0.868025567 |
| 0.245994101 | 0.402900084 |
| 0.101100928 | 0.211496246 |
| 0.16661719 | 0.305509772 |
| 0.019539222 | 0.058807856 |
| 0.004433063 | 0.0176625 |
| 0.708879881 | 0.814125883 |
| 0.469825164 | NA |
| 4.11E-10 | 7.68E-09 |
| NA | NA |
| NA | NA |
| 5.05E-06 | 4.91E-05 |
| 7.30E-09 | 1.17E-07 |
| 0.214435578 | 0.365339168 |
| 0.01086669 | 0.036844582 |
| 0.075799256 | 0.170849799 |
| 0.005370499 | 0.020670864 |
| 0.634244024 | 0.759670657 |
| NA | NA |
| 0.204361069 | 0.352861963 |
| 0.793689871 | NA |
| 8.34E-05 | 0.000596859 |
| 0.877183794 | 0.926433604 |
| 0.009548163 | 0.033153227 |
| 0.955792398 | NA |
| 0.650504938 | NA |
| 0.064373181 | 0.151180113 |
| 0.051327046 | 0.126612742 |
| 0.001016428 | 0.005179472 |
| 0.132974676 | 0.258943241 |
| 0.049881312 | 0.123520079 |
| 0.928004785 | 0.957035091 |
| 0.216420794 | 0.367709883 |
| 0.002577509 | 0.01132334 |
| 0.157510108 | NA |
| NA | NA |
| 0.043420848 | 0.110950404 |
| 0.746903277 | NA |
| 0.779092876 | 0.863795819 |
| 0.215036448 | 0.365932475 |
| 0.052195344 | 0.128236332 |

| | |
|---|---|
| 0.04010121 | 0.104395603 |
| 0.035444426 | 0.094664299 |
| 0.067223266 | 0.156269964 |
| 0.282476515 | 0.442889533 |
| 0.503548803 | 0.655617828 |
| 0.413576605 | 0.575770624 |
| 0.285315742 | 0.445872582 |
| 0.025338717 | 0.072445903 |
| 0.715264556 | NA |
| 0.08029759 | 0.178337523 |
| 0.015123646 | 0.048022014 |
| 0.239719289 | 0.395741331 |
| 0.013495911 | 0.04385388 |
| 0.759674447 | NA |
| 0.605524794 | 0.73820432 |
| 0.002878889 | 0.012384479 |
| 0.430368822 | 0.591604891 |
| 0.093684383 | 0.199809648 |
| NA | NA |
| 0.226935736 | 0.380138524 |
| 0.029755619 | 0.08244014 |
| 0.000807355 | 0.004260749 |
| 0.876992514 | 0.926329976 |
| 0.029323951 | 0.081502884 |
| 1.06E-20 | 4.72E-19 |
| 0.034872001 | 0.09337646 |
| 0.055049002 | 0.133783575 |
| 0.982996718 | NA |
| 0.000421206 | 0.002444657 |
| 0.49946781 | 0.652296322 |
| NA | NA |
| 0.000177338 | 0.001157827 |
| 0.712011519 | NA |
| 0.923262567 | NA |
| 0.900052437 | NA |
| 1.03E-05 | 9.31E-05 |
| 0.004200551 | 0.016888768 |
| 0.788448692 | 0.869873726 |
| 0.066943487 | 0.155771715 |
| 0.276263476 | 0.436107126 |
| 0.422057274 | 0.583278594 |
| 0.955792873 | NA |
| NA | NA |
| 0.089094467 | 0.192686977 |
| NA | NA |
| 0.154340447 | 0.289033695 |
| 0.462529676 | 0.621247749 |

| | |
|---|---|
| 0.000244973 | 0.001525599 |
| 0.007425719 | 0.026984268 |
| 1.81E-08 | 2.73E-07 |
| 0.586373156 | 0.723080005 |
| NA | NA |
| 0.648610918 | 0.770264094 |
| 0.665320878 | 0.782951031 |
| 0.93838321 | NA |
| 0.377168778 | 0.540054711 |
| 0.005092271 | 0.019817894 |
| 0.941520523 | 0.964703895 |
| 2.76E-05 | 0.000223944 |
| 0.074658541 | 0.168886391 |
| 0.059389497 | 0.141926586 |
| 0.001610838 | 0.007628563 |
| 0.253410971 | 0.411326237 |
| 0.323625864 | 0.486296895 |
| NA | NA |
| 0.839319108 | NA |
| 0.001647643 | 0.007778105 |
| 0.00728243 | 0.026540562 |
| 0.75341223 | 0.84541346 |
| 0.000593839 | 0.003282525 |
| 0.016114262 | 0.050493422 |
| 0.629343708 | 0.755989005 |
| 0.763995555 | NA |
| 0.171754052 | NA |
| 0.30080646 | 0.462211174 |
| 0.788906265 | 0.870055922 |
| 0.064948442 | 0.152320704 |
| NA | NA |
| 0.000237107 | 0.001482823 |
| 0.315105072 | 0.477472048 |
| 0.001828251 | 0.0084777 |
| 0.941278332 | NA |
| 7.01E-09 | 1.12E-07 |
| 0.107067547 | NA |
| 0.961416346 | 0.976514119 |
| 0.24269178 | NA |
| 0.162995757 | 0.300746125 |
| NA | NA |
| 0.66629613 | 0.783672506 |
| NA | NA |
| 0.895065287 | NA |
| 0.024331499 | 0.070215608 |
| NA | NA |
| NA | NA |

| | | |
|---|---|---|
| | 0.853595154 | NA |
| NA | | NA |
| | 0.827450996 | 0.895237779 |
| | 0.955792398 | NA |
| NA | | NA |
| | 0.039737647 | NA |
| | 0.539436434 | NA |
| | 0.839319108 | NA |
| NA | | NA |
| | 0.941278332 | NA |
| NA | | NA |
| | 0.924655456 | 0.954904945 |
| NA | | NA |
| | 0.105259437 | 0.217897471 |
| | 0.302225002 | NA |
| | 0.542784124 | 0.689583545 |
| | 0.2852949 | NA |
| | 0.05665376 | 0.13680357 |
| | 0.160359903 | 0.297110169 |
| | 0.908928276 | NA |
| | 0.908276342 | NA |
| | 0.311470131 | 0.473511091 |
| | 0.000777262 | 0.004124947 |
| | 0.925010181 | NA |
| | 0.000135367 | 0.000914306 |
| | 1.51E-07 | 1.97E-06 |
| NA | | NA |
| | 0.839319108 | NA |
| | 1.01E-05 | 9.14E-05 |
| | 0.141255271 | 0.271032484 |
| | 0.002033202 | 0.009276267 |
| | 0.729417018 | NA |
| NA | | NA |
| | 0.017108597 | 0.052975 |
| | 0.843788904 | NA |
| | 0.002385782 | 0.010612295 |
| | 0.646962081 | 0.769103558 |
| | 0.1108289 | 0.226354281 |
| | 0.037329077 | 0.09855532 |
| | 0.022989668 | 0.067041013 |
| NA | | NA |
| | 0.557531108 | 0.701317451 |
| | 0.447434497 | 0.607730256 |
| | 0.008486989 | 0.03006956 |
| | 0.146414244 | 0.278304386 |
| | 0.309111609 | 0.470697621 |
| | 0.892215567 | 0.935040806 |

| | | |
|---|---|---|
| | 0.61803259 | NA |
| | 0.000705198 | 0.003799256 |
| | 0.470254898 | 0.62807963 |
| | 0.153976031 | 0.288691413 |
| | 0.478622946 | 0.635405397 |
| | 0.075915123 | 0.17099761 |
| | 0.68395505 | 0.797115903 |
| NA | | NA |
| | 0.772336158 | 0.859386903 |
| | 3.70E-10 | 6.95E-09 |
| | 0.083367997 | NA |
| | 0.285852055 | 0.446509514 |
| | 0.439497255 | 0.60041129 |
| | 0.116246779 | 0.234523295 |
| | 0.000247919 | 0.001541928 |
| | 0.13657187 | 0.264283223 |
| | 0.386382735 | 0.54921958 |
| | 2.66E-11 | 5.76E-10 |
| | 0.016689841 | 0.051928314 |
| | 6.93E-09 | 1.11E-07 |
| | 0.003577747 | 0.014812973 |
| | 0.629982766 | 0.756391124 |
| | 1.03E-08 | 1.62E-07 |
| | 0.389893575 | 0.552461416 |
| | 0.15258314 | 0.286677459 |
| | 0.955792873 | NA |
| | 0.001662279 | 0.007832796 |
| | 0.435610559 | 0.596742592 |
| | 0.179922857 | 0.321945861 |
| | 0.000554797 | 0.003104205 |
| | 0.024582343 | 0.070784088 |
| | 0.062355861 | 0.147446275 |
| | 0.351268929 | 0.515109595 |
| NA | | NA |
| | 0.539708879 | 0.686886468 |
| | 0.051541602 | 0.126970938 |
| | 0.156929808 | 0.29268479 |
| | 0.117694784 | 0.236722263 |
| | 0.305286675 | 0.466922786 |
| | 0.637507285 | 0.761744398 |
| | 0.01309818 | 0.042801056 |
| | 0.918993213 | 0.951350632 |
| | 0.041971443 | 0.108152461 |
| NA | | NA |
| | 0.114204937 | 0.231384569 |
| | 0.139329274 | 0.268180122 |
| | 0.955791446 | NA |

| | | | |
|---|---|---|---|
| | 0.015849111 | | 0.049842299 |
| | 0.211007157 | | 0.361115354 |
| | 0.089730197 | | 0.193755912 |
| | 0.020530271 | | 0.061120889 |
| NA | | NA | |
| | 0.018998571 | | 0.057559538 |
| | 0.101273549 | | 0.211708504 |
| | 0.061988437 | | 0.146723229 |
| NA | | NA | |
| | 0.033541078 | | 0.090669476 |
| | 0.041544442 | | 0.107331034 |
| | 0.012540784 | | 0.04132573 |
| | 0.007876999 | | 0.028295852 |
| | 0.011408689 | | 0.038338222 |
| | 0.003570622 | | 0.014795949 |
| | 6.41E-05 | | 0.000473494 |
| | 0.002561429 | | 0.011268057 |
| | 6.66E-05 | | 0.000488886 |
| | 0.000914979 | | 0.00473545 |
| | 0.000188741 | | 0.0012201 |
| | 0.000577269 | | 0.003206572 |
| | 0.036300225 | | 0.096441339 |
| | 0.941276431 | NA | |
| | 0.955791922 | NA | |
| NA | | NA | |
| NA | | NA | |
| NA | | NA | |
| | 0.358385969 | | 0.521986466 |
| NA | | NA | |
| | 0.941277857 | NA | |
| | 0.382329453 | | 0.544910473 |
| NA | | NA | |
| | 0.373271173 | NA | |
| | 0.688181623 | | 0.800043909 |
| | 0.000607002 | | 0.003345117 |
| | 0.069677706 | | 0.160673199 |
| | 0.209575306 | | 0.359199838 |
| | 0.336838334 | | 0.500078095 |
| | 0.46741031 | | 0.625694339 |
| | 7.57E-21 | | 3.41E-19 |
| | 4.78E-06 | | 4.68E-05 |
| | 0.008173379 | | 0.029184334 |
| | 0.016370539 | | 0.051153022 |
| NA | | NA | |
| | 0.299745754 | | 0.461245697 |
| NA | | NA | |
| NA | | NA | |

| | | |
|---|---|---|
| | 0.979089689 | NA |
| NA | | NA |
| | 0.000219946 | 0.001392328 |
| | 0.056215713 | 0.136007968 |
| | 0.029411233 | 0.081702015 |
| | 0.367605256 | 0.530716839 |
| | 0.178005282 | 0.319475322 |
| | 0.697307639 | NA |
| | 0.96035716 | 0.975646469 |
| | 6.88E-23 | 3.50E-21 |
| | 0.050017953 | 0.123832673 |
| | 0.00801148 | 0.028688018 |
| | 0.06183845 | 0.146441029 |
| | 0.878528295 | 0.927099556 |
| | 0.890026919 | 0.933769151 |
| | 0.839318641 | NA |
| NA | | NA |
| NA | | NA |
| | 0.941277857 | NA |
| | 0.423914804 | 0.585369854 |
| | 0.698441549 | NA |
| | 0.469027426 | 0.627108796 |
| | 0.52698611 | 0.676044162 |
| NA | | NA |
| | 0.099802368 | 0.209294798 |
| | 0.296089972 | 0.457542915 |
| | 0.056611871 | 0.136744037 |
| | 0.000223632 | 0.001413032 |
| | 0.022528621 | 0.065924053 |
| | 0.762739603 | 0.852734995 |
| | 0.860720764 | 0.916679745 |
| | 0.003191614 | 0.013512415 |
| | 0.357793805 | 0.5213473 |
| NA | | NA |
| | 0.515582943 | 0.666627395 |
| | 0.000269181 | 0.001655567 |
| | 0.00547759 | 0.02098846 |
| NA | | NA |
| | 0.088869532 | 0.192437463 |
| | 0.955791922 | NA |
| | 0.419781177 | NA |
| | 0.67380855 | NA |
| | 0.143013743 | 0.273427363 |
| | 0.895164377 | 0.936820599 |
| NA | | NA |
| | 0.716973861 | 0.820097643 |
| | 0.024431211 | 0.070409455 |

| | |
|---|---|
| NA | NA |
| 0.000111351 | 0.000769785 |
| 0.839318641 | NA |
| 0.10931133 | 0.224009531 |
| 0.839319107 | NA |
| 0.647499676 | NA |
| 0.588379297 | NA |
| 0.501954369 | 0.654156695 |
| 0.986579384 | 0.991639555 |
| 0.490764282 | 0.645047538 |
| NA | NA |
| 0.381385099 | 0.543934815 |
| 0.042323538 | 0.108871161 |
| 0.000225635 | 0.001422669 |
| NA | NA |
| 0.318386106 | NA |
| 0.676593248 | NA |
| 4.45E-07 | 5.29E-06 |
| 0.002096749 | 0.009511497 |
| 0.609785273 | 0.741537274 |
| 0.277547835 | 0.43740936 |
| 0.739825989 | NA |
| 0.004539612 | 0.018023661 |
| 0.170678094 | NA |
| 0.080989186 | 0.179321359 |
| 7.90E-50 | 1.07E-47 |
| NA | NA |
| 0.926330958 | NA |
| NA | NA |
| 0.874352097 | 0.924906832 |
| 0.692898585 | 0.803173864 |
| 0.028703386 | 0.080187558 |
| 0.000149488 | 0.000996952 |
| 0.73316589 | 0.831664419 |
| 0.004585937 | 0.018177264 |
| NA | NA |
| 0.011922641 | 0.039695441 |
| 0.000210893 | 0.001342875 |
| 0.075069675 | 0.169574679 |
| 0.028992077 | 0.080729292 |
| 0.171627814 | 0.311790966 |
| 0.001522565 | 0.007262543 |
| 0.498281619 | NA |
| 0.002650326 | 0.011591889 |
| 0.800157022 | NA |
| NA | NA |
| 0.141400499 | 0.271223683 |

| | |
|---|---|
| 0.005169827 | 0.020080324 |
| 0.008311052 | 0.029591571 |
| 0.719778104 | 0.821725667 |
| 0.226306113 | 0.379333669 |
| 1.45E-12 | 3.58E-11 |
| NA | NA |
| 0.941275481 | NA |
| NA | NA |
| 0.055978655 | 0.135585244 |
| 0.925162031 | 0.955138003 |
| 9.71E-16 | 3.12E-14 |
| 7.34E-92 | 2.24E-89 |
| 0.839319108 | NA |
| 0.941277857 | NA |
| 0.834616295 | NA |
| 0.015926262 | 0.05005637 |
| 0.766668136 | 0.855078721 |
| 0.00977075 | 0.033773364 |
| 0.062336261 | 0.147414574 |
| 0.805124302 | 0.880868577 |
| 0.031234234 | 0.08562039 |
| 0.73622171 | 0.833647157 |
| NA | NA |
| 0.379975941 | NA |
| 9.75E-06 | 8.87E-05 |
| 1.28E-08 | 1.97E-07 |
| 0.942245039 | NA |
| 6.54E-05 | 0.000482014 |
| 0.268409925 | 0.428167887 |
| NA | NA |
| 0.523461673 | NA |
| 0.249625574 | 0.407069601 |
| 0.569901966 | 0.710794728 |
| 0.166086529 | 0.304891074 |
| 0.243103213 | 0.399577436 |
| 0.248930041 | 0.406269393 |
| NA | NA |
| 0.222964469 | 0.375371361 |
| 0.001501544 | 0.007180984 |
| 0.161092285 | 0.298082621 |
| 0.520588 | 0.67087361 |
| 0.464809123 | 0.623290847 |
| 0.660845339 | 0.779611457 |
| 0.667821383 | 0.784735757 |
| 0.007555058 | 0.027354046 |
| 0.49252751 | 0.646413842 |
| 0.312047225 | 0.474015448 |

| | | |
|---|---|---|
| | 0.330101234 | 0.493100889 |
| | 0.839319108 | NA |
| | 0.163968018 | 0.302095113 |
| | 0.981963082 | 0.98868314 |
| | 0.405157215 | NA |
| | 1.26E-05 | 0.000111347 |
| | 0.030341025 | 0.083711265 |
| | 4.24E-06 | 4.19E-05 |
| | 0.664857787 | 0.782560827 |
| | 0.406702187 | 0.568494565 |
| | 0.004291684 | 0.017188401 |
| | 2.78E-06 | 2.84E-05 |
| | 0.179529647 | 0.321338897 |
| | 0.240131206 | NA |
| | 0.016987301 | 0.052661075 |
| | 0.831413565 | NA |
| NA | | NA |
| NA | | NA |
| | 0.052585662 | 0.129035457 |
| | 0.83315501 | NA |
| | 0.312615024 | 0.474635631 |
| | 0.014521076 | 0.046461977 |
| | 0.924295044 | NA |
| | 0.844149115 | NA |
| NA | | NA |
| | 0.839319108 | NA |
| NA | | NA |
| NA | | NA |
| | 0.031035348 | 0.085192407 |
| NA | | NA |
| | 0.612903554 | 0.743661563 |
| | 0.003620239 | 0.014944317 |
| | 0.481980285 | 0.638225375 |
| | 0.004895809 | 0.019188217 |
| NA | | NA |
| NA | | NA |
| | 0.839318641 | NA |
| | 0.541310117 | 0.688462683 |
| | 0.145925693 | 0.277665948 |
| | 0.822050994 | 0.891707027 |
| | 0.069413333 | 0.160203254 |
| NA | | NA |
| NA | | NA |
| NA | | NA |
| | 0.402937653 | NA |
| | 0.22678826 | 0.379981287 |
| | 0.876683706 | 0.926202931 |

| | |
|---|---|
| 0.590849173 | 0.726752727 |
| NA | NA |
| 0.000497905 | 0.002827086 |
| 0.001126743 | 0.005653193 |
| 0.839319108 | NA |
| 0.873947307 | 0.924785321 |
| 0.277687211 | 0.437434077 |
| 0.941277857 | NA |
| 0.49990768 | 0.652704477 |
| 0.729243903 | 0.829416758 |
| 0.593913716 | 0.729015228 |
| 0.08095057 | 0.179252533 |
| 0.264203797 | 0.423187975 |
| NA | NA |
| 0.792316268 | NA |
| 0.17514352 | NA |
| 0.156604384 | 0.292215235 |
| 0.941279303 | NA |
| 0.303949908 | 0.465398244 |
| 0.955792398 | NA |
| 0.925250407 | 0.955182718 |
| 1.30E-07 | 1.71E-06 |
| 0.122989291 | 0.244744678 |
| 0.365115933 | 0.528425723 |
| 0.017614067 | 0.054166669 |
| 0.047022082 | 0.11797603 |
| 0.941278332 | NA |
| 5.10E-05 | 0.000385384 |
| 0.19912545 | 0.34653681 |
| NA | NA |
| 0.218312339 | 0.36958908 |
| NA | NA |
| 0.208797913 | 0.358210172 |
| 0.334213893 | 0.497495877 |
| 2.28E-09 | 3.87E-08 |
| 0.000166254 | 0.001096166 |
| 0.783658165 | 0.866967214 |
| 0.938572556 | NA |
| 0.000252128 | 0.001564427 |
| 0.920509814 | NA |
| 0.002311902 | 0.01034359 |
| NA | NA |
| 2.09E-12 | 5.08E-11 |
| 0.469272741 | 0.627295843 |
| NA | NA |
| 1.20E-22 | 6.00E-21 |
| 0.479357929 | 0.636061848 |

| | |
|---|---|
| 0.941277857 | NA |
| 0.953758092 | 0.971970805 |
| 0.110889183 | 0.226401126 |
| 0.503503031 | NA |
| 0.000708035 | 0.003812323 |
| NA | NA |
| 0.000739039 | 0.003957392 |
| 0.003943483 | 0.016012075 |
| 0.941279303 | NA |
| 0.989015128 | NA |
| 0.042061974 | 0.108327104 |
| 0.758395874 | NA |
| 0.422308277 | NA |
| 0.999955691 | 0.999955691 |
| 0.678831534 | 0.793086349 |
| 0.018306708 | 0.055892287 |
| 0.000771821 | 0.00410156 |
| 0.336084391 | NA |
| NA | NA |
| 0.08364922 | 0.183825938 |
| 0.376257912 | 0.539416131 |
| 0.149521436 | 0.282551092 |
| 0.708194269 | NA |
| 0.321023012 | 0.483821296 |
| 0.330204515 | 0.493193264 |
| 0.076900731 | 0.172613279 |
| 0.052086193 | 0.128073923 |
| 0.022821671 | 0.066642304 |
| 0.198171108 | 0.3452799 |
| 2.48E-05 | 0.000204496 |
| 0.088051349 | 0.191135267 |
| 3.67E-23 | 1.89E-21 |
| NA | NA |
| NA | NA |
| 0.730399633 | 0.830111521 |
| 0.200390854 | 0.347975713 |
| NA | NA |
| NA | NA |
| 0.941277857 | NA |
| NA | NA |
| NA | NA |
| NA | NA |
| NA | NA |
| 0.016897808 | 0.05244512 |
| 0.403241733 | 0.565250867 |
| 3.33E-07 | 4.06E-06 |
| 0.014482422 | 0.046369465 |

| | |
|---|---|
| NA | NA |
| 0.048188269 | 0.120253426 |
| NA | NA |
| 0.126138407 | 0.249343607 |
| 6.20E-08 | 8.59E-07 |
| 0.808255193 | 0.882400243 |
| 0.006904223 | 0.025399429 |
| 0.148690771 | 0.281534875 |
| 0.030576117 | 0.084203719 |
| 3.59E-09 | 5.96E-08 |
| 0.192992942 | 0.33873741 |
| 0.002910156 | 0.01250734 |
| 0.029413686 | 0.081702015 |
| 0.777502871 | 0.862997731 |
| NA | NA |
| 0.955791922 | NA |
| NA | NA |
| 0.941277857 | NA |
| 0.067722518 | 0.15713796 |
| 0.998476618 | NA |
| 0.839319108 | NA |
| NA | NA |
| 0.092746453 | 0.198467236 |
| 0.1997075 | 0.347183534 |
| 0.061167906 | 0.145257756 |
| 0.391843774 | 0.554183941 |
| 0.1714475 | 0.311582301 |
| 0.136654768 | 0.26438013 |
| 0.004619204 | 0.018290851 |
| 0.00867668 | 0.030619027 |
| 0.275225185 | 0.435102939 |
| 0.139498549 | 0.268332347 |
| 0.900438649 | 0.940018234 |
| 0.406335739 | NA |
| 0.029439618 | 0.081761507 |
| 0.480587551 | 0.637160636 |
| 0.189625537 | 0.334256612 |
| NA | NA |
| NA | NA |
| 0.033467795 | 0.090509304 |
| 0.004959535 | 0.019399647 |
| 0.941278332 | NA |
| 0.90731999 | 0.943748776 |
| 0.705325606 | 0.812065262 |
| 0.046267915 | 0.116535109 |
| 0.000121165 | 0.000829919 |
| NA | NA |

| | | |
|---|---|---|
| NA | 0.96681714 | 0.980159365 |
| | 0.001579497 | 0.007499538 |
| | 0.22825999 | 0.3817753 |
| | 0.296305295 | 0.45779322 |
| | 0.717470552 | 0.820340515 |
| | 0.449971879 | NA |
| | 0.188544202 | NA |
| | 0.14711573 | 0.279371997 |
| | 8.52E-15 | 2.53E-13 |
| NA | | NA |
| | 0.083571033 | 0.183738936 |
| | 0.949978359 | NA |
| | 0.839911896 | 0.903321135 |
| NA | | NA |
| | 0.896251637 | 0.937366265 |
| | 0.000298582 | 0.001816217 |
| | 0.000171292 | 0.001123783 |
| | 0.003699237 | 0.015209749 |
| | 0.017156292 | 0.053084602 |
| | 0.020482986 | 0.060995376 |
| | 0.847324423 | 0.908223719 |
| | 0.715732036 | NA |
| NA | | NA |
| | 0.772933943 | 0.859898977 |
| | 0.869654406 | NA |
| | 0.139574158 | 0.268439542 |
| | 0.005372655 | 0.020675818 |
| | 0.313994318 | 0.476152698 |
| | 0.150427627 | 0.283872067 |
| | 0.947601587 | NA |
| | 0.915756548 | NA |
| NA | | NA |
| NA | | NA |
| NA | | NA |
| | 0.839319108 | NA |
| | 0.955792873 | NA |
| | 0.020463362 | 0.06095219 |
| | 5.86E-45 | 7.08E-43 |
| | 0.006229554 | 0.023346851 |
| | 0.000926856 | 0.00477925 |
| | 1.19E-10 | 2.36E-09 |
| | 0.703932078 | 0.811026251 |
| | 0.901960544 | 0.940738118 |
| | 0.096171477 | 0.203690469 |
| | 0.127156513 | 0.250731449 |
| | 0.002502677 | 0.011047611 |
| NA | | NA |

| | | |
|---|---|---|
| 4.77E-05 | | 0.000363918 |
| 0.001380446 | | 0.006697385 |
| 6.09E-11 | | 1.26E-09 |
| 0.941278332 | NA | |
| 0.941278332 | NA | |
| NA | NA | |
| 0.941275481 | NA | |
| NA | NA | |
| NA | NA | |
| NA | NA | |
| NA | NA | |
| NA | NA | |
| NA | NA | |
| NA | NA | |
| 0.839319108 | NA | |
| 0.948205862 | NA | |
| NA | NA | |
| NA | NA | |
| NA | NA | |
| 0.71137011 | NA | |
| NA | NA | |
| NA | NA | |
| NA | NA | |
| NA | NA | |
| 0.542645608 | NA | |
| 0.941277857 | NA | |
| 0.083742318 | | 0.183962589 |
| 0.010673272 | | 0.036292173 |
| 0.196036626 | | 0.342614271 |
| 0.088927026 | | 0.192526931 |
| 0.275770429 | | 0.435588809 |
| 0.030894622 | | 0.084904029 |
| 0.386095979 | | 0.548943103 |
| 0.00614898 | | 0.023103081 |
| 0.26409615 | NA | |
| 0.941277857 | NA | |
| NA | NA | |
| NA | NA | |
| NA | NA | |
| NA | NA | |
| NA | NA | |
| 0.639584131 | | 0.763306332 |
| 0.00010086 | | 0.000706287 |
| 0.748393366 | NA | |
| NA | NA | |
| 2.00E-09 | | 3.42E-08 |
| 0.955792398 | NA | |

| | |
|---|---|
| 0.824760608 | 0.893568108 |
| NA | NA |
| 0.941278332 | NA |
| 0.000259153 | 0.00160148 |
| 0.18704112 | 0.330950991 |
| 0.441560162 | 0.602155092 |
| 0.822215708 | 0.891751494 |
| 0.325798391 | 0.488697587 |
| 0.955792873 | NA |
| 0.944609092 | 0.966578634 |
| NA | NA |
| NA | NA |
| 0.941278332 | NA |
| NA | NA |
| NA | NA |
| NA | NA |
| NA | NA |
| NA | NA |
| 0.000340695 | 0.002036983 |
| 0.792779811 | 0.872508672 |
| 0.172326238 | NA |
| 0.839319108 | NA |
| 0.267044564 | 0.426647517 |
| 0.409050103 | 0.571072235 |
| 0.751007805 | NA |
| 0.102266572 | 0.2133534 |
| 0.572529357 | 0.71242494 |
| 0.136112397 | 0.263629854 |
| 0.941278788 | NA |
| NA | NA |
| 6.14E-10 | 1.12E-08 |
| 0.530283873 | NA |
| 0.08865921 | 0.192156849 |
| 0.252691977 | 0.410551157 |
| 0.020370223 | 0.060770965 |
| 0.023272282 | 0.067717658 |
| 0.622352407 | 0.750759312 |
| 0.00043229 | 0.002499844 |
| 0.516869742 | NA |
| 0.523447224 | 0.6732824 |
| NA | NA |
| 0.955792873 | NA |
| NA | NA |
| NA | NA |
| NA | NA |
| NA | NA |
| 0.941278332 | NA |

| | | |
|---|---|---|
| | 1.15E-05 | 0.000102438 |
| NA | | NA |
| | 0.466590804 | NA |
| | 0.102165626 | 0.213161486 |
| | 0.93838321 | NA |
| | 0.375520259 | NA |
| | 0.824917228 | 0.893634157 |
| | 0.480377378 | 0.637059414 |
| | 0.058882858 | 0.140863392 |
| | 0.046535172 | 0.116999066 |
| | 0.006306954 | 0.023577526 |
| | 0.450639706 | 0.610583839 |
| | 0.000674274 | 0.003661663 |
| NA | | NA |
| | 0.057224484 | 0.137860037 |
| | 0.484255308 | NA |
| | 0.963652478 | 0.977867338 |
| | 0.695626711 | 0.805081541 |
| | 0.850336409 | NA |
| NA | | NA |
| NA | | NA |
| | 2.03E-19 | 8.28E-18 |
| NA | | NA |
| | 0.470924497 | 0.628515469 |
| | 0.471015477 | 0.628566404 |
| | 0.70490784 | NA |
| | 0.905490771 | NA |
| | 0.293551975 | 0.454763099 |
| NA | | NA |
| | 0.839318641 | NA |
| | 0.739496953 | 0.836211382 |
| | 0.077112877 | 0.172958979 |
| | 0.025428588 | 0.072607281 |
| | 0.103844502 | 0.215700047 |
| | 0.146875619 | 0.279039147 |
| | 0.187281565 | 0.331277929 |
| | 0.629010095 | 0.755802831 |
| | 0.408974992 | 0.571034361 |
| | 0.08891458 | 0.192517497 |
| NA | | NA |
| | 0.507683595 | NA |
| NA | | NA |
| NA | | NA |
| | 0.888002491 | NA |
| NA | | NA |
| | 0.088519591 | 0.191906668 |
| NA | | NA |

| | |
|---|---|
| NA | NA |
| 0.004517659 | 0.017951613 |
| 0.443050829 | 0.603618944 |
| NA | NA |
| 0.041231079 | 0.106683568 |
| 0.255607157 | 0.41364536 |
| 0.493161874 | 0.646841842 |
| 0.530967157 | NA |
| 0.036918555 | 0.097702406 |
| 0.839319108 | NA |
| 0.953255299 | NA |
| 0.107013963 | 0.220665888 |
| 0.396681312 | 0.559315716 |
| 0.004927215 | 0.019289074 |
| 0.120178669 | 0.240620178 |
| 0.825325614 | 0.893943129 |
| 0.000452492 | 0.002602759 |
| 0.691373385 | 0.801978899 |
| 0.01199356 | 0.039875781 |
| 0.487407938 | 0.642267826 |
| 0.941278332 | NA |
| 0.246222814 | 0.403178001 |
| 0.194417077 | 0.340533907 |
| NA | NA |
| 0.001832139 | 0.008492421 |
| 0.486329111 | NA |
| 0.383761806 | 0.546507981 |
| 0.938552414 | NA |
| NA | NA |
| NA | NA |
| 0.346867194 | 0.51074129 |
| 0.56003783 | 0.703356924 |
| 0.831929452 | NA |
| 0.534270056 | NA |
| 0.46442246 | NA |
| 0.839319108 | NA |
| 2.93E-08 | 4.27E-07 |
| NA | NA |
| 0.84401219 | NA |
| 0.853596091 | NA |
| NA | NA |
| NA | NA |
| NA | NA |
| 0.941278332 | NA |
| 0.952164477 | NA |
| 0.751597581 | NA |
| NA | NA |

| | |
|---|---|
| NA | NA |
| 0.941278332 | NA |
| NA | NA |
| 1.03E-60 | 1.86E-58 |
| 1.07E-180 | 8.76E-178 |
| 0.837259124 | NA |
| 5.84E-101 | 2.01E-98 |
| 2.86E-211 | 4.54E-208 |
| 0.172541683 | 0.312782722 |
| 0.612948498 | 0.743674184 |
| 0.278898119 | 0.438637044 |
| 0.903227318 | 0.941435305 |
| NA | NA |
| 0.955792398 | NA |
| 2.61E-06 | 2.70E-05 |
| NA | NA |
| 6.94E-06 | 6.53E-05 |
| 0.396089572 | 0.5588124 |
| NA | NA |
| NA | NA |
| NA | NA |
| 0.000582428 | 0.003229198 |
| 0.305804459 | 0.467426004 |
| 0.406386373 | 0.568234732 |
| 0.043926127 | 0.111905136 |
| 0.002687718 | 0.011729568 |
| NA | NA |
| NA | NA |
| 0.160959266 | 0.297935488 |
| 0.028206942 | 0.079032563 |
| 0.06460512 | 0.151664964 |
| 0.0274704 | 0.077460324 |
| 0.01324258 | 0.043213586 |
| 0.062731246 | 0.148098505 |
| 0.241606041 | NA |
| 0.563111645 | 0.705504968 |
| 0.010741598 | 0.036488013 |
| 0.161929123 | 0.299079458 |
| 0.002213674 | 0.009960279 |
| 0.066872017 | 0.155635842 |
| 0.673173565 | 0.788999447 |
| 0.020291403 | 0.060568845 |
| 0.328948537 | NA |
| NA | NA |
| NA | NA |
| 0.899553397 | NA |
| 0.212208871 | NA |

| | |
|---|---|
| NA | NA |
| 0.127135071 | 0.250730712 |
| 0.482462679 | NA |
| NA | NA |
| 0.821579542 | 0.891427098 |
| 1.44E-05 | 0.000125798 |
| 0.955791922 | NA |
| NA | NA |
| 8.61E-05 | 0.000613808 |
| 0.001799833 | 0.008370385 |
| 0.001275298 | 0.006267735 |
| 8.08E-05 | 0.000580375 |
| NA | NA |
| 0.955792873 | NA |
| 0.000290447 | 0.001773074 |
| NA | NA |
| NA | NA |
| 0.921493168 | NA |
| 0.409987546 | 0.571919725 |
| NA | NA |
| 0.17756764 | 0.319182263 |
| 0.948975296 | 0.968816796 |
| 0.579664225 | NA |
| 0.215273304 | 0.366230804 |
| 0.592757732 | 0.728234723 |
| 0.849597312 | 0.909682734 |
| 0.783972597 | 0.867199604 |
| 0.669477422 | 0.786132294 |
| 0.390497939 | NA |
| NA | NA |
| 0.939241089 | NA |
| 0.995865333 | 0.996917253 |
| 0.686935379 | 0.799102482 |
| 0.634780768 | NA |
| 0.003361929 | 0.014100716 |
| 0.876048126 | NA |
| 0.878916568 | 0.927385945 |
| 0.070846789 | 0.162618687 |
| 0.012948479 | 0.042393355 |
| NA | NA |
| 0.39144853 | 0.553906302 |
| 0.091137181 | 0.196063556 |
| 0.839318641 | NA |
| 0.002058228 | 0.009367102 |
| NA | NA |
| NA | NA |
| NA | NA |

| | |
|---|---|
| 0.839319108 | NA |
| 0.212643422 | 0.363208096 |
| NA | NA |
| 9.82E-05 | 0.00068992 |
| 0.778265992 | NA |
| 0.922849885 | NA |
| 0.019372246 | 0.058396429 |
| 0.401925774 | 0.564236937 |
| 0.031006161 | 0.085131925 |
| 0.165077895 | 0.303622909 |
| 0.864459726 | 0.91922223 |
| 0.853596091 | NA |
| NA | NA |
| 0.602882323 | 0.736145071 |
| 0.02567827 | 0.073196908 |
| 0.29732415 | 0.458735296 |
| 1.69E-20 | 7.41E-19 |
| NA | NA |
| 0.542047311 | NA |
| 0.001555569 | 0.007399212 |
| NA | NA |
| 0.400353521 | NA |
| 0.232212649 | 0.386566335 |
| 3.40E-11 | 7.26E-10 |
| NA | NA |
| 0.546628278 | NA |
| 0.960276057 | 0.975605716 |
| 0.472748249 | 0.630242737 |
| NA | NA |
| NA | NA |
| 1.64E-05 | 0.000141203 |
| 5.52E-10 | 1.02E-08 |
| 0.018605006 | 0.056643175 |
| NA | NA |
| NA | NA |
| 0.912035729 | 0.946889856 |
| 0.373827389 | 0.536984792 |
| 0.455147502 | 0.614869554 |
| 0.484680693 | 0.64037579 |
| 0.941277857 | NA |
| NA | NA |
| 0.377123372 | 0.540024697 |
| 0.839318641 | NA |
| 0.617993747 | NA |
| NA | NA |
| 0.924105347 | 0.954592539 |
| 0.987526972 | 0.992111978 |

| | | |
|---|---|---|
| | 0.253473861 | 0.41138651 |
| | 0.432787756 | NA |
| NA | | NA |
| | 0.839319108 | NA |
| | 0.605428681 | NA |
| | 0.33114425 | 0.494224695 |
| NA | | NA |
| | 0.59946117 | NA |
| | 1.98E-05 | 0.0001666 |
| | 0.025633125 | 0.073094482 |
| | 0.012066038 | 0.040055207 |
| | 0.493533727 | 0.646993147 |
| NA | | NA |
| NA | | NA |
| | 0.732377347 | 0.831176115 |
| | 0.213235351 | 0.363920785 |
| | 0.917582103 | 0.95044119 |
| | 0.317113511 | 0.479568955 |
| | 0.155901893 | 0.291242179 |
| | 0.910090002 | NA |
| NA | | NA |
| | 0.839318641 | NA |
| NA | | NA |
| NA | | NA |
| NA | | NA |
| NA | | NA |
| NA | | NA |
| NA | | NA |
| NA | | NA |
| | 0.955792398 | NA |
| NA | | NA |
| NA | | NA |
| NA | | NA |
| | 1.07E-05 | 9.57E-05 |
| | 0.009200601 | 0.032141141 |
| | 0.007251553 | 0.026452331 |
| | 0.040617296 | 0.105439293 |
| | 0.278339397 | 0.43798984 |
| | 0.811469192 | NA |
| | 0.812844687 | NA |
| | 0.315592691 | 0.478028334 |
| NA | | NA |
| | 0.000683408 | 0.00370113 |
| NA | | NA |
| | 0.632591582 | 0.758421641 |
| | 0.04433092 | 0.1126831 |
| NA | | NA |

| | | |
|---|---|---|
| | 0.044968081 | NA |
| | 0.779997632 | 0.864527319 |
| | 0.853595154 | NA |
| NA | | NA |
| NA | | NA |
| | 0.955792873 | NA |
| | 0.047827863 | 0.119579704 |
| | 0.01723565 | 0.053271387 |
| NA | | NA |
| NA | | NA |
| NA | | NA |
| | 0.19877373 | 0.346051222 |
| | 0.089979923 | 0.19420585 |
| | 0.538871523 | 0.68636008 |
| NA | | NA |
| | 0.55761467 | 0.701317451 |
| | 0.845000217 | 0.906712432 |
| | 0.068918989 | 0.159278548 |
| NA | | NA |
| | 0.908907149 | 0.944704653 |
| NA | | NA |
| | 0.839318641 | NA |
| | 0.081116291 | 0.179559042 |
| | 0.969742546 | 0.981982089 |
| | 0.990090771 | 0.99347163 |
| NA | | NA |
| | 0.937638937 | 0.963105753 |
| | 0.694130761 | 0.803858336 |
| | 0.919015422 | 0.951350632 |
| | 0.853595154 | NA |
| NA | | NA |
| NA | | NA |
| NA | | NA |
| NA | | NA |
| NA | | NA |
| | 9.40E-10 | 1.68E-08 |
| | 0.932644224 | NA |
| NA | | NA |
| NA | | NA |
| | 5.79E-05 | 0.000432228 |
| | 0.00010162 | 0.000710148 |
| | 0.88438961 | NA |
| NA | | NA |
| NA | | NA |
| NA | | NA |
| | 2.51E-06 | 2.60E-05 |
| | 0.870268939 | 0.92243237 |

| | |
|---|---|
| NA | NA |
| NA | NA |
| 1.63E-06 | 1.76E-05 |
| 0.655417156 | 0.775090135 |
| 0.103253283 | 0.214777971 |
| 0.006913788 | 0.025422829 |
| 0.831973572 | 0.898078683 |
| 0.617356485 | 0.746966198 |
| 0.126363889 | 0.249623478 |
| 0.48666523 | 0.64182213 |
| 0.752775715 | 0.844929151 |
| 0.69143582 | NA |
| NA | NA |
| 0.025812438 | 0.073535325 |
| 0.003844966 | 0.015708529 |
| 0.020961563 | 0.062109686 |
| 0.779257987 | 0.863898398 |
| 0.433308911 | 0.594547685 |
| 3.63E-05 | 0.000284814 |
| NA | NA |
| 0.941278788 | NA |
| NA | NA |
| 0.891969346 | 0.934936094 |
| 0.376745778 | 0.539727616 |
| 0.511052613 | 0.662619148 |
| 7.84E-13 | 1.99E-11 |
| NA | NA |
| 0.100667937 | 0.210794249 |
| 0.07383719 | 0.167521951 |
| 0.921383099 | NA |
| 0.839318641 | NA |
| 0.001018606 | 0.005186067 |
| 0.037183121 | NA |
| 0.483803728 | 0.639697482 |
| 0.952380021 | NA |
| 0.779078701 | NA |
| 9.61E-14 | 2.63E-12 |
| 0.733063054 | 0.831587399 |
| 0.61968884 | 0.748760535 |
| 0.280367934 | 0.440366738 |
| NA | NA |
| 0.839318641 | NA |
| 2.67E-43 | 3.10E-41 |
| 0.63865805 | 0.762662596 |
| 0.482476965 | 0.638598494 |
| 0.312645947 | 0.474652305 |
| 0.307525251 | 0.469032168 |

| | |
|---|---|
| 0.227133173 | 0.380371757 |
| 1.54E-06 | 1.66E-05 |
| 0.739991096 | NA |
| 9.25E-06 | 8.45E-05 |
| 0.768102123 | 0.856236733 |
| 0.306189198 | NA |
| 0.918368503 | NA |
| 0.914377546 | NA |
| NA | NA |
| 0.82074009 | 0.890678686 |
| NA | NA |
| NA | NA |
| NA | NA |
| NA | NA |
| 0.839319108 | NA |
| 0.996979947 | 0.997558905 |
| NA | NA |
| 0.955792398 | NA |
| 0.000786414 | 0.004169487 |
| NA | NA |
| 0.930758729 | 0.958753593 |
| 0.214351579 | 0.365318182 |
| 0.000113776 | 0.000784049 |
| 0.226887715 | 0.380117222 |
| 0.069321673 | 0.160044875 |
| 0.005202039 | 0.020165948 |
| 0.839318641 | NA |
| 0.631062358 | NA |
| 0.822151266 | NA |
| 0.698154614 | 0.80686944 |
| 0.941278332 | NA |
| NA | NA |
| 7.89E-19 | 3.10E-17 |
| NA | NA |
| NA | NA |
| 0.096746018 | 0.204488831 |
| 0.70347709 | 0.810844101 |
| 0.553825928 | NA |
| 0.017661359 | 0.054284049 |
| 0.969165048 | 0.981536315 |
| 0.075746203 | 0.170762561 |
| 0.941278838 | NA |
| 0.873177689 | 0.924280679 |
| 0.923509189 | 0.954466597 |
| 0.357140189 | 0.520718058 |
| 0.955792398 | NA |
| 0.148079894 | 0.280754949 |

| | | |
|---|---|---|
| | 3.02E-06 | 3.07E-05 |
| | 0.96722435 | 0.980329961 |
| | 0.080857117 | 0.179130044 |
| | 0.004976549 | 0.019456606 |
| | 0.053478956 | 0.130688513 |
| | 0.004427633 | 0.017643821 |
| | 0.304709557 | 0.466381382 |
| | 0.481087304 | 0.637487957 |
| | 8.12E-17 | 2.80E-15 |
| | 0.587063044 | NA |
| | 0.525866411 | 0.675044348 |
| | 0.28885402 | NA |
| | 0.640782619 | NA |
| | 0.955585653 | NA |
| | 0.217770815 | NA |
| | 0.955792873 | NA |
| NA | | NA |
| | 0.001727118 | 0.00809708 |
| NA | | NA |
| | 0.853595622 | NA |
| | 0.010398574 | 0.035535801 |
| | 0.941278332 | NA |
| | 0.764383825 | NA |
| | 0.236674089 | 0.392101111 |
| NA | | NA |
| | 0.352023653 | 0.51562315 |
| NA | | NA |
| NA | | NA |
| | 0.193186637 | 0.338902442 |
| NA | | NA |
| | 0.390691598 | 0.55319699 |
| NA | | NA |
| | 0.839318641 | NA |
| | 0.850608484 | 0.910431835 |
| | 0.941278788 | NA |
| | 0.000873159 | 0.004549676 |
| NA | | NA |
| | 0.801207117 | 0.878492596 |
| | 6.06E-14 | 1.68E-12 |
| | 2.60E-08 | 3.83E-07 |
| | 0.756853334 | NA |
| | 0.438189495 | NA |
| | 0.110884253 | 0.226401126 |
| | 0.5359355 | 0.684059243 |
| | 0.941277857 | NA |
| | 0.26034696 | 0.418899367 |
| NA | | NA |

| | | |
|---|---|---|
| | 0.29603767 | 0.457521501 |
| | 0.589635083 | 0.725709409 |
| NA | NA | |
| NA | NA | |
| | 0.582694895 | 0.720221546 |
| | 0.873743088 | 0.924714672 |
| | 0.941278332 | NA |
| NA | NA | |
| | 0.171564095 | 0.311722794 |
| | 7.12E-75 | 1.70E-72 |
| | 2.30E-05 | 0.000190654 |
| | 0.126855854 | 0.250262994 |
| | 0.838993826 | NA |
| | 0.001259939 | 0.006205061 |
| | 0.053258866 | 0.130405055 |
| | 0.731247391 | 0.830477641 |
| | 0.309862517 | NA |
| | 0.794266924 | 0.873822109 |
| | 0.956294952 | 0.973222697 |
| | 0.026087571 | 0.074159365 |
| | 0.695772268 | NA |
| | 0.250882167 | 0.408484061 |
| | 0.791252016 | 0.871750204 |
| NA | NA | |
| | 0.786252996 | 0.868628494 |
| NA | NA | |
| | 0.921978748 | 0.953547326 |
| | 0.668350078 | 0.785079631 |
| | 0.030127211 | 0.083295142 |
| | 0.624157117 | 0.752212036 |
| NA | NA | |
| NA | NA | |
| | 0.362481628 | NA |
| | 0.398739963 | 0.56155308 |
| | 0.039131383 | 0.102340972 |
| | 0.941276431 | NA |
| | 0.243818642 | 0.400288755 |
| | 0.061379523 | 0.145629525 |
| | 0.941278332 | NA |
| NA | NA | |
| | 0.856963873 | 0.914233074 |
| | 0.394963215 | 0.557520724 |
| | 0.839319574 | NA |
| | 0.550898295 | 0.696324212 |
| | 0.305894233 | 0.46750318 |
| | 0.307400916 | 0.46893268 |
| NA | NA | |

| | |
|---|---|
| NA | NA |
| 0.00093231 | 0.004804251 |
| NA | NA |
| 4.08E-05 | 0.000316658 |
| 9.30E-148 | 5.27E-145 |
| 0.760627917 | NA |
| 0.606143505 | 0.738807236 |
| 0.000348733 | 0.002073578 |
| 0.017923731 | 0.054947884 |
| 0.028922475 | 0.080619832 |
| NA | NA |
| 0.010416057 | 0.035575118 |
| 0.806497351 | 0.881532419 |
| 0.582422637 | 0.720034467 |
| 0.676235839 | 0.791324721 |
| 1.55E-17 | 5.61E-16 |
| 0.900622973 | 0.940137567 |
| 0.668502327 | 0.785219719 |
| 0.003292799 | 0.013868384 |
| 0.56006007 | NA |
| 0.576487038 | NA |
| 3.03E-13 | 7.94E-12 |
| 2.79E-162 | 2.01E-159 |
| 2.57E-15 | 7.99E-14 |
| 0.928412448 | NA |
| 1.31E-09 | 2.30E-08 |
| 0.165117279 | NA |
| 0.532530836 | NA |
| 0.66704116 | 0.784230075 |
| 4.72E-09 | 7.72E-08 |
| 0.943351682 | 0.965914856 |
| 0.14253356 | 0.272913185 |
| 0.955792398 | NA |
| 0.367560463 | 0.530716839 |
| 0.753866961 | 0.845834893 |
| 0.306731637 | 0.468301888 |
| 0.74303779 | 0.83852368 |
| 0.853595622 | NA |
| 0.000166454 | 0.001096885 |
| 0.381226823 | 0.543825835 |
| 0.970320416 | 0.982229107 |
| 0.293273568 | 0.45445947 |
| NA | NA |
| 0.955792398 | NA |
| 0.288180711 | 0.448817463 |
| 0.005356863 | 0.020628388 |
| 0.116854525 | 0.235350432 |

| | |
|---|---|
| 0.033863383 | 0.091271232 |
| 0.037877863 | 0.099742107 |
| 0.051402284 | 0.126732667 |
| 0.068375007 | 0.158298017 |
| 0.348614895 | 0.512525091 |
| 4.12E-10 | 7.69E-09 |
| 0.813515837 | 0.886151845 |
| 0.059893984 | 0.142788121 |
| 0.58394119 | 0.721125257 |
| 0.105763942 | 0.218675586 |
| NA | NA |
| NA | NA |
| NA | NA |
| 0.267702471 | NA |
| 1.22E-25 | 7.04E-24 |
| NA | NA |
| 0.470645532 | 0.628248834 |
| 0.006984824 | 0.025627515 |
| NA | NA |
| NA | NA |
| 0.01789605 | 0.054891982 |
| NA | NA |
| NA | NA |
| NA | NA |
| NA | NA |
| NA | NA |
| NA | NA |
| NA | NA |
| NA | NA |
| NA | NA |
| 0.921481694 | NA |
| 0.001301471 | 0.006371371 |
| 0.141633189 | NA |
| 0.008252187 | 0.029417187 |
| 0.790958851 | NA |
| NA | NA |
| NA | NA |
| NA | NA |
| NA | NA |
| NA | NA |
| NA | NA |
| NA | NA |
| NA | NA |
| 0.002441583 | 0.010818049 |
| 0.011111663 | 0.037520187 |
| 2.33E-06 | 2.43E-05 |
| NA | NA |

| | |
|---|---|
| 0.035699608 | 0.095228295 |
| 0.042418654 | 0.109009805 |
| 0.661789333 | NA |
| 0.011200855 | NA |
| 0.000653603 | 0.003559967 |
| NA | NA |
| 0.092373368 | NA |
| 0.551677292 | NA |
| NA | NA |
| 0.941277857 | NA |
| NA | NA |
| 0.941277857 | NA |
| NA | NA |
| 0.941277857 | NA |
| 0.708444157 | NA |
| NA | NA |
| NA | NA |
| 0.000500336 | 0.002838854 |
| 0.941278332 | NA |
| 0.839319108 | NA |
| NA | NA |
| NA | NA |
| NA | NA |
| NA | NA |
| NA | NA |
| NA | NA |
| NA | NA |
| 0.636896557 | 0.761245039 |
| NA | NA |
| NA | NA |
| 0.94608247 | NA |
| NA | NA |
| NA | NA |
| 0.941278332 | NA |
| 0.005729528 | 0.021802433 |
| 0.359889278 | NA |
| 0.941280787 | NA |
| 0.626484057 | NA |
| 0.420141146 | NA |
| 0.081182657 | NA |
| 0.396612505 | NA |
| 0.786733216 | NA |
| 3.31E-15 | 1.02E-13 |
| 0.220393668 | NA |
| 0.941279268 | NA |
| 0.923904649 | NA |
| 0.2749957 | NA |

| | | |
|---|---|---|
| | 0.941276431 | NA |
| | 0.937972656 | NA |
| | 0.693813816 | NA |
| | 0.669674127 | 0.78623601 |
| | 2.83E-08 | 4.13E-07 |
| | 0.000182415 | 0.001184353 |
| NA | | NA |
| | 0.007540852 | 0.027315076 |
| | 0.031207853 | 0.085578113 |
| NA | | NA |
| | 0.942583637 | NA |
| | 0.767923204 | NA |
| NA | | NA |
| NA | | NA |
| | 0.941277857 | NA |
| | 0.935650667 | NA |
| | 0.935503946 | NA |
| | 0.941278332 | NA |
| | 0.941278332 | NA |
| NA | | NA |
| | 0.941278332 | NA |
| | 0.318120138 | 0.480572197 |
| | 0.799533082 | NA |
| | 0.000962733 | 0.004939634 |
| | 0.128779827 | 0.253051882 |
| | 0.112133594 | 0.228217513 |
| | 0.233276843 | 0.387882393 |
| NA | | NA |
| | 4.63E-06 | 4.54E-05 |
| | 0.537073221 | 0.685033857 |
| | 0.134741748 | 0.261635102 |
| | 0.376583527 | 0.539641247 |
| | 0.034882032 | 0.093392814 |
| | 0.876109366 | 0.925901569 |
| | 0.839318641 | NA |
| NA | | NA |
| NA | | NA |
| | 0.788146828 | NA |
| NA | | NA |
| | 0.941277857 | NA |
| | 0.023154783 | 0.06745779 |
| | 2.18E-05 | 0.00018196 |
| | 0.140009711 | NA |
| | 0.624142788 | NA |
| | 0.919622047 | NA |
| | 0.950512078 | NA |
| | 0.914390259 | NA |

| | | |
|---|---|---|
| | 0.941278424 | NA |
| | 0.95456991 | NA |
| | 0.941278207 | NA |
| NA | | NA |
| | 0.955792873 | NA |
| NA | | NA |
| NA | | NA |
| NA | | NA |
| | 0.473289351 | NA |
| NA | | NA |
| NA | | NA |
| NA | | NA |
| NA | | NA |
| | 0.941277857 | NA |
| NA | | NA |
| | 0.941277857 | NA |
| NA | | NA |
| | 0.000954129 | 0.004901831 |
| | 0.420772627 | NA |
| | 0.626355361 | 0.753830727 |
| | 0.728836517 | NA |
| NA | | NA |
| | 0.398152403 | 0.560825157 |
| | 0.423359459 | 0.58470476 |
| NA | | NA |
| | 0.013042875 | 0.04265547 |
| | 0.000494948 | 0.002811637 |
| | 0.877940671 | 0.926725968 |
| | 0.842822009 | 0.905354701 |
| | 0.939610821 | NA |
| NA | | NA |
| | 0.74298716 | 0.83852368 |
| NA | | NA |
| | 0.941468986 | 0.964703895 |
| | 0.000331058 | 0.001985938 |
| | 0.941208171 | NA |
| | 0.132093961 | NA |
| | 0.891181173 | 0.934527019 |
| | 0.770409393 | 0.857884649 |
| | 0.043159717 | 0.110468375 |
| | 0.096605433 | 0.204300535 |
| | 0.086618574 | 0.188730803 |
| | 0.272684925 | 0.432581084 |
| | 7.09E-07 | 8.13E-06 |
| | 0.993327612 | 0.995209152 |
| | 0.301776571 | 0.463127477 |
| | 0.163832703 | 0.301976751 |

| | | |
|---|---|---|
| NA | | NA |
| | 0.006837332 | 0.025188387 |
| | 0.199647922 | 0.347183534 |
| | 6.49E-05 | 0.000478162 |
| | 0.101201654 | 0.211644056 |
| | 0.003758713 | 0.015409927 |
| | 0.180507803 | 0.322531846 |
| | 0.395772841 | NA |
| | 0.866999455 | 0.920524624 |
| | 0.277747299 | 0.437434077 |
| | 2.50E-12 | 6.01E-11 |
| | 0.941278332 | NA |
| | 0.043038855 | 0.110246538 |
| | 0.020238171 | 0.060447871 |
| | 0.014908017 | 0.047457618 |
| | 9.65E-11 | 1.94E-09 |
| | 4.36E-11 | 9.20E-10 |
| | 7.70E-27 | 4.71E-25 |
| | 1.25E-25 | 7.19E-24 |
| | 0.616346859 | NA |
| | 2.10E-10 | 4.06E-09 |
| | 0.005741475 | 0.021837422 |
| | 1.50E-10 | 2.95E-09 |
| | 2.06E-11 | 4.51E-10 |
| | 7.79E-13 | 1.98E-11 |
| | 9.67E-08 | 1.30E-06 |
| | 2.81E-10 | 5.35E-09 |
| | 0.062000298 | 0.146736711 |
| | 0.00199874 | 0.009164711 |
| NA | | NA |
| NA | | NA |
| | 0.941277857 | NA |
| | 0.955791922 | NA |
| | 0.955792398 | NA |
| | 1.84E-19 | 7.56E-18 |
| | 0.501863899 | 0.654102529 |
| | 3.37E-07 | 4.09E-06 |
| | 0.001912579 | 0.008818909 |
| | 0.179174107 | 0.320799015 |
| | 0.127668631 | 0.25152285 |
| | 0.359530234 | 0.523108908 |
| | 0.941278332 | NA |
| | 0.733799911 | 0.832164192 |
| | 0.809767542 | NA |
| | 0.62600446 | 0.753598935 |
| | 0.016646365 | 0.051833553 |
| | 0.001563437 | 0.007433666 |

| | | | |
|---|---|---|---|
| NA | | NA | |
| | 0.369840837 | | 0.533059926 |
| | 0.001359037 | | 0.006605636 |
| | 0.513051959 | | 0.664282495 |
| | 0.021422794 | | 0.063208795 |
| | 0.758487145 | NA | |
| | 0.955792398 | NA | |
| | 0.270823372 | | 0.430776389 |
| NA | | NA | |
| | 0.832162011 | NA | |
| | 0.631808673 | NA | |
| | 0.907106255 | NA | |
| | 0.148692333 | | 0.281534875 |
| | 0.074769913 | | 0.169057991 |
| | 0.68490145 | | 0.797633052 |
| | 2.17E-07 | | 2.75E-06 |
| | 0.916264098 | NA | |
| | 2.64E-11 | | 5.73E-10 |
| | 0.574808017 | NA | |
| NA | | NA | |
| NA | | NA | |
| | 0.476510314 | | 0.633524321 |
| | 0.016824165 | | 0.052250624 |
| NA | | NA | |
| | 0.212642699 | NA | |
| NA | | NA | |
| | 0.077605251 | NA | |
| | 0.663851495 | | 0.781956415 |
| NA | | NA | |
| | 0.531260224 | | 0.67996602 |
| | 0.408539526 | | 0.570638445 |
| | 0.839319574 | NA | |
| | 0.546143782 | NA | |
| | 0.28582276 | NA | |
| | 0.302899231 | NA | |
| | 0.746648309 | | 0.840668073 |
| | 0.396734338 | | 0.559324215 |
| | 0.608665996 | | 0.740895369 |
| | 0.152664665 | | 0.286752238 |
| | 0.705632596 | NA | |
| NA | | NA | |
| NA | | NA | |
| NA | | NA | |
| NA | | NA | |
| NA | | NA | |
| NA | | NA | |
| NA | | NA | |

| | |
|---|---|
| NA | NA |
| NA | NA |
| NA | NA |
| NA | NA |
| NA | NA |
| NA | NA |
| NA | NA |
| NA | NA |
| NA | NA |
| NA | NA |
| NA | NA |
| 0.731007615 | 0.830443073 |
| 0.676681116 | 0.791584276 |
| 0.307833448 | 0.469379123 |
| NA | NA |
| 0.941275956 | NA |
| 0.707151277 | 0.813239344 |
| 0.943923451 | 0.966167798 |
| 6.26E-06 | 5.96E-05 |
| 0.006364367 | 0.0237474 |
| 0.637493856 | 0.761744398 |
| 0.586958953 | 0.723508833 |
| 0.694638288 | NA |
| 0.941275956 | NA |
| 8.24E-06 | 7.61E-05 |
| 0.335947285 | 0.499139655 |
| 0.477473256 | NA |
| NA | NA |
| 0.832789929 | 0.898613783 |
| 0.258915527 | 0.41730142 |
| 0.323223287 | NA |
| 0.800209225 | NA |
| 0.516254276 | 0.667058482 |
| 0.494608778 | 0.647704563 |
| 2.45E-05 | 0.000201805 |
| 0.329314206 | 0.492326909 |
| 0.004194261 | 0.01686918 |
| 0.579639631 | 0.717488298 |
| 1.73E-15 | 5.47E-14 |
| 0.941277857 | NA |
| 0.894635019 | NA |
| 4.26E-06 | 4.22E-05 |
| 0.038754816 | 0.101513188 |
| 7.24E-39 | 7.12E-37 |
| NA | NA |
| 0.026988029 | 0.076381577 |
| 0.000188325 | 0.001217745 |

| | | | |
|---|---|---|---|
| | 0.65656001 | | 0.776056084 |
| | 0.897062103 | | 0.937872959 |
| | 0.191723455 | | 0.337006254 |
| NA | | NA | |
| | 0.376904764 | | 0.539833265 |
| | 0.817257543 | NA | |
| | 0.57036701 | | 0.711039416 |
| | 0.839319108 | NA | |
| | 0.010417987 | | 0.035576605 |
| | 0.696848688 | | 0.80579054 |
| | 0.597391594 | | 0.732000758 |
| | 6.50E-12 | | 1.51E-10 |
| | 0.146148154 | | 0.277978133 |
| NA | | NA | |
| | 0.488301749 | NA | |
| | 0.070985743 | | 0.162821881 |
| | 2.17E-08 | | 3.23E-07 |
| | 0.571638303 | | 0.711878559 |
| | 0.859992662 | | 0.916519447 |
| | 8.56E-27 | | 5.23E-25 |
| | 0.067303541 | | 0.156395528 |
| | 0.780251735 | NA | |
| | 0.839319574 | NA | |
| NA | | NA | |
| NA | | NA | |
| NA | | NA | |
| NA | | NA | |
| NA | | NA | |
| NA | | NA | |
| NA | | NA | |
| NA | | NA | |
| NA | | NA | |
| NA | | NA | |
| NA | | NA | |
| NA | | NA | |
| NA | | NA | |
| NA | | NA | |
| NA | | NA | |
| NA | | NA | |
| NA | | NA | |
| NA | | NA | |
| | 0.839318641 | NA | |
| | 6.06E-08 | | 8.39E-07 |
| NA | | NA | |
| | 2.37E-149 | | 1.41E-146 |
| | 0.005405625 | | 0.020765768 |
| | 6.44E-05 | | 0.000475161 |

| | | | |
|---|---|---|---|
| NA | | NA | |
| | 0.895572469 | | 0.936820599 |
| | 0.001868924 | | 0.00863937 |
| | 0.839319108 | NA | |
| NA | | NA | |
| NA | | NA | |
| NA | | NA | |
| NA | | NA | |
| NA | | NA | |
| NA | | NA | |
| NA | | NA | |
| NA | | NA | |
| | 0.690966234 | | 0.801792603 |
| NA | | NA | |
| | 0.755626499 | NA | |
| | 0.743511643 | NA | |
| NA | | NA | |
| | 0.839320041 | NA | |
| NA | | NA | |
| NA | | NA | |
| NA | | NA | |
| | 0.955792873 | NA | |
| | 0.953932387 | NA | |
| NA | | NA | |
| NA | | NA | |
| | 0.839319108 | NA | |
| NA | | NA | |
| NA | | NA | |
| NA | | NA | |
| | 0.955792873 | NA | |
| | 0.941278332 | NA | |
| NA | | NA | |
| | 0.004727715 | | 0.018664632 |
| | 0.67052885 | | 0.786722102 |
| | 0.008884268 | | 0.03123997 |
| | 0.388705346 | | 0.551500038 |
| | 0.211983161 | | 0.362420859 |
| NA | | NA | |
| | 9.28E-07 | | 1.04E-05 |
| | 0.019476844 | | 0.058644888 |
| NA | | NA | |
| | 0.739889323 | NA | |
| NA | | NA | |
| NA | | NA | |
| | 0.941275956 | NA | |
| | 0.305371124 | | 0.466954365 |
| | 0.212588442 | NA | |

| | |
|---|---|
| 0.936046576 | 0.962160157 |
| 0.12455405 | 0.247052958 |
| NA | NA |
| 0.043466123 | 0.111018419 |
| 0.524560949 | 0.674080713 |
| 0.70758666 | NA |
| 0.941277857 | NA |
| 0.047577559 | 0.119041422 |
| 0.166087625 | 0.304891074 |
| 0.938914 | 0.963601145 |
| 0.686802263 | 0.799025732 |
| 0.000126011 | 0.000859403 |
| 0.000280411 | 0.001718417 |
| 0.839318641 | NA |
| 0.476604523 | 0.633535398 |
| 0.169711995 | 0.30930206 |
| NA | NA |
| 0.602907823 | 0.736145071 |
| NA | NA |
| NA | NA |
| 0.791471477 | 0.871750204 |
| NA | NA |
| 0.941278332 | NA |
| 0.624142788 | NA |
| 0.941277857 | NA |
| NA | NA |
| NA | NA |
| 0.941278332 | NA |
| NA | NA |
| 0.955792873 | NA |
| 0.001184146 | 0.005892757 |
| 0.270473649 | 0.430536601 |
| 1.87E-12 | 4.57E-11 |
| 0.648839552 | 0.770412004 |
| 0.838333133 | NA |
| 0.002356215 | 0.010504818 |
| 0.269378257 | 0.429281017 |
| 0.157067698 | 0.292827238 |
| 0.148923218 | 0.281815108 |
| 0.043164139 | 0.110468375 |
| 0.332135343 | 0.495238078 |
| 1.27E-09 | 2.23E-08 |
| NA | NA |
| 0.055571324 | 0.134791467 |
| 0.839318641 | NA |
| 0.011928359 | 0.039703371 |
| 0.941277857 | NA |

| | |
|---|---|
| NA | NA |
| NA | NA |
| 0.645846934 | NA |
| NA | NA |
| NA | NA |
| NA | NA |
| NA | NA |
| 0.639777971 | NA |
| NA | NA |
| NA | NA |
| NA | NA |
| 0.941279303 | NA |
| NA | NA |
| NA | NA |
| NA | NA |
| NA | NA |
| NA | NA |
| NA | NA |
| NA | NA |
| 0.831413565 | NA |
| NA | NA |
| NA | NA |
| NA | NA |
| NA | NA |
| 0.839318641 | NA |
| NA | NA |
| NA | NA |
| NA | NA |
| 0.941277857 | NA |
| 0.941279051 | NA |
| NA | NA |
| NA | NA |
| NA | NA |
| NA | NA |
| NA | NA |
| NA | NA |
| 0.941281154 | NA |
| NA | NA |
| 0.832162011 | NA |
| 0.839319108 | NA |
| NA | NA |
| NA | NA |
| NA | NA |
| NA | NA |
| 0.825972908 | NA |
| 0.055232262 | 0.134133078 |
| NA | NA |

| | | | |
|---|---|---|---|
| | 0.941279303 | NA | |
| NA | | NA | |
| NA | | NA | |
| | 0.739991096 | NA | |
| NA | | NA | |
| NA | | NA | |
| NA | | NA | |
| NA | | NA | |
| NA | | NA | |
| | 0.348402291 | | 0.512399365 |
| | 0.941278332 | NA | |
| NA | | NA | |
| NA | | NA | |
| NA | | NA | |
| NA | | NA | |
| NA | | NA | |
| NA | | NA | |
| NA | | NA | |
| NA | | NA | |
| NA | | NA | |
| | 0.935650667 | NA | |
| NA | | NA | |
| NA | | NA | |
| NA | | NA | |
| | 0.941278332 | NA | |
| NA | | NA | |
| NA | | NA | |
| NA | | NA | |
| NA | | NA | |
| NA | | NA | |
| NA | | NA | |
| NA | | NA | |
| NA | | NA | |
| | 0.934047677 | NA | |
| | 0.839319574 | NA | |
| NA | | NA | |
| NA | | NA | |
| | 0.839318641 | NA | |
| | 0.941278332 | NA | |
| | 0.941278788 | NA | |
| NA | | NA | |
| | 0.941277857 | NA | |
| | 0.839319108 | NA | |
| NA | | NA | |
| NA | | NA | |
| NA | | NA | |
| NA | | NA | |

| | | | |
|---|---|---|---|
| | 0.941277857 | NA | |
| NA | | NA | |
| NA | | NA | |
| NA | | NA | |
| NA | | NA | |
| NA | | NA | |
| NA | | NA | |
| NA | | NA | |
| NA | | NA | |
| | 0.839318641 | NA | |
| NA | | NA | |
| | 0.941278332 | NA | |
| | 0.835561294 | NA | |
| NA | | NA | |
| NA | | NA | |
| | 0.941278332 | NA | |
| NA | | NA | |
| | 0.839319108 | NA | |
| NA | | NA | |
| NA | | NA | |
| NA | | NA | |
| NA | | NA | |
| NA | | NA | |
| NA | | NA | |
| NA | | NA | |
| NA | | NA | |
| NA | | NA | |
| | 0.838457485 | NA | |
| NA | | NA | |
| NA | | NA | |
| NA | | NA | |
| | 0.634372216 | | 0.75971383 |
| NA | | NA | |
| NA | | NA | |
| | 0.941277857 | NA | |
| NA | | NA | |
| | 0.941276431 | NA | |
| NA | | NA | |
| NA | | NA | |
| | 0.941277857 | NA | |
| NA | | NA | |
| NA | | NA | |
| NA | | NA | |
| NA | | NA | |
| NA | | NA | |
| NA | | NA | |
| NA | | NA | |

| | | |
|---|---|---|
| 0.685162694 | NA | |
| 0.703395612 | NA | |
| 0.941278332 | NA | |
| 0.928411644 | NA | |
| 0.928411644 | NA | |
| NA | NA | |
| 0.941278424 | NA | |
| NA | NA | |
| NA | NA | |
| 0.349178275 | | 0.513191806 |
| NA | NA | |
| 0.941279303 | NA | |
| NA | NA | |
| NA | NA | |
| NA | NA | |
| NA | NA | |
| NA | NA | |
| 0.941278332 | NA | |
| 0.941275481 | NA | |
| 0.645598806 | | 0.767952477 |
| NA | NA | |
| 0.773242253 | NA | |
| 0.955792398 | NA | |
| NA | NA | |
| NA | NA | |
| NA | NA | |
| 0.941279268 | NA | |
| NA | NA | |
| 0.941278332 | NA | |
| 0.941277857 | NA | |
| NA | NA | |
| NA | NA | |
| 0.941282915 | | 0.964703895 |
| NA | NA | |
| NA | NA | |
| NA | NA | |
| 0.512829654 | NA | |
| 0.955792873 | NA | |
| 0.941277857 | NA | |
| 0.581775751 | NA | |
| NA | NA | |
| 0.941275956 | NA | |
| 0.941280944 | NA | |
| 0.941277857 | NA | |
| NA | NA | |
| 0.874459303 | NA | |
| 0.941278332 | NA | |

| | |
|---|---|
| NA | NA |
| 0.024388048 | 0.070302086 |
| NA | NA |
| NA | NA |
| 0.839319108 | NA |
| NA | NA |
| 0.955792398 | NA |
| NA | NA |
| NA | NA |
| 0.614215097 | NA |
| NA | NA |
| 0.839320627 | NA |
| NA | NA |
| NA | NA |
| 0.222904666 | 0.375354602 |
| NA | NA |
| 0.821695621 | NA |
| 0.838457485 | NA |
| NA | NA |
| 0.941280558 | NA |
| NA | NA |
| 0.839321179 | NA |
| 0.821695621 | NA |
| NA | NA |
| 0.839318641 | NA |
| 0.7996816 | NA |
| 0.933117769 | NA |
| NA | NA |
| 0.647217777 | 0.769292261 |
| 0.907471713 | 0.943834027 |
| 0.968063599 | 0.980796398 |
| 0.941275956 | NA |
| NA | NA |
| 0.955792398 | NA |
| NA | NA |
| 0.275894439 | 0.435726855 |
| 0.514170061 | 0.665240505 |
| 0.841923686 | NA |
| 0.000674861 | 0.003664016 |
| 0.404381115 | 0.566414364 |
| 0.07428727 | 0.168270422 |
| 0.158838394 | 0.295088312 |
| 5.17E-19 | 2.06E-17 |
| 7.42E-21 | 3.35E-19 |
| 1.97E-05 | 0.000165902 |
| 0.188306723 | 0.332646328 |
| 0.572204955 | 0.712174354 |

| | |
|---|---|
| 0.254518747 | 0.412546261 |
| 7.74E-06 | 7.21E-05 |
| 0.026051394 | 0.074074217 |
| 0.252114215 | 0.409836261 |
| 0.537718222 | 0.685452502 |
| 0.386069013 | 0.548937554 |
| 0.001909229 | 0.008805169 |
| 0.944289243 | NA |
| 0.221045699 | 0.372958886 |
| 0.027914812 | 0.078393907 |
| 0.008516404 | 0.030157543 |
| 0.672730775 | NA |
| 0.005378912 | 0.020689862 |
| 0.007516538 | 0.027247735 |
| 0.004749752 | 0.01872402 |
| 0.084804779 | 0.185867711 |
| 0.274898079 | 0.434813608 |
| 8.09E-14 | 2.23E-12 |
| 0.01594535 | 0.050089906 |
| 0.096053407 | 0.203494722 |
| 0.024966147 | 0.071577898 |
| 6.64E-21 | 3.02E-19 |
| 0.920963372 | 0.952651559 |
| 0.575262228 | 0.71422417 |
| 0.001166071 | 0.005814963 |
| 0.62297176 | NA |
| 0.576279741 | 0.714928093 |
| 0.969020703 | 0.981431887 |
| 0.057953921 | 0.139091648 |
| 0.193425768 | 0.339196944 |
| 0.75678104 | NA |
| 0.941275956 | NA |
| 0.024456762 | 0.070466027 |
| 0.011735533 | 0.039220607 |
| 0.739992478 | NA |
| 1.19E-05 | 0.000105901 |
| 0.046886263 | 0.11776248 |
| 0.89655087 | NA |
| 0.06568625 | 0.153521613 |
| NA | NA |
| NA | NA |
| NA | NA |
| NA | NA |
| NA | NA |
| NA | NA |
| 0.476778891 | 0.633602986 |
| NA | NA |

| | |
|---|---|
| NA | NA |
| 0.111435825 | 0.227201246 |
| 0.372010217 | 0.535343845 |
| 0.523792079 | 0.673511106 |
| NA | NA |
| 0.283743457 | 0.444349087 |
| 0.853595622 | NA |
| NA | NA |
| NA | NA |
| NA | NA |
| NA | NA |
| 0.857854554 | NA |
| NA | NA |
| NA | NA |
| NA | NA |
| NA | NA |
| 0.593131403 | NA |
| 0.128868266 | 0.253162965 |
| NA | NA |
| NA | NA |
| 0.941075561 | NA |
| NA | NA |
| 0.896754649 | NA |
| 0.937000045 | 0.962629548 |
| 0.306101944 | 0.467760559 |
| 0.089912679 | 0.194095916 |
| NA | NA |
| 0.034824836 | 0.093271153 |
| 0.920351348 | 0.9524436 |
| 2.98E-12 | 7.12E-11 |
| 0.000216475 | 0.001374743 |
| 0.579200776 | NA |
| 0.045083196 | 0.114180702 |
| 0.023211088 | 0.067592576 |
| 1.11E-18 | 4.35E-17 |
| 3.12E-08 | 4.53E-07 |
| NA | NA |
| 0.122509679 | 0.244096382 |
| 1.29E-07 | 1.70E-06 |
| 0.633735058 | 0.759295372 |
| 0.853595622 | NA |
| 0.688097513 | NA |
| 0.098362862 | 0.207105344 |
| 0.580746692 | NA |
| NA | NA |
| 0.18974005 | 0.334309917 |
| 0.146047346 | 0.277830797 |

| | |
|---|---|
| NA | NA |
| 0.00078915 | 0.004179652 |
| 0.000888269 | 0.004613262 |
| 0.64176284 | 0.764985934 |
| 3.20E-05 | 0.000254706 |
| 0.009363403 | 0.032602064 |
| 0.032700335 | 0.08891174 |
| 0.00279953 | 0.012093362 |
| 0.515926695 | 0.666816203 |
| 0.344811492 | 0.508406315 |
| 2.24E-06 | 2.35E-05 |
| 1.33E-06 | 1.45E-05 |
| 0.107634232 | 0.221484395 |
| 0.023008267 | 0.067080201 |
| 0.154558028 | 0.289395074 |
| 0.633595638 | 0.759204761 |
| 0.009753909 | 0.033729849 |
| 0.952200245 | 0.970932242 |
| 0.014865358 | 0.04734715 |
| NA | NA |
| 0.672267128 | NA |
| 0.436369027 | NA |
| 0.867467864 | 0.920816576 |
| NA | NA |
| NA | NA |
| NA | NA |
| 0.941279268 | NA |
| NA | NA |
| 0.566901053 | NA |
| 0.306700596 | 0.468301888 |
| NA | NA |
| 0.205380889 | 0.35403215 |
| 0.716418217 | 0.819759372 |
| 0.021326626 | 0.062995327 |
| NA | NA |
| 0.33692741 | 0.500079298 |
| 0.040187764 | 0.104575179 |
| 0.00421763 | 0.016951709 |
| 0.839319108 | NA |
| 0.021626748 | 0.063723677 |
| 5.02E-05 | 0.000380588 |
| 0.334497321 | NA |
| 0.941277857 | NA |
| 0.396253347 | 0.5589259 |
| 0.615803493 | NA |
| 0.31827496 | 0.480775567 |
| 0.95456991 | NA |

| | | |
|---|---|---|
| NA | | NA |
| | 0.688501761 | NA |
| | 0.307158236 | 0.468812509 |
| | 1.27E-06 | 1.39E-05 |
| | 0.221920097 | 0.374168884 |
| NA | | NA |
| NA | | NA |
| | 0.052674406 | 0.129214 |
| | 0.061958371 | 0.146681236 |
| | 0.564738788 | 0.706620985 |
| | 0.006457236 | 0.02402028 |
| | 0.292011784 | 0.453257687 |
| | 0.000593432 | 0.003281038 |
| | 2.71E-06 | 2.79E-05 |
| | 0.15320697 | 0.287414027 |
| | 0.531546671 | NA |
| | 0.004512716 | 0.017940816 |
| | 0.033364668 | 0.090336233 |
| | 0.941277857 | NA |
| | 0.534931178 | 0.683033737 |
| | 0.940475126 | NA |
| | 0.000244248 | 0.001521881 |
| | 0.355947243 | 0.519582649 |
| | 0.010825189 | 0.03673003 |
| | 0.027372459 | 0.077249142 |
| | 0.670125748 | NA |
| NA | | NA |
| | 0.853806167 | 0.912048218 |
| | 0.47956728 | 0.636233231 |
| | 0.905257723 | 0.942519764 |
| NA | | NA |
| | 0.955791922 | NA |
| | 0.476844723 | NA |
| | 0.000300042 | 0.001822764 |
| | 0.128738614 | 0.253008737 |
| | 0.869101082 | 0.921575098 |
| | 0.334558688 | NA |
| | 0.205431747 | 0.354094173 |
| | 0.006317269 | 0.02361238 |
| | 0.906588518 | NA |
| | 0.060728021 | 0.144414863 |
| | 0.191756549 | 0.337039535 |
| | 0.008343867 | 0.029681769 |
| | 0.11260342 | 0.229075778 |
| | 0.019997213 | 0.059840793 |
| | 0.344100494 | 0.50770394 |
| | 0.239896315 | 0.395923734 |

| | | |
|---|---|---|
| | 0.731195786 | 0.830477641 |
| | 0.383866534 | NA |
| | 0.432547814 | NA |
| | 0.120215069 | 0.240672813 |
| | 0.828202129 | 0.89562173 |
| NA | | NA |
| | 0.016480275 | 0.051424163 |
| | 0.000492868 | 0.002801154 |
| | 0.035182347 | 0.09405933 |
| | 0.017925818 | 0.054947884 |
| | 4.40E-19 | 1.76E-17 |
| | 0.596365827 | 0.731120237 |
| | 0.080715818 | 0.178949135 |
| | 0.466248097 | 0.624374773 |
| | 0.001244214 | 0.006142873 |
| | 0.185275089 | NA |
| | 0.805187443 | 0.880868577 |
| NA | | NA |
| | 0.855721843 | NA |
| NA | | NA |
| | 1.66E-07 | 2.15E-06 |
| | 0.018778647 | 0.057055061 |
| | 4.97E-21 | 2.28E-19 |
| | 0.000236799 | 0.001481284 |
| | 1.80E-05 | 0.000152665 |
| | 0.474517858 | 0.632035482 |
| | 0.806334599 | 0.881435479 |
| | 0.186407154 | 0.330127257 |
| | 0.016326279 | 0.051037311 |
| | 0.948821832 | NA |
| | 1.99E-15 | 6.28E-14 |
| | 0.102420428 | 0.213591774 |
| | 4.60E-07 | 5.44E-06 |
| | 0.327800077 | 0.490525426 |
| | 0.231504393 | NA |
| | 0.021896665 | 0.064375547 |
| NA | | NA |
| | 0.129079027 | 0.253346995 |
| | 0.09076885 | 0.195483186 |
| | 0.865553264 | NA |
| | 0.001235262 | 0.006107543 |
| | 0.003171011 | 0.013444318 |
| NA | | NA |
| | 8.12E-10 | 1.46E-08 |
| NA | | NA |
| | 0.135345937 | NA |
| | 0.186034258 | 0.329585962 |

| | | | |
|---|---|---|---|
| NA | | NA | |
| | 0.904106384 | | 0.941735821 |
| | 6.81E-05 | | 0.000497747 |
| | 0.532352927 | | 0.680836641 |
| | 0.2457294 | | 0.402571509 |
| | 0.845835367 | | 0.907158717 |
| | 0.962819416 | | 0.977522084 |
| | 0.07865664 | | 0.175577731 |
| | 0.398873757 | | 0.561575367 |
| | 0.000759541 | | 0.00404805 |
| | 0.00620553 | | 0.023264139 |
| | 0.028916815 | NA | |
| | 0.915579684 | NA | |
| NA | | NA | |
| | 0.771607906 | NA | |
| | 0.941278788 | NA | |
| | 0.756306501 | NA | |
| | 0.012348547 | | 0.040820586 |
| | 0.08303983 | | 0.182729342 |
| | 0.298841664 | | 0.460509471 |
| NA | | NA | |
| | 0.941278332 | NA | |
| NA | | NA | |
| NA | | NA | |
| NA | | NA | |
| NA | | NA | |
| | 0.941278332 | NA | |
| NA | | NA | |
| NA | | NA | |
| NA | | NA | |
| NA | | NA | |
| NA | | NA | |
| NA | | NA | |
| NA | | NA | |
| NA | | NA | |
| NA | | NA | |
| NA | | NA | |
| NA | | NA | |
| | 0.839319108 | NA | |
| NA | | NA | |
| NA | | NA | |
| NA | | NA | |
| | 0.71780035 | NA | |
| | 0.153366829 | | 0.287662511 |
| | 0.839319108 | NA | |
| NA | | NA | |
| NA | | NA | |

| | |
|---|---|
| NA | NA |
| 0.613797171 | NA |
| 0.941275956 | NA |
| 0.570688953 | 0.7112545 |
| 0.019821068 | 0.059426929 |
| 1.92E-06 | 2.03E-05 |
| 2.69E-16 | 9.04E-15 |
| 0.831413565 | NA |
| 0.767520703 | 0.855748912 |
| 0.489929323 | 0.644341792 |
| 0.386281079 | 0.54914067 |
| 0.136198596 | 0.263741645 |
| NA | NA |
| 0.095759269 | 0.203106588 |
| 0.370920815 | 0.534195997 |
| 0.268152856 | 0.427929888 |
| 0.955791922 | NA |
| 0.911920286 | 0.946847262 |
| 0.003434035 | 0.014332263 |
| 0.409408361 | 0.57144194 |
| 0.912709487 | 0.94739633 |
| 0.716754769 | 0.819965251 |
| 0.84058606 | 0.903781811 |
| 1.07E-08 | 1.67E-07 |
| 0.931799938 | 0.959371849 |
| 0.018950814 | 0.05747233 |
| 0.487857535 | 0.642657697 |
| 1.30E-13 | 3.51E-12 |
| 0.844107149 | 0.906163487 |
| 0.289912744 | 0.450877298 |
| 0.594047282 | 0.729103976 |
| 0.955792398 | NA |
| 0.835047237 | 0.899970585 |
| 3.83E-05 | 0.000298576 |
| 0.000858537 | 0.004485272 |
| 0.99858453 | 0.998831218 |
| 0.814291008 | 0.886545425 |
| 0.941276431 | NA |
| 1.73E-11 | 3.82E-10 |
| 6.39E-06 | 6.07E-05 |
| NA | NA |
| 0.161567983 | 0.298620991 |
| NA | NA |
| NA | NA |
| NA | NA |
| 2.95E-05 | 0.000238094 |
| NA | NA |

| | | | |
|---|---|---|---|
| NA | | NA | |
| | 0.782204694 | | 0.866076017 |
| | 0.138676181 | | 0.267237735 |
| | 0.723448144 | NA | |
| | 0.839320041 | NA | |
| NA | | NA | |
| NA | | NA | |
| NA | | NA | |
| | 0.986112752 | NA | |
| | 0.005095482 | | 0.019823906 |
| | 0.281171898 | NA | |
| NA | | NA | |
| | 0.848602559 | | 0.908979709 |
| | 0.859173131 | | 0.916022897 |
| | 0.534136966 | NA | |
| NA | | NA | |
| NA | | NA | |
| | 0.804588217 | | 0.880558921 |
| | 0.191774691 | | 0.337046533 |
| | 0.000560534 | | 0.003130577 |
| | 0.589221877 | | 0.725350873 |
| | 0.877196627 | | 0.926433604 |
| | 9.30E-06 | | 8.49E-05 |
| | 0.883517302 | NA | |
| | 0.003199473 | | 0.013535971 |
| | 0.560246179 | | 0.70350723 |
| | 0.924127199 | NA | |
| NA | | NA | |
| NA | | NA | |
| | 0.146542278 | | 0.278482978 |
| | 0.941278332 | NA | |
| | 0.045457989 | | 0.114934253 |
| | 0.809061777 | | 0.882967072 |
| | 0.955792397 | NA | |
| | 0.059737889 | | 0.142544485 |
| | 0.080379371 | | 0.17845255 |
| | 0.437015214 | | 0.597944132 |
| NA | | NA | |
| | 0.897843632 | | 0.938413841 |
| | 0.468469335 | | 0.626573787 |
| | 0.381176473 | | 0.543825835 |
| | 0.277394036 | | 0.43728279 |
| | 0.001631901 | | 0.007716034 |
| NA | | NA | |
| NA | | NA | |
| | 0.904468064 | | 0.942027255 |
| | 0.941278332 | NA | |

| | |
|---|---|
| NA | NA |
| 0.155192098 | 0.290148638 |
| 0.256368395 | 0.414479452 |
| 0.00275684 | 0.011963734 |
| 0.853595622 | NA |
| 0.805130188 | 0.880868577 |
| 0.237694527 | 0.393189599 |
| 0.721613933 | NA |
| 0.27349547 | NA |
| 0.731633742 | 0.830757864 |
| NA | NA |
| NA | NA |
| 0.713105814 | 0.817559105 |
| 0.60428212 | 0.737245902 |
| NA | NA |
| NA | NA |
| 0.007639746 | 0.027601886 |
| NA | NA |
| NA | NA |
| 0.005701376 | 0.021716138 |
| 0.242009632 | 0.398268193 |
| 0.242026728 | NA |
| 1.06E-05 | 9.51E-05 |
| 2.73E-06 | 2.80E-05 |
| NA | NA |
| NA | NA |
| 0.077975272 | 0.174415825 |
| 0.000387909 | 0.002276382 |
| 1.82E-07 | 2.34E-06 |
| 0.248272139 | 0.405473683 |
| 0.793988952 | 0.873597053 |
| 0.059758585 | 0.142565283 |
| 0.076100027 | NA |
| 0.537288084 | 0.685138379 |
| 0.39422808 | 0.556747167 |
| 0.217965738 | 0.369254127 |
| 0.302714671 | NA |
| 0.169157701 | 0.308693676 |
| 0.727380904 | 0.828101606 |
| 4.54E-13 | 1.18E-11 |
| 0.266886906 | 0.426544813 |
| 0.204042294 | 0.352567309 |
| 0.012906423 | 0.042278928 |
| 0.853595154 | NA |
| NA | NA |
| NA | NA |
| NA | NA |

| | | |
|---|---|---|
| NA | | NA |
| NA | | NA |
| | 0.032539706 | 0.088555921 |
| | 0.291588604 | NA |
| | 0.04533057 | 0.114672998 |
| | 0.27093219 | 0.430805531 |
| | 0.379324534 | 0.542132974 |
| NA | | NA |
| | 0.000445574 | 0.002570419 |
| | 0.839319108 | NA |
| | 0.985739715 | 0.991027527 |
| | 0.226670579 | 0.37983759 |
| | 0.025625868 | 0.073082545 |
| | 0.000155616 | 0.001034049 |
| | 0.955792873 | NA |
| NA | | NA |
| | 0.107868413 | 0.221908727 |
| | 0.955792873 | NA |
| | 0.475000971 | 0.632341722 |
| | 0.350603628 | 0.514460733 |
| | 0.0029526 | 0.012655821 |
| | 0.004610052 | 0.018260685 |
| | 0.838510266 | NA |
| | 0.000105899 | 0.000735514 |
| | 0.024216104 | 0.069933477 |
| | 0.919271923 | 0.95153339 |
| | 0.536109274 | 0.68417593 |
| | 5.50E-07 | 6.43E-06 |
| | 0.014752598 | 0.047025758 |
| | 0.080245945 | 0.178272727 |
| | 0.168231233 | 0.307687091 |
| | 0.182284045 | 0.324706245 |
| | 0.941277857 | NA |
| | 0.924934574 | NA |
| | 0.542319124 | 0.689246911 |
| | 0.200825934 | 0.348400793 |
| | 1.08E-09 | 1.92E-08 |
| | 0.000896086 | 0.004651828 |
| | 0.188745433 | 0.333248186 |
| | 0.015020418 | 0.047757945 |
| | 0.768520482 | NA |
| | 0.00323126 | 0.013650406 |
| | 0.628634146 | NA |
| | 0.081299671 | 0.179858236 |
| | 0.430305426 | 0.591586076 |
| | 0.757538328 | NA |
| | 1.15E-09 | 2.02E-08 |

| | |
|---|---|
| 0.00158666 | 0.007524543 |
| 0.519128909 | 0.669754153 |
| 0.994512843 | 0.995988288 |
| 0.938047 | 0.963218063 |
| 0.002774845 | 0.012021635 |
| 0.000211778 | 0.001348151 |
| NA | NA |
| 0.858111099 | 0.915075826 |
| 4.16E-06 | 4.12E-05 |
| 0.019185368 | 0.057979955 |
| 0.440045307 | 0.600846627 |
| 0.159318092 | 0.295725589 |
| 0.091498307 | 0.196538463 |
| 0.886016551 | 0.931738356 |
| NA | NA |
| 0.088160425 | 0.19126738 |
| 0.710730653 | 0.815543123 |
| 0.607832902 | 0.740373509 |
| 0.040770698 | 0.105779828 |
| 0.43143047 | 0.592858845 |
| 1.06E-06 | 1.17E-05 |
| 0.203980737 | NA |
| 0.018203803 | 0.055663788 |
| 0.458213945 | 0.617786808 |
| 0.677303186 | 0.791961605 |
| NA | NA |
| 0.01809541 | 0.055370261 |
| NA | NA |
| 0.21403347 | 0.364877858 |
| 0.941276431 | NA |
| 0.001143675 | 0.005724871 |
| 0.291897054 | 0.453182029 |
| 0.081270139 | 0.179809615 |
| 0.573999132 | 0.71353814 |
| 0.3343262 | 0.497600814 |
| 0.009667981 | 0.033510598 |
| 0.434681962 | NA |
| 0.111300067 | 0.227046619 |
| 3.25E-09 | 5.41E-08 |
| 0.304290087 | 0.465889146 |
| 0.073568859 | 0.167120249 |
| 0.270969807 | 0.430807785 |
| 0.067428468 | 0.156609329 |
| 0.000901494 | 0.004674808 |
| 0.047422497 | 0.118715846 |
| NA | NA |
| 0.788554318 | 0.869909616 |

| | | |
|---|---|---|
| | 0.709386887 | NA |
| | 0.879782694 | NA |
| | 0.941278332 | NA |
| | 0.941275956 | NA |
| NA | | NA |
| NA | | NA |
| | 0.238804687 | 0.394614631 |
| | 0.156773594 | 0.292485115 |
| NA | | NA |
| | 0.018472058 | 0.056304831 |
| | 0.098577563 | 0.207384052 |
| NA | | NA |
| | 8.68E-06 | 7.97E-05 |
| NA | | NA |
| | 0.67380855 | NA |
| | 0.070872856 | 0.162626193 |
| | 0.215035268 | 0.365932475 |
| | 0.409021438 | NA |
| | 0.397168415 | 0.559737262 |
| | 0.000103339 | 0.000720255 |
| | 0.941278332 | NA |
| | 0.000219455 | 0.001389959 |
| | 0.000163024 | 0.001076667 |
| | 0.655116698 | NA |
| | 0.839319108 | NA |
| | 0.094051002 | 0.200345758 |
| | 0.153024143 | 0.287246108 |
| | 0.648507876 | NA |
| | 0.554743493 | 0.699549914 |
| | 0.391783096 | 0.554183941 |
| | 0.020350307 | 0.060721827 |
| | 0.914135133 | 0.948236923 |
| | 0.013589907 | 0.044069069 |
| NA | | NA |
| | 0.877816945 | 0.92667756 |
| | 1.87E-05 | 0.000158172 |
| | 0.001972807 | 0.009058017 |
| | 0.013460669 | 0.04377522 |
| NA | | NA |
| | 0.009636705 | 0.033407055 |
| | 0.014617032 | 0.046718758 |
| | 0.422989577 | 0.584352291 |
| | 0.010207757 | 0.034969061 |
| | 0.215957988 | 0.367106986 |
| | 0.000124481 | 0.000849942 |
| | 9.16E-06 | 8.38E-05 |
| | 0.282874832 | 0.443280449 |

| | |
|---|---|
| NA | NA |
| 0.266103999 | 0.425544704 |
| 0.994423848 | 0.995972078 |
| 0.000775901 | 0.004120483 |
| 0.096368393 | 0.20399862 |
| 0.879519381 | 0.927866593 |
| 0.217516948 | 0.368835107 |
| NA | NA |
| 0.005736886 | 0.021826943 |
| NA | NA |
| 0.012220239 | 0.040476779 |
| 0.00123191 | 0.0060935 |
| NA | NA |
| 1.01E-15 | 3.23E-14 |
| 0.000348899 | 0.00207405 |
| 0.012277715 | 0.04062758 |
| 0.488400206 | 0.643076933 |
| 0.839319108 | NA |
| 0.812967752 | 0.885838603 |
| 0.003776355 | 0.015468042 |
| 0.214796129 | 0.365680386 |
| 0.583646357 | 0.720873364 |
| 0.037429693 | 0.098758626 |
| 5.48E-26 | 3.22E-24 |
| 0.002316435 | 0.010354734 |
| 0.81308 | NA |
| 0.000311597 | 0.001881916 |
| 0.000630859 | 0.003453474 |
| 9.88E-09 | 1.55E-07 |
| 2.63E-112 | 1.03E-109 |
| 0.694766011 | 0.804437447 |
| NA | NA |
| 0.19530898 | 0.341618485 |
| 0.001774418 | 0.008281311 |
| 7.61E-08 | 1.04E-06 |
| 0.0842921 | 0.184982534 |
| 0.955792398 | NA |
| NA | NA |
| 0.82515277 | NA |
| NA | NA |
| 0.000146299 | 0.000977598 |
| 0.839319108 | NA |
| 1.04E-12 | 2.61E-11 |
| 0.09415403 | 0.20052382 |
| NA | NA |
| 0.00128043 | 0.006287762 |
| 0.050611226 | 0.125041354 |

| | |
|---|---|
| 0.067136416 | 0.156113812 |
| NA | NA |
| 0.049226039 | 0.122164339 |
| 0.712527963 | NA |
| 0.479711927 | 0.63638966 |
| 2.25E-73 | 5.14E-71 |
| 0.142499904 | NA |
| NA | NA |
| 0.941277857 | NA |
| 0.584588716 | 0.721625382 |
| 0.281481483 | 0.441678573 |
| 0.926159517 | NA |
| 0.03310336 | 0.089751245 |
| 0.380312759 | 0.542991079 |
| 0.292374878 | 0.453439429 |
| 0.368135061 | 0.531271239 |
| 0.093038038 | 0.198800088 |
| 4.99E-28 | 3.22E-26 |
| 0.878494502 | 0.927099556 |
| NA | NA |
| 3.13E-26 | 1.86E-24 |
| 5.38E-15 | 1.63E-13 |
| 0.090451892 | 0.194959335 |
| 0.720964594 | NA |
| 0.000245484 | 0.001527984 |
| 0.938893503 | NA |
| NA | NA |
| 0.928152905 | NA |
| 0.007345915 | 0.026734278 |
| 9.20E-09 | 1.45E-07 |
| 4.61E-16 | 1.53E-14 |
| NA | NA |
| NA | NA |
| NA | NA |
| NA | NA |
| NA | NA |
| NA | NA |
| NA | NA |
| NA | NA |
| NA | NA |
| NA | NA |
| 0.839319108 | NA |
| NA | NA |
| NA | NA |
| NA | NA |
| NA | NA |
| 0.92466283 | NA |

| | |
|---|---|
| NA | NA |
| NA | NA |
| 0.839318641 | NA |
| NA | NA |
| NA | NA |
| NA | NA |
| NA | NA |
| 0.372842195 | 0.536022585 |
| 0.444325217 | NA |
| 0.610050885 | 0.741784478 |
| 0.918340111 | NA |
| 0.365426211 | 0.52864977 |
| 1.36E-05 | 0.00011891 |
| 2.28E-11 | 4.97E-10 |
| 1.62E-73 | 3.75E-71 |
| 1.37E-35 | 1.20E-33 |
| NA | NA |
| 3.93E-21 | 1.81E-19 |
| 2.86E-11 | 6.15E-10 |
| 9.14E-07 | 1.02E-05 |
| 0.898259304 | 0.938641143 |
| 0.004946855 | 0.019356408 |
| 0.158845364 | NA |
| 0.92212779 | NA |
| 0.955792873 | NA |
| NA | NA |
| NA | NA |
| 1.13E-40 | 1.18E-38 |
| 0.368790282 | NA |
| 0.179014368 | 0.320633605 |
| 0.095828068 | 0.203180089 |
| NA | NA |
| 6.99E-06 | 6.57E-05 |
| 0.003367247 | 0.014112909 |
| 0.053513938 | 0.130760575 |
| 0.020690007 | 0.061483366 |
| 0.667925456 | 0.784735757 |
| 0.024740588 | 0.071094467 |
| 0.041186297 | 0.10663725 |
| 0.012730322 | 0.041822929 |
| 0.065160539 | 0.152589762 |
| 0.493603418 | 0.646993147 |
| 4.22E-24 | 2.26E-22 |
| 0.48267933 | 0.638724268 |
| 0.99172984 | 0.994333794 |
| 0.981342955 | 0.988464598 |
| 0.494753356 | 0.647786973 |

| | |
|---|---|
| NA | NA |
| 0.082753627 | 0.182194232 |
| 0.650017361 | 0.771271846 |
| 0.000427643 | 0.002476581 |
| 0.000127459 | 0.000867783 |
| 0.005553502 | 0.021234451 |
| 0.631871144 | 0.75791343 |
| NA | NA |
| 0.302404883 | 0.463748536 |
| 0.447952044 | 0.608153922 |
| 0.126300706 | 0.249602242 |
| 0.249933175 | 0.40748746 |
| 0.06301615 | 0.14863843 |
| 0.941277857 | NA |
| NA | NA |
| 0.552406218 | 0.697708932 |
| 0.295074024 | 0.456447125 |
| 0.239283071 | 0.39510341 |
| 1.78E-59 | 3.00E-57 |
| 8.68E-14 | 2.38E-12 |
| NA | NA |
| NA | NA |
| 0.941279303 | NA |
| 3.18E-16 | 1.06E-14 |
| 0.97309549 | 0.98375887 |
| 8.01E-31 | 5.76E-29 |
| 0.353947195 | 0.51758009 |
| 0.000391264 | 0.002293247 |
| 0.031725941 | 0.086688194 |
| 0.397268247 | NA |
| 6.60E-15 | 1.98E-13 |
| 0.002474358 | 0.010944928 |
| 0.853803533 | NA |
| 0.925454985 | NA |
| 0.046977582 | 0.117924532 |
| NA | NA |
| 0.044626516 | 0.113257052 |
| 0.328170429 | 0.491017887 |
| 0.535576014 | 0.683747065 |
| 1.25E-08 | 1.93E-07 |
| 0.763777612 | 0.853490059 |
| 0.047835909 | 0.119587261 |
| 0.839318641 | NA |
| 0.52152006 | 0.671638377 |
| 0.006065489 | 0.022850708 |
| 0.19022711 | 0.3349945 |
| NA | NA |

| | |
|---|---|
| 0.490488987 | 0.644888575 |
| NA | NA |
| 0.954259795 | NA |
| 0.882758955 | NA |
| 0.119637543 | 0.239758593 |
| 0.011756855 | 0.039262094 |
| 0.721623642 | 0.82332244 |
| 0.941277857 | NA |
| 0.493222754 | 0.646886048 |
| 0.389341922 | 0.551909739 |
| 0.790000192 | 0.870899197 |
| 0.415267696 | 0.577112262 |
| 0.751295529 | NA |
| NA | NA |
| 0.780780665 | 0.865183491 |
| 0.941278838 | NA |
| 0.666804601 | 0.784038093 |
| 0.953363385 | NA |
| NA | NA |
| 0.941278332 | NA |
| 2.93E-16 | 9.82E-15 |
| 0.120831138 | 0.241625524 |
| 0.98830424 | 0.992641469 |
| 0.427248176 | 0.588567314 |
| 7.68E-05 | 0.000553641 |
| NA | NA |
| 0.019597744 | 0.058941812 |
| 0.839318641 | NA |
| 0.739991096 | NA |
| NA | NA |
| 0.485893807 | NA |
| 0.24822396 | NA |
| 0.415976246 | NA |
| 0.711881532 | 0.816425444 |
| 0.956811732 | 0.973498882 |
| 0.638389735 | 0.76241608 |
| 0.724929736 | NA |
| 0.000178508 | 0.001163913 |
| 0.36411502 | 0.52755777 |
| 0.420654327 | 0.582211725 |
| NA | NA |
| 0.941277857 | NA |
| 0.236783024 | 0.392254283 |
| 1.88E-08 | 2.82E-07 |
| 0.28470204 | NA |
| 0.610426269 | 0.741975593 |
| 0.022943399 | 0.066932074 |

| | | |
|---|---|---|
| | 0.005902086 | 0.022348305 |
| | 0.819808217 | 0.890201991 |
| | 0.82479167 | 0.893568108 |
| | 0.014550205 | 0.046548921 |
| | 4.98E-05 | 0.000377733 |
| | 0.117758784 | 0.236830973 |
| | 0.202384762 | 0.350492004 |
| | 0.953798549 | NA |
| | 0.586996012 | 0.723508833 |
| | 0.000706312 | 0.003804399 |
| | 0.118715735 | 0.238292454 |
| NA | | NA |
| | 0.803888693 | NA |
| NA | | NA |
| | 9.15E-07 | 1.02E-05 |
| | 1.12E-05 | 1.00E-04 |
| | 0.234867508 | NA |
| | 0.029790818 | 0.082508849 |
| | 2.88E-07 | 3.56E-06 |
| | 0.918348326 | NA |
| | 0.49842988 | NA |
| | 2.00E-05 | 0.000167621 |
| NA | | NA |
| | 0.94127925 | NA |
| | 0.000100302 | 0.00070321 |
| | 0.839319574 | NA |
| | 0.019405516 | 0.058476573 |
| NA | | NA |
| | 1.58E-05 | 0.000136729 |
| | 0.447896101 | NA |
| NA | | NA |
| | 0.941278788 | NA |
| | 1.20E-28 | 7.95E-27 |
| | 0.839319574 | NA |
| | 2.47E-18 | 9.32E-17 |
| | 1.54E-20 | 6.77E-19 |
| | 0.635032494 | NA |
| | 2.45E-11 | 5.34E-10 |
| | 0.005786135 | 0.021979187 |
| | 0.277501962 | 0.437366025 |
| | 0.000622263 | 0.003411123 |
| | 1.01E-05 | 9.14E-05 |
| | 1.59E-29 | 1.09E-27 |
| | 5.24E-06 | 5.07E-05 |
| | 0.941278332 | NA |
| | 0.01291202 | 0.042291441 |
| | 0.379491873 | 0.542339572 |

| | |
|---|---|
| 0.000967985 | 0.004961234 |
| 9.41E-12 | 2.13E-10 |
| 7.04E-29 | 4.68E-27 |
| 6.70E-08 | 9.25E-07 |
| 6.92E-11 | 1.42E-09 |
| 2.84E-19 | 1.15E-17 |
| NA | NA |
| 0.941278332 | NA |
| 0.00773382 | 0.027886743 |
| 0.000240599 | 0.001501897 |
| 0.941277857 | NA |
| 0.839319108 | NA |
| 3.72E-08 | 5.33E-07 |
| NA | NA |
| NA | NA |
| 0.249018932 | NA |
| NA | NA |
| NA | NA |
| NA | NA |
| 0.474756595 | NA |
| NA | NA |
| NA | NA |
| NA | NA |
| 0.839320041 | NA |
| NA | NA |
| NA | NA |
| NA | NA |
| 0.218473018 | NA |
| 0.000139688 | 0.000938697 |
| 0.755994516 | NA |
| 0.608430836 | NA |
| NA | NA |
| 1.26E-07 | 1.67E-06 |
| 0.46988731 | NA |
| NA | NA |
| NA | NA |
| NA | NA |
| 0.839319108 | NA |
| NA | NA |
| 0.941275481 | NA |
| NA | NA |
| NA | NA |
| 0.787036781 | NA |
| NA | NA |
| NA | NA |
| 0.20665523 | 0.355629099 |
| 4.76E-15 | 1.45E-13 |

| | | | |
|---|---|---|---|
| | 0.817448418 | | 0.888605556 |
| | 0.166115774 | | 0.304919229 |
| | 6.07E-06 | | 5.80E-05 |
| | 0.000839143 | | 0.004400369 |
| | 0.362736294 | | 0.526432906 |
| | 0.651858658 | NA | |
| NA | | NA | |
| NA | | NA | |
| NA | | NA | |
| NA | | NA | |
| NA | | NA | |
| NA | | NA | |
| NA | | NA | |
| NA | | NA | |
| NA | | NA | |
| NA | | NA | |
| NA | | NA | |
| NA | | NA | |
| | 0.941278332 | NA | |
| NA | | NA | |
| NA | | NA | |
| NA | | NA | |
| NA | | NA | |
| NA | | NA | |
| NA | | NA | |
| NA | | NA | |
| NA | | NA | |
| NA | | NA | |
| NA | | NA | |
| NA | | NA | |
| | 0.941277857 | NA | |
| NA | | NA | |
| NA | | NA | |
| | 0.941277857 | NA | |
| NA | | NA | |
| NA | | NA | |
| NA | | NA | |
| NA | | NA | |
| NA | | NA | |
| NA | | NA | |
| NA | | NA | |
| NA | | NA | |
| | 0.166687836 | | 0.305553745 |
| NA | | NA | |
| | 0.839319108 | NA | |
| NA | | NA | |
| NA | | NA | |

| | | |
|---|---|---|
| | 0.286283055 | NA |
| | 0.955792398 | NA |
| | 3.76E-24 | 2.02E-22 |
| | 0.915881972 | NA |
| | 0.659227752 | NA |
| | 0.410809252 | NA |
| | 7.05E-06 | 6.61E-05 |
| | 0.446800685 | 0.607214223 |
| | 0.003375816 | 0.014143842 |
| | 0.011209745 | 0.03777635 |
| | 0.223217211 | 0.375664008 |
| | 0.757455185 | 0.848421097 |
| | 0.140168045 | 0.269338074 |
| | 0.555454473 | 0.699973458 |
| | 0.057831431 | 0.13885844 |
| | 0.00020049 | 0.001283505 |
| | 0.321405182 | 0.484274618 |
| | 8.00E-05 | 0.000575395 |
| | 0.647644754 | NA |
| | 0.111743474 | 0.227676611 |
| NA | | NA |
| | 0.012279555 | 0.04062802 |
| | 0.505210662 | NA |
| NA | | NA |
| | 0.255973176 | 0.413988824 |
| | 8.40E-15 | 2.49E-13 |
| | 0.47321694 | NA |
| | 0.158782839 | 0.295054192 |
| | 0.729744598 | NA |
| | 0.941279268 | NA |
| | 0.059440732 | 0.142034766 |
| | 0.048332431 | 0.120512101 |
| | 0.941277857 | NA |
| | 0.21570633 | 0.366731577 |
| | 0.839320041 | NA |
| | 0.839319109 | NA |
| | 0.955792398 | NA |
| NA | | NA |
| NA | | NA |
| | 0.878481972 | 0.927099556 |
| | 0.862672346 | 0.917823438 |
| | 0.122021317 | 0.243455275 |
| | 0.577114714 | 0.715702831 |
| | 5.46E-08 | 7.62E-07 |
| | 0.389188411 | 0.551790717 |
| | 0.037470212 | 0.098843621 |
| | 0.335279435 | 0.498489669 |

| | |
|---|---|
| 0.714015128 | 0.818089346 |
| 0.006422722 | 0.023945957 |
| 0.290480886 | 0.451561444 |
| 0.230724663 | 0.384619312 |
| 0.156280466 | 0.291816714 |
| 0.00061894 | 0.003398789 |
| 0.019864649 | 0.059518928 |
| 7.56E-33 | 5.92E-31 |
| 0.312582328 | 0.474635631 |
| 0.010633046 | 0.036176066 |
| 0.00569149 | 0.021688896 |
| 0.907350097 | 0.943748776 |
| 0.072897736 | 0.16592827 |
| 0.000109922 | 0.000761237 |
| NA | NA |
| 0.969295618 | NA |
| 1.28E-06 | 1.40E-05 |
| 0.669659632 | 0.78623601 |
| 0.440581336 | 0.601290214 |
| 0.868458974 | 0.921216246 |
| 0.801001929 | 0.878492596 |
| NA | NA |
| 0.596066917 | NA |
| 5.04E-08 | 7.05E-07 |
| 0.571555575 | NA |
| 0.256402773 | 0.414486471 |
| 0.839319108 | NA |
| NA | NA |
| NA | NA |
| 0.613829911 | 0.744153273 |
| 0.232638317 | 0.387085441 |
| 0.336863946 | 0.500078095 |
| 0.693314823 | NA |
| 0.001162161 | 0.005801544 |
| 0.721353511 | 0.823168869 |
| 0.521397153 | 0.671552762 |
| 1.70E-07 | 2.19E-06 |
| 0.013433304 | 0.043718894 |
| 0.111152054 | 0.226801646 |
| 0.692412725 | NA |
| NA | NA |
| 0.161862379 | 0.299002589 |
| 0.955791922 | NA |
| NA | NA |
| 0.041601234 | NA |
| NA | NA |
| 0.151906324 | 0.285846653 |

| | |
|---|---|
| 0.941278332 | NA |
| 0.147718417 | NA |
| 0.030981592 | 0.085089299 |
| NA | NA |
| 0.839319574 | NA |
| 0.267505098 | 0.427096615 |
| 0.027667682 | 0.077869951 |
| 0.02574645 | 0.0733649 |
| 0.000133632 | 0.000904641 |
| 0.013996536 | 0.04512333 |
| 0.001703132 | 0.008001964 |
| 0.000169992 | 0.001117415 |
| 8.10E-05 | 0.000580957 |
| NA | NA |
| 0.000299798 | 0.001821752 |
| 0.799586237 | 0.877620713 |
| 0.259118226 | 0.417543295 |
| 4.40E-12 | 1.03E-10 |
| 2.51E-06 | 2.60E-05 |
| 2.04E-10 | 3.95E-09 |
| 0.295354597 | 0.456752135 |
| NA | NA |
| NA | NA |
| 0.955792398 | NA |
| 0.002364499 | 0.010534381 |
| 0.853595154 | NA |
| NA | NA |
| 0.085202446 | 0.186486321 |
| 0.705632596 | NA |
| NA | NA |
| NA | NA |
| NA | NA |
| 0.185346339 | 0.328611812 |
| 0.799747637 | 0.877620713 |
| 0.774945345 | 0.861042344 |
| 0.899490324 | NA |
| 2.99E-13 | 7.86E-12 |
| 0.684608438 | 0.79740886 |
| 0.941278332 | NA |
| 0.542351976 | 0.689246911 |
| 0.446793246 | 0.607214223 |
| 0.558509983 | 0.702067843 |
| 0.008448621 | 0.029973777 |
| NA | NA |
| 0.895696894 | NA |
| 0.839319107 | NA |
| 0.653457494 | 0.774003044 |

| | | |
|---|---|---|
| NA | | NA |
| | 0.532819276 | NA |
| NA | | NA |
| NA | | NA |
| | 0.941279268 | NA |
| | 0.198009823 | 0.345167607 |
| | 0.052814362 | 0.129503189 |
| | 0.003466443 | 0.014443616 |
| | 0.004789036 | 0.01885668 |
| | 0.955792398 | NA |
| | 0.048962114 | 0.121686983 |
| | 0.534516507 | 0.682687374 |
| | 0.899764417 | 0.939595167 |
| | 0.003607189 | 0.014908546 |
| | 0.171894347 | 0.312055159 |
| | 0.120833065 | 0.241625524 |
| | 0.583283517 | 0.720557043 |
| | 0.709190636 | 0.814364894 |
| | 0.649475109 | 0.770749546 |
| | 0.203258484 | 0.351519178 |
| | 0.274190969 | 0.434073392 |
| | 0.40061956 | 0.563101117 |
| | 0.839318641 | NA |
| | 0.119440847 | 0.239465216 |
| | 0.184819279 | 0.327889572 |
| | 0.946922904 | 0.967625072 |
| | 8.56E-05 | 0.000611182 |
| | 0.424197706 | NA |
| | 0.231581853 | NA |
| | 0.839318641 | NA |
| | 0.49410139 | 0.647310858 |
| | 0.752864714 | 0.844989198 |
| | 0.574576868 | 0.713786167 |
| | 0.662782037 | 0.781005893 |
| | 0.955792398 | NA |
| | 0.577920756 | NA |
| | 0.274884903 | 0.434813608 |
| | 5.91E-06 | 5.67E-05 |
| | 0.390451828 | 0.553029754 |
| | 0.109009123 | 0.223617611 |
| | 0.300815605 | 0.462211174 |
| | 0.0009927 | 0.005078069 |
| | 0.839318641 | NA |
| | 0.322338792 | 0.485128544 |
| | 0.839318641 | NA |
| | 0.941275956 | NA |
| | 0.689373731 | NA |

| | |
|---|---|
| NA | NA |
| 0.000133772 | 0.000905326 |
| 0.000701208 | 0.003785473 |
| 0.486615491 | NA |
| 7.23E-08 | 9.93E-07 |
| 0.006985603 | 0.025627515 |
| NA | NA |
| 4.47E-14 | 1.26E-12 |
| 0.914954493 | NA |
| NA | NA |
| NA | NA |
| 0.378660279 | 0.541443708 |
| NA | NA |
| 0.34789336 | 0.511788579 |
| 0.953798549 | NA |
| 0.982333263 | 0.988869696 |
| 0.458503394 | NA |
| 0.129518636 | 0.253892116 |
| 3.55E-16 | 1.18E-14 |
| 0.800646606 | NA |
| 0.535874645 | 0.684054934 |
| 0.630444514 | NA |
| 0.955792873 | NA |
| 6.76E-06 | 6.39E-05 |
| 0.684805921 | NA |
| 0.734894808 | 0.832977211 |
| 0.028332243 | 0.079299629 |
| NA | NA |
| 0.56319071 | 0.705530167 |
| 0.370823886 | NA |
| NA | NA |
| NA | NA |
| NA | NA |
| NA | NA |
| NA | NA |
| NA | NA |
| NA | NA |
| NA | NA |
| NA | NA |
| 0.776964226 | NA |
| 0.001265096 | 0.006224023 |
| 0.055532872 | 0.13473939 |
| 0.016771157 | 0.052126807 |
| 0.003492527 | 0.014538146 |
| NA | NA |
| 0.259218252 | 0.417647928 |
| NA | NA |

| | |
|---|---|
| NA | NA |
| 0.012748226 | 0.041870193 |
| 0.457991906 | 0.617697379 |
| 0.16962246 | 0.309232168 |
| 5.64E-10 | 1.03E-08 |
| 0.762613297 | 0.852633834 |
| 0.069650228 | 0.160625398 |
| NA | NA |
| 0.495745204 | 0.648657432 |
| 0.808142355 | 0.882317524 |
| 0.861316963 | NA |
| 0.621731545 | 0.750200458 |
| 0.533627548 | 0.681881296 |
| 0.00155332 | 0.007391469 |
| 1.27E-25 | 7.27E-24 |
| 1.18E-06 | 1.29E-05 |
| NA | NA |
| 0.001080815 | 0.005458426 |
| 0.002430924 | 0.010776838 |
| 3.11E-10 | 5.89E-09 |
| 0.488027515 | NA |
| 2.65E-09 | 4.47E-08 |
| 1.66E-09 | 2.86E-08 |
| 0.273871332 | 0.433682751 |
| 0.277290387 | 0.437177306 |
| 3.26E-07 | 3.99E-06 |
| 0.273378886 | 0.433292143 |
| 0.955792398 | NA |
| 0.02412364 | 0.069734137 |
| 0.362168141 | 0.52585409 |
| 0.371560898 | 0.534831768 |
| 0.287264876 | 0.447830664 |
| 0.567120458 | 0.708774016 |
| NA | NA |
| NA | NA |
| 0.063850758 | 0.150234851 |
| NA | NA |
| 1.22E-08 | 1.88E-07 |
| 8.46E-06 | 7.80E-05 |
| 0.839322424 | NA |
| 0.941277857 | NA |
| 0.955792398 | NA |
| 7.13E-07 | 8.16E-06 |
| NA | NA |
| 0.002233659 | 0.010038812 |
| 0.34942689 | 0.51332554 |
| 0.608353112 | 0.740703911 |

| | | |
|---|---|---|
| | 0.44229423 | 0.602845288 |
| | 0.639362677 | 0.763156835 |
| | 0.016203858 | 0.050727899 |
| | 0.76746683 | NA |
| | 0.333313169 | 0.496558806 |
| | 0.027099192 | 0.076623303 |
| NA | | NA |
| | 0.817553865 | 0.888671704 |
| | 7.00E-12 | 1.62E-10 |
| | 0.011698922 | 0.039124266 |
| | 0.012218838 | 0.040476779 |
| | 0.455143577 | 0.614869554 |
| NA | | NA |
| | 0.000649431 | 0.003538863 |
| | 0.04818191 | 0.120250166 |
| | 0.37790614 | NA |
| | 0.898624174 | 0.938898757 |
| | 0.70275415 | NA |
| | 0.875738483 | 0.925643364 |
| | 5.46E-06 | 5.28E-05 |
| | 0.137854802 | 0.266114543 |
| | 0.941782916 | 0.964806618 |
| NA | | NA |
| | 0.243819275 | 0.400288755 |
| | 0.107025759 | 0.220671094 |
| | 6.21E-09 | 9.99E-08 |
| | 0.16660587 | 0.305509772 |
| | 1.84E-13 | 4.93E-12 |
| | 0.364174781 | 0.52755777 |
| | 0.453771562 | 0.613674472 |
| | 0.000189974 | 0.001226738 |
| | 2.49E-05 | 0.000205133 |
| | 0.342951801 | 0.506417195 |
| | 0.088373855 | 0.191660546 |
| | 0.015381229 | 0.048664604 |
| | 0.216795961 | 0.368189616 |
| | 0.297471457 | 0.45887334 |
| | 0.149135621 | 0.282172182 |
| | 0.577782469 | 0.716157805 |
| | 0.029187287 | 0.081196332 |
| | 0.75562621 | 0.847090008 |
| NA | | NA |
| | 0.991320506 | 0.994118841 |
| NA | | NA |
| | 0.955792873 | NA |
| NA | | NA |
| | 0.941278788 | NA |

| | |
|---|---|
| NA | NA |
| NA | NA |
| 0.941278207 | NA |
| NA | NA |
| NA | NA |
| NA | NA |
| NA | NA |
| NA | NA |
| NA | NA |
| NA | NA |
| NA | NA |
| NA | NA |
| NA | NA |
| NA | NA |
| NA | NA |
| NA | NA |
| 0.941279303 | NA |
| 0.922470252 | NA |
| NA | NA |
| 0.433395422 | 0.594608076 |
| 0.388870563 | 0.551561603 |
| 0.495805046 | NA |
| 0.447342113 | NA |
| 0.941278424 | NA |
| 0.095096654 | 0.202078602 |
| 0.024193018 | 0.06988704 |
| 0.107215102 | 0.220908401 |
| 0.972774691 | 0.983643451 |
| 0.000580843 | 0.003221159 |
| 0.707282732 | 0.813272637 |
| NA | NA |
| 0.009823249 | 0.033915429 |
| NA | NA |
| NA | NA |
| 0.955792873 | NA |
| 0.941278332 | NA |
| 0.839318641 | NA |
| NA | NA |
| NA | NA |
| 0.941275956 | NA |
| NA | NA |
| 0.86862908 | 0.921268575 |
| 0.026271768 | 0.074611697 |
| 0.073952024 | 0.167734433 |
| NA | NA |
| 0.420343523 | 0.581889993 |
| NA | NA |

| | | |
|---|---|---|
| | 0.820089927 | 0.89041969 |
| | 0.366174483 | 0.529346191 |
| | 0.000465764 | 0.002668778 |
| | 0.007691402 | 0.027759023 |
| | 3.37E-05 | 0.000266894 |
| | 0.057731914 | 0.138689446 |
| | 0.685869534 | NA |
| | 0.2705102 | 0.430545107 |
| | 0.554704257 | 0.699549914 |
| | 0.897568577 | NA |
| | 0.78534343 | 0.868025567 |
| | 0.955791922 | NA |
| NA | | NA |
| NA | | NA |
| NA | | NA |
| NA | | NA |
| NA | | NA |
| NA | | NA |
| | 0.839318641 | NA |
| NA | | NA |
| NA | | NA |
| | 0.853595622 | NA |
| | 0.941277857 | NA |
| NA | | NA |
| NA | | NA |
| | 0.952380021 | NA |
| | 0.941277857 | NA |
| | 0.475589961 | NA |
| NA | | NA |
| NA | | NA |
| | 0.955792398 | NA |
| NA | | NA |
| | 0.955792398 | NA |
| NA | | NA |
| NA | | NA |
| NA | | NA |
| | 0.728678906 | NA |
| | 0.793807415 | NA |
| | 0.896368437 | NA |
| NA | | NA |
| | 0.955792873 | NA |
| NA | | NA |
| NA | | NA |
| | 0.955792873 | NA |
| NA | | NA |
| NA | | NA |
| | 0.572194755 | NA |

| | | |
|---|---|---|
| | 0.702555031 | NA |
| NA | | NA |
| NA | | NA |
| NA | | NA |
| NA | | NA |
| NA | | NA |
| NA | | NA |
| NA | | NA |
| | 0.941277857 | NA |
| | 0.955792873 | NA |
| NA | | NA |
| NA | | NA |
| NA | | NA |
| NA | | NA |
| NA | | NA |
| NA | | NA |
| NA | | NA |
| NA | | NA |
| NA | | NA |
| | 0.941279303 | NA |
| | 0.440882514 | NA |
| NA | | NA |
| NA | | NA |
| NA | | NA |
| | 0.955792873 | NA |
| NA | | NA |
| NA | | NA |
| NA | | NA |
| NA | | NA |
| NA | | NA |
| | 0.941277857 | NA |
| NA | | NA |
| NA | | NA |
| NA | | NA |
| NA | | NA |
| NA | | NA |
| NA | | NA |
| | 0.839318641 | NA |
| NA | | NA |
| | 0.496739366 | | 0.649708232 |
| NA | | NA |
| NA | | NA |
| NA | | NA |
| NA | | NA |
| NA | | NA |
| NA | | NA |
| | 0.554138023 | | 0.699119751 |

| | |
|---|---|
| NA | NA |
| NA | NA |
| NA | NA |
| NA | NA |
| NA | NA |
| NA | NA |
| NA | NA |
| NA | NA |
| NA | NA |
| NA | NA |
| NA | NA |
| NA | NA |
| NA | NA |
| NA | NA |
| NA | NA |
| NA | NA |
| NA | NA |
| NA | NA |
| NA | NA |
| NA | NA |
| NA | NA |
| 0.529921126 | 0.678894592 |
| NA | NA |
| NA | NA |
| NA | NA |
| NA | NA |
| 0.533299718 | 0.681616287 |
| NA | NA |
| NA | NA |
| NA | NA |
| NA | NA |
| NA | NA |
| NA | NA |
| NA | NA |
| NA | NA |
| NA | NA |
| NA | NA |
| NA | NA |
| NA | NA |
| 0.955792873 | NA |
| NA | NA |
| NA | NA |
| NA | NA |
| 0.072787823 | 0.165720321 |
| NA | NA |
| 0.135489684 | 0.262680252 |
| 0.8718739 | NA |

| | |
|---|---|
| 0.28416278 | NA |
| 0.688877149 | 0.800461527 |
| 0.793294475 | NA |
| 0.25301104 | 0.410817161 |
| 0.331484766 | 0.494608852 |
| 0.03724496 | 0.098380261 |
| NA | NA |
| 0.267325608 | 0.426861948 |
| 2.88E-06 | 2.94E-05 |
| 5.42E-08 | 7.56E-07 |
| 0.042891092 | 0.109969125 |
| 0.028573933 | 0.079882164 |
| 6.26E-16 | 2.05E-14 |
| 4.72E-34 | 3.88E-32 |
| NA | NA |
| 0.034060096 | 0.091666486 |
| 0.941278332 | NA |
| 1.39E-11 | 3.10E-10 |
| 0.005997515 | 0.022651994 |
| 0.09837933 | 0.207105344 |
| 0.019843037 | 0.059484127 |
| 6.21E-05 | 0.000460304 |
| 0.004814315 | 0.018924909 |
| 0.027603543 | 0.077762994 |
| 0.092161101 | 0.197409874 |
| NA | NA |
| NA | NA |
| NA | NA |
| 0.416151574 | NA |
| 0.315888985 | 0.478385801 |
| 0.941279303 | NA |
| NA | NA |
| 0.007617422 | 0.027537952 |
| 0.053868045 | 0.131490842 |
| 0.514225662 | 0.665240505 |
| 0.390676336 | 0.55319699 |
| 0.90179821 | NA |
| NA | NA |
| 0.378254409 | NA |
| 0.441995844 | NA |
| NA | NA |
| NA | NA |
| NA | NA |
| NA | NA |
| NA | NA |
| NA | NA |
| NA | NA |

| | | |
|---|---|---|
| NA | NA | |
| 0.955792873 | NA | |
| 0.821695621 | NA | |
| NA | NA | |
| NA | NA | |
| NA | NA | |
| NA | NA | |
| 0.941275956 | NA | |
| NA | NA | |
| NA | NA | |
| NA | NA | |
| NA | NA | |
| NA | NA | |
| NA | NA | |
| NA | NA | |
| 0.56410312 | | 0.70628009 |
| NA | NA | |
| NA | NA | |
| NA | NA | |
| 0.955792873 | NA | |
| 0.743511643 | NA | |
| 0.92273787 | NA | |
| 0.67501279 | NA | |
| 0.699724571 | | 0.808252171 |
| NA | NA | |
| 0.975687806 | | 0.985207923 |
| NA | NA | |
| NA | NA | |
| 0.274607561 | | 0.434530739 |
| 0.818345676 | NA | |
| NA | NA | |
| NA | NA | |
| NA | NA | |
| NA | NA | |
| NA | NA | |
| NA | NA | |
| NA | NA | |
| NA | NA | |
| NA | NA | |
| NA | NA | |
| NA | NA | |
| NA | NA | |
| NA | NA | |
| NA | NA | |
| NA | NA | |
| NA | NA | |
| NA | NA | |

| | |
|---|---|
| NA | NA |
| NA | NA |
| NA | NA |
| NA | NA |
| NA | NA |
| NA | NA |
| NA | NA |
| NA | NA |
| NA | NA |
| NA | NA |
| NA | NA |
| NA | NA |
| NA | NA |
| NA | NA |
| NA | NA |
| NA | NA |
| NA | NA |
| NA | NA |
| NA | NA |
| NA | NA |
| NA | NA |
| NA | NA |
| 0.016053811 | 0.050345798 |
| NA | NA |
| NA | NA |
| NA | NA |
| NA | NA |
| NA | NA |
| NA | NA |
| NA | NA |
| NA | NA |
| NA | NA |
| NA | NA |
| NA | NA |
| NA | NA |
| NA | NA |
| NA | NA |
| NA | NA |
| NA | NA |
| NA | NA |
| NA | NA |
| NA | NA |
| NA | NA |
| NA | NA |
| NA | NA |
| NA | NA |
| NA | NA |

| | |
|---|---|
| NA | NA |
| NA | NA |
| NA | NA |
| NA | NA |
| 0.929914733 | NA |
| 0.839319416 | NA |
| 0.422863207 | NA |
| 0.096850434 | NA |
| NA | NA |
| 1.83E-23 | 9.51E-22 |
| 1.02E-10 | 2.03E-09 |
| 0.383987583 | 0.546696522 |
| 0.92050159 | NA |
| 1.27E-14 | 3.72E-13 |
| 0.189686112 | 0.334264369 |
| 0.39525955 | 0.557773629 |
| 0.399100652 | 0.561795122 |
| 1.25E-05 | 0.000110598 |
| 0.000807505 | 0.004260749 |
| 0.04790042 | 0.11968568 |
| 0.004973549 | 0.019448072 |
| 0.103240722 | 0.214777971 |
| NA | NA |
| 3.53E-25 | 1.99E-23 |
| 0.040338161 | 0.104840458 |
| 0.871755459 | 0.92337093 |
| 0.025043382 | 0.07175666 |
| 0.006425582 | 0.023949533 |
| 0.480547091 | 0.637160636 |
| 0.835281548 | 0.899994439 |
| 0.562012058 | 0.704757968 |
| NA | NA |
| 0.414495413 | 0.576476559 |
| 0.941278207 | NA |
| 0.3601161 | 0.523833247 |
| 0.41390565 | 0.576060243 |
| 0.941279303 | NA |
| NA | NA |
| 0.91732096 | NA |
| 0.659583309 | 0.778524506 |
| 0.760902806 | 0.851361284 |
| 4.58E-07 | 5.44E-06 |
| 0.342515459 | 0.505961208 |
| 0.016615299 | 0.051770826 |
| 0.157912101 | 0.293894547 |
| 0.052281544 | 0.128421601 |
| 0.166466081 | 0.305373847 |

| | |
|---|---|
| 0.031350491 | 0.085887707 |
| 0.003350474 | 0.014064935 |
| 0.60495738 | 0.737777095 |
| 0.015524578 | 0.049026934 |
| NA | NA |
| 0.016928902 | 0.052521078 |
| 0.004687398 | 0.018530052 |
| 0.68734002 | 0.799299743 |
| 0.138364403 | 0.266836243 |
| 0.149323266 | 0.282370094 |
| 0.462481537 | 0.621218145 |
| 0.545720925 | 0.692037915 |
| NA | NA |
| NA | NA |
| NA | NA |
| 0.5974232 | NA |
| NA | NA |
| NA | NA |
| NA | NA |
| NA | NA |
| NA | NA |
| NA | NA |
| NA | NA |
| NA | NA |
| NA | NA |
| NA | NA |
| 0.955792873 | NA |
| NA | NA |
| 0.941278332 | NA |
| NA | NA |
| NA | NA |
| 0.941278332 | NA |
| NA | NA |
| 0.838993826 | NA |
| NA | NA |
| NA | NA |
| NA | NA |
| 0.351936272 | 0.515621515 |
| 3.57E-12 | 8.48E-11 |
| 3.05E-09 | 5.11E-08 |
| 0.042345171 | 0.108891503 |
| 0.001387345 | 0.006720143 |
| 0.509590044 | 0.661283515 |
| 0.199046931 | 0.346425495 |
| 0.390974125 | 0.553465333 |
| 1.03E-06 | 1.14E-05 |
| 0.493898985 | 0.647237992 |

| | |
|---|---|
| 0.814156036 | 0.886443528 |
| 0.932142137 | NA |
| 0.934458219 | 0.960973711 |
| 0.259891536 | 0.418449459 |
| 0.002332203 | 0.01041484 |
| 0.152685659 | 0.286769042 |
| 0.408732111 | 0.57082918 |
| 0.007133393 | 0.026081263 |
| 0.975499124 | 0.985178051 |
| 0.029194957 | 0.081208176 |
| 1.65E-05 | 0.000141776 |
| 0.275538404 | 0.435443359 |
| 0.203486643 | 0.351811512 |
| 0.33042932 | 0.493374234 |
| NA | NA |
| 0.216834286 | 0.368227392 |
| 0.429168293 | 0.590466111 |
| 0.313167336 | 0.475116614 |
| 3.94E-20 | 1.70E-18 |
| NA | NA |
| 0.243305131 | 0.399830761 |
| 0.074424227 | 0.168500471 |
| 0.26554414 | NA |
| 0.487949709 | 0.642661519 |
| 0.161120651 | 0.298084865 |
| 0.317577291 | 0.480087309 |
| 0.997084403 | 0.997587346 |
| 0.841232181 | 0.904177393 |
| 0.907414906 | NA |
| 0.278247948 | 0.437937781 |
| 5.46E-05 | 0.000409885 |
| 0.933362696 | 0.960386545 |
| 0.165157043 | 0.303721544 |
| 0.321911867 | 0.484700548 |
| NA | NA |
| 0.45855693 | 0.618109189 |
| 0.039632967 | 0.103425489 |
| 0.484644504 | 0.64037579 |
| 0.994058414 | 0.99564797 |
| 0.67550328 | 0.790858573 |
| 0.000100895 | 0.000706324 |
| 7.15E-07 | 8.19E-06 |
| 0.756111545 | 0.84719517 |
| 0.108978088 | 0.223573217 |
| 2.24E-13 | 5.94E-12 |
| 0.008242142 | 0.029394585 |
| 2.68E-13 | 7.08E-12 |

| | | |
|---|---|---|
| | 0.89779942 | NA |
| | 0.013673386 | 0.044279448 |
| | 0.942268347 | 0.965098044 |
| | 0.401423801 | 0.563765005 |
| | 0.559492625 | 0.702857627 |
| | 0.000547127 | 0.00307067 |
| | 0.147177783 | 0.279422955 |
| NA | | NA |
| | 0.900135254 | NA |
| | 2.75E-28 | 1.80E-26 |
| | 0.350799607 | NA |
| | 4.17E-05 | 0.000322712 |
| | 0.131839943 | 0.257365235 |
| | 0.123683298 | 0.245673859 |
| | 0.040837453 | 0.105941479 |
| | 0.01219916 | 0.040423834 |
| NA | | NA |
| | 0.493786775 | 0.647162206 |
| | 0.654649504 | NA |
| | 0.839319574 | NA |
| | 0.667302144 | NA |
| NA | | NA |
| NA | | NA |
| NA | | NA |
| NA | | NA |
| NA | | NA |
| NA | | NA |
| NA | | NA |
| NA | | NA |
| NA | | NA |
| NA | | NA |
| NA | | NA |
| NA | | NA |
| NA | | NA |
| NA | | NA |
| NA | | NA |
| NA | | NA |
| NA | | NA |
| NA | | NA |
| NA | | NA |
| NA | | NA |
| NA | | NA |
| NA | | NA |
| NA | | NA |
| NA | | NA |
| NA | | NA |
| NA | | NA |

| | |
|---|---|
| NA | NA |
| NA | NA |
| NA | NA |
| NA | NA |
| NA | NA |
| NA | NA |
| NA | NA |
| NA | NA |
| NA | NA |
| NA | NA |
| NA | NA |
| NA | NA |
| NA | NA |
| NA | NA |
| NA | NA |
| NA | NA |
| NA | NA |
| NA | NA |
| NA | NA |
| NA | NA |
| NA | NA |
| NA | NA |
| NA | NA |
| NA | NA |
| NA | NA |
| NA | NA |
| NA | NA |
| NA | NA |
| NA | NA |
| NA | NA |
| NA | NA |
| NA | NA |
| NA | NA |
| NA | NA |
| NA | NA |
| NA | NA |
| NA | NA |
| NA | NA |
| NA | NA |
| NA | NA |
| NA | NA |
| NA | NA |
| NA | NA |
| NA | NA |
| NA | NA |
| NA | NA |
| NA | NA |

| | |
|---|---|
| NA | NA |
| NA | NA |
| NA | NA |
| NA | NA |
| NA | NA |
| NA | NA |
| NA | NA |
| NA | NA |
| NA | NA |
| NA | NA |
| NA | NA |
| NA | NA |
| NA | NA |
| NA | NA |
| NA | NA |
| NA | NA |
| NA | NA |
| NA | NA |
| NA | NA |
| NA | NA |
| NA | NA |
| NA | NA |
| NA | NA |
| NA | NA |
| NA | NA |
| NA | NA |
| NA | NA |
| NA | NA |
| NA | NA |
| NA | NA |
| NA | NA |
| NA | NA |
| 0.022519884 | 0.065906587 |
| 0.941278332 | NA |
| NA | NA |
| NA | NA |
| 0.955792873 | NA |
| NA | NA |
| NA | NA |
| NA | NA |
| NA | NA |
| NA | NA |
| NA | NA |
| NA | NA |
| NA | NA |
| NA | NA |
| NA | NA |

| | |
|---|---|
| NA | NA |
| NA | NA |
| 0.917521744 | 0.95042443 |
| NA | NA |
| NA | NA |
| NA | NA |
| NA | NA |
| NA | NA |
| 0.417304252 | 0.578860995 |
| 0.903362044 | 0.9415345 |
| 0.000780315 | 0.004138383 |
| 0.511214716 | 0.66274143 |
| 0.008749825 | 0.03085602 |
| 0.000259176 | 0.00160148 |
| 0.011148899 | 0.037613903 |
| 0.002715791 | 0.0118239 |
| 0.275376369 | 0.435226317 |
| 0.062583465 | 0.147811602 |
| NA | NA |
| NA | NA |
| 0.941277857 | NA |
| 0.58278589 | NA |
| 0.000360134 | 0.002132846 |
| 0.020299448 | 0.060585262 |
| 0.955792398 | NA |
| 8.10E-17 | 2.80E-15 |
| 0.400092471 | 0.562723875 |
| 0.019718413 | 0.059229893 |
| 2.81E-12 | 6.75E-11 |
| 0.941276431 | NA |
| 0.167723303 | 0.306946799 |
| 0.000107553 | 0.000745916 |
| 0.517767489 | 0.668505657 |
| 7.61E-06 | 7.09E-05 |
| NA | NA |
| 0.992264982 | NA |
| NA | NA |
| NA | NA |
| NA | NA |
| 4.18E-07 | 5.02E-06 |
| 0.024360085 | 0.070264027 |
| NA | NA |
| 1.00E-07 | 1.35E-06 |
| NA | NA |
| 0.955792398 | NA |
| 4.13E-07 | 4.96E-06 |
| 0.96390595 | 0.97808285 |

| | |
|---|---|
| 0.105493627 | 0.218249398 |
| 0.520034032 | 0.670380999 |
| NA | NA |
| 0.047881681 | 0.119674415 |
| 0.298273886 | NA |
| 0.005086398 | 0.019798274 |
| 0.029679687 | 0.082287235 |
| 0.493114809 | 0.646825735 |
| NA | NA |
| 0.242515633 | 0.398946513 |
| 0.331678094 | 0.494773252 |
| 0.506014192 | NA |
| 0.207054782 | 0.356066607 |
| 0.004311077 | 0.017251556 |
| 0.04758445 | 0.11904615 |
| 0.00991052 | 0.034132569 |
| 0.93912336 | 0.963699686 |
| 0.134841083 | 0.261742575 |
| 0.614997769 | NA |
| 0.004932079 | 0.01930494 |
| 2.10E-06 | 2.21E-05 |
| 0.955792398 | NA |
| 1.53E-08 | 2.33E-07 |
| 0.498476374 | 0.651478951 |
| 0.563021278 | NA |
| 0.800391721 | NA |
| 1.27E-05 | 0.000112011 |
| 0.910000593 | 0.945637814 |
| 0.815258863 | 0.88702503 |
| 0.792435537 | 0.872331452 |
| 0.002586137 | 0.0113527 |
| 0.111240288 | 0.226962226 |
| 0.403136119 | 0.56516939 |
| 9.28E-08 | 1.26E-06 |
| 0.782004189 | 0.865974863 |
| 2.70E-17 | 9.58E-16 |
| 0.617923477 | 0.747424261 |
| 1.06E-05 | 9.51E-05 |
| 4.49E-35 | 3.87E-33 |
| 0.004122124 | 0.016643731 |
| 0.076272999 | 0.171576403 |
| 0.131489852 | 0.256934691 |
| NA | NA |
| NA | NA |
| NA | NA |
| 0.354538104 | NA |
| 0.941278332 | NA |

| | | | |
|---|---|---|---|
| | 0.557014117 | | 0.701039024 |
| NA | | NA | |
| | 0.839319108 | NA | |
| NA | | NA | |
| NA | | NA | |
| | 0.941276431 | NA | |
| | 0.839319108 | NA | |
| NA | | NA | |
| NA | | NA | |
| NA | | NA | |
| NA | | NA | |
| NA | | NA | |
| | 0.786061823 | NA | |
| | 0.167090937 | | 0.306141868 |
| | 0.299076501 | | 0.460705725 |
| | 0.941277857 | NA | |
| | 0.917619186 | NA | |
| | 0.627141919 | | 0.754434042 |
| | 0.839319108 | NA | |
| | 0.955791446 | NA | |
| | 1.27E-07 | | 1.68E-06 |
| | 0.168767123 | | 0.308335513 |
| NA | | NA | |
| NA | | NA | |
| | 0.025684233 | | 0.073205138 |
| | 0.154095128 | | 0.288812524 |
| | 0.730476056 | | 0.830111521 |
| | 0.593518036 | | 0.728755032 |
| | 0.023146409 | | 0.067441649 |
| | 0.896897933 | | 0.937779975 |
| | 0.289090019 | | 0.44993923 |
| | 0.040314137 | | 0.104837712 |
| NA | | NA | |
| | 4.35E-06 | | 4.29E-05 |
| | 0.146342664 | | 0.278214703 |
| NA | | NA | |
| | 0.016065377 | | 0.050360608 |
| | 0.328965831 | | 0.491929679 |
| | 0.851821248 | | 0.911044421 |
| | 4.65E-07 | | 5.50E-06 |
| | 0.00503187 | | 0.019634191 |
| | 0.009393792 | | 0.032693527 |
| | 0.00521305 | | 0.020199529 |
| | 0.465874551 | | 0.6240787 |
| | 0.03454192 | | 0.092659392 |
| | 0.212509335 | | 0.363033062 |
| | 0.181780034 | | 0.324214576 |

| | | |
|---|---|---|
| | 0.302392665 | 0.463748536 |
| | 0.122523946 | 0.244104375 |
| | 5.32E-07 | 6.23E-06 |
| | 0.045629666 | 0.115258126 |
| | 0.910821506 | 0.94602573 |
| | 0.936551119 | 0.9623981 |
| | 0.375139536 | 0.538264876 |
| | 0.071132994 | 0.163049647 |
| | 0.134039346 | 0.260547034 |
| | 0.4153171 | 0.577147222 |
| | 0.012754561 | 0.041885219 |
| | 0.323417376 | 0.486083105 |
| | 0.001943247 | 0.008944723 |
| | 0.569649694 | 0.710736881 |
| | 0.568657781 | NA |
| NA | | NA |
| | 0.256557218 | 0.414679811 |
| | 0.004799142 | 0.018887099 |
| | 0.001450332 | 0.006971364 |
| | 0.183760748 | 0.326505921 |
| | 0.018472583 | 0.056304831 |
| | 0.468337298 | 0.626502802 |
| | 0.046736895 | 0.117469014 |
| | 0.761510396 | 0.851817685 |
| | 0.079425036 | 0.176911748 |
| | 8.03E-06 | 7.44E-05 |
| | 0.804200059 | 0.880395999 |
| | 1.24E-12 | 3.09E-11 |
| | 9.44E-27 | 5.75E-25 |
| | 0.079241379 | 0.17663451 |
| | 0.21070404 | 0.36075057 |
| | 0.016699319 | 0.051951013 |
| | 1.05E-11 | 2.38E-10 |
| | 0.569987722 | 0.71082719 |
| | 0.126303937 | NA |
| | 0.941278332 | NA |
| | 0.941278332 | NA |
| NA | | NA |
| | 0.311353803 | 0.473415307 |
| | 0.568326727 | NA |
| | 0.525899396 | 0.675050285 |
| | 0.774742687 | 0.860897546 |
| | 0.384539263 | 0.547351007 |
| | 0.019948658 | 0.059725531 |
| | 0.138682485 | 0.267237735 |
| | 0.978140929 | 0.986494311 |
| | 0.000142114 | 0.000952576 |

| | | |
|---|---|---|
| | 0.000209902 | 0.00133728 |
| | 9.02E-10 | 1.61E-08 |
| | 0.22955703 | 0.38329824 |
| | 0.201504675 | 0.349425489 |
| | 0.802369388 | 0.87912324 |
| | 0.435115825 | 0.59635245 |
| | 0.411681795 | 0.573770353 |
| | 0.131887718 | 0.257437384 |
| | 0.044920438 | 0.11384117 |
| | 5.97E-16 | 1.96E-14 |
| | 2.72E-07 | 3.38E-06 |
| | 0.899640297 | NA |
| | 0.27042407 | NA |
| NA | | NA |
| | 0.895132915 | 0.936820599 |
| NA | | NA |
| | 0.024340897 | 0.0702257 |
| | 0.885960202 | 0.931720264 |
| | 6.39E-06 | 6.07E-05 |
| | 0.328066881 | 0.490893814 |
| | 2.28E-05 | 0.000189038 |
| | 0.118702023 | 0.238285025 |
| NA | | NA |
| | 8.62E-10 | 1.54E-08 |
| | 0.849602856 | 0.909682734 |
| | 0.625560463 | 0.753331148 |
| | 0.112811464 | 0.229440183 |
| | 0.007249033 | 0.026447192 |
| | 0.000134259 | 0.000907591 |
| | 0.009018738 | 0.031642688 |
| | 0.864878264 | NA |
| | 0.003657333 | 0.015071301 |
| | 0.514884559 | NA |
| | 0.125227293 | NA |
| NA | | NA |
| | 0.075660049 | 0.17064915 |
| NA | | NA |
| | 0.001211296 | 0.006009018 |
| | 0.148608214 | 0.281509965 |
| | 0.072368698 | 0.165103015 |
| | 0.000346667 | 0.002062842 |
| | 0.016949293 | 0.052556939 |
| NA | | NA |
| NA | | NA |
| | 2.01E-09 | 3.43E-08 |
| | 6.80E-05 | 0.000497434 |
| | 0.955792398 | NA |

| | | |
|---|---|---|
| | 0.002831953 | 0.01221347 |
| | 0.033948478 | 0.09145916 |
| | 0.84530793 | 0.906845355 |
| | 0.03465002 | 0.09288656 |
| | 0.970411705 | 0.982229107 |
| | 0.043493564 | 0.111076588 |
| | 0.124821624 | 0.247418746 |
| | 0.500471177 | NA |
| | 0.277637175 | 0.437434077 |
| | 0.02266339 | 0.066236984 |
| NA | | NA |
| | 0.096867348 | 0.204672581 |
| | 0.267957789 | 0.42767594 |
| | 0.001826643 | 0.008471896 |
| | 0.110263913 | 0.225550159 |
| | 0.027023513 | 0.07646382 |
| | 0.536874159 | NA |
| | 0.590390988 | 0.726376837 |
| | 0.115115733 | 0.232754389 |
| | 0.941276431 | NA |
| | 4.67E-20 | 2.00E-18 |
| | 0.699275945 | NA |
| | 0.766545753 | 0.855022354 |
| | 0.193526562 | 0.339348699 |
| | 0.007107659 | 0.026005414 |
| NA | | NA |
| | 0.277885071 | 0.437533326 |
| | 0.000667027 | 0.003624957 |
| | 0.067303562 | 0.156395528 |
| | 0.131547053 | 0.256978723 |
| | 0.749908829 | NA |
| | 5.00E-21 | 2.29E-19 |
| | 1.88E-05 | 0.000158834 |
| | 0.281907237 | 0.442084336 |
| | 0.349351918 | 0.513256858 |
| | 0.17214101 | 0.312413292 |
| | 0.887575797 | 0.932978859 |
| | 0.293630884 | 0.454835501 |
| | 0.788146828 | NA |
| | 0.35766895 | 0.521229189 |
| | 0.000547766 | 0.003073532 |
| | 0.002588551 | 0.011357176 |
| | 9.47E-06 | 8.63E-05 |
| NA | | NA |
| | 0.550406443 | 0.69603518 |
| | 0.006064618 | 0.022850708 |
| | 0.001585326 | 0.007519717 |

| | | |
|---|---|---|
| | 0.953932387 | NA |
| NA | | NA |
| | 0.987362553 | 0.991988666 |
| | 0.047497283 | NA |
| | 1.43E-17 | 5.19E-16 |
| | 0.610939841 | 0.742227061 |
| | 0.454521243 | 0.614366148 |
| | 0.671277475 | NA |
| | 0.733909409 | 0.832172802 |
| | 0.768440697 | NA |
| | 0.594837891 | 0.729886147 |
| | 0.763228054 | 0.85297792 |
| NA | | NA |
| | 0.941278332 | NA |
| NA | | NA |
| | 0.830051232 | 0.896817041 |
| | 0.214012403 | 0.364877858 |
| | 0.022499597 | 0.065855313 |
| | 0.956298135 | NA |
| | 0.041195995 | 0.10665076 |
| | 0.1606621 | 0.297501113 |
| | 0.241413051 | 0.397637123 |
| | 0.369742217 | 0.532950055 |
| | 0.011837758 | 0.039468037 |
| | 0.075920619 | NA |
| | 0.956891194 | NA |
| | 5.87E-07 | 6.82E-06 |
| | 0.941279268 | NA |
| | 0.955791446 | NA |
| | 0.591099794 | 0.726910735 |
| | 0.941430486 | NA |
| | 0.248856128 | NA |
| | 0.169752584 | 0.309352347 |
| | 0.463999171 | 0.622659314 |
| | 0.537808348 | 0.685530675 |
| | 3.44E-12 | 8.19E-11 |
| | 0.680230048 | 0.794173006 |
| | 3.85E-13 | 1.00E-11 |
| NA | | NA |
| | 0.001664792 | 0.00783885 |
| | 0.01490215 | 0.047445288 |
| | 0.031208219 | 0.085578113 |
| | 0.844817288 | 0.906638762 |
| | 0.070829444 | 0.162618687 |
| | 0.020721247 | 0.061535511 |
| | 0.464123703 | 0.622666014 |
| | 0.488939955 | 0.643538421 |

| | | | |
|---|---|---|---|
| | 0.234790656 | | 0.38963168 |
| NA | | NA | |
| | 0.011189431 | | 0.037723913 |
| NA | | NA | |
| NA | | NA | |
| NA | | NA | |
| NA | | NA | |
| NA | | NA | |
| NA | | NA | |
| NA | | NA | |
| NA | | NA | |
| NA | | NA | |
| NA | | NA | |
| NA | | NA | |
| NA | | NA | |
| NA | | NA | |
| NA | | NA | |
| NA | | NA | |
| NA | | NA | |
| NA | | NA | |
| NA | | NA | |
| NA | | NA | |
| NA | | NA | |
| NA | | NA | |
| NA | | NA | |
| NA | | NA | |
| NA | | NA | |
| NA | | NA | |
| NA | | NA | |
| NA | | NA | |
| NA | | NA | |
| NA | | NA | |
| NA | | NA | |
| NA | | NA | |
| NA | | NA | |
| NA | | NA | |
| NA | | NA | |
| NA | | NA | |
| NA | | NA | |
| NA | | NA | |
| NA | | NA | |
| NA | | NA | |
| NA | | NA | |
| NA | | NA | |
| NA | | NA | |
| NA | | NA | |
| NA | | NA | |

| | |
|---|---|
| NA | NA |
| NA | NA |
| NA | NA |
| NA | NA |
| NA | NA |
| NA | NA |
| NA | NA |
| NA | NA |
| NA | NA |
| NA | NA |
| NA | NA |
| NA | NA |
| NA | NA |
| NA | NA |
| NA | NA |
| NA | NA |
| NA | NA |
| NA | NA |
| NA | NA |
| NA | NA |
| NA | NA |
| NA | NA |
| NA | NA |
| NA | NA |
| NA | NA |
| NA | NA |
| NA | NA |
| NA | NA |
| NA | NA |
| NA | NA |
| NA | NA |
| NA | NA |
| NA | NA |
| NA | NA |
| NA | NA |
| NA | NA |
| NA | NA |
| NA | NA |
| NA | NA |
| NA | NA |
| NA | NA |
| NA | NA |
| NA | NA |
| NA | NA |
| NA | NA |
| NA | NA |
| NA | NA |

| | | |
|---|---|---|
| NA | | NA |
| NA | | NA |
| NA | | NA |
| NA | | NA |
| NA | | NA |
| NA | | NA |
| NA | | NA |
| NA | | NA |
| NA | | NA |
| NA | | NA |
| NA | | NA |
| NA | | NA |
| | 0.955792873 | NA |
| NA | | NA |
| NA | | NA |
| NA | | NA |
| NA | | NA |
| NA | | NA |
| NA | | NA |
| NA | | NA |
| NA | | NA |
| NA | | NA |
| NA | | NA |
| NA | | NA |
| NA | | NA |
| NA | | NA |
| NA | | NA |
| NA | | NA |
| NA | | NA |
| NA | | NA |
| | 0.941278332 | NA |
| NA | | NA |
| NA | | NA |
| | 0.839319108 | NA |
| NA | | NA |
| | 1.63E-05 | 0.000140223 |
| | 0.523978306 | 0.673637877 |
| | 0.856771514 | NA |
| NA | | NA |
| | 0.254339449 | 0.41235526 |
| | 0.00985248 | 0.033981849 |
| | 0.796693189 | 0.875398914 |
| | 0.317490258 | 0.480047207 |
| | 0.160389655 | 0.29712966 |
| | 0.783554882 | 0.866925733 |
| | 0.941278332 | NA |
| | 0.005529259 | 0.021148551 |

| | |
|---|---|
| NA | NA |
| 4.79E-10 | 8.89E-09 |
| 0.542047311 | NA |
| NA | NA |
| 0.888657443 | NA |
| 0.10473128 | 0.216973968 |
| 0.687264961 | NA |
| 0.018284051 | 0.055841274 |
| 0.15080463 | 0.284448196 |
| 0.241574253 | 0.39781032 |
| 0.748393366 | NA |
| 0.369693095 | 0.532943798 |
| NA | NA |
| NA | NA |
| 4.64E-06 | 4.55E-05 |
| 1.76E-09 | 3.03E-08 |
| 0.054680515 | 0.133023877 |
| 0.731615005 | NA |
| 0.302278242 | 0.463666198 |
| 0.071623441 | 0.163873992 |
| 0.009718076 | 0.033654829 |
| 0.353540192 | 0.517143784 |
| NA | NA |
| 0.955791446 | NA |
| 0.481980553 | 0.638225375 |
| 0.966358451 | NA |
| 6.62E-05 | 0.000486495 |
| 0.059776703 | 0.142565641 |
| 0.464197919 | 0.622715373 |
| 0.001811584 | 0.008415169 |
| 0.003055507 | 0.013017217 |
| 0.436055419 | 0.597208998 |
| 0.788080684 | 0.869754548 |
| 0.777354769 | NA |
| 0.312683268 | 0.474678688 |
| 0.30699497 | 0.46864381 |
| 0.152309084 | 0.286378136 |
| 0.739991096 | NA |
| 0.406893612 | 0.568695346 |
| NA | NA |
| NA | NA |
| 0.538959458 | 0.686373089 |
| 0.873176765 | 0.924280679 |
| 0.003819186 | 0.015616608 |
| 0.07139006 | 0.163465725 |
| 0.353411541 | 0.517082708 |
| 0.058217831 | 0.139617211 |

| | |
|---|---|
| NA | NA |
| NA | NA |
| 0.728678906 | NA |
| NA | NA |
| 0.304671414 | NA |
| 0.13883745 | 0.267493037 |
| NA | NA |
| NA | NA |
| 0.000733061 | 0.003931535 |
| 0.444948931 | 0.605596664 |
| NA | NA |
| 1.42E-05 | 0.000124245 |
| 0.020392843 | 0.060803013 |
| 0.000870239 | 0.004536448 |
| NA | NA |
| NA | NA |
| NA | NA |
| 0.839319108 | NA |
| NA | NA |
| NA | NA |
| NA | NA |
| NA | NA |
| NA | NA |
| 0.941278788 | NA |
| NA | NA |
| NA | NA |
| NA | NA |
| NA | NA |
| NA | NA |
| NA | NA |
| NA | NA |
| NA | NA |
| NA | NA |
| 0.953336382 | NA |
| NA | NA |
| NA | NA |
| NA | NA |
| NA | NA |
| NA | NA |
| NA | NA |
| NA | NA |
| NA | NA |
| NA | NA |
| NA | NA |
| NA | NA |
| NA | NA |
| NA | NA |

| | |
|---|---|
| NA | NA |
| NA | NA |
| NA | NA |
| NA | NA |
| 0.698853538 | NA |
| 0.006431017 | 0.023962282 |
| 0.468207241 | 0.626434444 |
| 0.626506081 | 0.753935878 |
| 0.204829027 | NA |
| 0.000422294 | 0.002449781 |
| 0.001664786 | 0.00783885 |
| 0.034307988 | 0.092177304 |
| 0.849802318 | 0.909855372 |
| NA | NA |
| 0.853595154 | NA |
| 0.920968846 | 0.952651559 |
| 0.684635527 | NA |
| 0.839319108 | NA |
| NA | NA |
| 0.784161164 | NA |
| NA | NA |
| NA | NA |
| NA | NA |
| 0.15479187 | 0.2896051 |
| NA | NA |
| NA | NA |
| 0.728678906 | NA |
| NA | NA |
| 0.955792398 | NA |
| 0.001228169 | 0.006078785 |
| 0.839318641 | NA |
| NA | NA |
| 0.097802694 | 0.206224197 |
| 0.634914559 | NA |
| 0.95456991 | NA |
| NA | NA |
| 0.955792398 | NA |
| NA | NA |
| NA | NA |
| NA | NA |
| NA | NA |
| NA | NA |
| NA | NA |
| 0.853595622 | NA |
| NA | NA |
| NA | NA |
| NA | NA |

| | | |
|---|---|---|
| NA | NA | |
| NA | NA | |
| 0.706645317 | NA | |
| NA | NA | |
| 0.368175746 | | 0.531271239 |
| 0.495466152 | | 0.648399239 |
| 0.028288537 | | 0.079186612 |
| 0.805232791 | NA | |
| 0.146526356 | | 0.278474955 |
| 0.4979989 | NA | |
| NA | NA | |
| NA | NA | |
| 0.56124106 | | 0.70412501 |
| 0.001509757 | | 0.007209074 |
| NA | NA | |
| 0.750696126 | | 0.843499437 |
| 0.839319108 | NA | |
| 0.934588843 | NA | |
| 0.941278332 | NA | |
| 0.941278332 | NA | |
| NA | NA | |
| NA | NA | |
| NA | NA | |
| 0.707430518 | | 0.813403275 |
| 0.825067675 | | 0.893704414 |
| 0.636959113 | | 0.761280483 |
| NA | NA | |
| 0.531588591 | | 0.680225345 |
| NA | NA | |
| NA | NA | |
| NA | NA | |
| NA | NA | |
| NA | NA | |
| NA | NA | |
| 0.839318641 | NA | |
| 0.003453063 | | 0.014394011 |
| 0.165596925 | NA | |
| 0.365588999 | | 0.528788852 |
| 0.000574165 | | 0.003192307 |
| 0.76109692 | | 0.851498421 |
| 0.548489847 | | 0.694416527 |
| 0.00134788 | | 0.006562236 |
| NA | NA | |
| NA | NA | |
| NA | NA | |
| 0.853595154 | NA | |
| 0.472828156 | NA | |

| | | | |
|---|---|---|---|
| NA | | NA | |
| NA | | NA | |
| NA | | NA | |
| | 0.084578692 | | 0.18545758 |
| NA | | NA | |
| NA | | NA | |
| NA | | NA | |
| NA | | NA | |
| | 0.853595622 | NA | |
| NA | | NA | |
| | 0.697988805 | NA | |
| | 0.711727114 | | 0.816293007 |
| | 0.004644197 | | 0.018377586 |
| NA | | NA | |
| | 0.002373119 | | 0.010567816 |
| | 0.941278332 | NA | |
| | 8.06E-23 | | 4.07E-21 |
| | 0.059460083 | | 0.142066744 |
| | 0.810780549 | | 0.884102924 |
| | 0.667130643 | NA | |
| | 0.002481432 | | 0.010964405 |
| | 2.98E-07 | | 3.67E-06 |
| | 0.9228362 | NA | |
| | 0.099115402 | | 0.208176586 |
| | 0.056684786 | | 0.136836842 |
| | 0.094282646 | | 0.200743854 |
| | 0.0743189 | | 0.168326049 |
| | 0.14731447 | NA | |
| | 0.157009289 | | 0.292806001 |
| | 0.930143693 | | 0.958451889 |
| NA | | NA | |
| | 0.40039422 | | 0.563049937 |
| | 0.926045812 | NA | |
| NA | | NA | |
| | 0.921702854 | NA | |
| | 0.300039704 | | 0.46157866 |
| | 4.27E-06 | | 4.22E-05 |
| | 0.000361105 | | 0.002137 |
| | 0.000190146 | | 0.001227518 |
| | 0.000285765 | | 0.001746734 |
| | 0.821809878 | NA | |
| | 0.735804876 | NA | |
| NA | | NA | |
| | 0.941277857 | NA | |
| | 0.939587148 | NA | |
| | 0.941278332 | NA | |
| | 0.941278207 | NA | |

| | |
|---|---|
| 0.911087066 | NA |
| 0.941278332 | NA |
| 0.003937587 | 0.015999743 |
| NA | NA |
| NA | NA |
| 0.613323948 | NA |
| 0.941277857 | NA |
| NA | NA |
| NA | NA |
| NA | NA |
| 0.369643669 | 0.532904822 |
| 5.80E-08 | 8.06E-07 |
| 0.241804511 | 0.398079331 |
| NA | NA |
| 0.000678065 | 0.003676381 |
| 0.10407513 | 0.216047117 |
| NA | NA |
| 0.128045246 | 0.252087092 |
| NA | NA |
| NA | NA |
| NA | NA |
| NA | NA |
| 0.018860022 | 0.057265753 |
| 0.000144698 | 0.000967988 |
| 0.159935102 | 0.296662663 |
| 0.003026902 | 0.012924062 |
| 0.004671325 | 0.018478789 |
| 0.003921571 | 0.015958495 |
| 1.98E-06 | 2.09E-05 |
| 0.047092978 | 0.118102104 |
| 0.910850839 | 0.94602573 |
| 0.337282135 | 0.500529295 |
| 0.363740626 | 0.527404028 |
| 1.64E-08 | 2.49E-07 |
| 0.695599996 | 0.805081541 |
| 0.799283826 | NA |
| 0.000980626 | 0.005021701 |
| 2.24E-16 | 7.58E-15 |
| 0.624790632 | 0.752819904 |
| 0.01264825 | 0.041604981 |
| 0.029397255 | 0.081675437 |
| 0.686357885 | NA |
| NA | NA |
| 0.887059689 | 0.9326293 |
| 0.416842013 | 0.578440607 |
| 9.80E-14 | 2.68E-12 |
| 0.004894114 | 0.019184734 |

| | |
|---|---|
| 0.350278418 | 0.514078611 |
| 0.076199698 | 0.171476336 |
| 0.040211779 | 0.104614795 |
| NA | NA |
| 0.814508557 | 0.886610986 |
| 0.270445639 | 0.430536601 |
| 0.001445105 | 0.00695017 |
| 1.02E-05 | 9.18E-05 |
| 0.216535011 | 0.367825173 |
| 0.244021633 | 0.400455248 |
| 0.372982557 | 0.536191992 |
| 0.345858073 | 0.50959062 |
| 0.080872677 | 0.179130044 |
| 0.072398596 | 0.165107922 |
| 0.021868782 | 0.064325471 |
| 0.2078501 | 0.357098894 |
| 0.308982946 | NA |
| 2.13E-11 | 4.65E-10 |
| 0.066562572 | 0.155188788 |
| 0.025344722 | 0.072454368 |
| 1.44E-08 | 2.19E-07 |
| 0.039941933 | 0.104072019 |
| 0.452647904 | 0.612572516 |
| NA | NA |
| 0.002380551 | 0.010596946 |
| 0.698650219 | NA |
| 0.001234956 | 0.006107296 |
| 0.221769053 | NA |
| NA | NA |
| NA | NA |
| NA | NA |
| NA | NA |
| 2.08E-18 | 7.95E-17 |
| 0.000557893 | 0.003118594 |
| 0.989822602 | 0.993351574 |
| 0.338606426 | 0.5020876 |
| 0.640614549 | NA |
| 0.924372987 | 0.954663244 |
| 0.046451706 | 0.116866668 |
| 0.306441661 | 0.468073802 |
| 0.028977611 | 0.080716803 |
| 0.364755875 | 0.528065162 |
| 2.01E-08 | 3.00E-07 |
| 0.888917832 | 0.93342843 |
| NA | NA |
| 0.413480275 | NA |
| 0.192236718 | 0.337659092 |

| | | |
|---|---|---|
| | 4.38E-06 | 4.31E-05 |
| | 0.496783949 | NA |
| | 0.071588912 | 0.163826485 |
| | 0.178674939 | 0.320290774 |
| | 0.000446373 | 0.002574407 |
| | 0.271666328 | 0.431482846 |
| | 0.941278332 | NA |
| | 0.915068409 | NA |
| | 0.130585748 | 0.255629738 |
| | 0.043383299 | 0.110878265 |
| | 0.058762752 | 0.140640627 |
| | 8.01E-06 | 7.43E-05 |
| | 1.27E-07 | 1.67E-06 |
| NA | | NA |
| | 0.002031132 | 0.009270377 |
| | 0.451069353 | 0.611061625 |
| | 0.616480703 | NA |
| | 0.000680652 | 0.003687043 |
| | 3.17E-06 | 3.21E-05 |
| | 4.33E-29 | 2.91E-27 |
| | 0.839318641 | NA |
| | 0.985107596 | 0.990684934 |
| | 0.077297639 | 0.173258051 |
| | 0.12547751 | 0.248429187 |
| | 0.081540266 | 0.18025647 |
| | 0.468546896 | 0.626642311 |
| | 0.672504343 | NA |
| | 9.99E-11 | 2.00E-09 |
| | 0.592318491 | 0.727882943 |
| | 0.132970668 | 0.258943241 |
| | 0.144842313 | 0.276112185 |
| | 0.245489322 | 0.402281366 |
| | 0.925953722 | 0.955475364 |
| | 0.122741306 | 0.244455573 |
| | 4.95E-05 | 0.000375865 |
| | 0.007292219 | 0.0265681 |
| | 3.97E-05 | 0.000308516 |
| | 0.030835901 | 0.08477852 |
| NA | | NA |
| | 0.193409301 | 0.339196944 |
| | 0.018239238 | 0.055743496 |
| | 0.228027235 | 0.381493235 |
| | 0.840221195 | 0.903498752 |
| | 0.006201074 | 0.023254761 |
| | 0.284293948 | 0.444889189 |
| | 0.420804587 | 0.582325046 |
| NA | | NA |

| | |
|---|---|
| 0.484022098 | NA |
| 0.043717517 | 0.111494136 |
| 1.02E-07 | 1.36E-06 |
| 0.848500834 | 0.908952556 |
| 0.025295994 | 0.072349826 |
| 0.000369981 | 0.0021817 |
| 0.016054307 | 0.050345798 |
| 2.25E-07 | 2.84E-06 |
| 0.125165335 | 0.247893602 |
| 0.946935029 | 0.967625072 |
| NA | NA |
| 0.79747458 | NA |
| 3.89E-06 | 3.87E-05 |
| 0.10747685 | 0.221201858 |
| 0.58754757 | 0.723813843 |
| NA | NA |
| 0.019227902 | 0.058064264 |
| 0.269105168 | 0.428965026 |
| 2.58E-24 | 1.40E-22 |
| 0.463572544 | NA |
| 0.000140861 | 0.000945245 |
| 0.000109266 | 0.000757134 |
| 0.655386826 | 0.775090135 |
| 1.23E-05 | 0.000109109 |
| 0.140605819 | 0.270047584 |
| 0.124721559 | 0.247282178 |
| 0.941277857 | NA |
| 0.091758676 | 0.196846835 |
| 0.012349762 | 0.040820586 |
| 0.839318641 | NA |
| 2.30E-08 | 3.42E-07 |
| 7.47E-11 | 1.52E-09 |
| 0.033400176 | 0.090411805 |
| 0.721510618 | NA |
| 0.03368228 | 0.090937571 |
| NA | NA |
| 0.941277857 | NA |
| NA | NA |
| NA | NA |
| NA | NA |
| 0.837259124 | NA |
| 0.94127925 | NA |
| 0.780833705 | NA |
| NA | NA |
| NA | NA |
| NA | NA |
| NA | NA |

| | | |
|---|---|---|
| NA | NA | |
| NA | NA | |
| NA | NA | |
| NA | NA | |
| NA | NA | |
| NA | NA | |
| NA | NA | |
| NA | NA | |
| NA | NA | |
| NA | NA | |
| NA | NA | |
| NA | NA | |
| NA | NA | |
| NA | NA | |
| NA | NA | |
| NA | NA | |
| NA | NA | |
| NA | NA | |
| NA | NA | |
| NA | NA | |
| NA | NA | |
| NA | NA | |
| NA | NA | |
| NA | NA | |
| NA | NA | |
| NA | NA | |
| NA | NA | |
| NA | NA | |
| NA | NA | |
| 0.941278332 | NA | |
| 0.831499882 | NA | |
| 0.941278788 | NA | |
| 0.839319108 | NA | |
| NA | NA | |
| NA | NA | |
| NA | NA | |
| 0.839318641 | NA | |
| 0.493814536 | | 0.647162955 |
| 0.641670499 | NA | |
| 0.000161815 | | 0.001069272 |
| 0.941278332 | NA | |
| 0.216281208 | | 0.367551428 |
| 0.00775454 | | 0.027948752 |
| 0.089094589 | | 0.192686977 |
| 0.941277857 | NA | |
| 0.177233995 | | 0.318812239 |
| 0.007162186 | | 0.026170455 |

| | | | |
|---|---|---|---|
| NA | | NA | |
| | 0.274023017 | | 0.43386522 |
| | 0.296309652 | | 0.45779322 |
| | 0.955792873 | NA | |
| | 0.363954461 | | 0.52755777 |
| | 0.728577274 | | 0.829110414 |
| | 0.220714874 | | 0.372638348 |
| | 0.976374855 | | 0.985483601 |
| | 0.812998424 | NA | |
| | 0.975902858 | | 0.985350438 |
| | 0.649734015 | | 0.771012515 |
| | 0.000916218 | | 0.004739798 |
| | 0.653006857 | NA | |
| NA | | NA | |
| | 0.100756139 | | 0.210923274 |
| NA | | NA | |
| NA | | NA | |
| NA | | NA | |
| NA | | NA | |
| | 0.57858582 | | 0.716668386 |
| | 0.001099496 | | 0.005537497 |
| NA | | NA | |
| NA | | NA | |
| | 0.751182588 | | 0.843777629 |
| | 0.177654017 | | 0.319205926 |
| | 0.006567514 | | 0.024379285 |
| | 0.791947557 | NA | |
| | 0.000530208 | | 0.002984161 |
| | 0.00095574 | | 0.004908638 |
| | 6.98E-06 | | 6.56E-05 |
| NA | | NA | |
| | 0.001875352 | | 0.008665723 |
| | 6.96E-06 | | 6.55E-05 |
| | 0.877402602 | | 0.926494259 |
| | 0.645264599 | | 0.767737465 |
| | 0.603037594 | | 0.736190225 |
| | 0.159015192 | | 0.295324563 |
| | 0.239142018 | | 0.395052456 |
| NA | | NA | |
| | 0.910563393 | NA | |
| NA | | NA | |
| | 1.64E-09 | | 2.83E-08 |
| | 0.606581241 | | 0.739151523 |
| | 0.32279672 | | 0.485541427 |
| | 0.217378496 | | 0.368705406 |
| | 0.113300229 | | 0.230158919 |
| | 0.152862116 | | 0.286964594 |

| | |
|---|---|
| 0.721737919 | 0.823370684 |
| 0.655307898 | 0.775076456 |
| 0.270666071 | 0.430641298 |
| 0.602456465 | 0.735846752 |
| 0.365314667 | 0.52861514 |
| 0.903849498 | 0.941698566 |
| 0.945058913 | 0.966914213 |
| 0.993118754 | 0.995125561 |
| 0.176952458 | 0.318522454 |
| 0.209563155 | NA |
| 0.263101836 | 0.422019672 |
| 0.355966163 | NA |
| 0.839319108 | NA |
| 0.952811619 | NA |
| 0.152212459 | 0.286251396 |
| 0.44593404 | NA |
| 0.804019527 | 0.880356952 |
| 0.613705891 | NA |
| 0.776799055 | 0.862498069 |
| 0.818944694 | 0.889582038 |
| 0.429563592 | 0.590769691 |
| 0.635019898 | 0.760020207 |
| 0.863090034 | NA |
| 0.208371581 | 0.357705426 |
| NA | NA |
| 0.625174284 | 0.753056594 |
| 0.202159592 | 0.350229464 |
| 0.956016157 | 0.973105396 |
| 0.665706965 | 0.783218414 |
| 0.196191214 | 0.342859272 |
| 0.663097348 | NA |
| 0.372859328 | NA |
| 0.941275956 | NA |
| 0.381414343 | 0.54394393 |
| 0.807556402 | NA |
| 0.660589508 | NA |
| 0.955792873 | NA |
| NA | NA |
| NA | NA |
| 0.955792398 | NA |
| 0.854553095 | 0.91249811 |
| 0.791669975 | NA |
| NA | NA |
| NA | NA |
| NA | NA |
| 0.940363068 | NA |
| NA | NA |

| | | |
|---|---|---|
| NA | NA | |
| 0.955792398 | NA | |
| NA | NA | |
| 0.739991096 | NA | |
| 0.475435218 | | 0.632634449 |
| 0.571000547 | | 0.711456579 |
| 0.573435548 | | 0.71303484 |
| 0.93992387 | NA | |
| 0.332616682 | NA | |
| 0.580865915 | NA | |
| 0.465701976 | NA | |
| 0.871252751 | NA | |
| 0.437964828 | | 0.598830356 |
| 0.316048584 | NA | |
| 0.739991096 | NA | |
| 0.923227637 | | 0.954299919 |
| 0.955792398 | NA | |
| 0.839319108 | NA | |
| 0.171538476 | | 0.311700038 |
| 0.193518821 | NA | |
| 0.091784103 | | 0.196846835 |
| 0.74477664 | NA | |
| 0.943971627 | NA | |
| 0.955153912 | NA | |
| 0.52680736 | NA | |
| 0.258788838 | NA | |
| 0.826215771 | | 0.894418874 |
| 0.728580515 | | 0.829110414 |
| 0.724551561 | | 0.826065827 |
| 0.181955486 | | 0.324363727 |
| 0.580695167 | | 0.718421247 |
| 0.265722574 | NA | |
| 0.839319108 | NA | |
| 0.734737821 | NA | |
| 0.95195536 | NA | |
| 0.034106305 | | 0.091724862 |
| 0.887313045 | NA | |
| 0.88576845 | | 0.931642096 |
| 0.837390942 | NA | |
| 0.516512388 | | 0.667355762 |
| 0.062509019 | | 0.147705715 |
| 0.735211223 | NA | |
| 0.955791922 | NA | |
| 0.759609615 | NA | |
| 0.743665443 | NA | |
| 0.879528026 | | 0.927866593 |
| 0.699040713 | NA | |

| | | |
|---|---|---|
| 0.943971627 | NA | |
| 0.828287655 | NA | |
| 0.677454612 | NA | |
| 0.411022482 | NA | |
| NA | NA | |
| NA | NA | |
| 0.702535658 | NA | |
| 0.955792398 | NA | |
| 0.714677096 | NA | |
| NA | NA | |
| NA | NA | |
| NA | NA | |
| 0.839318641 | NA | |
| NA | NA | |
| NA | NA | |
| NA | NA | |
| NA | NA | |
| 0.725028456 | NA | |
| NA | NA | |
| NA | NA | |
| NA | NA | |
| NA | NA | |
| 0.547596417 | NA | |
| 0.941280349 | NA | |
| NA | NA | |
| NA | NA | |
| NA | NA | |
| 0.928601912 | NA | |
| 0.955868592 | | 0.973038415 |
| 0.369459787 | | 0.532736526 |
| 0.608602681 | NA | |
| 0.839319108 | NA | |
| 0.924895779 | NA | |
| 0.350832828 | NA | |
| 0.526821885 | | 0.675869914 |
| 0.739991096 | NA | |
| 0.592224698 | NA | |
| 0.804525138 | NA | |
| 0.082670212 | | 0.182044258 |
| 0.903892195 | | 0.941698566 |
| 0.708754351 | | 0.814125883 |
| 0.101629285 | | 0.212298853 |
| 0.479117158 | NA | |
| NA | NA | |
| 0.613971412 | NA | |
| 0.54076703 | NA | |
| NA | NA | |

| | | |
|---|---|---|
| 0.955541805 | NA | |
| 0.937256549 | NA | |
| 0.888032093 | NA | |
| 0.534855009 | NA | |
| 0.813398531 | NA | |
| 0.984367427 | NA | |
| 0.80483689 | NA | |
| 0.402275483 | | 0.564528097 |
| 0.95456991 | NA | |
| NA | NA | |
| 0.955791922 | NA | |
| NA | NA | |
| 0.039694417 | | 0.10356314 |
| 0.941159944 | NA | |
| 0.941277857 | NA | |
| 0.661285173 | NA | |
| NA | NA | |
| NA | NA | |
| NA | NA | |
| NA | NA | |
| NA | NA | |
| NA | NA | |
| 0.853595622 | NA | |
| 0.783815745 | NA | |
| NA | NA | |
| NA | NA | |
| 0.779617645 | NA | |
| NA | NA | |
| 0.598222039 | NA | |
| NA | NA | |
| NA | NA | |
| 0.955792873 | NA | |
| 0.839318641 | NA | |
| 0.853595154 | NA | |
| 0.26248862 | NA | |
| 0.658465057 | NA | |
| 0.942348831 | NA | |
| 0.941208171 | NA | |
| NA | NA | |
| 0.328974682 | NA | |
| 0.762851389 | NA | |
| 0.544259859 | NA | |
| NA | NA | |
| NA | NA | |
| 0.690128961 | NA | |
| 0.838197868 | NA | |
| 0.280351739 | NA | |

| | | |
|---|---|---|
| | 0.858235835 | NA |
| | 0.514258071 | 0.665240505 |
| NA | | NA |
| | 0.853595622 | NA |
| NA | | NA |
| | 0.839319108 | NA |
| NA | | NA |
| | 0.853595622 | NA |
| NA | | NA |
| | 0.955791922 | NA |
| | 0.955792398 | NA |
| | 0.941278332 | NA |
| | 0.839319574 | NA |
| | 0.901831873 | NA |
| | 0.614475241 | NA |
| | 0.575111999 | 0.714074899 |
| | 0.430963207 | 0.592319335 |
| | 0.393540959 | 0.55597471 |
| | 0.644985886 | 0.767595399 |
| NA | | NA |
| NA | | NA |
| | 0.626931626 | 0.754295434 |
| NA | | NA |
| | 2.64E-08 | 3.87E-07 |
| | 0.208491017 | 0.357812618 |
| | 0.414387163 | 0.576380135 |
| | 0.317842607 | NA |
| | 0.344078966 | NA |
| | 0.734749475 | 0.832944704 |
| | 0.817417153 | NA |
| | 0.698560799 | 0.807190003 |
| | 0.587364003 | NA |
| | 0.616994008 | 0.746603528 |
| | 0.726026327 | 0.827036474 |
| | 0.2184092 | NA |
| | 0.963320881 | 0.977822662 |
| | 0.686887999 | 0.79908642 |
| | 0.98702805 | 0.991861929 |
| | 0.420261506 | 0.5818511 |
| | 0.014899704 | 0.047445288 |
| | 0.167435811 | 0.306571765 |
| | 0.029036688 | 0.08082095 |
| | 0.361444737 | 0.525092019 |
| | 0.804804556 | 0.880692319 |
| | 0.965018269 | 0.978752549 |
| | 0.083128881 | 0.182868173 |
| | 0.010917089 | 0.036989119 |

| | |
|---|---|
| 0.854745547 | 0.91261431 |
| 0.286368858 | 0.447019384 |
| 0.043722639 | 0.111494136 |
| 0.028176943 | 0.078976402 |
| 0.322753731 | 0.485507446 |
| 0.52236547 | 0.672390001 |
| NA | NA |
| 0.300524779 | 0.46205492 |
| 0.10891249 | 0.223477487 |
| 0.6134385 | 0.744003219 |
| 0.260124983 | 0.418683719 |
| 0.613685501 | 0.744153273 |
| 0.253334116 | 0.41125758 |
| 0.086057032 | 0.187830612 |
| 3.26E-08 | 4.73E-07 |
| 0.254047865 | NA |
| 0.839318641 | NA |
| 0.474888582 | 0.632316963 |
| NA | NA |
| NA | NA |
| 0.026776725 | 0.075873764 |
| 0.414061778 | 0.576109104 |
| 0.015342069 | 0.048573016 |
| 0.21454968 | 0.365469941 |
| 0.273870989 | 0.433682751 |
| 8.22E-05 | 0.000589623 |
| 0.09170821 | 0.196778847 |
| 0.278876378 | 0.438637044 |
| 0.402733729 | 0.564771591 |
| 1.46E-07 | 1.91E-06 |
| NA | NA |
| 0.578073613 | 0.716366394 |
| 0.801692069 | 0.878782105 |
| 0.705482123 | 0.812065262 |
| 0.541916291 | 0.688959061 |
| 0.74627078 | NA |
| 0.511024065 | 0.662619148 |
| 0.003566097 | 0.014779774 |
| 0.000173756 | 0.001137753 |
| 5.53E-10 | 1.02E-08 |
| 0.882717314 | NA |
| 0.009976778 | 0.034325999 |
| NA | NA |
| 1.35E-18 | 5.21E-17 |
| NA | NA |
| 0.327435332 | NA |
| 0.651858658 | NA |

| | |
|---|---|
| 0.655222309 | 0.775013735 |
| 0.019540851 | 0.058807856 |
| 0.285155179 | 0.445672141 |
| 0.103178273 | 0.214728447 |
| 0.000345777 | 0.002058577 |
| 0.171410673 | 0.311562951 |
| 0.015793963 | 0.049725915 |
| 0.000507407 | 0.00287213 |
| 0.273406214 | 0.433292143 |
| 0.041223631 | 0.106680051 |
| 0.586636594 | 0.723328026 |
| 0.007621224 | 0.027547514 |
| NA | NA |
| 0.955792398 | NA |
| 0.22695086 | 0.380138524 |
| 0.661692076 | 0.78014028 |
| 0.501982907 | 0.65415807 |
| 0.74627078 | NA |
| 4.05E-14 | 1.15E-12 |
| NA | NA |
| NA | NA |
| 2.57E-10 | 4.93E-09 |
| NA | NA |
| NA | NA |
| 0.941275956 | NA |
| NA | NA |
| NA | NA |
| NA | NA |
| 0.890911405 | 0.934367626 |
| 0.400790595 | 0.563208038 |
| 0.839318641 | NA |
| 0.364183754 | 0.52755777 |
| 0.839318641 | NA |
| 0.955792398 | NA |
| NA | NA |
| 0.964844602 | NA |
| 0.889816242 | NA |
| 0.941278332 | NA |
| 2.34E-13 | 6.21E-12 |
| NA | NA |
| NA | NA |
| 0.010165428 | 0.034864194 |
| NA | NA |
| 4.44E-05 | 0.000340831 |
| NA | NA |
| 0.839319108 | NA |
| 0.941275956 | NA |

| | |
|---|---|
| 5.37E-05 | 0.000404111 |
| 0.691810231 | 0.802303017 |
| 0.347436874 | 0.511390296 |
| 0.504150438 | 0.656264081 |
| NA | NA |
| 0.955792873 | NA |
| 0.629372888 | 0.755989005 |
| 0.078704603 | 0.175651848 |
| NA | NA |
| NA | NA |
| 0.026719751 | 0.075742738 |
| 0.955792398 | NA |
| NA | NA |
| 0.281881956 | 0.442073818 |
| NA | NA |
| NA | NA |
| 0.822062318 | 0.891707027 |
| 0.839319108 | NA |
| 0.883505117 | NA |
| NA | NA |
| 8.78E-20 | 3.67E-18 |
| 0.14368241 | NA |
| 0.831929452 | NA |
| 1.11E-30 | 7.93E-29 |
| NA | NA |
| 0.000269369 | 0.001656298 |
| 0.592114819 | 0.727761843 |
| 0.868541038 | 0.921216246 |
| 0.81308 | NA |
| NA | NA |
| 0.549308442 | 0.694942039 |
| 0.206446117 | 0.355353665 |
| 0.087157948 | 0.189593628 |
| 0.517588415 | 0.668311565 |
| 0.086059141 | 0.187830612 |
| 0.673700915 | NA |
| 1.93E-07 | 2.47E-06 |
| 2.09E-42 | 2.37E-40 |
| 0.004408483 | 0.017585871 |
| 0.005483414 | 0.021003576 |
| 0.968942847 | 0.981394794 |
| 6.85E-08 | 9.44E-07 |
| 1.42E-08 | 2.17E-07 |
| NA | NA |
| 1.48E-05 | 0.000128947 |
| 0.1024257 | 0.213591774 |
| NA | NA |

| | |
|---|---|
| 0.001915857 | 0.008829571 |
| 0.022333167 | 0.065416445 |
| 0.00694411 | 0.025518559 |
| 0.023490328 | 0.068226576 |
| 0.001764088 | 0.008241182 |
| 0.018955161 | 0.057473979 |
| 0.572547851 | 0.71242494 |
| 4.18E-07 | 5.02E-06 |
| 0.79532565 | 0.874452825 |
| 0.748015676 | 0.841684363 |
| 0.886610634 | NA |
| 0.712544511 | 0.816954935 |
| 0.019189487 | 0.057981607 |
| 0.667823584 | NA |
| 0.223495553 | 0.375954713 |
| 7.21E-08 | 9.91E-07 |
| 0.880855459 | 0.928649583 |
| 0.942302759 | NA |
| 0.976695906 | 0.985724048 |
| 0.765355707 | 0.854255406 |
| NA | NA |
| 1.95E-07 | 2.49E-06 |
| 0.94412278 | 0.966330269 |
| 0.000403862 | 0.002354327 |
| 0.006454476 | 0.02401961 |
| 0.034478545 | 0.092541535 |
| NA | NA |
| 0.130805422 | 0.255917085 |
| 0.722659216 | 0.824224192 |
| 0.003047909 | 0.012994147 |
| 0.00279842 | 0.012090853 |
| 0.618593579 | 0.747919547 |
| 0.248581107 | 0.405783384 |
| 0.129352024 | 0.253652733 |
| 0.399804219 | 0.56246913 |
| 0.001916004 | 0.008829571 |
| 1.55E-28 | 1.02E-26 |
| 9.13E-07 | 1.02E-05 |
| 0.941278332 | NA |
| 0.000607058 | 0.003345117 |
| 0.689992455 | 0.801209895 |
| 0.628501034 | 0.755481581 |
| 0.115604155 | 0.233483847 |
| 9.84E-09 | 1.55E-07 |
| 0.984959776 | 0.990578129 |
| 0.048176638 | 0.120249616 |
| 0.893672581 | 0.935943802 |

| | |
|---|---|
| 0.874643226 | 0.925009245 |
| NA | NA |
| 0.555638011 | 0.700047424 |
| 0.006674882 | 0.024693122 |
| 0.165069904 | 0.303622909 |
| 0.113813192 | 0.230806986 |
| 0.003621238 | 0.014945847 |
| 1.53E-06 | 1.66E-05 |
| 0.897959346 | 0.938413841 |
| 0.004188102 | 0.0168558 |
| 0.365406834 | 0.52864977 |
| NA | NA |
| 0.001778071 | 0.008296733 |
| 0.853596091 | NA |
| 0.122814968 | 0.24452044 |
| NA | NA |
| NA | NA |
| 0.549985048 | 0.695576201 |
| 0.023437079 | 0.068105158 |
| 2.17E-21 | 1.01E-19 |
| 3.11E-09 | 5.20E-08 |
| NA | NA |
| 0.401554113 | 0.563881475 |
| 0.510561949 | 0.662075483 |
| 0.011364508 | 0.038227533 |
| 0.498347535 | NA |
| 0.001500459 | 0.007178677 |
| 0.874344983 | 0.924906832 |
| NA | NA |
| 0.644168085 | NA |
| NA | NA |
| 0.527551173 | 0.676649646 |
| 4.12E-07 | 4.96E-06 |
| 0.00876322 | 0.030891907 |
| 0.281480408 | 0.441678573 |
| 0.386317653 | NA |
| NA | NA |
| 0.443436613 | NA |
| 0.0299541 | 0.082915045 |
| NA | NA |
| 0.940983457 | NA |
| 0.011075337 | 0.037424072 |
| 0.000145992 | 0.000975864 |
| 0.283901507 | 0.444450387 |
| 9.24E-21 | 4.15E-19 |
| 0.529434329 | 0.678490061 |
| 0.188865707 | 0.333411078 |

| | |
|---|---|
| 0.065373177 | 0.152940077 |
| 0.461800579 | 0.620548601 |
| 0.433063922 | 0.59424579 |
| 5.04E-06 | 4.91E-05 |
| 0.653214481 | 0.773830932 |
| 0.526532867 | 0.675535538 |
| 0.720547752 | 0.822446535 |
| NA | NA |
| 1.79E-06 | 1.92E-05 |
| 0.451170752 | 0.611164205 |
| 2.55E-05 | 0.000209211 |
| 0.060707634 | 0.144395234 |
| 0.959632076 | 0.975242845 |
| 0.46921415 | 0.62725275 |
| 0.738338803 | NA |
| NA | NA |
| 0.688303299 | 0.800146282 |
| 0.941278332 | NA |
| 0.941277857 | NA |
| 0.853595622 | NA |
| NA | NA |
| NA | NA |
| NA | NA |
| NA | NA |
| NA | NA |
| NA | NA |
| NA | NA |
| NA | NA |
| NA | NA |
| NA | NA |
| NA | NA |
| NA | NA |
| NA | NA |
| NA | NA |
| NA | NA |
| NA | NA |
| NA | NA |
| NA | NA |
| NA | NA |
| NA | NA |
| NA | NA |
| NA | NA |
| NA | NA |
| NA | NA |
| NA | NA |
| NA | NA |
| NA | NA |

| | |
|---|---|
| NA | NA |
| NA | NA |
| NA | NA |
| 0.839319107 | NA |
| NA | NA |
| NA | NA |
| NA | NA |
| NA | NA |
| NA | NA |
| NA | NA |
| NA | NA |
| NA | NA |
| NA | NA |
| NA | NA |
| NA | NA |
| NA | NA |
| NA | NA |
| NA | NA |
| NA | NA |
| 0.878747746 | NA |
| NA | NA |
| NA | NA |
| NA | NA |
| NA | NA |
| NA | NA |
| NA | NA |
| NA | NA |
| NA | NA |
| NA | NA |
| 0.941278332 | NA |
| NA | NA |
| NA | NA |
| NA | NA |
| NA | NA |
| NA | NA |
| NA | NA |
| NA | NA |
| 0.000222531 | 0.001406822 |
| 0.941278332 | NA |
| 0.839319107 | NA |
| NA | NA |
| 0.984798628 | NA |
| 0.941277857 | NA |
| 0.673763827 | 0.789413075 |
| 0.559236117 | NA |
| 0.661238053 | 0.779765563 |
| 0.941277857 | NA |

| | |
|---|---|
| 5.13E-52 | 7.36E-50 |
| 0.839319108 | NA |
| 0.941278332 | NA |
| 0.40953057 | 0.571486851 |
| NA | NA |
| 0.545081774 | 0.691560783 |
| 0.941830289 | NA |
| 0.930977253 | 0.958895693 |
| 0.955791446 | NA |
| 0.955791922 | NA |
| 0.485012713 | 0.640530022 |
| 0.041089045 | 0.106466519 |
| 0.019849962 | 0.059489089 |
| 0.721489208 | NA |
| 0.246155509 | 0.403095523 |
| 0.151850346 | 0.285786506 |
| 0.554375796 | 0.699308533 |
| 0.826698785 | NA |
| 8.01E-05 | 0.000575818 |
| 0.00263508 | 0.011531581 |
| 0.045261487 | 0.114560415 |
| 0.338566558 | 0.50205976 |
| 0.955792873 | NA |
| 0.579402441 | 0.71730661 |
| 3.42E-06 | 3.44E-05 |
| 0.022627168 | 0.066147367 |
| NA | NA |
| 0.000138536 | 0.000933061 |
| 0.349980157 | 0.513799278 |
| 0.116629706 | 0.235046746 |
| 0.717069814 | 0.820167982 |
| 0.523682542 | 0.673469863 |
| 0.476763648 | 0.633602986 |
| 0.955792873 | NA |
| NA | NA |
| 0.193115608 | 0.338808306 |
| NA | NA |
| NA | NA |
| 0.738881751 | 0.835678947 |
| 0.041776522 | 0.107790221 |
| 0.071841977 | 0.164184612 |
| 0.065465976 | 0.153127078 |
| 0.001509514 | 0.007209074 |
| 0.941111787 | 0.964703895 |
| NA | NA |
| 0.009345166 | 0.032552851 |
| 0.071069191 | 0.162934779 |

| | |
|---|---|
| 2.03E-05 | 0.000170377 |
| 0.009891347 | 0.034078389 |
| 0.683071537 | 0.796437186 |
| 0.274222362 | 0.434094217 |
| 0.000160185 | 0.001058796 |
| NA | NA |
| 0.343216809 | 0.506777077 |
| 0.809894704 | 0.883463483 |
| 0.4728074 | NA |
| NA | NA |
| 0.178688772 | 0.320291448 |
| 0.283826137 | 0.444429079 |
| 0.0417206 | 0.107690051 |
| 0.002745473 | 0.011922595 |
| NA | NA |
| NA | NA |
| 0.83932147 | NA |
| 0.000203447 | 0.001300687 |
| 0.604842347 | NA |
| NA | NA |
| 0.430522344 | NA |
| NA | NA |
| 0.275088357 | 0.434965753 |
| 0.981922748 | 0.98868314 |
| 2.02E-16 | 6.83E-15 |
| 0.25453196 | 0.412546261 |
| 5.38E-05 | 0.00040455 |
| 0.177801023 | 0.319321374 |
| 0.098909358 | 0.207843253 |
| 0.012366696 | 0.040865208 |
| 0.001114888 | 0.005605525 |
| 0.002663494 | 0.011643066 |
| 0.078454166 | 0.175194981 |
| 0.221045753 | 0.372958886 |
| 0.038142499 | 0.10031393 |
| 0.063128567 | 0.148809792 |
| NA | NA |
| 0.106285615 | 0.219563461 |
| 0.774561545 | 0.860736444 |
| 0.020526834 | 0.061118303 |
| 0.002559795 | 0.011264233 |
| 0.288581774 | 0.449324484 |
| 0.020880932 | 0.061917024 |
| 1.12E-05 | 0.000100043 |
| 5.42E-13 | 1.39E-11 |
| 0.163886999 | 0.301996227 |
| 0.081437601 | 0.180062962 |

| | | |
|---|---|---|
| | 0.36797257 | 0.53107465 |
| | 4.94E-06 | 4.83E-05 |
| | 0.007849441 | 0.028229926 |
| | 0.092019735 | 0.197195798 |
| | 3.58E-05 | 0.000281759 |
| | 0.839319108 | NA |
| NA | | NA |
| | 1.79E-05 | 0.000152236 |
| | 0.08518824 | 0.186486321 |
| | 0.326917216 | 0.48969689 |
| | 0.018111818 | 0.055403866 |
| | 0.941276431 | NA |
| | 1.61E-05 | 0.000138582 |
| NA | | NA |
| | 0.916240375 | 0.949593024 |
| | 0.832703428 | NA |
| | 0.925062568 | 0.955076744 |
| | 5.10E-06 | 4.96E-05 |
| | 0.000555716 | 0.003108613 |
| | 0.095723406 | 0.203048615 |
| | 0.312598326 | 0.474635631 |
| | 3.63E-05 | 0.000284874 |
| NA | | NA |
| | 0.406291266 | 0.568153735 |
| NA | | NA |
| | 0.001438608 | 0.006923118 |
| | 0.903623355 | 0.941652848 |
| | 0.085212387 | 0.186486321 |
| | 0.037671298 | 0.099297037 |
| NA | | NA |
| | 0.014584435 | 0.046645891 |
| | 0.485444827 | NA |
| | 0.839319108 | NA |
| NA | | NA |
| | 0.799237614 | NA |
| | 0.375963517 | 0.539174769 |
| | 0.012550677 | 0.041346882 |
| | 0.657823382 | 0.777100741 |
| | 0.989887656 | 0.993351574 |
| | 0.187492396 | 0.331552303 |
| | 1.21E-05 | 0.000107637 |
| | 0.421284939 | 0.582810801 |
| | 0.646519826 | 0.768731386 |
| | 0.001480226 | 0.007093283 |
| | 0.652977679 | 0.773665923 |
| | 0.001220765 | 0.006050947 |
| | 0.530742416 | 0.679507879 |

| | | |
|---|---|---|
| | 0.145963937 | 0.277694319 |
| | 0.395148235 | 0.557715742 |
| | 0.283517786 | 0.444089955 |
| | 0.165790446 | 0.304556933 |
| | 2.74E-39 | 2.75E-37 |
| | 0.690978364 | 0.801792603 |
| | 0.60153315 | 0.735145412 |
| | 0.626490139 | 0.753935878 |
| | 0.941275956 | NA |
| | 0.733928723 | 0.832172802 |
| | 0.015834051 | 0.049832354 |
| | 0.544498218 | 0.691042149 |
| | 0.198065384 | 0.345171494 |
| | 0.392691544 | 0.555137146 |
| | 0.001079555 | 0.005454374 |
| | 0.144173829 | 0.275146366 |
| NA | | NA |
| | 1.48E-21 | 6.94E-20 |
| | 4.45E-15 | 1.37E-13 |
| | 0.099120463 | 0.208176586 |
| | 0.460729232 | 0.619773776 |
| | 0.197080646 | 0.343807793 |
| | 0.136203357 | 0.263741645 |
| | 0.027818093 | 0.078210045 |
| | 0.941275956 | NA |
| | 5.00E-09 | 8.16E-08 |
| | 0.307265971 | 0.468851733 |
| | 0.610002831 | NA |
| | 0.000182832 | 0.001186092 |
| | 0.293372853 | 0.454583728 |
| | 0.841679341 | NA |
| | 0.000609314 | 0.003354823 |
| | 0.813997823 | 0.886443528 |
| | 0.107120629 | 0.220809319 |
| | 0.605705656 | 0.738386992 |
| | 0.168259227 | 0.307714647 |
| | 0.310041154 | 0.471781073 |
| | 0.955791446 | NA |
| | 0.027175244 | 0.076819193 |
| | 0.287495723 | 0.448102494 |
| | 0.163550609 | 0.301559501 |
| | 0.262053854 | 0.420824068 |
| | 0.646153219 | 0.768447107 |
| | 8.81E-50 | 1.19E-47 |
| | 0.56722114 | 0.708862626 |
| | 0.445935434 | 0.606402228 |
| | 0.297011559 | 0.458550245 |

| | | | |
|---|---|---|---|
| NA | | NA | |
| | 0.442989081 | NA | |
| | 0.500001755 | NA | |
| | 0.028829222 | | 0.080444684 |
| | 0.000416725 | | 0.002422195 |
| | 0.115957966 | | 0.234036788 |
| | 4.77E-10 | | 8.86E-09 |
| | 0.168816109 | | 0.308377668 |
| | 0.149781071 | | 0.282966033 |
| NA | | NA | |
| NA | | NA | |
| NA | | NA | |
| NA | | NA | |
| NA | | NA | |
| NA | | NA | |
| | 6.58E-166 | | 5.05E-163 |
| | 0.30776912 | NA | |
| NA | | NA | |
| | 0.839319108 | NA | |
| NA | | NA | |
| | 0.941985068 | NA | |
| NA | | NA | |
| | 0.940528311 | NA | |
| | 0.55247782 | | 0.697708932 |
| | 0.335245693 | NA | |
| | 0.750444857 | NA | |
| NA | | NA | |
| NA | | NA | |
| NA | | NA | |
| | 0.048952871 | | 0.121676717 |
| NA | | NA | |
| NA | | NA | |
| | 0.941278332 | NA | |
| | 0.839319108 | NA | |
| NA | | NA | |
| NA | | NA | |
| NA | | NA | |
| NA | | NA | |
| NA | | NA | |
| | 0.898019658 | NA | |
| NA | | NA | |
| NA | | NA | |
| NA | | NA | |
| NA | | NA | |
| NA | | NA | |
| NA | | NA | |
| | 0.941278332 | NA | |

| | | |
|---|---|---|
| | 0.839318641 | NA |
| NA | | NA |
| | 0.941278332 | NA |
| | 0.941278207 | NA |
| NA | | NA |
| NA | | NA |
| NA | | NA |
| NA | | NA |
| NA | | NA |
| NA | | NA |
| | 0.955792398 | NA |
| NA | | NA |
| NA | | NA |
| NA | | NA |
| NA | | NA |
| NA | | NA |
| NA | | NA |
| NA | | NA |
| NA | | NA |
| NA | | NA |
| | 0.941277857 | NA |
| NA | | NA |
| NA | | NA |
| NA | | NA |
| NA | | NA |
| NA | | NA |
| | 0.941275956 | NA |
| NA | | NA |
| NA | | NA |
| NA | | NA |
| NA | | NA |
| NA | | NA |
| NA | | NA |
| NA | | NA |
| NA | | NA |
| NA | | NA |
| NA | | NA |
| NA | | NA |
| | 0.955792398 | NA |
| | 0.839319108 | NA |
| NA | | NA |
| NA | | NA |
| NA | | NA |
| NA | | NA |
| NA | | NA |
| NA | | NA |
| NA | | NA |

| | | |
|---|---|---|
| NA | NA | |
| NA | NA | |
| NA | NA | |
| NA | NA | |
| NA | NA | |
| 0.941277857 | NA | |
| NA | NA | |
| NA | NA | |
| NA | NA | |
| NA | NA | |
| 0.839318641 | NA | |
| 1.08E-06 | | 1.19E-05 |
| 0.000133282 | | 0.000902865 |
| NA | NA | |
| NA | NA | |
| NA | NA | |
| NA | NA | |
| NA | NA | |
| NA | NA | |
| NA | NA | |
| 0.534414242 | | 0.682660906 |
| NA | NA | |
| 0.955792398 | NA | |
| NA | NA | |
| 0.955792398 | NA | |
| NA | NA | |
| NA | NA | |
| NA | NA | |
| 0.94127925 | NA | |
| 0.941277857 | NA | |
| NA | NA | |
| NA | NA | |
| NA | NA | |
| NA | NA | |
| NA | NA | |
| NA | NA | |
| NA | NA | |
| 0.839318641 | NA | |
| NA | NA | |
| NA | NA | |
| NA | NA | |
| 0.955792398 | NA | |
| NA | NA | |
| NA | NA | |
| 0.676090387 | NA | |
| NA | NA | |
| NA | NA | |

| | | |
|---|---|---|
| | 0.839319108 | NA |
| | 0.437979186 | NA |
| | 6.69E-10 | 1.21E-08 |
| | 7.00E-72 | 1.54E-69 |
| NA | | NA |
| | 0.068121462 | 0.157910697 |
| NA | | NA |
| | 0.193798885 | NA |
| | 6.82E-26 | 3.98E-24 |
| | 0.839319108 | NA |
| | 1.75E-35 | 1.52E-33 |
| NA | | NA |
| | 4.73E-12 | 1.11E-10 |
| | 2.77E-11 | 5.97E-10 |
| | 3.27E-130 | 1.50E-127 |
| | 0.519927564 | NA |
| | 2.15E-49 | 2.90E-47 |
| | 0.364875997 | 0.528174815 |
| NA | | NA |
| NA | | NA |
| | 0.839319108 | NA |
| | 0.941277857 | NA |
| NA | | NA |
| | 0.444417298 | NA |
| | 0.484602853 | NA |
| NA | | NA |
| NA | | NA |
| | 6.38E-05 | 0.000471642 |
| NA | | NA |
| NA | | NA |
| NA | | NA |
| NA | | NA |
| NA | | NA |
| NA | | NA |
| NA | | NA |
| | 0.664002098 | 0.782095108 |
| NA | | NA |
| NA | | NA |
| NA | | NA |
| | 0.839318641 | NA |
| | 0.941277857 | NA |
| | 0.839319108 | NA |
| NA | | NA |
| | 0.834624451 | 0.899919872 |
| NA | | NA |
| | 0.941277857 | NA |
| | 0.839319107 | NA |

| | |
|---|---|
| NA | NA |
| NA | NA |
| NA | NA |
| NA | NA |
| NA | NA |
| 0.942543826 | NA |
| NA | NA |
| 0.622240336 | NA |
| 0.655290135 | NA |
| 0.926938668 | NA |
| 0.695704196 | NA |
| 0.954541598 | NA |
| NA | NA |
| NA | NA |
| NA | NA |
| 0.74297871 | 0.83852368 |
| 0.564659404 | 0.706620985 |
| 0.941278332 | NA |
| 7.24E-11 | 1.48E-09 |
| 0.839319108 | NA |
| 0.283831881 | 0.444429079 |
| 0.000559637 | 0.00312687 |
| 0.017895497 | 0.054891982 |
| 0.330733993 | 0.493767202 |
| 0.073542188 | 0.167091558 |
| 4.95E-12 | 1.16E-10 |
| 0.647001003 | 0.769111415 |
| 0.000477634 | 0.002728913 |
| 0.973022148 | 0.983726508 |
| 0.957691639 | 0.974061037 |
| 0.839318641 | NA |
| 0.222368444 | 0.374792076 |
| 0.365665706 | 0.528835528 |
| 5.21E-06 | 5.05E-05 |
| 0.933677794 | 0.960596449 |
| 0.571044176 | 0.711473696 |
| 0.018285308 | 0.055841274 |
| 0.804581005 | 0.880558921 |
| 0.030034936 | 0.083088278 |
| 3.30E-15 | 1.02E-13 |
| 2.03E-19 | 8.28E-18 |
| 0.79139344 | NA |
| 1.72E-05 | 0.000146632 |
| NA | NA |
| 0.042078314 | 0.108357463 |
| 0.941278332 | NA |
| NA | NA |

| | | | |
|---|---|---|---|
| | 8.67E-07 | | 9.75E-06 |
| | 0.189285271 | | 0.333894494 |
| NA | | NA | |
| | 0.139454824 | NA | |
| | 1.36E-14 | | 3.97E-13 |
| NA | | NA | |
| | 2.06E-107 | | 7.66E-105 |
| | 1.82E-08 | | 2.74E-07 |
| | 0.955791446 | NA | |
| NA | | NA | |
| NA | | NA | |
| NA | | NA | |
| NA | | NA | |
| | 0.941278332 | NA | |
| NA | | NA | |
| | 0.941277857 | NA | |
| NA | | NA | |
| | 0.853595154 | NA | |
| NA | | NA | |
| | 0.853596091 | NA | |
| NA | | NA | |
| | 0.955792873 | NA | |
| NA | | NA | |
| NA | | NA | |
| | 0.955792398 | NA | |
| NA | | NA | |
| NA | | NA | |
| NA | | NA | |
| NA | | NA | |
| NA | | NA | |
| | 0.955792873 | NA | |
| NA | | NA | |
| | 0.941277857 | NA | |
| | 0.94127925 | NA | |
| NA | | NA | |
| | 0.378229425 | NA | |
| NA | | NA | |
| | 0.941278332 | NA | |
| | 0.839319108 | NA | |
| | 0.758673704 | | 0.849426197 |
| | 0.70270036 | | 0.810252828 |
| | 0.45530321 | | 0.614942235 |
| NA | | NA | |
| | 0.002762529 | | 0.011985731 |
| NA | | NA | |
| NA | | NA | |
| NA | | NA | |

| | | | |
|---|---|---|---|
| NA | | NA | |
| | 9.31E-06 | | 8.50E-05 |
| | 0.853595622 | NA | |
| | 0.941277857 | NA | |
| NA | | NA | |
| NA | | NA | |
| | 0.95126354 | NA | |
| NA | | NA | |
| NA | | NA | |
| NA | | NA | |
| NA | | NA | |
| NA | | NA | |
| | 0.839318641 | NA | |
| NA | | NA | |
| NA | | NA | |
| NA | | NA | |
| NA | | NA | |
| | 0.941278424 | NA | |
| NA | | NA | |
| NA | | NA | |
| NA | | NA | |
| NA | | NA | |
| NA | | NA | |
| NA | | NA | |
| NA | | NA | |
| | 0.941278332 | NA | |
| NA | | NA | |
| NA | | NA | |
| NA | | NA | |
| NA | | NA | |
| NA | | NA | |
| NA | | NA | |
| NA | | NA | |
| NA | | NA | |
| | 0.941278332 | NA | |
| NA | | NA | |
| NA | | NA | |
| NA | | NA | |
| NA | | NA | |
| NA | | NA | |
| NA | | NA | |
| NA | | NA | |
| | 0.010123248 | | 0.034739556 |
| | 1.40E-07 | | 1.84E-06 |
| | 3.90E-05 | | 0.00030406 |
| | 0.56117657 | | 0.704091105 |
| | 1.87E-07 | | 2.40E-06 |

| | |
|---|---|
| 0.600905588 | 0.734592824 |
| 0.005947917 | 0.022486074 |
| 0.204719377 | 0.353278652 |
| 0.241293144 | 0.397484907 |
| 0.274934292 | 0.434813608 |
| 0.39981578 | 0.56246913 |
| 0.001439001 | 0.006923611 |
| NA | NA |
| 0.65478665 | 0.774613909 |
| 0.11037716 | 0.225712391 |
| 0.236990749 | 0.392434521 |
| 0.032640944 | 0.088780681 |
| 0.157935594 | 0.293894547 |
| 0.610179047 | 0.741788737 |
| 0.577652721 | 0.716095145 |
| 2.97E-06 | 3.03E-05 |
| 0.687089507 | 0.799229101 |
| 0.537176052 | 0.685091592 |
| 0.000879533 | 0.004574809 |
| 0.375409559 | 0.538510113 |
| 0.955469555 | NA |
| 0.041428395 | 0.107066088 |
| NA | NA |
| 0.727052235 | 0.827925428 |
| 0.991497397 | 0.994187598 |
| 0.118481199 | 0.237962155 |
| 6.68E-05 | 0.000490089 |
| 0.085496803 | 0.186936882 |
| 0.241532831 | 0.397769629 |
| 0.112676987 | 0.229186262 |
| 0.341928276 | 0.50531335 |
| NA | NA |
| 0.941278332 | NA |
| NA | NA |
| 0.846870508 | NA |
| 6.77E-06 | 6.39E-05 |
| 1.62E-13 | 4.35E-12 |
| 0.884366222 | 0.930962684 |
| 0.865655697 | 0.919767844 |
| 0.610562847 | 0.742065813 |
| 0.384978051 | 0.54778415 |
| 0.839319108 | NA |
| 0.021386623 | 0.06311772 |
| 0.173523957 | 0.314180528 |
| NA | NA |
| 0.000248121 | 0.001542383 |
| NA | NA |

| | | |
|---|---|---|
| | 0.941278332 | NA |
| | 0.000795159 | 0.00420773 |
| | 0.002509693 | 0.011074476 |
| | 0.014357946 | 0.046077608 |
| | 0.000178921 | 0.001165165 |
| | 0.537589982 | 0.685435866 |
| | 0.193412489 | 0.339196944 |
| | 0.806677938 | 0.881648833 |
| | 0.919010487 | NA |
| | 7.74E-06 | 7.21E-05 |
| | 0.286457872 | 0.44706011 |
| | 0.971495633 | 0.982809378 |
| NA | | NA |
| NA | | NA |
| | 0.427293638 | 0.588567314 |
| | 0.352022184 | 0.51562315 |
| | 0.716012842 | NA |
| NA | | NA |
| | 0.004749845 | 0.01872402 |
| | 0.941274531 | NA |
| NA | | NA |
| | 0.932343358 | NA |
| | 0.941278332 | NA |
| | 1.14E-10 | 2.27E-09 |
| | 0.153199336 | 0.287414027 |
| | 0.002750427 | 0.011941932 |
| | 0.000898439 | 0.004662013 |
| | 0.017804396 | 0.054653113 |
| | 0.258426669 | 0.416739267 |
| | 0.104663231 | 0.216870744 |
| | 2.05E-06 | 2.16E-05 |
| | 0.000786584 | 0.004169487 |
| | 0.00862588 | 0.030489038 |
| | 0.004581241 | 0.018161674 |
| NA | | NA |
| | 0.941277857 | NA |
| NA | | NA |
| | 0.553935418 | NA |
| | 0.65389284 | 0.774066813 |
| | 0.374794731 | NA |
| | 0.320951659 | NA |
| | 0.656348809 | 0.775844972 |
| | 0.001969364 | 0.009047442 |
| | 0.020986475 | 0.062152554 |
| | 1.27E-05 | 0.000112011 |
| | 0.168422349 | 0.307871036 |
| | 0.609454312 | 0.741490928 |

| | |
|---|---|
| 0.001491082 | 0.007138121 |
| NA | NA |
| NA | NA |
| 0.010870535 | 0.036852369 |
| NA | NA |
| 0.042008831 | 0.108225371 |
| NA | NA |
| 8.53E-05 | 0.000609322 |
| 1.48E-09 | 2.56E-08 |
| 0.750763007 | 0.843499437 |
| NA | NA |
| 0.424168033 | NA |
| 0.839319108 | NA |
| 0.799186804 | 0.877382689 |
| 1.89E-22 | 9.35E-21 |
| 0.401508134 | 0.563850174 |
| 4.01E-14 | 1.14E-12 |
| 0.775983087 | 0.861913735 |
| 0.850362178 | 0.910250071 |
| 7.87E-16 | 2.56E-14 |
| 0.836176408 | 0.900614094 |
| 0.064791949 | 0.151983638 |
| 0.931683819 | 0.959332854 |
| 0.029954898 | 0.082915045 |
| 2.58E-08 | 3.80E-07 |
| 0.005757856 | 0.021891946 |
| NA | NA |
| 0.886969051 | NA |
| NA | NA |
| 0.853595622 | NA |
| 0.94020055 | NA |
| 0.950446479 | NA |
| 0.299986579 | 0.461526763 |
| 0.839318641 | NA |
| 0.941277857 | NA |
| 0.303834011 | 0.465250716 |
| 0.000227519 | 0.001432271 |
| 0.791124991 | 0.871710058 |
| 0.275638944 | NA |
| 0.4760827 | 0.633163124 |
| 0.831521299 | 0.897834783 |
| 0.025058357 | 0.071777667 |
| 0.436493831 | 0.59750553 |
| 0.000123292 | 0.000843036 |
| 0.909461135 | NA |
| 0.000177356 | 0.001157827 |
| 0.055606303 | 0.13485042 |

| | |
|---|---|
| 3.16E-05 | 0.000252887 |
| 5.81E-16 | 1.91E-14 |
| 0.574524858 | NA |
| 0.844050799 | NA |
| 0.89581282 | NA |
| 0.263965168 | 0.423005381 |
| 0.154665672 | 0.289475803 |
| 0.601647754 | 0.735209972 |
| 0.461652209 | 0.62041928 |
| 0.336936883 | 0.500079298 |
| 0.085886264 | 0.187633304 |
| 0.047191075 | 0.118248443 |
| 0.941278332 | NA |
| 0.195298022 | 0.341618485 |
| 0.088078401 | 0.191159667 |
| 0.286610694 | 0.447176539 |
| 0.946080295 | 0.967272298 |
| NA | NA |
| 0.042799174 | 0.10976252 |
| 0.4643275 | 0.6228541 |
| NA | NA |
| 2.38E-24 | 1.29E-22 |
| 0.311697204 | 0.473695368 |
| 0.001181711 | 0.005883099 |
| 2.29E-06 | 2.40E-05 |
| 0.406499185 | 0.568277552 |
| 5.28E-09 | 8.59E-08 |
| 0.355893539 | 0.519582649 |
| 0.832265771 | 0.898271886 |
| 0.732619555 | 0.831242762 |
| 0.364580858 | 0.527948116 |
| 0.024948276 | 0.071544442 |
| 0.011046017 | 0.037340904 |
| 0.405907254 | 0.567920367 |
| 0.853596091 | NA |
| 0.862468543 | 0.91776092 |
| 0.659049448 | NA |
| 0.019370559 | 0.058396429 |
| 0.603816167 | 0.736872583 |
| 0.053191498 | 0.130253502 |
| NA | NA |
| 0.004601753 | 0.01823388 |
| 0.882172246 | 0.929502603 |
| 0.115241933 | 0.23291063 |
| NA | NA |
| 0.020446103 | 0.060918964 |
| 0.007443831 | 0.027033577 |

| | | |
|---|---|---|
| | 0.095922777 | 0.203338327 |
| | 0.617340265 | 0.746966198 |
| | 0.020150699 | 0.060231939 |
| | 0.032802638 | 0.089128811 |
| NA | | NA |
| | 0.298146661 | NA |
| | 0.344528608 | 0.508098786 |
| | 0.95182251 | NA |
| | 0.454566937 | 0.614366148 |
| | 0.455280785 | 0.614942235 |
| | 0.00068634 | 0.00371532 |
| | 0.940525674 | 0.964348759 |
| | 0.04151245 | 0.107271667 |
| NA | | NA |
| | 0.941277857 | NA |
| | 0.034886614 | 0.093394577 |
| NA | | NA |
| NA | | NA |
| | 0.970781505 | NA |
| NA | | NA |
| | 0.013223398 | 0.043156907 |
| | 0.000352319 | 0.002091769 |
| | 1.88E-35 | 1.64E-33 |
| | 9.95E-41 | 1.05E-38 |
| | 0.658498748 | 0.777652553 |
| | 0.014357165 | 0.046077608 |
| | 0.009123462 | 0.031944196 |
| | 0.000314603 | 0.001896704 |
| | 0.955792873 | NA |
| | 0.839319108 | NA |
| | 0.007738156 | 0.02789815 |
| | 0.443437398 | 0.604022945 |
| | 1.73E-18 | 6.67E-17 |
| NA | | NA |
| | 0.326541175 | 0.489405996 |
| | 0.048995614 | 0.121757528 |
| | 0.6405662 | NA |
| | 0.400787141 | 0.563208038 |
| | 0.553904228 | 0.698931152 |
| | 0.176771537 | 0.318375199 |
| | 0.003571832 | 0.014798391 |
| | 0.58427531 | 0.721417285 |
| | 0.003671972 | 0.015116766 |
| | 0.573657269 | 0.713236017 |
| | 0.019351692 | 0.058364036 |
| NA | | NA |
| | 0.079345625 | 0.176833762 |

| | |
|---|---|
| 0.767759895 | NA |
| 8.62E-12 | 1.96E-10 |
| 3.73E-10 | 7.01E-09 |
| 3.05E-15 | 9.45E-14 |
| 0.044240181 | 0.112524555 |
| 0.072097567 | 0.164626467 |
| 7.48E-07 | 8.52E-06 |
| 0.272341909 | 0.432209769 |
| 0.932664666 | 0.959844534 |
| 0.690655405 | 0.801588646 |
| 7.11E-07 | 8.15E-06 |
| 0.316272438 | 0.478699593 |
| 0.027840115 | 0.078250391 |
| 0.707523489 | 0.813437541 |
| 0.182072537 | 0.32448786 |
| 0.047885855 | 0.119674415 |
| 0.000180865 | 0.001176861 |
| 0.481919569 | 0.638221925 |
| 0.186604655 | 0.330375187 |
| 4.93E-06 | 4.81E-05 |
| 0.036808192 | 0.097475366 |
| 1.99E-06 | 2.10E-05 |
| 0.213649708 | 0.364484687 |
| 0.297233944 | 0.458715058 |
| 0.070238768 | 0.161606878 |
| 0.000619636 | 0.003401424 |
| 0.941279303 | NA |
| 0.853596091 | NA |
| 0.621925512 | 0.750358426 |
| 0.085703538 | 0.187285679 |
| 0.000452054 | 0.00260087 |
| 0.326780845 | 0.48963321 |
| 0.674994127 | 0.790543237 |
| 0.000480158 | 0.00274136 |
| 4.26E-27 | 2.63E-25 |
| 0.600334566 | NA |
| 0.172388254 | 0.31262269 |
| 0.419044237 | 0.580742974 |
| 0.149637815 | 0.282740278 |
| 0.003637907 | 0.014999041 |
| NA | NA |
| NA | NA |
| 0.941278332 | NA |
| NA | NA |
| 4.90E-07 | 5.77E-06 |
| 0.053416484 | 0.130618623 |
| 1.09E-33 | 8.85E-32 |

| | | | |
|---|---|---|---|
| NA | | NA | |
| | 0.000345616 | | 0.002058131 |
| | 0.70623323 | | 0.812504425 |
| | 0.609759839 | | 0.741537274 |
| | 1.02E-20 | | 4.57E-19 |
| | 0.544212839 | | 0.690882394 |
| | 0.996167178 | | 0.996963006 |
| | 0.501832748 | | 0.654102529 |
| | 0.003925724 | | 0.015964326 |
| | 0.839319107 | NA | |
| | 4.84E-53 | | 7.24E-51 |
| | 0.040358482 | | 0.10488067 |
| | 0.361196751 | | 0.524891938 |
| | 0.041364358 | | 0.106953055 |
| | 0.490507636 | | 0.644888575 |
| NA | | NA | |
| | 0.90894384 | | 0.944704653 |
| | 0.010431536 | | 0.035617763 |
| | 0.739885032 | | 0.836336778 |
| | 2.22E-06 | | 2.33E-05 |
| | 0.162894868 | | 0.300583273 |
| | 0.013643359 | | 0.044212286 |
| | 0.670927312 | NA | |
| NA | | NA | |
| | 0.937798132 | | 0.963153577 |
| | 0.283155009 | | 0.443661083 |
| | 0.913091329 | | 0.947567135 |
| | 0.938309692 | NA | |
| NA | | NA | |
| | 0.131511051 | | 0.256955019 |
| | 0.12507645 | | 0.247800039 |
| | 0.085226972 | | 0.186501092 |
| | 0.159052865 | | 0.295371482 |
| | 0.000203654 | | 0.001301306 |
| | 0.243632841 | | 0.400175894 |
| | 0.43534725 | | 0.596484875 |
| | 0.470004275 | | 0.627921068 |
| NA | | NA | |
| | 1.38E-10 | | 2.73E-09 |
| | 0.00675296 | | 0.024942051 |
| | 0.353505782 | | 0.517143784 |
| | 0.005678049 | | 0.021644607 |
| | 3.83E-05 | | 0.000298886 |
| | 0.263780991 | | 0.422824106 |
| | 0.101848049 | | 0.212648006 |
| | 0.301969372 | | 0.463289609 |
| | 0.003757206 | | 0.01540817 |

| | | |
|---|---|---|
| | 0.941278332 | NA |
| | 0.034503108 | 0.092586582 |
| | 0.193727221 | 0.339625493 |
| | 0.43163305 | 0.592931833 |
| | 0.481546488 | 0.63800554 |
| | 0.024659671 | 0.070921882 |
| | 0.7295792 | 0.82965275 |
| | 0.748393366 | NA |
| | 0.022885691 | 0.066780093 |
| | 1.21E-20 | 5.37E-19 |
| NA | | NA |
| | 2.34E-08 | 3.47E-07 |
| NA | | NA |
| | 3.54E-31 | 2.60E-29 |
| | 2.87E-34 | 2.39E-32 |
| | 1.27E-11 | 2.83E-10 |
| | 2.86E-05 | 0.000231722 |
| | 0.458714748 | 0.618146885 |
| | 1.65E-14 | 4.78E-13 |
| | 0.135467776 | 0.262659174 |
| | 0.52977214 | 0.678813333 |
| | 0.066474091 | 0.155028054 |
| NA | | NA |
| | 0.525445058 | 0.674768598 |
| | 0.708831757 | 0.814125883 |
| | 0.021488213 | 0.063378245 |
| NA | | NA |
| | 1.67E-05 | 0.00014315 |
| NA | | NA |
| | 0.054715537 | 0.133081873 |
| NA | | NA |
| | 0.949265502 | 0.968927376 |
| | 0.071000276 | 0.162839522 |
| NA | | NA |
| | 0.778522974 | 0.863592382 |
| NA | | NA |
| | 0.955792398 | NA |
| | 0.090334565 | 0.194825759 |
| | 0.773922576 | 0.860508485 |
| NA | | NA |
| | 0.003366308 | 0.014111457 |
| | 0.348929622 | 0.512921373 |
| | 0.765803506 | 0.854586046 |
| | 0.642951137 | NA |
| | 0.464819834 | NA |
| | 8.93E-05 | 0.000634804 |
| | 0.042916392 | 0.109992027 |

| | |
|---|---|
| NA | NA |
| 0.012086854 | 0.040101938 |
| 0.333768453 | 0.497019314 |
| 0.507486678 | 0.659093027 |
| 0.268372844 | 0.428146864 |
| 6.64E-08 | 9.16E-07 |
| 0.613031856 | NA |
| 0.941277857 | NA |
| NA | NA |
| 0.122962937 | 0.244733157 |
| 0.930961841 | 0.958895693 |
| 0.002210952 | 0.009951794 |
| NA | NA |
| 0.378222004 | 0.541061866 |
| 2.96E-05 | 0.000238866 |
| 0.003598785 | 0.014876395 |
| 0.272258015 | 0.432127426 |
| 0.11080924 | 0.226354281 |
| 0.707876809 | 0.813678645 |
| 0.020986248 | 0.062152554 |
| 0.250226413 | 0.407770031 |
| 0.000102755 | 0.000717028 |
| 0.286086346 | 0.446729662 |
| 0.656823497 | NA |
| 0.929475979 | NA |
| 0.855132678 | 0.912852312 |
| 0.123457411 | 0.245368511 |
| 0.184586879 | 0.327655821 |
| NA | NA |
| 0.05335388 | 0.130516861 |
| 0.228927502 | 0.382596012 |
| 2.26E-05 | 0.000187684 |
| 0.934170113 | 0.960895291 |
| 0.162738549 | 0.300341393 |
| 0.866999439 | 0.920524624 |
| 0.005693735 | 0.021693978 |
| 0.052028555 | 0.127958634 |
| 0.001154526 | 0.005768266 |
| 0.767474412 | NA |
| 0.941277857 | NA |
| 0.118424973 | 0.237889374 |
| 0.70816305 | 0.813813099 |
| 0.020614996 | 0.061327101 |
| 0.019980354 | 0.05979786 |
| 0.750126073 | 0.843051435 |
| 0.013902736 | 0.044887808 |
| 0.004172988 | 0.016814873 |

| | | |
|---|---|---|
| | 0.070164174 | 0.161497695 |
| | 0.942188863 | NA |
| | 0.804528531 | 0.880558921 |
| | 0.505987964 | 0.657720797 |
| | 0.053322504 | 0.13050717 |
| | 0.000196648 | 0.001261622 |
| NA | | NA |
| | 0.941281424 | NA |
| | 0.106546845 | 0.220007634 |
| | 0.955791922 | NA |
| | 0.734058779 | NA |
| NA | | NA |
| | 0.01371484 | 0.044383498 |
| | 0.511092553 | 0.662619148 |
| | 8.14E-05 | 0.000583807 |
| | 1.06E-24 | 5.93E-23 |
| NA | | NA |
| NA | | NA |
| | 0.439442356 | 0.60041129 |
| | 0.292274158 | 0.453408838 |
| | 0.798338933 | 0.876650672 |
| | 0.719404601 | 0.821417457 |
| NA | | NA |
| | 0.10447973 | 0.216641392 |
| | 0.041862184 | 0.107952732 |
| | 0.028866357 | 0.080536583 |
| | 0.007487475 | 0.027168213 |
| | 0.256437644 | NA |
| | 0.002585148 | 0.011350617 |
| | 0.35133775 | 0.515109595 |
| | 0.292470287 | 0.453539499 |
| NA | | NA |
| | 0.467594171 | 0.625860455 |
| | 4.30E-06 | 4.24E-05 |
| | 0.280186495 | 0.44015107 |
| | 1.21E-06 | 1.33E-05 |
| | 0.001863276 | 0.008619961 |
| | 0.996730645 | 0.997443045 |
| NA | | NA |
| NA | | NA |
| | 0.300572209 | 0.46205492 |
| | 0.387739293 | 0.550654573 |
| | 0.512771889 | 0.664140121 |
| | 0.151806571 | 0.28575774 |
| | 0.941278332 | NA |
| NA | | NA |
| NA | | NA |

| | |
|---|---|
| 0.252024233 | 0.409745956 |
| 1.09E-07 | 1.45E-06 |
| NA | NA |
| 0.002264357 | 0.010151861 |
| 0.048813921 | 0.121445485 |
| 0.061550681 | NA |
| 0.559345004 | 0.702728511 |
| NA | NA |
| 0.118472813 | 0.237962155 |
| 0.031500142 | 0.086229601 |
| 0.00094228 | 0.004850379 |
| 0.361676674 | 0.525332776 |
| 0.643541823 | 0.766323309 |
| NA | NA |
| 0.941277857 | NA |
| 0.955792398 | NA |
| 0.690457933 | NA |
| 0.847232876 | NA |
| 0.955792873 | NA |
| 0.922702563 | NA |
| 0.839318641 | NA |
| 0.839319574 | NA |
| 0.941279268 | NA |
| 0.955792398 | NA |
| 0.941275481 | NA |
| NA | NA |
| 0.864589717 | NA |
| NA | NA |
| 0.941275481 | NA |
| 0.499660115 | NA |
| 0.774487628 | NA |
| 0.755754584 | NA |
| 0.941278332 | NA |
| NA | NA |
| 0.290965325 | 0.452102601 |
| 0.542374365 | 0.689246911 |
| 1.52E-10 | 2.98E-09 |
| 0.009099291 | 0.031878322 |
| 0.039325747 | 0.102758966 |
| 0.147253605 | 0.279544608 |
| 0.13893358 | NA |
| 0.047974152 | 0.119832171 |
| 3.99E-06 | 3.96E-05 |
| 0.243386843 | 0.399927463 |
| 0.002000127 | 0.009169301 |
| 0.054589698 | 0.132843676 |
| 0.071870123 | 0.164233167 |

| | | |
|---|---|---|
| | 0.005297442 | 0.020455249 |
| | 1.03E-11 | 2.34E-10 |
| | 0.106481562 | NA |
| | 3.16E-20 | 1.37E-18 |
| | 2.59E-22 | 1.27E-20 |
| | 1.85E-108 | 6.99E-106 |
| NA | | NA |
| | 0.675748078 | NA |
| | 0.091954188 | 0.197108571 |
| | 0.007439508 | 0.027022001 |
| | 0.002798442 | 0.012090853 |
| | 0.378626672 | NA |
| NA | | NA |
| | 0.05341743 | 0.130618623 |
| | 3.33E-12 | 7.94E-11 |
| | 0.941277857 | NA |
| | 0.030474582 | 0.084001854 |
| | 0.840318812 | 0.903562901 |
| | 0.62907382 | 0.755803092 |
| | 0.600899828 | 0.734592824 |
| | 0.955792398 | NA |
| | 0.917015028 | 0.950147644 |
| | 9.32E-197 | 1.06E-193 |
| | 0.346846685 | NA |
| | 9.45E-15 | 2.78E-13 |
| | 0.000419255 | 0.002434524 |
| | 0.547826672 | 0.693804961 |
| | 0.005403145 | 0.02076294 |
| | 0.065136436 | 0.152578102 |
| | 0.007385597 | 0.026863077 |
| NA | | NA |
| | 0.859616793 | 0.916299154 |
| | 0.346783582 | 0.510649766 |
| | 0.461296423 | 0.620186255 |
| | 0.549134433 | 0.69480519 |
| | 2.99E-12 | 7.16E-11 |
| | 0.180497351 | 0.322531846 |
| | 9.13E-06 | 8.36E-05 |
| NA | | NA |
| | 4.47E-20 | 1.91E-18 |
| | 0.560577225 | 0.703718667 |
| | 0.278057994 | 0.43772066 |
| | 7.88E-06 | 7.32E-05 |
| | 0.046477602 | 0.116866668 |
| | 2.63E-05 | 0.000214843 |
| | 0.001354593 | 0.006589416 |
| | 0.367483476 | 0.530690553 |

| | | |
|---|---|---|
| | 0.481978238 | NA |
| | 0.427965993 | 0.589220648 |
| | 0.717972814 | NA |
| | 0.336920209 | 0.500079298 |
| | 0.616089936 | 0.745981842 |
| | 0.692058806 | 0.802420415 |
| | 0.099236043 | 0.208327421 |
| | 0.204328012 | 0.35283048 |
| | 0.082059083 | 0.181117423 |
| | 0.074008962 | NA |
| | 0.519212824 | 0.669754153 |
| | 0.559011113 | NA |
| | 8.05E-05 | 0.000579026 |
| | 0.483086474 | NA |
| NA | | NA |
| NA | | NA |
| NA | | NA |
| | 0.955791922 | NA |
| NA | | NA |
| NA | | NA |
| | 0.941277857 | NA |
| NA | | NA |
| NA | | NA |
| | 0.839319108 | NA |
| NA | | NA |
| | 0.955792398 | NA |
| | 0.263096576 | 0.422019672 |
| | 7.88E-05 | 0.000567744 |
| | 0.839319108 | NA |
| | 0.888756643 | NA |
| | 0.01712232 | 0.053003701 |
| | 2.90E-40 | 3.00E-38 |
| | 0.17260668 | NA |
| | 0.46125178 | 0.620186255 |
| | 0.257210849 | 0.41534777 |
| NA | | NA |
| | 5.23E-16 | 1.72E-14 |
| | 0.778132207 | NA |
| NA | | NA |
| | 0.839320041 | NA |
| | 0.737359007 | 0.834432515 |
| NA | | NA |
| | 0.006699458 | 0.024768638 |
| | 0.612521484 | NA |
| | 0.839319107 | NA |
| NA | | NA |
| | 0.071291823 | 0.163287912 |

| | | |
|---|---|---|
| | 0.284798697 | 0.445415507 |
| | 0.044404935 | 0.112835086 |
| | 0.17880717 | 0.320455408 |
| | 0.938802928 | 0.963601145 |
| | 0.001685511 | 0.007926997 |
| | 0.951145227 | NA |
| | 2.52E-07 | 3.15E-06 |
| | 0.670179572 | 0.786646324 |
| NA | | NA |
| | 0.941278332 | NA |
| NA | | NA |
| | 0.022285284 | 0.065300299 |
| | 0.955791922 | NA |
| | 0.508049739 | NA |
| NA | | NA |
| | 0.839318641 | NA |
| | 0.086781352 | 0.189033563 |
| | 0.060733251 | NA |
| | 0.941277857 | NA |
| NA | | NA |
| | 0.833801616 | NA |
| | 0.069782428 | 0.160883508 |
| | 0.030984433 | 0.085089299 |
| | 0.941198385 | NA |
| | 0.654357678 | 0.77439894 |
| | 0.106890757 | 0.220490135 |
| | 0.000310741 | 0.001877701 |
| | 0.134149584 | NA |
| NA | | NA |
| | 0.839318641 | NA |
| | 3.87E-06 | 3.86E-05 |
| | 0.505262121 | 0.65720799 |
| | 0.705423589 | 0.812065262 |
| | 0.505255595 | 0.65720799 |
| | 0.839318641 | NA |
| | 0.33556882 | 0.498733003 |
| | 0.941278424 | NA |
| | 0.008079432 | 0.028887885 |
| | 0.000270538 | 0.001662623 |
| | 0.941278332 | NA |
| NA | | NA |
| | 0.316959297 | 0.479396656 |
| | 0.941276431 | NA |
| | 0.853596091 | NA |
| | 0.112038755 | 0.228122022 |
| | 0.621657796 | 0.75018753 |
| | 0.377617707 | 0.540537446 |

| | | |
|---|---|---|
| | 0.941278332 | NA |
| | 0.030910714 | 0.084928642 |
| | 0.00437296 | 0.017475688 |
| NA | | NA |
| | 0.083181845 | 0.182967053 |
| NA | | NA |
| | 0.275767642 | 0.435588809 |
| | 0.170240014 | 0.309932141 |
| | 0.522104906 | 0.672173354 |
| | 0.000154965 | 0.001030298 |
| | 0.972295854 | 0.983304088 |
| | 0.089005305 | 0.192633286 |
| | 0.917322758 | NA |
| NA | | NA |
| NA | | NA |
| | 0.250073709 | 0.407663684 |
| NA | | NA |
| | 6.64E-05 | 0.000488075 |
| | 0.012496067 | 0.041206896 |
| | 0.549071892 | 0.69480519 |
| | 0.005590845 | 0.021370368 |
| | 3.37E-07 | 4.09E-06 |
| | 0.028525314 | 0.079777422 |
| | 0.043034445 | 0.110246538 |
| | 0.186029647 | NA |
| | 0.003012315 | 0.012874684 |
| | 0.00227257 | 0.010184847 |
| | 0.005449856 | 0.020898576 |
| | 0.941278332 | NA |
| | 0.439731174 | 0.600555509 |
| | 0.970881801 | 0.982438935 |
| | 0.44964055 | 0.609646238 |
| | 0.390404241 | 0.553029754 |
| | 3.67E-08 | 5.26E-07 |
| | 0.205454131 | 0.354107112 |
| | 5.46E-05 | 0.00040983 |
| | 0.888243689 | 0.933255773 |
| | 0.879049789 | NA |
| | 0.620372133 | NA |
| NA | | NA |
| | 0.638951193 | 0.762946033 |
| NA | | NA |
| | 0.322587652 | 0.485411006 |
| | 0.941277857 | NA |
| NA | | NA |
| | 0.839318641 | NA |
| | 0.118187586 | NA |

| | |
|---|---|
| 0.799305568 | NA |
| 0.61430391 | 0.744430877 |
| 3.69E-13 | 9.64E-12 |
| NA | NA |
| 0.145157751 | NA |
| NA | NA |
| 0.304261639 | NA |
| 0.191325153 | 0.336405504 |
| 7.31E-05 | 0.00053021 |
| 3.23E-05 | 0.000256781 |
| 0.820697573 | 0.890678686 |
| 0.961088062 | 0.976222338 |
| 0.266908062 | 0.426544813 |
| 0.006793025 | 0.025056189 |
| 0.007077594 | 0.025913074 |
| 0.293970433 | 0.455166979 |
| NA | NA |
| 0.122166152 | 0.243636258 |
| 0.161049922 | 0.298071675 |
| 0.332992754 | 0.496298906 |
| 0.941278332 | NA |
| 0.693146132 | 0.803382559 |
| 0.516622565 | 0.667442775 |
| 0.419036646 | 0.580742974 |
| 0.046779002 | 0.117537612 |
| 0.599063081 | NA |
| 0.993484923 | NA |
| NA | NA |
| NA | NA |
| NA | NA |
| 0.528059274 | 0.677165392 |
| NA | NA |
| NA | NA |
| 0.941277857 | NA |
| 0.570254958 | NA |
| NA | NA |
| NA | NA |
| NA | NA |
| NA | NA |
| 0.008301282 | 0.029565632 |
| 0.835025174 | NA |
| 0.306230083 | 0.46792633 |
| NA | NA |
| 0.433560237 | 0.59475539 |
| 0.336475532 | 0.499706159 |
| NA | NA |
| 0.839318641 | NA |

| | |
|---|---|
| NA | NA |
| 7.66E-06 | 7.14E-05 |
| 0.003118958 | 0.013240136 |
| 1.60E-05 | 0.000137721 |
| 0.338883364 | 0.50240435 |
| NA | NA |
| 5.46E-06 | 5.28E-05 |
| 0.002936915 | 0.012599693 |
| 0.777658416 | 0.863096307 |
| 0.642173668 | 0.765284664 |
| 6.73E-21 | 3.05E-19 |
| 3.10E-07 | 3.81E-06 |
| 0.614099514 | 0.744311876 |
| 1.68E-24 | 9.21E-23 |
| 0.003406208 | 0.014237155 |
| 0.520666018 | 0.670937825 |
| 0.037195153 | 0.098264633 |
| 0.043450673 | 0.110990869 |
| 0.000899179 | 0.004664832 |
| NA | NA |
| 0.494031933 | NA |
| NA | NA |
| 0.955792873 | NA |
| 1.39E-06 | 1.52E-05 |
| NA | NA |
| 9.57E-05 | 0.000674721 |
| 0.014438288 | NA |
| 0.992452696 | NA |
| 0.565290272 | NA |
| 0.000485584 | 0.002768887 |
| 0.00581858 | 0.022081287 |
| NA | NA |
| NA | NA |
| NA | NA |
| 0.947639761 | 0.968098269 |
| NA | NA |
| 0.554849982 | 0.699587007 |
| 0.701014614 | NA |
| 0.498309481 | 0.65133236 |
| 0.920932059 | NA |
| 0.674537207 | NA |
| 2.20E-05 | 0.000183551 |
| 0.955792398 | NA |
| 0.012597389 | 0.041477806 |
| NA | NA |
| NA | NA |
| 0.567355206 | 0.708955723 |

| | |
|---|---|
| 1.85E-06 | 1.97E-05 |
| 7.18E-12 | 1.65E-10 |
| 0.897685407 | NA |
| 0.955792873 | NA |
| 0.745981294 | 0.840278884 |
| NA | NA |
| 0.299181698 | 0.460749833 |
| 0.793764021 | NA |
| 0.003942431 | 0.016011753 |
| 0.956914524 | 0.973526307 |
| 0.175086948 | NA |
| 0.136200776 | 0.263741645 |
| 0.54739876 | 0.69355825 |
| 1.27E-05 | 0.00011261 |
| 0.019605496 | 0.058950229 |
| 1.84E-05 | 0.000155753 |
| 0.103794588 | 0.215675475 |
| NA | NA |
| 0.003806396 | 0.015569659 |
| 0.408768259 | 0.570842394 |
| 0.028229991 | 0.079059919 |
| 0.156874072 | 0.292626697 |
| 0.955670492 | 0.972955303 |
| 0.007072494 | 0.025902371 |
| 0.777064728 | 0.862672326 |
| 0.170588388 | 0.310376457 |
| 0.839320041 | NA |
| 0.178237415 | 0.319795488 |
| NA | NA |
| 0.853595622 | NA |
| 0.023041069 | 0.067159383 |
| 0.780065598 | NA |
| 0.00235377 | 0.01049933 |
| 0.02901613 | 0.080786282 |
| 0.968837231 | 0.981329578 |
| 0.420496157 | 0.582067434 |
| 0.15705188 | 0.292827238 |
| 0.024374853 | 0.070281073 |
| NA | NA |
| 0.009886253 | 0.034069277 |
| NA | NA |
| 0.581103892 | 0.718852182 |
| 0.020768916 | 0.06163862 |
| 0.624470027 | NA |
| 0.327741884 | 0.49050002 |
| 0.403296787 | NA |
| 0.101624909 | 0.212298853 |

| | |
|---|---|
| NA | NA |
| 0.151309039 | 0.285172704 |
| NA | NA |
| NA | NA |
| NA | NA |
| 0.715601778 | 0.819182883 |
| NA | NA |
| 0.391813946 | 0.554183941 |
| 0.941853951 | 0.964837862 |
| NA | NA |
| NA | NA |
| NA | NA |
| NA | NA |
| NA | NA |
| 0.020887787 | 0.061921922 |
| NA | NA |
| NA | NA |
| 0.654051421 | NA |
| 1.46E-06 | 1.58E-05 |
| 3.20E-05 | 0.000254706 |
| 0.866230384 | 0.920118511 |
| 0.941277857 | NA |
| NA | NA |
| NA | NA |
| NA | NA |
| 0.839319574 | NA |
| NA | NA |
| NA | NA |
| NA | NA |
| 0.750625527 | NA |
| 0.955791446 | NA |
| NA | NA |
| NA | NA |
| NA | NA |
| 0.683967488 | NA |
| NA | NA |
| NA | NA |
| NA | NA |
| NA | NA |
| NA | NA |
| NA | NA |
| NA | NA |
| NA | NA |
| 0.038856491 | 0.101733825 |
| NA | NA |
| NA | NA |
| NA | NA |

| | |
|---|---|
| NA | NA |
| 0.813076362 | 0.885916391 |
| 0.197053468 | 0.343802457 |
| 0.424656308 | 0.586019796 |
| 0.131233958 | 0.256544242 |
| 0.891068084 | 0.934449599 |
| 0.448255668 | 0.608427316 |
| 0.785603324 | 0.868223537 |
| 0.005851501 | 0.022192071 |
| NA | NA |
| 0.040135626 | 0.104462349 |
| 0.620605244 | 0.749411294 |
| 0.204113729 | 0.352588358 |
| 0.367899619 | NA |
| 0.567047348 | 0.70872057 |
| 0.808078659 | 0.882288452 |
| 0.955792398 | NA |
| 0.955792398 | NA |
| 0.396479807 | 0.559097835 |
| 0.300940774 | NA |
| 1.23E-22 | 6.13E-21 |
| 0.021350778 | 0.063043197 |
| 0.004399336 | 0.017554568 |
| NA | NA |
| NA | NA |
| NA | NA |
| NA | NA |
| NA | NA |
| NA | NA |
| NA | NA |
| NA | NA |
| NA | NA |
| 0.839320041 | NA |
| NA | NA |
| 0.839319108 | NA |
| NA | NA |
| NA | NA |
| NA | NA |
| NA | NA |
| NA | NA |
| NA | NA |
| NA | NA |
| NA | NA |
| NA | NA |
| NA | NA |
| NA | NA |
| NA | NA |

| | | |
|---|---|---|
| NA | NA | |
| NA | NA | |
| NA | NA | |
| NA | NA | |
| NA | NA | |
| NA | NA | |
| NA | NA | |
| NA | NA | |
| NA | NA | |
| NA | NA | |
| NA | NA | |
| NA | NA | |
| NA | NA | |
| NA | NA | |
| NA | NA | |
| NA | NA | |
| NA | NA | |
| NA | NA | |
| NA | NA | |
| NA | NA | |
| NA | NA | |
| NA | NA | |
| NA | NA | |
| NA | NA | |
| NA | NA | |
| NA | NA | |
| NA | NA | |
| NA | NA | |
| NA | NA | |
| NA | NA | |
| NA | NA | |
| NA | NA | |
| 0.839319107 | NA | |
| NA | NA | |
| 0.955792397 | NA | |
| NA | NA | |
| 0.941278332 | NA | |
| NA | NA | |
| 0.62609235 | | 0.753666622 |
| 0.941278332 | NA | |
| NA | NA | |
| 0.839318641 | NA | |
| 0.486167889 | NA | |
| 0.77998795 | NA | |
| 0.853595622 | NA | |
| NA | NA | |
| NA | NA | |

| | |
|---|---|
| NA | NA |
| NA | NA |
| NA | NA |
| 0.721600899 | 0.82332244 |
| 0.01982146 | 0.059426929 |
| 0.315311296 | 0.477723709 |
| 0.165543387 | 0.304243974 |
| 0.025977083 | 0.073915904 |
| 0.048283901 | 0.120454189 |
| 0.476316172 | 0.633402091 |
| NA | NA |
| 0.814558423 | NA |
| 0.002088265 | 0.009480239 |
| 0.822900215 | 0.892249951 |
| 0.185044128 | 0.328175344 |
| 0.215232164 | 0.366186989 |
| 1.75E-05 | 0.000148912 |
| 0.16189669 | 0.299042761 |
| 0.218405514 | 0.369709696 |
| 0.839320041 | NA |
| 0.497378358 | 0.65032958 |
| 0.904733585 | NA |
| 0.137744422 | 0.266034729 |
| NA | NA |
| 8.29E-16 | 2.68E-14 |
| 0.000635306 | 0.003472229 |
| 0.026849456 | 0.076070796 |
| 0.886783365 | NA |
| 0.229341357 | 0.383099374 |
| 1.09E-05 | 9.77E-05 |
| NA | NA |
| 0.284635348 | 0.44526878 |
| 0.29234001 | 0.453426099 |
| 0.398724572 | 0.56155308 |
| NA | NA |
| 0.263083401 | 0.422019672 |
| 0.457701163 | 0.617480196 |
| 0.170040295 | 0.309686976 |
| 0.700843313 | 0.808877387 |
| 0.54312029 | 0.689900157 |
| NA | NA |
| 2.61E-12 | 6.26E-11 |
| 0.505757877 | 0.65752944 |
| 0.913945927 | 0.948205952 |
| 0.143577944 | NA |
| 0.00164724 | 0.007777744 |
| 0.018258081 | 0.05579392 |

| | |
|---|---|
| 2.72E-06 | 2.79E-05 |
| 0.284884422 | 0.445432504 |
| 0.479493007 | 0.636170153 |
| NA | NA |
| 0.152060581 | 0.286069075 |
| 0.241794317 | 0.398079331 |
| NA | NA |
| 1.62E-05 | 0.000139771 |
| 0.544957839 | 0.69149624 |
| 4.64E-05 | 0.000354337 |
| 0.608708991 | 0.740909814 |
| 2.63E-31 | 1.94E-29 |
| 0.004148253 | 0.016729365 |
| NA | NA |
| 0.358219769 | 0.521820657 |
| 0.013711205 | 0.044377768 |
| 0.839319574 | NA |
| 0.021689316 | 0.06388431 |
| 0.402890337 | 0.564957924 |
| 0.004548023 | 0.018048022 |
| NA | NA |
| 0.568457954 | 0.70973665 |
| 0.228850446 | 0.38252095 |
| 0.001333037 | 0.006499733 |
| 0.046272301 | 0.116535109 |
| 0.000161958 | 0.00106992 |
| 0.22634346 | 0.379369553 |
| 0.287097107 | 0.447686409 |
| 0.90081492 | 0.940155956 |
| 0.027531653 | 0.077610433 |
| 0.357467731 | 0.521048832 |
| 0.002122559 | 0.009602958 |
| 0.017582875 | 0.054088279 |
| 0.010373142 | 0.035464165 |
| 1.79E-12 | 4.37E-11 |
| 0.941278332 | NA |
| 0.008106899 | 0.028973034 |
| 0.504775419 | 0.656898006 |
| NA | NA |
| NA | NA |
| 0.955792398 | NA |
| 8.09E-05 | 0.000580946 |
| 0.007803678 | 0.028091824 |
| 0.049367884 | 0.122464867 |
| 0.000451107 | 0.002596675 |
| 0.578485181 | 0.716655613 |
| 0.080290231 | 0.178337523 |

| | |
|---|---|
| 0.000921952 | 0.00476015 |
| 0.838882104 | NA |
| 0.808063896 | 0.882288452 |
| 0.115084844 | 0.232711703 |
| 2.76E-18 | 1.04E-16 |
| 0.641756927 | NA |
| 0.027316646 | 0.07713736 |
| 0.741815882 | 0.837564709 |
| NA | NA |
| NA | NA |
| 0.769641684 | 0.857314759 |
| NA | NA |
| NA | NA |
| 0.265064992 | 0.424196661 |
| 0.027632048 | 0.077821687 |
| 0.321060013 | 0.483835722 |
| NA | NA |
| 0.008009468 | 0.028685129 |
| 0.22090587 | 0.372828583 |
| 0.217775724 | 0.369117494 |
| 0.002704799 | 0.011786821 |
| 0.01619352 | 0.050702205 |
| 2.14E-19 | 8.72E-18 |
| NA | NA |
| 0.908830425 | NA |
| 0.953798549 | NA |
| 0.120919107 | 0.24171635 |
| 0.000191142 | 0.001232273 |
| 0.039628414 | 0.103424947 |
| NA | NA |
| 0.483948861 | 0.639799533 |
| 0.749657101 | 0.84281796 |
| 0.021270597 | 0.062868839 |
| 0.27701592 | 0.436889274 |
| 2.84E-09 | 4.77E-08 |
| 0.013333547 | 0.043450859 |
| 0.061866086 | 0.146491899 |
| 0.492218286 | 0.646173936 |
| 5.13E-05 | 0.000388157 |
| 9.60E-09 | 1.51E-07 |
| 0.325339754 | 0.488190169 |
| 0.925451256 | NA |
| 0.0057858 | 0.021979187 |
| 0.091185 | 0.196090073 |
| 0.000404417 | 0.002356986 |
| NA | NA |
| 0.60991817 | NA |

| | | |
|---|---|---|
| | 0.92069054 | NA |
| | 0.509822191 | 0.66146442 |
| | 0.554121083 | NA |
| | 0.057247167 | 0.137871081 |
| | 0.802275397 | 0.879097643 |
| | 0.001623607 | 0.007682919 |
| | 0.005101411 | 0.019837247 |
| NA | | NA |
| | 0.847499648 | NA |
| | 0.955792873 | NA |
| NA | | NA |
| NA | | NA |
| NA | | NA |
| | 0.839319574 | NA |
| | 0.078970625 | 0.176189127 |
| | 0.057925098 | 0.139041265 |
| | 0.138560009 | 0.267066591 |
| NA | | NA |
| | 0.122332568 | 0.243825137 |
| | 0.019754034 | 0.059299252 |
| | 0.346340166 | 0.510123058 |
| | 0.91631679 | 0.949630872 |
| | 0.674149292 | NA |
| NA | | NA |
| | 0.003323931 | 0.013970724 |
| NA | | NA |
| | 0.01891015 | 0.057374046 |
| | 5.36E-09 | 8.71E-08 |
| | 0.860964477 | 0.916857254 |
| | 0.901490533 | NA |
| | 0.839319574 | NA |
| | 0.913417426 | 0.947781577 |
| | 1.42E-06 | 1.54E-05 |
| | 0.320685859 | 0.483374379 |
| NA | | NA |
| | 0.178965554 | 0.320594424 |
| | 0.818477871 | 0.88927787 |
| NA | | NA |
| | 0.84377218 | 0.906007373 |
| NA | | NA |
| | 0.58306992 | 0.720497884 |
| | 1.15E-45 | 1.41E-43 |
| | 0.568129078 | 0.709438556 |
| | 0.525132542 | 0.674517315 |
| | 0.680493478 | 0.794365168 |
| | 8.86E-06 | 8.13E-05 |
| | 0.304729557 | 0.466382004 |

| | | |
|---|---|---|
| | 0.142843159 | 0.27328614 |
| | 0.620113018 | 0.748968897 |
| | 0.839319574 | NA |
| NA | | NA |
| NA | | NA |
| NA | | NA |
| | 0.630444514 | NA |
| | 0.941279303 | NA |
| | 0.012135252 | 0.040240076 |
| | 5.54E-05 | 0.000415172 |
| | 0.001355849 | 0.006592831 |
| | 0.435891607 | NA |
| | 0.928473254 | NA |
| | 0.49163756 | 0.645589606 |
| | 0.839320041 | NA |
| | 0.839323853 | NA |
| | 0.747503901 | 0.841412716 |
| | 0.116215259 | 0.234479578 |
| | 0.347428051 | 0.511390296 |
| | 0.92069054 | NA |
| | 0.556034768 | 0.700287822 |
| | 0.539582249 | NA |
| NA | | NA |
| | 0.073104892 | 0.166272589 |
| NA | | NA |
| NA | | NA |
| | 0.787269278 | NA |
| | 0.839318641 | NA |
| | 0.84920996 | NA |
| NA | | NA |
| NA | | NA |
| NA | | NA |
| | 0.955792873 | NA |
| | 0.943971627 | NA |
| NA | | NA |
| NA | | NA |
| NA | | NA |
| NA | | NA |
| | 1.82E-09 | 3.12E-08 |
| | 0.739991096 | NA |
| | 0.648112089 | NA |
| | 0.64318532 | 0.765991043 |
| | 0.003597787 | 0.014874853 |
| | 0.149614885 | NA |
| | 6.67E-12 | 1.54E-10 |
| | 0.001552445 | 0.007390257 |
| | 1.60E-57 | 2.57E-55 |

| | |
|---|---|
| 0.002121274 | 0.009600407 |
| NA | NA |
| 0.708019264 | NA |
| 0.170229769 | 0.309932141 |
| 0.190250333 | 0.335010611 |
| NA | NA |
| 0.216345804 | 0.367608712 |
| 0.048921002 | 0.121622904 |
| 0.839319108 | NA |
| 0.186032177 | 0.329585962 |
| 0.082418327 | 0.181741803 |
| 0.024959918 | 0.071569204 |
| 2.64E-05 | 0.000215646 |
| 0.000614619 | 0.003380123 |
| NA | NA |
| 0.316226435 | 0.478683643 |
| 0.989734257 | 0.993351574 |
| 0.799582457 | 0.877620713 |
| 0.081912562 | 0.180861114 |
| NA | NA |
| 0.629327856 | NA |
| NA | NA |
| 0.736737775 | 0.834046731 |
| 0.06993978 | 0.161152627 |
| 0.046179842 | 0.116376135 |
| 0.761533716 | 0.851817685 |
| 0.001730336 | 0.008107373 |
| 4.59E-16 | 1.52E-14 |
| 2.60E-17 | 9.25E-16 |
| 0.753235186 | 0.845318877 |
| 0.681723979 | NA |
| 0.006801769 | 0.025074274 |
| 0.000422807 | 0.002452159 |
| 0.034377719 | 0.092322967 |
| NA | NA |
| 0.718133377 | 0.820753416 |
| 0.000402081 | 0.002346818 |
| NA | NA |
| 0.735814446 | 0.833507461 |
| 0.057315931 | 0.137982583 |
| 0.690073013 | 0.801248259 |
| 0.68933486 | 0.800836994 |
| 0.217267212 | 0.368604775 |
| 0.672347642 | NA |
| 0.955792873 | NA |
| 0.138158503 | 0.266638181 |
| 0.181535869 | 0.323947905 |

| | |
|---|---|
| NA | NA |
| 0.000425538 | 0.002464992 |
| 0.012933065 | 0.042348717 |
| 0.005059584 | 0.019710019 |
| 0.881967302 | NA |
| 0.064257138 | 0.150952271 |
| 0.955792873 | NA |
| 0.941277857 | NA |
| NA | NA |
| 4.10E-05 | 0.000317803 |
| NA | NA |
| 0.853881291 | 0.912048218 |
| NA | NA |
| NA | NA |
| NA | NA |
| 0.001735521 | 0.008125268 |
| 0.008129908 | 0.029046543 |
| 0.790448062 | 0.87131222 |
| 0.021970758 | 0.064545542 |
| 0.322824344 | 0.485552292 |
| 0.256713042 | 0.41484716 |
| 0.490202061 | NA |
| 0.025060456 | 0.071777667 |
| 0.760285554 | NA |
| 0.008616965 | 0.030462054 |
| 0.170792818 | 0.310677161 |
| 0.936413394 | NA |
| 0.226555764 | 0.379698656 |
| 1.95E-08 | 2.92E-07 |
| 0.839319108 | NA |
| 0.134550276 | 0.261325075 |
| 0.043779046 | 0.111614059 |
| 0.191294254 | 0.336392422 |
| 0.941276431 | NA |
| 0.941277857 | NA |
| 0.839318641 | NA |
| NA | NA |
| NA | NA |
| 0.282789212 | 0.443208695 |
| 1.50E-13 | 4.05E-12 |
| 0.096575608 | 0.204291914 |
| 4.08E-16 | 1.36E-14 |
| 0.079359499 | 0.176834121 |
| NA | NA |
| 0.942087105 | 0.964993643 |
| 0.000296712 | 0.001806687 |
| 0.323071345 | 0.48577032 |

| | |
|---|---|
| NA | NA |
| 0.013065328 | 0.042711295 |
| 0.002839599 | 0.012244229 |
| NA | NA |
| NA | NA |
| NA | NA |
| 0.522394736 | 0.672390001 |
| 1.01E-09 | 1.80E-08 |
| 0.941278332 | NA |
| 0.381546943 | 0.544044286 |
| 0.006181037 | 0.023197893 |
| 0.007406149 | 0.026921373 |
| 0.000271459 | 0.001667424 |
| NA | NA |
| 0.941279268 | NA |
| 0.000216748 | 0.001376109 |
| NA | NA |
| 0.42498367 | NA |
| 0.592865426 | 0.728329438 |
| 5.00E-06 | 4.88E-05 |
| 0.656823497 | NA |
| 5.28E-209 | 7.85E-206 |
| 0.775747366 | NA |
| NA | NA |
| 0.016043576 | 0.050325409 |
| NA | NA |
| NA | NA |
| NA | NA |
| 0.056643875 | 0.136793579 |
| 7.37E-05 | 0.000534048 |
| NA | NA |
| 0.325187774 | NA |
| 0.875029985 | 0.92525621 |
| 0.424189132 | 0.585578803 |
| 0.538078435 | 0.685801494 |
| NA | NA |
| 1.78E-10 | 3.47E-09 |
| NA | NA |
| NA | NA |
| 0.839319574 | NA |
| NA | NA |
| NA | NA |
| NA | NA |
| NA | NA |
| 5.68E-09 | 9.21E-08 |
| NA | NA |
| 0.955792398 | NA |

| | | |
|---|---|---|
| | 0.968265853 | 0.980959557 |
| | 0.873022841 | 0.924280679 |
| | 0.914679494 | NA |
| | 0.877760873 | NA |
| | 1.17E-10 | 2.32E-09 |
| | 0.104307651 | 0.216397691 |
| NA | | NA |
| | 3.22E-05 | 0.000256369 |
| | 9.55E-05 | 0.000673668 |
| | 0.014590693 | 0.046655842 |
| NA | | NA |
| NA | | NA |
| NA | | NA |
| NA | | NA |
| | 0.000622697 | 0.003412715 |
| | 4.82E-06 | 4.71E-05 |
| | 0.887125346 | 0.932657133 |
| | 0.601315166 | NA |
| NA | | NA |
| | 0.588722496 | 0.724945212 |
| | 0.317693687 | 0.480145751 |
| | 0.129934306 | 0.254605774 |
| | 0.792365721 | NA |
| | 0.053999139 | 0.13172107 |
| | 0.839320041 | NA |
| | 0.895849763 | 0.937069482 |
| | 0.02549863 | 0.072780716 |
| | 0.955792398 | NA |
| NA | | NA |
| NA | | NA |
| | 9.83E-06 | 8.93E-05 |
| | 0.614284695 | 0.744430877 |
| | 0.865348054 | 0.919767844 |
| | 0.093496925 | 0.199515816 |
| NA | | NA |
| | 0.01499863 | 0.047714167 |
| | 0.376505948 | 0.53956256 |
| | 0.078433279 | 0.175194981 |
| | 0.067818995 | 0.157316804 |
| | 0.002682824 | 0.011712506 |
| | 0.027456564 | 0.077431415 |
| | 0.107404344 | 0.221201858 |
| | 0.94020055 | NA |
| | 0.030259321 | 0.083553638 |
| | 0.240807854 | NA |
| | 0.135526665 | NA |
| | 0.247844189 | 0.405038411 |

| | |
|---|---|
| 0.000796271 | 0.004212681 |
| 0.001487111 | 0.007121977 |
| 0.306907072 | 0.468539679 |
| NA | NA |
| 0.017319555 | 0.053495985 |
| 6.54E-11 | 1.35E-09 |
| 0.004899731 | 0.019197266 |
| 0.255424159 | 0.413494241 |
| 0.765831458 | 0.854586046 |
| 0.452370959 | 0.612371811 |
| 0.993805722 | 0.995520551 |
| 0.96415365 | 0.978200646 |
| 0.400620726 | 0.563101117 |
| 3.66E-08 | 5.25E-07 |
| 0.030148307 | 0.083325234 |
| 3.42E-05 | 0.000270458 |
| 2.97E-05 | 0.000239054 |
| 0.003559352 | 0.014759531 |
| 0.071171303 | 0.163121748 |
| 0.895081035 | NA |
| NA | NA |
| 0.024690806 | 0.070977122 |
| 0.004986943 | 0.01948444 |
| 0.116693048 | 0.235124687 |
| 0.884123828 | NA |
| 0.230315315 | 0.384177226 |
| 0.000103638 | 0.000722129 |
| 0.025113069 | 0.071901056 |
| 0.15132053 | 0.285172704 |
| 0.57795484 | 0.716296855 |
| 5.92E-06 | 5.67E-05 |
| 0.703855484 | 0.811016519 |
| 0.019647063 | 0.059038698 |
| NA | NA |
| 0.018904241 | 0.057363431 |
| 0.031878717 | 0.087025717 |
| 0.357045 | 0.5206928 |
| 0.086595163 | 0.188697067 |
| 0.03345499 | 0.090488146 |
| 0.204525932 | 0.353121009 |
| 0.041858803 | 0.107952732 |
| 0.746704675 | 0.840691769 |
| 0.019634106 | 0.059013891 |
| 0.003218251 | 0.01360583 |
| 0.302673771 | 0.464101082 |
| 0.13316332 | 0.259246961 |
| 0.122812873 | 0.24452044 |

| | | |
|---|---|---|
| | 1.35E-20 | 5.97E-19 |
| | 3.00E-05 | 0.000241413 |
| | 0.188045524 | 0.332406948 |
| | 0.029213259 | 0.081240098 |
| | 0.014473957 | 0.046348597 |
| | 0.181899966 | 0.324337627 |
| | 0.95655474 | NA |
| | 0.579456978 | 0.717336818 |
| | 0.070164021 | 0.161497695 |
| | 0.317947721 | 0.480372708 |
| | 2.07E-17 | 7.38E-16 |
| | 0.1574489 | 0.293243614 |
| | 0.471391336 | 0.628962194 |
| | 0.018555164 | 0.056513117 |
| | 0.941278207 | NA |
| | 6.60E-73 | 1.50E-70 |
| | 7.17E-12 | 1.65E-10 |
| | 6.26E-05 | 0.000463445 |
| | 0.053632795 | 0.130997208 |
| | 0.326123552 | 0.48909286 |
| | 0.030229456 | 0.083500233 |
| | 1.21E-06 | 1.33E-05 |
| NA | | NA |
| | 0.884305346 | 0.930962684 |
| | 0.259676581 | 0.418188226 |
| | 0.933094483 | NA |
| | 0.804544442 | NA |
| | 0.940475126 | NA |
| | 0.839318641 | NA |
| | 0.839319108 | NA |
| | 0.815215848 | 0.88702503 |
| | 0.001765062 | 0.008244116 |
| | 0.008814975 | 0.031023972 |
| | 0.463236281 | 0.621999042 |
| | 0.261793932 | 0.420602036 |
| | 0.680582596 | 0.794389424 |
| | 0.384261326 | 0.547020819 |
| | 0.03361146 | 0.090808169 |
| | 0.673544778 | 0.789272982 |
| | 0.000214833 | 0.001365407 |
| | 0.192585557 | 0.338122108 |
| | 0.003822984 | 0.015629453 |
| | 0.006448924 | 0.024006457 |
| | 0.022357001 | 0.065478201 |
| | 1.65E-137 | 8.01E-135 |
| | 2.03E-17 | 7.25E-16 |
| | 0.00119221 | 0.005929165 |

| | |
|---|---|
| 0.629042419 | 0.755803092 |
| 0.60848563 | 0.740751584 |
| 0.682048722 | 0.795556389 |
| 0.003112706 | 0.013220668 |
| 0.000147102 | 0.000981865 |
| 4.46E-06 | 4.39E-05 |
| 0.563412163 | NA |
| 1.49E-57 | 2.42E-55 |
| 0.151981618 | NA |
| 0.110349554 | 0.225686551 |
| 0.00042939 | 0.002485492 |
| 0.366552024 | 0.529666785 |
| 2.69E-05 | 0.000218427 |
| 0.800131796 | 0.87792085 |
| 0.239953077 | 0.395989956 |
| 0.055229337 | 0.134133078 |
| 1.10E-09 | 1.95E-08 |
| NA | NA |
| NA | NA |
| NA | NA |
| NA | NA |
| NA | NA |
| 1.19E-08 | 1.84E-07 |
| 0.634445456 | NA |
| 0.046765626 | 0.11751641 |
| 8.92E-07 | 1.00E-05 |
| 2.13E-10 | 4.11E-09 |
| 0.614108472 | NA |
| NA | NA |
| 0.040964371 | 0.106178151 |
| 0.490563842 | 0.644926847 |
| 0.042031059 | 0.108270915 |
| 1.04E-17 | 3.80E-16 |
| 0.560208076 | 0.703496498 |
| 0.064355788 | 0.151154181 |
| 0.941277857 | NA |
| 0.386657142 | 0.54949761 |
| 0.002326588 | 0.010393665 |
| NA | NA |
| 0.805943939 | 0.881129835 |
| 0.004922025 | 0.019271928 |
| 0.115722077 | 0.233642961 |
| 0.508827017 | 0.660437379 |
| 0.570533646 | 0.711172655 |
| 0.069868826 | 0.161020313 |
| 0.001935083 | 0.008910593 |
| 0.179666791 | 0.32156019 |

| | | |
|---|---|---|
| | 0.602725722 | 0.736073763 |
| | 0.709587801 | 0.814663749 |
| | 0.148452005 | 0.281348406 |
| NA | | NA |
| | 0.018611431 | 0.056655489 |
| NA | | NA |
| | 0.709823427 | 0.81481636 |
| NA | | NA |
| NA | | NA |
| | 0.922027496 | NA |
| | 0.093015135 | 0.198800088 |
| | 0.482416964 | 0.638554587 |
| | 7.13E-15 | 2.14E-13 |
| | 1.45E-05 | 0.000126218 |
| | 0.491434411 | 0.645465283 |
| | 0.17934469 | 0.321056131 |
| | 0.641823108 | 0.765019461 |
| | 0.79158977 | 0.871750204 |
| NA | | NA |
| | 0.567852248 | 0.709316149 |
| NA | | NA |
| NA | | NA |
| | 0.941277857 | NA |
| | 0.094021987 | 0.200332228 |
| NA | | NA |
| | 0.300511344 | 0.46205492 |
| | 0.160598217 | 0.297405957 |
| NA | | NA |
| | 0.823638073 | 0.89285054 |
| | 0.708753764 | 0.814125883 |
| | 0.654458048 | 0.77439894 |
| | 0.179144343 | 0.320784666 |
| NA | | NA |
| | 1.14E-12 | 2.84E-11 |
| | 0.900941072 | NA |
| NA | | NA |
| | 0.119316104 | 0.239295745 |
| NA | | NA |
| | 0.668182934 | 0.784922031 |
| | 0.814658946 | 0.886666616 |
| | 0.458555135 | 0.618109189 |
| | 0.429379314 | 0.590700241 |
| | 0.034526439 | 0.092638745 |
| | 0.104060242 | NA |
| NA | | NA |
| | 0.021263105 | 0.062854549 |
| | 0.600618703 | NA |

| | |
|---|---|
| 0.001740522 | 0.008147079 |
| 0.532867375 | 0.681239084 |
| 0.000499737 | 0.00283613 |
| 0.839319107 | NA |
| 0.931924112 | 0.959371849 |
| 0.35169962 | 0.515306516 |
| 0.289344995 | 0.450142374 |
| 0.955792398 | NA |
| 0.043229146 | 0.110603044 |
| 0.014682342 | 0.046870857 |
| NA | NA |
| 0.866395141 | 0.920211386 |
| 0.019763652 | 0.059299252 |
| 0.918725136 | NA |
| 1.20E-05 | 0.0001069 |
| 0.006575849 | 0.024398807 |
| 0.129271519 | 0.253536642 |
| NA | NA |
| 0.839319574 | NA |
| 0.000619013 | 0.003398789 |
| 0.613255755 | 0.743895295 |
| 0.65131295 | NA |
| NA | NA |
| 1.07E-06 | 1.18E-05 |
| NA | NA |
| NA | NA |
| 0.027614017 | 0.077783295 |
| NA | NA |
| 2.06E-09 | 3.52E-08 |
| NA | NA |
| NA | NA |
| 0.249530896 | 0.406970974 |
| 0.009572019 | 0.033226367 |
| 0.129251639 | 0.253518542 |
| 0.072637257 | 0.165493635 |
| 0.00833623 | 0.029663469 |
| 0.001770362 | 0.008265627 |
| 0.292854679 | NA |
| NA | NA |
| 0.000396646 | 0.002320219 |
| 0.203449942 | 0.35177361 |
| 0.941278332 | NA |
| 0.309001993 | 0.470560809 |
| 0.041371178 | 0.106953055 |
| 0.596448971 | 0.731146851 |
| 0.017960585 | 0.055033193 |
| 0.158549935 | 0.294800886 |

| | |
|---|---|
| 0.021168035 | 0.062620131 |
| 5.98E-07 | 6.94E-06 |
| 7.08E-17 | 2.46E-15 |
| NA | NA |
| NA | NA |
| 0.082514096 | 0.181818228 |
| 0.000183123 | 0.001187335 |
| 0.87482978 | NA |
| 0.093573473 | NA |
| 0.599156256 | 0.733144006 |
| 1.95E-05 | 0.00016383 |
| 4.59E-07 | 5.44E-06 |
| 0.000935025 | 0.004816157 |
| 0.046888288 | 0.11776248 |
| NA | NA |
| 0.16319626 | 0.301069398 |
| NA | NA |
| 0.01162281 | 0.038953271 |
| 4.93E-11 | 1.03E-09 |
| 0.721032868 | 0.822921329 |
| 0.399653789 | 0.562340948 |
| 0.571761704 | 0.711883249 |
| 0.005181451 | 0.020115624 |
| 0.907910675 | NA |
| 0.135377144 | 0.262547607 |
| 4.76E-06 | 4.66E-05 |
| 0.335082416 | 0.498321253 |
| 0.169121489 | 0.308674924 |
| NA | NA |
| 0.854004013 | 0.91208307 |
| 0.205258528 | 0.353872482 |
| 0.619158227 | 0.74838534 |
| 0.90773516 | NA |
| 0.376224412 | 0.539416131 |
| 0.448704306 | 0.60881295 |
| 0.890151523 | NA |
| NA | NA |
| 0.417270361 | NA |
| 0.51733586 | 0.668165833 |
| 0.210852927 | 0.36092926 |
| 0.217368447 | 0.368705406 |
| 0.814429895 | 0.886584171 |
| 0.981228087 | 0.98841216 |
| NA | NA |
| 0.004025783 | 0.016296718 |
| 0.775866844 | NA |
| 0.040254913 | 0.104708548 |

| | | | |
|---|---|---|---|
| | 0.004611388 | | 0.018262938 |
| | 0.000957855 | | 0.004917786 |
| | 0.308808665 | | 0.470326584 |
| | 0.011925512 | | 0.039699445 |
| | 0.001527773 | | 0.007284468 |
| | 0.080424048 | | 0.178528567 |
| | 0.036107592 | | 0.096079699 |
| | 0.538273787 | | 0.685954821 |
| | 0.46533858 | | 0.623753387 |
| | 0.002733674 | | 0.011882228 |
| | 0.108053824 | | 0.222117378 |
| | 5.57E-05 | | 0.00041667 |
| NA | | NA | |
| | 0.224644477 | | 0.377292397 |
| | 0.125831418 | | 0.248846487 |
| | 0.57912529 | | 0.71715 |
| | 0.007559581 | | 0.027362096 |
| | 0.000349391 | | 0.002076455 |
| NA | | NA | |
| | 0.108917237 | | 0.223477487 |
| | 0.046136835 | | 0.11629238 |
| | 0.938705482 | | 0.963601145 |
| | 0.008566887 | | 0.030312032 |
| | 0.01882225 | | 0.057172946 |
| | 0.373162102 | NA | |
| | 0.19476641 | | 0.340919995 |
| | 0.230098711 | | 0.383936457 |
| | 0.002010375 | | 0.0091968 |
| | 0.573873608 | NA | |
| | 0.366348176 | | 0.529565121 |
| | 0.636557484 | | 0.761067975 |
| NA | | NA | |
| | 0.30202522 | | 0.463344819 |
| | 0.123359743 | | 0.245256819 |
| | 0.835709626 | | 0.900202342 |
| | 0.968837183 | | 0.981329578 |
| | 0.124362617 | | 0.246782164 |
| | 0.48305283 | | 0.639040877 |
| NA | | NA | |
| NA | | NA | |
| NA | | NA | |
| | 0.077877393 | | 0.174230445 |
| | 0.012838713 | | 0.042127388 |
| | 0.839319108 | NA | |
| | 0.079635449 | | 0.17722609 |
| NA | | NA | |
| | 0.554872826 | | 0.699587007 |

| | | |
|---|---|---|
| | 0.839319107 | NA |
| | 0.067285757 | 0.156384688 |
| | 0.001838267 | 0.008515846 |
| | 0.788622918 | 0.869944974 |
| | 0.217151342 | NA |
| | 0.923654864 | 0.954564454 |
| | 0.267327854 | 0.426861948 |
| NA | | NA |
| | 0.239975711 | 0.395999853 |
| NA | | NA |
| | 0.839319108 | NA |
| | 0.284877316 | 0.445432504 |
| NA | | NA |
| | 0.636770987 | 0.76118881 |
| | 0.579725131 | NA |
| | 0.839318641 | NA |
| | 0.000286057 | 0.001748069 |
| | 0.847823672 | 0.908395855 |
| | 5.57E-11 | 1.16E-09 |
| | 0.000112665 | 0.00077797 |
| | 0.017484754 | 0.053866439 |
| | 0.853595622 | NA |
| NA | | NA |
| | 0.00280259 | 0.012102184 |
| | 0.269678252 | 0.429644026 |
| | 0.989008927 | NA |
| | 0.955791922 | NA |
| | 0.764896265 | NA |
| | 0.817199144 | NA |
| NA | | NA |
| NA | | NA |
| NA | | NA |
| | 0.137426118 | 0.265547692 |
| | 0.002281153 | 0.010221386 |
| | 0.941278332 | NA |
| NA | | NA |
| | 0.02182393 | 0.064225171 |
| | 0.839319108 | NA |
| | 0.902957639 | 0.941349627 |
| | 0.849978732 | NA |
| | 0.39641613 | 0.559041162 |
| | 0.002843885 | 0.012258267 |
| | 0.607671155 | 0.740244637 |
| NA | | NA |
| | 0.166730799 | 0.305576844 |
| | 0.021735894 | 0.063989827 |
| | 1.99E-05 | 0.000167489 |

| | | |
|---|---|---|
| | 0.741073673 | 0.83729508 |
| | 0.000895857 | 0.004651655 |
| | 1.97E-15 | 6.23E-14 |
| | 4.28E-18 | 1.59E-16 |
| | 0.599612951 | 0.733514286 |
| | 0.759911266 | NA |
| NA | | NA |
| | 0.003862881 | 0.015757378 |
| NA | | NA |
| | 0.85220602 | NA |
| | 0.974890765 | 0.984820897 |
| NA | | NA |
| | 0.221868596 | 0.374108552 |
| NA | | NA |
| | 7.99E-06 | 7.41E-05 |
| | 0.877603469 | 0.926657696 |
| | 0.839319108 | NA |
| | 0.780893052 | 0.865187191 |
| | 0.149342852 | 0.282381316 |
| NA | | NA |
| | 0.000110749 | 0.000766292 |
| | 0.009076567 | 0.031803392 |
| | 0.900936105 | NA |
| | 0.966382496 | 0.979969162 |
| | 0.062423858 | 0.147550725 |
| | 0.839319108 | NA |
| | 0.662848679 | NA |
| | 0.955792873 | NA |
| NA | | NA |
| | 0.044611894 | 0.113247766 |
| | 0.269702927 | 0.429654579 |
| | 0.347343662 | 0.511347962 |
| NA | | NA |
| | 0.599394565 | 0.733284819 |
| | 0.173460245 | 0.31411296 |
| | 0.023212407 | 0.067592576 |
| | 0.202519536 | 0.350623963 |
| | 0.576262226 | 0.714928093 |
| NA | | NA |
| | 0.032586282 | 0.088652266 |
| | 0.807556402 | NA |
| NA | | NA |
| | 0.884279636 | NA |
| | 0.096038448 | 0.203494722 |
| NA | | NA |
| | 0.460745232 | NA |
| | 0.662106071 | NA |

| | | | |
|---|---|---|---|
| NA | | NA | |
| | 0.046369787 | | 0.116737903 |
| | 0.011250233 | | 0.037891331 |
| | 0.000460126 | | 0.00264029 |
| | 0.778582419 | NA | |
| | 0.000392227 | | 0.002297756 |
| | 0.637581342 | | 0.761794644 |
| | 0.257168951 | | 0.415330124 |
| | 0.003505326 | | 0.014578678 |
| | 0.068699077 | NA | |
| | 0.007866062 | | 0.028269366 |
| | 9.11E-15 | | 2.69E-13 |
| | 1.07E-11 | | 2.41E-10 |
| | 0.63347378 | | 0.759182999 |
| | 0.092087603 | | 0.19728795 |
| | 0.784235247 | | 0.867327233 |
| | 0.955792398 | NA | |
| | 0.24866633 | | 0.405894664 |
| | 0.047334019 | | 0.118532268 |
| | 0.548273447 | | 0.694223034 |
| | 2.18E-20 | | 9.50E-19 |
| | 9.62E-234 | | 2.29E-230 |
| | 0.021864138 | | 0.064325471 |
| | 0.016921114 | | 0.052503763 |
| | 3.32E-05 | | 0.000263458 |
| | 8.82E-05 | | 0.000627572 |
| | 0.061791931 | | 0.146415288 |
| | 3.60E-05 | | 0.000283055 |
| NA | | NA | |
| | 0.821808679 | | 0.891558606 |
| | 0.1056668 | | 0.218531686 |
| | 0.051426761 | | 0.126779883 |
| | 0.108395059 | | 0.222623085 |
| | 0.839319108 | NA | |
| | 0.00425291 | | 0.017067571 |
| | 0.000185665 | | 0.001202176 |
| | 4.05E-06 | | 4.02E-05 |
| | 1.03E-05 | | 9.32E-05 |
| | 1.22E-25 | | 7.04E-24 |
| | 0.670122955 | NA | |
| | 0.52078043 | | 0.671012602 |
| | 0.001368267 | | 0.006643543 |
| | 0.109213278 | | 0.223959202 |
| | 0.912147115 | | 0.946958853 |
| | 0.000934981 | | 0.004816157 |
| | 0.591183134 | | 0.726975664 |
| | 0.844107925 | | 0.906163487 |

| | | |
|---|---|---|
| | 0.546721742 | 0.692921774 |
| | 1.87E-08 | 2.81E-07 |
| | 1.06E-06 | 1.17E-05 |
| | 0.096116221 | 0.203591555 |
| | 0.088508698 | 0.191906668 |
| | 0.183051563 | 0.325683458 |
| | 0.91775769 | NA |
| | 0.922282171 | 0.953653948 |
| | 0.983262957 | 0.989370601 |
| | 1.70E-06 | 1.83E-05 |
| | 0.295824214 | 0.457310381 |
| | 0.512905299 | 0.664175099 |
| | 0.407099665 | 0.568883122 |
| | 0.063246996 | NA |
| NA | | NA |
| NA | | NA |
| | 0.402980496 | 0.565005869 |
| | 0.001795541 | 0.008356954 |
| | 0.413981496 | 0.576109104 |
| | 0.002249615 | 0.010095274 |
| NA | | NA |
| NA | | NA |
| NA | | NA |
| | 0.839318641 | NA |
| | 0.184648234 | 0.327715849 |
| NA | | NA |
| NA | | NA |
| | 0.853596091 | NA |
| | 0.001016894 | 0.005179567 |
| | 0.849972972 | NA |
| | 0.021868822 | 0.064325471 |
| | 0.01738056 | 0.053628738 |
| | 0.033161875 | 0.089889415 |
| | 0.106746654 | 0.220303461 |
| NA | | NA |
| | 0.421899532 | 0.583193984 |
| | 0.941277857 | NA |
| NA | | NA |
| NA | | NA |
| | 5.91E-09 | 9.55E-08 |
| | 0.25070522 | 0.408354592 |
| | 1.66E-08 | 2.52E-07 |
| | 3.85E-32 | 2.94E-30 |
| | 4.30E-08 | 6.08E-07 |
| | 0.833801616 | NA |
| | 6.53E-06 | 6.19E-05 |
| | 0.493254175 | 0.646891612 |

| | |
|---|---|
| 0.955792873 | NA |
| 0.675334857 | NA |
| NA | NA |
| 0.941278788 | NA |
| 0.002610313 | 0.011442111 |
| 2.43E-08 | 3.59E-07 |
| 0.000576663 | 0.003204395 |
| 0.000250962 | 0.001558005 |
| NA | NA |
| 0.600766675 | 0.734592824 |
| 4.15E-07 | 4.99E-06 |
| 0.297168548 | 0.458643871 |
| NA | NA |
| 0.07201363 | 0.164482143 |
| 0.840890018 | 0.90397291 |
| 0.363271761 | 0.526910083 |
| NA | NA |
| 0.853595154 | NA |
| NA | NA |
| 0.955792398 | NA |
| NA | NA |
| NA | NA |
| NA | NA |
| 0.900439766 | NA |
| NA | NA |
| 0.500553137 | 0.653188912 |
| NA | NA |
| 0.839320041 | NA |
| 0.941275481 | NA |
| 0.941278788 | NA |
| NA | NA |
| 0.007207228 | 0.026318878 |
| 0.615780462 | 0.745817553 |
| 0.498896395 | 0.651822305 |
| 0.189213354 | 0.333826719 |
| 0.643010678 | 0.765821374 |
| 0.852724164 | 0.911466704 |
| 0.279534924 | 0.439338282 |
| 0.460943681 | 0.619889027 |
| 0.256859382 | 0.414970951 |
| 6.56E-09 | 1.05E-07 |
| 0.416503883 | 0.578155095 |
| 0.024616834 | 0.070866676 |
| 0.021726886 | 0.063979133 |
| 0.800935448 | 0.878492596 |
| 0.831743455 | 0.897952453 |
| 0.483817917 | 0.639697482 |

| | | |
|---|---|---|
| | 0.453906574 | 0.613717579 |
| | 1.05E-05 | 9.47E-05 |
| | 0.108193994 | 0.222290328 |
| | 0.21029302 | 0.3601849 |
| | 7.00E-08 | 9.63E-07 |
| | 0.724852663 | 0.826238017 |
| | 0.064147534 | 0.150724541 |
| | 0.044328226 | 0.1126831 |
| | 0.334355364 | NA |
| | 0.028962493 | 0.080684136 |
| | 0.131601122 | 0.257010456 |
| | 0.352136715 | 0.515724716 |
| | 0.633387322 | 0.759182999 |
| NA | NA | |
| | 0.962379624 | 0.977284005 |
| NA | NA | |
| | 0.741106875 | 0.83729508 |
| | 0.288905175 | 0.449739859 |
| | 0.975595054 | 0.985178051 |
| | 0.704658942 | 0.811627977 |
| | 0.085194448 | 0.186486321 |
| | 0.987792858 | 0.992253454 |
| | 0.175987249 | 0.317417813 |
| | 0.469522813 | 0.627489163 |
| | 0.002812152 | 0.012139072 |
| | 0.258212012 | 0.416534212 |
| | 0.555392787 | 0.69996077 |
| | 0.645439578 | 0.767868888 |
| | 0.634902962 | 0.76000605 |
| | 0.783168806 | 0.866703867 |
| | 0.745891329 | NA |
| | 0.008365656 | 0.02973262 |
| | 0.47656722 | 0.633524321 |
| | 0.817702872 | 0.888757758 |
| | 1.58E-70 | 3.38E-68 |
| | 0.000635822 | 0.003474252 |
| | 3.45E-05 | 0.000272461 |
| | 0.955792398 | NA |
| | 0.940685982 | NA |
| | 9.17E-09 | 1.45E-07 |
| NA | NA | |
| | 0.000672081 | 0.003650589 |
| | 0.000311301 | 0.001880603 |
| | 0.000788587 | 0.004177597 |
| NA | NA | |
| | 0.104124493 | 0.216099999 |
| | 0.166755508 | 0.305598599 |

| | | |
|---|---|---|
| | 0.764028074 | 0.8535741 |
| | 0.539163255 | 0.686485706 |
| | 0.005341314 | 0.020586401 |
| | 0.151767525 | 0.285753935 |
| | 0.623498358 | NA |
| | 0.406396935 | 0.568234732 |
| | 0.013410288 | 0.043669805 |
| | 2.41E-08 | 3.57E-07 |
| | 0.743511643 | NA |
| | 0.000417505 | 0.002426135 |
| | 0.799362055 | 0.877521245 |
| | 0.935677476 | NA |
| | 0.032191792 | 0.087709367 |
| | 0.168148875 | 0.307560096 |
| | 3.63E-05 | 0.000284874 |
| | 0.031027762 | 0.085181407 |
| | 0.08921991 | 0.192897838 |
| | 0.32870802 | 0.491629492 |
| | 0.009297668 | 0.032430115 |
| NA | | NA |
| | 0.946558437 | NA |
| | 0.182949633 | 0.325565658 |
| | 0.583187526 | 0.720526464 |
| | 0.07953271 | 0.177068335 |
| | 0.079942821 | 0.177703143 |
| | 8.09E-06 | 7.49E-05 |
| | 0.839319108 | NA |
| | 0.217100027 | 0.368495675 |
| NA | | NA |
| NA | | NA |
| | 0.743212514 | 0.83854495 |
| | 0.020130383 | 0.060193051 |
| | 0.80460217 | 0.880558921 |
| | 0.865547691 | 0.919767844 |
| | 0.865446654 | 0.919767844 |
| | 1.68E-06 | 1.81E-05 |
| | 0.036324892 | 0.096485711 |
| | 0.261702348 | 0.420483279 |
| | 3.17E-05 | 0.000252887 |
| | 0.941278332 | NA |
| | 0.941275956 | NA |
| | 0.4708189 | 0.628445013 |
| | 0.0991664 | 0.208217947 |
| | 0.207236128 | 0.356326976 |
| NA | | NA |
| | 0.823552641 | NA |
| | 0.887034996 | 0.9326293 |

| | |
|---|---|
| NA | NA |
| 0.750899265 | 0.843578813 |
| 5.95E-05 | 0.000442352 |
| 0.574474615 | 0.713786167 |
| 2.40E-10 | 4.62E-09 |
| 0.251268206 | 0.40896792 |
| 0.248892549 | 0.40623606 |
| 0.248503502 | 0.405740181 |
| 0.27825037 | 0.437937781 |
| 0.127231598 | 0.250815802 |
| 0.685221266 | 0.79781033 |
| 0.788226949 | 0.869831931 |
| NA | NA |
| 0.00119249 | 0.005929319 |
| 3.12E-06 | 3.16E-05 |
| 0.177846106 | 0.319321374 |
| 0.53915168 | 0.686485706 |
| 0.776590176 | 0.8623466 |
| 0.253987501 | 0.412037247 |
| 0.002647561 | 0.011581923 |
| NA | NA |
| 0.941277857 | NA |
| 0.011853433 | 0.039509232 |
| 0.96512443 | 0.978818513 |
| NA | NA |
| 0.000232159 | 0.001456468 |
| NA | NA |
| NA | NA |
| NA | NA |
| NA | NA |
| 0.02070356 | 0.061498333 |
| 2.17E-05 | 0.000181267 |
| 0.330258191 | 0.493206501 |
| 0.383497021 | 0.546253898 |
| 0.068782636 | 0.15907154 |
| 0.006091649 | 0.022923862 |
| 0.083050417 | 0.182729342 |
| 0.253791496 | 0.411803475 |
| 0.803544779 | 0.879967312 |
| 0.703466094 | NA |
| 0.019059657 | 0.057723001 |
| 4.40E-05 | 0.000338512 |
| 0.042966304 | 0.11009624 |
| 0.194926479 | 0.34117509 |
| 0.466062732 | 0.624196778 |
| 0.817460139 | NA |
| 0.000299591 | 0.001820954 |

| | | |
|---|---|---|
| NA | | NA |
| | 0.448718829 | 0.60881295 |
| | 0.695804868 | 0.805170278 |
| | 0.000484265 | 0.002762158 |
| | 0.070429785 | 0.161921158 |
| NA | | NA |
| | 0.798348036 | 0.876650672 |
| | 1.98E-31 | 1.47E-29 |
| | 0.141934728 | 0.272029181 |
| NA | | NA |
| | 0.000189936 | 0.001226738 |
| | 0.955792873 | NA |
| | 0.991119761 | 0.994085059 |
| | 0.202436794 | 0.350556607 |
| | 9.24E-08 | 1.25E-06 |
| | 0.441098885 | NA |
| NA | | NA |
| | 0.063967065 | 0.150452397 |
| | 0.616523147 | 0.746239537 |
| | 0.000117372 | 0.000806203 |
| NA | | NA |
| | 0.489165135 | 0.643583465 |
| | 8.43E-07 | 9.51E-06 |
| | 0.351639799 | 0.515282306 |
| | 8.50E-07 | 9.58E-06 |
| | 9.78E-16 | 3.14E-14 |
| | 0.002724668 | 0.01185604 |
| | 0.265558217 | 0.424871727 |
| NA | | NA |
| | 0.660301774 | 0.779201925 |
| | 0.199960703 | 0.347406179 |
| | 0.195429823 | 0.341779622 |
| NA | | NA |
| NA | | NA |
| | 0.570079683 | 0.710839312 |
| | 0.053334831 | 0.130516861 |
| | 0.468578741 | 0.626649691 |
| | 0.189397215 | 0.333973899 |
| | 0.052127987 | 0.128110516 |
| | 0.483156584 | NA |
| | 0.730957639 | NA |
| | 1.29E-05 | 0.000113453 |
| NA | | NA |
| | 0.941278332 | NA |
| | 0.209412421 | NA |
| NA | | NA |
| | 0.160881286 | 0.297837471 |

| | | |
|---|---|---|
| | 0.946099755 | 0.967272298 |
| | 0.37829802 | 0.541103347 |
| | 0.403210471 | 0.565240334 |
| | 0.038406904 | 0.10082289 |
| NA | | NA |
| | 7.31E-07 | 8.34E-06 |
| | 0.959689395 | 0.975259456 |
| | 0.201741288 | 0.349733877 |
| | 0.000831068 | 0.004365723 |
| | 0.941379378 | 0.964703895 |
| | 0.02851028 | 0.079751049 |
| NA | | NA |
| | 0.100221801 | 0.209970893 |
| NA | | NA |
| | 0.244201929 | 0.400668251 |
| | 0.89235891 | 0.935040806 |
| | 0.794880838 | NA |
| | 0.696611119 | NA |
| | 3.98E-18 | 1.48E-16 |
| NA | | NA |
| | 0.837581159 | 0.901433656 |
| | 0.041138911 | 0.106572526 |
| | 1.09E-06 | 1.20E-05 |
| | 0.094297786 | NA |
| NA | | NA |
| | 0.160076037 | 0.296772677 |
| NA | | NA |
| | 0.384781565 | NA |
| | 0.000797927 | 0.004219563 |
| | 0.407752181 | 0.569694612 |
| | 1.05E-06 | 1.17E-05 |
| | 0.17140699 | 0.311562951 |
| | 0.73202233 | 0.830985599 |
| | 0.121386958 | 0.242447963 |
| | 0.138350415 | 0.266836243 |
| | 0.653752872 | 0.774044961 |
| | 0.541695919 | 0.688789265 |
| | 0.182076497 | 0.32448786 |
| | 0.857880874 | NA |
| | 0.47163913 | 0.629222273 |
| | 0.408967758 | 0.571034361 |
| | 0.010015191 | 0.034433276 |
| | 0.000814981 | 0.004296387 |
| | 7.21E-12 | 1.66E-10 |
| | 0.058168066 | 0.139539431 |
| | 0.000158505 | 0.001049441 |
| NA | | NA |

| | | |
|---|---|---|
| | 2.18E-09 | 3.71E-08 |
| | 0.511702123 | NA |
| | 0.095928329 | 0.203338327 |
| NA | | NA |
| | 0.069479938 | 0.160325884 |
| | 0.001584641 | 0.007519717 |
| | 0.000565217 | 0.003149178 |
| | 0.4520103 | 0.611988007 |
| | 0.782704939 | 0.866267226 |
| | 0.995952046 | 0.996917253 |
| NA | | NA |
| | 0.214626973 | 0.365523127 |
| | 0.34050521 | 0.50405529 |
| | 0.43659756 | 0.597605627 |
| | 0.049291112 | 0.12228758 |
| | 0.656794882 | 0.776226666 |
| | 0.715511052 | NA |
| | 0.229505359 | 0.383238849 |
| | 0.527294169 | NA |
| NA | | NA |
| | 0.202279062 | 0.350385433 |
| | 0.603778302 | 0.736872583 |
| | 0.012481814 | 0.041177013 |
| NA | | NA |
| | 0.743511643 | NA |
| | 2.49E-74 | 5.86E-72 |
| | 0.839319108 | NA |
| NA | | NA |
| | 0.015274428 | 0.048397489 |
| | 6.80E-14 | 1.88E-12 |
| | 0.833801616 | NA |
| | 0.002586576 | 0.0113527 |
| | 0.768398692 | 0.856487107 |
| NA | | NA |
| | 0.470070836 | 0.627939484 |
| | 0.394099165 | NA |
| | 0.599114351 | 0.733130419 |
| NA | | NA |
| | 0.017626025 | 0.05419644 |
| | 0.955792397 | NA |
| | 0.631076715 | NA |
| | 0.719262098 | NA |
| | 0.884645759 | NA |
| | 0.186865673 | 0.33071431 |
| | 0.143163565 | 0.273613231 |
| | 0.001381255 | 0.006699943 |
| | 0.242854016 | 0.399309981 |

| | | |
|---|---|---|
| | 0.555063901 | 0.699694469 |
| | 0.8494572 | NA |
| | 0.143026681 | 0.273427363 |
| | 0.060467449 | 0.143881458 |
| NA | | NA |
| | 0.52170673 | NA |
| | 0.435086032 | NA |
| | 0.495645505 | NA |
| | 5.25E-09 | 8.55E-08 |
| | 0.702405612 | 0.81013027 |
| | 0.809126483 | NA |
| | 0.839319108 | NA |
| NA | | NA |
| | 0.207815978 | NA |
| | 1.94E-12 | 4.73E-11 |
| | 0.100667552 | 0.210794249 |
| | 0.116264874 | NA |
| | 0.862247703 | 0.917648997 |
| | 4.53E-15 | 1.39E-13 |
| | 0.065158811 | 0.152589762 |
| | 0.614897565 | NA |
| | 0.97451522 | 0.984554671 |
| | 0.047550829 | 0.118987051 |
| | 3.81E-36 | 3.40E-34 |
| | 6.70E-06 | 6.34E-05 |
| | 0.154045742 | 0.288753878 |
| | 0.242962002 | 0.39940622 |
| | 0.229646015 | 0.383393032 |
| | 0.157220062 | 0.292987364 |
| | 0.128687819 | 0.252996404 |
| | 2.53E-14 | 7.28E-13 |
| | 0.692232948 | NA |
| | 1.66E-08 | 2.52E-07 |
| | 0.001044384 | 0.005298046 |
| | 1.35E-08 | 2.08E-07 |
| | 0.870574482 | 0.922591889 |
| | 0.91073116 | NA |
| NA | | NA |
| | 6.13E-05 | 0.000454826 |
| | 0.079826782 | 0.177556963 |
| | 0.831929452 | NA |
| | 0.36210965 | 0.525801238 |
| | 0.2045896 | 0.353179697 |
| | 0.869175505 | 0.921601719 |
| | 0.796128334 | 0.874908486 |
| | 0.931999762 | 0.959371849 |
| | 0.040170363 | 0.104541328 |

| | | | |
|---|---|---|---|
| | 0.181715836 | | 0.32413072 |
| | 0.033379688 | | 0.090366621 |
| NA | | NA | |
| NA | | NA | |
| | 0.738587704 | | 0.835465475 |
| NA | | NA | |
| | 0.784461242 | | 0.867525274 |
| | 0.007023898 | | 0.025737799 |
| | 0.000408787 | | 0.002379737 |
| | 0.088737595 | | 0.192239214 |
| | 0.557355687 | | 0.701283505 |
| | 0.172522151 | | 0.312771136 |
| | 0.005160366 | | 0.020050121 |
| | 0.236938839 | | 0.392401142 |
| | 0.027359368 | | 0.077221354 |
| | 0.440597539 | | 0.601290214 |
| | 3.58E-06 | | 3.59E-05 |
| | 0.314437613 | | 0.476673082 |
| | 0.853694051 | | 0.91201193 |
| | 0.155395157 | | 0.29050546 |
| | 0.839319574 | NA | |
| | 0.174499101 | | 0.315394305 |
| | 0.000632916 | | 0.003463141 |
| NA | | NA | |
| | 0.069032724 | | 0.159494942 |
| | 2.76E-09 | | 4.64E-08 |
| | 0.518192254 | | 0.668908945 |
| | 0.225439727 | | 0.378334371 |
| NA | | NA | |
| NA | | NA | |
| NA | | NA | |
| NA | | NA | |
| NA | | NA | |
| NA | | NA | |
| NA | | NA | |
| NA | | NA | |
| NA | | NA | |
| NA | | NA | |
| NA | | NA | |
| NA | | NA | |
| NA | | NA | |
| NA | | NA | |
| NA | | NA | |
| NA | | NA | |
| NA | | NA | |
| NA | | NA | |
| NA | | NA | |

| | |
|---|---|
| NA | NA |
| NA | NA |
| NA | NA |
| NA | NA |
| NA | NA |
| NA | NA |
| NA | NA |
| NA | NA |
| NA | NA |
| NA | NA |
| NA | NA |
| NA | NA |
| NA | NA |
| NA | NA |
| NA | NA |
| NA | NA |
| NA | NA |
| NA | NA |
| NA | NA |
| NA | NA |
| NA | NA |
| NA | NA |
| NA | NA |
| NA | NA |
| NA | NA |
| NA | NA |
| NA | NA |
| NA | NA |
| NA | NA |
| NA | NA |
| NA | NA |
| NA | NA |
| 8.86E-07 | 9.95E-06 |
| 0.945970234 | 0.967272298 |
| 0.977986223 | 0.986440718 |
| 0.018863288 | 0.057268364 |
| 0.791409915 | 0.871750204 |
| 0.663426612 | NA |
| 0.803104406 | 0.879641576 |
| 0.839039497 | 0.902635847 |
| 0.74568045 | 0.840134719 |
| 0.463588084 | 0.622318608 |
| 0.003790498 | 0.015515294 |
| 0.003841074 | 0.015698017 |
| 0.441091195 | 0.601791392 |
| 0.176306566 | 0.317840558 |
| 0.30320065 | NA |

| | | |
|---|---|---|
| | 0.941278838 | NA |
| NA | | NA |
| | 0.051511823 | 0.126923853 |
| | 0.817237883 | 0.888457846 |
| | 0.746435768 | 0.84054706 |
| | 0.00524708 | 0.020304176 |
| | 0.005783105 | 0.021974692 |
| | 0.333143227 | 0.496409223 |
| | 0.950863585 | 0.970100945 |
| | 0.087791177 | 0.190671474 |
| | 0.142979614 | 0.27341257 |
| | 0.605280326 | 0.738028822 |
| | 0.893959798 | 0.936121034 |
| | 2.03E-05 | 0.000170141 |
| | 6.67E-05 | 0.000489315 |
| | 0.076301122 | 0.171623447 |
| | 0.10191065 | 0.212760047 |
| | 0.996013386 | 0.996917253 |
| | 0.235260303 | 0.390247804 |
| | 0.937404386 | NA |
| | 0.778208573 | NA |
| | 0.127350364 | 0.250986028 |
| | 0.000178186 | 0.001162287 |
| | 0.429091445 | 0.590428637 |
| | 0.109990716 | 0.2251468 |
| | 0.546272043 | 0.692499316 |
| | 0.369636085 | 0.532904822 |
| | 0.676197326 | 0.791324721 |
| | 0.001839108 | 0.008516431 |
| | 0.574434554 | 0.713786167 |
| | 2.50E-05 | 0.000205439 |
| NA | | NA |
| | 0.954720481 | NA |
| | 0.000506014 | 0.002865607 |
| | 0.56363559 | 0.70593845 |
| | 0.894331586 | 0.936363116 |
| | 0.113686163 | 0.230687329 |
| NA | | NA |
| | 0.044068899 | 0.112159698 |
| | 0.238062614 | 0.39368904 |
| | 0.668137638 | NA |
| | 0.839319574 | NA |
| | 0.188266318 | 0.332624325 |
| NA | | NA |
| NA | | NA |
| | 4.11E-05 | 0.000318437 |
| | 0.943117438 | NA |

| | | | |
|---|---|---|---|
| | 0.318048093 | NA | |
| | 0.909183738 | NA | |
| NA | | NA | |
| | 0.594279999 | NA | |
| | 0.952431307 | NA | |
| | 0.85444689 | NA | |
| | 0.00245243 | | 0.010855106 |
| | 0.090155395 | | 0.194531657 |
| | 0.018857083 | | 0.057264134 |
| | 0.895713045 | NA | |
| | 0.597072707 | NA | |
| | 0.918071596 | NA | |
| NA | | NA | |
| | 0.839319108 | NA | |
| | 0.92498945 | NA | |
| | 0.941502978 | NA | |
| NA | | NA | |
| NA | | NA | |
| NA | | NA | |
| NA | | NA | |
| NA | | NA | |
| | 0.027815652 | | 0.078210045 |
| | 0.959317227 | | 0.975112026 |
| | 0.206129032 | | 0.35492489 |
| | 7.55E-10 | | 1.36E-08 |
| | 0.012289397 | | 0.04064928 |
| | 0.686900553 | NA | |
| | 0.159993933 | | 0.296702423 |
| | 0.177588824 | | 0.319182263 |
| | 0.955792398 | NA | |
| | 0.529154773 | | 0.678314406 |
| | 1.01E-05 | | 9.17E-05 |
| | 0.095957135 | | 0.203363167 |
| NA | | NA | |
| | 2.87E-39 | | 2.87E-37 |
| | 1.16E-36 | | 1.07E-34 |
| | 1.38E-38 | | 1.33E-36 |
| | 0.29919122 | | 0.460749833 |
| NA | | NA | |
| NA | | NA | |
| | 0.00474758 | | 0.018721323 |
| | 0.096466003 | | 0.204132629 |
| | 1.33E-09 | | 2.33E-08 |
| | 0.094325463 | | 0.200801003 |
| NA | | NA | |
| | 1.18E-12 | | 2.95E-11 |
| | 0.934281796 | NA | |

| | | |
|---|---|---|
| | 0.941279303 | NA |
| | 7.59E-08 | 1.04E-06 |
| | 0.027539407 | 0.07761603 |
| | 0.909802956 | NA |
| NA | | NA |
| | 0.839319109 | NA |
| | 0.343776369 | NA |
| | 2.44E-07 | 3.05E-06 |
| | 0.789657476 | NA |
| | 3.28E-08 | 4.75E-07 |
| | 2.94E-35 | 2.55E-33 |
| | 0.242103322 | 0.39837838 |
| | 0.955791446 | NA |
| | 0.003666221 | 0.015102697 |
| | 0.015087543 | 0.047926559 |
| | 0.561123933 | NA |
| | 0.839319108 | NA |
| | 0.248528888 | 0.405753795 |
| | 6.42E-05 | 0.000474214 |
| | 0.008982659 | 0.031539348 |
| | 0.036325034 | 0.096485711 |
| NA | | NA |
| NA | | NA |
| | 0.251438201 | 0.409170229 |
| | 0.217170412 | 0.368588859 |
| NA | | NA |
| | 0.902983548 | NA |
| | 0.387099137 | NA |
| NA | | NA |
| NA | | NA |
| NA | | NA |
| | 0.448505844 | 0.608697462 |
| | 0.015397626 | 0.048697091 |
| | 0.124413261 | 0.246835413 |
| | 0.865574351 | 0.919767844 |
| NA | | NA |
| | 3.75E-06 | 3.75E-05 |
| NA | | NA |
| | 0.761622278 | 0.851817685 |
| | 0.001125412 | 0.005647703 |
| | 0.005381995 | 0.020698374 |
| | 0.004450068 | 0.017721351 |
| | 3.12E-07 | 3.83E-06 |
| | 0.812679666 | 0.885565234 |
| | 0.885623111 | 0.931530394 |
| | 0.004191683 | 0.01686166 |
| | 0.000408821 | 0.002379737 |

| | |
|---|---|
| 2.01E-10 | 3.90E-09 |
| 0.072758465 | 0.165703494 |
| NA | NA |
| 0.000874498 | 0.004552671 |
| 0.676920896 | 0.791715346 |
| 3.41E-05 | 0.000270069 |
| 0.920992565 | 0.952651559 |
| 0.470212745 | 0.628058573 |
| 0.881721835 | 0.929274887 |
| 0.091445624 | 0.196460757 |
| 2.78E-06 | 2.84E-05 |
| 0.604876783 | 0.737754403 |
| 0.23485379 | 0.389682113 |
| 0.955792398 | NA |
| 0.418507156 | 0.580123891 |
| 0.000173909 | 0.001138443 |
| 0.070089165 | 0.161371861 |
| 4.93E-05 | 0.000374669 |
| 0.907192016 | 0.943666829 |
| NA | NA |
| 0.055990888 | 0.135588067 |
| 0.703047262 | 0.810516852 |
| 0.955792398 | NA |
| 0.010073228 | 0.034602826 |
| 0.1159663 | 0.234036788 |
| 0.001423003 | 0.006859117 |
| 0.000736836 | 0.003947373 |
| 0.967053885 | 0.980280702 |
| 0.356524468 | 0.520192878 |
| 0.252777134 | 0.410605429 |
| 0.475278067 | 0.632517113 |
| 0.908808487 | NA |
| 0.474591498 | 0.632098195 |
| 0.100010195 | 0.209617732 |
| 0.841665239 | 0.904520318 |
| 0.941278332 | NA |
| 0.000402704 | 0.002349307 |
| 0.865589779 | 0.919767844 |
| NA | NA |
| 4.41E-07 | 5.26E-06 |
| 0.133933025 | 0.260404662 |
| 0.839320041 | NA |
| 0.883160997 | NA |
| 0.656443299 | NA |
| 0.292019637 | 0.453257687 |
| NA | NA |
| 0.934281796 | NA |

| | |
|---|---|
| 0.94020055 | NA |
| NA | NA |
| NA | NA |
| NA | NA |
| NA | NA |
| 0.955792873 | NA |
| NA | NA |
| NA | NA |
| NA | NA |
| NA | NA |
| NA | NA |
| 0.941278332 | NA |
| NA | NA |
| NA | NA |
| NA | NA |
| NA | NA |
| NA | NA |
| 0.941278332 | NA |
| NA | NA |
| NA | NA |
| NA | NA |
| NA | NA |
| NA | NA |
| NA | NA |
| NA | NA |
| NA | NA |
| 0.135678258 | 0.262960174 |
| 0.225520229 | 0.378442787 |
| 0.159973184 | NA |
| 0.835727577 | NA |
| NA | NA |
| NA | NA |
| NA | NA |
| 0.381057263 | 0.54376049 |
| NA | NA |
| NA | NA |
| NA | NA |
| 0.94127925 | NA |
| 0.955791446 | NA |
| NA | NA |
| NA | NA |
| NA | NA |
| NA | NA |
| NA | NA |
| NA | NA |
| 0.941278332 | NA |
| NA | NA |

| | |
|---|---|
| NA | NA |
| 0.525547108 | 0.674852842 |
| 0.041724126 | 0.107690051 |
| NA | NA |
| 0.034461576 | 0.09250642 |
| 0.003071528 | 0.013073518 |
| NA | NA |
| 0.011835088 | 0.039464663 |
| 0.001029163 | 0.005235334 |
| 0.00338271 | 0.014162753 |
| 0.643363261 | 0.766164624 |
| 0.627915604 | 0.755021328 |
| 0.030064113 | 0.083159329 |
| 0.44688165 | 0.607254912 |
| 8.56E-06 | 7.87E-05 |
| 3.72E-19 | 1.50E-17 |
| NA | NA |
| 0.896938159 | NA |
| 2.41E-06 | 2.51E-05 |
| 1.93E-05 | 0.000162487 |
| 0.000343811 | 0.002050977 |
| NA | NA |
| 0.350796489 | 0.514680271 |
| 0.044708168 | 0.113411895 |
| 0.424637208 | 0.586019796 |
| 0.727122217 | 0.827925808 |
| 2.31E-06 | 2.41E-05 |
| 0.171532197 | 0.311700038 |
| 1.54E-40 | 1.60E-38 |
| 0.969404899 | 0.981695686 |
| 0.635201847 | 0.760094348 |
| 0.699125643 | NA |
| 0.750461605 | 0.843325959 |
| 0.00035839 | 0.002124096 |
| 0.19570494 | 0.342160201 |
| 0.001651914 | 0.007792083 |
| NA | NA |
| 0.432148615 | 0.593354209 |
| 0.997014066 | 0.997558905 |
| 0.751746496 | 0.844145736 |
| 9.13E-05 | 0.000647304 |
| 0.47476691 | 0.632190321 |
| 4.02E-12 | 9.48E-11 |
| 0.334196049 | 0.497495877 |
| 0.000113074 | 0.000780256 |
| 0.764085074 | 0.853590585 |
| 0.033512763 | 0.090603225 |

| | | |
|---|---|---|
| | 0.058570481 | 0.140279189 |
| | 0.845314591 | 0.906845355 |
| | 0.001000396 | 0.00511194 |
| | 0.301416522 | 0.462739684 |
| | 0.683787103 | 0.797037249 |
| | 0.228422464 | 0.38201872 |
| | 1.70E-07 | 2.19E-06 |
| | 0.94127925 | NA |
| | 0.089262516 | 0.192954899 |
| | 5.59E-33 | 4.42E-31 |
| | 0.196474711 | 0.343174567 |
| | 0.715509863 | 0.819176213 |
| | 0.127824999 | 0.251736773 |
| | 3.04E-05 | 0.000243631 |
| | 0.420240035 | NA |
| | 0.672582567 | 0.788455982 |
| | 0.726991366 | 0.827895722 |
| | 0.661324652 | NA |
| | 0.007672527 | 0.027703503 |
| | 0.179152616 | 0.320784666 |
| | 0.2567456 | 0.414871606 |
| | 0.017379168 | 0.053628738 |
| NA | | NA |
| | 0.832717194 | NA |
| | 0.420088589 | 0.581740086 |
| | 0.387132268 | 0.549956565 |
| | 0.007487752 | 0.027168213 |
| | 0.009178325 | 0.032089086 |
| | 0.17744411 | 0.31903747 |
| | 0.163985136 | 0.302103267 |
| | 0.013932122 | 0.044960405 |
| | 0.484407906 | 0.640264159 |
| | 0.278286938 | 0.437965202 |
| | 3.81E-08 | 5.44E-07 |
| | 0.653830416 | 0.774059764 |
| | 0.783491778 | 0.866896211 |
| | 0.002665102 | 0.011647955 |
| | 0.031142263 | 0.085436624 |
| | 0.988167291 | 0.992545802 |
| NA | | NA |
| | 4.98E-21 | 2.28E-19 |
| | 0.31761081 | 0.480107487 |
| | 0.079230247 | 0.176626237 |
| | 0.905907505 | NA |
| | 0.955791446 | NA |
| | 0.177144509 | 0.318699442 |
| | 0.088835095 | 0.192380396 |

| | |
|---|---|
| 3.71E-05 | 0.000290058 |
| 0.056451946 | 0.136426969 |
| 0.082849786 | 0.182355335 |
| 0.181942098 | 0.324363727 |
| 0.50052276 | 0.653188912 |
| NA | NA |
| 0.154185245 | 0.288856135 |
| NA | NA |
| 0.0020798 | 0.00945082 |
| NA | NA |
| NA | NA |
| 0.196435909 | 0.343160917 |
| 0.028093671 | 0.078789391 |
| 0.021102779 | 0.062450373 |
| 0.024992816 | 0.071637644 |
| 0.724516911 | NA |
| 2.71E-07 | 3.37E-06 |
| 0.684388457 | 0.797279344 |
| NA | NA |
| 0.940475126 | NA |
| 0.095104802 | 0.202078602 |
| NA | NA |
| NA | NA |
| 0.231024981 | 0.384966158 |
| 0.891525237 | 0.934565988 |
| 0.109275348 | 0.223994239 |
| 0.534071615 | 0.682375467 |
| 0.746026507 | 0.840278884 |
| 0.774029225 | 0.860546668 |
| NA | NA |
| NA | NA |
| NA | NA |
| 0.015616229 | 0.049257552 |
| 0.897604142 | 0.938244864 |
| 0.468208153 | NA |
| 0.735819853 | 0.833507461 |
| 0.825465549 | 0.893990126 |
| 0.139965946 | 0.269079189 |
| 0.286632606 | 0.447176539 |
| 0.336753211 | 0.500056144 |
| 0.92730368 | 0.956477821 |
| 0.941277857 | NA |
| 0.056794349 | 0.137017951 |
| 0.312746966 | 0.474684561 |
| 0.0103575 | 0.035431047 |
| 0.229804802 | 0.383604313 |
| 0.585463345 | 0.722180687 |

| | |
|---|---|
| 9.67E-05 | 0.000680783 |
| 0.175924817 | 0.317344637 |
| 0.047181485 | 0.118243285 |
| NA | NA |
| NA | NA |
| 0.165409434 | 0.304044742 |
| 0.006120338 | 0.023009997 |
| 0.1502315 | 0.283569402 |
| 1.54E-05 | 0.000133717 |
| NA | NA |
| 1.50E-05 | 0.000130199 |
| 0.062729408 | NA |
| 0.052979072 | 0.129853554 |
| 0.032363166 | NA |
| 6.70E-05 | 0.000491409 |
| 0.134441298 | 0.261179544 |
| 0.777526424 | NA |
| 0.529684139 | 0.678737113 |
| 0.809854945 | 0.883463483 |
| 0.68730053 | 0.799299743 |
| 0.976143926 | 0.985459461 |
| 0.958374648 | 0.974464239 |
| 0.18118243 | NA |
| 1.93E-10 | 3.76E-09 |
| 0.132447487 | NA |
| 0.503571344 | 0.655617828 |
| 0.044371702 | 0.112762679 |
| 0.821695621 | NA |
| 0.261224818 | 0.419948636 |
| 0.189731609 | 0.334309917 |
| NA | NA |
| 9.28E-14 | 2.54E-12 |
| NA | NA |
| NA | NA |
| 0.033535027 | NA |
| NA | NA |
| 0.941277857 | NA |
| NA | NA |
| 0.093474139 | 0.199504256 |
| 0.000872198 | 0.004545665 |
| NA | NA |
| NA | NA |
| NA | NA |
| 0.644838193 | NA |
| NA | NA |
| NA | NA |
| 0.326812991 | 0.48963321 |

| | |
|---|---|
| 0.019249602 | 0.058115048 |
| 0.007417951 | 0.026960158 |
| 0.955792398 | NA |
| 0.682141412 | 0.795625516 |
| 0.94151042 | 0.964703895 |
| 0.006173264 | 0.023176031 |
| 0.218793525 | 0.370208537 |
| 0.769132337 | 0.857023917 |
| NA | NA |
| 0.00012087 | 0.000828156 |
| 0.000321945 | 0.00193802 |
| 0.000550543 | 0.003085084 |
| 0.008574722 | 0.030326232 |
| 0.336856926 | 0.500078095 |
| NA | NA |
| 0.955792398 | NA |
| 0.222613727 | 0.3750196 |
| 0.164977337 | 0.303578707 |
| NA | NA |
| NA | NA |
| 0.075858262 | 0.170934767 |
| 0.316297307 | 0.478699593 |
| 0.32654672 | 0.489405996 |
| 0.86547396 | 0.919767844 |
| NA | NA |
| NA | NA |
| NA | NA |
| NA | NA |
| NA | NA |
| 0.485017279 | NA |
| NA | NA |
| NA | NA |
| NA | NA |
| 0.518785422 | 0.669465062 |
| NA | NA |
| 0.955792398 | NA |
| 0.585592325 | 0.722264924 |
| NA | NA |
| NA | NA |
| NA | NA |
| 0.941277857 | NA |
| NA | NA |
| NA | NA |
| NA | NA |
| NA | NA |
| NA | NA |
| 0.941278424 | NA |

| | | |
|---|---|---|
| | 0.91534172 | NA |
| | 0.662120348 | NA |
| | 0.941279268 | NA |
| | 0.887921218 | 0.933164113 |
| | 0.019963649 | 0.059755379 |
| | 0.41774597 | NA |
| | 0.488319583 | 0.643036238 |
| | 0.263197616 | 0.422094416 |
| | 1.39E-05 | 0.000121463 |
| | 0.004383971 | 0.017510872 |
| | 0.941279303 | NA |
| NA | | NA |
| NA | | NA |
| | 1.06E-05 | 9.49E-05 |
| | 0.000331073 | 0.001985938 |
| | 0.51248259 | 0.663847033 |
| | 0.222650709 | 0.375028814 |
| | 0.0014997 | 0.007176492 |
| | 2.56E-11 | 5.57E-10 |
| | 0.00113037 | 0.005668999 |
| | 0.00356237 | 0.014766898 |
| | 6.01E-09 | 9.69E-08 |
| | 2.91E-05 | 0.000234575 |
| | 0.55466525 | 0.699549914 |
| | 0.204170331 | 0.352660539 |
| | 0.639006168 | 0.762946033 |
| | 5.81E-16 | 1.91E-14 |
| | 0.901929538 | NA |
| NA | | NA |
| | 0.464857901 | 0.623319687 |
| NA | | NA |
| NA | | NA |
| | 0.839318641 | NA |
| | 0.941278332 | NA |
| NA | | NA |
| | 0.549074006 | NA |
| | 0.955791922 | NA |
| | 0.929334299 | NA |
| NA | | NA |
| NA | | NA |
| | 0.853595622 | NA |
| NA | | NA |
| | 0.839318641 | NA |
| NA | | NA |
| NA | | NA |
| NA | | NA |
| NA | | NA |

| | | |
|---|---|---|
| NA | NA | |
| NA | NA | |
| NA | NA | |
| NA | NA | |
| NA | NA | |
| NA | NA | |
| 0.941275481 | NA | |
| NA | NA | |
| 0.839318641 | NA | |
| NA | NA | |
| 0.955792873 | NA | |
| 0.955791922 | NA | |
| NA | NA | |
| NA | NA | |
| NA | NA | |
| NA | NA | |
| NA | NA | |
| NA | NA | |
| NA | NA | |
| NA | NA | |
| NA | NA | |
| NA | NA | |
| NA | NA | |
| NA | NA | |
| NA | NA | |
| NA | NA | |
| NA | NA | |
| NA | NA | |
| 2.02E-55 | | 3.13E-53 |
| 5.27E-13 | | 1.36E-11 |
| 0.077473622 | NA | |
| 0.346820505 | NA | |
| 0.803875996 | NA | |
| 0.202915614 | NA | |
| 0.715555719 | NA | |
| 0.739991096 | NA | |
| NA | NA | |
| NA | NA | |
| NA | NA | |
| NA | NA | |
| NA | NA | |
| NA | NA | |
| NA | NA | |
| NA | NA | |
| NA | NA | |
| NA | NA | |
| NA | NA | |

| | | |
|---|---|---|
| NA | NA | |
| NA | NA | |
| NA | NA | |
| NA | NA | |
| NA | NA | |
| NA | NA | |
| NA | NA | |
| NA | NA | |
| NA | NA | |
| 0.941278332 | NA | |
| NA | NA | |
| NA | NA | |
| NA | NA | |
| 0.739991096 | NA | |
| 0.376807578 | | 0.539734848 |
| NA | NA | |
| NA | NA | |
| NA | NA | |
| NA | NA | |
| 0.955792873 | NA | |
| NA | NA | |
| NA | NA | |
| NA | NA | |
| NA | NA | |
| 0.941278332 | NA | |
| NA | NA | |
| NA | NA | |
| NA | NA | |
| NA | NA | |
| NA | NA | |
| 0.941278332 | NA | |
| NA | NA | |
| 0.839319107 | NA | |
| NA | NA | |
| 0.941278332 | NA | |
| 0.755626499 | NA | |
| NA | NA | |
| 0.941278332 | NA | |
| NA | NA | |
| NA | NA | |
| 0.839318641 | NA | |
| 0.755626499 | NA | |
| NA | NA | |
| NA | NA | |
| 0.982903359 | | 0.989178714 |
| 0.005597358 | | 0.021390067 |
| 9.37E-08 | | 1.26E-06 |

| | | | |
|---|---|---|---|
| | 0.777700466 | | 0.863096307 |
| NA | | NA | |
| NA | | NA | |
| | 0.001045246 | | 0.00530064 |
| | 0.942423631 | NA | |
| | 0.394826546 | NA | |
| | 0.053347038 | | 0.130516861 |
| NA | | NA | |
| NA | | NA | |
| NA | | NA | |
| | 0.372612677 | | 0.535822072 |
| | 0.14900846 | | 0.281954 |
| | 0.058264932 | | 0.13971609 |
| | 0.08616826 | | 0.187938874 |
| | 0.236401918 | | 0.391732 |
| NA | | NA | |
| | 0.741261802 | | 0.837390866 |
| | 0.741697849 | | 0.837510898 |
| | 0.364275024 | | 0.527657869 |
| | 0.273732472 | | 0.433520547 |
| NA | | NA | |
| | 0.077368186 | | 0.173351931 |
| | 0.862523133 | | 0.917777979 |
| NA | | NA | |
| | 0.056906065 | | 0.13723183 |
| NA | | NA | |
| | 0.096192614 | | 0.20371711 |
| | 0.016056466 | | 0.050345935 |
| NA | | NA | |
| | 0.631905808 | NA | |
| | 0.008314254 | | 0.029598545 |
| NA | | NA | |
| | 0.236915832 | | 0.392392362 |
| | 0.013669296 | | 0.044272224 |
| | 0.012361648 | | 0.040854199 |
| | 0.058180485 | | 0.139541708 |
| | 8.37E-07 | | 9.45E-06 |
| | 0.805839235 | | 0.881096305 |
| | 0.654874581 | | 0.774679428 |
| NA | | NA | |
| | 0.095975746 | | 0.2033845 |
| | 0.839319108 | NA | |
| | 0.955792873 | NA | |
| | 0.735462791 | | 0.833355932 |
| | 0.299263853 | | 0.460831872 |
| | 0.173663834 | | 0.314385958 |
| | 0.681127113 | | 0.794831962 |

| | |
|---|---|
| NA | NA |
| 0.021734326 | 0.063989827 |
| 0.207022795 | 0.356037324 |
| 0.1675159 | 0.306661549 |
| 0.835076557 | 0.899970585 |
| 0.771150455 | 0.85854919 |
| 0.002813218 | 0.012141473 |
| 0.287779418 | 0.448375852 |
| 2.07E-05 | 0.000173191 |
| 4.41E-05 | 0.000339053 |
| 0.383810589 | 0.546509909 |
| 0.026094714 | 0.074170813 |
| 0.715514388 | 0.819176213 |
| 0.123207686 | 0.244994933 |
| 0.958056098 | 0.974181957 |
| 0.438356797 | 0.599228605 |
| 0.000190618 | 0.001230231 |
| 0.935146874 | NA |
| 0.45454405 | 0.614366148 |
| 0.186133966 | 0.329713526 |
| NA | NA |
| 0.039790243 | 0.103756283 |
| 0.781377772 | NA |
| 0.661683423 | 0.78014028 |
| 0.000721651 | 0.003879117 |
| 0.008498464 | 0.030101255 |
| 0.727397625 | NA |
| 0.955792398 | NA |
| 0.538413041 | 0.686044283 |
| 0.075926786 | 0.171007699 |
| 0.117879292 | 0.236993235 |
| 0.519796731 | 0.670199414 |
| 1.05E-05 | 9.48E-05 |
| 0.130858859 | 0.255995936 |
| 0.955792398 | NA |
| 0.001634632 | 0.007725876 |
| NA | NA |
| 0.003069255 | 0.013068312 |
| 0.154230321 | 0.288917841 |
| 2.03E-07 | 2.59E-06 |
| 0.702703687 | 0.810252828 |
| 0.002225464 | 0.010005759 |
| NA | NA |
| 0.607644438 | 0.740244637 |
| 0.565201418 | 0.707081054 |
| 4.32E-14 | 1.22E-12 |
| 1.48E-17 | 5.36E-16 |

| | | |
|---|---|---|
| | 1.46E-22 | 7.26E-21 |
| | 1.28E-05 | 0.000113385 |
| | 9.65E-05 | 0.000679825 |
| | 1.12E-26 | 6.81E-25 |
| | 0.864985512 | 0.919740235 |
| | 0.006636831 | 0.024586749 |
| | 0.007562149 | 0.02736723 |
| | 0.753264994 | 0.845318877 |
| | 0.845082309 | NA |
| | 0.58763162 | 0.723857322 |
| | 7.19E-44 | 8.43E-42 |
| | 0.955792873 | NA |
| | 0.260923345 | NA |
| NA | | NA |
| | 0.195842931 | NA |
| | 0.633529806 | 0.759182999 |
| | 0.422023889 | 0.583266323 |
| | 0.04880187 | NA |
| | 0.222907205 | 0.375354602 |
| NA | | NA |
| | 0.070798955 | 0.162597137 |
| | 0.743511643 | NA |
| | 0.001305689 | 0.006385453 |
| | 0.196708718 | 0.343482596 |
| | 3.34E-05 | 0.000264966 |
| | 0.532664634 | 0.681053052 |
| | 2.45E-05 | 0.000201737 |
| | 0.006244978 | 0.023400971 |
| | 0.101992586 | 0.212893759 |
| | 0.157904434 | 0.293894547 |
| | 0.38447275 | 0.547289062 |
| | 0.000142332 | 0.000953771 |
| | 0.086101797 | 0.187861408 |
| | 6.83E-06 | 6.43E-05 |
| | 0.903857488 | NA |
| NA | | NA |
| | 0.918253994 | NA |
| | 0.001227127 | 0.006076155 |
| | 0.289419571 | 0.450216628 |
| | 0.943962327 | NA |
| | 0.515889518 | 0.666804361 |
| | 0.621964881 | NA |
| NA | | NA |
| | 4.94E-05 | 0.00037507 |
| | 0.839318641 | NA |
| | 0.839319108 | NA |
| | 0.159880064 | 0.296601034 |

| | |
|---|---|
| 0.00786022 | 0.028252636 |
| 0.002608514 | 0.011436331 |
| 0.500989323 | 0.653548115 |
| 0.978522934 | 0.986646267 |
| 3.44E-13 | 9.02E-12 |
| 0.393882773 | 0.556391558 |
| NA | NA |
| 0.000117808 | 0.000808786 |
| 0.007556412 | 0.027354784 |
| 0.171311358 | 0.311438305 |
| 0.941278332 | NA |
| 7.54E-09 | 1.20E-07 |
| NA | NA |
| NA | NA |
| 0.065123029 | 0.152578102 |
| 6.60E-05 | 0.000485406 |
| 0.839319108 | NA |
| 0.093229027 | 0.19914182 |
| 0.64607489 | NA |
| 0.0442637 | 0.112556945 |
| 9.09E-05 | 0.000644861 |
| 0.397136789 | 0.559725832 |
| NA | NA |
| 0.786425633 | NA |
| 0.347947512 | 0.511825393 |
| 0.079522531 | 0.177062235 |
| 0.044029997 | 0.112117978 |
| 0.364067589 | 0.52755777 |
| 0.492536822 | 0.646413842 |
| 0.955792873 | NA |
| 0.000170725 | 0.001120684 |
| 0.110728777 | 0.226248291 |
| 0.9997887 | 0.999872715 |
| 0.021881759 | 0.064347521 |
| 0.839724367 | 0.903250153 |
| 0.868203496 | 0.921186627 |
| 1.36E-11 | 3.02E-10 |
| 0.034539141 | 0.092659392 |
| 0.000608388 | 0.003350495 |
| 0.87646832 | 0.926140201 |
| 0.028420519 | 0.079528003 |
| 0.960981501 | 0.976155757 |
| 0.694103161 | 0.803858336 |
| 0.016065313 | 0.050360608 |
| 0.826066479 | NA |
| 0.002890456 | 0.012429745 |
| 0.002826386 | 0.01219167 |

| | |
|---|---|
| 0.141651641 | 0.271574092 |
| NA | NA |
| NA | NA |
| 0.70758666 | NA |
| 0.027157974 | 0.076780389 |
| 0.024915734 | 0.071492858 |
| 0.086437609 | 0.188455103 |
| 0.924309355 | NA |
| 0.405950713 | 0.567920367 |
| 0.033793791 | 0.091158231 |
| 0.871002132 | 0.922839651 |
| 0.353226493 | 0.516939067 |
| 0.381159982 | 0.543825835 |
| 0.010122593 | 0.034739556 |
| NA | NA |
| 0.00592344 | 0.022411393 |
| 0.683467258 | NA |
| NA | NA |
| 0.002028366 | 0.009264856 |
| NA | NA |
| 0.570862713 | 0.711393348 |
| NA | NA |
| 0.055500965 | 0.134689433 |
| 0.858854408 | 0.915798433 |
| 0.941278332 | NA |
| 0.547717604 | 0.693703741 |
| 8.96E-12 | 2.04E-10 |
| 0.356801142 | 0.520442504 |
| 0.001725736 | 0.008092314 |
| 0.076795578 | 0.172491114 |
| NA | NA |
| 0.754806586 | 0.846370488 |
| 0.788009767 | 0.869752305 |
| 0.150844263 | 0.284500408 |
| 0.449572759 | 0.609640068 |
| 0.630228727 | 0.756541464 |
| 0.464799308 | NA |
| 0.000764394 | 0.004069361 |
| 0.000562425 | 0.003139502 |
| 0.850967587 | 0.91048553 |
| 0.941277857 | NA |
| 1.44E-06 | 1.57E-05 |
| 8.81E-06 | 8.08E-05 |
| NA | NA |
| 0.089202264 | NA |
| 0.023777362 | 0.068900506 |
| 0.00377298 | 0.015459541 |

| | | |
|---|---|---|
| | 0.303439922 | 0.464796771 |
| | 0.057252472 | 0.137871081 |
| | 0.971418845 | NA |
| | 0.30342907 | 0.464796771 |
| | 0.812347483 | 0.88542389 |
| | 0.389003104 | 0.551626565 |
| | 0.001212814 | 0.006015292 |
| | 0.74627078 | NA |
| | 8.91E-12 | 2.03E-10 |
| | 0.639211747 | 0.763060957 |
| | 0.036976064 | 0.097832844 |
| | 0.089010952 | 0.192633286 |
| | 0.25875987 | 0.417135281 |
| | 0.56903721 | 0.710162735 |
| | 0.220149766 | 0.371842516 |
| NA | | NA |
| | 0.002128057 | 0.009624173 |
| | 0.910910965 | 0.946046898 |
| | 0.472209434 | NA |
| NA | | NA |
| | 0.107306331 | 0.221058099 |
| | 0.549001081 | NA |
| | 0.941277857 | NA |
| | 0.075811221 | 0.17086059 |
| | 1.32E-25 | 7.54E-24 |
| | 0.803579022 | 0.879967312 |
| | 0.100347478 | 0.210197189 |
| | 0.243790276 | 0.400288755 |
| | 0.020371882 | 0.060770965 |
| | 0.168340439 | 0.307815866 |
| | 0.209962716 | 0.359715869 |
| | 0.008149339 | 0.029107229 |
| | 0.862355155 | 0.91768129 |
| | 0.925865722 | NA |
| | 0.070233364 | 0.161606878 |
| | 0.500726946 | 0.65330827 |
| | 0.000329347 | 0.001977081 |
| | 0.000261455 | 0.001613469 |
| | 0.214401044 | 0.365339168 |
| | 0.011794347 | 0.039356377 |
| | 0.788749977 | 0.869975354 |
| | 0.517550762 | 0.668311565 |
| | 0.120917602 | 0.24171635 |
| | 0.990348631 | 0.993561791 |
| | 0.187362872 | 0.331397122 |
| | 0.493483056 | 0.646993147 |
| | 0.250445137 | 0.408042656 |

| | | |
|---|---|---|
| | 0.867919184 | 0.921060969 |
| | 0.839319108 | NA |
| | 0.743511643 | NA |
| | 9.71E-13 | 2.44E-11 |
| | 0.84465257 | NA |
| | 0.170185142 | 0.309879647 |
| | 0.048062514 | 0.120002514 |
| | 0.086043401 | 0.187830612 |
| | 0.098803274 | 0.207674038 |
| | 0.222842435 | 0.375298637 |
| | 0.378518828 | 0.541306486 |
| | 0.123169543 | 0.244994933 |
| | 0.693708837 | 0.803747042 |
| | 1.96E-10 | 3.82E-09 |
| | 0.506754451 | 0.65842947 |
| | 0.22491452 | 0.377671196 |
| | 0.183263673 | 0.325939024 |
| | 0.063890704 | 0.15031398 |
| | 0.280579934 | NA |
| | 0.912976426 | NA |
| | 0.304501192 | 0.466122412 |
| | 0.292068043 | 0.453290265 |
| | 0.365367781 | NA |
| | 0.000704693 | 0.003798256 |
| | 0.953477882 | 0.971893304 |
| | 3.30E-06 | 3.33E-05 |
| | 0.158808617 | 0.295079055 |
| | 0.000453215 | 0.002605588 |
| | 0.170861309 | 0.310730508 |
| | 0.806983902 | NA |
| | 2.92E-13 | 7.68E-12 |
| NA | | NA |
| | 4.51E-05 | 0.000345634 |
| | 0.002271692 | 0.010182828 |
| | 0.034423794 | 0.092415423 |
| | 2.09E-61 | 3.85E-59 |
| | 0.739991096 | NA |
| | 1.97E-20 | 8.61E-19 |
| | 0.000373728 | 0.002199672 |
| NA | | NA |
| | 0.853595622 | NA |
| | 0.941279268 | NA |
| | 1.09E-05 | 9.74E-05 |
| | 0.908095486 | 0.944328606 |
| | 0.078318661 | 0.1749874 |
| | 0.013213422 | 0.043130262 |
| | 0.135433956 | 0.262614991 |

| | |
|---|---|
| NA | NA |
| 0.0002718 | 0.001669085 |
| 0.274950828 | 0.434813608 |
| 0.883603679 | 0.930516537 |
| 0.057762241 | 0.138748295 |
| 0.014221201 | 0.045742301 |
| 0.083117985 | 0.182861104 |
| 0.771671307 | 0.858863398 |
| 0.851002923 | 0.91048553 |
| 0.307851335 | 0.469379123 |
| 0.034072199 | 0.091686392 |
| 0.216475857 | 0.367750935 |
| 0.029324876 | 0.081502884 |
| 4.17E-05 | 0.000322712 |
| 0.000308601 | 0.001867613 |
| 0.001807627 | 0.008400065 |
| NA | NA |
| 0.839319108 | NA |
| 0.629601093 | NA |
| 0.726714887 | NA |
| 0.839319108 | NA |
| 0.487687896 | 0.642530018 |
| 0.779513167 | NA |
| NA | NA |
| 0.670541605 | 0.786722102 |
| 0.069157268 | 0.159736176 |
| 0.154041187 | 0.288753878 |
| 0.093995547 | 0.200293823 |
| 1.24E-14 | 3.63E-13 |
| 0.327313041 | NA |
| 0.704915638 | NA |
| 5.39E-05 | 0.000405124 |
| 0.002042448 | 0.009307745 |
| 0.003853522 | 0.015734315 |
| 0.036844254 | 0.097535909 |
| NA | NA |
| 0.00402469 | 0.016296718 |
| NA | NA |
| 0.857970264 | NA |
| 0.346023532 | 0.509751322 |
| 0.489051674 | 0.643583465 |
| 0.195291579 | 0.341618485 |
| NA | NA |
| 0.000371565 | 0.002188922 |
| 0.557803593 | 0.701476257 |
| 0.955792873 | NA |
| 3.20E-07 | 3.92E-06 |

| | | |
|---|---|---|
| | 0.616554989 | NA |
| | 0.11703609 | 0.235616427 |
| | 0.001921188 | 0.00885003 |
| NA | | NA |
| | 0.849160616 | NA |
| | 0.008021199 | 0.028714179 |
| | 0.016438268 | 0.05129981 |
| | 0.004587556 | 0.018180654 |
| | 0.020944957 | 0.062075938 |
| | 0.773989752 | 0.860542978 |
| | 0.475323346 | NA |
| | 0.63421171 | 0.759670657 |
| | 0.389495813 | 0.55206213 |
| | 0.019160777 | 0.057938115 |
| | 0.146424915 | 0.278304386 |
| NA | | NA |
| NA | | NA |
| | 0.839318641 | NA |
| NA | | NA |
| | 0.798459083 | NA |
| | 0.242361511 | 0.398748095 |
| | 0.000301763 | 0.001830883 |
| | 0.301055733 | 0.462373819 |
| | 0.054649477 | 0.132975551 |
| | 0.388142431 | 0.551029831 |
| | 0.887786624 | 0.93306386 |
| | 0.683399668 | 0.796702693 |
| | 0.009842854 | 0.033978187 |
| | 0.000284916 | 0.001742438 |
| NA | | NA |
| | 0.123193126 | 0.244994933 |
| | 0.019147683 | 0.057924905 |
| | 0.905750601 | 0.94278549 |
| | 0.039116343 | 0.10231288 |
| | 0.000358488 | 0.002124096 |
| | 0.301785866 | 0.463127477 |
| NA | | NA |
| | 0.065317511 | 0.152854919 |
| NA | | NA |
| | 0.955963407 | 0.973093316 |
| | 0.013910335 | 0.044900161 |
| | 0.243426422 | 0.399947243 |
| | 0.017355694 | 0.0535937 |
| | 0.000166793 | 0.00109851 |
| | 3.84E-12 | 9.09E-11 |
| | 0.010663902 | 0.036270675 |
| | 0.018176527 | 0.055587524 |

| | | | |
|---|---|---|---|
| | 0.505350046 | | 0.657286436 |
| | 0.559189956 | | 0.702666895 |
| | 0.48919049 | | 0.643583465 |
| | 0.001221338 | | 0.006052527 |
| NA | | NA | |
| | 0.573311023 | | 0.712991743 |
| | 0.177182364 | | 0.318743454 |
| | 0.071233524 | | 0.163170084 |
| | 0.194319116 | | 0.340397705 |
| | 0.839320041 | NA | |
| | 0.801059594 | | 0.878492596 |
| | 1.67E-05 | | 0.000143266 |
| | 0.004026164 | | 0.016296718 |
| | 0.005523448 | | 0.02112972 |
| | 0.999036697 | | 0.999204617 |
| | 0.02261845 | | 0.066130002 |
| | 0.083802847 | | 0.184078569 |
| | 0.000396467 | | 0.002319743 |
| | 0.293565038 | | 0.454763099 |
| NA | | NA | |
| | 0.839318641 | NA | |
| | 5.71E-09 | | 9.26E-08 |
| NA | | NA | |
| | 0.039855784 | | 0.103889921 |
| NA | | NA | |
| | 4.45E-06 | | 4.38E-05 |
| | 0.631756409 | | 0.757871772 |
| | 6.54E-08 | | 9.03E-07 |
| | 0.133804592 | | 0.260226951 |
| | 0.007604001 | | 0.027506143 |
| | 0.520558845 | | 0.67087236 |
| NA | | NA | |
| | 0.628717668 | NA | |
| | 0.00017454 | | 0.001141944 |
| | 0.02532455 | | 0.072414097 |
| NA | | NA | |
| | 0.092569911 | | 0.198125081 |
| | 0.000324262 | | 0.001949501 |
| | 0.038798908 | | 0.101616594 |
| | 0.699968079 | | 0.808494212 |
| | 0.622331992 | | 0.750759312 |
| | 0.056691373 | | 0.136838866 |
| | 0.791000086 | | 0.871678506 |
| | 1.02E-08 | | 1.61E-07 |
| | 2.52E-07 | | 3.16E-06 |
| | 0.851707297 | | 0.911002778 |
| NA | | NA | |

| | |
|---|---|
| 0.077720747 | 0.173945389 |
| NA | NA |
| 6.72E-06 | 6.36E-05 |
| 0.673764363 | NA |
| 0.939680037 | NA |
| 0.940983457 | NA |
| 0.336796356 | 0.500078095 |
| 0.419494171 | 0.581153615 |
| 0.004038391 | 0.016338904 |
| 0.000237373 | 0.001483706 |
| NA | NA |
| NA | NA |
| NA | NA |
| 0.955792398 | NA |
| 0.877703711 | 0.926663806 |
| 0.085265809 | 0.186545368 |
| 0.864660582 | NA |
| 0.001135962 | 0.005691047 |
| 0.955791446 | NA |
| 1.13E-10 | 2.25E-09 |
| 0.897613296 | NA |
| NA | NA |
| 0.003444044 | 0.014363963 |
| 0.046125174 | 0.116275302 |
| 0.490595769 | NA |
| 1.45E-05 | 0.000126807 |
| 0.764251187 | 0.853630988 |
| 0.944289243 | NA |
| 0.016714768 | 0.051985484 |
| NA | NA |
| 0.955792398 | NA |
| 4.40E-07 | 5.25E-06 |
| 0.853595622 | NA |
| 0.301183182 | 0.462490658 |
| 0.305298686 | 0.466922786 |
| NA | NA |
| NA | NA |
| 0.955792398 | NA |
| NA | NA |
| 0.316660201 | 0.479126954 |
| 0.003113901 | 0.013223384 |
| 0.090133332 | 0.194501683 |
| 0.866692299 | 0.920403788 |
| 0.932798273 | 0.959940529 |
| NA | NA |
| 0.279231714 | 0.438996785 |
| 0.088039063 | 0.191126028 |

| | |
|---|---|
| 1.53E-16 | 5.19E-15 |
| 0.004853426 | 0.019046679 |
| NA | NA |
| 0.319954071 | 0.482638114 |
| 0.250863015 | 0.408484061 |
| NA | NA |
| 0.966720114 | 0.980144494 |
| 0.935876415 | 0.962068344 |
| 0.000429494 | 0.002485492 |
| 0.021435437 | 0.063230421 |
| NA | NA |
| 0.795915192 | 0.874871471 |
| NA | NA |
| 0.023302634 | 0.067788964 |
| 0.239235086 | 0.395092085 |
| 0.179217502 | 0.320852575 |
| NA | NA |
| 0.676259816 | 0.791324721 |
| 0.367277768 | 0.530457854 |
| 0.48016823 | NA |
| NA | NA |
| 0.908105935 | 0.944328606 |
| 0.936639337 | NA |
| 0.503947275 | 0.656071379 |
| 0.838993826 | NA |
| 0.925341217 | NA |
| 0.343499717 | 0.507037535 |
| 0.471903128 | 0.629405948 |
| 0.019757641 | 0.059299252 |
| 0.984241956 | 0.990065381 |
| 0.746505517 | 0.84054706 |
| 0.157085806 | 0.29283806 |
| 0.099613995 | 0.208991829 |
| 0.621398535 | NA |
| 0.659796076 | NA |
| 0.941278332 | NA |
| 0.839319108 | NA |
| 0.802647487 | NA |
| 3.56E-10 | 6.70E-09 |
| 0.021027717 | 0.062266947 |
| 0.776890031 | NA |
| 0.217269862 | 0.368604775 |
| 0.820623594 | NA |
| 0.906701153 | NA |
| 1.63E-05 | 0.000140223 |
| 0.82928908 | 0.896381426 |
| 0.212171536 | 0.362595303 |

| | |
|---|---|
| NA | NA |
| 0.75673401 | NA |
| 0.835127771 | NA |
| 0.839318641 | NA |
| 0.000146847 | 0.000980435 |
| 0.131819005 | 0.257349555 |
| 0.038531817 | 0.101017988 |
| 0.042150346 | 0.108519474 |
| 0.021400301 | 0.063150257 |
| NA | NA |
| 0.003108199 | 0.013206238 |
| NA | NA |
| 0.021942732 | 0.064487087 |
| 0.860539147 | 0.916609361 |
| 1.08E-07 | 1.44E-06 |
| 0.70865698 | 0.814125883 |
| NA | NA |
| 0.75127104 | NA |
| 0.941278332 | NA |
| 0.075589506 | 0.170538522 |
| 2.98E-05 | 0.000240211 |
| 0.003494698 | 0.01454464 |
| 0.209894917 | 0.359625598 |
| 0.002459527 | 0.010883372 |
| 0.03753468 | 0.098980773 |
| 0.003912605 | 0.015927453 |
| 0.690421481 | 0.801434387 |
| 0.114838923 | 0.232392114 |
| NA | NA |
| 1.45E-05 | 0.000126342 |
| 0.028909926 | 0.080603732 |
| 0.941277857 | NA |
| 0.955789428 | NA |
| 1.30E-10 | 2.58E-09 |
| 0.194633013 | 0.340786742 |
| 0.237248779 | 0.392697874 |
| 0.610293921 | 0.741890496 |
| 2.26E-05 | 0.000187636 |
| NA | NA |
| 0.072699104 | 0.165586992 |
| NA | NA |
| NA | NA |
| 0.839319108 | NA |
| NA | NA |
| NA | NA |
| NA | NA |
| 0.43827408 | 0.599149943 |

| | |
|---|---|
| NA | NA |
| 0.173478249 | NA |
| 0.584359518 | 0.721417285 |
| 0.005353549 | 0.020622296 |
| 0.002360464 | 0.010519335 |
| 0.068875718 | 0.159219031 |
| 0.941277857 | NA |
| NA | NA |
| 0.971070783 | 0.982586427 |
| 3.09E-08 | 4.49E-07 |
| 0.290393774 | 0.451525517 |
| 0.01201145 | 0.039924106 |
| 0.852099951 | 0.911172335 |
| 0.863364545 | NA |
| 0.027986859 | 0.078536101 |
| 2.56E-08 | 3.78E-07 |
| 0.214353152 | 0.365318182 |
| 0.374179748 | 0.537361311 |
| 0.024370381 | 0.070281073 |
| 0.853595622 | NA |
| 0.440931535 | 0.601673374 |
| 0.013277396 | 0.043313607 |
| 0.794990196 | NA |
| NA | NA |
| NA | NA |
| 0.000155849 | 0.001035021 |
| 0.003619399 | 0.014943441 |
| 0.292637712 | 0.453680897 |
| 0.010996136 | 0.037203985 |
| 0.941277857 | NA |
| 0.220343256 | 0.372142921 |
| NA | NA |
| NA | NA |
| NA | NA |
| NA | NA |
| NA | NA |
| NA | NA |
| NA | NA |
| 0.38504719 | NA |
| NA | NA |
| NA | NA |
| 0.941279803 | NA |
| 0.839319108 | NA |
| 0.411973993 | 0.574110362 |
| 0.65056996 | 0.77173517 |
| 0.708502718 | 0.814056764 |
| NA | NA |

| | | |
|---|---|---|
| | 0.475897701 | 0.633023196 |
| | 0.025819952 | 0.073541481 |
| | 0.044249186 | 0.112535434 |
| | 0.614334162 | NA |
| NA | | NA |
| | 0.995579151 | 0.996802697 |
| | 0.067677283 | 0.157064615 |
| | 0.830698075 | 0.897271536 |
| NA | | NA |
| | 0.839318641 | NA |
| | 0.063270881 | 0.149091525 |
| | 0.17688394 | 0.318495464 |
| | 0.24709981 | 0.404169164 |
| | 0.215196414 | 0.366178513 |
| | 0.248083764 | 0.405305083 |
| | 0.955792873 | NA |
| | 0.955792398 | NA |
| NA | | NA |
| NA | | NA |
| NA | | NA |
| | 0.321158546 | NA |
| | 0.839318641 | NA |
| | 0.517734507 | NA |
| | 0.941275956 | NA |
| NA | | NA |
| | 0.026499097 | 0.075167622 |
| | 0.005656538 | 0.021576429 |
| | 1.18E-13 | 3.20E-12 |
| | 0.52499383 | 0.674449691 |
| | 0.184127381 | 0.327084105 |
| | 2.63E-06 | 2.71E-05 |
| NA | | NA |
| | 0.93389938 | 0.960741368 |
| | 1.52E-13 | 4.10E-12 |
| | 0.941278207 | NA |
| | 0.084124793 | 0.184683482 |
| NA | | NA |
| | 0.390321252 | 0.552968659 |
| NA | | NA |
| | 0.853735184 | 0.912014938 |
| | 0.070500393 | 0.162067835 |
| | 0.730349419 | 0.830111521 |
| | 0.710285888 | 0.815150661 |
| | 0.462417268 | NA |
| NA | | NA |
| NA | | NA |
| | 0.683175831 | 0.796480756 |

| | | | |
|---|---|---|---|
| NA | | NA | |
| NA | | NA | |
| | 0.000431833 | | 0.002497812 |
| NA | | NA | |
| | 0.602478007 | | 0.735846752 |
| | 0.941278332 | NA | |
| | 0.926208399 | NA | |
| | 0.244666789 | NA | |
| | 0.734058779 | NA | |
| NA | | NA | |
| | 0.675884945 | | 0.791098983 |
| | 0.050923792 | | 0.1257092 |
| | 0.670825454 | | 0.786977547 |
| | 0.640580267 | NA | |
| | 0.296441255 | | 0.457907376 |
| | 0.721637981 | | 0.82332244 |
| | 0.396627312 | | 0.559272707 |
| | 0.90653467 | NA | |
| | 0.34997258 | | 0.513799278 |
| NA | | NA | |
| NA | | NA | |
| | 0.931428463 | NA | |
| | 0.196559769 | | 0.343276514 |
| | 0.016116231 | | 0.050493422 |
| | 0.361013343 | | 0.524689479 |
| | 0.93978302 | | 0.963757575 |
| | 0.533396317 | | 0.681695616 |
| | 0.12082079 | | 0.241625524 |
| | 0.901185089 | NA | |
| NA | | NA | |
| | 0.003951283 | | 0.016037173 |
| | 6.85E-05 | | 0.000499826 |
| | 0.014692634 | | 0.046891133 |
| | 0.139462099 | | 0.268305599 |
| | 0.739991096 | NA | |
| | 0.882952917 | NA | |
| | 0.946860719 | | 0.967625072 |
| | 0.908293155 | | 0.94444077 |
| | 0.015758692 | | 0.049634561 |
| | 0.011672705 | | 0.039076472 |
| | 0.019763926 | | 0.059299252 |
| NA | | NA | |
| NA | | NA | |
| | 0.751007199 | NA | |
| | 0.185566737 | | 0.328912667 |
| NA | | NA | |
| | 0.581394477 | NA | |

| | |
|---|---|
| NA | NA |
| 0.876780745 | 0.926202931 |
| NA | NA |
| NA | NA |
| 3.80E-13 | 9.92E-12 |
| NA | NA |
| NA | NA |
| 0.53529102 | 0.683419882 |
| 0.272110425 | 0.432044049 |
| 0.791059099 | NA |
| 0.011202069 | 0.037761175 |
| 0.553843713 | NA |
| 0.442059487 | 0.602594348 |
| 0.35147174 | 0.515109595 |
| 0.941183577 | 0.964703895 |
| 0.495559694 | NA |
| 0.84256253 | 0.905193905 |
| 0.018534243 | 0.056456622 |
| 0.714845536 | 0.818804305 |
| 0.143797285 | 0.274599742 |
| 0.329901221 | 0.492868062 |
| 0.35574216 | 0.519502723 |
| 0.062842033 | 0.148286515 |
| NA | NA |
| 0.853738599 | NA |
| 0.036296072 | 0.096441069 |
| 0.161057886 | 0.298071675 |
| 2.24E-05 | 0.000186532 |
| 0.208597966 | 0.357944544 |
| 0.195767199 | 0.342218777 |
| 0.000191471 | 0.001234062 |
| 0.661009296 | 0.779701886 |
| 0.005413609 | 0.020789727 |
| 0.927768867 | 0.95683325 |
| 0.016434137 | 0.05129981 |
| 0.799763531 | NA |
| NA | NA |
| 0.783997437 | NA |
| 0.884574912 | 0.930962684 |
| 0.439054691 | NA |
| 0.847698731 | NA |
| NA | NA |
| NA | NA |
| NA | NA |
| 0.941279303 | NA |
| NA | NA |
| NA | NA |

| | | | |
|---|---|---|---|
| NA | | NA | |
| | 0.288905237 | | 0.449739859 |
| | 1.15E-05 | | 0.000102027 |
| | 6.83E-09 | | 1.09E-07 |
| | 0.385688718 | | 0.548462353 |
| | 0.726152494 | | 0.827138288 |
| | 0.020056129 | | 0.059986929 |
| | 0.802000655 | | 0.879039399 |
| | 0.000255735 | | 0.001583099 |
| NA | | NA | |
| NA | | NA | |
| NA | | NA | |
| | 0.850989838 | | 0.91048553 |
| | 0.224342247 | | 0.377077478 |
| | 0.028692155 | | 0.080165592 |
| | 0.976887423 | | 0.985875533 |
| | 0.839319108 | NA | |
| NA | | NA | |
| NA | | NA | |
| | 0.324664342 | | 0.487611097 |
| | 0.001865746 | | 0.008629708 |
| | 0.001000338 | | 0.00511194 |
| | 0.074232822 | | 0.168163094 |
| | 0.134113655 | | 0.260606843 |
| | 8.29E-17 | | 2.85E-15 |
| | 0.953770009 | NA | |
| | 0.752065872 | NA | |
| | 0.697613061 | NA | |
| | 0.119053841 | | 0.238830132 |
| NA | | NA | |
| | 0.501573223 | | 0.653874786 |
| NA | | NA | |
| NA | | NA | |
| NA | | NA | |
| | 0.000306626 | | 0.001857075 |
| NA | | NA | |
| | 2.39E-15 | | 7.49E-14 |
| | 0.0240156 | | 0.069467816 |
| | 0.244610228 | NA | |
| | 0.92075031 | NA | |
| | 0.547382743 | NA | |
| NA | | NA | |
| | 0.955791922 | NA | |
| | 0.305386523 | NA | |
| NA | | NA | |
| | 2.00E-08 | | 2.99E-07 |
| | 0.653041268 | NA | |

| | | |
|---|---|---|
| | 0.198459143 | 0.345680535 |
| | 0.955792398 | NA |
| | 0.623281916 | 0.751385543 |
| | 0.728062495 | 0.828719024 |
| | 6.02E-08 | 8.35E-07 |
| | 0.793652823 | NA |
| NA | | NA |
| NA | | NA |
| | 0.479215912 | NA |
| | 0.290989821 | 0.452102601 |
| | 0.531125268 | NA |
| | 0.955791922 | NA |
| | 7.87E-36 | 6.99E-34 |
| | 0.001889372 | 0.008722743 |
| | 0.396013961 | NA |
| | 0.791710984 | 0.871750204 |
| | 0.604307385 | 0.737245902 |
| | 0.899547991 | 0.93958188 |
| | 0.119229037 | 0.23916143 |
| | 0.082250651 | 0.181472932 |
| NA | | NA |
| | 0.839319108 | NA |
| NA | | NA |
| NA | | NA |
| | 0.839319108 | NA |
| | 0.768552393 | 0.856578201 |
| | 0.74068855 | NA |
| | 0.955792873 | NA |
| | 0.695074144 | NA |
| | 0.934379906 | 0.960973711 |
| | 0.391288711 | 0.553778921 |
| NA | | NA |
| | 0.159794188 | 0.296470595 |
| | 0.239243014 | 0.395092085 |
| | 0.153096954 | 0.287337462 |
| NA | | NA |
| | 0.166664707 | 0.305549823 |
| | 0.006470474 | 0.024060338 |
| | 0.20824557 | 0.357571856 |
| | 0.936597705 | 0.9623981 |
| | 0.861253662 | 0.91708315 |
| | 0.839319108 | NA |
| | 0.839318641 | NA |
| | 0.345403215 | 0.509061583 |
| | 3.81E-08 | 5.45E-07 |
| NA | | NA |
| | 2.24E-28 | 1.47E-26 |

| | |
|---|---|
| NA | NA |
| 0.424897127 | NA |
| NA | NA |
| 0.904443126 | 0.942027255 |
| 0.839319574 | NA |
| 0.165624341 | 0.30432226 |
| NA | NA |
| 0.81924471 | 0.889826712 |
| NA | NA |
| NA | NA |
| 0.821581298 | NA |
| 0.955791446 | NA |
| 0.835058019 | 0.899970585 |
| NA | NA |
| NA | NA |
| NA | NA |
| NA | NA |
| NA | NA |
| 0.122607988 | 0.244230927 |
| 0.941278332 | NA |
| 0.892358641 | 0.935040806 |
| 0.069853218 | 0.160999931 |
| 0.839319108 | NA |
| 0.912969147 | NA |
| 0.010276367 | 0.035178783 |
| 0.245016506 | 0.401640474 |
| 0.006824695 | 0.025145727 |
| NA | NA |
| 0.002355374 | 0.010504518 |
| 0.018874802 | 0.057290931 |
| 0.055368817 | 0.134409851 |
| 0.184515079 | NA |
| 0.265505935 | NA |
| 0.978878876 | 0.986755675 |
| 0.095717618 | NA |
| 0.762989285 | 0.852875853 |
| 0.941278332 | NA |
| 0.033704502 | 0.09097693 |
| 9.31E-12 | 2.11E-10 |
| 0.716325178 | 0.819759372 |
| 0.139289628 | 0.268147179 |
| NA | NA |
| 0.812359253 | 0.88542389 |
| 3.64E-10 | 6.85E-09 |
| 0.955792398 | NA |
| NA | NA |
| 0.305246695 | 0.466903274 |

| | | |
|---|---|---|
| | 0.524540402 | 0.674080713 |
| | 0.003744689 | 0.01536478 |
| | 0.000571579 | 0.003179415 |
| | 0.941278332 | NA |
| | 6.47E-05 | 0.00047731 |
| | 0.578583671 | 0.716668386 |
| | 0.515887855 | 0.666804361 |
| NA | | NA |
| | 0.837694186 | 0.901514537 |
| | 0.019747261 | 0.059294096 |
| | 0.002873111 | 0.012366325 |
| | 0.33525156 | 0.498479363 |
| NA | | NA |
| | 0.115832225 | 0.233825683 |
| | 0.839319574 | NA |
| | 0.016031799 | 0.050301723 |
| | 0.893453094 | NA |
| | 0.526321081 | 0.67544586 |
| | 0.003241506 | 0.013687011 |
| | 0.31797943 | 0.480390123 |
| | 0.960804692 | 0.976017809 |
| | 0.014440172 | 0.046265309 |
| | 0.065258573 | 0.15274703 |
| | 0.533737984 | NA |
| | 0.007086993 | 0.025943495 |
| | 0.955791922 | NA |
| | 0.468384749 | 0.62651782 |
| | 0.941277857 | NA |
| | 0.065138591 | 0.152578102 |
| NA | | NA |
| NA | | NA |
| | 0.839319108 | NA |
| | 0.852729976 | NA |
| | 0.562545012 | NA |
| | 0.941278332 | NA |
| | 0.41806844 | NA |
| | 0.905922722 | 0.942923414 |
| | 0.021565349 | 0.06356175 |
| | 0.941275481 | NA |
| NA | | NA |
| | 0.59007316 | 0.7260984 |
| | 0.203085094 | 0.351295888 |
| | 8.55E-06 | 7.87E-05 |
| | 0.622551707 | NA |
| NA | | NA |
| | 0.99339223 | 0.995232 |
| | 7.43E-05 | 0.000538145 |

| | |
|---|---|
| 0.419228909 | 0.580887559 |
| 0.077226736 | 0.173149092 |
| NA | NA |
| NA | NA |
| 0.839320041 | NA |
| NA | NA |
| 0.917184594 | NA |
| 0.9127684 | NA |
| 0.923058171 | NA |
| NA | NA |
| NA | NA |
| 0.774260996 | 0.860603354 |
| NA | NA |
| NA | NA |
| 0.948858096 | 0.968760795 |
| 0.295964878 | 0.457468409 |
| 0.199882595 | 0.347321181 |
| 0.17759003 | 0.319182263 |
| 0.150008869 | 0.283194091 |
| 0.941277857 | NA |
| 0.839319574 | NA |
| 0.745499057 | NA |
| 0.107916248 | 0.221949584 |
| 0.788343313 | NA |
| 0.000217109 | 0.001377669 |
| 0.259021345 | 0.417443704 |
| 0.96706038 | 0.980280702 |
| NA | NA |
| 0.002795384 | 0.012084222 |
| 0.166047629 | 0.304887983 |
| 3.51E-05 | 0.000276547 |
| 0.000142867 | 0.000957086 |
| 0.333924526 | 0.497189514 |
| 0.004294754 | 0.017197801 |
| 0.566391179 | NA |
| NA | NA |
| NA | NA |
| 0.064138258 | 0.150717625 |
| NA | NA |
| 0.569038109 | NA |
| 0.941277857 | NA |
| NA | NA |
| NA | NA |
| NA | NA |
| 0.839319107 | NA |
| 0.175038288 | 0.316128791 |
| 0.82066222 | 0.890678686 |

| | | |
|---|---|---|
| | 0.152613668 | 0.286701699 |
| | 0.0381509 | 0.100316806 |
| | 0.978698598 | 0.986662209 |
| | 0.941276431 | NA |
| | 0.061796823 | 0.146415288 |
| | 0.190835409 | 0.335792445 |
| | 0.88858415 | 0.93342843 |
| | 0.52460714 | 0.674082225 |
| NA | | NA |
| | 0.112082247 | 0.228152026 |
| | 0.058541414 | 0.140223682 |
| | 0.000590308 | 0.003266041 |
| | 0.077982237 | 0.174415825 |
| | 0.021290553 | 0.062911162 |
| | 0.017323944 | 0.053502598 |
| | 0.422193993 | 0.583433664 |
| | 0.186908131 | 0.330740266 |
| | 0.871958525 | NA |
| NA | | NA |
| NA | | NA |
| | 0.65288389 | NA |
| NA | | NA |
| NA | | NA |
| | 0.739991096 | NA |
| | 0.321655999 | 0.484499183 |
| | 0.253226955 | NA |
| | 0.000505947 | 0.002865607 |
| | 0.363208971 | 0.526883223 |
| | 0.814691731 | NA |
| | 0.461404486 | 0.620261469 |
| NA | | NA |
| | 0.166271464 | 0.305099615 |
| | 0.941278332 | NA |
| | 0.839319574 | NA |
| | 0.03321061 | 0.089980529 |
| | 0.532438167 | 0.680836641 |
| | 0.43083208 | NA |
| | 0.093892767 | 0.200182339 |
| | 0.599082945 | NA |
| | 0.488650445 | NA |
| NA | | NA |
| | 0.941278788 | NA |
| | 0.019171167 | 0.057944396 |
| | 0.862809826 | 0.917839093 |
| | 0.6404377 | NA |
| | 0.730957639 | NA |
| | 2.16E-06 | 2.27E-05 |

| | | |
|---|---|---|
| | 0.941277857 | NA |
| | 0.80089593 | NA |
| NA | | NA |
| | 0.011641097 | 0.039003574 |
| | 0.000252548 | 0.001565808 |
| | 0.917818825 | NA |
| NA | | NA |
| NA | | NA |
| | 0.030352288 | 0.083722929 |
| | 0.028529763 | 0.079777422 |
| | 0.111739894 | 0.227676611 |
| NA | | NA |
| | 6.21E-05 | 0.0004604 |
| | 0.955664645 | 0.972955303 |
| | 0.175342105 | 0.316533393 |
| | 0.000249919 | 0.001551936 |
| | 0.941277857 | NA |
| | 0.738973018 | 0.835742458 |
| | 0.265755352 | 0.425101404 |
| | 0.99509094 | 0.996430566 |
| | 0.901975227 | NA |
| | 0.941275956 | NA |
| | 0.208870274 | 0.358237437 |
| | 0.850145647 | 0.910141082 |
| | 0.038532153 | 0.101017988 |
| | 0.454951435 | NA |
| | 0.000459955 | 0.002639945 |
| NA | | NA |
| | 0.92226807 | NA |
| | 0.671767839 | 0.787733674 |
| NA | | NA |
| | 0.616567798 | 0.746239537 |
| | 0.122910746 | 0.244649738 |
| | 0.941278332 | NA |
| | 0.000111482 | 0.000770471 |
| | 0.597178656 | 0.731822498 |
| | 6.11E-05 | 0.000453657 |
| | 0.142094731 | 0.272226239 |
| | 0.006089243 | 0.02291843 |
| | 0.776077797 | 0.861978709 |
| | 0.955792873 | NA |
| NA | | NA |
| NA | | NA |
| NA | | NA |
| NA | | NA |
| | 0.00961782 | 0.033361026 |
| | 0.299403452 | 0.460957372 |

| | |
|---|---|
| 0.257114923 | 0.415271046 |
| 0.894513362 | 0.936494702 |
| 0.7819393 | 0.865974863 |
| 0.700006698 | NA |
| 0.657666103 | 0.777093357 |
| 0.853595622 | NA |
| NA | NA |
| NA | NA |
| 0.002729395 | 0.011874441 |
| 4.33E-05 | 0.000333766 |
| 0.043988532 | NA |
| 0.639238088 | NA |
| 0.416370002 | 0.578104111 |
| NA | NA |
| 0.036563814 | 0.096968456 |
| NA | NA |
| 0.043428068 | 0.110956944 |
| NA | NA |
| 0.470413963 | 0.628186331 |
| 0.902855607 | 0.941349627 |
| NA | NA |
| NA | NA |
| 0.751265207 | NA |
| NA | NA |
| 1.89E-07 | 2.41E-06 |
| 0.861370197 | 0.917109919 |
| 0.83901926 | NA |
| 0.000286901 | 0.001752329 |
| 0.718944782 | 0.821089377 |
| 0.307164569 | 0.468812509 |
| 0.695947361 | 0.805261534 |
| 1.07E-38 | 1.04E-36 |
| 0.131975939 | 0.257567341 |
| 0.261417922 | 0.420168099 |
| 0.521035922 | 0.671232789 |
| 0.941278332 | NA |
| 0.240663118 | 0.396721624 |
| 0.470394182 | 0.628186331 |
| 0.085552685 | 0.187024708 |
| 1.90E-06 | 2.01E-05 |
| 0.036904663 | 0.097676503 |
| 0.20394416 | NA |
| 0.0007319 | 0.003927115 |
| 0.135841229 | 0.263190307 |
| 3.59E-90 | 1.08E-87 |
| 0.227473084 | 0.380860604 |
| 0.894747776 | NA |

| | | |
|---|---|---|
| | 0.025795819 | NA |
| | 0.102506703 | NA |
| NA | | NA |
| NA | | NA |
| NA | | NA |
| NA | | NA |
| | 0.005201257 | 0.020165948 |
| | 0.149874896 | 0.283053421 |
| | 0.754601615 | 0.846300129 |
| | 0.909745585 | NA |
| | 0.0013331 | 0.006499733 |
| | 0.380310773 | 0.542991079 |
| | 0.181622861 | 0.324013442 |
| | 0.210395441 | 0.360301624 |
| | 0.929996005 | 0.958402269 |
| | 0.024626206 | 0.070873907 |
| | 0.941275956 | NA |
| NA | | NA |
| NA | | NA |
| NA | | NA |
| | 0.839318641 | NA |
| NA | | NA |
| | 0.941278332 | NA |
| | 0.000448233 | 0.002582004 |
| | 0.839320041 | NA |
| | 0.193007564 | 0.338738095 |
| | 0.941278332 | NA |
| | 0.049016718 | 0.121797257 |
| | 0.31463061 | 0.476874541 |
| | 0.290703813 | NA |
| | 0.027998196 | 0.078558655 |
| | 0.941275481 | NA |
| NA | | NA |
| | 0.007095778 | 0.025971662 |
| | 0.906311961 | 0.943204814 |
| | 4.23E-05 | 0.00032636 |
| | 0.351442866 | 0.515109595 |
| | 0.55634696 | NA |
| | 0.000358137 | 0.002123664 |
| | 0.255356071 | 0.413468381 |
| | 0.222218208 | 0.374606684 |
| | 0.596251955 | 0.731120237 |
| | 0.694696796 | NA |
| | 0.222959909 | 0.375371361 |
| | 0.014730948 | 0.046985232 |
| | 0.702364436 | 0.810122034 |
| | 0.27710179 | 0.436995747 |

| | | |
|---|---|---|
| | 0.321578627 | 0.484443955 |
| | 0.760388458 | 0.850947478 |
| | 0.794877629 | 0.874089966 |
| | 4.63E-08 | 6.50E-07 |
| | 0.866062469 | 0.920022265 |
| | 0.098260678 | 0.206955195 |
| | 0.725135583 | 0.82635809 |
| | 0.77902631 | NA |
| | 0.012251323 | 0.040562801 |
| | 0.004721369 | 0.01864886 |
| NA | | NA |
| | 0.025428716 | 0.072607281 |
| | 0.55062273 | 0.696197727 |
| | 0.188813501 | 0.333343639 |
| | 0.364868269 | NA |
| | 0.550611635 | 0.696197727 |
| | 0.941278424 | NA |
| | 0.915816658 | NA |
| | 0.251983015 | 0.409734918 |
| | 0.484533596 | 0.640324879 |
| | 1.21E-206 | 1.70E-203 |
| | 0.098704099 | 0.20752186 |
| | 0.043876874 | 0.111815564 |
| | 0.000612234 | 0.003370119 |
| | 0.595393516 | NA |
| | 0.012505717 | 0.041233007 |
| | 0.941278332 | NA |
| | 0.71661376 | 0.819843341 |
| | 1.76E-05 | 0.000149908 |
| | 0.940685982 | NA |
| | 0.022832074 | 0.066656327 |
| | 0.007872494 | 0.028283941 |
| | 0.955792398 | NA |
| | 7.38E-06 | 6.90E-05 |
| | 0.411416035 | NA |
| NA | | NA |
| | 0.206963895 | 0.355961747 |
| | 0.072344621 | 0.165063906 |
| | 0.53886472 | 0.68636008 |
| | 6.02E-05 | 0.000447276 |
| | 0.598297247 | 0.73265814 |
| | 0.256381018 | 0.414479452 |
| | 0.742829353 | NA |
| | 0.000371508 | 0.002188922 |
| | 0.032852201 | 0.089232921 |
| | 2.28E-06 | 2.38E-05 |
| | 0.087310727 | 0.189873907 |

| | |
|---|---|
| 0.001418787 | 0.006844535 |
| 0.43895886 | 0.599913803 |
| 0.941278332 | NA |
| 9.39E-07 | 1.05E-05 |
| 0.017633509 | 0.054212448 |
| 0.56547294 | NA |
| 0.0302492 | 0.083535382 |
| 0.000494216 | 0.002808145 |
| 0.856591429 | 0.9138767 |
| 0.955792398 | NA |
| NA | NA |
| 4.21E-09 | 6.94E-08 |
| 0.001295466 | 0.006347281 |
| 0.020975426 | 0.062138544 |
| 0.77388967 | 0.860508485 |
| 0.094567177 | 0.20120568 |
| 4.38E-05 | 0.000337125 |
| 0.001324892 | 0.006463431 |
| 0.878239092 | 0.926958754 |
| 0.368424723 | 0.531533838 |
| 0.76323488 | 0.85297792 |
| 0.152517795 | 0.28664993 |
| 0.936872628 | 0.962580675 |
| NA | NA |
| NA | NA |
| NA | NA |
| 0.032869494 | 0.089258093 |
| 0.552641145 | 0.697860073 |
| 0.271123879 | 0.431023949 |
| 0.033336422 | 0.090290569 |
| 0.002249432 | 0.010095274 |
| NA | NA |
| 0.048588631 | 0.120998808 |
| 0.022731432 | 0.066403234 |
| 0.80399422 | NA |
| 0.00016461 | 0.001085935 |
| 0.839318641 | NA |
| 0.1546588 | 0.289475803 |
| 0.992949659 | 0.995039903 |
| 0.879091914 | NA |
| 0.037898529 | 0.099785486 |
| 0.34381288 | NA |
| 0.049719673 | 0.123196715 |
| 0.298115686 | NA |
| 0.725652862 | 0.826750978 |
| NA | NA |
| 0.122537983 | 0.244111908 |

| | |
|---|---|
| NA | NA |
| 2.86E-05 | 0.000231722 |
| 0.440955762 | NA |
| 0.003387992 | 0.01417988 |
| 1.27E-55 | 1.97E-53 |
| 6.20E-33 | 4.89E-31 |
| 0.59174828 | 0.727408393 |
| 0.014722395 | 0.04697352 |
| 0.367613164 | 0.530716839 |
| 0.100018006 | 0.209617732 |
| 0.001148468 | 0.005745237 |
| NA | NA |
| 0.853594685 | NA |
| 0.941278332 | NA |
| 0.853595154 | NA |
| NA | NA |
| NA | NA |
| NA | NA |
| 0.941277857 | NA |
| 0.941278332 | NA |
| NA | NA |
| NA | NA |
| 0.941277857 | NA |
| 0.631955598 | NA |
| 0.000295942 | 0.001803385 |
| 0.764313054 | NA |
| 0.938535472 | NA |
| 0.941278332 | NA |
| 0.838882104 | NA |
| 0.955791446 | NA |
| 0.936413394 | NA |
| NA | NA |
| 0.92069054 | NA |
| 0.945731973 | NA |
| 0.000358567 | 0.002124096 |
| 0.489069093 | 0.643583465 |
| 0.513285999 | 0.664449521 |
| 0.000208017 | 0.001326335 |
| 0.002062608 | 0.009379861 |
| 0.425630493 | NA |
| NA | NA |
| 0.01628866 | 0.050939783 |
| 0.668098601 | 0.7848617 |
| NA | NA |
| 0.078046902 | 0.174521749 |
| NA | NA |
| 0.343330997 | 0.506882793 |

| | |
|---|---|
| 0.635490228 | 0.760250222 |
| 0.86110685 | NA |
| 0.187958811 | 0.332303024 |
| NA | NA |
| 0.000196567 | 0.001261444 |
| 0.576897784 | NA |
| 0.652445508 | 0.773227843 |
| 0.9277323 | 0.95683325 |
| 1.75E-06 | 1.87E-05 |
| 0.955792398 | NA |
| 0.251260484 | 0.40896792 |
| 0.652787186 | NA |
| 0.015559964 | 0.049106107 |
| 0.088781719 | 0.192282299 |
| 0.10512012 | 0.217684801 |
| 0.769945577 | 0.857510576 |
| 0.503349242 | 0.655472079 |
| 0.854031683 | 0.91208307 |
| 0.846843839 | 0.907790356 |
| 0.705420328 | 0.812065262 |
| NA | NA |
| 0.477731677 | NA |
| 0.003592379 | 0.014857654 |
| 0.658210422 | NA |
| 0.437742401 | 0.598595004 |
| 0.65963915 | 0.778535779 |
| 0.793689871 | NA |
| 0.563885961 | NA |
| 0.023826398 | 0.068983813 |
| 0.001151323 | 0.005758306 |
| 7.43E-07 | 8.46E-06 |
| 0.108382194 | 0.222619345 |
| 0.17761682 | 0.319187282 |
| 0.089166273 | 0.192799385 |
| 0.955792398 | NA |
| 0.642909318 | NA |
| NA | NA |
| 0.587545261 | 0.723813843 |
| 0.129384815 | 0.253696133 |
| 0.955791446 | NA |
| 0.232787764 | 0.387279958 |
| 0.639277485 | NA |
| 0.553092566 | 0.69809679 |
| 0.003784384 | 0.015495597 |
| 1.70E-09 | 2.93E-08 |
| 1.21E-36 | 1.12E-34 |
| NA | NA |

| | | |
|---|---|---|
| | 0.839319108 | NA |
| | 0.229193754 | 0.382879684 |
| | 0.213047467 | 0.36374405 |
| | 0.469416069 | 0.627452198 |
| | 0.162891189 | 0.300583273 |
| | 0.493539776 | 0.646993147 |
| | 0.008255443 | 0.029424388 |
| | 0.014244896 | 0.045787577 |
| | 0.739991096 | NA |
| NA | | NA |
| | 0.000531278 | 0.002989475 |
| | 0.302912313 | 0.464377102 |
| | 0.00610434 | 0.022967988 |
| | 0.852262425 | 0.911218679 |
| | 0.019469933 | 0.058631495 |
| | 0.71112809 | 0.815773823 |
| NA | | NA |
| NA | | NA |
| NA | | NA |
| | 0.910533829 | 0.94586711 |
| NA | | NA |
| | 6.57E-08 | 9.08E-07 |
| | 0.644340304 | 0.76713625 |
| | 0.003619058 | 0.014943441 |
| | 0.207741971 | 0.356955385 |
| | 0.955792873 | NA |
| | 0.941277857 | NA |
| | 0.114931002 | 0.232519142 |
| | 0.024439855 | 0.070425839 |
| | 0.105412843 | 0.218133042 |
| | 0.620802252 | 0.749468748 |
| | 0.047595936 | NA |
| | 0.335061607 | 0.498321253 |
| | 0.839318641 | NA |
| | 0.6656049 | 0.783207822 |
| | 0.78569751 | 0.868287312 |
| | 0.88873844 | 0.93342843 |
| | 7.48E-11 | 1.52E-09 |
| | 9.91E-07 | 1.10E-05 |
| | 0.093714112 | 0.199855148 |
| | 0.000726341 | 0.003902567 |
| NA | | NA |
| | 0.129007574 | 0.253227634 |
| | 0.219279214 | 0.370803078 |
| | 0.228453679 | 0.38201872 |
| | 0.941278332 | NA |
| | 0.941277857 | NA |

| | |
|---|---|
| 2.29E-05 | 0.000189733 |
| 0.269633356 | 0.429601254 |
| NA | NA |
| NA | NA |
| NA | NA |
| NA | NA |
| 0.386195778 | 0.549052199 |
| NA | NA |
| 0.089782784 | 0.193834029 |
| 0.894616186 | NA |
| 0.026450135 | 0.075043674 |
| 2.32E-50 | 3.19E-48 |
| 0.186885016 | 0.330723952 |
| 0.955792398 | NA |
| NA | NA |
| 0.206507087 | 0.355432907 |
| NA | NA |
| 0.725478031 | 0.826607376 |
| 0.318657581 | 0.481078761 |
| NA | NA |
| 3.27E-11 | 6.98E-10 |
| 0.522449049 | 0.672398197 |
| 0.319107864 | 0.481605819 |
| NA | NA |
| 0.941278332 | NA |
| 0.685854386 | NA |
| 0.728678906 | NA |
| 0.091717693 | 0.196778847 |
| 0.7541385 | 0.846066498 |
| 0.000423374 | 0.002454845 |
| 0.786552437 | 0.868828713 |
| 4.15E-06 | 4.11E-05 |
| 0.77016954 | NA |
| 0.09736414 | 0.205539802 |
| 0.121250149 | 0.242259127 |
| 0.011463842 | 0.038480097 |
| 0.034375528 | 0.092322967 |
| 5.57E-08 | 7.76E-07 |
| 0.008345843 | 0.029684361 |
| 0.839319574 | NA |
| 0.736079242 | NA |
| 0.090258153 | 0.194718078 |
| 0.0002627 | 0.001619893 |
| 0.09752417 | 0.205749894 |
| 0.414991784 | 0.576829855 |
| 1.57E-42 | 1.79E-40 |
| 0.004385731 | 0.017514963 |

| | | |
|---|---|---|
| | 0.378206012 | 0.541061866 |
| | 0.016875534 | 0.052389653 |
| NA | | NA |
| | 0.373666148 | 0.536818062 |
| | 0.003054154 | 0.013013778 |
| | 2.29E-09 | 3.89E-08 |
| | 0.88998277 | 0.933769151 |
| | 0.224307226 | 0.37705571 |
| | 0.072541407 | 0.165328076 |
| | 0.248366531 | 0.405572185 |
| | 0.047111257 | 0.118117656 |
| | 0.126767902 | 0.250176116 |
| | 1.01E-09 | 1.80E-08 |
| | 0.002105835 | 0.00954362 |
| | 0.081537106 | NA |
| | 0.002479215 | 0.01095827 |
| | 0.001092225 | 0.005506703 |
| | 0.812169632 | 0.88537426 |
| | 0.216153363 | 0.367386629 |
| | 0.285936206 | 0.446578319 |
| | 0.001246196 | 0.00614883 |
| | 0.116832994 | 0.235331856 |
| | 0.009736501 | 0.033701428 |
| | 0.564244919 | 0.706292798 |
| | 0.062833047 | 0.14828001 |
| | 0.096935252 | 0.2047797 |
| | 0.714933291 | 0.818865415 |
| | 0.00200148 | 0.009171974 |
| | 0.77438617 | 0.860702293 |
| NA | | NA |
| | 0.910364749 | NA |
| | 0.677021642 | 0.791715346 |
| | 0.006118971 | 0.023009997 |
| | 0.460944475 | NA |
| | 0.001428432 | 0.006882157 |
| NA | | NA |
| NA | | NA |
| | 0.004907443 | 0.019221155 |
| | 0.637684151 | 0.761846856 |
| | 0.088155208 | 0.19126738 |
| | 0.767053257 | NA |
| | 0.057255763 | 0.137871081 |
| | 0.000534747 | 0.003006862 |
| | 0.094250462 | 0.20069328 |
| | 0.648261775 | 0.770105487 |
| | 0.0055876 | 0.021361398 |
| | 0.504364436 | 0.656506743 |

| | |
|---|---|
| 0.012348772 | 0.040820586 |
| 0.121948679 | 0.243345277 |
| 0.132479617 | 0.258243283 |
| 0.289486551 | 0.450291393 |
| 0.418149921 | 0.579763034 |
| 0.153839578 | 0.288503753 |
| 0.955792398 | NA |
| 0.033739549 | 0.091061202 |
| 0.041784339 | 0.107798703 |
| 0.108157933 | 0.222235422 |
| 0.006178533 | 0.023192151 |
| 6.33E-15 | 1.91E-13 |
| 0.257619218 | NA |
| 9.27E-39 | 9.08E-37 |
| 0.306530715 | 0.468115236 |
| 0.612493455 | 0.743447088 |
| 0.001759687 | 0.008225464 |
| 0.30641913 | 0.468073802 |
| 0.628868254 | 0.755696114 |
| 0.086475167 | 0.188504618 |
| 0.998466882 | 0.99876061 |
| 0.16388897 | 0.301996227 |
| 0.782652554 | 0.866267226 |
| 0.021385987 | 0.06311772 |
| 0.022316836 | 0.065376655 |
| 6.38E-05 | 0.000471371 |
| 1.05E-05 | 9.48E-05 |
| 0.091790733 | 0.196846835 |
| 0.222226972 | 0.374606684 |
| 0.006194388 | 0.023233348 |
| 1.65E-06 | 1.78E-05 |
| 7.08E-06 | 6.63E-05 |
| 0.954222681 | NA |
| 0.370716569 | 0.533998777 |
| 0.459705347 | 0.618816122 |
| 0.831595201 | 0.89785986 |
| 0.483587931 | 0.639571037 |
| 7.59E-05 | 0.000549088 |
| 0.479140316 | 0.635879442 |
| 0.022402622 | 0.06559567 |
| 0.00369584 | 0.015198407 |
| 0.017768055 | 0.05456267 |
| 0.47334311 | 0.630893278 |
| 1.29E-05 | 0.000114022 |
| 0.072771138 | 0.165703494 |
| 0.029204322 | 0.081224733 |
| 0.941278332 | NA |

| | |
|---|---|
| 0.786146446 | 0.868581799 |
| 0.086444535 | 0.188455103 |
| 2.29E-14 | 6.59E-13 |
| 0.435471966 | NA |
| 4.35E-13 | 1.13E-11 |
| 0.24909716 | NA |
| 2.88E-05 | 0.000232892 |
| 2.84E-32 | 2.18E-30 |
| 8.95E-20 | 3.73E-18 |
| 0.39402741 | 0.556562839 |
| 0.955792398 | NA |
| 0.167684512 | 0.306899404 |
| NA | NA |
| NA | NA |
| NA | NA |
| NA | NA |
| NA | NA |
| NA | NA |
| NA | NA |
| NA | NA |
| NA | NA |
| NA | NA |
| NA | NA |
| NA | NA |
| NA | NA |
| NA | NA |
| NA | NA |
| NA | NA |
| NA | NA |
| 0.997187557 | 0.997648616 |
| NA | NA |
| NA | NA |
| NA | NA |
| NA | NA |
| NA | NA |
| NA | NA |
| NA | NA |
| NA | NA |
| NA | NA |
| NA | NA |
| NA | NA |
| NA | NA |
| NA | NA |
| NA | NA |
| NA | NA |
| NA | NA |
| NA | NA |

| | |
|---|---|
| NA | NA |
| NA | NA |
| NA | NA |
| NA | NA |
| NA | NA |
| NA | NA |
| NA | NA |
| NA | NA |
| NA | NA |
| NA | NA |
| NA | NA |
| NA | NA |
| NA | NA |
| NA | NA |
| NA | NA |
| NA | NA |
| NA | NA |
| NA | NA |
| NA | NA |
| NA | NA |
| NA | NA |
| NA | NA |
| NA | NA |
| NA | NA |
| NA | NA |
| NA | NA |
| NA | NA |
| NA | NA |
| NA | NA |
| NA | NA |
| NA | NA |
| NA | NA |
| NA | NA |
| NA | NA |
| NA | NA |
| NA | NA |
| NA | NA |
| NA | NA |
| NA | NA |
| NA | NA |
| NA | NA |
| NA | NA |
| NA | NA |
| NA | NA |
| NA | NA |
| NA | NA |
| NA | NA |

| | | |
|---|---|---|
| NA | | NA |
| NA | | NA |
| NA | | NA |
| NA | | NA |
| NA | | NA |
| NA | | NA |
| NA | | NA |
| NA | | NA |
| NA | | NA |
| NA | | NA |
| NA | | NA |
| NA | | NA |
| NA | | NA |
| NA | | NA |
| NA | | NA |
| NA | | NA |
| NA | | NA |
| NA | | NA |
| NA | | NA |
| NA | | NA |
| NA | | NA |
| NA | | NA |
| NA | | NA |
| NA | | NA |
| NA | | NA |
| NA | | NA |
| | 0.941278332 | NA |
| NA | | NA |
| NA | | NA |
| | 0.955792873 | NA |
| NA | | NA |
| | 0.839319574 | NA |
| NA | | NA |
| NA | | NA |
| NA | | NA |
| NA | | NA |
| NA | | NA |
| NA | | NA |
| NA | | NA |
| NA | | NA |
| NA | | NA |
| NA | | NA |
| NA | | NA |
| NA | | NA |
| NA | | NA |
| NA | | NA |
| NA | | NA |

| | |
|---|---|
| NA | NA |
| NA | NA |
| NA | NA |
| NA | NA |
| NA | NA |
| NA | NA |
| NA | NA |
| NA | NA |
| NA | NA |
| NA | NA |
| NA | NA |
| NA | NA |
| NA | NA |
| NA | NA |
| NA | NA |
| NA | NA |
| NA | NA |
| NA | NA |
| NA | NA |
| NA | NA |
| NA | NA |
| NA | NA |
| NA | NA |
| NA | NA |
| NA | NA |
| NA | NA |
| NA | NA |
| NA | NA |
| NA | NA |
| NA | NA |
| NA | NA |
| NA | NA |
| NA | NA |
| NA | NA |
| NA | NA |
| NA | NA |
| NA | NA |
| NA | NA |
| NA | NA |
| NA | NA |
| NA | NA |
| NA | NA |
| NA | NA |
| NA | NA |
| NA | NA |
| NA | NA |
| NA | NA |

| | |
|---|---|
| NA | NA |
| NA | NA |
| NA | NA |
| NA | NA |
| NA | NA |
| NA | NA |
| NA | NA |
| NA | NA |
| NA | NA |
| NA | NA |
| NA | NA |
| NA | NA |
| NA | NA |
| NA | NA |
| NA | NA |
| NA | NA |
| NA | NA |
| NA | NA |
| NA | NA |
| NA | NA |
| NA | NA |
| NA | NA |
| NA | NA |
| NA | NA |
| NA | NA |
| NA | NA |
| NA | NA |
| NA | NA |
| NA | NA |
| NA | NA |
| NA | NA |
| NA | NA |
| NA | NA |
| NA | NA |
| NA | NA |
| NA | NA |
| NA | NA |
| NA | NA |
| NA | NA |
| NA | NA |
| NA | NA |
| NA | NA |
| NA | NA |
| NA | NA |
| NA | NA |
| NA | NA |
| NA | NA |

| | |
|---|---|
| NA | NA |
| NA | NA |
| NA | NA |
| NA | NA |
| NA | NA |
| NA | NA |
| NA | NA |
| NA | NA |
| NA | NA |
| NA | NA |
| NA | NA |
| 0.432633684 | 0.59382618 |
| NA | NA |
| 0.232473015 | 0.386891533 |
| 4.52E-07 | 5.37E-06 |
| 0.578423009 | 0.716629082 |
| 0.02769397 | 0.077925509 |
| 0.163525463 | 0.301536494 |
| 0.003107321 | 0.013204866 |
| 0.842049351 | 0.904803737 |
| 0.000107124 | 0.000743369 |
| 0.378915268 | 0.541721228 |
| 0.267036749 | NA |
| 0.112327924 | 0.228573929 |
| 0.839084951 | NA |
| 0.24439519 | 0.400880197 |
| 0.037270848 | 0.098426799 |
| 0.942499006 | NA |
| 0.138266316 | 0.26678136 |
| 0.941277857 | NA |
| 0.330855292 | 0.493917314 |
| 0.236126532 | 0.391302911 |
| 0.062809663 | 0.148239523 |
| NA | NA |
| 0.492429384 | 0.646344116 |
| NA | NA |
| 0.889426117 | 0.933455095 |
| 0.839863553 | 0.903318223 |
| 0.944436487 | NA |
| 0.743511643 | NA |
| 0.530852903 | NA |
| 0.564723438 | 0.706620985 |
| 0.001203926 | 0.005979934 |
| 0.889742173 | 0.933635021 |
| 0.39102338 | 0.55350214 |
| 0.000264836 | 0.001631796 |
| 0.001091028 | 0.005502352 |

| | |
|---|---|
| NA | NA |
| NA | NA |
| NA | NA |
| NA | NA |
| NA | NA |
| NA | NA |
| NA | NA |
| NA | NA |
| NA | NA |
| NA | NA |
| NA | NA |
| NA | NA |
| NA | NA |
| NA | NA |
| NA | NA |
| NA | NA |
| NA | NA |
| NA | NA |
| NA | NA |
| NA | NA |
| NA | NA |
| NA | NA |
| NA | NA |
| NA | NA |
| NA | NA |
| NA | NA |
| NA | NA |
| NA | NA |
| NA | NA |
| NA | NA |
| NA | NA |
| NA | NA |
| NA | NA |
| NA | NA |
| NA | NA |
| NA | NA |
| NA | NA |
| NA | NA |
| NA | NA |
| NA | NA |
| NA | NA |
| NA | NA |
| NA | NA |
| NA | NA |
| NA | NA |
| NA | NA |
| NA | NA |

| | |
|---|---|
| NA | NA |
| NA | NA |
| NA | NA |
| NA | NA |
| NA | NA |
| NA | NA |
| NA | NA |
| NA | NA |
| NA | NA |
| NA | NA |
| NA | NA |
| 0.659596891 | 0.778524506 |
| 0.941275481 | NA |
| 0.941275956 | NA |
| 0.003339496 | 0.014023764 |
| NA | NA |
| 0.185345535 | 0.328611812 |
| 0.44483775 | 0.605479964 |
| 0.278554384 | 0.438299163 |
| NA | NA |
| 0.189553477 | 0.334154336 |
| NA | NA |
| 0.054303282 | 0.132254856 |
| 0.600899186 | 0.734592824 |
| 0.443725804 | 0.604277468 |
| 0.290383463 | 0.451525517 |
| 0.857649346 | NA |
| NA | NA |
| 0.748393366 | NA |
| 0.07552557 | 0.17045379 |
| NA | NA |
| 0.83982623 | NA |
| 0.227687624 | NA |
| 0.839319108 | NA |
| 0.907021222 | NA |
| 0.164962923 | NA |
| 0.107832604 | 0.221854234 |
| 0.448755742 | 0.608828327 |
| 0.86853576 | 0.921216246 |
| 0.839318641 | NA |
| NA | NA |
| 0.892147254 | NA |
| 0.941278332 | NA |
| 0.257284578 | 0.41534777 |
| 0.840091215 | 0.903399796 |
| 0.74409079 | 0.839257403 |
| 0.001867853 | 0.008636098 |

| | | | |
|---|---|---|---|
| | 0.574525563 | | 0.713786167 |
| | 0.048811212 | | 0.121445485 |
| | 0.395621121 | | 0.558217672 |
| | 0.825858288 | | 0.894113212 |
| | 0.977022715 | | 0.985928463 |
| | 2.52E-06 | | 2.61E-05 |
| | 0.454765432 | | 0.614564193 |
| NA | | NA | |
| | 0.000648977 | | 0.003537201 |
| NA | | NA | |
| | 0.614172368 | | 0.744362274 |
| | 0.001393047 | | 0.006742032 |
| | 0.019164221 | | 0.057938115 |
| | 0.68860131 | | 0.800376082 |
| | 0.006609822 | | 0.024513398 |
| | 0.533305609 | | 0.681616287 |
| | 0.090355611 | | 0.194840031 |
| | 1.03E-05 | | 9.28E-05 |
| | 5.62E-06 | | 5.41E-05 |
| | 1.07E-06 | | 1.18E-05 |
| | 0.246995551 | | 0.404026397 |
| | 0.939489656 | | 0.963757575 |
| | 0.046673179 | | 0.11732126 |
| | 0.941278332 | NA | |
| | 0.499568093 | NA | |
| | 0.051387889 | | 0.126710301 |
| | 0.895203459 | NA | |
| | 0.130392632 | | 0.25531787 |
| NA | | NA | |
| | 0.299779861 | NA | |
| | 0.186529514 | | 0.330291287 |
| | 0.853595622 | NA | |
| | 0.258855664 | | 0.41723319 |
| | 0.429541907 | | 0.590769691 |
| | 0.839319108 | NA | |
| | 0.168110381 | | 0.307513317 |
| | 0.218530873 | | 0.369843546 |
| | 1.66E-27 | | 1.05E-25 |
| NA | | NA | |
| | 0.206354496 | | 0.355247339 |
| | 0.791189627 | NA | |
| | 0.4653984 | | 0.623798442 |
| | 8.20E-11 | | 1.66E-09 |
| | 0.304922396 | | 0.46655714 |
| | 0.013317999 | | 0.043418027 |
| | 0.372105092 | | 0.535415634 |
| | 0.018705641 | | 0.056884076 |

| | |
|---|---|
| 0.015864683 | 0.049875999 |
| 0.744872085 | 0.839740698 |
| NA | NA |
| NA | NA |
| NA | NA |
| NA | NA |
| NA | NA |
| NA | NA |
| 0.74627078 | NA |
| NA | NA |
| NA | NA |
| NA | NA |
| 0.955792873 | NA |
| 0.822151266 | NA |
| NA | NA |
| NA | NA |
| 0.225678917 | 0.378602311 |
| NA | NA |
| 0.706016082 | NA |
| 0.014382209 | 0.046135491 |
| 0.867467691 | NA |
| NA | NA |
| NA | NA |
| NA | NA |
| NA | NA |
| NA | NA |
| NA | NA |
| NA | NA |
| NA | NA |
| 0.923475547 | NA |
| NA | NA |
| NA | NA |
| NA | NA |
| NA | NA |
| 0.941278332 | NA |
| NA | NA |
| NA | NA |
| NA | NA |
| NA | NA |
| NA | NA |
| NA | NA |
| 3.40E-05 | 0.000269235 |
| 0.839318641 | NA |
| 5.07E-05 | 0.000383607 |
| 0.070224046 | 0.161606878 |
| 0.578995325 | NA |
| 0.866888433 | 0.920524624 |

| | | |
|---|---|---|
| | 0.87628572 | NA |
| | 0.739991096 | NA |
| | 7.06E-117 | 2.85E-114 |
| | 0.955792398 | NA |
| | 0.709950696 | NA |
| | 0.299682354 | 0.461207771 |
| | 0.66421716 | 0.782114139 |
| | 0.196231976 | 0.342880156 |
| | 4.82E-05 | 0.000367116 |
| | 0.478362419 | NA |
| | 0.277810645 | 0.437492379 |
| | 0.326485243 | 0.489405996 |
| | 0.450894789 | NA |
| | 0.260595437 | 0.419242486 |
| | 0.001037766 | 0.00527009 |
| | 0.021236431 | 0.062806849 |
| | 0.657817954 | 0.777100741 |
| | 0.839319574 | NA |
| | 0.883920612 | 0.930767935 |
| | 0.299389622 | 0.460957372 |
| | 0.888908943 | 0.93342843 |
| | 0.364746574 | 0.528065162 |
| NA | | NA |
| | 0.955792873 | NA |
| | 0.278090464 | NA |
| | 0.955792398 | NA |
| NA | | NA |
| NA | | NA |
| NA | | NA |
| NA | | NA |
| NA | | NA |
| NA | | NA |
| NA | | NA |
| NA | | NA |
| NA | | NA |
| NA | | NA |
| NA | | NA |
| NA | | NA |
| NA | | NA |
| | 0.834999579 | 0.899970585 |
| NA | | NA |
| | 0.30084876 | 0.462216768 |
| NA | | NA |
| | 0.955792398 | NA |
| | 0.861619314 | NA |
| | 0.214374551 | 0.365328494 |
| | 0.300669424 | 0.462099415 |

| | | |
|---|---|---|
| | 0.839319108 | NA |
| | 0.557870633 | 0.7015235 |
| | 0.676179866 | NA |
| | 4.25E-09 | 6.99E-08 |
| | 0.875267927 | 0.9253821 |
| | 0.839319108 | NA |
| NA | | NA |
| | 0.039226182 | 0.102555095 |
| | 0.935650667 | NA |
| | 0.024648164 | 0.070897353 |
| | 0.756079714 | 0.84719517 |
| | 0.660171324 | NA |
| NA | | NA |
| | 0.002305053 | 0.010318766 |
| NA | | NA |
| | 0.150902077 | 0.28456435 |
| | 0.570460482 | 0.711118697 |
| NA | | NA |
| | 1.75E-06 | 1.87E-05 |
| | 0.371232244 | 0.534547481 |
| | 0.853596091 | NA |
| | 0.947601587 | NA |
| | 0.884330429 | 0.930962684 |
| NA | | NA |
| | 1.69E-06 | 1.82E-05 |
| | 0.001049308 | 0.005316228 |
| NA | | NA |
| | 0.172350869 | 0.312579654 |
| NA | | NA |
| NA | | NA |
| NA | | NA |
| NA | | NA |
| NA | | NA |
| | 0.915308305 | NA |
| NA | | NA |
| NA | | NA |
| | 0.927535741 | NA |
| NA | | NA |
| NA | | NA |
| NA | | NA |
| NA | | NA |
| NA | | NA |
| | 0.716457703 | NA |
| NA | | NA |
| | 0.941278207 | NA |
| | 0.941277857 | NA |
| | 0.941277857 | NA |

| | | |
|---|---|---|
| | 0.204575212 | 0.353179697 |
| | 2.81E-07 | 3.48E-06 |
| | 5.80E-05 | 0.00043289 |
| | 0.051842811 | 0.127594104 |
| NA | | NA |
| | 0.955792398 | NA |
| | 0.611912509 | 0.742871648 |
| | 0.941278284 | NA |
| NA | | NA |
| NA | | NA |
| NA | | NA |
| NA | | NA |
| | 0.637227779 | 0.761563346 |
| NA | | NA |
| NA | | NA |
| | 0.127359434 | 0.250986028 |
| NA | | NA |
| NA | | NA |
| | 0.744481321 | 0.839498977 |
| | 0.996946363 | 0.997558905 |
| | 0.073607043 | 0.167154536 |
| | 0.036828667 | 0.097513031 |
| | 0.943535493 | NA |
| NA | | NA |
| NA | | NA |
| NA | | NA |
| NA | | NA |
| | 0.315588071 | NA |
| | 0.941277857 | NA |
| | 0.82739196 | NA |
| | 0.787267633 | 0.869255657 |
| | 0.000361417 | 0.002137828 |
| | 0.284650131 | 0.44526878 |
| NA | | NA |
| | 0.491410514 | 0.645465283 |
| | 0.739991096 | NA |
| | 0.986629365 | 0.991639555 |
| | 0.04254013 | 0.109274788 |
| | 0.839318641 | NA |
| NA | | NA |
| | 0.032304233 | 0.087982111 |
| | 0.431623249 | 0.592931833 |
| | 0.421510974 | 0.582963638 |
| | 0.871073592 | 0.922874283 |
| | 4.19E-06 | 4.15E-05 |
| | 0.000123961 | 0.000846877 |
| | 0.818277754 | 0.889101027 |

| | |
|---|---|
| 4.56E-08 | 6.42E-07 |
| 0.144155802 | 0.275134023 |
| NA | NA |
| NA | NA |
| 0.760678501 | NA |
| 0.033944409 | 0.09145855 |
| NA | NA |
| 3.15E-24 | 1.70E-22 |
| NA | NA |
| 0.215424117 | 0.366382625 |
| 3.55E-05 | 0.000279657 |
| 0.192913097 | 0.338622238 |
| 0.010800436 | 0.036651265 |
| 0.891370435 | 0.934565988 |
| 0.236505208 | NA |
| 0.139576833 | 0.268439542 |
| NA | NA |
| 0.280113826 | 0.440083782 |
| NA | NA |
| 0.839318641 | NA |
| 0.223441021 | NA |
| 0.12715446 | 0.250731449 |
| 0.610902077 | 0.742227061 |
| 0.937822002 | 0.963153577 |
| NA | NA |
| 0.330484876 | NA |
| NA | NA |
| 0.103838902 | 0.215700047 |
| 0.955792873 | NA |
| NA | NA |
| 0.715251169 | NA |
| NA | NA |
| NA | NA |
| NA | NA |
| NA | NA |
| NA | NA |
| NA | NA |
| NA | NA |
| NA | NA |
| NA | NA |
| 0.141498501 | 0.271324202 |
| 0.429578352 | NA |
| 0.955792873 | NA |
| 0.785811851 | 0.868333039 |
| NA | NA |
| 0.785559102 | NA |
| NA | NA |

| | | |
|---|---|---|
| NA | | NA |
| NA | | NA |
| NA | | NA |
| NA | | NA |
| NA | | NA |
| | 0.941278332 | NA |
| NA | | NA |
| NA | | NA |
| NA | | NA |
| NA | | NA |
| NA | | NA |
| NA | | NA |
| NA | | NA |
| NA | | NA |
| NA | | NA |
| NA | | NA |
| NA | | NA |
| NA | | NA |
| NA | | NA |
| NA | | NA |
| NA | | NA |
| NA | | NA |
| NA | | NA |
| NA | | NA |
| NA | | NA |
| NA | | NA |
| NA | | NA |
| NA | | NA |
| NA | | NA |
| NA | | NA |
| NA | | NA |
| NA | | NA |
| NA | | NA |
| NA | | NA |
| NA | | NA |
| NA | | NA |
| NA | | NA |
| NA | | NA |
| NA | | NA |
| NA | | NA |
| NA | | NA |
| NA | | NA |
| NA | | NA |
| NA | | NA |
| NA | | NA |
| NA | | NA |
| NA | | NA |

| | | | |
|---|---|---|---|
| NA | | NA | |
| NA | | NA | |
| NA | | NA | |
| NA | | NA | |
| NA | | NA | |
| NA | | NA | |
| NA | | NA | |
| | 0.461369029 | | 0.620248835 |
| NA | | NA | |
| NA | | NA | |
| NA | | NA | |
| NA | | NA | |
| NA | | NA | |
| NA | | NA | |
| NA | | NA | |
| NA | | NA | |
| NA | | NA | |
| NA | | NA | |
| NA | | NA | |
| NA | | NA | |
| NA | | NA | |
| NA | | NA | |
| NA | | NA | |
| NA | | NA | |
| NA | | NA | |
| NA | | NA | |
| NA | | NA | |
| NA | | NA | |
| NA | | NA | |
| NA | | NA | |
| NA | | NA | |
| NA | | NA | |
| NA | | NA | |
| NA | | NA | |
| NA | | NA | |
| NA | | NA | |
| NA | | NA | |
| NA | | NA | |
| NA | | NA | |
| NA | | NA | |
| NA | | NA | |
| NA | | NA | |
| NA | | NA | |
| NA | | NA | |
| NA | | NA | |
| NA | | NA | |
| NA | | NA | |

| | | | |
|---|---|---|---|
| NA | | NA | |
| | 0.154125564 | | 0.288812524 |
| NA | | NA | |
| NA | | NA | |
| NA | | NA | |
| NA | | NA | |
| NA | | NA | |
| NA | | NA | |
| NA | | NA | |
| NA | | NA | |
| NA | | NA | |
| NA | | NA | |
| NA | | NA | |
| NA | | NA | |
| NA | | NA | |
| NA | | NA | |
| NA | | NA | |
| NA | | NA | |
| NA | | NA | |
| NA | | NA | |
| NA | | NA | |
| NA | | NA | |
| NA | | NA | |
| | 0.941278332 | NA | |
| NA | | NA | |
| | 0.033420338 | | 0.090435527 |
| | 5.85E-19 | | 2.32E-17 |
| | 0.546158444 | | 0.692429066 |
| NA | | NA | |
| | 0.185974695 | NA | |
| NA | | NA | |
| | 0.437523791 | | 0.598364817 |
| | 0.391931158 | | 0.554259724 |
| | 0.631905808 | NA | |
| NA | | NA | |
| | 0.839319108 | NA | |
| NA | | NA | |
| | 0.001628787 | | 0.007702838 |
| | 0.41301589 | | 0.575191868 |
| | 0.001509662 | | 0.007209074 |
| | 0.557421247 | | 0.701317451 |
| | 0.022677244 | | 0.066269338 |
| | 0.704871192 | NA | |
| | 0.872249381 | | 0.923626646 |
| NA | | NA | |
| | 0.007458812 | | 0.027083852 |
| | 0.045629383 | | 0.115258126 |

| | |
|---|---|
| 0.067569289 | 0.156875168 |
| 0.142041323 | 0.272167733 |
| 0.000255078 | 0.001579437 |
| 0.971075105 | NA |
| 0.220409623 | 0.372228597 |
| NA | NA |
| 0.955792398 | NA |
| NA | NA |
| NA | NA |
| 0.000135874 | 0.00091669 |
| 0.619063371 | 0.748320477 |
| 0.459469916 | 0.61870915 |
| 0.033412181 | 0.090423733 |
| 0.482386214 | 0.638554587 |
| 0.371284803 | 0.534590822 |
| 0.250772535 | 0.40840834 |
| NA | NA |
| NA | NA |
| NA | NA |
| NA | NA |
| NA | NA |
| NA | NA |
| NA | NA |
| NA | NA |
| NA | NA |
| NA | NA |
| NA | NA |
| NA | NA |
| NA | NA |
| NA | NA |
| NA | NA |
| NA | NA |
| NA | NA |
| NA | NA |
| NA | NA |
| NA | NA |
| NA | NA |
| NA | NA |
| NA | NA |
| NA | NA |
| NA | NA |
| NA | NA |
| NA | NA |
| NA | NA |
| NA | NA |
| NA | NA |
| NA | NA |

| | | |
|---|---|---|
| NA | NA | |
| NA | NA | |
| NA | NA | |
| NA | NA | |
| 0.839318641 | NA | |
| NA | NA | |
| NA | NA | |
| NA | NA | |
| NA | NA | |
| 0.941278332 | NA | |
| NA | NA | |
| 0.059611918 | NA | |
| 0.004734834 | | 0.018680343 |
| 0.684286482 | | 0.797279344 |
| NA | NA | |
| 0.839319108 | NA | |
| 0.925579587 | NA | |
| NA | NA | |
| 0.776918731 | | 0.862550475 |
| 3.67E-19 | | 1.48E-17 |
| 0.718381738 | | 0.820958428 |
| 0.002002381 | | 0.009174336 |
| 0.184754611 | | 0.327831314 |
| 0.674980047 | | 0.790543237 |
| 0.75775842 | | 0.848601012 |
| 0.839319108 | NA | |
| NA | NA | |
| NA | NA | |
| 0.864210475 | NA | |
| NA | NA | |
| 0.880310464 | | 0.928321725 |
| NA | NA | |
| NA | NA | |
| NA | NA | |
| NA | NA | |
| 0.955792873 | NA | |
| NA | NA | |
| NA | NA | |
| 0.955792398 | NA | |
| NA | NA | |
| 0.955792873 | NA | |
| NA | NA | |
| NA | NA | |
| NA | NA | |
| NA | NA | |
| 0.033874789 | NA | |
| NA | NA | |

| | |
|---|---|
| NA | NA |
| NA | NA |
| NA | NA |
| NA | NA |
| 0.065209178 | 0.152674952 |
| NA | NA |
| NA | NA |
| NA | NA |
| 0.020190238 | 0.060327398 |
| NA | NA |
| NA | NA |
| NA | NA |
| 0.07146283 | 0.163616611 |
| 0.955792873 | NA |
| 0.451828122 | NA |
| 0.224570544 | 0.377286014 |
| 0.941278424 | NA |
| 0.228524776 | NA |
| 1.95E-07 | 2.49E-06 |
| 0.039493193 | 0.103128576 |
| 0.572195114 | 0.712174354 |
| 0.547691874 | 0.693703741 |
| NA | NA |
| 0.0014148 | 0.006834802 |
| NA | NA |
| NA | NA |
| NA | NA |
| NA | NA |
| NA | NA |
| NA | NA |
| 0.340769617 | 0.504321235 |
| NA | NA |
| NA | NA |
| 0.000158615 | 0.001049805 |
| NA | NA |
| 0.03026886 | 0.083570282 |
| NA | NA |
| NA | NA |
| NA | NA |
| NA | NA |
| NA | NA |
| NA | NA |
| NA | NA |
| NA | NA |
| 0.941278332 | NA |
| NA | NA |
| NA | NA |

| | | |
|---|---|---|
| | 0.941279268 | NA |
| | 0.944008553 | NA |
| | 0.803326486 | NA |
| NA | | NA |
| | 0.662686898 | NA |
| | 0.941276431 | NA |
| | 0.006808496 | 0.025093811 |
| | 0.099006963 | 0.207993268 |
| | 0.001095777 | 0.005523439 |
| | 2.41E-12 | 5.82E-11 |
| | 0.133455992 | 0.259646846 |
| | 2.17E-30 | 1.53E-28 |
| | 2.10E-05 | 0.00017539 |
| NA | | NA |
| | 0.028887594 | 0.080579224 |
| | 0.028428351 | NA |
| NA | | NA |
| NA | | NA |
| NA | | NA |
| | 7.49E-11 | 1.52E-09 |
| | 1.22E-58 | 2.03E-56 |
| | 8.82E-05 | 0.000627304 |
| | 0.000838143 | 0.004397017 |
| | 0.964081558 | 0.97817764 |
| | 0.049146329 | 0.122027075 |
| | 0.833944372 | 0.899431054 |
| | 0.910880429 | NA |
| | 0.092509867 | 0.198049995 |
| | 0.22524587 | NA |
| NA | | NA |
| | 0.982528086 | 0.988974438 |
| NA | | NA |
| NA | | NA |
| | 0.47778405 | NA |
| | 0.92663324 | NA |
| | 0.649010386 | 0.770576406 |
| NA | | NA |
| | 0.733851206 | 0.832164192 |
| NA | | NA |
| NA | | NA |
| | 4.34E-10 | 8.09E-09 |
| NA | | NA |
| | 0.839318641 | NA |
| NA | | NA |
| | 0.76631843 | NA |
| NA | | NA |
| | 0.214029052 | 0.364877858 |

| | | |
|---|---|---|
| | 0.941277857 | NA |
| | 0.88175842 | NA |
| | 0.941278332 | NA |
| NA | | NA |
| | 0.006269577 | 0.023463596 |
| | 0.01614606 | 0.050566909 |
| | 0.165745953 | 0.304498702 |
| | 0.020573702 | 0.061219558 |
| NA | | NA |
| | 0.080144964 | 0.178091991 |
| NA | | NA |
| | 0.435460814 | NA |
| NA | | NA |
| NA | | NA |
| | 0.003703277 | 0.01522373 |
| | 0.94517108 | 0.966925536 |
| NA | | NA |
| NA | | NA |
| | 0.839318641 | NA |
| NA | | NA |
| | 4.99E-06 | 4.87E-05 |
| | 0.266362589 | 0.425872394 |
| NA | | NA |
| | 0.853595154 | NA |
| NA | | NA |
| | 2.32E-11 | 5.07E-10 |
| NA | | NA |
| NA | | NA |
| NA | | NA |
| NA | | NA |
| NA | | NA |
| NA | | NA |
| | 0.941278332 | NA |
| | 0.092977821 | 0.198783625 |
| NA | | NA |
| | 0.919514217 | 0.951742798 |
| | 0.839319108 | NA |
| NA | | NA |
| | 0.007493867 | 0.027182112 |
| | 0.001574382 | 0.007478234 |
| | 0.980392286 | 0.98786289 |
| NA | | NA |
| NA | | NA |
| NA | | NA |
| NA | | NA |
| | 0.853596091 | NA |
| NA | | NA |

| | | |
|---|---|---|
| | 0.955792873 | NA |
| NA | | NA |
| | 0.109034429 | 0.223630977 |
| NA | | NA |
| | 0.176918278 | 0.318509012 |
| NA | | NA |
| NA | | NA |
| NA | | NA |
| | 0.005338436 | 0.020583039 |
| | 0.315525958 | 0.47795767 |
| NA | | NA |
| | 0.86731034 | NA |
| | 0.278635967 | 0.438398551 |
| | 0.46393267 | NA |
| NA | | NA |
| | 0.223986855 | 0.376586503 |
| | 0.035728372 | 0.095283665 |
| | 0.779617645 | NA |
| | 0.171442194 | 0.311582301 |
| | 0.367804384 | 0.530960695 |
| | 0.941278332 | NA |
| NA | | NA |
| NA | | NA |
| | 0.334785121 | NA |
| | 3.02E-05 | 0.00024256 |
| | 0.839319108 | NA |
| NA | | NA |
| | 0.455276939 | 0.614942235 |
| | 0.743511643 | NA |
| | 0.180065573 | 0.322007571 |
| NA | | NA |
| NA | | NA |
| NA | | NA |
| NA | | NA |
| | 0.946839907 | 0.967625072 |
| | 0.941278332 | NA |
| NA | | NA |
| NA | | NA |
| NA | | NA |
| NA | | NA |
| NA | | NA |
| NA | | NA |
| NA | | NA |
| NA | | NA |
| NA | | NA |
| NA | | NA |
| NA | | NA |

| | |
|---|---|
| NA | NA |
| NA | NA |
| NA | NA |
| NA | NA |
| NA | NA |
| NA | NA |
| NA | NA |
| NA | NA |
| NA | NA |
| NA | NA |
| 0.941277857 | NA |
| NA | NA |
| 0.021377024 | 0.063105037 |
| 0.641588395 | 0.764914359 |
| NA | NA |
| NA | NA |
| NA | NA |
| 0.007543724 | 0.027320436 |
| 0.390953134 | 0.553465333 |
| 0.109445766 | 0.224223458 |
| 0.477885423 | 0.634744913 |
| 0.941275956 | NA |
| 0.086056931 | 0.187830612 |
| NA | NA |
| 0.556706506 | NA |
| 0.005824764 | 0.022101125 |
| NA | NA |
| 0.00018233 | 0.001184142 |
| NA | NA |
| 0.060738946 | 0.144426412 |
| 0.029704838 | 0.082337784 |
| 0.301519342 | 0.462837829 |
| 0.465403084 | NA |
| 0.535970221 | 0.684066878 |
| 0.644437548 | 0.767213651 |
| NA | NA |
| 0.801029676 | NA |
| 0.036858996 | NA |
| 0.303149394 | 0.46460871 |
| 0.000920334 | 0.004752827 |
| 0.694614637 | 0.804340433 |
| 0.941281426 | NA |
| 0.00022181 | 0.001403381 |
| 0.941277857 | NA |
| NA | NA |
| NA | NA |
| 0.941277857 | NA |

| | |
|---|---|
| NA | NA |
| NA | NA |
| NA | NA |
| 0.001043311 | 0.005293729 |
| NA | NA |
| NA | NA |
| NA | NA |
| NA | NA |
| 0.941278332 | NA |
| 0.839318641 | NA |
| NA | NA |
| NA | NA |
| 0.887948318 | NA |
| 0.000193887 | 0.001247708 |
| 0.301883541 | 0.463247505 |
| 0.711184175 | 0.815773823 |
| NA | NA |
| 0.941276431 | NA |
| NA | NA |
| NA | NA |
| NA | NA |
| 0.000216936 | 0.001376939 |
| 0.00313037 | 0.013286208 |
| 0.941278332 | NA |
| 0.030173707 | 0.083355917 |
| 0.109951717 | 0.225124368 |
| NA | NA |
| 0.152720501 | 0.286811848 |
| NA | NA |
| 0.258018704 | 0.416250589 |
| 0.715227908 | NA |
| NA | NA |
| NA | NA |
| 0.895835233 | NA |
| NA | NA |
| 0.609413995 | 0.741490928 |
| 0.911096355 | 0.946156869 |
| NA | NA |
| NA | NA |
| 0.941275956 | NA |
| 0.941277857 | NA |
| 0.138669435 | 0.267237735 |
| 0.556359722 | 0.700585869 |
| 0.786135359 | 0.868581799 |
| 0.941275481 | NA |
| 0.249966808 | NA |
| 0.955791922 | NA |

| | | | |
|---|---|---|---|
| | 0.839319108 | NA | |
| | 0.096707521 | | 0.204480093 |
| | 0.828191776 | | 0.89562173 |
| | 0.727832274 | NA | |
| NA | | NA | |
| NA | | NA | |
| NA | | NA | |
| NA | | NA | |
| | 0.955792398 | NA | |
| NA | | NA | |
| NA | | NA | |
| NA | | NA | |
| NA | | NA | |
| NA | | NA | |
| NA | | NA | |
| | 0.048545779 | | 0.120917396 |
| NA | | NA | |
| NA | | NA | |
| NA | | NA | |
| NA | | NA | |
| NA | | NA | |
| NA | | NA | |
| NA | | NA | |
| | 0.284226464 | | 0.444812828 |
| | 0.291050704 | | 0.452133459 |
| NA | | NA | |
| NA | | NA | |
| NA | | NA | |
| NA | | NA | |
| NA | | NA | |
| NA | | NA | |
| NA | | NA | |
| NA | | NA | |
| NA | | NA | |
| NA | | NA | |
| | 0.740446682 | NA | |
| NA | | NA | |
| | 0.82736485 | | 0.895214936 |
| | 0.323460369 | NA | |
| | 0.839319108 | NA | |
| NA | | NA | |
| NA | | NA | |
| | 0.000183509 | | 0.00118951 |
| NA | | NA | |
| NA | | NA | |
| NA | | NA | |
| NA | | NA | |

| | |
|---|---|
| NA | NA |
| NA | NA |
| NA | NA |
| NA | NA |
| NA | NA |
| NA | NA |
| NA | NA |
| NA | NA |
| NA | NA |
| NA | NA |
| NA | NA |
| NA | NA |
| NA | NA |
| NA | NA |
| 0.748393366 | NA |
| NA | NA |
| NA | NA |
| NA | NA |
| NA | NA |
| NA | NA |
| NA | NA |
| NA | NA |
| NA | NA |
| NA | NA |
| NA | NA |
| NA | NA |
| NA | NA |
| NA | NA |
| NA | NA |
| 0.090437397 | NA |
| NA | NA |
| NA | NA |
| 0.839319574 | NA |
| 0.038392046 | 0.100815221 |
| NA | NA |
| NA | NA |
| 3.18E-07 | 3.89E-06 |
| NA | NA |
| 0.553798206 | NA |
| NA | NA |
| NA | NA |
| 0.48790741 | 0.642661519 |
| NA | NA |
| NA | NA |
| NA | NA |
| 0.297069016 | NA |
| 0.436617169 | 0.597605627 |

| | | | |
|---|---|---|---|
| | 0.632587615 | | 0.758421641 |
| | 0.790233216 | NA | |
| | 0.941275956 | NA | |
| | 0.551866766 | | 0.69717811 |
| | 0.636786471 | | 0.76118881 |
| NA | | NA | |
| NA | | NA | |
| NA | | NA | |
| | 0.938677532 | NA | |
| NA | | NA | |
| NA | | NA | |
| NA | | NA | |
| | 0.092900707 | | 0.198690145 |
| NA | | NA | |
| NA | | NA | |
| NA | | NA | |
| | 0.570911477 | | 0.711393348 |
| NA | | NA | |
| NA | | NA | |
| NA | | NA | |
| | 0.728678906 | NA | |
| NA | | NA | |
| NA | | NA | |
| | 0.941278332 | NA | |
| NA | | NA | |
| | 0.76259836 | NA | |
| | 0.839319108 | NA | |
| | 0.070300278 | | 0.161732658 |
| | 0.878312932 | | 0.926995584 |
| | 0.873399 | | 0.924442843 |
| | 0.376757178 | | 0.539727616 |
| | 0.03562287 | | 0.095067422 |
| | 0.139364883 | | 0.268186826 |
| NA | | NA | |
| NA | | NA | |
| NA | | NA | |
| | 0.748687767 | | 0.842225958 |
| | 0.017922552 | | 0.054947884 |
| NA | | NA | |
| | 0.061346694 | | 0.145566146 |
| | 0.617501707 | | 0.747046431 |
| | 0.03807769 | | 0.100201787 |
| | 3.81E-05 | | 0.000297417 |
| | 0.45607544 | NA | |
| | 0.220007054 | | 0.371680595 |
| NA | | NA | |
| | 0.680614382 | | 0.794389424 |

| | | |
|---|---|---|
| | 0.941277857 | NA |
| | 0.778265992 | NA |
| NA | | NA |
| | 0.109517676 | 0.22435147 |
| NA | | NA |
| | 0.329942329 | NA |
| | 0.877383683 | NA |
| NA | | NA |
| NA | | NA |
| | 0.362272765 | 0.525973914 |
| NA | | NA |
| | 0.382006564 | 0.544560115 |
| | 0.182662142 | 0.325258046 |
| | 0.292120672 | NA |
| | 5.85E-05 | 0.000435924 |
| | 0.947960636 | NA |
| | 0.714143889 | 0.818158107 |
| | 0.003421049 | 0.014285581 |
| | 0.578567798 | 0.716668386 |
| | 0.873868481 | 0.924785321 |
| NA | | NA |
| NA | | NA |
| | 0.943465155 | 0.965947936 |
| | 0.964815025 | 0.978629815 |
| | 0.839319108 | NA |
| NA | | NA |
| | 0.923889471 | 0.954592539 |
| | 0.000137154 | 0.00092506 |
| | 0.563762885 | 0.705986435 |
| | 0.000265335 | 0.001634027 |
| | 0.262837588 | 0.421794787 |
| NA | | NA |
| | 0.367739639 | NA |
| NA | | NA |
| | 0.774170251 | 0.860571326 |
| | 0.046926423 | 0.117820961 |
| | 0.075658231 | NA |
| | 0.402256385 | NA |
| | 0.68963014 | 0.800945477 |
| | 0.830380302 | NA |
| | 0.400450065 | 0.563095199 |
| | 1.11E-05 | 9.94E-05 |
| | 0.028616093 | 0.079971849 |
| | 0.652770835 | 0.773536365 |
| | 0.002162123 | 0.009758116 |
| | 0.256852767 | 0.414970951 |
| | 0.473797056 | 0.631216698 |

| | |
|---|---|
| 0.000325921 | 0.001958486 |
| NA | NA |
| 0.003339282 | 0.014023764 |
| 0.003185011 | 0.013491657 |
| 0.91058261 | 0.94586711 |
| 0.000988077 | 0.00505659 |
| 0.071517239 | 0.163693944 |
| 0.378391845 | 0.541189923 |
| 0.941278332 | NA |
| 9.00E-09 | 1.43E-07 |
| 0.000252453 | 0.001565631 |
| 0.782706437 | NA |
| 0.005942706 | 0.022473514 |
| 1.60E-07 | 2.08E-06 |
| 0.011314247 | 0.038081602 |
| 0.662203222 | NA |
| 0.030646735 | 0.084339646 |
| NA | NA |
| 0.955792398 | NA |
| 0.011660767 | 0.039063979 |
| 0.847791804 | 0.908395855 |
| 0.936900534 | 0.962580675 |
| 0.839318641 | NA |
| 0.008407621 | 0.029855021 |
| 0.002969617 | 0.01271274 |
| 2.52E-11 | 5.47E-10 |
| 0.007064704 | 0.025877823 |
| NA | NA |
| 0.000303908 | 0.001842019 |
| NA | NA |
| 3.29E-06 | 3.32E-05 |
| NA | NA |
| NA | NA |
| 0.091252829 | 0.196135077 |
| 0.481102268 | 0.637487957 |
| 0.002843228 | 0.012257654 |
| 0.801243415 | 0.878492596 |
| 0.955792398 | NA |
| 2.91E-06 | 2.97E-05 |
| 0.464810207 | 0.623290847 |
| 0.182087238 | 0.32448786 |
| 0.033117707 | 0.089779917 |
| 0.198529948 | 0.345753261 |
| 0.002004699 | 0.00918142 |
| 3.70E-05 | 0.000289291 |
| 0.941278332 | NA |
| NA | NA |

| | | | |
|---|---|---|---|
| | 0.225051888 | | 0.37781669 |
| NA | | NA | |
| NA | | NA | |
| NA | | NA | |
| NA | | NA | |
| NA | | NA | |
| NA | | NA | |
| NA | | NA | |
| | 0.393038001 | | 0.55552794 |
| | 0.941278332 | NA | |
| NA | | NA | |
| | 0.825159559 | NA | |
| | 0.560533652 | NA | |
| NA | | NA | |
| NA | | NA | |
| NA | | NA | |
| NA | | NA | |
| NA | | NA | |
| NA | | NA | |
| NA | | NA | |
| NA | | NA | |
| NA | | NA | |
| NA | | NA | |
| | 0.458806114 | | 0.618235004 |
| NA | | NA | |
| NA | | NA | |
| NA | | NA | |
| NA | | NA | |
| | 0.839318641 | NA | |
| | 0.955791922 | NA | |
| NA | | NA | |
| | 0.064052612 | | 0.150568463 |
| | 0.42058969 | NA | |
| NA | | NA | |
| | 0.044037937 | | 0.112117978 |
| | 0.328723778 | | 0.491629492 |
| | 1.31E-09 | | 2.29E-08 |
| | 0.853596091 | NA | |
| | 0.501979033 | NA | |
| | 0.031751792 | | 0.08672959 |
| | 0.941277857 | NA | |
| | 0.910567934 | | 0.94586711 |
| | 0.448099726 | | 0.608285028 |
| | 0.212758755 | | 0.36327718 |
| | 0.070235981 | | 0.161606878 |
| | 0.086015567 | | 0.187812359 |
| | 0.001566894 | | 0.007445639 |

| | | | |
|---|---|---|---|
| | 0.780881731 | | 0.865187191 |
| | 0.142259053 | | 0.272475254 |
| | 0.512049072 | | 0.663461731 |
| | 0.383222783 | | 0.545934203 |
| | 0.371564495 | | 0.534831768 |
| | 0.268760811 | | 0.428597748 |
| | 0.018683204 | | 0.056836111 |
| NA | | NA | |
| | 0.121307267 | | 0.242309129 |
| | 0.034006952 | | 0.091537522 |
| | 0.953770009 | NA | |
| | 0.782138158 | | 0.866042633 |
| NA | | NA | |
| NA | | NA | |
| NA | | NA | |
| NA | | NA | |
| | 0.844542648 | | 0.90649091 |
| NA | | NA | |
| NA | | NA | |
| | 0.839318641 | NA | |
| NA | | NA | |
| NA | | NA | |
| | 0.640874253 | NA | |
| NA | | NA | |
| | 0.941278332 | NA | |
| NA | | NA | |
| NA | | NA | |
| | 0.941277857 | NA | |
| | 0.955792398 | NA | |
| NA | | NA | |
| NA | | NA | |
| | 0.941278332 | NA | |
| NA | | NA | |
| | 0.699972323 | NA | |
| NA | | NA | |
| | 0.734918134 | | 0.832977211 |
| NA | | NA | |
| | 0.630770719 | NA | |
| | 1.07E-09 | | 1.89E-08 |
| | 0.931807121 | | 0.959371849 |
| NA | | NA | |
| | 0.911557252 | NA | |
| NA | | NA | |
| | 0.072146998 | | 0.164707736 |
| | 0.008317642 | | 0.029606177 |
| | 0.853595622 | NA | |
| NA | | NA | |

| | | |
|---|---|---|
| | 0.724271383 | NA |
| | 0.853595622 | NA |
| NA | | NA |
| NA | | NA |
| NA | | NA |
| | 0.791059099 | NA |
| NA | | NA |
| | 0.853595622 | NA |
| | 0.853596091 | NA |
| NA | | NA |
| | 0.941278332 | NA |
| NA | | NA |
| NA | | NA |
| NA | | NA |
| NA | | NA |
| | 5.42E-12 | 1.26E-10 |
| | 0.993848283 | 0.995521287 |
| NA | | NA |
| NA | | NA |
| NA | | NA |
| NA | | NA |
| | 0.941277857 | NA |
| | 0.249252733 | NA |
| NA | | NA |
| NA | | NA |
| | 0.007935845 | 0.028459993 |
| | 0.905610492 | 0.942722104 |
| | 0.230982664 | 0.384949543 |
| | 0.606173417 | NA |
| | 0.425693977 | NA |
| | 0.030327528 | 0.083683726 |
| NA | | NA |
| NA | | NA |
| | 0.548513785 | 0.694416527 |
| | 0.304750768 | 0.466384477 |
| NA | | NA |
| NA | | NA |
| | 0.687999471 | NA |
| | 0.020412429 | 0.060846167 |
| NA | | NA |
| | 0.839318641 | NA |
| | 0.000544838 | 0.003058967 |
| NA | | NA |
| NA | | NA |
| NA | | NA |
| NA | | NA |
| | 0.460765526 | 0.61978757 |

| | |
|---|---|
| NA | NA |
| NA | NA |
| NA | NA |
| NA | NA |
| 0.495301451 | 0.648219341 |
| 0.177466436 | 0.31903747 |
| NA | NA |
| 0.867643439 | NA |
| 0.941192604 | 0.964703895 |
| 0.390843635 | 0.553346431 |
| NA | NA |
| 0.642566655 | NA |
| NA | NA |
| NA | NA |
| 0.002555438 | 0.011247141 |
| 0.796862357 | 0.875503962 |
| 0.632461597 | 0.758414336 |
| 0.839319108 | NA |
| NA | NA |
| 0.983917399 | NA |
| NA | NA |
| 0.839320041 | NA |
| 0.700815618 | NA |
| 1.09E-19 | 4.53E-18 |
| NA | NA |
| 0.955792873 | NA |
| 0.455206658 | NA |
| 0.941277857 | NA |
| NA | NA |
| NA | NA |
| 0.895502608 | NA |
| NA | NA |
| NA | NA |
| NA | NA |
| NA | NA |
| NA | NA |
| 0.572585879 | 0.71242494 |
| NA | NA |
| 0.012787581 | 0.041987861 |
| 0.067088181 | 0.156016892 |
| 0.276361832 | 0.436233459 |
| 0.01683062 | 0.05226385 |
| NA | NA |
| NA | NA |
| NA | NA |
| 0.783997437 | NA |
| NA | NA |

| | | | |
|---|---|---|---|
| NA | | NA | |
| NA | | NA | |
| | 0.401353197 | | 0.563741795 |
| NA | | NA | |
| | 0.076050255 | | 0.171204782 |
| NA | | NA | |
| NA | | NA | |
| NA | | NA | |
| | 0.811511034 | NA | |
| | 0.839318641 | NA | |
| NA | | NA | |
| NA | | NA | |
| NA | | NA | |
| NA | | NA | |
| NA | | NA | |
| NA | | NA | |
| NA | | NA | |
| NA | | NA | |
| NA | | NA | |
| | 0.135055546 | NA | |
| NA | | NA | |
| NA | | NA | |
| NA | | NA | |
| NA | | NA | |
| NA | | NA | |
| | 0.014200738 | | 0.045695006 |
| NA | | NA | |
| NA | | NA | |
| NA | | NA | |
| | 0.955792398 | NA | |
| NA | | NA | |
| NA | | NA | |
| | 0.839318641 | NA | |
| | 0.671125131 | NA | |
| NA | | NA | |
| NA | | NA | |
| NA | | NA | |
| NA | | NA | |
| NA | | NA | |
| NA | | NA | |
| NA | | NA | |
| NA | | NA | |
| NA | | NA | |
| NA | | NA | |
| NA | | NA | |
| | 0.661866297 | NA | |
| NA | | NA | |

| | |
|---|---|
| NA | NA |
| 0.839318641 | NA |
| NA | NA |
| 0.708196277 | NA |
| 0.728879436 | NA |
| NA | NA |
| NA | NA |
| NA | NA |
| NA | NA |
| 0.412481655 | NA |
| NA | NA |
| NA | NA |
| NA | NA |
| NA | NA |
| 0.048566625 | 0.120956661 |
| 0.665745524 | 0.783218414 |
| NA | NA |
| 0.038949826 | 0.101922138 |
| NA | NA |
| 0.26892065 | 0.428781439 |
| 0.077095753 | 0.172958979 |
| NA | NA |
| NA | NA |
| 0.941275481 | NA |
| NA | NA |
| 0.955792398 | NA |
| NA | NA |
| 0.499615304 | NA |
| 0.33342091 | 0.496688228 |
| 0.000922478 | 0.004761834 |
| 0.557578115 | 0.701317451 |
| NA | NA |
| 0.921742362 | NA |
| NA | NA |
| 0.839319108 | NA |
| 0.300383465 | 0.461936638 |
| 0.018372004 | 0.056048504 |
| NA | NA |
| NA | NA |
| 0.004004774 | 0.016230482 |
| NA | NA |
| 0.370011016 | 0.533240628 |
| NA | NA |
| 0.941277857 | NA |
| 0.952268933 | NA |
| 0.941278332 | NA |
| 0.102279737 | 0.213362165 |

| | |
|---|---|
| 0.941278332 | NA |
| 0.33968032 | 0.503366391 |
| 0.227207592 | 0.380469615 |
| 0.653193515 | 0.773830932 |
| 0.853595154 | NA |
| 0.746399855 | NA |
| 0.505844756 | 0.657596115 |
| 0.941278332 | NA |
| 2.27E-05 | 0.000188997 |
| 0.140083469 | 0.269246137 |
| 0.955792873 | NA |
| 0.839319108 | NA |
| 0.743511643 | NA |
| 0.839318641 | NA |
| 0.02317767 | 0.067516205 |
| 0.92302243 | NA |
| NA | NA |
| NA | NA |
| 0.948037261 | 0.96821346 |
| 0.001282543 | 0.006294247 |
| NA | NA |
| 0.613936886 | 0.744228457 |
| 1.21E-41 | 1.33E-39 |
| NA | NA |
| 0.925128692 | NA |
| NA | NA |
| 0.003057488 | 0.013023322 |
| NA | NA |
| 0.839319574 | NA |
| 0.331818415 | 0.494951552 |
| 0.955792398 | NA |
| 0.011988824 | 0.039865604 |
| 0.044738155 | 0.11347587 |
| 0.757824894 | NA |
| 1.60E-05 | 0.000138082 |
| NA | NA |
| NA | NA |
| NA | NA |
| NA | NA |
| 0.203203699 | 0.351473321 |
| NA | NA |
| 0.000230823 | 0.001448856 |
| NA | NA |
| 0.838993826 | NA |
| 7.69E-08 | 1.05E-06 |
| NA | NA |
| 0.130053949 | NA |

| | | |
|---|---|---|
| | 0.016411199 | 0.051235919 |
| | 0.029518309 | 0.081897049 |
| | 0.140086722 | 0.269246137 |
| | 0.941277857 | NA |
| | 0.002432223 | 0.010780589 |
| | 0.941277857 | NA |
| | 0.00181939 | 0.008446483 |
| NA | | NA |
| | 0.538896747 | 0.68636008 |
| | 0.40011957 | 0.562723875 |
| | 0.501807179 | NA |
| | 0.931577835 | NA |
| | 0.797679751 | 0.876118935 |
| | 0.921453897 | NA |
| | 0.839319108 | NA |
| | 0.084918743 | 0.186082318 |
| | 0.486775671 | 0.641900641 |
| | 0.67608732 | 0.791278477 |
| | 0.486437616 | 0.641664161 |
| | 0.623068287 | 0.751385543 |
| NA | | NA |
| | 0.519801834 | 0.670199414 |
| NA | | NA |
| | 1.60E-05 | 0.000137815 |
| | 0.061510357 | 0.145863749 |
| | 0.881164011 | NA |
| NA | | NA |
| | 0.006006958 | 0.022680458 |
| | 0.007611479 | 0.027524829 |
| | 0.000247691 | 0.001540918 |
| | 0.001161668 | 0.0058003 |
| | 0.000906636 | 0.004696356 |
| | 0.783247691 | 0.866747014 |
| | 0.032130248 | 0.087551707 |
| | 0.94127925 | NA |
| | 0.055845595 | 0.135305054 |
| | 0.090500032 | 0.194998386 |
| NA | | NA |
| NA | | NA |
| NA | | NA |
| NA | | NA |
| | 0.07300615 | 0.166127378 |
| | 0.000342645 | 0.002045554 |
| | 0.907444822 | NA |
| NA | | NA |
| | 0.955792398 | NA |
| | 0.134783704 | 0.261673875 |

| | |
|---|---|
| 0.00652136 | 0.024223066 |
| NA | NA |
| 1.56E-41 | 1.69E-39 |
| 0.848397883 | 0.908883177 |
| 0.502129987 | 0.654242278 |
| 0.033959526 | 0.091478484 |
| 0.860474864 | 0.916609361 |
| 0.005605609 | 0.021416485 |
| 6.55E-10 | 1.19E-08 |
| 0.188280514 | 0.332624716 |
| 0.937966715 | 0.963177176 |
| 0.534362069 | NA |
| 0.01326262 | 0.043273049 |
| 0.00319955 | 0.013535971 |
| 0.134983269 | 0.261975846 |
| 0.839318641 | NA |
| 5.75E-14 | 1.60E-12 |
| 0.763243582 | 0.85297792 |
| NA | NA |
| 0.073438732 | NA |
| NA | NA |
| NA | NA |
| 0.944242109 | 0.966373729 |
| 0.722378157 | 0.824008053 |
| NA | NA |
| NA | NA |
| 0.000503107 | 0.00285254 |
| NA | NA |
| 0.195396156 | 0.341745852 |
| 0.748393366 | NA |
| 0.000701442 | 0.003785878 |
| 0.839318641 | NA |
| 0.000400016 | 0.002337058 |
| NA | NA |
| 0.137739227 | 0.266034729 |
| 0.752540247 | 0.844744527 |
| 0.94054529 | NA |
| 0.001268386 | 0.006238918 |
| 0.000159069 | 0.001052005 |
| NA | NA |
| NA | NA |
| NA | NA |
| NA | NA |
| NA | NA |
| 0.289352925 | 0.450142374 |
| NA | NA |
| NA | NA |

| | | | |
|---|---|---|---|
| NA | | NA | |
| NA | | NA | |
| NA | | NA | |
| NA | | NA | |
| NA | | NA | |
| NA | | NA | |
| NA | | NA | |
| NA | | NA | |
| NA | | NA | |
| NA | | NA | |
| NA | | NA | |
| | 0.382861522 | | 0.545609279 |
| | 0.941278332 | NA | |
| NA | | NA | |
| | 0.83901926 | NA | |
| | 0.870838969 | NA | |
| | 0.697545097 | | 0.806243306 |
| | 0.955792398 | NA | |
| | 0.11561384 | | 0.233483847 |
| | 0.874252224 | NA | |
| | 0.859762434 | NA | |
| | 0.555631916 | | 0.700047424 |
| | 0.063042771 | | 0.148671756 |
| | 0.276549522 | | 0.436298252 |
| NA | | NA | |
| NA | | NA | |
| | 0.12353975 | NA | |
| | 0.164267863 | | 0.302483651 |
| | 0.721342691 | NA | |
| | 0.096844929 | | 0.204661547 |
| NA | | NA | |
| | 0.118241088 | | 0.237571943 |
| NA | | NA | |
| | 0.128746439 | | 0.253008737 |
| NA | | NA | |
| | 0.128975635 | | 0.25318582 |
| | 0.184277487 | | 0.327227407 |
| NA | | NA | |
| NA | | NA | |
| | 0.555837257 | NA | |
| | 0.613864497 | NA | |
| | 0.71685749 | NA | |
| | 0.034600746 | | 0.092796277 |
| NA | | NA | |
| | 0.008676806 | | 0.030619027 |
| | 0.004221866 | | 0.016965871 |
| | 0.839319108 | NA | |

| | | | |
|---|---|---|---|
| | 0.598891728 | | 0.733026998 |
| | 0.033564893 | | 0.090713249 |
| | 0.174297851 | | 0.315102343 |
| | 0.236811112 | | 0.392262575 |
| NA | | NA | |
| | 0.049632343 | | 0.122996526 |
| | 0.699732277 | NA | |
| | 0.976356113 | NA | |
| | 0.839320041 | NA | |
| NA | | NA | |
| | 0.75589335 | NA | |
| | 0.387649941 | NA | |
| | 0.285043125 | | 0.445563569 |
| NA | | NA | |
| | 0.839318641 | NA | |
| | 0.002667183 | | 0.011654907 |
| | 0.552987946 | | 0.698038772 |
| | 0.292176447 | | 0.453357998 |
| NA | | NA | |
| | 0.743511643 | NA | |
| | 0.022908285 | NA | |
| | 2.11E-05 | | 0.000176312 |
| NA | | NA | |
| NA | | NA | |
| | 0.853595154 | NA | |
| NA | | NA | |
| | 0.000296325 | | 0.001805256 |
| | 0.311802877 | | 0.473765214 |
| | 0.127195054 | | 0.250786669 |
| | 0.065808015 | | 0.153760908 |
| | 0.076344837 | | 0.171689324 |
| | 0.204113404 | | 0.352588358 |
| | 0.013041299 | | 0.04265547 |
| NA | | NA | |
| | 0.898063058 | | 0.938477276 |
| | 0.918764101 | NA | |
| | 0.580432362 | | 0.718208103 |
| | 0.141834959 | | 0.271881751 |
| | 0.955792397 | NA | |
| NA | | NA | |
| | 0.626918656 | | 0.754295434 |
| | 0.151574461 | | 0.285560814 |
| | 0.129972285 | | 0.25465923 |
| | 0.963585911 | | 0.977841478 |
| | 0.154176347 | | 0.288856135 |
| | 0.047881959 | | 0.119674415 |
| | 0.001324642 | | 0.006463431 |

| | |
|---|---|
| 0.001538849 | 0.007332872 |
| 0.022267019 | 0.065273033 |
| 0.934276775 | 0.960921951 |
| 0.955792873 | NA |
| NA | NA |
| NA | NA |
| NA | NA |
| NA | NA |
| 0.491395733 | NA |
| 0.399767537 | 0.562467749 |
| 0.390506449 | 0.553065087 |
| NA | NA |
| 0.651466739 | 0.772452492 |
| NA | NA |
| 0.606004317 | 0.738675411 |
| 0.840788913 | 0.903905046 |
| 0.011385895 | 0.038294061 |
| 3.82E-31 | 2.79E-29 |
| NA | NA |
| 0.128467148 | 0.252729568 |
| 0.031611411 | 0.086484461 |
| 0.839318641 | NA |
| NA | NA |
| 0.889122431 | NA |
| NA | NA |
| NA | NA |
| NA | NA |
| 0.597252254 | NA |
| 0.608321552 | 0.740703376 |
| 0.941277857 | NA |
| 0.001698744 | 0.0079845 |
| 0.00045276 | 0.00260367 |
| 0.897402082 | NA |
| 0.765763569 | NA |
| 0.941278332 | NA |
| NA | NA |
| 0.558409149 | 0.701978164 |
| NA | NA |
| NA | NA |
| NA | NA |
| NA | NA |
| NA | NA |
| 0.683851058 | 0.797072763 |
| 0.022940786 | 0.066932074 |
| 7.41E-17 | 2.57E-15 |
| 8.37E-07 | 9.45E-06 |
| NA | NA |

| | |
|---|---|
| NA | NA |
| 3.87E-12 | 9.15E-11 |
| 0.839319108 | NA |
| NA | NA |
| NA | NA |
| 0.000268062 | 0.001649964 |
| NA | NA |
| NA | NA |
| 0.109020852 | 0.223622399 |
| 0.01940168 | 0.058476573 |
| 0.00367642 | 0.015131619 |
| NA | NA |
| NA | NA |
| NA | NA |
| NA | NA |
| 0.941278332 | NA |
| 0.009610669 | 0.033341079 |
| 0.72715692 | 0.827925808 |
| 0.925414408 | NA |
| 0.895163542 | NA |
| 0.977011145 | 0.985928463 |
| 0.542047311 | NA |
| NA | NA |
| 0.560490443 | 0.703665463 |
| NA | NA |
| NA | NA |
| 4.71E-11 | 9.87E-10 |
| NA | NA |
| 0.879117502 | NA |
| NA | NA |
| NA | NA |
| NA | NA |
| NA | NA |
| 0.941275956 | NA |
| 0.001324805 | 0.006463431 |
| NA | NA |
| NA | NA |
| 0.070380894 | 0.161855656 |
| 0.001946093 | 0.008956092 |
| NA | NA |
| NA | NA |
| NA | NA |
| 0.741771672 | 0.837554522 |
| NA | NA |
| 0.439598904 | 0.600443769 |
| 0.895572463 | 0.936820599 |
| NA | NA |

| | | | |
|---|---|---|---|
| | 0.001311436 | | 0.006410925 |
| NA | | NA | |
| | 0.941985068 | NA | |
| | 0.194246685 | NA | |
| | 0.277234122 | | 0.437146511 |
| | 0.626246427 | | 0.753775851 |
| | 0.257973303 | | 0.416205555 |
| NA | | NA | |
| NA | | NA | |
| NA | | NA | |
| NA | | NA | |
| NA | | NA | |
| NA | | NA | |
| NA | | NA | |
| | 0.838882104 | NA | |
| | 0.353370122 | | 0.517057972 |
| | 0.093984483 | | 0.200288179 |
| | 0.318363366 | | 0.480812253 |
| NA | | NA | |
| | 0.254193152 | | 0.412243084 |
| NA | | NA | |
| NA | | NA | |
| NA | | NA | |
| NA | | NA | |
| NA | | NA | |
| | 0.941278424 | NA | |
| | 0.0008441 | | 0.004422465 |
| | 0.16585958 | | 0.304660419 |
| | 0.758823415 | | 0.849513895 |
| | 0.941276431 | NA | |
| NA | | NA | |
| | 0.839318641 | NA | |
| NA | | NA | |
| NA | | NA | |
| NA | | NA | |
| NA | | NA | |
| | 0.104571476 | | 0.216760263 |
| | 0.955791446 | NA | |
| NA | | NA | |
| NA | | NA | |
| NA | | NA | |
| NA | | NA | |
| | 0.941278332 | NA | |
| NA | | NA | |
| NA | | NA | |
| NA | | NA | |
| NA | | NA | |

| | |
|---|---|
| NA | NA |
| NA | NA |
| NA | NA |
| NA | NA |
| NA | NA |
| 0.955792873 | NA |
| NA | NA |
| NA | NA |
| NA | NA |
| NA | NA |
| NA | NA |
| NA | NA |
| NA | NA |
| 0.712326933 | NA |
| NA | NA |
| NA | NA |
| NA | NA |
| NA | NA |
| NA | NA |
| NA | NA |
| NA | NA |
| NA | NA |
| NA | NA |
| NA | NA |
| NA | NA |
| NA | NA |
| NA | NA |
| 0.786986142 | 0.869176744 |
| 0.941278424 | NA |
| NA | NA |
| 0.651137077 | NA |
| 0.94010533 | NA |
| 0.735187901 | 0.833163949 |
| NA | NA |
| NA | NA |
| 0.239152628 | 0.395052456 |
| NA | NA |
| 0.934125979 | NA |
| 0.955792398 | NA |
| NA | NA |
| NA | NA |
| 0.118755604 | 0.238352382 |
| 0.955792873 | NA |
| NA | NA |
| NA | NA |
| NA | NA |
| 0.839319574 | NA |

| | | | |
|---|---|---|---|
| NA | | NA | |
| | 0.267228519 | NA | |
| | 0.831929452 | NA | |
| | 0.787313605 | NA | |
| | 0.010191893 | | 0.034936021 |
| NA | | NA | |
| NA | | NA | |
| NA | | NA | |
| | 0.839318641 | NA | |
| NA | | NA | |
| NA | | NA | |
| NA | | NA | |
| NA | | NA | |
| NA | | NA | |
| NA | | NA | |
| NA | | NA | |
| | 0.941278332 | NA | |
| NA | | NA | |
| NA | | NA | |
| NA | | NA | |
| NA | | NA | |
| NA | | NA | |
| NA | | NA | |
| NA | | NA | |
| NA | | NA | |
| | 0.839318641 | NA | |
| NA | | NA | |
| NA | | NA | |
| NA | | NA | |
| NA | | NA | |
| NA | | NA | |
| NA | | NA | |
| NA | | NA | |
| NA | | NA | |
| NA | | NA | |
| NA | | NA | |
| NA | | NA | |
| NA | | NA | |
| | 0.941275481 | NA | |
| NA | | NA | |
| NA | | NA | |
| NA | | NA | |
| NA | | NA | |
| NA | | NA | |
| NA | | NA | |
| NA | | NA | |
| NA | | NA | |

| | | | |
|---|---|---|---|
| NA | | NA | |
| NA | | NA | |
| | 0.955792398 | NA | |
| NA | | NA | |
| NA | | NA | |
| NA | | NA | |
| NA | | NA | |
| NA | | NA | |
| NA | | NA | |
| NA | | NA | |
| NA | | NA | |
| NA | | NA | |
| NA | | NA | |
| NA | | NA | |
| | 0.262919592 | NA | |
| | 0.949791594 | NA | |
| NA | | NA | |
| | 0.941277857 | NA | |
| | 0.303347593 | | 0.464773666 |
| NA | | NA | |
| NA | | NA | |
| NA | | NA | |
| NA | | NA | |
| NA | | NA | |
| NA | | NA | |
| NA | | NA | |
| NA | | NA | |
| NA | | NA | |
| NA | | NA | |
| NA | | NA | |
| | 0.254907541 | | 0.412938766 |
| NA | | NA | |
| NA | | NA | |
| | 0.678533033 | NA | |
| | 0.955792398 | NA | |
| NA | | NA | |
| NA | | NA | |
| NA | | NA | |
| NA | | NA | |
| NA | | NA | |
| NA | | NA | |
| NA | | NA | |
| NA | | NA | |
| NA | | NA | |
| NA | | NA | |
| NA | | NA | |
| NA | | NA | |

| | |
|---|---|
| NA | NA |
| NA | NA |
| NA | NA |
| NA | NA |
| 0.776133935 | NA |
| 0.15481371 | 0.289623197 |
| 0.038471403 | 0.100958801 |
| 0.774088837 | 0.860571326 |
| 0.066020696 | 0.154167038 |
| NA | NA |
| 0.691996526 | 0.802401778 |
| 0.666549431 | 0.783815501 |
| 0.839319574 | NA |
| 0.838051761 | 0.901858577 |
| NA | NA |
| 0.298076959 | 0.459539335 |
| 0.619015986 | 0.748320477 |
| 0.941280034 | NA |
| 0.010195534 | 0.034937243 |
| 0.000221372 | 0.00140098 |
| 0.005178703 | 0.020110569 |
| 1.27E-07 | 1.68E-06 |
| 0.0843026 | NA |
| 0.107434405 | 0.221201858 |
| 0.941275481 | NA |
| 0.133867651 | 0.260320085 |
| 1.35E-12 | 3.36E-11 |
| NA | NA |
| 0.122900712 | 0.244649738 |
| 1.34E-07 | 1.77E-06 |
| 0.01226844 | 0.040613827 |
| 0.664110551 | NA |
| 0.839318641 | NA |
| 0.130638914 | NA |
| 0.007896046 | 0.028351438 |
| 0.970192728 | 0.982200983 |
| 0.504870006 | NA |
| 0.259251805 | 0.417673715 |
| 1.16E-08 | 1.81E-07 |
| 0.692682485 | 0.80298527 |
| 0.496643046 | NA |
| 0.380430591 | 0.543126735 |
| NA | NA |
| NA | NA |
| 0.095910819 | 0.203337429 |
| 0.003025727 | 0.012922727 |
| 0.007138999 | 0.026097749 |

| | | |
|---|---|---|
| NA | NA | |
| NA | NA | |
| NA | NA | |
| 0.839318641 | NA | |
| 0.659550588 | | 0.778524506 |
| NA | NA | |
| NA | NA | |
| 0.839318641 | NA | |
| NA | NA | |
| NA | NA | |
| NA | NA | |
| NA | NA | |
| NA | NA | |
| NA | NA | |
| NA | NA | |
| NA | NA | |
| NA | NA | |
| NA | NA | |
| NA | NA | |
| NA | NA | |
| NA | NA | |
| 0.87313512 | | 0.924280679 |
| NA | NA | |
| 0.738325218 | NA | |
| 0.750895012 | NA | |
| 0.839318641 | NA | |
| NA | NA | |
| 0.00304451 | | 0.012981982 |
| 2.82E-06 | | 2.88E-05 |
| NA | NA | |
| NA | NA | |
| 0.365689069 | | 0.528837185 |
| 0.001017764 | | 0.005182887 |
| 0.252731019 | | 0.410566733 |
| 0.000574457 | | 0.003193187 |
| 0.080570022 | | 0.178792435 |
| 0.273355876 | | 0.433292143 |
| 0.16888172 | | 0.30842651 |
| 0.109088211 | NA | |
| 0.109194803 | | 0.223940609 |
| 0.036096622 | | 0.096062761 |
| NA | NA | |
| 0.907048939 | | 0.943641717 |
| 0.109322524 | | 0.224009531 |
| 0.596867963 | | 0.73158511 |
| 0.00231513 | | 0.010354376 |
| 0.889103932 | | 0.933455095 |

| | | |
|---|---|---|
| | 0.941277857 | NA |
| NA | | NA |
| NA | | NA |
| | 0.941278332 | NA |
| | 0.941277857 | NA |
| | 0.739991096 | NA |
| NA | | NA |
| NA | | NA |
| NA | | NA |
| NA | | NA |
| NA | | NA |
| NA | | NA |
| | 0.955791922 | NA |
| | 0.317521349 | NA |
| | 0.955792873 | NA |
| NA | | NA |
| NA | | NA |
| NA | | NA |
| NA | | NA |
| NA | | NA |
| | 0.473144974 | 0.630714602 |
| | 0.22692478 | NA |
| NA | | NA |
| | 0.066853813 | 0.155608688 |
| | 0.001729157 | 0.008103443 |
| | 1.26E-07 | 1.67E-06 |
| | 0.074044781 | 0.167896912 |
| NA | | NA |
| | 0.142209552 | 0.272424286 |
| | 0.002816091 | 0.012151667 |
| NA | | NA |
| NA | | NA |
| NA | | NA |
| | 0.955792873 | NA |
| | 0.791124563 | 0.871710058 |
| | 0.465637676 | 0.623968899 |
| | 0.044991295 | 0.113984333 |
| | 0.667295831 | 0.784328651 |
| | 0.908462139 | 0.944452264 |
| NA | | NA |
| | 0.455031733 | 0.614819783 |
| NA | | NA |
| NA | | NA |
| | 0.451234728 | NA |
| NA | | NA |
| NA | | NA |
| NA | | NA |

| | |
|---|---|
| NA | NA |
| NA | NA |
| NA | NA |
| 0.839318641 | NA |
| NA | NA |
| NA | NA |
| 0.068950743 | NA |
| 0.725259537 | 0.826437548 |
| NA | NA |
| NA | NA |
| NA | NA |
| NA | NA |
| NA | NA |
| NA | NA |
| NA | NA |
| NA | NA |
| NA | NA |
| 0.856368192 | 0.913720446 |
| 0.839319574 | NA |
| NA | NA |
| NA | NA |
| NA | NA |
| 0.001064592 | 0.005383346 |
| 0.600334566 | NA |
| NA | NA |
| NA | NA |
| NA | NA |
| 0.845923192 | 0.907209293 |
| 0.469868548 | 0.627798041 |
| NA | NA |
| NA | NA |
| 0.000926167 | 0.004776733 |
| 0.612666151 | 0.743559032 |
| 0.863982333 | NA |
| 0.941278332 | NA |
| 0.020277442 | 0.060534763 |
| 0.172400867 | 0.31262269 |
| 0.031133666 | 0.085422884 |
| NA | NA |
| 0.045179663 | 0.114388506 |
| 0.554791044 | NA |
| 0.955792398 | NA |
| 0.955791922 | NA |
| 0.739600213 | 0.836253113 |
| 0.030759406 | 0.084600806 |
| 0.733640093 | NA |
| 0.750146473 | 0.843051435 |

| | |
|---|---|
| 5.45E-07 | 6.37E-06 |
| NA | NA |
| NA | NA |
| 0.736071075 | 0.833647157 |
| 0.321510163 | 0.484371473 |
| NA | NA |
| 0.665442206 | NA |
| NA | NA |
| NA | NA |
| 0.713257135 | NA |
| 0.228222571 | 0.381766367 |
| 0.000661194 | 0.003597198 |
| 0.639959899 | 0.763639921 |
| NA | NA |
| NA | NA |
| NA | NA |
| 0.461116415 | 0.620084346 |
| 0.454494594 | 0.614366148 |
| NA | NA |
| 0.207638641 | 0.356814012 |
| 0.694485917 | NA |
| NA | NA |
| NA | NA |
| 0.839318641 | NA |
| 0.916080473 | NA |
| 0.913269402 | 0.947669296 |
| NA | NA |
| NA | NA |
| 0.465895909 | 0.6240787 |
| 0.054040394 | 0.131803408 |
| NA | NA |
| 0.297491605 | 0.458874679 |
| 0.551116025 | 0.696459896 |
| 0.011738217 | 0.039223108 |
| 0.002243505 | 0.010075455 |
| NA | NA |
| NA | NA |
| NA | NA |
| 0.839319108 | NA |
| 0.955791446 | NA |
| NA | NA |
| 0.172573461 | 0.312792683 |
| 0.048655142 | 0.121151762 |
| 0.230588452 | 0.384534879 |
| 0.941278332 | NA |
| 0.000173159 | 0.001134786 |
| 0.653810921 | 0.774059764 |

| | |
|---|---|
| 0.073742695 | 0.167371377 |
| 0.941278332 | NA |
| 0.571540585 | 0.711794109 |
| NA | NA |
| 0.875627664 | 0.925643364 |
| 0.017753001 | 0.054523476 |
| NA | NA |
| 0.557618384 | 0.701317451 |
| 0.839318641 | NA |
| 0.489362734 | 0.643774487 |
| 0.915561503 | NA |
| NA | NA |
| NA | NA |
| 0.839319108 | NA |
| NA | NA |
| 0.098384174 | 0.207105344 |
| 0.853595622 | NA |
| 0.010624914 | 0.036153567 |
| 0.255616627 | NA |
| NA | NA |
| 0.002870773 | 0.012358497 |
| 0.169520471 | 0.309185454 |
| 0.579495296 | 0.717346944 |
| 0.558212424 | 0.701804993 |
| 0.941278332 | NA |
| NA | NA |
| 0.941278332 | NA |
| 0.955792398 | NA |
| 0.419219611 | 0.580887559 |
| 0.003891478 | 0.015860438 |
| 0.593111289 | 0.728481083 |
| NA | NA |
| 0.955792398 | NA |
| 0.941275956 | NA |
| 0.941277857 | NA |
| 0.648331759 | 0.770116406 |
| NA | NA |
| NA | NA |
| NA | NA |
| 0.223509246 | 0.375954713 |
| NA | NA |
| 0.359781493 | NA |
| NA | NA |
| 0.941485052 | 0.964703895 |
| 0.281785028 | 0.441980048 |
| NA | NA |
| NA | NA |

| | |
|---|---|
| 2.88E-05 | 0.000232701 |
| 0.555494925 | 0.699978305 |
| 0.377814621 | 0.540656743 |
| 0.020811974 | 0.061758708 |
| 0.054690916 | 0.133035584 |
| 0.149352928 | 0.282381316 |
| 0.245732547 | 0.402571509 |
| 0.604319908 | 0.737245902 |
| 0.158283326 | NA |
| 0.123400486 | 0.245313704 |
| NA | NA |
| 0.781559182 | 0.865683449 |
| 0.285734369 | 0.446409547 |
| NA | NA |
| NA | NA |
| NA | NA |
| NA | NA |
| NA | NA |
| NA | NA |
| NA | NA |
| NA | NA |
| NA | NA |
| NA | NA |
| NA | NA |
| NA | NA |
| NA | NA |
| NA | NA |
| NA | NA |
| NA | NA |
| NA | NA |
| NA | NA |
| NA | NA |
| NA | NA |
| NA | NA |
| NA | NA |
| NA | NA |
| 0.955792873 | NA |
| NA | NA |
| NA | NA |
| NA | NA |
| NA | NA |
| NA | NA |
| NA | NA |
| NA | NA |
| NA | NA |
| NA | NA |
| NA | NA |

| | | | |
|---|---|---|---|
| NA | | NA | |
| NA | | NA | |
| NA | | NA | |
| NA | | NA | |
| NA | | NA | |
| NA | | NA | |
| NA | | NA | |
| NA | | NA | |
| NA | | NA | |
| NA | | NA | |
| NA | | NA | |
| NA | | NA | |
| NA | | NA | |
| NA | | NA | |
| NA | | NA | |
| NA | | NA | |
| | 0.272713791 | | 0.432598044 |
| | 0.845421691 | | 0.906886914 |
| | 0.908465329 | NA | |
| | 0.413561221 | | 0.575770624 |
| NA | | NA | |
| | 0.020148974 | | 0.060231939 |
| | 0.269348249 | | 0.429261937 |
| | 0.000746345 | | 0.003988439 |
| | 0.682547899 | | 0.796060618 |
| | 0.115387979 | | 0.233146407 |
| | 0.093882673 | | 0.200178749 |
| NA | | NA | |
| NA | | NA | |
| | 0.763974729 | | 0.853554567 |
| | 0.58812859 | NA | |
| | 0.941279803 | NA | |
| NA | | NA | |
| | 0.861129469 | NA | |
| | 0.006068944 | | 0.022857475 |
| NA | | NA | |
| | 0.000471398 | | 0.002698467 |
| | 0.00584197 | | 0.022159454 |
| NA | | NA | |
| NA | | NA | |
| | 0.642619647 | | 0.765585507 |
| | 3.51E-05 | | 0.000276547 |
| | 0.045178604 | | 0.114388506 |
| | 0.839319574 | NA | |
| | 3.62E-08 | | 5.21E-07 |
| | 0.052403625 | | 0.12864182 |
| | 0.747557982 | | 0.84141368 |

| | | |
|---|---|---|
| | 0.206772372 | 0.355735164 |
| | 0.941208171 | NA |
| | 0.888827393 | 0.93342843 |
| NA | | NA |
| | 0.040953015 | 0.106160277 |
| | 0.626380211 | NA |
| NA | | NA |
| NA | | NA |
| NA | | NA |
| | 2.25E-07 | 2.84E-06 |
| NA | | NA |
| | 0.6309611 | NA |
| | 0.931584637 | 0.959272227 |
| | 0.276488983 | 0.436289496 |
| NA | | NA |
| | 0.00914022 | 0.031988753 |
| | 0.307281495 | 0.468851733 |
| | 0.189068694 | 0.333659718 |
| | 0.743956064 | 0.839184979 |
| NA | | NA |
| | 0.815286387 | 0.88702503 |
| | 0.036586996 | 0.097019125 |
| | 0.070632129 | 0.16232364 |
| NA | | NA |
| | 0.000874299 | 0.004552631 |
| | 0.373523656 | 0.53676473 |
| | 0.123425011 | 0.245324602 |
| | 0.219151412 | 0.370682342 |
| | 0.721194359 | NA |
| NA | | NA |
| | 0.941278788 | NA |
| | 0.941277857 | NA |
| | 6.90E-05 | 0.000503209 |
| | 0.27690383 | NA |
| NA | | NA |
| | 0.941278332 | NA |
| | 0.012288895 | 0.04064928 |
| | 0.285837481 | 0.446509514 |
| | 0.202534648 | 0.350623963 |
| | 0.224443728 | 0.377168146 |
| | 0.379547741 | 0.542381878 |
| | 0.839319108 | NA |
| | 0.900002958 | 0.939761808 |
| | 0.244848291 | 0.401424371 |
| | 0.037459982 | 0.098827588 |
| | 0.665395446 | NA |
| | 0.284723158 | 0.445326627 |

| | | | |
|---|---|---|---|
| NA | | NA | |
| | 0.008477865 | | 0.030050655 |
| NA | | NA | |
| NA | | NA | |
| NA | | NA | |
| NA | | NA | |
| | 0.213779773 | | 0.364654304 |
| | 0.335600735 | | 0.498749295 |
| | 0.924559043 | NA | |
| | 0.522591485 | | 0.672545148 |
| NA | | NA | |
| NA | | NA | |
| NA | | NA | |
| NA | | NA | |
| NA | | NA | |
| NA | | NA | |
| NA | | NA | |
| NA | | NA | |
| NA | | NA | |
| NA | | NA | |
| NA | | NA | |
| NA | | NA | |
| NA | | NA | |
| NA | | NA | |
| NA | | NA | |
| NA | | NA | |
| NA | | NA | |
| NA | | NA | |
| NA | | NA | |
| NA | | NA | |
| | 0.978658652 | NA | |
| NA | | NA | |
| NA | | NA | |
| NA | | NA | |
| | 0.721484123 | | 0.823278446 |
| NA | | NA | |
| NA | | NA | |
| NA | | NA | |
| NA | | NA | |
| NA | | NA | |
| NA | | NA | |
| NA | | NA | |
| | 0.941278332 | NA | |
| NA | | NA | |
| | 0.900394534 | | 0.940018234 |
| | 0.955792398 | NA | |
| | 0.747653071 | NA | |

| | |
|---|---|
| 3.22E-20 | 1.40E-18 |
| NA | NA |
| 0.000189803 | 0.001226632 |
| NA | NA |
| 0.00271446 | 0.011820268 |
| NA | NA |
| 0.820623594 | NA |
| NA | NA |
| 0.015856205 | 0.049855932 |
| 0.015473133 | 0.048883927 |
| 0.027245671 | 0.076966916 |
| 0.479947121 | 0.636595229 |
| NA | NA |
| 0.568032912 | 0.709430128 |
| NA | NA |
| 0.594537428 | NA |
| 0.223461676 | 0.375942523 |
| NA | NA |
| 0.241241661 | 0.397455113 |
| NA | NA |
| 0.582712457 | NA |
| 0.27068482 | NA |
| 0.64327049 | NA |
| NA | NA |
| 0.014064384 | 0.045299115 |
| 0.16049221 | 0.297255901 |
| 0.897762655 | NA |
| NA | NA |
| NA | NA |
| 0.328066258 | NA |
| 0.941278332 | NA |
| NA | NA |
| 0.915077187 | NA |
| NA | NA |
| NA | NA |
| 0.649415093 | 0.770749546 |
| 0.002248434 | 0.010094733 |
| 0.366075321 | 0.529247931 |
| 0.061816344 | 0.146424602 |
| 0.941278284 | NA |
| 0.941278332 | NA |
| NA | NA |
| NA | NA |
| NA | NA |
| NA | NA |
| 0.839319108 | NA |
| NA | NA |

| | |
|---|---|
| NA | NA |
| NA | NA |
| 0.066414662 | 0.154934999 |
| 0.008199919 | 0.029270318 |
| NA | NA |
| 0.736139426 | 0.833647157 |
| 0.968547684 | 0.981203319 |
| 0.863192418 | 0.91816177 |
| 0.924039833 | 0.954592539 |
| 0.559752135 | 0.703109415 |
| 0.955639556 | 0.972955303 |
| 0.109666386 | 0.224611478 |
| NA | NA |
| NA | NA |
| 0.014280269 | 0.04589508 |
| NA | NA |
| NA | NA |
| 0.839319108 | NA |
| NA | NA |
| 0.017157375 | 0.053084602 |
| 0.07689512 | 0.172613279 |
| 1.74E-05 | 0.000148096 |
| 0.713301209 | NA |
| 0.000944682 | 0.00486064 |
| 0.839319107 | NA |
| 0.941277857 | NA |
| 0.727929945 | 0.828647403 |
| 0.508810217 | 0.660437379 |
| 0.160853676 | 0.297809521 |
| 0.098476747 | 0.207226907 |
| NA | NA |
| NA | NA |
| 0.790939096 | NA |
| 0.565355692 | 0.707165332 |
| 0.959158979 | 0.975065434 |
| 0.075026701 | 0.169493692 |
| 0.027329841 | 0.077156312 |
| 0.365228953 | 0.52852502 |
| 0.590128275 | NA |
| 0.955792398 | NA |
| NA | NA |
| 0.95182251 | NA |
| 0.713455398 | NA |
| 0.027080224 | 0.076587407 |
| 0.233938297 | 0.38856939 |
| 0.473283194 | 0.630849896 |
| NA | NA |

| | |
|---|---|
| NA | NA |
| NA | NA |
| NA | NA |
| 0.50227497 | 0.654323728 |
| NA | NA |
| 9.42E-05 | 0.00066613 |
| 0.634869658 | 0.760004406 |
| NA | NA |
| NA | NA |
| 0.839319108 | NA |
| 0.080847608 | 0.179130044 |
| 0.12894884 | NA |
| NA | NA |
| 0.305437043 | 0.466954365 |
| 0.955792873 | NA |
| NA | NA |
| NA | NA |
| NA | NA |
| NA | NA |
| NA | NA |
| NA | NA |
| NA | NA |
| 0.505120704 | 0.657167696 |
| 0.529599958 | 0.67866578 |
| NA | NA |
| 0.814434203 | 0.886584171 |
| 0.261193002 | 0.419948636 |
| 0.732849974 | 0.831464564 |
| 0.049156868 | 0.122030847 |
| 0.295553447 | 0.456980846 |
| 0.941278332 | NA |
| 0.05060058 | 0.125029528 |
| 0.715301883 | 0.819127853 |
| NA | NA |
| 0.237068027 | 0.392480571 |
| 0.416697011 | NA |
| 0.236039399 | 0.391286321 |
| 0.000110708 | 0.000766233 |
| 5.02E-05 | 0.000380588 |
| 0.707776586 | NA |
| 0.57242188 | NA |
| NA | NA |
| 0.655589897 | 0.775178873 |
| 0.373729113 | 0.536876115 |
| NA | NA |
| 0.741296019 | 0.837390866 |
| 0.823938988 | NA |

| | |
|---|---|
| NA | NA |
| NA | NA |
| NA | NA |
| 0.005767456 | 0.021918727 |
| NA | NA |
| 0.009255067 | 0.032300454 |
| 0.011246353 | 0.037883626 |
| 0.019345024 | 0.058358714 |
| 0.004378699 | 0.017495685 |
| 0.188137537 | 0.332495519 |
| 0.441015933 | NA |
| 0.001320551 | 0.006450187 |
| 0.132851404 | 0.258809061 |
| 0.672183795 | NA |
| 0.505164646 | NA |
| 0.019627103 | 0.059000292 |
| 0.901423019 | NA |
| NA | NA |
| 0.941277857 | NA |
| NA | NA |
| 0.839320041 | NA |
| 0.024055428 | 0.069562301 |
| 0.980153356 | 0.987685995 |
| 0.000178744 | 0.001164332 |
| 8.96E-11 | 1.81E-09 |
| 3.90E-11 | 8.27E-10 |
| 4.96E-42 | 5.49E-40 |
| 0.004179177 | 0.016831266 |
| 0.579758762 | NA |
| 0.768845362 | NA |
| 0.459207253 | NA |
| 0.687878118 | 0.799730131 |
| 0.955792397 | NA |
| NA | NA |
| NA | NA |
| NA | NA |
| 0.839319574 | NA |
| NA | NA |
| NA | NA |
| 0.730034537 | 0.830046911 |
| NA | NA |
| 0.600206867 | 0.734127638 |
| 0.759832911 | NA |
| 0.94127925 | NA |
| 0.37353882 | 0.53676473 |
| 0.611330216 | 0.742468556 |
| 0.155600058 | 0.29086567 |

| | | | |
|---|---|---|---|
| | 0.073899121 | | 0.167630505 |
| | 0.839318641 | NA | |
| | 0.732245526 | | 0.831135336 |
| NA | | NA | |
| NA | | NA | |
| | 0.047491839 | NA | |
| | 0.930430045 | | 0.958580961 |
| | 0.941277857 | NA | |
| | 0.790611578 | | 0.87134363 |
| | 0.643560903 | | 0.766323309 |
| NA | | NA | |
| NA | | NA | |
| NA | | NA | |
| NA | | NA | |
| | 0.839319108 | NA | |
| | 0.955792398 | NA | |
| | 0.701574326 | | 0.809446055 |
| NA | | NA | |
| | 0.797182307 | NA | |
| | 0.853595154 | NA | |
| | 0.564161056 | | 0.70628009 |
| NA | | NA | |
| NA | | NA | |
| | 0.167791992 | | 0.307046141 |
| | 0.008480514 | | 0.030055566 |
| | 0.466753896 | | 0.62494663 |
| | 0.410204306 | | 0.572171066 |
| | 0.002092091 | | 0.009493987 |
| | 0.285671314 | | 0.446340332 |
| | 0.228437992 | | 0.38201872 |
| NA | | NA | |
| NA | | NA | |
| NA | | NA | |
| NA | | NA | |
| | 0.485658279 | NA | |
| NA | | NA | |
| NA | | NA | |
| NA | | NA | |
| NA | | NA | |
| NA | | NA | |
| NA | | NA | |
| NA | | NA | |
| NA | | NA | |
| NA | | NA | |
| NA | | NA | |
| NA | | NA | |
| NA | | NA | |

| | | |
|---|---|---|
| NA | | NA |
| NA | | NA |
| NA | | NA |
| NA | | NA |
| NA | | NA |
| NA | | NA |
| NA | | NA |
| NA | | NA |
| NA | | NA |
| NA | | NA |
| NA | | NA |
| NA | | NA |
| NA | | NA |
| NA | | NA |
| NA | | NA |
| NA | | NA |
| NA | | NA |
| NA | | NA |
| NA | | NA |
| NA | | NA |
| NA | | NA |
| NA | | NA |
| NA | | NA |
| NA | | NA |
| NA | | NA |
| NA | | NA |
| NA | | NA |
| NA | | NA |
| NA | | NA |
| NA | | NA |
| NA | | NA |
| NA | | NA |
| NA | | NA |
| NA | | NA |
| NA | | NA |
| NA | | NA |
| NA | | NA |
| NA | | NA |
| NA | | NA |
| NA | | NA |
| NA | | NA |
| NA | | NA |
| NA | | NA |
| | 0.941278332 | NA |
| NA | | NA |
| NA | | NA |
| NA | | NA |

| | | |
|---|---|---|
| NA | | NA |
| NA | | NA |
| NA | | NA |
| NA | | NA |
| NA | | NA |
| NA | | NA |
| NA | | NA |
| NA | | NA |
| NA | | NA |
| NA | | NA |
| NA | | NA |
| NA | | NA |
| NA | | NA |
| NA | | NA |
| NA | | NA |
| NA | | NA |
| NA | | NA |
| NA | | NA |
| NA | | NA |
| NA | | NA |
| NA | | NA |
| NA | | NA |
| NA | | NA |
| NA | | NA |
| NA | | NA |
| NA | | NA |
| NA | | NA |
| NA | | NA |
| NA | | NA |
| NA | | NA |
| NA | | NA |
| NA | | NA |
| | 0.941278332 | NA |
| NA | | NA |
| NA | | NA |
| NA | | NA |
| NA | | NA |
| NA | | NA |
| NA | | NA |
| NA | | NA |
| NA | | NA |
| NA | | NA |
| NA | | NA |
| NA | | NA |
| NA | | NA |
| NA | | NA |
| NA | | NA |

| | | |
|---|---|---|
| | 0.839319108 | NA |
| NA | | NA |
| NA | | NA |
| NA | | NA |
| NA | | NA |
| NA | | NA |
| NA | | NA |
| NA | | NA |
| NA | | NA |
| NA | | NA |
| NA | | NA |
| NA | | NA |
| NA | | NA |
| | 0.799868295 | NA |
| NA | | NA |
| NA | | NA |
| NA | | NA |
| NA | | NA |
| NA | | NA |
| NA | | NA |
| NA | | NA |
| NA | | NA |
| NA | | NA |
| NA | | NA |
| NA | | NA |
| | 0.813126606 | NA |
| NA | | NA |
| NA | | NA |
| NA | | NA |
| NA | | NA |
| NA | | NA |
| NA | | NA |
| NA | | NA |
| NA | | NA |
| NA | | NA |
| NA | | NA |
| NA | | NA |
| NA | | NA |
| NA | | NA |
| NA | | NA |
| NA | | NA |
| NA | | NA |
| NA | | NA |
| NA | | NA |
| NA | | NA |
| NA | | NA |
| NA | | NA |

| | |
|---|---|
| NA | NA |
| NA | NA |
| NA | NA |
| NA | NA |
| NA | NA |
| NA | NA |
| NA | NA |
| NA | NA |
| NA | NA |
| NA | NA |
| NA | NA |
| NA | NA |
| NA | NA |
| NA | NA |
| NA | NA |
| NA | NA |
| NA | NA |
| NA | NA |
| NA | NA |
| NA | NA |
| NA | NA |
| NA | NA |
| NA | NA |
| NA | NA |
| NA | NA |
| NA | NA |
| NA | NA |
| NA | NA |
| NA | NA |
| NA | NA |
| NA | NA |
| NA | NA |
| NA | NA |
| NA | NA |
| NA | NA |
| NA | NA |
| NA | NA |
| NA | NA |
| NA | NA |
| NA | NA |
| NA | NA |
| NA | NA |
| NA | NA |
| NA | NA |
| NA | NA |
| NA | NA |
| NA | NA |

| | |
|---|---|
| NA | NA |
| NA | NA |
| NA | NA |
| 0.839319108 | NA |
| NA | NA |
| NA | NA |
| NA | NA |
| 0.24391625 | 0.400309906 |
| NA | NA |
| NA | NA |
| NA | NA |
| NA | NA |
| NA | NA |
| NA | NA |
| NA | NA |
| NA | NA |
| NA | NA |
| NA | NA |
| NA | NA |
| NA | NA |
| NA | NA |
| NA | NA |
| NA | NA |
| NA | NA |
| NA | NA |
| NA | NA |
| NA | NA |
| NA | NA |
| NA | NA |
| NA | NA |
| NA | NA |
| NA | NA |
| NA | NA |
| NA | NA |
| NA | NA |
| NA | NA |
| NA | NA |
| NA | NA |
| NA | NA |
| NA | NA |
| NA | NA |
| NA | NA |
| NA | NA |
| NA | NA |
| NA | NA |
| NA | NA |
| NA | NA |

| | | |
|---|---|---|
| NA | | NA |
| NA | | NA |
| NA | | NA |
| NA | | NA |
| NA | | NA |
| NA | | NA |
| NA | | NA |
| NA | | NA |
| NA | | NA |
| NA | | NA |
| NA | | NA |
| NA | | NA |
| NA | | NA |
| NA | | NA |
| NA | | NA |
| NA | | NA |
| NA | | NA |
| NA | | NA |
| NA | | NA |
| NA | | NA |
| NA | | NA |
| NA | | NA |
| NA | | NA |
| NA | | NA |
| NA | | NA |
| NA | | NA |
| NA | | NA |
| NA | | NA |
| NA | | NA |
| NA | | NA |
| NA | | NA |
| NA | | NA |
| NA | | NA |
| NA | | NA |
| NA | | NA |
| NA | | NA |
| NA | | NA |
| NA | | NA |
| NA | | NA |
| NA | | NA |
| NA | | NA |
| NA | | NA |
| NA | | NA |
| NA | | NA |
| | 0.955792398 | NA |
| NA | | NA |
| NA | | NA |

| | |
|---|---|
| NA | NA |
| NA | NA |
| NA | NA |
| NA | NA |
| NA | NA |
| NA | NA |
| NA | NA |
| NA | NA |
| NA | NA |
| NA | NA |
| NA | NA |
| NA | NA |
| NA | NA |
| NA | NA |
| NA | NA |
| NA | NA |
| NA | NA |
| NA | NA |
| NA | NA |
| NA | NA |
| NA | NA |
| NA | NA |
| NA | NA |
| NA | NA |
| NA | NA |
| NA | NA |
| NA | NA |
| NA | NA |
| NA | NA |
| NA | NA |
| NA | NA |
| NA | NA |
| NA | NA |
| NA | NA |
| NA | NA |
| NA | NA |
| NA | NA |
| NA | NA |
| NA | NA |
| NA | NA |
| NA | NA |
| NA | NA |
| NA | NA |
| NA | NA |
| NA | NA |
| NA | NA |
| NA | NA |

| | | |
|---|---|---|
| NA | | NA |
| NA | | NA |
| NA | | NA |
| NA | | NA |
| NA | | NA |
| NA | | NA |
| NA | | NA |
| NA | | NA |
| NA | | NA |
| NA | | NA |
| NA | | NA |
| NA | | NA |
| NA | | NA |
| NA | | NA |
| NA | | NA |
| NA | | NA |
| NA | | NA |
| NA | | NA |
| NA | | NA |
| NA | | NA |
| NA | | NA |
| NA | | NA |
| NA | | NA |
| NA | | NA |
| NA | | NA |
| NA | | NA |
| NA | | NA |
| NA | | NA |
| NA | | NA |
| NA | | NA |
| NA | | NA |
| NA | | NA |
| NA | | NA |
| NA | | NA |
| NA | | NA |
| NA | | NA |
| NA | | NA |
| NA | | NA |
| NA | | NA |
| NA | | NA |
| | 0.955792398 | NA |
| NA | | NA |
| NA | | NA |
| NA | | NA |
| NA | | NA |
| NA | | NA |
| NA | | NA |

| | |
|---|---|
| NA | NA |
| NA | NA |
| NA | NA |
| NA | NA |
| NA | NA |
| NA | NA |
| NA | NA |
| NA | NA |
| NA | NA |
| NA | NA |
| NA | NA |
| NA | NA |
| NA | NA |
| NA | NA |
| NA | NA |
| NA | NA |
| NA | NA |
| NA | NA |
| NA | NA |
| NA | NA |
| NA | NA |
| NA | NA |
| NA | NA |
| NA | NA |
| NA | NA |
| NA | NA |
| NA | NA |
| NA | NA |
| NA | NA |
| NA | NA |
| NA | NA |
| NA | NA |
| NA | NA |
| NA | NA |
| NA | NA |
| NA | NA |
| NA | NA |
| NA | NA |
| NA | NA |
| NA | NA |
| NA | NA |
| NA | NA |
| NA | NA |
| NA | NA |
| NA | NA |
| NA | NA |
| NA | NA |

| | |
|---|---|
| NA | NA |
| NA | NA |
| NA | NA |
| NA | NA |
| NA | NA |
| NA | NA |
| NA | NA |
| NA | NA |
| NA | NA |
| NA | NA |
| NA | NA |
| NA | NA |
| NA | NA |
| NA | NA |
| NA | NA |
| NA | NA |
| NA | NA |
| NA | NA |
| NA | NA |
| NA | NA |
| NA | NA |
| NA | NA |
| NA | NA |
| NA | NA |
| NA | NA |
| NA | NA |
| NA | NA |
| NA | NA |
| NA | NA |
| NA | NA |
| NA | NA |
| NA | NA |
| NA | NA |
| NA | NA |
| NA | NA |
| NA | NA |
| NA | NA |
| NA | NA |
| NA | NA |
| NA | NA |
| NA | NA |
| NA | NA |
| NA | NA |
| NA | NA |
| NA | NA |
| NA | NA |
| NA | NA |

| | | |
|---|---|---|
| | 0.955792398 | NA |
| NA | | NA |
| NA | | NA |
| NA | | NA |
| NA | | NA |
| NA | | NA |
| NA | | NA |
| NA | | NA |
| NA | | NA |
| NA | | NA |
| NA | | NA |
| NA | | NA |
| NA | | NA |
| NA | | NA |
| NA | | NA |
| NA | | NA |
| NA | | NA |
| NA | | NA |
| NA | | NA |
| NA | | NA |
| NA | | NA |
| NA | | NA |
| NA | | NA |
| NA | | NA |
| NA | | NA |
| NA | | NA |
| NA | | NA |
| NA | | NA |
| NA | | NA |
| NA | | NA |
| NA | | NA |
| NA | | NA |
| NA | | NA |
| NA | | NA |
| NA | | NA |
| NA | | NA |
| NA | | NA |
| NA | | NA |
| NA | | NA |
| NA | | NA |
| NA | | NA |
| NA | | NA |
| NA | | NA |
| NA | | NA |
| NA | | NA |
| NA | | NA |
| NA | | NA |

| | |
|---|---|
| NA | NA |
| NA | NA |
| NA | NA |
| NA | NA |
| NA | NA |
| NA | NA |
| NA | NA |
| NA | NA |
| NA | NA |
| NA | NA |
| NA | NA |
| NA | NA |
| NA | NA |
| NA | NA |
| NA | NA |
| NA | NA |
| NA | NA |
| NA | NA |
| NA | NA |
| NA | NA |
| NA | NA |
| NA | NA |
| NA | NA |
| NA | NA |
| NA | NA |
| NA | NA |
| NA | NA |
| NA | NA |
| NA | NA |
| NA | NA |
| NA | NA |
| NA | NA |
| NA | NA |
| NA | NA |
| NA | NA |
| NA | NA |
| NA | NA |
| NA | NA |
| NA | NA |
| NA | NA |
| NA | NA |
| NA | NA |
| NA | NA |
| NA | NA |
| NA | NA |
| NA | NA |
| NA | NA |

| | |
|---|---|
| NA | NA |
| NA | NA |
| NA | NA |
| NA | NA |
| NA | NA |
| NA | NA |
| NA | NA |
| NA | NA |
| NA | NA |
| NA | NA |
| NA | NA |
| NA | NA |
| NA | NA |
| NA | NA |
| NA | NA |
| NA | NA |
| NA | NA |
| NA | NA |
| NA | NA |
| NA | NA |
| NA | NA |
| NA | NA |
| NA | NA |
| NA | NA |
| NA | NA |
| NA | NA |
| NA | NA |
| NA | NA |
| NA | NA |
| NA | NA |
| NA | NA |
| NA | NA |
| NA | NA |
| NA | NA |
| NA | NA |
| NA | NA |
| NA | NA |
| NA | NA |
| NA | NA |
| NA | NA |
| NA | NA |
| NA | NA |
| NA | NA |
| NA | NA |
| NA | NA |
| NA | NA |
| NA | NA |

| | |
|---|---|
| NA | NA |
| NA | NA |
| NA | NA |
| NA | NA |
| NA | NA |
| NA | NA |
| NA | NA |
| NA | NA |
| NA | NA |
| NA | NA |
| NA | NA |
| NA | NA |
| NA | NA |
| NA | NA |
| NA | NA |
| NA | NA |
| NA | NA |
| NA | NA |
| NA | NA |
| NA | NA |
| NA | NA |
| NA | NA |
| NA | NA |
| NA | NA |
| NA | NA |
| NA | NA |
| NA | NA |
| NA | NA |
| NA | NA |
| NA | NA |
| NA | NA |
| NA | NA |
| NA | NA |
| NA | NA |
| NA | NA |
| NA | NA |
| NA | NA |
| NA | NA |
| NA | NA |
| NA | NA |
| NA | NA |
| NA | NA |
| NA | NA |
| NA | NA |
| NA | NA |
| NA | NA |
| NA | NA |

| | |
|---|---|
| NA | NA |
| NA | NA |
| NA | NA |
| NA | NA |
| NA | NA |
| NA | NA |
| NA | NA |
| NA | NA |
| NA | NA |
| NA | NA |
| NA | NA |
| NA | NA |
| NA | NA |
| NA | NA |
| 0.95509577 | 0.972692863 |
| NA | NA |
| NA | NA |
| NA | NA |
| NA | NA |
| NA | NA |
| NA | NA |
| NA | NA |
| NA | NA |
| NA | NA |
| NA | NA |
| NA | NA |
| NA | NA |
| NA | NA |
| NA | NA |
| 0.955792398 | NA |
| NA | NA |
| NA | NA |
| NA | NA |
| NA | NA |
| NA | NA |
| NA | NA |
| NA | NA |
| NA | NA |
| NA | NA |
| NA | NA |
| NA | NA |
| NA | NA |
| NA | NA |
| NA | NA |
| NA | NA |
| NA | NA |
| NA | NA |

| | | |
|---|---|---|
| NA | | NA |
| NA | | NA |
| NA | | NA |
| NA | | NA |
| NA | | NA |
| NA | | NA |
| NA | | NA |
| NA | | NA |
| NA | | NA |
| NA | | NA |
| NA | | NA |
| NA | | NA |
| NA | | NA |
| NA | | NA |
| NA | | NA |
| NA | | NA |
| NA | | NA |
| NA | | NA |
| | 0.955792398 | NA |
| NA | | NA |
| NA | | NA |
| NA | | NA |
| NA | | NA |
| NA | | NA |
| NA | | NA |
| NA | | NA |
| NA | | NA |
| NA | | NA |
| NA | | NA |
| NA | | NA |
| NA | | NA |
| NA | | NA |
| NA | | NA |
| NA | | NA |
| NA | | NA |
| NA | | NA |
| NA | | NA |
| NA | | NA |
| NA | | NA |
| NA | | NA |
| NA | | NA |
| NA | | NA |
| NA | | NA |
| NA | | NA |
| NA | | NA |
| NA | | NA |
| NA | | NA |

| | |
|---|---|
| NA | NA |
| NA | NA |
| NA | NA |
| NA | NA |
| NA | NA |
| NA | NA |
| NA | NA |
| NA | NA |
| NA | NA |
| NA | NA |
| NA | NA |
| NA | NA |
| NA | NA |
| NA | NA |
| NA | NA |
| NA | NA |
| NA | NA |
| NA | NA |
| NA | NA |
| NA | NA |
| NA | NA |
| NA | NA |
| NA | NA |
| NA | NA |
| NA | NA |
| NA | NA |
| NA | NA |
| NA | NA |
| NA | NA |
| NA | NA |
| NA | NA |
| NA | NA |
| NA | NA |
| NA | NA |
| NA | NA |
| NA | NA |
| NA | NA |
| NA | NA |
| NA | NA |
| NA | NA |
| NA | NA |
| NA | NA |
| NA | NA |
| NA | NA |
| NA | NA |
| NA | NA |
| NA | NA |

| | |
|---|---|
| NA | NA |
| NA | NA |
| NA | NA |
| NA | NA |
| NA | NA |
| NA | NA |
| NA | NA |
| NA | NA |
| NA | NA |
| NA | NA |
| NA | NA |
| NA | NA |
| NA | NA |
| NA | NA |
| NA | NA |
| NA | NA |
| NA | NA |
| NA | NA |
| NA | NA |
| NA | NA |
| NA | NA |
| NA | NA |
| NA | NA |
| NA | NA |
| NA | NA |
| NA | NA |
| NA | NA |
| NA | NA |
| NA | NA |
| NA | NA |
| NA | NA |
| NA | NA |
| NA | NA |
| NA | NA |
| NA | NA |
| NA | NA |
| NA | NA |
| NA | NA |
| NA | NA |
| NA | NA |
| NA | NA |
| NA | NA |
| NA | NA |
| NA | NA |
| NA | NA |
| NA | NA |
| NA | NA |

| | |
|---|---|
| NA | NA |
| NA | NA |
| NA | NA |
| NA | NA |
| NA | NA |
| NA | NA |
| NA | NA |
| NA | NA |
| NA | NA |
| NA | NA |
| NA | NA |
| NA | NA |
| NA | NA |
| NA | NA |
| NA | NA |
| NA | NA |
| NA | NA |
| NA | NA |
| NA | NA |
| NA | NA |
| NA | NA |
| NA | NA |
| NA | NA |
| NA | NA |
| NA | NA |
| NA | NA |
| NA | NA |
| NA | NA |
| NA | NA |
| NA | NA |
| NA | NA |
| NA | NA |
| NA | NA |
| NA | NA |
| NA | NA |
| NA | NA |
| NA | NA |
| NA | NA |
| NA | NA |
| NA | NA |
| NA | NA |
| NA | NA |
| NA | NA |
| NA | NA |
| NA | NA |
| NA | NA |
| NA | NA |

| | |
|---|---|
| NA | NA |
| NA | NA |
| NA | NA |
| NA | NA |
| NA | NA |
| NA | NA |
| NA | NA |
| NA | NA |
| NA | NA |
| NA | NA |
| NA | NA |
| NA | NA |
| NA | NA |
| NA | NA |
| NA | NA |
| NA | NA |
| NA | NA |
| NA | NA |
| NA | NA |
| NA | NA |
| NA | NA |
| NA | NA |
| NA | NA |
| NA | NA |
| NA | NA |
| NA | NA |
| NA | NA |
| NA | NA |
| NA | NA |
| NA | NA |
| NA | NA |
| NA | NA |
| NA | NA |
| NA | NA |
| NA | NA |
| NA | NA |
| NA | NA |
| NA | NA |
| NA | NA |
| NA | NA |
| NA | NA |
| NA | NA |
| NA | NA |
| NA | NA |
| NA | NA |
| NA | NA |
| NA | NA |

| | |
|---|---|
| NA | NA |
| NA | NA |
| NA | NA |
| NA | NA |
| NA | NA |
| NA | NA |
| NA | NA |
| NA | NA |
| NA | NA |
| NA | NA |
| NA | NA |
| NA | NA |
| NA | NA |
| NA | NA |
| NA | NA |
| NA | NA |
| NA | NA |
| NA | NA |
| NA | NA |
| NA | NA |
| NA | NA |
| NA | NA |
| NA | NA |
| NA | NA |
| NA | NA |
| NA | NA |
| NA | NA |
| NA | NA |
| NA | NA |
| NA | NA |
| NA | NA |
| NA | NA |
| NA | NA |
| NA | NA |
| NA | NA |
| NA | NA |
| NA | NA |
| NA | NA |
| NA | NA |
| NA | NA |
| NA | NA |
| NA | NA |
| NA | NA |
| NA | NA |
| NA | NA |
| NA | NA |
| NA | NA |

| | |
|---|---|
| NA | NA |
| NA | NA |
| NA | NA |
| NA | NA |
| NA | NA |
| NA | NA |
| NA | NA |
| NA | NA |
| NA | NA |
| NA | NA |
| NA | NA |
| NA | NA |
| NA | NA |
| NA | NA |
| NA | NA |
| NA | NA |
| NA | NA |
| NA | NA |
| NA | NA |
| NA | NA |
| NA | NA |
| NA | NA |
| NA | NA |
| NA | NA |
| NA | NA |
| NA | NA |
| NA | NA |
| NA | NA |
| NA | NA |
| NA | NA |
| NA | NA |
| NA | NA |
| NA | NA |
| NA | NA |
| NA | NA |
| NA | NA |
| NA | NA |
| NA | NA |
| NA | NA |
| NA | NA |
| NA | NA |
| NA | NA |
| NA | NA |
| NA | NA |
| NA | NA |
| NA | NA |
| NA | NA |

| | |
|---|---|
| NA | NA |
| NA | NA |
| NA | NA |
| NA | NA |
| NA | NA |
| NA | NA |
| NA | NA |
| NA | NA |
| NA | NA |
| NA | NA |
| NA | NA |
| NA | NA |
| NA | NA |
| NA | NA |
| NA | NA |
| NA | NA |
| NA | NA |
| NA | NA |
| NA | NA |
| NA | NA |
| NA | NA |
| NA | NA |
| NA | NA |
| NA | NA |
| NA | NA |
| NA | NA |
| NA | NA |
| NA | NA |
| NA | NA |
| NA | NA |
| NA | NA |
| NA | NA |
| NA | NA |
| NA | NA |
| NA | NA |
| NA | NA |
| NA | NA |
| NA | NA |
| NA | NA |
| NA | NA |
| NA | NA |
| NA | NA |
| NA | NA |
| NA | NA |
| NA | NA |
| NA | NA |
| NA | NA |

| | |
|---|---|
| NA | NA |
| NA | NA |
| NA | NA |
| NA | NA |
| NA | NA |
| NA | NA |
| NA | NA |
| NA | NA |
| NA | NA |
| NA | NA |
| NA | NA |
| NA | NA |
| NA | NA |
| NA | NA |
| NA | NA |
| NA | NA |
| NA | NA |
| NA | NA |
| NA | NA |
| NA | NA |
| NA | NA |
| NA | NA |
| NA | NA |
| NA | NA |
| NA | NA |
| NA | NA |
| NA | NA |
| NA | NA |
| NA | NA |
| NA | NA |
| NA | NA |
| NA | NA |
| NA | NA |
| NA | NA |
| NA | NA |
| NA | NA |
| NA | NA |
| NA | NA |
| NA | NA |
| NA | NA |
| NA | NA |
| NA | NA |
| NA | NA |
| NA | NA |
| NA | NA |
| NA | NA |
| NA | NA |

| | | |
|---|---|---|
| NA | | NA |
| NA | | NA |
| NA | | NA |
| NA | | NA |
| NA | | NA |
| NA | | NA |
| NA | | NA |
| NA | | NA |
| NA | | NA |
| NA | | NA |
| NA | | NA |
| NA | | NA |
| NA | | NA |
| NA | | NA |
| NA | | NA |
| | 0.955792398 | NA |
| NA | | NA |
| NA | | NA |
| NA | | NA |
| NA | | NA |
| NA | | NA |
| NA | | NA |
| NA | | NA |
| NA | | NA |
| NA | | NA |
| NA | | NA |
| NA | | NA |
| NA | | NA |
| NA | | NA |
| NA | | NA |
| NA | | NA |
| NA | | NA |
| NA | | NA |
| NA | | NA |
| NA | | NA |
| NA | | NA |
| NA | | NA |
| NA | | NA |
| NA | | NA |
| NA | | NA |
| NA | | NA |
| NA | | NA |
| NA | | NA |
| NA | | NA |
| NA | | NA |
| NA | | NA |
| NA | | NA |

| | |
|---|---|
| NA | NA |
| NA | NA |
| NA | NA |
| NA | NA |
| NA | NA |
| NA | NA |
| NA | NA |
| NA | NA |
| NA | NA |
| NA | NA |
| NA | NA |
| NA | NA |
| NA | NA |
| NA | NA |
| NA | NA |
| NA | NA |
| NA | NA |
| NA | NA |
| NA | NA |
| NA | NA |
| NA | NA |
| NA | NA |
| NA | NA |
| NA | NA |
| NA | NA |
| NA | NA |
| NA | NA |
| NA | NA |
| NA | NA |
| NA | NA |
| NA | NA |
| NA | NA |
| NA | NA |
| NA | NA |
| NA | NA |
| NA | NA |
| NA | NA |
| NA | NA |
| NA | NA |
| NA | NA |
| NA | NA |
| NA | NA |
| NA | NA |
| NA | NA |
| NA | NA |
| NA | NA |
| NA | NA |

| | |
|---|---|
| NA | NA |
| NA | NA |
| NA | NA |
| NA | NA |
| NA | NA |
| NA | NA |
| NA | NA |
| NA | NA |
| NA | NA |
| NA | NA |
| NA | NA |
| NA | NA |
| NA | NA |
| NA | NA |
| NA | NA |
| NA | NA |
| NA | NA |
| NA | NA |
| NA | NA |
| NA | NA |
| NA | NA |
| NA | NA |
| NA | NA |
| NA | NA |
| NA | NA |
| NA | NA |
| NA | NA |
| NA | NA |
| NA | NA |
| NA | NA |
| NA | NA |
| NA | NA |
| NA | NA |
| NA | NA |
| NA | NA |
| NA | NA |
| NA | NA |
| NA | NA |
| NA | NA |
| NA | NA |
| NA | NA |
| NA | NA |
| NA | NA |
| NA | NA |
| NA | NA |
| NA | NA |
| NA | NA |

| | |
|---|---|
| NA | NA |
| NA | NA |
| NA | NA |
| NA | NA |
| NA | NA |
| NA | NA |
| NA | NA |
| NA | NA |
| NA | NA |
| NA | NA |
| NA | NA |
| NA | NA |
| NA | NA |
| NA | NA |
| NA | NA |
| NA | NA |
| NA | NA |
| NA | NA |
| NA | NA |
| NA | NA |
| NA | NA |
| NA | NA |
| NA | NA |
| NA | NA |
| NA | NA |
| NA | NA |
| NA | NA |
| NA | NA |
| NA | NA |
| NA | NA |
| NA | NA |
| NA | NA |
| NA | NA |
| NA | NA |
| NA | NA |
| NA | NA |
| NA | NA |
| NA | NA |
| NA | NA |
| NA | NA |
| NA | NA |
| NA | NA |
| NA | NA |
| NA | NA |
| NA | NA |
| NA | NA |
| NA | NA |

| | |
|---|---|
| NA | NA |
| NA | NA |
| NA | NA |
| NA | NA |
| NA | NA |
| NA | NA |
| NA | NA |
| NA | NA |
| NA | NA |
| NA | NA |
| NA | NA |
| NA | NA |
| NA | NA |
| NA | NA |
| NA | NA |
| NA | NA |
| NA | NA |
| NA | NA |
| NA | NA |
| NA | NA |
| NA | NA |
| NA | NA |
| NA | NA |
| NA | NA |
| NA | NA |
| NA | NA |
| NA | NA |
| NA | NA |
| NA | NA |
| NA | NA |
| NA | NA |
| NA | NA |
| NA | NA |
| NA | NA |
| NA | NA |
| NA | NA |
| NA | NA |
| NA | NA |
| NA | NA |
| NA | NA |
| NA | NA |
| NA | NA |
| NA | NA |
| NA | NA |
| NA | NA |
| NA | NA |
| NA | NA |

| | |
|---|---|
| NA | NA |
| NA | NA |
| NA | NA |
| NA | NA |
| NA | NA |
| NA | NA |
| NA | NA |
| NA | NA |
| NA | NA |
| NA | NA |
| NA | NA |
| NA | NA |
| NA | NA |
| NA | NA |
| NA | NA |
| NA | NA |
| NA | NA |
| NA | NA |
| NA | NA |
| NA | NA |
| NA | NA |
| NA | NA |
| NA | NA |
| NA | NA |
| NA | NA |
| NA | NA |
| NA | NA |
| NA | NA |
| NA | NA |
| NA | NA |
| NA | NA |
| NA | NA |
| NA | NA |
| NA | NA |
| NA | NA |
| NA | NA |
| NA | NA |
| NA | NA |
| NA | NA |
| NA | NA |
| NA | NA |
| NA | NA |
| NA | NA |
| NA | NA |
| NA | NA |
| NA | NA |
| NA | NA |

| | |
|---|---|
| NA | NA |
| NA | NA |
| NA | NA |
| NA | NA |
| NA | NA |
| NA | NA |
| NA | NA |
| NA | NA |
| NA | NA |
| NA | NA |
| NA | NA |
| NA | NA |
| NA | NA |
| NA | NA |
| NA | NA |
| NA | NA |
| NA | NA |
| NA | NA |
| NA | NA |
| NA | NA |
| NA | NA |
| NA | NA |
| NA | NA |
| NA | NA |
| NA | NA |
| NA | NA |
| NA | NA |
| NA | NA |
| NA | NA |
| NA | NA |
| NA | NA |
| NA | NA |
| NA | NA |
| NA | NA |
| NA | NA |
| NA | NA |
| NA | NA |
| NA | NA |
| NA | NA |
| NA | NA |
| NA | NA |
| NA | NA |
| NA | NA |
| NA | NA |
| NA | NA |
| NA | NA |
| NA | NA |

| | | |
|---|---|---|
| NA | | NA |
| NA | | NA |
| NA | | NA |
| NA | | NA |
| NA | | NA |
| NA | | NA |
| NA | | NA |
| NA | | NA |
| NA | | NA |
| NA | | NA |
| NA | | NA |
| NA | | NA |
| NA | | NA |
| NA | | NA |
| NA | | NA |
| NA | | NA |
| NA | | NA |
| NA | | NA |
| NA | | NA |
| NA | | NA |
| NA | | NA |
| NA | | NA |
| NA | | NA |
| NA | | NA |
| NA | | NA |
| NA | | NA |
| NA | | NA |
| NA | | NA |
| NA | | NA |
| NA | | NA |
| NA | | NA |
| NA | | NA |
| NA | | NA |
| NA | | NA |
| NA | | NA |
| NA | | NA |
| NA | | NA |
| NA | | NA |
| | 0.941278332 | NA |
| NA | | NA |
| NA | | NA |
| NA | | NA |
| NA | | NA |
| NA | | NA |
| NA | | NA |
| NA | | NA |
| NA | | NA |

| | |
|---|---|
| NA | NA |
| NA | NA |
| NA | NA |
| NA | NA |
| NA | NA |
| NA | NA |
| NA | NA |
| NA | NA |
| NA | NA |
| NA | NA |
| NA | NA |
| NA | NA |
| NA | NA |
| NA | NA |
| NA | NA |
| NA | NA |
| NA | NA |
| NA | NA |
| NA | NA |
| NA | NA |
| NA | NA |
| NA | NA |
| NA | NA |
| NA | NA |
| NA | NA |
| NA | NA |
| 0.057519654 | 0.138375055 |
| 0.195772134 | NA |
| 0.207114579 | 0.356143709 |
| NA | NA |
| NA | NA |
